# Extended amygdala-parabrachial circuits alter threat assessment to regulate feeding

**DOI:** 10.1101/2020.03.03.975193

**Authors:** Dionnet L. Bhatti, Andrew T. Luskin, Christian E. Pedersen, Bernard Mulvey, Hannah Oden-Brunson, Kate Kimbell, Abbie Sawyer, Robert W. Gereau, Joseph D. Dougherty, Michael R. Bruchas

**Affiliations:** Department of Anesthesiology, Washington University School of Medicine, St. Louis, MO 63108, USA; Washington University Pain Center, Washington University School of Medicine, St. Louis, MO 63108, USA; Department of Biomedical Engineering, Washington University School of Medicine, St. Louis, MO, 63108 USA; Division of Biology and Biomedical Sciences, Washington University School of Medicine, St. Louis, MO, USA; Department of Genetics, Washington University School of Medicine, St. Louis, MO, USA; Department of Anesthesiology and Pain Medicine, University of Washington, Seattle, WA, USA; Center for Neurobiology of Addiction, Pain, and Emotion, University of Washington, Seattle, WA, USA; Graduate Program in Neuroscience, University of Washington, Seattle, WA, 98195; Department of Bioengineering, University of Washington, Seattle, WA, 98195; Department of Electrical Engineering, University of Washington, Seattle, WA, 98195; Program in Neuroscience, Harvard Medical School, Boston, MA 02115

**Author notes:** Corresponding author. (M.R.B.). these authors contributed equally (Dionnet L. Bhatti, Andrew T. Luskin).

**Keywords:** extended amygdala, bed nucleus of the stria terminalis, parabrachial nucleus, feeding, threat assessment, anxiety, motivation

## Abstract

An animal’s evolutionary success depends on the ability to seek and consume foods while avoiding environmental threats. However, how evolutionarily conserved threat detection circuits modulate feeding is unknown. In mammals, feeding and threat assessment are strongly influenced by the parabrachial nucleus (PBN), a structure that responds to threats and inhibits feeding. Here, we report that the PBN receives dense inputs from the bed nucleus of the stria terminalis (BNST), an extended amygdala structure that encodes affective information. Using a series of complementary approaches, we identify opposing BNST-PBN circuits that modulate a genetically-defined population of PBN neurons to control feeding. This previously unrecognized neural circuit integrates threat assessment with the intrinsic drive to eat.

## Introduction

All animals must successfully seek and consume food while avoiding environmental threats to survive. The internal state of an animal directly impacts the expression of risky behaviors, such as exploring a dangerous environment to obtain rewards and maintain homeostasis. Animals must adaptively prioritize certain behaviors to appropriately respond to their internal state (Alhadeff et al., 2018; Burnett et al., 2016). While many studies have explored the interaction of metabolic need states with behavior, how mammals integrate affective-threat assessment with internal need states remains largely unknown.

Several recent reports have found that in mammals, food consumption and threat assessment are heavily influenced by the parabrachial nucleus (PBN), a pontine structure that integrates visceral and sensory information to encode metabolic needs (Campos et al., 2016, 2018; Carter et al., 2013; Essner et al., 2017; Han et al., 2015; Mu et al., 2017; Roman et al., 2016; Ryan et al., 2017; Zséli et al., 2016). The amygdala and extended amygdala are evolutionarily conserved brain regions that encode and integrate valence, stress, and threat to alter behavioral states. Anatomical data suggest that the PBN receives input from several regions that may encode affective information, including the bed nucleus of the stria terminalis (BNST), a structure in the extended amygdala (Douglass et al., 2017; Kim et al., 2013; Mazzone et al., 2018; Zséli et al., 2016). However, the neural circuit mechanisms that underlie the integration of an animal’s own motivation to eat with internal affective states regarding environmental threats are still unknown. Here, we identified two previously unknown afferents from distinct populations within the BNST to the PBN that underlie the complex integration of threat-assessment and feeding signals to modulate PBN activity and ultimately regulate state-dependent feeding.

## Results

### Anatomical and Molecular Characterization of Opposing BNST-PBN Circuits

The BNST is a heterogeneous population comprised of glutamatergic, GABAergic, and peptidergic neurons (Daniel and Rainnie, 2016; Giardino et al., 2018; Gungor and Paré, 2016; Gungor et al., 2018; Jennings et al., 2013a, 2013b). To determine whether distinct neuronal circuits from the BNST innervate the PBN to alter feeding behaviors, we injected Cre-inducible anterograde AAVs expressing channelrhodopsin-2 with an eYFP reporter (AAV5-DIO-ChR2-eYFP) into the BNST of vesicular GABA transporter (vGAT)-Cre and vesicular glutamate transporter (vGLUT2)-Cre mice, which revealed robust axonal projections to the PBN from GABAergic (**Figures 1A and S1F-S1K**) and glutamatergic (**Figure 1D**) BNST neurons. To further substantiate our anterograde tracing findings, we used Cre-inducible retrograde adeno-associated viruses (retro-AAV2) expressing an enhanced yellow fluorescent protein (eYFP) reporter (AAV2retro-DIO-eYFP) (Tervo et al., 2016) into the PBN of vGAT-Cre or vGLUT2-Cre mice (**Figure S1A**). Retrograde tracing revealed populations of both GABAergic and glutamatergic BNST neurons that innervate the PBN (**Figure S1A**).

**Figure 1.**
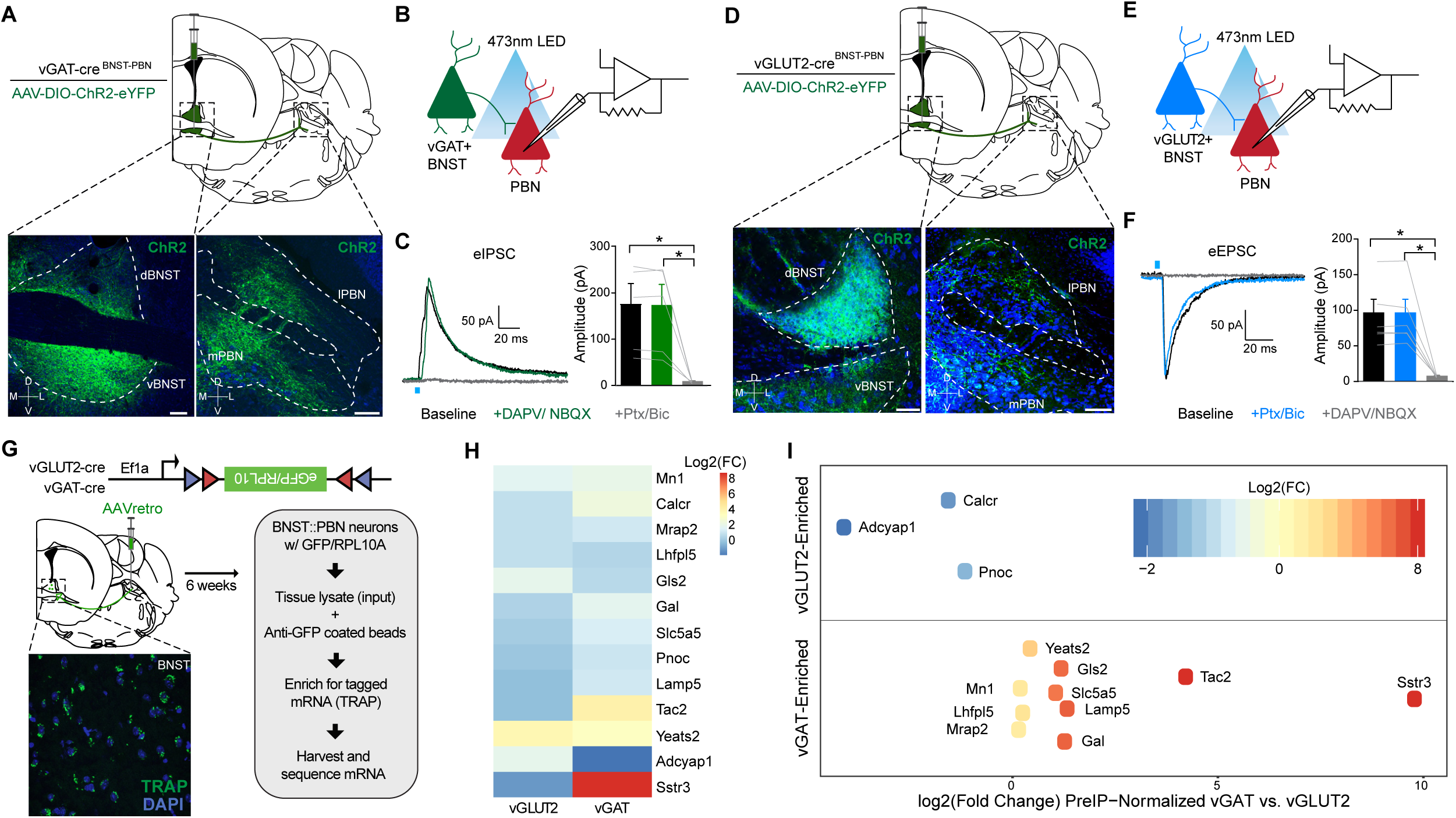
Anatomical and molecular characterization of opposing BNST-PBN circuits. **(A)** Schematic of viral injection and representative image depicting expression in BNST^vGAT^ soma and their terminals in the PBN (scale bar: 100 μm). Blue = Nissl. Green = eYFP. **(B)** Schematic of whole-cell patch clamp electrophysiology recordings of optically-evoked IPSCs. **(C)** Optogenetic activation of BNST^vGAT^ terminals elicit IPSCs in PBN neurons that are abolished by GABAA receptor antagonism (n=5 cells, 4 mice). **(D)** Schematic of viral injection and representative image depicting expression in BNST^vGLUT2^ soma and their terminals in the PBN (scale bar: 100 μm). Blue = Nissl. Green = eYFP. **(E)** Schematic of whole-cell patch clamp electrophysiology recordings of optically-evoked EPSCs. **(F)** Optogenetic activation of BNST^vGLUT2^ terminals elicit EPSCs in PBN neurons that are abolished by AMPA/NMDA receptor antagonism (n=6 cells, 5 mice). **(G)** Cartoon of injection of TRAP into PBN of vGLUT2-Cre or vGAT-Cre animals. Tagged mRNA was extracted from the BNST and sequenced. Inset: representative image of TRAP-GFP expression in BNST. **(H)** Heatmap of transcripts enriched in either vGAT or vGLUT2 projections from BNST to PBN over input homogenate (pre-IP) (n=3 vGLUT2-Cre samples; n=2 vGAT-Cre samples). **(I)** Transcripts enriched in either vGAT or vGLUT2 projections from BNST to PBN, after normalization to respective input (pre-IP) homogenates. Positive Log2(FC) values indicate transcript enrichment in BNST-PBN^vGAT^ neurons relative to BNST-PBN^vGLUT2^ neurons; negative Log2(FC) values indicate relative enrichment in BNST-PBN^vGLUT2^ neurons. \**p* < 0.05. Error bars indicate SEM. See also Figure S1 and S2.

We next assessed whether these distinct populations make functional monosynaptic connections to PBN neurons using *ex vivo* patch-clamp electrophysiology. We collected acute brain slices containing the PBN from either vGAT-Cre or vGLUT2-Cre mice expressing DIO-ChR2-eYFP in BNST-PBN projections. We optogenetically evoked post-synaptic currents in PBN neurons receiving BNST GABAergic (**Figure 1B**) or glutamatergic (**Figure 1E**) innervation during whole-cell patch clamp recording. Optogenetic activation of GABAergic BNST terminals in the PBN evoked IPSCs that were pharmacologically blocked using GABAA antagonists (**Figure 1C**), while optogenetic activation of glutamatergic BNST terminals in the PBN evoked EPSCs that were pharmacologically blocked using AMPAR/NMDAR antagonists (**Figure 1F**). Postsynaptic currents occurred <5ms after the light pulse, suggesting that both excitatory and inhibitory connections are monosynaptic (**Figure S1B-E**).

We further characterized the molecular expression profiles of these distinct inhibitory and excitatory BNST-PBN projections by using translating ribosome affinity purification (TRAP) to determine their translational signature (Doyle et al., 2008). We generated a new Cre-dependent TRAP construct within the retro-AAV2 vector (AAV2retro-DIO-TRAP) and injected it into the PBN of vGAT-Cre and vGLUT2-Cre mice to isolate RNA transcripts from each BNST-PBN projection population with genotype and projection specificity (Parker et al., 2019; Tervo et al., 2016). We then isolated and sequenced ribosome-bound mRNA from the BNST in each group (**Figures 1G, S2A, and S2B**). The vGAT and vGLUT2 projections are enriched in several genes of interest (**Figures 1H, 1I, and S2C-S2E**). BNST-PBN^vGAT^ neurons are enriched in the neuropeptide mRNAs *tachykinin 2* (*Tac2*) (Zelikowsky et al., 2018) and *galanin* (*Gal*), as well as a somatostatin receptor (*Sstr3*) and *melanocortin-2 receptor accessory protein 2* (*Mrap2*), which play important roles in limiting anxiety, responses and regulating feeding behavior (Ahrens et al., 2018; Asai et al., 2013; Bruschetta et al., 2018; Elsilä et al., 2018). BNST-PBN^vGLUT2^ neurons are enriched in *Adcyap1*, which encodes the neuropeptide PACAP, previously identified as a major regulator of stress responses in the BNST (Hammack et al., 2009; Roman et al., 2014), including in contexts of addiction and post-traumatic stress disorder (Miles et al., 2018; Ressler et al., 2011). Both BNST-PBN^vGAT^ and BNST-PBN^vGLUT2^ neurons express calcitonin receptor *(Calcr*), recently associated with the regulation of feeding (Cheng et al., 2020; Pan et al., 2018), and *nociceptin (Pnoc*), linked to motivated behaviors including feeding (Hardaway et al., 2019; Parker et al., 2019; Toll et al., 2016). We further validated these RNAseq findings by performing fluorescent in-situ hybridization experiments, which revelated co-expression of *Tac2, Sstr3*, and *Calcr* with vGAT and *Adcyap1* with vGLUT2 neurons within the BNST (**Figures S2F-S2I**).

### Feeding and Threat Differentially Engage Excitatory and Inhibitory BNST-PBN Circuits

To determine whether these distinct genetically-defined BNST-PBN circuits modulate feeding behavior, we used fiber photometry to monitor calcium-mediated fluorescence, a proxy for neuronal activity, of BNST-PBN terminals in freely-behaving mice (Gunaydin et al., 2014; Pologruto et al., 2004). To determine whether GABAergic BNST-PBN terminals are engaged during feeding behavior, we targeted AAVDJ-DIO-GCaMP6s to the BNST of vGAT-Cre mice and positioned optical fibers above the PBN for measurement of BNST-PBN GABAergic terminal calcium activity (**Figures 2A, 2B, and S3A, S3C, and S3E**). We found that BNST-PBN^vGAT^ GCaMP activity increased as an animal engaged in both sated and hedonic feeding (i.e. sucrose), as well as in other food seeking modalities including high-fat, food deprived, and normal chow within novel-anxiogenic environments (**Figures 2C-2K and S3G**). Conversely, to assess whether glutamatergic input to the PBN is altered during feeding behavior, we targeted AAVDJ-DIO-GCaMP6s to the BNST of vGLUT2-Cre mice and similarly positioned optical fibers above the PBN for measurement of BNST-PBN glutamatergic terminal calcium activity (**Figures 3A, 3B, and S3B, S3D, and S3F**). In these experiments, we found that BNST-PBN^vGLUT2^ GCaMP activity decreased when an animal engaged in these same feeding behaviors (**Figures 3C-3K**). Together, these data suggest a differential opposing role for GABAergic and glutamatergic BNST-PBN circuits in modulating feeding behavior.

**Figure 2.**
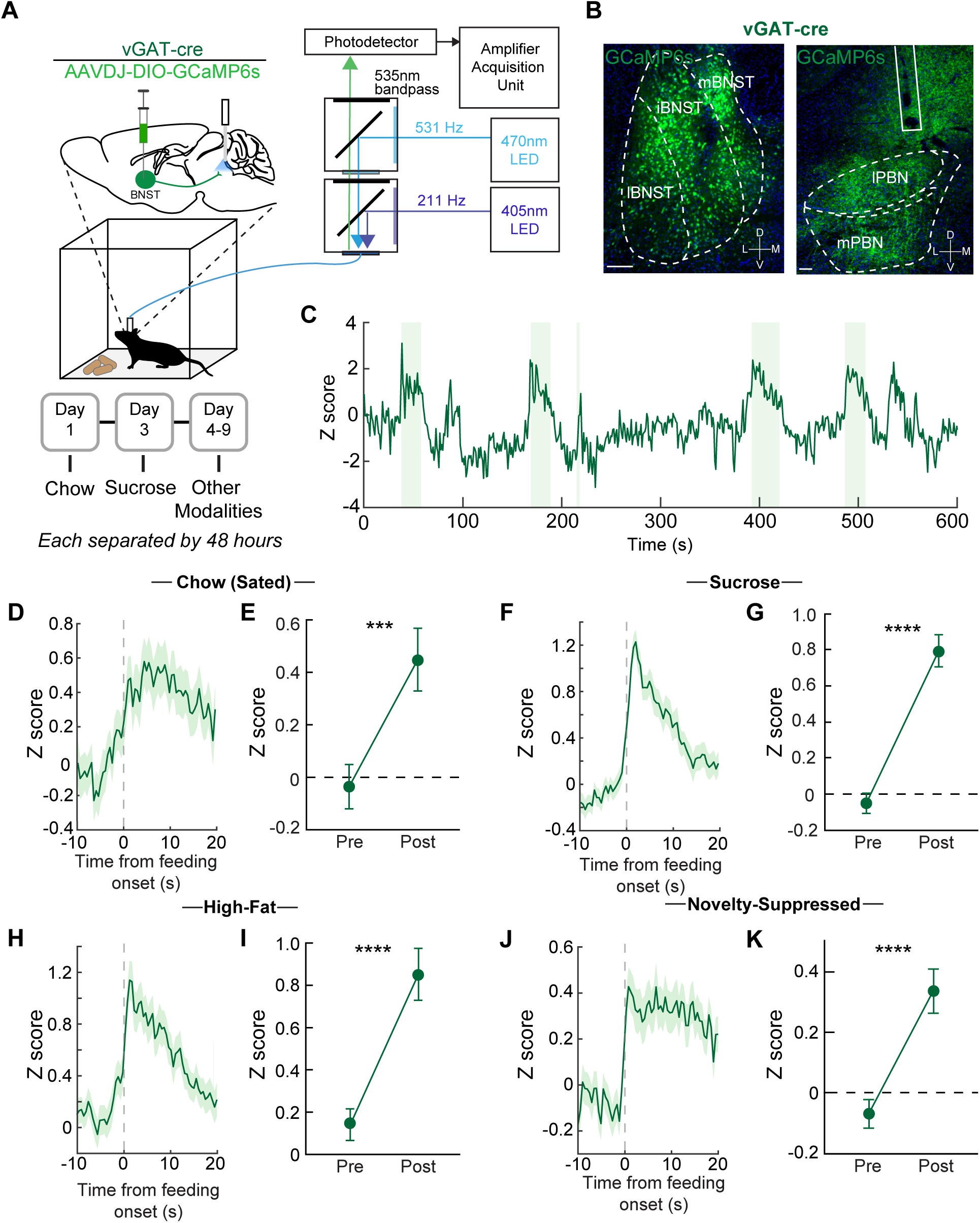
Feeding behavior engages an inhibitory BNST-PBN circuit. **(A)** Schematic of *in vivo* fiber photometry and behavior. **(B)** Representative GCaMP6s expression in the BNST and PBN of a vGAT-Cre mouse (scale bars: 100 μm BNST, 200 μm PBN). **(C)** Representative responses of BNST-PBN^vGAT^ terminals during food consumption trials (shaded areas represent periods of eating). **(D and E)** Average z-scored calcium response of BNST-PBN^vGAT^ terminals during consumption of normal chow under sated conditions (n=57 bouts; 7 mice) and averaged activity of 10-seconds pre- compared to post- consumption initiation over the testing period. **(F and G)** Average z-scored calcium response of BNST-PBN^vGAT^ terminals during consumption of sucrose (n=93 bouts; 7 mice) and averaged activity of 10-seconds pre- compared to post- consumption initiation over the testing period. **(H and I)** Average z-scored calcium response of BNST-PBN^vGAT^ terminals during consumption of high-fat (n=77 bouts; 7 mice) and averaged activity of 10-seconds pre- compared to post- consumption initiation over the testing period. **(J and K)** Average z-scored calcium response of BNST-PBN^vGAT^ terminals during consumption of normal chow under anxiogenic conditions (n=121 bouts; 7 mice) and averaged activity of 10-seconds pre- compared to post- consumption initiation over the testing period. \*\**p* < 0.01, \*\*\**p* < 0.001, \*\*\*\**p* < 0.0001. Error bars indicate SEM. See also Figure S3.

**Figure 3.**
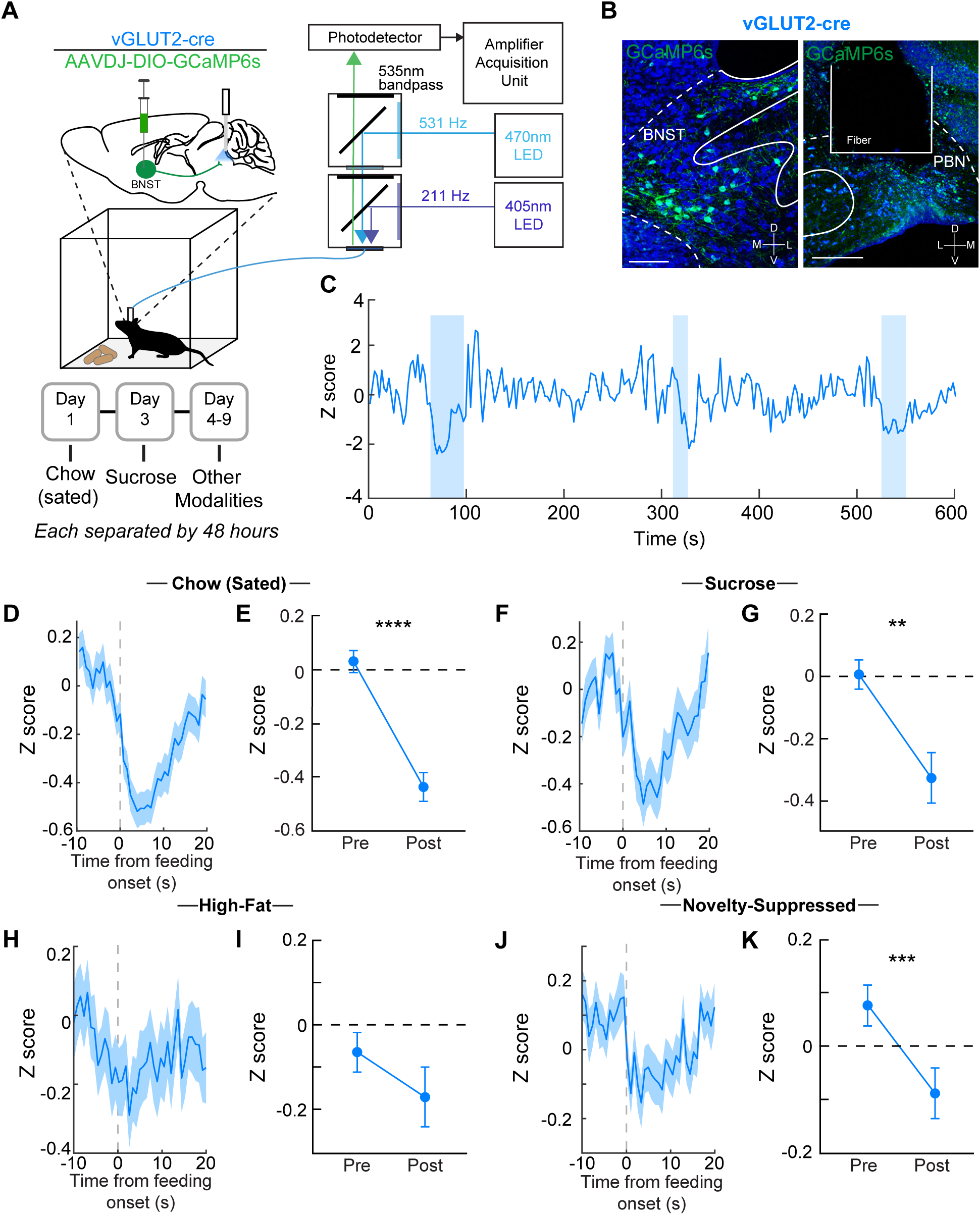
An excitatory BNST-PBN circuit is disengaged during feeding. **(A)** Schematic of *in vivo* fiber photometry and behavior. **(B)** Representative GCaMP6s expression in the BNST and PBN of a vGLUT2-Cre mouse (scale bars: 100 μm BNST, 200 μm PBN). **(C)** Representative responses of BNST-PBN^vGGLUT2^ terminals during food consumption trials (shaded areas represent periods of eating). **(D and E)** Average z-scored calcium response of BNST-PBN^vGLUT2^ terminals during consumption of normal chow under sated conditions (n=133 bouts; 6 mice) and averaged activity of 10-seconds pre- compared to post- consumption initiation over the testing period. **(F and G)** Average z-scored calcium response of BNST-PBN^vGLUT2^ terminals during consumption of sucrose (n=87 bouts; 6 mice) and averaged activity of 10-seconds pre- compared to post- consumption initiation over the testing period. **(H and I)** Average z-scored calcium response of BNST-PBN^vGLUT2^ terminals during consumption of high-fat (n=56 bouts; 6 mice) and averaged activity of 10-seconds pre- compared to post- consumption initiation over the testing period. **(J and K)** Average z-scored calcium response of BNST-PBN^vGLUT2^ terminals during consumption of normal chow under anxiogenic conditions (n=77 bouts; 6 mice) and averaged activity of 10-seconds pre- compared to post- consumption initiation over the testing period. \*\**p* < 0.01, \*\*\**p* < 0.001, \*\*\*\**p* < 0.0001. Error bars indicate SEM. See also Figure S3.

Food-seeking requires an alteration of threat assessment and anxiety-like behavior in order to adaptively seek out and consume food as necessary for survival. We therefore hypothesized that if these circuits indeed alter feeding behavior concurrently with threat assessment, GABAergic or glutamatergic BNST-PBN input may be differentially engaged during aversive threat stimuli, respectively. To assess this, we recorded from BNST-PBN terminals in vGAT-Cre and vGLUT-Cre mice during the presentation of multiple shock stimuli (**Figure 4A)**. When animals were presented with an aversive shock, BNST-PBN^vGAT^ calcium transient activity rapidly decreased in response (**Figures 4B-4D)**, while BNST-PBN^vGLUT2^ activity increased following shock (**Figures 4E-4F**). These observations indicate that inhibitory GABAergic BNST-PBN circuits may act to diminish threat signaling to engage and allow feeding, while excitatory glutamatergic BNST-PBN circuits could be recruited to enhance threat signaling and suppress feeding behaviors.

**Figure 4.**
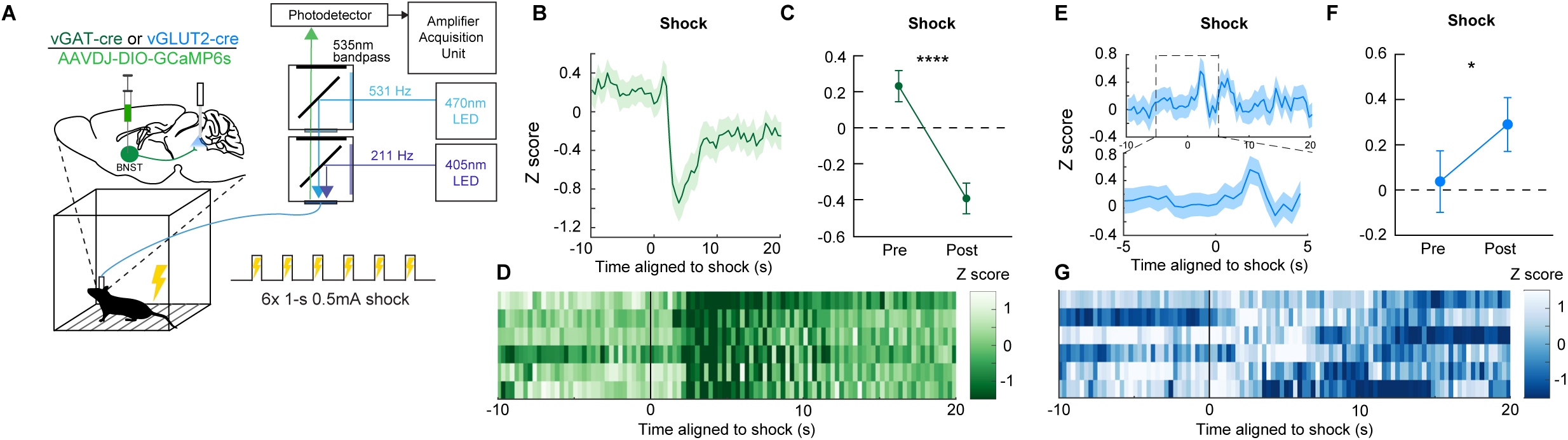
Distinct BNST-PBN circuits display opposing responses to aversive stimuli. **(A)** Schematic of *in vivo* fiber photometry and behavior. **(B)** Average z-scored calcium transient responses of BNST-PBN^vGAT^ terminals to an aversive shock (n=7 mice). **(C)** Mean z-scored calcium transient responses of BNST-PBN^vGAT^ terminals (n=7 mice) 10-seconds before and after the shock initiation. **(D)** Representative heatmap of BNST-PBN^vGAT^ terminal calcium transient activity of a single mouse during aversive shock presentation. **(E)** Average z-scored calcium transient responses of BNST-PBN^vGLUT2^ terminals to an aversive shock (n=6 mice). **(F)** Mean z-scored calcium transient responses of BNST-PBN^v GLUT2^ terminals (n=6 mice) 10-seconds before and after the shock initiation. **(G)** Representative heatmap of BNST-PBN^vGLUT2^ terminal calcium transient activity of a single mouse during aversive shock presentation. \*\**p* < 0.01, \*\*\**p* < 0.001, \*\*\*\**p* < 0.0001. Error bars indicate SEM. See also Figure S3.

### Excitatory and Inhibitory BNST-PBN Circuit Activation Drive Opposing Feeding Behavior

Since photometry measurements at the terminals revealed that these two opposing BNST-PBN projections are modulated during feeding and threat responses, we next utilized optogenetic approaches to assess whether manipulating neural circuit activity of BNST-PBN circuits alters feeding, threat, and affective valence, and to determine direct causality of this circuit in regulating behavior. We targeted AAV5-EF1α-DIO-ChR2 to the BNST of vGAT-Cre or vGLUT2-Cre mice and positioned optical fibers above the PBN for photo-stimulation of BNST-PBN GABAergic or glutamatergic terminals (**Figure 5A**). First, we measured food consumption with or without genetically defined neural circuit photo-activation while mice had free access to different foods (**Figure 5B**). Activation of ChR2 in BNST-PBN^vGAT^ neurons increased food consumption compared to control mice under sated conditions (**Figure 5C and S4B**). This increase also occurred with sucrose and high-fat foods (**Figures S4D-S4F**). Furthermore, photoactivation of BNST-PBN^vGAT^ signaling also increased consumption of less-palatable salt-enriched or bitter (i.e. quinine-enriched) foods (**Figure S4G-S4I**). Notably, this BNST-PBN^vGAT^-mediated increase in feeding was present when the animal was sated but was not evident after food deprivation (**Figure 5C, Figure S4C**). Together, these results suggest that the inhibitory BNST-PBN circuit is sufficient to drive feeding in sated states, regardless of food type, and likely overrides the valence of the food. In contrast, we saw a ceiling effect in the food-deprived state, suggesting that inhibitory BNST-PBN circuits are already engaged, consistent with our fiber photometry data (**Figure S3G)**. Additionally, optogenetic inhibition of the BNST-PBN^vGAT^ circuit using Arch3.0 revealed its necessity for hunger-driven feeding (**Figures S4P-S4R**). In contrast to activation of BNST-PBN^vGAT^, activation of BNST-PBN^vGLUT2^ circuits in a food-deprived state decreased consumption of normal chow (**Figures 5D and S5A-S5E)**, demonstrating that the excitatory BNST-PBN circuit is sufficient to reduce feeding when mice in a food-deprived state, suggesting binary and opposing BNST-PBN circuits.

**Figure 5.**
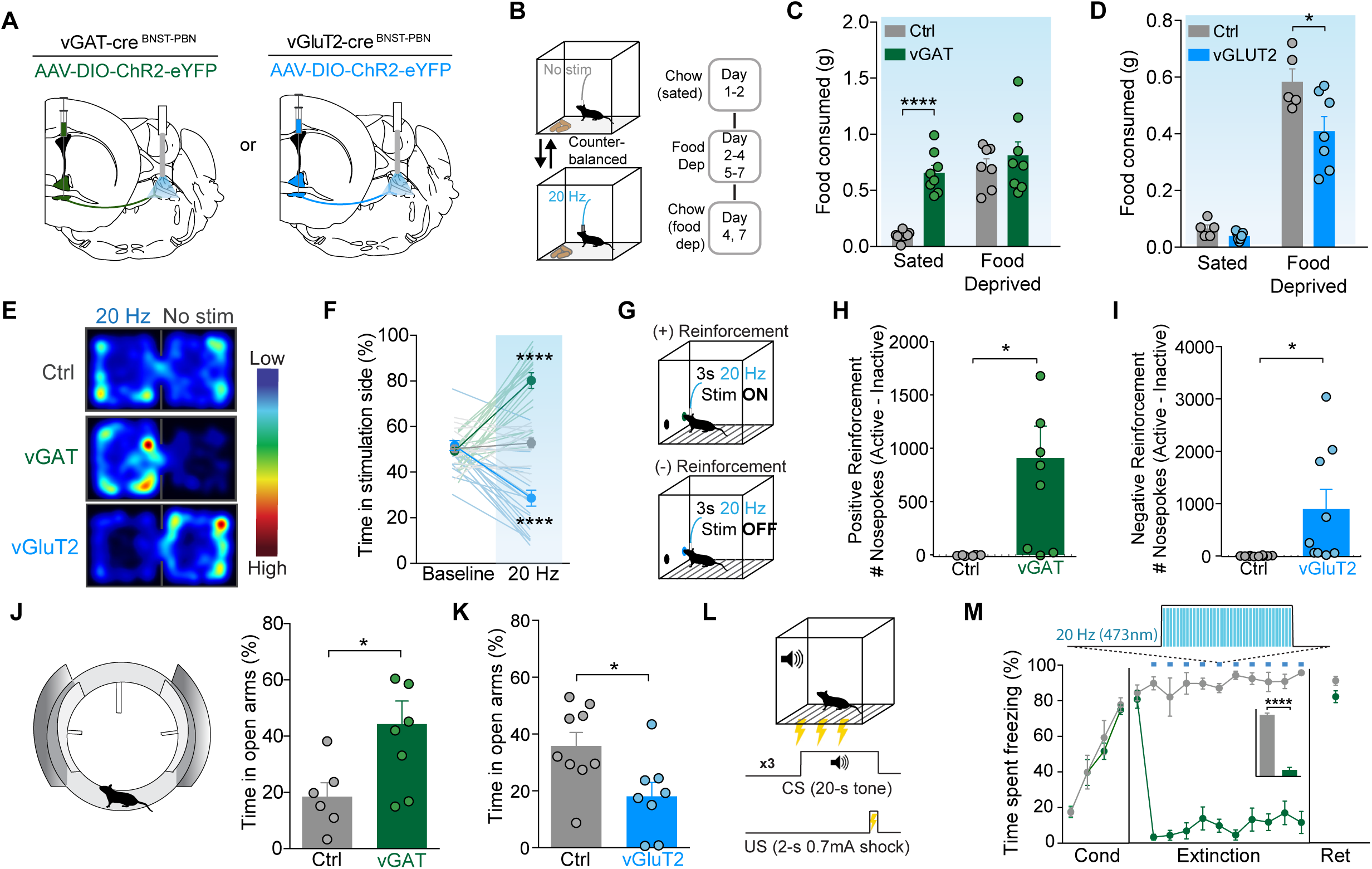
Distinct BNST-PBN circuits drive opposing feeding and affective behaviors. **(A)** Schematic of optogenetic approach to target BNST-PBN^vGAT^ and BNST-PBN^vGLUT2^. **(B)** Schematic of food consumption assay. **(C)** BNST-PBN^vGAT^ activation increases consumption of normal chow under sated conditions (ChR2, *n* = 8; Ctrl, *n* = 7). **(D)** BNST-PBN^vGLUT2^ activation decreases consumption of normal chow after food-deprivation (ChR2, *n* = 7; Ctrl, *n* = 5). **(E)** Representative heatmaps of time spent in the RTPP for Ctrl, BNST-PBN^vGAT^:ChR2, and BNST-PBN^vGLUT2^:ChR2 mice. **(F)** BNST-PBN^vGAT^ activation elicits a real-time place preference, while BNST-PBN^vGLUT2^ activation elicits a real-time place aversion compared to control mice (vGAT:ChR2, *n* = 18; vGLUT2:ChR2, n = 18; Ctrl, *n* = 25). **(G)** Schematic of positive (+) and negative (-) reinforcement tasks for assessment in BNST-PBN^vGAT^:ChR2 and BNST-PBN^vGLUT2^:ChR2, respectively. **(H)** BNST-PBN^vGAT^ activation is positively reinforcing in an operant-self stimulation task (ChR2, *n* = 8; Ctrl, *n* = 6). **(I)** BNST-PBN^vGLUT2^ activation is negatively reinforcing in an operant task to turn off optogentic stimulation (ChR2, *n* = 8; Ctrl, *n* = 6). **(J)** BNST-PBN^vGAT^ activation increases time spent in the open arms in the elevated zero maze (ChR2, *n* = 8; Ctrl, *n* = 6). **(K)** BNST-PBN^vGLUT2^ activation decreases time spent in the open arms in the elevated zero maze (ChR2, *n* = 9; Ctrl, *n* = 8). **(L)** Schematic of the fear conditioning protocol. **(M)** BNST-PBN^vGAT^ activation suppresses cued-defensive responses (i.e. freezing) after conditioning (ChR2, *n* =8; Ctrl, *n* = 6). \**p* < 0.05, \*\**p* < 0.01, \*\*\**p* < 0.001, \*\*\*\**p* < 0.0001. Error bars indicate SEM. See also Figure S4 and S5.

### Excitatory and Inhibitory BNST-PBN Circuit Activation Drive Opposing Affective States

Internal state (affective valence) alters an animal’s ability to assess potential threats and explore different environments, a requirement to locate and consume foods. Therefore, we sought to assess the affective valence of these distinct BNST-PBN circuits by subjecting mice to real-time-place-preference (RTPP) tests and operant-reinforcement assays (Al-Hasani et al., 2015; McCall et al., 2015; Seo et al., 2016). We injected AAV5-EF1α-DIO-ChR2 into the BNST of vGAT-Cre or vGLUT2-Cre mice, placing an optic fiber above the PBN, and tested whether stimulation of this BNST-PBN^vGAT^ pathway was reinforcing or aversive. In the RTPP assay, photoactivation of ChR2 in BNST-PBN^vGAT^ neurons resulted in a robust place preference for the photostimulation-paired side, while photoactivation of ChR2 in BNST-PBN^vGLUT2^ neurons resulted in a robust place aversion for the photostimulation-paired side, as compared to controls (**Figures 5E and 5F**). In an operant self-stimulation paradigm, photoactivation of BNST-PBN^vGAT^ neurons significantly increased the number of nosepokes for self-stimulation relative to controls (**Figure 5H**), demonstrating that BNST-PBN^vGAT^ stimulation is positively reinforcing. Conversely, photoactivation of BNST-PBN^vGLUT2^ neurons significantly increased the number of nosepokes mice performed to turn off photostimulation (**Figures 5I and S5F**), demonstrating that BNST-PBN^vGLUT2^ stimulation is negatively reinforcing. These data indicate that activation of GABAergic and glutamatergic BNST-PBN circuits have innate positive and negative affective valence, respectively (Namburi et al., 2016).

To assess whether BNST-PBN circuits modulate threat-assessment, we tested mice in the elevated zero maze (EZM), which measures anxiety-like behaviors and exploration in a novel brightly lit anxiogenic context (McCall et al., 2015). When EZM experiments were conducted in a brightly lit aversive setting, mice with photoactivation of BNST-PBN^vGAT^ neurons spent more time in the open arms compared to controls (**Figures 5J, S4K, and S4L**), consistent with increased exploratory drive. In contrast, mice with photoactivation of BNST-PBN^vGLUT2^ neurons showed decreased exploration of the open arms (**Figure 5K**), indicating that stimulation of this circuit is sufficient to reduce exploration of an anxiogenic environment. As a further test of whether BNST-PBN^vGAT^ activation can suppress defensive behaviors associated with threat, we subjected mice to threat-conditioning and measured their defensive response (freezing) to a conditioned stimulus. Mice were trained to associate a conditioned stimulus (tone) with a highly aversive unconditioned stimulus (0.7 mA, 2-s footshock). Photostimulation of BNST-PBN^vGAT^ neurons robustly reduced the defensive responses to the conditioned stimulus (**Figures 5L and 5M**). These data indicate that BNST-PBN circuits are sufficient and necessary for regulating opposing behavioral states including anxiety and threat within the context of feeding behavior.

### pDyn-expressing PBN neurons receive BNST input and alter feeding and affective behavior

We next aimed to determine the specific targets of BNST^vGAT^ and BNST^vGLUT2^ projections in the PBN. Previous studies have identified diverse neuronal populations within the primarily glutamatergic PBN (Carter et al., 2013; Geerling et al., 2015; Miller et al., 2011; Ryan et al., 2017). Here we examined dynorphin (pDyn) neurons, previously shown to regulate thermoregulation (Cintron-Colon et al., 2019; Geerling et al., 2015), and CGRP neurons (Campos et al., 2016; Carter et al., 2013; Han et al., 2015), shown to function as a general alarm system that responds to threats and inhibits feeding when activated. In-situ hybridization experiments demonstrated that pDyn neurons and CGRP neurons form genetically and anatomically distinct populations in the PBN (**Figures 6A and 6B**). Using a transynaptic rabies tracing method (Beier et al., 2015; Schwarz et al., 2015; Wickersham et al., 2007), we found that BNST neurons indeed form monosynaptic connections with pDyn-expressing neurons in the PBN (**Figures 6C-6E**) and may therefore be a critical downstream node for the behavioral effects of inhibitory and excitatory BNST-PBN inputs.

**Figure 6.**
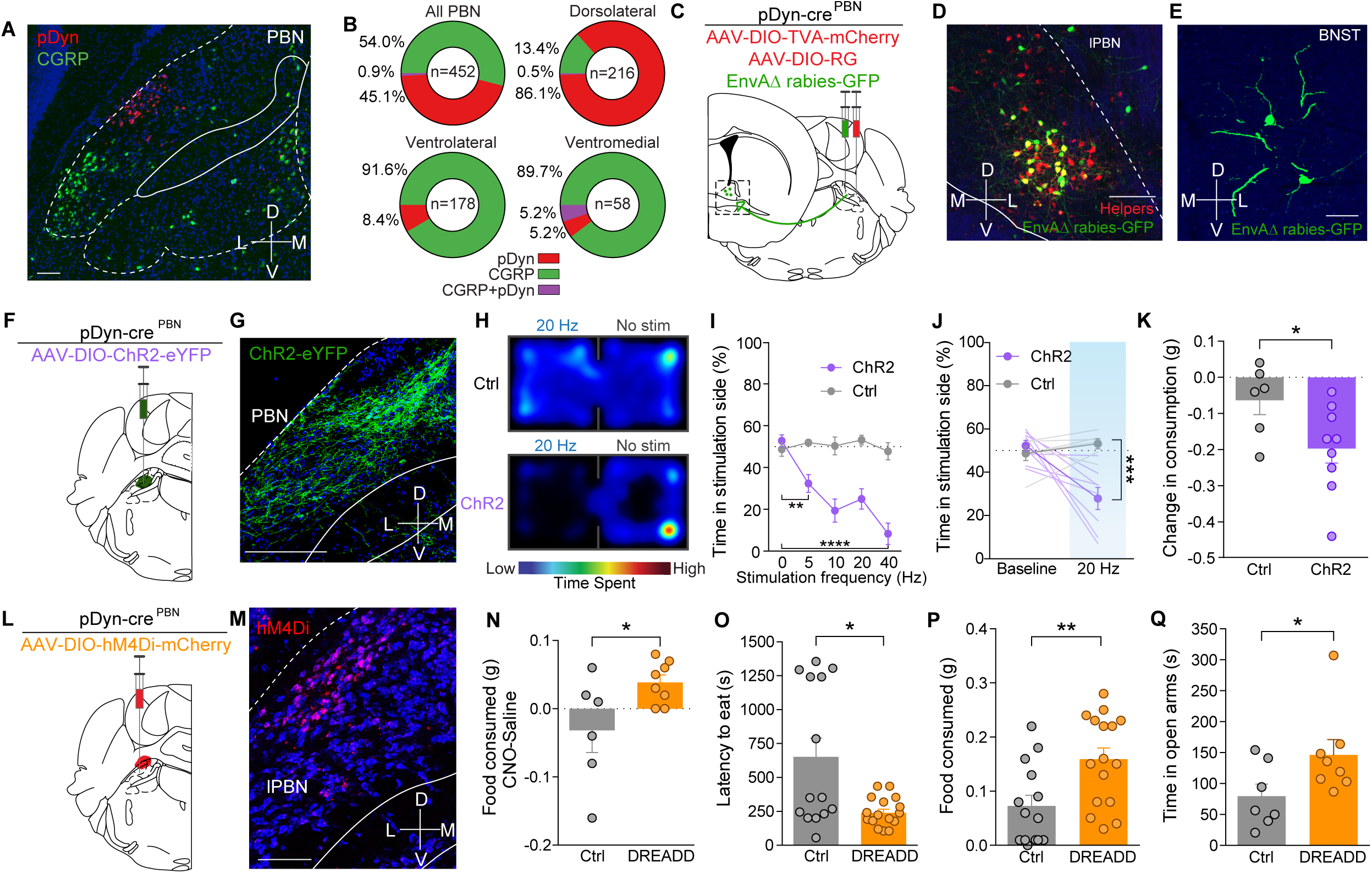
Dynorphin-expressing PBN neurons receive BNST input to bidirectionally modulate feeding and affective behaviors. **(A)** Fluorescent *in-situ* hybridization of pDyn and CGRP in the PBN (scale bar: 100μm). **(B)** pDyn and CGRP neurons are genetically and anatomically distinct populations in the PBN (n=452 cells). **(C)** Viral schematic for rabies tracing of monosynaptic inputs to PBN^pDyn^ neurons. **(D)** Starter cells in the PBN expressing TVA-mCherry and rabies-GFP (scale bar: 100 μm). **(E)** and their corresponding input cells in the BNST (scale bar: 50 μm). **(F)** Schematic of optogenetic approach to target PBN^pDyn^ neurons. **(G)** ChR2-eYFP expression in PBN^pDyn^ neurons (scale bar: 200 μm). **(H)** Representative heatmaps of time spent in the RTPP for Ctrl and PBN^pDyn^:ChR2 mice. **(I and J)** PBN^pDyn^ activation elicits a real-time place aversion in a **(I)** frequency-dependent manner including at **(J)** 20 Hz (ChR2, *n* = 8; Ctrl, *n* = 6). **(K)** PBN^pDyn^ activation decreases food consumption after food-deprivation (ChR2, *n* = 8; Ctrl, *n* = 6). **(L)** Schematic of chemogenetic approach to target PBN^pDyn^ neurons. **(K)** hM4Di -mCherry expression in PBN^pDyn^ neurons (scale bar: 100μm). **(N-Q)** Chemogenetic inhibition of PBN^pDyn^ neuron increases **(N)** food consumption, **(O)** decreases latency to eat and **(P)** while increasing food consumed in a novelty-suppressed feeding task and **(Q)** time spent in the open arms of elevated zero maze (DREADD, *n* = 7-8; Ctrl, *n* = 6-7). \**p* < 0.05, \*\**p* < 0.01, \*\*\**p* < 0.001, \*\*\*\**p* < 0.0001. Error bars indicate SEM. See also Figure S6.

We targeted dynorphin-expressing PBN neurons in pDyn-Cre mice with AAV5-EF1α-DIO-ChR2 and subjected the mice to a both feeding and affective behavioral tests (**Figures 6F and 6G**). Consistent with the effects of excitatory BNST-PBN activation, we found that photostimulation of PBN^pDyn^ produces a robust real-time place aversion (**Figures 6H-6J, S6A, and S6B**) and reduced feeding when animals were in a food-deprived state (**Figures 6K and S6C**). Additionally, inhibition of the broader PBN^vGLUT2^ population rapidly and reversibly increased food consumption (**Figures S6D and S6E)**. Likewise, chemogenetic inhibition of PBN^pDyn^ neurons using the hM4Di DREADD strategy (Armbruster et al., 2007) decreased the latency to consume food in an anxiogenic environment and increased overall food consumption in mice expressing the inhibitory DREADD when given CNO compared to saline, while control animals did not differ between treatments (**Figure 6L-6P**). Consistent with this finding, the inhibition of PBN^pDyn^ neurons produced anxiolytic behavior in an EZM test (**Figure 6Q**). These data reveal a previously unrecognized functional role of PBN^pDyn^ neurons as primary integrators of BNST afferents that coordinate feeding drive with threat assessment.

## Discussion

These studies indicate that discrete and opposing neural circuitry from the BNST to the PBN integrates internal states, threat assessment, and feeding behavior. The BNST has been shown to be involved in numerous functions related to threats and motivated behaviors (Lebow and Chen, 2016), and our findings demonstrate that the BNST conveys sensory and affective information to the PBN to alter behavior critical to an animal’s survival.

Recent studies have demonstrated BNST input to the PBN and revealed that BNST neurons may influence respiration and anxiety via increased metabolic activity in the parabrachial nucleus (Kim et al., 2013; Mazzone et al., 2018). Nonetheless, studies to date had not determined the identity of these projections. Employing the use of genetic targeting in our study, we unveil a separable role for discrete genetically-defined BNST-PBN projections. We also identify the complement of mRNAs actively translated by PBN-projecting BNST neurons in a cell-type specific manner, revealing several peptides and receptors that may act as neuromodulatory regulators influencing affect and feeding (**Figures 1G-1I**). Notably, these peptides and receptors provide an entryway to follow up studies and the development of new therapeutic strategies for treating feeding and affective diseases including obesity, generalized anxiety, and depression. Future studies employing Cre driver lines and pharmacological approaches will be critical to dissect the neuropeptide and receptor systems influencing these complex anxiety-like and feeding behaviors. In this study, we also probe the neural dynamics and functional roles of opposing BNST-PBN circuits using fiber photometry together with optogenetics. Our results reveal that the GABAergic BNST-PBN circuit is engaged by and drives feeding behavior and exploration, while the glutamatergic BNST-PBN circuit is disengaged during feeding and suppresses state-dependent feeding and exploration (**Figures 2-5**). Interestingly, the distinct circuits begin increasing or decreasing their activity prior to the consumption itself, suggesting that the circuit is not only encoding the consumption but the appetitive behavior as well. Furthermore, we demonstrate that these opposing inhibitory and excitatory circuits act to drive these behaviors partly via the attribution of valence inherent to the circuit itself (**Figure 5**). Further studies exploring both acute and learned threat detection with opposing feeding states will be needed to explore how these circuits adapt under various environmental influences. Here we demonstrate that activation of these circuits is sufficient to alter feeding, exploration, and threat assessment. Given that eating disorders are highly associated with varying affective states (Hardaway et al., 2015), further investigation of these circuits under varying affective states imposed by external adverse experiences such as pain or stress is necessary to fully understand the extent of their involvement.

The PBN consists of many cell-types that may mediate separable aspects of feeding and aversion (Campos et al., 2018; Fu et al., 2019; Park et al., 2020; Ryan et al., 2017). Of these cell-types, neurons expressing calcitonin gene-related peptide (CGRP) located in the vental/external lateral parabrachial (PBNel) thought to function as a general alarm system that sends aversive signals throughout the brain are of the most notable (Palmiter, 2018). However, our results demonstrate that GABAergic and glutamatergic BNST terminals are most dense in the dorsolateral and medial regions of the PBN and lacking in the ventrolateral region where CGRP neurons subside (**Figures 1A, 1D, 6A, 6B, and S1A**). This was both interesting and surprising to us because this reveals that although CGRP neurons mediate feeding and aversion, they are not the sole neuropeptide population in the PBN mediating these behaviors. In our studies, we have demonstrated that dynorphin-expressing neurons located in the dorsolateral PBN (PBN^pDyn^) are also critically poised as functional mediators of feeding and affective behaviors (Bhatti et al., 2018). Additionally, recent studies seem to suggest that PBN^pDyn^ may communicate with CGRP neurons in response to noxious pain stimuli (Chiang et al., 2019). Prior to this study, the primary function for PBN^pDyn^ was thought to be for thermoregulation (Cintron-Colon et al., 2019; Geerling et al., 2015). Altogether, these studies with our results suggest that a primary role for PBN^pDyn^ is to integrate internal states including threat assessment, anxiety, thermoregulation, or pain, in order to adaptively seek and consume or avoid food.

Here we present two opposing neural circuits from the BNST to the PBN that mediate threat-assessment, exploration, and feeding behaviors in part via their connections to dynorphin-expressing PBN neurons. Our results implicate these circuits, neuropeptide candidates, and both inhibitory and excitatory BNST neurons as critical integrators with the PBN for adaptive and affective behaviors necessary for survival. These findings may provide new avenues for the development of unique strategies to mitigate maladaptive behaviors related to eating patterns or affective state.

## Acknowledgments

We thank Lamley Lawson, Dylan Blumenthal, Michelle Chung, and Taylor Hobbs for animal colony maintenance; the Bruchas laboratory and UW NAPE Center for helpful discussions; Jordan G. McCall and Bryan A. Copits for assistance with electrophysiology. We thank Richard Palmiter (HHMI, UW) for thoughtful discussion and critiques regarding the manuscript.

## Funding

This study was supported by the National Institute on Mental Health (M.R.B., R01-MH112355), the National Institute on Drug Abuse (M.R.B., R01-DA033396), the National Institute of Neurological Disorders and Stroke (R.W.G., R56-NS048602, R01-NS106953), the National Institute of General Medical Sciences (A.T.L., T32GM008151), and National Institute of Mental Health (B.M., F30-MH116654, J.D.D., U01-MH109133). Sequencing of TRAP samples was made possible in part by Grant Number UL1-RR024992 from the NIH-National Center for Research Resources (NCRR).

## Author contributions

D.L.B., A.T.L., and M.R.B. conceptualized and designed the entire study. D.L.B., A.T.L., C.E.P., K.K., H.O.B., and A.S. performed stereotaxic surgeries, behavioral experiments, and histology. D.L.B. performed electrophysiological experiments. A.T.L. and C.E.P. performed fiber photometry experiments.

A.T.L. and B.M. performed RNA sequencing experiments. D.L.B., A.T.L., C.E.P, and B.M. performed statistical analysis. D.L.B., A.T.L., and M.R.B. made the figures. D.L.B., A.T.L., and M.R.B. wrote the manuscript with collective input from all authors. J.D.D. supervised RNA sequencing experiments. R.W.G. supervised electrophysiological experiments. M.R.B. supervised the entire study.

## Competing Interests

Although there are no studies we report here that are related, for full disclosure, Dr. Michael R. Bruchas is a co-founder and consultant for Neurolux, Inc, a neurotechnology company. J.D.D has previously received royalties related to the TRAP methodology.

## Data and materials

All data are available in the manuscript or the supplementary materials. RNA sequencing data is deposited in in the NCBI Gene Expression Omnibus (GEO) database with accession number GSE133484. We thank Chris Stander, K. Deisseroth, the Washington University HOPE Vector Core, and the University of North Carolina (UNC) Vector Core for viral constructs, prep and packaging.

## Methods

### Key Resources Table

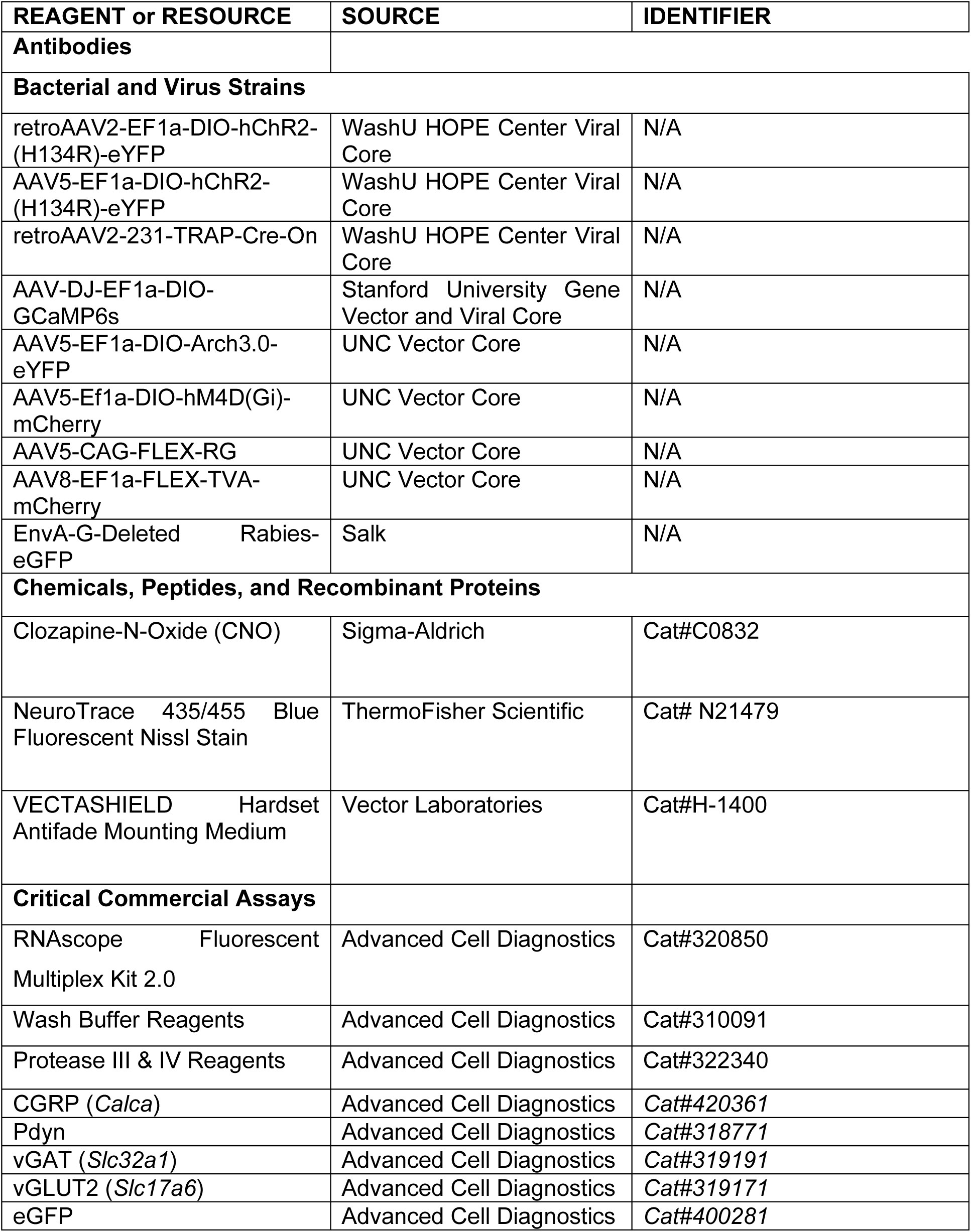
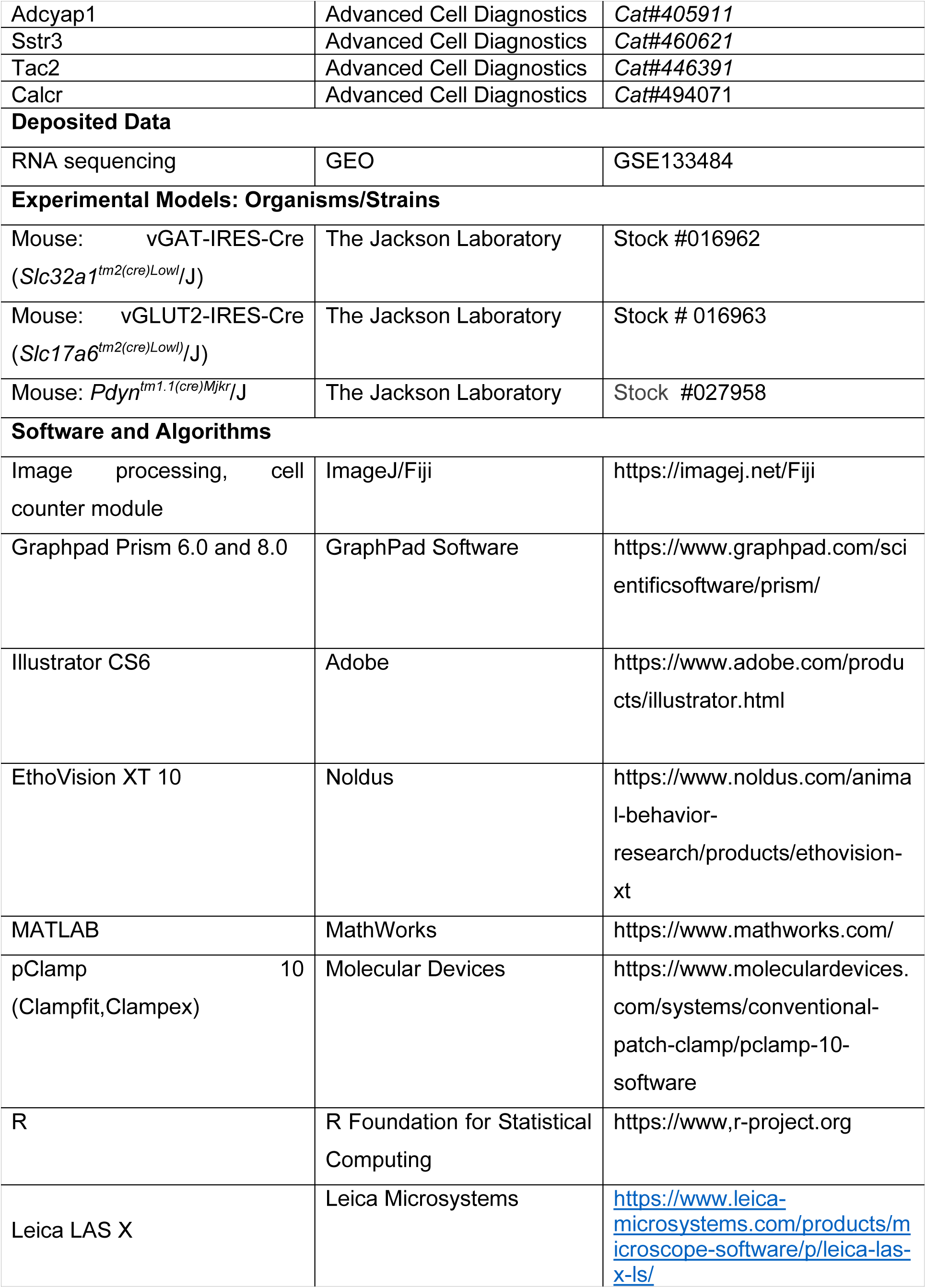
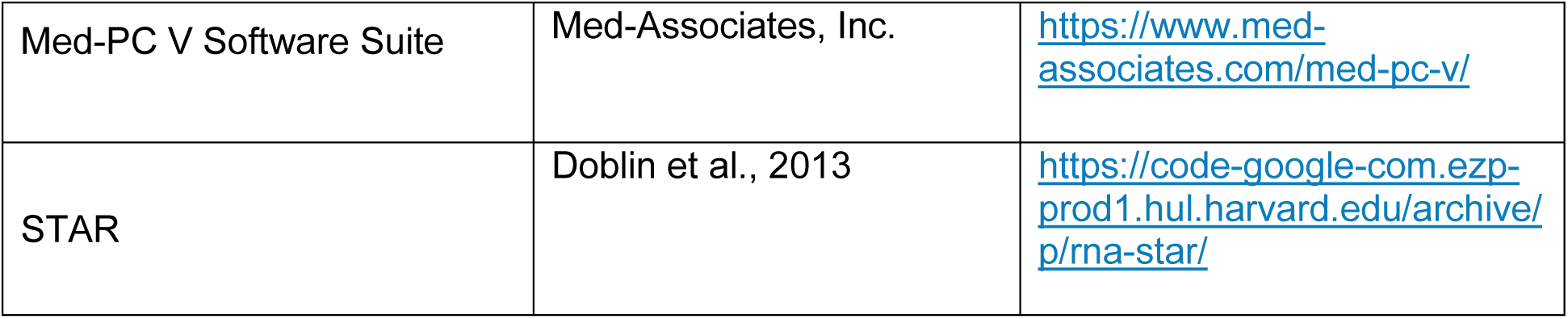

2 Key Resources Table
| REAGENT or RESOURCE | SOURCE | IDENTIFIER |
| --- | --- | --- |
| **Antibodies** |  |  |
| **Bacterial and Virus Strains** |  |  |
| retroAAV2-EF1a-DIO-hChR2-(H134R)-eYFP | WashU HOPE Center Viral Core | N/A |
| AAV5-EF1a-DIO-hChR2-(H134R)-eYFP | WashU HOPE Center Viral Core | N/A |
| retroAAV2-231-TRAP-Cre-On | WashU HOPE Center Viral Core | N/A |
| AAV-DJ-EF1a-DIO-GCaMP6s | Stanford University Gene Vector and Viral Core | N/A |
| AAV5-EF1a-DIO-Arch3.0-eYFP | UNC Vector Core | N/A |
| AAV5-Ef1a-DIO-hM4D(Gi)-mCherry | UNC Vector Core | N/A |
| AAV5-CAG-FLEX-RG | UNC Vector Core | N/A |
| AAV8-EF1a-FLEX-TVA-mCherry | UNC Vector Core | N/A |
| EnvA-G-Deleted Rabies-eGFP | Salk | N/A |
| **Chemicals, Peptides, and Recombinant Proteins** |  |  |
| Clozapine-N-Oxide (CNO) | Sigma-Aldrich | Cat#C0832 |
| NeuroTrace 435/455 Blue Fluorescent Nissl Stain | ThermoFisher Scientific | Cat# N21479 |
| VECTASHIELD Hardset Antifade Mounting Medium | Vector Laboratories | Cat#H-1400 |
| **Critical Commercial Assays** |  |  |
| RNAscope Fluorescent Multiplex Kit 2.0 | Advanced Cell Diagnostics | Cat#320850 |
| Wash Buffer Reagents | Advanced Cell Diagnostics | Cat#310091 |
| Protease III & IV Reagents | Advanced Cell Diagnostics | Cat#322340 |
| CGRP (*Calca*) | Advanced Cell Diagnostics | Cat#420361 |
| Pdyn | Advanced Cell Diagnostics | Cat#318771 |
| vGAT (*Slc32a1*) | Advanced Cell Diagnostics | Cat#319191 |
| vGLUT2 (*Slc17a6*) | Advanced Cell Diagnostics | Cat#319171 |
| eGFP | Advanced Cell Diagnostics | Cat#400281 |
| Adcyap1 | Advanced Cell Diagnostics | Cat#405911 |
| Sstr3 | Advanced Cell Diagnostics | Cat#460621 |
| Tac2 | Advanced Cell Diagnostics | Cat#446391 |
| Calcr | Advanced Cell Diagnostics | Cat#494071 |
| **Deposited Data** |  |  |
| RNA sequencing | GEO | GSE133484 |
| **Experimental Models: Organisms/Strains** |  |  |
| Mouse: vGAT-IRES-Cre<br>(*Slc32a1*<sup>tm2(cre)Lowl</sup>/J) | The Jackson Laboratory | Stock #016962 |
| Mouse: vGLUT2-IRES-Cre<br>(*Slc17a6*<sup>tm2(cre)Lowl</sup>/J) | The Jackson Laboratory | Stock # 016963 |
| Mouse: *Pdyn*<sup>tm1.1(cre)Mjkr</sup>/J | The Jackson Laboratory | Stock #027958 |
| **Software and Algorithms** |  |  |
| Image processing, cell counter module | ImageJ/Fiji | https://imagej.net/Fiji |
| Graphpad Prism 6.0 and 8.0 | GraphPad Software | https://www.graphpad.com/scientificsoftware/prism/ |
| Illustrator CS6 | Adobe | https://www.adobe.com/products/illustrator.html |
| EthoVision XT 10 | Noldus | https://www.noldus.com/animal-behavior-research/products/ethovision-xt |
| MATLAB | MathWorks | https://www.mathworks.com/ |
| pClamp 10<br>(Clampfit,Clampex) | Molecular Devices | https://www.moleculardevices.com/systems/conventional-patch-clamp/pclamp-10-software |
| R | R Foundation for Statistical Computing | https://www.r-project.org |
| Leica LAS X | Leica Microsystems | https://www.leica-microsystems.com/products/microscope-software/p/leica-las-x-ls/ |
| Med-PC V Software Suite | Med-Associates, Inc. | https://www.med-associates.com/med-pc-v/ |
| STAR | Doblin et al., 2013 | https://code-google-com.ezp-prod1.hul.harvard.edu/archive/p/rna-star/ |

### Contact for Reagent and Resource Sharing

Further information and requests for resources and reagents should be directed to and will be fulfilled by the Lead Contact, Dr. Michael Bruchas.

### Experimental Model and Subject Details

#### Animals

Adult male vGAT-IRES-Cre (*Slc32a1*^*tm2(cre)Lowl*^/J; Jackson Laboratory #016962) (Vong et al., 2011), vGLUT2-IRES-Cre (*Slc17a6*^*tm2(cre)Lowl)*^/J; Jackson Laboratory # 016963), and preprodynorphin(*Pdyn*)-IRES-Cre (*Pdyn*^*tm1.1(cre)Mjkr*^/J; Jackson Laboratory #027958) (Krashes et al., 2014) mice were group housed, given access to food and water *ad libitum*, and maintained on a 12:12 hr light:dark cycle (lights on at 7:00 AM). All animals were kept in an isolated and sound-attenuated holding facility within the lab and adjacent to behavior rooms one week prior to surgery, post-surgery, and throughout the duration of the behavioral assays to minimize stress. All procedures were approved by the Animal Care and Use Committee of Washington University and the University of Washington and conformed to US National Institutes of Health guidelines.

### Method Details Stereotaxic

#### Surgeries

After the animals were acclimated to the holding facility for at least seven days, the mice were anaesthetized in an induction chamber (2% isoflurane) and placed into a stereotaxic frame (Kopf Instruments, Model 1900) where they were maintained at 1-2% isoflurane. Mice were then injected using a blunt needle (86200, Hamilton) at a rate of 100 nL/min with 300-400 nL in the BNST (AP +0.14, ML ±0.9, DV -4.5) or PBN (AP -5.3, ML ±1.1, DV -3.5); unilateral for optogenetic activation, tracing, and fiber photometry experiments; bilateral for optogenetic or chemogenetic experiments. Needle was slowly removed from the brain 10 min after cessation of injection to allow for diffusion. For fiber photometry experiments, mice were also implanted with a 400-µm fiber optic (Doric Inc., MFC_400/430-0.48_MF2.5_FLT) in the same surgery. Mice recovered for at least 6 weeks prior to behavioral testing, permitting optimal expression of the virus. For optogenetic activation and fiber photometry experiments, five weeks after viral injection, intracranial optic fiber implants were directed above the PBN unilaterally (AP -5.3, ML ± 1.1, DV -3.0). For optogentic inhibition inhibition, fiber optics were placed bilaterally. The fiber optic implants were secured using two bone screws (CMA, 743102) and affixed with TitanBond (Horizon Dental Products) and dental cement (Lang Dental).

### Immunohistochemistry

Immunohistochemistry was performed as previously described (McCall et al., 2017). In brief, mice were anesthetized with sodium pentobarbital and transcardially perfused with 4% paraformaldehyde (PFA), post-fixed overnight in 4% PFA, and cryo-protected in 30% sucrose for at least 24 hours. Brains were then sectioned (30 µm) and placed in 0.1M PB until immunohistochemistry. Free-floating sections were washed in 0.1M PBS for 3 × 10 minutes intervals. Sections were then placed in blocking buffer (0.5% Triton X-100 and 5% natural goat serum in 0.1 M PBS) for 1 hr at room temperature. After blocking buffer, sections were incubated in NeuroTrace (1:400, 435/455 blue fluorescent Nissl stain, Invitrogen #N21479) for 1 hour, followed by 3 × 10 minute 0.1 MPBS then 3 × 10 minute 0.1 M PB washes. After immunostaining, sections were mounted and coverslipped with Vectashield HardSet mounting medium (Vector Laboratories) and imaged on a Leica TCS SPE confocal microscope. Animals that did not show targeted expression were excluded.

### Fluorescent In-Situ Hybridization (FISH)

Animals were anesthetized and rapidly decapitated. Brains were quickly removed and fresh frozen in dry ice, then stored at -80°C. Sections were cut at 20µm, mounted on slides, and stored at -80°C. Sections were fixed in 4% PFA for 15 min, dehydrated in serial ethanol concentrations (50%, 70%, and 100%) and processed with RNAscope (Advanced Cell Diagnostics, cat. No. 320293). Sections were hybridized with the probes listed below. Sections were then counterstained with DAPI and coverslipped. Confocal images were obtained on an Olympus FV3000RS microscope. Circular ROIs were drawn around each cell, and these ROIs were then used for co-expression analysis.

### Slice Electrophysiology

Acute brain slices were prepared using a protective cutting and recovery method (Ting et al., 2014). Anesthetized mice infected with AAV5-EF1a-DIO-hChR2-(H134R)-eYFP were transcadially perfused with NMDG-substituted aCSF containing (in mM) 93 NMDG, 2.5 KCl, 1.25 NaH2PO4, 30 NaHCO3, 20 HEPES, 25 glucose, 5 ascorbic acid, 2 thiourea, 3 Na-pyruvate, 12 N-acetyl-L-cysteine, 10 MgSO4, 0.5 CaCl2 (pH = 7.3-7.4). 200 mm thick coronal sections of the PBN were cut using a Vibratome VT1000s (Leica) and transferred to an oxygenated recovery chamber containing NMDG aCSF for 5–10 min at 32°C –34°C before being transferred to a holding chamber filled with modified aCSF containing (in mM) 92 NaCl, 2.5 KCl, 1.2 NaH2PO4, 30 NaHCO3, 20 HEPES, 25 glucose, 2 CaCl2, 2 MgCl2 (pH adjusted to 7.3-7.4 with NaOH), and Osm 290-310. Whole-cell patch-clamp recordings were made using fire-polished glass pipettes with a resistance of 3-5 MΩ filled with (in mM): 120 K+ gluconate, 5 NaCl, 2 MgCl2, 0.1 CaCl2, 10 HEPES, 1.1 EGTA, 4 Na2ATP, 0.4 Na2GTP, 15 phosphocreatine; pH adjusted to 7.3 with KOH, 291 mOsm. BNST axonal projections in the PBN were visualized through a 40x objective using IR-DIC microscopy on an Olympus BX51 microscope, and neurons with YFP+ axons in close proximity were identified using epifluorescent illumination. Recordings were made with Patchmaster software controlling a HEKA EPC10 amplifier. Following gigaseal formation and stable whole-cell access, currents elicited by 10 ms-pulse stimulation of ChR2-containing axonal terminals and isolated by blocking AMPA/KARs (10 µM NBQX, Abcam), NMDARs (50 µM D-APV, Abcam), GABAARs (100 µM picrotoxin and 50 µM bicuculline, Abcam) through bath application of the antagonists in aCSF solution. Neurons were voltage clamped at -70 mV for eEPSCs and 0 mV for eIPSCs. Input resistance was monitored to maintain cells with a stable Rs < 35 MΩ, and only these neurons were included in our analysis. eIPSC and eEPSC amplitudes were averaged across 5 sweeps per cell.

### Translating Ribosome Affinity Purification (TRAP)

In order to specifically capture transcribed RNA from GABAergic and glutamatergic neurons synapsing on the PBN, we generated an adenoviral vector encoding a floxed fusion GFP-ribosomal protein (RPL10A), and packaged it into the retrofecting AAV serotype rAAV2-retro (Tervo et al., 2016). This design restricts expression of the GFP•RPL10A fusion to a target (Cre-expressing) cell type, and further restricts expression to cells of that type synapsing onto the transduced region.

In order to selectively capture transcripts from those excitatory and inhibitory neurons in the BNST synapsing onto the PBN, we delivered the floxed eGFP•Rpl10a retro-AAV2 into the PBN of adult male mice ages 4m-10m expressing Cre recombinase under a glutamatergic (vGLUT2) or GABAergic (vGAT) neuron-specific promoter. At least 6 weeks were given for surgical recovery and transgene expression before mice were sacrificed for brain dissection, homogenization, and GFP-targeted immunoprecipitation.

In order to control for stress and circadian effects on BNST transcripts between replicates, mice to be sacrificed were always retrieved between 10AM and 12PM immediately before deep anesthetization by isoflurane, followed by rapid removal of the brain. A 1mm coronal slab of brain tissue was collected from AP ±0.5 mm, ML ±1.2mm, DV -3.8 to -4.8mm and microdissected in ice cold PBS containing RNAse inhibitors (rRNAsin, Superasin), 500 µM DTT, and cycloheximide (to arrest translation in progress to keep ribosomes associated with their RNAs for later capture). Three dissected BNSTs were pooled per replicate in order to minimize effects of dissection and incidental injection of structures adjacent to PBN. Pooled tissue was homogenized and processed using the standard TRAP protocol as previously described (Rieger et al., 2018). All RNA samples were quality checked for RINe (estimated RNA integrity number) on an Agilent Tapestation 4200 (Agilent Technologies, Santa Clara, California) using the High-Sensitivity RNA Assay Kit. All sequenced samples had a RINe of > 7.5 with total mass yielded per sample ranging from 2.5 ng to 250ng.

RNA samples were submitted to Washington University in St Louis’ Genome Technology Access Center for library preparation and mRNA sequencing. cDNAs were synthesized using polyA capture (Dynal mRNA Direct Kit, Life Technologies, Carlsbad, California). cDNAs were synthesized by polyA-targeted priming, followed by cDNA amplification using the Clontech SMARTer kit (Takara Bio, Mountain View, CA).

Single-read, 50bp RNA-seq of IP fractions and total homogenate from 5 replicates per cell type (20 samples total) was performed on a single lane of a HiSeq 3000 (Illumina, San Diego, California), yielding ∼350M reads, ranging from 10M to 25M per sample. Reads were processed using a standard pipeline of read trimming with Trimmomatic (Bolger et al., 2014), removal of rRNA reads using BowTie (Langmead et al., 2009), and alignment to mouse genome build 38.p5 along with gene-level feature counts using STAR aligner (Dobin et al., 2013).

Samplewise read counts were first subsetted to those reads with a normal distribution in order to perform downstream differential expression analysis, as recommended for analyses in EdgeR(McCarthy et al., 2012; Robinson et al., 2010). Gene counts were then quality-checked by hierarchical clustering to rule out batch effects and off-target cell enrichment. One replicate clustered separately from all other samples and was excluded. Additional quality checks included plotting relative enrichment / depletion of vGAT/vGLUT2-specific marker genes and that of off-target cell types (e.g. astrocytes and oligodendrocytes). Ultimately, 3 replicates for vGLUT2 neurons and 2 replicates for vGAT neurons were used for analysis.

Differential expression analysis was performed in EdgeR using the following model for single-cell type enriched transcripts, where group was vGLUT2 IP, vGLUT2 input, vGAT IP, or vGAT input:

Expression ∼ 0+group

Enrichment in cell types was then determined using EdgeR’s glmQLFtest function using contrasts of vGLUT2 Enriched = (vGLUT2 IP) – (vGLUT2 Input) and likewise for vGAT.

Differential expression analysis between cell types was performed using the same model as above but with a contrast of (vGLUT2-vGAT Differential Expression) = (vGLUT2 IP – vGLUT2 Input) – (vGAT IP – vGAT Input). This contrast thus regresses potential effects of dissection heterogeneity before comparing the target immunoprecipitated cell types. The supplementary data (**Table S2-S5**) contains differential expression results for all of the contrasts described above, as well as an additional contrast between immunopreciptates for vGLUT2 and vGAT without normalizing for input RNAs (that is, vGLUT2-vGAT differential expression, alternate calculation = vGLUT2 IP – vGAT IP). Raw data can be found in the NCBI Gene Expression Omnibus (GEO) database (GSE133484).

### Behavior

Behavioral assays were performed in sound attenuated rooms maintained at 23°C. Lighting was measured and stabilized at ∼100 lux for anxiety testing in vGAT-Cre experiments, ∼20 lux for anxiety testing in vGlut-Cre experiments, and ∼200 lux for place testing. All behavioral apparatuses were cleaned with 70% ethanol in between animals. Movements were video recorded and analyzed using Ethovision Software or recorded with Media Recorder.

#### Real-Time Place Preference (RTPP)

Mice were placed into a custom-made unbiased, balanced two-compartment conditioning apparatus (52.5 × 25.5 × 25.5 cm) as previously described (McCall et al., 2017) and allowed to freely roam the entire apparatus for 30 min. Entry into one compartment triggered constant photostimulation (0-40 Hz; 5-10 mW light power) while the animal remained in the light-paired chamber. Entry into the other chamber ended the photostimulation. The side paired with photostimulation was counterbalanced across mice. Time spent in each chamber and total distance traveled for the entire 30 min trial was measured using Ethovision 8.5 (Noldus). Data are expressed as mean +/- S.E.M percent time spent in photostimulation-paired chamber.

#### Operant Conditioning: Positive Reinforcement

For positive reinforcement induced by vGAT-containing BNST-PBN projections, mice with optical fibres implanted above the PBN were trained in one 1-hr session to nose poke on a fixed-ratio 1 schedule for optical self-stimulation (3-s, 20-Hz. ∼10 mW, 473nm) in a mouse operant chamber (17.8 × 15.2 × 18.4 cm; Med Associates) (Jennings et al., 2013). The following day, the mice were run again with the same conditions and the number of nosepokes recorded in the hour session was recorded.

#### Operant Conditioning: Negative Reinforcement

For negative reinforcement induced by vGLUT2-containing BNST-PBN projections, mice with optical fibres implanted above the PBN were trained in one 1-hr session to nose poke on a fixed-ratio 1 schedule to turn off optical self-stimulation in a mouse operant chamber (17.8 × 15.2 × 18.4 cm; Med Associates). Constant photostimulation (20-Hz, ∼10 mW, 473nm) began at the start of each session. Each nose poke resulted in the cessation of the photostimulation for 3-s. The following day, the mice were run again with the same conditions. The number of nosepokes recorded in the hour session was recorded. About 50% of Cre+ mice did not learn the operant task and thus were excluded from the results as non-responders (**Fig. S7F**). Non-responders were classified as animals whose difference of responses between active and inactive was < 5.

#### Threat Conditioning

Pavlovian threat conditioning was performed in Med Associates Fear Conditioning Chambers (NIR-022MD). This equipment consisted of a conductive grid floor inside a 29.53 cm L × 23.5 cm W × 20.96 cm H chamber inside of a soundproof box lit by an infrared light.

vGAT-Cre experiments: Mice were conditioned to four 20s tones co-terminating with 2-s 0.7mA shocks on Day 1. On Day 2, mice were placed back into the chamber. Eleven tone presentations in the absence of shock were presented. During the first tone, no optostimulation occurred. During the next ten tone presentations, 20 Hz blue light was delivered. On the third day, we assessed extinction recall by placing the mice into the chamber and presented the tone once. Freezing behavior was measured during all tone presentations.

All fiber photometry animals were conditioned to 6 20-second tones terminating with a 1-s 0.5mA shock.

#### Feeding Assays

Feeding assays were performed in a square plexiglass arena (27 cm × 27 cm). Chow, High-fat, sucrose, or salt pellets (Envigo) were placed in the corner of the arena. Mice were then introduced into the arena for 30 min. To record the amount of food consumed, pellets were measured before and after the assay. Baseline feeding and feeding whilst photostimulation (20 Hz; 5-10 mW light power) or photoinhibition (constant; ∼5 mW light power) were performed in a counterbalanced fashion across mice.

In addition to counterbalanced experiments across days, we also performed within-session manipulations to assess whether the effects are immediately reversible. Mice were introduced into the arena for 60 minutes: 20 min pre-stimulation, 20 min 20 Hz stimulation or constant inhibition, and 20 min post-stimulation. Food weight was measured at each 20 min point (**Fig. S5J, S7E, S8E**).

#### Elevated Zero Maze

EZM testing was performed as described previously (McCall et al., 2017), the EZM (Harvard Apparatus) was made of grey plastic, 200 cm in circumference, comprised of four 50 cm sections (two opened and two closed). The maze was elevated 50 cm above the floor and had a path width of 4 cm with a 0.5 cm lip on each open section. Mice were connected to fiber optic cables, positioned head first into a closed arm, and allowed to roam freely for 7 min. Animals received 20 Hz (10 ms pulse width) photostimulation (5-10 mW light power). Open arm time was the primary measure of anxiety-like behavior.

### Fiber Photometry

Fiber photometry recordings were performed as previously described (Parker et al., 2019). Briefly, an optic fiber was attached to the implanted fiber by a ferrule sleeve, then GCaMP6s was stimulated by two LEDs, a 531-Hz sinusoidal light (Thorlabs M470F3), bandpass filtered at 470 ± 20nm, and a 211-Hz sinusoidal light (Thorlabs M405FP1), bandpass filtered at 405 ± 10nm. (Filter cube: Doric FMC4; LED driver: DC4104). The 470 nm signal evokes Ca^2+^-dependent emission, while the 405 nm signal evokes Ca^2+^-independent isobestic control emission. Prior to recording, a 180s period of GCaMP6s excitation with both light channels was used to remove the majority of baseline drift. Laser intensity at the optic fiber tip was adjusted to ∼50 μW before each day of recording. GCaMP6s fluorescent signal was isolated by bandpass filtering (525 ± 25nm), transduced by a femtowatt silicon photoreceiver (Newport 2151), and recorded by a real-time processor (TDT RZ5P). The envelopes of 531 Hz and 211 Hz signals were extracted in real time by the TDT program Synapse at a sampling rate of 1017.25 Hz.

Fiber recordings were analyzed using custom MATLAB scripts available on Github. Motion artifacts were removed by subtracting the isobestic 405 nm signal from the 470 nm excitation signal. Baseline drift due to slow photobleaching artifacts was corrected by fitting a double exponential curve to the raw trace, then the photometry trace was z-scored relative to the mean and standard deviation of the signal. The mean z-score during the ten seconds preceding and following an event were compared using paired t tests.

### Statistical analyses

All summary data are expressed as mean ± SEM. Statistical significance was taken as *p < 0.05, **p < 0.01, ***p < 0.001, ****p < 0.0001, as determined by the Student’s t-test (paired and unpaired): One-Way Analysis of Variance (ANOVA) or One-Way Repeated Measures ANOVA, followed by Bonferroni post hoc tests as appropriate. Statistical analyses were performed in GraphPad Prism 6.0 or 8.0. All statistical information is listed in **Table S1.**

## Data and Code Availability

RNA sequencing data from Figure 1 have been deposited and are available from GEO (Accession: GSE133484). Custom MATLAB analysis code was created to appropriately organize, process, and combine photometry recording data with associated behavioral data. Analysis code for photometry from Figures 2, 3, and 4 is available online at https://www.github.com/BruchasLab. The full behavioral dataset supporting the current study are available from the corresponding author upon request.

