## Supplemental materials for "Extended amygdala-parabrachial circuits alter threat assessment to regulate feeding"

### SUPPLEMENTAL FIGURE 1

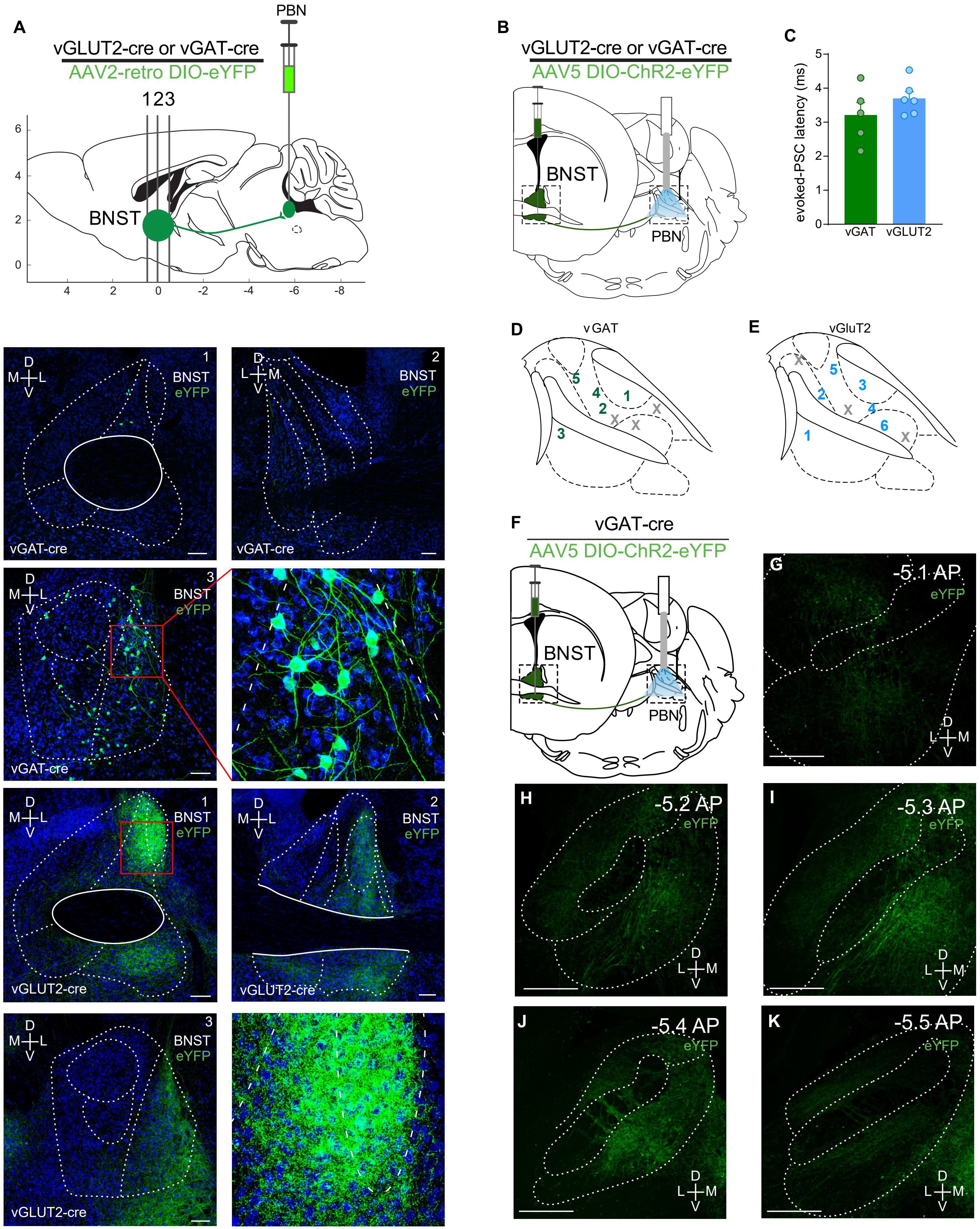

### SUPPLEMENTAL FIGURE 2

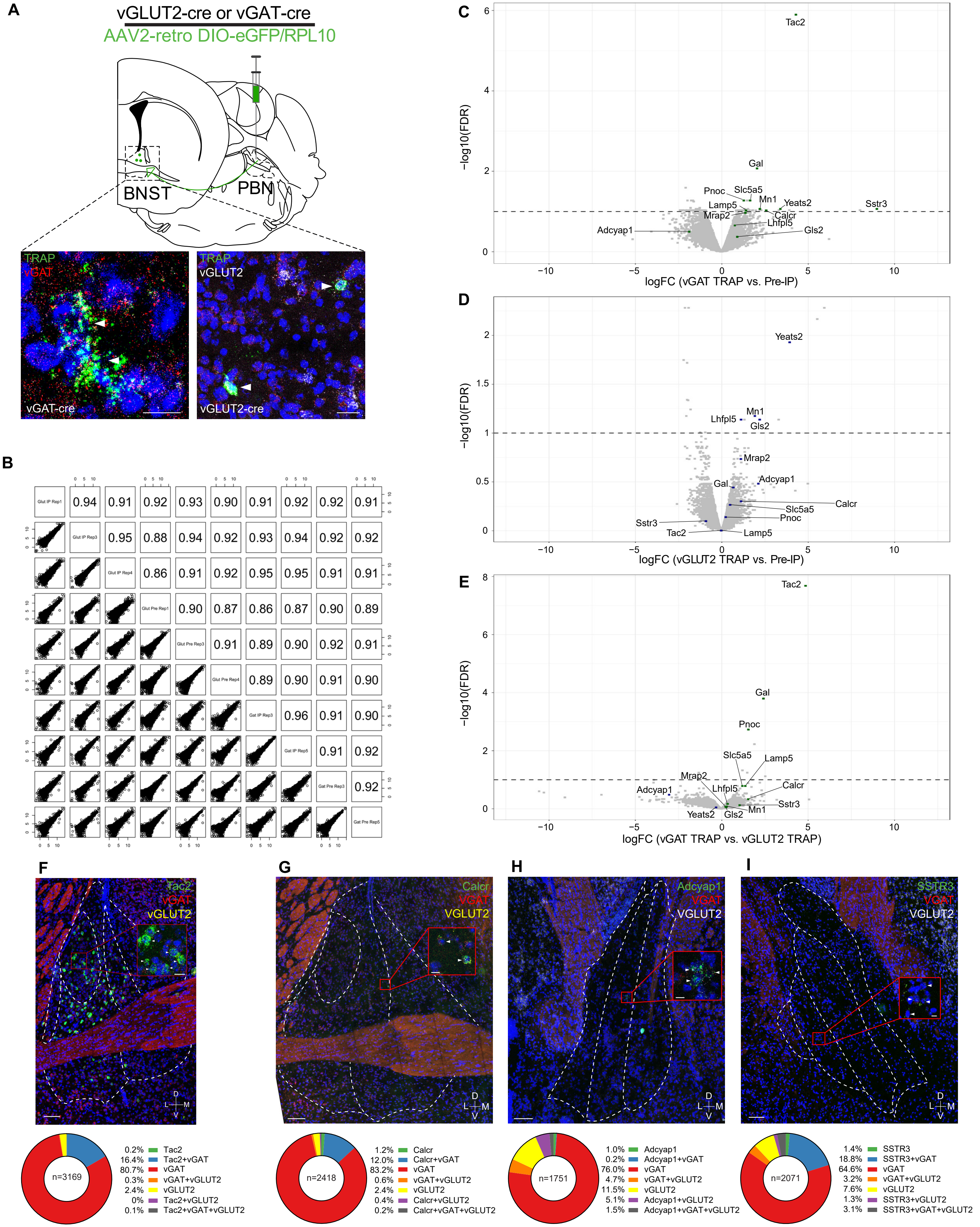

### SUPPLEMENTAL FIGURE 3

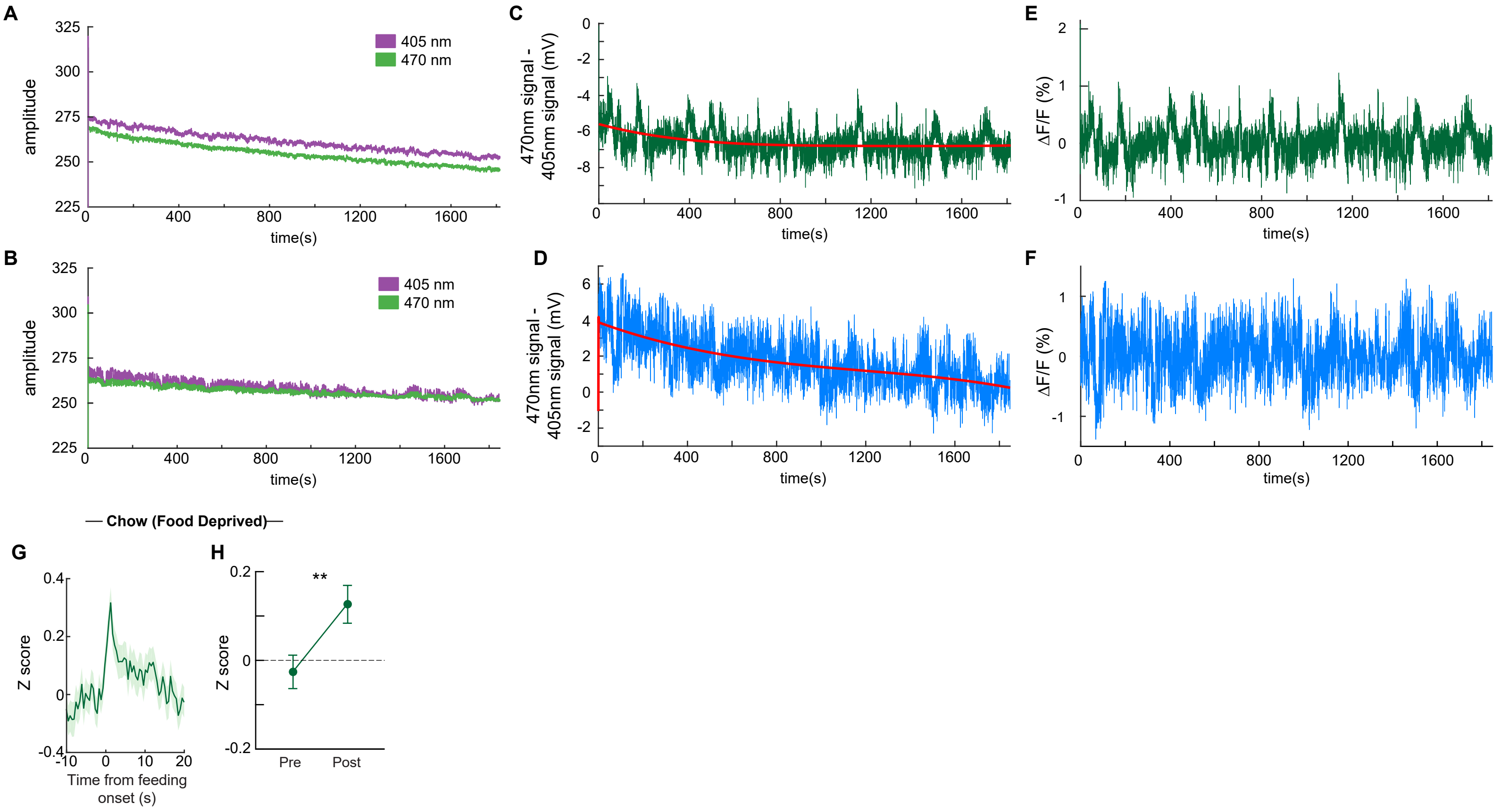

### SUPPLEMENTAL FIGURE 4

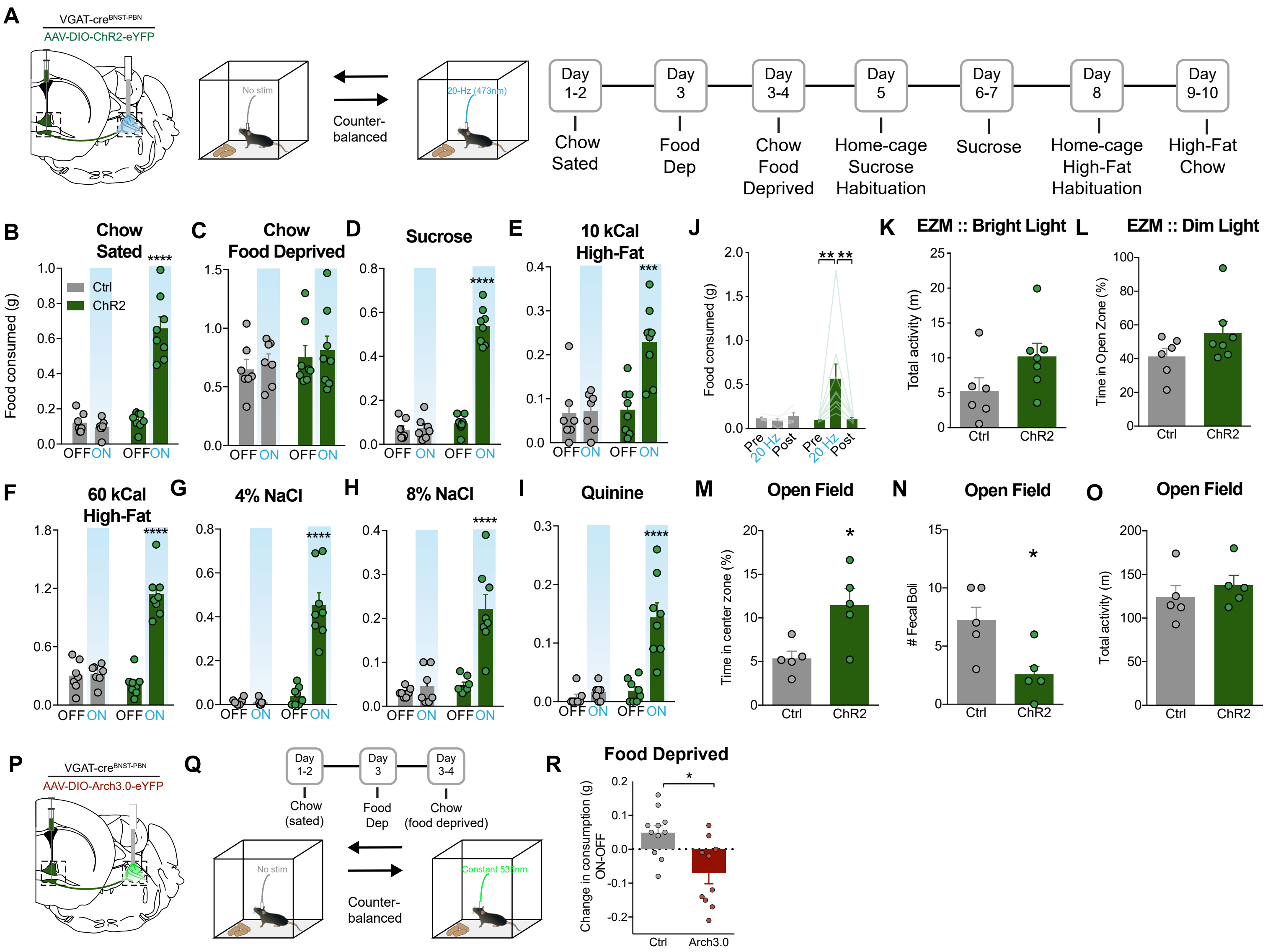

### SUPPLEMENTAL FIGURE 5

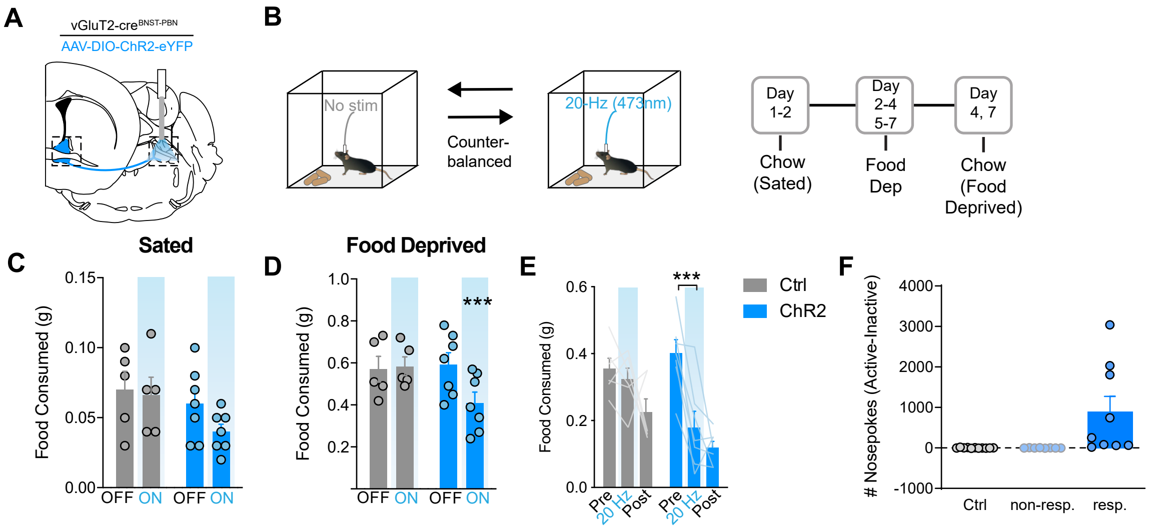

### SUPPLEMENTAL FIGURE 6

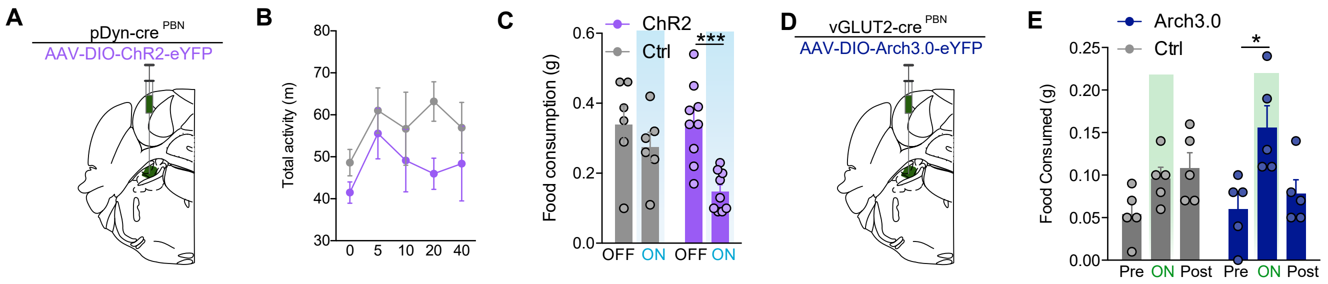

#### SUPPLEMENTARY MATERIALS

**Figure S1. Anatomical characterization of BNST-PBN projections. Related to Figure 1.**

**(A)** Viral and analysis schematic and images depicting retrogradely-labeled PBN-projecting $BNST^{vGAT}$ and $BNST^{vGLUT2}$ neurons across the A-P axis of the BNST (scale bar: 100μm).

**(B)** Viral schematic for electrophysiological recordings.

**(C)** Optically evoked IPSCs and EPSCs in the PBN have a latency <5 ms, suggesting a monosynaptic connection between the BNST-PBN.

**(D-E)** Numbers indicating optically responsive PBN neurons after **(D)** $BNST\text{-}PBN^{vGAT}$ or **(E)** $BNST\text{-}PBN^{vGLUT2}$ activation (x's represent non-responsive neurons).

**(F)** Viral schematic for anterograde tracing of $BNST\text{-}PBN^{vGAT}$.

**(G-K)** Terminal expression of ChR2-eYFP from $BNST^{vGAT}$ across the A-P axis of the BNST (scale bar: 200 μm).

**Figure S2. Validation of Cre-dependent retroTRAP. Related to Figure 1.**

**(A)** *In situ* hybridization images show coexpression of eGFP-TRAP with vGAT (*Slc31a1*) in BNST of vGAT-Cre animals (scale bar: 10 μm) and vGLUT2 (*Slc17a6*) in vGLUT2-Cre animals (scale bar: 20 μm) after injection of AAV2-retro-DIO-eGFP/RPL10 into PBN.

**(B)** Correlation of gene expression across all TRAP and input samples considered for analysis (n=3 vGLUT2-cre samples; n=2 vGAT-cre samples). Pearson correlation values are shown in the upper right-hand panel with sample identifiers along the diagonal.

(**C-E)** Volcano plot showing differential expression between **(C)** vGAT immunoprecipitated (IP) and pre-IP fraction, **(D)** vGLUT2 IP and pre-IP, and **(E)** vGAT IP and vGLUT2 IP. Each dot represents one gene; notable genes are labeled. X-axis shows log2(fold-change) of fold-change, with enrichment on right and depletion on left (**C** and **D**) and vGAT enrichment on right and vGLUT2 enrichment on left (**E**). Y-axis shows log of false discovery rate (FDR); area above horizontal line represents $p<0.05$

**(F-I)** Representative images for *in situ* hybridization of vGAT (*Slc31a1*) and vGLUT2 (*Slc17a6*) with **(F)** *Tac2*, **(G)** *Calcr*, **(H)** *Adcyap1*, and **(I)** *Sstr3* mRNA. Scale bars: 100 μm in main images; 10 μm in zoom-in.

**Figure S3. Monitoring BNST-PBN circuit activity. Related to Figure 2-4.**

**(A and B)** Transduced photon emissions in representative feeding session; excitation with 405 nm and 470 nm light in **(A)** vGAT and **(B)** vGLUT2 terminals**.**

**(C and D)** Subtracted signal (470 nm – 405 nm) in **(C)** vGAT and **(D)** vGLUT2 terminals in representative feeding session. Red line represents double exponential curve fit to signal.

**(E and F)** Baseline-corrected signal resulting from subtraction of double exponential curve in vGAT **(E)** and vGLUT2 **(F)** terminals in representative feeding session.

**(G and H)** Average z-scored calcium response of BNST-PBN$^{vGAT}$ terminals during consumption of normal chow after food deprivation (n=219 bouts; 7 mice) and averaged activity of 10-seconds pre- compared to post- consumption initiation over the testing period.

**Figure S4. An inhibitory BNST-PBN circuit that drives feeding. Related to Figure 5.**

**(A)** Schematic of optogenetic approach to target BNST-PBN$^{vGAT}$ and food consumption assays.

**(B-I)** If sated, BNST-PBN$^{vGAT}$ activation-elicited food consumption occurs regardless of the modality as shown by optogenetic activation compared to no-light within-subject controls as well as control mice.

**(J)** BNST-PBN$^{vGAT}$ activation-elicited food consumption is rapidly reversible.

**(K)** BNST-PBN$^{vGAT}$ activation does not significantly alter locomotion in the elevated zero maze, and **(L)** does not elicit anxiolysis when in a neutral environment.

**(M-O)** BNST-PBN$^{vGAT}$ activation **(M)** elicits anxiolysis, **(N)** suppresses novelty-induced fecal production, **(O)** but does not affect locomotion in the open field test.

**(P)** Schematic of optogenetic approach to target BNST-PBN$^{vGAT}$.

**(Q)** Schematic of food consumption assay.

**(R)** BNST-PBN$^{vGAT}$ inhibition suppresses food consumption of normal chow after food deprivation.

*$p < 0.05$, **$p < 0.01$, ***$p < 0.001$, ****$p < 0.0001$. Error bars indicate SEM.

**Figure S5. An excitatory BNST-PBN circuit that suppresses feeding. Related to Figure 5.**

**(A)** Schematic of optogenetic approach to target BNST-PBN$^{vGLUT2}$.

**(B)** Schematic of food consumption assays.

**(C-D)** BNST-PBN$^{vGLUT2}$ activation suppresses food consumption of normal chow after food deprivation but not in sated conditions.

**(E)** BNST-PBN$^{vGLUT2}$ activation rapidly elicits during activation and sustains food suppression after offset.

**(F)** BNST-PBN$^{vGLUT2}$ activation can guide negative-reinforcement learning in about 53% of mice determined to be ‘responders’.

*$p < 0.05$, **$p < 0.01$, ***$p < 0.001$, ****$p < 0.0001$. Error bars indicate SEM.

**Figure S6. PBN neurons receive input from the BNST to modulate feeding and affect. Related to Figure 6.**

**(A)** Schematic of optogenetic approach to target PBN$^{pDyn}$ with ChR2.

**(B)** PBN$^{pDyn}$ activation does not significantly alter locomotion at variable frequencies of stimulation.

**(C)** PBN$^{pDyn}$ activation suppresses food consumption of normal chow after food deprivation.

**(D)** Schematic of optogenetic approach to target PBN$^{vGLUT2}$ with Arch3.0.

**(E)** PBN$^{vGLUT2}$ inhibition elicits food consumption in a rapid and reversible manner.

*$p < 0.05$, ***$p < 0.001$. Error bars indicate SEM.

**Table S1.** Summary of statistical tests and results. **Related to all figures.**

**Table S2.** Differential gene expression levels between vGLUT2+ PBN-projecting BNST neurons and pre-IP background. **Related to Figure 1.**

**Table S3.** Differential gene expression levels between vGAT+ PBN-projecting BNST neurons and pre-IP background. **Related to Figure 1.**

**Table S4.** Differential gene expression levels between vGLUT2+ and vGAT+ PBN-projecting BNST neurons. **Related to Figure 1.**

**Table S5.** Differential gene expression levels between vGLUT2+ pre-IP and vGAT+ pre-IP background. **Related to Figure 1.**

### TABLE S1

| Figure | | Experimental Variables | Statistical Test | Results (Main Effect) | Post-hoc Test | Results (post-hoc) |
|---|---|---|---|---|---|---|
| 1 | C | Treatment (pharmacological agent) | Repeated measures one-way ANOVA | $F_{(1,4)}$=14.60; p=0.0185 | Tukey's multiple comparisons test | Baseline vs. DAPV/NBQX (p>0.9999). Baseline vs. Ptx/Bic (p=0.0382). DAPV/NBQX vs. Ptx/Bic (p=0.0420). n= 5. |
| | F | Treatment (pharmacological agent) | Repeated measures one-way ANOVA | $F_{(1,5)}$=23.93; p=0.0044 | Tukey's multiple comparisons test | Baseline vs. DAPV/NBQX (p=0.9976). Baseline vs. Ptx/Bic (p=0.0106). DAPV/NBQX vs. Ptx/Bic (p=0.0100). n= 6. |
| 2 | E | VGAT-cre :: 10s before/after feeding - chow (sated) | Paired two-tailed t-test | $t_{56}$=4.161; p=0.0001 | N/A | N/A |
| | G | VGAT-cre :: 10s before/after feeding - sucrose | Paired two-tailed t-test | $t_{92}$=9.676; p<0.0001 | N/A | N/A |
| | I | VGAT-cre :: 10s before/after feeding - high-fat | Paired two-tailed t-test | $t_{76}$=5.500; p<0.0001 | N/A | N/A |
| | K | VGAT-cre :: 10s before/after feeding - NSF | Paired two-tailed t-test | $t_{120}$=4.792; p<0.0001 | N/A | N/A |
| 3 | H | VGLUT2-cre :: 10s before/after feeding - chow (sated) | Paired two-tailed t-test | $t_{132}$=7.076; p<0.0001 | N/A | N/A |
| | I | VGLUT2-cre :: 10s before/after feeding - sucrose | Paired two-tailed t-test | $t_{87}$=3.312; p=0.0014 | N/A | N/A |
| | J | VGLUT2-cre :: 10s before/after feeding - high-fat | Paired two-tailed t-test | $t_{55}$=1.317; p=0.1932 | N/A | N/A |
| | K | VGLUT2-cre :: 10s before/after feeding - NSF | Paired two-tailed t-test | $t_{76}$=3.516; p=0.0007 | N/A | N/A |
| 4 | C | VGAT-cre :: 10s before/after shock | Paired two-tailed t-test | $t_{41}$=5.915; p<0.0001 | N/A | N/A |
| | F | VGLUT2-cre :: 2.5s before/after feeding | Paired two-tailed t-test | $t_{34}$=2.660; p=0.0118 | N/A | N/A |
| 5 | C | Chow: Ctrl [n=7] vs. ChR2 [n=8] | Unpaired two-tailed t-test | $t_{13}$=8.674; p<0.00001 | N/A | N/A |
| | | Food Dep:Ctrl [n=7] vs. ChR2 [n=8] | Unpaired two-tailed t-test | $t_{13}$=0.0324; p=0.9746 | N/A | N/A |
| | D | Chow: Ctrl [n=5] vs. ChR2 [n=7] | Unpaired two-tailed t-test | $t_{10}$=2.089; p=0.1223 | N/A | N/A |
| | | Food Dep: Ctrl [n=5] vs. ChR2 [n=7] | Unpaired two-tailed t-test | $t_{10}$=2.474; p=0.0329 | N/A | N/A |
| | F | Time [Baseline vs. 20 Hz] x Genotype [Ctrl, VGLUT-cre, VGAT-cre] | Repeated measures two-way ANOVA | F (2, 58) = 64.09; p<0.0001 | Bonferroni's multiple comparison test | Baseline: Ctrl [n=25] vs. vGAT [n=18], p>0.9999; Ctrl [n=25] vs. vGLUT2 [n=18], p>0.9999<br>20 Hz: Ctrl [n=25] vs. vGAT [n=18], p>0.0001; Ctrl [n=25] vs. vGLUT2 [n=18], p<0.0001 |
| | | Time [Baseline vs. 20 Hz] | | F (1, 58) = 3.563; p=0.0641 | | |
| | | Genotype [Ctrl, VGLUT-cre, VGAT-cre] | | F (2, 58) = 40.27; p<0.0001 | | |
| | | Subject | | F (58, 58) = 1.327; p=0.1422 | | |
| | H | Ctrl [n=6] vs. ChR2 [n=8] | Unpaired two-tailed t-test | $t_{12}$=2.811; p=0.0157 | N/A | N/A |
| | I | Ctrl [n=6] vs. ChR2 [n=8] | Unpaired two-tailed t-test | $t_{18}$=2.684; p=0.0151 | N/A | N/A |

| | | | | | | |
|---|---|---|---|---|---|---|
| | J | Ctrl [n=6] vs. ChR2 [n=8] | Unpaired two-tailed t-test | $t_{12}$=2.480; p=0.0290 | N/A | N/A |
| | K | Ctrl [n=9] vs. ChR2 [n=8] | Unpaired two-tailed t-test | $t_{15}$=2.608; p=0.0198 | N/A | N/A |
| | M | CS trails on Extinction day: Ctrl [n=6] vs. ChR2 [n=8] | Unpaired two-tailed t-test | $t_{12}$=16.54; p<0.0001 | N/A | N/A |
| 6 | I | Time [Frequency] x Genotype [Ctrl vs. ChR2] | Repeated measures two-way ANOVA | F (4, 48) = 7.725; p<0.0001 | Bonferroni's multiple comparison test | Ctrl [n=6] vs. ChR2 [n=8]: 0 Hz, p>0.9999; 5 Hz, p=0.008; 10 Hz, p=0.0156; 20 Hz, p=0.0063; 40 Hz, p=0.0043 |
| | | Time [Frequency] | | F (2.123, 25.47) = 7.919; p=0.0018 | | |
| | | Genotype [Ctrl vs. ChR2] | | F (1, 12) = 21.01; p=0.0006 | | |
| | | Subject | | F (12, 48) = 3.580; p=0.0008 | | |
| | J | Time [Baseline vs. 20 Hz] x Genotype [Ctrl vs. ChR2] | Repeated measures two-way ANOVA | F (1, 12) = 12.86; p=0.0037 | Bonferroni's multiple comparison test | Ctrl [n=6] vs. ChR2 [n=8]: Baseline, p=0.7518; 20 Hz, p=0.0001 |
| | | Time [Baseline vs. 20 Hz] | | F (1, 12) = 6.117; p=0.0293 | | |
| | | Genotype [Ctrl vs. ChR2] | | F (1, 12) = 10.11; p=0.0079 | | |
| | | Subject | | F (12, 12) = 0.7174; p=0.7130 | | |
| | K | Ctrl [n=6] vs. ChR2 [n=9] | Unpaired two-tailed t-test | $t_{13}$=2.225 p=0.0444 | N/A | N/A |
| | N | Ctrl [n=6] vs. DREADD [n=8] | Unpaired two-tailed t-test | $t_{12}$=2.306; p=0.0398 | N/A | N/A |
| | O | Ctrl [n=7] vs. DREADD [n=8] | Unpaired two-tailed t-test | $t_{13}$=2.287; p=0.0396 | N/A | N/A |
| | P | Ctrl [n=7] vs. DREADD [n=7] | Unpaired two-tailed t-test | $t_{13}$=3.600; p=0.0032 | N/A | N/A |
| | Q | Ctrl [n=7] vs. DREADD [n=7] | Unpaired two-tailed t-test | $t_{13}$=2.088 p=0.0570 | N/A | N/A |
| S3 | I | feeding - chow (food deprived) | Paired two-tailed t-test | $t_{218}$=3.006; p=0.0030 | N/A | N/A |
| | B | Time [OFF vs. ON] x Genotype [Ctrl vs. ChR2] | Repeated measures two-way ANOVA | F (1, 13) = 75.23; p<0.0001 | Bonferroni's multiple comparison test | Ctrl [n=7] vs. ChR2 [n=8]: OFF, p>0.9999;ON, p<0.0001 |
| | | Time [OFF vs. ON] | | F (1, 13) = 62.49; p<0.0001 | | |
| | | Genotype [Ctrl vs. ChR2] | | F (1, 13) = 43.07; p<0.0001 | | |
| | | Subject | | F (13, 13) = 1.926; p=0.1252 | | |
| | C | Time [OFF vs. ON] x Genotype [Ctrl vs. ChR2] | Repeated measures two-way ANOVA | F (1, 13) = 0.0003658; p=0.9850 | Bonferroni's multiple comparison test | Ctrl [n=7] vs. ChR2 [n=8]: OFF, p=0.9068; ON, p=0.9282 |
| | | Time [OFF vs. ON] | | F (1, 13) = 0.8081; p=0.3850 | | |
| | | Genotype [Ctrl vs. ChR2] | | F (1, 13) = 0.7293; p=0.4086 | | |
| | | Subject | | F (13, 13) = 3.456; p=0.0166 | | |
| | D | Time [OFF vs. ON] x Genotype [Ctrl vs. ChR2] | Repeated measures two-way ANOVA | F (1, 13) = 145.2; p<0.0001 | Bonferroni's multiple comparison test | Ctrl [n=7] vs. ChR2 [n=8]: OFF, p=0.6762; ON, p<0.0001 |
| | | Time [OFF vs. ON] | | F (1, 13) = 154.8; p<0.0001 | | |

| | | | | | | |
|---|---|---|---|---|---|---|
| | | Genotype [Ctrl vs. ChR2] | | F (1, 13) = 107.1; p<0.0001 | | |
| | | Subject | | F (13, 13) = 1.743; p=0.1644 | | |
| | E | Time [OFF vs. ON] x Genotype [Ctrl vs. ChR2] | Repeated measures two-way ANOVA | F (1, 13) = 12.53; p=0.0036 | Bonferroni's multiple comparison test | Ctrl [n=6] vs. ChR2 [n=8]: OFF, p>0.9999;ON, p=0.0002 |
| | | Time [OFF vs. ON] | | F (1, 13) = 14.00; p=0.0025 | | |
| | | Genotype [Ctrl vs. ChR2] | | F (1, 13) = 9.191; p=0.0096 | | |
| | | Subject | | F (13, 13) = 1.664; p=0.1850 | | |
| | F | Time [OFF vs. ON] x Genotype [Ctrl vs. ChR2] | Repeated measures two-way ANOVA | F (1, 13) = 78.71; p<0.0001 | Bonferroni's multiple comparison test | Ctrl [n=6] vs. ChR2 [n=8]: OFF, p=0.6268;ON, p<0.0001 |
| | | Time [OFF vs. ON] | | F (1, 13) = 85.31; p<0.0001 | | |
| | | Genotype [Ctrl vs. ChR2] | | F (1, 13) = 28.15; p=0.0001 | | |
| | | Subject | | F (13, 13) = 1.814; p=0.1478 | | |
| S4 | G | Time [OFF vs. ON] x Genotype [Ctrl vs. ChR2] | Repeated measures two-way ANOVA | F (1, 13) = 34.64; p<0.0001 | Bonferroni's multiple comparison test | Ctrl [n=7] vs. ChR2 [n=8]: OFF, p>0.9999;ON, p<0.0001 |
| | | Time [OFF vs. ON] | | F (1, 13) = 34.17; p<0.0001 | | |
| | | Genotype [Ctrl vs. ChR2] | | F (1, 13) = 65.66; p<0.0001 | | |
| | | Subject | | F (13, 13) = 0.6834; p=0.7490 | | |
| | H | Time [OFF vs. ON] x Genotype [Ctrl vs. ChR2] | Repeated measures two-way ANOVA | F (1, 13) = 20.48; p=0.0006 | Bonferroni's multiple comparison test | Ctrl [n=6] vs. ChR2 [n=8]: OFF, p>0.9999;ON, p<0.0001 |
| | | Time [OFF vs. ON] | | F (1, 13) = 28.54; p=0.0001 | | |
| | | Genotype [Ctrl vs. ChR2] | | F (1, 13) = 20.34; p=0.0006 | | |
| | | Subject | | F (13, 13) = 1.496; p=0.2389 | | |
| | I | Time [OFF vs. ON] x Genotype [Ctrl vs. ChR2] | Repeated measures two-way ANOVA | F (1, 13) = 20.01; p=0.0006 | Bonferroni's multiple comparison test | Ctrl [n=6] vs. ChR2 [n=8]: OFF, p>9999;ON, p<0.0001 |
| | | Time [OFF vs. ON] | | F (1, 13) = 26.33; p=0.0002 | | |
| | | Genotype [Ctrl vs. ChR2] | | F (1, 13) = 20.44; p=0.0006 | | |
| | | Subject | | F (13, 13) = 1.408; p=0.2732 | | |
| | J | Time [Pre vs. ON vs. Post] x Genotype [Ctrl vs. ChR2] | Repeated measures two-way ANOVA | F (2, 24) = 4.666; p=0.0194 | Bonferroni's multiple comparison test | Ctrl [n=5] vs. ChR2 [n=9]: Pre, p>0.9999; ON, p=0.0037; Post, p>0.9999 |
| | | Time [Pre vs. ON vs. Post] | | F (2, 24) = 3.431; p=0.0489 | | |
| | | Genotype [Ctrl vs. ChR2] | | F (1, 12) = 3.118; p=0.1028 | | |
| | | Subject | | F (12, 24) = 1.084; p=0.4143 | | |
| | K | Ctrl [n=6] vs. ChR2 [n=8] | Unpaired two-tailed t-test | $t_{12}$=1.418; p=0.1818 | N/A | N/A |
| | L | Ctrl [n=6] vs. ChR2 [n=7] | Unpaired two-tailed t-test | $t_{11}$=1.612; p=0.1352 | N/A | N/A |

| | | | | | | |
|---|---|---|---|---|---|---|
| | M | Ctrl [n=5] vs. ChR2 [n=5] | Unpaired two-tailed t-test | $t_8$=3.021; p=0.0165 | N/A | N/A |
| | N | Ctrl [n=5] vs. ChR2 [n=5] | Unpaired two-tailed t-test | $t_8$=2.799; p=0.0232 | N/A | N/A |
| | O | Ctrl [n=5] vs. ChR2 [n=5] | Unpaired two-tailed t-test | $t_8$=0.7674; p=0.4649 | N/A | N/A |
| | R | Ctrl [n=9] vs. Arch3.0 [n=10] | Unpaired two-tailed t-test | $t_{19}$=3.257; p=0.0041 | N/A | N/A |
| S5 | C | Time [OFF vs. ON] x Genotype [Ctrl vs. ChR2] | Repeated measures two-way ANOVA | F (1, 10) = 0.6161; p=0.4507 | Bonferroni's multiple comparison test | Ctrl [n=6] vs. ChR2 [n=8]: OFF, p>0.9999; ON, p=0.3190 |
| | | Time [OFF vs. ON] | | F (1, 10) = 1.386; p=0.2663 | | |
| | | Genotype [Ctrl vs. ChR2] | | F (1, 10) = 3.225; p=0.1027 | | |
| | | Subject | | F (10, 10) = 0.9670; p=0.5206 | | |
| | D | Time [OFF vs. ON] x Genotype [Ctrl vs. ChR2] | Repeated measures two-way ANOVA | F (1, 10) = 17.21; p=0.0020 | Bonferroni's multiple comparison test | Ctrl [n=6] vs. ChR2 [n=8]: OFF, p>0.9999;ON, p=0.0003 |
| | | Time [OFF vs. ON] | | F (1, 10) = 12.67; p=0.0052 | | |
| | | Genotype [Ctrl vs. ChR2] | | F (1, 10) = 1.036; p=0.3328 | | |
| | | Subject | | F (10, 10) = 9.796; p=0.0006 | | |
| | E | Time [Pre vs. ON vs. Post] x Genotype [Ctrl vs. ChR2] | Repeated measures two-way ANOVA | F (2, 22) = 4.085; p=0.0310 | Bonferroni's multiple comparison test | Ctrl [n=6] vs. ChR2 [n=7]: Pre, p>0.9999; ON, p=0.0276; Post, p=0.1499 |
| | | Time [Pre vs. ON vs. Post] | | F (1.699, 18.69) = 17.32; p<0.0001 | | |
| | | Genotype [Ctrl vs. ChR2] | | F (1, 11) = 4.369; p=0.0606 | | |
| | | Subject | | F (11, 22) = 1.277; p=0.2997 | | |
| S6 | C | Time [OFF vs. ON] x Genotype [Ctrl vs. ChR2] | Repeated measures two-way ANOVA | F (1, 13) = 4.948; p = 0.0444 | Bonferroni's multiple comparison test | ON vs. OFF: Ctrl [n=6], p=0.3914; ChR2 [n=9], p=0.0003 |
| | | Time [OFF vs. ON] | | F (1, 13) = 18.82; p =0.0008 | | |
| | | Genotype [Ctrl vs. ChR2] | | F (1, 13) = 1.817; p =0.2007 | | |
| | | Subject | | F (13, 13) = 2.248; p =0.0787 | | |
| | E | Time [Pre vs. ON vs. Post] x Genotype [Ctrl vs. Arch3.0] | Repeated measures two-way ANOVA | F (2, 16) = 2.919; p=0.0830 | Bonferroni's multiple comparison test | Ctrl [n=5] : Pre vs. Stim, p=0.3982; Stim vs. Post, p>0.9999; Arch3.0 [n=5] : Pre vs. |
| | | Time [Pre vs. ON vs. Post] | | F (2, 16) = 6.777; p=0.0074 | | |
| | | Genotype [Ctrl vs. Arch3.0] | | F (1, 8) = 0.8151; p=0.3930 | | |
| | | Subject | | F (8, 16) = 0.7539; p=0.6461 | | |
