## Supplemental Table 2 for "Extended amygdala-parabrachial circuits alter threat assessment to regulate feeding"

### TABLE S2

| ensembl_gene_id | external_gene_name | gene_biotype | log2 (fold change) | log2 (counts per million) | F-value | p-value | False discovery rate |
|---|---|---|---|---|---|---|---|
| ENSMUSG00000045471 | Hcrt | protein_coding | 5.961539218 | 10.01304624 | 92.819961 | 7.92E-07 | 0.005231196 |
| ENSMUSG00000109006 | B230209E15Rik | lincRNA | -1.889528185 | 6.39717013 | 56.907319 | 1.05E-06 | 0.005231196 |
| ENSMUSG00000019146 | Cacng2 | protein_coding | -2.103369497 | 5.129436191 | 54.01537 | 1.46E-06 | 0.005231196 |
| ENSMUSG00000017002 | Slpi | protein_coding | 5.557046254 | 2.919325958 | 49.486308 | 2.53E-06 | 0.006813198 |
| ENSMUSG00000041215 | Yeats2 | protein_coding | 3.958885353 | 5.218110279 | 49.185953 | 5.48E-06 | 0.01178542 |
| ENSMUSG00000064345 | mt-Nd2 | protein_coding | -2.18060363 | 10.31322046 | 41.528864 | 9.98E-06 | 0.017889353 |
| ENSMUSG00000064367 | mt-Nd5 | protein_coding | -1.977261229 | 9.814911841 | 39.410444 | 1.24E-05 | 0.019109762 |
| ENSMUSG00000064368 | mt-Nd6 | protein_coding | -1.978309921 | 8.052755857 | 34.148379 | 3.40E-05 | 0.045682316 |
| ENSMUSG00000064341 | mt-Nd1 | protein_coding | -1.923330515 | 12.02550762 | 32.021347 | 3.87E-05 | 0.046268722 |
| ENSMUSG00000064363 | mt-Nd4 | protein_coding | -1.77555009 | 10.34042568 | 30.470552 | 5.33E-05 | 0.057336021 |
| ENSMUSG00000064370 | mt-Cytb | protein_coding | -2.021979438 | 12.19260763 | 29.368155 | 7.01E-05 | 0.067018444 |
| ENSMUSG00000070576 | Mn1 | protein_coding | 1.938792132 | 8.317535647 | 29.884754 | 7.48E-05 | 0.067018444 |
| ENSMUSG00000030314 | Atg7 | protein_coding | 2.232503695 | 6.424224491 | 28.368315 | 0.0001059 | 0.072937956 |
| ENSMUSG00000037752 | Xkr8 | protein_coding | 3.322685943 | 2.798958383 | 25.67817 | 0.0001079 | 0.072937956 |
| ENSMUSG00000062252 | Lhfpl5 | protein_coding | 1.142537676 | 6.03676927 | 25.492658 | 0.000112 | 0.072937956 |
| ENSMUSG00000044005 | Gls2 | protein_coding | 2.218000516 | 3.5291459 | 25.425825 | 0.0001135 | 0.072937956 |
| ENSMUSG00000032604 | Qars | protein_coding | 1.336772871 | 5.644562786 | 25.092539 | 0.0001214 | 0.072937956 |
| ENSMUSG00000023064 | Sncg | protein_coding | 1.333912507 | 7.292200882 | 25.089922 | 0.0001221 | 0.072937956 |
| ENSMUSG00000028565 | Nfia | protein_coding | -1.229596859 | 6.998809161 | 23.392284 | 0.0001738 | 0.098344256 |
| ENSMUSG00000054162 | Spock3 | protein_coding | -0.930462652 | 7.329591639 | 23.120997 | 0.0001833 | 0.09855147 |
| ENSMUSG00000100826 | Snhg14 | processed_transcript | -1.156067892 | 6.081352785 | 22.453826 | 0.0002117 | 0.108425515 |
| ENSMUSG00000018451 | 6330403K07Rik | lincRNA | 0.932675329 | 10.97421339 | 21.814636 | 0.0002437 | 0.114655077 |
| ENSMUSG00000003410 | Elavl3 | protein_coding | -0.908156611 | 6.897769516 | 21.787066 | 0.0002452 | 0.114655077 |
| ENSMUSG00000092341 | Malat1 | lincRNA | -1.164653425 | 10.80327234 | 21.141829 | 0.0002834 | 0.124109993 |
| ENSMUSG00000042063 | Zfp386 | protein_coding | -1.450257622 | 5.182603598 | 21.062809 | 0.0002885 | 0.124109993 |
| ENSMUSG00000028161 | Ppp3ca | protein_coding | -0.82278238 | 9.571254196 | 20.444843 | 0.0003324 | 0.137465862 |
| ENSMUSG00000099696 | 2900052N01Rik | lincRNA | -1.115542686 | 5.699290823 | 20.079575 | 0.0003618 | 0.137495976 |
| ENSMUSG00000071267 | Zfp942 | protein_coding | -2.092440385 | 3.31002476 | 19.890467 | 0.0003782 | 0.137495976 |
| ENSMUSG00000063406 | Tmed5 | protein_coding | -1.431898104 | 5.217522506 | 19.83027 | 0.0003836 | 0.137495976 |
| ENSMUSG00000022091 | Sorbs3 | protein_coding | -1.324175157 | 4.802148404 | 19.829714 | 0.0003836 | 0.137495976 |
| ENSMUSG00000079184 | Mphosph8 | protein_coding | -0.875707485 | 6.514185979 | 19.386946 | 0.000426 | 0.142685713 |
| ENSMUSG00000071646 | Mta2 | protein_coding | 1.565862124 | 4.029628737 | 19.295658 | 0.0004354 | 0.142685713 |
| ENSMUSG00000029245 | Epha5 | protein_coding | -1.359040619 | 5.230722747 | 18.924697 | 0.0004759 | 0.142685713 |
| ENSMUSG00000100252 | Mir124-2hg | lincRNA | -1.183593651 | 5.424679353 | 18.895917 | 0.0004793 | 0.142685713 |
| ENSMUSG00000111080 | AC153899.1 | lincRNA | -1.098165359 | 5.911816474 | 18.850914 | 0.0004845 | 0.142685713 |
| ENSMUSG00000044308 | Ubr3 | protein_coding | 1.346556196 | 6.311988584 | 18.766554 | 0.0005009 | 0.142685713 |
| ENSMUSG00000059897 | Zfp930 | protein_coding | -1.981625328 | 3.991080406 | 18.698262 | 0.0005028 | 0.142685713 |
| ENSMUSG00000037204 | Atg101 | protein_coding | 0.8088889 | 6.702083432 | 18.686458 | 0.0005042 | 0.142685713 |
| ENSMUSG00000010554 | Mettl16 | protein_coding | 0.949897369 | 6.035521682 | 18.489153 | 0.0005291 | 0.145886395 |
| ENSMUSG00000022124 | Fbxl3 | protein_coding | -1.187532994 | 5.126446197 | 18.162058 | 0.0005735 | 0.154161439 |
| ENSMUSG00000020850 | Prpf8 | protein_coding | 0.931482094 | 6.328230262 | 18.059021 | 0.0005883 | 0.154289846 |
| ENSMUSG00000048495 | Tyw5 | protein_coding | -1.239983119 | 4.875694732 | 17.832764 | 0.0006224 | 0.156091012 |
| ENSMUSG00000035776 | Cd99l2 | protein_coding | 0.759867013 | 7.23010765 | 17.738974 | 0.0006372 | 0.156091012 |
| ENSMUSG00000066148 | Prpf4 | protein_coding | 1.354820105 | 4.626979966 | 17.729364 | 0.0006387 | 0.156091012 |
| ENSMUSG00000034647 | Ankrd12 | protein_coding | -1.241463168 | 6.328542907 | 17.529461 | 0.0006738 | 0.159748722 |
| ENSMUSG00000031246 | Sh3bgrl | protein_coding | -1.052090787 | 7.149897123 | 17.473743 | 0.0006834 | 0.159748722 |
| ENSMUSG00000031545 | Gpat4 | protein_coding | 1.112052633 | 6.547037953 | 16.893206 | 0.0007924 | 0.170835104 |
| ENSMUSG00000064339 | mt-Rnr2 | Mt_rRNA | -1.806071287 | 13.83599611 | 17.867885 | 0.0007945 | 0.170835104 |
| ENSMUSG00000021719 | Rgs7bp | protein_coding | -0.754433501 | 7.127353161 | 16.849876 | 0.0007988 | 0.170835104 |
| ENSMUSG00000024109 | Nrxn1 | protein_coding | -0.930722987 | 8.156305757 | 16.798593 | 0.0008094 | 0.170835104 |
| ENSMUSG00000028456 | Unc13b | protein_coding | -1.027348359 | 5.591983742 | 16.79468 | 0.0008102 | 0.170835104 |
| ENSMUSG00000038602 | Slc35f1 | protein_coding | -1.150474007 | 5.437663363 | 16.619345 | 0.0008479 | 0.17513299 |
| ENSMUSG00000056073 | Grik2 | protein_coding | -1.169247098 | 5.420068935 | 16.521358 | 0.0008698 | 0.17513299 |
| ENSMUSG00000038872 | Zfhx3 | protein_coding | -0.980141363 | 6.113214942 | 16.478966 | 0.0008795 | 0.17513299 |
| ENSMUSG00000025860 | Xiap | protein_coding | -0.894781849 | 6.136989412 | 16.371388 | 0.0009046 | 0.176855962 |
| ENSMUSG00000058589 | Anks1b | protein_coding | -0.691046937 | 7.130867999 | 16.295422 | 0.0009228 | 0.177194579 |
| ENSMUSG00000035847 | Ids | protein_coding | -0.880837505 | 6.718583316 | 16.071934 | 0.0009788 | 0.183637045 |
| ENSMUSG00000064351 | mt-Co1 | protein_coding | -1.536660754 | 11.50948277 | 16.662833 | 0.0010083 | 0.183637045 |
| ENSMUSG00000064337 | mt-Rnr1 | Mt_rRNA | -1.986141519 | 10.86487464 | 17.928622 | 0.0010244 | 0.183637045 |
| ENSMUSG00000018339 | Gpx3 | protein_coding | 0.944954563 | 9.533675749 | 15.899495 | 0.0010247 | 0.183637045 |
| ENSMUSG00000033629 | Hacd3 | protein_coding | 0.736983943 | 7.534286954 | 15.704409 | 0.0010795 | 0.18472957 |
| ENSMUSG00000042761 | Mrap2 | protein_coding | 1.131996216 | 5.658619907 | 15.647215 | 0.0010962 | 0.18472957 |
| ENSMUSG00000020462 | Cfap36 | protein_coding | -0.763190892 | 7.174457757 | 15.64252 | 0.0010975 | 0.18472957 |
| ENSMUSG00000020029 | Nudt4 | protein_coding | -0.858943942 | 6.206252499 | 15.635981 | 0.0010995 | 0.18472957 |
| ENSMUSG00000071669 | Snx29 | protein_coding | -1.862607417 | 3.290606917 | 15.567317 | 0.00112 | 0.185277692 |
| ENSMUSG00000060988 | Galnt13 | protein_coding | -1.173063033 | 5.405734543 | 15.464828 | 0.0011514 | 0.187585254 |
| ENSMUSG00000036198 | Arhgap36 | protein_coding | 1.154936923 | 5.900011865 | 15.167691 | 0.0012481 | 0.200316934 |
| ENSMUSG00000065979 | Cpped1 | protein_coding | -0.978373174 | 6.910962486 | 15.056047 | 0.0012868 | 0.202382216 |
| ENSMUSG00000050587 | Lrrc4c | protein_coding | -0.71648463 | 6.84101994 | 15.02275 | 0.0012986 | 0.202382216 |
| ENSMUSG00000050711 | Scg2 | protein_coding | 0.752704438 | 9.620289236 | 14.968671 | 0.0013181 | 0.202476295 |
| ENSMUSG00000030062 | Rpn1 | protein_coding | 0.694965372 | 6.965638885 | 14.823177 | 0.001372 | 0.207761233 |
| ENSMUSG00000042133 | Ppig | protein_coding | -0.911915843 | 6.474703558 | 14.738589 | 0.0014045 | 0.207761233 |
| ENSMUSG00000003279 | Dlgap1 | protein_coding | -0.839302299 | 6.176423712 | 14.677898 | 0.0014284 | 0.207761233 |
| ENSMUSG00000022089 | Bin3 | protein_coding | 1.480181584 | 3.915490179 | 14.674355 | 0.0014298 | 0.207761233 |

| ensembl_gene_id | external_gene_name | gene_biotype | log2 (fold change) | log2 (counts per million) | F-value | p-value | False discovery rate |
|---|---|---|---|---|---|---|---|
| ENSMUSG00000050148 | Ubqln2 | protein_coding | -0.726484904 | 6.796164998 | 14.438457 | 0.001527 | 0.218935225 |
| ENSMUSG00000027502 | Rtfdc1 | protein_coding | 0.798426468 | 7.356181442 | 14.333549 | 0.0015727 | 0.222511062 |
| ENSMUSG00000026384 | Ptpn4 | protein_coding | -0.919048291 | 5.596980002 | 14.2662 | 0.0016028 | 0.223825667 |
| ENSMUSG00000064372 | mt-Tp | Mt_tRNA | -1.51194388 | 4.066601396 | 14.098728 | 0.0016805 | 0.231670966 |
| ENSMUSG00000074746 | Pdzd8 | protein_coding | -0.8833236 | 5.42275172 | 14.02451 | 0.0017163 | 0.233610997 |
| ENSMUSG00000055373 | Fut9 | protein_coding | -1.06531092 | 5.562331203 | 13.803636 | 0.001828 | 0.245709036 |
| ENSMUSG00000021536 | Adcy2 | protein_coding | 1.133567898 | 5.192257329 | 13.558775 | 0.0019616 | 0.260407666 |
| ENSMUSG00000038871 | Bpgm | protein_coding | 0.704825736 | 7.396211372 | 13.442582 | 0.0020288 | 0.26373041 |
| ENSMUSG00000024293 | Esco1 | protein_coding | -1.10709552 | 4.417822002 | 13.430953 | 0.0020357 | 0.26373041 |
| ENSMUSG00000015837 | Sqstm1 | protein_coding | 0.753716548 | 8.745918331 | 13.169706 | 0.0021971 | 0.278629392 |
| ENSMUSG00000021423 | Ly86 | protein_coding | -1.41954538 | 4.544423758 | 13.161383 | 0.0022025 | 0.278629392 |
| ENSMUSG00000031425 | Plp1 | protein_coding | -0.807559671 | 10.61790917 | 13.101221 | 0.0022418 | 0.280304155 |
| ENSMUSG00000057982 | Zfp809 | protein_coding | -1.398217991 | 4.043580567 | 12.993569 | 0.0023141 | 0.284252356 |
| ENSMUSG00000034707 | Gns | protein_coding | 1.086417268 | 5.097607033 | 12.975902 | 0.0023263 | 0.284252356 |
| ENSMUSG00000025766 | D3Ertd751e | protein_coding | -1.328122746 | 4.628732869 | 12.892082 | 0.0023847 | 0.284538166 |
| ENSMUSG00000024150 | Mcfd2 | protein_coding | 0.690112939 | 7.672831725 | 12.849056 | 0.0024154 | 0.284538166 |
| ENSMUSG00000030095 | Tmem43 | protein_coding | 0.904586055 | 5.636489023 | 12.789455 | 0.0024586 | 0.284538166 |
| ENSMUSG00000039662 | Icmt | protein_coding | 1.15124117 | 4.319494949 | 12.707 | 0.0025199 | 0.284538166 |
| ENSMUSG00000046699 | Slitrk4 | protein_coding | -1.1475226 | 4.671661976 | 12.687489 | 0.0025346 | 0.284538166 |
| ENSMUSG00000028282 | Casp8ap2 | protein_coding | -1.395020657 | 3.979270051 | 12.661975 | 0.002554 | 0.284538166 |
| ENSMUSG00000003360 | Ddx23 | protein_coding | 1.465376287 | 3.709420481 | 12.64836 | 0.0025645 | 0.284538166 |
| ENSMUSG00000025607 | Copg2 | protein_coding | -0.89780249 | 7.752464664 | 12.641248 | 0.0025756 | 0.284538166 |
| ENSMUSG00000029126 | Nsg1 | protein_coding | 0.688748994 | 9.756698452 | 12.614361 | 0.0025907 | 0.284538166 |
| ENSMUSG00000043388 | Tmem130 | protein_coding | 0.796887028 | 9.615115595 | 12.596461 | 0.0026047 | 0.284538166 |
| ENSMUSG00000028176 | Lrrc7 | protein_coding | -1.389102799 | 4.433184604 | 12.573472 | 0.0026227 | 0.284538166 |
| ENSMUSG00000031931 | Ankrd49 | protein_coding | -1.042422102 | 4.469545221 | 12.527022 | 0.0026596 | 0.284538166 |
| ENSMUSG00000029193 | Cckar | protein_coding | 2.183339295 | 3.892365771 | 12.791567 | 0.0026752 | 0.284538166 |
| ENSMUSG00000023832 | Acat2 | protein_coding | 0.856398733 | 6.774164154 | 12.478151 | 0.0026991 | 0.284538166 |
| ENSMUSG00000025616 | Usp16 | protein_coding | -1.062012818 | 4.813608305 | 12.351764 | 0.0028042 | 0.288869471 |
| ENSMUSG00000042472 | Zfp410 | protein_coding | 1.013638593 | 5.151333987 | 12.315201 | 0.0028354 | 0.288869471 |
| ENSMUSG00000038174 | Fam126b | protein_coding | -0.760162189 | 6.424129411 | 12.309982 | 0.0028399 | 0.288869471 |
| ENSMUSG00000021555 | Naa35 | protein_coding | 0.703916683 | 6.4186992 | 12.301091 | 0.0028476 | 0.288869471 |
| ENSMUSG00000048965 | Mrgpre | protein_coding | 1.291099118 | 4.43760468 | 12.146978 | 0.0029844 | 0.296220378 |
| ENSMUSG00000034083 | Ccdc174 | protein_coding | -0.757923066 | 5.939229856 | 12.125562 | 0.003004 | 0.296220378 |
| ENSMUSG00000025272 | Tro | protein_coding | 0.876019412 | 7.082353565 | 12.104661 | 0.0030296 | 0.296220378 |
| ENSMUSG00000022759 | Lrrc74b | protein_coding | -1.754380456 | 3.16521939 | 12.089256 | 0.0030376 | 0.296220378 |
| ENSMUSG00000040483 | Xaf1 | protein_coding | -1.226450055 | 4.266145799 | 12.053258 | 0.0030713 | 0.296220378 |
| ENSMUSG00000062078 | Qk | protein_coding | -0.738567769 | 7.886346701 | 12.02422 | 0.0030988 | 0.296220378 |
| ENSMUSG00000022407 | Adsl | protein_coding | 0.899917049 | 6.061447949 | 12.009411 | 0.0031129 | 0.296220378 |
| ENSMUSG00000046138 | 9930021J03Rik | protein_coding | -1.153713038 | 5.848909932 | 11.957029 | 0.0031699 | 0.296929628 |
| ENSMUSG00000064037 | Gpn1 | protein_coding | 0.6680545 | 6.995836777 | 11.941269 | 0.0031788 | 0.296929628 |
| ENSMUSG00000009647 | Mcu | protein_coding | -0.788404164 | 6.260621807 | 11.896581 | 0.0032229 | 0.296929628 |
| ENSMUSG00000035199 | Arl6ip5 | protein_coding | 0.606150128 | 8.05023803 | 11.888674 | 0.0032308 | 0.296929628 |
| ENSMUSG00000038039 | Gcc2 | protein_coding | -0.8207073 | 5.948137453 | 11.80369 | 0.0033168 | 0.299848689 |
| ENSMUSG00000023960 | Enpp5 | protein_coding | 0.58339339 | 7.834927433 | 11.796334 | 0.0033244 | 0.299848689 |
| ENSMUSG00000054976 | Nyap2 | protein_coding | -1.128258746 | 4.313598731 | 11.768334 | 0.0033534 | 0.299848689 |
| ENSMUSG00000005936 | Kctd20 | protein_coding | 1.006606344 | 5.242159555 | 11.745145 | 0.0033776 | 0.299848689 |
| ENSMUSG00000032740 | Ccdc88a | protein_coding | -0.956299787 | 5.344034453 | 11.713444 | 0.003411 | 0.299848689 |
| ENSMUSG00000044528 | Tram1l1 | protein_coding | 0.631485178 | 6.654881795 | 11.695715 | 0.0034299 | 0.299848689 |
| ENSMUSG00000026456 | Cyb5r1 | protein_coding | 1.0200914 | 5.561723233 | 11.49058 | 0.0036569 | 0.315861417 |
| ENSMUSG00000036188 | Ankmy2 | protein_coding | 0.736401493 | 6.939125546 | 11.477605 | 0.0036718 | 0.315861417 |
| ENSMUSG00000025583 | Rptor | protein_coding | 0.957012875 | 4.777241995 | 11.398366 | 0.0037644 | 0.321258875 |
| ENSMUSG00000015176 | Nolc1 | protein_coding | 0.704097635 | 6.376195778 | 11.34965 | 0.0038227 | 0.323661386 |
| ENSMUSG00000056962 | Jmjd6 | protein_coding | 0.871535139 | 5.364287745 | 11.324808 | 0.0038528 | 0.323661386 |
| ENSMUSG00000043102 | Qrfp | protein_coding | 5.001459747 | 2.17994674 | 11.461943 | 0.0039735 | 0.329656574 |
| ENSMUSG00000004945 | Tmem242 | protein_coding | 0.650722855 | 6.377803356 | 11.217856 | 0.0039854 | 0.329656574 |
| ENSMUSG00000030083 | Abtb1 | protein_coding | 1.142043503 | 6.035089861 | 11.3004 | 0.0040239 | 0.330013136 |
| ENSMUSG00000087174 | 5530601H04Rik | processed_transcript | -0.881174295 | 5.187209242 | 11.125626 | 0.004104 | 0.330013136 |
| ENSMUSG00000053963 | Stum | protein_coding | -1.000043249 | 4.746367882 | 11.122797 | 0.0041077 | 0.330013136 |
| ENSMUSG00000024976 | Shoc2 | protein_coding | -0.897089603 | 8.430827714 | 11.125003 | 0.0041125 | 0.330013136 |
| ENSMUSG00000030036 | Mogs | protein_coding | 1.825851337 | 3.324228333 | 11.034651 | 0.0042249 | 0.330213317 |
| ENSMUSG00000071533 | Pcnp | protein_coding | -0.621021203 | 8.028674101 | 11.030542 | 0.0042305 | 0.330213317 |
| ENSMUSG00000027895 | Kcnc4 | protein_coding | -0.924728437 | 5.077709349 | 10.996105 | 0.0042773 | 0.330213317 |
| ENSMUSG00000030846 | Tial1 | protein_coding | -0.792183721 | 6.111302703 | 10.993509 | 0.0042809 | 0.330213317 |
| ENSMUSG00000024256 | Adcyap1 | protein_coding | 2.140956978 | 3.763489644 | 11.211114 | 0.0042921 | 0.330213317 |
| ENSMUSG00000028522 | Mier1 | protein_coding | -0.837422562 | 5.329494853 | 10.932481 | 0.0043655 | 0.330213317 |
| ENSMUSG00000061751 | Kalrn | protein_coding | 0.837155934 | 5.524769988 | 10.918336 | 0.0043854 | 0.330213317 |
| ENSMUSG00000002107 | Celf2 | protein_coding | -0.659496841 | 7.824730066 | 10.90344 | 0.0044064 | 0.330213317 |
| ENSMUSG00000027374 | Mrps5 | protein_coding | 0.783964886 | 5.855082703 | 10.894462 | 0.0044192 | 0.330213317 |
| ENSMUSG00000026830 | Ermn | protein_coding | -0.587458048 | 7.384948842 | 10.892399 | 0.0044221 | 0.330213317 |
| ENSMUSG00000032766 | Gng11 | protein_coding | -0.791903861 | 6.068835467 | 10.865059 | 0.0044612 | 0.33083421 |
| ENSMUSG00000052516 | Robo2 | protein_coding | -1.136343137 | 4.285642096 | 10.825826 | 0.0045179 | 0.331510404 |
| ENSMUSG00000047866 | Lonp2 | protein_coding | 0.731492223 | 6.782646359 | 10.816228 | 0.0045319 | 0.331510404 |
| ENSMUSG00000030067 | Foxp1 | protein_coding | -0.639541588 | 7.041363291 | 10.777859 | 0.0045885 | 0.332823167 |

| ensembl_gene_id | external_gene_name | gene_biotype | log2 (fold change) | log2 (counts per million) | F-value | p-value | False discovery rate |
|---|---|---|---|---|---|---|---|
| ENSMUSG00000074457 | S100a16 | protein_coding | -0.783097741 | 8.358842546 | 10.757056 | 0.0046194 | 0.332823167 |
| ENSMUSG00000060227 | Casc4 | protein_coding | -0.868586633 | 5.343589844 | 10.726228 | 0.0046658 | 0.332823167 |
| ENSMUSG00000057858 | Fam204a | protein_coding | -0.914813703 | 5.679382187 | 10.720976 | 0.0046737 | 0.332823167 |
| ENSMUSG00000039632 | Ccdc151 | protein_coding | 2.866858992 | 2.469709925 | 10.680891 | 0.0047348 | 0.33495941 |
| ENSMUSG00000052676 | Zmat1 | protein_coding | -1.216063983 | 4.490497069 | 10.629035 | 0.0048153 | 0.336237419 |
| ENSMUSG00000024526 | Cidea | protein_coding | 1.434209123 | 3.764927347 | 10.623835 | 0.0048235 | 0.336237419 |
| ENSMUSG00000027708 | Dcun1d1 | protein_coding | -0.873905624 | 5.159536717 | 10.605389 | 0.0048525 | 0.336237419 |
| ENSMUSG00000006010 | BC003331 | protein_coding | -0.805134942 | 5.896520046 | 10.557758 | 0.0049284 | 0.336237419 |
| ENSMUSG00000083282 | Ctsf | protein_coding | 1.096129376 | 5.385650902 | 10.527999 | 0.0049765 | 0.336237419 |
| ENSMUSG00000056260 | Lrif1 | protein_coding | -0.888335996 | 5.03457639 | 10.520577 | 0.0049886 | 0.336237419 |
| ENSMUSG00000031864 | Ints10 | protein_coding | 1.063781762 | 5.075568629 | 10.503921 | 0.0050159 | 0.336237419 |
| ENSMUSG00000041408 | Wapl | protein_coding | -0.627627622 | 6.719766122 | 10.467003 | 0.0050768 | 0.336237419 |
| ENSMUSG00000068151 | A230006K03Rik | lincRNA | -1.029948251 | 4.432775736 | 10.417378 | 0.0051601 | 0.336237419 |
| ENSMUSG00000036095 | Dgkb | protein_coding | -0.776945129 | 7.206414384 | 10.396576 | 0.0051955 | 0.336237419 |
| ENSMUSG00000062627 | Mysm1 | protein_coding | -0.862519985 | 5.109075998 | 10.374817 | 0.0052328 | 0.336237419 |
| ENSMUSG00000020570 | Sypl | protein_coding | -0.733026009 | 6.263794843 | 10.36755 | 0.0052453 | 0.336237419 |
| ENSMUSG00000037674 | Rfx7 | protein_coding | -1.221834823 | 3.692786048 | 10.361009 | 0.0052566 | 0.336237419 |
| ENSMUSG00000029998 | Pcyox1 | protein_coding | 0.720531084 | 6.168019748 | 10.358783 | 0.0052605 | 0.336237419 |
| ENSMUSG00000028961 | Pgd | protein_coding | 0.970728664 | 5.789461953 | 10.348876 | 0.0052777 | 0.336237419 |
| ENSMUSG00000038695 | Josd2 | protein_coding | -0.680824083 | 6.239917499 | 10.348508 | 0.0052783 | 0.336237419 |
| ENSMUSG00000026064 | Ptp4a1 | protein_coding | -1.100430759 | 4.085936593 | 10.313709 | 0.0053392 | 0.336237419 |
| ENSMUSG00000021500 | Ddx46 | protein_coding | 0.827624928 | 6.111591387 | 10.310086 | 0.0053456 | 0.336237419 |
| ENSMUSG00000026102 | Inpp1 | protein_coding | 0.784144683 | 5.806926255 | 10.296731 | 0.0053692 | 0.336237419 |
| ENSMUSG00000026739 | Bmi1 | protein_coding | -1.102534842 | 4.297099157 | 10.291584 | 0.0053783 | 0.336237419 |
| ENSMUSG00000048307 | Ankrd46 | protein_coding | 0.766469714 | 8.673423792 | 10.215619 | 0.0055151 | 0.342796509 |
| ENSMUSG00000000567 | Sox9 | protein_coding | -0.862077604 | 5.735015818 | 10.15317 | 0.0056305 | 0.346393985 |
| ENSMUSG00000019590 | Cyb561 | protein_coding | 0.703195588 | 7.175003432 | 10.131011 | 0.0056721 | 0.346393985 |
| ENSMUSG00000003226 | Ranbp2 | protein_coding | 1.287554213 | 3.727733926 | 10.125665 | 0.0056822 | 0.346393985 |
| ENSMUSG00000066392 | Nrxn3 | protein_coding | -0.665526808 | 7.907882488 | 10.107023 | 0.0057176 | 0.346393985 |
| ENSMUSG00000008575 | Nfib | protein_coding | -1.01718332 | 5.048399068 | 10.098372 | 0.005734 | 0.346393985 |
| ENSMUSG00000005892 | Trh | protein_coding | 1.022381946 | 7.312990643 | 10.330712 | 0.005781 | 0.347282171 |
| ENSMUSG00000022346 | Myc | protein_coding | 1.944470744 | 3.014558506 | 10.039534 | 0.0058476 | 0.348653361 |
| ENSMUSG00000047141 | Zfp654 | protein_coding | -1.253612856 | 3.831966821 | 10.028746 | 0.0058687 | 0.348653361 |
| ENSMUSG00000056394 | Lig1 | protein_coding | 1.118080294 | 4.074865149 | 9.9881587 | 0.0059489 | 0.351473577 |
| ENSMUSG00000026280 | Atg4b | protein_coding | 0.728492125 | 6.247778244 | 9.959332 | 0.0060066 | 0.352942481 |
| ENSMUSG00000015971 | Actr8 | protein_coding | 0.744763767 | 6.486960432 | 9.9045801 | 0.0061179 | 0.35727917 |
| ENSMUSG00000084957 | Bbip1 | protein_coding | -0.622724962 | 7.193835222 | 9.8763063 | 0.0061763 | 0.35727917 |
| ENSMUSG00000046574 | Prr12 | protein_coding | 1.324676015 | 4.780336741 | 9.8792549 | 0.00618 | 0.35727917 |
| ENSMUSG00000041119 | Pde9a | protein_coding | 0.970221148 | 5.060572639 | 9.8165352 | 0.0063018 | 0.359274276 |
| ENSMUSG00000028557 | Rnf11 | protein_coding | -0.561738509 | 7.771785049 | 9.8140261 | 0.0063071 | 0.359274276 |
| ENSMUSG00000046447 | Camk2n1 | protein_coding | -0.667068195 | 8.814693818 | 9.7603615 | 0.0064225 | 0.359274276 |
| ENSMUSG00000021130 | Galnt16 | protein_coding | 0.68811342 | 6.390037046 | 9.7340797 | 0.0064798 | 0.359274276 |
| ENSMUSG00000032911 | Cspg4 | protein_coding | 1.642365546 | 3.823510684 | 9.7238461 | 0.0065125 | 0.359274276 |
| ENSMUSG00000033186 | Mzt1 | protein_coding | -0.597319873 | 7.298392549 | 9.7002564 | 0.0065544 | 0.359274276 |
| ENSMUSG00000032328 | Tmem30a | protein_coding | -0.560579876 | 7.184056146 | 9.6999861 | 0.006555 | 0.359274276 |
| ENSMUSG00000000948 | Gm38393 | protein_coding | -0.96957517 | 4.792417404 | 9.6999343 | 0.0065552 | 0.359274276 |
| ENSMUSG00000025487 | Psmd13 | protein_coding | 0.590027787 | 7.360820135 | 9.6993332 | 0.0065565 | 0.359274276 |
| ENSMUSG00000026158 | Ogfrl1 | protein_coding | -0.800962807 | 6.023285108 | 9.6732131 | 0.0066148 | 0.359274276 |
| ENSMUSG00000059742 | Kcnh7 | protein_coding | -1.107450563 | 3.864897987 | 9.6554187 | 0.0066549 | 0.359274276 |
| ENSMUSG00000034850 | Tmem127 | protein_coding | 0.964349596 | 6.47245845 | 9.7208258 | 0.0067031 | 0.359274276 |
| ENSMUSG00000045767 | B230219D22Rik | protein_coding | -0.754266024 | 5.908005349 | 9.6296173 | 0.0067135 | 0.359274276 |
| ENSMUSG00000050192 | Eif5a2 | protein_coding | -0.798113579 | 5.390785133 | 9.6203592 | 0.0067347 | 0.359274276 |
| ENSMUSG00000069769 | Msi2 | protein_coding | -0.793809129 | 5.347387036 | 9.6056049 | 0.0067686 | 0.359274276 |
| ENSMUSG00000042182 | Bend6 | protein_coding | -0.614002617 | 7.256237673 | 9.6030252 | 0.0067745 | 0.359274276 |
| ENSMUSG00000040648 | Ppip5k2 | protein_coding | -0.990755498 | 4.174445386 | 9.5995447 | 0.0067825 | 0.359274276 |
| ENSMUSG00000002944 | Cd36 | protein_coding | 3.047955944 | 2.164736826 | 9.5835588 | 0.0068196 | 0.359464946 |
| ENSMUSG00000020780 | Srp68 | protein_coding | 0.646971848 | 6.480610392 | 9.5584018 | 0.0068783 | 0.3603004 |
| ENSMUSG00000025907 | Rb1cc1 | protein_coding | -0.812872672 | 5.252218328 | 9.5481392 | 0.0069024 | 0.3603004 |
| ENSMUSG00000024907 | Gal | protein_coding | 0.691517947 | 9.118265128 | 9.5238251 | 0.00696 | 0.361549035 |
| ENSMUSG00000027706 | Sec62 | protein_coding | -0.573098013 | 7.909474174 | 9.4851064 | 0.0070527 | 0.361958493 |
| ENSMUSG00000100837 | 1700063D05Rik | lincRNA | -1.405141098 | 3.539659676 | 9.4715549 | 0.0070855 | 0.361958493 |
| ENSMUSG00000001156 | Mxd1 | protein_coding | 1.122240478 | 3.812498202 | 9.4695542 | 0.0070904 | 0.361958493 |
| ENSMUSG00000066705 | Fxyd6 | protein_coding | 0.608841459 | 9.078928226 | 9.4645715 | 0.0071025 | 0.361958493 |
| ENSMUSG00000031532 | Saraf | protein_coding | 0.73591251 | 7.232649969 | 9.4312735 | 0.0071841 | 0.363361743 |
| ENSMUSG00000054894 | Atp5s | protein_coding | -0.75880971 | 5.555725959 | 9.4257729 | 0.0071976 | 0.363361743 |
| ENSMUSG00000031176 | Dynlt3 | protein_coding | -0.592165745 | 8.581354738 | 9.3484201 | 0.0073916 | 0.369950888 |
| ENSMUSG00000060002 | Chpt1 | protein_coding | -1.131590947 | 3.839606672 | 9.3463208 | 0.007397 | 0.369950888 |
| ENSMUSG00000020098 | Pcbd1 | protein_coding | 0.583783444 | 7.694197222 | 9.3034093 | 0.0075072 | 0.373526676 |
| ENSMUSG00000026721 | Rabgap1l | protein_coding | -0.673015033 | 6.152632248 | 9.2915832 | 0.0075379 | 0.373526676 |
| ENSMUSG00000104283 | Gm37459 | antisense_RNA | -1.051734549 | 4.559865584 | 9.2764356 | 0.0075775 | 0.373763965 |
| ENSMUSG00000021039 | Snw1 | protein_coding | 0.607979088 | 6.894242313 | 9.2506944 | 0.0076452 | 0.375383669 |
| ENSMUSG00000039539 | Sgcz | protein_coding | -1.308291388 | 4.028245037 | 9.2348438 | 0.0076873 | 0.375733062 |
| ENSMUSG00000040687 | Madd | protein_coding | 0.75164261 | 5.786761892 | 9.1931942 | 0.007799 | 0.375820349 |
| ENSMUSG00000066324 | Impad1 | protein_coding | -0.555133412 | 6.810191745 | 9.1854003 | 0.0078202 | 0.375820349 |

| ensembl_gene_id | external_gene_name | gene_biotype | log2 (fold change) | log2 (counts per million) | F-value | p-value | False discovery rate |
|---|---|---|---|---|---|---|---|
| ENSMUSG00000020176 | Grb10 | protein_coding | 0.829807564 | 5.982098793 | 9.1616449 | 0.0078849 | 0.375820349 |
| ENSMUSG00000037997 | Parp11 | protein_coding | 1.033542762 | 4.771992288 | 9.1573486 | 0.0078967 | 0.375820349 |
| ENSMUSG00000050875 | A730017C20Rik | protein_coding | 0.705383884 | 8.826962204 | 9.1372692 | 0.007952 | 0.375820349 |
| ENSMUSG00000063931 | Pepd | protein_coding | 0.826120347 | 6.764992624 | 9.1410033 | 0.0079532 | 0.375820349 |
| ENSMUSG00000097451 | Rian | processed_transcript | -0.74800825 | 5.684275856 | 9.1331362 | 0.0079635 | 0.375820349 |
| ENSMUSG00000021125 | Arg2 | protein_coding | 1.011874883 | 3.883070506 | 9.1312658 | 0.0079687 | 0.375820349 |
| ENSMUSG00000007872 | Id3 | protein_coding | -0.614749998 | 7.392787143 | 9.1079387 | 0.0080336 | 0.377230159 |
| ENSMUSG00000027499 | Pkia | protein_coding | -0.60210511 | 7.19892695 | 9.0752548 | 0.0081257 | 0.379039908 |
| ENSMUSG00000003746 | Man1a | protein_coding | -0.97205642 | 4.613703272 | 9.0682828 | 0.0081455 | 0.379039908 |
| ENSMUSG00000000838 | Fmr1 | protein_coding | -0.940998069 | 4.646406992 | 9.0485532 | 0.0082018 | 0.379039908 |
| ENSMUSG00000019986 | Ahi1 | protein_coding | 0.733358452 | 8.7779824 | 9.0445773 | 0.0082132 | 0.379039908 |
| ENSMUSG00000029635 | Cdk8 | protein_coding | -0.729385841 | 6.496972303 | 9.0033929 | 0.0083323 | 0.3804221 |
| ENSMUSG00000020580 | Rock2 | protein_coding | -0.643197462 | 6.259630581 | 8.9869658 | 0.0083804 | 0.3804221 |
| ENSMUSG00000006288 | Ttc5 | protein_coding | 0.606323881 | 6.543103202 | 8.9851543 | 0.0083857 | 0.3804221 |
| ENSMUSG00000025993 | Slc40a1 | protein_coding | 2.391633207 | 2.315840869 | 8.977701 | 0.0084076 | 0.3804221 |
| ENSMUSG00000029428 | Stx2 | protein_coding | -1.811664389 | 2.756617378 | 8.9735022 | 0.00842 | 0.3804221 |
| ENSMUSG00000025792 | Slc25a10 | protein_coding | -0.802150501 | 5.270183151 | 8.946472 | 0.0085002 | 0.380446146 |
| ENSMUSG00000032047 | Acat1 | protein_coding | 0.575058564 | 8.079841106 | 8.9448194 | 0.0085052 | 0.380446146 |
| ENSMUSG00000027206 | Cops2 | protein_coding | -0.671186023 | 7.228814751 | 8.9376175 | 0.0085267 | 0.380446146 |
| ENSMUSG00000038005 | Hpf1 | protein_coding | -0.635040772 | 5.994374637 | 8.9101022 | 0.0086095 | 0.382305872 |
| ENSMUSG00000071753 | C230004F18Rik | processed_transcript | -0.987956093 | 4.836514358 | 8.9002263 | 0.0086395 | 0.382305872 |
| ENSMUSG00000019923 | Zwint | protein_coding | 0.705933234 | 12.04740912 | 8.8721503 | 0.0087253 | 0.384228922 |
| ENSMUSG00000023495 | Pcbp4 | protein_coding | -0.948618476 | 5.32965233 | 8.8544334 | 0.0087799 | 0.384228922 |
| ENSMUSG00000028809 | Srrm1 | protein_coding | 0.573191636 | 7.215659472 | 8.8511374 | 0.0087901 | 0.384228922 |
| ENSMUSG00000031939 | Taf1d | protein_coding | -1.169577727 | 4.656538511 | 8.8332099 | 0.0088459 | 0.385100904 |
| ENSMUSG00000029250 | Polr2b | protein_coding | 0.774612334 | 5.613263363 | 8.8126233 | 0.0089104 | 0.386196192 |
| ENSMUSG00000086429 | Gt(ROSA)26Sor | antisense_RNA | -1.08300545 | 5.287847624 | 8.7909941 | 0.0089911 | 0.386196192 |
| ENSMUSG00000020777 | Acox1 | protein_coding | 0.692597282 | 6.792100537 | 8.7848516 | 0.0089983 | 0.386196192 |
| ENSMUSG00000068134 | Zfp120 | protein_coding | -1.127684027 | 4.44365837 | 8.7668752 | 0.0090557 | 0.386196192 |
| ENSMUSG00000032998 | Foxj3 | protein_coding | 0.842060316 | 5.043641327 | 8.7604515 | 0.0090764 | 0.386196192 |
| ENSMUSG00000032417 | Rwdd2a | protein_coding | 0.583397282 | 7.882959122 | 8.7302825 | 0.0091739 | 0.386196192 |
| ENSMUSG00000036242 | 3632451O06Rik | protein_coding | 0.907261454 | 4.742168553 | 8.7269984 | 0.0091846 | 0.386196192 |
| ENSMUSG00000055567 | Unc80 | protein_coding | -0.684169094 | 6.175450892 | 8.7242809 | 0.0091935 | 0.386196192 |
| ENSMUSG00000097875 | Gm26704 | lincRNA | -1.309159451 | 3.844733875 | 8.7173908 | 0.009216 | 0.386196192 |
| ENSMUSG00000025666 | Tmem47 | protein_coding | -0.676676003 | 6.171143608 | 8.7116966 | 0.0092346 | 0.386196192 |
| ENSMUSG00000022336 | Eif3e | protein_coding | -0.664206813 | 7.651434213 | 8.6946347 | 0.0092907 | 0.386196192 |
| ENSMUSG00000022696 | Sidt1 | protein_coding | 1.535659705 | 3.069529831 | 8.6770242 | 0.0093491 | 0.386196192 |
| ENSMUSG00000064115 | Cadm2 | protein_coding | -0.862329391 | 5.531232018 | 8.6486565 | 0.0094439 | 0.386196192 |
| ENSMUSG00000029093 | Sorcs2 | protein_coding | 1.391341543 | 3.639065966 | 8.64795 | 0.0094463 | 0.386196192 |
| ENSMUSG00000049775 | Tmsb4x | protein_coding | -0.825903618 | 11.71089182 | 8.6439607 | 0.0094724 | 0.386196192 |
| ENSMUSG00000063698 | Sfxn4 | protein_coding | 0.741270133 | 5.734147057 | 8.6364364 | 0.0094851 | 0.386196192 |
| ENSMUSG00000028576 | Ift74 | protein_coding | -0.693149 | 5.909639603 | 8.624587 | 0.0095252 | 0.386196192 |
| ENSMUSG00000061601 | Pclo | protein_coding | -0.93264961 | 5.385969975 | 8.6106646 | 0.0095726 | 0.386196192 |
| ENSMUSG00000044667 | Plppr4 | protein_coding | -0.616011589 | 6.517486683 | 8.6094364 | 0.0095768 | 0.386196192 |
| ENSMUSG00000074415 | 2610203C20Rik | processed_transcript | -0.772202224 | 5.094700376 | 8.6057735 | 0.0095894 | 0.386196192 |
| ENSMUSG00000024587 | Nars | protein_coding | 0.669056813 | 8.572052115 | 8.5875793 | 0.0096518 | 0.387261994 |
| ENSMUSG00000085181 | Gm12709 | antisense_RNA | -0.858408142 | 5.139537362 | 8.5371537 | 0.0098274 | 0.389414654 |
| ENSMUSG00000038503 | Mesd | protein_coding | 0.625317288 | 6.570156497 | 8.5345107 | 0.0098367 | 0.389414654 |
| ENSMUSG00000047213 | Ythdf3 | protein_coding | -0.984138731 | 4.409674026 | 8.5234115 | 0.0098759 | 0.389414654 |
| ENSMUSG00000018809 | Smyd4 | protein_coding | 1.151949182 | 4.364534529 | 8.5233089 | 0.0098763 | 0.389414654 |
| ENSMUSG00000018541 | Cwc25 | protein_coding | 0.876070295 | 4.840670748 | 8.5111512 | 0.0099194 | 0.389414654 |
| ENSMUSG00000068263 | Efcc1 | protein_coding | -1.17660593 | 3.664385717 | 8.5100213 | 0.0099234 | 0.389414654 |
| ENSMUSG00000064349 | mt-Tc | Mt_tRNA | -1.572720867 | 2.982255113 | 8.4991777 | 0.0099621 | 0.389414654 |
| ENSMUSG00000042157 | Sprr2i | protein_coding | 4.286261389 | 1.905783563 | 8.5094996 | 0.0099952 | 0.389414654 |
| ENSMUSG00000023017 | Asic1 | protein_coding | 0.796812685 | 5.062771338 | 8.4681458 | 0.0100736 | 0.389546889 |
| ENSMUSG00000000753 | Serpinf1 | protein_coding | 1.252389504 | 4.379507595 | 8.4689336 | 0.0100839 | 0.389546889 |
| ENSMUSG00000042901 | Aida | protein_coding | -0.736751844 | 5.122552065 | 8.4558306 | 0.0101183 | 0.389546889 |
| ENSMUSG00000030137 | Tuba8 | protein_coding | 1.8636079 | 3.06575667 | 8.4489146 | 0.0101435 | 0.389546889 |
| ENSMUSG00000026424 | Gpr37l1 | protein_coding | -0.749679673 | 6.583081309 | 8.4275909 | 0.0102216 | 0.389629484 |
| ENSMUSG00000017677 | Wsb1 | protein_coding | -1.133758417 | 4.565200691 | 8.4151551 | 0.0102675 | 0.389629484 |
| ENSMUSG00000040339 | Fam102b | protein_coding | -0.691040619 | 5.639958171 | 8.4118792 | 0.0102796 | 0.389629484 |
| ENSMUSG00000049300 | Prmt6 | protein_coding | -1.162763999 | 3.436988072 | 8.4089114 | 0.0102906 | 0.389629484 |
| ENSMUSG00000020111 | Micu1 | protein_coding | 0.601674088 | 7.00800492 | 8.3989436 | 0.0103276 | 0.389659402 |
| ENSMUSG00000086308 | G630016G05Rik | lincRNA | -1.704484743 | 2.940889393 | 8.3808506 | 0.0103952 | 0.390838479 |
| ENSMUSG00000091264 | Smim13 | protein_coding | -0.563221416 | 6.418624663 | 8.3619273 | 0.0104665 | 0.391116674 |
| ENSMUSG00000037857 | Nufip2 | protein_coding | -0.802987937 | 6.648928208 | 8.3590862 | 0.0104906 | 0.391116674 |
| ENSMUSG00000026383 | Epb41l5 | protein_coding | 1.187079103 | 3.350534585 | 8.3499756 | 0.0105117 | 0.391116674 |
| ENSMUSG00000064289 | Tank | protein_coding | -0.744615198 | 5.599823498 | 8.3137266 | 0.0106504 | 0.392140813 |
| ENSMUSG00000073982 | Rhog | protein_coding | -0.759124259 | 6.224358474 | 8.3087998 | 0.0106694 | 0.392140813 |
| ENSMUSG00000013465 | Nelfb | protein_coding | 0.619152834 | 5.997050314 | 8.3001156 | 0.0107031 | 0.392140813 |
| ENSMUSG00000054855 | Rnd1 | protein_coding | 0.946326385 | 5.994944056 | 8.3040605 | 0.0107371 | 0.392140813 |
| ENSMUSG00000022342 | Kcnv1 | protein_coding | -1.438841504 | 3.770552748 | 8.2923353 | 0.0107469 | 0.392140813 |
| ENSMUSG00000036499 | Eea1 | protein_coding | -0.999995118 | 4.669997963 | 8.285964 | 0.0107581 | 0.392140813 |
| ENSMUSG00000072704 | Smim10l1 | protein_coding | -0.764089536 | 8.020245595 | 8.2694221 | 0.0108365 | 0.393518325 |

| ensembl_gene_id | external_gene_name | gene_biotype | log2 (fold change) | log2 (counts per million) | F-value | p-value | False discovery rate |
|---|---|---|---|---|---|---|---|
| ENSMUSG00000020863 | Luc7l3 | protein_coding | -0.699094511 | 6.400903113 | 8.2576667 | 0.0108691 | 0.393518325 |
| ENSMUSG00000095567 | Noc2l | protein_coding | 0.953892261 | 4.321235495 | 8.243456 | 0.0109253 | 0.394010893 |
| ENSMUSG00000106106 | CT010467.1 | rRNA | -0.66134592 | 10.43028008 | 8.220998 | 0.0110148 | 0.394010893 |
| ENSMUSG00000025172 | Ankrd2 | protein_coding | 1.676593153 | 2.945274011 | 8.2201413 | 0.0110182 | 0.394010893 |
| ENSMUSG00000085925 | Rtl1 | protein_coding | -1.155091124 | 4.657096858 | 8.219303 | 0.0110354 | 0.394010893 |
| ENSMUSG00000016262 | Sertad4 | protein_coding | -1.04352753 | 4.413357749 | 8.2061116 | 0.0110746 | 0.394010893 |
| ENSMUSG00000051359 | Ncald | protein_coding | 0.647866534 | 8.885252362 | 8.1992006 | 0.0111025 | 0.394010893 |
| ENSMUSG00000046204 | Pnma2 | protein_coding | 0.619600501 | 7.583794278 | 8.1899381 | 0.01114 | 0.394041475 |
| ENSMUSG00000027677 | Ttc14 | protein_coding | -1.384730971 | 3.715954324 | 8.1599282 | 0.0112625 | 0.395764386 |
| ENSMUSG00000076617 | Ighm | IG_C_gene | 1.26327221 | 5.804644818 | 8.425443 | 0.0112983 | 0.395764386 |
| ENSMUSG00000021962 | Dcp1a | protein_coding | 1.148705299 | 3.482967852 | 8.1510319 | 0.0112991 | 0.395764386 |
| ENSMUSG00000109324 | Prmt1 | protein_coding | 0.569504528 | 7.313579922 | 8.1395064 | 0.0113468 | 0.396142454 |
| ENSMUSG00000027199 | Gatm | protein_coding | -0.657628509 | 8.833057624 | 8.1252329 | 0.0114061 | 0.396924327 |
| ENSMUSG00000066036 | Ubr4 | protein_coding | 0.87217798 | 5.281390209 | 8.1091751 | 0.0114732 | 0.397972968 |
| ENSMUSG00000027643 | Ghrh | protein_coding | 1.495124047 | 3.367773726 | 8.0992253 | 0.0115151 | 0.398139593 |
| ENSMUSG00000024477 | Pggt1b | protein_coding | -1.218676987 | 3.324418447 | 8.087313 | 0.0115654 | 0.398597396 |
| ENSMUSG00000033676 | Gabrb3 | protein_coding | -0.589205979 | 6.4470507 | 8.0514428 | 0.0117184 | 0.401097448 |
| ENSMUSG00000071302 | 2610044O15Rik8 | protein_coding | -0.839675886 | 4.952829646 | 8.0321311 | 0.0118017 | 0.401097448 |
| ENSMUSG00000044349 | Snhg11 | protein_coding | -0.864508024 | 7.403342186 | 8.1583983 | 0.0118158 | 0.401097448 |
| ENSMUSG00000045991 | Onecut2 | protein_coding | -0.880073909 | 4.74678289 | 8.0272828 | 0.0118227 | 0.401097448 |
| ENSMUSG00000026600 | Soat1 | protein_coding | 1.000782723 | 4.214065536 | 8.0268949 | 0.0118244 | 0.401097448 |
| ENSMUSG00000036503 | Rnf13 | protein_coding | -0.543755305 | 7.427291732 | 7.9761907 | 0.0120469 | 0.406702617 |
| ENSMUSG00000046844 | Vat1l | protein_coding | 0.646403836 | 7.407664796 | 7.9720487 | 0.0120653 | 0.406702617 |
| ENSMUSG00000005534 | Insr | protein_coding | -1.202552247 | 3.876891473 | 7.9286464 | 0.0122599 | 0.410521499 |
| ENSMUSG00000037366 | Pafah2 | protein_coding | 2.454207749 | 2.283650416 | 7.9279877 | 0.0122629 | 0.410521499 |
| ENSMUSG00000029771 | Irf5 | protein_coding | -1.763668788 | 2.881099797 | 7.9213115 | 0.0122931 | 0.410521499 |
| ENSMUSG00000031245 | Hmgn5 | protein_coding | -1.28470515 | 3.447118563 | 7.9126366 | 0.0123326 | 0.410563564 |
| ENSMUSG00000031422 | Morf4l2 | protein_coding | 0.620021484 | 8.396678086 | 7.8788275 | 0.0124877 | 0.41114836 |
| ENSMUSG00000026781 | Acbd5 | protein_coding | -0.775983589 | 5.32265234 | 7.875954 | 0.0125009 | 0.41114836 |
| ENSMUSG00000027835 | Pdcd10 | protein_coding | -0.645326882 | 6.234613368 | 7.864719 | 0.012553 | 0.41114836 |
| ENSMUSG00000039219 | Arid4b | protein_coding | -0.708033917 | 5.786670601 | 7.857042 | 0.0125888 | 0.41114836 |
| ENSMUSG00000025321 | Itgb8 | protein_coding | -1.399479465 | 3.993392886 | 7.8516391 | 0.0126288 | 0.41114836 |
| ENSMUSG00000040242 | Fgfr1op2 | protein_coding | -0.570078202 | 6.371582917 | 7.8426277 | 0.0126562 | 0.41114836 |
| ENSMUSG00000036775 | Decr2 | protein_coding | 0.563385147 | 6.638058043 | 7.8417084 | 0.0126605 | 0.41114836 |
| ENSMUSG00000026361 | Cdc73 | protein_coding | -0.831544598 | 4.876864289 | 7.833922 | 0.0126971 | 0.41114836 |
| ENSMUSG00000002064 | Sdf2 | protein_coding | 0.601101211 | 7.649452526 | 7.8223004 | 0.012752 | 0.41114836 |
| ENSMUSG00000059401 | Mamld1 | protein_coding | 0.993282928 | 4.822807118 | 7.8170747 | 0.0127767 | 0.41114836 |
| ENSMUSG00000028389 | Zfp37 | protein_coding | -0.892035553 | 4.774334343 | 7.8102767 | 0.012809 | 0.41114836 |
| ENSMUSG00000028409 | Smu1 | protein_coding | 0.588642601 | 6.920026931 | 7.7892606 | 0.0129094 | 0.41114836 |
| ENSMUSG00000028381 | Ugcg | protein_coding | -0.733285714 | 5.413771416 | 7.7876256 | 0.0129172 | 0.41114836 |
| ENSMUSG00000074733 | Zfp950 | protein_coding | -0.780505009 | 5.169887919 | 7.7801328 | 0.0129533 | 0.41114836 |
| ENSMUSG00000032042 | Srpr | protein_coding | 0.654743504 | 5.738268041 | 7.7785832 | 0.0129607 | 0.41114836 |
| ENSMUSG00000037236 | Matr3 | protein_coding | -0.54567777 | 8.047291825 | 7.7783418 | 0.0129619 | 0.41114836 |
| ENSMUSG00000006342 | Susd2 | protein_coding | 1.280141771 | 4.02023654 | 7.7647708 | 0.0130275 | 0.411717317 |
| ENSMUSG00000049184 | Purg | protein_coding | -0.827565015 | 4.934903238 | 7.7544498 | 0.0130777 | 0.411717317 |
| ENSMUSG00000021091 | Serpina3n | protein_coding | 0.82078989 | 4.794840131 | 7.7509516 | 0.0130947 | 0.411717317 |
| ENSMUSG00000034145 | Tmem63c | protein_coding | 1.309216033 | 3.439357991 | 7.7335855 | 0.0131797 | 0.41266057 |
| ENSMUSG00000095620 | 2010005H15Rik | protein_coding | 1.440945016 | 3.44310427 | 7.7291624 | 0.0132015 | 0.41266057 |
| ENSMUSG00000053510 | Nrd1 | protein_coding | 0.667306683 | 5.395380543 | 7.6813182 | 0.0134394 | 0.417434852 |
| ENSMUSG00000020792 | Exoc7 | protein_coding | 0.683133883 | 7.062563813 | 7.6778675 | 0.0134567 | 0.417434852 |
| ENSMUSG00000040759 | Cmtm5 | protein_coding | -0.749344746 | 7.823720568 | 7.6777816 | 0.013513 | 0.417434852 |
| ENSMUSG00000030654 | Arl6ip1 | protein_coding | 0.504132009 | 10.05420097 | 7.6634194 | 0.0135296 | 0.417434852 |
| ENSMUSG00000098087 | Gm17750 | lincRNA | -1.25155491 | 3.3932452 | 7.659732 | 0.0135483 | 0.417434852 |
| ENSMUSG00000028952 | Zbtb48 | protein_coding | 1.022779613 | 4.685661917 | 7.6359273 | 0.0136696 | 0.419223029 |
| ENSMUSG00000030286 | Emc3 | protein_coding | 0.665472559 | 6.178071852 | 7.6276546 | 0.013712 | 0.419223029 |
| ENSMUSG00000020362 | Cnot6 | protein_coding | -0.842587939 | 5.369454094 | 7.6254651 | 0.0137233 | 0.419223029 |
| ENSMUSG00000043498 | 9330132A10Rik | pseudogene | -1.120898008 | 3.451029808 | 7.5777428 | 0.0139713 | 0.424701956 |
| ENSMUSG00000032624 | Eml4 | protein_coding | 1.037709682 | 3.902866519 | 7.5757772 | 0.0139816 | 0.424701956 |
| ENSMUSG00000022962 | Gart | protein_coding | 1.050705231 | 4.633510336 | 7.5484837 | 0.0141259 | 0.424863291 |
| ENSMUSG00000028295 | Smim8 | protein_coding | -0.665867536 | 6.499060213 | 7.5445147 | 0.014147 | 0.424863291 |
| ENSMUSG00000014813 | Stc1 | protein_coding | 1.227321993 | 3.672506606 | 7.5439593 | 0.01415 | 0.424863291 |
| ENSMUSG00000035597 | Prpf39 | protein_coding | -0.979182385 | 4.118613969 | 7.5338388 | 0.014204 | 0.424863291 |
| ENSMUSG00000052942 | Glis3 | protein_coding | -1.251400693 | 3.356678143 | 7.5328425 | 0.0142093 | 0.424863291 |
| ENSMUSG00000000560 | Gabra2 | protein_coding | -1.136748128 | 6.153070811 | 7.7862935 | 0.0142322 | 0.424863291 |
| ENSMUSG00000004317 | Clcn5 | protein_coding | 1.006930679 | 4.298924838 | 7.522741 | 0.0142635 | 0.424863291 |
| ENSMUSG00000020571 | Pdia6 | protein_coding | 0.484870685 | 7.807010753 | 7.4993517 | 0.0143899 | 0.425847067 |
| ENSMUSG00000031979 | Cog2 | protein_coding | 0.895237912 | 4.659814181 | 7.4913101 | 0.0144336 | 0.425847067 |
| ENSMUSG00000004565 | Pnpla6 | protein_coding | 0.869967181 | 4.932264495 | 7.487533 | 0.0144542 | 0.425847067 |
| ENSMUSG00000020361 | Hspa4 | protein_coding | 0.725850868 | 5.412967386 | 7.4873996 | 0.014455 | 0.425847067 |
| ENSMUSG00000042607 | Asb4 | protein_coding | 1.246878338 | 4.45846655 | 7.47976 | 0.0145127 | 0.426379735 |
| ENSMUSG00000025645 | Ccdc51 | protein_coding | 1.227063903 | 3.507253039 | 7.4549324 | 0.0146334 | 0.428755011 |
| ENSMUSG00000034912 | Mdga2 | protein_coding | -0.992189915 | 5.501504469 | 7.4415081 | 0.014724 | 0.430092836 |
| ENSMUSG00000064373 | Selenop | protein_coding | -0.723647404 | 7.836042949 | 7.4261789 | 0.0148097 | 0.430092836 |
| ENSMUSG00000050945 | Zfp438 | protein_coding | 1.754979011 | 3.362616456 | 7.4263936 | 0.014851 | 0.430092836 |

| ensembl_gene_id | external_gene_name | gene_biotype | log2 (fold change) | log2 (counts per million) | F-value | p-value | False discovery rate |
|---|---|---|---|---|---|---|---|
| ENSMUSG00000097924 | A730020E08Rik | bidirectional_promoter | -1.713112047 | 2.845040836 | 7.413526 | 0.0148647 | 0.430092836 |
| ENSMUSG00000038070 | Cntln | protein_coding | -1.245240634 | 3.697882556 | 7.4065888 | 0.0149038 | 0.430092836 |
| ENSMUSG00000026824 | Kcnj3 | protein_coding | -0.806592495 | 4.63725668 | 7.3990822 | 0.0149463 | 0.430092836 |
| ENSMUSG00000000805 | Car4 | protein_coding | -1.155504115 | 3.783159165 | 7.3951841 | 0.0149684 | 0.430092836 |
| ENSMUSG00000021149 | Gtpbp4 | protein_coding | 0.59409922 | 6.593417034 | 7.389797 | 0.0149991 | 0.430092836 |
| ENSMUSG00000071252 | 2210408I21Rik | protein_coding | -0.82633912 | 6.947680985 | 7.4872012 | 0.0150409 | 0.430145606 |
| ENSMUSG00000024392 | Bag6 | protein_coding | -0.776351172 | 5.578006025 | 7.366144 | 0.0151344 | 0.430676043 |
| ENSMUSG00000030268 | Bcat1 | protein_coding | 0.753055466 | 7.583348572 | 7.3850898 | 0.0151516 | 0.430676043 |
| ENSMUSG00000029213 | Commd8 | protein_coding | -0.559791028 | 7.193414177 | 7.3582989 | 0.0151796 | 0.430676043 |
| ENSMUSG00000014349 | Ube2z | protein_coding | 0.6010716 | 6.026995833 | 7.3490203 | 0.0152333 | 0.43106102 |
| ENSMUSG00000019906 | Lin7a | protein_coding | -0.684229261 | 6.307667229 | 7.3322283 | 0.0153309 | 0.432685768 |
| ENSMUSG00000030059 | Tmf1 | protein_coding | -1.069779599 | 6.469920041 | 7.5430903 | 0.0155251 | 0.43537918 |
| ENSMUSG00000026986 | Hnmt | protein_coding | -0.522195915 | 6.539923914 | 7.2956517 | 0.0155461 | 0.43537918 |
| ENSMUSG00000053390 | Zfp952 | protein_coding | 1.036695287 | 4.308508244 | 7.2878558 | 0.0155924 | 0.43537918 |
| ENSMUSG00000043991 | Pura | protein_coding | -0.578682969 | 6.957728223 | 7.2855054 | 0.0156064 | 0.43537918 |
| ENSMUSG00000079084 | Ccdc82 | protein_coding | -0.994141075 | 4.377176218 | 7.2817518 | 0.0156288 | 0.43537918 |
| ENSMUSG00000063887 | Nlgn1 | protein_coding | -0.65504902 | 6.333251997 | 7.2136124 | 0.0160414 | 0.443722679 |
| ENSMUSG00000094483 | Purb | protein_coding | -0.49990011 | 8.418996873 | 7.2093884 | 0.0160674 | 0.443722679 |
| ENSMUSG00000002524 | Puf60 | protein_coding | 0.523092202 | 8.566875765 | 7.2019233 | 0.0161135 | 0.443722679 |
| ENSMUSG00000048978 | Nrsn1 | protein_coding | -0.497458755 | 9.176900978 | 7.2009609 | 0.0161194 | 0.443722679 |
| ENSMUSG00000032253 | Phip | protein_coding | -0.853411618 | 4.797293326 | 7.1764179 | 0.0162719 | 0.443722679 |
| ENSMUSG00000026864 | Hspa5 | protein_coding | 0.473629995 | 8.346771058 | 7.176181 | 0.0162734 | 0.443722679 |
| ENSMUSG00000041096 | Tspyl2 | protein_coding | 0.589242548 | 6.912719448 | 7.1675893 | 0.0163272 | 0.443722679 |
| ENSMUSG00000040297 | Suco | protein_coding | -0.82982817 | 4.478071577 | 7.1619432 | 0.0163626 | 0.443722679 |
| ENSMUSG00000018372 | Cep95 | protein_coding | -0.926680816 | 3.87349461 | 7.1618407 | 0.0163633 | 0.443722679 |
| ENSMUSG00000035027 | Map2k2 | protein_coding | -0.654771242 | 5.336854328 | 7.1590416 | 0.0163809 | 0.443722679 |
| ENSMUSG00000048304 | Slitrk3 | protein_coding | 1.001941995 | 4.373265654 | 7.1434805 | 0.0164792 | 0.443722679 |
| ENSMUSG00000064343 | mt-Tq | Mt_tRNA | -1.791832501 | 2.419131349 | 7.1354028 | 0.0165305 | 0.443722679 |
| ENSMUSG00000002307 | Daxx | protein_coding | 0.914531279 | 5.234720564 | 7.1306948 | 0.0165604 | 0.443722679 |
| ENSMUSG00000004789 | Dlst | protein_coding | 0.628336843 | 6.92417559 | 7.1283199 | 0.0165756 | 0.443722679 |
| ENSMUSG00000027963 | Extl2 | protein_coding | 0.54684443 | 6.745595982 | 7.1240748 | 0.0166027 | 0.443722679 |
| ENSMUSG00000034088 | Hdlbp | protein_coding | 0.621906701 | 6.210549682 | 7.1157329 | 0.0166561 | 0.443722679 |
| ENSMUSG00000027163 | Commd9 | protein_coding | 0.631271599 | 7.208490425 | 7.1137705 | 0.0166687 | 0.443722679 |
| ENSMUSG00000021302 | Ggps1 | protein_coding | -0.513495749 | 6.849266201 | 7.1134059 | 0.0166711 | 0.443722679 |
| ENSMUSG00000035781 | R3hdm4 | protein_coding | -0.610737949 | 7.261553499 | 7.0717956 | 0.0169408 | 0.447467182 |
| ENSMUSG00000022867 | Usp25 | protein_coding | -1.010591078 | 4.176363426 | 7.0699608 | 0.0169528 | 0.447467182 |
| ENSMUSG00000095362 | Gm14325 | protein_coding | -0.756550987 | 5.72133313 | 7.0625726 | 0.0170012 | 0.447467182 |
| ENSMUSG00000021549 | Rasa1 | protein_coding | -1.028022572 | 3.628780971 | 7.0597469 | 0.0170198 | 0.447467182 |
| ENSMUSG00000074968 | Ano3 | protein_coding | -0.961020136 | 5.341786411 | 7.0516775 | 0.0170903 | 0.447467182 |
| ENSMUSG00000114304 | AC165092.1 | lincRNA | 0.90163296 | 7.087818151 | 7.2094674 | 0.0171197 | 0.447467182 |
| ENSMUSG00000020594 | Pum2 | protein_coding | -0.518541728 | 7.253718798 | 7.0353496 | 0.0171811 | 0.447467182 |
| ENSMUSG00000017802 | Retreg3 | protein_coding | 0.984248491 | 3.661015455 | 7.0333268 | 0.0171946 | 0.447467182 |
| ENSMUSG00000027200 | Sema6d | protein_coding | -0.795238988 | 4.817091646 | 7.0324424 | 0.0172004 | 0.447467182 |
| ENSMUSG00000109865 | Hspa14 | protein_coding | 0.6709012 | 5.770359006 | 7.0219111 | 0.0172707 | 0.447467182 |
| ENSMUSG00000020152 | Actr2 | protein_coding | -0.52337655 | 9.058469489 | 7.0196377 | 0.0172859 | 0.447467182 |
| ENSMUSG00000042763 | Maneal | protein_coding | 0.99926939 | 4.41117024 | 7.007442 | 0.0173678 | 0.447467182 |
| ENSMUSG00000011877 | Git1 | protein_coding | -0.520400527 | 6.426263458 | 7.0027994 | 0.017399 | 0.447467182 |
| ENSMUSG00000029103 | Lrpap1 | protein_coding | 0.476587686 | 9.146954222 | 6.9950248 | 0.0174516 | 0.447467182 |
| ENSMUSG00000026841 | Fibcd1 | protein_coding | 1.140101931 | 3.830676593 | 6.994141 | 0.0174575 | 0.447467182 |
| ENSMUSG00000025823 | Pdia4 | protein_coding | 0.630583909 | 5.920511761 | 6.9877026 | 0.0175012 | 0.447467182 |
| ENSMUSG00000028020 | Glrb | protein_coding | -0.49539237 | 6.764037551 | 6.9691853 | 0.0176274 | 0.447467182 |
| ENSMUSG00000002997 | Prkar2b | protein_coding | -0.675093258 | 6.045390933 | 6.9657041 | 0.0176513 | 0.447467182 |
| ENSMUSG00000029629 | Phf14 | protein_coding | -0.859530531 | 4.886092772 | 6.9636171 | 0.0176656 | 0.447467182 |
| ENSMUSG00000022865 | Cxadr | protein_coding | -1.373789428 | 2.899589865 | 6.9539568 | 0.017732 | 0.447467182 |
| ENSMUSG00000027466 | Rbck1 | protein_coding | -0.646204335 | 6.158664113 | 6.9513234 | 0.0177502 | 0.447467182 |
| ENSMUSG00000049659 | Aftph | protein_coding | -0.661991622 | 5.669652592 | 6.950272 | 0.0177574 | 0.447467182 |
| ENSMUSG00000017858 | Ift52 | protein_coding | 0.655874202 | 6.539735167 | 6.9448357 | 0.017795 | 0.447467182 |
| ENSMUSG00000033061 | Resp18 | protein_coding | 0.548185695 | 9.264313916 | 6.9426004 | 0.0178105 | 0.447467182 |
| ENSMUSG00000042426 | Dhx29 | protein_coding | 0.867553418 | 3.953861934 | 6.9237251 | 0.0179418 | 0.448729514 |
| ENSMUSG00000001986 | Gria3 | protein_coding | -0.583236346 | 5.891999901 | 6.9191272 | 0.0179739 | 0.448729514 |
| ENSMUSG00000021548 | Ccnh | protein_coding | -0.50673842 | 7.14178214 | 6.9145164 | 0.0180062 | 0.448729514 |
| ENSMUSG00000034818 | Celf5 | protein_coding | -0.565260152 | 6.515981532 | 6.9114624 | 0.0180276 | 0.448729514 |
| ENSMUSG00000030061 | Uba3 | protein_coding | -0.539620927 | 6.785816306 | 6.8988285 | 0.0181166 | 0.449430488 |
| ENSMUSG00000024530 | Prelid3a | protein_coding | 0.574152375 | 6.054081993 | 6.8929041 | 0.0181585 | 0.449430488 |
| ENSMUSG00000034488 | Edil3 | protein_coding | -0.508744187 | 6.477288724 | 6.8897035 | 0.0181812 | 0.449430488 |
| ENSMUSG00000003037 | Rab8a | protein_coding | 0.506106299 | 7.495020344 | 6.8780574 | 0.018264 | 0.449920163 |
| ENSMUSG00000020230 | Prmt2 | protein_coding | 0.732413962 | 9.243598961 | 6.884296 | 0.0182847 | 0.449920163 |
| ENSMUSG00000027981 | Rnpc3 | protein_coding | -1.319960259 | 2.974047067 | 6.8611608 | 0.0183849 | 0.451352912 |
| ENSMUSG00000107872 | Gm44511 | protein_coding | -1.387422728 | 3.305188604 | 6.8312443 | 0.0186012 | 0.454801952 |
| ENSMUSG00000003411 | Rab3b | protein_coding | 0.503711884 | 8.485015231 | 6.8300385 | 0.01861 | 0.454801952 |
| ENSMUSG00000032469 | Dbr1 | protein_coding | 1.374658435 | 3.535123546 | 6.8207587 | 0.0186777 | 0.455421911 |
| ENSMUSG00000025086 | Trub1 | protein_coding | 0.762415847 | 5.321156264 | 6.7916373 | 0.018892 | 0.457884574 |
| ENSMUSG00000020326 | Ccng1 | protein_coding | -0.815150634 | 4.812654868 | 6.7827494 | 0.018958 | 0.457884574 |
| ENSMUSG00000036646 | Man1b1 | protein_coding | 0.694646257 | 5.165876527 | 6.7801246 | 0.0189775 | 0.457884574 |

| ensembl_gene_id | external_gene_name | gene_biotype | log2 (fold change) | log2 (counts per million) | F-value | p-value | False discovery rate |
|---|---|---|---|---|---|---|---|
| ENSMUSG00000001313 | Rnd2 | protein_coding | -0.595352817 | 7.485209106 | 6.778482 | 0.0189898 | 0.457884574 |
| ENSMUSG00000064061 | Dzip3 | protein_coding | -0.615287343 | 5.567912203 | 6.7782403 | 0.0189916 | 0.457884574 |
| ENSMUSG00000090015 | Gm15446 | protein_coding | -1.629348857 | 3.140716799 | 6.7260346 | 0.0193853 | 0.465298329 |
| ENSMUSG00000051504 | Siglech | protein_coding | -1.117284452 | 3.656645677 | 6.7215809 | 0.0194193 | 0.465298329 |
| ENSMUSG00000029787 | Avl9 | protein_coding | -0.823221742 | 4.490577068 | 6.7203276 | 0.0194289 | 0.465298329 |
| ENSMUSG00000062421 | Arf2 | protein_coding | 0.666170743 | 6.704694964 | 6.7023893 | 0.0195667 | 0.46755674 |
| ENSMUSG00000086405 | 9330198N18Rik | lincRNA | -1.424661593 | 2.958387547 | 6.6908674 | 0.0196558 | 0.468293117 |
| ENSMUSG00000025626 | Phf6 | protein_coding | -0.75823212 | 4.926960192 | 6.680526 | 0.0197361 | 0.468293117 |
| ENSMUSG00000000594 | Gm2a | protein_coding | -0.534971892 | 7.111728144 | 6.6696558 | 0.019821 | 0.468293117 |
| ENSMUSG00000044408 | Sptssa | protein_coding | -0.550157296 | 6.727464877 | 6.6585945 | 0.0199078 | 0.468293117 |
| ENSMUSG00000009687 | Fxyd5 | protein_coding | -1.86551804 | 2.35456016 | 6.6562489 | 0.0199262 | 0.468293117 |
| ENSMUSG00000027714 | Exosc9 | protein_coding | 0.538576052 | 6.244827269 | 6.6547713 | 0.0199379 | 0.468293117 |
| ENSMUSG00000060923 | Acyp2 | protein_coding | -0.478830135 | 7.248511188 | 6.6546573 | 0.0199388 | 0.468293117 |
| ENSMUSG00000081225 | Cyp2j12 | protein_coding | -0.728867579 | 4.902637011 | 6.6537549 | 0.0199459 | 0.468293117 |
| ENSMUSG00000016510 | Mtif3 | protein_coding | 0.718762396 | 5.049693734 | 6.6362513 | 0.0200844 | 0.469564832 |
| ENSMUSG00000027883 | Gpsm2 | protein_coding | -1.217530259 | 2.962965676 | 6.6257291 | 0.0201682 | 0.469564832 |
| ENSMUSG00000022536 | Glyr1 | protein_coding | 0.606684137 | 7.648161917 | 6.6241149 | 0.0201811 | 0.469564832 |
| ENSMUSG00000021032 | Ngb | protein_coding | 0.569022073 | 8.359207565 | 6.6229697 | 0.0201903 | 0.469564832 |
| ENSMUSG00000022119 | Rbm26 | protein_coding | -0.815594087 | 4.824486385 | 6.6194544 | 0.0202184 | 0.469564832 |
| ENSMUSG00000036840 | Siah1a | protein_coding | -1.031713339 | 3.974477674 | 6.6107316 | 0.0202884 | 0.46957239 |
| ENSMUSG00000024193 | Phf1 | protein_coding | -0.961193232 | 4.191752132 | 6.6066244 | 0.0203214 | 0.46957239 |
| ENSMUSG00000025512 | Chid1 | protein_coding | 0.552680853 | 6.30292777 | 6.6031119 | 0.0203497 | 0.46957239 |
| ENSMUSG00000018199 | Trove2 | protein_coding | -0.614582314 | 5.844787989 | 6.5826104 | 0.0205159 | 0.471312672 |
| ENSMUSG00000044950 | Pwwp2a | protein_coding | -0.892155961 | 4.588338942 | 6.5717735 | 0.0206043 | 0.471312672 |
| ENSMUSG00000020319 | Wdpcp | protein_coding | 1.134667328 | 3.361835588 | 6.5672311 | 0.0206415 | 0.471312672 |
| ENSMUSG00000022842 | Ece2 | protein_coding | 0.757735634 | 5.364220344 | 6.5670669 | 0.0206429 | 0.471312672 |
| ENSMUSG00000007989 | Fzd3 | protein_coding | -0.61046867 | 5.708642128 | 6.5668913 | 0.0206443 | 0.471312672 |
| ENSMUSG00000001380 | Hars | protein_coding | 0.541015234 | 8.238366538 | 6.5580378 | 0.0207171 | 0.471705191 |
| ENSMUSG00000036599 | Chst12 | protein_coding | 0.650951585 | 5.686100521 | 6.5541319 | 0.0207492 | 0.471705191 |
| ENSMUSG00000038518 | Jarid2 | protein_coding | 0.661179561 | 8.066763333 | 6.5430473 | 0.02086 | 0.47322212 |
| ENSMUSG00000034839 | Larp6 | protein_coding | 0.961237406 | 4.866273999 | 6.5095809 | 0.0211204 | 0.478121923 |
| ENSMUSG00000020464 | Pnpt1 | protein_coding | 1.199310848 | 3.13808061 | 6.5030411 | 0.0211755 | 0.478362735 |
| ENSMUSG00000021477 | Ctsl | protein_coding | 0.5124041 | 7.359115582 | 6.491916 | 0.0212697 | 0.47885631 |
| ENSMUSG00000038349 | Plcl1 | protein_coding | -0.737605252 | 4.898905079 | 6.4871542 | 0.0213101 | 0.47885631 |
| ENSMUSG00000025959 | Klf7 | protein_coding | -0.64521997 | 5.180760094 | 6.4844227 | 0.0213334 | 0.47885631 |
| ENSMUSG00000041632 | Mrps27 | protein_coding | 0.827175877 | 5.571094879 | 6.4773031 | 0.0213941 | 0.47885631 |
| ENSMUSG00000109088 | Gm44593 | lincRNA | -1.497685236 | 2.679908493 | 6.4529607 | 0.0216032 | 0.47885631 |
| ENSMUSG00000020994 | Pnn | protein_coding | -0.674684414 | 5.979140039 | 6.4491582 | 0.0216361 | 0.47885631 |
| ENSMUSG00000028772 | Zcchc17 | protein_coding | 0.436575809 | 8.941094121 | 6.4415011 | 0.0217024 | 0.47885631 |
| ENSMUSG00000021619 | Atg10 | protein_coding | -0.547126267 | 5.97382329 | 6.4397256 | 0.0217178 | 0.47885631 |
| ENSMUSG00000058704 | Memo1 | protein_coding | -0.875942424 | 4.05571093 | 6.439127 | 0.021723 | 0.47885631 |
| ENSMUSG00000020886 | Dlg4 | protein_coding | -0.553358714 | 5.726195246 | 6.4386952 | 0.0217268 | 0.47885631 |
| ENSMUSG00000029062 | Cdk11b | protein_coding | 0.68254932 | 4.978969342 | 6.4318645 | 0.0217863 | 0.47885631 |
| ENSMUSG00000006611 | Hfe | protein_coding | 1.60123358 | 2.95804863 | 6.4260118 | 0.0218374 | 0.47885631 |
| ENSMUSG00000035566 | Pcdh17 | protein_coding | -0.454193316 | 6.755909387 | 6.4230274 | 0.0218635 | 0.47885631 |
| ENSMUSG00000020321 | Mdh1 | protein_coding | 0.516777416 | 11.90593286 | 6.4227203 | 0.0218662 | 0.47885631 |
| ENSMUSG00000062070 | Pgk1 | protein_coding | 0.630282934 | 5.410924104 | 6.4217362 | 0.0218748 | 0.47885631 |
| ENSMUSG00000027175 | Tcp11l1 | protein_coding | -0.902756132 | 3.964658979 | 6.4177332 | 0.0219099 | 0.47885631 |
| ENSMUSG00000020385 | Clk4 | protein_coding | -0.712972033 | 5.68838557 | 6.3943816 | 0.022116 | 0.482008975 |
| ENSMUSG00000059119 | Nap1l4 | protein_coding | 0.673462667 | 8.400587276 | 6.3888407 | 0.0221848 | 0.482008975 |
| ENSMUSG00000026761 | Orc4 | protein_coding | -0.567786459 | 6.233257525 | 6.3862084 | 0.0221886 | 0.482008975 |
| ENSMUSG00000029863 | Casp2 | protein_coding | -1.353885997 | 2.862350715 | 6.3805829 | 0.0222388 | 0.482124704 |
| ENSMUSG00000078861 | Zfp931 | protein_coding | -0.969728786 | 5.300697294 | 6.3755608 | 0.022334 | 0.482253379 |
| ENSMUSG00000020091 | Eif4ebp2 | protein_coding | -0.739525853 | 4.73625418 | 6.3636322 | 0.0223907 | 0.482253379 |
| ENSMUSG00000060935 | Tmem263 | protein_coding | -0.522147738 | 7.272686587 | 6.3634831 | 0.0223921 | 0.482253379 |
| ENSMUSG00000020541 | Tom1l1 | protein_coding | -0.6667205 | 5.201902742 | 6.3599248 | 0.0224241 | 0.482253379 |
| ENSMUSG00000056763 | Cspp1 | protein_coding | -0.691655829 | 4.77568364 | 6.3444111 | 0.0225645 | 0.484050964 |
| ENSMUSG00000060261 | Gtf2i | protein_coding | 0.609289678 | 6.664395869 | 6.3393916 | 0.0226101 | 0.484050964 |
| ENSMUSG00000063142 | Kcnma1 | protein_coding | -0.5291265 | 6.222946109 | 6.3358045 | 0.0226428 | 0.484050964 |
| ENSMUSG00000100147 | 1700047M11Rik | lincRNA | -0.877619447 | 4.785041665 | 6.3307379 | 0.022689 | 0.484076985 |
| ENSMUSG00000010803 | Gabra1 | protein_coding | -0.481425902 | 7.603223781 | 6.3183868 | 0.0228022 | 0.485253714 |
| ENSMUSG00000078117 | Gm16485 | protein_coding | -1.943931694 | 2.420546414 | 6.3123967 | 0.0228573 | 0.485253714 |
| ENSMUSG00000027132 | Katnbl1 | protein_coding | -1.103587353 | 3.374344132 | 6.3026747 | 0.0229471 | 0.485253714 |
| ENSMUSG00000030870 | Ubfd1 | protein_coding | 0.521206355 | 6.262234854 | 6.3022287 | 0.0229512 | 0.485253714 |
| ENSMUSG00000075229 | Ccdc58 | protein_coding | -0.778525407 | 4.814919982 | 6.2985444 | 0.0229853 | 0.485253714 |
| ENSMUSG00000073131 | Vma21 | protein_coding | -0.512539927 | 6.259907015 | 6.2953544 | 0.0230149 | 0.485253714 |
| ENSMUSG00000026004 | Kansl1l | protein_coding | -1.329749197 | 3.449601336 | 6.2803356 | 0.0231549 | 0.486418427 |
| ENSMUSG00000028633 | Ctps | protein_coding | 0.694897115 | 5.140252002 | 6.2797204 | 0.0231606 | 0.486418427 |
| ENSMUSG00000000278 | Scpep1 | protein_coding | 1.188240475 | 3.180799759 | 6.2713299 | 0.0232393 | 0.487118539 |
| ENSMUSG00000031772 | Cntnap4 | protein_coding | 1.207149357 | 3.277471789 | 6.259516 | 0.0233505 | 0.487857436 |
| ENSMUSG00000032381 | Fam96a | protein_coding | -0.52059837 | 6.727894162 | 6.2538563 | 0.023404 | 0.487857436 |
| ENSMUSG00000028414 | Fktn | protein_coding | -0.651860413 | 4.980741172 | 6.2495633 | 0.0234447 | 0.487857436 |
| ENSMUSG00000027889 | Ampd2 | protein_coding | 0.72321542 | 6.126254103 | 6.2481076 | 0.0234787 | 0.487857436 |
| ENSMUSG00000021196 | Pfkp | protein_coding | 0.665165055 | 7.676349815 | 6.2335202 | 0.0236176 | 0.487857436 |

| ensembl_gene_id | external_gene_name | gene_biotype | log2 (fold change) | log2 (counts per million) | F-value | p-value | False discovery rate |
|---|---|---|---|---|---|---|---|
| ENSMUSG00000023104 | Rfc2 | protein_coding | 0.508315604 | 6.384480116 | 6.2256743 | 0.0236726 | 0.487857436 |
| ENSMUSG00000021716 | Srek1ip1 | protein_coding | -0.57841037 | 6.026478648 | 6.2131488 | 0.023793 | 0.487857436 |
| ENSMUSG00000021171 | Esyt2 | protein_coding | -0.844496703 | 4.758010707 | 6.2121617 | 0.0238026 | 0.487857436 |
| ENSMUSG00000021709 | Erbin | protein_coding | -0.803281638 | 4.967908066 | 6.208278 | 0.0238401 | 0.487857436 |
| ENSMUSG00000060882 | Kcnd2 | protein_coding | -0.505163789 | 6.631503232 | 6.2079764 | 0.023843 | 0.487857436 |
| ENSMUSG00000039431 | Mtmr7 | protein_coding | 0.612643735 | 6.083458304 | 6.2056432 | 0.0238656 | 0.487857436 |
| ENSMUSG00000028347 | Tmeff1 | protein_coding | -0.623974996 | 5.739402037 | 6.2011785 | 0.0239088 | 0.487857436 |
| ENSMUSG00000037286 | Stag1 | protein_coding | -0.707354237 | 5.854711725 | 6.1867943 | 0.0240488 | 0.487857436 |
| ENSMUSG00000042790 | Rnf214 | protein_coding | 0.622993088 | 5.510956496 | 6.1746999 | 0.0241673 | 0.487857436 |
| ENSMUSG00000033287 | Kctd17 | protein_coding | -0.678859053 | 5.488237808 | 6.1735569 | 0.0241785 | 0.487857436 |
| ENSMUSG00000063229 | Ldha | protein_coding | 0.468429852 | 8.55512679 | 6.1730058 | 0.0241839 | 0.487857436 |
| ENSMUSG00000020063 | Sirt1 | protein_coding | -1.308637219 | 2.913816698 | 6.170729 | 0.0242063 | 0.487857436 |
| ENSMUSG00000021939 | Ctsb | protein_coding | 0.485303844 | 10.06288877 | 6.1689794 | 0.0242236 | 0.487857436 |
| ENSMUSG00000046380 | Jrk | protein_coding | 1.329976538 | 3.524425754 | 6.1689502 | 0.0242238 | 0.487857436 |
| ENSMUSG00000041798 | Gck | protein_coding | 0.927243665 | 5.389062002 | 6.1694574 | 0.0242393 | 0.487857436 |
| ENSMUSG00000055963 | Triqk | protein_coding | -0.539357982 | 5.890380288 | 6.1614473 | 0.0242979 | 0.487857436 |
| ENSMUSG00000032580 | Rbm5 | protein_coding | 0.501499053 | 6.59973986 | 6.1613969 | 0.0242984 | 0.487857436 |
| ENSMUSG00000031297 | Slc7a3 | protein_coding | 0.953183864 | 4.918918982 | 6.1556235 | 0.024376 | 0.487857436 |
| ENSMUSG00000051285 | Pcmtd1 | protein_coding | -0.68511098 | 5.094840284 | 6.1451638 | 0.0244594 | 0.487857436 |
| ENSMUSG00000024800 | Rpp30 | protein_coding | -0.533146419 | 6.361405398 | 6.1450609 | 0.0244604 | 0.487857436 |
| ENSMUSG00000068040 | Tm9sf4 | protein_coding | 0.557935288 | 6.085282906 | 6.1426466 | 0.0244845 | 0.487857436 |
| ENSMUSG00000014873 | Surf2 | protein_coding | 0.582586236 | 6.407623706 | 6.1393404 | 0.0245175 | 0.487857436 |
| ENSMUSG00000035547 | Capn5 | protein_coding | 0.898027004 | 5.012830379 | 6.1339291 | 0.0245716 | 0.487857436 |
| ENSMUSG00000047085 | Lrrc4b | protein_coding | -0.778925168 | 4.831499957 | 6.1262756 | 0.0246483 | 0.487857436 |
| ENSMUSG00000004865 | Srpk1 | protein_coding | 0.589130523 | 6.884929363 | 6.1256411 | 0.0246547 | 0.487857436 |
| ENSMUSG00000029415 | Sdad1 | protein_coding | 1.035324727 | 4.236613644 | 6.1174775 | 0.0247369 | 0.487857436 |
| ENSMUSG00000027301 | Oxt | protein_coding | 1.955951726 | 4.513115714 | 6.5760713 | 0.0248466 | 0.487857436 |
| ENSMUSG00000027375 | Mal | protein_coding | -0.540558013 | 7.874490494 | 6.0907611 | 0.025008 | 0.487857436 |
| ENSMUSG00000066798 | Zbtb6 | protein_coding | -1.048013377 | 3.804663878 | 6.0853999 | 0.0250629 | 0.487857436 |
| ENSMUSG00000036572 | Upf3b | protein_coding | -0.636245665 | 5.476546506 | 6.0776458 | 0.0251424 | 0.487857436 |
| ENSMUSG00000052613 | Pcdh15 | protein_coding | -0.965469419 | 4.051659865 | 6.0749055 | 0.0251706 | 0.487857436 |
| ENSMUSG00000026603 | Smyd2 | protein_coding | 1.013678398 | 3.360309474 | 6.0735702 | 0.0251843 | 0.487857436 |
| ENSMUSG00000042066 | Tmcc2 | protein_coding | -0.648506994 | 6.075979298 | 6.0706424 | 0.0252145 | 0.487857436 |
| ENSMUSG00000027487 | Cdk5rap1 | protein_coding | 0.924106823 | 4.290621699 | 6.069826 | 0.0252229 | 0.487857436 |
| ENSMUSG00000052707 | Tnrc6a | protein_coding | 0.625450661 | 5.374586533 | 6.0644198 | 0.0252788 | 0.487857436 |
| ENSMUSG00000032034 | Kcnj5 | protein_coding | 1.795792317 | 2.643768098 | 6.0582985 | 0.0253422 | 0.487857436 |
| ENSMUSG00000015222 | Map2 | protein_coding | -0.447563392 | 8.291809547 | 6.0545991 | 0.0253806 | 0.487857436 |
| ENSMUSG00000075700 | Selenot | protein_coding | -0.684784199 | 5.747603591 | 6.0463404 | 0.0254665 | 0.487857436 |
| ENSMUSG00000027067 | Ssrp1 | protein_coding | 0.539159898 | 6.496944194 | 6.0449556 | 0.025481 | 0.487857436 |
| ENSMUSG00000053289 | Ddx10 | protein_coding | -0.816462134 | 5.026413245 | 6.0323112 | 0.0256133 | 0.487857436 |
| ENSMUSG00000058748 | Zfp958 | protein_coding | -1.120701865 | 3.542261307 | 6.0175259 | 0.0257691 | 0.487857436 |
| ENSMUSG00000090386 | Mir99ahg | processed_transcript | -0.991874815 | 4.000903403 | 6.0109307 | 0.0258389 | 0.487857436 |
| ENSMUSG00000027820 | Mme | protein_coding | -1.478486077 | 2.654254081 | 6.0106349 | 0.025842 | 0.487857436 |
| ENSMUSG00000026014 | Raph1 | protein_coding | -0.698219419 | 4.442371162 | 6.0076291 | 0.025874 | 0.487857436 |
| ENSMUSG00000026034 | Clk1 | protein_coding | -0.873682405 | 5.304603204 | 6.0082989 | 0.0258879 | 0.487857436 |
| ENSMUSG00000008855 | Hdac5 | protein_coding | -0.742502062 | 5.34606645 | 6.0051956 | 0.0258998 | 0.487857436 |
| ENSMUSG00000032115 | Hyou1 | protein_coding | 1.118961426 | 4.211311351 | 6.0043239 | 0.0259301 | 0.487857436 |
| ENSMUSG00000025764 | Jade1 | protein_coding | -0.567735327 | 5.582317761 | 6.0005707 | 0.025949 | 0.487857436 |
| ENSMUSG00000024479 | Mal2 | protein_coding | -0.612695558 | 5.853344285 | 5.9855127 | 0.0261101 | 0.487857436 |
| ENSMUSG00000045917 | Tmem268 | protein_coding | 0.908026467 | 4.102658642 | 5.98546 | 0.0261106 | 0.487857436 |
| ENSMUSG00000000766 | Oprm1 | protein_coding | -0.748693314 | 5.499168234 | 5.9833493 | 0.0261333 | 0.487857436 |
| ENSMUSG00000032306 | Mpi | protein_coding | 0.524341045 | 7.010687497 | 5.9777858 | 0.0261932 | 0.487857436 |
| ENSMUSG00000046096 | BC030336 | protein_coding | -0.620042483 | 5.853572325 | 5.9732733 | 0.0262418 | 0.487857436 |
| ENSMUSG00000062116 | Zfp954 | protein_coding | 0.860562262 | 5.134182017 | 5.9702493 | 0.0262745 | 0.487857436 |
| ENSMUSG00000020128 | Vps54 | protein_coding | -1.330668325 | 3.201127812 | 5.9666372 | 0.0263136 | 0.487857436 |
| ENSMUSG00000002413 | Braf | protein_coding | -0.597908555 | 6.161610104 | 5.9604413 | 0.0263807 | 0.487857436 |
| ENSMUSG00000030204 | Ddx47 | protein_coding | 0.52495461 | 6.951213676 | 5.9492604 | 0.0265025 | 0.487857436 |
| ENSMUSG00000038160 | Atg5 | protein_coding | 0.481240146 | 6.381312863 | 5.9460714 | 0.0265373 | 0.487857436 |
| ENSMUSG00000025791 | Pgm2 | protein_coding | 0.66451752 | 7.022991 | 5.9475551 | 0.0265424 | 0.487857436 |
| ENSMUSG00000024077 | Strn | protein_coding | -0.539387793 | 6.419242611 | 5.9450827 | 0.0265481 | 0.487857436 |
| ENSMUSG00000031027 | Stk33 | protein_coding | 1.262233085 | 3.030573786 | 5.9407799 | 0.0265953 | 0.487857436 |
| ENSMUSG00000018379 | Srsf1 | protein_coding | -0.565637495 | 5.664583714 | 5.9395228 | 0.026609 | 0.487857436 |
| ENSMUSG00000046314 | Stxbp6 | protein_coding | -0.563892702 | 5.801587031 | 5.9297563 | 0.0267164 | 0.487857436 |
| ENSMUSG00000020009 | Ifngr1 | protein_coding | 0.882247719 | 4.016042888 | 5.9289662 | 0.0267251 | 0.487857436 |
| ENSMUSG00000036975 | Tmem177 | protein_coding | 1.023521712 | 3.831928398 | 5.9232934 | 0.0267877 | 0.487857436 |
| ENSMUSG00000036257 | Pnpla8 | protein_coding | -0.512706075 | 6.179466432 | 5.9213102 | 0.0268097 | 0.487857436 |
| ENSMUSG00000056258 | Kcnq3 | protein_coding | -0.842094786 | 5.110848953 | 5.9178951 | 0.0268475 | 0.487857436 |
| ENSMUSG00000027965 | Olfm3 | protein_coding | -0.812989134 | 4.733912521 | 5.9168107 | 0.0268595 | 0.487857436 |
| ENSMUSG00000056895 | Hist3h2ba | protein_coding | -1.026537553 | 3.746305997 | 5.9125989 | 0.0269062 | 0.487857436 |
| ENSMUSG00000090641 | Zfp712 | protein_coding | -1.711952793 | 2.588525085 | 5.9125268 | 0.026907 | 0.487857436 |
| ENSMUSG00000041000 | Trim62 | protein_coding | 0.973502675 | 3.93206568 | 5.9117332 | 0.0269159 | 0.487857436 |
| ENSMUSG00000034610 | Zcchc11 | protein_coding | -0.928834359 | 4.181040372 | 5.9097872 | 0.0269375 | 0.487857436 |
| ENSMUSG00000028977 | Casz1 | protein_coding | 0.929308041 | 3.652476258 | 5.9080763 | 0.0269565 | 0.487857436 |
| ENSMUSG00000041377 | Ninj2 | protein_coding | -0.877032081 | 4.6352863 | 5.9052684 | 0.0269878 | 0.487857436 |

| ensembl_gene_id | external_gene_name | gene_biotype | log2 (fold change) | log2 (counts per million) | F-value | p-value | False discovery rate |
|---|---|---|---|---|---|---|---|
| ENSMUSG00000059975 | Zfp74 | protein_coding | -0.972370818 | 3.668341166 | 5.9023186 | 0.0270207 | 0.487857436 |
| ENSMUSG00000033819 | Ppp1r16a | protein_coding | -0.722452365 | 4.734216239 | 5.8981234 | 0.0270676 | 0.487857436 |
| ENSMUSG00000026127 | Imp4 | protein_coding | 0.709281746 | 4.426090176 | 5.8945013 | 0.0271081 | 0.487857436 |
| ENSMUSG00000033184 | Tmed7 | protein_coding | -0.806163231 | 5.589289033 | 5.8895059 | 0.0271857 | 0.487857436 |
| ENSMUSG00000022139 | Mbnl2 | protein_coding | -0.509821375 | 7.573170751 | 5.8870628 | 0.0271916 | 0.487857436 |
| ENSMUSG00000001999 | Blvra | protein_coding | 0.450275934 | 6.923639702 | 5.8816166 | 0.0272529 | 0.487857436 |
| ENSMUSG00000024359 | Hspa9 | protein_coding | 0.543180975 | 6.974329792 | 5.8748886 | 0.0273289 | 0.487857436 |
| ENSMUSG00000024188 | Luc7l | protein_coding | 0.4905703 | 6.515031364 | 5.8734239 | 0.0273455 | 0.487857436 |
| ENSMUSG00000000148 | Brat1 | protein_coding | -1.140774121 | 4.062798521 | 5.8737096 | 0.0273638 | 0.487857436 |
| ENSMUSG00000029430 | Ran | protein_coding | 0.488077609 | 8.036845059 | 5.8706246 | 0.0273772 | 0.487857436 |
| ENSMUSG00000010045 | Tmem115 | protein_coding | -0.621984255 | 5.388791014 | 5.8688194 | 0.0273976 | 0.487857436 |
| ENSMUSG00000014243 | Zswim7 | protein_coding | 0.622431157 | 5.390457261 | 5.8683359 | 0.0274031 | 0.487857436 |
| ENSMUSG00000039697 | Ncoa7 | protein_coding | -0.529737822 | 5.966730322 | 5.8572269 | 0.0275295 | 0.488546922 |
| ENSMUSG00000026872 | Zeb2 | protein_coding | -0.785678913 | 4.886389772 | 5.8569432 | 0.0275327 | 0.488546922 |
| ENSMUSG00000034931 | Dhx8 | protein_coding | 0.823740659 | 4.399326705 | 5.8459084 | 0.0276589 | 0.489849733 |
| ENSMUSG00000030091 | Nup210 | protein_coding | 1.656732715 | 2.789625532 | 5.8425677 | 0.0276973 | 0.489849733 |
| ENSMUSG00000025545 | Clybl | protein_coding | -0.461490807 | 7.084661771 | 5.8341444 | 0.0277942 | 0.490756904 |
| ENSMUSG00000072258 | Taf1a | protein_coding | 0.863008917 | 4.182932386 | 5.8271461 | 0.027875 | 0.49137724 |
| ENSMUSG00000033282 | Rpgrip1l | protein_coding | -0.937822174 | 4.09526138 | 5.8140082 | 0.0280275 | 0.492179093 |
| ENSMUSG00000015656 | Hspa8 | protein_coding | 0.746676062 | 8.910845924 | 5.8847439 | 0.0280547 | 0.492179093 |
| ENSMUSG00000028959 | Fastk | protein_coding | -0.545141357 | 6.605509675 | 5.811294 | 0.0280591 | 0.492179093 |
| ENSMUSG00000027665 | Pik3ca | protein_coding | -0.616788097 | 5.507951303 | 5.8052708 | 0.0281294 | 0.492179093 |
| ENSMUSG00000041168 | Lonp1 | protein_coding | 0.641117206 | 4.990670459 | 5.8035659 | 0.0281494 | 0.492179093 |
| ENSMUSG00000022978 | Mis18a | protein_coding | -0.857354153 | 3.910452885 | 5.7980626 | 0.0282138 | 0.492505566 |
| ENSMUSG00000044772 | Sntn | protein_coding | -1.181055747 | 3.33696982 | 5.7727769 | 0.0285123 | 0.495835093 |
| ENSMUSG00000056215 | Lrguk | protein_coding | 1.889638959 | 2.904019546 | 5.7731055 | 0.0285303 | 0.495835093 |
| ENSMUSG00000026491 | Ahctf1 | protein_coding | 1.147872443 | 3.264716847 | 5.7684716 | 0.0285634 | 0.495835093 |
| ENSMUSG00000031753 | Cog4 | protein_coding | 1.056391336 | 4.126550012 | 5.7651674 | 0.0286028 | 0.495835093 |
| ENSMUSG00000037958 | Nsrp1 | protein_coding | -0.603674408 | 6.112309632 | 5.7569172 | 0.0287013 | 0.495835093 |
| ENSMUSG00000036057 | Ptpn23 | protein_coding | -1.009736969 | 3.875969448 | 5.7562767 | 0.028709 | 0.495835093 |
| ENSMUSG00000021391 | Cenpp | protein_coding | -1.036580367 | 3.797325587 | 5.7538671 | 0.0287378 | 0.495835093 |
| ENSMUSG00000030613 | Ccdc90b | protein_coding | -0.656511609 | 5.437788353 | 5.7508932 | 0.0287735 | 0.495835093 |
| ENSMUSG00000048905 | 4930539E08Rik | protein_coding | 1.89171539 | 2.498641152 | 5.7461945 | 0.0288299 | 0.495877625 |
| ENSMUSG00000024335 | Brd2 | protein_coding | 0.482686298 | 6.404587949 | 5.7429716 | 0.0288687 | 0.495877625 |
| ENSMUSG00000101493 | 2810405F17Rik | lincRNA | -1.566645332 | 2.5205655 | 5.7391903 | 0.0289143 | 0.495877625 |
| ENSMUSG00000019828 | Grm1 | protein_coding | -0.619361671 | 5.464318844 | 5.7322788 | 0.0289978 | 0.496518328 |
| ENSMUSG00000049796 | Crh | protein_coding | -1.362581926 | 2.83028429 | 5.7266159 | 0.0290665 | 0.49690255 |
| ENSMUSG00000037363 | Letm2 | protein_coding | 1.114359995 | 3.508788132 | 5.7151335 | 0.0292062 | 0.498499484 |
| ENSMUSG00000045103 | Dmd | protein_coding | -0.667785617 | 4.828921698 | 5.7069835 | 0.0293059 | 0.499408085 |
| ENSMUSG00000061524 | Zic2 | protein_coding | -1.595937139 | 2.582367582 | 5.6882813 | 0.0295361 | 0.501300004 |
| ENSMUSG00000035011 | Zbtb7a | protein_coding | -0.740131521 | 5.645804851 | 5.6863839 | 0.0295596 | 0.501300004 |
| ENSMUSG00000021377 | Dek | protein_coding | -0.657439265 | 5.116669153 | 5.6852382 | 0.0295738 | 0.501300004 |
| ENSMUSG00000021518 | Ptdss1 | protein_coding | 1.061415622 | 4.259465503 | 5.676229 | 0.0297079 | 0.501300004 |
| ENSMUSG00000029482 | Aacs | protein_coding | 0.873132108 | 4.888742531 | 5.6733742 | 0.0297211 | 0.501300004 |
| ENSMUSG00000068739 | Sars | protein_coding | 0.549648372 | 8.568066418 | 5.6708883 | 0.0297521 | 0.501300004 |
| ENSMUSG00000023964 | Calcr | protein_coding | 1.143676207 | 3.543107807 | 5.6693043 | 0.0297719 | 0.501300004 |
| ENSMUSG00000110027 | C030029H02Rik | lincRNA | -0.782141922 | 4.059750618 | 5.6678594 | 0.0297899 | 0.501300004 |
| ENSMUSG00000016552 | Foxred2 | protein_coding | 1.432804675 | 2.864188343 | 5.6631588 | 0.0298487 | 0.501504053 |
| ENSMUSG00000038637 | Lrrc56 | protein_coding | 1.560789215 | 2.744091276 | 5.6569657 | 0.0299263 | 0.501982796 |
| ENSMUSG00000030917 | Tmem159 | protein_coding | 0.593251506 | 5.45839797 | 5.6451016 | 0.0300756 | 0.501982796 |
| ENSMUSG00000038342 | Mlxip | protein_coding | 1.64721731 | 2.475320782 | 5.6412992 | 0.0301237 | 0.501982796 |
| ENSMUSG00000028431 | Ikbkap | protein_coding | 0.631801316 | 5.093809927 | 5.6383636 | 0.0301608 | 0.501982796 |
| ENSMUSG00000061244 | Exoc5 | protein_coding | -0.713586613 | 4.714899718 | 5.6345405 | 0.0302093 | 0.501982796 |
| ENSMUSG00000085436 | Zfp335os | antisense_RNA | -1.045289017 | 3.328951338 | 5.6324515 | 0.0302358 | 0.501982796 |
| ENSMUSG00000054426 | A930005H10Rik | processed_transcript | -0.697716284 | 4.744549992 | 5.6293763 | 0.0302749 | 0.501982796 |
| ENSMUSG00000020300 | Cpeb4 | protein_coding | -0.635702414 | 5.245710737 | 5.6231298 | 0.0303545 | 0.501982796 |
| ENSMUSG00000057649 | Brd9 | protein_coding | 0.616345247 | 5.955178553 | 5.6208836 | 0.0303832 | 0.501982796 |
| ENSMUSG00000021379 | Id4 | protein_coding | -0.450766997 | 6.834048099 | 5.6144661 | 0.0304653 | 0.501982796 |
| ENSMUSG00000032727 | Mier3 | protein_coding | -0.614413109 | 5.296214816 | 5.614346 | 0.0304668 | 0.501982796 |
| ENSMUSG00000052852 | Reep1 | protein_coding | 0.556442069 | 6.004427529 | 5.6116452 | 0.0305015 | 0.501982796 |
| ENSMUSG00000032854 | Ugt8a | protein_coding | -0.56644706 | 5.687384174 | 5.610734 | 0.0305132 | 0.501982796 |
| ENSMUSG00000050538 | B230217C12Rik | protein_coding | -0.433316782 | 7.527299184 | 5.6066913 | 0.0305651 | 0.501982796 |
| ENSMUSG00000042670 | Immp1l | protein_coding | -0.619586847 | 5.770079126 | 5.601145 | 0.0306365 | 0.501982796 |
| ENSMUSG00000023046 | Igfbp6 | protein_coding | -0.955102038 | 4.68971003 | 5.5991007 | 0.0306855 | 0.501982796 |
| ENSMUSG00000029148 | Nrbp1 | protein_coding | 0.659268784 | 6.827885153 | 5.5915547 | 0.0307831 | 0.501982796 |
| ENSMUSG00000052298 | Cdc42se2 | protein_coding | 0.592726839 | 7.224808403 | 5.5901413 | 0.0308014 | 0.501982796 |
| ENSMUSG00000041141 | Pnmal1 | protein_coding | 0.607118486 | 7.956656141 | 5.5892876 | 0.0308125 | 0.501982796 |
| ENSMUSG00000026273 | Mterf4 | protein_coding | 0.600967806 | 5.478050371 | 5.5845869 | 0.0308509 | 0.501982796 |
| ENSMUSG00000063550 | Nup98 | protein_coding | 0.612437981 | 5.737977979 | 5.5840817 | 0.0308575 | 0.501982796 |
| ENSMUSG00000045414 | 1190002N15Rik | protein_coding | -0.91605343 | 3.857464086 | 5.5780046 | 0.0309366 | 0.502104401 |
| ENSMUSG00000067578 | Cbln4 | protein_coding | 1.029120523 | 4.364327424 | 5.5780768 | 0.0309584 | 0.502104401 |
| ENSMUSG00000004069 | Dnaja3 | protein_coding | 0.560819155 | 6.564819385 | 5.5714141 | 0.0310227 | 0.502390531 |
| ENSMUSG00000021057 | Akap5 | protein_coding | -0.525345908 | 6.766825998 | 5.5617947 | 0.0311489 | 0.503674882 |
| ENSMUSG00000032527 | Pccb | protein_coding | 0.682859472 | 5.911814377 | 5.5558096 | 0.0312277 | 0.504152143 |

| ensembl_gene_id | external_gene_name | gene_biotype | log2 (fold change) | log2 (counts per million) | F-value | p-value | False discovery rate |
|---|---|---|---|---|---|---|---|
| ENSMUSG00000039768 | Dnajc11 | protein_coding | 0.733536575 | 6.270257955 | 5.5587084 | 0.0312722 | 0.504152143 |
| ENSMUSG00000032235 | Ice2 | protein_coding | -1.342853267 | 2.950215542 | 5.5447345 | 0.0313741 | 0.504287236 |
| ENSMUSG00000001416 | Cct3 | protein_coding | 0.485395388 | 7.875360436 | 5.5447137 | 0.0313743 | 0.504287236 |
| ENSMUSG00000022494 | Shisa9 | protein_coding | -0.721097367 | 4.665970214 | 5.5386409 | 0.0314549 | 0.504828289 |
| ENSMUSG00000109156 | Gm45194 | protein_coding | -0.965467188 | 4.031249565 | 5.5333598 | 0.0315252 | 0.505202417 |
| ENSMUSG00000069520 | Tmem19 | protein_coding | 0.584595803 | 5.323552314 | 5.5278713 | 0.0315985 | 0.505270037 |
| ENSMUSG00000042595 | Fam199x | protein_coding | -1.02807663 | 3.526595474 | 5.5252992 | 0.0316329 | 0.505270037 |
| ENSMUSG00000052337 | Immt | protein_coding | 0.582062413 | 7.219270007 | 5.524203 | 0.0316704 | 0.505270037 |
| ENSMUSG00000030161 | Gabarapl1 | protein_coding | 0.487302935 | 10.32629476 | 5.5170663 | 0.0317433 | 0.505420604 |
| ENSMUSG00000027799 | Nbea | protein_coding | -0.519585543 | 6.13149796 | 5.5147889 | 0.0317739 | 0.505420604 |
| ENSMUSG00000002617 | Zfp40 | protein_coding | -1.145602476 | 3.713377652 | 5.5051213 | 0.0319042 | 0.506172661 |
| ENSMUSG00000022035 | Ccdc25 | protein_coding | 0.495243149 | 6.514769902 | 5.5030861 | 0.0319317 | 0.506172661 |
| ENSMUSG00000030060 | Hmces | protein_coding | 0.735826435 | 5.224205205 | 5.5008206 | 0.0319624 | 0.506172661 |
| ENSMUSG00000067787 | Blcap | protein_coding | 0.463100244 | 8.873666099 | 5.490546 | 0.0321018 | 0.506580782 |
| ENSMUSG00000015396 | Cd83 | protein_coding | 0.597307337 | 6.530667424 | 5.4879178 | 0.0321376 | 0.506580782 |
| ENSMUSG00000026000 | Lancl1 | protein_coding | 0.456861858 | 6.946182531 | 5.48604 | 0.0321632 | 0.506580782 |
| ENSMUSG00000025162 | Csnk1d | protein_coding | 0.548590613 | 6.384389758 | 5.482796 | 0.0322075 | 0.506580782 |
| ENSMUSG00000032340 | Neo1 | protein_coding | 0.806820541 | 4.574868656 | 5.4812706 | 0.0322283 | 0.506580782 |
| ENSMUSG00000021395 | Spin1 | protein_coding | -0.477567637 | 7.491724962 | 5.4781677 | 0.0322708 | 0.506580782 |
| ENSMUSG00000022253 | Nadk2 | protein_coding | -0.743323943 | 4.544090733 | 5.4679644 | 0.0324108 | 0.508037505 |
| ENSMUSG00000034255 | Arhgap27 | protein_coding | 0.68800689 | 5.421061186 | 5.4618316 | 0.0324953 | 0.50849674 |
| ENSMUSG00000022454 | Nell2 | protein_coding | 0.455707553 | 7.578508581 | 5.4563866 | 0.0325706 | 0.50849674 |
| ENSMUSG00000021930 | Spryd7 | protein_coding | -0.434312793 | 6.699836475 | 5.4498139 | 0.0326617 | 0.50849674 |
| ENSMUSG00000028354 | Fmn2 | protein_coding | -0.835008574 | 4.408030175 | 5.4479506 | 0.0326876 | 0.50849674 |
| ENSMUSG00000036282 | Naa30 | protein_coding | -0.887317035 | 4.880531943 | 5.4382168 | 0.0328464 | 0.50849674 |
| ENSMUSG00000026171 | Rnf25 | protein_coding | 0.574840548 | 6.474817037 | 5.4339508 | 0.0328828 | 0.50849674 |
| ENSMUSG00000020280 | Pus10 | protein_coding | -0.95895443 | 3.556230994 | 5.4332998 | 0.0328919 | 0.50849674 |
| ENSMUSG00000038538 | Ubn2 | protein_coding | -0.643909091 | 5.661004336 | 5.4304772 | 0.0329314 | 0.50849674 |
| ENSMUSG00000066306 | Numa1 | protein_coding | -0.74341325 | 4.956374096 | 5.4301189 | 0.0329364 | 0.50849674 |
| ENSMUSG00000034902 | Pip5k1c | protein_coding | -0.546571963 | 6.460003544 | 5.4262707 | 0.0329904 | 0.50849674 |
| ENSMUSG00000066613 | Zfp932 | protein_coding | -0.868846372 | 3.590341987 | 5.4218344 | 0.0330528 | 0.50849674 |
| ENSMUSG00000028549 | Itgb3bp | protein_coding | -0.736626574 | 4.194755483 | 5.4212378 | 0.0330612 | 0.50849674 |
| ENSMUSG00000044499 | Hs3st5 | protein_coding | -1.051260766 | 3.457253235 | 5.416718 | 0.0331249 | 0.50849674 |
| ENSMUSG00000098108 | Gm27008 | antisense_RNA | -1.30926758 | 3.144980135 | 5.4153183 | 0.0331446 | 0.50849674 |
| ENSMUSG00000042097 | Zfp239 | protein_coding | 0.536285099 | 6.28642213 | 5.4149763 | 0.0331495 | 0.50849674 |
| ENSMUSG00000045466 | Zfp956 | protein_coding | 0.933848037 | 4.374954297 | 5.3989373 | 0.0333769 | 0.510538797 |
| ENSMUSG00000031770 | Herpud1 | protein_coding | 0.505898378 | 6.77839581 | 5.3944616 | 0.0334406 | 0.510538797 |
| ENSMUSG00000063594 | Gng8 | protein_coding | 0.847509706 | 4.820213282 | 5.3923411 | 0.0334709 | 0.510538797 |
| ENSMUSG00000020024 | Cep83 | protein_coding | -0.871413482 | 4.004862667 | 5.3852302 | 0.0335726 | 0.510538797 |
| ENSMUSG00000063200 | Nol7 | protein_coding | -0.565577293 | 6.24434449 | 5.3847922 | 0.0335789 | 0.510538797 |
| ENSMUSG00000024614 | Tmx3 | protein_coding | -0.577793477 | 5.685585436 | 5.3718708 | 0.0337647 | 0.510538797 |
| ENSMUSG00000039166 | Akap7 | protein_coding | -0.530226257 | 6.00025257 | 5.3716922 | 0.0337672 | 0.510538797 |
| ENSMUSG00000045064 | Zc2hc1c | protein_coding | 1.34669796 | 3.334567697 | 5.3704297 | 0.0337855 | 0.510538797 |
| ENSMUSG00000032060 | Cryab | protein_coding | -0.495596703 | 9.198418857 | 5.3643004 | 0.0338741 | 0.510538797 |
| ENSMUSG00000073680 | Tmem88b | protein_coding | -0.734248945 | 5.823505136 | 5.3594411 | 0.033968 | 0.510538797 |
| ENSMUSG00000000167 | Pih1d2 | protein_coding | -0.883163297 | 3.671753911 | 5.3575 | 0.0339727 | 0.510538797 |
| ENSMUSG00000010362 | Rdm1 | protein_coding | -0.789909616 | 5.33457854 | 5.3571769 | 0.0339774 | 0.510538797 |
| ENSMUSG00000022636 | Alcam | protein_coding | -0.568131045 | 7.059554592 | 5.3571629 | 0.0339776 | 0.510538797 |
| ENSMUSG00000038011 | Dnah10 | protein_coding | -1.456232599 | 2.859504059 | 5.351441 | 0.0340608 | 0.510538797 |
| ENSMUSG00000029147 | Ppm1g | protein_coding | 0.635824361 | 5.239946908 | 5.3497253 | 0.0340858 | 0.510538797 |
| ENSMUSG00000030729 | Pgm2l1 | protein_coding | -0.539346048 | 6.782313586 | 5.3460591 | 0.0341393 | 0.510538797 |
| ENSMUSG00000041840 | Haus1 | protein_coding | -0.499128708 | 5.787045799 | 5.3447776 | 0.0341581 | 0.510538797 |
| ENSMUSG00000004187 | Kifc2 | protein_coding | -0.548729576 | 6.489650591 | 5.3414362 | 0.034207 | 0.510538797 |
| ENSMUSG00000028405 | Aco1 | protein_coding | 0.983615976 | 4.148423769 | 5.3397751 | 0.0342313 | 0.510538797 |
| ENSMUSG00000057060 | Slc35f3 | protein_coding | -1.027339822 | 3.816031417 | 5.3369237 | 0.0342731 | 0.510538797 |
| ENSMUSG00000029499 | Pxmp2 | protein_coding | -0.972522203 | 3.719381021 | 5.3364775 | 0.0342796 | 0.510538797 |
| ENSMUSG00000042579 | 4632404H12Rik | lincRNA | -0.87500615 | 3.727082463 | 5.3300486 | 0.0343742 | 0.511238292 |
| ENSMUSG00000067818 | Myl9 | protein_coding | -0.764247597 | 4.61961519 | 5.3147056 | 0.0346009 | 0.513766282 |
| ENSMUSG00000031327 | Chic1 | protein_coding | -0.594658341 | 5.439736471 | 5.3120278 | 0.0346407 | 0.513766282 |
| ENSMUSG00000033065 | Pfkm | protein_coding | 0.590593378 | 8.237464808 | 5.3098755 | 0.0346964 | 0.513766282 |
| ENSMUSG00000111375 | Btbd8 | protein_coding | 0.761795602 | 5.097936744 | 5.3047938 | 0.0347483 | 0.513766282 |
| ENSMUSG00000026675 | Hsd17b7 | protein_coding | 1.073532466 | 4.474511903 | 5.3086691 | 0.034783 | 0.513766282 |
| ENSMUSG00000039968 | Rsbn1l | protein_coding | -1.019742598 | 3.902598808 | 5.2917507 | 0.0349434 | 0.515427576 |
| ENSMUSG00000001025 | S100a6 | protein_coding | -0.784808708 | 5.360026536 | 5.2884846 | 0.0349925 | 0.515444001 |
| ENSMUSG00000028051 | Hcn3 | protein_coding | 1.032000681 | 3.320938554 | 5.2809771 | 0.0351055 | 0.516401814 |
| ENSMUSG00000042064 | Myo3b | protein_coding | -1.578485466 | 2.505294159 | 5.2766732 | 0.0351705 | 0.516651147 |
| ENSMUSG00000057182 | Scn3a | protein_coding | -0.574924604 | 5.406071106 | 5.2734293 | 0.0352196 | 0.51666632 |
| ENSMUSG00000054693 | Adam10 | protein_coding | -0.506060529 | 5.802093242 | 5.2685856 | 0.035293 | 0.517038216 |
| ENSMUSG00000046442 | Ppm1e | protein_coding | -0.590696552 | 5.906311378 | 5.2607261 | 0.0354126 | 0.517601337 |
| ENSMUSG00000010057 | Nprl2 | protein_coding | 0.600322362 | 6.55218835 | 5.255439 | 0.0354932 | 0.517601337 |
| ENSMUSG00000014763 | Fam120b | protein_coding | 0.527922912 | 6.325271253 | 5.2536403 | 0.0355207 | 0.517601337 |
| ENSMUSG00000092274 | Neat1 | lincRNA | -1.084921958 | 3.937045109 | 5.2549901 | 0.035524 | 0.517601337 |
| ENSMUSG00000053119 | Chmp3 | protein_coding | 0.452370543 | 7.855505258 | 5.2457478 | 0.0356417 | 0.518612732 |
| ENSMUSG00000064138 | Fam172a | protein_coding | -1.058986971 | 3.102864087 | 5.2374435 | 0.0357694 | 0.519768268 |

| ensembl_gene_id | external_gene_name | gene_biotype | log2 (fold change) | log2 (counts per million) | F-value | p-value | False discovery rate |
|---|---|---|---|---|---|---|---|
| ENSMUSG00000030145 | Zfp248 | protein_coding | -0.828550967 | 4.432422608 | 5.227184 | 0.035928 | 0.519872238 |
| ENSMUSG00000039585 | Myo9a | protein_coding | -1.366656019 | 2.871054901 | 5.2265537 | 0.0359377 | 0.519872238 |
| ENSMUSG00000021519 | Mterf3 | protein_coding | -0.558241137 | 5.873411583 | 5.2254249 | 0.0359552 | 0.519872238 |
| ENSMUSG00000037876 | Jmjd1c | protein_coding | 0.794649985 | 4.82454241 | 5.2244762 | 0.03597 | 0.519872238 |
| ENSMUSG00000007458 | M6pr | protein_coding | 0.487057477 | 7.138541407 | 5.2146433 | 0.0361229 | 0.521381792 |
| ENSMUSG00000058756 | Thra | protein_coding | -0.499393247 | 6.284864811 | 5.2111729 | 0.036177 | 0.521463503 |
| ENSMUSG00000032114 | Slc37a4 | protein_coding | 0.859273333 | 4.078771897 | 5.2056621 | 0.0362632 | 0.52200609 |
| ENSMUSG00000036093 | Arl5a | protein_coding | -0.729109322 | 4.645275477 | 5.201581 | 0.0363272 | 0.522027495 |
| ENSMUSG00000025474 | Tubgcp2 | protein_coding | 0.726227204 | 5.413787032 | 5.1993773 | 0.0363618 | 0.522027495 |
| ENSMUSG00000068856 | Sf3b4 | protein_coding | -0.811461038 | 4.936819582 | 5.1895283 | 0.0365169 | 0.523555358 |
| ENSMUSG00000034998 | Foxn2 | protein_coding | -0.745272546 | 4.48501583 | 5.1853952 | 0.0365822 | 0.523793472 |
| ENSMUSG00000038112 | AW551984 | protein_coding | 0.704899206 | 6.598811039 | 5.2182893 | 0.0366811 | 0.524511151 |
| ENSMUSG00000030805 | Stx4a | protein_coding | 0.554073264 | 6.131729887 | 5.1680002 | 0.0368586 | 0.525404165 |
| ENSMUSG00000036810 | Cnep1r1 | protein_coding | -0.636615875 | 5.715660577 | 5.1679315 | 0.0368597 | 0.525404165 |
| ENSMUSG00000006641 | Slc5a6 | protein_coding | 0.777242907 | 4.429235679 | 5.1645021 | 0.0369145 | 0.525404165 |
| ENSMUSG00000070056 | Mfhas1 | protein_coding | 0.678107616 | 4.650240112 | 5.1629681 | 0.036939 | 0.525404165 |
| ENSMUSG00000024563 | Smad2 | protein_coding | 0.761037904 | 4.776615974 | 5.1545203 | 0.0370745 | 0.526633976 |
| ENSMUSG00000031617 | Tmem184c | protein_coding | -0.80324244 | 4.827262625 | 5.1513015 | 0.0371262 | 0.526673399 |
| ENSMUSG00000001630 | Stk38l | protein_coding | 0.615794449 | 5.132107652 | 5.1474855 | 0.0371877 | 0.526687557 |
| ENSMUSG00000110185 | Igip | protein_coding | -0.718997326 | 5.364580679 | 5.1451615 | 0.0372252 | 0.526687557 |
| ENSMUSG00000095139 | Pou3f2 | protein_coding | -0.664576923 | 4.942613641 | 5.1403772 | 0.0373025 | 0.526687807 |
| ENSMUSG00000085829 | Gm4285 | antisense_RNA | -1.856582504 | 2.39625023 | 5.1376408 | 0.0373468 | 0.526687807 |
| ENSMUSG00000074519 | Zfp971 | protein_coding | -1.04099515 | 3.113243149 | 5.1360778 | 0.0373722 | 0.526687807 |
| ENSMUSG00000024268 | Celf4 | protein_coding | -0.598324716 | 9.063432229 | 5.1345091 | 0.037422 | 0.526699814 |
| ENSMUSG00000021870 | Slmap | protein_coding | -0.72744766 | 5.716524717 | 5.1284259 | 0.0375209 | 0.527332157 |
| ENSMUSG00000020598 | Nrcam | protein_coding | 0.528745371 | 5.843984081 | 5.1220986 | 0.0375996 | 0.527332157 |
| ENSMUSG00000067336 | Bmpr2 | protein_coding | -0.559779481 | 5.509089548 | 5.1171133 | 0.0376811 | 0.527332157 |
| ENSMUSG00000028444 | Cntfr | protein_coding | -0.978981682 | 3.91942776 | 5.1163756 | 0.0376932 | 0.527332157 |
| ENSMUSG00000021236 | Entpd5 | protein_coding | 0.75956694 | 4.612089594 | 5.1152215 | 0.0377121 | 0.527332157 |
| ENSMUSG00000089872 | Rps6kc1 | protein_coding | 0.648716513 | 4.90180226 | 5.1096052 | 0.0378043 | 0.527729246 |
| ENSMUSG00000027827 | Kcnab1 | protein_coding | -0.43925795 | 6.617753753 | 5.1075131 | 0.0378387 | 0.527729246 |
| ENSMUSG00000031539 | Ap3m2 | protein_coding | 0.592349806 | 7.416457161 | 5.1034651 | 0.0379298 | 0.528315402 |
| ENSMUSG00000004151 | Etv1 | protein_coding | -0.611282454 | 5.695036509 | 5.0949335 | 0.0380463 | 0.528639272 |
| ENSMUSG00000057594 | Arl16 | protein_coding | 0.448405549 | 6.821007018 | 5.094622 | 0.0380514 | 0.528639272 |
| ENSMUSG00000030512 | Snrpa1 | protein_coding | -0.423487731 | 6.516635699 | 5.0868258 | 0.0381807 | 0.529217611 |
| ENSMUSG00000043629 | 1700019D03Rik | protein_coding | -0.954323947 | 3.571831769 | 5.0851225 | 0.0382091 | 0.529217611 |
| ENSMUSG00000026787 | Gad2 | protein_coding | -0.479924369 | 8.99710077 | 5.0832223 | 0.0382407 | 0.529217611 |
| ENSMUSG00000074505 | Fat3 | protein_coding | -0.93503579 | 4.060654349 | 5.0754132 | 0.038371 | 0.529487473 |
| ENSMUSG00000040724 | Kcna2 | protein_coding | -0.688770058 | 5.230897557 | 5.0737118 | 0.0383994 | 0.529487473 |
| ENSMUSG00000113701 | AC121583.1 | transcribed_processe | -0.706822871 | 4.360972706 | 5.0669817 | 0.0385122 | 0.529487473 |
| ENSMUSG00000011306 | Sugp1 | protein_coding | 0.603999556 | 6.264808811 | 5.0651862 | 0.0385424 | 0.529487473 |
| ENSMUSG00000036822 | Topors | protein_coding | 1.1112128 | 4.026247427 | 5.0658973 | 0.0385551 | 0.529487473 |
| ENSMUSG00000017286 | Glod4 | protein_coding | 0.404004629 | 7.522430869 | 5.0618209 | 0.038599 | 0.529487473 |
| ENSMUSG00000029234 | Tmem165 | protein_coding | -1.203242487 | 3.647534826 | 5.0606186 | 0.0386439 | 0.529487473 |
| ENSMUSG00000019853 | Hebp2 | protein_coding | 0.786495677 | 3.909298826 | 5.0566723 | 0.0386857 | 0.529487473 |
| ENSMUSG00000024740 | Ddb1 | protein_coding | 0.655255432 | 7.069296325 | 5.0877694 | 0.038793 | 0.529487473 |
| ENSMUSG00000004698 | Hdac9 | protein_coding | -0.65820363 | 5.084762076 | 5.0430191 | 0.0389169 | 0.529487473 |
| ENSMUSG00000041679 | Lrrc29 | protein_coding | -1.306096654 | 2.848384261 | 5.0422302 | 0.0389303 | 0.529487473 |
| ENSMUSG00000033460 | Armcx1 | protein_coding | 0.539417609 | 6.736641341 | 5.0372762 | 0.0390147 | 0.529487473 |
| ENSMUSG00000052305 | Hbb-bs | protein_coding | -0.749234604 | 5.942453252 | 5.042301 | 0.0390191 | 0.529487473 |
| ENSMUSG00000048154 | Kmt2d | protein_coding | -0.77748862 | 6.139164003 | 5.0898321 | 0.0390324 | 0.529487473 |
| ENSMUSG00000037331 | Larp1 | protein_coding | 0.642005494 | 5.011395459 | 5.0357594 | 0.0390405 | 0.529487473 |
| ENSMUSG00000042742 | Bmt2 | protein_coding | -0.750083889 | 4.507488992 | 5.0331068 | 0.0390858 | 0.529487473 |
| ENSMUSG00000046793 | Gpr61 | protein_coding | 0.802682759 | 4.355467775 | 5.0324338 | 0.0390973 | 0.529487473 |
| ENSMUSG00000031570 | Plpp5 | protein_coding | -0.708214177 | 4.85331234 | 5.0128039 | 0.0394343 | 0.531365186 |
| ENSMUSG00000032745 | Gpbp1 | protein_coding | -0.547686152 | 6.97775992 | 5.0123413 | 0.0394423 | 0.531365186 |
| ENSMUSG00000028760 | Eif4g3 | protein_coding | -0.59058698 | 5.792021111 | 5.0101298 | 0.0394805 | 0.531365186 |
| ENSMUSG00000020921 | Tmem101 | protein_coding | 0.572267632 | 5.629054628 | 5.01009 | 0.0394811 | 0.531365186 |
| ENSMUSG00000097080 | 1700086O06Rik | lincRNA | -1.004471684 | 3.712583447 | 5.0098005 | 0.0394861 | 0.531365186 |
| ENSMUSG00000031311 | Nono | protein_coding | 0.696312782 | 6.397071582 | 5.0273733 | 0.0395916 | 0.531365186 |
| ENSMUSG00000008976 | Gabpa | protein_coding | -1.120852131 | 2.888751968 | 5.0030557 | 0.0396029 | 0.531365186 |
| ENSMUSG00000041720 | Pi4ka | protein_coding | -0.824062438 | 4.881560216 | 4.9999498 | 0.0396568 | 0.531365186 |
| ENSMUSG00000031647 | Mfap3l | protein_coding | -0.832386539 | 3.545629981 | 4.9960671 | 0.0397243 | 0.531365186 |
| ENSMUSG00000046873 | Mbtps2 | protein_coding | -0.899372931 | 3.778127387 | 4.9957062 | 0.0397306 | 0.531365186 |
| ENSMUSG00000049904 | Tmem17 | protein_coding | 0.759930119 | 3.952169071 | 4.9927982 | 0.0397812 | 0.531365186 |
| ENSMUSG00000047996 | Prrg1 | protein_coding | -0.953549163 | 3.350186742 | 4.990062 | 0.0398289 | 0.531365186 |
| ENSMUSG00000049583 | Grm5 | protein_coding | -0.604160875 | 7.011997702 | 4.9874676 | 0.0398992 | 0.531643032 |
| ENSMUSG00000090063 | Dlx6os1 | antisense_RNA | -1.010350348 | 4.559342189 | 4.9833584 | 0.0400369 | 0.532243722 |
| ENSMUSG00000034570 | Inpp5j | protein_coding | -1.138949977 | 3.112371007 | 4.9778188 | 0.0400433 | 0.532243722 |
| ENSMUSG00000010592 | Dazl | protein_coding | -2.569578893 | 2.350461149 | 5.0008339 | 0.0401469 | 0.532962185 |
| ENSMUSG00000097518 | Gm26694 | lincRNA | -0.797663839 | 4.020025057 | 4.9687503 | 0.0402029 | 0.5330476 |
| ENSMUSG00000003354 | Ccdc65 | protein_coding | 0.858091402 | 4.403472492 | 4.9612843 | 0.0403349 | 0.533816467 |
| ENSMUSG00000092083 | Kcnb2 | protein_coding | -0.961554602 | 3.41738996 | 4.9588818 | 0.0403774 | 0.533816467 |
| ENSMUSG00000022822 | Abcc5 | protein_coding | 0.463554681 | 6.360321778 | 4.9570565 | 0.0404098 | 0.533816467 |

| ensembl_gene_id | external_gene_name | gene_biotype | log2 (fold change) | log2 (counts per million) | F-value | p-value | False discovery rate |
|---|---|---|---|---|---|---|---|
| ENSMUSG00000027634 | Ndrg3 | protein_coding | 0.615697197 | 9.406259515 | 4.9555698 | 0.0404613 | 0.533840638 |
| ENSMUSG00000007021 | Syngr3 | protein_coding | 0.457836211 | 9.000692261 | 4.9328301 | 0.0408424 | 0.535936002 |
| ENSMUSG00000032077 | Bud13 | protein_coding | 1.178333318 | 4.458012439 | 4.9855594 | 0.0408779 | 0.535936002 |
| ENSMUSG00000021748 | Pdhb | protein_coding | 0.383604236 | 7.858428937 | 4.929614 | 0.0409002 | 0.535936002 |
| ENSMUSG00000042208 | 0610010F05Rik | protein_coding | -0.580544759 | 4.933790896 | 4.9240018 | 0.0410014 | 0.535936002 |
| ENSMUSG00000026888 | Grb14 | protein_coding | -0.633089728 | 6.202078848 | 4.9221657 | 0.0410598 | 0.535936002 |
| ENSMUSG00000042671 | Rgs8 | protein_coding | 0.623455549 | 6.430973914 | 4.917802 | 0.0411387 | 0.535936002 |
| ENSMUSG00000041837 | Pdcd7 | protein_coding | -0.874988819 | 4.364299716 | 4.9159722 | 0.0411466 | 0.535936002 |
| ENSMUSG00000057715 | A830018L16Rik | protein_coding | -0.700820377 | 4.937030067 | 4.915919 | 0.0411476 | 0.535936002 |
| ENSMUSG00000057554 | Lgals8 | protein_coding | 0.710425552 | 5.631530436 | 4.9142973 | 0.041177 | 0.535936002 |
| ENSMUSG00000055923 | Aasdh | protein_coding | 1.23881043 | 3.069737384 | 4.9133085 | 0.0411949 | 0.535936002 |
| ENSMUSG00000025745 | Hadha | protein_coding | 0.592334658 | 5.641655946 | 4.9122421 | 0.0412143 | 0.535936002 |
| ENSMUSG00000034152 | Exoc3 | protein_coding | 0.453550801 | 7.219471058 | 4.9120238 | 0.0412182 | 0.535936002 |
| ENSMUSG00000024982 | Zdhhc6 | protein_coding | 0.608740299 | 4.825078853 | 4.9092842 | 0.041268 | 0.535936002 |
| ENSMUSG00000105265 | Sox2ot | processed_transcript | -0.456041809 | 6.116986292 | 4.9014801 | 0.0414102 | 0.536310833 |
| ENSMUSG00000033628 | Pik3c3 | protein_coding | 0.63081778 | 5.531721256 | 4.899547 | 0.0414455 | 0.536310833 |
| ENSMUSG00000025892 | Gria4 | protein_coding | -0.623202619 | 5.401965913 | 4.8973728 | 0.0414853 | 0.536310833 |
| ENSMUSG00000048483 | Zdhhc22 | protein_coding | 1.072311783 | 7.562365395 | 5.2675011 | 0.041539 | 0.536310833 |
| ENSMUSG00000054196 | Cthrc1 | protein_coding | 0.527728663 | 5.38972456 | 4.8891083 | 0.0416368 | 0.536310833 |
| ENSMUSG00000046139 | Patl1 | protein_coding | 0.923723762 | 3.763403618 | 4.8889529 | 0.0416396 | 0.536310833 |
| ENSMUSG00000018347 | Zkscan6 | protein_coding | 1.675215185 | 3.488809905 | 4.9740507 | 0.0416625 | 0.536310833 |
| ENSMUSG00000020605 | Hs1bp3 | protein_coding | 0.862294176 | 3.979064474 | 4.8850122 | 0.0417121 | 0.536310833 |
| ENSMUSG00000051034 | Zfp11 | protein_coding | -1.162816496 | 3.349475125 | 4.881329 | 0.04178 | 0.536310833 |
| ENSMUSG00000026558 | Uck2 | protein_coding | 0.908306314 | 3.764361791 | 4.8804815 | 0.0417956 | 0.536310833 |
| ENSMUSG00000024378 | Stard4 | protein_coding | 0.945424519 | 4.103663219 | 4.8707228 | 0.0419762 | 0.537611973 |
| ENSMUSG00000045648 | Vwc2l | protein_coding | 0.767330011 | 3.978575379 | 4.8695982 | 0.041997 | 0.537611973 |
| ENSMUSG00000032101 | Ddx25 | protein_coding | 0.465171258 | 7.373943566 | 4.8639669 | 0.0421017 | 0.538310714 |
| ENSMUSG00000019817 | Plagl1 | protein_coding | -0.84660456 | 3.547039266 | 4.8571571 | 0.0422286 | 0.539292627 |
| ENSMUSG00000029098 | Acox3 | protein_coding | 0.935250723 | 3.777063605 | 4.8475959 | 0.0424076 | 0.540420372 |
| ENSMUSG00000026388 | 3110009E18Rik | protein_coding | -0.766157923 | 4.377404602 | 4.8470721 | 0.0424174 | 0.540420372 |
| ENSMUSG00000053950 | Adnp2 | protein_coding | 1.009073821 | 3.565987606 | 4.841757 | 0.0425174 | 0.540495268 |
| ENSMUSG00000048022 | Tmem229a | protein_coding | -0.852258796 | 4.868359132 | 4.8426102 | 0.0425268 | 0.540495268 |
| ENSMUSG00000027900 | Dram2 | protein_coding | -0.562399197 | 5.679446548 | 4.8341118 | 0.0426615 | 0.540495268 |
| ENSMUSG00000019791 | Hint3 | protein_coding | -0.772290428 | 4.098868876 | 4.8339873 | 0.0426639 | 0.540495268 |
| ENSMUSG00000074165 | Zfp788 | protein_coding | -0.774707351 | 4.507406213 | 4.8334182 | 0.0426746 | 0.540495268 |
| ENSMUSG00000033972 | Zfp944 | protein_coding | -0.805117848 | 3.843087249 | 4.8267625 | 0.0428007 | 0.541394346 |
| ENSMUSG00000026083 | Eif5b | protein_coding | -0.608506875 | 6.243742382 | 4.8243571 | 0.0428463 | 0.541394346 |
| ENSMUSG00000021537 | Cetn3 | protein_coding | -0.549378904 | 7.85686863 | 4.8206876 | 0.0429417 | 0.541809346 |
| ENSMUSG00000019796 | Lrp11 | protein_coding | 0.643034959 | 5.493221051 | 4.8156427 | 0.0430122 | 0.541809346 |
| ENSMUSG00000022203 | Efs | protein_coding | -1.120440583 | 2.928711855 | 4.8068945 | 0.0431795 | 0.541809346 |
| ENSMUSG00000041685 | Fcho2 | protein_coding | -0.819446228 | 3.901869086 | 4.8060422 | 0.0431958 | 0.541809346 |
| ENSMUSG00000020537 | Drg2 | protein_coding | 0.54850844 | 7.094896345 | 4.8059885 | 0.0432225 | 0.541809346 |
| ENSMUSG00000030770 | Parva | protein_coding | 0.553513882 | 5.503169889 | 4.7991651 | 0.0433279 | 0.541809346 |
| ENSMUSG00000003233 | Dvl3 | protein_coding | -0.934527434 | 4.062868915 | 4.7989744 | 0.0433315 | 0.541809346 |
| ENSMUSG00000004665 | Cnn2 | protein_coding | 1.54302102 | 2.957617468 | 4.7977786 | 0.0433545 | 0.541809346 |
| ENSMUSG00000022708 | Zbtb20 | protein_coding | -0.645179028 | 6.260583793 | 4.7989624 | 0.0433575 | 0.541809346 |
| ENSMUSG00000079083 | Jrkl | protein_coding | -1.186788812 | 2.992706332 | 4.7962993 | 0.043383 | 0.541809346 |
| ENSMUSG00000050394 | Armcx6 | protein_coding | 1.095115361 | 3.473449672 | 4.7894867 | 0.0435145 | 0.542042041 |
| ENSMUSG00000021596 | Mctp1 | protein_coding | -0.78427275 | 6.10910159 | 4.8398052 | 0.0435461 | 0.542042041 |
| ENSMUSG00000038628 | Polr3k | protein_coding | -0.39688531 | 7.76036592 | 4.785673 | 0.0435883 | 0.542042041 |
| ENSMUSG00000042258 | Isl1 | protein_coding | -0.459378149 | 6.31340539 | 4.7807198 | 0.0436844 | 0.542042041 |
| ENSMUSG00000061576 | Dpp6 | protein_coding | 0.457961545 | 7.247168666 | 4.7788142 | 0.0437214 | 0.542042041 |
| ENSMUSG00000032517 | Mobp | protein_coding | -0.426022242 | 7.82379215 | 4.7775247 | 0.0437465 | 0.542042041 |
| ENSMUSG00000039356 | Exosc2 | protein_coding | 0.887233803 | 3.985303725 | 4.7771131 | 0.0437545 | 0.542042041 |
| ENSMUSG00000033400 | Agl | protein_coding | 0.803333588 | 3.815284469 | 4.7694766 | 0.0439034 | 0.542252937 |
| ENSMUSG00000021097 | Clmn | protein_coding | -0.65812548 | 5.483942147 | 4.7685177 | 0.0439222 | 0.542252937 |
| ENSMUSG00000097350 | 4732491K20Rik | lincRNA | -0.65262618 | 4.549704638 | 4.7683259 | 0.0439259 | 0.542252937 |
| ENSMUSG00000026831 | 1700007K13Rik | protein_coding | -1.020692213 | 3.121678407 | 4.7652798 | 0.0439855 | 0.542252937 |
| ENSMUSG00000055531 | Cpsf6 | protein_coding | 0.57937861 | 5.774154191 | 4.7633292 | 0.0440237 | 0.542252937 |
| ENSMUSG00000033981 | Gria2 | protein_coding | -0.489362463 | 8.100468188 | 4.7587476 | 0.0441136 | 0.542455692 |
| ENSMUSG00000028034 | Fubp1 | protein_coding | 0.503689099 | 6.719120052 | 4.7560923 | 0.0441658 | 0.542455692 |
| ENSMUSG00000037001 | Zfp39 | protein_coding | -0.88622431 | 3.344581473 | 4.7495645 | 0.0442944 | 0.542455692 |
| ENSMUSG00000033948 | Zswim5 | protein_coding | 1.243016779 | 3.294930318 | 4.7491168 | 0.0443032 | 0.542455692 |
| ENSMUSG00000021577 | Sdha | protein_coding | 0.604828573 | 9.626183273 | 4.7501104 | 0.0443095 | 0.542455692 |
| ENSMUSG00000029192 | Tbc1d14 | protein_coding | 0.608786759 | 5.501214275 | 4.7463275 | 0.0443583 | 0.542455692 |
| ENSMUSG00000030411 | Nova2 | protein_coding | -0.634262734 | 5.176793 | 4.741499 | 0.0444539 | 0.542455692 |
| ENSMUSG00000042396 | Rbm7 | protein_coding | -0.541373436 | 5.8744885 | 4.7407491 | 0.0444688 | 0.542455692 |
| ENSMUSG00000085795 | Zfp703 | protein_coding | 1.265276446 | 2.951697461 | 4.7370873 | 0.0445415 | 0.542455692 |
| ENSMUSG00000058420 | Syt17 | protein_coding | 0.720680812 | 5.326430569 | 4.7328947 | 0.0446248 | 0.542455692 |
| ENSMUSG00000073856 | Iqck | protein_coding | -0.941732115 | 3.595928393 | 4.7318122 | 0.0446464 | 0.542455692 |
| ENSMUSG00000047423 | AI837181 | protein_coding | -0.509421761 | 6.38112456 | 4.7297168 | 0.0446881 | 0.542455692 |
| ENSMUSG00000039765 | Cc2d2a | protein_coding | 0.775410807 | 3.858945597 | 4.7288179 | 0.0447061 | 0.542455692 |
| ENSMUSG00000021038 | Vipas39 | protein_coding | 0.556621428 | 6.141613416 | 4.7267968 | 0.0447464 | 0.542455692 |
| ENSMUSG00000042505 | Sdhaf3 | protein_coding | -0.477997509 | 6.69178615 | 4.7200658 | 0.0448811 | 0.542645204 |

| ensembl_gene_id | external_gene_name | gene_biotype | log2 (fold change) | log2 (counts per million) | F-value | p-value | False discovery rate |
|---|---|---|---|---|---|---|---|
| ENSMUSG00000012422 | Tmem167 | protein_coding | -0.478791 | 7.382104096 | 4.7163391 | 0.0449558 | 0.542645204 |
| ENSMUSG00000002108 | Nr1h3 | protein_coding | -1.669549037 | 2.680416916 | 4.715456 | 0.0449736 | 0.542645204 |
| ENSMUSG00000024507 | Hsd17b4 | protein_coding | 0.540219192 | 5.464083413 | 4.7128908 | 0.0450251 | 0.542645204 |
| ENSMUSG00000058446 | Znrf2 | protein_coding | -0.877609366 | 3.726275141 | 4.708625 | 0.045111 | 0.542645204 |
| ENSMUSG00000031960 | Aars | protein_coding | 0.563953702 | 7.021415259 | 4.706038 | 0.0451892 | 0.542645204 |
| ENSMUSG00000037605 | Adgrl3 | protein_coding | -0.757227436 | 5.248406716 | 4.7038014 | 0.0452084 | 0.542645204 |
| ENSMUSG00000051579 | Tceal8 | protein_coding | -0.439638966 | 7.465920246 | 4.7033583 | 0.0452173 | 0.542645204 |
| ENSMUSG00000059895 | Ptp4a3 | protein_coding | -0.57515111 | 5.311808711 | 4.7032405 | 0.0452197 | 0.542645204 |
| ENSMUSG00000022538 | Lsg1 | protein_coding | 0.538185484 | 5.240216218 | 4.6991087 | 0.0453033 | 0.542645204 |
| ENSMUSG00000005804 | Bloc1s6 | protein_coding | 0.741846563 | 6.412023216 | 4.7462492 | 0.0453656 | 0.542645204 |
| ENSMUSG00000026347 | Tmem163 | protein_coding | 0.886963398 | 4.287206901 | 4.6959378 | 0.0453676 | 0.542645204 |
| ENSMUSG00000069601 | Ank3 | protein_coding | -0.442076445 | 6.505575988 | 4.6898418 | 0.0454915 | 0.543522529 |
| ENSMUSG00000048747 | E130114P18Rik | protein_coding | -0.94959833 | 4.640997814 | 4.6887166 | 0.0455812 | 0.543783492 |
| ENSMUSG00000020415 | Pttg1 | protein_coding | -0.36613959 | 7.486446678 | 4.683137 | 0.0456282 | 0.543783492 |
| ENSMUSG00000023572 | Ccndbp1 | protein_coding | 0.425591366 | 7.556442304 | 4.6813348 | 0.0456651 | 0.543783492 |
| ENSMUSG00000061755 | Bod1l | protein_coding | -0.68903703 | 5.3902274 | 4.6735241 | 0.0458251 | 0.544278469 |
| ENSMUSG00000040003 | Magi2 | protein_coding | -0.562483761 | 5.226439022 | 4.6731503 | 0.0458328 | 0.544278469 |
| ENSMUSG00000022533 | Atp13a3 | protein_coding | -0.804378153 | 4.077991839 | 4.6718982 | 0.0458585 | 0.544278469 |
| ENSMUSG00000016018 | Skiv2l2 | protein_coding | 0.607980097 | 5.557120922 | 4.6653411 | 0.0459934 | 0.544366824 |
| ENSMUSG00000043061 | Tmem18 | protein_coding | 0.667539986 | 4.516317477 | 4.6638494 | 0.0460242 | 0.544366824 |
| ENSMUSG00000026418 | Tnni1 | protein_coding | -0.587254142 | 4.722029144 | 4.6632308 | 0.046037 | 0.544366824 |
| ENSMUSG00000025873 | Faf2 | protein_coding | 0.544124884 | 5.906230298 | 4.6600048 | 0.0461036 | 0.544366824 |
| ENSMUSG00000042271 | Nxt2 | protein_coding | -0.560803777 | 6.680657324 | 4.6546103 | 0.0462415 | 0.544366824 |
| ENSMUSG00000027642 | Rpn2 | protein_coding | 0.36308477 | 7.653253879 | 4.6506918 | 0.0462966 | 0.544366824 |
| ENSMUSG00000029376 | Mthfd2l | protein_coding | -0.690967137 | 4.601838328 | 4.6489492 | 0.0463328 | 0.544366824 |
| ENSMUSG00000025428 | Atp5a1 | protein_coding | 0.537953435 | 10.5517582 | 4.637404 | 0.0465736 | 0.544366824 |
| ENSMUSG00000039607 | Rbms3 | protein_coding | -0.677174357 | 5.626298058 | 4.6368798 | 0.0465846 | 0.544366824 |
| ENSMUSG00000042390 | Gatad2b | protein_coding | -0.528026472 | 5.723837878 | 4.6367155 | 0.046588 | 0.544366824 |
| ENSMUSG00000027162 | Lin7c | protein_coding | -0.41931392 | 6.417872567 | 4.6359733 | 0.0466036 | 0.544366824 |
| ENSMUSG00000021906 | Oxnad1 | protein_coding | 0.699905135 | 4.883129261 | 4.635545 | 0.0466125 | 0.544366824 |
| ENSMUSG00000069072 | Slc7a14 | protein_coding | 0.486733325 | 6.089734951 | 4.6354295 | 0.0466149 | 0.544366824 |
| ENSMUSG00000037855 | Zfp365 | protein_coding | -0.401461845 | 7.027593811 | 4.6303641 | 0.0467212 | 0.544366824 |
| ENSMUSG00000053907 | Mat2a | protein_coding | 0.406661979 | 7.41689777 | 4.6300051 | 0.0467287 | 0.544366824 |
| ENSMUSG00000052738 | Suclg1 | protein_coding | 0.384097237 | 9.129198051 | 4.6277177 | 0.0467768 | 0.544366824 |
| ENSMUSG00000016409 | Nkap | protein_coding | 0.59043169 | 5.150353516 | 4.6249442 | 0.0468351 | 0.544366824 |
| ENSMUSG00000026782 | Abi2 | protein_coding | -0.436239753 | 7.19286247 | 4.6163628 | 0.0470162 | 0.544366824 |
| ENSMUSG00000021540 | Smad5 | protein_coding | 0.617229076 | 4.475325274 | 4.6024092 | 0.0473124 | 0.544366824 |
| ENSMUSG00000018189 | Uchl5 | protein_coding | -0.406657365 | 6.939610943 | 4.5955551 | 0.0474587 | 0.544366824 |
| ENSMUSG00000031169 | Porcn | protein_coding | 0.721700199 | 5.106679883 | 4.5949408 | 0.0474718 | 0.544366824 |
| ENSMUSG00000019948 | Actr6 | protein_coding | -0.460012373 | 6.397790488 | 4.5910605 | 0.0475549 | 0.544366824 |
| ENSMUSG00000021661 | Ankra2 | protein_coding | -0.546329585 | 5.411818967 | 4.586469 | 0.0476534 | 0.544366824 |
| ENSMUSG00000029088 | Kcnip4 | protein_coding | -0.517374841 | 5.534930803 | 4.5856829 | 0.0476703 | 0.544366824 |
| ENSMUSG00000029687 | Ezh2 | protein_coding | -1.398790195 | 3.123296772 | 4.586703 | 0.0476749 | 0.544366824 |
| ENSMUSG00000058624 | Gda | protein_coding | -0.491152359 | 6.490384451 | 4.5845827 | 0.0476939 | 0.544366824 |
| ENSMUSG00000023915 | Tnfrsf21 | protein_coding | 0.550458544 | 5.548326239 | 4.5836629 | 0.0477137 | 0.544366824 |
| ENSMUSG00000000125 | Wnt3 | protein_coding | 1.062967096 | 4.082509671 | 4.584697 | 0.047718 | 0.544366824 |
| ENSMUSG00000000792 | Slc5a5 | protein_coding | 0.509587496 | 6.950427196 | 4.5823627 | 0.0477417 | 0.544366824 |
| ENSMUSG00000049985 | Ankrd55 | protein_coding | 1.059053129 | 3.601224438 | 4.5809544 | 0.047772 | 0.544366824 |
| ENSMUSG00000032458 | Copb2 | protein_coding | 0.538466664 | 6.600300104 | 4.5797067 | 0.0477989 | 0.544366824 |
| ENSMUSG00000038535 | Zfp280d | protein_coding | -1.11803049 | 3.146602103 | 4.5762884 | 0.0478727 | 0.544366824 |
| ENSMUSG00000035399 | Oser1 | protein_coding | -0.726183349 | 3.785850373 | 4.5706863 | 0.0479938 | 0.544366824 |
| ENSMUSG00000035236 | Scai | protein_coding | -0.734760284 | 4.68379217 | 4.5655586 | 0.0481051 | 0.544366824 |
| ENSMUSG00000022761 | Lztr1 | protein_coding | 0.475899648 | 6.556894168 | 4.565227 | 0.0481123 | 0.544366824 |
| ENSMUSG00000031432 | Prps1 | protein_coding | 0.383986138 | 7.358218619 | 4.5632298 | 0.0481557 | 0.544366824 |
| ENSMUSG00000018042 | Cyb5r3 | protein_coding | 0.494143059 | 7.649355006 | 4.5625857 | 0.0481697 | 0.544366824 |
| ENSMUSG00000067150 | Xpo5 | protein_coding | 0.886366347 | 3.970514468 | 4.5598334 | 0.0482296 | 0.544366824 |
| ENSMUSG00000037257 | Aagab | protein_coding | 0.602067276 | 6.616531165 | 4.5582888 | 0.0482899 | 0.544366824 |
| ENSMUSG00000021772 | Nkiras1 | protein_coding | 0.374695787 | 8.258030439 | 4.5567503 | 0.0482968 | 0.544366824 |
| ENSMUSG00000035764 | Fbxo45 | protein_coding | -0.665267309 | 4.888335855 | 4.5566017 | 0.0483001 | 0.544366824 |
| ENSMUSG00000041506 | Rrp9 | protein_coding | -0.629807219 | 4.809833241 | 4.5559557 | 0.0483142 | 0.544366824 |
| ENSMUSG00000024581 | Napg | protein_coding | 0.439321579 | 7.799076739 | 4.5533665 | 0.0483707 | 0.544366824 |
| ENSMUSG00000028676 | Srsf10 | protein_coding | -0.516406906 | 5.481555705 | 4.5532736 | 0.0483727 | 0.544366824 |
| ENSMUSG00000019852 | Arfgef3 | protein_coding | -0.547934156 | 5.896840829 | 4.55236 | 0.0483927 | 0.544366824 |
| ENSMUSG00000032064 | Dixdc1 | protein_coding | -0.432804854 | 6.123348435 | 4.5505293 | 0.0484328 | 0.544366824 |
| ENSMUSG00000026277 | Stk25 | protein_coding | 0.445906578 | 7.586585753 | 4.5481852 | 0.0484841 | 0.544366824 |
| ENSMUSG00000042502 | Cd2bp2 | protein_coding | 0.540951634 | 6.020395609 | 4.546901 | 0.0485123 | 0.544366824 |
| ENSMUSG00000047466 | 8030462N17Rik | protein_coding | -1.029638011 | 3.532953595 | 4.5456748 | 0.0485392 | 0.544366824 |
| ENSMUSG00000046364 | Rpl27a | protein_coding | -0.498193213 | 6.757894458 | 4.541168 | 0.0486382 | 0.544366824 |
| ENSMUSG00000076498 | Trbc2 | TR_C_gene | 1.325582445 | 2.604877757 | 4.5405854 | 0.048651 | 0.544366824 |
| ENSMUSG00000029162 | Khk | protein_coding | 0.544672708 | 5.107388417 | 4.5380761 | 0.0487063 | 0.544366824 |
| ENSMUSG00000090112 | Shprh | protein_coding | -0.801414575 | 4.266955647 | 4.5375423 | 0.048718 | 0.544366824 |
| ENSMUSG00000024382 | Ercc3 | protein_coding | 0.697929748 | 4.83417328 | 4.5372749 | 0.0487239 | 0.544366824 |
| ENSMUSG00000006390 | Elovl1 | protein_coding | -0.645976823 | 4.990622228 | 4.53291 | 0.0488203 | 0.544366824 |
| ENSMUSG00000048410 | Zfp407 | protein_coding | -0.957755096 | 3.295447265 | 4.532134 | 0.0488374 | 0.544366824 |

| ensembl_gene_id | external_gene_name | gene_biotype | log2 (fold change) | log2 (counts per million) | F-value | p-value | False discovery rate |
|---|---|---|---|---|---|---|---|
| ENSMUSG00000020877 | Scrn2 | protein_coding | 1.123404466 | 3.29208002 | 4.5310603 | 0.0488612 | 0.544366824 |
| ENSMUSG00000026275 | Ppp1r7 | protein_coding | -0.407313259 | 7.815914867 | 4.5294279 | 0.0488973 | 0.544366824 |
| ENSMUSG00000044566 | Cage1 | protein_coding | -0.925278734 | 3.494822054 | 4.5260172 | 0.0489728 | 0.544366824 |
| ENSMUSG00000027254 | Map1a | protein_coding | -0.55905626 | 7.385341799 | 4.5259371 | 0.0490013 | 0.544366824 |
| ENSMUSG00000062931 | Zfp938 | protein_coding | -0.420820975 | 6.348800424 | 4.5224226 | 0.0490526 | 0.544366824 |
| ENSMUSG00000004264 | Phb2 | protein_coding | 0.891606998 | 3.591811995 | 4.5223876 | 0.0490534 | 0.544366824 |
| ENSMUSG00000047342 | Zfp286 | protein_coding | -0.62376441 | 5.013948161 | 4.5223027 | 0.0490553 | 0.544366824 |
| ENSMUSG00000029826 | Zc3hav1 | protein_coding | -0.922702272 | 3.925240295 | 4.5069727 | 0.0493973 | 0.547028615 |
| ENSMUSG00000028140 | Mrpl9 | protein_coding | 0.44597322 | 6.042274355 | 4.5066481 | 0.0494045 | 0.547028615 |
| ENSMUSG00000027429 | Sec23b | protein_coding | 0.66396538 | 5.954301935 | 4.5036759 | 0.049498 | 0.547028615 |
| ENSMUSG00000030615 | Tmem126a | protein_coding | -0.523479583 | 6.654889712 | 4.5022167 | 0.0495039 | 0.547028615 |
| ENSMUSG00000019087 | Atp6ap1 | protein_coding | 0.388432441 | 9.103616349 | 4.5001873 | 0.0495495 | 0.547028615 |
| ENSMUSG00000051413 | Plagl2 | protein_coding | 1.038245681 | 2.963117602 | 4.4916685 | 0.0497414 | 0.548266666 |
| ENSMUSG00000018848 | Rars | protein_coding | 0.490722883 | 6.479045316 | 4.4900419 | 0.0497782 | 0.548266666 |
| ENSMUSG00000035161 | Ints6 | protein_coding | -0.662185828 | 4.27250757 | 4.4873585 | 0.0498389 | 0.548266666 |
| ENSMUSG00000022779 | Top3b | protein_coding | 0.678663048 | 4.148305104 | 4.486177 | 0.0498656 | 0.548266666 |
| ENSMUSG00000041115 | Iqsec2 | protein_coding | 0.732546303 | 4.473498805 | 4.4833384 | 0.0499299 | 0.548413192 |
| ENSMUSG00000025217 | Btrc | protein_coding | 0.637159728 | 6.648808445 | 4.500198 | 0.0500606 | 0.548756211 |
| ENSMUSG00000024750 | Zfand5 | protein_coding | -0.479871709 | 6.018019954 | 4.4760574 | 0.0500954 | 0.548756211 |
| ENSMUSG00000020189 | Osbpl8 | protein_coding | -0.584630652 | 5.708584989 | 4.4752279 | 0.0501143 | 0.548756211 |
| ENSMUSG00000020211 | Sf3a2 | protein_coding | -0.809524771 | 3.820183956 | 4.4705544 | 0.0502208 | 0.549222631 |
| ENSMUSG00000005981 | Trap1 | protein_coding | 0.579455708 | 6.09934087 | 4.4667835 | 0.050307 | 0.549222631 |
| ENSMUSG00000020864 | Ankrd40 | protein_coding | -0.52764472 | 5.976955726 | 4.4666496 | 0.0503101 | 0.549222631 |
| ENSMUSG00000021816 | Ppp3cb | protein_coding | -0.430936123 | 7.762420812 | 4.4608295 | 0.0504434 | 0.550120053 |
| ENSMUSG00000058291 | Zfp68 | protein_coding | -0.545657209 | 5.270613456 | 4.4585546 | 0.0504957 | 0.550131833 |
| ENSMUSG00000021048 | Mthfd1 | protein_coding | 0.735649084 | 4.127595466 | 4.4546956 | 0.0505844 | 0.550541061 |
| ENSMUSG00000052534 | Pbx1 | protein_coding | -0.430320087 | 7.052924281 | 4.4481695 | 0.050735 | 0.55162093 |
| ENSMUSG00000097675 | 1700101I11Rik | processed_transcript | 1.163478441 | 2.873337155 | 4.4418328 | 0.0508816 | 0.552173322 |
| ENSMUSG00000036764 | Dnajc12 | protein_coding | 0.37996082 | 6.958223742 | 4.441537 | 0.0508885 | 0.552173322 |
| ENSMUSG00000022122 | Ednrb | protein_coding | -0.583601022 | 6.1008618 | 4.4356484 | 0.0510252 | 0.552503332 |
| ENSMUSG00000020387 | Jade2 | protein_coding | -0.519148601 | 5.322138839 | 4.4327831 | 0.0510919 | 0.552503332 |
| ENSMUSG00000032583 | Mon1a | protein_coding | -0.731173767 | 4.200117861 | 4.4302584 | 0.0511508 | 0.552503332 |
| ENSMUSG00000031901 | Dus2 | protein_coding | 0.6229364 | 5.148643558 | 4.4291076 | 0.0511776 | 0.552503332 |
| ENSMUSG00000029414 | Kntc1 | protein_coding | 1.730601667 | 2.226839101 | 4.4234927 | 0.0513088 | 0.552503332 |
| ENSMUSG00000013539 | Tango2 | protein_coding | 0.415622589 | 8.030168215 | 4.4188406 | 0.0514179 | 0.552503332 |
| ENSMUSG00000057156 | Homez | protein_coding | -0.820124765 | 3.484538364 | 4.4181856 | 0.0514333 | 0.552503332 |
| ENSMUSG00000049097 | Ankrd34a | protein_coding | -0.576965014 | 5.665853992 | 4.4164585 | 0.0514738 | 0.552503332 |
| ENSMUSG00000039988 | Ankrd13c | protein_coding | -0.487574418 | 5.982011039 | 4.415103 | 0.0515057 | 0.552503332 |
| ENSMUSG00000025939 | Ube2w | protein_coding | -0.393112048 | 8.205577493 | 4.4149142 | 0.0515101 | 0.552503332 |
| ENSMUSG00000003363 | Pld3 | protein_coding | 0.432017006 | 9.931445965 | 4.4134306 | 0.051545 | 0.552503332 |
| ENSMUSG00000054737 | Zfp182 | protein_coding | -1.42089385 | 2.357138798 | 4.4119728 | 0.0515793 | 0.552503332 |
| ENSMUSG00000065922 | n-R5-8s1 | rRNA | -0.745771032 | 5.306719367 | 4.4084198 | 0.0516903 | 0.552503332 |
| ENSMUSG00000038845 | Phb | protein_coding | 0.542314833 | 6.715993829 | 4.4069476 | 0.051725 | 0.552503332 |
| ENSMUSG00000044906 | 4930503L19Rik | protein_coding | -1.552892815 | 2.687705148 | 4.3980317 | 0.0519089 | 0.552503332 |
| ENSMUSG00000028530 | Jak1 | protein_coding | 0.41497648 | 6.645700066 | 4.3923019 | 0.052045 | 0.552503332 |
| ENSMUSG00000049119 | Fam110b | protein_coding | 0.441120822 | 6.340096032 | 4.391987 | 0.0520525 | 0.552503332 |
| ENSMUSG00000074782 | 4833422C13Rik | antisense_RNA | -0.793475023 | 4.016640669 | 4.3918423 | 0.052056 | 0.552503332 |
| ENSMUSG00000020297 | Nsg2 | protein_coding | 0.374010293 | 9.103249512 | 4.3915632 | 0.0520626 | 0.552503332 |
| ENSMUSG00000026181 | Ppm1f | protein_coding | 0.630063338 | 5.832316627 | 4.3911695 | 0.052072 | 0.552503332 |
| ENSMUSG00000026313 | Hdac4 | protein_coding | 0.56770711 | 5.850926802 | 4.3909206 | 0.0520779 | 0.552503332 |
| ENSMUSG00000095432 | Zfp748 | protein_coding | -0.866177398 | 3.508578526 | 4.3875817 | 0.0521575 | 0.552503332 |
| ENSMUSG00000021981 | Cab39l | protein_coding | -0.464656759 | 6.511037761 | 4.3858848 | 0.052198 | 0.552503332 |
| ENSMUSG00000008206 | Cers4 | protein_coding | 0.543942815 | 5.832784365 | 4.3853308 | 0.0522113 | 0.552503332 |
| ENSMUSG00000031555 | Adam9 | protein_coding | -0.818762854 | 3.504916784 | 4.3846993 | 0.0522264 | 0.552503332 |
| ENSMUSG00000055067 | Smyd3 | protein_coding | 0.771237663 | 6.563992212 | 4.4748713 | 0.0522548 | 0.552503332 |
| ENSMUSG00000024271 | Elp2 | protein_coding | 0.526209993 | 6.922050333 | 4.379198 | 0.0523836 | 0.553321585 |
| ENSMUSG00000026849 | Tor1a | protein_coding | 0.65396868 | 5.716089504 | 4.3676933 | 0.0526346 | 0.554727337 |
| ENSMUSG00000049957 | Ccdc137 | protein_coding | 0.587137994 | 4.829238784 | 4.3655191 | 0.052687 | 0.554727337 |
| ENSMUSG00000030980 | Knop1 | protein_coding | 0.519873245 | 5.433076153 | 4.3651848 | 0.0526951 | 0.554727337 |
| ENSMUSG00000039983 | Ccdc32 | protein_coding | 0.401302409 | 7.137188915 | 4.3637149 | 0.0527306 | 0.554727337 |
| ENSMUSG00000025268 | Maged2 | protein_coding | 0.575073164 | 7.775685718 | 4.3644617 | 0.0528591 | 0.554727337 |
| ENSMUSG00000039270 | Megf9 | protein_coding | -0.578606142 | 5.528848811 | 4.3576709 | 0.0528769 | 0.554727337 |
| ENSMUSG00000050914 | Ankrd37 | protein_coding | -0.584991128 | 4.687636827 | 4.3576313 | 0.0528778 | 0.554727337 |
| ENSMUSG00000104861 | 3110039M20Rik | lincRNA | -1.071176086 | 3.578127539 | 4.3499514 | 0.0530644 | 0.555719032 |
| ENSMUSG00000020315 | Sptbn1 | protein_coding | -0.518933688 | 7.420372831 | 4.3491974 | 0.0531101 | 0.555719032 |
| ENSMUSG00000032355 | Mlip | protein_coding | -0.758658945 | 4.229628769 | 4.3463643 | 0.0531518 | 0.555719032 |
| ENSMUSG00000055109 | Gm15155 | protein_coding | -1.562390334 | 2.188059548 | 4.3442113 | 0.0532044 | 0.555719032 |
| ENSMUSG00000014353 | Tmem87b | protein_coding | 0.696092944 | 5.005499003 | 4.3431294 | 0.0532308 | 0.555719032 |
| ENSMUSG00000085208 | Brip1os | antisense_RNA | 1.319290622 | 2.826125638 | 4.3309399 | 0.0535296 | 0.557265439 |
| ENSMUSG00000068732 | Tmem167b | protein_coding | 0.691759912 | 5.617912655 | 4.3301812 | 0.0535757 | 0.557265439 |
| ENSMUSG00000021712 | Trim23 | protein_coding | -0.725473238 | 5.5530116 | 4.3299646 | 0.053581 | 0.557265439 |
| ENSMUSG00000023235 | Ccl25 | protein_coding | -0.692984199 | 4.87212232 | 4.3276726 | 0.05361 | 0.557265439 |
| ENSMUSG00000006494 | Pdk1 | protein_coding | 0.89013586 | 4.046963452 | 4.3265342 | 0.053638 | 0.557265439 |
| ENSMUSG00000033713 | Foxn3 | protein_coding | -0.564532463 | 6.87413152 | 4.3223509 | 0.0537687 | 0.557645435 |

| ensembl_gene_id | external_gene_name | gene_biotype | log2 (fold change) | log2 (counts per million) | F-value | p-value | False discovery rate |
|---|---|---|---|---|---|---|---|
| ENSMUSG00000018583 | G3bp1 | protein_coding | 0.477273802 | 6.057077749 | 4.3179315 | 0.0538506 | 0.557645435 |
| ENSMUSG00000022329 | Stk3 | protein_coding | -0.86093489 | 4.112228634 | 4.3166908 | 0.0538813 | 0.557645435 |
| ENSMUSG00000021730 | Hcn1 | protein_coding | -0.619001348 | 4.651421315 | 4.3154086 | 0.0539131 | 0.557645435 |
| ENSMUSG00000069670 | Nkain2 | protein_coding | -0.772968575 | 5.140302414 | 4.315678 | 0.0539339 | 0.557645435 |
| ENSMUSG00000030942 | Thumpd1 | protein_coding | 0.677042991 | 5.519415912 | 4.3100002 | 0.0540474 | 0.558153318 |
| ENSMUSG00000036902 | Neto2 | protein_coding | -0.75122942 | 4.898348325 | 4.3066821 | 0.05413 | 0.558153318 |
| ENSMUSG00000039159 | Ube2h | protein_coding | -0.527943364 | 6.566999314 | 4.3054741 | 0.0541601 | 0.558153318 |
| ENSMUSG00000027630 | Tbl1xr1 | protein_coding | -0.566564694 | 5.024175438 | 4.3042482 | 0.0541907 | 0.558153318 |
| ENSMUSG00000031438 | Rnf128 | protein_coding | -0.772401857 | 4.235137812 | 4.3021305 | 0.0542435 | 0.558163091 |
| ENSMUSG00000057604 | Lmcd1 | protein_coding | -1.080429432 | 3.760950453 | 4.2960392 | 0.0544234 | 0.559253852 |
| ENSMUSG00000024480 | Ap3s1 | protein_coding | -0.880997561 | 3.781050752 | 4.2937396 | 0.0544535 | 0.559253852 |
| ENSMUSG00000032452 | Clstn2 | protein_coding | 0.68076198 | 4.601326026 | 4.2897143 | 0.0545546 | 0.559434826 |
| ENSMUSG00000024639 | Gnaq | protein_coding | -0.542808545 | 8.272937565 | 4.2899929 | 0.0545752 | 0.559434826 |
| ENSMUSG00000025232 | Hexa | protein_coding | 0.559766287 | 5.996006021 | 4.2857446 | 0.0546545 | 0.55971433 |
| ENSMUSG00000034867 | Ankrd27 | protein_coding | 0.836228957 | 4.345784258 | 4.2797523 | 0.0548057 | 0.560383973 |
| ENSMUSG00000056919 | Cep162 | protein_coding | 1.07500574 | 3.024399606 | 4.2790243 | 0.0548241 | 0.560383973 |
| ENSMUSG00000029560 | Snx8 | protein_coding | 0.895047935 | 4.573281168 | 4.2764773 | 0.0549162 | 0.560792148 |
| ENSMUSG00000021285 | Ppp1r13b | protein_coding | 0.97046803 | 3.25431394 | 4.2662133 | 0.0551492 | 0.562092013 |
| ENSMUSG00000026930 | Gpsm1 | protein_coding | -0.513729966 | 6.004158475 | 4.2649372 | 0.0551817 | 0.562092013 |
| ENSMUSG00000108350 | Gm44950 | TEC | 0.787686976 | 4.493757093 | 4.2640242 | 0.0552049 | 0.562092013 |
| ENSMUSG00000027805 | Pfn2 | protein_coding | 0.449995769 | 10.00691878 | 4.2621554 | 0.0552526 | 0.562092013 |
| ENSMUSG00000040415 | Dtx3 | protein_coding | -0.350126353 | 7.562950715 | 4.255231 | 0.0554296 | 0.56307549 |
| ENSMUSG00000041645 | Ddx24 | protein_coding | 0.418075039 | 8.638056338 | 4.2534912 | 0.0554742 | 0.56307549 |
| ENSMUSG00000031930 | Wwp2 | protein_coding | 0.703389752 | 5.607010301 | 4.2479085 | 0.0556453 | 0.56307549 |
| ENSMUSG00000040856 | Dlk1 | protein_coding | 0.425261278 | 6.572223561 | 4.2452535 | 0.0556858 | 0.56307549 |
| ENSMUSG00000022365 | Derl1 | protein_coding | 0.506983883 | 5.782097291 | 4.2446736 | 0.0557008 | 0.56307549 |
| ENSMUSG00000027953 | Slc50a1 | protein_coding | -0.433599969 | 5.91594276 | 4.2405193 | 0.0558079 | 0.56307549 |
| ENSMUSG00000033352 | Map2k4 | protein_coding | 0.405345954 | 7.402422423 | 4.23928 | 0.0558399 | 0.56307549 |
| ENSMUSG00000002015 | Bcap31 | protein_coding | 0.401897981 | 7.856845416 | 4.2362109 | 0.0559192 | 0.56307549 |
| ENSMUSG00000022051 | Bnip3l | protein_coding | -0.359872949 | 7.068522962 | 4.2361888 | 0.0559198 | 0.56307549 |
| ENSMUSG00000027439 | Gzf1 | protein_coding | 0.51754593 | 5.316569817 | 4.2334645 | 0.0559903 | 0.56307549 |
| ENSMUSG00000034681 | Rnps1 | protein_coding | 0.502188546 | 6.11300136 | 4.2330633 | 0.0560007 | 0.56307549 |
| ENSMUSG00000021286 | Zfyve21 | protein_coding | -0.627743537 | 5.29721669 | 4.23306 | 0.0560008 | 0.56307549 |
| ENSMUSG00000028914 | Casp9 | protein_coding | 1.643493915 | 2.50036652 | 4.2319342 | 0.05603 | 0.56307549 |
| ENSMUSG00000001964 | Emd | protein_coding | -0.650166664 | 5.350165237 | 4.2280498 | 0.0561309 | 0.56312069 |
| ENSMUSG00000051674 | Dcun1d4 | protein_coding | 0.489145022 | 6.531076334 | 4.2257144 | 0.0561916 | 0.56312069 |
| ENSMUSG00000055991 | Zkscan5 | protein_coding | 0.883336001 | 3.547191066 | 4.2216256 | 0.0562981 | 0.56312069 |
| ENSMUSG00000097412 | 1810014B01Rik | transcribed_unproces | -1.279096309 | 2.91370316 | 4.2207438 | 0.0563211 | 0.56312069 |
| ENSMUSG00000028407 | Toporsos | protein_coding | -0.844620853 | 3.603594937 | 4.2186222 | 0.0563765 | 0.56312069 |
| ENSMUSG00000033308 | Dpyd | protein_coding | 1.209217639 | 2.844302619 | 4.2172044 | 0.0564135 | 0.56312069 |
| ENSMUSG00000087620 | 5330434G04Rik | lincRNA | -0.782044754 | 4.666725903 | 4.216424 | 0.0564339 | 0.56312069 |
| ENSMUSG00000019210 | Atp6v1e1 | protein_coding | 0.371545473 | 10.1101088 | 4.2143342 | 0.0564886 | 0.56312069 |
| ENSMUSG00000030407 | Qpctl | protein_coding | 0.606609657 | 4.971526964 | 4.2084508 | 0.0566429 | 0.56312069 |
| ENSMUSG00000015597 | Zfp318 | protein_coding | 0.653359793 | 4.926958998 | 4.2077342 | 0.0566617 | 0.56312069 |
| ENSMUSG00000044550 | Tceal3 | protein_coding | 0.352778433 | 7.206967085 | 4.2066439 | 0.0566904 | 0.56312069 |
| ENSMUSG00000021065 | Fut8 | protein_coding | -0.615134798 | 4.914717648 | 4.2060175 | 0.0567069 | 0.56312069 |
| ENSMUSG00000028675 | Pnrc2 | protein_coding | -0.575207979 | 6.454867682 | 4.2015993 | 0.0568511 | 0.56312069 |
| ENSMUSG00000048175 | Asb8 | protein_coding | 0.465657908 | 7.244738122 | 4.1993548 | 0.0568824 | 0.56312069 |
| ENSMUSG00000019659 | Ccdc12 | protein_coding | -0.458057322 | 5.902746514 | 4.1985677 | 0.0569032 | 0.56312069 |
| ENSMUSG00000031767 | Nudt7 | protein_coding | 0.601287116 | 4.349246878 | 4.1981113 | 0.0569152 | 0.56312069 |
| ENSMUSG00000098973 | Mir6236 | miRNA | 0.788823784 | 8.725991134 | 4.3022881 | 0.0569248 | 0.56312069 |
| ENSMUSG00000068740 | Celsr2 | protein_coding | -0.607809501 | 5.933846653 | 4.1911045 | 0.0571007 | 0.563187453 |
| ENSMUSG00000030689 | Ino80e | protein_coding | -0.401787285 | 6.461476471 | 4.1904623 | 0.0571177 | 0.563187453 |
| ENSMUSG00000031577 | Tti2 | protein_coding | -0.863580192 | 3.608366245 | 4.1896351 | 0.0571396 | 0.563187453 |
| ENSMUSG00000014592 | Camta1 | protein_coding | -0.406242121 | 7.884971833 | 4.1881288 | 0.0571796 | 0.563187453 |
| ENSMUSG00000028954 | Nub1 | protein_coding | 0.476830862 | 6.897385071 | 4.1876097 | 0.0571934 | 0.563187453 |
| ENSMUSG00000008999 | Bmp7 | protein_coding | 1.118315202 | 3.158227919 | 4.1853982 | 0.0572522 | 0.563250451 |
| ENSMUSG00000071424 | Grid2 | protein_coding | -0.656725691 | 4.632350797 | 4.1819379 | 0.0573443 | 0.563640967 |
| ENSMUSG00000003234 | Abcf3 | protein_coding | 0.554910276 | 7.064597713 | 4.1787958 | 0.0574561 | 0.563702356 |
| ENSMUSG00000023988 | Bysl | protein_coding | 0.606133102 | 5.257597428 | 4.1768188 | 0.0574809 | 0.563702356 |
| ENSMUSG00000033871 | Ppargc1b | protein_coding | -0.937230517 | 3.937598235 | 4.1758113 | 0.0575078 | 0.563702356 |
| ENSMUSG00000007777 | 0610009B22Rik | protein_coding | -0.467076718 | 7.420469813 | 4.1721937 | 0.0576046 | 0.563840621 |
| ENSMUSG00000038248 | Sobp | protein_coding | -0.656803768 | 4.816461611 | 4.1699352 | 0.0576651 | 0.563840621 |
| ENSMUSG00000020076 | Ddx50 | protein_coding | -0.493448748 | 5.727135582 | 4.1679097 | 0.0577195 | 0.563840621 |
| ENSMUSG00000021665 | Hexb | protein_coding | 0.537002324 | 8.13720421 | 4.1681045 | 0.0577423 | 0.563840621 |
| ENSMUSG00000037419 | Endod1 | protein_coding | 0.453521384 | 6.359519041 | 4.1655054 | 0.0577841 | 0.563840621 |
| ENSMUSG00000019775 | Rgs17 | protein_coding | -0.427951244 | 7.274097657 | 4.1574872 | 0.0580001 | 0.565017249 |
| ENSMUSG00000037416 | Dmxl1 | protein_coding | -0.869714396 | 3.915605645 | 4.1571298 | 0.0580098 | 0.565017249 |
| ENSMUSG00000028351 | Brinp1 | protein_coding | 0.442402229 | 6.494578386 | 4.1492089 | 0.0582241 | 0.566592012 |
| ENSMUSG00000020312 | Shc2 | protein_coding | 0.921976713 | 3.328318316 | 4.143082 | 0.0583906 | 0.567085593 |
| ENSMUSG00000027405 | Nop56 | protein_coding | 0.456179764 | 7.276843481 | 4.1398402 | 0.0584789 | 0.567085593 |
| ENSMUSG00000022760 | Thap7 | protein_coding | -0.487639765 | 6.147698245 | 4.1397606 | 0.0584811 | 0.567085593 |
| ENSMUSG00000027858 | Tspan2 | protein_coding | -0.4561887 | 6.491047904 | 4.1395863 | 0.0584858 | 0.567085593 |
| ENSMUSG00000027109 | Sp3 | protein_coding | -0.615475574 | 4.960542373 | 4.1354033 | 0.0586 | 0.56735088 |

| ensembl_gene_id | external_gene_name | gene_biotype | log2 (fold change) | log2 (counts per million) | F-value | p-value | False discovery rate |
|---|---|---|---|---|---|---|---|
| ENSMUSG00000055401 | Fbxo6 | protein_coding | 0.517957946 | 5.581647577 | 4.1347189 | 0.0586187 | 0.56735088 |
| ENSMUSG00000005371 | Fbxo11 | protein_coding | -0.429645149 | 6.474362649 | 4.1269407 | 0.0588318 | 0.568560241 |
| ENSMUSG00000036528 | Ppfibp2 | protein_coding | 1.022596943 | 3.651401463 | 4.1251103 | 0.0588821 | 0.568560241 |
| ENSMUSG00000035595 | 1600002K03Rik | protein_coding | -0.839810375 | 3.89745917 | 4.1243747 | 0.0589023 | 0.568560241 |
| ENSMUSG00000072980 | Oip5 | protein_coding | -0.911819086 | 3.608398567 | 4.1187397 | 0.0590574 | 0.568941222 |
| ENSMUSG00000020134 | Peli1 | protein_coding | -0.624502232 | 5.027350715 | 4.1166046 | 0.0591163 | 0.568941222 |
| ENSMUSG00000004642 | Slbp | protein_coding | 0.52091713 | 6.246675971 | 4.1154192 | 0.0591491 | 0.568941222 |
| ENSMUSG00000057914 | Cacnb2 | protein_coding | -0.757792392 | 4.629845211 | 4.1125084 | 0.0592295 | 0.568941222 |
| ENSMUSG00000106981 | Gm45495 | lincRNA | 1.195627947 | 3.014933517 | 4.1110035 | 0.0592712 | 0.568941222 |
| ENSMUSG00000004530 | Coro1c | protein_coding | 0.701615391 | 6.0219746 | 4.1345301 | 0.0593424 | 0.568941222 |
| ENSMUSG00000025104 | Hdgfl3 | protein_coding | -0.495149275 | 6.422532832 | 4.1084295 | 0.0593425 | 0.568941222 |
| ENSMUSG00000038495 | Otud7b | protein_coding | -0.501112931 | 5.380455749 | 4.1057823 | 0.059416 | 0.568941222 |
| ENSMUSG00000022788 | Fgd4 | protein_coding | -0.702920881 | 3.77936611 | 4.1041196 | 0.0594622 | 0.568941222 |
| ENSMUSG00000030374 | Strn4 | protein_coding | -0.479630102 | 6.040028159 | 4.1027337 | 0.0595007 | 0.568941222 |
| ENSMUSG00000054013 | Tmem179 | protein_coding | 0.457201834 | 7.989735247 | 4.1019064 | 0.0595237 | 0.568941222 |
| ENSMUSG00000028015 | Ctso | protein_coding | -0.78673094 | 4.782366102 | 4.0999033 | 0.0596078 | 0.569002755 |
| ENSMUSG00000002320 | Tm9sf1 | protein_coding | 0.967045836 | 3.637929839 | 4.0977029 | 0.0596409 | 0.569002755 |
| ENSMUSG00000024845 | Tmem134 | processed_transcript | -0.82301434 | 3.994797671 | 4.0954869 | 0.0597027 | 0.569002755 |
| ENSMUSG00000068917 | Clk2 | protein_coding | 1.238486635 | 3.160121422 | 4.0940863 | 0.0597418 | 0.569002755 |
| ENSMUSG00000032553 | Srprb | protein_coding | 0.491869099 | 5.847270819 | 4.0774104 | 0.0602101 | 0.572954669 |
| ENSMUSG00000040667 | Nup88 | protein_coding | 0.534707915 | 6.057076029 | 4.0735055 | 0.0603203 | 0.573403919 |
| ENSMUSG00000028621 | Cyb5rl | protein_coding | 0.818963528 | 4.490891648 | 4.0719635 | 0.0603639 | 0.573403919 |
| ENSMUSG00000027210 | Meis2 | protein_coding | -0.472047015 | 7.258384013 | 4.0697936 | 0.0604253 | 0.57348076 |
| ENSMUSG00000025357 | Dgka | protein_coding | 1.535067113 | 2.571706719 | 4.0670364 | 0.0605035 | 0.573716126 |
| ENSMUSG00000038156 | Spon1 | protein_coding | 0.512976498 | 5.425957612 | 4.0650932 | 0.0605586 | 0.573733131 |
| ENSMUSG00000025203 | Scd2 | protein_coding | 0.44501666 | 8.816656589 | 4.0613562 | 0.0606649 | 0.57375847 |
| ENSMUSG00000023068 | Nus1 | protein_coding | -0.47954967 | 5.801789935 | 4.0578208 | 0.0607656 | 0.57375847 |
| ENSMUSG00000047603 | Zfp235 | protein_coding | -0.596469017 | 4.396018757 | 4.0547402 | 0.0608535 | 0.57375847 |
| ENSMUSG00000020648 | Dus4l | protein_coding | -1.296340539 | 2.939779055 | 4.0539469 | 0.0608761 | 0.57375847 |
| ENSMUSG00000000346 | Dazap2 | protein_coding | -0.391576469 | 8.570230436 | 4.0513133 | 0.0609514 | 0.57375847 |
| ENSMUSG00000034768 | Asb16 | protein_coding | -1.071960489 | 2.949665346 | 4.0499528 | 0.0609904 | 0.57375847 |
| ENSMUSG00000092536 | Gm20501 | bidirectional_promoter | -0.798986086 | 3.668349907 | 4.048317 | 0.0610372 | 0.57375847 |
| ENSMUSG00000005447 | Pafah1b3 | protein_coding | 0.44341647 | 7.042839988 | 4.047329 | 0.0610655 | 0.57375847 |
| ENSMUSG00000043962 | Thrap3 | protein_coding | 0.76847439 | 5.265451221 | 4.0466939 | 0.0611122 | 0.57375847 |
| ENSMUSG00000008958 | Vps72 | protein_coding | -0.415882293 | 6.102991075 | 4.0440676 | 0.0611591 | 0.57375847 |
| ENSMUSG00000094786 | Gm14403 | protein_coding | -0.905002687 | 3.592279995 | 4.0417801 | 0.0612249 | 0.57375847 |
| ENSMUSG00000039611 | Tmem246 | protein_coding | 0.476390758 | 6.775406665 | 4.0410359 | 0.0612463 | 0.57375847 |
| ENSMUSG00000062115 | Rai1 | protein_coding | 0.635371175 | 5.042439665 | 4.0407346 | 0.061255 | 0.57375847 |
| ENSMUSG00000022889 | Mrpl39 | protein_coding | 0.487440306 | 6.207912801 | 4.0388382 | 0.0613096 | 0.573770108 |
| ENSMUSG00000028909 | Ptpru | protein_coding | 1.085907008 | 3.507827806 | 4.0329836 | 0.0614785 | 0.574850758 |
| ENSMUSG00000063334 | Krr1 | protein_coding | -0.654244777 | 4.937537707 | 4.0276833 | 0.0616319 | 0.575135278 |
| ENSMUSG00000025337 | Sbds | protein_coding | 0.351124623 | 8.185856683 | 4.0243742 | 0.0617279 | 0.575135278 |
| ENSMUSG00000039316 | Rftn1 | protein_coding | 0.614925939 | 4.447628496 | 4.023077 | 0.0617656 | 0.575135278 |
| ENSMUSG00000030774 | Pak1 | protein_coding | 0.448129658 | 8.523614792 | 4.0226931 | 0.0617768 | 0.575135278 |
| ENSMUSG00000028540 | Dph2 | protein_coding | 0.75533157 | 4.539993622 | 4.0174482 | 0.0619294 | 0.575135278 |
| ENSMUSG00000069565 | Dazap1 | protein_coding | 0.929056617 | 3.087351188 | 4.0174363 | 0.0619298 | 0.575135278 |
| ENSMUSG00000031781 | Ciapin1 | protein_coding | 0.395666624 | 7.512321031 | 4.0168559 | 0.0619467 | 0.575135278 |
| ENSMUSG00000091476 | Gm16432 | protein_coding | -1.27540835 | 2.639064719 | 4.0166056 | 0.061954 | 0.575135278 |
| ENSMUSG00000021693 | Kif2a | protein_coding | -0.446662841 | 5.858300644 | 4.0123997 | 0.0620768 | 0.575135278 |
| ENSMUSG00000070822 | Zscan18 | protein_coding | 0.938763496 | 4.896560747 | 4.0489163 | 0.0621237 | 0.575135278 |
| ENSMUSG00000039354 | Smarcal1 | protein_coding | 0.827099876 | 4.604776588 | 4.0110611 | 0.0621445 | 0.575135278 |
| ENSMUSG00000025979 | Mob4 | protein_coding | -0.386230829 | 7.801513554 | 4.0098701 | 0.0621508 | 0.575135278 |
| ENSMUSG00000030682 | Cdipt | protein_coding | 0.444367304 | 6.957219419 | 3.9989647 | 0.0624709 | 0.577600976 |
| ENSMUSG00000041236 | Vps41 | protein_coding | 0.472426946 | 7.767959497 | 3.9951324 | 0.0625839 | 0.578148352 |
| ENSMUSG00000021957 | Tkt | protein_coding | 0.498770745 | 8.534920959 | 3.9930531 | 0.0626739 | 0.578444417 |
| ENSMUSG00000022829 | Stxbp5l | protein_coding | -0.802102246 | 3.902236103 | 3.9904066 | 0.0627235 | 0.578444417 |
| ENSMUSG00000028248 | Pnisr | protein_coding | -0.515773475 | 5.828569493 | 3.9859511 | 0.0628555 | 0.57916489 |
| ENSMUSG00000049728 | Zfp668 | protein_coding | 0.85589124 | 3.787904068 | 3.9812731 | 0.0629944 | 0.579432112 |
| ENSMUSG00000058672 | Tubb2a | protein_coding | 0.476712424 | 9.439810375 | 3.9795177 | 0.0630467 | 0.579432112 |
| ENSMUSG00000020085 | Aifm2 | protein_coding | 0.912758825 | 4.33042888 | 3.9789395 | 0.0630926 | 0.579432112 |
| ENSMUSG00000103103 | 4833445I07Rik | lincRNA | -1.169600976 | 2.77715913 | 3.9777246 | 0.0631001 | 0.579432112 |
| ENSMUSG00000028745 | Capzb | protein_coding | 0.384632914 | 8.992086046 | 3.9722214 | 0.0632643 | 0.579611864 |
| ENSMUSG00000038170 | Pde4dip | protein_coding | 0.440413591 | 6.043936981 | 3.9716084 | 0.0632826 | 0.579611864 |
| ENSMUSG00000039976 | Tbc1d16 | protein_coding | -0.642586048 | 4.131890637 | 3.9696857 | 0.0633401 | 0.579611864 |
| ENSMUSG00000073684 | Faap20 | protein_coding | 0.469651948 | 5.802245919 | 3.9687275 | 0.0633688 | 0.579611864 |
| ENSMUSG00000088595 | Gm24601 | rRNA | -0.712191387 | 4.51822652 | 3.9680477 | 0.0633892 | 0.579611864 |
| ENSMUSG00000037017 | Zscan21 | protein_coding | 0.721395519 | 4.961300475 | 3.9614185 | 0.0635881 | 0.580597993 |
| ENSMUSG00000057615 | Ldoc1 | protein_coding | 0.826423187 | 3.467957453 | 3.9608563 | 0.063605 | 0.580597993 |
| ENSMUSG00000027400 | Pdyn | protein_coding | 0.62854103 | 6.739999228 | 3.9945649 | 0.0637114 | 0.581075623 |
| ENSMUSG00000063018 | 2010204K13Rik | processed_transcript | 0.444079961 | 7.865832667 | 3.9533579 | 0.063831 | 0.581667427 |
| ENSMUSG00000060240 | Cend1 | protein_coding | 0.478436117 | 9.860547497 | 3.9515882 | 0.0638844 | 0.581667427 |
| ENSMUSG00000043372 | Hexim2 | protein_coding | 0.64662505 | 5.644263685 | 3.9489225 | 0.0639938 | 0.582170667 |
| ENSMUSG00000067028 | Cntnap5b | protein_coding | -1.106455738 | 2.91609367 | 3.9408756 | 0.0642092 | 0.583474856 |
| ENSMUSG00000037938 | Chchd5 | protein_coding | 0.393132471 | 6.292169243 | 3.9396752 | 0.0642457 | 0.583474856 |

| ensembl_gene_id | external_gene_name | gene_biotype | log2 (fold change) | log2 (counts per million) | F-value | p-value | False discovery rate |
|---|---|---|---|---|---|---|---|
| ENSMUSG00000031585 | Gtf2e2 | protein_coding | -0.437496505 | 5.868931169 | 3.9360568 | 0.0643559 | 0.583531049 |
| ENSMUSG00000022507 | 1810013L24Rik | protein_coding | -0.384376998 | 6.407993132 | 3.9359089 | 0.0643604 | 0.583531049 |
| ENSMUSG00000000826 | Dnajc5 | protein_coding | 0.390241307 | 8.25432264 | 3.9326477 | 0.06446 | 0.583941072 |
| ENSMUSG00000047671 | Tctex1d4 | protein_coding | -1.005252293 | 3.345862456 | 3.9293601 | 0.0645605 | 0.584359349 |
| ENSMUSG00000019027 | Dnah1 | protein_coding | 0.719571498 | 5.188644309 | 3.9283367 | 0.0646207 | 0.58441201 |
| ENSMUSG00000014932 | Yes1 | protein_coding | -1.432180913 | 2.407524458 | 3.9226652 | 0.0647657 | 0.585231992 |
| ENSMUSG00000039536 | Stau1 | protein_coding | 0.556750777 | 5.958777813 | 3.9175478 | 0.0649232 | 0.585540875 |
| ENSMUSG00000028811 | Yars | protein_coding | 0.514345332 | 7.068007096 | 3.9179881 | 0.0649385 | 0.585540875 |
| ENSMUSG00000029501 | Ankle2 | protein_coding | -0.646945369 | 4.398547778 | 3.9162455 | 0.0649633 | 0.585540875 |
| ENSMUSG00000040653 | Ppp1r14c | protein_coding | -0.858357575 | 3.440860724 | 3.9112835 | 0.0651165 | 0.585889097 |
| ENSMUSG00000078578 | Ube2d3 | protein_coding | -0.353186421 | 7.32672066 | 3.9085099 | 0.0652023 | 0.585889097 |
| ENSMUSG00000036611 | Eepd1 | protein_coding | 0.900027411 | 3.164771061 | 3.9083427 | 0.0652074 | 0.585889097 |
| ENSMUSG00000027800 | Tm4sf1 | protein_coding | -0.68848678 | 4.283675541 | 3.9079412 | 0.0652199 | 0.585889097 |
| ENSMUSG00000024146 | Cript | protein_coding | -0.407583385 | 8.259973186 | 3.9053039 | 0.0653016 | 0.586133691 |
| ENSMUSG00000040430 | Pitpnc1 | protein_coding | -0.467087036 | 7.311017842 | 3.9029778 | 0.0653738 | 0.586292225 |
| ENSMUSG00000026926 | Pmpca | protein_coding | 0.600949233 | 6.246171186 | 3.9011105 | 0.0654607 | 0.586582792 |
| ENSMUSG00000066456 | Hmgn3 | protein_coding | -0.482418965 | 7.04535961 | 3.8963067 | 0.0655813 | 0.586984742 |
| ENSMUSG00000035824 | Tk2 | protein_coding | 0.608957876 | 5.365178531 | 3.8936782 | 0.0656633 | 0.586984742 |
| ENSMUSG00000042744 | Hectd4 | protein_coding | -0.454444669 | 6.342138632 | 3.8902944 | 0.0657691 | 0.586984742 |
| ENSMUSG00000015090 | Ptgds | protein_coding | -0.575818124 | 9.368961098 | 3.9075738 | 0.0657777 | 0.586984742 |
| ENSMUSG00000009394 | Syn2 | protein_coding | 0.376511207 | 8.464177171 | 3.8899914 | 0.0657785 | 0.586984742 |
| ENSMUSG00000021268 | Meg3 | lincRNA | -0.958772058 | 7.508684122 | 4.1354904 | 0.0658834 | 0.587103411 |
| ENSMUSG00000023966 | Rsph9 | protein_coding | 0.38186595 | 6.664529236 | 3.8860409 | 0.0659022 | 0.587103411 |
| ENSMUSG00000024050 | Wiz | protein_coding | -0.843647513 | 3.622251832 | 3.8843386 | 0.0659556 | 0.587103411 |
| ENSMUSG00000035984 | Nme5 | protein_coding | -0.463061241 | 6.20005046 | 3.8813187 | 0.0660505 | 0.587427744 |
| ENSMUSG00000038319 | Kcnh2 | protein_coding | -1.072923035 | 2.905448428 | 3.8797015 | 0.0661013 | 0.587427744 |
| ENSMUSG00000078784 | 1810022K09Rik | protein_coding | -0.645541469 | 4.566680193 | 3.8754134 | 0.0662364 | 0.588030184 |
| ENSMUSG00000031153 | Gripap1 | protein_coding | 0.383903916 | 7.663267864 | 3.8733894 | 0.0663003 | 0.588030184 |
| ENSMUSG00000038742 | Angptl6 | protein_coding | 0.82144731 | 4.581476581 | 3.8732658 | 0.0663332 | 0.588030184 |
| ENSMUSG00000108112 | Gm45193 | processed_transcript | -0.86581684 | 3.974443416 | 3.8685516 | 0.0664532 | 0.588608841 |
| ENSMUSG00000022191 | Drosha | protein_coding | 0.464419835 | 6.414153116 | 3.8667937 | 0.0665089 | 0.588617123 |
| ENSMUSG00000029472 | Anapc5 | protein_coding | 0.457988183 | 7.048916681 | 3.8633523 | 0.066618 | 0.589098306 |
| ENSMUSG00000030738 | Eif3c | protein_coding | 0.426776033 | 7.789052658 | 3.8598215 | 0.0667302 | 0.589169551 |
| ENSMUSG00000036699 | Zcchc12 | protein_coding | 0.75459032 | 9.901316203 | 3.9431217 | 0.0667357 | 0.589169551 |
| ENSMUSG00000003847 | Nfat5 | protein_coding | -0.671040477 | 4.026495328 | 3.857026 | 0.0668192 | 0.589423353 |
| ENSMUSG00000034165 | Ccnd3 | protein_coding | -0.744843814 | 3.629713756 | 3.8502517 | 0.0670355 | 0.590251375 |
| ENSMUSG00000052997 | Uba2 | protein_coding | 0.38697954 | 6.448238628 | 3.849802 | 0.0670498 | 0.590251375 |
| ENSMUSG00000031255 | Sytl4 | protein_coding | -0.667841545 | 4.055596508 | 3.848899 | 0.0670787 | 0.590251375 |
| ENSMUSG00000050240 | Hic2 | protein_coding | 1.545911431 | 3.345082412 | 3.9015828 | 0.0671776 | 0.590251375 |
| ENSMUSG00000006418 | Rnf114 | protein_coding | 0.406792207 | 7.368095382 | 3.844107 | 0.0672323 | 0.590251375 |
| ENSMUSG00000019158 | Tmem160 | protein_coding | -0.75474842 | 4.02644066 | 3.8413522 | 0.0673208 | 0.590251375 |
| ENSMUSG00000000308 | Ckmt1 | protein_coding | 0.53147598 | 8.95899828 | 3.8418109 | 0.0673352 | 0.590251375 |
| ENSMUSG00000036834 | Plch1 | protein_coding | -1.13950303 | 3.273452275 | 3.8394369 | 0.0673824 | 0.590251375 |
| ENSMUSG00000059142 | Zfp945 | protein_coding | -0.622944356 | 4.555713559 | 3.8374406 | 0.0674466 | 0.590251375 |
| ENSMUSG00000023031 | Cela1 | protein_coding | -0.984771769 | 3.943138773 | 3.8378682 | 0.067462 | 0.590251375 |
| ENSMUSG00000001123 | Lgals9 | protein_coding | 0.946913406 | 3.411263067 | 3.8336609 | 0.0675685 | 0.590702665 |
| ENSMUSG00000049721 | Gal3st1 | protein_coding | -1.092756853 | 3.6048914 | 3.829526 | 0.0677313 | 0.591644786 |
| ENSMUSG00000026341 | Actr3 | protein_coding | -0.400445459 | 6.223189003 | 3.8225087 | 0.0679296 | 0.592728545 |
| ENSMUSG00000019761 | Krt10 | protein_coding | -0.553591396 | 5.047705005 | 3.8214012 | 0.0679656 | 0.592728545 |
| ENSMUSG00000019927 | Ube2d1 | protein_coding | -0.417552561 | 6.169708397 | 3.8077331 | 0.0684116 | 0.595923268 |
| ENSMUSG00000091474 | 2610021A01Rik | protein_coding | -0.875865011 | 3.469323336 | 3.8052081 | 0.0684943 | 0.595923268 |
| ENSMUSG00000026411 | Tmem9 | protein_coding | 0.365386542 | 7.866960137 | 3.8050899 | 0.0684982 | 0.595923268 |
| ENSMUSG00000039634 | Zfp189 | protein_coding | -0.963200485 | 3.583307246 | 3.801401 | 0.0686193 | 0.596494432 |
| ENSMUSG00000043183 | Simc1 | protein_coding | -0.886429101 | 3.341186191 | 3.7976032 | 0.0687443 | 0.59697318 |
| ENSMUSG00000050121 | Opalin | protein_coding | -0.384142015 | 7.410040088 | 3.7946712 | 0.0688409 | 0.59697318 |
| ENSMUSG00000022031 | Elp3 | protein_coding | 0.562064501 | 6.756834004 | 3.7984549 | 0.0688731 | 0.59697318 |
| ENSMUSG00000042042 | Csgalnact2 | protein_coding | -1.249210645 | 2.890965698 | 3.7929891 | 0.0688965 | 0.59697318 |
| ENSMUSG00000014905 | Dnajb9 | protein_coding | -0.61313694 | 5.484625982 | 3.7858998 | 0.0691311 | 0.5979572 |
| ENSMUSG00000004364 | Cul3 | protein_coding | -0.398906373 | 6.451955058 | 3.7852087 | 0.069154 | 0.5979572 |
| ENSMUSG00000022548 | Apod | protein_coding | -0.452818111 | 8.762868782 | 3.784519 | 0.0691769 | 0.5979572 |
| ENSMUSG00000019373 | Cops3 | protein_coding | 0.338925702 | 7.154969852 | 3.7826029 | 0.0692405 | 0.598026355 |
| ENSMUSG00000000194 | Gpr107 | protein_coding | 0.688905335 | 4.487791172 | 3.7802742 | 0.0693179 | 0.598167862 |
| ENSMUSG00000034156 | Tspoap1 | protein_coding | -0.761263571 | 4.054410639 | 3.7787646 | 0.0693681 | 0.598167862 |
| ENSMUSG00000039676 | Capsl | protein_coding | -0.897292253 | 4.066368369 | 3.7757447 | 0.0694981 | 0.59880819 |
| ENSMUSG00000074829 | 2010315B03Rik | protein_coding | -0.712706075 | 4.00988942 | 3.7645097 | 0.0698446 | 0.60023035 |
| ENSMUSG00000025139 | Tollip | protein_coding | 0.468823528 | 7.020217951 | 3.7641121 | 0.0698579 | 0.60023035 |
| ENSMUSG00000024993 | Fam45a | protein_coding | -0.431311723 | 6.807848676 | 3.7639433 | 0.0698636 | 0.60023035 |
| ENSMUSG00000055540 | Epha6 | protein_coding | -0.874692578 | 3.574033654 | 3.7580898 | 0.0700604 | 0.60023035 |
| ENSMUSG00000002308 | Cd320 | protein_coding | -0.637735256 | 4.901515219 | 3.7573644 | 0.0700848 | 0.60023035 |
| ENSMUSG00000038909 | Kat7 | protein_coding | 0.541572515 | 6.342927976 | 3.7582116 | 0.0700857 | 0.60023035 |
| ENSMUSG00000034612 | Chst11 | protein_coding | 0.662146303 | 3.991969294 | 3.7536196 | 0.0702112 | 0.60023035 |
| ENSMUSG00000000776 | Polr3d | protein_coding | 1.177792327 | 3.890417899 | 3.79103 | 0.0702548 | 0.60023035 |
| ENSMUSG00000044231 | Nhlrc1 | protein_coding | -0.832589441 | 3.692155523 | 3.7515334 | 0.0702816 | 0.60023035 |
| ENSMUSG00000047379 | B4gat1 | protein_coding | 0.499289165 | 7.438646218 | 3.7522001 | 0.0702885 | 0.60023035 |

| ensembl_gene_id | external_gene_name | gene_biotype | log2 (fold change) | log2 (counts per million) | F-value | p-value | False discovery rate |
|---|---|---|---|---|---|---|---|
| ENSMUSG00000029777 | Gars | protein_coding | 0.425159259 | 6.473768453 | 3.7478786 | 0.0704053 | 0.60023035 |
| ENSMUSG00000000751 | Rpa1 | protein_coding | 0.573926064 | 6.402588931 | 3.7455396 | 0.070514 | 0.60023035 |
| ENSMUSG00000028759 | Hp1bp3 | protein_coding | 0.427016074 | 7.591971683 | 3.7428517 | 0.0705758 | 0.60023035 |
| ENSMUSG00000027359 | Slc27a2 | protein_coding | 1.340174678 | 2.826299836 | 3.7427847 | 0.0705781 | 0.60023035 |
| ENSMUSG00000062761 | Zfp512 | protein_coding | 0.456372763 | 5.651797819 | 3.7421694 | 0.070599 | 0.60023035 |
| ENSMUSG00000015749 | Anp32e | protein_coding | 0.36302804 | 6.679126619 | 3.7403177 | 0.070662 | 0.60023035 |
| ENSMUSG00000030327 | Necap1 | protein_coding | 0.491571159 | 9.043497889 | 3.7403 | 0.070692 | 0.60023035 |
| ENSMUSG00000032492 | Pth1r | protein_coding | 0.969992404 | 3.535233835 | 3.7378691 | 0.0707453 | 0.60023035 |
| ENSMUSG00000058093 | Zfp729b | protein_coding | -1.388481368 | 2.732909415 | 3.7369158 | 0.0707778 | 0.60023035 |
| ENSMUSG00000042712 | Tceal9 | protein_coding | -0.429925656 | 9.303714622 | 3.7368665 | 0.0707795 | 0.60023035 |
| ENSMUSG00000002332 | Dhrs1 | protein_coding | 0.381841545 | 7.310267359 | 3.726281 | 0.0711414 | 0.602365822 |
| ENSMUSG00000014418 | Hps5 | protein_coding | -1.066750431 | 3.081212267 | 3.7262243 | 0.0711434 | 0.602365822 |
| ENSMUSG00000030357 | Fkbp4 | protein_coding | 0.372499092 | 7.238306529 | 3.7241002 | 0.0712162 | 0.602508509 |
| ENSMUSG00000040435 | Ppp1r15a | protein_coding | 0.727309112 | 4.902967388 | 3.7154738 | 0.0715131 | 0.604544727 |
| ENSMUSG00000064036 | Mro | protein_coding | -0.480899462 | 5.294121439 | 3.7134252 | 0.0715839 | 0.604667203 |
| ENSMUSG00000051606 | 2010001K21Rik | processed_transcript | -1.896631309 | 2.361829615 | 3.7086098 | 0.07178 | 0.604843039 |
| ENSMUSG00000079109 | Pms2 | protein_coding | 1.436000212 | 2.451916495 | 3.7055727 | 0.0718557 | 0.604843039 |
| ENSMUSG00000020926 | Adam11 | protein_coding | -0.565030324 | 4.884671681 | 3.7042313 | 0.0719023 | 0.604843039 |
| ENSMUSG00000024646 | Cyb5a | protein_coding | -0.433939874 | 8.28430428 | 3.7035032 | 0.0719276 | 0.604843039 |
| ENSMUSG00000037108 | Zcwpw1 | protein_coding | -0.780748824 | 3.452655585 | 3.7034269 | 0.0719302 | 0.604843039 |
| ENSMUSG00000026784 | Pdss1 | protein_coding | -0.882812861 | 3.362814986 | 3.7030653 | 0.0719428 | 0.604843039 |
| ENSMUSG00000010608 | Rbm25 | protein_coding | -0.380858056 | 6.623945702 | 3.7003459 | 0.0720373 | 0.604843039 |
| ENSMUSG00000019179 | Mdh2 | protein_coding | 0.426844075 | 10.78112278 | 3.6987756 | 0.072092 | 0.604843039 |
| ENSMUSG00000024891 | Slc29a2 | protein_coding | 0.658272402 | 4.347253447 | 3.6973234 | 0.0721426 | 0.604843039 |
| ENSMUSG00000029563 | Foxp2 | protein_coding | -0.586975248 | 5.536918771 | 3.6966188 | 0.0721672 | 0.604843039 |
| ENSMUSG00000022244 | Amacr | protein_coding | 0.782621923 | 4.10322344 | 3.6941504 | 0.0722533 | 0.605093366 |
| ENSMUSG00000052144 | Ppp4r2 | protein_coding | -0.535493308 | 4.914873638 | 3.6900915 | 0.0723952 | 0.60580999 |
| ENSMUSG00000026036 | Nif3l1 | protein_coding | 0.611158398 | 6.00136296 | 3.6877673 | 0.0725062 | 0.605820261 |
| ENSMUSG00000054640 | Slc8a1 | protein_coding | -0.963068823 | 3.27902357 | 3.6845179 | 0.0725906 | 0.605820261 |
| ENSMUSG00000022673 | Mcm4 | protein_coding | 0.874436278 | 3.356208428 | 3.6837322 | 0.0726182 | 0.605820261 |
| ENSMUSG00000030224 | Strap | protein_coding | -0.380145969 | 6.523884665 | 3.6816139 | 0.0726927 | 0.605820261 |
| ENSMUSG00000037134 | Prmt9 | protein_coding | 0.823053855 | 3.579216557 | 3.6802072 | 0.0727422 | 0.605820261 |
| ENSMUSG00000035575 | Utp6 | protein_coding | 0.625370907 | 4.648040619 | 3.6792268 | 0.0727767 | 0.605820261 |
| ENSMUSG00000037149 | Ddx1 | protein_coding | 0.369179532 | 7.647847429 | 3.6788254 | 0.0727908 | 0.605820261 |
| ENSMUSG00000069255 | Dusp22 | protein_coding | 0.569689173 | 5.260943692 | 3.6770873 | 0.0728521 | 0.605861186 |
| ENSMUSG00000022359 | Wdyhv1 | protein_coding | 0.435710493 | 5.77310825 | 3.6726688 | 0.0730081 | 0.606617126 |
| ENSMUSG00000097006 | 9530082P21Rik | lincRNA | 0.512271187 | 8.267374426 | 3.6721573 | 0.0730558 | 0.606617126 |
| ENSMUSG00000086155 | 9430041J12Rik | antisense_RNA | -1.555631552 | 2.106164436 | 3.6683083 | 0.0731624 | 0.606831141 |
| ENSMUSG00000030347 | D6Wsu163e | protein_coding | 0.489563224 | 5.091092312 | 3.6667022 | 0.0732194 | 0.606831141 |
| ENSMUSG00000029404 | Arl6ip4 | protein_coding | 0.332040686 | 7.407770941 | 3.665814 | 0.0732509 | 0.606831141 |
| ENSMUSG00000028132 | Tmem56 | protein_coding | -0.760218709 | 4.357039653 | 3.6618939 | 0.0733902 | 0.607330248 |
| ENSMUSG00000070639 | Lrrc8b | protein_coding | -0.636590629 | 4.292064938 | 3.6609402 | 0.0734241 | 0.607330248 |
| ENSMUSG00000022771 | Ppil2 | protein_coding | 0.463288134 | 6.472391962 | 3.6547722 | 0.073644 | 0.608680897 |
| ENSMUSG00000092035 | Peg10 | protein_coding | -0.583356787 | 4.61487077 | 3.6496448 | 0.0738274 | 0.609727942 |
| ENSMUSG00000037253 | Mex3c | protein_coding | -0.861084233 | 3.938099964 | 3.6468648 | 0.073927 | 0.610039078 |
| ENSMUSG00000038022 | Mindy4 | protein_coding | 0.988150573 | 3.076901349 | 3.6440207 | 0.0740291 | 0.610039078 |
| ENSMUSG00000026999 | Nup35 | protein_coding | -0.612864722 | 4.37657614 | 3.6438509 | 0.0740352 | 0.610039078 |
| ENSMUSG00000043441 | Gpr149 | protein_coding | -0.891100475 | 3.185985951 | 3.6394099 | 0.074195 | 0.610836031 |
| ENSMUSG00000097290 | 1300002E11Rik | lincRNA | -0.612426097 | 4.73278305 | 3.6365597 | 0.0742978 | 0.610836031 |
| ENSMUSG00000044986 | Tst | protein_coding | -0.586547858 | 6.522073545 | 3.6531949 | 0.0743024 | 0.610836031 |
| ENSMUSG00000028833 | Ncdn | protein_coding | -0.409910605 | 8.830321483 | 3.6273415 | 0.0746313 | 0.611749346 |
| ENSMUSG00000021737 | Psmd6 | protein_coding | 0.372654075 | 8.008013468 | 3.6269554 | 0.0746454 | 0.611749346 |
| ENSMUSG00000036241 | Ube2r2 | protein_coding | -0.369047767 | 6.414309663 | 3.6260082 | 0.0746797 | 0.611749346 |
| ENSMUSG00000036752 | Tubb4b | protein_coding | 0.439900239 | 7.668649757 | 3.6257519 | 0.074689 | 0.611749346 |
| ENSMUSG00000002326 | Gmpr2 | protein_coding | 0.568300347 | 5.289476233 | 3.6255069 | 0.0746979 | 0.611749346 |
| ENSMUSG00000038416 | Cdc16 | protein_coding | 0.570056184 | 5.149809649 | 3.6228532 | 0.0747944 | 0.61207303 |
| ENSMUSG00000042328 | Hps4 | protein_coding | 0.741364318 | 3.690043828 | 3.6204153 | 0.0748831 | 0.612333133 |
| ENSMUSG00000028289 | Epha7 | protein_coding | -0.51622902 | 5.22198334 | 3.6147947 | 0.0750881 | 0.613490561 |
| ENSMUSG00000044150 | A830080D01Rik | protein_coding | -1.038151503 | 3.460706837 | 3.6102083 | 0.075256 | 0.613490561 |
| ENSMUSG00000029778 | Adcyap1r1 | protein_coding | 0.435323805 | 6.178430116 | 3.6081863 | 0.0753301 | 0.613490561 |
| ENSMUSG00000030421 | Uri1 | protein_coding | -0.530110582 | 4.919120289 | 3.6075478 | 0.0753535 | 0.613490561 |
| ENSMUSG00000033355 | Rtp4 | protein_coding | -0.987090487 | 3.126391149 | 3.6069532 | 0.0753753 | 0.613490561 |
| ENSMUSG00000043411 | Usp48 | protein_coding | 0.588127502 | 4.625503793 | 3.6041524 | 0.0754782 | 0.613490561 |
| ENSMUSG00000001687 | Ubl3 | protein_coding | -0.353677565 | 8.338305244 | 3.6039596 | 0.0754853 | 0.613490561 |
| ENSMUSG00000043289 | Mei4 | protein_coding | -1.496960974 | 2.37639446 | 3.6033762 | 0.0755067 | 0.613490561 |
| ENSMUSG00000032898 | Fbxo21 | protein_coding | 0.458091586 | 6.329678203 | 3.6022876 | 0.0755468 | 0.613490561 |
| ENSMUSG00000059482 | 2610301B20Rik | protein_coding | -0.382897272 | 6.793202852 | 3.6000744 | 0.0756283 | 0.613490561 |
| ENSMUSG00000026510 | Trp53bp2 | protein_coding | -0.639524656 | 4.712471166 | 3.5994245 | 0.0756522 | 0.613490561 |
| ENSMUSG00000025393 | Atp5b | protein_coding | 0.463038118 | 10.30612259 | 3.5955359 | 0.0757957 | 0.613742029 |
| ENSMUSG00000049764 | Zfp280b | protein_coding | 0.832162918 | 3.189917158 | 3.5953784 | 0.0758016 | 0.613742029 |
| ENSMUSG00000071723 | Gspt2 | protein_coding | 0.499471855 | 5.012475864 | 3.59327 | 0.0758795 | 0.613742029 |
| ENSMUSG00000028842 | Ago3 | protein_coding | -0.786546895 | 4.372173985 | 3.5907294 | 0.0759736 | 0.613742029 |
| ENSMUSG00000030521 | Mphosph10 | protein_coding | -0.565776578 | 4.697835945 | 3.5901257 | 0.0759959 | 0.613742029 |
| ENSMUSG00000024958 | Gpr137 | protein_coding | -0.50055391 | 6.39624649 | 3.5867725 | 0.0761203 | 0.613742029 |

| ensembl_gene_id | external_gene_name | gene_biotype | log2 (fold change) | log2 (counts per million) | F-value | p-value | False discovery rate |
|---|---|---|---|---|---|---|---|
| ENSMUSG00000039480 | Nt5dc1 | protein_coding | 1.838075122 | 1.833752759 | 3.5862734 | 0.0761388 | 0.613742029 |
| ENSMUSG00000037111 | Setd7 | protein_coding | -0.535200584 | 4.984668212 | 3.5862462 | 0.0761399 | 0.613742029 |
| ENSMUSG00000020612 | Prkar1a | protein_coding | 0.611834534 | 9.700402142 | 3.6112943 | 0.0763727 | 0.613809872 |
| ENSMUSG00000040747 | Cd53 | protein_coding | -0.81897035 | 4.078831472 | 3.579511 | 0.0763905 | 0.613809872 |
| ENSMUSG00000026117 | Zap70 | protein_coding | 1.934242804 | 2.329439338 | 3.5770571 | 0.076512 | 0.613809872 |
| ENSMUSG00000006262 | Mob1b | protein_coding | -0.620633731 | 3.822096656 | 3.5755207 | 0.0765395 | 0.613809872 |
| ENSMUSG00000019471 | Cdc37 | protein_coding | 0.326915045 | 7.155075556 | 3.5754723 | 0.0765413 | 0.613809872 |
| ENSMUSG00000040929 | Rfx3 | protein_coding | -0.671238964 | 4.279747821 | 3.5751078 | 0.0765549 | 0.613809872 |
| ENSMUSG00000078877 | Gm14295 | protein_coding | -0.747277019 | 3.695255168 | 3.574464 | 0.076579 | 0.613809872 |
| ENSMUSG00000023827 | Agpat4 | protein_coding | 0.408198813 | 7.118018798 | 3.5725498 | 0.0766506 | 0.613809872 |
| ENSMUSG00000051354 | Samd3 | protein_coding | 1.209998374 | 3.681003651 | 3.6024724 | 0.0766974 | 0.613809872 |
| ENSMUSG00000024084 | Qpct | protein_coding | 0.402692491 | 6.503661628 | 3.5702715 | 0.0767359 | 0.613809872 |
| ENSMUSG00000034765 | Dusp5 | protein_coding | 1.36157467 | 2.497909975 | 3.5691978 | 0.0767762 | 0.613809872 |
| ENSMUSG00000048076 | Arf1 | protein_coding | 0.385575559 | 9.201643865 | 3.5668789 | 0.0768632 | 0.614049153 |
| ENSMUSG00000022602 | Arc | protein_coding | -0.648359412 | 5.510500588 | 3.5618383 | 0.0770828 | 0.615094743 |
| ENSMUSG00000026972 | Arrdc1 | protein_coding | 1.125465798 | 3.728158626 | 3.5806385 | 0.0771116 | 0.615094743 |
| ENSMUSG00000006711 | D130043K22Rik | protein_coding | 1.017445847 | 2.919162373 | 3.5588436 | 0.0771657 | 0.615094743 |
| ENSMUSG00000032606 | Nicn1 | protein_coding | 0.38757046 | 8.336555782 | 3.5528619 | 0.0773918 | 0.615443045 |
| ENSMUSG00000032249 | Anp32a | protein_coding | -0.383315402 | 6.482187072 | 3.5523675 | 0.0774105 | 0.615443045 |
| ENSMUSG00000022175 | Lrp10 | protein_coding | -0.850536371 | 3.736932533 | 3.5519969 | 0.0774246 | 0.615443045 |
| ENSMUSG00000031641 | Cbr4 | protein_coding | 0.543151761 | 5.240956239 | 3.5495283 | 0.0775181 | 0.615443045 |
| ENSMUSG00000073563 | Csnk1g3 | protein_coding | -0.512498445 | 5.131053326 | 3.5485999 | 0.0775534 | 0.615443045 |
| ENSMUSG00000028385 | Snx30 | protein_coding | 0.524168828 | 4.822640067 | 3.5474989 | 0.0775952 | 0.615443045 |
| ENSMUSG00000004446 | Bid | protein_coding | 0.592543347 | 5.242118399 | 3.5471073 | 0.07761 | 0.615443045 |
| ENSMUSG00000052833 | Sae1 | protein_coding | 0.482107651 | 7.901327901 | 3.5442961 | 0.0777469 | 0.615545874 |
| ENSMUSG00000028494 | Plin2 | protein_coding | 0.813607026 | 3.848423259 | 3.5402816 | 0.0778698 | 0.615545874 |
| ENSMUSG00000021189 | Atxn3 | protein_coding | 0.779283379 | 3.618118477 | 3.5396438 | 0.0778942 | 0.615545874 |
| ENSMUSG00000028882 | Ppp1r8 | protein_coding | 0.47781241 | 5.707958089 | 3.5356976 | 0.0780449 | 0.615545874 |
| ENSMUSG00000029723 | Tsc22d4 | protein_coding | -0.656176602 | 7.49352734 | 3.6033471 | 0.078057 | 0.615545874 |
| ENSMUSG00000000223 | Drp2 | protein_coding | 0.679871941 | 4.106157728 | 3.5340642 | 0.0781074 | 0.615545874 |
| ENSMUSG00000031633 | Slc25a4 | protein_coding | 0.37022756 | 11.73553787 | 3.5332878 | 0.0781371 | 0.615545874 |
| ENSMUSG00000071324 | Armc2 | protein_coding | 1.260302698 | 2.970125851 | 3.5332049 | 0.0781403 | 0.615545874 |
| ENSMUSG00000039715 | Wdr34 | protein_coding | 0.570808135 | 4.733743123 | 3.5329223 | 0.0781511 | 0.615545874 |
| ENSMUSG00000060703 | Cd302 | protein_coding | -0.64736773 | 5.575286645 | 3.532549 | 0.0781954 | 0.615545874 |
| ENSMUSG00000025582 | Nptx1 | protein_coding | 0.62877489 | 4.495206793 | 3.5300231 | 0.0782623 | 0.61562105 |
| ENSMUSG00000038324 | Trpc4ap | protein_coding | 0.3727743 | 6.560717276 | 3.5264721 | 0.0783986 | 0.616242982 |
| ENSMUSG00000022546 | Gpt | protein_coding | -0.783792578 | 4.057947977 | 3.5238801 | 0.0784983 | 0.616243562 |
| ENSMUSG00000005682 | Pan2 | protein_coding | 1.427118721 | 2.311549658 | 3.5234913 | 0.0785133 | 0.616243562 |
| ENSMUSG00000020706 | Ftsj3 | protein_coding | 0.720005149 | 4.824402061 | 3.5193869 | 0.0787016 | 0.616564702 |
| ENSMUSG00000049571 | Cfap46 | protein_coding | 1.076475779 | 3.510504677 | 3.5193675 | 0.0787024 | 0.616564702 |
| ENSMUSG00000022434 | Fam118a | protein_coding | 0.515866869 | 5.011475363 | 3.5179708 | 0.0787262 | 0.616564702 |
| ENSMUSG00000009739 | Pou6f1 | protein_coding | -0.573116818 | 4.692505443 | 3.514642 | 0.078855 | 0.61672197 |
| ENSMUSG00000024251 | Thada | protein_coding | -0.821248024 | 3.259667911 | 3.5139261 | 0.0788827 | 0.61672197 |
| ENSMUSG00000056820 | Tsnax | protein_coding | 0.434402032 | 6.500801362 | 3.5117144 | 0.0789684 | 0.61672197 |
| ENSMUSG00000041483 | Zfp281 | protein_coding | -0.630077534 | 4.817946654 | 3.5094428 | 0.0790565 | 0.61672197 |
| ENSMUSG00000022234 | Cct5 | protein_coding | 0.40226414 | 9.145480461 | 3.5082098 | 0.0791044 | 0.61672197 |
| ENSMUSG00000022574 | Naprt | protein_coding | -1.075355978 | 3.163063662 | 3.5057906 | 0.0791985 | 0.61672197 |
| ENSMUSG00000040428 | Plekha4 | protein_coding | -1.09563103 | 2.958626033 | 3.5051313 | 0.0792241 | 0.61672197 |
| ENSMUSG00000026019 | Wdr12 | protein_coding | 0.380789656 | 6.68670352 | 3.5050774 | 0.0792262 | 0.61672197 |
| ENSMUSG00000004791 | Pgf | protein_coding | 0.652950587 | 4.251531182 | 3.5034627 | 0.0792891 | 0.61672197 |
| ENSMUSG00000024782 | Ak3 | protein_coding | -0.996335357 | 2.964821122 | 3.5026733 | 0.0793199 | 0.61672197 |
| ENSMUSG00000024644 | Cndp2 | protein_coding | 0.577624528 | 6.333249275 | 3.5036093 | 0.0794275 | 0.617112972 |
| ENSMUSG00000029622 | Arpc1b | protein_coding | -0.610191788 | 4.752222559 | 3.496836 | 0.0795478 | 0.617169875 |
| ENSMUSG00000018507 | Trpv2 | protein_coding | 0.892832159 | 4.695499816 | 3.5250047 | 0.0795497 | 0.617169875 |
| ENSMUSG00000045211 | Nudt18 | protein_coding | 0.510349948 | 5.721261591 | 3.4922802 | 0.0797262 | 0.617794787 |
| ENSMUSG00000020661 | Dnmt3a | protein_coding | 0.417879307 | 5.956895013 | 3.4913148 | 0.079764 | 0.617794787 |
| ENSMUSG00000038708 | Golga4 | protein_coding | -0.477455597 | 5.144764638 | 3.4903333 | 0.0798026 | 0.617794787 |
| ENSMUSG00000018433 | Nol11 | protein_coding | 0.565926656 | 5.266250352 | 3.488301 | 0.0798824 | 0.61796788 |
| ENSMUSG00000054986 | Sec14l3 | protein_coding | 1.335779645 | 2.385090534 | 3.4852276 | 0.0800033 | 0.618151564 |
| ENSMUSG00000027255 | Arfgap2 | protein_coding | 0.717268392 | 4.615170239 | 3.4847752 | 0.0800211 | 0.618151564 |
| ENSMUSG00000026842 | Abl1 | protein_coding | 0.85518068 | 4.118710143 | 3.4809902 | 0.0802005 | 0.619092688 |
| ENSMUSG00000097545 | Mir124a-1hg | lincRNA | -0.867735346 | 3.619828558 | 3.4786203 | 0.0802639 | 0.619137883 |
| ENSMUSG00000029505 | Ep400 | protein_coding | 0.717711595 | 4.822313806 | 3.4726446 | 0.0805307 | 0.620750558 |
| ENSMUSG00000026385 | Dbi | protein_coding | -0.592741454 | 9.643997402 | 3.4996002 | 0.080608 | 0.620785491 |
| ENSMUSG00000017119 | Nbr1 | protein_coding | 0.445421478 | 6.019132822 | 3.4680839 | 0.0806817 | 0.620785491 |
| ENSMUSG00000032705 | Exd2 | protein_coding | -0.654916814 | 4.414390876 | 3.4661403 | 0.080759 | 0.620785491 |
| ENSMUSG00000023118 | Sympk | protein_coding | -0.665415967 | 4.058834445 | 3.4646716 | 0.0808175 | 0.620785491 |
| ENSMUSG00000069763 | Tmem100 | protein_coding | -0.658919963 | 5.158979621 | 3.4645104 | 0.0808239 | 0.620785491 |
| ENSMUSG00000021037 | Ahsa1 | protein_coding | 0.351289727 | 7.846547904 | 3.4610055 | 0.0809637 | 0.620993849 |
| ENSMUSG00000021136 | Smoc1 | protein_coding | 0.933594716 | 3.176641471 | 3.460935 | 0.0809666 | 0.620993849 |
| ENSMUSG00000006576 | Slc4a3 | protein_coding | -0.441613078 | 5.747084477 | 3.4591747 | 0.0810369 | 0.621090286 |
| ENSMUSG00000025278 | Flnb | protein_coding | -1.233789048 | 3.310572923 | 3.4613621 | 0.0811115 | 0.621219674 |
| ENSMUSG00000031337 | Mtm1 | protein_coding | -1.369973522 | 2.849705972 | 3.4556713 | 0.0811771 | 0.621279075 |
| ENSMUSG00000024240 | Epc1 | protein_coding | 0.576608297 | 4.111177096 | 3.4511989 | 0.0813565 | 0.622209094 |

| ensembl_gene_id | external_gene_name | gene_biotype | log2 (fold change) | log2 (counts per million) | F-value | p-value | False discovery rate |
|---|---|---|---|---|---|---|---|
| ENSMUSG00000109836 | Gm45884 | TEC | -0.726272325 | 3.596234696 | 3.4480141 | 0.0814845 | 0.622745272 |
| ENSMUSG00000041718 | Alg13 | protein_coding | -1.465592015 | 2.759858993 | 3.4423109 | 0.0817446 | 0.624087598 |
| ENSMUSG00000035910 | Dcdc2a | protein_coding | -0.696433022 | 4.129149183 | 3.4407791 | 0.0817762 | 0.624087598 |
| ENSMUSG00000019996 | Map7 | protein_coding | -0.800059634 | 3.207849493 | 3.436366 | 0.0819547 | 0.625006619 |
| ENSMUSG00000031285 | Dcx | protein_coding | -0.816520301 | 3.778773421 | 3.4322277 | 0.0821226 | 0.625783629 |
| ENSMUSG00000026489 | Coq8a | protein_coding | 0.539419415 | 5.611937598 | 3.4309861 | 0.082173 | 0.625783629 |
| ENSMUSG00000029212 | Gabrb1 | protein_coding | -0.423096418 | 6.368171514 | 3.4263445 | 0.0823619 | 0.626562516 |
| ENSMUSG00000074797 | Itpa | protein_coding | 0.371579296 | 6.312542748 | 3.4239231 | 0.0824606 | 0.626562516 |
| ENSMUSG00000058546 | Rpl23a | protein_coding | -0.828501866 | 3.615805118 | 3.4230076 | 0.082498 | 0.626562516 |
| ENSMUSG00000023923 | Tbc1d5 | protein_coding | 0.609226189 | 4.134017351 | 3.4227536 | 0.0825084 | 0.626562516 |
| ENSMUSG00000019936 | Epyc | protein_coding | -1.111827543 | 2.734746908 | 3.4158999 | 0.0827888 | 0.627475784 |
| ENSMUSG00000024947 | Men1 | protein_coding | 0.526414391 | 5.013387401 | 3.4156597 | 0.0827986 | 0.627475784 |
| ENSMUSG00000026924 | Sec16a | protein_coding | 0.646793656 | 3.965881154 | 3.4149996 | 0.0828257 | 0.627475784 |
| ENSMUSG00000022620 | Arsa | protein_coding | 0.664210691 | 4.693227459 | 3.4136756 | 0.08288 | 0.627475784 |
| ENSMUSG00000020238 | Ncln | protein_coding | -0.633809558 | 4.257319797 | 3.4110237 | 0.082989 | 0.627475784 |
| ENSMUSG00000024187 | Fam234a | protein_coding | -0.807716782 | 4.779985029 | 3.4146288 | 0.0830036 | 0.627475784 |
| ENSMUSG00000021076 | Actr10 | protein_coding | 0.363752699 | 8.860257403 | 3.4098536 | 0.0830371 | 0.627475784 |
| ENSMUSG00000038827 | Fam206a | protein_coding | -0.546814708 | 4.517711697 | 3.4077227 | 0.0831248 | 0.627697581 |
| ENSMUSG00000060314 | Zfp941 | protein_coding | 0.425926145 | 6.292525636 | 3.4052042 | 0.0832287 | 0.627983155 |
| ENSMUSG00000025915 | Sgk3 | protein_coding | -0.524574691 | 5.458691514 | 3.4039733 | 0.0832795 | 0.627983155 |
| ENSMUSG00000050967 | Creg2 | protein_coding | -0.617645446 | 4.896421896 | 3.4002452 | 0.0834335 | 0.628114867 |
| ENSMUSG00000032309 | Fbxo22 | protein_coding | 0.683046797 | 4.51369685 | 3.3961365 | 0.0836037 | 0.628114867 |
| ENSMUSG00000038544 | Inip | protein_coding | -0.48853618 | 5.42137329 | 3.3958827 | 0.0836143 | 0.628114867 |
| ENSMUSG00000024570 | Rbfa | protein_coding | 0.805222177 | 3.664244014 | 3.395539 | 0.0836285 | 0.628114867 |
| ENSMUSG00000039058 | Ak5 | protein_coding | 0.491011982 | 5.474999154 | 3.395106 | 0.0836465 | 0.628114867 |
| ENSMUSG00000048930 | Tada3 | protein_coding | -0.498517326 | 4.935587301 | 3.3927373 | 0.0837449 | 0.628114867 |
| ENSMUSG00000026925 | Inpp5e | protein_coding | 0.777840458 | 3.412562807 | 3.3922958 | 0.0837632 | 0.628114867 |
| ENSMUSG00000030110 | Ret | protein_coding | 0.771307205 | 3.503712827 | 3.3922713 | 0.0837642 | 0.628114867 |
| ENSMUSG00000113314 | AC122013.1 | lincRNA | 1.325818072 | 2.498443276 | 3.3907586 | 0.0838271 | 0.628148553 |
| ENSMUSG00000050783 | Htr1f | protein_coding | -1.545440948 | 2.268847071 | 3.3880591 | 0.0839395 | 0.628493197 |
| ENSMUSG00000027808 | Serp1 | protein_coding | -0.640221368 | 4.652541343 | 3.386848 | 0.08399 | 0.628493197 |
| ENSMUSG00000015568 | Lpl | protein_coding | -0.910095809 | 3.603553369 | 3.3813349 | 0.0842203 | 0.629094235 |
| ENSMUSG00000003948 | Mmd | protein_coding | -0.663471951 | 6.024027836 | 3.4095761 | 0.0842256 | 0.629094235 |
| ENSMUSG00000043881 | Kbtbd7 | protein_coding | -0.61892099 | 4.593530467 | 3.3807241 | 0.0842459 | 0.629094235 |
| ENSMUSG00000002222 | Rmnd5a | protein_coding | -0.491003734 | 5.427513093 | 3.3763899 | 0.0844275 | 0.629774749 |
| ENSMUSG00000015290 | Ubl4a | protein_coding | 0.388810154 | 7.399745294 | 3.3726295 | 0.0845854 | 0.629774749 |
| ENSMUSG00000040697 | Dnajc16 | protein_coding | 0.814240373 | 4.041502549 | 3.37262 | 0.0845858 | 0.629774749 |
| ENSMUSG00000110444 | Gm10033 | transcribed_unproces | -1.196561367 | 3.567408666 | 3.4006021 | 0.0845967 | 0.629774749 |
| ENSMUSG00000015968 | Cacna1d | protein_coding | -0.715929179 | 3.628391845 | 3.3715741 | 0.0846298 | 0.629774749 |
| ENSMUSG00000026775 | Yme1l1 | protein_coding | -0.483253381 | 6.837194338 | 3.3707259 | 0.084696 | 0.629831116 |
| ENSMUSG00000041935 | AW549877 | protein_coding | -0.484518235 | 5.234525089 | 3.3679913 | 0.0847807 | 0.629875411 |
| ENSMUSG00000025940 | Tmem70 | protein_coding | -0.551958271 | 4.937732559 | 3.3670822 | 0.0848191 | 0.629875411 |
| ENSMUSG00000022698 | Naa50 | protein_coding | -0.390974892 | 5.952970426 | 3.3639723 | 0.0849504 | 0.630367986 |
| ENSMUSG00000020650 | Bcap29 | protein_coding | -0.421979963 | 6.324469534 | 3.3621579 | 0.0850271 | 0.630367986 |
| ENSMUSG00000025151 | Maged1 | protein_coding | 0.518741384 | 10.0312578 | 3.3620709 | 0.0850613 | 0.630367986 |
| ENSMUSG00000066595 | Mfsd7b | protein_coding | -0.937028262 | 3.027479416 | 3.359386 | 0.0851445 | 0.630390665 |
| ENSMUSG00000003452 | Bicd1 | protein_coding | -0.450800582 | 5.506908073 | 3.3572795 | 0.0852338 | 0.630390665 |
| ENSMUSG00000097048 | 1600020E01Rik | antisense_RNA | -0.780104465 | 4.654607397 | 3.357148 | 0.0852699 | 0.630390665 |
| ENSMUSG00000035268 | Pkig | protein_coding | -0.369663543 | 7.141990585 | 3.3557483 | 0.0852988 | 0.630390665 |
| ENSMUSG00000005667 | Mthfd2 | protein_coding | 0.505947561 | 5.701857152 | 3.3541806 | 0.0853655 | 0.630426279 |
| ENSMUSG00000060397 | Zfp128 | protein_coding | -1.026379548 | 3.274873164 | 3.3525627 | 0.0854343 | 0.630426279 |
| ENSMUSG00000025492 | Ifitm3 | protein_coding | -0.679080542 | 4.155131299 | 3.3504824 | 0.0855228 | 0.630426279 |
| ENSMUSG00000029211 | Gabra4 | protein_coding | -0.528414193 | 6.717935929 | 3.3536695 | 0.0855506 | 0.630426279 |
| ENSMUSG00000054720 | Lrrc8c | protein_coding | -0.650981779 | 3.937073174 | 3.3469532 | 0.0856733 | 0.630426279 |
| ENSMUSG00000019864 | Rtn4ip1 | protein_coding | 0.623028786 | 4.815927102 | 3.3440553 | 0.0857972 | 0.630426279 |
| ENSMUSG00000071369 | Map3k5 | protein_coding | 0.596564641 | 4.299181997 | 3.3437537 | 0.0858101 | 0.630426279 |
| ENSMUSG00000032294 | Pkm | protein_coding | 0.511415781 | 10.22861395 | 3.3434428 | 0.0858539 | 0.630426279 |
| ENSMUSG00000036202 | Rif1 | protein_coding | -0.679964228 | 4.189042573 | 3.3416669 | 0.0858994 | 0.630426279 |
| ENSMUSG00000031785 | Adgrg1 | protein_coding | 0.417342871 | 6.230589893 | 3.3409507 | 0.0859301 | 0.630426279 |
| ENSMUSG00000028847 | Trappc3 | protein_coding | 0.319733337 | 7.886045481 | 3.3405186 | 0.0859486 | 0.630426279 |
| ENSMUSG00000024454 | Hdac3 | protein_coding | 0.372500734 | 7.052944055 | 3.3388125 | 0.0860217 | 0.630532713 |
| ENSMUSG00000031760 | Mt3 | protein_coding | -0.453728656 | 10.72742009 | 3.3364938 | 0.0861213 | 0.6308323 |
| ENSMUSG00000034453 | Polr3b | protein_coding | 0.757700879 | 3.459292314 | 3.3334646 | 0.0862515 | 0.631356197 |
| ENSMUSG00000053192 | Mllt11 | protein_coding | 0.508151846 | 6.47669126 | 3.3281635 | 0.0865105 | 0.632821459 |
| ENSMUSG00000030122 | Ptms | protein_coding | -0.502501407 | 6.578648641 | 3.3227246 | 0.0867457 | 0.633916775 |
| ENSMUSG00000028932 | Psmc2 | protein_coding | 0.357742551 | 8.499905704 | 3.3195311 | 0.0868536 | 0.633916775 |
| ENSMUSG00000027285 | Haus2 | protein_coding | -0.428063078 | 6.045529529 | 3.3179853 | 0.0869207 | 0.633916775 |
| ENSMUSG00000030291 | Med21 | protein_coding | -0.393127531 | 7.272504161 | 3.3176774 | 0.086934 | 0.633916775 |
| ENSMUSG00000059554 | Ccdc28a | protein_coding | 0.40212344 | 6.973733931 | 3.3171944 | 0.086955 | 0.633916775 |
| ENSMUSG00000070287 | Slc35g2 | protein_coding | 0.521835151 | 5.014595835 | 3.3147975 | 0.0870592 | 0.633974723 |
| ENSMUSG00000097431 | Gm26782 | lincRNA | -0.472735541 | 5.107348508 | 3.3142281 | 0.087084 | 0.633974723 |
| ENSMUSG00000031673 | Cdh11 | protein_coding | -0.575563209 | 4.760155225 | 3.3125464 | 0.0871572 | 0.633974723 |
| ENSMUSG00000001150 | Mcm3ap | protein_coding | 0.669111509 | 3.73704569 | 3.3115929 | 0.0871988 | 0.633974723 |
| ENSMUSG00000047446 | Arl4a | protein_coding | -0.603992224 | 6.549602201 | 3.3319011 | 0.087503 | 0.635756493 |

| ensembl_gene_id | external_gene_name | gene_biotype | log2 (fold change) | log2 (counts per million) | F-value | p-value | False discovery rate |
|---|---|---|---|---|---|---|---|
| ENSMUSG00000033880 | Lgals3bp | protein_coding | 0.924159307 | 4.716344849 | 3.3351835 | 0.087701 | 0.636764607 |
| ENSMUSG00000026679 | Enkur | protein_coding | -0.70041467 | 4.017114693 | 3.2964316 | 0.0878627 | 0.637269131 |
| ENSMUSG00000005378 | Wbscr22 | protein_coding | 0.383419009 | 6.300554617 | 3.2958343 | 0.087889 | 0.637269131 |
| ENSMUSG00000018143 | Mafk | protein_coding | 0.995567663 | 3.372681478 | 3.2888403 | 0.0881974 | 0.638386125 |
| ENSMUSG00000047935 | Gm5607 | sense_overlapping | -0.532870886 | 4.563505048 | 3.2884545 | 0.0882144 | 0.638386125 |
| ENSMUSG00000020766 | Galk1 | protein_coding | -0.697829293 | 4.268817448 | 3.2883025 | 0.0882211 | 0.638386125 |
| ENSMUSG00000026526 | Fh1 | protein_coding | 0.351679313 | 7.555256967 | 3.2864611 | 0.0883026 | 0.638545691 |
| ENSMUSG00000032024 | Clmp | protein_coding | -0.857967356 | 3.363255867 | 3.2828477 | 0.0884626 | 0.639273177 |
| ENSMUSG00000029661 | Col1a2 | protein_coding | -1.31355876 | 3.187938435 | 3.2906987 | 0.0887362 | 0.640819762 |
| ENSMUSG00000056413 | Adap1 | protein_coding | -0.469577262 | 6.48871669 | 3.272673 | 0.0889152 | 0.64100885 |
| ENSMUSG00000052033 | Pfdn4 | protein_coding | -0.457137734 | 7.429777655 | 3.2731138 | 0.0889262 | 0.64100885 |
| ENSMUSG00000024805 | Pcgf5 | protein_coding | -0.57894059 | 5.309325547 | 3.2720882 | 0.0889412 | 0.64100885 |
| ENSMUSG00000050377 | Il31ra | protein_coding | 0.641989888 | 5.067837887 | 3.267853 | 0.0891305 | 0.641942479 |
| ENSMUSG00000029823 | Luc7l2 | protein_coding | -0.473926458 | 5.357817522 | 3.2658572 | 0.0892198 | 0.642155863 |
| ENSMUSG00000020671 | Rab10 | protein_coding | -0.94447104 | 2.972997929 | 3.2623132 | 0.0893787 | 0.642214825 |
| ENSMUSG00000024926 | Kat5 | protein_coding | 0.469672492 | 6.447044186 | 3.260643 | 0.0894538 | 0.642214825 |
| ENSMUSG00000020817 | Rabep1 | protein_coding | -0.388475473 | 6.820638269 | 3.2602882 | 0.0894697 | 0.642214825 |
| ENSMUSG00000085408 | C530005A16Rik | processed_transcript | 0.880162373 | 3.387807311 | 3.2586594 | 0.0895429 | 0.642214825 |
| ENSMUSG00000028933 | Xrcc2 | protein_coding | 0.774480247 | 3.25736211 | 3.2579121 | 0.0895766 | 0.642214825 |
| ENSMUSG00000042312 | S100a13 | protein_coding | -0.649772137 | 6.416420582 | 3.2995311 | 0.0895874 | 0.642214825 |
| ENSMUSG00000030876 | Mettl9 | protein_coding | -0.58460555 | 5.042005265 | 3.254322 | 0.0897383 | 0.642214825 |
| ENSMUSG00000050334 | C130071C03Rik | lincRNA | -0.568690553 | 4.538321769 | 3.2504086 | 0.089915 | 0.642214825 |
| ENSMUSG00000068921 | Dap3 | protein_coding | 0.350402468 | 7.196531851 | 3.2487666 | 0.0899893 | 0.642214825 |
| ENSMUSG00000086171 | Pcsk2os1 | antisense_RNA | -1.166583678 | 2.510343127 | 3.248049 | 0.0900218 | 0.642214825 |
| ENSMUSG00000058076 | Sdhc | protein_coding | 0.341756228 | 9.022960936 | 3.2475334 | 0.0900451 | 0.642214825 |
| ENSMUSG00000038990 | Cables2 | protein_coding | 0.807423381 | 3.377384987 | 3.247347 | 0.0900535 | 0.642214825 |
| ENSMUSG00000085882 | 2610507I01Rik | processed_transcript | 0.889565954 | 3.233352781 | 3.2472292 | 0.0900589 | 0.642214825 |
| ENSMUSG00000069539 | Scyl2 | protein_coding | -1.038829072 | 2.866869818 | 3.2447701 | 0.0901703 | 0.642214825 |
| ENSMUSG00000036533 | Cdc42ep3 | protein_coding | -0.622602393 | 4.414159543 | 3.2441469 | 0.0901986 | 0.642214825 |
| ENSMUSG00000006273 | Atp6v1b2 | protein_coding | 0.388283703 | 9.505910288 | 3.2423645 | 0.0902795 | 0.642214825 |
| ENSMUSG00000044770 | Scml4 | protein_coding | 0.86524043 | 3.261107086 | 3.2415882 | 0.0903148 | 0.642214825 |
| ENSMUSG00000022523 | Fgf12 | protein_coding | -0.345706216 | 7.554931054 | 3.2399231 | 0.0903905 | 0.642214825 |
| ENSMUSG00000048988 | Elfn1 | protein_coding | -1.082231288 | 3.066622727 | 3.2372079 | 0.0905141 | 0.642214825 |
| ENSMUSG00000031551 | Ido1 | protein_coding | -0.751708763 | 3.973522036 | 3.2361942 | 0.0905603 | 0.642214825 |
| ENSMUSG00000028636 | Ppcs | protein_coding | 0.890151725 | 4.117834814 | 3.2362074 | 0.0906715 | 0.642214825 |
| ENSMUSG00000022420 | Dnal4 | protein_coding | -0.430441531 | 6.838531098 | 3.2302948 | 0.0908298 | 0.642214825 |
| ENSMUSG00000037594 | BC022687 | protein_coding | 0.88099416 | 3.430828508 | 3.2299152 | 0.0908471 | 0.642214825 |
| ENSMUSG00000031099 | Smarca1 | protein_coding | -0.538386624 | 5.353965234 | 3.2299008 | 0.0908478 | 0.642214825 |
| ENSMUSG00000053310 | Nrgn | protein_coding | -0.430676776 | 7.189110252 | 3.2271331 | 0.0909746 | 0.642214825 |
| ENSMUSG00000020601 | Trib2 | protein_coding | 0.470341197 | 5.577241253 | 3.2263724 | 0.0910094 | 0.642214825 |
| ENSMUSG00000071379 | Hpcal1 | protein_coding | 0.625379432 | 7.646691765 | 3.2829116 | 0.0910763 | 0.642214825 |
| ENSMUSG00000035228 | Ccdc106 | protein_coding | -0.563775046 | 5.471957661 | 3.2240312 | 0.0911169 | 0.642214825 |
| ENSMUSG00000002748 | Baz1b | protein_coding | 0.500928242 | 5.4255037 | 3.2216334 | 0.0912271 | 0.642214825 |
| ENSMUSG00000021974 | Fgf9 | protein_coding | -0.521435504 | 5.384579814 | 3.2203188 | 0.0912876 | 0.642214825 |
| ENSMUSG00000059923 | Grb2 | protein_coding | 0.317965659 | 6.953409347 | 3.2197975 | 0.0913116 | 0.642214825 |
| ENSMUSG00000049281 | Scn3b | protein_coding | 0.362732536 | 6.319144643 | 3.2196987 | 0.0913161 | 0.642214825 |
| ENSMUSG00000031729 | Ist1 | protein_coding | 0.596868282 | 5.561109839 | 3.2194716 | 0.0913574 | 0.642214825 |
| ENSMUSG00000020705 | Ddx42 | protein_coding | 0.421378073 | 5.811682995 | 3.2184909 | 0.0913717 | 0.642214825 |
| ENSMUSG00000106379 | Lhfpl3 | protein_coding | -0.418593168 | 6.027799078 | 3.216897 | 0.0914452 | 0.642214825 |
| ENSMUSG00000108456 | 4732496C06Rik | lincRNA | -0.858153326 | 3.32660542 | 3.2168667 | 0.0914466 | 0.642214825 |
| ENSMUSG00000020088 | Sar1a | protein_coding | 0.342497472 | 7.259950516 | 3.2148081 | 0.0915416 | 0.642214825 |
| ENSMUSG00000034551 | Hdx | protein_coding | -1.742684232 | 2.241436869 | 3.2136923 | 0.0915932 | 0.642214825 |
| ENSMUSG00000020608 | Smc6 | protein_coding | -0.444761681 | 5.399207557 | 3.2132152 | 0.0916152 | 0.642214825 |
| ENSMUSG00000021259 | Cyp46a1 | protein_coding | -0.560667625 | 5.252938892 | 3.2093468 | 0.0917942 | 0.642214825 |
| ENSMUSG00000040842 | Szrd1 | protein_coding | -0.517147905 | 5.368235244 | 3.2072055 | 0.0918935 | 0.642214825 |
| ENSMUSG00000083012 | Fam220a | protein_coding | -0.531853958 | 4.321405995 | 3.2056649 | 0.091965 | 0.642214825 |
| ENSMUSG00000020946 | Gosr2 | protein_coding | 0.414650715 | 6.890727503 | 3.2056059 | 0.0919677 | 0.642214825 |
| ENSMUSG00000020074 | Ccar1 | protein_coding | -0.47935264 | 5.314641543 | 3.2041108 | 0.0920372 | 0.642214825 |
| ENSMUSG00000071657 | Bscl2 | protein_coding | 0.342037275 | 7.871977081 | 3.200527 | 0.0922039 | 0.642214825 |
| ENSMUSG00000051890 | Klhdc1 | protein_coding | -0.568102673 | 4.471963416 | 3.1994946 | 0.092252 | 0.642214825 |
| ENSMUSG00000071076 | Jund | protein_coding | -0.546511766 | 4.43190559 | 3.1993174 | 0.0922603 | 0.642214825 |
| ENSMUSG00000060601 | Nr1h2 | protein_coding | -0.438227205 | 5.922656333 | 3.1992987 | 0.0922612 | 0.642214825 |
| ENSMUSG00000026490 | Cdc42bpa | protein_coding | -0.444646838 | 6.237104324 | 3.1960153 | 0.0924144 | 0.642214825 |
| ENSMUSG00000027406 | Idh3b | protein_coding | 0.351628164 | 9.161263292 | 3.1934924 | 0.0925323 | 0.642214825 |
| ENSMUSG00000033342 | Plppr5 | protein_coding | -0.549850621 | 4.639042414 | 3.1929168 | 0.0925592 | 0.642214825 |
| ENSMUSG00000021303 | Gng4 | protein_coding | -0.363907496 | 6.380812427 | 3.1925667 | 0.0925756 | 0.642214825 |
| ENSMUSG00000047388 | Atmin | protein_coding | 0.43554668 | 5.551702336 | 3.192245 | 0.0925906 | 0.642214825 |
| ENSMUSG00000005882 | Uqcc1 | protein_coding | 0.404482261 | 7.093360736 | 3.191162 | 0.0926413 | 0.642214825 |
| ENSMUSG00000080316 | Spaca6 | protein_coding | -0.709621294 | 3.488998129 | 3.1881509 | 0.0927825 | 0.642214825 |
| ENSMUSG00000023110 | Prmt5 | protein_coding | 0.361209875 | 6.795089092 | 3.1858281 | 0.0928916 | 0.642214825 |
| ENSMUSG00000078427 | Sarnp | protein_coding | -0.410890621 | 6.551563415 | 3.185189 | 0.0929216 | 0.642214825 |
| ENSMUSG00000021986 | Amer2 | protein_coding | -0.579162642 | 5.07212423 | 3.1847013 | 0.0929445 | 0.642214825 |
| ENSMUSG00000028953 | Abcf2 | protein_coding | 0.658881763 | 6.907046021 | 3.2407525 | 0.0929768 | 0.642214825 |
| ENSMUSG00000055817 | Mta3 | protein_coding | 0.617362134 | 4.98990718 | 3.1838745 | 0.0929834 | 0.642214825 |

| ensembl_gene_id | external_gene_name | gene_biotype | log2 (fold change) | log2 (counts per million) | F-value | p-value | False discovery rate |
|---|---|---|---|---|---|---|---|
| ENSMUSG00000038173 | Enpp6 | protein_coding | 1.096311122 | 3.093479553 | 3.1821625 | 0.093064 | 0.642214825 |
| ENSMUSG00000020627 | Klhl29 | protein_coding | -0.737947556 | 4.320473232 | 3.1821315 | 0.0930655 | 0.642214825 |
| ENSMUSG00000093483 | AA465934 | processed_transcript | -0.824211879 | 3.409305018 | 3.1784534 | 0.0932389 | 0.642214825 |
| ENSMUSG00000056296 | Synpr | protein_coding | -0.469744604 | 7.160050266 | 3.1789793 | 0.0932449 | 0.642214825 |
| ENSMUSG00000041817 | Fam169a | protein_coding | -0.539531739 | 5.870143142 | 3.177035 | 0.0933058 | 0.642214825 |
| ENSMUSG00000039347 | Atp6v0e2 | protein_coding | 0.364895929 | 10.37809293 | 3.1766425 | 0.0933244 | 0.642214825 |
| ENSMUSG00000020719 | Ddx5 | protein_coding | -0.349718442 | 8.230626734 | 3.1765056 | 0.0933308 | 0.642214825 |
| ENSMUSG00000068798 | Rap1a | protein_coding | -0.493581386 | 4.606959822 | 3.1758783 | 0.0933605 | 0.642214825 |
| ENSMUSG00000051527 | Usp29 | protein_coding | 0.50193725 | 5.338877038 | 3.1743546 | 0.0934326 | 0.642214825 |
| ENSMUSG00000063253 | Scoc | protein_coding | -0.353085538 | 7.516493739 | 3.1739347 | 0.0934524 | 0.642214825 |
| ENSMUSG00000024474 | Ik | protein_coding | 0.406172077 | 7.511914059 | 3.1727471 | 0.0935086 | 0.642214825 |
| ENSMUSG00000075334 | Rprm | protein_coding | 0.691705135 | 5.63172732 | 3.1962031 | 0.0935282 | 0.642214825 |
| ENSMUSG00000032316 | Clk3 | protein_coding | 0.498446076 | 5.255660421 | 3.1709859 | 0.0935921 | 0.642243705 |
| ENSMUSG00000068749 | Psma5 | protein_coding | 0.327117423 | 6.83374403 | 3.1678883 | 0.0937391 | 0.642817158 |
| ENSMUSG00000029038 | Ssu72 | protein_coding | 0.313687736 | 7.920760484 | 3.1652047 | 0.0938667 | 0.642817158 |
| ENSMUSG00000002881 | Nab1 | protein_coding | -0.665401401 | 4.198576992 | 3.1642965 | 0.0939099 | 0.642817158 |
| ENSMUSG00000083307 | AA414768 | processed_pseudoge | 0.930160356 | 2.909304346 | 3.1633199 | 0.0939564 | 0.642817158 |
| ENSMUSG00000055737 | Ghr | protein_coding | -0.832105653 | 2.985612569 | 3.1622517 | 0.0940073 | 0.642817158 |
| ENSMUSG00000038805 | Six3 | protein_coding | -0.58091719 | 5.097093561 | 3.1610284 | 0.0940656 | 0.642817158 |
| ENSMUSG00000024290 | Rock1 | protein_coding | -0.792450631 | 3.749513411 | 3.1604304 | 0.0940941 | 0.642817158 |
| ENSMUSG00000040354 | Mars | protein_coding | 0.68872837 | 4.800313522 | 3.1580134 | 0.0942404 | 0.642912636 |
| ENSMUSG00000068617 | Efcab1 | protein_coding | -0.44950812 | 5.490461706 | 3.1564815 | 0.0942828 | 0.642912636 |
| ENSMUSG00000021785 | Ngly1 | protein_coding | 0.618911343 | 4.401940673 | 3.156383 | 0.0942875 | 0.642912636 |
| ENSMUSG00000067629 | Syngap1 | protein_coding | -0.704498477 | 3.426343774 | 3.1534565 | 0.0944276 | 0.643061046 |
| ENSMUSG00000021669 | Col4a3bp | protein_coding | -0.508157738 | 4.648174305 | 3.1534206 | 0.0944293 | 0.643061046 |
| ENSMUSG00000097881 | Celrr | lincRNA | -1.36583562 | 2.872962828 | 3.1516331 | 0.0945459 | 0.643061046 |
| ENSMUSG00000042745 | Id1 | protein_coding | -0.669410297 | 3.894868094 | 3.1509355 | 0.0945485 | 0.643061046 |
| ENSMUSG00000043633 | Fam221b | protein_coding | -0.901319084 | 3.063791925 | 3.147521 | 0.0947125 | 0.643175072 |
| ENSMUSG00000033739 | Fkbpl | protein_coding | 0.527690527 | 5.200513763 | 3.1428005 | 0.0949398 | 0.643175072 |
| ENSMUSG00000024790 | Sac3d1 | protein_coding | -0.687250018 | 3.673260541 | 3.1400069 | 0.0950746 | 0.643175072 |
| ENSMUSG00000075415 | Fnbp1 | protein_coding | -0.423707535 | 6.990679437 | 3.1383952 | 0.0951525 | 0.643175072 |
| ENSMUSG00000078453 | Abracl | protein_coding | -0.631327954 | 4.923847524 | 3.1383839 | 0.095153 | 0.643175072 |
| ENSMUSG00000029674 | Limk1 | protein_coding | -1.302591632 | 2.789344449 | 3.13727 | 0.0952069 | 0.643175072 |
| ENSMUSG00000079235 | Ccdc13 | protein_coding | -0.784508114 | 3.461573038 | 3.1362714 | 0.0952553 | 0.643175072 |
| ENSMUSG00000031622 | Sin3b | protein_coding | 0.329991086 | 6.505434759 | 3.133835 | 0.0953733 | 0.643175072 |
| ENSMUSG00000052837 | Junb | protein_coding | -1.112130285 | 2.756355389 | 3.1331949 | 0.0954043 | 0.643175072 |
| ENSMUSG00000047417 | Rexo1 | protein_coding | -0.655840025 | 4.115535995 | 3.1324221 | 0.0954418 | 0.643175072 |
| ENSMUSG00000026889 | Rbm18 | protein_coding | -0.351015355 | 6.793040632 | 3.1323952 | 0.0954431 | 0.643175072 |
| ENSMUSG00000025909 | Sntg1 | protein_coding | -0.558299131 | 4.783253492 | 3.1312364 | 0.0954994 | 0.643175072 |
| ENSMUSG00000020108 | Ddit4 | protein_coding | 0.920562414 | 3.351009333 | 3.1308751 | 0.0955169 | 0.643175072 |
| ENSMUSG00000040813 | Tex264 | protein_coding | 0.347783753 | 6.528082836 | 3.130765 | 0.0955223 | 0.643175072 |
| ENSMUSG00000022016 | Akap11 | protein_coding | 0.56532502 | 6.148210347 | 3.1330524 | 0.0955594 | 0.643175072 |
| ENSMUSG00000036986 | Pml | protein_coding | 1.229297729 | 2.724934985 | 3.1287302 | 0.0956212 | 0.643175072 |
| ENSMUSG00000033933 | Vhl | protein_coding | 0.556156105 | 5.045703462 | 3.1278905 | 0.095662 | 0.643175072 |
| ENSMUSG00000000486 | 1-Sep | protein_coding | -0.967096185 | 2.941187693 | 3.1274285 | 0.0956845 | 0.643175072 |
| ENSMUSG00000058741 | Prr19 | protein_coding | 1.645356679 | 2.448788106 | 3.1253496 | 0.0958167 | 0.643175072 |
| ENSMUSG00000027425 | Kat14 | protein_coding | 0.56274036 | 4.035801704 | 3.1243155 | 0.0958362 | 0.643175072 |
| ENSMUSG00000028670 | Lypla2 | protein_coding | -0.372543071 | 6.34628073 | 3.1230689 | 0.095897 | 0.643175072 |
| ENSMUSG00000058388 | Phtf1 | protein_coding | 0.71024767 | 4.01175604 | 3.122941 | 0.0959032 | 0.643175072 |
| ENSMUSG00000020271 | Fbxw11 | protein_coding | 0.42534079 | 6.205915144 | 3.1221499 | 0.0959418 | 0.643175072 |
| ENSMUSG00000051232 | Tmem199 | protein_coding | 0.335917766 | 7.900208875 | 3.1209441 | 0.0960007 | 0.643175072 |
| ENSMUSG00000020130 | Tbc1d15 | protein_coding | 0.454168605 | 5.682578696 | 3.1178703 | 0.0961511 | 0.64336738 |
| ENSMUSG00000025198 | Erlin1 | protein_coding | -0.756599772 | 3.890880688 | 3.1176299 | 0.0961629 | 0.64336738 |
| ENSMUSG00000026887 | Mrrf | protein_coding | 0.483861611 | 5.462209478 | 3.115256 | 0.0962792 | 0.64336738 |
| ENSMUSG00000021098 | 4930447C04Rik | protein_coding | -1.045656373 | 2.70016333 | 3.1134141 | 0.0963695 | 0.64336738 |
| ENSMUSG00000044042 | Fmn1 | protein_coding | -0.757382606 | 3.170494221 | 3.112653 | 0.0964069 | 0.64336738 |
| ENSMUSG00000028799 | Zfp362 | protein_coding | -0.885117301 | 2.955070288 | 3.1119339 | 0.0964422 | 0.64336738 |
| ENSMUSG00000041775 | Mapk1ip1 | protein_coding | 0.494694258 | 5.904104184 | 3.111482 | 0.0964644 | 0.64336738 |
| ENSMUSG00000029775 | Klhdc10 | protein_coding | 0.428018898 | 5.376062898 | 3.1097398 | 0.0965501 | 0.64336738 |
| ENSMUSG00000040370 | Etfrf1 | protein_coding | -0.387553535 | 6.488541711 | 3.108768 | 0.0965979 | 0.64336738 |
| ENSMUSG00000042606 | Hirip3 | protein_coding | 0.624505926 | 4.380832311 | 3.1078391 | 0.0966437 | 0.64336738 |
| ENSMUSG00000028803 | Nipal3 | protein_coding | 0.505793098 | 5.249474232 | 3.1069473 | 0.0966876 | 0.64336738 |
| ENSMUSG00000022204 | Ngdn | protein_coding | 0.376280569 | 6.862385799 | 3.1025753 | 0.0969033 | 0.644404167 |
| ENSMUSG00000036160 | Surf6 | protein_coding | 0.68921764 | 4.498404518 | 3.1007777 | 0.0969922 | 0.644519411 |
| ENSMUSG00000028555 | Ttc39a | protein_coding | 1.29140256 | 2.707469355 | 3.0998008 | 0.0970405 | 0.644519411 |
| ENSMUSG00000026799 | Med27 | protein_coding | 0.403589181 | 5.612580468 | 3.0930916 | 0.0973733 | 0.646101414 |
| ENSMUSG00000022787 | Wdr53 | protein_coding | 0.522191338 | 4.781929122 | 3.0925763 | 0.0973989 | 0.646101414 |
| ENSMUSG00000025893 | Kbtbd3 | protein_coding | -0.498057382 | 5.278127537 | 3.0859218 | 0.0977304 | 0.647476537 |
| ENSMUSG00000020340 | Cyfip2 | protein_coding | 0.382175379 | 8.355776164 | 3.0844238 | 0.0978052 | 0.647476537 |
| ENSMUSG00000026245 | Farsb | protein_coding | 0.377318206 | 7.306000075 | 3.0828466 | 0.097884 | 0.647476537 |
| ENSMUSG00000097537 | 2610020C07Rik | lincRNA | -0.533927008 | 4.705334267 | 3.0827218 | 0.0978903 | 0.647476537 |
| ENSMUSG00000039463 | Slc9a8 | protein_coding | 0.966083963 | 4.368778682 | 3.1135789 | 0.0979073 | 0.647476537 |
| ENSMUSG00000020423 | Btg2 | protein_coding | 0.923191635 | 5.95360029 | 3.1842093 | 0.0982366 | 0.64802328 |
| ENSMUSG00000044629 | Cnrip1 | protein_coding | 0.313360376 | 7.54233379 | 3.0751879 | 0.0982679 | 0.64802328 |

| ensembl_gene_id | external_gene_name | gene_biotype | log2 (fold change) | log2 (counts per million) | F-value | p-value | False discovery rate |
|---|---|---|---|---|---|---|---|
| ENSMUSG00000041852 | Tcf20 | protein_coding | 0.532746283 | 5.792420449 | 3.0709904 | 0.0984791 | 0.64802328 |
| ENSMUSG00000039740 | Alg2 | protein_coding | 0.387533163 | 7.151120417 | 3.0702893 | 0.0985144 | 0.64802328 |
| ENSMUSG00000025613 | Cct8 | protein_coding | 0.443006983 | 8.639490942 | 3.0706575 | 0.0985269 | 0.64802328 |
| ENSMUSG00000020170 | Frs2 | protein_coding | 0.581488986 | 4.04767324 | 3.0693407 | 0.0985622 | 0.64802328 |
| ENSMUSG00000022621 | Rabl2 | protein_coding | 0.450430787 | 6.451840936 | 3.0666485 | 0.0986981 | 0.64802328 |
| ENSMUSG00000003559 | As3mt | protein_coding | 0.489268223 | 5.860821575 | 3.0657574 | 0.0987431 | 0.64802328 |
| ENSMUSG00000000631 | Myo18a | protein_coding | -0.496860655 | 5.756674525 | 3.0656816 | 0.0987469 | 0.64802328 |
| ENSMUSG00000021876 | Rnase4 | protein_coding | 0.643949936 | 4.138297218 | 3.0656735 | 0.0987473 | 0.64802328 |
| ENSMUSG00000031642 | Sh3rf1 | protein_coding | -0.464072086 | 4.948869836 | 3.0651544 | 0.0987735 | 0.64802328 |
| ENSMUSG00000021840 | Mapk1ip1l | protein_coding | -0.615179599 | 3.850109201 | 3.0636233 | 0.098851 | 0.64802328 |
| ENSMUSG00000063362 | Alg11 | protein_coding | 0.424402233 | 6.593971449 | 3.0633245 | 0.0988661 | 0.64802328 |
| ENSMUSG00000036249 | Rbm43 | protein_coding | 0.568842336 | 5.19496541 | 3.0631485 | 0.098875 | 0.64802328 |
| ENSMUSG00000028158 | Mttp | protein_coding | -0.761274018 | 3.913784523 | 3.0614461 | 0.0989612 | 0.64802328 |
| ENSMUSG00000041057 | Wdr43 | protein_coding | 0.450414142 | 5.112090577 | 3.0602668 | 0.099021 | 0.64802328 |
| ENSMUSG00000020740 | Gga3 | protein_coding | 0.460067818 | 5.906229905 | 3.0571969 | 0.0991768 | 0.64802328 |
| ENSMUSG00000006675 | P4htm | protein_coding | 0.40624192 | 6.530913841 | 3.0558557 | 0.0992449 | 0.64802328 |
| ENSMUSG00000039523 | Cep104 | protein_coding | -0.72601923 | 4.015936478 | 3.0534616 | 0.0993667 | 0.64802328 |
| ENSMUSG00000024486 | Hbegf | protein_coding | -0.594828118 | 4.366426274 | 3.053044 | 0.099388 | 0.64802328 |
| ENSMUSG00000025173 | Wdr45b | protein_coding | 0.35337666 | 6.486144189 | 3.0528988 | 0.0993954 | 0.64802328 |
| ENSMUSG00000017760 | Ctsa | protein_coding | 0.321089285 | 6.797332357 | 3.0526309 | 0.099409 | 0.64802328 |
| ENSMUSG00000031299 | Pdha1 | protein_coding | 0.662745621 | 7.285462606 | 3.1125878 | 0.0994199 | 0.64802328 |
| ENSMUSG00000036323 | Srp72 | protein_coding | 0.324165982 | 7.571869453 | 3.0516423 | 0.0994594 | 0.64802328 |
| ENSMUSG00000097709 | 2810429I04Rik | lincRNA | -0.808661294 | 3.477889688 | 3.0507698 | 0.0995039 | 0.64802328 |
| ENSMUSG00000021266 | Wars | protein_coding | 0.668448523 | 7.014786767 | 3.1089981 | 0.099597 | 0.64802328 |
| ENSMUSG00000036977 | Anapc10 | protein_coding | -0.519126799 | 5.966676204 | 3.0480133 | 0.0996757 | 0.64802328 |
| ENSMUSG00000020737 | Jpt1 | protein_coding | 0.326649955 | 8.50463391 | 3.0473716 | 0.0996773 | 0.64802328 |
| ENSMUSG00000023944 | Hsp90ab1 | protein_coding | 0.387318025 | 11.18772475 | 3.0425431 | 0.0999245 | 0.648867949 |
| ENSMUSG00000098649 | Gm27199 | antisense_RNA | 0.964530367 | 2.751579585 | 3.041851 | 0.0999599 | 0.648867949 |
| ENSMUSG00000027371 | Fahd2a | protein_coding | -0.388024039 | 6.322687499 | 3.0412978 | 0.0999883 | 0.648867949 |
| ENSMUSG00000033377 | Palmd | protein_coding | -0.928555374 | 3.561440039 | 3.038295 | 0.1001736 | 0.649344422 |
| ENSMUSG00000049411 | Tmem241 | protein_coding | 0.850232715 | 4.27090545 | 3.0404158 | 0.1001825 | 0.649344422 |
| ENSMUSG00000017491 | Rarb | protein_coding | -0.69070702 | 3.526248906 | 3.0331791 | 0.1004058 | 0.650134359 |
| ENSMUSG00000026737 | Pip4k2a | protein_coding | -0.406051333 | 5.635831493 | 3.0317868 | 0.1004776 | 0.650134359 |
| ENSMUSG00000063430 | Wscd2 | protein_coding | -0.742104486 | 3.351754842 | 3.030312 | 0.1005537 | 0.650134359 |
| ENSMUSG00000039716 | Dock3 | protein_coding | 0.649184657 | 3.952430827 | 3.0288364 | 0.1006299 | 0.650134359 |
| ENSMUSG00000037815 | Ctnna1 | protein_coding | 0.460044659 | 5.880425681 | 3.0282701 | 0.1006592 | 0.650134359 |
| ENSMUSG00000060424 | Pantr1 | lincRNA | -0.753484902 | 7.07974581 | 3.1330908 | 0.1006938 | 0.650134359 |
| ENSMUSG00000079467 | Gm14966 | antisense_RNA | -1.479874001 | 1.889569052 | 3.0269473 | 0.1007276 | 0.650134359 |
| ENSMUSG00000029201 | Ugdh | protein_coding | -0.554468537 | 4.528827159 | 3.0238063 | 0.1008903 | 0.650434094 |
| ENSMUSG00000027367 | Stard7 | protein_coding | 0.336665516 | 6.49919492 | 3.023398 | 0.1009115 | 0.650434094 |
| ENSMUSG00000022466 | Rpap3 | protein_coding | 0.43527829 | 5.866990366 | 3.0199295 | 0.1010915 | 0.650434094 |
| ENSMUSG00000027177 | Hipk3 | protein_coding | -0.512448516 | 5.20658307 | 3.0192403 | 0.1011273 | 0.650434094 |
| ENSMUSG00000017734 | Dbndd2 | protein_coding | -0.503494516 | 5.176144182 | 3.0189072 | 0.1011447 | 0.650434094 |
| ENSMUSG00000015143 | Actn1 | protein_coding | -0.503124827 | 4.509325244 | 3.0184886 | 0.1011664 | 0.650434094 |
| ENSMUSG00000049044 | Rapgef4 | protein_coding | -0.509224192 | 6.176587746 | 3.0180386 | 0.101221 | 0.650434094 |
| ENSMUSG00000044573 | Acp1 | protein_coding | -0.35382163 | 6.066362044 | 3.0167299 | 0.1012579 | 0.650434094 |
| ENSMUSG00000024524 | Gnal | protein_coding | -0.380065299 | 7.129695087 | 3.0138038 | 0.1014104 | 0.651024733 |
| ENSMUSG00000068115 | Ninl | protein_coding | -1.641500746 | 2.241747413 | 3.0117105 | 0.1015197 | 0.651337315 |
| ENSMUSG00000030603 | Psmc4 | protein_coding | 0.317984953 | 8.493986337 | 3.0078869 | 0.1017196 | 0.651716352 |
| ENSMUSG00000074682 | Zcchc3 | protein_coding | 0.769557293 | 3.928975272 | 3.007336 | 0.1017485 | 0.651716352 |
| ENSMUSG00000090626 | Tex9 | protein_coding | -0.738129034 | 3.860997671 | 3.0055767 | 0.1018407 | 0.651716352 |
| ENSMUSG00000097392 | D930016D06Rik | lincRNA | -1.155393928 | 2.721452666 | 3.0055547 | 0.1018418 | 0.651716352 |
| ENSMUSG00000004071 | Cdip1 | protein_coding | 0.410789992 | 10.04724437 | 3.0047377 | 0.1018847 | 0.651716352 |
| ENSMUSG00000022790 | Igsf11 | protein_coding | -0.834485703 | 3.397509605 | 3.0036082 | 0.1019439 | 0.651716352 |
| ENSMUSG00000034780 | B3galt1 | protein_coding | -0.459784714 | 5.905181875 | 3.000559 | 0.1021041 | 0.651716352 |
| ENSMUSG00000078862 | Gm14326 | protein_coding | -0.617035557 | 4.483002058 | 3.0003043 | 0.1021175 | 0.651716352 |
| ENSMUSG00000001518 | Itfg2 | protein_coding | -0.911323659 | 2.944837538 | 2.9981527 | 0.1022308 | 0.651716352 |
| ENSMUSG00000017978 | Cadps2 | protein_coding | 0.527712107 | 5.975778509 | 2.9956737 | 0.1023926 | 0.651716352 |
| ENSMUSG00000022951 | Rcan1 | protein_coding | 0.410030294 | 5.367254783 | 2.9949207 | 0.1024012 | 0.651716352 |
| ENSMUSG00000039661 | Dusp26 | protein_coding | 0.347092763 | 7.594853447 | 2.9945438 | 0.1024211 | 0.651716352 |
| ENSMUSG00000029782 | Tmem209 | protein_coding | 0.674285451 | 3.896533446 | 2.9937527 | 0.1024628 | 0.651716352 |
| ENSMUSG00000090125 | Pou3f1 | protein_coding | -0.675548864 | 3.977710097 | 2.9914912 | 0.1025823 | 0.651716352 |
| ENSMUSG00000073700 | Klhl21 | protein_coding | 0.809362772 | 3.31837598 | 2.9903893 | 0.1026406 | 0.651716352 |
| ENSMUSG00000035790 | Cep19 | protein_coding | 0.316250214 | 7.53305366 | 2.9902381 | 0.1026486 | 0.651716352 |
| ENSMUSG00000014426 | Map3k4 | protein_coding | 0.623923732 | 4.223032953 | 2.9886392 | 0.1027333 | 0.651716352 |
| ENSMUSG00000024787 | Snx15 | protein_coding | -0.339678441 | 6.245294448 | 2.9876096 | 0.1027878 | 0.651716352 |
| ENSMUSG00000035840 | Lysmd3 | protein_coding | -0.663851707 | 3.98831712 | 2.9838527 | 0.1029872 | 0.651716352 |
| ENSMUSG00000027247 | Arhgap1 | protein_coding | 0.508281943 | 6.71735667 | 2.9917924 | 0.1030154 | 0.651716352 |
| ENSMUSG00000036591 | Arhgap21 | protein_coding | -0.652973292 | 4.502984032 | 2.9829229 | 0.1030366 | 0.651716352 |
| ENSMUSG00000027222 | Pex16 | protein_coding | -0.415324871 | 6.157758827 | 2.9801154 | 0.103186 | 0.651716352 |
| ENSMUSG00000030982 | 9030624J02Rik | protein_coding | 0.606940936 | 6.165526476 | 3.0027345 | 0.1031982 | 0.651716352 |
| ENSMUSG00000039917 | Rhbdd2 | protein_coding | 0.943259814 | 6.527210063 | 3.1534064 | 0.1033663 | 0.651716352 |
| ENSMUSG00000007812 | Zfp655 | protein_coding | -0.451921133 | 5.639643847 | 2.9766167 | 0.1033726 | 0.651716352 |
| ENSMUSG00000014550 | Rbsn | protein_coding | 0.487055173 | 5.928368641 | 2.9748941 | 0.1034645 | 0.651716352 |

| ensembl_gene_id | external_gene_name | gene_biotype | log2 (fold change) | log2 (counts per million) | F-value | p-value | False discovery rate |
|---|---|---|---|---|---|---|---|
| ENSMUSG00000021087 | Rtn1 | protein_coding | 0.319691791 | 10.12214384 | 2.9741782 | 0.1035028 | 0.651716352 |
| ENSMUSG00000036570 | Fxyd1 | protein_coding | -0.693656154 | 6.27820526 | 3.0228175 | 0.1035986 | 0.651716352 |
| ENSMUSG00000019794 | Katna1 | protein_coding | -0.623820558 | 3.812878857 | 2.9722084 | 0.1036081 | 0.651716352 |
| ENSMUSG00000026878 | Rab14 | protein_coding | 0.339210953 | 7.850148033 | 2.9719287 | 0.1036231 | 0.651716352 |
| ENSMUSG00000024817 | Uhrf2 | protein_coding | -0.517017087 | 4.927797077 | 2.9694282 | 0.1037571 | 0.651716352 |
| ENSMUSG00000019124 | Scrn1 | protein_coding | 0.413343885 | 7.737756782 | 2.9692487 | 0.1037667 | 0.651716352 |
| ENSMUSG00000062937 | Mtap | protein_coding | 0.524990222 | 5.135717786 | 2.9674398 | 0.1038637 | 0.651716352 |
| ENSMUSG00000040351 | Ankib1 | protein_coding | -0.580711659 | 4.844242296 | 2.967208 | 0.1038762 | 0.651716352 |
| ENSMUSG00000048897 | Zfp710 | protein_coding | -0.55782079 | 4.031453304 | 2.966279 | 0.103926 | 0.651716352 |
| ENSMUSG00000048707 | Tprn | protein_coding | -0.620990239 | 4.150384956 | 2.9660825 | 0.1039366 | 0.651716352 |
| ENSMUSG00000031917 | Nip7 | protein_coding | 0.399473423 | 6.922530447 | 2.9629187 | 0.1041067 | 0.651716352 |
| ENSMUSG00000035067 | Xkr6 | protein_coding | -1.216874097 | 2.614266877 | 2.9628158 | 0.1041122 | 0.651716352 |
| ENSMUSG00000003039 | Fam32a | protein_coding | 0.304742963 | 8.117999032 | 2.9627157 | 0.1041176 | 0.651716352 |
| ENSMUSG00000070697 | Utp3 | protein_coding | 0.342852988 | 6.400541001 | 2.9613121 | 0.1041932 | 0.651716352 |
| ENSMUSG00000038296 | Galnt18 | protein_coding | 0.588553456 | 4.65128594 | 2.9593985 | 0.1042964 | 0.651716352 |
| ENSMUSG00000025477 | Inpp5a | protein_coding | 0.592484195 | 4.403235227 | 2.9582838 | 0.1043565 | 0.651716352 |
| ENSMUSG00000030610 | Det1 | protein_coding | 1.267365472 | 2.628626449 | 2.955262 | 0.1045198 | 0.651716352 |
| ENSMUSG00000028532 | Cachd1 | protein_coding | 0.712788482 | 3.756836825 | 2.9549619 | 0.104536 | 0.651716352 |
| ENSMUSG00000001911 | Nfix | protein_coding | -0.488567234 | 5.602617967 | 2.9546345 | 0.1045537 | 0.651716352 |
| ENSMUSG00000029817 | Tra2a | protein_coding | 0.522609059 | 4.997497304 | 2.9537735 | 0.1046003 | 0.651716352 |
| ENSMUSG00000029161 | Cgref1 | protein_coding | 0.518464934 | 5.114376841 | 2.9530203 | 0.1046411 | 0.651716352 |
| ENSMUSG00000051910 | Sox6 | protein_coding | -0.821860442 | 3.748186388 | 2.9526309 | 0.1046622 | 0.651716352 |
| ENSMUSG00000047714 | Ppp1r2 | protein_coding | -0.497531601 | 5.611116276 | 2.9514906 | 0.104724 | 0.651716352 |
| ENSMUSG00000026615 | Eprs | protein_coding | 0.421828722 | 6.566048291 | 2.9513438 | 0.1047319 | 0.651716352 |
| ENSMUSG00000073987 | Ggh | protein_coding | -0.519184492 | 5.054664238 | 2.9501734 | 0.1047954 | 0.651716352 |
| ENSMUSG00000031197 | Vbp1 | protein_coding | -0.313274288 | 7.867598489 | 2.9501622 | 0.104796 | 0.651716352 |
| ENSMUSG00000029004 | Kmt2e | protein_coding | -0.44810829 | 6.199021232 | 2.9497045 | 0.1048208 | 0.651716352 |
| ENSMUSG00000096257 | Ccer2 | protein_coding | 1.197362048 | 2.987224756 | 2.9487255 | 0.1049052 | 0.651716352 |
| ENSMUSG00000014554 | Dguok | protein_coding | -0.418958581 | 5.686858152 | 2.9480217 | 0.1049122 | 0.651716352 |
| ENSMUSG00000050071 | Bex1 | protein_coding | 0.331782295 | 8.358075561 | 2.9418319 | 0.1052491 | 0.652772477 |
| ENSMUSG00000032217 | Rnf111 | protein_coding | -0.54533439 | 4.463264588 | 2.941583 | 0.1052627 | 0.652772477 |
| ENSMUSG00000046240 | Hepacam | protein_coding | -0.719215311 | 4.693450718 | 2.9421255 | 0.1052643 | 0.652772477 |
| ENSMUSG00000042348 | Arl15 | protein_coding | -0.76897991 | 3.510940519 | 2.9404332 | 0.1053254 | 0.652774626 |
| ENSMUSG00000019894 | Slc6a15 | protein_coding | 0.443800748 | 5.604783337 | 2.9370177 | 0.105512 | 0.653398732 |
| ENSMUSG00000035171 | 1110059E24Rik | protein_coding | -0.35036661 | 7.010336727 | 2.9344932 | 0.1056502 | 0.653398732 |
| ENSMUSG00000025133 | Ints4 | protein_coding | 0.57786214 | 4.854161641 | 2.9307062 | 0.1058579 | 0.653398732 |
| ENSMUSG00000045441 | Gprin3 | protein_coding | -0.592413145 | 4.460587543 | 2.9288399 | 0.1059604 | 0.653398732 |
| ENSMUSG00000046269 | Usp27x | protein_coding | -0.64917017 | 3.983395356 | 2.9285163 | 0.1059782 | 0.653398732 |
| ENSMUSG00000050213 | Snip1 | protein_coding | 0.710338127 | 3.608189537 | 2.9276087 | 0.1060281 | 0.653398732 |
| ENSMUSG00000039963 | Ccdc40 | protein_coding | 0.817198379 | 3.778706287 | 2.9272202 | 0.1060495 | 0.653398732 |
| ENSMUSG00000028199 | Cryz | protein_coding | 0.569739894 | 5.182555888 | 2.9265511 | 0.1060863 | 0.653398732 |
| ENSMUSG00000025175 | Fn3k | protein_coding | -0.395126671 | 6.706386204 | 2.9261633 | 0.1061077 | 0.653398732 |
| ENSMUSG00000020124 | Usp15 | protein_coding | 0.383296993 | 5.88853095 | 2.9259168 | 0.1061212 | 0.653398732 |
| ENSMUSG00000027122 | Arl14ep | protein_coding | -0.440206188 | 6.619910862 | 2.9250054 | 0.1061714 | 0.653398732 |
| ENSMUSG00000029623 | Pdap1 | protein_coding | 0.330096622 | 6.886533944 | 2.924559 | 0.106196 | 0.653398732 |
| ENSMUSG00000021832 | Psmc6 | protein_coding | -0.329729994 | 7.89661194 | 2.9224092 | 0.1063146 | 0.653398732 |
| ENSMUSG00000057315 | Arhgap24 | protein_coding | 0.864420009 | 4.943632688 | 2.9563326 | 0.1063598 | 0.653398732 |
| ENSMUSG00000045589 | Frrs1l | protein_coding | -0.338527365 | 6.858846421 | 2.9206875 | 0.1064097 | 0.653398732 |
| ENSMUSG00000026662 | Sephs1 | protein_coding | 0.657209056 | 4.295783792 | 2.9203776 | 0.1064268 | 0.653398732 |
| ENSMUSG00000025408 | Ddit3 | protein_coding | -0.397781545 | 6.742839347 | 2.9197935 | 0.1064591 | 0.653398732 |
| ENSMUSG00000039137 | Whrn | protein_coding | -0.648108175 | 4.322699121 | 2.9175037 | 0.1065858 | 0.653803019 |
| ENSMUSG00000020928 | Higd1b | protein_coding | -1.030629001 | 2.886518888 | 2.9147936 | 0.1067359 | 0.653859877 |
| ENSMUSG00000033721 | Vav3 | protein_coding | -0.832193605 | 3.443028302 | 2.9140872 | 0.1067751 | 0.653859877 |
| ENSMUSG00000030718 | Ppme1 | protein_coding | 0.489561345 | 7.3473376 | 2.9222195 | 0.1071231 | 0.653859877 |
| ENSMUSG00000045790 | Ccdc149 | protein_coding | 0.552757719 | 4.636312051 | 2.9067799 | 0.1071814 | 0.653859877 |
| ENSMUSG00000036513 | Commd2 | protein_coding | -0.38092456 | 6.116908395 | 2.9067601 | 0.1071825 | 0.653859877 |
| ENSMUSG00000036502 | Tmem255a | protein_coding | 0.414736838 | 6.151645705 | 2.9066481 | 0.1071887 | 0.653859877 |
| ENSMUSG00000050379 | 6-Sep | protein_coding | 0.508415005 | 6.389171914 | 2.9073395 | 0.1072303 | 0.653859877 |
| ENSMUSG00000041477 | Dcp1b | protein_coding | -0.624870953 | 3.722479341 | 2.905899 | 0.1072305 | 0.653859877 |
| ENSMUSG00000040820 | Hlcs | protein_coding | -0.691607752 | 3.496479684 | 2.9057299 | 0.1072399 | 0.653859877 |
| ENSMUSG00000058318 | Phf21a | protein_coding | 0.525823879 | 4.990714443 | 2.9052362 | 0.1072675 | 0.653859877 |
| ENSMUSG00000013495 | Tmem175 | protein_coding | 0.54036244 | 5.834361965 | 2.9050934 | 0.1073067 | 0.653859877 |
| ENSMUSG00000019863 | Qrsl1 | protein_coding | 0.718915978 | 4.740587059 | 2.9047713 | 0.1073247 | 0.653859877 |
| ENSMUSG00000056023 | Gm9989 | processed_transcript | -0.839383325 | 3.638110107 | 2.902751 | 0.1074062 | 0.653985847 |
| ENSMUSG00000027998 | Plrg1 | protein_coding | 0.439804587 | 6.257843665 | 2.9000156 | 0.1075592 | 0.654475499 |
| ENSMUSG00000028613 | Lrp8 | protein_coding | -0.723731912 | 4.297031855 | 2.8986647 | 0.1076348 | 0.654475499 |
| ENSMUSG00000060181 | Slc35e3 | protein_coding | 0.751349566 | 3.131692399 | 2.8947479 | 0.1078545 | 0.654475499 |
| ENSMUSG00000022008 | Gpalpp1 | protein_coding | 0.525206093 | 4.981242341 | 2.8941094 | 0.1078903 | 0.654475499 |
| ENSMUSG00000078670 | Fam174b | protein_coding | 0.972085136 | 3.171188857 | 2.8938711 | 0.1079037 | 0.654475499 |
| ENSMUSG00000001467 | Cyp51 | protein_coding | 0.429371464 | 5.708776918 | 2.8932492 | 0.1079387 | 0.654475499 |
| ENSMUSG00000055421 | Pcdh9 | protein_coding | -0.358492754 | 6.98252191 | 2.8922328 | 0.1079958 | 0.654475499 |
| ENSMUSG00000045100 | Slc25a26 | protein_coding | -0.659959178 | 4.702150218 | 2.892019 | 0.1080078 | 0.654475499 |
| ENSMUSG00000025647 | Shisa5 | protein_coding | 0.379681542 | 7.52527952 | 2.8891502 | 0.1081693 | 0.654475499 |
| ENSMUSG00000024735 | Prpf19 | protein_coding | 0.417736541 | 5.857345862 | 2.888876 | 0.1081848 | 0.654475499 |

| ensembl_gene_id | external_gene_name | gene_biotype | log2 (fold change) | log2 (counts per million) | F-value | p-value | False discovery rate |
|---|---|---|---|---|---|---|---|
| ENSMUSG00000028256 | Odf2l | protein_coding | -0.966292394 | 3.174813505 | 2.8879226 | 0.1082385 | 0.654475499 |
| ENSMUSG00000026792 | Lrsam1 | protein_coding | 0.759766995 | 4.490933125 | 2.8878992 | 0.1082712 | 0.654475499 |
| ENSMUSG00000006315 | Tmem147 | protein_coding | 0.349898745 | 7.795517713 | 2.8872248 | 0.1082779 | 0.654475499 |
| ENSMUSG00000085396 | Firre | processed_transcript | -0.763244454 | 3.688050705 | 2.8845498 | 0.1084289 | 0.654772426 |
| ENSMUSG00000094870 | Zfp131 | protein_coding | -0.579683843 | 4.197405735 | 2.884198 | 0.1084488 | 0.654772426 |
| ENSMUSG00000039176 | Polg | protein_coding | 1.350125225 | 2.443151353 | 2.8819654 | 0.1085751 | 0.655166967 |
| ENSMUSG00000007659 | Bcl2l1 | protein_coding | 0.33576347 | 8.134873366 | 2.8797554 | 0.1087002 | 0.655484698 |
| ENSMUSG00000050530 | Fam171a1 | protein_coding | 0.424578071 | 5.133659877 | 2.8776483 | 0.1088197 | 0.655484698 |
| ENSMUSG00000027676 | Ccdc39 | protein_coding | -0.94348 | 3.229858903 | 2.8762117 | 0.1089013 | 0.655484698 |
| ENSMUSG00000039244 | E130309D02Rik | protein_coding | -0.629626087 | 3.943347011 | 2.875379 | 0.1089486 | 0.655484698 |
| ENSMUSG00000041669 | Prima1 | protein_coding | -1.255415 | 2.634618313 | 2.8750221 | 0.1089689 | 0.655484698 |
| ENSMUSG00000045639 | Zfp629 | protein_coding | -0.579261802 | 4.085104144 | 2.8745522 | 0.1089956 | 0.655484698 |
| ENSMUSG00000042750 | Bex2 | protein_coding | 0.329829395 | 10.3553781 | 2.8735181 | 0.1090544 | 0.655484698 |
| ENSMUSG00000020463 | Ppp4r3b | protein_coding | -0.463393801 | 5.015230073 | 2.8657559 | 0.1094971 | 0.657487872 |
| ENSMUSG00000024068 | Spast | protein_coding | -0.637089898 | 4.623684832 | 2.8633517 | 0.1096347 | 0.657487872 |
| ENSMUSG00000029397 | Rchy1 | protein_coding | -0.363697195 | 6.608216687 | 2.8624171 | 0.1096882 | 0.657487872 |
| ENSMUSG00000033904 | Ccp110 | protein_coding | -0.423181875 | 5.276146718 | 2.8602754 | 0.109811 | 0.657487872 |
| ENSMUSG00000035104 | Eva1a | protein_coding | -0.992312464 | 3.812427907 | 2.879928 | 0.1098309 | 0.657487872 |
| ENSMUSG00000041958 | Pigs | protein_coding | 0.495367256 | 5.845747439 | 2.8596697 | 0.1098458 | 0.657487872 |
| ENSMUSG00000063446 | Plppr1 | protein_coding | 0.471113923 | 4.933772752 | 2.8563529 | 0.1100363 | 0.657487872 |
| ENSMUSG00000034243 | Golgb1 | protein_coding | -0.469404028 | 5.276950784 | 2.8554763 | 0.1100868 | 0.657487872 |
| ENSMUSG00000027079 | Clp1 | protein_coding | 0.693069476 | 4.119925071 | 2.854398 | 0.1101488 | 0.657487872 |
| ENSMUSG00000038418 | Egr1 | protein_coding | -0.425769772 | 6.746067033 | 2.8539159 | 0.1101766 | 0.657487872 |
| ENSMUSG00000025785 | Exosc7 | protein_coding | 0.340869372 | 7.102316384 | 2.8530773 | 0.1102249 | 0.657487872 |
| ENSMUSG00000015023 | Ddx19a | protein_coding | 0.887707746 | 2.818227446 | 2.8514829 | 0.1103169 | 0.657487872 |
| ENSMUSG00000050751 | Pgbd5 | protein_coding | -0.350540169 | 6.575469217 | 2.8511635 | 0.1103353 | 0.657487872 |
| ENSMUSG00000025228 | Actr1a | protein_coding | 0.519400752 | 7.759377898 | 2.8693505 | 0.1105067 | 0.657487872 |
| ENSMUSG00000025007 | Aldh18a1 | protein_coding | 0.877295087 | 4.630399413 | 2.8743351 | 0.1106118 | 0.657487872 |
| ENSMUSG00000018217 | Pmp22 | protein_coding | -0.512832747 | 5.840557124 | 2.8460383 | 0.1106315 | 0.657487872 |
| ENSMUSG00000007617 | Homer1 | protein_coding | -0.47394891 | 5.929049542 | 2.8457733 | 0.1106468 | 0.657487872 |
| ENSMUSG00000026433 | Rab29 | protein_coding | -1.028167681 | 3.270456569 | 2.8434287 | 0.110814 | 0.657487872 |
| ENSMUSG00000020089 | Ppa1 | protein_coding | 0.381728602 | 7.898838711 | 2.8427818 | 0.1108202 | 0.657487872 |
| ENSMUSG00000026377 | Nifk | protein_coding | 0.390439877 | 5.969500023 | 2.8421755 | 0.1108554 | 0.657487872 |
| ENSMUSG00000061032 | Rrp1 | protein_coding | 0.380838706 | 8.565164267 | 2.8417394 | 0.1108807 | 0.657487872 |
| ENSMUSG00000022792 | Yars2 | protein_coding | 0.615657198 | 3.835039822 | 2.8406862 | 0.1109418 | 0.657487872 |
| ENSMUSG00000026627 | Tmem206 | protein_coding | 0.671819414 | 4.50691482 | 2.8400111 | 0.110981 | 0.657487872 |
| ENSMUSG00000021936 | Mapk8 | protein_coding | -0.445685565 | 5.316795201 | 2.8395056 | 0.1110104 | 0.657487872 |
| ENSMUSG00000040455 | Usp45 | protein_coding | -0.555692357 | 4.613921663 | 2.839285 | 0.1110232 | 0.657487872 |
| ENSMUSG00000001855 | Nup214 | protein_coding | 0.498629941 | 4.491393077 | 2.8388298 | 0.1110497 | 0.657487872 |
| ENSMUSG00000040867 | Begain | protein_coding | -0.512573612 | 4.989020161 | 2.8377933 | 0.11111 | 0.657487872 |
| ENSMUSG00000033444 | Specc1l | protein_coding | 0.512119854 | 4.588919484 | 2.8371889 | 0.1111452 | 0.657487872 |
| ENSMUSG00000030127 | Cops7a | protein_coding | 0.314424052 | 8.366976052 | 2.8369189 | 0.1111609 | 0.657487872 |
| ENSMUSG00000020114 | Cand1 | protein_coding | 0.375132797 | 6.489287104 | 2.8347625 | 0.1112865 | 0.657869059 |
| ENSMUSG00000010607 | Pigyl | protein_coding | -0.330268949 | 7.440696244 | 2.8319678 | 0.1114496 | 0.658360145 |
| ENSMUSG00000035297 | Cops4 | protein_coding | 0.30528641 | 8.429446949 | 2.8312409 | 0.111492 | 0.658360145 |
| ENSMUSG00000047766 | Lrrc49 | protein_coding | 0.407691866 | 6.572946789 | 2.8297104 | 0.1115815 | 0.658374391 |
| ENSMUSG00000028328 | Tmod1 | protein_coding | -0.534759429 | 4.726566109 | 2.8291052 | 0.1116169 | 0.658374391 |
| ENSMUSG00000012429 | Mplkip | protein_coding | -1.094555989 | 2.538806212 | 2.8244922 | 0.1118872 | 0.65949537 |
| ENSMUSG00000069662 | Marcks | protein_coding | -0.300561692 | 7.706302332 | 2.8233904 | 0.1119518 | 0.65949537 |
| ENSMUSG00000029406 | Pitpnm2 | protein_coding | -0.500284467 | 5.065491646 | 2.8214374 | 0.1120666 | 0.65949537 |
| ENSMUSG00000034813 | Grip1 | protein_coding | -0.49944191 | 4.547246663 | 2.8195819 | 0.1121757 | 0.65949537 |
| ENSMUSG00000051412 | Vamp7 | protein_coding | -0.393935624 | 6.712283676 | 2.8193485 | 0.1121895 | 0.65949537 |
| ENSMUSG00000032435 | Dync1li1 | protein_coding | 0.526754118 | 7.157449133 | 2.841967 | 0.1122262 | 0.65949537 |
| ENSMUSG00000027584 | Oprl1 | protein_coding | 0.425166221 | 6.424172654 | 2.8185537 | 0.1122363 | 0.65949537 |
| ENSMUSG00000058355 | Abce1 | protein_coding | 0.404771234 | 5.983356726 | 2.8166308 | 0.1123496 | 0.659800686 |
| ENSMUSG00000001833 | 7-Sep | protein_coding | -0.351090161 | 9.081041745 | 2.8145636 | 0.1124716 | 0.65984546 |
| ENSMUSG00000062098 | Btbd3 | protein_coding | 0.369060154 | 6.552591929 | 2.8144217 | 0.1124799 | 0.65984546 |
| ENSMUSG00000026333 | Gin1 | protein_coding | -0.65484477 | 4.444129451 | 2.8130003 | 0.1125639 | 0.659977978 |
| ENSMUSG00000039103 | Nexn | protein_coding | -0.725785519 | 4.231479967 | 2.8119488 | 0.1126261 | 0.659982625 |
| ENSMUSG00000027010 | Slc25a12 | protein_coding | 0.509592789 | 7.805494318 | 2.8309156 | 0.1127274 | 0.660216555 |
| ENSMUSG00000037722 | Gnpnat1 | protein_coding | -0.535029309 | 4.307557478 | 2.8070163 | 0.1129182 | 0.660373393 |
| ENSMUSG00000059436 | Max | protein_coding | -0.627163459 | 5.952105046 | 2.8263692 | 0.1129936 | 0.660373393 |
| ENSMUSG00000020255 | D10Wsu102e | protein_coding | 0.544688409 | 5.333206271 | 2.8054032 | 0.113014 | 0.660373393 |
| ENSMUSG00000023307 | 5-Mar | protein_coding | -0.48950487 | 5.014800071 | 2.8039944 | 0.1130977 | 0.660373393 |
| ENSMUSG00000031266 | Gla | protein_coding | 0.792291648 | 3.426680206 | 2.8036247 | 0.1131196 | 0.660373393 |
| ENSMUSG00000027652 | Ralgapb | protein_coding | 0.471524608 | 4.935392349 | 2.8035743 | 0.1131226 | 0.660373393 |
| ENSMUSG00000063320 | 1190007I07Rik | protein_coding | -0.505649293 | 4.499521849 | 2.8020788 | 0.1132116 | 0.660534095 |
| ENSMUSG00000020694 | Tlk2 | protein_coding | -0.444918567 | 5.276387066 | 2.8002606 | 0.1133199 | 0.660807216 |
| ENSMUSG00000038836 | Agbl3 | protein_coding | -0.450744141 | 5.249206097 | 2.7973783 | 0.1134917 | 0.661178843 |
| ENSMUSG00000052273 | Dnah3 | protein_coding | -1.113633203 | 2.341484185 | 2.7971299 | 0.1135066 | 0.661178843 |
| ENSMUSG00000025034 | Trim8 | protein_coding | -0.43360848 | 5.425123894 | 2.7947045 | 0.1136515 | 0.661664445 |
| ENSMUSG00000026587 | Astn1 | protein_coding | 0.391923492 | 6.196100042 | 2.7925437 | 0.1137807 | 0.661813412 |
| ENSMUSG00000024287 | Thoc1 | protein_coding | -0.374334002 | 5.986577429 | 2.7896936 | 0.1139515 | 0.661813412 |
| ENSMUSG00000032121 | Tmem218 | protein_coding | -0.437311432 | 5.524120132 | 2.7896033 | 0.1139569 | 0.661813412 |

| ensembl_gene_id | external_gene_name | gene_biotype | log2 (fold change) | log2 (counts per million) | F-value | p-value | False discovery rate |
|---|---|---|---|---|---|---|---|
| ENSMUSG00000057133 | Chd6 | protein_coding | 0.426218946 | 5.686323388 | 2.7888376 | 0.1140029 | 0.661813412 |
| ENSMUSG00000041079 | Rwdd2b | protein_coding | 0.551139353 | 4.046549818 | 2.7873997 | 0.1140892 | 0.661813412 |
| ENSMUSG00000063804 | Lin28b | protein_coding | -0.819666392 | 3.228588473 | 2.7861061 | 0.1141669 | 0.661813412 |
| ENSMUSG00000034777 | Vax2 | protein_coding | -1.093743303 | 2.562757672 | 2.7832229 | 0.1143404 | 0.661813412 |
| ENSMUSG00000033285 | Wdr3 | protein_coding | 1.412634131 | 2.434928954 | 2.7831773 | 0.1143432 | 0.661813412 |
| ENSMUSG00000004096 | Cwc15 | protein_coding | -0.348331099 | 8.159173265 | 2.78056 | 0.1145009 | 0.661813412 |
| ENSMUSG00000061666 | Gdpd1 | protein_coding | 0.316415522 | 6.787748303 | 2.7805392 | 0.1145022 | 0.661813412 |
| ENSMUSG00000021363 | Mak | protein_coding | 1.678273675 | 2.430673105 | 2.7803806 | 0.1145432 | 0.661813412 |
| ENSMUSG00000038957 | Edc3 | protein_coding | 0.515424616 | 5.546286261 | 2.7779788 | 0.1146568 | 0.661813412 |
| ENSMUSG00000029616 | Erp29 | protein_coding | 0.309888285 | 7.669565449 | 2.7778739 | 0.1146631 | 0.661813412 |
| ENSMUSG00000047414 | Flrt2 | protein_coding | -0.577516746 | 4.531359265 | 2.7775537 | 0.1146825 | 0.661813412 |
| ENSMUSG00000027712 | Anxa5 | protein_coding | 0.396986879 | 7.370112413 | 2.7758984 | 0.1147825 | 0.661813412 |
| ENSMUSG00000037234 | Hook3 | protein_coding | -0.62607341 | 4.477801318 | 2.7757214 | 0.1147933 | 0.661813412 |
| ENSMUSG00000031950 | Gabarapl2 | protein_coding | 0.326715987 | 6.722212429 | 2.7744896 | 0.1148678 | 0.661813412 |
| ENSMUSG00000034928 | Rnf44 | protein_coding | 0.442108096 | 5.358573775 | 2.7740088 | 0.1148969 | 0.661813412 |
| ENSMUSG00000063087 | Gm10125 | processed_transcript | -1.086182641 | 2.386144775 | 2.7738004 | 0.1149096 | 0.661813412 |
| ENSMUSG00000097527 | 1700112J16Rik | lincRNA | 1.011411047 | 2.740116355 | 2.7727341 | 0.1149742 | 0.661813412 |
| ENSMUSG00000004394 | Tmed4 | protein_coding | 0.312715908 | 8.130152195 | 2.7726279 | 0.1149806 | 0.661813412 |
| ENSMUSG00000028677 | Rnf220 | protein_coding | 0.319909754 | 6.594261474 | 2.7717959 | 0.1150311 | 0.661813412 |
| ENSMUSG00000027433 | Xrn2 | protein_coding | 0.399494659 | 5.935419471 | 2.7681161 | 0.1152546 | 0.662437968 |
| ENSMUSG00000030451 | Herc2 | protein_coding | -0.514965661 | 5.246005761 | 2.7669336 | 0.1153265 | 0.662437968 |
| ENSMUSG00000024276 | Zfp397 | protein_coding | -0.661109492 | 4.089525751 | 2.7668178 | 0.1153335 | 0.662437968 |
| ENSMUSG00000014077 | Chp1 | protein_coding | 0.371378015 | 7.876562314 | 2.7659549 | 0.1153861 | 0.662437968 |
| ENSMUSG00000039804 | Ncoa5 | protein_coding | 0.581469316 | 4.500782273 | 2.7646185 | 0.1154675 | 0.662551537 |
| ENSMUSG00000069919 | Hba-a1 | protein_coding | -0.744695051 | 4.437089775 | 2.7606109 | 0.1157434 | 0.663518436 |
| ENSMUSG00000031207 | Msn | protein_coding | 1.025382842 | 2.915599636 | 2.7587661 | 0.1158248 | 0.663518436 |
| ENSMUSG00000025200 | Cwf19l1 | protein_coding | 0.54119881 | 4.413403784 | 2.7587117 | 0.1158281 | 0.663518436 |
| ENSMUSG00000046985 | Tapt1 | protein_coding | 0.633961762 | 3.982898694 | 2.7566636 | 0.1159534 | 0.663518436 |
| ENSMUSG00000020986 | Sec23a | protein_coding | 0.761267835 | 4.69112239 | 2.7666329 | 0.1159819 | 0.663518436 |
| ENSMUSG00000029730 | Mcm7 | protein_coding | 0.804340714 | 3.548071418 | 2.7545129 | 0.1160853 | 0.663518436 |
| ENSMUSG00000038233 | Fam198a | protein_coding | 1.50108887 | 2.170744089 | 2.7544862 | 0.1160869 | 0.663518436 |
| ENSMUSG00000038127 | Ccdc50 | protein_coding | -0.387663947 | 5.518560666 | 2.7537898 | 0.1161296 | 0.663518436 |
| ENSMUSG00000006392 | Med8 | protein_coding | 0.368115307 | 6.431049831 | 2.7509849 | 0.1163019 | 0.664063078 |
| ENSMUSG00000030189 | Ybx3 | protein_coding | 0.64458611 | 3.853006636 | 2.7499457 | 0.1163658 | 0.664063078 |
| ENSMUSG00000036104 | Rab3gap1 | protein_coding | 0.431880019 | 6.15508334 | 2.7476644 | 0.1165062 | 0.664063078 |
| ENSMUSG00000040363 | Bcor | protein_coding | 0.466820199 | 4.932756 | 2.7471087 | 0.1165404 | 0.664063078 |
| ENSMUSG00000021775 | Nr1d2 | protein_coding | -0.417917427 | 5.451792979 | 2.7470681 | 0.1165429 | 0.664063078 |
| ENSMUSG00000094002 | Gm9866 | lincRNA | 0.325415887 | 6.629773502 | 2.7462155 | 0.1165955 | 0.664063078 |
| ENSMUSG00000001227 | Sema6b | protein_coding | -0.850848081 | 3.384376543 | 2.7440566 | 0.1167287 | 0.66446974 |
| ENSMUSG00000032216 | Nedd4 | protein_coding | 0.377480885 | 8.22302611 | 2.7423156 | 0.1168362 | 0.664730052 |
| ENSMUSG00000022237 | Ankrd33b | protein_coding | -1.075852357 | 3.099320655 | 2.7410317 | 0.1169471 | 0.665008847 |
| ENSMUSG00000098234 | Snhg6 | lincRNA | -0.69385548 | 5.926672747 | 2.777051 | 0.1171928 | 0.665235032 |
| ENSMUSG00000039795 | Zfand1 | protein_coding | -0.42806977 | 5.231060953 | 2.7360164 | 0.1172263 | 0.665235032 |
| ENSMUSG00000037007 | Zfp113 | protein_coding | -0.794090268 | 3.324830141 | 2.7359769 | 0.1172288 | 0.665235032 |
| ENSMUSG00000059991 | Nptx2 | protein_coding | 0.821619334 | 3.168592128 | 2.7358881 | 0.1172343 | 0.665235032 |
| ENSMUSG00000028030 | Tbck | protein_coding | 0.599125935 | 6.406016081 | 2.7650401 | 0.1173514 | 0.665548135 |
| ENSMUSG00000037463 | Fbxo27 | protein_coding | 1.033078869 | 2.454209972 | 2.7295684 | 0.1176273 | 0.665730841 |
| ENSMUSG00000055302 | Mrfap1 | protein_coding | -0.307079521 | 7.938451238 | 2.7285082 | 0.1176934 | 0.665730841 |
| ENSMUSG00000028607 | Cpt2 | protein_coding | 0.66107834 | 3.913188687 | 2.7268474 | 0.117797 | 0.665730841 |
| ENSMUSG00000027765 | P2ry1 | protein_coding | -0.827358804 | 3.143338903 | 2.7256715 | 0.1178704 | 0.665730841 |
| ENSMUSG00000026839 | Upp2 | protein_coding | -0.613103039 | 4.29042745 | 2.7251962 | 0.1179001 | 0.665730841 |
| ENSMUSG00000030435 | U2af2 | protein_coding | 0.690881256 | 3.440949511 | 2.7242234 | 0.1179609 | 0.665730841 |
| ENSMUSG00000069208 | Zfp825 | protein_coding | -0.647605059 | 3.554510274 | 2.7241553 | 0.1179652 | 0.665730841 |
| ENSMUSG00000022200 | Golph3 | protein_coding | -0.467467638 | 5.640855745 | 2.7229341 | 0.1180416 | 0.665730841 |
| ENSMUSG00000022212 | Cpne6 | protein_coding | 0.53404225 | 7.179004098 | 2.7534938 | 0.1180578 | 0.665730841 |
| ENSMUSG00000024431 | Nr3c1 | protein_coding | -0.327892019 | 6.420682602 | 2.7226594 | 0.1180588 | 0.665730841 |
| ENSMUSG00000087150 | BC064078 | transcribed_unproces | -0.88630787 | 3.367491776 | 2.7221444 | 0.118091 | 0.665730841 |
| ENSMUSG00000042045 | Sln | protein_coding | -0.800344828 | 6.315763863 | 2.8078904 | 0.1182208 | 0.665730841 |
| ENSMUSG00000061536 | Sec22c | protein_coding | 0.477313527 | 4.617431643 | 2.7187941 | 0.118301 | 0.665730841 |
| ENSMUSG00000005609 | Ctr9 | protein_coding | 0.472528809 | 5.359630466 | 2.7184272 | 0.118324 | 0.665730841 |
| ENSMUSG00000045409 | Trim39 | protein_coding | 0.732142033 | 3.885929486 | 2.7180306 | 0.1183489 | 0.665730841 |
| ENSMUSG00000097277 | 2900076A07Rik | lincRNA | -0.609641505 | 3.852739501 | 2.7172652 | 0.1183969 | 0.665730841 |
| ENSMUSG00000036845 | Lin37 | protein_coding | -0.320931625 | 6.935440291 | 2.7166422 | 0.1184361 | 0.665730841 |
| ENSMUSG00000097911 | Gm26691 | antisense_RNA | -0.952781967 | 2.904787901 | 2.7152912 | 0.118521 | 0.665821269 |
| ENSMUSG00000027456 | Sdcbp2 | protein_coding | -1.077140523 | 3.053528154 | 2.7144166 | 0.118576 | 0.665821269 |
| ENSMUSG00000028879 | Stx12 | protein_coding | 0.393137113 | 7.947623175 | 2.709685 | 0.1188741 | 0.666914548 |
| ENSMUSG00000030342 | Cd9 | protein_coding | -0.421710572 | 6.841465186 | 2.7098574 | 0.1188947 | 0.666914548 |
| ENSMUSG00000042851 | Zc3h6 | protein_coding | -0.727179969 | 3.161167287 | 2.7075962 | 0.1190061 | 0.667093919 |
| ENSMUSG00000040843 | Tiprl | protein_coding | 0.347018928 | 7.344423892 | 2.7068884 | 0.1190508 | 0.667093919 |
| ENSMUSG00000028136 | Snx27 | protein_coding | 0.375090028 | 6.71847641 | 2.7053785 | 0.1191463 | 0.667281346 |
| ENSMUSG00000038000 | Acd | protein_coding | 0.439851243 | 6.959643338 | 2.7047444 | 0.1192179 | 0.667334847 |
| ENSMUSG00000001750 | Tcirg1 | protein_coding | -0.954089948 | 2.990143412 | 2.7024965 | 0.1193288 | 0.667608277 |
| ENSMUSG00000025862 | Stag2 | protein_coding | -0.49702088 | 4.708151346 | 2.7005918 | 0.1194497 | 0.667723572 |
| ENSMUSG00000025283 | Sat1 | protein_coding | -0.365144452 | 6.838112687 | 2.7002142 | 0.1194736 | 0.667723572 |

| ensembl_gene_id | external_gene_name | gene_biotype | log2 (fold change) | log2 (counts per million) | F-value | p-value | False discovery rate |
|---|---|---|---|---|---|---|---|
| ENSMUSG00000007656 | Arpp19 | protein_coding | -0.416706019 | 9.503377922 | 2.6991102 | 0.1195438 | 0.66776839 |
| ENSMUSG00000028797 | Tmem234 | protein_coding | -0.287520045 | 7.596368689 | 2.6974861 | 0.119647 | 0.667998057 |
| ENSMUSG00000047648 | Fbxo30 | protein_coding | -0.953158377 | 2.810722436 | 2.6953252 | 0.1197845 | 0.668120665 |
| ENSMUSG00000033809 | Alg3 | protein_coding | 0.518954131 | 4.917568556 | 2.6944215 | 0.1198421 | 0.668120665 |
| ENSMUSG00000030301 | Ccdc91 | protein_coding | 0.402737332 | 5.880553558 | 2.6942136 | 0.1198554 | 0.668120665 |
| ENSMUSG00000066175 | 2510046G10Rik | antisense_RNA | -1.63762866 | 2.061809455 | 2.6885362 | 0.1202179 | 0.669343393 |
| ENSMUSG00000027165 | B230118H07Rik | protein_coding | -0.341640654 | 7.438964117 | 2.6877095 | 0.1202708 | 0.669343393 |
| ENSMUSG00000022044 | Stmn4 | protein_coding | 0.393140058 | 8.71982359 | 2.6873694 | 0.1202925 | 0.669343393 |
| ENSMUSG00000061130 | Ppm1b | protein_coding | -0.439407086 | 5.23265105 | 2.6859856 | 0.1203812 | 0.669343393 |
| ENSMUSG00000029034 | Ints11 | protein_coding | 0.877160709 | 3.329845632 | 2.6856802 | 0.1204007 | 0.669343393 |
| ENSMUSG00000020224 | Llph | protein_coding | -0.478457026 | 4.444461609 | 2.6839722 | 0.1205103 | 0.669343393 |
| ENSMUSG00000031387 | Renbp | protein_coding | 0.847997931 | 3.052426427 | 2.6834896 | 0.1205412 | 0.669343393 |
| ENSMUSG00000070866 | Zfp804a | protein_coding | -0.625666001 | 4.426000685 | 2.6799741 | 0.1207671 | 0.669343393 |
| ENSMUSG00000045201 | Lrrc3b | protein_coding | -0.475690376 | 4.919093809 | 2.6797245 | 0.1207831 | 0.669343393 |
| ENSMUSG00000073411 | H2-D1 | protein_coding | 0.863541405 | 4.646071214 | 2.7085401 | 0.120858 | 0.669343393 |
| ENSMUSG00000024059 | Clip4 | protein_coding | 0.402647267 | 5.871477717 | 2.6774623 | 0.1209288 | 0.669343393 |
| ENSMUSG00000031310 | Zmym3 | protein_coding | 0.32961274 | 6.639066481 | 2.6754008 | 0.1210617 | 0.669343393 |
| ENSMUSG00000103649 | Gm37768 | antisense_RNA | 1.00261442 | 2.671521566 | 2.6737825 | 0.1211661 | 0.669343393 |
| ENSMUSG00000092607 | Scnm1 | protein_coding | -0.325643045 | 7.329508744 | 2.6723753 | 0.121257 | 0.669343393 |
| ENSMUSG00000009076 | Zmat5 | protein_coding | -0.415904306 | 5.897945453 | 2.6705267 | 0.1213766 | 0.669343393 |
| ENSMUSG00000027852 | Nras | protein_coding | -0.292993295 | 7.041033276 | 2.6694249 | 0.1214479 | 0.669343393 |
| ENSMUSG00000021680 | Crhbp | protein_coding | 0.547544681 | 4.435789994 | 2.669312 | 0.1214552 | 0.669343393 |
| ENSMUSG00000027248 | Pdia3 | protein_coding | 0.398517373 | 6.219888738 | 2.6686411 | 0.1214987 | 0.669343393 |
| ENSMUSG00000002808 | Epdr1 | protein_coding | 0.373028592 | 6.191063094 | 2.6663867 | 0.1216449 | 0.669343393 |
| ENSMUSG00000030337 | Vamp1 | protein_coding | 0.442252481 | 7.451851913 | 2.6668854 | 0.121727 | 0.669343393 |
| ENSMUSG00000025008 | Tctn3 | protein_coding | 1.146166519 | 3.740737521 | 2.7026524 | 0.1218024 | 0.669343393 |
| ENSMUSG00000062995 | Ica1 | protein_coding | 0.476008877 | 6.464474724 | 2.6638307 | 0.1218424 | 0.669343393 |
| ENSMUSG00000039770 | Ypel5 | protein_coding | -0.291328409 | 8.954682556 | 2.6631302 | 0.1218564 | 0.669343393 |
| ENSMUSG00000004462 | Tbccd1 | protein_coding | 0.650901469 | 4.312858335 | 2.6617843 | 0.1219439 | 0.669343393 |
| ENSMUSG00000026803 | Ttf1 | protein_coding | -0.659651726 | 3.734458087 | 2.661581 | 0.1219572 | 0.669343393 |
| ENSMUSG00000014956 | Ppp1cb | protein_coding | -0.284248042 | 8.33464785 | 2.6602805 | 0.1220419 | 0.669343393 |
| ENSMUSG00000043259 | Fam13c | protein_coding | 0.386739573 | 5.533974813 | 2.6601507 | 0.1220503 | 0.669343393 |
| ENSMUSG00000050732 | Vamp8 | protein_coding | -0.693831369 | 5.692453053 | 2.6897824 | 0.1220503 | 0.669343393 |
| ENSMUSG00000044628 | Rnf208 | protein_coding | 0.386393134 | 6.932068985 | 2.6582641 | 0.1221733 | 0.669343393 |
| ENSMUSG00000041264 | Uspl1 | protein_coding | 0.444667359 | 5.374578541 | 2.6570391 | 0.1222532 | 0.669343393 |
| ENSMUSG00000035666 | Gtf3c4 | protein_coding | 0.696421504 | 3.779221143 | 2.6563022 | 0.1223013 | 0.669343393 |
| ENSMUSG00000028098 | Rnf115 | protein_coding | -0.380981168 | 5.421290025 | 2.6554456 | 0.1223573 | 0.669343393 |
| ENSMUSG00000024007 | Ppil1 | protein_coding | 0.338377893 | 6.774868429 | 2.6544989 | 0.1224192 | 0.669343393 |
| ENSMUSG00000043929 | Klhl15 | protein_coding | -0.678757015 | 4.742716628 | 2.6549795 | 0.1224193 | 0.669343393 |
| ENSMUSG00000022897 | Dyrk1a | protein_coding | -0.391679505 | 6.246967347 | 2.6539554 | 0.1224547 | 0.669343393 |
| ENSMUSG00000022762 | Ncam2 | protein_coding | -0.67987607 | 4.609970633 | 2.6541415 | 0.122474 | 0.669343393 |
| ENSMUSG00000101621 | Gm28905 | lincRNA | -1.225878706 | 2.29363569 | 2.6532685 | 0.1224997 | 0.669343393 |
| ENSMUSG00000044164 | Rnf182 | protein_coding | 0.894732825 | 3.699817746 | 2.6533078 | 0.1225286 | 0.669343393 |
| ENSMUSG00000017288 | Vps53 | protein_coding | 0.468752002 | 6.081862636 | 2.6507191 | 0.1226666 | 0.669343393 |
| ENSMUSG00000037960 | Card19 | protein_coding | -0.586634677 | 4.101817204 | 2.64926 | 0.1227623 | 0.669343393 |
| ENSMUSG00000070544 | Top1 | protein_coding | -0.400354823 | 6.256643158 | 2.6463775 | 0.1229516 | 0.669343393 |
| ENSMUSG00000029304 | Spp1 | protein_coding | -0.726615376 | 4.246992768 | 2.6468481 | 0.1229522 | 0.669343393 |
| ENSMUSG00000024370 | Cdc23 | protein_coding | 0.593772601 | 5.487269264 | 2.6461908 | 0.1229954 | 0.669343393 |
| ENSMUSG00000032375 | Aph1b | protein_coding | 0.542136911 | 4.121172696 | 2.6435963 | 0.1231346 | 0.669343393 |
| ENSMUSG00000019189 | Rnf145 | protein_coding | 0.4992225 | 5.142568878 | 2.6428406 | 0.1231843 | 0.669343393 |
| ENSMUSG00000056267 | Cep70 | protein_coding | -0.528084388 | 4.42457313 | 2.6422668 | 0.1232221 | 0.669343393 |
| ENSMUSG00000097616 | 1110019D14Rik | lincRNA | -0.819758126 | 3.245475465 | 2.6411053 | 0.1232987 | 0.669343393 |
| ENSMUSG00000024513 | Mbd2 | protein_coding | -0.643585707 | 3.827740814 | 2.6410407 | 0.123303 | 0.669343393 |
| ENSMUSG00000029096 | Htra3 | protein_coding | -0.68755557 | 3.500425552 | 2.6398689 | 0.1233803 | 0.669343393 |
| ENSMUSG00000031847 | 1700030J22Rik | protein_coding | -0.666933863 | 3.621338539 | 2.6370604 | 0.1235658 | 0.669343393 |
| ENSMUSG00000041084 | Ostc | protein_coding | 0.291790748 | 7.486781233 | 2.6370072 | 0.1235693 | 0.669343393 |
| ENSMUSG00000096472 | Cdkn2d | protein_coding | 0.286489239 | 8.239729479 | 2.6354812 | 0.1236703 | 0.669343393 |
| ENSMUSG00000048251 | Bcl11b | protein_coding | -0.392843768 | 6.005800764 | 2.6354704 | 0.123671 | 0.669343393 |
| ENSMUSG00000026163 | Sphkap | protein_coding | -0.363842453 | 5.666542243 | 2.632944 | 0.1238384 | 0.669343393 |
| ENSMUSG00000031870 | Pgr | protein_coding | -0.500763416 | 4.161279224 | 2.6320733 | 0.1238961 | 0.669343393 |
| ENSMUSG00000040548 | Tex2 | protein_coding | -0.34539539 | 6.582425221 | 2.6317876 | 0.1239151 | 0.669343393 |
| ENSMUSG00000090223 | Pcp4 | protein_coding | -0.46921849 | 8.449282519 | 2.6438944 | 0.1239184 | 0.669343393 |
| ENSMUSG00000022477 | Aco2 | protein_coding | 0.351218159 | 8.493533715 | 2.6296439 | 0.1240574 | 0.669343393 |
| ENSMUSG00000039130 | Zc3hc1 | protein_coding | 0.391234726 | 6.442612538 | 2.6276656 | 0.1241889 | 0.669343393 |
| ENSMUSG00000022120 | Rnf219 | protein_coding | 0.618269433 | 4.369410088 | 2.6275713 | 0.1241952 | 0.669343393 |
| ENSMUSG00000108621 | Gm37494 | lincRNA | -0.873136959 | 2.913498847 | 2.6271377 | 0.1242241 | 0.669343393 |
| ENSMUSG00000051346 | Spryd4 | protein_coding | 0.468295889 | 4.964114323 | 2.6271261 | 0.1242248 | 0.669343393 |
| ENSMUSG00000033773 | Rpap2 | protein_coding | 0.689773334 | 4.518197972 | 2.6270161 | 0.1242637 | 0.669343393 |
| ENSMUSG00000028433 | Ubap2 | protein_coding | 0.585700888 | 4.129500924 | 2.6258587 | 0.1243092 | 0.669343393 |
| ENSMUSG00000097316 | Gm10516 | antisense_RNA | -1.23191697 | 2.657396966 | 2.6244955 | 0.1244001 | 0.669343393 |
| ENSMUSG00000035704 | Alg8 | protein_coding | -0.807202773 | 3.612258277 | 2.62427 | 0.1244151 | 0.669343393 |
| ENSMUSG00000009907 | Vps4b | protein_coding | 0.41039152 | 5.311644651 | 2.6217663 | 0.1245822 | 0.669343393 |
| ENSMUSG00000026078 | Pdcl3 | protein_coding | -0.311003787 | 7.318520387 | 2.6203118 | 0.1246793 | 0.669343393 |
| ENSMUSG00000020435 | Osbp2 | protein_coding | -0.734837006 | 3.854302169 | 2.6193091 | 0.1247464 | 0.669343393 |

| ensembl_gene_id | external_gene_name | gene_biotype | log2 (fold change) | log2 (counts per million) | F-value | p-value | False discovery rate |
|---|---|---|---|---|---|---|---|
| ENSMUSG00000030180 | Kdm5a | protein_coding | -0.458497915 | 5.70130828 | 2.6188945 | 0.1247741 | 0.669343393 |
| ENSMUSG00000003382 | Etv3 | protein_coding | 0.558270438 | 3.984042676 | 2.6183723 | 0.1248091 | 0.669343393 |
| ENSMUSG00000026495 | Efcab2 | protein_coding | -0.467787731 | 5.377614684 | 2.6178788 | 0.1248421 | 0.669343393 |
| ENSMUSG00000023044 | Csad | protein_coding | 0.661536425 | 4.287033396 | 2.6170801 | 0.1248956 | 0.669343393 |
| ENSMUSG00000037977 | 6430571L13Rik | protein_coding | 0.857396191 | 3.498634964 | 2.617003 | 0.1249007 | 0.669343393 |
| ENSMUSG00000017801 | Mlx | protein_coding | -0.426234708 | 6.238427675 | 2.6168965 | 0.1249079 | 0.669343393 |
| ENSMUSG00000053477 | Tcf4 | protein_coding | -0.337775014 | 6.748185216 | 2.6160133 | 0.1249671 | 0.669343393 |
| ENSMUSG00000019699 | Akt3 | protein_coding | -0.456317584 | 5.479204504 | 2.6151992 | 0.1250217 | 0.669343393 |
| ENSMUSG00000019874 | Fabp7 | protein_coding | -0.658996567 | 6.935996241 | 2.6725335 | 0.1250523 | 0.669343393 |
| ENSMUSG00000030695 | Aldoa | protein_coding | 0.349166473 | 11.9914057 | 2.6122555 | 0.1252193 | 0.669343393 |
| ENSMUSG00000031201 | Brcc3 | protein_coding | -0.356689232 | 7.292192666 | 2.6121737 | 0.1252248 | 0.669343393 |
| ENSMUSG00000037971 | 1110032A03Rik | protein_coding | -0.286218343 | 7.593220878 | 2.6109794 | 0.1253051 | 0.669343393 |
| ENSMUSG00000029094 | Afap1 | protein_coding | -0.512097909 | 4.558819411 | 2.6099827 | 0.1253721 | 0.669343393 |
| ENSMUSG00000018474 | Chd3 | protein_coding | -0.324209555 | 6.673162646 | 2.6096726 | 0.125393 | 0.669343393 |
| ENSMUSG00000038352 | Arl5c | protein_coding | -0.903646735 | 4.748157757 | 2.6499559 | 0.1254022 | 0.669343393 |
| ENSMUSG00000034235 | Usp54 | protein_coding | -1.248717491 | 2.469394203 | 2.6093456 | 0.125415 | 0.669343393 |
| ENSMUSG00000023826 | Park2 | protein_coding | -0.513779945 | 4.804067349 | 2.6090046 | 0.1254379 | 0.669343393 |
| ENSMUSG00000021371 | Mcur1 | protein_coding | 0.827110918 | 3.262011593 | 2.607943 | 0.1255095 | 0.669343393 |
| ENSMUSG00000042797 | Aqp11 | protein_coding | 0.632627816 | 3.620436067 | 2.6073049 | 0.1255525 | 0.669343393 |
| ENSMUSG00000026469 | Xpr1 | protein_coding | -0.353606727 | 6.068042782 | 2.6052461 | 0.1256913 | 0.669751699 |
| ENSMUSG00000072769 | Gm10419 | lincRNA | -0.346447577 | 6.107039761 | 2.6017051 | 0.1259306 | 0.669858775 |
| ENSMUSG00000039943 | Plcb4 | protein_coding | -0.390480432 | 6.39625355 | 2.6012358 | 0.1259624 | 0.669858775 |
| ENSMUSG00000053985 | Zfp14 | protein_coding | 0.591480133 | 3.830076174 | 2.6008673 | 0.1259873 | 0.669858775 |
| ENSMUSG00000044716 | Dok7 | protein_coding | -0.815485471 | 3.466412865 | 2.6004017 | 0.1260189 | 0.669858775 |
| ENSMUSG00000068923 | Syt11 | protein_coding | 0.335296133 | 9.840075036 | 2.6003381 | 0.1260232 | 0.669858775 |
| ENSMUSG00000004221 | Ikbkg | protein_coding | 0.561932508 | 5.073525441 | 2.5994223 | 0.1260852 | 0.669858775 |
| ENSMUSG00000022679 | Mpv17l | protein_coding | -0.344748524 | 5.775120625 | 2.5979832 | 0.1261828 | 0.670046001 |
| ENSMUSG00000034342 | Cbl | protein_coding | -0.506543386 | 4.555585624 | 2.5967884 | 0.1262638 | 0.670145525 |
| ENSMUSG00000020150 | Gamt | protein_coding | -0.673634661 | 3.771287056 | 2.5940586 | 0.1264493 | 0.670798681 |
| ENSMUSG00000033295 | Ptprf | protein_coding | 0.380345427 | 5.670309118 | 2.5930713 | 0.1265164 | 0.670823993 |
| ENSMUSG00000040054 | Baz2a | protein_coding | 0.605742336 | 4.379592687 | 2.5900106 | 0.1267249 | 0.671598101 |
| ENSMUSG00000055862 | Izumo4 | protein_coding | 0.34906943 | 6.276003752 | 2.5873414 | 0.126907 | 0.671710999 |
| ENSMUSG00000033174 | Mgll | protein_coding | -0.411320092 | 6.842980407 | 2.5869959 | 0.1269621 | 0.671710999 |
| ENSMUSG00000027281 | Slx4ip | protein_coding | -0.662494341 | 3.290612586 | 2.585037 | 0.1270645 | 0.671710999 |
| ENSMUSG00000054667 | Irs4 | protein_coding | 0.601745134 | 4.720075632 | 2.5844903 | 0.1271019 | 0.671710999 |
| ENSMUSG00000001120 | Pcbp3 | protein_coding | 0.681885358 | 6.028750481 | 2.6234432 | 0.1271563 | 0.671710999 |
| ENSMUSG00000004788 | Eif2b2 | protein_coding | 0.341241405 | 6.234374159 | 2.5836604 | 0.1271587 | 0.671710999 |
| ENSMUSG00000005813 | Metap1 | protein_coding | 0.431146158 | 5.827359175 | 2.582478 | 0.1272397 | 0.671710999 |
| ENSMUSG00000070934 | Rraga | protein_coding | 0.447264424 | 6.785571419 | 2.5828467 | 0.1272459 | 0.671710999 |
| ENSMUSG00000028445 | Enho | protein_coding | -0.459072843 | 6.246349103 | 2.5774024 | 0.1276195 | 0.673022859 |
| ENSMUSG00000038485 | Socs7 | protein_coding | -0.456059935 | 5.370109791 | 2.5748554 | 0.1277632 | 0.673022859 |
| ENSMUSG00000039195 | 1110008P14Rik | protein_coding | 0.611902006 | 4.218693784 | 2.5747955 | 0.1277673 | 0.673022859 |
| ENSMUSG00000028229 | Rmdn1 | protein_coding | -0.537216802 | 4.287708467 | 2.5740797 | 0.1278166 | 0.673022859 |
| ENSMUSG00000023277 | Twf2 | protein_coding | -0.417281414 | 5.785012577 | 2.5736152 | 0.1278486 | 0.673022859 |
| ENSMUSG00000037852 | Cpe | protein_coding | 0.392613551 | 10.16661404 | 2.5733051 | 0.12787 | 0.673022859 |
| ENSMUSG00000019689 | Fmc1 | protein_coding | -0.344798678 | 7.439051326 | 2.5718444 | 0.1279707 | 0.67320937 |
| ENSMUSG00000068036 | Afdn | protein_coding | 0.507684566 | 5.070946904 | 2.5703871 | 0.1280713 | 0.67320937 |
| ENSMUSG00000025092 | Hspa12a | protein_coding | 0.361240136 | 5.960480157 | 2.5694557 | 0.1281356 | 0.67320937 |
| ENSMUSG00000055723 | Rras2 | protein_coding | 1.478794411 | 2.084447914 | 2.569163 | 0.1281558 | 0.67320937 |
| ENSMUSG00000027134 | Lpcat4 | protein_coding | 0.486328023 | 4.840404694 | 2.5662716 | 0.1283558 | 0.673654821 |
| ENSMUSG00000025130 | P4hb | protein_coding | 0.370270227 | 6.470404982 | 2.5661259 | 0.1283659 | 0.673654821 |
| ENSMUSG00000028671 | Gale | protein_coding | 0.570903963 | 4.816075524 | 2.5642774 | 0.128494 | 0.673926665 |
| ENSMUSG00000061119 | Prcp | protein_coding | 0.960483716 | 3.268052999 | 2.5640241 | 0.1285431 | 0.673926665 |
| ENSMUSG00000048728 | Zfp454 | protein_coding | -0.708224382 | 3.464650488 | 2.5605183 | 0.1287549 | 0.674442746 |
| ENSMUSG00000022617 | Chkb | protein_coding | -0.555924109 | 3.886913244 | 2.559938 | 0.1287953 | 0.674442746 |
| ENSMUSG00000029545 | Acads | protein_coding | -0.775771695 | 3.275808124 | 2.5591805 | 0.1288479 | 0.674442746 |
| ENSMUSG00000039911 | Spsb1 | protein_coding | 0.853729839 | 2.936041677 | 2.5579044 | 0.1289367 | 0.674442746 |
| ENSMUSG00000028063 | Lmna | protein_coding | 0.433581289 | 5.095959939 | 2.5576405 | 0.1289551 | 0.674442746 |
| ENSMUSG00000055670 | Zzef1 | protein_coding | -0.668215092 | 4.076059421 | 2.5549328 | 0.1291438 | 0.675101307 |
| ENSMUSG00000050244 | Heatr1 | protein_coding | -0.924530051 | 2.870058153 | 2.5537559 | 0.1292259 | 0.675202395 |
| ENSMUSG00000024095 | Hnrnpll | protein_coding | -0.536101287 | 5.037191559 | 2.550518 | 0.1294522 | 0.676056119 |
| ENSMUSG00000051224 | Tceanc | protein_coding | -1.073481951 | 2.64279464 | 2.5480262 | 0.1296267 | 0.676421124 |
| ENSMUSG00000038375 | Trp53inp2 | protein_coding | 0.42989408 | 5.434666145 | 2.547723 | 0.1296479 | 0.676421124 |
| ENSMUSG00000028863 | Meaf6 | protein_coding | 0.332910017 | 7.874483094 | 2.5439617 | 0.1299118 | 0.677426268 |
| ENSMUSG00000032050 | Rdx | protein_coding | 0.480026055 | 5.48007083 | 2.5418355 | 0.1300613 | 0.677426268 |
| ENSMUSG00000016349 | Eef1a2 | protein_coding | -0.493771441 | 5.491949823 | 2.5402692 | 0.1301716 | 0.677426268 |
| ENSMUSG00000030246 | Ldhb | protein_coding | 0.299474375 | 10.4757629 | 2.5401651 | 0.1301789 | 0.677426268 |
| ENSMUSG00000026843 | Fubp3 | protein_coding | -0.494412299 | 4.951118832 | 2.5395367 | 0.1302232 | 0.677426268 |
| ENSMUSG00000038806 | Sde2 | protein_coding | 0.324746311 | 6.14126393 | 2.5394566 | 0.1302288 | 0.677426268 |
| ENSMUSG00000028822 | Tmem50a | protein_coding | 0.289693763 | 8.468053722 | 2.5375976 | 0.1303599 | 0.677426268 |
| ENSMUSG00000037892 | Pcdh18 | protein_coding | -0.767582882 | 3.706613125 | 2.5370318 | 0.1303998 | 0.677426268 |
| ENSMUSG00000021476 | Habp4 | protein_coding | 0.354268656 | 6.94883417 | 2.5343346 | 0.1305903 | 0.677426268 |
| ENSMUSG00000030979 | Uros | protein_coding | 0.330123724 | 6.304832327 | 2.5337697 | 0.1306303 | 0.677426268 |
| ENSMUSG00000031485 | Prosc | protein_coding | 0.361002176 | 6.440097532 | 2.5336287 | 0.1306403 | 0.677426268 |

| ensembl_gene_id | external_gene_name | gene_biotype | log2 (fold change) | log2 (counts per million) | F-value | p-value | False discovery rate |
|---|---|---|---|---|---|---|---|
| ENSMUSG00000040520 | Manea | protein_coding | -0.575285532 | 4.031956797 | 2.5332998 | 0.1306635 | 0.677426268 |
| ENSMUSG00000018865 | Sult4a1 | protein_coding | 0.363240739 | 9.330066523 | 2.5317242 | 0.130775 | 0.677426268 |
| ENSMUSG00000031278 | Acsl4 | protein_coding | -0.401337393 | 6.224603325 | 2.5314492 | 0.1307945 | 0.677426268 |
| ENSMUSG00000061411 | Nol4l | protein_coding | 0.521926709 | 4.668333996 | 2.5313871 | 0.1307989 | 0.677426268 |
| ENSMUSG00000025722 | Wdr73 | protein_coding | 0.402412738 | 5.634725503 | 2.5306868 | 0.1308485 | 0.677426268 |
| ENSMUSG00000057219 | Armc7 | protein_coding | 0.542339443 | 4.448733052 | 2.5295328 | 0.1309304 | 0.677523645 |
| ENSMUSG00000032413 | Rasa2 | protein_coding | -0.533215055 | 4.582664916 | 2.5254097 | 0.1312232 | 0.67756906 |
| ENSMUSG00000039990 | Edrf1 | protein_coding | 0.869291791 | 3.266259873 | 2.5253392 | 0.1312282 | 0.67756906 |
| ENSMUSG00000021597 | Slf1 | protein_coding | -0.559896408 | 4.400162418 | 2.5242518 | 0.1313056 | 0.67756906 |
| ENSMUSG00000037217 | Syn1 | protein_coding | -0.37029208 | 6.267628301 | 2.5222795 | 0.131446 | 0.67756906 |
| ENSMUSG00000029878 | Dbpht2 | lincRNA | -0.584491122 | 8.515574096 | 2.5624663 | 0.1315025 | 0.67756906 |
| ENSMUSG00000058246 | Gm10037 | protein_coding | -0.531167018 | 4.268719226 | 2.5206589 | 0.1315616 | 0.67756906 |
| ENSMUSG00000038302 | Afg1l | protein_coding | 0.554448704 | 4.72901734 | 2.5199964 | 0.1316088 | 0.67756906 |
| ENSMUSG00000020635 | Fkbp1b | protein_coding | 0.563972637 | 4.312844413 | 2.5192517 | 0.131662 | 0.67756906 |
| ENSMUSG00000058317 | Ube2e2 | protein_coding | -0.318726662 | 6.828385677 | 2.5182034 | 0.1317369 | 0.67756906 |
| ENSMUSG00000021587 | Pcsk1 | protein_coding | 0.498514103 | 4.154065003 | 2.516861 | 0.1318329 | 0.67756906 |
| ENSMUSG00000019917 | 10-Sep | protein_coding | 0.819665402 | 3.195238907 | 2.5148668 | 0.1319756 | 0.67756906 |
| ENSMUSG00000047409 | Ctdspl | protein_coding | 0.453683264 | 4.700609006 | 2.5145682 | 0.1319969 | 0.67756906 |
| ENSMUSG00000025525 | Apool | protein_coding | -0.630401676 | 4.103545178 | 2.5140587 | 0.1320334 | 0.67756906 |
| ENSMUSG00000031996 | Aplp2 | protein_coding | 0.304489649 | 7.990537434 | 2.5140046 | 0.1320373 | 0.67756906 |
| ENSMUSG00000071359 | Tbpl1 | protein_coding | 0.392815211 | 6.377242744 | 2.5120418 | 0.1321781 | 0.67756906 |
| ENSMUSG00000000416 | Cttnbp2 | protein_coding | -0.52812849 | 5.046223057 | 2.5116967 | 0.1322028 | 0.67756906 |
| ENSMUSG00000059273 | Zc3h4 | protein_coding | -0.47121362 | 4.300804903 | 2.5109587 | 0.1322558 | 0.67756906 |
| ENSMUSG00000005483 | Dnajb1 | protein_coding | 0.336165093 | 6.685269881 | 2.5106661 | 0.1322768 | 0.67756906 |
| ENSMUSG00000027996 | Sfrp2 | protein_coding | 0.802979298 | 2.939593684 | 2.5099667 | 0.132327 | 0.67756906 |
| ENSMUSG00000063646 | Jakmip1 | protein_coding | 0.337491803 | 5.848733938 | 2.5089857 | 0.1323975 | 0.67756906 |
| ENSMUSG00000025375 | Aatk | protein_coding | -0.446104166 | 5.730584382 | 2.5089336 | 0.1324013 | 0.67756906 |
| ENSMUSG00000020166 | Cnot2 | protein_coding | 0.424380522 | 5.701456193 | 2.5085673 | 0.1324276 | 0.67756906 |
| ENSMUSG00000079435 | Rpl36a | protein_coding | -0.482338348 | 5.304824514 | 2.5084856 | 0.1324335 | 0.67756906 |
| ENSMUSG00000026594 | Ralgps2 | protein_coding | -0.502165677 | 4.878117885 | 2.508016 | 0.1324673 | 0.67756906 |
| ENSMUSG00000020218 | Wif1 | protein_coding | -0.973601188 | 3.057694366 | 2.5066556 | 0.1325652 | 0.67756906 |
| ENSMUSG00000058254 | Tspan7 | protein_coding | 0.302655054 | 11.05145679 | 2.5060608 | 0.132608 | 0.67756906 |
| ENSMUSG00000070394 | Tmem256 | protein_coding | -0.412940817 | 8.328333018 | 2.505294 | 0.1326948 | 0.67756906 |
| ENSMUSG00000087211 | Lhx1os | lincRNA | 0.502274383 | 4.070579514 | 2.504736 | 0.1327035 | 0.67756906 |
| ENSMUSG00000000325 | Arvcf | protein_coding | -0.638622522 | 3.429181469 | 2.5027732 | 0.1328451 | 0.677965659 |
| ENSMUSG00000038776 | Ephx1 | protein_coding | 0.558892382 | 4.720512754 | 2.5016444 | 0.1329266 | 0.677965659 |
| ENSMUSG00000026269 | Rnpepl1 | protein_coding | -0.663060937 | 3.453164609 | 2.5010394 | 0.1329703 | 0.677965659 |
| ENSMUSG00000034194 | R3hcc1 | protein_coding | 0.428538354 | 5.675439554 | 2.4985604 | 0.1331496 | 0.678230267 |
| ENSMUSG00000021592 | Arsk | protein_coding | -0.909302287 | 3.16525007 | 2.4974185 | 0.1332323 | 0.678230267 |
| ENSMUSG00000012117 | Dhdds | protein_coding | 0.372496188 | 6.15705973 | 2.497361 | 0.1332364 | 0.678230267 |
| ENSMUSG00000102252 | Snrpn | protein_coding | 0.43292788 | 5.158850636 | 2.4961326 | 0.1333255 | 0.678230267 |
| ENSMUSG00000022553 | Maf1 | protein_coding | -0.587783528 | 5.636313139 | 2.5047942 | 0.1333376 | 0.678230267 |
| ENSMUSG00000031023 | Akip1 | protein_coding | 0.635906659 | 3.610570956 | 2.4947673 | 0.1334245 | 0.678351616 |
| ENSMUSG00000097717 | 1700066J03Rik | lincRNA | -0.97372212 | 3.00396811 | 2.4925804 | 0.1335833 | 0.678531431 |
| ENSMUSG00000024002 | Brd4 | protein_coding | 0.355298506 | 6.56187823 | 2.4897347 | 0.1337903 | 0.678531431 |
| ENSMUSG00000019971 | Cep290 | protein_coding | -0.757492919 | 3.458790192 | 2.4891801 | 0.1338307 | 0.678531431 |
| ENSMUSG00000038886 | Man2a2 | protein_coding | 0.685132826 | 3.766803742 | 2.4890287 | 0.1338417 | 0.678531431 |
| ENSMUSG00000112121 | AC163903.1 | lincRNA | -1.028290772 | 2.695447453 | 2.4868739 | 0.1339988 | 0.678531431 |
| ENSMUSG00000056486 | Chn1 | protein_coding | 0.285477509 | 7.21260601 | 2.4867894 | 0.134005 | 0.678531431 |
| ENSMUSG00000020549 | Elac2 | protein_coding | 0.670919929 | 3.621125108 | 2.4852077 | 0.1341204 | 0.678531431 |
| ENSMUSG00000021290 | 2010107E04Rik | protein_coding | -0.407564961 | 9.27045432 | 2.4853313 | 0.1341429 | 0.678531431 |
| ENSMUSG00000057894 | Zfp329 | protein_coding | -0.478227588 | 5.127169057 | 2.4844369 | 0.1341767 | 0.678531431 |
| ENSMUSG00000049336 | Tenm2 | protein_coding | -0.491157822 | 5.487232192 | 2.4841131 | 0.1342004 | 0.678531431 |
| ENSMUSG00000034848 | Ttc21b | protein_coding | 1.026113021 | 2.747969089 | 2.4820522 | 0.1343511 | 0.678531431 |
| ENSMUSG00000051234 | Rnf7 | protein_coding | -0.274239175 | 7.855072363 | 2.4803427 | 0.1344763 | 0.678531431 |
| ENSMUSG00000086350 | B230369F24Rik | antisense_RNA | -0.618486201 | 3.605258578 | 2.4796569 | 0.1345265 | 0.678531431 |
| ENSMUSG00000063856 | Gpx1 | protein_coding | -0.289293214 | 9.114042541 | 2.4794331 | 0.1345429 | 0.678531431 |
| ENSMUSG00000031409 | Tceal6 | protein_coding | 0.311063826 | 6.238640912 | 2.478818 | 0.134588 | 0.678531431 |
| ENSMUSG00000025591 | Tma16 | protein_coding | -0.487946117 | 4.834137173 | 2.4786845 | 0.1345978 | 0.678531431 |
| ENSMUSG00000038506 | Dcun1d2 | protein_coding | 0.465856771 | 5.279136708 | 2.4786322 | 0.1346017 | 0.678531431 |
| ENSMUSG00000029436 | Mmp17 | protein_coding | 0.465958617 | 5.137459336 | 2.4782401 | 0.1346304 | 0.678531431 |
| ENSMUSG00000085282 | Gm15663 | processed_transcript | -0.579137866 | 3.665509334 | 2.4763744 | 0.1347674 | 0.678531431 |
| ENSMUSG00000022043 | Trim35 | protein_coding | 0.310631044 | 6.707283872 | 2.4761379 | 0.1347847 | 0.678531431 |
| ENSMUSG00000071103 | 1700029J07Rik | protein_coding | -1.57436432 | 2.093148936 | 2.4761345 | 0.134785 | 0.678531431 |
| ENSMUSG00000074918 | Inafm2 | protein_coding | -0.380838546 | 7.812709367 | 2.4735361 | 0.1349761 | 0.679142263 |
| ENSMUSG00000020910 | Adprm | protein_coding | -0.408389324 | 5.80374054 | 2.471933 | 0.1350941 | 0.679142263 |
| ENSMUSG00000052981 | Ube2ql1 | protein_coding | 0.344461136 | 6.85934638 | 2.4719098 | 0.1350958 | 0.679142263 |
| ENSMUSG00000057193 | Slc44a2 | protein_coding | 0.689554558 | 3.950299999 | 2.4696142 | 0.1352651 | 0.679374517 |
| ENSMUSG00000096910 | Zfp955b | protein_coding | -0.976682203 | 2.486901328 | 2.4695694 | 0.1352684 | 0.679374517 |
| ENSMUSG00000046062 | Ppp1r15b | protein_coding | 0.480370195 | 4.111451231 | 2.468553 | 0.1353434 | 0.67943397 |
| ENSMUSG00000041889 | Shisa4 | protein_coding | -0.305831535 | 6.555733141 | 2.4657426 | 0.1355511 | 0.68015915 |
| ENSMUSG00000029447 | Cct6a | processed_transcript | 0.301187072 | 8.111927685 | 2.464513 | 0.1356421 | 0.680298298 |
| ENSMUSG00000029752 | Asns | protein_coding | 0.523716453 | 8.118845226 | 2.4910501 | 0.1357423 | 0.680483652 |
| ENSMUSG00000018882 | Mrpl45 | protein_coding | 0.310570902 | 7.34872276 | 2.4609259 | 0.135908 | 0.680996565 |

| ensembl_gene_id | external_gene_name | gene_biotype | log2 (fold change) | log2 (counts per million) | F-value | p-value | False discovery rate |
|---|---|---|---|---|---|---|---|
| ENSMUSG00000026771 | Spopl | protein_coding | -0.708678336 | 3.182406445 | 2.4567172 | 0.1362208 | 0.682245913 |
| ENSMUSG00000041014 | Nrg3 | protein_coding | -0.517972045 | 4.681865863 | 2.4550728 | 0.1363432 | 0.682541231 |
| ENSMUSG00000033106 | Slc7a6os | protein_coding | 0.787611896 | 3.552645987 | 2.4531267 | 0.1364883 | 0.68294957 |
| ENSMUSG00000047789 | Slc38a9 | protein_coding | 0.540621672 | 3.844259702 | 2.4512093 | 0.1366314 | 0.683347774 |
| ENSMUSG00000024169 | Ift140 | protein_coding | 0.819288288 | 2.989982275 | 2.4502311 | 0.1367045 | 0.6833883 |
| ENSMUSG00000053801 | Grwd1 | protein_coding | 0.406897614 | 6.023791231 | 2.448052 | 0.1368675 | 0.6833883 |
| ENSMUSG00000041028 | Ghitm | protein_coding | 0.313635631 | 9.508313207 | 2.4471932 | 0.1369318 | 0.6833883 |
| ENSMUSG00000019303 | Psmc3ip | protein_coding | -0.599187984 | 3.827689081 | 2.4467955 | 0.1369616 | 0.6833883 |
| ENSMUSG00000044071 | Fam19a2 | protein_coding | -0.54518631 | 4.660714796 | 2.4460246 | 0.1370194 | 0.6833883 |
| ENSMUSG00000036545 | Adamts2 | protein_coding | 1.584377053 | 2.421073913 | 2.4475319 | 0.1370208 | 0.6833883 |
| ENSMUSG00000033006 | Sox10 | protein_coding | -0.489744085 | 5.169216814 | 2.4425173 | 0.1372825 | 0.684204972 |
| ENSMUSG00000025231 | Sufu | protein_coding | -0.85073506 | 3.201262282 | 2.4412715 | 0.1373762 | 0.684204972 |
| ENSMUSG00000042642 | Flad1 | protein_coding | 0.393357659 | 6.006308765 | 2.4412361 | 0.1373788 | 0.684204972 |
| ENSMUSG00000034706 | Dnaic2 | protein_coding | 0.607870555 | 3.603919497 | 2.4404347 | 0.1374391 | 0.684204972 |
| ENSMUSG00000028698 | Pik3r3 | protein_coding | 0.652600069 | 3.624051346 | 2.4390124 | 0.1375462 | 0.684293769 |
| ENSMUSG00000033510 | Otud7a | protein_coding | -0.975637229 | 2.772559865 | 2.4385071 | 0.1375842 | 0.684293769 |
| ENSMUSG00000003033 | Ap1m1 | protein_coding | 0.420163379 | 7.141286704 | 2.4375006 | 0.1376915 | 0.684324975 |
| ENSMUSG00000001016 | Ilf2 | protein_coding | 0.300016301 | 7.841223411 | 2.4361319 | 0.1377633 | 0.684324975 |
| ENSMUSG00000026834 | Acvr1c | protein_coding | -0.560870257 | 3.894645852 | 2.4358915 | 0.1377814 | 0.684324975 |
| ENSMUSG00000047368 | Abhd17b | protein_coding | -0.469020723 | 5.259547045 | 2.4320017 | 0.1380753 | 0.685468112 |
| ENSMUSG00000062981 | Mrpl42 | protein_coding | -0.376792665 | 8.561114125 | 2.4303345 | 0.1382015 | 0.685778038 |
| ENSMUSG00000037996 | Slc24a2 | protein_coding | -0.420241642 | 6.09735924 | 2.4283177 | 0.1383544 | 0.686143871 |
| ENSMUSG00000025137 | Pcyt2 | protein_coding | 0.362588486 | 7.272872055 | 2.4276785 | 0.1384029 | 0.686143871 |
| ENSMUSG00000004267 | Eno2 | protein_coding | 0.397346454 | 10.23750331 | 2.4250905 | 0.1385994 | 0.686801542 |
| ENSMUSG00000002733 | Plekha3 | protein_coding | -0.483336193 | 4.711575466 | 2.42313 | 0.1387485 | 0.687223749 |
| ENSMUSG00000020895 | Tmem107 | protein_coding | -0.404028049 | 5.722713389 | 2.4171778 | 0.1392024 | 0.688286303 |
| ENSMUSG00000024064 | Galnt14 | protein_coding | 0.557400385 | 3.847379419 | 2.4156209 | 0.1393214 | 0.688286303 |
| ENSMUSG00000042302 | Ehbp1 | protein_coding | -0.616620357 | 4.174940949 | 2.4151467 | 0.1393577 | 0.688286303 |
| ENSMUSG00000029343 | Crybb1 | protein_coding | -1.357076442 | 2.065168389 | 2.4150565 | 0.1393646 | 0.688286303 |
| ENSMUSG00000032641 | Gpr19 | protein_coding | 0.352611587 | 5.796899798 | 2.4144975 | 0.1394074 | 0.688286303 |
| ENSMUSG00000074863 | Platr25 | protein_coding | -1.021296792 | 2.442684069 | 2.4144336 | 0.1394123 | 0.688286303 |
| ENSMUSG00000030316 | Tamm41 | protein_coding | 0.68019409 | 3.63002157 | 2.4141538 | 0.1394337 | 0.688286303 |
| ENSMUSG00000052331 | Ankrd44 | protein_coding | -1.139672877 | 2.457030023 | 2.4134233 | 0.1394896 | 0.688286303 |
| ENSMUSG00000007827 | Ankrd26 | protein_coding | 0.611289448 | 4.709306544 | 2.4123729 | 0.1396016 | 0.688286303 |
| ENSMUSG00000056222 | Spock1 | protein_coding | 0.294660756 | 6.523959085 | 2.4107398 | 0.1396954 | 0.688286303 |
| ENSMUSG00000002058 | Unc119 | protein_coding | 0.538796542 | 4.628577171 | 2.4097305 | 0.1397729 | 0.688286303 |
| ENSMUSG00000022340 | Sybu | protein_coding | 0.295640716 | 6.519935658 | 2.40964 | 0.1397798 | 0.688286303 |
| ENSMUSG00000072501 | Phf20l1 | protein_coding | -0.524263862 | 4.447356078 | 2.4094403 | 0.1397952 | 0.688286303 |
| ENSMUSG00000049807 | Arhgap23 | protein_coding | -0.57336054 | 4.26667607 | 2.4060259 | 0.1400577 | 0.688992153 |
| ENSMUSG00000020400 | Tnip1 | protein_coding | 0.520326049 | 5.062295397 | 2.4059094 | 0.1400667 | 0.688992153 |
| ENSMUSG00000030701 | Plekhb1 | protein_coding | 0.347859682 | 9.101479327 | 2.4050183 | 0.1401353 | 0.689014543 |
| ENSMUSG00000018733 | Pex12 | protein_coding | 0.587963504 | 4.360905925 | 2.4018558 | 0.1403792 | 0.689258751 |
| ENSMUSG00000087148 | 1700030C14Rik | antisense_RNA | -1.211631488 | 2.385949082 | 2.4016077 | 0.1403983 | 0.689258751 |
| ENSMUSG00000027865 | Gdap2 | protein_coding | 0.548020291 | 5.846192724 | 2.4087694 | 0.140484 | 0.689258751 |
| ENSMUSG00000001300 | Efnb2 | protein_coding | -0.638534042 | 4.524014235 | 2.4001007 | 0.1405462 | 0.689258751 |
| ENSMUSG00000056917 | Sipa1 | protein_coding | 0.654258725 | 4.245049254 | 2.3991621 | 0.1405873 | 0.689258751 |
| ENSMUSG00000066607 | 6030419C18Rik | protein_coding | -0.921534626 | 2.612149884 | 2.3980439 | 0.1406738 | 0.689258751 |
| ENSMUSG00000075376 | Rc3h2 | protein_coding | -0.573380079 | 5.503473446 | 2.3995713 | 0.1407026 | 0.689258751 |
| ENSMUSG00000057363 | Uxs1 | protein_coding | -0.658196547 | 3.824973209 | 2.3962564 | 0.1408122 | 0.689258751 |
| ENSMUSG00000027663 | Zmat3 | protein_coding | 0.351979632 | 5.696992462 | 2.3958993 | 0.1408399 | 0.689258751 |
| ENSMUSG00000013707 | Tnfaip8l2 | protein_coding | -1.187053298 | 3.013897389 | 2.3982531 | 0.1408501 | 0.689258751 |
| ENSMUSG00000024261 | Syt4 | protein_coding | 0.309415633 | 8.374893494 | 2.3952525 | 0.1408901 | 0.689258751 |
| ENSMUSG00000057147 | Dph6 | protein_coding | 0.398932292 | 5.277548766 | 2.3942602 | 0.140967 | 0.689321711 |
| ENSMUSG00000052848 | C130026L21Rik | lincRNA | 0.48621107 | 4.471491748 | 2.3912371 | 0.1412018 | 0.690052954 |
| ENSMUSG00000031840 | Rab3a | protein_coding | 0.499914525 | 9.649739403 | 2.4072693 | 0.141258 | 0.690052954 |
| ENSMUSG00000048537 | Phldb1 | protein_coding | -0.460509962 | 4.846870434 | 2.3897227 | 0.1413196 | 0.690052954 |
| ENSMUSG00000003153 | Slc2a3 | protein_coding | 0.291322805 | 6.362208726 | 2.3886104 | 0.1414062 | 0.690052954 |
| ENSMUSG00000031889 | D230025D16Rik | protein_coding | 0.532242896 | 4.334105351 | 2.388101 | 0.1414459 | 0.690052954 |
| ENSMUSG00000073940 | Hbb-bt | protein_coding | -0.737943917 | 3.790202896 | 2.3873864 | 0.1415016 | 0.690052954 |
| ENSMUSG00000038344 | Txlng | protein_coding | -0.67187496 | 3.688826837 | 2.3854885 | 0.1416496 | 0.690461733 |
| ENSMUSG00000024941 | Scyl1 | protein_coding | 0.407485103 | 5.49164501 | 2.3823974 | 0.1418912 | 0.691325596 |
| ENSMUSG00000091985 | Gm17354 | lincRNA | 1.001786192 | 2.537460446 | 2.3814806 | 0.1419629 | 0.691361827 |
| ENSMUSG00000069917 | Hba-a2 | protein_coding | -0.706475866 | 3.810027504 | 2.3793619 | 0.1421288 | 0.691856572 |
| ENSMUSG00000038128 | Camk4 | protein_coding | -0.441337243 | 6.331167359 | 2.3762041 | 0.142408 | 0.692901857 |
| ENSMUSG00000028568 | Btf3l4 | protein_coding | -0.441303204 | 4.661104131 | 2.3746166 | 0.1425013 | 0.693020354 |
| ENSMUSG00000031672 | Got2 | protein_coding | 0.426897834 | 7.03732811 | 2.3749342 | 0.1425906 | 0.693020354 |
| ENSMUSG00000037010 | Apln | protein_coding | 0.728440055 | 3.299783963 | 2.3730355 | 0.1426257 | 0.693020354 |
| ENSMUSG00000034656 | Cacna1a | protein_coding | -0.81426696 | 3.40050763 | 2.3714618 | 0.1427496 | 0.693309239 |
| ENSMUSG00000076441 | Ass1 | protein_coding | 0.487123609 | 6.249848399 | 2.3724526 | 0.1429832 | 0.694130166 |
| ENSMUSG00000042210 | Abhd14a | protein_coding | -0.43483166 | 6.311576668 | 2.3674303 | 0.1430991 | 0.694379423 |
| ENSMUSG00000027291 | Vps39 | protein_coding | 0.55003709 | 5.778991377 | 2.3706391 | 0.143408 | 0.695175144 |
| ENSMUSG00000023175 | Bsg | protein_coding | 0.501419968 | 8.302447301 | 2.387077 | 0.1434342 | 0.695175144 |
| ENSMUSG00000031393 | Mecp2 | protein_coding | -0.387905027 | 5.871023054 | 2.3624966 | 0.1434581 | 0.695175144 |
| ENSMUSG00000063297 | Luzp2 | protein_coding | -0.426167308 | 6.133680057 | 2.361695 | 0.1435217 | 0.695175144 |

| ensembl_gene_id | external_gene_name | gene_biotype | log2 (fold change) | log2 (counts per million) | F-value | p-value | False discovery rate |
|---|---|---|---|---|---|---|---|
| ENSMUSG00000003657 | Calb2 | protein_coding | 0.478586205 | 10.09150668 | 2.3728053 | 0.1437871 | 0.695885053 |
| ENSMUSG00000027506 | Tpd52 | protein_coding | -0.397500786 | 6.305528105 | 2.3575309 | 0.1438524 | 0.695885053 |
| ENSMUSG00000030443 | Zfp583 | protein_coding | -0.620409044 | 3.713609321 | 2.356209 | 0.1439576 | 0.695885053 |
| ENSMUSG00000059439 | Bcas3 | protein_coding | 0.59421371 | 4.684346565 | 2.3547686 | 0.1440723 | 0.695885053 |
| ENSMUSG00000042213 | Zfand4 | protein_coding | -1.048328049 | 2.652720383 | 2.3540773 | 0.1441274 | 0.695885053 |
| ENSMUSG00000029434 | Vps33a | protein_coding | 0.489128971 | 5.761476132 | 2.3537483 | 0.1441851 | 0.695885053 |
| ENSMUSG00000032096 | Arcn1 | protein_coding | -0.36228783 | 5.576845699 | 2.353264 | 0.1441923 | 0.695885053 |
| ENSMUSG00000020402 | Vdac1 | protein_coding | 0.292246842 | 9.386725854 | 2.3522176 | 0.1442758 | 0.695885053 |
| ENSMUSG00000062234 | Gak | protein_coding | 0.388521249 | 5.490160043 | 2.3511634 | 0.14436 | 0.695885053 |
| ENSMUSG00000030704 | Rab6a | protein_coding | -0.279677042 | 8.306047026 | 2.3505446 | 0.1444094 | 0.695885053 |
| ENSMUSG00000018160 | Med1 | protein_coding | 0.410645689 | 6.664700456 | 2.3506073 | 0.1444358 | 0.695885053 |
| ENSMUSG00000021708 | Rasgrf2 | protein_coding | 0.390851749 | 6.114797076 | 2.350101 | 0.1444448 | 0.695885053 |
| ENSMUSG00000026202 | Tuba4a | protein_coding | 0.403388611 | 9.099061636 | 2.3490725 | 0.1445585 | 0.696028935 |
| ENSMUSG00000054387 | Mdm4 | protein_coding | -0.379851501 | 5.099620466 | 2.3467078 | 0.1447164 | 0.696028935 |
| ENSMUSG00000071708 | Sms | protein_coding | -0.339825695 | 6.018823401 | 2.3463067 | 0.1447485 | 0.696028935 |
| ENSMUSG00000024357 | Sil1 | protein_coding | 0.361838129 | 6.185084574 | 2.3457411 | 0.1447939 | 0.696028935 |
| ENSMUSG00000050608 | Minos1 | protein_coding | -0.346730648 | 9.318764556 | 2.3456852 | 0.1447984 | 0.696028935 |
| ENSMUSG00000098557 | Kctd12 | protein_coding | -0.498557477 | 4.537306075 | 2.343494 | 0.1449742 | 0.696481826 |
| ENSMUSG00000046032 | Snx12 | protein_coding | 0.286375722 | 6.618575788 | 2.3428752 | 0.1450239 | 0.696481826 |
| ENSMUSG00000022323 | Rida | protein_coding | -0.393852655 | 7.338296693 | 2.3411761 | 0.1451918 | 0.696481826 |
| ENSMUSG00000032733 | Snx33 | protein_coding | 0.808901191 | 3.184480107 | 2.3394477 | 0.1452995 | 0.696481826 |
| ENSMUSG00000007908 | Hmgcll1 | protein_coding | -0.582434684 | 4.293571825 | 2.338871 | 0.145346 | 0.696481826 |
| ENSMUSG00000031198 | Fundc2 | protein_coding | -0.306652553 | 7.531706821 | 2.3384256 | 0.1453818 | 0.696481826 |
| ENSMUSG00000042570 | Mier2 | protein_coding | -0.802223329 | 3.993611786 | 2.3420814 | 0.145399 | 0.696481826 |
| ENSMUSG00000020993 | Trappc6b | protein_coding | -0.402129512 | 5.564164391 | 2.3373133 | 0.1454715 | 0.696481826 |
| ENSMUSG00000026349 | Ccnt2 | protein_coding | -0.490782974 | 4.920341983 | 2.3370522 | 0.1454925 | 0.696481826 |
| ENSMUSG00000008658 | Rbfox1 | protein_coding | -0.28258982 | 8.260538718 | 2.3364603 | 0.1455403 | 0.696481826 |
| ENSMUSG00000067942 | Zfp160 | protein_coding | -0.617546711 | 3.655992398 | 2.3351674 | 0.1456446 | 0.696613911 |
| ENSMUSG00000020407 | Upp1 | protein_coding | -0.981233115 | 2.819865986 | 2.3345135 | 0.1456975 | 0.696613911 |
| ENSMUSG00000023452 | Pisd | protein_coding | 0.354266737 | 5.562063979 | 2.3335834 | 0.1457726 | 0.696663503 |
| ENSMUSG00000030432 | Rpl28 | protein_coding | 0.317578 | 6.389847888 | 2.3326953 | 0.1458444 | 0.696697061 |
| ENSMUSG00000037610 | Kcnmb2 | protein_coding | -1.060238612 | 2.971614265 | 2.3313397 | 0.1459855 | 0.697061317 |
| ENSMUSG00000025076 | Casp7 | protein_coding | -1.25249428 | 2.317533746 | 2.3280495 | 0.1462208 | 0.697104181 |
| ENSMUSG00000020364 | Zfp354a | protein_coding | -0.766431982 | 3.351531826 | 2.327629 | 0.1462549 | 0.697104181 |
| ENSMUSG00000038910 | Plcl2 | protein_coding | 0.513682719 | 5.087735322 | 2.3273129 | 0.1462806 | 0.697104181 |
| ENSMUSG00000022554 | Hgh1 | protein_coding | -0.905145926 | 2.639369943 | 2.3263823 | 0.1463561 | 0.697104181 |
| ENSMUSG00000078974 | Sec61g | protein_coding | -0.447533132 | 5.158550074 | 2.3260128 | 0.1463862 | 0.697104181 |
| ENSMUSG00000033751 | Gadd45gip1 | protein_coding | 0.970205009 | 3.098671897 | 2.3255512 | 0.1464237 | 0.697104181 |
| ENSMUSG00000060487 | Samd5 | protein_coding | -0.4884218 | 4.481120482 | 2.3252485 | 0.1464483 | 0.697104181 |
| ENSMUSG00000053574 | 4930563E22Rik | processed_transcript | 0.717604589 | 3.28713829 | 2.3232222 | 0.1466131 | 0.697463801 |
| ENSMUSG00000036908 | Unc93b1 | protein_coding | -1.136205195 | 3.136844393 | 2.3324308 | 0.146662 | 0.697463801 |
| ENSMUSG00000039713 | Plekhg5 | protein_coding | 0.52838036 | 4.772621883 | 2.3212317 | 0.1467753 | 0.697463801 |
| ENSMUSG00000045106 | Ccdc73 | protein_coding | -0.59120544 | 3.773213786 | 2.3211334 | 0.1467833 | 0.697463801 |
| ENSMUSG00000020225 | Tmbim4 | protein_coding | 0.315355565 | 6.932846328 | 2.3180825 | 0.1470323 | 0.698166209 |
| ENSMUSG00000025956 | Mettl21a | protein_coding | 0.47505673 | 4.838935925 | 2.3177312 | 0.147061 | 0.698166209 |
| ENSMUSG00000028243 | Ubxn2b | protein_coding | -0.402043133 | 5.753515982 | 2.3140367 | 0.1473632 | 0.69866459 |
| ENSMUSG00000025235 | Bbs4 | protein_coding | 0.449498733 | 6.460608854 | 2.3149982 | 0.1473985 | 0.69866459 |
| ENSMUSG00000056124 | B4galt6 | protein_coding | -0.319084981 | 6.352546689 | 2.3129164 | 0.147455 | 0.69866459 |
| ENSMUSG00000047155 | Cyp4x1 | protein_coding | -0.615207829 | 3.539105573 | 2.3122418 | 0.1475103 | 0.69866459 |
| ENSMUSG00000005481 | Ddx39 | protein_coding | 0.481695207 | 5.050695147 | 2.3119229 | 0.1475365 | 0.69866459 |
| ENSMUSG00000016252 | Atp5e | protein_coding | -0.331022575 | 9.763074852 | 2.3116879 | 0.1475558 | 0.69866459 |
| ENSMUSG00000067148 | Polr1c | protein_coding | 0.284060591 | 6.988972938 | 2.3105181 | 0.1476518 | 0.698776512 |
| ENSMUSG00000039206 | Daglb | protein_coding | 0.434158015 | 4.974757951 | 2.3098169 | 0.1477094 | 0.698776512 |
| ENSMUSG00000032526 | Ss18l2 | protein_coding | -0.281257763 | 8.073288735 | 2.3060994 | 0.1480152 | 0.699027295 |
| ENSMUSG00000040997 | Abhd4 | protein_coding | -0.313806577 | 6.797734357 | 2.3058002 | 0.1480398 | 0.699027295 |
| ENSMUSG00000032252 | Glce | protein_coding | -0.496312166 | 4.339791961 | 2.3042272 | 0.1481695 | 0.699027295 |
| ENSMUSG00000044847 | Lsm11 | protein_coding | 0.840865254 | 3.172878441 | 2.3037762 | 0.1482067 | 0.699027295 |
| ENSMUSG00000022305 | Lrp12 | protein_coding | -0.618360222 | 4.229442977 | 2.3035241 | 0.1482275 | 0.699027295 |
| ENSMUSG00000024383 | Map3k2 | protein_coding | -0.598182808 | 3.76212729 | 2.303477 | 0.1482314 | 0.699027295 |
| ENSMUSG00000021427 | Ssr1 | protein_coding | 0.315027048 | 8.64397154 | 2.3032353 | 0.1482513 | 0.699027295 |
| ENSMUSG00000023004 | Tuba1b | protein_coding | 0.297472415 | 7.738139621 | 2.3028577 | 0.1482825 | 0.699027295 |
| ENSMUSG00000055013 | Agap1 | protein_coding | -0.332610011 | 6.385730368 | 2.3018288 | 0.1483674 | 0.699080217 |
| ENSMUSG00000019432 | Ddx39b | protein_coding | 0.311034898 | 7.952588544 | 2.3011365 | 0.1484246 | 0.699080217 |
| ENSMUSG00000037126 | Psd | protein_coding | -0.464553972 | 5.103726207 | 2.300361 | 0.1484887 | 0.699080217 |
| ENSMUSG00000063626 | Unc5d | protein_coding | -0.439791988 | 5.112167552 | 2.2979641 | 0.1486871 | 0.699707668 |
| ENSMUSG00000048379 | Socs4 | protein_coding | -0.983592176 | 2.690753874 | 2.2969016 | 0.1487751 | 0.699815637 |
| ENSMUSG00000036907 | C1ql2 | protein_coding | 1.219977337 | 2.369910729 | 2.2951582 | 0.1489197 | 0.700189436 |
| ENSMUSG00000040612 | Ildr2 | protein_coding | -0.462319371 | 5.880273793 | 2.2936593 | 0.1490754 | 0.700615486 |
| ENSMUSG00000037685 | Atp8a1 | protein_coding | 0.650372961 | 3.502410305 | 2.2916303 | 0.1492128 | 0.700846581 |
| ENSMUSG00000017679 | Ttpal | protein_coding | 0.43727496 | 4.909761282 | 2.2911229 | 0.149255 | 0.700846581 |
| ENSMUSG00000011114 | Tbrg1 | protein_coding | 0.341062489 | 7.01306031 | 2.2894506 | 0.1493942 | 0.701152681 |
| ENSMUSG00000041380 | Htr2c | protein_coding | -0.381918196 | 5.492600064 | 2.2884005 | 0.1494817 | 0.701152681 |
| ENSMUSG00000029765 | Plxna4 | protein_coding | -0.509477216 | 4.416682776 | 2.2869385 | 0.1496036 | 0.701152681 |
| ENSMUSG00000057506 | Bloc1s2 | protein_coding | -0.398019396 | 5.078684327 | 2.286473 | 0.1496425 | 0.701152681 |

| ensembl_gene_id | external_gene_name | gene_biotype | log2 (fold change) | log2 (counts per million) | F-value | p-value | False discovery rate |
|---|---|---|---|---|---|---|---|
| ENSMUSG00000003464 | Pex19 | protein_coding | 0.346436328 | 7.878205664 | 2.2864288 | 0.1496462 | 0.701152681 |
| ENSMUSG00000034910 | Pygo1 | protein_coding | -0.480259793 | 4.763878088 | 2.2856299 | 0.1497129 | 0.701159708 |
| ENSMUSG00000006736 | Tspan31 | protein_coding | 0.299939829 | 6.733065879 | 2.2840258 | 0.1498469 | 0.701232202 |
| ENSMUSG00000039069 | Mtg2 | protein_coding | 0.524768001 | 4.653762963 | 2.283884 | 0.1498588 | 0.701232202 |
| ENSMUSG00000036686 | Cc2d1a | protein_coding | -0.573536998 | 4.848330308 | 2.2817203 | 0.1500399 | 0.701774052 |
| ENSMUSG00000031696 | Vps35 | protein_coding | 0.277638399 | 7.691568384 | 2.2794585 | 0.1502294 | 0.702287401 |
| ENSMUSG00000029326 | Enoph1 | protein_coding | 0.286281578 | 7.226127742 | 2.2781021 | 0.1503432 | 0.702287401 |
| ENSMUSG00000051319 | 1500011K16Rik | protein_coding | -0.298789768 | 7.289265781 | 2.2780746 | 0.1503455 | 0.702287401 |
| ENSMUSG00000036941 | Elac1 | protein_coding | 0.487078882 | 4.921002979 | 2.2753443 | 0.150575 | 0.702839901 |
| ENSMUSG00000050627 | Gpd1l | protein_coding | 0.468866298 | 5.678833813 | 2.2745142 | 0.1506448 | 0.702839901 |
| ENSMUSG00000027287 | Snap23 | protein_coding | -0.605202031 | 4.287965476 | 2.2736522 | 0.1507174 | 0.702839901 |
| ENSMUSG00000022160 | Mettl3 | protein_coding | 0.647417076 | 3.93605508 | 2.2734423 | 0.1507351 | 0.702839901 |
| ENSMUSG00000057455 | Rit2 | protein_coding | 0.322870737 | 9.683594199 | 2.2727826 | 0.1507906 | 0.702839901 |
| ENSMUSG00000019437 | Tlcd1 | protein_coding | -0.574817503 | 4.300287122 | 2.2719435 | 0.1508614 | 0.702864885 |
| ENSMUSG00000042148 | Cox10 | protein_coding | 0.405076523 | 5.255918177 | 2.2702482 | 0.1510044 | 0.703066301 |
| ENSMUSG00000037979 | Ccdc92 | protein_coding | -0.354528384 | 6.192813252 | 2.2688719 | 0.1511206 | 0.703066301 |
| ENSMUSG00000038007 | Acer2 | protein_coding | -0.511879357 | 4.297017974 | 2.2687852 | 0.1511279 | 0.703066301 |
| ENSMUSG00000031879 | Fam96b | protein_coding | -0.344765948 | 6.989638524 | 2.268333 | 0.1511661 | 0.703066301 |
| ENSMUSG00000021699 | Pde4d | protein_coding | -0.364780016 | 5.435247316 | 2.2636743 | 0.1515605 | 0.704412779 |
| ENSMUSG00000020818 | Mfsd11 | protein_coding | 0.397446865 | 5.251139658 | 2.2621184 | 0.1516925 | 0.704412779 |
| ENSMUSG00000070802 | Pnmal2 | protein_coding | 0.33747615 | 8.621589177 | 2.2612327 | 0.1517677 | 0.704412779 |
| ENSMUSG00000022816 | Fstl1 | protein_coding | -0.431292537 | 5.619074361 | 2.2612222 | 0.1517686 | 0.704412779 |
| ENSMUSG00000005716 | Pvalb | protein_coding | -0.537937236 | 7.802006038 | 2.2928313 | 0.1517832 | 0.704412779 |
| ENSMUSG00000034275 | Igsf9b | protein_coding | 0.587526148 | 4.741212726 | 2.2590458 | 0.1519848 | 0.704679123 |
| ENSMUSG00000055805 | Fmnl1 | protein_coding | -0.534249334 | 4.428708604 | 2.2584508 | 0.1520042 | 0.704679123 |
| ENSMUSG00000026452 | Syt2 | protein_coding | 0.498574334 | 4.272954533 | 2.2577442 | 0.1520643 | 0.704679123 |
| ENSMUSG00000027351 | Spred1 | protein_coding | -0.321570826 | 6.353192015 | 2.256663 | 0.1521564 | 0.704679123 |
| ENSMUSG00000034377 | Tulp4 | protein_coding | -0.356586863 | 5.511357516 | 2.2564397 | 0.1521754 | 0.704679123 |
| ENSMUSG00000027315 | Spint1 | protein_coding | 1.127910854 | 2.528566975 | 2.2553923 | 0.1522647 | 0.704679123 |
| ENSMUSG00000031060 | Rbm10 | protein_coding | 0.507738701 | 4.31472965 | 2.2549866 | 0.1522993 | 0.704679123 |
| ENSMUSG00000022404 | Slc25a17 | protein_coding | 0.302791718 | 7.268080084 | 2.2537749 | 0.1524027 | 0.704854208 |
| ENSMUSG00000036273 | Lrrk2 | protein_coding | -0.852587343 | 2.917905266 | 2.2511757 | 0.1526248 | 0.705451957 |
| ENSMUSG00000037071 | Scd1 | protein_coding | 0.455357941 | 5.692765077 | 2.2503359 | 0.1526966 | 0.705451957 |
| ENSMUSG00000024213 | Nudt3 | protein_coding | -0.299555835 | 8.135176112 | 2.2499604 | 0.1527287 | 0.705451957 |
| ENSMUSG00000022469 | Rapgef3 | protein_coding | 0.6216418 | 3.899572658 | 2.248006 | 0.1528961 | 0.705679301 |
| ENSMUSG00000079641 | Rpl39 | protein_coding | -0.447856491 | 6.23679866 | 2.247233 | 0.1529937 | 0.705679301 |
| ENSMUSG00000024664 | Fads3 | protein_coding | 0.762765423 | 4.01031925 | 2.2477926 | 0.1530281 | 0.705679301 |
| ENSMUSG00000039478 | Micu3 | protein_coding | -0.48912589 | 8.724196797 | 2.2686398 | 0.1530405 | 0.705679301 |
| ENSMUSG00000042857 | Gm9776 | bidirectional_promoter | -0.902978116 | 2.515001981 | 2.2446447 | 0.1531845 | 0.705826918 |
| ENSMUSG00000066026 | Dhrs3 | protein_coding | -0.426000501 | 5.994403766 | 2.244421 | 0.1532038 | 0.705826918 |
| ENSMUSG00000030306 | Tmtc1 | protein_coding | 0.458097741 | 4.54557722 | 2.2428897 | 0.1533354 | 0.706130839 |
| ENSMUSG00000052798 | Nup107 | protein_coding | 0.472634879 | 4.677629792 | 2.2399752 | 0.1535863 | 0.706983616 |
| ENSMUSG00000060373 | Hnrnpc | protein_coding | 0.269380953 | 7.866966741 | 2.2370579 | 0.153838 | 0.707144561 |
| ENSMUSG00000022247 | Brix1 | protein_coding | -0.295614387 | 6.667334184 | 2.2365294 | 0.1538836 | 0.707144561 |
| ENSMUSG00000033961 | Zfp446 | protein_coding | -0.665995759 | 3.355200966 | 2.2344182 | 0.1540661 | 0.707144561 |
| ENSMUSG00000015747 | Vps45 | protein_coding | 0.325958419 | 6.472370461 | 2.2341104 | 0.1540928 | 0.707144561 |
| ENSMUSG00000038738 | Shank1 | protein_coding | -0.497055361 | 5.505999891 | 2.2342349 | 0.1541133 | 0.707144561 |
| ENSMUSG00000030008 | Pradc1 | protein_coding | -0.787894805 | 3.403218735 | 2.2335458 | 0.1541416 | 0.707144561 |
| ENSMUSG00000026664 | Phyh | protein_coding | 0.25286599 | 8.053546593 | 2.2332004 | 0.1541716 | 0.707144561 |
| ENSMUSG00000024309 | Pfdn6 | protein_coding | 0.258595319 | 8.166682519 | 2.2325255 | 0.15423 | 0.707144561 |
| ENSMUSG00000031783 | Polr2c | protein_coding | 0.325574255 | 5.726039436 | 2.2321425 | 0.1542632 | 0.707144561 |
| ENSMUSG00000024491 | Rbm27 | protein_coding | -0.419387278 | 5.31306942 | 2.2319609 | 0.1542789 | 0.707144561 |
| ENSMUSG00000022639 | Dubr | lincRNA | -0.724712048 | 3.784583327 | 2.2310294 | 0.1543597 | 0.707213247 |
| ENSMUSG00000062797 | Hikeshi | protein_coding | -0.264083368 | 7.780947549 | 2.2295794 | 0.1544855 | 0.707251944 |
| ENSMUSG00000030042 | Pole4 | protein_coding | -0.374097503 | 5.649741215 | 2.2294159 | 0.1544997 | 0.707251944 |
| ENSMUSG00000027854 | Sike1 | protein_coding | -0.631353176 | 4.124325647 | 2.2283019 | 0.1545964 | 0.707264307 |
| ENSMUSG00000029073 | Cptp | protein_coding | 0.468828927 | 6.527474622 | 2.2378456 | 0.154634 | 0.707264307 |
| ENSMUSG00000031618 | Nr3c2 | protein_coding | -0.598374158 | 4.117893082 | 2.2271137 | 0.1546997 | 0.707264307 |
| ENSMUSG00000033029 | 1700088E04Rik | protein_coding | -0.577386727 | 3.823155226 | 2.2247084 | 0.154909 | 0.707466681 |
| ENSMUSG00000026755 | Arpc5l | protein_coding | -0.260089218 | 8.091165685 | 2.2245809 | 0.1549201 | 0.707466681 |
| ENSMUSG00000046814 | Gchfr | protein_coding | 0.646623247 | 3.452972226 | 2.2231458 | 0.1550452 | 0.707466681 |
| ENSMUSG00000021987 | Mtmr6 | protein_coding | 0.368942924 | 6.883620978 | 2.2215902 | 0.1551809 | 0.707466681 |
| ENSMUSG00000031590 | Frg1 | protein_coding | -0.293768836 | 7.061392723 | 2.2198308 | 0.1553345 | 0.707466681 |
| ENSMUSG00000042625 | Safb2 | protein_coding | 0.674311028 | 3.514037681 | 2.2195431 | 0.1553597 | 0.707466681 |
| ENSMUSG00000041362 | Shtn1 | protein_coding | -0.404377859 | 5.496170292 | 2.2190667 | 0.1554013 | 0.707466681 |
| ENSMUSG00000041245 | Wnk3 | protein_coding | -0.798584941 | 3.176383851 | 2.2184041 | 0.1554593 | 0.707466681 |
| ENSMUSG00000018931 | Natd1 | protein_coding | 0.943649967 | 2.723686703 | 2.218272 | 0.1554708 | 0.707466681 |
| ENSMUSG00000096370 | Gm21992 | protein_coding | -0.793898816 | 3.226450449 | 2.2182665 | 0.1554713 | 0.707466681 |
| ENSMUSG00000035725 | Prkx | protein_coding | 0.809518688 | 3.582011779 | 2.2184031 | 0.1554906 | 0.707466681 |
| ENSMUSG00000049225 | Pdp1 | protein_coding | -0.337165634 | 7.125775586 | 2.2172254 | 0.1555624 | 0.707466681 |
| ENSMUSG00000032368 | Zic1 | protein_coding | -0.354532265 | 6.105026732 | 2.2167133 | 0.1556072 | 0.707466681 |
| ENSMUSG00000078695 | Cisd3 | protein_coding | 0.281796617 | 6.448157169 | 2.2160523 | 0.1556652 | 0.707466681 |
| ENSMUSG00000026511 | Srp9 | protein_coding | 0.272001232 | 7.03976577 | 2.2147492 | 0.1557794 | 0.707466681 |
| ENSMUSG00000070306 | Ccdc153 | protein_coding | -0.470184171 | 5.855935236 | 2.2149086 | 0.1557966 | 0.707466681 |

| ensembl_gene_id | external_gene_name | gene_biotype | log2 (fold change) | log2 (counts per million) | F-value | p-value | False discovery rate |
|---|---|---|---|---|---|---|---|
| ENSMUSG00000022313 | Utp23 | protein_coding | 0.552092769 | 4.466096438 | 2.2134641 | 0.1558921 | 0.707560424 |
| ENSMUSG00000055639 | Dach1 | protein_coding | -0.55177021 | 3.902452405 | 2.2121323 | 0.1560091 | 0.707560424 |
| ENSMUSG00000041923 | Nol4 | protein_coding | -0.332808678 | 6.112688642 | 2.2120687 | 0.1560147 | 0.707560424 |
| ENSMUSG00000040606 | Kazn | protein_coding | -0.389974243 | 5.968817437 | 2.210687 | 0.1561361 | 0.707574772 |
| ENSMUSG00000032422 | Snx14 | protein_coding | 0.541236797 | 4.587299046 | 2.2105356 | 0.1561494 | 0.707574772 |
| ENSMUSG00000021752 | Kctd6 | protein_coding | 0.479925387 | 4.238287717 | 2.2077498 | 0.1563947 | 0.708354437 |
| ENSMUSG00000040536 | Necab1 | protein_coding | -0.552290277 | 5.67572211 | 2.2161218 | 0.1564532 | 0.708354437 |
| ENSMUSG00000022024 | Sugt1 | protein_coding | -0.269894086 | 7.235566152 | 2.2037123 | 0.156751 | 0.708637492 |
| ENSMUSG00000024220 | Zfp523 | protein_coding | -0.566306557 | 4.664263773 | 2.2017547 | 0.1569241 | 0.708637492 |
| ENSMUSG00000037062 | Sh3glb1 | protein_coding | -0.387756755 | 5.881245522 | 2.2016543 | 0.156933 | 0.708637492 |
| ENSMUSG00000035476 | Tab3 | protein_coding | -0.904642192 | 3.025920837 | 2.2011994 | 0.1569733 | 0.708637492 |
| ENSMUSG00000031959 | Wdr59 | protein_coding | -0.57643197 | 3.81467286 | 2.2008417 | 0.1570049 | 0.708637492 |
| ENSMUSG00000032625 | Thsd7a | protein_coding | -0.735163835 | 4.466602786 | 2.212969 | 0.1570713 | 0.708637492 |
| ENSMUSG00000036098 | Myrf | protein_coding | -0.500051487 | 4.412695229 | 2.1998105 | 0.1570963 | 0.708637492 |
| ENSMUSG00000037475 | Thoc2 | protein_coding | -0.531335239 | 4.708718785 | 2.198444 | 0.1572174 | 0.708637492 |
| ENSMUSG00000006335 | Tfpt | protein_coding | -0.342232095 | 5.883506186 | 2.1983922 | 0.157222 | 0.708637492 |
| ENSMUSG00000073557 | Ppp1r12b | protein_coding | -0.998317635 | 2.64936118 | 2.1980779 | 0.1572499 | 0.708637492 |
| ENSMUSG00000047635 | 2810006K23Rik | protein_coding | 0.431730507 | 5.227095628 | 2.1979717 | 0.1572593 | 0.708637492 |
| ENSMUSG00000031066 | Usp11 | protein_coding | 0.432235544 | 7.700349473 | 2.2092828 | 0.1573066 | 0.708637492 |
| ENSMUSG00000030243 | Recql | protein_coding | -0.652631597 | 3.821508345 | 2.1963023 | 0.1574075 | 0.708795265 |
| ENSMUSG00000033717 | Adra2a | protein_coding | 0.750232972 | 3.305469818 | 2.1936098 | 0.1576469 | 0.70957609 |
| ENSMUSG00000029821 | Dfna5 | protein_coding | -0.449405645 | 7.575944082 | 2.2054623 | 0.1578052 | 0.709799658 |
| ENSMUSG00000053553 | 3110082I17Rik | protein_coding | -0.478860025 | 4.48468091 | 2.1915697 | 0.1578286 | 0.709799658 |
| ENSMUSG00000015335 | Zdhhc12 | protein_coding | 0.581294192 | 4.070934075 | 2.1897179 | 0.1579937 | 0.710148356 |
| ENSMUSG00000000934 | Top1mt | protein_coding | 0.457970004 | 5.3546275 | 2.1891159 | 0.1580475 | 0.710148356 |
| ENSMUSG00000029223 | Uchl1 | protein_coding | 0.281222468 | 10.33781782 | 2.1880647 | 0.1581414 | 0.710148356 |
| ENSMUSG00000028149 | Rap1gds1 | protein_coding | 0.279604601 | 7.472948559 | 2.1866687 | 0.1582661 | 0.710148356 |
| ENSMUSG00000028639 | Ybx1 | protein_coding | -0.47907543 | 5.303736514 | 2.1852907 | 0.1583894 | 0.710148356 |
| ENSMUSG00000024083 | Pja2 | protein_coding | 0.338779867 | 9.105464165 | 2.185176 | 0.1583997 | 0.710148356 |
| ENSMUSG00000030726 | Pold3 | protein_coding | 0.505069497 | 5.231761122 | 2.1846148 | 0.1584499 | 0.710148356 |
| ENSMUSG00000020205 | Phlda1 | protein_coding | -0.566854046 | 4.042854638 | 2.1839499 | 0.1585095 | 0.710148356 |
| ENSMUSG00000025656 | Arhgef9 | protein_coding | -0.282378761 | 7.063057312 | 2.1836889 | 0.1585329 | 0.710148356 |
| ENSMUSG00000001663 | Gstt1 | protein_coding | 0.402822623 | 5.76642151 | 2.1825841 | 0.1586319 | 0.710148356 |
| ENSMUSG00000020640 | Itsn2 | protein_coding | -0.547860833 | 3.894198218 | 2.181243 | 0.1587522 | 0.710148356 |
| ENSMUSG00000038370 | Pcp4l1 | protein_coding | -0.408666843 | 8.590326869 | 2.1825917 | 0.1587526 | 0.710148356 |
| ENSMUSG00000027347 | Rasgrp1 | protein_coding | -0.31522551 | 6.741384254 | 2.1794724 | 0.1589113 | 0.710148356 |
| ENSMUSG00000085891 | Gm14634 | processed_transcript | 1.103905764 | 2.374153954 | 2.1788846 | 0.1589641 | 0.710148356 |
| ENSMUSG00000036949 | Slc39a12 | protein_coding | 0.562003386 | 4.149159605 | 2.1782772 | 0.1590187 | 0.710148356 |
| ENSMUSG00000022421 | Nptxr | protein_coding | 0.437808398 | 6.482046768 | 2.1776183 | 0.1591912 | 0.710148356 |
| ENSMUSG00000036620 | Mgat4b | protein_coding | -0.974744541 | 2.571468355 | 2.1760146 | 0.1592224 | 0.710148356 |
| ENSMUSG00000031226 | Pbdc1 | protein_coding | -0.293706489 | 6.387891772 | 2.1746144 | 0.1593486 | 0.710148356 |
| ENSMUSG00000031790 | Mmp15 | protein_coding | -0.559675708 | 3.531035954 | 2.1744385 | 0.1593645 | 0.710148356 |
| ENSMUSG00000031883 | Car7 | protein_coding | -0.616307115 | 3.743672053 | 2.1726684 | 0.1595242 | 0.710148356 |
| ENSMUSG00000028519 | Dab1 | protein_coding | -0.308374099 | 5.940746159 | 2.1721153 | 0.1595742 | 0.710148356 |
| ENSMUSG00000017390 | Aldoc | protein_coding | 0.29823047 | 10.98811899 | 2.1699454 | 0.1597704 | 0.710148356 |
| ENSMUSG00000029238 | Clock | protein_coding | -0.451518658 | 5.392085765 | 2.1693829 | 0.1598213 | 0.710148356 |
| ENSMUSG00000016150 | Tenm1 | protein_coding | -0.527677095 | 4.381687609 | 2.1689032 | 0.1598647 | 0.710148356 |
| ENSMUSG00000057421 | Las1l | protein_coding | 0.358750276 | 5.820323439 | 2.16888 | 0.1598668 | 0.710148356 |
| ENSMUSG00000020572 | Nampt | protein_coding | -0.38262647 | 4.97681015 | 2.1678595 | 0.1599592 | 0.710148356 |
| ENSMUSG00000006906 | Stambp | protein_coding | 0.320243353 | 6.173512635 | 2.1676868 | 0.1599749 | 0.710148356 |
| ENSMUSG00000036561 | Ppp6r2 | protein_coding | -0.543645025 | 4.045764098 | 2.1670111 | 0.1600361 | 0.710148356 |
| ENSMUSG00000028081 | Rps3a1 | protein_coding | -0.364128533 | 7.926400516 | 2.1659955 | 0.1601593 | 0.710148356 |
| ENSMUSG00000024180 | Tmem8 | protein_coding | -0.811512222 | 2.937878498 | 2.1654704 | 0.1601759 | 0.710148356 |
| ENSMUSG00000060675 | Pla2g16 | protein_coding | -0.447000439 | 6.167451262 | 2.1651556 | 0.1602356 | 0.710148356 |
| ENSMUSG00000057897 | Camk2b | protein_coding | 0.291900126 | 8.455445347 | 2.1642974 | 0.1602824 | 0.710148356 |
| ENSMUSG00000074282 | Zfp94 | protein_coding | 0.44011921 | 4.398316985 | 2.1635665 | 0.1603488 | 0.710148356 |
| ENSMUSG00000017943 | Gdap1l1 | protein_coding | 0.323489798 | 7.587768862 | 2.1631724 | 0.1603846 | 0.710148356 |
| ENSMUSG00000018040 | Rrp7a | protein_coding | 0.343897669 | 6.211056727 | 2.1631449 | 0.1603871 | 0.710148356 |
| ENSMUSG00000020827 | Mink1 | protein_coding | -0.411906496 | 4.869979965 | 2.1629459 | 0.1604052 | 0.710148356 |
| ENSMUSG00000020882 | Cacnb1 | protein_coding | -0.342227833 | 5.735647018 | 2.1628377 | 0.160415 | 0.710148356 |
| ENSMUSG00000005683 | Cs | protein_coding | 0.39269234 | 8.087842053 | 2.1628266 | 0.1604472 | 0.710148356 |
| ENSMUSG00000028232 | Tmem68 | protein_coding | -0.347716818 | 5.896256195 | 2.1621039 | 0.1604818 | 0.710148356 |
| ENSMUSG00000029388 | Eif2b1 | protein_coding | 0.273420244 | 6.75653842 | 2.1602005 | 0.160655 | 0.710512727 |
| ENSMUSG00000111977 | AC132265.1 | bidirectional_promoter | 0.820610096 | 3.175579365 | 2.1594171 | 0.1607264 | 0.710512727 |
| ENSMUSG00000086841 | 2410006H16Rik | processed_transcript | -0.32687308 | 7.585014337 | 2.1590231 | 0.1607623 | 0.710512727 |
| ENSMUSG00000025789 | St8sia2 | protein_coding | -0.970004691 | 2.608282858 | 2.1571607 | 0.1609322 | 0.710523808 |
| ENSMUSG00000013736 | Trnt1 | protein_coding | -0.417111908 | 6.412082725 | 2.1568063 | 0.1609957 | 0.710523808 |
| ENSMUSG00000027207 | Galk2 | protein_coding | 0.393232415 | 6.042863571 | 2.1559894 | 0.1610392 | 0.710523808 |
| ENSMUSG00000029101 | Rgs12 | protein_coding | 0.538635936 | 3.786857235 | 2.1550283 | 0.161127 | 0.710523808 |
| ENSMUSG00000032590 | Apeh | protein_coding | 0.512490729 | 4.439069016 | 2.1544494 | 0.16118 | 0.710523808 |
| ENSMUSG00000020774 | Aspa | protein_coding | -0.435074891 | 5.993678956 | 2.1537332 | 0.1612766 | 0.710523808 |
| ENSMUSG00000020077 | Srgn | protein_coding | -0.987129736 | 2.586494069 | 2.1521583 | 0.1613897 | 0.710523808 |
| ENSMUSG00000034675 | Dbn1 | protein_coding | -0.34899336 | 6.113673448 | 2.1520176 | 0.1614026 | 0.710523808 |
| ENSMUSG00000040451 | Sgms1 | protein_coding | -0.528632494 | 4.122359206 | 2.1520149 | 0.1614028 | 0.710523808 |

| ensembl_gene_id | external_gene_name | gene_biotype | log2 (fold change) | log2 (counts per million) | F-value | p-value | False discovery rate |
|---|---|---|---|---|---|---|---|
| ENSMUSG00000046768 | Rhoj | protein_coding | -0.97825691 | 2.680213226 | 2.1517663 | 0.1614256 | 0.710523808 |
| ENSMUSG00000027332 | Ivd | protein_coding | 0.417948186 | 6.87274443 | 2.1521631 | 0.161556 | 0.710800378 |
| ENSMUSG00000041815 | Poldip3 | protein_coding | -0.284944344 | 8.061233683 | 2.1496394 | 0.1616207 | 0.710800378 |
| ENSMUSG00000071550 | Cfap44 | protein_coding | -1.158204656 | 2.155988903 | 2.14805 | 0.1617666 | 0.711005013 |
| ENSMUSG00000036641 | Ccdc148 | protein_coding | -0.441632525 | 5.152694454 | 2.1476926 | 0.1617994 | 0.711005013 |
| ENSMUSG00000021470 | Ercc6l2 | protein_coding | -0.6450246 | 3.592632967 | 2.1469345 | 0.1618691 | 0.711020667 |
| ENSMUSG00000049680 | Urgcp | protein_coding | -0.467666759 | 4.460458606 | 2.144243 | 0.1621168 | 0.711083357 |
| ENSMUSG00000040464 | Gtpbp10 | protein_coding | -0.898332703 | 2.572768242 | 2.1436309 | 0.1621732 | 0.711083357 |
| ENSMUSG00000024966 | Stip1 | protein_coding | 0.363080708 | 8.728754636 | 2.143578 | 0.1621781 | 0.711083357 |
| ENSMUSG00000055409 | Nell1 | protein_coding | 0.314156538 | 5.860489754 | 2.1433792 | 0.1621964 | 0.711083357 |
| ENSMUSG00000031592 | Pcm1 | protein_coding | -0.447728979 | 5.501152417 | 2.1431881 | 0.162214 | 0.711083357 |
| ENSMUSG00000045007 | Tubg2 | protein_coding | 0.385319099 | 6.830641441 | 2.1399592 | 0.1625431 | 0.711497335 |
| ENSMUSG00000064202 | 4430402I18Rik | protein_coding | -0.818093983 | 3.057452635 | 2.1390414 | 0.1625969 | 0.711497335 |
| ENSMUSG00000000804 | Usp32 | protein_coding | -0.343646478 | 5.593076024 | 2.1387984 | 0.1626193 | 0.711497335 |
| ENSMUSG00000025475 | Adgra1 | protein_coding | 0.388358283 | 6.688221718 | 2.1382234 | 0.1627036 | 0.711497335 |
| ENSMUSG00000033021 | Gmppa | protein_coding | 0.446013328 | 5.686105646 | 2.1374922 | 0.1627402 | 0.711497335 |
| ENSMUSG00000062110 | Scfd2 | protein_coding | -0.602901053 | 4.006111283 | 2.1365608 | 0.1628264 | 0.711497335 |
| ENSMUSG00000021759 | Plpp1 | protein_coding | 0.422335552 | 4.696786493 | 2.1361406 | 0.1628653 | 0.711497335 |
| ENSMUSG00000062944 | 9130023H24Rik | protein_coding | -0.660430586 | 3.946583191 | 2.1353999 | 0.162934 | 0.711497335 |
| ENSMUSG00000024413 | Npc1 | protein_coding | 0.524882538 | 4.208555159 | 2.1348714 | 0.162983 | 0.711497335 |
| ENSMUSG00000041571 | Selenow | protein_coding | -0.364326666 | 12.37679339 | 2.1340713 | 0.1630572 | 0.711497335 |
| ENSMUSG00000055897 | Ppp4r1l-ps | transcribed_unproces | -0.636284414 | 3.477519279 | 2.133911 | 0.1630721 | 0.711497335 |
| ENSMUSG00000099689 | Zfp383 | protein_coding | -0.684224908 | 3.063736148 | 2.1323794 | 0.1632142 | 0.711497335 |
| ENSMUSG00000025037 | Maoa | protein_coding | 0.396949839 | 5.56564449 | 2.1323747 | 0.1632147 | 0.711497335 |
| ENSMUSG00000052102 | Gnpda1 | protein_coding | 0.454790851 | 5.444722279 | 2.1315427 | 0.163292 | 0.711497335 |
| ENSMUSG00000053581 | Zfand2a | protein_coding | 0.439066104 | 6.114160358 | 2.1309542 | 0.1633778 | 0.711497335 |
| ENSMUSG00000066441 | Rdh11 | protein_coding | 0.449421013 | 4.662928458 | 2.1304231 | 0.1633961 | 0.711497335 |
| ENSMUSG00000069049 | Eif2s3y | protein_coding | -0.484937693 | 5.038632738 | 2.1285737 | 0.1635682 | 0.711497335 |
| ENSMUSG00000048578 | Mlec | protein_coding | 0.36527381 | 5.514728006 | 2.1274611 | 0.1636719 | 0.711497335 |
| ENSMUSG00000080727 | C920021L13Rik | antisense_RNA | -0.877242996 | 2.533672648 | 2.1272707 | 0.1636896 | 0.711497335 |
| ENSMUSG00000001260 | Gabrg1 | protein_coding | -0.332528436 | 6.075152125 | 2.1271213 | 0.1637036 | 0.711497335 |
| ENSMUSG00000041836 | Ptpre | protein_coding | 0.498742546 | 4.376135082 | 2.1262899 | 0.1637811 | 0.711497335 |
| ENSMUSG00000024580 | Grpel2 | protein_coding | -0.383288097 | 5.73925752 | 2.1260594 | 0.1638026 | 0.711497335 |
| ENSMUSG00000057816 | 1700007G11Rik | protein_coding | -0.552597589 | 3.541309499 | 2.1257625 | 0.1638303 | 0.711497335 |
| ENSMUSG00000025089 | Gfra1 | protein_coding | 0.360820078 | 5.701463054 | 2.1227435 | 0.1641124 | 0.711549632 |
| ENSMUSG00000027465 | Tbc1d20 | protein_coding | -0.425301662 | 5.170926343 | 2.1222654 | 0.1641571 | 0.711549632 |
| ENSMUSG00000003355 | Fkbp11 | protein_coding | 0.736017043 | 3.242435879 | 2.1219746 | 0.1641843 | 0.711549632 |
| ENSMUSG00000001240 | Ramp2 | protein_coding | 0.958833751 | 2.70273353 | 2.1217602 | 0.1642044 | 0.711549632 |
| ENSMUSG00000027574 | Nkain4 | protein_coding | -0.576714441 | 6.610725985 | 2.1514017 | 0.1644741 | 0.711549632 |
| ENSMUSG00000079038 | D130040H23Rik | protein_coding | -1.055207222 | 2.546525431 | 2.1183973 | 0.1645194 | 0.711549632 |
| ENSMUSG00000043154 | Ppp2r3a | protein_coding | -0.46946116 | 4.875235619 | 2.116544 | 0.1646934 | 0.711549632 |
| ENSMUSG00000049950 | Rpp38 | protein_coding | -0.362139272 | 5.107521382 | 2.1165141 | 0.1646962 | 0.711549632 |
| ENSMUSG00000037608 | Bclaf1 | protein_coding | -0.325646068 | 7.408676833 | 2.1159103 | 0.164753 | 0.711549632 |
| ENSMUSG00000003949 | Hlf | protein_coding | -0.275132113 | 6.827037216 | 2.1158214 | 0.1647613 | 0.711549632 |
| ENSMUSG00000037098 | Rab11fip3 | protein_coding | -0.485337763 | 4.664841523 | 2.1152089 | 0.1648189 | 0.711549632 |
| ENSMUSG00000032178 | Ilf3 | protein_coding | 0.517452538 | 7.480326421 | 2.1430836 | 0.164872 | 0.711549632 |
| ENSMUSG00000102869 | 2900097C17Rik | lincRNA | 0.353720295 | 7.970443406 | 2.114049 | 0.164928 | 0.711549632 |
| ENSMUSG00000026317 | Cln8 | protein_coding | -0.465914788 | 5.354004225 | 2.1133449 | 0.1649943 | 0.711549632 |
| ENSMUSG00000015120 | Ube2i | protein_coding | 0.609956445 | 3.638052954 | 2.112871 | 0.1650389 | 0.711549632 |
| ENSMUSG00000024078 | Ttc27 | protein_coding | 0.738158933 | 4.069331207 | 2.1140368 | 0.1650741 | 0.711549632 |
| ENSMUSG00000024137 | E4f1 | protein_coding | 0.75500534 | 3.098904098 | 2.1108118 | 0.165233 | 0.711549632 |
| ENSMUSG00000022724 | Riox2 | protein_coding | 0.715375083 | 3.831922771 | 2.1105773 | 0.1652861 | 0.711549632 |
| ENSMUSG00000040097 | Flywch1 | protein_coding | 0.870473168 | 7.261020424 | 2.233312 | 0.1652983 | 0.711549632 |
| ENSMUSG00000048833 | Slc39a9 | protein_coding | 0.394929092 | 4.910425204 | 2.1099423 | 0.165315 | 0.711549632 |
| ENSMUSG00000055447 | Cd47 | protein_coding | -0.289419712 | 7.192665503 | 2.1098926 | 0.1653197 | 0.711549632 |
| ENSMUSG00000003541 | Ier3 | protein_coding | 0.399975089 | 4.830210852 | 2.1097339 | 0.1653347 | 0.711549632 |
| ENSMUSG00000064329 | Scn1a | protein_coding | 0.342780927 | 6.450607601 | 2.1091016 | 0.1653943 | 0.711549632 |
| ENSMUSG00000025027 | Xpnpep1 | protein_coding | 0.376431417 | 6.097822581 | 2.1074243 | 0.1655528 | 0.711549632 |
| ENSMUSG00000048895 | Cdk5r1 | protein_coding | -0.291241114 | 6.889457101 | 2.1071356 | 0.1655801 | 0.711549632 |
| ENSMUSG00000044018 | Mrpl50 | protein_coding | -0.298746019 | 7.294997489 | 2.1062559 | 0.1656633 | 0.711549632 |
| ENSMUSG00000026618 | Iars2 | protein_coding | 0.664809834 | 3.856479725 | 2.10489 | 0.1657926 | 0.711549632 |
| ENSMUSG00000030007 | Cct7 | protein_coding | 0.344214333 | 9.448790663 | 2.1041298 | 0.1658646 | 0.711549632 |
| ENSMUSG00000005732 | Ranbp1 | protein_coding | 0.300195785 | 7.170496367 | 2.1028106 | 0.1659897 | 0.711549632 |
| ENSMUSG00000025867 | Cplx2 | protein_coding | -0.285655563 | 8.226536737 | 2.1027204 | 0.1659983 | 0.711549632 |
| ENSMUSG00000020642 | Rnf144a | protein_coding | -0.642880344 | 3.501786462 | 2.1025462 | 0.1660148 | 0.711549632 |
| ENSMUSG00000019173 | Rab5c | protein_coding | -0.447675322 | 5.459684264 | 2.1025004 | 0.1660191 | 0.711549632 |
| ENSMUSG00000043831 | Lysmd4 | protein_coding | 0.283890326 | 6.736433411 | 2.1024276 | 0.166026 | 0.711549632 |
| ENSMUSG00000068876 | Cgn | protein_coding | 0.684357738 | 3.525450988 | 2.1003446 | 0.1662238 | 0.712113415 |
| ENSMUSG00000070319 | Eif3g | protein_coding | 0.257157579 | 7.844693706 | 2.0981713 | 0.1664305 | 0.712517157 |
| ENSMUSG00000026687 | Aldh9a1 | protein_coding | 0.362107359 | 6.127759362 | 2.0979599 | 0.1664506 | 0.712517157 |
| ENSMUSG00000037166 | Ppp1r14a | protein_coding | -0.67271511 | 3.985494082 | 2.0968466 | 0.1665876 | 0.712819775 |
| ENSMUSG00000020250 | Txnrd1 | protein_coding | 0.367337644 | 5.659791275 | 2.0935241 | 0.1668734 | 0.71375895 |
| ENSMUSG00000002741 | Ykt6 | protein_coding | 0.442726226 | 5.749168837 | 2.0927329 | 0.166949 | 0.713798191 |
| ENSMUSG00000000439 | Mkrn2 | protein_coding | 0.371459912 | 5.914693123 | 2.0902042 | 0.1671908 | 0.714423193 |

| ensembl_gene_id | external_gene_name | gene_biotype | log2 (fold change) | log2 (counts per million) | F-value | p-value | False discovery rate |
|---|---|---|---|---|---|---|---|
| ENSMUSG00000028121 | Bcar3 | protein_coding | 0.592011508 | 4.360199005 | 2.0898147 | 0.167228 | 0.714423193 |
| ENSMUSG00000042109 | Csdc2 | protein_coding | -0.310026638 | 8.009879572 | 2.0887805 | 0.1673271 | 0.714562388 |
| ENSMUSG00000005103 | Wdr1 | protein_coding | 0.429985117 | 7.835162807 | 2.0982869 | 0.1676197 | 0.715527857 |
| ENSMUSG00000037706 | Cd81 | protein_coding | 0.292553608 | 9.865788007 | 2.0848822 | 0.167701 | 0.715591023 |
| ENSMUSG00000029287 | Tgfbr3 | protein_coding | -1.140181758 | 2.925349882 | 2.0846258 | 0.1678382 | 0.715703404 |
| ENSMUSG00000040591 | 1110051M20Rik | protein_coding | 0.32586577 | 8.374392058 | 2.0821332 | 0.1679654 | 0.715703404 |
| ENSMUSG00000024131 | Slc3a1 | protein_coding | 0.293282548 | 7.649142147 | 2.0819658 | 0.1679815 | 0.715703404 |
| ENSMUSG00000027531 | Impa1 | protein_coding | 0.249174916 | 6.892489951 | 2.0818398 | 0.1679936 | 0.715703404 |
| ENSMUSG00000020629 | Adi1 | protein_coding | -0.355951712 | 6.30891027 | 2.0808923 | 0.1680849 | 0.715715057 |
| ENSMUSG00000020611 | Gna13 | protein_coding | 0.396748406 | 5.400920887 | 2.0801495 | 0.1681564 | 0.715715057 |
| ENSMUSG00000036815 | Dpp10 | protein_coding | -0.355625839 | 6.009901135 | 2.0790657 | 0.1682609 | 0.715715057 |
| ENSMUSG00000004895 | Prcc | protein_coding | 0.538993544 | 3.576970569 | 2.0790486 | 0.1682626 | 0.715715057 |
| ENSMUSG00000027828 | Ssr3 | protein_coding | -0.242885627 | 7.518890899 | 2.075935 | 0.1685633 | 0.71657387 |
| ENSMUSG00000003458 | Ncstn | protein_coding | 0.414409283 | 5.951255116 | 2.0752606 | 0.1686285 | 0.71657387 |
| ENSMUSG00000017831 | Rab5a | protein_coding | -0.525554275 | 5.016698875 | 2.0742426 | 0.168727 | 0.71657387 |
| ENSMUSG00000029455 | Aldh2 | protein_coding | 0.402106869 | 5.05775773 | 2.0742005 | 0.1687311 | 0.71657387 |
| ENSMUSG00000031928 | Mre11a | protein_coding | -0.684448448 | 3.140148877 | 2.0726516 | 0.1688811 | 0.716861161 |
| ENSMUSG00000029405 | G3bp2 | protein_coding | 0.380278459 | 7.275277119 | 2.0719618 | 0.1689788 | 0.716861161 |
| ENSMUSG00000026155 | Smap1 | protein_coding | 0.613223699 | 3.407709961 | 2.0709879 | 0.1690424 | 0.716861161 |
| ENSMUSG00000024082 | Ndufaf7 | protein_coding | 0.278150016 | 6.749561846 | 2.0702833 | 0.1691108 | 0.716861161 |
| ENSMUSG00000035382 | Pcsk7 | protein_coding | -0.725824351 | 3.056818934 | 2.070064 | 0.169132 | 0.716861161 |
| ENSMUSG00000053080 | 2700081O15Rik | protein_coding | 0.852589724 | 2.671389807 | 2.0679732 | 0.1693352 | 0.717214921 |
| ENSMUSG00000046598 | Bdh1 | protein_coding | 0.388878627 | 5.806172135 | 2.067832 | 0.1693489 | 0.717214921 |
| ENSMUSG00000052151 | Plpp2 | protein_coding | 0.458566907 | 4.438587461 | 2.0671424 | 0.169416 | 0.717216448 |
| ENSMUSG00000039230 | Tbcd | protein_coding | 0.437372759 | 4.638417579 | 2.066364 | 0.1694917 | 0.717249533 |
| ENSMUSG00000038976 | Ppp1r9b | protein_coding | -0.38241628 | 7.021831273 | 2.0652734 | 0.1696288 | 0.717249533 |
| ENSMUSG00000062794 | Zfp599 | protein_coding | -0.600094742 | 13.5488801 | 2.0899607 | 0.1696551 | 0.717249533 |
| ENSMUSG00000009894 | Snap47 | protein_coding | 0.383920755 | 10.21676749 | 2.0643877 | 0.1697151 | 0.717249533 |
| ENSMUSG00000032002 | Dcun1d5 | protein_coding | 0.320855759 | 5.856712684 | 2.0636382 | 0.1697573 | 0.717249533 |
| ENSMUSG00000032757 | Bet1 | protein_coding | -0.272863975 | 6.576747972 | 2.0619899 | 0.1699181 | 0.717439843 |
| ENSMUSG00000031841 | Cdh13 | protein_coding | 0.250255918 | 7.328179184 | 2.0609199 | 0.1700226 | 0.717439843 |
| ENSMUSG00000098332 | 2310009A05Rik | processed_transcript | -0.428654222 | 5.400427499 | 2.0601369 | 0.1700992 | 0.717439843 |
| ENSMUSG00000026110 | Mgat4a | protein_coding | -0.490494867 | 4.364640138 | 2.0599109 | 0.1701213 | 0.717439843 |
| ENSMUSG00000001127 | Araf | protein_coding | 0.263887219 | 9.716486823 | 2.059084 | 0.1702022 | 0.717439843 |
| ENSMUSG00000060981 | Hist1h4h | protein_coding | -1.085702267 | 2.313796193 | 2.057697 | 0.1703379 | 0.717439843 |
| ENSMUSG00000058503 | Fam133b | protein_coding | -0.3274808 | 6.062374776 | 2.057287 | 0.1703781 | 0.717439843 |
| ENSMUSG00000006673 | Qrich1 | protein_coding | 0.320337043 | 5.913338784 | 2.0566397 | 0.1704415 | 0.717439843 |
| ENSMUSG00000071860 | 2900055J20Rik | protein_coding | -0.580343363 | 3.835069793 | 2.0554745 | 0.1705558 | 0.717439843 |
| ENSMUSG00000030498 | Gas2 | protein_coding | -0.450906027 | 4.081728623 | 2.0548545 | 0.1706166 | 0.717439843 |
| ENSMUSG00000026556 | Vangl2 | protein_coding | 0.55617224 | 3.631927552 | 2.0545637 | 0.1706452 | 0.717439843 |
| ENSMUSG00000026116 | Tmem131 | protein_coding | -0.524627751 | 3.967843613 | 2.0544354 | 0.1706578 | 0.717439843 |
| ENSMUSG00000011257 | Pabpc4 | protein_coding | -0.656244453 | 3.198286458 | 2.0543138 | 0.1706697 | 0.717439843 |
| ENSMUSG00000044224 | Dnajc21 | protein_coding | -0.470370213 | 4.343740295 | 2.0531922 | 0.1707798 | 0.717622379 |
| ENSMUSG00000031446 | Cul4a | protein_coding | 0.562816409 | 4.612522061 | 2.0521205 | 0.1708852 | 0.717784545 |
| ENSMUSG00000017667 | Zfp334 | protein_coding | 0.422082807 | 4.632684367 | 2.0493741 | 0.1711555 | 0.718563562 |
| ENSMUSG00000046994 | Mars2 | protein_coding | -0.778317707 | 3.103361942 | 2.0479676 | 0.1712942 | 0.718563562 |
| ENSMUSG00000063015 | Ccni | protein_coding | -0.355379598 | 6.356034087 | 2.0477024 | 0.1713203 | 0.718563562 |
| ENSMUSG00000023328 | Ache | protein_coding | 0.305712295 | 6.236611995 | 2.0463443 | 0.1714543 | 0.718563562 |
| ENSMUSG00000097416 | Gm26670 | antisense_RNA | -0.997388127 | 2.352260028 | 2.04628 | 0.1714607 | 0.718563562 |
| ENSMUSG00000073427 | Gm4924 | protein_coding | 0.626351661 | 3.329425983 | 2.0451003 | 0.1715772 | 0.718563562 |
| ENSMUSG00000035505 | Cox18 | protein_coding | -0.57824557 | 3.559834666 | 2.0446607 | 0.1716207 | 0.718563562 |
| ENSMUSG00000002504 | Slc9a3r2 | protein_coding | -0.415389772 | 4.740477789 | 2.0439313 | 0.1716928 | 0.718563562 |
| ENSMUSG00000091537 | Tma7 | protein_coding | -0.269101611 | 7.239049652 | 2.0435453 | 0.171731 | 0.718563562 |
| ENSMUSG00000036111 | Lmo1 | protein_coding | 0.412707278 | 4.430360891 | 2.0426654 | 0.1718181 | 0.718563562 |
| ENSMUSG00000049755 | Zfp672 | protein_coding | 0.431436636 | 5.688238535 | 2.0423385 | 0.1718504 | 0.718563562 |
| ENSMUSG00000023990 | Tfeb | protein_coding | -1.230955506 | 2.436470241 | 2.0421153 | 0.1718725 | 0.718563562 |
| ENSMUSG00000026650 | Meig1 | protein_coding | -0.476799034 | 4.534744567 | 2.0386298 | 0.1722182 | 0.71884015 |
| ENSMUSG00000001098 | Kctd10 | protein_coding | 0.806725006 | 3.615844524 | 2.0387667 | 0.1722354 | 0.71884015 |
| ENSMUSG00000041650 | Pcca | protein_coding | -0.528211948 | 4.350311959 | 2.0383234 | 0.1722486 | 0.71884015 |
| ENSMUSG00000026797 | Stxbp1 | protein_coding | 0.417433467 | 9.183460742 | 2.0446181 | 0.1722499 | 0.71884015 |
| ENSMUSG00000048351 | Coa7 | protein_coding | 0.331714109 | 5.885505597 | 2.0376055 | 0.1723199 | 0.71884015 |
| ENSMUSG00000029838 | Ptn | protein_coding | -0.379416916 | 8.78621346 | 2.0362403 | 0.1724864 | 0.71884015 |
| ENSMUSG00000028106 | Rprd2 | protein_coding | -0.42781038 | 4.85400875 | 2.034664 | 0.1726125 | 0.71884015 |
| ENSMUSG00000029056 | Pank4 | protein_coding | 0.425935912 | 4.982483717 | 2.0335743 | 0.172721 | 0.71884015 |
| ENSMUSG00000020333 | Acsl6 | protein_coding | 0.415349281 | 5.723656483 | 2.0333154 | 0.1727468 | 0.71884015 |
| ENSMUSG00000069844 | Sco1 | protein_coding | 0.954379645 | 3.304374672 | 2.0365516 | 0.1727952 | 0.71884015 |
| ENSMUSG00000034317 | Trim59 | protein_coding | -0.709995472 | 4.038572998 | 2.0326081 | 0.1728481 | 0.71884015 |
| ENSMUSG00000029536 | Gatc | protein_coding | -0.303698742 | 5.99262506 | 2.031431 | 0.1729348 | 0.71884015 |
| ENSMUSG00000042078 | Svop | protein_coding | 0.266244049 | 7.24270854 | 2.030474 | 0.1730303 | 0.71884015 |
| ENSMUSG00000014039 | Prdm15 | protein_coding | 1.046417618 | 2.37350164 | 2.0282746 | 0.1732501 | 0.71884015 |
| ENSMUSG00000059921 | Unc5c | protein_coding | -0.588803359 | 4.057897879 | 2.0275402 | 0.1733236 | 0.71884015 |
| ENSMUSG00000015879 | Fam184b | protein_coding | 0.666604024 | 3.132677739 | 2.0271653 | 0.1733611 | 0.71884015 |
| ENSMUSG00000030282 | Cmas | protein_coding | 0.261477483 | 6.74470459 | 2.0268605 | 0.1733916 | 0.71884015 |
| ENSMUSG00000056938 | Acbd4 | protein_coding | -0.492043159 | 5.565065564 | 2.0271613 | 0.1733923 | 0.71884015 |

| ensembl_gene_id | external_gene_name | gene_biotype | log2 (fold change) | log2 (counts per million) | F-value | p-value | False discovery rate |
|---|---|---|---|---|---|---|---|
| ENSMUSG00000027748 | Trpc4 | protein_coding | -0.460354651 | 4.408237184 | 2.0252123 | 0.1735568 | 0.71884015 |
| ENSMUSG00000041556 | Fbxo2 | protein_coding | -0.460534115 | 4.92481782 | 2.0251624 | 0.1735618 | 0.71884015 |
| ENSMUSG00000004040 | Stat3 | protein_coding | 0.602874953 | 4.838964581 | 2.0260994 | 0.1735798 | 0.71884015 |
| ENSMUSG00000031026 | Trim66 | protein_coding | 0.572131338 | 3.792633113 | 2.0241536 | 0.1736629 | 0.71884015 |
| ENSMUSG00000051451 | Crebzf | protein_coding | -0.916026203 | 2.589739195 | 2.0239962 | 0.1736787 | 0.71884015 |
| ENSMUSG00000030380 | Mzf1 | protein_coding | -1.106073638 | 2.223890863 | 2.023665 | 0.173712 | 0.71884015 |
| ENSMUSG00000020520 | Galnt10 | protein_coding | -0.609244498 | 3.456535794 | 2.0236488 | 0.1737136 | 0.71884015 |
| ENSMUSG00000024816 | Frmd8 | protein_coding | -0.518692258 | 5.575737944 | 2.0247075 | 0.1737513 | 0.71884015 |
| ENSMUSG00000041670 | Rims1 | protein_coding | -0.313895429 | 6.013708503 | 2.02229 | 0.17385 | 0.71884015 |
| ENSMUSG00000030534 | Vps33b | protein_coding | 0.712295374 | 3.958415376 | 2.0225776 | 0.1738519 | 0.71884015 |
| ENSMUSG00000001576 | Ergic1 | protein_coding | -0.387862814 | 5.837601065 | 2.0214936 | 0.1739301 | 0.71884015 |
| ENSMUSG00000004936 | Map2k1 | protein_coding | 0.312353243 | 7.707290744 | 2.0213528 | 0.1739442 | 0.71884015 |
| ENSMUSG00000038456 | Dennd2a | protein_coding | -0.615842865 | 3.903286235 | 2.0202528 | 0.1740548 | 0.719020964 |
| ENSMUSG00000051671 | Coa6 | protein_coding | -0.314859767 | 6.509257988 | 2.0194372 | 0.1741369 | 0.719083776 |
| ENSMUSG00000028798 | Eif3i | protein_coding | 0.262425286 | 9.126720684 | 2.0177991 | 0.1743019 | 0.719488817 |
| ENSMUSG00000038855 | Itpkb | protein_coding | 0.539278891 | 4.182160553 | 2.0156247 | 0.1745212 | 0.720117663 |
| ENSMUSG00000108532 | Gm32647 | lincRNA | -1.11706369 | 2.491066607 | 2.0145364 | 0.1746311 | 0.720294755 |
| ENSMUSG00000035726 | Supt16 | protein_coding | 0.412276366 | 5.624381339 | 2.0093094 | 0.1751601 | 0.722199779 |
| ENSMUSG00000020623 | Map2k6 | protein_coding | 0.564653225 | 4.021855182 | 2.0084823 | 0.175244 | 0.722268751 |
| ENSMUSG00000015484 | Fam163a | protein_coding | -0.813665022 | 3.346538733 | 2.007569 | 0.1753367 | 0.722373896 |
| ENSMUSG00000073565 | Prr16 | protein_coding | -1.109883404 | 2.509511292 | 2.0066076 | 0.1754344 | 0.722499359 |
| ENSMUSG00000034042 | Gpbp1l1 | protein_coding | 0.574922865 | 3.696995942 | 2.0035294 | 0.1757475 | 0.722876339 |
| ENSMUSG00000074030 | Exoc8 | protein_coding | -0.422792401 | 4.648191444 | 2.0034992 | 0.1757505 | 0.722876339 |
| ENSMUSG00000040037 | Negr1 | protein_coding | -0.370285453 | 6.345821179 | 2.0034856 | 0.1757519 | 0.722876339 |
| ENSMUSG00000040720 | Virma | protein_coding | 0.484136767 | 4.250629407 | 2.0030645 | 0.1757948 | 0.722876339 |
| ENSMUSG00000027660 | Skil | protein_coding | -0.297094199 | 6.513864919 | 2.0015546 | 0.1759487 | 0.722975439 |
| ENSMUSG00000034636 | Zyg11b | protein_coding | 0.296509596 | 5.867173384 | 2.0014395 | 0.1759604 | 0.722975439 |
| ENSMUSG00000021245 | Mlh3 | protein_coding | 0.62797637 | 3.920570449 | 2.0006911 | 0.1760368 | 0.722975439 |
| ENSMUSG00000058571 | Gpc6 | protein_coding | -0.651762277 | 3.164254474 | 1.9986289 | 0.1762473 | 0.722975439 |
| ENSMUSG00000056214 | Pard6g | protein_coding | 0.737021106 | 3.154507109 | 1.9985884 | 0.1762515 | 0.722975439 |
| ENSMUSG00000034432 | Cops8 | protein_coding | 0.279825787 | 8.788019941 | 1.9980937 | 0.176302 | 0.722975439 |
| ENSMUSG00000054715 | Zscan22 | protein_coding | 0.621697962 | 4.282864331 | 1.9983497 | 0.1763066 | 0.722975439 |
| ENSMUSG00000028059 | Arhgef2 | protein_coding | -0.48023347 | 4.502508929 | 1.9975583 | 0.1763568 | 0.722975439 |
| ENSMUSG00000053024 | Cntn2 | protein_coding | 0.395358899 | 5.471520605 | 1.9961795 | 0.1764979 | 0.723149553 |
| ENSMUSG00000015889 | Lta4h | protein_coding | 0.324145848 | 6.221510192 | 1.9952776 | 0.1765902 | 0.723149553 |
| ENSMUSG00000030655 | Smg1 | protein_coding | -0.565603844 | 3.940697427 | 1.9951722 | 0.176601 | 0.723149553 |
| ENSMUSG00000028436 | Dcaf12 | protein_coding | 0.420325193 | 4.935003927 | 1.9938214 | 0.1767395 | 0.723405176 |
| ENSMUSG00000111088 | AC122428.3 | TEC | -0.8424816 | 3.354417989 | 1.99355 | 0.176798 | 0.723405176 |
| ENSMUSG00000025190 | Got1 | protein_coding | 0.367439489 | 9.60610963 | 1.9927655 | 0.1768785 | 0.723459201 |
| ENSMUSG00000020486 | 4-Sep | protein_coding | 0.294257993 | 8.938024311 | 1.9896983 | 0.1771629 | 0.724112933 |
| ENSMUSG00000035203 | Epn1 | protein_coding | -0.342778253 | 6.099293271 | 1.9896001 | 0.177173 | 0.724112933 |
| ENSMUSG00000069184 | Zfp72 | protein_coding | -0.723432968 | 3.149835557 | 1.9884343 | 0.1772929 | 0.724327912 |
| ENSMUSG00000035900 | Gramd4 | protein_coding | 0.61064939 | 3.297468782 | 1.9876942 | 0.1773692 | 0.724364036 |
| ENSMUSG00000041729 | Coro2b | protein_coding | 0.354740082 | 5.548722249 | 1.9858557 | 0.1775586 | 0.724708724 |
| ENSMUSG00000024120 | Lrpprc | protein_coding | 0.330046894 | 5.719920287 | 1.9855677 | 0.1775883 | 0.724708724 |
| ENSMUSG00000051695 | Pcbp1 | protein_coding | 0.257323995 | 6.761842645 | 1.9819193 | 0.1779652 | 0.72523861 |
| ENSMUSG00000028826 | Tmem57 | protein_coding | 0.425204672 | 5.475240805 | 1.9814934 | 0.1780093 | 0.72523861 |
| ENSMUSG00000027552 | E2f5 | protein_coding | -0.681384029 | 2.977907243 | 1.9800944 | 0.1781541 | 0.72523861 |
| ENSMUSG00000029125 | Stx18 | protein_coding | 0.371334087 | 5.479894927 | 1.9792814 | 0.1782383 | 0.72523861 |
| ENSMUSG00000035964 | Tmem59l | protein_coding | 0.284471679 | 9.53239765 | 1.9792435 | 0.1782422 | 0.72523861 |
| ENSMUSG00000032246 | Calml4 | protein_coding | -0.723420301 | 3.87623688 | 1.9795105 | 0.1782452 | 0.72523861 |
| ENSMUSG00000034473 | Sec22a | protein_coding | 0.297853566 | 6.196040873 | 1.978325 | 0.1783374 | 0.72523861 |
| ENSMUSG00000030888 | Rrp8 | protein_coding | -0.531835085 | 4.165826031 | 1.9781998 | 0.1783504 | 0.72523861 |
| ENSMUSG00000021282 | Eif5 | protein_coding | -0.261617556 | 6.971456499 | 1.9779355 | 0.1783778 | 0.72523861 |
| ENSMUSG00000040043 | Rbms2 | protein_coding | -0.578415287 | 3.612148986 | 1.9773449 | 0.1784391 | 0.72523861 |
| ENSMUSG00000010755 | Cars | protein_coding | 0.550815616 | 5.871903777 | 1.9918986 | 0.1784665 | 0.72523861 |
| ENSMUSG00000033684 | Qsox1 | protein_coding | 0.475046483 | 5.093274121 | 1.9755134 | 0.1786293 | 0.72523861 |
| ENSMUSG00000015341 | Golga7 | protein_coding | -0.320439489 | 5.737273515 | 1.9737701 | 0.1788106 | 0.72523861 |
| ENSMUSG00000032359 | Ctsh | protein_coding | -0.670350109 | 3.698743639 | 1.972457 | 0.1789473 | 0.72523861 |
| ENSMUSG00000051391 | Ywhag | protein_coding | 0.349551313 | 9.147746212 | 1.971485 | 0.1790485 | 0.72523861 |
| ENSMUSG00000044288 | Cnr1 | protein_coding | -0.329940494 | 6.287251988 | 1.971409 | 0.1790564 | 0.72523861 |
| ENSMUSG00000047804 | Akap10 | protein_coding | 0.901426224 | 2.627325968 | 1.9706228 | 0.1791384 | 0.72523861 |
| ENSMUSG00000025579 | Gaa | protein_coding | 0.281669954 | 9.350906561 | 1.9705778 | 0.1791431 | 0.72523861 |
| ENSMUSG00000002055 | Spag5 | protein_coding | -1.247867028 | 2.194848976 | 1.9704429 | 0.1791572 | 0.72523861 |
| ENSMUSG00000018012 | Rac3 | protein_coding | 0.484945572 | 4.290277025 | 1.9702532 | 0.179177 | 0.72523861 |
| ENSMUSG00000048118 | Arid4a | protein_coding | -0.350234821 | 6.112381513 | 1.9701227 | 0.1791906 | 0.72523861 |
| ENSMUSG00000048047 | Zbtb33 | protein_coding | -0.854371446 | 2.590932394 | 1.9690823 | 0.1792991 | 0.72523861 |
| ENSMUSG00000060860 | Ube2s | protein_coding | -0.863502932 | 2.896766933 | 1.9689026 | 0.1793179 | 0.72523861 |
| ENSMUSG00000022023 | Wbp4 | protein_coding | -0.255766878 | 6.75804316 | 1.9680841 | 0.1794034 | 0.72523861 |
| ENSMUSG00000031344 | Gabrq | protein_coding | 0.579208642 | 4.170744822 | 1.9676926 | 0.1794443 | 0.72523861 |
| ENSMUSG00000032288 | Imp3 | protein_coding | -0.484889237 | 4.824134145 | 1.9674298 | 0.1794718 | 0.72523861 |
| ENSMUSG00000071553 | Cpa2 | protein_coding | -0.445667165 | 4.811288757 | 1.9652127 | 0.1797037 | 0.725903048 |
| ENSMUSG00000020328 | Nudcd2 | protein_coding | -0.348570996 | 5.547238243 | 1.9641143 | 0.1798187 | 0.726094964 |
| ENSMUSG00000056131 | Pgm3 | protein_coding | 0.655610632 | 4.301884487 | 1.9625037 | 0.1800182 | 0.726375731 |

| ensembl_gene_id | external_gene_name | gene_biotype | log2 (fold change) | log2 (counts per million) | F-value | p-value | False discovery rate |
|---|---|---|---|---|---|---|---|
| ENSMUSG00000029219 | Slc10a4 | protein_coding | -0.586532387 | 3.807945829 | 1.9620446 | 0.1800358 | 0.726375731 |
| ENSMUSG00000036817 | Sun1 | protein_coding | 0.516914523 | 5.080491014 | 1.9611991 | 0.1801551 | 0.726375731 |
| ENSMUSG00000031144 | Syp | protein_coding | 0.445651646 | 6.858672657 | 1.9722229 | 0.1801585 | 0.726375731 |
| ENSMUSG00000015599 | Ttbk1 | protein_coding | -0.676853645 | 3.213615904 | 1.9586198 | 0.1803956 | 0.726395806 |
| ENSMUSG00000047284 | Neurl4 | protein_coding | -0.494847969 | 5.23873819 | 1.9586776 | 0.1804201 | 0.726395806 |
| ENSMUSG00000041112 | Elmo1 | protein_coding | -0.286477365 | 8.636083867 | 1.958367 | 0.1804222 | 0.726395806 |
| ENSMUSG00000028602 | Tnfrsf8 | protein_coding | 1.556551121 | 1.854737579 | 1.9580751 | 0.1804529 | 0.726395806 |
| ENSMUSG00000018765 | Fxr2 | protein_coding | 0.347808921 | 5.496278609 | 1.9563516 | 0.1806344 | 0.726395806 |
| ENSMUSG00000037493 | Cib2 | protein_coding | 0.375064986 | 5.405799481 | 1.9559249 | 0.1806794 | 0.726395806 |
| ENSMUSG00000012535 | Tnpo3 | protein_coding | 0.510314742 | 5.407096462 | 1.9559828 | 0.1807039 | 0.726395806 |
| ENSMUSG00000091705 | H2-Q2 | protein_coding | 0.95391239 | 2.492497233 | 1.9553049 | 0.1807447 | 0.726395806 |
| ENSMUSG00000056941 | Commd7 | protein_coding | 0.404159672 | 6.256936883 | 1.9546313 | 0.1808464 | 0.726395806 |
| ENSMUSG00000086968 | 4933431E20Rik | antisense_RNA | 0.58819896 | 4.024833679 | 1.9536467 | 0.1809197 | 0.726395806 |
| ENSMUSG00000021932 | Rnaseh2b | protein_coding | -0.379226581 | 5.32804282 | 1.9532631 | 0.1809602 | 0.726395806 |
| ENSMUSG00000008763 | Man1a2 | protein_coding | -0.430217373 | 5.403876338 | 1.9528057 | 0.1810085 | 0.726395806 |
| ENSMUSG00000031523 | Dlc1 | protein_coding | -0.680183772 | 3.484914263 | 1.9524923 | 0.1810416 | 0.726395806 |
| ENSMUSG00000019295 | Tmem129 | protein_coding | 0.407692085 | 5.344032066 | 1.9512335 | 0.1811747 | 0.726658627 |
| ENSMUSG00000071234 | Syndig1l | protein_coding | -0.49436466 | 5.554383131 | 1.9508659 | 0.1812442 | 0.726666116 |
| ENSMUSG00000026869 | Psmd5 | protein_coding | 0.496436391 | 6.091297663 | 1.9602485 | 0.1814097 | 0.72680373 |
| ENSMUSG00000067567 | Hdac8 | protein_coding | 0.876259265 | 3.922088003 | 1.9666324 | 0.1814137 | 0.72680373 |
| ENSMUSG00000032869 | Psmf1 | protein_coding | 0.370966586 | 6.785877124 | 1.9483407 | 0.1815116 | 0.726925085 |
| ENSMUSG00000042460 | C1galt1 | protein_coding | -0.970498993 | 2.674108706 | 1.9467124 | 0.1816537 | 0.727063544 |
| ENSMUSG00000045731 | Pnoc | protein_coding | 0.251423457 | 6.687693393 | 1.9463412 | 0.1816931 | 0.727063544 |
| ENSMUSG00000026915 | Strbp | protein_coding | -0.270592048 | 6.935576728 | 1.9447336 | 0.1818638 | 0.727063544 |
| ENSMUSG00000018481 | Appbp2 | protein_coding | 0.266192998 | 6.66161444 | 1.9446647 | 0.1818711 | 0.727063544 |
| ENSMUSG00000096433 | Zfp994 | protein_coding | -1.024084555 | 2.366309007 | 1.9445417 | 0.1818842 | 0.727063544 |
| ENSMUSG00000020733 | Slc9a3r1 | protein_coding | -0.464342108 | 4.971427444 | 1.9431085 | 0.1820366 | 0.727402374 |
| ENSMUSG00000057863 | Rpl36 | protein_coding | -0.33245434 | 5.553229542 | 1.9410913 | 0.1822514 | 0.727990082 |
| ENSMUSG00000031511 | Arhgef7 | protein_coding | 0.430411085 | 5.673300661 | 1.9398479 | 0.182384 | 0.728249025 |
| ENSMUSG00000030509 | Asb7 | protein_coding | -0.453473239 | 4.074378218 | 1.9386224 | 0.1825147 | 0.728500613 |
| ENSMUSG00000038671 | Arfrp1 | protein_coding | 0.279331026 | 6.881633397 | 1.9369547 | 0.1826928 | 0.728771803 |
| ENSMUSG00000059182 | Skap2 | protein_coding | 0.53277033 | 4.404641125 | 1.9352005 | 0.1828804 | 0.728771803 |
| ENSMUSG00000041423 | Paqr6 | protein_coding | -0.532628948 | 3.762613481 | 1.9350283 | 0.1828989 | 0.728771803 |
| ENSMUSG00000031256 | Cstf2 | protein_coding | 0.541576417 | 4.311827684 | 1.9341611 | 0.1829917 | 0.728771803 |
| ENSMUSG00000021810 | Ecd | protein_coding | 0.38488225 | 6.148060357 | 1.9341461 | 0.1829933 | 0.728771803 |
| ENSMUSG00000006732 | Mettl1 | protein_coding | 0.32645944 | 6.02174802 | 1.9332386 | 0.1830905 | 0.728771803 |
| ENSMUSG00000025782 | Taf3 | protein_coding | -0.407863015 | 4.81405377 | 1.9322978 | 0.1831914 | 0.728771803 |
| ENSMUSG00000053347 | Zfp943 | protein_coding | -0.735638338 | 4.009394861 | 1.937683 | 0.1831916 | 0.728771803 |
| ENSMUSG00000020912 | Krt12 | protein_coding | -0.943565943 | 2.457496946 | 1.9314602 | 0.1832813 | 0.728771803 |
| ENSMUSG00000062991 | Nrg1 | protein_coding | -0.367893337 | 5.108745726 | 1.9313584 | 0.1832922 | 0.728771803 |
| ENSMUSG00000026672 | Optn | protein_coding | 0.605355634 | 3.886804943 | 1.9310232 | 0.1833282 | 0.728771803 |
| ENSMUSG00000024151 | Msh2 | protein_coding | 0.535560433 | 5.128752653 | 1.9268356 | 0.1838088 | 0.730317484 |
| ENSMUSG00000028552 | Eps15 | protein_coding | -0.287571665 | 6.687543463 | 1.9261442 | 0.1838528 | 0.730317484 |
| ENSMUSG00000027546 | Atp9a | protein_coding | 0.270476042 | 7.260064044 | 1.9245864 | 0.1840207 | 0.730448274 |
| ENSMUSG00000045795 | Whamm | protein_coding | -0.953279337 | 2.505882743 | 1.9245783 | 0.1840216 | 0.730448274 |
| ENSMUSG00000037747 | Phyhipl | protein_coding | 0.299479088 | 7.142047207 | 1.9222964 | 0.1842679 | 0.730929757 |
| ENSMUSG00000053004 | Hrh1 | protein_coding | 0.455044799 | 4.110665507 | 1.9221951 | 0.1842789 | 0.730929757 |
| ENSMUSG00000037415 | Ranbp10 | protein_coding | -0.758138698 | 3.407233385 | 1.9202574 | 0.1844884 | 0.731490917 |
| ENSMUSG00000022054 | Nefm | protein_coding | 0.538564439 | 7.018960095 | 1.9461756 | 0.1846723 | 0.731950363 |
| ENSMUSG00000032370 | Lactb | protein_coding | 0.741218021 | 2.970459917 | 1.9161762 | 0.1849306 | 0.732361283 |
| ENSMUSG00000019297 | Nop9 | protein_coding | -0.889688901 | 2.89386203 | 1.9136095 | 0.1852094 | 0.732361283 |
| ENSMUSG00000072872 | Rybp | protein_coding | 0.883864136 | 2.674842252 | 1.9133146 | 0.1852414 | 0.732361283 |
| ENSMUSG00000016637 | Ift27 | protein_coding | -0.288278678 | 7.178958989 | 1.9122169 | 0.1853609 | 0.732361283 |
| ENSMUSG00000026504 | Sdccag8 | protein_coding | -0.601287083 | 4.75536557 | 1.9124003 | 0.1854516 | 0.732361283 |
| ENSMUSG00000032788 | Pdxk | protein_coding | 0.492703913 | 7.679197173 | 1.9342882 | 0.1855777 | 0.732361283 |
| ENSMUSG00000027253 | Lrp4 | protein_coding | 0.66280039 | 4.078801821 | 1.9090429 | 0.1857372 | 0.732361283 |
| ENSMUSG00000024197 | Plin3 | protein_coding | -0.42740768 | 5.200013871 | 1.9079928 | 0.1858213 | 0.732361283 |
| ENSMUSG00000105867 | Gm42517 | protein_coding | 0.748309365 | 2.859212297 | 1.9078308 | 0.185839 | 0.732361283 |
| ENSMUSG00000014602 | Kif1a | protein_coding | 0.279556376 | 8.857309886 | 1.9076756 | 0.185856 | 0.732361283 |
| ENSMUSG00000113429 | CT030140.1 | lincRNA | -0.740196333 | 2.950139412 | 1.9071462 | 0.1859138 | 0.732361283 |
| ENSMUSG00000098702 | 1500015A07Rik | antisense_RNA | -0.669296495 | 3.304849174 | 1.904418 | 0.1862121 | 0.732361283 |
| ENSMUSG00000015461 | Atf6b | protein_coding | -0.604807751 | 4.014220969 | 1.9041656 | 0.1862398 | 0.732361283 |
| ENSMUSG00000024799 | Tm7sf2 | protein_coding | 0.579666662 | 4.114726198 | 1.9038196 | 0.1862776 | 0.732361283 |
| ENSMUSG00000022378 | Fam49b | protein_coding | -0.269518676 | 6.70146057 | 1.9035663 | 0.1863054 | 0.732361283 |
| ENSMUSG00000036880 | Acaa2 | protein_coding | 0.332879666 | 5.883959439 | 1.9035541 | 0.1863067 | 0.732361283 |
| ENSMUSG00000034285 | Nipsnap1 | protein_coding | 0.245322597 | 8.165715942 | 1.9032614 | 0.1863388 | 0.732361283 |
| ENSMUSG00000002289 | Angptl4 | protein_coding | 0.887656166 | 2.480448562 | 1.9025083 | 0.1864213 | 0.732361283 |
| ENSMUSG00000055320 | Tead1 | protein_coding | -0.503935382 | 3.646694706 | 1.902359 | 0.1864377 | 0.732361283 |
| ENSMUSG00000009418 | Nav1 | protein_coding | -0.355343317 | 5.623536203 | 1.9020933 | 0.1864668 | 0.732361283 |
| ENSMUSG00000040412 | 5330417C22Rik | protein_coding | 0.476037937 | 4.990527782 | 1.9011886 | 0.186566 | 0.732361283 |
| ENSMUSG00000001419 | Mef2d | protein_coding | 0.321549644 | 5.675214612 | 1.9009542 | 0.1865918 | 0.732361283 |
| ENSMUSG00000047824 | Pygo2 | protein_coding | -0.458844794 | 4.81998272 | 1.8999716 | 0.1866996 | 0.732361283 |
| ENSMUSG00000037138 | Aff3 | protein_coding | 0.474939138 | 4.817062394 | 1.8993736 | 0.1867653 | 0.732361283 |
| ENSMUSG00000004056 | Akt2 | protein_coding | -0.389204446 | 4.845874673 | 1.8990801 | 0.1867976 | 0.732361283 |

| ensembl_gene_id | external_gene_name | gene_biotype | log2 (fold change) | log2 (counts per million) | F-value | p-value | False discovery rate |
|---|---|---|---|---|---|---|---|
| ENSMUSG00000031242 | 2610002M06Rik | protein_coding | -0.319061605 | 5.652742903 | 1.8984015 | 0.1868722 | 0.732361283 |
| ENSMUSG00000018428 | Akap1 | protein_coding | -0.644258759 | 3.861081418 | 1.8983781 | 0.1868747 | 0.732361283 |
| ENSMUSG00000051159 | Cited1 | protein_coding | 0.535372615 | 5.499660301 | 1.9053011 | 0.186891 | 0.732361283 |
| ENSMUSG00000090093 | Gm14399 | protein_coding | -0.845007809 | 2.701887579 | 1.897866 | 0.1869311 | 0.732361283 |
| ENSMUSG00000035845 | Alg12 | protein_coding | 0.61697934 | 3.806612406 | 1.8968557 | 0.1870422 | 0.732361283 |
| ENSMUSG00000026585 | Kifap3 | protein_coding | 0.278335612 | 7.567748367 | 1.8963278 | 0.1871004 | 0.732361283 |
| ENSMUSG00000034064 | Poglut1 | protein_coding | 0.353758759 | 6.366769192 | 1.8957358 | 0.1871656 | 0.732361283 |
| ENSMUSG00000024785 | Rcl1 | protein_coding | 0.539758576 | 4.641329716 | 1.8946088 | 0.1872898 | 0.732361283 |
| ENSMUSG00000001525 | Tubb5 | protein_coding | 0.352132726 | 10.28704004 | 1.8944577 | 0.1873064 | 0.732361283 |
| ENSMUSG00000011179 | Odc1 | protein_coding | 0.497941913 | 4.258614454 | 1.8942635 | 0.1873278 | 0.732361283 |
| ENSMUSG00000025533 | Asl | protein_coding | 0.27718514 | 6.434689736 | 1.8940551 | 0.1873508 | 0.732361283 |
| ENSMUSG00000046709 | Mapk10 | protein_coding | 0.311449185 | 8.463617021 | 1.89367 | 0.1873933 | 0.732361283 |
| ENSMUSG00000020441 | 2310033P09Rik | protein_coding | 0.542376837 | 3.652092943 | 1.8936205 | 0.1873988 | 0.732361283 |
| ENSMUSG00000056666 | Retsat | protein_coding | -0.604409191 | 3.650956387 | 1.8933178 | 0.1874322 | 0.732361283 |
| ENSMUSG00000036890 | Gtdc1 | protein_coding | -0.494115142 | 4.693488783 | 1.8923295 | 0.1875413 | 0.732521458 |
| ENSMUSG00000029202 | Pds5a | protein_coding | -0.566122902 | 3.485613207 | 1.8916579 | 0.1876155 | 0.732545156 |
| ENSMUSG00000068250 | Amn1 | protein_coding | -0.356730256 | 5.656343951 | 1.8891322 | 0.1878948 | 0.73291571 |
| ENSMUSG00000071655 | Ubxn1 | protein_coding | 0.259439276 | 6.632327151 | 1.8873069 | 0.1880971 | 0.73291571 |
| ENSMUSG00000021720 | Rnf180 | protein_coding | -0.4238724 | 4.500126184 | 1.8868852 | 0.1881438 | 0.73291571 |
| ENSMUSG00000030172 | Erc1 | protein_coding | -0.501100113 | 4.023491567 | 1.8865495 | 0.188181 | 0.73291571 |
| ENSMUSG00000059423 | Zfp933 | protein_coding | -0.766474094 | 3.282108402 | 1.8864164 | 0.1881958 | 0.73291571 |
| ENSMUSG00000071855 | Ccdc112 | protein_coding | -0.534112887 | 4.039742645 | 1.8863574 | 0.1882023 | 0.73291571 |
| ENSMUSG00000070738 | Dgkd | protein_coding | 0.51483382 | 4.513298388 | 1.8861142 | 0.1882293 | 0.73291571 |
| ENSMUSG00000037418 | Best1 | protein_coding | 1.020966912 | 2.896601473 | 1.8861503 | 0.1882557 | 0.73291571 |
| ENSMUSG00000030849 | Fgfr2 | protein_coding | -0.272494816 | 6.727220766 | 1.8835584 | 0.1885132 | 0.733357258 |
| ENSMUSG00000028345 | Tex10 | protein_coding | 0.513868025 | 4.033069419 | 1.8830948 | 0.1885647 | 0.733357258 |
| ENSMUSG00000030983 | Bccip | protein_coding | -0.247814462 | 7.669501024 | 1.8830142 | 0.1885737 | 0.733357258 |
| ENSMUSG00000024037 | Wdr4 | protein_coding | 0.39192234 | 4.752591358 | 1.8820351 | 0.1886826 | 0.733515555 |
| ENSMUSG00000022471 | Xrcc6 | protein_coding | 0.66749645 | 3.169702503 | 1.8810168 | 0.188796 | 0.733552709 |
| ENSMUSG00000022587 | Ly6e | protein_coding | 0.300320464 | 7.940055716 | 1.8807239 | 0.1888286 | 0.733552709 |
| ENSMUSG00000032786 | Alas1 | protein_coding | 0.456060634 | 6.36808781 | 1.8905245 | 0.1889169 | 0.733630554 |
| ENSMUSG00000031374 | Zfp92 | protein_coding | 0.825786104 | 2.641578821 | 1.8782862 | 0.1891004 | 0.733722848 |
| ENSMUSG00000085664 | Atxn7l1os2 | antisense_RNA | -0.811461569 | 2.93795205 | 1.8780277 | 0.1891293 | 0.733722848 |
| ENSMUSG00000046152 | Fut10 | protein_coding | 0.76155257 | 2.851395179 | 1.8778837 | 0.1891453 | 0.733722848 |
| ENSMUSG00000033454 | Zbtb1 | protein_coding | -0.869707723 | 2.644817207 | 1.8770287 | 0.1892408 | 0.733784811 |
| ENSMUSG00000074345 | Tnfaip8l3 | protein_coding | 0.74515419 | 3.454021263 | 1.8760319 | 0.1893522 | 0.733784811 |
| ENSMUSG00000042834 | Nrep | protein_coding | -0.322347261 | 6.433058837 | 1.8749991 | 0.1894677 | 0.733784811 |
| ENSMUSG00000020634 | Ubxn2a | protein_coding | -0.342769136 | 5.43505583 | 1.8746879 | 0.1895025 | 0.733784811 |
| ENSMUSG00000008384 | Sertad1 | protein_coding | -0.420367408 | 4.178920212 | 1.8745205 | 0.1895212 | 0.733784811 |
| ENSMUSG00000030315 | Vgll4 | protein_coding | 0.380789147 | 4.932348267 | 1.8737486 | 0.1896077 | 0.733784811 |
| ENSMUSG00000021484 | Lman2 | protein_coding | 0.416045721 | 5.333730486 | 1.8721724 | 0.1897843 | 0.733784811 |
| ENSMUSG00000038564 | Ift172 | protein_coding | 0.642202925 | 3.500283351 | 1.8721308 | 0.1897889 | 0.733784811 |
| ENSMUSG00000029502 | Golga3 | protein_coding | 0.401971826 | 5.084133763 | 1.871497 | 0.18986 | 0.733784811 |
| ENSMUSG00000005803 | Sqor | protein_coding | 0.572778935 | 3.767947818 | 1.8713815 | 0.1898729 | 0.733784811 |
| ENSMUSG00000022295 | Atp6v1c1 | protein_coding | 0.272806394 | 9.301728821 | 1.870978 | 0.1899182 | 0.733784811 |
| ENSMUSG00000053470 | Kdm3a | protein_coding | 0.4816338 | 4.227019001 | 1.8703521 | 0.1899885 | 0.733784811 |
| ENSMUSG00000049907 | Rasl11b | protein_coding | -0.448365616 | 4.109289556 | 1.8684453 | 0.1902027 | 0.733784811 |
| ENSMUSG00000033486 | Catsper2 | protein_coding | -0.655868859 | 3.225605393 | 1.8672324 | 0.1903391 | 0.733784811 |
| ENSMUSG00000040782 | Rfwd2 | protein_coding | -0.735584267 | 3.500210466 | 1.8667005 | 0.190399 | 0.733784811 |
| ENSMUSG00000051811 | Cox6b2 | protein_coding | -0.48035505 | 4.008247307 | 1.866121 | 0.1904642 | 0.733784811 |
| ENSMUSG00000018909 | Arrb1 | protein_coding | 0.285125778 | 6.224803928 | 1.8659566 | 0.1904827 | 0.733784811 |
| ENSMUSG00000060733 | Ipmk | protein_coding | -0.468680271 | 3.940777536 | 1.8654098 | 0.1905443 | 0.733784811 |
| ENSMUSG00000067873 | Htatsf1 | protein_coding | -0.33465148 | 5.837514164 | 1.8649777 | 0.190593 | 0.733784811 |
| ENSMUSG00000020334 | Slc22a4 | protein_coding | 0.724473547 | 3.121545902 | 1.864724 | 0.1906216 | 0.733784811 |
| ENSMUSG00000026764 | Kif5c | protein_coding | 0.320067585 | 7.28772482 | 1.8645435 | 0.1906419 | 0.733784811 |
| ENSMUSG00000031575 | Ash2l | protein_coding | 0.443755065 | 4.821138281 | 1.8643605 | 0.1906626 | 0.733784811 |
| ENSMUSG00000020524 | Gria1 | protein_coding | 0.25827551 | 8.10925162 | 1.8637328 | 0.1907334 | 0.73379468 |
| ENSMUSG00000030898 | Cckbr | protein_coding | 0.803757861 | 3.172540333 | 1.8613037 | 0.1910077 | 0.734409319 |
| ENSMUSG00000029684 | Wasl | protein_coding | -0.302962854 | 6.066731873 | 1.8611084 | 0.1910297 | 0.734409319 |
| ENSMUSG00000044881 | Coa4 | protein_coding | 0.431550579 | 4.985126148 | 1.8597681 | 0.1911813 | 0.734729364 |
| ENSMUSG00000038332 | Sesn1 | protein_coding | 0.365867489 | 5.305426125 | 1.8582827 | 0.1913495 | 0.734916532 |
| ENSMUSG00000032018 | Sc5d | protein_coding | 0.27167704 | 6.268769391 | 1.8578123 | 0.1914028 | 0.734916532 |
| ENSMUSG00000103428 | Gm20754 | lincRNA | -1.153044536 | 2.397947304 | 1.8573003 | 0.1914608 | 0.734916532 |
| ENSMUSG00000005698 | Ctcf | protein_coding | -0.332924991 | 6.059742147 | 1.8567566 | 0.1915224 | 0.734916532 |
| ENSMUSG00000030209 | Grin2b | protein_coding | -0.334157068 | 6.206190231 | 1.8563219 | 0.1915718 | 0.734916532 |
| ENSMUSG00000055150 | Zfp78 | protein_coding | -0.591907236 | 3.449240101 | 1.8553322 | 0.1916841 | 0.73508514 |
| ENSMUSG00000029610 | Aimp2 | protein_coding | 0.27603499 | 6.109981577 | 1.8539741 | 0.1918383 | 0.735106665 |
| ENSMUSG00000021809 | Nudt13 | protein_coding | 0.527029593 | 4.202712886 | 1.8530204 | 0.1919467 | 0.735106665 |
| ENSMUSG00000027550 | Lrrcc1 | protein_coding | -0.975388868 | 4.236047588 | 1.8868895 | 0.1919602 | 0.735106665 |
| ENSMUSG00000056698 | Elmod3 | protein_coding | -0.355342236 | 5.625959417 | 1.8521517 | 0.1920455 | 0.735106665 |
| ENSMUSG00000001089 | Luzp1 | protein_coding | 0.398006771 | 5.210286883 | 1.8520542 | 0.1920567 | 0.735106665 |
| ENSMUSG00000022995 | Enah | protein_coding | -0.277913054 | 6.376213725 | 1.8514995 | 0.1921198 | 0.735106665 |
| ENSMUSG00000026207 | Speg | protein_coding | -0.626771264 | 3.661828598 | 1.8508091 | 0.1921984 | 0.735106665 |
| ENSMUSG00000015709 | Arnt2 | protein_coding | 0.307190375 | 6.141255029 | 1.8503679 | 0.1922487 | 0.735106665 |

| ensembl_gene_id | external_gene_name | gene_biotype | log2 (fold change) | log2 (counts per million) | F-value | p-value | False discovery rate |
|---|---|---|---|---|---|---|---|
| ENSMUSG00000031885 | Cbfb | protein_coding | -0.455659102 | 4.202384283 | 1.8492418 | 0.192377 | 0.735106665 |
| ENSMUSG00000022339 | Ebag9 | protein_coding | -0.408075154 | 4.976404661 | 1.8467881 | 0.1926571 | 0.735106665 |
| ENSMUSG00000030643 | Rab30 | protein_coding | -0.439842364 | 5.044029253 | 1.846609 | 0.1926776 | 0.735106665 |
| ENSMUSG00000026353 | Ubxn4 | protein_coding | -0.30432684 | 5.908342931 | 1.8449072 | 0.1928722 | 0.735106665 |
| ENSMUSG00000035413 | Tmem98 | protein_coding | -0.360613958 | 5.736134457 | 1.8448795 | 0.1928753 | 0.735106665 |
| ENSMUSG00000022961 | Son | protein_coding | 0.317124015 | 7.369054877 | 1.8440592 | 0.1929692 | 0.735106665 |
| ENSMUSG00000040549 | Ckap5 | protein_coding | 0.275539513 | 6.407334351 | 1.842592 | 0.1931373 | 0.735106665 |
| ENSMUSG00000050373 | Snx21 | protein_coding | 0.471720822 | 4.355608397 | 1.8420894 | 0.1931949 | 0.735106665 |
| ENSMUSG00000026674 | Ddr2 | protein_coding | -0.718572558 | 2.860090175 | 1.8390701 | 0.1935414 | 0.735106665 |
| ENSMUSG00000030272 | Camk1 | protein_coding | 0.511492512 | 3.713426668 | 1.8386952 | 0.1935845 | 0.735106665 |
| ENSMUSG00000028138 | Adh5 | protein_coding | 0.249356322 | 7.589360747 | 1.8384083 | 0.1936175 | 0.735106665 |
| ENSMUSG00000028426 | Rad23b | protein_coding | -0.334933183 | 5.85898496 | 1.8383507 | 0.1936241 | 0.735106665 |
| ENSMUSG00000037916 | Ndufv1 | protein_coding | 0.286935897 | 9.217131399 | 1.8381626 | 0.1936458 | 0.735106665 |
| ENSMUSG00000078681 | Tm2d3 | protein_coding | 0.270911161 | 8.32003089 | 1.8380986 | 0.1936531 | 0.735106665 |
| ENSMUSG00000025370 | Cdh9 | protein_coding | 1.031425146 | 2.458546631 | 1.8380618 | 0.1936573 | 0.735106665 |
| ENSMUSG00000050234 | Gja4 | protein_coding | -0.996257948 | 2.662100191 | 1.8378658 | 0.1936799 | 0.735106665 |
| ENSMUSG00000022718 | Dgcr8 | protein_coding | 0.788536317 | 2.741823488 | 1.8377335 | 0.1936951 | 0.735106665 |
| ENSMUSG00000022661 | Cd200 | protein_coding | 0.299169234 | 8.070795326 | 1.83773 | 0.1936955 | 0.735106665 |
| ENSMUSG00000039759 | Thap3 | protein_coding | -0.309850624 | 6.023403703 | 1.8377057 | 0.1936983 | 0.735106665 |
| ENSMUSG00000020048 | Hsp90b1 | protein_coding | 0.242273267 | 7.071487555 | 1.8371567 | 0.1937615 | 0.735106665 |
| ENSMUSG00000036430 | Tbcc | protein_coding | -0.633473949 | 3.39072153 | 1.8368977 | 0.1937913 | 0.735106665 |
| ENSMUSG00000033943 | Mga | protein_coding | -0.428379633 | 5.35196869 | 1.8367865 | 0.1938041 | 0.735106665 |
| ENSMUSG00000026434 | Nucks1 | protein_coding | -0.280416507 | 7.003684231 | 1.8367443 | 0.1938089 | 0.735106665 |
| ENSMUSG00000106775 | C130093G08Rik | TEC | 0.767593048 | 4.64571464 | 1.8582335 | 0.193931 | 0.735116175 |
| ENSMUSG00000025371 | Chmp6 | protein_coding | 0.411728394 | 5.972846728 | 1.835797 | 0.1939482 | 0.735116175 |
| ENSMUSG00000014776 | Nol3 | protein_coding | 0.528225242 | 5.167329487 | 1.8337146 | 0.1941882 | 0.735766714 |
| ENSMUSG00000022462 | Slc38a2 | protein_coding | 0.363152191 | 5.304103256 | 1.8327687 | 0.1942672 | 0.735806867 |
| ENSMUSG00000031860 | Pbx4 | protein_coding | 0.459611821 | 4.024994436 | 1.8317656 | 0.1943831 | 0.735840773 |
| ENSMUSG00000086697 | 4933427G23Rik | transcribed_processe | 0.89514228 | 2.445828553 | 1.8312669 | 0.1944407 | 0.735840773 |
| ENSMUSG00000025262 | Fam120c | protein_coding | 0.847633122 | 3.31235211 | 1.8281247 | 0.1948345 | 0.735840773 |
| ENSMUSG00000043621 | Ubxn10 | protein_coding | -0.694866252 | 3.129834677 | 1.8277554 | 0.1948471 | 0.735840773 |
| ENSMUSG00000028901 | Gmeb1 | protein_coding | -0.548185913 | 4.083689922 | 1.8274426 | 0.1948834 | 0.735840773 |
| ENSMUSG00000037339 | Fam53a | protein_coding | -0.796603714 | 3.00363387 | 1.8264396 | 0.1949997 | 0.735840773 |
| ENSMUSG00000092329 | Gm20388 | protein_coding | -0.238104751 | 6.876614423 | 1.8262245 | 0.1950247 | 0.735840773 |
| ENSMUSG00000006435 | Neurl1a | protein_coding | -0.386598071 | 5.92054398 | 1.825961 | 0.1950552 | 0.735840773 |
| ENSMUSG00000020182 | Ddc | protein_coding | 0.350285788 | 5.347214123 | 1.825822 | 0.1950714 | 0.735840773 |
| ENSMUSG00000022412 | Mief1 | protein_coding | 0.477153639 | 5.00375469 | 1.8253957 | 0.1951209 | 0.735840773 |
| ENSMUSG00000021222 | Dcaf4 | protein_coding | 0.350974902 | 6.168689986 | 1.8249147 | 0.1951767 | 0.735840773 |
| ENSMUSG00000074736 | Syndig1 | protein_coding | -0.317999825 | 6.409774563 | 1.8245992 | 0.1952134 | 0.735840773 |
| ENSMUSG00000026283 | Ing5 | protein_coding | 0.397825684 | 4.677544366 | 1.8239506 | 0.1952887 | 0.735840773 |
| ENSMUSG00000047907 | Tshz2 | protein_coding | -0.431034255 | 4.067112296 | 1.8230423 | 0.1953943 | 0.735840773 |
| ENSMUSG00000032264 | Zw10 | protein_coding | -0.553584327 | 4.109129274 | 1.8224442 | 0.1954639 | 0.735840773 |
| ENSMUSG00000001773 | Folh1 | protein_coding | -0.814234807 | 2.654719932 | 1.8222348 | 0.1954883 | 0.735840773 |
| ENSMUSG00000029146 | Snx17 | protein_coding | 0.312474442 | 8.228083589 | 1.821667 | 0.1955544 | 0.735840773 |
| ENSMUSG00000042032 | Mat2b | protein_coding | 0.25487738 | 7.830334238 | 1.8206586 | 0.1956718 | 0.735840773 |
| ENSMUSG00000027882 | Stxbp3 | protein_coding | -0.434182702 | 4.760748714 | 1.8206131 | 0.1956771 | 0.735840773 |
| ENSMUSG00000025423 | Pias2 | protein_coding | -0.342806834 | 5.407554787 | 1.8200308 | 0.195745 | 0.735840773 |
| ENSMUSG00000033294 | Noc4l | protein_coding | -0.455985938 | 4.910377051 | 1.8194806 | 0.1958091 | 0.735840773 |
| ENSMUSG00000033983 | Coil | protein_coding | 0.432483482 | 4.528481232 | 1.8189694 | 0.1958687 | 0.735840773 |
| ENSMUSG00000029017 | Pmpcb | protein_coding | 0.260106217 | 7.100038551 | 1.8189364 | 0.1958726 | 0.735840773 |
| ENSMUSG00000001542 | Ell2 | protein_coding | -0.600819158 | 3.6254633 | 1.8185424 | 0.1959185 | 0.735840773 |
| ENSMUSG00000056153 | Socs6 | protein_coding | -0.452769007 | 4.079099879 | 1.8169787 | 0.1961011 | 0.736023726 |
| ENSMUSG00000058620 | Adra2b | protein_coding | -0.638261784 | 3.442978142 | 1.8164648 | 0.1961612 | 0.736023726 |
| ENSMUSG00000026192 | Atic | protein_coding | 0.424624467 | 5.524916594 | 1.8160559 | 0.196209 | 0.736023726 |
| ENSMUSG00000018974 | Sart3 | protein_coding | 0.482720805 | 4.178009768 | 1.8152999 | 0.1962974 | 0.736023726 |
| ENSMUSG00000020821 | Kif1c | protein_coding | -0.686781986 | 2.868441457 | 1.8151964 | 0.1963095 | 0.736023726 |
| ENSMUSG00000067860 | Zic3 | protein_coding | -0.463745648 | 3.910919561 | 1.8134443 | 0.1965146 | 0.736535923 |
| ENSMUSG00000052087 | Rgs14 | protein_coding | -0.40606087 | 5.188153035 | 1.8124485 | 0.1966313 | 0.736685802 |
| ENSMUSG00000038212 | Mfsd14b | protein_coding | -0.47737323 | 4.628872732 | 1.81167 | 0.1967226 | 0.736685802 |
| ENSMUSG00000029209 | Gnpda2 | protein_coding | 0.314331353 | 5.92283896 | 1.8109853 | 0.1968029 | 0.736685802 |
| ENSMUSG00000067931 | Zfp948 | protein_coding | -0.56955973 | 3.557058881 | 1.8107663 | 0.1968286 | 0.736685802 |
| ENSMUSG00000113216 | AL590503.1 | lincRNA | -0.564247753 | 4.160542779 | 1.8094555 | 0.1969826 | 0.73700545 |
| ENSMUSG00000024055 | Cyp4f13 | protein_coding | -0.928832272 | 2.949855005 | 1.8087897 | 0.1970608 | 0.73704175 |
| ENSMUSG00000027304 | Rtf1 | protein_coding | 0.40101023 | 5.315831213 | 1.8079738 | 0.1971568 | 0.737144208 |
| ENSMUSG00000017615 | Tnfaip1 | protein_coding | 0.358512477 | 6.625073356 | 1.807628 | 0.1972275 | 0.737152316 |
| ENSMUSG00000023456 | Tpi1 | protein_coding | 0.239175452 | 9.877387181 | 1.8064574 | 0.1973352 | 0.737221671 |
| ENSMUSG00000029070 | Mxra8 | protein_coding | 1.60418767 | 3.850094941 | 1.8994722 | 0.197574 | 0.737221671 |
| ENSMUSG00000024283 | Wac | protein_coding | -0.320809386 | 5.65565543 | 1.8041672 | 0.1976052 | 0.737221671 |
| ENSMUSG00000019139 | Isyna1 | protein_coding | 0.511956205 | 5.3840446 | 1.8041592 | 0.197767 | 0.737221671 |
| ENSMUSG00000020736 | Nt5c | protein_coding | 0.234702123 | 7.34385362 | 1.8022187 | 0.1978352 | 0.737221671 |
| ENSMUSG00000049536 | Tceal1 | protein_coding | -0.232801102 | 7.576970249 | 1.8020275 | 0.1978578 | 0.737221671 |
| ENSMUSG00000028187 | Rpf1 | protein_coding | -0.431513997 | 4.458672323 | 1.8015781 | 0.1979109 | 0.737221671 |
| ENSMUSG00000031787 | Katnb1 | protein_coding | 0.442571196 | 6.125970279 | 1.8041879 | 0.1981698 | 0.737221671 |
| ENSMUSG00000029484 | Anxa3 | protein_coding | -0.610843337 | 3.495999761 | 1.7993444 | 0.1981751 | 0.737221671 |

| ensembl_gene_id | external_gene_name | gene_biotype | log2 (fold change) | log2 (counts per million) | F-value | p-value | False discovery rate |
|---|---|---|---|---|---|---|---|
| ENSMUSG00000021767 | Kat6b | protein_coding | -0.468163845 | 4.30740422 | 1.7990956 | 0.1982046 | 0.737221671 |
| ENSMUSG00000040521 | Tsfm | protein_coding | 0.308415956 | 6.541211415 | 1.7985194 | 0.1982729 | 0.737221671 |
| ENSMUSG00000031600 | Vps37a | protein_coding | -0.33779008 | 5.384052491 | 1.7984872 | 0.1982767 | 0.737221671 |
| ENSMUSG00000048485 | Zbtb8b | protein_coding | 0.771980573 | 2.654409053 | 1.7976371 | 0.1983774 | 0.737221671 |
| ENSMUSG00000046753 | Ccdc66 | protein_coding | -0.681086808 | 3.093185447 | 1.7974021 | 0.1984053 | 0.737221671 |
| ENSMUSG00000001248 | Gramd1a | protein_coding | 0.523667789 | 4.273829548 | 1.7971372 | 0.1984367 | 0.737221671 |
| ENSMUSG00000035226 | Rims4 | protein_coding | 0.422084454 | 5.01318547 | 1.7969542 | 0.1984584 | 0.737221671 |
| ENSMUSG00000047434 | Xxylt1 | protein_coding | -0.542144381 | 3.392330409 | 1.796115 | 0.198558 | 0.737221671 |
| ENSMUSG00000035783 | Acta2 | protein_coding | -0.888614045 | 3.195979537 | 1.7960531 | 0.1985953 | 0.737221671 |
| ENSMUSG00000045078 | Rnf216 | protein_coding | 0.427269511 | 5.031410019 | 1.7957486 | 0.1986014 | 0.737221671 |
| ENSMUSG00000060568 | Fam78b | protein_coding | -0.612485238 | 3.485171774 | 1.7950501 | 0.1986844 | 0.737221671 |
| ENSMUSG00000006527 | Sfmbt1 | protein_coding | 0.420357488 | 4.659079906 | 1.7948005 | 0.198714 | 0.737221671 |
| ENSMUSG00000054723 | Vmac | protein_coding | 0.595985692 | 3.522435072 | 1.7940819 | 0.1987994 | 0.737221671 |
| ENSMUSG00000031985 | Gnpat | protein_coding | 0.393328206 | 5.414486197 | 1.7938844 | 0.1988229 | 0.737221671 |
| ENSMUSG00000020755 | Sap30bp | protein_coding | 0.239619141 | 6.926630139 | 1.791526 | 0.1991036 | 0.73800777 |
| ENSMUSG00000037851 | Iars | protein_coding | 0.452170549 | 4.834015032 | 1.7906699 | 0.1992055 | 0.738131362 |
| ENSMUSG00000027078 | Ube2l6 | protein_coding | 0.480626942 | 4.029431324 | 1.789757 | 0.1993144 | 0.738169996 |
| ENSMUSG00000049354 | Dcaf7 | protein_coding | 0.276410685 | 6.43481068 | 1.789431 | 0.1993533 | 0.738169996 |
| ENSMUSG00000038084 | Opa1 | protein_coding | 0.314041626 | 5.952056589 | 1.7884623 | 0.1994689 | 0.73819894 |
| ENSMUSG00000031765 | Mt1 | protein_coding | -0.515028215 | 10.16083644 | 1.8109966 | 0.1994984 | 0.73819894 |
| ENSMUSG00000008822 | Acyp1 | protein_coding | -0.278179157 | 6.363257055 | 1.7874309 | 0.199592 | 0.738291431 |
| ENSMUSG00000028411 | Aptx | protein_coding | 0.26238391 | 6.507321975 | 1.7866743 | 0.1996824 | 0.738371852 |
| ENSMUSG00000047786 | Lix1 | protein_coding | -0.416897673 | 5.070151099 | 1.7853369 | 0.1998424 | 0.73841325 |
| ENSMUSG00000030043 | Tacr1 | protein_coding | 0.642539243 | 3.236117114 | 1.7852253 | 0.1998558 | 0.73841325 |
| ENSMUSG00000031529 | Tnks | protein_coding | -0.345792447 | 5.229642601 | 1.7848585 | 0.1998997 | 0.73841325 |
| ENSMUSG00000027452 | Acss1 | protein_coding | 0.801601802 | 3.118755559 | 1.7830634 | 0.2001147 | 0.738953692 |
| ENSMUSG00000034993 | Vat1 | protein_coding | 0.319371473 | 6.200072375 | 1.7807274 | 0.2003949 | 0.739512047 |
| ENSMUSG00000068696 | Gpr88 | protein_coding | -0.711768905 | 6.335451933 | 1.8456951 | 0.200464 | 0.739512047 |
| ENSMUSG00000015291 | Gdi1 | protein_coding | 0.403661847 | 10.42006291 | 1.7842469 | 0.2005383 | 0.739512047 |
| ENSMUSG00000059474 | Mbtd1 | protein_coding | -0.528905549 | 3.747940993 | 1.7791215 | 0.2005879 | 0.739512047 |
| ENSMUSG00000029701 | Rbm28 | protein_coding | 0.398989249 | 5.289942024 | 1.7789034 | 0.2006141 | 0.739512047 |
| ENSMUSG00000055319 | Sec23ip | protein_coding | 0.398188745 | 5.61426494 | 1.7783678 | 0.2006785 | 0.739512047 |
| ENSMUSG00000069020 | Urm1 | protein_coding | -0.25301948 | 6.34336436 | 1.7769987 | 0.2008444 | 0.739869731 |
| ENSMUSG00000033554 | Dph5 | protein_coding | 0.332606185 | 5.479971465 | 1.7754379 | 0.2010314 | 0.740304897 |
| ENSMUSG00000029152 | Ociad1 | protein_coding | 0.273440443 | 8.870748976 | 1.7746891 | 0.2011217 | 0.74038388 |
| ENSMUSG00000034714 | Ttyh2 | protein_coding | -0.382737713 | 7.227593125 | 1.7795088 | 0.2013041 | 0.740589344 |
| ENSMUSG00000034040 | Wbscr17 | protein_coding | -0.464785096 | 4.346021108 | 1.7730855 | 0.2013152 | 0.740589344 |
| ENSMUSG00000057098 | Ebf1 | protein_coding | -0.470108174 | 3.995808102 | 1.7721516 | 0.2014281 | 0.740751036 |
| ENSMUSG00000097785 | B230217O12Rik | lincRNA | 0.415326817 | 5.528692352 | 1.7710961 | 0.2015557 | 0.740966949 |
| ENSMUSG00000075486 | Commd6 | protein_coding | -0.319333746 | 7.794029343 | 1.7698922 | 0.2017014 | 0.741094785 |
| ENSMUSG00000024277 | Mapre2 | protein_coding | 0.31276328 | 8.384898168 | 1.7696698 | 0.2017283 | 0.741094785 |
| ENSMUSG00000014867 | Surf4 | protein_coding | 0.368735577 | 5.453202944 | 1.7687749 | 0.2018367 | 0.741138655 |
| ENSMUSG00000029179 | Zcchc4 | protein_coding | -0.662226818 | 3.609632238 | 1.7684334 | 0.2018781 | 0.741138655 |
| ENSMUSG00000027525 | Phactr3 | protein_coding | -0.266631933 | 6.714599074 | 1.7678426 | 0.2019497 | 0.74114856 |
| ENSMUSG00000063785 | Utp14a | protein_coding | 0.711495078 | 3.571688047 | 1.7649208 | 0.2023343 | 0.741657995 |
| ENSMUSG00000050321 | Neto1 | protein_coding | -0.488921742 | 4.702651805 | 1.7645886 | 0.2023448 | 0.741657995 |
| ENSMUSG00000035234 | Abraxas1 | protein_coding | -0.749309756 | 2.869221752 | 1.7643838 | 0.2023697 | 0.741657995 |
| ENSMUSG00000032750 | Gab3 | protein_coding | -0.586142767 | 3.593346028 | 1.7630304 | 0.2025343 | 0.741657995 |
| ENSMUSG00000034024 | Cct2 | protein_coding | 0.254593339 | 8.002043955 | 1.7629199 | 0.2025478 | 0.741657995 |
| ENSMUSG00000056493 | Foxk1 | protein_coding | 0.564627545 | 3.329235493 | 1.7618818 | 0.2026742 | 0.741657995 |
| ENSMUSG00000025724 | Sec11a | protein_coding | -0.304045575 | 6.941212833 | 1.7609474 | 0.202788 | 0.741657995 |
| ENSMUSG00000016619 | Nup50 | protein_coding | 0.429912976 | 4.860087363 | 1.7595839 | 0.2029543 | 0.741657995 |
| ENSMUSG00000022231 | Sema5a | protein_coding | -0.477859306 | 4.641101121 | 1.7594367 | 0.2029723 | 0.741657995 |
| ENSMUSG00000054874 | Pcnx3 | protein_coding | 0.959547787 | 2.79137171 | 1.7593354 | 0.2029847 | 0.741657995 |
| ENSMUSG00000020709 | Adap2 | protein_coding | 0.88852188 | 2.499203079 | 1.7592266 | 0.2029979 | 0.741657995 |
| ENSMUSG00000027620 | Rbm39 | protein_coding | 0.263308154 | 7.1349074 | 1.7591257 | 0.2030103 | 0.741657995 |
| ENSMUSG00000033706 | Smyd5 | protein_coding | -0.283644517 | 6.38222208 | 1.7587061 | 0.2030615 | 0.741657995 |
| ENSMUSG00000021314 | Amph | protein_coding | 0.414062023 | 7.151884838 | 1.7679456 | 0.2031674 | 0.741657995 |
| ENSMUSG00000108778 | Gm20083 | lincRNA | -0.93301307 | 2.465986827 | 1.7571531 | 0.2032512 | 0.741657995 |
| ENSMUSG00000085741 | 5430405H02Rik | processed_transcript | -0.590920472 | 3.345413118 | 1.7570105 | 0.2032687 | 0.741657995 |
| ENSMUSG00000046722 | Cdc42se1 | protein_coding | 0.348770381 | 5.214550562 | 1.7563886 | 0.2033447 | 0.741657995 |
| ENSMUSG00000028455 | Stoml2 | protein_coding | 0.321467155 | 7.123481909 | 1.7556624 | 0.2034336 | 0.741657995 |
| ENSMUSG00000036052 | Dnajb5 | protein_coding | -0.426364818 | 5.281467032 | 1.7548933 | 0.2035277 | 0.741657995 |
| ENSMUSG00000037949 | Ano10 | protein_coding | 0.529905548 | 4.049037285 | 1.7544374 | 0.2035836 | 0.741657995 |
| ENSMUSG00000060450 | Rnf14 | protein_coding | 0.347609674 | 8.835925022 | 1.7535467 | 0.2037225 | 0.741657995 |
| ENSMUSG00000008489 | Elavl2 | protein_coding | -0.297494953 | 6.592545263 | 1.7531822 | 0.2037374 | 0.741657995 |
| ENSMUSG00000024422 | Dhx16 | protein_coding | -0.759268788 | 2.864319705 | 1.7525598 | 0.2038138 | 0.741657995 |
| ENSMUSG00000039278 | Pcsk1n | protein_coding | -0.372002612 | 5.94998412 | 1.7515279 | 0.2039404 | 0.741657995 |
| ENSMUSG00000050332 | Amer1 | protein_coding | 0.760883875 | 3.496551886 | 1.7500498 | 0.2041518 | 0.741657995 |
| ENSMUSG00000032397 | Tipin | protein_coding | -0.336930612 | 6.106471395 | 1.7491337 | 0.2042346 | 0.741657995 |
| ENSMUSG00000013236 | Ptprs | protein_coding | -0.300489742 | 7.138867334 | 1.7489841 | 0.204253 | 0.741657995 |
| ENSMUSG00000039652 | Cpeb3 | protein_coding | -0.406037041 | 5.090136494 | 1.7488109 | 0.2042743 | 0.741657995 |
| ENSMUSG00000097326 | A330048O09Rik | lincRNA | -0.71224476 | 3.431106651 | 1.7482097 | 0.2043483 | 0.741657995 |
| ENSMUSG00000085218 | BB218582 | processed_transcript | -0.724417449 | 2.970129781 | 1.7480204 | 0.2043716 | 0.741657995 |

| ensembl_gene_id | external_gene_name | gene_biotype | log2 (fold change) | log2 (counts per million) | F-value | p-value | False discovery rate |
|---|---|---|---|---|---|---|---|
| ENSMUSG00000042105 | Inpp5f | protein_coding | 0.325411806 | 8.659097926 | 1.7471408 | 0.20448 | 0.741657995 |
| ENSMUSG00000018387 | Shroom1 | protein_coding | 0.742571064 | 2.888356445 | 1.7467902 | 0.2045232 | 0.741657995 |
| ENSMUSG00000061603 | Akap6 | protein_coding | -0.320560304 | 5.891526183 | 1.7462438 | 0.2045905 | 0.741657995 |
| ENSMUSG00000020883 | Fbxl20 | protein_coding | -0.493465209 | 3.978598993 | 1.7462327 | 0.2045919 | 0.741657995 |
| ENSMUSG00000051934 | Spats2 | protein_coding | -0.379589716 | 4.648541293 | 1.7460805 | 0.2046106 | 0.741657995 |
| ENSMUSG00000041261 | Car8 | protein_coding | 0.518793277 | 4.349591531 | 1.7458656 | 0.2046371 | 0.741657995 |
| ENSMUSG00000089809 | Rasgef1b | protein_coding | 0.408450119 | 5.397372802 | 1.7454433 | 0.2046892 | 0.741657995 |
| ENSMUSG00000007944 | Ttc9b | protein_coding | -0.503502582 | 4.256573623 | 1.7447638 | 0.2047731 | 0.741657995 |
| ENSMUSG00000022674 | Ube2v2 | protein_coding | -0.38482456 | 4.830762363 | 1.7447201 | 0.2047784 | 0.741657995 |
| ENSMUSG00000085492 | Trmt61b | processed_transcript | -0.370996075 | 5.168343951 | 1.7440306 | 0.2048636 | 0.741666817 |
| ENSMUSG00000063600 | Egfem1 | protein_coding | -0.537538986 | 3.503946583 | 1.743459 | 0.2049342 | 0.741666817 |
| ENSMUSG00000002147 | Stat6 | protein_coding | 0.827407913 | 2.770354408 | 1.7427764 | 0.2050185 | 0.741666817 |
| ENSMUSG00000068114 | Ccdc134 | protein_coding | -0.695358742 | 3.203838353 | 1.741262 | 0.2052058 | 0.741666817 |
| ENSMUSG00000061718 | Ppp1r1b | protein_coding | -0.796374045 | 2.587541766 | 1.7412583 | 0.2052063 | 0.741666817 |
| ENSMUSG00000040618 | Pck2 | protein_coding | 0.391102377 | 5.48517705 | 1.7403338 | 0.2053207 | 0.741666817 |
| ENSMUSG00000003435 | Supt5 | protein_coding | 0.466422372 | 5.680561552 | 1.7410397 | 0.2053414 | 0.741666817 |
| ENSMUSG00000041120 | Nbl1 | protein_coding | -0.525963357 | 3.95341608 | 1.7400389 | 0.2053572 | 0.741666817 |
| ENSMUSG00000073542 | Cep76 | protein_coding | -0.726085468 | 2.759413771 | 1.7396806 | 0.2054016 | 0.741666817 |
| ENSMUSG00000109398 | Gm3854 | protein_coding | 0.809755539 | 3.106398389 | 1.7383146 | 0.205571 | 0.741801532 |
| ENSMUSG00000040423 | Rc3h1 | protein_coding | -0.365505301 | 4.843479795 | 1.7376885 | 0.2056486 | 0.741801532 |
| ENSMUSG00000041773 | Enc1 | protein_coding | 0.32747901 | 8.466463369 | 1.7362825 | 0.2058529 | 0.741801532 |
| ENSMUSG00000019820 | Utrn | protein_coding | -0.384121263 | 5.03436791 | 1.735902 | 0.2058705 | 0.741801532 |
| ENSMUSG00000036357 | Gpr101 | protein_coding | -0.418348757 | 4.869678402 | 1.7357964 | 0.2058836 | 0.741801532 |
| ENSMUSG00000002372 | Ranbp3 | protein_coding | -0.286739096 | 6.514298845 | 1.7357491 | 0.2058895 | 0.741801532 |
| ENSMUSG00000036087 | Slain2 | protein_coding | -0.838198883 | 3.484801061 | 1.7415114 | 0.2059333 | 0.741801532 |
| ENSMUSG00000017264 | Exosc10 | protein_coding | 0.417612676 | 5.261205355 | 1.7342057 | 0.2060814 | 0.741801532 |
| ENSMUSG00000029718 | Pcolce | protein_coding | 1.08264496 | 3.082834057 | 1.7436038 | 0.2061418 | 0.741801532 |
| ENSMUSG00000021983 | Atp8a2 | protein_coding | 0.468888492 | 4.255534243 | 1.7333481 | 0.2061881 | 0.741801532 |
| ENSMUSG00000062012 | Zfp13 | protein_coding | -0.374513433 | 5.172760166 | 1.7332705 | 0.2061978 | 0.741801532 |
| ENSMUSG00000055720 | Ubl7 | protein_coding | -0.27032852 | 7.95282427 | 1.7314827 | 0.2064205 | 0.742354538 |
| ENSMUSG00000001844 | Zdhhc4 | protein_coding | 0.340337578 | 5.724670165 | 1.7303306 | 0.2065643 | 0.74248471 |
| ENSMUSG00000019194 | Scn1b | protein_coding | -0.353631962 | 5.104356915 | 1.7300855 | 0.2065948 | 0.74248471 |
| ENSMUSG00000031540 | Kat6a | protein_coding | 0.487357682 | 4.057204075 | 1.7269983 | 0.2069806 | 0.743622671 |
| ENSMUSG00000047098 | Rnf31 | protein_coding | 0.550495064 | 4.120447558 | 1.72599 | 0.2071068 | 0.743827549 |
| ENSMUSG00000036333 | Kidins220 | protein_coding | 0.330620196 | 6.645574306 | 1.7247555 | 0.2072615 | 0.744134422 |
| ENSMUSG00000046688 | Tifa | protein_coding | -0.933661251 | 2.772496425 | 1.7235964 | 0.2074068 | 0.744194282 |
| ENSMUSG00000043251 | Exoc3l | protein_coding | 1.113881811 | 2.066138384 | 1.7234695 | 0.2074227 | 0.744194282 |
| ENSMUSG00000021811 | Dnajc9 | protein_coding | 0.373320175 | 6.019361797 | 1.7225155 | 0.2075424 | 0.744194282 |
| ENSMUSG00000000296 | Tpd52l1 | protein_coding | -0.233168508 | 7.58623538 | 1.7224157 | 0.207555 | 0.744194282 |
| ENSMUSG00000039108 | Lsm14b | protein_coding | -0.449532171 | 4.354056581 | 1.7215647 | 0.2076619 | 0.744303798 |
| ENSMUSG00000028167 | Bdh2 | protein_coding | -0.542558749 | 3.684436894 | 1.7209857 | 0.2077346 | 0.744303798 |
| ENSMUSG00000030226 | Lmo3 | protein_coding | -0.265337669 | 7.101001143 | 1.7205199 | 0.2077932 | 0.744303798 |
| ENSMUSG00000046785 | Epm2aip1 | protein_coding | -0.279065505 | 6.769392348 | 1.7196829 | 0.2078984 | 0.744432881 |
| ENSMUSG00000004127 | Trmt10a | protein_coding | -0.447097984 | 4.116830788 | 1.7175945 | 0.2081614 | 0.744908003 |
| ENSMUSG00000028397 | Kdm4c | protein_coding | -0.464685217 | 4.255247997 | 1.7171099 | 0.2082225 | 0.744908003 |
| ENSMUSG00000057234 | Mettl15 | protein_coding | -0.530712466 | 3.394642558 | 1.7168607 | 0.2082539 | 0.744908003 |
| ENSMUSG00000096054 | Syne1 | protein_coding | -0.304418413 | 6.143398477 | 1.7164029 | 0.2083116 | 0.744908003 |
| ENSMUSG00000021820 | Camk2g | protein_coding | 0.227499017 | 7.244062361 | 1.7155097 | 0.2084244 | 0.744908003 |
| ENSMUSG00000043079 | Synpo | protein_coding | -0.370377239 | 5.277590938 | 1.7150987 | 0.2084762 | 0.744908003 |
| ENSMUSG00000000581 | C1d | protein_coding | -0.284070491 | 9.177309877 | 1.714002 | 0.2086148 | 0.744908003 |
| ENSMUSG00000039879 | Heca | protein_coding | -0.467829768 | 3.829112711 | 1.7138106 | 0.208639 | 0.744908003 |
| ENSMUSG00000046637 | Ttc34 | protein_coding | -1.05899652 | 2.504701623 | 1.7135943 | 0.2086663 | 0.744908003 |
| ENSMUSG00000021012 | Zc3h14 | protein_coding | -0.445229047 | 4.454841109 | 1.713139 | 0.2087239 | 0.744908003 |
| ENSMUSG00000046079 | Lrrc8d | protein_coding | -0.396171029 | 4.44451498 | 1.7112114 | 0.2089678 | 0.745320555 |
| ENSMUSG00000102674 | 8030442B05Rik | lincRNA | -0.680176148 | 3.225545928 | 1.7111301 | 0.2089781 | 0.745320555 |
| ENSMUSG00000108776 | Gm45169 | lincRNA | -0.617715192 | 3.493116215 | 1.710417 | 0.2090684 | 0.74539553 |
| ENSMUSG00000020794 | Ube2g1 | protein_coding | -0.313616229 | 5.654516639 | 1.7078325 | 0.2093963 | 0.745961613 |
| ENSMUSG00000002341 | Ncan | protein_coding | 0.267593632 | 7.218162338 | 1.7071137 | 0.2094876 | 0.745961613 |
| ENSMUSG00000027180 | Fbxo3 | protein_coding | 0.274538588 | 7.1115003 | 1.7064226 | 0.2095754 | 0.745961613 |
| ENSMUSG00000020647 | Ncoa1 | protein_coding | -0.441610749 | 6.32571021 | 1.7153647 | 0.2095897 | 0.745961613 |
| ENSMUSG00000042185 | Nfrkb | protein_coding | 0.633843916 | 3.61727441 | 1.7054636 | 0.2096973 | 0.745961613 |
| ENSMUSG00000035048 | Anapc13 | protein_coding | -0.350767334 | 7.441151497 | 1.7057928 | 0.2097629 | 0.745961613 |
| ENSMUSG00000017009 | Sdc4 | protein_coding | -0.363966585 | 7.306298795 | 1.7069975 | 0.2098578 | 0.745961613 |
| ENSMUSG00000024325 | Ring1 | protein_coding | -0.777061593 | 2.785337148 | 1.7033011 | 0.2099727 | 0.745961613 |
| ENSMUSG00000047710 | Champ1 | protein_coding | 0.443750981 | 4.904989309 | 1.7032402 | 0.2099804 | 0.745961613 |
| ENSMUSG00000031775 | Pllp | protein_coding | -0.405551219 | 4.431402865 | 1.7028913 | 0.2100249 | 0.745961613 |
| ENSMUSG00000034667 | Xpot | protein_coding | 0.402868574 | 4.929697942 | 1.702749 | 0.210043 | 0.745961613 |
| ENSMUSG00000062961 | Ccdc177 | protein_coding | -0.530270945 | 3.767734098 | 1.7026183 | 0.2100597 | 0.745961613 |
| ENSMUSG00000032185 | Carm1 | protein_coding | -0.473646644 | 4.119040385 | 1.7013053 | 0.2102272 | 0.746020369 |
| ENSMUSG00000037788 | Vopp1 | protein_coding | 0.453989501 | 4.910781996 | 1.7009956 | 0.2102667 | 0.746020369 |
| ENSMUSG00000018401 | Mtmr4 | protein_coding | 0.383697232 | 5.241823159 | 1.7008523 | 0.210285 | 0.746020369 |
| ENSMUSG00000005069 | Pex5 | protein_coding | 0.435551109 | 4.408127076 | 1.7003138 | 0.2103537 | 0.746020369 |
| ENSMUSG00000059659 | Gm10069 | antisense_RNA | -0.691091566 | 3.082248651 | 1.6987076 | 0.210559 | 0.746502162 |
| ENSMUSG00000063160 | Numbl | protein_coding | 0.570752012 | 3.21841183 | 1.6977352 | 0.2106834 | 0.746519979 |

| ensembl_gene_id | external_gene_name | gene_biotype | log2 (fold change) | log2 (counts per million) | F-value | p-value | False discovery rate |
|---|---|---|---|---|---|---|---|
| ENSMUSG00000025425 | St8sia5 | protein_coding | 0.870469477 | 2.602638481 | 1.697583 | 0.2107029 | 0.746519979 |
| ENSMUSG00000038500 | Prr3 | protein_coding | 0.357991506 | 5.502780232 | 1.6969124 | 0.2107887 | 0.746578153 |
| ENSMUSG00000026176 | Ctdsp1 | protein_coding | -0.615685083 | 3.061700666 | 1.6959139 | 0.2109166 | 0.74667487 |
| ENSMUSG00000028247 | Coq3 | protein_coding | 0.326018515 | 6.189198552 | 1.6956152 | 0.2109549 | 0.74667487 |
| ENSMUSG00000029012 | Orc5 | protein_coding | 0.376951311 | 5.490140977 | 1.6946211 | 0.2110824 | 0.746835587 |
| ENSMUSG00000022763 | Aifm3 | protein_coding | 0.389810944 | 5.137190422 | 1.6936406 | 0.2112082 | 0.746835587 |
| ENSMUSG00000058325 | Dock1 | protein_coding | -0.487205565 | 4.34361608 | 1.6918787 | 0.2114346 | 0.746835587 |
| ENSMUSG00000034480 | Diaph2 | protein_coding | -0.521731654 | 4.128041007 | 1.6917784 | 0.2114475 | 0.746835587 |
| ENSMUSG00000038515 | Grtp1 | protein_coding | 0.674106235 | 3.133808473 | 1.6917643 | 0.2114493 | 0.746835587 |
| ENSMUSG00000000632 | Sez6 | protein_coding | -0.418919415 | 6.559710408 | 1.7003227 | 0.2114968 | 0.746835587 |
| ENSMUSG00000030069 | Prok2 | protein_coding | -0.533249249 | 3.528654604 | 1.6907121 | 0.2115846 | 0.746835587 |
| ENSMUSG00000020955 | Ap4s1 | protein_coding | -0.331006675 | 5.522627673 | 1.6902368 | 0.2116458 | 0.746835587 |
| ENSMUSG00000026238 | Ptma | protein_coding | -0.269543424 | 6.344461929 | 1.6898959 | 0.2116897 | 0.746835587 |
| ENSMUSG00000039033 | Tasp1 | protein_coding | -0.373820773 | 5.575886204 | 1.6898555 | 0.2116949 | 0.746835587 |
| ENSMUSG00000033712 | Ccar2 | protein_coding | 0.418871565 | 5.48746704 | 1.688797 | 0.2118312 | 0.746835652 |
| ENSMUSG00000018661 | Cog1 | protein_coding | 0.549925448 | 3.643888327 | 1.6885226 | 0.2118666 | 0.746835652 |
| ENSMUSG00000026288 | Inpp5d | protein_coding | 1.4406748 | 2.104741917 | 1.6884668 | 0.2119032 | 0.746835652 |
| ENSMUSG00000027316 | Gfra4 | protein_coding | -0.362819655 | 5.032923443 | 1.6874906 | 0.2119996 | 0.746930559 |
| ENSMUSG00000029403 | Cdkl2 | protein_coding | -0.359734567 | 5.228739217 | 1.6851523 | 0.2123015 | 0.746932526 |
| ENSMUSG00000005142 | Man2b1 | protein_coding | 0.415140728 | 4.325568572 | 1.6851409 | 0.212303 | 0.746932526 |
| ENSMUSG00000031154 | Otud5 | protein_coding | 0.404007686 | 5.270545097 | 1.6847763 | 0.2123501 | 0.746932526 |
| ENSMUSG00000056851 | Pcbp2 | protein_coding | -0.243965392 | 6.502847413 | 1.6846918 | 0.2123611 | 0.746932526 |
| ENSMUSG00000026374 | Tsn | protein_coding | 0.260041068 | 8.515474492 | 1.6843896 | 0.2124001 | 0.746932526 |
| ENSMUSG00000004366 | Sst | protein_coding | -0.357518896 | 8.91773104 | 1.6843627 | 0.2125107 | 0.746932526 |
| ENSMUSG00000037526 | Atg14 | protein_coding | 0.47660283 | 4.759605011 | 1.6833204 | 0.2125384 | 0.746932526 |
| ENSMUSG00000015757 | Ppil4 | protein_coding | 0.301421872 | 5.927004705 | 1.6831855 | 0.2125559 | 0.746932526 |
| ENSMUSG00000030471 | Zdhhc13 | protein_coding | 0.565090764 | 3.432563223 | 1.680853 | 0.212858 | 0.747749937 |
| ENSMUSG00000044328 | Trp53i13 | protein_coding | 0.509787148 | 3.714643279 | 1.6798554 | 0.2129874 | 0.747751663 |
| ENSMUSG00000031342 | Gpm6b | protein_coding | -0.282768557 | 8.856646969 | 1.67779 | 0.2132557 | 0.747751663 |
| ENSMUSG00000047731 | Wbp1l | protein_coding | -0.562653133 | 4.181229716 | 1.6775555 | 0.2132862 | 0.747751663 |
| ENSMUSG00000029512 | Ulk1 | protein_coding | 0.372594314 | 4.964379479 | 1.677308 | 0.2133183 | 0.747751663 |
| ENSMUSG00000092310 | Gm20509 | protein_coding | -0.592621167 | 3.420100164 | 1.6772455 | 0.2133265 | 0.747751663 |
| ENSMUSG00000022136 | Dnajc3 | protein_coding | 0.449996713 | 4.487794101 | 1.6761932 | 0.2134633 | 0.747751663 |
| ENSMUSG00000029249 | Rest | protein_coding | -1.069902675 | 2.127344686 | 1.6758922 | 0.2135025 | 0.747751663 |
| ENSMUSG00000028452 | Vcp | protein_coding | 0.806622692 | 2.751292978 | 1.6752968 | 0.21358 | 0.747751663 |
| ENSMUSG00000001445 | Mrpl10 | protein_coding | 0.298105489 | 6.93874724 | 1.6752705 | 0.2135835 | 0.747751663 |
| ENSMUSG00000003527 | Dgcr14 | protein_coding | 0.360461251 | 5.291420031 | 1.6748853 | 0.2136336 | 0.747751663 |
| ENSMUSG00000048249 | Crebrf | protein_coding | -0.264573093 | 6.30893955 | 1.6741006 | 0.2137359 | 0.747751663 |
| ENSMUSG00000028643 | Svbp | protein_coding | -0.332637515 | 5.704019942 | 1.6740596 | 0.2137412 | 0.747751663 |
| ENSMUSG00000054793 | Cadm4 | protein_coding | -0.589959828 | 4.443614982 | 1.6744696 | 0.2137947 | 0.747751663 |
| ENSMUSG00000041625 | Ggact | protein_coding | 0.359435355 | 5.141992694 | 1.6733627 | 0.2138321 | 0.747751663 |
| ENSMUSG00000039552 | Rsph4a | protein_coding | 0.947783498 | 2.769764578 | 1.6723723 | 0.2139613 | 0.7479513 |
| ENSMUSG00000025855 | Prkar1b | protein_coding | 0.379379697 | 10.12034065 | 1.6725534 | 0.2140952 | 0.7479513 |
| ENSMUSG00000003316 | Glg1 | protein_coding | 0.239158131 | 6.681295146 | 1.6713266 | 0.2140978 | 0.7479513 |
| ENSMUSG00000004947 | Dtx2 | protein_coding | 0.821291601 | 2.572193851 | 1.6704772 | 0.2142088 | 0.74805795 |
| ENSMUSG00000034640 | Tiparp | protein_coding | -0.526552578 | 4.451759757 | 1.6700284 | 0.2142675 | 0.74805795 |
| ENSMUSG00000035441 | Myo1d | protein_coding | -0.413704408 | 4.208263697 | 1.6692663 | 0.2143672 | 0.748091984 |
| ENSMUSG00000028708 | Mknk1 | protein_coding | 0.427859141 | 4.546987891 | 1.6687735 | 0.2144317 | 0.748091984 |
| ENSMUSG00000020704 | Asic2 | protein_coding | 0.281828014 | 5.867010755 | 1.667994 | 0.2145337 | 0.748091984 |
| ENSMUSG00000049800 | Sertad2 | protein_coding | -0.4522583 | 3.710481192 | 1.6678276 | 0.2145555 | 0.748091984 |
| ENSMUSG00000001323 | Srr | protein_coding | -0.359908473 | 4.668064243 | 1.6666933 | 0.2147042 | 0.748128944 |
| ENSMUSG00000031130 | Brs3 | protein_coding | 1.233196834 | 3.223265056 | 1.6886646 | 0.2147053 | 0.748128944 |
| ENSMUSG00000029810 | Tmem176b | protein_coding | 0.273062114 | 6.570724312 | 1.6659232 | 0.2148052 | 0.748173228 |
| ENSMUSG00000036782 | Klhl13 | protein_coding | -0.277889621 | 7.218380801 | 1.6648989 | 0.2149396 | 0.748173228 |
| ENSMUSG00000038816 | Ctnnal1 | protein_coding | -0.503460436 | 3.863867728 | 1.6642564 | 0.215024 | 0.748173228 |
| ENSMUSG00000034175 | Rhbdd3 | protein_coding | -0.397789015 | 4.91049185 | 1.6624517 | 0.2152612 | 0.748173228 |
| ENSMUSG00000056579 | Tug1 | processed_transcript | 0.441188384 | 6.521281845 | 1.672913 | 0.215394 | 0.748173228 |
| ENSMUSG00000021054 | Sgpp1 | protein_coding | -0.395659345 | 4.598487404 | 1.6613383 | 0.2154077 | 0.748173228 |
| ENSMUSG00000020246 | Hcfc2 | protein_coding | 0.427927264 | 4.458769391 | 1.6602853 | 0.2155464 | 0.748173228 |
| ENSMUSG00000046798 | Cldn12 | protein_coding | 0.404935802 | 4.887008155 | 1.660254 | 0.2155505 | 0.748173228 |
| ENSMUSG00000021376 | Tpmt | protein_coding | -0.441834361 | 3.970889341 | 1.660057 | 0.2155765 | 0.748173228 |
| ENSMUSG00000026970 | Rbms1 | protein_coding | -0.41866102 | 4.38989333 | 1.6584902 | 0.2157831 | 0.748173228 |
| ENSMUSG00000031533 | Mrps31 | protein_coding | 0.373880685 | 6.396049181 | 1.6584203 | 0.2158216 | 0.748173228 |
| ENSMUSG00000016181 | Diexf | protein_coding | 0.837572211 | 3.022252251 | 1.6578682 | 0.2158651 | 0.748173228 |
| ENSMUSG00000049624 | Slc17a5 | protein_coding | 0.453490087 | 4.188864397 | 1.6575638 | 0.2159053 | 0.748173228 |
| ENSMUSG00000003721 | Insig2 | protein_coding | 0.240935395 | 6.989022052 | 1.6570089 | 0.2159786 | 0.748173228 |
| ENSMUSG00000097059 | Fam120aos | lincRNA | -0.734506039 | 2.730748456 | 1.6563012 | 0.2160721 | 0.748173228 |
| ENSMUSG00000097393 | D030068K23Rik | lincRNA | -0.442349195 | 5.219533529 | 1.6562554 | 0.2160782 | 0.748173228 |
| ENSMUSG00000021190 | Lgmn | protein_coding | 0.297873546 | 7.624437304 | 1.6558925 | 0.2161262 | 0.748173228 |
| ENSMUSG00000019916 | P4ha1 | protein_coding | 0.453488522 | 4.435432049 | 1.655823 | 0.2161353 | 0.748173228 |
| ENSMUSG00000018820 | Zfyve27 | protein_coding | 0.349182198 | 5.296735211 | 1.6557723 | 0.216142 | 0.748173228 |
| ENSMUSG00000002010 | Idh3g | protein_coding | 0.238251225 | 8.415720531 | 1.6557452 | 0.2161456 | 0.748173228 |
| ENSMUSG00000062169 | Cnih4 | protein_coding | -0.319520597 | 5.796278403 | 1.6554919 | 0.2161791 | 0.748173228 |
| ENSMUSG00000062778 | Chia1 | protein_coding | 1.288049887 | 2.152560485 | 1.6548834 | 0.2162596 | 0.748211003 |

| ensembl_gene_id | external_gene_name | gene_biotype | log2 (fold change) | log2 (counts per million) | F-value | p-value | False discovery rate |
|---|---|---|---|---|---|---|---|
| ENSMUSG00000032478 | Nme6 | protein_coding | -0.33957341 | 5.070254869 | 1.6508034 | 0.2168004 | 0.749317668 |
| ENSMUSG00000036368 | Rmdn2 | protein_coding | -0.864762145 | 2.618125722 | 1.65069 | 0.2168154 | 0.749317668 |
| ENSMUSG00000039630 | Hnrnpu | protein_coding | -0.221778098 | 7.5492458 | 1.6502765 | 0.2168703 | 0.749317668 |
| ENSMUSG00000018774 | Cd68 | protein_coding | -0.442384963 | 5.002781027 | 1.6502505 | 0.2168738 | 0.749317668 |
| ENSMUSG00000068299 | Nat8f4 | protein_coding | 0.577396396 | 3.405434141 | 1.6498427 | 0.2169279 | 0.749317668 |
| ENSMUSG00000068735 | Trp53i11 | protein_coding | 0.341375602 | 5.634389852 | 1.6479522 | 0.2171792 | 0.749687803 |
| ENSMUSG00000026885 | Ttll11 | protein_coding | 0.451266466 | 4.24246534 | 1.6473911 | 0.2172539 | 0.749687803 |
| ENSMUSG00000060038 | Dhps | protein_coding | 0.407249559 | 5.142539167 | 1.6466059 | 0.2173584 | 0.749687803 |
| ENSMUSG00000018171 | Vmp1 | protein_coding | 0.245869009 | 7.624051019 | 1.6463456 | 0.2173931 | 0.749687803 |
| ENSMUSG00000024127 | Prepl | protein_coding | 0.751743954 | 7.106366955 | 1.7277257 | 0.2174491 | 0.749687803 |
| ENSMUSG00000000282 | Mnt | protein_coding | -0.402161451 | 4.597142169 | 1.6454805 | 0.2175084 | 0.749687803 |
| ENSMUSG00000015981 | Stk32c | protein_coding | -0.679915934 | 3.19143195 | 1.6453702 | 0.2175231 | 0.749687803 |
| ENSMUSG00000003814 | Calr | protein_coding | 0.246453965 | 8.158670102 | 1.6446391 | 0.2176206 | 0.749783485 |
| ENSMUSG00000031012 | Cask | protein_coding | 0.416711527 | 7.302078276 | 1.6569826 | 0.2177611 | 0.749860438 |
| ENSMUSG00000029407 | Uso1 | protein_coding | -0.342400145 | 5.557244643 | 1.6434266 | 0.2177824 | 0.749860438 |
| ENSMUSG00000028563 | Tm2d1 | protein_coding | -0.494725764 | 4.406926536 | 1.6414708 | 0.2180437 | 0.749884125 |
| ENSMUSG00000043670 | Diras1 | protein_coding | -0.500333944 | 5.99051786 | 1.6545467 | 0.2180824 | 0.749884125 |
| ENSMUSG00000027716 | Trpc3 | protein_coding | 0.839141717 | 2.94328663 | 1.6399871 | 0.2182422 | 0.749884125 |
| ENSMUSG00000057342 | Sphk2 | protein_coding | -0.429355688 | 4.929433609 | 1.6398177 | 0.2182649 | 0.749884125 |
| ENSMUSG00000020873 | Slc35b1 | protein_coding | 0.238408635 | 7.164479832 | 1.6392698 | 0.2183383 | 0.749884125 |
| ENSMUSG00000052681 | Rap1b | protein_coding | -0.351327064 | 5.520195589 | 1.6386009 | 0.2184279 | 0.749884125 |
| ENSMUSG00000017144 | Rnd3 | protein_coding | 0.403638427 | 5.151207108 | 1.6384537 | 0.2184476 | 0.749884125 |
| ENSMUSG00000033009 | Ogfod1 | protein_coding | 0.283577288 | 6.537881971 | 1.638399 | 0.218455 | 0.749884125 |
| ENSMUSG00000037754 | Ppp1r16b | protein_coding | -0.321890055 | 8.07945623 | 1.6382343 | 0.2185063 | 0.749884125 |
| ENSMUSG00000042444 | Mindy2 | protein_coding | -0.414500441 | 5.504832714 | 1.6379286 | 0.218518 | 0.749884125 |
| ENSMUSG00000032599 | Ip6k2 | protein_coding | 0.257025438 | 6.790141189 | 1.6371417 | 0.2186236 | 0.749884125 |
| ENSMUSG00000030168 | Adipor2 | protein_coding | 0.559382934 | 4.827109208 | 1.6408221 | 0.2186261 | 0.749884125 |
| ENSMUSG00000033416 | Gucd1 | protein_coding | 0.775052606 | 3.277746035 | 1.6339347 | 0.2190836 | 0.751213734 |
| ENSMUSG00000070476 | Fam217b | protein_coding | 0.284133879 | 5.854551818 | 1.6302371 | 0.2195525 | 0.752273726 |
| ENSMUSG00000020954 | Strn3 | protein_coding | -0.372660245 | 5.765313245 | 1.6301988 | 0.2195577 | 0.752273726 |
| ENSMUSG00000032423 | Syncrip | protein_coding | -0.534398551 | 3.30915469 | 1.6296716 | 0.2196289 | 0.752273726 |
| ENSMUSG00000043415 | Otud1 | protein_coding | -0.671992915 | 3.314810434 | 1.6293475 | 0.2196726 | 0.752273726 |
| ENSMUSG00000037679 | Inf2 | protein_coding | -0.414634756 | 4.847573096 | 1.6274684 | 0.2199265 | 0.752903342 |
| ENSMUSG00000036120 | Rfxank | protein_coding | -0.380519703 | 4.926668222 | 1.625669 | 0.2201699 | 0.753098322 |
| ENSMUSG00000044365 | Cxxc4 | protein_coding | -0.363920354 | 5.379609679 | 1.6255926 | 0.2201803 | 0.753098322 |
| ENSMUSG00000050029 | Rap2c | protein_coding | -0.445620109 | 4.081772515 | 1.6254946 | 0.2201935 | 0.753098322 |
| ENSMUSG00000018102 | Hist1h2bc | protein_coding | -0.460610587 | 5.968762913 | 1.6328467 | 0.2203312 | 0.753161677 |
| ENSMUSG00000026694 | Mettl13 | protein_coding | 0.568628329 | 3.717727346 | 1.6242649 | 0.2203601 | 0.753161677 |
| ENSMUSG00000008153 | Clstn3 | protein_coding | 0.272213888 | 8.246138266 | 1.6235122 | 0.2204622 | 0.753161677 |
| ENSMUSG00000031467 | Agpat5 | protein_coding | -0.357442981 | 5.353533336 | 1.6232908 | 0.2204922 | 0.753161677 |
| ENSMUSG00000047036 | Zfp445 | protein_coding | -0.407259882 | 4.849598076 | 1.6225473 | 0.2205931 | 0.753233288 |
| ENSMUSG00000042055 | Wdr11 | protein_coding | 0.384727466 | 4.757641252 | 1.6219775 | 0.2206705 | 0.753233288 |
| ENSMUSG00000037740 | Mrps26 | protein_coding | 0.352760574 | 4.850606707 | 1.6204195 | 0.2208822 | 0.753233288 |
| ENSMUSG00000025289 | Prdx4 | protein_coding | 0.341531537 | 5.927358822 | 1.6203778 | 0.2208878 | 0.753233288 |
| ENSMUSG00000026134 | Prim2 | protein_coding | 0.399125129 | 4.413397606 | 1.6180405 | 0.2212059 | 0.753233288 |
| ENSMUSG00000019505 | Ubb | protein_coding | 0.264481188 | 10.89795493 | 1.6176235 | 0.2212627 | 0.753233288 |
| ENSMUSG00000056537 | Rlim | protein_coding | 0.395933728 | 5.508096725 | 1.6172026 | 0.2213201 | 0.753233288 |
| ENSMUSG00000029022 | Miip | protein_coding | -0.522383987 | 3.589943655 | 1.616402 | 0.2214292 | 0.753233288 |
| ENSMUSG00000040111 | Gramd1b | protein_coding | -0.328739716 | 5.352504123 | 1.6162437 | 0.2214508 | 0.753233288 |
| ENSMUSG00000009535 | Rnmt | protein_coding | 0.29436304 | 6.789872447 | 1.6160792 | 0.2214733 | 0.753233288 |
| ENSMUSG00000031948 | Kars | protein_coding | 0.346092751 | 7.945595081 | 1.6162305 | 0.2215584 | 0.753233288 |
| ENSMUSG00000014748 | Tex261 | protein_coding | -0.379502621 | 5.957894525 | 1.615481 | 0.221584 | 0.753233288 |
| ENSMUSG00000021069 | Pygl | protein_coding | 0.939674965 | 2.657915326 | 1.6144942 | 0.2216896 | 0.753233288 |
| ENSMUSG00000041747 | Utp15 | protein_coding | 0.419333809 | 4.508978226 | 1.6141201 | 0.2217408 | 0.753233288 |
| ENSMUSG00000110080 | Gm6145 | processed_transcript | -0.996079097 | 3.08788904 | 1.6213326 | 0.2218173 | 0.753233288 |
| ENSMUSG00000022052 | Ppp2r2a | protein_coding | -0.329391988 | 6.583689077 | 1.6133447 | 0.2218467 | 0.753233288 |
| ENSMUSG00000047514 | Tspyl1 | protein_coding | -0.282001866 | 8.789173962 | 1.6123539 | 0.2219823 | 0.753233288 |
| ENSMUSG00000031723 | Txnl4b | protein_coding | 0.355702668 | 6.709429716 | 1.6114596 | 0.2221337 | 0.753233288 |
| ENSMUSG00000073155 | 1810058I24Rik | lincRNA | -0.23580457 | 6.7691686 | 1.6106731 | 0.2222124 | 0.753233288 |
| ENSMUSG00000027950 | Chrnb2 | protein_coding | 0.427945581 | 4.510799278 | 1.6102577 | 0.2222693 | 0.753233288 |
| ENSMUSG00000023942 | Slc29a1 | protein_coding | 0.433483719 | 4.982933342 | 1.6101415 | 0.2222852 | 0.753233288 |
| ENSMUSG00000050272 | Dscam | protein_coding | -0.399513921 | 4.946494286 | 1.6101254 | 0.2222875 | 0.753233288 |
| ENSMUSG00000070814 | 6330408A02Rik | protein_coding | 0.697260553 | 3.603768717 | 1.6099535 | 0.2223401 | 0.753233288 |
| ENSMUSG00000045519 | Zfp560 | protein_coding | -0.336299743 | 5.7476614 | 1.6092345 | 0.2224096 | 0.753233288 |
| ENSMUSG00000018923 | Med11 | protein_coding | 0.360051805 | 5.137953717 | 1.6086674 | 0.2224874 | 0.753233288 |
| ENSMUSG00000034403 | Pja1 | protein_coding | 0.34956377 | 8.745547709 | 1.6093271 | 0.2225025 | 0.753233288 |
| ENSMUSG00000075590 | Nrbp2 | protein_coding | 0.53089938 | 4.648332518 | 1.6087102 | 0.2225106 | 0.753233288 |
| ENSMUSG00000087184 | Gm11650 | processed_transcript | -0.864552599 | 2.571897185 | 1.606725 | 0.2227542 | 0.753233288 |
| ENSMUSG00000010453 | Kansl3 | protein_coding | 0.394233926 | 5.673572314 | 1.6059027 | 0.2228672 | 0.753233288 |
| ENSMUSG00000029121 | Crmp1 | protein_coding | 0.500535386 | 4.814668844 | 1.6058129 | 0.2229086 | 0.753233288 |
| ENSMUSG00000032356 | Rasgrf1 | protein_coding | 0.306947374 | 8.150316656 | 1.6047498 | 0.2230259 | 0.753233288 |
| ENSMUSG00000030120 | Mlf2 | protein_coding | -0.213671534 | 7.984126122 | 1.6040996 | 0.2231154 | 0.753233288 |
| ENSMUSG00000019132 | BC005537 | protein_coding | -0.264616664 | 6.782680237 | 1.6039575 | 0.223135 | 0.753233288 |
| ENSMUSG00000074649 | BC029722 | protein_coding | 0.499453012 | 4.083220011 | 1.6036817 | 0.223173 | 0.753233288 |

| ensembl_gene_id | external_gene_name | gene_biotype | log2 (fold change) | log2 (counts per million) | F-value | p-value | False discovery rate |
|---|---|---|---|---|---|---|---|
| ENSMUSG00000097121 | D130020L05Rik | antisense_RNA | -0.494298483 | 3.731361309 | 1.6034652 | 0.2232028 | 0.753233288 |
| ENSMUSG00000042293 | Gm5617 | protein_coding | -0.538071595 | 4.872114521 | 1.6042863 | 0.2232286 | 0.753233288 |
| ENSMUSG00000026249 | Serpine2 | protein_coding | 0.343842119 | 7.864507227 | 1.6031606 | 0.2233502 | 0.753233288 |
| ENSMUSG00000028560 | Usp1 | protein_coding | -0.65174987 | 3.026843183 | 1.6021127 | 0.2233893 | 0.753233288 |
| ENSMUSG00000027983 | Cyp2u1 | protein_coding | -0.461475114 | 4.690115659 | 1.602075 | 0.2233945 | 0.753233288 |
| ENSMUSG00000025968 | Ndufs1 | protein_coding | 0.420879752 | 5.702321585 | 1.6014636 | 0.2235078 | 0.753233288 |
| ENSMUSG00000022911 | Arl13b | protein_coding | -0.549322093 | 3.263438864 | 1.601229 | 0.2235112 | 0.753233288 |
| ENSMUSG00000028528 | Dnajc6 | protein_coding | 0.244280568 | 6.82525502 | 1.5997979 | 0.2237089 | 0.753233288 |
| ENSMUSG00000048271 | Rbm33 | protein_coding | -0.333863007 | 5.378286856 | 1.5995487 | 0.2237433 | 0.753233288 |
| ENSMUSG00000049550 | Clip1 | protein_coding | -0.287307468 | 6.116884207 | 1.5994577 | 0.2237559 | 0.753233288 |
| ENSMUSG00000000131 | Xpo6 | protein_coding | 0.535246558 | 5.281730071 | 1.6074746 | 0.223776 | 0.753233288 |
| ENSMUSG00000027523 | Gnas | protein_coding | -0.500341246 | 3.948568539 | 1.5988149 | 0.2238448 | 0.753233288 |
| ENSMUSG00000033055 | Ankrd54 | protein_coding | -0.516933579 | 3.488553445 | 1.5984256 | 0.2238986 | 0.753233288 |
| ENSMUSG00000029657 | Hsph1 | protein_coding | 0.252965854 | 7.818311086 | 1.5969142 | 0.2241079 | 0.753233288 |
| ENSMUSG00000029392 | Rilpl1 | protein_coding | 0.447142243 | 4.592945239 | 1.5966772 | 0.2241407 | 0.753233288 |
| ENSMUSG00000042046 | Dstyk | protein_coding | 0.341853411 | 5.169482576 | 1.5965659 | 0.2241561 | 0.753233288 |
| ENSMUSG00000018899 | Irf1 | protein_coding | 0.833906576 | 2.824912173 | 1.5962128 | 0.2242051 | 0.753233288 |
| ENSMUSG00000024218 | Taf11 | protein_coding | 0.240014038 | 6.62879639 | 1.5959326 | 0.2242439 | 0.753233288 |
| ENSMUSG00000064326 | Siva1 | protein_coding | -0.35935435 | 5.736593877 | 1.5958639 | 0.2242534 | 0.753233288 |
| ENSMUSG00000007950 | Abhd8 | protein_coding | -0.295539982 | 6.008048681 | 1.5955581 | 0.2242958 | 0.753233288 |
| ENSMUSG00000024963 | Dnajc4 | protein_coding | -0.320279654 | 6.0058377 | 1.594254 | 0.2244768 | 0.753605525 |
| ENSMUSG00000037709 | Fam13a | protein_coding | 0.484146888 | 3.768903823 | 1.592032 | 0.2247855 | 0.754406395 |
| ENSMUSG00000003031 | Cdkn1b | protein_coding | -0.442675549 | 5.148480046 | 1.5913603 | 0.2248789 | 0.754484475 |
| ENSMUSG00000022634 | Yaf2 | protein_coding | -0.256734011 | 6.893968802 | 1.5906859 | 0.2249728 | 0.754563956 |
| ENSMUSG00000029071 | Dvl1 | protein_coding | 0.415311842 | 4.1831954 | 1.5901215 | 0.2250513 | 0.75459217 |
| ENSMUSG00000021273 | Fdft1 | protein_coding | 0.280028738 | 7.190815928 | 1.5871648 | 0.2254636 | 0.755311188 |
| ENSMUSG00000042225 | Ammecr1 | protein_coding | -0.558600557 | 3.249757824 | 1.5862078 | 0.2255973 | 0.755311188 |
| ENSMUSG00000046432 | Bex3 | protein_coding | 0.259231583 | 9.760962919 | 1.5855327 | 0.2256916 | 0.755311188 |
| ENSMUSG00000029547 | Ints1 | protein_coding | 0.58546834 | 4.190595333 | 1.5838853 | 0.2259509 | 0.755311188 |
| ENSMUSG00000074555 | Gm10714 | antisense_RNA | -0.868297783 | 2.85445708 | 1.5830218 | 0.2260429 | 0.755311188 |
| ENSMUSG00000031661 | Nkd1 | protein_coding | -1.03475879 | 2.463176117 | 1.5815875 | 0.2262439 | 0.755311188 |
| ENSMUSG00000048240 | Gng7 | protein_coding | -0.308868154 | 8.62343048 | 1.5815662 | 0.2262469 | 0.755311188 |
| ENSMUSG00000029580 | Actb | protein_coding | 0.434558914 | 12.74089718 | 1.5886841 | 0.226304 | 0.755311188 |
| ENSMUSG00000039704 | Lmbrd2 | protein_coding | -0.930122812 | 2.339368941 | 1.5806915 | 0.2263696 | 0.755311188 |
| ENSMUSG00000084830 | Gm14539 | processed_pseudoge | -0.464675484 | 3.93243946 | 1.5803456 | 0.2264182 | 0.755311188 |
| ENSMUSG00000030591 | Psmd8 | protein_coding | 0.241685274 | 8.434043169 | 1.5802449 | 0.2264323 | 0.755311188 |
| ENSMUSG00000048794 | Cfap100 | protein_coding | -0.930424665 | 2.308252603 | 1.58001 | 0.2264653 | 0.755311188 |
| ENSMUSG00000036564 | Ndrg4 | protein_coding | 0.384014735 | 11.22154694 | 1.5807239 | 0.2265199 | 0.755311188 |
| ENSMUSG00000031148 | Gpkow | protein_coding | 0.435644843 | 5.156053859 | 1.5792234 | 0.2265758 | 0.755311188 |
| ENSMUSG00000020783 | Ncbp3 | protein_coding | 0.45272651 | 4.194204079 | 1.5791862 | 0.226581 | 0.755311188 |
| ENSMUSG00000028029 | Aimp1 | protein_coding | -0.28432316 | 7.773119981 | 1.5780364 | 0.2267426 | 0.755311188 |
| ENSMUSG00000020679 | Hnf1b | protein_coding | -0.578435893 | 3.596697642 | 1.5776317 | 0.2267995 | 0.755311188 |
| ENSMUSG00000002871 | Tpra1 | protein_coding | 0.350083874 | 5.906543138 | 1.5775549 | 0.2268103 | 0.755311188 |
| ENSMUSG00000032300 | 1700017B05Rik | protein_coding | 0.853276469 | 3.158139168 | 1.5771377 | 0.2268979 | 0.755311188 |
| ENSMUSG00000003423 | Pih1d1 | protein_coding | 0.371203978 | 5.332552352 | 1.5767493 | 0.2269237 | 0.755311188 |
| ENSMUSG00000079056 | Kcnip3 | protein_coding | -0.248659637 | 6.305152198 | 1.5761932 | 0.227002 | 0.755311188 |
| ENSMUSG00000028039 | Efna3 | protein_coding | -0.471703244 | 4.154770139 | 1.5755338 | 0.2270949 | 0.755311188 |
| ENSMUSG00000021079 | Timm9 | protein_coding | -0.291627928 | 7.126191626 | 1.5755056 | 0.2270988 | 0.755311188 |
| ENSMUSG00000041921 | Metap1d | protein_coding | -0.517934466 | 3.832582105 | 1.575148 | 0.2271492 | 0.755311188 |
| ENSMUSG00000029681 | Bcl7b | protein_coding | 0.31201394 | 6.484288191 | 1.5747639 | 0.2272034 | 0.755311188 |
| ENSMUSG00000024732 | Ccdc86 | protein_coding | -0.452619822 | 4.064614931 | 1.5742105 | 0.2272814 | 0.755311188 |
| ENSMUSG00000025227 | Mfsd13a | protein_coding | -0.377524838 | 4.583608346 | 1.5741767 | 0.2272862 | 0.755311188 |
| ENSMUSG00000001436 | Slc19a1 | protein_coding | -0.61746148 | 4.7001377 | 1.5810749 | 0.2274346 | 0.755311188 |
| ENSMUSG00000028483 | Snapc3 | protein_coding | -0.310709502 | 5.714900084 | 1.5724854 | 0.2275249 | 0.755311188 |
| ENSMUSG00000000568 | Hnrnpd | protein_coding | -0.279079397 | 6.889537929 | 1.5722997 | 0.2275512 | 0.755311188 |
| ENSMUSG00000040943 | Tet2 | protein_coding | 0.392640408 | 5.009119942 | 1.5721493 | 0.2275724 | 0.755311188 |
| ENSMUSG00000039234 | Sec24d | protein_coding | 0.63300583 | 3.237169065 | 1.5720995 | 0.2275795 | 0.755311188 |
| ENSMUSG00000023809 | Rps6ka2 | protein_coding | 0.478598933 | 4.225923846 | 1.5716166 | 0.2276477 | 0.755311188 |
| ENSMUSG00000037735 | 2810032G03Rik | lincRNA | 0.726922313 | 2.874450947 | 1.5714497 | 0.2276713 | 0.755311188 |
| ENSMUSG00000038530 | Rgs4 | protein_coding | 0.238047865 | 8.482805222 | 1.5709088 | 0.2277478 | 0.755311188 |
| ENSMUSG00000020078 | Vps26a | protein_coding | -0.283605174 | 6.546155547 | 1.5704306 | 0.2278154 | 0.755311188 |
| ENSMUSG00000029634 | Rnf6 | protein_coding | 0.351138694 | 7.05288655 | 1.571109 | 0.227874 | 0.755311188 |
| ENSMUSG00000004798 | Ulk2 | protein_coding | -0.384566403 | 5.441127595 | 1.5695504 | 0.22794 | 0.755311188 |
| ENSMUSG00000026219 | Trip12 | protein_coding | 0.270583702 | 6.114001535 | 1.5690898 | 0.2280052 | 0.755311188 |
| ENSMUSG00000032259 | Drd2 | protein_coding | -0.451571622 | 5.144395593 | 1.5687865 | 0.228077 | 0.755316256 |
| ENSMUSG00000067274 | Rplp0 | protein_coding | -0.225470607 | 8.351305797 | 1.5676998 | 0.2282022 | 0.755480361 |
| ENSMUSG00000049960 | Mrps16 | protein_coding | 0.239161169 | 6.888654785 | 1.5672425 | 0.2282671 | 0.755480361 |
| ENSMUSG00000048482 | Bdnf | protein_coding | 0.48995485 | 3.991353696 | 1.5653192 | 0.2285401 | 0.755813387 |
| ENSMUSG00000032602 | Slc25a20 | protein_coding | 0.373963672 | 4.862579175 | 1.5650448 | 0.2285791 | 0.755813387 |
| ENSMUSG00000035953 | Tmem55b | protein_coding | 0.351788189 | 5.558917797 | 1.565034 | 0.2285806 | 0.755813387 |
| ENSMUSG00000029594 | Rbm19 | protein_coding | -0.625326458 | 3.338040783 | 1.5645539 | 0.2286488 | 0.755813387 |
| ENSMUSG00000018412 | Kansl1 | protein_coding | -0.379513136 | 5.409290668 | 1.5631909 | 0.2288427 | 0.756040026 |
| ENSMUSG00000050796 | B3galt6 | protein_coding | 0.553660613 | 3.461346429 | 1.5630832 | 0.228858 | 0.756040026 |
| ENSMUSG00000029769 | Ccdc136 | protein_coding | -0.397091438 | 6.416344774 | 1.5654849 | 0.2292506 | 0.757104451 |

| ensembl_gene_id | external_gene_name | gene_biotype | log2 (fold change) | log2 (counts per million) | F-value | p-value | False discovery rate |
|---|---|---|---|---|---|---|---|
| ENSMUSG00000001143 | Lman2l | protein_coding | -0.341141207 | 5.726341757 | 1.5597462 | 0.2293336 | 0.757145854 |
| ENSMUSG00000052459 | Atp6v1a | protein_coding | 0.280710117 | 8.941268298 | 1.5584597 | 0.2295173 | 0.757221493 |
| ENSMUSG00000029054 | Gabrd | protein_coding | -0.417268072 | 5.524120706 | 1.5585154 | 0.229538 | 0.757221493 |
| ENSMUSG00000024241 | Sos1 | protein_coding | 0.566856776 | 3.380003633 | 1.5581063 | 0.2295678 | 0.757221493 |
| ENSMUSG00000109946 | D2Bwg1423e | protein_coding | 0.262847714 | 6.58515184 | 1.5560058 | 0.2298681 | 0.757897513 |
| ENSMUSG00000008129 | 4930432K21Rik | protein_coding | 0.882641007 | 2.80066687 | 1.5556154 | 0.229924 | 0.757897513 |
| ENSMUSG00000007836 | Hnrnpa0 | protein_coding | -0.277310911 | 7.269052776 | 1.5551674 | 0.2299882 | 0.757897513 |
| ENSMUSG00000027994 | Mcub | protein_coding | -0.715907383 | 3.54038633 | 1.5546929 | 0.2300849 | 0.757897513 |
| ENSMUSG00000031762 | Mt2 | protein_coding | -0.525518288 | 5.28709843 | 1.5620103 | 0.2301251 | 0.757897513 |
| ENSMUSG00000026142 | Rhbdd1 | protein_coding | 1.096126043 | 2.721468663 | 1.5527176 | 0.2304438 | 0.758498269 |
| ENSMUSG00000034187 | Nsf | protein_coding | 0.405330152 | 9.831542962 | 1.5597008 | 0.2305246 | 0.758498269 |
| ENSMUSG00000051177 | Plcb1 | protein_coding | -0.557771622 | 3.797051818 | 1.5508676 | 0.2306051 | 0.758498269 |
| ENSMUSG00000004980 | Hnrnpa2b1 | protein_coding | 0.222743111 | 8.948773507 | 1.5505119 | 0.2306563 | 0.758498269 |
| ENSMUSG00000030754 | Copb1 | protein_coding | 0.347150577 | 5.99782051 | 1.5494343 | 0.2308112 | 0.758498269 |
| ENSMUSG00000027568 | Ntsr1 | protein_coding | 0.806296131 | 2.674081521 | 1.5493507 | 0.2308233 | 0.758498269 |
| ENSMUSG00000072623 | Zfp9 | protein_coding | 0.583491454 | 4.1590308 | 1.549356 | 0.2308512 | 0.758498269 |
| ENSMUSG00000046727 | Cystm1 | protein_coding | -0.233029714 | 8.856223718 | 1.5489174 | 0.2308856 | 0.758498269 |
| ENSMUSG00000043587 | Pxylp1 | protein_coding | 0.498646287 | 3.931386906 | 1.5482069 | 0.2309879 | 0.758498269 |
| ENSMUSG00000020658 | Efr3b | protein_coding | 0.293820428 | 5.894749229 | 1.5473768 | 0.2311076 | 0.758498269 |
| ENSMUSG00000047126 | Cltc | protein_coding | -0.227331003 | 8.0806273 | 1.545906 | 0.2313197 | 0.758498269 |
| ENSMUSG00000034616 | Ssh3 | protein_coding | -0.651839181 | 3.420697223 | 1.5458984 | 0.2313208 | 0.758498269 |
| ENSMUSG00000006586 | Runx1t1 | protein_coding | -0.384973905 | 6.322733554 | 1.5452605 | 0.2315665 | 0.758498269 |
| ENSMUSG00000002731 | Prkra | protein_coding | 0.281026024 | 6.252492785 | 1.5434428 | 0.2316755 | 0.758498269 |
| ENSMUSG00000086363 | A330102I10Rik | lincRNA | -0.408439716 | 4.731109168 | 1.543431 | 0.2316772 | 0.758498269 |
| ENSMUSG00000030232 | Aebp2 | protein_coding | -0.348011021 | 5.211686 | 1.5424174 | 0.2318238 | 0.758498269 |
| ENSMUSG00000029703 | Lrwd1 | protein_coding | 0.621215857 | 3.433185979 | 1.5422403 | 0.2318494 | 0.758498269 |
| ENSMUSG00000041438 | Utp4 | protein_coding | 0.411640574 | 5.843235692 | 1.5398634 | 0.2322224 | 0.758498269 |
| ENSMUSG00000029471 | Camkk2 | protein_coding | -0.410071134 | 4.662695683 | 1.5395707 | 0.2322362 | 0.758498269 |
| ENSMUSG00000039656 | Rxrb | protein_coding | -0.351476639 | 5.001536529 | 1.5395185 | 0.2322438 | 0.758498269 |
| ENSMUSG00000009291 | Pttg1ip | protein_coding | 0.287091648 | 6.171831776 | 1.5389255 | 0.2323298 | 0.758498269 |
| ENSMUSG00000030752 | Kdm8 | protein_coding | 0.854924788 | 2.7555095 | 1.5388146 | 0.2323459 | 0.758498269 |
| ENSMUSG00000078813 | Leng1 | protein_coding | -0.783136532 | 2.817330814 | 1.538698 | 0.2323629 | 0.758498269 |
| ENSMUSG00000002210 | Smg9 | protein_coding | -0.428789213 | 4.2138044 | 1.5378022 | 0.2324929 | 0.758498269 |
| ENSMUSG00000025588 | Nat1 | protein_coding | -0.969498306 | 2.428411009 | 1.5375724 | 0.2325263 | 0.758498269 |
| ENSMUSG00000027775 | Mfsd1 | protein_coding | -0.555575963 | 3.452838777 | 1.5369026 | 0.2326236 | 0.758498269 |
| ENSMUSG00000053334 | Ficd | protein_coding | 0.613603616 | 3.187597239 | 1.53649 | 0.2326836 | 0.758498269 |
| ENSMUSG00000028692 | Akr1a1 | protein_coding | 0.250787808 | 10.57220494 | 1.5362728 | 0.2327152 | 0.758498269 |
| ENSMUSG00000019990 | Pde7b | protein_coding | -0.428066776 | 4.13932871 | 1.5362164 | 0.2327234 | 0.758498269 |
| ENSMUSG00000047879 | Usp14 | protein_coding | 0.285253246 | 7.679212796 | 1.5358254 | 0.2327803 | 0.758498269 |
| ENSMUSG00000023952 | Gtpbp2 | protein_coding | -0.453585593 | 4.165417954 | 1.5353538 | 0.2328489 | 0.758498269 |
| ENSMUSG00000097736 | 9530059O14Rik | lincRNA | -1.097627357 | 2.537022903 | 1.535441 | 0.2328648 | 0.758498269 |
| ENSMUSG00000050132 | Sarm1 | protein_coding | 0.618243785 | 3.056241751 | 1.5346323 | 0.232954 | 0.758498269 |
| ENSMUSG00000049252 | Lrp1b | protein_coding | -0.461407318 | 5.20458909 | 1.5348107 | 0.2329566 | 0.758498269 |
| ENSMUSG00000020962 | Gtf2a1 | protein_coding | -0.302123006 | 5.943670029 | 1.5341757 | 0.2330205 | 0.758498269 |
| ENSMUSG00000033813 | Tcea1 | protein_coding | -0.379092061 | 4.076756085 | 1.5340732 | 0.2330354 | 0.758498269 |
| ENSMUSG00000096210 | H1f0 | protein_coding | 0.274817883 | 5.834766493 | 1.5334484 | 0.2331265 | 0.758498269 |
| ENSMUSG00000078622 | Ccdc47 | protein_coding | 0.36457351 | 5.700669994 | 1.533275 | 0.2331517 | 0.758498269 |
| ENSMUSG00000003273 | Car11 | protein_coding | -0.392522638 | 5.660026022 | 1.5327454 | 0.233229 | 0.758498269 |
| ENSMUSG00000021036 | Sptlc2 | protein_coding | 0.544795439 | 3.764679548 | 1.5321704 | 0.2333128 | 0.758498269 |
| ENSMUSG00000034353 | Ramp1 | protein_coding | -0.424545388 | 6.466903309 | 1.5416144 | 0.2333672 | 0.758498269 |
| ENSMUSG00000000214 | Th | protein_coding | -0.431163234 | 5.627137338 | 1.5318668 | 0.2333857 | 0.758498269 |
| ENSMUSG00000038116 | Phf20 | protein_coding | 0.247899146 | 6.675524015 | 1.5312776 | 0.2334432 | 0.758498269 |
| ENSMUSG00000053841 | Txlna | protein_coding | 0.41741243 | 4.507375285 | 1.5312581 | 0.233446 | 0.758498269 |
| ENSMUSG00000004937 | Sgta | protein_coding | 0.372107779 | 8.72233234 | 1.5380825 | 0.2335608 | 0.758498269 |
| ENSMUSG00000090399 | Gm38399 | lincRNA | -0.677979709 | 2.784710787 | 1.5301844 | 0.2336029 | 0.758498269 |
| ENSMUSG00000044080 | S100a1 | protein_coding | -0.428380214 | 8.199146881 | 1.5462401 | 0.2336878 | 0.758498269 |
| ENSMUSG00000049305 | Ccdc71 | protein_coding | 0.376475629 | 5.392133224 | 1.5295598 | 0.2336942 | 0.758498269 |
| ENSMUSG00000005161 | Prdx2 | protein_coding | 0.217270983 | 9.48512235 | 1.5290831 | 0.2337639 | 0.758498269 |
| ENSMUSG00000040151 | Hs2st1 | protein_coding | -0.433303261 | 4.851491335 | 1.5277959 | 0.2339523 | 0.758880588 |
| ENSMUSG00000059834 | Sclt1 | protein_coding | -0.647792863 | 3.183665427 | 1.5259767 | 0.2342189 | 0.759216963 |
| ENSMUSG00000037386 | Rims2 | protein_coding | -0.252972821 | 6.12669985 | 1.5256326 | 0.2342694 | 0.759216963 |
| ENSMUSG00000000711 | Rab5b | protein_coding | 0.402941234 | 6.431023966 | 1.532884 | 0.234316 | 0.759216963 |
| ENSMUSG00000040183 | Ankrd6 | protein_coding | -0.360991627 | 5.466469459 | 1.5251619 | 0.2343384 | 0.759216963 |
| ENSMUSG00000027293 | Ehd4 | protein_coding | 0.512806155 | 3.606209092 | 1.5246022 | 0.2344206 | 0.75925439 |
| ENSMUSG00000024810 | Il33 | protein_coding | -0.280371648 | 8.05266783 | 1.5236439 | 0.2345613 | 0.759481521 |
| ENSMUSG00000037750 | Fam222b | protein_coding | 0.343847537 | 5.159409597 | 1.521965 | 0.2348082 | 0.760051927 |
| ENSMUSG00000055489 | Ano5 | protein_coding | -0.693749487 | 4.58816733 | 1.5359047 | 0.2350234 | 0.760367172 |
| ENSMUSG00000033047 | Eif3l | protein_coding | 0.245936939 | 7.38292893 | 1.5194297 | 0.2351816 | 0.760367172 |
| ENSMUSG00000000301 | Pemt | protein_coding | 0.568557014 | 3.320569159 | 1.5190444 | 0.2352384 | 0.760367172 |
| ENSMUSG00000001829 | Clpb | protein_coding | -0.294418224 | 5.379357391 | 1.5189917 | 0.2352462 | 0.760367172 |
| ENSMUSG00000051107 | Gm15440 | protein_coding | 0.437058753 | 3.974827409 | 1.5188795 | 0.2352628 | 0.760367172 |
| ENSMUSG00000055024 | Ep300 | protein_coding | -0.541437017 | 3.593335121 | 1.5176321 | 0.2354468 | 0.760367172 |
| ENSMUSG00000035131 | Brinp3 | protein_coding | 0.358291639 | 5.172183855 | 1.5170983 | 0.2355257 | 0.760367172 |
| ENSMUSG00000035493 | Tgfbi | protein_coding | -0.966955362 | 2.223095746 | 1.5169274 | 0.2355509 | 0.760367172 |

| ensembl_gene_id | external_gene_name | gene_biotype | log2 (fold change) | log2 (counts per million) | F-value | p-value | False discovery rate |
|---|---|---|---|---|---|---|---|
| ENSMUSG00000024533 | Spire1 | protein_coding | -0.342882791 | 5.694342266 | 1.5166675 | 0.2355893 | 0.760367172 |
| ENSMUSG00000020899 | Pfas | protein_coding | 0.374368337 | 5.876812559 | 1.5157802 | 0.235749 | 0.760367172 |
| ENSMUSG00000001100 | Poldip2 | protein_coding | 0.276465751 | 5.884132122 | 1.5154226 | 0.2357734 | 0.760367172 |
| ENSMUSG00000022141 | Nipbl | protein_coding | -0.559802145 | 4.416390134 | 1.5152696 | 0.2358245 | 0.760367172 |
| ENSMUSG00000016940 | Kctd2 | protein_coding | 0.371496426 | 5.36580741 | 1.5150747 | 0.2358248 | 0.760367172 |
| ENSMUSG00000022836 | Mylk | protein_coding | -0.831941097 | 2.678548687 | 1.5125285 | 0.236202 | 0.761092544 |
| ENSMUSG00000030801 | Kat8 | protein_coding | 0.350374568 | 4.772854295 | 1.5122651 | 0.236241 | 0.761092544 |
| ENSMUSG00000027667 | Zfp639 | protein_coding | -0.33184897 | 5.600760563 | 1.5119036 | 0.2362947 | 0.761092544 |
| ENSMUSG00000024238 | Zeb1 | protein_coding | -0.266284075 | 6.277915847 | 1.5116456 | 0.2363329 | 0.761092544 |
| ENSMUSG00000022750 | Klhl22 | protein_coding | 0.267448837 | 6.821613233 | 1.5098487 | 0.2365997 | 0.761184525 |
| ENSMUSG00000036777 | Anln | protein_coding | -0.361578607 | 4.705413853 | 1.5097183 | 0.2366191 | 0.761184525 |
| ENSMUSG00000085881 | Gm15912 | processed_transcript | -0.531090619 | 3.367241813 | 1.5090528 | 0.236718 | 0.761184525 |
| ENSMUSG00000041702 | Btbd7 | protein_coding | 0.521634677 | 3.842871206 | 1.5084791 | 0.2368034 | 0.761184525 |
| ENSMUSG00000025354 | Dnajc14 | protein_coding | 0.42285622 | 4.999085832 | 1.5073589 | 0.2369701 | 0.761184525 |
| ENSMUSG00000048429 | Timm29 | protein_coding | 0.368758256 | 5.196733817 | 1.5066625 | 0.2370738 | 0.761184525 |
| ENSMUSG00000002103 | Acp2 | protein_coding | 0.584657102 | 4.573478707 | 1.5111865 | 0.2371264 | 0.761184525 |
| ENSMUSG00000021901 | Bap1 | protein_coding | 0.229058758 | 6.542753954 | 1.5058212 | 0.2371992 | 0.761184525 |
| ENSMUSG00000050473 | Slc35d3 | protein_coding | -0.711340072 | 3.052097934 | 1.5052975 | 0.2372773 | 0.761184525 |
| ENSMUSG00000024515 | Smad4 | protein_coding | 0.368574991 | 4.955008203 | 1.5052926 | 0.237278 | 0.761184525 |
| ENSMUSG00000031835 | Mbtps1 | protein_coding | 0.327681105 | 5.428851872 | 1.5052855 | 0.2372791 | 0.761184525 |
| ENSMUSG00000105891 | A230001M10Rik | lincRNA | -0.536930057 | 3.815658261 | 1.5050334 | 0.2373167 | 0.761184525 |
| ENSMUSG00000010136 | Pifo | protein_coding | -0.392372533 | 4.485323866 | 1.5046686 | 0.2373711 | 0.761184525 |
| ENSMUSG00000044647 | Csrnp3 | protein_coding | -0.265815782 | 6.295928373 | 1.5043853 | 0.2374134 | 0.761184525 |
| ENSMUSG00000004508 | Gab2 | protein_coding | -0.436212509 | 3.938016817 | 1.5043188 | 0.2374233 | 0.761184525 |
| ENSMUSG00000021234 | Fam161b | protein_coding | 0.605850867 | 3.909700963 | 1.5032242 | 0.2376152 | 0.761529172 |
| ENSMUSG00000060534 | Dcc | protein_coding | -0.653799839 | 3.846926997 | 1.5033416 | 0.2376725 | 0.761529172 |
| ENSMUSG00000004347 | Pde1c | protein_coding | -0.391547714 | 5.082863341 | 1.5012939 | 0.2378754 | 0.761952402 |
| ENSMUSG00000041468 | Gpr12 | protein_coding | -0.412881681 | 4.7505083 | 1.5001393 | 0.2380483 | 0.762278995 |
| ENSMUSG00000033499 | Larp4b | protein_coding | -0.268087952 | 6.176354794 | 1.4994369 | 0.2381535 | 0.762289942 |
| ENSMUSG00000029504 | Ddx51 | protein_coding | 1.269335019 | 2.463090803 | 1.5030136 | 0.2381935 | 0.762289942 |
| ENSMUSG00000026247 | Ecel1 | protein_coding | 0.407559136 | 5.931074143 | 1.4991823 | 0.2382948 | 0.762387229 |
| ENSMUSG00000052684 | Jun | protein_coding | 0.533010797 | 3.651238088 | 1.4969365 | 0.2385286 | 0.762398392 |
| ENSMUSG00000021504 | B4galt7 | protein_coding | 0.406663075 | 4.868777202 | 1.4961747 | 0.238643 | 0.762398392 |
| ENSMUSG00000035572 | Dcaf10 | protein_coding | -0.527215537 | 3.641918906 | 1.4958066 | 0.2386984 | 0.762398392 |
| ENSMUSG00000111336 | AC160338.2 | processed_pseudoge | 0.756556214 | 2.495703663 | 1.495164 | 0.238795 | 0.762398392 |
| ENSMUSG00000027673 | Ndufb5 | protein_coding | 0.239273759 | 9.868636205 | 1.4950131 | 0.2388177 | 0.762398392 |
| ENSMUSG00000005871 | Apc | protein_coding | -0.360494555 | 5.403525281 | 1.4947528 | 0.2388568 | 0.762398392 |
| ENSMUSG00000006998 | Psmd2 | protein_coding | 0.303758452 | 7.72167652 | 1.4949163 | 0.2388606 | 0.762398392 |
| ENSMUSG00000024776 | Stambpl1 | protein_coding | 0.355397705 | 4.764129208 | 1.4943986 | 0.2389101 | 0.762398392 |
| ENSMUSG00000055403 | 4933427D06Rik | lincRNA | 0.749348334 | 3.222625254 | 1.4943588 | 0.2389444 | 0.762398392 |
| ENSMUSG00000074698 | Csnk2a1 | protein_coding | 0.307644718 | 5.274515442 | 1.4937533 | 0.2390073 | 0.762398392 |
| ENSMUSG00000014355 | Anapc1 | protein_coding | 0.463153835 | 4.217643733 | 1.4924137 | 0.2392091 | 0.762600108 |
| ENSMUSG00000011960 | Ccnt1 | protein_coding | 0.339048035 | 5.296576263 | 1.4919429 | 0.23928 | 0.762600108 |
| ENSMUSG00000031392 | Irak1 | protein_coding | -0.32491751 | 5.165498465 | 1.4919216 | 0.2392832 | 0.762600108 |
| ENSMUSG00000051483 | Cbr1 | protein_coding | 0.277699962 | 7.257001137 | 1.4910223 | 0.2394189 | 0.762720311 |
| ENSMUSG00000025373 | Rnf41 | protein_coding | 0.262962058 | 7.783121997 | 1.4907313 | 0.2394628 | 0.762720311 |
| ENSMUSG00000039157 | Fam102a | protein_coding | -0.257241519 | 6.538388626 | 1.4901089 | 0.2395568 | 0.762793684 |
| ENSMUSG00000032609 | Klhdc8b | protein_coding | 0.358623953 | 5.787174411 | 1.4893182 | 0.2396762 | 0.762794941 |
| ENSMUSG00000063919 | Srrm4 | protein_coding | 0.437210254 | 4.491326494 | 1.4888104 | 0.239753 | 0.762794941 |
| ENSMUSG00000066880 | Zfp617 | protein_coding | -0.401058586 | 3.954046888 | 1.4886979 | 0.23977 | 0.762794941 |
| ENSMUSG00000056755 | Grm7 | protein_coding | -0.375342176 | 5.387431684 | 1.487846 | 0.2398988 | 0.762964068 |
| ENSMUSG00000022680 | Pdxdc1 | protein_coding | 0.351354056 | 5.315056667 | 1.4856439 | 0.2402323 | 0.762964068 |
| ENSMUSG00000034274 | Thoc5 | protein_coding | 0.358087925 | 5.329991792 | 1.4848475 | 0.2403531 | 0.762964068 |
| ENSMUSG00000032062 | 2310030G06Rik | protein_coding | 0.588526883 | 3.466498892 | 1.4848048 | 0.2403596 | 0.762964068 |
| ENSMUSG00000085885 | Gm11906 | processed_transcript | -0.39643387 | 4.577046964 | 1.4839111 | 0.2404952 | 0.762964068 |
| ENSMUSG00000051951 | Xkr4 | protein_coding | -0.640587127 | 3.610802665 | 1.4838879 | 0.240527 | 0.762964068 |
| ENSMUSG00000036339 | Tmem260 | protein_coding | -0.563104966 | 3.371221441 | 1.4831408 | 0.2406121 | 0.762964068 |
| ENSMUSG00000024298 | Zfp871 | protein_coding | 0.404068866 | 4.790642478 | 1.4831074 | 0.2406172 | 0.762964068 |
| ENSMUSG00000043448 | Gjc2 | protein_coding | -0.968734389 | 2.313014584 | 1.4827692 | 0.2406686 | 0.762964068 |
| ENSMUSG00000049313 | Sorl1 | protein_coding | 0.276002583 | 6.405108232 | 1.4826571 | 0.2406856 | 0.762964068 |
| ENSMUSG00000027498 | Cstf1 | protein_coding | 0.326863582 | 5.412004816 | 1.4816406 | 0.2408401 | 0.762964068 |
| ENSMUSG00000014177 | Tvp23b | protein_coding | 0.258437195 | 6.339747396 | 1.481544 | 0.2408548 | 0.762964068 |
| ENSMUSG00000020283 | Pex13 | protein_coding | -0.421863979 | 4.641362991 | 1.4814347 | 0.2408714 | 0.762964068 |
| ENSMUSG00000089759 | 3632454L22Rik | processed_transcript | -0.767607381 | 2.614671891 | 1.4810189 | 0.2409347 | 0.762964068 |
| ENSMUSG00000079562 | Maea | protein_coding | 0.275634294 | 7.573083674 | 1.480461 | 0.2410196 | 0.762964068 |
| ENSMUSG00000056476 | Med12l | protein_coding | -0.548134695 | 3.301636444 | 1.4801199 | 0.2410715 | 0.762964068 |
| ENSMUSG00000087479 | Gm16835 | processed_transcript | 0.573621734 | 3.380178744 | 1.4790126 | 0.2412402 | 0.762964068 |
| ENSMUSG00000041375 | Ccdc9 | protein_coding | -0.588812921 | 3.067042876 | 1.4785158 | 0.2413159 | 0.762964068 |
| ENSMUSG00000011884 | Gltp | protein_coding | -0.371870161 | 6.231484029 | 1.4783164 | 0.2413746 | 0.762964068 |
| ENSMUSG00000043460 | Elfn2 | protein_coding | -0.598377318 | 3.91564847 | 1.4782731 | 0.2413812 | 0.762964068 |
| ENSMUSG00000032834 | Pwp2 | protein_coding | 0.520580273 | 4.081071624 | 1.4778083 | 0.2414238 | 0.762964068 |
| ENSMUSG00000028273 | Pdlim5 | protein_coding | 0.526249462 | 4.080537185 | 1.477752 | 0.2414324 | 0.762964068 |
| ENSMUSG00000051007 | Gatd1 | protein_coding | -0.270997785 | 7.261766072 | 1.4776034 | 0.2414551 | 0.762964068 |
| ENSMUSG00000042705 | Commd10 | protein_coding | -0.234520561 | 6.755597794 | 1.4768705 | 0.241567 | 0.763093278 |

| ensembl_gene_id | external_gene_name | gene_biotype | log2 (fold change) | log2 (counts per million) | F-value | p-value | False discovery rate |
|---|---|---|---|---|---|---|---|
| ENSMUSG00000047617 | Paxx | protein_coding | -0.228579283 | 7.046361588 | 1.4758576 | 0.2417217 | 0.763357737 |
| ENSMUSG00000024826 | Dpf2 | protein_coding | 0.284177283 | 6.602826591 | 1.4746143 | 0.2419117 | 0.763685103 |
| ENSMUSG00000043518 | Rai2 | protein_coding | 0.485807692 | 4.267342959 | 1.4740435 | 0.2419991 | 0.763685103 |
| ENSMUSG00000049517 | Rps23 | protein_coding | -0.579168943 | 3.084731653 | 1.4736394 | 0.2420609 | 0.763685103 |
| ENSMUSG00000041225 | Arhgap12 | protein_coding | 0.298097429 | 5.38148747 | 1.4733227 | 0.2421094 | 0.763685103 |
| ENSMUSG00000034354 | Mtmr3 | protein_coding | 0.405803277 | 5.133218822 | 1.47138 | 0.2424071 | 0.763868761 |
| ENSMUSG00000066129 | Kndc1 | protein_coding | -0.407835074 | 4.97263177 | 1.4710467 | 0.2424582 | 0.763868761 |
| ENSMUSG00000021271 | Zfp839 | protein_coding | 0.414471833 | 3.894737059 | 1.4709155 | 0.2424784 | 0.763868761 |
| ENSMUSG00000013822 | Elof1 | protein_coding | 0.219907565 | 8.163650204 | 1.4708214 | 0.2424928 | 0.763868761 |
| ENSMUSG00000071064 | Zfp827 | protein_coding | -0.399559263 | 4.348245981 | 1.4700925 | 0.2426047 | 0.763868761 |
| ENSMUSG00000028546 | Elavl4 | protein_coding | -0.303298561 | 7.37055083 | 1.469783 | 0.2426804 | 0.763868761 |
| ENSMUSG00000014075 | Tctex1d2 | protein_coding | -0.244765428 | 6.552979031 | 1.4695387 | 0.2426897 | 0.763868761 |
| ENSMUSG00000035596 | Mboat7 | protein_coding | 0.268867784 | 6.622442919 | 1.4692379 | 0.2427359 | 0.763868761 |
| ENSMUSG00000018209 | Stk4 | protein_coding | 0.377201403 | 4.73692849 | 1.4687331 | 0.2428135 | 0.76388931 |
| ENSMUSG00000034601 | 2700049A03Rik | protein_coding | 0.864576425 | 2.973556882 | 1.4680298 | 0.2429498 | 0.764022701 |
| ENSMUSG00000038774 | Ascc3 | protein_coding | 0.619605854 | 3.242007027 | 1.4671906 | 0.2430507 | 0.764022701 |
| ENSMUSG00000048874 | Phf3 | protein_coding | -0.3363805 | 5.110300952 | 1.4670716 | 0.2430691 | 0.764022701 |
| ENSMUSG00000029919 | Hpgds | protein_coding | -0.620056123 | 3.063518813 | 1.4655778 | 0.2432991 | 0.76452234 |
| ENSMUSG00000022623 | Shank3 | protein_coding | -0.655723062 | 2.872524293 | 1.4638966 | 0.2435584 | 0.764898973 |
| ENSMUSG00000027007 | Ssfa2 | protein_coding | -0.725122053 | 3.402114785 | 1.4636734 | 0.2436209 | 0.764898973 |
| ENSMUSG00000022681 | Ntan1 | protein_coding | -0.248802485 | 6.963301012 | 1.4631518 | 0.2436734 | 0.764898973 |
| ENSMUSG00000053453 | Thoc7 | protein_coding | -0.306491372 | 7.306993074 | 1.4627253 | 0.2437673 | 0.764898973 |
| ENSMUSG00000061374 | Fiz1 | protein_coding | -0.529075259 | 3.292743738 | 1.462496 | 0.2437746 | 0.764898973 |
| ENSMUSG00000024667 | Tmem216 | protein_coding | 0.324836578 | 5.050720056 | 1.4619126 | 0.2438648 | 0.764910729 |
| ENSMUSG00000038074 | Fkbp14 | protein_coding | 0.547465887 | 3.382830478 | 1.4615513 | 0.2439207 | 0.764910729 |
| ENSMUSG00000068184 | Ndufaf2 | protein_coding | 0.269254555 | 7.34723088 | 1.4603196 | 0.2441112 | 0.765285067 |
| ENSMUSG00000023021 | Cers5 | protein_coding | 0.35857459 | 5.106722379 | 1.4596869 | 0.2442091 | 0.765368985 |
| ENSMUSG00000027752 | Exosc8 | protein_coding | -0.280865767 | 5.884115606 | 1.4583017 | 0.2444238 | 0.765405998 |
| ENSMUSG00000048458 | Fam212b | protein_coding | -0.433157873 | 5.087369993 | 1.4580793 | 0.2444583 | 0.765405998 |
| ENSMUSG00000036594 | H2-Aa | protein_coding | -1.146413557 | 2.156284574 | 1.4578505 | 0.2444938 | 0.765405998 |
| ENSMUSG00000029199 | Lias | protein_coding | 0.247442671 | 6.878861536 | 1.4572777 | 0.2445826 | 0.765405998 |
| ENSMUSG00000026932 | Nacc2 | protein_coding | -0.331797104 | 6.10164232 | 1.4563548 | 0.2447259 | 0.765405998 |
| ENSMUSG00000009470 | Tnpo1 | protein_coding | -0.411968922 | 4.766316474 | 1.4562803 | 0.2447374 | 0.765405998 |
| ENSMUSG00000022307 | Oxr1 | protein_coding | -0.334734311 | 7.482523217 | 1.4579734 | 0.2448004 | 0.765405998 |
| ENSMUSG00000049323 | Smcr8 | protein_coding | 0.512568717 | 3.611316607 | 1.4554475 | 0.2448668 | 0.765405998 |
| ENSMUSG00000022324 | Matn2 | protein_coding | 0.616070763 | 3.743421757 | 1.4549374 | 0.2449742 | 0.765405998 |
| ENSMUSG00000028093 | Acp6 | protein_coding | 0.567734496 | 3.547751919 | 1.4539574 | 0.2450985 | 0.765405998 |
| ENSMUSG00000024302 | Dtna | protein_coding | 0.250501737 | 6.816373177 | 1.4539369 | 0.2451017 | 0.765405998 |
| ENSMUSG00000029119 | Man2b2 | protein_coding | -0.569569632 | 3.495648565 | 1.4538905 | 0.2451089 | 0.765405998 |
| ENSMUSG00000022668 | Gtpbp8 | protein_coding | -0.372377272 | 4.736968523 | 1.4536504 | 0.2451463 | 0.765405998 |
| ENSMUSG00000017132 | Cyth1 | protein_coding | -0.293162789 | 5.502105999 | 1.4518087 | 0.2454332 | 0.76599573 |
| ENSMUSG00000056515 | Rab31 | protein_coding | 0.251900341 | 6.519935439 | 1.4511285 | 0.2455392 | 0.76599573 |
| ENSMUSG00000008140 | Emc10 | protein_coding | -0.325612779 | 5.708759003 | 1.4501774 | 0.2456876 | 0.76599573 |
| ENSMUSG00000063273 | Naa15 | protein_coding | -0.377207762 | 5.168552921 | 1.4500504 | 0.2457074 | 0.76599573 |
| ENSMUSG00000024360 | Etf1 | protein_coding | -0.27528306 | 6.051335038 | 1.4499401 | 0.2457247 | 0.76599573 |
| ENSMUSG00000023707 | Ogfod2 | protein_coding | 0.349378689 | 5.971557086 | 1.4496971 | 0.2457626 | 0.76599573 |
| ENSMUSG00000067369 | Trmt2b | protein_coding | 0.469853923 | 4.47209768 | 1.448579 | 0.2459373 | 0.766317987 |
| ENSMUSG00000049516 | Spty2d1 | protein_coding | -0.454375313 | 3.873395935 | 1.4463054 | 0.246293 | 0.767203991 |
| ENSMUSG00000079593 | Gm14597 | protein_coding | 0.21383706 | 8.072834169 | 1.4445531 | 0.2465676 | 0.767836945 |
| ENSMUSG00000032224 | Fam81a | protein_coding | -0.340635112 | 8.374444457 | 1.4470733 | 0.2467018 | 0.767871253 |
| ENSMUSG00000024830 | Rps6kb2 | protein_coding | 0.901356826 | 3.145732886 | 1.4484859 | 0.2467214 | 0.767871253 |
| ENSMUSG00000025266 | Gnl3l | protein_coding | 0.448922204 | 8.742299325 | 1.4595921 | 0.2469468 | 0.76819849 |
| ENSMUSG00000022442 | Ttll1 | protein_coding | 0.342863287 | 7.433576324 | 1.4458302 | 0.2470813 | 0.76819849 |
| ENSMUSG00000038936 | Sccpdh | protein_coding | 0.257604303 | 7.117289642 | 1.4412185 | 0.2470912 | 0.76819849 |
| ENSMUSG00000018405 | Mrm1 | protein_coding | -0.584552373 | 3.018384625 | 1.4410844 | 0.2471123 | 0.76819849 |
| ENSMUSG00000031613 | Hpgd | protein_coding | -0.597215184 | 3.347126738 | 1.4388647 | 0.2474617 | 0.769023748 |
| ENSMUSG00000015829 | Tnr | protein_coding | -0.475770535 | 3.979884262 | 1.4384766 | 0.2475229 | 0.769023748 |
| ENSMUSG00000025825 | Iscu | protein_coding | 0.34746959 | 4.795017122 | 1.4380362 | 0.2475923 | 0.769023748 |
| ENSMUSG00000031913 | Vps4a | protein_coding | 0.501007816 | 5.448352667 | 1.4436413 | 0.2477945 | 0.769429605 |
| ENSMUSG00000086657 | Stamos | processed_transcript | -0.800671492 | 2.476351875 | 1.4362845 | 0.2478687 | 0.769437739 |
| ENSMUSG00000040599 | Mis12 | protein_coding | 0.397527192 | 4.846130486 | 1.4344488 | 0.2481588 | 0.769999308 |
| ENSMUSG00000025395 | Prim1 | protein_coding | 0.463995728 | 4.083517732 | 1.4332285 | 0.2483519 | 0.769999308 |
| ENSMUSG00000025586 | Cpeb1 | protein_coding | 0.468519178 | 5.172911502 | 1.433151 | 0.2484355 | 0.769999308 |
| ENSMUSG00000002957 | Ap2a2 | protein_coding | 0.305550618 | 7.263747972 | 1.4326863 | 0.2484657 | 0.769999308 |
| ENSMUSG00000031681 | Smad1 | protein_coding | 0.36478212 | 4.910375733 | 1.432505 | 0.2484665 | 0.769999308 |
| ENSMUSG00000032743 | D430042O09Rik | protein_coding | -0.767033209 | 2.810748478 | 1.4324242 | 0.2484793 | 0.769999308 |
| ENSMUSG00000038429 | Usp5 | protein_coding | 0.391924415 | 8.407859596 | 1.441311 | 0.248774 | 0.770142479 |
| ENSMUSG00000003420 | Fcgrt | protein_coding | -0.631148165 | 3.467224619 | 1.4300045 | 0.248863 | 0.770142479 |
| ENSMUSG00000037458 | Azin1 | protein_coding | 0.243680185 | 6.855944283 | 1.4299774 | 0.2488673 | 0.770142479 |
| ENSMUSG00000054469 | Lclat1 | protein_coding | 0.581173738 | 3.717781153 | 1.4298863 | 0.2488818 | 0.770142479 |
| ENSMUSG00000071547 | Nt5dc2 | protein_coding | 0.493825443 | 3.340278492 | 1.4298234 | 0.2488917 | 0.770142479 |
| ENSMUSG00000038175 | Mylip | protein_coding | -0.603854925 | 3.224945135 | 1.428371 | 0.2491225 | 0.770142479 |
| ENSMUSG00000003604 | Aven | protein_coding | -0.494514161 | 4.143550156 | 1.4282025 | 0.2491492 | 0.770142479 |
| ENSMUSG00000030759 | Far1 | protein_coding | -0.307712187 | 5.452490102 | 1.4281391 | 0.2491593 | 0.770142479 |

| ensembl_gene_id | external_gene_name | gene_biotype | log2 (fold change) | log2 (counts per million) | F-value | p-value | False discovery rate |
|---|---|---|---|---|---|---|---|
| ENSMUSG00000059974 | Ntm | protein_coding | 0.230116358 | 7.518673752 | 1.4275753 | 0.249249 | 0.770142479 |
| ENSMUSG00000055116 | Arntl | protein_coding | 0.454384712 | 3.893684507 | 1.4270045 | 0.2493398 | 0.770142479 |
| ENSMUSG00000001910 | Nacc1 | protein_coding | -0.378927337 | 4.97235212 | 1.4266642 | 0.249394 | 0.770142479 |
| ENSMUSG00000018446 | C1qbp | protein_coding | 0.357877471 | 5.541922467 | 1.4262056 | 0.249467 | 0.770142479 |
| ENSMUSG00000021023 | 1110008L16Rik | protein_coding | 0.5579774 | 3.348714097 | 1.4255854 | 0.2495658 | 0.770142479 |
| ENSMUSG00000040289 | Hey1 | protein_coding | 0.371636921 | 4.576717294 | 1.4252879 | 0.2496132 | 0.770142479 |
| ENSMUSG00000060579 | Fhit | protein_coding | -0.326712758 | 5.12763996 | 1.424996 | 0.2496597 | 0.770142479 |
| ENSMUSG00000013997 | Nit1 | protein_coding | 0.289609217 | 5.735274349 | 1.4249225 | 0.2496714 | 0.770142479 |
| ENSMUSG00000025255 | Zfhx4 | protein_coding | -0.367837558 | 4.583105789 | 1.4239486 | 0.2498267 | 0.770199517 |
| ENSMUSG00000087674 | 4930447M23Rik | lincRNA | -0.564979403 | 3.176993293 | 1.4232505 | 0.2499381 | 0.770199517 |
| ENSMUSG00000029108 | Pcdh7 | protein_coding | -0.307152878 | 5.97182435 | 1.4229908 | 0.2499796 | 0.770199517 |
| ENSMUSG00000023007 | Prpf40b | protein_coding | 0.41070936 | 5.256867665 | 1.4228478 | 0.2500024 | 0.770199517 |
| ENSMUSG00000026520 | Pycr2 | protein_coding | 0.468177277 | 5.07548035 | 1.4225545 | 0.2500771 | 0.770199517 |
| ENSMUSG00000033014 | Trim33 | protein_coding | -0.346926597 | 4.776449862 | 1.4216822 | 0.2501887 | 0.770199517 |
| ENSMUSG00000069495 | Epc2 | protein_coding | -0.413104561 | 4.693634474 | 1.4216658 | 0.2501913 | 0.770199517 |
| ENSMUSG00000046480 | Scn4b | protein_coding | 0.43819028 | 5.135063588 | 1.421345 | 0.2502704 | 0.770222558 |
| ENSMUSG00000034126 | Pomt2 | protein_coding | -0.933987609 | 2.751272529 | 1.420707 | 0.2503724 | 0.770265073 |
| ENSMUSG00000052423 | B4galt3 | protein_coding | -0.316115894 | 5.498150261 | 1.4199244 | 0.2504699 | 0.770265073 |
| ENSMUSG00000039057 | Myo16 | protein_coding | -0.77012943 | 2.929518244 | 1.4189947 | 0.2506187 | 0.770265073 |
| ENSMUSG00000028218 | Fam92a | protein_coding | -0.273888784 | 5.899350725 | 1.4179864 | 0.2507804 | 0.770265073 |
| ENSMUSG00000087461 | C230014O12Rik | lincRNA | 0.731252298 | 2.858043497 | 1.4178593 | 0.2508007 | 0.770265073 |
| ENSMUSG00000047554 | Tmem41b | protein_coding | -0.375364207 | 4.389520327 | 1.417762 | 0.2508164 | 0.770265073 |
| ENSMUSG00000056004 | 9330182L06Rik | protein_coding | -0.347737357 | 5.064102773 | 1.4175611 | 0.2508486 | 0.770265073 |
| ENSMUSG00000019768 | Esr1 | protein_coding | 0.682547351 | 3.539813594 | 1.4176801 | 0.2508573 | 0.770265073 |
| ENSMUSG00000034269 | Setd5 | protein_coding | 0.380816411 | 4.314935118 | 1.4162313 | 0.251062 | 0.770673604 |
| ENSMUSG00000044788 | Fads6 | protein_coding | 0.47135453 | 4.033293867 | 1.4153661 | 0.251201 | 0.770725077 |
| ENSMUSG00000039218 | Srrm2 | protein_coding | 0.299195011 | 6.124599516 | 1.4141114 | 0.2514027 | 0.770725077 |
| ENSMUSG00000032744 | Heyl | protein_coding | -0.574482896 | 3.977557591 | 1.4142157 | 0.2514137 | 0.770725077 |
| ENSMUSG00000027238 | Frmd5 | protein_coding | -0.393202324 | 4.258166199 | 1.4133077 | 0.2515321 | 0.770725077 |
| ENSMUSG00000033543 | Gtf2a2 | protein_coding | -0.241883359 | 7.381250823 | 1.4132688 | 0.2515383 | 0.770725077 |
| ENSMUSG00000031925 | Maml2 | protein_coding | -0.44254695 | 4.363464793 | 1.4131052 | 0.2515647 | 0.770725077 |
| ENSMUSG00000028902 | Sf3a3 | protein_coding | 0.249523616 | 6.182383047 | 1.4130068 | 0.2515805 | 0.770725077 |
| ENSMUSG00000055322 | Tns1 | protein_coding | 0.645179678 | 3.784379738 | 1.4116491 | 0.2519001 | 0.771480864 |
| ENSMUSG00000039828 | Wdr70 | protein_coding | -0.288169526 | 5.70048989 | 1.410586 | 0.2519707 | 0.771480864 |
| ENSMUSG00000024654 | Asrgl1 | protein_coding | 0.221985752 | 7.648444656 | 1.4087231 | 0.2522715 | 0.772182031 |
| ENSMUSG00000020902 | Ntn1 | protein_coding | -0.954896292 | 2.661444907 | 1.4083813 | 0.2523528 | 0.772211138 |
| ENSMUSG00000038028 | Tigar | protein_coding | 0.41522596 | 4.351674564 | 1.4075694 | 0.252458 | 0.772313333 |
| ENSMUSG00000049421 | Zfp260 | protein_coding | -0.336247725 | 5.497328134 | 1.4054317 | 0.2528041 | 0.772604239 |
| ENSMUSG00000006289 | Osgep | protein_coding | 0.432763929 | 4.89572541 | 1.405156 | 0.2528488 | 0.772604239 |
| ENSMUSG00000022048 | Dpysl2 | protein_coding | 0.415097728 | 11.92064053 | 1.4115417 | 0.2528914 | 0.772604239 |
| ENSMUSG00000086555 | Gm13446 | processed_transcript | 0.785967522 | 3.558892642 | 1.411403 | 0.2529138 | 0.772604239 |
| ENSMUSG00000001524 | Gtf2h4 | protein_coding | -0.340639266 | 5.099909183 | 1.4045759 | 0.2529429 | 0.772604239 |
| ENSMUSG00000028907 | Utp11 | protein_coding | -0.245636401 | 6.560606227 | 1.403923 | 0.2530488 | 0.772604239 |
| ENSMUSG00000030638 | Sh3gl3 | protein_coding | -0.386105552 | 5.200236941 | 1.4034874 | 0.2531195 | 0.772604239 |
| ENSMUSG00000035941 | Ibtk | protein_coding | 0.509867422 | 3.603984258 | 1.4032092 | 0.2531647 | 0.772604239 |
| ENSMUSG00000040112 | Mrps35 | protein_coding | 0.436950207 | 4.382661176 | 1.4028472 | 0.2532234 | 0.772604239 |
| ENSMUSG00000029697 | Fezf1 | protein_coding | 1.13419927 | 3.055115251 | 1.4172131 | 0.2532716 | 0.772604239 |
| ENSMUSG00000022425 | Enpp2 | protein_coding | 0.468832594 | 8.708527452 | 1.4213269 | 0.2534324 | 0.772647035 |
| ENSMUSG00000032939 | Nup93 | protein_coding | 0.372642514 | 4.417205493 | 1.4011584 | 0.2534979 | 0.772647035 |
| ENSMUSG00000000168 | Dlat | protein_coding | 0.26037201 | 6.283778534 | 1.4006973 | 0.2535729 | 0.772647035 |
| ENSMUSG00000022814 | Umps | protein_coding | 0.376229982 | 5.473291853 | 1.4005328 | 0.2535997 | 0.772647035 |
| ENSMUSG00000062257 | Opcml | protein_coding | -0.534768976 | 3.674555153 | 1.4002546 | 0.2536449 | 0.772647035 |
| ENSMUSG00000027596 | a | protein_coding | -0.678789741 | 2.687726089 | 1.3992191 | 0.2538136 | 0.772763604 |
| ENSMUSG00000028221 | Tmem55a | protein_coding | -0.217943133 | 7.032470232 | 1.3988562 | 0.2538727 | 0.772763604 |
| ENSMUSG00000024317 | Rnf138 | protein_coding | -0.374575542 | 4.513559622 | 1.398696 | 0.2538988 | 0.772763604 |
| ENSMUSG00000028100 | Nudt17 | protein_coding | 0.51361805 | 4.793880109 | 1.3985069 | 0.254062 | 0.773041461 |
| ENSMUSG00000060843 | Ctnna3 | protein_coding | -0.592941474 | 2.978928468 | 1.3967935 | 0.2542091 | 0.773270308 |
| ENSMUSG00000022099 | Dmtn | protein_coding | -0.236736978 | 7.798985178 | 1.3950471 | 0.2544944 | 0.773783131 |
| ENSMUSG00000091475 | 2810468N07Rik | processed_transcript | -0.247119319 | 6.296432369 | 1.3943479 | 0.2546088 | 0.773783131 |
| ENSMUSG00000071266 | Zfp946 | protein_coding | -0.70943174 | 2.83477977 | 1.3938727 | 0.2546865 | 0.773783131 |
| ENSMUSG00000028016 | Ints12 | protein_coding | -0.337421598 | 4.737708314 | 1.3936992 | 0.2547149 | 0.773783131 |
| ENSMUSG00000040044 | Orc3 | protein_coding | -0.331062936 | 6.001959922 | 1.3935607 | 0.2547375 | 0.773783131 |
| ENSMUSG00000019960 | Dusp6 | protein_coding | -0.353634983 | 5.189858179 | 1.3931215 | 0.2548095 | 0.773783131 |
| ENSMUSG00000030588 | Yif1b | protein_coding | -0.246543537 | 6.243423051 | 1.3926436 | 0.2548877 | 0.773802332 |
| ENSMUSG00000022351 | Sqle | protein_coding | 0.342916514 | 5.66798087 | 1.3911323 | 0.2551355 | 0.77384754 |
| ENSMUSG00000054459 | Vsnl1 | protein_coding | 0.261134823 | 10.53233629 | 1.3910233 | 0.2551533 | 0.77384754 |
| ENSMUSG00000003200 | Sh3gl1 | protein_coding | 0.479487434 | 4.02389838 | 1.3886232 | 0.2555475 | 0.77384754 |
| ENSMUSG00000036565 | Ttyh3 | protein_coding | -0.452321486 | 4.395065341 | 1.3884316 | 0.255579 | 0.77384754 |
| ENSMUSG00000029819 | Npy | protein_coding | -0.499275351 | 6.897415766 | 1.4150563 | 0.2556116 | 0.77384754 |
| ENSMUSG00000030264 | Thumpd3 | protein_coding | 0.356911123 | 4.766676556 | 1.3882208 | 0.2556136 | 0.77384754 |
| ENSMUSG00000043241 | Upf2 | protein_coding | -0.40202728 | 5.165862436 | 1.3881506 | 0.2556252 | 0.77384754 |
| ENSMUSG00000021374 | Nup153 | protein_coding | 0.505937531 | 3.758688117 | 1.3874197 | 0.2557454 | 0.77384754 |
| ENSMUSG00000016200 | Syt14 | protein_coding | -0.557845025 | 3.44593663 | 1.3869191 | 0.2558278 | 0.77384754 |
| ENSMUSG00000024665 | Fads2 | protein_coding | 0.442163433 | 4.030559259 | 1.3865963 | 0.2558809 | 0.77384754 |

| ensembl_gene_id | external_gene_name | gene_biotype | log2 (fold change) | log2 (counts per million) | F-value | p-value | False discovery rate |
|---|---|---|---|---|---|---|---|
| ENSMUSG00000016487 | Ppfibp1 | protein_coding | -0.588935017 | 3.142430266 | 1.3864909 | 0.2558983 | 0.77384754 |
| ENSMUSG00000028484 | Psip1 | protein_coding | -0.349658411 | 4.976152638 | 1.3863904 | 0.2559148 | 0.77384754 |
| ENSMUSG00000002968 | Med25 | protein_coding | -0.413961238 | 4.253980917 | 1.3858304 | 0.2560071 | 0.77384754 |
| ENSMUSG00000052581 | Lrrtm4 | protein_coding | -0.411651351 | 5.281600629 | 1.3859543 | 0.2560142 | 0.77384754 |
| ENSMUSG00000028766 | Alpl | protein_coding | -1.028546711 | 2.546182813 | 1.3857636 | 0.2560456 | 0.77384754 |
| ENSMUSG00000079317 | Trappc2 | protein_coding | -0.227976974 | 6.535422888 | 1.3850725 | 0.256132 | 0.77384754 |
| ENSMUSG00000034951 | Cog7 | protein_coding | 0.353020545 | 5.961421117 | 1.3842699 | 0.2562919 | 0.77384754 |
| ENSMUSG00000018442 | Derl2 | protein_coding | 0.300492013 | 5.580025005 | 1.3834232 | 0.2564041 | 0.77384754 |
| ENSMUSG00000058655 | Eif4b | protein_coding | 0.329687749 | 7.164427178 | 1.3850743 | 0.2564512 | 0.77384754 |
| ENSMUSG00000050052 | Tdrp | protein_coding | 0.410413737 | 4.529503358 | 1.3829601 | 0.2564806 | 0.77384754 |
| ENSMUSG00000031202 | Rab39b | protein_coding | -0.381847138 | 4.459768071 | 1.3827754 | 0.2565111 | 0.77384754 |
| ENSMUSG00000001783 | Rtcb | protein_coding | 0.298442943 | 6.885349732 | 1.3827016 | 0.2565508 | 0.77384754 |
| ENSMUSG00000006920 | Ezh1 | protein_coding | 0.243038704 | 6.198793368 | 1.3824924 | 0.2565578 | 0.77384754 |
| ENSMUSG00000039285 | Azi2 | protein_coding | -0.32323827 | 5.542731768 | 1.3808404 | 0.256831 | 0.773857816 |
| ENSMUSG00000030884 | Uqcrc2 | protein_coding | 0.2264528 | 8.887543791 | 1.3804951 | 0.2568882 | 0.773857816 |
| ENSMUSG00000066057 | Gm1976 | lincRNA | -0.631117745 | 3.030686164 | 1.3796956 | 0.2570205 | 0.773857816 |
| ENSMUSG00000026421 | Csrp1 | protein_coding | -0.234748674 | 8.291073707 | 1.3794623 | 0.2570592 | 0.773857816 |
| ENSMUSG00000042359 | Osbpl6 | protein_coding | 0.377024527 | 5.156949153 | 1.3788502 | 0.2571606 | 0.773857816 |
| ENSMUSG00000026893 | Gca | protein_coding | -0.443948698 | 4.339121022 | 1.3787303 | 0.2571805 | 0.773857816 |
| ENSMUSG00000069135 | Fgfr1op | protein_coding | 0.499487832 | 4.019923772 | 1.3778581 | 0.2573251 | 0.773857816 |
| ENSMUSG00000057719 | Sh3rf2 | protein_coding | -0.591765691 | 3.397565118 | 1.3772578 | 0.2574247 | 0.773857816 |
| ENSMUSG00000024312 | Wdr46 | protein_coding | 0.693549633 | 2.900480382 | 1.3772431 | 0.2574271 | 0.773857816 |
| ENSMUSG00000068391 | Chrac1 | protein_coding | 0.253966854 | 6.829943269 | 1.3771632 | 0.2574404 | 0.773857816 |
| ENSMUSG00000070047 | Fat1 | protein_coding | -0.574985627 | 3.598370895 | 1.3765952 | 0.2575347 | 0.773857816 |
| ENSMUSG00000021258 | Ccnk | protein_coding | 0.601186591 | 3.107589317 | 1.3762223 | 0.2575967 | 0.773857816 |
| ENSMUSG00000032212 | Sltm | protein_coding | -0.296355563 | 5.581230775 | 1.3747565 | 0.2578403 | 0.773857816 |
| ENSMUSG00000036327 | Qsox2 | protein_coding | 0.74889608 | 2.993632608 | 1.3746544 | 0.2578573 | 0.773857816 |
| ENSMUSG00000047412 | Zbtb44 | protein_coding | -0.559966222 | 3.245284772 | 1.3746304 | 0.2578613 | 0.773857816 |
| ENSMUSG00000026447 | Pik3c2b | protein_coding | 0.439688024 | 4.035664441 | 1.3742377 | 0.2579266 | 0.773857816 |
| ENSMUSG00000026965 | Anapc2 | protein_coding | 0.263257543 | 6.07619777 | 1.374048 | 0.2579582 | 0.773857816 |
| ENSMUSG00000023074 | Mospd1 | protein_coding | -0.301387696 | 5.229285499 | 1.3739491 | 0.2579746 | 0.773857816 |
| ENSMUSG00000005087 | Cd44 | protein_coding | -0.789516258 | 2.696449333 | 1.3731179 | 0.258113 | 0.773857816 |
| ENSMUSG00000018750 | Zbtb4 | protein_coding | -0.23421466 | 6.130943418 | 1.3729695 | 0.2581377 | 0.773857816 |
| ENSMUSG00000024227 | Pdzph1 | protein_coding | -0.509521088 | 4.119564187 | 1.3729319 | 0.258144 | 0.773857816 |
| ENSMUSG00000029875 | Ccdc184 | protein_coding | 0.373528111 | 6.12034788 | 1.3726834 | 0.2582852 | 0.773857816 |
| ENSMUSG00000054008 | Ndst1 | protein_coding | 0.525671606 | 4.169782026 | 1.3711798 | 0.2584636 | 0.773857816 |
| ENSMUSG00000049122 | Frmd3 | protein_coding | -0.69129038 | 3.188285896 | 1.3710111 | 0.2584643 | 0.773857816 |
| ENSMUSG00000019795 | Pcmt1 | protein_coding | 0.226912641 | 8.737572817 | 1.3709924 | 0.2584674 | 0.773857816 |
| ENSMUSG00000030978 | Rrm1 | protein_coding | 0.509635461 | 4.168672564 | 1.3703596 | 0.258573 | 0.773857816 |
| ENSMUSG00000031290 | Lrch2 | protein_coding | -0.402271292 | 3.809746072 | 1.3701029 | 0.2586158 | 0.773857816 |
| ENSMUSG00000036678 | Aaas | protein_coding | 0.359140218 | 5.155665353 | 1.3700728 | 0.2586209 | 0.773857816 |
| ENSMUSG00000039782 | Cpeb2 | protein_coding | -0.351661827 | 4.842610896 | 1.3699087 | 0.2586483 | 0.773857816 |
| ENSMUSG00000050069 | Grem2 | protein_coding | -0.528103569 | 3.546617738 | 1.3676142 | 0.2590319 | 0.774600376 |
| ENSMUSG00000019143 | Hars2 | protein_coding | 0.39982657 | 4.390408431 | 1.3672634 | 0.2590906 | 0.774600376 |
| ENSMUSG00000041530 | Ago1 | protein_coding | 0.398732128 | 4.381906915 | 1.3669528 | 0.2591426 | 0.774600376 |
| ENSMUSG00000079614 | Seh1l | protein_coding | 0.272949784 | 6.686796075 | 1.3659623 | 0.2593085 | 0.774600376 |
| ENSMUSG00000027187 | Cat | protein_coding | 0.34870907 | 5.842333269 | 1.3658774 | 0.2593227 | 0.774600376 |
| ENSMUSG00000038537 | Mc3r | protein_coding | 1.065714661 | 2.441413846 | 1.3660054 | 0.2593287 | 0.774600376 |
| ENSMUSG00000021103 | Mnat1 | protein_coding | -0.250787119 | 6.011421216 | 1.3645322 | 0.2595483 | 0.775023595 |
| ENSMUSG00000040771 | Oard1 | protein_coding | 0.249234779 | 6.457537831 | 1.3641376 | 0.2596145 | 0.775023595 |
| ENSMUSG00000022956 | Atp5o | protein_coding | 0.210569077 | 8.53452819 | 1.3636289 | 0.2596999 | 0.77506339 |
| ENSMUSG00000022208 | Jph4 | protein_coding | -0.259556588 | 6.716425145 | 1.3630856 | 0.2597912 | 0.775111371 |
| ENSMUSG00000058709 | Egln2 | protein_coding | -0.264132957 | 6.016477242 | 1.362675 | 0.2598602 | 0.775111371 |
| ENSMUSG00000026224 | 4933407L21Rik | protein_coding | -0.397685292 | 4.27700674 | 1.3609971 | 0.2601424 | 0.775315209 |
| ENSMUSG00000029196 | Tada2b | protein_coding | 0.48826663 | 3.571371922 | 1.359793 | 0.2603451 | 0.775315209 |
| ENSMUSG00000013622 | Atraid | protein_coding | -0.256772897 | 7.682198409 | 1.3587019 | 0.260529 | 0.775315209 |
| ENSMUSG00000032305 | Fam219b | protein_coding | 0.358793343 | 5.791319871 | 1.3582296 | 0.2606087 | 0.775315209 |
| ENSMUSG00000019792 | Trmt11 | protein_coding | -0.493602091 | 4.368417646 | 1.3577794 | 0.260712 | 0.775315209 |
| ENSMUSG00000022610 | Mapk12 | protein_coding | 0.959564161 | 2.362859331 | 1.3572387 | 0.2607759 | 0.775315209 |
| ENSMUSG00000003119 | Cdk12 | protein_coding | -0.430407635 | 4.176047042 | 1.356422 | 0.2609138 | 0.775315209 |
| ENSMUSG00000004099 | Dnmt1 | protein_coding | 0.371504203 | 4.668233799 | 1.3561665 | 0.260957 | 0.775315209 |
| ENSMUSG00000049176 | Frmpd4 | protein_coding | -0.333751521 | 5.270504808 | 1.3555764 | 0.2610568 | 0.775315209 |
| ENSMUSG00000041608 | Entpd3 | protein_coding | 0.58884668 | 3.918803749 | 1.3555214 | 0.2610934 | 0.775315209 |
| ENSMUSG00000021706 | Zfyve16 | protein_coding | -0.538623751 | 3.227398287 | 1.3550938 | 0.2611384 | 0.775315209 |
| ENSMUSG00000014599 | Csf1 | protein_coding | 0.592148622 | 2.841161253 | 1.3548262 | 0.2611836 | 0.775315209 |
| ENSMUSG00000036792 | Mbd5 | protein_coding | -0.330270672 | 4.79709081 | 1.3542044 | 0.2612889 | 0.775315209 |
| ENSMUSG00000017999 | Ddx27 | protein_coding | 0.405711901 | 4.486232458 | 1.3532576 | 0.2614492 | 0.775315209 |
| ENSMUSG00000046169 | Adamts6 | protein_coding | -0.552050684 | 3.154787281 | 1.3532369 | 0.2614527 | 0.775315209 |
| ENSMUSG00000026825 | Dnm1 | protein_coding | -0.264367349 | 6.683177301 | 1.352804 | 0.2615261 | 0.775315209 |
| ENSMUSG00000022629 | Kif21a | protein_coding | 0.259948226 | 7.104242049 | 1.3516334 | 0.2617246 | 0.775315209 |
| ENSMUSG00000027360 | Hdc | protein_coding | -0.55636731 | 3.202417784 | 1.3511867 | 0.2618004 | 0.775315209 |
| ENSMUSG00000015882 | Lcorl | protein_coding | -0.59848551 | 3.036228734 | 1.3509484 | 0.2618408 | 0.775315209 |
| ENSMUSG00000024045 | Akap8 | protein_coding | 0.344502901 | 4.802518512 | 1.3497635 | 0.2620421 | 0.775315209 |
| ENSMUSG00000001983 | Taco1 | protein_coding | 0.500658802 | 3.448332275 | 1.3486622 | 0.2622293 | 0.775315209 |

| ensembl_gene_id | external_gene_name | gene_biotype | log2 (fold change) | log2 (counts per million) | F-value | p-value | False discovery rate |
|---|---|---|---|---|---|---|---|
| ENSMUSG00000039050 | Osbpl2 | protein_coding | 0.288836409 | 5.793113036 | 1.3486455 | 0.2622321 | 0.775315209 |
| ENSMUSG00000017764 | Zswim1 | protein_coding | 0.370412972 | 5.908942062 | 1.3487328 | 0.2622445 | 0.775315209 |
| ENSMUSG00000056673 | Kdm5d | protein_coding | 0.59527973 | 2.893253497 | 1.3482913 | 0.2622924 | 0.775315209 |
| ENSMUSG00000041986 | Elmod1 | protein_coding | -0.406524899 | 6.195305318 | 1.3542085 | 0.262346 | 0.775315209 |
| ENSMUSG00000024424 | Ttc39c | protein_coding | -0.472831374 | 3.912295957 | 1.3476049 | 0.2624092 | 0.775315209 |
| ENSMUSG00000054226 | Tprkb | protein_coding | -0.206274214 | 7.424712086 | 1.3475657 | 0.2624159 | 0.775315209 |
| ENSMUSG00000025858 | Get4 | protein_coding | 0.419141755 | 3.690348332 | 1.3475093 | 0.2624255 | 0.775315209 |
| ENSMUSG00000033059 | Pygb | protein_coding | 0.339095325 | 6.971582932 | 1.3511275 | 0.2625061 | 0.775315209 |
| ENSMUSG00000032551 | 1110059G10Rik | protein_coding | -0.299649484 | 5.850311521 | 1.346991 | 0.2625137 | 0.775315209 |
| ENSMUSG00000074892 | B3galt5 | protein_coding | 0.348242501 | 4.952348581 | 1.3468515 | 0.2625375 | 0.775315209 |
| ENSMUSG00000020513 | Tubd1 | protein_coding | 0.815624888 | 2.857777701 | 1.3468207 | 0.2625427 | 0.775315209 |
| ENSMUSG00000002910 | Arrdc2 | protein_coding | 0.719884477 | 2.682520466 | 1.3466182 | 0.2625772 | 0.775315209 |
| ENSMUSG00000002984 | Tomm40 | protein_coding | -0.407154872 | 4.313692104 | 1.3465914 | 0.2625818 | 0.775315209 |
| ENSMUSG00000061143 | Maml3 | protein_coding | -0.581842923 | 3.033189784 | 1.3464172 | 0.2626115 | 0.775315209 |
| ENSMUSG00000020456 | Ogdh | protein_coding | 0.309122371 | 8.134128311 | 1.3463482 | 0.2626505 | 0.775315209 |
| ENSMUSG00000000171 | Sdhd | protein_coding | 0.218709123 | 9.071525417 | 1.3456239 | 0.2627467 | 0.775315209 |
| ENSMUSG00000055436 | Srsf11 | protein_coding | -0.300143604 | 5.796884674 | 1.3453681 | 0.2627904 | 0.775315209 |
| ENSMUSG00000007739 | Cct4 | protein_coding | 0.23022087 | 7.595742519 | 1.3447841 | 0.26289 | 0.775315209 |
| ENSMUSG00000108434 | Gm45768 | lincRNA | 0.72320972 | 3.28471313 | 1.3447644 | 0.2629206 | 0.775315209 |
| ENSMUSG00000045087 | S1pr5 | protein_coding | 0.73548191 | 2.742858132 | 1.3445636 | 0.2629276 | 0.775315209 |
| ENSMUSG00000023755 | Rhebl1 | protein_coding | 0.573025249 | 4.079893955 | 1.3441865 | 0.2630193 | 0.775315209 |
| ENSMUSG00000022132 | Cldn10 | protein_coding | -0.557107249 | 5.794280525 | 1.3585846 | 0.2631987 | 0.775315209 |
| ENSMUSG00000029757 | Dync1i1 | protein_coding | 0.282086797 | 7.483071199 | 1.3428813 | 0.263215 | 0.775315209 |
| ENSMUSG00000050229 | Pigm | protein_coding | 0.619683684 | 2.779106092 | 1.3427106 | 0.2632442 | 0.775315209 |
| ENSMUSG00000053929 | Cyhr1 | protein_coding | 0.218177635 | 6.914207001 | 1.342101 | 0.2633484 | 0.775315209 |
| ENSMUSG00000051343 | Rab11fip5 | protein_coding | 0.328485212 | 5.378459488 | 1.3415889 | 0.263436 | 0.775315209 |
| ENSMUSG00000050558 | Prokr2 | protein_coding | 0.815262536 | 2.716448841 | 1.3410063 | 0.2635358 | 0.775315209 |
| ENSMUSG00000097164 | Cep83os | bidirectional_promoter | -0.321563048 | 5.223949307 | 1.339971 | 0.2637131 | 0.775315209 |
| ENSMUSG00000015944 | Gatsl2 | protein_coding | -0.242453456 | 6.851639751 | 1.3397818 | 0.2637455 | 0.775315209 |
| ENSMUSG00000053838 | Nudcd3 | protein_coding | 0.30112421 | 7.133423902 | 1.3399108 | 0.2637506 | 0.775315209 |
| ENSMUSG00000059851 | Kmt5c | protein_coding | 0.656610695 | 3.019809196 | 1.3396698 | 0.2637647 | 0.775315209 |
| ENSMUSG00000035354 | Uvrag | protein_coding | 0.474501129 | 3.360669855 | 1.3393927 | 0.2638122 | 0.775315209 |
| ENSMUSG00000014633 | Cmc2 | protein_coding | -0.277089122 | 5.842276473 | 1.3393355 | 0.263822 | 0.775315209 |
| ENSMUSG00000066043 | Phactr4 | protein_coding | -0.660715134 | 2.921263514 | 1.3387641 | 0.26392 | 0.775354077 |
| ENSMUSG00000035329 | Fbxo33 | protein_coding | -0.325455386 | 5.444145011 | 1.3384177 | 0.2639795 | 0.775354077 |
| ENSMUSG00000036427 | Gpi1 | protein_coding | 0.307672883 | 9.013078224 | 1.3374108 | 0.2641795 | 0.775660502 |
| ENSMUSG00000005198 | Polr2a | protein_coding | -0.481701101 | 4.727635432 | 1.337128 | 0.2642281 | 0.775660502 |
| ENSMUSG00000031758 | Cdyl2 | protein_coding | 0.618627416 | 4.238801782 | 1.3418488 | 0.2644455 | 0.776086958 |
| ENSMUSG00000023186 | Vwa5a | protein_coding | 0.620611137 | 2.970044968 | 1.3338564 | 0.2647637 | 0.776808816 |
| ENSMUSG00000038425 | Poli | protein_coding | 0.528972825 | 3.520332659 | 1.3323556 | 0.2650224 | 0.776880452 |
| ENSMUSG00000024944 | Arl2 | protein_coding | 0.210009544 | 9.161043313 | 1.3323295 | 0.2650269 | 0.776880452 |
| ENSMUSG00000069516 | Lyz2 | protein_coding | -0.635703869 | 3.313845195 | 1.3319575 | 0.2650911 | 0.776880452 |
| ENSMUSG00000037355 | Uvssa | protein_coding | -0.704077065 | 2.934807809 | 1.3314118 | 0.2651853 | 0.776880452 |
| ENSMUSG00000062373 | Tmem65 | protein_coding | -0.297039456 | 5.907264928 | 1.3307847 | 0.2652936 | 0.776880452 |
| ENSMUSG00000042751 | Nmnat2 | protein_coding | 0.371521462 | 7.270128218 | 1.3395185 | 0.2654408 | 0.776880452 |
| ENSMUSG00000042541 | Sem1 | protein_coding | -0.289621958 | 9.098111375 | 1.3300373 | 0.2654498 | 0.776880452 |
| ENSMUSG00000025264 | Tsr2 | protein_coding | 0.237823225 | 7.826594417 | 1.3295983 | 0.2654986 | 0.776880452 |
| ENSMUSG00000025145 | Lrrc45 | protein_coding | 0.597737133 | 3.403219882 | 1.3295747 | 0.2655027 | 0.776880452 |
| ENSMUSG00000034254 | Agpat1 | protein_coding | -0.284145016 | 5.640121654 | 1.3293002 | 0.2655502 | 0.776880452 |
| ENSMUSG00000022800 | Fyttd1 | protein_coding | -0.258273947 | 6.968082929 | 1.3279545 | 0.2657831 | 0.776880452 |
| ENSMUSG00000028683 | Eif2b3 | protein_coding | 0.267414363 | 6.287355809 | 1.3277431 | 0.2658197 | 0.776880452 |
| ENSMUSG00000023050 | Map3k12 | protein_coding | 0.456930928 | 4.076085082 | 1.3275709 | 0.2658495 | 0.776880452 |
| ENSMUSG00000027519 | Rab22a | protein_coding | 0.269189621 | 5.427502453 | 1.3274117 | 0.2658771 | 0.776880452 |
| ENSMUSG00000053483 | Usp21 | protein_coding | -0.352835316 | 5.18299884 | 1.3274001 | 0.2658791 | 0.776880452 |
| ENSMUSG00000030207 | Fam234b | protein_coding | 0.313885969 | 6.781779912 | 1.3268268 | 0.2660055 | 0.776880452 |
| ENSMUSG00000058486 | Wdr91 | protein_coding | -0.492899942 | 4.671450756 | 1.3267369 | 0.2660211 | 0.776880452 |
| ENSMUSG00000002280 | Narfl | protein_coding | 0.532458175 | 3.723069689 | 1.3261914 | 0.2660886 | 0.776880452 |
| ENSMUSG00000063434 | Sorcs3 | protein_coding | -0.328736188 | 5.116461354 | 1.3250251 | 0.266291 | 0.777260241 |
| ENSMUSG00000036270 | Edc4 | protein_coding | -0.392658241 | 4.747502669 | 1.3244604 | 0.266389 | 0.777335471 |
| ENSMUSG00000048644 | Ctxn1 | protein_coding | -0.196050839 | 7.951408505 | 1.3237999 | 0.2665038 | 0.777341041 |
| ENSMUSG00000041153 | Osgin2 | protein_coding | -0.570880814 | 3.325612548 | 1.3236173 | 0.2665355 | 0.777341041 |
| ENSMUSG00000044708 | Kcnj10 | protein_coding | 0.405412889 | 4.889760016 | 1.3217883 | 0.2668537 | 0.777577606 |
| ENSMUSG00000020547 | Bzw2 | protein_coding | -0.25083196 | 6.436693677 | 1.3215637 | 0.2668928 | 0.777577606 |
| ENSMUSG00000045092 | S1pr1 | protein_coding | -0.356719036 | 5.733062734 | 1.3212591 | 0.2669458 | 0.777577606 |
| ENSMUSG00000047989 | Ino80c | protein_coding | 0.284277694 | 6.052578601 | 1.3211808 | 0.2669595 | 0.777577606 |
| ENSMUSG00000018008 | Cyth4 | protein_coding | 0.452408994 | 3.659424417 | 1.3210732 | 0.2669782 | 0.777577606 |
| ENSMUSG00000012114 | Med15 | protein_coding | -0.330442916 | 5.038707628 | 1.3201975 | 0.2671308 | 0.777595378 |
| ENSMUSG00000055022 | Cntn1 | protein_coding | -0.20865769 | 7.891555497 | 1.3201032 | 0.2671472 | 0.777595378 |
| ENSMUSG00000027500 | Stmn2 | protein_coding | 0.214090265 | 10.26118656 | 1.3196455 | 0.267227 | 0.777595378 |
| ENSMUSG00000025816 | Sec61a2 | protein_coding | 0.360432254 | 4.587601177 | 1.3193789 | 0.2672736 | 0.777595378 |
| ENSMUSG00000026516 | Nvl | protein_coding | 0.354344979 | 5.279776214 | 1.3181852 | 0.2674819 | 0.777753614 |
| ENSMUSG00000032086 | Bace1 | protein_coding | 0.301192205 | 5.419567206 | 1.3178512 | 0.2675402 | 0.777753614 |
| ENSMUSG00000062949 | Atp11c | protein_coding | -0.694442493 | 2.800095287 | 1.3173488 | 0.267628 | 0.777753614 |
| ENSMUSG00000021133 | Susd6 | protein_coding | 0.526388366 | 3.394380395 | 1.3169609 | 0.2676958 | 0.777753614 |

| ensembl_gene_id | external_gene_name | gene_biotype | log2 (fold change) | log2 (counts per million) | F-value | p-value | False discovery rate |
|---|---|---|---|---|---|---|---|
| ENSMUSG00000037989 | Wnk2 | protein_coding | -0.362314358 | 4.822597408 | 1.3164419 | 0.2677866 | 0.777753614 |
| ENSMUSG00000044442 | N6amt1 | protein_coding | -0.2194183 | 6.41823056 | 1.3162916 | 0.2678129 | 0.777753614 |
| ENSMUSG00000027324 | Rpusd2 | protein_coding | 0.662994902 | 2.66853841 | 1.3157949 | 0.2678998 | 0.777753614 |
| ENSMUSG00000044066 | Cep68 | protein_coding | 0.481328379 | 3.705952981 | 1.315756 | 0.2679066 | 0.777753614 |
| ENSMUSG00000031604 | Msmo1 | protein_coding | 0.211784728 | 7.175170499 | 1.3150164 | 0.268036 | 0.777867614 |
| ENSMUSG00000030652 | Coq7 | protein_coding | 0.197153216 | 7.84327492 | 1.3142284 | 0.2681741 | 0.777867614 |
| ENSMUSG00000038722 | Bud31 | protein_coding | -0.26690101 | 6.785825901 | 1.3135497 | 0.268293 | 0.777867614 |
| ENSMUSG00000008318 | Relt | protein_coding | -0.638226145 | 3.405330242 | 1.3135271 | 0.268297 | 0.777867614 |
| ENSMUSG00000061455 | Stx17 | protein_coding | 0.371185908 | 4.505653527 | 1.3134671 | 0.2683075 | 0.777867614 |
| ENSMUSG00000002718 | Cse1l | protein_coding | -0.279962329 | 5.626755388 | 1.3118503 | 0.2685912 | 0.778480239 |
| ENSMUSG00000066037 | Hnrnpr | protein_coding | 0.23928184 | 6.498009891 | 1.3104145 | 0.2688435 | 0.778909368 |
| ENSMUSG00000038085 | Cnbd2 | protein_coding | -0.510189656 | 3.309772347 | 1.3098364 | 0.2689452 | 0.778909368 |
| ENSMUSG00000022370 | Mrpl13 | protein_coding | -0.227377604 | 7.425329315 | 1.3086189 | 0.2691595 | 0.778909368 |
| ENSMUSG00000031513 | Leprotl1 | protein_coding | 0.317728481 | 6.383759213 | 1.3082356 | 0.2692539 | 0.778909368 |
| ENSMUSG00000021468 | Sptlc1 | protein_coding | 0.339617561 | 4.391982916 | 1.3077957 | 0.2693045 | 0.778909368 |
| ENSMUSG00000029559 | 2210016L21Rik | protein_coding | 0.227347402 | 8.426601944 | 1.3077849 | 0.2693064 | 0.778909368 |
| ENSMUSG00000063904 | 4930533O14Rik | protein_coding | 0.430897722 | 5.120789091 | 1.307695 | 0.2693492 | 0.778909368 |
| ENSMUSG00000020261 | Slc36a1 | protein_coding | 0.655669641 | 3.689174556 | 1.3090393 | 0.2693526 | 0.778909368 |
| ENSMUSG00000028854 | Slc9a1 | protein_coding | 0.387056836 | 4.565030444 | 1.3073036 | 0.2693912 | 0.778909368 |
| ENSMUSG00000047613 | A430005L14Rik | protein_coding | 0.200469644 | 7.307185625 | 1.3068901 | 0.2694642 | 0.778910759 |
| ENSMUSG00000027954 | Efna1 | protein_coding | -0.67888818 | 3.056885066 | 1.3048309 | 0.2698277 | 0.779751881 |
| ENSMUSG00000068284 | Usf3 | protein_coding | -0.529590435 | 3.504776773 | 1.3040646 | 0.2699631 | 0.779933668 |
| ENSMUSG00000048550 | Thnsl1 | protein_coding | 0.348711252 | 5.112092337 | 1.3027765 | 0.2701909 | 0.780382276 |
| ENSMUSG00000033808 | Tmem87a | protein_coding | -0.756643574 | 2.727547027 | 1.3014134 | 0.2704324 | 0.780618397 |
| ENSMUSG00000014769 | Psmb1 | protein_coding | 0.207887945 | 10.02840023 | 1.3011135 | 0.2704855 | 0.780618397 |
| ENSMUSG00000033099 | Nol12 | protein_coding | 0.201303053 | 6.686652263 | 1.3010855 | 0.2704905 | 0.780618397 |
| ENSMUSG00000028107 | Tars2 | protein_coding | -0.29905986 | 5.71337882 | 1.2996175 | 0.2707509 | 0.780903765 |
| ENSMUSG00000035032 | Nek11 | protein_coding | -0.787182929 | 2.447111599 | 1.2978896 | 0.2710578 | 0.780903765 |
| ENSMUSG00000045136 | Tubb2b | protein_coding | 0.284464307 | 5.423477762 | 1.2977909 | 0.2710753 | 0.780903765 |
| ENSMUSG00000039852 | Rere | protein_coding | -0.280500508 | 6.268421773 | 1.2971296 | 0.2711929 | 0.780903765 |
| ENSMUSG00000019810 | Fuca2 | protein_coding | 0.282111738 | 6.321853766 | 1.2969365 | 0.2712273 | 0.780903765 |
| ENSMUSG00000041660 | Bbox1 | protein_coding | 0.678289235 | 3.012470954 | 1.2968306 | 0.2712461 | 0.780903765 |
| ENSMUSG00000051721 | Wdcp | protein_coding | 0.501317835 | 4.719304903 | 1.2969101 | 0.2713758 | 0.780903765 |
| ENSMUSG00000062519 | Zfp398 | protein_coding | 0.503335929 | 3.625922666 | 1.2954852 | 0.2714856 | 0.780903765 |
| ENSMUSG00000033159 | Cnppd1 | protein_coding | -0.306955257 | 5.282652657 | 1.294972 | 0.2715771 | 0.780903765 |
| ENSMUSG00000058600 | Rpl30 | protein_coding | -0.308298852 | 6.950298277 | 1.2940171 | 0.2717741 | 0.780903765 |
| ENSMUSG00000029110 | Rnf4 | protein_coding | 0.261604957 | 6.827518734 | 1.2935912 | 0.2718233 | 0.780903765 |
| ENSMUSG00000022935 | Grik1 | protein_coding | 0.533197961 | 4.004442477 | 1.293727 | 0.2718259 | 0.780903765 |
| ENSMUSG00000034586 | Hid1 | protein_coding | 0.323037825 | 6.134864464 | 1.2933385 | 0.2718684 | 0.780903765 |
| ENSMUSG00000024990 | Rbp4 | protein_coding | -0.669898598 | 2.89401527 | 1.293291 | 0.2718769 | 0.780903765 |
| ENSMUSG00000031963 | Bmper | protein_coding | 0.6498106 | 2.732481153 | 1.2929049 | 0.2719458 | 0.780903765 |
| ENSMUSG00000031548 | Sfrp1 | protein_coding | 0.817855204 | 2.630183538 | 1.2924514 | 0.2720268 | 0.780903765 |
| ENSMUSG00000029729 | Zkscan1 | protein_coding | 0.294805035 | 5.761704343 | 1.292056 | 0.2720974 | 0.780903765 |
| ENSMUSG00000074748 | Atxn7l3b | protein_coding | -0.20098976 | 7.701684118 | 1.2912805 | 0.272236 | 0.780903765 |
| ENSMUSG00000029924 | Slc37a3 | protein_coding | 0.383157908 | 5.162331966 | 1.291229 | 0.2722452 | 0.780903765 |
| ENSMUSG00000029168 | Dpysl5 | protein_coding | 0.404386388 | 4.853573106 | 1.2910047 | 0.2722853 | 0.780903765 |
| ENSMUSG00000028890 | Mtf1 | protein_coding | -0.319969692 | 4.908268748 | 1.2904715 | 0.2723807 | 0.780903765 |
| ENSMUSG00000095325 | Zfp870 | protein_coding | 0.504616757 | 3.837029677 | 1.29039 | 0.2723953 | 0.780903765 |
| ENSMUSG00000021337 | Scgn | protein_coding | 0.621929449 | 3.085266767 | 1.290355 | 0.2724015 | 0.780903765 |
| ENSMUSG00000071660 | Ttc9c | protein_coding | 0.260144957 | 5.814119013 | 1.2899133 | 0.2724806 | 0.780903765 |
| ENSMUSG00000026335 | Pam | protein_coding | 0.276892703 | 6.194626399 | 1.2896222 | 0.2725327 | 0.780903765 |
| ENSMUSG00000008373 | Prpf31 | protein_coding | -0.218681922 | 6.59998186 | 1.2895726 | 0.2725416 | 0.780903765 |
| ENSMUSG00000029804 | Herc3 | protein_coding | 0.280506218 | 5.997854704 | 1.2892708 | 0.2725956 | 0.780903765 |
| ENSMUSG00000038879 | Nipal2 | protein_coding | 0.838489204 | 3.33169737 | 1.2962301 | 0.2726228 | 0.780903765 |
| ENSMUSG00000015668 | Pdzd11 | protein_coding | -0.204844169 | 7.534398526 | 1.2885908 | 0.2727175 | 0.780966978 |
| ENSMUSG00000031442 | Mcf2l | protein_coding | 0.354856247 | 4.428412457 | 1.2875512 | 0.2729039 | 0.781292713 |
| ENSMUSG00000030519 | Apba2 | protein_coding | 0.350332184 | 5.802000429 | 1.2872638 | 0.2729822 | 0.78130889 |
| ENSMUSG00000001962 | Fam50a | protein_coding | 0.516166865 | 4.370050922 | 1.2863082 | 0.2731537 | 0.78149547 |
| ENSMUSG00000032046 | Abhd12 | protein_coding | 0.228890786 | 6.967606994 | 1.285942 | 0.2731927 | 0.78149547 |
| ENSMUSG00000032531 | Amotl2 | protein_coding | 0.694281253 | 3.205997303 | 1.2852032 | 0.2733522 | 0.781576252 |
| ENSMUSG00000024350 | Dnajc18 | protein_coding | 0.275049645 | 7.567937018 | 1.2846529 | 0.2734244 | 0.781576252 |
| ENSMUSG00000033335 | Dnm2 | protein_coding | 0.609150248 | 4.189013686 | 1.2903541 | 0.273439 | 0.781576252 |
| ENSMUSG00000025982 | Sf3b1 | protein_coding | 0.20670707 | 7.046822769 | 1.2831453 | 0.2736957 | 0.781888226 |
| ENSMUSG00000025648 | Pfkfb4 | protein_coding | 0.590444986 | 3.353675628 | 1.2823033 | 0.2738474 | 0.781888226 |
| ENSMUSG00000090150 | Acad11 | protein_coding | 0.546964213 | 2.967308539 | 1.2822594 | 0.2738553 | 0.781888226 |
| ENSMUSG00000032366 | Tpm1 | protein_coding | -0.202574906 | 7.779207097 | 1.2821886 | 0.2738681 | 0.781888226 |
| ENSMUSG00000038481 | Cdk19 | protein_coding | -0.342730058 | 4.85149738 | 1.2819464 | 0.2739117 | 0.781888226 |
| ENSMUSG00000033128 | Gga1 | protein_coding | -0.361140557 | 4.297826087 | 1.281437 | 0.2740036 | 0.781942822 |
| ENSMUSG00000047719 | Ubiad1 | protein_coding | 0.517531355 | 3.207248485 | 1.2808576 | 0.2741081 | 0.782016025 |
| ENSMUSG00000020198 | Ap3d1 | protein_coding | 0.273906204 | 6.565677307 | 1.279362 | 0.2743782 | 0.782016025 |
| ENSMUSG00000023156 | Rpp14 | protein_coding | 0.219284664 | 6.519071113 | 1.2786721 | 0.2745029 | 0.782016025 |
| ENSMUSG00000039849 | Pcif1 | protein_coding | -0.247789661 | 6.315142522 | 1.2784057 | 0.274551 | 0.782016025 |
| ENSMUSG00000029486 | Mrpl1 | protein_coding | -0.302286296 | 5.919402217 | 1.2783579 | 0.2745597 | 0.782016025 |
| ENSMUSG00000054309 | Cpsf3 | protein_coding | 0.313196075 | 6.587729237 | 1.2782735 | 0.2746016 | 0.782016025 |

| ensembl_gene_id | external_gene_name | gene_biotype | log2 (fold change) | log2 (counts per million) | F-value | p-value | False discovery rate |
|---|---|---|---|---|---|---|---|
| ENSMUSG00000031231 | Cox7b | protein_coding | -0.273673246 | 9.469297054 | 1.2781257 | 0.2746017 | 0.782016025 |
| ENSMUSG00000043207 | Zmpste24 | protein_coding | -0.370676135 | 4.388046085 | 1.2780738 | 0.274611 | 0.782016025 |
| ENSMUSG00000109321 | A330076H08Rik | processed_transcript | -0.437548613 | 3.772797869 | 1.2772635 | 0.2747577 | 0.782028223 |
| ENSMUSG00000036298 | Slc2a13 | protein_coding | -0.373765139 | 5.510649288 | 1.2773937 | 0.2747608 | 0.782028223 |
| ENSMUSG00000033610 | Pank1 | protein_coding | 0.440825782 | 3.737443995 | 1.2760113 | 0.2749845 | 0.782457778 |
| ENSMUSG00000032376 | Usp3 | protein_coding | 0.356253831 | 4.465359218 | 1.2732111 | 0.2754925 | 0.783696044 |
| ENSMUSG00000021270 | Hsp90aa1 | protein_coding | 0.218306173 | 7.648787366 | 1.271819 | 0.2757456 | 0.784141479 |
| ENSMUSG00000021279 | Cdc42bpb | protein_coding | -0.504959767 | 3.217432205 | 1.2714971 | 0.2758041 | 0.784141479 |
| ENSMUSG00000027227 | Sord | protein_coding | 0.378168545 | 5.580487141 | 1.2710335 | 0.2759151 | 0.784141479 |
| ENSMUSG00000020178 | Adora2a | protein_coding | -0.486708615 | 4.517630999 | 1.2708924 | 0.2759408 | 0.784141479 |
| ENSMUSG00000031078 | Cttn | protein_coding | 0.320222666 | 6.768797952 | 1.2701627 | 0.2761438 | 0.784243119 |
| ENSMUSG00000021066 | Atl1 | protein_coding | 0.302156267 | 6.57713241 | 1.2690923 | 0.2762687 | 0.784243119 |
| ENSMUSG00000005533 | Igf1r | protein_coding | -0.40412179 | 4.398673351 | 1.2686163 | 0.2763289 | 0.784243119 |
| ENSMUSG00000026197 | Zfand2b | protein_coding | 0.276674616 | 5.663000998 | 1.2681359 | 0.2764166 | 0.784243119 |
| ENSMUSG00000040699 | Limd2 | protein_coding | 0.375696764 | 5.42800098 | 1.2679438 | 0.2764516 | 0.784243119 |
| ENSMUSG00000039953 | Clstn1 | protein_coding | 0.221718376 | 7.959736909 | 1.2677485 | 0.2764873 | 0.784243119 |
| ENSMUSG00000026708 | Cenpl | protein_coding | -0.768835172 | 2.71994695 | 1.2674047 | 0.2765501 | 0.784243119 |
| ENSMUSG00000022311 | Csmd3 | protein_coding | -0.45114778 | 4.866877705 | 1.2674957 | 0.27656 | 0.784243119 |
| ENSMUSG00000094707 | A830019P07Rik | lincRNA | -0.886728639 | 2.449636593 | 1.2665159 | 0.2767124 | 0.784443739 |
| ENSMUSG00000028696 | Ipp | protein_coding | 0.560426754 | 3.869324549 | 1.2663097 | 0.2767767 | 0.784443739 |
| ENSMUSG00000030401 | Rtn2 | protein_coding | -0.34322051 | 5.087100593 | 1.2654306 | 0.2769108 | 0.784617201 |
| ENSMUSG00000022789 | Dnm1l | protein_coding | -0.224578108 | 7.672574536 | 1.2643703 | 0.2771049 | 0.784707659 |
| ENSMUSG00000005267 | Zfp287 | protein_coding | -0.500641229 | 3.255660461 | 1.2637602 | 0.2772166 | 0.784707659 |
| ENSMUSG00000039046 | Usp6nl | protein_coding | 0.411170634 | 3.871457784 | 1.2631393 | 0.2773304 | 0.784707659 |
| ENSMUSG00000004567 | Mcoln1 | protein_coding | 0.387221302 | 4.725137671 | 1.2630827 | 0.2773408 | 0.784707659 |
| ENSMUSG00000045624 | Esf1 | protein_coding | -0.330560515 | 5.188180045 | 1.2628888 | 0.2773763 | 0.784707659 |
| ENSMUSG00000054814 | Usp46 | protein_coding | -0.29378339 | 6.444932736 | 1.2627657 | 0.2773989 | 0.784707659 |
| ENSMUSG00000025218 | Poll | protein_coding | -0.352553277 | 6.066550268 | 1.2626121 | 0.2774536 | 0.784707659 |
| ENSMUSG00000011658 | Fuz | protein_coding | 0.722274891 | 2.572785091 | 1.2618017 | 0.2775757 | 0.784789545 |
| ENSMUSG00000033278 | Ptprm | protein_coding | 0.412557765 | 4.481869238 | 1.261514 | 0.2776285 | 0.784789545 |
| ENSMUSG00000031399 | Fam3a | protein_coding | 0.311296186 | 5.110490453 | 1.261073 | 0.2777095 | 0.784812112 |
| ENSMUSG00000023845 | Lnpep | protein_coding | -0.937647314 | 2.367203165 | 1.2605458 | 0.2778063 | 0.784879498 |
| ENSMUSG00000049154 | Fam183b | protein_coding | 0.247966795 | 5.985182853 | 1.259341 | 0.2780278 | 0.784915295 |
| ENSMUSG00000046020 | Pofut1 | protein_coding | 0.683682939 | 2.626413497 | 1.259264 | 0.2780419 | 0.784915295 |
| ENSMUSG00000038665 | Dgki | protein_coding | -0.476014215 | 3.621750182 | 1.2587663 | 0.2781335 | 0.784915295 |
| ENSMUSG00000045095 | Magi1 | protein_coding | -0.260170973 | 6.208144879 | 1.2585999 | 0.2781641 | 0.784915295 |
| ENSMUSG00000040818 | Dennd6a | protein_coding | -0.427122451 | 3.729116788 | 1.2584921 | 0.278184 | 0.784915295 |
| ENSMUSG00000028630 | Dyrk2 | protein_coding | -0.427871858 | 3.865607839 | 1.257208 | 0.2784204 | 0.785187889 |
| ENSMUSG00000042111 | Ccdc115 | protein_coding | 0.295446913 | 5.416773609 | 1.2571745 | 0.2784266 | 0.785187889 |
| ENSMUSG00000054091 | 1810037I17Rik | protein_coding | -0.266730156 | 7.637371474 | 1.2567675 | 0.2785016 | 0.785193475 |
| ENSMUSG00000031111 | Igsf1 | protein_coding | 0.310890938 | 5.180803012 | 1.2550701 | 0.2788147 | 0.785759569 |
| ENSMUSG00000056832 | Ttc26 | protein_coding | 0.695623211 | 2.656063282 | 1.2547088 | 0.2788815 | 0.785759569 |
| ENSMUSG00000024069 | Slc30a6 | protein_coding | 0.421043051 | 4.55533128 | 1.2543317 | 0.2789511 | 0.785759569 |
| ENSMUSG00000042408 | Zmym6 | protein_coding | -0.628352903 | 2.98492359 | 1.2540958 | 0.2789947 | 0.785759569 |
| ENSMUSG00000024036 | Slc37a1 | protein_coding | 0.626017828 | 2.945747937 | 1.2532445 | 0.2791521 | 0.785996881 |
| ENSMUSG00000021607 | Mrpl36 | protein_coding | 0.191305776 | 7.686975598 | 1.2518066 | 0.2794181 | 0.786513256 |
| ENSMUSG00000058230 | Arhgap35 | protein_coding | 0.295072314 | 6.117530316 | 1.251463 | 0.2794817 | 0.786513256 |
| ENSMUSG00000090291 | Lrrc10b | protein_coding | -0.467775631 | 4.112851668 | 1.2497796 | 0.2797938 | 0.786644458 |
| ENSMUSG00000021798 | Ldb3 | protein_coding | 0.433055439 | 4.315403515 | 1.2482508 | 0.2800775 | 0.786644458 |
| ENSMUSG00000047797 | Gjb1 | protein_coding | -0.415447034 | 6.266397011 | 1.2567376 | 0.2801146 | 0.786644458 |
| ENSMUSG00000049858 | Suox | protein_coding | 0.560546943 | 3.829617284 | 1.2479908 | 0.2801522 | 0.786644458 |
| ENSMUSG00000046562 | Unc119b | protein_coding | 0.593633135 | 3.136478763 | 1.2476786 | 0.2801838 | 0.786644458 |
| ENSMUSG00000061393 | Acvr2b | protein_coding | 0.726073648 | 2.642841186 | 1.2474186 | 0.2802322 | 0.786644458 |
| ENSMUSG00000056770 | Setd3 | protein_coding | 0.22582381 | 6.51482753 | 1.2472952 | 0.2802551 | 0.786644458 |
| ENSMUSG00000027387 | Zc3h8 | protein_coding | -0.415567687 | 3.806664377 | 1.2470692 | 0.2802971 | 0.786644458 |
| ENSMUSG00000006740 | Kif5b | protein_coding | -0.265577559 | 6.115875941 | 1.2469666 | 0.2803162 | 0.786644458 |
| ENSMUSG00000005699 | Pard6a | protein_coding | -0.244832686 | 6.116825136 | 1.2467339 | 0.2803595 | 0.786644458 |
| ENSMUSG00000097055 | Gm4419 | lincRNA | -0.49024826 | 3.336072077 | 1.2466352 | 0.2803778 | 0.786644458 |
| ENSMUSG00000058298 | Mcm9 | protein_coding | -0.616140734 | 2.917679339 | 1.2464825 | 0.2804062 | 0.786644458 |
| ENSMUSG00000031924 | Cyb5b | protein_coding | 0.213216234 | 6.754261924 | 1.2455851 | 0.2805732 | 0.786721138 |
| ENSMUSG00000035620 | Ric8b | protein_coding | 0.296964425 | 5.915712258 | 1.2454995 | 0.2805892 | 0.786721138 |
| ENSMUSG00000035863 | Palm | protein_coding | -0.2895851 | 5.592000077 | 1.245032 | 0.2806763 | 0.786721138 |
| ENSMUSG00000020590 | Snx13 | protein_coding | -0.336118546 | 4.746669632 | 1.2442347 | 0.2808248 | 0.786721138 |
| ENSMUSG00000022742 | Cpox | protein_coding | -0.341606811 | 5.632461472 | 1.2440222 | 0.2808644 | 0.786721138 |
| ENSMUSG00000036959 | Bcorl1 | protein_coding | 0.454189359 | 3.776822993 | 1.2439788 | 0.2808725 | 0.786721138 |
| ENSMUSG00000027562 | Car2 | protein_coding | -0.232595569 | 10.41162244 | 1.2422119 | 0.2812022 | 0.786988621 |
| ENSMUSG00000028430 | Nol6 | protein_coding | 0.288122567 | 6.081971905 | 1.2416459 | 0.281308 | 0.786988621 |
| ENSMUSG00000029176 | Anapc4 | protein_coding | 0.255340192 | 6.037861913 | 1.2412954 | 0.2813735 | 0.786988621 |
| ENSMUSG00000085156 | Snhg15 | processed_transcript | -0.535241894 | 3.023525146 | 1.2410704 | 0.2814155 | 0.786988621 |
| ENSMUSG00000018572 | Phf23 | protein_coding | 0.275011532 | 5.843592551 | 1.2406058 | 0.2815024 | 0.786988621 |
| ENSMUSG00000024074 | Crim1 | protein_coding | -0.276873559 | 5.537589851 | 1.2399666 | 0.281622 | 0.786988621 |
| ENSMUSG00000032503 | Arpp21 | protein_coding | -0.348864014 | 10.18890769 | 1.244323 | 0.2817678 | 0.786988621 |
| ENSMUSG00000030619 | Eed | protein_coding | -0.432944938 | 4.505695003 | 1.2390137 | 0.2818004 | 0.786988621 |
| ENSMUSG00000022489 | Pde1b | protein_coding | -0.232093204 | 7.601957761 | 1.238967 | 0.2818091 | 0.786988621 |

| ensembl_gene_id | external_gene_name | gene_biotype | log2 (fold change) | log2 (counts per million) | F-value | p-value | False discovery rate |
|---|---|---|---|---|---|---|---|
| ENSMUSG00000079045 | Prox1os | antisense_RNA | -0.572656028 | 4.162944071 | 1.2417505 | 0.2818223 | 0.786988621 |
| ENSMUSG00000060961 | Slc4a4 | protein_coding | -0.382839358 | 5.376427232 | 1.239008 | 0.2818278 | 0.786988621 |
| ENSMUSG00000034349 | Smc4 | protein_coding | -0.799375293 | 2.374126917 | 1.2385162 | 0.2818936 | 0.786988621 |
| ENSMUSG00000049470 | Aff4 | protein_coding | 0.249178917 | 6.631452802 | 1.2382608 | 0.2819414 | 0.786988621 |
| ENSMUSG00000017376 | Nlk | protein_coding | -0.30673586 | 5.430241726 | 1.2376059 | 0.2820642 | 0.786988621 |
| ENSMUSG00000048000 | Gigyf2 | protein_coding | -0.232432879 | 6.073082081 | 1.2369083 | 0.282195 | 0.786988621 |
| ENSMUSG00000034110 | Kctd7 | protein_coding | 0.394696239 | 4.071132289 | 1.2366666 | 0.2822404 | 0.786988621 |
| ENSMUSG00000009293 | Ube2g2 | protein_coding | -0.253053864 | 6.617847669 | 1.2361357 | 0.28234 | 0.786988621 |
| ENSMUSG00000050310 | Rictor | protein_coding | -0.346842093 | 4.044007343 | 1.2361158 | 0.2823438 | 0.786988621 |
| ENSMUSG00000029203 | Ube2k | protein_coding | -0.225958791 | 7.384558187 | 1.2360369 | 0.2823586 | 0.786988621 |
| ENSMUSG00000042401 | Crtac1 | protein_coding | 0.464631369 | 4.769151047 | 1.2348572 | 0.2826065 | 0.787475526 |
| ENSMUSG00000027411 | Vps16 | protein_coding | 0.382339663 | 5.552159739 | 1.2342665 | 0.2827176 | 0.78758092 |
| ENSMUSG00000100164 | 2610306M01Rik | bidirectional_promoter | -0.295296911 | 5.269928062 | 1.232105 | 0.2830983 | 0.788437088 |
| ENSMUSG00000079557 | 2-Mar | protein_coding | -0.311256323 | 5.597889616 | 1.2309701 | 0.2833122 | 0.788817563 |
| ENSMUSG00000024065 | Ehd3 | protein_coding | 0.293029368 | 6.826757013 | 1.2307413 | 0.2833817 | 0.788817563 |
| ENSMUSG00000028358 | Zfp618 | protein_coding | 0.786204115 | 2.889924072 | 1.2294636 | 0.2835966 | 0.788817563 |
| ENSMUSG00000048782 | Insc | protein_coding | -0.829280767 | 2.653834911 | 1.2289963 | 0.2836849 | 0.788817563 |
| ENSMUSG00000059772 | Slx1b | protein_coding | -0.371232164 | 4.103154165 | 1.228085 | 0.2838572 | 0.788817563 |
| ENSMUSG00000029452 | Tmem116 | protein_coding | -0.510328659 | 3.387420825 | 1.2279928 | 0.2838746 | 0.788817563 |
| ENSMUSG00000035681 | Kcnc2 | protein_coding | -0.354405666 | 5.261467745 | 1.2266384 | 0.284131 | 0.788817563 |
| ENSMUSG00000057469 | E2f6 | protein_coding | 0.406035792 | 4.490085816 | 1.2264811 | 0.2841608 | 0.788817563 |
| ENSMUSG00000058260 | Serpina9 | protein_coding | -0.442216941 | 4.249092998 | 1.2264427 | 0.284168 | 0.788817563 |
| ENSMUSG00000056966 | Gjc3 | protein_coding | -0.297211665 | 5.573142038 | 1.2264379 | 0.2841689 | 0.788817563 |
| ENSMUSG00000047139 | Cd24a | protein_coding | 0.432165323 | 4.013841585 | 1.2258484 | 0.2842806 | 0.788817563 |
| ENSMUSG00000039221 | Rpl22l1 | protein_coding | -0.285676823 | 6.996212736 | 1.2257398 | 0.2843274 | 0.788817563 |
| ENSMUSG00000022913 | Psmg1 | protein_coding | 0.32822469 | 4.951692239 | 1.2254905 | 0.2843485 | 0.788817563 |
| ENSMUSG00000020841 | Cpd | protein_coding | 0.376454981 | 4.600038782 | 1.2248037 | 0.2844787 | 0.788817563 |
| ENSMUSG00000027016 | Zfp385b | protein_coding | -0.300000668 | 6.379825828 | 1.2247881 | 0.2844817 | 0.788817563 |
| ENSMUSG00000047921 | Trappc9 | protein_coding | 0.313773318 | 6.268041293 | 1.2248886 | 0.2844888 | 0.788817563 |
| ENSMUSG00000009640 | Dmap1 | protein_coding | 0.264911413 | 5.696855416 | 1.2242464 | 0.2845844 | 0.788817563 |
| ENSMUSG00000020687 | Cdc27 | protein_coding | -0.286189884 | 5.098841015 | 1.2229024 | 0.2848397 | 0.788817563 |
| ENSMUSG00000028271 | Gtf2b | protein_coding | 0.231847886 | 6.314934803 | 1.2218288 | 0.2850438 | 0.788817563 |
| ENSMUSG00000020409 | Slu7 | protein_coding | -0.233610542 | 7.116265078 | 1.2218095 | 0.2850475 | 0.788817563 |
| ENSMUSG00000039384 | Dusp10 | protein_coding | 0.56431757 | 4.084571816 | 1.2223017 | 0.2850942 | 0.788817563 |
| ENSMUSG00000070858 | Gm1673 | protein_coding | -0.478126723 | 3.401330592 | 1.2213394 | 0.2851369 | 0.788817563 |
| ENSMUSG00000000355 | Mcts1 | protein_coding | -0.220006089 | 7.797102568 | 1.2205145 | 0.2852939 | 0.788817563 |
| ENSMUSG00000022722 | Arl6 | protein_coding | -0.238883935 | 7.395387216 | 1.2198563 | 0.2854193 | 0.788817563 |
| ENSMUSG00000053062 | Jam2 | protein_coding | -0.380142806 | 3.995873277 | 1.2198482 | 0.2854208 | 0.788817563 |
| ENSMUSG00000053460 | Ggcx | protein_coding | 0.726251825 | 3.244690604 | 1.2198339 | 0.2854497 | 0.788817563 |
| ENSMUSG00000039450 | Dcxr | protein_coding | -0.440477653 | 4.156763777 | 1.2196838 | 0.2854522 | 0.788817563 |
| ENSMUSG00000020375 | Rufy1 | protein_coding | -0.317358457 | 4.966513053 | 1.219466 | 0.2854937 | 0.788817563 |
| ENSMUSG00000027984 | Hadh | protein_coding | 0.278900432 | 5.869148006 | 1.2192084 | 0.2855428 | 0.788817563 |
| ENSMUSG00000027305 | Ndufaf1 | protein_coding | -0.234916594 | 5.910435324 | 1.2182845 | 0.285719 | 0.788817563 |
| ENSMUSG00000006526 | Tmem110 | protein_coding | 0.605600774 | 3.326175884 | 1.2182118 | 0.2857329 | 0.788817563 |
| ENSMUSG00000067851 | Arfgef1 | protein_coding | -0.326515657 | 5.310221381 | 1.2179793 | 0.2857772 | 0.788817563 |
| ENSMUSG00000025352 | Gdf11 | protein_coding | 0.39164153 | 4.577487827 | 1.2177414 | 0.2858226 | 0.788817563 |
| ENSMUSG00000092368 | A930015D03Rik | antisense_RNA | -0.615321345 | 2.822723527 | 1.2173509 | 0.2858972 | 0.788817563 |
| ENSMUSG00000022545 | Ercc4 | protein_coding | 0.618990222 | 4.237128675 | 1.2235561 | 0.2860202 | 0.788817563 |
| ENSMUSG00000025213 | Kazald1 | protein_coding | 0.643536453 | 2.824612093 | 1.2161971 | 0.2861177 | 0.788817563 |
| ENSMUSG00000032238 | Rora | protein_coding | -0.335170457 | 6.226476772 | 1.2158768 | 0.286205 | 0.788817563 |
| ENSMUSG00000023106 | Denr | protein_coding | -0.210476153 | 7.38599362 | 1.2154364 | 0.2862631 | 0.788817563 |
| ENSMUSG00000004473 | Clec11a | protein_coding | -0.46279039 | 3.374650093 | 1.2145421 | 0.2864343 | 0.788817563 |
| ENSMUSG00000085913 | Gm15601 | antisense_RNA | 0.691388214 | 2.707428456 | 1.2141536 | 0.2865087 | 0.788817563 |
| ENSMUSG00000026425 | Srgap2 | protein_coding | -0.282898157 | 5.390911162 | 1.21415 | 0.2865094 | 0.788817563 |
| ENSMUSG00000003131 | Pafah1b2 | protein_coding | 0.220185292 | 7.544336209 | 1.2136914 | 0.2865972 | 0.788817563 |
| ENSMUSG00000026098 | Pms1 | protein_coding | -0.766820057 | 2.571576675 | 1.2129809 | 0.2867334 | 0.788817563 |
| ENSMUSG00000027002 | Nckap1 | protein_coding | -0.228718418 | 7.916296501 | 1.2129653 | 0.2867364 | 0.788817563 |
| ENSMUSG00000097585 | E230029C05Rik | processed_transcript | -0.481141526 | 3.481865834 | 1.2123618 | 0.2868522 | 0.788817563 |
| ENSMUSG00000028423 | Nfx1 | protein_coding | 0.278925326 | 6.440801969 | 1.2119595 | 0.2869293 | 0.788817563 |
| ENSMUSG00000026241 | Nppc | protein_coding | 0.547564645 | 3.262840325 | 1.2117163 | 0.286976 | 0.788817563 |
| ENSMUSG00000033671 | Cep350 | protein_coding | -0.654517908 | 2.879776363 | 1.211497 | 0.2870181 | 0.788817563 |
| ENSMUSG00000110332 | Gm45396 | protein_coding | -0.405466711 | 4.327348264 | 1.2111748 | 0.28708 | 0.788817563 |
| ENSMUSG00000061904 | Slc25a3 | protein_coding | 0.227178551 | 8.245816559 | 1.2109354 | 0.287126 | 0.788817563 |
| ENSMUSG00000091625 | Lsm5 | protein_coding | -0.574368882 | 3.039674698 | 1.2108597 | 0.2871405 | 0.788817563 |
| ENSMUSG00000073154 | 9330158H04Rik | lincRNA | 0.626546039 | 2.924406404 | 1.2108257 | 0.287147 | 0.788817563 |
| ENSMUSG00000020380 | Rad50 | protein_coding | 0.41263685 | 3.776395721 | 1.2104618 | 0.287217 | 0.788817563 |
| ENSMUSG00000109776 | Gm30400 | processed_transcript | -0.830536521 | 2.474284492 | 1.2103349 | 0.2872414 | 0.788817563 |
| ENSMUSG00000038215 | Cep44 | protein_coding | 0.638459333 | 3.199055763 | 1.2101881 | 0.2872696 | 0.788817563 |
| ENSMUSG00000043140 | Tmem186 | protein_coding | 0.390294774 | 5.17310996 | 1.2092012 | 0.2874593 | 0.789137171 |
| ENSMUSG00000009549 | Srp14 | protein_coding | 0.225458588 | 9.519047073 | 1.2078553 | 0.2877184 | 0.78951909 |
| ENSMUSG00000045980 | Tmem104 | protein_coding | 0.526704425 | 3.024620185 | 1.2075348 | 0.2877802 | 0.78951909 |
| ENSMUSG00000014074 | Rnf168 | protein_coding | 0.357099205 | 4.749605064 | 1.2073347 | 0.2878187 | 0.78951909 |
| ENSMUSG00000061898 | Rbak | protein_coding | 0.779652051 | 2.679502963 | 1.2068409 | 0.2879139 | 0.789578788 |
| ENSMUSG00000021731 | Mrps30 | protein_coding | -0.436300685 | 4.137049958 | 1.2064083 | 0.2879973 | 0.789606172 |

| ensembl_gene_id | external_gene_name | gene_biotype | log2 (fold change) | log2 (counts per million) | F-value | p-value | False discovery rate |
|---|---|---|---|---|---|---|---|
| ENSMUSG00000067071 | Hes6 | protein_coding | 0.48250424 | 3.233856932 | 1.2057421 | 0.2881259 | 0.789736005 |
| ENSMUSG00000033208 | S100b | protein_coding | -0.356918949 | 6.003589543 | 1.2058917 | 0.2881916 | 0.789736005 |
| ENSMUSG00000039286 | Fndc3b | protein_coding | -0.437592671 | 3.988063727 | 1.2042961 | 0.2884051 | 0.789978081 |
| ENSMUSG00000022564 | Grina | protein_coding | 0.214550554 | 8.836490859 | 1.2038111 | 0.2884989 | 0.789978081 |
| ENSMUSG00000021385 | Ippk | protein_coding | 0.424545846 | 4.175136875 | 1.2038036 | 0.2885003 | 0.789978081 |
| ENSMUSG00000025326 | Ube3a | protein_coding | -0.265460782 | 5.993583477 | 1.2022822 | 0.2887947 | 0.790204931 |
| ENSMUSG00000003038 | Hmgn2 | protein_coding | -0.274305239 | 5.561308851 | 1.2019609 | 0.2888569 | 0.790204931 |
| ENSMUSG00000013663 | Pten | protein_coding | -0.310624621 | 5.377594495 | 1.2019106 | 0.2888666 | 0.790204931 |
| ENSMUSG00000041781 | Cpsf2 | protein_coding | 0.262409072 | 5.601195972 | 1.2016879 | 0.2889098 | 0.790204931 |
| ENSMUSG00000039620 | 6430573F11Rik | protein_coding | 0.451157373 | 3.508994447 | 1.2013247 | 0.2889801 | 0.790204931 |
| ENSMUSG00000041638 | Gcn1l1 | protein_coding | -0.357197791 | 4.07384481 | 1.2010978 | 0.2890241 | 0.790204931 |
| ENSMUSG00000074657 | Kif5a | protein_coding | -0.27678594 | 8.897045906 | 1.1993111 | 0.2893966 | 0.790917199 |
| ENSMUSG00000059447 | Hadhb | protein_coding | 0.379131666 | 3.844665674 | 1.1984798 | 0.2895321 | 0.790917199 |
| ENSMUSG00000022010 | Tsc22d1 | protein_coding | -0.206503085 | 9.219451156 | 1.1984262 | 0.2895425 | 0.790917199 |
| ENSMUSG00000024889 | Rce1 | protein_coding | 0.357878591 | 4.453823746 | 1.1982393 | 0.2895788 | 0.790917199 |
| ENSMUSG00000035152 | Ap2b1 | protein_coding | -0.249905677 | 6.570604096 | 1.1973893 | 0.289744 | 0.791054151 |
| ENSMUSG00000026773 | Pfkfb3 | protein_coding | 0.554936165 | 3.827403228 | 1.197358 | 0.2897761 | 0.791054151 |
| ENSMUSG00000029922 | Mkrn1 | protein_coding | 0.217260781 | 6.384668255 | 1.1966971 | 0.2898787 | 0.791127432 |
| ENSMUSG00000040389 | Wdr47 | protein_coding | 0.258833082 | 6.156199779 | 1.1963304 | 0.2899501 | 0.791127432 |
| ENSMUSG00000034893 | Cog3 | protein_coding | 0.847009085 | 3.031055226 | 1.1974404 | 0.2902596 | 0.791526737 |
| ENSMUSG00000026707 | Nsun6 | protein_coding | 0.544728965 | 3.048525799 | 1.1942906 | 0.2903475 | 0.791526737 |
| ENSMUSG00000052374 | Actn2 | protein_coding | -0.754315696 | 3.558390301 | 1.2023022 | 0.2903679 | 0.791526737 |
| ENSMUSG00000028876 | Epha10 | protein_coding | -0.499562773 | 3.683833959 | 1.1940683 | 0.2903909 | 0.791526737 |
| ENSMUSG00000097195 | Snhg5 | processed_transcript | -0.378932743 | 4.857032958 | 1.1920499 | 0.2907849 | 0.791618987 |
| ENSMUSG00000032997 | Chpf | protein_coding | -0.346088057 | 5.077606247 | 1.1918407 | 0.2908258 | 0.791618987 |
| ENSMUSG00000034312 | Iqsec1 | protein_coding | -0.31935444 | 5.851191908 | 1.1917707 | 0.2908395 | 0.791618987 |
| ENSMUSG00000027679 | Dnajc19 | protein_coding | -0.2439886 | 6.113242534 | 1.1911401 | 0.2909628 | 0.791618987 |
| ENSMUSG00000024566 | Atp9b | protein_coding | 0.403943298 | 3.993432612 | 1.1910399 | 0.2909824 | 0.791618987 |
| ENSMUSG00000043572 | Pars2 | protein_coding | 0.371187496 | 3.973158608 | 1.190911 | 0.2910076 | 0.791618987 |
| ENSMUSG00000018196 | Glrx2 | protein_coding | -0.311718455 | 5.315577468 | 1.1907007 | 0.2910488 | 0.791618987 |
| ENSMUSG00000035069 | Oma1 | protein_coding | 0.475828585 | 3.725733586 | 1.1905492 | 0.2910784 | 0.791618987 |
| ENSMUSG00000078515 | Ddi2 | protein_coding | 0.316991935 | 5.409686099 | 1.1899869 | 0.2911884 | 0.791618987 |
| ENSMUSG00000097248 | Gm2694 | lincRNA | 0.426400985 | 4.397667393 | 1.1897199 | 0.2912407 | 0.791618987 |
| ENSMUSG00000024885 | Aldh3b1 | protein_coding | -0.461338444 | 3.921266022 | 1.1894991 | 0.291284 | 0.791618987 |
| ENSMUSG00000022437 | Samm50 | protein_coding | -0.263047448 | 6.467781763 | 1.1893758 | 0.2913081 | 0.791618987 |
| ENSMUSG00000002804 | Nudt14 | protein_coding | 0.259477505 | 5.569466343 | 1.188282 | 0.2915225 | 0.792001389 |
| ENSMUSG00000000861 | Bcl11a | protein_coding | -0.424121137 | 5.098905753 | 1.1868391 | 0.2918315 | 0.79233702 |
| ENSMUSG00000032295 | Man2c1 | protein_coding | -0.377168637 | 5.041759755 | 1.1857866 | 0.2920124 | 0.79233702 |
| ENSMUSG00000032507 | Fbxl2 | protein_coding | 0.347549646 | 5.618176864 | 1.1856012 | 0.2920488 | 0.79233702 |
| ENSMUSG00000066152 | Slc31a2 | protein_coding | 0.408224482 | 3.622977023 | 1.1851271 | 0.2921421 | 0.79233702 |
| ENSMUSG00000040557 | Mettl27 | protein_coding | -0.428831723 | 4.134411456 | 1.1850052 | 0.292166 | 0.79233702 |
| ENSMUSG00000071757 | Zhx2 | protein_coding | -0.534837691 | 3.222623385 | 1.1848501 | 0.2921965 | 0.79233702 |
| ENSMUSG00000036061 | Smug1 | protein_coding | 0.500550762 | 3.632797409 | 1.1844586 | 0.2922736 | 0.79233702 |
| ENSMUSG00000031990 | Jam3 | protein_coding | -0.316935069 | 4.968066662 | 1.1831032 | 0.2925404 | 0.79233702 |
| ENSMUSG00000022864 | D16Ertd472e | protein_coding | 0.8654552 | 2.559572326 | 1.1828816 | 0.2925841 | 0.79233702 |
| ENSMUSG00000056306 | Sertm1 | protein_coding | -0.529060995 | 3.222958381 | 1.1827214 | 0.2926157 | 0.79233702 |
| ENSMUSG00000021610 | Clptm1l | protein_coding | 0.346042926 | 5.774737366 | 1.1823378 | 0.2927171 | 0.79233702 |
| ENSMUSG00000060427 | Zfp868 | protein_coding | 0.457683073 | 4.786684045 | 1.1818731 | 0.2928087 | 0.79233702 |
| ENSMUSG00000093436 | Gm20646 | sense_overlapping | -0.420555142 | 3.643241201 | 1.1814949 | 0.2928576 | 0.79233702 |
| ENSMUSG00000022248 | Rad1 | protein_coding | 0.298198402 | 4.790968335 | 1.1813238 | 0.2928913 | 0.79233702 |
| ENSMUSG00000044627 | Swi5 | protein_coding | 0.245650978 | 9.426210742 | 1.1810697 | 0.2929415 | 0.79233702 |
| ENSMUSG00000002343 | Armc6 | protein_coding | 0.316426896 | 6.423998086 | 1.1810667 | 0.2929678 | 0.79233702 |
| ENSMUSG00000068457 | Uty | protein_coding | -0.470118081 | 3.355291775 | 1.1809289 | 0.2929693 | 0.79233702 |
| ENSMUSG00000029014 | Dnajc2 | protein_coding | -0.327231279 | 4.619513144 | 1.1807282 | 0.2930089 | 0.79233702 |
| ENSMUSG00000024063 | Lbh | protein_coding | 0.387431188 | 4.762518515 | 1.1799555 | 0.2931615 | 0.79233702 |
| ENSMUSG00000037703 | Lzts3 | protein_coding | -0.302207957 | 5.886241849 | 1.1790601 | 0.2933385 | 0.79233702 |
| ENSMUSG00000028976 | Slc2a5 | protein_coding | 1.106527868 | 2.785864423 | 1.1863099 | 0.2934774 | 0.79233702 |
| ENSMUSG00000097353 | A430046D13Rik | lincRNA | -0.397209733 | 3.66636268 | 1.1783516 | 0.2934787 | 0.79233702 |
| ENSMUSG00000042613 | Pbxip1 | protein_coding | -0.412850617 | 3.96877125 | 1.1783201 | 0.2934849 | 0.79233702 |
| ENSMUSG00000046671 | Mtfr1l | protein_coding | 0.207027517 | 7.657374727 | 1.1780585 | 0.2935367 | 0.79233702 |
| ENSMUSG00000032925 | Itgbl1 | protein_coding | -0.436750461 | 3.819230925 | 1.1779796 | 0.2935523 | 0.79233702 |
| ENSMUSG00000015476 | Prrt1 | protein_coding | -0.280451972 | 5.824490711 | 1.1779313 | 0.2935619 | 0.79233702 |
| ENSMUSG00000086496 | Gm14204 | processed_transcript | -0.358921882 | 5.415769256 | 1.1770508 | 0.2937362 | 0.792468983 |
| ENSMUSG00000001891 | Ugp2 | protein_coding | -0.218241174 | 6.183373685 | 1.1769402 | 0.2937581 | 0.792468983 |
| ENSMUSG00000036510 | Cdh8 | protein_coding | 0.435921074 | 4.200600969 | 1.1760876 | 0.2939271 | 0.792480363 |
| ENSMUSG00000038323 | 1700066M21Rik | protein_coding | -0.621053456 | 3.207639769 | 1.1759827 | 0.293948 | 0.792480363 |
| ENSMUSG00000037710 | Cisd1 | protein_coding | 0.295016055 | 8.734266486 | 1.1759334 | 0.2939835 | 0.792480363 |
| ENSMUSG00000028165 | Cisd2 | protein_coding | -0.242691026 | 6.175455156 | 1.174998 | 0.2941433 | 0.7927126 |
| ENSMUSG00000052921 | Arhgef15 | protein_coding | 0.269769359 | 6.334668996 | 1.1733899 | 0.2944628 | 0.793061093 |
| ENSMUSG00000034908 | Sidt2 | protein_coding | -0.369009654 | 4.407146496 | 1.173077 | 0.294525 | 0.793061093 |
| ENSMUSG00000031558 | Slit2 | protein_coding | -0.501161434 | 3.494476546 | 1.1730702 | 0.2945263 | 0.793061093 |
| ENSMUSG00000058761 | Rnf169 | protein_coding | 0.268659725 | 5.562580829 | 1.1728624 | 0.2945677 | 0.793061093 |
| ENSMUSG00000027423 | Snx5 | protein_coding | -0.246257571 | 6.114143889 | 1.1723202 | 0.2946755 | 0.793152923 |
| ENSMUSG00000029911 | Ssbp1 | protein_coding | -0.254885144 | 6.047762266 | 1.1707873 | 0.2949808 | 0.79358456 |

| ensembl_gene_id | external_gene_name | gene_biotype | log2 (fold change) | log2 (counts per million) | F-value | p-value | False discovery rate |
|---|---|---|---|---|---|---|---|
| ENSMUSG00000074886 | Grk6 | protein_coding | 0.347439427 | 4.896833007 | 1.1707736 | 0.2949835 | 0.79358456 |
| ENSMUSG00000019302 | Atp6v0a1 | protein_coding | 0.234141388 | 7.714074091 | 1.1698723 | 0.2951632 | 0.793755638 |
| ENSMUSG00000020284 | 1810043G02Rik | protein_coding | 0.249251681 | 6.869623216 | 1.1697141 | 0.2951947 | 0.793755638 |
| ENSMUSG00000046962 | Zbtb21 | protein_coding | -0.288348807 | 5.212693497 | 1.1683166 | 0.2954737 | 0.794307061 |
| ENSMUSG00000050947 | Amigo1 | protein_coding | 0.349264594 | 4.700277835 | 1.1665649 | 0.2958238 | 0.794957812 |
| ENSMUSG00000020577 | Tspan13 | protein_coding | 0.216423897 | 7.317139806 | 1.166366 | 0.2958636 | 0.794957812 |
| ENSMUSG00000032940 | Rbm11 | protein_coding | 0.377029621 | 4.260309132 | 1.1653002 | 0.2960769 | 0.795332341 |
| ENSMUSG00000029627 | Zkscan14 | protein_coding | 0.31321105 | 4.75515435 | 1.1648162 | 0.2961739 | 0.79539412 |
| ENSMUSG00000041341 | Atg2b | protein_coding | 0.427284018 | 3.948557634 | 1.1641436 | 0.2963087 | 0.795463759 |
| ENSMUSG00000060147 | Serpinb6a | protein_coding | 0.279132507 | 6.124552557 | 1.1625754 | 0.2966234 | 0.795463759 |
| ENSMUSG00000058013 | 11-Sep | protein_coding | 0.251768218 | 6.660549514 | 1.1623156 | 0.2966755 | 0.795463759 |
| ENSMUSG00000040128 | Pnrc1 | protein_coding | -0.258426165 | 5.564301215 | 1.1620228 | 0.2967343 | 0.795463759 |
| ENSMUSG00000028790 | Khdrbs1 | protein_coding | -0.293912865 | 5.447666043 | 1.1617035 | 0.2967985 | 0.795463759 |
| ENSMUSG00000002129 | Sf3a1 | protein_coding | 0.286262999 | 6.106236639 | 1.161444 | 0.2968506 | 0.795463759 |
| ENSMUSG00000026678 | Rgs5 | protein_coding | -0.440297304 | 4.246764995 | 1.1612666 | 0.2968863 | 0.795463759 |
| ENSMUSG00000030868 | Dctn5 | protein_coding | 0.24328504 | 5.84964316 | 1.1611909 | 0.2969015 | 0.795463759 |
| ENSMUSG00000028995 | Fam126a | protein_coding | -0.535654701 | 3.874948119 | 1.1606106 | 0.2970438 | 0.795463759 |
| ENSMUSG00000042757 | Tmem108 | protein_coding | -0.489699782 | 3.452302513 | 1.160056 | 0.2971298 | 0.795463759 |
| ENSMUSG00000097145 | 9230114K14Rik | processed_transcript | -0.696134381 | 2.639944073 | 1.1598331 | 0.2971747 | 0.795463759 |
| ENSMUSG00000048562 | Sp8 | protein_coding | -0.388748117 | 5.007978714 | 1.159258 | 0.2972905 | 0.795463759 |
| ENSMUSG00000001924 | Uba1 | protein_coding | 0.212949172 | 7.770458192 | 1.1591866 | 0.2973048 | 0.795463759 |
| ENSMUSG00000034593 | Myo5a | protein_coding | -0.268264475 | 7.076444363 | 1.1591496 | 0.2973379 | 0.795463759 |
| ENSMUSG00000002329 | Mdp1 | protein_coding | 0.191297018 | 7.037462489 | 1.1584666 | 0.2974499 | 0.795463759 |
| ENSMUSG00000066357 | Wdr6 | protein_coding | 0.570879052 | 7.794394425 | 1.1988001 | 0.2974599 | 0.795463759 |
| ENSMUSG00000022658 | Tagln3 | protein_coding | 0.208677899 | 10.66124834 | 1.1575846 | 0.2976278 | 0.795463759 |
| ENSMUSG00000048004 | Tmem196 | protein_coding | -0.424766626 | 4.160727152 | 1.1573853 | 0.297668 | 0.795463759 |
| ENSMUSG00000037049 | Smpd1 | protein_coding | 0.237886691 | 6.9351457 | 1.1570154 | 0.2977426 | 0.795463759 |
| ENSMUSG00000007411 | Mark3 | protein_coding | 0.404701947 | 4.971848746 | 1.1566688 | 0.2978381 | 0.795463759 |
| ENSMUSG00000058656 | Samd12 | protein_coding | -0.435028102 | 4.076291208 | 1.1563588 | 0.2978752 | 0.795463759 |
| ENSMUSG00000023206 | Il15ra | protein_coding | 0.524740979 | 3.391109557 | 1.1562978 | 0.2978875 | 0.795463759 |
| ENSMUSG00000027937 | Jtb | protein_coding | -0.329052879 | 5.714458474 | 1.155695 | 0.2980093 | 0.795463759 |
| ENSMUSG00000026248 | Mrpl44 | protein_coding | 0.210506331 | 7.060938854 | 1.1554979 | 0.2980491 | 0.795463759 |
| ENSMUSG00000032565 | Nudt16 | protein_coding | 0.354996045 | 4.08789586 | 1.1554972 | 0.2980492 | 0.795463759 |
| ENSMUSG00000070705 | Eid2b | protein_coding | 0.451472817 | 3.510431073 | 1.1549094 | 0.2981681 | 0.795583453 |
| ENSMUSG00000020491 | 2810021J22Rik | protein_coding | -0.46826247 | 3.813919539 | 1.1533867 | 0.2984762 | 0.796056514 |
| ENSMUSG00000037031 | Tspan15 | protein_coding | -0.416200331 | 4.183674783 | 1.1533016 | 0.2984934 | 0.796056514 |
| ENSMUSG00000040586 | Ofd1 | protein_coding | -0.703794874 | 2.848026959 | 1.1524315 | 0.2986697 | 0.796274902 |
| ENSMUSG00000071644 | Eef1g | protein_coding | 0.190212533 | 7.444402866 | 1.1520248 | 0.2987522 | 0.796274902 |
| ENSMUSG00000001376 | Vps50 | protein_coding | -0.343013161 | 5.090107022 | 1.1514654 | 0.2988656 | 0.796274902 |
| ENSMUSG00000031398 | Plxna3 | protein_coding | -0.640353756 | 2.688513231 | 1.1512178 | 0.2989159 | 0.796274902 |
| ENSMUSG00000022391 | Rangap1 | protein_coding | 0.221386944 | 7.905234947 | 1.1510714 | 0.2989456 | 0.796274902 |
| ENSMUSG00000021250 | Fos | protein_coding | 0.316085036 | 6.540849949 | 1.1512659 | 0.2990252 | 0.796289782 |
| ENSMUSG00000027933 | Ints3 | protein_coding | 0.334615282 | 4.900079861 | 1.1497611 | 0.2992116 | 0.79636299 |
| ENSMUSG00000016559 | H3f3b | protein_coding | 0.209818686 | 9.584166665 | 1.1496263 | 0.299239 | 0.79636299 |
| ENSMUSG00000030747 | Dgat2 | protein_coding | -0.28705607 | 5.811963899 | 1.1487193 | 0.2994234 | 0.79636299 |
| ENSMUSG00000073147 | 5031425E22Rik | lincRNA | -0.521466652 | 3.38855646 | 1.1485835 | 0.2994511 | 0.79636299 |
| ENSMUSG00000019782 | Rwdd1 | protein_coding | -0.384461374 | 4.027359334 | 1.1485383 | 0.2994602 | 0.79636299 |
| ENSMUSG00000025920 | Stau2 | protein_coding | -0.191081688 | 7.064248822 | 1.1483573 | 0.2994971 | 0.79636299 |
| ENSMUSG00000023030 | Slc11a2 | protein_coding | 0.428986713 | 4.586060056 | 1.147114 | 0.2997502 | 0.796644464 |
| ENSMUSG00000034245 | Hdac11 | protein_coding | 0.23075177 | 8.240556123 | 1.1471095 | 0.2997511 | 0.796644464 |
| ENSMUSG00000031536 | Polb | protein_coding | -0.183328228 | 7.048115771 | 1.1463175 | 0.2999125 | 0.796681428 |
| ENSMUSG00000039747 | Orai2 | protein_coding | 0.391294337 | 4.35907682 | 1.1452444 | 0.3001313 | 0.796681428 |
| ENSMUSG00000060371 | Caln1 | protein_coding | -0.301263138 | 6.668395795 | 1.1450934 | 0.3001876 | 0.796681428 |
| ENSMUSG00000045174 | Amer3 | protein_coding | 0.478375742 | 4.274527975 | 1.1449662 | 0.3002135 | 0.796681428 |
| ENSMUSG00000055761 | Nkain3 | protein_coding | -0.512121854 | 3.530744899 | 1.1447159 | 0.3002392 | 0.796681428 |
| ENSMUSG00000060260 | Pwwp2b | protein_coding | 0.439188392 | 4.46380528 | 1.1445625 | 0.3002705 | 0.796681428 |
| ENSMUSG00000027245 | Hypk | protein_coding | -0.370440706 | 4.01193018 | 1.1444982 | 0.3002836 | 0.796681428 |
| ENSMUSG00000036916 | Zfp280c | protein_coding | -0.549660032 | 3.532355404 | 1.1434196 | 0.300504 | 0.7968552 |
| ENSMUSG00000043301 | Kcnj6 | protein_coding | -0.580812484 | 2.906168264 | 1.1428411 | 0.3006222 | 0.7968552 |
| ENSMUSG00000015095 | Fbxw5 | protein_coding | 0.225407263 | 7.437499802 | 1.142733 | 0.3006444 | 0.7968552 |
| ENSMUSG00000052572 | Dlg2 | protein_coding | -0.233593531 | 6.437732956 | 1.1425638 | 0.300679 | 0.7968552 |
| ENSMUSG00000035629 | Rubcn | protein_coding | 0.331078774 | 5.172054063 | 1.1423649 | 0.3007197 | 0.7968552 |
| ENSMUSG00000002486 | Tchp | protein_coding | -0.571867075 | 2.932409377 | 1.141228 | 0.3009524 | 0.796983299 |
| ENSMUSG00000020941 | Map3k14 | protein_coding | -0.945932256 | 2.231274153 | 1.1411177 | 0.300975 | 0.796983299 |
| ENSMUSG00000045427 | Hnrnph2 | protein_coding | 0.279446525 | 5.821643913 | 1.1409076 | 0.301018 | 0.796983299 |
| ENSMUSG00000039987 | Phtf2 | protein_coding | -0.567906769 | 2.961224913 | 1.1406809 | 0.3010645 | 0.796983299 |
| ENSMUSG00000049076 | Acap2 | protein_coding | -0.352388739 | 4.383379589 | 1.1402408 | 0.3011547 | 0.797025859 |
| ENSMUSG00000024759 | Atl3 | protein_coding | -0.577350177 | 3.070248104 | 1.1397878 | 0.3012475 | 0.797075511 |
| ENSMUSG00000031224 | Magee2 | protein_coding | 0.374390175 | 5.034566242 | 1.1376387 | 0.3016887 | 0.797859835 |
| ENSMUSG00000067870 | Rpl31-ps8 | processed_pseudoge | -0.470964098 | 3.545993671 | 1.13712 | 0.3017954 | 0.797859835 |
| ENSMUSG00000017210 | Med24 | protein_coding | 0.354194417 | 5.606035835 | 1.1367992 | 0.3018867 | 0.797859835 |
| ENSMUSG00000054612 | Mgmt | protein_coding | 0.52693024 | 3.160805417 | 1.1364872 | 0.3019255 | 0.797859835 |
| ENSMUSG00000033658 | Ddx19b | protein_coding | 0.536688639 | 2.951968457 | 1.1357824 | 0.3020705 | 0.797859835 |
| ENSMUSG00000066442 | Mthfs | protein_coding | 0.56933604 | 2.857450493 | 1.1356986 | 0.3020878 | 0.797859835 |

| ensembl_gene_id | external_gene_name | gene_biotype | log2 (fold change) | log2 (counts per million) | F-value | p-value | False discovery rate |
|---|---|---|---|---|---|---|---|
| ENSMUSG00000021615 | Xrcc4 | protein_coding | -0.389868963 | 4.192460649 | 1.1356682 | 0.3020941 | 0.797859835 |
| ENSMUSG00000037818 | Abhd18 | protein_coding | -0.510484213 | 3.23669657 | 1.1352969 | 0.3021705 | 0.797859835 |
| ENSMUSG00000032470 | Mras | protein_coding | -0.217790194 | 6.058891247 | 1.134754 | 0.3022824 | 0.797859835 |
| ENSMUSG00000029265 | Dr1 | protein_coding | -0.403090989 | 4.416622029 | 1.1345703 | 0.3023202 | 0.797859835 |
| ENSMUSG00000090122 | Kcne1l | protein_coding | 0.87938382 | 2.519501753 | 1.133796 | 0.3024798 | 0.797859835 |
| ENSMUSG00000030629 | Zfand6 | protein_coding | -0.211557687 | 7.01366206 | 1.1322497 | 0.302799 | 0.797859835 |
| ENSMUSG00000000247 | Lhx2 | protein_coding | 0.702499724 | 3.10998722 | 1.1321492 | 0.302845 | 0.797859835 |
| ENSMUSG00000035372 | 1810055G02Rik | protein_coding | -0.372427018 | 4.514499838 | 1.1311343 | 0.3030295 | 0.797859835 |
| ENSMUSG00000034663 | Bmp2k | protein_coding | 0.647629732 | 2.757014937 | 1.1311105 | 0.3030344 | 0.797859835 |
| ENSMUSG00000045257 | Morn2 | protein_coding | -0.239938989 | 5.872086206 | 1.1306512 | 0.3031294 | 0.797859835 |
| ENSMUSG00000073792 | Alg6 | protein_coding | 0.409220213 | 3.70962646 | 1.130296 | 0.3032029 | 0.797859835 |
| ENSMUSG00000020277 | Pfkl | protein_coding | 0.323971225 | 7.5166223 | 1.1350433 | 0.3032037 | 0.797859835 |
| ENSMUSG00000031497 | Tnfsf13b | protein_coding | -0.614167571 | 3.010661282 | 1.129263 | 0.3034167 | 0.797859835 |
| ENSMUSG00000021671 | Poc5 | protein_coding | 0.340902177 | 4.472090136 | 1.1288344 | 0.3035055 | 0.797859835 |
| ENSMUSG00000049092 | Gpr137c | protein_coding | -0.64011763 | 2.982284169 | 1.1282979 | 0.3036167 | 0.797859835 |
| ENSMUSG00000026123 | Plekhb2 | protein_coding | 0.389699115 | 9.104807935 | 1.1387328 | 0.3036245 | 0.797859835 |
| ENSMUSG00000025261 | Huwe1 | protein_coding | -0.236017808 | 5.898105929 | 1.1281991 | 0.3036372 | 0.797859835 |
| ENSMUSG00000009575 | Cbx5 | protein_coding | -0.189785163 | 8.581273462 | 1.1279082 | 0.3036975 | 0.797859835 |
| ENSMUSG00000093738 | AI606473 | processed_transcript | -0.569701919 | 3.792607044 | 1.127564 | 0.3037941 | 0.797859835 |
| ENSMUSG00000039835 | Nhsl1 | protein_coding | 0.524500333 | 3.785744422 | 1.1274919 | 0.3038091 | 0.797859835 |
| ENSMUSG00000036376 | Abt1 | protein_coding | 0.276605458 | 6.069733086 | 1.1270129 | 0.3038833 | 0.797859835 |
| ENSMUSG00000074227 | Spint2 | protein_coding | 0.229200354 | 5.881870537 | 1.1269862 | 0.3038888 | 0.797859835 |
| ENSMUSG00000054934 | Kcnmb4 | protein_coding | -0.734228491 | 2.524912605 | 1.1264419 | 0.3040018 | 0.797859835 |
| ENSMUSG00000015711 | Prune1 | protein_coding | 0.39294124 | 5.363997029 | 1.1262894 | 0.3040587 | 0.797859835 |
| ENSMUSG00000032456 | Nmnat3 | protein_coding | -0.651124685 | 2.809270267 | 1.1257174 | 0.3041523 | 0.797859835 |
| ENSMUSG00000037224 | Zfyve28 | protein_coding | 0.368759418 | 3.891816337 | 1.1254788 | 0.3042019 | 0.797859835 |
| ENSMUSG00000026150 | Mff | protein_coding | -0.232009007 | 7.330525342 | 1.1249577 | 0.3043102 | 0.797859835 |
| ENSMUSG00000051067 | Lingo3 | protein_coding | 0.570743307 | 3.708246974 | 1.1250312 | 0.3043202 | 0.797859835 |
| ENSMUSG00000026554 | Dcaf8 | protein_coding | 0.277987468 | 7.101564655 | 1.1250033 | 0.304326 | 0.797859835 |
| ENSMUSG00000037649 | H2-DMa | protein_coding | -0.422956315 | 4.709291474 | 1.1247072 | 0.3043876 | 0.797859835 |
| ENSMUSG00000007987 | Ift22 | protein_coding | 0.190490919 | 8.388343311 | 1.1243806 | 0.3044303 | 0.797859835 |
| ENSMUSG00000007721 | Ccdc124 | protein_coding | -0.202532865 | 8.225386948 | 1.1240615 | 0.3044967 | 0.797859835 |
| ENSMUSG00000057666 | Gapdh | protein_coding | 0.365362974 | 4.429047748 | 1.1238685 | 0.3045369 | 0.797859835 |
| ENSMUSG00000036983 | Tfb1m | protein_coding | -0.421544798 | 3.668714948 | 1.12381 | 0.304549 | 0.797859835 |
| ENSMUSG00000025995 | Wdr75 | protein_coding | -0.248268548 | 5.615716546 | 1.1236319 | 0.3045861 | 0.797859835 |
| ENSMUSG00000027263 | Tubgcp4 | protein_coding | 0.531514199 | 3.016042096 | 1.1223151 | 0.3048605 | 0.797935784 |
| ENSMUSG00000041912 | Tdrkh | protein_coding | -0.246656592 | 5.926393016 | 1.1220911 | 0.3049072 | 0.797935784 |
| ENSMUSG00000022748 | Cmss1 | protein_coding | 0.32455774 | 5.107140504 | 1.1217662 | 0.3049749 | 0.797935784 |
| ENSMUSG00000025280 | Polr3a | protein_coding | 0.439742895 | 3.709208896 | 1.1216403 | 0.3050012 | 0.797935784 |
| ENSMUSG00000051705 | Senp8 | protein_coding | -0.367831146 | 4.582202443 | 1.1212429 | 0.3050841 | 0.797935784 |
| ENSMUSG00000027239 | Mdk | protein_coding | -0.573518072 | 4.884046734 | 1.1314003 | 0.3051286 | 0.797935784 |
| ENSMUSG00000038042 | Ptpdc1 | protein_coding | 0.418215945 | 4.040815844 | 1.1210012 | 0.3051346 | 0.797935784 |
| ENSMUSG00000070002 | Ell | protein_coding | -0.316915439 | 5.35422909 | 1.1202082 | 0.3053002 | 0.798057668 |
| ENSMUSG00000032415 | Ube2cbp | protein_coding | 0.437489037 | 3.39040437 | 1.1192839 | 0.3054933 | 0.798057668 |
| ENSMUSG00000032329 | Hmg20a | protein_coding | 0.26512868 | 6.190941691 | 1.1192234 | 0.305506 | 0.798057668 |
| ENSMUSG00000097219 | Gm26551 | lincRNA | -0.442662535 | 3.711355246 | 1.1191651 | 0.3055182 | 0.798057668 |
| ENSMUSG00000045210 | Vcpip1 | protein_coding | -0.259065126 | 5.468057633 | 1.119002 | 0.3055523 | 0.798057668 |
| ENSMUSG00000038848 | Ythdf1 | protein_coding | -0.350830981 | 5.274372104 | 1.1181692 | 0.3057265 | 0.798181429 |
| ENSMUSG00000036905 | C1qb | protein_coding | -0.465475944 | 6.327932409 | 1.1324544 | 0.3057481 | 0.798181429 |
| ENSMUSG00000050079 | Rspry1 | protein_coding | 0.320091179 | 4.873429617 | 1.1171879 | 0.305932 | 0.798282625 |
| ENSMUSG00000024493 | Lars | protein_coding | 0.294869855 | 5.57129113 | 1.1171718 | 0.3059353 | 0.798282625 |
| ENSMUSG00000032398 | Snapc5 | protein_coding | -0.252985891 | 6.535597671 | 1.11507 | 0.3063761 | 0.798421671 |
| ENSMUSG00000042816 | Gpr151 | protein_coding | 1.67089954 | 2.063609126 | 1.1251411 | 0.3064208 | 0.798421671 |
| ENSMUSG00000024560 | Cxxc1 | protein_coding | 0.495873738 | 4.553512164 | 1.1167292 | 0.3065399 | 0.798421671 |
| ENSMUSG00000062661 | Ncs1 | protein_coding | -0.25103707 | 10.21342779 | 1.1140077 | 0.3065992 | 0.798421671 |
| ENSMUSG00000028300 | 3110043O21Rik | protein_coding | 0.315796857 | 5.315404608 | 1.1137285 | 0.3066578 | 0.798421671 |
| ENSMUSG00000022332 | Khdrbs3 | protein_coding | -0.237290791 | 6.130064683 | 1.1132767 | 0.3067528 | 0.798421671 |
| ENSMUSG00000056602 | Fry | protein_coding | -0.272095512 | 5.901134414 | 1.1132569 | 0.306757 | 0.798421671 |
| ENSMUSG00000028126 | Pip5k1a | protein_coding | -0.274361383 | 5.848862134 | 1.1131905 | 0.306771 | 0.798421671 |
| ENSMUSG00000042331 | Specc1 | protein_coding | 0.363431284 | 5.358759164 | 1.1132985 | 0.3067733 | 0.798421671 |
| ENSMUSG00000062232 | Rapgef2 | protein_coding | -0.303492787 | 5.343580898 | 1.1131003 | 0.3067899 | 0.798421671 |
| ENSMUSG00000073838 | Tufm | protein_coding | 0.390649614 | 3.580335509 | 1.111657 | 0.3070937 | 0.798421671 |
| ENSMUSG00000008892 | Vdac3 | protein_coding | 0.186678309 | 7.591641338 | 1.1115731 | 0.3071113 | 0.798421671 |
| ENSMUSG00000024899 | Papss2 | protein_coding | -0.46274504 | 3.466976227 | 1.1114747 | 0.307132 | 0.798421671 |
| ENSMUSG00000109121 | Gm44644 | lincRNA | -0.542727479 | 3.242296823 | 1.11106 | 0.3072194 | 0.798421671 |
| ENSMUSG00000003469 | Phyhip | protein_coding | -0.21039535 | 8.650892577 | 1.1110136 | 0.3072292 | 0.798421671 |
| ENSMUSG00000044795 | Cyb5d1 | protein_coding | 0.651569716 | 3.322605512 | 1.1103445 | 0.3073953 | 0.798421671 |
| ENSMUSG00000005370 | Msh6 | protein_coding | 0.921209378 | 2.915605067 | 1.1148154 | 0.3074025 | 0.798421671 |
| ENSMUSG00000024425 | Ndfip1 | protein_coding | 0.202405987 | 6.807682116 | 1.1099618 | 0.3074509 | 0.798421671 |
| ENSMUSG00000036639 | Nudt1 | protein_coding | 0.332514061 | 4.102397497 | 1.1087887 | 0.3076985 | 0.798421671 |
| ENSMUSG00000039427 | Alg1 | protein_coding | 0.537156885 | 3.42903892 | 1.1084338 | 0.3077735 | 0.798421671 |
| ENSMUSG00000045854 | Lyrm2 | protein_coding | -0.277988015 | 6.16688225 | 1.1079916 | 0.3078669 | 0.798421671 |
| ENSMUSG00000044933 | Sstr3 | protein_coding | -0.890068491 | 1.991282608 | 1.1124948 | 0.3079043 | 0.798421671 |
| ENSMUSG00000033209 | Ttc28 | protein_coding | -0.616383408 | 2.80421923 | 1.107378 | 0.3079966 | 0.798421671 |

| ensembl_gene_id | external_gene_name | gene_biotype | log2 (fold change) | log2 (counts per million) | F-value | p-value | False discovery rate |
|---|---|---|---|---|---|---|---|
| ENSMUSG00000020990 | Cdkl1 | protein_coding | 0.365236888 | 4.588056942 | 1.1071801 | 0.3080384 | 0.798421671 |
| ENSMUSG00000032262 | Elovl4 | protein_coding | 0.241887114 | 5.962057517 | 1.1071392 | 0.3080471 | 0.798421671 |
| ENSMUSG00000066568 | Lsm14a | protein_coding | -0.251388451 | 5.43454583 | 1.1071072 | 0.3080539 | 0.798421671 |
| ENSMUSG00000001763 | Tspan33 | protein_coding | 0.511431921 | 3.839518618 | 1.107222 | 0.3080546 | 0.798421671 |
| ENSMUSG00000029127 | Zbtb49 | protein_coding | -1.186722395 | 2.185977151 | 1.1074723 | 0.3080677 | 0.798421671 |
| ENSMUSG00000046287 | Pnma3 | protein_coding | 0.444238116 | 4.538704125 | 1.1059048 | 0.3083333 | 0.798821305 |
| ENSMUSG00000032666 | 1700025G04Rik | protein_coding | -0.274803713 | 5.294701351 | 1.1052706 | 0.3084426 | 0.798821305 |
| ENSMUSG00000053134 | Supt7l | protein_coding | 0.323201021 | 4.77275833 | 1.104639 | 0.3085765 | 0.798821305 |
| ENSMUSG00000027296 | Itpka | protein_coding | -0.457012966 | 5.082096118 | 1.1088823 | 0.3085902 | 0.798821305 |
| ENSMUSG00000029576 | Radil | protein_coding | -0.505932242 | 3.178132561 | 1.1045597 | 0.3085933 | 0.798821305 |
| ENSMUSG00000000552 | Zfp385a | protein_coding | -0.403023307 | 4.803536637 | 1.1035995 | 0.308797 | 0.798935797 |
| ENSMUSG00000070509 | Rgma | protein_coding | -0.484842267 | 3.427260483 | 1.1030402 | 0.3089157 | 0.798935797 |
| ENSMUSG00000034949 | Zfr2 | protein_coding | -0.429016593 | 3.528318463 | 1.1029451 | 0.3089359 | 0.798935797 |
| ENSMUSG00000000399 | Ndufa9 | protein_coding | 0.1912337 | 9.01370572 | 1.1026181 | 0.3090053 | 0.798935797 |
| ENSMUSG00000028801 | Stpg1 | protein_coding | -0.55095433 | 3.122918865 | 1.1025441 | 0.309021 | 0.798935797 |
| ENSMUSG00000041763 | Tpp2 | protein_coding | 0.288867953 | 5.151239924 | 1.1018287 | 0.309173 | 0.798935797 |
| ENSMUSG00000028096 | Gpr89 | protein_coding | 0.312975105 | 4.975555012 | 1.1015251 | 0.3092376 | 0.798935797 |
| ENSMUSG00000027349 | Fam98b | protein_coding | 0.221194095 | 6.291079979 | 1.1003169 | 0.3094946 | 0.798935797 |
| ENSMUSG00000038866 | Zcchc2 | protein_coding | 0.519550521 | 2.898668744 | 1.1003054 | 0.3094971 | 0.798935797 |
| ENSMUSG00000025016 | Tm9sf3 | protein_coding | -0.272606246 | 6.079951102 | 1.1002522 | 0.3095084 | 0.798935797 |
| ENSMUSG00000025795 | Rassf3 | protein_coding | -0.577759011 | 2.979774299 | 1.0997671 | 0.3096117 | 0.798935797 |
| ENSMUSG00000013662 | Atad1 | protein_coding | -0.234414033 | 6.835617778 | 1.0995881 | 0.3096498 | 0.798935797 |
| ENSMUSG00000036478 | Btg1 | protein_coding | -0.343684236 | 5.003005241 | 1.0986801 | 0.3098433 | 0.798935797 |
| ENSMUSG00000031728 | Zfp821 | protein_coding | 0.378028086 | 4.6341549 | 1.0982898 | 0.3099265 | 0.798935797 |
| ENSMUSG00000011148 | Adssl1 | protein_coding | 0.300801389 | 4.719641668 | 1.097975 | 0.3099937 | 0.798935797 |
| ENSMUSG00000026496 | Parp1 | protein_coding | 0.273005514 | 5.257757209 | 1.0979009 | 0.3100095 | 0.798935797 |
| ENSMUSG00000003308 | Keap1 | protein_coding | 0.469177421 | 4.751555809 | 1.0997828 | 0.310082 | 0.798935797 |
| ENSMUSG00000043857 | Mgat5b | protein_coding | -0.365213908 | 4.474945001 | 1.0974247 | 0.3101111 | 0.798935797 |
| ENSMUSG00000032905 | Atg12 | protein_coding | -0.192577952 | 6.55226275 | 1.0971002 | 0.3101804 | 0.798935797 |
| ENSMUSG00000051323 | Pcdh19 | protein_coding | 0.279465147 | 5.411491613 | 1.0959622 | 0.3104235 | 0.798935797 |
| ENSMUSG00000031521 | Aga | protein_coding | 0.406967363 | 4.549416809 | 1.0958421 | 0.3104492 | 0.798935797 |
| ENSMUSG00000028128 | F3 | protein_coding | -0.378597281 | 5.787861744 | 1.0988081 | 0.3104522 | 0.798935797 |
| ENSMUSG00000054237 | Fra10ac1 | protein_coding | -0.268191459 | 5.396569907 | 1.0954124 | 0.3105411 | 0.798935797 |
| ENSMUSG00000066438 | Plekhd1 | protein_coding | 0.747280466 | 2.44947377 | 1.0952607 | 0.3105735 | 0.798935797 |
| ENSMUSG00000032652 | Crebl2 | protein_coding | 0.321371515 | 7.062361033 | 1.0991632 | 0.3107069 | 0.798935797 |
| ENSMUSG00000041769 | Ppp2r2d | protein_coding | 0.211807917 | 6.312592525 | 1.0944898 | 0.3107385 | 0.798935797 |
| ENSMUSG00000067288 | Rps28 | protein_coding | -0.368766698 | 4.775434699 | 1.0944129 | 0.3107549 | 0.798935797 |
| ENSMUSG00000028172 | Tacr3 | protein_coding | 0.574032774 | 2.731953969 | 1.0937878 | 0.3108888 | 0.798935797 |
| ENSMUSG00000051238 | Swsap1 | protein_coding | -0.562367263 | 2.90923937 | 1.0937716 | 0.3108922 | 0.798935797 |
| ENSMUSG00000058833 | 2810428I15Rik | protein_coding | -0.232733087 | 7.558352681 | 1.0932134 | 0.3110118 | 0.798935797 |
| ENSMUSG00000021102 | Glrx5 | protein_coding | -0.373486233 | 5.001211171 | 1.0914285 | 0.3113947 | 0.798935797 |
| ENSMUSG00000016918 | Sulf1 | protein_coding | 0.568174173 | 3.857128097 | 1.092953 | 0.3114575 | 0.798935797 |
| ENSMUSG00000032058 | Ppp2r1b | protein_coding | 0.345909994 | 4.65675267 | 1.0908419 | 0.3115206 | 0.798935797 |
| ENSMUSG00000070565 | Rasal2 | protein_coding | -0.331624033 | 5.085976539 | 1.090838 | 0.3115215 | 0.798935797 |
| ENSMUSG00000001052 | Sec24b | protein_coding | 0.286722001 | 4.881880454 | 1.0907693 | 0.3115362 | 0.798935797 |
| ENSMUSG00000050063 | Klk6 | protein_coding | -0.547155266 | 3.952359432 | 1.0911152 | 0.3115952 | 0.798935797 |
| ENSMUSG00000046679 | C87436 | protein_coding | 0.358054619 | 4.824337619 | 1.0904271 | 0.3116097 | 0.798935797 |
| ENSMUSG00000069045 | Ddx3y | protein_coding | -0.247816717 | 6.104745614 | 1.0903861 | 0.3116186 | 0.798935797 |
| ENSMUSG00000024841 | Eif1ad | protein_coding | -0.184917928 | 7.031500112 | 1.0901389 | 0.3116717 | 0.798935797 |
| ENSMUSG00000042558 | Adprhl2 | protein_coding | -0.441729248 | 3.937292687 | 1.0899766 | 0.3117066 | 0.798935797 |
| ENSMUSG00000027597 | Ahcy | protein_coding | 0.401660404 | 3.452462373 | 1.0897958 | 0.3117454 | 0.798935797 |
| ENSMUSG00000061474 | Mrps36 | protein_coding | -0.321226035 | 4.525805202 | 1.0896732 | 0.3117718 | 0.798935797 |
| ENSMUSG00000035376 | Hacd2 | protein_coding | -0.417068307 | 3.720250957 | 1.0893916 | 0.3118324 | 0.798935797 |
| ENSMUSG00000027361 | Gabpb1 | protein_coding | -0.366601926 | 4.01135822 | 1.0884843 | 0.3120276 | 0.799245562 |
| ENSMUSG00000057113 | Npm1 | protein_coding | -0.196442045 | 8.151725743 | 1.0876266 | 0.3122123 | 0.799271701 |
| ENSMUSG00000001053 | N4bp3 | protein_coding | -0.723827429 | 2.452162379 | 1.0868951 | 0.31237 | 0.799271701 |
| ENSMUSG00000038351 | Sgsm2 | protein_coding | 0.322827496 | 4.882327647 | 1.0866945 | 0.3124132 | 0.799271701 |
| ENSMUSG00000025420 | Katnal2 | protein_coding | 0.459906417 | 3.421721063 | 1.0861885 | 0.3125224 | 0.799271701 |
| ENSMUSG00000039910 | Cited2 | protein_coding | -0.310263135 | 4.336699034 | 1.0861013 | 0.3125412 | 0.799271701 |
| ENSMUSG00000028969 | Cdk5 | protein_coding | 0.195041315 | 8.546438567 | 1.0855578 | 0.3126585 | 0.799271701 |
| ENSMUSG00000047022 | Mipol1 | protein_coding | -0.387427823 | 4.213269487 | 1.0852385 | 0.3127275 | 0.799271701 |
| ENSMUSG00000058440 | Nrf1 | protein_coding | 0.295136388 | 4.588800568 | 1.0852224 | 0.3127309 | 0.799271701 |
| ENSMUSG00000021771 | Vdac2 | protein_coding | 0.204345662 | 9.130915207 | 1.0847453 | 0.312834 | 0.799271701 |
| ENSMUSG00000108414 | Snhg1 | processed_transcript | -0.323184582 | 4.599979832 | 1.0844235 | 0.3129035 | 0.799271701 |
| ENSMUSG00000029676 | Pot1a | protein_coding | -0.411038035 | 4.502546478 | 1.0838738 | 0.3130224 | 0.799271701 |
| ENSMUSG00000026826 | Nr4a2 | protein_coding | -0.611696134 | 2.621287073 | 1.083756 | 0.3130478 | 0.799271701 |
| ENSMUSG00000028793 | Rnf19b | protein_coding | -0.315974972 | 4.540440979 | 1.0836546 | 0.3130698 | 0.799271701 |
| ENSMUSG00000029836 | Cbx3 | protein_coding | -0.386223054 | 3.558430742 | 1.0836145 | 0.3130784 | 0.799271701 |
| ENSMUSG00000004880 | Lbr | protein_coding | 0.441609354 | 3.517130989 | 1.0832497 | 0.3131574 | 0.799283448 |
| ENSMUSG00000037072 | Selenof | protein_coding | -0.22342884 | 8.841278563 | 1.0826345 | 0.3132905 | 0.799301274 |
| ENSMUSG00000033728 | Lrrc14 | protein_coding | 0.359804562 | 4.191297207 | 1.0824235 | 0.3133362 | 0.799301274 |
| ENSMUSG00000045983 | Eif4g1 | protein_coding | 0.288945682 | 5.20967708 | 1.0820577 | 0.3134154 | 0.799301274 |
| ENSMUSG00000060166 | Zdhhc8 | protein_coding | -0.4121087 | 3.884171771 | 1.0817046 | 0.3134919 | 0.799301274 |
| ENSMUSG00000108030 | 9530062K07Rik | bidirectional_promoter | -0.811192722 | 3.071294766 | 1.0848977 | 0.3136154 | 0.799301274 |

| ensembl_gene_id | external_gene_name | gene_biotype | log2 (fold change) | log2 (counts per million) | F-value | p-value | False discovery rate |
|---|---|---|---|---|---|---|---|
| ENSMUSG00000024501 | Dpysl3 | protein_coding | 0.294406775 | 6.205575626 | 1.0806221 | 0.3137513 | 0.799301274 |
| ENSMUSG00000042616 | Oscp1 | protein_coding | 0.241501743 | 6.296014459 | 1.0803276 | 0.3137905 | 0.799301274 |
| ENSMUSG00000063632 | Sox11 | protein_coding | -0.249004856 | 6.041335213 | 1.0795627 | 0.3139565 | 0.799301274 |
| ENSMUSG00000031357 | Syap1 | protein_coding | -0.429429433 | 3.793191762 | 1.0794456 | 0.3139819 | 0.799301274 |
| ENSMUSG00000005299 | Letm1 | protein_coding | 0.271127682 | 5.787352324 | 1.0793031 | 0.3140129 | 0.799301274 |
| ENSMUSG00000074129 | Rpl13a | protein_coding | 0.271246008 | 5.367134901 | 1.0790938 | 0.3140583 | 0.799301274 |
| ENSMUSG00000025468 | Caly | protein_coding | 0.20565908 | 9.560052941 | 1.0787922 | 0.3141239 | 0.799301274 |
| ENSMUSG00000020422 | Tns3 | protein_coding | 0.487640062 | 3.221768076 | 1.0787286 | 0.3141377 | 0.799301274 |
| ENSMUSG00000036501 | Fam13b | protein_coding | 0.393536705 | 3.975044105 | 1.0780498 | 0.3142852 | 0.799301274 |
| ENSMUSG00000020235 | Fzr1 | protein_coding | -0.231837365 | 6.145964474 | 1.0778189 | 0.3143355 | 0.799301274 |
| ENSMUSG00000030815 | Phkg2 | protein_coding | 0.274106744 | 5.202607546 | 1.0777351 | 0.3143537 | 0.799301274 |
| ENSMUSG00000023330 | Dtwd1 | protein_coding | 0.246224688 | 5.508982348 | 1.0761774 | 0.3146928 | 0.799363258 |
| ENSMUSG00000039826 | Trub2 | protein_coding | 0.430624066 | 5.013049099 | 1.0765532 | 0.3147434 | 0.799363258 |
| ENSMUSG00000023723 | Mrps23 | protein_coding | 0.2038721 | 6.937090636 | 1.0734617 | 0.3152851 | 0.799363258 |
| ENSMUSG00000001383 | Zmat2 | protein_coding | 0.206526424 | 9.115223408 | 1.0730029 | 0.3153853 | 0.799363258 |
| ENSMUSG00000075706 | Gpx4 | protein_coding | -0.204072763 | 8.737888227 | 1.0723146 | 0.3155358 | 0.799363258 |
| ENSMUSG00000038967 | Pdk2 | protein_coding | 0.290833135 | 6.877811759 | 1.0720126 | 0.3156915 | 0.799363258 |
| ENSMUSG00000016554 | Eif3d | protein_coding | 0.202573719 | 6.399336327 | 1.0711285 | 0.3157953 | 0.799363258 |
| ENSMUSG00000021219 | Rgs6 | protein_coding | -0.412707184 | 4.009210378 | 1.0710291 | 0.315817 | 0.799363258 |
| ENSMUSG00000040424 | Hipk4 | protein_coding | 0.34323316 | 4.448475181 | 1.0709889 | 0.3158258 | 0.799363258 |
| ENSMUSG00000040990 | Sh3kbp1 | protein_coding | -0.252208733 | 5.563578914 | 1.0708181 | 0.3158632 | 0.799363258 |
| ENSMUSG00000030872 | Gga2 | protein_coding | 0.603522424 | 3.592282085 | 1.0710514 | 0.3159302 | 0.799363258 |
| ENSMUSG00000020566 | Atp6v1c2 | protein_coding | 0.454070636 | 3.341533126 | 1.0704668 | 0.3159402 | 0.799363258 |
| ENSMUSG00000034620 | Tmem5 | protein_coding | -0.324738001 | 4.559196807 | 1.0702418 | 0.3159895 | 0.799363258 |
| ENSMUSG00000041740 | Rnf10 | protein_coding | 0.255628953 | 5.532524674 | 1.0702123 | 0.3159959 | 0.799363258 |
| ENSMUSG00000042557 | Sin3a | protein_coding | -0.578087929 | 2.824291115 | 1.0701948 | 0.3159998 | 0.799363258 |
| ENSMUSG00000017485 | Top2b | protein_coding | -0.301478837 | 5.196539979 | 1.0700457 | 0.3160324 | 0.799363258 |
| ENSMUSG00000020743 | Mif4gd | protein_coding | -0.28579848 | 5.246410867 | 1.0698279 | 0.3160802 | 0.799363258 |
| ENSMUSG00000051427 | Ccdc157 | protein_coding | 0.54420109 | 2.943598465 | 1.0686496 | 0.3163386 | 0.799363258 |
| ENSMUSG00000026509 | Capn2 | protein_coding | -0.333167569 | 4.854152845 | 1.0682254 | 0.3164317 | 0.799363258 |
| ENSMUSG00000040506 | Ambra1 | protein_coding | 0.337311247 | 5.066546137 | 1.0677982 | 0.3165255 | 0.799363258 |
| ENSMUSG00000030058 | Copg1 | protein_coding | 0.254192283 | 7.934758162 | 1.0678519 | 0.3165383 | 0.799363258 |
| ENSMUSG00000074749 | Kiz | protein_coding | 0.403258151 | 4.178520559 | 1.0676142 | 0.3165659 | 0.799363258 |
| ENSMUSG00000018340 | Anxa6 | protein_coding | 0.251855384 | 6.697405451 | 1.066889 | 0.3167252 | 0.799363258 |
| ENSMUSG00000025192 | Entpd7 | protein_coding | 0.491109793 | 3.783401664 | 1.0665658 | 0.3167963 | 0.799363258 |
| ENSMUSG00000022841 | Ap2m1 | protein_coding | 0.242053204 | 7.793836002 | 1.0664719 | 0.3168169 | 0.799363258 |
| ENSMUSG00000022272 | Myo10 | protein_coding | -0.319073482 | 4.921313803 | 1.0660969 | 0.3168994 | 0.799363258 |
| ENSMUSG00000036466 | Megf11 | protein_coding | -0.325706325 | 4.842826329 | 1.0657489 | 0.316976 | 0.799363258 |
| ENSMUSG00000031921 | Terf2 | protein_coding | 0.31627655 | 4.788759732 | 1.0657405 | 0.3169778 | 0.799363258 |
| ENSMUSG00000045625 | Pigz | protein_coding | -0.317050969 | 4.635725031 | 1.0651401 | 0.31711 | 0.799363258 |
| ENSMUSG00000025220 | Mgea5 | protein_coding | 0.35623906 | 4.184251364 | 1.0644368 | 0.3172648 | 0.799363258 |
| ENSMUSG00000034825 | Nrip3 | protein_coding | 0.325502969 | 7.636221506 | 1.0701066 | 0.3172792 | 0.799363258 |
| ENSMUSG00000039194 | Rlbp1 | protein_coding | 0.494512134 | 4.133638272 | 1.0643481 | 0.317309 | 0.799363258 |
| ENSMUSG00000096943 | Gm26721 | lincRNA | 0.635860893 | 2.809165358 | 1.0636336 | 0.3174419 | 0.799363258 |
| ENSMUSG00000026279 | Thap4 | protein_coding | 0.283334243 | 4.762737754 | 1.0635867 | 0.3174522 | 0.799363258 |
| ENSMUSG00000024030 | Abcg1 | protein_coding | 0.243672065 | 7.836224138 | 1.0634544 | 0.3174814 | 0.799363258 |
| ENSMUSG00000032097 | Ddx6 | protein_coding | -0.201871187 | 6.223176284 | 1.0623499 | 0.3177251 | 0.799363258 |
| ENSMUSG00000089917 | Uckl1 | protein_coding | 0.323912577 | 5.065466382 | 1.0622844 | 0.3177396 | 0.799363258 |
| ENSMUSG00000017747 | Ghdc | protein_coding | 0.853037966 | 2.546931946 | 1.0619898 | 0.3178046 | 0.799363258 |
| ENSMUSG00000041777 | Cir1 | protein_coding | -0.491366398 | 2.987626848 | 1.0617687 | 0.3178535 | 0.799363258 |
| ENSMUSG00000038295 | Atg9b | protein_coding | 0.379738562 | 4.819288226 | 1.0617436 | 0.317859 | 0.799363258 |
| ENSMUSG00000014164 | Klhl3 | protein_coding | 0.32490362 | 6.45429583 | 1.0643172 | 0.3179179 | 0.799363258 |
| ENSMUSG00000027193 | Api5 | protein_coding | -0.242318166 | 5.813265692 | 1.0612522 | 0.3179676 | 0.799363258 |
| ENSMUSG00000031731 | Ap1g1 | protein_coding | 0.525328048 | 3.130153161 | 1.0612392 | 0.3179704 | 0.799363258 |
| ENSMUSG00000020891 | Alox8 | protein_coding | 0.759713655 | 2.964018178 | 1.0611589 | 0.3180127 | 0.799363258 |
| ENSMUSG00000000682 | Cd52 | protein_coding | -0.748758738 | 2.638470569 | 1.0610014 | 0.318023 | 0.799363258 |
| ENSMUSG00000036086 | Zranb3 | protein_coding | 0.738798964 | 2.837932009 | 1.0609115 | 0.3180429 | 0.799363258 |
| ENSMUSG00000030733 | Sh2b1 | protein_coding | 0.285923261 | 4.85364938 | 1.0606189 | 0.3181076 | 0.799363258 |
| ENSMUSG00000016541 | Atxn10 | protein_coding | 0.240280535 | 9.478745114 | 1.0605563 | 0.3181214 | 0.799363258 |
| ENSMUSG00000041415 | Dicer1 | protein_coding | 0.386007922 | 4.0657958 | 1.0602994 | 0.3181782 | 0.799363258 |
| ENSMUSG00000031483 | Erlin2 | protein_coding | 0.290751284 | 5.141166293 | 1.0592737 | 0.3184053 | 0.799363258 |
| ENSMUSG00000038570 | Saxo2 | protein_coding | -0.284545653 | 4.880712069 | 1.05888 | 0.3184924 | 0.799363258 |
| ENSMUSG00000031916 | Cog8 | protein_coding | -0.763693401 | 2.432140051 | 1.058757 | 0.3185197 | 0.799363258 |
| ENSMUSG00000031157 | Pqbp1 | protein_coding | 0.209262793 | 7.908240987 | 1.0586319 | 0.3185474 | 0.799363258 |
| ENSMUSG00000064225 | Paqr9 | protein_coding | 0.402693271 | 4.614825467 | 1.0585166 | 0.3185729 | 0.799363258 |
| ENSMUSG00000019966 | Kitl | protein_coding | -0.421373564 | 4.71695056 | 1.0576928 | 0.3187801 | 0.799363258 |
| ENSMUSG00000028756 | Pink1 | protein_coding | 0.223565709 | 7.55903131 | 1.0575797 | 0.3187806 | 0.799363258 |
| ENSMUSG00000046178 | Nxph1 | protein_coding | -0.253083707 | 5.835213888 | 1.05735 | 0.3188316 | 0.799363258 |
| ENSMUSG00000035202 | Lars2 | protein_coding | 0.250550885 | 9.543560959 | 1.0564206 | 0.3190379 | 0.799363258 |
| ENSMUSG00000063524 | Eno1 | protein_coding | 0.248816356 | 8.641938171 | 1.0563256 | 0.319059 | 0.799363258 |
| ENSMUSG00000112280 | AC153962.3 | processed_transcript | -0.387862847 | 3.952618323 | 1.0557272 | 0.3191919 | 0.799363258 |
| ENSMUSG00000003545 | Fosb | protein_coding | -0.51933093 | 3.305642416 | 1.0556759 | 0.3192033 | 0.799363258 |
| ENSMUSG00000028470 | Hint2 | protein_coding | 0.422984333 | 4.938380281 | 1.0560577 | 0.3192075 | 0.799363258 |
| ENSMUSG00000019888 | Mgat4c | protein_coding | -0.375608144 | 4.655284636 | 1.0556151 | 0.3192168 | 0.799363258 |

| ensembl_gene_id | external_gene_name | gene_biotype | log2 (fold change) | log2 (counts per million) | F-value | p-value | False discovery rate |
|---|---|---|---|---|---|---|---|
| ENSMUSG00000030287 | Itpr2 | protein_coding | -0.665824739 | 2.660606631 | 1.0544644 | 0.3194726 | 0.799363258 |
| ENSMUSG00000022843 | Clcn2 | protein_coding | 0.754883963 | 2.632605279 | 1.054427 | 0.319481 | 0.799363258 |
| ENSMUSG00000032297 | Celf6 | protein_coding | 0.329057921 | 5.126596465 | 1.0542731 | 0.3195152 | 0.799363258 |
| ENSMUSG00000029389 | Ddx55 | protein_coding | 0.278359609 | 5.187898048 | 1.0540658 | 0.3195613 | 0.799363258 |
| ENSMUSG00000037376 | Trmt6 | protein_coding | 0.293768916 | 5.655510675 | 1.0537554 | 0.3196304 | 0.799363258 |
| ENSMUSG00000014980 | Tsen15 | protein_coding | 0.28888387 | 4.869317829 | 1.0528931 | 0.3198225 | 0.799363258 |
| ENSMUSG00000026455 | Klhl12 | protein_coding | 0.411504539 | 4.628527613 | 1.0528058 | 0.3198664 | 0.799363258 |
| ENSMUSG00000036067 | Slc2a6 | protein_coding | 0.419733374 | 4.741101066 | 1.0527581 | 0.319877 | 0.799363258 |
| ENSMUSG00000038417 | Fig4 | protein_coding | 0.340363865 | 4.988663656 | 1.0522927 | 0.3199563 | 0.799363258 |
| ENSMUSG00000028753 | Vwa5b1 | protein_coding | 0.430791796 | 3.62853526 | 1.0515943 | 0.320112 | 0.799363258 |
| ENSMUSG00000053644 | Aldh7a1 | protein_coding | 0.238543065 | 5.997732818 | 1.0514876 | 0.3201358 | 0.799363258 |
| ENSMUSG00000079316 | Rab9 | protein_coding | -0.233976363 | 6.864754682 | 1.0510364 | 0.3202365 | 0.799363258 |
| ENSMUSG00000031104 | Rab33a | protein_coding | 0.217397033 | 6.802206864 | 1.050792 | 0.3202911 | 0.799363258 |
| ENSMUSG00000021890 | Eaf1 | protein_coding | -0.539018187 | 3.042197161 | 1.0504141 | 0.3203755 | 0.799363258 |
| ENSMUSG00000034413 | Neurl1b | protein_coding | -0.561373031 | 3.442703507 | 1.0501502 | 0.3204344 | 0.799363258 |
| ENSMUSG00000026576 | Atp1b1 | protein_coding | 0.244599871 | 8.629832626 | 1.0499175 | 0.3204864 | 0.799363258 |
| ENSMUSG00000000295 | Hddc2 | protein_coding | 0.230900813 | 7.101080233 | 1.0495675 | 0.3205646 | 0.799363258 |
| ENSMUSG00000038383 | Pigu | protein_coding | 0.311004933 | 5.496662148 | 1.0491108 | 0.3206667 | 0.799363258 |
| ENSMUSG00000027522 | Stx16 | protein_coding | 0.369312846 | 5.559585882 | 1.0493387 | 0.3207045 | 0.799363258 |
| ENSMUSG00000021025 | Nfkbia | protein_coding | 0.306523036 | 5.816459171 | 1.0485108 | 0.3208009 | 0.799363258 |
| ENSMUSG00000035851 | Ythdc1 | protein_coding | 0.28974316 | 5.114196526 | 1.048399 | 0.320826 | 0.799363258 |
| ENSMUSG00000025790 | Slco3a1 | protein_coding | -0.357376138 | 4.733153981 | 1.0482889 | 0.3208506 | 0.799363258 |
| ENSMUSG00000063296 | Tmem117 | protein_coding | 0.602103468 | 3.037931534 | 1.0481342 | 0.3208852 | 0.799363258 |
| ENSMUSG00000033214 | Slitrk5 | protein_coding | -0.300160935 | 4.334733499 | 1.0481105 | 0.3208905 | 0.799363258 |
| ENSMUSG00000030125 | Lrrc23 | protein_coding | -0.426650703 | 3.572157851 | 1.0479795 | 0.3209199 | 0.799363258 |
| ENSMUSG00000025868 | Higd2a | protein_coding | -0.18190538 | 8.450003104 | 1.0470482 | 0.3211285 | 0.799697598 |
| ENSMUSG00000044730 | 9930104L06Rik | processed_transcript | 0.512714249 | 3.161767278 | 1.0454933 | 0.3214771 | 0.800270436 |
| ENSMUSG00000005621 | Zfp592 | protein_coding | 0.470158042 | 3.739771541 | 1.0445775 | 0.3216827 | 0.800270436 |
| ENSMUSG00000025421 | Hdhd2 | protein_coding | 0.185511968 | 8.431281938 | 1.0443162 | 0.3217415 | 0.800270436 |
| ENSMUSG00000029076 | Sdf4 | protein_coding | 0.185339347 | 8.512245284 | 1.044268 | 0.3217523 | 0.800270436 |
| ENSMUSG00000007880 | Arid1a | protein_coding | -0.308459907 | 6.022103419 | 1.0441029 | 0.3218137 | 0.800270436 |
| ENSMUSG00000028581 | Laptm5 | protein_coding | -0.309837516 | 5.352613588 | 1.0439149 | 0.3218316 | 0.800270436 |
| ENSMUSG00000028580 | Pum1 | protein_coding | 0.289738853 | 6.789635868 | 1.044096 | 0.3218794 | 0.800270436 |
| ENSMUSG00000086477 | Gm15506 | processed_transcript | 0.553516634 | 2.935751709 | 1.0403069 | 0.322644 | 0.801847128 |
| ENSMUSG00000020135 | Apc2 | protein_coding | 0.411107352 | 4.679888566 | 1.0403316 | 0.3226627 | 0.801847128 |
| ENSMUSG00000022377 | Asap1 | protein_coding | -0.306613037 | 5.303138934 | 1.0372896 | 0.3233256 | 0.802860939 |
| ENSMUSG00000054871 | Tmem158 | protein_coding | -0.434949706 | 4.795902813 | 1.0373599 | 0.3233979 | 0.802860939 |
| ENSMUSG00000004054 | Map3k11 | protein_coding | -0.65628679 | 2.518704053 | 1.0367143 | 0.3234558 | 0.802860939 |
| ENSMUSG00000029598 | Plbd2 | protein_coding | -0.315310357 | 5.096875469 | 1.0365411 | 0.323495 | 0.802860939 |
| ENSMUSG00000034445 | Cyb561a3 | protein_coding | -0.689559014 | 2.512051655 | 1.0364142 | 0.3235237 | 0.802860939 |
| ENSMUSG00000074093 | Svip | protein_coding | -0.261761517 | 6.117589266 | 1.0358071 | 0.3236612 | 0.802860939 |
| ENSMUSG00000057378 | Ryr3 | protein_coding | -0.517446547 | 3.667007566 | 1.0356803 | 0.3237142 | 0.802860939 |
| ENSMUSG00000039210 | Gpatch2 | protein_coding | 0.760067369 | 2.641426006 | 1.0354546 | 0.3237411 | 0.802860939 |
| ENSMUSG00000029603 | Dtx1 | protein_coding | 0.286859944 | 5.30201011 | 1.035176 | 0.3238042 | 0.802860939 |
| ENSMUSG00000027864 | Ptgfrn | protein_coding | -0.431103907 | 4.140524915 | 1.0343793 | 0.3239849 | 0.802860939 |
| ENSMUSG00000028800 | Hdac1 | protein_coding | -0.371424446 | 3.739194439 | 1.0343561 | 0.3239902 | 0.802860939 |
| ENSMUSG00000032867 | Fbxw8 | protein_coding | -0.644417737 | 2.483064752 | 1.0343053 | 0.3240017 | 0.802860939 |
| ENSMUSG00000024844 | Banf1 | protein_coding | -0.231062757 | 7.963875365 | 1.0341307 | 0.3240413 | 0.802860939 |
| ENSMUSG00000024410 | 3110002H16Rik | protein_coding | 0.288755494 | 5.443924977 | 1.033215 | 0.3242492 | 0.803062949 |
| ENSMUSG00000021451 | Sema4d | protein_coding | 0.369449554 | 3.612911937 | 1.0329329 | 0.3243133 | 0.803062949 |
| ENSMUSG00000050830 | Vwc2 | protein_coding | -0.513690844 | 3.167405937 | 1.032785 | 0.3243469 | 0.803062949 |
| ENSMUSG00000020723 | Cacng4 | protein_coding | -0.306532615 | 5.752116786 | 1.031862 | 0.3245567 | 0.80333338 |
| ENSMUSG00000048100 | Taf13 | protein_coding | -0.303335754 | 5.28099885 | 1.0316474 | 0.3246056 | 0.80333338 |
| ENSMUSG00000025950 | Idh1 | protein_coding | 0.201234915 | 6.523686016 | 1.0313176 | 0.3246806 | 0.803334176 |
| ENSMUSG00000031210 | Gpr165 | protein_coding | 0.56415503 | 3.673757385 | 1.0293033 | 0.3251636 | 0.804344168 |
| ENSMUSG00000009079 | Ewsr1 | protein_coding | 0.245543374 | 5.932260034 | 1.0286249 | 0.3252942 | 0.804447944 |
| ENSMUSG00000028538 | St3gal3 | protein_coding | -0.472083973 | 3.241666298 | 1.0283575 | 0.3253552 | 0.804447944 |
| ENSMUSG00000090247 | Bloc1s1 | protein_coding | -0.283474837 | 5.050698492 | 1.0270464 | 0.3256546 | 0.804735496 |
| ENSMUSG00000026723 | Trdmt1 | protein_coding | -0.504452093 | 3.159473923 | 1.0269959 | 0.3256661 | 0.804735496 |
| ENSMUSG00000022094 | Slc39a14 | protein_coding | -0.583041561 | 3.064125829 | 1.0267876 | 0.3257137 | 0.804735496 |
| ENSMUSG00000025019 | Lcor | protein_coding | -0.332218927 | 4.180898007 | 1.0265379 | 0.3257708 | 0.804735496 |
| ENSMUSG00000069729 | Arid1b | protein_coding | -0.266484531 | 5.325468977 | 1.02599 | 0.3258961 | 0.804784422 |
| ENSMUSG00000046185 | Zfp84 | protein_coding | 0.534211549 | 3.379679398 | 1.0257968 | 0.3259403 | 0.804784422 |
| ENSMUSG00000017837 | Nkiras2 | protein_coding | -0.363109052 | 4.918589916 | 1.0245849 | 0.3262178 | 0.804866656 |
| ENSMUSG00000029191 | Rfc1 | protein_coding | 0.262910706 | 5.548123575 | 1.0243788 | 0.326265 | 0.804866656 |
| ENSMUSG00000000326 | Comt | protein_coding | 0.19366715 | 7.220056283 | 1.0240889 | 0.3263314 | 0.804866656 |
| ENSMUSG00000022021 | Diaph3 | protein_coding | -0.363143396 | 4.311244021 | 1.0238518 | 0.3263857 | 0.804866656 |
| ENSMUSG00000048799 | Cep120 | protein_coding | 0.304766866 | 4.952780151 | 1.0237317 | 0.3264133 | 0.804866656 |
| ENSMUSG00000031608 | Galnt7 | protein_coding | 0.524136873 | 3.425388949 | 1.0231648 | 0.3265433 | 0.804866656 |
| ENSMUSG00000031980 | Agt | protein_coding | -0.254491938 | 6.828260512 | 1.0220255 | 0.3268048 | 0.804866656 |
| ENSMUSG00000024070 | Prkd3 | protein_coding | -0.429253401 | 3.494450604 | 1.0219244 | 0.326828 | 0.804866656 |
| ENSMUSG00000025429 | Pstpip2 | protein_coding | -0.70037702 | 2.547898882 | 1.0216883 | 0.3268822 | 0.804866656 |
| ENSMUSG00000030662 | Ipo5 | protein_coding | 0.236671356 | 6.205584337 | 1.0215041 | 0.3269246 | 0.804866656 |
| ENSMUSG00000025170 | Rab40b | protein_coding | 0.388396111 | 4.990216302 | 1.0215229 | 0.3269444 | 0.804866656 |

| ensembl_gene_id | external_gene_name | gene_biotype | log2 (fold change) | log2 (counts per million) | F-value | p-value | False discovery rate |
|---|---|---|---|---|---|---|---|
| ENSMUSG00000020180 | Snrpd3 | protein_coding | 0.215301532 | 8.641579732 | 1.0211237 | 0.327012 | 0.804866656 |
| ENSMUSG00000040859 | Bsdc1 | protein_coding | 0.222055569 | 5.986133482 | 1.020891 | 0.3270655 | 0.804866656 |
| ENSMUSG00000045039 | Megf8 | protein_coding | -0.384289425 | 4.716612134 | 1.0206177 | 0.3271283 | 0.804866656 |
| ENSMUSG00000038257 | Glra3 | protein_coding | -0.459125929 | 3.159077533 | 1.0205994 | 0.3271325 | 0.804866656 |
| ENSMUSG00000036437 | Npy1r | protein_coding | 0.359522664 | 4.079519667 | 1.0199373 | 0.3272848 | 0.804866656 |
| ENSMUSG00000029521 | Chek2 | protein_coding | -0.494681892 | 3.049746519 | 1.0193714 | 0.3274151 | 0.804866656 |
| ENSMUSG00000040564 | Apoc1 | protein_coding | -0.499767397 | 3.600949766 | 1.0192729 | 0.3274378 | 0.804866656 |
| ENSMUSG00000053702 | Nebl | protein_coding | -0.283663325 | 4.805923661 | 1.0191991 | 0.3274548 | 0.804866656 |
| ENSMUSG00000040848 | Sft2d2 | protein_coding | -0.433845371 | 3.875666158 | 1.0181868 | 0.327688 | 0.804866656 |
| ENSMUSG00000079598 | Clec2l | protein_coding | 0.318745199 | 4.576287823 | 1.0180108 | 0.3277286 | 0.804866656 |
| ENSMUSG00000031391 | L1cam | protein_coding | 0.240704776 | 5.602604218 | 1.0179156 | 0.3277506 | 0.804866656 |
| ENSMUSG00000024579 | Pcyox1l | protein_coding | 0.366776066 | 3.862839148 | 1.0178134 | 0.3277741 | 0.804866656 |
| ENSMUSG00000028207 | Asph | protein_coding | -0.283778999 | 5.442789133 | 1.0176984 | 0.3278006 | 0.804866656 |
| ENSMUSG00000024743 | Syt7 | protein_coding | -0.259544785 | 5.813303596 | 1.0175067 | 0.3278449 | 0.804866656 |
| ENSMUSG00000031749 | St3gal2 | protein_coding | -0.278889578 | 5.079419444 | 1.0167722 | 0.3280144 | 0.80509896 |
| ENSMUSG00000031292 | Cdkl5 | protein_coding | -0.264818442 | 5.96809459 | 1.0163228 | 0.3281181 | 0.80516366 |
| ENSMUSG00000038679 | Trps1 | protein_coding | -0.325177252 | 4.421046374 | 1.0160096 | 0.3281905 | 0.80516366 |
| ENSMUSG00000021453 | Gadd45g | protein_coding | -0.324319385 | 5.83083445 | 1.0148398 | 0.3284849 | 0.805339948 |
| ENSMUSG00000085385 | Snhg17 | processed_transcript | -0.751251789 | 2.346316515 | 1.0147282 | 0.3284867 | 0.805339948 |
| ENSMUSG00000072825 | Cep170b | protein_coding | 0.243371064 | 6.618849116 | 1.0145588 | 0.3285259 | 0.805339948 |
| ENSMUSG00000047228 | A2ml1 | protein_coding | -0.247408128 | 5.711089496 | 1.0143065 | 0.3285843 | 0.805339948 |
| ENSMUSG00000020216 | Jsrp1 | protein_coding | -0.517415271 | 3.231822087 | 1.0140491 | 0.3286438 | 0.805339948 |
| ENSMUSG00000001285 | Myg1 | protein_coding | 0.215317996 | 7.240407338 | 1.0134443 | 0.3287839 | 0.805339948 |
| ENSMUSG00000046500 | Fam19a4 | protein_coding | -0.750383693 | 2.344135323 | 1.0130887 | 0.3288663 | 0.805339948 |
| ENSMUSG00000028886 | Eya3 | protein_coding | -0.564192521 | 3.124177612 | 1.0125663 | 0.3289873 | 0.805339948 |
| ENSMUSG00000069171 | Nr2f1 | protein_coding | -0.305799331 | 4.585679755 | 1.0122247 | 0.3290665 | 0.805339948 |
| ENSMUSG00000056870 | Gulp1 | protein_coding | -0.509250166 | 3.06054886 | 1.0121409 | 0.329086 | 0.805339948 |
| ENSMUSG00000024589 | Nedd4l | protein_coding | 0.249979931 | 6.077579682 | 1.01214 | 0.3290862 | 0.805339948 |
| ENSMUSG00000040957 | Cables1 | protein_coding | -0.454265318 | 3.644113282 | 1.010773 | 0.3294034 | 0.805719556 |
| ENSMUSG00000048218 | Amigo2 | protein_coding | 0.336040373 | 4.458766099 | 1.0103078 | 0.3295115 | 0.805719556 |
| ENSMUSG00000021701 | Plk2 | protein_coding | 0.198880328 | 7.13169582 | 1.0100701 | 0.3295667 | 0.805719556 |
| ENSMUSG00000021192 | Golga5 | protein_coding | 0.389592155 | 3.68689339 | 1.0097793 | 0.3296343 | 0.805719556 |
| ENSMUSG00000110841 | AC153368.1 | transcribed_processe | -0.336999849 | 4.00430989 | 1.0093189 | 0.3297414 | 0.805719556 |
| ENSMUSG00000013698 | Pea15a | protein_coding | 0.235375994 | 10.3333805 | 1.0086023 | 0.3299081 | 0.805719556 |
| ENSMUSG00000057103 | Nat8f1 | protein_coding | 0.321737527 | 4.708141597 | 1.0078978 | 0.3300721 | 0.805719556 |
| ENSMUSG00000029130 | Rnf32 | protein_coding | 0.308128274 | 4.393035802 | 1.0078546 | 0.3300821 | 0.805719556 |
| ENSMUSG00000047248 | C2cd3 | protein_coding | -0.926648731 | 2.391018639 | 1.0079168 | 0.3300916 | 0.805719556 |
| ENSMUSG00000070280 | Slc22a14 | protein_coding | 0.59199312 | 2.627265896 | 1.0074673 | 0.3301724 | 0.805719556 |
| ENSMUSG00000052040 | Klf13 | protein_coding | -0.260879816 | 8.319600223 | 1.0073563 | 0.3302222 | 0.805719556 |
| ENSMUSG00000037447 | Arid5a | protein_coding | -0.4451641 | 4.065854634 | 1.0066176 | 0.3303704 | 0.805719556 |
| ENSMUSG00000022012 | Enox1 | protein_coding | -0.392678854 | 4.283157381 | 1.0065265 | 0.3303917 | 0.805719556 |
| ENSMUSG00000041417 | Pik3r1 | protein_coding | -0.285942316 | 5.653165169 | 1.0065234 | 0.3303924 | 0.805719556 |
| ENSMUSG00000028651 | Ppie | protein_coding | 0.294272286 | 4.584048901 | 1.0063809 | 0.3304256 | 0.805719556 |
| ENSMUSG00000048310 | Pskh1 | protein_coding | 0.583013119 | 2.831746232 | 1.0055243 | 0.3306255 | 0.805719556 |
| ENSMUSG00000042770 | Hebp1 | protein_coding | -0.358787204 | 5.318491374 | 1.0056035 | 0.3306309 | 0.805719556 |
| ENSMUSG00000013787 | Ehmt2 | protein_coding | -0.280223595 | 5.583196149 | 1.0054431 | 0.3306445 | 0.805719556 |
| ENSMUSG00000035184 | Fam124a | protein_coding | 0.396592063 | 4.021235127 | 1.0051591 | 0.3307108 | 0.805719556 |
| ENSMUSG00000057093 | Zfp607b | protein_coding | 0.454992808 | 3.5890363 | 1.0048963 | 0.3307722 | 0.805719556 |
| ENSMUSG00000024304 | Cdh2 | protein_coding | 0.396986115 | 3.838995901 | 1.0047137 | 0.3308148 | 0.805719556 |
| ENSMUSG00000022223 | Sdr39u1 | protein_coding | 0.182976762 | 8.470249167 | 1.0041848 | 0.3309384 | 0.805838064 |
| ENSMUSG00000038398 | Upf3a | protein_coding | 0.344014839 | 4.345430702 | 1.0037776 | 0.3310336 | 0.805887388 |
| ENSMUSG00000035168 | Tanc1 | protein_coding | -0.385737654 | 3.707261857 | 1.0026883 | 0.3312885 | 0.806259941 |
| ENSMUSG00000029994 | Anxa4 | protein_coding | 0.612502929 | 2.78056413 | 1.0020574 | 0.3314363 | 0.806259941 |
| ENSMUSG00000108348 | Gm42372 | protein_coding | -0.226038084 | 6.623810946 | 1.0015763 | 0.331549 | 0.806259941 |
| ENSMUSG00000027195 | Hsd17b12 | protein_coding | 0.180056236 | 7.438420096 | 1.0015344 | 0.3315588 | 0.806259941 |
| ENSMUSG00000034066 | Farp2 | protein_coding | 0.540032871 | 2.859174941 | 1.000625 | 0.331772 | 0.806259941 |
| ENSMUSG00000030612 | Mrpl46 | protein_coding | 0.17753277 | 7.79607444 | 1.0004735 | 0.3318076 | 0.806259941 |
| ENSMUSG00000039914 | Coq10a | protein_coding | 0.321420358 | 5.652001736 | 0.9999415 | 0.3319563 | 0.806259941 |
| ENSMUSG00000020460 | Rps27a | protein_coding | 0.379535619 | 3.584484307 | 0.9997944 | 0.331967 | 0.806259941 |
| ENSMUSG00000060548 | Tnfrsf19 | protein_coding | 0.480639694 | 3.548605899 | 0.999793 | 0.3319673 | 0.806259941 |
| ENSMUSG00000021357 | Exoc2 | protein_coding | 0.377985683 | 4.732963836 | 0.9997668 | 0.3319735 | 0.806259941 |
| ENSMUSG00000025413 | Ttc4 | protein_coding | 0.278791448 | 5.924255249 | 0.999605 | 0.3320114 | 0.806259941 |
| ENSMUSG00000033326 | Kdm4a | protein_coding | 0.667248548 | 2.537096055 | 0.9987886 | 0.3322032 | 0.806543559 |
| ENSMUSG00000104939 | Gm33651 | lincRNA | -0.529606854 | 3.545577896 | 0.998374 | 0.3323245 | 0.806655897 |
| ENSMUSG00000050505 | Pcdh20 | protein_coding | -0.390226719 | 3.923139704 | 0.9977959 | 0.3324367 | 0.806746006 |
| ENSMUSG00000061371 | Zfp873 | protein_coding | -0.389396541 | 4.0858188 | 0.9969847 | 0.3326276 | 0.806771346 |
| ENSMUSG00000018861 | Fdxr | protein_coding | 0.394906958 | 3.581501674 | 0.9969253 | 0.3326416 | 0.806771346 |
| ENSMUSG00000073478 | D730003I15Rik | lincRNA | -0.401749763 | 4.288158583 | 0.9967952 | 0.3326722 | 0.806771346 |
| ENSMUSG00000022285 | Ywhaz | protein_coding | -0.199270431 | 10.25803832 | 0.9960176 | 0.3328554 | 0.807009912 |
| ENSMUSG00000039323 | Igfbp2 | protein_coding | -0.578550243 | 3.213648607 | 0.9957407 | 0.3329207 | 0.807009912 |
| ENSMUSG00000027088 | Phospho2 | protein_coding | -0.245978737 | 5.56835317 | 0.9950405 | 0.3330858 | 0.807071391 |
| ENSMUSG00000024798 | Htr7 | protein_coding | -0.37707125 | 4.349140398 | 0.9949966 | 0.3330961 | 0.807071391 |
| ENSMUSG00000074671 | Tspyl3 | protein_coding | 0.307024607 | 5.022841253 | 0.9943721 | 0.3332435 | 0.807114757 |
| ENSMUSG00000018415 | Gid4 | protein_coding | -0.278079981 | 5.948252585 | 0.9941604 | 0.3332935 | 0.807114757 |

| ensembl_gene_id | external_gene_name | gene_biotype | log2 (fold change) | log2 (counts per million) | F-value | p-value | False discovery rate |
|---|---|---|---|---|---|---|---|
| ENSMUSG00000056204 | Pgpep1 | protein_coding | 0.289393504 | 5.433848232 | 0.9936973 | 0.3334029 | 0.807114757 |
| ENSMUSG00000029596 | Sdsl | protein_coding | -0.635802986 | 2.705739573 | 0.993649 | 0.3334143 | 0.807114757 |
| ENSMUSG00000031109 | Enox2 | protein_coding | -0.315946298 | 4.325506399 | 0.9918365 | 0.3338429 | 0.807734214 |
| ENSMUSG00000029647 | Pan3 | protein_coding | -0.316933402 | 4.615411374 | 0.9917369 | 0.3338664 | 0.807734214 |
| ENSMUSG00000041684 | Bivm | protein_coding | 0.319814351 | 4.219318617 | 0.9915306 | 0.3339153 | 0.807734214 |
| ENSMUSG00000001440 | Kpnb1 | protein_coding | 0.212150193 | 6.058698675 | 0.9912967 | 0.3339706 | 0.807734214 |
| ENSMUSG00000057068 | Fam47e | protein_coding | -0.588607208 | 3.11634576 | 0.9905067 | 0.3341578 | 0.807777885 |
| ENSMUSG00000002778 | Kdelr1 | protein_coding | -0.321259843 | 5.216955976 | 0.9895368 | 0.3343877 | 0.807777885 |
| ENSMUSG00000035770 | Dync1li2 | protein_coding | 0.221167561 | 6.71919187 | 0.9895132 | 0.3343933 | 0.807777885 |
| ENSMUSG00000073609 | D2hgdh | protein_coding | 0.541829201 | 3.018181515 | 0.9894694 | 0.3344037 | 0.807777885 |
| ENSMUSG00000055240 | Zfp101 | protein_coding | -0.54673975 | 3.021592044 | 0.9893573 | 0.3344303 | 0.807777885 |
| ENSMUSG00000015002 | Efr3a | protein_coding | -0.228656478 | 5.889607461 | 0.9891291 | 0.3344845 | 0.807777885 |
| ENSMUSG00000018599 | Mief2 | protein_coding | -0.325254079 | 4.81715464 | 0.9890024 | 0.3345145 | 0.807777885 |
| ENSMUSG00000032279 | Idh3a | protein_coding | 0.216314027 | 9.358748694 | 0.9881785 | 0.3347102 | 0.807920008 |
| ENSMUSG00000061689 | Dlgap4 | protein_coding | -0.21425884 | 6.687834921 | 0.9881216 | 0.3347237 | 0.807920008 |
| ENSMUSG00000038014 | Fam120a | protein_coding | -0.297710369 | 5.576354021 | 0.9868692 | 0.3350214 | 0.808408865 |
| ENSMUSG00000032437 | Stt3b | protein_coding | 0.328216952 | 5.189159621 | 0.9863428 | 0.3351466 | 0.808408865 |
| ENSMUSG00000024604 | Rbm22 | protein_coding | 0.207025534 | 5.880171391 | 0.9863212 | 0.3351517 | 0.808408865 |
| ENSMUSG00000005338 | Cadm3 | protein_coding | -0.268439384 | 5.57829691 | 0.9858222 | 0.3352705 | 0.808514034 |
| ENSMUSG00000022793 | B4galt4 | protein_coding | 0.488666726 | 3.48912952 | 0.9852231 | 0.3354132 | 0.808551096 |
| ENSMUSG00000032251 | Irak1bp1 | protein_coding | -0.302287629 | 4.895445632 | 0.9851263 | 0.3354363 | 0.808551096 |
| ENSMUSG00000026904 | Slc4a10 | protein_coding | -0.281885704 | 5.330997393 | 0.9845885 | 0.3355645 | 0.808678802 |
| ENSMUSG00000086822 | 5330413P13Rik | lincRNA | -0.79819573 | 2.127317112 | 0.9833535 | 0.3358591 | 0.80896598 |
| ENSMUSG00000015697 | Setdb1 | protein_coding | 0.366629017 | 3.872200855 | 0.9831072 | 0.3359179 | 0.80896598 |
| ENSMUSG00000024792 | Zfpl1 | protein_coding | 0.513379989 | 3.797924973 | 0.9831578 | 0.3359295 | 0.80896598 |
| ENSMUSG00000027823 | Gmps | protein_coding | 0.276392256 | 7.305183761 | 0.9840195 | 0.3359846 | 0.80896598 |
| ENSMUSG00000089911 | Mfsd14a | protein_coding | -0.377470745 | 4.328370809 | 0.9818913 | 0.3362084 | 0.80921 |
| ENSMUSG00000074358 | Ccdc61 | protein_coding | 0.409774824 | 3.424313251 | 0.9812169 | 0.3363696 | 0.80921 |
| ENSMUSG00000021928 | Ebpl | protein_coding | 0.29862039 | 4.421422157 | 0.9808568 | 0.3364558 | 0.80921 |
| ENSMUSG00000055026 | Gabrg3 | protein_coding | -0.289811513 | 4.684126774 | 0.9808518 | 0.336457 | 0.80921 |
| ENSMUSG00000032560 | Dnajc13 | protein_coding | -0.38312028 | 4.102390063 | 0.9808101 | 0.336467 | 0.80921 |
| ENSMUSG00000031431 | Tsc22d3 | protein_coding | 0.198418225 | 8.258900288 | 0.9802499 | 0.3366011 | 0.80921 |
| ENSMUSG00000000339 | Rtca | protein_coding | 0.411684019 | 4.546336052 | 0.9800678 | 0.3366683 | 0.80921 |
| ENSMUSG00000022130 | Tgds | protein_coding | -0.69953981 | 2.404967611 | 0.9798871 | 0.336688 | 0.80921 |
| ENSMUSG00000056854 | Pou3f4 | protein_coding | 0.378533897 | 4.056695907 | 0.9782424 | 0.3370822 | 0.809885139 |
| ENSMUSG00000036948 | BC037034 | protein_coding | 0.405243068 | 3.583442074 | 0.9780871 | 0.3371195 | 0.809885139 |
| ENSMUSG00000029416 | Slc15a4 | protein_coding | -0.33213379 | 4.24000187 | 0.9774331 | 0.3372765 | 0.810081314 |
| ENSMUSG00000021520 | Uqcrb | protein_coding | -0.258070737 | 8.483374154 | 0.9769216 | 0.3374229 | 0.810090842 |
| ENSMUSG00000048216 | Gpr85 | protein_coding | 0.249553097 | 6.060611206 | 0.9767229 | 0.3374471 | 0.810090842 |
| ENSMUSG00000037369 | Kdm6a | protein_coding | -0.407785457 | 3.870583074 | 0.9759535 | 0.337632 | 0.810090842 |
| ENSMUSG00000035960 | Apex1 | protein_coding | 0.250547734 | 5.558135195 | 0.9758932 | 0.3376466 | 0.810090842 |
| ENSMUSG00000000869 | Il4 | protein_coding | 0.343982218 | 4.629032743 | 0.9758492 | 0.3376571 | 0.810090842 |
| ENSMUSG00000085316 | D330050G23Rik | lincRNA | -0.798401325 | 2.556195154 | 0.974857 | 0.3378959 | 0.810191945 |
| ENSMUSG00000049090 | Zadh2 | protein_coding | -0.364843033 | 3.667105702 | 0.97428 | 0.3380348 | 0.810191945 |
| ENSMUSG00000079658 | Eloc | protein_coding | -0.191713343 | 8.764212717 | 0.9742529 | 0.3380414 | 0.810191945 |
| ENSMUSG00000064043 | Trerf1 | processed_transcript | -0.307926381 | 4.211828547 | 0.9741093 | 0.338076 | 0.810191945 |
| ENSMUSG00000036934 | 4921524J17Rik | protein_coding | -0.265324054 | 5.390113529 | 0.9741092 | 0.338076 | 0.810191945 |
| ENSMUSG00000041879 | Ipo9 | protein_coding | -0.237600709 | 6.119711116 | 0.973699 | 0.3381748 | 0.810201007 |
| ENSMUSG00000025323 | Sp4 | protein_coding | -0.343886347 | 3.837975303 | 0.973228 | 0.3382884 | 0.810201007 |
| ENSMUSG00000024369 | Nelfe | protein_coding | 0.230041433 | 6.009581249 | 0.9727074 | 0.338414 | 0.810201007 |
| ENSMUSG00000036819 | Jmjd4 | protein_coding | 0.346008462 | 4.267717668 | 0.9726525 | 0.3384272 | 0.810201007 |
| ENSMUSG00000024897 | Apba1 | protein_coding | -0.273363793 | 5.352296882 | 0.9722745 | 0.3385184 | 0.810201007 |
| ENSMUSG00000068882 | Ssb | protein_coding | -0.221486622 | 7.356923846 | 0.9722189 | 0.3385319 | 0.810201007 |
| ENSMUSG00000032468 | Armc8 | protein_coding | -0.239944639 | 6.763155801 | 0.9705393 | 0.3389377 | 0.810414375 |
| ENSMUSG00000039530 | Tusc3 | protein_coding | 0.248955538 | 7.370832467 | 0.970296 | 0.33902 | 0.810414375 |
| ENSMUSG00000032754 | Slc8b1 | protein_coding | 0.60510823 | 2.605505296 | 0.969968 | 0.3390759 | 0.810414375 |
| ENSMUSG00000048997 | Atxn7l2 | protein_coding | 0.524002391 | 2.92928557 | 0.9687339 | 0.3393746 | 0.810414375 |
| ENSMUSG00000030088 | Aldh1l1 | protein_coding | 0.444205931 | 4.31170691 | 0.968792 | 0.3393841 | 0.810414375 |
| ENSMUSG00000060212 | Pcnx2 | protein_coding | 0.430670425 | 3.502275679 | 0.9682148 | 0.3395004 | 0.810414375 |
| ENSMUSG00000026211 | Obsl1 | protein_coding | -0.552957615 | 2.832879547 | 0.9682049 | 0.3395028 | 0.810414375 |
| ENSMUSG00000031538 | Plat | protein_coding | 0.389150599 | 4.335392688 | 0.9681738 | 0.3395104 | 0.810414375 |
| ENSMUSG00000022096 | Hr | protein_coding | -0.59704609 | 2.578028652 | 0.9676537 | 0.3396365 | 0.810414375 |
| ENSMUSG00000025821 | Zfp282 | protein_coding | -0.536128636 | 3.051024875 | 0.9672311 | 0.339739 | 0.810414375 |
| ENSMUSG00000031955 | Bcar1 | protein_coding | -0.351347458 | 4.683054308 | 0.9672276 | 0.3397398 | 0.810414375 |
| ENSMUSG00000037896 | Rcor1 | protein_coding | -0.395836534 | 4.042405478 | 0.9668419 | 0.3398334 | 0.810414375 |
| ENSMUSG00000024211 | Grm8 | protein_coding | 0.534249151 | 4.150109232 | 0.9704961 | 0.3398584 | 0.810414375 |
| ENSMUSG00000026807 | Ak8 | protein_coding | 0.654492221 | 3.527731632 | 0.9702242 | 0.3399241 | 0.810414375 |
| ENSMUSG00000022528 | Hes1 | protein_coding | -0.506062465 | 2.960252542 | 0.9660201 | 0.340033 | 0.810414375 |
| ENSMUSG00000033382 | Trappc8 | protein_coding | 0.413700573 | 3.871704066 | 0.9659442 | 0.3400514 | 0.810414375 |
| ENSMUSG00000045827 | Serpinb9 | protein_coding | 0.407158488 | 3.913009374 | 0.9655172 | 0.3401552 | 0.810414375 |
| ENSMUSG00000040022 | Rab11fip2 | protein_coding | -0.286617011 | 4.496235975 | 0.9650993 | 0.3402568 | 0.810414375 |
| ENSMUSG00000037172 | E330009J07Rik | protein_coding | -0.323927019 | 4.419615746 | 0.9647861 | 0.340333 | 0.810414375 |
| ENSMUSG00000000959 | Oxa1l | protein_coding | 0.228987823 | 6.370403043 | 0.9637504 | 0.340585 | 0.810414375 |
| ENSMUSG00000040383 | Aqr | protein_coding | 0.340483615 | 4.519892556 | 0.9637002 | 0.3405972 | 0.810414375 |

| ensembl_gene_id | external_gene_name | gene_biotype | log2 (fold change) | log2 (counts per million) | F-value | p-value | False discovery rate |
|---|---|---|---|---|---|---|---|
| ENSMUSG00000023830 | Igf2r | protein_coding | 0.450068005 | 3.655759802 | 0.9632618 | 0.340704 | 0.810414375 |
| ENSMUSG00000017314 | Mpp2 | protein_coding | -0.227973291 | 6.632723407 | 0.9631742 | 0.3407253 | 0.810414375 |
| ENSMUSG00000021978 | Extl3 | protein_coding | -0.29985485 | 5.404115986 | 0.9630885 | 0.3407462 | 0.810414375 |
| ENSMUSG00000059288 | Cdyl | protein_coding | -0.347393888 | 4.048326968 | 0.9630005 | 0.3407677 | 0.810414375 |
| ENSMUSG00000026944 | Abca2 | protein_coding | -0.264798955 | 6.357412888 | 0.9629103 | 0.3407896 | 0.810414375 |
| ENSMUSG00000028680 | Plk3 | protein_coding | 0.492242087 | 3.88673489 | 0.962918 | 0.3408112 | 0.810414375 |
| ENSMUSG00000110156 | Gm42067 | antisense_RNA | -0.453805422 | 3.177236938 | 0.9624561 | 0.3409003 | 0.810414375 |
| ENSMUSG00000079111 | Kdelr2 | protein_coding | 0.277133162 | 6.432431385 | 0.9622223 | 0.3409808 | 0.810414375 |
| ENSMUSG00000030087 | Klf15 | protein_coding | 0.456012225 | 3.61185396 | 0.9615377 | 0.3411244 | 0.810414375 |
| ENSMUSG00000022159 | Rab2b | protein_coding | 0.258524323 | 4.849298695 | 0.9612566 | 0.341193 | 0.810414375 |
| ENSMUSG00000040855 | Reps2 | protein_coding | -0.254266263 | 6.145058229 | 0.9611985 | 0.3412072 | 0.810414375 |
| ENSMUSG00000041354 | Rgl2 | protein_coding | -0.77856097 | 2.553597632 | 0.9609825 | 0.3412599 | 0.810414375 |
| ENSMUSG00000019558 | Slc6a8 | protein_coding | -0.268892554 | 5.170336647 | 0.9609538 | 0.3412669 | 0.810414375 |
| ENSMUSG00000041453 | Rpl21 | protein_coding | -0.251641701 | 5.190588427 | 0.9609285 | 0.3412731 | 0.810414375 |
| ENSMUSG00000008604 | Ubqln4 | protein_coding | -0.239270749 | 5.605717211 | 0.9603066 | 0.341425 | 0.810414375 |
| ENSMUSG00000029869 | Ephb6 | protein_coding | -0.421971634 | 4.24987142 | 0.9603035 | 0.3414258 | 0.810414375 |
| ENSMUSG00000022125 | Cln5 | protein_coding | -0.467476283 | 3.400346536 | 0.9598001 | 0.3415488 | 0.810414375 |
| ENSMUSG00000050199 | Lgr4 | protein_coding | -0.296389537 | 4.807990148 | 0.9597531 | 0.3415603 | 0.810414375 |
| ENSMUSG00000050565 | Tor1aip2 | protein_coding | -0.202591645 | 6.52170787 | 0.9587519 | 0.3418052 | 0.810782796 |
| ENSMUSG00000023284 | Zfp605 | protein_coding | -0.455879509 | 3.455143663 | 0.9585019 | 0.3418664 | 0.810782796 |
| ENSMUSG00000035247 | Hectd1 | protein_coding | 0.379477939 | 4.236072465 | 0.9563658 | 0.3423899 | 0.811382889 |
| ENSMUSG00000024854 | Pold4 | protein_coding | -0.252676792 | 5.691467525 | 0.9561448 | 0.3424441 | 0.811382889 |
| ENSMUSG00000021938 | Pspc1 | protein_coding | -0.353020288 | 4.338091377 | 0.9557512 | 0.3425407 | 0.811382889 |
| ENSMUSG00000060036 | Rpl3 | protein_coding | 0.232572661 | 7.69626689 | 0.9548466 | 0.3427629 | 0.811382889 |
| ENSMUSG00000020561 | Twistnb | protein_coding | -0.403505883 | 3.707681291 | 0.9546575 | 0.3428093 | 0.811382889 |
| ENSMUSG00000040414 | Slc25a28 | protein_coding | -0.389575672 | 4.194077487 | 0.954644 | 0.3428127 | 0.811382889 |
| ENSMUSG00000050954 | Zfp169 | protein_coding | -0.637086909 | 2.607587525 | 0.9537422 | 0.3430344 | 0.811382889 |
| ENSMUSG00000020079 | Supv3l1 | protein_coding | 0.621355144 | 3.067613605 | 0.9537242 | 0.3430388 | 0.811382889 |
| ENSMUSG00000022855 | Senp2 | protein_coding | 0.225110894 | 6.140582818 | 0.9535348 | 0.3430854 | 0.811382889 |
| ENSMUSG00000021643 | Serf1 | protein_coding | -0.334189036 | 4.613546209 | 0.9531266 | 0.3431859 | 0.811382889 |
| ENSMUSG00000034659 | Tmem109 | protein_coding | -0.25386527 | 6.521911539 | 0.9529928 | 0.3432188 | 0.811382889 |
| ENSMUSG00000022403 | St13 | protein_coding | -0.223524628 | 9.582884206 | 0.9528467 | 0.3432548 | 0.811382889 |
| ENSMUSG00000017778 | Cox7c | protein_coding | -0.373405707 | 3.971414894 | 0.9526871 | 0.3432941 | 0.811382889 |
| ENSMUSG00000064302 | Clasp1 | protein_coding | -0.289160554 | 5.440027815 | 0.952157 | 0.3434247 | 0.811382889 |
| ENSMUSG00000031323 | Dmrtc1a | protein_coding | 0.429289274 | 3.489503687 | 0.9521395 | 0.343429 | 0.811382889 |
| ENSMUSG00000028843 | Sh3bgrl3 | protein_coding | -0.192843026 | 8.961784637 | 0.9519566 | 0.343474 | 0.811382889 |
| ENSMUSG00000014496 | Ankrd28 | protein_coding | -0.416776815 | 3.551917331 | 0.9516282 | 0.343555 | 0.811382889 |
| ENSMUSG00000001985 | Grik3 | protein_coding | -0.315542516 | 4.438517667 | 0.9513525 | 0.343623 | 0.811382889 |
| ENSMUSG00000030276 | Ttll3 | protein_coding | -0.711937699 | 2.692870825 | 0.9512915 | 0.343638 | 0.811382889 |
| ENSMUSG00000020948 | Klhl28 | protein_coding | -0.266678752 | 4.843608937 | 0.9511636 | 0.3436696 | 0.811382889 |
| ENSMUSG00000041548 | Hspb8 | protein_coding | 0.316188493 | 4.908712143 | 0.9509306 | 0.343727 | 0.811382889 |
| ENSMUSG00000031565 | Fgfr1 | protein_coding | 0.254620875 | 5.45452829 | 0.9505327 | 0.3438252 | 0.811382889 |
| ENSMUSG00000051439 | Cd14 | protein_coding | -0.574909412 | 2.931237514 | 0.9503137 | 0.3438793 | 0.811382889 |
| ENSMUSG00000027751 | Supt20 | protein_coding | -0.39500524 | 4.469264427 | 0.9501068 | 0.3439304 | 0.811382889 |
| ENSMUSG00000061024 | Rrs1 | protein_coding | -0.417511577 | 3.855579368 | 0.949397 | 0.3441057 | 0.811441762 |
| ENSMUSG00000039456 | Morc3 | protein_coding | 0.415296104 | 4.556758285 | 0.9493417 | 0.3441426 | 0.811441762 |
| ENSMUSG00000023932 | Cdc5l | protein_coding | 0.244800655 | 5.338712987 | 0.9490577 | 0.3441895 | 0.811441762 |
| ENSMUSG00000018574 | Acadvl | protein_coding | 0.317586676 | 5.42607439 | 0.9486822 | 0.3442824 | 0.811441762 |
| ENSMUSG00000023089 | Ndufa5 | protein_coding | -0.279118071 | 10.15552811 | 0.9488205 | 0.3443326 | 0.811441762 |
| ENSMUSG00000097868 | A330069E16Rik | lincRNA | -0.427026061 | 4.657656107 | 0.9479703 | 0.3445429 | 0.811455784 |
| ENSMUSG00000040621 | Gemin8 | protein_coding | -0.341821038 | 3.708095508 | 0.9475512 | 0.3445622 | 0.811455784 |
| ENSMUSG00000038453 | Srcin1 | protein_coding | -0.236950498 | 5.644276852 | 0.9474557 | 0.3445858 | 0.811455784 |
| ENSMUSG00000040596 | Pogk | protein_coding | -0.313055252 | 4.689284965 | 0.9472353 | 0.3446404 | 0.811455784 |
| ENSMUSG00000034981 | Parm1 | protein_coding | 0.313487424 | 6.183718437 | 0.947667 | 0.3448127 | 0.811683603 |
| ENSMUSG00000023852 | Chd1 | protein_coding | -0.179068143 | 6.615970631 | 0.9454055 | 0.345094 | 0.812168004 |
| ENSMUSG00000048834 | Vstm2a | protein_coding | -0.197177213 | 6.941873829 | 0.9438445 | 0.3454816 | 0.812656754 |
| ENSMUSG00000085067 | Gm15631 | lincRNA | -0.346725037 | 5.229903911 | 0.9438716 | 0.345498 | 0.812656754 |
| ENSMUSG00000056771 | Gm10010 | lincRNA | 0.881081281 | 2.306700587 | 0.9436563 | 0.3455284 | 0.812656754 |
| ENSMUSG00000019856 | Fam184a | protein_coding | 0.400275146 | 3.47433877 | 0.9429924 | 0.3456935 | 0.812867214 |
| ENSMUSG00000007564 | Ppp2r1a | protein_coding | 0.220197285 | 9.864951749 | 0.9416718 | 0.3460222 | 0.812994012 |
| ENSMUSG00000040249 | Lrp1 | protein_coding | -0.269572859 | 5.675659049 | 0.9412407 | 0.3461296 | 0.812994012 |
| ENSMUSG00000079157 | Fam155a | protein_coding | -0.235033525 | 5.533616211 | 0.9407644 | 0.3462483 | 0.812994012 |
| ENSMUSG00000038141 | Tmem181a | protein_coding | 0.379447186 | 3.611116157 | 0.9405761 | 0.3462952 | 0.812994012 |
| ENSMUSG00000021794 | Glud1 | protein_coding | 0.237570019 | 6.788155854 | 0.9398941 | 0.3464653 | 0.812994012 |
| ENSMUSG00000014767 | Tbp | protein_coding | -0.261713272 | 5.374297906 | 0.9396314 | 0.3465309 | 0.812994012 |
| ENSMUSG00000033960 | Jcad | protein_coding | -0.440698622 | 4.348085359 | 0.9394048 | 0.3466106 | 0.812994012 |
| ENSMUSG00000060924 | Csmd1 | protein_coding | -0.427187385 | 4.672874283 | 0.9395055 | 0.3466704 | 0.812994012 |
| ENSMUSG00000034173 | Zbed5 | protein_coding | 0.441677754 | 3.221061799 | 0.938563 | 0.3467977 | 0.812994012 |
| ENSMUSG00000073616 | Cops9 | protein_coding | -0.260388278 | 8.663054637 | 0.938086 | 0.34694 | 0.812994012 |
| ENSMUSG00000006134 | Crkl | protein_coding | 0.329220553 | 3.826480559 | 0.9375197 | 0.3470585 | 0.812994012 |
| ENSMUSG00000038828 | Tmem214 | protein_coding | 0.321585489 | 4.820998701 | 0.9373159 | 0.3471095 | 0.812994012 |
| ENSMUSG00000027612 | Mmp24 | protein_coding | -0.508784768 | 3.06383252 | 0.9366727 | 0.3472705 | 0.812994012 |
| ENSMUSG00000026342 | Slc35f5 | protein_coding | 0.600862186 | 3.268655608 | 0.9366398 | 0.3473018 | 0.812994012 |
| ENSMUSG00000032011 | Thy1 | protein_coding | 0.240161237 | 7.286844666 | 0.9364685 | 0.3473447 | 0.812994012 |

| ensembl_gene_id | external_gene_name | gene_biotype | log2 (fold change) | log2 (counts per million) | F-value | p-value | False discovery rate |
|---|---|---|---|---|---|---|---|
| ENSMUSG00000011589 | Fsd1 | protein_coding | 0.247426812 | 7.226805919 | 0.9356276 | 0.3475554 | 0.812994012 |
| ENSMUSG00000032384 | Csnk1g1 | protein_coding | -0.277647759 | 5.506981906 | 0.9351845 | 0.3476434 | 0.812994012 |
| ENSMUSG00000032171 | Pin1 | protein_coding | -0.183681015 | 8.909003625 | 0.9346295 | 0.3477826 | 0.812994012 |
| ENSMUSG00000034412 | Tbc1d10a | protein_coding | 0.489898264 | 3.277671923 | 0.9342343 | 0.3478817 | 0.812994012 |
| ENSMUSG00000036820 | Amdhd2 | protein_coding | -0.303190432 | 5.454167321 | 0.9340888 | 0.3479183 | 0.812994012 |
| ENSMUSG00000025544 | Tm9sf2 | protein_coding | 0.353674178 | 5.955620366 | 0.9372204 | 0.348028 | 0.812994012 |
| ENSMUSG00000020628 | Trappc12 | protein_coding | -0.203156379 | 6.182638544 | 0.9334556 | 0.3480773 | 0.812994012 |
| ENSMUSG00000020923 | Ubtf | protein_coding | 0.232598662 | 6.328263927 | 0.9334554 | 0.3480773 | 0.812994012 |
| ENSMUSG00000029670 | Ing3 | protein_coding | -0.264537325 | 5.85921492 | 0.9332651 | 0.3481251 | 0.812994012 |
| ENSMUSG00000038280 | Ostm1 | protein_coding | -0.305067492 | 4.833108994 | 0.9332589 | 0.3481267 | 0.812994012 |
| ENSMUSG00000021892 | Sh3bp5 | protein_coding | 0.311506297 | 4.685489665 | 0.9329211 | 0.3482116 | 0.812994012 |
| ENSMUSG00000021185 | 9030617O03Rik | protein_coding | 0.503502714 | 4.068951454 | 0.9337821 | 0.3484327 | 0.812994012 |
| ENSMUSG00000027806 | Tsc22d2 | protein_coding | -0.313098419 | 4.615468649 | 0.9319165 | 0.3484642 | 0.812994012 |
| ENSMUSG00000072640 | Lyrm9 | protein_coding | -0.26001754 | 6.788462245 | 0.9318133 | 0.3485132 | 0.812994012 |
| ENSMUSG00000045284 | Dcaf12l1 | protein_coding | 0.217541519 | 6.420948291 | 0.9317043 | 0.3485176 | 0.812994012 |
| ENSMUSG00000033096 | Apmap | protein_coding | 0.216585886 | 5.88880109 | 0.9313511 | 0.3486065 | 0.812994012 |
| ENSMUSG00000026791 | Slc2a8 | protein_coding | 0.659440665 | 2.732372097 | 0.9312627 | 0.3486288 | 0.812994012 |
| ENSMUSG00000020083 | 2010107G23Rik | protein_coding | 0.199194072 | 6.60916479 | 0.9311332 | 0.3486614 | 0.812994012 |
| ENSMUSG00000024260 | Sap130 | protein_coding | -0.411454237 | 3.683428337 | 0.9300282 | 0.3489398 | 0.812994012 |
| ENSMUSG00000033931 | Rbm34 | protein_coding | -0.29742758 | 4.615637206 | 0.9298241 | 0.3489912 | 0.812994012 |
| ENSMUSG00000028673 | Fuca1 | protein_coding | 0.186538204 | 6.853897675 | 0.929801 | 0.348997 | 0.812994012 |
| ENSMUSG00000098659 | 1110015O18Rik | lincRNA | -0.618694795 | 2.736696433 | 0.9296111 | 0.3490449 | 0.812994012 |
| ENSMUSG00000038664 | Herc1 | protein_coding | 0.261748635 | 5.117133329 | 0.9295613 | 0.3490575 | 0.812994012 |
| ENSMUSG00000038206 | Fbxo8 | protein_coding | -0.309420374 | 4.913619376 | 0.9292231 | 0.3491428 | 0.812994012 |
| ENSMUSG00000023391 | Dlx2 | protein_coding | -0.585262534 | 2.893843547 | 0.9290291 | 0.3491918 | 0.812994012 |
| ENSMUSG00000038059 | Smim3 | protein_coding | 0.394223692 | 3.914729862 | 0.9286575 | 0.3492855 | 0.812994012 |
| ENSMUSG00000049940 | Pgrmc2 | protein_coding | -0.259197568 | 5.786279103 | 0.9286204 | 0.3492949 | 0.812994012 |
| ENSMUSG00000036781 | Rps27l | protein_coding | -0.249510341 | 6.179979603 | 0.9285804 | 0.349305 | 0.812994012 |
| ENSMUSG00000039308 | Ndst2 | protein_coding | 0.7655273 | 2.597608436 | 0.9285201 | 0.3493203 | 0.812994012 |
| ENSMUSG00000022103 | Gfra2 | protein_coding | 0.311436628 | 5.329226926 | 0.9281829 | 0.3494054 | 0.812994012 |
| ENSMUSG00000032382 | Snx1 | protein_coding | 0.194380283 | 7.24547317 | 0.9279012 | 0.3494766 | 0.812994012 |
| ENSMUSG00000038072 | Galnt11 | protein_coding | 0.419871592 | 3.718150019 | 0.9274683 | 0.349586 | 0.812994012 |
| ENSMUSG00000040883 | Tmem205 | protein_coding | -0.248295274 | 6.500854608 | 0.927334 | 0.3496199 | 0.812994012 |
| ENSMUSG00000068290 | Ddrgk1 | protein_coding | -0.211736752 | 7.561920124 | 0.9272305 | 0.3496461 | 0.812994012 |
| ENSMUSG00000079523 | Tmsb10 | protein_coding | -0.200522208 | 8.16261568 | 0.9270974 | 0.3496797 | 0.812994012 |
| ENSMUSG00000036779 | Papd5 | protein_coding | -0.340948939 | 4.076825432 | 0.9270804 | 0.349684 | 0.812994012 |
| ENSMUSG00000041697 | Cox6a1 | protein_coding | -0.290565582 | 9.755176941 | 0.9293514 | 0.3496968 | 0.812994012 |
| ENSMUSG00000015653 | Steap2 | protein_coding | 0.382156464 | 3.443828162 | 0.9268017 | 0.3497545 | 0.812994012 |
| ENSMUSG00000022474 | Pmm1 | protein_coding | 0.167300182 | 8.903386654 | 0.9258593 | 0.349993 | 0.813372512 |
| ENSMUSG00000049792 | Bag5 | protein_coding | 0.195577365 | 6.379971003 | 0.9253893 | 0.350112 | 0.813410298 |
| ENSMUSG00000039007 | Cpq | protein_coding | 0.339465038 | 4.352774455 | 0.9251978 | 0.3501605 | 0.813410298 |
| ENSMUSG00000025351 | Cd63 | protein_coding | -0.289459679 | 6.778323389 | 0.9261915 | 0.3505434 | 0.813842971 |
| ENSMUSG00000064179 | Tnnt1 | protein_coding | -0.596953962 | 4.643140755 | 0.9323702 | 0.3505826 | 0.813842971 |
| ENSMUSG00000015354 | Pcolce2 | protein_coding | -0.589824663 | 2.687559785 | 0.9234273 | 0.3506095 | 0.813842971 |
| ENSMUSG00000036062 | Phf24 | protein_coding | -0.209281599 | 6.335359416 | 0.9226514 | 0.3508066 | 0.813842971 |
| ENSMUSG00000036218 | Pdzrn4 | protein_coding | -0.360275673 | 3.917302898 | 0.9225776 | 0.3508253 | 0.813842971 |
| ENSMUSG00000026442 | Nfasc | protein_coding | 0.206657482 | 5.774130877 | 0.9222745 | 0.3509023 | 0.813842971 |
| ENSMUSG00000019907 | Ppp1r12a | protein_coding | -0.230463476 | 6.455738273 | 0.9218757 | 0.3510037 | 0.813842971 |
| ENSMUSG00000018585 | Atox1 | protein_coding | -0.194783987 | 8.537547706 | 0.9217512 | 0.3510354 | 0.813842971 |
| ENSMUSG00000069237 | Fam8a1 | protein_coding | -0.265405078 | 5.511208279 | 0.9215327 | 0.3510909 | 0.813842971 |
| ENSMUSG00000028559 | Osbpl9 | protein_coding | -0.213189777 | 5.932342339 | 0.9214826 | 0.3511036 | 0.813842971 |
| ENSMUSG00000047146 | Tet1 | protein_coding | -0.38151741 | 3.973923499 | 0.9210129 | 0.3512232 | 0.813880729 |
| ENSMUSG00000104597 | B230334C09Rik | antisense_RNA | -0.410061043 | 3.76481505 | 0.9208237 | 0.3512713 | 0.813880729 |
| ENSMUSG00000062729 | Ppox | protein_coding | -0.409470044 | 4.797051836 | 0.920694 | 0.3513875 | 0.813974567 |
| ENSMUSG00000051817 | Sox12 | protein_coding | -0.393669033 | 3.284725338 | 0.9198237 | 0.351526 | 0.814119917 |
| ENSMUSG00000029309 | Sparcl1 | protein_coding | -0.219135809 | 9.275472997 | 0.9189157 | 0.3517574 | 0.814442523 |
| ENSMUSG00000021466 | Ptch1 | protein_coding | -0.323744666 | 4.973170299 | 0.9186831 | 0.3518168 | 0.814442523 |
| ENSMUSG00000028896 | Rcc1 | protein_coding | 0.487415281 | 3.877977304 | 0.9181763 | 0.3519689 | 0.814591837 |
| ENSMUSG00000022390 | Zc3h7b | protein_coding | -0.306335148 | 5.210253924 | 0.9175778 | 0.3520989 | 0.814591837 |
| ENSMUSG00000021707 | Dhfr | protein_coding | -0.472773903 | 3.269350618 | 0.91754 | 0.3521085 | 0.814591837 |
| ENSMUSG00000031700 | Gpt2 | protein_coding | 0.321569443 | 4.450509533 | 0.9171955 | 0.3521965 | 0.814620186 |
| ENSMUSG00000033653 | Vps8 | protein_coding | 0.346067948 | 4.796670176 | 0.9162232 | 0.3524451 | 0.81501973 |
| ENSMUSG00000028617 | Lrrc42 | protein_coding | -0.270635213 | 4.923167692 | 0.9153892 | 0.3526585 | 0.815174182 |
| ENSMUSG00000042197 | Zfp451 | protein_coding | -0.235726625 | 5.442188885 | 0.9150832 | 0.3527368 | 0.815174182 |
| ENSMUSG00000034617 | Mtrr | protein_coding | 0.682124523 | 3.45573141 | 0.9196523 | 0.3527393 | 0.815174182 |
| ENSMUSG00000028597 | Gpx7 | protein_coding | 0.407722128 | 3.864937553 | 0.9132192 | 0.3532146 | 0.815902171 |
| ENSMUSG00000027419 | Pcsk2 | protein_coding | 0.20003485 | 6.944093578 | 0.9130845 | 0.3532491 | 0.815902171 |
| ENSMUSG00000025212 | Sfxn3 | protein_coding | -0.244892506 | 7.358371596 | 0.9128299 | 0.3533372 | 0.815902171 |
| ENSMUSG00000035878 | Hykk | protein_coding | 0.954032286 | 2.693860234 | 0.9161119 | 0.3533578 | 0.815902171 |
| ENSMUSG00000022428 | Cby1 | protein_coding | 0.190911746 | 7.321213859 | 0.9117874 | 0.3535822 | 0.816244969 |
| ENSMUSG00000046807 | Lrrc75b | protein_coding | 0.347832296 | 4.251037385 | 0.910926 | 0.3538036 | 0.816580865 |
| ENSMUSG00000018501 | Ncor1 | protein_coding | -0.220481333 | 5.848514274 | 0.9104313 | 0.3539309 | 0.816699271 |
| ENSMUSG00000000600 | Krit1 | protein_coding | -0.309788977 | 4.727434989 | 0.9096109 | 0.354142 | 0.816873183 |
| ENSMUSG00000036537 | Rnf113a1 | protein_coding | 0.289636515 | 5.094406699 | 0.9092632 | 0.3542316 | 0.816873183 |

| ensembl_gene_id | external_gene_name | gene_biotype | log2 (fold change) | log2 (counts per million) | F-value | p-value | False discovery rate |
|---|---|---|---|---|---|---|---|
| ENSMUSG00000005949 | Ctns | protein_coding | 0.469951854 | 3.660435599 | 0.9092534 | 0.3542341 | 0.816873183 |
| ENSMUSG00000046034 | Otulin | protein_coding | -0.42920741 | 3.192789776 | 0.9079815 | 0.3545621 | 0.817355002 |
| ENSMUSG00000034121 | Mks1 | protein_coding | -1.247814823 | 2.16448445 | 0.9114959 | 0.3545951 | 0.817355002 |
| ENSMUSG00000055917 | Zfp277 | protein_coding | -0.348804024 | 5.231259473 | 0.9055271 | 0.3552187 | 0.818372173 |
| ENSMUSG00000031987 | Egln1 | protein_coding | -0.253801543 | 5.548782701 | 0.905243 | 0.3552695 | 0.818372173 |
| ENSMUSG00000057236 | Rbbp4 | protein_coding | 0.237912315 | 7.282346161 | 0.9051908 | 0.3553057 | 0.818372173 |
| ENSMUSG00000038187 | Btbd10 | protein_coding | 0.244207798 | 5.697058457 | 0.9049676 | 0.3553408 | 0.818372173 |
| ENSMUSG00000032244 | Fem1b | protein_coding | 0.202536179 | 6.056165787 | 0.903788 | 0.3556463 | 0.818608362 |
| ENSMUSG00000062209 | Erbb4 | protein_coding | -0.330386783 | 4.623836947 | 0.9037022 | 0.3556685 | 0.818608362 |
| ENSMUSG00000053007 | Creb5 | protein_coding | -0.466214884 | 3.695559275 | 0.9029779 | 0.3558563 | 0.818608362 |
| ENSMUSG00000083695 | Rnf138rt1 | protein_coding | 0.507141938 | 3.793234133 | 0.9024958 | 0.356004 | 0.818608362 |
| ENSMUSG00000016344 | Ppdpf | protein_coding | -0.255088739 | 5.202265819 | 0.9009968 | 0.3563706 | 0.818608362 |
| ENSMUSG00000028654 | Mycl | protein_coding | 0.54238328 | 2.9758431 | 0.9009198 | 0.3563906 | 0.818608362 |
| ENSMUSG00000045691 | Thtpa | protein_coding | 0.367943809 | 5.335795227 | 0.9011582 | 0.35645 | 0.818608362 |
| ENSMUSG00000028646 | Rragc | protein_coding | 0.237107086 | 5.912414531 | 0.9006201 | 0.3564685 | 0.818608362 |
| ENSMUSG00000063714 | Sp3os | antisense_RNA | -0.406238021 | 3.530542364 | 0.9004922 | 0.3565017 | 0.818608362 |
| ENSMUSG00000112639 | AC134329.1 | lincRNA | -0.497894687 | 2.834979682 | 0.9003886 | 0.3565287 | 0.818608362 |
| ENSMUSG00000004896 | Rrnad1 | protein_coding | 0.377711771 | 3.883037571 | 0.9001403 | 0.3565933 | 0.818608362 |
| ENSMUSG00000020866 | Cacna1g | protein_coding | -0.531955641 | 3.129771186 | 0.8999188 | 0.3566509 | 0.818608362 |
| ENSMUSG00000057130 | Txnl4a | protein_coding | 0.220418485 | 6.757728205 | 0.8996278 | 0.3567266 | 0.818608362 |
| ENSMUSG00000016757 | Ttll12 | protein_coding | -0.465586417 | 3.701757573 | 0.8994543 | 0.3567718 | 0.818608362 |
| ENSMUSG00000006941 | Eif1b | protein_coding | 0.177323469 | 8.260969803 | 0.8993495 | 0.3567991 | 0.818608362 |
| ENSMUSG00000032498 | Mlh1 | protein_coding | -0.540831417 | 2.910645201 | 0.8992356 | 0.3568287 | 0.818608362 |
| ENSMUSG00000034903 | Cobll1 | protein_coding | -0.811687185 | 2.680774634 | 0.8985845 | 0.3570209 | 0.818608362 |
| ENSMUSG00000026817 | Ak1 | protein_coding | 0.207619674 | 9.070943259 | 0.898426 | 0.3570396 | 0.818608362 |
| ENSMUSG00000044122 | Proca1 | protein_coding | -0.358313146 | 3.87143142 | 0.8982512 | 0.3570852 | 0.818608362 |
| ENSMUSG00000029422 | Rsrc2 | protein_coding | -0.196468159 | 7.29315906 | 0.8981539 | 0.3571105 | 0.818608362 |
| ENSMUSG00000003872 | Lin7b | protein_coding | -0.200251553 | 6.138974701 | 0.8979779 | 0.3571564 | 0.818608362 |
| ENSMUSG00000045193 | Cirbp | protein_coding | 0.180989764 | 9.040230803 | 0.897942 | 0.3571658 | 0.818608362 |
| ENSMUSG00000039047 | Pigk | protein_coding | 0.22069364 | 6.552210073 | 0.897787 | 0.3572062 | 0.818608362 |
| ENSMUSG00000078640 | Gm11627 | protein_coding | -0.559692055 | 3.102746571 | 0.8969513 | 0.3574242 | 0.818608362 |
| ENSMUSG00000034295 | Fhod3 | protein_coding | 0.458729364 | 2.940591565 | 0.896777 | 0.3574697 | 0.818608362 |
| ENSMUSG00000029027 | Dffb | protein_coding | 0.731390989 | 2.44101323 | 0.8966887 | 0.3574928 | 0.818608362 |
| ENSMUSG00000042429 | Adora1 | protein_coding | -0.286858926 | 5.825502439 | 0.896538 | 0.3575321 | 0.818608362 |
| ENSMUSG00000040373 | Cacng5 | protein_coding | 0.339893971 | 4.872499501 | 0.8963177 | 0.3575896 | 0.818608362 |
| ENSMUSG00000025937 | Lactb2 | protein_coding | -0.410181977 | 3.577857389 | 0.8956649 | 0.3577602 | 0.818608362 |
| ENSMUSG00000022013 | Dnajc15 | protein_coding | -0.194512291 | 7.804727818 | 0.8956417 | 0.3577663 | 0.818608362 |
| ENSMUSG00000027339 | Rassf2 | protein_coding | 0.293920288 | 4.579458605 | 0.8954998 | 0.3578033 | 0.818608362 |
| ENSMUSG00000047712 | Ust | protein_coding | -0.48758427 | 3.075890735 | 0.894775 | 0.3579929 | 0.818691992 |
| ENSMUSG00000027438 | Napb | protein_coding | 0.216534618 | 8.827453936 | 0.8947622 | 0.3579962 | 0.818691992 |
| ENSMUSG00000049717 | Lig4 | protein_coding | -0.553340189 | 3.23115119 | 0.8944867 | 0.3580683 | 0.818691992 |
| ENSMUSG00000053025 | Sv2b | protein_coding | 0.267866688 | 5.410793827 | 0.8911143 | 0.3589524 | 0.820538944 |
| ENSMUSG00000031858 | Mau2 | protein_coding | -0.301572477 | 5.019563387 | 0.8901235 | 0.3592127 | 0.82056906 |
| ENSMUSG00000039579 | Grin3a | protein_coding | -0.225698748 | 5.533032447 | 0.8900605 | 0.3592293 | 0.82056906 |
| ENSMUSG00000026020 | Nop58 | protein_coding | 0.21503807 | 6.335977242 | 0.8897605 | 0.3593082 | 0.82056906 |
| ENSMUSG00000027737 | Slc7a11 | protein_coding | -0.4942828 | 3.218321286 | 0.8893079 | 0.3594272 | 0.82056906 |
| ENSMUSG00000024846 | Cst6 | protein_coding | -0.310135218 | 4.547371648 | 0.8892998 | 0.3594294 | 0.82056906 |
| ENSMUSG00000018651 | Tada2a | protein_coding | 0.332202614 | 5.035906239 | 0.8886309 | 0.3596055 | 0.82056906 |
| ENSMUSG00000006299 | Aamp | protein_coding | 0.228996341 | 9.100446211 | 0.8886023 | 0.359613 | 0.82056906 |
| ENSMUSG00000016319 | Slc25a5 | protein_coding | 0.163630103 | 9.055108358 | 0.888521 | 0.3596344 | 0.82056906 |
| ENSMUSG00000028044 | Cks1b | protein_coding | 0.420696541 | 3.204712317 | 0.8884528 | 0.3596524 | 0.82056906 |
| ENSMUSG00000039001 | Rps21 | protein_coding | -0.247965124 | 8.753982477 | 0.8877926 | 0.3598488 | 0.820772666 |
| ENSMUSG00000036745 | Ttll7 | protein_coding | -0.182836027 | 7.030682103 | 0.8873085 | 0.3599539 | 0.820772666 |
| ENSMUSG00000042572 | Ube2q1 | protein_coding | 0.247505864 | 5.737948593 | 0.8872452 | 0.3599706 | 0.820772666 |
| ENSMUSG00000028139 | Riiad1 | protein_coding | -0.280036261 | 5.2233371 | 0.8869132 | 0.3600582 | 0.820798324 |
| ENSMUSG00000071337 | Tia1 | protein_coding | -0.387715247 | 3.8261691 | 0.8865203 | 0.3601619 | 0.820860624 |
| ENSMUSG00000044014 | Npy5r | protein_coding | -0.637085767 | 2.983356318 | 0.8850014 | 0.360563 | 0.821600829 |
| ENSMUSG00000027639 | Samhd1 | protein_coding | 0.414600866 | 3.30342135 | 0.8840862 | 0.3608051 | 0.821978197 |
| ENSMUSG00000061559 | Wdr61 | protein_coding | 0.173950318 | 7.66152338 | 0.8835367 | 0.3609505 | 0.822135353 |
| ENSMUSG00000029581 | Fscn1 | protein_coding | -0.224097845 | 5.563897605 | 0.8830899 | 0.3610688 | 0.822205057 |
| ENSMUSG00000020455 | Trim11 | protein_coding | -0.423000074 | 3.555612013 | 0.8825972 | 0.3611994 | 0.822205057 |
| ENSMUSG00000022565 | Plec | protein_coding | -0.283477251 | 4.769189975 | 0.8824456 | 0.3612396 | 0.822205057 |
| ENSMUSG00000059534 | Uqcr10 | protein_coding | -0.224881632 | 10.21776316 | 0.8822667 | 0.361287 | 0.822205057 |
| ENSMUSG00000058135 | Gstm1 | protein_coding | -0.208086627 | 8.942497461 | 0.881848 | 0.361398 | 0.822283725 |
| ENSMUSG00000030469 | Zfp719 | protein_coding | -0.387792449 | 3.742661636 | 0.88138 | 0.3615222 | 0.822347257 |
| ENSMUSG00000020521 | Rnft1 | protein_coding | -0.32879738 | 4.037475334 | 0.8802393 | 0.3618251 | 0.822347257 |
| ENSMUSG00000040463 | Mybbp1a | protein_coding | -0.2859827 | 4.832715963 | 0.8801385 | 0.3618519 | 0.822347257 |
| ENSMUSG00000020431 | Adcy1 | protein_coding | -0.379362707 | 4.762574829 | 0.8800946 | 0.3618859 | 0.822347257 |
| ENSMUSG00000054792 | Klhl18 | protein_coding | 0.294242765 | 4.768380638 | 0.8795582 | 0.3620061 | 0.822347257 |
| ENSMUSG00000023094 | Msrb2 | protein_coding | 0.212480282 | 5.95553684 | 0.8795399 | 0.362011 | 0.822347257 |
| ENSMUSG00000020604 | Arsg | protein_coding | 0.397795242 | 4.808082423 | 0.8792848 | 0.36216 | 0.822347257 |
| ENSMUSG00000029020 | Mfn2 | protein_coding | 0.278633782 | 7.370764413 | 0.8819542 | 0.362237 | 0.822347257 |
| ENSMUSG00000014551 | Mrps25 | protein_coding | 0.172752871 | 7.710675343 | 0.8786336 | 0.3622521 | 0.822347257 |
| ENSMUSG00000024270 | Slc39a6 | protein_coding | 0.310077505 | 5.133146725 | 0.8786086 | 0.3622587 | 0.822347257 |

| ensembl_gene_id | external_gene_name | gene_biotype | log2 (fold change) | log2 (counts per million) | F-value | p-value | False discovery rate |
|---|---|---|---|---|---|---|---|
| ENSMUSG00000026858 | Miga2 | protein_coding | 0.260208309 | 5.634868065 | 0.8781754 | 0.362374 | 0.822347257 |
| ENSMUSG00000043164 | Tmem212 | protein_coding | -0.469781799 | 3.784289306 | 0.8781001 | 0.3624164 | 0.822347257 |
| ENSMUSG00000029290 | Zfp326 | protein_coding | -0.319801753 | 4.101239666 | 0.8778787 | 0.3624531 | 0.822347257 |
| ENSMUSG00000032333 | Stoml1 | protein_coding | 0.245531612 | 6.128324996 | 0.8775353 | 0.3625445 | 0.822347257 |
| ENSMUSG00000094152 | Slc6a16 | protein_coding | 0.429036417 | 3.775682246 | 0.8774282 | 0.3625731 | 0.822347257 |
| ENSMUSG00000034595 | Ppp1r18 | protein_coding | -0.635272398 | 2.624881968 | 0.876029 | 0.3629463 | 0.823020046 |
| ENSMUSG00000042035 | Igsf3 | protein_coding | 0.264342219 | 5.47660172 | 0.874885 | 0.3632518 | 0.823463025 |
| ENSMUSG00000030376 | Slc8a2 | protein_coding | -0.562741213 | 2.93078696 | 0.8747241 | 0.3632948 | 0.823463025 |
| ENSMUSG00000032872 | Cyb5r4 | protein_coding | 0.364189731 | 3.598388793 | 0.8739962 | 0.3634894 | 0.8237305 |
| ENSMUSG00000039419 | Cntnap2 | protein_coding | -0.180397606 | 7.646214872 | 0.8724596 | 0.3639007 | 0.823871204 |
| ENSMUSG00000036306 | Lzts1 | protein_coding | 0.196330325 | 8.608718317 | 0.8721712 | 0.363978 | 0.823871204 |
| ENSMUSG00000055541 | Lair1 | protein_coding | -0.475122343 | 3.076907307 | 0.8717961 | 0.3640786 | 0.823871204 |
| ENSMUSG00000058267 | Mrps14 | protein_coding | 0.177479443 | 8.757600693 | 0.8717842 | 0.3640818 | 0.823871204 |
| ENSMUSG00000038582 | Pptc7 | protein_coding | -0.330685183 | 4.48550398 | 0.8716659 | 0.3641135 | 0.823871204 |
| ENSMUSG00000036766 | Dner | protein_coding | -0.239313808 | 6.807641202 | 0.8716166 | 0.3641489 | 0.823871204 |
| ENSMUSG00000026657 | Frmd4a | protein_coding | -0.276249901 | 4.795458317 | 0.8714247 | 0.3641782 | 0.823871204 |
| ENSMUSG00000097587 | 4930578M01Rik | lincRNA | -0.459114536 | 3.936053073 | 0.871474 | 0.3641872 | 0.823871204 |
| ENSMUSG00000028527 | Ak4 | protein_coding | 0.286243854 | 4.634569626 | 0.8709871 | 0.3642956 | 0.823871204 |
| ENSMUSG00000056596 | Trnp1 | protein_coding | -0.277063779 | 5.774710519 | 0.8709048 | 0.3643176 | 0.823871204 |
| ENSMUSG00000035024 | Ncapd3 | protein_coding | 0.392757083 | 3.666744051 | 0.870279 | 0.3644857 | 0.824077861 |
| ENSMUSG00000030203 | Dusp16 | protein_coding | 0.459970914 | 3.462242681 | 0.8697394 | 0.3646306 | 0.824232237 |
| ENSMUSG00000028062 | Lamtor2 | protein_coding | 0.206838659 | 9.130326099 | 0.8673512 | 0.3652732 | 0.825016852 |
| ENSMUSG00000030302 | Atp2b2 | protein_coding | -0.196952146 | 8.150041567 | 0.8673301 | 0.3652789 | 0.825016852 |
| ENSMUSG00000104784 | Gm42984 | antisense_RNA | -0.618982838 | 2.66238835 | 0.8670598 | 0.3653517 | 0.825016852 |
| ENSMUSG00000027942 | 4933434E20Rik | protein_coding | -0.166377348 | 8.603960631 | 0.867046 | 0.3653554 | 0.825016852 |
| ENSMUSG00000043872 | Zmym1 | protein_coding | 0.439066177 | 3.116851037 | 0.867024 | 0.3653613 | 0.825016852 |
| ENSMUSG00000086370 | Ftx | lincRNA | -0.463004624 | 3.516053268 | 0.8663953 | 0.3655308 | 0.825161366 |
| ENSMUSG00000037902 | Sirpa | protein_coding | 0.194142995 | 6.931019803 | 0.8660089 | 0.365635 | 0.825161366 |
| ENSMUSG00000015733 | Capza2 | protein_coding | -0.173124223 | 9.262856819 | 0.865933 | 0.3656555 | 0.825161366 |
| ENSMUSG00000006235 | Epor | protein_coding | -0.47912508 | 2.937804514 | 0.8645647 | 0.366025 | 0.825673828 |
| ENSMUSG00000087478 | 4930506C21Rik | processed_transcript | -0.598088404 | 2.649936518 | 0.8642972 | 0.3660973 | 0.825673828 |
| ENSMUSG00000021916 | Glt8d1 | protein_coding | -0.389216809 | 4.0443837 | 0.8639533 | 0.3661902 | 0.825673828 |
| ENSMUSG00000023828 | Slc22a3 | protein_coding | 0.618836683 | 2.622116416 | 0.8637193 | 0.3662535 | 0.825673828 |
| ENSMUSG00000029571 | Tmem106b | protein_coding | -0.239588225 | 5.82573429 | 0.8636712 | 0.3662665 | 0.825673828 |
| ENSMUSG00000041881 | Ndufa7 | protein_coding | -0.218163934 | 9.535408849 | 0.8633126 | 0.3663636 | 0.825718343 |
| ENSMUSG00000013155 | Enkd1 | protein_coding | -0.379658228 | 3.709672084 | 0.8630306 | 0.3664399 | 0.825718343 |
| ENSMUSG00000035227 | Spcs2 | protein_coding | 0.192462869 | 8.275560973 | 0.8623209 | 0.366632 | 0.825978251 |
| ENSMUSG00000024870 | Rab1b | protein_coding | -0.190194521 | 8.0398978 | 0.8607371 | 0.3670614 | 0.826368279 |
| ENSMUSG00000057541 | Pus7 | protein_coding | 0.548206022 | 2.835257955 | 0.8605569 | 0.3671103 | 0.826368279 |
| ENSMUSG00000033157 | Abhd10 | protein_coding | -0.225591146 | 5.680328417 | 0.8602325 | 0.3671984 | 0.826368279 |
| ENSMUSG00000045932 | Ifit2 | protein_coding | -0.437159764 | 3.092937271 | 0.8600148 | 0.3672575 | 0.826368279 |
| ENSMUSG00000018995 | Nars2 | protein_coding | -0.378205591 | 3.687330034 | 0.8599386 | 0.3672782 | 0.826368279 |
| ENSMUSG00000063439 | B9d2 | protein_coding | -0.178809012 | 7.162228884 | 0.8598056 | 0.3673143 | 0.826368279 |
| ENSMUSG00000025404 | R3hdm2 | protein_coding | -0.219680375 | 5.633320548 | 0.8594319 | 0.3674158 | 0.826368279 |
| ENSMUSG00000037523 | Mavs | protein_coding | 0.382666101 | 4.380389987 | 0.8593088 | 0.3674492 | 0.826368279 |
| ENSMUSG00000023336 | Wfdc1 | protein_coding | 0.823739344 | 2.318903445 | 0.8589377 | 0.3675501 | 0.826368279 |
| ENSMUSG00000045608 | Dbx2 | protein_coding | -0.373376741 | 3.945594001 | 0.8588512 | 0.3675737 | 0.826368279 |
| ENSMUSG00000029910 | Mad2l1 | protein_coding | -0.345978896 | 4.158878799 | 0.8579819 | 0.3678101 | 0.826662972 |
| ENSMUSG00000032523 | Hhatl | protein_coding | -0.7115044 | 3.133468773 | 0.8608161 | 0.3678963 | 0.826662972 |
| ENSMUSG00000040482 | Dxo | protein_coding | -0.320414557 | 4.449983873 | 0.8575218 | 0.3679354 | 0.826662972 |
| ENSMUSG00000021703 | Serinc5 | protein_coding | -0.349561552 | 4.266735391 | 0.8563374 | 0.3682581 | 0.827080463 |
| ENSMUSG00000031358 | Msl3 | protein_coding | -0.304895229 | 4.63740218 | 0.8554123 | 0.3685104 | 0.827080463 |
| ENSMUSG00000042660 | Wdr55 | protein_coding | 0.267258407 | 4.827518776 | 0.8548207 | 0.3686719 | 0.827080463 |
| ENSMUSG00000066640 | Fbxl18 | protein_coding | 0.349503373 | 4.108800012 | 0.8548094 | 0.368675 | 0.827080463 |
| ENSMUSG00000022791 | Tnk2 | protein_coding | -0.284784883 | 5.128254215 | 0.8546268 | 0.3687249 | 0.827080463 |
| ENSMUSG00000051855 | Mest | protein_coding | -0.447589722 | 3.702198442 | 0.8546085 | 0.3687299 | 0.827080463 |
| ENSMUSG00000040029 | Ipo8 | protein_coding | 0.247494056 | 5.01949478 | 0.8543063 | 0.3688124 | 0.827080463 |
| ENSMUSG00000052105 | Mtcl1 | protein_coding | -0.423643965 | 3.993371631 | 0.8539445 | 0.3689113 | 0.827080463 |
| ENSMUSG00000059336 | Slc14a1 | protein_coding | 0.644054884 | 2.821442669 | 0.8534045 | 0.369059 | 0.827080463 |
| ENSMUSG00000021326 | Trim27 | protein_coding | 0.389826318 | 3.769012894 | 0.8533641 | 0.36907 | 0.827080463 |
| ENSMUSG00000074579 | Lekr1 | protein_coding | -0.454520226 | 3.043518446 | 0.853193 | 0.3691168 | 0.827080463 |
| ENSMUSG00000056749 | Nfil3 | protein_coding | -0.450434075 | 3.514816785 | 0.8529479 | 0.3691839 | 0.827080463 |
| ENSMUSG00000039145 | Camk1d | protein_coding | 0.211499645 | 7.706375568 | 0.852234 | 0.3693793 | 0.827080463 |
| ENSMUSG00000004035 | Gstm7 | protein_coding | -0.166629883 | 7.183323043 | 0.8521447 | 0.3694038 | 0.827080463 |
| ENSMUSG00000027618 | Nfs1 | protein_coding | 0.293638344 | 4.773280756 | 0.8520275 | 0.3694359 | 0.827080463 |
| ENSMUSG00000042992 | Borcs5 | protein_coding | 0.332890407 | 5.352748953 | 0.8519901 | 0.3694681 | 0.827080463 |
| ENSMUSG00000019977 | Hbs1l | protein_coding | 0.267887941 | 5.924672831 | 0.8518166 | 0.3694937 | 0.827080463 |
| ENSMUSG00000084159 | Gm12696 | processed_pseudoge | 0.472563066 | 2.892748836 | 0.8517727 | 0.3695057 | 0.827080463 |
| ENSMUSG00000025157 | Zdhhc16 | protein_coding | 0.346671711 | 3.534691485 | 0.8504488 | 0.3698687 | 0.827605504 |
| ENSMUSG00000031807 | Pgls | protein_coding | 0.314972841 | 4.35469058 | 0.8503559 | 0.3698942 | 0.827605504 |
| ENSMUSG00000002455 | Prpf6 | protein_coding | 0.263968811 | 6.141371447 | 0.8498143 | 0.3700648 | 0.827815054 |
| ENSMUSG00000019802 | Sec63 | protein_coding | -0.357115266 | 4.528129988 | 0.8484563 | 0.370416 | 0.828171032 |
| ENSMUSG00000001942 | Siae | protein_coding | 0.300816751 | 4.628530044 | 0.8479773 | 0.3705478 | 0.828171032 |
| ENSMUSG00000036632 | Alg5 | protein_coding | 0.314550364 | 5.020217067 | 0.8479703 | 0.3705497 | 0.828171032 |

| ensembl_gene_id | external_gene_name | gene_biotype | log2 (fold change) | log2 (counts per million) | F-value | p-value | False discovery rate |
|---|---|---|---|---|---|---|---|
| ENSMUSG00000035274 | Tpbg | protein_coding | 0.52545922 | 3.203808165 | 0.8478637 | 0.370579 | 0.828171032 |
| ENSMUSG00000001774 | Chordc1 | protein_coding | -0.173831877 | 6.496847039 | 0.8473096 | 0.3707315 | 0.828171032 |
| ENSMUSG00000041037 | Irgq | protein_coding | -0.286820362 | 4.537448549 | 0.8470821 | 0.3707942 | 0.828171032 |
| ENSMUSG00000027782 | Kpna4 | protein_coding | 0.217960396 | 5.463499398 | 0.8469201 | 0.3708388 | 0.828171032 |
| ENSMUSG00000108461 | AV356131 | bidirectional_promoter | -0.632840467 | 2.681893611 | 0.8469154 | 0.3708401 | 0.828171032 |
| ENSMUSG00000038366 | Lasp1 | protein_coding | 0.274649657 | 5.692131249 | 0.8465681 | 0.3709358 | 0.828212755 |
| ENSMUSG00000000374 | Trappc10 | protein_coding | -0.284528808 | 4.455311871 | 0.8453287 | 0.3712776 | 0.828665678 |
| ENSMUSG00000020175 | Rab36 | protein_coding | 0.234919084 | 6.417357746 | 0.8450947 | 0.3713421 | 0.828665678 |
| ENSMUSG00000029474 | Rnf34 | protein_coding | 0.217825302 | 6.829315571 | 0.8449943 | 0.3713698 | 0.828665678 |
| ENSMUSG00000078546 | Zfp995 | protein_coding | -0.504633288 | 2.667089842 | 0.8441022 | 0.3716162 | 0.829006402 |
| ENSMUSG00000045083 | Lingo2 | protein_coding | -0.261636911 | 4.931962584 | 0.8438832 | 0.3716767 | 0.829006402 |
| ENSMUSG00000060032 | H2afj | protein_coding | -0.503111136 | 2.916554437 | 0.8429866 | 0.3719247 | 0.829183682 |
| ENSMUSG00000040123 | Zmym5 | protein_coding | -0.241313892 | 5.017706106 | 0.8422436 | 0.3721303 | 0.829183682 |
| ENSMUSG00000041703 | Zic5 | protein_coding | -0.482399453 | 3.725062182 | 0.8422498 | 0.3721504 | 0.829183682 |
| ENSMUSG00000054263 | Lifr | protein_coding | -0.400794253 | 3.791130107 | 0.842064 | 0.37218 | 0.829183682 |
| ENSMUSG00000020745 | Pafah1b1 | protein_coding | -0.179457667 | 7.459927703 | 0.8416825 | 0.3722857 | 0.829183682 |
| ENSMUSG00000055943 | Emc7 | protein_coding | -0.167365084 | 7.536572216 | 0.84158 | 0.3723141 | 0.829183682 |
| ENSMUSG00000032480 | Dhx30 | protein_coding | 0.229289312 | 6.567190402 | 0.8415143 | 0.3723323 | 0.829183682 |
| ENSMUSG00000038115 | Ano2 | protein_coding | 0.615924591 | 3.114696155 | 0.8414458 | 0.3723731 | 0.829183682 |
| ENSMUSG00000054474 | Thnsl2 | protein_coding | 0.482245725 | 3.354334824 | 0.8405482 | 0.3726001 | 0.82921302 |
| ENSMUSG00000041939 | Mvk | protein_coding | -0.27633851 | 5.485362946 | 0.8404931 | 0.3726154 | 0.82921302 |
| ENSMUSG00000032405 | Pias1 | protein_coding | -0.218610629 | 7.093369764 | 0.8404851 | 0.3726176 | 0.82921302 |
| ENSMUSG00000027395 | Polr1b | protein_coding | 0.615608313 | 2.59214841 | 0.8400823 | 0.3727294 | 0.829290146 |
| ENSMUSG00000056268 | Dennd1b | protein_coding | -0.599960285 | 2.790479501 | 0.8393325 | 0.3729376 | 0.829401922 |
| ENSMUSG00000052698 | Tln2 | protein_coding | 0.250459455 | 5.70148989 | 0.8390741 | 0.3730094 | 0.829401922 |
| ENSMUSG00000062867 | Impdh2 | protein_coding | 0.341414637 | 4.062557865 | 0.839068 | 0.373011 | 0.829401922 |
| ENSMUSG00000030094 | Xpc | protein_coding | 0.385536263 | 3.523634083 | 0.838723 | 0.3731069 | 0.829443618 |
| ENSMUSG00000034390 | Cmip | protein_coding | 0.25051829 | 7.457598778 | 0.8383047 | 0.3733276 | 0.829676788 |
| ENSMUSG00000008036 | Ap2s1 | protein_coding | -0.167311608 | 8.101066345 | 0.8377909 | 0.3733661 | 0.829676788 |
| ENSMUSG00000047909 | Ankrd16 | protein_coding | 0.64989484 | 2.79214358 | 0.8371146 | 0.3735544 | 0.829699818 |
| ENSMUSG00000032434 | Cmtm6 | protein_coding | 0.613312288 | 2.968209498 | 0.8369492 | 0.3736004 | 0.829699818 |
| ENSMUSG00000086607 | 4930511M06Rik | processed_transcript | -0.47034804 | 3.102546833 | 0.8369221 | 0.373608 | 0.829699818 |
| ENSMUSG00000027809 | Etfdh | protein_coding | 0.28214588 | 4.954823637 | 0.8359549 | 0.3738775 | 0.830000808 |
| ENSMUSG00000025085 | Ablim1 | protein_coding | -0.269279111 | 5.591578019 | 0.8358817 | 0.3738979 | 0.830000808 |
| ENSMUSG00000019715 | Gle1 | protein_coding | 0.290078728 | 6.066302344 | 0.8351654 | 0.3741194 | 0.830321199 |
| ENSMUSG00000055334 | Snupn | protein_coding | 0.246341515 | 5.270604123 | 0.8342045 | 0.3743659 | 0.830599065 |
| ENSMUSG00000039601 | Rcan2 | protein_coding | 0.246241037 | 8.189909291 | 0.8338561 | 0.3745189 | 0.830599065 |
| ENSMUSG00000021728 | Emb | protein_coding | 0.254929773 | 5.713795227 | 0.8333686 | 0.3745995 | 0.830599065 |
| ENSMUSG00000071648 | Rom1 | protein_coding | 0.441069943 | 3.346982988 | 0.8333334 | 0.3746094 | 0.830599065 |
| ENSMUSG00000057265 | Bbof1 | protein_coding | -0.718633639 | 2.443925944 | 0.8332567 | 0.3746308 | 0.830599065 |
| ENSMUSG00000029636 | Wasf3 | protein_coding | -0.22578402 | 5.556638143 | 0.8323224 | 0.3748922 | 0.831007245 |
| ENSMUSG00000028778 | Hcrtr1 | protein_coding | 1.215965636 | 3.40245049 | 0.8597077 | 0.3752087 | 0.83140537 |
| ENSMUSG00000036873 | 2410004B18Rik | protein_coding | -0.364830613 | 4.247626815 | 0.8310103 | 0.3752598 | 0.83140537 |
| ENSMUSG00000030223 | Ptpro | protein_coding | 0.251710133 | 4.842292511 | 0.8305562 | 0.3753871 | 0.83140537 |
| ENSMUSG00000047658 | Gal3st3 | protein_coding | -0.537033762 | 3.135643891 | 0.830473 | 0.3754104 | 0.83140537 |
| ENSMUSG00000061313 | Ddhd2 | protein_coding | 0.345487857 | 7.095675097 | 0.8377828 | 0.3754811 | 0.83140537 |
| ENSMUSG00000003062 | Stard3nl | protein_coding | 0.172049547 | 7.038505656 | 0.8300263 | 0.3755357 | 0.83140537 |
| ENSMUSG00000034007 | Scaper | protein_coding | -0.230506184 | 5.731327711 | 0.829402 | 0.375711 | 0.83162217 |
| ENSMUSG00000017631 | Abr | protein_coding | 0.186884984 | 7.151394272 | 0.8287772 | 0.3758865 | 0.831839476 |
| ENSMUSG00000044952 | Kctd21 | protein_coding | 0.36086663 | 4.021924645 | 0.8275449 | 0.3762331 | 0.832025142 |
| ENSMUSG00000032897 | Nfyc | protein_coding | -0.301325037 | 4.118222849 | 0.8275275 | 0.376238 | 0.832025142 |
| ENSMUSG00000098112 | Bin2 | protein_coding | -0.837262899 | 2.31569101 | 0.8273951 | 0.3762752 | 0.832025142 |
| ENSMUSG00000052763 | Zfp212 | protein_coding | 0.407996288 | 3.652756293 | 0.8273783 | 0.3762799 | 0.832025142 |
| ENSMUSG00000074771 | Ankef1 | protein_coding | -0.606055655 | 2.771985152 | 0.8262026 | 0.3766111 | 0.832244199 |
| ENSMUSG00000024347 | Psd2 | protein_coding | 0.401738623 | 4.892576603 | 0.828032 | 0.3766484 | 0.832244199 |
| ENSMUSG00000039637 | Coro7 | protein_coding | -0.258611422 | 4.868359133 | 0.8259056 | 0.3766948 | 0.832244199 |
| ENSMUSG00000056310 | Tyw1 | protein_coding | 0.303658224 | 4.485944164 | 0.8253468 | 0.3768524 | 0.832244199 |
| ENSMUSG00000027599 | Armc1 | protein_coding | -0.199298432 | 7.614774014 | 0.8249328 | 0.3769692 | 0.832244199 |
| ENSMUSG00000036698 | Ago2 | protein_coding | 0.238517634 | 5.261314727 | 0.8248983 | 0.3769789 | 0.832244199 |
| ENSMUSG00000046434 | Hnrnpa1 | protein_coding | 0.308761846 | 3.84562228 | 0.8248874 | 0.376982 | 0.832244199 |
| ENSMUSG00000009207 | Lnpk | protein_coding | -0.234924624 | 5.532369626 | 0.8245699 | 0.3770716 | 0.832244199 |
| ENSMUSG00000031591 | Asah1 | protein_coding | 0.204087606 | 6.231860948 | 0.824366 | 0.3771292 | 0.832244199 |
| ENSMUSG00000020516 | Rps6kb1 | protein_coding | -0.281042959 | 5.715167585 | 0.8242817 | 0.377153 | 0.832244199 |
| ENSMUSG00000049643 | 2310022A10Rik | protein_coding | 0.224288661 | 5.683966829 | 0.8229665 | 0.3775247 | 0.83236519 |
| ENSMUSG00000040563 | Plppr2 | protein_coding | -0.292095914 | 5.355403126 | 0.8226232 | 0.3776218 | 0.83236519 |
| ENSMUSG00000026201 | Stk16 | protein_coding | -0.246641392 | 6.45245225 | 0.8224385 | 0.3776957 | 0.83236519 |
| ENSMUSG00000032067 | Pts | protein_coding | 0.219737822 | 7.21680756 | 0.8220322 | 0.3777891 | 0.83236519 |
| ENSMUSG00000054312 | Mrps21 | protein_coding | -0.227928141 | 7.97486002 | 0.8217626 | 0.377887 | 0.83236519 |
| ENSMUSG00000020803 | Txndc17 | protein_coding | 0.15816465 | 7.832674069 | 0.8214859 | 0.3779438 | 0.83236519 |
| ENSMUSG00000038372 | Gmds | protein_coding | -0.335640973 | 3.71384034 | 0.8212768 | 0.3780031 | 0.83236519 |
| ENSMUSG00000041130 | Zfp598 | protein_coding | -0.513919087 | 3.23157067 | 0.8211281 | 0.3780452 | 0.83236519 |
| ENSMUSG00000027569 | Mrgbp | protein_coding | -0.47695348 | 3.122264797 | 0.8208632 | 0.3781203 | 0.83236519 |
| ENSMUSG00000057572 | Zbtb8os | protein_coding | -0.291999896 | 4.87184687 | 0.8208208 | 0.3781323 | 0.83236519 |
| ENSMUSG00000011831 | Evi5 | protein_coding | -0.285192275 | 4.949072978 | 0.8208093 | 0.3781356 | 0.83236519 |

| ensembl_gene_id | external_gene_name | gene_biotype | log2 (fold change) | log2 (counts per million) | F-value | p-value | False discovery rate |
|---|---|---|---|---|---|---|---|
| ENSMUSG00000020467 | Efemp1 | protein_coding | -0.455202692 | 3.055751695 | 0.8207572 | 0.3781504 | 0.83236519 |
| ENSMUSG00000025869 | Nop16 | protein_coding | 0.171215667 | 7.109687438 | 0.8203939 | 0.3782534 | 0.83236519 |
| ENSMUSG00000049658 | Bdp1 | protein_coding | -0.359711387 | 3.753330249 | 0.8202594 | 0.3782915 | 0.83236519 |
| ENSMUSG00000028521 | Slc35d1 | protein_coding | -0.408001343 | 3.083161886 | 0.8195582 | 0.3784905 | 0.832512355 |
| ENSMUSG00000038235 | F11r | protein_coding | 0.431189718 | 3.493792847 | 0.819444 | 0.378523 | 0.832512355 |
| ENSMUSG00000032127 | Vps11 | protein_coding | 0.287747217 | 6.488605054 | 0.8216915 | 0.3786327 | 0.832512355 |
| ENSMUSG00000050973 | Gdpgp1 | protein_coding | 0.38682829 | 4.282659624 | 0.8184379 | 0.3788088 | 0.832512355 |
| ENSMUSG00000038650 | Rnh1 | protein_coding | 0.212526993 | 7.089744045 | 0.8183601 | 0.3788309 | 0.832512355 |
| ENSMUSG00000024999 | Noc3l | protein_coding | 0.529624667 | 2.803657768 | 0.8178147 | 0.378986 | 0.832512355 |
| ENSMUSG00000044502 | Bod1 | protein_coding | 0.280594839 | 5.512528293 | 0.8175432 | 0.3790632 | 0.832512355 |
| ENSMUSG00000020396 | Nefh | protein_coding | 0.25204653 | 5.388966316 | 0.8173797 | 0.3791098 | 0.832512355 |
| ENSMUSG00000070570 | Slc17a7 | protein_coding | -0.728619449 | 2.764082517 | 0.8174339 | 0.3791159 | 0.832512355 |
| ENSMUSG00000018239 | Zcchc10 | protein_coding | -0.231441893 | 5.560305535 | 0.8172994 | 0.3791326 | 0.832512355 |
| ENSMUSG00000031937 | Vstm5 | protein_coding | -0.324715256 | 4.592154197 | 0.81691 | 0.3792435 | 0.83258581 |
| ENSMUSG00000022110 | Sucla2 | protein_coding | 0.192760071 | 8.265718248 | 0.8158853 | 0.3795354 | 0.832694021 |
| ENSMUSG00000025817 | Nudt5 | protein_coding | 0.231838666 | 5.502467842 | 0.8155858 | 0.3796208 | 0.832694021 |
| ENSMUSG00000028393 | Alad | protein_coding | -0.174278451 | 6.709993692 | 0.8155186 | 0.37964 | 0.832694021 |
| ENSMUSG00000001909 | Trmt1 | protein_coding | -0.305118347 | 5.004510961 | 0.8154711 | 0.3796535 | 0.832694021 |
| ENSMUSG00000020366 | Mapk9 | protein_coding | 0.178500414 | 6.710022446 | 0.8153784 | 0.37968 | 0.832694021 |
| ENSMUSG00000032392 | Parp16 | protein_coding | -0.602131886 | 2.656371158 | 0.8146891 | 0.3798767 | 0.832926594 |
| ENSMUSG00000041607 | Mbp | protein_coding | -0.251742676 | 11.80476767 | 0.8145392 | 0.3799409 | 0.832926594 |
| ENSMUSG00000028869 | Gnl2 | protein_coding | 0.371908503 | 5.825993876 | 0.8181625 | 0.3801935 | 0.833071297 |
| ENSMUSG00000028999 | Rint1 | protein_coding | 0.301083449 | 3.966373281 | 0.8131852 | 0.3803063 | 0.833071297 |
| ENSMUSG00000019861 | Gopc | protein_coding | 0.249155362 | 4.726471528 | 0.8129354 | 0.3803778 | 0.833071297 |
| ENSMUSG00000036372 | Tmem258 | protein_coding | -0.26686084 | 7.304822578 | 0.815308 | 0.3805323 | 0.833071297 |
| ENSMUSG00000033319 | Fem1c | protein_coding | -0.332767958 | 3.986097007 | 0.8120954 | 0.3806182 | 0.833071297 |
| ENSMUSG00000030079 | Ruvbl1 | protein_coding | 0.260171451 | 5.052224155 | 0.8120585 | 0.3806287 | 0.833071297 |
| ENSMUSG00000072214 | 5-Sep | protein_coding | 0.195966751 | 7.012611073 | 0.811773 | 0.3807105 | 0.833071297 |
| ENSMUSG00000038502 | Ptov1 | protein_coding | 0.224008019 | 6.60639287 | 0.8108566 | 0.380973 | 0.833071297 |
| ENSMUSG00000031155 | Pim2 | protein_coding | 0.39068487 | 5.437067109 | 0.8143181 | 0.3810806 | 0.833071297 |
| ENSMUSG00000033020 | Polr2f | protein_coding | -0.205441318 | 7.719445482 | 0.8104699 | 0.3810839 | 0.833071297 |
| ENSMUSG00000030235 | Slco1c1 | protein_coding | 0.372597252 | 3.677615317 | 0.8104689 | 0.3810842 | 0.833071297 |
| ENSMUSG00000001054 | Rmnd5b | protein_coding | 0.233560035 | 5.63977067 | 0.8103811 | 0.3811094 | 0.833071297 |
| ENSMUSG00000019338 | Zfp687 | protein_coding | -0.329092456 | 3.918077177 | 0.8103551 | 0.3811169 | 0.833071297 |
| ENSMUSG00000045009 | Prrt3 | protein_coding | -0.558037225 | 2.80428381 | 0.810312 | 0.3811292 | 0.833071297 |
| ENSMUSG00000050855 | Zfp940 | protein_coding | -0.31327448 | 4.580419847 | 0.8101732 | 0.381169 | 0.833071297 |
| ENSMUSG00000050605 | Zfp61 | protein_coding | 0.338178003 | 4.759084114 | 0.8091649 | 0.3814585 | 0.833251581 |
| ENSMUSG00000035062 | Zc4h2 | protein_coding | -0.240008674 | 5.682403937 | 0.8091622 | 0.3814593 | 0.833251581 |
| ENSMUSG00000027971 | Ndst4 | protein_coding | 0.390064947 | 3.514078181 | 0.8090761 | 0.381484 | 0.833251581 |
| ENSMUSG00000054477 | Kcnn2 | protein_coding | -0.340372142 | 3.612487642 | 0.8087725 | 0.3815712 | 0.833272832 |
| ENSMUSG00000035529 | Prdm4 | protein_coding | 0.404658903 | 3.940585413 | 0.8080504 | 0.3817788 | 0.833434367 |
| ENSMUSG00000026820 | Ptges2 | protein_coding | -0.236609254 | 5.29739169 | 0.807976 | 0.3818002 | 0.833434367 |
| ENSMUSG00000026254 | Eif4e2 | protein_coding | 0.197229221 | 6.461110761 | 0.8071942 | 0.3820252 | 0.833756186 |
| ENSMUSG00000034532 | Fbxo16 | protein_coding | -0.26212814 | 5.05428201 | 0.8056959 | 0.3824569 | 0.834313953 |
| ENSMUSG00000019943 | Atp2b1 | protein_coding | -0.206305114 | 8.203782665 | 0.8056855 | 0.3824598 | 0.834313953 |
| ENSMUSG00000039738 | Slx4 | protein_coding | 0.50683628 | 2.92523349 | 0.8052694 | 0.3825799 | 0.834313953 |
| ENSMUSG00000051727 | Kctd14 | protein_coding | 0.751984679 | 2.14395777 | 0.8051073 | 0.3826266 | 0.834313953 |
| ENSMUSG00000030465 | Psd3 | protein_coding | -0.192148449 | 6.568826009 | 0.8046322 | 0.3827638 | 0.834313953 |
| ENSMUSG00000001415 | Smg5 | protein_coding | -0.265608281 | 4.749983484 | 0.8044693 | 0.3828108 | 0.834313953 |
| ENSMUSG00000025980 | Hspd1 | protein_coding | 0.183314291 | 7.043086391 | 0.804424 | 0.3828239 | 0.834313953 |
| ENSMUSG00000097166 | 9330179D12Rik | antisense_RNA | -0.431487765 | 3.657326077 | 0.8039369 | 0.3829646 | 0.834451469 |
| ENSMUSG00000020219 | Timm13 | protein_coding | -0.174534784 | 8.92112637 | 0.8034877 | 0.3830944 | 0.834565257 |
| ENSMUSG00000028373 | Astn2 | protein_coding | 0.270873124 | 5.382908985 | 0.8025533 | 0.3833647 | 0.834932978 |
| ENSMUSG00000027624 | Epb41l1 | protein_coding | 0.194354499 | 6.805994717 | 0.8023673 | 0.3834185 | 0.834932978 |
| ENSMUSG00000053293 | Pom121 | protein_coding | 0.395986633 | 4.121406808 | 0.8014931 | 0.3836716 | 0.835315079 |
| ENSMUSG00000036206 | Sh3bp4 | protein_coding | 0.424480402 | 3.751942726 | 0.8010852 | 0.3837898 | 0.835403258 |
| ENSMUSG00000040280 | Ndufa4l2 | protein_coding | -0.619344275 | 2.881207013 | 0.8000058 | 0.3841029 | 0.835691544 |
| ENSMUSG00000038888 | Ctu1 | protein_coding | -0.608768262 | 3.035940077 | 0.7995444 | 0.3842581 | 0.835691544 |
| ENSMUSG00000051185 | Fam174a | protein_coding | -0.257744766 | 5.226008723 | 0.7994591 | 0.3842616 | 0.835691544 |
| ENSMUSG00000031365 | Zfp275 | protein_coding | 0.307033541 | 4.555549452 | 0.7990993 | 0.3843661 | 0.835691544 |
| ENSMUSG00000034032 | Rpap1 | protein_coding | -0.400339864 | 3.352133866 | 0.7988824 | 0.3844291 | 0.835691544 |
| ENSMUSG00000015085 | Entpd2 | protein_coding | -0.475164367 | 2.870821829 | 0.7987077 | 0.3844799 | 0.835691544 |
| ENSMUSG00000036264 | Fstl4 | protein_coding | -0.487573543 | 3.592194445 | 0.79859 | 0.3845354 | 0.835691544 |
| ENSMUSG00000021478 | Drd1 | protein_coding | 0.305585947 | 5.486676655 | 0.7983263 | 0.384612 | 0.835691544 |
| ENSMUSG00000041695 | Kcnj2 | protein_coding | 0.899609869 | 2.236459618 | 0.7982929 | 0.3846217 | 0.835691544 |
| ENSMUSG00000033068 | Entpd6 | protein_coding | 0.179507718 | 6.758340762 | 0.7978746 | 0.3847221 | 0.835740849 |
| ENSMUSG00000087549 | Gm13427 | antisense_RNA | -0.380547798 | 6.941726025 | 0.8095195 | 0.3849898 | 0.836153468 |
| ENSMUSG00000041319 | Thoc6 | protein_coding | 0.4120896 | 4.222495636 | 0.7965641 | 0.3851249 | 0.836277885 |
| ENSMUSG00000016253 | Nelfcd | protein_coding | 0.251917569 | 6.811902554 | 0.7958033 | 0.3854025 | 0.836648108 |
| ENSMUSG00000027651 | Rprd1b | protein_coding | -0.32433845 | 4.07536025 | 0.7950873 | 0.3855343 | 0.836648108 |
| ENSMUSG00000039128 | Cdc123 | protein_coding | -0.179529981 | 8.75466243 | 0.7947274 | 0.3856394 | 0.836648108 |
| ENSMUSG00000038256 | Bcl9 | protein_coding | 0.353955203 | 4.243007085 | 0.7940423 | 0.3858394 | 0.836648108 |
| ENSMUSG00000024286 | Ccny | protein_coding | -0.334991565 | 5.07855017 | 0.7940812 | 0.3858493 | 0.836648108 |
| ENSMUSG00000027722 | Spata5 | protein_coding | 0.3845099 | 3.663025969 | 0.7939396 | 0.3858695 | 0.836648108 |

| ensembl_gene_id | external_gene_name | gene_biotype | log2 (fold change) | log2 (counts per million) | F-value | p-value | False discovery rate |
|---|---|---|---|---|---|---|---|
| ENSMUSG00000004994 | Ccdc130 | protein_coding | 0.24507065 | 5.078397919 | 0.7936974 | 0.3859403 | 0.836648108 |
| ENSMUSG00000001786 | Fbxo7 | protein_coding | 0.247591277 | 5.935291912 | 0.7936288 | 0.3859603 | 0.836648108 |
| ENSMUSG00000041297 | Cdk13 | protein_coding | 0.386516994 | 4.689720122 | 0.7937712 | 0.3859957 | 0.836648108 |
| ENSMUSG00000003929 | Zfp81 | protein_coding | -0.436187128 | 3.661235405 | 0.7926931 | 0.386234 | 0.836914212 |
| ENSMUSG00000044447 | Dock5 | protein_coding | -0.637338234 | 2.442328391 | 0.7920289 | 0.3864284 | 0.836914212 |
| ENSMUSG00000052928 | Ctif | protein_coding | 0.270748964 | 4.956431126 | 0.7919748 | 0.3864442 | 0.836914212 |
| ENSMUSG00000040624 | Plekhg1 | protein_coding | -0.380206127 | 3.843939393 | 0.7910548 | 0.3867138 | 0.836914212 |
| ENSMUSG00000030216 | Wbp11 | protein_coding | 0.186928785 | 6.936768788 | 0.7910267 | 0.3867221 | 0.836914212 |
| ENSMUSG00000017466 | Timp2 | protein_coding | 0.167514999 | 8.165904532 | 0.7907628 | 0.3867995 | 0.836914212 |
| ENSMUSG00000002844 | Adprh | protein_coding | 0.187464314 | 7.485039426 | 0.7904923 | 0.3868788 | 0.836914212 |
| ENSMUSG00000061286 | Exosc5 | protein_coding | -0.203537109 | 6.021888694 | 0.7903559 | 0.3869188 | 0.836914212 |
| ENSMUSG00000086316 | 2210013O21Rik | protein_coding | 0.150034329 | 7.655400986 | 0.7903076 | 0.386933 | 0.836914212 |
| ENSMUSG00000036158 | Prickle1 | protein_coding | 0.284299817 | 4.668239792 | 0.7902164 | 0.3869597 | 0.836914212 |
| ENSMUSG00000023072 | Cep89 | protein_coding | 0.377148555 | 3.758035032 | 0.7898852 | 0.387057 | 0.836914212 |
| ENSMUSG00000068762 | Gstm6 | protein_coding | 0.200589672 | 6.187682757 | 0.789381 | 0.387205 | 0.836914212 |
| ENSMUSG00000040225 | Prrc2c | protein_coding | 0.208649143 | 6.235597442 | 0.7893075 | 0.3872266 | 0.836914212 |
| ENSMUSG00000018417 | Myo1b | protein_coding | -0.307845816 | 3.79058272 | 0.7892432 | 0.3872455 | 0.836914212 |
| ENSMUSG00000029171 | Pgm1 | protein_coding | -0.556653317 | 2.853407313 | 0.7891057 | 0.3872859 | 0.836914212 |
| ENSMUSG00000043019 | Edem3 | protein_coding | -0.41738089 | 3.210635039 | 0.7885431 | 0.3874512 | 0.837103328 |
| ENSMUSG00000049971 | Glt1d1 | protein_coding | -0.455773098 | 3.186405473 | 0.78654 | 0.3880409 | 0.838200614 |
| ENSMUSG00000026439 | Rbbp5 | protein_coding | 0.253752747 | 5.231769556 | 0.7858325 | 0.3882495 | 0.838200614 |
| ENSMUSG00000037712 | Fermt2 | protein_coding | -0.239753354 | 5.541511576 | 0.785633 | 0.3883083 | 0.838200614 |
| ENSMUSG00000031453 | Rasa3 | protein_coding | 0.26095639 | 4.952652906 | 0.7855119 | 0.388344 | 0.838200614 |
| ENSMUSG00000021721 | Htr1a | protein_coding | 0.339372302 | 4.1537979 | 0.7852805 | 0.3884123 | 0.838200614 |
| ENSMUSG00000034071 | Zfp551 | protein_coding | 0.919190549 | 2.569405987 | 0.7879507 | 0.3884427 | 0.838200614 |
| ENSMUSG00000066415 | Msl2 | protein_coding | -0.356552532 | 4.015005728 | 0.7848514 | 0.388539 | 0.838200614 |
| ENSMUSG00000026766 | Mmadhc | protein_coding | 0.189438992 | 6.389745668 | 0.7844052 | 0.3886707 | 0.838200614 |
| ENSMUSG00000036606 | Plxnb2 | protein_coding | -0.418324784 | 3.938736052 | 0.784083 | 0.388787 | 0.838200614 |
| ENSMUSG00000062866 | Phactr2 | protein_coding | -0.242769178 | 5.45711007 | 0.783975 | 0.3887979 | 0.838200614 |
| ENSMUSG00000046854 | Pip5kl1 | protein_coding | 1.072782406 | 2.460729268 | 0.7879398 | 0.3889091 | 0.838200614 |
| ENSMUSG00000028648 | Ndufs5 | protein_coding | -0.175620603 | 7.458033273 | 0.7827052 | 0.3891734 | 0.838200614 |
| ENSMUSG00000060904 | Arl1 | protein_coding | -0.167769934 | 6.881057215 | 0.7825728 | 0.3892126 | 0.838200614 |
| ENSMUSG00000086272 | BC039966 | antisense_RNA | 0.453199981 | 3.834084902 | 0.7824156 | 0.3892802 | 0.838200614 |
| ENSMUSG00000029432 | Nipsnap2 | protein_coding | 0.233489052 | 8.179459623 | 0.7823267 | 0.3893065 | 0.838200614 |
| ENSMUSG00000060519 | Tor3a | protein_coding | 0.614329484 | 2.469766756 | 0.7821178 | 0.3893473 | 0.838200614 |
| ENSMUSG00000026749 | Nek6 | protein_coding | 0.323123059 | 4.331659509 | 0.7820524 | 0.3893667 | 0.838200614 |
| ENSMUSG00000030854 | Ptpn5 | protein_coding | -0.175046497 | 8.552007529 | 0.7818326 | 0.3894318 | 0.838200614 |
| ENSMUSG00000058056 | Palld | protein_coding | 0.47421073 | 3.232377554 | 0.7818044 | 0.3894402 | 0.838200614 |
| ENSMUSG00000022704 | Qtrt2 | protein_coding | 0.405893533 | 3.32029933 | 0.7809027 | 0.3897075 | 0.838582677 |
| ENSMUSG00000025040 | Fundc1 | protein_coding | -0.181320587 | 7.658821387 | 0.7805384 | 0.3898156 | 0.838582677 |
| ENSMUSG00000032228 | Tcf12 | protein_coding | -0.285982306 | 4.905642339 | 0.7804168 | 0.3898517 | 0.838582677 |
| ENSMUSG00000025158 | Rfng | protein_coding | 0.310627562 | 4.553115981 | 0.7791575 | 0.3902256 | 0.838636558 |
| ENSMUSG00000024943 | Smc5 | protein_coding | -0.372636624 | 4.320390629 | 0.7789926 | 0.3902746 | 0.838636558 |
| ENSMUSG00000031949 | Adat1 | protein_coding | -0.377018262 | 3.422585029 | 0.778898 | 0.3903028 | 0.838636558 |
| ENSMUSG00000039068 | Zzz3 | protein_coding | 0.290557942 | 4.790596783 | 0.7787106 | 0.3903585 | 0.838636558 |
| ENSMUSG00000024222 | Fkbp5 | protein_coding | 0.337149345 | 4.536240414 | 0.7780938 | 0.3905419 | 0.838636558 |
| ENSMUSG00000020918 | Kat2a | protein_coding | 0.22729432 | 5.123732653 | 0.7780333 | 0.3905599 | 0.838636558 |
| ENSMUSG00000048776 | Pthlh | protein_coding | 0.696172091 | 2.503974901 | 0.7779635 | 0.3905807 | 0.838636558 |
| ENSMUSG00000041774 | Ydjc | protein_coding | 0.457873304 | 3.276588869 | 0.7778139 | 0.3906252 | 0.838636558 |
| ENSMUSG00000033439 | Trmt13 | protein_coding | -0.509280455 | 2.783158122 | 0.7777097 | 0.3906563 | 0.838636558 |
| ENSMUSG00000000563 | Atp5f1 | protein_coding | 0.158951529 | 9.728584423 | 0.7777085 | 0.3906566 | 0.838636558 |
| ENSMUSG00000021288 | Klc1 | protein_coding | 0.199654821 | 7.094391431 | 0.7772735 | 0.3907861 | 0.838747158 |
| ENSMUSG00000005505 | Kbtbd4 | protein_coding | 0.390702639 | 3.955570353 | 0.7768159 | 0.3909225 | 0.838872341 |
| ENSMUSG00000056305 | Usp39 | protein_coding | 0.272504741 | 4.208385166 | 0.7761473 | 0.3911218 | 0.838941713 |
| ENSMUSG00000031095 | Cul4b | protein_coding | -0.376789008 | 3.859403634 | 0.7760529 | 0.3911499 | 0.838941713 |
| ENSMUSG00000084808 | 9430091E24Rik | processed_transcript | 0.366932939 | 3.72286899 | 0.7754764 | 0.3913219 | 0.838941713 |
| ENSMUSG00000031403 | Dkc1 | protein_coding | -0.210672326 | 5.864226478 | 0.7753338 | 0.3913645 | 0.838941713 |
| ENSMUSG00000009905 | Kdsr | protein_coding | 0.232054216 | 5.742732626 | 0.775166 | 0.3914146 | 0.838941713 |
| ENSMUSG00000003581 | Rnf215 | protein_coding | 0.341388688 | 3.454155927 | 0.7743785 | 0.3916498 | 0.838941713 |
| ENSMUSG00000050074 | Spink8 | protein_coding | -0.49414691 | 3.449041662 | 0.7744093 | 0.3916615 | 0.838941713 |
| ENSMUSG00000021114 | Atp6v1d | protein_coding | 0.159668786 | 9.903445503 | 0.7742049 | 0.3917017 | 0.838941713 |
| ENSMUSG00000028284 | Map3k7 | protein_coding | -0.327750536 | 3.835597982 | 0.7739709 | 0.3917716 | 0.838941713 |
| ENSMUSG00000032598 | Nckipsd | protein_coding | 0.223911015 | 5.747788628 | 0.7739077 | 0.3917905 | 0.838941713 |
| ENSMUSG00000021773 | Comtd1 | protein_coding | 0.467365394 | 3.079362413 | 0.7738325 | 0.391813 | 0.838941713 |
| ENSMUSG00000048148 | Nwd1 | protein_coding | -0.393676047 | 3.57062 | 0.7731688 | 0.3920116 | 0.839038711 |
| ENSMUSG00000027018 | Hat1 | protein_coding | -0.221120393 | 5.333454055 | 0.7730982 | 0.3920327 | 0.839038711 |
| ENSMUSG00000019814 | Ltv1 | protein_coding | 0.2960805 | 4.246042227 | 0.7723566 | 0.3922547 | 0.839038711 |
| ENSMUSG00000038811 | Gngt2 | protein_coding | -0.537569116 | 2.841497445 | 0.7723508 | 0.3922564 | 0.839038711 |
| ENSMUSG00000049553 | Polr1a | protein_coding | -0.39433015 | 3.578568344 | 0.7718511 | 0.3924062 | 0.839038711 |
| ENSMUSG00000015839 | Nfe2l2 | protein_coding | -0.357198738 | 4.716166321 | 0.7718996 | 0.3924125 | 0.839038711 |
| ENSMUSG00000026389 | Steap3 | protein_coding | -0.724862733 | 2.068060088 | 0.7742281 | 0.3925168 | 0.839038711 |
| ENSMUSG00000022312 | Eif3h | protein_coding | -0.164829104 | 8.916982619 | 0.7714054 | 0.3925398 | 0.839038711 |
| ENSMUSG00000026404 | Ddx59 | protein_coding | -0.439802315 | 2.920960976 | 0.7713361 | 0.3925606 | 0.839038711 |
| ENSMUSG00000053279 | Aldh1a1 | protein_coding | 0.333660391 | 4.881994317 | 0.7706377 | 0.3927909 | 0.839139389 |

| ensembl_gene_id | external_gene_name | gene_biotype | log2 (fold change) | log2 (counts per million) | F-value | p-value | False discovery rate |
|---|---|---|---|---|---|---|---|
| ENSMUSG00000075232 | Amd1 | protein_coding | -0.287253892 | 3.991844883 | 0.7701367 | 0.3929205 | 0.839139389 |
| ENSMUSG00000059552 | Trp53 | protein_coding | -0.236853251 | 5.633688644 | 0.7699295 | 0.3929827 | 0.839139389 |
| ENSMUSG00000058897 | Col25a1 | protein_coding | -0.281347633 | 4.848781409 | 0.7697886 | 0.393025 | 0.839139389 |
| ENSMUSG00000031134 | Rbmx | protein_coding | -0.22769492 | 6.32331531 | 0.7697799 | 0.3930276 | 0.839139389 |
| ENSMUSG00000024666 | Tmem138 | protein_coding | 0.277512388 | 5.474530119 | 0.7696193 | 0.3930759 | 0.839139389 |
| ENSMUSG00000022364 | Tbc1d31 | protein_coding | 0.664938383 | 2.316712019 | 0.7692187 | 0.3931963 | 0.839229814 |
| ENSMUSG00000093803 | Ppp2r3d | protein_coding | -0.358269942 | 3.402143341 | 0.7679142 | 0.3935887 | 0.839650105 |
| ENSMUSG00000020477 | Mrps24 | protein_coding | -0.313051476 | 4.17725496 | 0.7678038 | 0.393622 | 0.839650105 |
| ENSMUSG00000029413 | Naaa | protein_coding | 0.362240544 | 4.870785773 | 0.7678547 | 0.3936275 | 0.839650105 |
| ENSMUSG00000002266 | Zim1 | protein_coding | -0.400050863 | 3.127123016 | 0.7671834 | 0.3938089 | 0.83987042 |
| ENSMUSG00000020684 | Rasl10b | protein_coding | -0.336581759 | 5.728973139 | 0.7695964 | 0.3938897 | 0.8398762 |
| ENSMUSG00000066406 | Akap13 | protein_coding | -0.452647815 | 3.010877153 | 0.7662974 | 0.394076 | 0.840106815 |
| ENSMUSG00000023800 | Tiam2 | protein_coding | 0.334525664 | 4.999777228 | 0.7650691 | 0.3944675 | 0.84048878 |
| ENSMUSG00000025364 | Pa2g4 | protein_coding | 0.188220601 | 7.354736714 | 0.7649407 | 0.3944855 | 0.84048878 |
| ENSMUSG00000084215 | Gm11814 | processed_pseudoge | 0.487067952 | 3.054759638 | 0.7646429 | 0.3945755 | 0.84048878 |
| ENSMUSG00000020027 | Socs2 | protein_coding | -0.388077147 | 3.119462797 | 0.7645479 | 0.3946042 | 0.84048878 |
| ENSMUSG00000042354 | Gnl3 | protein_coding | 0.245714289 | 5.427918584 | 0.7642709 | 0.394688 | 0.84048878 |
| ENSMUSG00000040007 | Bahd1 | protein_coding | 0.48351554 | 3.270660705 | 0.7641091 | 0.3947369 | 0.84048878 |
| ENSMUSG00000090698 | Apold1 | protein_coding | -0.51645189 | 3.3684765 | 0.7639286 | 0.3948122 | 0.84048878 |
| ENSMUSG00000034442 | Trmt5 | protein_coding | -0.388782014 | 4.143886005 | 0.763703 | 0.3948804 | 0.84048878 |
| ENSMUSG00000020526 | Znhit3 | protein_coding | 0.165408939 | 7.147062682 | 0.7612128 | 0.3956142 | 0.841663633 |
| ENSMUSG00000085609 | 1700016P03Rik | processed_transcript | -0.403949585 | 3.263727687 | 0.760477 | 0.3958375 | 0.841663633 |
| ENSMUSG00000097729 | 2310015A10Rik | processed_transcript | -0.449116863 | 3.141519068 | 0.7600549 | 0.3959657 | 0.841663633 |
| ENSMUSG00000027893 | Ahcyl1 | protein_coding | -0.169458556 | 7.54395055 | 0.759969 | 0.3959919 | 0.841663633 |
| ENSMUSG00000024953 | Prdx5 | protein_coding | 0.160818062 | 10.25842496 | 0.7598019 | 0.3960426 | 0.841663633 |
| ENSMUSG00000028629 | Exo5 | protein_coding | -0.495015046 | 4.006934492 | 0.7616265 | 0.3962911 | 0.841663633 |
| ENSMUSG00000022416 | Cacna1i | protein_coding | 0.512713278 | 2.987274636 | 0.7584658 | 0.396449 | 0.841663633 |
| ENSMUSG00000030847 | Bag3 | protein_coding | 0.51026169 | 2.816494219 | 0.7580084 | 0.3965883 | 0.841663633 |
| ENSMUSG00000038034 | Igsf8 | protein_coding | 0.195923637 | 5.971416614 | 0.757481 | 0.3967489 | 0.841663633 |
| ENSMUSG00000002032 | Tmem25 | protein_coding | 0.238651646 | 6.284978924 | 0.7573309 | 0.3967947 | 0.841663633 |
| ENSMUSG00000022257 | Laptm4b | protein_coding | 0.205019921 | 6.147958706 | 0.7572939 | 0.396806 | 0.841663633 |
| ENSMUSG00000019772 | Vip | protein_coding | 0.430683309 | 3.329568435 | 0.7571519 | 0.3968493 | 0.841663633 |
| ENSMUSG00000040797 | Iqsec3 | protein_coding | -0.283968234 | 5.413933185 | 0.7565108 | 0.3970448 | 0.841663633 |
| ENSMUSG00000000197 | Nalcn | protein_coding | -0.275674368 | 5.587749691 | 0.7564546 | 0.3970619 | 0.841663633 |
| ENSMUSG00000021209 | Ppp4r4 | protein_coding | 0.299570537 | 4.206087628 | 0.7564072 | 0.3970764 | 0.841663633 |
| ENSMUSG00000040734 | Ppp1r13l | protein_coding | -0.577683241 | 2.762359993 | 0.7562791 | 0.3971155 | 0.841663633 |
| ENSMUSG00000032850 | Rnft2 | protein_coding | 0.293346655 | 4.516265606 | 0.7558808 | 0.3972371 | 0.841663633 |
| ENSMUSG00000007815 | Rhoa | protein_coding | 0.239545832 | 5.819720118 | 0.7557564 | 0.397275 | 0.841663633 |
| ENSMUSG00000029047 | Pex10 | protein_coding | 0.239355938 | 4.967391868 | 0.7552323 | 0.3974351 | 0.841663633 |
| ENSMUSG00000020811 | Wscd1 | protein_coding | 0.22185549 | 5.535521252 | 0.755024 | 0.3974988 | 0.841663633 |
| ENSMUSG00000018470 | Kcnab3 | protein_coding | -0.3344044 | 3.594478776 | 0.7547152 | 0.3975932 | 0.841663633 |
| ENSMUSG00000028226 | Mmp16 | protein_coding | -0.335966703 | 4.054769991 | 0.7545628 | 0.3976398 | 0.841663633 |
| ENSMUSG00000026753 | Ppp6c | protein_coding | 0.170392601 | 6.718598413 | 0.7543658 | 0.3977 | 0.841663633 |
| ENSMUSG00000042406 | Atf4 | protein_coding | -0.174273065 | 7.274473083 | 0.7540658 | 0.3977918 | 0.841663633 |
| ENSMUSG00000031149 | Praf2 | protein_coding | 0.201220426 | 6.38582939 | 0.7539897 | 0.3978151 | 0.841663633 |
| ENSMUSG00000026077 | Npas2 | protein_coding | 0.321654187 | 3.989062889 | 0.7539725 | 0.3978203 | 0.841663633 |
| ENSMUSG00000043065 | Spice1 | protein_coding | 0.410463087 | 3.476077722 | 0.7533122 | 0.3980225 | 0.841663633 |
| ENSMUSG00000030393 | Zik1 | protein_coding | -0.39441799 | 3.691285119 | 0.7532612 | 0.3980381 | 0.841663633 |
| ENSMUSG00000033579 | Fa2h | protein_coding | -0.27107389 | 4.900477609 | 0.7530349 | 0.3981074 | 0.841663633 |
| ENSMUSG00000001520 | Nrip2 | protein_coding | 0.243937501 | 6.737210411 | 0.7530108 | 0.3981354 | 0.841663633 |
| ENSMUSG00000030255 | Sspn | protein_coding | 0.575851546 | 2.634035781 | 0.75277 | 0.3981885 | 0.841663633 |
| ENSMUSG00000016257 | Prelid3b | protein_coding | 0.168948567 | 7.019083206 | 0.7524427 | 0.3982888 | 0.841663633 |
| ENSMUSG00000027447 | Cst3 | protein_coding | -0.339983396 | 11.15737554 | 0.7578275 | 0.3984034 | 0.841663633 |
| ENSMUSG00000038122 | Tbc1d32 | protein_coding | -0.528273366 | 2.883341405 | 0.7519812 | 0.3984304 | 0.841663633 |
| ENSMUSG00000033910 | Gucy1a3 | protein_coding | 0.155068157 | 6.664581123 | 0.7514541 | 0.3985921 | 0.841663633 |
| ENSMUSG00000051166 | Eml5 | protein_coding | -0.323100557 | 3.710090778 | 0.7513705 | 0.3986177 | 0.841663633 |
| ENSMUSG00000021044 | Adck1 | protein_coding | 0.274565119 | 5.485596723 | 0.7512987 | 0.3986398 | 0.841663633 |
| ENSMUSG00000025144 | Cenpx | protein_coding | -0.204862441 | 6.391339461 | 0.7511601 | 0.3986823 | 0.841663633 |
| ENSMUSG00000114362 | AC124433.1 | antisense_RNA | -0.756866807 | 2.15005337 | 0.751112 | 0.3986971 | 0.841663633 |
| ENSMUSG00000039771 | Polr2j | protein_coding | -0.191628351 | 7.22821898 | 0.7508787 | 0.3987687 | 0.841663633 |
| ENSMUSG00000020956 | Dtd2 | protein_coding | 0.429235042 | 5.50627464 | 0.7573534 | 0.3987897 | 0.841663633 |
| ENSMUSG00000023961 | Enpp4 | protein_coding | 0.258930574 | 5.192161877 | 0.75068 | 0.3988298 | 0.841663633 |
| ENSMUSG00000032615 | Nt5m | protein_coding | 0.200048433 | 6.081307107 | 0.7505837 | 0.3988593 | 0.841663633 |
| ENSMUSG00000020514 | Mrpl22 | protein_coding | 0.191037599 | 6.224653797 | 0.7505282 | 0.3988764 | 0.841663633 |
| ENSMUSG00000035891 | Cerk | protein_coding | 0.288653303 | 4.706977048 | 0.7498211 | 0.3990937 | 0.841945196 |
| ENSMUSG00000028402 | Mpdz | protein_coding | 0.353078251 | 3.322529361 | 0.7495847 | 0.3991664 | 0.841945196 |
| ENSMUSG00000029551 | Psmg3 | protein_coding | 0.483027131 | 2.713556447 | 0.7485259 | 0.3994923 | 0.842467255 |
| ENSMUSG00000036769 | Wdr44 | protein_coding | -0.394617398 | 4.062501324 | 0.7482719 | 0.399591 | 0.842490771 |
| ENSMUSG00000031610 | Scrg1 | protein_coding | -0.553137674 | 5.703641688 | 0.7673018 | 0.3997525 | 0.842490771 |
| ENSMUSG00000029772 | Ahcyl2 | protein_coding | -0.206389134 | 5.592556027 | 0.7475302 | 0.3997991 | 0.842490771 |
| ENSMUSG00000022515 | Anks3 | protein_coding | 0.348304033 | 4.971730314 | 0.7472962 | 0.3998918 | 0.842490771 |
| ENSMUSG00000034486 | Gbx2 | protein_coding | -0.667422708 | 3.344052447 | 0.7512745 | 0.3998952 | 0.842490771 |
| ENSMUSG00000020614 | Fam20a | protein_coding | -0.66140352 | 2.756788209 | 0.7467577 | 0.4000579 | 0.842615618 |
| ENSMUSG00000039737 | Prkrip1 | protein_coding | 0.193001169 | 6.35533797 | 0.7460331 | 0.4002611 | 0.842615618 |

| ensembl_gene_id | external_gene_name | gene_biotype | log2 (fold change) | log2 (counts per million) | F-value | p-value | False discovery rate |
|---|---|---|---|---|---|---|---|
| ENSMUSG00000064145 | Arih2 | protein_coding | 0.264714799 | 4.752227447 | 0.7460113 | 0.4002678 | 0.842615618 |
| ENSMUSG00000109336 | Samd4b | protein_coding | -0.28238109 | 5.297529667 | 0.7457943 | 0.4003348 | 0.842615618 |
| ENSMUSG00000003348 | Mob3a | protein_coding | 0.618439509 | 2.797709961 | 0.7457574 | 0.4003462 | 0.842615618 |
| ENSMUSG00000028249 | Sdcbp | protein_coding | 0.167001328 | 8.821410057 | 0.7454974 | 0.4004266 | 0.842619781 |
| ENSMUSG00000050619 | Zscan29 | protein_coding | -0.577264803 | 2.68258433 | 0.7440822 | 0.4008643 | 0.842848738 |
| ENSMUSG00000040865 | Ino80d | protein_coding | 0.331302297 | 3.558900076 | 0.7439982 | 0.4008903 | 0.842848738 |
| ENSMUSG00000059187 | Fam19a1 | protein_coding | 0.309172412 | 4.408622389 | 0.7439002 | 0.4009207 | 0.842848738 |
| ENSMUSG00000013701 | Timm23 | protein_coding | 0.400782857 | 3.06188704 | 0.7438936 | 0.4009227 | 0.842848738 |
| ENSMUSG00000044792 | Isca1 | protein_coding | 0.179093065 | 6.796535888 | 0.7437259 | 0.4009747 | 0.842848738 |
| ENSMUSG00000035696 | Rnf38 | protein_coding | 0.324078134 | 4.031856508 | 0.7435135 | 0.4010404 | 0.842848738 |
| ENSMUSG00000048332 | Lhfp | protein_coding | -0.458643717 | 3.587935089 | 0.7433589 | 0.4011088 | 0.842848738 |
| ENSMUSG00000022131 | Gpr180 | protein_coding | 0.369997991 | 3.381276287 | 0.7430504 | 0.4011839 | 0.842848738 |
| ENSMUSG00000026640 | Plxna2 | protein_coding | 0.280376649 | 4.949512398 | 0.7428578 | 0.4012436 | 0.842848738 |
| ENSMUSG00000004748 | Mtfp1 | protein_coding | 0.178295151 | 7.955389561 | 0.7426141 | 0.4013192 | 0.842848738 |
| ENSMUSG00000051113 | Fam71e1 | protein_coding | -0.556160094 | 2.910561462 | 0.7412614 | 0.401739 | 0.843404737 |
| ENSMUSG00000020812 | 1810032O08Rik | protein_coding | -0.481097179 | 2.780781175 | 0.7412016 | 0.4017576 | 0.843404737 |
| ENSMUSG00000030760 | Acer3 | protein_coding | -0.230896067 | 4.766273208 | 0.7410031 | 0.4018193 | 0.843404737 |
| ENSMUSG00000052139 | Babam2 | protein_coding | 0.190381152 | 6.603073595 | 0.7400986 | 0.4021004 | 0.843703817 |
| ENSMUSG00000027366 | Sppl2a | protein_coding | 0.25875084 | 4.958975193 | 0.74004 | 0.4021187 | 0.843703817 |
| ENSMUSG00000017221 | Psmd3 | protein_coding | -0.232499829 | 5.503592171 | 0.7394199 | 0.4023116 | 0.843789762 |
| ENSMUSG00000059714 | Flot1 | protein_coding | 0.204424136 | 6.609385333 | 0.7393797 | 0.4023241 | 0.843789762 |
| ENSMUSG00000015377 | Dennd6b | protein_coding | 0.384742525 | 3.564111178 | 0.7391519 | 0.402395 | 0.843789762 |
| ENSMUSG00000029624 | Ptcd1 | protein_coding | 0.36585419 | 3.945842654 | 0.7376758 | 0.4028551 | 0.844589731 |
| ENSMUSG00000028868 | Wasf2 | protein_coding | 0.260501644 | 4.571624132 | 0.7372039 | 0.4030023 | 0.844612597 |
| ENSMUSG00000029621 | Arpc1a | protein_coding | 0.147684042 | 9.187904977 | 0.7371336 | 0.4030243 | 0.844612597 |
| ENSMUSG00000039850 | Endov | protein_coding | -0.261552695 | 4.222574245 | 0.7368858 | 0.4031016 | 0.844612597 |
| ENSMUSG00000028104 | Polr3gl | protein_coding | -0.329902837 | 3.652093066 | 0.736577 | 0.4031981 | 0.844634312 |
| ENSMUSG00000028399 | Ptprd | protein_coding | -0.19858154 | 6.425603442 | 0.7363496 | 0.4032691 | 0.844634312 |
| ENSMUSG00000038335 | Tsr1 | protein_coding | -0.255597851 | 5.146566259 | 0.7357707 | 0.40345 | 0.844848676 |
| ENSMUSG00000064120 | Mocs1 | protein_coding | 0.732730462 | 2.373372985 | 0.7351286 | 0.4036508 | 0.845024554 |
| ENSMUSG00000029614 | Rpl6 | protein_coding | 0.16059392 | 6.90047633 | 0.7349997 | 0.4036912 | 0.845024554 |
| ENSMUSG00000020562 | Efcab10 | protein_coding | 0.221339235 | 5.368158711 | 0.7345856 | 0.4038208 | 0.845039504 |
| ENSMUSG00000027099 | Mtx2 | protein_coding | -0.264333174 | 5.076598187 | 0.7342776 | 0.4039172 | 0.845039504 |
| ENSMUSG00000026018 | Ica1l | protein_coding | -0.258362467 | 4.956794094 | 0.7340982 | 0.4039734 | 0.845039504 |
| ENSMUSG00000033066 | Gas7 | protein_coding | -0.246034283 | 6.236110823 | 0.7340377 | 0.4040127 | 0.845039504 |
| ENSMUSG00000049532 | Sall2 | protein_coding | 0.209319106 | 5.735768004 | 0.7334854 | 0.4041654 | 0.845144201 |
| ENSMUSG00000029283 | Cdc7 | protein_coding | -0.57861738 | 2.676501028 | 0.7332955 | 0.404225 | 0.845144201 |
| ENSMUSG00000036850 | Mrpl41 | protein_coding | 0.15201998 | 8.315883781 | 0.7329215 | 0.4043423 | 0.845144201 |
| ENSMUSG00000042492 | Tbc1d10b | protein_coding | -0.218182974 | 5.264162678 | 0.7326488 | 0.4044278 | 0.845144201 |
| ENSMUSG00000034108 | Ccs | protein_coding | 0.185721263 | 6.176152604 | 0.7324199 | 0.4044996 | 0.845144201 |
| ENSMUSG00000042688 | Mapk6 | protein_coding | 0.299358701 | 5.407855678 | 0.7323713 | 0.4045352 | 0.845144201 |
| ENSMUSG00000029090 | Adgra3 | protein_coding | -0.832993408 | 2.282222908 | 0.7321238 | 0.4046129 | 0.845144201 |
| ENSMUSG00000022529 | Zfp263 | protein_coding | -0.290810582 | 3.891581902 | 0.7305757 | 0.4050791 | 0.84595377 |
| ENSMUSG00000028937 | Acot7 | protein_coding | 0.216591888 | 10.6617693 | 0.7297984 | 0.4053238 | 0.846195454 |
| ENSMUSG00000038290 | Smg6 | protein_coding | 0.268473189 | 4.756265061 | 0.729708 | 0.4053523 | 0.846195454 |
| ENSMUSG00000049265 | Kcnk3 | protein_coding | -0.266566833 | 4.463893453 | 0.7290757 | 0.4055514 | 0.846446935 |
| ENSMUSG00000051065 | Mb21d2 | protein_coding | 0.253780067 | 4.829982531 | 0.7287036 | 0.4056687 | 0.846460564 |
| ENSMUSG00000052419 | 2610001J05Rik | protein_coding | -0.164396442 | 7.768716308 | 0.7285555 | 0.4057154 | 0.846460564 |
| ENSMUSG00000022899 | Slc15a2 | protein_coding | -0.424619491 | 3.892640066 | 0.7276303 | 0.4060275 | 0.846947408 |
| ENSMUSG00000061322 | Dnaic1 | protein_coding | -0.352085069 | 3.965382689 | 0.7272156 | 0.4061383 | 0.846982437 |
| ENSMUSG00000036748 | Cuedc2 | protein_coding | -0.151187383 | 8.642863803 | 0.726773 | 0.4062781 | 0.846982437 |
| ENSMUSG00000031239 | Itm2a | protein_coding | -0.234260231 | 5.464426864 | 0.7262402 | 0.4064465 | 0.846982437 |
| ENSMUSG00000017715 | Pgs1 | protein_coding | 0.278602835 | 4.039491949 | 0.726164 | 0.4064706 | 0.846982437 |
| ENSMUSG00000004383 | Large1 | protein_coding | 0.222102951 | 5.054282434 | 0.7258963 | 0.4065553 | 0.846982437 |
| ENSMUSG00000030727 | Rabep2 | protein_coding | -0.40622588 | 3.876506452 | 0.7259395 | 0.4065608 | 0.846982437 |
| ENSMUSG00000022983 | Scaf4 | protein_coding | -0.558370331 | 3.146398055 | 0.7258323 | 0.4065957 | 0.846982437 |
| ENSMUSG00000074781 | Ube2n | protein_coding | -0.185569381 | 6.876333105 | 0.7247065 | 0.4069319 | 0.84748101 |
| ENSMUSG00000040236 | Trappc5 | protein_coding | 0.178361167 | 6.273591413 | 0.7245148 | 0.4069926 | 0.84748101 |
| ENSMUSG00000028233 | Tgs1 | protein_coding | -0.202593845 | 5.482856638 | 0.723605 | 0.4072811 | 0.84789704 |
| ENSMUSG00000036106 | Prr5 | protein_coding | -0.421708973 | 3.567331264 | 0.7233791 | 0.4073527 | 0.84789704 |
| ENSMUSG00000006464 | Bbs1 | protein_coding | 0.450356531 | 3.40991514 | 0.7229201 | 0.4074984 | 0.84789704 |
| ENSMUSG00000008435 | Rdh13 | protein_coding | 0.205323282 | 6.406501572 | 0.7228904 | 0.4075078 | 0.84789704 |
| ENSMUSG00000042520 | Ubap2l | protein_coding | 0.344823599 | 4.45753949 | 0.7220912 | 0.4077617 | 0.848105893 |
| ENSMUSG00000114309 | AC163038.1 | bidirectional_promoter | -0.520638612 | 3.07934206 | 0.7220777 | 0.407766 | 0.848105893 |
| ENSMUSG00000026172 | Bcs1l | protein_coding | 0.206115032 | 6.012627498 | 0.7207371 | 0.4081923 | 0.848448463 |
| ENSMUSG00000041133 | Smc1a | protein_coding | 0.221826353 | 6.176882888 | 0.7207152 | 0.4081993 | 0.848448463 |
| ENSMUSG00000041907 | Gpr45 | protein_coding | -0.388030165 | 3.999234521 | 0.7205974 | 0.4082368 | 0.848448463 |
| ENSMUSG00000041020 | Map7d2 | protein_coding | -0.144784466 | 7.664329673 | 0.7204364 | 0.4082881 | 0.848448463 |
| ENSMUSG00000022751 | Nit2 | protein_coding | 0.204522366 | 5.828133495 | 0.7203198 | 0.4083252 | 0.848448463 |
| ENSMUSG00000028443 | Nudt2 | protein_coding | -0.17372072 | 6.671103307 | 0.7199673 | 0.4084375 | 0.848498935 |
| ENSMUSG00000015850 | Adamtsl4 | protein_coding | -0.624360686 | 2.53228929 | 0.7197482 | 0.4085073 | 0.848498935 |
| ENSMUSG00000006360 | Crip1 | protein_coding | -0.398387799 | 4.025108628 | 0.7192257 | 0.4086939 | 0.848710485 |
| ENSMUSG00000059213 | Ddn | protein_coding | -0.369756172 | 5.670490936 | 0.7232623 | 0.4089775 | 0.848710485 |
| ENSMUSG00000038046 | Mrm3 | protein_coding | 0.206445497 | 5.840858269 | 0.7181498 | 0.4090171 | 0.848710485 |

| ensembl_gene_id | external_gene_name | gene_biotype | log2 (fold change) | log2 (counts per million) | F-value | p-value | False discovery rate |
|---|---|---|---|---|---|---|---|
| ENSMUSG00000030881 | Arfip2 | protein_coding | 0.179120134 | 6.476021251 | 0.718146 | 0.4090184 | 0.848710485 |
| ENSMUSG00000022828 | Gtf2e1 | protein_coding | 0.330787964 | 3.880361525 | 0.717666 | 0.4091717 | 0.848710485 |
| ENSMUSG00000033845 | Mrpl15 | protein_coding | 0.150020254 | 7.412366499 | 0.717661 | 0.4091733 | 0.848710485 |
| ENSMUSG00000061979 | Rcc1l | protein_coding | 0.587865177 | 3.091089863 | 0.7176404 | 0.4091999 | 0.848710485 |
| ENSMUSG00000029458 | Brap | protein_coding | 0.240041791 | 4.55708519 | 0.7174503 | 0.4092406 | 0.848710485 |
| ENSMUSG00000026563 | Tada1 | protein_coding | 0.541323776 | 2.606274316 | 0.7171322 | 0.4093423 | 0.848757664 |
| ENSMUSG00000020186 | Csrp2 | protein_coding | -0.248795866 | 4.560132812 | 0.7167722 | 0.4094574 | 0.848832666 |
| ENSMUSG00000027030 | Stk39 | protein_coding | -0.215481719 | 5.963510753 | 0.7161703 | 0.4096499 | 0.848929742 |
| ENSMUSG00000022415 | Syngr1 | protein_coding | 0.145365959 | 7.596962092 | 0.7161323 | 0.4096621 | 0.848929742 |
| ENSMUSG00000038539 | Atf5 | protein_coding | -0.543727555 | 2.495895301 | 0.7151958 | 0.409962 | 0.849100183 |
| ENSMUSG00000036555 | Iqce | protein_coding | 0.336405938 | 4.140155216 | 0.7151716 | 0.4099697 | 0.849100183 |
| ENSMUSG00000058297 | Spock2 | protein_coding | 0.158955932 | 7.721449421 | 0.7151358 | 0.4099812 | 0.849100183 |
| ENSMUSG00000010095 | Slc3a2 | protein_coding | 0.270422943 | 5.734786904 | 0.7141198 | 0.410327 | 0.84923808 |
| ENSMUSG00000042178 | Armc5 | protein_coding | 0.497196416 | 3.018035505 | 0.713511 | 0.4105024 | 0.84923808 |
| ENSMUSG00000041258 | Zfp236 | protein_coding | -0.27508616 | 3.922609002 | 0.7134806 | 0.4105122 | 0.84923808 |
| ENSMUSG00000063787 | Chchd1 | protein_coding | -0.184536665 | 8.251797352 | 0.7134622 | 0.4105181 | 0.84923808 |
| ENSMUSG00000021831 | Ero1l | protein_coding | 0.524380945 | 3.134148566 | 0.7133623 | 0.4105502 | 0.84923808 |
| ENSMUSG00000030683 | Sez6l2 | protein_coding | -0.165289523 | 7.401941601 | 0.7127935 | 0.4107329 | 0.84923808 |
| ENSMUSG00000068580 | Zfyve19 | protein_coding | -0.202516557 | 6.224034757 | 0.7124417 | 0.410846 | 0.84923808 |
| ENSMUSG00000097929 | Tunar | protein_coding | -0.260358448 | 4.680065021 | 0.7124226 | 0.4108521 | 0.84923808 |
| ENSMUSG00000087574 | C030037D09Rik | lincRNA | -0.390743066 | 3.542319725 | 0.7118521 | 0.4110356 | 0.84923808 |
| ENSMUSG00000067279 | Ppp1r3c | protein_coding | 0.302927633 | 4.881953259 | 0.7118255 | 0.4110441 | 0.84923808 |
| ENSMUSG00000020022 | Ndufa12 | protein_coding | 0.155506296 | 8.523937266 | 0.7116854 | 0.4110892 | 0.84923808 |
| ENSMUSG00000039070 | Cpa4 | protein_coding | 0.72407286 | 2.618529772 | 0.7117033 | 0.4111034 | 0.84923808 |
| ENSMUSG00000062646 | Ganc | protein_coding | -0.302604239 | 3.615466265 | 0.7116257 | 0.4111084 | 0.84923808 |
| ENSMUSG00000013646 | Sh3bp5l | protein_coding | -0.213057754 | 5.210169796 | 0.7114857 | 0.4111535 | 0.84923808 |
| ENSMUSG00000026644 | Acbd7 | protein_coding | 0.397912539 | 3.201097795 | 0.7112142 | 0.4112409 | 0.849255522 |
| ENSMUSG00000026471 | Mr1 | protein_coding | -1.065597587 | 2.113181124 | 0.7132585 | 0.4113465 | 0.849293605 |
| ENSMUSG00000007653 | Gabrb2 | protein_coding | 0.376024763 | 4.267264857 | 0.7107222 | 0.4114193 | 0.849293605 |
| ENSMUSG00000053297 | AI854703 | protein_coding | 0.24291603 | 5.33239503 | 0.7100443 | 0.4116179 | 0.849293605 |
| ENSMUSG00000029175 | Slc35f6 | protein_coding | 0.251185328 | 5.03880609 | 0.7098501 | 0.4116805 | 0.849293605 |
| ENSMUSG00000025917 | Cops5 | protein_coding | 0.17170015 | 7.789224821 | 0.709605 | 0.4117595 | 0.849293605 |
| ENSMUSG00000024608 | Rps14 | protein_coding | -0.257851655 | 10.3060541 | 0.7107364 | 0.4119034 | 0.849293605 |
| ENSMUSG00000006932 | Ctnnb1 | protein_coding | 0.423914385 | 3.323877475 | 0.7091407 | 0.4119093 | 0.849293605 |
| ENSMUSG00000002052 | Supt6 | protein_coding | 0.200601472 | 6.11979242 | 0.7087935 | 0.4120214 | 0.849293605 |
| ENSMUSG00000061665 | Cd2ap | protein_coding | -0.583258926 | 2.505229017 | 0.708751 | 0.4120352 | 0.849293605 |
| ENSMUSG00000035829 | Ppp1r26 | protein_coding | 0.340928625 | 3.6414229 | 0.7085388 | 0.4121037 | 0.849293605 |
| ENSMUSG00000030029 | Lrig1 | protein_coding | 0.397492407 | 3.127578595 | 0.7082708 | 0.4121903 | 0.849293605 |
| ENSMUSG00000038437 | Mllt6 | protein_coding | -0.387645562 | 3.657007758 | 0.7082187 | 0.4122071 | 0.849293605 |
| ENSMUSG00000020457 | Drg1 | protein_coding | 0.16051925 | 8.483408787 | 0.7077148 | 0.41237 | 0.849336184 |
| ENSMUSG00000041570 | Camsap2 | protein_coding | 0.378622422 | 3.322878241 | 0.707666 | 0.4123858 | 0.849336184 |
| ENSMUSG00000025898 | Cwf19l2 | protein_coding | 0.300271987 | 4.25217616 | 0.7072434 | 0.4125225 | 0.849455024 |
| ENSMUSG00000071477 | Zfp777 | protein_coding | -0.359388126 | 3.500699086 | 0.7065719 | 0.4127398 | 0.849739816 |
| ENSMUSG00000021910 | Nisch | protein_coding | 0.156284831 | 7.4462223 | 0.7060901 | 0.4128958 | 0.849898362 |
| ENSMUSG00000024259 | Slc25a46 | protein_coding | -0.216150429 | 5.508648155 | 0.7055288 | 0.4130778 | 0.850044659 |
| ENSMUSG00000034801 | Sos2 | protein_coding | -0.242636017 | 4.382323573 | 0.7053831 | 0.413125 | 0.850044659 |
| ENSMUSG00000020701 | Tmem132e | protein_coding | 0.401672188 | 3.462792717 | 0.7049645 | 0.4132608 | 0.850060337 |
| ENSMUSG00000063428 | Ddo | protein_coding | 0.271394153 | 4.108612222 | 0.7048721 | 0.4132908 | 0.850060337 |
| ENSMUSG00000084349 | Rpl3-ps1 | processed_pseudoge | 0.441914521 | 3.374979959 | 0.7044272 | 0.4134352 | 0.850160312 |
| ENSMUSG00000059361 | Nrsn2 | protein_coding | 0.387196075 | 3.125861112 | 0.704049 | 0.413558 | 0.850160312 |
| ENSMUSG00000027810 | Eif2a | protein_coding | -0.159306065 | 6.400073667 | 0.7038472 | 0.4136235 | 0.850160312 |
| ENSMUSG00000075705 | Msrb1 | protein_coding | -0.23867488 | 4.944493655 | 0.7034797 | 0.4137429 | 0.850160312 |
| ENSMUSG00000041977 | Arhgef11 | protein_coding | -0.437822912 | 3.297365252 | 0.7034606 | 0.4137491 | 0.850160312 |
| ENSMUSG00000002031 | Ift46 | protein_coding | 0.195988531 | 7.052698323 | 0.7030915 | 0.4138691 | 0.850160312 |
| ENSMUSG00000028893 | Sesn2 | protein_coding | 0.433313458 | 3.422479289 | 0.7030187 | 0.4138928 | 0.850160312 |
| ENSMUSG00000029761 | Cald1 | protein_coding | -0.414489256 | 4.789121677 | 0.7058221 | 0.4139729 | 0.850162531 |
| ENSMUSG00000024420 | Zfp521 | protein_coding | 0.316003488 | 4.269042558 | 0.7019318 | 0.4142465 | 0.850422414 |
| ENSMUSG00000023192 | Grm2 | protein_coding | 0.541124776 | 3.081456582 | 0.701733 | 0.4143112 | 0.850422414 |
| ENSMUSG00000091994 | E130317F20Rik | bidirectional_promoter | -0.559132418 | 2.538111552 | 0.7016547 | 0.4143367 | 0.850422414 |
| ENSMUSG00000019809 | Pex3 | protein_coding | -0.271755086 | 5.017499214 | 0.7013859 | 0.4144243 | 0.850439867 |
| ENSMUSG00000069895 | Atxn1l | protein_coding | -0.464673503 | 3.812984742 | 0.7022845 | 0.414506 | 0.850445056 |
| ENSMUSG00000031683 | Lsm6 | protein_coding | 0.187064079 | 7.894789187 | 0.7003301 | 0.4147686 | 0.850751127 |
| ENSMUSG00000031576 | Kcnu1 | protein_coding | 0.639704178 | 2.930505954 | 0.7002534 | 0.4148134 | 0.850751127 |
| ENSMUSG00000087497 | 2810001G20Rik | antisense_RNA | -0.254639128 | 4.972236344 | 0.6998662 | 0.41492 | 0.850807484 |
| ENSMUSG00000020495 | Smg8 | protein_coding | -0.330584463 | 3.617206367 | 0.6996033 | 0.4150058 | 0.850821234 |
| ENSMUSG00000063568 | Jazf1 | protein_coding | -0.244812783 | 4.730185123 | 0.6984088 | 0.4153962 | 0.851301839 |
| ENSMUSG00000040565 | Btaf1 | protein_coding | -0.373301217 | 3.801692078 | 0.6982525 | 0.4154473 | 0.851301839 |
| ENSMUSG00000014846 | Tppp3 | protein_coding | 0.195729794 | 8.447401721 | 0.6981594 | 0.4154777 | 0.851301839 |
| ENSMUSG00000020917 | Acly | protein_coding | 0.209336792 | 7.114004496 | 0.6975606 | 0.4156934 | 0.851581541 |
| ENSMUSG00000042523 | Dnal1 | protein_coding | -0.239553818 | 5.155331092 | 0.6971816 | 0.4157978 | 0.851633103 |
| ENSMUSG00000113902 | AC121965.1 | bidirectional_promoter | -0.221769382 | 9.115190095 | 0.6966496 | 0.4159918 | 0.851749364 |
| ENSMUSG00000021311 | Mtr | protein_coding | -0.44819898 | 3.017225109 | 0.6965249 | 0.416013 | 0.851749364 |
| ENSMUSG00000039982 | Dtx4 | protein_coding | 0.293180918 | 4.140135314 | 0.6959094 | 0.4162148 | 0.852000372 |
| ENSMUSG00000021972 | Hmbox1 | protein_coding | 0.333340079 | 3.662169594 | 0.6952566 | 0.416429 | 0.852104068 |

| ensembl_gene_id | external_gene_name | gene_biotype | log2 (fold change) | log2 (counts per million) | F-value | p-value | False discovery rate |
|---|---|---|---|---|---|---|---|
| ENSMUSG00000050428 | Fbxo46 | protein_coding | -0.243410773 | 4.897242929 | 0.6951837 | 0.416453 | 0.852104068 |
| ENSMUSG00000060679 | Mrps9 | protein_coding | 0.179623547 | 6.640901099 | 0.6947953 | 0.4165805 | 0.852104068 |
| ENSMUSG00000023018 | Smarcd1 | protein_coding | -0.192220942 | 6.309771219 | 0.6947895 | 0.4165824 | 0.852104068 |
| ENSMUSG00000055553 | Kxd1 | protein_coding | 0.158850002 | 7.517410548 | 0.6942095 | 0.416773 | 0.852331788 |
| ENSMUSG00000046311 | Zfp62 | protein_coding | -0.34377849 | 3.823587933 | 0.6939045 | 0.4168733 | 0.852374764 |
| ENSMUSG00000039501 | Znfx1 | protein_coding | 0.330324689 | 3.843822906 | 0.6933255 | 0.4170638 | 0.852591028 |
| ENSMUSG00000039942 | Ptger4 | protein_coding | -0.320175612 | 4.093155842 | 0.693101 | 0.4171377 | 0.852591028 |
| ENSMUSG00000029141 | Slc4a1ap | protein_coding | 0.250288666 | 4.239304345 | 0.6927225 | 0.4172623 | 0.852683648 |
| ENSMUSG00000045259 | Klhdc9 | protein_coding | -0.288795781 | 4.511997835 | 0.6917431 | 0.417585 | 0.852965365 |
| ENSMUSG00000022076 | Klhl1 | protein_coding | 0.329504716 | 4.332187391 | 0.6916119 | 0.4176283 | 0.852965365 |
| ENSMUSG00000026790 | Odf2 | protein_coding | 0.249664146 | 5.846598924 | 0.6915821 | 0.4176381 | 0.852965365 |
| ENSMUSG00000025880 | Smad7 | protein_coding | -0.383000577 | 3.443489442 | 0.69126 | 0.4177444 | 0.852994047 |
| ENSMUSG00000020659 | Cbll1 | protein_coding | -0.28330638 | 4.67701495 | 0.6910377 | 0.4178177 | 0.852994047 |
| ENSMUSG00000028920 | Fbxo42 | protein_coding | 0.319001877 | 3.880096418 | 0.6908183 | 0.4178901 | 0.852994047 |
| ENSMUSG00000006050 | Sra1 | protein_coding | -0.169427219 | 7.857434219 | 0.6898788 | 0.4182005 | 0.853092495 |
| ENSMUSG00000048960 | Prex2 | protein_coding | -0.67454642 | 3.109215943 | 0.6923902 | 0.4182076 | 0.853092495 |
| ENSMUSG00000001105 | Ift20 | protein_coding | -0.145972896 | 7.772655175 | 0.6895959 | 0.4182939 | 0.853092495 |
| ENSMUSG00000080268 | Brms1 | protein_coding | -0.173428191 | 6.284589523 | 0.6893409 | 0.4183783 | 0.853092495 |
| ENSMUSG00000021400 | Wrnip1 | protein_coding | 0.234381152 | 5.408688002 | 0.6892927 | 0.4183942 | 0.853092495 |
| ENSMUSG00000057367 | Birc2 | protein_coding | 0.269299517 | 5.390641583 | 0.6890005 | 0.4184909 | 0.853092495 |
| ENSMUSG00000047123 | Ticam1 | protein_coding | -0.584272365 | 2.958278918 | 0.6890512 | 0.4184937 | 0.853092495 |
| ENSMUSG00000020799 | Tekt1 | protein_coding | 0.300612175 | 4.468957633 | 0.687318 | 0.4190481 | 0.853807396 |
| ENSMUSG00000022090 | Pdlim2 | protein_coding | -0.188076065 | 6.69297983 | 0.6872082 | 0.4190845 | 0.853807396 |
| ENSMUSG00000025193 | Cutc | protein_coding | -0.264036927 | 4.378400117 | 0.686903 | 0.4191857 | 0.853807396 |
| ENSMUSG00000029613 | Eif2ak1 | protein_coding | 0.253423652 | 4.672484427 | 0.6865111 | 0.4193157 | 0.853807396 |
| ENSMUSG00000014402 | Tsg101 | protein_coding | 0.154456296 | 6.732788674 | 0.6864163 | 0.4193472 | 0.853807396 |
| ENSMUSG00000026153 | Fam135a | protein_coding | -0.498688005 | 3.403982412 | 0.6863489 | 0.4193891 | 0.853807396 |
| ENSMUSG00000042548 | Asxl1 | protein_coding | 0.322874155 | 3.71268328 | 0.6862565 | 0.4194002 | 0.853807396 |
| ENSMUSG00000047003 | Zfp41 | protein_coding | 0.668832594 | 2.686236001 | 0.6853308 | 0.4197273 | 0.854191251 |
| ENSMUSG00000038132 | Rbm24 | protein_coding | 0.287716216 | 4.232803002 | 0.6845414 | 0.4199702 | 0.854191251 |
| ENSMUSG00000028318 | Polr1e | protein_coding | 0.298910183 | 4.451103162 | 0.6844977 | 0.4199847 | 0.854191251 |
| ENSMUSG00000026880 | Stom | protein_coding | 0.539065292 | 2.608445582 | 0.6844068 | 0.420015 | 0.854191251 |
| ENSMUSG00000056598 | Drc3 | protein_coding | 0.337148084 | 3.995195366 | 0.6841753 | 0.420092 | 0.854191251 |
| ENSMUSG00000069835 | Sat2 | protein_coding | 0.207067748 | 5.743220845 | 0.6840087 | 0.4201475 | 0.854191251 |
| ENSMUSG00000032112 | Trappc4 | protein_coding | 0.142949488 | 8.58449716 | 0.6839425 | 0.4201695 | 0.854191251 |
| ENSMUSG00000063870 | Chd4 | protein_coding | 0.176595079 | 6.028503614 | 0.683778 | 0.4202243 | 0.854191251 |
| ENSMUSG00000035139 | Secisbp2 | protein_coding | 0.269721815 | 4.315297583 | 0.6828066 | 0.4205479 | 0.854667208 |
| ENSMUSG00000039178 | Tbc1d19 | protein_coding | 0.196619356 | 5.520886412 | 0.6824452 | 0.4206685 | 0.854667208 |
| ENSMUSG00000020530 | Ggnbp2 | protein_coding | 0.159507007 | 7.034951486 | 0.68236 | 0.4206969 | 0.854667208 |
| ENSMUSG00000020459 | Mtif2 | protein_coding | -0.232035799 | 5.176760627 | 0.6817913 | 0.4208866 | 0.854850331 |
| ENSMUSG00000034755 | Pcdh11x | protein_coding | -0.237333617 | 5.168562025 | 0.6814773 | 0.4209915 | 0.854850331 |
| ENSMUSG00000097413 | A830052D11Rik | lincRNA | 0.543275612 | 2.599959074 | 0.6813754 | 0.4210255 | 0.854850331 |
| ENSMUSG00000031843 | Mphosph6 | protein_coding | -0.206626472 | 6.021009943 | 0.6808097 | 0.4212145 | 0.855072601 |
| ENSMUSG00000028780 | Sema3c | protein_coding | 0.627612617 | 2.825147627 | 0.6805765 | 0.4213119 | 0.855108954 |
| ENSMUSG00000029048 | Rer1 | protein_coding | -0.182024778 | 6.511122146 | 0.6798776 | 0.4215262 | 0.85538238 |
| ENSMUSG00000083844 | Ube2d-ps | transcribed_unproces | -0.194834927 | 5.765932777 | 0.6793276 | 0.4217103 | 0.855482751 |
| ENSMUSG00000038371 | Sbf2 | protein_coding | -0.261487108 | 4.293764089 | 0.6792545 | 0.4217348 | 0.855482751 |
| ENSMUSG00000037890 | Wdr19 | protein_coding | 0.404663998 | 3.403430134 | 0.6783516 | 0.4220373 | 0.855930961 |
| ENSMUSG00000028211 | Trp53inp1 | protein_coding | 0.367960398 | 3.28170641 | 0.6778341 | 0.4222109 | 0.855930961 |
| ENSMUSG00000036992 | Nxt1 | protein_coding | 0.246154053 | 4.667078708 | 0.6777363 | 0.4222436 | 0.855930961 |
| ENSMUSG00000032280 | Tle3 | protein_coding | 0.462511241 | 3.152596638 | 0.6775139 | 0.4223183 | 0.855930961 |
| ENSMUSG00000037270 | 4932438A13Rik | protein_coding | 0.311471952 | 4.134086467 | 0.6773002 | 0.42239 | 0.855930961 |
| ENSMUSG00000063531 | Sema3e | protein_coding | -0.455999499 | 2.783433438 | 0.6771712 | 0.4224333 | 0.855930961 |
| ENSMUSG00000026754 | Golga1 | protein_coding | 0.220352258 | 4.866080909 | 0.67674 | 0.4225781 | 0.856063031 |
| ENSMUSG00000039205 | Ciz1 | protein_coding | -0.305549574 | 4.429933006 | 0.6764426 | 0.422678 | 0.856104168 |
| ENSMUSG00000028184 | Adgrl2 | protein_coding | 0.250328343 | 4.233915542 | 0.6755515 | 0.4229777 | 0.856230127 |
| ENSMUSG00000021178 | Psmc1 | protein_coding | 0.150670022 | 9.082673382 | 0.6754138 | 0.423024 | 0.856230127 |
| ENSMUSG00000004558 | Ndrg2 | protein_coding | 0.184272692 | 9.746559123 | 0.6748961 | 0.4231982 | 0.856230127 |
| ENSMUSG00000002985 | Apoe | protein_coding | -0.386531092 | 7.517200894 | 0.6894556 | 0.4233355 | 0.856230127 |
| ENSMUSG00000021497 | Txndc15 | protein_coding | -0.184436366 | 6.261456886 | 0.6744014 | 0.4233649 | 0.856230127 |
| ENSMUSG00000039252 | Lgi2 | protein_coding | 0.245702531 | 4.613793116 | 0.674371 | 0.4233751 | 0.856230127 |
| ENSMUSG00000032855 | Pkd1 | protein_coding | -0.507952562 | 2.826235212 | 0.6743149 | 0.423394 | 0.856230127 |
| ENSMUSG00000022744 | Cldnd1 | protein_coding | -0.153603074 | 7.721319446 | 0.6741729 | 0.4234418 | 0.856230127 |
| ENSMUSG00000027012 | Dync1i2 | protein_coding | -0.156170007 | 7.194547716 | 0.6739043 | 0.4235324 | 0.856230127 |
| ENSMUSG00000021143 | Pacs2 | protein_coding | -0.224910369 | 5.061377253 | 0.6737069 | 0.4235989 | 0.856230127 |
| ENSMUSG00000030138 | Bms1 | protein_coding | 0.368818373 | 3.655076753 | 0.6736559 | 0.4236161 | 0.856230127 |
| ENSMUSG00000020142 | Slc1a4 | protein_coding | 0.265604511 | 4.882143168 | 0.6731698 | 0.4237801 | 0.856400539 |
| ENSMUSG00000017754 | Pltp | protein_coding | -0.29705999 | 4.931605448 | 0.6726351 | 0.4239606 | 0.856554555 |
| ENSMUSG00000050705 | 2310061I04Rik | protein_coding | 0.201742526 | 5.638533226 | 0.6723951 | 0.4240416 | 0.856554555 |
| ENSMUSG00000029145 | Eif2b4 | protein_coding | 0.211820644 | 5.925867278 | 0.6722363 | 0.4240953 | 0.856554555 |
| ENSMUSG00000026223 | Itm2c | protein_coding | 0.18254781 | 9.311222768 | 0.67125 | 0.4244286 | 0.85706688 |
| ENSMUSG00000059890 | Ube4a | protein_coding | -0.310791525 | 3.81874335 | 0.6708114 | 0.424577 | 0.857205594 |
| ENSMUSG00000061758 | Akr1b10 | protein_coding | 0.209352133 | 5.577998856 | 0.6698128 | 0.4249152 | 0.857727238 |
| ENSMUSG00000027215 | Cd82 | protein_coding | -0.212963105 | 6.399149265 | 0.6695214 | 0.425014 | 0.857765603 |

| ensembl_gene_id | external_gene_name | gene_biotype | log2 (fold change) | log2 (counts per million) | F-value | p-value | False discovery rate |
|---|---|---|---|---|---|---|---|
| ENSMUSG00000016831 | Tox4 | protein_coding | 0.21273218 | 6.124848092 | 0.6692071 | 0.4251205 | 0.857773248 |
| ENSMUSG00000033083 | Tbc1d4 | protein_coding | -0.382952066 | 3.271941174 | 0.6690397 | 0.4251773 | 0.857773248 |
| ENSMUSG00000026107 | Nabp1 | protein_coding | -0.360571149 | 3.476302538 | 0.6680973 | 0.4254971 | 0.858257441 |
| ENSMUSG00000029330 | Cds1 | protein_coding | 0.261164933 | 4.559815468 | 0.6674531 | 0.425716 | 0.858357103 |
| ENSMUSG00000073889 | Il11ra1 | protein_coding | 0.28784194 | 3.906025596 | 0.6672874 | 0.4257723 | 0.858357103 |
| ENSMUSG00000037300 | Ttc13 | protein_coding | 0.448436612 | 3.753155692 | 0.6672256 | 0.4258633 | 0.858357103 |
| ENSMUSG00000063281 | Zfp35 | protein_coding | -0.247750873 | 5.15766405 | 0.6668466 | 0.4259221 | 0.858357103 |
| ENSMUSG00000058587 | Tmod3 | protein_coding | -0.168506302 | 6.482394661 | 0.6667775 | 0.4259456 | 0.858357103 |
| ENSMUSG00000049091 | Sephs2 | protein_coding | 0.318276578 | 5.725464198 | 0.6686772 | 0.4260473 | 0.858401169 |
| ENSMUSG00000037625 | Cldn11 | protein_coding | -0.147761102 | 7.407840448 | 0.6661345 | 0.4261644 | 0.858444411 |
| ENSMUSG00000030551 | Nr2f2 | protein_coding | -0.277405423 | 6.558364138 | 0.6684242 | 0.4263722 | 0.858444411 |
| ENSMUSG00000003378 | Grik5 | protein_coding | -0.427506209 | 3.244811165 | 0.6650414 | 0.4265368 | 0.858444411 |
| ENSMUSG00000043456 | Zfp536 | protein_coding | -0.393663521 | 3.772374222 | 0.6649303 | 0.4265747 | 0.858444411 |
| ENSMUSG00000030533 | Unc45a | protein_coding | -0.276552016 | 4.486014819 | 0.6648844 | 0.4265903 | 0.858444411 |
| ENSMUSG00000051306 | Usp42 | protein_coding | 0.299230284 | 4.038454282 | 0.66475 | 0.4266362 | 0.858444411 |
| ENSMUSG00000024208 | Uqcc2 | protein_coding | -0.172173949 | 9.520094615 | 0.6644213 | 0.4267483 | 0.858444411 |
| ENSMUSG00000038020 | Rapgefl1 | protein_coding | 0.265688836 | 5.041302395 | 0.6642816 | 0.426796 | 0.858444411 |
| ENSMUSG00000040640 | Erc2 | protein_coding | -0.198959121 | 6.308424434 | 0.6642143 | 0.4268189 | 0.858444411 |
| ENSMUSG00000061589 | Dot1l | protein_coding | 0.355368314 | 3.989008239 | 0.664073 | 0.4268671 | 0.858444411 |
| ENSMUSG00000039477 | Tnrc18 | protein_coding | 0.32348949 | 3.305940623 | 0.6631593 | 0.4271792 | 0.858473971 |
| ENSMUSG00000030089 | Slc41a3 | protein_coding | 0.420116159 | 3.496801846 | 0.6631455 | 0.4271839 | 0.858473971 |
| ENSMUSG00000028583 | Pdpn | protein_coding | -0.348150464 | 4.183131412 | 0.663054 | 0.4272151 | 0.858473971 |
| ENSMUSG00000047606 | Ankrd34c | protein_coding | -0.377941227 | 3.019823095 | 0.6630184 | 0.4272273 | 0.858473971 |
| ENSMUSG00000097119 | B230354K17Rik | lincRNA | -0.389330366 | 3.440706363 | 0.6626536 | 0.427352 | 0.858473971 |
| ENSMUSG00000057335 | Cep170 | protein_coding | 0.217649447 | 5.24209487 | 0.6623216 | 0.4274656 | 0.858473971 |
| ENSMUSG00000057440 | Mpp7 | protein_coding | -0.395698184 | 3.335569132 | 0.6622482 | 0.4274907 | 0.858473971 |
| ENSMUSG00000066196 | Spag8 | protein_coding | 0.412733453 | 3.189612317 | 0.6620003 | 0.4275755 | 0.858473971 |
| ENSMUSG00000031068 | Glrx3 | protein_coding | -0.159380184 | 6.39204857 | 0.6619276 | 0.4276004 | 0.858473971 |
| ENSMUSG00000013593 | Ndufs2 | protein_coding | 0.162897702 | 8.539366537 | 0.6612729 | 0.4278245 | 0.85876367 |
| ENSMUSG00000026608 | Kctd3 | protein_coding | 0.288873288 | 4.791770273 | 0.6608692 | 0.4279628 | 0.858828148 |
| ENSMUSG00000020413 | Hus1 | protein_coding | 0.427055216 | 3.090170719 | 0.6607129 | 0.4280164 | 0.858828148 |
| ENSMUSG00000022517 | Mgrn1 | protein_coding | 0.354599831 | 6.009870723 | 0.6656318 | 0.428155 | 0.858945981 |
| ENSMUSG00000042439 | Zfp532 | protein_coding | -0.234375365 | 4.906140566 | 0.6598709 | 0.4283051 | 0.859000818 |
| ENSMUSG00000022914 | Brwd1 | protein_coding | 0.178084228 | 6.174824678 | 0.6597632 | 0.4283421 | 0.859000818 |
| ENSMUSG00000010307 | Tmem86a | protein_coding | -0.30499643 | 4.325601805 | 0.6586787 | 0.4287145 | 0.859381136 |
| ENSMUSG00000001767 | Crnkl1 | protein_coding | 0.278783781 | 4.060720349 | 0.6585403 | 0.4287621 | 0.859381136 |
| ENSMUSG00000063239 | Grm4 | protein_coding | -0.260122323 | 4.528026052 | 0.6584618 | 0.4287891 | 0.859381136 |
| ENSMUSG00000017493 | Igfbp4 | protein_coding | -0.397489463 | 4.478549762 | 0.6602118 | 0.4288514 | 0.859381136 |
| ENSMUSG00000026112 | Coa5 | protein_coding | -0.162570212 | 6.58948572 | 0.6579438 | 0.4289672 | 0.85938267 |
| ENSMUSG00000030254 | Rad18 | protein_coding | 0.38516153 | 3.375191328 | 0.6574738 | 0.4291289 | 0.85938267 |
| ENSMUSG00000025236 | Adpgk | protein_coding | 0.482404274 | 3.119365889 | 0.6570252 | 0.4292833 | 0.85938267 |
| ENSMUSG00000038646 | Fam103a1 | protein_coding | -0.159023608 | 6.473152165 | 0.6569625 | 0.4293049 | 0.85938267 |
| ENSMUSG00000096974 | Gm26881 | antisense_RNA | -0.405793292 | 2.940711554 | 0.6566499 | 0.4294126 | 0.85938267 |
| ENSMUSG00000024873 | Cnih2 | protein_coding | -0.275052094 | 5.668024471 | 0.6566101 | 0.4294454 | 0.85938267 |
| ENSMUSG00000046791 | Riox1 | protein_coding | 0.466969255 | 2.938538894 | 0.6564053 | 0.4294969 | 0.85938267 |
| ENSMUSG00000004562 | Arhgef40 | protein_coding | -0.306263102 | 5.044574257 | 0.6561173 | 0.4296152 | 0.85938267 |
| ENSMUSG00000095041 | AC149090.1 | protein_coding | -0.304137507 | 4.889587051 | 0.6560507 | 0.4296191 | 0.85938267 |
| ENSMUSG00000043510 | Hscb | protein_coding | 0.267396291 | 4.868453037 | 0.6559571 | 0.4296514 | 0.85938267 |
| ENSMUSG00000035151 | Elmod2 | protein_coding | 0.571791715 | 2.45865367 | 0.6554076 | 0.4298409 | 0.859555376 |
| ENSMUSG00000053931 | Cnn3 | protein_coding | 0.29355745 | 4.961678493 | 0.6548144 | 0.4300457 | 0.859555376 |
| ENSMUSG00000059208 | Hnrnpm | protein_coding | 0.20275439 | 6.265211582 | 0.6547865 | 0.4300553 | 0.859555376 |
| ENSMUSG00000040811 | Eml2 | protein_coding | -0.226390772 | 5.45460871 | 0.6546868 | 0.4300898 | 0.859555376 |
| ENSMUSG00000042590 | Ipo11 | protein_coding | 0.36043933 | 3.43643476 | 0.6543963 | 0.4301901 | 0.859555376 |
| ENSMUSG00000037260 | Hgsnat | protein_coding | 0.274140661 | 4.476158733 | 0.6543175 | 0.4302173 | 0.859555376 |
| ENSMUSG00000024856 | Cdk2ap2 | protein_coding | -0.196515724 | 5.554639604 | 0.6530081 | 0.4306702 | 0.86025686 |
| ENSMUSG00000097503 | 3110045C21Rik | antisense_RNA | -0.568080531 | 2.590778942 | 0.6528398 | 0.4307284 | 0.86025686 |
| ENSMUSG00000029233 | Srd5a3 | protein_coding | 0.409156931 | 3.477859204 | 0.6520825 | 0.4309908 | 0.860620922 |
| ENSMUSG00000109536 | 9330162G02Rik | TEC | -0.334479268 | 3.428943043 | 0.6514314 | 0.4312165 | 0.860884381 |
| ENSMUSG00000058444 | Map2k5 | protein_coding | -0.267329928 | 5.263199779 | 0.6512402 | 0.4312828 | 0.860884381 |
| ENSMUSG00000004113 | Cacna1b | protein_coding | 0.282105156 | 4.851315997 | 0.650647 | 0.4314887 | 0.861017732 |
| ENSMUSG00000027185 | Nat10 | protein_coding | 0.322451282 | 3.557744579 | 0.6505862 | 0.4315098 | 0.861017732 |
| ENSMUSG00000053754 | Chd8 | protein_coding | -0.443785143 | 3.055711784 | 0.6499091 | 0.431745 | 0.861186832 |
| ENSMUSG00000044786 | Zfp36 | protein_coding | -0.471666658 | 2.927527003 | 0.6498811 | 0.4317547 | 0.861186832 |
| ENSMUSG00000066235 | Pomgnt2 | protein_coding | 0.246453071 | 5.394538667 | 0.6493868 | 0.4319265 | 0.861258179 |
| ENSMUSG00000047044 | D030056L22Rik | protein_coding | -0.302886771 | 3.731754147 | 0.6493174 | 0.4319507 | 0.861258179 |
| ENSMUSG00000015312 | Gadd45b | protein_coding | -0.333742805 | 3.546688812 | 0.6489915 | 0.432064 | 0.861324527 |
| ENSMUSG00000047361 | Gm973 | protein_coding | 0.64403893 | 2.290108878 | 0.6481785 | 0.432347 | 0.861423681 |
| ENSMUSG00000029178 | Klf3 | protein_coding | -0.34894783 | 3.876269742 | 0.6477354 | 0.4325014 | 0.861423681 |
| ENSMUSG00000017686 | Rhot1 | protein_coding | -0.226558974 | 5.268118741 | 0.6476784 | 0.4325213 | 0.861423681 |
| ENSMUSG00000029328 | Hnrnpdl | protein_coding | -0.15300795 | 6.69906303 | 0.6476351 | 0.4325364 | 0.861423681 |
| ENSMUSG00000047067 | Dusp28 | protein_coding | 0.275539988 | 3.811464744 | 0.6469092 | 0.4327895 | 0.861423681 |
| ENSMUSG00000024527 | Afg3l2 | protein_coding | 0.238095122 | 5.199336318 | 0.6469032 | 0.4327916 | 0.861423681 |
| ENSMUSG00000087380 | 2210408F21Rik | processed_transcript | 0.206265613 | 4.930598638 | 0.6468935 | 0.4327949 | 0.861423681 |
| ENSMUSG00000033854 | Kcnk10 | protein_coding | -0.411667077 | 3.331571941 | 0.6468003 | 0.4328275 | 0.861423681 |

| ensembl_gene_id | external_gene_name | gene_biotype | log2 (fold change) | log2 (counts per million) | F-value | p-value | False discovery rate |
|---|---|---|---|---|---|---|---|
| ENSMUSG00000007029 | Vars | protein_coding | -0.401571092 | 3.074595686 | 0.6467793 | 0.4328348 | 0.861423681 |
| ENSMUSG00000024940 | Ltbp3 | protein_coding | 0.488172312 | 2.779765272 | 0.6464035 | 0.4329659 | 0.861482809 |
| ENSMUSG00000027613 | Eif6 | protein_coding | -0.154054687 | 6.846657421 | 0.64612 | 0.4330649 | 0.861482809 |
| ENSMUSG00000002059 | Rab34 | protein_coding | -0.343396946 | 3.398121639 | 0.6458363 | 0.433164 | 0.861482809 |
| ENSMUSG00000024824 | Rad9a | protein_coding | -0.285929456 | 4.462150438 | 0.6452275 | 0.4333767 | 0.861482809 |
| ENSMUSG00000038630 | Zkscan16 | protein_coding | 0.432768852 | 2.952419971 | 0.6451489 | 0.4334042 | 0.861482809 |
| ENSMUSG00000027787 | Nmd3 | protein_coding | -0.189415671 | 5.983076922 | 0.6448329 | 0.4335147 | 0.861482809 |
| ENSMUSG00000055204 | Ankrd17 | protein_coding | 0.174990394 | 6.665545523 | 0.6447921 | 0.433529 | 0.861482809 |
| ENSMUSG00000047507 | Baiap3 | protein_coding | -0.259291179 | 6.667296313 | 0.6468455 | 0.433545 | 0.861482809 |
| ENSMUSG00000026035 | Ppil3 | protein_coding | 0.206315415 | 6.186644103 | 0.6441355 | 0.4337588 | 0.861482809 |
| ENSMUSG00000041268 | Dmxl2 | protein_coding | 0.213649974 | 5.70629224 | 0.6441196 | 0.4337643 | 0.861482809 |
| ENSMUSG00000044783 | Hjurp | protein_coding | 0.242443328 | 4.757925192 | 0.6438074 | 0.4338736 | 0.861482809 |
| ENSMUSG00000033966 | Cdkl4 | protein_coding | 0.319000653 | 4.235279325 | 0.6436359 | 0.4339337 | 0.861482809 |
| ENSMUSG00000055214 | Pld5 | protein_coding | -0.215688256 | 5.240425973 | 0.6434644 | 0.4339938 | 0.861482809 |
| ENSMUSG00000034022 | Cpsf1 | protein_coding | 0.275337645 | 4.900114075 | 0.6433053 | 0.4340495 | 0.861482809 |
| ENSMUSG00000026626 | Ppp2r5a | protein_coding | -0.301715828 | 4.44567116 | 0.6430054 | 0.4341547 | 0.861482809 |
| ENSMUSG00000000876 | Pxmp4 | protein_coding | -0.208324735 | 5.15063373 | 0.6429259 | 0.4341825 | 0.861482809 |
| ENSMUSG00000057766 | Ankrd29 | protein_coding | -0.207441453 | 5.195263539 | 0.6426656 | 0.4342738 | 0.861482809 |
| ENSMUSG00000038079 | Tmem237 | protein_coding | 0.218743408 | 6.25793534 | 0.6423685 | 0.4343781 | 0.861482809 |
| ENSMUSG00000041124 | Msantd4 | protein_coding | 0.183916474 | 6.79558732 | 0.642192 | 0.43444 | 0.861482809 |
| ENSMUSG00000056553 | Ptprn2 | protein_coding | -0.337757458 | 3.91967776 | 0.6421157 | 0.4344668 | 0.861482809 |
| ENSMUSG00000047045 | Tmem164 | protein_coding | 0.329515365 | 3.627570521 | 0.6411625 | 0.4348015 | 0.861868369 |
| ENSMUSG00000036278 | Macrod1 | protein_coding | -0.247849553 | 5.257612405 | 0.6402729 | 0.4351144 | 0.861868369 |
| ENSMUSG00000056459 | Zbtb25 | protein_coding | -0.361368469 | 3.542563016 | 0.6398565 | 0.4352609 | 0.861868369 |
| ENSMUSG00000028998 | Tomm7 | protein_coding | -0.199965638 | 8.606368385 | 0.6398695 | 0.4352752 | 0.861868369 |
| ENSMUSG00000059811 | Atl2 | protein_coding | -0.22577762 | 4.999606337 | 0.6393669 | 0.4354333 | 0.861868369 |
| ENSMUSG00000017639 | Rab11fip4 | protein_coding | -0.218516957 | 7.063124303 | 0.6394297 | 0.4354796 | 0.861868369 |
| ENSMUSG00000033701 | Acbd6 | protein_coding | -0.21553206 | 5.386860559 | 0.6391824 | 0.4354983 | 0.861868369 |
| ENSMUSG00000024400 | Wdr33 | protein_coding | -0.170332106 | 6.93051735 | 0.6391097 | 0.4355239 | 0.861868369 |
| ENSMUSG00000026281 | Dtymk | protein_coding | -0.205533585 | 6.249767161 | 0.6389857 | 0.4355676 | 0.861868369 |
| ENSMUSG00000079427 | Mthfsl | protein_coding | -0.277624682 | 3.962724281 | 0.6386302 | 0.435693 | 0.861868369 |
| ENSMUSG00000000976 | Heatr6 | protein_coding | 0.258255004 | 4.608350955 | 0.6385187 | 0.4357322 | 0.861868369 |
| ENSMUSG00000029016 | Clcn6 | protein_coding | 0.198074925 | 5.548644841 | 0.6385145 | 0.4357337 | 0.861868369 |
| ENSMUSG00000038462 | Uqcrfs1 | protein_coding | 0.230479522 | 6.788603422 | 0.6390306 | 0.4357383 | 0.861868369 |
| ENSMUSG00000000794 | Kcnn3 | protein_coding | -0.395211973 | 3.752232933 | 0.6383235 | 0.4358199 | 0.861868369 |
| ENSMUSG00000028758 | Kif17 | protein_coding | 0.2523931 | 4.78534031 | 0.6381466 | 0.4358635 | 0.861868369 |
| ENSMUSG00000032648 | Pygm | protein_coding | 0.302144835 | 4.634923413 | 0.6376261 | 0.4360472 | 0.861955234 |
| ENSMUSG00000022560 | Slc52a2 | protein_coding | -0.308599105 | 4.264855553 | 0.6375679 | 0.4360677 | 0.861955234 |
| ENSMUSG00000044763 | Trmt10c | protein_coding | -0.280649799 | 4.636811629 | 0.6371103 | 0.4362293 | 0.86197061 |
| ENSMUSG00000062376 | Borcs7 | protein_coding | -0.147071991 | 7.115717169 | 0.6368324 | 0.4363275 | 0.86197061 |
| ENSMUSG00000025651 | Uqcrc1 | protein_coding | 0.234359071 | 9.629316734 | 0.6376345 | 0.436399 | 0.86197061 |
| ENSMUSG00000030397 | Mark4 | protein_coding | -0.280719854 | 4.220877947 | 0.6364504 | 0.4364626 | 0.86197061 |
| ENSMUSG00000027394 | Ttl | protein_coding | -0.225894891 | 4.823554834 | 0.6364115 | 0.4364763 | 0.86197061 |
| ENSMUSG00000022216 | Psme1 | protein_coding | 0.162261476 | 7.421260387 | 0.635931 | 0.4366463 | 0.86214791 |
| ENSMUSG00000070462 | Tlnrd1 | protein_coding | -0.319701393 | 4.426052836 | 0.6354892 | 0.4368026 | 0.862155501 |
| ENSMUSG00000034265 | Zdhhc14 | protein_coding | 0.314924802 | 4.272817134 | 0.6354671 | 0.4368105 | 0.862155501 |
| ENSMUSG00000054823 | Nsd3 | protein_coding | -0.196342487 | 6.613822115 | 0.6346339 | 0.4371056 | 0.862487346 |
| ENSMUSG00000032078 | Zpr1 | protein_coding | -0.310822775 | 4.798499595 | 0.6345925 | 0.437139 | 0.862487346 |
| ENSMUSG00000031954 | Cfdp1 | protein_coding | 0.137965093 | 8.68150032 | 0.6341994 | 0.4372596 | 0.86256704 |
| ENSMUSG00000002908 | Kcnn1 | protein_coding | -0.6144533 | 2.593229118 | 0.6339473 | 0.4373491 | 0.862585219 |
| ENSMUSG00000020728 | Cep112 | protein_coding | -0.425554609 | 2.978589964 | 0.6335382 | 0.4374942 | 0.862617017 |
| ENSMUSG00000020834 | Dhrs13 | protein_coding | 0.561612447 | 2.554185785 | 0.6334497 | 0.4375256 | 0.862617017 |
| ENSMUSG00000040822 | 1700123O20Rik | protein_coding | -0.168322159 | 6.609926099 | 0.6331553 | 0.4376301 | 0.862664884 |
| ENSMUSG00000020621 | Rdh14 | protein_coding | -0.227364293 | 5.326370321 | 0.6327866 | 0.4377611 | 0.862764868 |
| ENSMUSG00000059970 | Hspa2 | protein_coding | 0.275688689 | 5.644125734 | 0.6319148 | 0.4380897 | 0.863182695 |
| ENSMUSG00000045763 | Basp1 | protein_coding | -0.148590148 | 7.471415446 | 0.6315062 | 0.4382164 | 0.863182695 |
| ENSMUSG00000047832 | Cdca4 | protein_coding | -0.399786236 | 3.904284938 | 0.631289 | 0.4383124 | 0.863182695 |
| ENSMUSG00000056856 | Jakmip3 | protein_coding | -0.274554147 | 4.059510386 | 0.6311658 | 0.4383375 | 0.863182695 |
| ENSMUSG00000026049 | Tex30 | protein_coding | 0.262822018 | 4.010276521 | 0.631062 | 0.4383745 | 0.863182695 |
| ENSMUSG00000039000 | Ube3c | protein_coding | 0.209793969 | 5.459230692 | 0.6306626 | 0.4385167 | 0.863304727 |
| ENSMUSG00000048040 | Arxes2 | protein_coding | 0.176990037 | 6.371574271 | 0.6289848 | 0.4391151 | 0.864009491 |
| ENSMUSG00000073433 | Arhgdig | protein_coding | -0.212278428 | 5.627176082 | 0.6289837 | 0.4391155 | 0.864009491 |
| ENSMUSG00000054021 | Sirt5 | protein_coding | 0.241505723 | 5.581060247 | 0.6289392 | 0.4391314 | 0.864009491 |
| ENSMUSG00000023169 | Slc38a1 | protein_coding | 0.166771048 | 7.064916432 | 0.6285777 | 0.4392605 | 0.864009491 |
| ENSMUSG00000030530 | Furin | protein_coding | -0.419227797 | 2.998263109 | 0.628533 | 0.4392765 | 0.864009491 |
| ENSMUSG00000030717 | Nupr1 | protein_coding | -0.380172642 | 5.056776348 | 0.6314631 | 0.4393655 | 0.864026532 |
| ENSMUSG00000072875 | Gpr27 | protein_coding | -0.311518916 | 4.238632282 | 0.6278801 | 0.4395098 | 0.864081673 |
| ENSMUSG00000031903 | Pla2g15 | protein_coding | 0.249807533 | 4.10737651 | 0.6277558 | 0.4395542 | 0.864081673 |
| ENSMUSG00000075273 | Ttc30b | protein_coding | 0.248654556 | 4.086097015 | 0.6263611 | 0.4400535 | 0.864710154 |
| ENSMUSG00000024807 | Syvn1 | protein_coding | -0.263305344 | 4.645465583 | 0.6262817 | 0.4400819 | 0.864710154 |
| ENSMUSG00000048756 | Foxo3 | protein_coding | 0.249845904 | 5.828438081 | 0.6262407 | 0.4401152 | 0.864710154 |
| ENSMUSG00000021796 | Bmpr1a | protein_coding | -0.270498407 | 4.532161124 | 0.6247785 | 0.440621 | 0.865363254 |
| ENSMUSG00000084843 | B230312C02Rik | lincRNA | -0.391522436 | 3.027621948 | 0.6247646 | 0.440626 | 0.865363254 |
| ENSMUSG00000020823 | Sec14l1 | protein_coding | 0.216174213 | 5.454961313 | 0.6244451 | 0.4407407 | 0.865363254 |

| ensembl_gene_id | external_gene_name | gene_biotype | log2 (fold change) | log2 (counts per million) | F-value | p-value | False discovery rate |
|---|---|---|---|---|---|---|---|
| ENSMUSG00000098176 | Ccdc166 | protein_coding | 0.335322246 | 4.102231282 | 0.6240164 | 0.4408947 | 0.865363254 |
| ENSMUSG00000053730 | Tmem39b | protein_coding | 0.425846192 | 3.168198016 | 0.6239274 | 0.4409267 | 0.865363254 |
| ENSMUSG00000045251 | Zfp688 | protein_coding | -0.229745754 | 4.787095876 | 0.623917 | 0.4409305 | 0.865363254 |
| ENSMUSG00000068523 | Gng5 | protein_coding | -0.260794015 | 4.80179362 | 0.6232866 | 0.4411571 | 0.865525177 |
| ENSMUSG00000023249 | Parp3 | protein_coding | 0.453948229 | 2.909312316 | 0.6232398 | 0.441174 | 0.865525177 |
| ENSMUSG00000024642 | Tle4 | protein_coding | 0.231640001 | 5.585379741 | 0.6227942 | 0.4413343 | 0.865681808 |
| ENSMUSG00000036006 | Ripor2 | protein_coding | 0.237808412 | 5.540538924 | 0.621961 | 0.4416344 | 0.866091823 |
| ENSMUSG00000026796 | Fam129b | protein_coding | -0.399213776 | 2.90440795 | 0.6217523 | 0.4417096 | 0.866091823 |
| ENSMUSG00000002602 | Axl | protein_coding | 0.441972025 | 3.171210673 | 0.6215432 | 0.441785 | 0.866091823 |
| ENSMUSG00000031774 | Fam192a | protein_coding | 0.22166766 | 6.846178738 | 0.6211986 | 0.4419765 | 0.866237808 |
| ENSMUSG00000038702 | Dsel | protein_coding | -0.412649072 | 3.507317858 | 0.6208901 | 0.4420205 | 0.866237808 |
| ENSMUSG00000003660 | Snrnp200 | protein_coding | 0.22588274 | 5.519479476 | 0.6205339 | 0.4421491 | 0.86629347 |
| ENSMUSG00000029554 | Mad1l1 | protein_coding | 0.203892891 | 5.278598499 | 0.6201138 | 0.4423008 | 0.86629347 |
| ENSMUSG00000006442 | Srm | protein_coding | -0.231571367 | 5.229121767 | 0.6200267 | 0.4423323 | 0.86629347 |
| ENSMUSG00000020739 | Nup85 | protein_coding | 0.243555315 | 5.327087493 | 0.6195689 | 0.4424977 | 0.86629347 |
| ENSMUSG00000029267 | Mtf2 | protein_coding | 0.244738382 | 4.569289192 | 0.6195513 | 0.442504 | 0.86629347 |
| ENSMUSG00000048832 | Vps37c | protein_coding | 0.294663847 | 4.872612485 | 0.6194731 | 0.4425323 | 0.86629347 |
| ENSMUSG00000015843 | Rxrg | protein_coding | 0.160237904 | 6.702866231 | 0.6189011 | 0.4427392 | 0.866514067 |
| ENSMUSG00000022456 | 3-Sep | protein_coding | -0.156149354 | 8.375689342 | 0.6185436 | 0.4428685 | 0.866514067 |
| ENSMUSG00000093606 | B130034C11Rik | processed_transcript | -0.538561989 | 2.838125638 | 0.6184933 | 0.4428868 | 0.866514067 |
| ENSMUSG00000028706 | Nsun4 | protein_coding | 0.306702654 | 4.674578843 | 0.6177842 | 0.4431435 | 0.866779396 |
| ENSMUSG00000055202 | Zfp811 | protein_coding | -0.282757517 | 4.399401755 | 0.6175577 | 0.4432256 | 0.866779396 |
| ENSMUSG00000039754 | Alkbh4 | protein_coding | -0.341429383 | 4.552657248 | 0.6173113 | 0.4433334 | 0.866779396 |
| ENSMUSG00000045374 | Wdr81 | protein_coding | -0.492752928 | 2.921892044 | 0.6171844 | 0.443361 | 0.866779396 |
| ENSMUSG00000041488 | Stx3 | protein_coding | 0.287619315 | 3.858314056 | 0.6167434 | 0.4435209 | 0.866779396 |
| ENSMUSG00000019899 | Lama2 | protein_coding | -0.464059977 | 2.772551931 | 0.6164524 | 0.4436265 | 0.866779396 |
| ENSMUSG00000036053 | Fmnl2 | protein_coding | -0.291198562 | 4.242882389 | 0.6164307 | 0.4436344 | 0.866779396 |
| ENSMUSG00000026767 | Mindy3 | protein_coding | -0.17269995 | 6.240347076 | 0.6163058 | 0.4436797 | 0.866779396 |
| ENSMUSG00000042804 | Gpr153 | protein_coding | 0.322311479 | 3.871506597 | 0.6160002 | 0.4437907 | 0.866779396 |
| ENSMUSG00000049521 | Cdc42ep1 | protein_coding | 0.393708369 | 3.740462329 | 0.6156786 | 0.4439259 | 0.866779396 |
| ENSMUSG00000028033 | Kcnq5 | protein_coding | -0.252846902 | 4.809074714 | 0.6149334 | 0.4441784 | 0.866779396 |
| ENSMUSG00000034994 | Eef2 | protein_coding | 0.183997836 | 10.00423559 | 0.6149259 | 0.4441811 | 0.866779396 |
| ENSMUSG00000087143 | A830082K12Rik | antisense_RNA | -0.299990936 | 3.85780153 | 0.6148797 | 0.4441979 | 0.866779396 |
| ENSMUSG00000097310 | A930038B10Rik | lincRNA | 0.647336481 | 2.324610215 | 0.6148253 | 0.4442177 | 0.866779396 |
| ENSMUSG00000038267 | Slc22a23 | protein_coding | 0.238251655 | 4.568708152 | 0.6144672 | 0.444348 | 0.866779396 |
| ENSMUSG00000022820 | Ndufb4 | protein_coding | -0.207171552 | 5.958522135 | 0.6144174 | 0.4443661 | 0.866779396 |
| ENSMUSG00000005580 | Adcy9 | protein_coding | 0.380780511 | 4.012218975 | 0.6143949 | 0.4443927 | 0.866779396 |
| ENSMUSG00000066113 | Adamtsl1 | protein_coding | -0.612654183 | 3.038575232 | 0.6144813 | 0.4448119 | 0.867250381 |
| ENSMUSG00000031709 | Tbc1d9 | protein_coding | 0.223703352 | 4.883991709 | 0.6129085 | 0.4449158 | 0.867250381 |
| ENSMUSG00000009035 | Tmem184b | protein_coding | 0.303095612 | 4.128339592 | 0.6127516 | 0.444973 | 0.867250381 |
| ENSMUSG00000045435 | Tmem60 | protein_coding | -0.188887541 | 7.072074073 | 0.6126014 | 0.4450278 | 0.867250381 |
| ENSMUSG00000024098 | Twsg1 | protein_coding | -0.335072045 | 3.465440633 | 0.6122533 | 0.4451549 | 0.867250381 |
| ENSMUSG00000028741 | Mrto4 | protein_coding | 0.178136575 | 6.218993079 | 0.6121839 | 0.4451802 | 0.867250381 |
| ENSMUSG00000040213 | Kyat3 | protein_coding | 0.391663803 | 3.134046576 | 0.6121025 | 0.4452099 | 0.867250381 |
| ENSMUSG00000064254 | Ethe1 | protein_coding | 0.275023279 | 4.879331073 | 0.6119122 | 0.4452794 | 0.867250381 |
| ENSMUSG00000059486 | Kbtbd2 | protein_coding | -0.207539579 | 5.815664921 | 0.6116096 | 0.4453899 | 0.867308504 |
| ENSMUSG00000034098 | Fstl5 | protein_coding | 0.221142952 | 5.253125928 | 0.6108957 | 0.4456508 | 0.867457598 |
| ENSMUSG00000023367 | Tmem176a | protein_coding | 0.256746183 | 5.30402105 | 0.6108464 | 0.4456688 | 0.867457598 |
| ENSMUSG00000016528 | Mapkapk2 | protein_coding | 0.554720831 | 2.809409699 | 0.6104104 | 0.4458283 | 0.867457598 |
| ENSMUSG00000090553 | Snrpe | protein_coding | -0.199212456 | 7.173520655 | 0.6104586 | 0.445829 | 0.867457598 |
| ENSMUSG00000047394 | Odf3b | protein_coding | -0.380638703 | 3.508796897 | 0.610297 | 0.4458698 | 0.867457598 |
| ENSMUSG00000026809 | Spaca9 | protein_coding | -0.506378927 | 2.80355454 | 0.6100643 | 0.445955 | 0.867466287 |
| ENSMUSG00000030064 | Frmd4b | protein_coding | -0.361295058 | 3.473205466 | 0.6097279 | 0.4460781 | 0.867548982 |
| ENSMUSG00000044117 | 2900011O08Rik | protein_coding | 0.173666812 | 9.481265429 | 0.6092255 | 0.4462622 | 0.867749919 |
| ENSMUSG00000034781 | Gna11 | protein_coding | 0.179937243 | 8.147263377 | 0.6083379 | 0.4465877 | 0.868077671 |
| ENSMUSG00000057176 | Ccdc189 | protein_coding | -0.230221684 | 4.692951345 | 0.6083255 | 0.4465922 | 0.868077671 |
| ENSMUSG00000033793 | Atp6v1h | protein_coding | 0.17376053 | 8.77313999 | 0.608053 | 0.4466922 | 0.868115169 |
| ENSMUSG00000040265 | Dnm3 | protein_coding | 0.220605654 | 5.663832161 | 0.6076139 | 0.4468534 | 0.868186964 |
| ENSMUSG00000024952 | Rps6ka4 | protein_coding | -0.274995758 | 4.084557071 | 0.6075125 | 0.4468906 | 0.868186964 |
| ENSMUSG00000048029 | Eno4 | protein_coding | 0.398261314 | 3.208913066 | 0.607131 | 0.4470308 | 0.868279463 |
| ENSMUSG00000044250 | Pced1b | protein_coding | -0.402878002 | 2.992362881 | 0.6069434 | 0.4470997 | 0.868279463 |
| ENSMUSG00000026199 | Ankzf1 | protein_coding | 0.315390674 | 3.976778825 | 0.6065173 | 0.4472564 | 0.868426919 |
| ENSMUSG00000087178 | A230056P14Rik | processed_transcript | -0.464924674 | 3.420152517 | 0.6060109 | 0.447461 | 0.868449701 |
| ENSMUSG00000109554 | A230057D06Rik | lincRNA | -0.625672746 | 2.69635699 | 0.6058727 | 0.4475118 | 0.868449701 |
| ENSMUSG00000075595 | Zfp652 | protein_coding | 0.245381421 | 4.281857141 | 0.6057473 | 0.4475397 | 0.868449701 |
| ENSMUSG00000033790 | Tubgcp5 | protein_coding | 0.222988998 | 4.515937438 | 0.6053221 | 0.4476963 | 0.868449701 |
| ENSMUSG00000026688 | Mgst3 | protein_coding | 0.142053794 | 8.852648888 | 0.6052446 | 0.4477249 | 0.868449701 |
| ENSMUSG00000028291 | Akirin2 | protein_coding | -0.216157314 | 5.350126835 | 0.6051391 | 0.4477637 | 0.868449701 |
| ENSMUSG00000078676 | Casc3 | protein_coding | 0.23910583 | 4.850136266 | 0.6048025 | 0.4478878 | 0.868449701 |
| ENSMUSG00000031953 | Tmem170 | protein_coding | -0.301995475 | 4.277935296 | 0.6047308 | 0.4479143 | 0.868449701 |
| ENSMUSG00000071652 | Ints5 | protein_coding | 0.307102958 | 3.756312583 | 0.6040445 | 0.4481674 | 0.868506938 |
| ENSMUSG00000024831 | Ighmbp2 | protein_coding | -0.478918142 | 3.065338584 | 0.6039309 | 0.4482093 | 0.868506938 |
| ENSMUSG00000059323 | Tonsl | protein_coding | 0.142965199 | 9.574008816 | 0.6036214 | 0.4483235 | 0.868506938 |
| ENSMUSG00000029174 | Tbc1d1 | protein_coding | 0.393041782 | 3.007960314 | 0.6034978 | 0.4483692 | 0.868506938 |

| ensembl_gene_id | external_gene_name | gene_biotype | log2 (fold change) | log2 (counts per million) | F-value | p-value | False discovery rate |
|---|---|---|---|---|---|---|---|
| ENSMUSG00000019826 | Zbtb24 | protein_coding | -0.279233673 | 3.807397015 | 0.6033576 | 0.4484209 | 0.868506938 |
| ENSMUSG00000022358 | Fbxo32 | protein_coding | -0.380808465 | 3.737421218 | 0.6031354 | 0.448503 | 0.868506938 |
| ENSMUSG00000033557 | Fam20b | protein_coding | -0.223197413 | 5.503172762 | 0.6029293 | 0.4485792 | 0.868506938 |
| ENSMUSG00000055717 | Slain1 | protein_coding | -0.20601943 | 5.534441336 | 0.6029002 | 0.4485899 | 0.868506938 |
| ENSMUSG00000055675 | Kbtbd11 | protein_coding | 0.301987641 | 5.211612869 | 0.6027678 | 0.448705 | 0.868538371 |
| ENSMUSG00000030772 | Dkk3 | protein_coding | 0.263797945 | 5.609323161 | 0.6024685 | 0.4487677 | 0.868538371 |
| ENSMUSG00000028069 | Gpatch4 | protein_coding | -0.162622701 | 6.731263987 | 0.6018908 | 0.4489632 | 0.868680177 |
| ENSMUSG00000015766 | Eps8 | protein_coding | 0.298215477 | 4.559607813 | 0.6017846 | 0.4490026 | 0.868680177 |
| ENSMUSG00000022772 | Senp5 | protein_coding | -0.313394578 | 3.340804341 | 0.6005623 | 0.4494553 | 0.868740774 |
| ENSMUSG00000024245 | Tmem178 | protein_coding | -0.482392912 | 3.659488846 | 0.6023296 | 0.4495055 | 0.868740774 |
| ENSMUSG00000086629 | 2810403D21Rik | processed_transcript | -0.366947932 | 3.35964998 | 0.6003256 | 0.4495431 | 0.868740774 |
| ENSMUSG00000028878 | Fam76a | protein_coding | 0.224328138 | 5.551510911 | 0.6003189 | 0.4495456 | 0.868740774 |
| ENSMUSG00000042699 | Dhx9 | protein_coding | 0.262853468 | 4.608989933 | 0.6000094 | 0.4496604 | 0.868740774 |
| ENSMUSG00000052533 | Nup188 | protein_coding | 0.545507842 | 2.716644853 | 0.5996816 | 0.449782 | 0.868740774 |
| ENSMUSG00000074896 | Ifit3 | protein_coding | -0.384686518 | 3.128248239 | 0.5996634 | 0.4497888 | 0.868740774 |
| ENSMUSG00000036186 | Fam69b | protein_coding | -0.26872201 | 4.7655652 | 0.5994981 | 0.4498501 | 0.868740774 |
| ENSMUSG00000034557 | Zfyve9 | protein_coding | -0.203498641 | 5.23574843 | 0.5994891 | 0.4498535 | 0.868740774 |
| ENSMUSG00000020102 | Slc16a7 | protein_coding | -0.371735995 | 3.21491113 | 0.5994113 | 0.4498824 | 0.868740774 |
| ENSMUSG00000002012 | Pnck | protein_coding | 0.18844327 | 9.46024771 | 0.5991996 | 0.449961 | 0.868740774 |
| ENSMUSG00000022174 | Dad1 | protein_coding | 0.154792034 | 9.050245605 | 0.5990855 | 0.4500034 | 0.868740774 |
| ENSMUSG00000055305 | Zfp93 | protein_coding | 0.482435313 | 2.681189644 | 0.5986713 | 0.4501573 | 0.868881892 |
| ENSMUSG00000000142 | Axin2 | protein_coding | -0.352084576 | 3.539760292 | 0.597632 | 0.4505438 | 0.869238189 |
| ENSMUSG00000060992 | Copz1 | protein_coding | 0.149361225 | 8.602941946 | 0.5975599 | 0.4505707 | 0.869238189 |
| ENSMUSG00000036009 | Mettl25 | protein_coding | 0.398175072 | 2.995099745 | 0.597213 | 0.4506998 | 0.869238189 |
| ENSMUSG00000030213 | Atf7ip | protein_coding | 0.238032063 | 4.988359292 | 0.5970652 | 0.4507548 | 0.869238189 |
| ENSMUSG00000021460 | Auh | protein_coding | 0.221335688 | 5.498863594 | 0.5969058 | 0.4508142 | 0.869238189 |
| ENSMUSG00000045967 | Gpr158 | protein_coding | -0.215104371 | 5.215833255 | 0.5968719 | 0.4508269 | 0.869238189 |
| ENSMUSG00000031755 | Bbs2 | protein_coding | 0.301021007 | 4.505617287 | 0.5963748 | 0.4510121 | 0.869439475 |
| ENSMUSG00000027540 | Ptpn1 | protein_coding | -0.256773433 | 4.79176329 | 0.5961515 | 0.4510954 | 0.869444135 |
| ENSMUSG00000022476 | Polr3h | protein_coding | 0.182900702 | 6.335432108 | 0.5956293 | 0.4512902 | 0.869444513 |
| ENSMUSG00000027164 | Traf6 | protein_coding | -0.481981825 | 2.683158561 | 0.5954539 | 0.4513557 | 0.869444513 |
| ENSMUSG00000052214 | Opa3 | protein_coding | -0.156164844 | 6.439299332 | 0.5953347 | 0.4514002 | 0.869444513 |
| ENSMUSG00000045576 | St7l | protein_coding | 0.272795472 | 4.455683809 | 0.5951778 | 0.4514588 | 0.869444513 |
| ENSMUSG00000025287 | Acot9 | protein_coding | 0.206289181 | 5.547245871 | 0.5948391 | 0.4515853 | 0.869444513 |
| ENSMUSG00000047473 | Zfp30 | protein_coding | 0.22708088 | 4.606516946 | 0.5945261 | 0.4517022 | 0.869444513 |
| ENSMUSG00000028967 | Errfi1 | protein_coding | 0.286194323 | 4.431978901 | 0.5942553 | 0.4518035 | 0.869444513 |
| ENSMUSG00000037617 | Spag1 | protein_coding | 0.567810805 | 2.575763389 | 0.5940523 | 0.4518794 | 0.869444513 |
| ENSMUSG00000029465 | Arpc3 | protein_coding | -0.185950063 | 8.650120317 | 0.5940347 | 0.4518859 | 0.869444513 |
| ENSMUSG00000020163 | Uqcr11 | protein_coding | -0.193943475 | 9.283705867 | 0.5938442 | 0.4519572 | 0.869444513 |
| ENSMUSG00000028453 | Fancg | protein_coding | -0.279858256 | 3.980522971 | 0.5937699 | 0.451985 | 0.869444513 |
| ENSMUSG00000058392 | Rrp1b | protein_coding | 0.23294333 | 4.924290776 | 0.592443 | 0.4524819 | 0.869927952 |
| ENSMUSG00000028007 | Snx7 | protein_coding | -0.297685073 | 3.934943451 | 0.5922416 | 0.4525574 | 0.869927952 |
| ENSMUSG00000024835 | Coro1b | protein_coding | -0.150817213 | 6.337349066 | 0.5921846 | 0.4525788 | 0.869927952 |
| ENSMUSG00000038271 | Iffo1 | protein_coding | 0.299805173 | 4.008015608 | 0.5919799 | 0.4526555 | 0.869927952 |
| ENSMUSG00000024346 | Pfdn1 | protein_coding | 0.133027515 | 9.040204816 | 0.5917878 | 0.4527276 | 0.869927952 |
| ENSMUSG00000054452 | Aes | protein_coding | -0.1524847 | 7.901939294 | 0.5916968 | 0.4527617 | 0.869927952 |
| ENSMUSG00000056050 | Mia3 | protein_coding | -0.198044781 | 5.465371818 | 0.5915878 | 0.4528026 | 0.869927952 |
| ENSMUSG00000035206 | Sppl2b | protein_coding | 0.301396744 | 3.960929296 | 0.5906176 | 0.453167 | 0.870143121 |
| ENSMUSG00000028464 | Tpm2 | protein_coding | -0.30318136 | 4.452468545 | 0.5905656 | 0.4531865 | 0.870143121 |
| ENSMUSG00000021687 | Scamp1 | protein_coding | 0.1400469 | 7.292855386 | 0.5901156 | 0.4533556 | 0.870143121 |
| ENSMUSG00000024411 | Aqp4 | protein_coding | -0.237350634 | 5.041781743 | 0.5900722 | 0.453372 | 0.870143121 |
| ENSMUSG00000022375 | Lrrc6 | protein_coding | -0.433248421 | 3.852469351 | 0.5912356 | 0.4533937 | 0.870143121 |
| ENSMUSG00000021124 | Vti1b | protein_coding | 0.322041886 | 8.008059592 | 0.5982442 | 0.4534002 | 0.870143121 |
| ENSMUSG00000033917 | Gde1 | protein_coding | 0.237595996 | 5.794515996 | 0.5896313 | 0.4535558 | 0.870286469 |
| ENSMUSG00000075033 | Nxpe3 | protein_coding | 0.370533836 | 3.370713129 | 0.5893332 | 0.45365 | 0.870311926 |
| ENSMUSG00000035086 | Becn1 | protein_coding | 0.320528033 | 3.339431092 | 0.5885169 | 0.4539574 | 0.87049166 |
| ENSMUSG00000040021 | Lats1 | protein_coding | 0.365276413 | 3.305346806 | 0.5885054 | 0.4539618 | 0.87049166 |
| ENSMUSG00000002635 | Pdcd2l | protein_coding | 0.276220445 | 4.720181586 | 0.5884396 | 0.4539865 | 0.87049166 |
| ENSMUSG00000053768 | Chchd3 | protein_coding | -0.133443093 | 6.995113663 | 0.5877584 | 0.4542434 | 0.870658957 |
| ENSMUSG00000028329 | Xpa | protein_coding | -0.303833304 | 3.471350559 | 0.5874459 | 0.4543613 | 0.870658957 |
| ENSMUSG00000055538 | Zcchc24 | protein_coding | -0.24288414 | 5.011976266 | 0.5873722 | 0.4543891 | 0.870658957 |
| ENSMUSG00000022114 | Spry2 | protein_coding | -0.385654887 | 6.703076342 | 0.5998344 | 0.4543977 | 0.870658957 |
| ENSMUSG00000046441 | Cmtr2 | protein_coding | 0.465424267 | 3.056756815 | 0.5866931 | 0.4546455 | 0.870978648 |
| ENSMUSG00000085042 | Abhd11os | transcribed_unitary_p | 0.237078358 | 5.216477272 | 0.5863496 | 0.4547753 | 0.870999634 |
| ENSMUSG00000024122 | Pdpk1 | protein_coding | 0.165621559 | 6.553150578 | 0.5862354 | 0.4548185 | 0.870999634 |
| ENSMUSG00000043131 | Mob1a | protein_coding | 0.341337 | 3.664848471 | 0.5856669 | 0.4550334 | 0.87111368 |
| ENSMUSG00000003974 | Grm3 | protein_coding | 0.242842256 | 4.983939823 | 0.5856495 | 0.45504 | 0.87111368 |
| ENSMUSG00000022300 | Dcaf13 | protein_coding | -0.221237753 | 6.142582573 | 0.584828 | 0.4553689 | 0.871513007 |
| ENSMUSG00000025885 | Myo5b | protein_coding | -0.260420676 | 4.374766362 | 0.584327 | 0.4555408 | 0.871513007 |
| ENSMUSG00000062444 | Ap3b2 | protein_coding | 0.193877822 | 6.66180657 | 0.5842487 | 0.4555705 | 0.871513007 |
| ENSMUSG00000078908 | Mon1b | protein_coding | 0.575052214 | 2.834909832 | 0.5840223 | 0.4556741 | 0.871513007 |
| ENSMUSG00000079283 | 2310009B15Rik | protein_coding | -0.254957754 | 4.54184601 | 0.5839068 | 0.4557001 | 0.871513007 |
| ENSMUSG00000032187 | Smarca4 | protein_coding | 0.220774265 | 5.949081 | 0.5836915 | 0.4557817 | 0.871513007 |
| ENSMUSG00000073293 | Nudt10 | protein_coding | 0.264478067 | 5.579233833 | 0.5833269 | 0.4559379 | 0.871513007 |

| ensembl_gene_id | external_gene_name | gene_biotype | log2 (fold change) | log2 (counts per million) | F-value | p-value | False discovery rate |
|---|---|---|---|---|---|---|---|
| ENSMUSG00000018287 | Spag7 | protein_coding | 0.130351727 | 8.922908867 | 0.5827667 | 0.4561327 | 0.871513007 |
| ENSMUSG00000022860 | Chodl | protein_coding | -0.423922979 | 3.315417337 | 0.5825941 | 0.4561983 | 0.871513007 |
| ENSMUSG00000033316 | Galnt9 | protein_coding | -0.279004977 | 4.225352478 | 0.5823055 | 0.4563079 | 0.871513007 |
| ENSMUSG00000018537 | Pcgf2 | protein_coding | -0.351115218 | 3.225708111 | 0.5818228 | 0.4564914 | 0.871513007 |
| ENSMUSG00000033863 | Klf9 | protein_coding | -0.149773068 | 6.174847207 | 0.5817009 | 0.4565378 | 0.871513007 |
| ENSMUSG00000056999 | Ide | protein_coding | 0.221123321 | 4.665093056 | 0.5816707 | 0.4565493 | 0.871513007 |
| ENSMUSG00000035517 | Tdrd7 | protein_coding | -0.307200879 | 4.36072328 | 0.5814449 | 0.4566352 | 0.871513007 |
| ENSMUSG00000022817 | Itgb5 | protein_coding | -0.438382244 | 2.879385228 | 0.5812068 | 0.4567258 | 0.871513007 |
| ENSMUSG00000035232 | Pdk3 | protein_coding | 0.443544569 | 4.387157413 | 0.584855 | 0.456751 | 0.871513007 |
| ENSMUSG00000046593 | Tmem215 | protein_coding | -0.250752292 | 4.79170462 | 0.5808935 | 0.4568451 | 0.871513007 |
| ENSMUSG00000026571 | Dcaf6 | protein_coding | -0.228383191 | 4.34606051 | 0.5808886 | 0.4568469 | 0.871513007 |
| ENSMUSG00000037872 | Ackr1 | protein_coding | 0.255827783 | 4.243581478 | 0.5806661 | 0.4569317 | 0.871513007 |
| ENSMUSG00000059363 | Fxn | protein_coding | 0.420506777 | 2.881427468 | 0.5805646 | 0.4569703 | 0.871513007 |
| ENSMUSG00000029433 | Diablo | protein_coding | -0.201102964 | 4.82074795 | 0.5804687 | 0.4570069 | 0.871513007 |
| ENSMUSG00000028719 | Cmpk1 | protein_coding | -0.164389496 | 7.30232154 | 0.5804036 | 0.4570317 | 0.871513007 |
| ENSMUSG00000019945 | 1700040L02Rik | protein_coding | -0.315967996 | 3.569373878 | 0.5801622 | 0.4571237 | 0.871533871 |
| ENSMUSG00000050271 | Prag1 | protein_coding | 0.338800481 | 3.463287593 | 0.5793383 | 0.4574379 | 0.871978325 |
| ENSMUSG00000028865 | Cd164l2 | protein_coding | -0.260398041 | 3.901543846 | 0.5787995 | 0.4576436 | 0.872215748 |
| ENSMUSG00000026546 | Cfap45 | protein_coding | -0.337758393 | 3.53867362 | 0.5784284 | 0.4577853 | 0.872250902 |
| ENSMUSG00000071176 | Arhgef10 | protein_coding | -0.257280747 | 4.175205598 | 0.5783265 | 0.4578242 | 0.872250902 |
| ENSMUSG00000038077 | Kcna6 | protein_coding | -0.217703258 | 5.795123717 | 0.5774905 | 0.4581439 | 0.872499753 |
| ENSMUSG00000029466 | Anapc7 | protein_coding | 0.200808953 | 5.210385149 | 0.5774349 | 0.4581651 | 0.872499753 |
| ENSMUSG00000066189 | Cacng3 | protein_coding | 0.141588861 | 7.436770443 | 0.5769654 | 0.4583448 | 0.872499753 |
| ENSMUSG00000051652 | Lrrc3 | protein_coding | 0.587643368 | 3.470693892 | 0.5808727 | 0.4583712 | 0.872499753 |
| ENSMUSG00000021993 | Mipep | protein_coding | 0.252072277 | 4.298738987 | 0.5768548 | 0.4583872 | 0.872499753 |
| ENSMUSG00000033730 | Egr3 | protein_coding | -0.275164042 | 4.995065314 | 0.5766422 | 0.4584686 | 0.872499753 |
| ENSMUSG00000021483 | Cdk20 | protein_coding | 0.235681869 | 4.943556972 | 0.5763982 | 0.458562 | 0.872499753 |
| ENSMUSG00000050891 | Tatdn1 | protein_coding | -0.191227831 | 5.328396989 | 0.5762828 | 0.4586063 | 0.872499753 |
| ENSMUSG00000002475 | Abhd3 | protein_coding | 0.232726571 | 5.700202732 | 0.5760771 | 0.4586851 | 0.872499753 |
| ENSMUSG00000036026 | Tmem63b | protein_coding | -0.216273637 | 5.229959865 | 0.5752773 | 0.4589918 | 0.872728737 |
| ENSMUSG00000021243 | Fcf1 | protein_coding | -0.236098956 | 5.818224989 | 0.5752116 | 0.4590348 | 0.872728737 |
| ENSMUSG00000008540 | Mgst1 | protein_coding | -0.415119327 | 5.354484766 | 0.5806399 | 0.459049 | 0.872728737 |
| ENSMUSG00000040760 | Appl1 | protein_coding | 0.184023479 | 5.821678524 | 0.5746169 | 0.4592454 | 0.872812684 |
| ENSMUSG00000049556 | Lingo1 | protein_coding | 0.21836891 | 5.534299387 | 0.5745906 | 0.4592555 | 0.872812684 |
| ENSMUSG00000032612 | Usp4 | protein_coding | 0.16528785 | 6.156896791 | 0.57364 | 0.4596208 | 0.873179344 |
| ENSMUSG00000028467 | Gba2 | protein_coding | 0.18866389 | 5.388187898 | 0.5731143 | 0.4598231 | 0.873179344 |
| ENSMUSG00000022973 | Synj1 | protein_coding | 0.179008655 | 7.54218916 | 0.572944 | 0.4598886 | 0.873179344 |
| ENSMUSG00000068039 | Tcp1 | protein_coding | 0.163345666 | 8.356202542 | 0.5726765 | 0.4599917 | 0.873179344 |
| ENSMUSG00000021486 | Prelid1 | protein_coding | -0.155648429 | 7.383620694 | 0.5722599 | 0.4601521 | 0.873179344 |
| ENSMUSG00000023460 | Rab12 | protein_coding | 0.182802698 | 5.846956722 | 0.5722218 | 0.4601668 | 0.873179344 |
| ENSMUSG00000015536 | Mocs2 | protein_coding | -0.137303905 | 7.941274613 | 0.572023 | 0.4602434 | 0.873179344 |
| ENSMUSG00000019813 | Cep57l1 | protein_coding | -0.370026716 | 3.313911245 | 0.5717726 | 0.46034 | 0.873179344 |
| ENSMUSG00000068264 | Ap5s1 | protein_coding | -0.176656641 | 6.289212354 | 0.5712966 | 0.4605235 | 0.873179344 |
| ENSMUSG00000019718 | L3hypdh | protein_coding | 0.34270993 | 3.288056262 | 0.5712247 | 0.4605513 | 0.873179344 |
| ENSMUSG00000033364 | Usp37 | protein_coding | 0.314837847 | 5.195301526 | 0.5721487 | 0.4606454 | 0.873179344 |
| ENSMUSG00000018449 | Rpain | protein_coding | -0.187642585 | 5.882815107 | 0.5707613 | 0.4607302 | 0.873179344 |
| ENSMUSG00000028341 | Nr4a3 | protein_coding | -0.448084492 | 3.210663218 | 0.5705007 | 0.4608308 | 0.873179344 |
| ENSMUSG00000002345 | Borcs8 | protein_coding | 0.164179206 | 7.178621538 | 0.5704752 | 0.4608407 | 0.873179344 |
| ENSMUSG00000057054 | Inca1 | protein_coding | -0.280678559 | 3.848387541 | 0.5703088 | 0.460905 | 0.873179344 |
| ENSMUSG00000020451 | Limk2 | protein_coding | 0.214489159 | 6.679281051 | 0.5701963 | 0.4610125 | 0.873179344 |
| ENSMUSG00000030657 | Xylt1 | protein_coding | 0.43384128 | 3.242131493 | 0.5699132 | 0.4610579 | 0.873179344 |
| ENSMUSG00000005625 | Psmd4 | protein_coding | 0.142628713 | 8.643510323 | 0.5698015 | 0.461101 | 0.873179344 |
| ENSMUSG00000062006 | Rpl34 | protein_coding | -0.193254684 | 7.859558926 | 0.5696157 | 0.4611905 | 0.873179344 |
| ENSMUSG00000022261 | Sdc2 | protein_coding | 0.178322598 | 6.526697747 | 0.5691337 | 0.4613594 | 0.873179344 |
| ENSMUSG00000042680 | Garem1 | protein_coding | -0.322354781 | 3.877893025 | 0.5683766 | 0.4616525 | 0.873179344 |
| ENSMUSG00000006169 | Clint1 | protein_coding | 0.219419586 | 4.742445546 | 0.5681919 | 0.4617241 | 0.873179344 |
| ENSMUSG00000038526 | Car14 | protein_coding | -0.297585202 | 4.815138323 | 0.5680479 | 0.4617975 | 0.873179344 |
| ENSMUSG00000057637 | Prdm2 | protein_coding | 0.221675263 | 5.321888145 | 0.5678943 | 0.4618394 | 0.873179344 |
| ENSMUSG00000029767 | Calu | protein_coding | 0.211539617 | 5.515481613 | 0.5675839 | 0.4619598 | 0.873179344 |
| ENSMUSG00000057229 | Atp5sl | protein_coding | 0.169951973 | 6.897571732 | 0.5674986 | 0.4619929 | 0.873179344 |
| ENSMUSG00000035944 | Ttc38 | protein_coding | 0.343999921 | 3.406208913 | 0.5673273 | 0.4620594 | 0.873179344 |
| ENSMUSG00000028222 | Calb1 | protein_coding | -0.179729299 | 7.475597524 | 0.5673134 | 0.4620648 | 0.873179344 |
| ENSMUSG00000037936 | Scarb1 | protein_coding | 0.269495855 | 4.02383914 | 0.567211 | 0.4621045 | 0.873179344 |
| ENSMUSG00000022112 | Gpc5 | protein_coding | 0.274742528 | 4.158869144 | 0.5669521 | 0.462205 | 0.873179344 |
| ENSMUSG00000053317 | Sec61b | protein_coding | 0.150879284 | 6.311453429 | 0.5659946 | 0.4625769 | 0.873179344 |
| ENSMUSG00000022982 | Sod1 | protein_coding | 0.17967221 | 10.04575214 | 0.5659864 | 0.4625801 | 0.873179344 |
| ENSMUSG00000051615 | Rap2a | protein_coding | 0.1957475 | 5.783770997 | 0.5658693 | 0.4626256 | 0.873179344 |
| ENSMUSG00000021501 | Caml | protein_coding | -0.172721671 | 6.034386963 | 0.5656985 | 0.462692 | 0.873179344 |
| ENSMUSG00000042515 | Mum1l1 | protein_coding | 0.408318515 | 3.670943387 | 0.5657399 | 0.4626934 | 0.873179344 |
| ENSMUSG00000002781 | Tmem143 | protein_coding | -0.300900489 | 3.594656534 | 0.5656312 | 0.4627182 | 0.873179344 |
| ENSMUSG00000050890 | Pdik1l | protein_coding | -0.263133525 | 4.28095881 | 0.5655038 | 0.4627677 | 0.873179344 |
| ENSMUSG00000035325 | Sec31a | protein_coding | 0.1862441 | 6.04844774 | 0.5652892 | 0.4628512 | 0.873179344 |
| ENSMUSG00000027080 | Med19 | protein_coding | -0.179730257 | 6.069591869 | 0.5640628 | 0.4633288 | 0.873179344 |
| ENSMUSG00000021877 | Arf4 | protein_coding | 0.168114711 | 8.805973161 | 0.5640071 | 0.4633505 | 0.873179344 |

| ensembl_gene_id | external_gene_name | gene_biotype | log2 (fold change) | log2 (counts per million) | F-value | p-value | False discovery rate |
|---|---|---|---|---|---|---|---|
| ENSMUSG00000021576 | Pdcd6 | protein_coding | -0.142300233 | 7.915904575 | 0.5636166 | 0.4635028 | 0.873179344 |
| ENSMUSG00000062309 | Rpp25 | protein_coding | 0.252494033 | 4.69093654 | 0.5634544 | 0.4635661 | 0.873179344 |
| ENSMUSG00000008668 | Rps18 | protein_coding | -0.180281538 | 7.167909004 | 0.5633176 | 0.4636194 | 0.873179344 |
| ENSMUSG00000020290 | Xpo1 | protein_coding | -0.2215162 | 5.05689535 | 0.5632685 | 0.4636386 | 0.873179344 |
| ENSMUSG00000033906 | Zdhhc15 | protein_coding | -0.581506821 | 2.51327674 | 0.5632487 | 0.4636463 | 0.873179344 |
| ENSMUSG00000019230 | Lhx9 | protein_coding | 0.556239134 | 4.271214065 | 0.5704117 | 0.4638672 | 0.873179344 |
| ENSMUSG00000052544 | St6galnac3 | protein_coding | -0.30102801 | 3.529795754 | 0.5625837 | 0.4639059 | 0.873179344 |
| ENSMUSG00000062542 | Syt9 | protein_coding | 0.294482236 | 4.246402326 | 0.5622956 | 0.4640185 | 0.873179344 |
| ENSMUSG00000021647 | Cartpt | protein_coding | -0.230580101 | 6.853129418 | 0.5640267 | 0.4640216 | 0.873179344 |
| ENSMUSG00000034664 | Itga2b | protein_coding | 0.275544366 | 4.189217898 | 0.5621951 | 0.4640577 | 0.873179344 |
| ENSMUSG00000029319 | Coq2 | protein_coding | -0.213010191 | 4.903824083 | 0.5621518 | 0.4640746 | 0.873179344 |
| ENSMUSG00000049881 | 2810025M15Rik | transcribed_processe | -0.18089472 | 5.804134644 | 0.5620963 | 0.4640963 | 0.873179344 |
| ENSMUSG00000084898 | Gm12371 | lincRNA | -0.368260376 | 3.240525916 | 0.5620555 | 0.4641123 | 0.873179344 |
| ENSMUSG00000030538 | Cib1 | protein_coding | -0.260749901 | 4.759552767 | 0.5616737 | 0.4642616 | 0.873179344 |
| ENSMUSG00000041594 | Tmtc4 | protein_coding | 0.571759958 | 2.540256851 | 0.5616633 | 0.4642656 | 0.873179344 |
| ENSMUSG00000032418 | Me1 | protein_coding | 0.245148568 | 4.417696933 | 0.5610879 | 0.4644907 | 0.873179344 |
| ENSMUSG00000054893 | Zfp667 | protein_coding | 0.211360319 | 5.479913346 | 0.5609913 | 0.4645285 | 0.873179344 |
| ENSMUSG00000062184 | Hs6st2 | protein_coding | -0.195901573 | 5.469406566 | 0.5609286 | 0.4645531 | 0.873179344 |
| ENSMUSG00000028832 | Stmn1 | protein_coding | 0.144495876 | 7.60688748 | 0.5606935 | 0.4646451 | 0.873179344 |
| ENSMUSG00000035509 | Fbxl21 | protein_coding | 0.282613555 | 3.861763145 | 0.5606359 | 0.4646677 | 0.873179344 |
| ENSMUSG00000026113 | Inpp4a | protein_coding | 0.261617998 | 9.06552636 | 0.5638919 | 0.4647768 | 0.873179344 |
| ENSMUSG00000039519 | Cyp7b1 | protein_coding | -0.370268703 | 3.486738706 | 0.5601656 | 0.4648519 | 0.873179344 |
| ENSMUSG00000057409 | Zfp53 | protein_coding | -0.571984149 | 2.409120313 | 0.5601288 | 0.4648664 | 0.873179344 |
| ENSMUSG00000043987 | Cep164 | protein_coding | -0.443734924 | 2.793293955 | 0.5597859 | 0.4650008 | 0.873179344 |
| ENSMUSG00000022960 | Donson | protein_coding | -0.755781684 | 2.715215233 | 0.5613747 | 0.465275 | 0.873179344 |
| ENSMUSG00000039156 | Stim2 | protein_coding | 0.251551205 | 4.815811894 | 0.558741 | 0.4654108 | 0.873179344 |
| ENSMUSG00000051373 | Plpp7 | protein_coding | 0.224262679 | 4.706038046 | 0.5586572 | 0.4654437 | 0.873179344 |
| ENSMUSG00000032846 | Zswim6 | processed_transcript | -0.401957376 | 3.842680476 | 0.5585787 | 0.4655679 | 0.873179344 |
| ENSMUSG00000025871 | 4833439L19Rik | protein_coding | 0.187301528 | 8.268381697 | 0.5581887 | 0.4656452 | 0.873179344 |
| ENSMUSG00000040722 | Scamp5 | protein_coding | 0.138075716 | 7.379337979 | 0.5581158 | 0.4656564 | 0.873179344 |
| ENSMUSG00000026856 | Dolpp1 | protein_coding | 0.232158219 | 4.741127617 | 0.5578391 | 0.4657652 | 0.873179344 |
| ENSMUSG00000058070 | Eml1 | protein_coding | 0.35596792 | 4.21443811 | 0.5578644 | 0.4657726 | 0.873179344 |
| ENSMUSG00000002458 | Rgs19 | protein_coding | 0.176610755 | 5.987328493 | 0.5577301 | 0.465808 | 0.873179344 |
| ENSMUSG00000043535 | Setx | protein_coding | 0.251792888 | 4.54475546 | 0.5574593 | 0.4659145 | 0.873179344 |
| ENSMUSG00000027166 | Dnajc24 | protein_coding | -0.254211097 | 5.076899597 | 0.5574072 | 0.465935 | 0.873179344 |
| ENSMUSG00000039599 | Fam149b | protein_coding | -0.353798716 | 3.162414155 | 0.5572157 | 0.4660104 | 0.873179344 |
| ENSMUSG00000070498 | Tmem132b | protein_coding | -0.221785504 | 4.839907472 | 0.5567152 | 0.4662074 | 0.873179344 |
| ENSMUSG00000026312 | Cdh7 | protein_coding | -0.244158209 | 4.282803202 | 0.5566412 | 0.4662366 | 0.873179344 |
| ENSMUSG00000026701 | Prdx6 | protein_coding | -0.152732742 | 9.748615925 | 0.5564448 | 0.4663139 | 0.873179344 |
| ENSMUSG00000031479 | Vps36 | protein_coding | 0.202280571 | 5.319831092 | 0.5562607 | 0.4663865 | 0.873179344 |
| ENSMUSG00000038274 | Fau | protein_coding | 0.248703338 | 4.187082892 | 0.5559966 | 0.4664905 | 0.873179344 |
| ENSMUSG00000048106 | 4632415L05Rik | lincRNA | -0.333486698 | 3.111521231 | 0.5558649 | 0.4665425 | 0.873179344 |
| ENSMUSG00000024165 | Hn1l | protein_coding | 0.280180222 | 4.286297872 | 0.5557495 | 0.4665879 | 0.873179344 |
| ENSMUSG00000079242 | C730034F03Rik | protein_coding | -0.447585522 | 3.111918437 | 0.5556676 | 0.4666203 | 0.873179344 |
| ENSMUSG00000039328 | Rnf122 | protein_coding | -0.263898363 | 4.062670197 | 0.5553366 | 0.4667508 | 0.873179344 |
| ENSMUSG00000061132 | Blnk | protein_coding | -0.903635847 | 2.08573582 | 0.5560871 | 0.4667832 | 0.873179344 |
| ENSMUSG00000037737 | Actrt3 | protein_coding | 0.674722499 | 2.315384612 | 0.5552425 | 0.466788 | 0.873179344 |
| ENSMUSG00000032878 | Ccdc85a | protein_coding | -0.193974244 | 5.289002769 | 0.5552419 | 0.4667882 | 0.873179344 |
| ENSMUSG00000045071 | E130308A19Rik | protein_coding | -0.338174988 | 3.44162894 | 0.5550093 | 0.46688 | 0.873179344 |
| ENSMUSG00000027575 | Arfgap1 | protein_coding | 0.162704123 | 6.255567342 | 0.5549954 | 0.4668855 | 0.873179344 |
| ENSMUSG00000018476 | Kdm6b | protein_coding | 0.358784908 | 5.4998987 | 0.5587463 | 0.4668938 | 0.873179344 |
| ENSMUSG00000029712 | Actl6b | protein_coding | 0.174944735 | 6.161214538 | 0.5549092 | 0.4669195 | 0.873179344 |
| ENSMUSG00000074212 | Dnajb14 | protein_coding | 0.179438323 | 5.796248168 | 0.5545299 | 0.4670693 | 0.873179344 |
| ENSMUSG00000004633 | Chn2 | protein_coding | -0.534637456 | 2.262162866 | 0.5544549 | 0.467099 | 0.873179344 |
| ENSMUSG00000042569 | Dhrs7b | protein_coding | 0.198878654 | 5.481559796 | 0.5542935 | 0.4671627 | 0.873179344 |
| ENSMUSG00000029009 | Mthfr | protein_coding | 0.466854013 | 3.157893418 | 0.5537413 | 0.4673983 | 0.873467892 |
| ENSMUSG00000018858 | Mrpl58 | protein_coding | 0.135781586 | 7.702655499 | 0.5533709 | 0.4675275 | 0.873557513 |
| ENSMUSG00000033233 | Trim45 | protein_coding | 0.430864816 | 3.97313716 | 0.5544528 | 0.467772 | 0.873852349 |
| ENSMUSG00000099034 | 2810039B14Rik | transcribed_unitary_p | -0.340337576 | 3.229645535 | 0.5525322 | 0.4678596 | 0.873852349 |
| ENSMUSG00000024647 | Cbln2 | protein_coding | -0.233367282 | 4.755811278 | 0.5522566 | 0.4679688 | 0.873852349 |
| ENSMUSG00000000276 | Dgke | protein_coding | 0.246999135 | 4.35600267 | 0.5521509 | 0.4680106 | 0.873852349 |
| ENSMUSG00000031371 | Haus7 | protein_coding | 0.164399288 | 5.661302932 | 0.5515744 | 0.4682392 | 0.873852349 |
| ENSMUSG00000047656 | Trpt1 | protein_coding | 0.196415275 | 5.582264213 | 0.5512183 | 0.4683805 | 0.873852349 |
| ENSMUSG00000018761 | Mpdu1 | protein_coding | 0.157348374 | 6.826506948 | 0.5511588 | 0.4684041 | 0.873852349 |
| ENSMUSG00000047547 | Cltb | protein_coding | -0.177367025 | 7.68489647 | 0.5510324 | 0.4684543 | 0.873852349 |
| ENSMUSG00000019841 | Rev3l | protein_coding | -0.303918319 | 4.223288506 | 0.5509995 | 0.4684674 | 0.873852349 |
| ENSMUSG00000036879 | Phkb | protein_coding | 0.230542964 | 4.607449824 | 0.5508365 | 0.4685321 | 0.873852349 |
| ENSMUSG00000020160 | Meis1 | protein_coding | -0.519016453 | 3.539785769 | 0.5530061 | 0.4687236 | 0.873852349 |
| ENSMUSG00000070366 | Plpp4 | protein_coding | -0.307504943 | 4.061904239 | 0.5499017 | 0.4689036 | 0.873852349 |
| ENSMUSG00000007817 | Zmiz1 | protein_coding | 0.198657929 | 5.827892092 | 0.5496948 | 0.4689858 | 0.873852349 |
| ENSMUSG00000027014 | Cwc22 | protein_coding | 0.34753581 | 3.187849467 | 0.5493965 | 0.4691045 | 0.873852349 |
| ENSMUSG00000074578 | Zfas1 | processed_transcript | -0.198285168 | 7.938346674 | 0.5493533 | 0.4691389 | 0.873852349 |
| ENSMUSG00000033220 | Rac2 | protein_coding | -0.50284877 | 2.77756677 | 0.5492945 | 0.4691451 | 0.873852349 |
| ENSMUSG00000039395 | Mreg | protein_coding | -0.488818083 | 2.458589269 | 0.5492648 | 0.4691569 | 0.873852349 |

| ensembl_gene_id | external_gene_name | gene_biotype | log2 (fold change) | log2 (counts per million) | F-value | p-value | False discovery rate |
|---|---|---|---|---|---|---|---|
| ENSMUSG00000024827 | Gldc | protein_coding | 0.551361112 | 2.589045251 | 0.5488896 | 0.4693063 | 0.873852349 |
| ENSMUSG00000002846 | Timmdc1 | protein_coding | 0.280027675 | 5.077553258 | 0.5489181 | 0.4693122 | 0.873852349 |
| ENSMUSG00000018167 | Stard3 | protein_coding | -0.246593222 | 5.476916645 | 0.5488203 | 0.4693511 | 0.873852349 |
| ENSMUSG00000025290 | Rps24 | protein_coding | -0.193698116 | 8.061290589 | 0.5487179 | 0.4693919 | 0.873852349 |
| ENSMUSG00000035164 | Zc3h12c | protein_coding | -0.346051065 | 3.641309505 | 0.5478688 | 0.4697131 | 0.874298846 |
| ENSMUSG00000043885 | Slc36a4 | protein_coding | 0.222830241 | 5.17974341 | 0.5474468 | 0.4698814 | 0.874460786 |
| ENSMUSG00000022150 | Dab2 | protein_coding | -0.376214848 | 3.048865381 | 0.5470803 | 0.4700277 | 0.874581655 |
| ENSMUSG00000032386 | Trip4 | protein_coding | -0.203354914 | 4.950811401 | 0.5464415 | 0.4702828 | 0.874826678 |
| ENSMUSG00000025102 | 3110040N11Rik | protein_coding | 0.165501622 | 6.076094014 | 0.5462201 | 0.4703713 | 0.874826678 |
| ENSMUSG00000075227 | Znhit2 | protein_coding | -0.320737874 | 3.470282843 | 0.5460304 | 0.4704471 | 0.874826678 |
| ENSMUSG00000089665 | Fcor | protein_coding | -0.24672872 | 4.81290421 | 0.54583 | 0.4705273 | 0.874826678 |
| ENSMUSG00000022240 | Ctnnd2 | protein_coding | -0.13750214 | 7.273596759 | 0.5456375 | 0.4706043 | 0.874826678 |
| ENSMUSG00000026031 | Cflar | protein_coding | -0.324222922 | 3.42663038 | 0.5453513 | 0.4707188 | 0.874826678 |
| ENSMUSG00000014668 | Chfr | protein_coding | 0.663321145 | 2.161270994 | 0.5451851 | 0.4707853 | 0.874826678 |
| ENSMUSG00000049164 | Zfp518a | protein_coding | -0.378097856 | 2.82800712 | 0.5450694 | 0.4708316 | 0.874826678 |
| ENSMUSG00000086877 | A230072C01Rik | processed_transcript | -0.224896772 | 4.659431682 | 0.5448733 | 0.4709102 | 0.874826678 |
| ENSMUSG00000003809 | Gcdh | protein_coding | 0.272816526 | 4.636542732 | 0.5447166 | 0.4709729 | 0.874826678 |
| ENSMUSG00000034157 | Cipc | protein_coding | 0.172449136 | 6.034007373 | 0.5441029 | 0.4712188 | 0.875053246 |
| ENSMUSG00000039202 | Abhd2 | protein_coding | -0.253966683 | 4.758151013 | 0.5439377 | 0.4712851 | 0.875053246 |
| ENSMUSG00000036469 | 1-Mar | protein_coding | -0.19368622 | 4.910146209 | 0.5438033 | 0.471339 | 0.875053246 |
| ENSMUSG00000021495 | Fam193b | protein_coding | 0.425595977 | 3.323254284 | 0.5435002 | 0.4714706 | 0.875146404 |
| ENSMUSG00000030307 | Slc6a11 | protein_coding | 0.217283721 | 6.060838409 | 0.5429713 | 0.4716901 | 0.875328219 |
| ENSMUSG00000036896 | C1qc | protein_coding | -0.423969713 | 4.251901366 | 0.5453341 | 0.4717691 | 0.875328219 |
| ENSMUSG00000053799 | Exoc6 | protein_coding | 0.243604955 | 4.011580773 | 0.5426232 | 0.4718127 | 0.875328219 |
| ENSMUSG00000078671 | Chd2 | protein_coding | 0.220219882 | 4.680624463 | 0.5422911 | 0.4719462 | 0.875424818 |
| ENSMUSG00000030579 | Tyrobp | protein_coding | -0.318397973 | 5.019160979 | 0.5427968 | 0.4721801 | 0.875424939 |
| ENSMUSG00000064264 | Zfp428 | protein_coding | -0.31762518 | 3.237660318 | 0.5415534 | 0.4722429 | 0.875424939 |
| ENSMUSG00000052926 | Rnaseh2a | protein_coding | 0.181194803 | 5.930473243 | 0.54142 | 0.4722966 | 0.875424939 |
| ENSMUSG00000031349 | Nsdhl | protein_coding | 0.283526706 | 5.791566562 | 0.5430621 | 0.4723007 | 0.875424939 |
| ENSMUSG00000040877 | Wdr25 | protein_coding | 0.428260947 | 3.857686571 | 0.5427198 | 0.4724011 | 0.875424939 |
| ENSMUSG00000048497 | Mmgt2 | protein_coding | 0.344844208 | 3.594320019 | 0.5409984 | 0.4724663 | 0.875424939 |
| ENSMUSG00000024026 | Glo1 | protein_coding | 0.133308537 | 7.490100632 | 0.5408746 | 0.4725162 | 0.875424939 |
| ENSMUSG00000096916 | Zfp850 | protein_coding | -0.630147468 | 2.376784484 | 0.5404095 | 0.4727036 | 0.875621311 |
| ENSMUSG00000022604 | Cep97 | protein_coding | -0.277221185 | 3.975601437 | 0.5397602 | 0.4729654 | 0.875955395 |
| ENSMUSG00000042198 | Chchd7 | protein_coding | -0.207443225 | 5.275850413 | 0.538695 | 0.4733955 | 0.876342546 |
| ENSMUSG00000044167 | Foxo1 | protein_coding | 0.202293121 | 5.322789056 | 0.5386272 | 0.4734229 | 0.876342546 |
| ENSMUSG00000020329 | Polrmt | protein_coding | -0.426652192 | 3.309973205 | 0.5385768 | 0.4734432 | 0.876342546 |
| ENSMUSG00000054405 | Dnajc8 | protein_coding | 0.125313435 | 8.553718443 | 0.53809 | 0.47364 | 0.876342546 |
| ENSMUSG00000027244 | Atg13 | protein_coding | 0.189367903 | 6.63133618 | 0.537897 | 0.4737181 | 0.876342546 |
| ENSMUSG00000030697 | Ppp4c | protein_coding | 0.175321146 | 5.388606369 | 0.5376814 | 0.4738053 | 0.876342546 |
| ENSMUSG00000031360 | Ctps2 | protein_coding | 0.251202036 | 5.768637426 | 0.5378099 | 0.4738153 | 0.876342546 |
| ENSMUSG00000028416 | Bag1 | protein_coding | -0.162117909 | 6.204112354 | 0.5376293 | 0.4738264 | 0.876342546 |
| ENSMUSG00000025486 | Sirt3 | protein_coding | 0.148797704 | 7.720722773 | 0.5371066 | 0.474038 | 0.876541193 |
| ENSMUSG00000096225 | Lhx8 | protein_coding | -0.274266785 | 4.460769682 | 0.5365947 | 0.4742454 | 0.876541193 |
| ENSMUSG00000087651 | 1500009L16Rik | protein_coding | 0.151570807 | 6.322882621 | 0.5365743 | 0.4742537 | 0.876541193 |
| ENSMUSG00000021958 | Pinx1 | protein_coding | -0.223830024 | 4.210913605 | 0.5365589 | 0.4742599 | 0.876541193 |
| ENSMUSG00000040048 | Ndufb10 | protein_coding | -0.147600125 | 10.18834066 | 0.5361935 | 0.474408 | 0.876578179 |
| ENSMUSG00000028923 | Necap2 | protein_coding | -0.198467428 | 5.508739971 | 0.5358769 | 0.4745365 | 0.876578179 |
| ENSMUSG00000039179 | Tekt5 | protein_coding | 0.218019705 | 5.769587978 | 0.535336 | 0.474756 | 0.876578179 |
| ENSMUSG00000072494 | Ppp1r3e | protein_coding | -0.342177759 | 3.247803737 | 0.5352393 | 0.4747952 | 0.876578179 |
| ENSMUSG00000005575 | Ube2m | protein_coding | -0.168951472 | 6.758331894 | 0.5352273 | 0.4748001 | 0.876578179 |
| ENSMUSG00000038677 | Scube3 | protein_coding | -0.490255926 | 2.59670893 | 0.5350813 | 0.4748594 | 0.876578179 |
| ENSMUSG00000030447 | Cyfip1 | protein_coding | -0.208985464 | 5.532259004 | 0.5349954 | 0.4748943 | 0.876578179 |
| ENSMUSG00000042682 | Selenok | protein_coding | -0.164968079 | 9.326716745 | 0.5346091 | 0.4750513 | 0.876578179 |
| ENSMUSG00000029840 | Mtpn | protein_coding | -0.160247251 | 7.639171487 | 0.5345807 | 0.4750629 | 0.876578179 |
| ENSMUSG00000027674 | Pex5l | protein_coding | -0.33581581 | 3.225459184 | 0.5344129 | 0.475131 | 0.876578179 |
| ENSMUSG00000032570 | Atp2c1 | protein_coding | -0.214390943 | 5.012945453 | 0.5343008 | 0.4751766 | 0.876578179 |
| ENSMUSG00000047721 | Bola2 | protein_coding | -0.194971058 | 7.920219984 | 0.5339453 | 0.4753382 | 0.876601734 |
| ENSMUSG00000028889 | Yrdc | protein_coding | -0.289877068 | 4.415054992 | 0.5338686 | 0.4753524 | 0.876601734 |
| ENSMUSG00000032985 | 5730522E02Rik | protein_coding | -0.431545434 | 3.229163008 | 0.5333093 | 0.4755801 | 0.876762271 |
| ENSMUSG00000031378 | Abcd1 | protein_coding | 0.478086272 | 2.850826298 | 0.5332541 | 0.4756026 | 0.876762271 |
| ENSMUSG00000029007 | Agtrap | protein_coding | -0.263587424 | 4.356498875 | 0.5328861 | 0.4757525 | 0.876888312 |
| ENSMUSG00000027603 | Ggt7 | protein_coding | 0.172605373 | 7.073577587 | 0.531942 | 0.4761374 | 0.877224163 |
| ENSMUSG00000042050 | Wdr60 | protein_coding | -0.299979184 | 3.910344139 | 0.5317799 | 0.4762036 | 0.877224163 |
| ENSMUSG00000022840 | Adcy5 | protein_coding | -0.205921519 | 6.869010957 | 0.5319435 | 0.4762179 | 0.877224163 |
| ENSMUSG00000020149 | Rab1a | protein_coding | 0.154742927 | 8.128892063 | 0.5315558 | 0.4762951 | 0.877224163 |
| ENSMUSG00000026778 | Prkcq | protein_coding | -0.358081801 | 3.361686752 | 0.5314394 | 0.4763426 | 0.877224163 |
| ENSMUSG00000023939 | Mrpl14 | protein_coding | -0.143060007 | 7.523152473 | 0.5310302 | 0.4765097 | 0.877381703 |
| ENSMUSG00000041598 | Cdc42ep4 | protein_coding | 0.243385525 | 4.979978276 | 0.530597 | 0.4766868 | 0.877557431 |
| ENSMUSG00000035933 | Cog5 | protein_coding | -0.361107673 | 3.09656944 | 0.530066 | 0.4769039 | 0.877593031 |
| ENSMUSG00000021111 | Papola | protein_coding | -0.176778631 | 6.108188155 | 0.5300298 | 0.4769187 | 0.877593031 |
| ENSMUSG00000020859 | Spag9 | protein_coding | -0.156683841 | 6.745140716 | 0.5299511 | 0.476951 | 0.877593031 |
| ENSMUSG00000040272 | Accs | protein_coding | -0.667617014 | 2.430935469 | 0.5294318 | 0.4771804 | 0.87771131 |
| ENSMUSG00000003352 | Cacnb3 | protein_coding | -0.175177696 | 5.509101422 | 0.5293261 | 0.4772068 | 0.87771131 |

| ensembl_gene_id | external_gene_name | gene_biotype | log2 (fold change) | log2 (counts per million) | F-value | p-value | False discovery rate |
|---|---|---|---|---|---|---|---|
| ENSMUSG00000030231 | Plekha5 | protein_coding | -0.185904581 | 5.603126478 | 0.5286379 | 0.4774889 | 0.87771131 |
| ENSMUSG00000017897 | Eya2 | protein_coding | -0.329692884 | 3.05916692 | 0.5282242 | 0.4776585 | 0.87771131 |
| ENSMUSG00000040124 | Gorab | protein_coding | -0.231376444 | 4.695669483 | 0.5280641 | 0.4777242 | 0.87771131 |
| ENSMUSG00000021824 | Ap3m1 | protein_coding | -0.201556872 | 4.904119096 | 0.5279408 | 0.4777748 | 0.87771131 |
| ENSMUSG00000046027 | Stard5 | protein_coding | -0.241174357 | 4.094386568 | 0.5279358 | 0.4777768 | 0.87771131 |
| ENSMUSG00000021474 | Sfxn1 | protein_coding | -0.130237318 | 7.44795253 | 0.5278443 | 0.4778144 | 0.87771131 |
| ENSMUSG00000039474 | Wfs1 | protein_coding | -0.175777626 | 6.511998723 | 0.527786 | 0.4778383 | 0.87771131 |
| ENSMUSG00000029169 | Dhx15 | protein_coding | -0.197068938 | 6.032586644 | 0.5271743 | 0.4780895 | 0.87771131 |
| ENSMUSG00000079481 | Nhsl2 | protein_coding | -0.214022347 | 5.12842791 | 0.5269151 | 0.4781961 | 0.87771131 |
| ENSMUSG00000039717 | Ralyl | protein_coding | -0.138334658 | 7.948291506 | 0.5268464 | 0.4782243 | 0.87771131 |
| ENSMUSG00000038013 | Wipf2 | protein_coding | -0.151830473 | 6.463170301 | 0.526799 | 0.4782438 | 0.87771131 |
| ENSMUSG00000022389 | Tef | protein_coding | 0.123683619 | 7.112904169 | 0.5266087 | 0.4783221 | 0.87771131 |
| ENSMUSG00000031750 | Il34 | protein_coding | 0.197261966 | 5.729122828 | 0.5260732 | 0.4785423 | 0.87771131 |
| ENSMUSG00000034673 | Pbx2 | protein_coding | 0.234480693 | 4.721255609 | 0.5259054 | 0.4786114 | 0.87771131 |
| ENSMUSG00000025509 | Pnpla2 | protein_coding | 0.299431703 | 5.14181246 | 0.5264966 | 0.4786863 | 0.87771131 |
| ENSMUSG00000024430 | Cabyr | protein_coding | 0.503628005 | 2.451006749 | 0.5256448 | 0.4787187 | 0.87771131 |
| ENSMUSG00000024008 | Cpne5 | protein_coding | -0.194996885 | 7.202802253 | 0.5255595 | 0.4787706 | 0.87771131 |
| ENSMUSG00000020372 | Rack1 | protein_coding | -0.146164352 | 8.901453793 | 0.5254312 | 0.4788067 | 0.87771131 |
| ENSMUSG00000029227 | Fip1l1 | protein_coding | -0.312541858 | 4.152033898 | 0.5249741 | 0.478995 | 0.87771131 |
| ENSMUSG00000033323 | Ctdp1 | protein_coding | 0.60873416 | 2.510626562 | 0.5248298 | 0.4790545 | 0.87771131 |
| ENSMUSG00000029516 | Cit | protein_coding | 0.277189922 | 8.525516425 | 0.5280908 | 0.4793495 | 0.87771131 |
| ENSMUSG00000022749 | Tbc1d23 | protein_coding | 0.297353238 | 3.55848293 | 0.5240967 | 0.4793569 | 0.87771131 |
| ENSMUSG00000041544 | Disp3 | protein_coding | 0.396299749 | 3.256826416 | 0.523807 | 0.4794765 | 0.87771131 |
| ENSMUSG00000029632 | Ndufa4 | protein_coding | -0.179710338 | 11.03033526 | 0.523317 | 0.4796789 | 0.87771131 |
| ENSMUSG00000063275 | Hacd1 | protein_coding | -0.489915374 | 2.469023942 | 0.5232662 | 0.4796999 | 0.87771131 |
| ENSMUSG00000037197 | Rbm17 | protein_coding | 0.205745521 | 4.756584771 | 0.5232554 | 0.4797043 | 0.87771131 |
| ENSMUSG00000029535 | Triap1 | protein_coding | -0.161050494 | 6.337030239 | 0.5231231 | 0.479759 | 0.87771131 |
| ENSMUSG00000023945 | Slc5a7 | protein_coding | -0.333094733 | 3.343747888 | 0.5229075 | 0.4798482 | 0.87771131 |
| ENSMUSG00000028821 | Syf2 | protein_coding | -0.152199743 | 7.347891965 | 0.522836 | 0.4798777 | 0.87771131 |
| ENSMUSG00000033565 | Rbfox2 | protein_coding | -0.14101241 | 6.285508904 | 0.5226941 | 0.4799364 | 0.87771131 |
| ENSMUSG00000074923 | Pak6 | protein_coding | -0.219498764 | 4.58149921 | 0.5226599 | 0.4799505 | 0.87771131 |
| ENSMUSG00000032174 | Icam5 | protein_coding | -0.602959587 | 2.171267815 | 0.522659 | 0.4799509 | 0.87771131 |
| ENSMUSG00000048109 | Rbm15 | protein_coding | -0.330233179 | 5.00797031 | 0.5240452 | 0.480029 | 0.87771131 |
| ENSMUSG00000042589 | Cux2 | protein_coding | -0.345903776 | 5.710128165 | 0.5264272 | 0.48003 | 0.87771131 |
| ENSMUSG00000020388 | Pdlim4 | protein_coding | -0.339050337 | 3.714481648 | 0.5224549 | 0.4800354 | 0.87771131 |
| ENSMUSG00000035640 | Cbarp | protein_coding | -0.168827708 | 6.86161817 | 0.5212084 | 0.4805515 | 0.878457208 |
| ENSMUSG00000026260 | Ndufa10 | protein_coding | 0.206882906 | 8.476402603 | 0.5218949 | 0.4806067 | 0.878457208 |
| ENSMUSG00000028480 | Glipr2 | protein_coding | 0.483946091 | 2.854064298 | 0.5201395 | 0.4809949 | 0.879017441 |
| ENSMUSG00000002679 | Med6 | protein_coding | 0.198695882 | 5.727165433 | 0.5197062 | 0.4811748 | 0.879036049 |
| ENSMUSG00000021364 | Elovl2 | protein_coding | -0.609381524 | 2.667073904 | 0.5196908 | 0.481198 | 0.879036049 |
| ENSMUSG00000035284 | Vps13c | protein_coding | -0.240997363 | 4.709763561 | 0.5193393 | 0.4813273 | 0.879036049 |
| ENSMUSG00000032607 | Amt | protein_coding | 0.354289669 | 5.327476593 | 0.5227552 | 0.4813408 | 0.879036049 |
| ENSMUSG00000057177 | Gsk3a | protein_coding | -0.197173355 | 4.855785757 | 0.5191311 | 0.4814139 | 0.879036049 |
| ENSMUSG00000039096 | Rsad1 | protein_coding | 0.409901423 | 3.080677049 | 0.5186592 | 0.4816101 | 0.879245089 |
| ENSMUSG00000036438 | Calm2 | protein_coding | -0.165216422 | 12.2062153 | 0.5181997 | 0.4818014 | 0.879444966 |
| ENSMUSG00000020964 | Sel1l | protein_coding | 0.160515123 | 5.974239127 | 0.5179537 | 0.4819038 | 0.879482614 |
| ENSMUSG00000038884 | A230050P20Rik | protein_coding | -0.345915486 | 4.417019797 | 0.5182889 | 0.4820811 | 0.879656808 |
| ENSMUSG00000020482 | Ccdc117 | protein_coding | -0.36420026 | 3.228345873 | 0.5166299 | 0.4824557 | 0.880191004 |
| ENSMUSG00000056427 | Slit3 | protein_coding | -0.396569229 | 2.882208452 | 0.5163862 | 0.4825574 | 0.880221782 |
| ENSMUSG00000030410 | Dmwd | protein_coding | -0.175329394 | 6.561215865 | 0.5161973 | 0.4826363 | 0.880221782 |
| ENSMUSG00000047945 | Marcksl1 | protein_coding | -0.237733705 | 4.9562803 | 0.5154064 | 0.4829667 | 0.88055512 |
| ENSMUSG00000038861 | Pi4kb | protein_coding | 0.312712355 | 3.527739879 | 0.5153679 | 0.4829828 | 0.88055512 |
| ENSMUSG00000105879 | Gm6204 | processed_pseudoge | -0.27257988 | 3.852311768 | 0.5150289 | 0.4831246 | 0.880584051 |
| ENSMUSG00000006931 | P3h4 | protein_coding | 0.271393768 | 3.879283003 | 0.5149384 | 0.4831625 | 0.880584051 |
| ENSMUSG00000036966 | Spryd3 | protein_coding | 0.17698327 | 8.680516596 | 0.5142038 | 0.4834699 | 0.880906837 |
| ENSMUSG00000032369 | Plscr1 | protein_coding | 0.545837307 | 2.352033512 | 0.5141239 | 0.4835034 | 0.880906837 |
| ENSMUSG00000049526 | Tmem202 | protein_coding | -0.629061673 | 2.35832237 | 0.5137943 | 0.4836415 | 0.881009097 |
| ENSMUSG00000029669 | Tspan12 | protein_coding | -0.200106375 | 6.187878731 | 0.5131888 | 0.4838953 | 0.881322239 |
| ENSMUSG00000044636 | Csrnp2 | protein_coding | -0.279660724 | 4.041868163 | 0.5128937 | 0.4840191 | 0.881398423 |
| ENSMUSG00000068479 | Mfap1a | protein_coding | -0.23257258 | 4.009950903 | 0.5123407 | 0.4842513 | 0.881585434 |
| ENSMUSG00000058454 | Dhcr7 | protein_coding | -0.17339609 | 5.621171495 | 0.512135 | 0.4843377 | 0.881585434 |
| ENSMUSG00000025224 | Gbf1 | protein_coding | 0.20302733 | 5.734216641 | 0.5120634 | 0.4843678 | 0.881585434 |
| ENSMUSG00000024725 | Ostf1 | protein_coding | 0.340021414 | 3.297632081 | 0.5112473 | 0.4847108 | 0.881983111 |
| ENSMUSG00000031309 | Rps6ka3 | protein_coding | -0.302935937 | 3.483811245 | 0.510885 | 0.4848633 | 0.881983111 |
| ENSMUSG00000030731 | Syt3 | protein_coding | -0.209109914 | 4.880763602 | 0.5104861 | 0.4850313 | 0.881983111 |
| ENSMUSG00000038250 | Usp38 | protein_coding | -0.410083253 | 4.033244723 | 0.511947 | 0.4850589 | 0.881983111 |
| ENSMUSG00000072694 | 1500011B03Rik | protein_coding | 0.121629942 | 8.042027602 | 0.5104047 | 0.4850655 | 0.881983111 |
| ENSMUSG00000004637 | Wwox | protein_coding | 0.210151573 | 6.345004564 | 0.5104134 | 0.4850784 | 0.881983111 |
| ENSMUSG00000032840 | 2410131K14Rik | protein_coding | -0.183950452 | 5.490908084 | 0.5094259 | 0.4854781 | 0.882560579 |
| ENSMUSG00000046215 | Rprml | protein_coding | -0.366655767 | 3.693100388 | 0.5087964 | 0.4857603 | 0.882924295 |
| ENSMUSG00000003762 | Coq8b | protein_coding | -0.41560965 | 2.662124604 | 0.5081862 | 0.4860015 | 0.88321342 |
| ENSMUSG00000040794 | C1qtnf4 | protein_coding | 0.374969398 | 3.310333493 | 0.5079863 | 0.486086 | 0.883217723 |
| ENSMUSG00000020087 | Tysnd1 | protein_coding | 0.579589689 | 2.574259294 | 0.5075618 | 0.4862654 | 0.883252331 |
| ENSMUSG00000057069 | Ero1lb | protein_coding | 0.583673838 | 2.659979662 | 0.5074437 | 0.4863319 | 0.883252331 |

| ensembl_gene_id | external_gene_name | gene_biotype | log2 (fold change) | log2 (counts per million) | F-value | p-value | False discovery rate |
|---|---|---|---|---|---|---|---|
| ENSMUSG00000002812 | Flii | protein_coding | 0.263118181 | 5.044575425 | 0.5073971 | 0.4863516 | 0.883252331 |
| ENSMUSG00000045948 | Mrps12 | protein_coding | 0.12288413 | 7.801019874 | 0.5071392 | 0.4864443 | 0.883252331 |
| ENSMUSG00000093904 | Tomm20 | protein_coding | 0.145833603 | 6.853735272 | 0.5069702 | 0.4865158 | 0.883252331 |
| ENSMUSG00000051098 | Mblac2 | protein_coding | -0.229541166 | 5.543081478 | 0.5067764 | 0.4865979 | 0.883252331 |
| ENSMUSG00000033578 | Tmem35a | protein_coding | 0.146158204 | 6.739431291 | 0.5064697 | 0.4867278 | 0.883339013 |
| ENSMUSG00000032044 | Rpusd4 | protein_coding | 0.243772827 | 4.453725906 | 0.5056696 | 0.4870669 | 0.883755617 |
| ENSMUSG00000028032 | Papss1 | protein_coding | 0.135258871 | 6.706910355 | 0.5055406 | 0.4871217 | 0.883755617 |
| ENSMUSG00000064247 | Plcxd1 | protein_coding | 0.205995468 | 4.633945197 | 0.5050483 | 0.4873306 | 0.883985572 |
| ENSMUSG00000018634 | Crhr1 | protein_coding | 0.449825879 | 2.681756768 | 0.5047447 | 0.4874596 | 0.884070312 |
| ENSMUSG00000055653 | Gpc3 | protein_coding | 0.227134947 | 4.439939438 | 0.503943 | 0.4878003 | 0.884103897 |
| ENSMUSG00000030611 | Mrps11 | protein_coding | -0.137042052 | 7.327358573 | 0.5037646 | 0.4878762 | 0.884103897 |
| ENSMUSG00000021338 | Carmil1 | protein_coding | 0.302727905 | 3.396781773 | 0.5037008 | 0.4879033 | 0.884103897 |
| ENSMUSG00000020308 | Tpgs1 | protein_coding | 0.375235736 | 4.572860729 | 0.5058581 | 0.4879865 | 0.884103897 |
| ENSMUSG00000042605 | Atxn2 | protein_coding | -0.168196882 | 5.778144979 | 0.5033668 | 0.4880455 | 0.884103897 |
| ENSMUSG00000029089 | 5730480H06Rik | protein_coding | 0.338921354 | 3.863149373 | 0.5033286 | 0.4880781 | 0.884103897 |
| ENSMUSG00000021608 | Lpcat1 | protein_coding | -0.248281324 | 4.229477468 | 0.5025398 | 0.4883977 | 0.884103897 |
| ENSMUSG00000001506 | Col1a1 | protein_coding | -0.694809921 | 2.015746299 | 0.5025259 | 0.4884036 | 0.884103897 |
| ENSMUSG00000002825 | Qtrt1 | protein_coding | 0.194596621 | 5.198670867 | 0.5024761 | 0.4884249 | 0.884103897 |
| ENSMUSG00000032135 | Mcam | protein_coding | 0.549553689 | 2.90400085 | 0.5024767 | 0.488441 | 0.884103897 |
| ENSMUSG00000029095 | Ablim2 | protein_coding | -0.154604847 | 6.51450988 | 0.5023001 | 0.4884999 | 0.884103897 |
| ENSMUSG00000022092 | Ppp3cc | protein_coding | -0.384904539 | 3.041517414 | 0.5022416 | 0.4885248 | 0.884103897 |
| ENSMUSG00000066877 | Nck2 | protein_coding | -0.270352818 | 3.486586342 | 0.5021897 | 0.488547 | 0.884103897 |
| ENSMUSG00000062054 | Iah1 | protein_coding | -0.185094866 | 5.522841567 | 0.5015661 | 0.488813 | 0.88427825 |
| ENSMUSG00000053684 | BC048403 | protein_coding | -0.424154915 | 2.762959654 | 0.5015027 | 0.48884 | 0.88427825 |
| ENSMUSG00000032773 | Chrm1 | protein_coding | -0.242404278 | 5.403171046 | 0.501105 | 0.4890263 | 0.88427825 |
| ENSMUSG00000021134 | Srsf5 | protein_coding | 0.140418676 | 7.73424352 | 0.5010513 | 0.4890328 | 0.88427825 |
| ENSMUSG00000070737 | Tmem35b | protein_coding | 0.357341919 | 2.970141968 | 0.5009812 | 0.4890628 | 0.88427825 |
| ENSMUSG00000030386 | Zfp606 | protein_coding | 0.310049718 | 3.567408348 | 0.5002327 | 0.4893827 | 0.88427825 |
| ENSMUSG00000030720 | Cln3 | protein_coding | -0.306950653 | 4.129359323 | 0.5001856 | 0.4894029 | 0.88427825 |
| ENSMUSG00000084799 | Ino80dos | lincRNA | -0.372875967 | 2.857561626 | 0.5001357 | 0.4894242 | 0.88427825 |
| ENSMUSG00000006463 | Zdhhc24 | protein_coding | 0.289589404 | 3.984992583 | 0.5000269 | 0.4894707 | 0.88427825 |
| ENSMUSG00000005881 | Ergic3 | protein_coding | 0.121141877 | 8.70639423 | 0.500018 | 0.4894745 | 0.88427825 |
| ENSMUSG00000073424 | Cyp4f15 | protein_coding | 0.721092817 | 2.390839071 | 0.4998083 | 0.489628 | 0.88427825 |
| ENSMUSG00000039065 | Fam173b | protein_coding | 0.210181992 | 4.623880927 | 0.4995096 | 0.4896921 | 0.88427825 |
| ENSMUSG00000028577 | Plaa | protein_coding | -0.21245221 | 4.639243415 | 0.4994623 | 0.4897124 | 0.88427825 |
| ENSMUSG00000079330 | Lemd1 | protein_coding | 0.330944316 | 3.620997287 | 0.4990257 | 0.4898994 | 0.88437541 |
| ENSMUSG00000046897 | Zfp740 | protein_coding | 0.199140497 | 4.845189861 | 0.4988974 | 0.4899543 | 0.88437541 |
| ENSMUSG00000062929 | Cfl2 | protein_coding | -0.12969549 | 7.9738112 | 0.4986498 | 0.4900605 | 0.88437541 |
| ENSMUSG00000052889 | Prkcb | protein_coding | 0.174447807 | 8.464317642 | 0.498607 | 0.4900951 | 0.88437541 |
| ENSMUSG00000028479 | Gne | protein_coding | 0.216240911 | 4.514314055 | 0.498299 | 0.4902109 | 0.884435884 |
| ENSMUSG00000034105 | Tldc1 | protein_coding | 0.288706729 | 3.480613273 | 0.4976335 | 0.4904965 | 0.884783695 |
| ENSMUSG00000040729 | Cep126 | protein_coding | -0.485861462 | 3.213043619 | 0.4976281 | 0.4906606 | 0.884783695 |
| ENSMUSG00000034462 | Pkd2 | protein_coding | -0.39253838 | 2.742760216 | 0.4971798 | 0.4906914 | 0.884783695 |
| ENSMUSG00000034501 | Pcnx4 | protein_coding | 0.224568778 | 4.806627932 | 0.4969596 | 0.490786 | 0.884783695 |
| ENSMUSG00000022594 | Lynx1 | protein_coding | 0.153003956 | 7.52563258 | 0.4968823 | 0.4908192 | 0.884783695 |
| ENSMUSG00000049489 | Fam58b | protein_coding | 0.212686592 | 4.903408993 | 0.4967005 | 0.4908974 | 0.884783695 |
| ENSMUSG00000031458 | Coprs | protein_coding | 0.205253878 | 6.282476966 | 0.4957502 | 0.4913226 | 0.885174452 |
| ENSMUSG00000034300 | Fam53c | protein_coding | 0.218908486 | 5.380149082 | 0.495617 | 0.4913636 | 0.885174452 |
| ENSMUSG00000002459 | Rgs20 | protein_coding | 0.176455255 | 6.075147803 | 0.4956164 | 0.4913639 | 0.885174452 |
| ENSMUSG00000028111 | Ctsk | protein_coding | 0.578651611 | 3.124977368 | 0.4971995 | 0.4915167 | 0.885174452 |
| ENSMUSG00000057388 | Mrpl18 | protein_coding | -0.129878396 | 8.058049206 | 0.4951591 | 0.4915609 | 0.885174452 |
| ENSMUSG00000043964 | Orai3 | protein_coding | 0.302313496 | 3.812328674 | 0.4949206 | 0.4916637 | 0.885174452 |
| ENSMUSG00000021732 | Fgf10 | protein_coding | -0.494550126 | 2.872667115 | 0.4948588 | 0.4916904 | 0.885174452 |
| ENSMUSG00000039298 | Cdk5rap2 | protein_coding | 0.35631714 | 2.982512472 | 0.4946493 | 0.4917807 | 0.885188869 |
| ENSMUSG00000021514 | Zfp369 | protein_coding | -0.342525408 | 3.534012811 | 0.4938941 | 0.4921067 | 0.885478532 |
| ENSMUSG00000006304 | Arpc2 | protein_coding | 0.129492917 | 7.838878671 | 0.4938247 | 0.4921366 | 0.885478532 |
| ENSMUSG00000055435 | Maf | protein_coding | -0.328817599 | 3.232228002 | 0.4937041 | 0.4921887 | 0.885478532 |
| ENSMUSG00000020949 | Fkbp3 | protein_coding | -0.134998056 | 9.463429662 | 0.4931588 | 0.4924243 | 0.885700848 |
| ENSMUSG00000032126 | Hmbs | protein_coding | 0.31223713 | 3.707314394 | 0.493037 | 0.492477 | 0.885700848 |
| ENSMUSG00000026578 | Ccdc181 | protein_coding | 0.171447715 | 5.90773494 | 0.4926468 | 0.4926458 | 0.885739358 |
| ENSMUSG00000010277 | 2610507B11Rik | protein_coding | -0.221753378 | 5.370121584 | 0.4923902 | 0.4927568 | 0.885739358 |
| ENSMUSG00000030744 | Rps3 | protein_coding | 0.160379969 | 10.97915624 | 0.4922551 | 0.4928153 | 0.885739358 |
| ENSMUSG00000047221 | Fam185a | protein_coding | 0.377955361 | 3.551582404 | 0.4916711 | 0.4930844 | 0.885739358 |
| ENSMUSG00000032657 | Fam189b | protein_coding | -0.270697283 | 3.935314827 | 0.4915665 | 0.4931135 | 0.885739358 |
| ENSMUSG00000026960 | Arl6ip6 | protein_coding | -0.266598747 | 3.486995503 | 0.4915432 | 0.4931237 | 0.885739358 |
| ENSMUSG00000049112 | Oxtr | protein_coding | -0.43708257 | 2.82457283 | 0.490965 | 0.4933743 | 0.885739358 |
| ENSMUSG00000020359 | Phykpl | protein_coding | 0.281192967 | 3.634499591 | 0.4905795 | 0.4935416 | 0.885739358 |
| ENSMUSG00000022100 | Xpo7 | protein_coding | -0.183860676 | 5.314992387 | 0.4905584 | 0.4935508 | 0.885739358 |
| ENSMUSG00000001420 | Tmem79 | protein_coding | 0.430445948 | 2.657697046 | 0.490437 | 0.4936035 | 0.885739358 |
| ENSMUSG00000022195 | 6030458C11Rik | protein_coding | 0.261821105 | 5.026614836 | 0.490277 | 0.4936891 | 0.885739358 |
| ENSMUSG00000029063 | Nadk | protein_coding | 0.208497746 | 4.830288005 | 0.4898046 | 0.4938781 | 0.885739358 |
| ENSMUSG00000041870 | Ankrd13a | protein_coding | 0.262211838 | 3.997599059 | 0.4894692 | 0.4940239 | 0.885739358 |
| ENSMUSG00000058997 | Vwa8 | protein_coding | 0.364720898 | 4.08039628 | 0.4900431 | 0.494102 | 0.885739358 |
| ENSMUSG00000020593 | Lpin1 | protein_coding | 0.259554473 | 4.67513294 | 0.4889687 | 0.4942416 | 0.885739358 |

| ensembl_gene_id | external_gene_name | gene_biotype | log2 (fold change) | log2 (counts per million) | F-value | p-value | False discovery rate |
|---|---|---|---|---|---|---|---|
| ENSMUSG00000031711 | Zfp330 | protein_coding | 0.147360228 | 5.953053569 | 0.4889208 | 0.4942625 | 0.885739358 |
| ENSMUSG00000029053 | Prkcz | protein_coding | 0.137714117 | 7.089822294 | 0.4887916 | 0.4943187 | 0.885739358 |
| ENSMUSG00000054920 | Klhl5 | protein_coding | 0.212626036 | 4.89914094 | 0.4886445 | 0.4943827 | 0.885739358 |
| ENSMUSG00000047539 | Fbxo28 | protein_coding | -0.204934969 | 4.946111766 | 0.4886419 | 0.4943838 | 0.885739358 |
| ENSMUSG00000005413 | Hmox1 | protein_coding | 0.302422017 | 3.43502437 | 0.4877718 | 0.4947628 | 0.885739358 |
| ENSMUSG00000059866 | Tnip2 | protein_coding | -0.406394202 | 2.952295257 | 0.487634 | 0.4948229 | 0.885739358 |
| ENSMUSG00000031969 | Acad8 | protein_coding | 0.195136428 | 5.460269292 | 0.4873146 | 0.4949622 | 0.885739358 |
| ENSMUSG00000071650 | Ganab | protein_coding | 0.153804861 | 5.829963717 | 0.487095 | 0.495058 | 0.885739358 |
| ENSMUSG00000034006 | Pqlc1 | protein_coding | 0.224124316 | 5.558147472 | 0.4870922 | 0.4950592 | 0.885739358 |
| ENSMUSG00000004268 | Emg1 | protein_coding | -0.127087672 | 7.510535678 | 0.4869565 | 0.4951185 | 0.885739358 |
| ENSMUSG00000024556 | Me2 | protein_coding | 0.244398029 | 4.003990473 | 0.4868104 | 0.4951822 | 0.885739358 |
| ENSMUSG00000038025 | Phf2 | protein_coding | -0.37929022 | 3.106562759 | 0.4865566 | 0.495293 | 0.885739358 |
| ENSMUSG00000035285 | Nat14 | protein_coding | -0.178829521 | 6.095206322 | 0.4865241 | 0.4953072 | 0.885739358 |
| ENSMUSG00000029029 | Wrap73 | protein_coding | -0.170149711 | 5.753226431 | 0.4863134 | 0.4953993 | 0.885739358 |
| ENSMUSG00000027935 | Rab13 | protein_coding | -0.203525361 | 4.98848082 | 0.4860646 | 0.495508 | 0.885739358 |
| ENSMUSG00000019890 | Nts | protein_coding | -0.365559866 | 7.780597208 | 0.497645 | 0.4955222 | 0.885739358 |
| ENSMUSG00000060121 | Gemin2 | protein_coding | -0.201128291 | 4.848682691 | 0.4858663 | 0.4955947 | 0.885739358 |
| ENSMUSG00000029390 | Tmed2 | protein_coding | 0.322094376 | 3.063734541 | 0.4856733 | 0.495679 | 0.885739358 |
| ENSMUSG00000038860 | Garnl3 | protein_coding | 0.229152917 | 5.181698089 | 0.485594 | 0.4957137 | 0.885739358 |
| ENSMUSG00000000759 | Tubgcp3 | protein_coding | 0.220459822 | 4.827960238 | 0.4851798 | 0.4958949 | 0.885739358 |
| ENSMUSG00000022337 | Emc2 | protein_coding | -0.133409819 | 7.531188737 | 0.4848761 | 0.4960279 | 0.885739358 |
| ENSMUSG00000030265 | Kras | protein_coding | -0.199441755 | 5.529993642 | 0.4845241 | 0.4961821 | 0.885739358 |
| ENSMUSG00000020922 | Lsm12 | protein_coding | -0.187388648 | 6.049970308 | 0.4844648 | 0.496208 | 0.885739358 |
| ENSMUSG00000041623 | D11Wsu47e | protein_coding | 0.30886497 | 3.499927593 | 0.4842068 | 0.4963211 | 0.885739358 |
| ENSMUSG00000019843 | Fyn | protein_coding | 0.215940862 | 4.647893775 | 0.4839662 | 0.4964266 | 0.885739358 |
| ENSMUSG00000032582 | Rbm6 | protein_coding | -0.225590072 | 4.699735557 | 0.4837505 | 0.4965212 | 0.885739358 |
| ENSMUSG00000005514 | Por | protein_coding | 0.187687446 | 6.626504381 | 0.4837221 | 0.4965497 | 0.885739358 |
| ENSMUSG00000073888 | Ccl27a | protein_coding | -0.164325533 | 6.095853882 | 0.4835848 | 0.4965939 | 0.885739358 |
| ENSMUSG00000032009 | Sesn3 | protein_coding | -0.180628991 | 6.287416602 | 0.4829047 | 0.4968924 | 0.885739358 |
| ENSMUSG00000045733 | Sprn | protein_coding | 0.253085755 | 4.718273447 | 0.4826997 | 0.4969824 | 0.885739358 |
| ENSMUSG00000045176 | Borcs6 | protein_coding | 0.290412035 | 3.728893799 | 0.4826127 | 0.4970206 | 0.885739358 |
| ENSMUSG00000035954 | Dock4 | protein_coding | 0.253176628 | 4.222675166 | 0.4825162 | 0.4970631 | 0.885739358 |
| ENSMUSG00000020868 | Xylt2 | protein_coding | 0.267801875 | 3.705169652 | 0.4819468 | 0.4973134 | 0.885739358 |
| ENSMUSG00000054252 | Fgfr3 | protein_coding | -0.238159248 | 4.760512676 | 0.4814919 | 0.4975135 | 0.885739358 |
| ENSMUSG00000035126 | Wdr78 | protein_coding | 0.487003741 | 2.817560419 | 0.4814625 | 0.4975265 | 0.885739358 |
| ENSMUSG00000041733 | Coq5 | protein_coding | 0.145351337 | 6.582183234 | 0.4813034 | 0.4975965 | 0.885739358 |
| ENSMUSG00000024970 | AI846148 | protein_coding | 0.246646985 | 4.005249017 | 0.4812094 | 0.4976379 | 0.885739358 |
| ENSMUSG00000029577 | Ube3b | protein_coding | -0.203978272 | 4.872432996 | 0.4809909 | 0.4977342 | 0.885739358 |
| ENSMUSG00000025140 | Pycr1 | protein_coding | -0.337793611 | 3.771223146 | 0.4809309 | 0.4977606 | 0.885739358 |
| ENSMUSG00000068686 | Cd59b | protein_coding | 0.391619495 | 2.836200096 | 0.4808742 | 0.4977856 | 0.885739358 |
| ENSMUSG00000000120 | Ngfr | protein_coding | 0.397542423 | 2.71391218 | 0.4808576 | 0.4977929 | 0.885739358 |
| ENSMUSG00000027763 | Mbnl1 | protein_coding | -0.160011127 | 6.597685114 | 0.4807289 | 0.4978496 | 0.885739358 |
| ENSMUSG00000024561 | Mbd1 | protein_coding | 0.213419006 | 5.181178733 | 0.480684 | 0.4978694 | 0.885739358 |
| ENSMUSG00000024780 | Cdc37l1 | protein_coding | -0.138563977 | 6.340182071 | 0.4806786 | 0.4978717 | 0.885739358 |
| ENSMUSG00000091636 | Akain1 | protein_coding | 0.594919465 | 2.482868168 | 0.4804932 | 0.4979535 | 0.885739358 |
| ENSMUSG00000021710 | Nln | protein_coding | 0.241737322 | 4.26161093 | 0.4802989 | 0.4980391 | 0.885739358 |
| ENSMUSG00000039318 | Rab3gap2 | protein_coding | 0.281858378 | 4.163688123 | 0.4802185 | 0.4980746 | 0.885739358 |
| ENSMUSG00000070372 | Capza1 | protein_coding | 0.263947116 | 3.747598936 | 0.4802011 | 0.4980823 | 0.885739358 |
| ENSMUSG00000051111 | Sv2c | protein_coding | 0.228234032 | 4.970558256 | 0.4801676 | 0.498097 | 0.885739358 |
| ENSMUSG00000020900 | Myh10 | protein_coding | 0.178456002 | 6.443934694 | 0.4801124 | 0.4981214 | 0.885739358 |
| ENSMUSG00000028757 | Ddost | protein_coding | 0.225697629 | 5.732806411 | 0.4801468 | 0.4981222 | 0.885739358 |
| ENSMUSG00000030806 | Stx1b | protein_coding | 0.158257517 | 8.032936804 | 0.4799344 | 0.4981999 | 0.885739358 |
| ENSMUSG00000007039 | Ddah2 | protein_coding | -0.205418172 | 4.694998761 | 0.4798956 | 0.498217 | 0.885739358 |
| ENSMUSG00000039059 | Hrh3 | protein_coding | -0.267559672 | 3.775898326 | 0.4798709 | 0.4982279 | 0.885739358 |
| ENSMUSG00000041841 | Rpl37 | protein_coding | -0.213922102 | 8.0548986 | 0.481189 | 0.498306 | 0.885739358 |
| ENSMUSG00000038374 | Rbm8a | protein_coding | 0.131202076 | 7.190096563 | 0.4795385 | 0.4983746 | 0.885739358 |
| ENSMUSG00000020639 | Pfn4 | protein_coding | -0.266117134 | 4.264502937 | 0.479415 | 0.4984292 | 0.885739358 |
| ENSMUSG00000024959 | Bad | protein_coding | -0.130780217 | 7.611176989 | 0.4790574 | 0.4985871 | 0.885873633 |
| ENSMUSG00000021814 | Anxa7 | protein_coding | 0.165709682 | 6.427187044 | 0.4782158 | 0.4989592 | 0.886388301 |
| ENSMUSG00000059857 | Ntng1 | protein_coding | -0.271061845 | 5.235176209 | 0.4779704 | 0.4991423 | 0.886468667 |
| ENSMUSG00000029439 | Sfswap | protein_coding | 0.319418968 | 3.823565253 | 0.4777163 | 0.4991803 | 0.886468667 |
| ENSMUSG00000032180 | Tmed1 | protein_coding | 0.173151753 | 5.381310942 | 0.4775549 | 0.4992518 | 0.886468667 |
| ENSMUSG00000042532 | Golga7b | protein_coding | 0.168085226 | 7.965838073 | 0.4769971 | 0.4994989 | 0.886529675 |
| ENSMUSG00000031813 | Mvb12a | protein_coding | 0.241569311 | 4.721394405 | 0.4769087 | 0.4995381 | 0.886529675 |
| ENSMUSG00000002580 | Mien1 | protein_coding | -0.155277991 | 7.03482004 | 0.4768952 | 0.4995441 | 0.886529675 |
| ENSMUSG00000028637 | Ccdc30 | protein_coding | 0.186156481 | 5.308539103 | 0.4767331 | 0.4996159 | 0.886529675 |
| ENSMUSG00000024640 | Psat1 | protein_coding | 0.118761038 | 7.37906699 | 0.4762416 | 0.4998339 | 0.886587588 |
| ENSMUSG00000022551 | Cyc1 | protein_coding | -0.141354677 | 6.715909011 | 0.4760526 | 0.4999178 | 0.886587588 |
| ENSMUSG00000022292 | Rrm2b | protein_coding | -0.213212329 | 4.879641011 | 0.4759499 | 0.4999634 | 0.886587588 |
| ENSMUSG00000020536 | Llgl1 | protein_coding | 0.217342797 | 4.550861631 | 0.4757177 | 0.5000665 | 0.886587588 |
| ENSMUSG00000040761 | Spen | protein_coding | 0.313873303 | 4.045662314 | 0.4756859 | 0.5000966 | 0.886587588 |
| ENSMUSG00000028804 | Csmd2 | protein_coding | -0.371259362 | 3.077264071 | 0.4754956 | 0.5001652 | 0.886587588 |
| ENSMUSG00000021764 | Ndufs4 | protein_coding | -0.181262296 | 8.83922535 | 0.4753952 | 0.5002257 | 0.886587588 |
| ENSMUSG00000028906 | Epb41 | protein_coding | -0.198260616 | 4.773790182 | 0.4747572 | 0.5004935 | 0.886687624 |

| ensembl_gene_id | external_gene_name | gene_biotype | log2 (fold change) | log2 (counts per million) | F-value | p-value | False discovery rate |
|---|---|---|---|---|---|---|---|
| ENSMUSG00000038642 | Ctss | protein_coding | -0.256321683 | 6.462792108 | 0.4775347 | 0.5005166 | 0.886687624 |
| ENSMUSG00000040035 | Disp2 | protein_coding | 0.229035692 | 7.018399155 | 0.4767841 | 0.5005593 | 0.886687624 |
| ENSMUSG00000034560 | Washc4 | protein_coding | 0.28014753 | 4.050259752 | 0.4744217 | 0.5006427 | 0.886687624 |
| ENSMUSG00000070644 | Etnk2 | protein_coding | 0.448754006 | 2.743692198 | 0.4741231 | 0.5007757 | 0.886687624 |
| ENSMUSG00000034940 | Synrg | protein_coding | -0.1568819 | 6.096106445 | 0.4740887 | 0.500791 | 0.886687624 |
| ENSMUSG00000062040 | Zfp27 | protein_coding | 0.266819166 | 4.00792207 | 0.4739351 | 0.5008594 | 0.886687624 |
| ENSMUSG00000046532 | Ar | protein_coding | -0.183767128 | 5.82960088 | 0.4736632 | 0.5009805 | 0.886756089 |
| ENSMUSG00000028078 | Dclk2 | protein_coding | 0.183027992 | 4.942542086 | 0.4731513 | 0.5012087 | 0.88694001 |
| ENSMUSG00000018567 | Gabarap | protein_coding | -0.135406479 | 9.783309314 | 0.4730403 | 0.5012582 | 0.88694001 |
| ENSMUSG00000069094 | Pde7a | protein_coding | -0.341888362 | 3.208383452 | 0.4728751 | 0.5013318 | 0.88694001 |
| ENSMUSG00000061911 | Myt1l | protein_coding | -0.125207084 | 7.934148902 | 0.4722392 | 0.5016157 | 0.88728988 |
| ENSMUSG00000023266 | Frs3 | protein_coding | 0.236903782 | 4.120808735 | 0.4718778 | 0.5017771 | 0.88728988 |
| ENSMUSG00000027905 | Ddx20 | protein_coding | 0.375649257 | 3.076885073 | 0.4718511 | 0.5017891 | 0.88728988 |
| ENSMUSG00000034035 | Ccdc17 | protein_coding | -0.454599455 | 2.517724767 | 0.4716931 | 0.5018597 | 0.88728988 |
| ENSMUSG00000026623 | Lpgat1 | protein_coding | -0.137316048 | 6.42459055 | 0.4711074 | 0.5021216 | 0.887513982 |
| ENSMUSG00000015672 | Mrpl32 | protein_coding | -0.16456731 | 6.614622053 | 0.4710405 | 0.5021515 | 0.887513982 |
| ENSMUSG00000091864 | Gm17102 | lincRNA | -0.351935521 | 3.367061646 | 0.4702027 | 0.5025266 | 0.8878985 |
| ENSMUSG00000040447 | Spns2 | protein_coding | -0.338214528 | 4.353384316 | 0.4709022 | 0.5026163 | 0.8878985 |
| ENSMUSG00000030002 | Dusp11 | protein_coding | -0.247311383 | 3.916512805 | 0.4700014 | 0.5026168 | 0.8878985 |
| ENSMUSG00000023147 | Wrb | protein_coding | 0.123809038 | 8.827525129 | 0.4697726 | 0.5027193 | 0.887933785 |
| ENSMUSG00000039294 | BC017643 | protein_coding | 0.346786093 | 4.386000687 | 0.4706662 | 0.5028937 | 0.887987286 |
| ENSMUSG00000027472 | Pdrg1 | protein_coding | 0.132877406 | 7.819654886 | 0.4690588 | 0.5030395 | 0.887987286 |
| ENSMUSG00000037652 | Phc3 | protein_coding | -0.225910291 | 4.092420459 | 0.4690175 | 0.5030581 | 0.887987286 |
| ENSMUSG00000019969 | Psen1 | protein_coding | 0.175547537 | 5.820382091 | 0.4687432 | 0.5031812 | 0.887987286 |
| ENSMUSG00000021770 | Samd8 | protein_coding | -0.196491006 | 5.083330362 | 0.4685593 | 0.5032638 | 0.887987286 |
| ENSMUSG00000042323 | Pbrm1 | protein_coding | 0.179334228 | 5.235378555 | 0.4684207 | 0.503326 | 0.887987286 |
| ENSMUSG00000053646 | Plxnb1 | protein_coding | -0.520575294 | 3.351708276 | 0.4705546 | 0.5033277 | 0.887987286 |
| ENSMUSG00000035473 | Galm | protein_coding | 0.621379805 | 2.245934502 | 0.4680516 | 0.5034919 | 0.888021284 |
| ENSMUSG00000003438 | Timm50 | protein_coding | 0.211201547 | 4.759467136 | 0.4678054 | 0.5036026 | 0.888021284 |
| ENSMUSG00000021144 | Mta1 | protein_coding | -0.236836715 | 5.067801162 | 0.4676695 | 0.5036637 | 0.888021284 |
| ENSMUSG00000043940 | Wdfy3 | protein_coding | -0.213622373 | 5.725792591 | 0.4673602 | 0.5038029 | 0.888021284 |
| ENSMUSG00000063894 | Zkscan8 | protein_coding | 0.315836675 | 3.195151195 | 0.4665153 | 0.5041834 | 0.888021284 |
| ENSMUSG00000004626 | Stxbp2 | protein_coding | -0.25094391 | 4.167826793 | 0.4664891 | 0.5041952 | 0.888021284 |
| ENSMUSG00000024853 | Sf3b2 | protein_coding | 0.143944521 | 9.008018132 | 0.4663464 | 0.5042595 | 0.888021284 |
| ENSMUSG00000030424 | Zfp939 | transcribed_unproces | 0.322672566 | 3.067086335 | 0.4661009 | 0.5043702 | 0.888021284 |
| ENSMUSG00000038803 | Ost4 | protein_coding | -0.161092941 | 6.636590519 | 0.4658045 | 0.5045039 | 0.888021284 |
| ENSMUSG00000028608 | 0610037L13Rik | protein_coding | 0.135402011 | 6.864864031 | 0.4657696 | 0.5045196 | 0.888021284 |
| ENSMUSG00000031137 | Fgf13 | protein_coding | -0.184812835 | 6.550759424 | 0.4657718 | 0.5045343 | 0.888021284 |
| ENSMUSG00000008734 | Gprc5b | protein_coding | -0.253595271 | 4.896268762 | 0.4656504 | 0.5045734 | 0.888021284 |
| ENSMUSG00000057132 | Rpgrip1 | protein_coding | -0.342088165 | 3.430311073 | 0.4656449 | 0.5045759 | 0.888021284 |
| ENSMUSG00000003848 | Nob1 | protein_coding | 0.174804007 | 6.283997351 | 0.4655616 | 0.5046135 | 0.888021284 |
| ENSMUSG00000038894 | Irs2 | protein_coding | 0.202149458 | 5.25886737 | 0.4654985 | 0.5046419 | 0.888021284 |
| ENSMUSG00000037487 | Ubr5 | protein_coding | 0.182506311 | 5.318454565 | 0.4654402 | 0.5046683 | 0.888021284 |
| ENSMUSG00000032267 | Usp28 | protein_coding | -0.516594048 | 2.665603186 | 0.4650181 | 0.5048589 | 0.888211315 |
| ENSMUSG00000014606 | Slc25a11 | protein_coding | 0.129094623 | 8.391474826 | 0.4645621 | 0.5050649 | 0.88841651 |
| ENSMUSG00000108644 | Gm44920 | lincRNA | -0.500152049 | 2.522787418 | 0.4643944 | 0.5051408 | 0.88841651 |
| ENSMUSG00000031189 | Aff2 | protein_coding | -0.394966697 | 2.77269957 | 0.4639779 | 0.5053291 | 0.888439263 |
| ENSMUSG00000035674 | Ndufa3 | protein_coding | -0.189216929 | 10.02601084 | 0.4639596 | 0.505353 | 0.888439263 |
| ENSMUSG00000021990 | Spata13 | protein_coding | 0.272943063 | 4.088362386 | 0.4638178 | 0.5054016 | 0.888439263 |
| ENSMUSG00000031628 | Casp3 | protein_coding | -0.395427628 | 3.475953786 | 0.4636577 | 0.5054897 | 0.888448898 |
| ENSMUSG00000097040 | 2610316D01Rik | processed_transcript | -0.285245604 | 4.550037084 | 0.4632155 | 0.5056899 | 0.888490494 |
| ENSMUSG00000032589 | Bsn | protein_coding | 0.26244228 | 4.550189453 | 0.4629959 | 0.5057738 | 0.888490494 |
| ENSMUSG00000021254 | Gpatch2l | protein_coding | 0.357151693 | 3.772626543 | 0.4630157 | 0.5057804 | 0.888490494 |
| ENSMUSG00000031827 | Cotl1 | protein_coding | -0.145719419 | 5.986020748 | 0.4627622 | 0.5058797 | 0.888490494 |
| ENSMUSG00000041710 | Trpc5 | protein_coding | 0.355691093 | 3.258501206 | 0.462659 | 0.5059265 | 0.888490494 |
| ENSMUSG00000038766 | Gabpb2 | protein_coding | -0.431505165 | 2.501157165 | 0.4616426 | 0.5063877 | 0.888754563 |
| ENSMUSG00000026185 | Igfbp5 | protein_coding | -0.216356574 | 5.404350889 | 0.4616074 | 0.5064037 | 0.888754563 |
| ENSMUSG00000044807 | Zfp354c | protein_coding | 0.231170288 | 4.196527264 | 0.4614543 | 0.5064732 | 0.888754563 |
| ENSMUSG00000026553 | Copa | protein_coding | 0.179853264 | 6.898510448 | 0.4614025 | 0.5065124 | 0.888754563 |
| ENSMUSG00000034891 | Sncb | protein_coding | 0.166730544 | 9.225122231 | 0.4612208 | 0.5065793 | 0.888754563 |
| ENSMUSG00000048120 | Entpd1 | protein_coding | -0.285876859 | 3.685527013 | 0.46118 | 0.5065979 | 0.888754563 |
| ENSMUSG00000031402 | Mpp1 | protein_coding | 0.177836178 | 6.297692361 | 0.4610534 | 0.5066554 | 0.888754563 |
| ENSMUSG00000030739 | Myh14 | protein_coding | -0.295305246 | 3.342815442 | 0.4607007 | 0.5068158 | 0.888808033 |
| ENSMUSG00000019832 | Rab32 | protein_coding | -0.781351701 | 1.917970587 | 0.4606047 | 0.506875 | 0.888808033 |
| ENSMUSG00000028057 | Rit1 | protein_coding | 0.145885013 | 6.135760638 | 0.4598094 | 0.5072215 | 0.888808033 |
| ENSMUSG00000038065 | Mturn | protein_coding | 0.205675594 | 6.059530408 | 0.4598022 | 0.5072403 | 0.888808033 |
| ENSMUSG00000043190 | Rfesd | protein_coding | 0.183010275 | 5.511361375 | 0.4595817 | 0.5073252 | 0.888808033 |
| ENSMUSG00000054619 | Mettl7a1 | protein_coding | -0.316586725 | 3.794749842 | 0.4594474 | 0.5073864 | 0.888808033 |
| ENSMUSG00000033126 | Ybey | protein_coding | 0.268912479 | 4.195006508 | 0.4593043 | 0.5074516 | 0.888808033 |
| ENSMUSG00000009112 | Bcl2l13 | protein_coding | -0.266488819 | 4.002064157 | 0.4590594 | 0.5075632 | 0.888808033 |
| ENSMUSG00000034379 | Wdr5b | protein_coding | 0.328290219 | 3.381513871 | 0.4589991 | 0.5075908 | 0.888808033 |
| ENSMUSG00000030842 | Lamtor1 | protein_coding | -0.123643494 | 7.343612553 | 0.4589933 | 0.5075934 | 0.888808033 |
| ENSMUSG00000024191 | Bnip1 | protein_coding | -0.176727353 | 5.806986693 | 0.4589647 | 0.5076064 | 0.888808033 |
| ENSMUSG00000033054 | Npat | protein_coding | -0.225688917 | 4.384894328 | 0.4584535 | 0.5078397 | 0.888808033 |

| ensembl_gene_id | external_gene_name | gene_biotype | log2 (fold change) | log2 (counts per million) | F-value | p-value | False discovery rate |
|---|---|---|---|---|---|---|---|
| ENSMUSG00000039958 | Etfbkmt | protein_coding | -0.265835825 | 4.189351122 | 0.4575394 | 0.5082573 | 0.888808033 |
| ENSMUSG00000032714 | Syde1 | protein_coding | -0.589796843 | 2.675543045 | 0.4575977 | 0.5082871 | 0.888808033 |
| ENSMUSG00000042650 | Alkbh5 | protein_coding | -0.241828192 | 6.623995135 | 0.4600388 | 0.5083461 | 0.888808033 |
| ENSMUSG00000007038 | Neu1 | protein_coding | 0.192048013 | 5.20445601 | 0.4573026 | 0.5083656 | 0.888808033 |
| ENSMUSG00000109975 | Gm19196 | transcribed_unproces | -0.23389566 | 3.871161809 | 0.4570808 | 0.508467 | 0.888808033 |
| ENSMUSG00000003623 | Crot | protein_coding | 0.220372666 | 4.772818687 | 0.4564912 | 0.5087369 | 0.888808033 |
| ENSMUSG00000028709 | Mob3c | protein_coding | 0.652927308 | 2.685665053 | 0.4576843 | 0.5087931 | 0.888808033 |
| ENSMUSG00000044600 | Smim7 | protein_coding | -0.11623494 | 8.103929156 | 0.4563419 | 0.5088052 | 0.888808033 |
| ENSMUSG00000024414 | Mrpl27 | protein_coding | -0.132815409 | 7.467434165 | 0.4561086 | 0.5089121 | 0.888808033 |
| ENSMUSG00000047976 | Kcna1 | protein_coding | -0.237663481 | 4.522790484 | 0.4559864 | 0.5089681 | 0.888808033 |
| ENSMUSG00000000253 | Gmpr | protein_coding | -0.121007498 | 7.122825237 | 0.4559512 | 0.5089842 | 0.888808033 |
| ENSMUSG00000038026 | Kcnj9 | protein_coding | 0.341187525 | 4.986539953 | 0.4580333 | 0.5092002 | 0.888808033 |
| ENSMUSG00000021595 | Nsun2 | protein_coding | 0.19725407 | 5.686086394 | 0.4554676 | 0.5092059 | 0.888808033 |
| ENSMUSG00000030222 | Rerg | protein_coding | 0.170892509 | 5.623906843 | 0.4552356 | 0.5093124 | 0.888808033 |
| ENSMUSG00000024974 | Smc3 | protein_coding | -0.182006736 | 5.852107548 | 0.4551413 | 0.5093557 | 0.888808033 |
| ENSMUSG00000025207 | Sema4g | protein_coding | 0.408466877 | 3.063198317 | 0.454934 | 0.5094508 | 0.888808033 |
| ENSMUSG00000021668 | Polk | protein_coding | -0.326369282 | 3.087683292 | 0.4549131 | 0.5094604 | 0.888808033 |
| ENSMUSG00000061578 | Ksr2 | protein_coding | 0.221385653 | 4.148114964 | 0.4549091 | 0.5094622 | 0.888808033 |
| ENSMUSG00000023055 | Calcoco1 | protein_coding | -0.179561893 | 6.086909112 | 0.454449 | 0.5096735 | 0.888808033 |
| ENSMUSG00000042404 | Dennd4b | protein_coding | -0.216684373 | 4.202061161 | 0.4543319 | 0.5097274 | 0.888808033 |
| ENSMUSG00000027430 | Dtd1 | protein_coding | 0.117851538 | 8.941783108 | 0.4540016 | 0.5098792 | 0.888808033 |
| ENSMUSG00000041765 | Ubac2 | protein_coding | 0.179182776 | 5.337075299 | 0.4539387 | 0.5099081 | 0.888808033 |
| ENSMUSG00000085251 | Gm12326 | lincRNA | -0.425213691 | 3.29364466 | 0.4538678 | 0.5099562 | 0.888808033 |
| ENSMUSG00000022537 | Tmem44 | protein_coding | 0.297988462 | 3.575541263 | 0.4538001 | 0.5099718 | 0.888808033 |
| ENSMUSG00000021670 | Hmgcr | protein_coding | 0.156669521 | 5.731228708 | 0.4535842 | 0.5100711 | 0.888808033 |
| ENSMUSG00000067825 | Pex26 | protein_coding | -0.307191289 | 3.494029996 | 0.4533056 | 0.5101993 | 0.888808033 |
| ENSMUSG00000058638 | Zfp110 | protein_coding | 0.321447234 | 3.539110823 | 0.4531879 | 0.5102535 | 0.888808033 |
| ENSMUSG00000032562 | Gnai2 | protein_coding | -0.137262776 | 7.278666785 | 0.4530528 | 0.5103157 | 0.888808033 |
| ENSMUSG00000046982 | Tshz1 | protein_coding | -0.212859987 | 4.906798199 | 0.4526143 | 0.5105177 | 0.888808033 |
| ENSMUSG00000041229 | Phf8 | protein_coding | 0.224212289 | 4.269503119 | 0.4525522 | 0.5105463 | 0.888808033 |
| ENSMUSG00000030341 | Tnfrsf1a | protein_coding | 0.449350771 | 2.560831926 | 0.4523501 | 0.5106394 | 0.888808033 |
| ENSMUSG00000039542 | Ncam1 | protein_coding | 0.148848069 | 8.750398292 | 0.4523295 | 0.5106489 | 0.888808033 |
| ENSMUSG00000028992 | Nmnat1 | protein_coding | 0.273224089 | 3.606100699 | 0.4522057 | 0.510706 | 0.888808033 |
| ENSMUSG00000053332 | Gas5 | processed_transcript | -0.217278317 | 10.45470529 | 0.4534906 | 0.5107131 | 0.888808033 |
| ENSMUSG00000025665 | Rps6ka6 | protein_coding | -0.345673846 | 3.323199076 | 0.4520425 | 0.5107813 | 0.888808033 |
| ENSMUSG00000011958 | Bnip2 | protein_coding | -0.273355899 | 3.770114211 | 0.4519401 | 0.5108285 | 0.888808033 |
| ENSMUSG00000097536 | 2610037D02Rik | lincRNA | -0.477838605 | 2.550039944 | 0.4518337 | 0.5108776 | 0.888808033 |
| ENSMUSG00000018377 | Vezf1 | protein_coding | -0.186169978 | 5.292392694 | 0.4516739 | 0.5109513 | 0.888808033 |
| ENSMUSG00000034429 | Zfp707 | protein_coding | 0.273727287 | 4.951956453 | 0.4514993 | 0.5110474 | 0.888808033 |
| ENSMUSG00000039640 | Mrpl12 | protein_coding | 0.159554402 | 6.935011068 | 0.45141 | 0.5110731 | 0.888808033 |
| ENSMUSG00000039168 | Dap | protein_coding | 0.226282856 | 4.225448907 | 0.4513048 | 0.5111217 | 0.888808033 |
| ENSMUSG00000001424 | Snd1 | protein_coding | 0.167121446 | 5.71235447 | 0.4511684 | 0.5111847 | 0.888808033 |
| ENSMUSG00000016308 | Ube2a | protein_coding | -0.126906548 | 7.517151785 | 0.4510056 | 0.5112599 | 0.888808033 |
| ENSMUSG00000021235 | Coq6 | protein_coding | 0.226162538 | 5.401147564 | 0.4508871 | 0.5113147 | 0.888808033 |
| ENSMUSG00000060098 | Prmt7 | protein_coding | 0.292177007 | 5.276261344 | 0.4510181 | 0.5118511 | 0.889419502 |
| ENSMUSG00000029534 | St7 | protein_coding | -0.204650881 | 4.278496417 | 0.4497008 | 0.5118634 | 0.889419502 |
| ENSMUSG00000026470 | Stx6 | protein_coding | 0.187763319 | 5.734930447 | 0.4495902 | 0.5119146 | 0.889419502 |
| ENSMUSG00000037343 | Taf2 | protein_coding | 0.256270717 | 3.868903352 | 0.4493642 | 0.5120193 | 0.889457692 |
| ENSMUSG00000055371 | Stam2 | protein_coding | -0.216525268 | 4.296966598 | 0.4485487 | 0.5123973 | 0.889635204 |
| ENSMUSG00000004360 | 9330159F19Rik | protein_coding | -0.276996853 | 4.99647521 | 0.4486763 | 0.5124205 | 0.889635204 |
| ENSMUSG00000042363 | Lgalsl | protein_coding | -0.137060531 | 6.228389488 | 0.4484301 | 0.5124523 | 0.889635204 |
| ENSMUSG00000041346 | Wrap53 | protein_coding | -0.187937549 | 4.948486831 | 0.4484299 | 0.5124524 | 0.889635204 |
| ENSMUSG00000034037 | Fgd5 | protein_coding | -0.3289757 | 3.056742567 | 0.4476383 | 0.5128199 | 0.889997061 |
| ENSMUSG00000020156 | Mum1 | protein_coding | 0.217033718 | 5.315632701 | 0.4476243 | 0.5128264 | 0.889997061 |
| ENSMUSG00000035992 | Fnip1 | protein_coding | 0.205429344 | 4.657855697 | 0.4471675 | 0.5130386 | 0.890221815 |
| ENSMUSG00000021361 | Tmem14c | protein_coding | 0.15050125 | 6.955817858 | 0.4463981 | 0.5133966 | 0.890663946 |
| ENSMUSG00000032712 | 2810474O19Rik | protein_coding | -0.241787329 | 3.9110502 | 0.4458424 | 0.5136554 | 0.890663946 |
| ENSMUSG00000028114 | Mettl14 | protein_coding | 0.22316139 | 4.797595066 | 0.4456134 | 0.513762 | 0.890663946 |
| ENSMUSG00000032322 | Pstpip1 | protein_coding | 0.536349154 | 2.260251507 | 0.4455657 | 0.5137843 | 0.890663946 |
| ENSMUSG00000046157 | Tmem229b | protein_coding | 0.292227122 | 3.999142315 | 0.4455059 | 0.5138122 | 0.890663946 |
| ENSMUSG00000028848 | Gpn2 | protein_coding | -0.380622307 | 2.981742726 | 0.445411 | 0.5138564 | 0.890663946 |
| ENSMUSG00000043496 | Tril | protein_coding | -0.380201995 | 4.010766033 | 0.4465961 | 0.5138981 | 0.890663946 |
| ENSMUSG00000097638 | Carlr | lincRNA | -0.632795623 | 2.306244157 | 0.4450811 | 0.5140103 | 0.890663946 |
| ENSMUSG00000029003 | Mad2l2 | protein_coding | -0.195710198 | 5.318783669 | 0.4449279 | 0.5140817 | 0.890663946 |
| ENSMUSG00000058835 | Abi1 | protein_coding | -0.132756801 | 6.933388157 | 0.4448422 | 0.5141217 | 0.890663946 |
| ENSMUSG00000024797 | Vps51 | protein_coding | -0.223718458 | 4.582024347 | 0.4446535 | 0.5142098 | 0.890673011 |
| ENSMUSG00000028420 | Tmem38b | protein_coding | -0.368318311 | 3.341440637 | 0.4444537 | 0.5143031 | 0.890691132 |
| ENSMUSG00000031631 | Cfap97 | protein_coding | -0.179719271 | 5.248263559 | 0.444077 | 0.514479 | 0.890852336 |
| ENSMUSG00000037259 | Dzank1 | protein_coding | -0.257212016 | 4.315431019 | 0.4436804 | 0.5146644 | 0.890906108 |
| ENSMUSG00000079657 | Rab26 | protein_coding | 0.259123318 | 5.200088954 | 0.4435887 | 0.5147463 | 0.890906108 |
| ENSMUSG00000009630 | Ppp2cb | protein_coding | -0.15185145 | 6.220185394 | 0.4432617 | 0.5148603 | 0.890906108 |
| ENSMUSG00000039367 | Sec24c | protein_coding | -0.567789525 | 2.482691938 | 0.4430621 | 0.5149537 | 0.890906108 |
| ENSMUSG00000063882 | Uqcrh | protein_coding | -0.175085785 | 10.25289796 | 0.4430122 | 0.5149923 | 0.890906108 |
| ENSMUSG00000054945 | Gm9958 | protein_coding | -0.735650726 | 1.942477758 | 0.4428221 | 0.515066 | 0.890906108 |

| ensembl_gene_id | external_gene_name | gene_biotype | log2 (fold change) | log2 (counts per million) | F-value | p-value | False discovery rate |
|---|---|---|---|---|---|---|---|
| ENSMUSG00000075467 | Dnlz | protein_coding | -0.265391827 | 4.734856396 | 0.4428033 | 0.51509 | 0.890906108 |
| ENSMUSG00000032803 | Cdv3 | protein_coding | -0.161649039 | 5.753626047 | 0.4425589 | 0.5151893 | 0.89093445 |
| ENSMUSG00000074758 | Gm5535 | lincRNA | 0.398206535 | 2.770397321 | 0.4423569 | 0.5152839 | 0.890954809 |
| ENSMUSG00000060301 | 2610008E11Rik | protein_coding | -0.263071223 | 3.531408418 | 0.4418312 | 0.5155303 | 0.891187406 |
| ENSMUSG00000048349 | Pou4f1 | protein_coding | -1.509431382 | 1.427937485 | 0.4454288 | 0.515602 | 0.891187406 |
| ENSMUSG00000018849 | Wwc1 | protein_coding | 0.19232753 | 5.078103555 | 0.4413968 | 0.5157341 | 0.891187406 |
| ENSMUSG00000022205 | Sub1 | protein_coding | -0.141504994 | 10.23803381 | 0.4413605 | 0.5157512 | 0.891187406 |
| ENSMUSG00000034614 | Pik3ip1 | protein_coding | 0.25382624 | 4.147793106 | 0.4409109 | 0.5159623 | 0.891187406 |
| ENSMUSG00000036192 | Rorb | protein_coding | -0.366725792 | 5.784684724 | 0.4472365 | 0.5160241 | 0.891187406 |
| ENSMUSG00000002658 | Gtf2f1 | protein_coding | 0.159746782 | 6.596767364 | 0.440129 | 0.5163298 | 0.891187406 |
| ENSMUSG00000045761 | Togaram2 | protein_coding | 0.53067884 | 2.528370165 | 0.4398766 | 0.5164485 | 0.891187406 |
| ENSMUSG00000036636 | Clcn7 | protein_coding | 0.237928807 | 4.562534078 | 0.4398521 | 0.51646 | 0.891187406 |
| ENSMUSG00000073705 | Apitd1 | protein_coding | -0.218033504 | 4.583008262 | 0.4396282 | 0.5165654 | 0.891187406 |
| ENSMUSG00000001569 | Nom1 | protein_coding | 0.20974325 | 4.537747946 | 0.4395639 | 0.5165957 | 0.891187406 |
| ENSMUSG00000041959 | S100a10 | protein_coding | 0.144393369 | 6.685286142 | 0.4395398 | 0.516607 | 0.891187406 |
| ENSMUSG00000003072 | Atp5d | protein_coding | 0.3004428 | 8.518672717 | 0.4462865 | 0.5167819 | 0.891187406 |
| ENSMUSG00000045777 | Ifitm10 | protein_coding | 0.160720255 | 5.831106192 | 0.4390496 | 0.5168379 | 0.891187406 |
| ENSMUSG00000053094 | Tmem248 | protein_coding | 0.208571604 | 4.729966442 | 0.4390302 | 0.516847 | 0.891187406 |
| ENSMUSG00000028988 | Ctnnbip1 | protein_coding | 0.31090608 | 3.844294527 | 0.4387667 | 0.5169712 | 0.891187406 |
| ENSMUSG00000042709 | Atpaf2 | protein_coding | 0.179648767 | 5.953940537 | 0.4386615 | 0.5170208 | 0.891187406 |
| ENSMUSG00000003923 | Tfam | protein_coding | 0.167624058 | 5.862559057 | 0.4385752 | 0.5170615 | 0.891187406 |
| ENSMUSG00000045996 | Polr2k | protein_coding | 0.226307313 | 4.235161033 | 0.4384896 | 0.5171018 | 0.891187406 |
| ENSMUSG00000015087 | Rabl6 | protein_coding | 0.134872421 | 6.476375965 | 0.4383356 | 0.5171745 | 0.891187406 |
| ENSMUSG00000037486 | Asxl2 | protein_coding | -0.23128314 | 4.385081825 | 0.438313 | 0.5171851 | 0.891187406 |
| ENSMUSG00000031834 | Pik3r2 | protein_coding | -0.199487404 | 4.690326057 | 0.438193 | 0.5172418 | 0.891187406 |
| ENSMUSG00000021340 | Gpld1 | protein_coding | 0.281592403 | 3.527496448 | 0.4376816 | 0.5174831 | 0.891427087 |
| ENSMUSG00000020086 | H2afy2 | protein_coding | -0.465508523 | 2.575478223 | 0.4374708 | 0.5175827 | 0.891427087 |
| ENSMUSG00000055137 | Sugct | protein_coding | -0.336500198 | 2.948896426 | 0.4371261 | 0.5177456 | 0.891427087 |
| ENSMUSG00000021188 | Trip11 | protein_coding | 0.217042007 | 4.587727055 | 0.4371129 | 0.5177518 | 0.891427087 |
| ENSMUSG00000031029 | Eif3f | protein_coding | 0.162715023 | 6.454889722 | 0.4370207 | 0.5177954 | 0.891427087 |
| ENSMUSG00000025132 | Arhgdia | protein_coding | 0.138095389 | 9.882980384 | 0.43681 | 0.517895 | 0.891455923 |
| ENSMUSG00000035351 | Nup37 | protein_coding | -0.245423997 | 4.096007473 | 0.4364193 | 0.5180798 | 0.891631322 |
| ENSMUSG00000024962 | Vegfb | protein_coding | 0.190832186 | 5.333725805 | 0.4360084 | 0.5182744 | 0.891823354 |
| ENSMUSG00000029361 | Nos1 | protein_coding | -0.217022362 | 6.17370665 | 0.4356462 | 0.5187534 | 0.892503987 |
| ENSMUSG00000052430 | Bmpr1b | protein_coding | 0.342460568 | 3.043873098 | 0.4348237 | 0.5188359 | 0.892503987 |
| ENSMUSG00000030322 | Mbd4 | protein_coding | -0.33193567 | 3.104928882 | 0.4345557 | 0.5189631 | 0.892565105 |
| ENSMUSG00000025369 | Smarcc2 | protein_coding | -0.145011617 | 6.432757822 | 0.4343614 | 0.5190553 | 0.892565105 |
| ENSMUSG00000029675 | Eln | protein_coding | -0.607835845 | 3.211596894 | 0.4373232 | 0.5191204 | 0.892565105 |
| ENSMUSG00000030218 | Mgp | protein_coding | -0.535958855 | 2.11718742 | 0.4333674 | 0.5195277 | 0.893122465 |
| ENSMUSG00000041303 | Gtf3c3 | protein_coding | 0.178985875 | 5.121092947 | 0.4328179 | 0.5197891 | 0.893188979 |
| ENSMUSG00000017057 | Il13ra1 | protein_coding | 0.318626162 | 3.208087871 | 0.4324395 | 0.5199694 | 0.893188979 |
| ENSMUSG00000003199 | Mpnd | protein_coding | 0.174977214 | 5.730802558 | 0.4321052 | 0.5201286 | 0.893188979 |
| ENSMUSG00000038658 | Ric1 | protein_coding | -0.280544281 | 4.409328162 | 0.4321256 | 0.5201339 | 0.893188979 |
| ENSMUSG00000033361 | Prrg3 | protein_coding | 0.265102998 | 4.063571141 | 0.4319139 | 0.5202198 | 0.893188979 |
| ENSMUSG00000030751 | Psma1 | protein_coding | 0.126480135 | 8.507514574 | 0.4319113 | 0.5202211 | 0.893188979 |
| ENSMUSG00000054302 | Eapp | protein_coding | 0.131214521 | 7.104930646 | 0.4316302 | 0.5203551 | 0.893188979 |
| ENSMUSG00000062526 | Mppe1 | protein_coding | -0.386327771 | 2.980219548 | 0.4315167 | 0.5204093 | 0.893188979 |
| ENSMUSG00000036644 | Tbc1d9b | protein_coding | 0.194637596 | 5.50299736 | 0.4314843 | 0.5204247 | 0.893188979 |
| ENSMUSG00000022426 | Josd1 | protein_coding | -0.214689303 | 5.076976247 | 0.4310402 | 0.5206367 | 0.893188979 |
| ENSMUSG00000030750 | Nsmce1 | protein_coding | -0.139702935 | 6.353034097 | 0.4305686 | 0.520862 | 0.893188979 |
| ENSMUSG00000029781 | Fkbp9 | protein_coding | -0.366039222 | 3.42077307 | 0.4301867 | 0.5210595 | 0.893188979 |
| ENSMUSG00000033697 | Arhgap39 | protein_coding | -0.416677701 | 2.727258943 | 0.42976 | 0.5212486 | 0.893188979 |
| ENSMUSG00000075266 | Cenpw | protein_coding | -0.386232298 | 3.134142398 | 0.4293461 | 0.5214467 | 0.893188979 |
| ENSMUSG00000020430 | Pes1 | protein_coding | 0.214698932 | 5.057633856 | 0.4293237 | 0.5214574 | 0.893188979 |
| ENSMUSG00000018286 | Psmb6 | protein_coding | 0.131179411 | 9.771318944 | 0.4292732 | 0.5214816 | 0.893188979 |
| ENSMUSG00000027086 | Fastkd1 | protein_coding | 0.218246215 | 4.208332391 | 0.4292097 | 0.5215121 | 0.893188979 |
| ENSMUSG00000027350 | Chgb | protein_coding | 0.130330288 | 7.543770136 | 0.429114 | 0.5215579 | 0.893188979 |
| ENSMUSG00000017969 | Ptgis | protein_coding | 0.461395267 | 2.848786097 | 0.4290553 | 0.521586 | 0.893188979 |
| ENSMUSG00000026544 | Dusp23 | protein_coding | -0.285623793 | 3.664859156 | 0.4289927 | 0.521616 | 0.893188979 |
| ENSMUSG00000031343 | Gabra3 | protein_coding | 0.185712598 | 5.587913627 | 0.42872 | 0.5217467 | 0.893188979 |
| ENSMUSG00000022757 | Tfg | protein_coding | 0.182432657 | 6.460969884 | 0.4287387 | 0.5217527 | 0.893188979 |
| ENSMUSG00000023992 | Trem2 | protein_coding | -0.256621977 | 4.016970492 | 0.4285195 | 0.5218428 | 0.893188979 |
| ENSMUSG00000015804 | Med28 | protein_coding | 0.16370094 | 8.268673354 | 0.4284545 | 0.521889 | 0.893188979 |
| ENSMUSG00000026502 | Desi2 | protein_coding | -0.221599173 | 4.263983837 | 0.4281408 | 0.5220244 | 0.893188979 |
| ENSMUSG00000049882 | Vcpkmt | protein_coding | 0.341152317 | 2.982139181 | 0.428118 | 0.5220354 | 0.893188979 |
| ENSMUSG00000021090 | Lrrc9 | protein_coding | 0.468272907 | 2.667987494 | 0.4273758 | 0.5223917 | 0.893188979 |
| ENSMUSG00000044471 | Lncpint | lincRNA | -0.562870949 | 2.492313321 | 0.4270085 | 0.5225682 | 0.893188979 |
| ENSMUSG00000030077 | Chl1 | protein_coding | 0.160303174 | 6.308066585 | 0.4267949 | 0.5226709 | 0.893188979 |
| ENSMUSG00000031028 | Tub | protein_coding | -0.158193441 | 5.704199397 | 0.4267504 | 0.5226923 | 0.893188979 |
| ENSMUSG00000050390 | C77080 | protein_coding | -0.246758607 | 4.336900118 | 0.4267077 | 0.5227129 | 0.893188979 |
| ENSMUSG00000024104 | Washc2 | protein_coding | -0.219869954 | 5.126617546 | 0.426663 | 0.5227344 | 0.893188979 |
| ENSMUSG00000030199 | Etv6 | protein_coding | -0.426271897 | 3.028764161 | 0.426654 | 0.5227387 | 0.893188979 |
| ENSMUSG00000040138 | Ndp | protein_coding | -0.21718429 | 4.483514932 | 0.4266493 | 0.522741 | 0.893188979 |
| ENSMUSG00000006705 | Pknox1 | protein_coding | -0.2267677 | 4.007590986 | 0.4265306 | 0.5227981 | 0.893188979 |

| ensembl_gene_id | external_gene_name | gene_biotype | log2 (fold change) | log2 (counts per million) | F-value | p-value | False discovery rate |
|---|---|---|---|---|---|---|---|
| ENSMUSG00000028962 | Slc4a2 | protein_coding | -0.290737299 | 3.489299007 | 0.4265271 | 0.5227998 | 0.893188979 |
| ENSMUSG00000032890 | Rims3 | protein_coding | -0.236815036 | 7.353189369 | 0.4294449 | 0.5228166 | 0.893188979 |
| ENSMUSG00000001785 | Pwp1 | protein_coding | -0.156760264 | 5.304626079 | 0.4264545 | 0.5228347 | 0.893188979 |
| ENSMUSG00000039753 | Fbxl5 | protein_coding | -0.127223637 | 6.427888956 | 0.4263684 | 0.5228761 | 0.893188979 |
| ENSMUSG00000026473 | Glul | protein_coding | 0.131156168 | 8.303558326 | 0.4262339 | 0.5229409 | 0.893188979 |
| ENSMUSG00000026017 | Carf | protein_coding | -0.261805525 | 4.410506606 | 0.4261163 | 0.5229975 | 0.893188979 |
| ENSMUSG00000028690 | Mmachc | protein_coding | -0.219114931 | 4.731167092 | 0.4259967 | 0.5230551 | 0.893188979 |
| ENSMUSG00000027746 | Ufm1 | protein_coding | -0.146291603 | 6.165844384 | 0.4248845 | 0.5235912 | 0.893917157 |
| ENSMUSG00000034648 | Lrrn1 | protein_coding | -0.246416124 | 4.623744272 | 0.4247565 | 0.5236529 | 0.893917157 |
| ENSMUSG00000052253 | Zfp622 | protein_coding | -0.216575524 | 5.262507283 | 0.4245949 | 0.5237309 | 0.893917157 |
| ENSMUSG00000041199 | Rpusd1 | protein_coding | 0.127640584 | 6.885391051 | 0.4243123 | 0.5238673 | 0.894008156 |
| ENSMUSG00000021413 | Prpf4b | protein_coding | 0.178788046 | 5.784699678 | 0.4239919 | 0.5240221 | 0.894130404 |
| ENSMUSG00000031865 | Dctn1 | protein_coding | -0.166366482 | 6.87130974 | 0.4237348 | 0.5241612 | 0.894144923 |
| ENSMUSG00000034484 | Snx2 | protein_coding | -0.178661757 | 7.439300765 | 0.423777 | 0.5242056 | 0.894144923 |
| ENSMUSG00000026632 | Tatdn3 | protein_coding | -0.213087397 | 4.434896797 | 0.423439 | 0.5242894 | 0.894144923 |
| ENSMUSG00000028441 | 1110017D15Rik | protein_coding | -0.165689164 | 5.39075987 | 0.4232863 | 0.5243632 | 0.894144923 |
| ENSMUSG00000001034 | Mapk7 | protein_coding | 0.457675476 | 2.897732483 | 0.4228685 | 0.5245654 | 0.894274087 |
| ENSMUSG00000048603 | Gm9828 | bidirectional_promoter | -0.321680766 | 3.293833343 | 0.4227861 | 0.5246053 | 0.894274087 |
| ENSMUSG00000063760 | Rnf217 | protein_coding | 0.279290049 | 3.511575161 | 0.421964 | 0.5250036 | 0.89467035 |
| ENSMUSG00000005899 | Smpd4 | protein_coding | 0.298937713 | 3.595173308 | 0.4219629 | 0.5250042 | 0.89467035 |
| ENSMUSG00000021428 | Riok1 | protein_coding | 0.196624055 | 4.878698262 | 0.4213641 | 0.5252947 | 0.894971492 |
| ENSMUSG00000025083 | Afap1l2 | protein_coding | -0.523107102 | 2.378908864 | 0.4210291 | 0.5254573 | 0.894971492 |
| ENSMUSG00000029920 | Smarcad1 | protein_coding | -0.266451073 | 3.78108318 | 0.4209506 | 0.5254954 | 0.894971492 |
| ENSMUSG00000038576 | Susd4 | protein_coding | 0.141356462 | 7.183129061 | 0.4209127 | 0.5255138 | 0.894971492 |
| ENSMUSG00000062906 | Hdac10 | protein_coding | -0.441923977 | 3.334555254 | 0.4211315 | 0.5258046 | 0.895191696 |
| ENSMUSG00000033350 | Chst2 | protein_coding | -0.120270208 | 6.92829808 | 0.420304 | 0.5258096 | 0.895191696 |
| ENSMUSG00000027886 | 1700013F07Rik | protein_coding | 0.622038612 | 2.351329396 | 0.4193383 | 0.5262943 | 0.895875103 |
| ENSMUSG00000020576 | Nbas | protein_coding | 0.253818354 | 4.351054025 | 0.4183946 | 0.5267396 | 0.896300424 |
| ENSMUSG00000026641 | Usf1 | protein_coding | 0.138655757 | 6.31063339 | 0.4179165 | 0.5269728 | 0.896300424 |
| ENSMUSG00000021681 | Aggf1 | protein_coding | -0.195302983 | 4.869183155 | 0.4175644 | 0.5271448 | 0.896300424 |
| ENSMUSG00000029385 | Ccng2 | protein_coding | -0.233612338 | 4.500731803 | 0.4173601 | 0.5272446 | 0.896300424 |
| ENSMUSG00000026187 | Xrcc5 | protein_coding | -0.173470488 | 5.202809155 | 0.4173302 | 0.5272592 | 0.896300424 |
| ENSMUSG00000045665 | Mfsd5 | protein_coding | 0.185380083 | 4.838395001 | 0.4171096 | 0.5273671 | 0.896300424 |
| ENSMUSG00000034832 | Tet3 | protein_coding | 0.207608042 | 4.60386671 | 0.4170881 | 0.5273776 | 0.896300424 |
| ENSMUSG00000073725 | Lmbrd1 | protein_coding | 0.131901698 | 6.696811724 | 0.4170023 | 0.5274196 | 0.896300424 |
| ENSMUSG00000022559 | Fbxl6 | protein_coding | -0.329949923 | 3.018726712 | 0.4168284 | 0.5275046 | 0.896300424 |
| ENSMUSG00000025436 | Atp23 | protein_coding | -0.331946791 | 3.030248704 | 0.4167254 | 0.527555 | 0.896300424 |
| ENSMUSG00000074001 | Klhl40 | protein_coding | -0.562954268 | 2.161219232 | 0.4160728 | 0.5278744 | 0.896300424 |
| ENSMUSG00000026933 | Camsap1 | protein_coding | 0.180481242 | 5.571069479 | 0.4160034 | 0.5279084 | 0.896300424 |
| ENSMUSG00000044072 | Eml6 | protein_coding | -0.357395199 | 2.775507336 | 0.4158461 | 0.5279855 | 0.896300424 |
| ENSMUSG00000030616 | Sytl2 | protein_coding | 0.188523901 | 4.494897183 | 0.4157825 | 0.5280166 | 0.896300424 |
| ENSMUSG00000025381 | Cnpy2 | protein_coding | 0.121603079 | 7.272330767 | 0.4150825 | 0.5283598 | 0.896300424 |
| ENSMUSG00000030329 | Pianp | protein_coding | -0.158772144 | 6.795804784 | 0.4150138 | 0.5283935 | 0.896300424 |
| ENSMUSG00000030101 | Sumf1 | protein_coding | 0.306118611 | 4.039992714 | 0.4146236 | 0.5285997 | 0.896300424 |
| ENSMUSG00000046573 | Lyrm4 | protein_coding | 0.163851087 | 7.735868849 | 0.4145847 | 0.5286188 | 0.896300424 |
| ENSMUSG00000029602 | Rasal1 | protein_coding | 0.216076425 | 5.027993756 | 0.4144678 | 0.5286616 | 0.896300424 |
| ENSMUSG00000027646 | Src | protein_coding | -0.261470456 | 4.171514877 | 0.4142352 | 0.5287758 | 0.896300424 |
| ENSMUSG00000038279 | Nop2 | protein_coding | 0.222209404 | 5.250246065 | 0.4140745 | 0.5288548 | 0.896300424 |
| ENSMUSG00000044496 | 2510039O18Rik | protein_coding | 0.353437005 | 3.529021307 | 0.4138432 | 0.5289832 | 0.896300424 |
| ENSMUSG00000026227 | 2810459M11Rik | protein_coding | 0.533999187 | 2.526631939 | 0.4135091 | 0.5291328 | 0.896300424 |
| ENSMUSG00000023259 | Slc26a6 | protein_coding | -0.512730177 | 2.561602246 | 0.4134638 | 0.5291551 | 0.896300424 |
| ENSMUSG00000021028 | Mbip | protein_coding | -0.210992814 | 4.987960437 | 0.4134477 | 0.529163 | 0.896300424 |
| ENSMUSG00000033618 | Map3k13 | protein_coding | 0.323374105 | 2.837090241 | 0.4129193 | 0.5294231 | 0.896300424 |
| ENSMUSG00000029231 | Pdgfra | protein_coding | 0.355806461 | 3.623955412 | 0.4129032 | 0.5294456 | 0.896300424 |
| ENSMUSG00000097625 | Gm26561 | lincRNA | 0.164491491 | 6.104517801 | 0.4127228 | 0.5295198 | 0.896300424 |
| ENSMUSG00000022861 | Dgkg | protein_coding | 0.137280862 | 6.621568105 | 0.412689 | 0.5295365 | 0.896300424 |
| ENSMUSG00000030188 | Magohb | protein_coding | -0.195298829 | 5.418424642 | 0.412629 | 0.5295661 | 0.896300424 |
| ENSMUSG00000018648 | Dusp14 | protein_coding | 0.143160988 | 6.258316206 | 0.4125052 | 0.5296271 | 0.896300424 |
| ENSMUSG00000054057 | A930004D18Rik | protein_coding | -0.248410175 | 3.860349917 | 0.4124941 | 0.5296326 | 0.896300424 |
| ENSMUSG00000086290 | Snhg12 | lincRNA | -0.158677111 | 5.972992914 | 0.4121139 | 0.52982 | 0.896300424 |
| ENSMUSG00000032116 | Stt3a | protein_coding | 0.155220432 | 5.841603953 | 0.4120862 | 0.5298336 | 0.896300424 |
| ENSMUSG00000025609 | Mkln1 | protein_coding | 0.155036596 | 5.956628802 | 0.411686 | 0.530031 | 0.896300424 |
| ENSMUSG00000017417 | Plxdc1 | protein_coding | 0.238558338 | 3.993092198 | 0.4114664 | 0.5301395 | 0.896300424 |
| ENSMUSG00000036435 | Exoc1 | protein_coding | 0.193899447 | 5.289982686 | 0.4113747 | 0.5301847 | 0.896300424 |
| ENSMUSG00000068329 | Htra2 | protein_coding | 0.213080355 | 4.739776655 | 0.4110939 | 0.5303234 | 0.896300424 |
| ENSMUSG00000047675 | Rps8 | protein_coding | -0.166356692 | 6.217113011 | 0.4109522 | 0.5303934 | 0.896300424 |
| ENSMUSG00000020448 | Rnf185 | protein_coding | 0.20677564 | 5.802407097 | 0.4109192 | 0.5304243 | 0.896300424 |
| ENSMUSG00000067242 | Lgi1 | protein_coding | -0.213631542 | 4.060192259 | 0.4108271 | 0.5304552 | 0.896300424 |
| ENSMUSG00000026879 | Gsn | protein_coding | 0.130910181 | 6.200328088 | 0.4106237 | 0.5305558 | 0.896300424 |
| ENSMUSG00000029916 | Agk | protein_coding | -0.174540314 | 4.77577093 | 0.4105381 | 0.5305981 | 0.896300424 |
| ENSMUSG00000037646 | Vps13b | protein_coding | 0.322612224 | 3.46712772 | 0.4101209 | 0.5308044 | 0.896300424 |
| ENSMUSG00000024248 | Cox7a2l | protein_coding | 0.117091809 | 8.40287394 | 0.4099691 | 0.5308796 | 0.896300424 |
| ENSMUSG00000003068 | Stk11 | protein_coding | -0.189762216 | 5.19273725 | 0.4098868 | 0.5309203 | 0.896300424 |
| ENSMUSG00000025209 | Twnk | protein_coding | 0.332822927 | 3.574506826 | 0.4098669 | 0.5309302 | 0.896300424 |

| ensembl_gene_id | external_gene_name | gene_biotype | log2 (fold change) | log2 (counts per million) | F-value | p-value | False discovery rate |
|---|---|---|---|---|---|---|---|
| ENSMUSG00000036197 | Gxylt1 | protein_coding | -0.281844656 | 3.20701106 | 0.4097746 | 0.5309759 | 0.896300424 |
| ENSMUSG00000037926 | Ssh2 | protein_coding | -0.210311055 | 4.286228006 | 0.4095975 | 0.5310636 | 0.896300424 |
| ENSMUSG00000030539 | Sema4b | protein_coding | 0.345923856 | 2.720905013 | 0.4095285 | 0.5310977 | 0.896300424 |
| ENSMUSG00000013419 | Zfp651 | protein_coding | -0.24046891 | 4.179712255 | 0.4093199 | 0.5312011 | 0.896300424 |
| ENSMUSG00000087259 | 2610035D17Rik | lincRNA | -0.168252746 | 5.366582611 | 0.4091503 | 0.5312851 | 0.896300424 |
| ENSMUSG00000041220 | Elovl6 | protein_coding | -0.175299563 | 5.424769524 | 0.4087664 | 0.5314755 | 0.896300424 |
| ENSMUSG00000038784 | Cnot4 | protein_coding | -0.158687383 | 6.426545098 | 0.4086775 | 0.5315196 | 0.896300424 |
| ENSMUSG00000034930 | Rtkn | protein_coding | -0.204305627 | 4.703298896 | 0.4086264 | 0.5315449 | 0.896300424 |
| ENSMUSG00000062328 | Rpl17 | protein_coding | -0.280817015 | 3.274014848 | 0.4086197 | 0.5315483 | 0.896300424 |
| ENSMUSG00000061702 | Tmem91 | protein_coding | 0.1283621 | 7.338949389 | 0.4085679 | 0.5315739 | 0.896300424 |
| ENSMUSG00000059316 | Slc27a4 | protein_coding | 0.181724536 | 5.374491072 | 0.4084313 | 0.5316418 | 0.896300424 |
| ENSMUSG00000038696 | Mapkap1 | protein_coding | 0.191641771 | 5.030753625 | 0.4079985 | 0.5318566 | 0.896300424 |
| ENSMUSG00000072692 | Rpl37rt | transcribed_processe | -0.215343154 | 5.42589792 | 0.407941 | 0.5318997 | 0.896300424 |
| ENSMUSG00000021552 | Gkap1 | protein_coding | -0.299660209 | 3.370369014 | 0.4078735 | 0.5319187 | 0.896300424 |
| ENSMUSG00000028884 | Rpa2 | protein_coding | 0.197276081 | 5.220254905 | 0.4077802 | 0.531965 | 0.896300424 |
| ENSMUSG00000002043 | Trappc6a | protein_coding | 0.18236202 | 4.975157349 | 0.4076575 | 0.532026 | 0.896300424 |
| ENSMUSG00000068823 | Csde1 | protein_coding | -0.165522534 | 8.718473416 | 0.407466 | 0.5321357 | 0.896300424 |
| ENSMUSG00000031782 | Coq9 | protein_coding | 0.135145045 | 6.965553373 | 0.4073946 | 0.5321567 | 0.896300424 |
| ENSMUSG00000024273 | 2700062C07Rik | protein_coding | 0.16156038 | 5.358026639 | 0.4070664 | 0.5323199 | 0.896300424 |
| ENSMUSG00000051768 | Xrcc1 | protein_coding | 0.213332017 | 4.455173051 | 0.4070363 | 0.5323349 | 0.896300424 |
| ENSMUSG00000031546 | Gins4 | protein_coding | -0.166541587 | 4.980697551 | 0.4068639 | 0.5324207 | 0.896300424 |
| ENSMUSG00000075324 | Fign | protein_coding | -0.313174522 | 4.297662235 | 0.4073942 | 0.5324297 | 0.896300424 |
| ENSMUSG00000031487 | Brf2 | protein_coding | 0.18568146 | 5.21612389 | 0.4067226 | 0.532491 | 0.896300424 |
| ENSMUSG00000060152 | Pop5 | protein_coding | 0.19061279 | 5.033105139 | 0.4066795 | 0.5325125 | 0.896300424 |
| ENSMUSG00000031851 | Ntpcr | protein_coding | -0.131279026 | 6.280122565 | 0.4066128 | 0.5325457 | 0.896300424 |
| ENSMUSG00000030493 | Faap24 | protein_coding | -0.238308362 | 4.212610145 | 0.4060828 | 0.5328096 | 0.896502418 |
| ENSMUSG00000021606 | Ndufs6 | protein_coding | -0.136261044 | 8.767019294 | 0.4060371 | 0.5328324 | 0.896502418 |
| ENSMUSG00000058351 | Smim4 | protein_coding | -0.214126316 | 5.621326812 | 0.4047421 | 0.533493 | 0.897233936 |
| ENSMUSG00000037262 | Kin | protein_coding | 0.202855555 | 4.991138795 | 0.4044969 | 0.533601 | 0.897233936 |
| ENSMUSG00000064068 | Mtx1 | protein_coding | -0.13885309 | 5.985579184 | 0.4043905 | 0.5336542 | 0.897233936 |
| ENSMUSG00000062203 | Gspt1 | protein_coding | -0.347096946 | 2.725209974 | 0.4042315 | 0.5337337 | 0.897233936 |
| ENSMUSG00000023980 | Taf8 | protein_coding | 0.173768786 | 5.220399206 | 0.4035069 | 0.5340962 | 0.897233936 |
| ENSMUSG00000021420 | Fars2 | protein_coding | -0.151348563 | 5.774157552 | 0.4034345 | 0.5341324 | 0.897233936 |
| ENSMUSG00000020374 | Rasgef1c | protein_coding | 0.306952409 | 3.537564496 | 0.4033631 | 0.5341682 | 0.897233936 |
| ENSMUSG00000078485 | Plekhn1 | protein_coding | 0.512995925 | 3.134278056 | 0.4045506 | 0.5342338 | 0.897233936 |
| ENSMUSG00000040907 | Atp1a3 | protein_coding | 0.119330778 | 9.92749935 | 0.403149 | 0.5342754 | 0.897233936 |
| ENSMUSG00000041460 | Cacna2d4 | protein_coding | -0.441497176 | 2.942424916 | 0.4031289 | 0.5342854 | 0.897233936 |
| ENSMUSG00000014601 | Strip1 | protein_coding | -0.27395181 | 5.067694756 | 0.4039568 | 0.534314 | 0.897233936 |
| ENSMUSG00000066829 | Zfp810 | protein_coding | -0.353181107 | 3.844884453 | 0.4035267 | 0.534379 | 0.897233936 |
| ENSMUSG00000037669 | Ldah | protein_coding | 0.127166492 | 6.544297754 | 0.4028866 | 0.5344068 | 0.897233936 |
| ENSMUSG00000051403 | Ppp1r37 | protein_coding | 0.191095939 | 4.556805796 | 0.4027568 | 0.5344719 | 0.897233936 |
| ENSMUSG00000029524 | Sirt4 | protein_coding | -0.229371507 | 3.985969658 | 0.4023367 | 0.5346825 | 0.897233936 |
| ENSMUSG00000039531 | Zufsp | protein_coding | -0.228356989 | 3.712014587 | 0.40231 | 0.5346959 | 0.897233936 |
| ENSMUSG00000046402 | Rbp1 | protein_coding | -0.221116044 | 5.686358435 | 0.402173 | 0.5348171 | 0.897233936 |
| ENSMUSG00000042428 | Mgat3 | protein_coding | -0.160634031 | 5.993891387 | 0.4020684 | 0.5348171 | 0.897233936 |
| ENSMUSG00000041231 | Ublcp1 | protein_coding | 0.150152017 | 5.847100931 | 0.4019979 | 0.5348526 | 0.897233936 |
| ENSMUSG00000034203 | Chchd4 | protein_coding | 0.11658754 | 7.239234446 | 0.4016659 | 0.5350192 | 0.897234398 |
| ENSMUSG00000035293 | G2e3 | protein_coding | 0.437216137 | 2.867473345 | 0.4016649 | 0.5350197 | 0.897234398 |
| ENSMUSG00000057322 | Rpl38 | protein_coding | 0.213242077 | 4.578530249 | 0.4013716 | 0.5351671 | 0.897309224 |
| ENSMUSG00000035245 | Eogt | protein_coding | 0.360858732 | 2.911415084 | 0.4012439 | 0.5352312 | 0.897309224 |
| ENSMUSG00000028664 | Ephb2 | protein_coding | -0.3490398 | 2.859946521 | 0.4003659 | 0.5356728 | 0.897602244 |
| ENSMUSG00000053550 | Shisa7 | protein_coding | -0.225970235 | 4.438449316 | 0.4000474 | 0.5358332 | 0.897602244 |
| ENSMUSG00000051256 | Jagn1 | protein_coding | 0.131169561 | 6.482956711 | 0.3999504 | 0.535882 | 0.897602244 |
| ENSMUSG00000039621 | Prex1 | protein_coding | 0.291502696 | 4.392303219 | 0.3998714 | 0.5359741 | 0.897602244 |
| ENSMUSG00000041915 | Ammecr1l | protein_coding | 0.299223334 | 3.297961726 | 0.3996084 | 0.5360544 | 0.897602244 |
| ENSMUSG00000023904 | Hcfc1r1 | protein_coding | -0.11044243 | 8.646237457 | 0.3995762 | 0.5360706 | 0.897602244 |
| ENSMUSG00000022235 | Cmbl | protein_coding | 0.172095161 | 5.518609554 | 0.3993794 | 0.5361698 | 0.897602244 |
| ENSMUSG00000073702 | Rpl31 | protein_coding | -0.233533362 | 3.938345888 | 0.3991971 | 0.5362618 | 0.897602244 |
| ENSMUSG00000011254 | Thg1l | protein_coding | 0.195423473 | 5.010979857 | 0.3990789 | 0.5363214 | 0.897602244 |
| ENSMUSG00000046879 | Irgm1 | protein_coding | -0.302674638 | 4.267996404 | 0.3989296 | 0.5364489 | 0.897602244 |
| ENSMUSG00000022095 | Fam160b2 | protein_coding | -0.232015556 | 3.768285915 | 0.3985271 | 0.5365999 | 0.897602244 |
| ENSMUSG00000030045 | Mrpl19 | protein_coding | 0.114977228 | 7.065750676 | 0.3982421 | 0.5367439 | 0.897602244 |
| ENSMUSG00000074754 | Smim26 | protein_coding | -0.136881664 | 6.791577845 | 0.3981242 | 0.5368034 | 0.897602244 |
| ENSMUSG00000049657 | Zbtb5 | protein_coding | 0.348689188 | 3.629371052 | 0.3980433 | 0.5368587 | 0.897602244 |
| ENSMUSG00000051730 | Mettl5 | protein_coding | -0.133452101 | 6.12775311 | 0.3979936 | 0.5368694 | 0.897602244 |
| ENSMUSG00000028099 | Polr3c | protein_coding | 0.216337776 | 5.569447924 | 0.3976319 | 0.5370666 | 0.897602244 |
| ENSMUSG00000000184 | Ccnd2 | protein_coding | 0.190486145 | 5.165602179 | 0.3974418 | 0.5371485 | 0.897602244 |
| ENSMUSG00000046449 | C77370 | protein_coding | -0.214399255 | 4.756877016 | 0.3973228 | 0.5372087 | 0.897602244 |
| ENSMUSG00000035686 | Thrsp | protein_coding | -0.199403351 | 6.131810158 | 0.3974118 | 0.5372157 | 0.897602244 |
| ENSMUSG00000097554 | Gm26825 | lincRNA | 0.352245284 | 2.700900117 | 0.3970789 | 0.5373321 | 0.897602244 |
| ENSMUSG00000032281 | Acsbg1 | protein_coding | 0.136236016 | 7.206270198 | 0.3968895 | 0.5374281 | 0.897602244 |
| ENSMUSG00000040537 | Adam22 | protein_coding | 0.140206164 | 6.220758536 | 0.3968746 | 0.5374356 | 0.897602244 |
| ENSMUSG00000027566 | Psma7 | protein_coding | 0.127657546 | 9.496865509 | 0.3968173 | 0.5374646 | 0.897602244 |
| ENSMUSG00000029592 | Usp30 | protein_coding | -0.240541993 | 4.847948825 | 0.3967085 | 0.5375341 | 0.897602244 |

| ensembl_gene_id | external_gene_name | gene_biotype | log2 (fold change) | log2 (counts per million) | F-value | p-value | False discovery rate |
|---|---|---|---|---|---|---|---|
| ENSMUSG00000045482 | Trrap | protein_coding | 0.251659069 | 4.645502751 | 0.3966859 | 0.5375455 | 0.897602244 |
| ENSMUSG00000021431 | Snrnp48 | protein_coding | 0.216457403 | 4.203297408 | 0.3965968 | 0.5375763 | 0.897602244 |
| ENSMUSG00000085980 | Gm12408 | lincRNA | 0.367135951 | 2.757608371 | 0.3963153 | 0.537719 | 0.897701078 |
| ENSMUSG00000036315 | Znrd1 | protein_coding | -0.169593997 | 6.192882707 | 0.3959911 | 0.5378835 | 0.897733167 |
| ENSMUSG00000032816 | Igdcc4 | protein_coding | -0.318012356 | 3.276347996 | 0.3959482 | 0.5379052 | 0.897733167 |
| ENSMUSG00000043461 | Sptssb | protein_coding | 0.394777871 | 2.800323848 | 0.3955926 | 0.5380857 | 0.897815383 |
| ENSMUSG00000023092 | Fhl1 | protein_coding | -0.137508621 | 7.233668299 | 0.3954709 | 0.5381474 | 0.897815383 |
| ENSMUSG00000027999 | Pla2g12a | protein_coding | 0.185883911 | 4.434526774 | 0.3953576 | 0.538205 | 0.897815383 |
| ENSMUSG00000021685 | Otp | protein_coding | 0.39587562 | 2.728168457 | 0.3941315 | 0.5388284 | 0.898621713 |
| ENSMUSG00000005973 | Rcn1 | protein_coding | 0.1932836 | 5.132738455 | 0.3940304 | 0.5388798 | 0.898621713 |
| ENSMUSG00000031325 | 4930519F16Rik | transcribed_unitary_p | -0.402194596 | 2.588531123 | 0.3938215 | 0.5389862 | 0.898621713 |
| ENSMUSG00000007850 | Hnrnph1 | protein_coding | 0.125574232 | 6.886565233 | 0.3935545 | 0.5391222 | 0.898621713 |
| ENSMUSG00000020386 | Sar1b | protein_coding | 0.142499485 | 7.789736804 | 0.3933039 | 0.5392498 | 0.898621713 |
| ENSMUSG00000022335 | Zfat | protein_coding | -0.500314519 | 2.241048006 | 0.3931525 | 0.539327 | 0.898621713 |
| ENSMUSG00000015852 | Fcrls | protein_coding | 0.46229061 | 3.534348842 | 0.3952868 | 0.5393669 | 0.898621713 |
| ENSMUSG00000035713 | Usp35 | protein_coding | 0.300624739 | 3.237843402 | 0.3929642 | 0.5394231 | 0.898621713 |
| ENSMUSG00000003778 | Brd8 | protein_coding | 0.158403466 | 6.983895678 | 0.392958 | 0.5394404 | 0.898621713 |
| ENSMUSG00000027160 | Ccdc34 | protein_coding | -0.328678702 | 3.2779481 | 0.3926812 | 0.5395674 | 0.898673504 |
| ENSMUSG00000027168 | Pax6 | protein_coding | 0.244598436 | 3.913721954 | 0.3922254 | 0.5398001 | 0.898673504 |
| ENSMUSG00000055900 | Tmem69 | protein_coding | -0.252257093 | 3.362338281 | 0.3921663 | 0.5398303 | 0.898673504 |
| ENSMUSG00000043398 | Gpr135 | protein_coding | -0.315598252 | 3.060723677 | 0.392159 | 0.539834 | 0.898673504 |
| ENSMUSG00000005779 | Psmb4 | protein_coding | 0.125937644 | 7.730041608 | 0.3920068 | 0.5399118 | 0.898673504 |
| ENSMUSG00000037935 | Smarce1 | protein_coding | 0.175602982 | 5.526289286 | 0.3918656 | 0.5399839 | 0.898673504 |
| ENSMUSG00000026917 | Wdr5 | protein_coding | 0.267322989 | 3.894247691 | 0.3917235 | 0.5400566 | 0.898673504 |
| ENSMUSG00000009863 | Sdhb | protein_coding | 0.124566258 | 10.22007054 | 0.3913631 | 0.5402408 | 0.898841068 |
| ENSMUSG00000030335 | Mrpl51 | protein_coding | 0.11566144 | 6.91307832 | 0.3910746 | 0.5403884 | 0.898947537 |
| ENSMUSG00000020070 | Rufy2 | protein_coding | -0.189765952 | 5.062097477 | 0.3895565 | 0.5411664 | 0.899666525 |
| ENSMUSG00000027977 | Ndst3 | protein_coding | -0.386405753 | 2.657612592 | 0.3895226 | 0.5411838 | 0.899666525 |
| ENSMUSG00000074591 | Ankub1 | protein_coding | -0.445116696 | 2.561682603 | 0.3893023 | 0.5412969 | 0.899666525 |
| ENSMUSG00000022678 | Nde1 | protein_coding | -0.221709778 | 5.028134558 | 0.3892731 | 0.5413119 | 0.899666525 |
| ENSMUSG00000025189 | Cnnm1 | protein_coding | -0.139126606 | 5.847748916 | 0.3891614 | 0.5413692 | 0.899666525 |
| ENSMUSG00000035212 | Leprot | protein_coding | 0.187683948 | 5.555849346 | 0.3889893 | 0.5414576 | 0.899666525 |
| ENSMUSG00000033365 | Ipo13 | protein_coding | 0.183886668 | 5.310807978 | 0.3887478 | 0.5415817 | 0.899666525 |
| ENSMUSG00000107283 | Mpv17 | protein_coding | 0.124546854 | 6.948560434 | 0.388707 | 0.5416026 | 0.899666525 |
| ENSMUSG00000028191 | Bcl10 | protein_coding | -0.171992949 | 4.911238581 | 0.3886156 | 0.5416496 | 0.899666525 |
| ENSMUSG00000005204 | Senp3 | protein_coding | -0.184357272 | 5.074071555 | 0.3886007 | 0.5416573 | 0.899666525 |
| ENSMUSG00000030352 | Tspan9 | protein_coding | 0.293964699 | 3.740494061 | 0.3881883 | 0.5418694 | 0.899879725 |
| ENSMUSG00000041828 | Abca8a | protein_coding | -0.331007909 | 3.14266088 | 0.3876092 | 0.5421674 | 0.900167525 |
| ENSMUSG00000042447 | Mios | protein_coding | 0.274434868 | 3.605521355 | 0.3875264 | 0.5422101 | 0.900167525 |
| ENSMUSG00000020747 | Tmem94 | protein_coding | -0.183299472 | 4.770877502 | 0.3872984 | 0.5423275 | 0.900223462 |
| ENSMUSG00000063146 | Clip2 | protein_coding | -0.270728612 | 3.68875427 | 0.3867051 | 0.5426334 | 0.900332225 |
| ENSMUSG00000037400 | Atp11b | protein_coding | -0.373082998 | 3.520111788 | 0.3865574 | 0.5427851 | 0.900332225 |
| ENSMUSG00000039830 | Olig2 | protein_coding | -0.290785683 | 3.316571828 | 0.3863868 | 0.5427976 | 0.900332225 |
| ENSMUSG00000025427 | Rnf165 | protein_coding | 0.237720965 | 4.695807413 | 0.3860117 | 0.5429913 | 0.900332225 |
| ENSMUSG00000039860 | Srrm3 | protein_coding | -0.314808706 | 2.993255815 | 0.3860084 | 0.542993 | 0.900332225 |
| ENSMUSG00000026103 | Gls | protein_coding | -0.170367102 | 6.054018853 | 0.3859568 | 0.5430196 | 0.900332225 |
| ENSMUSG00000021193 | Pitrm1 | protein_coding | 0.23591404 | 4.532985356 | 0.3858026 | 0.5430993 | 0.900332225 |
| ENSMUSG00000025221 | Kcnip2 | protein_coding | -0.138429625 | 6.323218804 | 0.3857583 | 0.5431221 | 0.900332225 |
| ENSMUSG00000085438 | 1700020I14Rik | lincRNA | -0.117978487 | 6.815821602 | 0.3854826 | 0.5432647 | 0.900332225 |
| ENSMUSG00000021277 | Traf3 | protein_coding | 0.263029953 | 3.856128044 | 0.3854023 | 0.5433062 | 0.900332225 |
| ENSMUSG00000079610 | Ankrd39 | protein_coding | 0.236173218 | 3.616515507 | 0.385342 | 0.5433373 | 0.900332225 |
| ENSMUSG00000006423 | C330007P06Rik | protein_coding | 0.211555718 | 4.447876819 | 0.3852015 | 0.54341 | 0.900332225 |
| ENSMUSG00000061482 | Hist1h4d | protein_coding | 0.215759155 | 3.765522318 | 0.3849336 | 0.5435486 | 0.900332225 |
| ENSMUSG00000039683 | Sdk1 | protein_coding | 0.472683997 | 2.433209778 | 0.3847355 | 0.5436511 | 0.900332225 |
| ENSMUSG00000001039 | B9d1 | protein_coding | -0.154082245 | 5.906474746 | 0.3844674 | 0.5437899 | 0.900332225 |
| ENSMUSG00000026360 | Rgs2 | protein_coding | -0.173728742 | 7.852127661 | 0.3849539 | 0.5438037 | 0.900332225 |
| ENSMUSG00000034621 | Gpatch8 | protein_coding | -0.166327622 | 5.727575147 | 0.3844163 | 0.5438164 | 0.900332225 |
| ENSMUSG00000022314 | Rad21 | protein_coding | -0.156744059 | 5.920039703 | 0.3839979 | 0.5440332 | 0.900552457 |
| ENSMUSG00000020335 | Zfp354b | protein_coding | -0.29509509 | 3.46258822 | 0.3837373 | 0.5441683 | 0.900637543 |
| ENSMUSG00000022964 | Tmem50b | protein_coding | 0.145793234 | 7.304724141 | 0.3833873 | 0.5443499 | 0.900773367 |
| ENSMUSG00000097571 | Jpx | lincRNA | -0.389870239 | 2.721412026 | 0.3832562 | 0.5444179 | 0.900773367 |
| ENSMUSG00000022770 | Dlg1 | protein_coding | -0.163517466 | 5.375225563 | 0.3829882 | 0.544557 | 0.900857974 |
| ENSMUSG00000029608 | Rph3a | protein_coding | -0.184984554 | 7.193579012 | 0.3838394 | 0.5446592 | 0.900857974 |
| ENSMUSG00000001802 | Lrp3 | protein_coding | -0.236361736 | 4.149814786 | 0.3825597 | 0.5447796 | 0.900857974 |
| ENSMUSG00000021532 | Fastkd3 | protein_coding | -0.189564983 | 5.034490308 | 0.3824926 | 0.5448145 | 0.900857974 |
| ENSMUSG00000038126 | Mphosph9 | protein_coding | 0.249603917 | 3.540894495 | 0.3822171 | 0.5449578 | 0.900857974 |
| ENSMUSG00000031887 | Tradd | protein_coding | -0.382756824 | 2.985530773 | 0.3821902 | 0.5449717 | 0.900857974 |
| ENSMUSG00000079477 | Rab7 | protein_coding | 0.169587571 | 6.059395973 | 0.381905 | 0.5451201 | 0.900964769 |
| ENSMUSG00000021546 | Hnrnpk | protein_coding | 0.135842597 | 8.311756513 | 0.381479 | 0.5453419 | 0.901093935 |
| ENSMUSG00000018326 | Ywhab | protein_coding | 0.108830075 | 9.025884273 | 0.3814103 | 0.5453776 | 0.901093935 |
| ENSMUSG00000029759 | Pon3 | protein_coding | -0.614204518 | 2.371750793 | 0.3812988 | 0.5454497 | 0.901093935 |
| ENSMUSG00000010048 | Ifrd2 | protein_coding | -0.335391664 | 3.343851751 | 0.3804152 | 0.5458963 | 0.901323038 |
| ENSMUSG00000050856 | Atp5k | protein_coding | -0.169681668 | 9.580473233 | 0.3801449 | 0.5460514 | 0.901323038 |
| ENSMUSG00000027770 | Dhx36 | protein_coding | 0.16530238 | 6.075141452 | 0.3800386 | 0.5460929 | 0.901323038 |

| ensembl_gene_id | external_gene_name | gene_biotype | log2 (fold change) | log2 (counts per million) | F-value | p-value | False discovery rate |
|---|---|---|---|---|---|---|---|
| ENSMUSG00000041203 | Trir | protein_coding | 0.202979081 | 4.336467588 | 0.3799666 | 0.5461305 | 0.901323038 |
| ENSMUSG00000047842 | Diras2 | protein_coding | -0.187767521 | 5.535112795 | 0.3799362 | 0.5461464 | 0.901323038 |
| ENSMUSG00000084416 | Rpl10a-ps1 | processed_pseudoge | -0.298882214 | 2.886219556 | 0.3797099 | 0.5462646 | 0.901323038 |
| ENSMUSG00000061778 | Mospd2 | protein_coding | 0.245462422 | 3.429067216 | 0.3796133 | 0.5463151 | 0.901323038 |
| ENSMUSG00000071014 | Ndufb6 | protein_coding | 0.117063037 | 8.649696776 | 0.3795819 | 0.5463315 | 0.901323038 |
| ENSMUSG00000008226 | Scrn3 | protein_coding | 0.208643215 | 5.023483344 | 0.3795603 | 0.5463427 | 0.901323038 |
| ENSMUSG00000078348 | Sf3b5 | protein_coding | -0.121275745 | 8.019502921 | 0.3790288 | 0.5466206 | 0.901509468 |
| ENSMUSG00000025578 | Cbx8 | protein_coding | 0.309829919 | 3.14278901 | 0.3790235 | 0.5466234 | 0.901509468 |
| ENSMUSG00000037089 | Slc35b2 | protein_coding | 0.201168218 | 5.014499288 | 0.37856 | 0.546866 | 0.901694658 |
| ENSMUSG00000046997 | Spsb4 | protein_coding | 0.42978256 | 2.451580769 | 0.3784165 | 0.5469411 | 0.901694658 |
| ENSMUSG00000023034 | Nr4a1 | protein_coding | 0.149936503 | 5.870254993 | 0.3782404 | 0.5470333 | 0.901694658 |
| ENSMUSG00000029470 | P2rx4 | protein_coding | 0.325828651 | 3.457031536 | 0.3781682 | 0.5470711 | 0.901694658 |
| ENSMUSG00000031748 | Gnao1 | protein_coding | -0.123043677 | 8.517778331 | 0.3776097 | 0.5473639 | 0.902014551 |
| ENSMUSG00000057278 | Snrpg | protein_coding | -0.209085232 | 5.204688223 | 0.377478 | 0.547433 | 0.902014551 |
| ENSMUSG00000042608 | Stk40 | protein_coding | 0.290926 | 3.739329331 | 0.3771428 | 0.5476089 | 0.90206131 |
| ENSMUSG00000022037 | Clu | protein_coding | 0.143420746 | 9.99337249 | 0.3771042 | 0.5476291 | 0.90206131 |
| ENSMUSG00000001151 | Pcnt | protein_coding | -0.340070638 | 3.436726609 | 0.3766943 | 0.5478583 | 0.902126529 |
| ENSMUSG00000068663 | Clec16a | protein_coding | 0.30379213 | 4.878212848 | 0.37824 | 0.5478786 | 0.902126529 |
| ENSMUSG00000008301 | Phax | protein_coding | 0.104455038 | 7.659439916 | 0.3765084 | 0.5479421 | 0.902126529 |
| ENSMUSG00000040269 | Mrps28 | protein_coding | -0.118230462 | 6.852699931 | 0.3763031 | 0.54805 | 0.902126529 |
| ENSMUSG00000032353 | Tmed3 | protein_coding | 0.116438501 | 7.049319492 | 0.3760779 | 0.5481684 | 0.902126529 |
| ENSMUSG00000025810 | Nrp1 | protein_coding | 0.224254177 | 4.129597708 | 0.3759033 | 0.5482603 | 0.902126529 |
| ENSMUSG00000031833 | Mast3 | protein_coding | -0.181819485 | 5.532815314 | 0.375891 | 0.5482667 | 0.902126529 |
| ENSMUSG00000032449 | Slc25a36 | protein_coding | -0.15011208 | 5.766421646 | 0.3757519 | 0.5483399 | 0.902126529 |
| ENSMUSG00000002833 | Hdgfl2 | protein_coding | -0.189489552 | 5.880915318 | 0.3756026 | 0.5484323 | 0.902140549 |
| ENSMUSG00000026798 | Coq4 | protein_coding | -0.137733143 | 5.82078392 | 0.3753901 | 0.5485304 | 0.902153508 |
| ENSMUSG00000038876 | Rnf146 | protein_coding | -0.175455621 | 4.722159032 | 0.3752427 | 0.548608 | 0.902153508 |
| ENSMUSG00000026131 | Dst | protein_coding | -0.197888263 | 5.014893336 | 0.3740773 | 0.5492225 | 0.902601162 |
| ENSMUSG00000038762 | Abcf1 | protein_coding | 0.144974072 | 6.924859348 | 0.3740281 | 0.5492485 | 0.902601162 |
| ENSMUSG00000021534 | 1700001L19Rik | protein_coding | 0.151517202 | 6.731267775 | 0.3739033 | 0.5493144 | 0.902601162 |
| ENSMUSG00000058301 | Upf1 | protein_coding | -0.198848663 | 4.106842552 | 0.3736574 | 0.5494443 | 0.902601162 |
| ENSMUSG00000033685 | Ucp2 | protein_coding | 0.154291787 | 6.573361326 | 0.3734284 | 0.5495653 | 0.902601162 |
| ENSMUSG00000021573 | Tppp | protein_coding | 0.120923194 | 7.734587106 | 0.3731223 | 0.5497271 | 0.902601162 |
| ENSMUSG00000092094 | Zfp804b | protein_coding | -0.335489048 | 2.697701864 | 0.3731193 | 0.5497287 | 0.902601162 |
| ENSMUSG00000089687 | Rab42 | protein_coding | -0.353255691 | 2.882036493 | 0.37294 | 0.5498235 | 0.902601162 |
| ENSMUSG00000071253 | Slc25a16 | protein_coding | 0.193788999 | 4.729946211 | 0.3727364 | 0.5499312 | 0.902601162 |
| ENSMUSG00000023067 | Cdkn1a | protein_coding | -0.342336968 | 4.338982356 | 0.3742642 | 0.5499638 | 0.902601162 |
| ENSMUSG00000079508 | Apoo | protein_coding | -0.185673458 | 4.398444904 | 0.3726646 | 0.5499692 | 0.902601162 |
| ENSMUSG00000041552 | Ptchd1 | protein_coding | -0.267663536 | 3.196184494 | 0.3724848 | 0.5500644 | 0.902601162 |
| ENSMUSG00000027312 | Atrn | protein_coding | 0.216276654 | 5.29521595 | 0.372394 | 0.5501263 | 0.902601162 |
| ENSMUSG00000025142 | Aspscr1 | protein_coding | 0.288294527 | 3.519149347 | 0.37235 | 0.5501358 | 0.902601162 |
| ENSMUSG00000027288 | Zfp106 | protein_coding | -0.153236065 | 5.455991824 | 0.3723434 | 0.5501393 | 0.902601162 |
| ENSMUSG00000039372 | 4-Mar | protein_coding | 0.265327353 | 3.590618025 | 0.3718281 | 0.5504123 | 0.902834586 |
| ENSMUSG00000031385 | Plxnb3 | protein_coding | -0.401611926 | 2.818135004 | 0.371758 | 0.5504495 | 0.902834586 |
| ENSMUSG00000046707 | Csnk2a2 | protein_coding | 0.161860847 | 5.413038128 | 0.3710701 | 0.5508144 | 0.903027478 |
| ENSMUSG00000034973 | Dopey1 | protein_coding | -0.303669581 | 3.857142208 | 0.3710458 | 0.5508411 | 0.903027478 |
| ENSMUSG00000035765 | Dym | protein_coding | -0.146781047 | 5.538047024 | 0.3708913 | 0.5509094 | 0.903027478 |
| ENSMUSG00000021365 | Nedd9 | protein_coding | 0.439644787 | 2.474451511 | 0.3708625 | 0.5509247 | 0.903027478 |
| ENSMUSG00000020053 | Igf1 | protein_coding | -0.568092516 | 2.51168405 | 0.370771 | 0.550987 | 0.903027478 |
| ENSMUSG00000033615 | Cplx1 | protein_coding | -0.134668004 | 9.61942052 | 0.3705654 | 0.5510825 | 0.903046329 |
| ENSMUSG00000078607 | 1810010H24Rik | protein_coding | 0.552818065 | 2.574770644 | 0.3700523 | 0.551369 | 0.903378104 |
| ENSMUSG00000013921 | Clip3 | protein_coding | -0.138477793 | 8.694531453 | 0.3696605 | 0.5515637 | 0.903559437 |
| ENSMUSG00000057691 | Zfp746 | protein_coding | -0.23666521 | 3.760092009 | 0.3690351 | 0.5518967 | 0.903859537 |
| ENSMUSG00000048001 | Hes5 | protein_coding | -0.408968051 | 2.560221355 | 0.3690009 | 0.551915 | 0.903859537 |
| ENSMUSG00000022570 | Tsta3 | protein_coding | 0.112807665 | 6.72415395 | 0.3687036 | 0.5520734 | 0.903915114 |
| ENSMUSG00000028550 | Atg4c | protein_coding | -0.166115664 | 5.355859429 | 0.3685823 | 0.5521381 | 0.903915114 |
| ENSMUSG00000020485 | Supt4a | protein_coding | 0.117167769 | 6.644403082 | 0.3684643 | 0.5522011 | 0.903915114 |
| ENSMUSG00000022354 | Ndufb9 | protein_coding | -0.149870638 | 10.17307244 | 0.3683003 | 0.5522886 | 0.903920703 |
| ENSMUSG00000019232 | Etnppl | protein_coding | -0.152840439 | 5.211151314 | 0.3680048 | 0.5524463 | 0.904041295 |
| ENSMUSG00000024217 | Snrpc | protein_coding | 0.187370428 | 5.049865964 | 0.3677158 | 0.5526006 | 0.904135675 |
| ENSMUSG00000053641 | Dennd4a | protein_coding | -0.205761556 | 4.413775591 | 0.3675212 | 0.5527046 | 0.904135675 |
| ENSMUSG00000044676 | Zfp612 | protein_coding | -0.299431399 | 3.313462669 | 0.3674246 | 0.5527562 | 0.904135675 |
| ENSMUSG00000010721 | Lmbr1 | protein_coding | 0.167978876 | 4.76906295 | 0.366929 | 0.5530213 | 0.90441563 |
| ENSMUSG00000037771 | Slc32a1 | protein_coding | -0.299521788 | 4.616713118 | 0.3677824 | 0.5530956 | 0.90441563 |
| ENSMUSG00000040387 | Klhl32 | protein_coding | -0.198475324 | 4.658007617 | 0.3664139 | 0.5532969 | 0.904607278 |
| ENSMUSG00000021779 | Thrb | protein_coding | -0.200751926 | 5.798525101 | 0.3658561 | 0.5536095 | 0.904980668 |
| ENSMUSG00000021244 | Ylpm1 | protein_coding | -0.157725181 | 5.454123486 | 0.3654065 | 0.5538369 | 0.905137505 |
| ENSMUSG00000038668 | Lpar1 | protein_coding | 0.190247171 | 5.332223296 | 0.3653378 | 0.5538738 | 0.905137505 |
| ENSMUSG00000041930 | Fam222a | protein_coding | 0.495266315 | 2.58983844 | 0.3646165 | 0.5542611 | 0.905504087 |
| ENSMUSG00000027668 | Mfn1 | protein_coding | 0.229859033 | 4.57648401 | 0.3645996 | 0.5542702 | 0.905504087 |
| ENSMUSG00000036617 | Etl4 | protein_coding | -0.231448537 | 4.367446043 | 0.364294 | 0.5544344 | 0.905504087 |
| ENSMUSG00000044465 | Fam160a2 | protein_coding | -0.190386309 | 4.499321294 | 0.3641849 | 0.5544931 | 0.905504087 |
| ENSMUSG00000026750 | Psmb7 | protein_coding | 0.106484029 | 8.167252807 | 0.3641365 | 0.5545191 | 0.905504087 |
| ENSMUSG00000024294 | Mib1 | protein_coding | -0.15130232 | 6.045810957 | 0.3634915 | 0.5548662 | 0.905834081 |

| ensembl_gene_id | external_gene_name | gene_biotype | log2 (fold change) | log2 (counts per million) | F-value | p-value | False discovery rate |
|---|---|---|---|---|---|---|---|
| ENSMUSG00000004631 | Sgce | protein_coding | -0.134620927 | 5.822565911 | 0.3634479 | 0.5548897 | 0.905834081 |
| ENSMUSG00000009876 | Cox4i2 | protein_coding | -0.38157389 | 2.777996292 | 0.3632191 | 0.555013 | 0.905897751 |
| ENSMUSG00000000078 | Klf6 | protein_coding | -0.161120913 | 5.934176 | 0.3628532 | 0.5552101 | 0.906082067 |
| ENSMUSG00000025332 | Kdm5c | protein_coding | -0.206462924 | 4.416504247 | 0.3626272 | 0.555332 | 0.906143414 |
| ENSMUSG00000032184 | Lysmd2 | protein_coding | -0.120778844 | 6.451402441 | 0.3617909 | 0.5557834 | 0.906265612 |
| ENSMUSG00000040570 | Rundc3b | protein_coding | -0.185874605 | 5.015460729 | 0.3617423 | 0.5558097 | 0.906265612 |
| ENSMUSG00000020978 | Klhdc2 | protein_coding | 0.122904832 | 6.554160516 | 0.361693 | 0.5558363 | 0.906265612 |
| ENSMUSG00000028343 | Erp44 | protein_coding | -0.149149217 | 5.414849164 | 0.3616039 | 0.5558845 | 0.906265612 |
| ENSMUSG00000010911 | Apip | protein_coding | -0.141779692 | 5.986840214 | 0.3615647 | 0.5559056 | 0.906265612 |
| ENSMUSG00000025933 | Tmem14a | protein_coding | 0.126608724 | 6.769581659 | 0.3615519 | 0.5559126 | 0.906265612 |
| ENSMUSG00000074994 | Qser1 | protein_coding | 0.422624564 | 2.57466188 | 0.3612636 | 0.5560684 | 0.906382272 |
| ENSMUSG00000037306 | Man1c1 | protein_coding | -0.299957838 | 3.687824388 | 0.3605367 | 0.5564618 | 0.906827392 |
| ENSMUSG00000026730 | Pter | protein_coding | 0.303703725 | 4.465772053 | 0.361418 | 0.5565102 | 0.906827392 |
| ENSMUSG00000024135 | Srbd1 | protein_coding | 0.332682885 | 3.426965269 | 0.3602865 | 0.5566108 | 0.90685392 |
| ENSMUSG00000039410 | Prdm16 | protein_coding | -0.192516401 | 5.45472691 | 0.3600736 | 0.5567126 | 0.90688242 |
| ENSMUSG00000022577 | Ly6h | protein_coding | -0.12360193 | 6.972127586 | 0.3595129 | 0.5570167 | 0.907240275 |
| ENSMUSG00000022109 | Med4 | protein_coding | 0.131985377 | 6.378644572 | 0.3591113 | 0.5572347 | 0.907457877 |
| ENSMUSG00000059173 | Pde1a | protein_coding | -0.141592782 | 6.814877576 | 0.358711 | 0.5574521 | 0.9076745 |
| ENSMUSG00000031691 | Tnpo2 | protein_coding | 0.326712971 | 3.370183233 | 0.3585113 | 0.5575606 | 0.907713766 |
| ENSMUSG00000061461 | Smim20 | protein_coding | -0.108907005 | 7.151173044 | 0.3583455 | 0.5576508 | 0.907723084 |
| ENSMUSG00000037643 | Prkci | protein_coding | 0.260510809 | 4.427018228 | 0.3581659 | 0.5577619 | 0.9077666 |
| ENSMUSG00000035735 | Dagla | protein_coding | -0.267407669 | 3.67213159 | 0.357905 | 0.5578904 | 0.90782916 |
| ENSMUSG00000032299 | Commd4 | protein_coding | 0.124895889 | 7.677940495 | 0.3576154 | 0.558048 | 0.90782916 |
| ENSMUSG00000045690 | Wdr89 | protein_coding | -0.210005271 | 4.393169587 | 0.3575386 | 0.5580899 | 0.90782916 |
| ENSMUSG00000040688 | Tbl3 | protein_coding | -0.189526585 | 5.372885284 | 0.3574501 | 0.5581381 | 0.90782916 |
| ENSMUSG00000020038 | Cry1 | protein_coding | 0.342978143 | 3.987714875 | 0.3577384 | 0.5585032 | 0.907905605 |
| ENSMUSG00000014547 | Wdfy2 | protein_coding | -0.312889481 | 3.165943596 | 0.3565894 | 0.5586073 | 0.907905605 |
| ENSMUSG00000035383 | Pmch | protein_coding | -0.509789092 | 6.988678681 | 0.3729103 | 0.5586106 | 0.907905605 |
| ENSMUSG00000045962 | Wnk1 | protein_coding | -0.161286979 | 5.624088413 | 0.3564568 | 0.5586797 | 0.907905605 |
| ENSMUSG00000044229 | Nxpe4 | protein_coding | -0.344737081 | 3.301324228 | 0.3558765 | 0.5589965 | 0.907905605 |
| ENSMUSG00000007891 | Ctsd | protein_coding | 0.137885215 | 6.348899188 | 0.3555989 | 0.5591482 | 0.907905605 |
| ENSMUSG00000029575 | Mmab | protein_coding | -0.175311828 | 5.092512116 | 0.355576 | 0.5591607 | 0.907905605 |
| ENSMUSG00000007613 | Tgfbr1 | protein_coding | -0.458838342 | 3.187294785 | 0.3564809 | 0.5591875 | 0.907905605 |
| ENSMUSG00000031221 | Igbp1 | protein_coding | -0.171206511 | 6.132569825 | 0.3554908 | 0.5592208 | 0.907905605 |
| ENSMUSG00000013275 | Slc41a1 | protein_coding | -0.190806392 | 4.441277428 | 0.3549876 | 0.5594826 | 0.907905605 |
| ENSMUSG00000024395 | Lims2 | protein_coding | -0.176631046 | 5.325717595 | 0.3547017 | 0.5596391 | 0.907905605 |
| ENSMUSG00000053617 | Sh3pxd2a | protein_coding | -0.369407331 | 3.092813162 | 0.3542469 | 0.5598883 | 0.907905605 |
| ENSMUSG00000016256 | Ctsz | protein_coding | -0.162139847 | 6.410510985 | 0.3541956 | 0.5599298 | 0.907905605 |
| ENSMUSG00000033768 | Nrxn2 | protein_coding | -0.185692448 | 4.840511298 | 0.3540959 | 0.559971 | 0.907905605 |
| ENSMUSG00000079108 | Srp54b | protein_coding | -0.267654678 | 3.34042388 | 0.3537406 | 0.5601659 | 0.907905605 |
| ENSMUSG00000034757 | Tmub2 | protein_coding | -0.123250694 | 6.644310887 | 0.3536138 | 0.5602355 | 0.907905605 |
| ENSMUSG00000022949 | Clic6 | protein_coding | -0.358978582 | 4.046555903 | 0.3548453 | 0.5603749 | 0.907905605 |
| ENSMUSG00000040675 | Mthfd1l | protein_coding | 0.274049159 | 3.427437564 | 0.3531764 | 0.5604755 | 0.907905605 |
| ENSMUSG00000028518 | Prkaa2 | protein_coding | -0.195168304 | 4.330244399 | 0.3531527 | 0.5604886 | 0.907905605 |
| ENSMUSG00000020893 | Per1 | protein_coding | 0.225557223 | 4.241100739 | 0.3529008 | 0.560627 | 0.907905605 |
| ENSMUSG00000027829 | Ccnl1 | protein_coding | 0.24444493 | 3.801533746 | 0.3527941 | 0.5606856 | 0.907905605 |
| ENSMUSG00000024073 | Birc6 | protein_coding | -0.16656128 | 5.592319642 | 0.3525163 | 0.5608383 | 0.907905605 |
| ENSMUSG00000031754 | Nudt21 | protein_coding | 0.199122106 | 3.847981517 | 0.3524514 | 0.560874 | 0.907905605 |
| ENSMUSG00000052812 | Atad2b | protein_coding | 0.313673681 | 3.200018374 | 0.3524396 | 0.5608805 | 0.907905605 |
| ENSMUSG00000037275 | Gemin5 | protein_coding | 0.264189225 | 3.51674595 | 0.3522663 | 0.5609758 | 0.907905605 |
| ENSMUSG00000004849 | Ap1s1 | protein_coding | -0.114866378 | 9.61928205 | 0.3522491 | 0.5609853 | 0.907905605 |
| ENSMUSG00000026765 | Lypd6b | protein_coding | -0.205928435 | 4.229487079 | 0.3522023 | 0.5610111 | 0.907905605 |
| ENSMUSG00000009734 | Pou6f2 | protein_coding | -0.514942214 | 2.355691237 | 0.3518971 | 0.5611791 | 0.907905605 |
| ENSMUSG00000035179 | Ppp1r32 | protein_coding | 0.291177291 | 3.234266445 | 0.3516279 | 0.5613273 | 0.907905605 |
| ENSMUSG00000058793 | Cds2 | protein_coding | 0.115071801 | 7.027360207 | 0.3515469 | 0.5613719 | 0.907905605 |
| ENSMUSG00000025872 | Thoc3 | protein_coding | 0.17635603 | 5.108572263 | 0.351514 | 0.56139 | 0.907905605 |
| ENSMUSG00000024565 | Sall3 | protein_coding | -0.382992263 | 2.707476908 | 0.3514113 | 0.5614466 | 0.907905605 |
| ENSMUSG00000020840 | Blmh | protein_coding | 0.1200986 | 7.288699791 | 0.3512398 | 0.5615412 | 0.907905605 |
| ENSMUSG00000035310 | Lin54 | protein_coding | 0.313939972 | 2.820126201 | 0.3512203 | 0.5615519 | 0.907905605 |
| ENSMUSG00000011154 | Cfap161 | protein_coding | -0.20623456 | 4.412085731 | 0.3507366 | 0.5618187 | 0.907905605 |
| ENSMUSG00000051469 | Zfp24 | protein_coding | -0.158023209 | 5.445761587 | 0.3506169 | 0.5618848 | 0.907905605 |
| ENSMUSG00000028048 | Gba | protein_coding | 0.199104042 | 4.871983274 | 0.3505448 | 0.5619245 | 0.907905605 |
| ENSMUSG00000035437 | Rabgap1 | protein_coding | 0.313929817 | 3.37899824 | 0.3504734 | 0.561964 | 0.907905605 |
| ENSMUSG00000027881 | Prpf38b | protein_coding | -0.25626865 | 3.818091983 | 0.3503999 | 0.5620046 | 0.907905605 |
| ENSMUSG00000026074 | Map4k4 | protein_coding | -0.200118822 | 4.142938693 | 0.3503754 | 0.5620181 | 0.907905605 |
| ENSMUSG00000024397 | Aif1 | protein_coding | -0.284900886 | 4.614596879 | 0.3511436 | 0.5621107 | 0.907905605 |
| ENSMUSG00000056014 | A430033K04Rik | protein_coding | -0.217721539 | 4.071842789 | 0.35015 | 0.5621426 | 0.907905605 |
| ENSMUSG00000036613 | Tssc1 | protein_coding | 0.15145266 | 6.756816031 | 0.3501487 | 0.5621566 | 0.907905605 |
| ENSMUSG00000022571 | Pycrl | protein_coding | 0.121957927 | 6.757808045 | 0.3498073 | 0.562332 | 0.907905605 |
| ENSMUSG00000025875 | Tspan17 | protein_coding | -0.31958023 | 3.463116363 | 0.3497855 | 0.562344 | 0.907905605 |
| ENSMUSG00000034107 | Ano7 | protein_coding | -0.452743993 | 2.574093117 | 0.3495401 | 0.5624797 | 0.907905605 |
| ENSMUSG00000043284 | Tmem11 | protein_coding | -0.125187401 | 6.622788953 | 0.3495073 | 0.5624978 | 0.907905605 |
| ENSMUSG00000037536 | Fbxo34 | protein_coding | 0.223976783 | 3.509491537 | 0.3492194 | 0.5626571 | 0.907905605 |
| ENSMUSG00000036398 | Ppp1r11 | protein_coding | 0.1072921 | 9.340274285 | 0.3492042 | 0.5626655 | 0.907905605 |

| ensembl_gene_id | external_gene_name | gene_biotype | log2 (fold change) | log2 (counts per million) | F-value | p-value | False discovery rate |
|---|---|---|---|---|---|---|---|
| ENSMUSG00000001211 | Agpat3 | protein_coding | -0.182826105 | 5.813345945 | 0.3488272 | 0.5628876 | 0.907905605 |
| ENSMUSG00000055733 | Nap1l3 | protein_coding | -0.16230874 | 5.768470532 | 0.3487252 | 0.5629308 | 0.907905605 |
| ENSMUSG00000049892 | Rasd1 | protein_coding | 0.179677689 | 6.061518301 | 0.3487306 | 0.5629411 | 0.907905605 |
| ENSMUSG00000024176 | Sox8 | protein_coding | -0.201652146 | 4.254497487 | 0.3480643 | 0.5632972 | 0.907905605 |
| ENSMUSG00000026975 | Dph7 | protein_coding | 0.128225717 | 6.119075733 | 0.3476404 | 0.5635324 | 0.907905605 |
| ENSMUSG00000079020 | Slc45a4 | protein_coding | 0.269188771 | 3.704989403 | 0.3475788 | 0.5635666 | 0.907905605 |
| ENSMUSG00000024906 | Mus81 | protein_coding | 0.201644125 | 4.973403595 | 0.3475126 | 0.5636034 | 0.907905605 |
| ENSMUSG00000028978 | Nos3 | protein_coding | -0.288437109 | 3.220516984 | 0.3474708 | 0.5636265 | 0.907905605 |
| ENSMUSG00000023965 | Fbxl17 | protein_coding | -0.150466673 | 5.385805921 | 0.3472826 | 0.5637311 | 0.907905605 |
| ENSMUSG00000053111 | Fank1 | protein_coding | -0.236262018 | 4.313828583 | 0.3471327 | 0.5638144 | 0.907905605 |
| ENSMUSG00000021684 | Pde8b | protein_coding | 0.183942708 | 5.163452221 | 0.3467338 | 0.5640361 | 0.907905605 |
| ENSMUSG00000031229 | Atrx | protein_coding | -0.150230258 | 6.221157175 | 0.3463726 | 0.5642371 | 0.907905605 |
| ENSMUSG00000034723 | Tmx4 | protein_coding | -0.114627849 | 7.659285269 | 0.3463622 | 0.5642429 | 0.907905605 |
| ENSMUSG00000027628 | Aar2 | protein_coding | 0.134954992 | 6.484043331 | 0.3462641 | 0.5642975 | 0.907905605 |
| ENSMUSG00000026827 | Gpd2 | protein_coding | 0.119722058 | 6.542160794 | 0.34615 | 0.564361 | 0.907905605 |
| ENSMUSG00000034263 | Ints14 | protein_coding | -0.191014789 | 4.562746553 | 0.3458598 | 0.5645226 | 0.907905605 |
| ENSMUSG00000073468 | Sft2d1 | protein_coding | 0.188522072 | 4.717820799 | 0.3458595 | 0.5645228 | 0.907905605 |
| ENSMUSG00000031447 | Lamp1 | protein_coding | 0.245318375 | 3.552951563 | 0.3457973 | 0.5645574 | 0.907905605 |
| ENSMUSG00000050164 | Mchr1 | protein_coding | -0.233198554 | 4.577819094 | 0.3457862 | 0.5645636 | 0.907905605 |
| ENSMUSG00000031007 | Atp6ap2 | protein_coding | 0.107809072 | 8.573246147 | 0.3457304 | 0.5645947 | 0.907905605 |
| ENSMUSG00000044835 | Ankrd45 | protein_coding | -0.160458467 | 5.516952363 | 0.345408 | 0.5647744 | 0.907905605 |
| ENSMUSG00000023079 | Gtf2ird1 | protein_coding | 0.218130935 | 4.057280364 | 0.345327 | 0.5648196 | 0.907905605 |
| ENSMUSG00000043602 | Zfp3 | protein_coding | -0.224127549 | 3.811429342 | 0.3450191 | 0.5649913 | 0.907905605 |
| ENSMUSG00000035769 | Xylb | protein_coding | -0.290456643 | 3.277310768 | 0.344944 | 0.5650333 | 0.907905605 |
| ENSMUSG00000044348 | Slc25a53 | protein_coding | 0.174837365 | 4.529963874 | 0.344785 | 0.565122 | 0.907905605 |
| ENSMUSG00000052310 | Slc39a1 | protein_coding | -0.272984706 | 4.916541823 | 0.3453832 | 0.5652999 | 0.907905605 |
| ENSMUSG00000017692 | Rhbdl3 | protein_coding | -0.278811042 | 3.989578344 | 0.3443433 | 0.5653819 | 0.907905605 |
| ENSMUSG00000033228 | Scaf11 | protein_coding | 0.167133118 | 5.105056989 | 0.3442984 | 0.5653938 | 0.907905605 |
| ENSMUSG00000009406 | Elk1 | protein_coding | 0.206566109 | 4.138443608 | 0.3441981 | 0.5654499 | 0.907905605 |
| ENSMUSG00000073771 | Btbd19 | protein_coding | 0.295388409 | 3.129784981 | 0.344068 | 0.5655226 | 0.907905605 |
| ENSMUSG00000028005 | Gucy1b3 | protein_coding | 0.120392089 | 7.007594023 | 0.3440377 | 0.5655395 | 0.907905605 |
| ENSMUSG00000085389 | 1700003M07Rik | lincRNA | -0.261909541 | 3.252357839 | 0.3439392 | 0.5655946 | 0.907905605 |
| ENSMUSG00000028898 | Trnau1ap | protein_coding | -0.223439093 | 4.140814043 | 0.3438915 | 0.5656213 | 0.907905605 |
| ENSMUSG00000025978 | Rftn2 | protein_coding | -0.173875804 | 4.74961476 | 0.3438865 | 0.5656241 | 0.907905605 |
| ENSMUSG00000020802 | Ube2o | protein_coding | -0.171018307 | 4.962172103 | 0.3438163 | 0.5656633 | 0.907905605 |
| ENSMUSG00000059810 | Rgs3 | protein_coding | -0.263393796 | 3.626404954 | 0.3437715 | 0.5656884 | 0.907905605 |
| ENSMUSG00000026235 | Epha4 | protein_coding | -0.171948091 | 6.211637622 | 0.343567 | 0.565816 | 0.907905605 |
| ENSMUSG00000004892 | Bcan | protein_coding | 0.238932278 | 5.297750323 | 0.3440085 | 0.5659091 | 0.907905605 |
| ENSMUSG00000087590 | Epb41l4aos | lincRNA | -0.244823847 | 3.339230179 | 0.3433432 | 0.5659281 | 0.907905605 |
| ENSMUSG00000032118 | Fez1 | protein_coding | 0.133434705 | 9.195683626 | 0.3433031 | 0.5659506 | 0.907905605 |
| ENSMUSG00000020092 | Pald1 | protein_coding | 0.477700406 | 2.379014668 | 0.3432364 | 0.5659879 | 0.907905605 |
| ENSMUSG00000068859 | Sp9 | protein_coding | -0.214974385 | 3.855387841 | 0.3431734 | 0.5660232 | 0.907905605 |
| ENSMUSG00000024767 | Otub1 | protein_coding | 0.132907325 | 10.83118066 | 0.3430238 | 0.566107 | 0.907905605 |
| ENSMUSG00000037795 | N4bp2 | protein_coding | -0.355200378 | 2.840029752 | 0.342959 | 0.5661433 | 0.907905605 |
| ENSMUSG00000028108 | Ecm1 | protein_coding | 0.392687849 | 2.629739521 | 0.3427447 | 0.5662634 | 0.907905605 |
| ENSMUSG00000014837 | 4931428F04Rik | protein_coding | -0.245431635 | 5.338054235 | 0.3435288 | 0.5663342 | 0.907905605 |
| ENSMUSG00000050017 | Pitpnb | protein_coding | -0.134557381 | 6.832640067 | 0.3425318 | 0.5663827 | 0.907905605 |
| ENSMUSG00000058159 | T2 | processed_transcript | -0.429701817 | 2.287451488 | 0.341955 | 0.5667064 | 0.907905605 |
| ENSMUSG00000057738 | Sptan1 | protein_coding | 0.148031079 | 7.14596439 | 0.3418921 | 0.5667548 | 0.907905605 |
| ENSMUSG00000045594 | Glb1 | protein_coding | 0.250583744 | 5.061458579 | 0.3424278 | 0.5667762 | 0.907905605 |
| ENSMUSG00000045871 | Slitrk6 | protein_coding | -0.469412768 | 2.80035259 | 0.3418482 | 0.5667795 | 0.907905605 |
| ENSMUSG00000032330 | Cox7a2 | protein_coding | -0.144727875 | 10.09163773 | 0.341814 | 0.5667855 | 0.907905605 |
| ENSMUSG00000041420 | Meis3 | protein_coding | -0.190299756 | 4.46361733 | 0.3415826 | 0.5669155 | 0.907905605 |
| ENSMUSG00000047583 | Tyw3 | protein_coding | 0.174799878 | 4.365579562 | 0.341533 | 0.5669434 | 0.907905605 |
| ENSMUSG00000019433 | Gipc1 | protein_coding | -0.15729085 | 5.693327097 | 0.3414407 | 0.5669952 | 0.907905605 |
| ENSMUSG00000066735 | Vkorc1l1 | protein_coding | -0.17766965 | 4.973310593 | 0.3412855 | 0.5670825 | 0.907905605 |
| ENSMUSG00000063445 | Nmral1 | protein_coding | -0.12814618 | 6.160852941 | 0.3409579 | 0.5672667 | 0.907905605 |
| ENSMUSG00000047242 | Taf9b | protein_coding | -0.143117021 | 6.562037021 | 0.3407008 | 0.5674114 | 0.907905605 |
| ENSMUSG00000060538 | Tmem219 | protein_coding | 0.186482331 | 4.342771791 | 0.3404728 | 0.5675397 | 0.907905605 |
| ENSMUSG00000039568 | Ubald1 | protein_coding | 0.123535057 | 6.558367213 | 0.3404454 | 0.5675551 | 0.907905605 |
| ENSMUSG00000060477 | Irak2 | protein_coding | 0.247596796 | 3.333828828 | 0.3403346 | 0.5676175 | 0.907905605 |
| ENSMUSG00000035478 | Mbd3 | protein_coding | -0.191321845 | 5.413148314 | 0.3401563 | 0.567718 | 0.907905605 |
| ENSMUSG00000035486 | Plk5 | protein_coding | 0.288549912 | 3.165917379 | 0.3400825 | 0.5677595 | 0.907905605 |
| ENSMUSG00000002489 | Tiam1 | protein_coding | -0.163463074 | 5.244388358 | 0.3400492 | 0.5677783 | 0.907905605 |
| ENSMUSG00000022607 | Ptk2 | protein_coding | 0.147955868 | 5.103287679 | 0.3396673 | 0.5679936 | 0.907905605 |
| ENSMUSG00000046607 | Hrk | protein_coding | 0.187418035 | 4.154284163 | 0.3396152 | 0.568023 | 0.907905605 |
| ENSMUSG00000000085 | Scmh1 | protein_coding | -0.169508031 | 5.071774487 | 0.3394975 | 0.5680894 | 0.907905605 |
| ENSMUSG00000024998 | Plce1 | protein_coding | -0.291262853 | 2.957736556 | 0.3388123 | 0.5684762 | 0.907905605 |
| ENSMUSG00000036598 | Ccdc113 | protein_coding | -0.182199166 | 4.42532233 | 0.3385181 | 0.5686425 | 0.907905605 |
| ENSMUSG00000051339 | 2900026A02Rik | protein_coding | 0.226019201 | 4.055889803 | 0.3384218 | 0.5686969 | 0.907905605 |
| ENSMUSG00000029060 | Mib2 | protein_coding | 0.215775774 | 4.132679786 | 0.3384169 | 0.5686997 | 0.907905605 |
| ENSMUSG00000002028 | Kmt2a | protein_coding | -0.191415418 | 5.263140623 | 0.3384137 | 0.5687015 | 0.907905605 |
| ENSMUSG00000024308 | Tapbp | protein_coding | 0.164116596 | 4.960714711 | 0.3382469 | 0.5687958 | 0.907905605 |
| ENSMUSG00000062101 | Zfp119b | protein_coding | -0.45207 | 2.779244344 | 0.3382306 | 0.5688181 | 0.907905605 |

| ensembl_gene_id | external_gene_name | gene_biotype | log2 (fold change) | log2 (counts per million) | F-value | p-value | False discovery rate |
|---|---|---|---|---|---|---|---|
| ENSMUSG00000003849 | Nqo1 | protein_coding | 0.268901282 | 3.758894883 | 0.3381589 | 0.5688456 | 0.907905605 |
| ENSMUSG00000016481 | Cr1l | protein_coding | 0.219456547 | 3.992040996 | 0.3378187 | 0.5690381 | 0.907905605 |
| ENSMUSG00000037570 | Mcrs1 | protein_coding | -0.164717785 | 4.744328331 | 0.3376246 | 0.569148 | 0.907905605 |
| ENSMUSG00000038534 | Osbpl7 | protein_coding | 0.178789661 | 4.381384105 | 0.3375093 | 0.5692133 | 0.907905605 |
| ENSMUSG00000028164 | Manba | protein_coding | 0.352983243 | 2.762128959 | 0.3374245 | 0.5692613 | 0.907905605 |
| ENSMUSG00000032621 | Srek1 | protein_coding | -0.163383646 | 5.54780392 | 0.3374102 | 0.5692694 | 0.907905605 |
| ENSMUSG00000028434 | Epb41l4b | protein_coding | 0.168591511 | 4.797368241 | 0.3373419 | 0.5693081 | 0.907905605 |
| ENSMUSG00000042389 | Tsen2 | protein_coding | 0.156850636 | 5.267071006 | 0.3369271 | 0.5695432 | 0.907905605 |
| ENSMUSG00000079036 | Alkbh1 | protein_coding | 0.185281598 | 4.201292992 | 0.3366345 | 0.5697092 | 0.907905605 |
| ENSMUSG00000037847 | Nmrk1 | protein_coding | 0.271625838 | 3.704456929 | 0.3365459 | 0.5697595 | 0.907905605 |
| ENSMUSG00000044147 | Arf6 | protein_coding | 0.152574899 | 5.076528552 | 0.3364866 | 0.5697932 | 0.907905605 |
| ENSMUSG00000035615 | Frmpd1 | protein_coding | -0.53160922 | 2.221045393 | 0.3363844 | 0.5698512 | 0.907905605 |
| ENSMUSG00000022186 | Oxct1 | protein_coding | 0.107869278 | 8.154560547 | 0.3363039 | 0.5698969 | 0.907905605 |
| ENSMUSG00000025931 | Paqr8 | protein_coding | 0.208864197 | 4.716172339 | 0.3361618 | 0.5699775 | 0.907905605 |
| ENSMUSG00000034522 | Zfp395 | protein_coding | -0.228636196 | 3.499835818 | 0.3360707 | 0.5700293 | 0.907905605 |
| ENSMUSG00000039936 | Pik3cd | protein_coding | 0.393978365 | 2.828825237 | 0.3359212 | 0.5701142 | 0.907905605 |
| ENSMUSG00000093930 | Hmgcs1 | protein_coding | -0.137387983 | 7.080878806 | 0.3358047 | 0.5701804 | 0.907905605 |
| ENSMUSG00000035949 | Fbxw2 | protein_coding | -0.213262347 | 5.108968232 | 0.3357328 | 0.5702343 | 0.907905605 |
| ENSMUSG00000027698 | Nceh1 | protein_coding | 0.100388746 | 7.061547712 | 0.3355202 | 0.5703422 | 0.907905605 |
| ENSMUSG00000001158 | Snrnp27 | protein_coding | -0.133820439 | 8.072100003 | 0.3354178 | 0.5704004 | 0.907905605 |
| ENSMUSG00000028488 | Sh3gl2 | protein_coding | 0.121625487 | 7.615826739 | 0.3353944 | 0.5704137 | 0.907905605 |
| ENSMUSG00000039533 | Mmd2 | protein_coding | -0.153115091 | 7.968709712 | 0.3353924 | 0.5704278 | 0.907905605 |
| ENSMUSG00000017740 | Slc12a5 | protein_coding | 0.205510651 | 6.468301671 | 0.3360978 | 0.5708031 | 0.908259237 |
| ENSMUSG00000010175 | Prox1 | protein_coding | -0.248718491 | 4.413979625 | 0.3347053 | 0.5708189 | 0.908259237 |
| ENSMUSG00000006575 | Rundc3a | protein_coding | -0.136831304 | 6.008340285 | 0.3341334 | 0.5711319 | 0.908622787 |
| ENSMUSG00000033669 | Zfp7 | protein_coding | 0.275382817 | 3.10848337 | 0.3338668 | 0.5712839 | 0.908730206 |
| ENSMUSG00000027428 | Rbbp9 | protein_coding | -0.153012763 | 6.297842946 | 0.3332487 | 0.5716368 | 0.908730939 |
| ENSMUSG00000070371 | Prss36 | protein_coding | 0.247454711 | 4.542791938 | 0.3332639 | 0.5716411 | 0.908730939 |
| ENSMUSG00000045658 | Pid1 | protein_coding | -0.187962481 | 4.859347571 | 0.33323 | 0.5716475 | 0.908730939 |
| ENSMUSG00000029221 | Slc30a9 | protein_coding | -0.116522516 | 6.804376336 | 0.3330731 | 0.5717372 | 0.908730939 |
| ENSMUSG00000042529 | Kcnj12 | protein_coding | -0.293440931 | 3.044335872 | 0.3330087 | 0.571774 | 0.908730939 |
| ENSMUSG00000097789 | Gm2115 | protein_coding | -0.257728913 | 3.277335339 | 0.3329425 | 0.5718118 | 0.908730939 |
| ENSMUSG00000015755 | Tab2 | protein_coding | -0.19590029 | 5.182467895 | 0.3325778 | 0.5720204 | 0.908730939 |
| ENSMUSG00000028228 | Cpne3 | protein_coding | -0.314432118 | 3.502049258 | 0.3325855 | 0.5720289 | 0.908730939 |
| ENSMUSG00000052369 | Tmem106c | protein_coding | 0.104024699 | 7.361881729 | 0.3324669 | 0.5720838 | 0.908730939 |
| ENSMUSG00000024483 | Ankhd1 | protein_coding | -0.164856471 | 4.962412996 | 0.3321464 | 0.5722673 | 0.908730939 |
| ENSMUSG00000028080 | Lrba | protein_coding | -0.275100266 | 3.827978285 | 0.3317803 | 0.572477 | 0.908730939 |
| ENSMUSG00000049401 | Ogfr | protein_coding | 0.21799514 | 3.834392804 | 0.3316194 | 0.5725692 | 0.908730939 |
| ENSMUSG00000002930 | Ppp1r17 | protein_coding | 0.381142432 | 3.752463583 | 0.3328997 | 0.5726212 | 0.908730939 |
| ENSMUSG00000052593 | Adam17 | protein_coding | 0.466559013 | 2.509099436 | 0.3313397 | 0.5727295 | 0.908730939 |
| ENSMUSG00000044791 | Setd2 | protein_coding | -0.21407551 | 4.536648777 | 0.3312673 | 0.572771 | 0.908730939 |
| ENSMUSG00000036377 | C530008M17Rik | protein_coding | -0.18729517 | 4.964826786 | 0.3312194 | 0.5727985 | 0.908730939 |
| ENSMUSG00000039809 | Gabbr2 | protein_coding | 0.156373681 | 5.834550087 | 0.3310595 | 0.5728902 | 0.908730939 |
| ENSMUSG00000042207 | Kdm5b | protein_coding | -0.13950917 | 5.486204669 | 0.3310514 | 0.5728949 | 0.908730939 |
| ENSMUSG00000036667 | Tcaf1 | protein_coding | -0.133781624 | 7.530609626 | 0.3310455 | 0.5728983 | 0.908730939 |
| ENSMUSG00000032841 | Prr5l | protein_coding | -0.202867969 | 5.449710752 | 0.3304678 | 0.5732429 | 0.908730939 |
| ENSMUSG00000059013 | Sh2d3c | protein_coding | 0.232114081 | 4.324343061 | 0.3304211 | 0.5732568 | 0.908730939 |
| ENSMUSG00000042043 | Tbca | protein_coding | -0.14418446 | 7.893308059 | 0.3304186 | 0.5732711 | 0.908730939 |
| ENSMUSG00000053411 | Cbx7 | protein_coding | -0.156104683 | 5.132280649 | 0.3303352 | 0.5733062 | 0.908730939 |
| ENSMUSG00000029151 | Slc30a3 | protein_coding | -0.573950593 | 2.706361497 | 0.3314437 | 0.5733126 | 0.908730939 |
| ENSMUSG00000023805 | Synj2 | protein_coding | -0.184712854 | 4.969783912 | 0.3301643 | 0.5734044 | 0.908742418 |
| ENSMUSG00000032477 | Cdc25a | protein_coding | -0.363902527 | 3.205353042 | 0.3298393 | 0.5736041 | 0.908925054 |
| ENSMUSG00000029641 | Rasl11a | protein_coding | 0.319049417 | 3.948291261 | 0.3301697 | 0.5738329 | 0.909048581 |
| ENSMUSG00000026811 | St6galnac6 | protein_coding | -0.134125868 | 5.915697612 | 0.3293877 | 0.5738512 | 0.909048581 |
| ENSMUSG00000039725 | Trp53rka | protein_coding | 0.344027534 | 3.216718572 | 0.3290718 | 0.5740331 | 0.909202851 |
| ENSMUSG00000062151 | Unc13c | protein_coding | -0.175704352 | 6.355882941 | 0.3288655 | 0.5742208 | 0.909270161 |
| ENSMUSG00000041654 | Slc39a11 | protein_coding | 0.243099662 | 4.16824922 | 0.3287047 | 0.5742447 | 0.909270161 |
| ENSMUSG00000043257 | Pigv | protein_coding | 0.359827334 | 2.814847498 | 0.3282573 | 0.5745028 | 0.909519616 |
| ENSMUSG00000001794 | Capns1 | protein_coding | 0.106758137 | 7.23377314 | 0.3280067 | 0.5746475 | 0.909519616 |
| ENSMUSG00000000056 | Narf | protein_coding | 0.148135687 | 5.785371926 | 0.3278612 | 0.5747315 | 0.909519616 |
| ENSMUSG00000029387 | Gtf2h3 | protein_coding | 0.159280584 | 5.419838996 | 0.3278454 | 0.5747406 | 0.909519616 |
| ENSMUSG00000037470 | Uggt1 | protein_coding | 0.214014016 | 4.094917282 | 0.3275985 | 0.5748832 | 0.909611413 |
| ENSMUSG00000026090 | 2010300C02Rik | protein_coding | -0.149559437 | 5.642131526 | 0.3269197 | 0.5752758 | 0.909800658 |
| ENSMUSG00000031505 | Naxd | protein_coding | 0.118752146 | 6.902994987 | 0.3268034 | 0.5753431 | 0.909800658 |
| ENSMUSG00000028156 | Eif4e | protein_coding | -0.108933676 | 7.869576667 | 0.32677 | 0.5753624 | 0.909800658 |
| ENSMUSG00000023274 | Cd4 | protein_coding | -0.186561593 | 4.681910182 | 0.3267184 | 0.5753923 | 0.909800658 |
| ENSMUSG00000024777 | Ppp2r5b | protein_coding | -0.132620909 | 5.645437289 | 0.3266168 | 0.5754511 | 0.909800658 |
| ENSMUSG00000034308 | Sdr42e1 | protein_coding | 0.332192694 | 3.166429921 | 0.3265143 | 0.5755105 | 0.909800658 |
| ENSMUSG00000032594 | Ip6k1 | protein_coding | 0.1146255 | 7.034763964 | 0.3263261 | 0.5756195 | 0.909822533 |
| ENSMUSG00000045994 | B3gat1 | protein_coding | -0.177231316 | 4.885530652 | 0.3261983 | 0.5756935 | 0.909822533 |
| ENSMUSG00000034807 | Colgalt1 | protein_coding | -0.287301049 | 3.792657305 | 0.3258973 | 0.5758808 | 0.909984756 |
| ENSMUSG00000037351 | Actr1b | protein_coding | 0.137210189 | 8.367555443 | 0.3254769 | 0.5761119 | 0.9102162 |
| ENSMUSG00000022562 | Oplah | protein_coding | -0.366467514 | 2.775960112 | 0.3247167 | 0.5765535 | 0.910655103 |
| ENSMUSG00000035754 | Wdr18 | protein_coding | 0.119057753 | 8.189957898 | 0.3247071 | 0.5765591 | 0.910655103 |

| ensembl_gene_id | external_gene_name | gene_biotype | log2 (fold change) | log2 (counts per million) | F-value | p-value | False discovery rate |
|---|---|---|---|---|---|---|---|
| ENSMUSG00000028684 | Urod | protein_coding | 0.136010672 | 7.131386531 | 0.3242059 | 0.5768506 | 0.910872265 |
| ENSMUSG00000096141 | Dnah7a | protein_coding | 0.200744957 | 3.695228633 | 0.3240879 | 0.5769193 | 0.910872265 |
| ENSMUSG00000011096 | Akt1s1 | protein_coding | 0.337187223 | 2.861234585 | 0.3239938 | 0.576974 | 0.910872265 |
| ENSMUSG00000022516 | Nudt16l1 | protein_coding | -0.110120807 | 7.037987779 | 0.323872 | 0.577045 | 0.910872265 |
| ENSMUSG00000024592 | C330018D20Rik | protein_coding | -0.195300256 | 4.791125602 | 0.3237081 | 0.5771404 | 0.910872265 |
| ENSMUSG00000039183 | Nubp2 | protein_coding | -0.110194161 | 6.605297368 | 0.3235975 | 0.5772048 | 0.910872265 |
| ENSMUSG00000043909 | Trp53bp1 | protein_coding | -0.196999548 | 4.852830253 | 0.323072 | 0.5775112 | 0.910975821 |
| ENSMUSG00000031660 | Brd7 | protein_coding | -0.158173298 | 5.766305864 | 0.3229781 | 0.577566 | 0.910975821 |
| ENSMUSG00000019528 | Gyg | protein_coding | 0.132752047 | 6.934157067 | 0.3228218 | 0.5776572 | 0.910975821 |
| ENSMUSG00000020288 | Ahsa2 | protein_coding | -0.252147774 | 3.798943059 | 0.322808 | 0.5776652 | 0.910975821 |
| ENSMUSG00000026111 | Unc50 | protein_coding | -0.145899702 | 5.589462597 | 0.3225728 | 0.5778025 | 0.910975821 |
| ENSMUSG00000086003 | B230206L02Rik | processed_transcript | -0.200208544 | 4.057111651 | 0.3224089 | 0.5778982 | 0.910975821 |
| ENSMUSG00000019822 | Smpd2 | protein_coding | 0.228863041 | 4.065077862 | 0.3223234 | 0.5779482 | 0.910975821 |
| ENSMUSG00000019891 | Dcbld1 | protein_coding | 0.410155029 | 2.688547111 | 0.3223233 | 0.5779482 | 0.910975821 |
| ENSMUSG00000021807 | 2700060E02Rik | protein_coding | 0.117719071 | 6.484399322 | 0.3218994 | 0.578196 | 0.911232798 |
| ENSMUSG00000000881 | Dlg3 | protein_coding | 0.153376787 | 5.349493532 | 0.3217259 | 0.5782975 | 0.911259172 |
| ENSMUSG00000047747 | Rnf150 | protein_coding | -0.230815096 | 4.084373128 | 0.3213688 | 0.5785064 | 0.911279114 |
| ENSMUSG00000020132 | Rab21 | protein_coding | -0.108807401 | 6.693586283 | 0.3213201 | 0.5785349 | 0.911279114 |
| ENSMUSG00000040649 | Rimklb | protein_coding | 0.142240088 | 6.130343826 | 0.3212698 | 0.5785644 | 0.911279114 |
| ENSMUSG00000030795 | Fus | protein_coding | -0.158330376 | 5.029174441 | 0.3203968 | 0.5790759 | 0.911369989 |
| ENSMUSG00000031668 | Eif2ak3 | protein_coding | -0.344627852 | 3.672571185 | 0.3206142 | 0.5794386 | 0.911369989 |
| ENSMUSG00000023010 | Tmbim6 | protein_coding | 0.117128465 | 9.694523127 | 0.3194561 | 0.5796282 | 0.911369989 |
| ENSMUSG00000019872 | Smpdl3a | protein_coding | -0.188979135 | 4.455349923 | 0.3193168 | 0.5797101 | 0.911369989 |
| ENSMUSG00000026073 | Il1r2 | protein_coding | -0.540423264 | 2.054757729 | 0.3192923 | 0.5797245 | 0.911369989 |
| ENSMUSG00000061436 | Hipk2 | protein_coding | -0.261551884 | 4.56739696 | 0.3196901 | 0.5797293 | 0.911369989 |
| ENSMUSG00000054519 | Zfp867 | protein_coding | -0.223275352 | 3.704421995 | 0.3190864 | 0.5798455 | 0.911369989 |
| ENSMUSG00000025551 | Fgf14 | protein_coding | 0.182325066 | 5.814159515 | 0.318958 | 0.5799337 | 0.911369989 |
| ENSMUSG00000043531 | Sorcs1 | protein_coding | -0.21211348 | 3.709531103 | 0.3185951 | 0.5801346 | 0.911369989 |
| ENSMUSG00000020848 | Doc2b | protein_coding | 0.222750982 | 3.522851538 | 0.3185907 | 0.5801372 | 0.911369989 |
| ENSMUSG00000026188 | Tmem169 | protein_coding | -0.204160826 | 3.928872815 | 0.3185553 | 0.5801581 | 0.911369989 |
| ENSMUSG00000038563 | Efl1 | protein_coding | 0.161421246 | 4.51451738 | 0.3184984 | 0.5801916 | 0.911369989 |
| ENSMUSG00000022419 | Deptor | protein_coding | -0.26579724 | 4.525746661 | 0.3188984 | 0.5802116 | 0.911369989 |
| ENSMUSG00000040136 | Abcc8 | protein_coding | 0.73282481 | 2.080395375 | 0.3191953 | 0.5802704 | 0.911369989 |
| ENSMUSG00000029599 | Ddx54 | protein_coding | 0.281185098 | 3.394293668 | 0.3183041 | 0.580306 | 0.911369989 |
| ENSMUSG00000026466 | Tor1aip1 | protein_coding | -0.209460601 | 3.687747395 | 0.3180772 | 0.5804397 | 0.911369989 |
| ENSMUSG00000039959 | Hip1 | protein_coding | -0.167225616 | 4.775215907 | 0.3179876 | 0.5804925 | 0.911369989 |
| ENSMUSG00000003271 | Sult2b1 | protein_coding | -0.252786015 | 3.973833882 | 0.3177778 | 0.5806162 | 0.911369989 |
| ENSMUSG00000003526 | Prodh | protein_coding | -0.257921346 | 4.696585418 | 0.3181767 | 0.5807023 | 0.911369989 |
| ENSMUSG00000054676 | 1600014C10Rik | protein_coding | 0.184146229 | 4.517361593 | 0.3173542 | 0.5808662 | 0.911369989 |
| ENSMUSG00000044646 | Zbtb7c | protein_coding | 0.348358486 | 2.786383775 | 0.3173351 | 0.5808774 | 0.911369989 |
| ENSMUSG00000064105 | Cnnm2 | protein_coding | -0.288198881 | 3.6964863 | 0.3173105 | 0.5809045 | 0.911369989 |
| ENSMUSG00000038542 | Pcid2 | protein_coding | 0.290073534 | 3.686813196 | 0.3172877 | 0.580918 | 0.911369989 |
| ENSMUSG00000053070 | 9230110C19Rik | protein_coding | -0.205618795 | 4.620341163 | 0.3170325 | 0.5810561 | 0.911369989 |
| ENSMUSG00000027104 | Atf2 | protein_coding | -0.128350068 | 7.127111128 | 0.3170049 | 0.5810724 | 0.911369989 |
| ENSMUSG00000044968 | Napepld | protein_coding | 0.313373773 | 3.709934025 | 0.3170602 | 0.5810854 | 0.911369989 |
| ENSMUSG00000025935 | Tram1 | protein_coding | -0.171716761 | 5.064605052 | 0.3168855 | 0.581143 | 0.911369989 |
| ENSMUSG00000040945 | Rcc2 | protein_coding | -0.123729948 | 5.898254329 | 0.3165874 | 0.5813191 | 0.911369989 |
| ENSMUSG00000050144 | Slc25a44 | protein_coding | -0.118391915 | 7.042172236 | 0.3165043 | 0.5813683 | 0.911369989 |
| ENSMUSG00000017707 | Serinc3 | protein_coding | -0.137700324 | 6.07841394 | 0.3163729 | 0.581446 | 0.911369989 |
| ENSMUSG00000025555 | Farp1 | protein_coding | 0.199144077 | 4.541455002 | 0.3156821 | 0.5818549 | 0.911369989 |
| ENSMUSG00000020476 | Dbnl | protein_coding | -0.110491099 | 6.914025836 | 0.3156586 | 0.5818688 | 0.911369989 |
| ENSMUSG00000025178 | Pi4k2a | protein_coding | 0.148821812 | 6.30671777 | 0.3155362 | 0.5819413 | 0.911369989 |
| ENSMUSG00000048661 | Lemd3 | protein_coding | -0.202122393 | 3.832823905 | 0.3150589 | 0.5822243 | 0.911369989 |
| ENSMUSG00000000605 | Clcn4 | protein_coding | 0.108685603 | 7.351839235 | 0.314999 | 0.5822598 | 0.911369989 |
| ENSMUSG00000049744 | Arhgap15 | protein_coding | 0.404381395 | 2.51061778 | 0.3149963 | 0.5822614 | 0.911369989 |
| ENSMUSG00000028669 | Pithd1 | protein_coding | 0.114748846 | 6.480969073 | 0.3146689 | 0.5824557 | 0.911369989 |
| ENSMUSG00000029578 | Wipi2 | protein_coding | -0.111678311 | 6.48268592 | 0.3144549 | 0.5825827 | 0.911369989 |
| ENSMUSG00000024997 | Prdx3 | protein_coding | 0.097744712 | 8.346244959 | 0.314212 | 0.582727 | 0.911369989 |
| ENSMUSG00000025147 | Mob2 | protein_coding | 0.106689424 | 6.615878038 | 0.3141324 | 0.5827743 | 0.911369989 |
| ENSMUSG00000032265 | Fam46a | protein_coding | -0.212550745 | 3.586635144 | 0.314104 | 0.5827912 | 0.911369989 |
| ENSMUSG00000026883 | Dab2ip | protein_coding | 0.194628233 | 4.455905101 | 0.314075 | 0.5828084 | 0.911369989 |
| ENSMUSG00000046836 | Brox | protein_coding | 0.167774783 | 4.588628176 | 0.3139148 | 0.5829037 | 0.911369989 |
| ENSMUSG00000031021 | Tmem9b | protein_coding | 0.131971575 | 5.948648697 | 0.3137761 | 0.5829862 | 0.911369989 |
| ENSMUSG00000036046 | 5031439G07Rik | protein_coding | -0.154219282 | 5.875616793 | 0.3133617 | 0.5832327 | 0.911369989 |
| ENSMUSG00000064065 | Ipcef1 | protein_coding | -0.321286591 | 2.627359366 | 0.3133404 | 0.5832454 | 0.911369989 |
| ENSMUSG00000043391 | 2510009E07Rik | protein_coding | -0.147365441 | 5.119740428 | 0.3133164 | 0.5832597 | 0.911369989 |
| ENSMUSG00000057329 | Bcl2 | protein_coding | 0.247255671 | 3.884976949 | 0.3132519 | 0.5832981 | 0.911369989 |
| ENSMUSG00000020674 | Pxdn | protein_coding | 0.490777826 | 2.519421216 | 0.3129834 | 0.5834705 | 0.911369989 |
| ENSMUSG00000032532 | Cck | protein_coding | -0.295638446 | 3.604041295 | 0.3129581 | 0.5834856 | 0.911369989 |
| ENSMUSG00000021301 | Hecw1 | protein_coding | -0.16076288 | 5.212150038 | 0.3129072 | 0.5835034 | 0.911369989 |
| ENSMUSG00000034472 | Rasd2 | protein_coding | 0.131682388 | 6.691949808 | 0.3128732 | 0.5835237 | 0.911369989 |
| ENSMUSG00000097908 | 4933404O12Rik | lincRNA | -0.283384658 | 3.217324992 | 0.312863 | 0.5835298 | 0.911369989 |
| ENSMUSG00000021252 | Erg28 | protein_coding | 0.114035863 | 7.380017523 | 0.312704 | 0.5836245 | 0.911369989 |
| ENSMUSG00000052906 | Ubxn8 | protein_coding | -0.132009359 | 6.051718057 | 0.3126589 | 0.5836514 | 0.911369989 |

| ensembl_gene_id | external_gene_name | gene_biotype | log2 (fold change) | log2 (counts per million) | F-value | p-value | False discovery rate |
|---|---|---|---|---|---|---|---|
| ENSMUSG00000029207 | Apbb2 | protein_coding | 0.118046503 | 6.354913136 | 0.3126211 | 0.5836739 | 0.911369989 |
| ENSMUSG00000026027 | Stradb | protein_coding | 0.184051549 | 4.655099218 | 0.3125928 | 0.5836908 | 0.911369989 |
| ENSMUSG00000045268 | Zfp691 | protein_coding | -0.395478698 | 2.370564015 | 0.3125625 | 0.5837088 | 0.911369989 |
| ENSMUSG00000014195 | Dnajc7 | protein_coding | 0.115924769 | 6.780053273 | 0.3124904 | 0.5837518 | 0.911369989 |
| ENSMUSG00000034587 | 8430429K09Rik | processed_transcript | -0.213377158 | 4.422276254 | 0.3124649 | 0.583767 | 0.911369989 |
| ENSMUSG00000021969 | Zdhhc20 | protein_coding | -0.12454471 | 6.100182519 | 0.3124229 | 0.5837921 | 0.911369989 |
| ENSMUSG00000024099 | Ndufv2 | protein_coding | 0.11179586 | 10.3424644 | 0.3120868 | 0.5839927 | 0.911550741 |
| ENSMUSG00000019849 | Prep | protein_coding | 0.191794327 | 3.773300172 | 0.311554 | 0.5843108 | 0.911812965 |
| ENSMUSG00000028465 | Tln1 | protein_coding | 0.153049956 | 5.298574854 | 0.3115215 | 0.5843302 | 0.911812965 |
| ENSMUSG00000040652 | Oaz2 | protein_coding | 0.120187909 | 6.202725126 | 0.3111007 | 0.5845818 | 0.912073122 |
| ENSMUSG00000029066 | Mrpl20 | protein_coding | 0.114488591 | 8.825360776 | 0.3106459 | 0.584854 | 0.912218566 |
| ENSMUSG00000023026 | Dip2b | protein_coding | -0.156961872 | 5.396342755 | 0.310423 | 0.5849874 | 0.912218566 |
| ENSMUSG00000019806 | Aig1 | protein_coding | 0.160031231 | 6.224464513 | 0.3104296 | 0.5849959 | 0.912218566 |
| ENSMUSG00000059540 | Tcea2 | protein_coding | 0.223117085 | 4.56843756 | 0.3103781 | 0.5850143 | 0.912218566 |
| ENSMUSG00000021686 | Ap3b1 | protein_coding | 0.178684256 | 4.754380161 | 0.3100422 | 0.5852156 | 0.912258261 |
| ENSMUSG00000062352 | Itgb1bp1 | protein_coding | -0.138830616 | 6.483289988 | 0.3099535 | 0.5852688 | 0.912258261 |
| ENSMUSG00000030486 | Zfp108 | protein_coding | -0.374488374 | 2.501826416 | 0.309911 | 0.5852943 | 0.912258261 |
| ENSMUSG00000022500 | Litaf | protein_coding | -0.114910238 | 6.764656062 | 0.3093829 | 0.5856111 | 0.912525398 |
| ENSMUSG00000019883 | Echdc1 | protein_coding | -0.225059934 | 3.896634565 | 0.3093413 | 0.5856361 | 0.912525398 |
| ENSMUSG00000034059 | Ypel4 | protein_coding | 0.119880577 | 6.631333858 | 0.3092011 | 0.5857203 | 0.912525398 |
| ENSMUSG00000034226 | Rhov | protein_coding | 0.380006289 | 2.758434946 | 0.3089987 | 0.5858419 | 0.912582572 |
| ENSMUSG00000048921 | Zfp689 | protein_coding | -0.233525466 | 3.831255787 | 0.3088073 | 0.5859569 | 0.91262954 |
| ENSMUSG00000021218 | Gdi2 | protein_coding | 0.122236407 | 8.659815693 | 0.3083041 | 0.5862595 | 0.912843308 |
| ENSMUSG00000021112 | Mpp5 | protein_coding | -0.126535785 | 5.843263303 | 0.3082882 | 0.586269 | 0.912843308 |
| ENSMUSG00000028917 | Plekhm2 | protein_coding | -0.269740395 | 3.915621451 | 0.3077558 | 0.5866019 | 0.912843308 |
| ENSMUSG00000066760 | Psg16 | protein_coding | 0.460149898 | 2.924307113 | 0.3081335 | 0.5866937 | 0.912843308 |
| ENSMUSG00000024253 | Dync2li1 | protein_coding | 0.128237137 | 5.827767068 | 0.3075191 | 0.5867322 | 0.912843308 |
| ENSMUSG00000005362 | Crbn | protein_coding | -0.147440258 | 6.155560112 | 0.3073649 | 0.5868252 | 0.912843308 |
| ENSMUSG00000071719 | Tmem28 | protein_coding | 0.212438021 | 3.903333358 | 0.3071695 | 0.586943 | 0.912843308 |
| ENSMUSG00000024781 | Lipa | protein_coding | -0.153940676 | 5.126524875 | 0.3071082 | 0.58698 | 0.912843308 |
| ENSMUSG00000044991 | 1110034G24Rik | protein_coding | -0.161977521 | 5.267753978 | 0.3070565 | 0.5870112 | 0.912843308 |
| ENSMUSG00000042305 | Tmem183a | protein_coding | 0.150851164 | 5.834486723 | 0.3069039 | 0.5871033 | 0.912843308 |
| ENSMUSG00000026848 | Tor1b | protein_coding | 0.191140723 | 4.511419025 | 0.3068378 | 0.5871431 | 0.912843308 |
| ENSMUSG00000037656 | Slc20a2 | protein_coding | 0.188044373 | 4.918344931 | 0.3065871 | 0.5872945 | 0.912843308 |
| ENSMUSG00000091896 | Ube2d2a | protein_coding | -0.135099143 | 5.745542378 | 0.3064657 | 0.5873678 | 0.912843308 |
| ENSMUSG00000026229 | Psmd1 | protein_coding | -0.120024626 | 7.239032462 | 0.3063882 | 0.5874147 | 0.912843308 |
| ENSMUSG00000004393 | Ddx56 | protein_coding | 0.133435282 | 7.161439461 | 0.3061408 | 0.5875642 | 0.912843308 |
| ENSMUSG00000050821 | Fam131a | protein_coding | -0.102953159 | 6.95714607 | 0.3060633 | 0.587611 | 0.912843308 |
| ENSMUSG00000005469 | Prkaca | protein_coding | 0.12527286 | 7.918285294 | 0.305873 | 0.5877261 | 0.912843308 |
| ENSMUSG00000003575 | Crtc1 | protein_coding | -0.124992696 | 5.899694798 | 0.305772 | 0.5877872 | 0.912843308 |
| ENSMUSG00000029918 | Mrps33 | protein_coding | -0.134062961 | 6.964220476 | 0.3057109 | 0.5878242 | 0.912843308 |
| ENSMUSG00000061111 | Mcrip1 | protein_coding | -0.145763851 | 7.096390871 | 0.305577 | 0.5879175 | 0.912843308 |
| ENSMUSG00000024812 | Tjp2 | protein_coding | 0.227331236 | 3.754612464 | 0.3053544 | 0.58804 | 0.912843308 |
| ENSMUSG00000028035 | Dnajb4 | protein_coding | -0.124265116 | 6.584486295 | 0.3052999 | 0.5880729 | 0.912843308 |
| ENSMUSG00000028174 | Rpe65 | protein_coding | -0.680491237 | 2.05384736 | 0.3056042 | 0.588138 | 0.912843308 |
| ENSMUSG00000027298 | Tyro3 | protein_coding | -0.160472546 | 5.381614131 | 0.3050669 | 0.5882141 | 0.912843308 |
| ENSMUSG00000019998 | Stx7 | protein_coding | -0.11068621 | 7.730290443 | 0.3050631 | 0.5882164 | 0.912843308 |
| ENSMUSG00000055254 | Ntrk2 | protein_coding | 0.119225834 | 7.519889772 | 0.3048838 | 0.588325 | 0.912880113 |
| ENSMUSG00000032479 | Map4 | protein_coding | 0.134246268 | 8.692960752 | 0.3043524 | 0.5886474 | 0.912994513 |
| ENSMUSG00000029438 | Bcl7a | protein_coding | 0.17249854 | 5.314470997 | 0.3042762 | 0.5886936 | 0.912994513 |
| ENSMUSG00000026159 | Agfg1 | protein_coding | -0.147533182 | 5.152506144 | 0.3039787 | 0.5888742 | 0.912994513 |
| ENSMUSG00000025899 | Alkbh8 | protein_coding | -0.141318678 | 5.523790829 | 0.3039312 | 0.588903 | 0.912994513 |
| ENSMUSG00000045205 | Dpy19l4 | protein_coding | 0.205259549 | 3.949632797 | 0.3036355 | 0.5890827 | 0.912994513 |
| ENSMUSG00000026096 | Osgepl1 | protein_coding | 0.183032417 | 4.626322671 | 0.3035524 | 0.5891332 | 0.912994513 |
| ENSMUSG00000019362 | D8Ertd738e | protein_coding | 0.09953336 | 8.80868539 | 0.3035156 | 0.5891556 | 0.912994513 |
| ENSMUSG00000029461 | Fam168a | protein_coding | 0.133532413 | 7.503392389 | 0.303372 | 0.5892429 | 0.912994513 |
| ENSMUSG00000031010 | Usp9x | protein_coding | 0.182111987 | 4.380907038 | 0.3033712 | 0.5892434 | 0.912994513 |
| ENSMUSG00000053580 | Tanc2 | protein_coding | 0.137313994 | 5.970584882 | 0.3033392 | 0.5892629 | 0.912994513 |
| ENSMUSG00000046818 | Ddit4l | protein_coding | -0.192766812 | 4.730396204 | 0.3025702 | 0.589731 | 0.912994513 |
| ENSMUSG00000050840 | Cdh20 | protein_coding | 0.404843585 | 2.865321039 | 0.3025485 | 0.5897564 | 0.912994513 |
| ENSMUSG00000055629 | B4galnt4 | protein_coding | 0.146165023 | 5.690109937 | 0.3025028 | 0.589772 | 0.912994513 |
| ENSMUSG00000028439 | Fam219a | protein_coding | 0.323737155 | 2.820741868 | 0.3024793 | 0.5897863 | 0.912994513 |
| ENSMUSG00000037465 | Klf10 | protein_coding | 0.206142817 | 3.896625607 | 0.3024707 | 0.5897916 | 0.912994513 |
| ENSMUSG00000009073 | Nf2 | protein_coding | 0.167777918 | 4.799165403 | 0.3023661 | 0.5898553 | 0.912994513 |
| ENSMUSG00000059820 | AU019823 | protein_coding | -0.253279892 | 3.350332117 | 0.3022584 | 0.589921 | 0.912994513 |
| ENSMUSG00000025078 | Nhlrc2 | protein_coding | -0.315956192 | 2.748052881 | 0.3020083 | 0.5900734 | 0.912994513 |
| ENSMUSG00000024194 | Cuta | protein_coding | -0.09444788 | 7.473278638 | 0.3019998 | 0.5900786 | 0.912994513 |
| ENSMUSG00000024969 | Mark2 | protein_coding | -0.136756184 | 5.771430586 | 0.3019699 | 0.5900969 | 0.912994513 |
| ENSMUSG00000023938 | Aars2 | protein_coding | 0.373870913 | 2.543370175 | 0.3014093 | 0.5904391 | 0.913295291 |
| ENSMUSG00000054850 | Smim10l2a | protein_coding | 0.178300425 | 4.392290456 | 0.3011292 | 0.5906102 | 0.913295291 |
| ENSMUSG00000032959 | Pebp1 | protein_coding | 0.097319712 | 7.635538589 | 0.3010813 | 0.5906395 | 0.913295291 |
| ENSMUSG00000004846 | Plod3 | protein_coding | -0.258448735 | 3.690383343 | 0.3010742 | 0.5906438 | 0.913295291 |
| ENSMUSG00000041429 | Nthl1 | protein_coding | -0.277606856 | 3.525973992 | 0.3009561 | 0.590716 | 0.913295291 |
| ENSMUSG00000027221 | Chst1 | protein_coding | -0.119284241 | 6.988503255 | 0.3004581 | 0.5910206 | 0.913621322 |

| ensembl_gene_id | external_gene_name | gene_biotype | log2 (fold change) | log2 (counts per million) | F-value | p-value | False discovery rate |
|---|---|---|---|---|---|---|---|
| ENSMUSG00000020515 | Cnot8 | protein_coding | 0.131638454 | 6.864369852 | 0.3003336 | 0.5910968 | 0.913621322 |
| ENSMUSG00000034724 | Cnot6l | protein_coding | -0.156206074 | 5.221231872 | 0.2995547 | 0.591574 | 0.91388111 |
| ENSMUSG00000051989 | Smim11 | protein_coding | -0.172615676 | 5.259884456 | 0.2993511 | 0.5916989 | 0.91388111 |
| ENSMUSG00000002728 | Naa20 | protein_coding | 0.105160244 | 7.108867017 | 0.2991179 | 0.591842 | 0.91388111 |
| ENSMUSG00000021217 | Tshz3 | protein_coding | -0.265737807 | 2.952394091 | 0.2989112 | 0.5919689 | 0.91388111 |
| ENSMUSG00000038569 | Rad9b | protein_coding | 0.480427863 | 2.652886334 | 0.2989131 | 0.5919799 | 0.91388111 |
| ENSMUSG00000002233 | Rhoc | protein_coding | -0.232662723 | 4.492495405 | 0.2987432 | 0.5920842 | 0.91388111 |
| ENSMUSG00000083674 | Zfp133-ps | transcribed_unitary_p | -0.305255113 | 3.335927274 | 0.2985944 | 0.5921635 | 0.91388111 |
| ENSMUSG00000041801 | Phlda3 | protein_coding | 0.149286821 | 5.543786398 | 0.2985852 | 0.5921691 | 0.91388111 |
| ENSMUSG00000027680 | Fxr1 | protein_coding | -0.175484756 | 5.052254019 | 0.2985703 | 0.5921783 | 0.91388111 |
| ENSMUSG00000048878 | Hexim1 | protein_coding | -0.170023938 | 4.585483857 | 0.2983403 | 0.5923197 | 0.91388111 |
| ENSMUSG00000020844 | Nxn | protein_coding | -0.219137314 | 3.816429731 | 0.2982695 | 0.5923632 | 0.91388111 |
| ENSMUSG00000041426 | Hibch | protein_coding | -0.156589439 | 5.204312644 | 0.2982606 | 0.5923687 | 0.91388111 |
| ENSMUSG00000026857 | Ntmt1 | protein_coding | 0.106085855 | 7.068004118 | 0.2981283 | 0.5924501 | 0.91388111 |
| ENSMUSG00000038102 | Trappc11 | protein_coding | 0.160729485 | 4.883764086 | 0.2981208 | 0.5924547 | 0.91388111 |
| ENSMUSG00000000384 | Tbrg4 | protein_coding | -0.190043528 | 4.916363317 | 0.2976951 | 0.5927167 | 0.913890095 |
| ENSMUSG00000029359 | Tesc | protein_coding | -0.213170817 | 4.607347057 | 0.2976686 | 0.592733 | 0.913890095 |
| ENSMUSG00000025234 | Arih1 | protein_coding | -0.169336021 | 5.035524536 | 0.2975772 | 0.5927892 | 0.913890095 |
| ENSMUSG00000024231 | Cul2 | protein_coding | 0.160120861 | 5.685802949 | 0.2975299 | 0.5928184 | 0.913890095 |
| ENSMUSG00000001707 | Eef1e1 | protein_coding | -0.11367032 | 7.812687643 | 0.2973743 | 0.5929143 | 0.913890095 |
| ENSMUSG00000056201 | Cfl1 | protein_coding | -0.102963487 | 10.34545569 | 0.2972685 | 0.5929794 | 0.913890095 |
| ENSMUSG00000041560 | Nop53 | protein_coding | -0.166128625 | 6.640298729 | 0.2976253 | 0.5930687 | 0.913890095 |
| ENSMUSG00000004233 | Wars2 | protein_coding | 0.293485534 | 3.088499615 | 0.2970074 | 0.5931404 | 0.913890095 |
| ENSMUSG00000024712 | Rfk | protein_coding | -0.119327738 | 6.438470408 | 0.2965197 | 0.5934413 | 0.914203364 |
| ENSMUSG00000038172 | Ttc39b | protein_coding | -0.184298904 | 4.595200301 | 0.2964023 | 0.5935138 | 0.914203364 |
| ENSMUSG00000024269 | Tpgs2 | protein_coding | 0.106741045 | 6.612250471 | 0.2961108 | 0.5936938 | 0.914349734 |
| ENSMUSG00000037287 | Tbcel | protein_coding | 0.174704596 | 4.496771683 | 0.2956497 | 0.5939789 | 0.914440429 |
| ENSMUSG00000031482 | Slc25a15 | protein_coding | 0.252427473 | 3.020939249 | 0.2956241 | 0.5939947 | 0.914440429 |
| ENSMUSG00000049672 | Zbtb14 | protein_coding | 0.246441684 | 3.195792007 | 0.2956029 | 0.5940078 | 0.914440429 |
| ENSMUSG00000070643 | Sox13 | protein_coding | 0.406179837 | 2.646483976 | 0.2952973 | 0.5941969 | 0.914598303 |
| ENSMUSG00000052395 | Rft1 | protein_coding | 0.17550811 | 4.127595458 | 0.295148 | 0.5942894 | 0.914598303 |
| ENSMUSG00000071265 | 1700086L19Rik | transcribed_unitary_p | -0.131912736 | 5.636011337 | 0.2950249 | 0.5943656 | 0.914598303 |
| ENSMUSG00000073633 | Fbxo36 | protein_coding | -0.185060467 | 4.715544166 | 0.2945524 | 0.5946584 | 0.914886185 |
| ENSMUSG00000020630 | Rnaseh1 | protein_coding | -0.193257767 | 4.392954136 | 0.2944485 | 0.5947228 | 0.914886185 |
| ENSMUSG00000036955 | Kif1bp | protein_coding | -0.127565363 | 6.236492011 | 0.2941642 | 0.5948992 | 0.915026597 |
| ENSMUSG00000045404 | Kcnk13 | protein_coding | -0.31550691 | 3.372932918 | 0.2935912 | 0.595267 | 0.915175896 |
| ENSMUSG00000032702 | Kank1 | protein_coding | 0.219990185 | 3.697602007 | 0.2935296 | 0.5952933 | 0.915175896 |
| ENSMUSG00000020090 | Npffr1 | protein_coding | -0.279428815 | 3.041290467 | 0.293439 | 0.5953496 | 0.915175896 |
| ENSMUSG00000068742 | Cry2 | protein_coding | -0.137279456 | 5.848385492 | 0.2933405 | 0.5954108 | 0.915175896 |
| ENSMUSG00000030270 | Cpne9 | protein_coding | -0.37584707 | 5.478852177 | 0.2991376 | 0.5954218 | 0.915175896 |
| ENSMUSG00000036002 | Fam214b | protein_coding | -0.184252369 | 4.200672327 | 0.2929476 | 0.5956551 | 0.915226788 |
| ENSMUSG00000042506 | Usp22 | protein_coding | -0.142998211 | 8.288347525 | 0.292892 | 0.5957017 | 0.915226788 |
| ENSMUSG00000029165 | Agbl5 | protein_coding | -0.287024689 | 2.977751785 | 0.2926881 | 0.5958166 | 0.915226788 |
| ENSMUSG00000007837 | Prrg2 | protein_coding | -0.245410845 | 3.673805446 | 0.2925994 | 0.5958719 | 0.915226788 |
| ENSMUSG00000024500 | Ppp2r2b | protein_coding | -0.098398685 | 7.88228679 | 0.2925295 | 0.5959154 | 0.915226788 |
| ENSMUSG00000086191 | Zfp652os | antisense_RNA | -0.279035414 | 2.934209577 | 0.292449 | 0.5959656 | 0.915226788 |
| ENSMUSG00000015790 | Surf1 | protein_coding | -0.10495529 | 8.21324202 | 0.2921894 | 0.5961273 | 0.915344473 |
| ENSMUSG00000022468 | Endou | protein_coding | -0.573128531 | 3.049065569 | 0.2943466 | 0.5962353 | 0.91537952 |
| ENSMUSG00000020843 | Timm22 | protein_coding | 0.122111985 | 7.608335653 | 0.2916166 | 0.5964846 | 0.915631466 |
| ENSMUSG00000025138 | Sirt7 | protein_coding | 0.300846577 | 3.093681982 | 0.290963 | 0.5968927 | 0.916112934 |
| ENSMUSG00000097820 | E530011L22Rik | antisense_RNA | -0.372269835 | 2.661232199 | 0.2904073 | 0.5972403 | 0.916112934 |
| ENSMUSG00000057411 | Fam173a | protein_coding | 0.109364338 | 8.093885779 | 0.2903914 | 0.5972502 | 0.916112934 |
| ENSMUSG00000024387 | Csnk2b | protein_coding | -0.093471079 | 8.273179454 | 0.2903237 | 0.5972926 | 0.916112934 |
| ENSMUSG00000112343 | AC154734.1 | processed_transcript | -0.456528799 | 2.490998379 | 0.2902111 | 0.5973631 | 0.916112934 |
| ENSMUSG00000055044 | Pdlim1 | protein_coding | 0.352681417 | 2.578696262 | 0.2900159 | 0.5974853 | 0.916112934 |
| ENSMUSG00000034135 | Sik3 | protein_coding | 0.105890375 | 6.39715125 | 0.2898336 | 0.5975996 | 0.916112934 |
| ENSMUSG00000070733 | Fryl | protein_coding | 0.198393976 | 3.96075965 | 0.2897411 | 0.5976575 | 0.916112934 |
| ENSMUSG00000040785 | Ttc3 | protein_coding | 0.106953989 | 9.858794681 | 0.2895876 | 0.5977537 | 0.916112934 |
| ENSMUSG00000020775 | Mrpl38 | protein_coding | 0.166787163 | 6.000364874 | 0.2894919 | 0.5978256 | 0.916112934 |
| ENSMUSG00000023032 | Slc4a8 | protein_coding | 0.165712929 | 4.621661121 | 0.2894257 | 0.5978552 | 0.916112934 |
| ENSMUSG00000034800 | Zfp661 | protein_coding | 0.368367221 | 2.819067178 | 0.2891705 | 0.5980153 | 0.916112934 |
| ENSMUSG00000019710 | Mrpl24 | protein_coding | 0.105667532 | 6.704732725 | 0.2891043 | 0.5980568 | 0.916112934 |
| ENSMUSG00000022436 | Sh3bp1 | protein_coding | 0.107260977 | 7.128496643 | 0.2889312 | 0.5981655 | 0.916112934 |
| ENSMUSG00000036940 | Kdm1a | protein_coding | 0.241138877 | 3.272368553 | 0.2888935 | 0.5981892 | 0.916112934 |
| ENSMUSG00000027884 | Clcc1 | protein_coding | -0.16089075 | 4.867255731 | 0.2887452 | 0.5982823 | 0.916112934 |
| ENSMUSG00000078958 | Atp6ap1l | protein_coding | -0.245656291 | 3.491809677 | 0.2887439 | 0.5982831 | 0.916112934 |
| ENSMUSG00000029790 | Cep41 | protein_coding | -0.306858551 | 3.099118787 | 0.2886665 | 0.5983317 | 0.916112934 |
| ENSMUSG00000037280 | Galnt6 | protein_coding | 0.217096668 | 3.473773619 | 0.288404 | 0.5984967 | 0.916132278 |
| ENSMUSG00000030706 | Mrpl48 | protein_coding | 0.101618186 | 7.445085293 | 0.2883753 | 0.5985148 | 0.916132278 |
| ENSMUSG00000031714 | Gab1 | protein_coding | 0.175161729 | 4.408208257 | 0.2877405 | 0.5989141 | 0.916364219 |
| ENSMUSG00000034154 | Ino80 | protein_coding | 0.233694991 | 3.498932042 | 0.2876377 | 0.5989788 | 0.916364219 |
| ENSMUSG00000028133 | Rwdd3 | protein_coding | 0.157709434 | 4.843181953 | 0.2875343 | 0.5990439 | 0.916364219 |
| ENSMUSG00000003585 | Sec14l2 | protein_coding | 0.18873464 | 3.81467034 | 0.287461 | 0.5990901 | 0.916364219 |
| ENSMUSG00000034509 | Mad2l1bp | protein_coding | 0.14476687 | 5.073675527 | 0.2873906 | 0.5991344 | 0.916364219 |

| ensembl_gene_id | external_gene_name | gene_biotype | log2 (fold change) | log2 (counts per million) | F-value | p-value | False discovery rate |
|---|---|---|---|---|---|---|---|
| ENSMUSG00000042073 | Abhd14b | protein_coding | -0.233047127 | 3.539141639 | 0.2871744 | 0.5992707 | 0.916364219 |
| ENSMUSG00000019254 | Ppp1r12c | protein_coding | -0.173474027 | 4.829504984 | 0.2868645 | 0.5994661 | 0.916364219 |
| ENSMUSG00000039483 | Asb6 | protein_coding | 0.136945882 | 6.025324549 | 0.2868017 | 0.5995057 | 0.916364219 |
| ENSMUSG00000034109 | Golim4 | protein_coding | -0.446817706 | 2.425925254 | 0.2867547 | 0.5995354 | 0.916364219 |
| ENSMUSG00000038400 | Pmepa1 | protein_coding | -0.175605055 | 4.561801788 | 0.2865215 | 0.5996826 | 0.916364219 |
| ENSMUSG00000045034 | Ankrd34b | protein_coding | 0.223038751 | 3.798458628 | 0.2862316 | 0.5998656 | 0.916364219 |
| ENSMUSG00000037361 | Sf3b6 | protein_coding | -0.114915944 | 8.375932533 | 0.2861483 | 0.5999183 | 0.916364219 |
| ENSMUSG00000064210 | Ano6 | protein_coding | 0.234723043 | 3.670365067 | 0.286056 | 0.5999766 | 0.916364219 |
| ENSMUSG00000031594 | Fgl1 | protein_coding | -0.367712746 | 2.721858819 | 0.2858421 | 0.6001118 | 0.916364219 |
| ENSMUSG00000032285 | Dnaja4 | protein_coding | 0.309794252 | 2.787858811 | 0.2857954 | 0.6001413 | 0.916364219 |
| ENSMUSG00000028101 | Pias3 | protein_coding | 0.216637926 | 5.432729769 | 0.286306 | 0.6002523 | 0.916364219 |
| ENSMUSG00000039908 | Slc26a11 | protein_coding | 0.268474788 | 3.558136159 | 0.2855985 | 0.6002659 | 0.916364219 |
| ENSMUSG00000034210 | Efcab14 | protein_coding | -0.121229288 | 5.928916924 | 0.2855571 | 0.6002921 | 0.916364219 |
| ENSMUSG00000045573 | Penk | protein_coding | 0.172000812 | 10.40961684 | 0.2861573 | 0.6003716 | 0.916364219 |
| ENSMUSG00000047115 | Fam221a | protein_coding | -0.434909028 | 2.264758872 | 0.2853989 | 0.6003921 | 0.916364219 |
| ENSMUSG00000045038 | Prkce | protein_coding | -0.107686698 | 7.599534377 | 0.2852982 | 0.6004559 | 0.916364219 |
| ENSMUSG00000021013 | Ttc8 | protein_coding | -0.140469366 | 5.003376252 | 0.2851222 | 0.6005673 | 0.916383393 |
| ENSMUSG00000073423 | Zfp414 | protein_coding | -0.125694888 | 6.22489882 | 0.2850092 | 0.6006389 | 0.916383393 |
| ENSMUSG00000036099 | Vezt | protein_coding | 0.174733958 | 5.092441276 | 0.2845336 | 0.6009403 | 0.916493565 |
| ENSMUSG00000018921 | Pelp1 | protein_coding | -0.200781376 | 3.524670135 | 0.2844449 | 0.6009966 | 0.916493565 |
| ENSMUSG00000031139 | Mcf2 | protein_coding | -0.288963589 | 3.053798652 | 0.2842094 | 0.601146 | 0.916493565 |
| ENSMUSG00000002325 | Irf9 | protein_coding | 0.242361455 | 3.572885205 | 0.2841474 | 0.6011853 | 0.916493565 |
| ENSMUSG00000098097 | 6530403H02Rik | lincRNA | -0.314978352 | 2.770981704 | 0.2841446 | 0.6011871 | 0.916493565 |
| ENSMUSG00000030609 | Aen | protein_coding | 0.220721121 | 4.770893489 | 0.2839547 | 0.6013194 | 0.916493565 |
| ENSMUSG00000029649 | Pomp | protein_coding | 0.108534146 | 9.62776888 | 0.2836773 | 0.6014838 | 0.916493565 |
| ENSMUSG00000041147 | Brca2 | protein_coding | -0.263867922 | 3.318215813 | 0.2836357 | 0.6015103 | 0.916493565 |
| ENSMUSG00000023033 | Scn8a | protein_coding | -0.143204169 | 5.789346112 | 0.2835632 | 0.6015563 | 0.916493565 |
| ENSMUSG00000017776 | Crk | protein_coding | -0.13540642 | 6.282211737 | 0.2835521 | 0.6015634 | 0.916493565 |
| ENSMUSG00000096956 | Snhg18 | lincRNA | -0.356656936 | 2.863060401 | 0.2827793 | 0.602055 | 0.917112577 |
| ENSMUSG00000054733 | Msra | protein_coding | -0.13748062 | 5.175527977 | 0.2824289 | 0.6022782 | 0.917219373 |
| ENSMUSG00000041966 | Dcaf17 | protein_coding | -0.203693411 | 3.947191032 | 0.2824014 | 0.6022957 | 0.917219373 |
| ENSMUSG00000047187 | Rab2a | protein_coding | -0.094873066 | 8.097974959 | 0.28216 | 0.6024496 | 0.91727066 |
| ENSMUSG00000061360 | Phf5a | protein_coding | -0.126333145 | 6.590204573 | 0.2818522 | 0.6026459 | 0.91727066 |
| ENSMUSG00000033318 | Gstt2 | protein_coding | -0.229620595 | 3.994900866 | 0.2818247 | 0.6026634 | 0.91727066 |
| ENSMUSG00000050910 | Cdr2l | protein_coding | -0.178542164 | 5.033813653 | 0.281676 | 0.6027583 | 0.91727066 |
| ENSMUSG00000002006 | Pdzd4 | protein_coding | 0.202690661 | 4.355802485 | 0.2813577 | 0.6029616 | 0.91727066 |
| ENSMUSG00000032842 | Abcc10 | protein_coding | 0.396028467 | 2.920291954 | 0.2813529 | 0.6029764 | 0.91727066 |
| ENSMUSG00000050552 | Lamtor4 | protein_coding | -0.117091548 | 7.544119523 | 0.281302 | 0.6029972 | 0.91727066 |
| ENSMUSG00000032396 | Dis3l | protein_coding | 0.174159696 | 4.497877359 | 0.2812415 | 0.6030358 | 0.91727066 |
| ENSMUSG00000030417 | Pdcd5 | protein_coding | -0.207190697 | 4.30447361 | 0.2808981 | 0.6032553 | 0.91727066 |
| ENSMUSG00000020483 | Dynll2 | protein_coding | -0.151542639 | 8.058738353 | 0.2813141 | 0.6032886 | 0.91727066 |
| ENSMUSG00000024610 | Cd74 | protein_coding | -0.222677674 | 4.200338899 | 0.2805283 | 0.6034918 | 0.91727066 |
| ENSMUSG00000038644 | Pold1 | protein_coding | -0.388090042 | 2.400839149 | 0.2804277 | 0.6035563 | 0.91727066 |
| ENSMUSG00000036893 | Ehmt1 | protein_coding | -0.141140138 | 5.202402411 | 0.280411 | 0.6035669 | 0.91727066 |
| ENSMUSG00000029128 | Rab28 | protein_coding | -0.127873287 | 6.833190454 | 0.2801521 | 0.6037328 | 0.91727066 |
| ENSMUSG00000031730 | Dhodh | protein_coding | 0.124847263 | 6.121563963 | 0.2796923 | 0.6040274 | 0.91727066 |
| ENSMUSG00000074785 | Plxnc1 | protein_coding | 0.145422403 | 5.013057301 | 0.2796508 | 0.6040541 | 0.91727066 |
| ENSMUSG00000068706 | Gm10250 | protein_coding | -0.224366708 | 3.267852304 | 0.2796069 | 0.6040822 | 0.91727066 |
| ENSMUSG00000055128 | Cgrrf1 | protein_coding | 0.123110156 | 6.209589568 | 0.2795586 | 0.6041132 | 0.91727066 |
| ENSMUSG00000041235 | Chd7 | protein_coding | -0.212082059 | 3.645943808 | 0.2792241 | 0.6043278 | 0.91727066 |
| ENSMUSG00000003421 | Nosip | protein_coding | -0.099826755 | 7.377059727 | 0.2792055 | 0.6043397 | 0.91727066 |
| ENSMUSG00000066687 | Zbtb16 | protein_coding | -0.340603312 | 2.852946376 | 0.2791931 | 0.6043477 | 0.91727066 |
| ENSMUSG00000056679 | Gpr173 | protein_coding | 0.315681511 | 2.789059851 | 0.2789797 | 0.6044847 | 0.91727066 |
| ENSMUSG00000027495 | Fam210b | protein_coding | 0.165874014 | 4.945300352 | 0.2787174 | 0.6046533 | 0.91727066 |
| ENSMUSG00000036615 | Rfxap | protein_coding | -0.177537362 | 4.236576245 | 0.2785646 | 0.6047514 | 0.91727066 |
| ENSMUSG00000025035 | Arl3 | protein_coding | 0.10379555 | 9.134159162 | 0.2785644 | 0.6047516 | 0.91727066 |
| ENSMUSG00000023905 | Tnfrsf12a | protein_coding | -0.212872335 | 4.519644428 | 0.278492 | 0.6047981 | 0.91727066 |
| ENSMUSG00000034160 | Ogt | protein_coding | -0.158257108 | 5.72437829 | 0.2784843 | 0.6048031 | 0.91727066 |
| ENSMUSG00000003680 | Taf6l | protein_coding | -0.22033351 | 4.353004743 | 0.2783485 | 0.6048904 | 0.91727066 |
| ENSMUSG00000034111 | Tmed8 | protein_coding | 0.17242863 | 4.609132762 | 0.278269 | 0.6049416 | 0.91727066 |
| ENSMUSG00000023027 | Atf1 | protein_coding | -0.358312519 | 2.517143718 | 0.2781524 | 0.6050166 | 0.91727066 |
| ENSMUSG00000062270 | Morf4l1 | protein_coding | 0.108014717 | 6.593492639 | 0.2781127 | 0.6050421 | 0.91727066 |
| ENSMUSG00000059796 | Eif4a1 | protein_coding | 0.106353985 | 6.153011089 | 0.2780863 | 0.6050591 | 0.91727066 |
| ENSMUSG00000018565 | Elp5 | protein_coding | 0.097471875 | 8.18573126 | 0.277595 | 0.6053755 | 0.917605843 |
| ENSMUSG00000015214 | Mtmr1 | protein_coding | -0.182899053 | 4.185210537 | 0.277478 | 0.6054509 | 0.917605843 |
| ENSMUSG00000005150 | Wdr83 | protein_coding | 0.126663124 | 6.399766298 | 0.277205 | 0.6056269 | 0.917639745 |
| ENSMUSG00000060206 | Zfp462 | protein_coding | -0.185631441 | 4.066285577 | 0.2771786 | 0.6056439 | 0.917639745 |
| ENSMUSG00000027944 | Hax1 | protein_coding | 0.125843506 | 5.327219095 | 0.2769965 | 0.6057614 | 0.91768846 |
| ENSMUSG00000024937 | Ehbp1l1 | protein_coding | 0.133962093 | 5.694630554 | 0.2764673 | 0.6061031 | 0.91787125 |
| ENSMUSG00000049760 | 2410015M20Rik | protein_coding | 0.111186054 | 8.724607971 | 0.2764503 | 0.6061141 | 0.91787125 |
| ENSMUSG00000024101 | Washc1 | protein_coding | 0.144960234 | 5.043493892 | 0.2761917 | 0.6062812 | 0.91787125 |
| ENSMUSG00000030987 | Stim1 | protein_coding | 0.126291884 | 5.982933369 | 0.2761105 | 0.6063337 | 0.91787125 |
| ENSMUSG00000051950 | B3glct | protein_coding | 0.289701713 | 3.011145977 | 0.2761042 | 0.6063378 | 0.91787125 |
| ENSMUSG00000057551 | Zfp317 | protein_coding | -0.185640947 | 4.910413976 | 0.2759594 | 0.6064314 | 0.91787125 |

| ensembl_gene_id | external_gene_name | gene_biotype | log2 (fold change) | log2 (counts per million) | F-value | p-value | False discovery rate |
|---|---|---|---|---|---|---|---|
| ENSMUSG00000030677 | Kif22 | protein_coding | -0.212109214 | 3.465537622 | 0.2758849 | 0.6064796 | 0.91787125 |
| ENSMUSG00000018846 | Pank3 | protein_coding | -0.155424288 | 4.737048058 | 0.2753819 | 0.6068052 | 0.918187698 |
| ENSMUSG00000051736 | Fam229b | protein_coding | -0.135208107 | 5.959750319 | 0.275298 | 0.6068595 | 0.918187698 |
| ENSMUSG00000063410 | Stk24 | protein_coding | -0.180452202 | 3.879405367 | 0.275128 | 0.6069697 | 0.91822524 |
| ENSMUSG00000068099 | 1500009C09Rik | protein_coding | 0.11931835 | 5.961928734 | 0.2748637 | 0.607141 | 0.918266414 |
| ENSMUSG00000000884 | Gnb1l | protein_coding | 0.203899305 | 5.353469977 | 0.2750037 | 0.6071892 | 0.918266414 |
| ENSMUSG00000019951 | Uhrf1bp1l | protein_coding | -0.134235564 | 5.283490123 | 0.2746001 | 0.607312 | 0.918266414 |
| ENSMUSG00000078812 | Eif5a | protein_coding | 0.109939032 | 9.469711168 | 0.2745029 | 0.607375 | 0.918266414 |
| ENSMUSG00000038365 | Fbxo25 | protein_coding | 0.127541314 | 6.579606532 | 0.2744277 | 0.6074239 | 0.918266414 |
| ENSMUSG00000047084 | Ngrn | protein_coding | 0.125985255 | 6.385211894 | 0.2738702 | 0.607786 | 0.918569663 |
| ENSMUSG00000042873 | Lhfpl4 | protein_coding | 0.113831244 | 6.044644546 | 0.2737295 | 0.6078775 | 0.918569663 |
| ENSMUSG00000000420 | Galnt1 | protein_coding | 0.179970901 | 4.118701884 | 0.273629 | 0.6079428 | 0.918569663 |
| ENSMUSG00000030994 | D7Ertd443e | protein_coding | 0.190193485 | 4.277472516 | 0.2735931 | 0.6079662 | 0.918569663 |
| ENSMUSG00000042167 | Papd4 | protein_coding | -0.24371884 | 3.182694362 | 0.2734382 | 0.6080669 | 0.918592785 |
| ENSMUSG00000036214 | Znrd1as | protein_coding | -0.221490931 | 3.733411621 | 0.2732565 | 0.6081851 | 0.918593712 |
| ENSMUSG00000057561 | Eif1a | protein_coding | -0.108727694 | 7.456750366 | 0.2731747 | 0.6082384 | 0.918593712 |
| ENSMUSG00000041658 | Rragb | protein_coding | 0.382251257 | 5.215019542 | 0.2777534 | 0.6087157 | 0.918863136 |
| ENSMUSG00000020692 | Nle1 | protein_coding | -0.231544731 | 4.406804354 | 0.2724078 | 0.6087497 | 0.918863136 |
| ENSMUSG00000013089 | Etv5 | protein_coding | -0.180129475 | 4.202614693 | 0.2722755 | 0.6088244 | 0.918863136 |
| ENSMUSG00000031862 | Atp13a1 | protein_coding | 0.26157373 | 3.360370246 | 0.2722689 | 0.6088287 | 0.918863136 |
| ENSMUSG00000032293 | Ireb2 | protein_coding | -0.169000114 | 4.662883172 | 0.2722455 | 0.608844 | 0.918863136 |
| ENSMUSG00000085071 | Gm14066 | processed_transcript | -0.31288972 | 3.106832346 | 0.2719872 | 0.6090126 | 0.918988547 |
| ENSMUSG00000058975 | Kcnc1 | protein_coding | -0.143975557 | 5.32414464 | 0.2712126 | 0.6095187 | 0.919059219 |
| ENSMUSG00000046230 | Vps13a | protein_coding | -0.203977799 | 3.501451692 | 0.271046 | 0.6096277 | 0.919059219 |
| ENSMUSG00000026620 | Mark1 | protein_coding | 0.169803766 | 4.913049454 | 0.2709432 | 0.609695 | 0.919059219 |
| ENSMUSG00000027848 | Olfml3 | protein_coding | -0.18817458 | 4.462495399 | 0.2708535 | 0.6097537 | 0.919059219 |
| ENSMUSG00000026821 | Ralgds | protein_coding | -0.146097512 | 5.58292015 | 0.2706914 | 0.6098599 | 0.919059219 |
| ENSMUSG00000026634 | Angel2 | protein_coding | -0.129715811 | 5.422101788 | 0.2705873 | 0.609928 | 0.919059219 |
| ENSMUSG00000037190 | Cyb561d2 | protein_coding | 0.119817062 | 6.107873697 | 0.2704713 | 0.610004 | 0.919059219 |
| ENSMUSG00000005034 | Prkacb | protein_coding | 0.112924796 | 8.785418905 | 0.2704626 | 0.6100097 | 0.919059219 |
| ENSMUSG00000039473 | Ubn1 | protein_coding | -0.182294236 | 4.132008918 | 0.2703985 | 0.6100517 | 0.919059219 |
| ENSMUSG00000040356 | Skiv2l | protein_coding | -0.157988555 | 4.674149618 | 0.2703176 | 0.6101047 | 0.919059219 |
| ENSMUSG00000051586 | Mical3 | protein_coding | 0.218324274 | 4.129778632 | 0.2703033 | 0.6101141 | 0.919059219 |
| ENSMUSG00000090841 | Myl6 | protein_coding | 0.102567576 | 7.729844937 | 0.2702404 | 0.6101554 | 0.919059219 |
| ENSMUSG00000024462 | Gabbr1 | protein_coding | 0.118405384 | 7.898759309 | 0.2700292 | 0.6102938 | 0.919059219 |
| ENSMUSG00000001280 | Sp1 | protein_coding | 0.204479354 | 3.948194956 | 0.2700037 | 0.6103106 | 0.919059219 |
| ENSMUSG00000039114 | Nrn1 | protein_coding | 0.224747569 | 6.156109319 | 0.2715761 | 0.6103963 | 0.919059219 |
| ENSMUSG00000025857 | Dnaaf5 | protein_coding | -0.763002381 | 1.77534841 | 0.2705042 | 0.6104269 | 0.919059219 |
| ENSMUSG00000024039 | Cbs | protein_coding | 0.193860838 | 4.681258939 | 0.2694784 | 0.6106554 | 0.919143969 |
| ENSMUSG00000090546 | Cdr1 | protein_coding | -0.275762523 | 4.856864564 | 0.2707072 | 0.610781 | 0.919143969 |
| ENSMUSG00000078235 | Fam43b | protein_coding | 0.274578404 | 2.797049473 | 0.2692687 | 0.6107931 | 0.919143969 |
| ENSMUSG00000034220 | Gpc1 | protein_coding | 0.180223196 | 4.569326509 | 0.2692199 | 0.6108251 | 0.919143969 |
| ENSMUSG00000037426 | Depdc5 | protein_coding | 0.177773678 | 4.586188718 | 0.2690454 | 0.6109398 | 0.919174012 |
| ENSMUSG00000040188 | Scamp2 | protein_coding | 0.140881148 | 5.15352161 | 0.2689295 | 0.6110161 | 0.919174012 |
| ENSMUSG00000070436 | Serpinh1 | protein_coding | -0.279945266 | 3.617527688 | 0.2686351 | 0.6112211 | 0.919353879 |
| ENSMUSG00000040723 | Rcsd1 | protein_coding | -0.361297679 | 2.563251632 | 0.2683833 | 0.6113755 | 0.919457424 |
| ENSMUSG00000031626 | Sorbs2 | protein_coding | -0.159221057 | 4.848276821 | 0.2680588 | 0.6115892 | 0.91965026 |
| ENSMUSG00000070923 | Klhl9 | protein_coding | -0.105094531 | 8.232878104 | 0.2675349 | 0.6119347 | 0.919707586 |
| ENSMUSG00000045613 | Chrm2 | protein_coding | -0.177188328 | 4.4486178 | 0.2673236 | 0.6120741 | 0.919707586 |
| ENSMUSG00000005089 | Slc1a2 | protein_coding | 0.173096647 | 7.751621344 | 0.2686076 | 0.6121289 | 0.919707586 |
| ENSMUSG00000024038 | Ndufv3 | protein_coding | 0.108277805 | 9.184277111 | 0.267092 | 0.612227 | 0.919707586 |
| ENSMUSG00000020253 | Ppm1m | protein_coding | -0.190382554 | 4.064014177 | 0.2668663 | 0.6123762 | 0.919707586 |
| ENSMUSG00000049555 | Tmie | protein_coding | 0.192580677 | 4.130788794 | 0.2668588 | 0.6123811 | 0.919707586 |
| ENSMUSG00000039853 | Trim14 | protein_coding | -0.466213945 | 2.136635728 | 0.2667735 | 0.6124375 | 0.919707586 |
| ENSMUSG00000030086 | Chchd6 | protein_coding | -0.129214611 | 7.819536396 | 0.2667432 | 0.6124689 | 0.919707586 |
| ENSMUSG00000022765 | Snap29 | protein_coding | 0.184562791 | 4.01884437 | 0.2667083 | 0.6124806 | 0.919707586 |
| ENSMUSG00000000266 | Mid2 | protein_coding | -0.216200971 | 4.204205419 | 0.2667052 | 0.6124827 | 0.919707586 |
| ENSMUSG00000049932 | H2afx | protein_coding | 0.328727709 | 2.595392298 | 0.2665386 | 0.6125928 | 0.919744528 |
| ENSMUSG00000025237 | Parp6 | protein_coding | 0.115729188 | 5.991046716 | 0.2663332 | 0.6127287 | 0.919773249 |
| ENSMUSG00000021848 | Otx2 | protein_coding | -0.241578538 | 3.240621142 | 0.2662511 | 0.612783 | 0.919773249 |
| ENSMUSG00000003549 | Ercc1 | protein_coding | -0.107410509 | 6.386600643 | 0.2658998 | 0.6130156 | 0.919993998 |
| ENSMUSG00000024957 | Kcnk4 | protein_coding | -0.38200691 | 2.800689289 | 0.2650706 | 0.6135654 | 0.920690621 |
| ENSMUSG00000098188 | Sowahc | protein_coding | 0.252829377 | 3.065199733 | 0.2647199 | 0.6137983 | 0.920773015 |
| ENSMUSG00000062014 | Gmfb | protein_coding | -0.125378517 | 7.448109602 | 0.2646472 | 0.6138532 | 0.920773015 |
| ENSMUSG00000025762 | Larp1b | protein_coding | -0.266917322 | 3.278689258 | 0.2646011 | 0.6138772 | 0.920773015 |
| ENSMUSG00000039253 | Fn3krp | protein_coding | 0.36634787 | 3.490451767 | 0.2652728 | 0.6141201 | 0.921008777 |
| ENSMUSG00000061288 | Taok3 | protein_coding | -0.11610174 | 7.069856673 | 0.2640033 | 0.6142747 | 0.921112272 |
| ENSMUSG00000021975 | Ints9 | protein_coding | -0.177973254 | 4.656545547 | 0.263806 | 0.614406 | 0.921180681 |
| ENSMUSG00000073096 | Lrrc61 | protein_coding | -0.224354988 | 4.166503939 | 0.2636056 | 0.6145395 | 0.921252335 |
| ENSMUSG00000044364 | Tmem74b | protein_coding | -0.212152794 | 3.394795497 | 0.2632483 | 0.6147776 | 0.921352929 |
| ENSMUSG00000029802 | Abcg2 | protein_coding | 0.29246559 | 2.876853871 | 0.2631445 | 0.6148468 | 0.921352929 |
| ENSMUSG00000058706 | 0610030E20Rik | protein_coding | 0.301220836 | 2.85645559 | 0.2630809 | 0.6148892 | 0.921352929 |
| ENSMUSG00000010290 | AI597479 | protein_coding | 0.157154779 | 5.439774411 | 0.2629907 | 0.6149493 | 0.921352929 |
| ENSMUSG00000029617 | Ccz1 | protein_coding | 0.183908078 | 4.257759713 | 0.2623324 | 0.6153888 | 0.921778542 |

| ensembl_gene_id | external_gene_name | gene_biotype | log2 (fold change) | log2 (counts per million) | F-value | p-value | False discovery rate |
|---|---|---|---|---|---|---|---|
| ENSMUSG00000039623 | Ap5z1 | protein_coding | -0.266308188 | 3.859692997 | 0.2621028 | 0.6155535 | 0.921778542 |
| ENSMUSG00000097311 | Gm26871 | processed_transcript | -0.380069047 | 2.800488916 | 0.2618747 | 0.6156948 | 0.921778542 |
| ENSMUSG00000015092 | Edf1 | protein_coding | 0.105858208 | 7.716842567 | 0.2618516 | 0.6157102 | 0.921778542 |
| ENSMUSG00000020003 | Pex7 | protein_coding | -0.167946603 | 4.586048451 | 0.2618096 | 0.6157383 | 0.921778542 |
| ENSMUSG00000020412 | Ascc2 | protein_coding | 0.220969013 | 3.282315563 | 0.2617956 | 0.6157477 | 0.921778542 |
| ENSMUSG00000034748 | Sirt6 | protein_coding | -0.20228979 | 4.765305217 | 0.2615516 | 0.6159222 | 0.921911407 |
| ENSMUSG00000041355 | Ssr2 | protein_coding | -0.097769118 | 7.580759846 | 0.2612234 | 0.6161308 | 0.921980708 |
| ENSMUSG00000028982 | Slc25a33 | protein_coding | 0.206769007 | 4.497904763 | 0.2610913 | 0.6162193 | 0.921980708 |
| ENSMUSG00000032839 | Trpc1 | protein_coding | -0.240147418 | 3.585666934 | 0.2610816 | 0.6162257 | 0.921980708 |
| ENSMUSG00000028960 | Ube4b | protein_coding | 0.136621005 | 5.598998595 | 0.2609341 | 0.6163246 | 0.922000367 |
| ENSMUSG00000061838 | Suclg2 | protein_coding | 0.268341581 | 3.038395466 | 0.2605728 | 0.616567 | 0.92214088 |
| ENSMUSG00000005566 | Trim28 | protein_coding | -0.101403041 | 6.502968316 | 0.2605198 | 0.6166026 | 0.92214088 |
| ENSMUSG00000104674 | Gm42756 | antisense_RNA | -0.214741718 | 3.81264865 | 0.2604106 | 0.6166758 | 0.92214088 |
| ENSMUSG00000030096 | Slc6a6 | protein_coding | -0.130579784 | 5.537140466 | 0.2599057 | 0.6170149 | 0.922519681 |
| ENSMUSG00000061607 | Mdc1 | protein_coding | 0.462038939 | 2.20541745 | 0.2596194 | 0.6172074 | 0.922639138 |
| ENSMUSG00000018425 | Dhx40 | protein_coding | -0.149739231 | 4.628552036 | 0.2595317 | 0.6172664 | 0.922639138 |
| ENSMUSG00000019804 | Snx3 | protein_coding | 0.092557279 | 8.357148891 | 0.2592351 | 0.617466 | 0.922658799 |
| ENSMUSG00000029420 | Rimbp2 | protein_coding | -0.211464615 | 3.871527703 | 0.2591318 | 0.6175356 | 0.922658799 |
| ENSMUSG00000028127 | Abcd3 | protein_coding | -0.145205696 | 5.816527553 | 0.2588347 | 0.6177357 | 0.922658799 |
| ENSMUSG00000037999 | Arap2 | protein_coding | 0.159895551 | 4.659843276 | 0.2585684 | 0.6179152 | 0.922658799 |
| ENSMUSG00000026189 | Pecr | protein_coding | -0.278510936 | 2.841673926 | 0.2585209 | 0.6179472 | 0.922658799 |
| ENSMUSG00000040181 | Fmo1 | protein_coding | -0.255233461 | 3.594445406 | 0.2585096 | 0.6179548 | 0.922658799 |
| ENSMUSG00000028830 | AU040320 | protein_coding | 0.136121582 | 5.325684901 | 0.2582436 | 0.6181343 | 0.922658799 |
| ENSMUSG00000029059 | Fam213b | protein_coding | -0.127186582 | 9.193708674 | 0.2582147 | 0.6181538 | 0.922658799 |
| ENSMUSG00000048163 | Selplg | protein_coding | -0.215962673 | 4.155605552 | 0.2578175 | 0.618422 | 0.922658799 |
| ENSMUSG00000032120 | C2cd2l | protein_coding | -0.117797716 | 6.011683688 | 0.2576565 | 0.6185308 | 0.922658799 |
| ENSMUSG00000038181 | Chpf2 | protein_coding | -0.463129456 | 2.365429595 | 0.2576037 | 0.6185664 | 0.922658799 |
| ENSMUSG00000036371 | Serbp1 | protein_coding | -0.093762386 | 7.425135801 | 0.2575315 | 0.6186153 | 0.922658799 |
| ENSMUSG00000022561 | Gpaa1 | protein_coding | -0.156241252 | 5.135706021 | 0.2573204 | 0.618758 | 0.922658799 |
| ENSMUSG00000033063 | Cntnap3 | protein_coding | -0.323409994 | 2.680127865 | 0.2573074 | 0.6187668 | 0.922658799 |
| ENSMUSG00000038388 | Mpp6 | protein_coding | -0.159837342 | 4.397720151 | 0.2572817 | 0.6187842 | 0.922658799 |
| ENSMUSG00000030711 | Sult1a1 | protein_coding | 0.302192993 | 3.215347347 | 0.2572367 | 0.6188146 | 0.922658799 |
| ENSMUSG00000020638 | Cmpk2 | protein_coding | 0.13254319 | 5.485329651 | 0.2571814 | 0.618852 | 0.922658799 |
| ENSMUSG00000021096 | Ppm1a | protein_coding | -0.085820519 | 7.718985064 | 0.2570262 | 0.6189571 | 0.922658799 |
| ENSMUSG00000034158 | Lrrc58 | protein_coding | 0.146090559 | 5.653376863 | 0.2569844 | 0.6189853 | 0.922658799 |
| ENSMUSG00000075020 | Mir670hg | lincRNA | -0.232233401 | 4.707947878 | 0.2573007 | 0.6190556 | 0.922658799 |
| ENSMUSG00000024242 | Map4k3 | protein_coding | -0.161478438 | 5.042346066 | 0.2568424 | 0.6190815 | 0.922658799 |
| ENSMUSG00000031527 | Eri1 | protein_coding | -0.359221314 | 2.766883389 | 0.2566469 | 0.6192139 | 0.922679795 |
| ENSMUSG00000059146 | Ntrk3 | protein_coding | -0.115077327 | 6.111669655 | 0.2565682 | 0.6192672 | 0.922679795 |
| ENSMUSG00000029462 | Vps29 | protein_coding | 0.093413344 | 9.414165902 | 0.2561336 | 0.6195619 | 0.922821228 |
| ENSMUSG00000021711 | Trappc13 | protein_coding | 0.095718626 | 7.177096875 | 0.2559352 | 0.6196965 | 0.922821228 |
| ENSMUSG00000055676 | Gm5069 | transcribed_processe | -0.28233861 | 2.771218341 | 0.2551898 | 0.6202028 | 0.922821228 |
| ENSMUSG00000020708 | Psmc5 | protein_coding | 0.097734438 | 9.994707594 | 0.2549127 | 0.6203913 | 0.922821228 |
| ENSMUSG00000005225 | Plekha8 | protein_coding | 0.237797981 | 3.064504677 | 0.2546898 | 0.620543 | 0.922821228 |
| ENSMUSG00000062210 | Tnfaip8 | protein_coding | -0.207778451 | 4.226963676 | 0.2545944 | 0.620608 | 0.922821228 |
| ENSMUSG00000040410 | Fbxl4 | protein_coding | 0.148581304 | 4.835194913 | 0.2543624 | 0.620766 | 0.922821228 |
| ENSMUSG00000028469 | Npr2 | protein_coding | 0.246318436 | 3.538515871 | 0.2542385 | 0.6208505 | 0.922821228 |
| ENSMUSG00000030869 | Ndufab1 | protein_coding | 0.165737282 | 4.057246265 | 0.2541588 | 0.6209048 | 0.922821228 |
| ENSMUSG00000022376 | Adcy8 | protein_coding | 0.260336876 | 2.998032649 | 0.2541047 | 0.6209416 | 0.922821228 |
| ENSMUSG00000044813 | Shb | protein_coding | 0.312678084 | 3.022880809 | 0.2539848 | 0.6210234 | 0.922821228 |
| ENSMUSG00000021590 | Spata9 | protein_coding | -0.376115544 | 2.518650741 | 0.253826 | 0.6211317 | 0.922821228 |
| ENSMUSG00000033732 | Sf3b3 | protein_coding | -0.138280207 | 5.440594887 | 0.2537196 | 0.6212043 | 0.922821228 |
| ENSMUSG00000003868 | Ruvbl2 | protein_coding | 0.136986782 | 6.01352007 | 0.2536918 | 0.6212233 | 0.922821228 |
| ENSMUSG00000054752 | Fsd1l | protein_coding | -0.19704233 | 3.702558382 | 0.2536618 | 0.6212438 | 0.922821228 |
| ENSMUSG00000025439 | Clns1a | protein_coding | 0.091671898 | 7.542459201 | 0.2535017 | 0.621353 | 0.922821228 |
| ENSMUSG00000024429 | Gnl1 | protein_coding | 0.10354239 | 6.405451743 | 0.2530401 | 0.6216684 | 0.922821228 |
| ENSMUSG00000004568 | Arhgef18 | protein_coding | 0.248360426 | 3.849706534 | 0.2530044 | 0.6217039 | 0.922821228 |
| ENSMUSG00000027669 | Gnb4 | protein_coding | -0.109874886 | 6.130793214 | 0.2529777 | 0.6217111 | 0.922821228 |
| ENSMUSG00000026740 | Dnajc1 | protein_coding | -0.253153374 | 4.213419847 | 0.2531221 | 0.6218361 | 0.922821228 |
| ENSMUSG00000051344 | Plekhm3 | protein_coding | -0.1604913 | 4.396973582 | 0.2527235 | 0.6218849 | 0.922821228 |
| ENSMUSG00000052373 | Mpp3 | protein_coding | 0.165858916 | 5.007328505 | 0.2526544 | 0.6219322 | 0.922821228 |
| ENSMUSG00000032181 | Scg3 | protein_coding | 0.119769036 | 7.058210045 | 0.2525698 | 0.6219901 | 0.922821228 |
| ENSMUSG00000046111 | Cep295 | protein_coding | 0.262786431 | 2.984215369 | 0.2525465 | 0.622006 | 0.922821228 |
| ENSMUSG00000031568 | Rwdd4a | protein_coding | 0.148284487 | 4.71929133 | 0.2524177 | 0.6220942 | 0.922821228 |
| ENSMUSG00000087260 | Lamtor5 | protein_coding | -0.106531757 | 7.5296565 | 0.2523119 | 0.6221666 | 0.922821228 |
| ENSMUSG00000018669 | Cdk5rap3 | protein_coding | 0.129030961 | 6.585328663 | 0.2522605 | 0.6222019 | 0.922821228 |
| ENSMUSG00000027875 | Hmgcs2 | protein_coding | 0.164212848 | 4.074244339 | 0.2522534 | 0.6222067 | 0.922821228 |
| ENSMUSG00000038457 | Tmem255b | protein_coding | -0.486080472 | 2.342129433 | 0.2521605 | 0.6222814 | 0.922821228 |
| ENSMUSG00000020546 | Stxbp4 | protein_coding | -0.200378754 | 4.780325131 | 0.2520905 | 0.6223294 | 0.922821228 |
| ENSMUSG00000027544 | Nfatc2 | protein_coding | -0.200154333 | 4.198646993 | 0.252018 | 0.622368 | 0.922821228 |
| ENSMUSG00000039220 | Ppp1r10 | protein_coding | -0.169533848 | 5.239057029 | 0.2519019 | 0.6224476 | 0.922821228 |
| ENSMUSG00000048264 | Dip2c | protein_coding | 0.142798801 | 4.975060271 | 0.2517057 | 0.6225821 | 0.922821228 |
| ENSMUSG00000035772 | Mrps2 | protein_coding | 0.1273487 | 5.785814546 | 0.251641 | 0.6226265 | 0.922821228 |
| ENSMUSG00000002379 | Ndufa11 | protein_coding | -0.102653675 | 9.817108866 | 0.2516322 | 0.6226325 | 0.922821228 |

| ensembl_gene_id | external_gene_name | gene_biotype | log2 (fold change) | log2 (counts per million) | F-value | p-value | False discovery rate |
|---|---|---|---|---|---|---|---|
| ENSMUSG00000035045 | Zc3h12b | protein_coding | -0.159283356 | 4.20635853 | 0.2516181 | 0.6226422 | 0.922821228 |
| ENSMUSG00000030621 | Me3 | protein_coding | -0.202569582 | 4.22441266 | 0.2516131 | 0.6226456 | 0.922821228 |
| ENSMUSG00000023019 | Gpd1 | protein_coding | -0.202998285 | 3.716180756 | 0.2516048 | 0.6226513 | 0.922821228 |
| ENSMUSG00000021750 | Fam107a | protein_coding | -0.125017044 | 6.542335667 | 0.2510879 | 0.6230062 | 0.922821228 |
| ENSMUSG00000041313 | Slc7a1 | protein_coding | 0.185407045 | 3.810194806 | 0.2509148 | 0.6231251 | 0.922821228 |
| ENSMUSG00000021094 | Dhrs7 | protein_coding | 0.141655489 | 6.063722617 | 0.250856 | 0.6231656 | 0.922821228 |
| ENSMUSG00000021392 | Nol8 | protein_coding | 0.156835619 | 4.738627855 | 0.2506757 | 0.6232895 | 0.922821228 |
| ENSMUSG00000000916 | Nsun5 | protein_coding | -0.23128739 | 3.846835917 | 0.2506328 | 0.623319 | 0.922821228 |
| ENSMUSG00000023861 | Mpc1 | protein_coding | -0.099678694 | 6.224462493 | 0.250613 | 0.6233326 | 0.922821228 |
| ENSMUSG00000032123 | Dpagt1 | protein_coding | 0.284141676 | 3.034197488 | 0.2502948 | 0.6235515 | 0.922821228 |
| ENSMUSG00000028578 | Caap1 | protein_coding | -0.380278226 | 2.471576403 | 0.2502384 | 0.6235904 | 0.922821228 |
| ENSMUSG00000054418 | 2900041M22Rik | processed_transcript | 0.29478932 | 2.730016073 | 0.2501939 | 0.623621 | 0.922821228 |
| ENSMUSG00000079434 | Neu2 | protein_coding | -0.275623087 | 3.211015147 | 0.2500033 | 0.6237522 | 0.922821228 |
| ENSMUSG00000035078 | Mtmr9 | protein_coding | 0.128516877 | 5.870127254 | 0.2500008 | 0.623754 | 0.922821228 |
| ENSMUSG00000031451 | Gas6 | protein_coding | 0.291605098 | 2.816422772 | 0.2499024 | 0.6238217 | 0.922821228 |
| ENSMUSG00000027605 | Acss2 | protein_coding | 0.224186425 | 3.965329002 | 0.2498425 | 0.6238631 | 0.922821228 |
| ENSMUSG00000030605 | Mfge8 | protein_coding | -0.143062189 | 7.272475873 | 0.25023 | 0.6238759 | 0.922821228 |
| ENSMUSG00000032119 | Hinfp | protein_coding | -0.293623647 | 3.263631182 | 0.2497008 | 0.6239607 | 0.922821228 |
| ENSMUSG00000025630 | Hprt | protein_coding | 0.091125805 | 9.881867869 | 0.2494215 | 0.6241533 | 0.922821228 |
| ENSMUSG00000027778 | Ift80 | protein_coding | 0.205767421 | 3.804707773 | 0.2491495 | 0.624341 | 0.922821228 |
| ENSMUSG00000045215 | Asxl3 | protein_coding | -0.23169013 | 3.130231723 | 0.249093 | 0.62438 | 0.922821228 |
| ENSMUSG00000027533 | Fabp5 | protein_coding | 0.096590134 | 6.901610174 | 0.2489488 | 0.6244796 | 0.922821228 |
| ENSMUSG00000038690 | Atp5j2 | protein_coding | -0.122771294 | 9.890778011 | 0.248848 | 0.6245492 | 0.922821228 |
| ENSMUSG00000030839 | Sergef | protein_coding | 0.105747727 | 6.531287465 | 0.2487391 | 0.6246244 | 0.922821228 |
| ENSMUSG00000031839 | Hsbp1 | protein_coding | -0.102850445 | 10.62593126 | 0.248591 | 0.6247268 | 0.922821228 |
| ENSMUSG00000022614 | Lmf2 | protein_coding | -0.23592963 | 3.527637532 | 0.2485769 | 0.6247366 | 0.922821228 |
| ENSMUSG00000039257 | Vstm2b | protein_coding | -0.135347922 | 5.263091871 | 0.2485562 | 0.6247508 | 0.922821228 |
| ENSMUSG00000022453 | Naga | protein_coding | 0.155374982 | 5.324071958 | 0.2485303 | 0.6247688 | 0.922821228 |
| ENSMUSG00000041708 | Mpped1 | protein_coding | 0.127882 | 5.241067827 | 0.2482729 | 0.6249468 | 0.922872671 |
| ENSMUSG00000002797 | Ggct | protein_coding | -0.137274821 | 6.464061391 | 0.2482227 | 0.6249925 | 0.922872671 |
| ENSMUSG00000029426 | Scarb2 | protein_coding | 0.148765429 | 5.566327779 | 0.2481078 | 0.6250611 | 0.922872671 |
| ENSMUSG00000051518 | Rps19bp1 | protein_coding | 0.180720131 | 4.065044223 | 0.2477958 | 0.6252771 | 0.923064847 |
| ENSMUSG00000031644 | Nek1 | protein_coding | -0.253422888 | 3.354982581 | 0.2476413 | 0.6253841 | 0.923074393 |
| ENSMUSG00000027650 | Tti1 | protein_coding | -0.177408194 | 4.096851908 | 0.2475387 | 0.6254552 | 0.923074393 |
| ENSMUSG00000034248 | Slc25a37 | protein_coding | -0.301985272 | 2.954380726 | 0.2469707 | 0.6258492 | 0.923384858 |
| ENSMUSG00000042707 | Dnali1 | protein_coding | -0.272242088 | 3.274985517 | 0.246846 | 0.6259358 | 0.923384858 |
| ENSMUSG00000073481 | 2-Mar | protein_coding | 0.189713393 | 4.460534505 | 0.2467081 | 0.6260315 | 0.923384858 |
| ENSMUSG00000022635 | Zcrb1 | protein_coding | -0.141821427 | 5.522701848 | 0.2465319 | 0.6261539 | 0.923384858 |
| ENSMUSG00000028089 | Chd1l | protein_coding | 0.267976603 | 3.109815379 | 0.246529 | 0.626156 | 0.923384858 |
| ENSMUSG00000063535 | Zfp773 | protein_coding | 0.440949355 | 2.375556669 | 0.2464932 | 0.6261808 | 0.923384858 |
| ENSMUSG00000005886 | Ncoa2 | protein_coding | -0.16839451 | 4.780771823 | 0.2463149 | 0.6263048 | 0.923441013 |
| ENSMUSG00000052155 | Acvr2a | protein_coding | -0.191804712 | 3.817366856 | 0.245951 | 0.6265579 | 0.923641198 |
| ENSMUSG00000019494 | Cops6 | protein_coding | 0.0869249 | 9.482982842 | 0.2457987 | 0.626664 | 0.923641198 |
| ENSMUSG00000024594 | Prrc1 | protein_coding | -0.439757356 | 2.087124265 | 0.2457494 | 0.6266982 | 0.923641198 |
| ENSMUSG00000002190 | Clgn | protein_coding | -0.354111201 | 2.905563367 | 0.2451922 | 0.6270974 | 0.924102763 |
| ENSMUSG00000022848 | Dirc2 | protein_coding | -0.150362587 | 5.275796203 | 0.2449597 | 0.6272487 | 0.924199075 |
| ENSMUSG00000024758 | Rtn3 | protein_coding | 0.107168396 | 8.667423783 | 0.2446151 | 0.6274892 | 0.924302776 |
| ENSMUSG00000066510 | Ankdd1a | protein_coding | 0.282930988 | 3.148513677 | 0.2444678 | 0.6275921 | 0.924302776 |
| ENSMUSG00000032966 | Fkbp1a | protein_coding | -0.09684379 | 9.154511293 | 0.2443761 | 0.6276562 | 0.924302776 |
| ENSMUSG00000057181 | 5730455P16Rik | protein_coding | -0.116127719 | 6.975994754 | 0.2442611 | 0.6277365 | 0.924302776 |
| ENSMUSG00000085069 | Gm13111 | antisense_RNA | 0.336460776 | 2.502866299 | 0.2441826 | 0.6277915 | 0.924302776 |
| ENSMUSG00000037628 | Cdkn3 | protein_coding | -0.308156397 | 2.570795995 | 0.2441206 | 0.6278348 | 0.924302776 |
| ENSMUSG00000032802 | Srxn1 | protein_coding | 0.114904893 | 6.536059391 | 0.2439299 | 0.6279681 | 0.924349868 |
| ENSMUSG00000075702 | Selenom | protein_coding | -0.099895131 | 9.621468166 | 0.2437344 | 0.628105 | 0.924349868 |
| ENSMUSG00000046668 | Cxxc5 | protein_coding | 0.125569713 | 5.562815778 | 0.2437062 | 0.6281247 | 0.924349868 |
| ENSMUSG00000057530 | Ece1 | protein_coding | 0.132609876 | 5.311178642 | 0.2434932 | 0.6282738 | 0.924442849 |
| ENSMUSG00000032412 | Atp1b3 | protein_coding | 0.135537533 | 6.144788157 | 0.2432493 | 0.6284447 | 0.924567772 |
| ENSMUSG00000022210 | Dhrs4 | protein_coding | 0.17080511 | 4.737723596 | 0.2431071 | 0.6285443 | 0.924587878 |
| ENSMUSG00000061654 | Spry3 | protein_coding | 0.238821491 | 3.475000625 | 0.2427018 | 0.6288286 | 0.924827192 |
| ENSMUSG00000008333 | Snrpb2 | protein_coding | -0.120333807 | 6.805207652 | 0.2425623 | 0.6289265 | 0.924827192 |
| ENSMUSG00000025240 | Sacm1l | protein_coding | 0.293435098 | 3.773463814 | 0.2430414 | 0.6290085 | 0.924827192 |
| ENSMUSG00000113722 | AC140477.3 | lincRNA | 0.145694441 | 4.754180436 | 0.242385 | 0.6290511 | 0.924827192 |
| ENSMUSG00000022905 | Kpna1 | protein_coding | 0.104903788 | 6.03260121 | 0.2419285 | 0.6293719 | 0.925172345 |
| ENSMUSG00000061232 | H2-K1 | protein_coding | -0.262865059 | 3.521797605 | 0.2415476 | 0.6296506 | 0.925255811 |
| ENSMUSG00000054675 | Tmem119 | protein_coding | -0.277261684 | 3.114579512 | 0.2413679 | 0.6297664 | 0.925255811 |
| ENSMUSG00000059040 | Eno1b | protein_coding | 0.606221894 | 2.21628682 | 0.2417197 | 0.6299355 | 0.925255811 |
| ENSMUSG00000036854 | Hspb6 | protein_coding | -0.155954454 | 5.283787309 | 0.2410408 | 0.6299968 | 0.925255811 |
| ENSMUSG00000022394 | L3mbtl2 | protein_coding | -0.21162274 | 4.542254689 | 0.240946 | 0.6300744 | 0.925255811 |
| ENSMUSG00000072653 | Zfp783 | transcribed_unproces | 0.335004485 | 2.885958419 | 0.2405439 | 0.6303472 | 0.925255811 |
| ENSMUSG00000022040 | Ephx2 | protein_coding | 0.128276978 | 4.957748281 | 0.240428 | 0.630429 | 0.925255811 |
| ENSMUSG00000029478 | Ncor2 | protein_coding | 0.204438182 | 3.755960441 | 0.2404206 | 0.6304342 | 0.925255811 |
| ENSMUSG00000040651 | Fam208a | protein_coding | 0.228900057 | 3.492858956 | 0.2403295 | 0.6304986 | 0.925255811 |
| ENSMUSG00000026080 | Chst10 | protein_coding | -0.12097954 | 5.767554448 | 0.2403085 | 0.6305134 | 0.925255811 |
| ENSMUSG00000028312 | Smc2 | protein_coding | 0.321759989 | 2.554614109 | 0.2402962 | 0.6305221 | 0.925255811 |

| ensembl_gene_id | external_gene_name | gene_biotype | log2 (fold change) | log2 (counts per million) | F-value | p-value | False discovery rate |
|---|---|---|---|---|---|---|---|
| ENSMUSG00000107585 | 3300002P13Rik | lincRNA | -0.299443748 | 2.696054502 | 0.2401935 | 0.6305946 | 0.925255811 |
| ENSMUSG00000038150 | Ormdl3 | protein_coding | 0.110447513 | 6.097584382 | 0.2400402 | 0.6307029 | 0.925255811 |
| ENSMUSG00000054770 | Kctd18 | protein_coding | -0.236540349 | 3.477737419 | 0.2399312 | 0.63078 | 0.925255811 |
| ENSMUSG00000040767 | Snrnp25 | protein_coding | 0.101519447 | 6.89303129 | 0.239908 | 0.6307963 | 0.925255811 |
| ENSMUSG00000046922 | Gpr6 | protein_coding | 0.248208628 | 3.465183134 | 0.2397752 | 0.6308903 | 0.925255811 |
| ENSMUSG00000069581 | Tspear | protein_coding | -0.319355404 | 2.714406044 | 0.2397728 | 0.630892 | 0.925255811 |
| ENSMUSG00000038451 | Spsb2 | protein_coding | 0.177061693 | 3.76742242 | 0.2393174 | 0.6312142 | 0.925255811 |
| ENSMUSG00000000708 | Kat2b | protein_coding | -0.188835773 | 3.592394854 | 0.2392327 | 0.6312742 | 0.925255811 |
| ENSMUSG00000049422 | Chchd10 | protein_coding | -0.152201682 | 5.888721733 | 0.2390189 | 0.6314363 | 0.925255811 |
| ENSMUSG00000059495 | Arhgef12 | protein_coding | 0.12616752 | 5.470293133 | 0.2388342 | 0.6315565 | 0.925255811 |
| ENSMUSG00000002845 | Tmem39a | protein_coding | 0.274190182 | 3.255610213 | 0.2388176 | 0.6315683 | 0.925255811 |
| ENSMUSG00000029363 | Rfc5 | protein_coding | -0.301255876 | 2.78036276 | 0.2387645 | 0.6316059 | 0.925255811 |
| ENSMUSG00000051335 | Gfod1 | protein_coding | -0.178132663 | 3.964177096 | 0.2387144 | 0.6316414 | 0.925255811 |
| ENSMUSG00000084289 | Gm6977 | processed_pseudoge | -0.374931767 | 3.235061878 | 0.2393646 | 0.631714 | 0.925255811 |
| ENSMUSG00000032410 | Xrn1 | protein_coding | -0.189370968 | 4.021214016 | 0.2385632 | 0.6317487 | 0.925255811 |
| ENSMUSG00000017307 | Acot8 | protein_coding | 0.155996717 | 5.065814765 | 0.2383862 | 0.6318742 | 0.925255811 |
| ENSMUSG00000024975 | Pdcd4 | protein_coding | -0.173196058 | 7.436718025 | 0.2395557 | 0.6319578 | 0.925255811 |
| ENSMUSG00000015759 | Cnih1 | protein_coding | -0.094774598 | 7.541103114 | 0.238146 | 0.6320448 | 0.925255811 |
| ENSMUSG00000036833 | Pnpla7 | protein_coding | -0.330277476 | 2.789872672 | 0.2380382 | 0.6321213 | 0.925255811 |
| ENSMUSG00000024883 | Rin1 | protein_coding | 0.243142293 | 3.607593129 | 0.2380022 | 0.6321469 | 0.925255811 |
| ENSMUSG00000025026 | Add3 | protein_coding | 0.113665077 | 6.071688875 | 0.2377796 | 0.6323051 | 0.925255811 |
| ENSMUSG00000032308 | Ulk3 | protein_coding | -0.30798408 | 3.059243722 | 0.2377212 | 0.6323466 | 0.925255811 |
| ENSMUSG00000025466 | Fuom | protein_coding | -0.106559282 | 7.227118524 | 0.2375571 | 0.6324633 | 0.925255811 |
| ENSMUSG00000044857 | Lemd2 | protein_coding | -0.259090053 | 3.499210004 | 0.2374464 | 0.632542 | 0.925255811 |
| ENSMUSG00000024539 | Ptpn2 | protein_coding | -0.15576686 | 4.713151037 | 0.2373241 | 0.6326291 | 0.925255811 |
| ENSMUSG00000015776 | Med22 | protein_coding | -0.09928697 | 6.323748204 | 0.2372989 | 0.632647 | 0.925255811 |
| ENSMUSG00000022265 | Ank | protein_coding | 0.110819268 | 6.578249939 | 0.2370307 | 0.632838 | 0.925255811 |
| ENSMUSG00000027710 | Acad9 | protein_coding | 0.321821999 | 3.779285076 | 0.2378093 | 0.6330213 | 0.925255811 |
| ENSMUSG00000025066 | Sfr1 | protein_coding | -0.084317847 | 8.415339589 | 0.2366618 | 0.6331008 | 0.925255811 |
| ENSMUSG00000020785 | Camkk1 | protein_coding | 0.140885988 | 5.651688003 | 0.2366262 | 0.6331262 | 0.925255811 |
| ENSMUSG00000004591 | Pkn2 | protein_coding | -0.213014808 | 3.453411419 | 0.2365049 | 0.6332127 | 0.925255811 |
| ENSMUSG00000021996 | Esd | protein_coding | 0.089896344 | 8.145446247 | 0.2364637 | 0.6332421 | 0.925255811 |
| ENSMUSG00000005510 | Ndufs3 | protein_coding | 0.083623905 | 7.416447895 | 0.2362377 | 0.6334033 | 0.925255811 |
| ENSMUSG00000025911 | Adhfe1 | protein_coding | 0.188847444 | 3.611597344 | 0.2358508 | 0.6336796 | 0.925255811 |
| ENSMUSG00000059518 | Znhit1 | protein_coding | 0.095808402 | 7.934591468 | 0.2357865 | 0.6337256 | 0.925255811 |
| ENSMUSG00000020846 | Rflnb | protein_coding | 0.214513241 | 3.672663658 | 0.2356924 | 0.6337928 | 0.925255811 |
| ENSMUSG00000026482 | Rgl1 | protein_coding | -0.110459505 | 5.569146892 | 0.2351387 | 0.6341888 | 0.925255811 |
| ENSMUSG00000001988 | Npas1 | protein_coding | -0.287393147 | 3.054201747 | 0.2351276 | 0.6341968 | 0.925255811 |
| ENSMUSG00000023345 | Poc1a | protein_coding | -0.193163439 | 4.073827309 | 0.2349646 | 0.6343135 | 0.925255811 |
| ENSMUSG00000024593 | Megf10 | protein_coding | 0.306268776 | 2.874228502 | 0.2347174 | 0.6344906 | 0.925255811 |
| ENSMUSG00000001056 | Nhp2 | protein_coding | -0.105089017 | 7.596107777 | 0.234623 | 0.6345583 | 0.925255811 |
| ENSMUSG00000052299 | Ltn1 | protein_coding | 0.169810834 | 4.44658231 | 0.2344128 | 0.634709 | 0.925255811 |
| ENSMUSG00000038546 | Ranbp9 | protein_coding | -0.195483165 | 3.615199335 | 0.2342836 | 0.6348016 | 0.925255811 |
| ENSMUSG00000054199 | Gon4l | protein_coding | -0.250822791 | 4.01507341 | 0.2342381 | 0.6349613 | 0.925255811 |
| ENSMUSG00000042772 | Smg7 | protein_coding | 0.095114757 | 6.953742699 | 0.2339064 | 0.6350724 | 0.925255811 |
| ENSMUSG00000049321 | Zfp2 | protein_coding | -0.177930338 | 4.650543512 | 0.2338042 | 0.6351458 | 0.925255811 |
| ENSMUSG00000042104 | Uggt2 | protein_coding | -0.201522806 | 4.359631651 | 0.2336345 | 0.6352677 | 0.925255811 |
| ENSMUSG00000002102 | Psmc3 | protein_coding | 0.086456639 | 8.549169289 | 0.2333374 | 0.6354813 | 0.925255811 |
| ENSMUSG00000047030 | Spata2 | protein_coding | -0.187771747 | 4.479292543 | 0.2332991 | 0.6355089 | 0.925255811 |
| ENSMUSG00000062963 | Ufc1 | protein_coding | 0.092028091 | 8.435340463 | 0.2332815 | 0.6355215 | 0.925255811 |
| ENSMUSG00000046750 | Selenov | protein_coding | 0.150569139 | 4.165583799 | 0.2332495 | 0.6355446 | 0.925255811 |
| ENSMUSG00000037070 | Rbmxl1 | protein_coding | 0.157097048 | 4.031584788 | 0.2332338 | 0.6355559 | 0.925255811 |
| ENSMUSG00000035246 | Pcyt1b | protein_coding | 0.158029074 | 4.864259464 | 0.2330643 | 0.6356778 | 0.925255811 |
| ENSMUSG00000074220 | Zfp382 | protein_coding | 0.233415545 | 3.186077791 | 0.2330163 | 0.6357124 | 0.925255811 |
| ENSMUSG00000096847 | Tmem151b | protein_coding | -0.124976525 | 6.425331316 | 0.2327861 | 0.6358781 | 0.925255811 |
| ENSMUSG00000025794 | Rpl14 | protein_coding | -0.099564933 | 9.060645753 | 0.2327768 | 0.6358848 | 0.925255811 |
| ENSMUSG00000028403 | Zdhhc21 | protein_coding | -0.166051307 | 4.489614356 | 0.2326981 | 0.6359415 | 0.925255811 |
| ENSMUSG00000021967 | Mrpl57 | protein_coding | -0.116571202 | 6.480578311 | 0.2326342 | 0.6359876 | 0.925255811 |
| ENSMUSG00000032076 | Cadm1 | protein_coding | 0.095159298 | 7.275656087 | 0.2325549 | 0.6360447 | 0.925255811 |
| ENSMUSG00000014859 | E2f4 | protein_coding | -0.19525637 | 3.896729469 | 0.2325247 | 0.6360664 | 0.925255811 |
| ENSMUSG00000020770 | Unk | protein_coding | 0.315829414 | 3.006113874 | 0.2324804 | 0.6360984 | 0.925255811 |
| ENSMUSG00000037503 | Fam168b | protein_coding | 0.099412368 | 6.432525751 | 0.2320469 | 0.6364111 | 0.925255811 |
| ENSMUSG00000030779 | Rbbp6 | protein_coding | -0.112950523 | 5.982878442 | 0.2319751 | 0.6364629 | 0.925255811 |
| ENSMUSG00000035342 | Lzts2 | protein_coding | -0.297391449 | 3.030487529 | 0.2317334 | 0.6366374 | 0.925255811 |
| ENSMUSG00000030826 | Bcat2 | protein_coding | 0.18215344 | 4.651427735 | 0.2315997 | 0.636734 | 0.925255811 |
| ENSMUSG00000027520 | Zdbf2 | protein_coding | -0.179491686 | 4.992381259 | 0.2315148 | 0.6368059 | 0.925255811 |
| ENSMUSG00000021578 | Ccdc127 | protein_coding | 0.101100099 | 6.335168064 | 0.2312918 | 0.6369566 | 0.925255811 |
| ENSMUSG00000038552 | Fndc4 | protein_coding | -0.146995934 | 5.636499316 | 0.2312862 | 0.6369607 | 0.925255811 |
| ENSMUSG00000027739 | Rab33b | protein_coding | -0.124115246 | 5.633893349 | 0.2309867 | 0.6371774 | 0.925255811 |
| ENSMUSG00000002346 | Slc25a42 | protein_coding | -0.204401555 | 4.156457264 | 0.2309848 | 0.6371787 | 0.925255811 |
| ENSMUSG00000073207 | Ccdc160 | protein_coding | 0.255272713 | 3.255506659 | 0.2309165 | 0.6372282 | 0.925255811 |
| ENSMUSG00000020660 | Pomc | protein_coding | -0.188809319 | 4.148240725 | 0.2308432 | 0.6372812 | 0.925255811 |
| ENSMUSG00000034055 | Phka1 | protein_coding | 0.305157227 | 2.741177134 | 0.2308035 | 0.6373099 | 0.925255811 |
| ENSMUSG00000018919 | Tm4sf5 | protein_coding | -0.233841542 | 3.347728474 | 0.2307211 | 0.6373697 | 0.925255811 |

| ensembl_gene_id | external_gene_name | gene_biotype | log2 (fold change) | log2 (counts per million) | F-value | p-value | False discovery rate |
|---|---|---|---|---|---|---|---|
| ENSMUSG00000054604 | Cggbp1 | protein_coding | -0.112434701 | 5.712912326 | 0.2306181 | 0.6374443 | 0.925255811 |
| ENSMUSG00000031671 | Setd6 | protein_coding | 0.287542923 | 3.194264437 | 0.2306142 | 0.6374471 | 0.925255811 |
| ENSMUSG00000062908 | Acadm | protein_coding | -0.114229929 | 5.884754575 | 0.2304664 | 0.6375541 | 0.925255811 |
| ENSMUSG00000003444 | Med29 | protein_coding | 0.165905425 | 3.908901613 | 0.2304626 | 0.6375569 | 0.925255811 |
| ENSMUSG00000050103 | Agmo | protein_coding | -0.302405463 | 2.957131236 | 0.2303509 | 0.6376379 | 0.925255811 |
| ENSMUSG00000075316 | Scn9a | protein_coding | 0.181909348 | 3.5078237 | 0.2303408 | 0.6376452 | 0.925255811 |
| ENSMUSG00000005907 | Pex1 | protein_coding | 0.270379761 | 3.524648802 | 0.2302089 | 0.6377514 | 0.925255811 |
| ENSMUSG00000110834 | CAAA01180111.2 | lincRNA | -0.156063537 | 4.268034083 | 0.2301867 | 0.637757 | 0.925255811 |
| ENSMUSG00000000787 | Ddx3x | protein_coding | -0.100333036 | 6.582524005 | 0.2300296 | 0.637871 | 0.925255811 |
| ENSMUSG00000022201 | Zfr | protein_coding | -0.089322453 | 8.293012069 | 0.2299814 | 0.637906 | 0.925255811 |
| ENSMUSG00000020801 | Med31 | protein_coding | -0.126635244 | 6.786437743 | 0.2299433 | 0.637944 | 0.925255811 |
| ENSMUSG00000022338 | Eny2 | protein_coding | -0.094987292 | 7.138504758 | 0.2296957 | 0.6381133 | 0.925255811 |
| ENSMUSG00000057716 | Tmem178b | protein_coding | 0.139575142 | 5.542624092 | 0.2296324 | 0.6381593 | 0.925255811 |
| ENSMUSG00000052372 | Il1rapl1 | protein_coding | 0.140664534 | 7.527975867 | 0.2301287 | 0.6381819 | 0.925255811 |
| ENSMUSG00000067365 | Tmem128 | protein_coding | -0.111530062 | 6.726846509 | 0.2295689 | 0.6382054 | 0.925255811 |
| ENSMUSG00000024182 | Axin1 | protein_coding | -0.157355633 | 4.05334865 | 0.2295222 | 0.6382394 | 0.925255811 |
| ENSMUSG00000033938 | Ndufb7 | protein_coding | 0.099667178 | 9.54847961 | 0.2294959 | 0.6382585 | 0.925255811 |
| ENSMUSG00000071528 | Usmg5 | protein_coding | -0.122698159 | 7.503552794 | 0.2293462 | 0.6383778 | 0.925255811 |
| ENSMUSG00000105843 | Gm19439 | TEC | -0.358011983 | 4.624204269 | 0.2326768 | 0.6385586 | 0.925255811 |
| ENSMUSG00000039904 | Gpr37 | protein_coding | -0.129130811 | 6.394693251 | 0.2287114 | 0.6388353 | 0.925255811 |
| ENSMUSG00000028266 | Lmo4 | protein_coding | -0.103570003 | 6.567840984 | 0.228444 | 0.639024 | 0.925255811 |
| ENSMUSG00000073910 | Mob3b | protein_coding | -0.237455074 | 3.951102461 | 0.2284566 | 0.6390252 | 0.925255811 |
| ENSMUSG00000038369 | Ncoa6 | protein_coding | 0.185072364 | 4.480358588 | 0.2284254 | 0.6390375 | 0.925255811 |
| ENSMUSG00000090069 | E430024P14Rik | antisense_RNA | 0.32710511 | 3.360192038 | 0.2289307 | 0.6390491 | 0.925255811 |
| ENSMUSG00000079414 | Gm11110 | protein_coding | 0.361881663 | 2.612340391 | 0.2283695 | 0.6390783 | 0.925255811 |
| ENSMUSG00000040234 | Tm7sf3 | protein_coding | 0.154767112 | 4.603531011 | 0.2283149 | 0.6391181 | 0.925255811 |
| ENSMUSG00000020109 | Dnajb12 | protein_coding | -0.118038044 | 5.728433243 | 0.2280589 | 0.6393048 | 0.925255811 |
| ENSMUSG00000049299 | Trappc1 | protein_coding | -0.099260489 | 7.355933209 | 0.2279862 | 0.6393578 | 0.925255811 |
| ENSMUSG00000024858 | Grk2 | protein_coding | -0.111366271 | 6.36618441 | 0.2278182 | 0.6394804 | 0.925255811 |
| ENSMUSG00000022270 | Retreg1 | protein_coding | -0.111976753 | 5.980789228 | 0.2277155 | 0.6395554 | 0.925255811 |
| ENSMUSG00000017721 | Pigt | protein_coding | 0.180382368 | 4.034348605 | 0.2276333 | 0.6396155 | 0.925255811 |
| ENSMUSG00000028556 | Dock7 | protein_coding | -0.195233689 | 3.640016014 | 0.2275518 | 0.6396749 | 0.925255811 |
| ENSMUSG00000090523 | Gypc | protein_coding | 0.449613236 | 1.789287632 | 0.2274397 | 0.6397569 | 0.925255811 |
| ENSMUSG00000042962 | Gm5436 | processed_pseudoge | 0.176965218 | 3.503938644 | 0.227348 | 0.6398239 | 0.925255811 |
| ENSMUSG00000003644 | Rps6ka1 | protein_coding | 0.152264885 | 4.109013313 | 0.2273467 | 0.6398249 | 0.925255811 |
| ENSMUSG00000074582 | Arfgef2 | protein_coding | 0.208492133 | 4.268294228 | 0.2272175 | 0.6399297 | 0.925255811 |
| ENSMUSG00000031537 | Ikbkb | protein_coding | 0.143844088 | 4.460225865 | 0.2272006 | 0.6399317 | 0.925255811 |
| ENSMUSG00000041143 | Tmco4 | protein_coding | -0.339371905 | 2.414615072 | 0.2270354 | 0.6400525 | 0.925255811 |
| ENSMUSG00000042115 | Klhdc8a | protein_coding | -0.150741524 | 5.107572488 | 0.2269263 | 0.6401323 | 0.925255811 |
| ENSMUSG00000086938 | 4930481A15Rik | antisense_RNA | -0.161849852 | 3.896316853 | 0.2268551 | 0.6401844 | 0.925255811 |
| ENSMUSG00000011427 | Zfp790 | protein_coding | 0.228119947 | 3.195718129 | 0.2266947 | 0.6403018 | 0.925267854 |
| ENSMUSG00000026527 | Rgs7 | protein_coding | -0.12973154 | 5.555931476 | 0.2265239 | 0.6404269 | 0.925267854 |
| ENSMUSG00000034282 | Evpl | protein_coding | 0.208284802 | 3.819500099 | 0.2264912 | 0.6404509 | 0.925267854 |
| ENSMUSG00000037736 | Limch1 | protein_coding | -0.115689136 | 5.94337844 | 0.2261273 | 0.6407177 | 0.925426051 |
| ENSMUSG00000033161 | Atp1a1 | protein_coding | -0.167758705 | 4.929642025 | 0.2260643 | 0.6407639 | 0.925426051 |
| ENSMUSG00000032314 | Etfa | protein_coding | -0.087085649 | 6.758427469 | 0.2259897 | 0.6408186 | 0.925426051 |
| ENSMUSG00000031516 | Dctn6 | protein_coding | -0.094056568 | 8.032111938 | 0.2256616 | 0.6410594 | 0.925519638 |
| ENSMUSG00000026946 | Nmi | protein_coding | -0.219537331 | 3.25338944 | 0.225659 | 0.6410613 | 0.925519638 |
| ENSMUSG00000032399 | Rpl4 | protein_coding | 0.094920528 | 10.60178081 | 0.2255496 | 0.6411416 | 0.925519638 |
| ENSMUSG00000025277 | Abhd6 | protein_coding | -0.111917778 | 6.217601907 | 0.2250714 | 0.6414932 | 0.925534916 |
| ENSMUSG00000033530 | Ttc7b | protein_coding | 0.212738008 | 4.516190074 | 0.2251163 | 0.6414976 | 0.925534916 |
| ENSMUSG00000053398 | Phgdh | protein_coding | 0.131143173 | 5.139658463 | 0.2247314 | 0.6417433 | 0.925534916 |
| ENSMUSG00000029467 | Atp2a2 | protein_coding | 0.108486099 | 7.904779447 | 0.2245667 | 0.6418646 | 0.925534916 |
| ENSMUSG00000054364 | Rhob | protein_coding | -0.091783576 | 7.058097079 | 0.2243409 | 0.642031 | 0.925534916 |
| ENSMUSG00000033102 | Cdc14b | protein_coding | -0.199603582 | 3.484622043 | 0.224171 | 0.6421562 | 0.925534916 |
| ENSMUSG00000060143 | Gm10076 | protein_coding | -0.232859289 | 3.08257954 | 0.2240975 | 0.6422104 | 0.925534916 |
| ENSMUSG00000032504 | Pdcd6ip | protein_coding | -0.14243467 | 5.101560594 | 0.2239441 | 0.6423235 | 0.925534916 |
| ENSMUSG00000028391 | Wdr31 | protein_coding | 0.180917407 | 5.189647739 | 0.223933 | 0.642342 | 0.925534916 |
| ENSMUSG00000037148 | Arhgap10 | protein_coding | 0.361663724 | 2.855312343 | 0.2236409 | 0.6425576 | 0.925534916 |
| ENSMUSG00000028292 | Rars2 | protein_coding | 0.123214617 | 5.453790549 | 0.2236083 | 0.6425714 | 0.925534916 |
| ENSMUSG00000061315 | Naca | protein_coding | -0.105229297 | 6.815709084 | 0.2235919 | 0.6425835 | 0.925534916 |
| ENSMUSG00000041642 | Kif21b | protein_coding | 0.20802874 | 3.805694586 | 0.2234792 | 0.6426668 | 0.925534916 |
| ENSMUSG00000019868 | Vta1 | protein_coding | 0.091898423 | 6.910586962 | 0.223451 | 0.6426876 | 0.925534916 |
| ENSMUSG00000026457 | Adipor1 | protein_coding | 0.123658904 | 5.534447329 | 0.2232635 | 0.6428261 | 0.925534916 |
| ENSMUSG00000009281 | Rarres2 | protein_coding | -0.196391243 | 4.253690316 | 0.2232117 | 0.6428644 | 0.925534916 |
| ENSMUSG00000021948 | Prkcd | protein_coding | 0.215890313 | 5.889248203 | 0.2244438 | 0.6429555 | 0.925534916 |
| ENSMUSG00000026895 | Ndufa8 | protein_coding | 0.087696674 | 9.504452864 | 0.2230855 | 0.6429577 | 0.925534916 |
| ENSMUSG00000072915 | Gm12258 | protein_coding | -0.200937659 | 3.798291442 | 0.2230268 | 0.6430011 | 0.925534916 |
| ENSMUSG00000029364 | Wsb2 | protein_coding | 0.120985269 | 5.49505858 | 0.2230232 | 0.6430038 | 0.925534916 |
| ENSMUSG00000042216 | Sgsm1 | protein_coding | 0.13024902 | 6.090930906 | 0.2228182 | 0.6431555 | 0.925534916 |
| ENSMUSG00000085558 | 4930412C18Rik | processed_transcript | -0.208596605 | 3.611819385 | 0.2227272 | 0.6432229 | 0.925534916 |
| ENSMUSG00000026319 | 2310035C23Rik | protein_coding | -0.143793073 | 5.275347878 | 0.2226984 | 0.6432442 | 0.925534916 |
| ENSMUSG00000038056 | Kmt2c | protein_coding | 0.145603489 | 4.777930851 | 0.2226273 | 0.6432968 | 0.925534916 |
| ENSMUSG00000024925 | Rnaseh2c | protein_coding | 0.254921856 | 4.817628596 | 0.2237891 | 0.6433156 | 0.925534916 |

| ensembl_gene_id | external_gene_name | gene_biotype | log2 (fold change) | log2 (counts per million) | F-value | p-value | False discovery rate |
|---|---|---|---|---|---|---|---|
| ENSMUSG00000032030 | Cul5 | protein_coding | -0.123900469 | 5.90373286 | 0.2224632 | 0.6434183 | 0.925534916 |
| ENSMUSG00000057969 | Sema3b | protein_coding | 0.512697662 | 2.483645804 | 0.2229228 | 0.6434761 | 0.925534916 |
| ENSMUSG00000039089 | L3mbtl3 | protein_coding | -0.148823654 | 4.024540819 | 0.2221891 | 0.6436215 | 0.925550405 |
| ENSMUSG00000044734 | Serpinb1a | protein_coding | -0.121758661 | 5.844945195 | 0.2220849 | 0.6436988 | 0.925550405 |
| ENSMUSG00000068015 | Lrch1 | protein_coding | 0.274202294 | 3.917521528 | 0.2225589 | 0.6437451 | 0.925550405 |
| ENSMUSG00000038143 | Stox2 | protein_coding | -0.123249245 | 6.042121672 | 0.2218238 | 0.6438925 | 0.9255783 |
| ENSMUSG00000048108 | Tmem72 | protein_coding | -0.431931131 | 2.046472739 | 0.2215741 | 0.6440778 | 0.9255783 |
| ENSMUSG00000057531 | Dtnbp1 | protein_coding | 0.105800165 | 6.867280888 | 0.2215213 | 0.644117 | 0.9255783 |
| ENSMUSG00000027189 | Trim44 | protein_coding | -0.100876845 | 6.742039693 | 0.2214553 | 0.6441661 | 0.9255783 |
| ENSMUSG00000056185 | Snx32 | protein_coding | 0.093622198 | 7.503683569 | 0.2214164 | 0.6441949 | 0.9255783 |
| ENSMUSG00000045031 | Cetn4 | protein_coding | 0.177112933 | 3.838305497 | 0.2212682 | 0.6443051 | 0.925612869 |
| ENSMUSG00000037640 | Zfp60 | protein_coding | -0.154524113 | 4.904706894 | 0.2210561 | 0.6444628 | 0.925715756 |
| ENSMUSG00000037697 | Ddhd1 | protein_coding | -0.32539459 | 2.963253179 | 0.2205996 | 0.6448127 | 0.925929806 |
| ENSMUSG00000051331 | Cacna1c | protein_coding | -0.216829251 | 3.874655236 | 0.220345 | 0.6449922 | 0.925929806 |
| ENSMUSG00000027333 | Smox | protein_coding | -0.137608572 | 4.904103028 | 0.2203264 | 0.645006 | 0.925929806 |
| ENSMUSG00000034566 | Atp5h | protein_coding | -0.103674807 | 8.487546857 | 0.2201737 | 0.6451198 | 0.925929806 |
| ENSMUSG00000059995 | Atxn7l3 | protein_coding | 0.128083885 | 5.42908172 | 0.2201136 | 0.6451646 | 0.925929806 |
| ENSMUSG00000107476 | Zfp862-ps | transcribed_unitary_p | -0.268345162 | 2.780247633 | 0.2200602 | 0.6452045 | 0.925929806 |
| ENSMUSG00000025809 | Itgb1 | protein_coding | 0.167797357 | 4.437759961 | 0.2199544 | 0.6452834 | 0.925929806 |
| ENSMUSG00000028871 | Rspo1 | protein_coding | -0.28964443 | 3.104288689 | 0.2197537 | 0.6454331 | 0.925929806 |
| ENSMUSG00000039031 | Arhgap18 | protein_coding | -0.212907721 | 3.136624267 | 0.2197076 | 0.6454675 | 0.925929806 |
| ENSMUSG00000039886 | Tmem120a | protein_coding | 0.148856137 | 4.318940806 | 0.219602 | 0.6455464 | 0.925929806 |
| ENSMUSG00000031993 | Snx19 | protein_coding | 0.138315242 | 5.034120209 | 0.2195851 | 0.645559 | 0.925929806 |
| ENSMUSG00000035198 | Tubg1 | protein_coding | 0.135069217 | 6.697199905 | 0.2193416 | 0.6457885 | 0.926135421 |
| ENSMUSG00000025246 | Tbl1x | protein_coding | -0.111079853 | 6.155743411 | 0.2188956 | 0.6460744 | 0.926167478 |
| ENSMUSG00000039470 | Zdhhc2 | protein_coding | -0.141445785 | 5.22588489 | 0.2187964 | 0.6461486 | 0.926167478 |
| ENSMUSG00000066838 | Zfp772 | protein_coding | -0.23984759 | 3.219859574 | 0.2187666 | 0.6461709 | 0.926167478 |
| ENSMUSG00000048376 | F2r | protein_coding | -0.297538301 | 2.762143809 | 0.2187335 | 0.6461957 | 0.926167478 |
| ENSMUSG00000028277 | Ube2j1 | protein_coding | 0.114896089 | 6.01648963 | 0.218625 | 0.646277 | 0.926167478 |
| ENSMUSG00000006058 | Snf8 | protein_coding | -0.143939962 | 5.101828562 | 0.2185574 | 0.6463276 | 0.926167478 |
| ENSMUSG00000001552 | Jup | protein_coding | -0.145565204 | 4.976714421 | 0.2184004 | 0.6464452 | 0.926212558 |
| ENSMUSG00000052428 | Tmco1 | protein_coding | 0.098330738 | 7.076710616 | 0.2180719 | 0.6466914 | 0.926287421 |
| ENSMUSG00000021022 | Ppp2r3c | protein_coding | -0.139083201 | 4.722584434 | 0.2179922 | 0.6467512 | 0.926287421 |
| ENSMUSG00000029587 | Zfp12 | protein_coding | 0.340940146 | 3.779491193 | 0.2192724 | 0.6467798 | 0.926287421 |
| ENSMUSG00000050668 | Gpatch11 | protein_coding | 0.095898239 | 6.136313262 | 0.2178712 | 0.646842 | 0.926287421 |
| ENSMUSG00000030093 | Wnt7a | protein_coding | 0.273900485 | 2.849999121 | 0.2175342 | 0.647095 | 0.926526259 |
| ENSMUSG00000031652 | N4bp1 | protein_coding | -0.118847802 | 5.833230899 | 0.2171473 | 0.6473858 | 0.92665671 |
| ENSMUSG00000020100 | Slc29a3 | protein_coding | 0.24626065 | 3.130165832 | 0.2170886 | 0.6474299 | 0.92665671 |
| ENSMUSG00000066637 | Ttc32 | protein_coding | -0.308143527 | 2.762092343 | 0.2170586 | 0.6474524 | 0.92665671 |
| ENSMUSG00000024140 | Epas1 | protein_coding | -0.189215742 | 4.294059683 | 0.2169128 | 0.6475622 | 0.92665671 |
| ENSMUSG00000047013 | Fbxo41 | protein_coding | 0.156251074 | 5.231849758 | 0.2168399 | 0.647617 | 0.92665671 |
| ENSMUSG00000074575 | Kcng1 | protein_coding | -0.215126434 | 3.102649966 | 0.2166046 | 0.6477941 | 0.926786871 |
| ENSMUSG00000020133 | 2310011J03Rik | protein_coding | -0.09029571 | 6.388075915 | 0.216395 | 0.647952 | 0.926889427 |
| ENSMUSG00000042388 | Dlgap3 | protein_coding | 0.229859987 | 3.139058895 | 0.2162145 | 0.648088 | 0.926922135 |
| ENSMUSG00000039515 | Ptpa | protein_coding | 0.10254036 | 8.410019744 | 0.2158363 | 0.6483733 | 0.926922135 |
| ENSMUSG00000028782 | Adgrb2 | protein_coding | 0.233291434 | 3.140051897 | 0.2156828 | 0.6484892 | 0.926922135 |
| ENSMUSG00000022193 | Psmb5 | protein_coding | -0.087346903 | 6.843764382 | 0.2155974 | 0.6485537 | 0.926922135 |
| ENSMUSG00000025260 | Hsd17b10 | protein_coding | -0.085730421 | 7.110727815 | 0.2155806 | 0.6485664 | 0.926922135 |
| ENSMUSG00000062901 | Klhl24 | protein_coding | 0.136512832 | 5.141138767 | 0.2154016 | 0.6487016 | 0.926922135 |
| ENSMUSG00000069089 | Cdk7 | protein_coding | -0.124445466 | 5.207456483 | 0.2153235 | 0.6487606 | 0.926922135 |
| ENSMUSG00000037553 | Zdhhc18 | protein_coding | 0.181135523 | 4.081426624 | 0.2152413 | 0.6488228 | 0.926922135 |
| ENSMUSG00000056608 | Chd9 | protein_coding | -0.136395936 | 5.372321809 | 0.2152025 | 0.6488521 | 0.926922135 |
| ENSMUSG00000025534 | Gusb | protein_coding | 0.121236571 | 5.25349744 | 0.2151555 | 0.6488876 | 0.926922135 |
| ENSMUSG00000052794 | 1700030K09Rik | protein_coding | 0.176479437 | 4.025646093 | 0.2150837 | 0.6489419 | 0.926922135 |
| ENSMUSG00000020919 | Stat5b | protein_coding | 0.162873456 | 4.271669641 | 0.2149201 | 0.6490657 | 0.926922135 |
| ENSMUSG00000047454 | Gphn | protein_coding | -0.094827427 | 7.188718565 | 0.2148807 | 0.6490955 | 0.926922135 |
| ENSMUSG00000074182 | Znhit6 | protein_coding | -0.13278697 | 4.740682252 | 0.2142559 | 0.6495688 | 0.927435047 |
| ENSMUSG00000063317 | Usp31 | protein_coding | 0.132577352 | 5.091182228 | 0.2141789 | 0.6496272 | 0.927435047 |
| ENSMUSG00000006356 | Crip2 | protein_coding | -0.09042449 | 8.839673363 | 0.2138442 | 0.649881 | 0.927674325 |
| ENSMUSG00000002981 | Clptm1 | protein_coding | -0.108664838 | 5.646695765 | 0.2135052 | 0.6501384 | 0.927872954 |
| ENSMUSG00000033597 | Caskin1 | protein_coding | -0.156937274 | 5.013224174 | 0.2134338 | 0.6501928 | 0.927872954 |
| ENSMUSG00000020042 | Btbd11 | protein_coding | 0.143457246 | 5.056746215 | 0.2130924 | 0.6504523 | 0.927947158 |
| ENSMUSG00000030298 | Sec13 | protein_coding | 0.089750898 | 7.532545021 | 0.2128237 | 0.6506567 | 0.927947158 |
| ENSMUSG00000061731 | Ext1 | protein_coding | -0.154561696 | 4.01353138 | 0.2127809 | 0.6506893 | 0.927947158 |
| ENSMUSG00000068206 | Pick1 | protein_coding | 0.153061538 | 4.937320041 | 0.2127268 | 0.6507305 | 0.927947158 |
| ENSMUSG00000022498 | Txndc11 | protein_coding | 0.240659417 | 2.905277887 | 0.2126691 | 0.6507744 | 0.927947158 |
| ENSMUSG00000061981 | Flot2 | protein_coding | -0.10549334 | 6.29793369 | 0.2126469 | 0.6507914 | 0.927947158 |
| ENSMUSG00000003346 | Abhd17a | protein_coding | -0.162463162 | 4.933499331 | 0.2125715 | 0.6508488 | 0.927947158 |
| ENSMUSG00000042298 | Ttc19 | protein_coding | 0.127649914 | 5.377593297 | 0.2122082 | 0.6511257 | 0.928218832 |
| ENSMUSG00000022682 | Rrn3 | protein_coding | 0.164980076 | 7.475947597 | 0.2127685 | 0.6515974 | 0.928768164 |
| ENSMUSG00000022685 | Parn | protein_coding | 0.14605093 | 4.910161498 | 0.2109708 | 0.652071 | 0.929053985 |
| ENSMUSG00000027968 | Larp7 | protein_coding | 0.130289875 | 5.413909771 | 0.210736 | 0.6522508 | 0.929053985 |
| ENSMUSG00000037824 | Tspan14 | protein_coding | 0.14808618 | 4.513875636 | 0.210707 | 0.652273 | 0.929053985 |
| ENSMUSG00000022663 | Atg3 | protein_coding | -0.12525057 | 5.160404018 | 0.2104731 | 0.6524522 | 0.929053985 |

| ensembl_gene_id | external_gene_name | gene_biotype | log2 (fold change) | log2 (counts per million) | F-value | p-value | False discovery rate |
|---|---|---|---|---|---|---|---|
| ENSMUSG00000030788 | Rnf141 | protein_coding | -0.123665243 | 5.494603547 | 0.2104552 | 0.6524659 | 0.929053985 |
| ENSMUSG00000097068 | Gm26760 | lincRNA | -0.284574651 | 2.564848767 | 0.2103647 | 0.6525353 | 0.929053985 |
| ENSMUSG00000024436 | Mrps18b | protein_coding | 0.141174757 | 4.956158129 | 0.210335 | 0.6525581 | 0.929053985 |
| ENSMUSG00000049339 | Retreg2 | protein_coding | -0.099175525 | 6.910979518 | 0.2102533 | 0.6526207 | 0.929053985 |
| ENSMUSG00000034310 | Tmem132d | protein_coding | -0.242788409 | 2.98634704 | 0.2101212 | 0.652722 | 0.929053985 |
| ENSMUSG00000030032 | Wdr54 | protein_coding | -0.114976697 | 5.463490421 | 0.210113 | 0.6527283 | 0.929053985 |
| ENSMUSG00000010914 | Pdhx | protein_coding | 0.100783755 | 6.463653855 | 0.2100869 | 0.6527483 | 0.929053985 |
| ENSMUSG00000063019 | Manbal | protein_coding | 0.088459771 | 7.243091372 | 0.2099121 | 0.6528825 | 0.929121902 |
| ENSMUSG00000024132 | Eci1 | protein_coding | -0.169902638 | 5.003862725 | 0.2097865 | 0.6529888 | 0.929150283 |
| ENSMUSG00000041323 | Ak7 | protein_coding | -0.297780328 | 2.577050978 | 0.2096225 | 0.6531049 | 0.929192483 |
| ENSMUSG00000028837 | Psmb2 | protein_coding | 0.084459586 | 8.876893701 | 0.2093138 | 0.6533422 | 0.929250454 |
| ENSMUSG00000111947 | AC131120.1 | lincRNA | -0.34640731 | 2.78943609 | 0.20914 | 0.6534858 | 0.929250454 |
| ENSMUSG00000026942 | Traf2 | protein_coding | 0.157013085 | 4.716629366 | 0.2090335 | 0.6535578 | 0.929250454 |
| ENSMUSG00000033436 | Armcx2 | protein_coding | 0.107056259 | 6.195995287 | 0.2089117 | 0.6536516 | 0.929250454 |
| ENSMUSG00000031214 | Ophn1 | protein_coding | 0.144076144 | 4.328522202 | 0.2088334 | 0.6537118 | 0.929250454 |
| ENSMUSG00000034880 | Mrpl34 | protein_coding | 0.098351995 | 7.009545127 | 0.2082703 | 0.6541458 | 0.929250454 |
| ENSMUSG00000005893 | Nr2c2 | protein_coding | -0.147594501 | 4.941019602 | 0.2082107 | 0.6541918 | 0.929250454 |
| ENSMUSG00000009681 | Bcr | protein_coding | 0.097825321 | 6.228285443 | 0.2081523 | 0.6542369 | 0.929250454 |
| ENSMUSG00000029755 | Dlx5 | protein_coding | -0.118167248 | 5.722980699 | 0.2080053 | 0.6543504 | 0.929250454 |
| ENSMUSG00000019876 | Pkib | protein_coding | 0.12596531 | 4.949440417 | 0.2077072 | 0.6545805 | 0.929250454 |
| ENSMUSG00000028773 | Fabp3 | protein_coding | -0.093052554 | 6.611636062 | 0.2076741 | 0.6546061 | 0.929250454 |
| ENSMUSG00000019039 | Dalrd3 | protein_coding | -0.144561351 | 5.753914959 | 0.2074825 | 0.654764 | 0.929250454 |
| ENSMUSG00000097867 | Lppos | lincRNA | -0.269796516 | 2.724646276 | 0.2073754 | 0.6548371 | 0.929250454 |
| ENSMUSG00000074793 | Hspa12b | protein_coding | 0.25543907 | 2.950965648 | 0.207343 | 0.6548621 | 0.929250454 |
| ENSMUSG00000038347 | Tcte2 | protein_coding | -0.154337907 | 4.131523896 | 0.2073246 | 0.6548763 | 0.929250454 |
| ENSMUSG00000038622 | Med30 | protein_coding | -0.195704559 | 3.868644546 | 0.207226 | 0.6549526 | 0.929250454 |
| ENSMUSG00000038683 | Pak1ip1 | protein_coding | 0.104633882 | 7.238185548 | 0.2071901 | 0.6549804 | 0.929250454 |
| ENSMUSG00000057406 | Nsd2 | protein_coding | 0.114357122 | 5.332635908 | 0.2071179 | 0.6550362 | 0.929250454 |
| ENSMUSG00000028070 | Naxe | protein_coding | 0.11230923 | 7.422686091 | 0.2071259 | 0.6550399 | 0.929250454 |
| ENSMUSG00000059412 | Fxyd2 | protein_coding | -0.143059687 | 5.13876124 | 0.2070839 | 0.6550625 | 0.929250454 |
| ENSMUSG00000071037 | Camkmt | protein_coding | -0.170591965 | 4.431726802 | 0.2070759 | 0.6550688 | 0.929250454 |
| ENSMUSG00000026959 | Grin1 | protein_coding | -0.286176192 | 2.545413255 | 0.2070124 | 0.6551179 | 0.929250454 |
| ENSMUSG00000039660 | Spout1 | protein_coding | -0.117838386 | 5.892875434 | 0.2067984 | 0.6552836 | 0.929250454 |
| ENSMUSG00000027649 | Ctnnbl1 | protein_coding | 0.116940733 | 6.228980183 | 0.2067466 | 0.6553238 | 0.929250454 |
| ENSMUSG00000001674 | Ddx18 | protein_coding | 0.143553326 | 4.228747452 | 0.2067206 | 0.6553439 | 0.929250454 |
| ENSMUSG00000052387 | Trpm3 | protein_coding | -0.244831845 | 4.38765766 | 0.207266 | 0.655426 | 0.929250454 |
| ENSMUSG00000030763 | Lcmt1 | protein_coding | 0.083952174 | 7.912261747 | 0.2065464 | 0.6554789 | 0.929250454 |
| ENSMUSG00000009013 | Dynll1 | protein_coding | -0.114760214 | 10.1079042 | 0.2063424 | 0.6556371 | 0.929352193 |
| ENSMUSG00000056900 | Usp13 | protein_coding | -0.171488234 | 4.009411994 | 0.2059373 | 0.6559515 | 0.929675366 |
| ENSMUSG00000060681 | Slc9a6 | protein_coding | 0.107956518 | 5.87618612 | 0.2056041 | 0.6562104 | 0.929905741 |
| ENSMUSG00000026718 | Stam | protein_coding | -0.109253115 | 5.8683243 | 0.2054087 | 0.6563624 | 0.929905741 |
| ENSMUSG00000043702 | Pde12 | protein_coding | 0.323576724 | 2.91466877 | 0.2053284 | 0.6564347 | 0.929905741 |
| ENSMUSG00000039701 | Usp53 | protein_coding | -0.212907545 | 3.229360718 | 0.2051848 | 0.6565366 | 0.929905741 |
| ENSMUSG00000096995 | 2810029C07Rik | bidirectional_promoter | -0.181837681 | 3.791981806 | 0.2051341 | 0.6565761 | 0.929905741 |
| ENSMUSG00000067995 | Gtf2f2 | protein_coding | 0.112900431 | 5.655540052 | 0.2050611 | 0.656633 | 0.929905741 |
| ENSMUSG00000037434 | Slc30a1 | protein_coding | -0.207570223 | 3.463613346 | 0.2044515 | 0.657108 | 0.930193989 |
| ENSMUSG00000092060 | Bend4 | protein_coding | -0.205401675 | 4.913103167 | 0.2049375 | 0.6571087 | 0.930193989 |
| ENSMUSG00000037742 | Eef1a1 | protein_coding | 0.091092657 | 10.30163358 | 0.2044142 | 0.6571371 | 0.930193989 |
| ENSMUSG00000041193 | Pla2g5 | protein_coding | -0.274620559 | 2.690346538 | 0.2041497 | 0.6573436 | 0.930193989 |
| ENSMUSG00000016346 | Kcnq2 | protein_coding | 0.100563909 | 5.984644413 | 0.2041305 | 0.6573586 | 0.930193989 |
| ENSMUSG00000021660 | Btf3 | protein_coding | -0.103166128 | 6.743615765 | 0.2041081 | 0.6573761 | 0.930193989 |
| ENSMUSG00000031503 | Col4a2 | protein_coding | -0.324182563 | 2.444763669 | 0.2040236 | 0.657442 | 0.930193989 |
| ENSMUSG00000001517 | Foxm1 | protein_coding | -0.26432609 | 2.814224342 | 0.2036567 | 0.6577288 | 0.930477218 |
| ENSMUSG00000021147 | Wdr37 | protein_coding | -0.105696127 | 6.485592369 | 0.2034367 | 0.6579009 | 0.930514458 |
| ENSMUSG00000022451 | Twf1 | protein_coding | 0.287297142 | 2.529935583 | 0.2033453 | 0.6579724 | 0.930514458 |
| ENSMUSG00000094910 | D430019H16Rik | protein_coding | -0.154725219 | 5.818413957 | 0.2033509 | 0.6580147 | 0.930514458 |
| ENSMUSG00000028649 | Macf1 | protein_coding | -0.168196846 | 5.171982175 | 0.20291 | 0.658323 | 0.930636431 |
| ENSMUSG00000017418 | Arl5b | protein_coding | 0.234137137 | 3.134239237 | 0.2028614 | 0.6583514 | 0.930636431 |
| ENSMUSG00000040928 | S100pbp | protein_coding | -0.121095437 | 5.027817843 | 0.2028497 | 0.6583606 | 0.930636431 |
| ENSMUSG00000044709 | Gemin7 | protein_coding | -0.109238278 | 5.674839794 | 0.2023195 | 0.6587764 | 0.930853527 |
| ENSMUSG00000100417 | Gm28578 | lincRNA | -0.350040054 | 2.375551385 | 0.2022803 | 0.6588073 | 0.930853527 |
| ENSMUSG00000002324 | Rec8 | protein_coding | 0.253331607 | 3.110103132 | 0.2022354 | 0.6588425 | 0.930853527 |
| ENSMUSG00000029992 | Gfpt1 | protein_coding | -0.145832095 | 5.066774403 | 0.2022125 | 0.6588604 | 0.930853527 |
| ENSMUSG00000008475 | Arpc5 | protein_coding | -0.081568058 | 7.254298124 | 0.2019644 | 0.6590554 | 0.931006623 |
| ENSMUSG00000024352 | Spata24 | protein_coding | -0.153172277 | 4.421206474 | 0.2017859 | 0.6591957 | 0.931082528 |
| ENSMUSG00000038354 | Ankrd35 | protein_coding | 0.33531256 | 2.39288361 | 0.2015875 | 0.6593518 | 0.931167454 |
| ENSMUSG00000015474 | Ppt2 | protein_coding | 0.21754424 | 2.910200772 | 0.2013728 | 0.6595207 | 0.931167454 |
| ENSMUSG00000052336 | Cx3cr1 | protein_coding | 0.184348591 | 4.189887564 | 0.2010688 | 0.6597601 | 0.931167454 |
| ENSMUSG00000021846 | Peli2 | protein_coding | -0.160911214 | 3.828429506 | 0.2009589 | 0.6598467 | 0.931167454 |
| ENSMUSG00000004285 | Atp6v1f | protein_coding | 0.08713682 | 9.236162641 | 0.2008889 | 0.659902 | 0.931167454 |
| ENSMUSG00000038970 | Lmtk2 | protein_coding | 0.158531155 | 4.285647152 | 0.2006802 | 0.6600665 | 0.931167454 |
| ENSMUSG00000038252 | Ncapd2 | protein_coding | -0.391806836 | 2.557168031 | 0.2005713 | 0.6601622 | 0.931167454 |
| ENSMUSG00000025786 | Zdhhc3 | protein_coding | 0.156194775 | 5.349899634 | 0.200419 | 0.6602824 | 0.931167454 |
| ENSMUSG00000022525 | Hrasls | protein_coding | -0.113780186 | 6.23717462 | 0.200375 | 0.6603075 | 0.931167454 |

| ensembl_gene_id | external_gene_name | gene_biotype | log2 (fold change) | log2 (counts per million) | F-value | p-value | False discovery rate |
|---|---|---|---|---|---|---|---|
| ENSMUSG00000034574 | Daam1 | protein_coding | -0.149947136 | 4.86905192 | 0.200367 | 0.6603138 | 0.931167454 |
| ENSMUSG00000035849 | Krt222 | protein_coding | 0.113705766 | 5.881823538 | 0.2001051 | 0.6605207 | 0.931167454 |
| ENSMUSG00000048142 | Nat8l | protein_coding | 0.090794653 | 6.965309063 | 0.1998853 | 0.6606945 | 0.931167454 |
| ENSMUSG00000020716 | Nf1 | protein_coding | -0.135240278 | 5.142647013 | 0.1998362 | 0.6607332 | 0.931167454 |
| ENSMUSG00000025508 | Rplp2 | protein_coding | -0.09841396 | 8.84790649 | 0.1997876 | 0.6607717 | 0.931167454 |
| ENSMUSG00000043059 | Zfp513 | protein_coding | 0.360190745 | 2.252911747 | 0.1997743 | 0.6607822 | 0.931167454 |
| ENSMUSG00000021275 | Tecpr2 | protein_coding | 0.152317567 | 4.203637937 | 0.1997543 | 0.660798 | 0.931167454 |
| ENSMUSG00000037509 | Arhgef4 | protein_coding | -0.106962948 | 6.287495244 | 0.1996219 | 0.6609028 | 0.931167454 |
| ENSMUSG00000041308 | Sntb2 | protein_coding | 0.31106574 | 2.552772567 | 0.1996028 | 0.660918 | 0.931167454 |
| ENSMUSG00000085084 | 4930570G19Rik | lincRNA | -0.147211483 | 4.099744963 | 0.1995507 | 0.6609592 | 0.931167454 |
| ENSMUSG00000031657 | Heatr3 | protein_coding | -0.164257377 | 4.264892897 | 0.1992101 | 0.6612289 | 0.931167454 |
| ENSMUSG00000039018 | Mtg1 | protein_coding | 0.107373961 | 6.316391935 | 0.1990475 | 0.6613578 | 0.931167454 |
| ENSMUSG00000084786 | Ubl5 | protein_coding | -0.106512457 | 9.224991029 | 0.1988841 | 0.6614874 | 0.931167454 |
| ENSMUSG00000031786 | Drc7 | protein_coding | 0.326709596 | 3.185635984 | 0.1991722 | 0.6616097 | 0.931167454 |
| ENSMUSG00000009995 | Taz | protein_coding | 0.140871 | 4.485728644 | 0.198583 | 0.6617263 | 0.931167454 |
| ENSMUSG00000018548 | Trim37 | protein_coding | 0.084001232 | 6.692543958 | 0.1984758 | 0.6618115 | 0.931167454 |
| ENSMUSG00000042787 | Exog | protein_coding | 0.16000863 | 5.026639843 | 0.1984752 | 0.6618119 | 0.931167454 |
| ENSMUSG00000024079 | Eif2ak2 | protein_coding | -0.561638504 | 1.681537871 | 0.1984618 | 0.6618322 | 0.931167454 |
| ENSMUSG00000059920 | 4930453N24Rik | protein_coding | -0.142415289 | 4.780918003 | 0.1983265 | 0.6619301 | 0.931167454 |
| ENSMUSG00000002496 | Tsc2 | protein_coding | 0.155713951 | 4.776192968 | 0.1982981 | 0.6619526 | 0.931167454 |
| ENSMUSG00000032172 | Olfm2 | protein_coding | 0.090937631 | 6.814013233 | 0.1982874 | 0.6619611 | 0.931167454 |
| ENSMUSG00000012609 | Ttll5 | protein_coding | -0.158358155 | 5.153133943 | 0.1981256 | 0.6620898 | 0.931167454 |
| ENSMUSG00000022827 | Rabl3 | protein_coding | -0.166960403 | 4.725457774 | 0.1981195 | 0.6620946 | 0.931167454 |
| ENSMUSG00000072235 | Tuba1a | protein_coding | 0.107290308 | 9.916995326 | 0.1979756 | 0.662209 | 0.931167454 |
| ENSMUSG00000022781 | Pak2 | protein_coding | -0.13109291 | 5.053092836 | 0.1979647 | 0.6622177 | 0.931167454 |
| ENSMUSG00000024381 | Bin1 | protein_coding | -0.108062555 | 6.261509006 | 0.1977727 | 0.6623705 | 0.931167454 |
| ENSMUSG00000031934 | Panx1 | protein_coding | 0.191220814 | 3.856228081 | 0.1977178 | 0.6624142 | 0.931167454 |
| ENSMUSG00000020212 | Mdm1 | protein_coding | 0.190232739 | 3.198376845 | 0.1976604 | 0.6624599 | 0.931167454 |
| ENSMUSG00000053141 | Ptprt | protein_coding | -0.160908221 | 4.442839227 | 0.1975487 | 0.6625489 | 0.931170831 |
| ENSMUSG00000018821 | Avpi1 | protein_coding | -0.09296058 | 7.044228334 | 0.1971312 | 0.6628815 | 0.931199253 |
| ENSMUSG00000026463 | Atp2b4 | protein_coding | -0.104021222 | 6.025280533 | 0.1971178 | 0.6628922 | 0.931199253 |
| ENSMUSG00000067722 | BC003965 | protein_coding | 0.089342175 | 6.375988644 | 0.1971162 | 0.6628936 | 0.931199253 |
| ENSMUSG00000047193 | Dync2h1 | protein_coding | 0.13555522 | 5.172442651 | 0.1970887 | 0.6629155 | 0.931199253 |
| ENSMUSG00000032059 | Alg9 | protein_coding | 0.166168334 | 4.222902375 | 0.1964268 | 0.6634439 | 0.931616319 |
| ENSMUSG00000051223 | Bzw1 | protein_coding | 0.088531671 | 7.448089155 | 0.1964128 | 0.6634551 | 0.931616319 |
| ENSMUSG00000036155 | Mgat5 | protein_coding | -0.18977493 | 3.778046629 | 0.1963913 | 0.6634723 | 0.931616319 |
| ENSMUSG00000031375 | Bgn | protein_coding | 0.244336301 | 3.133467997 | 0.1962688 | 0.6635703 | 0.931632202 |
| ENSMUSG00000012483 | Rpa3 | protein_coding | -0.168697453 | 4.794647199 | 0.1961205 | 0.6636888 | 0.931676991 |
| ENSMUSG00000025777 | Gdap1 | protein_coding | -0.088335686 | 6.892165658 | 0.1959953 | 0.663789 | 0.931695998 |
| ENSMUSG00000035835 | Plppr3 | protein_coding | 0.12458868 | 5.406949151 | 0.1957603 | 0.6639771 | 0.93178503 |
| ENSMUSG00000086649 | Gm15286 | processed_transcript | -0.251340226 | 2.672795255 | 0.1956996 | 0.6640257 | 0.93178503 |
| ENSMUSG00000029480 | Dhx37 | protein_coding | 0.192866685 | 3.670939962 | 0.195099 | 0.6645073 | 0.932242013 |
| ENSMUSG00000071866 | Ppia | protein_coding | 0.081733857 | 7.158662897 | 0.1950306 | 0.6645621 | 0.932242013 |
| ENSMUSG00000031988 | Vps26b | protein_coding | 0.122503613 | 5.71486949 | 0.1949531 | 0.6646244 | 0.932242013 |
| ENSMUSG00000032965 | Ift57 | protein_coding | -0.241232759 | 3.069249186 | 0.1948218 | 0.6647298 | 0.932242013 |
| ENSMUSG00000021221 | Dpf3 | protein_coding | 0.211555077 | 3.478082958 | 0.1947018 | 0.6648262 | 0.932242013 |
| ENSMUSG00000019785 | Clvs2 | protein_coding | -0.111155982 | 5.983015569 | 0.1946454 | 0.6648716 | 0.932242013 |
| ENSMUSG00000030259 | Rassf8 | protein_coding | -0.180379172 | 3.946340098 | 0.1943054 | 0.6651449 | 0.932410614 |
| ENSMUSG00000025184 | R3hcc1l | protein_coding | 0.179738953 | 3.570688041 | 0.1940839 | 0.6653232 | 0.932410614 |
| ENSMUSG00000033373 | Fntb | protein_coding | 0.118420063 | 5.852446831 | 0.1940471 | 0.6653527 | 0.932410614 |
| ENSMUSG00000022812 | Gsk3b | protein_coding | -0.158651059 | 9.531558881 | 0.1950328 | 0.6653724 | 0.932410614 |
| ENSMUSG00000026812 | Tsc1 | protein_coding | -0.178282444 | 3.868228359 | 0.1939569 | 0.6654254 | 0.932410614 |
| ENSMUSG00000020107 | Anapc16 | protein_coding | 0.078484323 | 7.218092901 | 0.1936413 | 0.6656797 | 0.932543348 |
| ENSMUSG00000019975 | Ikbip | protein_coding | 0.185419925 | 3.845814942 | 0.1935439 | 0.6657582 | 0.932543348 |
| ENSMUSG00000055839 | Elob | protein_coding | 0.090884601 | 9.024725631 | 0.1934079 | 0.6658679 | 0.932543348 |
| ENSMUSG00000049511 | Htr1b | protein_coding | -0.212639608 | 3.790007403 | 0.193341 | 0.6659219 | 0.932543348 |
| ENSMUSG00000015165 | Hnrnpl | protein_coding | -0.094661052 | 6.560718103 | 0.193258 | 0.6659889 | 0.932543348 |
| ENSMUSG00000042453 | Reln | protein_coding | 0.183577169 | 4.225070867 | 0.1931941 | 0.6660404 | 0.932543348 |
| ENSMUSG00000031508 | Ankrd10 | protein_coding | 0.170878909 | 4.519871206 | 0.1930308 | 0.6661723 | 0.932606536 |
| ENSMUSG00000039361 | Picalm | protein_coding | -0.101042103 | 6.279522814 | 0.1928152 | 0.6663464 | 0.932655715 |
| ENSMUSG00000038736 | Nudcd1 | protein_coding | 0.166246041 | 4.382402137 | 0.1927726 | 0.6663809 | 0.932655715 |
| ENSMUSG00000035828 | Pim3 | protein_coding | 0.135385448 | 5.045713111 | 0.192598 | 0.666522 | 0.932731841 |
| ENSMUSG00000043432 | Leng9 | protein_coding | -0.147936088 | 4.373271908 | 0.1922528 | 0.6668014 | 0.932806167 |
| ENSMUSG00000038618 | Rassf7 | protein_coding | -0.232642762 | 3.373185711 | 0.1921265 | 0.6669036 | 0.932806167 |
| ENSMUSG00000068101 | Cenpm | protein_coding | 0.306199703 | 2.727712796 | 0.1920837 | 0.6669383 | 0.932806167 |
| ENSMUSG00000042604 | Kcna4 | protein_coding | -0.195178521 | 3.988612793 | 0.1920191 | 0.6669906 | 0.932806167 |
| ENSMUSG00000053166 | Cdh22 | protein_coding | -0.275499426 | 2.911345034 | 0.1919966 | 0.6670089 | 0.932806167 |
| ENSMUSG00000028437 | Ubap1 | protein_coding | 0.11736614 | 5.993346689 | 0.191769 | 0.6671934 | 0.932942856 |
| ENSMUSG00000059326 | Csf2ra | protein_coding | -0.258877275 | 2.661422324 | 0.1916546 | 0.6672862 | 0.932951241 |
| ENSMUSG00000020034 | Tcp11l2 | protein_coding | -0.176256803 | 5.714036698 | 0.1919693 | 0.6675017 | 0.933131239 |
| ENSMUSG00000026426 | Arl8a | protein_coding | 0.090593409 | 6.998271907 | 0.1911597 | 0.6676879 | 0.933270209 |
| ENSMUSG00000055692 | Tmem191c | protein_coding | -0.118394023 | 5.673885743 | 0.1910103 | 0.6678093 | 0.933318574 |
| ENSMUSG00000029538 | Srsf9 | protein_coding | -0.134075411 | 5.304814752 | 0.190663 | 0.6680917 | 0.933591976 |
| ENSMUSG00000037573 | Tob1 | protein_coding | -0.111992582 | 5.356894332 | 0.1905313 | 0.6681989 | 0.933593618 |

| ensembl_gene_id | external_gene_name | gene_biotype | log2 (fold change) | log2 (counts per million) | F-value | p-value | False discovery rate |
|---|---|---|---|---|---|---|---|
| ENSMUSG00000027357 | Crls1 | protein_coding | 0.117750728 | 4.835741882 | 0.1902299 | 0.6684443 | 0.933593618 |
| ENSMUSG00000061650 | Med9 | protein_coding | -0.164162492 | 4.35018074 | 0.1901109 | 0.6685413 | 0.933593618 |
| ENSMUSG00000033998 | Kcnk1 | protein_coding | 0.208820225 | 3.67952757 | 0.1898675 | 0.6687398 | 0.933593618 |
| ENSMUSG00000010505 | Myt1 | protein_coding | -0.273250413 | 2.486515823 | 0.1897501 | 0.6688356 | 0.933593618 |
| ENSMUSG00000025169 | Ogfod3 | protein_coding | -0.169167162 | 4.034516583 | 0.1896247 | 0.6689379 | 0.933593618 |
| ENSMUSG00000042632 | Pla2g6 | protein_coding | 0.120607704 | 5.294236621 | 0.189542 | 0.6690054 | 0.933593618 |
| ENSMUSG00000055053 | Nfic | protein_coding | -0.137534034 | 5.752961924 | 0.1894432 | 0.6690955 | 0.933593618 |
| ENSMUSG00000019080 | Mfsd3 | protein_coding | 0.174152417 | 4.22252676 | 0.1893829 | 0.6691353 | 0.933593618 |
| ENSMUSG00000022544 | Eef2kmt | protein_coding | 0.185675943 | 3.838473285 | 0.1893212 | 0.6691858 | 0.933593618 |
| ENSMUSG00000048720 | Tbc1d12 | protein_coding | -0.199605916 | 3.270310205 | 0.189299 | 0.6692039 | 0.933593618 |
| ENSMUSG00000052137 | Rbm12b2 | protein_coding | -0.228039508 | 3.454937519 | 0.1891969 | 0.6692874 | 0.933593618 |
| ENSMUSG00000032827 | Ppp1r9a | protein_coding | 0.112446898 | 7.15037439 | 0.1891868 | 0.669305 | 0.933593618 |
| ENSMUSG00000052962 | Mrpl35 | protein_coding | 0.075011818 | 7.399186012 | 0.1891502 | 0.6693255 | 0.933593618 |
| ENSMUSG00000063972 | Nr6a1 | protein_coding | 0.23276715 | 3.21914049 | 0.189065 | 0.6693952 | 0.933593618 |
| ENSMUSG00000038733 | Wdr26 | protein_coding | -0.12753436 | 6.151534961 | 0.1889317 | 0.6695136 | 0.933637634 |
| ENSMUSG00000038055 | Dexi | protein_coding | -0.158244493 | 4.185808353 | 0.1886155 | 0.669763 | 0.933864331 |
| ENSMUSG00000048416 | Mlf1 | protein_coding | -0.175973077 | 3.661680414 | 0.1884971 | 0.66986 | 0.933878485 |
| ENSMUSG00000020346 | Mgat1 | protein_coding | 0.120627871 | 5.171762214 | 0.1882585 | 0.6700556 | 0.933932049 |
| ENSMUSG00000000627 | Sema4f | protein_coding | 0.164524786 | 4.304261615 | 0.1880965 | 0.6701884 | 0.933932049 |
| ENSMUSG00000022550 | Adck5 | protein_coding | -0.18982306 | 4.420545386 | 0.1880763 | 0.6702143 | 0.933932049 |
| ENSMUSG00000042810 | Krba1 | protein_coding | -0.158610246 | 3.77363301 | 0.1879677 | 0.6702941 | 0.933932049 |
| ENSMUSG00000030516 | Tjp1 | protein_coding | -0.162936436 | 4.111698494 | 0.1879207 | 0.6703327 | 0.933932049 |
| ENSMUSG00000031065 | Cdk16 | protein_coding | -0.111261087 | 6.001277039 | 0.1874752 | 0.6706986 | 0.933969179 |
| ENSMUSG00000020982 | Nemf | protein_coding | 0.09079431 | 6.868900378 | 0.1873689 | 0.670786 | 0.933969179 |
| ENSMUSG00000112117 | RMST_1 | lincRNA | -0.176392341 | 3.455362308 | 0.1871763 | 0.6709444 | 0.933969179 |
| ENSMUSG00000045757 | Zfp764 | protein_coding | -0.461841263 | 2.131660146 | 0.1871657 | 0.6709624 | 0.933969179 |
| ENSMUSG00000021180 | Rps6ka5 | protein_coding | -0.208209697 | 4.180148764 | 0.1871737 | 0.6709703 | 0.933969179 |
| ENSMUSG00000066647 | Gm5113 | protein_coding | 0.119500572 | 4.778625466 | 0.1871344 | 0.6709789 | 0.933969179 |
| ENSMUSG00000040964 | Arhgef10l | protein_coding | -0.17213072 | 3.819511373 | 0.1870732 | 0.6710292 | 0.933969179 |
| ENSMUSG00000036402 | Gng12 | protein_coding | -0.140989458 | 4.976671488 | 0.1869095 | 0.6711639 | 0.933969179 |
| ENSMUSG00000017843 | Ppp2r5c | protein_coding | 0.089110903 | 7.854451144 | 0.1866193 | 0.671403 | 0.933969179 |
| ENSMUSG00000001334 | Fndc5 | protein_coding | -0.135577155 | 5.763481412 | 0.1865983 | 0.6714296 | 0.933969179 |
| ENSMUSG00000026414 | Tnnt2 | protein_coding | 0.179199861 | 3.784445905 | 0.1865553 | 0.6714558 | 0.933969179 |
| ENSMUSG00000052727 | Map1b | protein_coding | -0.18886171 | 9.563344979 | 0.1883921 | 0.6715378 | 0.933969179 |
| ENSMUSG00000054034 | Tceal5 | protein_coding | 0.088303384 | 6.457638063 | 0.1861357 | 0.6718019 | 0.933969179 |
| ENSMUSG00000010663 | Fads1 | protein_coding | 0.104449571 | 5.969591678 | 0.1861348 | 0.6718026 | 0.933969179 |
| ENSMUSG00000028004 | Npy2r | protein_coding | 0.242583687 | 4.064201765 | 0.1865315 | 0.6718354 | 0.933969179 |
| ENSMUSG00000030105 | Arl8b | protein_coding | -0.089945903 | 8.049807547 | 0.1860518 | 0.6718712 | 0.933969179 |
| ENSMUSG00000014232 | Cluap1 | protein_coding | -0.117377146 | 4.900014006 | 0.1860459 | 0.671876 | 0.933969179 |
| ENSMUSG00000040620 | Dhx33 | protein_coding | -0.235640091 | 2.79761533 | 0.1859893 | 0.6719228 | 0.933969179 |
| ENSMUSG00000030990 | Pgap2 | protein_coding | 0.126676037 | 4.838311353 | 0.185645 | 0.6722073 | 0.934077274 |
| ENSMUSG00000038902 | Pogz | protein_coding | -0.137650356 | 4.852381412 | 0.1856427 | 0.6722092 | 0.934077274 |
| ENSMUSG00000006818 | Sod2 | protein_coding | 0.086305745 | 7.601455699 | 0.1855798 | 0.6722611 | 0.934077274 |
| ENSMUSG00000037408 | Cnnm4 | protein_coding | 0.154807472 | 5.394827322 | 0.1853763 | 0.6724388 | 0.934203375 |
| ENSMUSG00000020744 | Slc25a19 | protein_coding | -0.108656953 | 5.895115586 | 0.1847527 | 0.672946 | 0.934787319 |
| ENSMUSG00000035834 | Polr3g | protein_coding | 0.411549255 | 1.979969554 | 0.1844415 | 0.6732042 | 0.935025123 |
| ENSMUSG00000022601 | Zbtb11 | protein_coding | 0.182370052 | 3.69462832 | 0.1842192 | 0.6733887 | 0.935129694 |
| ENSMUSG00000021650 | Ptcd2 | protein_coding | -0.09945419 | 6.435827119 | 0.1841414 | 0.6734534 | 0.935129694 |
| ENSMUSG00000021368 | Tbc1d7 | protein_coding | 0.090877444 | 6.389363671 | 0.1839275 | 0.6736311 | 0.935255739 |
| ENSMUSG00000029650 | Slc46a3 | protein_coding | 0.245363326 | 2.794712814 | 0.183668 | 0.6738469 | 0.935434548 |
| ENSMUSG00000027833 | Shox2 | protein_coding | -0.522392585 | 4.859036932 | 0.1906065 | 0.6741215 | 0.935532099 |
| ENSMUSG00000039512 | Uhrf1bp1 | protein_coding | 0.199354985 | 3.174498448 | 0.1832056 | 0.674232 | 0.935532099 |
| ENSMUSG00000030922 | Lyrm1 | protein_coding | -0.095869737 | 5.692454423 | 0.1830784 | 0.674338 | 0.935532099 |
| ENSMUSG00000036181 | Hist1h1c | protein_coding | 0.167829061 | 3.712771943 | 0.1829707 | 0.6744278 | 0.935532099 |
| ENSMUSG00000004110 | Cacna1e | protein_coding | 0.150208474 | 4.923582832 | 0.1829346 | 0.6744579 | 0.935532099 |
| ENSMUSG00000043192 | Gm1840 | processed_pseudoge | 0.381258861 | 2.559594542 | 0.1828395 | 0.6745464 | 0.935532099 |
| ENSMUSG00000059824 | Dbp | protein_coding | 0.164603466 | 3.487219614 | 0.1827293 | 0.6746292 | 0.935532099 |
| ENSMUSG00000029625 | Cpsf4 | protein_coding | 0.238933893 | 3.032354332 | 0.1826423 | 0.6747018 | 0.935532099 |
| ENSMUSG00000038299 | Wdr36 | protein_coding | 0.157837797 | 4.820468584 | 0.1824862 | 0.6748322 | 0.935532099 |
| ENSMUSG00000027422 | Rrbp1 | protein_coding | -0.155304865 | 3.931150346 | 0.1823949 | 0.6749084 | 0.935532099 |
| ENSMUSG00000037972 | Snn | protein_coding | 0.111098331 | 5.734400238 | 0.1823792 | 0.6749215 | 0.935532099 |
| ENSMUSG00000008690 | Ncaph2 | protein_coding | -0.10940668 | 7.207697295 | 0.1823427 | 0.6749612 | 0.935532099 |
| ENSMUSG00000014850 | Msh3 | protein_coding | -0.168607346 | 3.962347534 | 0.1819255 | 0.6753009 | 0.935882303 |
| ENSMUSG00000020681 | Ace | protein_coding | 0.22762693 | 3.3613673 | 0.1817347 | 0.6754606 | 0.935912385 |
| ENSMUSG00000017548 | Suz12 | protein_coding | -0.13295764 | 4.468041776 | 0.1816791 | 0.6755072 | 0.935912385 |
| ENSMUSG00000019845 | Tube1 | protein_coding | -0.327584785 | 2.32141918 | 0.1815877 | 0.6755837 | 0.935912385 |
| ENSMUSG00000091478 | Gm10039 | processed_pseudoge | 0.204321204 | 2.898952936 | 0.1813499 | 0.6757831 | 0.936053061 |
| ENSMUSG00000016382 | Pls3 | protein_coding | 0.092879421 | 6.281365613 | 0.1812315 | 0.6758823 | 0.936053061 |
| ENSMUSG00000061533 | Cep128 | protein_coding | -0.250079236 | 3.424717076 | 0.1810077 | 0.6760792 | 0.936053061 |
| ENSMUSG00000001175 | Calm1 | protein_coding | -0.086606247 | 12.85395451 | 0.1808868 | 0.6761716 | 0.936053061 |
| ENSMUSG00000023170 | Gps2 | protein_coding | 0.122020283 | 4.89222568 | 0.180836 | 0.6762142 | 0.936053061 |
| ENSMUSG00000052605 | Isoc2b | protein_coding | -0.154208626 | 4.50164993 | 0.1807979 | 0.6762462 | 0.936053061 |
| ENSMUSG00000002279 | Lmf1 | protein_coding | 0.163148813 | 4.875749006 | 0.1807076 | 0.6763312 | 0.936053061 |
| ENSMUSG00000031561 | Tenm3 | protein_coding | -0.197388676 | 3.863586045 | 0.1806366 | 0.6763817 | 0.936053061 |

| ensembl_gene_id | external_gene_name | gene_biotype | log2 (fold change) | log2 (counts per million) | F-value | p-value | False discovery rate |
|---|---|---|---|---|---|---|---|
| ENSMUSG00000026309 | Ilkap | protein_coding | 0.123365987 | 5.086444162 | 0.1803995 | 0.676581 | 0.936208468 |
| ENSMUSG00000007594 | Hapln4 | protein_coding | -0.131178102 | 5.143696213 | 0.1801297 | 0.6768081 | 0.936245585 |
| ENSMUSG00000024158 | Hagh | protein_coding | 0.081472531 | 8.304436477 | 0.1800997 | 0.6768333 | 0.936245585 |
| ENSMUSG00000038422 | Hdhd3 | protein_coding | 0.12850619 | 5.09188441 | 0.1800572 | 0.6768691 | 0.936245585 |
| ENSMUSG00000020014 | Cfap54 | protein_coding | -0.262722946 | 2.646951021 | 0.1798644 | 0.6770314 | 0.93634971 |
| ENSMUSG00000027384 | Ndufaf5 | protein_coding | 0.083476005 | 7.220808295 | 0.1794746 | 0.6773601 | 0.936578107 |
| ENSMUSG00000001289 | Pfdn5 | protein_coding | -0.10532644 | 9.841359291 | 0.1792092 | 0.677584 | 0.936578107 |
| ENSMUSG00000006715 | Gmnn | protein_coding | 0.187353967 | 4.321437448 | 0.1791734 | 0.6776233 | 0.936578107 |
| ENSMUSG00000002109 | Ddb2 | protein_coding | 0.261933472 | 2.829398467 | 0.1791528 | 0.6776317 | 0.936578107 |
| ENSMUSG00000019803 | Nr2e1 | protein_coding | -0.316867452 | 2.88878223 | 0.1791166 | 0.6776713 | 0.936578107 |
| ENSMUSG00000069744 | Psmb3 | protein_coding | 0.088111049 | 7.452217863 | 0.1790331 | 0.6777328 | 0.936578107 |
| ENSMUSG00000034467 | Dynlrb2 | protein_coding | -0.17684648 | 4.88891651 | 0.178938 | 0.6778461 | 0.936578107 |
| ENSMUSG00000032525 | Nktr | protein_coding | -0.188685844 | 4.031667074 | 0.1786035 | 0.6780959 | 0.936578107 |
| ENSMUSG00000026234 | Ncl | protein_coding | -0.084001266 | 8.209735329 | 0.1785875 | 0.6781095 | 0.936578107 |
| ENSMUSG00000025571 | Tnrc6c | protein_coding | 0.10401302 | 5.659112533 | 0.1784617 | 0.678216 | 0.936578107 |
| ENSMUSG00000016664 | Pacsin2 | protein_coding | -0.132563303 | 4.843475171 | 0.1783953 | 0.6782721 | 0.936578107 |
| ENSMUSG00000026991 | Pkp4 | protein_coding | 0.075754664 | 6.884406364 | 0.1782937 | 0.6783582 | 0.936578107 |
| ENSMUSG00000048486 | Fitm2 | protein_coding | 0.145463461 | 4.319730697 | 0.1779712 | 0.6786315 | 0.936578107 |
| ENSMUSG00000037152 | Ndufc1 | protein_coding | -0.097982267 | 9.412866579 | 0.1778257 | 0.6787548 | 0.936578107 |
| ENSMUSG00000047246 | Hist1h2be | protein_coding | -0.241500619 | 2.832654143 | 0.177705 | 0.6788572 | 0.936578107 |
| ENSMUSG00000027184 | Caprin1 | protein_coding | -0.093357613 | 6.502732555 | 0.1776729 | 0.6788845 | 0.936578107 |
| ENSMUSG00000003437 | Paf1 | protein_coding | 0.170454195 | 5.326865428 | 0.1779312 | 0.6788962 | 0.936578107 |
| ENSMUSG00000071072 | Ptges3 | protein_coding | 0.076093211 | 8.537608677 | 0.1776209 | 0.6789286 | 0.936578107 |
| ENSMUSG00000034430 | Zxdc | protein_coding | 0.158574952 | 3.944649992 | 0.1776027 | 0.6789441 | 0.936578107 |
| ENSMUSG00000020289 | Nprl3 | protein_coding | -0.114468392 | 5.770974429 | 0.1776019 | 0.6789447 | 0.936578107 |
| ENSMUSG00000027424 | Mgme1 | protein_coding | 0.14575532 | 3.799947461 | 0.1775066 | 0.6790257 | 0.936578107 |
| ENSMUSG00000030225 | Dera | protein_coding | -0.203372922 | 3.302145592 | 0.1769631 | 0.6794875 | 0.936601164 |
| ENSMUSG00000029173 | Sepsecs | protein_coding | 0.263746396 | 3.513726781 | 0.1770034 | 0.6796833 | 0.936601164 |
| ENSMUSG00000019726 | Lyst | protein_coding | -0.161750399 | 3.96989673 | 0.1766443 | 0.6797589 | 0.936601164 |
| ENSMUSG00000038612 | Mcl1 | protein_coding | 0.157381058 | 4.875636283 | 0.176607 | 0.6797907 | 0.936601164 |
| ENSMUSG00000031986 | Sprtn | protein_coding | 0.315214313 | 2.526641802 | 0.1765993 | 0.6797972 | 0.936601164 |
| ENSMUSG00000010025 | Aldh3a2 | protein_coding | 0.153592358 | 5.385840034 | 0.1765508 | 0.6798476 | 0.936601164 |
| ENSMUSG00000078307 | AI593442 | protein_coding | -0.129832831 | 8.096051681 | 0.1767994 | 0.6799983 | 0.936601164 |
| ENSMUSG00000032575 | Manf | protein_coding | 0.100956375 | 6.02686512 | 0.1760929 | 0.6802289 | 0.936601164 |
| ENSMUSG00000019054 | Fis1 | protein_coding | -0.085683934 | 10.28494368 | 0.1760249 | 0.6802869 | 0.936601164 |
| ENSMUSG00000016833 | Mrps18c | protein_coding | -0.113898138 | 7.215217141 | 0.1759651 | 0.6803706 | 0.936601164 |
| ENSMUSG00000020097 | Sgpl1 | protein_coding | 0.219782772 | 3.309907319 | 0.1757489 | 0.6805225 | 0.936601164 |
| ENSMUSG00000026339 | Ccdc93 | protein_coding | 0.230952189 | 2.93749674 | 0.1755425 | 0.6806989 | 0.936601164 |
| ENSMUSG00000037012 | Hk1 | protein_coding | 0.159120407 | 7.295175808 | 0.176428 | 0.6808167 | 0.936601164 |
| ENSMUSG00000018589 | Glra2 | protein_coding | 0.136209611 | 5.258017311 | 0.1751856 | 0.6810041 | 0.936601164 |
| ENSMUSG00000047777 | Phf13 | protein_coding | 0.170284364 | 3.441621594 | 0.1751838 | 0.6810056 | 0.936601164 |
| ENSMUSG00000024091 | Vapa | protein_coding | -0.072618198 | 8.549084186 | 0.1751657 | 0.6810211 | 0.936601164 |
| ENSMUSG00000021715 | Cwc27 | protein_coding | 0.185287325 | 3.686163553 | 0.1749919 | 0.6811699 | 0.936601164 |
| ENSMUSG00000063052 | Lrrc40 | protein_coding | -0.127683456 | 5.179749987 | 0.1748579 | 0.6812847 | 0.936601164 |
| ENSMUSG00000007041 | Clic1 | protein_coding | -0.146967584 | 4.372558021 | 0.1748345 | 0.6813048 | 0.936601164 |
| ENSMUSG00000020696 | Rffl | protein_coding | 0.222591223 | 3.31766545 | 0.1745422 | 0.6815553 | 0.936601164 |
| ENSMUSG00000024789 | Jak2 | protein_coding | -0.159240234 | 4.004079368 | 0.1745092 | 0.6815836 | 0.936601164 |
| ENSMUSG00000046093 | Hpcal4 | protein_coding | 0.081898049 | 8.117955572 | 0.1742973 | 0.6817654 | 0.936601164 |
| ENSMUSG00000030201 | Lrp6 | protein_coding | 0.168947677 | 3.641438725 | 0.1740672 | 0.681963 | 0.936601164 |
| ENSMUSG00000037251 | Pomk | protein_coding | 0.110188477 | 5.944052646 | 0.1740168 | 0.6820063 | 0.936601164 |
| ENSMUSG00000059456 | Ptk2b | protein_coding | -0.092615115 | 7.305700514 | 0.173849 | 0.6821505 | 0.936601164 |
| ENSMUSG00000031563 | Wwc2 | protein_coding | 0.147730377 | 4.512205655 | 0.173813 | 0.6821814 | 0.936601164 |
| ENSMUSG00000045664 | Cdc42ep2 | protein_coding | -0.128541133 | 5.455680769 | 0.1737917 | 0.6821998 | 0.936601164 |
| ENSMUSG00000026643 | Nmt2 | protein_coding | -0.108042803 | 8.632761936 | 0.1737795 | 0.6822192 | 0.936601164 |
| ENSMUSG00000001082 | Mfsd10 | protein_coding | 0.219534901 | 3.679602123 | 0.1737569 | 0.6822386 | 0.936601164 |
| ENSMUSG00000057778 | Cyb5d2 | protein_coding | -0.152136763 | 4.542652472 | 0.1737347 | 0.6822487 | 0.936601164 |
| ENSMUSG00000032564 | Cpne4 | protein_coding | -0.12194332 | 5.771340579 | 0.1736108 | 0.6823553 | 0.936601164 |
| ENSMUSG00000006651 | Aplp1 | protein_coding | -0.109974581 | 9.291884173 | 0.173429 | 0.6825207 | 0.936601164 |
| ENSMUSG00000111471 | AC122463.1 | antisense_RNA | -0.114047508 | 5.102127644 | 0.1733257 | 0.6826007 | 0.936601164 |
| ENSMUSG00000031808 | Slc27a1 | protein_coding | -0.139269029 | 5.635908775 | 0.1731357 | 0.6827734 | 0.936601164 |
| ENSMUSG00000031107 | Rbmx2 | protein_coding | 0.124064586 | 4.255340661 | 0.1730051 | 0.682877 | 0.936601164 |
| ENSMUSG00000050323 | Ndufaf6 | protein_coding | 0.258692452 | 2.951008596 | 0.1729735 | 0.6829042 | 0.936601164 |
| ENSMUSG00000055078 | Gabra5 | protein_coding | 0.208850902 | 3.08861533 | 0.1729183 | 0.6829518 | 0.936601164 |
| ENSMUSG00000049946 | BC030500 | protein_coding | 0.139229326 | 5.154877365 | 0.1727825 | 0.6830689 | 0.936601164 |
| ENSMUSG00000052146 | Rps10 | protein_coding | 0.112267395 | 5.612174742 | 0.1727805 | 0.6830707 | 0.936601164 |
| ENSMUSG00000022664 | Slc35a5 | protein_coding | -0.108289747 | 5.115606589 | 0.172579 | 0.6832447 | 0.936601164 |
| ENSMUSG00000025925 | Terf1 | protein_coding | -0.167701148 | 3.790794773 | 0.1724695 | 0.6833392 | 0.936601164 |
| ENSMUSG00000052920 | Prkg1 | protein_coding | -0.203564924 | 3.423851385 | 0.1724228 | 0.6833795 | 0.936601164 |
| ENSMUSG00000029348 | Asphd2 | protein_coding | -0.108641197 | 5.911058479 | 0.1724053 | 0.6833946 | 0.936601164 |
| ENSMUSG00000024927 | Rela | protein_coding | 0.133462757 | 5.16523938 | 0.1723129 | 0.6834745 | 0.936601164 |
| ENSMUSG00000074088 | Snrnp40 | protein_coding | 0.134197231 | 5.159838166 | 0.1722846 | 0.6834989 | 0.936601164 |
| ENSMUSG00000030103 | Bhlhe40 | protein_coding | -0.112781244 | 6.305069919 | 0.1721738 | 0.6835947 | 0.936601164 |
| ENSMUSG00000032425 | Zfp949 | protein_coding | 0.131349679 | 5.079971543 | 0.1720084 | 0.6837377 | 0.936601164 |
| ENSMUSG00000033965 | Slc16a2 | protein_coding | 0.220114382 | 3.931101434 | 0.1720946 | 0.68377 | 0.936601164 |

| ensembl_gene_id | external_gene_name | gene_biotype | log2 (fold change) | log2 (counts per million) | F-value | p-value | False discovery rate |
|---|---|---|---|---|---|---|---|
| ENSMUSG00000031168 | Ebp | protein_coding | -0.100019769 | 5.573726174 | 0.1718968 | 0.6838342 | 0.936601164 |
| ENSMUSG00000056367 | Actr3b | protein_coding | -0.093616632 | 6.426638227 | 0.1718132 | 0.6839066 | 0.936601164 |
| ENSMUSG00000003299 | Mrpl4 | protein_coding | -0.080781761 | 7.4879385 | 0.1716594 | 0.6840398 | 0.936601164 |
| ENSMUSG00000040543 | Pitpnm3 | protein_coding | -0.168675387 | 3.960726786 | 0.1715554 | 0.6841299 | 0.936601164 |
| ENSMUSG00000067367 | Lyar | protein_coding | -0.204171041 | 3.281575278 | 0.17148 | 0.6841953 | 0.936601164 |
| ENSMUSG00000015126 | Tsr3 | protein_coding | -0.129141766 | 4.581790649 | 0.1712293 | 0.6844126 | 0.936601164 |
| ENSMUSG00000041736 | Tspo | protein_coding | -0.181066071 | 3.492472395 | 0.1712215 | 0.6844194 | 0.936601164 |
| ENSMUSG00000053181 | A830005F24Rik | protein_coding | -0.244608366 | 3.006694887 | 0.1711136 | 0.684513 | 0.936601164 |
| ENSMUSG00000028789 | Azin2 | protein_coding | 0.132143404 | 4.471869449 | 0.170965 | 0.684642 | 0.936601164 |
| ENSMUSG00000034321 | Exosc1 | protein_coding | -0.102410163 | 6.37568515 | 0.170924 | 0.6846775 | 0.936601164 |
| ENSMUSG00000028796 | Phc2 | protein_coding | 0.108496341 | 5.428602898 | 0.1708084 | 0.684778 | 0.936601164 |
| ENSMUSG00000031556 | Tm2d2 | protein_coding | 0.087729268 | 7.927169006 | 0.1707843 | 0.6847989 | 0.936601164 |
| ENSMUSG00000063316 | Rpl27 | protein_coding | -0.153699573 | 3.778532114 | 0.1706822 | 0.6848876 | 0.936601164 |
| ENSMUSG00000038135 | Crygn | protein_coding | -0.2323718 | 3.045605241 | 0.1706596 | 0.6849072 | 0.936601164 |
| ENSMUSG00000019818 | Cd164 | protein_coding | -0.098390869 | 6.237024336 | 0.1706222 | 0.6849398 | 0.936601164 |
| ENSMUSG00000068748 | Ptprz1 | protein_coding | -0.175379624 | 5.973749057 | 0.1713407 | 0.6850713 | 0.936601164 |
| ENSMUSG00000038760 | Trhr | protein_coding | -0.364838383 | 2.638988981 | 0.1704782 | 0.6850738 | 0.936601164 |
| ENSMUSG00000006676 | Usp19 | protein_coding | 0.141595417 | 3.875609456 | 0.1704616 | 0.6850794 | 0.936601164 |
| ENSMUSG00000037151 | Lrrc20 | protein_coding | -0.138659812 | 4.799613182 | 0.1703083 | 0.6852128 | 0.936601164 |
| ENSMUSG00000038244 | Mical2 | protein_coding | 0.202464497 | 3.840897518 | 0.1701376 | 0.6853701 | 0.936601164 |
| ENSMUSG00000082536 | Gm13456 | processed_pseudoge | 0.287576692 | 2.286606122 | 0.1701129 | 0.6853828 | 0.936601164 |
| ENSMUSG00000061859 | Patj | protein_coding | 0.20339097 | 5.584489827 | 0.1708514 | 0.6855986 | 0.936601164 |
| ENSMUSG00000032497 | Lrrfip2 | protein_coding | 0.122976504 | 4.629103 | 0.16973 | 0.6857165 | 0.936601164 |
| ENSMUSG00000028986 | Klhl7 | protein_coding | 0.096227236 | 6.381126515 | 0.1697227 | 0.6857228 | 0.936601164 |
| ENSMUSG00000039488 | Cntn5 | protein_coding | -0.216703237 | 3.316897843 | 0.1696852 | 0.6857555 | 0.936601164 |
| ENSMUSG00000043004 | Gng2 | protein_coding | 0.090941171 | 7.764402545 | 0.1696787 | 0.6857612 | 0.936601164 |
| ENSMUSG00000032633 | Flcn | protein_coding | 0.147461678 | 5.481049435 | 0.1696197 | 0.6858215 | 0.936601164 |
| ENSMUSG00000038510 | Rpf2 | protein_coding | -0.0881173 | 6.323682725 | 0.1695635 | 0.6858617 | 0.936601164 |
| ENSMUSG00000020892 | Aloxe3 | protein_coding | -0.195863752 | 4.186535771 | 0.1695056 | 0.6859209 | 0.936601164 |
| ENSMUSG00000020137 | Thap2 | protein_coding | -0.17752497 | 3.538827538 | 0.1692923 | 0.6860983 | 0.936601164 |
| ENSMUSG00000003662 | Ciao1 | protein_coding | 0.096335465 | 6.247825886 | 0.1692173 | 0.6861639 | 0.936601164 |
| ENSMUSG00000034543 | Morc2a | protein_coding | 0.145333412 | 3.987447383 | 0.1691872 | 0.6861901 | 0.936601164 |
| ENSMUSG00000019261 | Map1s | protein_coding | -0.198809333 | 4.671903465 | 0.169547 | 0.6862173 | 0.936601164 |
| ENSMUSG00000032192 | Gnb5 | protein_coding | 0.074588304 | 7.735530839 | 0.1691432 | 0.6862286 | 0.936601164 |
| ENSMUSG00000018395 | Kif3a | protein_coding | 0.103637315 | 7.687993734 | 0.1690683 | 0.6863028 | 0.936601164 |
| ENSMUSG00000061759 | Armt1 | protein_coding | 0.127430759 | 4.666926094 | 0.1689941 | 0.6863589 | 0.936601164 |
| ENSMUSG00000031982 | Arv1 | protein_coding | -0.184108027 | 3.505064802 | 0.1684297 | 0.6868527 | 0.937156094 |
| ENSMUSG00000022018 | Rgcc | protein_coding | 0.249745058 | 3.124808967 | 0.1681857 | 0.6870665 | 0.937211079 |
| ENSMUSG00000053870 | Fpgt | protein_coding | -0.195898617 | 3.953645991 | 0.1681948 | 0.6870673 | 0.937211079 |
| ENSMUSG00000021113 | Snapc1 | protein_coding | 0.098306252 | 5.663202216 | 0.1680155 | 0.6872158 | 0.937294657 |
| ENSMUSG00000026854 | Usp20 | protein_coding | 0.10852047 | 5.97601616 | 0.1677757 | 0.6874262 | 0.937337983 |
| ENSMUSG00000026088 | Mitd1 | protein_coding | -0.148049415 | 4.400588202 | 0.1677348 | 0.6874621 | 0.937337983 |
| ENSMUSG00000051062 | Fbll1 | protein_coding | 0.121694556 | 5.381324325 | 0.1676813 | 0.687509 | 0.937337983 |
| ENSMUSG00000040462 | Os9 | protein_coding | 0.07632009 | 6.832922169 | 0.1669098 | 0.6881875 | 0.937476052 |
| ENSMUSG00000034216 | Vps18 | protein_coding | -0.172480776 | 5.153911227 | 0.1670924 | 0.688261 | 0.937476052 |
| ENSMUSG00000026393 | Nek7 | protein_coding | -0.120877975 | 6.044481512 | 0.1668315 | 0.6882651 | 0.937476052 |
| ENSMUSG00000004032 | Gstm5 | protein_coding | -0.094111144 | 9.655987292 | 0.166724 | 0.6883511 | 0.937476052 |
| ENSMUSG00000050965 | Prkca | protein_coding | -0.10441972 | 5.234489138 | 0.1666645 | 0.6884036 | 0.937476052 |
| ENSMUSG00000028857 | Tmem222 | protein_coding | 0.092471643 | 8.220295672 | 0.1665963 | 0.6884637 | 0.937476052 |
| ENSMUSG00000032763 | Ilvbl | protein_coding | 0.150639742 | 5.377961018 | 0.1665973 | 0.6884715 | 0.937476052 |
| ENSMUSG00000049130 | C5ar1 | protein_coding | -0.392234955 | 2.071131458 | 0.1665287 | 0.6885233 | 0.937476052 |
| ENSMUSG00000040710 | St8sia4 | protein_coding | -0.283374661 | 2.980059613 | 0.1665229 | 0.6885371 | 0.937476052 |
| ENSMUSG00000097039 | Pvt1 | lincRNA | 0.123221833 | 4.786443159 | 0.1662714 | 0.6887502 | 0.937476052 |
| ENSMUSG00000027797 | Dclk1 | protein_coding | 0.080829442 | 7.263284849 | 0.1660481 | 0.6889474 | 0.937476052 |
| ENSMUSG00000031938 | 4931406C07Rik | protein_coding | -0.186507493 | 3.821503406 | 0.1660253 | 0.6889675 | 0.937476052 |
| ENSMUSG00000026788 | Zbtb43 | protein_coding | 0.127005521 | 4.339041362 | 0.1660222 | 0.6889703 | 0.937476052 |
| ENSMUSG00000037784 | Dzip1l | protein_coding | 0.289416962 | 2.5623487 | 0.1657561 | 0.6892054 | 0.937476052 |
| ENSMUSG00000027087 | Itgav | protein_coding | -0.203025343 | 5.549576828 | 0.1666828 | 0.6892338 | 0.937476052 |
| ENSMUSG00000063480 | Snu13 | protein_coding | 0.105224724 | 5.062151622 | 0.1656958 | 0.6892587 | 0.937476052 |
| ENSMUSG00000008682 | Rpl10 | protein_coding | -0.196479807 | 3.033878391 | 0.1656817 | 0.6892712 | 0.937476052 |
| ENSMUSG00000031381 | Piga | protein_coding | 0.178656396 | 3.304020973 | 0.1656717 | 0.6892801 | 0.937476052 |
| ENSMUSG00000041046 | Ramp3 | protein_coding | -0.160329588 | 6.002881069 | 0.1662191 | 0.6893253 | 0.937476052 |
| ENSMUSG00000024773 | Atg2a | protein_coding | -0.196451057 | 3.524198162 | 0.165537 | 0.6893992 | 0.937476052 |
| ENSMUSG00000013523 | Bcas1 | protein_coding | 0.157848193 | 5.245615683 | 0.1654776 | 0.6894833 | 0.937476052 |
| ENSMUSG00000028603 | Scp2 | protein_coding | 0.222763232 | 3.075464304 | 0.1653626 | 0.6895536 | 0.937476052 |
| ENSMUSG00000036932 | Aifm1 | protein_coding | 0.122150485 | 5.357849465 | 0.1652927 | 0.6896155 | 0.937476052 |
| ENSMUSG00000027695 | Pld1 | protein_coding | 0.303809469 | 2.534102462 | 0.1651758 | 0.6897191 | 0.937480214 |
| ENSMUSG00000036422 | Pcdh8 | protein_coding | 0.163217701 | 4.455041384 | 0.1650924 | 0.6897929 | 0.937480214 |
| ENSMUSG00000021843 | Ktn1 | protein_coding | -0.110577158 | 5.683007978 | 0.1645666 | 0.6902593 | 0.937910629 |
| ENSMUSG00000056458 | Mok | protein_coding | -0.161262795 | 4.404013579 | 0.1644562 | 0.6903574 | 0.937910629 |
| ENSMUSG00000015305 | Sash1 | protein_coding | -0.157902113 | 4.177191885 | 0.1643248 | 0.6904741 | 0.937910629 |
| ENSMUSG00000043557 | Mdga1 | protein_coding | 0.215324605 | 3.316663069 | 0.1641388 | 0.6906394 | 0.937910629 |
| ENSMUSG00000091002 | Tcerg1l | protein_coding | -0.156436209 | 4.286745588 | 0.1640567 | 0.6907125 | 0.937910629 |
| ENSMUSG00000030584 | Dpf1 | protein_coding | -0.130333559 | 4.907281229 | 0.1640534 | 0.6907154 | 0.937910629 |

| ensembl_gene_id | external_gene_name | gene_biotype | log2 (fold change) | log2 (counts per million) | F-value | p-value | False discovery rate |
|---|---|---|---|---|---|---|---|
| ENSMUSG00000045312 | Lhfpl2 | protein_coding | 0.335863019 | 2.521567076 | 0.164048 | 0.6907202 | 0.937910629 |
| ENSMUSG00000029228 | Lnx1 | protein_coding | 0.177468323 | 4.27486884 | 0.1637882 | 0.6909601 | 0.938021346 |
| ENSMUSG00000027243 | Harbi1 | protein_coding | -0.178488126 | 4.173488341 | 0.1637563 | 0.6909798 | 0.938021346 |
| ENSMUSG00000056899 | Immp2l | protein_coding | -0.153123521 | 4.525311142 | 0.1636624 | 0.6910634 | 0.938021346 |
| ENSMUSG00000070780 | Rbm47 | protein_coding | -0.335190298 | 2.402039901 | 0.1635356 | 0.6911764 | 0.938056281 |
| ENSMUSG00000038984 | Tspyl5 | protein_coding | -0.106478076 | 6.739903537 | 0.1631537 | 0.6915256 | 0.938235075 |
| ENSMUSG00000026005 | Rpe | protein_coding | -0.16031136 | 4.424036185 | 0.1630121 | 0.6916433 | 0.938235075 |
| ENSMUSG00000052160 | Pld4 | protein_coding | -0.231905049 | 3.550459364 | 0.1629958 | 0.6916665 | 0.938235075 |
| ENSMUSG00000026484 | Rnf2 | protein_coding | 0.112186473 | 5.037563545 | 0.1629099 | 0.6917346 | 0.938235075 |
| ENSMUSG00000022400 | Rbx1 | protein_coding | 0.074796071 | 9.326508806 | 0.1628844 | 0.6917573 | 0.938235075 |
| ENSMUSG00000030134 | Rasgef1a | protein_coding | -0.102672997 | 7.583743779 | 0.1628002 | 0.6918412 | 0.938235075 |
| ENSMUSG00000001995 | Sipa1l2 | protein_coding | 0.183523454 | 3.881671861 | 0.1627036 | 0.6919189 | 0.938235075 |
| ENSMUSG00000038152 | 5033430I15Rik | lincRNA | -0.230758669 | 2.755846196 | 0.1622654 | 0.6923109 | 0.938336995 |
| ENSMUSG00000007338 | Mrpl49 | protein_coding | 0.075738786 | 7.222371625 | 0.1621968 | 0.6923723 | 0.938336995 |
| ENSMUSG00000035495 | Tstd2 | protein_coding | -0.175500267 | 3.222864669 | 0.162132 | 0.6924304 | 0.938336995 |
| ENSMUSG00000074024 | 4632427E13Rik | bidirectional_promoter | 0.4411959 | 2.528839498 | 0.1623471 | 0.6926663 | 0.938336995 |
| ENSMUSG00000047459 | Dynlrb1 | protein_coding | -0.088168489 | 10.28289572 | 0.1617648 | 0.6927594 | 0.938336995 |
| ENSMUSG00000025876 | Unc5a | protein_coding | -0.17684198 | 4.321028332 | 0.1617625 | 0.6927701 | 0.938336995 |
| ENSMUSG00000030098 | Grip2 | protein_coding | -0.24208374 | 3.227023866 | 0.1617174 | 0.692802 | 0.938336995 |
| ENSMUSG00000021177 | Tdp1 | protein_coding | -0.138238958 | 4.693445551 | 0.1616596 | 0.6928538 | 0.938336995 |
| ENSMUSG00000048058 | Ldlrad3 | protein_coding | 0.218882786 | 3.027808093 | 0.161524 | 0.6929755 | 0.938336995 |
| ENSMUSG00000084885 | 3010001F23Rik | processed_transcript | -0.214978171 | 2.823728381 | 0.1614148 | 0.6930736 | 0.938336995 |
| ENSMUSG00000058192 | Zfp846 | protein_coding | -0.16381928 | 4.151514707 | 0.1613783 | 0.6931063 | 0.938336995 |
| ENSMUSG00000028573 | Fggy | protein_coding | 0.120761688 | 5.243828792 | 0.1613751 | 0.6931092 | 0.938336995 |
| ENSMUSG00000015714 | Cers2 | protein_coding | -0.090014747 | 6.226918209 | 0.1612687 | 0.6932048 | 0.938336995 |
| ENSMUSG00000031875 | Cmtm3 | protein_coding | -0.183339676 | 3.327982326 | 0.161114 | 0.6933438 | 0.938336995 |
| ENSMUSG00000020713 | Gh | protein_coding | 0.292697461 | 3.159073707 | 0.1611902 | 0.6934937 | 0.938336995 |
| ENSMUSG00000016427 | Ndufa1 | protein_coding | -0.104686092 | 9.064829798 | 0.1602999 | 0.6940853 | 0.938336995 |
| ENSMUSG00000002949 | Timm44 | protein_coding | 0.100813531 | 6.385036777 | 0.1602619 | 0.694111 | 0.938336995 |
| ENSMUSG00000029640 | Usp12 | protein_coding | 0.208101984 | 3.012577258 | 0.1602571 | 0.6941154 | 0.938336995 |
| ENSMUSG00000063849 | Ppcdc | protein_coding | 0.105878062 | 5.543887898 | 0.1601012 | 0.694256 | 0.938336995 |
| ENSMUSG00000040102 | Klhl42 | protein_coding | 0.162421992 | 4.246095242 | 0.1600673 | 0.6942865 | 0.938336995 |
| ENSMUSG00000021645 | Smn1 | protein_coding | 0.101183142 | 5.4629213 | 0.1600527 | 0.6942997 | 0.938336995 |
| ENSMUSG00000005417 | Mprip | protein_coding | 0.111303051 | 5.350901569 | 0.1600086 | 0.6943395 | 0.938336995 |
| ENSMUSG00000043445 | Pgp | protein_coding | 0.128524046 | 4.355717368 | 0.1599919 | 0.6943547 | 0.938336995 |
| ENSMUSG00000007646 | Rad51c | protein_coding | -0.224660753 | 3.204790489 | 0.1596064 | 0.6947029 | 0.938336995 |
| ENSMUSG00000052752 | Traf7 | protein_coding | -0.107951954 | 4.874099156 | 0.1593705 | 0.6949162 | 0.938336995 |
| ENSMUSG00000034168 | Irf2bpl | protein_coding | -0.124718008 | 4.404039221 | 0.1593579 | 0.6949275 | 0.938336995 |
| ENSMUSG00000029642 | Polr1d | protein_coding | 0.082296953 | 6.929829932 | 0.1593402 | 0.6949436 | 0.938336995 |
| ENSMUSG00000023868 | Pde10a | protein_coding | -0.119453348 | 6.422342543 | 0.1590813 | 0.6952088 | 0.938336995 |
| ENSMUSG00000030505 | Prmt3 | protein_coding | 0.168563828 | 3.326721743 | 0.1589548 | 0.6952925 | 0.938336995 |
| ENSMUSG00000004929 | Thop1 | protein_coding | -0.175952404 | 4.018110319 | 0.1589058 | 0.6953369 | 0.938336995 |
| ENSMUSG00000039801 | 2410089E03Rik | protein_coding | -0.198327622 | 3.251676142 | 0.1589024 | 0.69534 | 0.938336995 |
| ENSMUSG00000030350 | Prmt8 | protein_coding | -0.09008361 | 6.015413048 | 0.1588497 | 0.6953877 | 0.938336995 |
| ENSMUSG00000041771 | Slc24a4 | protein_coding | -0.268250318 | 2.504585725 | 0.1586978 | 0.6955255 | 0.938336995 |
| ENSMUSG00000024843 | Chka | protein_coding | 0.142549006 | 4.607991209 | 0.1584522 | 0.6957483 | 0.938336995 |
| ENSMUSG00000057880 | Abat | protein_coding | 0.082747053 | 8.308179823 | 0.1583707 | 0.6958223 | 0.938336995 |
| ENSMUSG00000026568 | Mpc2 | protein_coding | -0.158128342 | 6.934030928 | 0.159443 | 0.6958266 | 0.938336995 |
| ENSMUSG00000028849 | Map7d1 | protein_coding | 0.083733639 | 6.582567633 | 0.1582843 | 0.6959008 | 0.938336995 |
| ENSMUSG00000030532 | Hddc3 | protein_coding | 0.156055674 | 5.144441123 | 0.158221 | 0.6959799 | 0.938336995 |
| ENSMUSG00000027509 | Rae1 | protein_coding | 0.122031682 | 4.86638759 | 0.1581348 | 0.6960366 | 0.938336995 |
| ENSMUSG00000069227 | Gprin1 | protein_coding | -0.147696806 | 3.98846622 | 0.1581231 | 0.6960473 | 0.938336995 |
| ENSMUSG00000046822 | Slc39a3 | protein_coding | 0.154212162 | 4.066614776 | 0.158062 | 0.6961028 | 0.938336995 |
| ENSMUSG00000036672 | Cenpt | protein_coding | 0.117505133 | 5.315053013 | 0.1578966 | 0.6962532 | 0.938336995 |
| ENSMUSG00000021666 | Gfm2 | protein_coding | -0.165901067 | 3.871473383 | 0.1578643 | 0.6962826 | 0.938336995 |
| ENSMUSG00000029028 | Lrrc47 | protein_coding | -0.242004453 | 2.925906026 | 0.1577704 | 0.696368 | 0.938336995 |
| ENSMUSG00000029776 | Hibadh | protein_coding | -0.097138485 | 6.256325144 | 0.1577656 | 0.6963724 | 0.938336995 |
| ENSMUSG00000061298 | Agbl4 | protein_coding | -0.141788161 | 4.51388219 | 0.1577306 | 0.6964042 | 0.938336995 |
| ENSMUSG00000022556 | Hsf1 | protein_coding | 0.143107354 | 4.899558544 | 0.1577021 | 0.6964302 | 0.938336995 |
| ENSMUSG00000023075 | Akirin1 | protein_coding | 0.150033969 | 4.188818032 | 0.1575674 | 0.6965528 | 0.938336995 |
| ENSMUSG00000063065 | Mapk3 | protein_coding | 0.071012651 | 7.28755616 | 0.1574641 | 0.6966469 | 0.938336995 |
| ENSMUSG00000071074 | Yipf3 | protein_coding | 0.121181738 | 5.260315035 | 0.1574461 | 0.6966633 | 0.938336995 |
| ENSMUSG00000026577 | Blzf1 | protein_coding | -0.114574419 | 5.141870769 | 0.1573333 | 0.6967662 | 0.938336995 |
| ENSMUSG00000027194 | Ttc17 | protein_coding | 0.137237146 | 3.704452578 | 0.1572919 | 0.6968038 | 0.938336995 |
| ENSMUSG00000041890 | Git2 | protein_coding | 0.162020909 | 4.123003064 | 0.1572158 | 0.6968732 | 0.938336995 |
| ENSMUSG00000030986 | Dhx32 | protein_coding | 0.12718766 | 4.597175037 | 0.1571614 | 0.6969229 | 0.938336995 |
| ENSMUSG00000021071 | Trim9 | protein_coding | 0.138163528 | 5.423972199 | 0.1570638 | 0.6970204 | 0.938336995 |
| ENSMUSG00000019831 | Wasf1 | protein_coding | -0.078442009 | 7.054750933 | 0.1570514 | 0.6970232 | 0.938336995 |
| ENSMUSG00000019897 | Ccdc59 | protein_coding | -0.076208668 | 7.036182763 | 0.15704 | 0.6970336 | 0.938336995 |
| ENSMUSG00000055932 | Fto | protein_coding | 0.148690919 | 8.114774257 | 0.157736 | 0.6972201 | 0.938336995 |
| ENSMUSG00000056167 | Cnot10 | protein_coding | -0.123934311 | 5.128757025 | 0.1568313 | 0.6972241 | 0.938336995 |
| ENSMUSG00000004961 | Syt5 | protein_coding | 0.070158089 | 8.28337684 | 0.1568002 | 0.6972525 | 0.938336995 |
| ENSMUSG00000056019 | Zfp709 | protein_coding | -0.23544049 | 2.776760095 | 0.1567295 | 0.6973171 | 0.938336995 |
| ENSMUSG00000031667 | Aktip | protein_coding | 0.084215471 | 8.165522375 | 0.1562219 | 0.6977812 | 0.93874064 |

| ensembl_gene_id | external_gene_name | gene_biotype | log2 (fold change) | log2 (counts per million) | F-value | p-value | False discovery rate |
|---|---|---|---|---|---|---|---|
| ENSMUSG00000022521 | Crebbp | protein_coding | -0.094944927 | 7.026471536 | 0.1562105 | 0.6977917 | 0.93874064 |
| ENSMUSG00000029253 | Cenpc1 | protein_coding | 0.139036095 | 4.863560757 | 0.1559522 | 0.6980282 | 0.938767076 |
| ENSMUSG00000028710 | Atpaf1 | protein_coding | -0.179250641 | 3.757950926 | 0.1558502 | 0.6981217 | 0.938767076 |
| ENSMUSG00000027076 | Timm10 | protein_coding | -0.089263508 | 7.283647256 | 0.1558241 | 0.6981456 | 0.938767076 |
| ENSMUSG00000020263 | Appl2 | protein_coding | -0.113550337 | 4.59630411 | 0.1557132 | 0.6982473 | 0.938767076 |
| ENSMUSG00000026890 | Lhx6 | protein_coding | -0.116790665 | 6.003669517 | 0.1557218 | 0.6982478 | 0.938767076 |
| ENSMUSG00000020591 | Ntsr2 | protein_coding | -0.177147189 | 5.596747364 | 0.1559514 | 0.6986921 | 0.939072265 |
| ENSMUSG00000022321 | Cdh10 | protein_coding | -0.134733685 | 4.203458714 | 0.155153 | 0.6987616 | 0.939072265 |
| ENSMUSG00000020532 | Acaca | protein_coding | -0.15174735 | 3.55394166 | 0.1551451 | 0.6987688 | 0.939072265 |
| ENSMUSG00000048279 | Sacs | protein_coding | 0.151073923 | 4.283864982 | 0.1550012 | 0.6989011 | 0.939072265 |
| ENSMUSG00000046550 | Spin2c | protein_coding | 0.120886073 | 5.370066525 | 0.1549331 | 0.6989637 | 0.939072265 |
| ENSMUSG00000055745 | Ldoc1l | protein_coding | 0.104760297 | 5.336780888 | 0.1548949 | 0.6989988 | 0.939072265 |
| ENSMUSG00000041975 | Mettl8 | protein_coding | -0.186802105 | 4.448576166 | 0.1545184 | 0.6995034 | 0.939597409 |
| ENSMUSG00000032383 | Ppib | protein_coding | 0.086290892 | 6.18592516 | 0.154237 | 0.6996048 | 0.939597409 |
| ENSMUSG00000046160 | Olig1 | protein_coding | -0.118537065 | 5.77983526 | 0.1541859 | 0.6996519 | 0.939597409 |
| ENSMUSG00000034826 | Nup54 | protein_coding | 0.190164594 | 3.960240692 | 0.1540295 | 0.6998046 | 0.939685112 |
| ENSMUSG00000021591 | Glrx | protein_coding | 0.089398235 | 6.682961607 | 0.1538464 | 0.6999652 | 0.939701346 |
| ENSMUSG00000024661 | Fth1 | protein_coding | -0.378965606 | 12.1728887 | 0.1596994 | 0.7000406 | 0.939701346 |
| ENSMUSG00000049922 | Slc35c1 | protein_coding | 0.154869169 | 4.063966979 | 0.1534506 | 0.700331 | 0.939701346 |
| ENSMUSG00000001017 | Chtop | protein_coding | -0.082779436 | 5.922652074 | 0.1534251 | 0.7003545 | 0.939701346 |
| ENSMUSG00000023473 | Celsr3 | protein_coding | -0.261844752 | 2.799257655 | 0.1532591 | 0.7005082 | 0.939701346 |
| ENSMUSG00000054414 | Slc30a7 | protein_coding | 0.204780232 | 3.354132197 | 0.1532176 | 0.7005465 | 0.939701346 |
| ENSMUSG00000002477 | Snrpd1 | protein_coding | -0.092382708 | 7.826227714 | 0.153202 | 0.700561 | 0.939701346 |
| ENSMUSG00000021488 | Nsd1 | protein_coding | 0.090712686 | 6.332371186 | 0.1531754 | 0.7005856 | 0.939701346 |
| ENSMUSG00000031627 | Irf2 | protein_coding | -0.100728016 | 4.944026901 | 0.1531565 | 0.7006032 | 0.939701346 |
| ENSMUSG00000003235 | Eif2b5 | protein_coding | 0.114185181 | 5.373775988 | 0.1529759 | 0.7007704 | 0.939738232 |
| ENSMUSG00000025495 | Ptdss2 | protein_coding | 0.125218299 | 5.097319969 | 0.15293 | 0.7008129 | 0.939738232 |
| ENSMUSG00000042535 | Gtpbp1 | protein_coding | 0.166560437 | 3.944895231 | 0.1528438 | 0.7008928 | 0.939738232 |
| ENSMUSG00000046567 | 4930430F08Rik | protein_coding | 0.124890395 | 4.367057172 | 0.1526295 | 0.7010916 | 0.939810471 |
| ENSMUSG00000062604 | Srpk2 | protein_coding | 0.082493487 | 7.969830073 | 0.1525683 | 0.7011484 | 0.939810471 |
| ENSMUSG00000053286 | Trmt1l | protein_coding | 0.164126419 | 3.963026734 | 0.152376 | 0.7013269 | 0.939810471 |
| ENSMUSG00000034192 | Lsm3 | protein_coding | -0.121345247 | 6.577785955 | 0.1525946 | 0.7013776 | 0.939810471 |
| ENSMUSG00000078435 | AU041133 | protein_coding | -0.226205839 | 2.924900475 | 0.1518716 | 0.7017957 | 0.939810471 |
| ENSMUSG00000029167 | Ppargc1a | protein_coding | -0.10468185 | 5.318008391 | 0.151783 | 0.7018781 | 0.939810471 |
| ENSMUSG00000074825 | Itpripl1 | protein_coding | -0.19616042 | 3.798788421 | 0.1517111 | 0.7019533 | 0.939810471 |
| ENSMUSG00000034889 | Cactin | protein_coding | 0.285899696 | 2.749299879 | 0.1515591 | 0.7020866 | 0.939810471 |
| ENSMUSG00000031557 | Plekha2 | protein_coding | 0.277559559 | 2.553370924 | 0.1515202 | 0.7021229 | 0.939810471 |
| ENSMUSG00000033970 | Rfc3 | protein_coding | 0.105114855 | 4.936916151 | 0.1515013 | 0.7021405 | 0.939810471 |
| ENSMUSG00000062198 | 2700097O09Rik | protein_coding | -0.130232542 | 4.295031524 | 0.1513682 | 0.7022646 | 0.939810471 |
| ENSMUSG00000020057 | Dram1 | protein_coding | 0.128130583 | 4.04375824 | 0.1512291 | 0.7023942 | 0.939810471 |
| ENSMUSG00000049134 | Nrap | protein_coding | 0.5933193 | 1.738939819 | 0.1513952 | 0.7025802 | 0.939810471 |
| ENSMUSG00000021003 | Galc | protein_coding | -0.198170952 | 3.282337328 | 0.1510077 | 0.7026008 | 0.939810471 |
| ENSMUSG00000037720 | Tmem33 | protein_coding | 0.101313223 | 6.126478516 | 0.1508894 | 0.7027113 | 0.939810471 |
| ENSMUSG00000051435 | Fhad1 | protein_coding | -0.097608579 | 5.190793183 | 0.1508787 | 0.7027212 | 0.939810471 |
| ENSMUSG00000059734 | Ndufs8 | protein_coding | 0.076879874 | 9.673630641 | 0.1507086 | 0.7028802 | 0.939810471 |
| ENSMUSG00000063260 | Syt10 | protein_coding | 0.145102494 | 3.929621671 | 0.1505256 | 0.7030513 | 0.939810471 |
| ENSMUSG00000021115 | Vrk1 | protein_coding | -0.099749842 | 5.201208222 | 0.1504551 | 0.7031172 | 0.939810471 |
| ENSMUSG00000047495 | Dlgap2 | protein_coding | -0.130693872 | 4.648677871 | 0.1504526 | 0.7031196 | 0.939810471 |
| ENSMUSG00000020778 | Ten1 | protein_coding | 0.071981564 | 7.058388038 | 0.1503829 | 0.7031847 | 0.939810471 |
| ENSMUSG00000032324 | Tspan3 | protein_coding | 0.099603966 | 7.434399191 | 0.150381 | 0.7031948 | 0.939810471 |
| ENSMUSG00000037110 | Ralgapa2 | protein_coding | -0.270253678 | 2.806873281 | 0.1502132 | 0.7033436 | 0.939810471 |
| ENSMUSG00000079197 | Psme2 | protein_coding | 0.080699356 | 6.052629701 | 0.1500978 | 0.7034517 | 0.939810471 |
| ENSMUSG00000078816 | Prkcg | protein_coding | -0.124875709 | 7.171257509 | 0.1506063 | 0.7034766 | 0.939810471 |
| ENSMUSG00000055760 | Gemin6 | protein_coding | -0.155095498 | 4.42354317 | 0.1499266 | 0.7036121 | 0.939810471 |
| ENSMUSG00000028899 | Taf12 | protein_coding | 0.100371849 | 6.018048433 | 0.1498074 | 0.7037239 | 0.939810471 |
| ENSMUSG00000020691 | Mettl2 | protein_coding | 0.160335063 | 4.728498721 | 0.1496428 | 0.7038866 | 0.939810471 |
| ENSMUSG00000010406 | Mrpl52 | protein_coding | 0.085992827 | 7.395968681 | 0.1495698 | 0.7039468 | 0.939810471 |
| ENSMUSG00000001901 | Kcnh6 | protein_coding | 0.237249681 | 2.875576545 | 0.1494252 | 0.7040826 | 0.939810471 |
| ENSMUSG00000066151 | Fkbp15 | protein_coding | 0.115250038 | 4.607863867 | 0.1493683 | 0.7041361 | 0.939810471 |
| ENSMUSG00000001305 | Rrp15 | protein_coding | -0.132006821 | 4.420875696 | 0.1492763 | 0.7042224 | 0.939810471 |
| ENSMUSG00000056069 | Fam105a | protein_coding | -0.18240689 | 3.276289901 | 0.1492577 | 0.7042399 | 0.939810471 |
| ENSMUSG00000061762 | Tac1 | protein_coding | -0.132761957 | 8.511015986 | 0.1499063 | 0.7042799 | 0.939810471 |
| ENSMUSG00000032946 | Rasgrp2 | protein_coding | -0.100978744 | 5.225065832 | 0.1487729 | 0.704696 | 0.939810471 |
| ENSMUSG00000028894 | Inpp5b | protein_coding | 0.141900633 | 4.148575402 | 0.1486871 | 0.7047769 | 0.939810471 |
| ENSMUSG00000027365 | Trpm7 | protein_coding | 0.139825532 | 3.924431 | 0.1486709 | 0.704792 | 0.939810471 |
| ENSMUSG00000005802 | Slc30a4 | protein_coding | 0.224244843 | 3.385005898 | 0.1485996 | 0.7048675 | 0.939810471 |
| ENSMUSG00000035093 | Secisbp2l | protein_coding | -0.098040215 | 6.285429562 | 0.1484587 | 0.704992 | 0.939810471 |
| ENSMUSG00000024011 | Pi16 | protein_coding | 0.280446256 | 3.090579411 | 0.1484108 | 0.7050764 | 0.939810471 |
| ENSMUSG00000031333 | Abcb7 | protein_coding | -0.13462688 | 4.066281437 | 0.1482198 | 0.7052174 | 0.939810471 |
| ENSMUSG00000030844 | Rgs10 | protein_coding | 0.074557876 | 8.342266456 | 0.147779 | 0.7056336 | 0.939810471 |
| ENSMUSG00000100618 | Gm29595 | lincRNA | -0.241578954 | 2.591720433 | 0.1477212 | 0.7056883 | 0.939810471 |
| ENSMUSG00000007610 | Gtpbp3 | protein_coding | 0.206830755 | 2.835083554 | 0.1476162 | 0.7057876 | 0.939810471 |
| ENSMUSG00000113170 | AC132344.1 | lincRNA | -0.137007507 | 3.751730224 | 0.1475766 | 0.7058251 | 0.939810471 |
| ENSMUSG00000031958 | Ldhd | protein_coding | 0.194596296 | 4.17706719 | 0.1477319 | 0.7058323 | 0.939810471 |

| ensembl_gene_id | external_gene_name | gene_biotype | log2 (fold change) | log2 (counts per million) | F-value | p-value | False discovery rate |
|---|---|---|---|---|---|---|---|
| ENSMUSG00000029735 | Tpk1 | protein_coding | -0.328931355 | 2.61527718 | 0.1474649 | 0.7059389 | 0.939810471 |
| ENSMUSG00000093674 | Rpl41 | protein_coding | -0.106410534 | 9.105067067 | 0.1471147 | 0.706292 | 0.939810471 |
| ENSMUSG00000005823 | Gpr108 | protein_coding | 0.108854282 | 5.589183926 | 0.1470699 | 0.7063049 | 0.939810471 |
| ENSMUSG00000039672 | Kcne2 | protein_coding | 0.389339641 | 3.019242124 | 0.1483494 | 0.7063253 | 0.939810471 |
| ENSMUSG00000035649 | Zcchc7 | protein_coding | 0.118034519 | 4.421059595 | 0.1468629 | 0.7065012 | 0.939810471 |
| ENSMUSG00000030822 | Prr14 | protein_coding | -0.124635973 | 5.12615451 | 0.146861 | 0.706503 | 0.939810471 |
| ENSMUSG00000049811 | Fam161a | protein_coding | 0.196774455 | 2.959912832 | 0.1468351 | 0.7065275 | 0.939810471 |
| ENSMUSG00000012296 | Tjap1 | protein_coding | -0.279952986 | 2.745423782 | 0.1467953 | 0.7065653 | 0.939810471 |
| ENSMUSG00000024687 | Osbp | protein_coding | -0.146135037 | 4.320719299 | 0.1467006 | 0.7066552 | 0.939810471 |
| ENSMUSG00000037428 | Vgf | protein_coding | -0.160616687 | 4.032652325 | 0.1465447 | 0.7068033 | 0.939810471 |
| ENSMUSG00000054256 | Msi1 | protein_coding | 0.163315816 | 4.022097204 | 0.1465348 | 0.7068127 | 0.939810471 |
| ENSMUSG00000034336 | Ina | protein_coding | 0.072026804 | 7.552055652 | 0.1465311 | 0.7068161 | 0.939810471 |
| ENSMUSG00000020032 | Nuak1 | protein_coding | 0.123171629 | 4.065982924 | 0.1465036 | 0.7068422 | 0.939810471 |
| ENSMUSG00000022858 | Tra2b | protein_coding | -0.085859618 | 6.079785217 | 0.1464597 | 0.706884 | 0.939810471 |
| ENSMUSG00000020171 | Yeats4 | protein_coding | -0.139869323 | 4.308757763 | 0.1464046 | 0.7069364 | 0.939810471 |
| ENSMUSG00000039254 | Pomt1 | protein_coding | -0.218100951 | 2.894868136 | 0.1464022 | 0.7069386 | 0.939810471 |
| ENSMUSG00000078566 | Bnip3 | protein_coding | 0.10641086 | 5.546691871 | 0.1463641 | 0.7069749 | 0.939810471 |
| ENSMUSG00000070880 | Gad1 | protein_coding | -0.076740507 | 8.867156089 | 0.1460635 | 0.7072607 | 0.939810471 |
| ENSMUSG00000066892 | Fbxl12 | protein_coding | 0.164742979 | 3.706845045 | 0.1458488 | 0.7074652 | 0.939810471 |
| ENSMUSG00000050854 | Tmem125 | protein_coding | -0.193215704 | 3.590188888 | 0.1458179 | 0.7074945 | 0.939810471 |
| ENSMUSG00000067925 | Cxx1a | protein_coding | 0.134981401 | 3.974048044 | 0.1457819 | 0.7075289 | 0.939810471 |
| ENSMUSG00000060716 | Plekhh1 | protein_coding | 0.209837869 | 3.439004716 | 0.1457613 | 0.7075566 | 0.939810471 |
| ENSMUSG00000026021 | Sumo1 | protein_coding | -0.073217899 | 8.260689262 | 0.1457035 | 0.7076036 | 0.939810471 |
| ENSMUSG00000027956 | Tmem144 | protein_coding | -0.164839829 | 3.501868317 | 0.1455862 | 0.7077154 | 0.939810471 |
| ENSMUSG00000033569 | Adgrb3 | protein_coding | -0.118593588 | 5.961160046 | 0.1455894 | 0.7077204 | 0.939810471 |
| ENSMUSG00000002342 | Tmem161a | protein_coding | 0.138204809 | 4.850548156 | 0.1454176 | 0.7078762 | 0.939810471 |
| ENSMUSG00000025485 | Ric8a | protein_coding | 0.10633703 | 5.790532906 | 0.1452206 | 0.7080642 | 0.939810471 |
| ENSMUSG00000032373 | Car12 | protein_coding | 0.246745718 | 3.89923021 | 0.1458004 | 0.7081501 | 0.939810471 |
| ENSMUSG00000039131 | Gipc2 | protein_coding | 0.156007673 | 3.787851571 | 0.1450156 | 0.70826 | 0.939810471 |
| ENSMUSG00000021481 | Zfp346 | protein_coding | -0.131841966 | 4.227970794 | 0.1449912 | 0.7082833 | 0.939810471 |
| ENSMUSG00000054423 | Cadps | protein_coding | -0.072022903 | 7.090859496 | 0.1448935 | 0.7083767 | 0.939810471 |
| ENSMUSG00000041459 | Tardbp | protein_coding | 0.083652389 | 6.137299951 | 0.1448553 | 0.7084132 | 0.939810471 |
| ENSMUSG00000053091 | Lins1 | protein_coding | 0.192865772 | 3.081821021 | 0.1446296 | 0.7086291 | 0.939810471 |
| ENSMUSG00000023977 | Ubr2 | protein_coding | 0.154984338 | 3.830373238 | 0.1445617 | 0.7086942 | 0.939810471 |
| ENSMUSG00000028060 | 2810403A07Rik | protein_coding | 0.128333316 | 4.220048193 | 0.1445044 | 0.708749 | 0.939810471 |
| ENSMUSG00000057895 | Zfp105 | protein_coding | -0.229966999 | 2.794361976 | 0.1444683 | 0.7087835 | 0.939810471 |
| ENSMUSG00000045822 | Zswim3 | protein_coding | 0.168441718 | 3.408235743 | 0.1443848 | 0.7088635 | 0.939810471 |
| ENSMUSG00000037266 | Rsrp1 | protein_coding | 0.247232942 | 5.794475185 | 0.1469205 | 0.7089554 | 0.939810471 |
| ENSMUSG00000001036 | Epn2 | protein_coding | 0.081586281 | 6.300248836 | 0.1442392 | 0.7090031 | 0.939810471 |
| ENSMUSG00000060938 | Rpl26 | protein_coding | -0.098261821 | 8.23068949 | 0.144176 | 0.7090718 | 0.939810471 |
| ENSMUSG00000006021 | Kptn | protein_coding | 0.137352073 | 5.224893488 | 0.1441599 | 0.7090872 | 0.939810471 |
| ENSMUSG00000026094 | Stk17b | protein_coding | -0.411979395 | 2.255065074 | 0.1441801 | 0.7091029 | 0.939810471 |
| ENSMUSG00000044352 | Sowaha | protein_coding | -0.155103667 | 5.361425796 | 0.1443682 | 0.7091726 | 0.939810471 |
| ENSMUSG00000028066 | Pmf1 | protein_coding | -0.117942966 | 4.495058868 | 0.1439487 | 0.7092818 | 0.939810471 |
| ENSMUSG00000043153 | Ispd | protein_coding | -0.143725146 | 4.227464938 | 0.1438239 | 0.7094015 | 0.939810471 |
| ENSMUSG00000018678 | Sp2 | protein_coding | -0.31907776 | 2.484319972 | 0.1437531 | 0.7094695 | 0.939810471 |
| ENSMUSG00000021000 | Ctage5 | protein_coding | -0.148630575 | 4.251313208 | 0.1436184 | 0.7095989 | 0.939810471 |
| ENSMUSG00000025576 | Rbfox3 | protein_coding | -0.075177977 | 6.697180347 | 0.1435363 | 0.7096778 | 0.939810471 |
| ENSMUSG00000054455 | Vapb | protein_coding | 0.103909378 | 5.438720093 | 0.143505 | 0.7097079 | 0.939810471 |
| ENSMUSG00000016921 | Srsf6 | protein_coding | 0.127930468 | 5.095380755 | 0.1433668 | 0.7098408 | 0.939810471 |
| ENSMUSG00000087075 | A230065H16Rik | protein_coding | -0.090047573 | 6.340546695 | 0.1433639 | 0.7098436 | 0.939810471 |
| ENSMUSG00000033707 | Lrrc24 | protein_coding | -0.152090864 | 4.087936264 | 0.1433625 | 0.709845 | 0.939810471 |
| ENSMUSG00000071042 | Rasgrp3 | protein_coding | -0.199265731 | 3.738488421 | 0.1432176 | 0.7099924 | 0.939810471 |
| ENSMUSG00000039680 | Mrps6 | protein_coding | -0.104979645 | 6.241458897 | 0.1430444 | 0.7101511 | 0.939810471 |
| ENSMUSG00000033526 | Ppip5k1 | protein_coding | 0.118455856 | 4.9857567 | 0.1430231 | 0.7101717 | 0.939810471 |
| ENSMUSG00000052584 | Serp2 | protein_coding | -0.075617394 | 6.425509516 | 0.1430149 | 0.7101796 | 0.939810471 |
| ENSMUSG00000027551 | Zfp64 | protein_coding | -0.130648714 | 5.124007071 | 0.1430133 | 0.7101811 | 0.939810471 |
| ENSMUSG00000019055 | Plod1 | protein_coding | -0.152959685 | 3.98051833 | 0.1430092 | 0.710185 | 0.939810471 |
| ENSMUSG00000003863 | Ppfia3 | protein_coding | 0.111641155 | 4.668904969 | 0.1429833 | 0.71021 | 0.939810471 |
| ENSMUSG00000038473 | Nos1ap | protein_coding | -0.097928595 | 5.935121701 | 0.1429016 | 0.7102887 | 0.939810471 |
| ENSMUSG00000042694 | Stn1 | protein_coding | 0.112598604 | 4.440789371 | 0.1427976 | 0.7103889 | 0.939810471 |
| ENSMUSG00000008683 | Rps15a | protein_coding | -0.094074486 | 7.305842225 | 0.1427898 | 0.7104044 | 0.939810471 |
| ENSMUSG00000024878 | Cbwd1 | protein_coding | 0.107613752 | 4.8534708 | 0.1426759 | 0.7105063 | 0.939810471 |
| ENSMUSG00000051497 | Kcnj16 | protein_coding | -0.179219625 | 3.231320837 | 0.14258 | 0.7105989 | 0.939810471 |
| ENSMUSG00000033781 | Asb13 | protein_coding | -0.090845203 | 5.697185403 | 0.1424186 | 0.7107547 | 0.939810471 |
| ENSMUSG00000041609 | Bicdl1 | protein_coding | 0.152632927 | 4.482857563 | 0.1424073 | 0.7107655 | 0.939810471 |
| ENSMUSG00000003534 | Ddr1 | protein_coding | 0.12132037 | 4.699261605 | 0.1423479 | 0.7108229 | 0.939810471 |
| ENSMUSG00000027282 | Mtch2 | protein_coding | 0.06400947 | 7.647824628 | 0.1421777 | 0.7109873 | 0.939912287 |
| ENSMUSG00000030500 | Slc17a6 | protein_coding | 0.346917754 | 6.462160859 | 0.1474288 | 0.7111523 | 0.940014803 |
| ENSMUSG00000026213 | Stk11ip | protein_coding | -0.192186297 | 3.316187816 | 0.1418841 | 0.7112713 | 0.940056534 |
| ENSMUSG00000003418 | St8sia6 | protein_coding | -0.253866795 | 2.527975686 | 0.1417733 | 0.7113785 | 0.94007129 |
| ENSMUSG00000006024 | Napa | protein_coding | 0.078384186 | 9.942773001 | 0.1416022 | 0.7115442 | 0.94007129 |
| ENSMUSG00000029135 | Fosl2 | protein_coding | -0.200193252 | 3.461338492 | 0.1415602 | 0.7115849 | 0.94007129 |
| ENSMUSG00000049739 | Zfp646 | protein_coding | 0.175110354 | 3.184393989 | 0.1415 | 0.7116433 | 0.94007129 |

| ensembl_gene_id | external_gene_name | gene_biotype | log2 (fold change) | log2 (counts per million) | F-value | p-value | False discovery rate |
|---|---|---|---|---|---|---|---|
| ENSMUSG00000024172 | St6gal2 | protein_coding | -0.254415464 | 2.881335926 | 0.1414213 | 0.7117196 | 0.94007129 |
| ENSMUSG00000041329 | Atp1b2 | protein_coding | 0.098681692 | 7.359503043 | 0.1412754 | 0.711869 | 0.940153164 |
| ENSMUSG00000110723 | AC153912.1 | protein_coding | -0.231182774 | 2.861603887 | 0.1407688 | 0.712353 | 0.940420026 |
| ENSMUSG00000032312 | Csk | protein_coding | 0.137634908 | 4.611753996 | 0.1407638 | 0.7123578 | 0.940420026 |
| ENSMUSG00000071054 | Safb | protein_coding | -0.120392495 | 4.510427583 | 0.1407274 | 0.7123932 | 0.940420026 |
| ENSMUSG00000002319 | Ipo4 | protein_coding | 0.102640491 | 6.218564489 | 0.1406188 | 0.7124988 | 0.940420026 |
| ENSMUSG00000027274 | Mkks | protein_coding | -0.088494287 | 7.441326678 | 0.14057 | 0.7125463 | 0.940420026 |
| ENSMUSG00000040850 | Psme4 | protein_coding | -0.120981317 | 4.122706037 | 0.1403061 | 0.7128032 | 0.940420026 |
| ENSMUSG00000040472 | Rabggta | protein_coding | -0.088325133 | 6.178422067 | 0.1402629 | 0.7128453 | 0.940420026 |
| ENSMUSG00000033075 | Senp1 | protein_coding | -0.203471709 | 3.370551733 | 0.1402607 | 0.7128474 | 0.940420026 |
| ENSMUSG00000062591 | Tubb4a | protein_coding | -0.112406693 | 11.07819344 | 0.1402793 | 0.7128581 | 0.940420026 |
| ENSMUSG00000005078 | Jkamp | protein_coding | 0.068754259 | 7.001393794 | 0.1400268 | 0.7130753 | 0.940545769 |
| ENSMUSG00000110277 | Gm45871 | protein_coding | -0.19128166 | 3.264618075 | 0.1398792 | 0.7132193 | 0.940545769 |
| ENSMUSG00000024170 | Telo2 | protein_coding | -0.144179516 | 4.060941039 | 0.1398083 | 0.7132885 | 0.940545769 |
| ENSMUSG00000030092 | Cntn6 | protein_coding | -0.141872826 | 4.10731142 | 0.1396642 | 0.7134291 | 0.940545769 |
| ENSMUSG00000055065 | Ddx17 | protein_coding | -0.109442345 | 5.506969951 | 0.1394922 | 0.7135972 | 0.940545769 |
| ENSMUSG00000030880 | Polr3e | protein_coding | -0.168341046 | 4.903822005 | 0.1397741 | 0.7136098 | 0.940545769 |
| ENSMUSG00000028013 | Ppa2 | protein_coding | 0.085924856 | 6.441135138 | 0.1394302 | 0.7136578 | 0.940545769 |
| ENSMUSG00000025558 | Dock9 | protein_coding | 0.148590452 | 4.382703992 | 0.1392547 | 0.7138293 | 0.940545769 |
| ENSMUSG00000018076 | Med13l | protein_coding | 0.109634222 | 5.03335934 | 0.1392287 | 0.7138548 | 0.940545769 |
| ENSMUSG00000021186 | Fbln5 | protein_coding | -0.295447573 | 2.269903079 | 0.1391686 | 0.7139136 | 0.940545769 |
| ENSMUSG00000038095 | Sbno1 | protein_coding | 0.090508795 | 6.455760382 | 0.1391666 | 0.7139156 | 0.940545769 |
| ENSMUSG00000031379 | Pir | protein_coding | -0.178924126 | 3.804556121 | 0.1389901 | 0.7140883 | 0.940658082 |
| ENSMUSG00000045107 | Saysd1 | protein_coding | -0.101446178 | 5.068478322 | 0.1388562 | 0.7142195 | 0.940715648 |
| ENSMUSG00000073678 | Pgap1 | protein_coding | -0.114898931 | 4.58660552 | 0.1384053 | 0.7146617 | 0.941182731 |
| ENSMUSG00000064246 | Chil1 | protein_coding | 0.153892741 | 4.459438099 | 0.1382135 | 0.7148579 | 0.941272403 |
| ENSMUSG00000020023 | Tmcc3 | protein_coding | -0.331938371 | 2.391541803 | 0.1381577 | 0.7149048 | 0.941272403 |
| ENSMUSG00000010097 | Nxf1 | protein_coding | 0.064571187 | 7.710549011 | 0.1380433 | 0.7150172 | 0.941305122 |
| ENSMUSG00000003345 | Csnk1g2 | protein_coding | -0.126857745 | 5.019597493 | 0.1378693 | 0.7151884 | 0.941385922 |
| ENSMUSG00000021917 | Spcs1 | protein_coding | -0.090596328 | 8.833726053 | 0.137723 | 0.7153324 | 0.941385922 |
| ENSMUSG00000022505 | Emp2 | protein_coding | -0.210613242 | 3.382255567 | 0.1377219 | 0.7153412 | 0.941385922 |
| ENSMUSG00000041353 | Tmem29 | protein_coding | 0.0751656 | 6.878532586 | 0.1375868 | 0.7154665 | 0.941435489 |
| ENSMUSG00000026204 | Ptprn | protein_coding | 0.081743065 | 7.339596512 | 0.1372854 | 0.7157635 | 0.941580221 |
| ENSMUSG00000028517 | Plpp3 | protein_coding | -0.118549352 | 6.071146533 | 0.1371248 | 0.7159504 | 0.941580221 |
| ENSMUSG00000022353 | Mtss1 | protein_coding | -0.09465908 | 5.152988467 | 0.1370764 | 0.7159697 | 0.941580221 |
| ENSMUSG00000026614 | Slc30a10 | protein_coding | -0.216765966 | 3.199459821 | 0.1370738 | 0.7159722 | 0.941580221 |
| ENSMUSG00000079179 | Rab10os | antisense_RNA | 0.103043421 | 6.313609855 | 0.1370022 | 0.7160508 | 0.941580221 |
| ENSMUSG00000042978 | Sbk1 | protein_coding | -0.147019193 | 4.205062604 | 0.1368679 | 0.7161756 | 0.941580221 |
| ENSMUSG00000046341 | Gm11223 | processed_pseudoge | 0.160265447 | 3.20080586 | 0.1367612 | 0.716281 | 0.941580221 |
| ENSMUSG00000039901 | 9130011E15Rik | protein_coding | -0.159262063 | 3.936812485 | 0.1366033 | 0.7164371 | 0.941580221 |
| ENSMUSG00000032667 | Pon2 | protein_coding | -0.110598601 | 5.017661309 | 0.1364768 | 0.7165622 | 0.941580221 |
| ENSMUSG00000050846 | Zfp623 | protein_coding | 0.122900888 | 4.333014828 | 0.1363753 | 0.7166626 | 0.941580221 |
| ENSMUSG00000024511 | Rab27b | protein_coding | -0.108024792 | 6.756997328 | 0.1364653 | 0.716706 | 0.941580221 |
| ENSMUSG00000044876 | Zfp444 | protein_coding | 0.118442057 | 5.120859974 | 0.1363046 | 0.7167327 | 0.941580221 |
| ENSMUSG00000041360 | Pum3 | protein_coding | -0.093465887 | 5.666340357 | 0.1362966 | 0.7167406 | 0.941580221 |
| ENSMUSG00000021919 | Chat | protein_coding | 0.202869011 | 3.470664016 | 0.1362131 | 0.7168311 | 0.941580221 |
| ENSMUSG00000073639 | Rab18 | protein_coding | 0.077115555 | 8.258163609 | 0.1361215 | 0.716914 | 0.941580221 |
| ENSMUSG00000030630 | Fah | protein_coding | 0.098761829 | 5.376407871 | 0.1360575 | 0.7169775 | 0.941580221 |
| ENSMUSG00000032320 | Rcn2 | protein_coding | 0.108077338 | 6.618201067 | 0.1359671 | 0.7171089 | 0.941637783 |
| ENSMUSG00000005125 | Ndrg1 | protein_coding | -0.101143592 | 5.389788109 | 0.135685 | 0.717347 | 0.94177514 |
| ENSMUSG00000019738 | Polr2i | protein_coding | -0.090583683 | 6.638654352 | 0.1355901 | 0.7174412 | 0.94177514 |
| ENSMUSG00000040018 | Cox15 | protein_coding | 0.169881249 | 3.750012357 | 0.1353586 | 0.7176713 | 0.94177514 |
| ENSMUSG00000029860 | Zyx | protein_coding | -0.126227557 | 4.171575472 | 0.1352871 | 0.7177423 | 0.94177514 |
| ENSMUSG00000040824 | Snrpd2 | protein_coding | -0.08761115 | 7.163819088 | 0.1352002 | 0.7178287 | 0.94177514 |
| ENSMUSG00000047153 | Khnyn | protein_coding | 0.262811843 | 2.890848492 | 0.1351477 | 0.7178888 | 0.94177514 |
| ENSMUSG00000052364 | B630019K06Rik | protein_coding | 0.102268973 | 5.775078088 | 0.1351269 | 0.7179017 | 0.94177514 |
| ENSMUSG00000020940 | 1700023F06Rik | protein_coding | -0.114438649 | 4.456806218 | 0.1351144 | 0.7179141 | 0.94177514 |
| ENSMUSG00000036550 | Cnot1 | protein_coding | 0.118648728 | 4.584060115 | 0.1349404 | 0.7180874 | 0.941887481 |
| ENSMUSG00000045160 | Bola3 | protein_coding | 0.12616068 | 4.126265745 | 0.1347099 | 0.7183171 | 0.942014539 |
| ENSMUSG00000063873 | Slc24a3 | protein_coding | -0.10107152 | 6.203920954 | 0.1345952 | 0.7184314 | 0.942014539 |
| ENSMUSG00000007207 | Stx1a | protein_coding | 0.116942302 | 7.031155404 | 0.1349678 | 0.7184471 | 0.942014539 |
| ENSMUSG00000005262 | Ufd1 | protein_coding | -0.09006511 | 7.572505418 | 0.1344161 | 0.7186179 | 0.942123631 |
| ENSMUSG00000026663 | Atf6 | protein_coding | -0.120481536 | 4.068017822 | 0.1343071 | 0.718719 | 0.942141279 |
| ENSMUSG00000021226 | Acot2 | protein_coding | 0.129238825 | 5.08055963 | 0.1341949 | 0.7188309 | 0.942173215 |
| ENSMUSG00000032518 | Rpsa | protein_coding | 0.138278275 | 3.711441235 | 0.1340683 | 0.7189575 | 0.942178339 |
| ENSMUSG00000024991 | Eif3a | protein_coding | -0.104663794 | 6.49308973 | 0.1338457 | 0.7191878 | 0.942178339 |
| ENSMUSG00000113679 | AC159633.2 | processed_transcript | -0.151659532 | 3.571613855 | 0.133821 | 0.7192048 | 0.942178339 |
| ENSMUSG00000002820 | Atg4d | protein_coding | -0.17150621 | 3.81406085 | 0.1337282 | 0.7192977 | 0.942178339 |
| ENSMUSG00000061808 | Ttr | protein_coding | -0.33958401 | 11.33229296 | 0.1387586 | 0.7193323 | 0.942178339 |
| ENSMUSG00000041540 | Sox5 | protein_coding | 0.132915567 | 4.527138898 | 0.1336654 | 0.7193606 | 0.942178339 |
| ENSMUSG00000029106 | Add1 | protein_coding | -0.068496269 | 7.534351231 | 0.1334959 | 0.7195303 | 0.942253564 |
| ENSMUSG00000020790 | Ankfy1 | protein_coding | 0.132602469 | 4.178651392 | 0.1332603 | 0.7197665 | 0.942253564 |
| ENSMUSG00000029131 | Dnajb6 | protein_coding | -0.119956405 | 4.287201928 | 0.1332456 | 0.7197812 | 0.942253564 |
| ENSMUSG00000028656 | Cap1 | protein_coding | 0.091516762 | 6.506580688 | 0.1332216 | 0.7198054 | 0.942253564 |

| ensembl_gene_id | external_gene_name | gene_biotype | log2 (fold change) | log2 (counts per million) | F-value | p-value | False discovery rate |
|---|---|---|---|---|---|---|---|
| ENSMUSG00000013150 | Gfod2 | protein_coding | -0.07068016 | 6.447896045 | 0.13297 | 0.7200579 | 0.942253564 |
| ENSMUSG00000039703 | Nploc4 | protein_coding | -0.102885743 | 4.976776944 | 0.1329139 | 0.7201143 | 0.942253564 |
| ENSMUSG00000046312 | AI464131 | protein_coding | -0.233988417 | 2.567405261 | 0.1329095 | 0.7201187 | 0.942253564 |
| ENSMUSG00000005501 | Usp40 | protein_coding | 0.192352309 | 2.889453536 | 0.1327635 | 0.7202654 | 0.942253564 |
| ENSMUSG00000050556 | Kcnb1 | protein_coding | -0.119265165 | 5.253583299 | 0.1326527 | 0.7203768 | 0.942253564 |
| ENSMUSG00000033799 | Fam208b | protein_coding | -0.278705895 | 2.470023968 | 0.1324987 | 0.7205317 | 0.942253564 |
| ENSMUSG00000003099 | Ppp5c | protein_coding | 0.09123101 | 7.03799125 | 0.13244 | 0.7205985 | 0.942253564 |
| ENSMUSG00000061136 | Prpf40a | protein_coding | -0.105464694 | 5.872851531 | 0.1324173 | 0.7206136 | 0.942253564 |
| ENSMUSG00000057729 | Prtn3 | protein_coding | 0.342458582 | 2.570975663 | 0.1324027 | 0.7206695 | 0.942253564 |
| ENSMUSG00000039458 | Mtmr12 | protein_coding | -0.154332874 | 3.255889402 | 0.1323289 | 0.7207026 | 0.942253564 |
| ENSMUSG00000015961 | Adss | protein_coding | 0.083788136 | 5.890868046 | 0.1322994 | 0.7207324 | 0.942253564 |
| ENSMUSG00000078161 | Erich3 | protein_coding | 0.150092939 | 4.031065691 | 0.1321921 | 0.7208404 | 0.942280247 |
| ENSMUSG00000034799 | Unc13a | protein_coding | 0.139040216 | 3.774578642 | 0.1318153 | 0.7212205 | 0.942315154 |
| ENSMUSG00000020151 | Ptprr | protein_coding | 0.100984978 | 4.876245521 | 0.1317295 | 0.7213071 | 0.942315154 |
| ENSMUSG00000010936 | Vac14 | protein_coding | -0.115508863 | 4.58232797 | 0.1316012 | 0.7214367 | 0.942315154 |
| ENSMUSG00000030166 | Rad52 | protein_coding | -0.110546012 | 4.579699803 | 0.1314996 | 0.7215393 | 0.942315154 |
| ENSMUSG00000024457 | Trim26 | protein_coding | 0.119659777 | 4.668947304 | 0.1314957 | 0.7215432 | 0.942315154 |
| ENSMUSG00000004100 | Ppan | protein_coding | -0.164504324 | 3.711390121 | 0.1314538 | 0.7215856 | 0.942315154 |
| ENSMUSG00000037664 | Cdkn1c | protein_coding | 0.279353783 | 2.328990306 | 0.1314371 | 0.7216025 | 0.942315154 |
| ENSMUSG00000035049 | Rrp12 | protein_coding | 0.120921932 | 4.313849598 | 0.1313659 | 0.7216744 | 0.942315154 |
| ENSMUSG00000038807 | Rap1gap2 | protein_coding | -0.110036383 | 5.432078849 | 0.1311568 | 0.721886 | 0.942315154 |
| ENSMUSG00000071691 | Gm960 | protein_coding | 0.308053606 | 2.546245427 | 0.1311631 | 0.7218872 | 0.942315154 |
| ENSMUSG00000045294 | Insig1 | protein_coding | -0.102006675 | 5.877428147 | 0.1311095 | 0.7219339 | 0.942315154 |
| ENSMUSG00000025889 | Snca | protein_coding | -0.066432622 | 9.380044398 | 0.1310739 | 0.7219699 | 0.942315154 |
| ENSMUSG00000032826 | Ank2 | protein_coding | 0.083908064 | 6.614685541 | 0.1309364 | 0.7221092 | 0.942315154 |
| ENSMUSG00000040550 | Otud6b | protein_coding | -0.100640363 | 5.995032259 | 0.1309229 | 0.7221229 | 0.942315154 |
| ENSMUSG00000001138 | Cnnm3 | protein_coding | 0.139988928 | 4.023338172 | 0.1308649 | 0.7221816 | 0.942315154 |
| ENSMUSG00000070939 | Tgfbrap1 | protein_coding | -0.121429833 | 4.076641298 | 0.1304277 | 0.7226252 | 0.942779472 |
| ENSMUSG00000002014 | Ssr4 | protein_coding | 0.08761759 | 7.614212691 | 0.1302629 | 0.7227926 | 0.942828525 |
| ENSMUSG00000005312 | Ubqln1 | protein_coding | -0.065409009 | 7.206045527 | 0.1302181 | 0.7228381 | 0.942828525 |
| ENSMUSG00000020953 | Coch | protein_coding | 0.147169147 | 3.620251044 | 0.1300811 | 0.7229773 | 0.942864012 |
| ENSMUSG00000032402 | Smad3 | protein_coding | -0.131826487 | 4.130400917 | 0.1299573 | 0.7231033 | 0.942864012 |
| ENSMUSG00000075284 | Wipf1 | protein_coding | -0.168693952 | 3.142378486 | 0.1298921 | 0.7231696 | 0.942864012 |
| ENSMUSG00000015342 | Xk | protein_coding | 0.312823336 | 3.146942023 | 0.1304022 | 0.7232519 | 0.942864012 |
| ENSMUSG00000068220 | Lgals1 | protein_coding | -0.104846477 | 5.289753324 | 0.1297352 | 0.7233294 | 0.942864012 |
| ENSMUSG00000038759 | Nup205 | protein_coding | -0.176248969 | 3.095527149 | 0.1296743 | 0.7233914 | 0.942864012 |
| ENSMUSG00000092416 | Zfp141 | protein_coding | -0.178648307 | 3.008401772 | 0.1294473 | 0.7236228 | 0.943031022 |
| ENSMUSG00000018169 | Mfng | protein_coding | -0.1548832 | 3.752551007 | 0.1293765 | 0.723695 | 0.943031022 |
| ENSMUSG00000022179 | 4931414P19Rik | protein_coding | 0.239424812 | 2.714418146 | 0.1292788 | 0.7237947 | 0.943046679 |
| ENSMUSG00000074909 | Ranbp6 | protein_coding | 0.1204453 | 5.08870337 | 0.1291387 | 0.7239377 | 0.943085679 |
| ENSMUSG00000067430 | Zfp763 | protein_coding | 0.130847181 | 4.197765692 | 0.1290777 | 0.724 | 0.943085679 |
| ENSMUSG00000057230 | Aak1 | protein_coding | 0.071403148 | 6.234859702 | 0.128886 | 0.7241959 | 0.943226549 |
| ENSMUSG00000028413 | B4galt1 | protein_coding | -0.187429858 | 2.837557655 | 0.1287776 | 0.7243067 | 0.943234405 |
| ENSMUSG00000090061 | Nwd2 | protein_coding | -0.263746287 | 2.857913007 | 0.1286873 | 0.7244066 | 0.943234405 |
| ENSMUSG00000017291 | Taok1 | protein_coding | 0.126224336 | 3.751936543 | 0.1285595 | 0.7245299 | 0.943234405 |
| ENSMUSG00000066551 | Hmgb1 | protein_coding | -0.066972823 | 6.525799609 | 0.1285371 | 0.7245528 | 0.943234405 |
| ENSMUSG00000019428 | Fkbp8 | protein_coding | -0.06524976 | 8.438425054 | 0.1281856 | 0.7249129 | 0.943589011 |
| ENSMUSG00000037531 | Mrpl47 | protein_coding | -0.094992013 | 5.937371966 | 0.1276278 | 0.7254857 | 0.944061122 |
| ENSMUSG00000025410 | Dctn2 | protein_coding | 0.085380889 | 8.007037595 | 0.1275123 | 0.7256044 | 0.944061122 |
| ENSMUSG00000019578 | Ubxn6 | protein_coding | 0.076932594 | 6.444794577 | 0.1274497 | 0.7256688 | 0.944061122 |
| ENSMUSG00000001441 | Npepps | protein_coding | -0.067523053 | 6.845432145 | 0.127339 | 0.7257828 | 0.944061122 |
| ENSMUSG00000031932 | Gpr83 | protein_coding | 0.112754249 | 5.762851993 | 0.1272951 | 0.7258355 | 0.944061122 |
| ENSMUSG00000037085 | Trmt12 | protein_coding | 0.104295999 | 5.502149929 | 0.1272562 | 0.725868 | 0.944061122 |
| ENSMUSG00000005354 | Txn2 | protein_coding | -0.069276201 | 9.312066701 | 0.1271157 | 0.7260127 | 0.944061122 |
| ENSMUSG00000032883 | Acsl3 | protein_coding | -0.097997819 | 5.964220527 | 0.1270434 | 0.7260872 | 0.944061122 |
| ENSMUSG00000070000 | Fcho1 | protein_coding | -0.150018771 | 3.832681954 | 0.1270269 | 0.7261042 | 0.944061122 |
| ENSMUSG00000020752 | Recql5 | protein_coding | -0.157120956 | 3.326451752 | 0.1269175 | 0.726217 | 0.944061122 |
| ENSMUSG00000038633 | Degs1 | protein_coding | -0.062450166 | 7.198477339 | 0.1268932 | 0.7262421 | 0.944061122 |
| ENSMUSG00000030583 | Sipa1l3 | protein_coding | -0.18858535 | 3.080284127 | 0.1268088 | 0.7263292 | 0.944061122 |
| ENSMUSG00000022283 | Pabpc1 | protein_coding | -0.096411225 | 5.356366693 | 0.1265844 | 0.7265607 | 0.9442254 |
| ENSMUSG00000033237 | Arid2 | protein_coding | 0.115035123 | 4.539216334 | 0.1265038 | 0.726644 | 0.9442254 |
| ENSMUSG00000034075 | Zdhhc5 | protein_coding | 0.146949959 | 3.375278264 | 0.1264312 | 0.726719 | 0.9442254 |
| ENSMUSG00000009555 | Cdk9 | protein_coding | -0.123489148 | 4.755560864 | 0.1263126 | 0.7268416 | 0.944265626 |
| ENSMUSG00000032407 | U2surp | protein_coding | -0.073402738 | 6.560169412 | 0.1260605 | 0.7271024 | 0.944265626 |
| ENSMUSG00000022940 | Pigp | protein_coding | 0.098624641 | 5.775707716 | 0.1259329 | 0.7272347 | 0.944265626 |
| ENSMUSG00000078794 | Dact3 | protein_coding | -0.120131276 | 4.872785388 | 0.1259214 | 0.7272465 | 0.944265626 |
| ENSMUSG00000052557 | Gan | protein_coding | 0.159574786 | 3.380614522 | 0.1256816 | 0.727495 | 0.944265626 |
| ENSMUSG00000017167 | Cntnap1 | protein_coding | -0.116321876 | 4.798864447 | 0.1253541 | 0.7278349 | 0.944265626 |
| ENSMUSG00000061086 | Myl4 | protein_coding | 0.190838565 | 2.909886367 | 0.125322 | 0.7278683 | 0.944265626 |
| ENSMUSG00000037242 | Clic4 | protein_coding | -0.11258091 | 5.37808778 | 0.1252832 | 0.7279085 | 0.944265626 |
| ENSMUSG00000031156 | Slc35a2 | protein_coding | -0.091890915 | 5.421110488 | 0.1252704 | 0.7279219 | 0.944265626 |
| ENSMUSG00000040466 | Blvrb | protein_coding | -0.086157335 | 5.167728801 | 0.1252308 | 0.727963 | 0.944265626 |
| ENSMUSG00000026974 | Zmynd19 | protein_coding | -0.123357639 | 3.884006846 | 0.1251695 | 0.7280267 | 0.944265626 |
| ENSMUSG00000046861 | Hectd3 | protein_coding | 0.135427017 | 4.242507124 | 0.125066 | 0.7281343 | 0.944265626 |

| ensembl_gene_id | external_gene_name | gene_biotype | log2 (fold change) | log2 (counts per million) | F-value | p-value | False discovery rate |
|---|---|---|---|---|---|---|---|
| ENSMUSG00000064215 | Ifi27 | protein_coding | -0.074958337 | 6.399140843 | 0.1250573 | 0.7281434 | 0.944265626 |
| ENSMUSG00000022752 | Tomm70a | protein_coding | 0.069135263 | 7.013920706 | 0.1249941 | 0.7282091 | 0.944265626 |
| ENSMUSG00000030905 | Crym | protein_coding | -0.133966095 | 7.711231282 | 0.1256128 | 0.7282904 | 0.944265626 |
| ENSMUSG00000028634 | Hivep3 | protein_coding | -0.133645691 | 4.62086736 | 0.1248693 | 0.7283389 | 0.944265626 |
| ENSMUSG00000030678 | Maz | protein_coding | 0.128455456 | 4.644659329 | 0.1248175 | 0.7283928 | 0.944265626 |
| ENSMUSG00000039682 | Lap3 | protein_coding | -0.077222541 | 6.696552077 | 0.1247554 | 0.7284575 | 0.944265626 |
| ENSMUSG00000059689 | Zfp637 | protein_coding | 0.062741616 | 7.198845951 | 0.1245294 | 0.7286929 | 0.944265626 |
| ENSMUSG00000020153 | Ndufs7 | protein_coding | 0.068000906 | 9.745717511 | 0.1244324 | 0.7287941 | 0.944265626 |
| ENSMUSG00000061613 | U2af1 | protein_coding | 0.11217158 | 4.06321943 | 0.1244014 | 0.7288264 | 0.944265626 |
| ENSMUSG00000054400 | Cklf | protein_coding | -0.229560387 | 2.540702746 | 0.1242507 | 0.7289837 | 0.944265626 |
| ENSMUSG00000022965 | Ifngr2 | protein_coding | 0.093686472 | 5.18798113 | 0.1242024 | 0.7290341 | 0.944265626 |
| ENSMUSG00000027346 | Gpcpd1 | protein_coding | -0.09844103 | 4.817931451 | 0.1241022 | 0.7291387 | 0.944265626 |
| ENSMUSG00000022885 | St6gal1 | protein_coding | -0.16955698 | 3.565684672 | 0.1240258 | 0.7292185 | 0.944265626 |
| ENSMUSG00000043671 | Dpy19l3 | protein_coding | 0.139459057 | 4.19884492 | 0.1239921 | 0.7292538 | 0.944265626 |
| ENSMUSG00000037692 | Ahdc1 | protein_coding | 0.155192677 | 3.881753752 | 0.1239296 | 0.7293191 | 0.944265626 |
| ENSMUSG00000052512 | Nav2 | protein_coding | -0.112647649 | 5.38433187 | 0.1238805 | 0.7293704 | 0.944265626 |
| ENSMUSG00000029156 | Sgcb | protein_coding | -0.108487313 | 5.265246916 | 0.1237597 | 0.7294967 | 0.944265626 |
| ENSMUSG00000046909 | Tefm | protein_coding | 0.0993015 | 5.180163379 | 0.1237413 | 0.7295159 | 0.944265626 |
| ENSMUSG00000032131 | Abcg4 | protein_coding | 0.100258581 | 5.149193701 | 0.1236828 | 0.7295772 | 0.944265626 |
| ENSMUSG00000035278 | Plekhj1 | protein_coding | -0.071712663 | 6.151296234 | 0.1236679 | 0.7295928 | 0.944265626 |
| ENSMUSG00000024862 | Klc2 | protein_coding | -0.128060904 | 5.55964312 | 0.1236424 | 0.7296478 | 0.944265626 |
| ENSMUSG00000048706 | Lurap1l | protein_coding | -0.117142311 | 4.247336239 | 0.1231593 | 0.7301256 | 0.944770282 |
| ENSMUSG00000078652 | Psme3 | protein_coding | -0.071473329 | 6.240512895 | 0.1229972 | 0.7302957 | 0.944770954 |
| ENSMUSG00000001521 | Tulp3 | protein_coding | -0.168329934 | 3.3495638 | 0.1229913 | 0.7303019 | 0.944770954 |
| ENSMUSG00000041328 | Pcf11 | protein_coding | 0.148268851 | 3.966971098 | 0.1227346 | 0.7305715 | 0.944899946 |
| ENSMUSG00000022142 | Nup155 | protein_coding | -0.174950161 | 2.915053981 | 0.1227291 | 0.7305773 | 0.944899946 |
| ENSMUSG00000039428 | Tmem135 | protein_coding | 0.098169539 | 5.142708133 | 0.1225042 | 0.7308138 | 0.945006298 |
| ENSMUSG00000033916 | Chmp2a | protein_coding | -0.06113822 | 9.030813832 | 0.122474 | 0.7308456 | 0.945006298 |
| ENSMUSG00000037622 | Wdtc1 | protein_coding | -0.10852686 | 4.902716224 | 0.1224002 | 0.7309232 | 0.945006298 |
| ENSMUSG00000039745 | Htatip2 | protein_coding | 0.132113449 | 4.621718336 | 0.1221938 | 0.7311406 | 0.945173727 |
| ENSMUSG00000036450 | Hif1an | protein_coding | 0.132903308 | 4.210286783 | 0.1219039 | 0.7314463 | 0.945176903 |
| ENSMUSG00000024142 | Mlst8 | protein_coding | 0.142543583 | 4.379486391 | 0.1218683 | 0.7314838 | 0.945176903 |
| ENSMUSG00000040452 | Cdh12 | protein_coding | -0.201087903 | 3.24618239 | 0.1217896 | 0.7315668 | 0.945176903 |
| ENSMUSG00000040033 | Stat2 | protein_coding | -0.242836598 | 2.603208494 | 0.1217778 | 0.7315793 | 0.945176903 |
| ENSMUSG00000014226 | Cacybp | protein_coding | -0.076547162 | 7.549578527 | 0.1216932 | 0.7316687 | 0.945176903 |
| ENSMUSG00000027993 | Trim2 | protein_coding | -0.064537514 | 7.573465912 | 0.1216517 | 0.7317125 | 0.945176903 |
| ENSMUSG00000029659 | Medag | protein_coding | 0.300216429 | 2.621532017 | 0.1216152 | 0.7317584 | 0.945176903 |
| ENSMUSG00000027894 | Slc6a17 | protein_coding | 0.081438491 | 6.121089919 | 0.1213734 | 0.7320066 | 0.945302565 |
| ENSMUSG00000027890 | Gstm4 | protein_coding | -0.096510023 | 4.802565336 | 0.1213499 | 0.7320315 | 0.945302565 |
| ENSMUSG00000029153 | Ociad2 | protein_coding | -0.058880465 | 7.775326845 | 0.1211025 | 0.7322933 | 0.945339727 |
| ENSMUSG00000010797 | Wnt2 | protein_coding | 0.189802718 | 3.163097649 | 0.1210121 | 0.732389 | 0.945339727 |
| ENSMUSG00000017412 | Cacnb4 | protein_coding | -0.082136702 | 5.601837244 | 0.1208966 | 0.7325115 | 0.945339727 |
| ENSMUSG00000005357 | Slc1a6 | protein_coding | 0.130165286 | 4.15953709 | 0.1206942 | 0.732726 | 0.945339727 |
| ENSMUSG00000028468 | Rgp1 | protein_coding | -0.162730454 | 3.156965295 | 0.1206606 | 0.7327618 | 0.945339727 |
| ENSMUSG00000032186 | Tmod2 | protein_coding | -0.065359369 | 8.400191226 | 0.120655 | 0.7327676 | 0.945339727 |
| ENSMUSG00000040952 | Rps19 | protein_coding | -0.086893817 | 8.554367268 | 0.1206025 | 0.7328307 | 0.945339727 |
| ENSMUSG00000036882 | Arhgap33 | protein_coding | 0.10045776 | 5.61869211 | 0.1202925 | 0.7331527 | 0.945339727 |
| ENSMUSG00000029246 | Ppat | protein_coding | 0.169437471 | 4.337523247 | 0.1203877 | 0.7331899 | 0.945339727 |
| ENSMUSG00000060012 | Kif13b | protein_coding | -0.142035364 | 3.707502773 | 0.1202125 | 0.7332377 | 0.945339727 |
| ENSMUSG00000043542 | Zc2hc1a | protein_coding | 0.066360071 | 7.167439517 | 0.1201806 | 0.7332716 | 0.945339727 |
| ENSMUSG00000058881 | Zfp516 | protein_coding | 0.247631552 | 2.882451343 | 0.1201209 | 0.7333424 | 0.945339727 |
| ENSMUSG00000021713 | Ppwd1 | protein_coding | 0.298708087 | 2.292183636 | 0.1201046 | 0.7333524 | 0.945339727 |
| ENSMUSG00000071645 | Tut1 | protein_coding | 0.107123743 | 5.181577807 | 0.1199426 | 0.7335249 | 0.945339727 |
| ENSMUSG00000031167 | Rbm3 | protein_coding | -0.109714808 | 4.087314132 | 0.1198813 | 0.7335902 | 0.945339727 |
| ENSMUSG00000059248 | 9-Sep | protein_coding | -0.158530151 | 5.15528298 | 0.1201976 | 0.7336167 | 0.945339727 |
| ENSMUSG00000078713 | Tomm5 | protein_coding | -0.071880662 | 6.628975089 | 0.1197607 | 0.7337187 | 0.945339727 |
| ENSMUSG00000025503 | Taldo1 | protein_coding | -0.061127975 | 8.388558491 | 0.1197339 | 0.7337472 | 0.945339727 |
| ENSMUSG00000021156 | Zmynd11 | protein_coding | -0.119931774 | 4.511812128 | 0.1196594 | 0.7338267 | 0.945339727 |
| ENSMUSG00000060073 | Psma3 | protein_coding | 0.078823448 | 6.12581061 | 0.1195992 | 0.7338908 | 0.945339727 |
| ENSMUSG00000021366 | Hivep1 | protein_coding | -0.251471585 | 2.754575673 | 0.1194045 | 0.7340986 | 0.945339727 |
| ENSMUSG00000036019 | Tmtc2 | protein_coding | 0.149675781 | 3.387662185 | 0.1193918 | 0.7341121 | 0.945339727 |
| ENSMUSG00000068551 | Zfp467 | protein_coding | 0.122125387 | 4.430571319 | 0.1193522 | 0.7341544 | 0.945339727 |
| ENSMUSG00000044221 | Grsf1 | protein_coding | -0.072678534 | 6.971257597 | 0.1193374 | 0.7341702 | 0.945339727 |
| ENSMUSG00000019579 | Mydgf | protein_coding | 0.068379254 | 7.3175133 | 0.1190996 | 0.7344243 | 0.945553645 |
| ENSMUSG00000086859 | Snhg20 | lincRNA | -0.14810043 | 3.822661677 | 0.1189856 | 0.7345461 | 0.9455973 |
| ENSMUSG00000030030 | 1700003E16Rik | protein_coding | -0.097341548 | 5.293670112 | 0.1188328 | 0.7347097 | 0.945694624 |
| ENSMUSG00000040681 | Hmgn1 | protein_coding | -0.104102229 | 4.808654613 | 0.1186027 | 0.734956 | 0.945898504 |
| ENSMUSG00000028567 | Txndc12 | protein_coding | 0.076169934 | 6.104744527 | 0.1184188 | 0.7351532 | 0.946030409 |
| ENSMUSG00000028672 | Hmgcl | protein_coding | 0.070633689 | 7.215860882 | 0.1183323 | 0.7352459 | 0.946030409 |
| ENSMUSG00000020741 | Cluh | protein_coding | 0.090856522 | 5.834445549 | 0.1178362 | 0.7357788 | 0.946030409 |
| ENSMUSG00000024201 | Kdm4b | protein_coding | -0.175512115 | 4.745665976 | 0.1182309 | 0.7357932 | 0.946030409 |
| ENSMUSG00000000902 | Smarcb1 | protein_coding | 0.07911056 | 7.733330558 | 0.1176915 | 0.7359344 | 0.946030409 |
| ENSMUSG00000025279 | Dnase1l3 | protein_coding | -0.364296783 | 1.981800295 | 0.1176295 | 0.7360012 | 0.946030409 |
| ENSMUSG00000002900 | Lamb1 | protein_coding | 0.167096083 | 3.348845359 | 0.1174659 | 0.7361774 | 0.946030409 |

| ensembl_gene_id | external_gene_name | gene_biotype | log2 (fold change) | log2 (counts per million) | F-value | p-value | False discovery rate |
|---|---|---|---|---|---|---|---|
| ENSMUSG00000020458 | Rtn4 | protein_coding | -0.07325687 | 8.419615773 | 0.1174371 | 0.7362084 | 0.946030409 |
| ENSMUSG00000026514 | Cnih3 | protein_coding | -0.111440625 | 4.499758929 | 0.1172168 | 0.7364459 | 0.946030409 |
| ENSMUSG00000035762 | Tmem161b | protein_coding | -0.10362063 | 4.604566558 | 0.117192 | 0.7364726 | 0.946030409 |
| ENSMUSG00000045598 | Zfp553 | protein_coding | -0.133701352 | 3.911544166 | 0.1170568 | 0.7366185 | 0.946030409 |
| ENSMUSG00000090733 | Rps27 | protein_coding | -0.114212754 | 4.099858358 | 0.1170421 | 0.7366344 | 0.946030409 |
| ENSMUSG00000092203 | 1110038B12Rik | processed_transcript | -0.090463019 | 5.013220904 | 0.1169022 | 0.7367854 | 0.946030409 |
| ENSMUSG00000023911 | Flywch2 | protein_coding | 0.081474282 | 5.777949659 | 0.116897 | 0.7367911 | 0.946030409 |
| ENSMUSG00000034731 | Dgkh | protein_coding | 0.160029982 | 3.239338009 | 0.1168914 | 0.7367971 | 0.946030409 |
| ENSMUSG00000041035 | Gm17018 | protein_coding | 0.063691095 | 8.814747378 | 0.1168863 | 0.7368026 | 0.946030409 |
| ENSMUSG00000048915 | Efna5 | protein_coding | -0.130797658 | 6.530370452 | 0.1174148 | 0.7368456 | 0.946030409 |
| ENSMUSG00000027488 | Snta1 | protein_coding | 0.113482094 | 4.297580333 | 0.1168235 | 0.7368705 | 0.946030409 |
| ENSMUSG00000084910 | C630043F03Rik | processed_transcript | -0.138049683 | 3.895247764 | 0.1166022 | 0.7371097 | 0.946030409 |
| ENSMUSG00000097287 | D130017N08Rik | lincRNA | 0.133823613 | 3.860943306 | 0.1165669 | 0.737148 | 0.946030409 |
| ENSMUSG00000021690 | Jmy | protein_coding | 0.124349209 | 4.531544115 | 0.1165635 | 0.7371516 | 0.946030409 |
| ENSMUSG00000035632 | Cnot3 | protein_coding | -0.127758055 | 3.568040834 | 0.116535 | 0.7371824 | 0.946030409 |
| ENSMUSG00000030304 | Ergic2 | protein_coding | -0.069900004 | 6.167755877 | 0.1165244 | 0.7371939 | 0.946030409 |
| ENSMUSG00000021432 | Slc35b3 | protein_coding | -0.180902117 | 3.149178668 | 0.1164795 | 0.7372425 | 0.946030409 |
| ENSMUSG00000028331 | Trmo | protein_coding | 0.292444872 | 3.10188917 | 0.1167636 | 0.7373706 | 0.946030409 |
| ENSMUSG00000031314 | Taf1 | protein_coding | -0.106130867 | 5.39495505 | 0.1163128 | 0.737423 | 0.946030409 |
| ENSMUSG00000028622 | Mrpl37 | protein_coding | -0.078277495 | 6.302739605 | 0.1160244 | 0.7377356 | 0.946030409 |
| ENSMUSG00000046841 | Ckap4 | protein_coding | 0.114596887 | 4.205987618 | 0.1157926 | 0.7379872 | 0.946030409 |
| ENSMUSG00000050312 | Nsun3 | protein_coding | 0.13805628 | 3.963659659 | 0.1157802 | 0.7380007 | 0.946030409 |
| ENSMUSG00000053253 | Ndfip2 | protein_coding | -0.079701959 | 5.975936153 | 0.1157097 | 0.7380772 | 0.946030409 |
| ENSMUSG00000021982 | Cdadc1 | protein_coding | 0.073508903 | 6.603513022 | 0.115709 | 0.7380781 | 0.946030409 |
| ENSMUSG00000028252 | Ccnc | protein_coding | -0.080629236 | 7.047618896 | 0.1156663 | 0.7381244 | 0.946030409 |
| ENSMUSG00000027834 | Serpini1 | protein_coding | 0.070878381 | 6.329136714 | 0.1156037 | 0.7381924 | 0.946030409 |
| ENSMUSG00000063765 | Chadl | protein_coding | -0.137838892 | 3.341909579 | 0.1155963 | 0.7382005 | 0.946030409 |
| ENSMUSG00000025921 | Rdh10 | protein_coding | -0.208073063 | 2.745578129 | 0.115583 | 0.7382149 | 0.946030409 |
| ENSMUSG00000066621 | Tecpr1 | protein_coding | -0.103754692 | 5.740775339 | 0.1155032 | 0.7383017 | 0.946030409 |
| ENSMUSG00000001666 | Ddt | protein_coding | -0.089008022 | 5.151267093 | 0.1154922 | 0.7383137 | 0.946030409 |
| ENSMUSG00000035614 | Togaram1 | protein_coding | -0.117506851 | 4.576164842 | 0.1153335 | 0.7384864 | 0.946109826 |
| ENSMUSG00000005615 | Pcyt1a | protein_coding | 0.114623556 | 5.212254866 | 0.1152294 | 0.7385996 | 0.946109826 |
| ENSMUSG00000073733 | Rsg1 | protein_coding | -0.26320027 | 2.950123744 | 0.1152198 | 0.7386442 | 0.946109826 |
| ENSMUSG00000030707 | Coro1a | protein_coding | -0.067468065 | 7.390618051 | 0.1150985 | 0.7387422 | 0.946109826 |
| ENSMUSG00000051401 | Kctd16 | protein_coding | -0.190510991 | 2.779442668 | 0.1149527 | 0.7389012 | 0.946109826 |
| ENSMUSG00000029686 | Cul1 | protein_coding | -0.074119241 | 6.502790169 | 0.1149505 | 0.7389036 | 0.946109826 |
| ENSMUSG00000025971 | Maip1 | protein_coding | -0.09302966 | 4.989493481 | 0.1145588 | 0.7393311 | 0.94642629 |
| ENSMUSG00000030669 | Calca | protein_coding | -0.318346147 | 3.089485496 | 0.115047 | 0.7394048 | 0.94642629 |
| ENSMUSG00000023015 | Racgap1 | protein_coding | -0.133148731 | 3.922597179 | 0.1143982 | 0.7395066 | 0.94642629 |
| ENSMUSG00000078572 | Ndufaf8 | protein_coding | 0.087200726 | 6.474336998 | 0.1143852 | 0.7395209 | 0.94642629 |
| ENSMUSG00000021803 | Cdhr1 | protein_coding | -0.217653962 | 2.655449623 | 0.1143212 | 0.7395908 | 0.94642629 |
| ENSMUSG00000032913 | Lrig2 | protein_coding | 0.114451057 | 3.960089999 | 0.1141759 | 0.7397499 | 0.946501173 |
| ENSMUSG00000031166 | Wdr13 | protein_coding | 0.069350318 | 5.967335504 | 0.1141026 | 0.7398301 | 0.946501173 |
| ENSMUSG00000045333 | Zfp423 | protein_coding | 0.124459678 | 4.549716161 | 0.1140265 | 0.7399134 | 0.946501173 |
| ENSMUSG00000015224 | Cyp2j9 | protein_coding | -0.117804765 | 4.778190594 | 0.1137745 | 0.7401897 | 0.946535073 |
| ENSMUSG00000036356 | Csgalnact1 | protein_coding | 0.177000042 | 3.439311521 | 0.1135671 | 0.7404172 | 0.946535073 |
| ENSMUSG00000025059 | Gk | protein_coding | -0.11216052 | 4.091340859 | 0.1134732 | 0.7405204 | 0.946535073 |
| ENSMUSG00000022808 | Snx4 | protein_coding | 0.093763742 | 5.947934191 | 0.1134301 | 0.7405677 | 0.946535073 |
| ENSMUSG00000052560 | Cpne8 | protein_coding | -0.094726831 | 5.446482968 | 0.1134288 | 0.7405691 | 0.946535073 |
| ENSMUSG00000030541 | Idh2 | protein_coding | 0.098194421 | 4.56323447 | 0.11342 | 0.7405788 | 0.946535073 |
| ENSMUSG00000012848 | Rps5 | protein_coding | -0.078345351 | 10.12689909 | 0.1133511 | 0.7406545 | 0.946535073 |
| ENSMUSG00000006411 | Nectin4 | protein_coding | 0.254303495 | 3.299685051 | 0.1135263 | 0.7408233 | 0.946535073 |
| ENSMUSG00000044367 | Slc16a13 | protein_coding | 0.358386954 | 2.677996523 | 0.1134281 | 0.7409986 | 0.946535073 |
| ENSMUSG00000021384 | Susd3 | protein_coding | 0.217055212 | 2.919735398 | 0.1129561 | 0.7410891 | 0.946535073 |
| ENSMUSG00000071573 | Rnls | protein_coding | -0.153186775 | 3.221765086 | 0.1128953 | 0.741156 | 0.946535073 |
| ENSMUSG00000045128 | Rpl18a | protein_coding | -0.080509825 | 8.363054074 | 0.1127388 | 0.7413285 | 0.946535073 |
| ENSMUSG00000041216 | Clvs1 | protein_coding | -0.124096022 | 4.45779578 | 0.1126749 | 0.7413989 | 0.946535073 |
| ENSMUSG00000028958 | Tmub1 | protein_coding | -0.069726079 | 6.454854854 | 0.1126019 | 0.7414794 | 0.946535073 |
| ENSMUSG00000007867 | Ift43 | protein_coding | -0.083669002 | 5.788973458 | 0.1125039 | 0.7415875 | 0.946535073 |
| ENSMUSG00000014786 | Slc9a5 | protein_coding | 0.15151692 | 3.415460126 | 0.1124501 | 0.7416469 | 0.946535073 |
| ENSMUSG00000022306 | Zfpm2 | protein_coding | -0.241556894 | 2.911789258 | 0.1120453 | 0.7421013 | 0.946535073 |
| ENSMUSG00000053040 | Aph1c | protein_coding | -0.14863573 | 3.38231935 | 0.1119747 | 0.7421724 | 0.946535073 |
| ENSMUSG00000029780 | Nt5c3 | protein_coding | 0.097102393 | 5.587108197 | 0.1119229 | 0.7422297 | 0.946535073 |
| ENSMUSG00000015488 | Cacfd1 | protein_coding | -0.120021384 | 4.85552178 | 0.1118519 | 0.7423083 | 0.946535073 |
| ENSMUSG00000042726 | Trafd1 | protein_coding | 0.119766425 | 4.927957212 | 0.1118097 | 0.742355 | 0.946535073 |
| ENSMUSG00000021248 | Tmed10 | protein_coding | 0.067337539 | 6.826569887 | 0.1117638 | 0.7424059 | 0.946535073 |
| ENSMUSG00000030245 | Golt1b | protein_coding | 0.074194263 | 7.497290907 | 0.1117045 | 0.7424716 | 0.946535073 |
| ENSMUSG00000044149 | Nkrf | protein_coding | -0.116925504 | 4.948426562 | 0.1116749 | 0.7425043 | 0.946535073 |
| ENSMUSG00000037214 | Thap1 | protein_coding | 0.115444167 | 3.795572481 | 0.111597 | 0.7425907 | 0.946535073 |
| ENSMUSG00000019082 | Slc25a22 | protein_coding | -0.097851324 | 7.807801147 | 0.111756 | 0.7426693 | 0.946535073 |
| ENSMUSG00000021276 | Cinp | protein_coding | 0.064532924 | 6.88534505 | 0.1115218 | 0.7426741 | 0.946535073 |
| ENSMUSG00000034371 | Tkfc | protein_coding | 0.127027492 | 3.857554488 | 0.1114811 | 0.7427192 | 0.946535073 |
| ENSMUSG00000025223 | Ldb1 | protein_coding | 0.094345785 | 5.148441697 | 0.1113477 | 0.7428673 | 0.946535073 |
| ENSMUSG00000031534 | Smim19 | protein_coding | 0.107217166 | 4.746452526 | 0.1113321 | 0.7428846 | 0.946535073 |

| ensembl_gene_id | external_gene_name | gene_biotype | log2 (fold change) | log2 (counts per million) | F-value | p-value | False discovery rate |
|---|---|---|---|---|---|---|---|
| ENSMUSG00000030075 | Cntn3 | protein_coding | -0.148366126 | 3.320332865 | 0.1113028 | 0.7429171 | 0.946535073 |
| ENSMUSG00000063800 | Prpf38a | protein_coding | 0.091180495 | 6.011208155 | 0.111251 | 0.7429746 | 0.946535073 |
| ENSMUSG00000039844 | Rapgef1 | protein_coding | -0.111608183 | 4.591278604 | 0.1111462 | 0.7430911 | 0.946535073 |
| ENSMUSG00000021629 | Slc30a5 | protein_coding | 0.116132834 | 4.547295748 | 0.1109163 | 0.7433467 | 0.946535073 |
| ENSMUSG00000055041 | Commd5 | protein_coding | 0.075358558 | 6.244817319 | 0.1108106 | 0.7434643 | 0.946535073 |
| ENSMUSG00000062393 | Dgkk | protein_coding | -0.116824786 | 4.040301523 | 0.1107278 | 0.7435565 | 0.946535073 |
| ENSMUSG00000089832 | Shkbp1 | protein_coding | -0.274208 | 2.76160648 | 0.1105906 | 0.7437163 | 0.946535073 |
| ENSMUSG00000097156 | Gm3764 | lincRNA | -0.101220378 | 5.037532287 | 0.1105393 | 0.7437665 | 0.946535073 |
| ENSMUSG00000018363 | Smurf2 | protein_coding | -0.11495457 | 4.560869623 | 0.1104769 | 0.7438361 | 0.946535073 |
| ENSMUSG00000064325 | Hhip | protein_coding | -0.195256586 | 2.593913966 | 0.1103968 | 0.7439253 | 0.946535073 |
| ENSMUSG00000024900 | Cpt1a | protein_coding | 0.269706921 | 2.519854994 | 0.1103639 | 0.743962 | 0.946535073 |
| ENSMUSG00000021116 | Eif2s1 | protein_coding | -0.061047188 | 6.639158588 | 0.1103317 | 0.7439979 | 0.946535073 |
| ENSMUSG00000015812 | Gnrh1 | protein_coding | 0.16991655 | 5.32651247 | 0.1109312 | 0.7440066 | 0.946535073 |
| ENSMUSG00000020589 | Fam49a | protein_coding | -0.167845321 | 3.25155899 | 0.1103008 | 0.7440324 | 0.946535073 |
| ENSMUSG00000060279 | Ap2a1 | protein_coding | -0.241749987 | 4.119981076 | 0.1111234 | 0.7440339 | 0.946535073 |
| ENSMUSG00000026240 | Cops7b | protein_coding | 0.104963563 | 4.814157823 | 0.1102244 | 0.7441177 | 0.946535073 |
| ENSMUSG00000040502 | 9-Mar | protein_coding | -0.164950959 | 3.218846446 | 0.1101782 | 0.7441693 | 0.946535073 |
| ENSMUSG00000023011 | Faim2 | protein_coding | 0.081330203 | 9.330157212 | 0.1100383 | 0.7443255 | 0.946535073 |
| ENSMUSG00000034708 | Grn | protein_coding | 0.078070895 | 5.539982496 | 0.1100244 | 0.7443411 | 0.946535073 |
| ENSMUSG00000020523 | Fam114a2 | protein_coding | 0.078465715 | 6.023009542 | 0.1099024 | 0.7444775 | 0.946535073 |
| ENSMUSG00000027598 | Itch | protein_coding | 0.097545746 | 4.714672349 | 0.109872 | 0.7445114 | 0.946535073 |
| ENSMUSG00000002428 | Hltf | protein_coding | 0.104056056 | 4.522616814 | 0.1097275 | 0.7446732 | 0.946535073 |
| ENSMUSG00000026109 | Tmeff2 | protein_coding | -0.072014251 | 7.305166309 | 0.1097187 | 0.744683 | 0.946535073 |
| ENSMUSG00000039233 | Tbce | protein_coding | 0.11093712 | 5.077393966 | 0.1096694 | 0.7447381 | 0.946535073 |
| ENSMUSG00000055835 | Zfp1 | protein_coding | -0.120069961 | 4.897413214 | 0.1096309 | 0.7447813 | 0.946535073 |
| ENSMUSG00000054099 | Slc25a40 | protein_coding | -0.150371434 | 3.363760727 | 0.1094693 | 0.7449623 | 0.946636522 |
| ENSMUSG00000020898 | Ctc1 | protein_coding | -0.116566421 | 4.549825919 | 0.1094025 | 0.7450372 | 0.946636522 |
| ENSMUSG00000039263 | Npepl1 | protein_coding | -0.36900762 | 2.256624466 | 0.1093839 | 0.7451894 | 0.946718031 |
| ENSMUSG00000038773 | Kdm3b | protein_coding | -0.108253714 | 3.911557916 | 0.1091426 | 0.7453287 | 0.946783172 |
| ENSMUSG00000002017 | Fam98a | protein_coding | -0.105287751 | 4.176499394 | 0.1087954 | 0.7457188 | 0.947166803 |
| ENSMUSG00000018796 | Acsl1 | protein_coding | 0.112469905 | 4.888774052 | 0.108645 | 0.745888 | 0.947192338 |
| ENSMUSG00000028729 | Ebna1bp2 | protein_coding | 0.072147453 | 6.966038955 | 0.1084997 | 0.7460516 | 0.947192338 |
| ENSMUSG00000006373 | Pgrmc1 | protein_coding | 0.063661243 | 9.279987835 | 0.1084625 | 0.7460935 | 0.947192338 |
| ENSMUSG00000056342 | Usp34 | protein_coding | -0.079533116 | 5.862036858 | 0.1083899 | 0.7461753 | 0.947192338 |
| ENSMUSG00000057101 | Zfp180 | protein_coding | 0.090650206 | 5.044179791 | 0.1083729 | 0.7461944 | 0.947192338 |
| ENSMUSG00000024754 | Tmem2 | protein_coding | 0.182125018 | 3.101771873 | 0.1081836 | 0.7464079 | 0.947192338 |
| ENSMUSG00000005687 | Bcas2 | protein_coding | -0.065402474 | 8.263035608 | 0.1080356 | 0.7465749 | 0.947192338 |
| ENSMUSG00000097102 | 2310069G16Rik | lincRNA | -0.187184109 | 3.187040995 | 0.1079894 | 0.746627 | 0.947192338 |
| ENSMUSG00000051978 | Erich1 | protein_coding | -0.118693283 | 4.202474229 | 0.1079778 | 0.7466401 | 0.947192338 |
| ENSMUSG00000024085 | Man2a1 | protein_coding | -0.293045527 | 2.203001725 | 0.1077984 | 0.7468428 | 0.947192338 |
| ENSMUSG00000020889 | Nr1d1 | protein_coding | -0.098562425 | 6.574850309 | 0.1078946 | 0.7468521 | 0.947192338 |
| ENSMUSG00000037461 | Ints7 | protein_coding | -0.098031218 | 5.262373825 | 0.1077656 | 0.7468799 | 0.947192338 |
| ENSMUSG00000010392 | Gosr1 | protein_coding | 0.110630872 | 5.264274629 | 0.1077619 | 0.746884 | 0.947192338 |
| ENSMUSG00000042410 | Agps | protein_coding | -0.107816475 | 4.159991566 | 0.1076621 | 0.746997 | 0.94719745 |
| ENSMUSG00000049878 | Rlf | protein_coding | -0.132190986 | 3.754233877 | 0.1076026 | 0.7470642 | 0.94719745 |
| ENSMUSG00000071662 | Polr2g | protein_coding | 0.070640589 | 8.576597452 | 0.1074983 | 0.7471822 | 0.947226831 |
| ENSMUSG00000035798 | Zdhhc17 | protein_coding | 0.082125282 | 5.272496121 | 0.1074265 | 0.7472636 | 0.947226831 |
| ENSMUSG00000021215 | Net1 | protein_coding | 0.199661574 | 2.936039139 | 0.1071189 | 0.7476122 | 0.947453724 |
| ENSMUSG00000075271 | Ttc30a1 | protein_coding | -0.20544904 | 2.943196988 | 0.1071131 | 0.7476188 | 0.947453724 |
| ENSMUSG00000033595 | Lgi3 | protein_coding | 0.080002715 | 5.970040407 | 0.1069675 | 0.747784 | 0.947551386 |
| ENSMUSG00000030527 | Crtc3 | protein_coding | 0.192161615 | 3.106435 | 0.1062236 | 0.7486301 | 0.948327257 |
| ENSMUSG00000024909 | Efemp2 | protein_coding | 0.221391846 | 4.049244831 | 0.1067795 | 0.7486603 | 0.948327257 |
| ENSMUSG00000079055 | Slc8a3 | protein_coding | 0.117734425 | 4.086510186 | 0.1061966 | 0.7486608 | 0.948327257 |
| ENSMUSG00000040669 | Phc1 | protein_coding | -0.129349861 | 4.314819874 | 0.1059802 | 0.7489076 | 0.94852805 |
| ENSMUSG00000026478 | Lamc1 | protein_coding | 0.163974865 | 3.023260391 | 0.1058128 | 0.7490988 | 0.948578005 |
| ENSMUSG00000048078 | Tenm4 | protein_coding | 0.126341368 | 4.19453274 | 0.1057347 | 0.7491879 | 0.948578005 |
| ENSMUSG00000039686 | Zer1 | protein_coding | -0.14848789 | 5.816096672 | 0.1062934 | 0.7492117 | 0.948578005 |
| ENSMUSG00000030659 | Nucb2 | protein_coding | -0.081835732 | 5.219319898 | 0.1054778 | 0.7494816 | 0.948650558 |
| ENSMUSG00000024384 | Iws1 | protein_coding | -0.128025485 | 3.589956295 | 0.1054417 | 0.749523 | 0.948650558 |
| ENSMUSG00000040365 | Trim41 | protein_coding | 0.141408333 | 3.930646536 | 0.1054324 | 0.7495336 | 0.948650558 |
| ENSMUSG00000024033 | Rsph1 | protein_coding | -0.078265341 | 5.553946355 | 0.1052788 | 0.7497094 | 0.948761152 |
| ENSMUSG00000031703 | Itfg1 | protein_coding | -0.087224326 | 6.88123795 | 0.1051658 | 0.7498456 | 0.948761152 |
| ENSMUSG00000046613 | Vwa5b2 | protein_coding | 0.189611108 | 2.903223785 | 0.1051093 | 0.7499036 | 0.948761152 |
| ENSMUSG00000052917 | Senp7 | protein_coding | -0.081528648 | 5.549636345 | 0.105048 | 0.7499739 | 0.948761152 |
| ENSMUSG00000010064 | Slc38a3 | protein_coding | -0.122566526 | 4.503629925 | 0.1048663 | 0.7501824 | 0.948913191 |
| ENSMUSG00000039629 | Strip2 | protein_coding | -0.134924207 | 4.260741772 | 0.1047872 | 0.7502731 | 0.948916389 |
| ENSMUSG00000015652 | Steap1 | protein_coding | -0.420768692 | 2.66212385 | 0.1053562 | 0.7505294 | 0.949037215 |
| ENSMUSG00000045464 | 2810002D19Rik | processed_transcript | -0.148409404 | 4.38772 | 0.1045303 | 0.7505834 | 0.949037215 |
| ENSMUSG00000020585 | Laptm4a | protein_coding | -0.070758266 | 8.703104482 | 0.1044737 | 0.7506335 | 0.949037215 |
| ENSMUSG00000044134 | Fam109a | protein_coding | 0.30265117 | 2.104216316 | 0.1043651 | 0.7507583 | 0.9490771 |
| ENSMUSG00000005873 | Reep5 | protein_coding | 0.06085219 | 8.580468128 | 0.1042929 | 0.7508415 | 0.9490771 |
| ENSMUSG00000084883 | Ccdc85c | protein_coding | -0.13646323 | 5.217603897 | 0.1041586 | 0.7512428 | 0.949370194 |
| ENSMUSG00000029279 | Brdt | protein_coding | -0.13901793 | 3.734904963 | 0.1039384 | 0.75125 | 0.949370194 |
| ENSMUSG00000033763 | Mtss1l | protein_coding | 0.07933186 | 5.913357951 | 0.1038497 | 0.7513524 | 0.949388012 |

| ensembl_gene_id | external_gene_name | gene_biotype | log2 (fold change) | log2 (counts per million) | F-value | p-value | False discovery rate |
|---|---|---|---|---|---|---|---|
| ENSMUSG00000027203 | Dut | protein_coding | -0.102200636 | 4.373505902 | 0.1036434 | 0.7515905 | 0.949546425 |
| ENSMUSG00000040936 | Ulk4 | protein_coding | 0.128575418 | 3.71758564 | 0.1035689 | 0.7516766 | 0.949546425 |
| ENSMUSG00000042507 | Elmsan1 | protein_coding | 0.18084045 | 2.772001058 | 0.1035117 | 0.7517426 | 0.949546425 |
| ENSMUSG00000040666 | Sh3bgr | protein_coding | -0.294678232 | 2.521521413 | 0.1033257 | 0.7519645 | 0.949715133 |
| ENSMUSG00000027167 | Elp4 | protein_coding | -0.10725976 | 3.932947935 | 0.1031963 | 0.7521076 | 0.949775844 |
| ENSMUSG00000020220 | Vps13d | protein_coding | -0.129340098 | 4.030778721 | 0.1030925 | 0.7522278 | 0.949775844 |
| ENSMUSG00000020522 | Mfap3 | protein_coding | 0.106098333 | 4.986215384 | 0.1030496 | 0.7522776 | 0.949775844 |
| ENSMUSG00000035969 | Rusc2 | protein_coding | 0.10731158 | 4.488377439 | 0.1029569 | 0.7523851 | 0.949800029 |
| ENSMUSG00000038178 | Slc43a2 | protein_coding | -0.109290179 | 4.133100289 | 0.1026631 | 0.752726 | 0.950118868 |
| ENSMUSG00000064030 | Pym1 | protein_coding | 0.097001552 | 4.753457262 | 0.1025719 | 0.7528319 | 0.95014099 |
| ENSMUSG00000074738 | Fndc10 | protein_coding | -0.113378092 | 4.143632959 | 0.1024658 | 0.7529552 | 0.950185114 |
| ENSMUSG00000030714 | Sgf29 | protein_coding | -0.09446244 | 5.825382955 | 0.1023447 | 0.7530961 | 0.950212867 |
| ENSMUSG00000050730 | Arhgap42 | protein_coding | -0.307845087 | 2.211125599 | 0.1020685 | 0.7534176 | 0.950212867 |
| ENSMUSG00000006241 | Ccdc159 | protein_coding | 0.139233501 | 3.402527741 | 0.102033 | 0.753459 | 0.950212867 |
| ENSMUSG00000035248 | Zcchc6 | protein_coding | -0.1508678 | 3.076259494 | 0.1020307 | 0.7534617 | 0.950212867 |
| ENSMUSG00000062296 | Trank1 | protein_coding | 0.125823365 | 4.826616183 | 0.1020253 | 0.7534747 | 0.950212867 |
| ENSMUSG00000032579 | Hemk1 | protein_coding | 0.105624745 | 4.990889154 | 0.1019208 | 0.7535898 | 0.950212867 |
| ENSMUSG00000028974 | Dffa | protein_coding | -0.085219846 | 5.691760046 | 0.1018446 | 0.7536786 | 0.950212867 |
| ENSMUSG00000020814 | Mxra7 | protein_coding | -0.121079186 | 3.90545563 | 0.1018265 | 0.7536998 | 0.950212867 |
| ENSMUSG00000108608 | Gm6916 | transcribed_processe | -0.608958546 | 2.572950252 | 0.1038356 | 0.7537725 | 0.950212867 |
| ENSMUSG00000017929 | B4galt5 | protein_coding | 0.095061188 | 4.77261447 | 0.1016651 | 0.7538881 | 0.950247207 |
| ENSMUSG00000020305 | Asb3 | protein_coding | -0.091676028 | 5.64929895 | 0.1015582 | 0.754013 | 0.950262697 |
| ENSMUSG00000013076 | Amotl1 | protein_coding | 0.151899222 | 7.001349671 | 0.1029714 | 0.7540771 | 0.950262697 |
| ENSMUSG00000056216 | Cebpg | protein_coding | -0.094597388 | 4.363496584 | 0.1013809 | 0.7542202 | 0.95033158 |
| ENSMUSG00000026743 | Mllt10 | protein_coding | 0.097004783 | 4.706873075 | 0.1012879 | 0.7543291 | 0.950357401 |
| ENSMUSG00000037032 | Apbb1 | protein_coding | 0.077414663 | 9.16330702 | 0.1011128 | 0.754534 | 0.950504236 |
| ENSMUSG00000005672 | Kit | protein_coding | -0.169126481 | 3.267229625 | 0.101027 | 0.7546346 | 0.950519564 |
| ENSMUSG00000024081 | Cebpz | protein_coding | -0.090399737 | 5.211813581 | 0.1009467 | 0.7547286 | 0.95052663 |
| ENSMUSG00000029999 | Tgfa | protein_coding | -0.091743835 | 4.891510745 | 0.1007421 | 0.7549686 | 0.950583314 |
| ENSMUSG00000059263 | Usp47 | protein_coding | -0.080850252 | 5.505801422 | 0.1005827 | 0.7551557 | 0.950583314 |
| ENSMUSG00000000827 | Tpd52l2 | protein_coding | 0.061574968 | 7.251869599 | 0.1005648 | 0.7551767 | 0.950583314 |
| ENSMUSG00000021131 | Erh | protein_coding | 0.120579252 | 3.646676036 | 0.1005403 | 0.7552055 | 0.950583314 |
| ENSMUSG00000025813 | Homer2 | protein_coding | 0.11249433 | 4.084011976 | 0.1005318 | 0.7552156 | 0.950583314 |
| ENSMUSG00000033487 | Fndc3a | protein_coding | 0.077938534 | 5.590292676 | 0.1004529 | 0.7553083 | 0.95058869 |
| ENSMUSG00000025739 | Gng13 | protein_coding | 0.104225545 | 4.249985549 | 0.1002815 | 0.7555098 | 0.950631903 |
| ENSMUSG00000101585 | 1600010M07Rik | antisense_RNA | -0.193812279 | 2.830697685 | 0.1002734 | 0.7555194 | 0.950631903 |
| ENSMUSG00000078619 | Smarcd2 | protein_coding | 0.259971599 | 2.691362152 | 0.1000525 | 0.7557861 | 0.950684252 |
| ENSMUSG00000020484 | Xbp1 | protein_coding | 0.090335017 | 5.883448601 | 0.1000302 | 0.7558058 | 0.950684252 |
| ENSMUSG00000035621 | Midn | protein_coding | -0.148616289 | 2.965152088 | 0.0998885 | 0.7559727 | 0.950684252 |
| ENSMUSG00000023087 | Noct | protein_coding | -0.091360634 | 4.931298851 | 0.0998832 | 0.755979 | 0.950684252 |
| ENSMUSG00000034341 | Wbp2 | protein_coding | 0.084053886 | 10.2406158 | 0.099858 | 0.7560153 | 0.950684252 |
| ENSMUSG00000036580 | Spg20 | protein_coding | 0.089249213 | 5.385816295 | 0.0997878 | 0.7560915 | 0.950684252 |
| ENSMUSG00000075296 | Aldh3b2 | protein_coding | 0.092484994 | 5.345022307 | 0.0996066 | 0.7563053 | 0.950841958 |
| ENSMUSG00000031295 | Phka2 | protein_coding | -0.186463492 | 3.009510972 | 0.0995088 | 0.7564208 | 0.950875993 |
| ENSMUSG00000071415 | Rpl23 | protein_coding | -0.080441866 | 7.520585782 | 0.0993955 | 0.7565614 | 0.950941561 |
| ENSMUSG00000047181 | Samd14 | protein_coding | 0.087455561 | 5.095027515 | 0.0990664 | 0.7569443 | 0.951022678 |
| ENSMUSG00000020184 | Mdm2 | protein_coding | -0.093928774 | 4.96848928 | 0.0989161 | 0.7571223 | 0.951022678 |
| ENSMUSG00000022106 | Rcbtb2 | protein_coding | 0.166365824 | 3.564056249 | 0.0987827 | 0.7572871 | 0.951022678 |
| ENSMUSG00000015671 | Psma2 | protein_coding | 0.06634352 | 8.831763041 | 0.0987401 | 0.7573311 | 0.951022678 |
| ENSMUSG00000034000 | Neu4 | protein_coding | -0.113217141 | 4.023571202 | 0.0986867 | 0.7573945 | 0.951022678 |
| ENSMUSG00000038587 | Akap12 | protein_coding | -0.13935125 | 3.967296444 | 0.0986663 | 0.7574187 | 0.951022678 |
| ENSMUSG00000022387 | Brd1 | protein_coding | -0.098181392 | 4.943559841 | 0.0986614 | 0.7574245 | 0.951022678 |
| ENSMUSG00000037533 | Rapgef6 | protein_coding | 0.095722486 | 5.157853183 | 0.0985988 | 0.7574988 | 0.951022678 |
| ENSMUSG00000039262 | Prrc2b | protein_coding | 0.08421736 | 5.773422834 | 0.0985518 | 0.7575547 | 0.951022678 |
| ENSMUSG00000040118 | Cacna2d1 | protein_coding | -0.074320541 | 5.728301998 | 0.0984194 | 0.757712 | 0.951022678 |
| ENSMUSG00000023912 | Slc25a27 | protein_coding | 0.08427896 | 4.990672139 | 0.0982118 | 0.7579589 | 0.951022678 |
| ENSMUSG00000029446 | Psph | protein_coding | 0.085082031 | 5.792015047 | 0.0981314 | 0.7580547 | 0.951022678 |
| ENSMUSG00000025949 | Pikfyve | protein_coding | -0.107802116 | 4.167803261 | 0.0981212 | 0.7580668 | 0.951022678 |
| ENSMUSG00000041762 | Gpr155 | protein_coding | -0.131470449 | 4.032145918 | 0.0980168 | 0.7581911 | 0.951022678 |
| ENSMUSG00000040025 | Ythdf2 | protein_coding | -0.066735391 | 6.242873669 | 0.0979864 | 0.7582274 | 0.951022678 |
| ENSMUSG00000035242 | Oaz1 | protein_coding | -0.058202832 | 8.286674294 | 0.0978856 | 0.7583475 | 0.951022678 |
| ENSMUSG00000015357 | Clpx | protein_coding | 0.084600722 | 5.129757506 | 0.0978721 | 0.7583637 | 0.951022678 |
| ENSMUSG00000036432 | Siah2 | protein_coding | 0.150187512 | 3.735798607 | 0.0978019 | 0.7584474 | 0.951022678 |
| ENSMUSG00000021327 | Zkscan3 | protein_coding | -0.118697305 | 3.967106536 | 0.0977933 | 0.7584576 | 0.951022678 |
| ENSMUSG00000028691 | Prdx1 | protein_coding | -0.064824302 | 9.35749687 | 0.0977781 | 0.7584757 | 0.951022678 |
| ENSMUSG00000020183 | Cpm | protein_coding | 0.136120744 | 3.680920737 | 0.0977718 | 0.7584833 | 0.951022678 |
| ENSMUSG00000024558 | Mapk4 | protein_coding | -0.093087341 | 4.869485368 | 0.0976148 | 0.7586707 | 0.951129245 |
| ENSMUSG00000030556 | Lrrc28 | protein_coding | 0.06908981 | 5.877708572 | 0.0974803 | 0.7588313 | 0.951129245 |
| ENSMUSG00000025156 | Gps1 | protein_coding | -0.072139286 | 5.477462297 | 0.097459 | 0.7588568 | 0.951129245 |
| ENSMUSG00000039043 | Arpin | protein_coding | 0.100675443 | 5.12965249 | 0.0974044 | 0.7589221 | 0.951129245 |
| ENSMUSG00000050931 | Sgms2 | protein_coding | -0.278029007 | 2.269271934 | 0.0972392 | 0.7591196 | 0.951265969 |
| ENSMUSG00000021108 | Prkch | protein_coding | -0.123516079 | 4.471156913 | 0.0969728 | 0.7594387 | 0.951554958 |
| ENSMUSG00000029185 | Fam114a1 | protein_coding | 0.313291865 | 2.987001015 | 0.0973378 | 0.7595854 | 0.951627867 |
| ENSMUSG00000058594 | Fbxo18 | protein_coding | -0.074258862 | 5.709384351 | 0.0967705 | 0.7596814 | 0.951637215 |

| ensembl_gene_id | external_gene_name | gene_biotype | log2 (fold change) | log2 (counts per million) | F-value | p-value | False discovery rate |
|---|---|---|---|---|---|---|---|
| ENSMUSG00000024896 | Minpp1 | protein_coding | 0.10444792 | 4.77867413 | 0.0965928 | 0.7598947 | 0.951735657 |
| ENSMUSG00000000127 | Fer | protein_coding | 0.104812087 | 4.309187938 | 0.0965539 | 0.7599414 | 0.951735657 |
| ENSMUSG00000028990 | Lzic | protein_coding | -0.116843915 | 4.126036374 | 0.0964274 | 0.7600935 | 0.951735657 |
| ENSMUSG00000032536 | Trak1 | protein_coding | 0.073983749 | 5.859126942 | 0.0964103 | 0.760114 | 0.951735657 |
| ENSMUSG00000033540 | Idua | protein_coding | -0.112138934 | 4.047153992 | 0.0961469 | 0.760431 | 0.951873199 |
| ENSMUSG00000033769 | Exoc6b | protein_coding | -0.0848158 | 5.277434501 | 0.0961234 | 0.7604593 | 0.951873199 |
| ENSMUSG00000074259 | Gramd2 | protein_coding | 0.252673742 | 2.559464492 | 0.0960983 | 0.7604894 | 0.951873199 |
| ENSMUSG00000022064 | Pibf1 | protein_coding | -0.153919391 | 3.199530729 | 0.0957958 | 0.7608542 | 0.952058156 |
| ENSMUSG00000053825 | Ppfia2 | protein_coding | -0.06729478 | 6.398884823 | 0.0956217 | 0.7610644 | 0.952058156 |
| ENSMUSG00000074305 | Peak1 | protein_coding | 0.151652338 | 3.140702015 | 0.0955234 | 0.761183 | 0.952058156 |
| ENSMUSG00000020099 | Unc5b | protein_coding | -0.117779508 | 4.237964072 | 0.0954849 | 0.7612296 | 0.952058156 |
| ENSMUSG00000038312 | Edem2 | protein_coding | 0.155407451 | 4.458289532 | 0.0956818 | 0.7612412 | 0.952058156 |
| ENSMUSG00000002661 | Alkbh7 | protein_coding | -0.063287506 | 6.488680113 | 0.0953534 | 0.7613887 | 0.952058156 |
| ENSMUSG00000044285 | Gm1821 | processed_pseudoge | 0.313438763 | 5.447086433 | 0.0982924 | 0.7616577 | 0.952058156 |
| ENSMUSG00000020376 | Rnf130 | protein_coding | -0.12666729 | 3.970070322 | 0.0950794 | 0.7617204 | 0.952058156 |
| ENSMUSG00000024212 | Mllt1 | protein_coding | 0.11985155 | 4.066042031 | 0.0950077 | 0.7618072 | 0.952058156 |
| ENSMUSG00000003344 | Btbd2 | protein_coding | -0.068717308 | 6.043958302 | 0.0949671 | 0.7618565 | 0.952058156 |
| ENSMUSG00000007589 | Tinf2 | protein_coding | -0.133158854 | 3.47534983 | 0.0948699 | 0.7619744 | 0.952058156 |
| ENSMUSG00000097486 | Gm17733 | bidirectional_promoter | -0.334810981 | 2.426052377 | 0.0949708 | 0.7620854 | 0.952058156 |
| ENSMUSG00000022350 | Washc5 | protein_coding | 0.095161829 | 4.438826356 | 0.09473 | 0.7621442 | 0.952058156 |
| ENSMUSG00000050002 | Idnk | protein_coding | -0.065703188 | 7.140977413 | 0.0946889 | 0.7621941 | 0.952058156 |
| ENSMUSG00000008305 | Tle1 | protein_coding | -0.097290501 | 4.754083591 | 0.0946406 | 0.7622527 | 0.952058156 |
| ENSMUSG00000018800 | Abca5 | protein_coding | 0.108362621 | 3.999379252 | 0.0945992 | 0.762303 | 0.952058156 |
| ENSMUSG00000034424 | Gcsh | protein_coding | 0.058628267 | 7.694820109 | 0.0945521 | 0.7623602 | 0.952058156 |
| ENSMUSG00000065990 | Aurkaip1 | protein_coding | 0.059619915 | 8.863702639 | 0.094415 | 0.7625269 | 0.952058156 |
| ENSMUSG00000076928 | Trac | TR_C_gene | -0.284592408 | 2.465869828 | 0.0942108 | 0.7627819 | 0.952058156 |
| ENSMUSG00000076439 | Mog | protein_coding | -0.064253438 | 7.849745662 | 0.0941839 | 0.7628082 | 0.952058156 |
| ENSMUSG00000031429 | Psmd10 | protein_coding | 0.062426017 | 6.94919868 | 0.0940929 | 0.762919 | 0.952058156 |
| ENSMUSG00000036169 | Sostdc1 | protein_coding | -0.161062166 | 5.072898366 | 0.0945211 | 0.7629433 | 0.952058156 |
| ENSMUSG00000007670 | Khsrp | protein_coding | 0.08154748 | 5.071241666 | 0.0939457 | 0.7630985 | 0.952058156 |
| ENSMUSG00000032537 | Ephb1 | protein_coding | -0.213987779 | 2.574719015 | 0.0939046 | 0.7631487 | 0.952058156 |
| ENSMUSG00000039191 | Rbpj | protein_coding | 0.137265741 | 3.246021187 | 0.0938963 | 0.7631588 | 0.952058156 |
| ENSMUSG00000020656 | Grhl1 | protein_coding | -0.177309468 | 3.637464104 | 0.0939032 | 0.7631845 | 0.952058156 |
| ENSMUSG00000009216 | Fam163b | protein_coding | -0.108662433 | 4.936179858 | 0.093847 | 0.763219 | 0.952058156 |
| ENSMUSG00000005470 | Asf1b | protein_coding | 0.364045341 | 2.029709115 | 0.0938514 | 0.7632366 | 0.952058156 |
| ENSMUSG00000052609 | Plekhg3 | protein_coding | -0.136045081 | 3.484572256 | 0.0937624 | 0.7633222 | 0.952058156 |
| ENSMUSG00000030237 | Slco1a4 | protein_coding | 0.233979459 | 2.316407609 | 0.0935989 | 0.7635219 | 0.952058156 |
| ENSMUSG00000051022 | Hs3st1 | protein_coding | -0.10207047 | 4.575954066 | 0.0935229 | 0.7636147 | 0.952058156 |
| ENSMUSG00000005897 | Nr2c1 | protein_coding | -0.182117482 | 2.943919074 | 0.0934525 | 0.7637009 | 0.952058156 |
| ENSMUSG00000028964 | Park7 | protein_coding | 0.055240085 | 10.18779602 | 0.0934284 | 0.7637303 | 0.952058156 |
| ENSMUSG00000087403 | Kantr | protein_coding | -0.08444533 | 6.665898781 | 0.0933059 | 0.7638867 | 0.952058156 |
| ENSMUSG00000008859 | Rala | protein_coding | 0.078530381 | 5.393500259 | 0.0932818 | 0.7639097 | 0.952058156 |
| ENSMUSG00000034923 | Ly6g6f | protein_coding | 0.164133784 | 3.087477309 | 0.0931664 | 0.7640511 | 0.952058156 |
| ENSMUSG00000029518 | Rab35 | protein_coding | -0.07841236 | 5.597696146 | 0.0931085 | 0.7641221 | 0.952058156 |
| ENSMUSG00000046658 | Zfp316 | protein_coding | -0.185913513 | 2.786277545 | 0.093108 | 0.7641227 | 0.952058156 |
| ENSMUSG00000028454 | Pigo | protein_coding | 0.167729472 | 3.200219405 | 0.0930676 | 0.7641722 | 0.952058156 |
| ENSMUSG00000006127 | Inpp5k | protein_coding | 0.096715081 | 5.897336812 | 0.0930675 | 0.7641787 | 0.952058156 |
| ENSMUSG00000078789 | Dph1 | protein_coding | -0.145630625 | 3.712056252 | 0.092739 | 0.7645755 | 0.952088495 |
| ENSMUSG00000029550 | Sppl3 | protein_coding | -0.096677152 | 4.9790143 | 0.0927191 | 0.7645999 | 0.952088495 |
| ENSMUSG00000031024 | St5 | protein_coding | 0.149136345 | 3.050117708 | 0.0927056 | 0.7646165 | 0.952088495 |
| ENSMUSG00000042010 | Acacb | protein_coding | 0.148896314 | 4.116900543 | 0.0926763 | 0.7646588 | 0.952088495 |
| ENSMUSG00000048490 | Nrip1 | protein_coding | -0.100789239 | 4.388132433 | 0.0926673 | 0.7646635 | 0.952088495 |
| ENSMUSG00000027475 | Kif3b | protein_coding | -0.088797115 | 4.750649199 | 0.09259 | 0.7647586 | 0.952088495 |
| ENSMUSG00000042684 | Npl | protein_coding | 0.17581462 | 3.115437471 | 0.0925246 | 0.764839 | 0.952088495 |
| ENSMUSG00000047844 | Bex4 | protein_coding | 0.078365732 | 5.16928169 | 0.0923797 | 0.7650172 | 0.952088495 |
| ENSMUSG00000037966 | Ninj1 | protein_coding | 0.094905174 | 4.936713085 | 0.0923067 | 0.7651071 | 0.952088495 |
| ENSMUSG00000041040 | Fam117b | protein_coding | -0.195592929 | 2.756767461 | 0.092266 | 0.7651573 | 0.952088495 |
| ENSMUSG00000042719 | Naa25 | protein_coding | 0.137936381 | 3.837378938 | 0.0922499 | 0.765177 | 0.952088495 |
| ENSMUSG00000030545 | Pex11a | protein_coding | 0.12769559 | 4.157662437 | 0.0917055 | 0.7658489 | 0.952647164 |
| ENSMUSG00000024403 | Atp6v1g2 | protein_coding | 0.059947163 | 10.46978886 | 0.0916955 | 0.7658612 | 0.952647164 |
| ENSMUSG00000067604 | Nms | protein_coding | 0.186642096 | 2.993198315 | 0.0916707 | 0.7658918 | 0.952647164 |
| ENSMUSG00000056383 | AI987944 | protein_coding | -0.184430899 | 3.222489646 | 0.0914518 | 0.7661689 | 0.952795388 |
| ENSMUSG00000025006 | Sorbs1 | protein_coding | -0.067361358 | 6.236332475 | 0.0914221 | 0.7661993 | 0.952795388 |
| ENSMUSG00000079426 | Arpc4 | protein_coding | -0.061040615 | 8.860484356 | 0.0912474 | 0.7664157 | 0.952795388 |
| ENSMUSG00000048787 | Dcun1d3 | protein_coding | -0.120504197 | 3.568965369 | 0.0912228 | 0.7664463 | 0.952795388 |
| ENSMUSG00000029344 | Tpst2 | protein_coding | 0.130312487 | 3.349081953 | 0.0911977 | 0.7664774 | 0.952795388 |
| ENSMUSG00000038712 | Mindy1 | protein_coding | -0.114193812 | 4.8086224 | 0.0911451 | 0.7665426 | 0.952795388 |
| ENSMUSG00000003955 | Fam162a | protein_coding | 0.081383521 | 6.252638151 | 0.0910078 | 0.7667129 | 0.952896921 |
| ENSMUSG00000035443 | Thyn1 | protein_coding | 0.055228117 | 7.727956212 | 0.0909028 | 0.7668432 | 0.952926756 |
| ENSMUSG00000024875 | Yif1a | protein_coding | -0.066041096 | 6.495635627 | 0.0905973 | 0.7672231 | 0.952926756 |
| ENSMUSG00000020037 | Rfx4 | protein_coding | -0.103078718 | 3.874908467 | 0.0905705 | 0.7672563 | 0.952926756 |
| ENSMUSG00000021929 | Kpna3 | protein_coding | -0.064313211 | 6.483431306 | 0.0905192 | 0.7673203 | 0.952926756 |
| ENSMUSG00000036707 | Cab39 | protein_coding | 0.073277688 | 6.547629035 | 0.0903679 | 0.7675086 | 0.952926756 |
| ENSMUSG00000055660 | Mettl4 | protein_coding | 0.185748181 | 2.489487307 | 0.0903495 | 0.7675315 | 0.952926756 |

| ensembl_gene_id | external_gene_name | gene_biotype | log2 (fold change) | log2 (counts per million) | F-value | p-value | False discovery rate |
|---|---|---|---|---|---|---|---|
| ENSMUSG00000034898 | Filip1 | protein_coding | 0.11072823 | 4.020894749 | 0.0902448 | 0.767662 | 0.952926756 |
| ENSMUSG00000039693 | Msantd3 | protein_coding | 0.118415454 | 3.699884254 | 0.0901208 | 0.7678167 | 0.952926756 |
| ENSMUSG00000025041 | Nt5c2 | protein_coding | 0.081632831 | 5.858135852 | 0.0900792 | 0.7678687 | 0.952926756 |
| ENSMUSG00000011158 | Brf1 | protein_coding | -0.118217517 | 3.943897732 | 0.0900685 | 0.767882 | 0.952926756 |
| ENSMUSG00000059355 | Wdr83os | protein_coding | -0.059389146 | 6.405259769 | 0.089981 | 0.7679913 | 0.952926756 |
| ENSMUSG00000025451 | Paip1 | protein_coding | -0.113597935 | 8.314837663 | 0.090391 | 0.7680856 | 0.952926756 |
| ENSMUSG00000026087 | Mrpl30 | protein_coding | -0.062922646 | 8.558953754 | 0.0898263 | 0.7681845 | 0.952926756 |
| ENSMUSG00000046010 | Zfp830 | protein_coding | 0.095529268 | 4.747410748 | 0.0898226 | 0.7681892 | 0.952926756 |
| ENSMUSG00000029861 | Fam131b | protein_coding | -0.133293206 | 3.633777531 | 0.0897013 | 0.7683408 | 0.952926756 |
| ENSMUSG00000050812 | AI314180 | protein_coding | 0.08046282 | 5.528441864 | 0.0896078 | 0.7684578 | 0.952926756 |
| ENSMUSG00000033735 | Spr | protein_coding | 0.119359376 | 5.109772652 | 0.089419 | 0.7687168 | 0.952926756 |
| ENSMUSG00000046876 | Atxn1 | protein_coding | 0.109563123 | 6.992813897 | 0.0897737 | 0.7688092 | 0.952926756 |
| ENSMUSG00000057788 | Ddx49 | protein_coding | -0.104929732 | 5.217780257 | 0.0893049 | 0.7688373 | 0.952926756 |
| ENSMUSG00000021418 | Rpp40 | protein_coding | -0.114678973 | 3.772606096 | 0.0892793 | 0.7688694 | 0.952926756 |
| ENSMUSG00000031983 | 2310022B05Rik | protein_coding | 0.09601788 | 4.985790845 | 0.089171 | 0.7690052 | 0.952926756 |
| ENSMUSG00000037239 | Spred3 | protein_coding | -0.153438856 | 2.938059179 | 0.0891135 | 0.7690774 | 0.952926756 |
| ENSMUSG00000050908 | Tvp23a | protein_coding | -0.113942901 | 3.957495416 | 0.0890099 | 0.7692076 | 0.952926756 |
| ENSMUSG00000028309 | Rnf20 | protein_coding | 0.078033764 | 5.622701641 | 0.0889604 | 0.7692698 | 0.952926756 |
| ENSMUSG00000027489 | Necab3 | protein_coding | -0.151696015 | 5.198566932 | 0.0893312 | 0.7693333 | 0.952926756 |
| ENSMUSG00000028948 | Nol9 | protein_coding | 0.151554746 | 3.071242924 | 0.0888782 | 0.7693731 | 0.952926756 |
| ENSMUSG00000028926 | Cdk14 | protein_coding | 0.088781325 | 7.358194696 | 0.0890461 | 0.7694021 | 0.952926756 |
| ENSMUSG00000003161 | Sri | protein_coding | 0.054328683 | 7.586269207 | 0.08884 | 0.7694211 | 0.952926756 |
| ENSMUSG00000037990 | Sh3rf3 | protein_coding | 0.11213679 | 3.771848816 | 0.08882 | 0.7694462 | 0.952926756 |
| ENSMUSG00000025437 | Usp33 | protein_coding | 0.16271842 | 3.952541438 | 0.0887666 | 0.7696711 | 0.952926756 |
| ENSMUSG00000006057 | Atp5g1 | protein_coding | -0.063426535 | 5.899637889 | 0.0885807 | 0.7697474 | 0.952926756 |
| ENSMUSG00000018841 | Rad51d | protein_coding | -0.086476621 | 5.734623597 | 0.08854 | 0.7697988 | 0.952926756 |
| ENSMUSG00000026209 | Dnpep | protein_coding | 0.066384471 | 6.566438219 | 0.0885249 | 0.7698177 | 0.952926756 |
| ENSMUSG00000026494 | Kif26b | protein_coding | -0.13905106 | 3.373505931 | 0.088496 | 0.7698542 | 0.952926756 |
| ENSMUSG00000079003 | Samd1 | protein_coding | -0.1669579 | 3.283322186 | 0.0884839 | 0.7698694 | 0.952926756 |
| ENSMUSG00000045969 | Ing1 | protein_coding | 0.069718519 | 5.778528828 | 0.088438 | 0.7699272 | 0.952926756 |
| ENSMUSG00000042229 | Rabif | protein_coding | -0.060224442 | 6.905860524 | 0.0883425 | 0.7700477 | 0.952958109 |
| ENSMUSG00000078700 | D030028A08Rik | processed_transcript | 0.210999638 | 2.384144999 | 0.0882091 | 0.7702161 | 0.952958109 |
| ENSMUSG00000026637 | Traf5 | protein_coding | -0.147046802 | 2.904134733 | 0.0881907 | 0.7702392 | 0.952958109 |
| ENSMUSG00000021696 | Elovl7 | protein_coding | -0.148169259 | 3.896187709 | 0.0881099 | 0.7703475 | 0.952958109 |
| ENSMUSG00000058153 | Sez6l | protein_coding | 0.066875574 | 5.904705611 | 0.0880503 | 0.7704166 | 0.952958109 |
| ENSMUSG00000044477 | Zfand3 | protein_coding | -0.092580726 | 4.897481481 | 0.0879906 | 0.7704922 | 0.952958109 |
| ENSMUSG00000030020 | Prickle2 | protein_coding | 0.072284057 | 5.767254028 | 0.08782 | 0.7707079 | 0.952958109 |
| ENSMUSG00000027342 | Pcna | protein_coding | 0.076080184 | 5.227734481 | 0.0877914 | 0.770744 | 0.952958109 |
| ENSMUSG00000032549 | Rab6b | protein_coding | -0.055273332 | 8.76489705 | 0.0876988 | 0.7708613 | 0.952958109 |
| ENSMUSG00000055491 | Pprc1 | protein_coding | 0.149602196 | 2.946576924 | 0.0876798 | 0.7708854 | 0.952958109 |
| ENSMUSG00000000134 | Tfe3 | protein_coding | 0.094959022 | 4.709438247 | 0.0874642 | 0.7711586 | 0.952958109 |
| ENSMUSG00000028657 | Ppt1 | protein_coding | -0.058133825 | 7.447321557 | 0.0874318 | 0.7711998 | 0.952958109 |
| ENSMUSG00000026655 | Fam107b | protein_coding | -0.230564344 | 3.145436871 | 0.0876186 | 0.7712008 | 0.952958109 |
| ENSMUSG00000054074 | Skida1 | protein_coding | -0.193599808 | 2.673199539 | 0.087279 | 0.7713937 | 0.952958109 |
| ENSMUSG00000048911 | Rnf24 | protein_coding | 0.165241921 | 3.282949124 | 0.087178 | 0.771522 | 0.952958109 |
| ENSMUSG00000014503 | Pkd2l2 | protein_coding | -0.069517673 | 5.546689647 | 0.0871119 | 0.771606 | 0.952958109 |
| ENSMUSG00000000378 | Ccm2 | protein_coding | 0.065545803 | 5.708089225 | 0.0870233 | 0.7717187 | 0.952958109 |
| ENSMUSG00000029716 | Tfr2 | protein_coding | -0.116847834 | 4.00157809 | 0.0870152 | 0.771729 | 0.952958109 |
| ENSMUSG00000060962 | Dmkn | protein_coding | -0.141863764 | 4.473782362 | 0.0871154 | 0.7717577 | 0.952958109 |
| ENSMUSG00000020988 | L2hgdh | protein_coding | -0.113126649 | 3.620809624 | 0.0869498 | 0.7718122 | 0.952958109 |
| ENSMUSG00000034842 | Art3 | protein_coding | -0.261972631 | 2.4869802 | 0.0869534 | 0.7718136 | 0.952958109 |
| ENSMUSG00000051502 | Ufsp1 | protein_coding | 0.111739235 | 3.98069418 | 0.0867722 | 0.7720382 | 0.953056879 |
| ENSMUSG00000031320 | Rps4x | protein_coding | -0.072870735 | 6.337535257 | 0.0867466 | 0.7720709 | 0.953056879 |
| ENSMUSG00000016386 | Mpped2 | protein_coding | -0.082656102 | 5.370563768 | 0.0865147 | 0.7723666 | 0.953096957 |
| ENSMUSG00000002980 | Bcam | protein_coding | 0.219516686 | 3.430448586 | 0.0867569 | 0.7724287 | 0.953096957 |
| ENSMUSG00000034525 | Ice1 | protein_coding | 0.103629467 | 4.221830255 | 0.0864294 | 0.7724754 | 0.953096957 |
| ENSMUSG00000029726 | Mepce | protein_coding | 0.078219871 | 5.396212388 | 0.0863906 | 0.7725249 | 0.953096957 |
| ENSMUSG00000079018 | Ly6c1 | protein_coding | -0.235692676 | 2.631200017 | 0.086347 | 0.7725807 | 0.953096957 |
| ENSMUSG00000061046 | Haghl | protein_coding | -0.056280239 | 7.77156325 | 0.0862495 | 0.7727052 | 0.953096957 |
| ENSMUSG00000040174 | Alkbh3 | protein_coding | 0.160306791 | 2.89348082 | 0.086197 | 0.7727723 | 0.953096957 |
| ENSMUSG00000020432 | Tcn2 | protein_coding | 0.130989146 | 4.297761331 | 0.0861021 | 0.7728998 | 0.953096957 |
| ENSMUSG00000022856 | Tmem41a | protein_coding | -0.090790997 | 4.42514257 | 0.0860472 | 0.7729639 | 0.953096957 |
| ENSMUSG00000054277 | Arfgap3 | protein_coding | -0.088214538 | 5.491942616 | 0.086027 | 0.7729897 | 0.953096957 |
| ENSMUSG00000021895 | Arhgef3 | protein_coding | 0.079617026 | 5.025681311 | 0.0858489 | 0.7732179 | 0.953268949 |
| ENSMUSG00000030279 | C2cd5 | protein_coding | -0.090767348 | 4.359382349 | 0.0856336 | 0.7734939 | 0.953478314 |
| ENSMUSG00000027589 | Pcmtd2 | protein_coding | 0.077698717 | 5.765213064 | 0.0855782 | 0.773565 | 0.953478314 |
| ENSMUSG00000053414 | Hunk | protein_coding | -0.091642427 | 4.479379812 | 0.08542 | 0.7737682 | 0.953486596 |
| ENSMUSG00000030016 | Zfp638 | protein_coding | -0.091728299 | 5.660084374 | 0.0853703 | 0.773832 | 0.953486596 |
| ENSMUSG00000066571 | 4931406P16Rik | protein_coding | -0.105185998 | 3.750215517 | 0.0853658 | 0.7738378 | 0.953486596 |
| ENSMUSG00000058441 | Panx2 | protein_coding | -0.10897177 | 4.069168426 | 0.08514 | 0.7741283 | 0.953735244 |
| ENSMUSG00000027220 | Syt13 | protein_coding | 0.060883693 | 6.202347776 | 0.0847657 | 0.7746105 | 0.954129524 |
| ENSMUSG00000024535 | Snx24 | protein_coding | -0.110241805 | 4.484463318 | 0.0846884 | 0.7747104 | 0.954129524 |
| ENSMUSG00000052748 | Swt1 | protein_coding | -0.106995946 | 4.030191547 | 0.0845759 | 0.7748556 | 0.954129524 |
| ENSMUSG00000044337 | Ackr3 | protein_coding | -0.240272283 | 2.357359568 | 0.0845587 | 0.7748779 | 0.954129524 |

| ensembl_gene_id | external_gene_name | gene_biotype | log2 (fold change) | log2 (counts per million) | F-value | p-value | False discovery rate |
|---|---|---|---|---|---|---|---|
| ENSMUSG00000021700 | Rab3c | protein_coding | -0.08899718 | 8.36536048 | 0.0846421 | 0.7750038 | 0.954129524 |
| ENSMUSG00000097560 | Gm26904 | lincRNA | -0.154223162 | 3.585983041 | 0.0844608 | 0.7750103 | 0.954129524 |
| ENSMUSG00000075703 | Selenoi | protein_coding | 0.111397235 | 5.201008099 | 0.0844151 | 0.7750694 | 0.954129524 |
| ENSMUSG00000060512 | 0610040J01Rik | protein_coding | -0.281675218 | 2.003837371 | 0.0842387 | 0.7752917 | 0.954293857 |
| ENSMUSG00000032475 | Nck1 | protein_coding | 0.076687969 | 5.326819621 | 0.0838802 | 0.7757564 | 0.954727915 |
| ENSMUSG00000053046 | Brsk2 | protein_coding | 0.07938708 | 5.330846141 | 0.0838218 | 0.7758321 | 0.954727915 |
| ENSMUSG00000024603 | Dctn4 | protein_coding | 0.088007618 | 7.585918399 | 0.0839405 | 0.7759107 | 0.954727915 |
| ENSMUSG00000070284 | Gmppb | protein_coding | 0.102232628 | 4.677332472 | 0.0836391 | 0.7760695 | 0.954803471 |
| ENSMUSG00000048234 | Rnf149 | protein_coding | 0.165210878 | 3.818176469 | 0.0836447 | 0.7762292 | 0.954803471 |
| ENSMUSG00000038286 | Bphl | protein_coding | 0.052833037 | 6.518229383 | 0.0835091 | 0.7762384 | 0.954803471 |
| ENSMUSG00000026810 | Dpm2 | protein_coding | -0.048846766 | 7.574163744 | 0.083356 | 0.7764376 | 0.954858063 |
| ENSMUSG00000056116 | H2-T22 | protein_coding | 0.11433885 | 4.006138147 | 0.0833385 | 0.7764604 | 0.954858063 |
| ENSMUSG00000074247 | Dda1 | protein_coding | 0.085511853 | 5.191936078 | 0.0831242 | 0.7767398 | 0.955008946 |
| ENSMUSG00000039307 | Hexdc | protein_coding | -0.114751825 | 3.88887667 | 0.0831081 | 0.7767607 | 0.955008946 |
| ENSMUSG00000046947 | Adck2 | protein_coding | -0.182336646 | 3.16324353 | 0.0829687 | 0.7769486 | 0.955130653 |
| ENSMUSG00000034936 | Arl4d | protein_coding | 0.155253798 | 4.098473613 | 0.0829178 | 0.7772401 | 0.955379896 |
| ENSMUSG00000040713 | Creg1 | protein_coding | 0.110394139 | 4.165173706 | 0.0825946 | 0.7774316 | 0.955459595 |
| ENSMUSG00000035919 | Bbs9 | protein_coding | -0.092758095 | 4.403336251 | 0.0825555 | 0.7774827 | 0.955459595 |
| ENSMUSG00000019920 | Lims1 | protein_coding | -0.095825533 | 4.735409188 | 0.0824721 | 0.777592 | 0.955484662 |
| ENSMUSG00000041343 | Ankrd42 | protein_coding | -0.141784985 | 3.169587784 | 0.0822918 | 0.7778281 | 0.955641687 |
| ENSMUSG00000041528 | Rnf123 | protein_coding | 0.248189198 | 2.638792401 | 0.0822385 | 0.777904 | 0.955641687 |
| ENSMUSG00000035559 | Mpv17l2 | protein_coding | -0.1587541 | 3.124717197 | 0.0821413 | 0.7780256 | 0.955641687 |
| ENSMUSG00000033991 | Ttc37 | protein_coding | 0.141129806 | 3.353452265 | 0.0819984 | 0.7782132 | 0.955641687 |
| ENSMUSG00000020677 | Ddx52 | protein_coding | 0.069168452 | 5.736859524 | 0.0819123 | 0.7783264 | 0.955641687 |
| ENSMUSG00000019370 | Calm3 | protein_coding | -0.053538166 | 9.637650955 | 0.0818301 | 0.7784345 | 0.955641687 |
| ENSMUSG00000033419 | Snap91 | protein_coding | 0.057544466 | 7.467942265 | 0.081743 | 0.7785491 | 0.955641687 |
| ENSMUSG00000049493 | Pls1 | protein_coding | -0.111959147 | 4.987409284 | 0.0817013 | 0.7786099 | 0.955641687 |
| ENSMUSG00000031284 | Pak3 | protein_coding | -0.053087255 | 7.575495308 | 0.0814716 | 0.7789066 | 0.955641687 |
| ENSMUSG00000032902 | Slc16a1 | protein_coding | 0.090977742 | 4.132903036 | 0.0813507 | 0.779066 | 0.955641687 |
| ENSMUSG00000022022 | Mtrf1 | protein_coding | -0.107684776 | 4.828859959 | 0.0811605 | 0.7793171 | 0.955641687 |
| ENSMUSG00000061118 | Dnajc30 | protein_coding | 0.07500125 | 5.483751363 | 0.0811103 | 0.7793834 | 0.955641687 |
| ENSMUSG00000037514 | Pank2 | protein_coding | 0.105278796 | 4.32589004 | 0.0810064 | 0.7795208 | 0.955641687 |
| ENSMUSG00000026836 | Acvr1 | protein_coding | 0.113744822 | 3.561656619 | 0.0809614 | 0.7795803 | 0.955641687 |
| ENSMUSG00000042734 | Ttc9 | protein_coding | 0.08406713 | 5.301513792 | 0.0807831 | 0.7798163 | 0.955641687 |
| ENSMUSG00000015149 | Sirt2 | protein_coding | 0.051852677 | 8.34260309 | 0.0807195 | 0.7799006 | 0.955641687 |
| ENSMUSG00000091811 | Inafm1 | protein_coding | 0.106576134 | 4.222060814 | 0.0805675 | 0.780102 | 0.955641687 |
| ENSMUSG00000025531 | Chm | protein_coding | -0.093808664 | 4.489383728 | 0.0805595 | 0.7801127 | 0.955641687 |
| ENSMUSG00000002983 | Relb | protein_coding | -0.191004917 | 2.645367851 | 0.0804991 | 0.7801928 | 0.955641687 |
| ENSMUSG00000035401 | Emsy | protein_coding | -0.093617952 | 4.647654301 | 0.080403 | 0.7803203 | 0.955641687 |
| ENSMUSG00000047888 | Tnrc6b | protein_coding | 0.080272416 | 4.947256467 | 0.0803813 | 0.7803492 | 0.955641687 |
| ENSMUSG00000042496 | Prdm10 | protein_coding | 0.138405287 | 3.93350366 | 0.0802471 | 0.7805334 | 0.955641687 |
| ENSMUSG00000031791 | Tmem38a | protein_coding | 0.052171741 | 7.106816414 | 0.0802177 | 0.7805665 | 0.955641687 |
| ENSMUSG00000024914 | Drap1 | protein_coding | 0.05512248 | 6.442584975 | 0.0802014 | 0.7805883 | 0.955641687 |
| ENSMUSG00000029528 | Pxn | protein_coding | -0.194182336 | 2.558738636 | 0.0801624 | 0.78064 | 0.955641687 |
| ENSMUSG00000032307 | Ube2q2 | protein_coding | -0.07198985 | 5.230685422 | 0.0801127 | 0.7807062 | 0.955641687 |
| ENSMUSG00000003380 | Rabac1 | protein_coding | -0.072990667 | 8.737440246 | 0.0800563 | 0.7807871 | 0.955641687 |
| ENSMUSG00000041997 | Tlk1 | protein_coding | -0.069307512 | 5.412870422 | 0.0800449 | 0.7807964 | 0.955641687 |
| ENSMUSG00000078786 | BC024978 | protein_coding | -0.105686132 | 4.375174023 | 0.0799745 | 0.7808901 | 0.955641687 |
| ENSMUSG00000061477 | Rps7 | protein_coding | 0.076254458 | 5.938776465 | 0.0798784 | 0.7810181 | 0.955641687 |
| ENSMUSG00000026851 | BC005624 | protein_coding | 0.053459901 | 7.926438961 | 0.0798572 | 0.7810465 | 0.955641687 |
| ENSMUSG00000039041 | Adrm1 | protein_coding | -0.097159732 | 4.483912201 | 0.0798045 | 0.7811167 | 0.955641687 |
| ENSMUSG00000022947 | Cbr3 | protein_coding | -0.144020096 | 3.36984053 | 0.0797757 | 0.7811551 | 0.955641687 |
| ENSMUSG00000041396 | Mettl18 | protein_coding | 0.088255867 | 4.638651767 | 0.0797267 | 0.7812204 | 0.955641687 |
| ENSMUSG00000039842 | Mcph1 | protein_coding | 0.098675512 | 4.052234881 | 0.0796966 | 0.7812605 | 0.955641687 |
| ENSMUSG00000040270 | Bach2 | protein_coding | 0.137023898 | 3.30761333 | 0.079653 | 0.7813187 | 0.955641687 |
| ENSMUSG00000063659 | Zbtb18 | protein_coding | -0.102682685 | 3.932208791 | 0.0796447 | 0.7813298 | 0.955641687 |
| ENSMUSG00000055707 | Klhl26 | protein_coding | 0.111071828 | 5.529094558 | 0.0797987 | 0.7813369 | 0.955641687 |
| ENSMUSG00000102215 | Gm2464 | lincRNA | 0.092897966 | 4.146585718 | 0.0795839 | 0.781411 | 0.955641687 |
| ENSMUSG00000031158 | Timm17b | protein_coding | 0.065184726 | 6.777739209 | 0.0795594 | 0.7814437 | 0.955641687 |
| ENSMUSG00000001627 | Ifrd1 | protein_coding | -0.145608101 | 3.542411069 | 0.0795531 | 0.7814521 | 0.955641687 |
| ENSMUSG00000032336 | Nptn | protein_coding | 0.053739509 | 8.112727407 | 0.0795529 | 0.7814524 | 0.955641687 |
| ENSMUSG00000032485 | Scap | protein_coding | -0.095392294 | 4.527274056 | 0.0794713 | 0.7815613 | 0.95566627 |
| ENSMUSG00000097062 | Gm17586 | lincRNA | -0.123426209 | 3.253532886 | 0.0793394 | 0.7817377 | 0.955773227 |
| ENSMUSG00000021774 | Ube2e1 | protein_coding | 0.06839882 | 5.5198214 | 0.0792668 | 0.7818348 | 0.955783213 |
| ENSMUSG00000085793 | Lin52 | protein_coding | 0.068837892 | 5.72416178 | 0.0790261 | 0.7821572 | 0.955938966 |
| ENSMUSG00000023236 | Scg5 | protein_coding | 0.058364579 | 9.418817556 | 0.0789255 | 0.7822921 | 0.955938966 |
| ENSMUSG00000040321 | Zfp770 | protein_coding | 0.13651948 | 3.689128903 | 0.0789059 | 0.7823183 | 0.955938966 |
| ENSMUSG00000033389 | Arhgap44 | protein_coding | 0.08431676 | 8.415104961 | 0.0790152 | 0.7823969 | 0.955938966 |
| ENSMUSG00000027560 | Dok5 | protein_coding | 0.073634014 | 5.700823367 | 0.0788071 | 0.7824509 | 0.955938966 |
| ENSMUSG00000036751 | Cox6b1 | protein_coding | -0.063084429 | 10.25496019 | 0.0786952 | 0.7826011 | 0.955938966 |
| ENSMUSG00000048755 | Mcat | protein_coding | -0.171239968 | 3.814426052 | 0.0787816 | 0.7827242 | 0.955938966 |
| ENSMUSG00000020610 | Amz2 | protein_coding | 0.063151495 | 6.09536867 | 0.0785124 | 0.7828469 | 0.955938966 |
| ENSMUSG00000062075 | Lmnb2 | protein_coding | 0.141871584 | 3.295389858 | 0.0783173 | 0.7831095 | 0.955938966 |
| ENSMUSG00000022433 | Csnk1e | protein_coding | 0.088059461 | 5.460540342 | 0.0783066 | 0.7831239 | 0.955938966 |

| ensembl_gene_id | external_gene_name | gene_biotype | log2 (fold change) | log2 (counts per million) | F-value | p-value | False discovery rate |
|---|---|---|---|---|---|---|---|
| ENSMUSG00000020644 | Id2 | protein_coding | -0.0550524 | 8.614376753 | 0.0782612 | 0.783185 | 0.955938966 |
| ENSMUSG00000042121 | Ssh1 | protein_coding | -0.083840828 | 4.392451141 | 0.0782296 | 0.7832276 | 0.955938966 |
| ENSMUSG00000063457 | Rps15 | protein_coding | -0.067749446 | 8.929819648 | 0.078216 | 0.7832459 | 0.955938966 |
| ENSMUSG00000015189 | Casd1 | protein_coding | -0.073968925 | 5.565796682 | 0.0781884 | 0.7832832 | 0.955938966 |
| ENSMUSG00000041241 | Mul1 | protein_coding | 0.052587504 | 7.04934122 | 0.0781791 | 0.7832957 | 0.955938966 |
| ENSMUSG00000045534 | Kcna5 | protein_coding | 0.115675212 | 3.796053901 | 0.0777685 | 0.78385 | 0.95644131 |
| ENSMUSG00000036285 | Noa1 | protein_coding | -0.127525726 | 3.49091992 | 0.0777425 | 0.7838852 | 0.95644131 |
| ENSMUSG00000034687 | Fras1 | protein_coding | 0.267518014 | 2.484782701 | 0.0776713 | 0.7840025 | 0.956464948 |
| ENSMUSG00000021662 | Arhgef28 | protein_coding | 0.119613895 | 4.535902081 | 0.0775466 | 0.784156 | 0.956464948 |
| ENSMUSG00000030688 | Stard10 | protein_coding | -0.087635343 | 4.304015813 | 0.0774824 | 0.7842373 | 0.956464948 |
| ENSMUSG00000030802 | Bckdk | protein_coding | 0.077189984 | 6.812872469 | 0.0774696 | 0.7842603 | 0.956464948 |
| ENSMUSG00000004934 | Pias4 | protein_coding | 0.257840213 | 2.134810029 | 0.0772998 | 0.7844847 | 0.956499166 |
| ENSMUSG00000020528 | Prpsap2 | protein_coding | -0.083311618 | 5.112066053 | 0.077276 | 0.784517 | 0.956499166 |
| ENSMUSG00000043733 | Ptpn11 | protein_coding | 0.088583962 | 5.782791783 | 0.0772232 | 0.7845945 | 0.956499166 |
| ENSMUSG00000026860 | Sh3glb2 | protein_coding | 0.06162791 | 8.027200636 | 0.0771823 | 0.7846442 | 0.956499166 |
| ENSMUSG00000026547 | Tagln2 | protein_coding | 0.098777579 | 4.606795362 | 0.0770466 | 0.7848285 | 0.956615411 |
| ENSMUSG00000053716 | Dusp7 | protein_coding | 0.09694486 | 4.516290361 | 0.076971 | 0.7849313 | 0.956632182 |
| ENSMUSG00000047767 | Atg16l2 | protein_coding | -0.12654034 | 4.074407515 | 0.0768597 | 0.7850883 | 0.956640597 |
| ENSMUSG00000035637 | Grhpr | protein_coding | 0.058979342 | 7.164007776 | 0.0768351 | 0.7851161 | 0.956640597 |
| ENSMUSG00000020721 | Helz | protein_coding | 0.067252907 | 5.362374502 | 0.0766777 | 0.7853305 | 0.95679344 |
| ENSMUSG00000028261 | Ndufaf4 | protein_coding | -0.059814873 | 6.343887909 | 0.0765248 | 0.7855388 | 0.95693878 |
| ENSMUSG00000024548 | Setbp1 | protein_coding | 0.132850321 | 3.466152354 | 0.0763798 | 0.7857367 | 0.957071475 |
| ENSMUSG00000058064 | Gm10036 | protein_coding | 0.295788445 | 2.964323997 | 0.0765966 | 0.7859955 | 0.957081005 |
| ENSMUSG00000031302 | Nlgn3 | protein_coding | 0.056694961 | 6.648636821 | 0.0761869 | 0.7860003 | 0.957081005 |
| ENSMUSG00000058488 | Kl | protein_coding | -0.294579188 | 3.713946412 | 0.0774841 | 0.7860116 | 0.957081005 |
| ENSMUSG00000020131 | Pcsk4 | protein_coding | -0.137899218 | 3.616326002 | 0.0760218 | 0.7862261 | 0.957233874 |
| ENSMUSG00000060743 | H3f3a | protein_coding | 0.092110199 | 3.826424919 | 0.0756865 | 0.7866856 | 0.957324537 |
| ENSMUSG00000028333 | Anp32b | protein_coding | -0.080166186 | 5.070327518 | 0.0756668 | 0.7867125 | 0.957324537 |
| ENSMUSG00000026344 | Lypd1 | protein_coding | 0.155177794 | 2.678574672 | 0.075666 | 0.7867136 | 0.957324537 |
| ENSMUSG00000006728 | Cdk4 | protein_coding | 0.053387666 | 7.036813166 | 0.0756486 | 0.7867376 | 0.957324537 |
| ENSMUSG00000027459 | Fam110a | protein_coding | -0.107820085 | 5.031921134 | 0.075637 | 0.7867592 | 0.957324537 |
| ENSMUSG00000032026 | Rexo2 | protein_coding | -0.067467244 | 5.948169437 | 0.0755778 | 0.7868348 | 0.957324537 |
| ENSMUSG00000042737 | Dpm3 | protein_coding | -0.066293584 | 6.893419205 | 0.0751357 | 0.7874427 | 0.957866962 |
| ENSMUSG00000040006 | Ginm1 | protein_coding | 0.080103388 | 5.331043321 | 0.0751241 | 0.7874588 | 0.957866962 |
| ENSMUSG00000031370 | Zrsr2 | protein_coding | -0.057878232 | 6.645237934 | 0.0750487 | 0.7875626 | 0.957884966 |
| ENSMUSG00000024446 | Rpp21 | protein_coding | -0.047821387 | 7.644382785 | 0.074934 | 0.7877208 | 0.957963453 |
| ENSMUSG00000027378 | Nphp1 | protein_coding | -0.119276794 | 3.735808116 | 0.0748727 | 0.7878053 | 0.957963453 |
| ENSMUSG00000028064 | Sema4a | protein_coding | 0.111485437 | 4.49220998 | 0.0747547 | 0.7879682 | 0.958024008 |
| ENSMUSG00000021646 | Mccc2 | protein_coding | 0.09277802 | 5.453756839 | 0.0746787 | 0.7880789 | 0.958024008 |
| ENSMUSG00000009733 | Tfcp2 | protein_coding | -0.089005396 | 4.626596057 | 0.0745024 | 0.788317 | 0.958024008 |
| ENSMUSG00000034932 | Mrpl54 | protein_coding | 0.105075308 | 6.746762172 | 0.0748742 | 0.7883511 | 0.958024008 |
| ENSMUSG00000031951 | Tmem231 | protein_coding | 0.100385073 | 4.565835644 | 0.0744556 | 0.7883818 | 0.958024008 |
| ENSMUSG00000035934 | Pknox2 | protein_coding | 0.076600771 | 5.223197322 | 0.0744499 | 0.7883897 | 0.958024008 |
| ENSMUSG00000073643 | Wdfy1 | protein_coding | 0.211031899 | 4.899426387 | 0.0754433 | 0.7886964 | 0.958138088 |
| ENSMUSG00000029123 | Stk32b | protein_coding | 0.137992557 | 4.482213564 | 0.0743769 | 0.7887089 | 0.958138088 |
| ENSMUSG00000037933 | Bicd2 | protein_coding | 0.086687314 | 4.4609733 | 0.0741724 | 0.7887741 | 0.958138088 |
| ENSMUSG00000038406 | Scaf1 | protein_coding | -0.093941374 | 4.706982245 | 0.0740189 | 0.7889873 | 0.958138088 |
| ENSMUSG00000028703 | Lrrc41 | protein_coding | 0.118205273 | 3.908571667 | 0.073915 | 0.7891316 | 0.958138088 |
| ENSMUSG00000032462 | Pik3cb | protein_coding | 0.157783274 | 3.052079524 | 0.0739122 | 0.7891354 | 0.958138088 |
| ENSMUSG00000022490 | Ppp1r1a | protein_coding | -0.075965616 | 5.34981323 | 0.0739111 | 0.789137 | 0.958138088 |
| ENSMUSG00000038068 | Rnf144b | protein_coding | 0.117679795 | 3.326195331 | 0.0738003 | 0.789291 | 0.958138088 |
| ENSMUSG00000000532 | Acvr1b | protein_coding | -0.097007312 | 4.535971057 | 0.0736427 | 0.7895103 | 0.958138088 |
| ENSMUSG00000027133 | Nop10 | protein_coding | -0.066884231 | 9.048247012 | 0.0736357 | 0.78952 | 0.958138088 |
| ENSMUSG00000056091 | St3gal5 | protein_coding | 0.083000594 | 4.89099431 | 0.073615 | 0.7895488 | 0.958138088 |
| ENSMUSG00000028991 | Mtor | protein_coding | 0.087248525 | 4.714338801 | 0.073576 | 0.7896032 | 0.958138088 |
| ENSMUSG00000027951 | Adar | protein_coding | 0.194272088 | 3.419891742 | 0.0736849 | 0.7897381 | 0.958138088 |
| ENSMUSG00000037058 | Paip2 | protein_coding | -0.046759977 | 7.838311285 | 0.0734778 | 0.7897401 | 0.958138088 |
| ENSMUSG00000042747 | Krtcap2 | protein_coding | 0.057244838 | 7.698861476 | 0.0734204 | 0.7898201 | 0.958138088 |
| ENSMUSG00000021747 | 4930452B06Rik | protein_coding | -0.094289685 | 3.836289135 | 0.0731954 | 0.7901342 | 0.958307268 |
| ENSMUSG00000068966 | Zbtb34 | protein_coding | -0.150893714 | 2.946997983 | 0.0731927 | 0.7901378 | 0.958307268 |
| ENSMUSG00000025208 | Mrpl43 | protein_coding | -0.05284099 | 7.39468562 | 0.0730468 | 0.7903418 | 0.95841581 |
| ENSMUSG00000004031 | Brinp2 | protein_coding | -0.080266331 | 4.603007488 | 0.0729865 | 0.7904262 | 0.95841581 |
| ENSMUSG00000070953 | Rabepk | protein_coding | -0.053214145 | 6.597109698 | 0.0729375 | 0.7904947 | 0.95841581 |
| ENSMUSG00000032352 | Lrrc1 | protein_coding | -0.122018514 | 3.422985212 | 0.072814 | 0.7906676 | 0.958418223 |
| ENSMUSG00000024622 | Hmgxb3 | protein_coding | -0.134146977 | 3.1985186 | 0.0727789 | 0.7907168 | 0.958418223 |
| ENSMUSG00000035378 | Shq1 | protein_coding | 0.150972507 | 2.995015407 | 0.0724829 | 0.7911321 | 0.958418223 |
| ENSMUSG00000030894 | Tpp1 | protein_coding | 0.080251047 | 6.126023615 | 0.0724695 | 0.7911564 | 0.958418223 |
| ENSMUSG00000035439 | Haus8 | protein_coding | 0.074012231 | 5.555559497 | 0.0724109 | 0.7912333 | 0.958418223 |
| ENSMUSG00000078716 | Tmem8b | protein_coding | -0.103969493 | 4.758628436 | 0.0722657 | 0.7914374 | 0.958418223 |
| ENSMUSG00000022892 | App | protein_coding | 0.055125527 | 9.150916813 | 0.072189 | 0.7915453 | 0.958418223 |
| ENSMUSG00000020810 | Cygb | protein_coding | -0.052046795 | 7.364913844 | 0.0721734 | 0.7915673 | 0.958418223 |
| ENSMUSG00000023353 | Agap3 | protein_coding | -0.06218949 | 5.872220697 | 0.0721017 | 0.7916682 | 0.958418223 |
| ENSMUSG00000074971 | Fibin | protein_coding | -0.160703958 | 2.923401288 | 0.072075 | 0.7917057 | 0.958418223 |
| ENSMUSG00000032680 | 6820408C15Rik | protein_coding | -0.11764306 | 4.215739749 | 0.072013 | 0.7917987 | 0.958418223 |

| ensembl_gene_id | external_gene_name | gene_biotype | log2 (fold change) | log2 (counts per million) | F-value | p-value | False discovery rate |
|---|---|---|---|---|---|---|---|
| ENSMUSG00000041712 | Ubr7 | protein_coding | 0.088413913 | 4.846411082 | 0.0719915 | 0.7918235 | 0.958418223 |
| ENSMUSG00000097023 | Mir9-3hg | lincRNA | -0.11127724 | 3.721432233 | 0.0717861 | 0.7921132 | 0.958418223 |
| ENSMUSG00000031133 | Arhgef6 | protein_coding | -0.166926413 | 2.854956478 | 0.0717702 | 0.7921356 | 0.958418223 |
| ENSMUSG00000034653 | Ythdc2 | protein_coding | -0.119268035 | 3.328496467 | 0.0717487 | 0.792166 | 0.958418223 |
| ENSMUSG00000021047 | Nova1 | protein_coding | -0.061111332 | 6.313334427 | 0.0717133 | 0.7922159 | 0.958418223 |
| ENSMUSG00000027573 | Gid8 | protein_coding | -0.07351816 | 5.507733195 | 0.0716928 | 0.7922449 | 0.958418223 |
| ENSMUSG00000033885 | Pxk | protein_coding | 0.085640853 | 5.072323438 | 0.071593 | 0.7923859 | 0.958418223 |
| ENSMUSG00000040359 | Ufl1 | protein_coding | -0.128749436 | 2.978506641 | 0.0715699 | 0.7924186 | 0.958418223 |
| ENSMUSG00000084403 | Rps15a-ps8 | processed_pseudoge | 0.462214341 | 1.126234636 | 0.0715077 | 0.7925065 | 0.958418223 |
| ENSMUSG00000027581 | Stmn3 | protein_coding | 0.053368155 | 11.21256015 | 0.0714537 | 0.792583 | 0.958418223 |
| ENSMUSG00000039913 | Pak7 | protein_coding | -0.078255363 | 4.788729336 | 0.0713138 | 0.7927811 | 0.958418223 |
| ENSMUSG00000032534 | Cep63 | protein_coding | 0.084960809 | 4.898120913 | 0.0713089 | 0.792788 | 0.958418223 |
| ENSMUSG00000058006 | Mdn1 | protein_coding | 0.158088942 | 3.343406399 | 0.0712646 | 0.7928563 | 0.958418223 |
| ENSMUSG00000027822 | Slc33a1 | protein_coding | -0.088976502 | 4.249301208 | 0.0712013 | 0.7929405 | 0.958418223 |
| ENSMUSG00000049148 | Plcxd3 | protein_coding | -0.085048738 | 5.864056823 | 0.0711641 | 0.7929987 | 0.958418223 |
| ENSMUSG00000025532 | Crcp | protein_coding | 0.052578398 | 7.105217199 | 0.0711352 | 0.7930343 | 0.958418223 |
| ENSMUSG00000040712 | Camta2 | protein_coding | 0.093669445 | 6.010244327 | 0.0712499 | 0.7930718 | 0.958418223 |
| ENSMUSG00000038594 | Cep85l | protein_coding | -0.176577768 | 3.084781103 | 0.0711058 | 0.7930815 | 0.958418223 |
| ENSMUSG00000043463 | Rab9b | protein_coding | 0.073895237 | 5.675280788 | 0.07093 | 0.7933256 | 0.958479067 |
| ENSMUSG00000064293 | Cntn4 | protein_coding | -0.088619261 | 4.833247032 | 0.0708977 | 0.7933715 | 0.958479067 |
| ENSMUSG00000028851 | Nudc | protein_coding | 0.098962179 | 5.935962443 | 0.071028 | 0.7933993 | 0.958479067 |
| ENSMUSG00000048581 | E130311K13Rik | protein_coding | 0.189617752 | 2.635295738 | 0.070752 | 0.7935786 | 0.958493273 |
| ENSMUSG00000027784 | Ppm1l | protein_coding | -0.072003413 | 5.153257303 | 0.0707445 | 0.7935893 | 0.958493273 |
| ENSMUSG00000030814 | Bcl7c | protein_coding | 0.15806008 | 2.647911309 | 0.0705823 | 0.7938203 | 0.958586284 |
| ENSMUSG00000033949 | Trim36 | protein_coding | 0.120769794 | 3.119246863 | 0.0704928 | 0.7939479 | 0.958586284 |
| ENSMUSG00000054728 | Phactr1 | protein_coding | -0.048779754 | 8.711495992 | 0.0704136 | 0.7940607 | 0.958586284 |
| ENSMUSG00000081534 | Slc48a1 | protein_coding | -0.065742214 | 7.330296116 | 0.0703757 | 0.7941203 | 0.958586284 |
| ENSMUSG00000026037 | Orc2 | protein_coding | 0.211108426 | 2.489259787 | 0.0703184 | 0.7941966 | 0.958586284 |
| ENSMUSG00000004044 | Cavin1 | protein_coding | 0.070752991 | 5.39230093 | 0.0702948 | 0.7942302 | 0.958586284 |
| ENSMUSG00000044912 | Syt16 | protein_coding | 0.083361098 | 4.870033124 | 0.0701719 | 0.7944057 | 0.958586284 |
| ENSMUSG00000063172 | Hspb11 | protein_coding | -0.063387908 | 5.895855498 | 0.0701326 | 0.794462 | 0.958586284 |
| ENSMUSG00000049670 | Morn4 | protein_coding | 0.046234834 | 7.601645334 | 0.0700428 | 0.7945903 | 0.958586284 |
| ENSMUSG00000035885 | Cox8a | protein_coding | -0.093781876 | 9.631013762 | 0.0703375 | 0.7946003 | 0.958586284 |
| ENSMUSG00000056121 | Fez2 | protein_coding | 0.077032074 | 4.617112887 | 0.0699606 | 0.7947079 | 0.958586284 |
| ENSMUSG00000038718 | Pbx3 | protein_coding | -0.084411767 | 5.39424119 | 0.0698969 | 0.7947992 | 0.958586284 |
| ENSMUSG00000037894 | H2afz | protein_coding | 0.084341233 | 3.996993766 | 0.0698404 | 0.7948801 | 0.958586284 |
| ENSMUSG00000003528 | Slc25a1 | protein_coding | 0.095306085 | 5.133553603 | 0.069792 | 0.7949549 | 0.958586284 |
| ENSMUSG00000032420 | Nt5e | protein_coding | 0.175856346 | 2.641088691 | 0.0697391 | 0.7950254 | 0.958586284 |
| ENSMUSG00000045180 | Shroom2 | protein_coding | 0.065994207 | 6.313908682 | 0.0696344 | 0.7951755 | 0.958586284 |
| ENSMUSG00000064181 | Rab3ip | protein_coding | 0.081722186 | 4.458143797 | 0.0695612 | 0.7952806 | 0.958586284 |
| ENSMUSG00000010376 | Nedd8 | protein_coding | 0.054738803 | 9.396075068 | 0.069508 | 0.7953569 | 0.958586284 |
| ENSMUSG00000060572 | Mfap2 | protein_coding | -0.100484963 | 4.044965662 | 0.0695058 | 0.7953601 | 0.958586284 |
| ENSMUSG00000053137 | Mapk11 | protein_coding | -0.108104568 | 3.555196215 | 0.0693992 | 0.7955132 | 0.958632626 |
| ENSMUSG00000039199 | Zdhhc1 | protein_coding | -0.155410999 | 2.754886922 | 0.0693088 | 0.7956432 | 0.958632626 |
| ENSMUSG00000032083 | Apoa1 | protein_coding | -0.127632742 | 3.085571856 | 0.069293 | 0.795666 | 0.958632626 |
| ENSMUSG00000021585 | Cast | protein_coding | 0.133295717 | 3.004113261 | 0.0691452 | 0.7958787 | 0.958781543 |
| ENSMUSG00000069206 | Zfp874a | protein_coding | -0.10983426 | 3.615797756 | 0.0690562 | 0.796007 | 0.958828592 |
| ENSMUSG00000020697 | Lig3 | protein_coding | 0.161561213 | 2.791081646 | 0.068953 | 0.7961558 | 0.958900419 |
| ENSMUSG00000091512 | Lamtor3 | protein_coding | -0.055607132 | 6.806980112 | 0.0687116 | 0.7965044 | 0.959212896 |
| ENSMUSG00000027176 | Cstf3 | protein_coding | -0.057717198 | 6.278474817 | 0.0685285 | 0.7967692 | 0.959291896 |
| ENSMUSG00000022711 | Pmm2 | protein_coding | 0.071436323 | 5.048380884 | 0.0684952 | 0.7968173 | 0.959291896 |
| ENSMUSG00000019978 | Epb41l2 | protein_coding | 0.084996882 | 4.435299091 | 0.0684301 | 0.7969117 | 0.959291896 |
| ENSMUSG00000045867 | Cradd | protein_coding | -0.072431473 | 4.529806171 | 0.0683795 | 0.796985 | 0.959291896 |
| ENSMUSG00000072964 | Bhlhb9 | protein_coding | -0.064769537 | 5.937865122 | 0.0682847 | 0.7971224 | 0.959291896 |
| ENSMUSG00000025986 | Slc39a10 | protein_coding | -0.058333431 | 5.661117132 | 0.0681908 | 0.7972586 | 0.959291896 |
| ENSMUSG00000037773 | Pced1a | protein_coding | 0.12753471 | 3.918021502 | 0.0681621 | 0.7973056 | 0.959291896 |
| ENSMUSG00000003813 | Rad23a | protein_coding | 0.051736806 | 8.108862216 | 0.0680599 | 0.7974486 | 0.959291896 |
| ENSMUSG00000036825 | Ssx2ip | protein_coding | 0.064263968 | 5.533175361 | 0.0680547 | 0.7974561 | 0.959291896 |
| ENSMUSG00000033740 | St18 | protein_coding | 0.091516414 | 3.954821333 | 0.0680506 | 0.7974621 | 0.959291896 |
| ENSMUSG00000050299 | Gm9843 | protein_coding | -0.079038735 | 4.56489152 | 0.0678396 | 0.7977689 | 0.959382911 |
| ENSMUSG00000025024 | Smndc1 | protein_coding | -0.081398869 | 4.371692579 | 0.0678182 | 0.7978002 | 0.959382911 |
| ENSMUSG00000025094 | Slc18a2 | protein_coding | 0.145306634 | 3.023974236 | 0.0678089 | 0.7978136 | 0.959382911 |
| ENSMUSG00000027692 | Tnik | protein_coding | -0.091631395 | 5.67519799 | 0.0677666 | 0.7978947 | 0.959382911 |
| ENSMUSG00000073176 | Zfp449 | protein_coding | 0.100592738 | 3.378219399 | 0.0675825 | 0.7981434 | 0.959477822 |
| ENSMUSG00000057841 | Rpl32 | protein_coding | -0.062861286 | 9.869372076 | 0.0674333 | 0.7983612 | 0.959477822 |
| ENSMUSG00000004843 | Chmp2b | protein_coding | -0.073511376 | 5.617063188 | 0.0674246 | 0.7983739 | 0.959477822 |
| ENSMUSG00000052656 | Rnf103 | protein_coding | 0.125575726 | 4.638591862 | 0.0675652 | 0.7983759 | 0.959477822 |
| ENSMUSG00000028755 | Cda | protein_coding | 0.09012653 | 4.950900767 | 0.0670772 | 0.7988818 | 0.959477822 |
| ENSMUSG00000034845 | Plvap | protein_coding | 0.091905511 | 3.980863032 | 0.0670318 | 0.7989481 | 0.959477822 |
| ENSMUSG00000063047 | Zfp780b | protein_coding | -0.072939335 | 4.690286311 | 0.0669918 | 0.7990068 | 0.959477822 |
| ENSMUSG00000016526 | Dyrk3 | protein_coding | 0.205455681 | 2.25077061 | 0.0669095 | 0.7991274 | 0.959477822 |
| ENSMUSG00000035469 | Rcbtb1 | protein_coding | -0.083105858 | 5.131448079 | 0.0668591 | 0.7992013 | 0.959477822 |
| ENSMUSG00000003184 | Irf3 | protein_coding | 0.129015128 | 3.253010263 | 0.0667926 | 0.7992989 | 0.959477822 |
| ENSMUSG00000015846 | Rxra | protein_coding | -0.071486256 | 4.58347294 | 0.0666886 | 0.7994514 | 0.959477822 |

| ensembl_gene_id | external_gene_name | gene_biotype | log2 (fold change) | log2 (counts per million) | F-value | p-value | False discovery rate |
|---|---|---|---|---|---|---|---|
| ENSMUSG00000000305 | Cdh4 | protein_coding | -0.066431914 | 5.535970234 | 0.0665462 | 0.7996607 | 0.959477822 |
| ENSMUSG00000022684 | Bfar | protein_coding | 0.063354921 | 5.791579699 | 0.0665246 | 0.7996925 | 0.959477822 |
| ENSMUSG00000022641 | Bbx | protein_coding | 0.07748581 | 5.438121776 | 0.0664924 | 0.7997399 | 0.959477822 |
| ENSMUSG00000026709 | Dars2 | protein_coding | 0.091975509 | 4.442531 | 0.0664659 | 0.7997789 | 0.959477822 |
| ENSMUSG00000063801 | Ap3s2 | protein_coding | 0.051712619 | 6.881998965 | 0.0664114 | 0.7998591 | 0.959477822 |
| ENSMUSG00000028861 | Mrps15 | protein_coding | -0.064945907 | 6.071185528 | 0.0663382 | 0.7999668 | 0.959477822 |
| ENSMUSG00000037994 | Slc9b2 | protein_coding | -0.157782327 | 2.693653155 | 0.0662771 | 0.8000568 | 0.959477822 |
| ENSMUSG00000026639 | Lamb3 | protein_coding | 0.156578864 | 3.199071581 | 0.0662112 | 0.8001539 | 0.959477822 |
| ENSMUSG00000026048 | Ercc5 | protein_coding | 0.117353434 | 3.90961275 | 0.0661129 | 0.800299 | 0.959477822 |
| ENSMUSG00000030177 | Ccdc77 | protein_coding | -0.187338064 | 2.536813618 | 0.0661111 | 0.8003016 | 0.959477822 |
| ENSMUSG00000021639 | Gtf2h2 | protein_coding | -0.060800145 | 5.981032414 | 0.0660722 | 0.8003589 | 0.959477822 |
| ENSMUSG00000027932 | Slc27a3 | protein_coding | -0.189018069 | 2.496547743 | 0.0659801 | 0.800495 | 0.959477822 |
| ENSMUSG00000025159 | Mms19 | protein_coding | 0.092695152 | 4.342493088 | 0.0659796 | 0.8004956 | 0.959477822 |
| ENSMUSG00000001018 | Snapin | protein_coding | 0.079596497 | 6.521484819 | 0.0660131 | 0.800581 | 0.959477822 |
| ENSMUSG00000024576 | Csnk1a1 | protein_coding | -0.061230886 | 7.22639045 | 0.0658854 | 0.8006349 | 0.959477822 |
| ENSMUSG00000024924 | Vldlr | protein_coding | -0.086645484 | 5.759781989 | 0.0658495 | 0.8006933 | 0.959477822 |
| ENSMUSG00000040385 | Ppp1ca | protein_coding | 0.047157474 | 9.192560825 | 0.0657717 | 0.800803 | 0.959477822 |
| ENSMUSG00000037013 | Ss18 | protein_coding | -0.101736661 | 4.372798749 | 0.065733 | 0.8008603 | 0.959477822 |
| ENSMUSG00000020832 | Eral1 | protein_coding | -0.080538623 | 5.595521897 | 0.0656608 | 0.8009671 | 0.959477822 |
| ENSMUSG00000025964 | Adam23 | protein_coding | 0.058040802 | 6.261421837 | 0.0656605 | 0.8009676 | 0.959477822 |
| ENSMUSG00000026833 | Olfm1 | protein_coding | -0.050922693 | 7.687292613 | 0.0656319 | 0.80101 | 0.959477822 |
| ENSMUSG00000042292 | Mkl1 | protein_coding | -0.078441782 | 4.91846856 | 0.0655032 | 0.8012008 | 0.959477822 |
| ENSMUSG00000015542 | Nat9 | protein_coding | 0.068273569 | 5.966383651 | 0.065452 | 0.8012766 | 0.959477822 |
| ENSMUSG00000038844 | Kif16b | protein_coding | 0.130840357 | 3.107738631 | 0.0654405 | 0.8012938 | 0.959477822 |
| ENSMUSG00000028466 | Creb3 | protein_coding | 0.050402595 | 7.342910945 | 0.0653421 | 0.8014397 | 0.959477822 |
| ENSMUSG00000061080 | Lsamp | protein_coding | -0.043451561 | 7.72007222 | 0.0653245 | 0.8014658 | 0.959477822 |
| ENSMUSG00000020807 | 4933427D14Rik | protein_coding | -0.102901261 | 3.610852911 | 0.0653007 | 0.8015011 | 0.959477822 |
| ENSMUSG00000033417 | Cacul1 | protein_coding | 0.074469267 | 5.058586534 | 0.0652747 | 0.8015398 | 0.959477822 |
| ENSMUSG00000048285 | Frmd6 | protein_coding | 0.17376714 | 2.624950927 | 0.0652727 | 0.8015428 | 0.959477822 |
| ENSMUSG00000021947 | Cryl1 | protein_coding | 0.067277964 | 5.011775904 | 0.0650923 | 0.801811 | 0.959692102 |
| ENSMUSG00000044317 | Gpr4 | protein_coding | -0.200985542 | 2.413926807 | 0.0648192 | 0.8022177 | 0.959739512 |
| ENSMUSG00000010660 | Plcd1 | protein_coding | 0.182999273 | 3.05950199 | 0.0647897 | 0.8022808 | 0.959739512 |
| ENSMUSG00000059669 | Taf1b | protein_coding | 0.085224921 | 4.718518786 | 0.0647694 | 0.802292 | 0.959739512 |
| ENSMUSG00000026203 | Dnajb2 | protein_coding | -0.048359393 | 6.674401916 | 0.0647398 | 0.8023361 | 0.959739512 |
| ENSMUSG00000018293 | Pfn1 | protein_coding | -0.056872965 | 6.820414673 | 0.0647387 | 0.8023378 | 0.959739512 |
| ENSMUSG00000031129 | Slc9a9 | protein_coding | -0.096587621 | 3.493578143 | 0.0646541 | 0.802464 | 0.959739512 |
| ENSMUSG00000021541 | Trpc7 | protein_coding | -0.074898462 | 5.12309614 | 0.064611 | 0.8025284 | 0.959739512 |
| ENSMUSG00000010086 | Rnf112 | protein_coding | -0.097964187 | 4.615824824 | 0.06449 | 0.8027092 | 0.959739512 |
| ENSMUSG00000029463 | Fam216a | protein_coding | 0.097034872 | 4.12607671 | 0.0644162 | 0.8028197 | 0.959739512 |
| ENSMUSG00000099964 | Gm29514 | antisense_RNA | 0.089357655 | 3.670833723 | 0.0643949 | 0.8028515 | 0.959739512 |
| ENSMUSG00000022637 | Cblb | protein_coding | 0.161583477 | 3.005102669 | 0.0642837 | 0.803018 | 0.959739512 |
| ENSMUSG00000097207 | 6030443J06Rik | lincRNA | -0.120814769 | 3.104366737 | 0.0642516 | 0.8030661 | 0.959739512 |
| ENSMUSG00000025538 | Sumf2 | protein_coding | 0.097182862 | 5.000445832 | 0.0641505 | 0.8032228 | 0.959739512 |
| ENSMUSG00000051726 | Kcnf1 | protein_coding | -0.102155839 | 4.390670137 | 0.0640835 | 0.8033181 | 0.959739512 |
| ENSMUSG00000074218 | Cox7a1 | protein_coding | -0.090853697 | 3.850723237 | 0.0640813 | 0.8033213 | 0.959739512 |
| ENSMUSG00000031174 | Rpgr | protein_coding | -0.076525747 | 4.631533362 | 0.0640674 | 0.8033422 | 0.959739512 |
| ENSMUSG00000036721 | Zscan12 | protein_coding | 0.11368125 | 3.671863815 | 0.0640503 | 0.8033679 | 0.959739512 |
| ENSMUSG00000049327 | Kmt5a | protein_coding | -0.085582935 | 4.106663803 | 0.0638363 | 0.8036893 | 0.959958664 |
| ENSMUSG00000048583 | Igf2 | protein_coding | -0.147527699 | 4.068313374 | 0.0640109 | 0.8037299 | 0.959958664 |
| ENSMUSG00000020358 | Hnrnpab | protein_coding | 0.15248583 | 2.762757059 | 0.0636289 | 0.8040013 | 0.960045201 |
| ENSMUSG00000021179 | Nrde2 | protein_coding | 0.108235535 | 4.271587522 | 0.0635975 | 0.8040486 | 0.960045201 |
| ENSMUSG00000035958 | Tdp2 | protein_coding | -0.093676604 | 3.702396804 | 0.0635264 | 0.8041558 | 0.960045201 |
| ENSMUSG00000037868 | Egr2 | protein_coding | 0.164663466 | 2.595432528 | 0.0635239 | 0.8041595 | 0.960045201 |
| ENSMUSG00000028945 | Rheb | protein_coding | 0.049263466 | 6.803813429 | 0.0632899 | 0.8045126 | 0.960360125 |
| ENSMUSG00000011832 | Evi5l | protein_coding | 0.066263734 | 4.973127487 | 0.0630109 | 0.8049344 | 0.960495961 |
| ENSMUSG00000019929 | Dcn | protein_coding | -0.118098365 | 3.742444667 | 0.0629383 | 0.8050443 | 0.960495961 |
| ENSMUSG00000014504 | Srp19 | protein_coding | 0.056559036 | 8.272780773 | 0.0628926 | 0.8051135 | 0.960495961 |
| ENSMUSG00000020849 | Ywhae | protein_coding | 0.053056977 | 11.16070672 | 0.0628684 | 0.8051503 | 0.960495961 |
| ENSMUSG00000067424 | Zfp563 | protein_coding | -0.091907565 | 3.802605472 | 0.0628057 | 0.8052453 | 0.960495961 |
| ENSMUSG00000087679 | Tmem250-ps | lincRNA | 0.062557335 | 9.365723529 | 0.0627804 | 0.8052837 | 0.960495961 |
| ENSMUSG00000035051 | Dhx57 | protein_coding | 0.088592125 | 4.112709344 | 0.0627561 | 0.8053206 | 0.960495961 |
| ENSMUSG00000085336 | Gm11732 | processed_transcript | 0.147937394 | 3.231971655 | 0.0626877 | 0.8054243 | 0.960495961 |
| ENSMUSG00000006005 | Tpr | protein_coding | 0.071716863 | 5.924069098 | 0.0626838 | 0.8054303 | 0.960495961 |
| ENSMUSG00000095930 | Nim1k | protein_coding | 0.065989089 | 5.445212786 | 0.0624894 | 0.8057256 | 0.960544055 |
| ENSMUSG00000021007 | Spata7 | protein_coding | -0.073352597 | 5.025590544 | 0.0624457 | 0.8057921 | 0.960544055 |
| ENSMUSG00000000983 | Wfdc18 | protein_coding | -0.115098272 | 3.370804042 | 0.0624306 | 0.8058151 | 0.960544055 |
| ENSMUSG00000042156 | Dzip1 | protein_coding | 0.075574011 | 5.309869513 | 0.0624221 | 0.8058279 | 0.960544055 |
| ENSMUSG00000066643 | Wdr35 | protein_coding | 0.074335711 | 5.363477915 | 0.0622846 | 0.8060372 | 0.960687041 |
| ENSMUSG00000029722 | Agfg2 | protein_coding | -0.072548743 | 5.019407381 | 0.0621209 | 0.8062867 | 0.960712687 |
| ENSMUSG00000039197 | Adk | protein_coding | 0.044709873 | 7.492440639 | 0.0621042 | 0.8063123 | 0.960712687 |
| ENSMUSG00000022452 | Smdt1 | protein_coding | -0.052598122 | 8.365848453 | 0.0620675 | 0.8063682 | 0.960712687 |
| ENSMUSG00000103183 | Gm37090 | TEC | 0.249760686 | 2.365240019 | 0.0620382 | 0.806418 | 0.960712687 |
| ENSMUSG00000107002 | 0610012G03Rik | protein_coding | 0.086464862 | 4.20281554 | 0.0619127 | 0.8066045 | 0.960712687 |
| ENSMUSG00000029673 | Auts2 | protein_coding | -0.047478592 | 6.258686383 | 0.0618283 | 0.8067336 | 0.960712687 |

| ensembl_gene_id | external_gene_name | gene_biotype | log2 (fold change) | log2 (counts per million) | F-value | p-value | False discovery rate |
|---|---|---|---|---|---|---|---|
| ENSMUSG00000032463 | Faim | protein_coding | -0.075525689 | 4.670042742 | 0.0616864 | 0.8069507 | 0.960712687 |
| ENSMUSG00000062372 | Otof | protein_coding | -0.132813081 | 3.280649575 | 0.0616827 | 0.8069564 | 0.960712687 |
| ENSMUSG00000031701 | Dnaja2 | protein_coding | -0.053460378 | 7.840731032 | 0.0616762 | 0.8069664 | 0.960712687 |
| ENSMUSG00000045237 | 1110012L19Rik | protein_coding | 0.05426887 | 5.830513754 | 0.0616238 | 0.8070466 | 0.960712687 |
| ENSMUSG00000054280 | Prr14l | protein_coding | -0.129517628 | 3.099167122 | 0.0616085 | 0.8070701 | 0.960712687 |
| ENSMUSG00000034602 | Mon2 | protein_coding | 0.083594988 | 4.907468043 | 0.0615688 | 0.8071309 | 0.960712687 |
| ENSMUSG00000018666 | Cbx1 | protein_coding | -0.050165971 | 6.193500757 | 0.0613496 | 0.8074671 | 0.960858843 |
| ENSMUSG00000022000 | Zc3h13 | protein_coding | 0.069405744 | 5.432840269 | 0.0613396 | 0.8074825 | 0.960858843 |
| ENSMUSG00000007036 | Abhd16a | protein_coding | -0.062846169 | 5.766196043 | 0.0612893 | 0.8075597 | 0.960858843 |
| ENSMUSG00000026519 | Tmem63a | protein_coding | 0.094012289 | 4.576866555 | 0.0611669 | 0.8077478 | 0.960858843 |
| ENSMUSG00000074622 | Mafb | protein_coding | 0.180522092 | 2.802937455 | 0.0610922 | 0.8078628 | 0.960858843 |
| ENSMUSG00000031352 | Hccs | protein_coding | -0.094917138 | 4.604801972 | 0.061083 | 0.807877 | 0.960858843 |
| ENSMUSG00000087598 | Zfp111 | protein_coding | -0.131043723 | 2.732039677 | 0.0610356 | 0.8079499 | 0.960858843 |
| ENSMUSG00000059003 | Grin2a | protein_coding | 0.101967882 | 4.417501533 | 0.0610235 | 0.8079685 | 0.960858843 |
| ENSMUSG00000047496 | Rnf152 | protein_coding | 0.107183672 | 3.959589499 | 0.0608848 | 0.8081823 | 0.961006774 |
| ENSMUSG00000071649 | B3gat3 | protein_coding | 0.051142605 | 6.366993141 | 0.0607788 | 0.8083457 | 0.961094771 |
| ENSMUSG00000018334 | Ksr1 | protein_coding | -0.117168215 | 3.4061664 | 0.0606327 | 0.8085714 | 0.961256809 |
| ENSMUSG00000026437 | Cdk18 | protein_coding | 0.091282766 | 5.932591511 | 0.0606567 | 0.80873 | 0.961306148 |
| ENSMUSG00000031837 | Necab2 | protein_coding | 0.060728451 | 6.648110719 | 0.0604902 | 0.8087917 | 0.961306148 |
| ENSMUSG00000036167 | Pphln1 | protein_coding | -0.06936314 | 5.291290637 | 0.0603649 | 0.8089855 | 0.961411852 |
| ENSMUSG00000097915 | A330009N23Rik | lincRNA | 0.118744672 | 4.085278012 | 0.0602528 | 0.8091643 | 0.961411852 |
| ENSMUSG00000054027 | Nt5dc3 | protein_coding | -0.086869626 | 4.951660942 | 0.060238 | 0.8091823 | 0.961411852 |
| ENSMUSG00000054611 | Kdm2a | protein_coding | -0.068943678 | 4.948489974 | 0.0600442 | 0.809483 | 0.961411852 |
| ENSMUSG00000002205 | Vrk3 | protein_coding | -0.076924417 | 5.766803364 | 0.0600449 | 0.8094869 | 0.961411852 |
| ENSMUSG00000004897 | Hdgf | protein_coding | 0.044777972 | 7.34081843 | 0.0599751 | 0.8095903 | 0.961411852 |
| ENSMUSG00000050663 | Trhde | protein_coding | 0.182539306 | 2.690121764 | 0.0599565 | 0.8096193 | 0.961411852 |
| ENSMUSG00000027300 | Ubox5 | protein_coding | 0.078910087 | 4.192564187 | 0.0598286 | 0.8098181 | 0.961411852 |
| ENSMUSG00000031367 | Ap1s2 | protein_coding | -0.064782647 | 6.820562455 | 0.0597997 | 0.8098681 | 0.961411852 |
| ENSMUSG00000034361 | Cpne2 | protein_coding | -0.06965333 | 5.225440251 | 0.0597061 | 0.8100088 | 0.961411852 |
| ENSMUSG00000029334 | Prkg2 | protein_coding | -0.131753551 | 3.431655454 | 0.0597016 | 0.8100158 | 0.961411852 |
| ENSMUSG00000053333 | Dis3l2 | protein_coding | 0.142291181 | 2.901809079 | 0.0596709 | 0.8100637 | 0.961411852 |
| ENSMUSG00000111966 | AL589661.1 | processed_pseudoge | -0.115362004 | 3.01471568 | 0.0596562 | 0.8100866 | 0.961411852 |
| ENSMUSG00000031878 | Nae1 | protein_coding | -0.055869451 | 6.584111225 | 0.0596269 | 0.8101323 | 0.961411852 |
| ENSMUSG00000005442 | Cic | protein_coding | 0.089697515 | 3.706908666 | 0.059529 | 0.810285 | 0.961422451 |
| ENSMUSG00000036040 | Adamtsl2 | protein_coding | -0.106642697 | 3.44761555 | 0.059458 | 0.8103959 | 0.961422451 |
| ENSMUSG00000015243 | Abca1 | protein_coding | -0.11323504 | 3.073190224 | 0.0594492 | 0.8104095 | 0.961422451 |
| ENSMUSG00000049690 | Nckap5 | protein_coding | -0.204601014 | 2.575471611 | 0.0593784 | 0.8105251 | 0.961453581 |
| ENSMUSG00000021589 | Rhobtb3 | protein_coding | 0.098965609 | 3.888652789 | 0.0592899 | 0.8106584 | 0.961505564 |
| ENSMUSG00000024404 | Riok3 | protein_coding | -0.086164927 | 4.928221499 | 0.0591758 | 0.810837 | 0.961528603 |
| ENSMUSG00000032936 | Camkv | protein_coding | 0.051188349 | 8.008859896 | 0.0590401 | 0.8110495 | 0.961528603 |
| ENSMUSG00000039648 | Kyat1 | protein_coding | -0.103388441 | 4.543496185 | 0.0590008 | 0.8111161 | 0.961528603 |
| ENSMUSG00000020427 | Igfbp3 | protein_coding | 0.146000084 | 2.704675004 | 0.0589814 | 0.8111415 | 0.961528603 |
| ENSMUSG00000045252 | Zfp574 | protein_coding | -0.092419968 | 4.009958688 | 0.0588722 | 0.8113128 | 0.961528603 |
| ENSMUSG00000051246 | Msantd1 | protein_coding | 0.129784591 | 3.377604844 | 0.0588295 | 0.8113798 | 0.961528603 |
| ENSMUSG00000024902 | Mrpl11 | protein_coding | 0.04475791 | 7.861113835 | 0.058824 | 0.8113884 | 0.961528603 |
| ENSMUSG00000035557 | Krt17 | protein_coding | 0.140484666 | 2.761113525 | 0.0587723 | 0.8114697 | 0.961528603 |
| ENSMUSG00000040055 | Gjb6 | protein_coding | -0.08370369 | 5.585771799 | 0.0587672 | 0.8114826 | 0.961528603 |
| ENSMUSG00000022111 | Uchl3 | protein_coding | 0.055888023 | 6.159675216 | 0.0586138 | 0.8117188 | 0.961702522 |
| ENSMUSG00000026289 | Atg16l1 | protein_coding | -0.094318874 | 4.945084035 | 0.0584774 | 0.8119385 | 0.96185686 |
| ENSMUSG00000044674 | Fzd1 | protein_coding | -0.195959333 | 2.389092387 | 0.0582986 | 0.8122154 | 0.962078877 |
| ENSMUSG00000028334 | Nans | protein_coding | 0.102292962 | 3.834587374 | 0.0581324 | 0.8124778 | 0.962283751 |
| ENSMUSG00000074890 | Lcmt2 | protein_coding | 0.139314556 | 3.090681703 | 0.0579359 | 0.8127886 | 0.962545792 |
| ENSMUSG00000073418 | C4b | protein_coding | 0.212077616 | 2.333341342 | 0.0577236 | 0.8131249 | 0.962594159 |
| ENSMUSG00000022555 | Dgat1 | protein_coding | 0.117658307 | 3.529617965 | 0.0576884 | 0.8131808 | 0.962594159 |
| ENSMUSG00000015478 | Rnf5 | protein_coding | -0.045316499 | 6.738153462 | 0.0575806 | 0.8133518 | 0.962594159 |
| ENSMUSG00000024613 | Tcof1 | protein_coding | -0.082220809 | 4.44965161 | 0.0575716 | 0.8133661 | 0.962594159 |
| ENSMUSG00000022881 | Rfc4 | protein_coding | 0.131727997 | 4.263347673 | 0.0577136 | 0.8133842 | 0.962594159 |
| ENSMUSG00000067377 | Tspan6 | protein_coding | 0.051154717 | 6.552946552 | 0.057533 | 0.8134275 | 0.962594159 |
| ENSMUSG00000022019 | Tdrd3 | protein_coding | 0.071117829 | 4.73038923 | 0.057515 | 0.8134561 | 0.962594159 |
| ENSMUSG00000022255 | Mtdh | protein_coding | 0.082649642 | 4.957280925 | 0.0573253 | 0.8137577 | 0.96284518 |
| ENSMUSG00000071679 | Zcchc16 | protein_coding | 0.145161806 | 2.834051564 | 0.0570098 | 0.8142608 | 0.962952571 |
| ENSMUSG00000025155 | Dus1l | protein_coding | 0.09185318 | 4.475230336 | 0.0569907 | 0.8142912 | 0.962952571 |
| ENSMUSG00000028944 | Prkag2 | protein_coding | -0.055621413 | 5.982183124 | 0.0569607 | 0.8143392 | 0.962952571 |
| ENSMUSG00000025465 | Echs1 | protein_coding | 0.041614729 | 7.770651461 | 0.056951 | 0.8143547 | 0.962952571 |
| ENSMUSG00000010066 | Cacna2d2 | protein_coding | 0.089717147 | 4.557096645 | 0.0569312 | 0.8143863 | 0.962952571 |
| ENSMUSG00000032193 | Ldlr | protein_coding | -0.136298667 | 3.684855885 | 0.0568943 | 0.8144937 | 0.962952571 |
| ENSMUSG00000035585 | Tsen34 | protein_coding | -0.087969404 | 4.467188683 | 0.0568451 | 0.8145239 | 0.962952571 |
| ENSMUSG00000029111 | Nelfa | protein_coding | 0.082765206 | 5.752998423 | 0.0568305 | 0.8145649 | 0.962952571 |
| ENSMUSG00000025511 | Tspan4 | protein_coding | -0.090283422 | 4.134808784 | 0.056672 | 0.8148008 | 0.963125542 |
| ENSMUSG00000026311 | Asb1 | protein_coding | 0.080928261 | 4.529768471 | 0.0565512 | 0.8149944 | 0.963248549 |
| ENSMUSG00000026914 | Psmd14 | protein_coding | 0.048158857 | 8.344813924 | 0.0564788 | 0.8151105 | 0.963279839 |
| ENSMUSG00000027259 | Adal | protein_coding | 0.079724534 | 4.295735333 | 0.0563316 | 0.8153469 | 0.963444013 |
| ENSMUSG00000021792 | Fam213a | protein_coding | 0.052193857 | 7.844558865 | 0.0562808 | 0.8154286 | 0.963444013 |
| ENSMUSG00000060090 | Rp2 | protein_coding | -0.153549611 | 2.647798057 | 0.0560375 | 0.8158201 | 0.963622575 |

| ensembl_gene_id | external_gene_name | gene_biotype | log2 (fold change) | log2 (counts per million) | F-value | p-value | False discovery rate |
|---|---|---|---|---|---|---|---|
| ENSMUSG00000024985 | Tcf7l2 | protein_coding | -0.213346492 | 8.165222098 | 0.057982 | 0.8158578 | 0.963622575 |
| ENSMUSG00000015094 | Npdc1 | protein_coding | 0.064384499 | 6.540215449 | 0.0559617 | 0.815947 | 0.963622575 |
| ENSMUSG00000025658 | Cnksr2 | protein_coding | 0.077761255 | 5.375949277 | 0.0559453 | 0.8159686 | 0.963622575 |
| ENSMUSG00000031072 | Oraov1 | protein_coding | -0.062102932 | 5.316259961 | 0.0559086 | 0.8160278 | 0.963622575 |
| ENSMUSG00000030047 | Arhgap25 | protein_coding | -0.196590913 | 2.611281715 | 0.0557906 | 0.8162232 | 0.963654138 |
| ENSMUSG00000034345 | Gtf2h5 | protein_coding | -0.043970291 | 8.783717357 | 0.0556998 | 0.8163651 | 0.963654138 |
| ENSMUSG00000061273 | Mmgt1 | protein_coding | 0.074194234 | 4.400901462 | 0.0556343 | 0.816471 | 0.963654138 |
| ENSMUSG00000024154 | Gtf2a1l | protein_coding | 0.163912297 | 2.657533945 | 0.055584 | 0.8165523 | 0.963654138 |
| ENSMUSG00000036023 | Parp2 | protein_coding | 0.055946674 | 5.474415451 | 0.0555193 | 0.8166571 | 0.963654138 |
| ENSMUSG00000025202 | Scd3 | protein_coding | -0.087261104 | 4.335583189 | 0.0555174 | 0.8166601 | 0.963654138 |
| ENSMUSG00000032640 | Chsy1 | protein_coding | 0.086510042 | 3.73724095 | 0.0554683 | 0.8167396 | 0.963654138 |
| ENSMUSG00000055200 | Sertad3 | protein_coding | 0.127097176 | 3.194498465 | 0.0553585 | 0.8169175 | 0.963654138 |
| ENSMUSG00000015522 | Arnt | protein_coding | 0.100764111 | 3.379198166 | 0.0553159 | 0.8169867 | 0.963654138 |
| ENSMUSG00000071172 | Srsf3 | protein_coding | -0.046381389 | 7.133198226 | 0.0551865 | 0.8171967 | 0.963654138 |
| ENSMUSG00000032586 | Traip | protein_coding | 0.072295185 | 4.521045935 | 0.0551467 | 0.8172613 | 0.963654138 |
| ENSMUSG00000048371 | Pdp2 | protein_coding | 0.100177093 | 5.615301896 | 0.0553203 | 0.8172718 | 0.963654138 |
| ENSMUSG00000021018 | Polr2h | protein_coding | 0.050800215 | 6.12571714 | 0.0551084 | 0.8173236 | 0.963654138 |
| ENSMUSG00000056158 | Car10 | protein_coding | 0.078856124 | 6.29530553 | 0.055178 | 0.8173968 | 0.963654138 |
| ENSMUSG00000029651 | Mtus2 | protein_coding | 0.056142761 | 5.4084515 | 0.0549289 | 0.8176156 | 0.963654138 |
| ENSMUSG00000030811 | Fbxl19 | protein_coding | -0.216855109 | 2.807103069 | 0.0549461 | 0.8177735 | 0.963654138 |
| ENSMUSG00000024955 | Esrra | protein_coding | 0.171165396 | 2.598698475 | 0.0547962 | 0.8178318 | 0.963654138 |
| ENSMUSG00000024764 | Naa40 | protein_coding | 0.101135742 | 3.508491131 | 0.0547773 | 0.8178626 | 0.963654138 |
| ENSMUSG00000025967 | Eef1b2 | protein_coding | -0.053516223 | 9.00386979 | 0.0547641 | 0.8178842 | 0.963654138 |
| ENSMUSG00000038205 | Prkab2 | protein_coding | -0.166395261 | 2.740696418 | 0.0546857 | 0.818012 | 0.963654138 |
| ENSMUSG00000024983 | Vti1a | protein_coding | 0.076779752 | 5.623948769 | 0.054678 | 0.8180294 | 0.963654138 |
| ENSMUSG00000057963 | Itpk1 | protein_coding | -0.05879512 | 5.643085714 | 0.0545989 | 0.8181538 | 0.963654138 |
| ENSMUSG00000033900 | Map9 | protein_coding | 0.05584051 | 6.439453988 | 0.0545694 | 0.818202 | 0.963654138 |
| ENSMUSG00000023921 | Mut | protein_coding | -0.068086941 | 5.10118876 | 0.0544594 | 0.8183818 | 0.963654138 |
| ENSMUSG00000006476 | Nsmf | protein_coding | -0.076935328 | 6.355059657 | 0.0545678 | 0.8183897 | 0.963654138 |
| ENSMUSG00000029464 | Gpn3 | protein_coding | -0.070383033 | 5.759942848 | 0.0544259 | 0.8184366 | 0.963654138 |
| ENSMUSG00000005610 | Eif4g2 | protein_coding | -0.055868034 | 8.797451722 | 0.0544029 | 0.8184742 | 0.963654138 |
| ENSMUSG00000037843 | Vstm2l | protein_coding | -0.145578553 | 2.609806328 | 0.0542961 | 0.818649 | 0.963701093 |
| ENSMUSG00000039633 | Lonrf1 | protein_coding | 0.113271973 | 3.347394235 | 0.0542691 | 0.8186933 | 0.963701093 |
| ENSMUSG00000067370 | B3galt4 | protein_coding | -0.139564783 | 3.63878684 | 0.0542302 | 0.8188786 | 0.963722695 |
| ENSMUSG00000009828 | Ick | protein_coding | 0.114144916 | 3.469201804 | 0.054132 | 0.8189182 | 0.963722695 |
| ENSMUSG00000061479 | Snrpa | protein_coding | -0.065118743 | 6.147516646 | 0.054094 | 0.8189806 | 0.963722695 |
| ENSMUSG00000022992 | Kansl2 | protein_coding | -0.133562046 | 2.796807331 | 0.0539246 | 0.8192587 | 0.963944548 |
| ENSMUSG00000027457 | Snph | protein_coding | 0.05494765 | 6.359652237 | 0.0537825 | 0.8194927 | 0.964114338 |
| ENSMUSG00000020519 | Sap30l | protein_coding | 0.127625892 | 2.941717624 | 0.0536759 | 0.8196683 | 0.964215374 |
| ENSMUSG00000021791 | Dydc2 | protein_coding | -0.126557349 | 2.879453552 | 0.0534315 | 0.8200716 | 0.964584269 |
| ENSMUSG00000025060 | Slk | protein_coding | -0.094230107 | 4.682045416 | 0.0533663 | 0.820184 | 0.964611053 |
| ENSMUSG00000039086 | Ss18l1 | protein_coding | -0.067489097 | 4.984078631 | 0.0532772 | 0.8203267 | 0.964673376 |
| ENSMUSG00000060771 | Tsga10 | protein_coding | -0.12271252 | 3.247839513 | 0.0531676 | 0.8205082 | 0.964781268 |
| ENSMUSG00000044475 | Ascc1 | protein_coding | 0.044691297 | 6.756643452 | 0.0530525 | 0.820699 | 0.964900062 |
| ENSMUSG00000015575 | Atp6v0e | protein_coding | 0.06264875 | 7.084785121 | 0.0527336 | 0.8212458 | 0.965346265 |
| ENSMUSG00000030257 | Srgap3 | protein_coding | -0.055512706 | 6.169413465 | 0.0527159 | 0.821258 | 0.965346265 |
| ENSMUSG00000087165 | 2010001A14Rik | processed_transcript | 0.081298124 | 4.217475595 | 0.0525008 | 0.8216164 | 0.965423018 |
| ENSMUSG00000026003 | Acadl | protein_coding | -0.07654093 | 5.020147576 | 0.0524833 | 0.8216456 | 0.965423018 |
| ENSMUSG00000032215 | Rsl24d1 | protein_coding | 0.046049748 | 6.40011498 | 0.0524376 | 0.8217219 | 0.965423018 |
| ENSMUSG00000074872 | Ctxn2 | protein_coding | -0.050289103 | 8.340177171 | 0.0523227 | 0.8219136 | 0.965423018 |
| ENSMUSG00000014856 | Tmem208 | protein_coding | 0.049704859 | 6.816027486 | 0.0522856 | 0.8219756 | 0.965423018 |
| ENSMUSG00000039110 | Mycbpap | protein_coding | -0.134197564 | 2.801243388 | 0.0522804 | 0.8219843 | 0.965423018 |
| ENSMUSG00000087128 | Gm12655 | processed_transcript | 0.157527334 | 2.679446956 | 0.0522574 | 0.8220227 | 0.965423018 |
| ENSMUSG00000025010 | Ccnj | protein_coding | -0.127334735 | 3.045179781 | 0.0522462 | 0.8220416 | 0.965423018 |
| ENSMUSG00000010047 | Hyal2 | protein_coding | 0.080211406 | 4.857066359 | 0.052144 | 0.8222124 | 0.96547052 |
| ENSMUSG00000003134 | Tbc1d8 | protein_coding | -0.096649005 | 3.701743693 | 0.0520806 | 0.8223185 | 0.96547052 |
| ENSMUSG00000036790 | Slitrk2 | protein_coding | -0.091851926 | 3.373395421 | 0.052061 | 0.8223514 | 0.96547052 |
| ENSMUSG00000024006 | Stk38 | protein_coding | 0.089333254 | 3.52532164 | 0.0519651 | 0.8225122 | 0.965553903 |
| ENSMUSG00000039126 | Prune2 | protein_coding | -0.101666056 | 8.868720627 | 0.0524461 | 0.8226396 | 0.965558107 |
| ENSMUSG00000032875 | Arhgef17 | protein_coding | -0.047270759 | 6.310172175 | 0.0518227 | 0.822751 | 0.965558107 |
| ENSMUSG00000009030 | Pdcl | protein_coding | 0.065184209 | 5.005014653 | 0.0518024 | 0.8227852 | 0.965558107 |
| ENSMUSG00000011752 | Pgam1 | protein_coding | 0.079754926 | 3.725204375 | 0.0517273 | 0.8229114 | 0.965600847 |
| ENSMUSG00000033429 | Mcee | protein_coding | -0.052094812 | 6.651822384 | 0.0515098 | 0.8232773 | 0.96582423 |
| ENSMUSG00000019518 | Ap4m1 | protein_coding | -0.123001328 | 2.850539078 | 0.0513645 | 0.8235222 | 0.96582423 |
| ENSMUSG00000034729 | Mrps10 | protein_coding | 0.048764981 | 6.322368147 | 0.0512172 | 0.8237709 | 0.96582423 |
| ENSMUSG00000043300 | B3galnt1 | protein_coding | -0.053422049 | 6.205817635 | 0.0511827 | 0.8238292 | 0.96582423 |
| ENSMUSG00000028949 | Smarcd3 | protein_coding | -0.087291019 | 3.796072841 | 0.0511685 | 0.8238531 | 0.96582423 |
| ENSMUSG00000031659 | Adcy7 | protein_coding | -0.112063565 | 3.286394113 | 0.0511667 | 0.8238562 | 0.96582423 |
| ENSMUSG00000102437 | Gm38048 | sense_overlapping | -0.124474871 | 3.008105649 | 0.0511615 | 0.8238651 | 0.96582423 |
| ENSMUSG00000022464 | Slc38a4 | protein_coding | -0.15226114 | 2.634927549 | 0.051145 | 0.8238929 | 0.96582423 |
| ENSMUSG00000008393 | Carhsp1 | protein_coding | 0.048517196 | 6.747320037 | 0.0511348 | 0.8239101 | 0.96582423 |
| ENSMUSG00000045348 | Nyap1 | protein_coding | -0.116151604 | 3.3711116 | 0.0509177 | 0.8242778 | 0.966149859 |
| ENSMUSG00000062190 | Lancl2 | protein_coding | 0.160081269 | 3.077568906 | 0.0507873 | 0.8245035 | 0.966208167 |
| ENSMUSG00000018736 | Ndel1 | protein_coding | 0.048601945 | 6.757699419 | 0.050763 | 0.8245401 | 0.966208167 |

| ensembl_gene_id | external_gene_name | gene_biotype | log2 (fold change) | log2 (counts per million) | F-value | p-value | False discovery rate |
|---|---|---|---|---|---|---|---|
| ENSMUSG00000036390 | Gadd45a | protein_coding | 0.119399948 | 2.947059194 | 0.0507294 | 0.8245971 | 0.966208167 |
| ENSMUSG00000026179 | Pnkd | protein_coding | -0.04451421 | 8.05180221 | 0.0504404 | 0.8250886 | 0.966451952 |
| ENSMUSG00000020847 | Rph3al | protein_coding | -0.111967943 | 3.492750588 | 0.0504211 | 0.8251214 | 0.966451952 |
| ENSMUSG00000045636 | Mtus1 | protein_coding | -0.069494439 | 5.259558151 | 0.0504118 | 0.8251374 | 0.966451952 |
| ENSMUSG00000020257 | Wdr82 | protein_coding | -0.04979092 | 6.41093458 | 0.0503778 | 0.8251953 | 0.966451952 |
| ENSMUSG00000030757 | Zkscan2 | protein_coding | 0.148444514 | 2.533081149 | 0.050343 | 0.8252545 | 0.966451952 |
| ENSMUSG00000071454 | Dtnb | protein_coding | 0.060943529 | 4.995225655 | 0.0500571 | 0.8257428 | 0.966687228 |
| ENSMUSG00000018697 | Aatf | protein_coding | 0.108021344 | 3.967412359 | 0.0500517 | 0.8257566 | 0.966687228 |
| ENSMUSG00000002625 | Akap8l | protein_coding | 0.056507131 | 5.114808277 | 0.0500287 | 0.8257913 | 0.966687228 |
| ENSMUSG00000057522 | Spop | protein_coding | -0.0507968 | 8.272857424 | 0.0500148 | 0.825815 | 0.966687228 |
| ENSMUSG00000034259 | Exosc4 | protein_coding | -0.050413068 | 6.135253347 | 0.0499519 | 0.8259228 | 0.966708184 |
| ENSMUSG00000002799 | Jag2 | protein_coding | -0.16335332 | 2.957578976 | 0.0498304 | 0.8261355 | 0.966741523 |
| ENSMUSG00000026649 | Cfap126 | protein_coding | -0.08306773 | 4.686892357 | 0.0496855 | 0.8263795 | 0.966741523 |
| ENSMUSG00000028771 | Ptpn12 | protein_coding | -0.096693572 | 4.087179303 | 0.0495901 | 0.8265433 | 0.966741523 |
| ENSMUSG00000038991 | Txndc5 | protein_coding | 0.075750538 | 5.050696154 | 0.0495011 | 0.8266963 | 0.966741523 |
| ENSMUSG00000028910 | Mecr | protein_coding | 0.06851689 | 5.23956045 | 0.0494821 | 0.826729 | 0.966741523 |
| ENSMUSG00000033004 | Mycbp2 | protein_coding | 0.063042087 | 6.269808914 | 0.049436 | 0.8268128 | 0.966741523 |
| ENSMUSG00000079017 | Ifi27l2a | protein_coding | -0.157895087 | 2.489332967 | 0.0493705 | 0.8269211 | 0.966741523 |
| ENSMUSG00000024067 | Dpy30 | protein_coding | -0.046268927 | 8.230608169 | 0.0493 | 0.8270426 | 0.966741523 |
| ENSMUSG00000031355 | Arhgap6 | protein_coding | 0.070964305 | 4.359166471 | 0.0491979 | 0.8272187 | 0.966741523 |
| ENSMUSG00000021027 | Ralgapa1 | protein_coding | -0.060654114 | 5.251444168 | 0.0491856 | 0.82724 | 0.966741523 |
| ENSMUSG00000029708 | Gcc1 | protein_coding | 0.08136791 | 3.788800149 | 0.0491019 | 0.8273844 | 0.966741523 |
| ENSMUSG00000020267 | Hint1 | protein_coding | 0.044519733 | 10.43714047 | 0.0490423 | 0.8274875 | 0.966741523 |
| ENSMUSG00000031059 | Ndufb11 | protein_coding | 0.050886582 | 9.367674835 | 0.0489055 | 0.8277241 | 0.966741523 |
| ENSMUSG00000024855 | Pacs1 | protein_coding | -0.061338172 | 5.570134098 | 0.0488992 | 0.827735 | 0.966741523 |
| ENSMUSG00000097466 | D430036J16Rik | antisense_RNA | 0.110268876 | 3.323470962 | 0.0488683 | 0.8277885 | 0.966741523 |
| ENSMUSG00000075410 | Prcd | protein_coding | 0.148632214 | 3.463079104 | 0.0489541 | 0.8278149 | 0.966741523 |
| ENSMUSG00000051048 | P4ha3 | protein_coding | -0.170137564 | 2.603268759 | 0.0488497 | 0.8278208 | 0.966741523 |
| ENSMUSG00000078970 | Wdr92 | protein_coding | -0.085899468 | 4.645180636 | 0.0488467 | 0.8278259 | 0.966741523 |
| ENSMUSG00000062202 | Btbd9 | protein_coding | -0.066039277 | 5.808401651 | 0.0487491 | 0.8279951 | 0.966741523 |
| ENSMUSG00000018509 | Cenpv | protein_coding | 0.074958434 | 4.041840797 | 0.048721 | 0.8280438 | 0.966741523 |
| ENSMUSG00000041439 | Mfsd6 | protein_coding | 0.055577975 | 5.493051545 | 0.0487071 | 0.8280679 | 0.966741523 |
| ENSMUSG00000029001 | Fbxo44 | protein_coding | -0.037770556 | 7.966833578 | 0.0486852 | 0.8281058 | 0.966741523 |
| ENSMUSG00000025384 | Faap100 | protein_coding | -0.099656028 | 4.090612436 | 0.0485852 | 0.828284 | 0.966741523 |
| ENSMUSG00000021905 | Dph3 | protein_coding | -0.052168419 | 6.193024074 | 0.0484784 | 0.8284651 | 0.966741523 |
| ENSMUSG00000047428 | Dlk2 | protein_coding | -0.114472224 | 3.196595907 | 0.0484677 | 0.8284837 | 0.966741523 |
| ENSMUSG00000051864 | Tbc1d22a | protein_coding | 0.075616285 | 4.965717648 | 0.0484383 | 0.8285349 | 0.966741523 |
| ENSMUSG00000017404 | Rpl19 | protein_coding | 0.087236704 | 3.484945015 | 0.0484264 | 0.8285556 | 0.966741523 |
| ENSMUSG00000030753 | Thap12 | protein_coding | -0.063939975 | 4.885944996 | 0.0484259 | 0.8285564 | 0.966741523 |
| ENSMUSG00000033488 | BC026585 | protein_coding | 0.091022167 | 4.166418027 | 0.0484247 | 0.8285585 | 0.966741523 |
| ENSMUSG00000001700 | Gramd3 | protein_coding | -0.102524883 | 3.769375656 | 0.0483695 | 0.8286547 | 0.966748837 |
| ENSMUSG00000061740 | Cyp2d22 | protein_coding | -0.104958828 | 3.458521513 | 0.0482437 | 0.8288739 | 0.966831432 |
| ENSMUSG00000004561 | Mettl17 | protein_coding | 0.073336099 | 4.410173875 | 0.0482256 | 0.8289053 | 0.966831432 |
| ENSMUSG00000050822 | Slc29a4 | protein_coding | 0.097473956 | 3.279897872 | 0.0480374 | 0.829234 | 0.967109872 |
| ENSMUSG00000029569 | Tmem168 | protein_coding | -0.128137313 | 3.063842105 | 0.0479846 | 0.8293263 | 0.967112659 |
| ENSMUSG00000096146 | Kcnj11 | protein_coding | 0.089818984 | 3.954955941 | 0.0478046 | 0.8296412 | 0.967209511 |
| ENSMUSG00000036054 | Sugp2 | protein_coding | 0.057659203 | 4.989714878 | 0.0477443 | 0.829747 | 0.967209511 |
| ENSMUSG00000039678 | Tbc1d13 | protein_coding | -0.091079136 | 3.440285826 | 0.0476448 | 0.8299215 | 0.967209511 |
| ENSMUSG00000044562 | Rasip1 | protein_coding | -0.139815252 | 3.036528528 | 0.0475001 | 0.8301757 | 0.967209511 |
| ENSMUSG00000028700 | Pomgnt1 | protein_coding | -0.069046437 | 5.107725066 | 0.0473915 | 0.8303666 | 0.967209511 |
| ENSMUSG00000074607 | Tox2 | protein_coding | 0.08955685 | 5.997810365 | 0.0475942 | 0.8303715 | 0.967209511 |
| ENSMUSG00000040532 | Abhd11 | protein_coding | 0.04615151 | 6.511618318 | 0.0472092 | 0.8306878 | 0.967209511 |
| ENSMUSG00000036578 | Fxyd7 | protein_coding | -0.041142252 | 7.396395722 | 0.0471481 | 0.8307956 | 0.967209511 |
| ENSMUSG00000050447 | Lypd6 | protein_coding | -0.077523133 | 4.566478356 | 0.0471327 | 0.8308228 | 0.967209511 |
| ENSMUSG00000025372 | Baiap2 | protein_coding | 0.048945965 | 7.550732892 | 0.047086 | 0.8309053 | 0.967209511 |
| ENSMUSG00000044864 | Ankrd50 | protein_coding | -0.075540554 | 4.326761059 | 0.0470519 | 0.8309655 | 0.967209511 |
| ENSMUSG00000073002 | Vamp5 | protein_coding | -0.118951542 | 3.080028733 | 0.0470374 | 0.8309911 | 0.967209511 |
| ENSMUSG00000002395 | Use1 | protein_coding | 0.043730127 | 8.642709291 | 0.046918 | 0.8312021 | 0.967209511 |
| ENSMUSG00000026135 | Zfp142 | protein_coding | 0.169674057 | 2.531789091 | 0.0468964 | 0.8312405 | 0.967209511 |
| ENSMUSG00000051790 | Nlgn2 | protein_coding | 0.04889796 | 6.043379295 | 0.046853 | 0.8313172 | 0.967209511 |
| ENSMUSG00000039840 | Epg5 | protein_coding | 0.130617006 | 3.388539015 | 0.0467959 | 0.8314227 | 0.967209511 |
| ENSMUSG00000087197 | Gm13780 | lincRNA | -0.100917478 | 3.154261479 | 0.0466507 | 0.8316758 | 0.967209511 |
| ENSMUSG00000021068 | Nin | protein_coding | -0.101952659 | 3.182937395 | 0.0466007 | 0.8317645 | 0.967209511 |
| ENSMUSG00000024695 | Zfp91 | protein_coding | -0.050227701 | 5.647609796 | 0.0465942 | 0.831776 | 0.967209511 |
| ENSMUSG00000003154 | Foxj2 | protein_coding | -0.084551389 | 3.932937719 | 0.046475 | 0.8319879 | 0.967209511 |
| ENSMUSG00000034252 | Senp6 | protein_coding | 0.065867015 | 5.136205662 | 0.046431 | 0.8320661 | 0.967209511 |
| ENSMUSG00000028293 | Slc35a1 | protein_coding | 0.060170765 | 5.190766832 | 0.0464125 | 0.8320991 | 0.967209511 |
| ENSMUSG00000024537 | Psmg2 | protein_coding | -0.044867366 | 6.137538241 | 0.0463617 | 0.8321895 | 0.967209511 |
| ENSMUSG00000069814 | Ccdc92b | protein_coding | 0.075950956 | 4.020722003 | 0.0462877 | 0.8323213 | 0.967209511 |
| ENSMUSG00000070348 | Ccnd1 | protein_coding | -0.16734115 | 2.568116953 | 0.0462486 | 0.832391 | 0.967209511 |
| ENSMUSG00000072437 | Nanos1 | protein_coding | -0.069998497 | 4.553238287 | 0.0462062 | 0.8324665 | 0.967209511 |
| ENSMUSG00000010517 | Faf1 | protein_coding | 0.069353227 | 5.091484325 | 0.0461658 | 0.8325386 | 0.967209511 |
| ENSMUSG00000038489 | Polr2l | protein_coding | -0.05844802 | 4.985064175 | 0.0461492 | 0.8325684 | 0.967209511 |
| ENSMUSG00000020903 | Stx8 | protein_coding | -0.04032181 | 6.872106506 | 0.0461466 | 0.8325728 | 0.967209511 |

| ensembl_gene_id | external_gene_name | gene_biotype | log2 (fold change) | log2 (counts per million) | F-value | p-value | False discovery rate |
|---|---|---|---|---|---|---|---|
| ENSMUSG00000059713 | Rcan3 | protein_coding | 0.051309141 | 5.330877705 | 0.046077 | 0.8326971 | 0.967209511 |
| ENSMUSG00000048796 | Cyb561d1 | protein_coding | 0.098540058 | 3.55124083 | 0.0460302 | 0.8327808 | 0.967209511 |
| ENSMUSG00000017421 | Zfp207 | protein_coding | -0.044743714 | 7.026281644 | 0.0460183 | 0.8328021 | 0.967209511 |
| ENSMUSG00000012076 | Brms1l | protein_coding | -0.051614509 | 5.794557637 | 0.0460051 | 0.8328256 | 0.967209511 |
| ENSMUSG00000000088 | Cox5a | protein_coding | 0.06241863 | 4.60852763 | 0.0459944 | 0.8328448 | 0.967209511 |
| ENSMUSG00000020381 | Mrnip | protein_coding | 0.06100591 | 5.218172705 | 0.0459345 | 0.8329518 | 0.967209511 |
| ENSMUSG00000022948 | Setd4 | protein_coding | 0.153258751 | 3.210991503 | 0.0458965 | 0.8331313 | 0.967209511 |
| ENSMUSG00000004270 | Lpcat3 | protein_coding | -0.088301175 | 4.037494218 | 0.0458032 | 0.833187 | 0.967209511 |
| ENSMUSG00000001366 | Fbxo9 | protein_coding | 0.039727774 | 6.406147436 | 0.0456829 | 0.8334027 | 0.967209511 |
| ENSMUSG00000044469 | Tnfaip8l1 | protein_coding | 0.104625746 | 3.351556379 | 0.045644 | 0.8334725 | 0.967209511 |
| ENSMUSG00000073102 | Drc1 | protein_coding | 0.062930392 | 4.882209668 | 0.0456118 | 0.8335303 | 0.967209511 |
| ENSMUSG00000041278 | Ttc1 | protein_coding | -0.036483571 | 7.870327927 | 0.0454619 | 0.8337998 | 0.967209511 |
| ENSMUSG00000032679 | Cd59a | protein_coding | 0.06477821 | 5.657192249 | 0.0454489 | 0.8338231 | 0.967209511 |
| ENSMUSG00000015217 | Hmgb3 | protein_coding | -0.077537697 | 3.751086296 | 0.0453985 | 0.8339138 | 0.967209511 |
| ENSMUSG00000038121 | Fam210a | protein_coding | 0.060460463 | 5.42499516 | 0.0453798 | 0.8339474 | 0.967209511 |
| ENSMUSG00000004933 | Matk | protein_coding | -0.048235013 | 7.258529824 | 0.0453662 | 0.8339719 | 0.967209511 |
| ENSMUSG00000026356 | Dars | protein_coding | -0.058419846 | 6.447264805 | 0.045294 | 0.8341063 | 0.967209511 |
| ENSMUSG00000019878 | Hsf2 | protein_coding | 0.083919358 | 5.061571402 | 0.0452846 | 0.8341232 | 0.967209511 |
| ENSMUSG00000047897 | Ripply2 | protein_coding | -0.111722731 | 2.911430618 | 0.0452798 | 0.8341277 | 0.967209511 |
| ENSMUSG00000052551 | Adarb2 | protein_coding | 0.069178951 | 4.072963191 | 0.0452434 | 0.8341933 | 0.967209511 |
| ENSMUSG00000044715 | Gskip | protein_coding | -0.039123054 | 7.505913195 | 0.04523 | 0.8342174 | 0.967209511 |
| ENSMUSG00000035967 | Ints6l | protein_coding | 0.130281548 | 2.801393177 | 0.0451928 | 0.8342845 | 0.967209511 |
| ENSMUSG00000059278 | Naa38 | protein_coding | -0.050377955 | 7.56424352 | 0.0451643 | 0.8343359 | 0.967209511 |
| ENSMUSG00000070695 | Cntnap5a | protein_coding | 0.071345943 | 4.667547053 | 0.0451035 | 0.8344458 | 0.967209511 |
| ENSMUSG00000016946 | Kctd5 | protein_coding | 0.157148326 | 3.874375361 | 0.0452808 | 0.8345464 | 0.967209511 |
| ENSMUSG00000024851 | Pitpnm1 | protein_coding | -0.053866283 | 5.789043041 | 0.0450088 | 0.8346169 | 0.967209511 |
| ENSMUSG00000026958 | Dpp7 | protein_coding | -0.068361048 | 4.544340655 | 0.044998 | 0.8346364 | 0.967209511 |
| ENSMUSG00000019777 | Hdac2 | protein_coding | 0.069491484 | 5.824625068 | 0.0449952 | 0.8346458 | 0.967209511 |
| ENSMUSG00000024597 | Slc12a2 | protein_coding | -0.064570737 | 5.596955192 | 0.0449552 | 0.8347137 | 0.967209511 |
| ENSMUSG00000049804 | Armcx4 | protein_coding | -0.095999366 | 4.416621524 | 0.0449562 | 0.8347163 | 0.967209511 |
| ENSMUSG00000048701 | Ccdc6 | protein_coding | 0.043361836 | 6.963461909 | 0.0449017 | 0.8348106 | 0.967214512 |
| ENSMUSG00000052446 | Zfp961 | protein_coding | 0.075600234 | 4.061898998 | 0.0448419 | 0.8349189 | 0.967235863 |
| ENSMUSG00000016179 | Camk1g | protein_coding | 0.05424999 | 6.34527043 | 0.0445923 | 0.8353717 | 0.967656178 |
| ENSMUSG00000039164 | Naif1 | protein_coding | -0.104781567 | 3.153482618 | 0.0441428 | 0.8361908 | 0.968492789 |
| ENSMUSG00000021241 | Isca2 | protein_coding | -0.035758253 | 8.298252088 | 0.0440084 | 0.8364364 | 0.968492789 |
| ENSMUSG00000008090 | Fgfrl1 | protein_coding | -0.165003263 | 2.626197143 | 0.0439283 | 0.8365831 | 0.968492789 |
| ENSMUSG00000053093 | Myh7 | protein_coding | 0.11042719 | 2.921770434 | 0.0439068 | 0.8366225 | 0.968492789 |
| ENSMUSG00000004018 | Fancl | protein_coding | -0.065912098 | 4.841292438 | 0.0438738 | 0.836683 | 0.968492789 |
| ENSMUSG00000024544 | Ldlrad4 | protein_coding | 0.097598776 | 4.545320812 | 0.0438591 | 0.8367904 | 0.968492789 |
| ENSMUSG00000042380 | Smim12 | protein_coding | -0.043205817 | 7.240841027 | 0.0438021 | 0.8368144 | 0.968492789 |
| ENSMUSG00000039887 | Alg14 | protein_coding | 0.048790476 | 5.90807092 | 0.0437719 | 0.8368698 | 0.968492789 |
| ENSMUSG00000034518 | Hmgxb4 | protein_coding | 0.070755824 | 3.850290875 | 0.0437529 | 0.8369046 | 0.968492789 |
| ENSMUSG00000000538 | Tom1l2 | protein_coding | 0.052594569 | 7.247091143 | 0.0436885 | 0.8370271 | 0.968501883 |
| ENSMUSG00000031776 | Arl2bp | protein_coding | -0.042421079 | 6.444304096 | 0.0436505 | 0.8370926 | 0.968501883 |
| ENSMUSG00000030423 | Pop4 | protein_coding | -0.050730792 | 6.037266014 | 0.0435885 | 0.8372067 | 0.968529683 |
| ENSMUSG00000026806 | Ddx31 | protein_coding | 0.134518689 | 2.812605396 | 0.0435376 | 0.8373002 | 0.968533696 |
| ENSMUSG00000034163 | Zfc3h1 | protein_coding | -0.129014062 | 2.682463866 | 0.0434264 | 0.8375051 | 0.968604317 |
| ENSMUSG00000057835 | Zfp119a | protein_coding | 0.126279407 | 3.001432612 | 0.0433509 | 0.8376441 | 0.968604317 |
| ENSMUSG00000022180 | Slc7a8 | protein_coding | 0.060040423 | 4.673576006 | 0.0433264 | 0.8376893 | 0.968604317 |
| ENSMUSG00000035314 | Gdpd5 | protein_coding | -0.081640768 | 3.631621812 | 0.0433089 | 0.8377216 | 0.968604317 |
| ENSMUSG00000040896 | Kcnd3 | protein_coding | 0.083433447 | 4.172259452 | 0.0431645 | 0.8379882 | 0.96873878 |
| ENSMUSG00000022241 | Tars | protein_coding | 0.062546128 | 5.555735571 | 0.0430615 | 0.8381787 | 0.96873878 |
| ENSMUSG00000032258 | Lca5 | protein_coding | 0.206507916 | 2.502113867 | 0.0430798 | 0.8382208 | 0.96873878 |
| ENSMUSG00000035397 | Klf16 | protein_coding | 0.059731052 | 5.288354694 | 0.0429898 | 0.8383115 | 0.96873878 |
| ENSMUSG00000062580 | Timm17a | protein_coding | 0.036060145 | 9.022382621 | 0.0429727 | 0.8383431 | 0.96873878 |
| ENSMUSG00000040276 | Pacsin1 | protein_coding | 0.042541108 | 7.277581248 | 0.0429537 | 0.8383784 | 0.96873878 |
| ENSMUSG00000059005 | Hnrnpa3 | protein_coding | -0.066306913 | 3.884910127 | 0.0425217 | 0.8391809 | 0.969561834 |
| ENSMUSG00000025521 | Tmem192 | protein_coding | 0.083403498 | 3.808437796 | 0.0423932 | 0.8394205 | 0.969689443 |
| ENSMUSG00000048039 | Isg20l2 | protein_coding | -0.086116765 | 3.647626949 | 0.0423409 | 0.8395181 | 0.969689443 |
| ENSMUSG00000038982 | Bloc1s5 | protein_coding | -0.050761685 | 5.813263215 | 0.0421879 | 0.8398041 | 0.969689443 |
| ENSMUSG00000029049 | Morn1 | protein_coding | -0.103523702 | 3.800028213 | 0.0421396 | 0.8398986 | 0.969689443 |
| ENSMUSG00000030598 | Fbxo17 | protein_coding | -0.085590739 | 4.047245717 | 0.0421211 | 0.839929 | 0.969689443 |
| ENSMUSG00000026443 | Lrrn2 | protein_coding | 0.048821072 | 5.853491411 | 0.0421207 | 0.8399298 | 0.969689443 |
| ENSMUSG00000030082 | Sec61a1 | protein_coding | 0.056538233 | 4.915062763 | 0.0420866 | 0.8399937 | 0.969689443 |
| ENSMUSG00000038555 | Reep2 | protein_coding | 0.045967889 | 7.814522904 | 0.0420764 | 0.8400128 | 0.969689443 |
| ENSMUSG00000039714 | Cplx3 | protein_coding | 0.143644407 | 3.717022402 | 0.0421033 | 0.8402918 | 0.969817805 |
| ENSMUSG00000032482 | Cspg5 | protein_coding | 0.049946154 | 7.199515376 | 0.0419209 | 0.8403043 | 0.969817805 |
| ENSMUSG00000020876 | Snx11 | protein_coding | -0.085809735 | 4.596379397 | 0.041708 | 0.8407086 | 0.970103055 |
| ENSMUSG00000024911 | Fibp | protein_coding | 0.042132626 | 6.886098027 | 0.0416657 | 0.840784 | 0.970103055 |
| ENSMUSG00000001270 | Ckb | protein_coding | -0.051664131 | 10.89649594 | 0.0416185 | 0.8408728 | 0.970103055 |
| ENSMUSG00000028163 | Nfkb1 | protein_coding | -0.09762059 | 3.062743816 | 0.0415915 | 0.8409237 | 0.970103055 |
| ENSMUSG00000027671 | Actl6a | protein_coding | 0.068323579 | 4.085812605 | 0.041549 | 0.8410038 | 0.970103055 |
| ENSMUSG00000040481 | Bptf | protein_coding | 0.049867787 | 6.101804355 | 0.0414479 | 0.8411946 | 0.970103055 |
| ENSMUSG00000018427 | Ypel2 | protein_coding | -0.039473254 | 6.565486051 | 0.0414412 | 0.8412073 | 0.970103055 |

| ensembl_gene_id | external_gene_name | gene_biotype | log2 (fold change) | log2 (counts per million) | F-value | p-value | False discovery rate |
|---|---|---|---|---|---|---|---|
| ENSMUSG00000074170 | Plekhf1 | protein_coding | -0.098792219 | 3.88026458 | 0.0413664 | 0.8413527 | 0.970103055 |
| ENSMUSG00000039787 | Cercam | protein_coding | 0.192348425 | 3.018022681 | 0.0415458 | 0.8413634 | 0.970103055 |
| ENSMUSG00000046058 | Eid2 | protein_coding | 0.068794594 | 4.640219936 | 0.0412778 | 0.841516 | 0.970174952 |
| ENSMUSG00000070699 | Sars2 | protein_coding | -0.084345984 | 4.124511531 | 0.0412162 | 0.8416326 | 0.970205363 |
| ENSMUSG00000028931 | Kcnab2 | protein_coding | -0.040345998 | 7.385451449 | 0.0409927 | 0.8420564 | 0.970249901 |
| ENSMUSG00000027811 | 4930579G24Rik | protein_coding | -0.142612719 | 2.607655136 | 0.0409733 | 0.8420933 | 0.970249901 |
| ENSMUSG00000031517 | Gpm6a | protein_coding | -0.036992218 | 9.910279319 | 0.0409256 | 0.8421839 | 0.970249901 |
| ENSMUSG00000085622 | 3110056K07Rik | processed_transcript | 0.086824253 | 4.492442256 | 0.0409232 | 0.8421926 | 0.970249901 |
| ENSMUSG00000024163 | Mapk8ip3 | protein_coding | -0.03717923 | 7.311188667 | 0.0408825 | 0.8422657 | 0.970249901 |
| ENSMUSG00000015994 | Fnta | protein_coding | -0.057778301 | 5.94889738 | 0.0408734 | 0.8422832 | 0.970249901 |
| ENSMUSG00000021900 | Btd | protein_coding | 0.073301902 | 4.185095894 | 0.040863 | 0.8423029 | 0.970249901 |
| ENSMUSG00000028049 | Scamp3 | protein_coding | 0.042633748 | 6.211898574 | 0.0407719 | 0.8424763 | 0.970271109 |
| ENSMUSG00000027091 | Zc3h15 | protein_coding | -0.050673697 | 6.253885872 | 0.0406264 | 0.8427536 | 0.970271109 |
| ENSMUSG00000027878 | Notch2 | protein_coding | -0.106780264 | 3.242617903 | 0.040623 | 0.8427601 | 0.970271109 |
| ENSMUSG00000027936 | Crtc2 | protein_coding | -0.111103673 | 3.167573966 | 0.0406219 | 0.8427621 | 0.970271109 |
| ENSMUSG00000033623 | Pcgf3 | protein_coding | 0.075190713 | 4.33184919 | 0.0406165 | 0.8427725 | 0.970271109 |
| ENSMUSG00000039100 | 6-Mar | protein_coding | -0.048459798 | 6.267039497 | 0.0404772 | 0.8430385 | 0.970349113 |
| ENSMUSG00000046756 | Mrps7 | protein_coding | -0.040411241 | 6.693885339 | 0.0403006 | 0.8433764 | 0.970349113 |
| ENSMUSG00000041891 | Lman1 | protein_coding | 0.088015053 | 3.955768525 | 0.0402966 | 0.8433841 | 0.970349113 |
| ENSMUSG00000037364 | Srrt | protein_coding | -0.050791481 | 5.570223714 | 0.0402846 | 0.843407 | 0.970349113 |
| ENSMUSG00000022280 | Rnf19a | protein_coding | -0.070044758 | 4.079640863 | 0.0402798 | 0.8434162 | 0.970349113 |
| ENSMUSG00000021373 | Cap2 | protein_coding | 0.042836593 | 7.068830187 | 0.0402603 | 0.8434536 | 0.970349113 |
| ENSMUSG00000023262 | Acy1 | protein_coding | -0.06882178 | 4.934477265 | 0.0402393 | 0.8434938 | 0.970349113 |
| ENSMUSG00000022427 | Tomm22 | protein_coding | 0.035705308 | 7.662045527 | 0.040155 | 0.8436556 | 0.970349113 |
| ENSMUSG00000024908 | Ppp6r3 | protein_coding | -0.068395868 | 4.469791417 | 0.0401462 | 0.8436725 | 0.970349113 |
| ENSMUSG00000043419 | Chd3os | protein_coding | -0.035339573 | 8.809065314 | 0.0400623 | 0.8438336 | 0.970349113 |
| ENSMUSG00000057134 | Ado | protein_coding | 0.052604123 | 8.137691572 | 0.0400281 | 0.8439034 | 0.970349113 |
| ENSMUSG00000020894 | Vamp2 | protein_coding | -0.038922053 | 9.219983842 | 0.0399701 | 0.8440109 | 0.970349113 |
| ENSMUSG00000053063 | Clec12a | protein_coding | 0.189451188 | 2.502526598 | 0.0399286 | 0.8441055 | 0.970349113 |
| ENSMUSG00000020075 | Ddx21 | protein_coding | -0.071513975 | 3.940671259 | 0.0399016 | 0.8441428 | 0.970349113 |
| ENSMUSG00000054942 | Miga1 | protein_coding | -0.051214114 | 5.453316848 | 0.0398602 | 0.8442224 | 0.970349113 |
| ENSMUSG00000022671 | Mzt2 | protein_coding | -0.040849321 | 6.378649776 | 0.0398282 | 0.844284 | 0.970349113 |
| ENSMUSG00000024867 | Pip5k1b | protein_coding | -0.055053887 | 5.233252831 | 0.039624 | 0.8446783 | 0.970580578 |
| ENSMUSG00000058407 | Txndc9 | protein_coding | -0.037821766 | 7.320790892 | 0.0396059 | 0.8447133 | 0.970580578 |
| ENSMUSG00000038291 | Snx25 | protein_coding | 0.071430705 | 4.481618643 | 0.0395432 | 0.8448345 | 0.970580578 |
| ENSMUSG00000005681 | Apoa2 | protein_coding | 0.084528148 | 3.382292466 | 0.0395213 | 0.8448769 | 0.970580578 |
| ENSMUSG00000072946 | Ptgr2 | protein_coding | -0.068591101 | 5.058398107 | 0.0394743 | 0.8449678 | 0.970580578 |
| ENSMUSG00000025962 | Fastkd2 | protein_coding | -0.060333538 | 5.107570736 | 0.0394008 | 0.8451103 | 0.970580578 |
| ENSMUSG00000096935 | 1700113A16Rik | antisense_RNA | -0.148955953 | 2.436714362 | 0.0393972 | 0.8451173 | 0.970580578 |
| ENSMUSG00000038525 | Armc10 | protein_coding | -0.097250627 | 3.275490753 | 0.0392904 | 0.8453245 | 0.970625165 |
| ENSMUSG00000035215 | Lsm7 | protein_coding | -0.075950844 | 3.812510282 | 0.0392523 | 0.8453983 | 0.970625165 |
| ENSMUSG00000031227 | Magee1 | protein_coding | -0.035918855 | 6.897244192 | 0.0392376 | 0.8454269 | 0.970625165 |
| ENSMUSG00000031796 | Cfap20 | protein_coding | 0.058733862 | 6.061019757 | 0.0391793 | 0.8455442 | 0.970626789 |
| ENSMUSG00000026696 | Vamp4 | protein_coding | 0.044428146 | 7.902477043 | 0.0390703 | 0.8457523 | 0.970626789 |
| ENSMUSG00000032239 | Rp9 | protein_coding | 0.058574723 | 4.929225882 | 0.0390566 | 0.845779 | 0.970626789 |
| ENSMUSG00000038916 | Soga3 | protein_coding | 0.060764124 | 5.687258894 | 0.0390513 | 0.8457894 | 0.970626789 |
| ENSMUSG00000061950 | Ppp4r1 | protein_coding | 0.10212773 | 3.309485181 | 0.0389696 | 0.8459484 | 0.970698915 |
| ENSMUSG00000033632 | AW554918 | protein_coding | 0.067945012 | 3.894266693 | 0.0388732 | 0.8461365 | 0.970698915 |
| ENSMUSG00000028610 | Dmrtb1 | protein_coding | 0.149562396 | 3.322500982 | 0.0389399 | 0.8462498 | 0.970698915 |
| ENSMUSG00000026727 | Rsu1 | protein_coding | 0.057059828 | 5.540845219 | 0.0388018 | 0.846276 | 0.970698915 |
| ENSMUSG00000001173 | Ocrl | protein_coding | 0.075376922 | 5.090427923 | 0.0387897 | 0.8463036 | 0.970698915 |
| ENSMUSG00000037492 | Zmat4 | protein_coding | 0.066992088 | 5.848493678 | 0.0387251 | 0.8464441 | 0.970756544 |
| ENSMUSG00000074576 | Mocs3 | protein_coding | -0.157004733 | 2.224110288 | 0.0386493 | 0.8465743 | 0.970794856 |
| ENSMUSG00000017428 | Psmd11 | protein_coding | -0.037304946 | 7.521194612 | 0.0386065 | 0.8466581 | 0.970794856 |
| ENSMUSG00000038467 | Chmp4b | protein_coding | -0.057900385 | 5.568378355 | 0.0385279 | 0.8468121 | 0.970867905 |
| ENSMUSG00000022849 | Hspbap1 | protein_coding | 0.115178446 | 3.208849033 | 0.0384659 | 0.8469337 | 0.970903854 |
| ENSMUSG00000006599 | Gtf2h1 | protein_coding | 0.067009946 | 4.52194204 | 0.0383349 | 0.8471911 | 0.971095443 |
| ENSMUSG00000030691 | Fchsd2 | protein_coding | -0.106348725 | 3.146760616 | 0.0382361 | 0.8473856 | 0.971155606 |
| ENSMUSG00000028234 | Rps20 | protein_coding | 0.049388268 | 7.008257084 | 0.0382184 | 0.8474243 | 0.971155606 |
| ENSMUSG00000012126 | Ubxn11 | protein_coding | -0.042550912 | 5.614019991 | 0.0380558 | 0.8477409 | 0.971287763 |
| ENSMUSG00000032601 | Prkar2a | protein_coding | -0.041884773 | 5.871923199 | 0.0380428 | 0.8477666 | 0.971287763 |
| ENSMUSG00000071659 | Hnrnpul2 | protein_coding | -0.053778419 | 5.875006162 | 0.0379977 | 0.8478556 | 0.971287763 |
| ENSMUSG00000103034 | Gm8797 | protein_coding | -0.092561031 | 2.947511634 | 0.0379748 | 0.8479009 | 0.971287763 |
| ENSMUSG00000073415 | Gm10501 | antisense_RNA | 0.112469137 | 2.689804278 | 0.0378846 | 0.8480793 | 0.971388634 |
| ENSMUSG00000033940 | Brk1 | protein_coding | 0.034581804 | 9.750060747 | 0.0377872 | 0.8482721 | 0.971505987 |
| ENSMUSG00000043668 | Tox3 | protein_coding | 0.088357772 | 3.709354237 | 0.0376914 | 0.8484619 | 0.971537838 |
| ENSMUSG00000034709 | Ppp1r21 | protein_coding | 0.048916687 | 5.805066777 | 0.037682 | 0.8484806 | 0.971537838 |
| ENSMUSG00000006395 | Hyi | polymorphic_pseudog | 0.100473646 | 2.861671572 | 0.0375344 | 0.8487737 | 0.971769965 |
| ENSMUSG00000001473 | Tubb6 | protein_coding | 0.119617848 | 2.699897975 | 0.0373897 | 0.8490617 | 0.971996259 |
| ENSMUSG00000023791 | Pigx | protein_coding | 0.088133743 | 4.053541785 | 0.0372635 | 0.8493172 | 0.972185236 |
| ENSMUSG00000038390 | Gpr162 | protein_coding | -0.069710794 | 4.424986241 | 0.0371411 | 0.8495576 | 0.972350939 |
| ENSMUSG00000032020 | Ubash3b | protein_coding | -0.073706342 | 6.093082325 | 0.0372175 | 0.8496428 | 0.972350939 |
| ENSMUSG00000037353 | Letmd1 | protein_coding | 0.054572941 | 5.638732501 | 0.0369533 | 0.8499335 | 0.97242798 |
| ENSMUSG00000020190 | Mknk2 | protein_coding | 0.080818724 | 3.544430512 | 0.0369486 | 0.8499429 | 0.97242798 |

| ensembl_gene_id | external_gene_name | gene_biotype | log2 (fold change) | log2 (counts per million) | F-value | p-value | False discovery rate |
|---|---|---|---|---|---|---|---|
| ENSMUSG00000054836 | Elp6 | protein_coding | -0.046809146 | 5.750769458 | 0.0368765 | 0.8500875 | 0.97242798 |
| ENSMUSG00000043091 | Tuba1c | protein_coding | 0.268274119 | 1.936353087 | 0.0368975 | 0.8501966 | 0.97242798 |
| ENSMUSG00000022802 | Lmln | protein_coding | -0.138903029 | 2.455071012 | 0.0368039 | 0.8502334 | 0.97242798 |
| ENSMUSG00000057110 | Cntrl | protein_coding | -0.155772415 | 2.464555657 | 0.0367942 | 0.8502528 | 0.97242798 |
| ENSMUSG00000021550 | 2210016F16Rik | protein_coding | 0.083268696 | 3.754677248 | 0.0367242 | 0.8503934 | 0.972485402 |
| ENSMUSG00000025134 | Alyref | protein_coding | 0.064919915 | 4.548844003 | 0.0365752 | 0.8506934 | 0.972716069 |
| ENSMUSG00000067194 | Eif1ax | protein_coding | -0.036932799 | 7.481456596 | 0.0364854 | 0.8508743 | 0.972716069 |
| ENSMUSG00000022747 | St3gal6 | protein_coding | -0.082491995 | 3.685599646 | 0.0364519 | 0.8509419 | 0.972716069 |
| ENSMUSG00000019461 | Plscr3 | protein_coding | 0.063608444 | 4.880976774 | 0.0364239 | 0.8509986 | 0.972716069 |
| ENSMUSG00000069631 | Strada | protein_coding | 0.070044606 | 4.689361972 | 0.0363997 | 0.8510474 | 0.972716069 |
| ENSMUSG00000002550 | Uck1 | protein_coding | 0.086713957 | 3.433031781 | 0.036301 | 0.8512469 | 0.972762174 |
| ENSMUSG00000038648 | Creb3l2 | protein_coding | -0.163612937 | 2.243356346 | 0.0362902 | 0.8512687 | 0.972762174 |
| ENSMUSG00000063382 | Bcl9l | protein_coding | -0.082148639 | 4.306735591 | 0.0362238 | 0.851407 | 0.972816827 |
| ENSMUSG00000028412 | Slc44a1 | protein_coding | -0.054290002 | 5.617019208 | 0.0358934 | 0.8520739 | 0.973047598 |
| ENSMUSG00000032777 | Gtf3c1 | protein_coding | 0.062519338 | 5.043845491 | 0.035871 | 0.8521195 | 0.973047598 |
| ENSMUSG00000031583 | Wrn | protein_coding | 0.102749042 | 3.024138021 | 0.035777 | 0.8523109 | 0.973047598 |
| ENSMUSG00000020758 | Itgb4 | protein_coding | -0.081162353 | 3.371077176 | 0.0357329 | 0.8524006 | 0.973047598 |
| ENSMUSG00000030313 | Dennd5b | protein_coding | 0.048175248 | 4.922465207 | 0.0357302 | 0.8524062 | 0.973047598 |
| ENSMUSG00000050786 | Ccdc126 | protein_coding | -0.076754763 | 3.822217161 | 0.0357047 | 0.8524583 | 0.973047598 |
| ENSMUSG00000059981 | Taok2 | protein_coding | -0.067878986 | 4.662937901 | 0.0357 | 0.8524679 | 0.973047598 |
| ENSMUSG00000014959 | Gorasp2 | protein_coding | 0.037017425 | 6.525024979 | 0.0356986 | 0.8524708 | 0.973047598 |
| ENSMUSG00000027273 | Snap25 | protein_coding | -0.049717586 | 12.17103416 | 0.0356743 | 0.8525202 | 0.973047598 |
| ENSMUSG00000003500 | Impdh1 | protein_coding | -0.083256755 | 4.04516045 | 0.0356671 | 0.852535 | 0.973047598 |
| ENSMUSG00000050022 | Amz1 | protein_coding | 0.06065235 | 4.858056728 | 0.0356164 | 0.8526385 | 0.973047598 |
| ENSMUSG00000030835 | Nomo1 | protein_coding | 0.042232422 | 6.283509334 | 0.0355011 | 0.8528743 | 0.973047598 |
| ENSMUSG00000030878 | Cdr2 | protein_coding | -0.080969219 | 3.563215937 | 0.0354687 | 0.8529407 | 0.973047598 |
| ENSMUSG00000024472 | Dcp2 | protein_coding | 0.063147783 | 4.493941162 | 0.0354346 | 0.8530103 | 0.973047598 |
| ENSMUSG00000044155 | Lsm8 | protein_coding | -0.051815503 | 6.069492638 | 0.0354292 | 0.8530216 | 0.973047598 |
| ENSMUSG00000022619 | Mapk8ip2 | protein_coding | 0.047546755 | 6.320424847 | 0.035412 | 0.8530568 | 0.973047598 |
| ENSMUSG00000044700 | Tmem201 | protein_coding | 0.063136244 | 4.577270721 | 0.0352509 | 0.8533873 | 0.973321319 |
| ENSMUSG00000008136 | Fhl2 | protein_coding | 0.125944684 | 3.259917399 | 0.0351532 | 0.8536257 | 0.973490029 |
| ENSMUSG00000045302 | Preb | protein_coding | -0.046815616 | 5.968168153 | 0.0349763 | 0.8539523 | 0.973749988 |
| ENSMUSG00000031820 | Babam1 | protein_coding | 0.040212569 | 7.129763984 | 0.0349363 | 0.8540348 | 0.973749988 |
| ENSMUSG00000021262 | Evl | protein_coding | 0.085460743 | 3.886538547 | 0.0348443 | 0.8542247 | 0.973810637 |
| ENSMUSG00000027286 | Lrrc57 | protein_coding | 0.048071707 | 5.643362814 | 0.0347895 | 0.8543381 | 0.973810637 |
| ENSMUSG00000031090 | Nadsyn1 | protein_coding | -0.124530462 | 2.574717907 | 0.034779 | 0.8543597 | 0.973810637 |
| ENSMUSG00000036990 | Otud4 | protein_coding | 0.073246164 | 4.057387711 | 0.0346076 | 0.8547148 | 0.974007564 |
| ENSMUSG00000022884 | Eif4a2 | protein_coding | -0.040754039 | 10.53247066 | 0.034605 | 0.8547202 | 0.974007564 |
| ENSMUSG00000049100 | Pcdh10 | protein_coding | -0.073209484 | 5.010605996 | 0.0345368 | 0.8548653 | 0.974007564 |
| ENSMUSG00000007570 | Fance | protein_coding | 0.127554203 | 2.876004176 | 0.0345208 | 0.8548948 | 0.974007564 |
| ENSMUSG00000035671 | Zswim4 | protein_coding | 0.065154578 | 3.825529101 | 0.0344654 | 0.8550099 | 0.974035572 |
| ENSMUSG00000028693 | Nasp | protein_coding | -0.064529089 | 4.348576499 | 0.0343529 | 0.8552438 | 0.974198782 |
| ENSMUSG00000038145 | Snrk | protein_coding | 0.064316556 | 4.663585786 | 0.0342119 | 0.8555376 | 0.974336522 |
| ENSMUSG00000032735 | Ablim3 | protein_coding | -0.0593125 | 4.937772889 | 0.0342079 | 0.8555459 | 0.974336522 |
| ENSMUSG00000047881 | Rell1 | protein_coding | -0.14571466 | 2.650813724 | 0.0339456 | 0.8560941 | 0.974488225 |
| ENSMUSG00000031935 | Med17 | protein_coding | 0.084389996 | 3.988971524 | 0.0339362 | 0.8561139 | 0.974488225 |
| ENSMUSG00000032350 | Gclc | protein_coding | -0.050734202 | 5.5127045 | 0.0339125 | 0.8561636 | 0.974488225 |
| ENSMUSG00000039952 | Dag1 | protein_coding | -0.055218944 | 4.91240035 | 0.0338144 | 0.8563693 | 0.974488225 |
| ENSMUSG00000019763 | Rmnd1 | protein_coding | 0.061309077 | 3.99486324 | 0.0337596 | 0.8564843 | 0.974488225 |
| ENSMUSG00000036225 | Kctd1 | protein_coding | 0.067455812 | 5.301943962 | 0.033759 | 0.8564892 | 0.974488225 |
| ENSMUSG00000019961 | Tmpo | protein_coding | 0.053712715 | 4.458552505 | 0.0337552 | 0.8564936 | 0.974488225 |
| ENSMUSG00000004356 | Utp20 | protein_coding | -0.139087901 | 2.419253035 | 0.0337219 | 0.8565635 | 0.974488225 |
| ENSMUSG00000022197 | Pdzd2 | protein_coding | 0.081951567 | 3.694721318 | 0.0335599 | 0.8569043 | 0.974488225 |
| ENSMUSG00000002660 | Clpp | protein_coding | -0.032986519 | 8.997684785 | 0.0335592 | 0.8569059 | 0.974488225 |
| ENSMUSG00000027997 | Casp6 | protein_coding | -0.106595305 | 2.845442278 | 0.0335484 | 0.8569286 | 0.974488225 |
| ENSMUSG00000030256 | Bhlhe41 | protein_coding | -0.105428638 | 7.981567153 | 0.0341425 | 0.8570212 | 0.974488225 |
| ENSMUSG00000025510 | Cd151 | protein_coding | -0.091680865 | 3.943163194 | 0.0335025 | 0.8570291 | 0.974488225 |
| ENSMUSG00000041936 | Agrn | protein_coding | -0.066361683 | 4.319190457 | 0.0334785 | 0.857076 | 0.974488225 |
| ENSMUSG00000032291 | Crabp1 | protein_coding | -0.071857416 | 4.617059342 | 0.0334588 | 0.8571177 | 0.974488225 |
| ENSMUSG00000039546 | Ajap1 | protein_coding | -0.04307041 | 5.796713387 | 0.0334533 | 0.8571291 | 0.974488225 |
| ENSMUSG00000027341 | Tmem230 | protein_coding | 0.058834448 | 4.33229292 | 0.033263 | 0.8575313 | 0.974842384 |
| ENSMUSG00000035064 | Eef2k | protein_coding | 0.066639905 | 3.847565914 | 0.0331479 | 0.8577752 | 0.975016565 |
| ENSMUSG00000034189 | Hsdl1 | protein_coding | 0.043193287 | 6.774531571 | 0.0330079 | 0.8580723 | 0.975108269 |
| ENSMUSG00000040659 | Efhd2 | protein_coding | 0.040757902 | 6.788462381 | 0.0329476 | 0.8582005 | 0.975108269 |
| ENSMUSG00000006215 | Zbtb17 | protein_coding | -0.085304656 | 4.625695513 | 0.0329736 | 0.8582438 | 0.975108269 |
| ENSMUSG00000043913 | Ccdc60 | protein_coding | -0.139956029 | 2.315341886 | 0.0328383 | 0.8584332 | 0.975108269 |
| ENSMUSG00000020732 | Rab37 | protein_coding | -0.152521783 | 4.52075295 | 0.0333356 | 0.8584723 | 0.975108269 |
| ENSMUSG00000037606 | Osbpl5 | protein_coding | -0.063381129 | 4.151074454 | 0.032744 | 0.8586343 | 0.975108269 |
| ENSMUSG00000030663 | 1110004F10Rik | protein_coding | 0.033068019 | 8.6347426 | 0.0327238 | 0.8586774 | 0.975108269 |
| ENSMUSG00000063808 | Gpatch1 | protein_coding | -0.048712642 | 5.479376929 | 0.0326604 | 0.8588129 | 0.975108269 |
| ENSMUSG00000045287 | Rtn4rl1 | protein_coding | -0.064603694 | 3.935299572 | 0.032646 | 0.8588435 | 0.975108269 |
| ENSMUSG00000026575 | Nme7 | protein_coding | -0.062653883 | 4.734404409 | 0.0325657 | 0.8590152 | 0.975108269 |
| ENSMUSG00000021375 | Kif13a | protein_coding | 0.07595927 | 3.691015578 | 0.0325236 | 0.8591053 | 0.975108269 |
| ENSMUSG00000061983 | Rps12 | protein_coding | -0.084730015 | 3.176514267 | 0.0325029 | 0.8591497 | 0.975108269 |

| ensembl_gene_id | external_gene_name | gene_biotype | log2 (fold change) | log2 (counts per million) | F-value | p-value | False discovery rate |
|---|---|---|---|---|---|---|---|
| ENSMUSG00000059183 | Mtfmt | protein_coding | -0.064899367 | 4.232497729 | 0.0325004 | 0.859155 | 0.975108269 |
| ENSMUSG00000008167 | Fbxw9 | protein_coding | -0.073641527 | 4.700894323 | 0.0324705 | 0.8592228 | 0.975108269 |
| ENSMUSG00000029763 | Exoc4 | protein_coding | 0.045740979 | 5.708463691 | 0.0324304 | 0.8593051 | 0.975108269 |
| ENSMUSG00000054509 | Parp4 | protein_coding | -0.123273042 | 2.787169199 | 0.0323894 | 0.8593932 | 0.975108269 |
| ENSMUSG00000001864 | Aif1l | protein_coding | -0.052636819 | 5.280336379 | 0.0323595 | 0.8594573 | 0.975108269 |
| ENSMUSG00000028614 | Ndc1 | protein_coding | 0.114952354 | 3.068110825 | 0.0323385 | 0.8595023 | 0.975108269 |
| ENSMUSG00000042246 | Tmc7 | protein_coding | -0.110074515 | 3.010522308 | 0.032297 | 0.8595915 | 0.975108269 |
| ENSMUSG00000038822 | Hace1 | protein_coding | 0.064732864 | 4.195022349 | 0.0322391 | 0.8597159 | 0.975108269 |
| ENSMUSG00000022707 | Gbe1 | protein_coding | 0.081393199 | 3.4421343 | 0.0322185 | 0.8597602 | 0.975108269 |
| ENSMUSG00000029311 | Hsd17b11 | protein_coding | -0.098520946 | 3.207451278 | 0.0320687 | 0.8600829 | 0.97518081 |
| ENSMUSG00000021493 | Pdlim7 | protein_coding | -0.034056101 | 6.876904623 | 0.0318803 | 0.8604899 | 0.97518081 |
| ENSMUSG00000030057 | Cnbp | protein_coding | 0.033009518 | 10.42446665 | 0.031754 | 0.8607635 | 0.97518081 |
| ENSMUSG00000005360 | Slc1a3 | protein_coding | -0.042226791 | 7.808350894 | 0.0317333 | 0.8608083 | 0.97518081 |
| ENSMUSG00000006276 | Eps15l1 | protein_coding | 0.047495754 | 5.558917699 | 0.0317259 | 0.8608245 | 0.97518081 |
| ENSMUSG00000031697 | Orc6 | protein_coding | -0.047990133 | 6.86565501 | 0.0316927 | 0.8609 | 0.97518081 |
| ENSMUSG00000040459 | Arglu1 | protein_coding | 0.042183226 | 7.490944732 | 0.031681 | 0.8609219 | 0.97518081 |
| ENSMUSG00000018677 | Slc25a39 | protein_coding | 0.035176784 | 6.66780126 | 0.0315586 | 0.8611878 | 0.97518081 |
| ENSMUSG00000037519 | Ppfia1 | protein_coding | -0.071274962 | 3.816122203 | 0.0314778 | 0.8613637 | 0.97518081 |
| ENSMUSG00000021945 | Zmym2 | protein_coding | -0.048289491 | 5.437061708 | 0.0314729 | 0.8613742 | 0.97518081 |
| ENSMUSG00000002068 | Ccne1 | protein_coding | -0.078905961 | 3.513192971 | 0.031376 | 0.8615854 | 0.97518081 |
| ENSMUSG00000028398 | Tmem261 | protein_coding | 0.03757573 | 6.682360992 | 0.0313734 | 0.8615913 | 0.97518081 |
| ENSMUSG00000009090 | Ap1b1 | protein_coding | 0.04434053 | 7.029784393 | 0.0313533 | 0.8616387 | 0.97518081 |
| ENSMUSG00000030816 | Rnf40 | protein_coding | -0.052299849 | 4.378118553 | 0.0313432 | 0.861657 | 0.97518081 |
| ENSMUSG00000029312 | Klhl8 | protein_coding | -0.056085183 | 4.255544535 | 0.0313259 | 0.8616949 | 0.97518081 |
| ENSMUSG00000020657 | Dnajc27 | protein_coding | 0.03384923 | 6.523434476 | 0.0313233 | 0.8617004 | 0.97518081 |
| ENSMUSG00000059878 | Zfp422 | protein_coding | -0.051446431 | 5.255728432 | 0.0312846 | 0.8617849 | 0.97518081 |
| ENSMUSG00000019952 | Poc1b | protein_coding | 0.126201089 | 2.66438283 | 0.0312494 | 0.8618619 | 0.97518081 |
| ENSMUSG00000020564 | Atxn7l1 | protein_coding | 0.045171577 | 6.226863511 | 0.0312141 | 0.8619391 | 0.97518081 |
| ENSMUSG00000039967 | Zfp292 | protein_coding | -0.06389661 | 4.74236974 | 0.0312135 | 0.8619405 | 0.97518081 |
| ENSMUSG00000058400 | Qrfpr | protein_coding | -0.09986494 | 2.91734529 | 0.0311838 | 0.8620054 | 0.97518081 |
| ENSMUSG00000021559 | Dapk1 | protein_coding | -0.046602331 | 5.818781129 | 0.0311356 | 0.862111 | 0.97518081 |
| ENSMUSG00000016495 | Plgrkt | protein_coding | -0.050929213 | 5.452036591 | 0.0311235 | 0.8621374 | 0.97518081 |
| ENSMUSG00000023999 | Kif6 | protein_coding | -0.134612127 | 2.428414314 | 0.0311173 | 0.862151 | 0.97518081 |
| ENSMUSG00000026305 | Lrrfip1 | protein_coding | 0.059428485 | 4.908103017 | 0.0311053 | 0.8621773 | 0.97518081 |
| ENSMUSG00000006800 | Sulf2 | protein_coding | 0.054034366 | 5.070675358 | 0.0310622 | 0.8622718 | 0.97518081 |
| ENSMUSG00000017176 | Nt5c3b | protein_coding | 0.034852078 | 6.755125826 | 0.0310618 | 0.8622728 | 0.97518081 |
| ENSMUSG00000030284 | Creld1 | protein_coding | 0.042942957 | 6.220781924 | 0.0309366 | 0.8625475 | 0.975363232 |
| ENSMUSG00000023088 | Abcc1 | protein_coding | 0.178485277 | 2.924059921 | 0.0310442 | 0.8626155 | 0.975363232 |
| ENSMUSG00000036112 | Metap2 | protein_coding | 0.037530397 | 7.489610262 | 0.0307772 | 0.8628983 | 0.975467137 |
| ENSMUSG00000028820 | Sfpq | protein_coding | 0.042113213 | 5.63432158 | 0.0307765 | 0.8628997 | 0.975467137 |
| ENSMUSG00000029182 | 1700001C02Rik | protein_coding | -0.080427047 | 3.55603683 | 0.0307317 | 0.8629985 | 0.975467137 |
| ENSMUSG00000047260 | Emc6 | protein_coding | -0.06249511 | 4.534435682 | 0.0306992 | 0.8630702 | 0.975467137 |
| ENSMUSG00000033379 | Atp6v0b | protein_coding | 0.037498501 | 8.426530837 | 0.0305837 | 0.8633253 | 0.975652804 |
| ENSMUSG00000071654 | Uqcc3 | protein_coding | -0.041307081 | 6.871597521 | 0.0304426 | 0.8636375 | 0.975837001 |
| ENSMUSG00000043421 | Hilpda | protein_coding | -0.052109882 | 4.813175792 | 0.0304131 | 0.8637028 | 0.975837001 |
| ENSMUSG00000032555 | Topbp1 | protein_coding | 0.144927574 | 2.651248365 | 0.0303815 | 0.8637765 | 0.975837001 |
| ENSMUSG00000028838 | Extl1 | protein_coding | 0.053568427 | 4.424538236 | 0.0303351 | 0.8638759 | 0.975837001 |
| ENSMUSG00000035805 | Mlc1 | protein_coding | 0.07179399 | 6.046922441 | 0.0304243 | 0.8639791 | 0.975837001 |
| ENSMUSG00000040282 | BC052040 | protein_coding | -0.058715248 | 4.305137647 | 0.0302636 | 0.8640347 | 0.975837001 |
| ENSMUSG00000095845 | Gm5741 | protein_coding | -0.496336755 | 2.704893232 | 0.0311555 | 0.8643435 | 0.975837001 |
| ENSMUSG00000032932 | Hspa13 | protein_coding | 0.046419461 | 5.451743719 | 0.0301006 | 0.8643976 | 0.975837001 |
| ENSMUSG00000086382 | Chrna1os | antisense_RNA | -0.100256361 | 2.764194342 | 0.0300535 | 0.8645025 | 0.975837001 |
| ENSMUSG00000022438 | Parvb | protein_coding | -0.065132896 | 3.865977338 | 0.0300529 | 0.8645039 | 0.975837001 |
| ENSMUSG00000063802 | Hspbp1 | protein_coding | -0.038261381 | 7.8338441 | 0.0300243 | 0.8645677 | 0.975837001 |
| ENSMUSG00000054484 | Tmem62 | protein_coding | 0.064395939 | 4.323852073 | 0.0300117 | 0.8645958 | 0.975837001 |
| ENSMUSG00000021010 | Npas3 | protein_coding | 0.125987587 | 2.326806981 | 0.0299531 | 0.8647267 | 0.975837001 |
| ENSMUSG00000052031 | Tagap1 | protein_coding | 0.053111298 | 5.618157086 | 0.0299182 | 0.8648045 | 0.975837001 |
| ENSMUSG00000031376 | Atp2b3 | protein_coding | -0.073744004 | 3.234894051 | 0.0298941 | 0.8648585 | 0.975837001 |
| ENSMUSG00000032261 | Sh3bgrl2 | protein_coding | 0.068061083 | 5.201580126 | 0.0298632 | 0.8649402 | 0.975837001 |
| ENSMUSG00000026867 | Gapvd1 | protein_coding | -0.043263995 | 5.793790567 | 0.0297697 | 0.8651371 | 0.975877381 |
| ENSMUSG00000018707 | Dync1h1 | protein_coding | -0.053294152 | 5.922824251 | 0.0297426 | 0.8652013 | 0.975877381 |
| ENSMUSG00000031751 | Amfr | protein_coding | -0.048573302 | 6.103461629 | 0.0296361 | 0.8654369 | 0.975877381 |
| ENSMUSG00000021494 | Ddx41 | protein_coding | -0.046147827 | 5.333514977 | 0.0295931 | 0.8655334 | 0.975877381 |
| ENSMUSG00000037541 | Shank2 | protein_coding | 0.060093289 | 4.28895981 | 0.0295475 | 0.8656361 | 0.975877381 |
| ENSMUSG00000033306 | Lpp | protein_coding | 0.071485503 | 3.491464405 | 0.0295381 | 0.8656571 | 0.975877381 |
| ENSMUSG00000020929 | Eftud2 | protein_coding | 0.050474802 | 5.697167108 | 0.0295247 | 0.8656873 | 0.975877381 |
| ENSMUSG00000019970 | Sgk1 | protein_coding | 0.054206171 | 7.726937866 | 0.0295998 | 0.8657021 | 0.975877381 |
| ENSMUSG00000053925 | Gm9929 | processed_transcript | -0.095806111 | 2.965741851 | 0.0294767 | 0.8657955 | 0.975880433 |
| ENSMUSG00000054648 | Zfp869 | protein_coding | -0.077991086 | 3.365947779 | 0.0293991 | 0.8659705 | 0.975975326 |
| ENSMUSG00000100594 | 2810414N06Rik | lincRNA | -0.083938113 | 3.632830618 | 0.0293205 | 0.8661479 | 0.976072977 |
| ENSMUSG00000024376 | Epb41l4a | protein_coding | -0.080254053 | 3.386500114 | 0.0292442 | 0.8663202 | 0.976080511 |
| ENSMUSG00000022075 | Rhobtb2 | protein_coding | -0.047591506 | 5.279647874 | 0.0292052 | 0.8664086 | 0.976080511 |
| ENSMUSG00000036452 | Arhgap26 | protein_coding | -0.047219562 | 5.249432187 | 0.0291835 | 0.8664576 | 0.976080511 |
| ENSMUSG00000020925 | Ccdc43 | protein_coding | 0.041498137 | 5.857816366 | 0.029157 | 0.8665177 | 0.976080511 |

| ensembl_gene_id | external_gene_name | gene_biotype | log2 (fold change) | log2 (counts per million) | F-value | p-value | False discovery rate |
|---|---|---|---|---|---|---|---|
| ENSMUSG00000005986 | Ankrd13d | protein_coding | 0.056165907 | 5.30003387 | 0.0290622 | 0.8667326 | 0.97615987 |
| ENSMUSG00000036968 | Cnpy4 | protein_coding | 0.052237199 | 5.693023427 | 0.0290159 | 0.8668379 | 0.97615987 |
| ENSMUSG00000041444 | Arhgap32 | protein_coding | -0.045573244 | 4.925505771 | 0.0290059 | 0.8668605 | 0.97615987 |
| ENSMUSG00000022386 | Trmu | protein_coding | 0.055287533 | 5.064305744 | 0.0289412 | 0.8670075 | 0.976190998 |
| ENSMUSG00000024258 | Polr2d | protein_coding | -0.031810246 | 7.113924232 | 0.028906 | 0.8670877 | 0.976190998 |
| ENSMUSG00000078941 | Ak6 | protein_coding | 0.046252265 | 5.52430676 | 0.0288722 | 0.8671645 | 0.976190998 |
| ENSMUSG00000028152 | Tspan5 | protein_coding | 0.035846444 | 6.044116231 | 0.0288341 | 0.8672512 | 0.976190998 |
| ENSMUSG00000027327 | 1700037H04Rik | protein_coding | 0.033373637 | 9.438842398 | 0.0285188 | 0.8679718 | 0.976765464 |
| ENSMUSG00000025793 | Hgs | protein_coding | 0.042318779 | 5.683935347 | 0.0285046 | 0.8680044 | 0.976765464 |
| ENSMUSG00000062760 | 1810041L15Rik | protein_coding | 0.057274369 | 5.549646128 | 0.0284061 | 0.8682338 | 0.976765464 |
| ENSMUSG00000055850 | Rnf181 | protein_coding | 0.030093425 | 7.940958936 | 0.0283759 | 0.8682998 | 0.976765464 |
| ENSMUSG00000027985 | Lef1 | protein_coding | 0.171390613 | 3.205573211 | 0.0285665 | 0.8684001 | 0.976765464 |
| ENSMUSG00000063450 | Syne2 | protein_coding | 0.101043439 | 3.309267407 | 0.0283174 | 0.8684377 | 0.976765464 |
| ENSMUSG00000003873 | Bax | protein_coding | 0.035084054 | 6.257187138 | 0.0282898 | 0.8684976 | 0.976765464 |
| ENSMUSG00000022475 | Hdac7 | protein_coding | -0.07174327 | 4.279906756 | 0.0282687 | 0.8685461 | 0.976765464 |
| ENSMUSG00000036972 | Zic4 | protein_coding | 0.062118249 | 5.174582636 | 0.0281905 | 0.8687297 | 0.976765464 |
| ENSMUSG00000031574 | Star | protein_coding | -0.093419802 | 2.702912723 | 0.0281163 | 0.8688975 | 0.976765464 |
| ENSMUSG00000028744 | Pqlc2 | protein_coding | -0.061744456 | 4.249229138 | 0.0281112 | 0.8689093 | 0.976765464 |
| ENSMUSG00000029064 | Gnb1 | protein_coding | -0.053803542 | 12.19785351 | 0.0281131 | 0.8689536 | 0.976765464 |
| ENSMUSG00000039016 | Timm8b | protein_coding | 0.04502608 | 9.334659322 | 0.0280499 | 0.8690542 | 0.976765464 |
| ENSMUSG00000020411 | Nipal4 | protein_coding | 0.104492965 | 2.931015967 | 0.0280275 | 0.8691027 | 0.976765464 |
| ENSMUSG00000005054 | Cstb | protein_coding | 0.035655022 | 8.031710396 | 0.0280116 | 0.8691395 | 0.976765464 |
| ENSMUSG00000024393 | Prrc2a | protein_coding | -0.052244488 | 4.643013439 | 0.0279116 | 0.8693709 | 0.976765464 |
| ENSMUSG00000041757 | Plekha6 | protein_coding | 0.053946424 | 4.590886825 | 0.0279114 | 0.8693713 | 0.976765464 |
| ENSMUSG00000046417 | Lrrc75a | protein_coding | -0.124926626 | 2.392586218 | 0.0279005 | 0.8693967 | 0.976765464 |
| ENSMUSG00000032012 | Nectin1 | protein_coding | 0.071647679 | 4.533618981 | 0.0277735 | 0.8696946 | 0.976998138 |
| ENSMUSG00000019577 | Pdk4 | protein_coding | -0.118659404 | 2.415017495 | 0.0277121 | 0.8698341 | 0.97705272 |
| ENSMUSG00000087687 | Pet100 | protein_coding | -0.044945416 | 6.789267763 | 0.0276373 | 0.8700114 | 0.977149329 |
| ENSMUSG00000022025 | Cnmd | protein_coding | 0.164942327 | 2.401022234 | 0.0275793 | 0.8701518 | 0.977149329 |
| ENSMUSG00000055407 | Map6 | protein_coding | 0.050579869 | 5.469052074 | 0.0275247 | 0.8702705 | 0.977149329 |
| ENSMUSG00000032540 | Abhd5 | protein_coding | 0.066039158 | 3.705300625 | 0.0275192 | 0.8702836 | 0.977149329 |
| ENSMUSG00000033478 | Fam160b1 | protein_coding | 0.065105882 | 3.699283433 | 0.0274656 | 0.8704087 | 0.977187744 |
| ENSMUSG00000060803 | Gstp1 | protein_coding | 0.030149453 | 8.774563866 | 0.0274147 | 0.8705277 | 0.977219372 |
| ENSMUSG00000034544 | Rsrc1 | protein_coding | 0.037506472 | 6.519231588 | 0.0273084 | 0.8707765 | 0.977314766 |
| ENSMUSG00000036860 | Mrpl55 | protein_coding | 0.031872621 | 7.83164215 | 0.0272757 | 0.8708531 | 0.977314766 |
| ENSMUSG00000027804 | Ppid | protein_coding | 0.03000738 | 6.992843414 | 0.0272619 | 0.8708853 | 0.977314766 |
| ENSMUSG00000038533 | Cbfa2t2 | protein_coding | 0.059599985 | 6.28692202 | 0.0271759 | 0.8712896 | 0.97763451 |
| ENSMUSG00000030614 | Tmem126b | protein_coding | 0.033546143 | 7.484926532 | 0.026989 | 0.8715271 | 0.97763451 |
| ENSMUSG00000019984 | Med23 | protein_coding | 0.059189647 | 3.592601219 | 0.0269695 | 0.8715731 | 0.97763451 |
| ENSMUSG00000038780 | Smurf1 | protein_coding | 0.056489349 | 3.930851603 | 0.0269668 | 0.8715795 | 0.97763451 |
| ENSMUSG00000026705 | Klhl20 | protein_coding | -0.054842623 | 3.967137319 | 0.0269476 | 0.8716249 | 0.97763451 |
| ENSMUSG00000021994 | Wnt5a | protein_coding | -0.073960021 | 3.22810331 | 0.0267982 | 0.8719778 | 0.977788104 |
| ENSMUSG00000035277 | Arx | protein_coding | -0.080178057 | 3.12653984 | 0.0267346 | 0.8721284 | 0.977788104 |
| ENSMUSG00000023806 | Rsph3b | protein_coding | 0.084513478 | 2.821497084 | 0.0266966 | 0.8722184 | 0.977788104 |
| ENSMUSG00000006930 | Hap1 | protein_coding | 0.034099915 | 7.80132675 | 0.0266523 | 0.8723235 | 0.977788104 |
| ENSMUSG00000025153 | Fasn | protein_coding | 0.046716113 | 6.044390897 | 0.0266008 | 0.8724458 | 0.977788104 |
| ENSMUSG00000002265 | Peg3 | protein_coding | 0.03693125 | 8.475969773 | 0.0265241 | 0.8726279 | 0.977788104 |
| ENSMUSG00000038260 | Trpm4 | protein_coding | 0.053468065 | 4.247393398 | 0.0265112 | 0.8726587 | 0.977788104 |
| ENSMUSG00000026255 | Efhd1 | protein_coding | -0.063898606 | 4.00969525 | 0.0264539 | 0.8727952 | 0.977788104 |
| ENSMUSG00000046862 | Pramef8 | protein_coding | 0.063518312 | 4.561603272 | 0.0264513 | 0.8728013 | 0.977788104 |
| ENSMUSG00000057375 | Yipf1 | protein_coding | 0.034159925 | 6.431579522 | 0.0264167 | 0.8728839 | 0.977788104 |
| ENSMUSG00000100691 | 2010320M18Rik | lincRNA | -0.037332103 | 5.567277269 | 0.0264097 | 0.8729005 | 0.977788104 |
| ENSMUSG00000038705 | Gmeb2 | protein_coding | -0.073718846 | 3.403363273 | 0.0263916 | 0.8729435 | 0.977788104 |
| ENSMUSG00000023571 | C1qtnf12 | protein_coding | -0.099940682 | 3.022558791 | 0.0263737 | 0.8729862 | 0.977788104 |
| ENSMUSG00000034190 | Chmp7 | protein_coding | -0.042655139 | 6.459667764 | 0.0263386 | 0.8730701 | 0.977788104 |
| ENSMUSG00000074064 | Mlycd | protein_coding | 0.063300563 | 4.413333649 | 0.0263043 | 0.8731519 | 0.977788104 |
| ENSMUSG00000043716 | Rpl7 | protein_coding | 0.037979891 | 8.566012322 | 0.0262772 | 0.8732167 | 0.977788104 |
| ENSMUSG00000005674 | Tomm40l | protein_coding | 0.048955998 | 5.84229215 | 0.0262152 | 0.8733651 | 0.977794006 |
| ENSMUSG00000059277 | R74862 | processed_transcript | -0.096092249 | 2.99845578 | 0.026199 | 0.8734038 | 0.977794006 |
| ENSMUSG00000034820 | Cpsf7 | protein_coding | 0.133641376 | 2.343303687 | 0.0260729 | 0.873706 | 0.978029136 |
| ENSMUSG00000021754 | Map3k1 | protein_coding | -0.063191131 | 3.794129042 | 0.0260039 | 0.8738716 | 0.978029136 |
| ENSMUSG00000031171 | Ftsj1 | protein_coding | -0.04324899 | 5.746934887 | 0.0259976 | 0.8738867 | 0.978029136 |
| ENSMUSG00000031902 | Nfatc3 | protein_coding | 0.074918641 | 3.247578072 | 0.0259349 | 0.8740376 | 0.978034286 |
| ENSMUSG00000052456 | Asna1 | protein_coding | -0.029222177 | 8.054059734 | 0.0259201 | 0.8740732 | 0.978034286 |
| ENSMUSG00000032459 | Mrps22 | protein_coding | 0.050328301 | 5.211931843 | 0.0258798 | 0.8741703 | 0.978041171 |
| ENSMUSG00000042350 | Arel1 | protein_coding | 0.062018482 | 4.131847776 | 0.0255457 | 0.8749777 | 0.978593307 |
| ENSMUSG00000045875 | Adra1a | protein_coding | -0.071157225 | 3.405582033 | 0.0255419 | 0.874987 | 0.978593307 |
| ENSMUSG00000029427 | Zcchc8 | protein_coding | -0.064957759 | 3.828438522 | 0.0254753 | 0.8751486 | 0.978593307 |
| ENSMUSG00000028223 | Decr1 | protein_coding | 0.048006221 | 5.156132739 | 0.0254589 | 0.8751884 | 0.978593307 |
| ENSMUSG00000090110 | Cmc4 | protein_coding | -0.057818669 | 3.930517205 | 0.0254548 | 0.8751983 | 0.978593307 |
| ENSMUSG00000028478 | Clta | protein_coding | -0.037500105 | 6.887830154 | 0.0254477 | 0.8752157 | 0.978593307 |
| ENSMUSG00000024772 | Ehd1 | protein_coding | 0.044648415 | 4.516743433 | 0.0254006 | 0.8753303 | 0.978593307 |
| ENSMUSG00000029386 | Tctn2 | processed_transcript | -0.04747782 | 4.322253723 | 0.0253752 | 0.8753919 | 0.978593307 |
| ENSMUSG00000027577 | Chrna4 | protein_coding | 0.089008762 | 4.640889132 | 0.0253062 | 0.8758338 | 0.978927392 |

| ensembl_gene_id | external_gene_name | gene_biotype | log2 (fold change) | log2 (counts per million) | F-value | p-value | False discovery rate |
|---|---|---|---|---|---|---|---|
| ENSMUSG00000033423 | Eri3 | protein_coding | 0.027576284 | 8.52696204 | 0.025078 | 0.8761172 | 0.978927392 |
| ENSMUSG00000032968 | Inha | protein_coding | -0.1147426 | 2.437307666 | 0.0250747 | 0.8761255 | 0.978927392 |
| ENSMUSG00000003810 | Mast2 | protein_coding | -0.040297693 | 5.41546014 | 0.0250449 | 0.8761984 | 0.978927392 |
| ENSMUSG00000023919 | Cenpq | protein_coding | -0.087438748 | 3.48608627 | 0.0250116 | 0.8762832 | 0.978927392 |
| ENSMUSG00000035530 | Eif1 | protein_coding | -0.037653992 | 9.399045328 | 0.0249705 | 0.8763807 | 0.978927392 |
| ENSMUSG00000036022 | Fam122b | protein_coding | -0.073827985 | 3.233161775 | 0.0248968 | 0.8765618 | 0.978927392 |
| ENSMUSG00000051041 | Olfml1 | protein_coding | 0.050935523 | 5.013972872 | 0.0248225 | 0.8767446 | 0.978927392 |
| ENSMUSG00000028394 | Pole3 | protein_coding | -0.031914395 | 6.041170929 | 0.0247688 | 0.8768766 | 0.978927392 |
| ENSMUSG00000073062 | Zxdb | protein_coding | -0.076896126 | 3.790263018 | 0.0247064 | 0.8770336 | 0.978927392 |
| ENSMUSG00000056692 | D17Wsu92e | protein_coding | 0.031224306 | 6.607723939 | 0.0247044 | 0.8770355 | 0.978927392 |
| ENSMUSG00000028572 | Hook1 | protein_coding | -0.062369777 | 3.667438773 | 0.0246709 | 0.8771181 | 0.978927392 |
| ENSMUSG00000027309 | 4930402H24Rik | protein_coding | 0.040156465 | 5.915773539 | 0.0246432 | 0.8771865 | 0.978927392 |
| ENSMUSG00000063889 | Crem | protein_coding | 0.043707377 | 5.032857539 | 0.024565 | 0.8773799 | 0.978927392 |
| ENSMUSG00000004902 | Slc25a18 | protein_coding | -0.040159479 | 6.351347819 | 0.02453 | 0.8774665 | 0.978927392 |
| ENSMUSG00000031708 | Tecr | protein_coding | 0.031517739 | 10.67126226 | 0.0244911 | 0.8775628 | 0.978927392 |
| ENSMUSG00000019188 | H13 | protein_coding | -0.059847357 | 5.09278234 | 0.0244685 | 0.8776221 | 0.978927392 |
| ENSMUSG00000032051 | Fdx1 | protein_coding | 0.065434211 | 3.43317196 | 0.0244579 | 0.8776452 | 0.978927392 |
| ENSMUSG00000035561 | Aldh1b1 | protein_coding | -0.051567603 | 4.460342344 | 0.0244472 | 0.8776717 | 0.978927392 |
| ENSMUSG00000020607 | Fam84a | protein_coding | -0.031406323 | 7.303704423 | 0.02444 | 0.8776895 | 0.978927392 |
| ENSMUSG00000054408 | Spcs3 | protein_coding | -0.03625837 | 5.892872514 | 0.0244187 | 0.8777425 | 0.978927392 |
| ENSMUSG00000022797 | Tfrc | protein_coding | 0.051155518 | 4.404060591 | 0.0243668 | 0.8778714 | 0.978927392 |
| ENSMUSG00000050541 | Adra1b | protein_coding | -0.115005539 | 2.48256062 | 0.0243326 | 0.8779562 | 0.978927392 |
| ENSMUSG00000060671 | Atp8b2 | protein_coding | 0.069447506 | 3.471157215 | 0.0243068 | 0.8780204 | 0.978927392 |
| ENSMUSG00000097213 | Gm26745 | lincRNA | -0.089481457 | 2.703866839 | 0.0242826 | 0.8780806 | 0.978927392 |
| ENSMUSG00000039671 | Zmynd8 | protein_coding | 0.03077702 | 6.581414069 | 0.0242732 | 0.878104 | 0.978927392 |
| ENSMUSG00000027427 | Polr3f | protein_coding | -0.053052742 | 4.529543924 | 0.0242499 | 0.8781622 | 0.978927392 |
| ENSMUSG00000037373 | Ctbp1 | protein_coding | 0.033459254 | 6.682706994 | 0.0241751 | 0.8783486 | 0.978927392 |
| ENSMUSG00000058799 | Nap1l1 | protein_coding | -0.02938858 | 6.744221483 | 0.0241381 | 0.8784409 | 0.978927392 |
| ENSMUSG00000028322 | Exosc3 | protein_coding | 0.073333799 | 3.460895537 | 0.0241155 | 0.8784973 | 0.978927392 |
| ENSMUSG00000000340 | Dbt | protein_coding | -0.082407191 | 3.982134135 | 0.0241331 | 0.8785129 | 0.978927392 |
| ENSMUSG00000018559 | Ctdnep1 | protein_coding | 0.042728202 | 5.025039211 | 0.0238369 | 0.8791952 | 0.97933313 |
| ENSMUSG00000029561 | Oasl2 | protein_coding | 0.189785975 | 3.05310636 | 0.0241308 | 0.8792117 | 0.97933313 |
| ENSMUSG00000061306 | Slc38a10 | protein_coding | 0.055716007 | 5.193890745 | 0.0238052 | 0.8792781 | 0.97933313 |
| ENSMUSG00000020196 | Cabin1 | protein_coding | 0.05692428 | 5.004642735 | 0.0237064 | 0.8795269 | 0.97933313 |
| ENSMUSG00000059586 | Nsmce2 | protein_coding | -0.036792229 | 6.325690648 | 0.0236859 | 0.8795754 | 0.97933313 |
| ENSMUSG00000006369 | Fbln1 | protein_coding | -0.087641867 | 3.155322925 | 0.0236828 | 0.879583 | 0.97933313 |
| ENSMUSG00000031665 | Sall1 | protein_coding | 0.081135883 | 3.208844785 | 0.0236812 | 0.8795872 | 0.97933313 |
| ENSMUSG00000086040 | Wipf3 | protein_coding | 0.094894431 | 2.898000175 | 0.0236739 | 0.8796056 | 0.97933313 |
| ENSMUSG00000023927 | Satb1 | protein_coding | 0.054328599 | 3.755547552 | 0.0235552 | 0.8799054 | 0.979500942 |
| ENSMUSG00000086851 | A430108G06Rik | processed_transcript | -0.172240046 | 2.553515066 | 0.023577 | 0.8800383 | 0.979500942 |
| ENSMUSG00000059839 | Zfp874b | protein_coding | 0.081349723 | 3.056650539 | 0.0234077 | 0.8802787 | 0.979500942 |
| ENSMUSG00000004319 | Clcn3 | protein_coding | 0.028438746 | 7.199467733 | 0.0233968 | 0.8803063 | 0.979500942 |
| ENSMUSG00000026566 | Mpzl1 | protein_coding | 0.08306668 | 3.369692735 | 0.0233797 | 0.8803497 | 0.979500942 |
| ENSMUSG00000034858 | Fam214a | protein_coding | -0.0527805 | 4.271265925 | 0.0233494 | 0.8804266 | 0.979500942 |
| ENSMUSG00000042487 | Leo1 | protein_coding | -0.048160012 | 6.18371895 | 0.0233476 | 0.8804426 | 0.979500942 |
| ENSMUSG00000020354 | Sgcd | protein_coding | -0.097140094 | 2.665280259 | 0.0233264 | 0.8804851 | 0.979500942 |
| ENSMUSG00000035864 | Syt1 | protein_coding | 0.030379747 | 9.368887695 | 0.0232174 | 0.8807624 | 0.979598941 |
| ENSMUSG00000028137 | Celf3 | protein_coding | -0.050522078 | 5.27909203 | 0.0231802 | 0.8808571 | 0.979598941 |
| ENSMUSG00000022972 | 1110004E09Rik | protein_coding | 0.058492601 | 6.017678376 | 0.023216 | 0.8809525 | 0.979598941 |
| ENSMUSG00000104864 | Gm43655 | processed_pseudoge | -0.078202676 | 2.811837455 | 0.0230986 | 0.8810654 | 0.979598941 |
| ENSMUSG00000039449 | Prpf18 | protein_coding | -0.04294305 | 5.607846361 | 0.0230662 | 0.8811482 | 0.979598941 |
| ENSMUSG00000001229 | Dpp9 | protein_coding | 0.049363418 | 5.095200661 | 0.0230467 | 0.881198 | 0.979598941 |
| ENSMUSG00000044894 | Uqcrq | protein_coding | -0.033530178 | 10.39844636 | 0.0229931 | 0.881335 | 0.979598941 |
| ENSMUSG00000020069 | Hnrnph3 | protein_coding | 0.053318455 | 4.443107926 | 0.022838 | 0.8817328 | 0.979598941 |
| ENSMUSG00000030310 | Slc6a1 | protein_coding | -0.036934889 | 7.111988007 | 0.0228174 | 0.8817857 | 0.979598941 |
| ENSMUSG00000037103 | Dcaf15 | protein_coding | -0.083457932 | 3.497198755 | 0.0228143 | 0.8817966 | 0.979598941 |
| ENSMUSG00000000915 | Hip1r | protein_coding | -0.057997486 | 4.0628248 | 0.0228124 | 0.8817986 | 0.979598941 |
| ENSMUSG00000025854 | Fam20c | protein_coding | 0.065424471 | 4.759858499 | 0.0228115 | 0.8818154 | 0.979598941 |
| ENSMUSG00000039414 | Heatr5b | protein_coding | 0.064477972 | 3.54909801 | 0.0227731 | 0.8818994 | 0.979598941 |
| ENSMUSG00000027313 | Chac1 | protein_coding | 0.05267375 | 4.619900276 | 0.0227612 | 0.8819302 | 0.979598941 |
| ENSMUSG00000026956 | Uap1l1 | protein_coding | 0.072606331 | 3.614291088 | 0.0227543 | 0.881948 | 0.979598941 |
| ENSMUSG00000029247 | Paics | protein_coding | 0.028209396 | 7.582638901 | 0.0227221 | 0.8820308 | 0.979598941 |
| ENSMUSG00000057497 | Fam136a | protein_coding | -0.030177376 | 6.854044648 | 0.0225929 | 0.8823639 | 0.979867711 |
| ENSMUSG00000018965 | Ywhah | protein_coding | 0.027423294 | 8.259081373 | 0.0225355 | 0.8825125 | 0.979893878 |
| ENSMUSG00000042766 | Trim46 | protein_coding | -0.035285959 | 5.439437472 | 0.0225133 | 0.8825697 | 0.979893878 |
| ENSMUSG00000033124 | Atg9a | protein_coding | 0.103056203 | 3.263364428 | 0.0224641 | 0.8827583 | 0.980002016 |
| ENSMUSG00000018593 | Sparc | protein_coding | -0.053696431 | 9.204871844 | 0.0223472 | 0.8832574 | 0.980454879 |
| ENSMUSG00000034789 | Rab24 | protein_coding | 0.058923245 | 4.431594642 | 0.0221137 | 0.8836083 | 0.980606681 |
| ENSMUSG00000023991 | Foxp4 | protein_coding | 0.069740814 | 3.535904737 | 0.0220785 | 0.8837004 | 0.980606681 |
| ENSMUSG00000062691 | Cebpzos | protein_coding | -0.042412713 | 4.80714867 | 0.0220506 | 0.8837734 | 0.980606681 |
| ENSMUSG00000015942 | Gtf2ird2 | protein_coding | 0.062599843 | 3.549050209 | 0.0219797 | 0.883959 | 0.980606681 |
| ENSMUSG00000054910 | 4931415C17Rik | processed_transcript | 0.108401157 | 2.736321017 | 0.0219537 | 0.8840269 | 0.980606681 |
| ENSMUSG00000079442 | St6galnac4 | protein_coding | -0.058812321 | 4.861569362 | 0.0219237 | 0.8841086 | 0.980606681 |
| ENSMUSG00000027455 | Nsfl1c | protein_coding | 0.028829769 | 7.593696853 | 0.0218604 | 0.8842717 | 0.980606681 |

| ensembl_gene_id | external_gene_name | gene_biotype | log2 (fold change) | log2 (counts per million) | F-value | p-value | False discovery rate |
|---|---|---|---|---|---|---|---|
| ENSMUSG00000000489 | Pdgfb | protein_coding | -0.069446782 | 3.367482814 | 0.0217735 | 0.8845004 | 0.980606681 |
| ENSMUSG00000027479 | Mapre1 | protein_coding | 0.027942787 | 7.318382081 | 0.0217475 | 0.8845687 | 0.980606681 |
| ENSMUSG00000031584 | Gsr | protein_coding | 0.039171373 | 5.228194358 | 0.0216771 | 0.8847542 | 0.980606681 |
| ENSMUSG00000024072 | Yipf4 | protein_coding | 0.044092576 | 5.55137118 | 0.0216731 | 0.8847647 | 0.980606681 |
| ENSMUSG00000028779 | Pef1 | protein_coding | -0.029813878 | 7.551714064 | 0.0216696 | 0.8847741 | 0.980606681 |
| ENSMUSG00000030536 | Iqgap1 | protein_coding | -0.075571281 | 3.601172273 | 0.0216511 | 0.8848228 | 0.980606681 |
| ENSMUSG00000040888 | Gfer | protein_coding | 0.042834031 | 4.47667064 | 0.0216114 | 0.8849277 | 0.980606681 |
| ENSMUSG00000059291 | Rpl11 | protein_coding | 0.044525355 | 4.430452932 | 0.0215897 | 0.8849849 | 0.980606681 |
| ENSMUSG00000087370 | Tmem170b | protein_coding | -0.047003381 | 5.335219299 | 0.0215891 | 0.8849867 | 0.980606681 |
| ENSMUSG00000078429 | Ctdsp2 | protein_coding | -0.051908071 | 4.81346619 | 0.0215867 | 0.884993 | 0.980606681 |
| ENSMUSG00000040260 | Daam2 | protein_coding | 0.049796177 | 4.34735342 | 0.0215706 | 0.8850356 | 0.980606681 |
| ENSMUSG00000033216 | Eefsec | protein_coding | -0.055377303 | 4.607461669 | 0.0213763 | 0.8855505 | 0.980855221 |
| ENSMUSG00000030035 | Wbp1 | protein_coding | 0.058294612 | 4.942696081 | 0.0213618 | 0.885592 | 0.980855221 |
| ENSMUSG00000040731 | Eif4h | protein_coding | -0.0288107 | 8.360380443 | 0.0213583 | 0.8855984 | 0.980855221 |
| ENSMUSG00000018398 | 8-Sep | protein_coding | 0.026353081 | 6.969690706 | 0.0213484 | 0.8856248 | 0.980855221 |
| ENSMUSG00000028180 | Zranb2 | protein_coding | 0.028042359 | 7.576783338 | 0.0212641 | 0.885849 | 0.980926463 |
| ENSMUSG00000039154 | Shd | protein_coding | -0.039014346 | 6.023462431 | 0.0212557 | 0.8858716 | 0.980926463 |
| ENSMUSG00000044934 | Zfp367 | protein_coding | 0.048252262 | 4.7212702 | 0.0211531 | 0.8861451 | 0.981126372 |
| ENSMUSG00000042446 | Zmym4 | protein_coding | 0.095383298 | 2.574492042 | 0.0210994 | 0.8862888 | 0.981126372 |
| ENSMUSG00000030647 | Ndufc2 | protein_coding | -0.033941619 | 9.492654567 | 0.0210683 | 0.8863719 | 0.981126372 |
| ENSMUSG00000047371 | Zfp768 | protein_coding | -0.109784229 | 2.477287658 | 0.0210184 | 0.8865055 | 0.981126372 |
| ENSMUSG00000041132 | N4bp2l1 | protein_coding | 0.041987147 | 5.07898896 | 0.0210024 | 0.8865485 | 0.981126372 |
| ENSMUSG00000047230 | Cldn2 | protein_coding | 0.186056797 | 1.917464903 | 0.0211031 | 0.8866417 | 0.981126372 |
| ENSMUSG00000025314 | Ptprj | protein_coding | 0.085405173 | 2.738237893 | 0.0209301 | 0.8867423 | 0.981126372 |
| ENSMUSG00000031605 | Klhl2 | protein_coding | -0.033862326 | 6.092562795 | 0.0208336 | 0.887002 | 0.981126372 |
| ENSMUSG00000039208 | Metrnl | protein_coding | 0.091685322 | 2.675614047 | 0.0208292 | 0.8870137 | 0.981126372 |
| ENSMUSG00000055866 | Per2 | protein_coding | 0.084987193 | 3.069343328 | 0.020751 | 0.8872246 | 0.981126372 |
| ENSMUSG00000024012 | Mtch1 | protein_coding | 0.027765348 | 9.233100313 | 0.020729 | 0.887284 | 0.981126372 |
| ENSMUSG00000029655 | N4bp2l2 | protein_coding | 0.049406874 | 5.050342002 | 0.0207188 | 0.8873114 | 0.981126372 |
| ENSMUSG00000013833 | Med16 | protein_coding | 0.048123116 | 3.914232501 | 0.0207032 | 0.8873535 | 0.981126372 |
| ENSMUSG00000104960 | Snhg8 | processed_transcript | -0.051800878 | 6.492113512 | 0.020675 | 0.8876346 | 0.981126372 |
| ENSMUSG00000033955 | Tnks1bp1 | protein_coding | -0.104678219 | 2.644913488 | 0.0205857 | 0.8876713 | 0.981126372 |
| ENSMUSG00000039563 | 2210406O10Rik | lincRNA | -0.064614991 | 3.185292419 | 0.0205628 | 0.8877334 | 0.981126372 |
| ENSMUSG00000037174 | Elf2 | protein_coding | 0.054582203 | 3.947263548 | 0.0205613 | 0.8877373 | 0.981126372 |
| ENSMUSG00000027860 | Vangl1 | protein_coding | 0.105889814 | 3.138916828 | 0.0205853 | 0.8877461 | 0.981126372 |
| ENSMUSG00000025958 | Creb1 | protein_coding | -0.04448424 | 5.110658551 | 0.0205435 | 0.8877857 | 0.981126372 |
| ENSMUSG00000030000 | Add2 | protein_coding | 0.03041089 | 7.219291826 | 0.0204977 | 0.8879101 | 0.98116296 |
| ENSMUSG00000036391 | Sec24a | protein_coding | -0.055659266 | 3.856373162 | 0.0204594 | 0.8880141 | 0.981177084 |
| ENSMUSG00000058886 | Deaf1 | protein_coding | -0.049659861 | 4.746996997 | 0.0203695 | 0.8882584 | 0.981269836 |
| ENSMUSG00000059493 | Nhs | protein_coding | -0.061446485 | 3.366856046 | 0.0203389 | 0.8883418 | 0.981269836 |
| ENSMUSG00000058740 | Kcnt1 | protein_coding | -0.055993503 | 3.781215717 | 0.0203279 | 0.8883718 | 0.981269836 |
| ENSMUSG00000034290 | Nek9 | protein_coding | -0.061873204 | 3.854974297 | 0.020287 | 0.8884834 | 0.981275222 |
| ENSMUSG00000039824 | Myl6b | protein_coding | -0.073700835 | 2.879614197 | 0.0202592 | 0.8885592 | 0.981275222 |
| ENSMUSG00000006471 | Ndor1 | protein_coding | 0.067879009 | 3.286308488 | 0.0201896 | 0.8887494 | 0.981384501 |
| ENSMUSG00000022217 | Emc9 | protein_coding | 0.025863889 | 6.944502775 | 0.0201311 | 0.8889098 | 0.981455738 |
| ENSMUSG00000109284 | B230311B06Rik | lincRNA | -0.067342487 | 3.309155098 | 0.0200645 | 0.8890923 | 0.981455738 |
| ENSMUSG00000044098 | Rsbn1 | protein_coding | -0.07911365 | 2.860981529 | 0.0200347 | 0.8891742 | 0.981455738 |
| ENSMUSG00000020262 | Adarb1 | protein_coding | 0.046145287 | 5.996686249 | 0.0200332 | 0.8891887 | 0.981455738 |
| ENSMUSG00000062031 | Pgghg | protein_coding | -0.137111681 | 2.076154502 | 0.0199997 | 0.8892703 | 0.981455738 |
| ENSMUSG00000021725 | Parp8 | protein_coding | -0.06207482 | 3.485293454 | 0.019903 | 0.8895363 | 0.981541912 |
| ENSMUSG00000020248 | Nfyb | protein_coding | 0.038238881 | 5.796719178 | 0.0198496 | 0.8896837 | 0.981541912 |
| ENSMUSG00000056708 | Ier5 | protein_coding | -0.077841511 | 3.078491672 | 0.0198366 | 0.8897196 | 0.981541912 |
| ENSMUSG00000002963 | Pnkp | protein_coding | -0.038908931 | 4.95504095 | 0.0198178 | 0.8897715 | 0.981541912 |
| ENSMUSG00000029600 | Rita1 | protein_coding | 0.035099439 | 5.501720603 | 0.0198057 | 0.8898048 | 0.981541912 |
| ENSMUSG00000034793 | G6pc3 | protein_coding | -0.03871484 | 6.389852792 | 0.0197244 | 0.8900326 | 0.98169249 |
| ENSMUSG00000047363 | Cstad | protein_coding | -0.057889269 | 3.822675004 | 0.0196621 | 0.8902022 | 0.981778869 |
| ENSMUSG00000047250 | Ptgs1 | protein_coding | -0.055160601 | 3.772499142 | 0.0195471 | 0.8905215 | 0.981988992 |
| ENSMUSG00000037624 | Kcnk2 | protein_coding | -0.041881454 | 5.383507972 | 0.0195278 | 0.8905753 | 0.981988992 |
| ENSMUSG00000018411 | Mapt | protein_coding | -0.035838002 | 8.897666719 | 0.0194282 | 0.8908557 | 0.981994552 |
| ENSMUSG00000018906 | P4ha2 | protein_coding | -0.060288501 | 4.267600162 | 0.0193988 | 0.8909349 | 0.981994552 |
| ENSMUSG00000021738 | Atxn7 | protein_coding | -0.061204873 | 3.312715562 | 0.0193537 | 0.8910609 | 0.981994552 |
| ENSMUSG00000079339 | Ifit1bl1 | protein_coding | 0.168442974 | 2.273117401 | 0.0193865 | 0.8910778 | 0.981994552 |
| ENSMUSG00000047259 | Mc4r | protein_coding | -0.064747704 | 3.845417213 | 0.0193296 | 0.8911283 | 0.981994552 |
| ENSMUSG00000033392 | Clasp2 | protein_coding | 0.038602445 | 7.111265289 | 0.0193349 | 0.8911283 | 0.981994552 |
| ENSMUSG00000020869 | Lrrc59 | protein_coding | -0.036850472 | 5.356798914 | 0.0192454 | 0.891364 | 0.982153644 |
| ENSMUSG00000005102 | Eif2ak4 | protein_coding | -0.060401888 | 3.634315011 | 0.0191634 | 0.891594 | 0.982160207 |
| ENSMUSG00000068373 | D430041D05Rik | protein_coding | 0.044131859 | 4.776379239 | 0.0190956 | 0.8917848 | 0.982160207 |
| ENSMUSG00000045973 | Slc25a51 | protein_coding | -0.032297509 | 6.216469553 | 0.0190289 | 0.8919726 | 0.982160207 |
| ENSMUSG00000036352 | Ubac1 | protein_coding | -0.035960959 | 5.215568652 | 0.019008 | 0.8920315 | 0.982160207 |
| ENSMUSG00000032232 | Cgnl1 | protein_coding | -0.134860158 | 1.933576761 | 0.0189132 | 0.8922995 | 0.982160207 |
| ENSMUSG00000042807 | Hecw2 | protein_coding | -0.050374932 | 4.125532092 | 0.0188655 | 0.8924343 | 0.982160207 |
| ENSMUSG00000028382 | Ptbp3 | protein_coding | 0.053044076 | 3.931620566 | 0.0188525 | 0.8924711 | 0.982160207 |
| ENSMUSG00000022463 | Srebf2 | protein_coding | 0.03110875 | 5.999905884 | 0.0188294 | 0.8925366 | 0.982160207 |
| ENSMUSG00000031732 | Phlpp2 | protein_coding | 0.049185768 | 4.01577232 | 0.0186355 | 0.8930876 | 0.982160207 |

| ensembl_gene_id | external_gene_name | gene_biotype | log2 (fold change) | log2 (counts per million) | F-value | p-value | False discovery rate |
|---|---|---|---|---|---|---|---|
| ENSMUSG00000024948 | Map4k2 | protein_coding | -0.066448647 | 4.428703934 | 0.0186425 | 0.8931378 | 0.982160207 |
| ENSMUSG00000029166 | Mapre3 | protein_coding | -0.026986564 | 6.801726034 | 0.0185915 | 0.8932132 | 0.982160207 |
| ENSMUSG00000040972 | Igsf21 | protein_coding | 0.069169105 | 4.150300322 | 0.0185767 | 0.8933073 | 0.982160207 |
| ENSMUSG00000052914 | Cyp2j6 | protein_coding | -0.063133657 | 4.464166553 | 0.0185516 | 0.8933419 | 0.982160207 |
| ENSMUSG00000028541 | B4galt2 | protein_coding | 0.055852746 | 3.494569312 | 0.0185141 | 0.8934341 | 0.982160207 |
| ENSMUSG00000089889 | 0610040B10Rik | antisense_RNA | -0.05504472 | 3.573180574 | 0.0184819 | 0.8935263 | 0.982160207 |
| ENSMUSG00000081552 | Gm12328 | processed_pseudoge | 0.077832596 | 2.681845599 | 0.0183522 | 0.8938981 | 0.982160207 |
| ENSMUSG00000029068 | Ccnl2 | protein_coding | 0.049724085 | 3.8551514 | 0.0183251 | 0.893976 | 0.982160207 |
| ENSMUSG00000031562 | Dctd | protein_coding | -0.056635577 | 4.074161882 | 0.0183028 | 0.89404 | 0.982160207 |
| ENSMUSG00000052026 | Slc6a7 | protein_coding | 0.057857585 | 3.676002892 | 0.0182989 | 0.8940513 | 0.982160207 |
| ENSMUSG00000006333 | Rps9 | protein_coding | 0.027926228 | 9.87165058 | 0.0182632 | 0.8941541 | 0.982160207 |
| ENSMUSG00000021203 | Otub2 | protein_coding | 0.088225486 | 3.546582868 | 0.0182546 | 0.8942843 | 0.982160207 |
| ENSMUSG00000025743 | Sdc3 | protein_coding | -0.06180688 | 3.225559943 | 0.01818 | 0.8943939 | 0.982160207 |
| ENSMUSG00000026728 | Vim | protein_coding | 0.055548453 | 4.265749196 | 0.0181796 | 0.8943951 | 0.982160207 |
| ENSMUSG00000064284 | Cdpf1 | protein_coding | 0.033190823 | 6.128265536 | 0.0181722 | 0.8944164 | 0.982160207 |
| ENSMUSG00000036275 | 9530068E07Rik | protein_coding | -0.032064025 | 6.025936184 | 0.0181043 | 0.8946126 | 0.982160207 |
| ENSMUSG00000021061 | Sptb | protein_coding | -0.065275153 | 3.921841883 | 0.018081 | 0.8946826 | 0.982160207 |
| ENSMUSG00000078570 | 1110065P20Rik | protein_coding | -0.089031382 | 2.752640373 | 0.0180618 | 0.8947355 | 0.982160207 |
| ENSMUSG00000021830 | Txndc16 | protein_coding | 0.035013333 | 5.510028045 | 0.0179854 | 0.8949571 | 0.982160207 |
| ENSMUSG00000024319 | Vps52 | protein_coding | 0.0386286 | 5.761971041 | 0.0179835 | 0.8949627 | 0.982160207 |
| ENSMUSG00000019647 | Sema6a | protein_coding | -0.046055052 | 4.673341871 | 0.0179702 | 0.8950011 | 0.982160207 |
| ENSMUSG00000039217 | Il18 | protein_coding | -0.047300614 | 5.980826172 | 0.0179912 | 0.8950385 | 0.982160207 |
| ENSMUSG00000019947 | Arid5b | protein_coding | 0.039331208 | 5.545585943 | 0.0179357 | 0.8951013 | 0.982160207 |
| ENSMUSG00000004207 | Psap | protein_coding | 0.02791524 | 9.011552708 | 0.0178935 | 0.8952239 | 0.982160207 |
| ENSMUSG00000004931 | Apba3 | protein_coding | 0.079646389 | 2.61079719 | 0.0178919 | 0.8952287 | 0.982160207 |
| ENSMUSG00000057614 | Gnai1 | protein_coding | -0.025190916 | 6.97412425 | 0.0178765 | 0.8952734 | 0.982160207 |
| ENSMUSG00000017344 | Vtn | protein_coding | -0.045277678 | 4.202792313 | 0.0178761 | 0.8952746 | 0.982160207 |
| ENSMUSG00000028743 | Akr7a5 | protein_coding | 0.049243099 | 3.887303776 | 0.0178605 | 0.8953201 | 0.982160207 |
| ENSMUSG00000036898 | Zfp157 | protein_coding | -0.044195452 | 4.376131062 | 0.017782 | 0.895549 | 0.982160207 |
| ENSMUSG00000074194 | Zfp791 | protein_coding | -0.059176324 | 3.226863141 | 0.0177419 | 0.895666 | 0.982160207 |
| ENSMUSG00000025340 | Rabgef1 | protein_coding | 0.034716818 | 5.840807769 | 0.0177019 | 0.8957829 | 0.982160207 |
| ENSMUSG00000096188 | Cmtm4 | protein_coding | 0.050770994 | 4.718665735 | 0.0176897 | 0.8958186 | 0.982160207 |
| ENSMUSG00000020599 | Rgs9 | protein_coding | -0.030826506 | 6.232323517 | 0.0176796 | 0.8958483 | 0.982160207 |
| ENSMUSG00000036634 | Mag | protein_coding | 0.025809303 | 7.307725008 | 0.0176768 | 0.8958565 | 0.982160207 |
| ENSMUSG00000039275 | Foxk2 | protein_coding | 0.040068516 | 5.098792257 | 0.017671 | 0.8958735 | 0.982160207 |
| ENSMUSG00000047261 | Gap43 | protein_coding | -0.026266719 | 11.00325107 | 0.0176599 | 0.8959061 | 0.982160207 |
| ENSMUSG00000044551 | 9930012K11Rik | protein_coding | 0.101502814 | 2.885556435 | 0.0176412 | 0.8959633 | 0.982160207 |
| ENSMUSG00000009927 | Rps25 | protein_coding | -0.031726162 | 6.467116773 | 0.0176343 | 0.8959811 | 0.982160207 |
| ENSMUSG00000026278 | Bok | protein_coding | 0.035551004 | 5.664162849 | 0.017577 | 0.8961491 | 0.982160207 |
| ENSMUSG00000085971 | Gm15411 | processed_transcript | 0.102076652 | 2.348571445 | 0.017567 | 0.8961785 | 0.982160207 |
| ENSMUSG00000022414 | Tab1 | protein_coding | 0.050908825 | 5.193363112 | 0.0174732 | 0.8964568 | 0.982160207 |
| ENSMUSG00000031112 | Stk26 | protein_coding | 0.099596432 | 2.454219891 | 0.0174415 | 0.8965476 | 0.982160207 |
| ENSMUSG00000029120 | Ppp2r2c | protein_coding | -0.0243736 | 7.304229255 | 0.0173545 | 0.8968044 | 0.982160207 |
| ENSMUSG00000001755 | Coasy | protein_coding | 0.042069491 | 5.350843705 | 0.0173207 | 0.8969043 | 0.982160207 |
| ENSMUSG00000098923 | Tmem185b | protein_coding | 0.044596599 | 4.597714848 | 0.0173174 | 0.8969141 | 0.982160207 |
| ENSMUSG00000038446 | Cdc40 | protein_coding | 0.047697033 | 5.089583073 | 0.0173175 | 0.8969163 | 0.982160207 |
| ENSMUSG00000025856 | Pdgfa | protein_coding | -0.035376663 | 5.019665444 | 0.0172898 | 0.8969957 | 0.982160207 |
| ENSMUSG00000029084 | Cd38 | protein_coding | 0.065658242 | 3.063188531 | 0.017276 | 0.8970366 | 0.982160207 |
| ENSMUSG00000003518 | Dusp3 | protein_coding | 0.036395239 | 8.317145625 | 0.0172697 | 0.8970694 | 0.982160207 |
| ENSMUSG00000001729 | Akt1 | protein_coding | 0.02500209 | 6.455986 | 0.0172294 | 0.8971747 | 0.982160207 |
| ENSMUSG00000020181 | Nav3 | protein_coding | 0.036573306 | 5.39169619 | 0.0172195 | 0.8972041 | 0.982160207 |
| ENSMUSG00000001157 | Gmcl1 | protein_coding | 0.054921019 | 4.452459458 | 0.0172129 | 0.8972264 | 0.982160207 |
| ENSMUSG00000100975 | Gm28875 | lincRNA | -0.083027832 | 3.551389565 | 0.0172391 | 0.8972446 | 0.982160207 |
| ENSMUSG00000037791 | Phf12 | protein_coding | 0.049949344 | 4.091898863 | 0.0171992 | 0.8972644 | 0.982160207 |
| ENSMUSG00000033596 | Rfwd3 | protein_coding | 0.063951792 | 3.150216625 | 0.0171498 | 0.897411 | 0.982160207 |
| ENSMUSG00000053898 | Ech1 | protein_coding | 0.02392994 | 7.728086844 | 0.0171196 | 0.897501 | 0.982160207 |
| ENSMUSG00000025402 | Nab2 | protein_coding | -0.060082182 | 3.529868613 | 0.0170693 | 0.8976506 | 0.982160207 |
| ENSMUSG00000042303 | Sgsm3 | protein_coding | 0.037929849 | 6.572978962 | 0.0170406 | 0.8977457 | 0.982160207 |
| ENSMUSG00000024423 | Impact | protein_coding | 0.036472994 | 10.99090957 | 0.017037 | 0.8977496 | 0.982160207 |
| ENSMUSG00000038520 | Tbc1d17 | protein_coding | 0.03240923 | 6.289337849 | 0.0170305 | 0.8977664 | 0.982160207 |
| ENSMUSG00000060126 | Tpt1 | protein_coding | -0.052489891 | 3.675308604 | 0.0169802 | 0.8979166 | 0.982160207 |
| ENSMUSG00000061458 | Nol10 | protein_coding | -0.046327573 | 4.249977819 | 0.016973 | 0.8979381 | 0.982160207 |
| ENSMUSG00000063235 | Ptpmt1 | processed_transcript | -0.046810449 | 3.817350706 | 0.0169411 | 0.8980335 | 0.982160207 |
| ENSMUSG00000007892 | Rplp1 | protein_coding | -0.032896229 | 9.342253724 | 0.016938 | 0.8980453 | 0.982160207 |
| ENSMUSG00000075704 | Txnrd2 | protein_coding | -0.056365788 | 3.669811175 | 0.016902 | 0.8981504 | 0.982160207 |
| ENSMUSG00000033022 | Cdo1 | protein_coding | -0.034577306 | 5.595285487 | 0.0168787 | 0.8982204 | 0.982160207 |
| ENSMUSG00000029198 | Grpel1 | protein_coding | 0.024520003 | 8.453275361 | 0.0168169 | 0.8984058 | 0.982186242 |
| ENSMUSG00000074461 | Gm10699 | TEC | -0.028760737 | 5.997095982 | 0.0167684 | 0.8985513 | 0.982186242 |
| ENSMUSG00000025742 | Prps2 | protein_coding | 0.041010619 | 4.834751372 | 0.0167481 | 0.8986126 | 0.982186242 |
| ENSMUSG00000034875 | Nudt19 | protein_coding | 0.022994958 | 6.684341927 | 0.0167206 | 0.8986952 | 0.982186242 |
| ENSMUSG00000020952 | Scfd1 | protein_coding | -0.026578823 | 6.245587856 | 0.0166785 | 0.898822 | 0.982186242 |
| ENSMUSG00000025757 | Hspa4l | protein_coding | 0.031388135 | 6.93759472 | 0.0166095 | 0.8990303 | 0.982186242 |
| ENSMUSG00000071398 | 2410004P03Rik | protein_coding | 0.062020248 | 3.660535499 | 0.0166048 | 0.8990445 | 0.982186242 |
| ENSMUSG00000033883 | D3Ertd254e | protein_coding | -0.044281184 | 4.246837968 | 0.0165742 | 0.8991371 | 0.982186242 |

| ensembl_gene_id | external_gene_name | gene_biotype | log2 (fold change) | log2 (counts per million) | F-value | p-value | False discovery rate |
|---|---|---|---|---|---|---|---|
| ENSMUSG00000034591 | Slc41a2 | protein_coding | 0.044779734 | 3.769568938 | 0.0165338 | 0.8992593 | 0.982186242 |
| ENSMUSG00000042662 | Dusp15 | protein_coding | -0.066538638 | 2.943083448 | 0.0165286 | 0.8992752 | 0.982186242 |
| ENSMUSG00000042249 | Grk3 | protein_coding | -0.052453899 | 4.939557764 | 0.0165134 | 0.8993239 | 0.982186242 |
| ENSMUSG00000025905 | Oprk1 | protein_coding | -0.05113602 | 4.134242713 | 0.0165017 | 0.8993566 | 0.982186242 |
| ENSMUSG00000015937 | H2afy | protein_coding | 0.052941301 | 4.298275487 | 0.016477 | 0.8994316 | 0.982186242 |
| ENSMUSG00000064127 | Med14 | protein_coding | -0.046898277 | 4.923458654 | 0.0163426 | 0.8998404 | 0.982335258 |
| ENSMUSG00000003402 | Prkcsh | protein_coding | 0.047524254 | 5.289178934 | 0.0163402 | 0.89985 | 0.982335258 |
| ENSMUSG00000024427 | Spry4 | protein_coding | 0.042312451 | 4.300934509 | 0.0162979 | 0.8999765 | 0.982335258 |
| ENSMUSG00000023020 | Cox14 | protein_coding | -0.030060075 | 8.847201036 | 0.0162784 | 0.9000359 | 0.982335258 |
| ENSMUSG00000024534 | Sncaip | protein_coding | 0.094313383 | 2.571521674 | 0.016271 | 0.9000585 | 0.982335258 |
| ENSMUSG00000035173 | Ccdc186 | protein_coding | -0.038343373 | 5.177513361 | 0.0162227 | 0.9002063 | 0.982335258 |
| ENSMUSG00000024847 | Aip | protein_coding | 0.024754254 | 8.863937582 | 0.0161194 | 0.9005226 | 0.982335258 |
| ENSMUSG00000041272 | Tox | protein_coding | 0.074980487 | 6.058983612 | 0.0163538 | 0.9005662 | 0.982335258 |
| ENSMUSG00000049038 | Mterf2 | protein_coding | 0.067537591 | 3.173659394 | 0.0161049 | 0.9005673 | 0.982335258 |
| ENSMUSG00000003808 | Farsa | protein_coding | -0.04009675 | 5.889888517 | 0.0160783 | 0.9006515 | 0.982335258 |
| ENSMUSG00000021198 | Unc79 | protein_coding | -0.054158466 | 3.788694174 | 0.0160438 | 0.9007549 | 0.982335258 |
| ENSMUSG00000018166 | Erbb3 | protein_coding | 0.096206649 | 2.661932717 | 0.0160128 | 0.9008503 | 0.982335258 |
| ENSMUSG00000026596 | Abl2 | protein_coding | 0.040390249 | 4.873128831 | 0.0159668 | 0.9009921 | 0.982335258 |
| ENSMUSG00000020273 | Papolg | protein_coding | 0.045561533 | 4.466994118 | 0.0159535 | 0.9010329 | 0.982335258 |
| ENSMUSG00000055681 | Cope | protein_coding | -0.022176912 | 9.279655957 | 0.0159505 | 0.9010422 | 0.982335258 |
| ENSMUSG00000036180 | Gatad2a | protein_coding | 0.034157387 | 4.89311502 | 0.0158784 | 0.9012647 | 0.982335258 |
| ENSMUSG00000063236 | 1110038F14Rik | protein_coding | 0.03313843 | 5.627800157 | 0.015877 | 0.9012692 | 0.982335258 |
| ENSMUSG00000020260 | Pofut2 | protein_coding | -0.042326217 | 5.643181593 | 0.0158308 | 0.9014147 | 0.982335258 |
| ENSMUSG00000039148 | Sart1 | protein_coding | -0.052932646 | 4.561525235 | 0.0157938 | 0.9015292 | 0.982335258 |
| ENSMUSG00000019066 | Rab3d | protein_coding | 0.116492303 | 2.520365655 | 0.0157821 | 0.9015656 | 0.982335258 |
| ENSMUSG00000024234 | Mtpap | protein_coding | -0.035825928 | 5.238940671 | 0.0157722 | 0.9015937 | 0.982335258 |
| ENSMUSG00000020776 | Fbf1 | protein_coding | 0.049302491 | 3.362885389 | 0.0157532 | 0.9016527 | 0.982335258 |
| ENSMUSG00000018378 | Cuedc1 | protein_coding | 0.047928424 | 3.943362675 | 0.0157479 | 0.9016692 | 0.982335258 |
| ENSMUSG00000087269 | D330023K18Rik | processed_transcript | 0.032414501 | 5.379798181 | 0.0157009 | 0.9018152 | 0.982393717 |
| ENSMUSG00000003135 | Cnot11 | protein_coding | -0.058875409 | 2.972213039 | 0.0156719 | 0.9019056 | 0.982393717 |
| ENSMUSG00000075289 | Carns1 | protein_coding | 0.059422723 | 3.162253939 | 0.0156137 | 0.9020867 | 0.982491484 |
| ENSMUSG00000020829 | Slc46a1 | protein_coding | 0.032172659 | 5.153076633 | 0.0154484 | 0.9026037 | 0.982876129 |
| ENSMUSG00000014294 | Ndufa2 | protein_coding | -0.03044907 | 9.642070955 | 0.0154423 | 0.9026227 | 0.982876129 |
| ENSMUSG00000036036 | Zfp57 | protein_coding | -0.125684972 | 2.74899328 | 0.015429 | 0.902816 | 0.982933126 |
| ENSMUSG00000015806 | Qdpr | protein_coding | 0.024628244 | 6.779004221 | 0.0153513 | 0.9029084 | 0.982933126 |
| ENSMUSG00000019790 | Stxbp5 | protein_coding | 0.034363104 | 5.111077448 | 0.0153383 | 0.9029493 | 0.982933126 |
| ENSMUSG00000040433 | Zbtb38 | protein_coding | -0.037265081 | 7.245201503 | 0.0152892 | 0.9032002 | 0.983106774 |
| ENSMUSG00000032724 | Abtb2 | protein_coding | 0.09270663 | 2.470819993 | 0.0152246 | 0.9033078 | 0.983124331 |
| ENSMUSG00000025505 | Tmem80 | protein_coding | 0.032615195 | 6.001325798 | 0.0151725 | 0.9034725 | 0.983204107 |
| ENSMUSG00000055725 | Paqr3 | protein_coding | -0.072764778 | 5.331199923 | 0.0152638 | 0.9036073 | 0.983251337 |
| ENSMUSG00000028668 | Eloa | protein_coding | -0.061199375 | 3.051228497 | 0.0150676 | 0.903805 | 0.983305286 |
| ENSMUSG00000020538 | Srebf1 | protein_coding | -0.051405772 | 4.174964171 | 0.0150298 | 0.903925 | 0.983305286 |
| ENSMUSG00000024617 | Camk2a | protein_coding | 0.03187746 | 7.991297627 | 0.0150286 | 0.9039313 | 0.983305286 |
| ENSMUSG00000020368 | Canx | protein_coding | -0.021343074 | 9.236388523 | 0.0149486 | 0.9041837 | 0.983480437 |
| ENSMUSG00000024668 | Sdhaf2 | protein_coding | -0.022661647 | 6.812997948 | 0.0148405 | 0.904529 | 0.983656294 |
| ENSMUSG00000014245 | Pigl | protein_coding | -0.069706116 | 2.891857036 | 0.0148172 | 0.9046035 | 0.983656294 |
| ENSMUSG00000025403 | Shmt2 | protein_coding | 0.05210945 | 4.19618175 | 0.0148121 | 0.9046198 | 0.983656294 |
| ENSMUSG00000037235 | Mxd4 | protein_coding | -0.044554749 | 4.331217165 | 0.0147373 | 0.9048597 | 0.983775855 |
| ENSMUSG00000024096 | Ralbp1 | protein_coding | 0.028201003 | 5.912335662 | 0.0147101 | 0.904947 | 0.983775855 |
| ENSMUSG00000027742 | Cog6 | protein_coding | 0.035691509 | 5.242379059 | 0.0146738 | 0.9050639 | 0.983775855 |
| ENSMUSG00000015289 | Lage3 | protein_coding | -0.033303465 | 5.787581657 | 0.0146461 | 0.9051529 | 0.983775855 |
| ENSMUSG00000020436 | Gabrg2 | protein_coding | -0.042389487 | 3.993808384 | 0.0146147 | 0.9052543 | 0.983775855 |
| ENSMUSG00000019467 | Arhgef25 | protein_coding | 0.03884777 | 4.862343665 | 0.0145889 | 0.9053374 | 0.983775855 |
| ENSMUSG00000041205 | Map6d1 | protein_coding | 0.041707962 | 4.165930521 | 0.0145787 | 0.9053702 | 0.983775855 |
| ENSMUSG00000032177 | Pde4a | protein_coding | 0.023749317 | 6.706429982 | 0.014534 | 0.9055148 | 0.983833592 |
| ENSMUSG00000062127 | Cttnbp2nl | protein_coding | 0.045429545 | 3.730858449 | 0.0144775 | 0.9056978 | 0.983892813 |
| ENSMUSG00000048222 | Mfap1b | protein_coding | 0.033853742 | 4.988416576 | 0.0144362 | 0.9058315 | 0.983892813 |
| ENSMUSG00000055782 | Abcd2 | protein_coding | -0.058220735 | 3.920591442 | 0.0144233 | 0.9058759 | 0.983892813 |
| ENSMUSG00000036885 | Arhgef26 | protein_coding | -0.058847029 | 3.349895326 | 0.014378 | 0.9060206 | 0.983892813 |
| ENSMUSG00000036534 | Slc38a7 | protein_coding | 0.063498753 | 3.731879295 | 0.0143382 | 0.9061525 | 0.983892813 |
| ENSMUSG00000059708 | Akap17b | protein_coding | 0.042605605 | 3.972768792 | 0.0143296 | 0.9061782 | 0.983892813 |
| ENSMUSG00000020121 | Srgap1 | protein_coding | -0.050213228 | 4.487032466 | 0.0143206 | 0.9062098 | 0.983892813 |
| ENSMUSG00000032199 | Polr2m | protein_coding | -0.023604294 | 6.88665138 | 0.0142829 | 0.9063303 | 0.983924241 |
| ENSMUSG00000067847 | Romo1 | protein_coding | 0.024656212 | 8.20766826 | 0.0142433 | 0.9064595 | 0.983965213 |
| ENSMUSG00000022176 | Rem2 | protein_coding | 0.101682465 | 2.818293959 | 0.0141416 | 0.9068411 | 0.983975468 |
| ENSMUSG00000006731 | B4galnt1 | protein_coding | 0.048628547 | 3.284752277 | 0.0140953 | 0.9069447 | 0.983975468 |
| ENSMUSG00000058715 | Fcer1g | protein_coding | -0.051715625 | 4.652242436 | 0.0140676 | 0.9070441 | 0.983975468 |
| ENSMUSG00000022526 | Zfp251 | protein_coding | 0.039008458 | 4.270325311 | 0.0140445 | 0.9071114 | 0.983975468 |
| ENSMUSG00000015501 | Hivep2 | protein_coding | 0.079591749 | 2.894831611 | 0.0140426 | 0.9071177 | 0.983975468 |
| ENSMUSG00000030782 | Tgfb1i1 | protein_coding | 0.04346995 | 4.468546827 | 0.0140368 | 0.9071368 | 0.983975468 |
| ENSMUSG00000021963 | Sap18 | protein_coding | 0.021806376 | 6.985810543 | 0.0140057 | 0.9072393 | 0.983975468 |
| ENSMUSG00000041577 | Prelp | protein_coding | -0.093641152 | 2.359007947 | 0.0139919 | 0.9072847 | 0.983975468 |
| ENSMUSG00000025499 | Hras | protein_coding | 0.029428682 | 6.458936108 | 0.0139895 | 0.9072925 | 0.983975468 |
| ENSMUSG00000028936 | Rpl22 | protein_coding | 0.026907722 | 7.214161766 | 0.0139017 | 0.9075827 | 0.984094729 |

| ensembl_gene_id | external_gene_name | gene_biotype | log2 (fold change) | log2 (counts per million) | F-value | p-value | False discovery rate |
|---|---|---|---|---|---|---|---|
| ENSMUSG00000004085 | Map3k20 | protein_coding | -0.074501737 | 3.11867091 | 0.0138694 | 0.9076895 | 0.984094729 |
| ENSMUSG00000026987 | Baz2b | protein_coding | -0.032252029 | 4.862876181 | 0.0137972 | 0.907929 | 0.984094729 |
| ENSMUSG00000026458 | Ppfia4 | protein_coding | 0.033793018 | 5.510509538 | 0.0137971 | 0.9079294 | 0.984094729 |
| ENSMUSG00000038497 | Tmco3 | protein_coding | -0.038508766 | 5.312085953 | 0.0137597 | 0.9080536 | 0.984094729 |
| ENSMUSG00000044148 | 1810030O07Rik | protein_coding | -0.029580552 | 5.633291775 | 0.0137209 | 0.9081826 | 0.984094729 |
| ENSMUSG00000022185 | Acin1 | protein_coding | -0.040920227 | 3.949173567 | 0.0136916 | 0.9082803 | 0.984094729 |
| ENSMUSG00000048807 | Slc35e4 | protein_coding | -0.042170309 | 4.387235814 | 0.013661 | 0.9083823 | 0.984094729 |
| ENSMUSG00000033060 | Lmo7 | protein_coding | 0.052194398 | 3.790753184 | 0.0136599 | 0.908386 | 0.984094729 |
| ENSMUSG00000045659 | Plekha7 | protein_coding | -0.061331434 | 3.350783315 | 0.0136449 | 0.908436 | 0.984094729 |
| ENSMUSG00000034575 | Papd7 | protein_coding | -0.03547338 | 4.694228269 | 0.0135596 | 0.9087213 | 0.984094729 |
| ENSMUSG00000030560 | Ctsc | protein_coding | -0.097018649 | 2.630697253 | 0.0135229 | 0.9088466 | 0.984094729 |
| ENSMUSG00000039735 | Fnbp1l | protein_coding | -0.033407624 | 4.974653609 | 0.013501 | 0.9089179 | 0.984094729 |
| ENSMUSG00000090946 | Ccdc71l | protein_coding | -0.080469144 | 2.466316339 | 0.0134816 | 0.908983 | 0.984094729 |
| ENSMUSG00000029440 | Psmd9 | protein_coding | -0.025519142 | 6.265659844 | 0.0133859 | 0.9093049 | 0.984094729 |
| ENSMUSG00000018698 | Lhx1 | protein_coding | -0.074568166 | 2.559117788 | 0.0133826 | 0.9093163 | 0.984094729 |
| ENSMUSG00000021918 | Nek4 | protein_coding | -0.042647703 | 4.061334243 | 0.0133507 | 0.9094238 | 0.984094729 |
| ENSMUSG00000058569 | Tmed9 | protein_coding | 0.035772703 | 8.118184108 | 0.0133763 | 0.9094333 | 0.984094729 |
| ENSMUSG00000010067 | Rassf1 | protein_coding | 0.03688516 | 4.843780091 | 0.013345 | 0.909443 | 0.984094729 |
| ENSMUSG00000021417 | Eci2 | protein_coding | 0.033747165 | 5.131709028 | 0.013301 | 0.9095918 | 0.984094729 |
| ENSMUSG00000023043 | Krt18 | protein_coding | -0.072949534 | 3.282090587 | 0.0132971 | 0.9096072 | 0.984094729 |
| ENSMUSG00000040040 | Ift88 | protein_coding | 0.043438981 | 4.567561074 | 0.0132207 | 0.9098639 | 0.984094729 |
| ENSMUSG00000036442 | Thap11 | protein_coding | 0.040387422 | 4.678581292 | 0.0132065 | 0.909912 | 0.984094729 |
| ENSMUSG00000073755 | 5730409E04Rik | protein_coding | -0.029075778 | 5.339607032 | 0.0131515 | 0.910099 | 0.984094729 |
| ENSMUSG00000026028 | Trak2 | protein_coding | 0.037667764 | 5.226720733 | 0.0131336 | 0.9101599 | 0.984094729 |
| ENSMUSG00000023150 | Ivns1abp | protein_coding | -0.022510492 | 6.256843816 | 0.0131277 | 0.9101799 | 0.984094729 |
| ENSMUSG00000028194 | Ddah1 | protein_coding | 0.027048069 | 7.196137723 | 0.0131155 | 0.9102214 | 0.984094729 |
| ENSMUSG00000026058 | Khdrbs2 | protein_coding | 0.03266474 | 4.980876755 | 0.0131015 | 0.9102692 | 0.984094729 |
| ENSMUSG00000005656 | Snx6 | protein_coding | -0.023523485 | 7.146636293 | 0.0131004 | 0.910273 | 0.984094729 |
| ENSMUSG00000063145 | Bbs5 | protein_coding | -0.036632835 | 4.886398385 | 0.0130955 | 0.9102897 | 0.984094729 |
| ENSMUSG00000003992 | Ssbp2 | protein_coding | -0.020707368 | 7.282676456 | 0.0130577 | 0.9104185 | 0.984094729 |
| ENSMUSG00000024456 | Diaph1 | protein_coding | -0.043752105 | 4.603002717 | 0.0130463 | 0.9104577 | 0.984094729 |
| ENSMUSG00000022672 | Prkdc | protein_coding | -0.094681333 | 2.64344309 | 0.0130342 | 0.9105011 | 0.984094729 |
| ENSMUSG00000051355 | Commd1 | protein_coding | -0.042207274 | 3.530138769 | 0.0129781 | 0.9106909 | 0.984094729 |
| ENSMUSG00000019235 | Rps6kl1 | protein_coding | -0.031203614 | 5.743658113 | 0.0129696 | 0.91072 | 0.984094729 |
| ENSMUSG00000025478 | Dpysl4 | protein_coding | 0.037382399 | 5.052584115 | 0.0129549 | 0.9107704 | 0.984094729 |
| ENSMUSG00000032087 | Dscaml1 | protein_coding | -0.064755311 | 3.56028053 | 0.0129502 | 0.9107887 | 0.984094729 |
| ENSMUSG00000054051 | Ercc6 | protein_coding | 0.079703556 | 2.562444111 | 0.012869 | 0.9110652 | 0.984203568 |
| ENSMUSG00000050315 | Synpo2 | protein_coding | 0.061810108 | 4.186734964 | 0.0128926 | 0.9110725 | 0.984203568 |
| ENSMUSG00000059498 | Fcgr3 | protein_coding | 0.056334688 | 4.39498967 | 0.0128618 | 0.9111664 | 0.984206162 |
| ENSMUSG00000037993 | Dhx38 | protein_coding | 0.046229838 | 4.389647886 | 0.0127479 | 0.9114828 | 0.984305913 |
| ENSMUSG00000113029 | AC122190.1 | antisense_RNA | -0.056752981 | 3.301354643 | 0.0127227 | 0.9115699 | 0.984305913 |
| ENSMUSG00000037349 | Nudt22 | protein_coding | 0.03246274 | 4.836511979 | 0.0126744 | 0.9117373 | 0.984305913 |
| ENSMUSG00000023909 | Paqr4 | protein_coding | -0.04166959 | 4.688858335 | 0.0126426 | 0.9118474 | 0.984305913 |
| ENSMUSG00000027340 | Slc23a2 | protein_coding | -0.034197586 | 5.141563334 | 0.0126295 | 0.9118929 | 0.984305913 |
| ENSMUSG00000034958 | Atcay | protein_coding | -0.034268735 | 6.108660041 | 0.0125924 | 0.9120242 | 0.984305913 |
| ENSMUSG00000060376 | Bckdha | protein_coding | 0.054444557 | 3.830387795 | 0.012592 | 0.9120257 | 0.984305913 |
| ENSMUSG00000021140 | Pcnx | protein_coding | 0.052864141 | 3.71304744 | 0.0125699 | 0.9121001 | 0.984305913 |
| ENSMUSG00000029864 | Gstk1 | protein_coding | -0.056805421 | 4.460523854 | 0.0125751 | 0.9121693 | 0.984305913 |
| ENSMUSG00000052949 | Rnf157 | protein_coding | 0.021204383 | 6.718553286 | 0.0125087 | 0.9123135 | 0.984305913 |
| ENSMUSG00000055493 | Epm2a | protein_coding | 0.03914155 | 4.133105188 | 0.0124683 | 0.9124546 | 0.984305913 |
| ENSMUSG00000053819 | Camk2d | protein_coding | 0.021797817 | 7.235924366 | 0.0124481 | 0.9125254 | 0.984305913 |
| ENSMUSG00000035007 | Rundc1 | protein_coding | 0.046288742 | 4.464754984 | 0.0124406 | 0.9125537 | 0.984305913 |
| ENSMUSG00000006412 | Pfdn2 | protein_coding | 0.023618924 | 8.392453779 | 0.0124391 | 0.9125569 | 0.984305913 |
| ENSMUSG00000023913 | Pla2g7 | protein_coding | 0.025103807 | 7.018557148 | 0.0124143 | 0.9126437 | 0.984305913 |
| ENSMUSG00000071661 | Zbtb3 | protein_coding | -0.057281484 | 3.124408689 | 0.0123915 | 0.9127234 | 0.984305913 |
| ENSMUSG00000031578 | Mak16 | protein_coding | 0.034402943 | 4.675581515 | 0.0123554 | 0.9128502 | 0.98434403 |
| ENSMUSG00000042675 | Ypel3 | protein_coding | 0.027615797 | 6.043903776 | 0.0121836 | 0.9134556 | 0.984486284 |
| ENSMUSG00000029310 | Nudt9 | protein_coding | -0.029561262 | 5.61468914 | 0.0121655 | 0.9135194 | 0.984486284 |
| ENSMUSG00000029401 | Rilpl2 | protein_coding | 0.042119643 | 4.513481641 | 0.0121654 | 0.91352 | 0.984486284 |
| ENSMUSG00000037104 | Socs5 | protein_coding | -0.025705512 | 5.970340127 | 0.0121007 | 0.9137493 | 0.984486284 |
| ENSMUSG00000022656 | Nectin3 | protein_coding | 0.064686656 | 3.045823265 | 0.0120692 | 0.913861 | 0.984486284 |
| ENSMUSG00000036299 | BC031181 | protein_coding | -0.020746693 | 9.008096586 | 0.0120647 | 0.913877 | 0.984486284 |
| ENSMUSG00000034744 | Nagk | protein_coding | -0.022755273 | 7.326640144 | 0.0120384 | 0.9139704 | 0.984486284 |
| ENSMUSG00000022967 | Ifnar1 | protein_coding | -0.04138117 | 3.699766087 | 0.0120336 | 0.9139876 | 0.984486284 |
| ENSMUSG00000000441 | Raf1 | protein_coding | 0.06643287 | 2.610805292 | 0.0120317 | 0.9139943 | 0.984486284 |
| ENSMUSG00000071451 | Psmg4 | protein_coding | 0.031218601 | 5.430961972 | 0.0120185 | 0.9140416 | 0.984486284 |
| ENSMUSG00000003208 | Ccdc94 | protein_coding | -0.052799173 | 3.964551078 | 0.0119681 | 0.9142233 | 0.984486284 |
| ENSMUSG00000024097 | Srsf7 | protein_coding | -0.022887922 | 8.150589319 | 0.0119483 | 0.914292 | 0.984486284 |
| ENSMUSG00000006699 | Cdc42 | protein_coding | 0.019095318 | 9.861276353 | 0.0119329 | 0.914347 | 0.984486284 |
| ENSMUSG00000039246 | Lyplal1 | protein_coding | -0.034098752 | 4.955391354 | 0.0118921 | 0.9144927 | 0.984486284 |
| ENSMUSG00000027952 | Pmvk | protein_coding | 0.020317502 | 8.011103667 | 0.0118868 | 0.9145118 | 0.984486284 |
| ENSMUSG00000018770 | Atp5g3 | protein_coding | 0.021323668 | 10.45908229 | 0.0118732 | 0.9145603 | 0.984486284 |
| ENSMUSG00000043683 | Fem1a | protein_coding | -0.035122436 | 4.919768379 | 0.0118542 | 0.9146284 | 0.984486284 |
| ENSMUSG00000022946 | Dopey2 | protein_coding | 0.03934339 | 4.038381557 | 0.0118537 | 0.9146301 | 0.984486284 |

| ensembl_gene_id | external_gene_name | gene_biotype | log2 (fold change) | log2 (counts per million) | F-value | p-value | False discovery rate |
|---|---|---|---|---|---|---|---|
| ENSMUSG00000022710 | Usp7 | protein_coding | -0.025427852 | 5.755904599 | 0.0117679 | 0.9149386 | 0.984719711 |
| ENSMUSG00000028525 | Pde4b | protein_coding | 0.023559242 | 7.408735093 | 0.0117232 | 0.9150995 | 0.98473107 |
| ENSMUSG00000038611 | Phrf1 | protein_coding | 0.036576454 | 4.427450614 | 0.0117141 | 0.9151323 | 0.98473107 |
| ENSMUSG00000048170 | Mcmbp | protein_coding | -0.029821184 | 5.869126509 | 0.0116707 | 0.915289 | 0.984736716 |
| ENSMUSG00000029050 | Ski | protein_coding | 0.035470943 | 5.481192407 | 0.0116552 | 0.915345 | 0.984736716 |
| ENSMUSG00000006095 | Tbcb | protein_coding | -0.019364873 | 9.009886997 | 0.0116263 | 0.9154498 | 0.984736716 |
| ENSMUSG00000017670 | Elmo2 | protein_coding | -0.028014222 | 6.908676478 | 0.0115488 | 0.9157331 | 0.984736716 |
| ENSMUSG00000062397 | Zfp706 | protein_coding | -0.021636467 | 8.195450523 | 0.0115359 | 0.9157777 | 0.984736716 |
| ENSMUSG00000040738 | Ints8 | protein_coding | -0.045425168 | 3.724447073 | 0.0114883 | 0.9159509 | 0.984736716 |
| ENSMUSG00000001847 | Rac1 | protein_coding | -0.021765483 | 6.536388108 | 0.0114794 | 0.9159834 | 0.984736716 |
| ENSMUSG00000028524 | Sgip1 | protein_coding | -0.031205577 | 4.884116224 | 0.0114682 | 0.9160242 | 0.984736716 |
| ENSMUSG00000003824 | Syce2 | protein_coding | -0.059373208 | 3.052079266 | 0.0114191 | 0.9162033 | 0.984736716 |
| ENSMUSG00000010110 | Stx5a | protein_coding | -0.022960377 | 6.442202614 | 0.0113966 | 0.9162858 | 0.984736716 |
| ENSMUSG00000040028 | Elavl1 | protein_coding | 0.026395465 | 5.450016364 | 0.0113918 | 0.9163034 | 0.984736716 |
| ENSMUSG00000015745 | Plekho1 | protein_coding | 0.056616227 | 3.719621721 | 0.0113895 | 0.916314 | 0.984736716 |
| ENSMUSG00000070730 | Rmdn3 | protein_coding | 0.038088485 | 4.036268146 | 0.0113714 | 0.9163778 | 0.984736716 |
| ENSMUSG00000026565 | Pou2f1 | protein_coding | -0.057442366 | 3.145914919 | 0.0113478 | 0.9164646 | 0.984736716 |
| ENSMUSG00000051510 | Mafg | protein_coding | 0.025853826 | 5.839539025 | 0.0113041 | 0.916625 | 0.984736716 |
| ENSMUSG00000026610 | Esrrg | protein_coding | -0.035812576 | 4.177014255 | 0.0112819 | 0.9167065 | 0.984736716 |
| ENSMUSG00000031634 | Ufsp2 | protein_coding | -0.019008512 | 7.256195566 | 0.0112274 | 0.9169069 | 0.984736716 |
| ENSMUSG00000056429 | Tgoln1 | protein_coding | 0.020326803 | 6.553580373 | 0.0112176 | 0.9169432 | 0.984736716 |
| ENSMUSG00000066149 | Cdc26 | protein_coding | -0.026700806 | 5.822246247 | 0.0111899 | 0.9170452 | 0.984736716 |
| ENSMUSG00000030518 | Fam189a1 | protein_coding | 0.052187659 | 4.028151011 | 0.0111887 | 0.9170521 | 0.984736716 |
| ENSMUSG00000022508 | Bcl6 | protein_coding | -0.058495974 | 3.048045473 | 0.0111736 | 0.9171055 | 0.984736716 |
| ENSMUSG00000021891 | Mettl6 | protein_coding | -0.029529096 | 5.307868571 | 0.0111483 | 0.9171992 | 0.984736716 |
| ENSMUSG00000020646 | Mboat2 | protein_coding | 0.032170216 | 4.425745491 | 0.0111114 | 0.9173356 | 0.984736716 |
| ENSMUSG00000018547 | Pip4k2b | protein_coding | 0.038215416 | 4.264914796 | 0.0111078 | 0.9173491 | 0.984736716 |
| ENSMUSG00000027188 | Pamr1 | protein_coding | 0.064354117 | 2.718552927 | 0.0110868 | 0.917427 | 0.984736716 |
| ENSMUSG00000027198 | Ext2 | protein_coding | 0.044329209 | 3.945682192 | 0.011045 | 0.9175819 | 0.984804739 |
| ENSMUSG00000027397 | Slc20a1 | protein_coding | -0.057019009 | 2.862421355 | 0.0108824 | 0.9181887 | 0.98500499 |
| ENSMUSG00000010609 | Psen2 | protein_coding | 0.033251906 | 4.709192614 | 0.0108674 | 0.9182449 | 0.98500499 |
| ENSMUSG00000026220 | Slc16a14 | protein_coding | -0.086173298 | 3.227649234 | 0.0108319 | 0.9185428 | 0.98500499 |
| ENSMUSG00000024442 | 0610009O20Rik | protein_coding | -0.023413243 | 5.770186713 | 0.010764 | 0.9186331 | 0.98500499 |
| ENSMUSG00000033249 | Hsf4 | protein_coding | -0.061180771 | 2.823140125 | 0.0107599 | 0.9186487 | 0.98500499 |
| ENSMUSG00000032263 | Bckdhb | protein_coding | -0.037523379 | 3.918022141 | 0.0107584 | 0.9186542 | 0.98500499 |
| ENSMUSG00000032870 | Smap2 | protein_coding | 0.017629592 | 7.112393624 | 0.0107507 | 0.9186834 | 0.98500499 |
| ENSMUSG00000078451 | Ppil6 | protein_coding | 0.058024881 | 2.86356817 | 0.0107252 | 0.9187795 | 0.98500499 |
| ENSMUSG00000037787 | Apopt1 | protein_coding | -0.019032719 | 6.798427294 | 0.01072 | 0.9187989 | 0.98500499 |
| ENSMUSG00000053226 | Dand5 | protein_coding | 0.031435785 | 4.903094634 | 0.0106955 | 0.9188915 | 0.98500499 |
| ENSMUSG00000068328 | Aup1 | protein_coding | 0.040052355 | 6.236428025 | 0.0106898 | 0.9190765 | 0.98500499 |
| ENSMUSG00000038486 | Sv2a | protein_coding | -0.018901258 | 8.389974562 | 0.0106311 | 0.9191353 | 0.98500499 |
| ENSMUSG00000016206 | H2-M3 | protein_coding | 0.058554743 | 3.008335465 | 0.0105931 | 0.9192795 | 0.98500499 |
| ENSMUSG00000022890 | Atp5j | protein_coding | -0.022401014 | 10.19202003 | 0.0105836 | 0.9193154 | 0.98500499 |
| ENSMUSG00000075318 | Scn2a | protein_coding | -0.019084624 | 6.959248327 | 0.0105662 | 0.9193813 | 0.98500499 |
| ENSMUSG00000044033 | Ccdc141 | protein_coding | 0.059214293 | 2.953889257 | 0.0105555 | 0.9194222 | 0.98500499 |
| ENSMUSG00000036918 | Ttc7 | protein_coding | -0.079752602 | 2.377443965 | 0.0105378 | 0.9194896 | 0.98500499 |
| ENSMUSG00000052712 | BC004004 | protein_coding | 0.018573552 | 6.97331961 | 0.010502 | 0.9196258 | 0.98500499 |
| ENSMUSG00000039067 | Psmd7 | protein_coding | -0.017968898 | 8.601291436 | 0.010492 | 0.9196639 | 0.98500499 |
| ENSMUSG00000040396 | Abhd13 | protein_coding | 0.039604861 | 3.6968648 | 0.0104696 | 0.9197494 | 0.98500499 |
| ENSMUSG00000022199 | Slc22a17 | protein_coding | -0.021417632 | 8.546739703 | 0.010465 | 0.919767 | 0.98500499 |
| ENSMUSG00000032301 | Psma4 | protein_coding | 0.021815053 | 7.577675327 | 0.0104606 | 0.9197838 | 0.98500499 |
| ENSMUSG00000022450 | Ndufa6 | protein_coding | -0.022747459 | 10.01225996 | 0.0104145 | 0.9199602 | 0.985095764 |
| ENSMUSG00000007783 | Cpt1c | protein_coding | -0.024127248 | 6.709707017 | 0.0103517 | 0.9202008 | 0.985182015 |
| ENSMUSG00000025239 | Limd1 | protein_coding | -0.037546477 | 4.056360226 | 0.0103229 | 0.9203118 | 0.985182015 |
| ENSMUSG00000004535 | Tax1bp1 | protein_coding | 0.017583623 | 7.530073695 | 0.0102912 | 0.9204336 | 0.985182015 |
| ENSMUSG00000028619 | Tceanc2 | protein_coding | 0.042257395 | 3.57439876 | 0.0102689 | 0.9205196 | 0.985182015 |
| ENSMUSG00000024502 | Jakmip2 | protein_coding | -0.02234562 | 6.198559828 | 0.0102636 | 0.9205401 | 0.985182015 |
| ENSMUSG00000043639 | Rbm20 | protein_coding | -0.068617452 | 2.429036329 | 0.0102505 | 0.9205904 | 0.985182015 |
| ENSMUSG00000028383 | Hsdl2 | protein_coding | 0.023762522 | 5.550690746 | 0.0101959 | 0.9208017 | 0.985310083 |
| ENSMUSG00000032298 | Neil1 | protein_coding | -0.069851778 | 3.125484663 | 0.0101702 | 0.9209032 | 0.985320616 |
| ENSMUSG00000054679 | Srsf12 | protein_coding | 0.0369694 | 5.326645041 | 0.0101444 | 0.921003 | 0.985329303 |
| ENSMUSG00000026853 | Crat | protein_coding | -0.03316219 | 4.648621722 | 0.0100685 | 0.9212963 | 0.985444782 |
| ENSMUSG00000058966 | Fam57b | protein_coding | -0.04738181 | 4.642708723 | 0.0100777 | 0.9213228 | 0.985444782 |
| ENSMUSG00000022151 | Ttc33 | protein_coding | -0.029020697 | 4.688553336 | 0.010044 | 0.9213918 | 0.985444782 |
| ENSMUSG00000030604 | Zfp626 | protein_coding | -0.061872483 | 2.699631647 | 0.0099899 | 0.921603 | 0.985444782 |
| ENSMUSG00000020453 | Patz1 | protein_coding | 0.048445971 | 3.463945877 | 0.0099725 | 0.9216711 | 0.985444782 |
| ENSMUSG00000017868 | Sgk2 | protein_coding | 0.062538304 | 3.413828535 | 0.0099578 | 0.9217393 | 0.985444782 |
| ENSMUSG00000027330 | Cdc25b | protein_coding | 0.05451252 | 3.054587964 | 0.0099482 | 0.921766 | 0.985444782 |
| ENSMUSG00000087006 | Gm13889 | protein_coding | -0.041314784 | 3.586865845 | 0.0099283 | 0.921844 | 0.985444782 |
| ENSMUSG00000034343 | Ube2f | protein_coding | 0.020929801 | 6.296843582 | 0.0098909 | 0.921991 | 0.985469737 |
| ENSMUSG00000038482 | Tfdp1 | protein_coding | -0.033470978 | 4.656460446 | 0.0098322 | 0.9222219 | 0.985469737 |
| ENSMUSG00000097383 | 1500026H17Rik | bidirectional_promoter | 0.034276511 | 3.728712492 | 0.0098051 | 0.9223289 | 0.985469737 |
| ENSMUSG00000032109 | Nlrx1 | protein_coding | 0.094985517 | 2.261595834 | 0.0097997 | 0.9223503 | 0.985469737 |
| ENSMUSG00000053475 | Tnfaip6 | protein_coding | -0.029814509 | 5.348703418 | 0.0097788 | 0.9224325 | 0.985469737 |

| ensembl_gene_id | external_gene_name | gene_biotype | log2 (fold change) | log2 (counts per million) | F-value | p-value | False discovery rate |
|---|---|---|---|---|---|---|---|
| ENSMUSG00000039375 | Wdr17 | protein_coding | 0.035409085 | 4.796422682 | 0.0097681 | 0.922475 | 0.985469737 |
| ENSMUSG00000008429 | Herpud2 | protein_coding | 0.03900429 | 4.95469928 | 0.00976 | 0.9225089 | 0.985469737 |
| ENSMUSG00000024829 | Mrpl21 | protein_coding | -0.018011853 | 7.109861112 | 0.0096595 | 0.9229055 | 0.985613416 |
| ENSMUSG00000037119 | Fam91a1 | protein_coding | 0.031122978 | 4.981655417 | 0.0096542 | 0.9229268 | 0.985613416 |
| ENSMUSG00000030275 | Etnk1 | protein_coding | -0.072399384 | 2.811776747 | 0.009654 | 0.9229292 | 0.985613416 |
| ENSMUSG00000042272 | Sestd1 | protein_coding | -0.044216973 | 3.482998893 | 0.0096333 | 0.9230101 | 0.985613416 |
| ENSMUSG00000028173 | Wls | protein_coding | 0.030757959 | 4.860362235 | 0.0095984 | 0.923149 | 0.985663893 |
| ENSMUSG00000016024 | Lbp | protein_coding | 0.07662549 | 2.970601539 | 0.0095718 | 0.9232924 | 0.985699554 |
| ENSMUSG00000040407 | Akap9 | protein_coding | -0.024247063 | 5.75019992 | 0.0095145 | 0.9234846 | 0.985699554 |
| ENSMUSG00000031347 | Cetn2 | protein_coding | 0.022112323 | 7.865143906 | 0.0094199 | 0.9238646 | 0.985699554 |
| ENSMUSG00000004460 | Dnajb11 | protein_coding | -0.029204016 | 4.93697652 | 0.0094176 | 0.9238738 | 0.985699554 |
| ENSMUSG00000031386 | Hcfc1 | protein_coding | -0.035237618 | 5.197956462 | 0.0094047 | 0.923928 | 0.985699554 |
| ENSMUSG00000045671 | Spred2 | protein_coding | 0.038189938 | 4.025440524 | 0.0093841 | 0.924009 | 0.985699554 |
| ENSMUSG00000035158 | Mitf | protein_coding | 0.068926798 | 2.723060343 | 0.0093777 | 0.9240347 | 0.985699554 |
| ENSMUSG00000055239 | Kcmf1 | protein_coding | 0.026985141 | 5.772593006 | 0.0093674 | 0.9240763 | 0.985699554 |
| ENSMUSG00000041617 | Ccdc74a | protein_coding | -0.044030735 | 3.585730457 | 0.009365 | 0.9240859 | 0.985699554 |
| ENSMUSG00000021911 | Parg | protein_coding | -0.032448301 | 4.457585054 | 0.0093618 | 0.9240991 | 0.985699554 |
| ENSMUSG00000018459 | Slc13a3 | protein_coding | 0.07544617 | 3.046341568 | 0.0093347 | 0.9242622 | 0.9857625 |
| ENSMUSG00000015943 | Bola1 | protein_coding | -0.018682346 | 6.70976838 | 0.0092682 | 0.9244781 | 0.9857625 |
| ENSMUSG00000026988 | Wdsub1 | protein_coding | -0.039641777 | 3.950590657 | 0.0092652 | 0.9244902 | 0.9857625 |
| ENSMUSG00000052488 | Cherp | protein_coding | -0.055475508 | 4.63832913 | 0.0092973 | 0.9245248 | 0.9857625 |
| ENSMUSG00000029993 | Nfu1 | protein_coding | 0.017670452 | 8.380902641 | 0.0092315 | 0.9246271 | 0.985773914 |
| ENSMUSG00000025241 | Fyco1 | protein_coding | 0.050156492 | 3.763028103 | 0.0091041 | 0.9251494 | 0.986232968 |
| ENSMUSG00000040524 | Zfp609 | protein_coding | 0.038569152 | 3.534331021 | 0.0090714 | 0.9252815 | 0.986275965 |
| ENSMUSG00000067199 | Frat1 | protein_coding | -0.028727282 | 7.740331066 | 0.0089947 | 0.9256708 | 0.986593185 |
| ENSMUSG00000019731 | Slc35e1 | protein_coding | -0.039792158 | 4.019405831 | 0.0089392 | 0.9258264 | 0.986661198 |
| ENSMUSG00000074060 | Fbxw15 | protein_coding | 0.043865866 | 3.577313877 | 0.0088868 | 0.9260435 | 0.986789392 |
| ENSMUSG00000030303 | Far2 | protein_coding | 0.047819665 | 4.74885854 | 0.0088876 | 0.9261549 | 0.986789392 |
| ENSMUSG00000096199 | Ptrhd1 | protein_coding | 0.02591734 | 5.25331376 | 0.0088438 | 0.926222 | 0.986789392 |
| ENSMUSG00000033411 | Ctdspl2 | protein_coding | 0.033764173 | 4.635409623 | 0.0088133 | 0.9263489 | 0.98682683 |
| ENSMUSG00000015980 | Lrrc27 | protein_coding | 0.037113693 | 4.809992088 | 0.0087714 | 0.9265254 | 0.986917003 |
| ENSMUSG00000002416 | Ndufb2 | protein_coding | 0.019605714 | 6.208844997 | 0.0087317 | 0.9266898 | 0.986994375 |
| ENSMUSG00000045410 | Akr1e1 | protein_coding | 0.019056713 | 6.841923166 | 0.0086877 | 0.9268738 | 0.987034505 |
| ENSMUSG00000032081 | Apoc3 | protein_coding | 0.049063237 | 3.179063449 | 0.0086018 | 0.9272352 | 0.987034505 |
| ENSMUSG00000041164 | Zmiz2 | protein_coding | 0.020671826 | 6.669697262 | 0.0085906 | 0.9272825 | 0.987034505 |
| ENSMUSG00000024190 | Dusp1 | protein_coding | 0.032073126 | 4.473585659 | 0.0085825 | 0.9273166 | 0.987034505 |
| ENSMUSG00000030741 | Spns1 | protein_coding | -0.026446453 | 5.835921161 | 0.0085219 | 0.9275732 | 0.987034505 |
| ENSMUSG00000021745 | Ptprg | protein_coding | -0.038239823 | 4.195560502 | 0.0085021 | 0.9276568 | 0.987034505 |
| ENSMUSG00000026918 | Brd3 | protein_coding | 0.019140915 | 6.624802106 | 0.0084942 | 0.9276906 | 0.987034505 |
| ENSMUSG00000039670 | Oxld1 | protein_coding | 0.061744279 | 2.546771813 | 0.0084676 | 0.9278034 | 0.987034505 |
| ENSMUSG00000026977 | 7-Mar | protein_coding | -0.023256515 | 5.295410902 | 0.0084667 | 0.9278071 | 0.987034505 |
| ENSMUSG00000034201 | Gas2l1 | protein_coding | 0.048727268 | 3.767728582 | 0.0084251 | 0.9279862 | 0.987034505 |
| ENSMUSG00000087618 | Gm14114 | lincRNA | -0.059145146 | 2.454754012 | 0.0084067 | 0.9280629 | 0.987034505 |
| ENSMUSG00000009070 | Rsph14 | protein_coding | 0.046603437 | 3.214988781 | 0.0083541 | 0.9282877 | 0.987034505 |
| ENSMUSG00000036622 | Atp13a2 | protein_coding | 0.030933701 | 5.209432498 | 0.0083509 | 0.9283012 | 0.987034505 |
| ENSMUSG00000037455 | Slc18b1 | protein_coding | 0.044633372 | 3.00630673 | 0.0083407 | 0.9283447 | 0.987034505 |
| ENSMUSG00000032637 | Atxn2l | protein_coding | -0.043049375 | 3.237501483 | 0.0082954 | 0.928539 | 0.987034505 |
| ENSMUSG00000063511 | Snrnp70 | protein_coding | -0.026958614 | 4.871768543 | 0.0082887 | 0.9285679 | 0.987034505 |
| ENSMUSG00000043384 | Gprasp1 | protein_coding | -0.051554487 | 3.13095495 | 0.0082862 | 0.9285785 | 0.987034505 |
| ENSMUSG00000040490 | Lrfn2 | protein_coding | -0.045016802 | 2.951923989 | 0.0082791 | 0.9286091 | 0.987034505 |
| ENSMUSG00000061207 | Stk19 | protein_coding | -0.01913151 | 6.162909754 | 0.0082743 | 0.9286298 | 0.987034505 |
| ENSMUSG00000032411 | Tfdp2 | protein_coding | -0.031913398 | 4.105395577 | 0.0082636 | 0.928676 | 0.987034505 |
| ENSMUSG00000006717 | Acot13 | protein_coding | 0.018573822 | 8.428009377 | 0.0082475 | 0.928745 | 0.987034505 |
| ENSMUSG00000040390 | Map3k10 | protein_coding | -0.061771221 | 3.157233329 | 0.0082434 | 0.9287648 | 0.987034505 |
| ENSMUSG00000021991 | Cacna2d3 | protein_coding | 0.03145117 | 5.559323087 | 0.0081932 | 0.9289812 | 0.987034505 |
| ENSMUSG00000021902 | Phf7 | protein_coding | 0.086347011 | 2.481842382 | 0.0081932 | 0.9289841 | 0.987034505 |
| ENSMUSG00000005583 | Mef2c | protein_coding | 0.023213992 | 6.856169045 | 0.0081837 | 0.9290222 | 0.987034505 |
| ENSMUSG00000021287 | Xrcc3 | protein_coding | -0.053925214 | 2.800430518 | 0.0081543 | 0.9291478 | 0.987070411 |
| ENSMUSG00000021852 | Slc35f4 | protein_coding | 0.037424948 | 3.69587323 | 0.0080789 | 0.929475 | 0.987235016 |
| ENSMUSG00000038403 | Hfe2 | protein_coding | -0.055497545 | 2.767876911 | 0.0080689 | 0.9295188 | 0.987235016 |
| ENSMUSG00000020064 | Herc4 | protein_coding | 0.038804834 | 3.692384802 | 0.0080552 | 0.9295782 | 0.987235016 |
| ENSMUSG00000055762 | Eef1d | protein_coding | 0.018241035 | 6.868658105 | 0.0080099 | 0.9297763 | 0.987323027 |
| ENSMUSG00000034211 | Mrps17 | protein_coding | -0.017331743 | 7.895572155 | 0.0079731 | 0.9299369 | 0.987323027 |
| ENSMUSG00000060657 | Marf1 | protein_coding | -0.051137926 | 7.253309898 | 0.0081149 | 0.9299634 | 0.987323027 |
| ENSMUSG00000033918 | Parl | protein_coding | -0.022333877 | 6.333580154 | 0.0078942 | 0.9302838 | 0.987323027 |
| ENSMUSG00000063354 | Slc39a4 | protein_coding | -0.169546944 | 1.36567172 | 0.0079097 | 0.9304463 | 0.987323027 |
| ENSMUSG00000037905 | Bri3bp | protein_coding | -0.020997585 | 5.625456159 | 0.0078472 | 0.9304909 | 0.987323027 |
| ENSMUSG00000053519 | Kcnip1 | protein_coding | -0.019883749 | 6.198908436 | 0.0078439 | 0.9305055 | 0.987323027 |
| ENSMUSG00000038615 | Nfe2l1 | protein_coding | 0.016273442 | 6.872197934 | 0.0078301 | 0.9305664 | 0.987323027 |
| ENSMUSG00000059602 | Syn3 | protein_coding | 0.041123577 | 3.615109331 | 0.007808 | 0.9306643 | 0.987323027 |
| ENSMUSG00000053436 | Mapk14 | protein_coding | 0.026225767 | 5.388833924 | 0.0077905 | 0.930742 | 0.987323027 |
| ENSMUSG00000041073 | Nacad | protein_coding | 0.027757433 | 4.855918819 | 0.0077537 | 0.9309052 | 0.987323027 |
| ENSMUSG00000077450 | Rab11b | protein_coding | 0.019404702 | 7.827942109 | 0.0077475 | 0.9309327 | 0.987323027 |
| ENSMUSG00000030660 | Pik3c2a | protein_coding | 0.053696311 | 2.710355144 | 0.0077444 | 0.9309465 | 0.987323027 |

| ensembl_gene_id | external_gene_name | gene_biotype | log2 (fold change) | log2 (counts per million) | F-value | p-value | False discovery rate |
|---|---|---|---|---|---|---|---|
| ENSMUSG00000025981 | Coq10b | protein_coding | 0.02600872 | 5.300111345 | 0.0077294 | 0.931013 | 0.987323027 |
| ENSMUSG00000074794 | Arrdc3 | protein_coding | 0.021879551 | 5.529296633 | 0.0077238 | 0.9310384 | 0.987323027 |
| ENSMUSG00000025050 | Pcgf6 | protein_coding | 0.059653715 | 2.536865775 | 0.0076714 | 0.9312716 | 0.987378004 |
| ENSMUSG00000029695 | Aass | protein_coding | 0.08728103 | 1.638383169 | 0.0076444 | 0.9313924 | 0.987378004 |
| ENSMUSG00000021024 | Psma6 | protein_coding | 0.017106423 | 9.087785455 | 0.0076213 | 0.9314962 | 0.987378004 |
| ENSMUSG00000028618 | Tmem59 | protein_coding | 0.01625501 | 9.205034219 | 0.0076196 | 0.9315039 | 0.987378004 |
| ENSMUSG00000021973 | Micu2 | protein_coding | 0.031383712 | 4.362806374 | 0.0076094 | 0.9315493 | 0.987378004 |
| ENSMUSG00000022443 | Myh9 | protein_coding | -0.035512512 | 3.503120734 | 0.007515 | 0.9319741 | 0.987459232 |
| ENSMUSG00000021040 | Slirp | protein_coding | -0.028632164 | 4.494074905 | 0.0075013 | 0.9320362 | 0.987459232 |
| ENSMUSG00000063049 | Ing2 | protein_coding | 0.033478288 | 4.181450923 | 0.0075007 | 0.9320389 | 0.987459232 |
| ENSMUSG00000039954 | Stk32a | protein_coding | -0.058461311 | 3.783346815 | 0.0075114 | 0.9321271 | 0.987459232 |
| ENSMUSG00000068205 | Macrod2 | protein_coding | 0.025793713 | 5.345425192 | 0.0074711 | 0.9321725 | 0.987459232 |
| ENSMUSG00000020525 | Ppm1d | protein_coding | 0.041250638 | 3.356789693 | 0.0074429 | 0.9323002 | 0.987459232 |
| ENSMUSG00000049106 | Dcaf5 | protein_coding | -0.0203688 | 5.500459631 | 0.0074284 | 0.9323661 | 0.987459232 |
| ENSMUSG00000055296 | Tmem245 | protein_coding | 0.03208307 | 4.413912717 | 0.0073944 | 0.9325209 | 0.987459232 |
| ENSMUSG00000021782 | Dlg5 | protein_coding | -0.035829391 | 3.431185895 | 0.007391 | 0.9325364 | 0.987459232 |
| ENSMUSG00000028274 | Rngtt | protein_coding | 0.028596378 | 5.010765698 | 0.0073754 | 0.9326072 | 0.987459232 |
| ENSMUSG00000033430 | Terf2ip | protein_coding | -0.023277324 | 5.577778088 | 0.0073668 | 0.9326464 | 0.987459232 |
| ENSMUSG00000051029 | Serpinb1b | protein_coding | 0.027064626 | 4.661446212 | 0.0073489 | 0.9327279 | 0.987459232 |
| ENSMUSG00000041992 | Rapgef5 | protein_coding | 0.023351517 | 5.380962526 | 0.0072212 | 0.9333138 | 0.987889088 |
| ENSMUSG00000029705 | Cux1 | protein_coding | 0.020645497 | 6.038084386 | 0.0072162 | 0.9333369 | 0.987889088 |
| ENSMUSG00000033007 | Asic4 | protein_coding | -0.080169878 | 2.312963814 | 0.0071963 | 0.93343 | 0.987889088 |
| ENSMUSG00000032036 | Kirrel3 | protein_coding | 0.023947902 | 5.479960075 | 0.0071805 | 0.9335014 | 0.987889088 |
| ENSMUSG00000002578 | Ikzf4 | protein_coding | 0.029453242 | 4.382670822 | 0.0071571 | 0.9336096 | 0.987906355 |
| ENSMUSG00000032220 | Myo1e | protein_coding | 0.048243815 | 2.891451718 | 0.0070922 | 0.9339106 | 0.988094012 |
| ENSMUSG00000032519 | Slc25a38 | protein_coding | 0.025259128 | 4.960370605 | 0.0070792 | 0.9339708 | 0.988094012 |
| ENSMUSG00000025781 | Atp5c1 | protein_coding | 0.01554053 | 10.3395293 | 0.0070454 | 0.9341284 | 0.988163548 |
| ENSMUSG00000019877 | Serinc1 | protein_coding | 0.014739598 | 9.253247427 | 0.0069877 | 0.9343978 | 0.988182514 |
| ENSMUSG00000073676 | Hspe1 | protein_coding | -0.017463196 | 7.452301563 | 0.0069826 | 0.9344219 | 0.988182514 |
| ENSMUSG00000048546 | Tob2 | protein_coding | -0.023024207 | 4.887642111 | 0.0069674 | 0.9344932 | 0.988182514 |
| ENSMUSG00000037957 | Wdr20 | protein_coding | -0.030162706 | 4.071077667 | 0.0069391 | 0.9346262 | 0.988182514 |
| ENSMUSG00000060402 | Chst8 | protein_coding | -0.046590063 | 3.486511991 | 0.0068871 | 0.9348726 | 0.988182514 |
| ENSMUSG00000004667 | Polr2e | protein_coding | 0.015456909 | 8.368268103 | 0.006874 | 0.9349328 | 0.988182514 |
| ENSMUSG00000025204 | Ndufb8 | protein_coding | -0.017775013 | 9.711001295 | 0.0068595 | 0.9350012 | 0.988182514 |
| ENSMUSG00000042462 | Dctpp1 | protein_coding | 0.024840456 | 5.573848654 | 0.0068439 | 0.9350749 | 0.988182514 |
| ENSMUSG00000014313 | Cox6c | protein_coding | 0.020644996 | 10.12988312 | 0.0068437 | 0.9350758 | 0.988182514 |
| ENSMUSG00000022557 | Bop1 | protein_coding | 0.028329751 | 5.784828613 | 0.0068399 | 0.9350998 | 0.988182514 |
| ENSMUSG00000042826 | Fgf11 | protein_coding | -0.037624031 | 3.445099074 | 0.0068198 | 0.935189 | 0.988182514 |
| ENSMUSG00000001768 | Rin2 | protein_coding | 0.028970966 | 4.377031292 | 0.0068071 | 0.9352491 | 0.988182514 |
| ENSMUSG00000005268 | Prlr | protein_coding | -0.047187379 | 3.490090336 | 0.0067396 | 0.9355721 | 0.9882664 |
| ENSMUSG00000081051 | Gm15427 | processed_pseudoge | -0.042266231 | 2.95974261 | 0.006733 | 0.935602 | 0.9882664 |
| ENSMUSG00000024892 | Pcx | protein_coding | -0.021347942 | 5.608768198 | 0.0067325 | 0.9356042 | 0.9882664 |
| ENSMUSG00000024583 | Txnl1 | protein_coding | 0.016075074 | 7.176357436 | 0.0066956 | 0.9357805 | 0.988268735 |
| ENSMUSG00000006763 | Saal1 | protein_coding | -0.027405384 | 4.029950953 | 0.0066776 | 0.9358669 | 0.988268735 |
| ENSMUSG00000005846 | Rsl1d1 | protein_coding | 0.017582434 | 7.142042132 | 0.0066744 | 0.9358821 | 0.988268735 |
| ENSMUSG00000035189 | Ano4 | protein_coding | 0.0271844 | 4.021598688 | 0.0066145 | 0.9361698 | 0.988355621 |
| ENSMUSG00000030761 | Myo7a | protein_coding | -0.058223365 | 2.559450715 | 0.0065992 | 0.9362434 | 0.988355621 |
| ENSMUSG00000070426 | Rnf121 | protein_coding | 0.017919323 | 5.735341833 | 0.0065981 | 0.9362484 | 0.988355621 |
| ENSMUSG00000039345 | Mettl22 | protein_coding | -0.017537261 | 6.464124396 | 0.0065706 | 0.9363812 | 0.988355621 |
| ENSMUSG00000020015 | Cdk17 | protein_coding | 0.021656605 | 6.730462063 | 0.0065621 | 0.936424 | 0.988355621 |
| ENSMUSG00000029189 | Sel1l3 | protein_coding | 0.047716894 | 2.90428747 | 0.0065421 | 0.9365191 | 0.988358981 |
| ENSMUSG00000047694 | Yipf6 | protein_coding | 0.032738815 | 4.256792494 | 0.0064597 | 0.9369195 | 0.988503712 |
| ENSMUSG00000020664 | Dld | protein_coding | -0.020318054 | 7.390086463 | 0.0064522 | 0.9369573 | 0.988503712 |
| ENSMUSG00000033307 | Mif | protein_coding | -0.01512893 | 7.599309128 | 0.0064302 | 0.9370632 | 0.988503712 |
| ENSMUSG00000050188 | Lsm10 | protein_coding | -0.020972321 | 5.930104451 | 0.0064088 | 0.9371679 | 0.988503712 |
| ENSMUSG00000061518 | Cox5b | protein_coding | 0.020116248 | 8.013800626 | 0.0063625 | 0.9373961 | 0.988503712 |
| ENSMUSG00000032202 | Rab27a | protein_coding | -0.035793041 | 3.891648485 | 0.0063467 | 0.9374722 | 0.988503712 |
| ENSMUSG00000042831 | Alkbh6 | protein_coding | -0.024223693 | 7.43084631 | 0.0063484 | 0.9375258 | 0.988503712 |
| ENSMUSG00000020471 | Pold2 | protein_coding | 0.023501598 | 5.39565547 | 0.0062952 | 0.9377257 | 0.988503712 |
| ENSMUSG00000036097 | Slf2 | protein_coding | -0.022271784 | 5.110730074 | 0.006281 | 0.9377959 | 0.988503712 |
| ENSMUSG00000040250 | Ints13 | protein_coding | 0.030943608 | 3.385165044 | 0.0062774 | 0.937814 | 0.988503712 |
| ENSMUSG00000019889 | Ptprk | protein_coding | 0.039981292 | 3.102611883 | 0.006261 | 0.9378947 | 0.988503712 |
| ENSMUSG00000034278 | Dnajc17 | protein_coding | -0.025851154 | 4.320435729 | 0.006252 | 0.9379395 | 0.988503712 |
| ENSMUSG00000023935 | Spats1 | protein_coding | -0.045851112 | 2.793312523 | 0.0062422 | 0.9379879 | 0.988503712 |
| ENSMUSG00000003970 | Rpl8 | protein_coding | -0.016281837 | 10.42760603 | 0.0062392 | 0.9380027 | 0.988503712 |
| ENSMUSG00000021704 | Mtx3 | protein_coding | 0.025905083 | 4.285084892 | 0.0062245 | 0.9380755 | 0.988503712 |
| ENSMUSG00000021819 | Zswim8 | protein_coding | -0.025426263 | 4.343796839 | 0.0062142 | 0.9381271 | 0.988503712 |
| ENSMUSG00000022769 | Sdf2l1 | protein_coding | -0.046540631 | 2.504649534 | 0.0061844 | 0.9382749 | 0.988562662 |
| ENSMUSG00000044340 | Phlpp1 | protein_coding | -0.021182602 | 5.578806587 | 0.0061513 | 0.9384404 | 0.988640085 |
| ENSMUSG00000007603 | Dus3l | protein_coding | -0.018841638 | 6.250032188 | 0.0061262 | 0.9385655 | 0.988674989 |
| ENSMUSG00000037499 | Nenf | protein_coding | -0.019705846 | 5.674159859 | 0.0060732 | 0.9388315 | 0.988858401 |
| ENSMUSG00000033909 | Usp36 | protein_coding | -0.025657286 | 4.481393143 | 0.0060203 | 0.9390978 | 0.98897582 |
| ENSMUSG00000038914 | Dido1 | protein_coding | 0.021427638 | 4.996768125 | 0.0060145 | 0.939127 | 0.98897582 |
| ENSMUSG00000009292 | Trpm2 | protein_coding | -0.037895937 | 3.85111753 | 0.0059531 | 0.9394393 | 0.989207839 |

| ensembl_gene_id | external_gene_name | gene_biotype | log2 (fold change) | log2 (counts per million) | F-value | p-value | False discovery rate |
|---|---|---|---|---|---|---|---|
| ENSMUSG00000004768 | Rab23 | protein_coding | 0.031967141 | 4.504457909 | 0.005909 | 0.9396639 | 0.9892581 |
| ENSMUSG00000027602 | Map1lc3a | protein_coding | -0.014180303 | 8.415411366 | 0.0058919 | 0.9397496 | 0.9892581 |
| ENSMUSG00000046330 | Rpl37a | protein_coding | 0.025000873 | 7.908050191 | 0.0058779 | 0.9399141 | 0.9892581 |
| ENSMUSG00000070044 | Fam149a | protein_coding | 0.025610289 | 4.159628302 | 0.0058287 | 0.9400728 | 0.9892581 |
| ENSMUSG00000042628 | Zfyve1 | protein_coding | 0.023729971 | 4.594394296 | 0.0058277 | 0.940078 | 0.9892581 |
| ENSMUSG00000060288 | Ppih | protein_coding | 0.035567423 | 3.363588497 | 0.0057393 | 0.9405329 | 0.9892581 |
| ENSMUSG00000031353 | Rbbp7 | protein_coding | 0.016216293 | 7.023521773 | 0.0056796 | 0.9408424 | 0.9892581 |
| ENSMUSG00000032040 | Dcps | protein_coding | -0.023870879 | 4.857427172 | 0.005665 | 0.9409185 | 0.9892581 |
| ENSMUSG00000054708 | Ankrd24 | protein_coding | 0.0305547 | 3.893537642 | 0.0056494 | 0.9409998 | 0.9892581 |
| ENSMUSG00000032571 | Pik3r4 | protein_coding | -0.030413247 | 4.210568491 | 0.0056375 | 0.941062 | 0.9892581 |
| ENSMUSG00000048481 | Mypop | protein_coding | 0.025970248 | 3.951406015 | 0.0056255 | 0.9411247 | 0.9892581 |
| ENSMUSG00000029270 | Fam69a | protein_coding | 0.018996437 | 5.543227714 | 0.0056011 | 0.941252 | 0.9892581 |
| ENSMUSG00000029036 | Atad3a | protein_coding | -0.033748099 | 3.588426218 | 0.0055891 | 0.941315 | 0.9892581 |
| ENSMUSG00000006378 | Gcat | protein_coding | 0.023754032 | 5.257505579 | 0.0055816 | 0.9413542 | 0.9892581 |
| ENSMUSG00000028645 | Slc2a1 | protein_coding | 0.036172564 | 3.207892489 | 0.0055797 | 0.9413644 | 0.9892581 |
| ENSMUSG00000075012 | Fjx1 | protein_coding | -0.043519197 | 3.278786675 | 0.0055733 | 0.9413979 | 0.9892581 |
| ENSMUSG00000015112 | Slc25a13 | protein_coding | -0.037518959 | 3.432724529 | 0.0055561 | 0.941488 | 0.9892581 |
| ENSMUSG00000021482 | Aaed1 | protein_coding | -0.026491423 | 4.010025099 | 0.0055537 | 0.9415009 | 0.9892581 |
| ENSMUSG00000052406 | Rexo4 | protein_coding | 0.023908838 | 5.485134177 | 0.0055519 | 0.9415099 | 0.9892581 |
| ENSMUSG00000002996 | Hbp1 | protein_coding | -0.029017583 | 3.632420087 | 0.0055498 | 0.9415214 | 0.9892581 |
| ENSMUSG00000086075 | Gm15728 | lincRNA | 0.038200339 | 3.171824099 | 0.005536 | 0.9415935 | 0.9892581 |
| ENSMUSG00000027257 | Pacsin3 | protein_coding | -0.028515252 | 4.267205859 | 0.0055341 | 0.9416038 | 0.9892581 |
| ENSMUSG00000035142 | Nubpl | protein_coding | -0.035009391 | 3.289785091 | 0.0055284 | 0.941634 | 0.9892581 |
| ENSMUSG00000023025 | Larp4 | protein_coding | -0.021101284 | 4.523088923 | 0.0055168 | 0.941695 | 0.9892581 |
| ENSMUSG00000021313 | Ryr2 | protein_coding | 0.035966304 | 4.29272841 | 0.0054968 | 0.9418182 | 0.989290908 |
| ENSMUSG00000038619 | Ensa | protein_coding | 0.014283325 | 9.758406998 | 0.0053939 | 0.9423468 | 0.989749515 |
| ENSMUSG00000041949 | Tango6 | protein_coding | -0.026728186 | 3.55573917 | 0.0053453 | 0.9426068 | 0.989911555 |
| ENSMUSG00000017781 | Pitpna | protein_coding | 0.0134132 | 7.228051855 | 0.0053165 | 0.942761 | 0.989911555 |
| ENSMUSG00000027849 | Syt6 | protein_coding | -0.033527081 | 4.318855024 | 0.0053138 | 0.9427773 | 0.989911555 |
| ENSMUSG00000086119 | Gm2415 | antisense_RNA | -0.045083684 | 2.633322965 | 0.0052851 | 0.9429301 | 0.989975372 |
| ENSMUSG00000087200 | Lrp8os3 | bidirectional_promoter | -0.030524406 | 3.219258825 | 0.0052014 | 0.943383 | 0.990351653 |
| ENSMUSG00000048355 | Arxes1 | protein_coding | -0.024557881 | 4.788125196 | 0.0051849 | 0.9434727 | 0.990351653 |
| ENSMUSG00000020173 | Cobl | protein_coding | 0.022309665 | 4.979195501 | 0.0051452 | 0.9436893 | 0.990482309 |
| ENSMUSG00000039461 | Tcta | protein_coding | -0.016955685 | 6.791598511 | 0.005094 | 0.9439693 | 0.990517541 |
| ENSMUSG00000027303 | Ptpra | protein_coding | 0.021817454 | 4.260738267 | 0.0050929 | 0.9439755 | 0.990517541 |
| ENSMUSG00000021557 | Agtpbp1 | protein_coding | -0.016553407 | 6.05202309 | 0.0050815 | 0.9440382 | 0.990517541 |
| ENSMUSG00000036980 | Taf6 | protein_coding | -0.026513492 | 5.728070284 | 0.0050753 | 0.9441118 | 0.990517541 |
| ENSMUSG00000094344 | Gm11942 | processed_pseudoge | 0.098710539 | 1.972589933 | 0.0050608 | 0.9442194 | 0.990517541 |
| ENSMUSG00000063358 | Mapk1 | protein_coding | 0.016403872 | 8.06731653 | 0.0050384 | 0.9442756 | 0.990517541 |
| ENSMUSG00000028496 | Mllt3 | protein_coding | 0.018615021 | 5.335618792 | 0.0050107 | 0.9444287 | 0.990581528 |
| ENSMUSG00000029229 | Chic2 | protein_coding | -0.030707638 | 3.422371939 | 0.004944 | 0.9447992 | 0.990739867 |
| ENSMUSG00000029291 | Rufy3 | protein_coding | -0.014147028 | 8.129277051 | 0.0049236 | 0.9449127 | 0.990739867 |
| ENSMUSG00000021884 | Hacl1 | protein_coding | 0.030408989 | 3.644691839 | 0.0049145 | 0.9449639 | 0.990739867 |
| ENSMUSG00000050953 | Gja1 | protein_coding | -0.017632146 | 7.008087782 | 0.0048817 | 0.9451489 | 0.990739867 |
| ENSMUSG00000040420 | Cdh18 | protein_coding | -0.027777244 | 3.766926113 | 0.0048753 | 0.9451832 | 0.990739867 |
| ENSMUSG00000067924 | Cxx1b | protein_coding | 0.018432338 | 4.928086095 | 0.0048732 | 0.945195 | 0.990739867 |
| ENSMUSG00000102805 | Gm37240 | protein_coding | 0.033336794 | 3.117351977 | 0.0048656 | 0.9452378 | 0.990739867 |
| ENSMUSG00000028785 | Hpca | protein_coding | -0.015336166 | 9.923674582 | 0.0048452 | 0.9453523 | 0.990739867 |
| ENSMUSG00000050963 | Kcns2 | protein_coding | 0.045705853 | 2.687444094 | 0.0048352 | 0.9454089 | 0.990739867 |
| ENSMUSG00000062753 | AI413582 | protein_coding | 0.013892016 | 7.381961825 | 0.0048127 | 0.9455358 | 0.990773101 |
| ENSMUSG00000097574 | C920006O11Rik | lincRNA | -0.026603221 | 3.671111301 | 0.0047969 | 0.9456249 | 0.990773101 |
| ENSMUSG00000020115 | Tbk1 | protein_coding | -0.022484507 | 4.763770872 | 0.004734 | 0.9459824 | 0.990812401 |
| ENSMUSG00000002732 | Fkbp7 | protein_coding | -0.040319478 | 2.801679118 | 0.0047273 | 0.9460205 | 0.990812401 |
| ENSMUSG00000039838 | Slc45a1 | protein_coding | 0.020340624 | 6.048189818 | 0.0047196 | 0.9460657 | 0.990812401 |
| ENSMUSG00000028792 | Ak2 | protein_coding | -0.01878183 | 5.347829628 | 0.0046939 | 0.9462109 | 0.990812401 |
| ENSMUSG00000040374 | Pex2 | protein_coding | 0.013926397 | 6.584902452 | 0.0046911 | 0.946227 | 0.990812401 |
| ENSMUSG00000036529 | Sbf1 | protein_coding | 0.018978474 | 5.353329883 | 0.0046778 | 0.9463032 | 0.990812401 |
| ENSMUSG00000014361 | Mertk | protein_coding | -0.047672201 | 2.70795975 | 0.0046585 | 0.9464138 | 0.990812401 |
| ENSMUSG00000020307 | Cdc34 | protein_coding | -0.023402789 | 4.500428073 | 0.0046466 | 0.9464821 | 0.990812401 |
| ENSMUSG00000037525 | Bcdin3d | protein_coding | 0.031733509 | 4.013761634 | 0.004641 | 0.9465157 | 0.990812401 |
| ENSMUSG00000026966 | Ssna1 | protein_coding | -0.013170236 | 7.620011426 | 0.0045803 | 0.9468651 | 0.990812401 |
| ENSMUSG00000001507 | Itga3 | protein_coding | -0.027345984 | 3.834342419 | 0.0045709 | 0.9469194 | 0.990812401 |
| ENSMUSG00000044030 | Irf2bp1 | protein_coding | 0.02157207 | 4.346356885 | 0.004563 | 0.9469654 | 0.990812401 |
| ENSMUSG00000037443 | Cep85 | protein_coding | 0.027629565 | 3.550220314 | 0.0045337 | 0.9471355 | 0.990812401 |
| ENSMUSG00000034751 | Mast4 | protein_coding | -0.028494998 | 3.656852043 | 0.0045296 | 0.9471595 | 0.990812401 |
| ENSMUSG00000048772 | Tmem53 | protein_coding | 0.023440652 | 4.221569174 | 0.0045261 | 0.9471797 | 0.990812401 |
| ENSMUSG00000031161 | Hdac6 | protein_coding | -0.023989454 | 4.879867729 | 0.0045147 | 0.947246 | 0.990812401 |
| ENSMUSG00000026670 | Uap1 | protein_coding | 0.019626605 | 5.067554371 | 0.004505 | 0.9473027 | 0.990812401 |
| ENSMUSG00000058690 | Ccser2 | protein_coding | 0.016523532 | 5.493256827 | 0.0044912 | 0.9473835 | 0.990812401 |
| ENSMUSG00000025470 | Zfp511 | protein_coding | 0.020606112 | 4.322987393 | 0.0044861 | 0.9474131 | 0.990812401 |
| ENSMUSG00000025759 | Mfsd8 | protein_coding | -0.052840542 | 2.851632995 | 0.0044326 | 0.9477289 | 0.990826949 |
| ENSMUSG00000030823 | 9130019O22Rik | protein_coding | 0.067076905 | 1.986000247 | 0.0044277 | 0.9477563 | 0.990826949 |
| ENSMUSG00000005374 | Tbl2 | protein_coding | 0.028595897 | 3.460875466 | 0.0044067 | 0.9478801 | 0.990826949 |
| ENSMUSG00000073139 | Tmem185a | protein_coding | -0.015566623 | 5.789414709 | 0.0044038 | 0.9478971 | 0.990826949 |

| ensembl_gene_id | external_gene_name | gene_biotype | log2 (fold change) | log2 (counts per million) | F-value | p-value | False discovery rate |
|---|---|---|---|---|---|---|---|
| ENSMUSG00000047205 | Dusp18 | protein_coding | -0.022493506 | 4.869226442 | 0.0043925 | 0.947964 | 0.990826949 |
| ENSMUSG00000061013 | Mkx | protein_coding | 0.047619899 | 2.553565789 | 0.0043898 | 0.9479799 | 0.990826949 |
| ENSMUSG00000028410 | Dnaja1 | protein_coding | 0.011802849 | 7.796491924 | 0.0043587 | 0.9481642 | 0.990838342 |
| ENSMUSG00000031715 | Smarca5 | protein_coding | 0.018818349 | 5.237627904 | 0.0043569 | 0.9481751 | 0.990838342 |
| ENSMUSG00000058402 | Zfp420 | protein_coding | -0.023275851 | 3.993449678 | 0.0043296 | 0.9483373 | 0.99086401 |
| ENSMUSG00000112390 | CT025654.1 | processed_transcript | -0.037612769 | 3.127733802 | 0.0043218 | 0.9483839 | 0.99086401 |
| ENSMUSG00000028582 | Cc2d1b | protein_coding | -0.032507688 | 3.403675715 | 0.004244 | 0.9488496 | 0.991195415 |
| ENSMUSG00000041961 | Znrf3 | protein_coding | -0.021356949 | 5.062820556 | 0.0042184 | 0.9490038 | 0.991195415 |
| ENSMUSG00000005148 | Klf5 | protein_coding | -0.028885279 | 3.456679139 | 0.004211 | 0.9490487 | 0.991195415 |
| ENSMUSG00000041895 | Wipi1 | protein_coding | 0.035646835 | 4.434203153 | 0.0042137 | 0.9491113 | 0.991195415 |
| ENSMUSG00000037348 | Paqr7 | protein_coding | -0.015980325 | 6.590218466 | 0.0041722 | 0.9492833 | 0.991195415 |
| ENSMUSG00000065954 | Tacc1 | protein_coding | -0.014776947 | 6.421678638 | 0.0041662 | 0.9493199 | 0.991195415 |
| ENSMUSG00000028653 | Trit1 | protein_coding | -0.021802591 | 4.177133886 | 0.0041619 | 0.9493464 | 0.991195415 |
| ENSMUSG00000028419 | Chmp5 | protein_coding | 0.011688617 | 8.829841351 | 0.0041388 | 0.9494867 | 0.991245678 |
| ENSMUSG00000021871 | Pnp | protein_coding | -0.035396787 | 2.807099839 | 0.0040555 | 0.9499969 | 0.991654074 |
| ENSMUSG00000022774 | Ncbp2 | protein_coding | -0.011165331 | 7.268332187 | 0.0040449 | 0.9500623 | 0.991654074 |
| ENSMUSG00000065126 | Snord104 | snoRNA | -0.032348828 | 3.124400437 | 0.0040147 | 0.9502486 | 0.991716596 |
| ENSMUSG00000025025 | Mxi1 | protein_coding | -0.017655343 | 5.435271417 | 0.0040053 | 0.9503067 | 0.991716596 |
| ENSMUSG00000021194 | Chga | protein_coding | -0.012790503 | 7.63615483 | 0.0039725 | 0.9505106 | 0.991753775 |
| ENSMUSG00000085139 | A730046J19Rik | protein_coding | -0.050964736 | 2.279239719 | 0.0039531 | 0.9506312 | 0.991753775 |
| ENSMUSG00000029402 | Snrnp35 | protein_coding | -0.018459468 | 5.151248136 | 0.0039512 | 0.9506431 | 0.991753775 |
| ENSMUSG00000017561 | Crlf3 | protein_coding | -0.030395108 | 3.450944053 | 0.0039403 | 0.9507112 | 0.991753775 |
| ENSMUSG00000020474 | Polm | protein_coding | 0.036311851 | 3.367098616 | 0.0039027 | 0.9509476 | 0.991762737 |
| ENSMUSG00000022108 | Itm2b | protein_coding | 0.011686283 | 8.615978133 | 0.0038997 | 0.9509654 | 0.991762737 |
| ENSMUSG00000036887 | C1qa | protein_coding | 0.031656548 | 3.806809501 | 0.0038873 | 0.9510446 | 0.991762737 |
| ENSMUSG00000112964 | CT009718.1 | lincRNA | 0.031155143 | 3.165644942 | 0.0038271 | 0.9514232 | 0.991762737 |
| ENSMUSG00000018322 | Tomm34 | protein_coding | 0.013015577 | 7.916772627 | 0.0038149 | 0.9515005 | 0.991762737 |
| ENSMUSG00000023286 | Ube2j2 | protein_coding | -0.019834094 | 4.369163291 | 0.003805 | 0.9515639 | 0.991762737 |
| ENSMUSG00000029672 | Fam3c | protein_coding | -0.015902136 | 5.096168148 | 0.0037979 | 0.9516085 | 0.991762737 |
| ENSMUSG00000026173 | Plcd4 | protein_coding | 0.026963399 | 3.898973072 | 0.0037651 | 0.9518178 | 0.991762737 |
| ENSMUSG00000024811 | Tnks2 | protein_coding | -0.020771174 | 5.379517139 | 0.0037443 | 0.9519521 | 0.991762737 |
| ENSMUSG00000025878 | Uimc1 | protein_coding | -0.019448678 | 4.440030561 | 0.0037414 | 0.9519693 | 0.991762737 |
| ENSMUSG00000021508 | Cxcl14 | protein_coding | 0.027644856 | 4.651294608 | 0.003739 | 0.9520068 | 0.991762737 |
| ENSMUSG00000044533 | Rps2 | protein_coding | 0.021328327 | 4.659442267 | 0.0037317 | 0.952032 | 0.991762737 |
| ENSMUSG00000024054 | Smchd1 | protein_coding | -0.027830226 | 3.379495894 | 0.0037276 | 0.9520579 | 0.991762737 |
| ENSMUSG00000026819 | Slc25a25 | protein_coding | 0.014243072 | 5.904882908 | 0.0037131 | 0.9521512 | 0.991762737 |
| ENSMUSG00000009731 | Kcnd1 | protein_coding | -0.056420544 | 2.365475782 | 0.0036918 | 0.9522901 | 0.991762737 |
| ENSMUSG00000029366 | Dck | protein_coding | 0.019960412 | 4.599533717 | 0.0036705 | 0.9524262 | 0.991762737 |
| ENSMUSG00000040918 | Slc19a2 | protein_coding | -0.031257088 | 3.250004622 | 0.0036672 | 0.9524475 | 0.991762737 |
| ENSMUSG00000003934 | Efnb3 | protein_coding | -0.012517921 | 7.187374307 | 0.0036607 | 0.9524896 | 0.991762737 |
| ENSMUSG00000063063 | Ctnna2 | protein_coding | -0.012645869 | 6.555486821 | 0.0036535 | 0.9525365 | 0.991762737 |
| ENSMUSG00000030269 | Mtmr14 | protein_coding | -0.029990074 | 2.959908207 | 0.0036492 | 0.9525645 | 0.991762737 |
| ENSMUSG00000030515 | Tarsl2 | protein_coding | 0.030903699 | 2.91404841 | 0.0036177 | 0.9527693 | 0.991879963 |
| ENSMUSG00000036693 | Nop14 | protein_coding | 0.018844418 | 5.11560451 | 0.0035964 | 0.9529085 | 0.99192884 |
| ENSMUSG00000039163 | Cmc1 | protein_coding | -0.020587349 | 4.134331646 | 0.0035544 | 0.9531837 | 0.992119298 |
| ENSMUSG00000021264 | Yy1 | protein_coding | 0.021772843 | 4.865081392 | 0.0035327 | 0.953327 | 0.992172412 |
| ENSMUSG00000037325 | Bbs7 | protein_coding | -0.019113845 | 4.537572092 | 0.0035045 | 0.9535131 | 0.992270094 |
| ENSMUSG00000087056 | Gm14004 | processed_transcript | -0.020392669 | 5.122867176 | 0.0034583 | 0.9538203 | 0.99241243 |
| ENSMUSG00000038517 | Tbkbp1 | protein_coding | 0.033188495 | 3.123857561 | 0.0034462 | 0.9539008 | 0.99241243 |
| ENSMUSG00000002546 | Golga2 | protein_coding | -0.017572282 | 5.61340806 | 0.0034311 | 0.9540021 | 0.99241243 |
| ENSMUSG00000026617 | Bpnt1 | protein_coding | -0.011794697 | 6.995437354 | 0.0034285 | 0.9540191 | 0.99241243 |
| ENSMUSG00000019808 | Adat2 | protein_coding | -0.030226822 | 2.972440111 | 0.0033908 | 0.9542724 | 0.992457521 |
| ENSMUSG00000048281 | Dleu7 | protein_coding | 0.021645992 | 3.777676594 | 0.0033827 | 0.9543268 | 0.992457521 |
| ENSMUSG00000062488 | Ifit3b | protein_coding | 0.036136834 | 2.6873316 | 0.0033615 | 0.9544702 | 0.992457521 |
| ENSMUSG00000037730 | Mynn | protein_coding | -0.019695636 | 4.006506073 | 0.0033591 | 0.9544867 | 0.992457521 |
| ENSMUSG00000045438 | Cox19 | protein_coding | 0.012134562 | 7.156107596 | 0.0033536 | 0.9545239 | 0.992457521 |
| ENSMUSG00000086943 | 4732414G09Rik | lincRNA | 0.051667309 | 2.822618559 | 0.0033363 | 0.9546856 | 0.992529692 |
| ENSMUSG00000047649 | Cd3eap | protein_coding | 0.013460835 | 6.083496836 | 0.0032716 | 0.9550822 | 0.992813714 |
| ENSMUSG00000071604 | Fam189a2 | protein_coding | -0.039243135 | 2.759313938 | 0.0032521 | 0.9552162 | 0.992813714 |
| ENSMUSG00000022552 | Sharpin | protein_coding | -0.023274361 | 5.492261759 | 0.0032557 | 0.9552358 | 0.992813714 |
| ENSMUSG00000038740 | Mvb12b | protein_coding | -0.018367297 | 4.934710172 | 0.0032141 | 0.9554788 | 0.992892309 |
| ENSMUSG00000046329 | Slc25a23 | protein_coding | -0.020327085 | 3.69950055 | 0.0032116 | 0.9554961 | 0.992892309 |
| ENSMUSG00000032388 | Spg21 | protein_coding | 0.015922808 | 5.239548365 | 0.0031726 | 0.9557664 | 0.993076946 |
| ENSMUSG00000013160 | Atp6v0d1 | protein_coding | 0.009681737 | 8.515038157 | 0.0031594 | 0.9558585 | 0.993076946 |
| ENSMUSG00000058174 | Gm5148 | protein_coding | 0.018981125 | 4.361021659 | 0.0031339 | 0.956037 | 0.99316642 |
| ENSMUSG00000034639 | Setmar | protein_coding | -0.033000307 | 2.899282814 | 0.0030825 | 0.9563982 | 0.993346416 |
| ENSMUSG00000027108 | Ola1 | protein_coding | -0.010486576 | 7.789073664 | 0.003058 | 0.9565722 | 0.993346416 |
| ENSMUSG00000040302 | Rbm48 | protein_coding | -0.01737056 | 4.46789051 | 0.0030502 | 0.9566275 | 0.993346416 |
| ENSMUSG00000063694 | Cycs | protein_coding | -0.01461716 | 5.154449995 | 0.003044 | 0.9566711 | 0.993346416 |
| ENSMUSG00000050737 | Ptges | protein_coding | -0.03254552 | 2.854792656 | 0.0030244 | 0.9568109 | 0.993346416 |
| ENSMUSG00000039105 | Atp6v1g1 | protein_coding | 0.009756944 | 8.419649537 | 0.0030183 | 0.9568546 | 0.993346416 |
| ENSMUSG00000043162 | Pyurf | protein_coding | -0.026988866 | 5.055327612 | 0.0030279 | 0.9568569 | 0.993346416 |
| ENSMUSG00000037940 | Inpp4b | protein_coding | 0.021852414 | 4.371399073 | 0.0029458 | 0.9573754 | 0.993354318 |
| ENSMUSG00000075478 | Slitrk1 | protein_coding | -0.018082723 | 4.949897252 | 0.0029028 | 0.9576871 | 0.993354318 |

| ensembl_gene_id | external_gene_name | gene_biotype | log2 (fold change) | log2 (counts per million) | F-value | p-value | False discovery rate |
|---|---|---|---|---|---|---|---|
| ENSMUSG00000044017 | Adgrd1 | protein_coding | -0.025545683 | 2.95835921 | 0.0029001 | 0.9577065 | 0.993354318 |
| ENSMUSG00000041592 | Sdk2 | protein_coding | 0.029705213 | 3.169684741 | 0.0028964 | 0.9577336 | 0.993354318 |
| ENSMUSG00000023873 | 1700010I14Rik | protein_coding | 0.038520747 | 2.972098165 | 0.0028844 | 0.9578224 | 0.993354318 |
| ENSMUSG00000019854 | Reps1 | protein_coding | -0.025730993 | 4.639470301 | 0.0028792 | 0.9579006 | 0.993354318 |
| ENSMUSG00000049751 | Rpl36al | protein_coding | 0.025198201 | 3.012442042 | 0.0028652 | 0.9579616 | 0.993354318 |
| ENSMUSG00000052133 | Sema5b | protein_coding | 0.030317249 | 2.685225902 | 0.002851 | 0.958066 | 0.993354318 |
| ENSMUSG00000063796 | Slc22a8 | protein_coding | -0.045195528 | 2.651915332 | 0.0028367 | 0.9581725 | 0.993354318 |
| ENSMUSG00000011837 | Snapc2 | protein_coding | 0.015697569 | 5.29531142 | 0.0028359 | 0.9581769 | 0.993354318 |
| ENSMUSG00000006191 | Cdkal1 | protein_coding | -0.01734089 | 4.641648167 | 0.0028309 | 0.9582143 | 0.993354318 |
| ENSMUSG00000040147 | Maob | protein_coding | -0.020630681 | 6.05615431 | 0.0028145 | 0.9583996 | 0.993354318 |
| ENSMUSG00000025422 | Agap2 | protein_coding | 0.024293172 | 3.675226007 | 0.0027952 | 0.9584781 | 0.993354318 |
| ENSMUSG00000048758 | Rpl29 | protein_coding | -0.012823437 | 5.486489102 | 0.0027937 | 0.9584895 | 0.993354318 |
| ENSMUSG00000037210 | Fam193a | protein_coding | -0.018588324 | 4.910194379 | 0.0027929 | 0.9584953 | 0.993354318 |
| ENSMUSG00000038459 | Abhd17c | protein_coding | 0.01870994 | 4.349748203 | 0.0027914 | 0.9585062 | 0.993354318 |
| ENSMUSG00000017664 | Slc35c2 | protein_coding | -0.011852565 | 6.684281518 | 0.0027853 | 0.9585518 | 0.993354318 |
| ENSMUSG00000025597 | Klhl4 | protein_coding | 0.018780161 | 3.970025512 | 0.0027573 | 0.9587598 | 0.993354318 |
| ENSMUSG00000029345 | Tfip11 | protein_coding | 0.022828454 | 4.892480515 | 0.0027493 | 0.9588412 | 0.993354318 |
| ENSMUSG00000030595 | Nfkbib | protein_coding | 0.015687128 | 5.707432319 | 0.0027083 | 0.9591281 | 0.993354318 |
| ENSMUSG00000026154 | Sdhaf4 | protein_coding | 0.011935734 | 8.414364103 | 0.0027029 | 0.9591687 | 0.993354318 |
| ENSMUSG00000060216 | Arrb2 | protein_coding | 0.018997433 | 4.17512589 | 0.0026981 | 0.9592049 | 0.993354318 |
| ENSMUSG00000068874 | Selenbp1 | protein_coding | -0.021093833 | 3.323188015 | 0.0026925 | 0.9592474 | 0.993354318 |
| ENSMUSG00000039239 | Tgfb2 | protein_coding | -0.019431819 | 4.117506558 | 0.0026637 | 0.9594658 | 0.993354318 |
| ENSMUSG00000036291 | Ap5m1 | protein_coding | -0.02117207 | 3.98614786 | 0.0026633 | 0.9594686 | 0.993354318 |
| ENSMUSG00000003865 | Gys1 | protein_coding | -0.020132585 | 4.209137295 | 0.0026544 | 0.959536 | 0.993354318 |
| ENSMUSG00000024528 | Srfbp1 | protein_coding | -0.017438347 | 4.73445239 | 0.0026491 | 0.9595768 | 0.993354318 |
| ENSMUSG00000038069 | Cdkn2aip | protein_coding | -0.024088503 | 3.533200707 | 0.0026482 | 0.9595833 | 0.993354318 |
| ENSMUSG00000031848 | Lsm4 | protein_coding | 0.028522253 | 4.547718722 | 0.0026413 | 0.9597174 | 0.993354318 |
| ENSMUSG00000052397 | Ezr | protein_coding | 0.018708854 | 4.466363293 | 0.0026294 | 0.959727 | 0.993354318 |
| ENSMUSG00000028973 | Abcb8 | protein_coding | -0.011737823 | 6.129467242 | 0.0026293 | 0.9597283 | 0.993354318 |
| ENSMUSG00000016477 | E2f3 | protein_coding | 0.021439393 | 3.467306497 | 0.0025983 | 0.9599655 | 0.993504268 |
| ENSMUSG00000022721 | Trmt2a | protein_coding | 0.01813389 | 5.02992924 | 0.0025616 | 0.9602489 | 0.993623219 |
| ENSMUSG00000007415 | Gatad1 | protein_coding | 0.011678876 | 7.549900113 | 0.0025531 | 0.9603152 | 0.993623219 |
| ENSMUSG00000033545 | Znrf1 | protein_coding | -0.013564859 | 5.488763128 | 0.0025415 | 0.9604051 | 0.993623219 |
| ENSMUSG00000021257 | Angel1 | protein_coding | -0.031297153 | 3.090701824 | 0.0025358 | 0.9604501 | 0.993623219 |
| ENSMUSG00000024498 | Tcerg1 | protein_coding | -0.014662728 | 5.115507489 | 0.0025222 | 0.9605559 | 0.99363709 |
| ENSMUSG00000032500 | Dclk3 | protein_coding | 0.020741883 | 4.730840745 | 0.0024781 | 0.9609031 | 0.99370596 |
| ENSMUSG00000037060 | Cavin3 | protein_coding | -0.013780321 | 5.077329017 | 0.0024763 | 0.9609163 | 0.99370596 |
| ENSMUSG00000027582 | Zgpat | protein_coding | 0.037446061 | 2.583425405 | 0.0024701 | 0.9609653 | 0.99370596 |
| ENSMUSG00000100599 | 1700120C14Rik | processed_transcript | -0.032359541 | 2.57138685 | 0.0024667 | 0.9609921 | 0.99370596 |
| ENSMUSG00000075701 | Selenos | protein_coding | -0.01250059 | 7.051738636 | 0.0024441 | 0.9611714 | 0.993795813 |
| ENSMUSG00000026427 | Eif2d | protein_coding | -0.013603546 | 5.489170157 | 0.0024263 | 0.9613127 | 0.993845192 |
| ENSMUSG00000038412 | Higd1a | protein_coding | 0.009478657 | 8.510514161 | 0.0024075 | 0.9614625 | 0.993845192 |
| ENSMUSG00000024588 | Fech | protein_coding | -0.017039967 | 5.324261847 | 0.0024034 | 0.9614965 | 0.993845192 |
| ENSMUSG00000021051 | Ppp2r5e | protein_coding | -0.011121716 | 5.915104441 | 0.0023844 | 0.9616478 | 0.993906088 |
| ENSMUSG00000026748 | Plxdc2 | protein_coding | -0.017317314 | 4.74135173 | 0.0023688 | 0.9617729 | 0.993939804 |
| ENSMUSG00000047238 | Mageh1 | protein_coding | -0.010258145 | 6.191350289 | 0.002349 | 0.9619332 | 0.994010013 |
| ENSMUSG00000030102 | Itpr1 | protein_coding | 0.015804437 | 6.325948694 | 0.0023287 | 0.9621329 | 0.994120794 |
| ENSMUSG00000037221 | Mospd3 | protein_coding | -0.02715442 | 4.650157983 | 0.0022979 | 0.9624311 | 0.994153135 |
| ENSMUSG00000038975 | Rabggtb | protein_coding | 0.00811345 | 7.224132248 | 0.0022834 | 0.9624678 | 0.994153135 |
| ENSMUSG00000017188 | Coa3 | protein_coding | -0.009142961 | 9.831433548 | 0.0022802 | 0.9624942 | 0.994153135 |
| ENSMUSG00000021224 | Numb | protein_coding | 0.018314515 | 4.281775495 | 0.0022754 | 0.962534 | 0.994153135 |
| ENSMUSG00000039706 | Ldb2 | protein_coding | -0.010841702 | 6.280817788 | 0.0022505 | 0.9627388 | 0.994269127 |
| ENSMUSG00000036957 | Lrfn3 | protein_coding | -0.036697109 | 2.342207843 | 0.0022012 | 0.9631495 | 0.994488996 |
| ENSMUSG00000094626 | Cecr6 | protein_coding | -0.029316095 | 3.036081441 | 0.0021736 | 0.9633804 | 0.994488996 |
| ENSMUSG00000021676 | Iqgap2 | protein_coding | -0.026186338 | 3.054315862 | 0.0021696 | 0.9634146 | 0.994488996 |
| ENSMUSG00000015247 | Nipsnap3b | protein_coding | -0.01065891 | 6.272602591 | 0.0021644 | 0.9634584 | 0.994488996 |
| ENSMUSG00000012405 | Rpl15 | protein_coding | 0.009057717 | 6.940425499 | 0.0021548 | 0.9635391 | 0.994488996 |
| ENSMUSG00000026051 | 1500015O10Rik | protein_coding | -0.025923029 | 3.884914148 | 0.0021563 | 0.9635582 | 0.994488996 |
| ENSMUSG00000068922 | Msto1 | protein_coding | -0.013010215 | 4.987511977 | 0.0021366 | 0.9636937 | 0.994488996 |
| ENSMUSG00000022895 | Ets2 | protein_coding | -0.01646578 | 5.444192148 | 0.0021221 | 0.9638181 | 0.994488996 |
| ENSMUSG00000020720 | Psmd12 | protein_coding | -0.008790863 | 8.112982504 | 0.0021192 | 0.9638419 | 0.994488996 |
| ENSMUSG00000020496 | Rnf187 | protein_coding | 0.009594679 | 8.020059388 | 0.0021151 | 0.9638765 | 0.994488996 |
| ENSMUSG00000021242 | Npc2 | protein_coding | 0.00890534 | 7.42587383 | 0.0020929 | 0.9640668 | 0.99449713 |
| ENSMUSG00000022807 | Osbpl11 | protein_coding | -0.018761682 | 3.436729191 | 0.0020926 | 0.9640694 | 0.99449713 |
| ENSMUSG00000063884 | Ptcd3 | protein_coding | -0.013012942 | 5.521540319 | 0.0020642 | 0.9643137 | 0.99461131 |
| ENSMUSG00000043252 | Tmem64 | protein_coding | 0.016773255 | 4.869118991 | 0.0020582 | 0.9643651 | 0.99461131 |
| ENSMUSG00000037022 | Mmaa | protein_coding | 0.014177074 | 4.865514758 | 0.0020382 | 0.9645386 | 0.994694863 |
| ENSMUSG00000072949 | Acot1 | protein_coding | 0.025351629 | 2.791619419 | 0.0020124 | 0.9647636 | 0.994831518 |
| ENSMUSG00000066042 | Med18 | protein_coding | 0.013702585 | 4.473760807 | 0.0019962 | 0.9649058 | 0.994882766 |
| ENSMUSG00000042129 | Rassf4 | protein_coding | 0.018755155 | 3.448326088 | 0.0019721 | 0.9651183 | 0.994997925 |
| ENSMUSG00000049866 | Arl4c | protein_coding | 0.011914898 | 5.539892239 | 0.0019597 | 0.9652276 | 0.994997925 |
| ENSMUSG00000015363 | Trabd | protein_coding | 0.016054294 | 4.444487945 | 0.0019521 | 0.9652951 | 0.994997925 |
| ENSMUSG00000030649 | Anapc15 | protein_coding | -0.017898184 | 3.334285731 | 0.0019394 | 0.9654087 | 0.995009427 |
| ENSMUSG00000036835 | Psenen | protein_coding | 0.011727866 | 4.977596897 | 0.0019301 | 0.9654913 | 0.995009427 |

| ensembl_gene_id | external_gene_name | gene_biotype | log2 (fold change) | log2 (counts per million) | F-value | p-value | False discovery rate |
|---|---|---|---|---|---|---|---|
| ENSMUSG00000017453 | Pipox | protein_coding | -0.029671052 | 2.536586131 | 0.0019147 | 0.9656292 | 0.995056131 |
| ENSMUSG00000042743 | Sgtb | protein_coding | 0.012437327 | 5.508618151 | 0.001899 | 0.9657708 | 0.995106644 |
| ENSMUSG00000025894 | Aasdhppt | protein_coding | -0.009730855 | 6.465648914 | 0.0018411 | 0.9662956 | 0.995293344 |
| ENSMUSG00000053565 | Eif3k | protein_coding | -0.008672854 | 9.322931709 | 0.0018246 | 0.9664471 | 0.995293344 |
| ENSMUSG00000030271 | Ogg1 | protein_coding | -0.014893268 | 3.96687806 | 0.0018138 | 0.9665468 | 0.995293344 |
| ENSMUSG00000035898 | Uba6 | protein_coding | -0.017760942 | 4.259359949 | 0.0017881 | 0.9667841 | 0.995293344 |
| ENSMUSG00000020669 | Sh3yl1 | protein_coding | -0.018246419 | 4.384485981 | 0.0017791 | 0.9668683 | 0.995293344 |
| ENSMUSG00000026239 | Pde6d | protein_coding | -0.01152837 | 4.990558208 | 0.0017749 | 0.9669068 | 0.995293344 |
| ENSMUSG00000018932 | Map2k3 | protein_coding | 0.021911473 | 2.856853344 | 0.0017701 | 0.9669516 | 0.995293344 |
| ENSMUSG00000052302 | Tbc1d30 | protein_coding | -0.010691086 | 5.205387389 | 0.0017622 | 0.9670255 | 0.995293344 |
| ENSMUSG00000025580 | Eif4a3 | protein_coding | 0.008444175 | 6.742675217 | 0.0017604 | 0.9670425 | 0.995293344 |
| ENSMUSG00000034292 | Traf3ip1 | protein_coding | 0.019008569 | 3.585416351 | 0.0017495 | 0.9671443 | 0.995293344 |
| ENSMUSG00000009356 | Lpo | protein_coding | 0.04819463 | 2.031690728 | 0.0017493 | 0.9671473 | 0.995293344 |
| ENSMUSG00000035653 | Lrfn5 | protein_coding | -0.012612135 | 5.506144873 | 0.0017449 | 0.967188 | 0.995293344 |
| ENSMUSG00000078137 | Ankrd63 | protein_coding | 0.016049196 | 4.836188436 | 0.0017444 | 0.9671935 | 0.995293344 |
| ENSMUSG00000022811 | Zfp148 | protein_coding | 0.008499835 | 6.010483517 | 0.0017364 | 0.9672677 | 0.995293344 |
| ENSMUSG00000027201 | Myef2 | protein_coding | 0.025459742 | 2.690352553 | 0.0017245 | 0.9673795 | 0.995293344 |
| ENSMUSG00000048920 | Fkrp | protein_coding | -0.012408875 | 4.62235271 | 0.001716 | 0.9674608 | 0.995293344 |
| ENSMUSG00000086401 | Gm15559 | antisense_RNA | 0.02758183 | 3.268404032 | 0.0017083 | 0.9675381 | 0.995293344 |
| ENSMUSG00000015932 | Dstn | protein_coding | 0.008041365 | 8.557411813 | 0.0016994 | 0.967618 | 0.995293344 |
| ENSMUSG00000072582 | Ptrh2 | protein_coding | 0.009376254 | 6.181312669 | 0.0016699 | 0.9679002 | 0.99534094 |
| ENSMUSG00000027011 | Ube2e3 | protein_coding | -0.007305993 | 6.868104404 | 0.0016651 | 0.9679465 | 0.99534094 |
| ENSMUSG00000027508 | Pag1 | protein_coding | -0.021380664 | 3.019349105 | 0.0016626 | 0.9679706 | 0.99534094 |
| ENSMUSG00000032349 | Elovl5 | protein_coding | -0.009372426 | 5.954478647 | 0.0016559 | 0.9680346 | 0.99534094 |
| ENSMUSG00000059939 | 9430015G10Rik | protein_coding | 0.015644193 | 3.558765264 | 0.0016385 | 0.9682028 | 0.995368456 |
| ENSMUSG00000048388 | Fam171b | protein_coding | -0.007935112 | 7.116365488 | 0.0016192 | 0.9683907 | 0.995368456 |
| ENSMUSG00000070282 | 3000002C10Rik | transcribed_processe | -0.034861832 | 2.154410955 | 0.0016163 | 0.9684192 | 0.995368456 |
| ENSMUSG00000034613 | Ppm1h | protein_coding | -0.015018519 | 7.551319464 | 0.0016094 | 0.9685641 | 0.995368456 |
| ENSMUSG00000028609 | Magoh | protein_coding | -0.009617101 | 6.642206271 | 0.0015992 | 0.968587 | 0.995368456 |
| ENSMUSG00000019837 | Gtf3c6 | protein_coding | -0.008716243 | 6.844510486 | 0.0015961 | 0.9686167 | 0.995368456 |
| ENSMUSG00000038305 | Spats2l | protein_coding | 0.014682197 | 4.865048663 | 0.0015647 | 0.9689275 | 0.995548552 |
| ENSMUSG00000069678 | Pcgf1 | protein_coding | 0.010041468 | 5.263945238 | 0.0015597 | 0.9689771 | 0.995548552 |
| ENSMUSG00000061887 | Ssbp3 | protein_coding | 0.009757258 | 8.011994118 | 0.001536 | 0.9692141 | 0.995549476 |
| ENSMUSG00000079215 | Zfp664 | protein_coding | 0.011160621 | 4.983712089 | 0.0015154 | 0.9694206 | 0.995549476 |
| ENSMUSG00000020231 | Dip2a | protein_coding | 0.013596595 | 4.17885028 | 0.0015134 | 0.9694405 | 0.995549476 |
| ENSMUSG00000020668 | Kif3c | protein_coding | 0.029445911 | 2.57057705 | 0.0015106 | 0.9694683 | 0.995549476 |
| ENSMUSG00000022558 | Mroh1 | protein_coding | 0.012731097 | 4.884030128 | 0.0015098 | 0.969477 | 0.995549476 |
| ENSMUSG00000049232 | Tigd2 | protein_coding | 0.016772701 | 4.020976715 | 0.0014984 | 0.9695923 | 0.995549476 |
| ENSMUSG00000025374 | Nabp2 | protein_coding | 0.011466466 | 5.12100011 | 0.0014938 | 0.9696385 | 0.995549476 |
| ENSMUSG00000032038 | St3gal4 | protein_coding | -0.012084787 | 4.551804892 | 0.0014836 | 0.9697426 | 0.995549476 |
| ENSMUSG00000026259 | Ngef | protein_coding | -0.013608083 | 6.559742315 | 0.0014817 | 0.9698113 | 0.995549476 |
| ENSMUSG00000025400 | Tac2 | protein_coding | -0.008232328 | 9.178170504 | 0.0014427 | 0.9701625 | 0.995735534 |
| ENSMUSG00000036712 | Cyld | protein_coding | 0.007751074 | 6.611239859 | 0.0014398 | 0.9701925 | 0.995735534 |
| ENSMUSG00000024044 | Epb41l3 | protein_coding | 0.008985366 | 7.654101313 | 0.0014237 | 0.9703594 | 0.995735534 |
| ENSMUSG00000042425 | Frmpd3 | protein_coding | -0.023616295 | 2.928074851 | 0.0014234 | 0.9703629 | 0.995735534 |
| ENSMUSG00000024663 | Rab3il1 | protein_coding | 0.018093953 | 3.605714295 | 0.0014045 | 0.97056 | 0.995842677 |
| ENSMUSG00000025969 | Nrp2 | protein_coding | 0.01399688 | 4.422069096 | 0.0013928 | 0.9706829 | 0.995854123 |
| ENSMUSG00000022783 | Spag6l | protein_coding | 0.015246395 | 3.789404451 | 0.0013761 | 0.9708593 | 0.995854123 |
| ENSMUSG00000025575 | Cant1 | protein_coding | 0.013882962 | 4.199755079 | 0.0013744 | 0.9708767 | 0.995854123 |
| ENSMUSG00000031792 | Usb1 | protein_coding | -0.012480563 | 4.800074597 | 0.0013683 | 0.9709416 | 0.995854123 |
| ENSMUSG00000045896 | Paip2b | protein_coding | -0.008768138 | 5.696696941 | 0.0013383 | 0.9712612 | 0.99608697 |
| ENSMUSG00000020475 | Pgam2 | protein_coding | 0.011602989 | 4.342649937 | 0.0013275 | 0.9713782 | 0.996111934 |
| ENSMUSG00000026032 | Ndufb3 | protein_coding | -0.009278652 | 8.401086337 | 0.0012942 | 0.9717396 | 0.996200435 |
| ENSMUSG00000038954 | Supt3 | protein_coding | -0.011657533 | 4.465072397 | 0.0012667 | 0.9720409 | 0.996200435 |
| ENSMUSG00000023272 | Creld2 | protein_coding | -0.022921007 | 3.29677908 | 0.0012668 | 0.9720424 | 0.996200435 |
| ENSMUSG00000028514 | Usp24 | protein_coding | 0.019823884 | 3.687224854 | 0.0012624 | 0.9720912 | 0.996200435 |
| ENSMUSG00000038002 | Cramp1l | protein_coding | 0.015874567 | 3.327462723 | 0.0012544 | 0.9721773 | 0.996200435 |
| ENSMUSG00000004771 | Rab11a | protein_coding | 0.006372547 | 6.915055833 | 0.0012542 | 0.9721784 | 0.996200435 |
| ENSMUSG00000032342 | Mto1 | protein_coding | 0.014063672 | 3.61128198 | 0.0012459 | 0.9722715 | 0.996200435 |
| ENSMUSG00000021120 | Pigh | protein_coding | 0.019644866 | 3.330616155 | 0.0012378 | 0.9723619 | 0.996200435 |
| ENSMUSG00000027655 | Dhx35 | protein_coding | 0.014994708 | 4.10466349 | 0.0012161 | 0.9726051 | 0.996200435 |
| ENSMUSG00000027678 | Ncoa3 | protein_coding | -0.012784677 | 4.86053662 | 0.0012101 | 0.9726725 | 0.996200435 |
| ENSMUSG00000018740 | Slc25a35 | protein_coding | -0.024877047 | 2.782725625 | 0.0012047 | 0.9727333 | 0.996200435 |
| ENSMUSG00000024683 | Mrpl16 | protein_coding | 0.007455224 | 6.373796248 | 0.0012006 | 0.9727794 | 0.996200435 |
| ENSMUSG00000022401 | Xpnpep3 | protein_coding | 0.015418661 | 3.690002345 | 0.0011996 | 0.9727907 | 0.996200435 |
| ENSMUSG00000078439 | Smim24 | protein_coding | -0.02100869 | 2.738258183 | 0.0011838 | 0.9729706 | 0.996200435 |
| ENSMUSG00000024145 | Pigf | protein_coding | 0.01233045 | 5.052367098 | 0.0011786 | 0.9730298 | 0.996200435 |
| ENSMUSG00000032673 | Prorsd1 | protein_coding | -0.010513951 | 4.986987756 | 0.0011574 | 0.9732737 | 0.996200435 |
| ENSMUSG00000040774 | Cept1 | protein_coding | -0.006994491 | 6.258797858 | 0.0011518 | 0.9733385 | 0.996200435 |
| ENSMUSG00000046324 | Ermp1 | protein_coding | -0.014227687 | 4.23663807 | 0.0011311 | 0.9735787 | 0.996200435 |
| ENSMUSG00000071856 | Mcc | protein_coding | 0.031510176 | 2.394947163 | 0.0011294 | 0.9736 | 0.996200435 |
| ENSMUSG00000005045 | Chd5 | protein_coding | -0.008275547 | 6.187593633 | 0.001126 | 0.9736389 | 0.996200435 |
| ENSMUSG00000042211 | Fbxo38 | protein_coding | -0.008785961 | 5.164586381 | 0.0011209 | 0.9736988 | 0.996200435 |
| ENSMUSG00000021694 | Ercc8 | protein_coding | 0.014098766 | 4.122049179 | 0.0011201 | 0.9737072 | 0.996200435 |

| ensembl_gene_id | external_gene_name | gene_biotype | log2 (fold change) | log2 (counts per million) | F-value | p-value | False discovery rate |
|---|---|---|---|---|---|---|---|
| ENSMUSG00000019732 | Calr3 | protein_coding | -0.027636366 | 3.324885192 | 0.0011236 | 0.9737241 | 0.996200435 |
| ENSMUSG00000032547 | Ryk | protein_coding | 0.015641997 | 3.831005932 | 0.0011032 | 0.9739067 | 0.996200435 |
| ENSMUSG00000037965 | Zc3h7a | protein_coding | -0.009047909 | 6.426642557 | 0.0010951 | 0.9740035 | 0.996200435 |
| ENSMUSG00000070283 | Ndufaf3 | protein_coding | 0.006513586 | 8.101443415 | 0.0010913 | 0.974048 | 0.996200435 |
| ENSMUSG00000006403 | Adamts4 | protein_coding | -0.016919322 | 3.873262184 | 0.00109 | 0.9740639 | 0.996200435 |
| ENSMUSG00000003731 | Kpna6 | protein_coding | -0.009673806 | 4.824788122 | 0.0010825 | 0.9741524 | 0.996200435 |
| ENSMUSG00000052713 | Zfp608 | protein_coding | -0.01215025 | 4.004855985 | 0.001071 | 0.9742902 | 0.996200435 |
| ENSMUSG00000051351 | Zfp46 | protein_coding | -0.014272489 | 3.520899645 | 0.0010668 | 0.9743409 | 0.996200435 |
| ENSMUSG00000060739 | Nsa2 | protein_coding | 0.012554969 | 3.599785677 | 0.0010653 | 0.9743585 | 0.996200435 |
| ENSMUSG00000096914 | Galntl6 | protein_coding | 0.008980408 | 5.198842689 | 0.0010563 | 0.974467 | 0.996200435 |
| ENSMUSG00000037278 | Tmem97 | protein_coding | -0.009802633 | 4.261964516 | 0.0010507 | 0.9745346 | 0.996200435 |
| ENSMUSG00000029371 | Cxcl5 | protein_coding | 0.034106467 | 2.469770503 | 0.0010462 | 0.9746144 | 0.996200435 |
| ENSMUSG00000025269 | Apex2 | protein_coding | -0.015844933 | 3.326395185 | 0.0010305 | 0.9747806 | 0.996275649 |
| ENSMUSG00000024130 | Abca3 | protein_coding | 0.018278542 | 2.720833177 | 0.0010147 | 0.9749749 | 0.99637953 |
| ENSMUSG00000038764 | Ptpn3 | protein_coding | 0.017752723 | 4.327253601 | 0.0010038 | 0.9751482 | 0.996461951 |
| ENSMUSG00000024181 | Mrpl28 | protein_coding | 0.005794275 | 8.842048939 | 0.0009789 | 0.9754199 | 0.996529254 |
| ENSMUSG00000031105 | Slc25a14 | protein_coding | -0.006036767 | 7.199902593 | 0.0009748 | 0.9754714 | 0.996529254 |
| ENSMUSG00000068747 | Sort1 | protein_coding | 0.009282588 | 5.841445731 | 0.0009732 | 0.9754921 | 0.996529254 |
| ENSMUSG00000050989 | Selenon | protein_coding | 0.016872792 | 3.32866713 | 0.000953 | 0.9757478 | 0.996695707 |
| ENSMUSG00000041351 | Rap1gap | protein_coding | -0.008641985 | 6.109828099 | 0.0009455 | 0.9758428 | 0.996698073 |
| ENSMUSG00000020700 | Map3k3 | protein_coding | -0.01536955 | 3.35893511 | 0.000929 | 0.9760501 | 0.996813748 |
| ENSMUSG00000008035 | Mid1ip1 | protein_coding | 0.008676428 | 6.306994888 | 0.0009223 | 0.9761414 | 0.996813748 |
| ENSMUSG00000046572 | Zfp518b | protein_coding | -0.022268308 | 2.427074338 | 0.0009097 | 0.9763042 | 0.9968377 |
| ENSMUSG00000034021 | Pds5b | protein_coding | 0.006538778 | 6.033313339 | 0.0009062 | 0.9763503 | 0.9968377 |
| ENSMUSG00000037196 | Pacrg | protein_coding | 0.006240014 | 6.015750497 | 0.0008806 | 0.9766859 | 0.996839304 |
| ENSMUSG00000076432 | Ywhaq | protein_coding | 0.005096014 | 8.08258164 | 0.0008788 | 0.9767102 | 0.996839304 |
| ENSMUSG00000027879 | Sec22b | protein_coding | 0.005775812 | 7.907287196 | 0.000875 | 0.9767614 | 0.996839304 |
| ENSMUSG00000025377 | Tepsin | protein_coding | 0.016563637 | 2.691341674 | 0.0008679 | 0.9768551 | 0.996839304 |
| ENSMUSG00000028053 | Ash1l | protein_coding | -0.007289643 | 6.306372161 | 0.0008641 | 0.9769058 | 0.996839304 |
| ENSMUSG00000042700 | Sipa1l1 | protein_coding | -0.008235437 | 6.147735034 | 0.0008543 | 0.9770367 | 0.996839304 |
| ENSMUSG00000024901 | Peli3 | protein_coding | -0.010571262 | 3.903887518 | 0.0008461 | 0.9771482 | 0.996839304 |
| ENSMUSG00000042215 | Bag2 | protein_coding | -0.009399495 | 4.929207901 | 0.0008446 | 0.9771681 | 0.996839304 |
| ENSMUSG00000093460 | Six3os1 | processed_transcript | -0.013864394 | 3.60449471 | 0.0008429 | 0.9771905 | 0.996839304 |
| ENSMUSG00000032908 | Sgpp2 | protein_coding | 0.032269868 | 2.288440636 | 0.0008301 | 0.977379 | 0.996839304 |
| ENSMUSG00000101930 | Gm5441 | antisense_RNA | -0.006100656 | 6.242236178 | 0.0008278 | 0.9773957 | 0.996839304 |
| ENSMUSG00000074925 | Ptar1 | protein_coding | -0.030168798 | 2.381256175 | 0.0008236 | 0.9774643 | 0.996839304 |
| ENSMUSG00000022677 | Fopnl | protein_coding | 0.005477069 | 8.132747093 | 0.0008118 | 0.977616 | 0.996899446 |
| ENSMUSG00000027962 | Vcam1 | protein_coding | 0.014092721 | 3.937435322 | 0.0008013 | 0.9777607 | 0.996938516 |
| ENSMUSG00000026667 | Uhmk1 | protein_coding | 0.00573502 | 6.742017928 | 0.0007797 | 0.9780619 | 0.996938516 |
| ENSMUSG00000038608 | Dock10 | protein_coding | 0.010710945 | 4.563913217 | 0.0007754 | 0.9781222 | 0.996938516 |
| ENSMUSG00000062822 | 4833420G17Rik | protein_coding | 0.007935422 | 5.393469938 | 0.0007623 | 0.9783082 | 0.996938516 |
| ENSMUSG00000034926 | Dhcr24 | protein_coding | -0.01445581 | 3.241174384 | 0.0007565 | 0.978391 | 0.996938516 |
| ENSMUSG00000033964 | Zbtb41 | protein_coding | 0.012916424 | 3.457689117 | 0.0007553 | 0.9784084 | 0.996938516 |
| ENSMUSG00000028224 | Nbn | protein_coding | 0.013010509 | 3.964025293 | 0.000748 | 0.9785139 | 0.996938516 |
| ENSMUSG00000035392 | Dennd1a | protein_coding | -0.00838197 | 4.580423137 | 0.0007477 | 0.9785171 | 0.996938516 |
| ENSMUSG00000078591 | Hs3st4 | protein_coding | -0.007305218 | 5.317790841 | 0.0007437 | 0.9785747 | 0.996938516 |
| ENSMUSG00000025812 | Pard3 | protein_coding | -0.008265686 | 4.74976956 | 0.0007432 | 0.9785814 | 0.996938516 |
| ENSMUSG00000043262 | Uevld | protein_coding | -0.016920834 | 2.707248116 | 0.0007296 | 0.9787788 | 0.996940755 |
| ENSMUSG00000007050 | Lsm2 | protein_coding | 0.012630203 | 3.413094897 | 0.0007222 | 0.9788872 | 0.996940755 |
| ENSMUSG00000037296 | Lsm1 | protein_coding | -0.007490994 | 6.027523324 | 0.0006919 | 0.9793348 | 0.996940755 |
| ENSMUSG00000025350 | Rdh5 | protein_coding | -0.017310044 | 3.235796726 | 0.0006895 | 0.979372 | 0.996940755 |
| ENSMUSG00000023963 | Cyp39a1 | protein_coding | -0.026771171 | 2.887792642 | 0.0006919 | 0.9793791 | 0.996940755 |
| ENSMUSG00000032348 | Gsta4 | protein_coding | 0.005061884 | 7.262439524 | 0.0006827 | 0.9794714 | 0.996940755 |
| ENSMUSG00000059070 | Rpl18 | protein_coding | -0.00625139 | 6.174859405 | 0.0006773 | 0.9795526 | 0.996940755 |
| ENSMUSG00000023967 | Mrps18a | protein_coding | -0.006474945 | 5.128822715 | 0.0006762 | 0.9795699 | 0.996940755 |
| ENSMUSG00000051853 | Arf3 | protein_coding | -0.005261843 | 9.245091916 | 0.0006619 | 0.9797869 | 0.996940755 |
| ENSMUSG00000066900 | Suds3 | protein_coding | 0.006203568 | 5.491229556 | 0.0006585 | 0.9798386 | 0.996940755 |
| ENSMUSG00000028655 | Mfsd2a | protein_coding | 0.02819325 | 2.405620596 | 0.0006591 | 0.9798498 | 0.996940755 |
| ENSMUSG00000054863 | Fam19a5 | protein_coding | 0.005846025 | 6.30314532 | 0.0006558 | 0.979881 | 0.996940755 |
| ENSMUSG00000095105 | Edaradd | protein_coding | 0.031968307 | 1.826958411 | 0.0006485 | 0.9799935 | 0.996940755 |
| ENSMUSG00000024921 | Smarca2 | protein_coding | 0.005897938 | 6.0614837 | 0.0006449 | 0.9800489 | 0.996940755 |
| ENSMUSG00000001827 | Folr1 | protein_coding | -0.023338764 | 3.469133099 | 0.00065 | 0.980089 | 0.996940755 |
| ENSMUSG00000020828 | Pld2 | protein_coding | -0.010620346 | 3.411782976 | 0.0006326 | 0.9802394 | 0.996940755 |
| ENSMUSG00000028367 | Txn1 | protein_coding | -0.00482681 | 8.31226565 | 0.0006297 | 0.980284 | 0.996940755 |
| ENSMUSG00000038301 | Snx10 | protein_coding | -0.00566262 | 8.472617737 | 0.0006195 | 0.9804454 | 0.996940755 |
| ENSMUSG00000022883 | Robo1 | protein_coding | 0.008658661 | 3.967815851 | 0.0006129 | 0.9805501 | 0.996940755 |
| ENSMUSG00000015748 | Prpf3 | protein_coding | 0.009213679 | 4.036123094 | 0.0005987 | 0.9807755 | 0.996940755 |
| ENSMUSG00000006522 | Itih3 | protein_coding | 0.009418159 | 5.66703038 | 0.0005965 | 0.9808305 | 0.996940755 |
| ENSMUSG00000022263 | Trio | protein_coding | -0.006601133 | 5.447010341 | 0.0005872 | 0.980962 | 0.996940755 |
| ENSMUSG00000006307 | Kmt2b | protein_coding | -0.014489511 | 3.089273277 | 0.0005756 | 0.9811501 | 0.996940755 |
| ENSMUSG00000060475 | Wtap | protein_coding | 0.011509294 | 5.476082691 | 0.0005781 | 0.9811511 | 0.996940755 |
| ENSMUSG00000021033 | Gstz1 | protein_coding | 0.004581858 | 7.07085356 | 0.0005583 | 0.9814354 | 0.996940755 |
| ENSMUSG00000033256 | Shf | protein_coding | -0.013853069 | 2.677663578 | 0.0005395 | 0.9817512 | 0.996940755 |
| ENSMUSG00000071341 | Egr4 | protein_coding | 0.022733092 | 1.664414579 | 0.0005278 | 0.9819497 | 0.996940755 |

| ensembl_gene_id | external_gene_name | gene_biotype | log2 (fold change) | log2 (counts per million) | F-value | p-value | False discovery rate |
|---|---|---|---|---|---|---|---|
| ENSMUSG00000024516 | Sec11c | protein_coding | -0.005803358 | 7.690142537 | 0.0005247 | 0.9820042 | 0.996940755 |
| ENSMUSG00000042228 | Lyn | protein_coding | 0.018265396 | 2.375049205 | 0.0005205 | 0.9820752 | 0.996940755 |
| ENSMUSG00000051154 | Commd3 | protein_coding | 0.004259087 | 8.214268243 | 0.0005114 | 0.9822326 | 0.996940755 |
| ENSMUSG00000012640 | Zfp715 | protein_coding | 0.009274684 | 3.863644095 | 0.0005069 | 0.9823101 | 0.996940755 |
| ENSMUSG00000027006 | Dnajc10 | protein_coding | -0.0047535 | 5.867635521 | 0.0004939 | 0.982539 | 0.996940755 |
| ENSMUSG00000040725 | Hnrnpul1 | protein_coding | -0.007335497 | 5.585243096 | 0.0004939 | 0.9825392 | 0.996940755 |
| ENSMUSG00000034247 | Plekhm1 | protein_coding | -0.008508628 | 3.720036447 | 0.0004887 | 0.9826318 | 0.996940755 |
| ENSMUSG00000026894 | Morn5 | protein_coding | -0.011164222 | 3.188588865 | 0.0004663 | 0.9830337 | 0.996940755 |
| ENSMUSG00000008200 | Fnbp4 | protein_coding | 0.007777575 | 3.868926687 | 0.0004653 | 0.9830529 | 0.996940755 |
| ENSMUSG00000046711 | Hmga1 | protein_coding | -0.008039513 | 3.853199304 | 0.0004522 | 0.9832914 | 0.996940755 |
| ENSMUSG00000040746 | Rnf167 | protein_coding | -0.004201665 | 7.602093668 | 0.0004497 | 0.9833382 | 0.996940755 |
| ENSMUSG00000041078 | Grid1 | protein_coding | 0.005412051 | 5.258684022 | 0.0004468 | 0.9833921 | 0.996940755 |
| ENSMUSG00000056629 | Fkbp2 | protein_coding | -0.004528618 | 9.108188008 | 0.0004442 | 0.9834402 | 0.996940755 |
| ENSMUSG00000028010 | Gar1 | protein_coding | -0.015890981 | 2.31814993 | 0.0004417 | 0.9834879 | 0.996940755 |
| ENSMUSG00000060950 | Trmt61a | protein_coding | 0.009130053 | 3.441033552 | 0.0004358 | 0.9835983 | 0.996940755 |
| ENSMUSG00000050088 | 1600012H06Rik | protein_coding | -0.01956061 | 2.783557498 | 0.0004348 | 0.9836337 | 0.996940755 |
| ENSMUSG00000025135 | Anapc11 | protein_coding | 0.003653931 | 9.282356923 | 0.0004289 | 0.9837292 | 0.996940755 |
| ENSMUSG00000035521 | Gnptg | protein_coding | 0.003899344 | 7.368568313 | 0.0004274 | 0.9837566 | 0.996940755 |
| ENSMUSG00000028134 | Ptbp2 | protein_coding | 0.005022314 | 6.239387828 | 0.0004269 | 0.9837656 | 0.996940755 |
| ENSMUSG00000027270 | Lamp5 | protein_coding | -0.005592763 | 7.01568764 | 0.0004235 | 0.9838314 | 0.996940755 |
| ENSMUSG00000046201 | Scaf8 | protein_coding | -0.007162744 | 4.444221024 | 0.0004195 | 0.983907 | 0.996940755 |
| ENSMUSG00000002608 | Ccdc97 | protein_coding | 0.006473441 | 5.891737021 | 0.0004155 | 0.9839858 | 0.996940755 |
| ENSMUSG00000033577 | Myo6 | protein_coding | 0.005659415 | 5.153348015 | 0.0004128 | 0.9840359 | 0.996940755 |
| ENSMUSG00000078350 | Smim1 | protein_coding | 0.005397 | 5.456210545 | 0.0004094 | 0.9841017 | 0.996940755 |
| ENSMUSG00000033272 | Slc35a4 | protein_coding | -0.00495097 | 6.143286942 | 0.0003983 | 0.9843196 | 0.996940755 |
| ENSMUSG00000021240 | Abcd4 | protein_coding | 0.012189513 | 2.905909648 | 0.0003872 | 0.9845399 | 0.996940755 |
| ENSMUSG00000032481 | Smarcc1 | protein_coding | -0.005106795 | 5.557874425 | 0.0003863 | 0.9845584 | 0.996940755 |
| ENSMUSG00000020056 | Washc3 | protein_coding | -0.004618033 | 6.423032868 | 0.0003845 | 0.9845927 | 0.996940755 |
| ENSMUSG00000037907 | Ankrd13b | protein_coding | 0.010296796 | 3.239422366 | 0.0003803 | 0.9846777 | 0.996940755 |
| ENSMUSG00000062647 | Rpl7a | protein_coding | -0.003934295 | 6.067649454 | 0.0003755 | 0.9847742 | 0.996940755 |
| ENSMUSG00000031093 | Dock11 | protein_coding | -0.015366127 | 2.307063063 | 0.0003667 | 0.9849538 | 0.996940755 |
| ENSMUSG00000037563 | Rps16 | protein_coding | 0.010314429 | 2.816220096 | 0.0003642 | 0.9850065 | 0.996940755 |
| ENSMUSG00000036989 | Trim3 | protein_coding | -0.004308183 | 6.15959805 | 0.0003565 | 0.9851653 | 0.996940755 |
| ENSMUSG00000027601 | Mtfr1 | protein_coding | 0.007814131 | 3.960358547 | 0.0003379 | 0.9855564 | 0.996940755 |
| ENSMUSG00000050721 | Plekho2 | protein_coding | 0.006441983 | 4.310670604 | 0.0003332 | 0.9856573 | 0.996940755 |
| ENSMUSG00000025702 | 8-Mar | protein_coding | -0.007357289 | 4.250494278 | 0.000331 | 0.985705 | 0.996940755 |
| ENSMUSG00000024112 | Cacna1h | protein_coding | -0.013898523 | 2.669602741 | 0.0003251 | 0.9858325 | 0.996940755 |
| ENSMUSG00000005506 | Celf1 | protein_coding | 0.003475741 | 6.52656855 | 0.0003199 | 0.9859469 | 0.996940755 |
| ENSMUSG00000035901 | Dennd5a | protein_coding | 0.003657063 | 6.827601737 | 0.0003169 | 0.9860123 | 0.996940755 |
| ENSMUSG00000031872 | Bean1 | protein_coding | -0.010853063 | 2.534409631 | 0.0003082 | 0.9862062 | 0.996940755 |
| ENSMUSG00000042385 | Gzmk | protein_coding | -0.015433491 | 2.587285795 | 0.0003081 | 0.9862096 | 0.996940755 |
| ENSMUSG00000027534 | Snx16 | protein_coding | 0.009457313 | 3.755280104 | 0.0003078 | 0.9862162 | 0.996940755 |
| ENSMUSG00000046182 | Gsg1l | protein_coding | -0.008393897 | 3.759179788 | 0.0002921 | 0.9865717 | 0.996940755 |
| ENSMUSG00000020120 | Plek | protein_coding | -0.00714916 | 3.509583045 | 0.0002893 | 0.9866361 | 0.996940755 |
| ENSMUSG00000073236 | 2500004C02Rik | processed_transcript | -0.005271158 | 4.325522771 | 0.0002867 | 0.9866951 | 0.996940755 |
| ENSMUSG00000036943 | Rab8b | protein_coding | 0.005584459 | 4.885630103 | 0.0002798 | 0.986856 | 0.996940755 |
| ENSMUSG00000006782 | Cnp | protein_coding | -0.004601722 | 9.811024305 | 0.0002771 | 0.9869211 | 0.996940755 |
| ENSMUSG00000039578 | Ccser1 | protein_coding | -0.005572962 | 4.043679451 | 0.000276 | 0.986946 | 0.996940755 |
| ENSMUSG00000029833 | Trim24 | protein_coding | -0.005865157 | 3.973895543 | 0.0002749 | 0.9869719 | 0.996940755 |
| ENSMUSG00000001418 | Glmp | protein_coding | -0.003682779 | 6.461224664 | 0.0002743 | 0.986988 | 0.996940755 |
| ENSMUSG00000027404 | Snrpb | protein_coding | 0.003651247 | 7.354725503 | 0.0002715 | 0.987054 | 0.996940755 |
| ENSMUSG00000034118 | Tpst1 | protein_coding | -0.005925895 | 4.475285247 | 0.0002711 | 0.9870642 | 0.996940755 |
| ENSMUSG00000021951 | Eef1akmt1 | protein_coding | 0.002857208 | 7.540472218 | 0.0002709 | 0.9870687 | 0.996940755 |
| ENSMUSG00000028145 | Them4 | protein_coding | -0.003774874 | 6.586980391 | 0.0002707 | 0.9870733 | 0.996940755 |
| ENSMUSG00000071073 | Lrrc73 | protein_coding | -0.00616755 | 3.816430592 | 0.0002643 | 0.9872253 | 0.996940755 |
| ENSMUSG00000004451 | Ralb | protein_coding | -0.004492183 | 4.742937865 | 0.0002574 | 0.9873949 | 0.996940755 |
| ENSMUSG00000034324 | Tmem132c | protein_coding | -0.010840065 | 2.717999183 | 0.0002522 | 0.9875214 | 0.996940755 |
| ENSMUSG00000090266 | Mettl23 | protein_coding | 0.005714185 | 4.816072364 | 0.0002399 | 0.9878302 | 0.996940755 |
| ENSMUSG00000031922 | Cep57 | protein_coding | 0.003929771 | 5.132438851 | 0.0002398 | 0.9878324 | 0.996940755 |
| ENSMUSG00000029725 | Ppp1r35 | protein_coding | -0.009292201 | 2.750171303 | 0.0002386 | 0.9878625 | 0.996940755 |
| ENSMUSG00000025362 | Rps26 | protein_coding | -0.004134941 | 7.665758863 | 0.0002362 | 0.9879262 | 0.996940755 |
| ENSMUSG00000038240 | Pdss2 | protein_coding | -0.004485644 | 4.928223451 | 0.0002349 | 0.9879584 | 0.996940755 |
| ENSMUSG00000027035 | Cers6 | protein_coding | -0.00507293 | 4.366479279 | 0.0002256 | 0.9881988 | 0.996940755 |
| ENSMUSG00000063077 | Kif1b | protein_coding | 0.002841389 | 7.914263442 | 0.0002234 | 0.9882561 | 0.996940755 |
| ENSMUSG00000061451 | Tmem151a | protein_coding | -0.003856913 | 5.821230767 | 0.0002198 | 0.9883501 | 0.996940755 |
| ENSMUSG00000056211 | R3hdm1 | protein_coding | -0.002704868 | 6.656809729 | 0.0002131 | 0.9885311 | 0.996940755 |
| ENSMUSG00000078584 | AU022252 | protein_coding | 0.004295843 | 4.762003024 | 0.0002111 | 0.9885836 | 0.996940755 |
| ENSMUSG00000043923 | Ccdc84 | protein_coding | -0.012512963 | 2.271907717 | 0.0002105 | 0.9885999 | 0.996940755 |
| ENSMUSG00000046519 | Golph3l | protein_coding | -0.005312593 | 4.820095839 | 0.0002092 | 0.9886359 | 0.996940755 |
| ENSMUSG00000049439 | Cyp20a1 | protein_coding | -0.005293274 | 4.164389823 | 0.0002089 | 0.9886445 | 0.996940755 |
| ENSMUSG00000089945 | Pakap | protein_coding | -0.007368814 | 3.012438572 | 0.000204 | 0.9887786 | 0.996940755 |
| ENSMUSG00000044252 | Osbpl1a | protein_coding | 0.002792821 | 6.867745508 | 0.0002008 | 0.9888658 | 0.996940755 |
| ENSMUSG00000030838 | Ush1c | protein_coding | 0.014834956 | 2.411824723 | 0.0001934 | 0.9890785 | 0.996940755 |
| ENSMUSG00000008348 | Ubc | protein_coding | 0.003266715 | 8.069522431 | 0.0001919 | 0.9891156 | 0.996940755 |

| ensembl_gene_id | external_gene_name | gene_biotype | log2 (fold change) | log2 (counts per million) | F-value | p-value | False discovery rate |
|---|---|---|---|---|---|---|---|
| ENSMUSG00000031778 | Cx3cl1 | protein_coding | 0.002809071 | 7.131790328 | 0.0001906 | 0.9891513 | 0.996940755 |
| ENSMUSG00000038368 | Focad | protein_coding | -0.007728578 | 2.994303433 | 0.000188 | 0.9892253 | 0.996940755 |
| ENSMUSG00000017400 | Stac2 | protein_coding | 0.005783532 | 4.566754954 | 0.0001875 | 0.9892413 | 0.996940755 |
| ENSMUSG00000021692 | Dimt1 | protein_coding | 0.004626059 | 3.968309027 | 0.0001791 | 0.9894855 | 0.996940755 |
| ENSMUSG00000035642 | Aamdc | protein_coding | -0.002826614 | 7.16647457 | 0.0001763 | 0.9895682 | 0.996940755 |
| ENSMUSG00000101892 | 9130401M01Rik | protein_coding | -0.003760265 | 4.625813844 | 0.0001684 | 0.9898046 | 0.996940755 |
| ENSMUSG00000036894 | Rap2b | protein_coding | -0.003562474 | 6.864916024 | 0.000164 | 0.9899399 | 0.996940755 |
| ENSMUSG00000028082 | Sh3d19 | protein_coding | -0.005738858 | 3.122541729 | 0.0001621 | 0.9899958 | 0.996940755 |
| ENSMUSG00000035601 | Trmt10b | protein_coding | 0.002801548 | 5.834775679 | 0.0001592 | 0.9900874 | 0.996940755 |
| ENSMUSG00000024621 | Csf1r | protein_coding | -0.004522643 | 4.449327018 | 0.0001586 | 0.9901061 | 0.996940755 |
| ENSMUSG00000054428 | Atpif1 | protein_coding | 0.00305753 | 9.284073866 | 0.0001562 | 0.9901784 | 0.996940755 |
| ENSMUSG00000034171 | Faah | protein_coding | -0.004353711 | 4.293621047 | 0.000156 | 0.9901871 | 0.996940755 |
| ENSMUSG00000034449 | Dhrs11 | protein_coding | 0.005976051 | 3.101245341 | 0.0001535 | 0.9902665 | 0.996940755 |
| ENSMUSG00000034327 | Kctd9 | protein_coding | 0.004668193 | 3.86237387 | 0.000152 | 0.9903136 | 0.996940755 |
| ENSMUSG00000043311 | D17H6S53E | protein_coding | -0.00604104 | 5.399066154 | 0.0001517 | 0.9903442 | 0.996940755 |
| ENSMUSG00000042423 | Fbrs | protein_coding | 0.005651618 | 3.257937636 | 0.0001501 | 0.9903731 | 0.996940755 |
| ENSMUSG00000041126 | H2afv | protein_coding | -0.002916942 | 5.493399484 | 0.00015 | 0.990376 | 0.996940755 |
| ENSMUSG00000032231 | Anxa2 | protein_coding | -0.004181309 | 4.781141088 | 0.0001481 | 0.9904389 | 0.996940755 |
| ENSMUSG00000039349 | C130074G19Rik | protein_coding | -0.006918603 | 3.535374711 | 0.0001439 | 0.9905746 | 0.996940755 |
| ENSMUSG00000054766 | Set | protein_coding | -0.002811663 | 8.405427607 | 0.0001436 | 0.990585 | 0.996940755 |
| ENSMUSG00000042364 | Snx18 | protein_coding | -0.005380662 | 3.823215141 | 0.0001426 | 0.9906185 | 0.996940755 |
| ENSMUSG00000073664 | Nbeal1 | protein_coding | -0.00526802 | 3.613402452 | 0.0001402 | 0.9906953 | 0.996940755 |
| ENSMUSG00000002221 | Paxip1 | protein_coding | -0.004031884 | 4.563424626 | 0.0001394 | 0.9907223 | 0.996940755 |
| ENSMUSG00000028883 | Sema3a | protein_coding | 0.005970135 | 3.199347871 | 0.0001379 | 0.9907722 | 0.996940755 |
| ENSMUSG00000024766 | Lipo3 | protein_coding | -0.005506701 | 3.2692085 | 0.0001364 | 0.9908223 | 0.996940755 |
| ENSMUSG00000040189 | Ccdc114 | protein_coding | 0.015207968 | 2.03408562 | 0.0001365 | 0.9908303 | 0.996940755 |
| ENSMUSG00000033720 | Sfxn5 | protein_coding | -0.003721778 | 8.774874846 | 0.0001361 | 0.9908435 | 0.996940755 |
| ENSMUSG00000070687 | Htr1d | protein_coding | 0.011002322 | 2.709892867 | 0.0001325 | 0.9909645 | 0.996940755 |
| ENSMUSG00000042079 | Hnrnpf | protein_coding | 0.005497875 | 2.968149836 | 0.0001302 | 0.9910354 | 0.996940755 |
| ENSMUSG00000044748 | Defb1 | protein_coding | -0.011480579 | 3.007935018 | 0.0001282 | 0.9911253 | 0.996940755 |
| ENSMUSG00000038949 | Cnst | protein_coding | 0.004459024 | 4.301464429 | 0.0001244 | 0.9912352 | 0.996940755 |
| ENSMUSG00000031232 | Magt1 | protein_coding | 0.004893813 | 3.95752326 | 0.000124 | 0.9912519 | 0.996940755 |
| ENSMUSG00000056076 | Eif3b | protein_coding | -0.002636929 | 5.88964188 | 0.0001235 | 0.9912674 | 0.996940755 |
| ENSMUSG00000048814 | Lonrf2 | protein_coding | 0.003209483 | 6.857009959 | 0.0001232 | 0.9912817 | 0.996940755 |
| ENSMUSG00000041794 | Myrip | protein_coding | -0.00480891 | 3.511552876 | 0.000122 | 0.9913215 | 0.996940755 |
| ENSMUSG00000037395 | Rcor3 | protein_coding | 0.00469167 | 3.439315693 | 0.0001192 | 0.9914229 | 0.996940755 |
| ENSMUSG00000028461 | Ccdc107 | protein_coding | 0.005603603 | 5.916558428 | 0.0001175 | 0.9915211 | 0.996940755 |
| ENSMUSG00000040209 | Zfp704 | protein_coding | 0.006408311 | 2.930957915 | 0.0001158 | 0.9915446 | 0.996940755 |
| ENSMUSG00000022009 | Nufip1 | protein_coding | -0.003871396 | 3.825513868 | 0.0001133 | 0.991636 | 0.996940755 |
| ENSMUSG00000062590 | Armc9 | protein_coding | 0.002791638 | 5.167580818 | 0.0001082 | 0.9918254 | 0.996940755 |
| ENSMUSG00000019470 | Xab2 | protein_coding | 0.003847934 | 5.777824972 | 0.0001048 | 0.9919659 | 0.996940755 |
| ENSMUSG00000043323 | Fbrsl1 | protein_coding | -0.004662128 | 3.680549112 | 0.0001042 | 0.9919784 | 0.996940755 |
| ENSMUSG00000027774 | Gfm1 | protein_coding | -0.003166563 | 4.742264326 | 0.0001035 | 0.9920061 | 0.996940755 |
| ENSMUSG00000053329 | D10Jhu81e | protein_coding | 0.002862401 | 5.636115395 | 0.0001016 | 0.9920783 | 0.996940755 |
| ENSMUSG00000021621 | Zcchc9 | protein_coding | 0.003510398 | 4.912392051 | 9.87E-05 | 0.9921945 | 0.996940755 |
| ENSMUSG00000027272 | Ubr1 | protein_coding | 0.003454859 | 3.851006777 | 9.82E-05 | 0.9922124 | 0.996940755 |
| ENSMUSG00000028278 | Rragd | protein_coding | -0.002162321 | 6.228608853 | 8.69E-05 | 0.9926736 | 0.997204829 |
| ENSMUSG00000048970 | C1galt1c1 | protein_coding | -0.002663003 | 5.775163264 | 8.66E-05 | 0.9926881 | 0.997204829 |
| ENSMUSG00000025199 | Chuk | protein_coding | -0.005774621 | 3.326487007 | 8.51E-05 | 0.9927534 | 0.997204829 |
| ENSMUSG00000048330 | Ric3 | protein_coding | -0.002791729 | 5.306662517 | 7.95E-05 | 0.9929948 | 0.997273454 |
| ENSMUSG00000024487 | Yipf5 | protein_coding | -0.002096448 | 5.843376927 | 7.92E-05 | 0.9930072 | 0.997273454 |
| ENSMUSG00000020268 | Lyrm7 | protein_coding | 0.001925649 | 5.825074949 | 7.33E-05 | 0.9932739 | 0.997358106 |
| ENSMUSG00000042155 | Klhl23 | protein_coding | -0.002461587 | 5.279083359 | 7.32E-05 | 0.993277 | 0.997358106 |
| ENSMUSG00000029822 | Osbpl3 | protein_coding | -0.004937455 | 2.998039118 | 6.99E-05 | 0.993431 | 0.99741954 |
| ENSMUSG00000040479 | Dgkz | protein_coding | -0.002188719 | 6.007074155 | 6.78E-05 | 0.9935324 | 0.997422919 |
| ENSMUSG00000041298 | Katnal1 | protein_coding | 0.001986674 | 6.237262926 | 6.56E-05 | 0.9936375 | 0.997422919 |
| ENSMUSG00000003269 | Cyth2 | protein_coding | 0.002530335 | 5.170669015 | 6.36E-05 | 0.9937334 | 0.997422919 |
| ENSMUSG00000054808 | Actn4 | protein_coding | 0.002767501 | 4.526153361 | 6.15E-05 | 0.993837 | 0.997422919 |
| ENSMUSG00000051557 | Pusl1 | protein_coding | 0.003304578 | 4.982828726 | 5.79E-05 | 0.9940225 | 0.997422919 |
| ENSMUSG00000021756 | Il6st | protein_coding | 0.002728769 | 4.07293837 | 5.75E-05 | 0.9940405 | 0.997422919 |
| ENSMUSG00000054079 | Utp18 | protein_coding | 0.002389896 | 4.102257036 | 5.59E-05 | 0.9941267 | 0.997422919 |
| ENSMUSG00000025537 | Phkg1 | protein_coding | 0.003681307 | 3.493481303 | 5.49E-05 | 0.9941764 | 0.997422919 |
| ENSMUSG00000022214 | Dcaf11 | protein_coding | 0.002203796 | 5.684362016 | 5.20E-05 | 0.9943314 | 0.997449493 |
| ENSMUSG00000024949 | Sf1 | protein_coding | 0.001599416 | 5.964313519 | 5.10E-05 | 0.9943884 | 0.997449493 |
| ENSMUSG00000024219 | Anks1 | protein_coding | 0.003092024 | 3.484281808 | 4.90E-05 | 0.9944999 | 0.99746832 |
| ENSMUSG00000027940 | Tpm3 | protein_coding | 0.001298224 | 6.75232842 | 4.55E-05 | 0.9946972 | 0.997573168 |
| ENSMUSG00000054843 | Atrnl1 | protein_coding | 0.001968117 | 4.743823263 | 4.09E-05 | 0.994975 | 0.997758664 |
| ENSMUSG00000033170 | Card10 | protein_coding | -0.007477538 | 2.435956805 | 3.89E-05 | 0.9951086 | 0.997799633 |
| ENSMUSG00000025584 | Pde8a | protein_coding | -0.00225792 | 4.010309423 | 2.97E-05 | 0.9957143 | 0.998167505 |
| ENSMUSG00000021265 | Slc25a29 | protein_coding | -0.00249234 | 3.224824225 | 2.87E-05 | 0.9957872 | 0.998167505 |
| ENSMUSG00000020444 | Guk1 | protein_coding | -0.001650809 | 7.484912509 | 2.83E-05 | 0.995827 | 0.998167505 |
| ENSMUSG00000113771 | AC160108.1 | antisense_RNA | -0.002919803 | 3.105544482 | 2.79E-05 | 0.9958468 | 0.998167505 |
| ENSMUSG00000029469 | Ift81 | protein_coding | 0.001390971 | 5.347854405 | 2.42E-05 | 0.996137 | 0.998304324 |
| ENSMUSG00000029033 | Acap3 | protein_coding | -0.001403433 | 4.663657039 | 2.29E-05 | 0.9962439 | 0.998304324 |

| ensembl_gene_id | external_gene_name | gene_biotype | log2 (fold change) | log2 (counts per million) | F-value | p-value | False discovery rate |
|---|---|---|---|---|---|---|---|
| ENSMUSG00000021072 | Tmx1 | protein_coding | 0.001005343 | 7.163245087 | 2.26E-05 | 0.9962672 | 0.998304324 |
| ENSMUSG00000060735 | Rxfp3 | protein_coding | -0.003293389 | 2.619317142 | 2.15E-05 | 0.9963547 | 0.998304324 |
| ENSMUSG00000021635 | Rad17 | protein_coding | -0.001407839 | 4.620835006 | 1.98E-05 | 0.9965014 | 0.998308502 |
| ENSMUSG00000022488 | Nckap1l | protein_coding | 0.004661864 | 2.372215698 | 1.94E-05 | 0.9965445 | 0.998308502 |
| ENSMUSG00000016534 | Lamp2 | protein_coding | 0.001028557 | 6.080716892 | 1.81E-05 | 0.996658 | 0.99832914 |
| ENSMUSG00000018001 | Cyth3 | protein_coding | -0.00156761 | 3.947395335 | 1.65E-05 | 0.9968063 | 0.998363153 |
| ENSMUSG00000052539 | Magi3 | protein_coding | 0.002431308 | 2.938889907 | 1.53E-05 | 0.9969224 | 0.998363153 |
| ENSMUSG00000037887 | Dusp8 | protein_coding | -0.000949414 | 8.228678214 | 1.49E-05 | 0.9969705 | 0.998363153 |
| ENSMUSG00000000409 | Lck | protein_coding | -0.001805957 | 3.777898361 | 1.30E-05 | 0.9971656 | 0.998465519 |
| ENSMUSG00000037653 | Kctd8 | protein_coding | -0.001275432 | 3.702296862 | 9.00E-06 | 0.9976425 | 0.998850034 |
| ENSMUSG00000030126 | Tmcc1 | protein_coding | -0.000798714 | 5.601854664 | 7.43E-06 | 0.9978583 | 0.998956031 |
| ENSMUSG00000034758 | Tle6 | protein_coding | -0.002677254 | 2.579938696 | 6.19E-06 | 0.998047 | 0.998956031 |
| ENSMUSG00000044768 | D1Ertd622e | protein_coding | 0.000818647 | 4.60672709 | 5.98E-06 | 0.9980781 | 0.998956031 |
| ENSMUSG00000040560 | Wdr7 | protein_coding | 0.000527098 | 6.03494408 | 5.45E-06 | 0.9981661 | 0.998956031 |
| ENSMUSG00000030323 | Ift122 | protein_coding | 0.000721189 | 4.801334896 | 5.17E-06 | 0.9982128 | 0.998956031 |
| ENSMUSG00000031918 | Mtmr2 | protein_coding | 0.000556745 | 5.617265635 | 3.57E-06 | 0.9985144 | 0.999164834 |
| ENSMUSG00000048503 | Tmem136 | protein_coding | 0.000624803 | 3.52924235 | 2.62E-06 | 0.9987274 | 0.999220164 |
| ENSMUSG00000028788 | Ptp4a2 | protein_coding | 0.00036265 | 8.424714195 | 2.51E-06 | 0.9987555 | 0.999220164 |
| ENSMUSG00000044988 | Ucn3 | protein_coding | -0.000717506 | 3.261059349 | 1.70E-06 | 0.9989753 | 0.999239745 |
| ENSMUSG00000053774 | Ubxn7 | protein_coding | -0.000648986 | 3.209179744 | 1.50E-06 | 0.9990377 | 0.999239745 |
| ENSMUSG00000038831 | Ralgps1 | protein_coding | 0.000449359 | 4.464031335 | 1.30E-06 | 0.9991037 | 0.999239745 |
| ENSMUSG00000024601 | Isoc1 | protein_coding | -0.000346891 | 5.166928495 | 1.18E-06 | 0.9991468 | 0.999239745 |
| ENSMUSG00000040446 | Rprd1a | protein_coding | 0.000154568 | 3.754154012 | 1.41E-07 | 0.9997047 | 0.999704748 |
