## Supplemental Table 3 for "Extended amygdala-parabrachial circuits alter threat assessment to regulate feeding"

### TABLE S3

| ensembl_gene_id | external_gene_name | gene_biotype | log2 (fold change) | log2 (counts per million) | F-value | p-value | False discovery rate |
|---|---|---|---|---|---|---|---|
| ENSMUSG00000025400 | Tac2 | protein_coding | 4.317350176 | 9.178170504 | 204.70745 | 1.18E-10 | 1.27E-06 |
| ENSMUSG00000024907 | Gal | protein_coding | 2.070327984 | 9.118265128 | 53.348064 | 1.58E-06 | 0.008491817 |
| ENSMUSG00000092341 | Malat1 | lincRNA | -2.057219772 | 10.80327234 | 41.763497 | 7.15E-06 | 0.025634954 |
| ENSMUSG00000045731 | Pnoc | protein_coding | 1.298918771 | 6.687693393 | 33.930631 | 2.39E-05 | 0.053225749 |
| ENSMUSG00000100826 | Snhg14 | processed_transcript | -1.739780789 | 6.081352785 | 32.576294 | 3.01E-05 | 0.053225749 |
| ENSMUSG00000003872 | Lin7b | protein_coding | 1.500839601 | 6.138974701 | 31.903893 | 3.38E-05 | 0.053225749 |
| ENSMUSG00000000792 | Slc5a5 | protein_coding | 1.670215405 | 6.950427196 | 31.382555 | 3.70E-05 | 0.053225749 |
| ENSMUSG00000030844 | Rgs10 | protein_coding | 1.3140955 | 8.342266456 | 29.590401 | 5.10E-05 | 0.053225749 |
| ENSMUSG00000055373 | Fut9 | protein_coding | -1.955408953 | 5.562331203 | 29.452536 | 5.23E-05 | 0.053225749 |
| ENSMUSG00000028565 | Nfia | protein_coding | -1.716571336 | 6.998809161 | 29.425274 | 5.29E-05 | 0.053225749 |
| ENSMUSG00000109006 | B230209E15Rik | lincRNA | -1.644759053 | 6.39717013 | 29.236356 | 5.44E-05 | 0.053225749 |
| ENSMUSG00000040118 | Cacna2d1 | protein_coding | -1.59394463 | 5.728301998 | 28.450811 | 6.30E-05 | 0.056462121 |
| ENSMUSG00000079184 | Mphosph8 | protein_coding | -1.24952905 | 6.514185979 | 25.833597 | 0.0001046 | 0.081568287 |
| ENSMUSG00000022124 | Fbxl3 | protein_coding | -1.775415979 | 5.126446197 | 25.75666 | 0.0001062 | 0.081568287 |
| ENSMUSG00000025860 | Xiap | protein_coding | -1.368899662 | 6.136989412 | 24.884381 | 0.0001267 | 0.086552793 |
| ENSMUSG00000074746 | Pdzd8 | protein_coding | -1.468923938 | 5.42275172 | 24.782076 | 0.0001294 | 0.086552793 |
| ENSMUSG00000070576 | Mn1 | protein_coding | 2.237579835 | 8.317535647 | 25.912731 | 0.0001502 | 0.086552793 |
| ENSMUSG00000044933 | Sstr3 | protein_coding | 9.004482197 | 1.991282608 | 24.884287 | 0.0001526 | 0.086552793 |
| ENSMUSG00000025647 | Shisa5 | protein_coding | 1.354057615 | 7.52527952 | 23.763806 | 0.0001599 | 0.086552793 |
| ENSMUSG00000062078 | Qk | protein_coding | -1.278564241 | 7.886346701 | 23.433525 | 0.0001715 | 0.086552793 |
| ENSMUSG00000031425 | Plp1 | protein_coding | -1.332503708 | 10.61790917 | 23.272066 | 0.0001775 | 0.086552793 |
| ENSMUSG00000041215 | Yeats2 | protein_coding | 3.420966882 | 5.218110279 | 25.063191 | 0.000181 | 0.086552793 |
| ENSMUSG00000038518 | Jarid2 | protein_coding | 1.550129637 | 8.066763333 | 23.095487 | 0.0001851 | 0.086552793 |
| ENSMUSG00000061518 | Cox5b | protein_coding | 1.511173851 | 8.013800626 | 22.785761 | 0.0001979 | 0.088661854 |
| ENSMUSG00000029404 | Arl6ip4 | protein_coding | 1.00721221 | 7.407770941 | 22.08369 | 0.0002297 | 0.092475219 |
| ENSMUSG00000008036 | Ap2s1 | protein_coding | 1.061898421 | 8.101066345 | 22.018751 | 0.000233 | 0.092475219 |
| ENSMUSG00000069769 | Msi2 | protein_coding | -1.472499447 | 5.347387036 | 21.233165 | 0.0002776 | 0.092475219 |
| ENSMUSG00000019590 | Cyb561 | protein_coding | 1.257087477 | 7.175003432 | 21.066553 | 0.0002883 | 0.092475219 |
| ENSMUSG00000036357 | Gpr101 | protein_coding | 1.828855749 | 4.869678402 | 21.016188 | 0.0002916 | 0.092475219 |
| ENSMUSG00000019146 | Cacng2 | protein_coding | -1.576058507 | 5.129436191 | 20.968935 | 0.0002948 | 0.092475219 |
| ENSMUSG00000021377 | Dek | protein_coding | -1.582879977 | 5.116669153 | 20.941042 | 0.0002966 | 0.092475219 |
| ENSMUSG00000054013 | Tmem179 | protein_coding | 1.283870784 | 7.989735247 | 20.937738 | 0.0002969 | 0.092475219 |
| ENSMUSG00000025862 | Stag2 | protein_coding | -1.765124207 | 4.708151346 | 20.901083 | 0.0002994 | 0.092475219 |
| ENSMUSG00000032740 | Ccdc88a | protein_coding | -1.594640571 | 5.344034453 | 20.848513 | 0.000303 | 0.092475219 |
| ENSMUSG00000062115 | Rai1 | protein_coding | 1.830881744 | 5.042439665 | 20.806492 | 0.0003059 | 0.092475219 |
| ENSMUSG00000000838 | Fmr1 | protein_coding | -1.769840481 | 4.646406992 | 20.300109 | 0.0003437 | 0.092475219 |
| ENSMUSG00000046314 | Stxbp6 | protein_coding | -1.297535719 | 5.801587031 | 20.192799 | 0.0003524 | 0.092475219 |
| ENSMUSG00000033629 | Hacd3 | protein_coding | 1.021759277 | 7.534286954 | 19.839607 | 0.0003827 | 0.092475219 |
| ENSMUSG00000041966 | Dcaf17 | protein_coding | -2.204339836 | 3.947191032 | 19.791935 | 0.000387 | 0.092475219 |
| ENSMUSG00000034083 | Ccdc174 | protein_coding | -1.195652296 | 5.939229856 | 19.731307 | 0.0003926 | 0.092475219 |
| ENSMUSG00000018451 | 6330403K07Rik | lincRNA | 1.088150099 | 10.97421339 | 19.676777 | 0.0003977 | 0.092475219 |
| ENSMUSG00000021719 | Rgs7bp | protein_coding | -1.006219794 | 7.127353161 | 19.674279 | 0.0003979 | 0.092475219 |
| ENSMUSG00000022577 | Ly6h | protein_coding | 1.137285814 | 6.972127586 | 19.617719 | 0.0004033 | 0.092475219 |
| ENSMUSG00000030062 | Rpn1 | protein_coding | 0.986128133 | 6.965638885 | 19.537809 | 0.000411 | 0.092475219 |
| ENSMUSG00000026389 | Steap3 | protein_coding | 8.025859631 | 2.068060088 | 20.047619 | 0.0004222 | 0.092475219 |
| ENSMUSG00000064345 | mt-Nd2 | protein_coding | -1.831842214 | 10.31322046 | 20.00769 | 0.0004251 | 0.092475219 |
| ENSMUSG00000004748 | Mtfp1 | protein_coding | 1.128189693 | 7.955389561 | 19.362685 | 0.0004284 | 0.092475219 |
| ENSMUSG00000064341 | mt-Nd1 | protein_coding | -1.836476383 | 12.02550762 | 19.572329 | 0.0004476 | 0.092475219 |
| ENSMUSG00000026830 | Ermn | protein_coding | -0.962971481 | 7.384948842 | 19.162354 | 0.0004495 | 0.092475219 |
| ENSMUSG00000030067 | Foxp1 | protein_coding | -1.056832474 | 7.041363291 | 19.158571 | 0.0004499 | 0.092475219 |
| ENSMUSG00000026411 | Tmem9 | protein_coding | 1.012642544 | 7.866960137 | 19.135648 | 0.0004524 | 0.092475219 |
| ENSMUSG00000019768 | Esr1 | protein_coding | 3.332245148 | 3.539813594 | 18.852166 | 0.0004861 | 0.092475219 |
| ENSMUSG00000032766 | Gng11 | protein_coding | -1.296340546 | 6.068835467 | 18.815724 | 0.0004887 | 0.092475219 |
| ENSMUSG00000017002 | Slpi | protein_coding | 3.937964928 | 2.919325958 | 18.807807 | 0.0004896 | 0.092475219 |
| ENSMUSG00000028673 | Fuca1 | protein_coding | 1.040939165 | 6.853897675 | 18.747627 | 0.0004968 | 0.092475219 |
| ENSMUSG00000043498 | 9330132A10Rik | pseudogene | -2.245618901 | 3.451029808 | 18.628658 | 0.0005114 | 0.092475219 |
| ENSMUSG00000022119 | Rbm26 | protein_coding | -1.722147788 | 4.824486385 | 18.46914 | 0.0005317 | 0.092475219 |
| ENSMUSG00000028522 | Mier1 | protein_coding | -1.354398637 | 5.329494853 | 18.453901 | 0.0005337 | 0.092475219 |
| ENSMUSG00000027162 | Lin7c | protein_coding | -1.035053378 | 6.417872567 | 18.441226 | 0.0005354 | 0.092475219 |
| ENSMUSG00000020152 | Actr2 | protein_coding | -1.046794809 | 9.058469489 | 18.407168 | 0.0005399 | 0.092475219 |
| ENSMUSG00000018169 | Mfng | protein_coding | 2.322698596 | 3.752551007 | 18.397999 | 0.0005411 | 0.092475219 |
| ENSMUSG00000017978 | Cadps2 | protein_coding | 1.629521017 | 5.975778509 | 18.35629 | 0.0005485 | 0.092475219 |
| ENSMUSG00000030172 | Erc1 | protein_coding | -2.016580457 | 4.023491567 | 18.306161 | 0.0005534 | 0.092475219 |
| ENSMUSG00000062627 | Mysm1 | protein_coding | -1.422591657 | 5.109075998 | 18.286252 | 0.0005561 | 0.092475219 |
| ENSMUSG00000031309 | Rps6ka3 | protein_coding | -2.382183181 | 3.483811245 | 18.265535 | 0.000559 | 0.092475219 |
| ENSMUSG00000064367 | mt-Nd5 | protein_coding | -1.641900113 | 9.814911841 | 18.512087 | 0.0005794 | 0.09379567 |
| ENSMUSG00000105265 | Sox2ot | processed_transcript | -1.085309243 | 6.116986292 | 18.085606 | 0.0005844 | 0.09379567 |
| ENSMUSG00000023964 | Calcr | protein_coding | 2.597572177 | 3.543107807 | 18.006916 | 0.000596 | 0.094034455 |
| ENSMUSG00000036578 | Fxyd7 | protein_coding | 0.990143897 | 7.396395722 | 17.926774 | 0.000608 | 0.094034455 |
| ENSMUSG00000027270 | Lamp5 | protein_coding | 1.433714553 | 7.01568764 | 17.898563 | 0.0006143 | 0.094034455 |
| ENSMUSG00000027858 | Tspan2 | protein_coding | -1.178242952 | 6.491047904 | 17.840131 | 0.0006212 | 0.094034455 |
| ENSMUSG00000000567 | Sox9 | protein_coding | -1.420847891 | 5.735015818 | 17.71759 | 0.0006406 | 0.094034455 |
| ENSMUSG00000060988 | Galnt13 | protein_coding | -1.55844823 | 5.405734543 | 17.603947 | 0.0006592 | 0.094034455 |
| ENSMUSG00000063887 | Nlgn1 | protein_coding | -1.2721091 | 6.333251997 | 17.598571 | 0.00066 | 0.094034455 |
| ENSMUSG00000031529 | Tnks | protein_coding | -1.357796671 | 5.229642601 | 17.560442 | 0.0006664 | 0.094034455 |
| ENSMUSG00000064368 | mt-Nd6 | protein_coding | -1.763850021 | 8.052755857 | 18.370499 | 0.0006673 | 0.094034455 |
| ENSMUSG00000058589 | Anks1b | protein_coding | -0.882511208 | 7.130867999 | 17.500508 | 0.0006766 | 0.094034455 |
| ENSMUSG00000015396 | Cd83 | protein_coding | 1.322233502 | 6.530667424 | 17.397763 | 0.0006944 | 0.094034455 |

| ensembl_gene_id | external_gene_name | gene_biotype | log2 (fold change) | log2 (counts per million) | F-value | p-value | False discovery rate |
|---|---|---|---|---|---|---|---|
| ENSMUSG00000021665 | Hexb | protein_coding | 1.363121881 | 8.13720421 | 17.384253 | 0.000699 | 0.094034455 |
| ENSMUSG00000099696 | 2900052N01Rik | lincRNA | -1.277969311 | 5.699290823 | 17.317837 | 0.0007086 | 0.094034455 |
| ENSMUSG00000034998 | Foxn2 | protein_coding | -1.734341231 | 4.48501583 | 17.279872 | 0.0007155 | 0.094034455 |
| ENSMUSG00000069045 | Ddx3y | protein_coding | -1.22503769 | 6.104745614 | 17.186103 | 0.0007328 | 0.094034455 |
| ENSMUSG00000028176 | Lrrc7 | protein_coding | -2.079992028 | 4.433184604 | 17.17121 | 0.0007356 | 0.094034455 |
| ENSMUSG00000006941 | Eif1b | protein_coding | 0.957165628 | 8.260969803 | 17.142018 | 0.0007411 | 0.094034455 |
| ENSMUSG00000031297 | Slc7a3 | protein_coding | 2.002048087 | 4.918918982 | 17.106955 | 0.0007501 | 0.094034455 |
| ENSMUSG00000046574 | Prr12 | protein_coding | 2.228508264 | 4.780336741 | 17.050916 | 0.000761 | 0.094034455 |
| ENSMUSG00000050121 | Opalin | protein_coding | -1.008640938 | 7.410040088 | 17.036041 | 0.0007614 | 0.094034455 |
| ENSMUSG00000018012 | Rac3 | protein_coding | 1.793489296 | 4.290277025 | 16.970532 | 0.0007744 | 0.094034455 |
| ENSMUSG00000010803 | Gabra1 | protein_coding | -0.974062629 | 7.603223781 | 16.950756 | 0.0007783 | 0.094034455 |
| ENSMUSG00000109324 | Prmt1 | protein_coding | 1.012605049 | 7.313579922 | 16.905186 | 0.0007875 | 0.094086243 |
| ENSMUSG00000048644 | Ctxn1 | protein_coding | 0.862032067 | 7.951408505 | 16.715452 | 0.000827 | 0.094376567 |
| ENSMUSG00000064370 | mt-Cytb | protein_coding | -1.892851143 | 12.19260763 | 17.305854 | 0.0008312 | 0.094376567 |
| ENSMUSG00000002107 | Celf2 | protein_coding | -1.004998397 | 7.824730066 | 16.629581 | 0.0008457 | 0.094376567 |
| ENSMUSG00000025907 | Rb1cc1 | protein_coding | -1.332090092 | 5.252218328 | 16.61346 | 0.0008492 | 0.094376567 |
| ENSMUSG00000024293 | Esco1 | protein_coding | -1.523828937 | 4.417822002 | 16.588503 | 0.0008547 | 0.094376567 |
| ENSMUSG00000025607 | Copg2 | protein_coding | -1.270920666 | 7.752464664 | 16.597644 | 0.0008553 | 0.094376567 |
| ENSMUSG00000051504 | Siglech | protein_coding | -2.285304265 | 3.656645677 | 16.569327 | 0.000859 | 0.094376567 |
| ENSMUSG00000057130 | Txnl4a | protein_coding | 1.175640302 | 6.757728205 | 16.561714 | 0.0008607 | 0.094376567 |
| ENSMUSG00000029245 | Epha5 | protein_coding | -1.571031128 | 5.230722747 | 16.525409 | 0.0008689 | 0.094376567 |
| ENSMUSG00000021379 | Id4 | protein_coding | -0.95227093 | 6.834048099 | 16.343119 | 0.0009113 | 0.097994341 |
| ENSMUSG00000035199 | Arl6ip5 | protein_coding | 0.873074151 | 8.05023803 | 16.298427 | 0.0009221 | 0.098168982 |
| ENSMUSG00000053414 | Hunk | protein_coding | 1.579838105 | 4.479379812 | 16.178194 | 0.0009517 | 0.100331913 |
| ENSMUSG00000021091 | Serpina3n | protein_coding | 1.443982517 | 4.794840131 | 15.976849 | 0.0010038 | 0.104795432 |
| ENSMUSG00000026384 | Ptpn4 | protein_coding | -1.197067677 | 5.596980002 | 15.863141 | 0.0010346 | 0.106426806 |
| ENSMUSG00000029088 | Kcnip4 | protein_coding | -1.195925766 | 5.534930803 | 15.803769 | 0.0010512 | 0.106426806 |
| ENSMUSG00000013539 | Tango2 | protein_coding | 0.968854951 | 8.030168215 | 15.781662 | 0.0010574 | 0.106426806 |
| ENSMUSG00000006010 | BC003331 | protein_coding | -1.220076645 | 5.896520046 | 15.775874 | 0.001059 | 0.106426806 |
| ENSMUSG00000032253 | Phip | protein_coding | -1.606243635 | 4.797293326 | 15.714481 | 0.0010766 | 0.106858837 |
| ENSMUSG00000063406 | Tmed5 | protein_coding | -1.562391303 | 5.217522506 | 15.66072 | 0.0010922 | 0.106858837 |
| ENSMUSG00000028345 | Tex10 | protein_coding | 1.932683749 | 4.033069419 | 15.657532 | 0.0010931 | 0.106858837 |
| ENSMUSG00000045302 | Preb | protein_coding | 1.229274799 | 5.968168153 | 15.563661 | 0.0011211 | 0.10791039 |
| ENSMUSG00000020042 | Btbd11 | protein_coding | 1.535326682 | 5.056746215 | 15.500386 | 0.0011404 | 0.10791039 |
| ENSMUSG00000114304 | AC165092.1 | lincRNA | 1.68427431 | 7.087818151 | 16.164365 | 0.0011414 | 0.10791039 |
| ENSMUSG00000042761 | Mrap2 | protein_coding | 1.391563138 | 5.658619907 | 15.451347 | 0.0011556 | 0.10791039 |
| ENSMUSG00000060002 | Chpt1 | protein_coding | -1.841190692 | 3.839606672 | 15.443666 | 0.001158 | 0.10791039 |
| ENSMUSG00000039539 | Sgcz | protein_coding | -2.13563711 | 4.028245037 | 15.337147 | 0.0011919 | 0.10791039 |
| ENSMUSG00000022500 | Litaf | protein_coding | -1.002933687 | 6.764656062 | 15.333488 | 0.0011931 | 0.10791039 |
| ENSMUSG00000024109 | Nrxn1 | protein_coding | -1.092888284 | 8.156305757 | 15.331225 | 0.0011938 | 0.10791039 |
| ENSMUSG00000031229 | Atrx | protein_coding | -1.249034384 | 6.221157175 | 15.32993 | 0.0011942 | 0.10791039 |
| ENSMUSG00000025666 | Tmem47 | protein_coding | -1.109975661 | 6.171143608 | 15.122963 | 0.0012635 | 0.112030916 |
| ENSMUSG00000051285 | Pcmtd1 | protein_coding | -1.335546869 | 5.094840284 | 15.070305 | 0.0012818 | 0.112030916 |
| ENSMUSG00000053477 | Tcf4 | protein_coding | -1.002479582 | 6.748185216 | 15.044571 | 0.0012909 | 0.112030916 |
| ENSMUSG00000036503 | Rnf13 | protein_coding | -0.921646519 | 7.427291732 | 15.026054 | 0.0012975 | 0.112030916 |
| ENSMUSG00000049184 | Purg | protein_coding | -1.44150686 | 4.934903238 | 15.021285 | 0.0012992 | 0.112030916 |
| ENSMUSG00000034647 | Ankrd12 | protein_coding | -1.424906669 | 6.328542907 | 15.02239 | 0.0013023 | 0.112030916 |
| ENSMUSG00000021098 | 4930447C04Rik | protein_coding | -3.029063122 | 2.70016333 | 14.977875 | 0.0013148 | 0.112202614 |
| ENSMUSG00000020863 | Luc7l3 | protein_coding | -1.165700816 | 6.400903113 | 14.921189 | 0.0013354 | 0.112444692 |
| ENSMUSG00000029484 | Anxa3 | protein_coding | -2.264497175 | 3.495999761 | 14.898887 | 0.0013436 | 0.112444692 |
| ENSMUSG00000064363 | mt-Nd4 | protein_coding | -1.526395752 | 10.34042568 | 15.232131 | 0.001349 | 0.112444692 |
| ENSMUSG00000035835 | Plppr3 | protein_coding | 1.356293227 | 5.406949151 | 14.778743 | 0.001389 | 0.114129168 |
| ENSMUSG00000043388 | Tmem130 | protein_coding | 1.061867446 | 9.615115595 | 14.775061 | 0.0013904 | 0.114129168 |
| ENSMUSG00000028389 | Zfp37 | protein_coding | -1.531558449 | 4.774334343 | 14.682796 | 0.0014264 | 0.114664043 |
| ENSMUSG00000021032 | Ngb | protein_coding | 1.044779325 | 8.359207565 | 14.679277 | 0.0014278 | 0.114664043 |
| ENSMUSG00000042078 | Svop | protein_coding | 0.882524198 | 7.24270854 | 14.676545 | 0.0014289 | 0.114664043 |
| ENSMUSG00000036111 | Lmo1 | protein_coding | 1.376447629 | 4.430360891 | 14.583188 | 0.0014665 | 0.116809824 |
| ENSMUSG00000029126 | Nsg1 | protein_coding | 0.906362066 | 9.756698452 | 14.462314 | 0.0015169 | 0.119650549 |
| ENSMUSG00000044667 | Plppr4 | protein_coding | -0.986356645 | 6.517486683 | 14.436319 | 0.0015279 | 0.119650549 |
| ENSMUSG00000030682 | Cdipt | protein_coding | 1.04440447 | 6.957219419 | 14.4186 | 0.0015356 | 0.119650549 |
| ENSMUSG00000079657 | Rab26 | protein_coding | 1.870556705 | 5.200088954 | 14.307039 | 0.0015949 | 0.121124117 |
| ENSMUSG00000026679 | Enkur | protein_coding | -1.857049267 | 4.017114693 | 14.246864 | 0.0016115 | 0.121124117 |
| ENSMUSG00000023150 | Ivns1abp | protein_coding | -0.925715408 | 6.256843816 | 14.234575 | 0.0016171 | 0.121124117 |
| ENSMUSG00000074863 | Platr25 | protein_coding | -3.342637114 | 2.442684069 | 14.183024 | 0.0016409 | 0.121124117 |
| ENSMUSG00000024976 | Shoc2 | protein_coding | -1.250568865 | 8.430827714 | 14.174544 | 0.001649 | 0.121124117 |
| ENSMUSG00000032854 | Ugt8a | protein_coding | -1.118439455 | 5.687384174 | 14.155848 | 0.0016535 | 0.121124117 |
| ENSMUSG00000038776 | Ephx1 | protein_coding | 1.686189128 | 4.720512754 | 14.146586 | 0.0016579 | 0.121124117 |
| ENSMUSG00000054162 | Spock3 | protein_coding | -0.893189223 | 7.329591639 | 14.143107 | 0.0016595 | 0.121124117 |
| ENSMUSG00000035202 | Lars2 | protein_coding | 1.136233994 | 9.543560959 | 14.141282 | 0.0016603 | 0.121124117 |
| ENSMUSG00000022454 | Nell2 | protein_coding | 0.903256212 | 7.578508581 | 14.126945 | 0.0016671 | 0.121124117 |
| ENSMUSG00000032368 | Zic1 | protein_coding | -1.107825068 | 6.105026732 | 14.023121 | 0.001717 | 0.123909823 |
| ENSMUSG00000039103 | Nexn | protein_coding | -2.078525872 | 4.231479967 | 13.982402 | 0.001737 | 0.124146111 |
| ENSMUSG00000042133 | Ppig | protein_coding | -1.094900345 | 6.474703558 | 13.969592 | 0.0017433 | 0.124146111 |
| ENSMUSG00000050377 | Il31ra | protein_coding | 1.655869722 | 5.067837887 | 13.896984 | 0.0017798 | 0.125911159 |
| ENSMUSG00000028456 | Unc13b | protein_coding | -1.155625773 | 5.591983742 | 13.810712 | 0.0018243 | 0.128214981 |
| ENSMUSG00000018199 | Trove2 | protein_coding | -1.097183094 | 5.844787989 | 13.614606 | 0.0019302 | 0.130159644 |
| ENSMUSG00000027677 | Ttc14 | protein_coding | -2.30481976 | 3.715954324 | 13.592664 | 0.0019425 | 0.130159644 |
| ENSMUSG00000098973 | Mir6236 | miRNA | 1.812760303 | 8.725991134 | 14.433133 | 0.0019444 | 0.130159644 |

| ensembl_gene_id | external_gene_name | gene_biotype | log2 (fold change) | log2 (counts per million) | F-value | p-value | False discovery rate |
|---|---|---|---|---|---|---|---|
| ENSMUSG00000034145 | Tmem63c | protein_coding | 2.227372268 | 3.439357991 | 13.574521 | 0.0019527 | 0.130159644 |
| ENSMUSG00000015837 | Sqstm1 | protein_coding | 0.9404254 | 8.745918331 | 13.566678 | 0.0019571 | 0.130159644 |
| ENSMUSG00000007038 | Neu1 | protein_coding | 1.306660888 | 5.20445601 | 13.565347 | 0.0019579 | 0.130159644 |
| ENSMUSG00000100147 | 1700047M11Rik | lincRNA | -1.616197656 | 4.785041665 | 13.559939 | 0.0019609 | 0.130159644 |
| ENSMUSG00000039886 | Tmem120a | protein_coding | 1.491664596 | 4.318940806 | 13.541765 | 0.0019713 | 0.130159644 |
| ENSMUSG00000032417 | Rwdd2a | protein_coding | 0.892712999 | 7.882959122 | 13.53406 | 0.0019757 | 0.130159644 |
| ENSMUSG00000074738 | Fndc10 | protein_coding | 1.65875172 | 4.143632959 | 13.503171 | 0.0019934 | 0.130159644 |
| ENSMUSG00000110027 | C030029H02Rik | lincRNA | -1.523520552 | 4.059750618 | 13.489723 | 0.0020012 | 0.130159644 |
| ENSMUSG00000060487 | Samd5 | protein_coding | -1.468263334 | 4.481120482 | 13.453822 | 0.0020222 | 0.130159644 |
| ENSMUSG00000050875 | A730017C20Rik | protein_coding | 1.054137418 | 8.826962204 | 13.443556 | 0.0020282 | 0.130159644 |
| ENSMUSG00000028576 | Ift74 | protein_coding | -1.065047058 | 5.909639603 | 13.424148 | 0.0020397 | 0.130159644 |
| ENSMUSG00000020940 | 1700023F06Rik | protein_coding | 1.428170637 | 4.456806218 | 13.419527 | 0.0020425 | 0.130159644 |
| ENSMUSG00000026223 | Itm2c | protein_coding | 1.009568671 | 9.311222768 | 13.403138 | 0.0020522 | 0.130159644 |
| ENSMUSG00000045414 | 1190002N15Rik | protein_coding | -1.785904715 | 3.857464086 | 13.372344 | 0.0020707 | 0.130159644 |
| ENSMUSG00000040929 | Rfx3 | protein_coding | -1.642615065 | 4.279747821 | 13.353826 | 0.0020819 | 0.130159644 |
| ENSMUSG00000026377 | Nifk | protein_coding | 1.050685171 | 5.969500023 | 13.353709 | 0.002082 | 0.130159644 |
| ENSMUSG00000020634 | Ubxn2a | protein_coding | -1.131573548 | 5.43505583 | 13.309092 | 0.0021093 | 0.130900779 |
| ENSMUSG00000003949 | Hlf | protein_coding | -0.853582804 | 6.827037216 | 13.294648 | 0.0021182 | 0.130900779 |
| ENSMUSG00000020098 | Pcbd1 | protein_coding | 0.8534166 | 7.694197222 | 13.149723 | 0.0022101 | 0.135342324 |
| ENSMUSG00000025915 | Sgk3 | protein_coding | -1.285724874 | 5.458691514 | 13.123135 | 0.0022274 | 0.135342324 |
| ENSMUSG00000037628 | Cdkn3 | protein_coding | 3.017039517 | 2.570795995 | 13.092473 | 0.0022476 | 0.135342324 |
| ENSMUSG00000027287 | Snap23 | protein_coding | -1.8471285 | 4.287965476 | 13.086138 | 0.0022518 | 0.135342324 |
| ENSMUSG00000056073 | Grik2 | protein_coding | -1.283734263 | 5.420068935 | 13.080011 | 0.0022559 | 0.135342324 |
| ENSMUSG00000033068 | Entpd6 | protein_coding | 0.895985628 | 6.758340762 | 13.065433 | 0.0022656 | 0.135342324 |
| ENSMUSG00000027347 | Rasgrp1 | protein_coding | -0.957857886 | 6.741384254 | 13.042962 | 0.0022806 | 0.135459494 |
| ENSMUSG00000020775 | Mrpl38 | protein_coding | 1.408967147 | 6.000364874 | 13.032781 | 0.0022927 | 0.135459494 |
| ENSMUSG00000038039 | Gcc2 | protein_coding | -1.060570304 | 5.948137453 | 12.991591 | 0.0023155 | 0.135459494 |
| ENSMUSG00000007989 | Fzd3 | protein_coding | -1.063473587 | 5.708642128 | 12.96864 | 0.0023313 | 0.135459494 |
| ENSMUSG00000021025 | Nfkbia | protein_coding | 1.352613268 | 5.816459171 | 12.964283 | 0.0023343 | 0.135459494 |
| ENSMUSG00000046447 | Camk2n1 | protein_coding | -0.946315091 | 8.814693818 | 12.951502 | 0.0023431 | 0.135459494 |
| ENSMUSG00000037966 | Ninj1 | protein_coding | 1.417282566 | 4.936713085 | 12.914306 | 0.0023691 | 0.135891945 |
| ENSMUSG00000042063 | Zfp386 | protein_coding | -1.384960792 | 5.182603598 | 12.90465 | 0.0023759 | 0.135891945 |
| ENSMUSG00000039607 | Rbms3 | protein_coding | -1.406051912 | 5.626298058 | 12.849329 | 0.0024152 | 0.136460872 |
| ENSMUSG00000034173 | Zbed5 | protein_coding | 2.146702794 | 3.221061799 | 12.842072 | 0.0024204 | 0.136460872 |
| ENSMUSG00000020571 | Pdia6 | protein_coding | 0.779708742 | 7.807010753 | 12.83727 | 0.0024239 | 0.136460872 |
| ENSMUSG00000061601 | Pclo | protein_coding | -1.414404841 | 5.385969975 | 12.776122 | 0.0024684 | 0.137110361 |
| ENSMUSG00000007021 | Syngr3 | protein_coding | 0.907197984 | 9.000692261 | 12.758476 | 0.0024814 | 0.137110361 |
| ENSMUSG00000018339 | Gpx3 | protein_coding | 1.037468573 | 9.533675749 | 12.732072 | 0.0025011 | 0.137110361 |
| ENSMUSG00000038174 | Fam126b | protein_coding | -0.953285479 | 6.424129411 | 12.718251 | 0.0025114 | 0.137110361 |
| ENSMUSG00000029563 | Foxp2 | protein_coding | -1.362388452 | 5.536918771 | 12.715037 | 0.0025138 | 0.137110361 |
| ENSMUSG00000026781 | Acbd5 | protein_coding | -1.229507968 | 5.32265234 | 12.704721 | 0.0025216 | 0.137110361 |
| ENSMUSG00000027327 | 1700037H04Rik | protein_coding | 0.869048555 | 9.438842398 | 12.700613 | 0.0025247 | 0.137110361 |
| ENSMUSG00000020133 | 2310011J03Rik | protein_coding | 0.853672723 | 6.388075915 | 12.643543 | 0.0025682 | 0.137802309 |
| ENSMUSG00000010608 | Rbm25 | protein_coding | -0.868859045 | 6.623945702 | 12.634953 | 0.0025748 | 0.137802309 |
| ENSMUSG00000018865 | Sult4a1 | protein_coding | 1.002497434 | 9.330066523 | 12.633552 | 0.0025759 | 0.137802309 |
| ENSMUSG00000021759 | Plpp1 | protein_coding | 1.281020247 | 4.696786493 | 12.602748 | 0.0025998 | 0.13839264 |
| ENSMUSG00000021993 | Mipep | protein_coding | 1.489104052 | 4.298738987 | 12.50407 | 0.002678 | 0.14123642 |
| ENSMUSG00000030314 | Atg7 | protein_coding | 1.854585443 | 6.424224491 | 13.229905 | 0.0026795 | 0.14123642 |
| ENSMUSG00000098557 | Kctd12 | protein_coding | -1.449752792 | 4.537306075 | 12.439832 | 0.0027304 | 0.141457651 |
| ENSMUSG00000038602 | Slc35f1 | protein_coding | -1.219763193 | 5.437663363 | 12.438286 | 0.0027317 | 0.141457651 |
| ENSMUSG00000014956 | Ppp1cb | protein_coding | -0.75709424 | 8.33464785 | 12.432219 | 0.0027367 | 0.141457651 |
| ENSMUSG00000064339 | mt-Rnr2 | Mt_rRNA | -1.892117993 | 13.83599611 | 13.001276 | 0.0027461 | 0.141457651 |
| ENSMUSG00000052372 | Il1rapl1 | protein_coding | 1.298742664 | 7.527975867 | 12.681432 | 0.0027494 | 0.141457651 |
| ENSMUSG00000026915 | Strbp | protein_coding | -0.844725373 | 6.935576728 | 12.355499 | 0.002801 | 0.143106928 |
| ENSMUSG00000035964 | Tmem59l | protein_coding | 0.876133626 | 9.53239765 | 12.347122 | 0.0028081 | 0.143106928 |
| ENSMUSG00000042380 | Smim12 | protein_coding | 0.892815327 | 7.240841027 | 12.23356 | 0.0029067 | 0.146840523 |
| ENSMUSG00000064351 | mt-Co1 | protein_coding | -1.64532156 | 11.50948277 | 12.653332 | 0.0029232 | 0.146840523 |
| ENSMUSG00000002617 | Zfp40 | protein_coding | -2.18623706 | 3.713377652 | 12.211051 | 0.0029267 | 0.146840523 |
| ENSMUSG00000003363 | Pld3 | protein_coding | 0.884495199 | 9.931445965 | 12.191606 | 0.0029441 | 0.146840523 |
| ENSMUSG00000004961 | Syt5 | protein_coding | 0.76233231 | 8.28337684 | 12.174815 | 0.0029592 | 0.146840523 |
| ENSMUSG00000031903 | Pla2g15 | protein_coding | 1.395565289 | 4.10737651 | 12.17025 | 0.0029633 | 0.146840523 |
| ENSMUSG00000037999 | Arap2 | protein_coding | -1.374485112 | 4.659843276 | 12.131429 | 0.0029986 | 0.147910435 |
| ENSMUSG00000031327 | Chic1 | protein_coding | -1.112254135 | 5.439736471 | 12.058169 | 0.0030667 | 0.149958199 |
| ENSMUSG00000027799 | Nbea | protein_coding | -0.949340018 | 6.13149796 | 12.056685 | 0.0030681 | 0.149958199 |
| ENSMUSG00000027602 | Map1lc3a | protein_coding | 0.789547739 | 8.415411366 | 12.02343 | 0.0030995 | 0.150101985 |
| ENSMUSG00000037710 | Cisd1 | protein_coding | 1.170790127 | 8.734266486 | 12.020859 | 0.0031084 | 0.150101985 |
| ENSMUSG00000031545 | Gpat4 | protein_coding | 1.151425619 | 6.547037953 | 12.016147 | 0.0031129 | 0.150101985 |
| ENSMUSG00000026469 | Xpr1 | protein_coding | -0.937898965 | 6.068042782 | 11.934178 | 0.0031858 | 0.151269232 |
| ENSMUSG00000046699 | Slitrk4 | protein_coding | -1.380020894 | 4.671661976 | 11.880977 | 0.0032385 | 0.151269232 |
| ENSMUSG00000048118 | Arid4a | protein_coding | -1.065287002 | 6.112381513 | 11.879821 | 0.0032396 | 0.151269232 |
| ENSMUSG00000025393 | Atp5b | protein_coding | 1.039434046 | 10.30612259 | 11.868181 | 0.0032513 | 0.151269232 |
| ENSMUSG00000004945 | Tmem242 | protein_coding | 0.822107949 | 6.377803356 | 11.844147 | 0.0032755 | 0.151269232 |
| ENSMUSG00000027568 | Ntsr1 | protein_coding | 2.983237325 | 2.674081521 | 11.83721 | 0.0032826 | 0.151269232 |
| ENSMUSG00000034681 | Rnps1 | protein_coding | 1.039507059 | 6.11300136 | 11.829115 | 0.0032908 | 0.151269232 |
| ENSMUSG00000074733 | Zfp950 | protein_coding | -1.194506587 | 5.169887919 | 11.822651 | 0.0032974 | 0.151269232 |
| ENSMUSG00000031782 | Coq9 | protein_coding | 0.901778725 | 6.965553373 | 11.816332 | 0.0033039 | 0.151269232 |
| ENSMUSG00000068580 | Zfyve19 | protein_coding | 1.021071857 | 6.224034757 | 11.794799 | 0.003326 | 0.151269232 |
| ENSMUSG00000050587 | Lrrc4c | protein_coding | -0.779340165 | 6.84101994 | 11.778674 | 0.0033426 | 0.151269232 |

| ensembl_gene_id | external_gene_name | gene_biotype | log2 (fold change) | log2 (counts per million) | F-value | p-value | False discovery rate |
|---|---|---|---|---|---|---|---|
| ENSMUSG00000049744 | Arhgap15 | protein_coding | -3.359138797 | 2.51061778 | 11.768177 | 0.0033535 | 0.151269232 |
| ENSMUSG00000041408 | Wapl | protein_coding | -0.817394063 | 6.719766122 | 11.756999 | 0.0033652 | 0.151269232 |
| ENSMUSG00000005447 | Pafah1b3 | protein_coding | 0.930532992 | 7.042839988 | 11.755466 | 0.0033668 | 0.151269232 |
| ENSMUSG00000037400 | Atp11b | protein_coding | -2.767546658 | 3.520111788 | 11.777341 | 0.0033802 | 0.151269232 |
| ENSMUSG00000042750 | Bex2 | protein_coding | 0.820740148 | 10.3553781 | 11.729394 | 0.0033941 | 0.151269232 |
| ENSMUSG00000041958 | Pigs | protein_coding | 1.250039138 | 5.845747439 | 11.721236 | 0.0034028 | 0.151269232 |
| ENSMUSG00000030069 | Prok2 | protein_coding | 1.734452559 | 3.528654604 | 11.711251 | 0.0034133 | 0.151269232 |
| ENSMUSG00000038538 | Ubn2 | protein_coding | -1.178670309 | 5.661004336 | 11.704211 | 0.0034208 | 0.151269232 |
| ENSMUSG00000026721 | Rabgap1l | protein_coding | -0.934204271 | 6.152632248 | 11.679032 | 0.0034477 | 0.151269232 |
| ENSMUSG00000019132 | BC005537 | protein_coding | -0.884511518 | 6.782680237 | 11.661721 | 0.0034664 | 0.151269232 |
| ENSMUSG00000046598 | Bdh1 | protein_coding | 1.146535491 | 5.806172135 | 11.649738 | 0.0034793 | 0.151269232 |
| ENSMUSG00000059288 | Cdyl | protein_coding | -1.528857533 | 4.048326968 | 11.644792 | 0.0034847 | 0.151269232 |
| ENSMUSG00000023707 | Ogfod2 | protein_coding | 1.235322535 | 5.971557086 | 11.621275 | 0.0035103 | 0.151269232 |
| ENSMUSG00000028857 | Tmem222 | protein_coding | 0.954459723 | 8.220295672 | 11.619619 | 0.0035122 | 0.151269232 |
| ENSMUSG00000029823 | Luc7l2 | protein_coding | -1.111810166 | 5.357817522 | 11.619224 | 0.0035126 | 0.151269232 |
| ENSMUSG00000045980 | Tmem104 | protein_coding | 2.105332905 | 3.024620185 | 11.615292 | 0.0035169 | 0.151269232 |
| ENSMUSG00000037236 | Matr3 | protein_coding | -0.819803836 | 8.047291825 | 11.591526 | 0.0035431 | 0.151788829 |
| ENSMUSG00000020003 | Pex7 | protein_coding | -1.40353648 | 4.586048451 | 11.577759 | 0.0035584 | 0.151838352 |
| ENSMUSG00000032959 | Pebp1 | protein_coding | 0.742042712 | 7.635538589 | 11.534706 | 0.0036066 | 0.153010689 |
| ENSMUSG00000024952 | Rps6ka4 | protein_coding | 1.529785898 | 4.084557071 | 11.507013 | 0.0036381 | 0.153010689 |
| ENSMUSG00000026171 | Rnf25 | protein_coding | 1.032278855 | 6.474817037 | 11.505174 | 0.0036402 | 0.153010689 |
| ENSMUSG00000060301 | 2610008E11Rik | protein_coding | -1.698775082 | 3.531408418 | 11.488316 | 0.0036595 | 0.153010689 |
| ENSMUSG00000042208 | 0610010F05Rik | protein_coding | -1.096091727 | 4.933790896 | 11.478876 | 0.0036703 | 0.153010689 |
| ENSMUSG00000025423 | Pias2 | protein_coding | -1.066330647 | 5.407554787 | 11.478082 | 0.0036712 | 0.153010689 |
| ENSMUSG00000046178 | Nxph1 | protein_coding | -1.035766038 | 5.835213888 | 11.436539 | 0.0037195 | 0.153387715 |
| ENSMUSG00000036907 | C1ql2 | protein_coding | 3.801647285 | 2.369910729 | 11.430019 | 0.0037271 | 0.153387715 |
| ENSMUSG00000075334 | Rprm | protein_coding | 1.661803813 | 5.63172732 | 11.654636 | 0.0037301 | 0.153387715 |
| ENSMUSG00000036533 | Cdc42ep3 | protein_coding | -1.481984159 | 4.414159543 | 11.421291 | 0.0037373 | 0.153387715 |
| ENSMUSG00000001833 | 7-Sep | protein_coding | -0.869008948 | 9.081041745 | 11.348037 | 0.0038246 | 0.15571177 |
| ENSMUSG00000049583 | Grm5 | protein_coding | -1.129948911 | 7.011997702 | 11.328378 | 0.0038558 | 0.15571177 |
| ENSMUSG00000035596 | Mboat7 | protein_coding | 0.925329859 | 6.622442919 | 11.321964 | 0.0038562 | 0.15571177 |
| ENSMUSG00000028455 | Stoml2 | protein_coding | 1.01165624 | 7.123481909 | 11.318878 | 0.00386 | 0.15571177 |
| ENSMUSG00000035992 | Fnip1 | protein_coding | -1.292968906 | 4.657855697 | 11.313644 | 0.0038664 | 0.15571177 |
| ENSMUSG00000063550 | Nup98 | protein_coding | 1.084911355 | 5.737977979 | 11.288392 | 0.0038974 | 0.156374306 |
| ENSMUSG00000061576 | Dpp6 | protein_coding | 0.867521924 | 7.247168666 | 11.273504 | 0.0039158 | 0.156528699 |
| ENSMUSG00000044628 | Rnf208 | protein_coding | 0.98339872 | 6.932068985 | 11.253533 | 0.0039406 | 0.156699391 |
| ENSMUSG00000006315 | Tmem147 | protein_coding | 0.850907971 | 7.795517713 | 11.244817 | 0.0039515 | 0.156699391 |
| ENSMUSG00000019936 | Epyc | protein_coding | -2.606742937 | 2.734746908 | 11.235054 | 0.0039638 | 0.156699391 |
| ENSMUSG00000094483 | Purb | protein_coding | -0.765040697 | 8.418996873 | 11.163306 | 0.0040551 | 0.159064293 |
| ENSMUSG00000015804 | Med28 | protein_coding | 1.035113997 | 8.268673354 | 11.166992 | 0.004058 | 0.159064293 |
| ENSMUSG00000007041 | Clic1 | protein_coding | -1.498911315 | 4.372558021 | 11.153349 | 0.004068 | 0.159064293 |
| ENSMUSG00000025739 | Gng13 | protein_coding | 1.375900014 | 4.249985549 | 11.14092 | 0.0040841 | 0.159116066 |
| ENSMUSG00000042444 | Mindy2 | protein_coding | -1.349847996 | 5.504832714 | 11.111385 | 0.0041227 | 0.159799065 |
| ENSMUSG00000035772 | Mrps2 | protein_coding | 1.052121008 | 5.785814546 | 11.102902 | 0.0041338 | 0.159799065 |
| ENSMUSG00000039219 | Arid4b | protein_coding | -1.039619438 | 5.786670601 | 11.093545 | 0.0041462 | 0.159799065 |
| ENSMUSG00000024926 | Kat5 | protein_coding | 1.071867143 | 6.447044186 | 11.070485 | 0.0041768 | 0.160404298 |
| ENSMUSG00000069601 | Ank3 | protein_coding | -0.838799421 | 6.505575988 | 11.043735 | 0.0042126 | 0.161204953 |
| ENSMUSG00000038587 | Akap12 | protein_coding | -1.894791186 | 3.967296444 | 11.012914 | 0.0042544 | 0.161788116 |
| ENSMUSG00000022548 | Apod | protein_coding | -0.953505986 | 8.762868782 | 11.010269 | 0.004258 | 0.161788116 |
| ENSMUSG00000031807 | Pgls | protein_coding | 1.412919436 | 4.35469058 | 10.993989 | 0.0042802 | 0.162060932 |
| ENSMUSG00000020608 | Smc6 | protein_coding | -1.017267502 | 5.399207557 | 10.976568 | 0.0043042 | 0.162251351 |
| ENSMUSG00000037605 | Adgrl3 | protein_coding | -1.450383272 | 5.248406716 | 10.963109 | 0.0043228 | 0.162251351 |
| ENSMUSG00000007036 | Abhd16a | protein_coding | 1.046404676 | 5.766196043 | 10.95553 | 0.0043333 | 0.162251351 |
| ENSMUSG00000023927 | Satb1 | protein_coding | -1.482996748 | 3.755547552 | 10.938459 | 0.0043571 | 0.162251351 |
| ENSMUSG00000036641 | Ccdc148 | protein_coding | -1.238033669 | 5.152694454 | 10.902545 | 0.0044077 | 0.162251351 |
| ENSMUSG00000028863 | Meaf6 | protein_coding | 0.848958044 | 7.874483094 | 10.881726 | 0.0044373 | 0.162251351 |
| ENSMUSG00000025208 | Mrpl43 | protein_coding | 0.794894641 | 7.39468562 | 10.865598 | 0.0044604 | 0.162251351 |
| ENSMUSG00000085251 | Gm12326 | lincRNA | -2.837371478 | 3.29364466 | 10.86148 | 0.0044744 | 0.162251351 |
| ENSMUSG00000039145 | Camk1d | protein_coding | 0.931739749 | 7.706375568 | 10.853882 | 0.0044773 | 0.162251351 |
| ENSMUSG00000034932 | Mrpl54 | protein_coding | 1.646069911 | 6.746762172 | 11.417764 | 0.0044794 | 0.162251351 |
| ENSMUSG00000046432 | Bex3 | protein_coding | 0.835802478 | 9.760962919 | 10.850015 | 0.0044828 | 0.162251351 |
| ENSMUSG00000060935 | Tmem263 | protein_coding | -0.840888761 | 7.272686587 | 10.849768 | 0.0044832 | 0.162251351 |
| ENSMUSG00000026739 | Bmi1 | protein_coding | -1.405178821 | 4.297099157 | 10.849234 | 0.004484 | 0.162251351 |
| ENSMUSG00000032172 | Olfm2 | protein_coding | 0.827352008 | 6.814013233 | 10.831679 | 0.0045094 | 0.162251351 |
| ENSMUSG00000007458 | M6pr | protein_coding | 0.864241093 | 7.138541407 | 10.830183 | 0.0045116 | 0.162251351 |
| ENSMUSG00000028414 | Fktn | protein_coding | -1.062801934 | 4.980741172 | 10.813378 | 0.0045361 | 0.162589586 |
| ENSMUSG00000045589 | Frrs1l | protein_coding | -0.802027335 | 6.858846421 | 10.781041 | 0.0045837 | 0.163198427 |
| ENSMUSG00000020774 | Aspa | protein_coding | -1.214132587 | 5.993678956 | 10.774783 | 0.0046013 | 0.163198427 |
| ENSMUSG00000024810 | Il33 | protein_coding | -0.920767192 | 8.05266783 | 10.757312 | 0.004619 | 0.163198427 |
| ENSMUSG00000046311 | Zfp62 | protein_coding | -1.707045386 | 3.823587933 | 10.749749 | 0.0046304 | 0.163198427 |
| ENSMUSG00000036437 | Npy1r | protein_coding | 1.455778103 | 4.079519667 | 10.724516 | 0.0046683 | 0.163198427 |
| ENSMUSG00000026687 | Aldh9a1 | protein_coding | 1.015505131 | 6.127759362 | 10.723902 | 0.0046693 | 0.163198427 |
| ENSMUSG00000031458 | Coprs | protein_coding | 1.192591907 | 6.282476966 | 10.720209 | 0.0046832 | 0.163198427 |
| ENSMUSG00000027122 | Arl14ep | protein_coding | -1.04320253 | 6.619910862 | 10.706606 | 0.0046955 | 0.163198427 |
| ENSMUSG00000026678 | Rgs5 | protein_coding | -1.697307592 | 4.246764995 | 10.693351 | 0.0047157 | 0.163198427 |
| ENSMUSG00000047228 | A2ml1 | protein_coding | 0.981893365 | 5.711089496 | 10.691611 | 0.0047184 | 0.163198427 |
| ENSMUSG00000064337 | mt-Rnr1 | Mt_rRNA | -1.962644068 | 10.86487464 | 11.683781 | 0.0047201 | 0.163198427 |
| ENSMUSG00000057219 | Armc7 | protein_coding | 1.395449859 | 4.448733052 | 10.65753 | 0.0047709 | 0.164078841 |

| ensembl_gene_id | external_gene_name | gene_biotype | log2 (fold change) | log2 (counts per million) | F-value | p-value | False discovery rate |
|---|---|---|---|---|---|---|---|
| ENSMUSG00000072704 | Smim10l1 | protein_coding | -1.06885121 | 8.020245595 | 10.658957 | 0.0047771 | 0.164078841 |
| ENSMUSG00000020156 | Mum1 | protein_coding | 1.325094872 | 5.315632701 | 10.636016 | 0.0048044 | 0.164078841 |
| ENSMUSG00000042742 | Bmt2 | protein_coding | -1.359388082 | 4.507488992 | 10.629898 | 0.004814 | 0.164078841 |
| ENSMUSG00000031292 | Cdkl5 | protein_coding | -1.066668321 | 5.96809459 | 10.623428 | 0.0048241 | 0.164078841 |
| ENSMUSG00000079593 | Gm14597 | protein_coding | 0.712920361 | 8.072834169 | 10.615175 | 0.0048371 | 0.164078841 |
| ENSMUSG00000006675 | P4htm | protein_coding | 0.933830253 | 6.530913841 | 10.597735 | 0.0048646 | 0.164494296 |
| ENSMUSG00000026834 | Acvr1c | protein_coding | -1.467328945 | 3.894645852 | 10.555377 | 0.0049323 | 0.166046244 |
| ENSMUSG00000013465 | Nelfb | protein_coding | 0.860022231 | 5.997050314 | 10.549707 | 0.0049414 | 0.166046244 |
| ENSMUSG00000029060 | Mib2 | protein_coding | 1.533919384 | 4.132679786 | 10.527017 | 0.0049781 | 0.166760003 |
| ENSMUSG00000048965 | Mrgpre | protein_coding | 1.481278694 | 4.43760468 | 10.507391 | 0.0050102 | 0.167311975 |
| ENSMUSG00000015224 | Cyp2j9 | protein_coding | -1.424071381 | 4.778190594 | 10.495085 | 0.0050304 | 0.167466709 |
| ENSMUSG00000045777 | Ifitm10 | protein_coding | 0.967068135 | 5.831106192 | 10.467947 | 0.0050753 | 0.16815468 |
| ENSMUSG00000046160 | Olig1 | protein_coding | -1.22021328 | 5.77983526 | 10.463708 | 0.0050823 | 0.16815468 |
| ENSMUSG00000026872 | Zeb2 | protein_coding | -1.316132886 | 4.886389772 | 10.451667 | 0.0051024 | 0.168301679 |
| ENSMUSG00000056153 | Socs6 | protein_coding | -1.347711962 | 4.079099879 | 10.436205 | 0.0051284 | 0.168639828 |
| ENSMUSG00000045948 | Mrps12 | protein_coding | 0.685330704 | 7.801019874 | 10.420495 | 0.0051549 | 0.168685263 |
| ENSMUSG00000039740 | Alg2 | protein_coding | 0.879765097 | 7.151120417 | 10.407613 | 0.0051767 | 0.168685263 |
| ENSMUSG00000059119 | Nap1l4 | protein_coding | 1.060175647 | 8.400587276 | 10.412783 | 0.0051768 | 0.168685263 |
| ENSMUSG00000067942 | Zfp160 | protein_coding | -1.656728262 | 3.655992398 | 10.398261 | 0.0051926 | 0.168690081 |
| ENSMUSG00000039347 | Atp6v0e2 | protein_coding | 0.811661983 | 10.37809293 | 10.371532 | 0.0052385 | 0.16966628 |
| ENSMUSG00000025869 | Nop16 | protein_coding | 0.74792468 | 7.109687438 | 10.344968 | 0.0052845 | 0.170408001 |
| ENSMUSG00000035236 | Scai | protein_coding | -1.37881004 | 4.68379217 | 10.340033 | 0.0052931 | 0.170408001 |
| ENSMUSG00000041199 | Rpusd1 | protein_coding | 0.774236466 | 6.885391051 | 10.309125 | 0.0053473 | 0.170831901 |
| ENSMUSG00000049336 | Tenm2 | protein_coding | -1.246149398 | 5.487232192 | 10.290746 | 0.0053798 | 0.170831901 |
| ENSMUSG00000025155 | Dus1l | protein_coding | 1.567496542 | 4.475230336 | 10.285651 | 0.0053888 | 0.170831901 |
| ENSMUSG00000024735 | Prpf19 | protein_coding | 0.976313747 | 5.857345862 | 10.275048 | 0.0054077 | 0.170831901 |
| ENSMUSG00000028266 | Lmo4 | protein_coding | -0.863346321 | 6.567840984 | 10.272995 | 0.0054114 | 0.170831901 |
| ENSMUSG00000031613 | Hpgd | protein_coding | -2.062329617 | 3.347126738 | 10.265146 | 0.0054255 | 0.170831901 |
| ENSMUSG00000004035 | Gstm7 | protein_coding | -0.712713688 | 7.183323043 | 10.264817 | 0.0054261 | 0.170831901 |
| ENSMUSG00000025766 | D3Ertd751e | protein_coding | -1.446624605 | 4.628732869 | 10.260347 | 0.0054341 | 0.170831901 |
| ENSMUSG00000081225 | Cyp2j12 | protein_coding | -1.11770311 | 4.902637011 | 10.251936 | 0.0054492 | 0.170831901 |
| ENSMUSG00000045103 | Dmd | protein_coding | -1.114380195 | 4.828921698 | 10.22137 | 0.0055046 | 0.172066762 |
| ENSMUSG00000074457 | S100a16 | protein_coding | -0.937206196 | 8.358842546 | 10.197962 | 0.0055475 | 0.172594666 |
| ENSMUSG00000031878 | Nae1 | protein_coding | -0.902263115 | 6.584111225 | 10.186711 | 0.0055682 | 0.172594666 |
| ENSMUSG00000042097 | Zfp239 | protein_coding | 0.907065135 | 6.28642213 | 10.185922 | 0.0055696 | 0.172594666 |
| ENSMUSG00000066324 | Impad1 | protein_coding | -0.717481461 | 6.810191745 | 10.161821 | 0.0056144 | 0.173480438 |
| ENSMUSG00000022564 | Grina | protein_coding | 0.766496028 | 8.836490859 | 10.133199 | 0.005668 | 0.174613568 |
| ENSMUSG00000039850 | Endov | protein_coding | 1.217764508 | 4.222574245 | 10.116668 | 0.0056992 | 0.174613568 |
| ENSMUSG00000028161 | Ppp3ca | protein_coding | -0.707862092 | 9.571254196 | 10.116398 | 0.0056997 | 0.174613568 |
| ENSMUSG00000017485 | Top2b | protein_coding | -1.159445544 | 5.196539979 | 10.10026 | 0.0057304 | 0.175055093 |
| ENSMUSG00000028653 | Trit1 | protein_coding | 1.355455978 | 4.177133886 | 10.086972 | 0.0057558 | 0.175328401 |
| ENSMUSG00000033949 | Trim36 | protein_coding | -1.828386098 | 3.119246863 | 10.078563 | 0.005772 | 0.175328401 |
| ENSMUSG00000032727 | Mier3 | protein_coding | -1.013029096 | 5.296214816 | 10.069342 | 0.0057898 | 0.175372647 |
| ENSMUSG00000054196 | Cthrc1 | protein_coding | 0.936742923 | 5.38972456 | 10.054654 | 0.0058182 | 0.175738644 |
| ENSMUSG00000071644 | Eef1g | protein_coding | 0.690259947 | 7.444402866 | 10.03555 | 0.0058554 | 0.176109803 |
| ENSMUSG00000026158 | Ogfrl1 | protein_coding | -1.003170253 | 6.023285108 | 10.031545 | 0.0058632 | 0.176109803 |
| ENSMUSG00000037857 | Nufip2 | protein_coding | -1.082145262 | 6.648928208 | 9.9878151 | 0.0059592 | 0.177776506 |
| ENSMUSG00000025959 | Klf7 | protein_coding | -0.988651018 | 5.180760094 | 9.9815521 | 0.005962 | 0.177776506 |
| ENSMUSG00000021811 | Dnajc9 | protein_coding | 1.117595077 | 6.019361797 | 9.9702569 | 0.0059846 | 0.177776506 |
| ENSMUSG00000097625 | Gm26561 | lincRNA | 0.999101966 | 6.104517801 | 9.9610559 | 0.0060031 | 0.177776506 |
| ENSMUSG00000027210 | Meis2 | protein_coding | -0.913991274 | 7.258384013 | 9.9605822 | 0.006004 | 0.177776506 |
| ENSMUSG00000035285 | Nat14 | protein_coding | 1.005435209 | 6.095206322 | 9.9536911 | 0.0060179 | 0.177776506 |
| ENSMUSG00000085609 | 1700016P03Rik | processed_transcript | -1.892305609 | 3.263727687 | 9.9190347 | 0.0060883 | 0.178207196 |
| ENSMUSG00000042744 | Hectd4 | protein_coding | -0.896043089 | 6.342138632 | 9.9063149 | 0.0061143 | 0.178207196 |
| ENSMUSG00000046442 | Ppm1e | protein_coding | -1.002352014 | 5.906311378 | 9.8913446 | 0.0061451 | 0.178207196 |
| ENSMUSG00000014776 | Nol3 | protein_coding | 1.54268065 | 5.167329487 | 9.8950833 | 0.0061473 | 0.178207196 |
| ENSMUSG00000030516 | Tjp1 | protein_coding | -1.498667849 | 4.111698494 | 9.8778196 | 0.0061731 | 0.178207196 |
| ENSMUSG00000063889 | Crem | protein_coding | -1.092436726 | 5.032857539 | 9.8708355 | 0.0061876 | 0.178207196 |
| ENSMUSG00000074227 | Spint2 | protein_coding | 0.84175075 | 5.881870537 | 9.870184 | 0.006189 | 0.178207196 |
| ENSMUSG00000028082 | Sh3d19 | protein_coding | -1.817355991 | 3.122541729 | 9.8598584 | 0.0062105 | 0.178207196 |
| ENSMUSG00000066798 | Zbtb6 | protein_coding | -1.665018392 | 3.804663878 | 9.851437 | 0.0062282 | 0.178207196 |
| ENSMUSG00000028232 | Tmem68 | protein_coding | -0.91790383 | 5.896256195 | 9.8434143 | 0.006245 | 0.178207196 |
| ENSMUSG00000029838 | Ptn | protein_coding | -1.031926173 | 8.78621346 | 9.8429177 | 0.006256 | 0.178207196 |
| ENSMUSG00000027199 | Gatm | protein_coding | -0.890015702 | 8.833057624 | 9.8323609 | 0.0062683 | 0.178207196 |
| ENSMUSG00000045665 | Mfsd5 | protein_coding | 1.12488134 | 4.838395001 | 9.8238438 | 0.0062863 | 0.178207196 |
| ENSMUSG00000051412 | Vamp7 | protein_coding | -0.906793142 | 6.712283676 | 9.8205593 | 0.0062933 | 0.178207196 |
| ENSMUSG00000063297 | Luzp2 | protein_coding | -1.079747405 | 6.133680057 | 9.8190013 | 0.0062966 | 0.178207196 |
| ENSMUSG00000030161 | Gabarapl1 | protein_coding | 0.799324912 | 10.32629476 | 9.8184925 | 0.0062977 | 0.178207196 |
| ENSMUSG00000030729 | Pgm2l1 | protein_coding | -0.899159275 | 6.782313586 | 9.794727 | 0.0063483 | 0.179169879 |
| ENSMUSG00000000560 | Gabra2 | protein_coding | -1.611262828 | 6.153070811 | 10.196907 | 0.0063681 | 0.179256087 |
| ENSMUSG00000074129 | Rpl13a | protein_coding | 1.010514261 | 5.367134901 | 9.7717866 | 0.0063977 | 0.179620136 |
| ENSMUSG00000015092 | Edf1 | protein_coding | 0.797056135 | 7.716842567 | 9.7402642 | 0.0064662 | 0.18107166 |
| ENSMUSG00000031342 | Gpm6b | protein_coding | -0.840267158 | 8.856646969 | 9.7319473 | 0.0064845 | 0.181110287 |
| ENSMUSG00000003352 | Cacnb3 | protein_coding | 0.932609016 | 5.509101422 | 9.7125151 | 0.0065273 | 0.181428945 |
| ENSMUSG00000035754 | Wdr18 | protein_coding | 0.801937826 | 8.189957898 | 9.6973356 | 0.0065609 | 0.181428945 |
| ENSMUSG00000043372 | Hexim2 | protein_coding | 1.252540128 | 5.644263685 | 9.6924959 | 0.006582 | 0.181428945 |
| ENSMUSG00000024925 | Rnaseh2c | protein_coding | 2.227939295 | 4.817628596 | 10.091889 | 0.0065821 | 0.181428945 |
| ENSMUSG00000031232 | Magt1 | protein_coding | -1.752374296 | 3.95752326 | 9.6853079 | 0.0065877 | 0.181428945 |

| ensembl_gene_id | external_gene_name | gene_biotype | log2 (fold change) | log2 (counts per million) | F-value | p-value | False discovery rate |
|---|---|---|---|---|---|---|---|
| ENSMUSG00000027778 | Ift80 | protein_coding | -1.611773167 | 3.804707773 | 9.6773785 | 0.0066055 | 0.181428945 |
| ENSMUSG00000026418 | Tnni1 | protein_coding | -1.043974188 | 4.722029144 | 9.6735869 | 0.006614 | 0.181428945 |
| ENSMUSG00000022021 | Diaph3 | protein_coding | 1.400263004 | 4.311244021 | 9.646799 | 0.0066744 | 0.182620718 |
| ENSMUSG00000026986 | Hnmt | protein_coding | -0.737776269 | 6.539923914 | 9.6063151 | 0.0067669 | 0.184569141 |
| ENSMUSG00000034777 | Vax2 | protein_coding | -2.622489964 | 2.562757672 | 9.6006672 | 0.00678 | 0.184569141 |
| ENSMUSG00000030020 | Prickle2 | protein_coding | -0.942419631 | 5.767254028 | 9.5616118 | 0.0068708 | 0.185330788 |
| ENSMUSG00000042258 | Isl1 | protein_coding | 0.798846425 | 6.31340539 | 9.5598782 | 0.0068748 | 0.185330788 |
| ENSMUSG00000003279 | Dlgap1 | protein_coding | -0.83139483 | 6.176423712 | 9.5597123 | 0.0068752 | 0.185330788 |
| ENSMUSG00000031149 | Praf2 | protein_coding | 0.885741328 | 6.38582939 | 9.5590151 | 0.0068769 | 0.185330788 |
| ENSMUSG00000097392 | D930016D06Rik | lincRNA | -2.684372177 | 2.721452666 | 9.4949152 | 0.0070291 | 0.188263863 |
| ENSMUSG00000027706 | Sec62 | protein_coding | -0.703796131 | 7.909474174 | 9.493073 | 0.0070335 | 0.188263863 |
| ENSMUSG00000019804 | Snx3 | protein_coding | 0.689366577 | 8.357148891 | 9.4869616 | 0.0070483 | 0.188263863 |
| ENSMUSG00000066705 | Fxyd6 | protein_coding | 0.747833119 | 9.078928226 | 9.4838652 | 0.0070557 | 0.188263863 |
| ENSMUSG00000022051 | Bnip3l | protein_coding | -0.661615773 | 7.068522962 | 9.4480955 | 0.0071427 | 0.190113113 |
| ENSMUSG00000023068 | Nus1 | protein_coding | -0.906473407 | 5.801789935 | 9.4281585 | 0.0071917 | 0.19094498 |
| ENSMUSG00000039640 | Mrpl12 | protein_coding | 0.90061351 | 6.935011068 | 9.4086182 | 0.0072401 | 0.191756728 |
| ENSMUSG00000025283 | Sat1 | protein_coding | -0.842341243 | 6.838112687 | 9.3955258 | 0.0072728 | 0.192147939 |
| ENSMUSG00000020315 | Sptbn1 | protein_coding | -0.941841791 | 7.420372831 | 9.3725349 | 0.0073415 | 0.193305754 |
| ENSMUSG00000031949 | Adat1 | protein_coding | 1.679760542 | 3.422585029 | 9.354277 | 0.0073767 | 0.193305754 |
| ENSMUSG00000049076 | Acap2 | protein_coding | -1.27051256 | 4.383379589 | 9.3473345 | 0.0073944 | 0.193305754 |
| ENSMUSG00000027309 | 4930402H24Rik | protein_coding | -0.972127545 | 5.915773539 | 9.3394843 | 0.0074144 | 0.193305754 |
| ENSMUSG00000030204 | Ddx47 | protein_coding | 0.808380993 | 6.951213676 | 9.3364765 | 0.0074221 | 0.193305754 |
| ENSMUSG00000024805 | Pcgf5 | protein_coding | -1.211347372 | 5.309325547 | 9.3355456 | 0.0074245 | 0.193305754 |
| ENSMUSG00000042607 | Asb4 | protein_coding | 1.728006483 | 4.45846655 | 9.3185804 | 0.0074791 | 0.193818049 |
| ENSMUSG00000019923 | Zwint | protein_coding | 0.88835395 | 12.04740912 | 9.3138588 | 0.0074802 | 0.193818049 |
| ENSMUSG00000040624 | Plekhg1 | protein_coding | -1.656283696 | 3.843939393 | 9.3018616 | 0.0075112 | 0.194154244 |
| ENSMUSG00000037204 | Atg101 | protein_coding | 0.697378606 | 6.702083432 | 9.2876322 | 0.0075482 | 0.194571779 |
| ENSMUSG00000002416 | Ndufb2 | protein_coding | 0.791126033 | 6.208844997 | 9.281752 | 0.0075636 | 0.194571779 |
| ENSMUSG00000002524 | Puf60 | protein_coding | 0.729049681 | 8.566875765 | 9.2621182 | 0.0076151 | 0.195429165 |
| ENSMUSG00000034891 | Sncb | protein_coding | 0.920981767 | 9.225122231 | 9.2309123 | 0.0076977 | 0.19708068 |
| ENSMUSG00000038555 | Reep2 | protein_coding | 0.839030925 | 7.814522904 | 9.2163646 | 0.0077366 | 0.197307766 |
| ENSMUSG00000038248 | Sobp | protein_coding | -1.210498998 | 4.816461611 | 9.1976052 | 0.0077871 | 0.197307766 |
| ENSMUSG00000033184 | Tmed7 | protein_coding | -1.253583511 | 5.589289033 | 9.1963671 | 0.0078018 | 0.197307766 |
| ENSMUSG00000048251 | Bcl11b | protein_coding | -0.911222192 | 6.005800764 | 9.186534 | 0.0078171 | 0.197307766 |
| ENSMUSG00000019990 | Pde7b | protein_coding | -1.322208087 | 4.13932871 | 9.1823219 | 0.0078285 | 0.197307766 |
| ENSMUSG00000040370 | Etfrf1 | protein_coding | -0.822847544 | 6.488541711 | 9.1804204 | 0.0078337 | 0.197307766 |
| ENSMUSG00000032911 | Cspg4 | protein_coding | 1.997321298 | 3.823510684 | 9.1826347 | 0.0078391 | 0.197307766 |
| ENSMUSG00000052572 | Dlg2 | protein_coding | -0.820720803 | 6.437732956 | 9.165587 | 0.0078741 | 0.197307766 |
| ENSMUSG00000020415 | Pttg1 | protein_coding | -0.629390347 | 7.486446678 | 9.1603124 | 0.0078886 | 0.197307766 |
| ENSMUSG00000026641 | Usf1 | protein_coding | 0.80147772 | 6.31063339 | 9.142703 | 0.007937 | 0.197307766 |
| ENSMUSG00000028399 | Ptprd | protein_coding | -0.868315106 | 6.425603442 | 9.1414199 | 0.0079406 | 0.197307766 |
| ENSMUSG00000028444 | Cntfr | protein_coding | -1.664333336 | 3.91942776 | 9.1397868 | 0.0079451 | 0.197307766 |
| ENSMUSG00000046138 | 9930021J03Rik | protein_coding | -1.246547374 | 5.848909932 | 9.1439132 | 0.0079452 | 0.197307766 |
| ENSMUSG00000033307 | Mif | protein_coding | 0.701404683 | 7.599309128 | 9.132048 | 0.0079665 | 0.197381854 |
| ENSMUSG00000085181 | Gm12709 | antisense_RNA | -1.092077271 | 5.139537362 | 9.1176174 | 0.0080066 | 0.1974709 |
| ENSMUSG00000032328 | Tmem30a | protein_coding | -0.668164089 | 7.184056146 | 9.1157868 | 0.0080117 | 0.1974709 |
| ENSMUSG00000046844 | Vat1l | protein_coding | 0.850745758 | 7.407664796 | 9.1109623 | 0.0080252 | 0.1974709 |
| ENSMUSG00000061731 | Ext1 | protein_coding | -1.263114847 | 4.01353138 | 9.0763258 | 0.0081227 | 0.199175867 |
| ENSMUSG00000027502 | Rtfdc1 | protein_coding | 0.778747141 | 7.356181442 | 9.0678389 | 0.0081467 | 0.199175867 |
| ENSMUSG00000028743 | Akr7a5 | protein_coding | 1.404065971 | 3.887303776 | 9.0633265 | 0.0081596 | 0.199175867 |
| ENSMUSG00000006262 | Mob1b | protein_coding | -1.239958495 | 3.822096656 | 9.0601744 | 0.0081686 | 0.199175867 |
| ENSMUSG00000036198 | Arhgap36 | protein_coding | 1.089992068 | 5.900011865 | 9.047485 | 0.0082048 | 0.199183179 |
| ENSMUSG00000023175 | Bsg | protein_coding | 1.231996008 | 8.302447301 | 9.2998107 | 0.0082059 | 0.199183179 |
| ENSMUSG00000029998 | Pcyox1 | protein_coding | 0.827756188 | 6.168019748 | 9.0132567 | 0.0083036 | 0.201100638 |
| ENSMUSG00000036646 | Man1b1 | protein_coding | 0.991772377 | 5.165876527 | 8.9803007 | 0.0084 | 0.202977498 |
| ENSMUSG00000030613 | Ccdc90b | protein_coding | -1.011845378 | 5.437788353 | 8.9671621 | 0.0084388 | 0.203457243 |
| ENSMUSG00000024500 | Ppp2r2b | protein_coding | 0.669827256 | 7.88228679 | 8.9527447 | 0.0084815 | 0.203983589 |
| ENSMUSG00000067787 | Blcap | protein_coding | 0.726305124 | 8.873666099 | 8.9470429 | 0.0084985 | 0.203983589 |
| ENSMUSG00000104597 | B230334C09Rik | antisense_RNA | -1.628316157 | 3.76481505 | 8.923109 | 0.0085703 | 0.204736877 |
| ENSMUSG00000022594 | Lynx1 | protein_coding | 0.798494453 | 7.52563258 | 8.9162241 | 0.008591 | 0.204736877 |
| ENSMUSG00000101621 | Gm28905 | lincRNA | -2.972155854 | 2.29363569 | 8.90647 | 0.0086205 | 0.204736877 |
| ENSMUSG00000085282 | Gm15663 | processed_transcript | 1.374683553 | 3.665509334 | 8.9059643 | 0.0086221 | 0.204736877 |
| ENSMUSG00000069729 | Arid1b | protein_coding | -0.972252192 | 5.325468977 | 8.8964646 | 0.0086509 | 0.204736877 |
| ENSMUSG00000020131 | Pcsk4 | protein_coding | 1.952699502 | 3.616326002 | 8.88934 | 0.0086726 | 0.204736877 |
| ENSMUSG00000030654 | Arl6ip1 | protein_coding | 0.666260488 | 10.05420097 | 8.8877055 | 0.0086776 | 0.204736877 |
| ENSMUSG00000029775 | Klhdc10 | protein_coding | 0.896881062 | 5.376062898 | 8.8861986 | 0.0086822 | 0.204736877 |
| ENSMUSG00000022076 | Klhl1 | protein_coding | 1.475167082 | 4.332187391 | 8.85448 | 0.0087798 | 0.20596458 |
| ENSMUSG00000041935 | AW549877 | protein_coding | -0.972891088 | 5.234525089 | 8.8418505 | 0.008819 | 0.20596458 |
| ENSMUSG00000024597 | Slc12a2 | protein_coding | -1.138355477 | 5.596955192 | 8.8382626 | 0.0088301 | 0.20596458 |
| ENSMUSG00000040813 | Tex264 | protein_coding | 0.718782886 | 6.528082836 | 8.8282945 | 0.0088613 | 0.20596458 |
| ENSMUSG00000025470 | Zfp511 | protein_coding | 1.138498222 | 4.322987393 | 8.8245651 | 0.0088729 | 0.20596458 |
| ENSMUSG00000036499 | Eea1 | protein_coding | -1.280912187 | 4.669997963 | 8.8225581 | 0.0088792 | 0.20596458 |
| ENSMUSG00000022759 | Lrrc74b | protein_coding | -1.853973929 | 3.16521939 | 8.8173262 | 0.0088956 | 0.20596458 |
| ENSMUSG00000006740 | Kif5b | protein_coding | -0.874260474 | 6.115875941 | 8.7940772 | 0.008969 | 0.20596458 |
| ENSMUSG00000032503 | Arpp21 | protein_coding | -1.159063973 | 10.18890769 | 8.9314176 | 0.008986 | 0.20596458 |
| ENSMUSG00000064372 | mt-Tp | Mt_tRNA | -1.479481926 | 4.066601396 | 8.7842769 | 0.0090002 | 0.20596458 |
| ENSMUSG00000030059 | Tmf1 | protein_coding | -1.462046521 | 6.469920041 | 9.1218083 | 0.0090055 | 0.20596458 |
| ENSMUSG00000035561 | Aldh1b1 | protein_coding | 1.215426735 | 4.460342344 | 8.7797281 | 0.0090146 | 0.20596458 |

| ensembl_gene_id | external_gene_name | gene_biotype | log2 (fold change) | log2 (counts per million) | F-value | p-value | False discovery rate |
|---|---|---|---|---|---|---|---|
| ENSMUSG00000033510 | Otud7a | protein_coding | -2.379725429 | 2.772559865 | 8.7796181 | 0.009015 | 0.20596458 |
| ENSMUSG00000028093 | Acp6 | protein_coding | 1.778033772 | 3.547751919 | 8.7767547 | 0.0090241 | 0.20596458 |
| ENSMUSG00000039842 | Mcph1 | protein_coding | 1.298001364 | 4.052234881 | 8.7746053 | 0.009031 | 0.20596458 |
| ENSMUSG00000066613 | Zfp932 | protein_coding | -1.366285519 | 3.590341987 | 8.7715547 | 0.0090408 | 0.20596458 |
| ENSMUSG00000010045 | Tmem115 | protein_coding | 0.940605776 | 5.388791014 | 8.7512745 | 0.0091059 | 0.207010425 |
| ENSMUSG00000064115 | Cadm2 | protein_coding | -1.075217686 | 5.531232018 | 8.7339042 | 0.0091622 | 0.207393476 |
| ENSMUSG00000033809 | Alg3 | protein_coding | 1.162851421 | 4.917568556 | 8.7247413 | 0.009192 | 0.207393476 |
| ENSMUSG00000049252 | Lrp1b | protein_coding | -1.377260463 | 5.20458909 | 8.72048 | 0.0092184 | 0.207393476 |
| ENSMUSG00000071042 | Rasgrp3 | protein_coding | -1.993888784 | 3.738488421 | 8.7203538 | 0.0092188 | 0.207393476 |
| ENSMUSG00000071252 | 2210408I21Rik | protein_coding | -1.110208918 | 6.947680985 | 8.8532676 | 0.0092312 | 0.207393476 |
| ENSMUSG00000020230 | Prmt2 | protein_coding | 1.014473728 | 9.243598961 | 8.7243992 | 0.0092385 | 0.207393476 |
| ENSMUSG00000021423 | Ly86 | protein_coding | -1.426851543 | 4.544423758 | 8.69601 | 0.0092862 | 0.207656499 |
| ENSMUSG00000028289 | Epha7 | protein_coding | -0.997270274 | 5.22198334 | 8.6952138 | 0.0092888 | 0.207656499 |
| ENSMUSG00000024227 | Pdzph1 | protein_coding | -1.634595693 | 4.119564187 | 8.6850688 | 0.0093224 | 0.20790993 |
| ENSMUSG00000029772 | Ahcyl2 | protein_coding | -0.871324443 | 5.592556027 | 8.6801053 | 0.0093388 | 0.20790993 |
| ENSMUSG00000032060 | Cryab | protein_coding | -0.774082436 | 9.198418857 | 8.654893 | 0.009423 | 0.208996337 |
| ENSMUSG00000026275 | Ppp1r7 | protein_coding | -0.692587855 | 7.815914867 | 8.6538378 | 0.0094265 | 0.208996337 |
| ENSMUSG00000021774 | Ube2e1 | protein_coding | 0.885548353 | 5.5198214 | 8.6410557 | 0.0094695 | 0.20918413 |
| ENSMUSG00000024287 | Thoc1 | protein_coding | -0.813207085 | 5.986577429 | 8.6344004 | 0.009492 | 0.20918413 |
| ENSMUSG00000015090 | Ptgds | protein_coding | -1.0615663 | 9.368961098 | 8.7053827 | 0.0094933 | 0.20918413 |
| ENSMUSG00000069516 | Lyz2 | protein_coding | -2.097493325 | 3.313845195 | 8.595799 | 0.0096236 | 0.211542556 |
| ENSMUSG00000032413 | Rasa2 | protein_coding | -1.23235326 | 4.582664916 | 8.5910992 | 0.0096397 | 0.211542556 |
| ENSMUSG00000026247 | Ecel1 | protein_coding | 1.20447716 | 5.931074143 | 8.5924473 | 0.0096817 | 0.212031028 |
| ENSMUSG00000021277 | Traf3 | protein_coding | 1.578110794 | 3.856128044 | 8.566088 | 0.0097262 | 0.212573602 |
| ENSMUSG00000021608 | Lpcat1 | protein_coding | 1.289098448 | 4.229477468 | 8.5491496 | 0.0097853 | 0.213178908 |
| ENSMUSG00000025616 | Usp16 | protein_coding | -1.086549187 | 4.813608305 | 8.5467943 | 0.0097936 | 0.213178908 |
| ENSMUSG00000029430 | Ran | protein_coding | 0.723117014 | 8.036845059 | 8.5382017 | 0.0098237 | 0.213403488 |
| ENSMUSG00000032553 | Srprb | protein_coding | 0.87517551 | 5.847270819 | 8.5229561 | 0.0098775 | 0.214139121 |
| ENSMUSG00000027884 | Clcc1 | protein_coding | 1.094523912 | 4.867255731 | 8.5133233 | 0.0099117 | 0.21430609 |
| ENSMUSG00000055567 | Unc80 | protein_coding | -0.833455597 | 6.175450892 | 8.5062558 | 0.0099368 | 0.21430609 |
| ENSMUSG00000043991 | Pura | protein_coding | -0.769004439 | 6.957728223 | 8.5039559 | 0.009945 | 0.21430609 |
| ENSMUSG00000031647 | Mfap3l | protein_coding | -1.384976767 | 3.545629981 | 8.4967411 | 0.0099708 | 0.214431585 |
| ENSMUSG00000109121 | Gm44644 | lincRNA | -1.888570868 | 3.242296823 | 8.489022 | 0.0099984 | 0.214597032 |
| ENSMUSG00000050334 | C130071C03Rik | lincRNA | -1.152220004 | 4.538321769 | 8.4652954 | 0.010084 | 0.216001785 |
| ENSMUSG00000040759 | Cmtm5 | protein_coding | -0.969255479 | 7.823720568 | 8.4703601 | 0.0101134 | 0.216201673 |
| ENSMUSG00000030847 | Bag3 | protein_coding | 2.221737577 | 2.816494219 | 8.4345964 | 0.0101959 | 0.217532284 |
| ENSMUSG00000031422 | Morf4l2 | protein_coding | 0.786806454 | 8.396678086 | 8.4179062 | 0.0102573 | 0.217569707 |
| ENSMUSG00000018906 | P4ha2 | protein_coding | 1.57023546 | 4.267600162 | 8.4136799 | 0.0102729 | 0.217569707 |
| ENSMUSG00000030086 | Chchd6 | protein_coding | 0.896337849 | 7.819536396 | 8.4101288 | 0.0102994 | 0.217569707 |
| ENSMUSG00000039943 | Plcb4 | protein_coding | -0.867495074 | 6.39625355 | 8.4061855 | 0.0103007 | 0.217569707 |
| ENSMUSG00000018931 | Natd1 | protein_coding | 2.419155265 | 2.723686703 | 8.4036718 | 0.01031 | 0.217569707 |
| ENSMUSG00000003119 | Cdk12 | protein_coding | -1.343423273 | 4.176047042 | 8.4012527 | 0.010319 | 0.217569707 |
| ENSMUSG00000038481 | Cdk19 | protein_coding | -1.097643964 | 4.85149738 | 8.3867271 | 0.0103732 | 0.218284009 |
| ENSMUSG00000007721 | Ccdc124 | protein_coding | 0.679882545 | 8.225386948 | 8.3716816 | 0.0104297 | 0.219043418 |
| ENSMUSG00000036572 | Upf3b | protein_coding | -0.923870852 | 5.476546506 | 8.3641568 | 0.010458 | 0.219190826 |
| ENSMUSG00000015222 | Map2 | protein_coding | -0.646081983 | 8.291809547 | 8.3590191 | 0.0104775 | 0.219190826 |
| ENSMUSG00000022090 | Pdlim2 | protein_coding | -0.810228356 | 6.69297983 | 8.3417853 | 0.0105429 | 0.220131571 |
| ENSMUSG00000042447 | Mios | protein_coding | 1.638882758 | 3.605521355 | 8.3358322 | 0.0105656 | 0.220178273 |
| ENSMUSG00000063047 | Zfp780b | protein_coding | -1.010034231 | 4.690286311 | 8.3136206 | 0.0106508 | 0.220690112 |
| ENSMUSG00000048497 | Mmgt2 | protein_coding | 1.747334036 | 3.594320019 | 8.3067728 | 0.0106773 | 0.220690112 |
| ENSMUSG00000031144 | Syp | protein_coding | 1.14392563 | 6.858672657 | 8.4437728 | 0.0106795 | 0.220690112 |
| ENSMUSG00000047205 | Dusp18 | protein_coding | 1.229878973 | 4.869226442 | 8.3036672 | 0.0106893 | 0.220690112 |
| ENSMUSG00000028581 | Laptm5 | protein_coding | -1.085818989 | 5.352613588 | 8.3002759 | 0.0107024 | 0.220690112 |
| ENSMUSG00000030759 | Far1 | protein_coding | -0.91740264 | 5.452490102 | 8.2974705 | 0.0107133 | 0.220690112 |
| ENSMUSG00000078681 | Tm2d3 | protein_coding | 0.707042158 | 8.32003089 | 8.2785053 | 0.0107872 | 0.220929312 |
| ENSMUSG00000074782 | 4833422C13Rik | antisense_RNA | -1.38067867 | 4.016640669 | 8.2747991 | 0.0108017 | 0.220929312 |
| ENSMUSG00000111080 | AC153899.1 | lincRNA | -0.887683005 | 5.911816474 | 8.2727832 | 0.0108096 | 0.220929312 |
| ENSMUSG00000021716 | Srek1ip1 | protein_coding | -0.823717657 | 6.026478648 | 8.2719618 | 0.0108128 | 0.220929312 |
| ENSMUSG00000020850 | Prpf8 | protein_coding | 0.774062066 | 6.328230262 | 8.2681844 | 0.0108277 | 0.220929312 |
| ENSMUSG00000042390 | Gatad2b | protein_coding | -0.869265824 | 5.723837878 | 8.2448499 | 0.0109198 | 0.222032428 |
| ENSMUSG00000029152 | Ociad1 | protein_coding | 0.725050802 | 8.870748976 | 8.2440275 | 0.010923 | 0.222032428 |
| ENSMUSG00000104674 | Gm42756 | antisense_RNA | -1.509328853 | 3.81264865 | 8.2203807 | 0.0110173 | 0.223189423 |
| ENSMUSG00000042302 | Ehbp1 | protein_coding | -1.427432417 | 4.174940949 | 8.2087276 | 0.0110641 | 0.223189423 |
| ENSMUSG00000031210 | Gpr165 | protein_coding | 2.016717514 | 3.673757385 | 8.2100221 | 0.0110727 | 0.223189423 |
| ENSMUSG00000032042 | Srpr | protein_coding | 0.828353347 | 5.738268041 | 8.206101 | 0.0110747 | 0.223189423 |
| ENSMUSG00000054976 | Nyap2 | protein_coding | -1.152717512 | 4.313598731 | 8.1934209 | 0.0111259 | 0.223189423 |
| ENSMUSG00000030231 | Plekha5 | protein_coding | -0.906800216 | 5.603126478 | 8.1917803 | 0.0111325 | 0.223189423 |
| ENSMUSG00000027499 | Pkia | protein_coding | -0.70214014 | 7.19892695 | 8.1898525 | 0.0111404 | 0.223189423 |
| ENSMUSG00000020644 | Id2 | protein_coding | -0.693018542 | 8.614376753 | 8.1685612 | 0.0112271 | 0.223189423 |
| ENSMUSG00000011958 | Bnip2 | protein_coding | -1.487314818 | 3.770114211 | 8.1671755 | 0.0112328 | 0.223189423 |
| ENSMUSG00000039910 | Cited2 | protein_coding | -1.057750785 | 4.336699034 | 8.1662958 | 0.0112364 | 0.223189423 |
| ENSMUSG00000034488 | Edil3 | protein_coding | -0.682799625 | 6.477288724 | 8.1565649 | 0.0112764 | 0.223189423 |
| ENSMUSG00000032249 | Anp32a | protein_coding | -0.714847654 | 6.482187072 | 8.1514509 | 0.0112974 | 0.223189423 |
| ENSMUSG00000041220 | Elovl6 | protein_coding | -0.968377649 | 5.424769524 | 8.148804 | 0.0113083 | 0.223189423 |
| ENSMUSG00000028974 | Dffa | protein_coding | 0.942840869 | 5.691760046 | 8.1418023 | 0.0113373 | 0.223189423 |
| ENSMUSG00000056966 | Gjc3 | protein_coding | -0.94758316 | 5.573142038 | 8.1405853 | 0.0113423 | 0.223189423 |
| ENSMUSG00000003308 | Keap1 | protein_coding | 1.627480326 | 4.751555809 | 8.2044009 | 0.0113456 | 0.223189423 |
| ENSMUSG00000034845 | Plvap | protein_coding | 1.258241659 | 3.980863032 | 8.135908 | 0.0113617 | 0.223189423 |

| ensembl_gene_id | external_gene_name | gene_biotype | log2 (fold change) | log2 (counts per million) | F-value | p-value | False discovery rate |
|---|---|---|---|---|---|---|---|
| ENSMUSG00000062252 | Lhfpl5 | protein_coding | 0.784030457 | 6.03676927 | 8.1304346 | 0.0113844 | 0.223189423 |
| ENSMUSG00000021108 | Prkch | protein_coding | -1.430664446 | 4.471156913 | 8.1230261 | 0.0114153 | 0.223189423 |
| ENSMUSG00000095362 | Gm14325 | protein_coding | -1.001140082 | 5.72133313 | 8.1215624 | 0.0114214 | 0.223189423 |
| ENSMUSG00000018379 | Srsf1 | protein_coding | -0.815504579 | 5.664583714 | 8.1097635 | 0.0114708 | 0.223189423 |
| ENSMUSG00000040037 | Negr1 | protein_coding | -0.921893964 | 6.345821179 | 8.1052262 | 0.0114898 | 0.223189423 |
| ENSMUSG00000007872 | Id3 | protein_coding | -0.714148904 | 7.392787143 | 8.1049088 | 0.0114911 | 0.223189423 |
| ENSMUSG00000020448 | Rnf185 | protein_coding | 1.14525095 | 5.802407097 | 8.1050183 | 0.0115048 | 0.223189423 |
| ENSMUSG00000041560 | Nop53 | protein_coding | 1.087243062 | 6.640298729 | 8.1853616 | 0.0115155 | 0.223189423 |
| ENSMUSG00000062753 | AI413582 | protein_coding | 0.70083565 | 7.381961825 | 8.0944304 | 0.0115353 | 0.223189423 |
| ENSMUSG00000025059 | Gk | protein_coding | -1.193199811 | 4.091340859 | 8.0917962 | 0.0115464 | 0.223189423 |
| ENSMUSG00000037773 | Pced1a | protein_coding | 1.761332914 | 3.918021502 | 8.0916689 | 0.0115611 | 0.223189423 |
| ENSMUSG00000027787 | Nmd3 | protein_coding | -0.827963945 | 5.983076922 | 8.0751282 | 0.0116171 | 0.223441297 |
| ENSMUSG00000097006 | 9530082P21Rik | lincRNA | 0.937015656 | 8.267374426 | 8.0759114 | 0.011628 | 0.223441297 |
| ENSMUSG00000000708 | Kat2b | protein_coding | -1.394926292 | 3.592394854 | 8.0695513 | 0.0116408 | 0.223441297 |
| ENSMUSG00000047656 | Trpt1 | protein_coding | 0.924717204 | 5.582264213 | 8.059948 | 0.0116819 | 0.223441297 |
| ENSMUSG00000037031 | Tspan15 | protein_coding | -1.390641993 | 4.183674783 | 8.0597533 | 0.0116827 | 0.223441297 |
| ENSMUSG00000024530 | Prelid3a | protein_coding | 0.764160879 | 6.054081993 | 8.0559961 | 0.0116988 | 0.223441297 |
| ENSMUSG00000048978 | Nrsn1 | protein_coding | -0.645368554 | 9.176900978 | 8.0468122 | 0.0117383 | 0.223797621 |
| ENSMUSG00000034560 | Washc4 | protein_coding | -1.444184323 | 4.050259752 | 8.0380473 | 0.0117761 | 0.223975408 |
| ENSMUSG00000045007 | Tubg2 | protein_coding | 0.920358498 | 6.830641441 | 8.03831 | 0.0117893 | 0.223975408 |
| ENSMUSG00000039968 | Rsbn1l | protein_coding | -1.572593623 | 3.902598808 | 8.0296831 | 0.0118123 | 0.224017275 |
| ENSMUSG00000032238 | Rora | protein_coding | -1.066354937 | 6.226476772 | 8.0170821 | 0.0118815 | 0.22485113 |
| ENSMUSG00000064373 | Selenop | protein_coding | -0.925541853 | 7.836042949 | 8.0132734 | 0.0118981 | 0.22485113 |
| ENSMUSG00000026456 | Cyb5r1 | protein_coding | 1.041781626 | 5.561723233 | 8.0002947 | 0.0119405 | 0.225257415 |
| ENSMUSG00000011831 | Evi5 | protein_coding | -1.111178141 | 4.949072978 | 7.9927405 | 0.0119738 | 0.225488489 |
| ENSMUSG00000034171 | Faah | protein_coding | 1.256234575 | 4.293621047 | 7.9809381 | 0.0120259 | 0.225964211 |
| ENSMUSG00000031755 | Bbs2 | protein_coding | 1.38948107 | 4.505617287 | 7.9703787 | 0.0120727 | 0.225964211 |
| ENSMUSG00000058254 | Tspan7 | protein_coding | 0.663254414 | 11.05145679 | 7.9677697 | 0.0120843 | 0.225964211 |
| ENSMUSG00000023919 | Cenpq | protein_coding | -1.988616496 | 3.48608627 | 7.9644407 | 0.0121137 | 0.225964211 |
| ENSMUSG00000043602 | Zfp3 | protein_coding | 1.364236297 | 3.811429342 | 7.9600005 | 0.012119 | 0.225964211 |
| ENSMUSG00000027109 | Sp3 | protein_coding | -1.065239104 | 4.960542373 | 7.9563711 | 0.0121352 | 0.225964211 |
| ENSMUSG00000029182 | 1700001C02Rik | protein_coding | -1.638969197 | 3.55603683 | 7.9517816 | 0.0121557 | 0.225964211 |
| ENSMUSG00000109088 | Gm44593 | lincRNA | -2.097835012 | 2.679908493 | 7.9492338 | 0.0121671 | 0.225964211 |
| ENSMUSG00000024940 | Ltbp3 | protein_coding | 2.248236126 | 2.779765272 | 7.9399464 | 0.0122089 | 0.226285394 |
| ENSMUSG00000055538 | Zcchc24 | protein_coding | -1.117395908 | 5.011976266 | 7.9360313 | 0.0122265 | 0.226285394 |
| ENSMUSG00000025162 | Csnk1d | protein_coding | 0.812905502 | 6.384389758 | 7.9263519 | 0.0122703 | 0.226455289 |
| ENSMUSG00000022139 | Mbnl2 | protein_coding | -0.728819629 | 7.573170751 | 7.9246846 | 0.0122778 | 0.226455289 |
| ENSMUSG00000026078 | Pdcl3 | protein_coding | -0.665118302 | 7.318520387 | 7.9187066 | 0.0123049 | 0.226566978 |
| ENSMUSG00000056215 | Lrguk | protein_coding | 2.927677058 | 2.904019546 | 7.9000386 | 0.0124048 | 0.227271429 |
| ENSMUSG00000085925 | Rtl1 | protein_coding | -1.405455769 | 4.657096858 | 7.8998439 | 0.0124057 | 0.227271429 |
| ENSMUSG00000057863 | Rpl36 | protein_coding | 0.830040578 | 5.553229542 | 7.8942262 | 0.0124167 | 0.227271429 |
| ENSMUSG00000027562 | Car2 | protein_coding | -0.721230597 | 10.41162244 | 7.8861383 | 0.0124539 | 0.227271429 |
| ENSMUSG00000034254 | Agpat1 | protein_coding | 0.855532837 | 5.640121654 | 7.8811774 | 0.0124768 | 0.227271429 |
| ENSMUSG00000070866 | Zfp804a | protein_coding | -1.350300949 | 4.426000685 | 7.8796487 | 0.0124839 | 0.227271429 |
| ENSMUSG00000023277 | Twf2 | protein_coding | 0.901727978 | 5.785012577 | 7.8780701 | 0.0124912 | 0.227271429 |
| ENSMUSG00000029211 | Gabra4 | protein_coding | -1.00544764 | 6.717935929 | 7.8892406 | 0.0125187 | 0.227388559 |
| ENSMUSG00000020189 | Osbpl8 | protein_coding | -0.961744199 | 5.708584989 | 7.8609097 | 0.0125708 | 0.227948242 |
| ENSMUSG00000008575 | Nfib | protein_coding | -1.122355775 | 5.048399068 | 7.8510133 | 0.0126169 | 0.228400212 |
| ENSMUSG00000029103 | Lrpap1 | protein_coding | 0.618612228 | 9.146954222 | 7.8322862 | 0.0127048 | 0.229604695 |
| ENSMUSG00000073131 | Vma21 | protein_coding | -0.702431008 | 6.259907015 | 7.8177416 | 0.0127735 | 0.230182127 |
| ENSMUSG00000022762 | Ncam2 | protein_coding | -1.46543412 | 4.609970633 | 7.8190898 | 0.0127821 | 0.230182127 |
| ENSMUSG00000031600 | Vps37a | protein_coding | -0.870805372 | 5.384052491 | 7.8082881 | 0.0128185 | 0.230182127 |
| ENSMUSG00000039197 | Adk | protein_coding | -0.61741218 | 7.492440639 | 7.8074628 | 0.0128224 | 0.230182127 |
| ENSMUSG00000042595 | Fam199x | protein_coding | -1.508080962 | 3.526595474 | 7.7934763 | 0.0128892 | 0.230533425 |
| ENSMUSG00000063142 | Kcnma1 | protein_coding | -0.724468047 | 6.222946109 | 7.791584 | 0.0128982 | 0.230533425 |
| ENSMUSG00000038679 | Trps1 | protein_coding | -1.139025275 | 4.421046374 | 7.7871501 | 0.0129195 | 0.230533425 |
| ENSMUSG00000048833 | Slc39a9 | protein_coding | 0.94367519 | 4.910425204 | 7.7854451 | 0.0129277 | 0.230533425 |
| ENSMUSG00000022466 | Rpap3 | protein_coding | 0.863719789 | 5.866990366 | 7.773866 | 0.0129835 | 0.231144893 |
| ENSMUSG00000023026 | Dip2b | protein_coding | -0.979257403 | 5.396342755 | 7.7489447 | 0.0131045 | 0.232621552 |
| ENSMUSG00000059742 | Kcnh7 | protein_coding | -1.23906555 | 3.864897987 | 7.7440551 | 0.0131284 | 0.232621552 |
| ENSMUSG00000042215 | Bag2 | protein_coding | 1.11516994 | 4.929207901 | 7.7418796 | 0.013139 | 0.232621552 |
| ENSMUSG00000025142 | Aspscr1 | protein_coding | 1.680932531 | 3.519149347 | 7.7390331 | 0.013153 | 0.232621552 |
| ENSMUSG00000025104 | Hdgfl3 | protein_coding | -0.837703121 | 6.422532832 | 7.7262551 | 0.0132158 | 0.233234495 |
| ENSMUSG00000046402 | Rbp1 | protein_coding | -1.219178445 | 5.686358435 | 7.7277038 | 0.013264 | 0.233234495 |
| ENSMUSG00000028053 | Ash1l | protein_coding | -0.851415058 | 6.306372161 | 7.7151627 | 0.0132706 | 0.233234495 |
| ENSMUSG00000025326 | Ube3a | protein_coding | -0.830727809 | 5.993583477 | 7.7124913 | 0.0132838 | 0.233234495 |
| ENSMUSG00000059355 | Wdr83os | protein_coding | 0.676110226 | 6.405259769 | 7.7100157 | 0.0132961 | 0.233234495 |
| ENSMUSG00000026255 | Efhd1 | protein_coding | -1.375995945 | 4.00969525 | 7.7045312 | 0.0133233 | 0.233299219 |
| ENSMUSG00000071533 | Pcnp | protein_coding | -0.635861579 | 8.028674101 | 7.6999141 | 0.0133463 | 0.233299219 |
| ENSMUSG00000101493 | 2810405F17Rik | lincRNA | -2.304962512 | 2.5205655 | 7.6931735 | 0.01338 | 0.233299219 |
| ENSMUSG00000041483 | Zfp281 | protein_coding | -1.162307282 | 4.817946654 | 7.6896467 | 0.0133976 | 0.233299219 |
| ENSMUSG00000028772 | Zcchc17 | protein_coding | 0.584917541 | 8.941094121 | 7.6844204 | 0.0134238 | 0.233299219 |
| ENSMUSG00000006418 | Rnf114 | protein_coding | 0.706981625 | 7.368095382 | 7.6831924 | 0.0134299 | 0.233299219 |
| ENSMUSG00000036181 | Hist1h1c | protein_coding | -1.403767102 | 3.712771943 | 7.6760844 | 0.0134657 | 0.233542625 |
| ENSMUSG00000086363 | A330102I10Rik | lincRNA | 1.141214538 | 4.731109168 | 7.6711543 | 0.0134905 | 0.233596785 |
| ENSMUSG00000024098 | Twsg1 | protein_coding | -1.517183074 | 3.465440633 | 7.6586833 | 0.0135536 | 0.234201766 |
| ENSMUSG00000031990 | Jam3 | protein_coding | -1.001188029 | 4.968066662 | 7.655646 | 0.013569 | 0.234201766 |
| ENSMUSG00000062209 | Erbb4 | protein_coding | -1.201447948 | 4.623836947 | 7.6336251 | 0.0136814 | 0.235276672 |

| ensembl_gene_id | external_gene_name | gene_biotype | log2 (fold change) | log2 (counts per million) | F-value | p-value | False discovery rate |
|---|---|---|---|---|---|---|---|
| ENSMUSG00000026843 | Fubp3 | protein_coding | -1.065109578 | 4.951118832 | 7.6298817 | 0.0137006 | 0.235276672 |
| ENSMUSG00000067931 | Zfp948 | protein_coding | -1.485649497 | 3.557058881 | 7.6264531 | 0.0137182 | 0.235276672 |
| ENSMUSG00000030465 | Psd3 | protein_coding | -0.730140591 | 6.568826009 | 7.6255255 | 0.013723 | 0.235276672 |
| ENSMUSG00000043629 | 1700019D03Rik | protein_coding | -1.442693883 | 3.571831769 | 7.6186927 | 0.0137582 | 0.235276672 |
| ENSMUSG00000017943 | Gdap1l1 | protein_coding | 0.746569816 | 7.587768862 | 7.6178394 | 0.0137626 | 0.235276672 |
| ENSMUSG00000041840 | Haus1 | protein_coding | -0.734076778 | 5.787045799 | 7.6070489 | 0.0138184 | 0.235742491 |
| ENSMUSG00000025439 | Clns1a | protein_coding | 0.617425882 | 7.542459201 | 7.6041046 | 0.0138337 | 0.235742491 |
| ENSMUSG00000041377 | Ninj2 | protein_coding | -1.233700675 | 4.6352863 | 7.58294 | 0.0139441 | 0.237247435 |
| ENSMUSG00000015290 | Ubl4a | protein_coding | 0.717545645 | 7.399745294 | 7.578103 | 0.0139694 | 0.23730354 |
| ENSMUSG00000006930 | Hap1 | protein_coding | 0.70719583 | 7.80132675 | 7.567422 | 0.0140256 | 0.237882413 |
| ENSMUSG00000025323 | Sp4 | protein_coding | -1.213251211 | 3.837975303 | 7.562339 | 0.0140525 | 0.237962229 |
| ENSMUSG00000033061 | Resp18 | protein_coding | 0.701664889 | 9.264313916 | 7.5559562 | 0.0140862 | 0.238159206 |
| ENSMUSG00000020564 | Atxn7l1 | protein_coding | 0.866139814 | 6.226863511 | 7.5426424 | 0.014157 | 0.238979814 |
| ENSMUSG00000025656 | Arhgef9 | protein_coding | -0.646261745 | 7.063057312 | 7.5338142 | 0.0142041 | 0.239268569 |
| ENSMUSG00000034910 | Pygo1 | protein_coding | -1.087205997 | 4.763878088 | 7.5311122 | 0.0142186 | 0.239268569 |
| ENSMUSG00000028347 | Tmeff1 | protein_coding | -0.848481782 | 5.739402037 | 7.5219069 | 0.014268 | 0.239724805 |
| ENSMUSG00000053004 | Hrh1 | protein_coding | 1.109242208 | 4.110665507 | 7.5121534 | 0.0143206 | 0.239883164 |
| ENSMUSG00000030629 | Zfand6 | protein_coding | -0.670799022 | 7.01366206 | 7.5070234 | 0.0143483 | 0.239883164 |
| ENSMUSG00000026361 | Cdc73 | protein_coding | -1.010904618 | 4.876864289 | 7.5064581 | 0.0143514 | 0.239883164 |
| ENSMUSG00000027375 | Mal | protein_coding | -0.738359134 | 7.874490494 | 7.5036328 | 0.0143667 | 0.239883164 |
| ENSMUSG00000025579 | Gaa | protein_coding | 0.675406841 | 9.350906561 | 7.4977089 | 0.0143988 | 0.240005579 |
| ENSMUSG00000030315 | Vgll4 | protein_coding | 0.947325345 | 4.932348267 | 7.4940639 | 0.0144186 | 0.240005579 |
| ENSMUSG00000039345 | Mettl22 | protein_coding | 0.728051352 | 6.464124396 | 7.4812093 | 0.0144888 | 0.240559912 |
| ENSMUSG00000056870 | Gulp1 | protein_coding | -1.765672648 | 3.06054886 | 7.4797679 | 0.0144967 | 0.240559912 |
| ENSMUSG00000026203 | Dnajb2 | protein_coding | 0.640193525 | 6.674401916 | 7.4670967 | 0.0145663 | 0.241342053 |
| ENSMUSG00000003072 | Atp5d | protein_coding | 1.61275813 | 8.518672717 | 8.1256177 | 0.0146435 | 0.242247812 |
| ENSMUSG00000027889 | Ampd2 | protein_coding | 0.971636155 | 6.126254103 | 7.4452255 | 0.0147033 | 0.242749551 |
| ENSMUSG00000020892 | Aloxe3 | protein_coding | 1.623164546 | 4.186535771 | 7.4301632 | 0.0147874 | 0.242749551 |
| ENSMUSG00000035649 | Zcchc7 | protein_coding | -1.044791964 | 4.421059595 | 7.4269069 | 0.0147895 | 0.242749551 |
| ENSMUSG00000041360 | Pum3 | protein_coding | 0.855006704 | 5.666340357 | 7.4264554 | 0.014792 | 0.242749551 |
| ENSMUSG00000028261 | Ndufaf4 | protein_coding | -0.7260189 | 6.343887909 | 7.4184233 | 0.0148371 | 0.242749551 |
| ENSMUSG00000031673 | Cdh11 | protein_coding | -1.067776643 | 4.760155225 | 7.4180323 | 0.0148393 | 0.242749551 |
| ENSMUSG00000012126 | Ubxn11 | protein_coding | 0.734269071 | 5.614019991 | 7.4089462 | 0.0148905 | 0.242749551 |
| ENSMUSG00000020132 | Rab21 | protein_coding | -0.644147128 | 6.693586283 | 7.4086128 | 0.0148924 | 0.242749551 |
| ENSMUSG00000059273 | Zc3h4 | protein_coding | -1.010376101 | 4.300804903 | 7.4074884 | 0.0148987 | 0.242749551 |
| ENSMUSG00000040483 | Xaf1 | protein_coding | -1.182952151 | 4.266145799 | 7.4073443 | 0.0148995 | 0.242749551 |
| ENSMUSG00000024610 | Cd74 | protein_coding | -1.476412968 | 4.200338899 | 7.3896287 | 0.015 | 0.244016823 |
| ENSMUSG00000029073 | Cptp | protein_coding | 1.055678325 | 6.527474622 | 7.4636277 | 0.0150309 | 0.244149694 |
| ENSMUSG00000022604 | Cep97 | protein_coding | -1.286355016 | 3.975601437 | 7.3744098 | 0.0150869 | 0.24469066 |
| ENSMUSG00000024074 | Crim1 | protein_coding | -0.83362163 | 5.537589851 | 7.3411104 | 0.0152792 | 0.246014104 |
| ENSMUSG00000005804 | Bloc1s6 | protein_coding | 1.147280442 | 6.412023216 | 7.4458781 | 0.0153117 | 0.246014104 |
| ENSMUSG00000000902 | Smarcb1 | protein_coding | 0.770434434 | 7.733330558 | 7.3307371 | 0.0153396 | 0.246014104 |
| ENSMUSG00000044252 | Osbpl1a | protein_coding | -0.658156164 | 6.867745508 | 7.3299381 | 0.0153443 | 0.246014104 |
| ENSMUSG00000029571 | Tmem106b | protein_coding | -0.862203329 | 5.82573429 | 7.3280534 | 0.0153553 | 0.246014104 |
| ENSMUSG00000007029 | Vars | protein_coding | 1.772380339 | 3.074595686 | 7.3245701 | 0.0153757 | 0.246014104 |
| ENSMUSG00000047379 | B4gat1 | protein_coding | 0.859620737 | 7.438646218 | 7.3269132 | 0.0153784 | 0.246014104 |
| ENSMUSG00000024395 | Lims2 | protein_coding | -0.995396196 | 5.325717595 | 7.3194852 | 0.0154055 | 0.246014104 |
| ENSMUSG00000055435 | Maf | protein_coding | -1.662739467 | 3.232228002 | 7.3174071 | 0.0154177 | 0.246014104 |
| ENSMUSG00000067851 | Arfgef1 | protein_coding | -0.994967174 | 5.310221381 | 7.3098792 | 0.015462 | 0.246014104 |
| ENSMUSG00000062794 | Zfp599 | protein_coding | -1.41754393 | 13.5488801 | 7.5341935 | 0.0154705 | 0.246014104 |
| ENSMUSG00000033096 | Apmap | protein_coding | 0.74874571 | 5.88880109 | 7.3075912 | 0.0154755 | 0.246014104 |
| ENSMUSG00000004642 | Slbp | protein_coding | 0.85494678 | 6.246675971 | 7.3067934 | 0.0154802 | 0.246014104 |
| ENSMUSG00000042831 | Alkbh6 | protein_coding | 1.025899766 | 7.43084631 | 7.4140626 | 0.0154935 | 0.246014104 |
| ENSMUSG00000027805 | Pfn2 | protein_coding | 0.72362152 | 10.00691878 | 7.299844 | 0.0155213 | 0.246014104 |
| ENSMUSG00000025958 | Creb1 | protein_coding | -1.039276196 | 5.110658551 | 7.2975925 | 0.0155346 | 0.246014104 |
| ENSMUSG00000024411 | Aqp4 | protein_coding | -1.047431239 | 5.041781743 | 7.2872086 | 0.0155963 | 0.246627484 |
| ENSMUSG00000056260 | Lrif1 | protein_coding | -0.903716511 | 5.03457639 | 7.2716002 | 0.0156895 | 0.247737289 |
| ENSMUSG00000035776 | Cd99l2 | protein_coding | 0.594773107 | 7.23010765 | 7.2674654 | 0.0157143 | 0.247765076 |
| ENSMUSG00000036745 | Ttll7 | protein_coding | -0.644874069 | 7.030682103 | 7.2611152 | 0.0157525 | 0.24800337 |
| ENSMUSG00000029629 | Phf14 | protein_coding | -1.084082658 | 4.886092772 | 7.2536812 | 0.0157973 | 0.248345508 |
| ENSMUSG00000065979 | Cpped1 | protein_coding | -0.830348208 | 6.910962486 | 7.2335336 | 0.0159195 | 0.249466109 |
| ENSMUSG00000074060 | Fbxw15 | protein_coding | -1.575800034 | 3.577313877 | 7.2325171 | 0.0159257 | 0.249466109 |
| ENSMUSG00000039058 | Ak5 | protein_coding | 0.884953627 | 5.474999154 | 7.2273747 | 0.0159571 | 0.249466109 |
| ENSMUSG00000062908 | Acadm | protein_coding | -0.791652229 | 5.884754575 | 7.222144 | 0.0159891 | 0.249466109 |
| ENSMUSG00000004446 | Bid | protein_coding | 1.044566196 | 5.242118399 | 7.2203592 | 0.016 | 0.249466109 |
| ENSMUSG00000032827 | Ppp1r9a | protein_coding | -0.859863392 | 7.15037439 | 7.2218338 | 0.0160078 | 0.249466109 |
| ENSMUSG00000067847 | Romo1 | protein_coding | 0.682929487 | 8.20766826 | 7.2123827 | 0.016049 | 0.249479888 |
| ENSMUSG00000042055 | Wdr11 | protein_coding | 1.017838335 | 4.757641252 | 7.2102001 | 0.0160624 | 0.249479888 |
| ENSMUSG00000021087 | Rtn1 | protein_coding | 0.611327112 | 10.12214384 | 7.2074307 | 0.0160795 | 0.249479888 |
| ENSMUSG00000056962 | Jmjd6 | protein_coding | 0.851866692 | 5.364287745 | 7.2038665 | 0.0161015 | 0.249479888 |
| ENSMUSG00000026826 | Nr4a2 | protein_coding | -2.024193334 | 2.621287073 | 7.1951259 | 0.0161555 | 0.24995728 |
| ENSMUSG00000085336 | Gm11732 | processed_transcript | 1.965243138 | 3.231971655 | 7.1778335 | 0.0162631 | 0.251259856 |
| ENSMUSG00000040339 | Fam102b | protein_coding | -0.787264728 | 5.639958171 | 7.1626053 | 0.0163585 | 0.252001907 |
| ENSMUSG00000109836 | Gm45884 | TEC | -1.301652157 | 3.596234696 | 7.1509658 | 0.0164318 | 0.252001907 |
| ENSMUSG00000035372 | 1810055G02Rik | protein_coding | 1.179366094 | 4.514499838 | 7.1489817 | 0.0164444 | 0.252001907 |
| ENSMUSG00000044349 | Snhg11 | protein_coding | -1.007592084 | 7.403342186 | 7.2516877 | 0.0164611 | 0.252001907 |
| ENSMUSG00000002059 | Rab34 | protein_coding | -1.450916996 | 3.398121639 | 7.1455214 | 0.0164663 | 0.252001907 |
| ENSMUSG00000062210 | Tnfaip8 | protein_coding | -1.388351123 | 4.226963676 | 7.1261082 | 0.0165897 | 0.252001907 |

| ensembl_gene_id | external_gene_name | gene_biotype | log2 (fold change) | log2 (counts per million) | F-value | p-value | False discovery rate |
|---|---|---|---|---|---|---|---|
| ENSMUSG00000028033 | Kcnq5 | protein_coding | -1.074694218 | 4.809074714 | 7.1203286 | 0.0166267 | 0.252001907 |
| ENSMUSG00000047368 | Abhd17b | protein_coding | -0.993066597 | 5.259547045 | 7.1192663 | 0.0166335 | 0.252001907 |
| ENSMUSG00000031409 | Tceal6 | protein_coding | 0.648128052 | 6.238640912 | 7.1183414 | 0.0166394 | 0.252001907 |
| ENSMUSG00000018166 | Erbb3 | protein_coding | -2.676993016 | 2.661932717 | 7.117611 | 0.0166441 | 0.252001907 |
| ENSMUSG00000058799 | Nap1l1 | protein_coding | -0.622138681 | 6.744221483 | 7.1159165 | 0.0166549 | 0.252001907 |
| ENSMUSG00000055862 | Izumo4 | protein_coding | 0.711905984 | 6.276003752 | 7.1158772 | 0.0166552 | 0.252001907 |
| ENSMUSG00000046027 | Stard5 | protein_coding | -1.105276747 | 4.094386568 | 7.1157144 | 0.0166562 | 0.252001907 |
| ENSMUSG00000020024 | Cep83 | protein_coding | -1.243927827 | 4.004862667 | 7.1155337 | 0.0166574 | 0.252001907 |
| ENSMUSG00000031860 | Pbx4 | protein_coding | 1.138179282 | 4.024994436 | 7.1147182 | 0.0166626 | 0.252001907 |
| ENSMUSG00000000214 | Th | protein_coding | 1.140804242 | 5.627137338 | 7.1088035 | 0.0167178 | 0.252103234 |
| ENSMUSG00000038805 | Six3 | protein_coding | -1.080759255 | 5.097093561 | 7.1016384 | 0.0167468 | 0.252103234 |
| ENSMUSG00000033669 | Zfp7 | protein_coding | 1.603072644 | 3.10848337 | 7.1009786 | 0.0167511 | 0.252103234 |
| ENSMUSG00000000568 | Hnrnpd | protein_coding | -0.731099995 | 6.889537929 | 7.099118 | 0.0167631 | 0.252103234 |
| ENSMUSG00000026857 | Ntmt1 | protein_coding | 0.63673479 | 7.068004118 | 7.0943585 | 0.0167939 | 0.252103823 |
| ENSMUSG00000023087 | Noct | protein_coding | -0.956855035 | 4.931298851 | 7.0918684 | 0.01681 | 0.252103823 |
| ENSMUSG00000085438 | 1700020I14Rik | lincRNA | -0.624731669 | 6.815821602 | 7.0843888 | 0.0168586 | 0.252480183 |
| ENSMUSG00000022536 | Glyr1 | protein_coding | 0.770792334 | 7.648161917 | 7.0719884 | 0.0169395 | 0.253126003 |
| ENSMUSG00000034610 | Zcchc11 | protein_coding | -1.257022521 | 4.181040372 | 7.0705645 | 0.0169488 | 0.253126003 |
| ENSMUSG00000040640 | Erc2 | protein_coding | -0.802842234 | 6.308424434 | 7.0607279 | 0.0170133 | 0.25334132 |
| ENSMUSG00000021288 | Klc1 | protein_coding | 0.7467366 | 7.094391431 | 7.0605404 | 0.0170146 | 0.25334132 |
| ENSMUSG00000040407 | Akap9 | protein_coding | -0.8227556 | 5.75019992 | 7.0494585 | 0.0170876 | 0.25334132 |
| ENSMUSG00000021703 | Serinc5 | protein_coding | -1.264147675 | 4.266735391 | 7.0458755 | 0.0171113 | 0.25334132 |
| ENSMUSG00000060227 | Casc4 | protein_coding | -0.863582774 | 5.343589844 | 7.0360239 | 0.0171766 | 0.25334132 |
| ENSMUSG00000014592 | Camta1 | protein_coding | -0.646682107 | 7.884971833 | 7.0359502 | 0.0171771 | 0.25334132 |
| ENSMUSG00000020451 | Limk2 | protein_coding | 0.93321919 | 6.679281051 | 7.0441527 | 0.0171859 | 0.25334132 |
| ENSMUSG00000021876 | Rnase4 | protein_coding | -1.22173228 | 4.138297218 | 7.0329918 | 0.0171968 | 0.25334132 |
| ENSMUSG00000041975 | Mettl8 | protein_coding | 1.622038547 | 4.448576166 | 7.0799066 | 0.0172148 | 0.25334132 |
| ENSMUSG00000057411 | Fam173a | protein_coding | 0.661908072 | 8.093885779 | 7.0293437 | 0.0172211 | 0.25334132 |
| ENSMUSG00000037752 | Xkr8 | protein_coding | 2.024942674 | 2.798958383 | 7.0271029 | 0.017236 | 0.25334132 |
| ENSMUSG00000031285 | Dcx | protein_coding | -1.490134818 | 3.778773421 | 7.0256129 | 0.017246 | 0.25334132 |
| ENSMUSG00000036769 | Wdr44 | protein_coding | -1.5373155 | 4.062501324 | 7.0242062 | 0.0172728 | 0.253377598 |
| ENSMUSG00000039086 | Ss18l1 | protein_coding | 0.965223132 | 4.984078631 | 7.0153242 | 0.0173148 | 0.253377598 |
| ENSMUSG00000005813 | Metap1 | protein_coding | 0.877452453 | 5.827359175 | 7.0146808 | 0.0173191 | 0.253377598 |
| ENSMUSG00000030337 | Vamp1 | protein_coding | 0.883065658 | 7.451851913 | 7.0120967 | 0.0174 | 0.25406958 |
| ENSMUSG00000045827 | Serpinb9 | protein_coding | 1.385103623 | 3.913009374 | 7.0006295 | 0.0174137 | 0.25406958 |
| ENSMUSG00000035597 | Prpf39 | protein_coding | -1.162392358 | 4.118613969 | 6.9905624 | 0.0174818 | 0.254717657 |
| ENSMUSG00000110332 | Gm45396 | protein_coding | -1.233384853 | 4.327348264 | 6.9793336 | 0.0175581 | 0.255181108 |
| ENSMUSG00000028431 | Ikbkap | protein_coding | 0.873856929 | 5.093809927 | 6.9761417 | 0.0175799 | 0.255181108 |
| ENSMUSG00000016386 | Mpped2 | protein_coding | -0.926040003 | 5.370563768 | 6.9754223 | 0.0175848 | 0.255181108 |
| ENSMUSG00000009647 | Mcu | protein_coding | -0.740648053 | 6.260621807 | 6.9529753 | 0.0177388 | 0.256940991 |
| ENSMUSG00000044550 | Tceal3 | protein_coding | 0.55613096 | 7.206967085 | 6.9507896 | 0.0177539 | 0.256940991 |
| ENSMUSG00000026201 | Stk16 | protein_coding | 0.883674972 | 6.45245225 | 6.9390514 | 0.0178528 | 0.257042328 |
| ENSMUSG00000028480 | Glipr2 | protein_coding | -2.278038701 | 2.854064298 | 6.9360808 | 0.0178557 | 0.257042328 |
| ENSMUSG00000043913 | Ccdc60 | protein_coding | -2.64857093 | 2.315341886 | 6.9319868 | 0.0178842 | 0.257042328 |
| ENSMUSG00000026743 | Mllt10 | protein_coding | -0.998328261 | 4.706873075 | 6.9310129 | 0.0178909 | 0.257042328 |
| ENSMUSG00000078429 | Ctdsp2 | protein_coding | -1.165554912 | 4.81346619 | 6.9286561 | 0.0179073 | 0.257042328 |
| ENSMUSG00000024065 | Ehd3 | protein_coding | 0.856212241 | 6.826757013 | 6.9232047 | 0.0179632 | 0.257042328 |
| ENSMUSG00000036916 | Zfp280c | protein_coding | -1.727654996 | 3.532355404 | 6.9183667 | 0.0179792 | 0.257042328 |
| ENSMUSG00000027088 | Phospho2 | protein_coding | 0.806590111 | 5.56835317 | 6.9149664 | 0.018003 | 0.257042328 |
| ENSMUSG00000020737 | Jpt1 | protein_coding | 0.604192426 | 8.50463391 | 6.9147477 | 0.0180046 | 0.257042328 |
| ENSMUSG00000021820 | Camk2g | protein_coding | 0.561340213 | 7.244062361 | 6.9085416 | 0.0180482 | 0.257042328 |
| ENSMUSG00000030538 | Cib1 | protein_coding | -1.139331953 | 4.759552767 | 6.9079051 | 0.0180526 | 0.257042328 |
| ENSMUSG00000002032 | Tmem25 | protein_coding | 0.888154898 | 6.284978924 | 6.9053255 | 0.0180708 | 0.257042328 |
| ENSMUSG00000040297 | Suco | protein_coding | -1.006008787 | 4.478071577 | 6.9045583 | 0.0180762 | 0.257042328 |
| ENSMUSG00000036106 | Prr5 | protein_coding | 1.69970728 | 3.567331264 | 6.9008984 | 0.018102 | 0.257042328 |
| ENSMUSG00000024835 | Coro1b | protein_coding | 0.637555574 | 6.337349066 | 6.8984309 | 0.0181194 | 0.257042328 |
| ENSMUSG00000068457 | Uty | protein_coding | -1.420927434 | 3.355291775 | 6.8882165 | 0.0181917 | 0.257694623 |
| ENSMUSG00000033981 | Gria2 | protein_coding | -0.723657409 | 8.100468188 | 6.8811325 | 0.0182421 | 0.257694623 |
| ENSMUSG00000039983 | Ccdc32 | protein_coding | 0.618525982 | 7.137188915 | 6.8765471 | 0.0182748 | 0.257694623 |
| ENSMUSG00000024477 | Pggt1b | protein_coding | -1.397013543 | 3.324418447 | 6.8756071 | 0.0182815 | 0.257694623 |
| ENSMUSG00000028048 | Gba | protein_coding | 1.091189687 | 4.871983274 | 6.8730043 | 0.0183 | 0.257694623 |
| ENSMUSG00000057858 | Fam204a | protein_coding | -0.900582866 | 5.679382187 | 6.8681219 | 0.018335 | 0.257694623 |
| ENSMUSG00000033632 | AW554918 | protein_coding | -1.130487448 | 3.894266693 | 6.8671945 | 0.0183416 | 0.257694623 |
| ENSMUSG00000028906 | Epb41 | protein_coding | -0.938408318 | 4.773790182 | 6.8650292 | 0.0183571 | 0.257694623 |
| ENSMUSG00000024234 | Mtpap | protein_coding | -0.924477131 | 5.238940671 | 6.8557828 | 0.0184235 | 0.258289968 |
| ENSMUSG00000045441 | Gprin3 | protein_coding | -1.12677676 | 4.460587543 | 6.8512168 | 0.0184565 | 0.258414367 |
| ENSMUSG00000041540 | Sox5 | protein_coding | -1.187203603 | 4.527138898 | 6.830795 | 0.0186044 | 0.260147757 |
| ENSMUSG00000033099 | Nol12 | protein_coding | 0.566477533 | 6.686652263 | 6.8192006 | 0.0186891 | 0.260186736 |
| ENSMUSG00000061751 | Kalrn | protein_coding | 0.825056481 | 5.524769988 | 6.8179176 | 0.0186985 | 0.260186736 |
| ENSMUSG00000038845 | Phb | protein_coding | 0.82857197 | 6.715993829 | 6.8181009 | 0.0187153 | 0.260186736 |
| ENSMUSG00000020137 | Thap2 | protein_coding | -1.422841133 | 3.538827538 | 6.814381 | 0.0187244 | 0.260186736 |
| ENSMUSG00000035824 | Tk2 | protein_coding | 0.994800051 | 5.365178531 | 6.8136722 | 0.0187296 | 0.260186736 |
| ENSMUSG00000024621 | Csf1r | protein_coding | -1.170034678 | 4.449327018 | 6.8085992 | 0.0187668 | 0.260186736 |
| ENSMUSG00000109946 | D2Bwg1423e | protein_coding | 0.676560776 | 6.58515184 | 6.8072705 | 0.0187766 | 0.260186736 |
| ENSMUSG00000062542 | Syt9 | protein_coding | 1.285393136 | 4.246402326 | 6.7987514 | 0.0188394 | 0.260464081 |
| ENSMUSG00000020521 | Rnft1 | protein_coding | -1.139884452 | 4.037475334 | 6.7938672 | 0.0188755 | 0.260464081 |
| ENSMUSG00000034850 | Tmem127 | protein_coding | 0.993963185 | 6.47245845 | 6.8400907 | 0.0188791 | 0.260464081 |
| ENSMUSG00000049957 | Ccdc137 | protein_coding | 0.902981632 | 4.829238784 | 6.7883688 | 0.0189163 | 0.260464081 |

| ensembl_gene_id | external_gene_name | gene_biotype | log2 (fold change) | log2 (counts per million) | F-value | p-value | False discovery rate |
|---|---|---|---|---|---|---|---|
| ENSMUSG00000064289 | Tank | protein_coding | -0.826471351 | 5.599823498 | 6.7822047 | 0.0189621 | 0.260464081 |
| ENSMUSG00000026879 | Gsn | protein_coding | -0.655715809 | 6.200328088 | 6.7806217 | 0.0189738 | 0.260464081 |
| ENSMUSG00000003099 | Ppp5c | protein_coding | 0.806721134 | 7.03799125 | 6.7824156 | 0.0189787 | 0.260464081 |
| ENSMUSG00000024309 | Pfdn6 | protein_coding | 0.553338814 | 8.166682519 | 6.7783942 | 0.0189904 | 0.260464081 |
| ENSMUSG00000028602 | Tnfrsf8 | protein_coding | 4.092462125 | 1.854737579 | 6.7720368 | 0.0190379 | 0.260464081 |
| ENSMUSG00000047388 | Atmin | protein_coding | 0.782393654 | 5.551702336 | 6.769109 | 0.0190598 | 0.260464081 |
| ENSMUSG00000060923 | Acyp2 | protein_coding | -0.592472738 | 7.248511188 | 6.7666766 | 0.019078 | 0.260464081 |
| ENSMUSG00000005371 | Fbxo11 | protein_coding | -0.678474509 | 6.474362649 | 6.765436 | 0.0190873 | 0.260464081 |
| ENSMUSG00000037646 | Vps13b | protein_coding | -1.69169084 | 3.46712772 | 6.7584132 | 0.01914 | 0.260764882 |
| ENSMUSG00000039246 | Lyplal1 | protein_coding | 1.007701441 | 4.955391354 | 6.7559575 | 0.0191585 | 0.260764882 |
| ENSMUSG00000022811 | Zfp148 | protein_coding | -0.656281732 | 6.010483517 | 6.7528265 | 0.0191821 | 0.260764882 |
| ENSMUSG00000020463 | Ppp4r3b | protein_coding | -0.879257926 | 5.015230073 | 6.7464167 | 0.0192305 | 0.260997693 |
| ENSMUSG00000050953 | Gja1 | protein_coding | -0.81200613 | 7.008087782 | 6.7446983 | 0.0192619 | 0.260997693 |
| ENSMUSG00000080268 | Brms1 | protein_coding | 0.669768372 | 6.284589523 | 6.7409281 | 0.019272 | 0.260997693 |
| ENSMUSG00000020083 | 2010107G23Rik | protein_coding | 0.659496608 | 6.60916479 | 6.7367207 | 0.019304 | 0.261101134 |
| ENSMUSG00000020340 | Cyfip2 | protein_coding | 0.694243229 | 8.355776164 | 6.7273736 | 0.0193751 | 0.261505256 |
| ENSMUSG00000015981 | Stk32c | protein_coding | -1.74030347 | 3.19143195 | 6.7264051 | 0.0193825 | 0.261505256 |
| ENSMUSG00000025789 | St8sia2 | protein_coding | -2.193986459 | 2.608282858 | 6.718423 | 0.0194435 | 0.261936732 |
| ENSMUSG00000028518 | Prkaa2 | protein_coding | -1.057358919 | 4.330244399 | 6.7158524 | 0.0194632 | 0.261936732 |
| ENSMUSG00000060036 | Rpl3 | protein_coding | 0.759493585 | 7.69626689 | 6.7088683 | 0.0195168 | 0.262330038 |
| ENSMUSG00000037625 | Cldn11 | protein_coding | -0.577371818 | 7.407840448 | 6.6944453 | 0.0196281 | 0.263496251 |
| ENSMUSG00000063884 | Ptcd3 | protein_coding | 0.926149669 | 5.521540319 | 6.6820251 | 0.0197245 | 0.264460183 |
| ENSMUSG00000027215 | Cd82 | protein_coding | -0.831457902 | 6.399149265 | 6.6722502 | 0.0198007 | 0.265151886 |
| ENSMUSG00000049643 | 2310022A10Rik | protein_coding | 0.785782275 | 5.683966829 | 6.6674078 | 0.0198386 | 0.265328954 |
| ENSMUSG00000037996 | Slc24a2 | protein_coding | -0.860163748 | 6.09735924 | 6.6602226 | 0.019895 | 0.265353633 |
| ENSMUSG00000054051 | Ercc6 | protein_coding | 2.442638105 | 2.562444111 | 6.6583721 | 0.0199095 | 0.265353633 |
| ENSMUSG00000018677 | Slc25a39 | protein_coding | 0.628689841 | 6.66780126 | 6.6577443 | 0.0199145 | 0.265353633 |
| ENSMUSG00000096054 | Syne1 | protein_coding | -0.744618081 | 6.143398477 | 6.6529703 | 0.0199521 | 0.26542898 |
| ENSMUSG00000039988 | Ankrd13c | protein_coding | -0.736422783 | 5.982011039 | 6.6471794 | 0.0199978 | 0.26542898 |
| ENSMUSG00000053094 | Tmem248 | protein_coding | 1.011973925 | 4.729966442 | 6.645647 | 0.0200099 | 0.26542898 |
| ENSMUSG00000034880 | Mrpl34 | protein_coding | 0.684046503 | 7.009545127 | 6.6414894 | 0.0200429 | 0.26542898 |
| ENSMUSG00000026970 | Rbms1 | protein_coding | -1.053344968 | 4.38989333 | 6.6414018 | 0.0200436 | 0.26542898 |
| ENSMUSG00000089872 | Rps6kc1 | protein_coding | 0.912225309 | 4.90180226 | 6.6340766 | 0.0201017 | 0.265800698 |
| ENSMUSG00000068617 | Efcab1 | protein_coding | -0.799785774 | 5.490461706 | 6.6288955 | 0.020143 | 0.265800698 |
| ENSMUSG00000019907 | Ppp1r12a | protein_coding | -0.762589692 | 6.455738273 | 6.6285439 | 0.0201458 | 0.265800698 |
| ENSMUSG00000030043 | Tacr1 | protein_coding | 1.532117073 | 3.236117114 | 6.622417 | 0.0201947 | 0.266119643 |
| ENSMUSG00000038836 | Agbl3 | protein_coding | -0.854695077 | 5.249206097 | 6.6013694 | 0.0203638 | 0.267849365 |
| ENSMUSG00000013593 | Ndufs2 | protein_coding | 0.632898544 | 8.539366537 | 6.5986948 | 0.0203854 | 0.267849365 |
| ENSMUSG00000024150 | Mcfd2 | protein_coding | 0.605209223 | 7.672831725 | 6.596804 | 0.0204007 | 0.267849365 |
| ENSMUSG00000090112 | Shprh | protein_coding | -1.208006482 | 4.266955647 | 6.5860556 | 0.0204878 | 0.267965172 |
| ENSMUSG00000042460 | C1galt1 | protein_coding | -2.296306893 | 2.674108706 | 6.5834852 | 0.0205088 | 0.267965172 |
| ENSMUSG00000000127 | Fer | protein_coding | -1.080872959 | 4.309187938 | 6.582137 | 0.0205197 | 0.267965172 |
| ENSMUSG00000030591 | Psmd8 | protein_coding | 0.606134649 | 8.434043169 | 6.581 | 0.020529 | 0.267965172 |
| ENSMUSG00000026098 | Pms1 | protein_coding | -2.307669203 | 2.571576675 | 6.5762729 | 0.0205675 | 0.267965172 |
| ENSMUSG00000071379 | Hpcal1 | protein_coding | 1.111989484 | 7.646691765 | 6.7777463 | 0.020576 | 0.267965172 |
| ENSMUSG00000022708 | Zbtb20 | protein_coding | -0.934606495 | 6.260583793 | 6.5757807 | 0.0205905 | 0.267965172 |
| ENSMUSG00000020441 | 2310033P09Rik | protein_coding | 1.272938104 | 3.652092943 | 6.5693245 | 0.0206244 | 0.267965172 |
| ENSMUSG00000102215 | Gm2464 | lincRNA | 1.04899861 | 4.146585718 | 6.5627103 | 0.0206786 | 0.267965172 |
| ENSMUSG00000097545 | Mir124a-1hg | lincRNA | -1.494208103 | 3.619828558 | 6.560013 | 0.0207008 | 0.267965172 |
| ENSMUSG00000028670 | Lypla2 | protein_coding | 0.663553222 | 6.34628073 | 6.557944 | 0.0207178 | 0.267965172 |
| ENSMUSG00000028393 | Alad | protein_coding | 0.60785453 | 6.709993692 | 6.557325 | 0.0207229 | 0.267965172 |
| ENSMUSG00000038046 | Mrm3 | protein_coding | 0.770717415 | 5.840858269 | 6.5543839 | 0.0207472 | 0.267965172 |
| ENSMUSG00000008489 | Elavl2 | protein_coding | -0.707986376 | 6.592545263 | 6.5530223 | 0.0207584 | 0.267965172 |
| ENSMUSG00000104283 | Gm37459 | antisense_RNA | -1.090711114 | 4.559865584 | 6.5476069 | 0.0208031 | 0.268139532 |
| ENSMUSG00000026824 | Kcnj3 | protein_coding | -0.929040754 | 4.63725668 | 6.5448109 | 0.0208263 | 0.268139532 |
| ENSMUSG00000027166 | Dnajc24 | protein_coding | -1.084711601 | 5.076899597 | 6.5349337 | 0.0209083 | 0.268139532 |
| ENSMUSG00000026860 | Sh3glb2 | protein_coding | 0.69803896 | 8.027200636 | 6.5339998 | 0.020916 | 0.268139532 |
| ENSMUSG00000038235 | F11r | protein_coding | -1.528670573 | 3.493792847 | 6.5320008 | 0.0209327 | 0.268139532 |
| ENSMUSG00000040022 | Rab11fip2 | protein_coding | -0.924663614 | 4.496235975 | 6.529703 | 0.0209518 | 0.268139532 |
| ENSMUSG00000032123 | Dpagt1 | protein_coding | 1.864626572 | 3.034197488 | 6.519746 | 0.0210351 | 0.268139532 |
| ENSMUSG00000024082 | Ndufaf7 | protein_coding | 0.606906798 | 6.749561846 | 6.517295 | 0.0210556 | 0.268139532 |
| ENSMUSG00000054474 | Thnsl2 | protein_coding | 1.687945859 | 3.354334824 | 6.5134618 | 0.0210878 | 0.268139532 |
| ENSMUSG00000028070 | Naxe | protein_coding | 0.777529597 | 7.422686091 | 6.5148741 | 0.0210951 | 0.268139532 |
| ENSMUSG00000036169 | Sostdc1 | protein_coding | 1.76540387 | 5.072898366 | 6.7106047 | 0.0211036 | 0.268139532 |
| ENSMUSG00000039976 | Tbc1d16 | protein_coding | -1.021050315 | 4.131890637 | 6.5091834 | 0.0211238 | 0.268139532 |
| ENSMUSG00000026181 | Ppm1f | protein_coding | 0.947697259 | 5.832316627 | 6.5080742 | 0.0211331 | 0.268139532 |
| ENSMUSG00000038544 | Inip | protein_coding | -0.835096717 | 5.42137329 | 6.5071769 | 0.0211407 | 0.268139532 |
| ENSMUSG00000018415 | Gid4 | protein_coding | -0.884166162 | 5.948252585 | 6.5065509 | 0.0211459 | 0.268139532 |
| ENSMUSG00000027459 | Fam110a | protein_coding | 1.24919525 | 5.031921134 | 6.5020345 | 0.0212033 | 0.26835373 |
| ENSMUSG00000025979 | Mob4 | protein_coding | -0.604003786 | 7.801513554 | 6.4958297 | 0.0212365 | 0.26835373 |
| ENSMUSG00000071855 | Ccdc112 | protein_coding | -1.248859346 | 4.039742645 | 6.494515 | 0.0212477 | 0.26835373 |
| ENSMUSG00000002413 | Braf | protein_coding | -0.76965578 | 6.161610104 | 6.4927461 | 0.0212627 | 0.26835373 |
| ENSMUSG00000003746 | Man1a | protein_coding | -1.01598068 | 4.613703272 | 6.4863806 | 0.0213167 | 0.2687207 |
| ENSMUSG00000019775 | Rgs17 | protein_coding | -0.655226752 | 7.274097657 | 6.4809809 | 0.0213627 | 0.26888278 |
| ENSMUSG00000004394 | Tmed4 | protein_coding | 0.58697167 | 8.130152195 | 6.4766081 | 0.0214 | 0.26888278 |
| ENSMUSG00000029919 | Hpgds | protein_coding | -1.655098575 | 3.063518813 | 6.4733313 | 0.0214281 | 0.26888278 |
| ENSMUSG00000059436 | Max | protein_coding | -1.187662896 | 5.952105046 | 6.5615697 | 0.0214296 | 0.26888278 |
| ENSMUSG00000035919 | Bbs9 | protein_coding | 1.025437262 | 4.403336251 | 6.4573192 | 0.0215656 | 0.269783093 |

| ensembl_gene_id | external_gene_name | gene_biotype | log2 (fold change) | log2 (counts per million) | F-value | p-value | False discovery rate |
|---|---|---|---|---|---|---|---|
| ENSMUSG00000057342 | Sphk2 | protein_coding | 1.066226619 | 4.929433609 | 6.4559395 | 0.0215775 | 0.269783093 |
| ENSMUSG00000111947 | AC131120.1 | lincRNA | -2.508167378 | 2.78943609 | 6.4571292 | 0.0215866 | 0.269783093 |
| ENSMUSG00000086657 | Stamos | processed_transcript | -2.188728565 | 2.476351875 | 6.4523723 | 0.0216083 | 0.269783093 |
| ENSMUSG00000027708 | Dcun1d1 | protein_coding | -0.836561095 | 5.159536717 | 6.4502082 | 0.021627 | 0.269783093 |
| ENSMUSG00000042396 | Rbm7 | protein_coding | -0.780059519 | 5.8744885 | 6.4473283 | 0.0216519 | 0.269783093 |
| ENSMUSG00000032640 | Chsy1 | protein_coding | -1.168015213 | 3.73724095 | 6.4397492 | 0.0217176 | 0.270289067 |
| ENSMUSG00000044042 | Fmn1 | protein_coding | -1.36294516 | 3.170494221 | 6.43493 | 0.0217596 | 0.270375863 |
| ENSMUSG00000053801 | Grwd1 | protein_coding | 0.811327536 | 6.023791231 | 6.4331693 | 0.0217749 | 0.270375863 |
| ENSMUSG00000031485 | Prosc | protein_coding | 0.707896231 | 6.440097532 | 6.4097109 | 0.0219805 | 0.272331376 |
| ENSMUSG00000032826 | Ank2 | protein_coding | -0.726098133 | 6.614685541 | 6.4074968 | 0.022 | 0.272331376 |
| ENSMUSG00000014551 | Mrps25 | protein_coding | 0.572932741 | 7.710675343 | 6.4065457 | 0.0220084 | 0.272331376 |
| ENSMUSG00000038495 | Otud7b | protein_coding | -0.773620008 | 5.380455749 | 6.4002834 | 0.0220637 | 0.272435373 |
| ENSMUSG00000032306 | Mpi | protein_coding | 0.665188276 | 7.010687497 | 6.3881013 | 0.0221718 | 0.272435373 |
| ENSMUSG00000026754 | Golga1 | protein_coding | -0.834352129 | 4.866080909 | 6.3853073 | 0.0221967 | 0.272435373 |
| ENSMUSG00000048495 | Tyw5 | protein_coding | -0.908335542 | 4.875694732 | 6.382948 | 0.0222177 | 0.272435373 |
| ENSMUSG00000022091 | Sorbs3 | protein_coding | -0.915357694 | 4.802148404 | 6.3823248 | 0.0222233 | 0.272435373 |
| ENSMUSG00000020134 | Peli1 | protein_coding | -0.968524816 | 5.027350715 | 6.3807557 | 0.0222373 | 0.272435373 |
| ENSMUSG00000034525 | Ice1 | protein_coding | -1.112195176 | 4.221830255 | 6.3799407 | 0.0222445 | 0.272435373 |
| ENSMUSG00000029840 | Mtpn | protein_coding | -0.68271273 | 7.639171487 | 6.3795191 | 0.0222483 | 0.272435373 |
| ENSMUSG00000028677 | Rnf220 | protein_coding | 0.597538484 | 6.594261474 | 6.3787233 | 0.0222554 | 0.272435373 |
| ENSMUSG00000035954 | Dock4 | protein_coding | -1.155533245 | 4.222675166 | 6.3770773 | 0.0222701 | 0.272435373 |
| ENSMUSG00000022619 | Mapk8ip2 | protein_coding | -0.788317917 | 6.320424847 | 6.3692481 | 0.0223403 | 0.272982892 |
| ENSMUSG00000036972 | Zic4 | protein_coding | -1.167163317 | 5.174582636 | 6.3673333 | 0.0223772 | 0.27312349 |
| ENSMUSG00000021549 | Rasa1 | protein_coding | -1.212300547 | 3.628780971 | 6.3610914 | 0.0224136 | 0.2732581 |
| ENSMUSG00000005823 | Gpr108 | protein_coding | 0.88348737 | 5.589183926 | 6.3558809 | 0.0224606 | 0.273520975 |
| ENSMUSG00000060012 | Kif13b | protein_coding | -1.30376658 | 3.707502773 | 6.3483237 | 0.022529 | 0.273714571 |
| ENSMUSG00000069670 | Nkain2 | protein_coding | -1.164914737 | 5.140302414 | 6.3493884 | 0.0225391 | 0.273714571 |
| ENSMUSG00000031902 | Nfatc3 | protein_coding | -1.485271687 | 3.247578072 | 6.3456885 | 0.0225529 | 0.273714571 |
| ENSMUSG00000037416 | Dmxl1 | protein_coding | -1.338330634 | 3.915605645 | 6.3404762 | 0.0226002 | 0.273979996 |
| ENSMUSG00000072501 | Phf20l1 | protein_coding | -1.068924065 | 4.447356078 | 6.3339266 | 0.0226599 | 0.274192493 |
| ENSMUSG00000037795 | N4bp2 | protein_coding | -1.983952079 | 2.840029752 | 6.3314894 | 0.0226821 | 0.274192493 |
| ENSMUSG00000020459 | Mtif2 | protein_coding | 0.876663882 | 5.176760627 | 6.3272624 | 0.0227208 | 0.274192493 |
| ENSMUSG00000019943 | Atp2b1 | protein_coding | -0.712321821 | 8.203782665 | 6.326709 | 0.0227258 | 0.274192493 |
| ENSMUSG00000037958 | Nsrp1 | protein_coding | -0.776227441 | 6.112309632 | 6.3245892 | 0.0227453 | 0.274192493 |
| ENSMUSG00000030126 | Tmcc1 | protein_coding | -0.912423885 | 5.601854664 | 6.3193823 | 0.022793 | 0.27446058 |
| ENSMUSG00000031503 | Col4a2 | protein_coding | 2.37899158 | 2.444763669 | 6.3079634 | 0.0228982 | 0.275417953 |
| ENSMUSG00000034757 | Tmub2 | protein_coding | 0.640139305 | 6.644310887 | 6.305194 | 0.0229237 | 0.275417953 |
| ENSMUSG00000032396 | Dis3l | protein_coding | 1.023716102 | 4.497877359 | 6.2974313 | 0.0229956 | 0.275578425 |
| ENSMUSG00000039701 | Usp53 | protein_coding | -1.496462529 | 3.229360718 | 6.293694 | 0.0230303 | 0.275578425 |
| ENSMUSG00000020300 | Cpeb4 | protein_coding | -0.831210586 | 5.245710737 | 6.2931671 | 0.0230352 | 0.275578425 |
| ENSMUSG00000036225 | Kctd1 | protein_coding | -1.153115281 | 5.301943962 | 6.2948456 | 0.0230396 | 0.275578425 |
| ENSMUSG00000028106 | Rprd2 | protein_coding | -0.938795594 | 4.85400875 | 6.2873939 | 0.023089 | 0.275861978 |
| ENSMUSG00000031626 | Sorbs2 | protein_coding | -0.961860422 | 4.848276821 | 6.2836364 | 0.023124 | 0.275971195 |
| ENSMUSG00000030421 | Uri1 | protein_coding | -0.86314709 | 4.919120289 | 6.2807903 | 0.0231506 | 0.275971195 |
| ENSMUSG00000031770 | Herpud1 | protein_coding | 0.670174219 | 6.77839581 | 6.2752889 | 0.0232021 | 0.275971195 |
| ENSMUSG00000090386 | Mir99ahg | processed_transcript | -1.253638717 | 4.000903403 | 6.2737827 | 0.0232162 | 0.275971195 |
| ENSMUSG00000005897 | Nr2c1 | protein_coding | -1.903916452 | 2.943919074 | 6.2726958 | 0.0232264 | 0.275971195 |
| ENSMUSG00000001248 | Gramd1a | protein_coding | 1.218322656 | 4.273829548 | 6.2666383 | 0.0232834 | 0.276342223 |
| ENSMUSG00000108456 | 4732496C06Rik | lincRNA | -1.508965441 | 3.32660542 | 6.2587088 | 0.0233581 | 0.276923866 |
| ENSMUSG00000050394 | Armcx6 | protein_coding | 1.546171161 | 3.473449672 | 6.2501264 | 0.0234394 | 0.277456252 |
| ENSMUSG00000029617 | Ccz1 | protein_coding | -1.124011946 | 4.257759713 | 6.248515 | 0.0234546 | 0.277456252 |
| ENSMUSG00000097537 | 2610020C07Rik | lincRNA | -0.942057889 | 4.705334267 | 6.2451609 | 0.0234865 | 0.277527786 |
| ENSMUSG00000044950 | Pwwp2a | protein_coding | -1.077440136 | 4.588338942 | 6.2380931 | 0.0235538 | 0.277892985 |
| ENSMUSG00000030095 | Tmem43 | protein_coding | 0.769798848 | 5.636489023 | 6.2364888 | 0.0235691 | 0.277892985 |
| ENSMUSG00000040563 | Plppr2 | protein_coding | 1.004463583 | 5.355403126 | 6.2327439 | 0.0236049 | 0.278009888 |
| ENSMUSG00000029616 | Erp29 | protein_coding | 0.56971736 | 7.669565449 | 6.2286483 | 0.023644 | 0.278166787 |
| ENSMUSG00000030512 | Snrpa1 | protein_coding | -0.576231806 | 6.516635699 | 6.2240921 | 0.0236877 | 0.278210143 |
| ENSMUSG00000086822 | 5330413P13Rik | lincRNA | -2.619923279 | 2.127317112 | 6.2207074 | 0.0237202 | 0.278210143 |
| ENSMUSG00000038500 | Prr3 | protein_coding | 0.842280741 | 5.502780232 | 6.2201764 | 0.0237254 | 0.278210143 |
| ENSMUSG00000020658 | Efr3b | protein_coding | 0.730260693 | 5.894749229 | 6.2107223 | 0.0238165 | 0.27877639 |
| ENSMUSG00000041733 | Coq5 | protein_coding | 0.64302005 | 6.582183234 | 6.209281 | 0.0238304 | 0.27877639 |
| ENSMUSG00000059897 | Zfp930 | protein_coding | -1.379279439 | 3.991080406 | 6.2071059 | 0.0238514 | 0.27877639 |
| ENSMUSG00000045624 | Esf1 | protein_coding | -0.910041696 | 5.188180045 | 6.2032448 | 0.0238888 | 0.278910203 |
| ENSMUSG00000025492 | Ifitm3 | protein_coding | -1.169740194 | 4.155131299 | 6.2004741 | 0.0239157 | 0.278921075 |
| ENSMUSG00000027811 | 4930579G24Rik | protein_coding | 2.353424117 | 2.607655136 | 6.1974692 | 0.0239449 | 0.278958843 |
| ENSMUSG00000040648 | Ppip5k2 | protein_coding | -0.972898273 | 4.174445386 | 6.1835153 | 0.0240809 | 0.280240117 |
| ENSMUSG00000003316 | Glg1 | protein_coding | 0.565927465 | 6.681295146 | 6.1796174 | 0.0241191 | 0.280380772 |
| ENSMUSG00000029559 | 2210016L21Rik | protein_coding | 0.607005237 | 8.426601944 | 6.1750832 | 0.0241635 | 0.280594457 |
| ENSMUSG00000046312 | AI464131 | protein_coding | -2.076253602 | 2.567405261 | 6.1715531 | 0.0241982 | 0.280694194 |
| ENSMUSG00000111471 | AC122463.1 | antisense_RNA | -0.846562431 | 5.102127644 | 6.1674834 | 0.0242383 | 0.2808522 |
| ENSMUSG00000023004 | Tuba1b | protein_coding | 0.597337108 | 7.738139621 | 6.1644505 | 0.0242682 | 0.2808522 |
| ENSMUSG00000001964 | Emd | protein_coding | -0.971555768 | 5.350165237 | 6.1608142 | 0.0243041 | 0.2808522 |
| ENSMUSG00000044167 | Foxo1 | protein_coding | -0.851518016 | 5.322789056 | 6.157808 | 0.0243339 | 0.2808522 |
| ENSMUSG00000055202 | Zfp811 | protein_coding | -1.105321009 | 4.399401755 | 6.1555708 | 0.024356 | 0.2808522 |
| ENSMUSG00000041740 | Rnf10 | protein_coding | 0.755279581 | 5.532524674 | 6.1523892 | 0.0243876 | 0.2808522 |
| ENSMUSG00000001025 | S100a6 | protein_coding | -1.053342922 | 5.360026536 | 6.1516724 | 0.0243947 | 0.2808522 |
| ENSMUSG00000054843 | Atrnl1 | protein_coding | -0.945446894 | 4.743823263 | 6.1379859 | 0.024531 | 0.281531072 |
| ENSMUSG00000026649 | Cfap126 | protein_coding | -1.155656411 | 4.686892357 | 6.1348978 | 0.0245619 | 0.281531072 |

| ensembl_gene_id | external_gene_name | gene_biotype | log2 (fold change) | log2 (counts per million) | F-value | p-value | False discovery rate |
|---|---|---|---|---|---|---|---|
| ENSMUSG00000023939 | Mrpl14 | protein_coding | 0.597668416 | 7.523152473 | 6.1330238 | 0.0245806 | 0.281531072 |
| ENSMUSG00000032712 | 2810474O19Rik | protein_coding | -1.119841742 | 3.9110502 | 6.132223 | 0.0245886 | 0.281531072 |
| ENSMUSG00000001380 | Hars | protein_coding | 0.641274305 | 8.238366538 | 6.1303128 | 0.0246078 | 0.281531072 |
| ENSMUSG00000056698 | Elmod3 | protein_coding | -0.795625157 | 5.625959417 | 6.1300184 | 0.0246107 | 0.281531072 |
| ENSMUSG00000064037 | Gpn1 | protein_coding | 0.585805947 | 6.995836777 | 6.1239693 | 0.0246715 | 0.281685589 |
| ENSMUSG00000032534 | Cep63 | protein_coding | 0.974769672 | 4.898120913 | 6.1234571 | 0.0246766 | 0.281685589 |
| ENSMUSG00000046096 | BC030336 | protein_coding | -0.773622989 | 5.853572325 | 6.1198056 | 0.0247134 | 0.281806069 |
| ENSMUSG00000029014 | Dnajc2 | protein_coding | -0.921678628 | 4.619513144 | 6.1140248 | 0.0247717 | 0.281817568 |
| ENSMUSG00000027546 | Atp9a | protein_coding | 0.592449143 | 7.260064044 | 6.1130051 | 0.024782 | 0.281817568 |
| ENSMUSG00000057594 | Arl16 | protein_coding | 0.601573894 | 6.821007018 | 6.1119171 | 0.024793 | 0.281817568 |
| ENSMUSG00000053192 | Mllt11 | protein_coding | 0.846894314 | 6.47669126 | 6.1068274 | 0.0248652 | 0.282339803 |
| ENSMUSG00000002985 | Apoe | protein_coding | -1.510403736 | 7.517200894 | 6.6563891 | 0.024894 | 0.282368059 |
| ENSMUSG00000034243 | Golgb1 | protein_coding | -0.846932122 | 5.276950784 | 6.096601 | 0.0249485 | 0.282582663 |
| ENSMUSG00000039270 | Megf9 | protein_coding | -0.845089504 | 5.528848811 | 6.0949354 | 0.0249655 | 0.282582663 |
| ENSMUSG00000005732 | Ranbp1 | protein_coding | 0.628651348 | 7.170496367 | 6.0882011 | 0.0250342 | 0.283062978 |
| ENSMUSG00000069763 | Tmem100 | protein_coding | -1.082527491 | 5.158979621 | 6.0788827 | 0.0251297 | 0.283844353 |
| ENSMUSG00000044465 | Fam160a2 | protein_coding | -0.969810117 | 4.499321294 | 6.0714759 | 0.0252059 | 0.284406396 |
| ENSMUSG00000038644 | Pold1 | protein_coding | 2.387951835 | 2.400839149 | 6.0665865 | 0.0252564 | 0.284676952 |
| ENSMUSG00000011306 | Sugp1 | protein_coding | 0.814336897 | 6.264808811 | 6.0616153 | 0.0253078 | 0.284957772 |
| ENSMUSG00000025209 | Twnk | protein_coding | -1.622892819 | 3.574506826 | 6.0529476 | 0.0253977 | 0.28567127 |
| ENSMUSG00000009112 | Bcl2l13 | protein_coding | 1.215917719 | 4.002064157 | 6.0489533 | 0.0254393 | 0.285675199 |
| ENSMUSG00000028540 | Dph2 | protein_coding | 1.154100313 | 4.539993622 | 6.0478096 | 0.0254512 | 0.285675199 |
| ENSMUSG00000018042 | Cyb5r3 | protein_coding | 0.698359004 | 7.649355006 | 6.038616 | 0.0255472 | 0.286453988 |
| ENSMUSG00000015214 | Mtmr1 | protein_coding | -1.064721111 | 4.185210537 | 6.0322134 | 0.0256143 | 0.286863632 |
| ENSMUSG00000057229 | Atp5sl | protein_coding | 0.681857908 | 6.897571732 | 6.0290333 | 0.0256478 | 0.286863632 |
| ENSMUSG00000032366 | Tpm1 | protein_coding | -0.540703216 | 7.779207097 | 6.0255189 | 0.0256847 | 0.286863632 |
| ENSMUSG00000029524 | Sirt4 | protein_coding | -1.110351694 | 3.985969658 | 6.0249747 | 0.0256905 | 0.286863632 |
| ENSMUSG00000053841 | Txlna | protein_coding | -1.032021032 | 4.507375285 | 6.0192016 | 0.0257514 | 0.287245364 |
| ENSMUSG00000027374 | Mrps5 | protein_coding | 0.712973833 | 5.855082703 | 6.0164334 | 0.0257806 | 0.287273747 |
| ENSMUSG00000016179 | Camk1g | protein_coding | 0.775199903 | 6.34527043 | 6.0101949 | 0.0258467 | 0.28771194 |
| ENSMUSG00000063015 | Ccni | protein_coding | -0.752678462 | 6.356034087 | 6.0066368 | 0.0258845 | 0.28783448 |
| ENSMUSG00000031837 | Necab2 | protein_coding | -0.749293323 | 6.648110719 | 5.9995954 | 0.0259594 | 0.288186588 |
| ENSMUSG00000023988 | Bysl | protein_coding | 0.894227209 | 5.257597428 | 5.9986275 | 0.0259698 | 0.288186588 |
| ENSMUSG00000030093 | Wnt7a | protein_coding | -1.846391677 | 2.849999121 | 5.9924756 | 0.0260355 | 0.288617937 |
| ENSMUSG00000036502 | Tmem255a | protein_coding | 0.732254099 | 6.151645705 | 5.9837558 | 0.0261289 | 0.289044872 |
| ENSMUSG00000093483 | AA465934 | processed_transcript | -1.419557964 | 3.409305018 | 5.9795655 | 0.026174 | 0.289044872 |
| ENSMUSG00000019087 | Atp6ap1 | protein_coding | 0.549330762 | 9.103616349 | 5.9794423 | 0.0261753 | 0.289044872 |
| ENSMUSG00000052298 | Cdc42se2 | protein_coding | 0.754065491 | 7.224808403 | 5.9796668 | 0.026194 | 0.289044872 |
| ENSMUSG00000054612 | Mgmt | protein_coding | 1.503671634 | 3.160805417 | 5.9763721 | 0.0262084 | 0.289044872 |
| ENSMUSG00000020580 | Rock2 | protein_coding | -0.64494355 | 6.259630581 | 5.9714481 | 0.0262615 | 0.289334138 |
| ENSMUSG00000097675 | 1700101I11Rik | processed_transcript | 1.679328314 | 2.873337155 | 5.9646199 | 0.0263354 | 0.289459039 |
| ENSMUSG00000041229 | Phf8 | protein_coding | -1.005809711 | 4.269503119 | 5.9626685 | 0.0263566 | 0.289459039 |
| ENSMUSG00000032839 | Trpc1 | protein_coding | -1.466891894 | 3.585666934 | 5.9544515 | 0.0264459 | 0.289459039 |
| ENSMUSG00000048100 | Taf13 | protein_coding | -0.901850226 | 5.28099885 | 5.9531484 | 0.0264601 | 0.289459039 |
| ENSMUSG00000003410 | Elavl3 | protein_coding | -0.57858549 | 6.897769516 | 5.9524383 | 0.0264678 | 0.289459039 |
| ENSMUSG00000019906 | Lin7a | protein_coding | -0.755510537 | 6.307667229 | 5.9487171 | 0.0265084 | 0.289459039 |
| ENSMUSG00000034480 | Diaph2 | protein_coding | -1.233991671 | 4.128041007 | 5.9469948 | 0.0265272 | 0.289459039 |
| ENSMUSG00000058135 | Gstm1 | protein_coding | -0.664665704 | 8.942497461 | 5.9455071 | 0.0265435 | 0.289459039 |
| ENSMUSG00000054752 | Fsd1l | protein_coding | -1.19080576 | 3.702558382 | 5.9440598 | 0.0265593 | 0.289459039 |
| ENSMUSG00000055717 | Slain1 | protein_coding | -0.803703149 | 5.534441336 | 5.9424396 | 0.0265771 | 0.289459039 |
| ENSMUSG00000055421 | Pcdh9 | protein_coding | -0.63277117 | 6.98252191 | 5.9420354 | 0.0265815 | 0.289459039 |
| ENSMUSG00000036617 | Etl4 | protein_coding | -1.171136051 | 4.367446043 | 5.9387294 | 0.0266177 | 0.289459039 |
| ENSMUSG00000030209 | Grin2b | protein_coding | -0.738209286 | 6.206190231 | 5.9356146 | 0.0266519 | 0.289459039 |
| ENSMUSG00000030655 | Smg1 | protein_coding | -1.226591147 | 3.940697427 | 5.9350143 | 0.0266585 | 0.289459039 |
| ENSMUSG00000028847 | Trappc3 | protein_coding | 0.522878418 | 7.886045481 | 5.9312453 | 0.0267 | 0.289459039 |
| ENSMUSG00000028622 | Mrpl37 | protein_coding | 0.688269203 | 6.302739605 | 5.9309228 | 0.0267036 | 0.289459039 |
| ENSMUSG00000022587 | Ly6e | protein_coding | 0.653932508 | 7.940055716 | 5.9146464 | 0.0268835 | 0.291116069 |
| ENSMUSG00000044768 | D1Ertd622e | protein_coding | -1.008498239 | 4.60672709 | 5.9094447 | 0.0269413 | 0.291448449 |
| ENSMUSG00000078671 | Chd2 | protein_coding | -0.901187933 | 4.680624463 | 5.8980298 | 0.0270686 | 0.292531687 |
| ENSMUSG00000039195 | 1110008P14Rik | protein_coding | 1.149659835 | 4.218693784 | 5.8949982 | 0.0271026 | 0.292604354 |
| ENSMUSG00000047935 | Gm5607 | sense_overlapping | -0.884969162 | 4.563505048 | 5.880804 | 0.0272621 | 0.293789833 |
| ENSMUSG00000022617 | Chkb | protein_coding | 1.045047372 | 3.886913244 | 5.8803685 | 0.027267 | 0.293789833 |
| ENSMUSG00000026260 | Ndufa10 | protein_coding | 0.859494811 | 8.476402603 | 5.9010072 | 0.0274708 | 0.294804485 |
| ENSMUSG00000013150 | Gfod2 | protein_coding | 0.577403132 | 6.447896045 | 5.8616075 | 0.0274796 | 0.294804485 |
| ENSMUSG00000002981 | Clptm1 | protein_coding | 0.706518269 | 5.646695765 | 5.8576407 | 0.0275248 | 0.294804485 |
| ENSMUSG00000073565 | Prr16 | protein_coding | -2.463158466 | 2.509511292 | 5.8555368 | 0.0275488 | 0.294804485 |
| ENSMUSG00000071477 | Zfp777 | protein_coding | 1.311660404 | 3.500699086 | 5.8553945 | 0.0275504 | 0.294804485 |
| ENSMUSG00000067430 | Zfp763 | protein_coding | -1.100402603 | 4.197765692 | 5.8552255 | 0.0275523 | 0.294804485 |
| ENSMUSG00000024785 | Rcl1 | protein_coding | 1.213933556 | 4.641329716 | 5.8547323 | 0.027558 | 0.294804485 |
| ENSMUSG00000027200 | Sema6d | protein_coding | -0.894204928 | 4.817091646 | 5.8504621 | 0.0276068 | 0.294804485 |
| ENSMUSG00000041417 | Pik3r1 | protein_coding | -0.852344817 | 5.653165169 | 5.8503601 | 0.0276079 | 0.294804485 |
| ENSMUSG00000003299 | Mrpl4 | protein_coding | 0.579042149 | 7.4879385 | 5.8436824 | 0.0276845 | 0.295161437 |
| ENSMUSG00000032735 | Ablim3 | protein_coding | 0.961927933 | 4.937772889 | 5.8426548 | 0.0276963 | 0.295161437 |
| ENSMUSG00000030127 | Cops7a | protein_coding | 0.553560075 | 8.366976052 | 5.8363057 | 0.0277693 | 0.295646649 |
| ENSMUSG00000006373 | Pgrmc1 | protein_coding | 0.573379441 | 9.279987835 | 5.8252766 | 0.0278967 | 0.295712742 |
| ENSMUSG00000020719 | Ddx5 | protein_coding | -0.581882322 | 8.230626734 | 5.8205392 | 0.0279516 | 0.295712742 |
| ENSMUSG00000047996 | Prrg1 | protein_coding | -1.271256752 | 3.350186742 | 5.8203392 | 0.0279539 | 0.295712742 |
| ENSMUSG00000021395 | Spin1 | protein_coding | -0.604031944 | 7.491724962 | 5.8197537 | 0.0279607 | 0.295712742 |

| ensembl_gene_id | external_gene_name | gene_biotype | log2 (fold change) | log2 (counts per million) | F-value | p-value | False discovery rate |
|---|---|---|---|---|---|---|---|
| ENSMUSG00000025813 | Homer2 | protein_coding | -1.065324881 | 4.084011976 | 5.8196596 | 0.0279618 | 0.295712742 |
| ENSMUSG00000033721 | Vav3 | protein_coding | -1.474593148 | 3.443028302 | 5.8160449 | 0.0280038 | 0.295712742 |
| ENSMUSG00000060240 | Cend1 | protein_coding | 0.71287792 | 9.860547497 | 5.8159605 | 0.0280048 | 0.295712742 |
| ENSMUSG00000074505 | Fat3 | protein_coding | -1.230750942 | 4.060654349 | 5.8144729 | 0.0280221 | 0.295712742 |
| ENSMUSG00000040363 | Bcor | protein_coding | 0.843597872 | 4.932756 | 5.8143946 | 0.028023 | 0.295712742 |
| ENSMUSG00000089945 | Pakap | protein_coding | -1.560383815 | 3.012438572 | 5.8071874 | 0.028107 | 0.29606055 |
| ENSMUSG00000030521 | Mphosph10 | protein_coding | -0.882136769 | 4.697835945 | 5.8068457 | 0.028111 | 0.29606055 |
| ENSMUSG00000020362 | Cnot6 | protein_coding | -0.909880606 | 5.369454094 | 5.8022098 | 0.0281652 | 0.296341294 |
| ENSMUSG00000028412 | Slc44a1 | protein_coding | -0.856015666 | 5.617019208 | 5.7986448 | 0.028207 | 0.296380657 |
| ENSMUSG00000044080 | S100a1 | protein_coding | -1.041077424 | 8.199146881 | 5.9573531 | 0.0282241 | 0.296380657 |
| ENSMUSG00000041014 | Nrg3 | protein_coding | -0.988839606 | 4.681865863 | 5.7890281 | 0.0283201 | 0.296695135 |
| ENSMUSG00000032745 | Gpbp1 | protein_coding | -0.722782758 | 6.97775992 | 5.787247 | 0.028341 | 0.296695135 |
| ENSMUSG00000036095 | Dgkb | protein_coding | -0.710802735 | 7.206414384 | 5.7854239 | 0.0283626 | 0.296695135 |
| ENSMUSG00000025264 | Tsr2 | protein_coding | 0.608845412 | 7.826594417 | 5.7849418 | 0.0283682 | 0.296695135 |
| ENSMUSG00000050965 | Prkca | protein_coding | -0.761567675 | 5.234489138 | 5.7813586 | 0.0284106 | 0.296695135 |
| ENSMUSG00000061032 | Rrp1 | protein_coding | 0.667324318 | 8.565164267 | 5.7805955 | 0.0284196 | 0.296695135 |
| ENSMUSG00000034210 | Efcab14 | protein_coding | -0.675666918 | 5.928916924 | 5.7781855 | 0.0284481 | 0.296704865 |
| ENSMUSG00000039031 | Arhgap18 | protein_coding | -1.39990095 | 3.136624267 | 5.7711792 | 0.0285312 | 0.29707452 |
| ENSMUSG00000030342 | Cd9 | protein_coding | -0.762510162 | 6.841465186 | 5.7723864 | 0.0285388 | 0.29707452 |
| ENSMUSG00000029053 | Prkcz | protein_coding | 0.58222991 | 7.089822294 | 5.7623952 | 0.0286358 | 0.297550106 |
| ENSMUSG00000033943 | Mga | protein_coding | -0.940864675 | 5.35196869 | 5.7620587 | 0.0286399 | 0.297550106 |
| ENSMUSG00000055707 | Klhl26 | protein_coding | 1.190662259 | 5.529094558 | 5.8233721 | 0.0287129 | 0.298021463 |
| ENSMUSG00000049119 | Fam110b | protein_coding | 0.619707883 | 6.340096032 | 5.7498046 | 0.0287865 | 0.298176057 |
| ENSMUSG00000063659 | Zbtb18 | protein_coding | -1.098269932 | 3.932208791 | 5.7496422 | 0.0287885 | 0.298176057 |
| ENSMUSG00000051256 | Jagn1 | protein_coding | 0.612665124 | 6.482956711 | 5.7472687 | 0.028817 | 0.298176057 |
| ENSMUSG00000035885 | Cox8a | protein_coding | 1.063274633 | 9.631013762 | 5.8916916 | 0.0288501 | 0.298176057 |
| ENSMUSG00000021027 | Ralgapa1 | protein_coding | -0.81104608 | 5.251444168 | 5.735551 | 0.0289582 | 0.298176057 |
| ENSMUSG00000004535 | Tax1bp1 | protein_coding | -0.510204272 | 7.530073695 | 5.7344598 | 0.0289714 | 0.298176057 |
| ENSMUSG00000001773 | Folh1 | protein_coding | -1.820074354 | 2.654719932 | 5.73431 | 0.0289732 | 0.298176057 |
| ENSMUSG00000026034 | Clk1 | protein_coding | -1.065334682 | 5.304603204 | 5.734428 | 0.0289939 | 0.298176057 |
| ENSMUSG00000004071 | Cdip1 | protein_coding | 0.697014633 | 10.04724437 | 5.7322952 | 0.0289976 | 0.298176057 |
| ENSMUSG00000041685 | Fcho2 | protein_coding | -1.112312701 | 3.901869086 | 5.7298906 | 0.0290267 | 0.298176057 |
| ENSMUSG00000005881 | Ergic3 | protein_coding | 0.503280312 | 8.70639423 | 5.7291618 | 0.0290356 | 0.298176057 |
| ENSMUSG00000067377 | Tspan6 | protein_coding | 0.627745664 | 6.552946552 | 5.7271001 | 0.0290606 | 0.298176057 |
| ENSMUSG00000025508 | Rplp2 | protein_coding | 0.647915175 | 8.84790649 | 5.7239588 | 0.0290987 | 0.298183511 |
| ENSMUSG00000052748 | Swt1 | protein_coding | -1.106230842 | 4.030191547 | 5.7206474 | 0.029139 | 0.298183511 |
| ENSMUSG00000050192 | Eif5a2 | protein_coding | -0.753293979 | 5.390785133 | 5.7198951 | 0.0291482 | 0.298183511 |
| ENSMUSG00000021057 | Akap5 | protein_coding | -0.65642889 | 6.766825998 | 5.7153972 | 0.029203 | 0.298183511 |
| ENSMUSG00000007739 | Cct4 | protein_coding | 0.584027297 | 7.595742519 | 5.7149296 | 0.0292087 | 0.298183511 |
| ENSMUSG00000031960 | Aars | protein_coding | 0.765663414 | 7.021415259 | 5.7151894 | 0.0292277 | 0.298183511 |
| ENSMUSG00000014077 | Chp1 | protein_coding | 0.655207697 | 7.876562314 | 5.7032634 | 0.0293515 | 0.299147347 |
| ENSMUSG00000060961 | Slc4a4 | protein_coding | -1.02736106 | 5.376427232 | 5.702931 | 0.0293778 | 0.299147347 |
| ENSMUSG00000033208 | S100b | protein_coding | -0.964914006 | 6.003589543 | 5.6943597 | 0.0295419 | 0.30043296 |
| ENSMUSG00000031622 | Sin3b | protein_coding | 0.546829378 | 6.505434759 | 5.6861973 | 0.0295619 | 0.30043296 |
| ENSMUSG00000018501 | Ncor1 | protein_coding | -0.679672809 | 5.848514274 | 5.6841 | 0.0295879 | 0.30043296 |
| ENSMUSG00000036478 | Btg1 | protein_coding | -0.970915479 | 5.003005241 | 5.6815753 | 0.0296192 | 0.300467055 |
| ENSMUSG00000054387 | Mdm4 | protein_coding | -0.72868202 | 5.099620466 | 5.6755495 | 0.029694 | 0.300820974 |
| ENSMUSG00000074872 | Ctxn2 | protein_coding | 0.643691553 | 8.340177171 | 5.6709941 | 0.0297508 | 0.300820974 |
| ENSMUSG00000032293 | Ireb2 | protein_coding | -0.95328354 | 4.662883172 | 5.6689512 | 0.0297763 | 0.300820974 |
| ENSMUSG00000036934 | 4921524J17Rik | protein_coding | -0.792247497 | 5.390113529 | 5.6685764 | 0.0297809 | 0.300820974 |
| ENSMUSG00000002968 | Med25 | protein_coding | 1.058429921 | 4.253980917 | 5.6675345 | 0.0297939 | 0.300820974 |
| ENSMUSG00000053730 | Tmem39b | protein_coding | 1.641422094 | 3.168198016 | 5.6644694 | 0.0298323 | 0.300925202 |
| ENSMUSG00000032936 | Camkv | protein_coding | 0.618223741 | 8.008859896 | 5.6606976 | 0.0298795 | 0.301119041 |
| ENSMUSG00000074212 | Dnajb14 | protein_coding | -0.708430083 | 5.796248168 | 5.6552517 | 0.0299478 | 0.301245066 |
| ENSMUSG00000024841 | Eif1ad | protein_coding | 0.517426115 | 7.031500112 | 5.6534994 | 0.0299698 | 0.301245066 |
| ENSMUSG00000028059 | Arhgef2 | protein_coding | -1.010286458 | 4.502508929 | 5.6509427 | 0.030002 | 0.301245066 |
| ENSMUSG00000037787 | Apopt1 | protein_coding | 0.537712421 | 6.798427294 | 5.6507803 | 0.030004 | 0.301245066 |
| ENSMUSG00000068859 | Sp9 | protein_coding | -1.096421976 | 3.855387841 | 5.645241 | 0.0300739 | 0.301447484 |
| ENSMUSG00000042312 | S100a13 | protein_coding | -1.063963161 | 6.416420582 | 5.7540899 | 0.0300803 | 0.301447484 |
| ENSMUSG00000045767 | B230219D22Rik | protein_coding | -0.709803103 | 5.908005349 | 5.6390392 | 0.0301523 | 0.30188785 |
| ENSMUSG00000030357 | Fkbp4 | protein_coding | 0.561945325 | 7.238306529 | 5.6272108 | 0.0303025 | 0.301974025 |
| ENSMUSG00000024947 | Men1 | protein_coding | 0.839512864 | 5.013387401 | 5.6264668 | 0.030312 | 0.301974025 |
| ENSMUSG00000031202 | Rab39b | protein_coding | -0.948303141 | 4.459768071 | 5.6242861 | 0.0303398 | 0.301974025 |
| ENSMUSG00000031617 | Tmem184c | protein_coding | -1.04344959 | 4.827262625 | 5.6239473 | 0.0303441 | 0.301974025 |
| ENSMUSG00000035521 | Gnptg | protein_coding | 0.550152736 | 7.368568313 | 5.6239076 | 0.0303446 | 0.301974025 |
| ENSMUSG00000003970 | Rpl8 | protein_coding | 0.600747065 | 10.42760603 | 5.6222384 | 0.0303659 | 0.301974025 |
| ENSMUSG00000026202 | Tuba4a | protein_coding | 0.767262483 | 9.099061636 | 5.6210791 | 0.0304032 | 0.301974025 |
| ENSMUSG00000025468 | Caly | protein_coding | 0.57654537 | 9.560052941 | 5.6192909 | 0.0304035 | 0.301974025 |
| ENSMUSG00000067722 | BC003965 | protein_coding | 0.586532048 | 6.375988644 | 5.6185009 | 0.0304136 | 0.301974025 |
| ENSMUSG00000017664 | Slc35c2 | protein_coding | 0.654324441 | 6.684281518 | 5.6132149 | 0.0304813 | 0.302016312 |
| ENSMUSG00000060371 | Caln1 | protein_coding | -0.824755273 | 6.668395795 | 5.6139865 | 0.030494 | 0.302016312 |
| ENSMUSG00000021939 | Ctsb | protein_coding | 0.567378495 | 10.06288877 | 5.6115906 | 0.0305022 | 0.302016312 |
| ENSMUSG00000020422 | Tns3 | protein_coding | -1.403302882 | 3.221768076 | 5.6025701 | 0.0306182 | 0.302885958 |
| ENSMUSG00000005779 | Psmb4 | protein_coding | 0.58529855 | 7.730041608 | 5.5948694 | 0.0307176 | 0.303362838 |
| ENSMUSG00000018750 | Zbtb4 | protein_coding | -0.581722534 | 6.130943418 | 5.5921186 | 0.0307532 | 0.303362838 |
| ENSMUSG00000031246 | Sh3bgrl | protein_coding | -0.726128427 | 7.149897123 | 5.5913235 | 0.0307861 | 0.303362838 |
| ENSMUSG00000052584 | Serp2 | protein_coding | 0.584026248 | 6.425509516 | 5.5890741 | 0.0307927 | 0.303362838 |
| ENSMUSG00000019303 | Psmc3ip | protein_coding | -1.138870751 | 3.827689081 | 5.5874637 | 0.0308136 | 0.303362838 |

| ensembl_gene_id | external_gene_name | gene_biotype | log2 (fold change) | log2 (counts per million) | F-value | p-value | False discovery rate |
|---|---|---|---|---|---|---|---|
| ENSMUSG00000024007 | Ppil1 | protein_coding | 0.601889977 | 6.774868429 | 5.5857633 | 0.0308356 | 0.303362838 |
| ENSMUSG00000029466 | Anapc7 | protein_coding | 0.772305099 | 5.210385149 | 5.579052 | 0.030923 | 0.303944025 |
| ENSMUSG00000032480 | Dhx30 | protein_coding | 0.729655494 | 6.567190402 | 5.5730822 | 0.0310009 | 0.304431717 |
| ENSMUSG00000078789 | Dph1 | protein_coding | 1.444129263 | 3.712056252 | 5.5644857 | 0.0311135 | 0.305258842 |
| ENSMUSG00000039578 | Ccser1 | protein_coding | -0.983374027 | 4.043679451 | 5.5553232 | 0.0312341 | 0.306162219 |
| ENSMUSG00000068882 | Ssb | protein_coding | -0.65063252 | 7.356923846 | 5.5513471 | 0.0312866 | 0.306397335 |
| ENSMUSG00000047250 | Ptgs1 | protein_coding | 1.154248223 | 3.772499142 | 5.5483688 | 0.0313259 | 0.306402865 |
| ENSMUSG00000024064 | Galnt14 | protein_coding | 1.044348659 | 3.847379419 | 5.5469953 | 0.0313441 | 0.306402865 |
| ENSMUSG00000031133 | Arhgef6 | protein_coding | -1.854842947 | 2.854956478 | 5.5430725 | 0.0313961 | 0.306632292 |
| ENSMUSG00000057649 | Brd9 | protein_coding | 0.750873857 | 5.955178553 | 5.5368411 | 0.0314789 | 0.307074109 |
| ENSMUSG00000048022 | Tmem229a | protein_coding | -1.129807033 | 4.868359132 | 5.5354975 | 0.0315196 | 0.307074109 |
| ENSMUSG00000057234 | Mettl15 | protein_coding | -1.192086882 | 3.394642558 | 5.5332273 | 0.031527 | 0.307074109 |
| ENSMUSG00000039515 | Ptpa | protein_coding | 0.638457766 | 8.410019744 | 5.5292716 | 0.0315798 | 0.307309772 |
| ENSMUSG00000037855 | Zfp365 | protein_coding | -0.539343195 | 7.027593811 | 5.5242011 | 0.0316476 | 0.307369536 |
| ENSMUSG00000046240 | Hepacam | protein_coding | -1.229441967 | 4.693450718 | 5.525008 | 0.0316596 | 0.307369536 |
| ENSMUSG00000072214 | 5-Sep | protein_coding | 0.628764944 | 7.012611073 | 5.5175046 | 0.0317374 | 0.307369536 |
| ENSMUSG00000025151 | Maged1 | protein_coding | 0.816867487 | 10.0312578 | 5.5157811 | 0.0317834 | 0.307369536 |
| ENSMUSG00000029309 | Sparcl1 | protein_coding | -0.660261741 | 9.275472997 | 5.5132243 | 0.0317949 | 0.307369536 |
| ENSMUSG00000063229 | Ldha | protein_coding | 0.542659163 | 8.55512679 | 5.5131731 | 0.0317956 | 0.307369536 |
| ENSMUSG00000033713 | Foxn3 | protein_coding | -0.784301259 | 6.87413152 | 5.511719 | 0.0318381 | 0.307369536 |
| ENSMUSG00000109536 | 9330162G02Rik | TEC | -1.219872406 | 3.428943043 | 5.5098368 | 0.0318405 | 0.307369536 |
| ENSMUSG00000039768 | Dnajc11 | protein_coding | 0.896734336 | 6.270257955 | 5.5158281 | 0.0318432 | 0.307369536 |
| ENSMUSG00000030782 | Tgfb1i1 | protein_coding | 1.066867558 | 4.468546827 | 5.5052528 | 0.0319024 | 0.307665109 |
| ENSMUSG00000051579 | Tceal8 | protein_coding | -0.58435257 | 7.465920246 | 5.4970967 | 0.0320128 | 0.308453336 |
| ENSMUSG00000036275 | 9530068E07Rik | protein_coding | 0.690629047 | 6.025936184 | 5.490019 | 0.032109 | 0.309103056 |
| ENSMUSG00000079562 | Maea | protein_coding | 0.652390706 | 7.573083674 | 5.4852416 | 0.0321741 | 0.30938355 |
| ENSMUSG00000027582 | Zgpat | protein_coding | 2.337268012 | 2.583425405 | 5.4821015 | 0.032217 | 0.30938355 |
| ENSMUSG00000063253 | Scoc | protein_coding | -0.56948222 | 7.516493739 | 5.4815275 | 0.0322248 | 0.30938355 |
| ENSMUSG00000025239 | Limd1 | protein_coding | -1.084066551 | 4.056360226 | 5.4783963 | 0.0322677 | 0.30938355 |
| ENSMUSG00000026527 | Rgs7 | protein_coding | -0.789341902 | 5.555931476 | 5.4773493 | 0.032282 | 0.30938355 |
| ENSMUSG00000051111 | Sv2c | protein_coding | 0.954307589 | 4.970558256 | 5.4735634 | 0.0323339 | 0.309605023 |
| ENSMUSG00000032925 | Itgbl1 | protein_coding | -1.170398225 | 3.819230925 | 5.4712166 | 0.0323661 | 0.309637834 |
| ENSMUSG00000050705 | 2310061I04Rik | protein_coding | 0.71090937 | 5.638533226 | 5.4669143 | 0.0324253 | 0.309700855 |
| ENSMUSG00000028745 | Capzb | protein_coding | 0.55356142 | 8.992086046 | 5.4665493 | 0.0324303 | 0.309700855 |
| ENSMUSG00000002031 | Ift46 | protein_coding | 0.671417261 | 7.052698323 | 5.4586778 | 0.0325389 | 0.310289012 |
| ENSMUSG00000024194 | Cuta | protein_coding | 0.492832823 | 7.473278638 | 5.4535436 | 0.03261 | 0.310289012 |
| ENSMUSG00000007603 | Dus3l | protein_coding | 0.692330357 | 6.250032188 | 5.4529667 | 0.032618 | 0.310289012 |
| ENSMUSG00000061244 | Exoc5 | protein_coding | -0.865430075 | 4.714899718 | 5.4529543 | 0.0326181 | 0.310289012 |
| ENSMUSG00000021814 | Anxa7 | protein_coding | 0.690781719 | 6.427187044 | 5.4516522 | 0.0326362 | 0.310289012 |
| ENSMUSG00000031833 | Mast3 | protein_coding | -0.859557167 | 5.532815314 | 5.4474148 | 0.032695 | 0.310308792 |
| ENSMUSG00000048720 | Tbc1d12 | protein_coding | -1.343368779 | 3.270310205 | 5.4469548 | 0.0327014 | 0.310308792 |
| ENSMUSG00000064254 | Ethe1 | protein_coding | 1.018909002 | 4.879331073 | 5.4405874 | 0.0327901 | 0.310308792 |
| ENSMUSG00000020946 | Gosr2 | protein_coding | 0.664127889 | 6.890727503 | 5.4395221 | 0.0328049 | 0.310308792 |
| ENSMUSG00000021188 | Trip11 | protein_coding | -0.956204075 | 4.587727055 | 5.4389468 | 0.032813 | 0.310308792 |
| ENSMUSG00000030839 | Sergef | protein_coding | 0.607603648 | 6.531287465 | 5.4369693 | 0.0328406 | 0.310308792 |
| ENSMUSG00000037486 | Asxl2 | protein_coding | -1.018964333 | 4.385081825 | 5.4357366 | 0.0328578 | 0.310308792 |
| ENSMUSG00000001750 | Tcirg1 | protein_coding | 1.697061549 | 2.990143412 | 5.4348342 | 0.0328704 | 0.310308792 |
| ENSMUSG00000026701 | Prdx6 | protein_coding | -0.586470895 | 9.748615925 | 5.4312011 | 0.0329213 | 0.310308792 |
| ENSMUSG00000018661 | Cog1 | protein_coding | 1.233949215 | 3.643888327 | 5.4285543 | 0.0329584 | 0.310308792 |
| ENSMUSG00000017764 | Zswim1 | protein_coding | 0.917957464 | 5.908942062 | 5.4279308 | 0.0329904 | 0.310308792 |
| ENSMUSG00000078427 | Sarnp | protein_coding | -0.659568294 | 6.551563415 | 5.4243383 | 0.0330176 | 0.310308792 |
| ENSMUSG00000029334 | Prkg2 | protein_coding | -1.658299752 | 3.431655454 | 5.4224999 | 0.0330434 | 0.310308792 |
| ENSMUSG00000104939 | Gm33651 | lincRNA | -1.578994016 | 3.545577896 | 5.4239749 | 0.0330459 | 0.310308792 |
| ENSMUSG00000008958 | Vps72 | protein_coding | 0.590781646 | 6.102991075 | 5.4190432 | 0.0330921 | 0.310308792 |
| ENSMUSG00000048895 | Cdk5r1 | protein_coding | -0.576438461 | 6.889457101 | 5.4184831 | 0.0331 | 0.310308792 |
| ENSMUSG00000059401 | Mamld1 | protein_coding | 1.021471609 | 4.822807118 | 5.415034 | 0.0331487 | 0.3104943 |
| ENSMUSG00000069049 | Eif2s3y | protein_coding | -0.955175234 | 5.038632738 | 5.410461 | 0.0332133 | 0.310829082 |
| ENSMUSG00000020248 | Nfyb | protein_coding | -0.780053473 | 5.796719178 | 5.4010583 | 0.0333467 | 0.311472423 |
| ENSMUSG00000022865 | Cxadr | protein_coding | -1.517676234 | 2.899589865 | 5.3985619 | 0.0333822 | 0.311472423 |
| ENSMUSG00000027883 | Gpsm2 | protein_coding | -1.368618765 | 2.962965676 | 5.3980305 | 0.0333898 | 0.311472423 |
| ENSMUSG00000015839 | Nfe2l2 | protein_coding | -1.184080527 | 4.716166321 | 5.3990992 | 0.0333979 | 0.311472423 |
| ENSMUSG00000033918 | Parl | protein_coding | 0.722289749 | 6.333580154 | 5.3942584 | 0.0334435 | 0.311554929 |
| ENSMUSG00000025234 | Arih1 | protein_coding | -0.893192069 | 5.035524536 | 5.3927744 | 0.0334647 | 0.311554929 |
| ENSMUSG00000030619 | Eed | protein_coding | -1.127040071 | 4.505695003 | 5.3903918 | 0.0334987 | 0.311601999 |
| ENSMUSG00000041444 | Arhgap32 | protein_coding | 0.776132249 | 4.925505771 | 5.3876865 | 0.0335374 | 0.31169229 |
| ENSMUSG00000031885 | Cbfb | protein_coding | -0.958602235 | 4.202384283 | 5.3776726 | 0.0336811 | 0.312644393 |
| ENSMUSG00000010025 | Aldh3a2 | protein_coding | 1.059765446 | 5.385840034 | 5.3781257 | 0.033698 | 0.312644393 |
| ENSMUSG00000096370 | Gm21992 | protein_coding | -1.557524944 | 3.226450449 | 5.373384 | 0.0337428 | 0.312649899 |
| ENSMUSG00000030870 | Ubfd1 | protein_coding | 0.588572712 | 6.262234854 | 5.3724183 | 0.0337568 | 0.312649899 |
| ENSMUSG00000005142 | Man2b1 | protein_coding | 0.921422879 | 4.325568572 | 5.3649334 | 0.0338649 | 0.313359163 |
| ENSMUSG00000029614 | Rpl6 | protein_coding | 0.532500197 | 6.90047633 | 5.3630872 | 0.0338916 | 0.313359163 |
| ENSMUSG00000043881 | Kbtbd7 | protein_coding | -0.969015933 | 4.593530467 | 5.3562556 | 0.0339908 | 0.313922397 |
| ENSMUSG00000030994 | D7Ertd443e | protein_coding | -1.058748861 | 4.277472516 | 5.3515816 | 0.0340588 | 0.313922397 |
| ENSMUSG00000068040 | Tm9sf4 | protein_coding | 0.641270563 | 6.085282906 | 5.3507772 | 0.0340705 | 0.313922397 |
| ENSMUSG00000020471 | Pold2 | protein_coding | 0.846320575 | 5.39565547 | 5.347998 | 0.034111 | 0.313922397 |
| ENSMUSG00000058454 | Dhcr7 | protein_coding | 0.691244387 | 5.621171495 | 5.3463283 | 0.0341354 | 0.313922397 |
| ENSMUSG00000022742 | Cpox | protein_coding | -0.877806428 | 5.632461472 | 5.3457067 | 0.0341445 | 0.313922397 |
| ENSMUSG00000032038 | St3gal4 | protein_coding | 0.901836689 | 4.551804892 | 5.3448571 | 0.0341569 | 0.313922397 |

| ensembl_gene_id | external_gene_name | gene_biotype | log2 (fold change) | log2 (counts per million) | F-value | p-value | False discovery rate |
|---|---|---|---|---|---|---|---|
| ENSMUSG00000019874 | Fabp7 | protein_coding | -1.176956614 | 6.935996241 | 5.5401748 | 0.0342265 | 0.314293033 |
| ENSMUSG00000014767 | Tbp | protein_coding | 0.772289517 | 5.374297906 | 5.3340777 | 0.0343149 | 0.314836162 |
| ENSMUSG00000078117 | Gm16485 | protein_coding | 2.278594887 | 2.420546414 | 5.3205822 | 0.0345139 | 0.316391815 |
| ENSMUSG00000028174 | Rpe65 | protein_coding | -4.046132764 | 2.05384736 | 5.3493705 | 0.0345691 | 0.3166286 |
| ENSMUSG00000108778 | Gm20083 | lincRNA | -2.106705135 | 2.465986827 | 5.3125447 | 0.034633 | 0.31694355 |
| ENSMUSG00000034349 | Smc4 | protein_coding | -2.121773816 | 2.374126917 | 5.310341 | 0.0346657 | 0.316973471 |
| ENSMUSG00000024256 | Adcyap1 | protein_coding | -1.853901344 | 3.763489644 | 5.3688611 | 0.0347212 | 0.317210902 |
| ENSMUSG00000108461 | AV356131 | bidirectional_promote | 2.044779135 | 2.681893611 | 5.2987648 | 0.0348384 | 0.317815699 |
| ENSMUSG00000039202 | Abhd2 | protein_coding | -1.00466075 | 4.758151013 | 5.2982187 | 0.0348465 | 0.317815699 |
| ENSMUSG00000097875 | Gm26704 | lincRNA | -1.243650837 | 3.844733875 | 5.2961115 | 0.0348781 | 0.31783374 |
| ENSMUSG00000067578 | Cbln4 | protein_coding | 1.25901056 | 4.364327424 | 5.2948846 | 0.0349202 | 0.317948295 |
| ENSMUSG00000066026 | Dhrs3 | protein_coding | -0.809870239 | 5.994403766 | 5.2865234 | 0.035022 | 0.318605119 |
| ENSMUSG00000107872 | Gm44511 | protein_coding | -1.520979259 | 3.305188604 | 5.2834984 | 0.0350675 | 0.318749769 |
| ENSMUSG00000046862 | Pramef8 | protein_coding | 1.123701509 | 4.561603272 | 5.2805639 | 0.0351118 | 0.318868685 |
| ENSMUSG00000048142 | Nat8l | protein_coding | 0.574213054 | 6.965309063 | 5.2771517 | 0.0351633 | 0.318868685 |
| ENSMUSG00000019841 | Rev3l | protein_coding | -1.166241878 | 4.223288506 | 5.2767371 | 0.0351696 | 0.318868685 |
| ENSMUSG00000020572 | Nampt | protein_coding | -0.741599406 | 4.97681015 | 5.274498 | 0.0352034 | 0.31890686 |
| ENSMUSG00000027351 | Spred1 | protein_coding | -0.604962066 | 6.353192015 | 5.2688645 | 0.0352888 | 0.319411217 |
| ENSMUSG00000040455 | Usp45 | protein_coding | -0.932885542 | 4.613921663 | 5.263837 | 0.0353652 | 0.319833501 |
| ENSMUSG00000017776 | Crk | protein_coding | -0.721120012 | 6.282211737 | 5.2589803 | 0.0354392 | 0.319933472 |
| ENSMUSG00000006299 | Aamp | protein_coding | 0.685120823 | 9.100446211 | 5.2584146 | 0.0354478 | 0.319933472 |
| ENSMUSG00000059183 | Mtfmt | protein_coding | 1.033606976 | 4.232497729 | 5.2569905 | 0.0354695 | 0.319933472 |
| ENSMUSG00000024750 | Zfand5 | protein_coding | -0.640659363 | 6.018019954 | 5.2553063 | 0.0354953 | 0.319933472 |
| ENSMUSG00000061702 | Tmem91 | protein_coding | 0.564465197 | 7.338949389 | 5.2471409 | 0.0356203 | 0.320738318 |
| ENSMUSG00000021619 | Atg10 | protein_coding | -0.605126543 | 5.97382329 | 5.2455814 | 0.0356442 | 0.320738318 |
| ENSMUSG00000034818 | Celf5 | protein_coding | -0.606102896 | 6.515981532 | 5.2435569 | 0.0356753 | 0.320749731 |
| ENSMUSG00000022323 | Rida | protein_coding | -0.726655448 | 7.338296693 | 5.243011 | 0.0357077 | 0.320772425 |
| ENSMUSG00000039278 | Pcsk1n | protein_coding | 0.795045941 | 5.94998412 | 5.2354014 | 0.0358009 | 0.320927763 |
| ENSMUSG00000062931 | Zfp938 | protein_coding | -0.555826437 | 6.348800424 | 5.2346925 | 0.0358119 | 0.320927763 |
| ENSMUSG00000025408 | Ddit3 | protein_coding | -0.655738848 | 6.742839347 | 5.234521 | 0.0358145 | 0.320927763 |
| ENSMUSG00000041660 | Bbox1 | protein_coding | -1.722780133 | 3.012470954 | 5.2287678 | 0.0359034 | 0.321456898 |
| ENSMUSG00000055214 | Pld5 | protein_coding | -0.760078741 | 5.240425973 | 5.2257293 | 0.0359505 | 0.321610616 |
| ENSMUSG00000057715 | A830018L16Rik | protein_coding | -0.891078822 | 4.937030067 | 5.2227948 | 0.0359961 | 0.321750273 |
| ENSMUSG00000006342 | Susd2 | protein_coding | 1.283671928 | 4.02023654 | 5.2154336 | 0.0361106 | 0.322505793 |
| ENSMUSG00000061524 | Zic2 | protein_coding | -1.928171494 | 2.582367582 | 5.2120255 | 0.0361637 | 0.322587468 |
| ENSMUSG00000026761 | Orc4 | protein_coding | -0.629747819 | 6.233257525 | 5.2110019 | 0.0361797 | 0.322587468 |
| ENSMUSG00000033730 | Egr3 | protein_coding | -1.047173854 | 4.995065314 | 5.2052439 | 0.0362698 | 0.322909836 |
| ENSMUSG00000028799 | Zfp362 | protein_coding | -1.418534395 | 2.955070288 | 5.2048515 | 0.0362759 | 0.322909836 |
| ENSMUSG00000015377 | Dennd6b | protein_coding | 1.283681671 | 3.564111178 | 5.2023704 | 0.0363148 | 0.322988694 |
| ENSMUSG00000044447 | Dock5 | protein_coding | -2.092295209 | 2.442328391 | 5.1997355 | 0.0363562 | 0.323089337 |
| ENSMUSG00000001525 | Tubb5 | protein_coding | 0.717121209 | 10.28704004 | 5.1978022 | 0.0363866 | 0.323092308 |
| ENSMUSG00000047866 | Lonp2 | protein_coding | 0.620942672 | 6.782646359 | 5.1870538 | 0.036556 | 0.32387861 |
| ENSMUSG00000034445 | Cyb561a3 | protein_coding | 1.974809751 | 2.512051655 | 5.1860957 | 0.0365712 | 0.32387861 |
| ENSMUSG00000059834 | Sclt1 | protein_coding | -1.509105997 | 3.183665427 | 5.1850296 | 0.036588 | 0.32387861 |
| ENSMUSG00000032212 | Sltm | protein_coding | -0.710887647 | 5.581230775 | 5.1827767 | 0.0366237 | 0.32387861 |
| ENSMUSG00000026526 | Fh1 | protein_coding | 0.542285866 | 7.555256967 | 5.182649 | 0.0366257 | 0.32387861 |
| ENSMUSG00000004100 | Ppan | protein_coding | 1.298655806 | 3.711390121 | 5.1795012 | 0.0366756 | 0.324053445 |
| ENSMUSG00000034867 | Ankrd27 | protein_coding | 1.142071239 | 4.345784258 | 5.1707636 | 0.0368146 | 0.324649679 |
| ENSMUSG00000036810 | Cnep1r1 | protein_coding | -0.787622892 | 5.715660577 | 5.1707255 | 0.0368152 | 0.324649679 |
| ENSMUSG00000073982 | Rhog | protein_coding | -0.734134549 | 6.224358474 | 5.1678143 | 0.0368616 | 0.324649679 |
| ENSMUSG00000052581 | Lrrtm4 | protein_coding | -0.993862111 | 5.281600629 | 5.166316 | 0.0369098 | 0.324649679 |
| ENSMUSG00000022208 | Jph4 | protein_coding | -0.622223108 | 6.716425145 | 5.1595978 | 0.036993 | 0.324649679 |
| ENSMUSG00000011884 | Gltp | protein_coding | -0.86191842 | 6.231484029 | 5.1610552 | 0.0369939 | 0.324649679 |
| ENSMUSG00000005936 | Kctd20 | protein_coding | 0.820834392 | 5.242159555 | 5.1576502 | 0.0370242 | 0.324649679 |
| ENSMUSG00000113701 | AC121583.1 | transcribed_processe | -0.883786609 | 4.360972706 | 5.1563524 | 0.0370451 | 0.324649679 |
| ENSMUSG00000067336 | Bmpr2 | protein_coding | -0.692413301 | 5.509089548 | 5.1543028 | 0.037078 | 0.324649679 |
| ENSMUSG00000021012 | Zc3h14 | protein_coding | -0.960366265 | 4.454841109 | 5.1524558 | 0.0371077 | 0.324649679 |
| ENSMUSG00000001552 | Jup | protein_coding | -0.874839771 | 4.976714421 | 5.1515727 | 0.0371219 | 0.324649679 |
| ENSMUSG00000028692 | Akr1a1 | protein_coding | 0.563649547 | 10.57220494 | 5.1472604 | 0.0371913 | 0.324649679 |
| ENSMUSG00000026510 | Trp53bp2 | protein_coding | -0.947570678 | 4.712471166 | 5.1456964 | 0.0372166 | 0.324649679 |
| ENSMUSG00000025474 | Tubgcp2 | protein_coding | 0.89116572 | 5.413787032 | 5.1444496 | 0.0372367 | 0.324649679 |
| ENSMUSG00000026159 | Agfg1 | protein_coding | -0.751702095 | 5.152506144 | 5.1439546 | 0.0372447 | 0.324649679 |
| ENSMUSG00000039361 | Picalm | protein_coding | -0.64576472 | 6.279522814 | 5.142224 | 0.0372726 | 0.324649679 |
| ENSMUSG00000039942 | Ptger4 | protein_coding | -1.086181722 | 4.093155842 | 5.141955 | 0.037277 | 0.324649679 |
| ENSMUSG00000026740 | Dnajc1 | protein_coding | -1.447190048 | 4.213419847 | 5.1700724 | 0.0373178 | 0.324649679 |
| ENSMUSG00000001763 | Tspan33 | protein_coding | 1.375466383 | 3.839518618 | 5.1408212 | 0.0373197 | 0.324649679 |
| ENSMUSG00000075318 | Scn2a | protein_coding | -0.517619248 | 6.959248327 | 5.1376335 | 0.0373469 | 0.324649679 |
| ENSMUSG00000029577 | Ube3b | protein_coding | 0.831316515 | 4.872432996 | 5.1310615 | 0.0374536 | 0.325313856 |
| ENSMUSG00000021451 | Sema4d | protein_coding | -1.022648372 | 3.612911937 | 5.1260445 | 0.0375353 | 0.325561807 |
| ENSMUSG00000029759 | Pon3 | protein_coding | -3.015023796 | 2.371750793 | 5.1270864 | 0.0375427 | 0.325561807 |
| ENSMUSG00000029501 | Ankle2 | protein_coding | -0.917230975 | 4.398547778 | 5.1219981 | 0.0376013 | 0.325802454 |
| ENSMUSG00000021986 | Amer2 | protein_coding | -0.907284821 | 5.07212423 | 5.1201755 | 0.037631 | 0.325802454 |
| ENSMUSG00000029810 | Tmem176b | protein_coding | -0.58804734 | 6.570724312 | 5.1096939 | 0.0378028 | 0.327026328 |
| ENSMUSG00000098097 | 6530403H02Rik | lincRNA | -1.720113333 | 2.770981704 | 5.1045927 | 0.0378867 | 0.32726461 |
| ENSMUSG00000033295 | Ptprf | protein_coding | 0.654855003 | 5.670309118 | 5.1024549 | 0.037922 | 0.32726461 |
| ENSMUSG00000059003 | Grin2a | protein_coding | -1.160847706 | 4.417501533 | 5.1014435 | 0.0379387 | 0.32726461 |
| ENSMUSG00000024781 | Lipa | protein_coding | 0.777127028 | 5.126524875 | 5.0992419 | 0.037975 | 0.32726461 |
| ENSMUSG00000022112 | Gpc5 | protein_coding | -1.03263891 | 4.158869144 | 5.0987862 | 0.0379825 | 0.32726461 |

| ensembl_gene_id | external_gene_name | gene_biotype | log2 (fold change) | log2 (counts per million) | F-value | p-value | False discovery rate |
|---|---|---|---|---|---|---|---|
| ENSMUSG00000022099 | Dmtn | protein_coding | 0.556141266 | 7.798985178 | 5.0951611 | 0.0380425 | 0.327274085 |
| ENSMUSG00000020994 | Pnn | protein_coding | -0.735059466 | 5.979140039 | 5.0907048 | 0.0381163 | 0.327274085 |
| ENSMUSG00000029472 | Anapc5 | protein_coding | 0.646469125 | 7.048916681 | 5.090353 | 0.0381222 | 0.327274085 |
| ENSMUSG00000032333 | Stoml1 | protein_coding | 0.728135613 | 6.128324996 | 5.0894242 | 0.0381376 | 0.327274085 |
| ENSMUSG00000026385 | Dbi | protein_coding | -0.884546695 | 9.643997402 | 5.1460487 | 0.0381572 | 0.327274085 |
| ENSMUSG00000044117 | 2900011O08Rik | protein_coding | 0.616695556 | 9.481265429 | 5.0876975 | 0.0381663 | 0.327274085 |
| ENSMUSG00000039759 | Thap3 | protein_coding | 0.634500422 | 6.023403703 | 5.0846397 | 0.0382171 | 0.327420892 |
| ENSMUSG00000027291 | Vps39 | protein_coding | 0.999233756 | 5.778991377 | 5.1098495 | 0.0382635 | 0.327420892 |
| ENSMUSG00000042670 | Immp1l | protein_coding | -0.727182606 | 5.770079126 | 5.0799448 | 0.0382953 | 0.327420892 |
| ENSMUSG00000024238 | Zeb1 | protein_coding | -0.600305034 | 6.277915847 | 5.0762981 | 0.0383562 | 0.327420892 |
| ENSMUSG00000049960 | Mrps16 | protein_coding | 0.529119348 | 6.888654785 | 5.07606 | 0.0383602 | 0.327420892 |
| ENSMUSG00000070733 | Fryl | protein_coding | -1.044819863 | 3.96075965 | 5.0751856 | 0.0383748 | 0.327420892 |
| ENSMUSG00000038648 | Creb3l2 | protein_coding | -2.516369106 | 2.243356346 | 5.0738856 | 0.0383965 | 0.327420892 |
| ENSMUSG00000026634 | Angel2 | protein_coding | -0.692891265 | 5.422101788 | 5.0684949 | 0.0384868 | 0.327930911 |
| ENSMUSG00000026895 | Ndufa8 | protein_coding | 0.512993529 | 9.504452864 | 5.0630985 | 0.0385775 | 0.328334148 |
| ENSMUSG00000097023 | Mir9-3hg | lincRNA | -1.170222341 | 3.721432233 | 5.0620439 | 0.0385952 | 0.328334148 |
| ENSMUSG00000056596 | Trnp1 | protein_coding | -0.824829906 | 5.774710519 | 5.0541545 | 0.0387283 | 0.329029071 |
| ENSMUSG00000000600 | Krit1 | protein_coding | -0.914322979 | 4.727434989 | 5.0535719 | 0.0387381 | 0.329029071 |
| ENSMUSG00000025609 | Mkln1 | protein_coding | -0.669300655 | 5.956628802 | 5.0502035 | 0.0387951 | 0.32925297 |
| ENSMUSG00000002845 | Tmem39a | protein_coding | 1.603357233 | 3.255610213 | 5.0457976 | 0.0388698 | 0.32938753 |
| ENSMUSG00000036777 | Anln | protein_coding | -0.825441763 | 4.705413853 | 5.0449061 | 0.0388849 | 0.32938753 |
| ENSMUSG00000027466 | Rbck1 | protein_coding | 0.67858391 | 6.158664113 | 5.0438493 | 0.0389028 | 0.32938753 |
| ENSMUSG00000038526 | Car14 | protein_coding | -1.109155095 | 4.815138323 | 5.0402108 | 0.0389894 | 0.329467769 |
| ENSMUSG00000044071 | Fam19a2 | protein_coding | -0.965477066 | 4.660714796 | 5.0379051 | 0.0390039 | 0.329467769 |
| ENSMUSG00000021478 | Drd1 | protein_coding | -0.959412022 | 5.486676655 | 5.0393421 | 0.0390042 | 0.329467769 |
| ENSMUSG00000057738 | Sptan1 | protein_coding | -0.700882552 | 7.14596439 | 5.0372549 | 0.0390398 | 0.329509287 |
| ENSMUSG00000026490 | Cdc42bpa | protein_coding | -0.686848485 | 6.237104324 | 5.0329235 | 0.0390889 | 0.329665188 |
| ENSMUSG00000044647 | Csrnp3 | protein_coding | -0.598163415 | 6.295928373 | 5.030682 | 0.0391272 | 0.329690698 |
| ENSMUSG00000030061 | Uba3 | protein_coding | -0.566207715 | 6.785816306 | 5.0238306 | 0.0392446 | 0.329690698 |
| ENSMUSG00000005716 | Pvalb | protein_coding | -0.995710755 | 7.802006038 | 5.1490622 | 0.0392519 | 0.329690698 |
| ENSMUSG00000018322 | Tomm34 | protein_coding | 0.580181076 | 7.916772627 | 5.0213803 | 0.0392866 | 0.329690698 |
| ENSMUSG00000003581 | Rnf215 | protein_coding | 1.088145978 | 3.454155927 | 5.0199608 | 0.039311 | 0.329690698 |
| ENSMUSG00000035179 | Ppp1r32 | protein_coding | 1.393026075 | 3.234266445 | 5.0198574 | 0.0393128 | 0.329690698 |
| ENSMUSG00000110185 | Igip | protein_coding | -0.873730194 | 5.364580679 | 5.01846 | 0.0393368 | 0.329690698 |
| ENSMUSG00000048000 | Gigyf2 | protein_coding | -0.576731915 | 6.073082081 | 5.0179827 | 0.039345 | 0.329690698 |
| ENSMUSG00000038371 | Sbf2 | protein_coding | -0.884906595 | 4.293764089 | 5.0135715 | 0.039421 | 0.329690698 |
| ENSMUSG00000027206 | Cops2 | protein_coding | -0.615386182 | 7.228814751 | 5.0115013 | 0.0394568 | 0.329690698 |
| ENSMUSG00000006641 | Slc5a6 | protein_coding | 0.941377207 | 4.429235679 | 5.0113668 | 0.0394591 | 0.329690698 |
| ENSMUSG00000096472 | Cdkn2d | protein_coding | 0.484568012 | 8.239729479 | 5.0113217 | 0.0394599 | 0.329690698 |
| ENSMUSG00000004667 | Polr2e | protein_coding | 0.512300834 | 8.368268103 | 5.0084352 | 0.0395097 | 0.329851112 |
| ENSMUSG00000003528 | Slc25a1 | protein_coding | 0.9981993 | 5.133553603 | 5.0015123 | 0.0396546 | 0.330771021 |
| ENSMUSG00000035228 | Ccdc106 | protein_coding | 0.866666565 | 5.471957661 | 4.9983813 | 0.039684 | 0.330771021 |
| ENSMUSG00000026344 | Lypd1 | protein_coding | 1.566122913 | 2.678574672 | 4.9967604 | 0.0397122 | 0.330771021 |
| ENSMUSG00000038070 | Cntln | protein_coding | -1.270033155 | 3.697882556 | 4.9947802 | 0.0397467 | 0.330801806 |
| ENSMUSG00000035164 | Zc3h12c | protein_coding | -1.31413915 | 3.641309505 | 4.9899755 | 0.0398304 | 0.331242518 |
| ENSMUSG00000001445 | Mrpl10 | protein_coding | 0.631662796 | 6.93874724 | 4.98574 | 0.0399044 | 0.331601507 |
| ENSMUSG00000048040 | Arxes2 | protein_coding | 0.614526092 | 6.371574271 | 4.9808755 | 0.0399896 | 0.331967433 |
| ENSMUSG00000072694 | 1500011B03Rik | protein_coding | 0.465778655 | 8.042027602 | 4.9731915 | 0.0401246 | 0.331967433 |
| ENSMUSG00000006920 | Ezh1 | protein_coding | 0.5681777 | 6.198793368 | 4.9728982 | 0.0401298 | 0.331967433 |
| ENSMUSG00000023913 | Pla2g7 | protein_coding | -0.619971354 | 7.018557148 | 4.9718562 | 0.0401481 | 0.331967433 |
| ENSMUSG00000061118 | Dnajc30 | protein_coding | 0.723237663 | 5.483751363 | 4.9690573 | 0.0401975 | 0.331967433 |
| ENSMUSG00000021376 | Tpmt | protein_coding | -0.945605121 | 3.970889341 | 4.967606 | 0.0402231 | 0.331967433 |
| ENSMUSG00000027827 | Kcnab1 | protein_coding | -0.532023275 | 6.617753753 | 4.966445 | 0.0402436 | 0.331967433 |
| ENSMUSG00000027698 | Nceh1 | protein_coding | 0.473972493 | 7.061547712 | 4.9660535 | 0.0402505 | 0.331967433 |
| ENSMUSG00000041236 | Vps41 | protein_coding | 0.646566217 | 7.767959497 | 4.9657745 | 0.0402554 | 0.331967433 |
| ENSMUSG00000068220 | Lgals1 | protein_coding | -0.812835019 | 5.289753324 | 4.9633806 | 0.0402977 | 0.331967433 |
| ENSMUSG00000068523 | Gng5 | protein_coding | -0.921694413 | 4.80179362 | 4.9633235 | 0.0402988 | 0.331967433 |
| ENSMUSG00000020541 | Tom1l1 | protein_coding | -0.726289913 | 5.201902742 | 4.9604231 | 0.0403501 | 0.331967433 |
| ENSMUSG00000055639 | Dach1 | protein_coding | -1.032225412 | 3.902452405 | 4.9570274 | 0.0404103 | 0.331967433 |
| ENSMUSG00000086003 | B230206L02Rik | processed_transcript | -0.975314938 | 4.057111651 | 4.9562437 | 0.0404242 | 0.331967433 |
| ENSMUSG00000037251 | Pomk | protein_coding | 0.725398364 | 5.944052646 | 4.956238 | 0.0404243 | 0.331967433 |
| ENSMUSG00000029076 | Sdf4 | protein_coding | 0.495722214 | 8.512245284 | 4.9548236 | 0.0404495 | 0.331967433 |
| ENSMUSG00000027406 | Idh3b | protein_coding | 0.537161285 | 9.161263292 | 4.9527577 | 0.0404862 | 0.331967433 |
| ENSMUSG00000021870 | Slmap | protein_coding | -0.8839845 | 5.716524717 | 4.9531572 | 0.0405042 | 0.331967433 |
| ENSMUSG00000068739 | Sars | protein_coding | 0.629431281 | 8.568066418 | 4.9473889 | 0.0405818 | 0.332150532 |
| ENSMUSG00000030376 | Slc8a2 | protein_coding | -1.682354412 | 2.93078696 | 4.9465259 | 0.0405972 | 0.332150532 |
| ENSMUSG00000011589 | Fsd1 | protein_coding | 0.699391499 | 7.226805919 | 4.9464088 | 0.0406244 | 0.332150532 |
| ENSMUSG00000035530 | Eif1 | protein_coding | 0.651805412 | 9.399045328 | 4.9408327 | 0.0406989 | 0.332150532 |
| ENSMUSG00000032840 | 2410131K14Rik | protein_coding | 0.705528661 | 5.490908084 | 4.9386617 | 0.0407378 | 0.332150532 |
| ENSMUSG00000032459 | Mrps22 | protein_coding | 0.862284101 | 5.211931843 | 4.9384105 | 0.0407423 | 0.332150532 |
| ENSMUSG00000033006 | Sox10 | protein_coding | -0.858570893 | 5.169216814 | 4.9363567 | 0.0407791 | 0.332150532 |
| ENSMUSG00000059811 | Atl2 | protein_coding | -0.775200558 | 4.999606337 | 4.9358466 | 0.0407883 | 0.332150532 |
| ENSMUSG00000041815 | Poldip3 | protein_coding | -0.529994498 | 8.061233683 | 4.9337212 | 0.0408264 | 0.332150532 |
| ENSMUSG00000029675 | Eln | protein_coding | -2.695254425 | 3.211596894 | 5.0720771 | 0.0408354 | 0.332150532 |
| ENSMUSG00000021918 | Nek4 | protein_coding | 1.029716847 | 4.061334243 | 4.9294044 | 0.040904 | 0.332347392 |
| ENSMUSG00000038206 | Fbxo8 | protein_coding | -0.87912413 | 4.913619376 | 4.9283732 | 0.0409226 | 0.332347392 |
| ENSMUSG00000028104 | Polr3gl | protein_coding | 1.067955925 | 3.652093066 | 4.9267228 | 0.0409523 | 0.332347392 |
| ENSMUSG00000071014 | Ndufb6 | protein_coding | 0.517338097 | 8.649696776 | 4.9176 | 0.0411171 | 0.333114815 |

| ensembl_gene_id | external_gene_name | gene_biotype | log2 (fold change) | log2 (counts per million) | F-value | p-value | False discovery rate |
|---|---|---|---|---|---|---|---|
| ENSMUSG00000022272 | Myo10 | protein_coding | -0.853905351 | 4.921313803 | 4.9153175 | 0.0411585 | 0.333114815 |
| ENSMUSG00000022750 | Klhl22 | protein_coding | 0.593437636 | 6.821613233 | 4.9139545 | 0.0411832 | 0.333114815 |
| ENSMUSG00000062691 | Cebpzos | protein_coding | 0.784838102 | 4.80714867 | 4.9111225 | 0.0412346 | 0.333114815 |
| ENSMUSG00000025200 | Cwf19l1 | protein_coding | 0.89672248 | 4.413403784 | 4.9106519 | 0.0412432 | 0.333114815 |
| ENSMUSG00000027177 | Hipk3 | protein_coding | -0.807487281 | 5.20658307 | 4.9102195 | 0.041251 | 0.333114815 |
| ENSMUSG00000097585 | E230029C05Rik | processed_transcript | -1.206797182 | 3.481865834 | 4.9065617 | 0.0413176 | 0.333114815 |
| ENSMUSG00000070953 | Rabepk | protein_coding | 0.53749759 | 6.597109698 | 4.9062125 | 0.0413239 | 0.333114815 |
| ENSMUSG00000029999 | Tgfa | protein_coding | -0.797912244 | 4.891510745 | 4.9055741 | 0.0413355 | 0.333114815 |
| ENSMUSG00000050321 | Neto1 | protein_coding | -1.008657745 | 4.702651805 | 4.904415 | 0.0413567 | 0.333114815 |
| ENSMUSG00000052698 | Tln2 | protein_coding | -0.747764379 | 5.70148989 | 4.8993947 | 0.0414483 | 0.33338788 |
| ENSMUSG00000021548 | Ccnh | protein_coding | -0.523441346 | 7.14178214 | 4.8977274 | 0.0414788 | 0.33338788 |
| ENSMUSG00000059316 | Slc27a4 | protein_coding | 0.776697765 | 5.374491072 | 4.8952432 | 0.0415243 | 0.33338788 |
| ENSMUSG00000030443 | Zfp583 | protein_coding | 1.12107327 | 3.713609321 | 4.8947578 | 0.0415331 | 0.33338788 |
| ENSMUSG00000059714 | Flot1 | protein_coding | 0.64797029 | 6.609385333 | 4.8940787 | 0.0415456 | 0.33338788 |
| ENSMUSG00000033623 | Pcgf3 | protein_coding | -1.050500453 | 4.33184919 | 4.8856901 | 0.0416996 | 0.334374577 |
| ENSMUSG00000035632 | Cnot3 | protein_coding | 1.028725037 | 3.568040834 | 4.8817677 | 0.0417719 | 0.334688415 |
| ENSMUSG00000044066 | Cep68 | protein_coding | 1.177020419 | 3.705952981 | 4.8786906 | 0.0418287 | 0.334688415 |
| ENSMUSG00000020562 | Efcab10 | protein_coding | 0.702866144 | 5.368158711 | 4.8778533 | 0.0418442 | 0.334688415 |
| ENSMUSG00000024937 | Ehbp1l1 | protein_coding | 0.693214541 | 5.694630554 | 4.8756338 | 0.0418852 | 0.334688415 |
| ENSMUSG00000037351 | Actr1b | protein_coding | 0.652905333 | 8.367555443 | 4.8739448 | 0.0419165 | 0.334688415 |
| ENSMUSG00000002732 | Fkbp7 | protein_coding | -1.645822581 | 2.801679118 | 4.8734547 | 0.0419255 | 0.334688415 |
| ENSMUSG00000029250 | Polr2b | protein_coding | 0.709591858 | 5.613263363 | 4.8709858 | 0.0419713 | 0.334786783 |
| ENSMUSG00000067818 | Myl9 | protein_coding | -0.905585225 | 4.61961519 | 4.8680037 | 0.0420266 | 0.334786783 |
| ENSMUSG00000050587 | Ak1 | protein_coding | 0.593614183 | 9.070943259 | 4.8663529 | 0.0420573 | 0.334786783 |
| ENSMUSG00000042228 | Lyn | protein_coding | -2.308652334 | 2.375049205 | 4.8647951 | 0.0420863 | 0.334786783 |
| ENSMUSG00000059498 | Fcgr3 | protein_coding | -1.388155211 | 4.39498967 | 4.9071798 | 0.0421562 | 0.334786783 |
| ENSMUSG00000049225 | Pdp1 | protein_coding | -0.615626657 | 7.125775586 | 4.8592704 | 0.0421892 | 0.334786783 |
| ENSMUSG00000048772 | Tmem53 | protein_coding | 0.955112959 | 4.221569174 | 4.8586778 | 0.0422002 | 0.334786783 |
| ENSMUSG00000036098 | Myrf | protein_coding | -0.924293837 | 4.412695229 | 4.8572606 | 0.0422267 | 0.334786783 |
| ENSMUSG00000052305 | Hbb-bs | protein_coding | -0.910093195 | 5.942453252 | 4.8621827 | 0.0422274 | 0.334786783 |
| ENSMUSG00000106379 | Lhfpl3 | protein_coding | -0.632045097 | 6.027799078 | 4.8560556 | 0.0422492 | 0.334786783 |
| ENSMUSG00000066456 | Hmgn3 | protein_coding | -0.663621486 | 7.04535961 | 4.8516914 | 0.0423308 | 0.335067477 |
| ENSMUSG00000048416 | Mlf1 | protein_coding | -1.114861448 | 3.661680414 | 4.8481829 | 0.0423966 | 0.335067477 |
| ENSMUSG00000087549 | Gm13427 | antisense_RNA | -1.188194315 | 6.941726025 | 5.087897 | 0.042398 | 0.335067477 |
| ENSMUSG00000041241 | Mul1 | protein_coding | 0.508017763 | 7.04934122 | 4.8450261 | 0.0424559 | 0.335067477 |
| ENSMUSG00000026414 | Tnnt2 | protein_coding | 1.150559933 | 3.784445905 | 4.8427602 | 0.0424985 | 0.335067477 |
| ENSMUSG00000023011 | Faim2 | protein_coding | 0.663443121 | 9.330157212 | 4.8412367 | 0.0425271 | 0.335067477 |
| ENSMUSG00000006736 | Tspan31 | protein_coding | 0.536738302 | 6.733065879 | 4.8374449 | 0.0425986 | 0.335067477 |
| ENSMUSG00000037286 | Stag1 | protein_coding | -0.772088257 | 5.854711725 | 4.8372186 | 0.0426029 | 0.335067477 |
| ENSMUSG00000059708 | Akap17b | protein_coding | -0.973517436 | 3.972768792 | 4.8369054 | 0.0426088 | 0.335067477 |
| ENSMUSG00000071267 | Zfp942 | protein_coding | -1.232848214 | 3.31002476 | 4.8363726 | 0.0426188 | 0.335067477 |
| ENSMUSG00000068798 | Rap1a | protein_coding | -0.753408939 | 4.606959822 | 4.8347439 | 0.0426496 | 0.335067477 |
| ENSMUSG00000053574 | 4930563E22Rik | processed_transcript | -1.269221937 | 3.28713829 | 4.83427 | 0.0426585 | 0.335067477 |
| ENSMUSG00000040181 | Fmo1 | protein_coding | -1.38775344 | 3.594445406 | 4.8294839 | 0.0427491 | 0.33553358 |
| ENSMUSG00000024479 | Mal2 | protein_coding | -0.67711601 | 5.853344285 | 4.8268597 | 0.0427988 | 0.335678954 |
| ENSMUSG00000025255 | Zfhx4 | protein_coding | -0.833558131 | 4.583105789 | 4.820225 | 0.0429249 | 0.336422243 |
| ENSMUSG00000046982 | Tshz1 | protein_coding | -0.862914404 | 4.906798199 | 4.817199 | 0.0429825 | 0.336448097 |
| ENSMUSG00000022751 | Nit2 | protein_coding | 0.651145859 | 5.828133495 | 4.8167667 | 0.0429908 | 0.336448097 |
| ENSMUSG00000017288 | Vps53 | protein_coding | 0.776605145 | 6.081862636 | 4.8109598 | 0.0431016 | 0.336619097 |
| ENSMUSG00000036699 | Zcchc12 | protein_coding | 1.038611038 | 9.901316203 | 4.9331274 | 0.0431133 | 0.336619097 |
| ENSMUSG00000002222 | Rmnd5a | protein_coding | -0.723864315 | 5.427513093 | 4.8095245 | 0.0431291 | 0.336619097 |
| ENSMUSG00000005378 | Wbscr22 | protein_coding | 0.567741323 | 6.300554617 | 4.8043481 | 0.0432283 | 0.336619097 |
| ENSMUSG00000036036 | Zfp57 | protein_coding | 2.991165793 | 2.74899328 | 4.8852987 | 0.0432534 | 0.336619097 |
| ENSMUSG00000020954 | Strn3 | protein_coding | -0.790638183 | 5.765313245 | 4.8022717 | 0.0432681 | 0.336619097 |
| ENSMUSG00000025584 | Pde8a | protein_coding | -1.133502198 | 4.010309423 | 4.8016948 | 0.0432792 | 0.336619097 |
| ENSMUSG00000025094 | Slc18a2 | protein_coding | -1.53015978 | 3.023974236 | 4.8016927 | 0.0432793 | 0.336619097 |
| ENSMUSG00000039630 | Hnrnpu | protein_coding | -0.465128906 | 7.5492458 | 4.8005873 | 0.0433005 | 0.336619097 |
| ENSMUSG00000005681 | Apoa2 | protein_coding | 1.157291929 | 3.382292466 | 4.7943982 | 0.0434197 | 0.336619097 |
| ENSMUSG00000034595 | Ppp1r18 | protein_coding | -1.888467492 | 2.624881968 | 4.7914262 | 0.043477 | 0.336619097 |
| ENSMUSG00000002997 | Prkar2b | protein_coding | -0.689995768 | 6.045390933 | 4.7844699 | 0.0436117 | 0.336619097 |
| ENSMUSG00000036202 | Rif1 | protein_coding | -1.014861939 | 4.189042573 | 4.7793139 | 0.0437117 | 0.336619097 |
| ENSMUSG00000057982 | Zfp809 | protein_coding | -1.040333044 | 4.043580567 | 4.7773247 | 0.0437504 | 0.336619097 |
| ENSMUSG00000022704 | Qtrt2 | protein_coding | 1.269072072 | 3.32029933 | 4.7772583 | 0.0437517 | 0.336619097 |
| ENSMUSG00000044709 | Gemin7 | protein_coding | 0.654694566 | 5.674839794 | 4.7760196 | 0.0437758 | 0.336619097 |
| ENSMUSG00000020635 | Fkbp1b | protein_coding | 0.971580526 | 4.312844413 | 4.7742856 | 0.0438096 | 0.336619097 |
| ENSMUSG00000046822 | Slc39a3 | protein_coding | -1.056979513 | 4.066614776 | 4.7729183 | 0.0438362 | 0.336619097 |
| ENSMUSG00000046785 | Epm2aip1 | protein_coding | -0.571713438 | 6.769392348 | 4.7697719 | 0.0438977 | 0.336619097 |
| ENSMUSG00000026864 | Hspa5 | protein_coding | 0.473091458 | 8.346771058 | 4.7691888 | 0.043909 | 0.336619097 |
| ENSMUSG00000032842 | Abcc10 | protein_coding | 2.1228723 | 2.920291954 | 4.7703863 | 0.0439114 | 0.336619097 |
| ENSMUSG00000024870 | Rab1b | protein_coding | 0.550335651 | 8.0398978 | 4.7690298 | 0.0439121 | 0.336619097 |
| ENSMUSG00000022674 | Ube2v2 | protein_coding | -0.786652197 | 4.830762363 | 4.7681993 | 0.0439284 | 0.336619097 |
| ENSMUSG00000037386 | Rims2 | protein_coding | -0.551434787 | 6.12669985 | 4.7667266 | 0.0439572 | 0.336619097 |
| ENSMUSG00000059182 | Skap2 | protein_coding | -1.031412377 | 4.404641125 | 4.7658904 | 0.0439735 | 0.336619097 |
| ENSMUSG00000059070 | Rpl18 | protein_coding | 0.645399388 | 6.174859405 | 4.7641318 | 0.044008 | 0.336619097 |
| ENSMUSG00000025428 | Atp5a1 | protein_coding | 0.668775451 | 10.5517582 | 4.7626841 | 0.0440363 | 0.336619097 |
| ENSMUSG00000023832 | Acat2 | protein_coding | 0.646356023 | 6.774164154 | 4.7622646 | 0.0440446 | 0.336619097 |
| ENSMUSG00000022285 | Ywhaz | protein_coding | -0.53496896 | 10.25803832 | 4.759904 | 0.0440909 | 0.336619097 |
| ENSMUSG00000036905 | C1qb | protein_coding | -1.206869372 | 6.327932409 | 4.9195665 | 0.0440939 | 0.336619097 |

| ensembl_gene_id | external_gene_name | gene_biotype | log2 (fold change) | log2 (counts per million) | F-value | p-value | False discovery rate |
|---|---|---|---|---|---|---|---|
| ENSMUSG00000029502 | Golga3 | protein_coding | 0.792227516 | 5.084133763 | 4.7586265 | 0.044116 | 0.336619097 |
| ENSMUSG00000027660 | Skil | protein_coding | -0.565419774 | 6.513864919 | 4.7584282 | 0.0441199 | 0.336619097 |
| ENSMUSG00000049985 | Ankrd55 | protein_coding | 1.322370855 | 3.601224438 | 4.7550186 | 0.0441869 | 0.336619097 |
| ENSMUSG00000031762 | Mt2 | protein_coding | -1.154338361 | 5.28709843 | 4.8052897 | 0.0442073 | 0.336619097 |
| ENSMUSG00000015656 | Hspa8 | protein_coding | 0.827198643 | 8.910845924 | 4.8034012 | 0.0442438 | 0.336619097 |
| ENSMUSG00000054855 | Rnd1 | protein_coding | 0.873965351 | 5.994944056 | 4.7559402 | 0.0442628 | 0.336619097 |
| ENSMUSG00000025981 | Coq10b | protein_coding | -0.801162844 | 5.300111345 | 4.7510936 | 0.0442642 | 0.336619097 |
| ENSMUSG00000047747 | Rnf150 | protein_coding | -1.099573717 | 4.084373128 | 4.750218 | 0.0442815 | 0.336619097 |
| ENSMUSG00000041096 | Tspyl2 | protein_coding | 0.587363999 | 6.912719448 | 4.7487792 | 0.0443099 | 0.336619097 |
| ENSMUSG00000027722 | Spata5 | protein_coding | 1.178796855 | 3.663025969 | 4.7476809 | 0.0443316 | 0.336619097 |
| ENSMUSG00000035851 | Ythdc1 | protein_coding | -0.760216615 | 5.114196526 | 4.7474508 | 0.0443361 | 0.336619097 |
| ENSMUSG00000003235 | Eif2b5 | protein_coding | 0.785063331 | 5.373775988 | 4.7472913 | 0.0443393 | 0.336619097 |
| ENSMUSG00000032517 | Mobp | protein_coding | -0.521558778 | 7.82379215 | 4.7463088 | 0.0443587 | 0.336619097 |
| ENSMUSG00000025899 | Alkbh8 | protein_coding | -0.687532371 | 5.523790829 | 4.7445429 | 0.0443936 | 0.336646562 |
| ENSMUSG00000029635 | Cdk8 | protein_coding | 0.649098012 | 6.496972303 | 4.7419578 | 0.0444448 | 0.336797217 |
| ENSMUSG00000031130 | Brs3 | protein_coding | 2.614344983 | 3.223265056 | 4.8684871 | 0.0445011 | 0.336986067 |
| ENSMUSG00000029038 | Ssu72 | protein_coding | 0.4704172 | 7.920760484 | 4.7298982 | 0.0446845 | 0.337408901 |
| ENSMUSG00000035847 | Ids | protein_coding | -0.585081557 | 6.718583316 | 4.7295898 | 0.0446907 | 0.337408901 |
| ENSMUSG00000037703 | Lzts3 | protein_coding | -0.748678657 | 5.886241849 | 4.72886 | 0.0447052 | 0.337408901 |
| ENSMUSG00000027163 | Commd9 | protein_coding | 0.629938981 | 7.208490425 | 4.7286672 | 0.0447091 | 0.337408901 |
| ENSMUSG00000030846 | Tial1 | protein_coding | -0.635811082 | 6.111302703 | 4.7264015 | 0.0447543 | 0.337408901 |
| ENSMUSG00000006732 | Mettl1 | protein_coding | 0.62595115 | 6.02174802 | 4.7256406 | 0.0447695 | 0.337408901 |
| ENSMUSG00000028969 | Cdk5 | protein_coding | 0.499422354 | 8.546438567 | 4.7239915 | 0.0448025 | 0.337408901 |
| ENSMUSG00000030881 | Arfip2 | protein_coding | 0.564634415 | 6.476021251 | 4.7237177 | 0.044808 | 0.337408901 |
| ENSMUSG00000033960 | Jcad | protein_coding | -1.241797965 | 4.348085359 | 4.720823 | 0.0448919 | 0.337804334 |
| ENSMUSG00000034714 | Ttyh2 | protein_coding | -0.773274927 | 7.227593125 | 4.7500745 | 0.0449453 | 0.337970118 |
| ENSMUSG00000062101 | Zfp119b | protein_coding | -2.145001381 | 2.779244344 | 4.7163978 | 0.0449806 | 0.337999215 |
| ENSMUSG00000043964 | Orai3 | protein_coding | 1.177615938 | 3.812328674 | 4.7116855 | 0.0450494 | 0.338172122 |
| ENSMUSG00000031375 | Bgn | protein_coding | -1.493637799 | 3.133467997 | 4.710832 | 0.0450666 | 0.338172122 |
| ENSMUSG00000025223 | Ldb1 | protein_coding | 0.761150147 | 5.148441697 | 4.7081779 | 0.04512 | 0.338337319 |
| ENSMUSG00000026547 | Tagln2 | protein_coding | -0.959329828 | 4.606795362 | 4.7044237 | 0.0451958 | 0.338669327 |
| ENSMUSG00000040855 | Reps2 | protein_coding | -0.694497145 | 6.145058229 | 4.6985475 | 0.0453147 | 0.339323804 |
| ENSMUSG00000005871 | Apc | protein_coding | -0.793738594 | 5.403525281 | 4.6962599 | 0.0453611 | 0.339434773 |
| ENSMUSG00000029121 | Crmp1 | protein_coding | 1.058013601 | 4.814668844 | 4.6951066 | 0.0454106 | 0.339568773 |
| ENSMUSG00000022797 | Tfrc | protein_coding | -0.881982555 | 4.404060591 | 4.6918972 | 0.0454497 | 0.339625169 |
| ENSMUSG00000040351 | Ankib1 | protein_coding | -0.908502003 | 4.844242296 | 4.6898454 | 0.0454914 | 0.339701036 |
| ENSMUSG00000079111 | Kdelr2 | protein_coding | 0.75646418 | 6.432431385 | 4.6839132 | 0.0456385 | 0.340119865 |
| ENSMUSG00000022311 | Csmd3 | protein_coding | -1.085140716 | 4.866877705 | 4.6838993 | 0.0456388 | 0.340119865 |
| ENSMUSG00000086496 | Gm14204 | processed_transcript | 0.877753282 | 5.415769256 | 4.6824427 | 0.0456424 | 0.340119865 |
| ENSMUSG00000028556 | Dock7 | protein_coding | -1.107411787 | 3.640016014 | 4.6799873 | 0.0456926 | 0.340233848 |
| ENSMUSG00000054519 | Zfp867 | protein_coding | -1.069744474 | 3.704421995 | 4.6786012 | 0.045721 | 0.340233848 |
| ENSMUSG00000019975 | Ikbip | protein_coding | -1.135037075 | 3.845814942 | 4.6734205 | 0.0458272 | 0.340384011 |
| ENSMUSG00000020431 | Adcy1 | protein_coding | -1.08873508 | 4.762574829 | 4.672799 | 0.0458661 | 0.340384011 |
| ENSMUSG00000036398 | Ppp1r11 | protein_coding | 0.481463471 | 9.340274285 | 4.6675419 | 0.0459481 | 0.340384011 |
| ENSMUSG00000042389 | Tsen2 | protein_coding | 0.722257498 | 5.267071006 | 4.6669647 | 0.04596 | 0.340384011 |
| ENSMUSG00000016344 | Ppdpf | protein_coding | 0.715780628 | 5.202265819 | 4.6633216 | 0.0460351 | 0.340384011 |
| ENSMUSG00000001864 | Aif1l | protein_coding | -0.786027056 | 5.280336379 | 4.6632946 | 0.0460357 | 0.340384011 |
| ENSMUSG00000075702 | Selenom | protein_coding | 0.536544162 | 9.621468166 | 4.6626385 | 0.0460492 | 0.340384011 |
| ENSMUSG00000040771 | Oard1 | protein_coding | -0.567255701 | 6.457537831 | 4.6608286 | 0.0460866 | 0.340384011 |
| ENSMUSG00000047259 | Mc4r | protein_coding | 1.239804355 | 3.845417213 | 4.6605838 | 0.0460916 | 0.340384011 |
| ENSMUSG00000096914 | Galntl6 | protein_coding | -0.738194763 | 5.198842689 | 4.6604501 | 0.0460944 | 0.340384011 |
| ENSMUSG00000058672 | Tubb2a | protein_coding | 0.632805461 | 9.439810375 | 4.6600477 | 0.0461027 | 0.340384011 |
| ENSMUSG00000019948 | Actr6 | protein_coding | -0.570060092 | 6.397790488 | 4.6582282 | 0.0461404 | 0.340384011 |
| ENSMUSG00000026434 | Nucks1 | protein_coding | -0.549958668 | 7.003684231 | 4.6576322 | 0.0461527 | 0.340384011 |
| ENSMUSG00000025228 | Actr1a | protein_coding | 0.81860322 | 7.759377898 | 4.7054771 | 0.0461863 | 0.340398117 |
| ENSMUSG00000061298 | Agbl4 | protein_coding | 0.95612258 | 4.51388219 | 4.6535153 | 0.046238 | 0.340546044 |
| ENSMUSG00000032231 | Anxa2 | protein_coding | -0.929663916 | 4.781141088 | 4.6498171 | 0.0463148 | 0.340878131 |
| ENSMUSG00000030064 | Frmd4b | protein_coding | -1.257280638 | 3.473205466 | 4.6476519 | 0.0463598 | 0.340883444 |
| ENSMUSG00000052369 | Tmem106c | protein_coding | 0.477543697 | 7.361881729 | 4.6466386 | 0.0463809 | 0.340883444 |
| ENSMUSG00000022658 | Tagln3 | protein_coding | 0.513041688 | 10.66124834 | 4.645212 | 0.0464106 | 0.340883444 |
| ENSMUSG00000000308 | Ckmt1 | protein_coding | 0.716953253 | 8.95899828 | 4.6443018 | 0.0464559 | 0.340982847 |
| ENSMUSG00000057894 | Zfp329 | protein_coding | -0.805757323 | 5.127169057 | 4.6408869 | 0.0465008 | 0.341080071 |
| ENSMUSG00000046607 | Hrk | protein_coding | -0.855702478 | 4.154284163 | 4.6365326 | 0.0465919 | 0.341514781 |
| ENSMUSG00000001783 | Rtcb | protein_coding | 0.671656853 | 6.885349732 | 4.6340918 | 0.0466693 | 0.341528987 |
| ENSMUSG00000003037 | Rab8a | protein_coding | 0.508493041 | 7.495020344 | 4.631216 | 0.0467033 | 0.341528987 |
| ENSMUSG00000029283 | Cdc7 | protein_coding | 1.851409608 | 2.676501028 | 4.6304421 | 0.0467195 | 0.341528987 |
| ENSMUSG00000021596 | Mctp1 | protein_coding | -0.950792879 | 6.10910159 | 4.6793689 | 0.0467208 | 0.341528987 |
| ENSMUSG00000022031 | Elp3 | protein_coding | 0.762812984 | 6.756834004 | 4.6331788 | 0.0468032 | 0.341898852 |
| ENSMUSG00000002881 | Nab1 | protein_coding | -1.001188417 | 4.198576992 | 4.6203379 | 0.0469322 | 0.342579878 |
| ENSMUSG00000032590 | Apeh | protein_coding | 0.933119939 | 4.439069016 | 4.6190135 | 0.0469602 | 0.342579878 |
| ENSMUSG00000020780 | Srp68 | protein_coding | 0.551337216 | 6.480610392 | 4.610917 | 0.0471315 | 0.343437214 |
| ENSMUSG00000025373 | Rnf41 | protein_coding | 0.567984778 | 7.783121997 | 4.6090012 | 0.0471722 | 0.343437214 |
| ENSMUSG00000038530 | Rgs4 | protein_coding | -0.500775835 | 8.482805222 | 4.6089389 | 0.0471735 | 0.343437214 |
| ENSMUSG00000021587 | Pcsk1 | protein_coding | -0.844935903 | 4.154065003 | 4.6045818 | 0.0472661 | 0.343780879 |
| ENSMUSG00000019173 | Rab5c | protein_coding | -0.814783311 | 5.459684264 | 4.6011367 | 0.0473395 | 0.343780879 |
| ENSMUSG00000001524 | Gtf2h4 | protein_coding | 0.759938708 | 5.099909183 | 4.6005067 | 0.0473529 | 0.343780879 |
| ENSMUSG00000039611 | Tmem246 | protein_coding | 0.62337393 | 6.775406665 | 4.5994206 | 0.0473761 | 0.343780879 |
| ENSMUSG00000040410 | Fbxl4 | protein_coding | 0.786020574 | 4.835194913 | 4.5984749 | 0.0473963 | 0.343780879 |

| ensembl_gene_id | external_gene_name | gene_biotype | log2 (fold change) | log2 (counts per million) | F-value | p-value | False discovery rate |
|---|---|---|---|---|---|---|---|
| ENSMUSG00000006095 | Tbcb | protein_coding | 0.472723418 | 9.009886997 | 4.5977137 | 0.0474125 | 0.343780879 |
| ENSMUSG00000015095 | Fbxw5 | protein_coding | 0.554514216 | 7.437499802 | 4.5862409 | 0.0476583 | 0.345272243 |
| ENSMUSG00000018669 | Cdk5rap3 | protein_coding | 0.677415614 | 6.585328663 | 4.5844008 | 0.0476978 | 0.345272243 |
| ENSMUSG00000050732 | Vamp8 | protein_coding | -1.143643659 | 5.692453053 | 4.6580434 | 0.0477429 | 0.345272243 |
| ENSMUSG00000025940 | Tmem70 | protein_coding | -0.793376688 | 4.937732559 | 4.5801904 | 0.0477885 | 0.345272243 |
| ENSMUSG00000024797 | Vps51 | protein_coding | 0.894544431 | 4.582024347 | 4.5786086 | 0.0478226 | 0.345272243 |
| ENSMUSG00000053475 | Tnfaip6 | protein_coding | -0.799055254 | 5.348703418 | 4.5778644 | 0.0478387 | 0.345272243 |
| ENSMUSG00000039568 | Ubald1 | protein_coding | 0.557771428 | 6.558367213 | 4.5756385 | 0.0478867 | 0.345272243 |
| ENSMUSG00000028633 | Ctps | protein_coding | 0.730218229 | 5.140252002 | 4.5745365 | 0.0479105 | 0.345272243 |
| ENSMUSG00000048905 | 4930539E08Rik | protein_coding | 2.11741512 | 2.498641152 | 4.5739119 | 0.047924 | 0.345272243 |
| ENSMUSG00000028755 | Cda | protein_coding | 0.915642039 | 4.950900767 | 4.5723236 | 0.0479584 | 0.345272243 |
| ENSMUSG00000037017 | Zscan21 | protein_coding | 0.95742836 | 4.961300475 | 4.569938 | 0.0480101 | 0.345272243 |
| ENSMUSG00000024382 | Ercc3 | protein_coding | 0.859863651 | 4.83417328 | 4.5665291 | 0.048084 | 0.345272243 |
| ENSMUSG00000036752 | Tubb4b | protein_coding | 0.605194682 | 7.668649757 | 4.5644949 | 0.0481282 | 0.345272243 |
| ENSMUSG00000032384 | Csnk1g1 | protein_coding | -0.76130533 | 5.506981906 | 4.5621309 | 0.0481796 | 0.345272243 |
| ENSMUSG00000037119 | Fam91a1 | protein_coding | -0.838175171 | 4.981655417 | 4.5609879 | 0.0482045 | 0.345272243 |
| ENSMUSG00000078812 | Eif5a | protein_coding | 0.550523768 | 9.469711168 | 4.5602054 | 0.0482215 | 0.345272243 |
| ENSMUSG00000036241 | Ube2r2 | protein_coding | -0.508129597 | 6.414309663 | 4.5587699 | 0.0482528 | 0.345272243 |
| ENSMUSG00000002068 | Ccne1 | protein_coding | 1.187992939 | 3.513192971 | 4.5578504 | 0.0482728 | 0.345272243 |
| ENSMUSG00000051323 | Pcdh19 | protein_coding | -0.704661287 | 5.411491613 | 4.5558707 | 0.048316 | 0.345272243 |
| ENSMUSG00000018040 | Rrp7a | protein_coding | 0.611491438 | 6.211056727 | 4.5558622 | 0.0483162 | 0.345272243 |
| ENSMUSG00000028759 | Hp1bp3 | protein_coding | 0.579291044 | 7.591971683 | 4.5551459 | 0.0483318 | 0.345272243 |
| ENSMUSG00000040612 | Ildr2 | protein_coding | -0.803405546 | 5.880273793 | 4.5562233 | 0.0483349 | 0.345272243 |
| ENSMUSG00000025525 | Apool | protein_coding | -1.047929326 | 4.103545178 | 4.5522409 | 0.0483953 | 0.345272243 |
| ENSMUSG00000072769 | Gm10419 | lincRNA | -0.564477158 | 6.107039761 | 4.5514487 | 0.0484127 | 0.345272243 |
| ENSMUSG00000040280 | Ndufa4l2 | protein_coding | -1.866784369 | 2.881207013 | 4.5510694 | 0.048421 | 0.345272243 |
| ENSMUSG00000067604 | Nms | protein_coding | -1.663145697 | 2.993198315 | 4.5490806 | 0.0484645 | 0.345353746 |
| ENSMUSG00000040097 | Flywch1 | protein_coding | 1.634361274 | 7.261020424 | 5.0124283 | 0.0485651 | 0.345398184 |
| ENSMUSG00000037742 | Eef1a1 | protein_coding | 0.52740602 | 10.30163358 | 4.5437415 | 0.0485816 | 0.345398184 |
| ENSMUSG00000034744 | Nagk | protein_coding | 0.543462496 | 7.326640144 | 4.5421924 | 0.0486157 | 0.345398184 |
| ENSMUSG00000032399 | Rpl4 | protein_coding | 0.523042181 | 10.60178081 | 4.541747 | 0.0486255 | 0.345398184 |
| ENSMUSG00000026966 | Ssna1 | protein_coding | 0.509284059 | 7.620011426 | 4.5405107 | 0.0486527 | 0.345398184 |
| ENSMUSG00000029627 | Zkscan14 | protein_coding | 0.760815762 | 4.75515435 | 4.5400202 | 0.0486635 | 0.345398184 |
| ENSMUSG00000029757 | Dync1i1 | protein_coding | 0.636824889 | 7.483071199 | 4.5332194 | 0.0488134 | 0.346234056 |
| ENSMUSG00000038000 | Acd | protein_coding | 0.698911676 | 6.959643338 | 4.5297561 | 0.0489167 | 0.34639822 |
| ENSMUSG00000025658 | Cnksr2 | protein_coding | -0.870124179 | 5.375949277 | 4.5272058 | 0.0489465 | 0.34639822 |
| ENSMUSG00000015942 | Gtf2ird2 | protein_coding | 1.122643902 | 3.549050209 | 4.5266751 | 0.0489582 | 0.34639822 |
| ENSMUSG00000019828 | Grm1 | protein_coding | -0.677143286 | 5.464318844 | 4.526351 | 0.0489654 | 0.34639822 |
| ENSMUSG00000005360 | Slc1a3 | protein_coding | -0.621233157 | 7.808350894 | 4.5161512 | 0.0491922 | 0.347255747 |
| ENSMUSG00000038664 | Herc1 | protein_coding | -0.716970174 | 5.117133329 | 4.5152431 | 0.0492124 | 0.347255747 |
| ENSMUSG00000048175 | Asb8 | protein_coding | 0.592032452 | 7.244738122 | 4.5143417 | 0.0492325 | 0.347255747 |
| ENSMUSG00000108776 | Gm45169 | lincRNA | -1.239770439 | 3.493116215 | 4.5135483 | 0.0492502 | 0.347255747 |
| ENSMUSG00000033379 | Atp6v0b | protein_coding | 0.559613276 | 8.426530837 | 4.5130349 | 0.0492617 | 0.347255747 |
| ENSMUSG00000044528 | Tram1l1 | protein_coding | 0.480008098 | 6.654881795 | 4.5116113 | 0.0492935 | 0.347255747 |
| ENSMUSG00000045268 | Zfp691 | protein_coding | 1.975365435 | 2.370564015 | 4.5094407 | 0.049342 | 0.347255747 |
| ENSMUSG00000023094 | Msrb2 | protein_coding | 0.591361829 | 5.95553684 | 4.5093074 | 0.049345 | 0.347255747 |
| ENSMUSG00000034259 | Exosc4 | protein_coding | 0.58852735 | 6.135253347 | 4.5000937 | 0.0495516 | 0.347944489 |
| ENSMUSG00000031134 | Rbmx | protein_coding | -0.679161079 | 6.32331531 | 4.4990812 | 0.0495744 | 0.347944489 |
| ENSMUSG00000055137 | Sugct | protein_coding | -1.364222526 | 2.948896426 | 4.4984737 | 0.049588 | 0.347944489 |
| ENSMUSG00000020810 | Cygb | protein_coding | 0.504668406 | 7.364913844 | 4.4980662 | 0.0495972 | 0.347944489 |
| ENSMUSG00000048218 | Amigo2 | protein_coding | 0.871558479 | 4.458766099 | 4.4976966 | 0.0496055 | 0.347944489 |
| ENSMUSG00000070287 | Slc35g2 | protein_coding | 0.753154442 | 5.014595835 | 4.4962983 | 0.049637 | 0.347944489 |
| ENSMUSG00000053007 | Creb5 | protein_coding | -1.294780877 | 3.695559275 | 4.491731 | 0.04974 | 0.348228978 |
| ENSMUSG00000041670 | Rims1 | protein_coding | -0.575638965 | 6.013708503 | 4.4893063 | 0.0497948 | 0.348228978 |
| ENSMUSG00000031532 | Saraf | protein_coding | 0.622564128 | 7.232649969 | 4.4890269 | 0.0498011 | 0.348228978 |
| ENSMUSG00000020474 | Polm | protein_coding | 1.561368969 | 3.367098616 | 4.4881549 | 0.0498477 | 0.348228978 |
| ENSMUSG00000043391 | 2510009E07Rik | protein_coding | -0.69266971 | 5.119740428 | 4.4849533 | 0.0498933 | 0.348228978 |
| ENSMUSG00000003271 | Sult2b1 | protein_coding | 1.159314945 | 3.973833882 | 4.4816557 | 0.0499681 | 0.348228978 |
| ENSMUSG00000065990 | Aurkaip1 | protein_coding | 0.504298905 | 8.863702639 | 4.4807252 | 0.0499892 | 0.348228978 |
| ENSMUSG00000025651 | Uqcrc1 | protein_coding | 0.76693494 | 9.629316734 | 4.5031586 | 0.0499897 | 0.348228978 |
| ENSMUSG00000034245 | Hdac11 | protein_coding | 0.560053378 | 8.240556123 | 4.4805124 | 0.0499941 | 0.348228978 |
| ENSMUSG00000001285 | Myg1 | protein_coding | 0.555613317 | 7.240407338 | 4.4793118 | 0.0500213 | 0.348228978 |
| ENSMUSG00000020267 | Hint1 | protein_coding | 0.522478705 | 10.43714047 | 4.4785267 | 0.0500392 | 0.348228978 |
| ENSMUSG00000083282 | Ctsf | protein_coding | 0.880317903 | 5.385650902 | 4.4773385 | 0.0500662 | 0.348228978 |
| ENSMUSG00000007837 | Prrg2 | protein_coding | -1.189678945 | 3.673805446 | 4.4699075 | 0.0502356 | 0.349181289 |
| ENSMUSG00000039294 | BC017643 | protein_coding | 1.334786701 | 4.386000687 | 4.503783 | 0.0504462 | 0.350418843 |
| ENSMUSG00000024290 | Rock1 | protein_coding | -1.186160667 | 3.749513411 | 4.4541443 | 0.0505971 | 0.351232011 |
| ENSMUSG00000028977 | Casz1 | protein_coding | 0.998952167 | 3.652476258 | 4.4501115 | 0.0506901 | 0.351232011 |
| ENSMUSG00000028222 | Calb1 | protein_coding | -0.619826996 | 7.475597524 | 4.44852 | 0.0507269 | 0.351232011 |
| ENSMUSG00000032309 | Fbxo22 | protein_coding | 0.962752744 | 4.51369685 | 4.4477845 | 0.0507439 | 0.351232011 |
| ENSMUSG00000055839 | Elob | protein_coding | 0.535285401 | 9.024725631 | 4.4472471 | 0.0507563 | 0.351232011 |
| ENSMUSG00000022949 | Clic6 | protein_coding | -1.702066242 | 4.046555903 | 4.5189794 | 0.0507593 | 0.351232011 |
| ENSMUSG00000051497 | Kcnj16 | protein_coding | -1.23839781 | 3.231320837 | 4.4417634 | 0.0508832 | 0.351863244 |
| ENSMUSG00000070923 | Klhl9 | protein_coding | -0.52616939 | 8.232878104 | 4.438784 | 0.0509524 | 0.352114798 |
| ENSMUSG00000034007 | Scaper | protein_coding | -0.657125794 | 5.731327711 | 4.434341 | 0.0510556 | 0.35222807 |
| ENSMUSG00000027581 | Stmn3 | protein_coding | 0.516185816 | 11.21256015 | 4.4330079 | 0.0510867 | 0.35222807 |
| ENSMUSG00000019852 | Arfgef3 | protein_coding | -0.665986864 | 5.896840829 | 4.431991 | 0.0511104 | 0.35222807 |
| ENSMUSG00000002012 | Pnck | protein_coding | 0.629690505 | 9.46024771 | 4.4297675 | 0.0511622 | 0.35222807 |

| ensembl_gene_id | external_gene_name | gene_biotype | log2 (fold change) | log2 (counts per million) | F-value | p-value | False discovery rate |
|---|---|---|---|---|---|---|---|
| ENSMUSG00000028766 | Alpl | protein_coding | 2.570719099 | 2.546182813 | 4.4289078 | 0.0512094 | 0.35222807 |
| ENSMUSG00000039967 | Zfp292 | protein_coding | -0.943713695 | 4.74236974 | 4.4276052 | 0.0512127 | 0.35222807 |
| ENSMUSG00000041328 | Pcf11 | protein_coding | -1.112694786 | 3.966971098 | 4.4269268 | 0.0512285 | 0.35222807 |
| ENSMUSG00000037847 | Nmrk1 | protein_coding | -1.219174943 | 3.704456929 | 4.4268302 | 0.0512308 | 0.35222807 |
| ENSMUSG00000030102 | Itpr1 | protein_coding | -0.861780108 | 6.325948694 | 4.4677641 | 0.0512669 | 0.352251122 |
| ENSMUSG00000059482 | 2610301B20Rik | protein_coding | -0.520871593 | 6.793202852 | 4.4210688 | 0.0513656 | 0.35250953 |
| ENSMUSG00000071669 | Snx29 | protein_coding | -1.211335175 | 3.290606917 | 4.4206837 | 0.0513746 | 0.35250953 |
| ENSMUSG00000021730 | Hcn1 | protein_coding | -0.774757923 | 4.651421315 | 4.4170424 | 0.0514601 | 0.35250953 |
| ENSMUSG00000002458 | Rgs19 | protein_coding | 0.609000686 | 5.987328493 | 4.4163053 | 0.0514774 | 0.35250953 |
| ENSMUSG00000037938 | Chchd5 | protein_coding | 0.51058797 | 6.292169243 | 4.4149572 | 0.0515091 | 0.35250953 |
| ENSMUSG00000035382 | Pcsk7 | protein_coding | 1.330031789 | 3.056818934 | 4.4119164 | 0.0515807 | 0.35250953 |
| ENSMUSG00000000346 | Dazap2 | protein_coding | -0.501261022 | 8.570230436 | 4.4115713 | 0.0515888 | 0.35250953 |
| ENSMUSG00000038418 | Egr1 | protein_coding | -0.654235812 | 6.746067033 | 4.4107945 | 0.0516071 | 0.35250953 |
| ENSMUSG00000059474 | Mbtd1 | protein_coding | -1.050787336 | 3.747940993 | 4.4087238 | 0.0516559 | 0.35250953 |
| ENSMUSG00000026796 | Fam129b | protein_coding | 1.338906639 | 2.90440795 | 4.4079406 | 0.0516744 | 0.35250953 |
| ENSMUSG00000039713 | Plekhg5 | protein_coding | 0.899562272 | 4.772621883 | 4.4074358 | 0.0516863 | 0.35250953 |
| ENSMUSG00000033382 | Trappc8 | protein_coding | -1.103241579 | 3.871704066 | 4.4069452 | 0.0516979 | 0.35250953 |
| ENSMUSG00000020260 | Pofut2 | protein_coding | 0.876378643 | 5.643181593 | 4.4036106 | 0.0518039 | 0.352851475 |
| ENSMUSG00000038708 | Golga4 | protein_coding | -0.660128119 | 5.144764638 | 4.3989501 | 0.0518871 | 0.352851475 |
| ENSMUSG00000062647 | Rpl7a | protein_coding | 0.523101129 | 6.067649454 | 4.3975646 | 0.05192 | 0.352851475 |
| ENSMUSG00000026277 | Stk25 | protein_coding | 0.538677737 | 7.586585753 | 4.3970052 | 0.0519333 | 0.352851475 |
| ENSMUSG00000024811 | Tnks2 | protein_coding | -0.88335598 | 5.379517139 | 4.3975229 | 0.0519469 | 0.352851475 |
| ENSMUSG00000009734 | Pou6f2 | protein_coding | -2.305050615 | 2.355691237 | 4.3962098 | 0.0519521 | 0.352851475 |
| ENSMUSG00000034160 | Ogt | protein_coding | -0.783710314 | 5.72437829 | 4.3947791 | 0.0519861 | 0.352851475 |
| ENSMUSG00000097911 | Gm26691 | antisense_RNA | -1.511719032 | 2.904787901 | 4.3937515 | 0.0520106 | 0.352851475 |
| ENSMUSG00000025092 | Hspa12a | protein_coding | 0.582438042 | 5.960480157 | 4.3912125 | 0.052071 | 0.353038652 |
| ENSMUSG00000060098 | Prmt7 | protein_coding | 1.131209202 | 5.276261344 | 4.4257108 | 0.0523091 | 0.354142573 |
| ENSMUSG00000009640 | Dmap1 | protein_coding | 0.615587307 | 5.696855416 | 4.3810491 | 0.0523137 | 0.354142573 |
| ENSMUSG00000028080 | Lrba | protein_coding | -1.246057398 | 3.827978285 | 4.3792598 | 0.0523565 | 0.354142573 |
| ENSMUSG00000022982 | Sod1 | protein_coding | 0.614341372 | 10.04575214 | 4.3788834 | 0.0523655 | 0.354142573 |
| ENSMUSG00000024911 | Fibp | protein_coding | 0.531489485 | 6.886098027 | 4.3754768 | 0.0524473 | 0.354443547 |
| ENSMUSG00000034424 | Gcsh | protein_coding | 0.490229225 | 7.694820109 | 4.373273 | 0.0525002 | 0.354443547 |
| ENSMUSG00000022507 | 1810013L24Rik | protein_coding | -0.498838771 | 6.407993132 | 4.3729108 | 0.0525089 | 0.354443547 |
| ENSMUSG00000022174 | Dad1 | protein_coding | 0.513404768 | 9.050245605 | 4.3709685 | 0.0525557 | 0.354536426 |
| ENSMUSG00000025362 | Rps26 | protein_coding | 0.692898854 | 7.665758863 | 4.3691815 | 0.052698 | 0.355273426 |
| ENSMUSG00000030704 | Rab6a | protein_coding | -0.467575289 | 8.306047026 | 4.3613544 | 0.0527877 | 0.355525012 |
| ENSMUSG00000049760 | 2410015M20Rik | protein_coding | 0.542414979 | 8.724607971 | 4.3607875 | 0.0528014 | 0.355525012 |
| ENSMUSG00000008976 | Gabpa | protein_coding | -1.303330514 | 2.888751968 | 4.3528198 | 0.0529946 | 0.356602892 |
| ENSMUSG00000000374 | Trappc10 | protein_coding | -0.80138774 | 4.455311871 | 4.3495636 | 0.0530739 | 0.356862512 |
| ENSMUSG00000038517 | Tbkbp1 | protein_coding | -1.483415749 | 3.123857561 | 4.3485065 | 0.0530996 | 0.356862512 |
| ENSMUSG00000032744 | Heyl | protein_coding | -1.242194937 | 3.977557591 | 4.3463701 | 0.0531791 | 0.357081671 |
| ENSMUSG00000022565 | Plec | protein_coding | -0.777445041 | 4.769189975 | 4.3444461 | 0.0531986 | 0.357081671 |
| ENSMUSG00000052613 | Pcdh15 | protein_coding | -1.013854629 | 4.051659865 | 4.3406529 | 0.0532913 | 0.357260559 |
| ENSMUSG00000004151 | Etv1 | protein_coding | -0.694614716 | 5.695036509 | 4.3400673 | 0.0533057 | 0.357260559 |
| ENSMUSG00000058325 | Dock1 | protein_coding | -0.973335743 | 4.34361608 | 4.3376995 | 0.0533636 | 0.357260559 |
| ENSMUSG00000068328 | Aup1 | protein_coding | 1.011393869 | 6.236428025 | 4.4253651 | 0.0534117 | 0.357260559 |
| ENSMUSG00000078435 | AU041133 | protein_coding | -1.519658995 | 2.924900475 | 4.3356818 | 0.0534131 | 0.357260559 |
| ENSMUSG00000071657 | Bscl2 | protein_coding | 0.487570417 | 7.871977081 | 4.3352121 | 0.0534246 | 0.357260559 |
| ENSMUSG00000039208 | Metrnl | protein_coding | 1.71405928 | 2.675614047 | 4.3327352 | 0.0534854 | 0.35727165 |
| ENSMUSG00000058833 | 2810428I15Rik | protein_coding | 0.569472979 | 7.558352681 | 4.3289253 | 0.0535791 | 0.35727165 |
| ENSMUSG00000022428 | Cby1 | protein_coding | 0.510860375 | 7.321213859 | 4.3286693 | 0.0535854 | 0.35727165 |
| ENSMUSG00000100252 | Mir124-2hg | lincRNA | -0.691214019 | 5.424679353 | 4.3284223 | 0.0535915 | 0.35727165 |
| ENSMUSG00000017831 | Rab5a | protein_coding | -0.943092769 | 5.016698875 | 4.328386 | 0.0535924 | 0.35727165 |
| ENSMUSG00000054640 | Slc8a1 | protein_coding | -1.310301827 | 3.27902357 | 4.3245667 | 0.0536866 | 0.357600144 |
| ENSMUSG00000008763 | Man1a2 | protein_coding | -0.793066608 | 5.403876338 | 4.3220608 | 0.0537484 | 0.357600144 |
| ENSMUSG00000039206 | Daglb | protein_coding | 0.732625823 | 4.974757951 | 4.3219942 | 0.0537501 | 0.357600144 |
| ENSMUSG00000031604 | Msmo1 | protein_coding | -0.471480557 | 7.175170499 | 4.3198686 | 0.0538026 | 0.357600144 |
| ENSMUSG00000031539 | Ap3m2 | protein_coding | 0.668435963 | 7.416457161 | 4.3203869 | 0.0538173 | 0.357600144 |
| ENSMUSG00000031983 | 2310022B05Rik | protein_coding | -0.83052236 | 4.985790845 | 4.3183091 | 0.0538412 | 0.357600144 |
| ENSMUSG00000025475 | Adgra1 | protein_coding | 0.678535542 | 6.688221718 | 4.3180472 | 0.0538752 | 0.357604906 |
| ENSMUSG00000011658 | Fuz | protein_coding | 1.684806231 | 2.572785091 | 4.3137934 | 0.0539532 | 0.357901457 |
| ENSMUSG00000004565 | Pnpla6 | protein_coding | 0.806599984 | 4.932264495 | 4.3095958 | 0.0540574 | 0.358372194 |
| ENSMUSG00000028098 | Rnf115 | protein_coding | 0.599401427 | 5.421290025 | 4.3025658 | 0.0542326 | 0.359066237 |
| ENSMUSG00000027010 | Slc25a12 | protein_coding | 0.77672973 | 7.805494318 | 4.3454695 | 0.0542371 | 0.359066237 |
| ENSMUSG00000042042 | Csgalnact2 | protein_coding | -1.650487172 | 2.890965698 | 4.3013776 | 0.0542623 | 0.359066237 |
| ENSMUSG00000044807 | Zfp354c | protein_coding | -0.864263984 | 4.196527264 | 4.2947845 | 0.0544273 | 0.359936649 |
| ENSMUSG00000006390 | Elovl1 | protein_coding | -0.775068964 | 4.990622228 | 4.2921405 | 0.0544937 | 0.360039357 |
| ENSMUSG00000061762 | Tac1 | protein_coding | -0.887902584 | 8.511015986 | 4.379526 | 0.0545098 | 0.360039357 |
| ENSMUSG00000040620 | Dhx33 | protein_coding | -1.430680331 | 2.79761533 | 4.289115 | 0.0545697 | 0.360213508 |
| ENSMUSG00000028381 | Ugcg | protein_coding | -0.670690519 | 5.413771416 | 4.2876482 | 0.0546066 | 0.360235963 |
| ENSMUSG00000033060 | Lmo7 | protein_coding | -1.192777056 | 3.790753184 | 4.2816512 | 0.0547578 | 0.360996852 |
| ENSMUSG00000025855 | Prkar1b | protein_coding | 0.746807829 | 10.12034065 | 4.2862792 | 0.0547891 | 0.360996852 |
| ENSMUSG00000048834 | Vstm2a | protein_coding | -0.515686557 | 6.941873829 | 4.2781006 | 0.0548475 | 0.361083776 |
| ENSMUSG00000053581 | Zfand2a | protein_coding | 0.767928829 | 6.114160358 | 4.2774959 | 0.0548904 | 0.361083776 |
| ENSMUSG00000020364 | Zfp354a | protein_coding | -1.31292243 | 3.351531826 | 4.2758498 | 0.0549045 | 0.361083776 |
| ENSMUSG00000037174 | Elf2 | protein_coding | -0.98092345 | 3.947263548 | 4.2745825 | 0.0549366 | 0.361083776 |
| ENSMUSG00000025823 | Pdia4 | protein_coding | 0.605887311 | 5.920511761 | 4.2711384 | 0.055024 | 0.361421757 |
| ENSMUSG00000029223 | Uchl1 | protein_coding | 0.481877635 | 10.33781782 | 4.2699073 | 0.0550552 | 0.361421757 |

| ensembl_gene_id | external_gene_name | gene_biotype | log2 (fold change) | log2 (counts per million) | F-value | p-value | False discovery rate |
|---|---|---|---|---|---|---|---|
| ENSMUSG00000004031 | Brinp2 | protein_coding | 0.761741247 | 4.603007488 | 4.266256 | 0.0551481 | 0.361680921 |
| ENSMUSG00000000266 | Mid2 | protein_coding | -1.069809043 | 4.204205419 | 4.2638791 | 0.0552086 | 0.361680921 |
| ENSMUSG00000066568 | Lsm14a | protein_coding | -0.609774201 | 5.43454583 | 4.2626954 | 0.0552388 | 0.361680921 |
| ENSMUSG00000020077 | Srgn | protein_coding | -1.753093117 | 2.586494069 | 4.2617099 | 0.055264 | 0.361680921 |
| ENSMUSG00000025193 | Cutc | protein_coding | -0.810732286 | 4.378400117 | 4.2608877 | 0.055285 | 0.361680921 |
| ENSMUSG00000037049 | Smpd1 | protein_coding | 0.561062002 | 6.9351457 | 4.258342 | 0.05535 | 0.361680921 |
| ENSMUSG00000057841 | Rpl32 | protein_coding | 0.61407403 | 9.869372076 | 4.2577675 | 0.0553647 | 0.361680921 |
| ENSMUSG00000066441 | Rdh11 | protein_coding | -0.781665714 | 4.662928458 | 4.2566825 | 0.0553925 | 0.361680921 |
| ENSMUSG00000042271 | Nxt2 | protein_coding | -0.657782972 | 6.680657324 | 4.2575713 | 0.0553974 | 0.361680921 |
| ENSMUSG00000028648 | Ndufs5 | protein_coding | 0.502912443 | 7.458033273 | 4.2527732 | 0.0554926 | 0.362082477 |
| ENSMUSG00000028411 | Aptx | protein_coding | 0.496649325 | 6.507321975 | 4.2462151 | 0.0556611 | 0.362633363 |
| ENSMUSG00000030313 | Dennd5b | protein_coding | -0.648446099 | 4.922465207 | 4.245004 | 0.0556923 | 0.362633363 |
| ENSMUSG00000057098 | Ebf1 | protein_coding | 0.893656975 | 3.995808102 | 4.2447665 | 0.0556984 | 0.362633363 |
| ENSMUSG00000026674 | Ddr2 | protein_coding | -1.386676368 | 2.860090175 | 4.2434346 | 0.0557327 | 0.362633363 |
| ENSMUSG00000019986 | Ahi1 | protein_coding | 0.61447692 | 8.7779824 | 4.2429318 | 0.0557456 | 0.362633363 |
| ENSMUSG00000021534 | 1700001L19Rik | protein_coding | 0.626726997 | 6.731267775 | 4.2369678 | 0.0558997 | 0.363415322 |
| ENSMUSG00000038766 | Gabpb2 | protein_coding | -1.644402628 | 2.501157165 | 4.2320368 | 0.0560274 | 0.364025466 |
| ENSMUSG00000009394 | Syn2 | protein_coding | 0.481160446 | 8.464177171 | 4.2288374 | 0.0561104 | 0.36434486 |
| ENSMUSG00000013822 | Elof1 | protein_coding | 0.457268131 | 8.163650204 | 4.2245132 | 0.0562229 | 0.364445458 |
| ENSMUSG00000026782 | Abi2 | protein_coding | -0.513274825 | 7.19286247 | 4.2243655 | 0.0562267 | 0.364445458 |
| ENSMUSG00000027189 | Trim44 | protein_coding | -0.542779111 | 6.742039693 | 4.2243325 | 0.0562276 | 0.364445458 |
| ENSMUSG00000054723 | Vmac | protein_coding | -1.122821505 | 3.522435072 | 4.2206888 | 0.0563225 | 0.364777876 |
| ENSMUSG00000051910 | Sox6 | protein_coding | -1.222868788 | 3.748186388 | 4.2197632 | 0.0563467 | 0.364777876 |
| ENSMUSG00000067279 | Ppp1r3c | protein_coding | -0.912964699 | 4.881953259 | 4.2159103 | 0.0564474 | 0.365196946 |
| ENSMUSG00000034593 | Myo5a | protein_coding | -0.630383952 | 7.076444363 | 4.215753 | 0.0564794 | 0.365196946 |
| ENSMUSG00000040774 | Cept1 | protein_coding | 0.521912081 | 6.258797858 | 4.2086371 | 0.056638 | 0.366002794 |
| ENSMUSG00000025531 | Chm | protein_coding | -0.84576646 | 4.489383728 | 4.2020849 | 0.0568104 | 0.366855524 |
| ENSMUSG00000041708 | Mpped1 | protein_coding | 0.647019013 | 5.241067827 | 4.1961421 | 0.0569673 | 0.366855524 |
| ENSMUSG00000030980 | Knop1 | protein_coding | 0.627745759 | 5.433076153 | 4.1960522 | 0.0569697 | 0.366855524 |
| ENSMUSG00000028278 | Rragd | protein_coding | -0.587296326 | 6.228608853 | 4.1958418 | 0.0569752 | 0.366855524 |
| ENSMUSG00000022489 | Pde1b | protein_coding | -0.525727942 | 7.601957761 | 4.1940524 | 0.0570226 | 0.366855524 |
| ENSMUSG00000045967 | Gpr158 | protein_coding | -0.707092703 | 5.215833255 | 4.1938349 | 0.0570283 | 0.366855524 |
| ENSMUSG00000021775 | Nr1d2 | protein_coding | -0.63725503 | 5.451792979 | 4.1933995 | 0.0570399 | 0.366855524 |
| ENSMUSG00000040138 | Ndp | protein_coding | -0.851392439 | 4.483514932 | 4.1932839 | 0.0570429 | 0.366855524 |
| ENSMUSG00000024944 | Arl2 | protein_coding | 0.456629926 | 9.161043313 | 4.1866798 | 0.0572181 | 0.367252993 |
| ENSMUSG00000045237 | 1110012L19Rik | protein_coding | 0.5513648 | 5.830513754 | 4.1854856 | 0.0572499 | 0.367252993 |
| ENSMUSG00000019820 | Utrn | protein_coding | -0.738082798 | 5.03436791 | 4.1854837 | 0.0572499 | 0.367252993 |
| ENSMUSG00000011960 | Ccnt1 | protein_coding | 0.700867277 | 5.296576263 | 4.1836589 | 0.0572985 | 0.367252993 |
| ENSMUSG00000025576 | Rbfox3 | protein_coding | 0.500276248 | 6.697180347 | 4.1833578 | 0.0573065 | 0.367252993 |
| ENSMUSG00000001520 | Nrip2 | protein_coding | 0.704939544 | 6.737210411 | 4.1842899 | 0.0573096 | 0.367252993 |
| ENSMUSG00000031667 | Aktip | protein_coding | 0.535601561 | 8.165522375 | 4.1786308 | 0.0574325 | 0.367480628 |
| ENSMUSG00000073433 | Arhgdig | protein_coding | 0.671450114 | 5.627176082 | 4.1785007 | 0.057436 | 0.367480628 |
| ENSMUSG00000021969 | Zdhhc20 | protein_coding | -0.560054792 | 6.100182519 | 4.1771817 | 0.0574712 | 0.367480628 |
| ENSMUSG00000042707 | Dnali1 | protein_coding | -1.405256264 | 3.274985517 | 4.1767824 | 0.0574819 | 0.367480628 |
| ENSMUSG00000037493 | Cib2 | protein_coding | 0.672135188 | 5.405799481 | 4.1750555 | 0.057528 | 0.367557218 |
| ENSMUSG00000025545 | Clybl | protein_coding | -0.478755809 | 7.084661771 | 4.1721488 | 0.0576058 | 0.367759341 |
| ENSMUSG00000059456 | Ptk2b | protein_coding | -0.559695704 | 7.305700514 | 4.1713188 | 0.0576281 | 0.367759341 |
| ENSMUSG00000044986 | Tst | protein_coding | -0.775828791 | 6.522073545 | 4.1912934 | 0.0576624 | 0.36776012 |
| ENSMUSG00000028292 | Rars2 | protein_coding | -0.656300751 | 5.453790549 | 4.166374 | 0.0577607 | 0.368169127 |
| ENSMUSG00000051502 | Ufsp1 | protein_coding | 0.969757091 | 3.98069418 | 4.1613724 | 0.0578953 | 0.368808329 |
| ENSMUSG00000022698 | Naa50 | protein_coding | -0.534966096 | 5.952970426 | 4.1573846 | 0.0580029 | 0.369274758 |
| ENSMUSG00000035681 | Kcnc2 | protein_coding | -0.804703367 | 5.261467745 | 4.1518851 | 0.0581516 | 0.370002508 |
| ENSMUSG00000020834 | Dhrs13 | protein_coding | 1.837892414 | 2.554185785 | 4.1446611 | 0.0583476 | 0.370813103 |
| ENSMUSG00000015747 | Vps45 | protein_coding | 0.546350739 | 6.472370461 | 4.1446488 | 0.058348 | 0.370813103 |
| ENSMUSG00000021621 | Zcchc9 | protein_coding | 0.89535251 | 4.912392051 | 4.1428515 | 0.0583969 | 0.370904599 |
| ENSMUSG00000027665 | Pik3ca | protein_coding | -0.64050667 | 5.507951303 | 4.1401882 | 0.0584694 | 0.371146097 |
| ENSMUSG00000057378 | Ryr3 | protein_coding | -1.31126218 | 3.667007566 | 4.1388929 | 0.0585328 | 0.371329623 |
| ENSMUSG00000024357 | Sil1 | protein_coding | 0.589831792 | 6.185084574 | 4.1354099 | 0.0585998 | 0.371535218 |
| ENSMUSG00000007892 | Rplp1 | protein_coding | 0.63174811 | 9.342253724 | 4.1315015 | 0.0587349 | 0.372012617 |
| ENSMUSG00000025920 | Stau2 | protein_coding | -0.445050137 | 7.064248822 | 4.127912 | 0.0588051 | 0.372012617 |
| ENSMUSG00000037979 | Ccdc92 | protein_coding | 0.58820341 | 6.192813252 | 4.1270233 | 0.0588295 | 0.372012617 |
| ENSMUSG00000007836 | Hnrnpa0 | protein_coding | -0.555331996 | 7.269052776 | 4.1263959 | 0.0588467 | 0.372012617 |
| ENSMUSG00000034371 | Tkfc | protein_coding | 0.957970013 | 3.857554488 | 4.1263467 | 0.0588481 | 0.372012617 |
| ENSMUSG00000059493 | Nhs | protein_coding | -1.08484407 | 3.366856046 | 4.1243564 | 0.0589028 | 0.372139556 |
| ENSMUSG00000090015 | Gm15446 | protein_coding | -1.563377518 | 3.140716799 | 4.1230536 | 0.0589386 | 0.372147261 |
| ENSMUSG00000063457 | Rps15 | protein_coding | 0.604375664 | 8.929819648 | 4.1190086 | 0.05905 | 0.372631875 |
| ENSMUSG00000028741 | Mrto4 | protein_coding | 0.567304827 | 6.218993079 | 4.1176081 | 0.0590886 | 0.372656945 |
| ENSMUSG00000030137 | Tuba8 | protein_coding | 1.568905977 | 3.06575667 | 4.1160419 | 0.0591319 | 0.372711024 |
| ENSMUSG00000021282 | Eif5 | protein_coding | -0.464280617 | 6.971456499 | 4.1141194 | 0.059185 | 0.372827306 |
| ENSMUSG00000031633 | Slc25a4 | protein_coding | 0.489582793 | 11.73553787 | 4.1105716 | 0.0592832 | 0.372891958 |
| ENSMUSG00000059554 | Ccdc28a | protein_coding | 0.548808304 | 6.973733931 | 4.1094788 | 0.0593134 | 0.372891958 |
| ENSMUSG00000031714 | Gab1 | protein_coding | -0.823444877 | 4.408208257 | 4.1061604 | 0.0594055 | 0.372891958 |
| ENSMUSG00000034413 | Neurl1b | protein_coding | -1.401260425 | 3.442703507 | 4.1054267 | 0.0594259 | 0.372891958 |
| ENSMUSG00000038642 | Ctss | protein_coding | -0.942363497 | 6.462792108 | 4.1845632 | 0.0594884 | 0.372891958 |
| ENSMUSG00000028923 | Necap2 | protein_coding | -0.677605017 | 5.508739971 | 4.1029579 | 0.0594945 | 0.372891958 |
| ENSMUSG00000100837 | 1700063D05Rik | lincRNA | -1.129251425 | 3.539659676 | 4.1025883 | 0.0595048 | 0.372891958 |
| ENSMUSG00000048240 | Gng7 | protein_coding | -0.611578216 | 8.62343048 | 4.1007565 | 0.0595558 | 0.372891958 |
| ENSMUSG00000029833 | Trim24 | protein_coding | -0.884453038 | 3.973895543 | 4.1005936 | 0.0595603 | 0.372891958 |

| ensembl_gene_id | external_gene_name | gene_biotype | log2 (fold change) | log2 (counts per million) | F-value | p-value | False discovery rate |
|---|---|---|---|---|---|---|---|
| ENSMUSG00000028430 | Nol6 | protein_coding | 0.645146578 | 6.081971905 | 4.0999971 | 0.0595769 | 0.372891958 |
| ENSMUSG00000020873 | Slc35b1 | protein_coding | 0.462616348 | 7.164479832 | 4.0962443 | 0.0596816 | 0.372891958 |
| ENSMUSG00000052560 | Cpne8 | protein_coding | -0.703670689 | 5.446482968 | 4.0952856 | 0.0597083 | 0.372891958 |
| ENSMUSG00000050786 | Ccdc126 | protein_coding | 1.027445294 | 3.822217161 | 4.0949548 | 0.0597176 | 0.372891958 |
| ENSMUSG00000071064 | Zfp827 | protein_coding | -0.834182846 | 4.348245981 | 4.0922216 | 0.059794 | 0.372891958 |
| ENSMUSG00000036840 | Siah1a | protein_coding | -0.993203999 | 3.974477674 | 4.0919129 | 0.0598026 | 0.372891958 |
| ENSMUSG00000019984 | Med23 | protein_coding | -0.912863488 | 3.592601219 | 4.0911004 | 0.0598254 | 0.372891958 |
| ENSMUSG00000031715 | Smarca5 | protein_coding | -0.714264903 | 5.237627904 | 4.0903054 | 0.0598476 | 0.372891958 |
| ENSMUSG00000034021 | Pds5b | protein_coding | -0.542380416 | 6.033313339 | 4.0882204 | 0.0599061 | 0.372891958 |
| ENSMUSG00000061046 | Haghl | protein_coding | 0.475451229 | 7.77156325 | 4.0881121 | 0.0599091 | 0.372891958 |
| ENSMUSG00000092416 | Zfp141 | protein_coding | -1.260008088 | 3.008401772 | 4.0873994 | 0.0599291 | 0.372891958 |
| ENSMUSG00000069662 | Marcks | protein_coding | -0.444063308 | 7.706302332 | 4.084139 | 0.0600206 | 0.372891958 |
| ENSMUSG00000027405 | Nop56 | protein_coding | 0.555092363 | 7.276843481 | 4.0838833 | 0.0600278 | 0.372891958 |
| ENSMUSG00000028527 | Ak4 | protein_coding | 0.765060257 | 4.634569626 | 4.0836844 | 0.0600334 | 0.372891958 |
| ENSMUSG00000060568 | Fam78b | protein_coding | -1.145693992 | 3.485171774 | 4.0830267 | 0.0600519 | 0.372891958 |
| ENSMUSG00000064349 | mt-Tc | Mt_tRNA | -1.346748511 | 2.982255113 | 4.0826601 | 0.0600622 | 0.372891958 |
| ENSMUSG00000028636 | Ppcs | protein_coding | 1.242806934 | 4.117834814 | 4.0831068 | 0.0601526 | 0.37323765 |
| ENSMUSG00000021803 | Cdhr1 | protein_coding | -1.646822987 | 2.655449623 | 4.07731 | 0.0602129 | 0.37331376 |
| ENSMUSG00000056383 | AI987944 | protein_coding | -1.545945564 | 3.222489646 | 4.077556 | 0.0602343 | 0.37331376 |
| ENSMUSG00000025173 | Wdr45b | protein_coding | 0.501296855 | 6.486144189 | 4.0700692 | 0.0604175 | 0.374233752 |
| ENSMUSG00000056214 | Pard6g | protein_coding | 1.324076311 | 3.154507109 | 4.0652341 | 0.0605546 | 0.3747365 |
| ENSMUSG00000041362 | Shtn1 | protein_coding | -0.674604587 | 5.496170292 | 4.0636528 | 0.0605996 | 0.3747365 |
| ENSMUSG00000057193 | Slc44a2 | protein_coding | -1.109813832 | 3.950299999 | 4.0629375 | 0.0606199 | 0.3747365 |
| ENSMUSG00000061207 | Stk19 | protein_coding | 0.521282947 | 6.162909754 | 4.0599613 | 0.0607046 | 0.3747365 |
| ENSMUSG00000033233 | Trim45 | protein_coding | 1.483108084 | 3.97313716 | 4.0978204 | 0.0607348 | 0.3747365 |
| ENSMUSG00000027900 | Dram2 | protein_coding | -0.632533611 | 5.679446548 | 4.0578686 | 0.0607642 | 0.3747365 |
| ENSMUSG00000057440 | Mpp7 | protein_coding | -1.211097702 | 3.335569132 | 4.0574914 | 0.060775 | 0.3747365 |
| ENSMUSG00000031592 | Pcm1 | protein_coding | -0.762397257 | 5.501152417 | 4.0574022 | 0.0607775 | 0.3747365 |
| ENSMUSG00000042675 | Ypel3 | protein_coding | 0.622270682 | 6.043903776 | 4.0554617 | 0.0608329 | 0.374862918 |
| ENSMUSG00000028300 | 3110043O21Rik | protein_coding | 0.743651383 | 5.315404608 | 4.050788 | 0.0609665 | 0.37538936 |
| ENSMUSG00000001988 | Npas1 | protein_coding | 1.464541559 | 3.054201747 | 4.0492392 | 0.0610108 | 0.37538936 |
| ENSMUSG00000035441 | Myo1d | protein_coding | -0.804046343 | 4.208263697 | 4.0488123 | 0.061023 | 0.37538936 |
| ENSMUSG00000030707 | Coro1a | protein_coding | 0.491094179 | 7.390618051 | 4.0429946 | 0.06119 | 0.37620115 |
| ENSMUSG00000060534 | Dcc | protein_coding | -1.336457311 | 3.846926997 | 4.0449523 | 0.0612372 | 0.376276128 |
| ENSMUSG00000031242 | 2610002M06Rik | protein_coding | -0.572360609 | 5.652742903 | 4.0375657 | 0.0613462 | 0.376731123 |
| ENSMUSG00000064181 | Rab3ip | protein_coding | 0.770076037 | 4.458143797 | 4.0343466 | 0.0614391 | 0.376984585 |
| ENSMUSG00000021338 | Carmil1 | protein_coding | -1.074070911 | 3.396781773 | 4.0327016 | 0.0614867 | 0.376984585 |
| ENSMUSG00000034175 | Rhbdd3 | protein_coding | 0.75986392 | 4.91049185 | 4.0324927 | 0.0614927 | 0.376984585 |
| ENSMUSG00000037960 | Card19 | protein_coding | -0.89620293 | 4.101817204 | 4.0264736 | 0.061667 | 0.377759754 |
| ENSMUSG00000039244 | E130309D02Rik | protein_coding | 0.918838774 | 3.943347011 | 4.0246821 | 0.061719 | 0.377759754 |
| ENSMUSG00000052151 | Plpp2 | protein_coding | 0.786911298 | 4.438587461 | 4.0233452 | 0.0617578 | 0.377759754 |
| ENSMUSG00000040591 | 1110051M20Rik | protein_coding | 0.555632635 | 8.374392058 | 4.022114 | 0.0617936 | 0.377759754 |
| ENSMUSG00000029535 | Triap1 | protein_coding | 0.549210009 | 6.337030239 | 4.0220725 | 0.0617948 | 0.377759754 |
| ENSMUSG00000041346 | Wrap53 | protein_coding | -0.692078765 | 4.948486831 | 4.0177863 | 0.0619196 | 0.37820298 |
| ENSMUSG00000045867 | Cradd | protein_coding | 0.685552513 | 4.529806171 | 4.017166 | 0.0619376 | 0.37820298 |
| ENSMUSG00000022560 | Slc52a2 | protein_coding | 0.948428459 | 4.264855553 | 4.0155888 | 0.0619836 | 0.37826906 |
| ENSMUSG00000015980 | Lrrc27 | protein_coding | 0.979785419 | 4.809992088 | 4.0147382 | 0.062037 | 0.378276172 |
| ENSMUSG00000021843 | Ktn1 | protein_coding | -0.677675896 | 5.683007978 | 4.0124914 | 0.0620741 | 0.378276172 |
| ENSMUSG00000025132 | Arhgdia | protein_coding | 0.513816363 | 9.882980384 | 4.0119356 | 0.0620903 | 0.378276172 |
| ENSMUSG00000002103 | Acp2 | protein_coding | 1.180833398 | 4.573478707 | 4.0332734 | 0.0621975 | 0.37855816 |
| ENSMUSG00000063800 | Prpf38a | protein_coding | -0.67545093 | 6.011208155 | 4.0079487 | 0.062207 | 0.37855816 |
| ENSMUSG00000031671 | Setd6 | protein_coding | 1.505028559 | 3.194264437 | 4.0055858 | 0.0622763 | 0.378609403 |
| ENSMUSG00000002332 | Dhrs1 | protein_coding | 0.485285813 | 7.310267359 | 4.0037269 | 0.0623309 | 0.378609403 |
| ENSMUSG00000033938 | Ndufb7 | protein_coding | 0.511148531 | 9.54847961 | 4.0033095 | 0.0623431 | 0.378609403 |
| ENSMUSG00000040043 | Rbms2 | protein_coding | -1.016231167 | 3.612148986 | 4.0028619 | 0.0623563 | 0.378609403 |
| ENSMUSG00000039477 | Tnrc18 | protein_coding | 0.996436376 | 3.305940623 | 4.0008756 | 0.0624147 | 0.378750101 |
| ENSMUSG00000018800 | Abca5 | protein_coding | -0.885201052 | 3.999379252 | 3.9996508 | 0.0624507 | 0.378755059 |
| ENSMUSG00000031227 | Magee1 | protein_coding | 0.445877094 | 6.897244192 | 3.9957305 | 0.0625663 | 0.379241793 |
| ENSMUSG00000064247 | Plcxd1 | protein_coding | 0.711301259 | 4.633945197 | 3.9885322 | 0.062779 | 0.380233144 |
| ENSMUSG00000045594 | Glb1 | protein_coding | 1.064578747 | 5.061458579 | 4.0113585 | 0.0628365 | 0.380233144 |
| ENSMUSG00000078640 | Gm11627 | protein_coding | -1.491708293 | 3.102746571 | 3.9865403 | 0.062838 | 0.380233144 |
| ENSMUSG00000050071 | Bex1 | protein_coding | 0.473315188 | 8.358075561 | 3.9801126 | 0.063029 | 0.380233144 |
| ENSMUSG00000024335 | Brd2 | protein_coding | 0.494145036 | 6.404587949 | 3.9789269 | 0.0630643 | 0.380233144 |
| ENSMUSG00000108621 | Gm37494 | lincRNA | -1.354306418 | 2.913498847 | 3.9787884 | 0.0630684 | 0.380233144 |
| ENSMUSG00000070306 | Ccdc153 | protein_coding | -0.782547959 | 5.855935236 | 3.9792792 | 0.0630824 | 0.380233144 |
| ENSMUSG00000026888 | Grb14 | protein_coding | -0.701008199 | 6.202078848 | 3.9783491 | 0.0631101 | 0.380233144 |
| ENSMUSG00000035139 | Secisbp2 | protein_coding | 0.823736348 | 4.315297583 | 3.9773238 | 0.063112 | 0.380233144 |
| ENSMUSG00000003662 | Ciao1 | protein_coding | 0.574348504 | 6.247825886 | 3.9765109 | 0.0631362 | 0.380233144 |
| ENSMUSG00000031865 | Dctn1 | protein_coding | 0.628916217 | 6.87130974 | 3.9765677 | 0.0631632 | 0.380233144 |
| ENSMUSG00000037652 | Phc3 | protein_coding | -0.824392763 | 4.092420459 | 3.9751462 | 0.0631769 | 0.380233144 |
| ENSMUSG00000061755 | Bod1l | protein_coding | -0.780701874 | 5.3902274 | 3.9737282 | 0.0632193 | 0.380233144 |
| ENSMUSG00000029771 | Irf5 | protein_coding | -1.519504413 | 2.881099797 | 3.973541 | 0.0632249 | 0.380233144 |
| ENSMUSG00000004788 | Eif2b2 | protein_coding | 0.519712285 | 6.234374159 | 3.9692494 | 0.0633532 | 0.380488804 |
| ENSMUSG00000033272 | Slc35a4 | protein_coding | 0.611144708 | 6.143286942 | 3.9689924 | 0.0633609 | 0.380488804 |
| ENSMUSG00000018736 | Ndel1 | protein_coding | 0.52973256 | 6.757699419 | 3.96765 | 0.0634011 | 0.380488804 |
| ENSMUSG00000058624 | Gda | protein_coding | -0.560985684 | 6.490384451 | 3.9671712 | 0.0634154 | 0.380488804 |
| ENSMUSG00000029234 | Tmem165 | protein_coding | -1.351951622 | 3.647534826 | 3.9671658 | 0.0634443 | 0.380488804 |
| ENSMUSG00000041997 | Tlk1 | protein_coding | -0.601082114 | 5.412870422 | 3.9634186 | 0.063528 | 0.380741929 |

| ensembl_gene_id | external_gene_name | gene_biotype | log2 (fold change) | log2 (counts per million) | F-value | p-value | False discovery rate |
|---|---|---|---|---|---|---|---|
| ENSMUSG00000062760 | 1810041L15Rik | protein_coding | -0.835096575 | 5.549646128 | 3.9633987 | 0.0635573 | 0.380741929 |
| ENSMUSG00000048249 | Crebrf | protein_coding | -0.501346451 | 6.30893955 | 3.9600854 | 0.0636282 | 0.380850498 |
| ENSMUSG00000030763 | Lcmt1 | protein_coding | 0.450963739 | 7.912261747 | 3.9594841 | 0.0636463 | 0.380850498 |
| ENSMUSG00000030301 | Ccdc91 | protein_coding | 0.599188963 | 5.880553558 | 3.9558133 | 0.0637569 | 0.381300056 |
| ENSMUSG00000025871 | 4833439L19Rik | protein_coding | 0.612914699 | 8.268381697 | 3.9524732 | 0.0638864 | 0.381862662 |
| ENSMUSG00000049755 | Zfp672 | protein_coding | 0.736855499 | 5.688238535 | 3.9462565 | 0.0640458 | 0.382602661 |
| ENSMUSG00000038497 | Tmco3 | protein_coding | 0.807920464 | 5.312085953 | 3.9445713 | 0.0640969 | 0.38260702 |
| ENSMUSG00000028572 | Hook1 | protein_coding | -0.986748451 | 3.667438773 | 3.9438867 | 0.0641177 | 0.38260702 |
| ENSMUSG00000036023 | Parp2 | protein_coding | -0.581125283 | 5.474415451 | 3.9403973 | 0.0642238 | 0.382946997 |
| ENSMUSG00000022885 | St6gal1 | protein_coding | -1.203804297 | 3.565684672 | 3.9396686 | 0.0642459 | 0.382946997 |
| ENSMUSG00000059277 | R74862 | processed_transcript | -1.469021322 | 2.99845578 | 3.9359076 | 0.0643605 | 0.383417297 |
| ENSMUSG00000005667 | Mthfd2 | protein_coding | 0.673847401 | 5.701857152 | 3.9337301 | 0.0644269 | 0.383490072 |
| ENSMUSG00000005533 | Igf1r | protein_coding | -0.885194405 | 4.398673351 | 3.9331698 | 0.064444 | 0.383490072 |
| ENSMUSG00000015002 | Efr3a | protein_coding | -0.560530249 | 5.889607461 | 3.9299334 | 0.0645429 | 0.383563705 |
| ENSMUSG00000022438 | Parvb | protein_coding | -0.92022866 | 3.865977338 | 3.9299139 | 0.0645435 | 0.383563705 |
| ENSMUSG00000058402 | Zfp420 | protein_coding | -0.866352435 | 3.993449678 | 3.9260761 | 0.0646611 | 0.383563705 |
| ENSMUSG00000024999 | Noc3l | protein_coding | -1.474172999 | 2.803657768 | 3.9246419 | 0.0647051 | 0.383563705 |
| ENSMUSG00000021130 | Galnt16 | protein_coding | 0.535410689 | 6.390037046 | 3.922916 | 0.064758 | 0.383563705 |
| ENSMUSG00000091264 | Smim13 | protein_coding | -0.473409406 | 6.418624663 | 3.9217314 | 0.0647944 | 0.383563705 |
| ENSMUSG00000002320 | Tm9sf1 | protein_coding | 1.161362438 | 3.637929839 | 3.9183939 | 0.0648971 | 0.383563705 |
| ENSMUSG00000070802 | Pnmal2 | protein_coding | 0.544911762 | 8.621589177 | 3.9180887 | 0.0649065 | 0.383563705 |
| ENSMUSG00000031176 | Dynlt3 | protein_coding | -0.468893098 | 8.581354738 | 3.9179532 | 0.0649107 | 0.383563705 |
| ENSMUSG00000018761 | Mpdu1 | protein_coding | 0.515925189 | 6.826506948 | 3.9166235 | 0.0649516 | 0.383563705 |
| ENSMUSG00000022391 | Rangap1 | protein_coding | 0.501195363 | 7.905234947 | 3.914665 | 0.065012 | 0.383563705 |
| ENSMUSG00000034220 | Gpc1 | protein_coding | 0.853957041 | 4.569326509 | 3.913344 | 0.0650528 | 0.383563705 |
| ENSMUSG00000037296 | Lsm1 | protein_coding | 0.69654448 | 6.027523324 | 3.9131146 | 0.0650599 | 0.383563705 |
| ENSMUSG00000022474 | Pmm1 | protein_coding | 0.421926387 | 8.903386654 | 3.9121577 | 0.0650894 | 0.383563705 |
| ENSMUSG00000050945 | Zfp438 | protein_coding | 1.611186689 | 3.362616456 | 3.9149607 | 0.0651079 | 0.383563705 |
| ENSMUSG00000022956 | Atp5o | protein_coding | 0.437415802 | 8.53452819 | 3.911411 | 0.0651125 | 0.383563705 |
| ENSMUSG00000028952 | Zbtb48 | protein_coding | 0.899825562 | 4.685661917 | 3.9112189 | 0.0651185 | 0.383563705 |
| ENSMUSG00000003360 | Ddx23 | protein_coding | 1.000125612 | 3.709420481 | 3.9108857 | 0.0651288 | 0.383563705 |
| ENSMUSG00000015714 | Cers2 | protein_coding | -0.548744655 | 6.226918209 | 3.9107119 | 0.0651341 | 0.383563705 |
| ENSMUSG00000079084 | Ccdc82 | protein_coding | -0.89628854 | 4.377176218 | 3.9089955 | 0.0651872 | 0.383666276 |
| ENSMUSG00000048310 | Pskh1 | protein_coding | 1.472589615 | 2.831746232 | 3.9066127 | 0.065261 | 0.383890501 |
| ENSMUSG00000053111 | Fank1 | protein_coding | 0.983675902 | 4.313828583 | 3.9043939 | 0.0653298 | 0.38394279 |
| ENSMUSG00000014932 | Yes1 | protein_coding | -1.777396511 | 2.407524458 | 3.9040235 | 0.0653413 | 0.38394279 |
| ENSMUSG00000033706 | Smyd5 | protein_coding | 0.51829321 | 6.38222208 | 3.901088 | 0.0654325 | 0.384086303 |
| ENSMUSG00000036298 | Slc2a13 | protein_coding | -0.811814571 | 5.510649288 | 3.9018682 | 0.0654372 | 0.384086303 |
| ENSMUSG00000034659 | Tmem109 | protein_coding | 0.632079362 | 6.521911539 | 3.8989906 | 0.0654977 | 0.38423202 |
| ENSMUSG00000037062 | Sh3glb1 | protein_coding | -0.63724762 | 5.881245522 | 3.8974012 | 0.0655472 | 0.384306885 |
| ENSMUSG00000030718 | Ppme1 | protein_coding | 0.696695691 | 7.3473376 | 3.9195825 | 0.0655975 | 0.384306885 |
| ENSMUSG00000026836 | Acvr1 | protein_coding | -0.987802729 | 3.561656619 | 3.8951398 | 0.0656177 | 0.384306885 |
| ENSMUSG00000043162 | Pyurf | protein_coding | 1.230267698 | 5.055327612 | 3.9503957 | 0.0657868 | 0.384878085 |
| ENSMUSG00000021796 | Bmpr1a | protein_coding | -0.83611763 | 4.532161124 | 3.8885016 | 0.0658252 | 0.384878085 |
| ENSMUSG00000053119 | Chmp3 | protein_coding | 0.477179761 | 7.855505258 | 3.884791 | 0.0659414 | 0.384878085 |
| ENSMUSG00000029787 | Avl9 | protein_coding | -0.772232211 | 4.490577768 | 3.8844724 | 0.0659514 | 0.384878085 |
| ENSMUSG00000058446 | Znrf2 | protein_coding | -0.982031966 | 3.726275141 | 3.8831154 | 0.065994 | 0.384878085 |
| ENSMUSG00000056367 | Actr3b | protein_coding | -0.54804964 | 6.426638227 | 3.8823309 | 0.0660187 | 0.384878085 |
| ENSMUSG00000101892 | 9130401M01Rik | protein_coding | 0.707345615 | 4.625813844 | 3.8821695 | 0.0660237 | 0.384878085 |
| ENSMUSG00000022390 | Zc3h7b | protein_coding | -0.780976264 | 5.210253924 | 3.8805942 | 0.0660733 | 0.384878085 |
| ENSMUSG00000108112 | Gm45193 | processed_transcript | -1.082704193 | 3.974443416 | 3.8804528 | 0.0660777 | 0.384878085 |
| ENSMUSG00000040003 | Magi2 | protein_coding | -0.631404525 | 5.226439022 | 3.8762862 | 0.0662089 | 0.384878085 |
| ENSMUSG00000047153 | Khnyn | protein_coding | 1.798938915 | 2.890848492 | 3.875211 | 0.0662718 | 0.384878085 |
| ENSMUSG00000038457 | Tmem255b | protein_coding | -2.472011675 | 2.342129433 | 3.8747629 | 0.0662859 | 0.384878085 |
| ENSMUSG00000042246 | Tmc7 | protein_coding | -1.538131009 | 3.010522308 | 3.8736497 | 0.066292 | 0.384878085 |
| ENSMUSG00000028063 | Lmna | protein_coding | 0.661070398 | 5.095959939 | 3.8731319 | 0.0663084 | 0.384878085 |
| ENSMUSG00000078861 | Zfp931 | protein_coding | -0.92674803 | 5.300697294 | 3.8745789 | 0.0663369 | 0.384878085 |
| ENSMUSG00000047371 | Zfp768 | protein_coding | -1.919290016 | 2.477287658 | 3.87134 | 0.066365 | 0.384878085 |
| ENSMUSG00000004451 | Ralb | protein_coding | 0.678436462 | 4.742937865 | 3.8699426 | 0.0664092 | 0.384878085 |
| ENSMUSG00000024420 | Zfp521 | protein_coding | 0.918223768 | 4.269042558 | 3.8692412 | 0.0664314 | 0.384878085 |
| ENSMUSG00000024457 | Trim26 | protein_coding | 0.806724651 | 4.668947304 | 3.8689228 | 0.0664414 | 0.384878085 |
| ENSMUSG00000024914 | Drap1 | protein_coding | 0.470424021 | 6.442584975 | 3.8686783 | 0.0664492 | 0.384878085 |
| ENSMUSG00000039168 | Dap | protein_coding | -0.824746253 | 4.225448907 | 3.8681185 | 0.0664669 | 0.384878085 |
| ENSMUSG00000087574 | C030037D09Rik | lincRNA | -1.141374961 | 3.542319725 | 3.8660984 | 0.0665309 | 0.384986697 |
| ENSMUSG00000062372 | Otof | protein_coding | 1.307907293 | 3.280649575 | 3.8652669 | 0.0665573 | 0.384986697 |
| ENSMUSG00000027835 | Pdcd10 | protein_coding | -0.55501239 | 6.234613368 | 3.863203 | 0.0666228 | 0.385050245 |
| ENSMUSG00000055493 | Epm2a | protein_coding | -0.855395658 | 4.133105188 | 3.8616971 | 0.0666706 | 0.385050245 |
| ENSMUSG00000027133 | Nop10 | protein_coding | 0.595300244 | 9.048247012 | 3.861537 | 0.0666757 | 0.385050245 |
| ENSMUSG00000029821 | Dfna5 | protein_coding | -0.734132879 | 7.575944082 | 3.8913414 | 0.0668601 | 0.385739458 |
| ENSMUSG00000031245 | Hmgn5 | protein_coding | -1.092610026 | 3.447118563 | 3.8555342 | 0.0668668 | 0.385739458 |
| ENSMUSG00000036053 | Fmnl2 | protein_coding | -0.911892932 | 4.242882389 | 3.8542446 | 0.0669079 | 0.38576981 |
| ENSMUSG00000051469 | Zfp24 | protein_coding | -0.647160839 | 5.445761587 | 3.8474168 | 0.0671262 | 0.386598796 |
| ENSMUSG00000043164 | Tmem212 | protein_coding | -1.241899146 | 3.784289306 | 3.8474535 | 0.0671541 | 0.386598796 |
| ENSMUSG00000043411 | Usp48 | protein_coding | -0.750103535 | 4.625503793 | 3.846376 | 0.0671595 | 0.386598796 |
| ENSMUSG00000074591 | Ankub1 | protein_coding | -1.802993122 | 2.561682603 | 3.8451702 | 0.0671982 | 0.386614372 |
| ENSMUSG00000021266 | Wars | protein_coding | 0.924290025 | 7.014786767 | 3.9305398 | 0.0673313 | 0.387114267 |
| ENSMUSG00000020687 | Cdc27 | protein_coding | -0.626370841 | 5.098841015 | 3.8402229 | 0.0673571 | 0.387114267 |
| ENSMUSG00000039830 | Olig2 | protein_coding | -1.145619091 | 3.316571828 | 3.8355128 | 0.0675088 | 0.387625422 |

| ensembl_gene_id | external_gene_name | gene_biotype | log2 (fold change) | log2 (counts per million) | F-value | p-value | False discovery rate |
|---|---|---|---|---|---|---|---|
| ENSMUSG00000043831 | Lysmd4 | protein_coding | 0.470143738 | 6.736433411 | 3.8352227 | 0.0675181 | 0.387625422 |
| ENSMUSG00000038369 | Ncoa6 | protein_coding | -0.945092252 | 4.480358588 | 3.8328279 | 0.0675954 | 0.387862049 |
| ENSMUSG00000038005 | Hpf1 | protein_coding | -0.51063285 | 5.994374637 | 3.8300266 | 0.067686 | 0.387933464 |
| ENSMUSG00000024142 | Mlst8 | protein_coding | 0.991112937 | 4.379486391 | 3.8292861 | 0.0677099 | 0.387933464 |
| ENSMUSG00000006356 | Crip2 | protein_coding | 0.469672486 | 8.839673363 | 3.8275269 | 0.0677669 | 0.387933464 |
| ENSMUSG00000022680 | Pdxdc1 | protein_coding | 0.694439816 | 5.315056667 | 3.8252587 | 0.0678404 | 0.387933464 |
| ENSMUSG00000006676 | Usp19 | protein_coding | 0.837984864 | 3.875609456 | 3.8244537 | 0.0678665 | 0.387933464 |
| ENSMUSG00000097326 | A330048O09Rik | lincRNA | -1.341986592 | 3.431106651 | 3.8234484 | 0.0678991 | 0.387933464 |
| ENSMUSG00000061119 | Prcp | protein_coding | -1.452435373 | 3.268052999 | 3.824302 | 0.0679006 | 0.387933464 |
| ENSMUSG00000042472 | Zfp410 | protein_coding | 0.690486905 | 5.151333987 | 3.8228754 | 0.0679177 | 0.387933464 |
| ENSMUSG00000026885 | Ttll11 | protein_coding | 0.85047778 | 4.24246534 | 3.8224189 | 0.0679326 | 0.387933464 |
| ENSMUSG00000026849 | Tor1a | protein_coding | 0.750980356 | 5.716089504 | 3.8206734 | 0.0679893 | 0.388051335 |
| ENSMUSG00000052942 | Glis3 | protein_coding | -1.107773791 | 3.356678143 | 3.819035 | 0.0680426 | 0.38814951 |
| ENSMUSG00000042323 | Pbrm1 | protein_coding | -0.632048331 | 5.235378555 | 3.8154462 | 0.0681595 | 0.388597065 |
| ENSMUSG00000027893 | Ahcyl1 | protein_coding | -0.467441255 | 7.54395055 | 3.8142301 | 0.0681992 | 0.388597065 |
| ENSMUSG00000073147 | 5031425E22Rik | lincRNA | -1.16962028 | 3.38855646 | 3.8122225 | 0.0682647 | 0.388597065 |
| ENSMUSG00000036466 | Megf11 | protein_coding | -0.760675602 | 4.842826329 | 3.8121957 | 0.0682656 | 0.388597065 |
| ENSMUSG00000027605 | Acss2 | protein_coding | 1.100019919 | 3.965329002 | 3.8093326 | 0.0683592 | 0.38892408 |
| ENSMUSG00000019578 | Ubxn6 | protein_coding | 0.519368052 | 6.444794577 | 3.8047091 | 0.0685107 | 0.389292459 |
| ENSMUSG00000038352 | Arl5c | protein_coding | -1.368752239 | 4.748157757 | 3.8818277 | 0.0685134 | 0.389292459 |
| ENSMUSG00000037608 | Bclaf1 | protein_coding | -0.536577315 | 7.408676833 | 3.8040427 | 0.0685326 | 0.389292459 |
| ENSMUSG00000069495 | Epc2 | protein_coding | -0.840251344 | 4.693634474 | 3.8028381 | 0.0685721 | 0.389311437 |
| ENSMUSG00000028936 | Rpl22 | protein_coding | 0.547745072 | 7.214161766 | 3.7986268 | 0.0687106 | 0.389891746 |
| ENSMUSG00000026090 | 2010300C02Rik | protein_coding | -0.633885627 | 5.642131526 | 3.7952319 | 0.0688224 | 0.390320556 |
| ENSMUSG00000010047 | Hyal2 | protein_coding | 0.844658973 | 4.857066359 | 3.7922301 | 0.0689215 | 0.390475121 |
| ENSMUSG00000016206 | H2-M3 | protein_coding | 1.406906726 | 3.008335465 | 3.7922063 | 0.0689223 | 0.390475121 |
| ENSMUSG00000071547 | Nt5dc2 | protein_coding | 0.986084273 | 3.340278492 | 3.7906305 | 0.0689744 | 0.390564435 |
| ENSMUSG00000057719 | Sh3rf2 | protein_coding | -1.25498576 | 3.397565118 | 3.7892062 | 0.0690215 | 0.390625543 |
| ENSMUSG00000020238 | Ncln | protein_coding | 0.82988122 | 4.257319797 | 3.7857056 | 0.0691375 | 0.39107602 |
| ENSMUSG00000025354 | Dnajc14 | protein_coding | 0.827540686 | 4.999085832 | 3.7817021 | 0.0692704 | 0.391536847 |
| ENSMUSG00000092310 | Gm20509 | protein_coding | -1.115210321 | 3.420100164 | 3.7793025 | 0.0693502 | 0.391536847 |
| ENSMUSG00000040021 | Lats1 | protein_coding | -1.157746557 | 3.305346806 | 3.7786738 | 0.0693711 | 0.391536847 |
| ENSMUSG00000073910 | Mob3b | protein_coding | 1.221122956 | 3.951102461 | 3.7794623 | 0.0693742 | 0.391536847 |
| ENSMUSG00000033186 | Mzt1 | protein_coding | -0.45666398 | 7.298392549 | 3.7777762 | 0.069401 | 0.391536847 |
| ENSMUSG00000036990 | Otud4 | protein_coding | -0.952466917 | 4.057387711 | 3.7713039 | 0.069617 | 0.392549365 |
| ENSMUSG00000037364 | Srrt | protein_coding | 0.604108521 | 5.570223714 | 3.7693627 | 0.0696819 | 0.392709555 |
| ENSMUSG00000048154 | Kmt2d | protein_coding | -0.829093406 | 6.139164003 | 3.7996957 | 0.0697472 | 0.392810536 |
| ENSMUSG00000027438 | Napb | protein_coding | 0.545570594 | 8.827453936 | 3.7666463 | 0.0697729 | 0.392810536 |
| ENSMUSG00000026482 | Rgl1 | protein_coding | 0.544619353 | 5.569146892 | 3.7585633 | 0.0700445 | 0.393903258 |
| ENSMUSG00000030207 | Fam234b | protein_coding | 0.650838587 | 6.781779912 | 3.7591833 | 0.070053 | 0.393903258 |
| ENSMUSG00000070639 | Lrrc8b | protein_coding | -0.800330532 | 4.292064938 | 3.7569919 | 0.0700974 | 0.393903258 |
| ENSMUSG00000020088 | Sar1a | protein_coding | 0.454587911 | 7.259950516 | 3.756513 | 0.0701135 | 0.393903258 |
| ENSMUSG00000019802 | Sec63 | protein_coding | -0.931974027 | 4.528129988 | 3.7545868 | 0.0701785 | 0.394062365 |
| ENSMUSG00000029560 | Snx8 | protein_coding | 1.037423854 | 4.573281168 | 3.7538206 | 0.0702338 | 0.394166818 |
| ENSMUSG00000034636 | Zyg11b | protein_coding | -0.498502567 | 5.867173384 | 3.7502034 | 0.0703266 | 0.394296816 |
| ENSMUSG00000021198 | Unc79 | protein_coding | -1.039276674 | 3.788694174 | 3.7500595 | 0.0703315 | 0.394296816 |
| ENSMUSG00000031781 | Ciapin1 | protein_coding | 0.468302257 | 7.512321031 | 3.7479598 | 0.0704026 | 0.394296816 |
| ENSMUSG00000037001 | Zfp39 | protein_coding | -0.970828582 | 3.344581473 | 3.7459884 | 0.0704694 | 0.394296816 |
| ENSMUSG00000043004 | Gng2 | protein_coding | 0.524677272 | 7.764402545 | 3.7459488 | 0.0704707 | 0.394296816 |
| ENSMUSG00000038126 | Mphosph9 | protein_coding | -0.982175958 | 3.540894495 | 3.7457656 | 0.0704769 | 0.394296816 |
| ENSMUSG00000051855 | Mest | protein_coding | -1.177402987 | 3.702198442 | 3.7430817 | 0.070568 | 0.394469036 |
| ENSMUSG00000036570 | Fxyd1 | protein_coding | -0.968360127 | 6.27820526 | 3.8177254 | 0.0705811 | 0.394469036 |
| ENSMUSG00000026036 | Nif3l1 | protein_coding | 0.757272919 | 6.00136296 | 3.7410932 | 0.070665 | 0.394614305 |
| ENSMUSG00000008153 | Clstn3 | protein_coding | 0.50685664 | 8.246138266 | 3.739232 | 0.0706989 | 0.394614305 |
| ENSMUSG00000031078 | Cttn | protein_coding | 0.676403428 | 6.768797952 | 3.741841 | 0.0707172 | 0.394614305 |
| ENSMUSG00000038173 | Enpp6 | protein_coding | -1.486070446 | 3.093479553 | 3.7366629 | 0.0707865 | 0.394711965 |
| ENSMUSG00000021222 | Dcaf4 | protein_coding | 0.617282072 | 6.168689986 | 3.7360224 | 0.0708083 | 0.394711965 |
| ENSMUSG00000036067 | Slc2a6 | protein_coding | 0.971333512 | 4.741101066 | 3.7358156 | 0.0708448 | 0.394711965 |
| ENSMUSG00000025512 | Chid1 | protein_coding | 0.508418108 | 6.30292777 | 3.7328166 | 0.0709177 | 0.394913641 |
| ENSMUSG00000000804 | Usp32 | protein_coding | -0.558978051 | 5.593076024 | 3.7314924 | 0.070963 | 0.394961125 |
| ENSMUSG00000063626 | Unc5d | protein_coding | -0.687884362 | 5.112167552 | 3.7220186 | 0.0712878 | 0.396416785 |
| ENSMUSG00000024993 | Fam45a | protein_coding | -0.52596337 | 6.807848676 | 3.7209388 | 0.0713249 | 0.396416785 |
| ENSMUSG00000057729 | Prtn3 | protein_coding | 2.383426589 | 2.570975663 | 3.7252424 | 0.0713351 | 0.396416785 |
| ENSMUSG00000025889 | Snca | protein_coding | 0.434167926 | 9.380044398 | 3.7180967 | 0.0714227 | 0.396562575 |
| ENSMUSG00000020594 | Pum2 | protein_coding | -0.462449519 | 7.253718798 | 3.7175673 | 0.071441 | 0.396562575 |
| ENSMUSG00000024943 | Smc5 | protein_coding | -1.013030853 | 4.320390629 | 3.7154532 | 0.0715139 | 0.396562575 |
| ENSMUSG00000033423 | Eri3 | protein_coding | 0.411600101 | 8.52696204 | 3.7134688 | 0.0715824 | 0.396562575 |
| ENSMUSG00000020464 | Pnpt1 | protein_coding | -1.117249615 | 3.13808061 | 3.7120531 | 0.0716313 | 0.396562575 |
| ENSMUSG00000033237 | Arid2 | protein_coding | -0.77178619 | 4.539216334 | 3.7093661 | 0.0717242 | 0.396562575 |
| ENSMUSG00000010554 | Mettl16 | protein_coding | 0.518449109 | 6.035521682 | 3.708902 | 0.0717403 | 0.396562575 |
| ENSMUSG00000024576 | Csnk1a1 | protein_coding | -0.56603529 | 7.22639045 | 3.7080221 | 0.0717708 | 0.396562575 |
| ENSMUSG00000020674 | Pxdn | protein_coding | 2.231494703 | 2.519421216 | 3.7087806 | 0.071774 | 0.396562575 |
| ENSMUSG00000026958 | Dpp7 | protein_coding | 0.76995063 | 4.544340655 | 3.7071473 | 0.0718011 | 0.396562575 |
| ENSMUSG00000029598 | Plbd2 | protein_coding | 0.736162841 | 5.096875469 | 3.7067027 | 0.0718165 | 0.396562575 |
| ENSMUSG00000002635 | Pdcd2l | protein_coding | 0.863193774 | 4.720181586 | 3.7065472 | 0.0718219 | 0.396562575 |
| ENSMUSG00000021693 | Kif2a | protein_coding | -0.527063574 | 5.858300644 | 3.7055655 | 0.071856 | 0.396562575 |
| ENSMUSG00000021287 | Xrcc3 | protein_coding | 1.428381549 | 2.800430518 | 3.7049404 | 0.0718777 | 0.396562575 |
| ENSMUSG00000047710 | Champ1 | protein_coding | 0.810736958 | 4.904989309 | 3.7021875 | 0.0719733 | 0.396594403 |

| ensembl_gene_id | external_gene_name | gene_biotype | log2 (fold change) | log2 (counts per million) | F-value | p-value | False discovery rate |
|---|---|---|---|---|---|---|---|
| ENSMUSG00000038671 | Arfrp1 | protein_coding | 0.47320027 | 6.881633397 | 3.702054 | 0.0719779 | 0.396594403 |
| ENSMUSG00000025034 | Trim8 | protein_coding | -0.615935023 | 5.425123894 | 3.7011644 | 0.0720089 | 0.396594403 |
| ENSMUSG00000032673 | Prorsd1 | protein_coding | -0.73267461 | 4.986987756 | 3.6971611 | 0.0721483 | 0.396594403 |
| ENSMUSG00000022792 | Yars2 | protein_coding | 0.872427347 | 3.835039822 | 3.6971265 | 0.0721495 | 0.396594403 |
| ENSMUSG00000034875 | Nudt19 | protein_coding | 0.42012031 | 6.684341927 | 3.6958049 | 0.0721956 | 0.396594403 |
| ENSMUSG00000009281 | Rarres2 | protein_coding | -1.008723281 | 4.253690316 | 3.6949016 | 0.0722271 | 0.396594403 |
| ENSMUSG00000028639 | Ybx1 | protein_coding | -0.775805981 | 5.303736514 | 3.6933816 | 0.0722802 | 0.396594403 |
| ENSMUSG00000031197 | Vbp1 | protein_coding | -0.429863501 | 7.867598489 | 3.6921888 | 0.0723219 | 0.396594403 |
| ENSMUSG00000028559 | Osbpl9 | protein_coding | 0.527807672 | 5.932342339 | 3.6914469 | 0.0723478 | 0.396594403 |
| ENSMUSG00000063931 | Pepd | protein_coding | 0.641917681 | 6.764992624 | 3.6918448 | 0.0723635 | 0.396594403 |
| ENSMUSG00000038742 | Angptl6 | protein_coding | 0.97952602 | 4.581476581 | 3.6880851 | 0.0724951 | 0.396594403 |
| ENSMUSG00000024446 | Rpp21 | protein_coding | 0.411521835 | 7.644382785 | 3.6866532 | 0.0725157 | 0.396594403 |
| ENSMUSG00000036249 | Rbm43 | protein_coding | 0.771110809 | 5.19496541 | 3.6858948 | 0.0725423 | 0.396594403 |
| ENSMUSG00000040724 | Kcna2 | protein_coding | -0.721072368 | 5.230897557 | 3.6846989 | 0.0725843 | 0.396594403 |
| ENSMUSG00000102869 | 2900097C17Rik | lincRNA | 0.573380676 | 7.970443406 | 3.6846648 | 0.0725855 | 0.396594403 |
| ENSMUSG00000017667 | Zfp334 | protein_coding | -0.700265684 | 4.632684367 | 3.6841152 | 0.0726048 | 0.396594403 |
| ENSMUSG00000021967 | Mrpl57 | protein_coding | 0.573254566 | 6.480578311 | 3.6837851 | 0.0726164 | 0.396594403 |
| ENSMUSG00000057816 | 1700007G11Rik | protein_coding | -0.909184242 | 3.541309499 | 3.6828949 | 0.0726476 | 0.396594403 |
| ENSMUSG00000037369 | Kdm6a | protein_coding | -0.989526528 | 3.870583074 | 3.6828478 | 0.0726493 | 0.396594403 |
| ENSMUSG00000023266 | Frs3 | protein_coding | 0.819090118 | 4.120808735 | 3.6826014 | 0.072658 | 0.396594403 |
| ENSMUSG00000029146 | Snx17 | protein_coding | 0.544773541 | 8.228083589 | 3.6759041 | 0.0728938 | 0.397564963 |
| ENSMUSG00000039672 | Kcne2 | protein_coding | -5.570421388 | 3.019242124 | 3.8048918 | 0.0729235 | 0.397564963 |
| ENSMUSG00000046204 | Pnma2 | protein_coding | 0.507632075 | 7.583794278 | 3.6744068 | 0.0729467 | 0.397564963 |
| ENSMUSG00000097709 | 2810429I04Rik | lincRNA | -1.094497974 | 3.477889688 | 3.6733025 | 0.0729857 | 0.397576064 |
| ENSMUSG00000066392 | Nrxn3 | protein_coding | -0.490654171 | 7.907882488 | 3.6722057 | 0.0730245 | 0.397585861 |
| ENSMUSG00000009216 | Fam163b | protein_coding | 0.838797111 | 4.936179858 | 3.6695873 | 0.0731171 | 0.397888878 |
| ENSMUSG00000053565 | Eif3k | protein_coding | 0.477210603 | 9.322931709 | 3.6664784 | 0.0732273 | 0.398286969 |
| ENSMUSG00000007207 | Stx1a | protein_coding | 0.754342192 | 7.031155404 | 3.7053964 | 0.0733366 | 0.398384982 |
| ENSMUSG00000029819 | Npy | protein_coding | -1.014982929 | 6.897415766 | 3.797629 | 0.0733672 | 0.398384982 |
| ENSMUSG00000020284 | 1810043G02Rik | protein_coding | 0.540725058 | 6.869623216 | 3.6624751 | 0.0733695 | 0.398384982 |
| ENSMUSG00000000934 | Top1mt | protein_coding | 0.7269174 | 5.3546275 | 3.6609137 | 0.073425 | 0.398384982 |
| ENSMUSG00000030744 | Rps3 | protein_coding | 0.537043988 | 10.97915624 | 3.6600772 | 0.0734548 | 0.398384982 |
| ENSMUSG00000022307 | Oxr1 | protein_coding | -0.652817319 | 7.482523217 | 3.6672463 | 0.0735441 | 0.398384982 |
| ENSMUSG00000021707 | Dhfr | protein_coding | 1.190302067 | 3.269350618 | 3.6570129 | 0.073564 | 0.398384982 |
| ENSMUSG00000052337 | Immt | protein_coding | 0.580774634 | 7.219270007 | 3.6576495 | 0.073571 | 0.398384982 |
| ENSMUSG00000042111 | Ccdc115 | protein_coding | 0.619580623 | 5.416773609 | 3.6552738 | 0.0736261 | 0.398384982 |
| ENSMUSG00000036819 | Jmjd4 | protein_coding | 0.833849475 | 4.267717668 | 3.6546479 | 0.0736484 | 0.398384982 |
| ENSMUSG00000035032 | Nek11 | protein_coding | -1.648297079 | 2.447111599 | 3.6545235 | 0.0736529 | 0.398384982 |
| ENSMUSG00000071649 | B3gat3 | protein_coding | 0.487598991 | 6.366993141 | 3.6519137 | 0.0737462 | 0.398492199 |
| ENSMUSG00000068205 | Macrod2 | protein_coding | -0.706818607 | 5.345425192 | 3.6518771 | 0.0737475 | 0.398492199 |
| ENSMUSG00000024146 | Cript | protein_coding | -0.483071053 | 8.259973186 | 3.6506555 | 0.0737912 | 0.398492199 |
| ENSMUSG00000018405 | Mrm1 | protein_coding | 1.155374585 | 3.018384625 | 3.6490962 | 0.073847 | 0.398492199 |
| ENSMUSG00000027820 | Mme | protein_coding | -1.457498436 | 2.654254081 | 3.6487904 | 0.073858 | 0.398492199 |
| ENSMUSG00000090399 | Gm38399 | lincRNA | -1.307739236 | 2.784710787 | 3.6443339 | 0.0740179 | 0.399110031 |
| ENSMUSG00000024077 | Strn | protein_coding | -0.518871714 | 6.419242611 | 3.6430333 | 0.0740646 | 0.399110031 |
| ENSMUSG00000035891 | Cerk | protein_coding | -0.784760896 | 4.706977048 | 3.6423545 | 0.074089 | 0.399110031 |
| ENSMUSG00000024924 | Vldlr | protein_coding | -0.800264216 | 5.759781989 | 3.6422935 | 0.074121 | 0.399110031 |
| ENSMUSG00000028173 | Wls | protein_coding | -0.74265242 | 4.860362235 | 3.6387246 | 0.0742197 | 0.399441861 |
| ENSMUSG00000047045 | Tmem164 | protein_coding | -0.983958989 | 3.627570521 | 3.635086 | 0.074351 | 0.399857387 |
| ENSMUSG00000042216 | Sgsm1 | protein_coding | 0.650075297 | 6.090930906 | 3.634524 | 0.0743713 | 0.399857387 |
| ENSMUSG00000079523 | Tmsb10 | protein_coding | 0.486841844 | 8.16261568 | 3.6290617 | 0.074569 | 0.400719726 |
| ENSMUSG00000020794 | Ube2g1 | protein_coding | -0.565117863 | 5.654516639 | 3.6261016 | 0.0746763 | 0.4009302 |
| ENSMUSG00000026433 | Rab29 | protein_coding | -1.433121248 | 3.270456569 | 3.6254952 | 0.0747281 | 0.4009302 |
| ENSMUSG00000044734 | Serpinb1a | protein_coding | -0.610262865 | 5.844945195 | 3.6223045 | 0.0748143 | 0.4009302 |
| ENSMUSG00000028164 | Manba | protein_coding | 1.472785755 | 2.762128959 | 3.6212827 | 0.0748515 | 0.4009302 |
| ENSMUSG00000034248 | Slc25a37 | protein_coding | 1.496537417 | 2.954380726 | 3.6210025 | 0.0748617 | 0.4009302 |
| ENSMUSG00000036975 | Tmem177 | protein_coding | 0.985423407 | 3.831928398 | 3.6209857 | 0.0748623 | 0.4009302 |
| ENSMUSG00000044229 | Nxpe4 | protein_coding | -1.393390928 | 3.301324228 | 3.6197964 | 0.0749057 | 0.4009302 |
| ENSMUSG00000034135 | Sik3 | protein_coding | -0.460487792 | 6.39715125 | 3.6197753 | 0.0749064 | 0.4009302 |
| ENSMUSG00000019428 | Fkbp8 | protein_coding | 0.425554433 | 8.438425054 | 3.6187407 | 0.0749441 | 0.400932442 |
| ENSMUSG00000024219 | Anks1 | protein_coding | 1.073861758 | 3.484281808 | 3.6169751 | 0.0750085 | 0.401077437 |
| ENSMUSG00000078439 | Smim24 | protein_coding | 1.447720116 | 2.738258183 | 3.6157855 | 0.075052 | 0.40109085 |
| ENSMUSG00000033285 | Wdr3 | protein_coding | 2.034499657 | 2.434928954 | 3.6141258 | 0.0751126 | 0.40109085 |
| ENSMUSG00000042515 | Mum1l1 | protein_coding | -1.275995233 | 3.670943387 | 3.6146583 | 0.0751229 | 0.40109085 |
| ENSMUSG00000028552 | Eps15 | protein_coding | -0.483917214 | 6.687543463 | 3.611956 | 0.075192 | 0.401160286 |
| ENSMUSG00000019179 | Mdh2 | protein_coding | 0.516993859 | 10.78112278 | 3.6114478 | 0.0752106 | 0.401160286 |
| ENSMUSG00000028248 | Pnisr | protein_coding | -0.604442887 | 5.828569493 | 3.6101574 | 0.0752578 | 0.401213335 |
| ENSMUSG00000027935 | Rab13 | protein_coding | 0.683748083 | 4.98848082 | 3.6082849 | 0.0753265 | 0.401380253 |
| ENSMUSG00000098176 | Ccdc166 | protein_coding | 1.013065739 | 4.102231282 | 3.6029516 | 0.0755224 | 0.401948414 |
| ENSMUSG00000020716 | Nf1 | protein_coding | -0.71017426 | 5.142647013 | 3.6023398 | 0.0755449 | 0.401948414 |
| ENSMUSG00000031381 | Piga | protein_coding | -1.03702544 | 3.304020973 | 3.6023301 | 0.0755452 | 0.401948414 |
| ENSMUSG00000048004 | Tmem196 | protein_coding | -0.92149118 | 4.160727152 | 3.6006633 | 0.0756066 | 0.402075974 |
| ENSMUSG00000062991 | Nrg1 | protein_coding | -0.620026904 | 5.108745726 | 3.5978544 | 0.0757101 | 0.402124318 |
| ENSMUSG00000037916 | Ndufv1 | protein_coding | 0.49247752 | 9.217131399 | 3.5974529 | 0.075725 | 0.402124318 |
| ENSMUSG00000041439 | Mfsd6 | protein_coding | 0.590590965 | 5.493051545 | 3.597374 | 0.0757279 | 0.402124318 |
| ENSMUSG00000034361 | Cpne2 | protein_coding | 0.669920644 | 5.225440251 | 3.594738 | 0.0758252 | 0.402442564 |
| ENSMUSG00000033965 | Slc16a2 | protein_coding | -1.271532381 | 3.931101434 | 3.6031142 | 0.0758752 | 0.402469799 |
| ENSMUSG00000024397 | Aif1 | protein_coding | -1.149527472 | 4.614596879 | 3.6224845 | 0.075963 | 0.402469799 |

| ensembl_gene_id | external_gene_name | gene_biotype | log2 (fold change) | log2 (counts per million) | F-value | p-value | False discovery rate |
|---|---|---|---|---|---|---|---|
| ENSMUSG00000040564 | Apoc1 | protein_coding | -1.17299214 | 3.600949766 | 3.5907617 | 0.0759724 | 0.402469799 |
| ENSMUSG00000039349 | C130074G19Rik | protein_coding | -1.369176833 | 3.535374711 | 3.5913601 | 0.0759801 | 0.402469799 |
| ENSMUSG00000030350 | Prmt8 | protein_coding | -0.528289442 | 6.015413048 | 3.5877228 | 0.076085 | 0.402505626 |
| ENSMUSG00000055541 | Lair1 | protein_coding | -1.20247867 | 3.076907307 | 3.5874268 | 0.076096 | 0.402505626 |
| ENSMUSG00000034586 | Hid1 | protein_coding | 0.663316797 | 6.134864464 | 3.5872146 | 0.0761039 | 0.402505626 |
| ENSMUSG00000085069 | Gm13111 | antisense_RNA | -1.63015046 | 2.502866299 | 3.5863349 | 0.0761366 | 0.402505626 |
| ENSMUSG00000034664 | Itga2b | protein_coding | 0.861719777 | 4.189217898 | 3.581939 | 0.0763 | 0.403171627 |
| ENSMUSG00000027881 | Prpf38b | protein_coding | -1.027108091 | 3.818091983 | 3.5803425 | 0.0763595 | 0.403230427 |
| ENSMUSG00000049044 | Rapgef4 | protein_coding | -0.688029526 | 6.176587746 | 3.5804297 | 0.0763862 | 0.403230427 |
| ENSMUSG00000024197 | Plin3 | protein_coding | -0.725245076 | 5.200013871 | 3.576639 | 0.0764977 | 0.403620958 |
| ENSMUSG00000063317 | Usp31 | protein_coding | -0.66907566 | 5.091182228 | 3.5755552 | 0.0765382 | 0.403636533 |
| ENSMUSG00000036022 | Fam122b | protein_coding | -1.106786664 | 3.233161775 | 3.5690399 | 0.0767821 | 0.404228243 |
| ENSMUSG00000020785 | Camkk1 | protein_coding | 0.675728693 | 5.651688003 | 3.5689962 | 0.0767837 | 0.404228243 |
| ENSMUSG00000040359 | Ufl1 | protein_coding | -1.131247393 | 2.978506641 | 3.568861 | 0.0767888 | 0.404228243 |
| ENSMUSG00000031483 | Erlin2 | protein_coding | 0.6604231 | 5.141166293 | 3.5679069 | 0.0768246 | 0.404228243 |
| ENSMUSG00000033411 | Ctdspl2 | protein_coding | -0.842848636 | 4.635409623 | 3.5672517 | 0.0768492 | 0.404228243 |
| ENSMUSG00000041329 | Atp1b2 | protein_coding | -0.611293233 | 7.359503043 | 3.5672355 | 0.0768798 | 0.404228243 |
| ENSMUSG00000042388 | Dlgap3 | protein_coding | -1.177953873 | 3.139058895 | 3.5652664 | 0.0769238 | 0.404228243 |
| ENSMUSG00000034255 | Arhgap27 | protein_coding | 0.678991647 | 5.421061186 | 3.5637252 | 0.0769818 | 0.404228243 |
| ENSMUSG00000043284 | Tmem11 | protein_coding | 0.490924114 | 6.622788953 | 3.5635412 | 0.0769887 | 0.404228243 |
| ENSMUSG00000040729 | Cep126 | protein_coding | -1.667660343 | 3.213043619 | 3.5693523 | 0.0770673 | 0.40444359 |
| ENSMUSG00000070319 | Eif3g | protein_coding | 0.410498153 | 7.844693706 | 3.5576791 | 0.0772097 | 0.404992903 |
| ENSMUSG00000032523 | Hhatl | protein_coding | -1.840557798 | 3.133468773 | 3.5861011 | 0.0773044 | 0.40521013 |
| ENSMUSG00000056553 | Ptprn2 | protein_coding | -1.002754585 | 3.91967776 | 3.554589 | 0.0773264 | 0.40521013 |
| ENSMUSG00000024807 | Syvn1 | protein_coding | 0.780880143 | 4.645465583 | 3.5516929 | 0.0774361 | 0.405466371 |
| ENSMUSG00000056222 | Spock1 | protein_coding | 0.439060298 | 6.523959085 | 3.550871 | 0.0774672 | 0.405466371 |
| ENSMUSG00000017009 | Sdc4 | protein_coding | -0.6480956 | 7.306298795 | 3.5596909 | 0.077494 | 0.405466371 |
| ENSMUSG00000048047 | Zbtb33 | protein_coding | -1.428150218 | 2.590932394 | 3.5493166 | 0.0775262 | 0.405466371 |
| ENSMUSG00000022450 | Ndufa6 | protein_coding | 0.515340706 | 10.01225996 | 3.5436004 | 0.0777434 | 0.406404826 |
| ENSMUSG00000027952 | Pmvk | protein_coding | 0.430555892 | 8.011103667 | 3.5425328 | 0.077784 | 0.406419753 |
| ENSMUSG00000020455 | Trim11 | protein_coding | 1.066049861 | 3.555612013 | 3.536663 | 0.078008 | 0.407138153 |
| ENSMUSG00000040565 | Btaf1 | protein_coding | -1.064797143 | 3.801692078 | 3.5358655 | 0.0780385 | 0.407138153 |
| ENSMUSG00000028690 | Mmachc | protein_coding | 0.779159463 | 4.731167092 | 3.5354047 | 0.0780561 | 0.407138153 |
| ENSMUSG00000026219 | Trip12 | protein_coding | -0.499896113 | 6.114001535 | 3.5339919 | 0.0781102 | 0.407138153 |
| ENSMUSG00000039620 | 6430573F11Rik | protein_coding | -0.96305551 | 3.508994447 | 3.5339738 | 0.0781109 | 0.407138153 |
| ENSMUSG00000027255 | Arfgap2 | protein_coding | 0.888529707 | 4.615170239 | 3.5300011 | 0.0782631 | 0.407491225 |
| ENSMUSG00000030213 | Atf7ip | protein_coding | -0.714464123 | 4.988359292 | 3.5293879 | 0.0782866 | 0.407491225 |
| ENSMUSG00000060512 | 0610040J01Rik | protein_coding | -2.394464252 | 2.003837371 | 3.524971 | 0.0784564 | 0.407491225 |
| ENSMUSG00000070730 | Rmdn3 | protein_coding | 0.829009612 | 4.036268146 | 3.5241116 | 0.0784894 | 0.407491225 |
| ENSMUSG00000028223 | Decr1 | protein_coding | -0.699595507 | 5.156132739 | 3.5232875 | 0.0785212 | 0.407491225 |
| ENSMUSG00000033075 | Senp1 | protein_coding | -1.279977857 | 3.370551733 | 3.5212012 | 0.0786016 | 0.407491225 |
| ENSMUSG00000013833 | Med16 | protein_coding | 0.797476427 | 3.914232501 | 3.5167609 | 0.078773 | 0.407491225 |
| ENSMUSG00000049299 | Trappc1 | protein_coding | 0.478770412 | 7.355933209 | 3.5159709 | 0.0788036 | 0.407491225 |
| ENSMUSG00000024614 | Tmx3 | protein_coding | -0.57498659 | 5.685585436 | 3.5157483 | 0.0788122 | 0.407491225 |
| ENSMUSG00000020015 | Cdk17 | protein_coding | -0.619258769 | 6.730462063 | 3.5163162 | 0.0788203 | 0.407491225 |
| ENSMUSG00000028861 | Mrps15 | protein_coding | 0.5851955 | 6.071185528 | 3.5147377 | 0.0788513 | 0.407491225 |
| ENSMUSG00000027603 | Ggt7 | protein_coding | 0.545170468 | 7.073577587 | 3.5128537 | 0.0789242 | 0.407491225 |
| ENSMUSG00000030268 | Bcat1 | protein_coding | 0.635571669 | 7.583348572 | 3.5188096 | 0.0789259 | 0.407491225 |
| ENSMUSG00000051107 | Gm15440 | protein_coding | -0.820657795 | 3.974827409 | 3.5113093 | 0.0789841 | 0.407491225 |
| ENSMUSG00000026192 | Atic | protein_coding | 0.728624521 | 5.524916594 | 3.5102581 | 0.0790249 | 0.407491225 |
| ENSMUSG00000031783 | Polr2c | protein_coding | 0.500523116 | 5.726039436 | 3.5102039 | 0.079027 | 0.407491225 |
| ENSMUSG00000048307 | Ankrd46 | protein_coding | 0.548552812 | 8.673423792 | 3.5094272 | 0.0790571 | 0.407491225 |
| ENSMUSG00000056602 | Fry | protein_coding | -0.59751243 | 5.901134414 | 3.5088011 | 0.0790814 | 0.407491225 |
| ENSMUSG00000086316 | 2210013O21Rik | protein_coding | 0.387487581 | 7.655400986 | 3.5079052 | 0.0791162 | 0.407491225 |
| ENSMUSG00000037965 | Zc3h7a | protein_coding | 0.630556881 | 6.426642557 | 3.5074488 | 0.0791641 | 0.407491225 |
| ENSMUSG00000042363 | Lgalsl | protein_coding | -0.471990018 | 6.228389488 | 3.5064409 | 0.0791732 | 0.407491225 |
| ENSMUSG00000020962 | Gtf2a1 | protein_coding | -0.563171652 | 5.943670029 | 3.5063345 | 0.0791773 | 0.407491225 |
| ENSMUSG00000027080 | Med19 | protein_coding | 0.552666608 | 6.069591869 | 3.5061594 | 0.0791841 | 0.407491225 |
| ENSMUSG00000009927 | Rps25 | protein_coding | 0.550885196 | 6.467116773 | 3.5056534 | 0.0792038 | 0.407491225 |
| ENSMUSG00000036160 | Surf6 | protein_coding | 0.908863287 | 4.498404518 | 3.5055566 | 0.0792076 | 0.407491225 |
| ENSMUSG00000049097 | Ankrd34a | protein_coding | 0.632719027 | 5.665853992 | 3.5018695 | 0.0793512 | 0.407491225 |
| ENSMUSG00000001844 | Zdhhc4 | protein_coding | 0.59553777 | 5.724670165 | 3.5011558 | 0.079379 | 0.407491225 |
| ENSMUSG00000038806 | Sde2 | protein_coding | 0.467724962 | 6.14126393 | 3.4999191 | 0.0794273 | 0.407491225 |
| ENSMUSG00000027630 | Tbl1xr1 | protein_coding | -0.629720682 | 5.024175438 | 3.4997934 | 0.0794322 | 0.407491225 |
| ENSMUSG00000006728 | Cdk4 | protein_coding | 0.445743868 | 7.036813166 | 3.4988218 | 0.0794702 | 0.407491225 |
| ENSMUSG00000064036 | Mro | protein_coding | -0.574196769 | 5.294121439 | 3.4972881 | 0.0795301 | 0.407491225 |
| ENSMUSG00000041891 | Lman1 | protein_coding | 1.044918647 | 3.955768525 | 3.4967759 | 0.0795501 | 0.407491225 |
| ENSMUSG00000030060 | Hmces | protein_coding | 0.722433932 | 5.224205205 | 3.4962205 | 0.0795718 | 0.407491225 |
| ENSMUSG00000051951 | Xkr4 | protein_coding | -1.223687175 | 3.610802665 | 3.4966937 | 0.0795835 | 0.407491225 |
| ENSMUSG00000053137 | Mapk11 | protein_coding | 0.955514706 | 3.555196215 | 3.4955004 | 0.0796 | 0.407491225 |
| ENSMUSG00000057110 | Cntrl | protein_coding | -1.95449294 | 2.464555657 | 3.4952793 | 0.0796087 | 0.407491225 |
| ENSMUSG00000016554 | Eif3d | protein_coding | 0.449088041 | 6.399336327 | 3.4951094 | 0.0796153 | 0.407491225 |
| ENSMUSG00000003423 | Pih1d1 | protein_coding | 0.68092501 | 5.332552352 | 3.4950252 | 0.0796186 | 0.407491225 |
| ENSMUSG00000019977 | Hbs1l | protein_coding | 0.668390484 | 5.924672831 | 3.492731 | 0.0797085 | 0.407652043 |
| ENSMUSG00000024982 | Zdhhc6 | protein_coding | 0.631486586 | 4.825078853 | 3.4922881 | 0.0797259 | 0.407652043 |
| ENSMUSG00000037926 | Ssh2 | protein_coding | -0.763289107 | 4.286228006 | 3.4882471 | 0.0798845 | 0.407734428 |
| ENSMUSG00000046994 | Mars2 | protein_coding | -1.268279356 | 3.103361942 | 3.4874092 | 0.0799174 | 0.407734428 |
| ENSMUSG00000045083 | Lingo2 | protein_coding | -0.654249633 | 4.931962584 | 3.4858814 | 0.0799776 | 0.407734428 |

| ensembl_gene_id | external_gene_name | gene_biotype | log2 (fold change) | log2 (counts per million) | F-value | p-value | False discovery rate |
|---|---|---|---|---|---|---|---|
| ENSMUSG00000059412 | Fxyd2 | protein_coding | 0.722356039 | 5.13876124 | 3.4854693 | 0.0799938 | 0.407734428 |
| ENSMUSG00000039179 | Tekt5 | protein_coding | 0.682098458 | 5.769587978 | 3.4851298 | 0.0800071 | 0.407734428 |
| ENSMUSG00000047454 | Gphn | protein_coding | 0.469273276 | 7.188718565 | 3.4841629 | 0.0800452 | 0.407734428 |
| ENSMUSG00000020085 | Aifm2 | protein_coding | 1.051207997 | 4.33042888 | 3.4844856 | 0.0800627 | 0.407734428 |
| ENSMUSG00000025791 | Pgm2 | protein_coding | 0.623640654 | 7.022991 | 3.4843754 | 0.080067 | 0.407734428 |
| ENSMUSG00000022760 | Thap7 | protein_coding | 0.550278633 | 6.147698245 | 3.4827428 | 0.0801012 | 0.407734428 |
| ENSMUSG00000005078 | Jkamp | protein_coding | 0.421516532 | 7.001393794 | 3.4822366 | 0.0801212 | 0.407734428 |
| ENSMUSG00000054850 | Smim10l2a | protein_coding | 0.743484881 | 4.392290456 | 3.4797522 | 0.0802192 | 0.40804035 |
| ENSMUSG00000036860 | Mrpl55 | protein_coding | 0.442102511 | 7.83164215 | 3.4783451 | 0.0802748 | 0.408080535 |
| ENSMUSG00000061136 | Prpf40a | protein_coding | -0.667298519 | 5.872851531 | 3.4760824 | 0.0803643 | 0.408080535 |
| ENSMUSG00000064343 | mt-Tq | Mt_tRNA | -1.530888065 | 2.419131349 | 3.476036 | 0.0803662 | 0.408080535 |
| ENSMUSG00000019996 | Map7 | protein_coding | -1.009674372 | 3.207849493 | 3.4757137 | 0.0803789 | 0.408080535 |
| ENSMUSG00000028013 | Ppa2 | protein_coding | 0.528037089 | 6.441135138 | 3.4731571 | 0.0804802 | 0.408264355 |
| ENSMUSG00000028180 | Zranb2 | protein_coding | -0.440304149 | 7.576783338 | 3.4714233 | 0.080549 | 0.408264355 |
| ENSMUSG00000073676 | Hspe1 | protein_coding | 0.478067514 | 7.452301563 | 3.4706108 | 0.0805813 | 0.408264355 |
| ENSMUSG00000021069 | Pygl | protein_coding | -1.737721589 | 2.657915326 | 3.4697693 | 0.0806147 | 0.408264355 |
| ENSMUSG00000036264 | Fstl4 | protein_coding | -1.273046104 | 3.592194445 | 3.4702503 | 0.0806258 | 0.408264355 |
| ENSMUSG00000054057 | A930004D18Rik | protein_coding | 0.893957421 | 3.860349917 | 3.4686043 | 0.080661 | 0.408264355 |
| ENSMUSG00000035011 | Zbtb7a | protein_coding | -0.709751078 | 5.645804851 | 3.4681032 | 0.0806809 | 0.408264355 |
| ENSMUSG00000030075 | Cntn3 | protein_coding | -1.027970947 | 3.320332865 | 3.4665473 | 0.0807428 | 0.40838541 |
| ENSMUSG00000021537 | Cetn3 | protein_coding | -0.571090367 | 7.85686863 | 3.464314 | 0.080862 | 0.408618899 |
| ENSMUSG00000006435 | Neurl1a | protein_coding | -0.656972269 | 5.92054398 | 3.4615725 | 0.0809411 | 0.408618899 |
| ENSMUSG00000025272 | Tro | protein_coding | 0.572948478 | 7.082353565 | 3.4618338 | 0.0809609 | 0.408618899 |
| ENSMUSG00000049807 | Arhgap23 | protein_coding | -0.850778065 | 4.26667607 | 3.4606862 | 0.0809765 | 0.408618899 |
| ENSMUSG00000008658 | Rbfox1 | protein_coding | 0.421752668 | 8.260538718 | 3.4606242 | 0.080979 | 0.408618899 |
| ENSMUSG00000027366 | Sppl2a | protein_coding | -0.693308736 | 4.958975193 | 3.4593544 | 0.0810297 | 0.408683131 |
| ENSMUSG00000021592 | Arsk | protein_coding | -1.306045935 | 3.16525007 | 3.4562982 | 0.081152 | 0.409107906 |
| ENSMUSG00000020744 | Slc25a19 | protein_coding | 0.576678101 | 5.895115586 | 3.4531501 | 0.0812781 | 0.409551951 |
| ENSMUSG00000063796 | Slc22a8 | protein_coding | 2.119115931 | 2.651915332 | 3.4520037 | 0.0813544 | 0.409560334 |
| ENSMUSG00000029376 | Mthfd2l | protein_coding | -0.735617999 | 4.601838328 | 3.4512108 | 0.081356 | 0.409560334 |
| ENSMUSG00000020692 | Nle1 | protein_coding | 1.037399578 | 4.406804354 | 3.4508354 | 0.0814013 | 0.409596759 |
| ENSMUSG00000008683 | Rps15a | protein_coding | 0.568787174 | 7.305842225 | 3.4497896 | 0.0814433 | 0.409616547 |
| ENSMUSG00000023990 | Tfeb | protein_coding | -2.015181039 | 2.436470241 | 3.4448607 | 0.0816115 | 0.410270313 |
| ENSMUSG00000035725 | Prkx | protein_coding | -1.254513641 | 3.582011779 | 3.4425706 | 0.0817341 | 0.41069485 |
| ENSMUSG00000042745 | Id1 | protein_coding | -0.89011437 | 3.894868094 | 3.437595 | 0.081905 | 0.411062044 |
| ENSMUSG00000045733 | Sprn | protein_coding | -0.836151555 | 4.718273447 | 3.4372256 | 0.0819199 | 0.411062044 |
| ENSMUSG00000031157 | Pqbp1 | protein_coding | 0.462403189 | 7.908240987 | 3.4366651 | 0.0819426 | 0.411062044 |
| ENSMUSG00000006494 | Pdk1 | protein_coding | 0.982941781 | 4.046963452 | 3.4362335 | 0.0819601 | 0.411062044 |
| ENSMUSG00000040536 | Necab1 | protein_coding | -0.850554441 | 5.67572211 | 3.4538563 | 0.0820196 | 0.411168743 |
| ENSMUSG00000022840 | Adcy5 | protein_coding | -0.646049635 | 6.869010957 | 3.4341013 | 0.0821916 | 0.411576887 |
| ENSMUSG00000042763 | Maneal | protein_coding | 0.856713253 | 4.41117024 | 3.4302985 | 0.082201 | 0.411576887 |
| ENSMUSG00000055963 | Triqk | protein_coding | -0.493152643 | 5.890380288 | 3.4299322 | 0.0822159 | 0.411576887 |
| ENSMUSG00000046793 | Gpr61 | protein_coding | 0.819341608 | 4.355467775 | 3.428812 | 0.0822614 | 0.411588384 |
| ENSMUSG00000002658 | Gtf2f1 | protein_coding | 0.549140131 | 6.596767364 | 3.4271227 | 0.0823302 | 0.411588384 |
| ENSMUSG00000030802 | Bckdk | protein_coding | 0.632481155 | 6.812872469 | 3.4277984 | 0.082333 | 0.411588384 |
| ENSMUSG00000032407 | U2surp | protein_coding | -0.470969544 | 6.560169412 | 3.4217938 | 0.0825476 | 0.412444058 |
| ENSMUSG00000087165 | 2010001A14Rik | processed_transcript | -0.813271537 | 4.217475595 | 3.4204093 | 0.0826042 | 0.412444058 |
| ENSMUSG00000031723 | Txnl4b | protein_coding | 0.636987445 | 6.709429716 | 3.4207827 | 0.0826192 | 0.412444058 |
| ENSMUSG00000007589 | Tinf2 | protein_coding | 1.009365136 | 3.47534983 | 3.4175264 | 0.0827221 | 0.412766197 |
| ENSMUSG00000001891 | Ugp2 | protein_coding | -0.458509902 | 6.183373685 | 3.4155568 | 0.0828029 | 0.41284069 |
| ENSMUSG00000020014 | Cfap54 | protein_coding | -1.436635536 | 2.646951021 | 3.4152886 | 0.0828139 | 0.41284069 |
| ENSMUSG00000046873 | Mbtps2 | protein_coding | -0.919253246 | 3.778127387 | 3.4136857 | 0.0828796 | 0.412977099 |
| ENSMUSG00000035376 | Hacd2 | protein_coding | -0.919322548 | 3.720250957 | 3.4114406 | 0.0829718 | 0.413245138 |
| ENSMUSG00000009555 | Cdk9 | protein_coding | 0.793462805 | 4.755560864 | 3.4103075 | 0.0830184 | 0.413285771 |
| ENSMUSG00000025937 | Lactb2 | protein_coding | -0.991361809 | 3.577857389 | 3.4060457 | 0.0831939 | 0.413700885 |
| ENSMUSG00000058006 | Mdn1 | protein_coding | 1.407121885 | 3.343406399 | 3.4061959 | 0.0832181 | 0.413700885 |
| ENSMUSG00000040521 | Tsfm | protein_coding | 0.522142202 | 6.541211415 | 3.4054168 | 0.0832199 | 0.413700885 |
| ENSMUSG00000032411 | Tfdp2 | protein_coding | -0.811225831 | 4.105395577 | 3.4045484 | 0.0832557 | 0.413700885 |
| ENSMUSG00000053062 | Jam2 | protein_coding | -0.793938181 | 3.995873277 | 3.4025807 | 0.083337 | 0.413809508 |
| ENSMUSG00000031403 | Dkc1 | protein_coding | -0.54323927 | 5.864226478 | 3.4012954 | 0.0833901 | 0.413809508 |
| ENSMUSG00000040747 | Cd53 | protein_coding | -0.990784987 | 4.078831472 | 3.4005258 | 0.0834219 | 0.413809508 |
| ENSMUSG00000024854 | Pold4 | protein_coding | -0.586617714 | 5.691467525 | 3.3994276 | 0.0834674 | 0.413809508 |
| ENSMUSG00000022339 | Ebag9 | protein_coding | -0.685060065 | 4.976404661 | 3.3993492 | 0.0834706 | 0.413809508 |
| ENSMUSG00000030519 | Apba2 | protein_coding | 0.701258258 | 5.802000429 | 3.3991686 | 0.0835085 | 0.413809508 |
| ENSMUSG00000022533 | Atp13a3 | protein_coding | -0.848158164 | 4.077991839 | 3.3974553 | 0.0835491 | 0.413819926 |
| ENSMUSG00000010505 | Myt1 | protein_coding | 1.499367102 | 2.486515823 | 3.3925018 | 0.0837546 | 0.414557369 |
| ENSMUSG00000029569 | Tmem168 | protein_coding | -1.347802022 | 3.063842105 | 3.3920108 | 0.0837751 | 0.414557369 |
| ENSMUSG00000020290 | Xpo1 | protein_coding | -0.670770602 | 5.05689535 | 3.3890732 | 0.0838973 | 0.414971279 |
| ENSMUSG00000039693 | Msantd3 | protein_coding | -0.902346581 | 3.699884254 | 3.3877151 | 0.0839539 | 0.415060235 |
| ENSMUSG00000034343 | Ube2f | protein_coding | 0.47692905 | 6.296843582 | 3.3863051 | 0.0840127 | 0.415098025 |
| ENSMUSG00000054364 | Rhob | protein_coding | -0.438499018 | 7.058097079 | 3.385425 | 0.0840494 | 0.415098025 |
| ENSMUSG00000024483 | Ankhd1 | protein_coding | -0.650901408 | 4.962412996 | 3.3847557 | 0.0840773 | 0.415098025 |
| ENSMUSG00000046593 | Tmem215 | protein_coding | 0.737174314 | 4.79170462 | 3.3800309 | 0.0842749 | 0.415576491 |
| ENSMUSG00000034152 | Exoc3 | protein_coding | 0.46151423 | 7.219471058 | 3.3794608 | 0.0842987 | 0.415576491 |
| ENSMUSG00000014355 | Anapc1 | protein_coding | 0.878516996 | 4.217643733 | 3.3788446 | 0.0843246 | 0.415576491 |
| ENSMUSG00000021196 | Pfkp | protein_coding | 0.599853781 | 7.676349815 | 3.379469 | 0.0843288 | 0.415576491 |
| ENSMUSG00000028226 | Mmp16 | protein_coding | -0.876222137 | 4.054769991 | 3.3773445 | 0.0843874 | 0.415674813 |
| ENSMUSG00000024248 | Cox7a2l | protein_coding | 0.412148691 | 8.40287394 | 3.3762345 | 0.084434 | 0.415713724 |

| ensembl_gene_id | external_gene_name | gene_biotype | log2 (fold change) | log2 (counts per million) | F-value | p-value | False discovery rate |
|---|---|---|---|---|---|---|---|
| ENSMUSG00000037876 | Jmjd1c | protein_coding | -0.786143171 | 4.82454241 | 3.3741166 | 0.0845229 | 0.415833382 |
| ENSMUSG00000055447 | Cd47 | protein_coding | -0.450344374 | 7.192665503 | 3.3738139 | 0.0845356 | 0.415833382 |
| ENSMUSG00000005267 | Zfp287 | protein_coding | -1.008522774 | 3.255660461 | 3.3720151 | 0.0846113 | 0.416015099 |
| ENSMUSG00000018507 | Trpv2 | protein_coding | 1.075540367 | 4.695499816 | 3.3970782 | 0.0846711 | 0.416119207 |
| ENSMUSG00000047261 | Gap43 | protein_coding | 0.445155842 | 11.00325107 | 3.3684421 | 0.0847617 | 0.416374094 |
| ENSMUSG00000000326 | Comt | protein_coding | 0.431650182 | 7.220056283 | 3.3672907 | 0.0848103 | 0.416422382 |
| ENSMUSG00000040124 | Gorab | protein_coding | 0.728535991 | 4.695669483 | 3.3663619 | 0.0848495 | 0.416424646 |
| ENSMUSG00000038816 | Ctnnal1 | protein_coding | -0.893192459 | 3.863867728 | 3.3646033 | 0.0849237 | 0.416598972 |
| ENSMUSG00000053297 | AI854703 | protein_coding | 0.64779374 | 5.33239503 | 3.361891 | 0.0850384 | 0.416971347 |
| ENSMUSG00000052155 | Acvr2a | protein_coding | -0.875443384 | 3.817366856 | 3.3601312 | 0.0851129 | 0.417146474 |
| ENSMUSG00000022814 | Umps | protein_coding | 0.719405459 | 5.473291853 | 3.3580359 | 0.0852017 | 0.417391524 |
| ENSMUSG00000058076 | Sdhc | protein_coding | 0.4257629 | 9.022960936 | 3.3565395 | 0.0852652 | 0.41751237 |
| ENSMUSG00000044788 | Fads6 | protein_coding | -0.901052224 | 4.033293867 | 3.3542839 | 0.0853611 | 0.417581649 |
| ENSMUSG00000029463 | Fam216a | protein_coding | 0.870737204 | 4.12607671 | 3.3539996 | 0.0853731 | 0.417581649 |
| ENSMUSG00000055653 | Gpc3 | protein_coding | 0.725143264 | 4.439939438 | 3.3530967 | 0.0854115 | 0.417581649 |
| ENSMUSG00000006005 | Tpr | protein_coding | -0.646732544 | 5.924069098 | 3.3522303 | 0.0854484 | 0.417581649 |
| ENSMUSG00000004929 | Thop1 | protein_coding | 1.018542759 | 4.018110319 | 3.3516393 | 0.0854736 | 0.417581649 |
| ENSMUSG00000085980 | Gm12408 | lincRNA | 1.319486384 | 2.757608371 | 3.3480183 | 0.0856279 | 0.418020244 |
| ENSMUSG00000005413 | Hmox1 | protein_coding | 0.988805457 | 3.43502437 | 3.3477089 | 0.0856411 | 0.418020244 |
| ENSMUSG00000032294 | Pkm | protein_coding | 0.627431202 | 10.22861395 | 3.3462219 | 0.0857351 | 0.418191854 |
| ENSMUSG00000031370 | Zrsr2 | protein_coding | 0.47538017 | 6.645237934 | 3.3450643 | 0.085754 | 0.418191854 |
| ENSMUSG00000031523 | Dlc1 | protein_coding | -1.123709466 | 3.484914263 | 3.3427034 | 0.085855 | 0.418282194 |
| ENSMUSG00000029536 | Gatc | protein_coding | 0.479551703 | 5.99262506 | 3.3418536 | 0.0858914 | 0.418282194 |
| ENSMUSG00000027078 | Ube2l6 | protein_coding | 0.812561251 | 4.029431324 | 3.3412898 | 0.0859155 | 0.418282194 |
| ENSMUSG00000028064 | Sema4a | protein_coding | 0.919308784 | 4.49220998 | 3.340368 | 0.085955 | 0.418282194 |
| ENSMUSG00000021072 | Tmx1 | protein_coding | 0.475621718 | 7.163245087 | 3.3400874 | 0.085967 | 0.418282194 |
| ENSMUSG00000020537 | Drg2 | protein_coding | 0.559650849 | 7.094896345 | 3.3398346 | 0.0860084 | 0.418294164 |
| ENSMUSG00000036402 | Gng12 | protein_coding | -0.740397603 | 4.976671488 | 3.3372124 | 0.0860904 | 0.418503586 |
| ENSMUSG00000032352 | Lrrc1 | protein_coding | -1.033191528 | 3.422985212 | 3.3360804 | 0.086139 | 0.418550733 |
| ENSMUSG00000033066 | Gas7 | protein_coding | -0.64868957 | 6.236110823 | 3.335765 | 0.0861831 | 0.418575785 |
| ENSMUSG00000017344 | Vtn | protein_coding | -0.768128345 | 4.202792313 | 3.3339737 | 0.0862296 | 0.418612511 |
| ENSMUSG00000100594 | 2810414N06Rik | lincRNA | -1.11166026 | 3.632830618 | 3.3315676 | 0.0863332 | 0.418740309 |
| ENSMUSG00000047213 | Ythdf3 | protein_coding | -0.759827162 | 4.409674026 | 3.3299801 | 0.0864016 | 0.418740309 |
| ENSMUSG00000026248 | Mrpl44 | protein_coding | 0.438674319 | 7.060938854 | 3.3295175 | 0.0864215 | 0.418740309 |
| ENSMUSG00000006273 | Atp6v1b2 | protein_coding | 0.482414525 | 9.505910288 | 3.3294765 | 0.0864233 | 0.418740309 |
| ENSMUSG00000022231 | Sema5a | protein_coding | -0.819348145 | 4.641101121 | 3.3288435 | 0.0864506 | 0.418740309 |
| ENSMUSG00000087590 | Epb41l4aos | lincRNA | 0.956124086 | 3.339230179 | 3.3277581 | 0.0864975 | 0.418778607 |
| ENSMUSG00000086370 | Ftx | lincRNA | -1.159550129 | 3.516053268 | 3.3218498 | 0.086753 | 0.418983185 |
| ENSMUSG00000048379 | Socs4 | protein_coding | -1.484312039 | 2.690753874 | 3.3212544 | 0.0867788 | 0.418983185 |
| ENSMUSG00000015243 | Abca1 | protein_coding | -1.060940827 | 3.073190224 | 3.3208808 | 0.086795 | 0.418983185 |
| ENSMUSG00000030310 | Slc6a1 | protein_coding | -0.549195036 | 7.111988007 | 3.3192943 | 0.0868638 | 0.418983185 |
| ENSMUSG00000031901 | Dus2 | protein_coding | 0.658141579 | 5.148643558 | 3.31902 | 0.0868757 | 0.418983185 |
| ENSMUSG00000022601 | Zbtb11 | protein_coding | 0.966630166 | 3.69462832 | 3.3184618 | 0.0869 | 0.418983185 |
| ENSMUSG00000034118 | Tpst1 | protein_coding | -0.811479346 | 4.475285247 | 3.3164499 | 0.0869874 | 0.418983185 |
| ENSMUSG00000051166 | Eml5 | protein_coding | -0.844303589 | 3.710090778 | 3.3156279 | 0.0870231 | 0.418983185 |
| ENSMUSG00000030660 | Pik3c2a | protein_coding | -1.414910679 | 2.710355144 | 3.3152125 | 0.0870412 | 0.418983185 |
| ENSMUSG00000061461 | Smim20 | protein_coding | -0.406806391 | 7.151173044 | 3.3151944 | 0.087042 | 0.418983185 |
| ENSMUSG00000020848 | Doc2b | protein_coding | 0.895277764 | 3.522851538 | 3.3139114 | 0.0870978 | 0.418983185 |
| ENSMUSG00000000078 | Klf6 | protein_coding | -0.601528506 | 5.934176 | 3.3131277 | 0.0871319 | 0.418983185 |
| ENSMUSG00000025351 | Cd63 | protein_coding | -0.678656022 | 6.778323389 | 3.3321851 | 0.0871513 | 0.418983185 |
| ENSMUSG00000040548 | Tex2 | protein_coding | 0.475993314 | 6.582425221 | 3.3119141 | 0.0871848 | 0.418983185 |
| ENSMUSG00000026339 | Ccdc93 | protein_coding | -1.253713158 | 2.93749674 | 3.3119091 | 0.087185 | 0.418983185 |
| ENSMUSG00000034000 | Neu4 | protein_coding | -0.803366794 | 4.023571202 | 3.3100922 | 0.0872642 | 0.418983185 |
| ENSMUSG00000073139 | Tmem185a | protein_coding | 0.524595688 | 5.789414709 | 3.3096413 | 0.0872839 | 0.418983185 |
| ENSMUSG00000031937 | Vstm5 | protein_coding | 0.80468813 | 4.592154197 | 3.3083631 | 0.0873397 | 0.418983185 |
| ENSMUSG00000020176 | Grb10 | protein_coding | 0.610647596 | 5.982098793 | 3.3074842 | 0.0873781 | 0.418983185 |
| ENSMUSG00000043301 | Kcnj6 | protein_coding | -1.233396719 | 2.906168264 | 3.3043966 | 0.0875132 | 0.418983185 |
| ENSMUSG00000054720 | Lrrc8c | protein_coding | -0.803828123 | 3.937073174 | 3.3034359 | 0.0875552 | 0.418983185 |
| ENSMUSG00000049521 | Cdc42ep1 | protein_coding | -1.151090179 | 3.740462329 | 3.3026776 | 0.0876191 | 0.418983185 |
| ENSMUSG00000032702 | Kank1 | protein_coding | -0.930787595 | 3.697602007 | 3.3018358 | 0.0876254 | 0.418983185 |
| ENSMUSG00000070697 | Utp3 | protein_coding | 0.444512534 | 6.400541001 | 3.3011849 | 0.0876539 | 0.418983185 |
| ENSMUSG00000038347 | Tcte2 | protein_coding | -0.76352719 | 4.131523896 | 3.3008361 | 0.0876692 | 0.418983185 |
| ENSMUSG00000097881 | Celrr | lincRNA | -1.751883572 | 2.872962828 | 3.301434 | 0.0876736 | 0.418983185 |
| ENSMUSG00000027346 | Gpcpd1 | protein_coding | -0.629988763 | 4.817931451 | 3.300545 | 0.087682 | 0.418983185 |
| ENSMUSG00000003814 | Calr | protein_coding | 0.42826663 | 8.158670102 | 3.3005064 | 0.0876837 | 0.418983185 |
| ENSMUSG00000068299 | Nat8f4 | protein_coding | -1.011871648 | 3.405434141 | 3.299624 | 0.0877224 | 0.418983185 |
| ENSMUSG00000024847 | Aip | protein_coding | 0.434408398 | 8.863937582 | 3.298637 | 0.0877658 | 0.418983185 |
| ENSMUSG00000016487 | Ppfibp1 | protein_coding | -1.132859923 | 3.142430266 | 3.298204 | 0.0877848 | 0.418983185 |
| ENSMUSG00000037669 | Ldah | protein_coding | -0.447464823 | 6.544297754 | 3.2980719 | 0.0877906 | 0.418983185 |
| ENSMUSG00000048874 | Phf3 | protein_coding | -0.622236422 | 5.110300952 | 3.2972764 | 0.0878255 | 0.418983185 |
| ENSMUSG00000038976 | Ppp1r9b | protein_coding | -0.594777679 | 7.021831273 | 3.2965921 | 0.0878863 | 0.419086885 |
| ENSMUSG00000035849 | Krt222 | protein_coding | -0.567365894 | 5.881823538 | 3.2874093 | 0.0882606 | 0.420685455 |
| ENSMUSG00000041710 | Trpc5 | protein_coding | 1.167893888 | 3.258501206 | 3.2861791 | 0.088315 | 0.420758365 |
| ENSMUSG00000021476 | Habp4 | protein_coding | 0.495288691 | 6.94883417 | 3.2822933 | 0.0884872 | 0.421367792 |
| ENSMUSG00000029826 | Zc3hav1 | protein_coding | -0.975182722 | 3.925240295 | 3.2815241 | 0.0885213 | 0.421367792 |
| ENSMUSG00000031820 | Babam1 | protein_coding | 0.477788457 | 7.129763984 | 3.2793743 | 0.0886168 | 0.42163557 |
| ENSMUSG00000034640 | Tiparp | protein_coding | -0.919351526 | 4.451759757 | 3.2736186 | 0.088873 | 0.422501088 |
| ENSMUSG00000003420 | Fcgrt | protein_coding | -1.189815161 | 3.467224619 | 3.273522 | 0.0888773 | 0.422501088 |

| ensembl_gene_id | external_gene_name | gene_biotype | log2 (fold change) | log2 (counts per million) | F-value | p-value | False discovery rate |
|---|---|---|---|---|---|---|---|
| ENSMUSG00000053825 | Ppfia2 | protein_coding | -0.484923166 | 6.398884823 | 3.2724649 | 0.0889244 | 0.422538422 |
| ENSMUSG00000037475 | Thoc2 | protein_coding | -0.802257663 | 4.708718785 | 3.2676979 | 0.0891374 | 0.423199961 |
| ENSMUSG00000001416 | Cct3 | protein_coding | 0.456413585 | 7.875360436 | 3.2670888 | 0.0891647 | 0.423199961 |
| ENSMUSG00000037348 | Paqr7 | protein_coding | 0.550300082 | 6.590218466 | 3.2667081 | 0.0891817 | 0.423199961 |
| ENSMUSG00000029344 | Tpst2 | protein_coding | -0.976013618 | 3.349081953 | 3.265537 | 0.0892342 | 0.423262052 |
| ENSMUSG00000024403 | Atp6v1g2 | protein_coding | 0.438814022 | 10.46978886 | 3.2633716 | 0.0893312 | 0.423535681 |
| ENSMUSG00000033964 | Zbtb41 | protein_coding | -1.054466678 | 3.457689117 | 3.2600343 | 0.0894811 | 0.423932646 |
| ENSMUSG00000020743 | Mif4gd | protein_coding | -0.615059227 | 5.246410867 | 3.2597513 | 0.0894938 | 0.423932646 |
| ENSMUSG00000038056 | Kmt2c | protein_coding | -0.690901956 | 4.777930851 | 3.258211 | 0.0895631 | 0.424074002 |
| ENSMUSG00000064210 | Ano6 | protein_coding | -0.986155276 | 3.670365067 | 3.2561851 | 0.0896543 | 0.42431905 |
| ENSMUSG00000033813 | Tcea1 | protein_coding | -0.683057498 | 4.076756085 | 3.2543118 | 0.0897388 | 0.424381005 |
| ENSMUSG00000025140 | Pycr1 | protein_coding | -1.092012364 | 3.771223146 | 3.2539532 | 0.0897549 | 0.424381005 |
| ENSMUSG00000068696 | Gpr88 | protein_coding | -1.218169103 | 6.335451933 | 3.4347045 | 0.0897858 | 0.424381005 |
| ENSMUSG00000022386 | Trmu | protein_coding | 0.72260427 | 5.064305744 | 3.2523835 | 0.0898258 | 0.424383406 |
| ENSMUSG00000073680 | Tmem88b | protein_coding | -0.705279509 | 5.823505136 | 3.2512423 | 0.089908 | 0.424585515 |
| ENSMUSG00000033671 | Cep350 | protein_coding | -1.355966706 | 2.879776363 | 3.2488034 | 0.0899876 | 0.424774696 |
| ENSMUSG00000036368 | Rmdn2 | protein_coding | -1.507972452 | 2.618125722 | 3.2461981 | 0.0901056 | 0.425011113 |
| ENSMUSG00000032316 | Clk3 | protein_coding | 0.617419544 | 5.255660421 | 3.245952 | 0.0901167 | 0.425011113 |
| ENSMUSG00000020078 | Vps26a | protein_coding | -0.50027364 | 6.546155547 | 3.2414954 | 0.090319 | 0.425693883 |
| ENSMUSG00000029863 | Casp2 | protein_coding | -1.202538761 | 2.862350715 | 3.2396765 | 0.0904017 | 0.425693883 |
| ENSMUSG00000063077 | Kif1b | protein_coding | -0.420255276 | 7.914263442 | 3.239163 | 0.0904251 | 0.425693883 |
| ENSMUSG00000038172 | Ttc39b | protein_coding | -0.754781136 | 4.595200301 | 3.2388725 | 0.0904383 | 0.425693883 |
| ENSMUSG00000056486 | Chn1 | protein_coding | 0.399604433 | 7.21260601 | 3.2378209 | 0.0904862 | 0.425693883 |
| ENSMUSG00000049327 | Kmt5a | protein_coding | -0.757767305 | 4.106663803 | 3.2375389 | 0.090499 | 0.425693883 |
| ENSMUSG00000028549 | Itgb3bp | protein_coding | -0.696210393 | 4.194755483 | 3.2326197 | 0.0907235 | 0.426562951 |
| ENSMUSG00000071553 | Cpa2 | protein_coding | 0.700730894 | 4.811288757 | 3.2272375 | 0.0909698 | 0.427428395 |
| ENSMUSG00000022337 | Emc2 | protein_coding | -0.42322709 | 7.531188737 | 3.2268609 | 0.090987 | 0.427428395 |
| ENSMUSG00000031278 | Acsl4 | protein_coding | -0.556946736 | 6.224603325 | 3.2246017 | 0.0910907 | 0.427628387 |
| ENSMUSG00000090063 | Dlx6os1 | antisense_RNA | -0.999233355 | 4.559342189 | 3.2242771 | 0.0912175 | 0.427628387 |
| ENSMUSG00000060090 | Rp2 | protein_coding | -1.452293648 | 2.647798057 | 3.2218294 | 0.0912181 | 0.427628387 |
| ENSMUSG00000002345 | Borcs8 | protein_coding | 0.479249686 | 7.178621538 | 3.2215005 | 0.0912332 | 0.427628387 |
| ENSMUSG00000030030 | 1700003E16Rik | protein_coding | 0.62654638 | 5.293670112 | 3.2203342 | 0.0912868 | 0.427628387 |
| ENSMUSG00000014349 | Ube2z | protein_coding | 0.489181632 | 6.026995833 | 3.2200483 | 0.0913 | 0.427628387 |
| ENSMUSG00000029388 | Eif2b1 | protein_coding | 0.410024347 | 6.75653842 | 3.2198751 | 0.091308 | 0.427628387 |
| ENSMUSG00000075706 | Gpx4 | protein_coding | 0.4336649 | 8.737888227 | 3.2167438 | 0.0914523 | 0.428021194 |
| ENSMUSG00000044005 | Gls2 | protein_coding | 0.925894043 | 3.5291459 | 3.2157371 | 0.0914987 | 0.428021194 |
| ENSMUSG00000033972 | Zfp944 | protein_coding | -0.807101058 | 3.843087249 | 3.2131379 | 0.0916188 | 0.428021194 |
| ENSMUSG00000048106 | 4632415L05Rik | lincRNA | -0.999001774 | 3.111521231 | 3.2127703 | 0.0916358 | 0.428021194 |
| ENSMUSG00000037224 | Zfyve28 | protein_coding | -0.766196381 | 3.891816337 | 3.2117736 | 0.0916819 | 0.428021194 |
| ENSMUSG00000048728 | Zfp454 | protein_coding | 0.988609313 | 3.464650488 | 3.2114883 | 0.0916951 | 0.428021194 |
| ENSMUSG00000020009 | Ifngr1 | protein_coding | 0.801941266 | 4.016042888 | 3.2105201 | 0.0917399 | 0.428021194 |
| ENSMUSG00000019080 | Mfsd3 | protein_coding | 0.884881018 | 4.22252676 | 3.2104246 | 0.0917443 | 0.428021194 |
| ENSMUSG00000047547 | Cltb | protein_coding | 0.527268147 | 7.68489647 | 3.2102995 | 0.0917501 | 0.428021194 |
| ENSMUSG00000032217 | Rnf111 | protein_coding | -0.703943212 | 4.463264588 | 3.2085357 | 0.0918318 | 0.428095826 |
| ENSMUSG00000032288 | Imp3 | protein_coding | 0.760273925 | 4.824134145 | 3.2082356 | 0.0918457 | 0.428095826 |
| ENSMUSG00000034724 | Cnot6l | protein_coding | -0.632293656 | 5.221231872 | 3.2065071 | 0.0919259 | 0.428283888 |
| ENSMUSG00000078652 | Psme3 | protein_coding | -0.448714415 | 6.240512895 | 3.2055309 | 0.0919712 | 0.42830946 |
| ENSMUSG00000024901 | Peli3 | protein_coding | 0.808940687 | 3.903887518 | 3.2036551 | 0.0920584 | 0.428394492 |
| ENSMUSG00000005583 | Mef2c | protein_coding | -0.566415229 | 6.856169045 | 3.2040861 | 0.0920692 | 0.428394492 |
| ENSMUSG00000022130 | Tgds | protein_coding | -1.594034823 | 2.404967611 | 3.202336 | 0.0921197 | 0.428444366 |
| ENSMUSG00000027763 | Mbnl1 | protein_coding | -0.508196886 | 6.597685114 | 3.2012034 | 0.0921724 | 0.428504177 |
| ENSMUSG00000026224 | 4933407L21Rik | protein_coding | -0.745992293 | 4.27700674 | 3.1968913 | 0.0923734 | 0.429158734 |
| ENSMUSG00000024429 | Gnl1 | protein_coding | 0.453162981 | 6.405451743 | 3.1944938 | 0.0924854 | 0.429158734 |
| ENSMUSG00000031765 | Mt1 | protein_coding | -0.849687921 | 10.16083644 | 3.2544746 | 0.0924938 | 0.429158734 |
| ENSMUSG00000023456 | Tpi1 | protein_coding | 0.389844121 | 9.877387181 | 3.1939228 | 0.0925121 | 0.429158734 |
| ENSMUSG00000060538 | Tmem219 | protein_coding | 0.705451682 | 4.342771791 | 3.193909 | 0.0925128 | 0.429158734 |
| ENSMUSG00000031156 | Slc35a2 | protein_coding | 0.573671036 | 5.421110488 | 3.190619 | 0.0926668 | 0.429687724 |
| ENSMUSG00000026004 | Kansl1l | protein_coding | -1.17203401 | 3.449601336 | 3.1863132 | 0.0928688 | 0.430438776 |
| ENSMUSG00000068134 | Zfp120 | protein_coding | -0.831591237 | 4.44365837 | 3.1852638 | 0.0929181 | 0.430450168 |
| ENSMUSG00000040842 | Szrd1 | protein_coding | -0.635762704 | 5.368235244 | 3.1845575 | 0.0929513 | 0.430450168 |
| ENSMUSG00000047766 | Lrrc49 | protein_coding | 0.53081569 | 6.572946789 | 3.1832893 | 0.093011 | 0.430541037 |
| ENSMUSG00000036882 | Arhgap33 | protein_coding | -0.637140073 | 5.61869211 | 3.1778366 | 0.093268 | 0.431001429 |
| ENSMUSG00000087679 | Tmem250-ps | lincRNA | 0.546714736 | 9.365723529 | 3.1769875 | 0.0933081 | 0.431001429 |
| ENSMUSG00000035969 | Rusc2 | protein_coding | -0.732808257 | 4.488377439 | 3.1768673 | 0.0933138 | 0.431001429 |
| ENSMUSG00000026083 | Eif5b | protein_coding | -0.606193228 | 6.243742382 | 3.1751945 | 0.0933928 | 0.431001429 |
| ENSMUSG00000040652 | Oaz2 | protein_coding | 0.472291422 | 6.202725126 | 3.1749688 | 0.0934035 | 0.431001429 |
| ENSMUSG00000027519 | Rab22a | protein_coding | -0.51103398 | 5.427502453 | 3.1744995 | 0.0934257 | 0.431001429 |
| ENSMUSG00000041828 | Abca8a | protein_coding | 1.171810909 | 3.14266088 | 3.1744726 | 0.093427 | 0.431001429 |
| ENSMUSG00000038451 | Spsb2 | protein_coding | -0.802529925 | 3.76742242 | 3.1736064 | 0.093468 | 0.431001429 |
| ENSMUSG00000022442 | Ttll1 | protein_coding | 0.625224571 | 7.433576324 | 3.1901478 | 0.0934758 | 0.431001429 |
| ENSMUSG00000052331 | Ankrd44 | protein_coding | -1.631727798 | 2.457030023 | 3.1721436 | 0.0935372 | 0.431001429 |
| ENSMUSG00000022475 | Hdac7 | protein_coding | 0.936933134 | 4.279906756 | 3.1718464 | 0.0935513 | 0.431001429 |
| ENSMUSG00000036257 | Pnpla8 | protein_coding | -0.460218219 | 6.179466432 | 3.1702474 | 0.0936271 | 0.43116593 |
| ENSMUSG00000024664 | Fads3 | protein_coding | 1.120978458 | 4.01031925 | 3.1693439 | 0.0937823 | 0.43122367 |
| ENSMUSG00000059866 | Tnip2 | protein_coding | -1.286066976 | 2.952295257 | 3.1656747 | 0.0938443 | 0.43122367 |
| ENSMUSG00000027864 | Ptgfrn | protein_coding | -0.939384429 | 4.140524915 | 3.1654216 | 0.0938563 | 0.43122367 |
| ENSMUSG00000040659 | Efhd2 | protein_coding | 0.490685431 | 6.788462381 | 3.1652973 | 0.0938623 | 0.43122367 |
| ENSMUSG00000061758 | Akr1b10 | protein_coding | -0.559741219 | 5.577998856 | 3.1650192 | 0.0938755 | 0.43122367 |

| ensembl_gene_id | external_gene_name | gene_biotype | log2 (fold change) | log2 (counts per million) | F-value | p-value | False discovery rate |
|---|---|---|---|---|---|---|---|
| ENSMUSG00000029446 | Psph | protein_coding | -0.596079236 | 5.792015047 | 3.1649187 | 0.0938803 | 0.43122367 |
| ENSMUSG00000033540 | Idua | protein_coding | -0.793549869 | 4.047153992 | 3.1601547 | 0.0941073 | 0.432081828 |
| ENSMUSG00000021607 | Mrpl36 | protein_coding | 0.372537599 | 7.686975598 | 3.1585883 | 0.0941821 | 0.432234855 |
| ENSMUSG00000113029 | AC122190.1 | antisense_RNA | -1.133901975 | 3.301354643 | 3.1569689 | 0.0942595 | 0.432234855 |
| ENSMUSG00000027584 | Oprl1 | protein_coding | 0.551952264 | 6.424172654 | 3.1569324 | 0.0942612 | 0.432234855 |
| ENSMUSG00000052981 | Ube2ql1 | protein_coding | 0.477190642 | 6.85934638 | 3.1534617 | 0.0944273 | 0.432812014 |
| ENSMUSG00000032802 | Srxn1 | protein_coding | 0.506781019 | 6.536059391 | 3.1505861 | 0.0945652 | 0.43312028 |
| ENSMUSG00000034912 | Mdga2 | protein_coding | -0.797241674 | 5.501504469 | 3.1505324 | 0.0945987 | 0.43312028 |
| ENSMUSG00000048076 | Arf1 | protein_coding | 0.443892114 | 9.201643865 | 3.1482477 | 0.0946775 | 0.43312028 |
| ENSMUSG00000032081 | Apoc3 | protein_coding | 1.143704611 | 3.179063449 | 3.1482239 | 0.0946787 | 0.43312028 |
| ENSMUSG00000030604 | Zfp626 | protein_coding | -1.374223787 | 2.699631647 | 3.1478642 | 0.094696 | 0.43312028 |
| ENSMUSG00000062866 | Phactr2 | protein_coding | -0.595609099 | 5.45711007 | 3.1434936 | 0.0949064 | 0.433898028 |
| ENSMUSG00000022561 | Gpaa1 | protein_coding | 0.676531388 | 5.135706021 | 3.1416899 | 0.0949934 | 0.434111152 |
| ENSMUSG00000070047 | Fat1 | protein_coding | -1.083638132 | 3.598370895 | 3.1362202 | 0.0952577 | 0.435134397 |
| ENSMUSG00000025375 | Aatk | protein_coding | -0.613086615 | 5.730584382 | 3.1273659 | 0.0956876 | 0.436912243 |
| ENSMUSG00000024498 | Tcerg1 | protein_coding | -0.636144933 | 5.115507489 | 3.1232137 | 0.0958899 | 0.437519222 |
| ENSMUSG00000026235 | Epha4 | protein_coding | -0.641155064 | 6.211637622 | 3.1230273 | 0.09593 | 0.437519222 |
| ENSMUSG00000047141 | Zfp654 | protein_coding | -0.857172491 | 3.831966821 | 3.1221353 | 0.0959426 | 0.437519222 |
| ENSMUSG00000030411 | Nova2 | protein_coding | -0.63178816 | 5.176793 | 3.1158998 | 0.0962476 | 0.438533054 |
| ENSMUSG00000026342 | Slc35f5 | protein_coding | 1.395670969 | 3.268655608 | 3.115938 | 0.0962767 | 0.438533054 |
| ENSMUSG00000075704 | Txnrd2 | protein_coding | 0.959071742 | 3.669811175 | 3.1150917 | 0.0962872 | 0.438533054 |
| ENSMUSG00000004698 | Hdac9 | protein_coding | -0.637947329 | 5.084762076 | 3.1130575 | 0.096387 | 0.43880178 |
| ENSMUSG00000041355 | Ssr2 | protein_coding | 0.413923323 | 7.580759846 | 3.1111997 | 0.0964783 | 0.439031402 |
| ENSMUSG00000046671 | Mtfr1l | protein_coding | 0.412709415 | 7.657374727 | 3.1097608 | 0.0965491 | 0.439051635 |
| ENSMUSG00000032298 | Neil1 | protein_coding | -1.521125557 | 3.125484663 | 3.1100789 | 0.0965644 | 0.439051635 |
| ENSMUSG00000026443 | Lrrn2 | protein_coding | 0.517233414 | 5.853491411 | 3.1068925 | 0.0966903 | 0.439433414 |
| ENSMUSG00000058013 | 11-Sep | protein_coding | -0.506048727 | 6.660549514 | 3.1055468 | 0.0967566 | 0.439433414 |
| ENSMUSG00000022321 | Cdh10 | protein_coding | -0.740664442 | 4.203458714 | 3.1043914 | 0.0968136 | 0.439433414 |
| ENSMUSG00000044364 | Tmem74b | protein_coding | 0.901609142 | 3.394795497 | 3.1036076 | 0.0968523 | 0.439433414 |
| ENSMUSG00000052430 | Bmpr1b | protein_coding | -1.14703228 | 3.043873098 | 3.1029015 | 0.0968872 | 0.439433414 |
| ENSMUSG00000026005 | Rpe | protein_coding | -0.860120825 | 4.424036185 | 3.1027728 | 0.0968936 | 0.439433414 |
| ENSMUSG00000020263 | Appl2 | protein_coding | -0.633580593 | 4.59630411 | 3.0964716 | 0.0972055 | 0.440125583 |
| ENSMUSG00000038909 | Kat7 | protein_coding | 0.605958065 | 6.342927976 | 3.0966558 | 0.0972274 | 0.440125583 |
| ENSMUSG00000021711 | Trappc13 | protein_coding | -0.408606956 | 7.177096875 | 3.0943221 | 0.0973122 | 0.440125583 |
| ENSMUSG00000020570 | Sypl | protein_coding | -0.492744743 | 6.263794843 | 3.0934087 | 0.0973575 | 0.440125583 |
| ENSMUSG00000022972 | 1110004E09Rik | protein_coding | 0.848184317 | 6.017678376 | 3.1312584 | 0.0973797 | 0.440125583 |
| ENSMUSG00000030605 | Mfge8 | protein_coding | -0.621186609 | 7.272475873 | 3.1078628 | 0.0974334 | 0.440125583 |
| ENSMUSG00000020706 | Ftsj3 | protein_coding | 0.833605026 | 4.824402061 | 3.0924341 | 0.097437 | 0.440125583 |
| ENSMUSG00000044786 | Zfp36 | protein_coding | 1.300885023 | 2.927527003 | 3.0908789 | 0.0974833 | 0.440125583 |
| ENSMUSG00000075415 | Fnbp1 | protein_coding | -0.516186334 | 6.990679437 | 3.0902556 | 0.0975144 | 0.440125583 |
| ENSMUSG00000066877 | Nck2 | protein_coding | -0.833306085 | 3.486586342 | 3.0891504 | 0.0975694 | 0.440125583 |
| ENSMUSG00000002996 | Hbp1 | protein_coding | 0.856290097 | 3.632420087 | 3.0876724 | 0.097643 | 0.440125583 |
| ENSMUSG00000034024 | Cct2 | protein_coding | 0.413038166 | 8.002043955 | 3.0876044 | 0.0976464 | 0.440125583 |
| ENSMUSG00000047507 | Baiap3 | protein_coding | 0.698889714 | 6.667296313 | 3.1093115 | 0.0977729 | 0.440125583 |
| ENSMUSG00000024565 | Sall3 | protein_coding | -1.451920227 | 2.707476908 | 3.0850194 | 0.0977754 | 0.440125583 |
| ENSMUSG00000031979 | Cog2 | protein_coding | 0.704416447 | 4.659814181 | 3.08425 | 0.0978139 | 0.440125583 |
| ENSMUSG00000018585 | Atox1 | protein_coding | 0.437109268 | 8.537547706 | 3.0838663 | 0.097833 | 0.440125583 |
| ENSMUSG00000032018 | Sc5d | protein_coding | 0.43030457 | 6.268769391 | 3.0828831 | 0.0978822 | 0.440125583 |
| ENSMUSG00000035934 | Pknox2 | protein_coding | 0.608038132 | 5.223197322 | 3.0824915 | 0.0979018 | 0.440125583 |
| ENSMUSG00000052917 | Senp7 | protein_coding | -0.54482162 | 5.549636345 | 3.0820923 | 0.0979218 | 0.440125583 |
| ENSMUSG00000027933 | Ints3 | protein_coding | 0.673708949 | 4.900079861 | 3.0812461 | 0.0979641 | 0.440125583 |
| ENSMUSG00000026511 | Srp9 | protein_coding | 0.39426542 | 7.03976577 | 3.0812245 | 0.0979652 | 0.440125583 |
| ENSMUSG00000025505 | Tmem80 | protein_coding | 0.572407019 | 6.001325798 | 3.0809656 | 0.0979781 | 0.440125583 |
| ENSMUSG00000045664 | Cdc42ep2 | protein_coding | -0.667903943 | 5.455680769 | 3.0807767 | 0.0979876 | 0.440125583 |
| ENSMUSG00000038181 | Chpf2 | protein_coding | 2.050288174 | 2.365429595 | 3.0745537 | 0.0982998 | 0.441009858 |
| ENSMUSG00000036966 | Spryd3 | protein_coding | 0.531208744 | 8.680516596 | 3.0744887 | 0.098303 | 0.441009858 |
| ENSMUSG00000067028 | Cntnap5b | protein_coding | -1.208884871 | 2.91609367 | 3.0737366 | 0.0983409 | 0.441009858 |
| ENSMUSG00000026771 | Spopl | protein_coding | -0.98123928 | 3.182406445 | 3.0726509 | 0.0983955 | 0.441009858 |
| ENSMUSG00000028470 | Hint2 | protein_coding | 0.924018342 | 4.938380281 | 3.07433 | 0.0984239 | 0.441009858 |
| ENSMUSG00000040997 | Abhd4 | protein_coding | -0.445291631 | 6.797734357 | 3.0719539 | 0.0984305 | 0.441009858 |
| ENSMUSG00000040599 | Mis12 | protein_coding | 0.721565022 | 4.846130486 | 3.069941 | 0.0985319 | 0.44127392 |
| ENSMUSG00000020696 | Rffl | protein_coding | 1.196857699 | 3.31766545 | 3.0691551 | 0.0985716 | 0.44127392 |
| ENSMUSG00000022517 | Mgrn1 | protein_coding | 0.951451828 | 6.009870723 | 3.1265099 | 0.0986631 | 0.44145944 |
| ENSMUSG00000025777 | Gdap1 | protein_coding | -0.429738561 | 6.892165658 | 3.0667068 | 0.0986951 | 0.44145944 |
| ENSMUSG00000019772 | Vip | protein_coding | 1.035038084 | 3.329568435 | 3.0656848 | 0.0987467 | 0.441506729 |
| ENSMUSG00000011877 | Git1 | protein_coding | -0.423161541 | 6.426263458 | 3.0644646 | 0.0988084 | 0.441598899 |
| ENSMUSG00000021268 | Meg3 | lincRNA | -1.048141545 | 7.508684122 | 3.2268093 | 0.0988984 | 0.441817185 |
| ENSMUSG00000020366 | Mapk9 | protein_coding | 0.42552047 | 6.710022446 | 3.0612509 | 0.0989711 | 0.441958563 |
| ENSMUSG00000047583 | Tyw3 | protein_coding | -0.642398046 | 4.365579562 | 3.0585898 | 0.099106 | 0.442078173 |
| ENSMUSG00000046756 | Mrps7 | protein_coding | 0.431832852 | 6.693885339 | 3.0579075 | 0.0991407 | 0.442078173 |
| ENSMUSG00000085622 | 3110056K07Rik | processed_transcript | -0.931859798 | 4.492442256 | 3.0584571 | 0.0991439 | 0.442078173 |
| ENSMUSG00000033931 | Rbm34 | protein_coding | -0.664318824 | 4.615637206 | 3.0572107 | 0.0991761 | 0.442078173 |
| ENSMUSG00000035378 | Shq1 | protein_coding | 1.251708858 | 2.995015407 | 3.0566718 | 0.0992034 | 0.442078173 |
| ENSMUSG00000032500 | Dclk3 | protein_coding | -0.909932527 | 4.730840745 | 3.0561334 | 0.0992619 | 0.442155413 |
| ENSMUSG00000060675 | Pla2g16 | protein_coding | -0.657018976 | 6.167451262 | 3.0542781 | 0.0993563 | 0.442370592 |
| ENSMUSG00000000325 | Arvcf | protein_coding | -0.868917851 | 3.429181469 | 3.0522566 | 0.0994281 | 0.442370592 |
| ENSMUSG00000060924 | Csmd1 | protein_coding | -0.960802171 | 4.672874283 | 3.0548963 | 0.099439 | 0.442370592 |
| ENSMUSG00000024581 | Napg | protein_coding | 0.44091821 | 7.799076739 | 3.0511247 | 0.0994858 | 0.442370592 |

| ensembl_gene_id | external_gene_name | gene_biotype | log2 (fold change) | log2 (counts per million) | F-value | p-value | False discovery rate |
|---|---|---|---|---|---|---|---|
| ENSMUSG00000019579 | Mydgf | protein_coding | 0.424535469 | 7.3175133 | 3.0500057 | 0.0995428 | 0.442370592 |
| ENSMUSG00000091705 | H2-Q2 | protein_coding | 1.495567302 | 2.492497233 | 3.0497276 | 0.099557 | 0.442370592 |
| ENSMUSG00000017734 | Dbndd2 | protein_coding | -0.627647927 | 5.176144182 | 3.0477087 | 0.0996601 | 0.442645735 |
| ENSMUSG00000097587 | 4930578M01Rik | lincRNA | -1.075903016 | 3.936053073 | 3.045956 | 0.0997808 | 0.442998839 |
| ENSMUSG00000034765 | Dusp5 | protein_coding | 1.612382832 | 2.497909975 | 3.0425778 | 0.0999227 | 0.443085229 |
| ENSMUSG00000029209 | Gnpda2 | protein_coding | -0.500794077 | 5.92283896 | 3.042541 | 0.0999246 | 0.443085229 |
| ENSMUSG00000036093 | Arl5a | protein_coding | -0.680344635 | 4.645275477 | 3.0422845 | 0.0999377 | 0.443085229 |
| ENSMUSG00000029108 | Pcdh7 | protein_coding | -0.55206407 | 5.97182435 | 3.0417501 | 0.0999651 | 0.443085229 |
| ENSMUSG00000021690 | Jmy | protein_coding | -0.790961933 | 4.531544115 | 3.0393547 | 0.100088 | 0.443447271 |
| ENSMUSG00000036893 | Ehmt1 | protein_coding | 0.574259079 | 5.202402411 | 3.0380589 | 0.1001546 | 0.44355945 |
| ENSMUSG00000019528 | Gyg | protein_coding | 0.500238838 | 6.934157067 | 3.0366036 | 0.1002294 | 0.443708078 |
| ENSMUSG00000031980 | Agt | protein_coding | -0.539648188 | 6.828260512 | 3.0351601 | 0.1003037 | 0.443854208 |
| ENSMUSG00000063018 | 2010204K13Rik | processed_transcript | 0.476715083 | 7.865832667 | 3.0328144 | 0.1004246 | 0.443960858 |
| ENSMUSG00000096210 | H1f0 | protein_coding | 0.477152706 | 5.834766493 | 3.0326765 | 0.1004317 | 0.443960858 |
| ENSMUSG00000020409 | Slu7 | protein_coding | -0.451589934 | 7.116265078 | 3.0322887 | 0.1004517 | 0.443960858 |
| ENSMUSG00000001517 | Foxm1 | protein_coding | 1.271551268 | 2.814224342 | 3.0303359 | 0.1005524 | 0.443982438 |
| ENSMUSG00000038872 | Zfhx3 | protein_coding | -0.511785233 | 6.113214942 | 3.0301669 | 0.1005612 | 0.443982438 |
| ENSMUSG00000048562 | Sp8 | protein_coding | -0.786411276 | 5.007978714 | 3.028922 | 0.1006255 | 0.443982438 |
| ENSMUSG00000053289 | Ddx10 | protein_coding | -0.712880437 | 5.026413245 | 3.0283622 | 0.1006544 | 0.443982438 |
| ENSMUSG00000034126 | Pomt2 | protein_coding | 1.733636192 | 2.751272529 | 3.0279358 | 0.1007076 | 0.443982438 |
| ENSMUSG00000002028 | Kmt2a | protein_coding | -0.708358656 | 5.263140623 | 3.0265184 | 0.1007498 | 0.443982438 |
| ENSMUSG00000022858 | Tra2b | protein_coding | -0.481065353 | 6.079785217 | 3.0263851 | 0.1007567 | 0.443982438 |
| ENSMUSG00000028669 | Pithd1 | protein_coding | 0.436949134 | 6.480969073 | 3.0250336 | 0.1008267 | 0.443982438 |
| ENSMUSG00000028729 | Ebna1bp2 | protein_coding | 0.467809923 | 6.966038955 | 3.0250051 | 0.1008282 | 0.443982438 |
| ENSMUSG00000021171 | Esyt2 | protein_coding | -0.725123822 | 4.758010707 | 3.0237497 | 0.1008932 | 0.444087075 |
| ENSMUSG00000043670 | Diras1 | protein_coding | 0.840938599 | 5.99051786 | 3.0585312 | 0.1009979 | 0.444365837 |
| ENSMUSG00000034781 | Gna11 | protein_coding | 0.492113874 | 8.147263377 | 3.0178779 | 0.1011982 | 0.445065069 |
| ENSMUSG00000021669 | Col4a3bp | protein_coding | -0.610130509 | 4.648174305 | 3.0164058 | 0.1012748 | 0.445220008 |
| ENSMUSG00000001018 | Snapin | protein_coding | 0.669574667 | 6.521484819 | 3.0294052 | 0.1013947 | 0.445482943 |
| ENSMUSG00000015120 | Ube2i | protein_coding | 0.903116367 | 3.638052954 | 3.0136299 | 0.1014195 | 0.445482943 |
| ENSMUSG00000025371 | Chmp6 | protein_coding | 0.648846485 | 5.972846728 | 3.0134718 | 0.1014589 | 0.445482943 |
| ENSMUSG00000094002 | Gm9866 | lincRNA | 0.416993011 | 6.629773502 | 3.0113339 | 0.1015394 | 0.44565422 |
| ENSMUSG00000030878 | Cdr2 | protein_coding | 0.918539472 | 3.563215937 | 3.0078338 | 0.1017224 | 0.445838244 |
| ENSMUSG00000020705 | Ddx42 | protein_coding | 0.502403954 | 5.811682995 | 3.0076631 | 0.1017314 | 0.445838244 |
| ENSMUSG00000023088 | Abcc1 | protein_coding | 2.374416492 | 2.924059921 | 3.0586926 | 0.1017531 | 0.445838244 |
| ENSMUSG00000019966 | Kitl | protein_coding | -0.883311529 | 4.71695056 | 3.0071553 | 0.1017891 | 0.445838244 |
| ENSMUSG00000035086 | Becn1 | protein_coding | 0.892442483 | 3.339431092 | 3.0055227 | 0.1018435 | 0.445838244 |
| ENSMUSG00000029804 | Herc3 | protein_coding | 0.525912567 | 5.997854704 | 3.0049922 | 0.1018713 | 0.445838244 |
| ENSMUSG00000055436 | Srsf11 | protein_coding | -0.553974166 | 5.796884674 | 3.0046286 | 0.1018904 | 0.445838244 |
| ENSMUSG00000031618 | Nr3c2 | protein_coding | -0.865407901 | 4.117893082 | 3.0032212 | 0.1019643 | 0.445838244 |
| ENSMUSG00000044894 | Uqcrq | protein_coding | 0.47034025 | 10.39844636 | 3.0028451 | 0.101984 | 0.445838244 |
| ENSMUSG00000012405 | Rpl15 | protein_coding | 0.415426743 | 6.940425499 | 3.0025919 | 0.1019973 | 0.445838244 |
| ENSMUSG00000021710 | Nln | protein_coding | 0.752877867 | 4.26161093 | 3.0013292 | 0.1020636 | 0.445838244 |
| ENSMUSG00000041126 | H2afv | protein_coding | -0.511597112 | 5.493399484 | 3.0007669 | 0.1020932 | 0.445838244 |
| ENSMUSG00000008668 | Rps18 | protein_coding | 0.51124757 | 7.167909004 | 2.9999891 | 0.1021341 | 0.445838244 |
| ENSMUSG00000036890 | Gtdc1 | protein_coding | -0.766660007 | 4.693488783 | 2.9993259 | 0.102169 | 0.445838244 |
| ENSMUSG00000032583 | Mon1a | protein_coding | 0.7441869 | 4.200117861 | 2.9971084 | 0.1022858 | 0.445838244 |
| ENSMUSG00000021493 | Pdlim7 | protein_coding | 0.405124812 | 6.876904623 | 2.9967077 | 0.1023069 | 0.445838244 |
| ENSMUSG00000022639 | Dubr | lincRNA | -1.042783293 | 3.784583327 | 2.9965464 | 0.1023154 | 0.445838244 |
| ENSMUSG00000051950 | B3glct | protein_coding | -1.189683789 | 3.011145977 | 2.9959002 | 0.1023495 | 0.445838244 |
| ENSMUSG00000024208 | Uqcc2 | protein_coding | 0.448536358 | 9.520094615 | 2.9947249 | 0.1024115 | 0.445838244 |
| ENSMUSG00000016541 | Atxn10 | protein_coding | 0.495478889 | 9.478745114 | 2.9942822 | 0.1024349 | 0.445838244 |
| ENSMUSG00000020530 | Ggnbp2 | protein_coding | -0.410304976 | 7.034951486 | 2.9939577 | 0.102452 | 0.445838244 |
| ENSMUSG00000063531 | Sema3e | protein_coding | -1.205928097 | 2.783433438 | 2.9921742 | 0.1025462 | 0.446067761 |
| ENSMUSG00000061603 | Akap6 | protein_coding | -0.517559007 | 5.891526183 | 2.9893767 | 0.1026942 | 0.446167089 |
| ENSMUSG00000042978 | Sbk1 | protein_coding | -0.859606489 | 4.205062604 | 2.9882582 | 0.1027535 | 0.446167089 |
| ENSMUSG00000041645 | Ddx24 | protein_coding | 0.429210133 | 8.638056338 | 2.987847 | 0.1027753 | 0.446167089 |
| ENSMUSG00000037343 | Taf2 | protein_coding | 0.820394798 | 3.868903352 | 2.9874247 | 0.1027977 | 0.446167089 |
| ENSMUSG00000004937 | Sgta | protein_coding | 0.639273228 | 8.72233234 | 3.0101701 | 0.1028081 | 0.446167089 |
| ENSMUSG00000064138 | Fam172a | protein_coding | -0.995674349 | 3.102864087 | 2.9865876 | 0.102842 | 0.446167089 |
| ENSMUSG00000023852 | Chd1 | protein_coding | -0.391054499 | 6.615970631 | 2.9862585 | 0.1028595 | 0.446167089 |
| ENSMUSG00000021003 | Galc | protein_coding | 1.128527634 | 3.282337328 | 2.9829047 | 0.1030376 | 0.446759465 |
| ENSMUSG00000039488 | Cntn5 | protein_coding | -1.131608454 | 3.316897843 | 2.9800576 | 0.1031891 | 0.446761829 |
| ENSMUSG00000022842 | Ece2 | protein_coding | 0.625702809 | 5.364220344 | 2.9798212 | 0.1032017 | 0.446761829 |
| ENSMUSG00000059586 | Nsmce2 | protein_coding | 0.507521984 | 6.325690648 | 2.9794691 | 0.1032205 | 0.446761829 |
| ENSMUSG00000047115 | Fam221a | protein_coding | 1.777841244 | 2.264758872 | 2.9790911 | 0.1032406 | 0.446761829 |
| ENSMUSG00000040907 | Atp1a3 | protein_coding | 0.397896649 | 9.92749935 | 2.9789916 | 0.1032459 | 0.446761829 |
| ENSMUSG00000025137 | Pcyt2 | protein_coding | 0.491552419 | 7.272872055 | 2.976849 | 0.1033602 | 0.447076319 |
| ENSMUSG00000021486 | Prelid1 | protein_coding | 0.435148883 | 7.383620694 | 2.9751241 | 0.1034522 | 0.447150433 |
| ENSMUSG00000031333 | Abcb7 | protein_coding | -0.751583683 | 4.066281437 | 2.9749705 | 0.1034605 | 0.447150433 |
| ENSMUSG00000029001 | Fbxo44 | protein_coding | 0.361961382 | 7.966833578 | 2.9715347 | 0.1036442 | 0.447764609 |
| ENSMUSG00000029924 | Slc37a3 | protein_coding | 0.714904844 | 5.162331966 | 2.9683949 | 0.1038125 | 0.448311399 |
| ENSMUSG00000037253 | Mex3c | protein_coding | -0.97197104 | 3.938099964 | 2.9668085 | 0.1038976 | 0.44849822 |
| ENSMUSG00000019845 | Tube1 | protein_coding | -1.664857421 | 2.32141918 | 2.9656833 | 0.103958 | 0.44849822 |
| ENSMUSG00000112117 | RMST_1 | lincRNA | -0.86542776 | 3.455362308 | 2.9645736 | 0.1040177 | 0.44849822 |
| ENSMUSG00000029364 | Wsb2 | protein_coding | 0.544510562 | 5.49505858 | 2.964483 | 0.1040226 | 0.44849822 |
| ENSMUSG00000022427 | Tomm22 | protein_coding | 0.376398598 | 7.662045527 | 2.9622256 | 0.104144 | 0.448678097 |
| ENSMUSG00000020549 | Elac2 | protein_coding | 0.905193925 | 3.621125108 | 2.9614365 | 0.1041865 | 0.448678097 |

| ensembl_gene_id | external_gene_name | gene_biotype | log2 (fold change) | log2 (counts per million) | F-value | p-value | False discovery rate |
|---|---|---|---|---|---|---|---|
| ENSMUSG00000028798 | Eif3i | protein_coding | 0.389664093 | 9.126720684 | 2.9613821 | 0.1041895 | 0.448678097 |
| ENSMUSG00000021066 | Atl1 | protein_coding | 0.568279472 | 6.57713241 | 2.9606064 | 0.1042625 | 0.448779901 |
| ENSMUSG00000056854 | Pou3f4 | protein_coding | -0.823058687 | 4.056695907 | 2.9578155 | 0.1043818 | 0.448779901 |
| ENSMUSG00000026708 | Cenpl | protein_coding | 1.481638011 | 2.71994695 | 2.9575454 | 0.1043964 | 0.448779901 |
| ENSMUSG00000057604 | Lmcd1 | protein_coding | -1.104370027 | 3.760950453 | 2.9569563 | 0.1044595 | 0.448779901 |
| ENSMUSG00000033256 | Shf | protein_coding | -1.281607603 | 2.677663578 | 2.9549645 | 0.1045359 | 0.448779901 |
| ENSMUSG00000002343 | Armc6 | protein_coding | 0.613873202 | 6.423998086 | 2.9552051 | 0.1045541 | 0.448779901 |
| ENSMUSG00000032178 | Ilf3 | protein_coding | 0.753375831 | 7.480326421 | 3.0041591 | 0.1045743 | 0.448779901 |
| ENSMUSG00000039431 | Mtmr7 | protein_coding | 0.518067699 | 6.083458304 | 2.9533378 | 0.1046239 | 0.448779901 |
| ENSMUSG00000025231 | Sufu | protein_coding | -1.16894945 | 3.201262282 | 2.9530548 | 0.1046392 | 0.448779901 |
| ENSMUSG00000030110 | Ret | protein_coding | 0.893315091 | 3.503712827 | 2.9527332 | 0.1046566 | 0.448779901 |
| ENSMUSG00000030216 | Wbp11 | protein_coding | 0.443295373 | 6.936768788 | 2.9517615 | 0.1047093 | 0.448779901 |
| ENSMUSG00000055660 | Mettl4 | protein_coding | -1.327400803 | 2.489487307 | 2.9516762 | 0.1047139 | 0.448779901 |
| ENSMUSG00000017868 | Sgk2 | protein_coding | -1.353984167 | 3.413828535 | 2.9525245 | 0.1048355 | 0.448994062 |
| ENSMUSG00000025816 | Sec61a2 | protein_coding | 0.669114609 | 4.587601177 | 2.9492153 | 0.1048474 | 0.448994062 |
| ENSMUSG00000036591 | Arhgap21 | protein_coding | -0.806374582 | 4.502984032 | 2.94579 | 0.1050335 | 0.449511338 |
| ENSMUSG00000021124 | Vti1b | protein_coding | 0.900743871 | 8.008059592 | 3.0460763 | 0.1050518 | 0.449511338 |
| ENSMUSG00000034462 | Pkd2 | protein_coding | -1.195728018 | 2.742760216 | 2.9413859 | 0.1052734 | 0.449988606 |
| ENSMUSG00000020107 | Anapc16 | protein_coding | 0.376042791 | 7.218092901 | 2.9402518 | 0.1053353 | 0.449988606 |
| ENSMUSG00000015812 | Gnrh1 | protein_coding | -1.101234727 | 5.32651247 | 2.996516 | 0.1053445 | 0.449988606 |
| ENSMUSG00000002871 | Tpra1 | protein_coding | 0.586025926 | 5.906543138 | 2.9398097 | 0.1053594 | 0.449988606 |
| ENSMUSG00000020354 | Sgcd | protein_coding | -1.38760808 | 2.665280259 | 2.9388519 | 0.1054117 | 0.449988606 |
| ENSMUSG00000021824 | Ap3m1 | protein_coding | -0.587142529 | 4.904119096 | 2.9388028 | 0.1054144 | 0.449988606 |
| ENSMUSG00000022841 | Ap2m1 | protein_coding | 0.492847276 | 7.793836002 | 2.9373088 | 0.1054961 | 0.450158479 |
| ENSMUSG00000031887 | Tradd | protein_coding | 1.344227035 | 2.985530773 | 2.9353142 | 0.1056052 | 0.450404642 |
| ENSMUSG00000057068 | Fam47e | protein_coding | -1.339026002 | 3.11634576 | 2.9342594 | 0.105663 | 0.450404642 |
| ENSMUSG00000005986 | Ankrd13d | protein_coding | 0.695388496 | 5.30003387 | 2.9339593 | 0.1056794 | 0.450404642 |
| ENSMUSG00000038570 | Saxo2 | protein_coding | 0.583611872 | 4.880712069 | 2.9305262 | 0.1058678 | 0.451028514 |
| ENSMUSG00000031453 | Rasa3 | protein_coding | 0.622710459 | 4.952652906 | 2.9277682 | 0.1060193 | 0.451495406 |
| ENSMUSG00000096433 | Zfp994 | protein_coding | -1.555084009 | 2.366309007 | 2.9245567 | 0.1061962 | 0.451912293 |
| ENSMUSG00000076439 | Mog | protein_coding | -0.440199035 | 7.849745662 | 2.9244639 | 0.1062013 | 0.451912293 |
| ENSMUSG00000001506 | Col1a1 | protein_coding | -2.174141389 | 2.015746299 | 2.9230437 | 0.1062796 | 0.452066667 |
| ENSMUSG00000028228 | Cpne3 | protein_coding | -1.166881134 | 3.502049258 | 2.92001 | 0.1064784 | 0.452599001 |
| ENSMUSG00000049796 | Crh | protein_coding | 1.170053351 | 2.83028429 | 2.9192538 | 0.1064889 | 0.452599001 |
| ENSMUSG00000000632 | Sez6 | protein_coding | -0.681132617 | 6.559710408 | 2.9397485 | 0.1065745 | 0.452631197 |
| ENSMUSG00000022452 | Smdt1 | protein_coding | 0.44311656 | 8.365848453 | 2.9175952 | 0.1065807 | 0.452631197 |
| ENSMUSG00000025078 | Nhlrc2 | protein_coding | -1.234283673 | 2.748052881 | 2.9166683 | 0.106632 | 0.452670395 |
| ENSMUSG00000046079 | Lrrc8d | protein_coding | -0.641573669 | 4.44451498 | 2.9142885 | 0.1067639 | 0.453051515 |
| ENSMUSG00000027676 | Ccdc39 | protein_coding | -1.17111327 | 3.229858903 | 2.9124963 | 0.1068634 | 0.453061201 |
| ENSMUSG00000041079 | Rwdd2b | protein_coding | -0.697503743 | 4.046549818 | 2.9122526 | 0.1068769 | 0.453061201 |
| ENSMUSG00000041429 | Nthl1 | protein_coding | 1.076882599 | 3.525973992 | 2.9109799 | 0.1069476 | 0.453061201 |
| ENSMUSG00000022476 | Polr3h | protein_coding | 0.497518319 | 6.335432108 | 2.9107345 | 0.1069613 | 0.453061201 |
| ENSMUSG00000030979 | Uros | protein_coding | 0.43348558 | 6.304832327 | 2.910454 | 0.1069769 | 0.453061201 |
| ENSMUSG00000028433 | Ubap2 | protein_coding | -0.771431098 | 4.129500924 | 2.9082922 | 0.1070972 | 0.453392058 |
| ENSMUSG00000032228 | Tcf12 | protein_coding | -0.682796607 | 4.905642339 | 2.9057238 | 0.1072403 | 0.453819256 |
| ENSMUSG00000071302 | 2610044O15Rik8 | protein_coding | -0.617529335 | 4.952829646 | 2.9037421 | 0.1073509 | 0.4541085 |
| ENSMUSG00000038552 | Fndc4 | protein_coding | -0.641812414 | 5.636499316 | 2.901756 | 0.1074618 | 0.454399111 |
| ENSMUSG00000051557 | Pusl1 | protein_coding | 0.913769051 | 4.982828726 | 2.9130623 | 0.1076303 | 0.454932568 |
| ENSMUSG00000005892 | Trh | protein_coding | -0.663043332 | 7.312990643 | 2.9256678 | 0.1077358 | 0.455199452 |
| ENSMUSG00000004789 | Dlst | protein_coding | 0.489983092 | 6.92417559 | 2.8950482 | 0.1078376 | 0.455450792 |
| ENSMUSG00000086429 | Gt(ROSA)26Sor | antisense_RNA | -0.760840056 | 5.287847624 | 2.8923244 | 0.108022 | 0.455618446 |
| ENSMUSG00000056050 | Mia3 | protein_coding | -0.539956314 | 5.465371818 | 2.8906889 | 0.1080827 | 0.455618446 |
| ENSMUSG00000063145 | Bbs5 | protein_coding | -0.672430099 | 4.886398385 | 2.8904177 | 0.1080979 | 0.455618446 |
| ENSMUSG00000021779 | Thrb | protein_coding | -0.695574099 | 5.798525101 | 2.8909601 | 0.1080987 | 0.455618446 |
| ENSMUSG00000038034 | Igsf8 | protein_coding | 0.472064116 | 5.971416614 | 2.890203 | 0.10811 | 0.455618446 |
| ENSMUSG00000038489 | Polr2l | protein_coding | 0.571023322 | 4.985064175 | 2.889821 | 0.1081315 | 0.455618446 |
| ENSMUSG00000068206 | Pick1 | protein_coding | 0.696733687 | 4.937320041 | 2.8884092 | 0.1082111 | 0.455674841 |
| ENSMUSG00000042406 | Atf4 | protein_coding | -0.418858857 | 7.274473083 | 2.8880795 | 0.1082297 | 0.455674841 |
| ENSMUSG00000092035 | Peg10 | protein_coding | -0.636352029 | 4.61487077 | 2.885799 | 0.1083583 | 0.456038025 |
| ENSMUSG00000031665 | Sall1 | protein_coding | -1.118081889 | 3.208844785 | 2.8837885 | 0.1084719 | 0.456299284 |
| ENSMUSG00000029004 | Kmt2e | protein_coding | -0.543169947 | 6.199021232 | 2.8831986 | 0.1085053 | 0.456299284 |
| ENSMUSG00000019970 | Sgk1 | protein_coding | -0.662275375 | 7.726937866 | 2.908733 | 0.1087256 | 0.457046857 |
| ENSMUSG00000074305 | Peak1 | protein_coding | -1.035405692 | 3.140702015 | 2.8782858 | 0.1087835 | 0.457111951 |
| ENSMUSG00000034075 | Zdhhc5 | protein_coding | -0.863261465 | 3.375278264 | 2.8769219 | 0.108861 | 0.457235978 |
| ENSMUSG00000027293 | Ehd4 | protein_coding | 0.881632224 | 3.606209092 | 2.8762677 | 0.1088981 | 0.457235978 |
| ENSMUSG00000040896 | Kcnd3 | protein_coding | -0.845433377 | 4.172259452 | 2.8750405 | 0.1089678 | 0.457350169 |
| ENSMUSG00000023572 | Ccndbp1 | protein_coding | 0.408087302 | 7.556442304 | 2.8718697 | 0.1091483 | 0.457928675 |
| ENSMUSG00000050530 | Fam171a1 | protein_coding | -0.522999618 | 5.133659877 | 2.8685379 | 0.1093382 | 0.45854676 |
| ENSMUSG00000068151 | A230006K03Rik | lincRNA | -0.659295714 | 4.432775736 | 2.8650458 | 0.1095378 | 0.459028382 |
| ENSMUSG00000047767 | Atg16l2 | protein_coding | 0.957690934 | 4.074407515 | 2.8650071 | 0.1095713 | 0.459028382 |
| ENSMUSG00000041132 | N4bp2l1 | protein_coding | -0.606145419 | 5.07898896 | 2.8640092 | 0.1095971 | 0.459028382 |
| ENSMUSG00000029755 | Dlx5 | protein_coding | 0.538342205 | 5.722980699 | 2.8630072 | 0.1096544 | 0.459028382 |
| ENSMUSG00000005514 | Por | protein_coding | 0.561499337 | 6.626504381 | 2.862489 | 0.1097155 | 0.459028382 |
| ENSMUSG00000049130 | C5ar1 | protein_coding | -2.085220622 | 2.071131458 | 2.8613963 | 0.1097467 | 0.459028382 |
| ENSMUSG00000021936 | Mapk8 | protein_coding | -0.550345001 | 5.316795201 | 2.8613067 | 0.1097519 | 0.459028382 |
| ENSMUSG00000021700 | Rab3c | protein_coding | -0.638902038 | 8.36536048 | 2.8808047 | 0.1098552 | 0.459137899 |
| ENSMUSG00000038349 | Plcl1 | protein_coding | -0.601214997 | 4.898905079 | 2.8593617 | 0.1098635 | 0.459137899 |
| ENSMUSG00000026249 | Serpine2 | protein_coding | 0.564204374 | 7.864507227 | 2.8597027 | 0.1099579 | 0.459214411 |

| ensembl_gene_id | external_gene_name | gene_biotype | log2 (fold change) | log2 (counts per million) | F-value | p-value | False discovery rate |
|---|---|---|---|---|---|---|---|
| ENSMUSG00000039853 | Trim14 | protein_coding | -1.975984629 | 2.136635728 | 2.8575559 | 0.1099672 | 0.459214411 |
| ENSMUSG00000020219 | Timm13 | protein_coding | 0.403665894 | 8.92112637 | 2.8561018 | 0.1100508 | 0.459385137 |
| ENSMUSG00000048485 | Zbtb8b | protein_coding | 1.204254275 | 2.654409053 | 2.8529214 | 0.1102339 | 0.45964063 |
| ENSMUSG00000096956 | Snhg18 | lincRNA | -1.448731163 | 2.863060401 | 2.8524966 | 0.1102584 | 0.45964063 |
| ENSMUSG00000068922 | Msto1 | protein_coding | 0.582073591 | 4.987511977 | 2.8521988 | 0.1102756 | 0.45964063 |
| ENSMUSG00000034154 | Ino80 | protein_coding | -0.92578667 | 3.498932042 | 2.8520704 | 0.110283 | 0.45964063 |
| ENSMUSG00000029655 | N4bp2l2 | protein_coding | -0.717561912 | 5.050342002 | 2.8511568 | 0.1103357 | 0.459656343 |
| ENSMUSG00000024589 | Nedd4l | protein_coding | -0.517179531 | 6.077579682 | 2.8505232 | 0.1103722 | 0.459656343 |
| ENSMUSG00000002846 | Timmdc1 | protein_coding | 0.792149385 | 5.077553258 | 2.8497229 | 0.1104498 | 0.459714552 |
| ENSMUSG00000020109 | Dnajb12 | protein_coding | 0.514096467 | 5.728433243 | 2.8469164 | 0.1105807 | 0.459714552 |
| ENSMUSG00000027695 | Pld1 | protein_coding | -1.592951453 | 2.534102462 | 2.8466912 | 0.1105937 | 0.459714552 |
| ENSMUSG00000026975 | Dph7 | protein_coding | 0.451883382 | 6.119075733 | 2.8463919 | 0.110611 | 0.459714552 |
| ENSMUSG00000054252 | Fgfr3 | protein_coding | -0.715145872 | 4.760512676 | 2.8454531 | 0.1106654 | 0.459714552 |
| ENSMUSG00000031792 | Usb1 | protein_coding | 0.704005883 | 4.800074597 | 2.844867 | 0.1106993 | 0.459714552 |
| ENSMUSG00000039770 | Ypel5 | protein_coding | -0.369051408 | 8.954682556 | 2.8446716 | 0.1107106 | 0.459714552 |
| ENSMUSG00000022346 | Myc | protein_coding | 1.239833248 | 3.014558506 | 2.8443678 | 0.1107282 | 0.459714552 |
| ENSMUSG00000049624 | Slc17a5 | protein_coding | 0.728458887 | 4.188864397 | 2.8423315 | 0.1108463 | 0.459936959 |
| ENSMUSG00000020467 | Efemp1 | protein_coding | -1.052681551 | 3.055751695 | 2.8416699 | 0.1108847 | 0.459936959 |
| ENSMUSG00000022342 | Kcnv1 | protein_coding | -1.069491936 | 3.770552748 | 2.8417724 | 0.1109101 | 0.459936959 |
| ENSMUSG00000049571 | Cfap46 | protein_coding | 1.176608788 | 3.510504677 | 2.8397933 | 0.1110251 | 0.460073789 |
| ENSMUSG00000023020 | Cox14 | protein_coding | 0.487447392 | 8.847201036 | 2.8389433 | 0.1110431 | 0.460073789 |
| ENSMUSG00000005469 | Prkaca | protein_coding | 0.46870995 | 7.918285294 | 2.8384554 | 0.1110715 | 0.460073789 |
| ENSMUSG00000004356 | Utp20 | protein_coding | 1.612105464 | 2.419253035 | 2.8369132 | 0.1111612 | 0.460268189 |
| ENSMUSG00000025534 | Gusb | protein_coding | 0.542316726 | 5.25349744 | 2.8355708 | 0.1112394 | 0.460414645 |
| ENSMUSG00000026694 | Mettl13 | protein_coding | -0.932838591 | 3.717727346 | 2.8326948 | 0.1114071 | 0.460931435 |
| ENSMUSG00000049090 | Zadh2 | protein_coding | -0.780713948 | 3.667105702 | 2.829738 | 0.1115799 | 0.461324502 |
| ENSMUSG00000022711 | Pmm2 | protein_coding | -0.566495337 | 5.048380884 | 2.8291429 | 0.1116147 | 0.461324502 |
| ENSMUSG00000022663 | Atg3 | protein_coding | -0.567447159 | 5.160404018 | 2.8284243 | 0.1116567 | 0.461324502 |
| ENSMUSG00000037363 | Letm2 | protein_coding | 0.966055752 | 3.508788132 | 2.8272556 | 0.1117252 | 0.461324502 |
| ENSMUSG00000057181 | 5730455P16Rik | protein_coding | -0.485070882 | 6.975994754 | 2.8259984 | 0.1117988 | 0.461324502 |
| ENSMUSG00000086859 | Snhg20 | lincRNA | 0.895131997 | 3.822661677 | 2.8251915 | 0.1118462 | 0.461324502 |
| ENSMUSG00000006471 | Ndor1 | protein_coding | 1.007090674 | 3.286308488 | 2.8246391 | 0.1118786 | 0.461324502 |
| ENSMUSG00000035401 | Emsy | protein_coding | -0.684561424 | 4.647654301 | 2.8231323 | 0.111967 | 0.461324502 |
| ENSMUSG00000035486 | Plk5 | protein_coding | 1.042232441 | 3.165917379 | 2.8227728 | 0.1119881 | 0.461324502 |
| ENSMUSG00000042737 | Dpm3 | protein_coding | 0.499362722 | 6.893419205 | 2.8226128 | 0.1119975 | 0.461324502 |
| ENSMUSG00000068101 | Cenpm | protein_coding | 1.470501733 | 2.727712796 | 2.822304 | 0.1120157 | 0.461324502 |
| ENSMUSG00000054418 | 2900041M22Rik | processed_transcript | 1.217807829 | 2.730016073 | 2.8222819 | 0.112017 | 0.461324502 |
| ENSMUSG00000033769 | Exoc6b | protein_coding | 0.566360216 | 5.277434501 | 2.8202679 | 0.1121354 | 0.461635358 |
| ENSMUSG00000044730 | 9930104L06Rik | processed_transcript | -1.040187375 | 3.161767278 | 2.8183883 | 0.112246 | 0.461913998 |
| ENSMUSG00000070705 | Eid2b | protein_coding | 0.867763211 | 3.510431073 | 2.8132867 | 0.112547 | 0.462473717 |
| ENSMUSG00000041078 | Grid1 | protein_coding | 0.529954495 | 5.258684022 | 2.8130132 | 0.1125631 | 0.462473717 |
| ENSMUSG00000036376 | Abt1 | protein_coding | 0.537159797 | 6.069733086 | 2.8127649 | 0.1125778 | 0.462473717 |
| ENSMUSG00000026393 | Nek7 | protein_coding | -0.612444671 | 6.044481512 | 2.8120899 | 0.1126491 | 0.462473717 |
| ENSMUSG00000041774 | Ydjc | protein_coding | 1.078599456 | 3.276588869 | 2.8107925 | 0.1126945 | 0.462473717 |
| ENSMUSG00000015757 | Ppil4 | protein_coding | -0.479243674 | 5.927004705 | 2.8103524 | 0.1127205 | 0.462473717 |
| ENSMUSG00000073987 | Ggh | protein_coding | -0.626052257 | 5.054664238 | 2.8102992 | 0.1127237 | 0.462473717 |
| ENSMUSG00000010797 | Wnt2 | protein_coding | -1.147561966 | 3.163097649 | 2.8102585 | 0.1127261 | 0.462473717 |
| ENSMUSG00000034614 | Pik3ip1 | protein_coding | -0.793744451 | 4.147793106 | 2.8093175 | 0.1127818 | 0.462510958 |
| ENSMUSG00000031266 | Gla | protein_coding | 0.97858024 | 3.426680206 | 2.8077121 | 0.112877 | 0.462510958 |
| ENSMUSG00000109336 | Samd4b | protein_coding | -0.677274036 | 5.297529667 | 2.807496 | 0.1128898 | 0.462510958 |
| ENSMUSG00000038344 | Txlng | protein_coding | -0.910535239 | 3.688826837 | 2.807202 | 0.1129072 | 0.462510958 |
| ENSMUSG00000010064 | Slc38a3 | protein_coding | -0.796646846 | 4.503629925 | 2.8041335 | 0.1130894 | 0.462916215 |
| ENSMUSG00000071753 | C230004F18Rik | processed_transcript | -0.67613907 | 4.836514358 | 2.8040181 | 0.1130963 | 0.462916215 |
| ENSMUSG00000022009 | Nufip1 | protein_coding | 0.757339586 | 3.825513868 | 2.8033616 | 0.1131353 | 0.462916215 |
| ENSMUSG00000019715 | Gle1 | protein_coding | 0.654630378 | 6.066302344 | 2.8005219 | 0.1133357 | 0.46331229 |
| ENSMUSG00000041781 | Cpsf2 | protein_coding | 0.494616669 | 5.601195972 | 2.7986991 | 0.1134129 | 0.46331229 |
| ENSMUSG00000061859 | Patj | protein_coding | 1.025611187 | 5.584489827 | 2.8501309 | 0.1134391 | 0.46331229 |
| ENSMUSG00000020430 | Pes1 | protein_coding | 0.677770665 | 5.057633856 | 2.7976132 | 0.1134777 | 0.46331229 |
| ENSMUSG00000052146 | Rps10 | protein_coding | 0.556436263 | 5.612174742 | 2.797409 | 0.1134899 | 0.46331229 |
| ENSMUSG00000018417 | Myo1b | protein_coding | -0.727622772 | 3.79058272 | 2.796695 | 0.1135325 | 0.46331229 |
| ENSMUSG00000050017 | Pitpnb | protein_coding | -0.471868992 | 6.832640067 | 2.7966755 | 0.1135337 | 0.46331229 |
| ENSMUSG00000034390 | Cmip | protein_coding | 0.562272041 | 7.457598778 | 2.7973917 | 0.1136413 | 0.463562971 |
| ENSMUSG00000040123 | Zmym5 | protein_coding | -0.543720521 | 5.017706106 | 2.7936892 | 0.1137122 | 0.463562971 |
| ENSMUSG00000048039 | Isg20l2 | protein_coding | 0.878248844 | 3.647626949 | 2.7934841 | 0.1137245 | 0.463562971 |
| ENSMUSG00000026917 | Wdr5 | protein_coding | 0.886917171 | 3.894247691 | 2.790638 | 0.1138949 | 0.464081814 |
| ENSMUSG00000026755 | Arpc5l | protein_coding | 0.357157943 | 8.091165685 | 2.788557 | 0.1140197 | 0.464320423 |
| ENSMUSG00000020841 | Cpd | protein_coding | -0.701201254 | 4.600038782 | 2.7882221 | 0.1140398 | 0.464320423 |
| ENSMUSG00000020912 | Krt12 | protein_coding | -1.411802944 | 2.457496946 | 2.7863682 | 0.1141512 | 0.464597921 |
| ENSMUSG00000029725 | Ppp1r35 | protein_coding | 1.256801618 | 2.750171303 | 2.7847163 | 0.1142505 | 0.464772332 |
| ENSMUSG00000037754 | Ppp1r16b | protein_coding | -0.517332684 | 8.07945623 | 2.7847406 | 0.1142805 | 0.464772332 |
| ENSMUSG00000049305 | Ccdc71 | protein_coding | 0.62617709 | 5.392133224 | 2.7825915 | 0.1143785 | 0.464947719 |
| ENSMUSG00000053226 | Dand5 | protein_coding | 0.624944062 | 4.903094634 | 2.7820405 | 0.1144117 | 0.464947719 |
| ENSMUSG00000022064 | Pibf1 | protein_coding | -1.031696762 | 3.199530729 | 2.7813492 | 0.1144533 | 0.464947719 |
| ENSMUSG00000034667 | Xpot | protein_coding | -0.636420442 | 4.929697942 | 2.7789457 | 0.1145984 | 0.465335941 |
| ENSMUSG00000028580 | Pum1 | protein_coding | 0.58270285 | 6.789635868 | 2.7799208 | 0.1146537 | 0.465335941 |
| ENSMUSG00000021676 | Iqgap2 | protein_coding | -1.187565447 | 3.054315862 | 2.7769753 | 0.1147174 | 0.465335941 |
| ENSMUSG00000041084 | Ostc | protein_coding | 0.36698484 | 7.486781233 | 2.7768903 | 0.1147226 | 0.465335941 |
| ENSMUSG00000042045 | Sln | protein_coding | -1.006115457 | 6.315763863 | 2.8664512 | 0.1148032 | 0.465335941 |

| ensembl_gene_id | external_gene_name | gene_biotype | log2 (fold change) | log2 (counts per million) | F-value | p-value | False discovery rate |
|---|---|---|---|---|---|---|---|
| ENSMUSG00000020111 | Micu1 | protein_coding | 0.423441381 | 7.00800492 | 2.7754688 | 0.1148085 | 0.465335941 |
| ENSMUSG00000031376 | Atp2b3 | protein_coding | 0.884653486 | 3.234894051 | 2.7743501 | 0.1148763 | 0.465435001 |
| ENSMUSG00000039911 | Spsb1 | protein_coding | -1.106535902 | 2.936041677 | 2.772529 | 0.1149866 | 0.46570664 |
| ENSMUSG00000028792 | Ak2 | protein_coding | 0.564268585 | 5.347829628 | 2.7705328 | 0.1151077 | 0.46602163 |
| ENSMUSG00000062203 | Gspt1 | protein_coding | -1.12513467 | 2.725209974 | 2.7690124 | 0.1152001 | 0.466219973 |
| ENSMUSG00000021709 | Erbin | protein_coding | -0.665134636 | 4.967908066 | 2.7659823 | 0.1153844 | 0.46675452 |
| ENSMUSG00000009356 | Lpo | protein_coding | -2.470933085 | 2.031690728 | 2.7659306 | 0.115419 | 0.46675452 |
| ENSMUSG00000022671 | Mzt2 | protein_coding | 0.417383686 | 6.378649776 | 2.7601507 | 0.1157401 | 0.467162655 |
| ENSMUSG00000026930 | Gpsm1 | protein_coding | -0.50720264 | 6.004158475 | 2.759729 | 0.1157659 | 0.467162655 |
| ENSMUSG00000039239 | Tgfb2 | protein_coding | -0.779417298 | 4.117506558 | 2.7584966 | 0.1158412 | 0.467162655 |
| ENSMUSG00000044408 | Sptssa | protein_coding | -0.435500773 | 6.727464877 | 2.7583203 | 0.115852 | 0.467162655 |
| ENSMUSG00000015733 | Capza2 | protein_coding | -0.378912365 | 9.262856819 | 2.7574416 | 0.1159058 | 0.467162655 |
| ENSMUSG00000041594 | Tmtc4 | protein_coding | 1.587377116 | 2.540256851 | 2.7573768 | 0.1159098 | 0.467162655 |
| ENSMUSG00000030045 | Mrpl19 | protein_coding | 0.371262549 | 7.065750676 | 2.7572781 | 0.1159158 | 0.467162655 |
| ENSMUSG00000026319 | 2310035C23Rik | protein_coding | -0.624935815 | 5.275347878 | 2.7571768 | 0.115922 | 0.467162655 |
| ENSMUSG00000020186 | Csrp2 | protein_coding | 0.603382642 | 4.560132812 | 2.7568833 | 0.11594 | 0.467162655 |
| ENSMUSG00000003435 | Supt5 | protein_coding | 0.723165139 | 5.680561552 | 2.7574322 | 0.1160207 | 0.467162655 |
| ENSMUSG00000030876 | Mettl9 | protein_coding | -0.666962797 | 5.042005265 | 2.7555531 | 0.1160215 | 0.467162655 |
| ENSMUSG00000024646 | Cyb5a | protein_coding | -0.45894868 | 8.28430428 | 2.7552307 | 0.1160412 | 0.467162655 |
| ENSMUSG00000071073 | Lrrc73 | protein_coding | 0.781745312 | 3.816430592 | 2.7537471 | 0.1161322 | 0.467327552 |
| ENSMUSG00000036594 | H2-Aa | protein_coding | -2.200454103 | 2.156284574 | 2.7525111 | 0.1162081 | 0.467327552 |
| ENSMUSG00000000416 | Cttnbp2 | protein_coding | -0.68778156 | 5.046223057 | 2.7509981 | 0.116301 | 0.467327552 |
| ENSMUSG00000054693 | Adam10 | protein_coding | -0.449894195 | 5.802093242 | 2.7509926 | 0.1163014 | 0.467327552 |
| ENSMUSG00000037012 | Hk1 | protein_coding | 0.781982806 | 7.295175808 | 2.800739 | 0.1163398 | 0.467327552 |
| ENSMUSG00000090150 | Acad11 | protein_coding | -0.9885206 | 2.967308539 | 2.7503164 | 0.116343 | 0.467327552 |
| ENSMUSG00000028937 | Acot7 | protein_coding | 0.515921687 | 10.6617693 | 2.7488944 | 0.1164304 | 0.467464379 |
| ENSMUSG00000048911 | Rnf24 | protein_coding | 1.175794784 | 3.282949124 | 2.7482365 | 0.1164709 | 0.467464379 |
| ENSMUSG00000015843 | Rxrg | protein_coding | 0.415607073 | 6.702866231 | 2.7476439 | 0.1165074 | 0.467464379 |
| ENSMUSG00000003200 | Sh3gl1 | protein_coding | 0.843338682 | 4.02389838 | 2.7442896 | 0.1167143 | 0.468119605 |
| ENSMUSG00000028445 | Enho | protein_coding | -0.583754571 | 6.246349103 | 2.7429384 | 0.1168292 | 0.468353104 |
| ENSMUSG00000024346 | Pfdn1 | protein_coding | 0.351047604 | 9.040204816 | 2.7419371 | 0.1168596 | 0.468353104 |
| ENSMUSG00000071337 | Tia1 | protein_coding | -0.85211356 | 3.8261691 | 2.7401374 | 0.1169709 | 0.468539874 |
| ENSMUSG00000015943 | Bola1 | protein_coding | 0.394336665 | 6.70976838 | 2.7390635 | 0.1170374 | 0.468539874 |
| ENSMUSG00000062590 | Armc9 | protein_coding | 0.547938543 | 5.167580818 | 2.7388992 | 0.1170476 | 0.468539874 |
| ENSMUSG00000004530 | Coro1c | protein_coding | 0.703238543 | 6.0219746 | 2.751368 | 0.1170805 | 0.468539874 |
| ENSMUSG00000021149 | Gtpbp4 | protein_coding | 0.442151396 | 6.593417034 | 2.734794 | 0.1173022 | 0.46916257 |
| ENSMUSG00000034473 | Sec22a | protein_coding | 0.429767044 | 6.196040873 | 2.7344535 | 0.1173234 | 0.46916257 |
| ENSMUSG00000034432 | Cops8 | protein_coding | 0.401050581 | 8.788019941 | 2.7331175 | 0.1174064 | 0.469320044 |
| ENSMUSG00000052681 | Rap1b | protein_coding | -0.560172035 | 5.520195589 | 2.7308076 | 0.1175501 | 0.469368078 |
| ENSMUSG00000003380 | Rabac1 | protein_coding | 0.523325147 | 8.737440246 | 2.7269059 | 0.1178248 | 0.469368078 |
| ENSMUSG00000022314 | Rad21 | protein_coding | -0.51586028 | 5.920039703 | 2.7262017 | 0.1178373 | 0.469368078 |
| ENSMUSG00000040462 | Os9 | protein_coding | 0.378328552 | 6.832922169 | 2.7257927 | 0.1178628 | 0.469368078 |
| ENSMUSG00000036820 | Amdhd2 | protein_coding | 0.636127228 | 5.454167321 | 2.7252906 | 0.1178942 | 0.469368078 |
| ENSMUSG00000002015 | Bcap31 | protein_coding | 0.394622585 | 7.856845416 | 2.7252397 | 0.1178974 | 0.469368078 |
| ENSMUSG00000020444 | Guk1 | protein_coding | 0.63588872 | 7.484912509 | 2.749308 | 0.1179731 | 0.469368078 |
| ENSMUSG00000051041 | Olfml1 | protein_coding | -0.65824914 | 5.013972872 | 2.7235799 | 0.1180012 | 0.469368078 |
| ENSMUSG00000075376 | Rc3h2 | protein_coding | -0.756759547 | 5.503473446 | 2.7245488 | 0.1180876 | 0.469368078 |
| ENSMUSG00000026117 | Zap70 | protein_coding | 2.099680158 | 2.329439338 | 2.7219644 | 0.1181337 | 0.469368078 |
| ENSMUSG00000029467 | Atp2a2 | protein_coding | -0.464157338 | 7.904779447 | 2.72081 | 0.1181746 | 0.469368078 |
| ENSMUSG00000036825 | Ssx2ip | protein_coding | 0.501959048 | 5.533175361 | 2.7205581 | 0.1181904 | 0.469368078 |
| ENSMUSG00000024431 | Nr3c1 | protein_coding | -0.403752917 | 6.420682602 | 2.7201509 | 0.1182159 | 0.469368078 |
| ENSMUSG00000043154 | Ppp2r3a | protein_coding | -0.657885522 | 4.875235619 | 2.7195899 | 0.118251 | 0.469368078 |
| ENSMUSG00000039016 | Timm8b | protein_coding | 0.544721581 | 9.334659322 | 2.719712 | 0.1182749 | 0.469368078 |
| ENSMUSG00000051232 | Tmem199 | protein_coding | 0.384002034 | 7.900208875 | 2.7191736 | 0.1182772 | 0.469368078 |
| ENSMUSG00000030978 | Rrm1 | protein_coding | -0.888486842 | 4.168672564 | 2.7186212 | 0.1183118 | 0.469368078 |
| ENSMUSG00000026185 | Igfbp5 | protein_coding | -0.64915702 | 5.404350889 | 2.7176534 | 0.1183726 | 0.469368078 |
| ENSMUSG00000066235 | Pomgnt2 | protein_coding | 0.620221195 | 5.394538667 | 2.7173536 | 0.1183914 | 0.469368078 |
| ENSMUSG00000003153 | Slc2a3 | protein_coding | 0.381218855 | 6.362208726 | 2.717311 | 0.1183941 | 0.469368078 |
| ENSMUSG00000034708 | Grn | protein_coding | 0.476247455 | 5.539982496 | 2.7171982 | 0.1184011 | 0.469368078 |
| ENSMUSG00000097248 | Gm2694 | lincRNA | -0.803643962 | 4.397667393 | 2.7169853 | 0.1184145 | 0.469368078 |
| ENSMUSG00000059142 | Zfp945 | protein_coding | -0.647334911 | 4.555713559 | 2.7166745 | 0.118434 | 0.469368078 |
| ENSMUSG00000036833 | Pnpla7 | protein_coding | -1.401029886 | 2.789872672 | 2.7161658 | 0.118466 | 0.469368078 |
| ENSMUSG00000034430 | Zxdc | protein_coding | 0.770025743 | 3.944649992 | 2.714971 | 0.1185411 | 0.469407752 |
| ENSMUSG00000014496 | Ankrd28 | protein_coding | -0.877281839 | 3.551917331 | 2.7143751 | 0.1185786 | 0.469407752 |
| ENSMUSG00000030695 | Aldoa | protein_coding | 0.43618143 | 11.9914057 | 2.7139243 | 0.118607 | 0.469407752 |
| ENSMUSG00000040025 | Ythdf2 | protein_coding | -0.433136115 | 6.242873669 | 2.7131378 | 0.1186565 | 0.469429444 |
| ENSMUSG00000067369 | Trmt2b | protein_coding | 0.801284772 | 4.47209768 | 2.7124506 | 0.1186998 | 0.469429444 |
| ENSMUSG00000023960 | Enpp5 | protein_coding | 0.34235759 | 7.834927433 | 2.7117435 | 0.1187443 | 0.469433005 |
| ENSMUSG00000001300 | Efnb2 | protein_coding | -0.844780518 | 4.524014235 | 2.7110041 | 0.1188224 | 0.469498851 |
| ENSMUSG00000024767 | Otub1 | protein_coding | 0.458423288 | 10.83118066 | 2.7100945 | 0.1188483 | 0.469498851 |
| ENSMUSG00000018698 | Lhx1 | protein_coding | -1.329848348 | 2.559117788 | 2.7089991 | 0.1189174 | 0.469516846 |
| ENSMUSG00000028886 | Eya3 | protein_coding | 1.141103845 | 3.124177612 | 2.7086389 | 0.1189402 | 0.469516846 |
| ENSMUSG00000028790 | Khdrbs1 | protein_coding | -0.553893781 | 5.447666043 | 2.7073611 | 0.1190209 | 0.469534949 |
| ENSMUSG00000028367 | Txn1 | protein_coding | 0.388172473 | 8.31226565 | 2.7068693 | 0.119052 | 0.469534949 |
| ENSMUSG00000021468 | Sptlc1 | protein_coding | 0.609129332 | 4.391982916 | 2.7062219 | 0.1190929 | 0.469534949 |
| ENSMUSG00000026211 | Obsl1 | protein_coding | 1.149611156 | 2.832879547 | 2.7047202 | 0.119188 | 0.469534949 |
| ENSMUSG00000055320 | Tead1 | protein_coding | -0.75330741 | 3.646694706 | 2.7041217 | 0.1192259 | 0.469534949 |
| ENSMUSG00000045991 | Onecut2 | protein_coding | -0.625276431 | 4.74678289 | 2.7036701 | 0.1192545 | 0.469534949 |

| ensembl_gene_id | external_gene_name | gene_biotype | log2 (fold change) | log2 (counts per million) | F-value | p-value | False discovery rate |
|---|---|---|---|---|---|---|---|
| ENSMUSG00000020074 | Ccar1 | protein_coding | -0.540582081 | 5.314641543 | 2.7032268 | 0.1192826 | 0.469534949 |
| ENSMUSG00000030868 | Dctn5 | protein_coding | 0.45570763 | 5.84964316 | 2.7030446 | 0.1192941 | 0.469534949 |
| ENSMUSG00000028995 | Fam126a | protein_coding | -1.02980144 | 3.874948119 | 2.7011053 | 0.1194486 | 0.469547822 |
| ENSMUSG00000029265 | Dr1 | protein_coding | -0.77040178 | 4.416622029 | 2.7003158 | 0.1194672 | 0.469547822 |
| ENSMUSG00000039782 | Cpeb2 | protein_coding | -0.609399525 | 4.842610896 | 2.7000759 | 0.1194824 | 0.469547822 |
| ENSMUSG00000039108 | Lsm14b | protein_coding | -0.696522622 | 4.354056581 | 2.6998882 | 0.1194943 | 0.469547822 |
| ENSMUSG00000041141 | Pnmal1 | protein_coding | 0.516462321 | 7.956656141 | 2.7000476 | 0.1195157 | 0.469547822 |
| ENSMUSG00000028933 | Xrcc2 | protein_coding | 0.870349404 | 3.25736211 | 2.6980327 | 0.1196122 | 0.469726013 |
| ENSMUSG00000026617 | Bpnt1 | protein_coding | 0.406495982 | 6.995437354 | 2.695866 | 0.1197501 | 0.469726013 |
| ENSMUSG00000069814 | Ccdc92b | protein_coding | -0.716989489 | 4.020722003 | 2.6955987 | 0.1197671 | 0.469726013 |
| ENSMUSG00000034472 | Rasd2 | protein_coding | -0.477372195 | 6.691949808 | 2.695386 | 0.1197807 | 0.469726013 |
| ENSMUSG00000017417 | Plxdc1 | protein_coding | 0.755737395 | 3.993092198 | 2.6947739 | 0.1198197 | 0.469726013 |
| ENSMUSG00000003348 | Mob3a | protein_coding | 1.48843121 | 2.797709961 | 2.694719 | 0.1198232 | 0.469726013 |
| ENSMUSG00000044566 | Cage1 | protein_coding | -0.87346966 | 3.494822054 | 2.6931728 | 0.1199217 | 0.469727772 |
| ENSMUSG00000023966 | Rsph9 | protein_coding | 0.389458732 | 6.664529236 | 2.692843 | 0.1199428 | 0.469727772 |
| ENSMUSG00000057963 | Itpk1 | protein_coding | 0.508663901 | 5.643085714 | 2.6926565 | 0.1199547 | 0.469727772 |
| ENSMUSG00000080316 | Spaca6 | protein_coding | -0.807350345 | 3.488998129 | 2.6906002 | 0.1200859 | 0.470070662 |
| ENSMUSG00000009406 | Elk1 | protein_coding | 0.719496713 | 4.138443608 | 2.6890304 | 0.1201863 | 0.470227493 |
| ENSMUSG00000022829 | Stxbp5l | protein_coding | -0.823034775 | 3.902236103 | 2.6883442 | 0.1202302 | 0.470227493 |
| ENSMUSG00000028896 | Rcc1 | protein_coding | 1.035675564 | 3.877977304 | 2.6884138 | 0.1202572 | 0.470227493 |
| ENSMUSG00000097785 | B230217O12Rik | lincRNA | 0.627021776 | 5.528692352 | 2.6868096 | 0.1203284 | 0.470334857 |
| ENSMUSG00000028706 | Nsun4 | protein_coding | 0.78757792 | 4.674578843 | 2.6826454 | 0.1205954 | 0.471095022 |
| ENSMUSG00000097451 | Rian | processed_transcript | -0.497805289 | 5.684275856 | 2.6824107 | 0.1206105 | 0.471095022 |
| ENSMUSG00000009894 | Snap47 | protein_coding | 0.536426436 | 10.21676749 | 2.6798898 | 0.120804 | 0.471679546 |
| ENSMUSG00000037070 | Rbmxl1 | protein_coding | 0.65912778 | 4.031584788 | 2.6783566 | 0.1208712 | 0.471770548 |
| ENSMUSG00000047446 | Arl4a | protein_coding | -0.668529879 | 6.549602201 | 2.6941351 | 0.1210804 | 0.472415581 |
| ENSMUSG00000024012 | Mtch1 | protein_coding | 0.386857274 | 9.233100313 | 2.6741803 | 0.1211405 | 0.472478526 |
| ENSMUSG00000029148 | Nrbp1 | protein_coding | 0.557264633 | 6.827885153 | 2.6732333 | 0.1212331 | 0.472668421 |
| ENSMUSG00000010406 | Mrpl52 | protein_coding | 0.446287942 | 7.395968681 | 2.6703976 | 0.121385 | 0.473088977 |
| ENSMUSG00000040952 | Rps19 | protein_coding | 0.502113138 | 8.554367268 | 2.6701074 | 0.1214352 | 0.473113436 |
| ENSMUSG00000025645 | Ccdc51 | protein_coding | -0.885992056 | 3.507253039 | 2.6668071 | 0.1216176 | 0.473521809 |
| ENSMUSG00000024427 | Spry4 | protein_coding | -0.67297306 | 4.300934509 | 2.6666449 | 0.1216281 | 0.473521809 |
| ENSMUSG00000021906 | Oxnad1 | protein_coding | 0.654257797 | 4.883129261 | 2.665298 | 0.1217155 | 0.473690632 |
| ENSMUSG00000029066 | Mrpl20 | protein_coding | 0.41125098 | 8.825360776 | 2.6634701 | 0.1218343 | 0.4738349 |
| ENSMUSG00000005225 | Plekha8 | protein_coding | -0.949178724 | 3.064504677 | 2.6623979 | 0.121904 | 0.4738349 |
| ENSMUSG00000068284 | Usf3 | protein_coding | -0.940949604 | 3.504776773 | 2.6623926 | 0.1219044 | 0.4738349 |
| ENSMUSG00000028035 | Dnajb4 | protein_coding | -0.451507746 | 6.584486295 | 2.6620159 | 0.1219289 | 0.4738349 |
| ENSMUSG00000024891 | Slc29a2 | protein_coding | 0.682954585 | 4.347253447 | 2.6578571 | 0.1221998 | 0.474716365 |
| ENSMUSG00000017969 | Ptgis | protein_coding | 1.43101117 | 2.848786097 | 2.6570197 | 0.1222545 | 0.474757152 |
| ENSMUSG00000030265 | Kras | protein_coding | -0.573844833 | 5.529993642 | 2.6557233 | 0.1223392 | 0.474914422 |
| ENSMUSG00000028483 | Snapc3 | protein_coding | 0.497439374 | 5.714900084 | 2.6543412 | 0.1224295 | 0.474947024 |
| ENSMUSG00000029033 | Acap3 | protein_coding | 0.59505951 | 4.663657039 | 2.6539213 | 0.122457 | 0.474947024 |
| ENSMUSG00000033294 | Noc4l | protein_coding | 0.672936708 | 4.910377051 | 2.6535681 | 0.1224801 | 0.474947024 |
| ENSMUSG00000042157 | Sprr2i | protein_coding | -2.844003221 | 1.905783563 | 2.6544181 | 0.1225933 | 0.475214855 |
| ENSMUSG00000041697 | Cox6a1 | protein_coding | 0.604177316 | 9.755176941 | 2.6618639 | 0.1227421 | 0.475478661 |
| ENSMUSG00000020650 | Bcap29 | protein_coding | -0.459357425 | 6.324469534 | 2.6494503 | 0.1227498 | 0.475478661 |
| ENSMUSG00000021938 | Pspc1 | protein_coding | -0.725415603 | 4.338091377 | 2.6485702 | 0.1228076 | 0.475492017 |
| ENSMUSG00000056019 | Zfp709 | protein_coding | -1.201640335 | 2.776760095 | 2.6480502 | 0.1228417 | 0.475492017 |
| ENSMUSG00000030091 | Nup210 | protein_coding | 1.378562536 | 2.789625532 | 2.6466814 | 0.1229316 | 0.475610434 |
| ENSMUSG00000053253 | Ndfip2 | protein_coding | -0.469347721 | 5.975936153 | 2.646238 | 0.1229608 | 0.475610434 |
| ENSMUSG00000029212 | Gabrb1 | protein_coding | -0.456418817 | 6.368171514 | 2.6455279 | 0.1230074 | 0.47561992 |
| ENSMUSG00000049659 | Aftph | protein_coding | -0.503691612 | 5.669652592 | 2.6418512 | 0.1232495 | 0.476231948 |
| ENSMUSG00000024442 | 0610009O20Rik | protein_coding | -0.449832563 | 5.770186713 | 2.6417787 | 0.1232543 | 0.476231948 |
| ENSMUSG00000021112 | Mpp5 | protein_coding | -0.454884169 | 5.843263303 | 2.6373797 | 0.1235447 | 0.47718258 |
| ENSMUSG00000043251 | Exoc3l | protein_coding | 1.714326531 | 2.066138384 | 2.634937 | 0.1237063 | 0.477325686 |
| ENSMUSG00000022436 | Sh3bp1 | protein_coding | 0.398327773 | 7.128496643 | 2.6342874 | 0.1237494 | 0.477325686 |
| ENSMUSG00000038388 | Mpp6 | protein_coding | -0.631591888 | 4.397720151 | 2.634239 | 0.1237526 | 0.477325686 |
| ENSMUSG00000049489 | Fam58b | protein_coding | 0.604262404 | 4.903408993 | 2.6341368 | 0.1237593 | 0.477325686 |
| ENSMUSG00000023104 | Rfc2 | protein_coding | 0.404906993 | 6.384480116 | 2.6326971 | 0.1238547 | 0.477522453 |
| ENSMUSG00000006241 | Ccdc159 | protein_coding | 0.870740926 | 3.402527741 | 2.630406 | 0.1240068 | 0.477795745 |
| ENSMUSG00000025949 | Pikfyve | protein_coding | -0.70183531 | 4.167803261 | 2.6302662 | 0.1240161 | 0.477795745 |
| ENSMUSG00000028383 | Hsdl2 | protein_coding | -0.470449834 | 5.550690746 | 2.629621 | 0.1240589 | 0.477795745 |
| ENSMUSG00000003437 | Paf1 | protein_coding | 0.818254625 | 5.326865428 | 2.6402744 | 0.124157 | 0.477959239 |
| ENSMUSG00000057322 | Rpl38 | protein_coding | 0.675419183 | 4.578530249 | 2.6276454 | 0.1241903 | 0.477959239 |
| ENSMUSG00000041380 | Htr2c | protein_coding | -0.507479637 | 5.492600064 | 2.626919 | 0.1242386 | 0.477974189 |
| ENSMUSG00000051065 | Mb21d2 | protein_coding | 0.594401301 | 4.829982531 | 2.6255944 | 0.1243268 | 0.478008033 |
| ENSMUSG00000048581 | E130311K13Rik | protein_coding | 1.502362649 | 2.635295738 | 2.6254518 | 0.1243363 | 0.478008033 |
| ENSMUSG00000079426 | Arpc4 | protein_coding | 0.401543193 | 8.860484356 | 2.6236621 | 0.1244556 | 0.478295702 |
| ENSMUSG00000032121 | Tmem218 | protein_coding | -0.521433125 | 5.524120132 | 2.6212602 | 0.124616 | 0.478502586 |
| ENSMUSG00000036054 | Sugp2 | protein_coding | 0.531220784 | 4.989714878 | 2.6209631 | 0.1246358 | 0.478502586 |
| ENSMUSG00000037487 | Ubr5 | protein_coding | -0.534023985 | 5.318454565 | 2.6208561 | 0.124643 | 0.478502586 |
| ENSMUSG00000046753 | Ccdc66 | protein_coding | -1.019985695 | 3.093185447 | 2.62011 | 0.1246928 | 0.47852319 |
| ENSMUSG00000021097 | Clmn | protein_coding | -0.599784268 | 5.483942147 | 2.6175832 | 0.1248619 | 0.479001031 |
| ENSMUSG00000034799 | Unc13a | protein_coding | -0.767888177 | 3.774578642 | 2.6158887 | 0.1249754 | 0.479152722 |
| ENSMUSG00000020462 | Cfap36 | protein_coding | -0.38100632 | 7.174457757 | 2.615663 | 0.1249906 | 0.479152722 |
| ENSMUSG00000057363 | Uxs1 | protein_coding | -0.848686696 | 3.824973209 | 2.6147825 | 0.1250496 | 0.479208263 |
| ENSMUSG00000002820 | Atg4d | protein_coding | -0.955265121 | 3.81406085 | 2.6123463 | 0.1252132 | 0.479664114 |
| ENSMUSG00000024829 | Mrpl21 | protein_coding | 0.363387988 | 7.109861112 | 2.6115727 | 0.1252652 | 0.479692402 |

| ensembl_gene_id | external_gene_name | gene_biotype | log2 (fold change) | log2 (counts per million) | F-value | p-value | False discovery rate |
|---|---|---|---|---|---|---|---|
| ENSMUSG00000020153 | Ndufs7 | protein_coding | 0.381961944 | 9.745717511 | 2.6103113 | 0.12535 | 0.479788914 |
| ENSMUSG00000050240 | Hic2 | protein_coding | 1.580852764 | 3.345082412 | 2.6381337 | 0.125408 | 0.479788914 |
| ENSMUSG00000017264 | Exosc10 | protein_coding | 0.631881927 | 5.261205355 | 2.6092084 | 0.1254242 | 0.479788914 |
| ENSMUSG00000059278 | Naa38 | protein_coding | 0.469975801 | 7.56424352 | 2.6057753 | 0.1256556 | 0.480503163 |
| ENSMUSG00000000827 | Tpd52l2 | protein_coding | 0.384881332 | 7.251869599 | 2.604092 | 0.1257693 | 0.480766799 |
| ENSMUSG00000038150 | Ormdl3 | protein_coding | 0.447387318 | 6.097584382 | 2.6025889 | 0.1258709 | 0.480982339 |
| ENSMUSG00000039738 | Slx4 | protein_coding | 1.136954364 | 2.92523349 | 2.6013787 | 0.1259527 | 0.480982339 |
| ENSMUSG00000020823 | Sec14l1 | protein_coding | -0.547043341 | 5.454961313 | 2.6002117 | 0.1260317 | 0.480982339 |
| ENSMUSG00000027752 | Exosc8 | protein_coding | 0.461865903 | 5.884115606 | 2.6000838 | 0.1260404 | 0.480982339 |
| ENSMUSG00000037679 | Inf2 | protein_coding | -0.651220698 | 4.847573096 | 2.5999521 | 0.1260493 | 0.480982339 |
| ENSMUSG00000025745 | Hadha | protein_coding | 0.529205292 | 5.641655946 | 2.5978822 | 0.1261896 | 0.481346878 |
| ENSMUSG00000032604 | Qars | protein_coding | 0.516269955 | 5.644562786 | 2.5923866 | 0.126563 | 0.482568517 |
| ENSMUSG00000024513 | Mbd2 | protein_coding | -0.794027763 | 3.827740814 | 2.5918488 | 0.1265996 | 0.482568517 |
| ENSMUSG00000038668 | Lpar1 | protein_coding | -0.626179618 | 5.332223296 | 2.5905774 | 0.1266862 | 0.482727524 |
| ENSMUSG00000027238 | Frmd5 | protein_coding | 0.663518031 | 4.258166199 | 2.5894492 | 0.1267632 | 0.482849509 |
| ENSMUSG00000029141 | Slc4a1ap | protein_coding | -0.599127844 | 4.239304345 | 2.58566 | 0.1270219 | 0.482878472 |
| ENSMUSG00000018340 | Anxa6 | protein_coding | 0.480889338 | 6.697405451 | 2.5838039 | 0.1271489 | 0.482878472 |
| ENSMUSG00000033444 | Specc1l | protein_coding | 0.605633702 | 4.588919484 | 2.5835687 | 0.127165 | 0.482878472 |
| ENSMUSG00000033845 | Mrpl15 | protein_coding | 0.349400458 | 7.412366499 | 2.5831263 | 0.1271952 | 0.482878472 |
| ENSMUSG00000074785 | Plxnc1 | protein_coding | -0.541864822 | 5.013057301 | 2.582739 | 0.1272218 | 0.482878472 |
| ENSMUSG00000048782 | Insc | protein_coding | -1.506656846 | 2.653834911 | 2.582713 | 0.1272236 | 0.482878472 |
| ENSMUSG00000041852 | Tcf20 | protein_coding | 0.60098939 | 5.792420449 | 2.5826491 | 0.1272279 | 0.482878472 |
| ENSMUSG00000085882 | 2610507I01Rik | processed_transcript | 0.983350952 | 3.233352781 | 2.58248 | 0.1272395 | 0.482878472 |
| ENSMUSG00000084786 | Ubl5 | protein_coding | 0.471122644 | 9.224991029 | 2.5824454 | 0.1272419 | 0.482878472 |
| ENSMUSG00000083307 | AA414768 | processed_pseudoge | -1.026752753 | 2.909304346 | 2.580755 | 0.1273578 | 0.482878472 |
| ENSMUSG00000030225 | Dera | protein_coding | -0.96413692 | 3.302145592 | 2.5802678 | 0.1273912 | 0.482878472 |
| ENSMUSG00000032398 | Snapc5 | protein_coding | -0.473260469 | 6.535597671 | 2.5784816 | 0.1275138 | 0.482878472 |
| ENSMUSG00000005674 | Tomm40l | protein_coding | 0.597014677 | 5.84229215 | 2.5784516 | 0.1275159 | 0.482878472 |
| ENSMUSG00000044835 | Ankrd45 | protein_coding | -0.540415908 | 5.516952363 | 2.5782951 | 0.1275266 | 0.482878472 |
| ENSMUSG00000024683 | Mrpl16 | protein_coding | 0.425160212 | 6.373796248 | 2.5778081 | 0.1275601 | 0.482878472 |
| ENSMUSG00000036611 | Eepd1 | protein_coding | 0.921916581 | 3.164771061 | 2.5776488 | 0.127571 | 0.482878472 |
| ENSMUSG00000030774 | Pak1 | protein_coding | 0.439435495 | 8.523614792 | 2.5774733 | 0.1275831 | 0.482878472 |
| ENSMUSG00000026313 | Hdac4 | protein_coding | 0.535402452 | 5.850926802 | 2.575115 | 0.1277453 | 0.482878472 |
| ENSMUSG00000030659 | Nucb2 | protein_coding | 0.500787452 | 5.219319898 | 2.5745059 | 0.1277872 | 0.482878472 |
| ENSMUSG00000037007 | Zfp113 | protein_coding | -0.949799655 | 3.324830141 | 2.5733497 | 0.1278669 | 0.482878472 |
| ENSMUSG00000039697 | Ncoa7 | protein_coding | -0.431936979 | 5.966730322 | 2.5720201 | 0.1279586 | 0.482878472 |
| ENSMUSG00000035769 | Xylb | protein_coding | 0.991332175 | 3.277310768 | 2.5718401 | 0.127971 | 0.482878472 |
| ENSMUSG00000002324 | Rec8 | protein_coding | 1.145231446 | 3.110103132 | 2.5717128 | 0.1279798 | 0.482878472 |
| ENSMUSG00000028436 | Dcaf12 | protein_coding | -0.586084172 | 4.935003927 | 2.5714963 | 0.1279947 | 0.482878472 |
| ENSMUSG00000062901 | Klhl24 | protein_coding | -0.58356602 | 5.141138767 | 2.5711749 | 0.1280169 | 0.482878472 |
| ENSMUSG00000036188 | Ankmy2 | protein_coding | 0.425869597 | 6.939125546 | 2.5710513 | 0.1280254 | 0.482878472 |
| ENSMUSG00000063354 | Slc39a4 | protein_coding | 6.236370232 | 1.36567172 | 2.630258 | 0.1280966 | 0.482878472 |
| ENSMUSG00000038539 | Atf5 | protein_coding | -1.316598704 | 2.495895301 | 2.569962 | 0.1281006 | 0.482878472 |
| ENSMUSG00000027639 | Samhd1 | protein_coding | -0.875766432 | 3.30342135 | 2.569609 | 0.128125 | 0.482878472 |
| ENSMUSG00000048970 | C1galt1c1 | protein_coding | -0.563683343 | 5.775163264 | 2.567927 | 0.1282413 | 0.482878472 |
| ENSMUSG00000033597 | Caskin1 | protein_coding | 0.677394395 | 5.013224174 | 2.567616 | 0.1282628 | 0.482878472 |
| ENSMUSG00000019189 | Rnf145 | protein_coding | 0.605492068 | 5.142568878 | 2.5674803 | 0.1282722 | 0.482878472 |
| ENSMUSG00000044551 | 9930012K11Rik | protein_coding | 1.561215601 | 2.885556435 | 2.5659374 | 0.1284105 | 0.482878472 |
| ENSMUSG00000058709 | Egln2 | protein_coding | 0.445764919 | 6.016477242 | 2.56544 | 0.1284134 | 0.482878472 |
| ENSMUSG00000029554 | Mad1l1 | protein_coding | 0.508796874 | 5.278598499 | 2.5652489 | 0.1284267 | 0.482878472 |
| ENSMUSG00000027618 | Nfs1 | protein_coding | 0.627322604 | 4.773280756 | 2.5644719 | 0.1284805 | 0.482878472 |
| ENSMUSG00000001036 | Epn2 | protein_coding | -0.424250842 | 6.300248836 | 2.563961 | 0.1285159 | 0.482878472 |
| ENSMUSG00000018634 | Crhr1 | protein_coding | 1.259937386 | 2.681756768 | 2.5635666 | 0.1285433 | 0.482878472 |
| ENSMUSG00000032946 | Rasgrp2 | protein_coding | 0.516577365 | 5.225065832 | 2.563187 | 0.1285696 | 0.482878472 |
| ENSMUSG00000026131 | Dst | protein_coding | -0.642873985 | 5.014893336 | 2.5629745 | 0.1285844 | 0.482878472 |
| ENSMUSG00000021302 | Ggps1 | protein_coding | -0.377659015 | 6.849266201 | 2.5625773 | 0.1286119 | 0.482878472 |
| ENSMUSG00000047554 | Tmem41b | protein_coding | -0.622177285 | 4.389520327 | 2.5617584 | 0.1286688 | 0.482923306 |
| ENSMUSG00000024843 | Chka | protein_coding | 0.712795189 | 4.607991209 | 2.5584231 | 0.1289006 | 0.483544866 |
| ENSMUSG00000026809 | Spaca9 | protein_coding | -1.303959119 | 2.80355454 | 2.5571823 | 0.128987 | 0.483544866 |
| ENSMUSG00000068748 | Ptprz1 | protein_coding | -0.847104246 | 5.973749057 | 2.5956617 | 0.1290266 | 0.483544866 |
| ENSMUSG00000054986 | Sec14l3 | protein_coding | -1.400643453 | 2.385090534 | 2.5551166 | 0.129131 | 0.483544866 |
| ENSMUSG00000029782 | Tmem209 | protein_coding | 0.769768198 | 3.896533446 | 2.5544686 | 0.1291762 | 0.483544866 |
| ENSMUSG00000032202 | Rab27a | protein_coding | 0.896813432 | 3.891648485 | 2.5537371 | 0.1292273 | 0.483544866 |
| ENSMUSG00000031835 | Mbtps1 | protein_coding | 0.525253278 | 5.428851872 | 2.5535384 | 0.1292411 | 0.483544866 |
| ENSMUSG00000042557 | Sin3a | protein_coding | -1.116851777 | 2.824291115 | 2.5529492 | 0.1292823 | 0.483544866 |
| ENSMUSG00000052763 | Zfp212 | protein_coding | 0.886831417 | 3.652756293 | 2.5527561 | 0.1292958 | 0.483544866 |
| ENSMUSG00000018217 | Pmp22 | protein_coding | -0.598259558 | 5.840557124 | 2.5523547 | 0.1293238 | 0.483544866 |
| ENSMUSG00000006711 | D130043K22Rik | protein_coding | 1.074259397 | 2.919162373 | 2.5522798 | 0.129329 | 0.483544866 |
| ENSMUSG00000034639 | Setmar | protein_coding | 1.190524591 | 2.899282814 | 2.5509268 | 0.1294236 | 0.483691234 |
| ENSMUSG00000024424 | Ttc39c | protein_coding | 0.806937138 | 3.912295957 | 2.5504331 | 0.1294581 | 0.483691234 |
| ENSMUSG00000020561 | Twistnb | protein_coding | -0.807945572 | 3.707681291 | 2.5492961 | 0.1295377 | 0.48381991 |
| ENSMUSG00000026109 | Tmeff2 | protein_coding | 0.426616561 | 7.305166309 | 2.5481644 | 0.129617 | 0.48381991 |
| ENSMUSG00000008855 | Hdac5 | protein_coding | -0.594103983 | 5.34606645 | 2.5472502 | 0.129681 | 0.48381991 |
| ENSMUSG00000005882 | Uqcc1 | protein_coding | 0.44333613 | 7.093360736 | 2.5469757 | 0.1297003 | 0.48381991 |
| ENSMUSG00000035277 | Arx | protein_coding | -0.969338779 | 3.12653984 | 2.5467296 | 0.1297175 | 0.48381991 |
| ENSMUSG00000036613 | Tssc1 | protein_coding | 0.501822669 | 6.756816031 | 2.5463838 | 0.1297733 | 0.483831217 |
| ENSMUSG00000024400 | Wdr33 | protein_coding | -0.417072201 | 6.93051735 | 2.5452207 | 0.1298234 | 0.483831217 |
| ENSMUSG00000044442 | N6amt1 | protein_coding | -0.374707568 | 6.41823056 | 2.5430781 | 0.1299739 | 0.483831217 |

| ensembl_gene_id | external_gene_name | gene_biotype | log2 (fold change) | log2 (counts per million) | F-value | p-value | False discovery rate |
|---|---|---|---|---|---|---|---|
| ENSMUSG00000030727 | Rabep2 | protein_coding | -0.938006323 | 3.876506452 | 2.5434188 | 0.12998 | 0.483831217 |
| ENSMUSG00000027282 | Mtch2 | protein_coding | 0.332083919 | 7.647824628 | 2.5424645 | 0.1300171 | 0.483831217 |
| ENSMUSG00000041642 | Kif21b | protein_coding | 0.873325581 | 3.805694586 | 2.5422009 | 0.1300356 | 0.483831217 |
| ENSMUSG00000023025 | Larp4 | protein_coding | -0.563160662 | 4.523088923 | 2.5419086 | 0.1300562 | 0.483831217 |
| ENSMUSG00000006611 | Hfe | protein_coding | -1.225988695 | 2.95804863 | 2.5415622 | 0.1300805 | 0.483831217 |
| ENSMUSG00000012296 | Tjap1 | protein_coding | 1.489370369 | 2.745423782 | 2.5397929 | 0.1302051 | 0.484127098 |
| ENSMUSG00000084957 | Bbip1 | protein_coding | -0.38608771 | 7.193835222 | 2.5391087 | 0.1302533 | 0.484127224 |
| ENSMUSG00000079610 | Ankrd39 | protein_coding | 0.749214133 | 3.616515507 | 2.5385148 | 0.1302952 | 0.484127224 |
| ENSMUSG00000021699 | Pde4d | protein_coding | -0.475236393 | 5.435247316 | 2.5375103 | 0.130366 | 0.484223159 |
| ENSMUSG00000020178 | Adora2a | protein_coding | -0.868826349 | 4.517630999 | 2.537123 | 0.1304249 | 0.484274401 |
| ENSMUSG00000053580 | Tanc2 | protein_coding | -0.48936943 | 5.970584882 | 2.5343618 | 0.1305884 | 0.484564475 |
| ENSMUSG00000028675 | Pnrc2 | protein_coding | -0.548138427 | 6.454867682 | 2.5342756 | 0.130626 | 0.484564475 |
| ENSMUSG00000027411 | Vps16 | protein_coding | 0.674046949 | 5.552159739 | 2.5339189 | 0.1306512 | 0.484564475 |
| ENSMUSG00000020612 | Prkar1a | protein_coding | 0.630039281 | 9.700402142 | 2.550092 | 0.1307029 | 0.484564475 |
| ENSMUSG00000041817 | Fam169a | protein_coding | -0.591827464 | 5.870143142 | 2.5323842 | 0.1307283 | 0.484564475 |
| ENSMUSG00000098188 | Sowahc | protein_coding | -0.965953075 | 3.065199733 | 2.5310896 | 0.13082 | 0.484706405 |
| ENSMUSG00000070002 | Ell | protein_coding | -0.586005865 | 5.35422909 | 2.5299611 | 0.1309 | 0.484706405 |
| ENSMUSG00000032425 | Zfp949 | protein_coding | 0.621920541 | 5.079971543 | 2.529935 | 0.1309018 | 0.484706405 |
| ENSMUSG00000078908 | Mon1b | protein_coding | 1.5080184 | 2.834909832 | 2.5291133 | 0.1309916 | 0.484719608 |
| ENSMUSG00000031546 | Gins4 | protein_coding | 0.511576215 | 4.980697551 | 2.5278621 | 0.1310489 | 0.484719608 |
| ENSMUSG00000029165 | Agbl5 | protein_coding | 1.050541246 | 2.977751785 | 2.5263106 | 0.1311591 | 0.484719608 |
| ENSMUSG00000041912 | Tdrkh | protein_coding | -0.453997485 | 5.926393016 | 2.5261078 | 0.1311735 | 0.484719608 |
| ENSMUSG00000046818 | Ddit4l | protein_coding | -0.688495412 | 4.730396204 | 2.5259156 | 0.1311872 | 0.484719608 |
| ENSMUSG00000027829 | Ccnl1 | protein_coding | -0.819155561 | 3.801533746 | 2.5256601 | 0.1312054 | 0.484719608 |
| ENSMUSG00000029864 | Gstk1 | protein_coding | -1.009338307 | 4.460523854 | 2.5427066 | 0.1312209 | 0.484719608 |
| ENSMUSG00000079435 | Rpl36a | protein_coding | 0.593903673 | 5.304824514 | 2.5236724 | 0.1313468 | 0.485017921 |
| ENSMUSG00000022946 | Dopey2 | protein_coding | 0.71232661 | 4.038381557 | 2.5207491 | 0.1315551 | 0.485539259 |
| ENSMUSG00000032297 | Celf6 | protein_coding | 0.626847917 | 5.126596465 | 2.5204245 | 0.1315783 | 0.485539259 |
| ENSMUSG00000072692 | Rpl37rt | transcribed_processe | 0.65976314 | 5.42589792 | 2.5190649 | 0.1317068 | 0.485846908 |
| ENSMUSG00000053702 | Nebl | protein_coding | -0.550157984 | 4.805923661 | 2.5174736 | 0.131789 | 0.485983409 |
| ENSMUSG00000052456 | Asna1 | protein_coding | 0.353127518 | 8.054059734 | 2.5153541 | 0.1319407 | 0.486062895 |
| ENSMUSG00000028871 | Rspo1 | protein_coding | 1.274591951 | 3.104288689 | 2.5145515 | 0.1319981 | 0.486062895 |
| ENSMUSG00000034994 | Eef2 | protein_coding | 0.456490891 | 10.00423559 | 2.5141451 | 0.1320273 | 0.486062895 |
| ENSMUSG00000078591 | Hs3st4 | protein_coding | 0.524163066 | 5.317790841 | 2.5112466 | 0.1322351 | 0.486062895 |
| ENSMUSG00000022335 | Zfat | protein_coding | 1.612645508 | 2.241048006 | 2.5106203 | 0.1322801 | 0.486062895 |
| ENSMUSG00000035765 | Dym | protein_coding | 0.471224178 | 5.538047024 | 2.5105155 | 0.1322876 | 0.486062895 |
| ENSMUSG00000058260 | Serpina9 | protein_coding | -0.806932561 | 4.249092998 | 2.5096218 | 0.1323518 | 0.486062895 |
| ENSMUSG00000042671 | Rgs8 | protein_coding | 0.546573551 | 6.430973914 | 2.5098308 | 0.1323683 | 0.486062895 |
| ENSMUSG00000045039 | Megf8 | protein_coding | -0.742579664 | 4.716612134 | 2.5093728 | 0.1323697 | 0.486062895 |
| ENSMUSG00000022623 | Shank3 | protein_coding | -1.07855514 | 2.872524293 | 2.5088508 | 0.1324072 | 0.486062895 |
| ENSMUSG00000035278 | Plekhj1 | protein_coding | 0.397005642 | 6.151296234 | 2.5081879 | 0.1324549 | 0.486062895 |
| ENSMUSG00000075289 | Carns1 | protein_coding | 0.942221409 | 3.162253939 | 2.5079319 | 0.1324733 | 0.486062895 |
| ENSMUSG00000005886 | Ncoa2 | protein_coding | -0.67096209 | 4.780771823 | 2.5075662 | 0.1324996 | 0.486062895 |
| ENSMUSG00000025597 | Klhl4 | protein_coding | -0.699647462 | 3.970025512 | 2.5073951 | 0.1325119 | 0.486062895 |
| ENSMUSG00000044772 | Sntn | protein_coding | -0.963642406 | 3.33696982 | 2.5071287 | 0.1325311 | 0.486062895 |
| ENSMUSG00000060450 | Rnf14 | protein_coding | 0.510039213 | 8.835925022 | 2.5075285 | 0.1325338 | 0.486062895 |
| ENSMUSG00000044231 | Nhlrc1 | protein_coding | -0.83672651 | 3.692155523 | 2.5061855 | 0.132599 | 0.486136148 |
| ENSMUSG00000019971 | Cep290 | protein_coding | -0.945026551 | 3.458790192 | 2.5053785 | 0.1326572 | 0.48615901 |
| ENSMUSG00000020120 | Plek | protein_coding | -0.820614314 | 3.509583045 | 2.504844 | 0.1326957 | 0.48615901 |
| ENSMUSG00000020700 | Map3k3 | protein_coding | 1.024551226 | 3.35893511 | 2.5040886 | 0.1327502 | 0.486192926 |
| ENSMUSG00000018770 | Atp5g3 | protein_coding | 0.379661617 | 10.45908229 | 2.5018918 | 0.1329087 | 0.486488702 |
| ENSMUSG00000029474 | Rnf34 | protein_coding | 0.460556768 | 6.829315571 | 2.5014083 | 0.1329436 | 0.486488702 |
| ENSMUSG00000034993 | Vat1 | protein_coding | 0.464786076 | 6.200072375 | 2.5008988 | 0.1329805 | 0.486488702 |
| ENSMUSG00000022336 | Eif3e | protein_coding | -0.435632451 | 7.651434213 | 2.500464 | 0.1330119 | 0.486488702 |
| ENSMUSG00000001909 | Trmt1 | protein_coding | 0.662255153 | 5.004510961 | 2.4983286 | 0.1331664 | 0.486846759 |
| ENSMUSG00000070644 | Etnk2 | protein_coding | 1.284279127 | 2.743692198 | 2.4972727 | 0.1332428 | 0.486846759 |
| ENSMUSG00000030016 | Zfp638 | protein_coding | -0.614679343 | 5.660084374 | 2.497223 | 0.1332464 | 0.486846759 |
| ENSMUSG00000022529 | Zfp263 | protein_coding | 0.673981229 | 3.891581902 | 2.4959559 | 0.1333383 | 0.486846759 |
| ENSMUSG00000029390 | Tmed2 | protein_coding | -0.904075118 | 3.063734541 | 2.493909 | 0.1334868 | 0.486846759 |
| ENSMUSG00000024639 | Gnaq | protein_coding | -0.506823885 | 8.272937565 | 2.4942663 | 0.1334924 | 0.486846759 |
| ENSMUSG00000024474 | Ik | protein_coding | 0.441683556 | 7.511914059 | 2.4937567 | 0.1334979 | 0.486846759 |
| ENSMUSG00000033059 | Pygb | protein_coding | 0.56749896 | 6.971582932 | 2.502602 | 0.1336608 | 0.486846759 |
| ENSMUSG00000023046 | Igfbp6 | protein_coding | -0.789911449 | 4.68971003 | 2.4902133 | 0.133787 | 0.486846759 |
| ENSMUSG00000035958 | Tdp2 | protein_coding | -0.72410554 | 3.702396804 | 2.4895012 | 0.1338073 | 0.486846759 |
| ENSMUSG00000014226 | Cacybp | protein_coding | -0.425920016 | 7.549578527 | 2.4891158 | 0.1338354 | 0.486846759 |
| ENSMUSG00000048483 | Zdhhc22 | protein_coding | 0.920424529 | 7.562365395 | 2.6033083 | 0.1338583 | 0.486846759 |
| ENSMUSG00000052539 | Magi3 | protein_coding | -1.202285009 | 2.938889907 | 2.4881553 | 0.1339054 | 0.486846759 |
| ENSMUSG00000050148 | Ubqln2 | protein_coding | -0.368564651 | 6.796164998 | 2.4878942 | 0.1339244 | 0.486846759 |
| ENSMUSG00000058159 | T2 | processed_transcript | 1.44826548 | 2.287451488 | 2.4867352 | 0.1340089 | 0.486846759 |
| ENSMUSG00000021071 | Trim9 | protein_coding | -0.682410838 | 5.423972199 | 2.4865858 | 0.1340513 | 0.486846759 |
| ENSMUSG00000055078 | Gabra5 | protein_coding | 0.975928077 | 3.08861533 | 2.4860659 | 0.1340578 | 0.486846759 |
| ENSMUSG00000015305 | Sash1 | protein_coding | -0.771505496 | 4.177191885 | 2.4859633 | 0.1340652 | 0.486846759 |
| ENSMUSG00000044629 | Cnrip1 | protein_coding | 0.345045171 | 7.54233379 | 2.4859421 | 0.1340668 | 0.486846759 |
| ENSMUSG00000029528 | Pxn | protein_coding | -1.377403018 | 2.558738636 | 2.4851144 | 0.1341272 | 0.486846759 |
| ENSMUSG00000003444 | Med29 | protein_coding | 0.671772839 | 3.908901613 | 2.4846956 | 0.1341578 | 0.486846759 |
| ENSMUSG00000030327 | Necap1 | protein_coding | 0.490694661 | 9.043497889 | 2.4842461 | 0.1342221 | 0.486846759 |
| ENSMUSG00000005089 | Slc1a2 | protein_coding | -0.654367262 | 7.751621344 | 2.5178379 | 0.1342478 | 0.486846759 |
| ENSMUSG00000020142 | Slc1a4 | protein_coding | 0.636851361 | 4.882143168 | 2.4824543 | 0.1343217 | 0.486846759 |

| ensembl_gene_id | external_gene_name | gene_biotype | log2 (fold change) | log2 (counts per million) | F-value | p-value | False discovery rate |
|---|---|---|---|---|---|---|---|
| ENSMUSG00000033526 | Ppip5k1 | protein_coding | 0.608411358 | 4.9857567 | 2.4818235 | 0.1343678 | 0.486846759 |
| ENSMUSG00000005802 | Slc30a4 | protein_coding | -1.145954698 | 3.385005898 | 2.4821234 | 0.1343774 | 0.486846759 |
| ENSMUSG00000041679 | Lrrc29 | protein_coding | -1.116695319 | 2.848384261 | 2.4811398 | 0.1344179 | 0.486846759 |
| ENSMUSG00000036902 | Neto2 | protein_coding | -0.702720143 | 4.898348325 | 2.4808604 | 0.1344383 | 0.486846759 |
| ENSMUSG00000052889 | Prkcb | protein_coding | -0.478322704 | 8.464317642 | 2.4810253 | 0.1344578 | 0.486846759 |
| ENSMUSG00000031392 | Irak1 | protein_coding | -0.51654342 | 5.165498465 | 2.479983 | 0.1345026 | 0.486846759 |
| ENSMUSG00000024764 | Naa40 | protein_coding | -0.845748032 | 3.508491131 | 2.479837 | 0.1345133 | 0.486846759 |
| ENSMUSG00000056310 | Tyw1 | protein_coding | 0.647858683 | 4.485944164 | 2.4786509 | 0.1346003 | 0.48692667 |
| ENSMUSG00000030788 | Rnf141 | protein_coding | -0.523931683 | 5.494603547 | 2.4783008 | 0.134626 | 0.48692667 |
| ENSMUSG00000050914 | Ankrd37 | protein_coding | -0.54417596 | 4.687636827 | 2.4771143 | 0.134713 | 0.487077772 |
| ENSMUSG00000074579 | Lekr1 | protein_coding | -0.956871436 | 3.043518446 | 2.4754301 | 0.1348368 | 0.48736124 |
| ENSMUSG00000026880 | Stom | protein_coding | 1.285122363 | 2.608445582 | 2.4740278 | 0.1349399 | 0.487370966 |
| ENSMUSG00000020627 | Klhl29 | protein_coding | -0.803285984 | 4.320473232 | 2.473612 | 0.1349705 | 0.487370966 |
| ENSMUSG00000041771 | Slc24a4 | protein_coding | -1.331078361 | 2.504585725 | 2.4735448 | 0.1349754 | 0.487370966 |
| ENSMUSG00000021957 | Tkt | protein_coding | 0.480585157 | 8.534920959 | 2.4725523 | 0.13508 | 0.487461289 |
| ENSMUSG00000000384 | Tbrg4 | protein_coding | 0.677529933 | 4.916363317 | 2.4719737 | 0.1350911 | 0.487461289 |
| ENSMUSG00000021504 | B4galt7 | protein_coding | 0.641922863 | 4.868777202 | 2.4698795 | 0.1352455 | 0.487561188 |
| ENSMUSG00000029729 | Zkscan1 | protein_coding | -0.501361827 | 5.761704343 | 2.4698433 | 0.1352482 | 0.487561188 |
| ENSMUSG00000004473 | Clec11a | protein_coding | 0.825868507 | 3.374650093 | 2.4688317 | 0.1353228 | 0.487561188 |
| ENSMUSG00000038128 | Camk4 | protein_coding | -0.554588485 | 6.331167359 | 2.4689267 | 0.1353473 | 0.487561188 |
| ENSMUSG00000042439 | Zfp532 | protein_coding | -0.559874772 | 4.906140566 | 2.4676915 | 0.135407 | 0.487561188 |
| ENSMUSG00000058355 | Abce1 | protein_coding | -0.466003106 | 5.983356726 | 2.4674332 | 0.1354261 | 0.487561188 |
| ENSMUSG00000004558 | Ndrg2 | protein_coding | -0.432197301 | 9.746559123 | 2.466468 | 0.1354975 | 0.487561188 |
| ENSMUSG00000022552 | Sharpin | protein_coding | 0.799669637 | 5.492261759 | 2.4826723 | 0.135526 | 0.487561188 |
| ENSMUSG00000073733 | Rsg1 | protein_coding | 1.541033979 | 2.950123744 | 2.4675728 | 0.1355665 | 0.487561188 |
| ENSMUSG00000034252 | Senp6 | protein_coding | -0.591318969 | 5.136205662 | 2.4649343 | 0.1356109 | 0.487561188 |
| ENSMUSG00000040321 | Zfp770 | protein_coding | -0.949582818 | 3.689128903 | 2.4648447 | 0.1356175 | 0.487561188 |
| ENSMUSG00000058835 | Abi1 | protein_coding | -0.384249584 | 6.933388157 | 2.4637478 | 0.1356988 | 0.487576012 |
| ENSMUSG00000031982 | Arv1 | protein_coding | 0.867258964 | 3.505064802 | 2.4635643 | 0.1357124 | 0.487576012 |
| ENSMUSG00000035829 | Ppp1r26 | protein_coding | 0.785641271 | 3.6414229 | 2.4622045 | 0.1358131 | 0.487744169 |
| ENSMUSG00000038453 | Srcin1 | protein_coding | -0.47133636 | 5.644276852 | 2.4617091 | 0.1358499 | 0.487744169 |
| ENSMUSG00000041841 | Rpl37 | protein_coding | 0.597021141 | 8.0548986 | 2.4784266 | 0.1359144 | 0.487813052 |
| ENSMUSG00000027007 | Ssfa2 | protein_coding | -1.195465517 | 3.402114785 | 2.4597308 | 0.1360282 | 0.487994285 |
| ENSMUSG00000034275 | Igsf9b | protein_coding | 0.765380342 | 4.741212726 | 2.4593607 | 0.1360557 | 0.487994285 |
| ENSMUSG00000019471 | Cdc37 | protein_coding | 0.33213628 | 7.155075556 | 2.4580003 | 0.1361253 | 0.488060435 |
| ENSMUSG00000039483 | Asb6 | protein_coding | 0.492399848 | 6.025324549 | 2.457232 | 0.1361825 | 0.488060435 |
| ENSMUSG00000032621 | Srek1 | protein_coding | -0.545632834 | 5.54780392 | 2.4568579 | 0.1362103 | 0.488060435 |
| ENSMUSG00000029580 | Actb | protein_coding | 0.665394061 | 12.74089718 | 2.4705745 | 0.13634 | 0.488362401 |
| ENSMUSG00000036019 | Tmtc2 | protein_coding | 0.843547312 | 3.387662185 | 2.4542965 | 0.1364011 | 0.488408888 |
| ENSMUSG00000036214 | Znrd1as | protein_coding | -0.821050704 | 3.733411621 | 2.4537231 | 0.1364438 | 0.488408888 |
| ENSMUSG00000030145 | Zfp248 | protein_coding | -0.697021759 | 4.432422608 | 2.4529864 | 0.1364988 | 0.488442997 |
| ENSMUSG00000021715 | Cwc27 | protein_coding | -0.85977309 | 3.686163553 | 2.4495932 | 0.1367522 | 0.488575125 |
| ENSMUSG00000022185 | Acin1 | protein_coding | -0.678712506 | 3.949173567 | 2.4495221 | 0.1367575 | 0.488575125 |
| ENSMUSG00000037740 | Mrps26 | protein_coding | 0.534510465 | 4.850606707 | 2.4477401 | 0.1368908 | 0.488575125 |
| ENSMUSG00000040423 | Rc3h1 | protein_coding | -0.53434475 | 4.843479795 | 2.4464935 | 0.1369842 | 0.488575125 |
| ENSMUSG00000002190 | Clgn | protein_coding | -1.412145735 | 2.905563367 | 2.4468142 | 0.1369917 | 0.488575125 |
| ENSMUSG00000032224 | Fam81a | protein_coding | -0.544779106 | 8.374444457 | 2.4531481 | 0.1370814 | 0.488575125 |
| ENSMUSG00000059291 | Rpl11 | protein_coding | 0.582054818 | 4.430452932 | 2.444894 | 0.1371041 | 0.488575125 |
| ENSMUSG00000015312 | Gadd45b | protein_coding | 0.812557989 | 3.546688812 | 2.4439045 | 0.1371784 | 0.488575125 |
| ENSMUSG00000028149 | Rap1gds1 | protein_coding | 0.362423441 | 7.472948559 | 2.4437102 | 0.137193 | 0.488575125 |
| ENSMUSG00000093674 | Rpl41 | protein_coding | 0.532824125 | 9.105067067 | 2.4439261 | 0.1372912 | 0.488575125 |
| ENSMUSG00000015568 | Lpl | protein_coding | -0.964193482 | 3.603553369 | 2.4423471 | 0.1372953 | 0.488575125 |
| ENSMUSG00000029178 | Klf3 | protein_coding | -0.841425511 | 3.876269742 | 2.4422244 | 0.1373045 | 0.488575125 |
| ENSMUSG00000026773 | Pfkfb3 | protein_coding | 0.98670207 | 3.827403228 | 2.4425655 | 0.1373104 | 0.488575125 |
| ENSMUSG00000021782 | Dlg5 | protein_coding | -0.807543687 | 3.431185895 | 2.4409742 | 0.1373985 | 0.488575125 |
| ENSMUSG00000005899 | Smpd4 | protein_coding | -0.899615004 | 3.595173308 | 2.4406085 | 0.137426 | 0.488575125 |
| ENSMUSG00000030177 | Ccdc77 | protein_coding | -1.449519781 | 2.536813618 | 2.4405562 | 0.13743 | 0.488575125 |
| ENSMUSG00000025588 | Nat1 | protein_coding | -1.547201717 | 2.428411009 | 2.4404552 | 0.1374376 | 0.488575125 |
| ENSMUSG00000028434 | Epb41l4b | protein_coding | -0.558541855 | 4.797368241 | 2.4400801 | 0.1374658 | 0.488575125 |
| ENSMUSG00000074671 | Tspyl3 | protein_coding | 0.59239885 | 5.022841253 | 2.4393718 | 0.1375191 | 0.488575125 |
| ENSMUSG00000042272 | Sestd1 | protein_coding | -0.87150661 | 3.482998893 | 2.4392816 | 0.1375259 | 0.488575125 |
| ENSMUSG00000027550 | Lrrcc1 | protein_coding | -1.383069715 | 4.236047588 | 2.4922807 | 0.13757 | 0.488575125 |
| ENSMUSG00000024772 | Ehd1 | protein_coding | -0.539528159 | 4.516743433 | 2.4374821 | 0.1376615 | 0.488575125 |
| ENSMUSG00000031111 | Igsf1 | protein_coding | 0.534148296 | 5.180803012 | 2.4370873 | 0.1376912 | 0.488575125 |
| ENSMUSG00000059689 | Zfp637 | protein_coding | 0.340715264 | 7.198845951 | 2.4365937 | 0.1377284 | 0.488575125 |
| ENSMUSG00000026977 | 7-Mar | protein_coding | -0.487024743 | 5.295410902 | 2.4363377 | 0.1377478 | 0.488575125 |
| ENSMUSG00000018572 | Phf23 | protein_coding | 0.473216248 | 5.843592551 | 2.4362768 | 0.1377523 | 0.488575125 |
| ENSMUSG00000087211 | Lhx1os | lincRNA | -0.615009455 | 4.070579514 | 2.4361427 | 0.1377625 | 0.488575125 |
| ENSMUSG00000041992 | Rapgef5 | protein_coding | -0.530434513 | 5.380962526 | 2.4353261 | 0.1378241 | 0.488592831 |
| ENSMUSG00000015749 | Anp32e | protein_coding | -0.360189269 | 6.679126619 | 2.4348726 | 0.1378583 | 0.488592831 |
| ENSMUSG00000025278 | Flnb | protein_coding | -1.310218928 | 3.310572923 | 2.4364951 | 0.1379046 | 0.488595737 |
| ENSMUSG00000004393 | Ddx56 | protein_coding | 0.461623419 | 7.161439461 | 2.4320631 | 0.1380707 | 0.488884071 |
| ENSMUSG00000024038 | Ndufv3 | protein_coding | 0.400809522 | 9.184277111 | 2.4319808 | 0.1380769 | 0.488884071 |
| ENSMUSG00000032193 | Ldlr | protein_coding | 1.111839251 | 3.684855885 | 2.4315994 | 0.1384178 | 0.4897469 |
| ENSMUSG00000001827 | Folr1 | protein_coding | -5.132266092 | 3.469133099 | 2.5273589 | 0.1385169 | 0.4897469 |
| ENSMUSG00000028630 | Dyrk2 | protein_coding | 0.744352883 | 3.865607839 | 2.425926 | 0.1385359 | 0.4897469 |
| ENSMUSG00000028351 | Brinp1 | protein_coding | 0.415006814 | 6.494578386 | 2.4253723 | 0.138578 | 0.4897469 |
| ENSMUSG00000060166 | Zdhhc8 | protein_coding | 0.767430172 | 3.884171771 | 2.4253458 | 0.13858 | 0.4897469 |

| ensembl_gene_id | external_gene_name | gene_biotype | log2 (fold change) | log2 (counts per million) | F-value | p-value | False discovery rate |
|---|---|---|---|---|---|---|---|
| ENSMUSG00000113722 | AC140477.3 | lincRNA | 0.567726735 | 4.754180436 | 2.4251634 | 0.1385939 | 0.4897469 |
| ENSMUSG00000003199 | Mpnd | protein_coding | 0.509757242 | 5.730802558 | 2.4233616 | 0.1387309 | 0.489921634 |
| ENSMUSG00000054414 | Slc30a7 | protein_coding | -1.000309179 | 3.354132197 | 2.423315 | 0.1387344 | 0.489921634 |
| ENSMUSG00000032519 | Slc25a38 | protein_coding | 0.583042744 | 4.960370605 | 2.4207028 | 0.1389334 | 0.490463113 |
| ENSMUSG00000027674 | Pex5l | protein_coding | -0.881156901 | 3.225459184 | 2.4199983 | 0.1389871 | 0.490491726 |
| ENSMUSG00000020225 | Tmbim4 | protein_coding | 0.395004373 | 6.932846328 | 2.4184193 | 0.1391076 | 0.490749106 |
| ENSMUSG00000049354 | Dcaf7 | protein_coding | -0.393964336 | 6.43481068 | 2.417209 | 0.1392 | 0.490749106 |
| ENSMUSG00000010914 | Pdhx | protein_coding | 0.420642882 | 6.463653855 | 2.4168659 | 0.1392262 | 0.490749106 |
| ENSMUSG00000020149 | Rab1a | protein_coding | 0.405305209 | 8.128892063 | 2.416652 | 0.1392426 | 0.490749106 |
| ENSMUSG00000035246 | Pcyt1b | protein_coding | -0.627048916 | 4.864259464 | 2.4154037 | 0.139338 | 0.490924634 |
| ENSMUSG00000038576 | Susd4 | protein_coding | 0.415079851 | 7.183129061 | 2.4130412 | 0.1395189 | 0.49140088 |
| ENSMUSG00000022176 | Rem2 | protein_coding | -1.691142451 | 2.818293959 | 2.4192968 | 0.139678 | 0.491800241 |
| ENSMUSG00000031358 | Msl3 | protein_coding | 0.630145188 | 4.63740218 | 2.4101433 | 0.1397412 | 0.491861486 |
| ENSMUSG00000102805 | Gm37240 | protein_coding | -0.922031762 | 3.117351977 | 2.4092882 | 0.1398068 | 0.491931585 |
| ENSMUSG00000052833 | Sae1 | protein_coding | 0.486738574 | 7.901327901 | 2.4080544 | 0.1399331 | 0.492085448 |
| ENSMUSG00000028560 | Usp1 | protein_coding | -0.998858959 | 3.026843183 | 2.4074312 | 0.1399496 | 0.492085448 |
| ENSMUSG00000024962 | Vegfb | protein_coding | 0.552974354 | 5.333725805 | 2.4069335 | 0.1399879 | 0.492085448 |
| ENSMUSG00000032135 | Mcam | protein_coding | -1.506809405 | 2.90400085 | 2.4059357 | 0.1400961 | 0.492197282 |
| ENSMUSG00000038831 | Ralgps1 | protein_coding | -0.756535568 | 4.464031335 | 2.4053097 | 0.1401128 | 0.492197282 |
| ENSMUSG00000032757 | Bet1 | protein_coding | -0.363074058 | 6.576747972 | 2.4043958 | 0.1401833 | 0.492197282 |
| ENSMUSG00000025178 | Pi4k2a | protein_coding | 0.505596947 | 6.30671777 | 2.4041427 | 0.1402028 | 0.492197282 |
| ENSMUSG00000109156 | Gm45194 | protein_coding | -0.791545492 | 4.031249565 | 2.4006983 | 0.1404686 | 0.49285815 |
| ENSMUSG00000037071 | Scd1 | protein_coding | -0.579820251 | 5.692765077 | 2.4005156 | 0.1404827 | 0.49285815 |
| ENSMUSG00000024383 | Map3k2 | protein_coding | -0.755841976 | 3.76212729 | 2.3991263 | 0.1405901 | 0.49307406 |
| ENSMUSG00000037697 | Ddhd1 | protein_coding | -1.344654968 | 2.963253179 | 2.3977268 | 0.1407298 | 0.493117492 |
| ENSMUSG00000038546 | Ranbp9 | protein_coding | -0.771671528 | 3.615199335 | 2.3969665 | 0.1407572 | 0.493117492 |
| ENSMUSG00000063765 | Chadl | protein_coding | 0.781362754 | 3.341909579 | 2.3966489 | 0.1407818 | 0.493117492 |
| ENSMUSG00000006998 | Psmd2 | protein_coding | 0.472005928 | 7.72167652 | 2.3969693 | 0.1407885 | 0.493117492 |
| ENSMUSG00000029020 | Mfn2 | protein_coding | 0.566741363 | 7.370764413 | 2.4110874 | 0.1408874 | 0.493117492 |
| ENSMUSG00000019210 | Atp6v1e1 | protein_coding | 0.342984093 | 10.1101088 | 2.3948363 | 0.1409223 | 0.493117492 |
| ENSMUSG00000027330 | Cdc25b | protein_coding | -1.052802865 | 3.054587964 | 2.3932324 | 0.1410468 | 0.493117492 |
| ENSMUSG00000025153 | Fasn | protein_coding | 0.54805922 | 6.044390897 | 2.3930078 | 0.1410642 | 0.493117492 |
| ENSMUSG00000027167 | Elp4 | protein_coding | -0.638653587 | 3.932947935 | 2.3929364 | 0.1410698 | 0.493117492 |
| ENSMUSG00000030880 | Polr3e | protein_coding | -0.870569689 | 4.903822005 | 2.4077101 | 0.1410725 | 0.493117492 |
| ENSMUSG00000027533 | Fabp5 | protein_coding | 0.367596449 | 6.901610174 | 2.3924585 | 0.1411069 | 0.493117492 |
| ENSMUSG00000053898 | Ech1 | protein_coding | -0.347301613 | 7.728086844 | 2.3916584 | 0.1411691 | 0.493174489 |
| ENSMUSG00000044477 | Zfand3 | protein_coding | 0.601175128 | 4.897481481 | 2.3907967 | 0.1412361 | 0.493248304 |
| ENSMUSG00000085385 | Snhg17 | processed_transcript | -1.440262747 | 2.346316515 | 2.3895002 | 0.1413369 | 0.493440328 |
| ENSMUSG00000014813 | Stc1 | protein_coding | 0.841338939 | 3.672506606 | 2.3875934 | 0.1414855 | 0.493798258 |
| ENSMUSG00000024120 | Lrpprc | protein_coding | -0.445723017 | 5.719920287 | 2.3870053 | 0.1415313 | 0.493798258 |
| ENSMUSG00000027823 | Gmps | protein_coding | -0.52913663 | 7.305183761 | 2.388814 | 0.1417683 | 0.494464558 |
| ENSMUSG00000028060 | 2810403A07Rik | protein_coding | -0.646221059 | 4.220048193 | 2.3826743 | 0.1418695 | 0.494657191 |
| ENSMUSG00000022802 | Lmln | protein_coding | 1.426374763 | 2.455071012 | 2.3818676 | 0.1419326 | 0.494716773 |
| ENSMUSG00000079477 | Rab7 | protein_coding | 0.520787083 | 6.059395973 | 2.3797979 | 0.1420946 | 0.495121117 |
| ENSMUSG00000029670 | Ing3 | protein_coding | -0.519851305 | 5.85921492 | 2.3769839 | 0.1423153 | 0.495729453 |
| ENSMUSG00000057914 | Cacnb2 | protein_coding | -0.709973407 | 4.629845211 | 2.3759917 | 0.1423932 | 0.495840217 |
| ENSMUSG00000042726 | Trafd1 | protein_coding | -0.682991725 | 4.927957212 | 2.3748344 | 0.1424842 | 0.495996278 |
| ENSMUSG00000022621 | Rabl2 | protein_coding | 0.485031571 | 6.451840936 | 2.3727847 | 0.1426454 | 0.496396863 |
| ENSMUSG00000024883 | Rin1 | protein_coding | -0.959310975 | 3.607593129 | 2.3684096 | 0.1429904 | 0.49731257 |
| ENSMUSG00000020526 | Znhit3 | protein_coding | 0.357617842 | 7.147062682 | 2.3682739 | 0.1430011 | 0.49731257 |
| ENSMUSG00000110723 | AC153912.1 | protein_coding | -1.184647332 | 2.861603887 | 2.3670968 | 0.143094 | 0.497475031 |
| ENSMUSG00000006411 | Nectin4 | protein_coding | 1.458164613 | 3.299685051 | 2.3844635 | 0.1433375 | 0.498160308 |
| ENSMUSG00000041323 | Ak7 | protein_coding | -1.251099914 | 2.577050978 | 2.3622401 | 0.1434785 | 0.498483809 |
| ENSMUSG00000057329 | Bcl2 | protein_coding | -0.850999598 | 3.884976949 | 2.3615618 | 0.1435323 | 0.498483809 |
| ENSMUSG00000026577 | Blzf1 | protein_coding | -0.54655593 | 5.141870769 | 2.3604855 | 0.1436177 | 0.498483809 |
| ENSMUSG00000038160 | Atg5 | protein_coding | 0.371406506 | 6.381312863 | 2.3602156 | 0.1436391 | 0.498483809 |
| ENSMUSG00000060703 | Cd302 | protein_coding | -0.654308735 | 5.575286645 | 2.360289 | 0.1436647 | 0.498483809 |
| ENSMUSG00000021681 | Aggf1 | protein_coding | -0.573190788 | 4.869183155 | 2.3593296 | 0.1437095 | 0.498483809 |
| ENSMUSG00000024182 | Axin1 | protein_coding | -0.622295194 | 4.05334865 | 2.3586322 | 0.1437649 | 0.498483809 |
| ENSMUSG00000002844 | Adprh | protein_coding | 0.397197963 | 7.485039426 | 2.3581722 | 0.1438014 | 0.498483809 |
| ENSMUSG00000000171 | Sdhd | protein_coding | 0.354884606 | 9.071525417 | 2.3570326 | 0.1438921 | 0.498637223 |
| ENSMUSG00000041423 | Paqr6 | protein_coding | -0.723747205 | 3.762613481 | 2.3517126 | 0.1443161 | 0.499561383 |
| ENSMUSG00000024231 | Cul2 | protein_coding | -0.55443067 | 5.685802949 | 2.3516439 | 0.1443216 | 0.499561383 |
| ENSMUSG00000030432 | Rpl28 | protein_coding | 0.390072182 | 6.389847888 | 2.351551 | 0.144329 | 0.499561383 |
| ENSMUSG00000030272 | Camk1 | protein_coding | 0.713428459 | 3.713426668 | 2.351356 | 0.1443446 | 0.499561383 |
| ENSMUSG00000038736 | Nudcd1 | protein_coding | -0.715142826 | 4.382402137 | 2.3501537 | 0.1444406 | 0.49965062 |
| ENSMUSG00000040883 | Tmem205 | protein_coding | 0.486878466 | 6.500854608 | 2.3498703 | 0.1444633 | 0.49965062 |
| ENSMUSG00000037972 | Snn | protein_coding | -0.491635508 | 5.734400238 | 2.348965 | 0.1445357 | 0.499731189 |
| ENSMUSG00000086629 | 2810403D21Rik | processed_transcript | -0.896747598 | 3.35964998 | 2.348417 | 0.1445795 | 0.499731189 |
| ENSMUSG00000032381 | Fam96a | protein_coding | -0.391131399 | 6.727894162 | 2.3469509 | 0.1446969 | 0.499976226 |
| ENSMUSG00000114362 | AC124433.1 | antisense_RNA | -1.697484586 | 2.15005337 | 2.3462685 | 0.1447516 | 0.500004486 |
| ENSMUSG00000040260 | Daam2 | protein_coding | -0.645939936 | 4.34735342 | 2.3446023 | 0.1448852 | 0.50017565 |
| ENSMUSG00000007617 | Homer1 | protein_coding | -0.531985362 | 5.929049542 | 2.3421221 | 0.1450844 | 0.50017565 |
| ENSMUSG00000036442 | Thap11 | protein_coding | 0.66515192 | 4.678581292 | 2.3418362 | 0.1451074 | 0.50017565 |
| ENSMUSG00000040621 | Gemin8 | protein_coding | -0.665079653 | 3.708095508 | 2.3413363 | 0.1451475 | 0.50017565 |
| ENSMUSG00000004994 | Ccdc130 | protein_coding | 0.517374684 | 5.078397919 | 2.3409822 | 0.145176 | 0.50017565 |
| ENSMUSG00000000753 | Serpinf1 | protein_coding | 0.798661031 | 4.379507595 | 2.3412277 | 0.1451877 | 0.50017565 |
| ENSMUSG00000056267 | Cep70 | protein_coding | -0.61502064 | 4.42457313 | 2.3407199 | 0.1451971 | 0.50017565 |

| ensembl_gene_id | external_gene_name | gene_biotype | log2 (fold change) | log2 (counts per million) | F-value | p-value | False discovery rate |
|---|---|---|---|---|---|---|---|
| ENSMUSG00000036834 | Plch1 | protein_coding | 1.110750268 | 3.273452275 | 2.3406516 | 0.1452026 | 0.50017565 |
| ENSMUSG00000003344 | Btbd2 | protein_coding | 0.419201118 | 6.043958302 | 2.3404382 | 0.1452198 | 0.50017565 |
| ENSMUSG00000054021 | Sirt5 | protein_coding | 0.572968722 | 5.581060247 | 2.3398316 | 0.1452686 | 0.500183582 |
| ENSMUSG00000019494 | Cops6 | protein_coding | 0.328598228 | 9.482982842 | 2.3365844 | 0.1455303 | 0.50086143 |
| ENSMUSG00000036186 | Fam69b | protein_coding | 0.657253329 | 4.7655652 | 2.3362329 | 0.1455586 | 0.50086143 |
| ENSMUSG00000030579 | Tyrobp | protein_coding | -0.818723088 | 5.019160979 | 2.34496 | 0.1456572 | 0.501040193 |
| ENSMUSG00000028343 | Erp44 | protein_coding | -0.465761867 | 5.414849164 | 2.3338548 | 0.1457507 | 0.50113362 |
| ENSMUSG00000030223 | Ptpro | protein_coding | 0.518829129 | 4.842292511 | 2.3333505 | 0.1457914 | 0.50113362 |
| ENSMUSG00000028132 | Tmem56 | protein_coding | -0.753652264 | 4.357039653 | 2.3329459 | 0.1458242 | 0.50113362 |
| ENSMUSG00000038121 | Fam210a | protein_coding | -0.535323361 | 5.42499516 | 2.330434 | 0.1460275 | 0.501570727 |
| ENSMUSG00000086382 | Chrna1os | antisense_RNA | -1.090266358 | 2.764194342 | 2.3301658 | 0.1460492 | 0.501570727 |
| ENSMUSG00000028382 | Ptbp3 | protein_coding | -0.728582386 | 3.931620566 | 2.3296465 | 0.1460913 | 0.501570727 |
| ENSMUSG00000060733 | Ipmk | protein_coding | -0.646026422 | 3.940777536 | 2.3280612 | 0.1462198 | 0.501851888 |
| ENSMUSG00000024668 | Sdhaf2 | protein_coding | 0.34831115 | 6.812997948 | 2.3264583 | 0.14635 | 0.502029435 |
| ENSMUSG00000036792 | Mbd5 | protein_coding | -0.53291031 | 4.79709081 | 2.3262739 | 0.1463649 | 0.502029435 |
| ENSMUSG00000029161 | Cgref1 | protein_coding | 0.565213635 | 5.114376841 | 2.3243617 | 0.1465204 | 0.502402334 |
| ENSMUSG00000040936 | Ulk4 | protein_coding | -0.758745875 | 3.71758564 | 2.3205425 | 0.1468315 | 0.503271079 |
| ENSMUSG00000049672 | Zbtb14 | protein_coding | 0.860647257 | 3.195792007 | 2.3199272 | 0.1468816 | 0.503271079 |
| ENSMUSG00000038127 | Ccdc50 | protein_coding | -0.437389196 | 5.518560666 | 2.3195289 | 0.1469142 | 0.503271079 |
| ENSMUSG00000035900 | Gramd4 | protein_coding | -0.816351143 | 3.297468782 | 2.3183983 | 0.1470065 | 0.503359797 |
| ENSMUSG00000016150 | Tenm1 | protein_coding | -0.675697016 | 4.381687609 | 2.317129 | 0.1471102 | 0.503359797 |
| ENSMUSG00000022636 | Alcam | protein_coding | -0.458590596 | 7.059554592 | 2.316986 | 0.1471219 | 0.503359797 |
| ENSMUSG00000001630 | Stk38l | protein_coding | -0.506548458 | 5.132107652 | 2.3166415 | 0.14715 | 0.503359797 |
| ENSMUSG00000039001 | Rps21 | protein_coding | 0.491544657 | 8.753982477 | 2.3158029 | 0.14725 | 0.503359797 |
| ENSMUSG00000029992 | Gfpt1 | protein_coding | -0.608287401 | 5.066774403 | 2.3154142 | 0.1472504 | 0.503359797 |
| ENSMUSG00000027808 | Serp1 | protein_coding | -0.656827091 | 4.652541343 | 2.3149184 | 0.147291 | 0.503359797 |
| ENSMUSG00000036537 | Rnf113a1 | protein_coding | 0.569286505 | 5.094406699 | 2.314631 | 0.1473145 | 0.503359797 |
| ENSMUSG00000110834 | CAAA01180111.2 | lincRNA | 0.607380789 | 4.268034083 | 2.3135608 | 0.1474022 | 0.503376077 |
| ENSMUSG00000032589 | Bsn | protein_coding | -0.726678161 | 4.550189453 | 2.3129638 | 0.1474511 | 0.503376077 |
| ENSMUSG00000016481 | Cr1l | protein_coding | -0.709326351 | 3.992040996 | 2.3125733 | 0.1474831 | 0.503376077 |
| ENSMUSG00000079045 | Prox1os | antisense_RNA | -0.969790181 | 4.162944071 | 2.3200579 | 0.1475066 | 0.503376077 |
| ENSMUSG00000032186 | Tmod2 | protein_coding | -0.35081451 | 8.400191226 | 2.3098561 | 0.1477062 | 0.503777415 |
| ENSMUSG00000024387 | Csnk2b | protein_coding | 0.323179285 | 8.273179454 | 2.3096369 | 0.1477242 | 0.503777415 |
| ENSMUSG00000041607 | Mbp | protein_coding | -0.520268365 | 11.80476767 | 2.3093259 | 0.1477811 | 0.503777415 |
| ENSMUSG00000022488 | Nckap1l | protein_coding | -2.084724791 | 2.372215698 | 2.3163229 | 0.1478116 | 0.503777415 |
| ENSMUSG00000024885 | Aldh3b1 | protein_coding | -0.800765802 | 3.921266022 | 2.3068969 | 0.1479495 | 0.50393604 |
| ENSMUSG00000021193 | Pitrm1 | protein_coding | 0.713627111 | 4.532985356 | 2.3059824 | 0.1480248 | 0.50393604 |
| ENSMUSG00000025422 | Agap2 | protein_coding | -0.873598402 | 3.675226007 | 2.3058961 | 0.1480319 | 0.50393604 |
| ENSMUSG00000014633 | Cmc2 | protein_coding | 0.446484319 | 5.842276473 | 2.3054371 | 0.1480697 | 0.50393604 |
| ENSMUSG00000027848 | Olfml3 | protein_coding | -0.675023535 | 4.462495399 | 2.3051617 | 0.1480924 | 0.50393604 |
| ENSMUSG00000024158 | Hagh | protein_coding | 0.357400328 | 8.304436477 | 2.302617 | 0.1483023 | 0.504490635 |
| ENSMUSG00000032599 | Ip6k2 | protein_coding | 0.373019022 | 6.790141189 | 2.3002658 | 0.1484966 | 0.504689497 |
| ENSMUSG00000008734 | Gprc5b | protein_coding | -0.693048113 | 4.896268762 | 2.3002307 | 0.1484995 | 0.504689497 |
| ENSMUSG00000008206 | Cers4 | protein_coding | 0.481746167 | 5.832784365 | 2.3001471 | 0.1485064 | 0.504689497 |
| ENSMUSG00000053141 | Ptprt | protein_coding | -0.685223918 | 4.442839227 | 2.2992279 | 0.1485824 | 0.504689497 |
| ENSMUSG00000033543 | Gtf2a2 | protein_coding | -0.378143392 | 7.381250823 | 2.2984 | 0.148651 | 0.504689497 |
| ENSMUSG00000029922 | Mkrn1 | protein_coding | 0.370768156 | 6.384668255 | 2.2982107 | 0.1486666 | 0.504689497 |
| ENSMUSG00000041836 | Ptpre | protein_coding | 0.648499536 | 4.376135082 | 2.2977351 | 0.148706 | 0.504689497 |
| ENSMUSG00000001768 | Rin2 | protein_coding | -0.663017556 | 4.377031292 | 2.2973523 | 0.1487377 | 0.504689497 |
| ENSMUSG00000022407 | Adsl | protein_coding | 0.478875982 | 6.061447949 | 2.2967181 | 0.1487903 | 0.504689497 |
| ENSMUSG00000034551 | Hdx | protein_coding | -1.833611024 | 2.241436869 | 2.2952775 | 0.1489098 | 0.504689497 |
| ENSMUSG00000024998 | Plce1 | protein_coding | -0.942009742 | 2.957736556 | 2.2951452 | 0.1489207 | 0.504689497 |
| ENSMUSG00000070858 | Gm1673 | protein_coding | 0.814365421 | 3.401330592 | 2.295106 | 0.148924 | 0.504689497 |
| ENSMUSG00000045659 | Plekha7 | protein_coding | -0.998984637 | 3.350783315 | 2.2901111 | 0.1493392 | 0.505887337 |
| ENSMUSG00000034903 | Cobll1 | protein_coding | -1.657010385 | 2.680774634 | 2.2900987 | 0.1493716 | 0.505887337 |
| ENSMUSG00000035696 | Rnf38 | protein_coding | -0.706915252 | 4.031856508 | 2.2882841 | 0.1494914 | 0.506085867 |
| ENSMUSG00000059323 | Tonsl | protein_coding | 0.341173725 | 9.574008816 | 2.2874478 | 0.1495611 | 0.506085867 |
| ENSMUSG00000032171 | Pin1 | protein_coding | 0.352268012 | 8.909003625 | 2.2873254 | 0.1495714 | 0.506085867 |
| ENSMUSG00000018474 | Chd3 | protein_coding | -0.372962321 | 6.673162646 | 2.2842271 | 0.1498301 | 0.506801876 |
| ENSMUSG00000003868 | Ruvbl2 | protein_coding | 0.504965666 | 6.01352007 | 2.2834304 | 0.1498967 | 0.506867761 |
| ENSMUSG00000048758 | Rpl29 | protein_coding | 0.450062969 | 5.486489102 | 2.2805549 | 0.1501375 | 0.50742948 |
| ENSMUSG00000032867 | Fbxw8 | protein_coding | -1.186940435 | 2.483064752 | 2.2803195 | 0.1501572 | 0.50742948 |
| ENSMUSG00000026312 | Cdh7 | protein_coding | -0.612099857 | 4.282803202 | 2.2785247 | 0.1503078 | 0.50748607 |
| ENSMUSG00000028960 | Ube4b | protein_coding | -0.498464858 | 5.598998595 | 2.2785246 | 0.1503078 | 0.50748607 |
| ENSMUSG00000068039 | Tcp1 | protein_coding | 0.399482478 | 8.356202542 | 2.2776954 | 0.1503774 | 0.50748607 |
| ENSMUSG00000001054 | Rmnd5b | protein_coding | 0.481812194 | 5.63977067 | 2.2774553 | 0.1503975 | 0.50748607 |
| ENSMUSG00000030505 | Prmt3 | protein_coding | 0.801488641 | 3.326721743 | 2.2770794 | 0.1504291 | 0.50748607 |
| ENSMUSG00000025135 | Anapc11 | protein_coding | 0.326418327 | 9.282356923 | 2.2766541 | 0.1504649 | 0.50748607 |
| ENSMUSG00000021809 | Nudt13 | protein_coding | 0.721611784 | 4.202712886 | 2.2760935 | 0.150512 | 0.50748607 |
| ENSMUSG00000097536 | 2610037D02Rik | lincRNA | -1.336086003 | 2.550039944 | 2.2751162 | 0.1505942 | 0.50748607 |
| ENSMUSG00000001687 | Ubl3 | protein_coding | -0.344456335 | 8.338305244 | 2.2745364 | 0.1506429 | 0.50748607 |
| ENSMUSG00000024587 | Nars | protein_coding | 0.42070556 | 8.572052115 | 2.2745011 | 0.1506459 | 0.50748607 |
| ENSMUSG00000030982 | 9030624J02Rik | protein_coding | 0.649778008 | 6.165526476 | 2.2881914 | 0.1507129 | 0.507552682 |
| ENSMUSG00000019232 | Etnppl | protein_coding | -0.468157496 | 5.211151314 | 2.272866 | 0.1507836 | 0.50763182 |
| ENSMUSG00000015755 | Tab2 | protein_coding | -0.634890096 | 5.182467895 | 2.2721032 | 0.1508479 | 0.50766086 |
| ENSMUSG00000022216 | Psme1 | protein_coding | 0.375976164 | 7.421260387 | 2.2709346 | 0.1509464 | 0.50766086 |
| ENSMUSG00000051413 | Plagl2 | protein_coding | 0.912807873 | 2.963117602 | 2.2702133 | 0.1510073 | 0.50766086 |
| ENSMUSG00000035354 | Uvrag | protein_coding | -0.770045845 | 3.360669855 | 2.2699332 | 0.1510309 | 0.50766086 |

| ensembl_gene_id | external_gene_name | gene_biotype | log2 (fold change) | log2 (counts per million) | F-value | p-value | False discovery rate |
|---|---|---|---|---|---|---|---|
| ENSMUSG00000093930 | Hmgcs1 | protein_coding | -0.438987578 | 7.080878806 | 2.2697949 | 0.1510426 | 0.50766086 |
| ENSMUSG00000048583 | Igf2 | protein_coding | -1.140464019 | 4.068313374 | 2.2911891 | 0.1510755 | 0.50766086 |
| ENSMUSG00000019139 | Isyna1 | protein_coding | 0.714225479 | 5.3840446 | 2.2672588 | 0.1514247 | 0.508675474 |
| ENSMUSG00000089917 | Uckl1 | protein_coding | 0.581847774 | 5.065466382 | 2.2642704 | 0.15151 | 0.508802788 |
| ENSMUSG00000040102 | Klhl42 | protein_coding | 0.755813724 | 4.246095242 | 2.2619167 | 0.1517096 | 0.509294003 |
| ENSMUSG00000003527 | Dgcr14 | protein_coding | 0.514878262 | 5.291420031 | 2.260881 | 0.1517975 | 0.509294003 |
| ENSMUSG00000029111 | Nelfa | protein_coding | 0.641993333 | 5.752998423 | 2.2613726 | 0.1518695 | 0.509294003 |
| ENSMUSG00000027552 | E2f5 | protein_coding | -0.90257458 | 2.977907243 | 2.2595666 | 0.1519093 | 0.509294003 |
| ENSMUSG00000042249 | Grk3 | protein_coding | -0.756981526 | 4.939557764 | 2.2598017 | 0.1519206 | 0.509294003 |
| ENSMUSG00000050002 | Idnk | protein_coding | 0.393604467 | 7.140977413 | 2.2588372 | 0.1519713 | 0.509294003 |
| ENSMUSG00000021340 | Gpld1 | protein_coding | 0.793808245 | 3.527496448 | 2.2586435 | 0.1519878 | 0.509294003 |
| ENSMUSG00000039262 | Prrc2b | protein_coding | 0.498750785 | 5.773422834 | 2.2574758 | 0.1520872 | 0.509313294 |
| ENSMUSG00000050963 | Kcns2 | protein_coding | 1.242197005 | 2.687444094 | 2.2574629 | 0.1520883 | 0.509313294 |
| ENSMUSG00000056069 | Fam105a | protein_coding | 0.891513515 | 3.276289901 | 2.2552372 | 0.1522779 | 0.509345685 |
| ENSMUSG00000019235 | Rps6kl1 | protein_coding | 0.505764025 | 5.743658113 | 2.2550426 | 0.1522945 | 0.509345685 |
| ENSMUSG00000022376 | Adcy8 | protein_coding | 0.965291175 | 2.998032649 | 2.2546666 | 0.1523266 | 0.509345685 |
| ENSMUSG00000026748 | Plxdc2 | protein_coding | -0.663364391 | 4.74135173 | 2.2544829 | 0.1523423 | 0.509345685 |
| ENSMUSG00000040687 | Madd | protein_coding | 0.455253964 | 5.786761892 | 2.2542408 | 0.1523629 | 0.509345685 |
| ENSMUSG00000027544 | Nfatc2 | protein_coding | 0.741681654 | 4.198646993 | 2.2540157 | 0.1523821 | 0.509345685 |
| ENSMUSG00000029249 | Rest | protein_coding | -1.559077557 | 2.127344686 | 2.252542 | 0.152508 | 0.509366981 |
| ENSMUSG00000043448 | Gjc2 | protein_coding | -1.486433781 | 2.313014584 | 2.2524645 | 0.1525146 | 0.509366981 |
| ENSMUSG00000025757 | Hspa4l | protein_coding | -0.449597779 | 6.93759472 | 2.2516088 | 0.1525877 | 0.509366981 |
| ENSMUSG00000063802 | Hspbp1 | protein_coding | 0.406504209 | 7.8338441 | 2.2513294 | 0.1526116 | 0.509366981 |
| ENSMUSG00000022234 | Cct5 | protein_coding | 0.394693402 | 9.145480461 | 2.2511688 | 0.1526254 | 0.509366981 |
| ENSMUSG00000020089 | Ppa1 | protein_coding | 0.416605209 | 7.898838711 | 2.2499058 | 0.1527334 | 0.509414147 |
| ENSMUSG00000042532 | Golga7b | protein_coding | 0.447935768 | 7.965838073 | 2.2498962 | 0.1527342 | 0.509414147 |
| ENSMUSG00000097789 | Gm2115 | protein_coding | -0.835330467 | 3.277335339 | 2.2486887 | 0.1528376 | 0.509600947 |
| ENSMUSG00000058897 | Col25a1 | protein_coding | 0.594734087 | 4.848781409 | 2.247432 | 0.1529453 | 0.509801974 |
| ENSMUSG00000033083 | Tbc1d4 | protein_coding | -0.867603829 | 3.271941174 | 2.2461706 | 0.1530535 | 0.509874209 |
| ENSMUSG00000071662 | Polr2g | protein_coding | 0.396074609 | 8.576597452 | 2.2455607 | 0.1531059 | 0.509874209 |
| ENSMUSG00000037126 | Psd | protein_coding | -0.568038721 | 5.103726207 | 2.2452525 | 0.1531323 | 0.509874209 |
| ENSMUSG00000047789 | Slc38a9 | protein_coding | 0.644648138 | 3.844259702 | 2.2449692 | 0.1531567 | 0.509874209 |
| ENSMUSG00000021996 | Esd | protein_coding | 0.339708081 | 8.145446247 | 2.2438102 | 0.1532563 | 0.51004782 |
| ENSMUSG00000029695 | Aass | protein_coding | -2.018244152 | 1.638383169 | 2.2431304 | 0.1533147 | 0.510084454 |
| ENSMUSG00000026709 | Dars2 | protein_coding | 0.659250016 | 4.442531 | 2.2407527 | 0.1535193 | 0.5104973 |
| ENSMUSG00000001924 | Uba1 | protein_coding | 0.363020622 | 7.770458192 | 2.2392846 | 0.1536459 | 0.5104973 |
| ENSMUSG00000063785 | Utp14a | protein_coding | -0.981106916 | 3.571688047 | 2.2392708 | 0.1536783 | 0.5104973 |
| ENSMUSG00000024393 | Prrc2a | protein_coding | 0.576463072 | 4.643013439 | 2.2387011 | 0.1536962 | 0.5104973 |
| ENSMUSG00000030092 | Cntn6 | protein_coding | -0.699269536 | 4.10731142 | 2.2385604 | 0.1537083 | 0.5104973 |
| ENSMUSG00000032551 | 1110059G10Rik | protein_coding | -0.475245858 | 5.850311521 | 2.2383827 | 0.1537236 | 0.5104973 |
| ENSMUSG00000035686 | Thrsp | protein_coding | -0.585740528 | 6.131810158 | 2.2385122 | 0.1538261 | 0.510679788 |
| ENSMUSG00000029166 | Mapre3 | protein_coding | 0.363271311 | 6.801726034 | 2.235235 | 0.1539955 | 0.510797054 |
| ENSMUSG00000024069 | Slc30a6 | protein_coding | -0.690025718 | 4.55533128 | 2.234592 | 0.1540511 | 0.510797054 |
| ENSMUSG00000038390 | Gpr162 | protein_coding | 0.671841277 | 4.424986241 | 2.2342064 | 0.1540845 | 0.510797054 |
| ENSMUSG00000030805 | Stx4a | protein_coding | 0.447586622 | 6.131729887 | 2.2341958 | 0.1540854 | 0.510797054 |
| ENSMUSG00000031129 | Slc9a9 | protein_coding | -0.702192673 | 3.493578143 | 2.233168 | 0.1541744 | 0.510797054 |
| ENSMUSG00000024526 | Cidea | protein_coding | -0.79663336 | 3.764927347 | 2.2331355 | 0.1541772 | 0.510797054 |
| ENSMUSG00000000884 | Gnb1l | protein_coding | 0.718940627 | 5.353469977 | 2.2365606 | 0.1542559 | 0.510797054 |
| ENSMUSG00000072915 | Gm12258 | protein_coding | -0.790465992 | 3.798291442 | 2.2320067 | 0.1542749 | 0.510797054 |
| ENSMUSG00000017561 | Crlf3 | protein_coding | -0.913700648 | 3.450944053 | 2.2318454 | 0.1542889 | 0.510797054 |
| ENSMUSG00000003234 | Abcf3 | protein_coding | 0.496352028 | 7.064597713 | 2.2310572 | 0.1543885 | 0.510969387 |
| ENSMUSG00000042185 | Nfrkb | protein_coding | -0.888750432 | 3.61727441 | 2.2284147 | 0.1545866 | 0.511417868 |
| ENSMUSG00000006392 | Med8 | protein_coding | -0.406154809 | 6.431049831 | 2.2280403 | 0.1546191 | 0.511417868 |
| ENSMUSG00000034687 | Fras1 | protein_coding | -1.819609922 | 2.484782701 | 2.2282107 | 0.1547179 | 0.511587132 |
| ENSMUSG00000061371 | Zfp873 | protein_coding | -0.719438912 | 4.0858188 | 2.2262373 | 0.1547759 | 0.511609906 |
| ENSMUSG00000087370 | Tmem170b | protein_coding | -0.58896151 | 5.335219299 | 2.2243757 | 0.154938 | 0.511609906 |
| ENSMUSG00000015937 | H2afy | protein_coding | 0.760503622 | 4.298275487 | 2.2235108 | 0.1550133 | 0.511609906 |
| ENSMUSG00000066829 | Zfp810 | protein_coding | -1.034848458 | 3.844884453 | 2.230427 | 0.155045 | 0.511609906 |
| ENSMUSG00000031948 | Kars | protein_coding | 0.497726877 | 7.945595081 | 2.2236141 | 0.1551179 | 0.511609906 |
| ENSMUSG00000029415 | Sdad1 | protein_coding | 0.761608933 | 4.236613644 | 2.2222537 | 0.155123 | 0.511609906 |
| ENSMUSG00000022634 | Yaf2 | protein_coding | -0.372480401 | 6.893968802 | 2.221878 | 0.1551557 | 0.511609906 |
| ENSMUSG00000020257 | Wdr82 | protein_coding | -0.406648069 | 6.41093458 | 2.2212128 | 0.1552138 | 0.511609906 |
| ENSMUSG00000041040 | Fam117b | protein_coding | -1.212484962 | 2.756767461 | 2.220517 | 0.1552746 | 0.511609906 |
| ENSMUSG00000022202 | Sugt1 | protein_coding | -0.33241554 | 7.235566152 | 2.2202189 | 0.1553006 | 0.511609906 |
| ENSMUSG00000020684 | Rasl10b | protein_coding | -0.710777304 | 5.728973139 | 2.233931 | 0.155319 | 0.511609906 |
| ENSMUSG00000037266 | Rsrp1 | protein_coding | -1.243288118 | 5.794475185 | 2.3348987 | 0.1553312 | 0.511609906 |
| ENSMUSG00000031708 | Tecr | protein_coding | 0.367960432 | 10.67126226 | 2.2196755 | 0.1553481 | 0.511609906 |
| ENSMUSG00000051705 | Senp8 | protein_coding | -0.64009156 | 4.582202443 | 2.2191862 | 0.1553909 | 0.511609906 |
| ENSMUSG00000052676 | Zmat1 | protein_coding | -0.674292922 | 4.490497069 | 2.2168699 | 0.1555935 | 0.512120398 |
| ENSMUSG00000034354 | Mtmr3 | protein_coding | 0.615497483 | 5.133218822 | 2.2138548 | 0.1558578 | 0.512800949 |
| ENSMUSG00000031284 | Pak3 | protein_coding | -0.339199697 | 7.575495308 | 2.2134236 | 0.1558957 | 0.512800949 |
| ENSMUSG00000063632 | Sox11 | protein_coding | -0.439738479 | 6.041335213 | 2.2127778 | 0.1559524 | 0.512830571 |
| ENSMUSG00000010755 | Cars | protein_coding | 0.716244485 | 5.871903777 | 2.2291417 | 0.1560805 | 0.513095007 |
| ENSMUSG00000030407 | Qpctl | protein_coding | 0.539352278 | 4.971526964 | 2.2105161 | 0.1561512 | 0.513112723 |
| ENSMUSG00000021772 | Nkiras1 | protein_coding | 0.319499982 | 8.258030439 | 2.2096776 | 0.1562249 | 0.513112723 |
| ENSMUSG00000038324 | Trpc4ap | protein_coding | 0.361993821 | 6.560717276 | 2.2096307 | 0.1562291 | 0.513112723 |
| ENSMUSG00000039546 | Ajap1 | protein_coding | -0.432002111 | 5.796713387 | 2.2087311 | 0.1563083 | 0.51321609 |
| ENSMUSG00000021728 | Emb | protein_coding | 0.510471081 | 5.713795227 | 2.2066468 | 0.1564919 | 0.513662346 |

| ensembl_gene_id | external_gene_name | gene_biotype | log2 (fold change) | log2 (counts per million) | F-value | p-value | False discovery rate |
|---|---|---|---|---|---|---|---|
| ENSMUSG00000022477 | Aco2 | protein_coding | 0.394089342 | 8.493533715 | 2.2051671 | 0.1566225 | 0.513934034 |
| ENSMUSG00000035953 | Tmem55b | protein_coding | 0.511949979 | 5.558917797 | 2.2041123 | 0.1567157 | 0.514082803 |
| ENSMUSG00000032376 | Usp3 | protein_coding | -0.584591157 | 4.465359218 | 2.2032501 | 0.1567919 | 0.514116224 |
| ENSMUSG00000029673 | Auts2 | protein_coding | -0.349277751 | 6.258686383 | 2.2029152 | 0.1568215 | 0.514116224 |
| ENSMUSG00000033595 | Lgi3 | protein_coding | 0.44736945 | 5.970040407 | 2.2020733 | 0.1568959 | 0.514203572 |
| ENSMUSG00000019831 | Wasf1 | protein_coding | -0.361458592 | 7.054750933 | 2.2003901 | 0.1570449 | 0.514481814 |
| ENSMUSG00000049721 | Gal3st1 | protein_coding | -1.017004053 | 3.6048914 | 2.2003856 | 0.1570765 | 0.514481814 |
| ENSMUSG00000020642 | Rnf144a | protein_coding | -0.811925844 | 3.501786462 | 2.198973 | 0.1571705 | 0.514632908 |
| ENSMUSG00000030226 | Lmo3 | protein_coding | -0.368027105 | 7.101001143 | 2.1951288 | 0.1575118 | 0.515446549 |
| ENSMUSG00000037110 | Ralgapa2 | protein_coding | -1.29221492 | 2.806873281 | 2.1947866 | 0.1575422 | 0.515446549 |
| ENSMUSG00000024050 | Wiz | protein_coding | -0.777994847 | 3.622251832 | 2.1940165 | 0.1576107 | 0.515446549 |
| ENSMUSG00000030282 | Cmas | protein_coding | 0.333922716 | 6.74470459 | 2.1934306 | 0.1576629 | 0.515446549 |
| ENSMUSG00000047412 | Zbtb44 | protein_coding | -0.871386285 | 3.245284772 | 2.1930891 | 0.1576933 | 0.515446549 |
| ENSMUSG00000024953 | Prdx5 | protein_coding | 0.334896505 | 10.25842496 | 2.1925907 | 0.1577376 | 0.515446549 |
| ENSMUSG00000021250 | Fos | protein_coding | 0.538626143 | 6.540849949 | 2.1936508 | 0.157789 | 0.515446549 |
| ENSMUSG00000032878 | Ccdc85a | protein_coding | -0.473794956 | 5.289002769 | 2.1917974 | 0.1578083 | 0.515446549 |
| ENSMUSG00000038564 | Ift172 | protein_coding | 0.865050757 | 3.500283351 | 2.191325 | 0.1578504 | 0.515446549 |
| ENSMUSG00000029348 | Asphd2 | protein_coding | -0.476512462 | 5.911058479 | 2.1902953 | 0.1579422 | 0.515589786 |
| ENSMUSG00000017677 | Wsb1 | protein_coding | -0.714929638 | 4.565200691 | 2.1879746 | 0.1581494 | 0.515751887 |
| ENSMUSG00000031028 | Tub | protein_coding | 0.441097145 | 5.704199397 | 2.1878164 | 0.1581635 | 0.515751887 |
| ENSMUSG00000029123 | Stk32b | protein_coding | 0.914734992 | 4.482213564 | 2.2009962 | 0.1581997 | 0.515751887 |
| ENSMUSG00000032383 | Ppib | protein_coding | 0.399119293 | 6.18592516 | 2.1872182 | 0.158217 | 0.515751887 |
| ENSMUSG00000086350 | B230369F24Rik | antisense_RNA | -0.714134155 | 3.605258578 | 2.1870537 | 0.1582317 | 0.515751887 |
| ENSMUSG00000037234 | Hook3 | protein_coding | -0.68460907 | 4.477801318 | 2.1855533 | 0.1583659 | 0.516032923 |
| ENSMUSG00000020736 | Nt5c | protein_coding | 0.316589029 | 7.34385362 | 2.1835334 | 0.1585468 | 0.516166696 |
| ENSMUSG00000102437 | Gm38048 | sense_overlapping | 1.029781697 | 3.008105649 | 2.1821032 | 0.158675 | 0.516166696 |
| ENSMUSG00000033063 | Cntnap3 | protein_coding | -1.220353623 | 2.680127865 | 2.1819576 | 0.1586881 | 0.516166696 |
| ENSMUSG00000011752 | Pgam1 | protein_coding | 0.641090421 | 3.725204375 | 2.1812395 | 0.1587526 | 0.516166696 |
| ENSMUSG00000006818 | Sod2 | protein_coding | 0.363386784 | 7.601455699 | 2.1806185 | 0.1588083 | 0.516166696 |
| ENSMUSG00000020926 | Adam11 | protein_coding | 0.533586455 | 4.884671681 | 2.1805785 | 0.1588119 | 0.516166696 |
| ENSMUSG00000030869 | Ndufab1 | protein_coding | 0.601024455 | 4.057246265 | 2.1805564 | 0.1588139 | 0.516166696 |
| ENSMUSG00000033880 | Lgals3bp | protein_coding | 0.920702274 | 4.716344849 | 2.1974089 | 0.1588636 | 0.516166696 |
| ENSMUSG00000021147 | Wdr37 | protein_coding | -0.425145969 | 6.485592369 | 2.1795072 | 0.1589081 | 0.516166696 |
| ENSMUSG00000074030 | Exoc8 | protein_coding | 0.54397227 | 4.648191444 | 2.1789149 | 0.1589614 | 0.516166696 |
| ENSMUSG00000033712 | Ccar2 | protein_coding | 0.585951603 | 5.48746704 | 2.1786918 | 0.1589814 | 0.516166696 |
| ENSMUSG00000020921 | Tmem101 | protein_coding | 0.460915302 | 5.629054628 | 2.1786747 | 0.158983 | 0.516166696 |
| ENSMUSG00000003585 | Sec14l2 | protein_coding | -0.645414647 | 3.81467034 | 2.17692 | 0.1591409 | 0.516420483 |
| ENSMUSG00000039795 | Zfand1 | protein_coding | -0.469498611 | 5.231060953 | 2.1767386 | 0.1591572 | 0.516420483 |
| ENSMUSG00000028779 | Pef1 | protein_coding | 0.366532265 | 7.551714064 | 2.1758405 | 0.1592381 | 0.516527096 |
| ENSMUSG00000032846 | Zswim6 | processed_transcript | -0.997065356 | 3.842680476 | 2.1732792 | 0.159636 | 0.517564353 |
| ENSMUSG00000062012 | Zfp13 | protein_coding | 0.51634136 | 5.172760166 | 2.1712306 | 0.1596541 | 0.517564353 |
| ENSMUSG00000026017 | Carf | protein_coding | -0.726834 | 4.410506606 | 2.1700585 | 0.1597601 | 0.517751856 |
| ENSMUSG00000024054 | Smchd1 | protein_coding | -0.834284915 | 3.379495894 | 2.1676093 | 0.1599819 | 0.51831435 |
| ENSMUSG00000037514 | Pank2 | protein_coding | -0.675062862 | 4.32589004 | 2.1659386 | 0.1601334 | 0.518578082 |
| ENSMUSG00000021361 | Tmem14c | protein_coding | 0.406908208 | 6.955817858 | 2.1652698 | 0.1601941 | 0.518578082 |
| ENSMUSG00000022142 | Nup155 | protein_coding | -0.916024224 | 2.915053981 | 2.1651167 | 0.160208 | 0.518578082 |
| ENSMUSG00000032030 | Cul5 | protein_coding | -0.475420207 | 5.90373286 | 2.1638935 | 0.1603191 | 0.518781494 |
| ENSMUSG00000020694 | Tlk2 | protein_coding | -0.4802719 | 5.276387066 | 2.1606003 | 0.1606186 | 0.519095252 |
| ENSMUSG00000055900 | Tmem69 | protein_coding | 0.743132816 | 3.362338281 | 2.1605164 | 0.1606263 | 0.519095252 |
| ENSMUSG00000001260 | Gabrg1 | protein_coding | -0.410089381 | 6.075152125 | 2.1602908 | 0.1606468 | 0.519095252 |
| ENSMUSG00000004285 | Atp6v1f | protein_coding | 0.350348302 | 9.236162641 | 2.1600265 | 0.1606709 | 0.519095252 |
| ENSMUSG00000049536 | Tceal1 | protein_coding | -0.312248809 | 7.576970249 | 2.1598145 | 0.1606902 | 0.519095252 |
| ENSMUSG00000027834 | Serpini1 | protein_coding | -0.3772795 | 6.329136714 | 2.1596448 | 0.1607057 | 0.519095252 |
| ENSMUSG00000005103 | Wdr1 | protein_coding | 0.536462404 | 7.835162807 | 2.1720538 | 0.1607861 | 0.5191992 |
| ENSMUSG00000030509 | Asb7 | protein_coding | -0.591584043 | 4.074378218 | 2.1579098 | 0.1608639 | 0.519294268 |
| ENSMUSG00000028034 | Fubp1 | protein_coding | 0.414720184 | 6.719120052 | 2.1561656 | 0.1610231 | 0.519557373 |
| ENSMUSG00000022338 | Eny2 | protein_coding | -0.357552233 | 7.138504758 | 2.1559588 | 0.161042 | 0.519557373 |
| ENSMUSG00000047003 | Zfp41 | protein_coding | 1.516241399 | 2.686236001 | 2.155188 | 0.1611436 | 0.519729055 |
| ENSMUSG00000009470 | Tnpo1 | protein_coding | -0.624614671 | 4.766316474 | 2.152697 | 0.1613404 | 0.520207753 |
| ENSMUSG00000050379 | 6-Sep | protein_coding | 0.536688079 | 6.389171914 | 2.1521792 | 0.1614673 | 0.520321469 |
| ENSMUSG00000045107 | Saysd1 | protein_coding | 0.491846885 | 5.068478322 | 2.1512558 | 0.1614724 | 0.520321469 |
| ENSMUSG00000062421 | Arf2 | protein_coding | 0.46153329 | 6.704694964 | 2.1498128 | 0.1616047 | 0.520539722 |
| ENSMUSG00000013698 | Pea15a | protein_coding | 0.421303911 | 10.3333805 | 2.1491015 | 0.16167 | 0.520539722 |
| ENSMUSG00000028020 | Glrb | protein_coding | -0.337430642 | 6.764037551 | 2.1484352 | 0.1617312 | 0.520539722 |
| ENSMUSG00000042700 | Sipa1l1 | protein_coding | -0.510585485 | 6.147735034 | 2.1476936 | 0.1617993 | 0.520539722 |
| ENSMUSG00000020346 | Mgat1 | protein_coding | -0.504681477 | 5.171762214 | 2.1476463 | 0.1618037 | 0.520539722 |
| ENSMUSG00000040433 | Zbtb38 | protein_coding | -0.545520797 | 7.245201503 | 2.1600533 | 0.1618739 | 0.520539722 |
| ENSMUSG00000033228 | Scaf11 | protein_coding | -0.517649097 | 5.105056989 | 2.1462685 | 0.1619304 | 0.520539722 |
| ENSMUSG00000056258 | Kcnq3 | protein_coding | -0.622023094 | 5.110848953 | 2.1460509 | 0.1619504 | 0.520539722 |
| ENSMUSG00000004069 | Dnaja3 | protein_coding | 0.426230593 | 6.564819385 | 2.1452684 | 0.1620224 | 0.520539722 |
| ENSMUSG00000034235 | Usp54 | protein_coding | -1.389560941 | 2.469394203 | 2.1445258 | 0.1620908 | 0.520539722 |
| ENSMUSG00000037355 | Uvssa | protein_coding | -1.121728655 | 2.934807809 | 2.1442385 | 0.1621172 | 0.520539722 |
| ENSMUSG00000029592 | Usp30 | protein_coding | -0.689626688 | 4.847948825 | 2.1444257 | 0.1621311 | 0.520539722 |
| ENSMUSG00000045795 | Whamm | protein_coding | -1.239626477 | 2.505882743 | 2.1423594 | 0.1622904 | 0.520539722 |
| ENSMUSG00000022788 | Fgd4 | protein_coding | -0.62713387 | 3.77936611 | 2.141506 | 0.1623692 | 0.520539722 |
| ENSMUSG00000000167 | Pih1d2 | protein_coding | -0.68622047 | 3.671753911 | 2.1414968 | 0.16237 | 0.520539722 |
| ENSMUSG00000002957 | Ap2a2 | protein_coding | 0.459118915 | 7.263747972 | 2.1416378 | 0.1623881 | 0.520539722 |
| ENSMUSG00000059213 | Ddn | protein_coding | -0.798643978 | 5.670490936 | 2.1680007 | 0.162401 | 0.520539722 |

| ensembl_gene_id | external_gene_name | gene_biotype | log2 (fold change) | log2 (counts per million) | F-value | p-value | False discovery rate |
|---|---|---|---|---|---|---|---|
| ENSMUSG00000010057 | Nprl2 | protein_coding | 0.467202581 | 6.55218835 | 2.1410477 | 0.1624115 | 0.520539722 |
| ENSMUSG00000021647 | Cartpt | protein_coding | 0.553573415 | 6.853129418 | 2.1523844 | 0.1625739 | 0.520885269 |
| ENSMUSG00000024527 | Afg3l2 | protein_coding | 0.532914978 | 5.199336318 | 2.1388324 | 0.1626162 | 0.520885269 |
| ENSMUSG00000061654 | Spry3 | protein_coding | 0.886438738 | 3.475000625 | 2.1360797 | 0.162871 | 0.520903298 |
| ENSMUSG00000031139 | Mcf2 | protein_coding | 0.982502219 | 3.053798652 | 2.1360621 | 0.1628726 | 0.520903298 |
| ENSMUSG00000022217 | Emc9 | protein_coding | 0.326469665 | 6.944502775 | 2.1356546 | 0.1629104 | 0.520903298 |
| ENSMUSG00000027304 | Rtf1 | protein_coding | -0.534529216 | 5.315831213 | 2.1355958 | 0.1629158 | 0.520903298 |
| ENSMUSG00000041205 | Map6d1 | protein_coding | -0.625403925 | 4.165930521 | 2.1355534 | 0.1629197 | 0.520903298 |
| ENSMUSG00000020163 | Uqcr11 | protein_coding | 0.451257133 | 9.283705867 | 2.1351949 | 0.162953 | 0.520903298 |
| ENSMUSG00000028403 | Zdhhc21 | protein_coding | -0.621189677 | 4.489614356 | 2.135015 | 0.1629696 | 0.520903298 |
| ENSMUSG00000053317 | Sec61b | protein_coding | 0.359627238 | 6.311453429 | 2.1345868 | 0.1630094 | 0.520903298 |
| ENSMUSG00000027221 | Chst1 | protein_coding | 0.391120148 | 6.988503255 | 2.1322706 | 0.1632243 | 0.52114557 |
| ENSMUSG00000018167 | Stard3 | protein_coding | 0.600308249 | 5.476916645 | 2.1324314 | 0.1632405 | 0.52114557 |
| ENSMUSG00000085396 | Firre | processed_transcript | -0.814304284 | 3.688050705 | 2.1318182 | 0.1632664 | 0.52114557 |
| ENSMUSG00000026727 | Rsu1 | protein_coding | -0.522154696 | 5.540845219 | 2.1316821 | 0.163279 | 0.52114557 |
| ENSMUSG00000021488 | Nsd1 | protein_coding | -0.415909509 | 6.332371186 | 2.1306339 | 0.1633765 | 0.521238091 |
| ENSMUSG00000010607 | Pigyl | protein_coding | 0.350836891 | 7.440696244 | 2.1303276 | 0.163405 | 0.521238091 |
| ENSMUSG00000028557 | Rnf11 | protein_coding | -0.319995222 | 7.771785049 | 2.1295178 | 0.1634803 | 0.521323824 |
| ENSMUSG00000021646 | Mccc2 | protein_coding | 0.611714465 | 5.453756839 | 2.1288913 | 0.1635697 | 0.521454182 |
| ENSMUSG00000057406 | Nsd2 | protein_coding | -0.452805341 | 5.332635908 | 2.1257506 | 0.1638314 | 0.52213377 |
| ENSMUSG00000028234 | Rps20 | protein_coding | 0.452715024 | 7.008257084 | 2.1241133 | 0.1640154 | 0.522262202 |
| ENSMUSG00000010136 | Pifo | protein_coding | -0.580706304 | 4.485323866 | 2.1233438 | 0.1640562 | 0.522262202 |
| ENSMUSG00000056459 | Zbtb25 | protein_coding | -0.829104722 | 3.542563016 | 2.1228283 | 0.1641044 | 0.522262202 |
| ENSMUSG00000009630 | Ppp2cb | protein_coding | -0.408308131 | 6.220185394 | 2.1219464 | 0.1641869 | 0.522262202 |
| ENSMUSG00000045763 | Basp1 | protein_coding | -0.334124294 | 7.471415446 | 2.1218082 | 0.1641999 | 0.522262202 |
| ENSMUSG00000042719 | Naa25 | protein_coding | 0.810373992 | 3.837378938 | 2.1216859 | 0.1642113 | 0.522262202 |
| ENSMUSG00000032560 | Dnajc13 | protein_coding | -0.707473621 | 4.102390063 | 2.1216816 | 0.1642117 | 0.522262202 |
| ENSMUSG00000063049 | Ing2 | protein_coding | -0.699330978 | 4.181450923 | 2.1206331 | 0.1643099 | 0.522386003 |
| ENSMUSG00000091864 | Gm17102 | lincRNA | -0.919334804 | 3.367061646 | 2.1202283 | 0.1643478 | 0.522386003 |
| ENSMUSG00000020289 | Nprl3 | protein_coding | 0.486416777 | 5.770974429 | 2.1193158 | 0.1644333 | 0.522503381 |
| ENSMUSG00000006021 | Kptn | protein_coding | 0.648887139 | 5.224893488 | 2.1176835 | 0.1646175 | 0.522933991 |
| ENSMUSG00000030717 | Nupr1 | protein_coding | -0.869597896 | 5.056776348 | 2.1369657 | 0.1646749 | 0.522961858 |
| ENSMUSG00000002147 | Stat6 | protein_coding | -1.132388339 | 2.770354408 | 2.1147597 | 0.1648611 | 0.523241294 |
| ENSMUSG00000039621 | Prex1 | protein_coding | -0.838926299 | 4.392303219 | 2.1157188 | 0.1648837 | 0.523241294 |
| ENSMUSG00000003039 | Fam32a | protein_coding | 0.315239445 | 8.117999032 | 2.1139877 | 0.1649338 | 0.523241294 |
| ENSMUSG00000046658 | Zfp316 | protein_coding | -1.101051894 | 2.786277545 | 2.1137354 | 0.1649575 | 0.523241294 |
| ENSMUSG00000029674 | Limk1 | protein_coding | -1.316326003 | 2.789344449 | 2.1128568 | 0.1650402 | 0.523349321 |
| ENSMUSG00000028441 | 1110017D15Rik | protein_coding | -0.456659864 | 5.39075987 | 2.1116309 | 0.1651557 | 0.523490635 |
| ENSMUSG00000033014 | Trim33 | protein_coding | -0.525908682 | 4.776449862 | 2.1113505 | 0.1651822 | 0.523490635 |
| ENSMUSG00000027596 | a | protein_coding | -1.033778161 | 2.687726089 | 2.1104215 | 0.1652698 | 0.523596131 |
| ENSMUSG00000015522 | Arnt | protein_coding | 0.780481129 | 3.379198166 | 2.1092178 | 0.1653834 | 0.523596131 |
| ENSMUSG00000020361 | Hspa4 | protein_coding | 0.472769377 | 5.412967386 | 2.1090384 | 0.1654003 | 0.523596131 |
| ENSMUSG00000052516 | Robo2 | protein_coding | -0.61110411 | 4.285642096 | 2.1089335 | 0.1654102 | 0.523596131 |
| ENSMUSG00000030612 | Mrpl46 | protein_coding | 0.315801309 | 7.79607444 | 2.1058759 | 0.1656993 | 0.52379473 |
| ENSMUSG00000011837 | Snapc2 | protein_coding | 0.528301749 | 5.29531142 | 2.105709 | 0.1657151 | 0.52379473 |
| ENSMUSG00000000787 | Ddx3x | protein_coding | -0.373696627 | 6.582524005 | 2.1056539 | 0.1657203 | 0.52379473 |
| ENSMUSG00000028397 | Kdm4c | protein_coding | -0.636687759 | 4.255247997 | 2.105601 | 0.1657253 | 0.52379473 |
| ENSMUSG00000059991 | Nptx2 | protein_coding | 0.884841044 | 3.168592128 | 2.1055802 | 0.1657273 | 0.52379473 |
| ENSMUSG00000028953 | Abcf2 | protein_coding | 0.654764562 | 6.907046021 | 2.1329432 | 0.1658083 | 0.52379473 |
| ENSMUSG00000039838 | Slc45a1 | protein_coding | 0.526974039 | 6.048189818 | 2.1049923 | 0.1658139 | 0.52379473 |
| ENSMUSG00000068329 | Htra2 | protein_coding | 0.593009566 | 4.739776655 | 2.1040623 | 0.165871 | 0.523821246 |
| ENSMUSG00000032492 | Pth1r | protein_coding | 0.88554578 | 3.535233835 | 2.1030093 | 0.1659709 | 0.523982592 |
| ENSMUSG00000039660 | Spout1 | protein_coding | 0.461819417 | 5.892875434 | 2.1022759 | 0.1660404 | 0.524048381 |
| ENSMUSG00000024533 | Spire1 | protein_coding | -0.49752593 | 5.694342266 | 2.0998733 | 0.1662686 | 0.524201148 |
| ENSMUSG00000074968 | Ano3 | protein_coding | -0.642180822 | 5.341786411 | 2.0992595 | 0.1663579 | 0.524201148 |
| ENSMUSG00000026107 | Nabp1 | protein_coding | -0.786107531 | 3.476302538 | 2.0984593 | 0.1664031 | 0.524201148 |
| ENSMUSG00000063260 | Syt10 | protein_coding | -0.669311797 | 3.929621671 | 2.0981902 | 0.1664287 | 0.524201148 |
| ENSMUSG00000018593 | Sparc | protein_coding | -0.644249496 | 9.204871844 | 2.1261342 | 0.1664408 | 0.524201148 |
| ENSMUSG00000029385 | Ccng2 | protein_coding | -0.649166763 | 4.500731803 | 2.0974056 | 0.1665033 | 0.524201148 |
| ENSMUSG00000020536 | Llgl1 | protein_coding | 0.566365741 | 4.550861631 | 2.0968797 | 0.1665534 | 0.524201148 |
| ENSMUSG00000035152 | Ap2b1 | protein_coding | -0.406953785 | 6.570604096 | 2.0964383 | 0.1665955 | 0.524201148 |
| ENSMUSG00000041552 | Ptchd1 | protein_coding | -0.783329821 | 3.196184494 | 2.0954563 | 0.1666891 | 0.524201148 |
| ENSMUSG00000016181 | Diexf | protein_coding | -1.157594204 | 3.022252251 | 2.0947493 | 0.1667565 | 0.524201148 |
| ENSMUSG00000019849 | Prep | protein_coding | -0.616579253 | 3.773300172 | 2.0946558 | 0.1667654 | 0.524201148 |
| ENSMUSG00000027828 | Ssr3 | protein_coding | -0.299216929 | 7.518890899 | 2.0946371 | 0.1667672 | 0.524201148 |
| ENSMUSG00000033904 | Ccp110 | protein_coding | -0.445329465 | 5.276146718 | 2.0945842 | 0.1667722 | 0.524201148 |
| ENSMUSG00000018102 | Hist1h2bc | protein_coding | -0.648632512 | 5.968762913 | 2.1069349 | 0.1668001 | 0.524201148 |
| ENSMUSG00000034820 | Cpsf7 | protein_coding | 1.516659486 | 2.343303687 | 2.0936645 | 0.16686 | 0.524201148 |
| ENSMUSG00000032872 | Cyb5r4 | protein_coding | -0.697552491 | 3.598388793 | 2.0917148 | 0.1670463 | 0.524201148 |
| ENSMUSG00000051331 | Cacna1c | protein_coding | -0.82841085 | 3.874655236 | 2.0916264 | 0.1670547 | 0.524201148 |
| ENSMUSG00000054893 | Zfp667 | protein_coding | 0.504980241 | 5.479913346 | 2.091435 | 0.167073 | 0.524201148 |
| ENSMUSG00000037523 | Mavs | protein_coding | 0.736443679 | 4.380389987 | 2.0912584 | 0.1670899 | 0.524201148 |
| ENSMUSG00000053091 | Lins1 | protein_coding | 0.91968433 | 3.081821021 | 2.0912572 | 0.16709 | 0.524201148 |
| ENSMUSG00000039682 | Lap3 | protein_coding | -0.388695334 | 6.696552077 | 2.0905711 | 0.1671557 | 0.524201148 |
| ENSMUSG00000024191 | Bnip1 | protein_coding | 0.463342093 | 5.806986693 | 2.0905119 | 0.1671613 | 0.524201148 |
| ENSMUSG00000069678 | Pcgf1 | protein_coding | 0.452368541 | 5.263945238 | 2.0862434 | 0.1675704 | 0.525269939 |
| ENSMUSG00000079283 | 2310009B15Rik | protein_coding | 0.595319636 | 4.54184601 | 2.085936 | 0.1675998 | 0.525269939 |
| ENSMUSG00000025583 | Rptor | protein_coding | 0.495004348 | 4.777241995 | 2.0828251 | 0.1678988 | 0.525750425 |

| ensembl_gene_id | external_gene_name | gene_biotype | log2 (fold change) | log2 (counts per million) | F-value | p-value | False discovery rate |
|---|---|---|---|---|---|---|---|
| ENSMUSG00000029036 | Atad3a | protein_coding | 0.814746999 | 3.588426218 | 2.0824603 | 0.1679339 | 0.525750425 |
| ENSMUSG00000074001 | Klhl40 | protein_coding | -1.593473416 | 2.161219232 | 2.0824295 | 0.1679369 | 0.525750425 |
| ENSMUSG00000022964 | Tmem50b | protein_coding | 0.416603485 | 7.304724141 | 2.0823062 | 0.1679487 | 0.525750425 |
| ENSMUSG00000027523 | Gnas | protein_coding | 0.716467581 | 3.948568539 | 2.0802433 | 0.1681474 | 0.526219151 |
| ENSMUSG00000039678 | Tbc1d13 | protein_coding | -0.743452317 | 3.440285826 | 2.0781898 | 0.1683455 | 0.526514684 |
| ENSMUSG00000026603 | Smyd2 | protein_coding | 0.721552767 | 3.360309474 | 2.0780068 | 0.1683631 | 0.526514684 |
| ENSMUSG00000003604 | Aven | protein_coding | -0.735019045 | 4.143550156 | 2.0757123 | 0.1685848 | 0.526514684 |
| ENSMUSG00000097867 | Lppos | lincRNA | -1.058381127 | 2.724646276 | 2.075528 | 0.1686026 | 0.526514684 |
| ENSMUSG00000038112 | AW551984 | protein_coding | 0.544340827 | 6.598811039 | 2.0836448 | 0.1686091 | 0.526514684 |
| ENSMUSG00000021048 | Mthfd1 | protein_coding | 0.622391426 | 4.127595466 | 2.0751037 | 0.1686437 | 0.526514684 |
| ENSMUSG00000031060 | Rbm10 | protein_coding | 0.602026369 | 4.31472965 | 2.074899 | 0.1686635 | 0.526514684 |
| ENSMUSG00000019897 | Ccdc59 | protein_coding | -0.339742568 | 7.036182763 | 2.0747975 | 0.1686733 | 0.526514684 |
| ENSMUSG00000022656 | Nectin3 | protein_coding | -1.051958434 | 3.045823265 | 2.0746859 | 0.1686841 | 0.526514684 |
| ENSMUSG00000021846 | Peli2 | protein_coding | -0.651339104 | 3.828429506 | 2.0741961 | 0.1687315 | 0.526514684 |
| ENSMUSG00000057897 | Camk2b | protein_coding | 0.350244643 | 8.455445347 | 2.0725043 | 0.1688953 | 0.526873093 |
| ENSMUSG00000062944 | 9130023H24Rik | protein_coding | -0.807139414 | 3.946583191 | 2.0715355 | 0.1689893 | 0.526952284 |
| ENSMUSG00000028821 | Syf2 | protein_coding | -0.37188432 | 7.347891965 | 2.0711906 | 0.1690227 | 0.526952284 |
| ENSMUSG00000025931 | Paqr8 | protein_coding | -0.644938481 | 4.716172339 | 2.0696329 | 0.1691739 | 0.526952284 |
| ENSMUSG00000027601 | Mtfr1 | protein_coding | -0.761339626 | 3.960358547 | 2.069187 | 0.1692172 | 0.526952284 |
| ENSMUSG00000038095 | Sbno1 | protein_coding | -0.430181423 | 6.455760382 | 2.069027 | 0.1692327 | 0.526952284 |
| ENSMUSG00000053460 | Ggcx | protein_coding | 1.183198277 | 3.244690604 | 2.0691699 | 0.1692498 | 0.526952284 |
| ENSMUSG00000035575 | Utp6 | protein_coding | 0.579087583 | 4.648040619 | 2.0684319 | 0.1692906 | 0.526952284 |
| ENSMUSG00000036615 | Rfxap | protein_coding | 0.602225749 | 4.236576245 | 2.0678925 | 0.169343 | 0.526952284 |
| ENSMUSG00000045962 | Wnk1 | protein_coding | -0.480280109 | 5.624088413 | 2.067452 | 0.1693858 | 0.526952284 |
| ENSMUSG00000020297 | Nsg2 | protein_coding | 0.314257808 | 9.103249512 | 2.0669497 | 0.1694347 | 0.526952284 |
| ENSMUSG00000031171 | Ftsj1 | protein_coding | 0.473345908 | 5.746934887 | 2.0666921 | 0.1694598 | 0.526952284 |
| ENSMUSG00000016346 | Kcnq2 | protein_coding | 0.393602436 | 5.984644413 | 2.0652913 | 0.1695962 | 0.527223902 |
| ENSMUSG00000008373 | Prpf31 | protein_coding | 0.33949836 | 6.59998186 | 2.0622819 | 0.1698896 | 0.527983563 |
| ENSMUSG00000029765 | Plxna4 | protein_coding | -0.59534242 | 4.416682776 | 2.0612352 | 0.1699918 | 0.527991041 |
| ENSMUSG00000009876 | Cox4i2 | protein_coding | -1.146292966 | 2.777996292 | 2.0612348 | 0.1699919 | 0.527991041 |
| ENSMUSG00000033618 | Map3k13 | protein_coding | -0.899036825 | 2.837090241 | 2.0607491 | 0.1700393 | 0.527991041 |
| ENSMUSG00000033740 | St18 | protein_coding | -0.622556244 | 3.954821333 | 2.0596283 | 0.1701489 | 0.52817876 |
| ENSMUSG00000006527 | Sfmbt1 | protein_coding | -0.555246234 | 4.659079906 | 2.0579597 | 0.1703122 | 0.528533104 |
| ENSMUSG00000020799 | Tekt1 | protein_coding | -0.645494359 | 4.468957633 | 2.056507 | 0.1704545 | 0.528822159 |
| ENSMUSG00000030603 | Psmc4 | protein_coding | 0.322026647 | 8.493986337 | 2.0558505 | 0.1705189 | 0.528869259 |
| ENSMUSG00000071176 | Arhgef10 | protein_coding | -0.603616169 | 4.175205598 | 2.0539807 | 0.1707024 | 0.52913021 |
| ENSMUSG00000021244 | Ylpm1 | protein_coding | 0.462890658 | 5.454123486 | 2.0530147 | 0.1707973 | 0.52913021 |
| ENSMUSG00000064179 | Tnnt1 | protein_coding | -1.102068436 | 4.643140755 | 2.0836988 | 0.1708149 | 0.52913021 |
| ENSMUSG00000061130 | Ppm1b | protein_coding | -0.473741546 | 5.23265105 | 2.0527085 | 0.1708274 | 0.52913021 |
| ENSMUSG00000038914 | Dido1 | protein_coding | -0.486184004 | 4.996768125 | 2.0518057 | 0.1709162 | 0.52913021 |
| ENSMUSG00000020986 | Sec23a | protein_coding | 0.81076508 | 4.69112239 | 2.0571809 | 0.1710141 | 0.52913021 |
| ENSMUSG00000026887 | Mrrf | protein_coding | 0.481283955 | 5.462209478 | 2.0504999 | 0.1710446 | 0.52913021 |
| ENSMUSG00000033174 | Mgll | protein_coding | -0.452002708 | 6.842980407 | 2.050538 | 0.1710717 | 0.52913021 |
| ENSMUSG00000087174 | 5530601H04Rik | processed_transcript | -0.461503533 | 5.187209242 | 2.0499818 | 0.1710957 | 0.52913021 |
| ENSMUSG00000049555 | Tmie | protein_coding | 0.654087022 | 4.130788794 | 2.0496059 | 0.1711327 | 0.52913021 |
| ENSMUSG00000031066 | Usp11 | protein_coding | 0.511007903 | 7.700349473 | 2.0600444 | 0.1711443 | 0.52913021 |
| ENSMUSG00000054091 | 1810037I17Rik | protein_coding | -0.418315247 | 7.637371474 | 2.0485877 | 0.171233 | 0.529237605 |
| ENSMUSG00000068686 | Cd59b | protein_coding | 1.000950612 | 2.836200096 | 2.0481366 | 0.1712775 | 0.529237605 |
| ENSMUSG00000024276 | Zfp397 | protein_coding | -0.700987154 | 4.089525751 | 2.0460434 | 0.1714841 | 0.529723664 |
| ENSMUSG00000045573 | Penk | protein_coding | 0.566499773 | 10.40961684 | 2.0562884 | 0.1716721 | 0.530152367 |
| ENSMUSG00000020758 | Itgb4 | protein_coding | -0.765660621 | 3.371077176 | 2.0430672 | 0.1717783 | 0.530245404 |
| ENSMUSG00000028354 | Fmn2 | protein_coding | -0.624843017 | 4.408030175 | 2.042758 | 0.1718089 | 0.530245404 |
| ENSMUSG00000025268 | Maged2 | protein_coding | 0.480941183 | 7.775685718 | 2.0440118 | 0.1718502 | 0.530245404 |
| ENSMUSG00000021635 | Rad17 | protein_coding | -0.558156197 | 4.620835006 | 2.0415299 | 0.1719305 | 0.530341086 |
| ENSMUSG00000059208 | Hnrnpm | protein_coding | 0.440804236 | 6.265211582 | 2.0403526 | 0.1720472 | 0.53054885 |
| ENSMUSG00000041120 | Nbl1 | protein_coding | -0.716674018 | 3.95341608 | 2.0386237 | 0.1722188 | 0.530912018 |
| ENSMUSG00000013419 | Zfp651 | protein_coding | -0.657744758 | 4.179712255 | 2.0377819 | 0.1723024 | 0.530912018 |
| ENSMUSG00000027575 | Arfgap1 | protein_coding | 0.381969455 | 6.255567342 | 2.0374073 | 0.1723396 | 0.530912018 |
| ENSMUSG00000025821 | Zfp282 | protein_coding | 0.974112673 | 3.051024875 | 2.037177 | 0.1723625 | 0.530912018 |
| ENSMUSG00000030905 | Crym | protein_coding | -0.671286929 | 7.711231282 | 2.0669358 | 0.1724321 | 0.530974264 |
| ENSMUSG00000051234 | Rnf7 | protein_coding | -0.304342281 | 7.855072363 | 2.0342554 | 0.1726532 | 0.531311254 |
| ENSMUSG00000030447 | Cyfip1 | protein_coding | -0.503202633 | 5.532259004 | 2.0342231 | 0.1726564 | 0.531311254 |
| ENSMUSG00000027993 | Trim2 | protein_coding | -0.32378302 | 7.573465912 | 2.0333616 | 0.1727422 | 0.531311254 |
| ENSMUSG00000033249 | Hsf4 | protein_coding | 1.048875917 | 2.823140125 | 2.0329936 | 0.1727789 | 0.531311254 |
| ENSMUSG00000032609 | Klhdc8b | protein_coding | 0.515711272 | 5.787174411 | 2.0328966 | 0.1727886 | 0.531311254 |
| ENSMUSG00000031557 | Plekha2 | protein_coding | -1.278708054 | 2.553370924 | 2.0313347 | 0.1729444 | 0.531638309 |
| ENSMUSG00000038485 | Socs7 | protein_coding | -0.498774339 | 5.370109791 | 2.0297384 | 0.1731038 | 0.531939301 |
| ENSMUSG00000031357 | Syap1 | protein_coding | -0.728075217 | 3.793191762 | 2.0293637 | 0.1731412 | 0.531939301 |
| ENSMUSG00000017404 | Rpl19 | protein_coding | 0.700391612 | 3.484945015 | 2.0273805 | 0.1733396 | 0.532396621 |
| ENSMUSG00000058400 | Qrfpr | protein_coding | 0.98457897 | 2.91734529 | 2.0260708 | 0.1734707 | 0.53247543 |
| ENSMUSG00000022434 | Fam118a | protein_coding | 0.481367033 | 5.011475363 | 2.0254886 | 0.1735291 | 0.53247543 |
| ENSMUSG00000024271 | Elp2 | protein_coding | 0.438511302 | 6.922050333 | 2.0257614 | 0.1735307 | 0.53247543 |
| ENSMUSG00000020253 | Ppm1m | protein_coding | 0.63984485 | 4.064014177 | 2.025147 | 0.1735633 | 0.53247543 |
| ENSMUSG00000026037 | Orc2 | protein_coding | 1.447264774 | 2.489259787 | 2.0230253 | 0.1737762 | 0.532904604 |
| ENSMUSG00000022951 | Rcan1 | protein_coding | 0.413558129 | 5.367254783 | 2.0227649 | 0.1738023 | 0.532904604 |
| ENSMUSG00000029634 | Rnf6 | protein_coding | 0.489690386 | 7.05288655 | 2.0229395 | 0.1739499 | 0.533171123 |
| ENSMUSG00000041303 | Gtf3c3 | protein_coding | -0.476555796 | 5.121092947 | 2.0204287 | 0.1740371 | 0.533171123 |
| ENSMUSG00000028649 | Macf1 | protein_coding | -0.658227101 | 5.171982175 | 2.0204981 | 0.1740609 | 0.533171123 |

| ensembl_gene_id | external_gene_name | gene_biotype | log2 (fold change) | log2 (counts per million) | F-value | p-value | False discovery rate |
|---|---|---|---|---|---|---|---|
| ENSMUSG00000002660 | Clpp | protein_coding | 0.313725657 | 8.997684785 | 2.0199273 | 0.1740876 | 0.533171123 |
| ENSMUSG00000047786 | Lix1 | protein_coding | -0.545765936 | 5.070151099 | 2.0190386 | 0.174177 | 0.533293195 |
| ENSMUSG00000063511 | Snrnp70 | protein_coding | -0.517732268 | 4.871768543 | 2.0180358 | 0.1742781 | 0.53345059 |
| ENSMUSG00000028619 | Tceanc2 | protein_coding | 0.732463774 | 3.57439876 | 2.0168507 | 0.1743975 | 0.533643008 |
| ENSMUSG00000026883 | Dab2ip | protein_coding | -0.609500755 | 4.455905101 | 2.0164278 | 0.1744402 | 0.533643008 |
| ENSMUSG00000038257 | Glra3 | protein_coding | -0.798814353 | 3.159077533 | 2.0153463 | 0.1745493 | 0.533738211 |
| ENSMUSG00000028461 | Ccdc107 | protein_coding | 0.921656403 | 5.916558428 | 2.067868 | 0.1746048 | 0.533738211 |
| ENSMUSG00000024360 | Etf1 | protein_coding | -0.400053391 | 6.051335038 | 2.0141354 | 0.1746716 | 0.533738211 |
| ENSMUSG00000025337 | Sbds | protein_coding | 0.304294097 | 8.185856683 | 2.013931 | 0.1746923 | 0.533738211 |
| ENSMUSG00000021910 | Nisch | protein_coding | 0.324009793 | 7.4462223 | 2.0135868 | 0.1747271 | 0.533738211 |
| ENSMUSG00000039714 | Cplx3 | protein_coding | -1.223549149 | 3.717022402 | 2.037415 | 0.1747691 | 0.533738211 |
| ENSMUSG00000020733 | Slc9a3r1 | protein_coding | 0.586860043 | 4.971427444 | 2.0125168 | 0.1748353 | 0.533746966 |
| ENSMUSG00000032036 | Kirrel3 | protein_coding | -0.494319356 | 5.479960075 | 2.0114165 | 0.1749466 | 0.533746966 |
| ENSMUSG00000050908 | Tvp23a | protein_coding | 0.666412686 | 3.957495416 | 2.0110244 | 0.1749863 | 0.533746966 |
| ENSMUSG00000021286 | Zfyve21 | protein_coding | -0.531848359 | 5.29721669 | 2.0101733 | 0.1750726 | 0.533746966 |
| ENSMUSG00000021279 | Cdc42bpb | protein_coding | -0.784959653 | 3.217432205 | 2.0101245 | 0.1750775 | 0.533746966 |
| ENSMUSG00000052921 | Arhgef15 | protein_coding | 0.433307765 | 6.334668996 | 2.0099263 | 0.1750976 | 0.533746966 |
| ENSMUSG00000038954 | Supt3 | protein_coding | 0.574871299 | 4.465072397 | 2.0097109 | 0.1751194 | 0.533746966 |
| ENSMUSG00000059923 | Grb2 | protein_coding | 0.307902911 | 6.953409347 | 2.0090634 | 0.1751851 | 0.533795778 |
| ENSMUSG00000040249 | Lrp1 | protein_coding | -0.485058768 | 5.675659049 | 2.0075268 | 0.175341 | 0.534119522 |
| ENSMUSG00000058407 | Txndc9 | protein_coding | -0.33043873 | 7.320790892 | 2.006436 | 0.1754518 | 0.53430567 |
| ENSMUSG00000008305 | Tle1 | protein_coding | -0.551356011 | 4.754083591 | 2.0050627 | 0.1755914 | 0.534487924 |
| ENSMUSG00000062006 | Rpl34 | protein_coding | 0.445009516 | 7.859558926 | 2.0049806 | 0.1756305 | 0.534487924 |
| ENSMUSG00000026102 | Inpp1 | protein_coding | 0.422055698 | 5.806926255 | 2.0032667 | 0.1757742 | 0.534487924 |
| ENSMUSG00000071650 | Ganab | protein_coding | 0.383847321 | 5.829963717 | 2.002772 | 0.1758246 | 0.534487924 |
| ENSMUSG00000032047 | Acat1 | protein_coding | 0.332748017 | 8.079841106 | 2.002752 | 0.1758266 | 0.534487924 |
| ENSMUSG00000029162 | Khk | protein_coding | 0.440982933 | 5.107388417 | 2.002198 | 0.1758831 | 0.534487924 |
| ENSMUSG00000078451 | Ppil6 | protein_coding | -0.988360263 | 2.86356817 | 2.0021228 | 0.1758908 | 0.534487924 |
| ENSMUSG00000033658 | Ddx19b | protein_coding | 0.879984398 | 2.951968457 | 2.0019409 | 0.1759093 | 0.534487924 |
| ENSMUSG00000024073 | Birc6 | protein_coding | -0.490024281 | 5.592319642 | 1.9995666 | 0.1761516 | 0.53507278 |
| ENSMUSG00000027430 | Dtd1 | protein_coding | 0.303024736 | 8.941783108 | 1.998087 | 0.1763027 | 0.535366684 |
| ENSMUSG00000027673 | Ndufb5 | protein_coding | 0.338933512 | 9.868636205 | 1.9976454 | 0.1763479 | 0.535366684 |
| ENSMUSG00000018372 | Cep95 | protein_coding | -0.595881209 | 3.87349461 | 1.9970254 | 0.1764113 | 0.535408038 |
| ENSMUSG00000057182 | Scn3a | protein_coding | -0.434330193 | 5.406071106 | 1.9962649 | 0.1764891 | 0.535493088 |
| ENSMUSG00000096146 | Kcnj11 | protein_coding | 0.719673774 | 3.954955941 | 1.9953425 | 0.1765836 | 0.535515157 |
| ENSMUSG00000017188 | Coa3 | protein_coding | 0.331577738 | 9.831433548 | 1.9952212 | 0.176596 | 0.535515157 |
| ENSMUSG00000036817 | Sun1 | protein_coding | 0.643052822 | 5.080491014 | 1.9941143 | 0.1767401 | 0.535801143 |
| ENSMUSG00000039316 | Rftn1 | protein_coding | -0.531243693 | 4.447628496 | 1.9926612 | 0.1768585 | 0.535986594 |
| ENSMUSG00000042453 | Reln | protein_coding | -0.742697783 | 4.225070867 | 1.99191 | 0.1769356 | 0.535986594 |
| ENSMUSG00000013707 | Tnfaip8l2 | protein_coding | -1.343303055 | 3.013897389 | 1.9935925 | 0.1769508 | 0.535986594 |
| ENSMUSG00000025025 | Mxi1 | protein_coding | -0.486050207 | 5.435271417 | 1.9911339 | 0.1770153 | 0.536030865 |
| ENSMUSG00000049940 | Pgrmc2 | protein_coding | -0.467039854 | 5.786279103 | 1.9902819 | 0.1771029 | 0.536034095 |
| ENSMUSG00000020220 | Vps13d | protein_coding | 0.706679614 | 4.030778721 | 1.9885809 | 0.1772779 | 0.536034095 |
| ENSMUSG00000026349 | Ccnt2 | protein_coding | -0.560423281 | 4.920341983 | 1.9884077 | 0.1772957 | 0.536034095 |
| ENSMUSG00000001120 | Pcbp3 | protein_coding | 0.741959677 | 6.028750481 | 2.0138246 | 0.1773238 | 0.536034095 |
| ENSMUSG00000043419 | Chd3os | protein_coding | 0.305089765 | 8.809065314 | 1.9876266 | 0.1773761 | 0.536034095 |
| ENSMUSG00000022696 | Sidt1 | protein_coding | 0.905587294 | 3.069529831 | 1.9867974 | 0.1774616 | 0.536034095 |
| ENSMUSG00000052406 | Rexo4 | protein_coding | 0.556863728 | 5.485134177 | 1.9865375 | 0.1774883 | 0.536034095 |
| ENSMUSG00000078695 | Cisd3 | protein_coding | 0.326990722 | 6.448157169 | 1.9862605 | 0.1775169 | 0.536034095 |
| ENSMUSG00000012848 | Rps5 | protein_coding | 0.402278257 | 10.12689909 | 1.9861054 | 0.1775329 | 0.536034095 |
| ENSMUSG00000053929 | Cyhr1 | protein_coding | 0.325712882 | 6.914207001 | 1.9860147 | 0.1775422 | 0.536034095 |
| ENSMUSG00000034570 | Inpp5j | protein_coding | -0.886734854 | 3.112371007 | 1.9852544 | 0.1776207 | 0.536034095 |
| ENSMUSG00000013089 | Etv5 | protein_coding | -0.607973785 | 4.202614693 | 1.9851466 | 0.1776318 | 0.536034095 |
| ENSMUSG00000040734 | Ppp1r13l | protein_coding | -1.173374227 | 2.762359993 | 1.9848304 | 0.1776644 | 0.536034095 |
| ENSMUSG00000000247 | Lhx2 | protein_coding | -1.152397712 | 3.10998722 | 1.9831546 | 0.1778682 | 0.536457874 |
| ENSMUSG00000071074 | Yipf3 | protein_coding | 0.532098833 | 5.260315035 | 1.9822524 | 0.1779308 | 0.536457874 |
| ENSMUSG00000038301 | Snx10 | protein_coding | -0.392953898 | 8.472617737 | 1.9820223 | 0.1779545 | 0.536457874 |
| ENSMUSG00000029359 | Tesc | protein_coding | -0.68499282 | 4.607347057 | 1.9777871 | 0.1783932 | 0.537375326 |
| ENSMUSG00000042606 | Hirip3 | protein_coding | 0.609608148 | 4.380832311 | 1.9776945 | 0.1784028 | 0.537375326 |
| ENSMUSG00000020359 | Phykpl | protein_coding | 0.698653774 | 3.634499591 | 1.9773694 | 0.1784366 | 0.537375326 |
| ENSMUSG00000020840 | Blmh | protein_coding | 0.349493661 | 7.288699791 | 1.9767162 | 0.1785044 | 0.537375326 |
| ENSMUSG00000045608 | Dbx2 | protein_coding | -0.704282268 | 3.945594001 | 1.976674 | 0.1785088 | 0.537375326 |
| ENSMUSG00000061232 | H2-K1 | protein_coding | -0.939581003 | 3.521797605 | 1.9756376 | 0.1786471 | 0.537578747 |
| ENSMUSG00000010277 | 2610507B11Rik | protein_coding | 0.548122577 | 5.370121584 | 1.9743776 | 0.1787474 | 0.537578747 |
| ENSMUSG00000001755 | Coasy | protein_coding | 0.553937452 | 5.350843705 | 1.9737024 | 0.1788176 | 0.537578747 |
| ENSMUSG00000062151 | Unc13c | protein_coding | -0.532081502 | 6.355882941 | 1.9750918 | 0.1788374 | 0.537578747 |
| ENSMUSG00000028811 | Yars | protein_coding | 0.446902113 | 7.068007096 | 1.9732458 | 0.1788958 | 0.537578747 |
| ENSMUSG00000079467 | Gm14966 | antisense_RNA | -1.488364753 | 1.889569052 | 1.9727854 | 0.1789131 | 0.537578747 |
| ENSMUSG00000036751 | Cox6b1 | protein_coding | 0.387403467 | 10.25496019 | 1.9726585 | 0.1789263 | 0.537578747 |
| ENSMUSG00000030199 | Etv6 | protein_coding | 1.173704697 | 3.028764161 | 1.9716911 | 0.1790271 | 0.537731254 |
| ENSMUSG00000027950 | Chrnb2 | protein_coding | 0.585421513 | 4.510799278 | 1.9704501 | 0.1791564 | 0.537736989 |
| ENSMUSG00000026280 | Atg4b | protein_coding | 0.394959756 | 6.247778244 | 1.969958 | 0.1792078 | 0.537736989 |
| ENSMUSG00000019853 | Hebp2 | protein_coding | 0.601981836 | 3.909298826 | 1.9695062 | 0.1792549 | 0.537736989 |
| ENSMUSG00000061666 | Gdpd1 | protein_coding | 0.32674279 | 6.787748303 | 1.9689814 | 0.1793097 | 0.537736989 |
| ENSMUSG00000038774 | Ascc3 | protein_coding | -0.886769851 | 3.242007027 | 1.9688833 | 0.1793199 | 0.537736989 |
| ENSMUSG00000053838 | Nudcd3 | protein_coding | 0.447775251 | 7.133423902 | 1.9690896 | 0.179329 | 0.537736989 |
| ENSMUSG00000022545 | Ercc4 | protein_coding | 0.971643913 | 4.237128675 | 1.9804541 | 0.1796448 | 0.538533877 |
| ENSMUSG00000042743 | Sgtb | protein_coding | -0.492476737 | 5.508618151 | 1.9636863 | 0.1798636 | 0.538600836 |

| ensembl_gene_id | external_gene_name | gene_biotype | log2 (fold change) | log2 (counts per million) | F-value | p-value | False discovery rate |
|---|---|---|---|---|---|---|---|
| ENSMUSG00000110444 | Gm10033 | transcribed_unproces | -1.116432539 | 3.567408666 | 1.9748095 | 0.1798896 | 0.538600836 |
| ENSMUSG00000018189 | Uchl5 | protein_coding | -0.325608705 | 6.939610943 | 1.9630852 | 0.1799266 | 0.538600836 |
| ENSMUSG00000052534 | Pbx1 | protein_coding | -0.350412345 | 7.052924281 | 1.9623001 | 0.180009 | 0.538600836 |
| ENSMUSG00000075486 | Commd6 | protein_coding | -0.412596379 | 7.794029343 | 1.9622624 | 0.1800129 | 0.538600836 |
| ENSMUSG00000026851 | BC005624 | protein_coding | 0.324728693 | 7.926438961 | 1.9605186 | 0.180196 | 0.538600836 |
| ENSMUSG00000020212 | Mdm1 | protein_coding | -0.746825356 | 3.198376845 | 1.960137 | 0.1802361 | 0.538600836 |
| ENSMUSG00000069919 | Hba-a1 | protein_coding | -0.776351279 | 4.437089775 | 1.9603436 | 0.180245 | 0.538600836 |
| ENSMUSG00000026790 | Odf2 | protein_coding | 0.516813973 | 5.846598924 | 1.9600377 | 0.1802465 | 0.538600836 |
| ENSMUSG00000063019 | Manbal | protein_coding | 0.331572487 | 7.243091372 | 1.9600258 | 0.1802478 | 0.538600836 |
| ENSMUSG00000020022 | Ndufa12 | protein_coding | 0.3163952 | 8.523937266 | 1.9599438 | 0.1802564 | 0.538600836 |
| ENSMUSG00000036896 | C1qc | protein_coding | -1.018457842 | 4.251901366 | 1.9776486 | 0.1802682 | 0.538600836 |
| ENSMUSG00000060216 | Arrb2 | protein_coding | 0.63198825 | 4.17512589 | 1.956599 | 0.1806083 | 0.539397301 |
| ENSMUSG00000027668 | Mfn1 | protein_coding | -0.664208598 | 4.57648401 | 1.9563446 | 0.1806351 | 0.539397301 |
| ENSMUSG00000032625 | Thsd7a | protein_coding | -0.85859747 | 4.466602786 | 1.9659802 | 0.1807372 | 0.539533294 |
| ENSMUSG00000041654 | Slc39a11 | protein_coding | 0.730947237 | 4.16824922 | 1.9548384 | 0.1807939 | 0.539533294 |
| ENSMUSG00000063200 | Nol7 | protein_coding | -0.417927562 | 6.24434449 | 1.9540994 | 0.1808719 | 0.539533294 |
| ENSMUSG00000040797 | Iqsec3 | protein_coding | -0.562325395 | 5.413933185 | 1.95371 | 0.180913 | 0.539533294 |
| ENSMUSG00000035383 | Pmch | protein_coding | -1.513458746 | 6.988678681 | 2.1539234 | 0.1809316 | 0.539533294 |
| ENSMUSG00000046711 | Hmga1 | protein_coding | 0.652629081 | 3.853199304 | 1.9529561 | 0.1809926 | 0.539534372 |
| ENSMUSG00000063434 | Sorcs3 | protein_coding | 0.492412834 | 5.116461354 | 1.952581 | 0.1810323 | 0.539534372 |
| ENSMUSG00000022883 | Robo1 | protein_coding | 0.610137582 | 3.967815851 | 1.9510319 | 0.181196 | 0.539872823 |
| ENSMUSG00000002205 | Vrk3 | protein_coding | 0.538526895 | 5.766803364 | 1.9496885 | 0.1813688 | 0.540237808 |
| ENSMUSG00000042747 | Krtcap2 | protein_coding | 0.361715352 | 7.698861476 | 1.9473921 | 0.1815816 | 0.540721878 |
| ENSMUSG00000022757 | Tfg | protein_coding | 0.479249882 | 6.460969884 | 1.9461204 | 0.1817471 | 0.540880485 |
| ENSMUSG00000004947 | Dtx2 | protein_coding | 1.106412594 | 2.572193851 | 1.9448086 | 0.1818558 | 0.540880485 |
| ENSMUSG00000054871 | Tmem158 | protein_coding | -0.743428898 | 4.795902813 | 1.9457179 | 0.1818703 | 0.540880485 |
| ENSMUSG00000035845 | Alg12 | protein_coding | 0.771525354 | 3.806612406 | 1.9444959 | 0.1818891 | 0.540880485 |
| ENSMUSG00000020064 | Herc4 | protein_coding | -0.751678019 | 3.692384802 | 1.9440714 | 0.1819342 | 0.540880485 |
| ENSMUSG00000029822 | Osbpl3 | protein_coding | -1.036028156 | 2.998039118 | 1.9437792 | 0.1819653 | 0.540880485 |
| ENSMUSG00000000682 | Cd52 | protein_coding | -1.28877466 | 2.638470569 | 1.9430677 | 0.182041 | 0.540880485 |
| ENSMUSG00000062232 | Rapgef2 | protein_coding | -0.494656404 | 5.343580898 | 1.9426153 | 0.1820891 | 0.540880485 |
| ENSMUSG00000056124 | B4galt6 | protein_coding | -0.359268613 | 6.352546689 | 1.9420615 | 0.1821481 | 0.540880485 |
| ENSMUSG00000035901 | Dennd5a | protein_coding | -0.351634585 | 6.827601737 | 1.9419275 | 0.1821623 | 0.540880485 |
| ENSMUSG00000034832 | Tet3 | protein_coding | 0.556699316 | 4.60386671 | 1.9413576 | 0.182223 | 0.540880485 |
| ENSMUSG00000048578 | Mlec | protein_coding | 0.42817937 | 5.514728006 | 1.9410113 | 0.1822599 | 0.540880485 |
| ENSMUSG00000022122 | Ednrb | protein_coding | -0.473717795 | 6.1008618 | 1.9406629 | 0.1822971 | 0.540880485 |
| ENSMUSG00000031536 | Polb | protein_coding | -0.292592141 | 7.048115771 | 1.940269 | 0.182339 | 0.540880485 |
| ENSMUSG00000018412 | Kansl1 | protein_coding | -0.522580551 | 5.409290668 | 1.9394331 | 0.1824282 | 0.540995702 |
| ENSMUSG00000009575 | Cbx5 | protein_coding | -0.304862387 | 8.581273462 | 1.9381385 | 0.1825664 | 0.541031142 |
| ENSMUSG00000062393 | Dgkk | protein_coding | 0.605496163 | 4.040301523 | 1.9380125 | 0.1825798 | 0.541031142 |
| ENSMUSG00000036693 | Nop14 | protein_coding | 0.540054309 | 5.11560451 | 1.9377788 | 0.1826048 | 0.541031142 |
| ENSMUSG00000037994 | Slc9b2 | protein_coding | 1.062687328 | 2.693653155 | 1.9368467 | 0.1827044 | 0.541031142 |
| ENSMUSG00000048216 | Gpr85 | protein_coding | 0.431837062 | 6.060611206 | 1.9365652 | 0.1827345 | 0.541031142 |
| ENSMUSG00000042298 | Ttc19 | protein_coding | -0.475992788 | 5.377593297 | 1.9354963 | 0.1828488 | 0.541031142 |
| ENSMUSG00000029686 | Cul1 | protein_coding | -0.374852149 | 6.502790169 | 1.9353305 | 0.1828665 | 0.541031142 |
| ENSMUSG00000020923 | Ubtf | protein_coding | 0.414223351 | 6.328263927 | 1.9351278 | 0.1828882 | 0.541031142 |
| ENSMUSG00000034105 | Tldc1 | protein_coding | 0.702926126 | 3.480613273 | 1.9347541 | 0.1829282 | 0.541031142 |
| ENSMUSG00000023911 | Flywch2 | protein_coding | 0.40656233 | 5.777949659 | 1.9346133 | 0.1829433 | 0.541031142 |
| ENSMUSG00000018377 | Vezf1 | protein_coding | -0.474220251 | 5.292392694 | 1.9335601 | 0.1830561 | 0.54114786 |
| ENSMUSG00000042401 | Crtac1 | protein_coding | -0.720531445 | 4.769151047 | 1.933336 | 0.1831106 | 0.54114786 |
| ENSMUSG00000025204 | Ndufb8 | protein_coding | 0.365929608 | 9.711001295 | 1.9327198 | 0.1831462 | 0.54114786 |
| ENSMUSG00000047606 | Ankrd34c | protein_coding | 0.81233523 | 3.019823095 | 1.9323663 | 0.1831841 | 0.54114786 |
| ENSMUSG00000020919 | Stat5b | protein_coding | -0.607321737 | 4.271669641 | 1.9306943 | 0.1833635 | 0.541529092 |
| ENSMUSG00000007987 | Ift22 | protein_coding | 0.30574384 | 8.388343311 | 1.9291791 | 0.1835262 | 0.541860977 |
| ENSMUSG00000036469 | 1-Mar | protein_coding | -0.446470365 | 4.910146209 | 1.9274574 | 0.1837114 | 0.541953577 |
| ENSMUSG00000048222 | Mfap1b | protein_coding | -0.482646705 | 4.988416576 | 1.9274146 | 0.183716 | 0.541953577 |
| ENSMUSG00000004633 | Chn2 | protein_coding | -1.236539824 | 2.262162866 | 1.9271009 | 0.1837498 | 0.541953577 |
| ENSMUSG00000039904 | Gpr37 | protein_coding | -0.462144704 | 6.394693251 | 1.9271781 | 0.1837592 | 0.541953577 |
| ENSMUSG00000029455 | Aldh2 | protein_coding | 0.47961496 | 5.05775773 | 1.9241552 | 0.1840672 | 0.542486928 |
| ENSMUSG00000091474 | 2610021A01Rik | protein_coding | -0.76008162 | 3.469323336 | 1.9240601 | 0.1840775 | 0.542486928 |
| ENSMUSG00000049401 | Ogfr | protein_coding | -0.644715694 | 3.834392804 | 1.9239314 | 0.1840914 | 0.542486928 |
| ENSMUSG00000038486 | Sv2a | protein_coding | 0.311337624 | 8.389974562 | 1.9190169 | 0.1846226 | 0.543903344 |
| ENSMUSG00000033054 | Npat | protein_coding | -0.574323685 | 4.384894328 | 1.9180114 | 0.1847316 | 0.544075203 |
| ENSMUSG00000025792 | Slc25a10 | protein_coding | 0.451815362 | 5.270183151 | 1.9170066 | 0.1848405 | 0.54421882 |
| ENSMUSG00000062040 | Zfp27 | protein_coding | 0.669618865 | 4.00792207 | 1.9166281 | 0.1848816 | 0.54421882 |
| ENSMUSG00000037257 | Aagab | protein_coding | 0.477617068 | 6.616531165 | 1.9159638 | 0.1849841 | 0.544371624 |
| ENSMUSG00000038633 | Degs1 | protein_coding | 0.29808002 | 7.198477339 | 1.9143721 | 0.1851265 | 0.544417779 |
| ENSMUSG00000027968 | Larp7 | protein_coding | -0.482566285 | 5.413909771 | 1.9142658 | 0.185138 | 0.544417779 |
| ENSMUSG00000029145 | Eif2b4 | protein_coding | 0.438953674 | 5.925867278 | 1.912734 | 0.1853046 | 0.544417779 |
| ENSMUSG00000039114 | Nrn1 | protein_coding | -0.744851184 | 6.156109319 | 1.9401159 | 0.1853151 | 0.544417779 |
| ENSMUSG00000020778 | Ten1 | protein_coding | 0.314531589 | 7.058388038 | 1.9125355 | 0.1853262 | 0.544417779 |
| ENSMUSG00000059839 | Zfp874b | protein_coding | -0.917518265 | 3.056650539 | 1.9117617 | 0.1854104 | 0.544417779 |
| ENSMUSG00000060843 | Ctnna3 | protein_coding | -0.866388341 | 2.978928468 | 1.9113805 | 0.1854519 | 0.544417779 |
| ENSMUSG00000022781 | Pak2 | protein_coding | -0.502237445 | 5.053092836 | 1.9109794 | 0.1854956 | 0.544417779 |
| ENSMUSG00000073176 | Zfp449 | protein_coding | -0.658815339 | 3.378219399 | 1.9109504 | 0.1854988 | 0.544417779 |
| ENSMUSG00000029624 | Ptcd1 | protein_coding | 0.728798727 | 3.945842654 | 1.9105552 | 0.1855418 | 0.544417779 |
| ENSMUSG00000030987 | Stim1 | protein_coding | 0.409443277 | 5.982933369 | 1.9099213 | 0.1856109 | 0.544417779 |
| ENSMUSG00000045128 | Rpl18a | protein_coding | 0.406579498 | 8.363054074 | 1.9092796 | 0.1856809 | 0.544417779 |

| ensembl_gene_id | external_gene_name | gene_biotype | log2 (fold change) | log2 (counts per million) | F-value | p-value | False discovery rate |
|---|---|---|---|---|---|---|---|
| ENSMUSG00000022554 | Hgh1 | protein_coding | -1.010106681 | 2.639369943 | 1.9091081 | 0.1856996 | 0.544417779 |
| ENSMUSG00000032114 | Slc37a4 | protein_coding | -0.634516036 | 4.078771897 | 1.9090259 | 0.1857086 | 0.544417779 |
| ENSMUSG00000034403 | Pja1 | protein_coding | 0.46664445 | 8.745547709 | 1.9088724 | 0.1858359 | 0.544642594 |
| ENSMUSG00000047342 | Zfp286 | protein_coding | -0.495137113 | 5.013948161 | 1.9068131 | 0.1859502 | 0.544689797 |
| ENSMUSG00000026179 | Pnkd | protein_coding | 0.335486303 | 8.05180221 | 1.9066034 | 0.1859731 | 0.544689797 |
| ENSMUSG00000024759 | Atl3 | protein_coding | -0.921472388 | 3.070248104 | 1.9063207 | 0.186004 | 0.544689797 |
| ENSMUSG00000018143 | Mafk | protein_coding | 0.935219531 | 3.372681478 | 1.9049632 | 0.1861525 | 0.544976125 |
| ENSMUSG00000041986 | Elmod1 | protein_coding | -0.596445919 | 6.195305318 | 1.914835 | 0.1862557 | 0.545016326 |
| ENSMUSG00000062526 | Mppe1 | protein_coding | 1.023149233 | 2.980219548 | 1.9039117 | 0.1862676 | 0.545016326 |
| ENSMUSG00000020844 | Nxn | protein_coding | -0.681636351 | 3.816429731 | 1.9033125 | 0.1863332 | 0.545060039 |
| ENSMUSG00000097164 | Cep83os | bidirectional_promote | -0.470412868 | 5.223949307 | 1.9012232 | 0.1865622 | 0.545581677 |
| ENSMUSG00000029676 | Pot1a | protein_coding | 0.681216759 | 4.502546478 | 1.9006616 | 0.1866239 | 0.545613534 |
| ENSMUSG00000041073 | Nacad | protein_coding | 0.534408622 | 4.855918819 | 1.8990174 | 0.1868045 | 0.545845518 |
| ENSMUSG00000038894 | Irs2 | protein_coding | 0.503638679 | 5.25886737 | 1.8990148 | 0.1868048 | 0.545845518 |
| ENSMUSG00000001542 | Ell2 | protein_coding | -0.76530386 | 3.6254633 | 1.8962214 | 0.1871121 | 0.546492137 |
| ENSMUSG00000042066 | Tmcc2 | protein_coding | -0.445059136 | 6.075979298 | 1.89604 | 0.1871321 | 0.546492137 |
| ENSMUSG00000021497 | Txndc15 | protein_coding | 0.379966056 | 6.261456886 | 1.8952083 | 0.1872237 | 0.546492137 |
| ENSMUSG00000042548 | Asxl1 | protein_coding | -0.665018793 | 3.71268328 | 1.895157 | 0.1872293 | 0.546492137 |
| ENSMUSG00000044149 | Nkrf | protein_coding | 0.595819262 | 4.948426562 | 1.894468 | 0.1873053 | 0.546517881 |
| ENSMUSG00000021120 | Pigh | protein_coding | 0.953792002 | 3.330616155 | 1.8939662 | 0.1873606 | 0.546517881 |
| ENSMUSG00000026335 | Pam | protein_coding | 0.416837794 | 6.194626399 | 1.8936943 | 0.1873906 | 0.546517881 |
| ENSMUSG00000036880 | Acaa2 | protein_coding | -0.41113291 | 5.883959439 | 1.8919766 | 0.1875803 | 0.546843623 |
| ENSMUSG00000027423 | Snx5 | protein_coding | -0.386017446 | 6.114143889 | 1.8917616 | 0.187604 | 0.546843623 |
| ENSMUSG00000039879 | Heca | protein_coding | -0.603710153 | 3.829112711 | 1.8909171 | 0.1876974 | 0.54684573 |
| ENSMUSG00000026810 | Dpm2 | protein_coding | 0.285131433 | 7.574163744 | 1.8906109 | 0.1877312 | 0.54684573 |
| ENSMUSG00000030760 | Acer3 | protein_coding | -0.454778734 | 4.766273208 | 1.890375 | 0.1877573 | 0.54684573 |
| ENSMUSG00000031347 | Cetn2 | protein_coding | -0.384346109 | 7.865143906 | 1.8894441 | 0.1878603 | 0.546997544 |
| ENSMUSG00000003575 | Crtc1 | protein_coding | 0.382692827 | 5.899694798 | 1.8888901 | 0.1879216 | 0.547028008 |
| ENSMUSG00000032002 | Dcun1d5 | protein_coding | 0.376746848 | 5.856712684 | 1.8876279 | 0.1880615 | 0.547188298 |
| ENSMUSG00000031590 | Frg1 | protein_coding | -0.331821405 | 7.061392723 | 1.8874327 | 0.1880831 | 0.547188298 |
| ENSMUSG00000028517 | Plpp3 | protein_coding | -0.544541969 | 6.071146533 | 1.8880107 | 0.1881294 | 0.547188298 |
| ENSMUSG00000016534 | Lamp2 | protein_coding | -0.41030114 | 6.080716892 | 1.8859007 | 0.188253 | 0.54739987 |
| ENSMUSG00000025876 | Unc5a | protein_coding | 0.750211254 | 4.321028332 | 1.8846871 | 0.1884181 | 0.547621696 |
| ENSMUSG00000075229 | Ccdc58 | protein_coding | -0.523549562 | 4.814919982 | 1.8842142 | 0.1884403 | 0.547621696 |
| ENSMUSG00000024359 | Hspa9 | protein_coding | 0.378178644 | 6.974329792 | 1.8831561 | 0.1885579 | 0.547621696 |
| ENSMUSG00000026048 | Ercc5 | protein_coding | -0.776428004 | 3.90961275 | 1.8830587 | 0.1885687 | 0.547621696 |
| ENSMUSG00000049751 | Rpl36al | protein_coding | 0.799913449 | 3.012442042 | 1.8825085 | 0.1886299 | 0.547621696 |
| ENSMUSG00000021276 | Cinp | protein_coding | 0.325575389 | 6.88534505 | 1.8824641 | 0.1886349 | 0.547621696 |
| ENSMUSG00000039298 | Cdk5rap2 | protein_coding | -0.878338783 | 2.982512472 | 1.8806024 | 0.1888421 | 0.547875223 |
| ENSMUSG00000045087 | S1pr5 | protein_coding | -1.078670339 | 2.742858132 | 1.8803959 | 0.1888651 | 0.547875223 |
| ENSMUSG00000018589 | Glra2 | protein_coding | 0.549833348 | 5.258017311 | 1.8799224 | 0.1889179 | 0.547875223 |
| ENSMUSG00000029238 | Clock | protein_coding | -0.518295789 | 5.392085765 | 1.8794168 | 0.1889743 | 0.547875223 |
| ENSMUSG00000042225 | Ammecr1 | protein_coding | -0.755510516 | 3.249757824 | 1.8793928 | 0.188977 | 0.547875223 |
| ENSMUSG00000036955 | Kif1bp | protein_coding | -0.396906041 | 6.236492011 | 1.8786969 | 0.1890546 | 0.547897888 |
| ENSMUSG00000038319 | Kcnh2 | protein_coding | 0.91749887 | 2.905448428 | 1.8784092 | 0.1890867 | 0.547897888 |
| ENSMUSG00000053411 | Cbx7 | protein_coding | -0.45719101 | 5.132280649 | 1.8765159 | 0.1892981 | 0.548362731 |
| ENSMUSG00000029427 | Zcchc8 | protein_coding | 0.699746536 | 3.828438522 | 1.8751661 | 0.189449 | 0.548652099 |
| ENSMUSG00000031889 | D230025D16Rik | protein_coding | 0.579412509 | 4.334105351 | 1.8741554 | 0.1895621 | 0.548765539 |
| ENSMUSG00000001911 | Nfix | protein_coding | -0.479951762 | 5.602617967 | 1.873904 | 0.1895903 | 0.548765539 |
| ENSMUSG00000029861 | Fam131b | protein_coding | -0.756911383 | 3.633777531 | 1.8734327 | 0.189643 | 0.548770593 |
| ENSMUSG00000037736 | Limch1 | protein_coding | -0.412008221 | 5.94337844 | 1.8718333 | 0.1898223 | 0.549141495 |
| ENSMUSG00000029084 | Cd38 | protein_coding | -0.852623883 | 3.063188531 | 1.8689653 | 0.1901442 | 0.549858664 |
| ENSMUSG00000019194 | Scn1b | protein_coding | 0.449908252 | 5.104356915 | 1.8684714 | 0.1901997 | 0.549858664 |
| ENSMUSG00000032754 | Slc8b1 | protein_coding | 1.057455657 | 2.605505296 | 1.8682593 | 0.1902236 | 0.549858664 |
| ENSMUSG00000066735 | Vkorc1l1 | protein_coding | -0.513000966 | 4.973310593 | 1.8675842 | 0.1902995 | 0.549930288 |
| ENSMUSG00000061740 | Cyp2d22 | protein_coding | -0.818907394 | 3.458521513 | 1.8671035 | 0.1903536 | 0.549938807 |
| ENSMUSG00000014867 | Surf4 | protein_coding | 0.464981816 | 5.453202944 | 1.8633972 | 0.1907712 | 0.550852738 |
| ENSMUSG00000036192 | Rorb | protein_coding | -0.945759537 | 5.784684724 | 1.9175101 | 0.1907724 | 0.550852738 |
| ENSMUSG00000030269 | Mtmr14 | protein_coding | 0.845433666 | 2.959908207 | 1.8629253 | 0.1908245 | 0.550855301 |
| ENSMUSG00000027367 | Stard7 | protein_coding | 0.324261569 | 6.49919492 | 1.862044 | 0.190924 | 0.550994656 |
| ENSMUSG00000029434 | Vps33a | protein_coding | 0.536973796 | 5.761476132 | 1.8615589 | 0.1910091 | 0.551092215 |
| ENSMUSG00000022521 | Crebbp | protein_coding | -0.403322058 | 7.026471536 | 1.8605842 | 0.191089 | 0.55117493 |
| ENSMUSG00000028453 | Fancg | protein_coding | -0.606465162 | 3.980522971 | 1.8591835 | 0.1912475 | 0.551248841 |
| ENSMUSG00000025967 | Eef1b2 | protein_coding | 0.382448221 | 9.00386979 | 1.8589208 | 0.1912772 | 0.551248841 |
| ENSMUSG00000029681 | Bcl7b | protein_coding | 0.416785238 | 6.484288191 | 1.8583821 | 0.1913382 | 0.551248841 |
| ENSMUSG00000035509 | Fbxl21 | protein_coding | 0.641606271 | 3.861763145 | 1.8581928 | 0.1913597 | 0.551248841 |
| ENSMUSG00000032024 | Clmp | protein_coding | 0.80289319 | 3.363255867 | 1.8580931 | 0.191371 | 0.551248841 |
| ENSMUSG00000020130 | Tbc1d15 | protein_coding | -0.430231812 | 5.682578696 | 1.8574934 | 0.1914389 | 0.551295495 |
| ENSMUSG00000051730 | Mettl5 | protein_coding | -0.354639234 | 6.12775311 | 1.8569183 | 0.1915041 | 0.551295495 |
| ENSMUSG00000025921 | Rdh10 | protein_coding | -1.042933104 | 2.745578129 | 1.8552463 | 0.1916938 | 0.551295495 |
| ENSMUSG00000056755 | Grm7 | protein_coding | -0.516317859 | 5.387431684 | 1.8549071 | 0.1917323 | 0.551295495 |
| ENSMUSG00000028322 | Exosc3 | protein_coding | -0.801914917 | 3.460895537 | 1.854895 | 0.1917337 | 0.551295495 |
| ENSMUSG00000021028 | Mbip | protein_coding | -0.550083393 | 4.987960437 | 1.8548771 | 0.1917357 | 0.551295495 |
| ENSMUSG00000025374 | Nabp2 | protein_coding | 0.497754961 | 5.12100011 | 1.853799 | 0.1918582 | 0.551295495 |
| ENSMUSG00000027642 | Rpn2 | protein_coding | 0.280819861 | 7.653253879 | 1.8536119 | 0.1918795 | 0.551295495 |
| ENSMUSG00000027680 | Fxr1 | protein_coding | -0.537194229 | 5.052254019 | 1.8535908 | 0.1918819 | 0.551295495 |
| ENSMUSG00000026797 | Stxbp1 | protein_coding | 0.487786531 | 9.183460742 | 1.858833 | 0.1918998 | 0.551295495 |
| ENSMUSG00000024392 | Bag6 | protein_coding | -0.475885346 | 5.578006025 | 1.8521275 | 0.1920483 | 0.551401473 |

| ensembl_gene_id | external_gene_name | gene_biotype | log2 (fold change) | log2 (counts per million) | F-value | p-value | False discovery rate |
|---|---|---|---|---|---|---|---|
| ENSMUSG00000028822 | Tmem50a | protein_coding | 0.303147462 | 8.468053722 | 1.8515775 | 0.1921109 | 0.551401473 |
| ENSMUSG00000034616 | Ssh3 | protein_coding | 0.88426245 | 3.420697223 | 1.8511953 | 0.1921544 | 0.551401473 |
| ENSMUSG00000009863 | Sdhb | protein_coding | 0.332140802 | 10.22007054 | 1.8511656 | 0.1921578 | 0.551401473 |
| ENSMUSG00000026553 | Copa | protein_coding | 0.442834573 | 6.898510448 | 1.8511207 | 0.1921931 | 0.551401473 |
| ENSMUSG00000028243 | Ubxn2b | protein_coding | -0.443018423 | 5.753515982 | 1.849072 | 0.1923964 | 0.551837435 |
| ENSMUSG00000010609 | Psen2 | protein_coding | 0.53188327 | 4.709192614 | 1.8478346 | 0.1925376 | 0.551863392 |
| ENSMUSG00000039662 | Icmt | protein_coding | -0.536896461 | 4.319494949 | 1.8471519 | 0.1926155 | 0.551863392 |
| ENSMUSG00000038068 | Rnf144b | protein_coding | -0.729503214 | 3.326195331 | 1.8469908 | 0.192634 | 0.551863392 |
| ENSMUSG00000040464 | Gtpbp10 | protein_coding | -1.037488303 | 2.572768242 | 1.8465311 | 0.1926865 | 0.551863392 |
| ENSMUSG00000031299 | Pdha1 | protein_coding | 0.62984745 | 7.285462606 | 1.8714827 | 0.1927975 | 0.551863392 |
| ENSMUSG00000061013 | Mkx | protein_coding | -1.226609211 | 2.553565789 | 1.8454145 | 0.1928141 | 0.551863392 |
| ENSMUSG00000030533 | Unc45a | protein_coding | 0.569011148 | 4.486014819 | 1.8453764 | 0.1928185 | 0.551863392 |
| ENSMUSG00000113314 | AC122013.1 | lincRNA | 1.19811349 | 2.498443276 | 1.84526 | 0.1928318 | 0.551863392 |
| ENSMUSG00000023019 | Gpd1 | protein_coding | -0.688200897 | 3.716180756 | 1.8449492 | 0.1928674 | 0.551863392 |
| ENSMUSG00000022253 | Nadk2 | protein_coding | -0.531543445 | 4.544090733 | 1.8438439 | 0.1929939 | 0.551906785 |
| ENSMUSG00000020889 | Nr1d1 | protein_coding | -0.500406205 | 6.574850309 | 1.8479605 | 0.1930419 | 0.551906785 |
| ENSMUSG00000060735 | Rxfp3 | protein_coding | -1.183889743 | 2.619317142 | 1.8426477 | 0.1931309 | 0.551906785 |
| ENSMUSG00000054459 | Vsnl1 | protein_coding | 0.368314642 | 10.53233629 | 1.8421964 | 0.1931826 | 0.551906785 |
| ENSMUSG00000031153 | Gripap1 | protein_coding | 0.324227347 | 7.663267864 | 1.8421899 | 0.1931834 | 0.551906785 |
| ENSMUSG00000023330 | Dtwd1 | protein_coding | 0.394914808 | 5.508982348 | 1.8421005 | 0.1931936 | 0.551906785 |
| ENSMUSG00000045252 | Zfp574 | protein_coding | 0.645550098 | 4.009958688 | 1.8413479 | 0.1932799 | 0.551906785 |
| ENSMUSG00000020811 | Wscd1 | protein_coding | -0.426451445 | 5.535521252 | 1.8412328 | 0.1932931 | 0.551906785 |
| ENSMUSG00000009733 | Tfcp2 | protein_coding | 0.548991388 | 4.626596057 | 1.8405161 | 0.1933754 | 0.551995028 |
| ENSMUSG00000026447 | Pik3c2b | protein_coding | -0.630576639 | 4.035664441 | 1.8392813 | 0.1935172 | 0.552253211 |
| ENSMUSG00000021033 | Gstz1 | protein_coding | 0.322560285 | 7.07085356 | 1.8381695 | 0.193645 | 0.55247128 |
| ENSMUSG00000022103 | Gfra2 | protein_coding | 0.539721611 | 5.329226926 | 1.8372526 | 0.1937504 | 0.552594927 |
| ENSMUSG00000037594 | BC022687 | protein_coding | 0.81713795 | 3.430828508 | 1.8368994 | 0.1937911 | 0.552594927 |
| ENSMUSG00000038365 | Fbxo25 | protein_coding | 0.404722345 | 6.579606532 | 1.8353108 | 0.193974 | 0.552853471 |
| ENSMUSG00000031367 | Ap1s2 | protein_coding | 0.441005596 | 6.820562455 | 1.835481 | 0.1939846 | 0.552853471 |
| ENSMUSG00000039704 | Lmbrd2 | protein_coding | -1.244576677 | 2.339368941 | 1.834683 | 0.1940464 | 0.552863064 |
| ENSMUSG00000031672 | Got2 | protein_coding | 0.459769123 | 7.03732811 | 1.8346394 | 0.194161 | 0.552863064 |
| ENSMUSG00000027854 | Sike1 | protein_coding | -0.706255032 | 4.124325647 | 1.8331664 | 0.1942213 | 0.552863064 |
| ENSMUSG00000027384 | Ndufaf5 | protein_coding | 0.327138836 | 7.220808295 | 1.8330992 | 0.1942291 | 0.552863064 |
| ENSMUSG00000042410 | Agps | protein_coding | -0.5533328 | 4.159991566 | 1.8329613 | 0.194245 | 0.552863064 |
| ENSMUSG00000002550 | Uck1 | protein_coding | 0.771331204 | 3.433031781 | 1.8304769 | 0.1945321 | 0.553443016 |
| ENSMUSG00000002058 | Unc119 | protein_coding | 0.576984626 | 4.628577171 | 1.8295929 | 0.1946343 | 0.553443016 |
| ENSMUSG00000014873 | Surf2 | protein_coding | 0.388389304 | 6.407623706 | 1.8293131 | 0.1946667 | 0.553443016 |
| ENSMUSG00000021577 | Sdha | protein_coding | 0.459076134 | 9.626183273 | 1.8294969 | 0.1946756 | 0.553443016 |
| ENSMUSG00000041415 | Dicer1 | protein_coding | 0.632386476 | 4.0657958 | 1.8289729 | 0.1947061 | 0.553443016 |
| ENSMUSG00000019039 | Dalrd3 | protein_coding | 0.527232645 | 5.753914959 | 1.8283985 | 0.1948028 | 0.553458069 |
| ENSMUSG00000050796 | B3galt6 | protein_coding | 0.739638555 | 3.461346429 | 1.8280384 | 0.1948144 | 0.553458069 |
| ENSMUSG00000033319 | Fem1c | protein_coding | -0.616694006 | 3.986097007 | 1.8269398 | 0.1949417 | 0.553575848 |
| ENSMUSG00000029551 | Psmg3 | protein_coding | 0.929932945 | 2.713556447 | 1.8264894 | 0.1949939 | 0.553575848 |
| ENSMUSG00000058690 | Ccser2 | protein_coding | -0.411131752 | 5.493256827 | 1.8263487 | 0.1950103 | 0.553575848 |
| ENSMUSG00000040018 | Cox15 | protein_coding | 0.773790425 | 3.750012357 | 1.8258129 | 0.1950724 | 0.553606175 |
| ENSMUSG00000025041 | Nt5c2 | protein_coding | 0.45338794 | 5.858135852 | 1.8251173 | 0.1951532 | 0.553689234 |
| ENSMUSG00000078572 | Ndufaf8 | protein_coding | 0.428762748 | 6.474336998 | 1.8236522 | 0.1953234 | 0.554025987 |
| ENSMUSG00000030652 | Coq7 | protein_coding | 0.2845935 | 7.84327492 | 1.8227094 | 0.195433 | 0.554190785 |
| ENSMUSG00000022403 | St13 | protein_coding | -0.378971868 | 9.582884206 | 1.8216655 | 0.1955545 | 0.554389089 |
| ENSMUSG00000034501 | Pcnx4 | protein_coding | 0.53285437 | 4.806627932 | 1.8210934 | 0.1956211 | 0.554431773 |
| ENSMUSG00000047719 | Ubiad1 | protein_coding | 0.772178141 | 3.207248485 | 1.8203639 | 0.1957061 | 0.554526503 |
| ENSMUSG00000038286 | Bphl | protein_coding | 0.303193805 | 6.518229383 | 1.819047 | 0.1958597 | 0.554737071 |
| ENSMUSG00000047193 | Dync2h1 | protein_coding | -0.50876289 | 5.172442651 | 1.8188417 | 0.1958836 | 0.554737071 |
| ENSMUSG00000090093 | Gm14399 | protein_coding | -1.014703052 | 2.701887579 | 1.8172807 | 0.1960658 | 0.554979866 |
| ENSMUSG00000049553 | Polr1a | protein_coding | 0.756805643 | 3.578568344 | 1.8164971 | 0.1961574 | 0.554979866 |
| ENSMUSG00000018858 | Mrpl58 | protein_coding | 0.301547124 | 7.702655499 | 1.8162778 | 0.196183 | 0.554979866 |
| ENSMUSG00000028341 | Nr4a3 | protein_coding | -1.012371098 | 3.210663218 | 1.8155118 | 0.1962726 | 0.554979866 |
| ENSMUSG00000025266 | Gnl3l | protein_coding | 0.618870821 | 8.742299325 | 1.840443 | 0.1963079 | 0.554979866 |
| ENSMUSG00000032788 | Pdxk | protein_coding | 0.589402661 | 7.679197173 | 1.8373493 | 0.1963082 | 0.554979866 |
| ENSMUSG00000038370 | Pcp4l1 | protein_coding | 0.457964581 | 8.590326869 | 1.8160174 | 0.1963306 | 0.554979866 |
| ENSMUSG00000036282 | Naa30 | protein_coding | -0.631936746 | 4.880531943 | 1.8138541 | 0.1964967 | 0.555210778 |
| ENSMUSG00000037640 | Zfp60 | protein_coding | -0.547657677 | 4.904706894 | 1.8130339 | 0.1965627 | 0.555210778 |
| ENSMUSG00000038279 | Nop2 | protein_coding | 0.568362683 | 5.250246065 | 1.8129952 | 0.1965672 | 0.555210778 |
| ENSMUSG00000020899 | Pfas | protein_coding | 0.503958205 | 5.876812559 | 1.8116153 | 0.1967591 | 0.5554906 |
| ENSMUSG00000058600 | Rpl30 | protein_coding | 0.448219136 | 6.950298277 | 1.8115254 | 0.1967696 | 0.5554906 |
| ENSMUSG00000074358 | Ccdc61 | protein_coding | 0.688303869 | 3.424313251 | 1.810061 | 0.1969115 | 0.555648028 |
| ENSMUSG00000001383 | Zmat2 | protein_coding | 0.328684815 | 9.115223408 | 1.8090465 | 0.1970306 | 0.555648028 |
| ENSMUSG00000065126 | Snord104 | snoRNA | 0.854398221 | 3.124400437 | 1.8088838 | 0.1970498 | 0.555648028 |
| ENSMUSG00000028530 | Jak1 | protein_coding | 0.327471456 | 6.645700066 | 1.8087832 | 0.1970616 | 0.555648028 |
| ENSMUSG00000030826 | Bcat2 | protein_coding | 0.626789623 | 4.651427735 | 1.8085947 | 0.1970838 | 0.555648028 |
| ENSMUSG00000024974 | Smc3 | protein_coding | -0.448605668 | 5.852107548 | 1.8069781 | 0.1972739 | 0.555725188 |
| ENSMUSG00000029405 | G3bp2 | protein_coding | 0.435068636 | 7.275277119 | 1.8067025 | 0.1973364 | 0.555725188 |
| ENSMUSG00000012117 | Dhdds | protein_coding | 0.387868818 | 6.15705973 | 1.8063004 | 0.1973537 | 0.555725188 |
| ENSMUSG00000028567 | Txndc12 | protein_coding | 0.365691021 | 6.104744527 | 1.8059607 | 0.1973938 | 0.555725188 |
| ENSMUSG00000050315 | Synpo2 | protein_coding | -0.913496302 | 4.186734964 | 1.8158503 | 0.1973978 | 0.555725188 |
| ENSMUSG00000031090 | Nadsyn1 | protein_coding | -1.109219981 | 2.574717907 | 1.8054764 | 0.1974508 | 0.555725188 |
| ENSMUSG00000061778 | Mospd2 | protein_coding | -0.66395382 | 3.429067216 | 1.8052891 | 0.1974729 | 0.555725188 |
| ENSMUSG00000041147 | Brca2 | protein_coding | -0.826350036 | 3.318215813 | 1.8047251 | 0.1975394 | 0.555735846 |

| ensembl_gene_id | external_gene_name | gene_biotype | log2 (fold change) | log2 (counts per million) | F-value | p-value | False discovery rate |
|---|---|---|---|---|---|---|---|
| ENSMUSG00000032750 | Gab3 | protein_coding | -0.72648818 | 3.593346028 | 1.8041911 | 0.1976024 | 0.555735846 |
| ENSMUSG00000030609 | Aen | protein_coding | 0.681498952 | 4.770893489 | 1.8041967 | 0.1976317 | 0.555735846 |
| ENSMUSG00000051224 | Tceanc | protein_coding | -1.12771093 | 2.64279464 | 1.8030997 | 0.1977312 | 0.555753596 |
| ENSMUSG00000015291 | Gdi1 | protein_coding | 0.498024359 | 10.42006291 | 1.8075841 | 0.1977699 | 0.555753596 |
| ENSMUSG00000033933 | Vhl | protein_coding | 0.521610719 | 5.045703462 | 1.8025753 | 0.1977931 | 0.555753596 |
| ENSMUSG00000014550 | Rbsn | protein_coding | 0.465288974 | 5.928368641 | 1.8002722 | 0.1980653 | 0.55597261 |
| ENSMUSG00000020770 | Unk | protein_coding | -1.093566324 | 3.006113874 | 1.8001971 | 0.1980742 | 0.55597261 |
| ENSMUSG00000032667 | Pon2 | protein_coding | -0.498401272 | 5.017661309 | 1.7990913 | 0.1982051 | 0.55597261 |
| ENSMUSG00000054792 | Klhl18 | protein_coding | 0.519341347 | 4.768380638 | 1.7990766 | 0.1982069 | 0.55597261 |
| ENSMUSG00000042992 | Borcs5 | protein_coding | 0.59637717 | 5.352748953 | 1.79925 | 0.1982163 | 0.55597261 |
| ENSMUSG00000045205 | Dpy19l4 | protein_coding | -0.615970548 | 3.949632797 | 1.7987828 | 0.1982416 | 0.55597261 |
| ENSMUSG00000022025 | Cnmd | protein_coding | -1.686355002 | 2.401022234 | 1.7992354 | 0.1982648 | 0.55597261 |
| ENSMUSG00000078350 | Smim1 | protein_coding | 0.44015818 | 5.456210545 | 1.7980805 | 0.1983249 | 0.55597261 |
| ENSMUSG00000040472 | Rabggta | protein_coding | 0.38850151 | 6.178422067 | 1.7975656 | 0.1983859 | 0.55597261 |
| ENSMUSG00000074758 | Gm5535 | lincRNA | -0.996271906 | 2.770397321 | 1.7975472 | 0.1983881 | 0.55597261 |
| ENSMUSG00000027895 | Kcnc4 | protein_coding | -0.456359577 | 5.077709349 | 1.7970472 | 0.1984474 | 0.555976484 |
| ENSMUSG00000039354 | Smarcal1 | protein_coding | 0.680272942 | 4.604776588 | 1.7969164 | 0.1984929 | 0.555976484 |
| ENSMUSG00000020114 | Cand1 | protein_coding | -0.366664975 | 6.489287104 | 1.7959823 | 0.1985737 | 0.555994009 |
| ENSMUSG00000060771 | Tsga10 | protein_coding | -0.895776964 | 3.247839513 | 1.7957395 | 0.1986025 | 0.555994009 |
| ENSMUSG00000020205 | Phlda1 | protein_coding | -0.640293778 | 4.042854638 | 1.7944139 | 0.19876 | 0.556047722 |
| ENSMUSG00000041638 | Gcn1l1 | protein_coding | -0.532710272 | 4.07384481 | 1.7941204 | 0.1987949 | 0.556047722 |
| ENSMUSG00000014294 | Ndufa2 | protein_coding | 0.402661742 | 9.642070955 | 1.7940839 | 0.1987992 | 0.556047722 |
| ENSMUSG00000044988 | Ucn3 | protein_coding | -0.918338403 | 3.261059349 | 1.7934719 | 0.198872 | 0.556047722 |
| ENSMUSG00000000276 | Dgke | protein_coding | -0.556161296 | 4.35600267 | 1.7934021 | 0.1988803 | 0.556047722 |
| ENSMUSG00000027164 | Traf6 | protein_coding | 1.055342964 | 2.683158561 | 1.7929249 | 0.198937 | 0.556061854 |
| ENSMUSG00000030058 | Copg1 | protein_coding | 0.403567334 | 7.934758162 | 1.7926969 | 0.1989941 | 0.556076912 |
| ENSMUSG00000019795 | Pcmt1 | protein_coding | 0.317891345 | 8.737572817 | 1.7909491 | 0.1991723 | 0.556195961 |
| ENSMUSG00000039395 | Mreg | protein_coding | 1.102910493 | 2.458589269 | 1.7907833 | 0.199192 | 0.556195961 |
| ENSMUSG00000041143 | Tmco4 | protein_coding | 1.195602466 | 2.414615072 | 1.7904233 | 0.1992349 | 0.556195961 |
| ENSMUSG00000032348 | Gsta4 | protein_coding | 0.318006081 | 7.262439524 | 1.7903503 | 0.1992436 | 0.556195961 |
| ENSMUSG00000074736 | Syndig1 | protein_coding | 0.38648064 | 6.409774563 | 1.7873424 | 0.1996026 | 0.556998857 |
| ENSMUSG00000026587 | Astn1 | protein_coding | -0.385756888 | 6.196100042 | 1.7870725 | 0.1996349 | 0.556998857 |
| ENSMUSG00000068823 | Csde1 | protein_coding | -0.425090651 | 8.718473416 | 1.7852741 | 0.1998799 | 0.557506132 |
| ENSMUSG00000044224 | Dnajc21 | protein_coding | -0.537736433 | 4.343740295 | 1.784306 | 0.1999658 | 0.557506132 |
| ENSMUSG00000056268 | Dennd1b | protein_coding | -1.09381245 | 2.790479501 | 1.7842526 | 0.1999722 | 0.557506132 |
| ENSMUSG00000028708 | Mknk1 | protein_coding | 0.544582163 | 4.546987891 | 1.7829526 | 0.200128 | 0.557795733 |
| ENSMUSG00000051435 | Fhad1 | protein_coding | -0.413733184 | 5.190793183 | 1.7820939 | 0.2002309 | 0.557831024 |
| ENSMUSG00000043259 | Fam13c | protein_coding | 0.387879363 | 5.533974813 | 1.7819819 | 0.2002444 | 0.557831024 |
| ENSMUSG00000030530 | Furin | protein_coding | -0.884008454 | 2.998263109 | 1.7805132 | 0.2004207 | 0.558177491 |
| ENSMUSG00000078348 | Sf3b5 | protein_coding | 0.322189752 | 8.019502921 | 1.7795173 | 0.2005403 | 0.558231032 |
| ENSMUSG00000034453 | Polr3b | protein_coding | 0.684387256 | 3.459292314 | 1.7794146 | 0.2005527 | 0.558231032 |
| ENSMUSG00000055540 | Epha6 | protein_coding | -0.741193394 | 3.574033654 | 1.7782827 | 0.2006888 | 0.558231032 |
| ENSMUSG00000016308 | Ube2a | protein_coding | 0.309243433 | 7.517151785 | 1.778194 | 0.2006994 | 0.558231032 |
| ENSMUSG00000014606 | Slc25a11 | protein_coding | 0.309711098 | 8.391474826 | 1.7781443 | 0.2007054 | 0.558231032 |
| ENSMUSG00000032606 | Nicn1 | protein_coding | 0.335517772 | 8.336555782 | 1.7776823 | 0.200761 | 0.558231032 |
| ENSMUSG00000005268 | Prlr | protein_coding | 0.946805438 | 3.490090336 | 1.7775796 | 0.2008033 | 0.558231032 |
| ENSMUSG00000067288 | Rps28 | protein_coding | 0.578715929 | 4.775434699 | 1.7758561 | 0.200981 | 0.558580573 |
| ENSMUSG00000058748 | Zfp958 | protein_coding | -0.739727239 | 3.542261307 | 1.775139 | 0.2010674 | 0.558676444 |
| ENSMUSG00000020170 | Frs2 | protein_coding | 0.547176704 | 4.04767324 | 1.774487 | 0.2011461 | 0.55868064 |
| ENSMUSG00000031753 | Cog4 | protein_coding | 0.712187393 | 4.126550012 | 1.7742651 | 0.2011728 | 0.55868064 |
| ENSMUSG00000028967 | Errfi1 | protein_coding | -0.607335798 | 4.431978901 | 1.773472 | 0.2012686 | 0.558715677 |
| ENSMUSG00000026150 | Mff | protein_coding | -0.35763458 | 7.330525342 | 1.772968 | 0.2013294 | 0.558715677 |
| ENSMUSG00000069184 | Zfp72 | protein_coding | -0.852894189 | 3.149835557 | 1.7725504 | 0.2013799 | 0.558715677 |
| ENSMUSG00000028464 | Tpm2 | protein_coding | -0.650204012 | 4.452468545 | 1.7724394 | 0.2013933 | 0.558715677 |
| ENSMUSG00000021391 | Cenpp | protein_coding | -0.70954997 | 3.797325587 | 1.7708276 | 0.2015882 | 0.558945312 |
| ENSMUSG00000027087 | Itgav | protein_coding | -0.832845119 | 5.549576828 | 1.7948442 | 0.2016145 | 0.558945312 |
| ENSMUSG00000071234 | Syndig1l | protein_coding | -0.583514924 | 5.554383131 | 1.7707123 | 0.201632 | 0.558945312 |
| ENSMUSG00000102674 | 8030442B05Rik | lincRNA | -0.861042605 | 3.225545928 | 1.7687354 | 0.2018415 | 0.559381873 |
| ENSMUSG00000074282 | Zfp94 | protein_coding | 0.496407871 | 4.398316985 | 1.7681494 | 0.2019125 | 0.559434505 |
| ENSMUSG00000023915 | Tnfrsf21 | protein_coding | 0.419759954 | 5.548326239 | 1.7670574 | 0.202045 | 0.559626695 |
| ENSMUSG00000021606 | Ndufs6 | protein_coding | 0.348482615 | 8.767019294 | 1.7667195 | 0.202086 | 0.559626695 |
| ENSMUSG00000046854 | Pip5kl1 | protein_coding | 2.004842066 | 2.460729268 | 1.7809508 | 0.2021839 | 0.55973258 |
| ENSMUSG00000024101 | Washc1 | protein_coding | 0.450813969 | 5.043493892 | 1.7655471 | 0.2022283 | 0.55973258 |
| ENSMUSG00000038400 | Pmepa1 | protein_coding | -0.54257142 | 4.561801788 | 1.7649645 | 0.2022991 | 0.559784436 |
| ENSMUSG00000039164 | Naif1 | protein_coding | 0.813083643 | 3.153482618 | 1.7642369 | 0.2023876 | 0.559785558 |
| ENSMUSG00000028683 | Eif2b3 | protein_coding | 0.377718197 | 6.287355809 | 1.7637567 | 0.2024459 | 0.559785558 |
| ENSMUSG00000013155 | Enkd1 | protein_coding | 0.670715862 | 3.709672084 | 1.7636766 | 0.2024557 | 0.559785558 |
| ENSMUSG00000063428 | Ddo | protein_coding | -0.532790248 | 4.108612222 | 1.7609107 | 0.2027925 | 0.560531938 |
| ENSMUSG00000020283 | Pex13 | protein_coding | -0.572225973 | 4.641362991 | 1.7606041 | 0.2028299 | 0.560531938 |
| ENSMUSG00000040524 | Zfp609 | protein_coding | -0.664989288 | 3.534331021 | 1.7598493 | 0.2029219 | 0.560642264 |
| ENSMUSG00000032564 | Cpne4 | protein_coding | -0.477958961 | 5.771340579 | 1.7576317 | 0.2031927 | 0.561246193 |
| ENSMUSG00000046269 | Usp27x | protein_coding | 0.627905136 | 3.983395356 | 1.7564155 | 0.2033414 | 0.561302787 |
| ENSMUSG00000022095 | Fam160b2 | protein_coding | -0.606629766 | 3.768285915 | 1.756018 | 0.2033901 | 0.561302787 |
| ENSMUSG00000023259 | Slc26a6 | protein_coding | 1.328057176 | 2.561602246 | 1.7555695 | 0.2034449 | 0.561302787 |
| ENSMUSG00000051451 | Crebzf | protein_coding | -1.059646993 | 2.589739195 | 1.7552345 | 0.203486 | 0.561302787 |
| ENSMUSG00000022555 | Dgat1 | protein_coding | 0.802738435 | 3.529617965 | 1.754404 | 0.2035877 | 0.561302787 |
| ENSMUSG00000039676 | Capsl | protein_coding | -0.753161643 | 4.066368369 | 1.754082 | 0.2036569 | 0.561302787 |
| ENSMUSG00000005198 | Polr2a | protein_coding | -0.680306612 | 4.727635432 | 1.7538532 | 0.203685 | 0.561302787 |

| ensembl_gene_id | external_gene_name | gene_biotype | log2 (fold change) | log2 (counts per million) | F-value | p-value | False discovery rate |
|---|---|---|---|---|---|---|---|
| ENSMUSG00000029422 | Rsrc2 | protein_coding | -0.337365027 | 7.29315906 | 1.7535075 | 0.2036975 | 0.561302787 |
| ENSMUSG00000027634 | Ndrg3 | protein_coding | 0.447757619 | 9.406259515 | 1.753028 | 0.2037861 | 0.561302787 |
| ENSMUSG00000028532 | Cachd1 | protein_coding | 0.673557195 | 3.756836825 | 1.7524701 | 0.2038248 | 0.561302787 |
| ENSMUSG00000036327 | Qsox2 | protein_coding | 1.07611287 | 2.993632608 | 1.7522882 | 0.2038471 | 0.561302787 |
| ENSMUSG00000090626 | Tex9 | protein_coding | -0.691256545 | 3.860997671 | 1.7521702 | 0.2038616 | 0.561302787 |
| ENSMUSG00000005125 | Ndrg1 | protein_coding | -0.447648598 | 5.389788109 | 1.7519237 | 0.2038918 | 0.561302787 |
| ENSMUSG00000042105 | Inpp5f | protein_coding | 0.399174022 | 8.659097926 | 1.7507294 | 0.2040385 | 0.561461311 |
| ENSMUSG00000034429 | Zfp707 | protein_coding | 0.664767409 | 4.951956453 | 1.7508343 | 0.2040554 | 0.561461311 |
| ENSMUSG00000077450 | Rab11b | protein_coding | 0.357605864 | 7.827942109 | 1.7501796 | 0.204106 | 0.561461311 |
| ENSMUSG00000085913 | Gm15601 | antisense_RNA | -1.018152439 | 2.707428456 | 1.749136 | 0.2042344 | 0.561535017 |
| ENSMUSG00000053644 | Aldh7a1 | protein_coding | -0.377125107 | 5.997732818 | 1.7491122 | 0.2042373 | 0.561535017 |
| ENSMUSG00000020728 | Cep112 | protein_coding | 0.879301068 | 2.978589964 | 1.748563 | 0.2043049 | 0.561577213 |
| ENSMUSG00000036087 | Slain2 | protein_coding | -1.06082921 | 3.484801061 | 1.7542526 | 0.204366 | 0.561601857 |
| ENSMUSG00000025144 | Cenpx | protein_coding | 0.383271897 | 6.391339461 | 1.7453226 | 0.2047041 | 0.562270144 |
| ENSMUSG00000020456 | Ogdh | protein_coding | 0.431741615 | 8.134128311 | 1.7450983 | 0.2047615 | 0.562270144 |
| ENSMUSG00000030649 | Anapc15 | protein_coding | 0.663216293 | 3.334285731 | 1.7448201 | 0.2047661 | 0.562270144 |
| ENSMUSG00000085829 | Gm4285 | antisense_RNA | -1.291140754 | 2.39625023 | 1.7440736 | 0.2048583 | 0.562379604 |
| ENSMUSG00000019916 | P4ha1 | protein_coding | 0.575617776 | 4.435432049 | 1.7425569 | 0.2050457 | 0.562662272 |
| ENSMUSG00000097059 | Fam120aos | lincRNA | -0.924668626 | 2.730748456 | 1.7420661 | 0.2051063 | 0.562662272 |
| ENSMUSG00000069917 | Hba-a2 | protein_coding | -0.755901577 | 3.810027504 | 1.7417238 | 0.2051487 | 0.562662272 |
| ENSMUSG00000050711 | Scg2 | protein_coding | 0.313074875 | 9.620289236 | 1.7415471 | 0.2051705 | 0.562662272 |
| ENSMUSG00000086290 | Snhg12 | lincRNA | 0.399701918 | 5.972992914 | 1.7410372 | 0.2052336 | 0.562691796 |
| ENSMUSG00000042305 | Tmem183a | protein_coding | 0.442729707 | 5.834486723 | 1.7398812 | 0.2053768 | 0.562940718 |
| ENSMUSG00000029659 | Medag | protein_coding | -1.423443508 | 2.621532017 | 1.7381832 | 0.205617 | 0.563171845 |
| ENSMUSG00000026425 | Srgap2 | protein_coding | -0.416387883 | 5.390911162 | 1.7376464 | 0.2056539 | 0.563171845 |
| ENSMUSG00000035242 | Oaz1 | protein_coding | 0.300609414 | 8.286674294 | 1.7374373 | 0.2056798 | 0.563171845 |
| ENSMUSG00000042660 | Wdr55 | protein_coding | 0.467783353 | 4.827518776 | 1.7374294 | 0.2056808 | 0.563171845 |
| ENSMUSG00000026234 | Ncl | protein_coding | -0.321349445 | 8.209735329 | 1.7370897 | 0.205723 | 0.563171845 |
| ENSMUSG00000061436 | Hipk2 | protein_coding | -0.754751508 | 4.56739696 | 1.7397795 | 0.2059509 | 0.563652449 |
| ENSMUSG00000022679 | Mpv17l | protein_coding | -0.444906093 | 5.775120625 | 1.7337089 | 0.2061432 | 0.564035109 |
| ENSMUSG00000020514 | Mrpl22 | protein_coding | 0.355955151 | 6.224653797 | 1.7328469 | 0.2062505 | 0.564185213 |
| ENSMUSG00000029231 | Pdgfra | protein_coding | -0.907435097 | 3.623955412 | 1.7326064 | 0.2063102 | 0.564204871 |
| ENSMUSG00000085885 | Gm11906 | processed_transcript | -0.529957467 | 4.577046964 | 1.7316416 | 0.2064007 | 0.564308941 |
| ENSMUSG00000051427 | Ccdc157 | protein_coding | -0.854667045 | 2.943598465 | 1.7289018 | 0.2067426 | 0.564942101 |
| ENSMUSG00000006586 | Runx1t1 | protein_coding | -0.501895706 | 6.322733554 | 1.7301263 | 0.2067489 | 0.564942101 |
| ENSMUSG00000039959 | Hip1 | protein_coding | -0.48482825 | 4.775215907 | 1.7285234 | 0.2067899 | 0.564942101 |
| ENSMUSG00000028229 | Rmdn1 | protein_coding | -0.541994791 | 4.287708467 | 1.7271017 | 0.2069677 | 0.565112393 |
| ENSMUSG00000026035 | Ppil3 | protein_coding | 0.41498662 | 6.186644103 | 1.7265104 | 0.2070417 | 0.565112393 |
| ENSMUSG00000053347 | Zfp943 | protein_coding | -0.860880363 | 4.009394861 | 1.7304128 | 0.2071147 | 0.565112393 |
| ENSMUSG00000032312 | Csk | protein_coding | 0.596162876 | 4.611753996 | 1.7258207 | 0.207128 | 0.565112393 |
| ENSMUSG00000052336 | Cx3cr1 | protein_coding | 0.677228546 | 4.189887564 | 1.7256133 | 0.207154 | 0.565112393 |
| ENSMUSG00000020591 | Ntsr2 | protein_coding | -0.733034655 | 5.596747364 | 1.7436915 | 0.2072503 | 0.565112393 |
| ENSMUSG00000072258 | Taf1a | protein_coding | 0.575385123 | 4.182932386 | 1.7242242 | 0.2073281 | 0.565112393 |
| ENSMUSG00000041773 | Enc1 | protein_coding | 0.399875907 | 8.466463369 | 1.7242067 | 0.2073599 | 0.565112393 |
| ENSMUSG00000043131 | Mob1a | protein_coding | -0.728064798 | 3.664848471 | 1.7239159 | 0.2073667 | 0.565112393 |
| ENSMUSG00000031583 | Wrn | protein_coding | -0.893281828 | 3.024138021 | 1.7236441 | 0.2074008 | 0.565112393 |
| ENSMUSG00000071723 | Gspt2 | protein_coding | 0.423117412 | 5.012475864 | 1.7234086 | 0.2074304 | 0.565112393 |
| ENSMUSG00000023806 | Rsph3b | protein_coding | 0.8351499 | 2.821497084 | 1.7225576 | 0.2075372 | 0.565260126 |
| ENSMUSG00000024773 | Atg2a | protein_coding | 0.792851999 | 3.524198162 | 1.7216195 | 0.207655 | 0.565437793 |
| ENSMUSG00000028004 | Npy2r | protein_coding | 0.902958847 | 4.064201765 | 1.7298078 | 0.2077793 | 0.565606365 |
| ENSMUSG00000004865 | Srpk1 | protein_coding | 0.382540041 | 6.884929363 | 1.7202901 | 0.2078221 | 0.565606365 |
| ENSMUSG00000030047 | Arhgap25 | protein_coding | -1.389557829 | 2.611281715 | 1.7188901 | 0.2080279 | 0.565678764 |
| ENSMUSG00000022132 | Cldn10 | protein_coding | -0.781045269 | 5.794280525 | 1.7417342 | 0.2080297 | 0.565678764 |
| ENSMUSG00000093436 | Gm20646 | sense_overlapping | -0.629247464 | 3.643241201 | 1.7185728 | 0.2080382 | 0.565678764 |
| ENSMUSG00000018565 | Elp5 | protein_coding | 0.297249163 | 8.18573126 | 1.717697 | 0.2081485 | 0.565678764 |
| ENSMUSG00000048796 | Cyb561d1 | protein_coding | 0.747678868 | 3.55124083 | 1.7176797 | 0.2081507 | 0.565678764 |
| ENSMUSG00000047363 | Cstad | protein_coding | 0.671787485 | 3.822675004 | 1.7174773 | 0.2081762 | 0.565678764 |
| ENSMUSG00000021756 | Il6st | protein_coding | -0.590658355 | 4.07293837 | 1.7171541 | 0.2082169 | 0.565678764 |
| ENSMUSG00000055013 | Agap1 | protein_coding | -0.352380686 | 6.385730368 | 1.7145396 | 0.2085468 | 0.566431993 |
| ENSMUSG00000073423 | Zfp414 | protein_coding | 0.377579781 | 6.22489882 | 1.7132803 | 0.208706 | 0.566721133 |
| ENSMUSG00000042292 | Mkl1 | protein_coding | -0.497205427 | 4.91846856 | 1.712267 | 0.2088342 | 0.566782083 |
| ENSMUSG00000055733 | Nap1l3 | protein_coding | -0.442206391 | 5.768470532 | 1.7121442 | 0.2088497 | 0.566782083 |
| ENSMUSG00000028931 | Kcnab2 | protein_coding | 0.320281487 | 7.385451449 | 1.7118529 | 0.2088866 | 0.566782083 |
| ENSMUSG00000087618 | Gm14114 | lincRNA | -1.046829089 | 2.454754012 | 1.7103311 | 0.2090793 | 0.567161949 |
| ENSMUSG00000013160 | Atp6v0d1 | protein_coding | 0.27589272 | 8.515038157 | 1.7064044 | 0.2095777 | 0.568166832 |
| ENSMUSG00000024404 | Riok3 | protein_coding | -0.569860658 | 4.928221499 | 1.7057958 | 0.2096551 | 0.568166832 |
| ENSMUSG00000025133 | Ints4 | protein_coding | 0.543791529 | 4.854161641 | 1.705591 | 0.2096811 | 0.568166832 |
| ENSMUSG00000025364 | Pa2g4 | protein_coding | 0.344319796 | 7.354736714 | 1.7054803 | 0.2096952 | 0.568166832 |
| ENSMUSG00000045983 | Eif4g1 | protein_coding | 0.44766638 | 5.20967708 | 1.705333 | 0.209714 | 0.568166832 |
| ENSMUSG00000027956 | Tmem144 | protein_coding | -0.697036951 | 3.501868317 | 1.7039019 | 0.2098961 | 0.568517139 |
| ENSMUSG00000020747 | Tmem94 | protein_coding | -0.47442953 | 4.770877502 | 1.7033219 | 0.20997 | 0.568574038 |
| ENSMUSG00000074657 | Kif5a | protein_coding | -0.404671639 | 8.897045906 | 1.7027962 | 0.2100666 | 0.568615988 |
| ENSMUSG00000058291 | Zfp68 | protein_coding | -0.414146402 | 5.270613456 | 1.7023707 | 0.2100913 | 0.568615988 |
| ENSMUSG00000096916 | Zfp850 | protein_coding | 1.401298325 | 2.376784484 | 1.7012787 | 0.2102306 | 0.5688424 |
| ENSMUSG00000029253 | Cenpc1 | protein_coding | -0.566615955 | 4.863560757 | 1.7008857 | 0.2102807 | 0.5688424 |
| ENSMUSG00000036120 | Rfxank | protein_coding | -0.476508123 | 4.926668222 | 1.700216 | 0.2103662 | 0.568930611 |
| ENSMUSG00000029447 | Cct6a | processed_transcript | 0.306414135 | 8.111927685 | 1.6975314 | 0.2107095 | 0.569715627 |
| ENSMUSG00000042810 | Krba1 | protein_coding | 0.593306673 | 3.77363301 | 1.696262 | 0.210872 | 0.569848614 |

| ensembl_gene_id | external_gene_name | gene_biotype | log2 (fold change) | log2 (counts per million) | F-value | p-value | False discovery rate |
|---|---|---|---|---|---|---|---|
| ENSMUSG00000028167 | Bdh2 | protein_coding | -0.666985977 | 3.684436894 | 1.6962289 | 0.2108763 | 0.569848614 |
| ENSMUSG00000022791 | Tnk2 | protein_coding | -0.496978366 | 5.128254215 | 1.6957118 | 0.2109425 | 0.569848614 |
| ENSMUSG00000025321 | Itgb8 | protein_coding | -0.802637035 | 3.993392886 | 1.6957228 | 0.2109706 | 0.569848614 |
| ENSMUSG00000023206 | Il15ra | protein_coding | 0.78507961 | 3.391109557 | 1.694722 | 0.2110695 | 0.569972336 |
| ENSMUSG00000021047 | Nova1 | protein_coding | -0.364199765 | 6.313334427 | 1.6942942 | 0.2111243 | 0.569977415 |
| ENSMUSG00000028803 | Nipal3 | protein_coding | 0.457433306 | 5.249474232 | 1.6930181 | 0.2112882 | 0.570185985 |
| ENSMUSG00000018334 | Ksr1 | protein_coding | -0.772295704 | 3.4061664 | 1.6919607 | 0.211424 | 0.570185985 |
| ENSMUSG00000062234 | Gak | protein_coding | -0.405036567 | 5.490160043 | 1.6917044 | 0.211457 | 0.570185985 |
| ENSMUSG00000027394 | Ttl | protein_coding | 0.456617606 | 4.823554834 | 1.6916756 | 0.2114607 | 0.570185985 |
| ENSMUSG00000024472 | Dcp2 | protein_coding | -0.532176255 | 4.493941162 | 1.6914309 | 0.2114921 | 0.570185985 |
| ENSMUSG00000026019 | Wdr12 | protein_coding | -0.323638089 | 6.68670352 | 1.6912162 | 0.2115198 | 0.570185985 |
| ENSMUSG00000030588 | Yif1b | protein_coding | 0.333159125 | 6.243423051 | 1.6886756 | 0.2118468 | 0.570886448 |
| ENSMUSG00000029228 | Lnx1 | protein_coding | 0.705394033 | 4.27486884 | 1.6886023 | 0.2118858 | 0.570886448 |
| ENSMUSG00000023156 | Rpp14 | protein_coding | 0.308778442 | 6.519071113 | 1.6865675 | 0.2121188 | 0.57117059 |
| ENSMUSG00000055128 | Cgrrf1 | protein_coding | -0.369892131 | 6.209589568 | 1.6864432 | 0.2121348 | 0.57117059 |
| ENSMUSG00000048776 | Pthlh | protein_coding | -1.275159151 | 2.503974901 | 1.6863209 | 0.2121506 | 0.57117059 |
| ENSMUSG00000026341 | Actr3 | protein_coding | -0.326820322 | 6.223189003 | 1.6854349 | 0.212265 | 0.571173223 |
| ENSMUSG00000025905 | Oprk1 | protein_coding | 0.644772888 | 4.134242713 | 1.6850817 | 0.2123107 | 0.571173223 |
| ENSMUSG00000089759 | 3632454L22Rik | processed_transcript | -1.008766823 | 2.614671891 | 1.6847916 | 0.2123482 | 0.571173223 |
| ENSMUSG00000036918 | Ttc7 | protein_coding | -1.249140723 | 2.377443965 | 1.6846687 | 0.212364 | 0.571173223 |
| ENSMUSG00000078877 | Gm14295 | protein_coding | -0.634027633 | 3.695255168 | 1.6834673 | 0.2125194 | 0.571275707 |
| ENSMUSG00000057236 | Rbbp4 | protein_coding | 0.398535621 | 7.282346161 | 1.683089 | 0.2125978 | 0.571275707 |
| ENSMUSG00000033439 | Trmt13 | protein_coding | -0.928871703 | 2.783158122 | 1.6827699 | 0.2126097 | 0.571275707 |
| ENSMUSG00000031104 | Rab33a | protein_coding | 0.336503178 | 6.802206864 | 1.6827316 | 0.2126147 | 0.571275707 |
| ENSMUSG00000028578 | Caap1 | protein_coding | -1.229375233 | 2.471576403 | 1.6821982 | 0.2126837 | 0.571318527 |
| ENSMUSG00000026749 | Nek6 | protein_coding | 0.588100452 | 4.331659509 | 1.6814624 | 0.2127791 | 0.571431857 |
| ENSMUSG00000026778 | Prkcq | protein_coding | -0.784842638 | 3.361686752 | 1.6802312 | 0.2129387 | 0.571717797 |
| ENSMUSG00000026452 | Syt2 | protein_coding | 0.525257738 | 4.272954533 | 1.6797677 | 0.2129988 | 0.57173651 |
| ENSMUSG00000039220 | Ppp1r10 | protein_coding | -0.538334729 | 5.239057029 | 1.6784932 | 0.2131643 | 0.571884046 |
| ENSMUSG00000032009 | Sesn3 | protein_coding | -0.415817948 | 6.287416602 | 1.6777462 | 0.2132614 | 0.571884046 |
| ENSMUSG00000073940 | Hbb-bt | protein_coding | -0.772154108 | 3.790202896 | 1.6773348 | 0.2133149 | 0.571884046 |
| ENSMUSG00000049176 | Frmpd4 | protein_coding | -0.457688049 | 5.270504808 | 1.6771235 | 0.2133423 | 0.571884046 |
| ENSMUSG00000038417 | Fig4 | protein_coding | -0.531912002 | 4.988663656 | 1.6769324 | 0.2133672 | 0.571884046 |
| ENSMUSG00000020691 | Mettl2 | protein_coding | 0.659270393 | 4.728498721 | 1.6762993 | 0.213479 | 0.571884046 |
| ENSMUSG00000021773 | Comtd1 | protein_coding | 0.84942375 | 3.079362413 | 1.6758936 | 0.2135023 | 0.571884046 |
| ENSMUSG00000026858 | Miga2 | protein_coding | 0.441173584 | 5.634868065 | 1.6756304 | 0.2135366 | 0.571884046 |
| ENSMUSG00000047635 | 2810006K23Rik | protein_coding | 0.462049573 | 5.227095628 | 1.6754237 | 0.2135635 | 0.571884046 |
| ENSMUSG00000011832 | Evi5l | protein_coding | -0.420005274 | 4.973127487 | 1.6746577 | 0.2136633 | 0.571884046 |
| ENSMUSG00000026667 | Uhmk1 | protein_coding | 0.326671702 | 6.742017928 | 1.6746114 | 0.2136693 | 0.571884046 |
| ENSMUSG00000034377 | Tulp4 | protein_coding | -0.377783396 | 5.511357516 | 1.6740369 | 0.2137442 | 0.571884046 |
| ENSMUSG00000020918 | Kat2a | protein_coding | 0.40858848 | 5.123732653 | 1.6738942 | 0.2137628 | 0.571884046 |
| ENSMUSG00000029094 | Afap1 | protein_coding | 0.506761834 | 4.558819411 | 1.6734909 | 0.2138154 | 0.571884046 |
| ENSMUSG00000031570 | Plpp5 | protein_coding | 0.503202914 | 4.85331234 | 1.6732134 | 0.2138516 | 0.571884046 |
| ENSMUSG00000046573 | Lyrm4 | protein_coding | 0.404206014 | 7.735868849 | 1.6726601 | 0.2139531 | 0.572013454 |
| ENSMUSG00000009207 | Lnpk | protein_coding | -0.411295797 | 5.532369626 | 1.6703932 | 0.2142198 | 0.572584055 |
| ENSMUSG00000027378 | Nphp1 | protein_coding | 0.704021735 | 3.735808116 | 1.6697242 | 0.2143073 | 0.572605771 |
| ENSMUSG00000022635 | Zcrb1 | protein_coding | -0.456912759 | 5.522701848 | 1.6695166 | 0.2143344 | 0.572605771 |
| ENSMUSG00000026707 | Nsun6 | protein_coding | -0.790447461 | 3.048525799 | 1.669007 | 0.2144011 | 0.572641621 |
| ENSMUSG00000057093 | Zfp607b | protein_coding | 0.720957976 | 3.5890363 | 1.6682753 | 0.2144969 | 0.572707936 |
| ENSMUSG00000001700 | Gramd3 | protein_coding | -0.752290322 | 3.769375656 | 1.6672325 | 0.2146335 | 0.572707936 |
| ENSMUSG00000063646 | Jakmip1 | protein_coding | 0.338853793 | 5.848733938 | 1.6666942 | 0.2147041 | 0.572707936 |
| ENSMUSG00000035268 | Pkig | protein_coding | -0.32036639 | 7.141990585 | 1.6666408 | 0.2147111 | 0.572707936 |
| ENSMUSG00000048897 | Zfp710 | protein_coding | -0.519642071 | 4.031453304 | 1.665869 | 0.2148123 | 0.572707936 |
| ENSMUSG00000021037 | Ahsa1 | protein_coding | 0.298222344 | 7.846547904 | 1.6651041 | 0.2149127 | 0.572707936 |
| ENSMUSG00000040877 | Wdr25 | protein_coding | 0.915803362 | 3.857686571 | 1.6729805 | 0.2149575 | 0.572707936 |
| ENSMUSG00000041341 | Atg2b | protein_coding | 0.630223465 | 3.948557634 | 1.664677 | 0.2149687 | 0.572707936 |
| ENSMUSG00000073002 | Vamp5 | protein_coding | -0.87725111 | 3.080028733 | 1.6646386 | 0.2149738 | 0.572707936 |
| ENSMUSG00000109284 | B230311B06Rik | lincRNA | -0.762478536 | 3.309155098 | 1.6643305 | 0.2150142 | 0.572707936 |
| ENSMUSG00000032034 | Kcnj5 | protein_coding | 1.129411797 | 2.643768098 | 1.6643289 | 0.2150144 | 0.572707936 |
| ENSMUSG00000038677 | Scube3 | protein_coding | -1.07366809 | 2.59670893 | 1.6636386 | 0.2151051 | 0.572707936 |
| ENSMUSG00000020829 | Slc46a1 | protein_coding | 0.408093485 | 5.153076633 | 1.6635381 | 0.2151183 | 0.572707936 |
| ENSMUSG00000021190 | Lgmn | protein_coding | 0.365757458 | 7.624437304 | 1.660824 | 0.2154754 | 0.573015946 |
| ENSMUSG00000024135 | Srbd1 | protein_coding | -0.889860834 | 3.426965269 | 1.6609756 | 0.2154848 | 0.573015946 |
| ENSMUSG00000014602 | Kif1a | protein_coding | 0.319582953 | 8.857309886 | 1.6606178 | 0.2155026 | 0.573015946 |
| ENSMUSG00000024240 | Epc1 | protein_coding | 0.493423344 | 4.111177096 | 1.6605245 | 0.2155149 | 0.573015946 |
| ENSMUSG00000036898 | Zfp157 | protein_coding | -0.525183466 | 4.376131062 | 1.6600478 | 0.2155777 | 0.573015946 |
| ENSMUSG00000051007 | Gatd1 | protein_coding | 0.351603451 | 7.261766072 | 1.6599705 | 0.2155879 | 0.573015946 |
| ENSMUSG00000030335 | Mrpl51 | protein_coding | 0.292178399 | 6.91307832 | 1.6587099 | 0.2157541 | 0.573015946 |
| ENSMUSG00000029304 | Spp1 | protein_coding | -0.717492064 | 4.246992768 | 1.6589109 | 0.2157569 | 0.573015946 |
| ENSMUSG00000031841 | Cdh13 | protein_coding | 0.275431616 | 7.328179184 | 1.658265 | 0.2158128 | 0.573015946 |
| ENSMUSG00000026003 | Acadl | protein_coding | -0.532160469 | 5.020147576 | 1.6581832 | 0.2158236 | 0.573015946 |
| ENSMUSG00000021958 | Pinx1 | protein_coding | -0.485522441 | 4.210913605 | 1.6581771 | 0.2158244 | 0.573015946 |
| ENSMUSG00000007653 | Gabrb2 | protein_coding | -0.70095012 | 4.267264857 | 1.6575866 | 0.2159316 | 0.573015946 |
| ENSMUSG00000048747 | E130114P18Rik | protein_coding | -0.699075037 | 4.640997814 | 1.6577707 | 0.215953 | 0.573015946 |
| ENSMUSG00000025586 | Cpeb1 | protein_coding | 0.620313388 | 5.172911502 | 1.6575655 | 0.2159801 | 0.573015946 |
| ENSMUSG00000027646 | Src | protein_coding | -0.643644112 | 4.171514877 | 1.6546545 | 0.2162899 | 0.573696482 |
| ENSMUSG00000020255 | D10Wsu102e | protein_coding | 0.514892124 | 5.333206271 | 1.6539671 | 0.2163809 | 0.573796289 |
| ENSMUSG00000033499 | Larp4b | protein_coding | -0.346913332 | 6.176354794 | 1.6534594 | 0.2164481 | 0.573833076 |

| ensembl_gene_id | external_gene_name | gene_biotype | log2 (fold change) | log2 (counts per million) | F-value | p-value | False discovery rate |
|---|---|---|---|---|---|---|---|
| ENSMUSG00000021189 | Atxn3 | protein_coding | 0.656708361 | 3.618118477 | 1.652906 | 0.2165215 | 0.57383736 |
| ENSMUSG00000027620 | Rbm39 | protein_coding | -0.314849468 | 7.1349074 | 1.6526418 | 0.2165565 | 0.57383736 |
| ENSMUSG00000020876 | Snx11 | protein_coding | 0.663783531 | 4.596379397 | 1.6522728 | 0.2166347 | 0.573903168 |
| ENSMUSG00000044147 | Arf6 | protein_coding | 0.415977088 | 5.076528552 | 1.6514036 | 0.2167207 | 0.573989553 |
| ENSMUSG00000021481 | Zfp346 | protein_coding | -0.548716813 | 4.227970794 | 1.6506313 | 0.2168232 | 0.574055692 |
| ENSMUSG00000021680 | Crhbp | protein_coding | -0.52918104 | 4.435789994 | 1.6502771 | 0.2168702 | 0.574055692 |
| ENSMUSG00000032965 | Ift57 | protein_coding | -0.871002439 | 3.069249186 | 1.6500091 | 0.2169058 | 0.574055692 |
| ENSMUSG00000032402 | Smad3 | protein_coding | -0.58464745 | 4.130400917 | 1.6485463 | 0.2171002 | 0.574270088 |
| ENSMUSG00000029153 | Ociad2 | protein_coding | -0.266477649 | 7.775326845 | 1.6482576 | 0.2171386 | 0.574270088 |
| ENSMUSG00000051177 | Plcb1 | protein_coding | -0.715489986 | 3.797051818 | 1.6477999 | 0.2171995 | 0.574270088 |
| ENSMUSG00000096943 | Gm26721 | lincRNA | -0.982734393 | 2.809165358 | 1.6477928 | 0.2172005 | 0.574270088 |
| ENSMUSG00000044475 | Ascc1 | protein_coding | -0.305274562 | 6.756643452 | 1.6466465 | 0.217353 | 0.574532257 |
| ENSMUSG00000026245 | Farsb | protein_coding | 0.337457713 | 7.306000075 | 1.6447819 | 0.2176015 | 0.575047786 |
| ENSMUSG00000071037 | Camkmt | protein_coding | -0.597799176 | 4.431726802 | 1.6436565 | 0.2177517 | 0.575084168 |
| ENSMUSG00000030125 | Lrrc23 | protein_coding | 0.659489806 | 3.572157851 | 1.6435735 | 0.2177628 | 0.575084168 |
| ENSMUSG00000022528 | Hes1 | protein_coding | -0.83192753 | 2.960252542 | 1.6431131 | 0.2178243 | 0.575084168 |
| ENSMUSG00000020590 | Snx13 | protein_coding | -0.475721843 | 4.746669632 | 1.6430758 | 0.2178292 | 0.575084168 |
| ENSMUSG00000014905 | Dnajb9 | protein_coding | -0.496458975 | 5.484625982 | 1.6375884 | 0.2185637 | 0.576881465 |
| ENSMUSG00000045038 | Prkce | protein_coding | -0.316530518 | 7.599534377 | 1.6363745 | 0.2187266 | 0.577169759 |
| ENSMUSG00000037306 | Man1c1 | protein_coding | -0.782388779 | 3.687824388 | 1.6359098 | 0.218789 | 0.577192776 |
| ENSMUSG00000054619 | Mettl7a1 | protein_coding | -0.742876105 | 3.794749842 | 1.635225 | 0.218881 | 0.577224959 |
| ENSMUSG00000005483 | Dnajb1 | protein_coding | 0.335026229 | 6.685269881 | 1.6348992 | 0.2189248 | 0.577224959 |
| ENSMUSG00000024493 | Lars | protein_coding | -0.437523522 | 5.57129113 | 1.6346205 | 0.2189622 | 0.577224959 |
| ENSMUSG00000053181 | A830005F24Rik | protein_coding | -0.932185152 | 3.006694887 | 1.6334928 | 0.2191139 | 0.577380029 |
| ENSMUSG00000031787 | Katnb1 | protein_coding | 0.515309079 | 6.125970279 | 1.6374909 | 0.2191284 | 0.577380029 |
| ENSMUSG00000027300 | Ubox5 | protein_coding | 0.511417079 | 4.192564187 | 1.6329658 | 0.2191848 | 0.577387146 |
| ENSMUSG00000032532 | Cck | protein_coding | -0.840076499 | 3.604041295 | 1.6326296 | 0.2192593 | 0.5774418 |
| ENSMUSG00000101585 | 1600010M07Rik | antisense_RNA | -0.972760613 | 2.830697685 | 1.6309734 | 0.2194532 | 0.577610685 |
| ENSMUSG00000021972 | Hmbox1 | protein_coding | -0.633091506 | 3.662169594 | 1.6308388 | 0.2194714 | 0.577610685 |
| ENSMUSG00000071679 | Zcchc16 | protein_coding | -0.944236519 | 2.834051564 | 1.6294378 | 0.2196604 | 0.577610685 |
| ENSMUSG00000026279 | Thap4 | protein_coding | 0.431597414 | 4.762737754 | 1.6290958 | 0.2197066 | 0.577610685 |
| ENSMUSG00000028607 | Cpt2 | protein_coding | 0.640331827 | 3.913188687 | 1.628277 | 0.2198172 | 0.577610685 |
| ENSMUSG00000033883 | D3Ertd254e | protein_coding | -0.5396822 | 4.246837968 | 1.628213 | 0.2198258 | 0.577610685 |
| ENSMUSG00000032586 | Traip | protein_coding | 0.483119135 | 4.521045935 | 1.6277338 | 0.2198906 | 0.577610685 |
| ENSMUSG00000026123 | Plekhb2 | protein_coding | 0.574987116 | 9.104807935 | 1.6462655 | 0.2198957 | 0.577610685 |
| ENSMUSG00000034203 | Chchd4 | protein_coding | 0.287688055 | 7.239234446 | 1.6273641 | 0.2199406 | 0.577610685 |
| ENSMUSG00000046836 | Brox | protein_coding | 0.470812481 | 4.588628176 | 1.6265657 | 0.2200486 | 0.577610685 |
| ENSMUSG00000049100 | Pcdh10 | protein_coding | -0.622185782 | 5.010605996 | 1.6267171 | 0.2200572 | 0.577610685 |
| ENSMUSG00000058388 | Phtf1 | protein_coding | 0.636174996 | 4.01175604 | 1.6264742 | 0.220061 | 0.577610685 |
| ENSMUSG00000020668 | Kif3c | protein_coding | 1.202294404 | 2.57057705 | 1.626471 | 0.2200614 | 0.577610685 |
| ENSMUSG00000004347 | Pde1c | protein_coding | -0.501781929 | 5.082863341 | 1.6263672 | 0.2200754 | 0.577610685 |
| ENSMUSG00000059810 | Rgs3 | protein_coding | 0.72224509 | 3.626404954 | 1.6240005 | 0.220396 | 0.578248473 |
| ENSMUSG00000023991 | Foxp4 | protein_coding | 0.737334822 | 3.535904737 | 1.6235413 | 0.2204583 | 0.578248473 |
| ENSMUSG00000052848 | C130026L21Rik | lincRNA | 0.489127204 | 4.471491748 | 1.6226788 | 0.2205753 | 0.578248473 |
| ENSMUSG00000024695 | Zfp91 | protein_coding | -0.36577937 | 5.647609796 | 1.622277 | 0.2206298 | 0.578248473 |
| ENSMUSG00000028525 | Pde4b | protein_coding | -0.340296037 | 7.408735093 | 1.6220626 | 0.2206589 | 0.578248473 |
| ENSMUSG00000032011 | Thy1 | protein_coding | 0.3878454 | 7.286844666 | 1.6219761 | 0.2206998 | 0.578248473 |
| ENSMUSG00000030237 | Slco1a4 | protein_coding | -1.253177724 | 2.316407609 | 1.6213705 | 0.2207529 | 0.578248473 |
| ENSMUSG00000021712 | Trim23 | protein_coding | -0.546143205 | 5.5530116 | 1.6213593 | 0.2207835 | 0.578248473 |
| ENSMUSG00000040822 | 1700123O20Rik | protein_coding | -0.330350148 | 6.609926099 | 1.6210063 | 0.2208024 | 0.578248473 |
| ENSMUSG00000098332 | 2310009A05Rik | processed_transcript | -0.465908177 | 5.400427499 | 1.6198868 | 0.2209546 | 0.578506178 |
| ENSMUSG00000063804 | Lin28b | protein_coding | -0.771305596 | 3.228588473 | 1.6184062 | 0.2211561 | 0.578892783 |
| ENSMUSG00000028089 | Chd1l | protein_coding | 0.852971657 | 3.109815379 | 1.6175159 | 0.2212774 | 0.579069285 |
| ENSMUSG00000042129 | Rassf4 | protein_coding | -0.666827025 | 3.448326088 | 1.6158133 | 0.2215096 | 0.579535843 |
| ENSMUSG00000000489 | Pdgfb | protein_coding | 0.747953922 | 3.367482814 | 1.6145947 | 0.2216759 | 0.579830012 |
| ENSMUSG00000019124 | Scrn1 | protein_coding | 0.373573815 | 7.737756782 | 1.6141132 | 0.2217417 | 0.579861024 |
| ENSMUSG00000014599 | Csf1 | protein_coding | 0.801520225 | 2.841161253 | 1.6133952 | 0.2218398 | 0.579976599 |
| ENSMUSG00000028820 | Sfpq | protein_coding | -0.375125523 | 5.63432158 | 1.6128619 | 0.2219128 | 0.580026251 |
| ENSMUSG00000046093 | Hpcal4 | protein_coding | 0.305821176 | 8.117955572 | 1.6112877 | 0.2221282 | 0.580448252 |
| ENSMUSG00000000711 | Rab5b | protein_coding | 0.507371274 | 6.431023966 | 1.6170026 | 0.2224742 | 0.581119365 |
| ENSMUSG00000038059 | Smim3 | protein_coding | -0.645499393 | 3.914729862 | 1.6086259 | 0.2224931 | 0.581119365 |
| ENSMUSG00000068762 | Gstm6 | protein_coding | 0.350932326 | 6.187682757 | 1.6059758 | 0.2228572 | 0.581920493 |
| ENSMUSG00000015971 | Actr8 | protein_coding | 0.365716008 | 6.486960432 | 1.6056058 | 0.2229081 | 0.581920493 |
| ENSMUSG00000030917 | Tmem159 | protein_coding | 0.386033066 | 5.45839797 | 1.6051054 | 0.2229769 | 0.581958962 |
| ENSMUSG00000073792 | Alg6 | protein_coding | -0.613220761 | 3.70962646 | 1.6042404 | 0.223096 | 0.582128454 |
| ENSMUSG00000028394 | Pole3 | protein_coding | 0.316134969 | 6.041170929 | 1.603701 | 0.2231703 | 0.582181053 |
| ENSMUSG00000021597 | Slf1 | protein_coding | -0.551634274 | 4.400162418 | 1.6021556 | 0.2233834 | 0.582595491 |
| ENSMUSG00000030304 | Ergic2 | protein_coding | -0.317433784 | 6.167755877 | 1.6014699 | 0.223478 | 0.58264101 |
| ENSMUSG00000004187 | Kifc2 | protein_coding | -0.368917833 | 6.489650591 | 1.6012438 | 0.2235092 | 0.58264101 |
| ENSMUSG00000000594 | Gm2a | protein_coding | -0.321937822 | 7.111728144 | 1.5993994 | 0.223764 | 0.583163805 |
| ENSMUSG00000056493 | Foxk1 | protein_coding | -0.663275279 | 3.329235493 | 1.5977895 | 0.2239867 | 0.583602792 |
| ENSMUSG00000007815 | Rhoa | protein_coding | 0.430168439 | 5.819720118 | 1.5971339 | 0.2240774 | 0.583697839 |
| ENSMUSG00000024277 | Mapre2 | protein_coding | 0.364227722 | 8.384898168 | 1.5965083 | 0.2241641 | 0.583710991 |
| ENSMUSG00000026463 | Atp2b4 | protein_coding | -0.364348114 | 6.025280533 | 1.595762 | 0.2242675 | 0.583710991 |
| ENSMUSG00000028291 | Akirin2 | protein_coding | -0.432611552 | 5.350126835 | 1.5955775 | 0.2242931 | 0.583710991 |
| ENSMUSG00000040675 | Mthfd1l | protein_coding | 0.721306601 | 3.427437564 | 1.5953062 | 0.2243308 | 0.583710991 |
| ENSMUSG00000024268 | Celf4 | protein_coding | -0.407787735 | 9.063432229 | 1.5953482 | 0.2243539 | 0.583710991 |
| ENSMUSG00000033909 | Usp36 | protein_coding | -0.514979546 | 4.481393143 | 1.5938963 | 0.2245264 | 0.58389344 |

| ensembl_gene_id | external_gene_name | gene_biotype | log2 (fold change) | log2 (counts per million) | F-value | p-value | False discovery rate |
|---|---|---|---|---|---|---|---|
| ENSMUSG00000002661 | Alkbh7 | protein_coding | 0.317502336 | 6.488680113 | 1.5932984 | 0.2246095 | 0.58389344 |
| ENSMUSG00000033004 | Mycbp2 | protein_coding | -0.441825121 | 6.269808914 | 1.593306 | 0.2246374 | 0.58389344 |
| ENSMUSG00000053063 | Clec12a | protein_coding | -1.51763859 | 2.502526598 | 1.5938275 | 0.2246412 | 0.58389344 |
| ENSMUSG00000052551 | Adarb2 | protein_coding | -0.505980098 | 4.072963191 | 1.5917984 | 0.224818 | 0.584035605 |
| ENSMUSG00000038690 | Atp5j2 | protein_coding | 0.380753688 | 9.890778011 | 1.5914148 | 0.2248713 | 0.584035605 |
| ENSMUSG00000030595 | Nfkbib | protein_coding | 0.46761165 | 5.707432319 | 1.5908344 | 0.2249521 | 0.584035605 |
| ENSMUSG00000056673 | Kdm5d | protein_coding | 0.807331126 | 2.893253497 | 1.5906266 | 0.224981 | 0.584035605 |
| ENSMUSG00000045180 | Shroom2 | protein_coding | 0.387679702 | 6.313908682 | 1.5903488 | 0.2250197 | 0.584035605 |
| ENSMUSG00000002379 | Ndufa11 | protein_coding | 0.316319032 | 9.817108866 | 1.5900238 | 0.225065 | 0.584035605 |
| ENSMUSG00000037239 | Spred3 | protein_coding | 0.805126634 | 2.938059179 | 1.5895364 | 0.2251329 | 0.584035605 |
| ENSMUSG00000036333 | Kidins220 | protein_coding | -0.391005136 | 6.645574306 | 1.589476 | 0.2251413 | 0.584035605 |
| ENSMUSG00000036427 | Gpi1 | protein_coding | 0.411103931 | 9.013078224 | 1.5892767 | 0.225198 | 0.584035605 |
| ENSMUSG00000024055 | Cyp4f13 | protein_coding | 1.103105113 | 2.949855005 | 1.5887743 | 0.2252391 | 0.584035605 |
| ENSMUSG00000033557 | Fam20b | protein_coding | 0.446392479 | 5.503172762 | 1.5874353 | 0.2254259 | 0.584379035 |
| ENSMUSG00000029642 | Polr1d | protein_coding | 0.318175999 | 6.929829932 | 1.5861325 | 0.2256078 | 0.584635189 |
| ENSMUSG00000070283 | Ndufaf3 | protein_coding | 0.30442658 | 8.101443415 | 1.5859491 | 0.2256334 | 0.584635189 |
| ENSMUSG00000022899 | Slc15a2 | protein_coding | -0.779037495 | 3.892640066 | 1.583993 | 0.2259359 | 0.585120597 |
| ENSMUSG00000019917 | 10-Sep | protein_coding | -0.804292848 | 3.195238907 | 1.5823461 | 0.2261376 | 0.585120597 |
| ENSMUSG00000039952 | Dag1 | protein_coding | -0.467835524 | 4.91240035 | 1.5820428 | 0.2261801 | 0.585120597 |
| ENSMUSG00000025010 | Ccnj | protein_coding | 0.8636099 | 3.045179781 | 1.5815623 | 0.2262475 | 0.585120597 |
| ENSMUSG00000019302 | Atp6v0a1 | protein_coding | 0.33381779 | 7.714074091 | 1.5812087 | 0.2262971 | 0.585120597 |
| ENSMUSG00000030541 | Idh2 | protein_coding | 0.459104552 | 4.56323447 | 1.5811574 | 0.2263043 | 0.585120597 |
| ENSMUSG00000022570 | Tsta3 | protein_coding | 0.286138952 | 6.72415395 | 1.5811446 | 0.2263061 | 0.585120597 |
| ENSMUSG00000020792 | Exoc7 | protein_coding | 0.378326042 | 7.062563813 | 1.5810142 | 0.2263244 | 0.585120597 |
| ENSMUSG00000040356 | Skiv2l | protein_coding | 0.472903854 | 4.674149618 | 1.5807948 | 0.2263551 | 0.585120597 |
| ENSMUSG00000034958 | Atcay | protein_coding | 0.470803019 | 6.108660041 | 1.5809311 | 0.2263649 | 0.585120597 |
| ENSMUSG00000028007 | Snx7 | protein_coding | -0.600165627 | 3.934943451 | 1.5802359 | 0.2264336 | 0.585157457 |
| ENSMUSG00000030738 | Eif3c | protein_coding | 0.334686604 | 7.789052658 | 1.5797571 | 0.2265008 | 0.58519055 |
| ENSMUSG00000029790 | Cep41 | protein_coding | 0.902149128 | 3.099118787 | 1.5772418 | 0.2268544 | 0.585675938 |
| ENSMUSG00000000627 | Sema4f | protein_coding | 0.586718958 | 4.304261615 | 1.5771555 | 0.2268665 | 0.585675938 |
| ENSMUSG00000029602 | Rasal1 | protein_coding | 0.518094439 | 5.027993756 | 1.5771495 | 0.2268674 | 0.585675938 |
| ENSMUSG00000024188 | Luc7l | protein_coding | 0.311954808 | 6.515031364 | 1.5768712 | 0.2269065 | 0.585675938 |
| ENSMUSG00000033863 | Klf9 | protein_coding | -0.304233761 | 6.174847207 | 1.5763495 | 0.22698 | 0.585724912 |
| ENSMUSG00000056895 | Hist3h2ba | protein_coding | 0.648111605 | 3.746305997 | 1.5758662 | 0.227048 | 0.585759995 |
| ENSMUSG00000038884 | A230050P20Rik | protein_coding | 0.748175853 | 4.417019797 | 1.5786709 | 0.2272013 | 0.586014655 |
| ENSMUSG00000047284 | Neurl4 | protein_coding | -0.544982381 | 5.23873819 | 1.5727221 | 0.2275203 | 0.586696922 |
| ENSMUSG00000023328 | Ache | protein_coding | 0.329105369 | 6.236611995 | 1.5720546 | 0.2275858 | 0.586725034 |
| ENSMUSG00000023030 | Slc11a2 | protein_coding | 0.616656793 | 4.586060056 | 1.5708838 | 0.2277513 | 0.58690958 |
| ENSMUSG00000063160 | Numbl | protein_coding | 0.681698978 | 3.21841183 | 1.570776 | 0.2277665 | 0.58690958 |
| ENSMUSG00000049321 | Zfp2 | protein_coding | 0.568498471 | 4.650543512 | 1.5700584 | 0.2278681 | 0.586956994 |
| ENSMUSG00000026110 | Mgat4a | protein_coding | -0.529419379 | 4.364640138 | 1.5698744 | 0.2278941 | 0.586956994 |
| ENSMUSG00000026784 | Pdss1 | protein_coding | -0.712098608 | 3.362814986 | 1.5689555 | 0.2280242 | 0.587151506 |
| ENSMUSG00000007564 | Ppp2r1a | protein_coding | 0.348304128 | 9.864951749 | 1.5684043 | 0.2281023 | 0.587211978 |
| ENSMUSG00000027316 | Gfra4 | protein_coding | 0.431015167 | 5.032923443 | 1.5679707 | 0.2281638 | 0.587229605 |
| ENSMUSG00000039745 | Htatip2 | protein_coding | 0.575172121 | 4.621718336 | 1.5668392 | 0.2283243 | 0.587501999 |
| ENSMUSG00000031224 | Magee2 | protein_coding | 0.540193776 | 5.034566242 | 1.5661079 | 0.2284281 | 0.587522338 |
| ENSMUSG00000058392 | Rrp1b | protein_coding | 0.467196066 | 4.924290776 | 1.5658857 | 0.2284596 | 0.587522338 |
| ENSMUSG00000098112 | Bin2 | protein_coding | -1.438136003 | 2.31569101 | 1.5656289 | 0.2284961 | 0.587522338 |
| ENSMUSG00000047067 | Dusp28 | protein_coding | 0.52992953 | 3.811464744 | 1.5624602 | 0.2289467 | 0.588404283 |
| ENSMUSG00000045215 | Asxl3 | protein_coding | -0.720391433 | 3.130231723 | 1.5624474 | 0.2289485 | 0.588404283 |
| ENSMUSG00000003929 | Zfp81 | protein_coding | -0.763716329 | 3.661235405 | 1.5619494 | 0.2290195 | 0.588445935 |
| ENSMUSG00000025410 | Dctn2 | protein_coding | 0.366629515 | 8.007037595 | 1.56093 | 0.2291647 | 0.58846643 |
| ENSMUSG00000025280 | Polr3a | protein_coding | 0.642761402 | 3.709208896 | 1.5607209 | 0.2291945 | 0.58846643 |
| ENSMUSG00000028959 | Fastk | protein_coding | 0.346070054 | 6.605509675 | 1.5599934 | 0.2292983 | 0.58846643 |
| ENSMUSG00000054667 | Irs4 | protein_coding | 0.569839944 | 4.720075632 | 1.5597653 | 0.2293308 | 0.58846643 |
| ENSMUSG00000087259 | 2610035D17Rik | lincRNA | -0.40355162 | 5.366582611 | 1.5595944 | 0.2293552 | 0.58846643 |
| ENSMUSG00000061024 | Rrs1 | protein_coding | -0.660286021 | 3.855579368 | 1.5592359 | 0.2294064 | 0.58846643 |
| ENSMUSG00000041308 | Sntb2 | protein_coding | -1.102807507 | 2.552772567 | 1.5592072 | 0.2294105 | 0.58846643 |
| ENSMUSG00000014837 | 4931428F04Rik | protein_coding | 0.642726303 | 5.338054235 | 1.5664833 | 0.2294902 | 0.588530353 |
| ENSMUSG00000058706 | 0610030E20Rik | protein_coding | 0.919448598 | 2.85645559 | 1.5579988 | 0.2295831 | 0.588628349 |
| ENSMUSG00000040794 | C1qtnf4 | protein_coding | -0.822901518 | 3.310333493 | 1.557217 | 0.2296949 | 0.588774471 |
| ENSMUSG00000071253 | Slc25a16 | protein_coding | -0.487112003 | 4.729946211 | 1.5554074 | 0.2299538 | 0.589297755 |
| ENSMUSG00000058655 | Eif4b | protein_coding | 0.428824641 | 7.164427178 | 1.5570522 | 0.2300517 | 0.589408216 |
| ENSMUSG00000005505 | Kbtbd4 | protein_coding | 0.680018458 | 3.955570353 | 1.5538058 | 0.2301833 | 0.5894666 |
| ENSMUSG00000035493 | Tgfbi | protein_coding | -1.252149942 | 2.223095746 | 1.5538 | 0.2301842 | 0.5894666 |
| ENSMUSG00000031105 | Slc25a14 | protein_coding | 0.295528745 | 7.199902593 | 1.5528093 | 0.2303263 | 0.589690073 |
| ENSMUSG00000079215 | Zfp664 | protein_coding | -0.440679082 | 4.983712089 | 1.5512717 | 0.230547 | 0.590057235 |
| ENSMUSG00000019818 | Cd164 | protein_coding | 0.365429405 | 6.237024336 | 1.5501373 | 0.2307101 | 0.590057235 |
| ENSMUSG00000027185 | Nat10 | protein_coding | 0.618005264 | 3.557744579 | 1.5501307 | 0.2307111 | 0.590057235 |
| ENSMUSG00000015597 | Zfp318 | protein_coding | 0.486495835 | 4.926958998 | 1.5499585 | 0.2307358 | 0.590057235 |
| ENSMUSG00000021482 | Aaed1 | protein_coding | -0.547908652 | 4.010025099 | 1.5493815 | 0.2308189 | 0.590057235 |
| ENSMUSG00000021951 | Eef1akmt1 | protein_coding | 0.264715363 | 7.540472218 | 1.5493278 | 0.2308266 | 0.590057235 |
| ENSMUSG00000106981 | Gm45495 | lincRNA | -0.89687708 | 3.014933517 | 1.5491387 | 0.2308538 | 0.590057235 |
| ENSMUSG00000033487 | Fndc3a | protein_coding | -0.375608041 | 5.590292676 | 1.5480422 | 0.2310117 | 0.590320464 |
| ENSMUSG00000026473 | Glul | protein_coding | -0.306585277 | 8.303558326 | 1.5473574 | 0.2311104 | 0.59043233 |
| ENSMUSG00000027520 | Zdbf2 | protein_coding | -0.572217053 | 4.992381259 | 1.5466743 | 0.2312375 | 0.590535477 |
| ENSMUSG00000018547 | Pip4k2b | protein_coding | 0.55957069 | 4.264914796 | 1.5463157 | 0.2312606 | 0.590535477 |
| ENSMUSG00000030302 | Atp2b2 | protein_coding | -0.322522911 | 8.150041567 | 1.5447856 | 0.2314814 | 0.590959099 |

| ensembl_gene_id | external_gene_name | gene_biotype | log2 (fold change) | log2 (counts per million) | F-value | p-value | False discovery rate |
|---|---|---|---|---|---|---|---|
| ENSMUSG00000040651 | Fam208a | protein_coding | -0.710355009 | 3.492858956 | 1.5434765 | 0.2316706 | 0.591301706 |
| ENSMUSG00000028778 | Hcrtr1 | protein_coding | 2.030919386 | 3.40245049 | 1.614241 | 0.2320703 | 0.592141533 |
| ENSMUSG00000059326 | Csf2ra | protein_coding | 0.922760404 | 2.661422324 | 1.5404428 | 0.2321098 | 0.592141533 |
| ENSMUSG00000048078 | Tenm4 | protein_coding | 0.603156814 | 4.19453274 | 1.5390552 | 0.232311 | 0.592246756 |
| ENSMUSG00000042010 | Acacb | protein_coding | 0.74773292 | 4.116900543 | 1.538907 | 0.2323611 | 0.592246756 |
| ENSMUSG00000040055 | Gjb6 | protein_coding | -0.528654897 | 5.585771799 | 1.5384911 | 0.2324215 | 0.592246756 |
| ENSMUSG00000051391 | Ywhag | protein_coding | 0.378365104 | 9.147746212 | 1.5381515 | 0.2324422 | 0.592246756 |
| ENSMUSG00000056305 | Usp39 | protein_coding | 0.472730933 | 4.208385166 | 1.5381382 | 0.2324441 | 0.592246756 |
| ENSMUSG00000022433 | Csnk1e | protein_coding | -0.480212172 | 5.460540342 | 1.5378495 | 0.2324861 | 0.592246756 |
| ENSMUSG00000079339 | Ifit1bl1 | protein_coding | 1.896724159 | 2.273117401 | 1.5451674 | 0.2325366 | 0.592246756 |
| ENSMUSG00000015889 | Lta4h | protein_coding | 0.349519541 | 6.221510192 | 1.5370462 | 0.2326028 | 0.592275083 |
| ENSMUSG00000044308 | Ubr3 | protein_coding | 0.469058355 | 6.311988584 | 1.5366278 | 0.2327676 | 0.592554424 |
| ENSMUSG00000054199 | Gon4l | protein_coding | -0.811214042 | 4.01507341 | 1.5379002 | 0.2328227 | 0.592554424 |
| ENSMUSG00000029406 | Pitpnm2 | protein_coding | -0.45486308 | 5.065491646 | 1.5350501 | 0.2328931 | 0.592593428 |
| ENSMUSG00000027936 | Crtc2 | protein_coding | 0.857205491 | 3.167573966 | 1.5345437 | 0.2329669 | 0.592612192 |
| ENSMUSG00000027852 | Nras | protein_coding | -0.272929497 | 7.041033276 | 1.5337758 | 0.2330787 | 0.592612192 |
| ENSMUSG00000047921 | Trappc9 | protein_coding | 0.431342521 | 6.268041293 | 1.5337189 | 0.2331156 | 0.592612192 |
| ENSMUSG00000041438 | Utp4 | protein_coding | 0.503893495 | 5.843235692 | 1.5336824 | 0.2331209 | 0.592612192 |
| ENSMUSG00000015745 | Plekho1 | protein_coding | -0.809853446 | 3.719621721 | 1.5327437 | 0.2332578 | 0.592819913 |
| ENSMUSG00000041319 | Thoc6 | protein_coding | 0.705784028 | 4.222495636 | 1.5313102 | 0.233467 | 0.593211371 |
| ENSMUSG00000039754 | Alkbh4 | protein_coding | 0.664655443 | 4.552657248 | 1.5302727 | 0.2336185 | 0.593456222 |
| ENSMUSG00000036667 | Tcaf1 | protein_coding | -0.352748861 | 7.530609626 | 1.529542 | 0.2336968 | 0.593514771 |
| ENSMUSG00000035864 | Syt1 | protein_coding | -0.302292811 | 9.368887695 | 1.5283707 | 0.2338681 | 0.593809717 |
| ENSMUSG00000050229 | Pigm | protein_coding | -0.815168711 | 2.779106092 | 1.527504 | 0.233995 | 0.593991671 |
| ENSMUSG00000002395 | Use1 | protein_coding | 0.305882184 | 8.642709291 | 1.5266494 | 0.2341203 | 0.594169289 |
| ENSMUSG00000028944 | Prkag2 | protein_coding | 0.354439884 | 5.982183124 | 1.525969 | 0.23422 | 0.594196401 |
| ENSMUSG00000026792 | Lrsam1 | protein_coding | 0.680382905 | 4.490933125 | 1.5260174 | 0.2342415 | 0.594196401 |
| ENSMUSG00000031775 | Pllp | protein_coding | 0.480522975 | 4.431402865 | 1.5232631 | 0.2346173 | 0.594802988 |
| ENSMUSG00000039963 | Ccdc40 | protein_coding | 0.718764393 | 3.778706287 | 1.5231628 | 0.234632 | 0.594802988 |
| ENSMUSG00000025537 | Phkg1 | protein_coding | 0.769644645 | 3.493481303 | 1.5230642 | 0.2346465 | 0.594802988 |
| ENSMUSG00000027983 | Cyp2u1 | protein_coding | 0.559007537 | 4.690115659 | 1.5213709 | 0.2348956 | 0.595294075 |
| ENSMUSG00000041870 | Ankrd13a | protein_coding | -0.578354987 | 3.997599059 | 1.5202754 | 0.235057 | 0.595527963 |
| ENSMUSG00000078619 | Smarcd2 | protein_coding | -1.258878275 | 2.691362152 | 1.5201857 | 0.2350987 | 0.595527963 |
| ENSMUSG00000087620 | 5330434G04Rik | lincRNA | -0.575157779 | 4.666725903 | 1.5180931 | 0.2353788 | 0.596097039 |
| ENSMUSG00000072653 | Zfp783 | transcribed_unproces | 1.048076276 | 2.885958419 | 1.5174084 | 0.2354799 | 0.59621264 |
| ENSMUSG00000043061 | Tmem18 | protein_coding | 0.468368492 | 4.516317477 | 1.5161936 | 0.2356594 | 0.59652662 |
| ENSMUSG00000037720 | Tmem33 | protein_coding | -0.395183563 | 6.126478516 | 1.5155011 | 0.2357618 | 0.596571238 |
| ENSMUSG00000066196 | Spag8 | protein_coding | 0.779938743 | 3.189612317 | 1.5148364 | 0.2358601 | 0.596571238 |
| ENSMUSG00000032112 | Trappc4 | protein_coding | 0.260654546 | 8.58449716 | 1.5139562 | 0.2359904 | 0.596571238 |
| ENSMUSG00000040520 | Manea | protein_coding | -0.554337839 | 4.031956797 | 1.5138384 | 0.2360079 | 0.596571238 |
| ENSMUSG00000027456 | Sdcbp2 | protein_coding | -0.994189236 | 3.053528154 | 1.5137984 | 0.2360138 | 0.596571238 |
| ENSMUSG00000037949 | Ano10 | protein_coding | 0.613226196 | 4.049037285 | 1.5132588 | 0.2360937 | 0.596571238 |
| ENSMUSG00000078570 | 1110065P20Rik | protein_coding | 1.021656056 | 2.752640373 | 1.5130841 | 0.2361196 | 0.596571238 |
| ENSMUSG00000026869 | Psmd5 | protein_coding | -0.540301234 | 6.091297663 | 1.5202522 | 0.2361645 | 0.596571238 |
| ENSMUSG00000052144 | Ppp4r2 | protein_coding | -0.423560904 | 4.914873638 | 1.51234 | 0.2362299 | 0.596571238 |
| ENSMUSG00000002459 | Rgs20 | protein_coding | -0.379837218 | 6.075147803 | 1.5118518 | 0.2363023 | 0.596571238 |
| ENSMUSG00000030393 | Zik1 | protein_coding | -0.694692488 | 3.691285119 | 1.511697 | 0.2363253 | 0.596571238 |
| ENSMUSG00000047123 | Ticam1 | protein_coding | -1.078009505 | 2.958278918 | 1.5117712 | 0.2363427 | 0.596571238 |
| ENSMUSG00000055022 | Cntn1 | protein_coding | -0.273790892 | 7.891555497 | 1.5085888 | 0.2367871 | 0.597552501 |
| ENSMUSG00000040928 | S100pbp | protein_coding | -0.407709914 | 5.027817843 | 1.5075694 | 0.2369388 | 0.597795045 |
| ENSMUSG00000090733 | Rps27 | protein_coding | 0.505450482 | 4.099858358 | 1.5067714 | 0.2370576 | 0.597893929 |
| ENSMUSG00000020334 | Slc22a4 | protein_coding | 0.799926558 | 3.121545902 | 1.5065592 | 0.2370892 | 0.597893929 |
| ENSMUSG00000014504 | Srp19 | protein_coding | 0.339576867 | 8.272780773 | 1.5058392 | 0.2371965 | 0.597893929 |
| ENSMUSG00000047714 | Ppp1r2 | protein_coding | -0.437629838 | 5.611116276 | 1.5051328 | 0.2373019 | 0.597893929 |
| ENSMUSG00000020978 | Klhdc2 | protein_coding | 0.308331002 | 6.554160516 | 1.5051103 | 0.2373052 | 0.597893929 |
| ENSMUSG00000005506 | Celf1 | protein_coding | -0.293793191 | 6.52656855 | 1.5050677 | 0.2373116 | 0.597893929 |
| ENSMUSG00000021661 | Ankra2 | protein_coding | -0.382985396 | 5.411818967 | 1.5043996 | 0.2374113 | 0.598004981 |
| ENSMUSG00000045996 | Polr2k | protein_coding | 0.516440067 | 4.235161033 | 1.5035677 | 0.2375355 | 0.598177723 |
| ENSMUSG00000027220 | Syt13 | protein_coding | 0.314682241 | 6.202347776 | 1.5021304 | 0.2377503 | 0.598509298 |
| ENSMUSG00000033306 | Lpp | protein_coding | -0.631933997 | 3.491464405 | 1.5019419 | 0.2377785 | 0.598509298 |
| ENSMUSG00000027222 | Pex16 | protein_coding | 0.361288488 | 6.157758827 | 1.5007555 | 0.237956 | 0.598623337 |
| ENSMUSG00000027782 | Kpna4 | protein_coding | -0.357359962 | 5.463499398 | 1.5006216 | 0.237976 | 0.598623337 |
| ENSMUSG00000020413 | Hus1 | protein_coding | -0.792306079 | 3.090170719 | 1.5005231 | 0.2379908 | 0.598623337 |
| ENSMUSG00000064225 | Paqr9 | protein_coding | 0.58930532 | 4.614825467 | 1.4989141 | 0.2382319 | 0.598829295 |
| ENSMUSG00000003955 | Fam162a | protein_coding | 0.406368025 | 6.252638151 | 1.4988695 | 0.2382385 | 0.598829295 |
| ENSMUSG00000023452 | Pisd | protein_coding | 0.347796734 | 5.562063979 | 1.4988616 | 0.2382397 | 0.598829295 |
| ENSMUSG00000017721 | Pigt | protein_coding | 0.567468121 | 4.034348605 | 1.4978788 | 0.2383871 | 0.598895244 |
| ENSMUSG00000022223 | Sdr39u1 | protein_coding | 0.273691019 | 8.470249167 | 1.4976156 | 0.2384266 | 0.598895244 |
| ENSMUSG00000050552 | Lamtor4 | protein_coding | 0.331565874 | 7.544119523 | 1.4975729 | 0.238433 | 0.598895244 |
| ENSMUSG00000028426 | Rad23b | protein_coding | -0.371630084 | 5.85898496 | 1.4968844 | 0.2385364 | 0.599014975 |
| ENSMUSG00000007891 | Ctsd | protein_coding | 0.347014188 | 6.348899188 | 1.4958286 | 0.2386951 | 0.599250307 |
| ENSMUSG00000029534 | St7 | protein_coding | 0.461851124 | 4.278496417 | 1.495519 | 0.2387416 | 0.599250307 |
| ENSMUSG00000032350 | Gclc | protein_coding | -0.413996771 | 5.5127045 | 1.4950705 | 0.238809 | 0.599279703 |
| ENSMUSG00000027712 | Anxa5 | protein_coding | -0.357019301 | 7.370112413 | 1.4940198 | 0.2389671 | 0.599536517 |
| ENSMUSG00000019373 | Cops3 | protein_coding | 0.261085325 | 7.154969852 | 1.4932482 | 0.2390833 | 0.599688123 |
| ENSMUSG00000029640 | Usp12 | protein_coding | 0.806026094 | 3.012577258 | 1.4920583 | 0.2392626 | 0.599997948 |
| ENSMUSG00000000253 | Gmpr | protein_coding | 0.268587392 | 7.122825237 | 1.4913478 | 0.2393698 | 0.60012673 |
| ENSMUSG00000038594 | Cep85l | protein_coding | -1.006834104 | 3.084781103 | 1.4901764 | 0.2395749 | 0.600430672 |

| ensembl_gene_id | external_gene_name | gene_biotype | log2 (fold change) | log2 (counts per million) | F-value | p-value | False discovery rate |
|---|---|---|---|---|---|---|---|
| ENSMUSG00000072980 | Oip5 | protein_coding | -0.677374516 | 3.608398567 | 1.4896919 | 0.2396198 | 0.600430672 |
| ENSMUSG00000039706 | Ldb2 | protein_coding | -0.343582094 | 6.280817788 | 1.488809 | 0.2397532 | 0.600430672 |
| ENSMUSG00000026944 | Abca2 | protein_coding | -0.405006514 | 6.357412888 | 1.4883876 | 0.2398169 | 0.600430672 |
| ENSMUSG00000097207 | 6030443J06Rik | lincRNA | -0.720348485 | 3.104366737 | 1.4883417 | 0.2398239 | 0.600430672 |
| ENSMUSG00000053963 | Stum | protein_coding | -0.446666301 | 4.746367882 | 1.4883272 | 0.2398261 | 0.600430672 |
| ENSMUSG00000040722 | Scamp5 | protein_coding | 0.276686874 | 7.379337979 | 1.4873898 | 0.2399679 | 0.600504488 |
| ENSMUSG00000020864 | Ankrd40 | protein_coding | -0.37429389 | 5.976955726 | 1.4873038 | 0.2399809 | 0.600504488 |
| ENSMUSG00000031399 | Fam3a | protein_coding | 0.413106709 | 5.110490453 | 1.4863315 | 0.2401281 | 0.600504488 |
| ENSMUSG00000026028 | Trak2 | protein_coding | -0.49665399 | 5.226720733 | 1.4856345 | 0.2402338 | 0.600504488 |
| ENSMUSG00000030264 | Thumpd3 | protein_coding | -0.453527954 | 4.766676556 | 1.4851511 | 0.240307 | 0.600504488 |
| ENSMUSG00000003974 | Grm3 | protein_coding | -0.480054494 | 4.983939823 | 1.4850235 | 0.2403264 | 0.600504488 |
| ENSMUSG00000028221 | Tmem55a | protein_coding | -0.275746685 | 7.032470232 | 1.4846201 | 0.2403876 | 0.600504488 |
| ENSMUSG00000018651 | Tada2a | protein_coding | 0.527275269 | 5.035906239 | 1.4844358 | 0.2404155 | 0.600504488 |
| ENSMUSG00000037722 | Gnpnat1 | protein_coding | -0.475658749 | 4.307557478 | 1.4844321 | 0.2404161 | 0.600504488 |
| ENSMUSG00000004798 | Ulk2 | protein_coding | -0.461460839 | 5.441127595 | 1.4843164 | 0.2404337 | 0.600504488 |
| ENSMUSG00000042569 | Dhrs7b | protein_coding | 0.400603442 | 5.481559796 | 1.4837203 | 0.2405241 | 0.600504488 |
| ENSMUSG00000075700 | Selenot | protein_coding | -0.415279652 | 5.747603591 | 1.48371 | 0.2405257 | 0.600504488 |
| ENSMUSG00000037217 | Syn1 | protein_coding | -0.348343825 | 6.267628301 | 1.4830565 | 0.2406249 | 0.600612779 |
| ENSMUSG00000058441 | Panx2 | protein_coding | -0.561554061 | 4.069168426 | 1.4818841 | 0.2408031 | 0.600777293 |
| ENSMUSG00000024858 | Grk2 | protein_coding | 0.348193337 | 6.36618441 | 1.4816505 | 0.2408386 | 0.600777293 |
| ENSMUSG00000033361 | Prrg3 | protein_coding | -0.59995826 | 4.063571141 | 1.4815201 | 0.2408584 | 0.600777293 |
| ENSMUSG00000020671 | Rab10 | protein_coding | -0.785854764 | 2.972997929 | 1.4808991 | 0.2409529 | 0.600803942 |
| ENSMUSG00000062867 | Impdh2 | protein_coding | 0.555190965 | 4.062557865 | 1.4807154 | 0.2409809 | 0.600803942 |
| ENSMUSG00000028909 | Ptpru | protein_coding | 0.797919777 | 3.507827806 | 1.4800106 | 0.2410882 | 0.600932129 |
| ENSMUSG00000095325 | Zfp870 | protein_coding | -0.668442295 | 3.837029677 | 1.4775995 | 0.2414557 | 0.601611112 |
| ENSMUSG00000058317 | Ube2e2 | protein_coding | 0.299180647 | 6.828385677 | 1.4774895 | 0.2414725 | 0.601611112 |
| ENSMUSG00000015094 | Npdc1 | protein_coding | 0.406473007 | 6.540215449 | 1.4769841 | 0.2415778 | 0.601640874 |
| ENSMUSG00000025487 | Psmd13 | protein_coding | 0.281236656 | 7.360820135 | 1.4766385 | 0.2416024 | 0.601640874 |
| ENSMUSG00000000948 | Gm38393 | protein_coding | -0.461337346 | 4.792417404 | 1.4761602 | 0.2416754 | 0.601640874 |
| ENSMUSG00000018076 | Med13l | protein_coding | -0.443208021 | 5.03335934 | 1.4759456 | 0.2417082 | 0.601640874 |
| ENSMUSG00000035863 | Palm | protein_coding | 0.389533244 | 5.592000077 | 1.4746359 | 0.2419084 | 0.601894224 |
| ENSMUSG00000024033 | Rsph1 | protein_coding | -0.362219187 | 5.553946355 | 1.4744408 | 0.2419383 | 0.601894224 |
| ENSMUSG00000085558 | 4930412C18Rik | processed_transcript | -0.658525418 | 3.611819385 | 1.4740252 | 0.2420019 | 0.601894224 |
| ENSMUSG00000027014 | Cwc22 | protein_coding | -0.70032975 | 3.187849467 | 1.4726994 | 0.2422049 | 0.601894224 |
| ENSMUSG00000042350 | Arel1 | protein_coding | 0.585181624 | 4.131847776 | 1.4725711 | 0.2422245 | 0.601894224 |
| ENSMUSG00000052139 | Babam2 | protein_coding | 0.329354818 | 6.603073595 | 1.4723102 | 0.2422645 | 0.601894224 |
| ENSMUSG00000002728 | Naa20 | protein_coding | 0.286420594 | 7.108867017 | 1.4722649 | 0.2422714 | 0.601894224 |
| ENSMUSG00000006289 | Osgep | protein_coding | 0.555152247 | 4.89572541 | 1.4717686 | 0.2423475 | 0.601894224 |
| ENSMUSG00000021337 | Scgn | protein_coding | -0.808002088 | 3.085266767 | 1.4705886 | 0.2425285 | 0.601894224 |
| ENSMUSG00000024325 | Ring1 | protein_coding | 0.891993244 | 2.785337148 | 1.4705292 | 0.2425376 | 0.601894224 |
| ENSMUSG00000024899 | Papss2 | protein_coding | -0.652257223 | 3.466976227 | 1.4704702 | 0.2425467 | 0.601894224 |
| ENSMUSG00000041623 | D11Wsu47e | protein_coding | 0.661509713 | 3.499927593 | 1.470443 | 0.2425509 | 0.601894224 |
| ENSMUSG00000042043 | Tbca | protein_coding | 0.373155291 | 7.893308059 | 1.4705098 | 0.2425688 | 0.601894224 |
| ENSMUSG00000047888 | Tnrc6b | protein_coding | -0.424468182 | 4.947256467 | 1.4701643 | 0.2425937 | 0.601894224 |
| ENSMUSG00000033998 | Kcnk1 | protein_coding | 0.723075719 | 3.67952757 | 1.4686619 | 0.2428244 | 0.602238248 |
| ENSMUSG00000019810 | Fuca2 | protein_coding | 0.367590839 | 6.321853766 | 1.4685326 | 0.2428443 | 0.602238248 |
| ENSMUSG00000025857 | Dnaaf5 | protein_coding | -2.526177206 | 1.77534841 | 1.4740732 | 0.2430867 | 0.602441769 |
| ENSMUSG00000026347 | Tmem163 | protein_coding | -0.605951004 | 4.287206901 | 1.4657134 | 0.2432782 | 0.602441769 |
| ENSMUSG00000066148 | Prpf4 | protein_coding | 0.468726418 | 4.626979966 | 1.4656735 | 0.2432844 | 0.602441769 |
| ENSMUSG00000007659 | Bcl2l1 | protein_coding | 0.293400612 | 8.134873366 | 1.4654739 | 0.2433151 | 0.602441769 |
| ENSMUSG00000097431 | Gm26782 | lincRNA | 0.386329037 | 5.107348508 | 1.4654094 | 0.2433251 | 0.602441769 |
| ENSMUSG00000050069 | Grem2 | protein_coding | 0.675953991 | 3.546617738 | 1.4652174 | 0.2433547 | 0.602441769 |
| ENSMUSG00000014850 | Msh3 | protein_coding | -0.589958761 | 3.962347534 | 1.465129 | 0.2433683 | 0.602441769 |
| ENSMUSG00000030224 | Strap | protein_coding | -0.294122267 | 6.523884665 | 1.4650881 | 0.2433746 | 0.602441769 |
| ENSMUSG00000033460 | Armcx1 | protein_coding | 0.355731541 | 6.736641341 | 1.4636108 | 0.2436025 | 0.602867101 |
| ENSMUSG00000031508 | Ankrd10 | protein_coding | -0.581606026 | 4.519871206 | 1.4627352 | 0.2437377 | 0.603062911 |
| ENSMUSG00000055407 | Map6 | protein_coding | -0.45384643 | 5.469052074 | 1.4613278 | 0.2439552 | 0.603462247 |
| ENSMUSG00000009090 | Ap1b1 | protein_coding | 0.370953285 | 7.029784393 | 1.4582752 | 0.244456 | 0.603928434 |
| ENSMUSG00000024169 | Ift140 | protein_coding | 0.782302083 | 2.989982275 | 1.4579155 | 0.2444837 | 0.603928434 |
| ENSMUSG00000044150 | A830080D01Rik | protein_coding | -0.824980302 | 3.460706837 | 1.4576078 | 0.2445314 | 0.603928434 |
| ENSMUSG00000021518 | Ptdss1 | protein_coding | 0.652039166 | 4.259465503 | 1.457441 | 0.2445853 | 0.603928434 |
| ENSMUSG00000053166 | Cdh22 | protein_coding | -0.945778828 | 2.911345034 | 1.4567287 | 0.2446678 | 0.603928434 |
| ENSMUSG00000026657 | Frmd4a | protein_coding | -0.441952451 | 4.795458317 | 1.456577 | 0.2446914 | 0.603928434 |
| ENSMUSG00000037606 | Osbpl5 | protein_coding | 0.524322776 | 4.151074454 | 1.4565476 | 0.2446959 | 0.603928434 |
| ENSMUSG00000031343 | Gabra3 | protein_coding | 0.420810246 | 5.587913627 | 1.4565395 | 0.2446972 | 0.603928434 |
| ENSMUSG00000078958 | Atp6ap1l | protein_coding | -0.695047725 | 3.491809677 | 1.4565369 | 0.2446976 | 0.603928434 |
| ENSMUSG00000037111 | Setd7 | protein_coding | -0.420692035 | 4.984668212 | 1.4562394 | 0.2447438 | 0.603928434 |
| ENSMUSG00000032083 | Apoa1 | protein_coding | 0.715938231 | 3.085571856 | 1.4561256 | 0.2447615 | 0.603928434 |
| ENSMUSG00000028684 | Urod | protein_coding | 0.353341998 | 7.131386531 | 1.4554919 | 0.2448599 | 0.60403274 |
| ENSMUSG00000024948 | Map4k2 | protein_coding | 0.734544974 | 4.428703934 | 1.4590138 | 0.2450297 | 0.604303643 |
| ENSMUSG00000021686 | Ap3b1 | protein_coding | 0.479818599 | 4.754380161 | 1.4538234 | 0.2451194 | 0.604303643 |
| ENSMUSG00000055681 | Cope | protein_coding | 0.259496287 | 9.279655957 | 1.4537016 | 0.2451383 | 0.604303643 |
| ENSMUSG00000024725 | Ostf1 | protein_coding | -0.728229758 | 3.297632081 | 1.4519079 | 0.2454177 | 0.604853674 |
| ENSMUSG00000021314 | Amph | protein_coding | 0.461109087 | 7.151884838 | 1.4562771 | 0.2458944 | 0.605472263 |
| ENSMUSG00000051510 | Mafg | protein_coding | -0.358218053 | 5.839539025 | 1.4486641 | 0.245924 | 0.605472263 |
| ENSMUSG00000032485 | Scap | protein_coding | 0.505205827 | 4.527274056 | 1.4485792 | 0.2459372 | 0.605472263 |
| ENSMUSG00000034709 | Ppp1r21 | protein_coding | -0.372693669 | 5.805066777 | 1.447588 | 0.2460922 | 0.605472263 |
| ENSMUSG00000028066 | Pmf1 | protein_coding | 0.458656122 | 4.495058868 | 1.4471728 | 0.2461572 | 0.605472263 |

| ensembl_gene_id | external_gene_name | gene_biotype | log2 (fold change) | log2 (counts per million) | F-value | p-value | False discovery rate |
|---|---|---|---|---|---|---|---|
| ENSMUSG00000057147 | Dph6 | protein_coding | 0.381084465 | 5.277548766 | 1.4470176 | 0.2461815 | 0.605472263 |
| ENSMUSG00000025420 | Katnal2 | protein_coding | -0.655130058 | 3.421721063 | 1.4468348 | 0.2462101 | 0.605472263 |
| ENSMUSG00000030284 | Creld1 | protein_coding | 0.359904623 | 6.220781924 | 1.4466077 | 0.2462456 | 0.605472263 |
| ENSMUSG00000038957 | Edc3 | protein_coding | 0.455389751 | 5.546286261 | 1.4458694 | 0.2463612 | 0.605472263 |
| ENSMUSG00000021791 | Dydc2 | protein_coding | -0.820934473 | 2.879453552 | 1.4455417 | 0.2464126 | 0.605472263 |
| ENSMUSG00000047496 | Rnf152 | protein_coding | -0.65125466 | 3.959589499 | 1.4451702 | 0.2464708 | 0.605472263 |
| ENSMUSG00000020180 | Snrpd3 | protein_coding | 0.314015513 | 8.641579732 | 1.4449532 | 0.2465048 | 0.605472263 |
| ENSMUSG00000071645 | Tut1 | protein_coding | 0.456805668 | 5.181577807 | 1.4447339 | 0.2465392 | 0.605472263 |
| ENSMUSG00000020273 | Papolg | protein_coding | -0.537016273 | 4.466994118 | 1.4447294 | 0.2465399 | 0.605472263 |
| ENSMUSG00000021748 | Pdhb | protein_coding | 0.254256288 | 7.858428937 | 1.4446146 | 0.2465579 | 0.605472263 |
| ENSMUSG00000026904 | Slc4a10 | protein_coding | -0.42287545 | 5.330997393 | 1.44454 | 0.2465696 | 0.605472263 |
| ENSMUSG00000028785 | Hpca | protein_coding | 0.324632176 | 9.923674582 | 1.4436437 | 0.2467102 | 0.605560473 |
| ENSMUSG00000029290 | Zfp326 | protein_coding | -0.512439418 | 4.101239666 | 1.4435931 | 0.2467182 | 0.605560473 |
| ENSMUSG00000046709 | Mapk10 | protein_coding | 0.333160728 | 8.463617021 | 1.4430523 | 0.2468031 | 0.605562862 |
| ENSMUSG00000047648 | Fbxo30 | protein_coding | -0.862030084 | 2.810722436 | 1.4428694 | 0.2468318 | 0.605562862 |
| ENSMUSG00000069835 | Sat2 | protein_coding | 0.368763742 | 5.743220845 | 1.4420476 | 0.2469609 | 0.605741387 |
| ENSMUSG00000002280 | Narfl | protein_coding | 0.687972804 | 3.723069689 | 1.4403187 | 0.2472328 | 0.606203102 |
| ENSMUSG00000024647 | Cbln2 | protein_coding | -0.461317281 | 4.755811278 | 1.4401338 | 0.2472619 | 0.606203102 |
| ENSMUSG00000024548 | Setbp1 | protein_coding | -0.71334597 | 3.466152354 | 1.4392739 | 0.2473973 | 0.606369736 |
| ENSMUSG00000105867 | Gm42517 | protein_coding | -0.803960594 | 2.859212297 | 1.4389578 | 0.2474471 | 0.606369736 |
| ENSMUSG00000031256 | Cstf2 | protein_coding | -0.571100663 | 4.311827684 | 1.4386282 | 0.247499 | 0.606369736 |
| ENSMUSG00000002797 | Ggct | protein_coding | 0.405646965 | 6.464061391 | 1.437976 | 0.2476298 | 0.606503563 |
| ENSMUSG00000002014 | Ssr4 | protein_coding | 0.35704311 | 7.614212691 | 1.4375663 | 0.2476664 | 0.606503563 |
| ENSMUSG00000033389 | Arhgap44 | protein_coding | 0.442219142 | 8.415104961 | 1.4438622 | 0.2477599 | 0.606594403 |
| ENSMUSG00000071528 | Usmg5 | protein_coding | 0.377229476 | 7.503552794 | 1.4367712 | 0.2478198 | 0.606600867 |
| ENSMUSG00000071415 | Rpl23 | protein_coding | 0.375295245 | 7.520585782 | 1.4357451 | 0.2479818 | 0.606600867 |
| ENSMUSG00000041035 | Gm17018 | protein_coding | 0.273592545 | 8.814747378 | 1.435564 | 0.2479825 | 0.606600867 |
| ENSMUSG00000032258 | Lca5 | protein_coding | -1.480564183 | 2.502113867 | 1.4384401 | 0.2480306 | 0.606600867 |
| ENSMUSG00000045875 | Adra1a | protein_coding | -0.658076371 | 3.405582033 | 1.4346688 | 0.248124 | 0.606600867 |
| ENSMUSG00000028478 | Clta | protein_coding | 0.346400997 | 6.887830154 | 1.434617 | 0.2481322 | 0.606600867 |
| ENSMUSG00000052609 | Plekhg3 | protein_coding | -0.66307723 | 3.484572256 | 1.4344573 | 0.2481575 | 0.606600867 |
| ENSMUSG00000079038 | D130040H23Rik | protein_coding | -1.077926681 | 2.546525431 | 1.4331346 | 0.2483668 | 0.606974512 |
| ENSMUSG00000039585 | Myo9a | protein_coding | -0.867199332 | 2.871054901 | 1.4315993 | 0.24861 | 0.607121157 |
| ENSMUSG00000032641 | Gpr19 | protein_coding | 0.331798188 | 5.796899798 | 1.4310264 | 0.2487008 | 0.607121157 |
| ENSMUSG00000025933 | Tmem14a | protein_coding | 0.308793924 | 6.769581659 | 1.4308552 | 0.248728 | 0.607121157 |
| ENSMUSG00000048482 | Bdnf | protein_coding | -0.584493272 | 3.991353696 | 1.4308177 | 0.2487339 | 0.607121157 |
| ENSMUSG00000058741 | Prr19 | protein_coding | 1.352404057 | 2.448788106 | 1.4308848 | 0.2487512 | 0.607121157 |
| ENSMUSG00000031996 | Aplp2 | protein_coding | -0.281784923 | 7.990537434 | 1.4306185 | 0.2487655 | 0.607121157 |
| ENSMUSG00000022195 | 6030458C11Rik | protein_coding | 0.551706347 | 5.026614836 | 1.4299371 | 0.2489016 | 0.607315298 |
| ENSMUSG00000038412 | Higd1a | protein_coding | -0.283189965 | 8.510514161 | 1.4291312 | 0.2490017 | 0.607421679 |
| ENSMUSG00000052926 | Rnaseh2a | protein_coding | 0.361247923 | 5.930473243 | 1.4282114 | 0.2491478 | 0.607640432 |
| ENSMUSG00000034467 | Dynlrb2 | protein_coding | -0.622592655 | 4.88891651 | 1.4277849 | 0.2493169 | 0.607794821 |
| ENSMUSG00000028651 | Ppie | protein_coding | 0.430914641 | 4.584048901 | 1.4271026 | 0.2493242 | 0.607794821 |
| ENSMUSG00000037461 | Ints7 | protein_coding | 0.438106064 | 5.262373825 | 1.4248077 | 0.2496897 | 0.608547939 |
| ENSMUSG00000036943 | Rab8b | protein_coding | -0.49170611 | 4.885630103 | 1.4238388 | 0.2498442 | 0.608724548 |
| ENSMUSG00000029328 | Hnrnpdl | protein_coding | -0.279479578 | 6.69906303 | 1.4236436 | 0.2498754 | 0.608724548 |
| ENSMUSG00000022602 | Arc | protein_coding | -0.507189596 | 5.510500588 | 1.4228755 | 0.2500259 | 0.608787866 |
| ENSMUSG00000005683 | Cs | protein_coding | 0.390205172 | 8.087842053 | 1.4219817 | 0.2501686 | 0.608787866 |
| ENSMUSG00000036534 | Slc38a7 | protein_coding | 0.775127218 | 3.731879295 | 1.4218772 | 0.2501853 | 0.608787866 |
| ENSMUSG00000082536 | Gm13456 | processed_pseudoge | 1.029018567 | 2.286606122 | 1.4216364 | 0.250196 | 0.608787866 |
| ENSMUSG00000000399 | Ndufa9 | protein_coding | 0.266055415 | 9.01370572 | 1.4216335 | 0.2501964 | 0.608787866 |
| ENSMUSG00000056004 | 9330182L06Rik | protein_coding | -0.429693108 | 5.064102773 | 1.421174 | 0.2502699 | 0.608787866 |
| ENSMUSG00000057969 | Sema3b | protein_coding | 1.637054214 | 2.483645804 | 1.4276857 | 0.2503109 | 0.608787866 |
| ENSMUSG00000005621 | Zfp592 | protein_coding | -0.680135847 | 3.739771541 | 1.4204142 | 0.2503915 | 0.608787866 |
| ENSMUSG00000020483 | Dynll2 | protein_coding | -0.418543821 | 8.058738353 | 1.4247378 | 0.2504109 | 0.608787866 |
| ENSMUSG00000048546 | Tob2 | protein_coding | -0.409446685 | 4.887642111 | 1.4195794 | 0.2505251 | 0.608927743 |
| ENSMUSG00000025892 | Gria4 | protein_coding | -0.412142689 | 5.401965913 | 1.4190346 | 0.2506124 | 0.609002195 |
| ENSMUSG00000021552 | Gkap1 | protein_coding | -0.691039958 | 3.370369014 | 1.4184793 | 0.2507013 | 0.609017521 |
| ENSMUSG00000039007 | Cpq | protein_coding | -0.526002772 | 4.352774455 | 1.4181496 | 0.2507542 | 0.609017521 |
| ENSMUSG00000022201 | Zfr | protein_coding | -0.272147718 | 8.293012069 | 1.4179351 | 0.2507886 | 0.609017521 |
| ENSMUSG00000021221 | Dpf3 | protein_coding | -0.702335321 | 3.478082958 | 1.4166739 | 0.2509909 | 0.609074859 |
| ENSMUSG00000020029 | Nudt4 | protein_coding | -0.315422969 | 6.206252499 | 1.4166317 | 0.2509977 | 0.609074859 |
| ENSMUSG00000054942 | Miga1 | protein_coding | -0.375546947 | 5.453316848 | 1.4160738 | 0.2510873 | 0.609074859 |
| ENSMUSG00000022463 | Srebf2 | protein_coding | -0.332366131 | 5.999905884 | 1.4157215 | 0.2511439 | 0.609074859 |
| ENSMUSG00000027997 | Casp6 | protein_coding | -0.859696493 | 2.845442278 | 1.4154108 | 0.2511938 | 0.609074859 |
| ENSMUSG00000029246 | Ppat | protein_coding | 0.721931101 | 4.337523247 | 1.4186833 | 0.2511963 | 0.609074859 |
| ENSMUSG00000019432 | Ddx39b | protein_coding | 0.298763558 | 7.952588544 | 1.4153181 | 0.2512087 | 0.609074859 |
| ENSMUSG00000045100 | Slc25a26 | protein_coding | 0.566036606 | 4.702150218 | 1.4144216 | 0.2513528 | 0.609279682 |
| ENSMUSG00000020368 | Canx | protein_coding | -0.254420447 | 9.236388523 | 1.4130254 | 0.2515775 | 0.609279682 |
| ENSMUSG00000031558 | Slit2 | protein_coding | -0.683945454 | 3.494476546 | 1.4129577 | 0.2515884 | 0.609279682 |
| ENSMUSG00000051022 | Hs3st1 | protein_coding | 0.495553087 | 4.575954066 | 1.4127893 | 0.2516155 | 0.609279682 |
| ENSMUSG00000083695 | Rnf138rt1 | protein_coding | 0.783204668 | 3.793234133 | 1.4127092 | 0.2516562 | 0.609279682 |
| ENSMUSG00000018428 | Akap1 | protein_coding | -0.684634041 | 3.861081418 | 1.4119965 | 0.2517432 | 0.609279682 |
| ENSMUSG00000066687 | Zbtb16 | protein_coding | -0.949196848 | 2.852946376 | 1.4118474 | 0.2517673 | 0.609279682 |
| ENSMUSG00000001039 | B9d1 | protein_coding | 0.361675947 | 5.906474746 | 1.411646 | 0.2517997 | 0.609279682 |
| ENSMUSG00000032077 | Bud13 | protein_coding | 0.758782276 | 4.458012439 | 1.4175653 | 0.2519245 | 0.609279682 |
| ENSMUSG00000024298 | Zfp871 | protein_coding | -0.485742028 | 4.790642478 | 1.4108008 | 0.251936 | 0.609279682 |
| ENSMUSG00000022668 | Gtpbp8 | protein_coding | 0.451948732 | 4.736968523 | 1.4106109 | 0.2519667 | 0.609279682 |

| ensembl_gene_id | external_gene_name | gene_biotype | log2 (fold change) | log2 (counts per million) | F-value | p-value | False discovery rate |
|---|---|---|---|---|---|---|---|
| ENSMUSG00000029056 | Pank4 | protein_coding | 0.434166776 | 4.982483717 | 1.4105711 | 0.2519731 | 0.609279682 |
| ENSMUSG00000003233 | Dvl3 | protein_coding | -0.618444116 | 4.062868915 | 1.4098278 | 0.2520931 | 0.609395237 |
| ENSMUSG00000090223 | Pcp4 | protein_coding | -0.42049864 | 8.449282519 | 1.4139655 | 0.2521342 | 0.609395237 |
| ENSMUSG00000025429 | Pstpip2 | protein_coding | -1.012039377 | 2.547898882 | 1.4083274 | 0.2523355 | 0.609744555 |
| ENSMUSG00000042772 | Smg7 | protein_coding | -0.286839873 | 6.953742699 | 1.4079455 | 0.2523972 | 0.609756743 |
| ENSMUSG00000033578 | Tmem35a | protein_coding | 0.298503807 | 6.739431291 | 1.4075331 | 0.2524639 | 0.60978089 |
| ENSMUSG00000020411 | Nipal4 | protein_coding | -0.910182277 | 2.931015967 | 1.4066765 | 0.2526025 | 0.609939747 |
| ENSMUSG00000032369 | Plscr1 | protein_coding | -1.110570832 | 2.352033512 | 1.4064257 | 0.2526431 | 0.609939747 |
| ENSMUSG00000032705 | Exd2 | protein_coding | -0.509304736 | 4.414390876 | 1.4058188 | 0.2527414 | 0.610040067 |
| ENSMUSG00000024782 | Ak3 | protein_coding | -0.776169309 | 2.964821122 | 1.405109 | 0.2528565 | 0.610180758 |
| ENSMUSG00000020385 | Clk4 | protein_coding | -0.410464131 | 5.68838557 | 1.4043042 | 0.252987 | 0.610298181 |
| ENSMUSG00000028138 | Adh5 | protein_coding | 0.267168703 | 7.589360747 | 1.4039207 | 0.2530492 | 0.610298181 |
| ENSMUSG00000033697 | Arhgap39 | protein_coding | -0.940694833 | 2.727258943 | 1.403488 | 0.2531194 | 0.610298181 |
| ENSMUSG00000039982 | Dtx4 | protein_coding | 0.513665429 | 4.140135314 | 1.4034094 | 0.2531321 | 0.610298181 |
| ENSMUSG00000005470 | Asf1b | protein_coding | 1.795477293 | 2.029709115 | 1.4024891 | 0.2533822 | 0.610727616 |
| ENSMUSG00000074247 | Dda1 | protein_coding | 0.434724039 | 5.191936078 | 1.4016138 | 0.2534238 | 0.610727616 |
| ENSMUSG00000029781 | Fkbp9 | protein_coding | 0.841848433 | 3.42077307 | 1.4013944 | 0.2534872 | 0.610743403 |
| ENSMUSG00000004933 | Matk | protein_coding | 0.328937822 | 7.258529824 | 1.4000574 | 0.253677 | 0.610803544 |
| ENSMUSG00000107283 | Mpv17 | protein_coding | 0.289836426 | 6.948560434 | 1.3999479 | 0.2536949 | 0.610803544 |
| ENSMUSG00000004626 | Stxbp2 | protein_coding | -0.538375369 | 4.167826793 | 1.3995899 | 0.2537532 | 0.610803544 |
| ENSMUSG00000044098 | Rsbn1 | protein_coding | -0.826392386 | 2.860981529 | 1.3995792 | 0.2537549 | 0.610803544 |
| ENSMUSG00000036639 | Nudt1 | protein_coding | 0.458679451 | 4.102397497 | 1.3993258 | 0.2537962 | 0.610803544 |
| ENSMUSG00000063063 | Ctnna2 | protein_coding | -0.303947852 | 6.555486821 | 1.3988047 | 0.2538811 | 0.610832938 |
| ENSMUSG00000054874 | Pcnx3 | protein_coding | 1.054238769 | 2.79137171 | 1.3984784 | 0.2539343 | 0.610832938 |
| ENSMUSG00000006333 | Rps9 | protein_coding | 0.29955437 | 9.87165058 | 1.3981914 | 0.2539811 | 0.610832938 |
| ENSMUSG00000040269 | Mrps28 | protein_coding | 0.279612678 | 6.852699931 | 1.3978568 | 0.2540356 | 0.610832938 |
| ENSMUSG00000051811 | Cox6b2 | protein_coding | -0.519417271 | 4.008247307 | 1.3961658 | 0.2543116 | 0.610880057 |
| ENSMUSG00000047044 | D030056L22Rik | protein_coding | -0.545126916 | 3.731754147 | 1.395943 | 0.254348 | 0.610880057 |
| ENSMUSG00000011427 | Zfp790 | protein_coding | 0.709781536 | 3.195718129 | 1.3957945 | 0.2543723 | 0.610880057 |
| ENSMUSG00000034522 | Zfp395 | protein_coding | -0.570041538 | 3.499835818 | 1.3953908 | 0.2544382 | 0.610880057 |
| ENSMUSG00000087478 | 4930506C21Rik | processed_transcript | -0.936880401 | 2.649936518 | 1.3951077 | 0.2544845 | 0.610880057 |
| ENSMUSG00000028876 | Epha10 | protein_coding | 0.664099425 | 3.683833959 | 1.3950058 | 0.2545012 | 0.610880057 |
| ENSMUSG00000044365 | Cxxc4 | protein_coding | -0.414704097 | 5.379609679 | 1.3949435 | 0.2545113 | 0.610880057 |
| ENSMUSG00000028756 | Pink1 | protein_coding | 0.314980523 | 7.55903131 | 1.3947724 | 0.2545393 | 0.610880057 |
| ENSMUSG00000085891 | Gm14634 | processed_transcript | -1.092665122 | 2.374153954 | 1.3944405 | 0.2545936 | 0.610880057 |
| ENSMUSG00000042207 | Kdm5b | protein_coding | -0.353540577 | 5.486204669 | 1.3942589 | 0.2546233 | 0.610880057 |
| ENSMUSG00000050188 | Lsm10 | protein_coding | 0.380061447 | 5.930104451 | 1.3936918 | 0.2547161 | 0.610966376 |
| ENSMUSG00000047459 | Dynlrb1 | protein_coding | 0.317025642 | 10.28289572 | 1.3916784 | 0.2550459 | 0.611047956 |
| ENSMUSG00000018599 | Mief2 | protein_coding | 0.477745179 | 4.81715464 | 1.3914889 | 0.255077 | 0.611047956 |
| ENSMUSG00000027313 | Chac1 | protein_coding | 0.503716754 | 4.619900276 | 1.3914223 | 0.2550879 | 0.611047956 |
| ENSMUSG00000004561 | Mettl17 | protein_coding | 0.480541288 | 4.410173875 | 1.3912361 | 0.2551184 | 0.611047956 |
| ENSMUSG00000033910 | Gucy1a3 | protein_coding | -0.259700011 | 6.664581123 | 1.3911412 | 0.255134 | 0.611047956 |
| ENSMUSG00000023460 | Rab12 | protein_coding | 0.35185368 | 5.846956722 | 1.3909469 | 0.2551659 | 0.611047956 |
| ENSMUSG00000032905 | Atg12 | protein_coding | -0.266168157 | 6.55226275 | 1.3909061 | 0.2551726 | 0.611047956 |
| ENSMUSG00000035898 | Uba6 | protein_coding | -0.617052393 | 4.259359949 | 1.3901879 | 0.2552904 | 0.611047956 |
| ENSMUSG00000058246 | Gm10037 | protein_coding | 0.484479471 | 4.268719226 | 1.3900321 | 0.255316 | 0.611047956 |
| ENSMUSG00000031189 | Aff2 | protein_coding | -0.846574864 | 2.77269957 | 1.3898834 | 0.2553404 | 0.611047956 |
| ENSMUSG00000034780 | B3galt1 | protein_coding | -0.384133486 | 5.905181875 | 1.3896235 | 0.2553831 | 0.611047956 |
| ENSMUSG00000028961 | Pgd | protein_coding | 0.432552898 | 5.789461953 | 1.3890691 | 0.2554742 | 0.611047956 |
| ENSMUSG00000017747 | Ghdc | protein_coding | -1.203315769 | 2.546931946 | 1.3888147 | 0.255516 | 0.611047956 |
| ENSMUSG00000037072 | Selenof | protein_coding | -0.310236523 | 8.841278563 | 1.3886341 | 0.2555457 | 0.611047956 |
| ENSMUSG00000019843 | Fyn | protein_coding | 0.449648752 | 4.647893775 | 1.3873104 | 0.2557634 | 0.61126701 |
| ENSMUSG00000038383 | Pigu | protein_coding | 0.437375491 | 5.496662148 | 1.3870745 | 0.2558022 | 0.61126701 |
| ENSMUSG00000064043 | Trerf1 | processed_transcript | -0.45834822 | 4.211828547 | 1.3868068 | 0.2558463 | 0.61126701 |
| ENSMUSG00000045092 | S1pr1 | protein_coding | -0.452558929 | 5.733062734 | 1.386695 | 0.2558647 | 0.61126701 |
| ENSMUSG00000015994 | Fnta | protein_coding | 0.414771681 | 5.94889738 | 1.3861603 | 0.2559527 | 0.611341551 |
| ENSMUSG00000070738 | Dgkd | protein_coding | 0.542509782 | 4.513298388 | 1.3856045 | 0.2560443 | 0.611424441 |
| ENSMUSG00000023032 | Slc4a8 | protein_coding | 0.445965079 | 4.621661121 | 1.3846389 | 0.2562035 | 0.611668772 |
| ENSMUSG00000036158 | Prickle1 | protein_coding | -0.465341304 | 4.668239792 | 1.3828343 | 0.2565013 | 0.612243951 |
| ENSMUSG00000047084 | Ngrn | protein_coding | 0.348280588 | 6.385211894 | 1.3817187 | 0.2566857 | 0.612289045 |
| ENSMUSG00000026568 | Mpc2 | protein_coding | 0.582741205 | 6.934030928 | 1.4010282 | 0.2566888 | 0.612289045 |
| ENSMUSG00000024593 | Megf10 | protein_coding | -0.925086995 | 2.874228502 | 1.3816864 | 0.2566911 | 0.612289045 |
| ENSMUSG00000078137 | Ankrd63 | protein_coding | -0.562428146 | 4.836188436 | 1.3803539 | 0.256939 | 0.612725658 |
| ENSMUSG00000033021 | Gmppa | protein_coding | 0.438901182 | 5.686105646 | 1.3798915 | 0.2569881 | 0.612725658 |
| ENSMUSG00000070280 | Slc22a14 | protein_coding | -0.854805839 | 2.627265896 | 1.3788806 | 0.2571555 | 0.612989044 |
| ENSMUSG00000016510 | Mtif3 | protein_coding | 0.400927502 | 5.049693734 | 1.3781147 | 0.2572825 | 0.613155835 |
| ENSMUSG00000060882 | Kcnd2 | protein_coding | -0.291813558 | 6.631503232 | 1.3771169 | 0.2574481 | 0.613414434 |
| ENSMUSG00000029817 | Tra2a | protein_coding | -0.439350838 | 4.997497304 | 1.3765776 | 0.2575376 | 0.613491848 |
| ENSMUSG00000029407 | Uso1 | protein_coding | -0.385018709 | 5.557244643 | 1.3748433 | 0.2578259 | 0.614042389 |
| ENSMUSG00000023284 | Zfp605 | protein_coding | -0.674552785 | 3.455143663 | 1.3737701 | 0.2580044 | 0.614092527 |
| ENSMUSG00000000296 | Tpd52l1 | protein_coding | 0.255454013 | 7.58623538 | 1.3737021 | 0.2580157 | 0.614092527 |
| ENSMUSG00000021877 | Arf4 | protein_coding | 0.321818571 | 8.805973161 | 1.3736872 | 0.2580182 | 0.614092527 |
| ENSMUSG00000031095 | Cul4b | protein_coding | -0.627205368 | 3.859403634 | 1.3720946 | 0.2582835 | 0.614466154 |
| ENSMUSG00000023827 | Agpat4 | protein_coding | 0.309426133 | 7.118018798 | 1.3720588 | 0.2582895 | 0.614466154 |
| ENSMUSG00000039059 | Hrh3 | protein_coding | 0.567487667 | 3.775898326 | 1.3714092 | 0.2583978 | 0.614563368 |
| ENSMUSG00000002210 | Smg9 | protein_coding | -0.496143015 | 4.2138044 | 1.3711284 | 0.2584447 | 0.614563368 |
| ENSMUSG00000097416 | Gm26670 | antisense_RNA | -1.012988933 | 2.352260028 | 1.3694427 | 0.2587261 | 0.615096627 |
| ENSMUSG00000020495 | Smg8 | protein_coding | 0.573558364 | 3.617206367 | 1.3678197 | 0.2589975 | 0.615605651 |

| ensembl_gene_id | external_gene_name | gene_biotype | log2 (fold change) | log2 (counts per million) | F-value | p-value | False discovery rate |
|---|---|---|---|---|---|---|---|
| ENSMUSG00000039530 | Tusc3 | protein_coding | 0.363128858 | 7.370832467 | 1.3674648 | 0.2590843 | 0.615623817 |
| ENSMUSG00000025357 | Dgka | protein_coding | 1.085725943 | 2.571706719 | 1.3666795 | 0.2591884 | 0.615623817 |
| ENSMUSG00000038354 | Ankrd35 | protein_coding | 1.119521262 | 2.39288361 | 1.3665987 | 0.2592019 | 0.615623817 |
| ENSMUSG00000030323 | Ift122 | protein_coding | -0.458892135 | 4.801334896 | 1.366406 | 0.2592342 | 0.615623817 |
| ENSMUSG00000004568 | Arhgef18 | protein_coding | -0.715141489 | 3.849706534 | 1.3661586 | 0.259303 | 0.615623817 |
| ENSMUSG00000025978 | Rftn2 | protein_coding | -0.426837669 | 4.74961476 | 1.3657229 | 0.2593486 | 0.615623817 |
| ENSMUSG00000045136 | Tubb2b | protein_coding | 0.357895649 | 5.423477762 | 1.3648078 | 0.2595021 | 0.615852099 |
| ENSMUSG00000038233 | Fam198a | protein_coding | 1.291979343 | 2.170744089 | 1.3640533 | 0.2596287 | 0.615904232 |
| ENSMUSG00000021240 | Abcd4 | protein_coding | 0.912571095 | 2.905909648 | 1.3639858 | 0.25964 | 0.615904232 |
| ENSMUSG00000071550 | Cfap44 | protein_coding | 1.184880706 | 2.155988903 | 1.3634013 | 0.2597381 | 0.615904232 |
| ENSMUSG00000033628 | Pik3c3 | protein_coding | 0.407630443 | 5.531721256 | 1.363312 | 0.2597532 | 0.615904232 |
| ENSMUSG00000026031 | Cflar | protein_coding | -0.640596923 | 3.42663038 | 1.3619075 | 0.2599892 | 0.61604277 |
| ENSMUSG00000022329 | Stk3 | protein_coding | -0.593503354 | 4.112228634 | 1.3618674 | 0.2599959 | 0.61604277 |
| ENSMUSG00000028030 | Tbck | protein_coding | 0.518292092 | 6.406016081 | 1.3712538 | 0.2600892 | 0.61604277 |
| ENSMUSG00000047804 | Akap10 | protein_coding | -0.918306151 | 2.627325968 | 1.3612992 | 0.2600915 | 0.61604277 |
| ENSMUSG00000020638 | Cmpk2 | protein_coding | 0.374617858 | 5.485329651 | 1.3612605 | 0.260098 | 0.61604277 |
| ENSMUSG00000049232 | Tigd2 | protein_coding | 0.631915009 | 4.020976715 | 1.3605833 | 0.260212 | 0.616176982 |
| ENSMUSG00000022787 | Wdr53 | protein_coding | 0.424277667 | 4.781929122 | 1.3600915 | 0.2602948 | 0.616237394 |
| ENSMUSG00000033220 | Rac2 | protein_coding | -0.970606004 | 2.77756677 | 1.3590164 | 0.260476 | 0.616530536 |
| ENSMUSG00000041609 | Bicdl1 | protein_coding | 0.582813619 | 4.482857563 | 1.3584408 | 0.260573 | 0.616624521 |
| ENSMUSG00000021364 | Elovl2 | protein_coding | -1.218844637 | 2.667073904 | 1.3572617 | 0.2607993 | 0.616832149 |
| ENSMUSG00000022962 | Gart | protein_coding | 0.544646719 | 4.633510336 | 1.3569468 | 0.2608252 | 0.616832149 |
| ENSMUSG00000024851 | Pitpnm1 | protein_coding | 0.362998424 | 5.789043041 | 1.3565362 | 0.2608945 | 0.616832149 |
| ENSMUSG00000109554 | A230057D06Rik | lincRNA | -1.153141695 | 2.69635699 | 1.356249 | 0.2609704 | 0.616832149 |
| ENSMUSG00000038762 | Abcf1 | protein_coding | 0.33943697 | 6.924859348 | 1.355876 | 0.2610061 | 0.616832149 |
| ENSMUSG00000026496 | Parp1 | protein_coding | 0.374867374 | 5.257757209 | 1.3557289 | 0.261031 | 0.616832149 |
| ENSMUSG00000020522 | Mfap3 | protein_coding | 0.473995969 | 4.986215384 | 1.3555435 | 0.2610623 | 0.616832149 |
| ENSMUSG00000020917 | Acly | protein_coding | 0.358055889 | 7.114004496 | 1.3541783 | 0.2613206 | 0.617094696 |
| ENSMUSG00000047514 | Tspyl1 | protein_coding | -0.316748605 | 8.789173962 | 1.3539251 | 0.2613361 | 0.617094696 |
| ENSMUSG00000059857 | Ntng1 | protein_coding | -0.565280041 | 5.235176209 | 1.3546233 | 0.2613456 | 0.617094696 |
| ENSMUSG00000033773 | Rpap2 | protein_coding | 0.604339281 | 4.518197972 | 1.3535957 | 0.2614192 | 0.617132655 |
| ENSMUSG00000030007 | Cct7 | protein_coding | 0.338039051 | 9.448790663 | 1.3530967 | 0.2614765 | 0.617132655 |
| ENSMUSG00000040035 | Disp2 | protein_coding | 0.47495073 | 7.018399155 | 1.360774 | 0.2618404 | 0.617722068 |
| ENSMUSG00000057497 | Fam136a | protein_coding | 0.286249346 | 6.854044648 | 1.3507607 | 0.2618727 | 0.617722068 |
| ENSMUSG00000015711 | Prune1 | protein_coding | 0.527212233 | 5.363997029 | 1.350285 | 0.2619807 | 0.617722068 |
| ENSMUSG00000028889 | Yrdc | protein_coding | -0.570987015 | 4.415054992 | 1.3498608 | 0.2620255 | 0.617722068 |
| ENSMUSG00000025016 | Tm9sf3 | protein_coding | -0.371349929 | 6.079951102 | 1.349845 | 0.2620282 | 0.617722068 |
| ENSMUSG00000033352 | Map2k4 | protein_coding | 0.279639534 | 7.402422423 | 1.3495939 | 0.2620709 | 0.617722068 |
| ENSMUSG00000022035 | Ccdc25 | protein_coding | 0.30024045 | 6.514769902 | 1.3485146 | 0.2622544 | 0.617757598 |
| ENSMUSG00000036948 | BC037034 | protein_coding | 0.581778276 | 3.583442074 | 1.3484863 | 0.2622592 | 0.617757598 |
| ENSMUSG00000098702 | 1500015A07Rik | antisense_RNA | 0.697582482 | 3.304849174 | 1.3483348 | 0.262285 | 0.617757598 |
| ENSMUSG00000034602 | Mon2 | protein_coding | -0.483926511 | 4.907468043 | 1.348154 | 0.2623157 | 0.617757598 |
| ENSMUSG00000050079 | Rspry1 | protein_coding | 0.434598151 | 4.873429617 | 1.3475963 | 0.2624107 | 0.617845807 |
| ENSMUSG00000010110 | Stx5a | protein_coding | 0.306559974 | 6.442202614 | 1.3470172 | 0.2625093 | 0.617942713 |
| ENSMUSG00000028224 | Nbn | protein_coding | -0.682136879 | 3.964025293 | 1.3460838 | 0.2626956 | 0.618034818 |
| ENSMUSG00000024370 | Cdc23 | protein_coding | 0.518038451 | 5.487269264 | 1.3460834 | 0.2626956 | 0.618034818 |
| ENSMUSG00000029017 | Pmpcb | protein_coding | 0.274097065 | 7.100038551 | 1.345626 | 0.2627464 | 0.618034818 |
| ENSMUSG00000027247 | Arhgap1 | protein_coding | 0.416856022 | 6.71735667 | 1.347741 | 0.2627783 | 0.618034818 |
| ENSMUSG00000029054 | Gabrd | protein_coding | 0.481978885 | 5.524120706 | 1.3435707 | 0.2631244 | 0.618606875 |
| ENSMUSG00000044471 | Lncpint | lincRNA | 1.275918543 | 2.492313321 | 1.3429125 | 0.2632097 | 0.618606875 |
| ENSMUSG00000024165 | Hn1l | protein_coding | 0.536055968 | 4.286297872 | 1.3428916 | 0.2632132 | 0.618606875 |
| ENSMUSG00000034449 | Dhrs11 | protein_coding | 0.695535754 | 3.101245341 | 1.3425494 | 0.2632717 | 0.618606875 |
| ENSMUSG00000038608 | Dock10 | protein_coding | -0.552490169 | 4.563913217 | 1.3423304 | 0.2633092 | 0.618606875 |
| ENSMUSG00000034071 | Zfp551 | protein_coding | 1.518880497 | 2.569405987 | 1.3472009 | 0.2635336 | 0.618998864 |
| ENSMUSG00000050890 | Pdik1l | protein_coding | -0.499751801 | 4.28095881 | 1.3403016 | 0.2636565 | 0.619152182 |
| ENSMUSG00000041343 | Ankrd42 | protein_coding | 0.720509691 | 3.169587784 | 1.3398883 | 0.2637273 | 0.619183255 |
| ENSMUSG00000022822 | Abcc5 | protein_coding | 0.295142219 | 6.360321778 | 1.3392974 | 0.2638286 | 0.619237479 |
| ENSMUSG00000042046 | Dstyk | protein_coding | 0.385811977 | 5.169482576 | 1.3390818 | 0.2638655 | 0.619237479 |
| ENSMUSG00000016253 | Nelfcd | protein_coding | 0.400216347 | 6.811902554 | 1.3392762 | 0.263931 | 0.619256036 |
| ENSMUSG00000040188 | Scamp2 | protein_coding | -0.385104407 | 5.15352161 | 1.3380728 | 0.2640387 | 0.619366172 |
| ENSMUSG00000010376 | Nedd8 | protein_coding | 0.294321534 | 9.396075068 | 1.3373074 | 0.2641701 | 0.619366172 |
| ENSMUSG00000024570 | Rbfa | protein_coding | 0.614241215 | 3.664244014 | 1.3372522 | 0.2641796 | 0.619366172 |
| ENSMUSG00000020745 | Pafah1b1 | protein_coding | -0.277519725 | 7.459927703 | 1.3368215 | 0.2642536 | 0.619366172 |
| ENSMUSG00000016252 | Atp5e | protein_coding | 0.308240917 | 9.763074852 | 1.3365683 | 0.2642971 | 0.619366172 |
| ENSMUSG00000030259 | Rassf8 | protein_coding | -0.584090817 | 3.946340098 | 1.336213 | 0.2643581 | 0.619366172 |
| ENSMUSG00000039680 | Mrps6 | protein_coding | 0.395820572 | 6.241458897 | 1.3356979 | 0.2644467 | 0.619366172 |
| ENSMUSG00000032232 | Cgnl1 | protein_coding | -1.451340933 | 1.933576761 | 1.3355058 | 0.2644798 | 0.619366172 |
| ENSMUSG00000021650 | Ptcd2 | protein_coding | 0.328418908 | 6.435827119 | 1.3354093 | 0.2644964 | 0.619366172 |
| ENSMUSG00000032902 | Slc16a1 | protein_coding | -0.456176553 | 4.132903036 | 1.3348081 | 0.2645998 | 0.619473533 |
| ENSMUSG00000039990 | Edrf1 | protein_coding | 0.779371326 | 3.266259873 | 1.3339183 | 0.264753 | 0.619571373 |
| ENSMUSG00000111375 | Btbd8 | protein_coding | -0.467330644 | 5.097936744 | 1.3338962 | 0.2647569 | 0.619571373 |
| ENSMUSG00000024312 | Wdr46 | protein_coding | 0.844130166 | 2.900480382 | 1.3327144 | 0.2649605 | 0.61989939 |
| ENSMUSG00000049946 | BC030500 | protein_coding | 0.476100402 | 5.154877365 | 1.3324142 | 0.2650123 | 0.61989939 |
| ENSMUSG00000006731 | B4galnt1 | protein_coding | -0.58244273 | 3.284752277 | 1.331916 | 0.2650983 | 0.619965599 |
| ENSMUSG00000006360 | Crip1 | protein_coding | -0.678641383 | 4.025108628 | 1.3304167 | 0.2653843 | 0.620499521 |
| ENSMUSG00000062604 | Srpk2 | protein_coding | -0.298740517 | 7.969830073 | 1.3294711 | 0.2655206 | 0.62068334 |
| ENSMUSG00000005682 | Pan2 | protein_coding | -1.060956165 | 2.311549658 | 1.3281287 | 0.2657529 | 0.6210913 |
| ENSMUSG00000046876 | Atxn1 | protein_coding | -0.521820532 | 6.992813897 | 1.3412437 | 0.2659365 | 0.621235156 |

| ensembl_gene_id | external_gene_name | gene_biotype | log2 (fold change) | log2 (counts per million) | F-value | p-value | False discovery rate |
|---|---|---|---|---|---|---|---|
| ENSMUSG00000023944 | Hsp90ab1 | protein_coding | 0.313084303 | 11.18772475 | 1.3267273 | 0.2659957 | 0.621235156 |
| ENSMUSG00000023027 | Atf1 | protein_coding | -0.968519116 | 2.517143718 | 1.3263441 | 0.2660621 | 0.621235156 |
| ENSMUSG00000024511 | Rab27b | protein_coding | -0.412850212 | 6.756997328 | 1.3288281 | 0.2660915 | 0.621235156 |
| ENSMUSG00000024131 | Slc3a1 | protein_coding | 0.286674328 | 7.649142147 | 1.3257159 | 0.2661711 | 0.621235156 |
| ENSMUSG00000020859 | Spag9 | protein_coding | -0.304649703 | 6.745140716 | 1.3242983 | 0.2664172 | 0.621235156 |
| ENSMUSG00000063065 | Mapk3 | protein_coding | 0.252637604 | 7.28755616 | 1.323295 | 0.2665915 | 0.621235156 |
| ENSMUSG00000050891 | Tatdn1 | protein_coding | -0.355102783 | 5.328396989 | 1.3227723 | 0.2666824 | 0.621235156 |
| ENSMUSG00000048304 | Slitrk3 | protein_coding | 0.532969836 | 4.373265654 | 1.3221873 | 0.2667842 | 0.621235156 |
| ENSMUSG00000015143 | Actn1 | protein_coding | -0.409394475 | 4.509325244 | 1.3220879 | 0.2668015 | 0.621235156 |
| ENSMUSG00000025421 | Hdhd2 | protein_coding | 0.255756151 | 8.431281938 | 1.3220539 | 0.2668074 | 0.621235156 |
| ENSMUSG00000020605 | Hs1bp3 | protein_coding | 0.549367946 | 3.979064474 | 1.3220276 | 0.266812 | 0.621235156 |
| ENSMUSG00000002496 | Tsc2 | protein_coding | 0.499321308 | 4.776192968 | 1.3219805 | 0.2668202 | 0.621235156 |
| ENSMUSG00000022111 | Uchl3 | protein_coding | 0.325743538 | 6.159675216 | 1.3218817 | 0.2668374 | 0.621235156 |
| ENSMUSG00000047797 | Gjb1 | protein_coding | -0.525680216 | 6.266397011 | 1.3312088 | 0.2668664 | 0.621235156 |
| ENSMUSG00000021194 | Chga | protein_coding | 0.286000712 | 7.63615483 | 1.3216661 | 0.2668749 | 0.621235156 |
| ENSMUSG00000031951 | Tmem231 | protein_coding | 0.523503538 | 4.565835644 | 1.3216466 | 0.2668783 | 0.621235156 |
| ENSMUSG00000026889 | Rbm18 | protein_coding | -0.279479076 | 6.793040632 | 1.3215135 | 0.2669015 | 0.621235156 |
| ENSMUSG00000003873 | Bax | protein_coding | 0.294829429 | 6.257187138 | 1.3214524 | 0.2669122 | 0.621235156 |
| ENSMUSG00000031561 | Tenm3 | protein_coding | -0.663623886 | 3.863586045 | 1.3201349 | 0.2671417 | 0.621634878 |
| ENSMUSG00000067995 | Gtf2f2 | protein_coding | -0.352405412 | 5.655540052 | 1.3195175 | 0.2672494 | 0.621741838 |
| ENSMUSG00000041164 | Zmiz2 | protein_coding | 0.314627188 | 6.669697262 | 1.3192083 | 0.2673033 | 0.621741838 |
| ENSMUSG00000026764 | Kif5c | protein_coding | -0.330354444 | 7.28772482 | 1.3169403 | 0.2676994 | 0.622528543 |
| ENSMUSG00000001157 | Gmcl1 | protein_coding | -0.591278862 | 4.452459458 | 1.3165266 | 0.2677988 | 0.622551799 |
| ENSMUSG00000040463 | Mybbp1a | protein_coding | 0.431140975 | 4.832715963 | 1.3162209 | 0.2678252 | 0.622551799 |
| ENSMUSG00000042807 | Hecw2 | protein_coding | -0.528493018 | 4.125532092 | 1.3151265 | 0.2680168 | 0.622827469 |
| ENSMUSG00000046500 | Fam19a4 | protein_coding | -1.056287735 | 2.344135323 | 1.3145904 | 0.2681107 | 0.622827469 |
| ENSMUSG00000002963 | Pnkp | protein_coding | 0.39025916 | 4.95504095 | 1.3145509 | 0.2681176 | 0.622827469 |
| ENSMUSG00000097929 | Tunar | protein_coding | 0.434119713 | 4.680065021 | 1.3132876 | 0.268339 | 0.62310028 |
| ENSMUSG00000044952 | Kctd21 | protein_coding | 0.556335532 | 4.021924645 | 1.3129835 | 0.2683924 | 0.62310028 |
| ENSMUSG00000020640 | Itsn2 | protein_coding | -0.526279203 | 3.894198218 | 1.3128894 | 0.2684089 | 0.62310028 |
| ENSMUSG00000026112 | Coa5 | protein_coding | 0.281696014 | 6.58948572 | 1.3105322 | 0.2688229 | 0.623627231 |
| ENSMUSG00000040354 | Mars | protein_coding | 0.544358195 | 4.800313522 | 1.3106698 | 0.2688256 | 0.623627231 |
| ENSMUSG00000041556 | Fbxo2 | protein_coding | 0.455681773 | 4.92481782 | 1.3102469 | 0.268873 | 0.623627231 |
| ENSMUSG00000009828 | Ick | protein_coding | -0.697014461 | 3.469201804 | 1.3100889 | 0.2689008 | 0.623627231 |
| ENSMUSG00000033530 | Ttc7b | protein_coding | 0.634924109 | 4.516190074 | 1.3105036 | 0.2689258 | 0.623627231 |
| ENSMUSG00000000420 | Galnt1 | protein_coding | -0.486541463 | 4.118701884 | 1.3093763 | 0.2690262 | 0.623725366 |
| ENSMUSG00000009681 | Bcr | protein_coding | -0.302333712 | 6.228285443 | 1.3068877 | 0.2694646 | 0.624231652 |
| ENSMUSG00000020520 | Galnt10 | protein_coding | -0.604939697 | 3.456535794 | 1.3067469 | 0.2694894 | 0.624231652 |
| ENSMUSG00000024176 | Sox8 | protein_coding | -0.488887518 | 4.254497487 | 1.3066529 | 0.269506 | 0.624231652 |
| ENSMUSG00000028333 | Anp32b | protein_coding | -0.413749891 | 5.070327518 | 1.3063048 | 0.2695674 | 0.624231652 |
| ENSMUSG00000029561 | Oasl2 | protein_coding | 1.800780155 | 3.05310636 | 1.3436514 | 0.2696009 | 0.624231652 |
| ENSMUSG00000037443 | Cep85 | protein_coding | -0.582330786 | 3.550220314 | 1.3057436 | 0.2696665 | 0.624231652 |
| ENSMUSG00000019362 | D8Ertd738e | protein_coding | 0.252984611 | 8.80868539 | 1.3052431 | 0.2697548 | 0.624231652 |
| ENSMUSG00000018427 | Ypel2 | protein_coding | -0.272328458 | 6.565486051 | 1.3039098 | 0.2699904 | 0.624231652 |
| ENSMUSG00000048458 | Fam212b | protein_coding | -0.510019674 | 5.087369993 | 1.3037265 | 0.2700229 | 0.624231652 |
| ENSMUSG00000028465 | Tln1 | protein_coding | 0.38558577 | 5.298574854 | 1.3035478 | 0.2700545 | 0.624231652 |
| ENSMUSG00000018286 | Psmb6 | protein_coding | 0.280173319 | 9.771318944 | 1.3035195 | 0.2700595 | 0.624231652 |
| ENSMUSG00000038291 | Snx25 | protein_coding | -0.507639405 | 4.481618643 | 1.3034818 | 0.2700661 | 0.624231652 |
| ENSMUSG00000004849 | Ap1s1 | protein_coding | 0.270807101 | 9.61928205 | 1.3034607 | 0.2700699 | 0.624231652 |
| ENSMUSG00000024127 | Prepl | protein_coding | 0.819308159 | 7.106366955 | 1.3595675 | 0.2701148 | 0.624231652 |
| ENSMUSG00000007908 | Hmgcll1 | protein_coding | -0.535287174 | 4.293571825 | 1.3030927 | 0.270135 | 0.624231652 |
| ENSMUSG00000055917 | Zfp277 | protein_coding | -0.512924302 | 5.231259473 | 1.3028391 | 0.2702068 | 0.624231652 |
| ENSMUSG00000020740 | Gga3 | protein_coding | 0.367412751 | 5.906229905 | 1.3025479 | 0.2702314 | 0.624231652 |
| ENSMUSG00000046215 | Rprml | protein_coding | 0.728717451 | 3.693100388 | 1.3015047 | 0.270443 | 0.624476454 |
| ENSMUSG00000029669 | Tspan12 | protein_coding | -0.39116947 | 6.187878731 | 1.3010808 | 0.2704913 | 0.624476454 |
| ENSMUSG00000029802 | Abcg2 | protein_coding | 0.802481526 | 2.876853871 | 1.3009663 | 0.2705116 | 0.624476454 |
| ENSMUSG00000112639 | AC134329.1 | lincRNA | -0.739939262 | 2.834979682 | 1.2995359 | 0.2707653 | 0.624928023 |
| ENSMUSG00000020803 | Txndc17 | protein_coding | 0.243372222 | 7.832674069 | 1.2970952 | 0.271199 | 0.62506613 |
| ENSMUSG00000031732 | Phlpp2 | protein_coding | 0.511844413 | 4.01577232 | 1.2968904 | 0.2712355 | 0.62506613 |
| ENSMUSG00000003808 | Farsa | protein_coding | 0.443706428 | 5.889888517 | 1.2969726 | 0.2712477 | 0.62506613 |
| ENSMUSG00000026825 | Dnm1 | protein_coding | -0.317517036 | 6.683177301 | 1.2965189 | 0.2713016 | 0.62506613 |
| ENSMUSG00000032575 | Manf | protein_coding | 0.338658577 | 6.02686512 | 1.2959628 | 0.2714006 | 0.62506613 |
| ENSMUSG00000025241 | Fyco1 | protein_coding | -0.76217976 | 3.763028103 | 1.2959424 | 0.271431 | 0.62506613 |
| ENSMUSG00000034951 | Cog7 | protein_coding | 0.418614718 | 5.961421117 | 1.2958239 | 0.2714521 | 0.62506613 |
| ENSMUSG00000003534 | Ddr1 | protein_coding | -0.451671189 | 4.699261605 | 1.2956468 | 0.2714568 | 0.62506613 |
| ENSMUSG00000027965 | Olfm3 | protein_coding | -0.464344451 | 4.733912521 | 1.295534 | 0.2714769 | 0.62506613 |
| ENSMUSG00000024317 | Rnf138 | protein_coding | -0.444732403 | 4.513559622 | 1.2955064 | 0.2714818 | 0.62506613 |
| ENSMUSG00000024853 | Sf3b2 | protein_coding | 0.29401562 | 9.008018132 | 1.2951104 | 0.2715524 | 0.62506613 |
| ENSMUSG00000051934 | Spats2 | protein_coding | -0.404086552 | 4.648541293 | 1.2949009 | 0.2715897 | 0.62506613 |
| ENSMUSG00000039804 | Ncoa5 | protein_coding | 0.492435376 | 4.500782273 | 1.2946082 | 0.2716419 | 0.62506613 |
| ENSMUSG00000036634 | Mag | protein_coding | -0.271553721 | 7.307725008 | 1.2945254 | 0.2716567 | 0.62506613 |
| ENSMUSG00000027679 | Dnajc19 | protein_coding | 0.312950417 | 6.113242534 | 1.2942986 | 0.2716971 | 0.62506613 |
| ENSMUSG00000028407 | Toporsos | protein_coding | -0.571822835 | 3.603594937 | 1.2932398 | 0.271886 | 0.625240452 |
| ENSMUSG00000034587 | 8430429K09Rik | processed_transcript | 0.536401107 | 4.422276254 | 1.293222 | 0.2718892 | 0.625240452 |
| ENSMUSG00000041720 | Pi4ka | protein_coding | 0.514376246 | 4.881560216 | 1.2921098 | 0.2720878 | 0.625563412 |
| ENSMUSG00000033318 | Gstt2 | protein_coding | 0.606993661 | 3.994900866 | 1.2915475 | 0.2721883 | 0.625650107 |
| ENSMUSG00000044847 | Lsm11 | protein_coding | 0.778026654 | 3.172878441 | 1.29108 | 0.2722719 | 0.625650107 |
| ENSMUSG00000032386 | Trip4 | protein_coding | -0.382736088 | 4.950811401 | 1.2909223 | 0.2723001 | 0.625650107 |

| ensembl_gene_id | external_gene_name | gene_biotype | log2 (fold change) | log2 (counts per million) | F-value | p-value | False discovery rate |
|---|---|---|---|---|---|---|---|
| ENSMUSG00000024941 | Scyl1 | protein_coding | 0.367390597 | 5.49164501 | 1.2902165 | 0.2724263 | 0.62572533 |
| ENSMUSG00000026856 | Dolpp1 | protein_coding | 0.432606027 | 4.741127617 | 1.290041 | 0.2724577 | 0.62572533 |
| ENSMUSG00000006335 | Tfpt | protein_coding | 0.322040903 | 5.883506186 | 1.2897197 | 0.2725153 | 0.62572533 |
| ENSMUSG00000026424 | Gpr37l1 | protein_coding | -0.359184257 | 6.583081309 | 1.2894388 | 0.2725656 | 0.62572533 |
| ENSMUSG00000039217 | Il18 | protein_coding | -0.495489813 | 5.980826172 | 1.293967 | 0.2727316 | 0.62584805 |
| ENSMUSG00000054302 | Eapp | protein_coding | -0.278475634 | 7.104930646 | 1.2884907 | 0.2727354 | 0.62584805 |
| ENSMUSG00000046324 | Ermp1 | protein_coding | -0.597455741 | 4.23663807 | 1.2873249 | 0.2729445 | 0.626079346 |
| ENSMUSG00000021094 | Dhrs7 | protein_coding | -0.395243123 | 6.063722617 | 1.2872792 | 0.2729527 | 0.626079346 |
| ENSMUSG00000026239 | Pde6d | protein_coding | -0.380992342 | 4.990558208 | 1.2868272 | 0.2730338 | 0.626088842 |
| ENSMUSG00000039096 | Rsad1 | protein_coding | 0.805711403 | 3.080677049 | 1.2863271 | 0.2731236 | 0.626088842 |
| ENSMUSG00000069844 | Sco1 | protein_coding | -0.935707315 | 3.304374672 | 1.2880757 | 0.2731315 | 0.626088842 |
| ENSMUSG00000029034 | Ints11 | protein_coding | 0.756676761 | 3.329845632 | 1.2849216 | 0.2733761 | 0.626351185 |
| ENSMUSG00000004896 | Rrnad1 | protein_coding | 0.556349801 | 3.883037571 | 1.2848556 | 0.273388 | 0.626351185 |
| ENSMUSG00000056608 | Chd9 | protein_coding | -0.410872178 | 5.372321809 | 1.2845109 | 0.27345 | 0.626351185 |
| ENSMUSG00000002475 | Abhd3 | protein_coding | -0.428699496 | 5.700202732 | 1.2841268 | 0.2735191 | 0.626351185 |
| ENSMUSG00000031112 | Stk26 | protein_coding | -1.073270318 | 2.454219891 | 1.2835743 | 0.2736185 | 0.626351185 |
| ENSMUSG00000020380 | Rad50 | protein_coding | -0.524272561 | 3.776395721 | 1.283434 | 0.2736438 | 0.626351185 |
| ENSMUSG00000048787 | Dcun1d3 | protein_coding | -0.56128854 | 3.568965369 | 1.283379 | 0.2736537 | 0.626351185 |
| ENSMUSG00000020532 | Acaca | protein_coding | -0.540929021 | 3.55394166 | 1.2827257 | 0.2737713 | 0.626487121 |
| ENSMUSG00000054400 | Cklf | protein_coding | -0.904010063 | 2.540702746 | 1.2822744 | 0.2738526 | 0.626539843 |
| ENSMUSG00000017119 | Nbr1 | protein_coding | -0.333851662 | 6.019132822 | 1.2817001 | 0.2739561 | 0.62664335 |
| ENSMUSG00000013495 | Tmem175 | protein_coding | 0.438167761 | 5.834361965 | 1.2800872 | 0.2742739 | 0.627236697 |
| ENSMUSG00000026273 | Mterf4 | protein_coding | 0.351354212 | 5.478050371 | 1.2785329 | 0.274528 | 0.627456343 |
| ENSMUSG00000007670 | Khsrp | protein_coding | -0.369924679 | 5.071241666 | 1.2784673 | 0.2745399 | 0.627456343 |
| ENSMUSG00000039629 | Strip2 | protein_coding | -0.58825913 | 4.260741772 | 1.2784391 | 0.274545 | 0.627456343 |
| ENSMUSG00000002546 | Golga2 | protein_coding | -0.417526693 | 5.61340806 | 1.2773484 | 0.2747423 | 0.627613597 |
| ENSMUSG00000020891 | Alox8 | protein_coding | 1.031765925 | 2.964018178 | 1.2773429 | 0.27477 | 0.627613597 |
| ENSMUSG00000073838 | Tufm | protein_coding | 0.512801181 | 3.580335509 | 1.2770684 | 0.274793 | 0.627613597 |
| ENSMUSG00000034926 | Dhcr24 | protein_coding | 0.735231829 | 3.241174384 | 1.2764941 | 0.274897 | 0.627613597 |
| ENSMUSG00000055745 | Ldoc1l | protein_coding | 0.370334343 | 5.336780888 | 1.2764465 | 0.2749056 | 0.627613597 |
| ENSMUSG00000014786 | Slc9a5 | protein_coding | 0.637241916 | 3.415460126 | 1.2760867 | 0.2749708 | 0.627629149 |
| ENSMUSG00000022498 | Txndc11 | protein_coding | 0.741688835 | 2.905277887 | 1.2752881 | 0.2751156 | 0.627826305 |
| ENSMUSG00000059820 | AU019823 | protein_coding | -0.649060295 | 3.350332117 | 1.2745718 | 0.2752455 | 0.627856544 |
| ENSMUSG00000021265 | Slc25a29 | protein_coding | 0.6512146 | 3.224824225 | 1.2745713 | 0.2752456 | 0.627856544 |
| ENSMUSG00000111966 | AL589661.1 | processed_pseudoge | 0.658171689 | 3.01471568 | 1.2736672 | 0.2754097 | 0.628097629 |
| ENSMUSG00000028538 | St3gal3 | protein_coding | -0.649196444 | 3.241666298 | 1.2732436 | 0.2754866 | 0.628139866 |
| ENSMUSG00000041468 | Gpr12 | protein_coding | -0.4693021 | 4.7505083 | 1.2722829 | 0.2756612 | 0.628284381 |
| ENSMUSG00000019978 | Epb41l2 | protein_coding | -0.454698889 | 4.435299091 | 1.2722519 | 0.2756669 | 0.628284381 |
| ENSMUSG00000007656 | Arpp19 | protein_coding | -0.350308772 | 9.503377922 | 1.2716693 | 0.2757728 | 0.628392665 |
| ENSMUSG00000042502 | Cd2bp2 | protein_coding | 0.350152909 | 6.020395609 | 1.2705744 | 0.2759721 | 0.628713528 |
| ENSMUSG00000004637 | Wwox | protein_coding | 0.406881011 | 6.345004564 | 1.2693916 | 0.2762142 | 0.629109967 |
| ENSMUSG00000024059 | Clip4 | protein_coding | -0.341036795 | 5.871477717 | 1.2685465 | 0.2763417 | 0.629109967 |
| ENSMUSG00000024454 | Hdac3 | protein_coding | 0.281161839 | 7.052944055 | 1.2681305 | 0.2764176 | 0.629109967 |
| ENSMUSG00000028207 | Asph | protein_coding | -0.389178481 | 5.442789133 | 1.2681139 | 0.2764206 | 0.629109967 |
| ENSMUSG00000031584 | Gsr | protein_coding | -0.370346377 | 5.228194358 | 1.2679168 | 0.2764566 | 0.629109967 |
| ENSMUSG00000049038 | Mterf2 | protein_coding | -0.740010567 | 3.173659394 | 1.2666393 | 0.2766899 | 0.629109967 |
| ENSMUSG00000062995 | Ica1 | protein_coding | 0.401829074 | 6.464474724 | 1.2667408 | 0.2766979 | 0.629109967 |
| ENSMUSG00000069744 | Psmb3 | protein_coding | 0.287392542 | 7.452217863 | 1.2665184 | 0.276712 | 0.629109967 |
| ENSMUSG00000037366 | Pafah2 | protein_coding | 1.168128278 | 2.283650416 | 1.2657991 | 0.2768435 | 0.629109967 |
| ENSMUSG00000043909 | Trp53bp1 | protein_coding | -0.478358639 | 4.852830253 | 1.265086 | 0.2769739 | 0.629109967 |
| ENSMUSG00000022400 | Rbx1 | protein_coding | 0.255473676 | 9.326508806 | 1.2650632 | 0.2769781 | 0.629109967 |
| ENSMUSG00000027628 | Aar2 | protein_coding | 0.316696072 | 6.484043331 | 1.2650584 | 0.2769789 | 0.629109967 |
| ENSMUSG00000112280 | AC153962.3 | processed_transcript | -0.524838112 | 3.952618323 | 1.2648044 | 0.2770254 | 0.629109967 |
| ENSMUSG00000035798 | Zdhhc17 | protein_coding | -0.347207569 | 5.272496121 | 1.2646334 | 0.2770567 | 0.629109967 |
| ENSMUSG00000074896 | Ifit3 | protein_coding | -0.698065729 | 3.128248239 | 1.2635032 | 0.2772637 | 0.629109967 |
| ENSMUSG00000021368 | Tbc1d7 | protein_coding | 0.291782197 | 6.389363671 | 1.2634188 | 0.2772792 | 0.629109967 |
| ENSMUSG00000020993 | Trappc6b | protein_coding | -0.363105308 | 5.564164391 | 1.2630844 | 0.2773405 | 0.629109967 |
| ENSMUSG00000024425 | Ndfip1 | protein_coding | -0.26549613 | 6.807682116 | 1.2630472 | 0.2773473 | 0.629109967 |
| ENSMUSG00000037499 | Nenf | protein_coding | 0.34946283 | 5.674159859 | 1.2627352 | 0.2774045 | 0.629109967 |
| ENSMUSG00000026240 | Cops7b | protein_coding | 0.438925568 | 4.814157823 | 1.2626063 | 0.2774281 | 0.629109967 |
| ENSMUSG00000031750 | Il34 | protein_coding | 0.374286604 | 5.729122828 | 1.2624152 | 0.2774632 | 0.629109967 |
| ENSMUSG00000044367 | Slc16a13 | protein_coding | 1.513137921 | 2.677996523 | 1.2711157 | 0.2774887 | 0.629109967 |
| ENSMUSG00000033781 | Asb13 | protein_coding | -0.33166635 | 5.697185403 | 1.2622595 | 0.2774917 | 0.629109967 |
| ENSMUSG00000026833 | Olfm1 | protein_coding | 0.274114444 | 7.687292613 | 1.2611355 | 0.277698 | 0.629444923 |
| ENSMUSG00000043885 | Slc36a4 | protein_coding | 0.415831936 | 5.17974341 | 1.2586262 | 0.2781593 | 0.63035759 |
| ENSMUSG00000031928 | Mre11a | protein_coding | -0.655660292 | 3.140148877 | 1.2576702 | 0.2783353 | 0.630399842 |
| ENSMUSG00000012114 | Med15 | protein_coding | 0.397151838 | 5.038707628 | 1.2573918 | 0.2783866 | 0.630399842 |
| ENSMUSG00000021810 | Ecd | protein_coding | 0.381102274 | 6.148060357 | 1.2573347 | 0.2783971 | 0.630399842 |
| ENSMUSG00000024957 | Kcnk4 | protein_coding | 1.0322902 | 2.800689289 | 1.2571555 | 0.2784301 | 0.630399842 |
| ENSMUSG00000023967 | Mrps18a | protein_coding | 0.342971954 | 5.128822715 | 1.2569334 | 0.2784711 | 0.630399842 |
| ENSMUSG00000073678 | Pgap1 | protein_coding | -0.427293747 | 4.58660552 | 1.2565652 | 0.2785389 | 0.630420756 |
| ENSMUSG00000027067 | Ssrp1 | protein_coding | 0.301223433 | 6.496944194 | 1.2560614 | 0.2786318 | 0.630498311 |
| ENSMUSG00000038665 | Dgki | protein_coding | -0.588471114 | 3.621750182 | 1.2556679 | 0.2787044 | 0.630529908 |
| ENSMUSG00000032299 | Commd4 | protein_coding | 0.286612997 | 7.677940495 | 1.2548776 | 0.2788503 | 0.630727226 |
| ENSMUSG00000027748 | Trpc4 | protein_coding | 0.446977046 | 4.408237184 | 1.2533107 | 0.2791398 | 0.631191926 |
| ENSMUSG00000042962 | Gm5436 | processed_pseudoge | 0.509327511 | 3.503938644 | 1.2531305 | 0.2791731 | 0.631191926 |
| ENSMUSG00000027165 | B230118H07Rik | protein_coding | -0.285692196 | 7.438964117 | 1.2526288 | 0.279266 | 0.631205494 |
| ENSMUSG00000044469 | Tnfaip8l1 | protein_coding | -0.672552801 | 3.351556379 | 1.2524634 | 0.2792965 | 0.631205494 |

| ensembl_gene_id | external_gene_name | gene_biotype | log2 (fold change) | log2 (counts per million) | F-value | p-value | False discovery rate |
|---|---|---|---|---|---|---|---|
| ENSMUSG00000040033 | Stat2 | protein_coding | -0.957357121 | 2.603208494 | 1.2516923 | 0.2794393 | 0.631248048 |
| ENSMUSG00000005161 | Prdx2 | protein_coding | 0.240786794 | 9.48512235 | 1.2515118 | 0.2794727 | 0.631248048 |
| ENSMUSG00000022453 | Naga | protein_coding | 0.429415944 | 5.324071958 | 1.2514104 | 0.2794915 | 0.631248048 |
| ENSMUSG00000075410 | Prcd | protein_coding | 0.949187521 | 3.463079104 | 1.255573 | 0.2797475 | 0.63162708 |
| ENSMUSG00000020484 | Xbp1 | protein_coding | -0.39250872 | 5.883448601 | 1.2496241 | 0.2798226 | 0.63162708 |
| ENSMUSG00000027227 | Sord | protein_coding | 0.459567279 | 5.580487141 | 1.2493358 | 0.2799025 | 0.63162708 |
| ENSMUSG00000029878 | Dbpht2 | lincRNA | -0.502122143 | 8.515574096 | 1.262118 | 0.2799432 | 0.63162708 |
| ENSMUSG00000039958 | Etfbkmt | protein_coding | -0.540148175 | 4.189351122 | 1.2486315 | 0.2800068 | 0.63162708 |
| ENSMUSG00000024560 | Cxxc1 | protein_coding | 0.652545391 | 4.553512164 | 1.251495 | 0.2800141 | 0.63162708 |
| ENSMUSG00000096847 | Tmem151b | protein_coding | 0.355281831 | 6.425331316 | 1.2481293 | 0.2801001 | 0.63162708 |
| ENSMUSG00000047126 | Cltc | protein_coding | -0.250473984 | 8.0806273 | 1.2478007 | 0.2801611 | 0.63162708 |
| ENSMUSG00000041028 | Ghitm | protein_coding | 0.274303627 | 9.508313207 | 1.2474879 | 0.2802193 | 0.63162708 |
| ENSMUSG00000026074 | Map4k4 | protein_coding | -0.468481199 | 4.142938693 | 1.2471484 | 0.2802824 | 0.63162708 |
| ENSMUSG00000061477 | Rps7 | protein_coding | 0.370919547 | 5.938776465 | 1.2467415 | 0.2803581 | 0.63162708 |
| ENSMUSG00000059518 | Znhit1 | protein_coding | 0.270112825 | 7.934591468 | 1.2466144 | 0.2803817 | 0.63162708 |
| ENSMUSG00000031930 | Wwp2 | protein_coding | 0.467199598 | 5.607010301 | 1.2465176 | 0.2804261 | 0.63162708 |
| ENSMUSG00000005204 | Senp3 | protein_coding | -0.406131167 | 5.074071555 | 1.2460699 | 0.280483 | 0.63162708 |
| ENSMUSG00000074682 | Zcchc3 | protein_coding | 0.612614325 | 3.928975272 | 1.2457616 | 0.2805404 | 0.63162708 |
| ENSMUSG00000037824 | Tspan14 | protein_coding | -0.442420149 | 4.513875636 | 1.2452645 | 0.280633 | 0.631703208 |
| ENSMUSG00000027598 | Itch | protein_coding | -0.404947736 | 4.714672349 | 1.2443113 | 0.2808106 | 0.631891699 |
| ENSMUSG00000022210 | Dhrs4 | protein_coding | -0.477868839 | 4.737723596 | 1.2438437 | 0.2808977 | 0.631891699 |
| ENSMUSG00000027962 | Vcam1 | protein_coding | -0.697597216 | 3.937435322 | 1.2438012 | 0.280932 | 0.631891699 |
| ENSMUSG00000025235 | Bbs4 | protein_coding | 0.403458005 | 6.460608854 | 1.2438399 | 0.2809943 | 0.631891699 |
| ENSMUSG00000055897 | Ppp4r1l-ps | transcribed_unproces | -0.600019037 | 3.477519279 | 1.243239 | 0.2810105 | 0.631891699 |
| ENSMUSG00000035726 | Supt16 | protein_coding | -0.396572889 | 5.624381339 | 1.2427008 | 0.281111 | 0.631943955 |
| ENSMUSG00000060803 | Gstp1 | protein_coding | 0.248738249 | 8.774563866 | 1.2424847 | 0.2811513 | 0.631943955 |
| ENSMUSG00000022884 | Eif4a2 | protein_coding | 0.299262361 | 10.53247066 | 1.2418316 | 0.2812733 | 0.632086006 |
| ENSMUSG00000030255 | Sspn | protein_coding | -0.903603973 | 2.634035781 | 1.2400447 | 0.2816074 | 0.632507075 |
| ENSMUSG00000021704 | Mtx3 | protein_coding | -0.447880757 | 4.285084892 | 1.2399028 | 0.2816339 | 0.632507075 |
| ENSMUSG00000000085 | Scmh1 | protein_coding | -0.401470333 | 5.071774487 | 1.2398521 | 0.2816434 | 0.632507075 |
| ENSMUSG00000020964 | Sel1l | protein_coding | -0.3056402 | 5.974239127 | 1.2395714 | 0.2816959 | 0.632507075 |
| ENSMUSG00000022685 | Parn | protein_coding | 0.43634898 | 4.910161498 | 1.2384717 | 0.2819019 | 0.632718424 |
| ENSMUSG00000035173 | Ccdc186 | protein_coding | -0.411324738 | 5.177513361 | 1.2384404 | 0.2819077 | 0.632718424 |
| ENSMUSG00000069089 | Cdk7 | protein_coding | -0.367613279 | 5.207456483 | 1.2370159 | 0.2821748 | 0.633185748 |
| ENSMUSG00000022721 | Trmt2a | protein_coding | 0.490283124 | 5.02992924 | 1.2361429 | 0.2823387 | 0.633421254 |
| ENSMUSG00000036270 | Edc4 | protein_coding | -0.467658882 | 4.747502669 | 1.2353976 | 0.2824787 | 0.633603098 |
| ENSMUSG00000037997 | Parp11 | protein_coding | 0.462043281 | 4.771992288 | 1.2343067 | 0.2826838 | 0.633804905 |
| ENSMUSG00000046330 | Rpl37a | protein_coding | 0.446567338 | 7.908050191 | 1.2427542 | 0.2827008 | 0.633804905 |
| ENSMUSG00000003469 | Phyhip | protein_coding | 0.271772441 | 8.650892577 | 1.2339786 | 0.2827455 | 0.633804905 |
| ENSMUSG00000022008 | Gpalpp1 | protein_coding | -0.420674639 | 4.981242341 | 1.2334592 | 0.2828432 | 0.633871616 |
| ENSMUSG00000032050 | Rdx | protein_coding | -0.411579261 | 5.48007083 | 1.2330718 | 0.2829161 | 0.633871616 |
| ENSMUSG00000021982 | Cdadc1 | protein_coding | 0.29410908 | 6.603513022 | 1.2327637 | 0.2829742 | 0.633871616 |
| ENSMUSG00000031986 | Sprtn | protein_coding | -1.036808996 | 2.526641802 | 1.2325679 | 0.283011 | 0.633871616 |
| ENSMUSG00000024792 | Zfpl1 | protein_coding | 0.727196737 | 3.797924973 | 1.2299827 | 0.2835248 | 0.634890152 |
| ENSMUSG00000035671 | Zswim4 | protein_coding | 0.486452912 | 3.825529101 | 1.2276561 | 0.2839383 | 0.635560887 |
| ENSMUSG00000036764 | Dnajc12 | protein_coding | -0.244464577 | 6.958223742 | 1.2275351 | 0.2839612 | 0.635560887 |
| ENSMUSG00000023999 | Kif6 | protein_coding | -1.065006314 | 2.428414314 | 1.2272426 | 0.2840166 | 0.635560887 |
| ENSMUSG00000059534 | Uqcr10 | protein_coding | 0.325013146 | 10.21776316 | 1.2270091 | 0.2840608 | 0.635560887 |
| ENSMUSG00000032263 | Bckdhb | protein_coding | 0.493204163 | 3.918022141 | 1.2262173 | 0.2842107 | 0.635764073 |
| ENSMUSG00000049091 | Sephs2 | protein_coding | 0.533336622 | 5.725464198 | 1.2306865 | 0.2843826 | 0.635841907 |
| ENSMUSG00000040272 | Accs | protein_coding | 1.292228305 | 2.430935469 | 1.2252661 | 0.2844172 | 0.635841907 |
| ENSMUSG00000070695 | Cntnap5a | protein_coding | -0.459116047 | 4.667547053 | 1.2249888 | 0.2844436 | 0.635841907 |
| ENSMUSG00000026489 | Coq8a | protein_coding | 0.396200585 | 5.611937598 | 1.224578 | 0.2845215 | 0.635841907 |
| ENSMUSG00000025213 | Kazald1 | protein_coding | 0.808314026 | 2.824612093 | 1.2244744 | 0.2845412 | 0.635841907 |
| ENSMUSG00000025555 | Farp1 | protein_coding | -0.484615963 | 4.541455002 | 1.2237387 | 0.2846808 | 0.63584237 |
| ENSMUSG00000052928 | Ctif | protein_coding | 0.413289085 | 4.956431126 | 1.2236039 | 0.2847064 | 0.63584237 |
| ENSMUSG00000002129 | Sf3a1 | protein_coding | 0.361206742 | 6.106236639 | 1.2234227 | 0.2847408 | 0.63584237 |
| ENSMUSG00000040044 | Orc3 | protein_coding | -0.380352354 | 6.001959922 | 1.2226868 | 0.2848806 | 0.63584237 |
| ENSMUSG00000036940 | Kdm1a | protein_coding | 0.617690264 | 3.272368553 | 1.2226256 | 0.2848923 | 0.63584237 |
| ENSMUSG00000026442 | Nfasc | protein_coding | 0.292799787 | 5.774130877 | 1.2226051 | 0.2848962 | 0.63584237 |
| ENSMUSG00000031840 | Rab3a | protein_coding | 0.437141657 | 9.649739403 | 1.2273464 | 0.2850771 | 0.636114063 |
| ENSMUSG00000020523 | Fam114a2 | protein_coding | 0.322046831 | 6.023009542 | 1.2213113 | 0.2851422 | 0.636127454 |
| ENSMUSG00000001424 | Snd1 | protein_coding | 0.336943992 | 5.71235447 | 1.2209989 | 0.2852017 | 0.636128125 |
| ENSMUSG00000055200 | Sertad3 | protein_coding | -0.740025095 | 3.194498465 | 1.2204918 | 0.2852982 | 0.636211532 |
| ENSMUSG00000053483 | Usp21 | protein_coding | 0.417081339 | 5.18299884 | 1.2190982 | 0.2855638 | 0.636651285 |
| ENSMUSG00000055489 | Ano5 | protein_coding | -0.762635635 | 4.58816733 | 1.2297343 | 0.2856139 | 0.636651285 |
| ENSMUSG00000039419 | Cntnap2 | protein_coding | 0.261171545 | 7.646214872 | 1.2181872 | 0.2857376 | 0.636719711 |
| ENSMUSG00000022295 | Atp6v1c1 | protein_coding | 0.269601026 | 9.301728821 | 1.218054 | 0.285763 | 0.636719711 |
| ENSMUSG00000019082 | Slc25a22 | protein_coding | 0.397207 | 7.807801147 | 1.2223956 | 0.2858881 | 0.636866627 |
| ENSMUSG00000021945 | Zmym2 | protein_coding | -0.371102858 | 5.437061708 | 1.2168416 | 0.2859945 | 0.636971576 |
| ENSMUSG00000001270 | Ckb | protein_coding | 0.342426459 | 10.89649594 | 1.2156144 | 0.2862291 | 0.637362064 |
| ENSMUSG00000021114 | Atp6v1d | protein_coding | 0.245037693 | 9.903445503 | 1.2146189 | 0.2864196 | 0.637654206 |
| ENSMUSG00000035666 | Gtf3c4 | protein_coding | 0.57857364 | 3.779221143 | 1.212413 | 0.2868423 | 0.638463174 |
| ENSMUSG00000068551 | Zfp467 | protein_coding | -0.480463827 | 4.430571319 | 1.2118695 | 0.2869466 | 0.63856312 |
| ENSMUSG00000005656 | Snx6 | protein_coding | -0.277724864 | 7.146636293 | 1.2109631 | 0.2871206 | 0.638712853 |
| ENSMUSG00000066042 | Med18 | protein_coding | 0.416676324 | 4.473760807 | 1.2109003 | 0.2871327 | 0.638712853 |
| ENSMUSG00000021917 | Spcs1 | protein_coding | 0.329398982 | 8.833726053 | 1.2105868 | 0.2871929 | 0.638714732 |
| ENSMUSG00000031574 | Star | protein_coding | -0.758421354 | 2.702912723 | 1.2101251 | 0.2872817 | 0.638779951 |

| ensembl_gene_id | external_gene_name | gene_biotype | log2 (fold change) | log2 (counts per million) | F-value | p-value | False discovery rate |
|---|---|---|---|---|---|---|---|
| ENSMUSG00000029287 | Tgfbr3 | protein_coding | -1.073328267 | 2.925349882 | 1.2100543 | 0.28739 | 0.638888713 |
| ENSMUSG00000038084 | Opa1 | protein_coding | -0.31692189 | 5.952056589 | 1.2086881 | 0.2875581 | 0.639130201 |
| ENSMUSG00000041037 | Irgq | protein_coding | -0.419084717 | 4.537448549 | 1.2076116 | 0.2877654 | 0.639458821 |
| ENSMUSG00000015023 | Ddx19a | protein_coding | 0.703680235 | 2.818227446 | 1.2071282 | 0.2878585 | 0.639533654 |
| ENSMUSG00000040532 | Abhd11 | protein_coding | 0.285995035 | 6.511618318 | 1.2056146 | 0.2881505 | 0.639923101 |
| ENSMUSG00000021670 | Hmgcr | protein_coding | -0.313626285 | 5.731228708 | 1.2056023 | 0.2881529 | 0.639923101 |
| ENSMUSG00000030706 | Mrpl48 | protein_coding | 0.254824728 | 7.445085293 | 1.2045251 | 0.2883609 | 0.640166193 |
| ENSMUSG00000047842 | Diras2 | protein_coding | -0.411824841 | 5.535112795 | 1.204419 | 0.2883814 | 0.640166193 |
| ENSMUSG00000040128 | Pnrc1 | protein_coding | -0.323588831 | 5.564301215 | 1.202155 | 0.2888193 | 0.641005966 |
| ENSMUSG00000063172 | Hspb11 | protein_coding | -0.322218408 | 5.895855498 | 1.201727 | 0.2889022 | 0.641057623 |
| ENSMUSG00000003033 | Ap1m1 | protein_coding | 0.360687144 | 7.141286704 | 1.2005232 | 0.2891614 | 0.641221405 |
| ENSMUSG00000078786 | BC024978 | protein_coding | -0.501917132 | 4.375174023 | 1.1997095 | 0.2892933 | 0.641221405 |
| ENSMUSG00000056537 | Rlim | protein_coding | -0.418476414 | 5.508096725 | 1.1994457 | 0.2893445 | 0.641221405 |
| ENSMUSG00000024220 | Zfp523 | protein_coding | 0.520158273 | 4.664263773 | 1.1994062 | 0.2893522 | 0.641221405 |
| ENSMUSG00000064293 | Cntn4 | protein_coding | -0.451620401 | 4.833247032 | 1.1992554 | 0.2893815 | 0.641221405 |
| ENSMUSG00000040653 | Ppp1r14c | protein_coding | 0.587898046 | 3.440860724 | 1.1990207 | 0.289427 | 0.641221405 |
| ENSMUSG00000031938 | 4931406C07Rik | protein_coding | -0.618389061 | 3.821503406 | 1.198809 | 0.2894681 | 0.641221405 |
| ENSMUSG00000023192 | Grm2 | protein_coding | 0.887358017 | 3.081456582 | 1.1984238 | 0.289543 | 0.641221405 |
| ENSMUSG00000078816 | Prkcg | protein_coding | -0.434624205 | 7.171257509 | 1.2063854 | 0.2895812 | 0.641221405 |
| ENSMUSG00000039050 | Osbpl2 | protein_coding | 0.335087281 | 5.793113036 | 1.1979391 | 0.2896372 | 0.641221405 |
| ENSMUSG00000004366 | Sst | protein_coding | -0.369575995 | 8.91773104 | 1.198338 | 0.289654 | 0.641221405 |
| ENSMUSG00000034040 | Wbscr17 | protein_coding | -0.470348573 | 4.346021108 | 1.1973268 | 0.2897562 | 0.641221405 |
| ENSMUSG00000036908 | Unc93b1 | protein_coding | -1.002343294 | 3.136844393 | 1.2006698 | 0.2897658 | 0.641221405 |
| ENSMUSG00000036062 | Phf24 | protein_coding | -0.293595916 | 6.335359416 | 1.1966129 | 0.2898951 | 0.641221405 |
| ENSMUSG00000039661 | Dusp26 | protein_coding | 0.268712669 | 7.594853447 | 1.1965806 | 0.2899014 | 0.641221405 |
| ENSMUSG00000022614 | Lmf2 | protein_coding | 0.63984167 | 3.527637532 | 1.196433 | 0.2899301 | 0.641221405 |
| ENSMUSG00000018919 | Tm4sf5 | protein_coding | 0.655330141 | 3.347728474 | 1.1959078 | 0.2900324 | 0.641315631 |
| ENSMUSG00000033022 | Cdo1 | protein_coding | 0.359261153 | 5.595285487 | 1.1946719 | 0.2902732 | 0.641716129 |
| ENSMUSG00000022544 | Eef2kmt | protein_coding | -0.580853682 | 3.838473285 | 1.1938351 | 0.2904363 | 0.641944921 |
| ENSMUSG00000020623 | Map2k6 | protein_coding | -0.541550174 | 4.021855182 | 1.191353 | 0.2909211 | 0.64287097 |
| ENSMUSG00000026064 | Ptp4a1 | protein_coding | -0.455854954 | 4.085936593 | 1.1910782 | 0.2909749 | 0.64287097 |
| ENSMUSG00000003623 | Crot | protein_coding | -0.440497234 | 4.772818687 | 1.1904655 | 0.2910948 | 0.642987365 |
| ENSMUSG00000029389 | Ddx55 | protein_coding | 0.363214647 | 5.187898048 | 1.1901978 | 0.2911472 | 0.642987365 |
| ENSMUSG00000031168 | Ebp | protein_coding | 0.322785136 | 5.573726174 | 1.1898744 | 0.2912105 | 0.642995154 |
| ENSMUSG00000013236 | Ptprs | protein_coding | -0.304268503 | 7.138867334 | 1.1892696 | 0.2913289 | 0.643124622 |
| ENSMUSG00000032291 | Crabp1 | protein_coding | -0.528914021 | 4.617059342 | 1.1889019 | 0.291401 | 0.643151651 |
| ENSMUSG00000091985 | Gm17354 | lincRNA | -0.878402949 | 2.537460446 | 1.1885014 | 0.2914795 | 0.643192883 |
| ENSMUSG00000039826 | Trub2 | protein_coding | 0.557919174 | 5.013049099 | 1.1876637 | 0.2917823 | 0.643729084 |
| ENSMUSG00000022240 | Ctnnd2 | protein_coding | -0.249159693 | 7.273596759 | 1.1857791 | 0.2920139 | 0.64410772 |
| ENSMUSG00000042035 | Igsf3 | protein_coding | 0.377911922 | 5.47660172 | 1.1850085 | 0.2921654 | 0.644309751 |
| ENSMUSG00000022377 | Asap1 | protein_coding | -0.40338085 | 5.303138934 | 1.1843924 | 0.2922866 | 0.64432076 |
| ENSMUSG00000063882 | Uqcrh | protein_coding | 0.350955567 | 10.25289796 | 1.1842289 | 0.2923446 | 0.64432076 |
| ENSMUSG00000031337 | Mtm1 | protein_coding | -0.989538089 | 2.849705972 | 1.1840695 | 0.2923501 | 0.64432076 |
| ENSMUSG00000037573 | Tob1 | protein_coding | -0.346708776 | 5.356894332 | 1.1834652 | 0.2924691 | 0.644450946 |
| ENSMUSG00000051029 | Serpinb1b | protein_coding | 0.426164898 | 4.661446212 | 1.1827773 | 0.2926047 | 0.644617498 |
| ENSMUSG00000019864 | Rtn4ip1 | protein_coding | 0.45230946 | 4.815927102 | 1.1815518 | 0.2928463 | 0.644896687 |
| ENSMUSG00000019969 | Psen1 | protein_coding | 0.341983514 | 5.820382091 | 1.1808549 | 0.2929839 | 0.644896687 |
| ENSMUSG00000019230 | Lhx9 | protein_coding | -1.012952092 | 4.271214065 | 1.2032788 | 0.2930235 | 0.644896687 |
| ENSMUSG00000056342 | Usp34 | protein_coding | -0.323060925 | 5.862036858 | 1.180626 | 0.2930291 | 0.644896687 |
| ENSMUSG00000032570 | Atp2c1 | protein_coding | -0.39465661 | 5.012945453 | 1.1806149 | 0.2930313 | 0.644896687 |
| ENSMUSG00000031931 | Ankrd49 | protein_coding | -0.395643974 | 4.469545221 | 1.1801604 | 0.293121 | 0.644962215 |
| ENSMUSG00000032329 | Hmg20a | protein_coding | 0.334346389 | 6.190941691 | 1.1789261 | 0.293365 | 0.645334854 |
| ENSMUSG00000026404 | Ddx59 | protein_coding | -0.673579901 | 2.920960976 | 1.1786966 | 0.2934104 | 0.645334854 |
| ENSMUSG00000004221 | Ikbkg | protein_coding | 0.465186005 | 5.073525441 | 1.1759859 | 0.2939473 | 0.64638355 |
| ENSMUSG00000033653 | Vps8 | protein_coding | 0.481887543 | 4.796670176 | 1.1741898 | 0.2943038 | 0.646969497 |
| ENSMUSG00000053025 | Sv2b | protein_coding | -0.379541937 | 5.410793827 | 1.1740372 | 0.2943341 | 0.646969497 |
| ENSMUSG00000049907 | Rasl11b | protein_coding | 0.439017347 | 4.109289556 | 1.1734292 | 0.294455 | 0.647102821 |
| ENSMUSG00000021392 | Nol8 | protein_coding | -0.417119139 | 4.738627855 | 1.1729356 | 0.2945531 | 0.647186225 |
| ENSMUSG00000034310 | Tmem132d | protein_coding | -0.704725684 | 2.98634704 | 1.1720289 | 0.2947335 | 0.647267617 |
| ENSMUSG00000019785 | Clvs2 | protein_coding | -0.334736467 | 5.983015569 | 1.1719278 | 0.2947536 | 0.647267617 |
| ENSMUSG00000037892 | Pcdh18 | protein_coding | -0.638498864 | 3.706613125 | 1.1718419 | 0.2947707 | 0.647267617 |
| ENSMUSG00000039519 | Cyp7b1 | protein_coding | -0.663453085 | 3.486738706 | 1.1706323 | 0.2950117 | 0.647655051 |
| ENSMUSG00000071172 | Srsf3 | protein_coding | -0.262271196 | 7.133198226 | 1.1701136 | 0.2951151 | 0.647655051 |
| ENSMUSG00000061950 | Ppp4r1 | protein_coding | 0.698163361 | 3.309485181 | 1.1697914 | 0.2951793 | 0.647655051 |
| ENSMUSG00000079317 | Trappc2 | protein_coding | -0.256574859 | 6.535422888 | 1.1697474 | 0.2951881 | 0.647655051 |
| ENSMUSG00000015668 | Pdzd11 | protein_coding | -0.239153398 | 7.534398526 | 1.1691381 | 0.2953096 | 0.647751248 |
| ENSMUSG00000055805 | Fmnl1 | protein_coding | -0.47612051 | 4.428708604 | 1.1689239 | 0.2953524 | 0.647751248 |
| ENSMUSG00000050390 | C77080 | protein_coding | -0.506371099 | 4.336900118 | 1.1680531 | 0.2955263 | 0.647769565 |
| ENSMUSG00000029213 | Commd8 | protein_coding | -0.272669925 | 7.193414177 | 1.168027 | 0.2955315 | 0.647769565 |
| ENSMUSG00000017412 | Cacnb4 | protein_coding | -0.312989656 | 5.601837244 | 1.1677943 | 0.295578 | 0.647769565 |
| ENSMUSG00000019188 | H13 | protein_coding | 0.508813784 | 5.09278234 | 1.167804 | 0.2956017 | 0.647769565 |
| ENSMUSG00000022637 | Cblb | protein_coding | -0.855103984 | 3.005102669 | 1.1665472 | 0.2958273 | 0.648049854 |
| ENSMUSG00000071369 | Map3k5 | protein_coding | -0.431173859 | 4.299181997 | 1.1664331 | 0.2958502 | 0.648049854 |
| ENSMUSG00000078713 | Tomm5 | protein_coding | 0.274816343 | 6.628975089 | 1.1657503 | 0.2959868 | 0.648074048 |
| ENSMUSG00000032187 | Smarca4 | protein_coding | 0.384016404 | 5.949081 | 1.1651157 | 0.2961139 | 0.648074048 |
| ENSMUSG00000052738 | Suclg1 | protein_coding | 0.235794934 | 9.129198051 | 1.1646699 | 0.2962032 | 0.648074048 |
| ENSMUSG00000016946 | Kctd5 | protein_coding | -0.990014069 | 3.874375361 | 1.1771145 | 0.2962187 | 0.648074048 |
| ENSMUSG00000029319 | Coq2 | protein_coding | -0.376597884 | 4.903824083 | 1.1643393 | 0.2962695 | 0.648074048 |

| ensembl_gene_id | external_gene_name | gene_biotype | log2 (fold change) | log2 (counts per million) | F-value | p-value | False discovery rate |
|---|---|---|---|---|---|---|---|
| ENSMUSG00000018171 | Vmp1 | protein_coding | 0.253610992 | 7.624051019 | 1.164119 | 0.2963137 | 0.648074048 |
| ENSMUSG00000005575 | Ube2m | protein_coding | 0.305982717 | 6.758331894 | 1.1640995 | 0.2963176 | 0.648074048 |
| ENSMUSG00000067860 | Zic3 | protein_coding | -0.458278999 | 3.910919561 | 1.1639708 | 0.2963434 | 0.648074048 |
| ENSMUSG00000022556 | Hsf1 | protein_coding | 0.479572688 | 4.899558544 | 1.1632625 | 0.2964854 | 0.648252925 |
| ENSMUSG00000028139 | Riiad1 | protein_coding | -0.395534057 | 5.2233371 | 1.1621305 | 0.2967127 | 0.648495958 |
| ENSMUSG00000053716 | Dusp7 | protein_coding | -0.465104503 | 4.516290361 | 1.1621081 | 0.2967172 | 0.648495958 |
| ENSMUSG00000000916 | Nsun5 | protein_coding | 0.609759419 | 3.846835917 | 1.1609401 | 0.296952 | 0.648877135 |
| ENSMUSG00000036790 | Slitrk2 | protein_coding | 0.541441197 | 3.373395421 | 1.1606296 | 0.2970144 | 0.648881721 |
| ENSMUSG00000044792 | Isca1 | protein_coding | 0.274394102 | 6.796535888 | 1.1594472 | 0.2972524 | 0.64911466 |
| ENSMUSG00000034187 | Nsf | protein_coding | 0.429063104 | 9.831542962 | 1.1645163 | 0.2972895 | 0.64911466 |
| ENSMUSG00000020053 | Igf1 | protein_coding | -1.239319836 | 2.51168405 | 1.1592243 | 0.2973228 | 0.64911466 |
| ENSMUSG00000020277 | Pfkl | protein_coding | 0.402528962 | 7.5166223 | 1.1638364 | 0.2973625 | 0.64911466 |
| ENSMUSG00000041801 | Phlda3 | protein_coding | 0.360452747 | 5.543786398 | 1.158386 | 0.2974662 | 0.649209199 |
| ENSMUSG00000001017 | Chtop | protein_coding | -0.280678185 | 5.922652074 | 1.1567914 | 0.2977878 | 0.6496337 |
| ENSMUSG00000021143 | Pacs2 | protein_coding | 0.364668768 | 5.061377253 | 1.1567662 | 0.2977929 | 0.6496337 |
| ENSMUSG00000040824 | Snrpd2 | protein_coding | 0.314393918 | 7.163819088 | 1.156401 | 0.2978666 | 0.6496337 |
| ENSMUSG00000041703 | Zic5 | protein_coding | -0.706703713 | 3.725062182 | 1.1563508 | 0.2979023 | 0.6496337 |
| ENSMUSG00000015085 | Entpd2 | protein_coding | -0.704329016 | 2.870821829 | 1.1552937 | 0.2980904 | 0.649636398 |
| ENSMUSG00000028454 | Pigo | protein_coding | 0.734081132 | 3.200219405 | 1.1552136 | 0.2981066 | 0.649636398 |
| ENSMUSG00000058571 | Gpc6 | protein_coding | -0.620046372 | 3.164254474 | 1.155031 | 0.2981435 | 0.649636398 |
| ENSMUSG00000005625 | Psmd4 | protein_coding | 0.248712889 | 8.643510323 | 1.1546803 | 0.2982144 | 0.649636398 |
| ENSMUSG00000063275 | Hacd1 | protein_coding | -0.900382194 | 2.469023942 | 1.154382 | 0.2982747 | 0.649636398 |
| ENSMUSG00000031731 | Ap1g1 | protein_coding | 0.681072096 | 3.130153161 | 1.1543084 | 0.2982896 | 0.649636398 |
| ENSMUSG00000027350 | Chgb | protein_coding | 0.262300039 | 7.543770136 | 1.1537437 | 0.2984039 | 0.649636398 |
| ENSMUSG00000040506 | Ambra1 | protein_coding | -0.430523415 | 5.066546137 | 1.1535211 | 0.298449 | 0.649636398 |
| ENSMUSG00000007827 | Ankrd26 | protein_coding | -0.518778351 | 4.709306544 | 1.1530483 | 0.2985702 | 0.649636398 |
| ENSMUSG00000025381 | Cnpy2 | protein_coding | 0.248230009 | 7.272330767 | 1.1523773 | 0.2986807 | 0.649636398 |
| ENSMUSG00000024906 | Mus81 | protein_coding | 0.451687142 | 4.973403595 | 1.1522377 | 0.298709 | 0.649636398 |
| ENSMUSG00000033906 | Zdhhc15 | protein_coding | -1.030042994 | 2.51327674 | 1.1519658 | 0.2987641 | 0.649636398 |
| ENSMUSG00000031954 | Cfdp1 | protein_coding | 0.227826334 | 8.68150032 | 1.151921 | 0.2987732 | 0.649636398 |
| ENSMUSG00000014763 | Fam120b | protein_coding | 0.30299193 | 6.325271253 | 1.1518327 | 0.2987911 | 0.649636398 |
| ENSMUSG00000027447 | Cst3 | protein_coding | -0.516588305 | 11.15737554 | 1.1625472 | 0.2988098 | 0.649636398 |
| ENSMUSG00000028385 | Snx30 | protein_coding | -0.365650898 | 4.822640067 | 1.1510095 | 0.2989581 | 0.649645611 |
| ENSMUSG00000035069 | Oma1 | protein_coding | -0.573211595 | 3.725733586 | 1.1508686 | 0.2989867 | 0.649645611 |
| ENSMUSG00000015961 | Adss | protein_coding | -0.303994674 | 5.890868046 | 1.1503761 | 0.2990867 | 0.649645611 |
| ENSMUSG00000025190 | Got1 | protein_coding | 0.341854177 | 9.60610963 | 1.1502202 | 0.2991439 | 0.649645611 |
| ENSMUSG00000046230 | Vps13a | protein_coding | -0.522712134 | 3.501451692 | 1.1500107 | 0.2991609 | 0.649645611 |
| ENSMUSG00000028423 | Nfx1 | protein_coding | 0.334124335 | 6.440801969 | 1.1495934 | 0.2992457 | 0.649645611 |
| ENSMUSG00000050856 | Atp5k | protein_coding | 0.361833055 | 9.580473233 | 1.1495715 | 0.2992756 | 0.649645611 |
| ENSMUSG00000003208 | Ccdc94 | protein_coding | 0.648834648 | 3.964551078 | 1.1494019 | 0.2993101 | 0.649645611 |
| ENSMUSG00000040697 | Dnajc16 | protein_coding | 0.589286632 | 4.041502549 | 1.1490422 | 0.2993578 | 0.649645611 |
| ENSMUSG00000004364 | Cul3 | protein_coding | -0.269920274 | 6.451955058 | 1.1483948 | 0.2994894 | 0.649679015 |
| ENSMUSG00000028838 | Extl1 | protein_coding | 0.404847115 | 4.424538236 | 1.148117 | 0.299546 | 0.649679015 |
| ENSMUSG00000025437 | Usp33 | protein_coding | 0.734175558 | 3.952541438 | 1.151274 | 0.2995544 | 0.649679015 |
| ENSMUSG00000022214 | Dcaf11 | protein_coding | 0.402498575 | 5.684362016 | 1.1471206 | 0.2997488 | 0.649969557 |
| ENSMUSG00000035984 | Nme5 | protein_coding | -0.309196437 | 6.20005046 | 1.1465618 | 0.2998627 | 0.650085349 |
| ENSMUSG00000018583 | G3bp1 | protein_coding | 0.301429017 | 6.057077749 | 1.145919 | 0.2999937 | 0.650238331 |
| ENSMUSG00000021591 | Glrx | protein_coding | 0.298273374 | 6.682961607 | 1.145267 | 0.3001267 | 0.650308539 |
| ENSMUSG00000026073 | Il1r2 | protein_coding | 1.27551109 | 2.054757729 | 1.1449124 | 0.3001991 | 0.650308539 |
| ENSMUSG00000059878 | Zfp422 | protein_coding | 0.383379584 | 5.255728432 | 1.1448709 | 0.3002075 | 0.650308539 |
| ENSMUSG00000038132 | Rbm24 | protein_coding | 0.463472583 | 4.232803002 | 1.1445544 | 0.3002722 | 0.650317532 |
| ENSMUSG00000021065 | Fut8 | protein_coding | -0.393079351 | 4.914717648 | 1.1433905 | 0.3005099 | 0.65070142 |
| ENSMUSG00000105879 | Gm6204 | processed_pseudoge | 0.499202144 | 3.852311768 | 1.1429381 | 0.3006024 | 0.650770634 |
| ENSMUSG00000064030 | Pym1 | protein_coding | 0.398097756 | 4.753457262 | 1.1425872 | 0.3006742 | 0.650794997 |
| ENSMUSG00000047181 | Samd14 | protein_coding | 0.365966591 | 5.095027515 | 1.1419221 | 0.3008103 | 0.650828127 |
| ENSMUSG00000019558 | Slc6a8 | protein_coding | -0.361096678 | 5.170336647 | 1.1415562 | 0.3008852 | 0.650828127 |
| ENSMUSG00000056771 | Gm10010 | lincRNA | 1.190027644 | 2.306700587 | 1.1412961 | 0.3009384 | 0.650828127 |
| ENSMUSG00000012483 | Rpa3 | protein_coding | -0.504284894 | 4.794647199 | 1.1412225 | 0.3009535 | 0.650828127 |
| ENSMUSG00000024766 | Lipo3 | protein_coding | -0.629084282 | 3.2692085 | 1.1406056 | 0.3010799 | 0.650828127 |
| ENSMUSG00000019832 | Rab32 | protein_coding | -1.534870785 | 1.917970587 | 1.140711 | 0.3010837 | 0.650828127 |
| ENSMUSG00000019790 | Stxbp5 | protein_coding | -0.366546707 | 5.111077448 | 1.1404432 | 0.3011132 | 0.650828127 |
| ENSMUSG00000035559 | Mpv17l2 | protein_coding | 0.731620445 | 3.124717197 | 1.139963 | 0.3012116 | 0.65089161 |
| ENSMUSG00000041915 | Ammecr1l | protein_coding | 0.626344909 | 3.297961726 | 1.139572 | 0.3012918 | 0.65089161 |
| ENSMUSG00000048707 | Tprn | protein_coding | -0.472635433 | 4.150384956 | 1.138964 | 0.3014166 | 0.65089161 |
| ENSMUSG00000035198 | Tubg1 | protein_coding | 0.376703123 | 6.697199905 | 1.1392077 | 0.3014851 | 0.65089161 |
| ENSMUSG00000059423 | Zfp933 | protein_coding | -0.733519464 | 3.282108402 | 1.1386108 | 0.3014891 | 0.65089161 |
| ENSMUSG00000028893 | Sesn2 | protein_coding | 0.685859709 | 3.422479289 | 1.1383022 | 0.3015524 | 0.65089161 |
| ENSMUSG00000079316 | Rab9 | protein_coding | -0.29855491 | 6.864754682 | 1.1377683 | 0.3016621 | 0.65089161 |
| ENSMUSG00000041923 | Nol4 | protein_coding | -0.29339386 | 6.112688642 | 1.1377611 | 0.3016636 | 0.65089161 |
| ENSMUSG00000052423 | B4galt3 | protein_coding | 0.346266226 | 5.498150261 | 1.1376456 | 0.3016873 | 0.65089161 |
| ENSMUSG00000025810 | Nrp1 | protein_coding | -0.487495833 | 4.129597708 | 1.1368553 | 0.3018498 | 0.650982415 |
| ENSMUSG00000109776 | Gm30400 | processed_transcript | -0.98355421 | 2.474284492 | 1.1366172 | 0.3018988 | 0.650982415 |
| ENSMUSG00000022191 | Drosha | protein_coding | 0.309423282 | 6.414153116 | 1.1365576 | 0.301911 | 0.650982415 |
| ENSMUSG00000021420 | Fars2 | protein_coding | 0.311659321 | 5.774157552 | 1.1355492 | 0.3021186 | 0.651201134 |
| ENSMUSG00000041617 | Ccdc74a | protein_coding | -0.596936743 | 3.585730457 | 1.1353897 | 0.3021514 | 0.651201134 |
| ENSMUSG00000056629 | Fkbp2 | protein_coding | 0.280658434 | 9.108188008 | 1.1351822 | 0.3021941 | 0.651201134 |
| ENSMUSG00000014846 | Tppp3 | protein_coding | 0.305849475 | 8.447401721 | 1.13454 | 0.3023265 | 0.65124368 |
| ENSMUSG00000050022 | Amz1 | protein_coding | 0.422705532 | 4.858056728 | 1.1344984 | 0.302335 | 0.65124368 |

| ensembl_gene_id | external_gene_name | gene_biotype | log2 (fold change) | log2 (counts per million) | F-value | p-value | False discovery rate |
|---|---|---|---|---|---|---|---|
| ENSMUSG00000025499 | Hras | protein_coding | 0.325111334 | 6.458936108 | 1.1337926 | 0.3024805 | 0.651426636 |
| ENSMUSG00000022044 | Stmn4 | protein_coding | 0.312681344 | 8.71982359 | 1.1329428 | 0.3026559 | 0.651559095 |
| ENSMUSG00000040888 | Gfer | protein_coding | 0.380841563 | 4.47667064 | 1.1325071 | 0.3027458 | 0.651559095 |
| ENSMUSG00000024308 | Tapbp | protein_coding | 0.369926071 | 4.960714711 | 1.1323906 | 0.3027699 | 0.651559095 |
| ENSMUSG00000024251 | Thada | protein_coding | -0.569914151 | 3.259667911 | 1.1323202 | 0.3027844 | 0.651559095 |
| ENSMUSG00000028452 | Vcp | protein_coding | 0.818897068 | 2.751292978 | 1.1319237 | 0.3028663 | 0.651604964 |
| ENSMUSG00000070814 | 6330408A02Rik | protein_coding | 0.722354109 | 3.603768717 | 1.1304846 | 0.3031891 | 0.652168966 |
| ENSMUSG00000019467 | Arhgef25 | protein_coding | -0.41961342 | 4.862343665 | 1.1297989 | 0.3033058 | 0.652289347 |
| ENSMUSG00000026466 | Tor1aip1 | protein_coding | -0.486817234 | 3.687747395 | 1.1280613 | 0.3036658 | 0.652808841 |
| ENSMUSG00000022620 | Arsa | protein_coding | 0.465184062 | 4.693227459 | 1.128047 | 0.3036687 | 0.652808841 |
| ENSMUSG00000021578 | Ccdc127 | protein_coding | 0.27442441 | 6.335168064 | 1.1270026 | 0.3038854 | 0.652992403 |
| ENSMUSG00000094707 | A830019P07Rik | lincRNA | -1.034690188 | 2.449636593 | 1.1260482 | 0.3040836 | 0.652992403 |
| ENSMUSG00000032571 | Pik3r4 | protein_coding | 0.530520362 | 4.210568491 | 1.1251725 | 0.3042656 | 0.652992403 |
| ENSMUSG00000005615 | Pcyt1a | protein_coding | -0.439735922 | 5.212254866 | 1.1250496 | 0.3042911 | 0.652992403 |
| ENSMUSG00000029029 | Wrap73 | protein_coding | 0.318015144 | 5.753226431 | 1.1248232 | 0.3043382 | 0.652992403 |
| ENSMUSG00000043671 | Dpy19l3 | protein_coding | 0.524555046 | 4.19884492 | 1.1243714 | 0.3044322 | 0.652992403 |
| ENSMUSG00000027806 | Tsc22d2 | protein_coding | -0.427023674 | 4.615468649 | 1.1242474 | 0.304458 | 0.652992403 |
| ENSMUSG00000086649 | Gm15286 | processed_transcript | 0.743917423 | 2.672795255 | 1.1240864 | 0.3044915 | 0.652992403 |
| ENSMUSG00000085084 | 4930570G19Rik | lincRNA | -0.433192296 | 4.099744963 | 1.1238768 | 0.3045351 | 0.652992403 |
| ENSMUSG00000075478 | Slitrk1 | protein_coding | 0.438368094 | 4.949897252 | 1.1236691 | 0.3045784 | 0.652992403 |
| ENSMUSG00000003354 | Ccdc65 | protein_coding | 0.497802253 | 4.403472492 | 1.1236486 | 0.3045827 | 0.652992403 |
| ENSMUSG00000021484 | Lman2 | protein_coding | 0.395372251 | 5.333730486 | 1.1234905 | 0.3046156 | 0.652992403 |
| ENSMUSG00000001729 | Akt1 | protein_coding | 0.247425913 | 6.455986 | 1.1234443 | 0.3046252 | 0.652992403 |
| ENSMUSG00000035093 | Secisbp2l | protein_coding | -0.332558853 | 6.285429562 | 1.1233197 | 0.3046511 | 0.652992403 |
| ENSMUSG00000027599 | Armc1 | protein_coding | 0.285510844 | 7.614774014 | 1.1231386 | 0.3046889 | 0.652992403 |
| ENSMUSG00000021848 | Otx2 | protein_coding | -0.640574082 | 3.240621142 | 1.1226225 | 0.3047964 | 0.652992403 |
| ENSMUSG00000033579 | Fa2h | protein_coding | -0.405740265 | 4.900477609 | 1.1224064 | 0.3048414 | 0.652992403 |
| ENSMUSG00000002580 | Mien1 | protein_coding | 0.291972489 | 7.03482004 | 1.1220779 | 0.3049099 | 0.652992403 |
| ENSMUSG00000029198 | Grpel1 | protein_coding | 0.24543853 | 8.453275361 | 1.122018 | 0.3049224 | 0.652992403 |
| ENSMUSG00000030120 | Mlf2 | protein_coding | -0.219009923 | 7.984126122 | 1.1217748 | 0.3049732 | 0.652992403 |
| ENSMUSG00000004843 | Chmp2b | protein_coding | -0.370356393 | 5.617063188 | 1.1215053 | 0.3050294 | 0.652992403 |
| ENSMUSG00000028334 | Nans | protein_coding | 0.54815002 | 3.834587374 | 1.1203015 | 0.3052807 | 0.653106115 |
| ENSMUSG00000027774 | Gfm1 | protein_coding | -0.406619753 | 4.742264326 | 1.1200538 | 0.3053324 | 0.653106115 |
| ENSMUSG00000024078 | Ttc27 | protein_coding | 0.660737027 | 4.069331207 | 1.1203395 | 0.3053904 | 0.653106115 |
| ENSMUSG00000031813 | Mvb12a | protein_coding | 0.458485405 | 4.721394405 | 1.1195604 | 0.3054355 | 0.653106115 |
| ENSMUSG00000020585 | Laptm4a | protein_coding | -0.284108304 | 8.703104482 | 1.1195225 | 0.3054434 | 0.653106115 |
| ENSMUSG00000038429 | Usp5 | protein_coding | 0.424452781 | 8.407859596 | 1.1267329 | 0.3054733 | 0.653106115 |
| ENSMUSG00000067629 | Syngap1 | protein_coding | -0.525184817 | 3.426343774 | 1.1191288 | 0.3055258 | 0.653106115 |
| ENSMUSG00000023050 | Map3k12 | protein_coding | -0.512380243 | 4.076085082 | 1.1188963 | 0.3055744 | 0.653106115 |
| ENSMUSG00000048351 | Coa7 | protein_coding | 0.30125341 | 5.885505597 | 1.1186345 | 0.3056291 | 0.653106115 |
| ENSMUSG00000047248 | C2cd3 | protein_coding | -1.209091168 | 2.391018639 | 1.1182852 | 0.3057274 | 0.653139541 |
| ENSMUSG00000020814 | Mxra7 | protein_coding | -0.495237566 | 3.90545563 | 1.1179793 | 0.3057662 | 0.653139541 |
| ENSMUSG00000020476 | Dbnl | protein_coding | 0.255211927 | 6.914025836 | 1.1171597 | 0.3059379 | 0.653356867 |
| ENSMUSG00000035615 | Frmpd1 | protein_coding | 1.203035636 | 2.221045393 | 1.1169133 | 0.3059895 | 0.653356867 |
| ENSMUSG00000026926 | Pmpca | protein_coding | 0.394353779 | 6.246171186 | 1.1158621 | 0.306235 | 0.65367292 |
| ENSMUSG00000020326 | Ccng1 | protein_coding | -0.405013384 | 4.812654868 | 1.1156274 | 0.3062591 | 0.65367292 |
| ENSMUSG00000046139 | Patl1 | protein_coding | -0.533790631 | 3.763403618 | 1.1140482 | 0.3065907 | 0.6540031 |
| ENSMUSG00000000295 | Hddc2 | protein_coding | 0.291852097 | 7.101080233 | 1.1136043 | 0.306684 | 0.6540031 |
| ENSMUSG00000020524 | Gria1 | protein_coding | 0.244601545 | 8.10925162 | 1.1132343 | 0.3067617 | 0.6540031 |
| ENSMUSG00000004096 | Cwc15 | protein_coding | -0.269985786 | 8.159173265 | 1.1131985 | 0.3067693 | 0.6540031 |
| ENSMUSG00000041453 | Rpl21 | protein_coding | 0.3323505 | 5.190588427 | 1.1130378 | 0.3068031 | 0.6540031 |
| ENSMUSG00000029471 | Camkk2 | protein_coding | -0.431877428 | 4.662695683 | 1.1130363 | 0.3068034 | 0.6540031 |
| ENSMUSG00000021022 | Ppp2r3c | protein_coding | 0.386626484 | 4.722584434 | 1.1127507 | 0.3068635 | 0.6540031 |
| ENSMUSG00000044122 | Proca1 | protein_coding | -0.495463745 | 3.87143142 | 1.1121409 | 0.3069918 | 0.6540031 |
| ENSMUSG00000043940 | Wdfy3 | protein_coding | -0.406017529 | 5.725792591 | 1.1120973 | 0.307001 | 0.6540031 |
| ENSMUSG00000028101 | Pias3 | protein_coding | 0.524558231 | 5.432729769 | 1.115989 | 0.3071027 | 0.6540031 |
| ENSMUSG00000042229 | Rabif | protein_coding | 0.261845911 | 6.905860524 | 1.110569 | 0.3073229 | 0.6540031 |
| ENSMUSG00000043496 | Tril | protein_coding | -0.750047388 | 4.010766033 | 1.1151525 | 0.3073318 | 0.6540031 |
| ENSMUSG00000042628 | Zfyve1 | protein_coding | 0.40522647 | 4.594394296 | 1.1104405 | 0.30735 | 0.6540031 |
| ENSMUSG00000067870 | Rpl31-ps8 | processed_pseudoge | 0.574010448 | 3.545993671 | 1.1103712 | 0.3073646 | 0.6540031 |
| ENSMUSG00000063146 | Clip2 | protein_coding | 0.567612633 | 3.68875427 | 1.1101685 | 0.3074073 | 0.6540031 |
| ENSMUSG00000044763 | Trmt10c | protein_coding | -0.458916192 | 4.636811629 | 1.1100867 | 0.3074246 | 0.6540031 |
| ENSMUSG00000024095 | Hnrnpll | protein_coding | -0.435450999 | 5.037191559 | 1.1099769 | 0.3074478 | 0.6540031 |
| ENSMUSG00000036057 | Ptpn23 | protein_coding | -0.540376211 | 3.875969448 | 1.1086256 | 0.3077329 | 0.6544536 |
| ENSMUSG00000041420 | Meis3 | protein_coding | 0.424870084 | 4.46361733 | 1.1083969 | 0.3077813 | 0.6544536 |
| ENSMUSG00000037022 | Mmaa | protein_coding | -0.406617936 | 4.865514758 | 1.1079876 | 0.3078677 | 0.654508049 |
| ENSMUSG00000074415 | 2610203C20Rik | processed_transcript | -0.339655607 | 5.094700376 | 1.1072769 | 0.308018 | 0.654523493 |
| ENSMUSG00000040112 | Mrps35 | protein_coding | 0.47798464 | 4.382661176 | 1.1069486 | 0.3080874 | 0.654523493 |
| ENSMUSG00000033488 | BC026585 | protein_coding | -0.542242703 | 4.166418027 | 1.1066985 | 0.3081403 | 0.654523493 |
| ENSMUSG00000048332 | Lhfp | protein_coding | -0.693134807 | 3.587935089 | 1.1064338 | 0.3082214 | 0.654523493 |
| ENSMUSG00000023904 | Hcfc1r1 | protein_coding | 0.225133191 | 8.646237457 | 1.1062157 | 0.3082425 | 0.654523493 |
| ENSMUSG00000050751 | Pgbd5 | protein_coding | -0.26828377 | 6.575469217 | 1.1058765 | 0.3083143 | 0.654523493 |
| ENSMUSG00000006057 | Atp5g1 | protein_coding | 0.275139065 | 5.899637889 | 1.1058121 | 0.3083279 | 0.654523493 |
| ENSMUSG00000096910 | Zfp955b | protein_coding | -0.799015612 | 2.486901328 | 1.10538 | 0.3084194 | 0.654523493 |
| ENSMUSG00000075324 | Fign | protein_coding | -0.638582862 | 4.297662235 | 1.1075857 | 0.3084228 | 0.654523493 |
| ENSMUSG00000049728 | Zfp668 | protein_coding | 0.551253056 | 3.787904068 | 1.1044486 | 0.3086169 | 0.654806042 |
| ENSMUSG00000097616 | 1110019D14Rik | lincRNA | -0.64947194 | 3.245475465 | 1.1041052 | 0.3086897 | 0.65483131 |
| ENSMUSG00000035642 | Aamdc | protein_coding | 0.274751696 | 7.16647457 | 1.1037451 | 0.3087661 | 0.654848801 |

| ensembl_gene_id | external_gene_name | gene_biotype | log2 (fold change) | log2 (counts per million) | F-value | p-value | False discovery rate |
|---|---|---|---|---|---|---|---|
| ENSMUSG00000042680 | Garem1 | protein_coding | -0.552981021 | 3.877893025 | 1.1034923 | 0.3088197 | 0.654848801 |
| ENSMUSG00000022389 | Tef | protein_coding | -0.219842932 | 7.112904169 | 1.1027048 | 0.3089869 | 0.655000082 |
| ENSMUSG00000022247 | Brix1 | protein_coding | -0.254320566 | 6.667334184 | 1.1022907 | 0.3090749 | 0.655000082 |
| ENSMUSG00000004267 | Eno2 | protein_coding | 0.327852451 | 10.23750331 | 1.1017086 | 0.3091986 | 0.655000082 |
| ENSMUSG00000021816 | Ppp3cb | protein_coding | -0.262312322 | 7.762420812 | 1.1016131 | 0.3092189 | 0.655000082 |
| ENSMUSG00000028613 | Lrp8 | protein_coding | -0.549058209 | 4.297031855 | 1.1015183 | 0.309239 | 0.655000082 |
| ENSMUSG00000055403 | 4933427D06Rik | lincRNA | -0.781229005 | 3.222625254 | 1.1015535 | 0.3092565 | 0.655000082 |
| ENSMUSG00000097868 | A330069E16Rik | lincRNA | -0.57104004 | 4.657656107 | 1.1011499 | 0.3094081 | 0.655077439 |
| ENSMUSG00000022515 | Anks3 | protein_coding | 0.523080565 | 4.971730314 | 1.1006719 | 0.309444 | 0.655077439 |
| ENSMUSG00000048997 | Atxn7l2 | protein_coding | -0.689405546 | 2.92928557 | 1.1004052 | 0.3094758 | 0.655077439 |
| ENSMUSG00000028111 | Ctsk | protein_coding | -1.069630016 | 3.124977368 | 1.104451 | 0.3098952 | 0.65571077 |
| ENSMUSG00000109865 | Hspa14 | protein_coding | 0.324728725 | 5.770359006 | 1.0982977 | 0.3099248 | 0.65571077 |
| ENSMUSG00000003382 | Etv3 | protein_coding | -0.448261013 | 3.984042676 | 1.0981425 | 0.309958 | 0.65571077 |
| ENSMUSG00000054477 | Kcnn2 | protein_coding | -0.487985298 | 3.612487642 | 1.0969257 | 0.3102177 | 0.655830439 |
| ENSMUSG00000050821 | Fam131a | protein_coding | 0.239483642 | 6.95714607 | 1.0965412 | 0.3102998 | 0.655830439 |
| ENSMUSG00000066129 | Kndc1 | protein_coding | -0.431884137 | 4.97263177 | 1.0964053 | 0.3103288 | 0.655830439 |
| ENSMUSG00000059013 | Sh2d3c | protein_coding | 0.521913286 | 4.324343061 | 1.0963116 | 0.3103488 | 0.655830439 |
| ENSMUSG00000026020 | Nop58 | protein_coding | -0.292571978 | 6.335977242 | 1.0962475 | 0.3103625 | 0.655830439 |
| ENSMUSG00000025290 | Rps24 | protein_coding | 0.335709586 | 8.061290589 | 1.0962802 | 0.3103805 | 0.655830439 |
| ENSMUSG00000023274 | Cd4 | protein_coding | 0.425630982 | 4.681910182 | 1.0945543 | 0.3107246 | 0.656071807 |
| ENSMUSG00000039801 | 2410089E03Rik | protein_coding | 0.658258877 | 3.251676142 | 1.0941755 | 0.3108057 | 0.656071807 |
| ENSMUSG00000052373 | Mpp3 | protein_coding | 0.423350257 | 5.007328505 | 1.0939424 | 0.3108556 | 0.656071807 |
| ENSMUSG00000025724 | Sec11a | protein_coding | 0.293607483 | 6.941212833 | 1.0939333 | 0.3108576 | 0.656071807 |
| ENSMUSG00000079157 | Fam155a | protein_coding | -0.311198981 | 5.533616211 | 1.093797 | 0.3108868 | 0.656071807 |
| ENSMUSG00000031934 | Panx1 | protein_coding | 0.564042465 | 3.856228081 | 1.0934855 | 0.3109535 | 0.656071807 |
| ENSMUSG00000054708 | Ankrd24 | protein_coding | 0.523978816 | 3.893537642 | 1.09346 | 0.310959 | 0.656071807 |
| ENSMUSG00000035517 | Tdrd7 | protein_coding | 0.530427308 | 4.36072328 | 1.0933489 | 0.3109828 | 0.656071807 |
| ENSMUSG00000058715 | Fcer1g | protein_coding | -0.570861105 | 4.652242436 | 1.0934221 | 0.3110576 | 0.656100863 |
| ENSMUSG00000000125 | Wnt3 | protein_coding | -0.630573637 | 4.082509671 | 1.0921829 | 0.3112577 | 0.656394101 |
| ENSMUSG00000038506 | Dcun1d2 | protein_coding | 0.378689328 | 5.279136708 | 1.0907633 | 0.3115375 | 0.656550201 |
| ENSMUSG00000032097 | Ddx6 | protein_coding | -0.251535988 | 6.223176284 | 1.0901794 | 0.311663 | 0.656550201 |
| ENSMUSG00000022094 | Slc39a14 | protein_coding | -0.738696023 | 3.064125829 | 1.0900549 | 0.3116898 | 0.656550201 |
| ENSMUSG00000026618 | Iars2 | protein_coding | 0.589536934 | 3.856479725 | 1.0895653 | 0.311795 | 0.656550201 |
| ENSMUSG00000001985 | Grik3 | protein_coding | -0.412178827 | 4.438517667 | 1.088789 | 0.311962 | 0.656550201 |
| ENSMUSG00000069171 | Nr2f1 | protein_coding | -0.387197439 | 4.585679755 | 1.0887878 | 0.3119623 | 0.656550201 |
| ENSMUSG00000048537 | Phldb1 | protein_coding | -0.384371045 | 4.846870434 | 1.0886767 | 0.3119862 | 0.656550201 |
| ENSMUSG00000035637 | Grhpr | protein_coding | 0.271752584 | 7.164007776 | 1.0884642 | 0.3120319 | 0.656550201 |
| ENSMUSG00000020515 | Cnot8 | protein_coding | 0.307687284 | 6.864369852 | 1.0878999 | 0.3121535 | 0.656550201 |
| ENSMUSG00000018293 | Pfn1 | protein_coding | 0.286636609 | 6.820414673 | 1.0878571 | 0.3121627 | 0.656550201 |
| ENSMUSG00000074578 | Zfas1 | processed_transcript | 0.342493524 | 7.938346674 | 1.0879385 | 0.31217 | 0.656550201 |
| ENSMUSG00000029048 | Rer1 | protein_coding | 0.283138146 | 6.511122146 | 1.0876861 | 0.3121995 | 0.656550201 |
| ENSMUSG00000044795 | Cyb5d1 | protein_coding | 0.793449865 | 3.322605512 | 1.0873852 | 0.3122892 | 0.656550201 |
| ENSMUSG00000020321 | Mdh1 | protein_coding | 0.259851274 | 11.90593286 | 1.0868464 | 0.3123805 | 0.656550201 |
| ENSMUSG00000021385 | Ippk | protein_coding | 0.499830394 | 4.175136875 | 1.0861349 | 0.312534 | 0.656550201 |
| ENSMUSG00000020599 | Rgs9 | protein_coding | -0.298573014 | 6.232323517 | 1.0860038 | 0.3125623 | 0.656550201 |
| ENSMUSG00000055531 | Cpsf6 | protein_coding | 0.339605052 | 5.774154191 | 1.0859713 | 0.3125693 | 0.656550201 |
| ENSMUSG00000030201 | Lrp6 | protein_coding | -0.529651913 | 3.641438725 | 1.085737 | 0.3126198 | 0.656550201 |
| ENSMUSG00000022125 | Cln5 | protein_coding | -0.612638659 | 3.400346536 | 1.0856702 | 0.3126342 | 0.656550201 |
| ENSMUSG00000002325 | Irf9 | protein_coding | 0.591640231 | 3.572885205 | 1.0853781 | 0.3126973 | 0.656550201 |
| ENSMUSG00000044148 | 1810030O07Rik | protein_coding | 0.322869663 | 5.633291775 | 1.085343 | 0.3127049 | 0.656550201 |
| ENSMUSG00000023367 | Tmem176a | protein_coding | -0.423306584 | 5.30402105 | 1.0851884 | 0.3127383 | 0.656550201 |
| ENSMUSG00000061273 | Mmgt1 | protein_coding | -0.403068389 | 4.400901462 | 1.0849754 | 0.3127843 | 0.656550201 |
| ENSMUSG00000033436 | Armcx2 | protein_coding | 0.299459851 | 6.195995287 | 1.0849163 | 0.312797 | 0.656550201 |
| ENSMUSG00000029047 | Pex10 | protein_coding | 0.352257055 | 4.967391868 | 1.0843154 | 0.3129269 | 0.656637009 |
| ENSMUSG00000068250 | Amn1 | protein_coding | -0.330823738 | 5.656343951 | 1.0841598 | 0.3129605 | 0.656637009 |
| ENSMUSG00000035790 | Cep19 | protein_coding | 0.233105542 | 7.53305366 | 1.0837113 | 0.3130575 | 0.656653979 |
| ENSMUSG00000037730 | Mynn | protein_coding | -0.436936459 | 4.006506073 | 1.0835575 | 0.3130908 | 0.656653979 |
| ENSMUSG00000061393 | Acvr2b | protein_coding | 0.842167955 | 2.642841186 | 1.0829875 | 0.3132141 | 0.656755101 |
| ENSMUSG00000000131 | Xpo6 | protein_coding | 0.540720425 | 5.281730071 | 1.087144 | 0.313278 | 0.656755101 |
| ENSMUSG00000025939 | Ube2w | protein_coding | -0.238217541 | 8.205577493 | 1.0824881 | 0.3133222 | 0.656755101 |
| ENSMUSG00000002980 | Bcam | protein_coding | 0.953641803 | 3.430448586 | 1.0890776 | 0.3134079 | 0.656806663 |
| ENSMUSG00000000826 | Dnajc5 | protein_coding | 0.250596841 | 8.25432264 | 1.081631 | 0.3135079 | 0.656888157 |
| ENSMUSG00000055850 | Rnf181 | protein_coding | -0.22763553 | 7.940958936 | 1.0806029 | 0.3137307 | 0.657009419 |
| ENSMUSG00000029661 | Col1a2 | protein_coding | -0.924985413 | 3.187938435 | 1.0828366 | 0.3137488 | 0.657009419 |
| ENSMUSG00000031479 | Vps36 | protein_coding | 0.347495669 | 5.319831092 | 1.0803599 | 0.3137835 | 0.657009419 |
| ENSMUSG00000024283 | Wac | protein_coding | -0.305665816 | 5.65565543 | 1.080237 | 0.3138101 | 0.657009419 |
| ENSMUSG00000036894 | Rap2b | protein_coding | 0.355895447 | 6.864916024 | 1.0799767 | 0.3139566 | 0.657024268 |
| ENSMUSG00000034006 | Pqlc1 | protein_coding | 0.410662362 | 5.558147472 | 1.0792597 | 0.3140223 | 0.657024268 |
| ENSMUSG00000053040 | Aph1c | protein_coding | 0.570738195 | 3.38231935 | 1.0791368 | 0.314049 | 0.657024268 |
| ENSMUSG00000036306 | Lzts1 | protein_coding | 0.267784297 | 8.608718317 | 1.0790786 | 0.3140616 | 0.657024268 |
| ENSMUSG00000024072 | Yipf4 | protein_coding | -0.383725157 | 5.55137118 | 1.0786067 | 0.3141642 | 0.65711094 |
| ENSMUSG00000038260 | Trpm4 | protein_coding | 0.421088146 | 4.247393398 | 1.0782311 | 0.3142458 | 0.657153888 |
| ENSMUSG00000050473 | Slc35d3 | protein_coding | -0.747674681 | 3.052097934 | 1.0776109 | 0.3143807 | 0.65730814 |
| ENSMUSG00000020335 | Zfp354b | protein_coding | -0.607246804 | 3.46258822 | 1.0755689 | 0.3148253 | 0.658007974 |
| ENSMUSG00000007050 | Lsm2 | protein_coding | 0.606215456 | 3.413094897 | 1.0755117 | 0.3148378 | 0.658007974 |
| ENSMUSG00000050063 | Klk6 | protein_coding | -0.670663683 | 3.952359432 | 1.0745811 | 0.3151731 | 0.658456534 |
| ENSMUSG00000035478 | Mbd3 | protein_coding | -0.418471993 | 5.413148314 | 1.0739664 | 0.3151749 | 0.658456534 |
| ENSMUSG00000048481 | Mypop | protein_coding | -0.439086559 | 3.951406015 | 1.0735503 | 0.3152658 | 0.658508158 |

| ensembl_gene_id | external_gene_name | gene_biotype | log2 (fold change) | log2 (counts per million) | F-value | p-value | False discovery rate |
|---|---|---|---|---|---|---|---|
| ENSMUSG00000029089 | 5730480H06Rik | protein_coding | 0.614342193 | 3.863149373 | 1.0734054 | 0.3153221 | 0.658508158 |
| ENSMUSG00000031533 | Mrps31 | protein_coding | 0.368642379 | 6.396049181 | 1.0728838 | 0.315436 | 0.658514514 |
| ENSMUSG00000026058 | Khdrbs2 | protein_coding | -0.366802756 | 4.980876755 | 1.0724599 | 0.315504 | 0.658514514 |
| ENSMUSG00000022052 | Ppp2r2a | protein_coding | -0.330518609 | 6.583689077 | 1.0722177 | 0.315557 | 0.658514514 |
| ENSMUSG00000039275 | Foxk2 | protein_coding | -0.384502314 | 5.098792257 | 1.0721577 | 0.3155701 | 0.658514514 |
| ENSMUSG00000044716 | Dok7 | protein_coding | 0.63739899 | 3.466412865 | 1.0705069 | 0.3159314 | 0.659140542 |
| ENSMUSG00000041313 | Slc7a1 | protein_coding | 0.4793883 | 3.810194806 | 1.0690158 | 0.3162582 | 0.65959003 |
| ENSMUSG00000032115 | Hyou1 | protein_coding | 0.570227558 | 4.211311351 | 1.0690766 | 0.3162695 | 0.65959003 |
| ENSMUSG00000027479 | Mapre1 | protein_coding | -0.240461555 | 7.318382081 | 1.0681849 | 0.3164406 | 0.659818751 |
| ENSMUSG00000027879 | Sec22b | protein_coding | -0.247484872 | 7.907287196 | 1.0671923 | 0.3166586 | 0.659890336 |
| ENSMUSG00000031959 | Wdr59 | protein_coding | -0.493959289 | 3.81467286 | 1.066377 | 0.3168378 | 0.659890336 |
| ENSMUSG00000056023 | Gm9989 | processed_transcript | -0.625605646 | 3.638110107 | 1.0662044 | 0.3168757 | 0.659890336 |
| ENSMUSG00000001418 | Glmp | protein_coding | 0.281987658 | 6.461224664 | 1.0659048 | 0.3169417 | 0.659890336 |
| ENSMUSG00000047242 | Taf9b | protein_coding | -0.310745448 | 6.562037021 | 1.0654304 | 0.317046 | 0.659890336 |
| ENSMUSG00000038014 | Fam120a | protein_coding | -0.382158512 | 5.576354021 | 1.0651723 | 0.3171029 | 0.659890336 |
| ENSMUSG00000045973 | Slc25a51 | protein_coding | 0.297780343 | 6.216469553 | 1.0650365 | 0.3171328 | 0.659890336 |
| ENSMUSG00000056899 | Immp2l | protein_coding | -0.479632981 | 4.525311142 | 1.0648322 | 0.3171778 | 0.659890336 |
| ENSMUSG00000031311 | Nono | protein_coding | 0.391876114 | 6.397071582 | 1.0662745 | 0.3172642 | 0.659890336 |
| ENSMUSG00000043557 | Mdga1 | protein_coding | -0.678178609 | 3.316663069 | 1.0643538 | 0.3172831 | 0.659890336 |
| ENSMUSG00000033454 | Zbtb1 | protein_coding | -0.808869352 | 2.644817207 | 1.0641916 | 0.3173189 | 0.659890336 |
| ENSMUSG00000034064 | Poglut1 | protein_coding | 0.324609624 | 6.366769192 | 1.0641858 | 0.3173201 | 0.659890336 |
| ENSMUSG00000042616 | Oscp1 | protein_coding | 0.293577078 | 6.296014459 | 1.0640827 | 0.3173429 | 0.659890336 |
| ENSMUSG00000045034 | Ankrd34b | protein_coding | -0.52941066 | 3.798458628 | 1.0638392 | 0.3173966 | 0.659890336 |
| ENSMUSG00000000194 | Gpr107 | protein_coding | 0.449106492 | 4.487791172 | 1.0638297 | 0.3173987 | 0.659890336 |
| ENSMUSG00000017167 | Cntnap1 | protein_coding | 0.419553845 | 4.798864447 | 1.0635661 | 0.3174568 | 0.659890336 |
| ENSMUSG00000021313 | Ryr2 | protein_coding | -0.625424955 | 4.29272841 | 1.0631673 | 0.31784 | 0.660390626 |
| ENSMUSG00000078670 | Fam174b | protein_coding | 0.718324516 | 3.171188857 | 1.0616937 | 0.31787 | 0.660390626 |
| ENSMUSG00000053134 | Supt7l | protein_coding | 0.389935343 | 4.77275833 | 1.0616408 | 0.3178817 | 0.660390626 |
| ENSMUSG00000033214 | Slitrk5 | protein_coding | -0.372179868 | 4.334733499 | 1.0612668 | 0.3179643 | 0.660434741 |
| ENSMUSG00000025147 | Mob2 | protein_coding | 0.240245063 | 6.615878038 | 1.0609861 | 0.3180264 | 0.660436037 |
| ENSMUSG00000047098 | Rnf31 | protein_coding | 0.532544646 | 4.120447558 | 1.0604158 | 0.3181525 | 0.66053833 |
| ENSMUSG00000063594 | Gng8 | protein_coding | -0.463325697 | 4.820213282 | 1.0602078 | 0.3181985 | 0.66053833 |
| ENSMUSG00000047721 | Bola2 | protein_coding | 0.336786925 | 7.920219984 | 1.0596835 | 0.3183391 | 0.660553653 |
| ENSMUSG00000022948 | Setd4 | protein_coding | 0.916068313 | 3.210991503 | 1.0622715 | 0.3183686 | 0.660553653 |
| ENSMUSG00000026819 | Slc25a25 | protein_coding | 0.295735391 | 5.904882908 | 1.0593417 | 0.3183902 | 0.660553653 |
| ENSMUSG00000058503 | Fam133b | protein_coding | -0.288367092 | 6.062374776 | 1.0586312 | 0.3185476 | 0.660709605 |
| ENSMUSG00000037259 | Dzank1 | protein_coding | -0.489203307 | 4.315431019 | 1.0582162 | 0.3186395 | 0.660709605 |
| ENSMUSG00000037085 | Trmt12 | protein_coding | 0.369770111 | 5.502149929 | 1.0580646 | 0.3186731 | 0.660709605 |
| ENSMUSG00000022553 | Maf1 | protein_coding | -0.469520812 | 5.636313139 | 1.0601372 | 0.3187111 | 0.660709605 |
| ENSMUSG00000028780 | Sema3c | protein_coding | -0.973125532 | 2.825147627 | 1.0574409 | 0.3188359 | 0.660827768 |
| ENSMUSG00000001995 | Sipa1l2 | protein_coding | 0.580509444 | 3.881671861 | 1.0570821 | 0.318891 | 0.660827768 |
| ENSMUSG00000018160 | Med1 | protein_coding | -0.337188922 | 6.664700456 | 1.0554833 | 0.3192706 | 0.66148679 |
| ENSMUSG00000020423 | Btg2 | protein_coding | 0.660525909 | 5.95360029 | 1.0735809 | 0.3194574 | 0.661726761 |
| ENSMUSG00000026705 | Klhl20 | protein_coding | -0.425866546 | 3.967137319 | 1.0539234 | 0.319593 | 0.661726761 |
| ENSMUSG00000049154 | Fam183b | protein_coding | -0.279343356 | 5.985182853 | 1.0536781 | 0.3196476 | 0.661726761 |
| ENSMUSG00000022557 | Bop1 | protein_coding | 0.433702341 | 5.784828613 | 1.0539911 | 0.3196669 | 0.661726761 |
| ENSMUSG00000032064 | Dixdc1 | protein_coding | -0.255818042 | 6.123348435 | 1.0534694 | 0.3196941 | 0.661726761 |
| ENSMUSG00000020669 | Sh3yl1 | protein_coding | 0.547059134 | 4.384485981 | 1.0521788 | 0.3200061 | 0.662032315 |
| ENSMUSG00000073154 | 9330158H04Rik | lincRNA | -0.716164361 | 2.924406404 | 1.0519931 | 0.3200231 | 0.662032315 |
| ENSMUSG00000064202 | 4430402I18Rik | protein_coding | -0.7065458 | 3.057452635 | 1.0519407 | 0.3200348 | 0.662032315 |
| ENSMUSG00000040007 | Bahd1 | protein_coding | -0.69906588 | 3.270660705 | 1.0517021 | 0.320088 | 0.662032315 |
| ENSMUSG00000060938 | Rpl26 | protein_coding | 0.325290931 | 8.23068949 | 1.049933 | 0.3205074 | 0.662772236 |
| ENSMUSG00000024644 | Cndp2 | protein_coding | 0.385540917 | 6.333924575 | 1.0497789 | 0.3206343 | 0.662831671 |
| ENSMUSG00000008540 | Mgst1 | protein_coding | -0.695765753 | 5.354484766 | 1.0622042 | 0.3206608 | 0.662831671 |
| ENSMUSG00000024151 | Msh2 | protein_coding | 0.486938867 | 5.128752653 | 1.0489772 | 0.320721 | 0.662831671 |
| ENSMUSG00000024193 | Phf1 | protein_coding | -0.466703186 | 4.191752132 | 1.0476354 | 0.3209969 | 0.663274353 |
| ENSMUSG00000024304 | Cdh2 | protein_coding | -0.496184129 | 3.838995901 | 1.046711 | 0.321204 | 0.663509201 |
| ENSMUSG00000032215 | Rsl24d1 | protein_coding | 0.252415232 | 6.40011498 | 1.046571 | 0.3212354 | 0.663509201 |
| ENSMUSG00000029920 | Smarcad1 | protein_coding | -0.523669303 | 3.78108318 | 1.0463022 | 0.3212957 | 0.663509201 |
| ENSMUSG00000054452 | Aes | protein_coding | 0.248577619 | 7.901939294 | 1.0456919 | 0.3214326 | 0.663664451 |
| ENSMUSG00000022774 | Ncbp2 | protein_coding | 0.220202773 | 7.268332187 | 1.0451322 | 0.3215582 | 0.663796342 |
| ENSMUSG00000041695 | Kcnj2 | protein_coding | 1.321709802 | 2.236459618 | 1.0448557 | 0.3216446 | 0.663847357 |
| ENSMUSG00000024687 | Osbp | protein_coding | 0.485013771 | 4.320719299 | 1.043766 | 0.3218651 | 0.664174873 |
| ENSMUSG00000038055 | Dexi | protein_coding | -0.458775694 | 4.185808353 | 1.042607 | 0.3221258 | 0.66451202 |
| ENSMUSG00000030235 | Slco1c1 | protein_coding | 0.541239843 | 3.677615317 | 1.0424519 | 0.3221607 | 0.66451202 |
| ENSMUSG00000019791 | Hint3 | protein_coding | -0.438755162 | 4.098868876 | 1.0422157 | 0.3222139 | 0.66451202 |
| ENSMUSG00000026753 | Ppp6c | protein_coding | -0.245825204 | 6.718598413 | 1.0418451 | 0.3222973 | 0.664556653 |
| ENSMUSG00000014601 | Strip1 | protein_coding | 0.544744582 | 5.067694756 | 1.0447393 | 0.3223927 | 0.664625971 |
| ENSMUSG00000026867 | Gapvd1 | protein_coding | -0.315052541 | 5.793790567 | 1.0405703 | 0.3225846 | 0.664816202 |
| ENSMUSG00000026803 | Ttf1 | protein_coding | 0.513637842 | 3.734458087 | 1.0404636 | 0.3226087 | 0.664816202 |
| ENSMUSG00000028468 | Rgp1 | protein_coding | -0.587968384 | 3.156965295 | 1.0400534 | 0.3227012 | 0.664879484 |
| ENSMUSG00000051098 | Mblac2 | protein_coding | 0.404366886 | 5.543081478 | 1.0396315 | 0.3227964 | 0.66494819 |
| ENSMUSG00000039533 | Mmd2 | protein_coding | -0.330795212 | 7.968709712 | 1.0394534 | 0.3228609 | 0.664953697 |
| ENSMUSG00000058488 | Kl | protein_coding | -1.565428104 | 3.713946412 | 1.0732516 | 0.3229294 | 0.664967322 |
| ENSMUSG00000004771 | Rab11a | protein_coding | -0.225254558 | 6.915055833 | 1.0386701 | 0.3230135 | 0.665013243 |
| ENSMUSG00000013736 | Trnt1 | protein_coding | -0.35440551 | 6.412082725 | 1.0382311 | 0.323137 | 0.665140116 |
| ENSMUSG00000026999 | Nup35 | protein_coding | -0.401781004 | 4.37657614 | 1.0371706 | 0.3233525 | 0.665395601 |
| ENSMUSG00000022235 | Cmbl | protein_coding | -0.338846511 | 5.518609554 | 1.0370276 | 0.3233849 | 0.665395601 |

| ensembl_gene_id | external_gene_name | gene_biotype | log2 (fold change) | log2 (counts per million) | F-value | p-value | False discovery rate |
|---|---|---|---|---|---|---|---|
| ENSMUSG00000026014 | Raph1 | protein_coding | -0.356906552 | 4.442371162 | 1.036747 | 0.3234484 | 0.665398949 |
| ENSMUSG00000021830 | Txndc16 | protein_coding | -0.32744321 | 5.510028045 | 1.0363333 | 0.323542 | 0.665464305 |
| ENSMUSG00000027254 | Map1a | protein_coding | -0.327239252 | 7.385341799 | 1.0360662 | 0.3236268 | 0.665511364 |
| ENSMUSG00000097560 | Gm26904 | lincRNA | 0.673987784 | 3.585983041 | 1.0346859 | 0.3239396 | 0.66597235 |
| ENSMUSG00000058997 | Vwa8 | protein_coding | -0.65795696 | 4.08039628 | 1.0365935 | 0.3239748 | 0.66597235 |
| ENSMUSG00000025169 | Ogfod3 | protein_coding | -0.493415221 | 4.034516583 | 1.0333269 | 0.3242238 | 0.666049826 |
| ENSMUSG00000025478 | Dpysl4 | protein_coding | 0.409581051 | 5.052584115 | 1.03296 | 0.3243071 | 0.666049826 |
| ENSMUSG00000024544 | Ldlrad4 | protein_coding | 0.586953091 | 4.545320812 | 1.0349748 | 0.3243079 | 0.666049826 |
| ENSMUSG00000001289 | Pfdn5 | protein_coding | 0.309956723 | 9.841359291 | 1.03267 | 0.324373 | 0.666049826 |
| ENSMUSG00000024827 | Gldc | protein_coding | -0.935611069 | 2.589045251 | 1.0325984 | 0.3243893 | 0.666049826 |
| ENSMUSG00000000197 | Nalcn | protein_coding | -0.395258007 | 5.587749691 | 1.0322642 | 0.3244653 | 0.666049826 |
| ENSMUSG00000032184 | Lysmd2 | protein_coding | -0.249927231 | 6.451402441 | 1.0319348 | 0.3245402 | 0.666049826 |
| ENSMUSG00000009293 | Ube2g2 | protein_coding | 0.283673757 | 6.617847669 | 1.0318097 | 0.3245686 | 0.666049826 |
| ENSMUSG00000008822 | Acyp1 | protein_coding | -0.259502349 | 6.363257055 | 1.0318037 | 0.32457 | 0.666049826 |
| ENSMUSG00000078622 | Ccdc47 | protein_coding | 0.366733411 | 5.700669994 | 1.0310678 | 0.3247374 | 0.666191236 |
| ENSMUSG00000032850 | Rnft2 | protein_coding | 0.423189171 | 4.516265606 | 1.0309564 | 0.3247628 | 0.666191236 |
| ENSMUSG00000036564 | Ndrg4 | protein_coding | 0.379796719 | 11.22154694 | 1.0306208 | 0.324969 | 0.666406461 |
| ENSMUSG00000020777 | Acox1 | protein_coding | 0.290262353 | 6.792100537 | 1.0299515 | 0.3249917 | 0.666406461 |
| ENSMUSG00000057101 | Zfp180 | protein_coding | -0.34491974 | 5.044179791 | 1.0279321 | 0.3254523 | 0.667223678 |
| ENSMUSG00000013662 | Atad1 | protein_coding | -0.278212032 | 6.835617778 | 1.0273221 | 0.3255916 | 0.667307291 |
| ENSMUSG00000034290 | Nek9 | protein_coding | 0.55070427 | 3.854974297 | 1.0269946 | 0.3256664 | 0.667307291 |
| ENSMUSG00000029420 | Rimbp2 | protein_coding | -0.520961302 | 3.871527703 | 1.0269386 | 0.3256792 | 0.667307291 |
| ENSMUSG00000042572 | Ube2q1 | protein_coding | 0.327764695 | 5.737948593 | 1.025815 | 0.3259361 | 0.667471064 |
| ENSMUSG00000052102 | Gnpda1 | protein_coding | 0.385608046 | 5.444722279 | 1.0257941 | 0.3259409 | 0.667471064 |
| ENSMUSG00000087687 | Pet100 | protein_coding | 0.336427049 | 6.789267763 | 1.0254062 | 0.3260538 | 0.667471064 |
| ENSMUSG00000036052 | Dnajb5 | protein_coding | 0.402061914 | 5.281467032 | 1.0250819 | 0.3261039 | 0.667471064 |
| ENSMUSG00000022724 | Riox2 | protein_coding | 0.612988381 | 3.831922771 | 1.0249386 | 0.3261609 | 0.667471064 |
| ENSMUSG00000024896 | Minpp1 | protein_coding | 0.421926393 | 4.77867413 | 1.0248089 | 0.3261665 | 0.667471064 |
| ENSMUSG00000030189 | Ybx3 | protein_coding | -0.48788012 | 3.853006636 | 1.0246886 | 0.326194 | 0.667471064 |
| ENSMUSG00000028410 | Dnaja1 | protein_coding | -0.221955448 | 7.796491924 | 1.0243235 | 0.3262777 | 0.667471064 |
| ENSMUSG00000063904 | 4930533O14Rik | protein_coding | 0.468622941 | 5.120789091 | 1.0241456 | 0.3263425 | 0.667471064 |
| ENSMUSG00000031758 | Cdyl2 | protein_coding | -0.664797465 | 4.238801782 | 1.0279704 | 0.3263799 | 0.667471064 |
| ENSMUSG00000025782 | Taf3 | protein_coding | -0.364828941 | 4.81405377 | 1.0233675 | 0.3264968 | 0.667583211 |
| ENSMUSG00000024867 | Pip5k1b | protein_coding | -0.344706697 | 5.233252831 | 1.0206953 | 0.3271105 | 0.668594456 |
| ENSMUSG00000006763 | Saal1 | protein_coding | 0.419273811 | 4.029950953 | 1.0204725 | 0.3271617 | 0.668594456 |
| ENSMUSG00000028081 | Rps3a1 | protein_coding | 0.306073933 | 7.926400516 | 1.020487 | 0.3271825 | 0.668594456 |
| ENSMUSG00000062444 | Ap3b2 | protein_coding | 0.314281092 | 6.66180657 | 1.0198047 | 0.3273154 | 0.668594456 |
| ENSMUSG00000022490 | Ppp1r1a | protein_coding | 0.348235263 | 5.34981323 | 1.0197872 | 0.3273194 | 0.668594456 |
| ENSMUSG00000030000 | Add2 | protein_coding | 0.263181158 | 7.219291826 | 1.0195915 | 0.3273644 | 0.668594456 |
| ENSMUSG00000028156 | Eif4e | protein_coding | 0.235831424 | 7.869576667 | 1.0190458 | 0.3274901 | 0.668724112 |
| ENSMUSG00000040818 | Dennd6a | protein_coding | -0.468168158 | 3.729116788 | 1.0183805 | 0.3276434 | 0.668910092 |
| ENSMUSG00000002718 | Cse1l | protein_coding | 0.303863861 | 5.626755388 | 1.0166035 | 0.3280533 | 0.669619846 |
| ENSMUSG00000023923 | Tbc1d5 | protein_coding | 0.41017412 | 4.134017351 | 1.0161156 | 0.328166 | 0.669642706 |
| ENSMUSG00000079442 | St6galnac4 | protein_coding | 0.49265448 | 4.861569362 | 1.0159982 | 0.3282172 | 0.669642706 |
| ENSMUSG00000029469 | Ift81 | protein_coding | -0.351221395 | 5.347854405 | 1.0157462 | 0.3282513 | 0.669642706 |
| ENSMUSG00000060181 | Slc35e3 | protein_coding | 0.543557641 | 3.131692399 | 1.015314 | 0.3283512 | 0.669719443 |
| ENSMUSG00000043398 | Gpr135 | protein_coding | 0.626023234 | 3.060723677 | 1.014491 | 0.3285416 | 0.669836347 |
| ENSMUSG00000035284 | Vps13c | protein_coding | -0.414582138 | 4.709763561 | 1.0143715 | 0.3285692 | 0.669836347 |
| ENSMUSG00000034940 | Synrg | protein_coding | -0.28212749 | 6.096106445 | 1.0142583 | 0.3285954 | 0.669836347 |
| ENSMUSG00000032607 | Amt | protein_coding | 0.612533113 | 5.327476593 | 1.0223698 | 0.3287795 | 0.67008451 |
| ENSMUSG00000034265 | Zdhhc14 | protein_coding | 0.491279041 | 4.272817134 | 1.0128432 | 0.3289232 | 0.670154267 |
| ENSMUSG00000024594 | Prrc1 | protein_coding | -1.107101529 | 2.087124265 | 1.0127776 | 0.3289384 | 0.670154267 |
| ENSMUSG00000073664 | Nbeal1 | protein_coding | -0.552796889 | 3.613402452 | 1.0124394 | 0.3290168 | 0.670187007 |
| ENSMUSG00000025246 | Tbl1x | protein_coding | -0.293397065 | 6.155743411 | 1.0115729 | 0.3292177 | 0.6704694 |
| ENSMUSG00000037108 | Zcwpw1 | protein_coding | 0.502269059 | 3.452655585 | 1.011204 | 0.3293034 | 0.67051676 |
| ENSMUSG00000039648 | Kyat1 | protein_coding | -0.523102254 | 4.543496185 | 1.0105645 | 0.3294758 | 0.670740926 |
| ENSMUSG00000059920 | 4930453N24Rik | protein_coding | -0.398839459 | 4.780918003 | 1.0095781 | 0.3296811 | 0.671031722 |
| ENSMUSG00000070436 | Serpinh1 | protein_coding | -0.678842446 | 3.617527688 | 1.0088957 | 0.3298637 | 0.671202843 |
| ENSMUSG00000052374 | Actn2 | protein_coding | 0.884402496 | 3.558390301 | 1.0149652 | 0.32989 | 0.671202843 |
| ENSMUSG00000020121 | Srgap1 | protein_coding | -0.519969189 | 4.487032466 | 1.0083087 | 0.3300004 | 0.671300405 |
| ENSMUSG00000006403 | Adamts4 | protein_coding | -0.648814157 | 3.873262184 | 1.007864 | 0.3301039 | 0.671375205 |
| ENSMUSG00000024213 | Nudt3 | protein_coding | -0.24563631 | 8.135176112 | 1.0075118 | 0.330162 | 0.671375205 |
| ENSMUSG00000008999 | Bmp7 | protein_coding | -0.667869689 | 3.158227919 | 1.0068745 | 0.3303105 | 0.671521913 |
| ENSMUSG00000090061 | Nwd2 | protein_coding | -0.92719789 | 2.857913007 | 1.0066537 | 0.3303859 | 0.671521913 |
| ENSMUSG00000003458 | Ncstn | protein_coding | 0.353981029 | 5.951255116 | 1.0061677 | 0.3304754 | 0.671521913 |
| ENSMUSG00000058318 | Phf21a | protein_coding | -0.378256375 | 4.990714443 | 1.0061009 | 0.3304909 | 0.671521913 |
| ENSMUSG00000038683 | Pak1ip1 | protein_coding | 0.282588512 | 7.238185548 | 1.0058632 | 0.3305464 | 0.671521913 |
| ENSMUSG00000031325 | 4930519F16Rik | transcribed_unitary_p | -0.802204563 | 2.588531123 | 1.0052662 | 0.3306858 | 0.671607448 |
| ENSMUSG00000038374 | Rbm8a | protein_coding | 0.233155505 | 7.190096563 | 1.0051254 | 0.3307187 | 0.671607448 |
| ENSMUSG00000022849 | Hspbap1 | protein_coding | -0.732451189 | 3.208849033 | 1.0048804 | 0.3307759 | 0.671607448 |
| ENSMUSG00000107002 | 0610012G03Rik | protein_coding | 0.42852542 | 4.20281554 | 1.0041998 | 0.3309349 | 0.671803521 |
| ENSMUSG00000043140 | Tmem186 | protein_coding | 0.437415532 | 5.17310996 | 1.0034979 | 0.331099 | 0.671921648 |
| ENSMUSG00000097166 | 9330179D12Rik | antisense_RNA | -0.598926695 | 3.657326077 | 1.0034165 | 0.3311181 | 0.671921648 |
| ENSMUSG00000042359 | Osbpl6 | protein_coding | -0.394105398 | 5.156949153 | 1.0030695 | 0.3311993 | 0.671959609 |
| ENSMUSG00000021374 | Nup153 | protein_coding | -0.535596101 | 3.758688117 | 1.0026517 | 0.3312971 | 0.672031231 |
| ENSMUSG00000048550 | Thnsl1 | protein_coding | -0.374183046 | 5.112092337 | 1.0016335 | 0.3315356 | 0.672233751 |
| ENSMUSG00000051817 | Sox12 | protein_coding | 0.508761201 | 3.284725338 | 1.0014668 | 0.3315746 | 0.672233751 |
| ENSMUSG00000037221 | Mospd3 | protein_coding | 0.709560974 | 4.650157983 | 1.0101736 | 0.3315845 | 0.672233751 |

| ensembl_gene_id | external_gene_name | gene_biotype | log2 (fold change) | log2 (counts per million) | F-value | p-value | False discovery rate |
|---|---|---|---|---|---|---|---|
| ENSMUSG00000044348 | Slc25a53 | protein_coding | -0.365092612 | 4.529963874 | 1.0007622 | 0.3317399 | 0.672421985 |
| ENSMUSG00000020387 | Jade2 | protein_coding | -0.30301584 | 5.322138839 | 1.0001714 | 0.3318785 | 0.672576194 |
| ENSMUSG00000029012 | Orc5 | protein_coding | -0.354456767 | 5.490140977 | 0.9996863 | 0.3319924 | 0.672680215 |
| ENSMUSG00000029068 | Ccnl2 | protein_coding | 0.460603569 | 3.8551514 | 0.9990045 | 0.3321525 | 0.672748101 |
| ENSMUSG00000037172 | E330009J07Rik | protein_coding | -0.409481113 | 4.419615746 | 0.9987729 | 0.3322069 | 0.672748101 |
| ENSMUSG00000057060 | Slc35f3 | protein_coding | 0.55330366 | 3.816031417 | 0.9985295 | 0.3322641 | 0.672748101 |
| ENSMUSG00000028277 | Ube2j1 | protein_coding | 0.302102366 | 6.01648963 | 0.9984785 | 0.3322761 | 0.672748101 |
| ENSMUSG00000008475 | Arpc5 | protein_coding | 0.222664919 | 7.254298124 | 0.997389 | 0.3325324 | 0.673140241 |
| ENSMUSG00000031023 | Akip1 | protein_coding | -0.494263385 | 3.610570956 | 0.9967457 | 0.3326839 | 0.673269189 |
| ENSMUSG00000037032 | Apbb1 | protein_coding | 0.297909706 | 9.16330702 | 0.9965866 | 0.3327213 | 0.673269189 |
| ENSMUSG00000028145 | Them4 | protein_coding | 0.281089213 | 6.586980391 | 0.9959456 | 0.3328724 | 0.67339147 |
| ENSMUSG00000027239 | Mdk | protein_coding | -0.667097267 | 4.884046734 | 1.0044741 | 0.332907 | 0.67339147 |
| ENSMUSG00000014748 | Tex261 | protein_coding | -0.366573173 | 5.957894525 | 0.9952039 | 0.333071 | 0.673596565 |
| ENSMUSG00000025261 | Huwe1 | protein_coding | -0.272684722 | 5.898105929 | 0.9927491 | 0.333627 | 0.674528195 |
| ENSMUSG00000034110 | Kctd7 | protein_coding | -0.434338011 | 4.071132289 | 0.9922522 | 0.3337445 | 0.674528195 |
| ENSMUSG00000030551 | Nr2f2 | protein_coding | 0.416647231 | 6.558364138 | 0.9974095 | 0.3337457 | 0.674528195 |
| ENSMUSG00000026563 | Tada1 | protein_coding | 0.783987846 | 2.606274316 | 0.9920569 | 0.3337907 | 0.674528195 |
| ENSMUSG00000028920 | Fbxo42 | protein_coding | 0.475387321 | 3.880096418 | 0.9918259 | 0.3338454 | 0.674528195 |
| ENSMUSG00000027865 | Gdap2 | protein_coding | 0.432417659 | 5.846192724 | 0.99279 | 0.3340988 | 0.674804214 |
| ENSMUSG00000029312 | Klhl8 | protein_coding | 0.391373633 | 4.255544535 | 0.9906695 | 0.3341192 | 0.674804214 |
| ENSMUSG00000020948 | Klhl28 | protein_coding | -0.333586021 | 4.843608937 | 0.9904542 | 0.3341702 | 0.674804214 |
| ENSMUSG00000031255 | Sytl4 | protein_coding | 0.411113517 | 4.055596508 | 0.9892317 | 0.3344601 | 0.675262795 |
| ENSMUSG00000029587 | Zfp12 | protein_coding | 0.914624793 | 3.779491193 | 0.9987153 | 0.3345308 | 0.675278623 |
| ENSMUSG00000002341 | Ncan | protein_coding | -0.249672279 | 7.218162338 | 0.9882168 | 0.3347011 | 0.675380441 |
| ENSMUSG00000032246 | Calml4 | protein_coding | -0.669760048 | 3.87623688 | 0.9880405 | 0.3347667 | 0.675380441 |
| ENSMUSG00000015790 | Surf1 | protein_coding | 0.236597772 | 8.21324202 | 0.9879282 | 0.3347696 | 0.675380441 |
| ENSMUSG00000008136 | Fhl2 | protein_coding | 0.835164697 | 3.259917399 | 0.9882184 | 0.3349358 | 0.675524811 |
| ENSMUSG00000029916 | Agk | protein_coding | -0.332064312 | 4.77577093 | 0.9870986 | 0.3349668 | 0.675524811 |
| ENSMUSG00000038860 | Garnl3 | protein_coding | 0.401723364 | 5.181698089 | 0.9867857 | 0.3350412 | 0.675548159 |
| ENSMUSG00000068735 | Trp53i11 | protein_coding | 0.325334161 | 5.634389852 | 0.9859708 | 0.3352352 | 0.675777773 |
| ENSMUSG00000090841 | Myl6 | protein_coding | 0.240126519 | 7.729844937 | 0.9857791 | 0.3352808 | 0.675777773 |
| ENSMUSG00000031563 | Wwc2 | protein_coding | -0.431339247 | 4.512205655 | 0.985368 | 0.3353787 | 0.675848417 |
| ENSMUSG00000059921 | Unc5c | protein_coding | -0.513719699 | 4.057897879 | 0.9849383 | 0.3354811 | 0.675928062 |
| ENSMUSG00000096257 | Ccer2 | protein_coding | 0.8299683 | 2.987224756 | 0.9839092 | 0.3357501 | 0.67634343 |
| ENSMUSG00000028998 | Tomm7 | protein_coding | 0.303866861 | 8.606368385 | 0.9833392 | 0.3358861 | 0.676474453 |
| ENSMUSG00000034923 | Ly6g6f | protein_coding | -0.652006431 | 3.087477309 | 0.9827551 | 0.3360019 | 0.676474453 |
| ENSMUSG00000059890 | Ube4a | protein_coding | -0.463562674 | 3.81874335 | 0.9827469 | 0.3360039 | 0.676474453 |
| ENSMUSG00000024800 | Rpp30 | protein_coding | -0.2610153 | 6.361405398 | 0.9815498 | 0.33629 | 0.676889296 |
| ENSMUSG00000031099 | Smarca1 | protein_coding | -0.364284842 | 5.353965234 | 0.9813581 | 0.3363359 | 0.676889296 |
| ENSMUSG00000026974 | Zmynd19 | protein_coding | 0.427735559 | 3.884006846 | 0.9794754 | 0.3367866 | 0.677617614 |
| ENSMUSG00000021096 | Ppm1a | protein_coding | -0.205378982 | 7.718985064 | 0.9793201 | 0.3368238 | 0.677617614 |
| ENSMUSG00000040782 | Rfwd2 | protein_coding | -0.657045813 | 3.500210466 | 0.9780999 | 0.3371164 | 0.67778947 |
| ENSMUSG00000015748 | Prpf3 | protein_coding | 0.460824123 | 4.036123094 | 0.977848 | 0.3371769 | 0.67778947 |
| ENSMUSG00000038074 | Fkbp14 | protein_coding | 0.552576837 | 3.382830478 | 0.9777232 | 0.3372068 | 0.67778947 |
| ENSMUSG00000038446 | Cdc40 | protein_coding | -0.443246274 | 5.089583073 | 0.9775121 | 0.3372811 | 0.67778947 |
| ENSMUSG00000051062 | Fbll1 | protein_coding | 0.360853075 | 5.381324325 | 0.9773701 | 0.3372916 | 0.67778947 |
| ENSMUSG00000046768 | Rhoj | protein_coding | -0.797259519 | 2.680213226 | 0.9770894 | 0.337359 | 0.67778947 |
| ENSMUSG00000043190 | Rfesd | protein_coding | -0.32721728 | 5.511361375 | 0.9766438 | 0.3374661 | 0.67778947 |
| ENSMUSG00000020231 | Dip2a | protein_coding | -0.426006167 | 4.17885028 | 0.9766325 | 0.3374688 | 0.67778947 |
| ENSMUSG00000042293 | Gm5617 | protein_coding | 0.517859989 | 4.872114521 | 0.9770702 | 0.3374765 | 0.67778947 |
| ENSMUSG00000055302 | Mrfap1 | protein_coding | -0.22500642 | 7.938451238 | 0.9760977 | 0.3375974 | 0.677861344 |
| ENSMUSG00000045210 | Vcpip1 | protein_coding | -0.296341543 | 5.468057633 | 0.9759272 | 0.3376384 | 0.677861344 |
| ENSMUSG00000074649 | BC029722 | protein_coding | 0.484918041 | 4.083220011 | 0.9750235 | 0.3378558 | 0.678171266 |
| ENSMUSG00000030842 | Lamtor1 | protein_coding | 0.220729399 | 7.343612553 | 0.974333 | 0.3380221 | 0.678378407 |
| ENSMUSG00000043587 | Pxylp1 | protein_coding | 0.488546327 | 3.931386906 | 0.9735764 | 0.3382044 | 0.678602134 |
| ENSMUSG00000066189 | Cacng3 | protein_coding | 0.225432436 | 7.436770443 | 0.9733467 | 0.3382598 | 0.678602134 |
| ENSMUSG00000031644 | Nek1 | protein_coding | -0.625162973 | 3.354982581 | 0.9723911 | 0.3384903 | 0.678708904 |
| ENSMUSG00000022255 | Mtdh | protein_coding | -0.422081634 | 4.957280925 | 0.9723255 | 0.3385061 | 0.678708904 |
| ENSMUSG00000097290 | 1300002E11Rik | lincRNA | -0.389847928 | 4.73278305 | 0.9723201 | 0.3385074 | 0.678708904 |
| ENSMUSG00000022574 | Naprt | protein_coding | -0.688891514 | 3.163063662 | 0.9720797 | 0.3385655 | 0.678708904 |
| ENSMUSG00000074622 | Mafb | protein_coding | -0.896477436 | 2.802937455 | 0.9716986 | 0.3386575 | 0.678766835 |
| ENSMUSG00000018841 | Rad51d | protein_coding | 0.351292141 | 5.734623597 | 0.9707634 | 0.3388835 | 0.6789566 |
| ENSMUSG00000045690 | Wdr89 | protein_coding | 0.426631902 | 4.393169587 | 0.9706499 | 0.3389109 | 0.6789566 |
| ENSMUSG00000056116 | H2-T22 | protein_coding | 0.482414085 | 4.006138147 | 0.9705231 | 0.3389416 | 0.6789566 |
| ENSMUSG00000023861 | Mpc1 | protein_coding | -0.241384157 | 6.224462493 | 0.9695158 | 0.3391853 | 0.679318223 |
| ENSMUSG00000024269 | Tpgs2 | protein_coding | 0.236692583 | 6.612250471 | 0.9692093 | 0.3392595 | 0.6793403 |
| ENSMUSG00000063316 | Rpl27 | protein_coding | 0.45328047 | 3.778532114 | 0.9685867 | 0.3394103 | 0.679515763 |
| ENSMUSG00000021559 | Dapk1 | protein_coding | -0.320668259 | 5.818781129 | 0.9675974 | 0.3396501 | 0.679869289 |
| ENSMUSG00000035614 | Togaram1 | protein_coding | 0.424423008 | 4.576164842 | 0.9671429 | 0.3397604 | 0.679940475 |
| ENSMUSG00000038116 | Phf20 | protein_coding | 0.241479286 | 6.675524015 | 0.9669296 | 0.3398122 | 0.679940475 |
| ENSMUSG00000029482 | Aacs | protein_coding | 0.440488906 | 4.888742531 | 0.9658771 | 0.3400677 | 0.680289447 |
| ENSMUSG00000032435 | Dync1li1 | protein_coding | 0.377385782 | 7.157449133 | 0.9698836 | 0.3401131 | 0.680289447 |
| ENSMUSG00000114309 | AC163038.1 | bidirectional_promote | 0.732149442 | 3.07934206 | 0.9651939 | 0.3402338 | 0.680378998 |
| ENSMUSG00000022538 | Lsg1 | protein_coding | -0.298203101 | 5.240216218 | 0.9648423 | 0.3403193 | 0.680378998 |
| ENSMUSG00000042303 | Sgsm3 | protein_coding | 0.350826263 | 6.572978962 | 0.9649186 | 0.3403859 | 0.680378998 |
| ENSMUSG00000096995 | 2810029C07Rik | bidirectional_promote | 0.489554869 | 3.791981806 | 0.9644655 | 0.340411 | 0.680378998 |
| ENSMUSG00000024780 | Cdc37l1 | protein_coding | -0.241252854 | 6.340182071 | 0.9638217 | 0.3405676 | 0.680565651 |
| ENSMUSG00000039070 | Cpa4 | protein_coding | -1.043997719 | 2.618529772 | 0.9635709 | 0.3406521 | 0.680608009 |

| ensembl_gene_id | external_gene_name | gene_biotype | log2 (fold change) | log2 (counts per million) | F-value | p-value | False discovery rate |
|---|---|---|---|---|---|---|---|
| ENSMUSG00000056394 | Lig1 | protein_coding | -0.417913988 | 4.074865149 | 0.9623684 | 0.3409217 | 0.681020154 |
| ENSMUSG00000037138 | Aff3 | protein_coding | -0.415722839 | 4.817062394 | 0.9620846 | 0.340991 | 0.681031915 |
| ENSMUSG00000038323 | 1700066M21Rik | protein_coding | -0.694781704 | 3.207639769 | 0.9612456 | 0.3411957 | 0.681314202 |
| ENSMUSG00000014554 | Dguok | protein_coding | 0.293011094 | 5.686858152 | 0.96094 | 0.3412703 | 0.681336703 |
| ENSMUSG00000021311 | Mtr | protein_coding | -0.650221899 | 3.017225109 | 0.9602825 | 0.3414309 | 0.681530781 |
| ENSMUSG00000023868 | Pde10a | protein_coding | -0.363265335 | 6.422342543 | 0.9602958 | 0.3415126 | 0.681567409 |
| ENSMUSG00000038462 | Uqcrfs1 | protein_coding | -0.347532823 | 6.788603422 | 0.9602471 | 0.3416714 | 0.681610056 |
| ENSMUSG00000005949 | Ctns | protein_coding | -0.596078932 | 3.660435599 | 0.9590031 | 0.3417437 | 0.681610056 |
| ENSMUSG00000035227 | Spcs2 | protein_coding | 0.248725075 | 8.275560973 | 0.9588775 | 0.3417744 | 0.681610056 |
| ENSMUSG00000015289 | Lage3 | protein_coding | -0.33123829 | 5.787581657 | 0.9587896 | 0.341796 | 0.681610056 |
| ENSMUSG00000006442 | Srm | protein_coding | 0.354046801 | 5.229121767 | 0.9585651 | 0.3418509 | 0.681610056 |
| ENSMUSG00000038733 | Wdr26 | protein_coding | -0.352952734 | 6.151534961 | 0.9583161 | 0.3419352 | 0.681651751 |
| ENSMUSG00000047777 | Phf13 | protein_coding | 0.492007359 | 3.441621594 | 0.9577458 | 0.3420515 | 0.681757222 |
| ENSMUSG00000014039 | Prdm15 | protein_coding | 0.890962237 | 2.37350164 | 0.9570432 | 0.3422238 | 0.681893868 |
| ENSMUSG00000005354 | Txn2 | protein_coding | 0.232908314 | 9.312066701 | 0.9569486 | 0.3422469 | 0.681893868 |
| ENSMUSG00000008435 | Rdh13 | protein_coding | 0.289764935 | 6.406501572 | 0.9566169 | 0.3423283 | 0.681929586 |
| ENSMUSG00000001082 | Mfsd10 | protein_coding | 0.628735179 | 3.679602123 | 0.9562035 | 0.342453 | 0.681962908 |
| ENSMUSG00000059824 | Dbp | protein_coding | -0.466620545 | 3.487219614 | 0.9560318 | 0.3424718 | 0.681962908 |
| ENSMUSG00000027589 | Pcmtd2 | protein_coding | -0.320185262 | 5.765213064 | 0.955753 | 0.3425403 | 0.681972862 |
| ENSMUSG00000048429 | Timm29 | protein_coding | 0.361110825 | 5.196733817 | 0.9553383 | 0.3426421 | 0.682049301 |
| ENSMUSG00000031696 | Vps35 | protein_coding | 0.220325692 | 7.691568384 | 0.954481 | 0.3428527 | 0.682342297 |
| ENSMUSG00000019470 | Xab2 | protein_coding | 0.451571915 | 5.777824972 | 0.9569544 | 0.3431495 | 0.682582312 |
| ENSMUSG00000047675 | Rps8 | protein_coding | 0.310825996 | 6.217113011 | 0.9532463 | 0.3431564 | 0.682582312 |
| ENSMUSG00000014418 | Hps5 | protein_coding | -0.661029104 | 3.081212267 | 0.9531278 | 0.3431856 | 0.682582312 |
| ENSMUSG00000029062 | Cdk11b | protein_coding | -0.321579516 | 4.978969342 | 0.9529586 | 0.3432272 | 0.682582312 |
| ENSMUSG00000033885 | Pxk | protein_coding | 0.38422687 | 5.072323438 | 0.9523672 | 0.3433729 | 0.682745662 |
| ENSMUSG00000049670 | Morn4 | protein_coding | 0.209008478 | 7.601645334 | 0.9519343 | 0.3434796 | 0.68278086 |
| ENSMUSG00000019808 | Adat2 | protein_coding | -0.612433293 | 2.972440111 | 0.95178 | 0.3435176 | 0.68278086 |
| ENSMUSG00000025971 | Maip1 | protein_coding | -0.328339288 | 4.989493481 | 0.9503009 | 0.3438824 | 0.683379774 |
| ENSMUSG00000051615 | Rap2a | protein_coding | 0.311417463 | 5.783770997 | 0.9493694 | 0.3441125 | 0.683562145 |
| ENSMUSG00000022141 | Nipbl | protein_coding | -0.547243061 | 4.416390134 | 0.9494095 | 0.3441258 | 0.683562145 |
| ENSMUSG00000058093 | Zfp729b | protein_coding | -0.865074604 | 2.732909415 | 0.9491573 | 0.3441649 | 0.683562145 |
| ENSMUSG00000061578 | Ksr2 | protein_coding | -0.395326 | 4.148114964 | 0.9476798 | 0.3445304 | 0.684161616 |
| ENSMUSG00000018326 | Ywhab | protein_coding | 0.210145299 | 9.025884273 | 0.9465501 | 0.3448102 | 0.684552714 |
| ENSMUSG00000021987 | Mtmr6 | protein_coding | 0.295511345 | 6.883620978 | 0.946269 | 0.3448799 | 0.684552714 |
| ENSMUSG00000025982 | Sf3b1 | protein_coding | -0.217744762 | 7.046822769 | 0.9459709 | 0.3449537 | 0.684552714 |
| ENSMUSG00000030035 | Wbp1 | protein_coding | 0.478948875 | 4.942696081 | 0.9456978 | 0.3450447 | 0.684552714 |
| ENSMUSG00000029016 | Clcn6 | protein_coding | 0.296670984 | 5.548644841 | 0.9455413 | 0.3450603 | 0.684552714 |
| ENSMUSG00000041426 | Hibch | protein_coding | -0.341628745 | 5.204312644 | 0.945073 | 0.3451765 | 0.684552714 |
| ENSMUSG00000023800 | Tiam2 | protein_coding | -0.461940998 | 4.999777228 | 0.9449576 | 0.3452283 | 0.684552714 |
| ENSMUSG00000024608 | Rps14 | protein_coding | 0.364829523 | 10.3060541 | 0.9470985 | 0.3452663 | 0.684552714 |
| ENSMUSG00000032714 | Syde1 | protein_coding | -1.083633198 | 2.675543045 | 0.9446079 | 0.3453762 | 0.684552714 |
| ENSMUSG00000090266 | Mettl23 | protein_coding | 0.443238808 | 4.816072364 | 0.9438976 | 0.3454684 | 0.684552714 |
| ENSMUSG00000044708 | Kcnj10 | protein_coding | -0.421835733 | 4.889760016 | 0.9433961 | 0.3455931 | 0.684552714 |
| ENSMUSG00000085067 | Gm15631 | lincRNA | -0.426579757 | 5.229903911 | 0.943367 | 0.3456235 | 0.684552714 |
| ENSMUSG00000021177 | Tdp1 | protein_coding | 0.410756747 | 4.693445551 | 0.9432563 | 0.3456278 | 0.684552714 |
| ENSMUSG00000021257 | Angel1 | protein_coding | -0.741419901 | 3.090701824 | 0.9432179 | 0.3456374 | 0.684552714 |
| ENSMUSG00000040265 | Dnm3 | protein_coding | 0.338005469 | 5.663832161 | 0.9430375 | 0.3456823 | 0.684552714 |
| ENSMUSG00000035529 | Prdm4 | protein_coding | 0.538884171 | 3.940585413 | 0.9419202 | 0.3459603 | 0.684809053 |
| ENSMUSG00000016427 | Ndufa1 | protein_coding | 0.31111487 | 9.064829798 | 0.9414858 | 0.3460916 | 0.684809053 |
| ENSMUSG00000040389 | Wdr47 | protein_coding | 0.282650287 | 6.156199779 | 0.9408681 | 0.3462224 | 0.684809053 |
| ENSMUSG00000062778 | Chia1 | protein_coding | 1.197045515 | 2.152560485 | 0.9401699 | 0.3463965 | 0.684809053 |
| ENSMUSG00000028964 | Park7 | protein_coding | 0.214710085 | 10.18779602 | 0.9401526 | 0.3464008 | 0.684809053 |
| ENSMUSG00000025665 | Rps6ka6 | protein_coding | 0.613496502 | 3.323199076 | 0.9400358 | 0.34643 | 0.684809053 |
| ENSMUSG00000050668 | Gpatch11 | protein_coding | 0.244333682 | 6.136313262 | 0.9398989 | 0.3464641 | 0.684809053 |
| ENSMUSG00000009292 | Trpm2 | protein_coding | 0.590049676 | 3.85111753 | 0.9397229 | 0.3465312 | 0.684809053 |
| ENSMUSG00000060548 | Tnfrsf19 | protein_coding | -0.573889677 | 3.548605899 | 0.9395978 | 0.3465393 | 0.684809053 |
| ENSMUSG00000052812 | Atad2b | protein_coding | -0.630116217 | 3.200018374 | 0.9394979 | 0.3465642 | 0.684809053 |
| ENSMUSG00000028865 | Cd164l2 | protein_coding | -0.406867167 | 3.901543846 | 0.9394404 | 0.3465786 | 0.684809053 |
| ENSMUSG00000038002 | Cramp1l | protein_coding | 0.542691527 | 3.327462723 | 0.9393335 | 0.3466053 | 0.684809053 |
| ENSMUSG00000009035 | Tmem184b | protein_coding | -0.46380392 | 4.128339592 | 0.9391435 | 0.3466527 | 0.684809053 |
| ENSMUSG00000025964 | Adam23 | protein_coding | 0.269704336 | 6.261421837 | 0.9389409 | 0.3467033 | 0.684809053 |
| ENSMUSG00000036941 | Elac1 | protein_coding | 0.386435305 | 4.921002979 | 0.9382192 | 0.3468836 | 0.684939063 |
| ENSMUSG00000041881 | Ndufa7 | protein_coding | 0.278557944 | 9.535408849 | 0.9375856 | 0.347042 | 0.684939063 |
| ENSMUSG00000039018 | Mtg1 | protein_coding | 0.285036732 | 6.316391935 | 0.9374394 | 0.3470786 | 0.684939063 |
| ENSMUSG00000038803 | Ost4 | protein_coding | 0.280566836 | 6.636590519 | 0.9372377 | 0.3471291 | 0.684939063 |
| ENSMUSG00000047187 | Rab2a | protein_coding | -0.211976096 | 8.097974959 | 0.9370491 | 0.3471763 | 0.684939063 |
| ENSMUSG00000041798 | Gck | protein_coding | 0.438824049 | 5.389062002 | 0.936991 | 0.3472139 | 0.684939063 |
| ENSMUSG00000034591 | Slc41a2 | protein_coding | 0.418501114 | 3.769568938 | 0.9363651 | 0.3473475 | 0.684939063 |
| ENSMUSG00000063524 | Eno1 | protein_coding | 0.286931985 | 8.641938171 | 0.9360425 | 0.3474283 | 0.684939063 |
| ENSMUSG00000044636 | Csrnp2 | protein_coding | -0.466511708 | 4.041868163 | 0.935868 | 0.3474721 | 0.684939063 |
| ENSMUSG00000015575 | Atp6v0e | protein_coding | -0.32515004 | 7.084785121 | 0.935936 | 0.3475389 | 0.684939063 |
| ENSMUSG00000028842 | Ago3 | protein_coding | -0.502573481 | 4.372173985 | 0.9355062 | 0.3475627 | 0.684939063 |
| ENSMUSG00000032777 | Gtf3c1 | protein_coding | 0.394164689 | 5.043845491 | 0.9354712 | 0.3475715 | 0.684939063 |
| ENSMUSG00000052387 | Trpm3 | protein_coding | -0.652227557 | 4.38765766 | 0.9400981 | 0.3475972 | 0.684939063 |
| ENSMUSG00000014769 | Psmb1 | protein_coding | 0.215805186 | 10.02840023 | 0.9346299 | 0.3477825 | 0.68494103 |
| ENSMUSG00000027285 | Haus2 | protein_coding | -0.278249531 | 6.045529529 | 0.9345433 | 0.3478042 | 0.68494103 |
| ENSMUSG00000059363 | Fxn | protein_coding | -0.665202966 | 2.881427468 | 0.9344791 | 0.3478203 | 0.68494103 |

| ensembl_gene_id | external_gene_name | gene_biotype | log2 (fold change) | log2 (counts per million) | F-value | p-value | False discovery rate |
|---|---|---|---|---|---|---|---|
| ENSMUSG00000057541 | Pus7 | protein_coding | -0.708446531 | 2.835257955 | 0.9341597 | 0.3479005 | 0.68494103 |
| ENSMUSG00000052752 | Traf7 | protein_coding | -0.320165563 | 4.874099156 | 0.9340612 | 0.3479252 | 0.68494103 |
| ENSMUSG00000038650 | Rnh1 | protein_coding | 0.278353737 | 7.089744045 | 0.9338415 | 0.3479804 | 0.68494103 |
| ENSMUSG00000108350 | Gm44950 | TEC | 0.452798307 | 4.493757093 | 0.9331663 | 0.34815 | 0.685149443 |
| ENSMUSG00000021990 | Spata13 | protein_coding | 0.48517947 | 4.088362386 | 0.9324601 | 0.3483275 | 0.685359724 |
| ENSMUSG00000029428 | Stx2 | protein_coding | -0.692297025 | 2.756617378 | 0.9320358 | 0.3484342 | 0.685359724 |
| ENSMUSG00000004018 | Fancl | protein_coding | 0.372104825 | 4.841292438 | 0.9319345 | 0.3484597 | 0.685359724 |
| ENSMUSG00000097277 | 2900076A07Rik | lincRNA | -0.443047259 | 3.852739501 | 0.9317202 | 0.3485136 | 0.685359724 |
| ENSMUSG00000031201 | Brcc3 | protein_coding | -0.261147497 | 7.292192666 | 0.9314392 | 0.3485843 | 0.685359724 |
| ENSMUSG00000031921 | Terf2 | protein_coding | 0.363431613 | 4.788759732 | 0.9312211 | 0.3486392 | 0.685359724 |
| ENSMUSG00000026854 | Usp20 | protein_coding | 0.313107464 | 5.97601616 | 0.9308703 | 0.3487276 | 0.685408087 |
| ENSMUSG00000024036 | Slc37a1 | protein_coding | -0.662690925 | 2.945747937 | 0.9299196 | 0.3489671 | 0.68557499 |
| ENSMUSG00000034673 | Pbx2 | protein_coding | 0.386123494 | 4.721255609 | 0.9298031 | 0.3489965 | 0.68557499 |
| ENSMUSG00000039512 | Uhrf1bp1 | protein_coding | 0.5621416 | 3.174498448 | 0.9297744 | 0.3490038 | 0.68557499 |
| ENSMUSG00000032422 | Snx14 | protein_coding | -0.432456785 | 4.587299046 | 0.9291783 | 0.3491541 | 0.685626802 |
| ENSMUSG00000066057 | Gm1976 | lincRNA | -0.636599571 | 3.030686164 | 0.9288251 | 0.3492432 | 0.685626802 |
| ENSMUSG00000076498 | Trbc2 | TR_C_gene | -0.733788257 | 2.604877757 | 0.9287165 | 0.3492707 | 0.685626802 |
| ENSMUSG00000000305 | Cdh4 | protein_coding | 0.30378706 | 5.535970234 | 0.928537 | 0.349316 | 0.685626802 |
| ENSMUSG00000028657 | Ppt1 | protein_coding | 0.232396709 | 7.447321557 | 0.9284064 | 0.3493489 | 0.685626802 |
| ENSMUSG00000068917 | Clk2 | protein_coding | 0.729104679 | 3.160121422 | 0.9280996 | 0.3494264 | 0.685653751 |
| ENSMUSG00000042757 | Tmem108 | protein_coding | 0.55071324 | 3.452302513 | 0.9272732 | 0.3496353 | 0.685832271 |
| ENSMUSG00000052794 | 1700030K09Rik | protein_coding | 0.449180639 | 4.025646093 | 0.9272348 | 0.349645 | 0.685832271 |
| ENSMUSG00000023495 | Pcbp4 | protein_coding | 0.375594368 | 5.32965233 | 0.9264567 | 0.3498418 | 0.685897462 |
| ENSMUSG00000058298 | Mcm9 | protein_coding | -0.655240107 | 2.917679339 | 0.9264323 | 0.349848 | 0.685897462 |
| ENSMUSG00000060073 | Psma3 | protein_coding | 0.269708752 | 6.12581061 | 0.9261479 | 0.3499199 | 0.685897462 |
| ENSMUSG00000090125 | Pou3f1 | protein_coding | -0.467460215 | 3.977710097 | 0.9260948 | 0.3499334 | 0.685897462 |
| ENSMUSG00000030527 | Crtc3 | protein_coding | 0.702862001 | 3.106435 | 0.9256949 | 0.3500346 | 0.685927089 |
| ENSMUSG00000004567 | Mcoln1 | protein_coding | 0.406868626 | 4.725137671 | 0.9253986 | 0.3501097 | 0.685927089 |
| ENSMUSG00000055239 | Kcmf1 | protein_coding | -0.3293414 | 5.772593006 | 0.9248498 | 0.3502487 | 0.685927089 |
| ENSMUSG00000022961 | Son | protein_coding | -0.275742941 | 7.369054877 | 0.9243012 | 0.3503878 | 0.685927089 |
| ENSMUSG00000039205 | Ciz1 | protein_coding | 0.441758485 | 4.429933006 | 0.9241986 | 0.3504138 | 0.685927089 |
| ENSMUSG00000040435 | Ppp1r15a | protein_coding | 0.44433512 | 4.902967388 | 0.9241088 | 0.3504366 | 0.685927089 |
| ENSMUSG00000071652 | Ints5 | protein_coding | -0.465875731 | 3.756312583 | 0.9240539 | 0.3504505 | 0.685927089 |
| ENSMUSG00000028514 | Usp24 | protein_coding | -0.668506582 | 3.687224854 | 0.9239843 | 0.3505515 | 0.685927089 |
| ENSMUSG00000102252 | Snrpn | protein_coding | 0.321294346 | 5.158850636 | 0.923489 | 0.3505939 | 0.685927089 |
| ENSMUSG00000019792 | Trmt11 | protein_coding | -0.500025034 | 4.368417646 | 0.9234446 | 0.3506281 | 0.685927089 |
| ENSMUSG00000046434 | Hnrnpa1 | protein_coding | 0.400994981 | 3.84562228 | 0.9231528 | 0.3506792 | 0.685927089 |
| ENSMUSG00000057375 | Yipf1 | protein_coding | 0.247341664 | 6.431579522 | 0.9229821 | 0.3507226 | 0.685927089 |
| ENSMUSG00000001156 | Mxd1 | protein_coding | -0.429658932 | 3.812498202 | 0.922221 | 0.3509159 | 0.685927089 |
| ENSMUSG00000034648 | Lrrn1 | protein_coding | -0.448407663 | 4.623744272 | 0.9221791 | 0.3509266 | 0.685927089 |
| ENSMUSG00000025203 | Scd2 | protein_coding | -0.259631142 | 8.816656589 | 0.9221241 | 0.3509406 | 0.685927089 |
| ENSMUSG00000079109 | Pms2 | protein_coding | 0.866338356 | 2.451916495 | 0.9217626 | 0.3510325 | 0.685927089 |
| ENSMUSG00000026626 | Ppp2r5a | protein_coding | -0.44829616 | 4.44567116 | 0.9215661 | 0.3510824 | 0.685927089 |
| ENSMUSG00000039835 | Nhsl1 | protein_coding | 0.590117152 | 3.785744422 | 0.9216 | 0.3510967 | 0.685927089 |
| ENSMUSG00000023873 | 1700010I14Rik | protein_coding | 0.855622957 | 2.972098165 | 0.9206406 | 0.3513408 | 0.686279337 |
| ENSMUSG00000064120 | Mocs1 | protein_coding | 1.014932275 | 2.373372985 | 0.919859 | 0.351517 | 0.686498735 |
| ENSMUSG00000047414 | Flrt2 | protein_coding | -0.413120751 | 4.531359265 | 0.9195462 | 0.3515967 | 0.686529695 |
| ENSMUSG00000024218 | Taf11 | protein_coding | 0.222864077 | 6.62879639 | 0.9184843 | 0.3518675 | 0.686933708 |
| ENSMUSG00000024350 | Dnajc18 | protein_coding | 0.284726994 | 7.567937018 | 0.9176986 | 0.352068 | 0.687195922 |
| ENSMUSG00000058569 | Tmed9 | protein_coding | -0.364412417 | 8.118184108 | 0.9211568 | 0.3521296 | 0.687195922 |
| ENSMUSG00000021732 | Fgf10 | protein_coding | -0.835777864 | 2.872667115 | 0.9171218 | 0.3522154 | 0.687238575 |
| ENSMUSG00000030731 | Syt3 | protein_coding | 0.344424764 | 4.880763602 | 0.916689 | 0.352326 | 0.68732966 |
| ENSMUSG00000020079 | Supv3l1 | protein_coding | 0.747654672 | 3.067613605 | 0.9161643 | 0.3524601 | 0.687455675 |
| ENSMUSG00000037103 | Dcaf15 | protein_coding | 0.661372687 | 3.497198755 | 0.9160145 | 0.3525213 | 0.687455675 |
| ENSMUSG00000033685 | Ucp2 | protein_coding | -0.297131646 | 6.573361326 | 0.9154446 | 0.3526443 | 0.687455675 |
| ENSMUSG00000032359 | Ctsh | protein_coding | -0.569622233 | 3.698743639 | 0.9154368 | 0.3526463 | 0.687455675 |
| ENSMUSG00000032330 | Cox7a2 | protein_coding | 0.29013997 | 10.09163773 | 0.9145267 | 0.3528793 | 0.687752456 |
| ENSMUSG00000010911 | Apip | protein_coding | 0.276768201 | 5.986840214 | 0.9142286 | 0.3529557 | 0.687752456 |
| ENSMUSG00000054737 | Zfp182 | protein_coding | -0.785671323 | 2.357138798 | 0.9140933 | 0.3529904 | 0.687752456 |
| ENSMUSG00000027833 | Shox2 | protein_coding | -1.480813013 | 4.859036932 | 0.9736394 | 0.3530838 | 0.687779919 |
| ENSMUSG00000002949 | Timm44 | protein_coding | 0.296290181 | 6.385036777 | 0.9135394 | 0.3531324 | 0.687779919 |
| ENSMUSG00000040767 | Snrnp25 | protein_coding | 0.242948015 | 6.89303129 | 0.9129904 | 0.3532733 | 0.687867614 |
| ENSMUSG00000008167 | Fbxw9 | protein_coding | 0.477726627 | 4.700894323 | 0.9128693 | 0.3533271 | 0.687867614 |
| ENSMUSG00000024422 | Dhx16 | protein_coding | 0.68050646 | 2.864319705 | 0.9123717 | 0.3534321 | 0.687867614 |
| ENSMUSG00000056941 | Commd7 | protein_coding | 0.339098173 | 6.256936883 | 0.9121352 | 0.3535156 | 0.687867614 |
| ENSMUSG00000035314 | Gdpd5 | protein_coding | 0.464711585 | 3.631621812 | 0.9119128 | 0.35355 | 0.687867614 |
| ENSMUSG00000021738 | Atxn7 | protein_coding | -0.51979205 | 3.312715562 | 0.9118688 | 0.3535613 | 0.687867614 |
| ENSMUSG00000050538 | B230217C12Rik | protein_coding | -0.21342223 | 7.527299184 | 0.9107749 | 0.3538425 | 0.688154553 |
| ENSMUSG00000057615 | Ldoc1 | protein_coding | 0.481655337 | 3.467957453 | 0.9106695 | 0.3538696 | 0.688154553 |
| ENSMUSG00000028069 | Gpatch4 | protein_coding | 0.245358879 | 6.731263987 | 0.9105484 | 0.3539007 | 0.688154553 |
| ENSMUSG00000039110 | Mycbpap | protein_coding | -0.691468091 | 2.801243388 | 0.9102766 | 0.3539707 | 0.688166082 |
| ENSMUSG00000090641 | Zfp712 | protein_coding | -0.809064879 | 2.588525085 | 0.9093681 | 0.3542046 | 0.688496383 |
| ENSMUSG00000021301 | Hecw1 | protein_coding | -0.336854453 | 5.212150038 | 0.9088272 | 0.354344 | 0.688642828 |
| ENSMUSG00000021994 | Wnt5a | protein_coding | 0.547545177 | 3.22810331 | 0.9084051 | 0.3544528 | 0.688656222 |
| ENSMUSG00000037990 | Sh3rf3 | protein_coding | -0.445092439 | 3.771848816 | 0.9083036 | 0.354479 | 0.688656222 |
| ENSMUSG00000027298 | Tyro3 | protein_coding | -0.342488423 | 5.381614131 | 0.9078638 | 0.3545924 | 0.688725672 |
| ENSMUSG00000036775 | Decr2 | protein_coding | 0.234062755 | 6.638058043 | 0.9074313 | 0.354704 | 0.688725672 |
| ENSMUSG00000005893 | Nr2c2 | protein_coding | -0.380608199 | 4.941019602 | 0.9071858 | 0.3547674 | 0.688725672 |

| ensembl_gene_id | external_gene_name | gene_biotype | log2 (fold change) | log2 (counts per million) | F-value | p-value | False discovery rate |
|---|---|---|---|---|---|---|---|
| ENSMUSG00000042182 | Bend6 | protein_coding | -0.230259894 | 7.256237673 | 0.9071134 | 0.3547861 | 0.688725672 |
| ENSMUSG00000095139 | Pou3f2 | protein_coding | -0.342665184 | 4.942613641 | 0.9069242 | 0.354835 | 0.688725672 |
| ENSMUSG00000022364 | Tbc1d31 | protein_coding | -0.896327555 | 2.316712019 | 0.9062657 | 0.3550051 | 0.688838653 |
| ENSMUSG00000042901 | Aida | protein_coding | -0.294173416 | 5.122552065 | 0.9060101 | 0.3550711 | 0.688838653 |
| ENSMUSG00000028760 | Eif4g3 | protein_coding | -0.308132493 | 5.792021111 | 0.9059552 | 0.3550853 | 0.688838653 |
| ENSMUSG00000021023 | 1110008L16Rik | protein_coding | 0.548741236 | 3.348714097 | 0.9052426 | 0.3552696 | 0.689071874 |
| ENSMUSG00000029723 | Tsc22d4 | protein_coding | -0.404741681 | 7.49352734 | 0.9116754 | 0.3555509 | 0.68949297 |
| ENSMUSG00000048794 | Cfap100 | protein_coding | -0.857315439 | 2.308252603 | 0.9036051 | 0.3556937 | 0.68964549 |
| ENSMUSG00000024799 | Tm7sf2 | protein_coding | 0.492842325 | 4.114726198 | 0.9026667 | 0.355937 | 0.689992869 |
| ENSMUSG00000020739 | Nup85 | protein_coding | 0.362388788 | 5.327087493 | 0.9023676 | 0.3560146 | 0.690018916 |
| ENSMUSG00000029632 | Ndufa4 | protein_coding | 0.288972022 | 11.03033526 | 0.9010594 | 0.3563543 | 0.690552857 |
| ENSMUSG00000040383 | Aqr | protein_coding | 0.405646712 | 4.519892556 | 0.900524 | 0.3564935 | 0.69069812 |
| ENSMUSG00000028637 | Ccdc30 | protein_coding | 0.314633186 | 5.308539103 | 0.9001267 | 0.3565968 | 0.690773806 |
| ENSMUSG00000025139 | Tollip | protein_coding | 0.280848443 | 7.020217951 | 0.8988949 | 0.3569174 | 0.691164379 |
| ENSMUSG00000039461 | Tcta | protein_coding | 0.276775412 | 6.791598511 | 0.8988583 | 0.356927 | 0.691164379 |
| ENSMUSG00000066357 | Wdr6 | protein_coding | 0.614632938 | 7.794394425 | 0.9255068 | 0.3571503 | 0.691397394 |
| ENSMUSG00000009076 | Zmat5 | protein_coding | -0.29583706 | 5.897945453 | 0.897793 | 0.3572046 | 0.691397394 |
| ENSMUSG00000030341 | Tnfrsf1a | protein_coding | 0.794395896 | 2.560831926 | 0.8976565 | 0.3572402 | 0.691397394 |
| ENSMUSG00000027809 | Etfdh | protein_coding | 0.360977127 | 4.954823637 | 0.8968463 | 0.3574516 | 0.69146932 |
| ENSMUSG00000055026 | Gabrg3 | protein_coding | 0.343540814 | 4.684126774 | 0.8961988 | 0.3576207 | 0.69146932 |
| ENSMUSG00000022159 | Rab2b | protein_coding | 0.306556314 | 4.849298695 | 0.8961467 | 0.3576343 | 0.69146932 |
| ENSMUSG00000045284 | Dcaf12l1 | protein_coding | 0.262678029 | 6.420948291 | 0.8954341 | 0.3578205 | 0.69146932 |
| ENSMUSG00000034731 | Dgkh | protein_coding | -0.546021898 | 3.239338009 | 0.895405 | 0.3578281 | 0.69146932 |
| ENSMUSG00000026187 | Xrcc5 | protein_coding | 0.312580871 | 5.202809155 | 0.8952491 | 0.3578689 | 0.69146932 |
| ENSMUSG00000026471 | Mr1 | protein_coding | -1.479785897 | 2.113181124 | 0.8982783 | 0.3579409 | 0.69146932 |
| ENSMUSG00000045466 | Zfp956 | protein_coding | 0.463821437 | 4.374954297 | 0.8949675 | 0.3579425 | 0.69146932 |
| ENSMUSG00000029345 | Tfip11 | protein_coding | 0.508049438 | 4.892480515 | 0.8964788 | 0.3580049 | 0.69146932 |
| ENSMUSG00000040236 | Trappc5 | protein_coding | 0.243373029 | 6.273591413 | 0.8946523 | 0.358025 | 0.69146932 |
| ENSMUSG00000025702 | 8-Mar | protein_coding | -0.473263475 | 4.250494278 | 0.8946348 | 0.3580296 | 0.69146932 |
| ENSMUSG00000021236 | Entpd5 | protein_coding | 0.387576762 | 4.612089594 | 0.8945604 | 0.358049 | 0.69146932 |
| ENSMUSG00000042834 | Nrep | protein_coding | -0.273603881 | 6.433058837 | 0.8942581 | 0.3581281 | 0.691497913 |
| ENSMUSG00000043668 | Tox3 | protein_coding | -0.532125643 | 3.709354237 | 0.8938613 | 0.358232 | 0.691574296 |
| ENSMUSG00000074771 | Ankef1 | protein_coding | -0.770300028 | 2.771985152 | 0.8930463 | 0.3584455 | 0.691626938 |
| ENSMUSG00000074582 | Arfgef2 | protein_coding | 0.513050013 | 4.268294228 | 0.8930428 | 0.358469 | 0.691626938 |
| ENSMUSG00000028062 | Lamtor2 | protein_coding | 0.25711404 | 9.130326099 | 0.892888 | 0.358487 | 0.691626938 |
| ENSMUSG00000097040 | 2610316D01Rik | processed_transcript | 0.488749856 | 4.550037084 | 0.8928478 | 0.3585201 | 0.691626938 |
| ENSMUSG00000018848 | Rars | protein_coding | 0.267537691 | 6.479045316 | 0.89253 | 0.3585809 | 0.691626938 |
| ENSMUSG00000025867 | Cplx2 | protein_coding | -0.228090941 | 8.226536737 | 0.8921277 | 0.3586864 | 0.691706395 |
| ENSMUSG00000024732 | Ccdc86 | protein_coding | 0.422908246 | 4.064614931 | 0.8912502 | 0.3589167 | 0.691956539 |
| ENSMUSG00000037674 | Rfx7 | protein_coding | -0.439804127 | 3.692786048 | 0.8911432 | 0.3589448 | 0.691956539 |
| ENSMUSG00000034189 | Hsdl1 | protein_coding | -0.275148448 | 6.774531571 | 0.8906981 | 0.3590617 | 0.692057871 |
| ENSMUSG00000111336 | AC160338.2 | processed_pseudoge | 0.716624566 | 2.495703663 | 0.8898242 | 0.3592914 | 0.692198092 |
| ENSMUSG00000031374 | Zfp92 | protein_coding | 0.697973861 | 2.641578821 | 0.8895506 | 0.3593634 | 0.692198092 |
| ENSMUSG00000000142 | Axin2 | protein_coding | -0.529772141 | 3.539760292 | 0.8893008 | 0.3594291 | 0.692198092 |
| ENSMUSG00000003452 | Bicd1 | protein_coding | -0.285223528 | 5.506908073 | 0.8891071 | 0.3594801 | 0.692198092 |
| ENSMUSG00000000776 | Polr3d | protein_coding | -0.707761132 | 3.890417899 | 0.8927633 | 0.3595132 | 0.692198092 |
| ENSMUSG00000021794 | Glud1 | protein_coding | -0.283923869 | 6.788155854 | 0.8889528 | 0.3595207 | 0.692198092 |
| ENSMUSG00000075266 | Cenpw | protein_coding | -0.695234759 | 3.134142398 | 0.8884056 | 0.3596648 | 0.692351509 |
| ENSMUSG00000018809 | Smyd4 | protein_coding | 0.448514898 | 4.364534529 | 0.8875864 | 0.3598806 | 0.69264303 |
| ENSMUSG00000031872 | Bean1 | protein_coding | 0.708755174 | 2.534409631 | 0.8858524 | 0.3603382 | 0.69337936 |
| ENSMUSG00000038077 | Kcna6 | protein_coding | 0.331670667 | 5.795123717 | 0.8856406 | 0.3603941 | 0.69337936 |
| ENSMUSG00000062963 | Ufc1 | protein_coding | 0.219663384 | 8.435340463 | 0.8850105 | 0.3605606 | 0.69337936 |
| ENSMUSG00000028719 | Cmpk1 | protein_coding | -0.249126378 | 7.30232154 | 0.8850079 | 0.3605613 | 0.69337936 |
| ENSMUSG00000025269 | Apex2 | protein_coding | 0.571024939 | 3.326395185 | 0.8849159 | 0.3605856 | 0.69337936 |
| ENSMUSG00000054509 | Parp4 | protein_coding | -0.802221253 | 2.787169199 | 0.8842832 | 0.360753 | 0.693502054 |
| ENSMUSG00000053080 | 2700081O15Rik | protein_coding | -0.682866455 | 2.671389807 | 0.8839655 | 0.360837 | 0.693502054 |
| ENSMUSG00000018965 | Ywhah | protein_coding | 0.210614216 | 8.259081373 | 0.8839432 | 0.3608429 | 0.693502054 |
| ENSMUSG00000033350 | Chst2 | protein_coding | 0.214259193 | 6.92829808 | 0.8831006 | 0.361066 | 0.693687628 |
| ENSMUSG00000037692 | Ahdc1 | protein_coding | -0.510155322 | 3.881753752 | 0.8828495 | 0.3611325 | 0.693687628 |
| ENSMUSG00000027424 | Mgme1 | protein_coding | 0.399407822 | 3.799947461 | 0.8828477 | 0.361133 | 0.693687628 |
| ENSMUSG00000054894 | Atp5s | protein_coding | 0.284018367 | 5.555725959 | 0.8823362 | 0.3612686 | 0.693824033 |
| ENSMUSG00000032192 | Gnb5 | protein_coding | -0.208829516 | 7.735530839 | 0.8816068 | 0.361462 | 0.693987597 |
| ENSMUSG00000029610 | Aimp2 | protein_coding | 0.233048399 | 6.109981577 | 0.8815284 | 0.3614828 | 0.693987597 |
| ENSMUSG00000024112 | Cacna1h | protein_coding | 0.908011291 | 2.669602741 | 0.8805582 | 0.3617403 | 0.694358078 |
| ENSMUSG00000020956 | Dtd2 | protein_coding | 0.582032595 | 5.50627464 | 0.8883332 | 0.361862 | 0.694451543 |
| ENSMUSG00000086075 | Gm15728 | lincRNA | 0.596748014 | 3.171824099 | 0.8797309 | 0.3619602 | 0.694451543 |
| ENSMUSG00000021745 | Ptprg | protein_coding | -0.482300615 | 4.195560502 | 0.8793798 | 0.3620535 | 0.694451543 |
| ENSMUSG00000046962 | Zbtb21 | protein_coding | -0.308663924 | 5.212693497 | 0.8789006 | 0.362181 | 0.694451543 |
| ENSMUSG00000044533 | Rps2 | protein_coding | 0.404632367 | 4.659442267 | 0.8787907 | 0.3622103 | 0.694451543 |
| ENSMUSG00000021040 | Slirp | protein_coding | 0.381053016 | 4.494074905 | 0.8786495 | 0.3622478 | 0.694451543 |
| ENSMUSG00000039637 | Coro7 | protein_coding | 0.331042873 | 4.868359133 | 0.8783796 | 0.3623197 | 0.694451543 |
| ENSMUSG00000038844 | Kif16b | protein_coding | -0.590570355 | 3.107738631 | 0.8782568 | 0.3623524 | 0.694451543 |
| ENSMUSG00000029440 | Psmd9 | protein_coding | 0.254150388 | 6.265659844 | 0.8781895 | 0.3623703 | 0.694451543 |
| ENSMUSG00000032305 | Fam219b | protein_coding | 0.355751332 | 5.791319871 | 0.8773433 | 0.3625957 | 0.694759741 |
| ENSMUSG00000029199 | Lias | protein_coding | 0.235204162 | 6.878861536 | 0.8768809 | 0.362719 | 0.694872095 |
| ENSMUSG00000022747 | St3gal6 | protein_coding | -0.498286665 | 3.685599646 | 0.8762378 | 0.3628905 | 0.695059297 |
| ENSMUSG00000040136 | Abcc8 | protein_coding | 1.533085808 | 2.080395375 | 0.8792797 | 0.362946 | 0.695059297 |
| ENSMUSG00000013646 | Sh3bp5l | protein_coding | 0.29041229 | 5.210169796 | 0.8756182 | 0.3630559 | 0.695146043 |

| ensembl_gene_id | external_gene_name | gene_biotype | log2 (fold change) | log2 (counts per million) | F-value | p-value | False discovery rate |
|---|---|---|---|---|---|---|---|
| ENSMUSG00000028676 | Srsf10 | protein_coding | -0.278814564 | 5.481555705 | 0.8753556 | 0.363126 | 0.695156529 |
| ENSMUSG00000047881 | Rell1 | protein_coding | -0.923739433 | 2.650813724 | 0.8749767 | 0.3632272 | 0.695226524 |
| ENSMUSG00000042590 | Ipo11 | protein_coding | 0.521221351 | 3.43643476 | 0.8746613 | 0.3633116 | 0.695264126 |
| ENSMUSG00000040964 | Arhgef10l | protein_coding | -0.459734188 | 3.819511373 | 0.8741833 | 0.3634394 | 0.695384937 |
| ENSMUSG00000067274 | Rplp0 | protein_coding | 0.206160103 | 8.351305797 | 0.8737402 | 0.3635579 | 0.69546423 |
| ENSMUSG00000010362 | Rdm1 | protein_coding | -0.391678736 | 5.33457854 | 0.8735448 | 0.3636101 | 0.69546423 |
| ENSMUSG00000064061 | Dzip3 | protein_coding | -0.270895556 | 5.567912203 | 0.8722831 | 0.363948 | 0.695786886 |
| ENSMUSG00000020990 | Cdkl1 | protein_coding | 0.396175732 | 4.588056942 | 0.8721597 | 0.3639811 | 0.695786886 |
| ENSMUSG00000032320 | Rcn2 | protein_coding | -0.338183444 | 6.618201067 | 0.8725575 | 0.3639938 | 0.695786886 |
| ENSMUSG00000028932 | Psmc2 | protein_coding | 0.224410239 | 8.499905704 | 0.8719486 | 0.3640377 | 0.695786886 |
| ENSMUSG00000028973 | Abcb8 | protein_coding | -0.261941529 | 6.129467242 | 0.8707611 | 0.3643562 | 0.696271973 |
| ENSMUSG00000031610 | Scrg1 | protein_coding | -0.73870741 | 5.703641688 | 0.8946557 | 0.3645061 | 0.696434661 |
| ENSMUSG00000039887 | Alg14 | protein_coding | -0.267024512 | 5.90807092 | 0.8696674 | 0.36465 | 0.696585721 |
| ENSMUSG00000019947 | Arid5b | protein_coding | -0.338626402 | 5.545585943 | 0.8689834 | 0.3648338 | 0.696813168 |
| ENSMUSG00000034931 | Dhx8 | protein_coding | 0.389322535 | 4.399326705 | 0.8685362 | 0.3649541 | 0.696835273 |
| ENSMUSG00000029361 | Nos1 | protein_coding | 0.375981945 | 6.17370665 | 0.8701422 | 0.364975 | 0.696835273 |
| ENSMUSG00000041702 | Btbd7 | protein_coding | 0.489729915 | 3.842871206 | 0.8678394 | 0.3651417 | 0.697029705 |
| ENSMUSG00000034898 | Filip1 | protein_coding | -0.427906303 | 4.020894749 | 0.867457 | 0.3652447 | 0.697053246 |
| ENSMUSG00000028986 | Klhl7 | protein_coding | -0.267343016 | 6.381126515 | 0.8673123 | 0.3652836 | 0.697053246 |
| ENSMUSG00000050088 | 1600012H06Rik | protein_coding | 1.093219322 | 2.783557498 | 0.8689437 | 0.365705 | 0.697647737 |
| ENSMUSG00000054426 | A930005H10Rik | processed_transcript | -0.335059547 | 4.744549992 | 0.8656757 | 0.3657249 | 0.697647737 |
| ENSMUSG00000027663 | Zmat3 | protein_coding | -0.258901267 | 5.696992462 | 0.8640367 | 0.3661677 | 0.698281404 |
| ENSMUSG00000026600 | Soat1 | protein_coding | 0.407594064 | 4.214065536 | 0.8639653 | 0.366187 | 0.698281404 |
| ENSMUSG00000046807 | Lrrc75b | protein_coding | 0.414865621 | 4.251037385 | 0.8635299 | 0.3663048 | 0.698312254 |
| ENSMUSG00000026173 | Plcd4 | protein_coding | 0.50259131 | 3.898973072 | 0.8632018 | 0.3663935 | 0.698312254 |
| ENSMUSG00000113902 | AC121965.1 | bidirectional_promote | 0.302531503 | 9.115190095 | 0.8630883 | 0.3664464 | 0.698312254 |
| ENSMUSG00000022752 | Tomm70a | protein_coding | -0.22325211 | 7.013920706 | 0.8626687 | 0.3665378 | 0.698312254 |
| ENSMUSG00000002944 | Cd36 | protein_coding | -1.042899068 | 2.164736826 | 0.8626615 | 0.3665398 | 0.698312254 |
| ENSMUSG00000028128 | F3 | protein_coding | -0.414359433 | 5.787861744 | 0.864554 | 0.3665928 | 0.698312254 |
| ENSMUSG00000022516 | Nudt16l1 | protein_coding | 0.220355074 | 7.037987779 | 0.8618122 | 0.3667698 | 0.698443442 |
| ENSMUSG00000047361 | Gm973 | protein_coding | -0.924910071 | 2.290108878 | 0.861732 | 0.3667916 | 0.698443442 |
| ENSMUSG00000074519 | Zfp971 | protein_coding | -0.519812527 | 3.113243149 | 0.8614159 | 0.3668773 | 0.698482924 |
| ENSMUSG00000061613 | U2af1 | protein_coding | 0.363000215 | 4.06321943 | 0.8602212 | 0.3672014 | 0.698976277 |
| ENSMUSG00000048878 | Hexim1 | protein_coding | -0.357475441 | 4.585483857 | 0.8598988 | 0.367289 | 0.699019162 |
| ENSMUSG00000034285 | Nipsnap1 | protein_coding | 0.201740311 | 8.165715942 | 0.859259 | 0.3674628 | 0.699162111 |
| ENSMUSG00000059146 | Ntrk3 | protein_coding | -0.259229067 | 6.111669655 | 0.8591437 | 0.3674941 | 0.699162111 |
| ENSMUSG00000049775 | Tmsb4x | protein_coding | -0.317001237 | 11.71089182 | 0.8586694 | 0.3676452 | 0.699210725 |
| ENSMUSG00000018932 | Map2k3 | protein_coding | -0.594542253 | 2.856853344 | 0.8582329 | 0.3677418 | 0.699210725 |
| ENSMUSG00000056708 | Ier5 | protein_coding | -0.652188033 | 3.078491672 | 0.8581262 | 0.3677708 | 0.699210725 |
| ENSMUSG00000045409 | Trim39 | protein_coding | -0.50586141 | 3.885929486 | 0.8580416 | 0.3677939 | 0.699210725 |
| ENSMUSG00000071266 | Zfp946 | protein_coding | -0.67953101 | 2.83477977 | 0.8578545 | 0.3678448 | 0.699210725 |
| ENSMUSG00000022722 | Arl6 | protein_coding | -0.245454628 | 7.395387216 | 0.857411 | 0.3679655 | 0.699243159 |
| ENSMUSG00000060261 | Gtf2i | protein_coding | 0.274505046 | 6.664395869 | 0.8573141 | 0.3679919 | 0.699243159 |
| ENSMUSG00000024242 | Map4k3 | protein_coding | -0.363716317 | 5.042346066 | 0.8561093 | 0.3683203 | 0.699561812 |
| ENSMUSG00000018846 | Pank3 | protein_coding | -0.338016912 | 4.737048058 | 0.8559951 | 0.3683514 | 0.699561812 |
| ENSMUSG00000022359 | Wdyhv1 | protein_coding | 0.257772739 | 5.77310825 | 0.855836 | 0.3683948 | 0.699561812 |
| ENSMUSG00000002109 | Ddb2 | protein_coding | -0.708984434 | 2.829398467 | 0.8557442 | 0.3684198 | 0.699561812 |
| ENSMUSG00000030002 | Dusp11 | protein_coding | -0.415700485 | 3.916512805 | 0.8547079 | 0.3687027 | 0.699770733 |
| ENSMUSG00000051339 | 2900026A02Rik | protein_coding | -0.439325513 | 4.055889803 | 0.854455 | 0.3687718 | 0.699770733 |
| ENSMUSG00000031691 | Tnpo2 | protein_coding | 0.627226834 | 3.370183233 | 0.8543645 | 0.3687965 | 0.699770733 |
| ENSMUSG00000000378 | Ccm2 | protein_coding | 0.252675429 | 5.708089225 | 0.8542993 | 0.3688143 | 0.699770733 |
| ENSMUSG00000024068 | Spast | protein_coding | -0.429105476 | 4.623684832 | 0.8539783 | 0.3689021 | 0.699770733 |
| ENSMUSG00000024259 | Slc25a46 | protein_coding | -0.292529725 | 5.508648155 | 0.853741 | 0.368967 | 0.699770733 |
| ENSMUSG00000031221 | Igbp1 | protein_coding | 0.32639309 | 6.132569825 | 0.8534211 | 0.3690764 | 0.699770733 |
| ENSMUSG00000028691 | Prdx1 | protein_coding | -0.234738953 | 9.35749687 | 0.8532013 | 0.3691146 | 0.699770733 |
| ENSMUSG00000047539 | Fbxo28 | protein_coding | -0.332930642 | 4.946111766 | 0.8530987 | 0.3691426 | 0.699770733 |
| ENSMUSG00000006906 | Stambp | protein_coding | 0.246330461 | 6.173512635 | 0.8528805 | 0.3692023 | 0.699770733 |
| ENSMUSG00000027193 | Api5 | protein_coding | -0.267703771 | 5.813265692 | 0.8522709 | 0.3693692 | 0.699770733 |
| ENSMUSG00000028884 | Rpa2 | protein_coding | 0.351381588 | 5.220254905 | 0.8522095 | 0.369386 | 0.699770733 |
| ENSMUSG00000072825 | Cep170b | protein_coding | 0.273577416 | 6.618849116 | 0.8521381 | 0.3694056 | 0.699770733 |
| ENSMUSG00000038335 | Tsr1 | protein_coding | -0.338368476 | 5.146566259 | 0.852009 | 0.3694409 | 0.699770733 |
| ENSMUSG00000055053 | Nfic | protein_coding | 0.358972308 | 5.752961924 | 0.851238 | 0.3696742 | 0.700011651 |
| ENSMUSG00000019806 | Aig1 | protein_coding | -0.325317798 | 6.224464513 | 0.85115 | 0.3696983 | 0.700011651 |
| ENSMUSG00000025395 | Prim1 | protein_coding | -0.439677368 | 4.083517732 | 0.8499934 | 0.3699937 | 0.700447509 |
| ENSMUSG00000030814 | Bcl7c | protein_coding | 0.681370231 | 2.647911309 | 0.8494953 | 0.3701305 | 0.700488496 |
| ENSMUSG00000025006 | Sorbs1 | protein_coding | -0.252169513 | 6.236332475 | 0.8494402 | 0.3701456 | 0.700488496 |
| ENSMUSG00000022175 | Lrp10 | protein_coding | -0.511068132 | 3.736932533 | 0.8484998 | 0.370404 | 0.700854237 |
| ENSMUSG00000028488 | Sh3gl2 | protein_coding | 0.237351525 | 7.615826739 | 0.8480253 | 0.3705346 | 0.700977873 |
| ENSMUSG00000026696 | Vamp4 | protein_coding | 0.253674514 | 7.902477043 | 0.8469839 | 0.3708212 | 0.701396781 |
| ENSMUSG00000023723 | Mrps23 | protein_coding | 0.221616876 | 6.937090636 | 0.8456316 | 0.371194 | 0.701978355 |
| ENSMUSG00000052727 | Map1b | protein_coding | -0.494885359 | 9.563344979 | 0.8584534 | 0.371438 | 0.702316246 |
| ENSMUSG00000025511 | Tspan4 | protein_coding | 0.428248875 | 4.134808784 | 0.8440933 | 0.3716187 | 0.702442409 |
| ENSMUSG00000027303 | Ptpra | protein_coding | 0.346623382 | 4.260738267 | 0.844033 | 0.3716353 | 0.702442409 |
| ENSMUSG00000038156 | Spon1 | protein_coding | -0.287043268 | 5.425957612 | 0.8434204 | 0.3718047 | 0.702458322 |
| ENSMUSG00000055932 | Fto | protein_coding | 0.423865442 | 8.114774257 | 0.850972 | 0.3718552 | 0.702458322 |
| ENSMUSG00000023021 | Cers5 | protein_coding | 0.335921739 | 5.106722379 | 0.8431595 | 0.3718768 | 0.702458322 |
| ENSMUSG00000006191 | Cdkal1 | protein_coding | -0.369711836 | 4.641648167 | 0.8430574 | 0.3719051 | 0.702458322 |
| ENSMUSG00000032353 | Tmed3 | protein_coding | 0.213645281 | 7.049319492 | 0.842707 | 0.372002 | 0.702518045 |

| ensembl_gene_id | external_gene_name | gene_biotype | log2 (fold change) | log2 (counts per million) | F-value | p-value | False discovery rate |
|---|---|---|---|---|---|---|---|
| ENSMUSG00000026502 | Desi2 | protein_coding | -0.381839818 | 4.263983837 | 0.8419161 | 0.372221 | 0.702645542 |
| ENSMUSG00000024369 | Nelfe | protein_coding | 0.262139439 | 6.009581249 | 0.8417806 | 0.3722585 | 0.702645542 |
| ENSMUSG00000020388 | Pdlim4 | protein_coding | -0.533132551 | 3.714481648 | 0.8415885 | 0.3723117 | 0.702645542 |
| ENSMUSG00000027716 | Trpc3 | protein_coding | -0.738042728 | 2.94328663 | 0.8415193 | 0.3723309 | 0.702645542 |
| ENSMUSG00000108434 | Gm45768 | lincRNA | -0.706366638 | 3.28471313 | 0.8407248 | 0.372573 | 0.702744341 |
| ENSMUSG00000043518 | Rai2 | protein_coding | -0.450791324 | 4.267342959 | 0.8404336 | 0.3726319 | 0.702744341 |
| ENSMUSG00000036285 | Noa1 | protein_coding | -0.517757991 | 3.49091992 | 0.8404153 | 0.372637 | 0.702744341 |
| ENSMUSG00000034949 | Zfr2 | protein_coding | 0.463026687 | 3.528318463 | 0.8403331 | 0.3726598 | 0.702744341 |
| ENSMUSG00000001986 | Gria3 | protein_coding | -0.248765923 | 5.891999901 | 0.8401521 | 0.37271 | 0.702744341 |
| ENSMUSG00000029649 | Pomp | protein_coding | 0.228798624 | 9.62776888 | 0.8395438 | 0.3728789 | 0.7028755 |
| ENSMUSG00000022960 | Donson | protein_coding | -1.159823929 | 2.715215233 | 0.843457 | 0.3729165 | 0.7028755 |
| ENSMUSG00000027714 | Exosc9 | protein_coding | 0.23397407 | 6.244827269 | 0.8390076 | 0.3730278 | 0.7028755 |
| ENSMUSG00000037503 | Fam168b | protein_coding | 0.232501751 | 6.432525751 | 0.83896 | 0.3730411 | 0.7028755 |
| ENSMUSG00000010307 | Tmem86a | protein_coding | 0.429342655 | 4.325601805 | 0.8386841 | 0.3731177 | 0.702896801 |
| ENSMUSG00000031634 | Ufsp2 | protein_coding | 0.201502144 | 7.256195566 | 0.83805 | 0.373294 | 0.703105775 |
| ENSMUSG00000046922 | Gpr6 | protein_coding | -0.580007058 | 3.465183134 | 0.8374366 | 0.3734647 | 0.703304042 |
| ENSMUSG00000045064 | Zc2hc1c | protein_coding | 0.640281742 | 3.334567697 | 0.8371276 | 0.3735507 | 0.703342877 |
| ENSMUSG00000041736 | Tspo | protein_coding | -0.504452762 | 3.492472395 | 0.8365285 | 0.3737176 | 0.703410771 |
| ENSMUSG00000033728 | Lrrc14 | protein_coding | 0.396148148 | 4.191297207 | 0.8364917 | 0.3737279 | 0.703410771 |
| ENSMUSG00000026821 | Ralgds | protein_coding | -0.316458086 | 5.58292015 | 0.8362937 | 0.3737831 | 0.703410771 |
| ENSMUSG00000035595 | 1600002K03Rik | protein_coding | -0.463056031 | 3.89745917 | 0.835918 | 0.3738878 | 0.703484737 |
| ENSMUSG00000039068 | Zzz3 | protein_coding | -0.371458372 | 4.790596783 | 0.8349635 | 0.374154 | 0.703642306 |
| ENSMUSG00000028851 | Nudc | protein_coding | 0.421555714 | 5.935962443 | 0.8375356 | 0.3742813 | 0.703642306 |
| ENSMUSG00000001910 | Nacc1 | protein_coding | 0.357661465 | 4.97235212 | 0.8344852 | 0.3742876 | 0.703642306 |
| ENSMUSG00000032652 | Crebl2 | protein_coding | 0.344881482 | 7.062361033 | 0.8372087 | 0.3743721 | 0.703642306 |
| ENSMUSG00000035960 | Apex1 | protein_coding | 0.284123382 | 5.558135195 | 0.8341486 | 0.3743816 | 0.703642306 |
| ENSMUSG00000015806 | Qdpr | protein_coding | -0.223732261 | 6.779004221 | 0.8340638 | 0.3744053 | 0.703642306 |
| ENSMUSG00000026207 | Speg | protein_coding | 0.532907528 | 3.661828598 | 0.8335709 | 0.374543 | 0.703642306 |
| ENSMUSG00000028793 | Rnf19b | protein_coding | -0.33749609 | 4.540440979 | 0.833416 | 0.3745863 | 0.703642306 |
| ENSMUSG00000027263 | Tubgcp4 | protein_coding | 0.567680449 | 3.016042096 | 0.8333241 | 0.374612 | 0.703642306 |
| ENSMUSG00000039047 | Pigk | protein_coding | 0.260969397 | 6.552210073 | 0.8330987 | 0.374675 | 0.703642306 |
| ENSMUSG00000001786 | Fbxo7 | protein_coding | -0.312258217 | 5.935291912 | 0.8328354 | 0.3747487 | 0.703642306 |
| ENSMUSG00000005148 | Klf5 | protein_coding | 0.506058429 | 3.456679139 | 0.8328065 | 0.3747568 | 0.703642306 |
| ENSMUSG00000039456 | Morc3 | protein_coding | -0.479853972 | 4.556758285 | 0.8317048 | 0.3750869 | 0.704139177 |
| ENSMUSG00000003660 | Snrnp200 | protein_coding | -0.321959032 | 5.519479476 | 0.8311366 | 0.3752244 | 0.704274349 |
| ENSMUSG00000019945 | 1700040L02Rik | protein_coding | -0.475248259 | 3.569373878 | 0.8294256 | 0.3757044 | 0.704931089 |
| ENSMUSG00000031808 | Slc27a1 | protein_coding | -0.37478837 | 5.635908775 | 0.8291028 | 0.3758167 | 0.704931089 |
| ENSMUSG00000017631 | Abr | protein_coding | 0.229430274 | 7.151394272 | 0.8289068 | 0.3758501 | 0.704931089 |
| ENSMUSG00000027971 | Ndst4 | protein_coding | -0.48556386 | 3.514078181 | 0.8288358 | 0.3758701 | 0.704931089 |
| ENSMUSG00000062961 | Ccdc177 | protein_coding | -0.457722922 | 3.767734098 | 0.8287219 | 0.3759021 | 0.704931089 |
| ENSMUSG00000002319 | Ipo4 | protein_coding | 0.304767588 | 6.218564489 | 0.8279756 | 0.3761119 | 0.705191169 |
| ENSMUSG00000005370 | Msh6 | protein_coding | -0.967738193 | 2.915605067 | 0.83075 | 0.3761733 | 0.705191169 |
| ENSMUSG00000035168 | Tanc1 | protein_coding | -0.433680595 | 3.707261857 | 0.827317 | 0.3762972 | 0.705191169 |
| ENSMUSG00000018765 | Fxr2 | protein_coding | 0.277679416 | 5.496278609 | 0.8272962 | 0.3763031 | 0.705191169 |
| ENSMUSG00000038533 | Cbfa2t2 | protein_coding | 0.405701551 | 6.28692202 | 0.8315934 | 0.3764141 | 0.705276254 |
| ENSMUSG00000022770 | Dlg1 | protein_coding | -0.296276596 | 5.375225563 | 0.8253368 | 0.3768552 | 0.705964395 |
| ENSMUSG00000022312 | Eif3h | protein_coding | 0.208859526 | 8.916982619 | 0.8251332 | 0.3769126 | 0.705964395 |
| ENSMUSG00000025533 | Asl | protein_coding | -0.223573803 | 6.434689736 | 0.8246293 | 0.3770548 | 0.706107767 |
| ENSMUSG00000043289 | Mei4 | protein_coding | -0.868601305 | 2.37639446 | 0.8238259 | 0.3772818 | 0.706409675 |
| ENSMUSG00000025037 | Maoa | protein_coding | 0.302810098 | 5.56564449 | 0.8229933 | 0.3775171 | 0.706616428 |
| ENSMUSG00000049300 | Prmt6 | protein_coding | -0.442584703 | 3.436988072 | 0.8229704 | 0.3775236 | 0.706616428 |
| ENSMUSG00000006463 | Zdhhc24 | protein_coding | 0.459606871 | 3.984992583 | 0.8223723 | 0.3776928 | 0.706745708 |
| ENSMUSG00000061286 | Exosc5 | protein_coding | 0.254919309 | 6.021888694 | 0.8222617 | 0.3777241 | 0.706745708 |
| ENSMUSG00000033486 | Catsper2 | protein_coding | -0.533975807 | 3.225605393 | 0.8196127 | 0.3784751 | 0.708027534 |
| ENSMUSG00000075316 | Scn9a | protein_coding | -0.423749285 | 3.5078237 | 0.8182009 | 0.3788762 | 0.708654598 |
| ENSMUSG00000031993 | Snx19 | protein_coding | 0.328804133 | 5.034120209 | 0.8176564 | 0.379031 | 0.708820991 |
| ENSMUSG00000053931 | Cnn3 | protein_coding | 0.410719964 | 4.961678493 | 0.8170289 | 0.3792096 | 0.709031682 |
| ENSMUSG00000001441 | Npepps | protein_coding | -0.209900268 | 6.845432145 | 0.8156684 | 0.3795973 | 0.709475761 |
| ENSMUSG00000055991 | Zkscan5 | protein_coding | 0.471359711 | 3.547191066 | 0.8155239 | 0.3796385 | 0.709475761 |
| ENSMUSG00000021209 | Ppp4r4 | protein_coding | 0.385733949 | 4.206087628 | 0.8155008 | 0.3796451 | 0.709475761 |
| ENSMUSG00000074575 | Kcng1 | protein_coding | -0.516840228 | 3.102649966 | 0.8145152 | 0.3799263 | 0.709877953 |
| ENSMUSG00000000409 | Lck | protein_coding | 0.55224017 | 3.777898361 | 0.8142537 | 0.3800225 | 0.709934243 |
| ENSMUSG00000001151 | Pcnt | protein_coding | -0.614759627 | 3.436726609 | 0.8139132 | 0.3801197 | 0.709992603 |
| ENSMUSG00000051890 | Klhdc1 | protein_coding | -0.355471768 | 4.471963416 | 0.8132647 | 0.3802836 | 0.710175348 |
| ENSMUSG00000044072 | Eml6 | protein_coding | 0.621480067 | 2.775507336 | 0.8130005 | 0.3803591 | 0.710193067 |
| ENSMUSG00000005102 | Eif2ak4 | protein_coding | -0.485523716 | 3.634315011 | 0.8120813 | 0.3806222 | 0.710346541 |
| ENSMUSG00000025582 | Nptx1 | protein_coding | 0.370131202 | 4.495206793 | 0.8120679 | 0.380626 | 0.710346541 |
| ENSMUSG00000031778 | Cx3cl1 | protein_coding | -0.225126704 | 7.131790328 | 0.8120207 | 0.3806395 | 0.710346541 |
| ENSMUSG00000051306 | Usp42 | protein_coding | 0.408115799 | 4.038454282 | 0.8114946 | 0.3807902 | 0.710473851 |
| ENSMUSG00000037541 | Shank2 | protein_coding | -0.394992858 | 4.28895981 | 0.8113212 | 0.3808399 | 0.710473851 |
| ENSMUSG00000094626 | Cecr6 | protein_coding | 0.702420155 | 3.036081441 | 0.8109283 | 0.3809525 | 0.710491262 |
| ENSMUSG00000062646 | Ganc | protein_coding | -0.397600571 | 3.615466265 | 0.8107163 | 0.3810133 | 0.710491262 |
| ENSMUSG00000025485 | Ric8a | protein_coding | 0.308378466 | 5.790532906 | 0.8104556 | 0.381088 | 0.710491262 |
| ENSMUSG00000039148 | Sart1 | protein_coding | 0.468011811 | 4.561525235 | 0.8101416 | 0.3811995 | 0.710491262 |
| ENSMUSG00000032340 | Neo1 | protein_coding | 0.380590689 | 4.574868656 | 0.8098964 | 0.3812485 | 0.710491262 |
| ENSMUSG00000037349 | Nudt22 | protein_coding | 0.318304984 | 4.836511979 | 0.8097066 | 0.381303 | 0.710491262 |
| ENSMUSG00000039725 | Trp53rka | protein_coding | 0.679578913 | 3.216718572 | 0.8093993 | 0.3813912 | 0.710491262 |
| ENSMUSG00000084830 | Gm14539 | processed_pseudoge | 0.408161053 | 3.93243946 | 0.8093868 | 0.3813948 | 0.710491262 |

| ensembl_gene_id | external_gene_name | gene_biotype | log2 (fold change) | log2 (counts per million) | F-value | p-value | False discovery rate |
|---|---|---|---|---|---|---|---|
| ENSMUSG00000049411 | Tmem241 | protein_coding | 0.534078133 | 4.27090545 | 0.8094866 | 0.3814686 | 0.710491262 |
| ENSMUSG00000027739 | Rab33b | protein_coding | -0.286991596 | 5.633893349 | 0.8089858 | 0.38151 | 0.710491262 |
| ENSMUSG00000054823 | Nsd3 | protein_coding | 0.272345971 | 6.613822115 | 0.8084659 | 0.3816594 | 0.710646426 |
| ENSMUSG00000021713 | Ppwd1 | protein_coding | -0.964571091 | 2.292183636 | 0.8074164 | 0.3819612 | 0.710958465 |
| ENSMUSG00000042766 | Trim46 | protein_coding | 0.260244675 | 5.439437472 | 0.8069337 | 0.3821002 | 0.710958465 |
| ENSMUSG00000042520 | Ubap2l | protein_coding | -0.448357696 | 4.45753949 | 0.8068242 | 0.3821317 | 0.710958465 |
| ENSMUSG00000039157 | Fam102a | protein_coding | -0.23221838 | 6.538388626 | 0.8065341 | 0.3822153 | 0.710958465 |
| ENSMUSG00000028954 | Nub1 | protein_coding | 0.256034338 | 6.897385071 | 0.8064186 | 0.3822485 | 0.710958465 |
| ENSMUSG00000008892 | Vdac3 | protein_coding | 0.194763617 | 7.591641338 | 0.8055158 | 0.3825088 | 0.710958465 |
| ENSMUSG00000042541 | Sem1 | protein_coding | 0.276076242 | 9.098111375 | 0.8055256 | 0.3825273 | 0.710958465 |
| ENSMUSG00000068114 | Ccdc134 | protein_coding | -0.585718963 | 3.203838353 | 0.8054051 | 0.3825407 | 0.710958465 |
| ENSMUSG00000028645 | Slc2a1 | protein_coding | 0.53818214 | 3.207892489 | 0.8053518 | 0.3825561 | 0.710958465 |
| ENSMUSG00000006276 | Eps15l1 | protein_coding | 0.294015591 | 5.558917699 | 0.8050895 | 0.3826318 | 0.710958465 |
| ENSMUSG00000024667 | Tmem216 | protein_coding | 0.294354642 | 5.050720056 | 0.8046938 | 0.382746 | 0.710958465 |
| ENSMUSG00000032322 | Pstpip1 | protein_coding | 0.88520568 | 2.260251507 | 0.8046931 | 0.3827462 | 0.710958465 |
| ENSMUSG00000024604 | Rbm22 | protein_coding | -0.229497693 | 5.880171391 | 0.8042079 | 0.3828863 | 0.710958465 |
| ENSMUSG00000030203 | Dusp16 | protein_coding | -0.545685528 | 3.462242681 | 0.804131 | 0.3829085 | 0.710958465 |
| ENSMUSG00000078862 | Gm14326 | protein_coding | -0.393206523 | 4.483002058 | 0.8038194 | 0.3829985 | 0.710958465 |
| ENSMUSG00000045534 | Kcna5 | protein_coding | -0.466691857 | 3.796053901 | 0.8037337 | 0.3830233 | 0.710958465 |
| ENSMUSG00000044562 | Rasip1 | protein_coding | 0.714817797 | 3.036528528 | 0.8036746 | 0.3830404 | 0.710958465 |
| ENSMUSG00000030303 | Far2 | protein_coding | 0.564563277 | 4.74885854 | 0.8080341 | 0.3830813 | 0.710958465 |
| ENSMUSG00000028409 | Smu1 | protein_coding | 0.23113835 | 6.920026931 | 0.8035265 | 0.3830832 | 0.710958465 |
| ENSMUSG00000041907 | Gpr45 | protein_coding | -0.505760877 | 3.999234521 | 0.8026111 | 0.3833479 | 0.71132706 |
| ENSMUSG00000038302 | Afg1l | protein_coding | 0.383447016 | 4.72901734 | 0.8016122 | 0.3836371 | 0.711647189 |
| ENSMUSG00000025722 | Wdr73 | protein_coding | 0.277103128 | 5.634725503 | 0.801558 | 0.3836528 | 0.711647189 |
| ENSMUSG00000031349 | Nsdhl | protein_coding | 0.422164496 | 5.791566562 | 0.8036777 | 0.3838653 | 0.71166935 |
| ENSMUSG00000072640 | Lyrm9 | protein_coding | 0.295593899 | 6.788462245 | 0.8008523 | 0.3838786 | 0.71166935 |
| ENSMUSG00000019295 | Tmem129 | protein_coding | 0.320163852 | 5.344032066 | 0.8007467 | 0.383888 | 0.71166935 |
| ENSMUSG00000046727 | Cystm1 | protein_coding | -0.205261348 | 8.856223718 | 0.8006035 | 0.3839295 | 0.71166935 |
| ENSMUSG00000022199 | Slc22a17 | protein_coding | 0.22952079 | 8.546739703 | 0.7998982 | 0.3841341 | 0.711925955 |
| ENSMUSG00000012609 | Ttll5 | protein_coding | 0.389936315 | 5.153133943 | 0.7989228 | 0.3844174 | 0.712218167 |
| ENSMUSG00000021219 | Rgs6 | protein_coding | 0.445882641 | 4.009210378 | 0.7986873 | 0.3844858 | 0.712218167 |
| ENSMUSG00000029071 | Dvl1 | protein_coding | 0.36404714 | 4.1831954 | 0.7986712 | 0.3844905 | 0.712218167 |
| ENSMUSG00000067924 | Cxx1b | protein_coding | 0.290941964 | 4.928086095 | 0.7979434 | 0.3847021 | 0.712257853 |
| ENSMUSG00000004902 | Slc25a18 | protein_coding | -0.281890354 | 6.351347819 | 0.7978207 | 0.3847378 | 0.712257853 |
| ENSMUSG00000010936 | Vac14 | protein_coding | 0.352010637 | 4.58232797 | 0.7977062 | 0.3847712 | 0.712257853 |
| ENSMUSG00000039917 | Rhbdd2 | protein_coding | 0.592716194 | 6.527210063 | 0.819186 | 0.3847878 | 0.712257853 |
| ENSMUSG00000032481 | Smarcc1 | protein_coding | 0.284780804 | 5.557874425 | 0.7974589 | 0.3848431 | 0.712257853 |
| ENSMUSG00000038256 | Bcl9 | protein_coding | 0.435689616 | 4.243007085 | 0.7967177 | 0.3850589 | 0.712432315 |
| ENSMUSG00000021156 | Zmynd11 | protein_coding | -0.381397878 | 4.511812128 | 0.7966801 | 0.3850699 | 0.712432315 |
| ENSMUSG00000073418 | C4b | protein_coding | 0.99283789 | 2.333341342 | 0.7953647 | 0.3854534 | 0.712823432 |
| ENSMUSG00000061143 | Maml3 | protein_coding | 0.558477259 | 3.033189784 | 0.7947582 | 0.3856304 | 0.712823432 |
| ENSMUSG00000024661 | Fth1 | protein_coding | 1.077832544 | 12.1728887 | 0.8432437 | 0.3857014 | 0.712823432 |
| ENSMUSG00000003721 | Insig2 | protein_coding | 0.204787361 | 6.989022052 | 0.7940996 | 0.3858227 | 0.712823432 |
| ENSMUSG00000001467 | Cyp51 | protein_coding | 0.275603415 | 5.708776918 | 0.7940728 | 0.3858305 | 0.712823432 |
| ENSMUSG00000036959 | Bcorl1 | protein_coding | -0.445025711 | 3.776822993 | 0.7940542 | 0.385836 | 0.712823432 |
| ENSMUSG00000008333 | Snrpb2 | protein_coding | -0.267468074 | 6.805207652 | 0.7938966 | 0.385882 | 0.712823432 |
| ENSMUSG00000027981 | Rnpc3 | protein_coding | -0.551493689 | 2.974047067 | 0.7934036 | 0.3860261 | 0.712823432 |
| ENSMUSG00000005580 | Adcy9 | protein_coding | 0.535726819 | 4.012218975 | 0.7934577 | 0.3860315 | 0.712823432 |
| ENSMUSG00000043702 | Pde12 | protein_coding | 0.809452977 | 2.91466877 | 0.793439 | 0.386037 | 0.712823432 |
| ENSMUSG00000020790 | Ankfy1 | protein_coding | -0.401792772 | 4.178651392 | 0.7931333 | 0.3861052 | 0.712823432 |
| ENSMUSG00000055553 | Kxd1 | protein_coding | 0.207766033 | 7.517410548 | 0.7928928 | 0.3861755 | 0.712823432 |
| ENSMUSG00000029708 | Gcc1 | protein_coding | -0.403646379 | 3.788800149 | 0.7926216 | 0.3862549 | 0.712823432 |
| ENSMUSG00000038866 | Zcchc2 | protein_coding | 0.551261077 | 2.898668744 | 0.792612 | 0.3862577 | 0.712823432 |
| ENSMUSG00000035399 | Oser1 | protein_coding | -0.370836114 | 3.785850373 | 0.7925506 | 0.3862757 | 0.712823432 |
| ENSMUSG00000074218 | Cox7a1 | protein_coding | 0.394694923 | 3.850723237 | 0.7919483 | 0.386452 | 0.71282418 |
| ENSMUSG00000023961 | Enpp4 | protein_coding | -0.328633338 | 5.192161877 | 0.7917314 | 0.3865156 | 0.71282418 |
| ENSMUSG00000031922 | Cep57 | protein_coding | -0.277595934 | 5.132438851 | 0.7910676 | 0.3867101 | 0.71282418 |
| ENSMUSG00000005312 | Ubqln1 | protein_coding | 0.197822539 | 7.206045527 | 0.7909435 | 0.3867465 | 0.71282418 |
| ENSMUSG00000036206 | Sh3bp4 | protein_coding | -0.521597368 | 3.751942726 | 0.7909124 | 0.3867556 | 0.71282418 |
| ENSMUSG00000031862 | Atp13a1 | protein_coding | 0.562090053 | 3.360370246 | 0.7907629 | 0.3867994 | 0.71282418 |
| ENSMUSG00000029608 | Rph3a | protein_coding | -0.326122163 | 7.193579012 | 0.7931703 | 0.3869144 | 0.71282418 |
| ENSMUSG00000022771 | Ppil2 | protein_coding | 0.26335459 | 6.472391962 | 0.7902888 | 0.3869385 | 0.71282418 |
| ENSMUSG00000032966 | Fkbp1a | protein_coding | -0.213489938 | 9.154511293 | 0.7902565 | 0.386948 | 0.71282418 |
| ENSMUSG00000040183 | Ankrd6 | protein_coding | 0.317822006 | 5.466469459 | 0.7895254 | 0.3871626 | 0.71282418 |
| ENSMUSG00000048285 | Frmd6 | protein_coding | 0.759977042 | 2.624950927 | 0.7893767 | 0.3872063 | 0.71282418 |
| ENSMUSG00000018008 | Cyth4 | protein_coding | 0.42959491 | 3.659424417 | 0.7890404 | 0.3873051 | 0.71282418 |
| ENSMUSG00000040481 | Bptf | protein_coding | -0.267603703 | 6.101804355 | 0.788904 | 0.3873452 | 0.71282418 |
| ENSMUSG00000026623 | Lpgat1 | protein_coding | -0.218054612 | 6.42459055 | 0.7888885 | 0.3873497 | 0.71282418 |
| ENSMUSG00000022710 | Usp7 | protein_coding | -0.255787482 | 5.755904599 | 0.7888477 | 0.3873617 | 0.71282418 |
| ENSMUSG00000021470 | Ercc6l2 | protein_coding | -0.482958994 | 3.592632967 | 0.78863 | 0.3874257 | 0.71282418 |
| ENSMUSG00000020701 | Tmem132e | protein_coding | 0.531762232 | 3.462792717 | 0.7885314 | 0.3874547 | 0.71282418 |
| ENSMUSG00000034793 | G6pc3 | protein_coding | 0.300134789 | 6.389852792 | 0.7884004 | 0.3875143 | 0.71282418 |
| ENSMUSG00000042529 | Kcnj12 | protein_coding | -0.553709146 | 3.044335872 | 0.7878027 | 0.387669 | 0.71282418 |
| ENSMUSG00000041669 | Prima1 | protein_coding | -0.800375437 | 2.634618313 | 0.7877386 | 0.3876879 | 0.71282418 |
| ENSMUSG00000031969 | Acad8 | protein_coding | 0.307020416 | 5.460269292 | 0.7876435 | 0.3877159 | 0.71282418 |
| ENSMUSG00000014232 | Cluap1 | protein_coding | -0.297116899 | 4.900014006 | 0.7875804 | 0.3877345 | 0.71282418 |
| ENSMUSG00000032131 | Abcg4 | protein_coding | -0.308383333 | 5.149193701 | 0.7865458 | 0.3880391 | 0.713262383 |

| ensembl_gene_id | external_gene_name | gene_biotype | log2 (fold change) | log2 (counts per million) | F-value | p-value | False discovery rate |
|---|---|---|---|---|---|---|---|
| ENSMUSG00000038991 | Txndc5 | protein_coding | 0.371131928 | 5.050696154 | 0.7849349 | 0.3885143 | 0.714013738 |
| ENSMUSG00000033342 | Plppr5 | protein_coding | -0.335053381 | 4.639042414 | 0.784494 | 0.3886445 | 0.714097579 |
| ENSMUSG00000005846 | Rsl1d1 | protein_coding | 0.233722895 | 7.142042132 | 0.7843307 | 0.3886927 | 0.714097579 |
| ENSMUSG00000039318 | Rab3gap2 | protein_coding | 0.444853181 | 4.163688123 | 0.783867 | 0.3888298 | 0.714227276 |
| ENSMUSG00000017418 | Arl5b | protein_coding | -0.564705037 | 3.134239237 | 0.7831811 | 0.3890326 | 0.714316885 |
| ENSMUSG00000020547 | Bzw2 | protein_coding | 0.237021774 | 6.436693677 | 0.7831352 | 0.3890462 | 0.714316885 |
| ENSMUSG00000026509 | Capn2 | protein_coding | -0.3526907 | 4.854152845 | 0.7828603 | 0.3891275 | 0.714316885 |
| ENSMUSG00000047613 | A430005L14Rik | protein_coding | 0.189934613 | 7.307185625 | 0.7825945 | 0.3892062 | 0.714316885 |
| ENSMUSG00000037361 | Sf3b6 | protein_coding | -0.232938033 | 8.375932533 | 0.7825793 | 0.3892107 | 0.714316885 |
| ENSMUSG00000035949 | Fbxw2 | protein_coding | -0.399914626 | 5.108968232 | 0.7823215 | 0.389308 | 0.714339542 |
| ENSMUSG00000026639 | Lamb3 | protein_coding | -0.670078176 | 3.199071581 | 0.7820889 | 0.3893559 | 0.714339542 |
| ENSMUSG00000091896 | Ube2d2a | protein_coding | -0.265313561 | 5.745542378 | 0.7815668 | 0.3895106 | 0.714446916 |
| ENSMUSG00000020847 | Rph3al | protein_coding | 0.54119448 | 3.492750588 | 0.7808344 | 0.3897277 | 0.714446916 |
| ENSMUSG00000037214 | Thap1 | protein_coding | -0.379655329 | 3.795572481 | 0.7806315 | 0.3897879 | 0.714446916 |
| ENSMUSG00000029227 | Fip1l1 | protein_coding | -0.464694223 | 4.152033898 | 0.7805102 | 0.3898239 | 0.714446916 |
| ENSMUSG00000020235 | Fzr1 | protein_coding | 0.242149747 | 6.145964474 | 0.780285 | 0.3898908 | 0.714446916 |
| ENSMUSG00000037706 | Cd81 | protein_coding | 0.219144614 | 9.865788007 | 0.7802754 | 0.3898936 | 0.714446916 |
| ENSMUSG00000027086 | Fastkd1 | protein_coding | 0.365215768 | 4.208332391 | 0.7800884 | 0.3899491 | 0.714446916 |
| ENSMUSG00000020982 | Nemf | protein_coding | -0.2275791 | 6.868900378 | 0.7800287 | 0.3899668 | 0.714446916 |
| ENSMUSG00000030471 | Zdhhc13 | protein_coding | -0.473606795 | 3.432563223 | 0.779869 | 0.3900143 | 0.714446916 |
| ENSMUSG00000018476 | Kdm6b | protein_coding | 0.526830872 | 5.4998987 | 0.7857144 | 0.3900788 | 0.714446916 |
| ENSMUSG00000049739 | Zfp646 | protein_coding | -0.505218502 | 3.184393989 | 0.7789076 | 0.3902999 | 0.714730041 |
| ENSMUSG00000018861 | Fdxr | protein_coding | 0.430727413 | 3.581501674 | 0.778216 | 0.3905056 | 0.714984916 |
| ENSMUSG00000027002 | Nckap1 | protein_coding | -0.224738004 | 7.916296501 | 0.777516 | 0.3907139 | 0.71524462 |
| ENSMUSG00000035640 | Cbarp | protein_coding | 0.252714258 | 6.86161817 | 0.7766236 | 0.3909798 | 0.7154827 |
| ENSMUSG00000068876 | Cgn | protein_coding | 0.510655818 | 3.525450988 | 0.7764246 | 0.3910391 | 0.7154827 |
| ENSMUSG00000069227 | Gprin1 | protein_coding | -0.402655297 | 3.98846622 | 0.7762877 | 0.3910799 | 0.7154827 |
| ENSMUSG00000003500 | Impdh1 | protein_coding | 0.484191695 | 4.04516045 | 0.7761093 | 0.3911331 | 0.7154827 |
| ENSMUSG00000018433 | Nol11 | protein_coding | 0.327642609 | 5.266250352 | 0.7759632 | 0.3911767 | 0.7154827 |
| ENSMUSG00000037993 | Dhx38 | protein_coding | 0.446657371 | 4.389647886 | 0.7752486 | 0.3913899 | 0.715552902 |
| ENSMUSG00000035027 | Map2k2 | protein_coding | 0.263128543 | 5.336854328 | 0.775202 | 0.3914038 | 0.715552902 |
| ENSMUSG00000016559 | H3f3b | protein_coding | 0.211056099 | 9.584166665 | 0.7751212 | 0.391428 | 0.715552902 |
| ENSMUSG00000023945 | Slc5a7 | protein_coding | -0.501605685 | 3.343747888 | 0.7749428 | 0.3914812 | 0.715552902 |
| ENSMUSG00000049106 | Dcaf5 | protein_coding | 0.256571261 | 5.500459631 | 0.774504 | 0.3916123 | 0.715634515 |
| ENSMUSG00000057788 | Ddx49 | protein_coding | 0.379230985 | 5.217780257 | 0.774147 | 0.391719 | 0.715634515 |
| ENSMUSG00000021013 | Ttc8 | protein_coding | -0.28519732 | 5.003376252 | 0.7739769 | 0.3917698 | 0.715634515 |
| ENSMUSG00000025794 | Rpl14 | protein_coding | 0.222466061 | 9.060645753 | 0.7739025 | 0.3917921 | 0.715634515 |
| ENSMUSG00000022180 | Slc7a8 | protein_coding | -0.313786589 | 4.673576006 | 0.7735585 | 0.391895 | 0.715700824 |
| ENSMUSG00000039194 | Rlbp1 | protein_coding | 0.523531135 | 4.133638272 | 0.7722312 | 0.3923132 | 0.716342895 |
| ENSMUSG00000055676 | Gm5069 | transcribed_processe | 0.611146827 | 2.771218341 | 0.7714569 | 0.3925243 | 0.716546869 |
| ENSMUSG00000100599 | 1700120C14Rik | processed_transcript | -0.706575095 | 2.57138685 | 0.7712578 | 0.392584 | 0.716546869 |
| ENSMUSG00000039057 | Myo16 | protein_coding | -0.700577634 | 2.929518244 | 0.7706609 | 0.3927631 | 0.716546869 |
| ENSMUSG00000022551 | Cyc1 | protein_coding | 0.221014013 | 6.715909011 | 0.7706262 | 0.3927735 | 0.716546869 |
| ENSMUSG00000042570 | Mier2 | protein_coding | 0.572889012 | 3.993611786 | 0.7710328 | 0.3928583 | 0.716546869 |
| ENSMUSG00000042734 | Ttc9 | protein_coding | -0.319321684 | 5.301513792 | 0.7700383 | 0.39295 | 0.716546869 |
| ENSMUSG00000108030 | 9530062K07Rik | bidirectional_promote | -0.842411944 | 3.071294766 | 0.7721855 | 0.3929912 | 0.716546869 |
| ENSMUSG00000022913 | Psmg1 | protein_coding | 0.318484641 | 4.951692239 | 0.7698197 | 0.3930157 | 0.716546869 |
| ENSMUSG00000003865 | Gys1 | protein_coding | -0.425521677 | 4.209137295 | 0.7694206 | 0.3931356 | 0.716546869 |
| ENSMUSG00000079641 | Rpl39 | protein_coding | 0.321205994 | 6.23679866 | 0.7694109 | 0.3931593 | 0.716546869 |
| ENSMUSG00000002017 | Fam98a | protein_coding | -0.343581929 | 4.176499394 | 0.769012 | 0.3932584 | 0.716546869 |
| ENSMUSG00000022881 | Rfc4 | protein_coding | -0.597336372 | 4.263347673 | 0.7724205 | 0.3932673 | 0.716546869 |
| ENSMUSG00000034889 | Cactin | protein_coding | 0.795159756 | 2.749299879 | 0.7689033 | 0.3932911 | 0.716546869 |
| ENSMUSG00000026494 | Kif26b | protein_coding | -0.503165006 | 3.373505931 | 0.7682539 | 0.3934865 | 0.716781311 |
| ENSMUSG00000026615 | Eprs | protein_coding | -0.263730286 | 6.566048291 | 0.7678118 | 0.3936196 | 0.716900182 |
| ENSMUSG00000022817 | Itgb5 | protein_coding | -0.62441374 | 2.879385228 | 0.7675943 | 0.3936851 | 0.716900182 |
| ENSMUSG00000004040 | Stat3 | protein_coding | -0.456235728 | 4.838964581 | 0.7669693 | 0.3939489 | 0.717156547 |
| ENSMUSG00000019952 | Poc1b | protein_coding | -0.781613489 | 2.66438283 | 0.7665058 | 0.3940131 | 0.717156547 |
| ENSMUSG00000025332 | Kdm5c | protein_coding | -0.373209427 | 4.416504247 | 0.7662907 | 0.394078 | 0.717156547 |
| ENSMUSG00000043415 | Otud1 | protein_coding | 0.582729304 | 3.314810434 | 0.7660239 | 0.3941585 | 0.717156547 |
| ENSMUSG00000028945 | Rheb | protein_coding | -0.210925397 | 6.803813429 | 0.7657279 | 0.3942478 | 0.717156547 |
| ENSMUSG00000032174 | Icam5 | protein_coding | -0.917487328 | 2.171267815 | 0.7657109 | 0.3942529 | 0.717156547 |
| ENSMUSG00000011154 | Cfap161 | protein_coding | -0.372226597 | 4.412085731 | 0.7652299 | 0.3943982 | 0.717156547 |
| ENSMUSG00000025868 | Higd2a | protein_coding | 0.190367469 | 8.450003104 | 0.7648101 | 0.394525 | 0.717156547 |
| ENSMUSG00000043241 | Upf2 | protein_coding | -0.367834383 | 5.165862436 | 0.7647379 | 0.3945468 | 0.717156547 |
| ENSMUSG00000029130 | Rnf32 | protein_coding | -0.330797525 | 4.393035802 | 0.7647298 | 0.3945492 | 0.717156547 |
| ENSMUSG00000030435 | U2af2 | protein_coding | 0.452649731 | 3.440949511 | 0.7646959 | 0.3945595 | 0.717156547 |
| ENSMUSG00000009013 | Dynll1 | protein_coding | 0.270722587 | 10.1079042 | 0.7644538 | 0.3946327 | 0.717168311 |
| ENSMUSG00000037152 | Ndufc1 | protein_coding | 0.248797629 | 9.412866579 | 0.7633229 | 0.3949747 | 0.717668682 |
| ENSMUSG00000020723 | Cacng4 | protein_coding | -0.324068482 | 5.752116786 | 0.7629152 | 0.3950982 | 0.717771673 |
| ENSMUSG00000006235 | Epor | protein_coding | -0.554673369 | 2.937804514 | 0.7621417 | 0.3953325 | 0.717981992 |
| ENSMUSG00000030232 | Aebp2 | protein_coding | -0.30130135 | 5.211686 | 0.7620923 | 0.3953475 | 0.717981992 |
| ENSMUSG00000024104 | Washc2 | protein_coding | -0.361575496 | 5.126617546 | 0.7613691 | 0.3955668 | 0.718150593 |
| ENSMUSG00000098087 | Gm17750 | lincRNA | -0.483210196 | 3.3932452 | 0.7613456 | 0.3955739 | 0.718150593 |
| ENSMUSG00000029505 | Ep400 | protein_coding | -0.410502396 | 4.822313806 | 0.7599893 | 0.3960064 | 0.718798785 |
| ENSMUSG00000085156 | Snhg15 | processed_transcript | -0.512877422 | 3.023525146 | 0.7597296 | 0.3960646 | 0.718798785 |
| ENSMUSG00000020486 | 4-Sep | protein_coding | -0.222570261 | 8.938024311 | 0.7587071 | 0.3963756 | 0.719186725 |
| ENSMUSG00000004934 | Pias4 | protein_coding | 1.072799872 | 2.134810029 | 0.7585869 | 0.3964121 | 0.719186725 |
| ENSMUSG00000028773 | Fabp3 | protein_coding | 0.217884687 | 6.611636062 | 0.7581427 | 0.3965474 | 0.719272455 |

| ensembl_gene_id | external_gene_name | gene_biotype | log2 (fold change) | log2 (counts per million) | F-value | p-value | False discovery rate |
|---|---|---|---|---|---|---|---|
| ENSMUSG00000040859 | Bsdc1 | protein_coding | 0.234929684 | 5.986133482 | 0.7578728 | 0.3966296 | 0.719272455 |
| ENSMUSG00000018559 | Ctdnep1 | protein_coding | 0.297104366 | 5.025039211 | 0.7577727 | 0.3966601 | 0.719272455 |
| ENSMUSG00000034192 | Lsm3 | protein_coding | -0.332042325 | 6.577785955 | 0.7593086 | 0.396824 | 0.719357872 |
| ENSMUSG00000063052 | Lrrc40 | protein_coding | -0.327304923 | 5.179749987 | 0.757108 | 0.3968626 | 0.719357872 |
| ENSMUSG00000021514 | Zfp369 | protein_coding | -0.523029305 | 3.534012811 | 0.7567886 | 0.39696 | 0.719357872 |
| ENSMUSG00000030486 | Zfp108 | protein_coding | -0.721607773 | 2.501826416 | 0.7566957 | 0.3969884 | 0.719357872 |
| ENSMUSG00000001123 | Lgals9 | protein_coding | 0.517977015 | 3.411263067 | 0.756521 | 0.3970417 | 0.719357872 |
| ENSMUSG00000042787 | Exog | protein_coding | 0.382696598 | 5.026639843 | 0.7562355 | 0.3971288 | 0.719372768 |
| ENSMUSG00000030741 | Spns1 | protein_coding | 0.307104386 | 5.835921161 | 0.7560556 | 0.3971837 | 0.719372768 |
| ENSMUSG00000060143 | Gm10076 | protein_coding | 0.527131194 | 3.08257954 | 0.7557119 | 0.3972886 | 0.719441665 |
| ENSMUSG00000034059 | Ypel4 | protein_coding | 0.229344403 | 6.631333858 | 0.7554501 | 0.3973686 | 0.719465283 |
| ENSMUSG00000025192 | Entpd7 | protein_coding | 0.505739356 | 3.783401664 | 0.755099 | 0.3974759 | 0.719538385 |
| ENSMUSG00000063535 | Zfp773 | protein_coding | -0.951472304 | 2.375556669 | 0.7548717 | 0.3975453 | 0.719542971 |
| ENSMUSG00000020271 | Fbxw11 | protein_coding | 0.256298616 | 6.205915144 | 0.7538653 | 0.3978531 | 0.719594691 |
| ENSMUSG00000030630 | Fah | protein_coding | 0.285660629 | 5.376407871 | 0.7538081 | 0.3978707 | 0.719594691 |
| ENSMUSG00000038611 | Phrf1 | protein_coding | 0.362517734 | 4.427450614 | 0.7536546 | 0.3979176 | 0.719594691 |
| ENSMUSG00000025538 | Sumf2 | protein_coding | 0.407504536 | 5.000445832 | 0.7536839 | 0.3979293 | 0.719594691 |
| ENSMUSG00000028328 | Tmod1 | protein_coding | -0.342476412 | 4.726566109 | 0.7535939 | 0.3979362 | 0.719594691 |
| ENSMUSG00000035828 | Pim3 | protein_coding | -0.329662607 | 5.045713111 | 0.7532459 | 0.3980428 | 0.719594691 |
| ENSMUSG00000040414 | Slc25a28 | protein_coding | -0.425693564 | 4.194077487 | 0.753197 | 0.3980577 | 0.719594691 |
| ENSMUSG00000022890 | Atp5j | protein_coding | 0.231546967 | 10.19202003 | 0.7528778 | 0.3981555 | 0.719594691 |
| ENSMUSG00000027360 | Hdc | protein_coding | 0.522481664 | 3.202417784 | 0.7528104 | 0.3981762 | 0.719594691 |
| ENSMUSG00000016526 | Dyrk3 | protein_coding | 0.859595626 | 2.25077061 | 0.752498 | 0.3982719 | 0.719646769 |
| ENSMUSG00000047989 | Ino80c | protein_coding | -0.263214031 | 6.052578601 | 0.751733 | 0.3985065 | 0.719840063 |
| ENSMUSG00000025893 | Kbtbd3 | protein_coding | -0.302862663 | 5.278127537 | 0.7517125 | 0.3985128 | 0.719840063 |
| ENSMUSG00000049556 | Lingo1 | protein_coding | 0.306972636 | 5.534299387 | 0.7513526 | 0.3986232 | 0.71988559 |
| ENSMUSG00000063296 | Tmem117 | protein_coding | -0.627188636 | 3.037931534 | 0.750807 | 0.3987907 | 0.71988559 |
| ENSMUSG00000079056 | Kcnip3 | protein_coding | -0.210919943 | 6.305152198 | 0.7507625 | 0.3988044 | 0.71988559 |
| ENSMUSG00000034226 | Rhov | protein_coding | 0.73542145 | 2.758434946 | 0.7507581 | 0.3988058 | 0.71988559 |
| ENSMUSG00000024079 | Eif2ak2 | protein_coding | -1.357509333 | 1.681537871 | 0.7503973 | 0.3989372 | 0.719974507 |
| ENSMUSG00000027243 | Harbi1 | protein_coding | 0.46751144 | 4.173488341 | 0.7501619 | 0.3989889 | 0.719974507 |
| ENSMUSG00000021501 | Caml | protein_coding | 0.244857668 | 6.034386963 | 0.7496771 | 0.399138 | 0.720065384 |
| ENSMUSG00000015766 | Eps8 | protein_coding | 0.413631086 | 4.559607813 | 0.7493954 | 0.3992246 | 0.720065384 |
| ENSMUSG00000075020 | Mir670hg | lincRNA | -0.492701517 | 4.707947878 | 0.7510504 | 0.3992402 | 0.720065384 |
| ENSMUSG00000025236 | Adpgk | protein_coding | 0.638080501 | 3.119365889 | 0.7490609 | 0.3993276 | 0.720102197 |
| ENSMUSG00000062761 | Zfp512 | protein_coding | 0.250034579 | 5.651797819 | 0.7482496 | 0.3995774 | 0.720431875 |
| ENSMUSG00000066442 | Mthfs | protein_coding | 0.567355596 | 2.857450493 | 0.7480146 | 0.3996498 | 0.720441621 |
| ENSMUSG00000002428 | Hltf | protein_coding | -0.335647343 | 4.522616814 | 0.746737 | 0.4000438 | 0.72103094 |
| ENSMUSG00000000439 | Mkrn2 | protein_coding | 0.272615738 | 5.914693123 | 0.7464294 | 0.4001387 | 0.721081232 |
| ENSMUSG00000048330 | Ric3 | protein_coding | 0.332531749 | 5.306662517 | 0.7455252 | 0.400418 | 0.721463573 |
| ENSMUSG00000097145 | 9230114K14Rik | processed_transcript | -0.691290258 | 2.639944073 | 0.7444546 | 0.4007491 | 0.721659603 |
| ENSMUSG00000025613 | Cct8 | protein_coding | 0.266865624 | 8.639490942 | 0.7443463 | 0.400803 | 0.721659603 |
| ENSMUSG00000008604 | Ubqln4 | protein_coding | -0.259151167 | 5.605717211 | 0.7439157 | 0.4009159 | 0.721659603 |
| ENSMUSG00000041765 | Ubac2 | protein_coding | 0.282289541 | 5.337075299 | 0.7436898 | 0.4009858 | 0.721659603 |
| ENSMUSG00000032181 | Scg3 | protein_coding | -0.252393236 | 7.058210045 | 0.7434149 | 0.401071 | 0.721659603 |
| ENSMUSG00000021706 | Zfyve16 | protein_coding | -0.491506857 | 3.227398287 | 0.7433506 | 0.4010909 | 0.721659603 |
| ENSMUSG00000027678 | Ncoa3 | protein_coding | 0.394480322 | 4.86053662 | 0.7433237 | 0.4010992 | 0.721659603 |
| ENSMUSG00000003402 | Prkcsh | protein_coding | 0.394076212 | 5.289178934 | 0.7433154 | 0.4011222 | 0.721659603 |
| ENSMUSG00000039159 | Ube2h | protein_coding | -0.268612807 | 6.566999314 | 0.7432219 | 0.4011308 | 0.721659603 |
| ENSMUSG00000040451 | Sgms1 | protein_coding | -0.384634755 | 4.122359206 | 0.7429767 | 0.4012068 | 0.721675572 |
| ENSMUSG00000029432 | Nipsnap2 | protein_coding | 0.279100979 | 8.179459623 | 0.7424248 | 0.4013983 | 0.721727898 |
| ENSMUSG00000043716 | Rpl7 | protein_coding | 0.247529824 | 8.566012322 | 0.742247 | 0.4014331 | 0.721727898 |
| ENSMUSG00000045636 | Mtus1 | protein_coding | -0.32929017 | 5.259558151 | 0.7421783 | 0.4014544 | 0.721727898 |
| ENSMUSG00000034484 | Snx2 | protein_coding | 0.289591552 | 7.439300765 | 0.7423702 | 0.4015044 | 0.721727898 |
| ENSMUSG00000070044 | Fam149a | protein_coding | 0.358354466 | 4.159628302 | 0.7416526 | 0.4016176 | 0.721810729 |
| ENSMUSG00000036529 | Sbf1 | protein_coding | 0.294196403 | 5.353329883 | 0.7412809 | 0.401733 | 0.721897497 |
| ENSMUSG00000017493 | Igfbp4 | protein_coding | -0.523751032 | 4.478549762 | 0.7428154 | 0.4019645 | 0.722009996 |
| ENSMUSG00000030079 | Ruvbl1 | protein_coding | 0.304525538 | 5.052224155 | 0.7404907 | 0.4019785 | 0.722009996 |
| ENSMUSG00000029125 | Stx18 | protein_coding | 0.278917288 | 5.479894927 | 0.7404312 | 0.401997 | 0.722009996 |
| ENSMUSG00000025402 | Nab2 | protein_coding | 0.49555694 | 3.529868613 | 0.7392917 | 0.4023515 | 0.722526074 |
| ENSMUSG00000024067 | Dpy30 | protein_coding | 0.219519598 | 8.230608169 | 0.7387138 | 0.4025315 | 0.72272853 |
| ENSMUSG00000040712 | Camta2 | protein_coding | 0.37108229 | 6.010244327 | 0.7408835 | 0.4025988 | 0.72272864 |
| ENSMUSG00000005054 | Cstb | protein_coding | 0.22434438 | 8.031710396 | 0.7382649 | 0.4026714 | 0.722738362 |
| ENSMUSG00000022110 | Sucla2 | protein_coding | 0.224754873 | 8.265718248 | 0.7377524 | 0.4028312 | 0.722904512 |
| ENSMUSG00000031729 | Ist1 | protein_coding | 0.349888416 | 5.561109839 | 0.7371559 | 0.4030377 | 0.723154338 |
| ENSMUSG00000029135 | Fosl2 | protein_coding | -0.581059186 | 3.461338492 | 0.736649 | 0.4031756 | 0.723281075 |
| ENSMUSG00000039089 | L3mbtl3 | protein_coding | 0.335809767 | 4.024540819 | 0.7361281 | 0.4033383 | 0.723452343 |
| ENSMUSG00000024084 | Qpct | protein_coding | -0.224003431 | 6.503661628 | 0.7356397 | 0.403491 | 0.723520462 |
| ENSMUSG00000050605 | Zfp61 | protein_coding | -0.396560655 | 4.759084114 | 0.735495 | 0.4035362 | 0.723520462 |
| ENSMUSG00000021902 | Phf7 | protein_coding | 1.041567107 | 2.481842382 | 0.7355271 | 0.4035781 | 0.723520462 |
| ENSMUSG00000019878 | Hsf2 | protein_coding | 0.418648958 | 5.061571402 | 0.7349763 | 0.4037188 | 0.723615601 |
| ENSMUSG00000050310 | Rictor | protein_coding | -0.331704733 | 4.044007343 | 0.7347612 | 0.4037658 | 0.723615601 |
| ENSMUSG00000043183 | Simc1 | protein_coding | 0.479092641 | 3.341186191 | 0.7340856 | 0.4039774 | 0.723874099 |
| ENSMUSG00000064246 | Chil1 | protein_coding | -0.440085876 | 4.459438099 | 0.7336362 | 0.4041385 | 0.724042149 |
| ENSMUSG00000026241 | Nppc | protein_coding | 0.525583968 | 3.262840325 | 0.7329812 | 0.4043235 | 0.724253031 |
| ENSMUSG00000025035 | Arl3 | protein_coding | 0.206235018 | 9.134159162 | 0.7324847 | 0.4044793 | 0.724411395 |
| ENSMUSG00000030094 | Xpc | protein_coding | -0.445262251 | 3.523634083 | 0.7321316 | 0.4045902 | 0.724451369 |
| ENSMUSG00000040225 | Prrc2c | protein_coding | -0.246384337 | 6.235597442 | 0.7317855 | 0.4046989 | 0.724451369 |

| ensembl_gene_id | external_gene_name | gene_biotype | log2 (fold change) | log2 (counts per million) | F-value | p-value | False discovery rate |
|---|---|---|---|---|---|---|---|
| ENSMUSG00000051343 | Rab11fip5 | protein_coding | 0.297516475 | 5.378459488 | 0.7316182 | 0.4047514 | 0.724451369 |
| ENSMUSG00000092536 | Gm20501 | bidirectional_promote | 0.421063216 | 3.668349907 | 0.7315555 | 0.4047711 | 0.724451369 |
| ENSMUSG00000074994 | Qser1 | protein_coding | 0.768060952 | 2.57466188 | 0.7311691 | 0.4048925 | 0.724545634 |
| ENSMUSG00000027035 | Cers6 | protein_coding | 0.356744366 | 4.366479279 | 0.7308629 | 0.4049888 | 0.724545634 |
| ENSMUSG00000038332 | Sesn1 | protein_coding | -0.283671301 | 5.305426125 | 0.7306807 | 0.4050461 | 0.724545634 |
| ENSMUSG00000033983 | Coil | protein_coding | 0.337673002 | 4.528481232 | 0.7304408 | 0.4051216 | 0.724545634 |
| ENSMUSG00000003923 | Tfam | protein_coding | 0.265602568 | 5.862559057 | 0.7303165 | 0.4051607 | 0.724545634 |
| ENSMUSG00000029761 | Cald1 | protein_coding | -0.521231199 | 4.789121677 | 0.7324313 | 0.4055066 | 0.724971226 |
| ENSMUSG00000024181 | Mrpl28 | protein_coding | 0.193728115 | 8.842048939 | 0.7288822 | 0.4056124 | 0.724971226 |
| ENSMUSG00000063856 | Gpx1 | protein_coding | 0.19198569 | 9.114042541 | 0.7282863 | 0.4058003 | 0.724971226 |
| ENSMUSG00000073888 | Ccl27a | protein_coding | 0.247426827 | 6.095853882 | 0.728286 | 0.4058004 | 0.724971226 |
| ENSMUSG00000020381 | Mrnip | protein_coding | 0.297650185 | 5.218172705 | 0.7282473 | 0.4058126 | 0.724971226 |
| ENSMUSG00000038861 | Pi4kb | protein_coding | 0.455495598 | 3.527739879 | 0.7281145 | 0.4058545 | 0.724971226 |
| ENSMUSG00000033216 | Eefsec | protein_coding | 0.396708283 | 4.607461669 | 0.7279567 | 0.4059043 | 0.724971226 |
| ENSMUSG00000050271 | Prag1 | protein_coding | -0.467374806 | 3.463287593 | 0.7278498 | 0.405938 | 0.724971226 |
| ENSMUSG00000018567 | Gabarap | protein_coding | 0.205731662 | 9.783309314 | 0.727435 | 0.406069 | 0.725029837 |
| ENSMUSG00000023912 | Slc25a27 | protein_coding | 0.282009818 | 4.990672139 | 0.7273186 | 0.4061057 | 0.725029837 |
| ENSMUSG00000031916 | Cog8 | protein_coding | -0.782715202 | 2.432140051 | 0.727052 | 0.4061899 | 0.72505982 |
| ENSMUSG00000038764 | Ptpn3 | protein_coding | -0.5927876 | 4.327253601 | 0.7295793 | 0.406621 | 0.72570885 |
| ENSMUSG00000022016 | Akap11 | protein_coding | -0.334424921 | 6.148210347 | 0.7251101 | 0.4069006 | 0.726053557 |
| ENSMUSG00000050855 | Zfp940 | protein_coding | 0.363922406 | 4.580419847 | 0.7246519 | 0.4069492 | 0.726053557 |
| ENSMUSG00000028402 | Mpdz | protein_coding | -0.429636403 | 3.322529361 | 0.7240468 | 0.407141 | 0.726275198 |
| ENSMUSG00000038305 | Spats2l | protein_coding | -0.388940799 | 4.865048663 | 0.7234657 | 0.4073253 | 0.726479123 |
| ENSMUSG00000074749 | Kiz | protein_coding | -0.406976965 | 4.178520559 | 0.7232604 | 0.4073904 | 0.726479123 |
| ENSMUSG00000051978 | Erich1 | protein_coding | 0.378901615 | 4.202474229 | 0.7230221 | 0.407466 | 0.726493508 |
| ENSMUSG00000000168 | Dlat | protein_coding | -0.2304712 | 6.283778534 | 0.7223597 | 0.4076764 | 0.726716858 |
| ENSMUSG00000015944 | Gatsl2 | protein_coding | -0.218420534 | 6.851639751 | 0.722202 | 0.4077265 | 0.726716858 |
| ENSMUSG00000022100 | Xpo7 | protein_coding | -0.27433302 | 5.314992387 | 0.7215354 | 0.4079384 | 0.726974039 |
| ENSMUSG00000033676 | Gabrb3 | protein_coding | -0.216453084 | 6.4470507 | 0.7210218 | 0.4081017 | 0.72714463 |
| ENSMUSG00000048108 | Tmem72 | protein_coding | 1.432028883 | 2.046472739 | 0.7204058 | 0.4082978 | 0.727373464 |
| ENSMUSG00000044881 | Coa4 | protein_coding | 0.326590114 | 4.985126148 | 0.7198762 | 0.4084665 | 0.727383951 |
| ENSMUSG00000031917 | Nip7 | protein_coding | 0.240855422 | 6.922530447 | 0.7198712 | 0.4084681 | 0.727383951 |
| ENSMUSG00000016024 | Lbp | protein_coding | -0.857565006 | 2.970601539 | 0.7209049 | 0.4085186 | 0.727383951 |
| ENSMUSG00000094910 | D430019H16Rik | protein_coding | -0.35872126 | 5.818413957 | 0.7198397 | 0.4085743 | 0.727383951 |
| ENSMUSG00000027746 | Ufm1 | protein_coding | -0.23332278 | 6.165844384 | 0.7189153 | 0.4087728 | 0.727617003 |
| ENSMUSG00000048376 | F2r | protein_coding | -0.671898905 | 2.762143809 | 0.7176811 | 0.4091668 | 0.728178742 |
| ENSMUSG00000027613 | Eif6 | protein_coding | 0.198688628 | 6.846657421 | 0.7175026 | 0.4092239 | 0.728178742 |
| ENSMUSG00000078546 | Zfp995 | protein_coding | -0.573630218 | 2.667089842 | 0.7166104 | 0.4095091 | 0.728565785 |
| ENSMUSG00000032388 | Spg21 | protein_coding | -0.294758699 | 5.239548365 | 0.7163621 | 0.4095886 | 0.728586565 |
| ENSMUSG00000028854 | Slc9a1 | protein_coding | 0.353061627 | 4.565030444 | 0.7153731 | 0.4099052 | 0.728924007 |
| ENSMUSG00000022150 | Dab2 | protein_coding | -0.545857964 | 3.048865381 | 0.7153461 | 0.4099138 | 0.728924007 |
| ENSMUSG00000022351 | Sqle | protein_coding | 0.301724244 | 5.66798087 | 0.7142786 | 0.4102561 | 0.729411969 |
| ENSMUSG00000012076 | Brms1l | protein_coding | -0.250190202 | 5.794557637 | 0.7140302 | 0.4103358 | 0.729433027 |
| ENSMUSG00000023942 | Slc29a1 | protein_coding | 0.353271297 | 4.982933342 | 0.7135814 | 0.4104798 | 0.729568505 |
| ENSMUSG00000060147 | Serpinb6a | protein_coding | -0.268642152 | 6.124552557 | 0.7129105 | 0.4106953 | 0.729743882 |
| ENSMUSG00000035007 | Rundc1 | protein_coding | -0.434890648 | 4.464754984 | 0.7129138 | 0.4107142 | 0.729743882 |
| ENSMUSG00000050428 | Fbxo46 | protein_coding | -0.3049156 | 4.897242929 | 0.7121867 | 0.410928 | 0.730003041 |
| ENSMUSG00000018442 | Derl2 | protein_coding | 0.265464114 | 5.580025005 | 0.7119709 | 0.4109974 | 0.730005756 |
| ENSMUSG00000032940 | Rbm11 | protein_coding | 0.361943214 | 4.260309132 | 0.7116648 | 0.4110958 | 0.730060061 |
| ENSMUSG00000053398 | Phgdh | protein_coding | -0.287101385 | 5.139658463 | 0.7113429 | 0.4111995 | 0.730123478 |
| ENSMUSG00000028519 | Dab1 | protein_coding | 0.216484138 | 5.940746159 | 0.7111235 | 0.4112701 | 0.730128369 |
| ENSMUSG00000040761 | Spen | protein_coding | 0.474448731 | 4.045662314 | 0.7103129 | 0.4115512 | 0.730346648 |
| ENSMUSG00000048960 | Prex2 | protein_coding | -0.84748319 | 3.109215943 | 0.712862 | 0.4115602 | 0.730346648 |
| ENSMUSG00000031027 | Stk33 | protein_coding | 0.530930049 | 3.030573786 | 0.7101095 | 0.4115968 | 0.730346648 |
| ENSMUSG00000027104 | Atf2 | protein_coding | -0.2356581 | 7.127111128 | 0.7091593 | 0.4119033 | 0.730634074 |
| ENSMUSG00000044250 | Pced1b | protein_coding | -0.535781527 | 2.992362881 | 0.7090964 | 0.4119237 | 0.730634074 |
| ENSMUSG00000046897 | Zfp740 | protein_coding | 0.293285941 | 4.845189861 | 0.7082905 | 0.4121839 | 0.730634074 |
| ENSMUSG00000031169 | Porcn | protein_coding | 0.346206265 | 5.106679883 | 0.708266 | 0.4121918 | 0.730634074 |
| ENSMUSG00000020817 | Rabep1 | protein_coding | -0.222280492 | 6.820638269 | 0.7079659 | 0.4122888 | 0.730634074 |
| ENSMUSG00000047022 | Mipol1 | protein_coding | -0.385602503 | 4.213269487 | 0.7077991 | 0.4123427 | 0.730634074 |
| ENSMUSG00000019505 | Ubb | protein_coding | 0.21418073 | 10.89795493 | 0.7075032 | 0.4124384 | 0.730634074 |
| ENSMUSG00000025427 | Rnf165 | protein_coding | -0.395199857 | 4.695807413 | 0.7073556 | 0.4124862 | 0.730634074 |
| ENSMUSG00000035310 | Lin54 | protein_coding | -0.551191289 | 2.820126201 | 0.7070916 | 0.4125716 | 0.730634074 |
| ENSMUSG00000039328 | Rnf122 | protein_coding | -0.369030835 | 4.062670197 | 0.7070784 | 0.4125759 | 0.730634074 |
| ENSMUSG00000025793 | Hgs | protein_coding | 0.258493474 | 5.683935347 | 0.7069282 | 0.4126245 | 0.730634074 |
| ENSMUSG00000098108 | Gm27008 | antisense_RNA | -0.56694385 | 3.144980135 | 0.7068624 | 0.4126458 | 0.730634074 |
| ENSMUSG00000029095 | Ablim2 | protein_coding | 0.225797261 | 6.51450988 | 0.7066999 | 0.4126984 | 0.730634074 |
| ENSMUSG00000068290 | Ddrgk1 | protein_coding | 0.226812971 | 7.561920124 | 0.7060717 | 0.4129018 | 0.730634074 |
| ENSMUSG00000058704 | Memo1 | protein_coding | -0.355215224 | 4.05571093 | 0.7059617 | 0.4129375 | 0.730634074 |
| ENSMUSG00000014668 | Chfr | protein_coding | 0.941954371 | 2.161270994 | 0.7058463 | 0.4129749 | 0.730634074 |
| ENSMUSG00000021271 | Zfp839 | protein_coding | -0.352521773 | 3.894737059 | 0.705829 | 0.4129805 | 0.730634074 |
| ENSMUSG00000042523 | Dnal1 | protein_coding | -0.298069001 | 5.155331092 | 0.7058091 | 0.4129869 | 0.730634074 |
| ENSMUSG00000073096 | Lrrc61 | protein_coding | -0.449994481 | 4.166503939 | 0.7056151 | 0.4130498 | 0.730634074 |
| ENSMUSG00000073557 | Ppp1r12b | protein_coding | -0.690209004 | 2.64936118 | 0.7022769 | 0.4141342 | 0.732162895 |
| ENSMUSG00000043535 | Setx | protein_coding | -0.349250204 | 4.54475546 | 0.7022688 | 0.4141368 | 0.732162895 |
| ENSMUSG00000029599 | Ddx54 | protein_coding | 0.517832763 | 3.394293668 | 0.7019927 | 0.4142267 | 0.732162895 |
| ENSMUSG00000003378 | Grik5 | protein_coding | 0.551966099 | 3.244811165 | 0.7019103 | 0.4142535 | 0.732162895 |
| ENSMUSG00000022947 | Cbr3 | protein_coding | -0.535146819 | 3.36984053 | 0.7019071 | 0.4142545 | 0.732162895 |

| ensembl_gene_id | external_gene_name | gene_biotype | log2 (fold change) | log2 (counts per million) | F-value | p-value | False discovery rate |
|---|---|---|---|---|---|---|---|
| ENSMUSG00000029003 | Mad2l2 | protein_coding | 0.30095448 | 5.318783669 | 0.7015153 | 0.4143822 | 0.7322681 |
| ENSMUSG00000028005 | Gucy1b3 | protein_coding | -0.211169869 | 7.007594023 | 0.7007724 | 0.4146243 | 0.732559631 |
| ENSMUSG00000021258 | Ccnk | protein_coding | -0.525429366 | 3.107589317 | 0.7005912 | 0.4146834 | 0.732559631 |
| ENSMUSG00000034317 | Trim59 | protein_coding | -0.517863007 | 4.038572998 | 0.6999732 | 0.4149048 | 0.732704471 |
| ENSMUSG00000066415 | Msl2 | protein_coding | -0.417731652 | 4.015005728 | 0.6996128 | 0.4150027 | 0.732704471 |
| ENSMUSG00000022200 | Golph3 | protein_coding | -0.291251409 | 5.640855745 | 0.6995985 | 0.4150074 | 0.732704471 |
| ENSMUSG00000064284 | Cdpf1 | protein_coding | 0.253143716 | 6.128265536 | 0.6994937 | 0.4150416 | 0.732704471 |
| ENSMUSG00000046287 | Pnma3 | protein_coding | -0.431110078 | 4.538704125 | 0.6992506 | 0.4151408 | 0.732704471 |
| ENSMUSG00000051483 | Cbr1 | protein_coding | 0.232900939 | 7.257001137 | 0.6990878 | 0.4151742 | 0.732704471 |
| ENSMUSG00000063873 | Slc24a3 | protein_coding | -0.283256251 | 6.203920954 | 0.6986412 | 0.4153202 | 0.732841794 |
| ENSMUSG00000057531 | Dtnbp1 | protein_coding | 0.230523916 | 6.867280888 | 0.69831 | 0.4154285 | 0.732912623 |
| ENSMUSG00000036992 | Nxt1 | protein_coding | -0.306093604 | 4.667078708 | 0.6970796 | 0.4158312 | 0.733256208 |
| ENSMUSG00000021991 | Cacna2d3 | protein_coding | -0.358547526 | 5.559323087 | 0.6971037 | 0.4158431 | 0.733256208 |
| ENSMUSG00000019158 | Tmem160 | protein_coding | 0.393412246 | 4.02644066 | 0.6969624 | 0.4158696 | 0.733256208 |
| ENSMUSG00000005087 | Cd44 | protein_coding | -0.696010339 | 2.696449333 | 0.6968278 | 0.4159137 | 0.733256208 |
| ENSMUSG00000027404 | Snrpb | protein_coding | 0.227045442 | 7.354725503 | 0.6966738 | 0.4159642 | 0.733256208 |
| ENSMUSG00000020832 | Eral1 | protein_coding | 0.322217988 | 5.595521897 | 0.6961681 | 0.41613 | 0.733280274 |
| ENSMUSG00000037843 | Vstm2l | protein_coding | -0.650075377 | 2.609806328 | 0.6958882 | 0.4162218 | 0.733280274 |
| ENSMUSG00000001942 | Siae | protein_coding | 0.334773515 | 4.628530044 | 0.6954916 | 0.4163519 | 0.733280274 |
| ENSMUSG00000020843 | Timm22 | protein_coding | 0.231133713 | 7.608335653 | 0.6951453 | 0.4164656 | 0.733280274 |
| ENSMUSG00000031365 | Zfp275 | protein_coding | -0.355887396 | 4.555549452 | 0.6947474 | 0.4165963 | 0.733280274 |
| ENSMUSG00000037887 | Dusp8 | protein_coding | 0.251732328 | 8.228678214 | 0.6946842 | 0.4166367 | 0.733280274 |
| ENSMUSG00000060475 | Wtap | protein_coding | 0.498252494 | 5.476082691 | 0.6995895 | 0.4166951 | 0.733280274 |
| ENSMUSG00000023072 | Cep89 | protein_coding | 0.438492605 | 3.758035032 | 0.6942593 | 0.4167567 | 0.733280274 |
| ENSMUSG00000045251 | Zfp688 | protein_coding | 0.297928246 | 4.787095876 | 0.6941958 | 0.4167775 | 0.733280274 |
| ENSMUSG00000024245 | Tmem178 | protein_coding | -0.655676414 | 3.659488846 | 0.6963649 | 0.4168295 | 0.733280274 |
| ENSMUSG00000036002 | Fam214b | protein_coding | 0.348233144 | 4.200672327 | 0.694028 | 0.4168327 | 0.733280274 |
| ENSMUSG00000036104 | Rab3gap1 | protein_coding | 0.266438537 | 6.15508334 | 0.694017 | 0.4168363 | 0.733280274 |
| ENSMUSG00000072872 | Rybp | protein_coding | 0.649607629 | 2.674842252 | 0.6938957 | 0.4168762 | 0.733280274 |
| ENSMUSG00000029063 | Nadk | protein_coding | 0.306386892 | 4.830288005 | 0.6935664 | 0.4169845 | 0.733280274 |
| ENSMUSG00000035248 | Zcchc6 | protein_coding | -0.489813837 | 3.076259494 | 0.6935171 | 0.4170007 | 0.733280274 |
| ENSMUSG00000025377 | Tepsin | protein_coding | 0.582825866 | 2.691341674 | 0.6929495 | 0.4171875 | 0.733488832 |
| ENSMUSG00000018347 | Zkscan6 | protein_coding | 0.744600741 | 3.488809905 | 0.6958334 | 0.4174575 | 0.733841662 |
| ENSMUSG00000057103 | Nat8f1 | protein_coding | -0.32655529 | 4.708141597 | 0.6914695 | 0.4176753 | 0.733841662 |
| ENSMUSG00000042079 | Hnrnpf | protein_coding | -0.497270186 | 2.968149836 | 0.6912218 | 0.417757 | 0.733841662 |
| ENSMUSG00000022537 | Tmem44 | protein_coding | -0.461362017 | 3.575541263 | 0.6910845 | 0.4178023 | 0.733841662 |
| ENSMUSG00000047246 | Hist1h2be | protein_coding | -0.591557059 | 2.832654143 | 0.6909741 | 0.4178387 | 0.733841662 |
| ENSMUSG00000049881 | 2810025M15Rik | transcribed_processe | -0.246533264 | 5.804134644 | 0.6908385 | 0.4178835 | 0.733841662 |
| ENSMUSG00000028898 | Trnau1ap | protein_coding | 0.39204813 | 4.140814043 | 0.6905771 | 0.4179698 | 0.733841662 |
| ENSMUSG00000044030 | Irf2bp1 | protein_coding | 0.328089829 | 4.346356885 | 0.6904311 | 0.418018 | 0.733841662 |
| ENSMUSG00000040945 | Rcc2 | protein_coding | 0.224168391 | 5.898254329 | 0.6903548 | 0.4180432 | 0.733841662 |
| ENSMUSG00000038936 | Sccpdh | protein_coding | 0.218480016 | 7.117289642 | 0.6901405 | 0.418114 | 0.733841662 |
| ENSMUSG00000006464 | Bbs1 | protein_coding | -0.543829433 | 3.40991514 | 0.6899038 | 0.4181922 | 0.733841662 |
| ENSMUSG00000032612 | Usp4 | protein_coding | 0.22283524 | 6.156896791 | 0.6898328 | 0.4182156 | 0.733841662 |
| ENSMUSG00000087143 | A830082K12Rik | antisense_RNA | -0.390610253 | 3.85780153 | 0.6894854 | 0.4183305 | 0.733841662 |
| ENSMUSG00000037533 | Rapgef6 | protein_coding | 0.313117976 | 5.157853183 | 0.6894456 | 0.4183437 | 0.733841662 |
| ENSMUSG00000070348 | Ccnd1 | protein_coding | -0.802224672 | 2.568116953 | 0.688654 | 0.4186055 | 0.73396536 |
| ENSMUSG00000021273 | Fdft1 | protein_coding | 0.225935178 | 7.190815928 | 0.6882187 | 0.4187496 | 0.73396536 |
| ENSMUSG00000026737 | Pip4k2a | protein_coding | -0.23754242 | 5.635831493 | 0.6881579 | 0.4187698 | 0.73396536 |
| ENSMUSG00000031790 | Mmp15 | protein_coding | -0.388378106 | 3.531035954 | 0.6879314 | 0.4188448 | 0.73396536 |
| ENSMUSG00000030134 | Rasgef1a | protein_coding | -0.259246284 | 7.583743779 | 0.6878415 | 0.4188942 | 0.73396536 |
| ENSMUSG00000005150 | Wdr83 | protein_coding | 0.24377513 | 6.399766298 | 0.687612 | 0.4189506 | 0.73396536 |
| ENSMUSG00000024376 | Epb41l4a | protein_coding | 0.486432796 | 3.386500114 | 0.6874594 | 0.4190012 | 0.73396536 |
| ENSMUSG00000029151 | Slc30a3 | protein_coding | 1.05512542 | 2.706361497 | 0.6903262 | 0.4190308 | 0.73396536 |
| ENSMUSG00000028140 | Mrpl9 | protein_coding | 0.213351415 | 6.042274355 | 0.68729 | 0.4190574 | 0.73396536 |
| ENSMUSG00000063698 | Sfxn4 | protein_coding | -0.255365344 | 5.734147057 | 0.6871159 | 0.4191151 | 0.73396536 |
| ENSMUSG00000062397 | Zfp706 | protein_coding | 0.204934765 | 8.195450523 | 0.686801 | 0.4192196 | 0.73396536 |
| ENSMUSG00000052713 | Zfp608 | protein_coding | -0.378876442 | 4.004855985 | 0.6867596 | 0.4192333 | 0.73396536 |
| ENSMUSG00000049112 | Oxtr | protein_coding | 0.640739931 | 2.82457283 | 0.6862574 | 0.4193999 | 0.73413764 |
| ENSMUSG00000030556 | Lrrc28 | protein_coding | 0.225377998 | 5.877708572 | 0.6860228 | 0.4194778 | 0.734154446 |
| ENSMUSG00000031534 | Smim19 | protein_coding | -0.327939767 | 4.746452526 | 0.685083 | 0.4197901 | 0.734340064 |
| ENSMUSG00000030352 | Tspan9 | protein_coding | -0.479937891 | 3.740494061 | 0.6850804 | 0.419791 | 0.734340064 |
| ENSMUSG00000029910 | Mad2l1 | protein_coding | 0.382150342 | 4.158878799 | 0.6849408 | 0.4198374 | 0.734340064 |
| ENSMUSG00000036377 | C530008M17Rik | protein_coding | -0.331040974 | 4.964826786 | 0.6848816 | 0.419857 | 0.734340064 |
| ENSMUSG00000031985 | Gnpat | protein_coding | 0.298884935 | 5.414486197 | 0.6846554 | 0.4199323 | 0.734352221 |
| ENSMUSG00000038695 | Josd2 | protein_coding | -0.213358026 | 6.239917499 | 0.6842701 | 0.4200605 | 0.734456928 |
| ENSMUSG00000024561 | Mbd1 | protein_coding | 0.312067196 | 5.181178733 | 0.6840214 | 0.4201432 | 0.734482241 |
| ENSMUSG00000021930 | Spryd7 | protein_coding | -0.188828082 | 6.699836475 | 0.6834526 | 0.4203327 | 0.734565589 |
| ENSMUSG00000032773 | Chrm1 | protein_coding | -0.347779351 | 5.403171046 | 0.6833433 | 0.4203886 | 0.734565589 |
| ENSMUSG00000041353 | Tmem29 | protein_coding | 0.205230164 | 6.878532586 | 0.6830749 | 0.4204585 | 0.734565589 |
| ENSMUSG00000031577 | Tti2 | protein_coding | 0.425336032 | 3.608366245 | 0.6830579 | 0.4204642 | 0.734565589 |
| ENSMUSG00000068115 | Ninl | protein_coding | 0.98077088 | 2.241747413 | 0.6823422 | 0.4207028 | 0.734863088 |
| ENSMUSG00000044627 | Swi5 | protein_coding | 0.228576962 | 9.426210742 | 0.6816807 | 0.4209236 | 0.735129318 |
| ENSMUSG00000030386 | Zfp606 | protein_coding | -0.45244274 | 3.567408348 | 0.6802891 | 0.4213886 | 0.735739067 |
| ENSMUSG00000028753 | Vwa5b1 | protein_coding | -0.43063888 | 3.62853526 | 0.6802263 | 0.4214096 | 0.735739067 |
| ENSMUSG00000033701 | Acbd6 | protein_coding | 0.27299588 | 5.386860559 | 0.6799897 | 0.4214887 | 0.735757789 |
| ENSMUSG00000062184 | Hs6st2 | protein_coding | -0.265043933 | 5.469406566 | 0.6794809 | 0.421659 | 0.735866396 |
| ENSMUSG00000042210 | Abhd14a | protein_coding | 0.284938419 | 6.311576668 | 0.679453 | 0.4216878 | 0.735866396 |

| ensembl_gene_id | external_gene_name | gene_biotype | log2 (fold change) | log2 (counts per million) | F-value | p-value | False discovery rate |
|---|---|---|---|---|---|---|---|
| ENSMUSG00000032044 | Rpusd4 | protein_coding | 0.347325849 | 4.453725906 | 0.6789211 | 0.4218464 | 0.735887237 |
| ENSMUSG00000022682 | Rrn3 | protein_coding | 0.3632373 | 7.475947597 | 0.6840521 | 0.4218862 | 0.735887237 |
| ENSMUSG00000032540 | Abhd5 | protein_coding | -0.406482543 | 3.705300625 | 0.6784412 | 0.4220073 | 0.735887237 |
| ENSMUSG00000026504 | Sdccag8 | protein_coding | -0.435638127 | 4.75536557 | 0.6786144 | 0.4220199 | 0.735887237 |
| ENSMUSG00000020246 | Hcfc2 | protein_coding | -0.336944477 | 4.458769391 | 0.6783379 | 0.4220419 | 0.735887237 |
| ENSMUSG00000039901 | 9130011E15Rik | protein_coding | 0.434496224 | 3.936812485 | 0.6779516 | 0.4221714 | 0.735993733 |
| ENSMUSG00000031641 | Cbr4 | protein_coding | -0.290084615 | 5.240956239 | 0.6764713 | 0.4226684 | 0.736240659 |
| ENSMUSG00000041650 | Pcca | protein_coding | -0.377273548 | 4.350311959 | 0.6764191 | 0.4226859 | 0.736240659 |
| ENSMUSG00000078453 | Abracl | protein_coding | 0.359785755 | 4.923847524 | 0.6762865 | 0.4227305 | 0.736240659 |
| ENSMUSG00000049658 | Bdp1 | protein_coding | -0.414281321 | 3.753330249 | 0.6759315 | 0.4228499 | 0.736240659 |
| ENSMUSG00000063694 | Cycs | protein_coding | -0.266829192 | 5.154449995 | 0.6758372 | 0.4228816 | 0.736240659 |
| ENSMUSG00000026383 | Epb41l5 | protein_coding | -0.412447901 | 3.350534585 | 0.6758158 | 0.4228887 | 0.736240659 |
| ENSMUSG00000021660 | Btf3 | protein_coding | 0.230034592 | 6.743615765 | 0.6757688 | 0.4229046 | 0.736240659 |
| ENSMUSG00000052419 | 2610001J05Rik | protein_coding | -0.194153918 | 7.768716308 | 0.6757289 | 0.422918 | 0.736240659 |
| ENSMUSG00000024528 | Srfbp1 | protein_coding | -0.341314671 | 4.73445239 | 0.6756953 | 0.4229293 | 0.736240659 |
| ENSMUSG00000109321 | A330076H08Rik | processed_transcript | -0.396420918 | 3.772797869 | 0.6751975 | 0.4230968 | 0.736413025 |
| ENSMUSG00000068264 | Ap5s1 | protein_coding | 0.235375809 | 6.289212354 | 0.6742044 | 0.4234312 | 0.736550104 |
| ENSMUSG00000074220 | Zfp382 | protein_coding | -0.483000565 | 3.186077791 | 0.6740965 | 0.4234676 | 0.736550104 |
| ENSMUSG00000024959 | Bad | protein_coding | 0.190079015 | 7.611176989 | 0.673712 | 0.4235972 | 0.736550104 |
| ENSMUSG00000017400 | Stac2 | protein_coding | 0.435688552 | 4.566754954 | 0.6737308 | 0.4236102 | 0.736550104 |
| ENSMUSG00000030669 | Calca | protein_coding | -0.962718335 | 3.089485496 | 0.6785328 | 0.4236331 | 0.736550104 |
| ENSMUSG00000024273 | 2700062C07Rik | protein_coding | -0.255622711 | 5.358026639 | 0.6735864 | 0.4236396 | 0.736550104 |
| ENSMUSG00000039478 | Micu3 | protein_coding | -0.32692946 | 8.724196797 | 0.6770432 | 0.423655 | 0.736550104 |
| ENSMUSG00000027428 | Rbbp9 | protein_coding | -0.267038834 | 6.297842946 | 0.6732161 | 0.4237645 | 0.736621316 |
| ENSMUSG00000104960 | Snhg8 | processed_transcript | 0.365093729 | 6.492113512 | 0.6769058 | 0.4238957 | 0.736730388 |
| ENSMUSG00000024556 | Me2 | protein_coding | 0.354658399 | 4.003990473 | 0.6725151 | 0.4240011 | 0.736751223 |
| ENSMUSG00000027577 | Chrna4 | protein_coding | 0.565008205 | 4.640889132 | 0.6773007 | 0.4240448 | 0.736751223 |
| ENSMUSG00000021180 | Rps6ka5 | protein_coding | -0.492292265 | 4.180148764 | 0.6722867 | 0.4241277 | 0.736776259 |
| ENSMUSG00000026238 | Ptma | protein_coding | -0.208576539 | 6.344461929 | 0.6717302 | 0.4242663 | 0.736887916 |
| ENSMUSG00000032067 | Pts | protein_coding | -0.243406723 | 7.21680756 | 0.6714681 | 0.4243549 | 0.736887916 |
| ENSMUSG00000037395 | Rcor3 | protein_coding | 0.438398683 | 3.439315693 | 0.6710647 | 0.4244913 | 0.736887916 |
| ENSMUSG00000024099 | Ndufv2 | protein_coding | 0.200777068 | 10.3424644 | 0.6706082 | 0.4246458 | 0.736887916 |
| ENSMUSG00000013523 | Bcas1 | protein_coding | -0.392620027 | 5.245615683 | 0.6706675 | 0.4246959 | 0.736887916 |
| ENSMUSG00000031344 | Gabrq | protein_coding | 0.409142569 | 4.170744822 | 0.6704473 | 0.4247003 | 0.736887916 |
| ENSMUSG00000025340 | Rabgef1 | protein_coding | 0.262873781 | 5.840807769 | 0.6704141 | 0.4247115 | 0.736887916 |
| ENSMUSG00000034543 | Morc2a | protein_coding | 0.356383702 | 3.987447383 | 0.6702834 | 0.4247558 | 0.736887916 |
| ENSMUSG00000028029 | Aimp1 | protein_coding | -0.2269523 | 7.773119981 | 0.6701271 | 0.4248087 | 0.736887916 |
| ENSMUSG00000035505 | Cox18 | protein_coding | 0.4096054 | 3.559834666 | 0.6697788 | 0.4249267 | 0.736973726 |
| ENSMUSG00000046034 | Otulin | protein_coding | 0.45384517 | 3.192789776 | 0.6686622 | 0.4253054 | 0.737511451 |
| ENSMUSG00000033323 | Ctdp1 | protein_coding | -0.845747096 | 2.510626562 | 0.6683867 | 0.4253989 | 0.737554634 |
| ENSMUSG00000016382 | Pls3 | protein_coding | -0.218901739 | 6.281365613 | 0.668098 | 0.4254969 | 0.737582936 |
| ENSMUSG00000010290 | AI597479 | protein_coding | 0.308360285 | 5.439774411 | 0.6679346 | 0.4255524 | 0.737582936 |
| ENSMUSG00000027452 | Acss1 | protein_coding | -0.604932767 | 3.118755559 | 0.6674064 | 0.4257318 | 0.737728135 |
| ENSMUSG00000030689 | Ino80e | protein_coding | 0.196473532 | 6.461476471 | 0.6671959 | 0.4258034 | 0.737728135 |
| ENSMUSG00000002983 | Relb | protein_coding | 0.684677966 | 2.645367851 | 0.6670824 | 0.425842 | 0.737728135 |
| ENSMUSG00000043460 | Elfn2 | protein_coding | 0.495417604 | 3.91564847 | 0.6668226 | 0.4259496 | 0.737795714 |
| ENSMUSG00000023186 | Vwa5a | protein_coding | -0.545655266 | 2.970044968 | 0.666294 | 0.4261102 | 0.737954998 |
| ENSMUSG00000062190 | Lancl2 | protein_coding | 0.715734455 | 3.077568906 | 0.6660313 | 0.4262188 | 0.73802434 |
| ENSMUSG00000027784 | Ppm1l | protein_coding | -0.271192871 | 5.153257303 | 0.665691 | 0.4263155 | 0.738072794 |
| ENSMUSG00000031026 | Trim66 | protein_coding | -0.401304861 | 3.792633113 | 0.6649898 | 0.4265544 | 0.738278102 |
| ENSMUSG00000002608 | Ccdc97 | protein_coding | 0.3183681 | 5.891737021 | 0.6649965 | 0.4265714 | 0.738278102 |
| ENSMUSG00000024261 | Syt4 | protein_coding | 0.199669207 | 8.374893494 | 0.6643706 | 0.4267656 | 0.7384954 |
| ENSMUSG00000105891 | A230001M10Rik | lincRNA | -0.43773343 | 3.815658261 | 0.6637838 | 0.4269658 | 0.738723028 |
| ENSMUSG00000041921 | Metap1d | protein_coding | 0.41380178 | 3.832582105 | 0.6629731 | 0.4272428 | 0.739016555 |
| ENSMUSG00000031521 | Aga | protein_coding | -0.388891549 | 4.549416809 | 0.6628359 | 0.4272897 | 0.739016555 |
| ENSMUSG00000029735 | Tpk1 | protein_coding | -0.860378044 | 2.61527718 | 0.66274 | 0.4273417 | 0.739016555 |
| ENSMUSG00000049764 | Zfp280b | protein_coding | -0.437334213 | 3.189917158 | 0.6621991 | 0.4275075 | 0.739184398 |
| ENSMUSG00000026798 | Coq4 | protein_coding | 0.224640091 | 5.82078392 | 0.6617943 | 0.427646 | 0.739186037 |
| ENSMUSG00000032336 | Nptn | protein_coding | 0.190190742 | 8.112727407 | 0.6617336 | 0.4276668 | 0.739186037 |
| ENSMUSG00000031913 | Vps4a | protein_coding | 0.42319074 | 5.448352667 | 0.6637698 | 0.4277146 | 0.739186037 |
| ENSMUSG00000061479 | Snrpa | protein_coding | 0.278852617 | 6.147516646 | 0.65993 | 0.4282849 | 0.739998539 |
| ENSMUSG00000023010 | Tmbim6 | protein_coding | 0.206249665 | 9.694523127 | 0.6598205 | 0.4283224 | 0.739998539 |
| ENSMUSG00000031155 | Pim2 | protein_coding | 0.436799246 | 5.437067109 | 0.6621756 | 0.4284964 | 0.740006864 |
| ENSMUSG00000040596 | Pogk | protein_coding | -0.32222279 | 4.689284965 | 0.6592887 | 0.428505 | 0.740006864 |
| ENSMUSG00000049493 | Pls1 | protein_coding | -0.394088712 | 4.987409284 | 0.6589562 | 0.4286383 | 0.740006864 |
| ENSMUSG00000024966 | Stip1 | protein_coding | 0.246265878 | 8.728754636 | 0.6588093 | 0.4286696 | 0.740006864 |
| ENSMUSG00000038811 | Gngt2 | protein_coding | -0.613328142 | 2.841497445 | 0.6588044 | 0.4286713 | 0.740006864 |
| ENSMUSG00000029461 | Fam168a | protein_coding | 0.241405482 | 7.503392389 | 0.6579177 | 0.4289761 | 0.740368763 |
| ENSMUSG00000071604 | Fam189a2 | protein_coding | 0.689019098 | 2.759313938 | 0.6575962 | 0.4290868 | 0.740368763 |
| ENSMUSG00000097736 | 9530059O14Rik | lincRNA | -0.948348451 | 2.537022903 | 0.6574659 | 0.4291507 | 0.740368763 |
| ENSMUSG00000026643 | Nmt2 | protein_coding | -0.257518865 | 8.632761936 | 0.6572423 | 0.4292277 | 0.740368763 |
| ENSMUSG00000047146 | Tet1 | protein_coding | -0.400800417 | 3.973923499 | 0.656966 | 0.4293037 | 0.740368763 |
| ENSMUSG00000026189 | Pecr | protein_coding | -0.546694761 | 2.841673926 | 0.6568274 | 0.4293514 | 0.740368763 |
| ENSMUSG00000033793 | Atp6v1h | protein_coding | 0.221300222 | 8.77313999 | 0.656794 | 0.4293629 | 0.740368763 |
| ENSMUSG00000028809 | Srrm1 | protein_coding | 0.191189961 | 7.215659472 | 0.6560838 | 0.4296077 | 0.74067202 |
| ENSMUSG00000003464 | Pex19 | protein_coding | 0.22701058 | 7.878205664 | 0.6554454 | 0.4298279 | 0.740827912 |
| ENSMUSG00000022248 | Rad1 | protein_coding | -0.273619371 | 4.790968335 | 0.6554026 | 0.4298426 | 0.740827912 |
| ENSMUSG00000001576 | Ergic1 | protein_coding | -0.270911364 | 5.837601065 | 0.6552226 | 0.4299048 | 0.740827912 |

| ensembl_gene_id | external_gene_name | gene_biotype | log2 (fold change) | log2 (counts per million) | F-value | p-value | False discovery rate |
|---|---|---|---|---|---|---|---|
| ENSMUSG00000036748 | Cuedc2 | protein_coding | 0.175787751 | 8.642863803 | 0.6545476 | 0.4301378 | 0.741002762 |
| ENSMUSG00000062198 | 2700097O09Rik | protein_coding | -0.332764555 | 4.295031524 | 0.6542769 | 0.4302314 | 0.741002762 |
| ENSMUSG00000059974 | Ntm | protein_coding | -0.191069068 | 7.518673752 | 0.6542719 | 0.4302331 | 0.741002762 |
| ENSMUSG00000079108 | Srp54b | protein_coding | -0.450343471 | 3.34042388 | 0.6541307 | 0.4302819 | 0.741002762 |
| ENSMUSG00000062110 | Scfd2 | protein_coding | 0.406967912 | 4.006111283 | 0.6535262 | 0.4304909 | 0.741243998 |
| ENSMUSG00000056211 | R3hdm1 | protein_coding | -0.18421568 | 6.656809729 | 0.6532191 | 0.4305972 | 0.741257578 |
| ENSMUSG00000022012 | Enox1 | protein_coding | -0.392833366 | 4.283157381 | 0.653105 | 0.4306367 | 0.741257578 |
| ENSMUSG00000022914 | Brwd1 | protein_coding | -0.217265344 | 6.174824678 | 0.6528213 | 0.4307348 | 0.741307904 |
| ENSMUSG00000056900 | Usp13 | protein_coding | 0.380226655 | 4.009411994 | 0.6524527 | 0.4308625 | 0.741350433 |
| ENSMUSG00000046572 | Zfp518b | protein_coding | -0.735333355 | 2.427074338 | 0.6523518 | 0.4308974 | 0.741350433 |
| ENSMUSG00000038537 | Mc3r | protein_coding | 0.904554568 | 2.441413846 | 0.6521875 | 0.4309734 | 0.74136248 |
| ENSMUSG00000066306 | Numa1 | protein_coding | -0.314910906 | 4.956374096 | 0.6512001 | 0.4312967 | 0.741800032 |
| ENSMUSG00000025968 | Ndufs1 | protein_coding | 0.329327695 | 5.702321585 | 0.6505864 | 0.4315287 | 0.742005965 |
| ENSMUSG00000015653 | Steap2 | protein_coding | 0.39728749 | 3.443828162 | 0.6504575 | 0.4315545 | 0.742005965 |
| ENSMUSG00000022120 | Rnf219 | protein_coding | 0.3808823 | 4.369410088 | 0.6496272 | 0.431843 | 0.742225043 |
| ENSMUSG00000027953 | Slc50a1 | protein_coding | 0.207888447 | 5.91594276 | 0.649602 | 0.4318517 | 0.742225043 |
| ENSMUSG00000033209 | Ttc28 | protein_coding | 0.584917913 | 2.80421923 | 0.6491927 | 0.431994 | 0.742225043 |
| ENSMUSG00000034247 | Plekhm1 | protein_coding | -0.384377355 | 3.720036447 | 0.6491675 | 0.4320028 | 0.742225043 |
| ENSMUSG00000040667 | Nup88 | protein_coding | 0.261383571 | 6.057076029 | 0.6490979 | 0.432027 | 0.742225043 |
| ENSMUSG00000038020 | Rapgefl1 | protein_coding | 0.323239521 | 5.041302395 | 0.6488215 | 0.4321232 | 0.74226629 |
| ENSMUSG00000071359 | Tbpl1 | protein_coding | 0.24488907 | 6.377242744 | 0.6486321 | 0.4321891 | 0.74226629 |
| ENSMUSG00000031627 | Irf2 | protein_coding | -0.25661951 | 4.944026901 | 0.648044 | 0.4323939 | 0.74228328 |
| ENSMUSG00000046185 | Zfp84 | protein_coding | -0.521088999 | 3.379679398 | 0.6479199 | 0.4324371 | 0.74228328 |
| ENSMUSG00000029090 | Adgra3 | protein_coding | 0.964864729 | 2.282222908 | 0.6478656 | 0.432475 | 0.74228328 |
| ENSMUSG00000025218 | Poll | protein_coding | 0.310203866 | 6.066550268 | 0.6478652 | 0.4324751 | 0.74228328 |
| ENSMUSG00000039263 | Npepl1 | protein_coding | -1.125475786 | 2.256624466 | 0.6484524 | 0.4326114 | 0.742316142 |
| ENSMUSG00000020953 | Coch | protein_coding | -0.409257694 | 3.620251044 | 0.6473598 | 0.4326323 | 0.742316142 |
| ENSMUSG00000046329 | Slc25a23 | protein_coding | -0.356518437 | 3.69950055 | 0.6470275 | 0.4327482 | 0.742396523 |
| ENSMUSG00000006522 | Itih3 | protein_coding | -0.381369543 | 5.66703038 | 0.6484873 | 0.4329739 | 0.742665244 |
| ENSMUSG00000001334 | Fndc5 | protein_coding | -0.309242423 | 5.763481412 | 0.6455411 | 0.433286 | 0.743053331 |
| ENSMUSG00000009549 | Srp14 | protein_coding | 0.201838419 | 9.519047073 | 0.6453373 | 0.4333384 | 0.743053331 |
| ENSMUSG00000034156 | Tspoap1 | protein_coding | -0.387783758 | 4.054410639 | 0.6449604 | 0.4334701 | 0.743135022 |
| ENSMUSG00000072875 | Gpr27 | protein_coding | -0.390243153 | 4.238632282 | 0.6447831 | 0.4335321 | 0.743135022 |
| ENSMUSG00000020436 | Gabrg2 | protein_coding | 0.346556136 | 3.993808384 | 0.6445703 | 0.4336066 | 0.743135022 |
| ENSMUSG00000073889 | Il11ra1 | protein_coding | 0.347917946 | 3.906025596 | 0.6444107 | 0.4336624 | 0.743135022 |
| ENSMUSG00000066643 | Wdr35 | protein_coding | 0.294235185 | 5.363477915 | 0.6434667 | 0.433993 | 0.743582944 |
| ENSMUSG00000038871 | Bpgm | protein_coding | 0.188017014 | 7.396211372 | 0.6430002 | 0.4341565 | 0.743645435 |
| ENSMUSG00000022641 | Bbx | protein_coding | -0.296763574 | 5.438121776 | 0.6429681 | 0.4341678 | 0.743645435 |
| ENSMUSG00000036422 | Pcdh8 | protein_coding | -0.39768013 | 4.455041384 | 0.6422041 | 0.4344358 | 0.743985954 |
| ENSMUSG00000020713 | Gh | protein_coding | -0.733324187 | 3.159073707 | 0.6431402 | 0.4345886 | 0.744129168 |
| ENSMUSG00000034107 | Ano7 | protein_coding | -0.754165579 | 2.574093117 | 0.6415416 | 0.4346683 | 0.744147228 |
| ENSMUSG00000060679 | Mrps9 | protein_coding | -0.211458391 | 6.640901099 | 0.6413276 | 0.4347435 | 0.744157488 |
| ENSMUSG00000049516 | Spty2d1 | protein_coding | -0.372982918 | 3.873395935 | 0.6408848 | 0.4348992 | 0.744305366 |
| ENSMUSG00000015176 | Nolc1 | protein_coding | 0.203926782 | 6.376195778 | 0.6406335 | 0.4349875 | 0.744338172 |
| ENSMUSG00000012535 | Tnpo3 | protein_coding | 0.357646921 | 5.407096462 | 0.6403286 | 0.4351136 | 0.744435411 |
| ENSMUSG00000022212 | Cpne6 | protein_coding | 0.31547299 | 7.179004098 | 0.6430555 | 0.4352825 | 0.744605892 |
| ENSMUSG00000034336 | Ina | protein_coding | 0.184367115 | 7.552055652 | 0.6393457 | 0.4354408 | 0.744716438 |
| ENSMUSG00000004460 | Dnajb11 | protein_coding | 0.295607397 | 4.93697652 | 0.6392184 | 0.4354856 | 0.744716438 |
| ENSMUSG00000031575 | Ash2l | protein_coding | -0.319560782 | 4.821138281 | 0.6388835 | 0.4356036 | 0.744799805 |
| ENSMUSG00000024347 | Psd2 | protein_coding | 0.435876473 | 4.892576603 | 0.6398236 | 0.4357523 | 0.744935509 |
| ENSMUSG00000049550 | Clip1 | protein_coding | 0.222889602 | 6.116884207 | 0.6377405 | 0.4360068 | 0.745171385 |
| ENSMUSG00000058192 | Zfp846 | protein_coding | -0.402540455 | 4.151514707 | 0.637678 | 0.4360289 | 0.745171385 |
| ENSMUSG00000026839 | Upp2 | protein_coding | -0.364053815 | 4.29042745 | 0.6373995 | 0.4361272 | 0.745221048 |
| ENSMUSG00000016200 | Syt14 | protein_coding | -0.465962344 | 3.44593663 | 0.6370812 | 0.4362397 | 0.745258013 |
| ENSMUSG00000060572 | Mfap2 | protein_coding | 0.374428584 | 4.044965662 | 0.6367048 | 0.4363726 | 0.745258013 |
| ENSMUSG00000007415 | Gatad1 | protein_coding | 0.226329866 | 7.549900113 | 0.6366762 | 0.4363828 | 0.745258013 |
| ENSMUSG00000032452 | Clstn2 | protein_coding | -0.323931304 | 4.601326026 | 0.6365536 | 0.4364261 | 0.745258013 |
| ENSMUSG00000004768 | Rab23 | protein_coding | -0.407945026 | 4.504457909 | 0.6358557 | 0.4366917 | 0.745593174 |
| ENSMUSG00000024603 | Dctn4 | protein_coding | 0.298083289 | 7.585918399 | 0.6374486 | 0.4368373 | 0.745662374 |
| ENSMUSG00000025066 | Sfr1 | protein_coding | 0.169271318 | 8.415339589 | 0.6352964 | 0.4368709 | 0.745662374 |
| ENSMUSG00000056917 | Sipa1 | protein_coding | 0.41042667 | 4.245049254 | 0.6348373 | 0.4370335 | 0.745821534 |
| ENSMUSG00000028312 | Smc2 | protein_coding | -0.637567576 | 2.554614109 | 0.6336942 | 0.4374389 | 0.746385946 |
| ENSMUSG00000043252 | Tmem64 | protein_coding | 0.364647466 | 4.869118991 | 0.6335132 | 0.4375031 | 0.746385946 |
| ENSMUSG00000025544 | Tm9sf2 | protein_coding | -0.357681213 | 5.955620366 | 0.6351046 | 0.4376647 | 0.746532461 |
| ENSMUSG00000022793 | B4galt4 | protein_coding | -0.486607186 | 3.48912952 | 0.6328803 | 0.4377278 | 0.746532461 |
| ENSMUSG00000086697 | 4933427G23Rik | transcribed_processe | -0.646698715 | 2.445828553 | 0.6326478 | 0.4378104 | 0.746554925 |
| ENSMUSG00000038712 | Mindy1 | protein_coding | 0.372167437 | 4.8086224 | 0.6324286 | 0.4378883 | 0.746569364 |
| ENSMUSG00000022204 | Ngdn | protein_coding | -0.208202565 | 6.862385799 | 0.6319079 | 0.4380734 | 0.74674409 |
| ENSMUSG00000037376 | Trmt6 | protein_coding | -0.279298281 | 5.655510675 | 0.6317498 | 0.4381297 | 0.74674409 |
| ENSMUSG00000034292 | Traf3ip1 | protein_coding | -0.443629901 | 3.585416351 | 0.6310466 | 0.43838 | 0.746949955 |
| ENSMUSG00000048279 | Sacs | protein_coding | -0.382361454 | 4.283864982 | 0.6310201 | 0.4383894 | 0.746949955 |
| ENSMUSG00000026439 | Rbbp5 | protein_coding | 0.28026166 | 5.231769556 | 0.6305261 | 0.4385654 | 0.747027591 |
| ENSMUSG00000026356 | Dars | protein_coding | -0.268581411 | 6.447264805 | 0.6305545 | 0.4385739 | 0.747027591 |
| ENSMUSG00000024845 | Tmem134 | processed_transcript | 0.397448203 | 3.994797671 | 0.6297556 | 0.4388401 | 0.747362556 |
| ENSMUSG00000001855 | Nup214 | protein_coding | -0.286975173 | 4.491393077 | 0.6285775 | 0.4392606 | 0.74770292 |
| ENSMUSG00000022270 | Retreg1 | protein_coding | -0.229250986 | 5.980789228 | 0.6285358 | 0.4392755 | 0.74770292 |
| ENSMUSG00000047428 | Dlk2 | protein_coding | 0.50933089 | 3.196595907 | 0.6282079 | 0.4393926 | 0.74770292 |
| ENSMUSG00000014313 | Cox6c | protein_coding | 0.242424063 | 10.12988312 | 0.6281419 | 0.4394162 | 0.74770292 |

| ensembl_gene_id | external_gene_name | gene_biotype | log2 (fold change) | log2 (counts per million) | F-value | p-value | False discovery rate |
|---|---|---|---|---|---|---|---|
| ENSMUSG00000022415 | Syngr1 | protein_coding | 0.166861394 | 7.596962092 | 0.6280719 | 0.4394412 | 0.74770292 |
| ENSMUSG00000090553 | Snrpe | protein_coding | 0.247837675 | 7.173520655 | 0.6280795 | 0.4394571 | 0.74770292 |
| ENSMUSG00000025503 | Taldo1 | protein_coding | -0.171523173 | 8.388558491 | 0.6276926 | 0.4395769 | 0.747788329 |
| ENSMUSG00000031851 | Ntpcr | protein_coding | 0.199638709 | 6.280122565 | 0.6274955 | 0.4396474 | 0.747789946 |
| ENSMUSG00000086119 | Gm2415 | antisense_RNA | -0.605924185 | 2.633322965 | 0.6270794 | 0.4397962 | 0.747892984 |
| ENSMUSG00000040412 | 5330417C22Rik | protein_coding | 0.337185669 | 4.990527782 | 0.6269375 | 0.439847 | 0.747892984 |
| ENSMUSG00000001211 | Agpat3 | protein_coding | 0.302156432 | 5.813345945 | 0.6266944 | 0.4399527 | 0.747954325 |
| ENSMUSG00000051727 | Kctd14 | protein_coding | -0.815951698 | 2.14395777 | 0.6260827 | 0.4401532 | 0.748097707 |
| ENSMUSG00000056185 | Snx32 | protein_coding | 0.192822374 | 7.503683569 | 0.6259435 | 0.4402031 | 0.748097707 |
| ENSMUSG00000027886 | 1700013F07Rik | protein_coding | -0.938668962 | 2.351329396 | 0.6258764 | 0.4402457 | 0.748097707 |
| ENSMUSG00000019961 | Tmpo | protein_coding | -0.28572937 | 4.458552505 | 0.6253596 | 0.4404125 | 0.748160783 |
| ENSMUSG00000044288 | Cnr1 | protein_coding | 0.227318828 | 6.287251988 | 0.6251959 | 0.4404712 | 0.748160783 |
| ENSMUSG00000048915 | Efna5 | protein_coding | -0.371285362 | 6.530370452 | 0.6295798 | 0.4404916 | 0.748160783 |
| ENSMUSG00000071072 | Ptges3 | protein_coding | -0.174922362 | 8.537608677 | 0.6248865 | 0.4405822 | 0.748196578 |
| ENSMUSG00000028273 | Pdlim5 | protein_coding | -0.420267391 | 4.080537185 | 0.6245448 | 0.4407049 | 0.748225853 |
| ENSMUSG00000030772 | Dkk3 | protein_coding | 0.331628432 | 5.609323161 | 0.6245026 | 0.4407386 | 0.748225853 |
| ENSMUSG00000053617 | Sh3pxd2a | protein_coding | -0.613171904 | 3.092813162 | 0.6239075 | 0.4409339 | 0.748346651 |
| ENSMUSG00000036026 | Tmem63b | protein_coding | 0.276837393 | 5.229959865 | 0.6238118 | 0.4409683 | 0.748346651 |
| ENSMUSG00000027301 | Oxt | protein_coding | -0.785458707 | 4.513115714 | 0.6348303 | 0.4411109 | 0.748346651 |
| ENSMUSG00000051335 | Gfod1 | protein_coding | -0.357337867 | 3.964177096 | 0.6232388 | 0.4411743 | 0.748346651 |
| ENSMUSG00000028758 | Kif17 | protein_coding | 0.306582448 | 4.78534031 | 0.6231419 | 0.4412092 | 0.748346651 |
| ENSMUSG00000028804 | Csmd2 | protein_coding | 0.524848888 | 3.077264071 | 0.623056 | 0.4412401 | 0.748346651 |
| ENSMUSG00000033991 | Ttc37 | protein_coding | -0.473978704 | 3.353452265 | 0.622898 | 0.441297 | 0.748346651 |
| ENSMUSG00000062116 | Zfp954 | protein_coding | 0.337088835 | 5.134182017 | 0.6224809 | 0.4414471 | 0.748363674 |
| ENSMUSG00000040972 | Igsf21 | protein_coding | -0.496372084 | 4.150300322 | 0.6233807 | 0.4414735 | 0.748363674 |
| ENSMUSG00000051736 | Fam229b | protein_coding | -0.249535351 | 5.959750319 | 0.6222902 | 0.4415158 | 0.748363674 |
| ENSMUSG00000037415 | Ranbp10 | protein_coding | 0.53479569 | 3.407233385 | 0.6210196 | 0.4419738 | 0.748971595 |
| ENSMUSG00000025369 | Smarcc2 | protein_coding | -0.212685388 | 6.432757822 | 0.6207319 | 0.4420776 | 0.748971595 |
| ENSMUSG00000039041 | Adrm1 | protein_coding | 0.334373442 | 4.483912201 | 0.6207159 | 0.4420834 | 0.748971595 |
| ENSMUSG00000061983 | Rps12 | protein_coding | 0.457623752 | 3.176514267 | 0.6205219 | 0.4421534 | 0.748972202 |
| ENSMUSG00000022856 | Tmem41a | protein_coding | 0.30132777 | 4.42514257 | 0.6196089 | 0.4424832 | 0.74941281 |
| ENSMUSG00000029167 | Ppargc1a | protein_coding | -0.260120993 | 5.318008391 | 0.6190168 | 0.4426973 | 0.749657346 |
| ENSMUSG00000036086 | Zranb3 | protein_coding | 0.691227537 | 2.837932009 | 0.6187796 | 0.4427832 | 0.749684653 |
| ENSMUSG00000052914 | Cyp2j6 | protein_coding | -0.449444931 | 4.464166553 | 0.6186665 | 0.442923 | 0.749803414 |
| ENSMUSG00000009291 | Pttg1ip | protein_coding | 0.224454599 | 6.171831776 | 0.6176381 | 0.4431965 | 0.750106466 |
| ENSMUSG00000007613 | Tgfbr1 | protein_coding | 0.764424172 | 3.187294785 | 0.6191655 | 0.4433591 | 0.750106466 |
| ENSMUSG00000057666 | Gapdh | protein_coding | 0.333398949 | 4.429047748 | 0.6171477 | 0.4433743 | 0.750106466 |
| ENSMUSG00000078607 | 1810010H24Rik | protein_coding | -0.88422691 | 2.574770644 | 0.6169932 | 0.4434487 | 0.750106466 |
| ENSMUSG00000003062 | Stard3nl | protein_coding | 0.181908087 | 7.038505656 | 0.6169122 | 0.4434597 | 0.750106466 |
| ENSMUSG00000047603 | Zfp235 | protein_coding | -0.284519236 | 4.396018757 | 0.6167443 | 0.4435206 | 0.750106466 |
| ENSMUSG00000096974 | Gm26881 | antisense_RNA | -0.485123207 | 2.940711554 | 0.616547 | 0.4435922 | 0.750109591 |
| ENSMUSG00000026842 | Abl1 | protein_coding | -0.441268039 | 4.118710143 | 0.6161552 | 0.4437528 | 0.750263221 |
| ENSMUSG00000022332 | Khdrbs3 | protein_coding | 0.216549084 | 6.130064683 | 0.6156699 | 0.4439107 | 0.750336161 |
| ENSMUSG00000030222 | Rerg | protein_coding | 0.245076503 | 5.623906843 | 0.6156015 | 0.4439355 | 0.750336161 |
| ENSMUSG00000064329 | Scn1a | protein_coding | -0.226974926 | 6.450607601 | 0.6147178 | 0.4442568 | 0.750692314 |
| ENSMUSG00000040713 | Creg1 | protein_coding | -0.373445008 | 4.165173706 | 0.614327 | 0.444399 | 0.750692314 |
| ENSMUSG00000026614 | Slc30a10 | protein_coding | -0.564431768 | 3.199459821 | 0.6142933 | 0.4444113 | 0.750692314 |
| ENSMUSG00000023921 | Mut | protein_coding | -0.280690797 | 5.10118876 | 0.6139451 | 0.4445381 | 0.750692314 |
| ENSMUSG00000031605 | Klhl2 | protein_coding | -0.226837445 | 6.092562795 | 0.6138812 | 0.4445613 | 0.750692314 |
| ENSMUSG00000020593 | Lpin1 | protein_coding | 0.357799271 | 4.67513294 | 0.6138136 | 0.4445859 | 0.750692314 |
| ENSMUSG00000042156 | Dzip1 | protein_coding | -0.291519349 | 5.309869513 | 0.6136792 | 0.4446349 | 0.750692314 |
| ENSMUSG00000040550 | Otud6b | protein_coding | -0.267423517 | 5.995032259 | 0.6131323 | 0.4448342 | 0.750858215 |
| ENSMUSG00000035674 | Ndufa3 | protein_coding | 0.266455573 | 10.02601084 | 0.6127408 | 0.4449954 | 0.750858215 |
| ENSMUSG00000027371 | Fahd2a | protein_coding | 0.213267905 | 6.322687499 | 0.612615 | 0.4450229 | 0.750858215 |
| ENSMUSG00000096935 | 1700113A16Rik | antisense_RNA | -0.717329787 | 2.436714362 | 0.6124231 | 0.4450929 | 0.750858215 |
| ENSMUSG00000038011 | Dnah10 | protein_coding | -0.59505841 | 2.859504059 | 0.6124117 | 0.445097 | 0.750858215 |
| ENSMUSG00000026965 | Anapc2 | protein_coding | -0.215467236 | 6.07619777 | 0.612123 | 0.4452024 | 0.750858215 |
| ENSMUSG00000062373 | Tmem65 | protein_coding | -0.247454628 | 5.907264928 | 0.6118966 | 0.4452851 | 0.750858215 |
| ENSMUSG00000039552 | Rsph4a | protein_coding | 0.695511831 | 2.769764578 | 0.6118782 | 0.4452918 | 0.750858215 |
| ENSMUSG00000074829 | 2010315B03Rik | protein_coding | -0.35217971 | 4.00988942 | 0.6114186 | 0.4454597 | 0.750910565 |
| ENSMUSG00000069520 | Tmem19 | protein_coding | 0.23670736 | 5.323552314 | 0.6114108 | 0.4454625 | 0.750910565 |
| ENSMUSG00000055923 | Aasdh | protein_coding | 0.529494348 | 3.069737384 | 0.6105795 | 0.4457664 | 0.751305092 |
| ENSMUSG00000051674 | Dcun1d4 | protein_coding | 0.227831355 | 6.531076334 | 0.6100964 | 0.4459432 | 0.751485271 |
| ENSMUSG00000027569 | Mrgbp | protein_coding | -0.505873315 | 3.122264797 | 0.6097021 | 0.4460876 | 0.751604316 |
| ENSMUSG00000023826 | Park2 | protein_coding | -0.306389699 | 4.804067349 | 0.6089051 | 0.4463796 | 0.751604316 |
| ENSMUSG00000026455 | Klhl12 | protein_coding | 0.388008654 | 4.628527613 | 0.6088675 | 0.4464117 | 0.751604316 |
| ENSMUSG00000035601 | Trmt10b | protein_coding | -0.212700334 | 5.834775679 | 0.6087297 | 0.4464439 | 0.751604316 |
| ENSMUSG00000044876 | Zfp444 | protein_coding | 0.308156613 | 5.120859974 | 0.6085843 | 0.4464973 | 0.751604316 |
| ENSMUSG00000017670 | Elmo2 | protein_coding | 0.249986807 | 6.908676478 | 0.6084573 | 0.4465621 | 0.751604316 |
| ENSMUSG00000053684 | BC048403 | protein_coding | 0.574489098 | 2.762959654 | 0.6083862 | 0.4465699 | 0.751604316 |
| ENSMUSG00000038773 | Kdm3b | protein_coding | 0.31681743 | 3.911557916 | 0.6080212 | 0.4467039 | 0.751604316 |
| ENSMUSG00000044014 | Npy5r | protein_coding | 0.647054974 | 2.983356318 | 0.6078946 | 0.4467503 | 0.751604316 |
| ENSMUSG00000034216 | Vps18 | protein_coding | 0.403747737 | 5.153911227 | 0.6091987 | 0.4467627 | 0.751604316 |
| ENSMUSG00000027649 | Ctnnbl1 | protein_coding | 0.245900511 | 6.228980183 | 0.6078063 | 0.4467827 | 0.751604316 |
| ENSMUSG00000039735 | Fnbp1l | protein_coding | -0.277647429 | 4.974653609 | 0.6071441 | 0.447026 | 0.751736639 |
| ENSMUSG00000083674 | Zfp133-ps | transcribed_unitary_p | -0.531191266 | 3.335927274 | 0.6070465 | 0.4470619 | 0.751736639 |
| ENSMUSG00000027244 | Atg13 | protein_coding | 0.246912146 | 6.63133618 | 0.6070213 | 0.4470711 | 0.751736639 |
| ENSMUSG00000027160 | Ccdc34 | protein_coding | 0.513610667 | 3.2779481 | 0.6067879 | 0.4471569 | 0.751763346 |

| ensembl_gene_id | external_gene_name | gene_biotype | log2 (fold change) | log2 (counts per million) | F-value | p-value | False discovery rate |
|---|---|---|---|---|---|---|---|
| ENSMUSG00000015536 | Mocs2 | protein_coding | -0.173348042 | 7.941274613 | 0.6064617 | 0.4472769 | 0.751766254 |
| ENSMUSG00000000794 | Kcnn3 | protein_coding | -0.475315617 | 3.752232933 | 0.6064526 | 0.4472985 | 0.751766254 |
| ENSMUSG00000053286 | Trmt1l | protein_coding | -0.403660321 | 3.963026734 | 0.6054386 | 0.4476534 | 0.752245218 |
| ENSMUSG00000024862 | Klc2 | protein_coding | 0.347766686 | 5.55964312 | 0.6053486 | 0.4477567 | 0.752250945 |
| ENSMUSG00000036339 | Tmem260 | protein_coding | 0.452153876 | 3.371221441 | 0.6050496 | 0.4477967 | 0.752250945 |
| ENSMUSG00000034758 | Tle6 | protein_coding | 1.026629903 | 2.579938696 | 0.6066197 | 0.4481288 | 0.752570863 |
| ENSMUSG00000027624 | Epb41l1 | protein_coding | -0.206221499 | 6.805994717 | 0.6040363 | 0.4481704 | 0.752570863 |
| ENSMUSG00000026546 | Cfap45 | protein_coding | 0.424460143 | 3.53867362 | 0.6036615 | 0.4483087 | 0.752570863 |
| ENSMUSG00000027770 | Dhx36 | protein_coding | -0.256445354 | 6.075141452 | 0.6036249 | 0.4483222 | 0.752570863 |
| ENSMUSG00000022965 | Ifngr2 | protein_coding | 0.253600581 | 5.18798113 | 0.6035846 | 0.4483371 | 0.752570863 |
| ENSMUSG00000042148 | Cox10 | protein_coding | 0.256201215 | 5.255918177 | 0.6033792 | 0.448413 | 0.752580747 |
| ENSMUSG00000090110 | Cmc4 | protein_coding | -0.346794195 | 3.930517205 | 0.6026984 | 0.4486645 | 0.752885384 |
| ENSMUSG00000024384 | Iws1 | protein_coding | -0.378568969 | 3.589956295 | 0.6023746 | 0.4487843 | 0.752913517 |
| ENSMUSG00000028049 | Scamp3 | protein_coding | 0.200322841 | 6.211898574 | 0.6017214 | 0.4490259 | 0.752913517 |
| ENSMUSG00000093904 | Tomm20 | protein_coding | -0.195004622 | 6.853735272 | 0.6016063 | 0.4490685 | 0.752913517 |
| ENSMUSG00000034341 | Wbp2 | protein_coding | 0.252809845 | 10.2406158 | 0.6015323 | 0.4491141 | 0.752913517 |
| ENSMUSG00000029363 | Rfc5 | protein_coding | -0.597365433 | 2.78036276 | 0.6014832 | 0.4491141 | 0.752913517 |
| ENSMUSG00000037818 | Abhd18 | protein_coding | -0.460955834 | 3.23669657 | 0.601454 | 0.4491249 | 0.752913517 |
| ENSMUSG00000097820 | E530011L22Rik | antisense_RNA | 0.664882526 | 2.661232199 | 0.6008155 | 0.4493614 | 0.752913517 |
| ENSMUSG00000039954 | Stk32a | protein_coding | -0.648970013 | 3.783346815 | 0.6045862 | 0.4494045 | 0.752913517 |
| ENSMUSG00000006782 | Cnp | protein_coding | -0.262626424 | 9.811024305 | 0.600451 | 0.4495147 | 0.752913517 |
| ENSMUSG00000039828 | Wdr70 | protein_coding | -0.230822485 | 5.70048989 | 0.6002644 | 0.4495658 | 0.752913517 |
| ENSMUSG00000021432 | Slc35b3 | protein_coding | 0.505706471 | 3.149178668 | 0.5996839 | 0.4497811 | 0.752913517 |
| ENSMUSG00000079235 | Ccdc13 | protein_coding | -0.420112198 | 3.461573038 | 0.5995241 | 0.4498405 | 0.752913517 |
| ENSMUSG00000020482 | Ccdc117 | protein_coding | -0.484985271 | 3.228345873 | 0.5994144 | 0.4498812 | 0.752913517 |
| ENSMUSG00000020224 | Llph | protein_coding | -0.276257194 | 4.444461609 | 0.5994107 | 0.4498826 | 0.752913517 |
| ENSMUSG00000052906 | Ubxn8 | protein_coding | -0.224577555 | 6.051718057 | 0.5993099 | 0.44992 | 0.752913517 |
| ENSMUSG00000038079 | Tmem237 | protein_coding | 0.259346691 | 6.25793534 | 0.5992831 | 0.44993 | 0.752913517 |
| ENSMUSG00000036550 | Cnot1 | protein_coding | 0.310627969 | 4.584060115 | 0.5992675 | 0.4499358 | 0.752913517 |
| ENSMUSG00000038372 | Gmds | protein_coding | 0.354510697 | 3.71384034 | 0.5990772 | 0.4500064 | 0.752913517 |
| ENSMUSG00000034755 | Pcdh11x | protein_coding | -0.272842417 | 5.168562025 | 0.5989577 | 0.4500508 | 0.752913517 |
| ENSMUSG00000008682 | Rpl10 | protein_coding | -0.459878315 | 3.033878391 | 0.5986949 | 0.4501485 | 0.752913517 |
| ENSMUSG00000019647 | Sema6a | protein_coding | -0.329311892 | 4.673341871 | 0.5986863 | 0.4501517 | 0.752913517 |
| ENSMUSG00000044770 | Scml4 | protein_coding | 0.453395504 | 3.261107086 | 0.5981103 | 0.4503658 | 0.753154558 |
| ENSMUSG00000020435 | Osbp2 | protein_coding | -0.432969694 | 3.854302169 | 0.5974051 | 0.4506283 | 0.753476258 |
| ENSMUSG00000040430 | Pitpnc1 | protein_coding | -0.223588155 | 7.311017842 | 0.5970651 | 0.4507549 | 0.753515356 |
| ENSMUSG00000022000 | Zc3h13 | protein_coding | -0.266013553 | 5.432840269 | 0.596966 | 0.4507918 | 0.753515356 |
| ENSMUSG00000041351 | Rap1gap | protein_coding | 0.26847687 | 6.109828099 | 0.5962788 | 0.4510479 | 0.753751214 |
| ENSMUSG00000023909 | Paqr4 | protein_coding | 0.354047805 | 4.688858335 | 0.5960489 | 0.4511337 | 0.753751214 |
| ENSMUSG00000023169 | Slc38a1 | protein_coding | 0.199403605 | 7.064916432 | 0.5960233 | 0.4511432 | 0.753751214 |
| ENSMUSG00000019889 | Ptprk | protein_coding | -0.488005691 | 3.102611883 | 0.5954111 | 0.4513716 | 0.754015728 |
| ENSMUSG00000020576 | Nbas | protein_coding | -0.371832535 | 4.351054025 | 0.5950132 | 0.4515202 | 0.75414682 |
| ENSMUSG00000040174 | Alkbh3 | protein_coding | 0.517105294 | 2.89348082 | 0.5939024 | 0.4519354 | 0.75471972 |
| ENSMUSG00000019066 | Rab3d | protein_coding | 0.905456971 | 2.520365655 | 0.5937684 | 0.4520036 | 0.75471972 |
| ENSMUSG00000034379 | Wdr5b | protein_coding | 0.45815945 | 3.381513871 | 0.5932323 | 0.4521862 | 0.754771355 |
| ENSMUSG00000061455 | Stx17 | protein_coding | -0.305746961 | 4.505653527 | 0.5928352 | 0.4523349 | 0.754771355 |
| ENSMUSG00000039536 | Stau1 | protein_coding | 0.26569136 | 5.958777813 | 0.592561 | 0.4524377 | 0.754771355 |
| ENSMUSG00000052798 | Nup107 | protein_coding | 0.297263355 | 4.677629792 | 0.5925084 | 0.4524574 | 0.754771355 |
| ENSMUSG00000086171 | Pcsk2os1 | antisense_RNA | -0.607650964 | 2.510343127 | 0.5924424 | 0.4524821 | 0.754771355 |
| ENSMUSG00000017740 | Slc12a5 | protein_coding | -0.336506254 | 6.468301671 | 0.5950783 | 0.4525924 | 0.754771355 |
| ENSMUSG00000081534 | Slc48a1 | protein_coding | 0.234827592 | 7.330296116 | 0.5917308 | 0.452767 | 0.754771355 |
| ENSMUSG00000024798 | Htr7 | protein_coding | 0.355045093 | 4.349140398 | 0.5916736 | 0.4527704 | 0.754771355 |
| ENSMUSG00000057614 | Gnai1 | protein_coding | -0.178114961 | 6.97412425 | 0.5916332 | 0.4527856 | 0.754771355 |
| ENSMUSG00000036555 | Iqce | protein_coding | 0.379381675 | 4.140155216 | 0.5914801 | 0.4528431 | 0.754771355 |
| ENSMUSG00000030380 | Mzf1 | protein_coding | 0.746094845 | 2.223890863 | 0.591375 | 0.4528825 | 0.754771355 |
| ENSMUSG00000073542 | Cep76 | protein_coding | 0.520495356 | 2.759413771 | 0.5912266 | 0.4529382 | 0.754771355 |
| ENSMUSG00000027194 | Ttc17 | protein_coding | -0.327700745 | 3.704452578 | 0.5911643 | 0.4529616 | 0.754771355 |
| ENSMUSG00000086040 | Wipf3 | protein_coding | -0.582747901 | 2.898000175 | 0.5910162 | 0.4530172 | 0.754771355 |
| ENSMUSG00000032565 | Nudt16 | protein_coding | 0.31316333 | 4.08789586 | 0.5899952 | 0.4534009 | 0.754988295 |
| ENSMUSG00000026259 | Ngef | protein_coding | -0.33575326 | 6.559742315 | 0.5930421 | 0.4534237 | 0.754988295 |
| ENSMUSG00000038375 | Trp53inp2 | protein_coding | -0.253257662 | 5.434666145 | 0.5896633 | 0.4535257 | 0.754988295 |
| ENSMUSG00000039067 | Psmd7 | protein_coding | 0.165147445 | 8.601291436 | 0.5896629 | 0.4535259 | 0.754988295 |
| ENSMUSG00000058793 | Cds2 | protein_coding | -0.182981441 | 7.027360207 | 0.5895908 | 0.453553 | 0.754988295 |
| ENSMUSG00000022018 | Rgcc | protein_coding | -0.58394018 | 3.124808967 | 0.5895491 | 0.4535687 | 0.754988295 |
| ENSMUSG00000028958 | Tmub1 | protein_coding | 0.195111405 | 6.454854854 | 0.5892751 | 0.4536718 | 0.755043077 |
| ENSMUSG00000027312 | Atrn | protein_coding | -0.336882047 | 5.29521595 | 0.5884747 | 0.4539913 | 0.755437672 |
| ENSMUSG00000034042 | Gpbp1l1 | protein_coding | 0.382815759 | 3.696995942 | 0.5880968 | 0.4541158 | 0.755437672 |
| ENSMUSG00000022075 | Rhobtb2 | protein_coding | -0.263222457 | 5.279647874 | 0.5880864 | 0.4541197 | 0.755437672 |
| ENSMUSG00000002307 | Daxx | protein_coding | 0.31840207 | 5.234720564 | 0.5871719 | 0.4544647 | 0.755781703 |
| ENSMUSG00000038178 | Slc43a2 | protein_coding | 0.324253056 | 4.133100289 | 0.5869218 | 0.4545591 | 0.755781703 |
| ENSMUSG00000039470 | Zdhhc2 | protein_coding | -0.284350878 | 5.22588489 | 0.5868494 | 0.4545865 | 0.755781703 |
| ENSMUSG00000020091 | Eif4ebp2 | protein_coding | -0.27434774 | 4.73625418 | 0.5866172 | 0.4546742 | 0.755781703 |
| ENSMUSG00000027560 | Dok5 | protein_coding | 0.2458242 | 5.700823367 | 0.5864727 | 0.4547288 | 0.755781703 |
| ENSMUSG00000006476 | Nsmf | protein_coding | -0.310402694 | 6.355059657 | 0.5881842 | 0.4547779 | 0.755781703 |
| ENSMUSG00000047085 | Lrrc4b | protein_coding | -0.29503353 | 4.831499957 | 0.5860747 | 0.4548792 | 0.755781703 |
| ENSMUSG00000029438 | Bcl7a | protein_coding | -0.294005922 | 5.314470997 | 0.5860494 | 0.4548888 | 0.755781703 |
| ENSMUSG00000019888 | Mgat4c | protein_coding | -0.345033566 | 4.655284636 | 0.5856245 | 0.4550495 | 0.755829744 |
| ENSMUSG00000024740 | Ddb1 | protein_coding | 0.272009556 | 7.069296325 | 0.5867276 | 0.4550894 | 0.755829744 |

| ensembl_gene_id | external_gene_name | gene_biotype | log2 (fold change) | log2 (counts per million) | F-value | p-value | False discovery rate |
|---|---|---|---|---|---|---|---|
| ENSMUSG00000030532 | Hddc3 | protein_coding | 0.372494368 | 5.144441123 | 0.5855364 | 0.4551286 | 0.755829744 |
| ENSMUSG00000037656 | Slc20a2 | protein_coding | -0.319047525 | 4.918344931 | 0.5848944 | 0.4553258 | 0.756040564 |
| ENSMUSG00000021671 | Poc5 | protein_coding | 0.301919068 | 4.472090136 | 0.5843494 | 0.4555323 | 0.756223643 |
| ENSMUSG00000030401 | Rtn2 | protein_coding | 0.286898465 | 5.087100593 | 0.5840221 | 0.4556564 | 0.756223643 |
| ENSMUSG00000017707 | Serinc3 | protein_coding | 0.226933251 | 6.07841394 | 0.5837868 | 0.4557456 | 0.756223643 |
| ENSMUSG00000068036 | Afdn | protein_coding | -0.297317156 | 5.070946904 | 0.5834154 | 0.4558865 | 0.756223643 |
| ENSMUSG00000040843 | Tiprl | protein_coding | 0.197297429 | 7.344423892 | 0.5833961 | 0.4558938 | 0.756223643 |
| ENSMUSG00000001098 | Kctd10 | protein_coding | 0.528281831 | 3.615844524 | 0.5833921 | 0.4559132 | 0.756223643 |
| ENSMUSG00000032998 | Foxj3 | protein_coding | 0.265350643 | 5.043641327 | 0.583305 | 0.4559284 | 0.756223643 |
| ENSMUSG00000019813 | Cep57l1 | protein_coding | 0.465545758 | 3.313911245 | 0.5829361 | 0.4560684 | 0.756230036 |
| ENSMUSG00000030410 | Dmwd | protein_coding | 0.228842123 | 6.561215865 | 0.5829243 | 0.4560729 | 0.756230036 |
| ENSMUSG00000027797 | Dclk1 | protein_coding | -0.186076017 | 7.263284849 | 0.5826861 | 0.4561633 | 0.756263409 |
| ENSMUSG00000033009 | Ogfod1 | protein_coding | 0.206901834 | 6.537881971 | 0.5819852 | 0.4564297 | 0.756588323 |
| ENSMUSG00000036850 | Mrpl41 | protein_coding | 0.165851258 | 8.315883781 | 0.5815952 | 0.456578 | 0.756645281 |
| ENSMUSG00000063358 | Mapk1 | protein_coding | -0.216229689 | 8.06731653 | 0.5815248 | 0.4566048 | 0.756645281 |
| ENSMUSG00000034973 | Dopey1 | protein_coding | -0.472438074 | 3.857142208 | 0.5812118 | 0.4567417 | 0.756755568 |
| ENSMUSG00000037013 | Ss18 | protein_coding | -0.373312021 | 4.372798749 | 0.5806581 | 0.4569347 | 0.75680202 |
| ENSMUSG00000078566 | Bnip3 | protein_coding | -0.260246835 | 5.546691871 | 0.5804241 | 0.4570239 | 0.75680202 |
| ENSMUSG00000030838 | Ush1c | protein_coding | 0.973602436 | 2.411824723 | 0.581293 | 0.4570297 | 0.75680202 |
| ENSMUSG00000031548 | Sfrp1 | protein_coding | -0.689124522 | 2.630183538 | 0.5803409 | 0.4570556 | 0.75680202 |
| ENSMUSG00000066151 | Fkbp15 | protein_coding | -0.279329825 | 4.607863867 | 0.5800143 | 0.4571801 | 0.75680202 |
| ENSMUSG00000017390 | Aldoc | protein_coding | 0.188737296 | 10.98811899 | 0.5799829 | 0.457192 | 0.75680202 |
| ENSMUSG00000029613 | Eif2ak1 | protein_coding | 0.289026669 | 4.672484427 | 0.5795841 | 0.4573441 | 0.756913531 |
| ENSMUSG00000052544 | St6galnac3 | protein_coding | -0.379645108 | 3.529795754 | 0.5794023 | 0.4574135 | 0.756913531 |
| ENSMUSG00000068732 | Tmem167b | protein_coding | 0.309413215 | 5.617912655 | 0.5792993 | 0.4574706 | 0.756913531 |
| ENSMUSG00000032883 | Acsl3 | protein_coding | -0.258128224 | 5.964220527 | 0.5786654 | 0.4576948 | 0.757135584 |
| ENSMUSG00000027187 | Cat | protein_coding | 0.279239064 | 5.842333269 | 0.5785323 | 0.4577456 | 0.757135584 |
| ENSMUSG00000021068 | Nin | protein_coding | 0.449212408 | 3.182937395 | 0.5781755 | 0.4578819 | 0.757244618 |
| ENSMUSG00000039100 | 6-Mar | protein_coding | -0.225500627 | 6.267039497 | 0.5777154 | 0.4580578 | 0.757318791 |
| ENSMUSG00000038637 | Lrrc56 | protein_coding | 0.597752411 | 2.744091276 | 0.5775117 | 0.4581357 | 0.757318791 |
| ENSMUSG00000075033 | Nxpe3 | protein_coding | 0.453608343 | 3.370713129 | 0.5772321 | 0.4582427 | 0.757318791 |
| ENSMUSG00000006717 | Acot13 | protein_coding | 0.1904454 | 8.428009377 | 0.5772208 | 0.4582471 | 0.757318791 |
| ENSMUSG00000005034 | Prkacb | protein_coding | 0.202166724 | 8.785418905 | 0.5770621 | 0.4583078 | 0.757318791 |
| ENSMUSG00000024855 | Pacs1 | protein_coding | 0.258875252 | 5.570134098 | 0.5765384 | 0.4585084 | 0.757318791 |
| ENSMUSG00000048921 | Zfp689 | protein_coding | 0.395524057 | 3.831255787 | 0.5764244 | 0.458552 | 0.757318791 |
| ENSMUSG00000043445 | Pgp | protein_coding | 0.299738118 | 4.355717368 | 0.576048 | 0.4586963 | 0.757318791 |
| ENSMUSG00000026554 | Dcaf8 | protein_coding | 0.243748665 | 7.101564655 | 0.5759797 | 0.4587402 | 0.757318791 |
| ENSMUSG00000032418 | Me1 | protein_coding | -0.305381526 | 4.417696933 | 0.5759114 | 0.4587486 | 0.757318791 |
| ENSMUSG00000022292 | Rrm2b | protein_coding | -0.288771523 | 4.879641011 | 0.5758639 | 0.4587668 | 0.757318791 |
| ENSMUSG00000021589 | Rhobtb3 | protein_coding | -0.377111551 | 3.888652789 | 0.5758092 | 0.4587878 | 0.757318791 |
| ENSMUSG00000002010 | Idh3g | protein_coding | 0.171965638 | 8.415720531 | 0.5756515 | 0.4588483 | 0.757318791 |
| ENSMUSG00000071654 | Uqcc3 | protein_coding | -0.220327895 | 6.871597521 | 0.5754833 | 0.4589128 | 0.757318791 |
| ENSMUSG00000016921 | Srsf6 | protein_coding | -0.316081045 | 5.095380755 | 0.5752983 | 0.4589838 | 0.757319743 |
| ENSMUSG00000022261 | Sdc2 | protein_coding | -0.220301462 | 6.526697747 | 0.5747871 | 0.45918 | 0.757407792 |
| ENSMUSG00000026788 | Zbtb43 | protein_coding | -0.294186444 | 4.339041362 | 0.5745875 | 0.4592567 | 0.757407792 |
| ENSMUSG00000061315 | Naca | protein_coding | 0.206825145 | 6.815709084 | 0.5745374 | 0.4592759 | 0.757407792 |
| ENSMUSG00000028419 | Chmp5 | protein_coding | 0.168703348 | 8.829841351 | 0.5742675 | 0.4593796 | 0.757407792 |
| ENSMUSG00000002804 | Nudt14 | protein_coding | 0.220667585 | 5.569466343 | 0.5742421 | 0.4593893 | 0.757407792 |
| ENSMUSG00000046750 | Selenov | protein_coding | 0.289971674 | 4.165583799 | 0.5734781 | 0.4596831 | 0.75757938 |
| ENSMUSG00000024414 | Mrpl27 | protein_coding | 0.182466236 | 7.467434165 | 0.5732689 | 0.4597636 | 0.75757938 |
| ENSMUSG00000056201 | Cfl1 | protein_coding | 0.17508585 | 10.34545569 | 0.5728034 | 0.4599428 | 0.75757938 |
| ENSMUSG00000017291 | Taok1 | protein_coding | -0.327957381 | 3.751936543 | 0.5724865 | 0.4600648 | 0.75757938 |
| ENSMUSG00000037531 | Mrpl47 | protein_coding | 0.247665849 | 5.937371966 | 0.5722562 | 0.4601536 | 0.75757938 |
| ENSMUSG00000038295 | Atg9b | protein_coding | 0.339879284 | 4.819288226 | 0.5721234 | 0.4602047 | 0.75757938 |
| ENSMUSG00000032101 | Ddx25 | protein_coding | 0.195212274 | 7.373943566 | 0.5720121 | 0.4602476 | 0.75757938 |
| ENSMUSG00000035783 | Acta2 | protein_coding | -0.620161429 | 3.195979537 | 0.5719647 | 0.4602835 | 0.75757938 |
| ENSMUSG00000037492 | Zmat4 | protein_coding | -0.316215323 | 5.848493678 | 0.5719382 | 0.4603563 | 0.75757938 |
| ENSMUSG00000059173 | Pde1a | protein_coding | -0.218983759 | 6.814877576 | 0.571682 | 0.4603749 | 0.75757938 |
| ENSMUSG00000028833 | Ncdn | protein_coding | 0.199264199 | 8.830321483 | 0.5716793 | 0.4603759 | 0.75757938 |
| ENSMUSG00000028546 | Elavl4 | protein_coding | -0.231526146 | 7.37055083 | 0.5715 | 0.4604627 | 0.75757938 |
| ENSMUSG00000044857 | Lemd2 | protein_coding | -0.504259069 | 3.499210004 | 0.5711494 | 0.4605804 | 0.75757938 |
| ENSMUSG00000005338 | Cadm3 | protein_coding | -0.250847942 | 5.57829691 | 0.5710892 | 0.4606036 | 0.75757938 |
| ENSMUSG00000098659 | 1110015O18Rik | lincRNA | -0.590910432 | 2.736696433 | 0.5709429 | 0.4606601 | 0.75757938 |
| ENSMUSG00000110080 | Gm6145 | processed_transcript | -0.722155263 | 3.08788904 | 0.5725365 | 0.4606885 | 0.75757938 |
| ENSMUSG00000027566 | Psma7 | protein_coding | 0.187608017 | 9.496865509 | 0.5708625 | 0.4606911 | 0.75757938 |
| ENSMUSG00000031432 | Prps1 | protein_coding | 0.166206649 | 7.358218619 | 0.5703763 | 0.4608789 | 0.757772287 |
| ENSMUSG00000029267 | Mtf2 | protein_coding | 0.289526801 | 4.569289192 | 0.569916 | 0.4610568 | 0.757948878 |
| ENSMUSG00000030500 | Slc17a6 | protein_coding | -0.863156853 | 6.462160859 | 0.5996996 | 0.461282 | 0.758203133 |
| ENSMUSG00000022468 | Endou | protein_coding | -0.99348303 | 3.049065569 | 0.5738086 | 0.4616846 | 0.758748917 |
| ENSMUSG00000074793 | Hspa12b | protein_coding | 0.525634206 | 2.950965648 | 0.5676583 | 0.461931 | 0.758932149 |
| ENSMUSG00000097717 | 1700066J03Rik | lincRNA | -0.569768238 | 3.00396811 | 0.5675137 | 0.461987 | 0.758932149 |
| ENSMUSG00000004462 | Tbccd1 | protein_coding | 0.36870779 | 4.312858335 | 0.5673377 | 0.4620553 | 0.758932149 |
| ENSMUSG00000021131 | Erh | protein_coding | 0.354012621 | 3.646676036 | 0.5669863 | 0.4621917 | 0.758932149 |
| ENSMUSG00000041658 | Rragb | protein_coding | 0.692402641 | 5.215019542 | 0.5806013 | 0.4622149 | 0.758932149 |
| ENSMUSG00000033569 | Adgrb3 | protein_coding | -0.288230336 | 5.961160046 | 0.5666639 | 0.4623344 | 0.758932149 |
| ENSMUSG00000029007 | Agtrap | protein_coding | 0.33754331 | 4.356498875 | 0.5665994 | 0.4623419 | 0.758932149 |
| ENSMUSG00000042385 | Gzmk | protein_coding | -0.818051776 | 2.587285795 | 0.5665214 | 0.4623898 | 0.758932149 |
| ENSMUSG00000032463 | Faim | protein_coding | 0.287091632 | 4.670042742 | 0.5662242 | 0.4624876 | 0.758932149 |

| ensembl_gene_id | external_gene_name | gene_biotype | log2 (fold change) | log2 (counts per million) | F-value | p-value | False discovery rate |
|---|---|---|---|---|---|---|---|
| ENSMUSG00000087056 | Gm14004 | processed_transcript | 0.319647948 | 5.122867176 | 0.5657456 | 0.4626737 | 0.758932149 |
| ENSMUSG00000063273 | Naa15 | protein_coding | -0.291162834 | 5.168552921 | 0.5657416 | 0.4626753 | 0.758932149 |
| ENSMUSG00000027333 | Smox | protein_coding | -0.272301211 | 4.904103028 | 0.5653934 | 0.4628107 | 0.758932149 |
| ENSMUSG00000038013 | Wipf2 | protein_coding | -0.192867762 | 6.463170301 | 0.5651678 | 0.4628985 | 0.758932149 |
| ENSMUSG00000030029 | Lrig1 | protein_coding | 0.439794567 | 3.127578595 | 0.5650748 | 0.4629347 | 0.758932149 |
| ENSMUSG00000029366 | Dck | protein_coding | -0.304949867 | 4.599533717 | 0.564943 | 0.462986 | 0.758932149 |
| ENSMUSG00000042816 | Gpr151 | protein_coding | -1.456020704 | 2.063609126 | 0.5685128 | 0.4630996 | 0.758932149 |
| ENSMUSG00000018287 | Spag7 | protein_coding | 0.157160528 | 8.922908867 | 0.5646479 | 0.4631009 | 0.758932149 |
| ENSMUSG00000052273 | Dnah3 | protein_coding | -0.625478047 | 2.341484185 | 0.5646013 | 0.463119 | 0.758932149 |
| ENSMUSG00000039285 | Azi2 | protein_coding | -0.253459928 | 5.542731768 | 0.5645549 | 0.4631371 | 0.758932149 |
| ENSMUSG00000028848 | Gpn2 | protein_coding | -0.529947563 | 2.981742726 | 0.5638382 | 0.4634164 | 0.75927124 |
| ENSMUSG00000039913 | Pak7 | protein_coding | -0.270994289 | 4.788729336 | 0.5636616 | 0.4634852 | 0.75927124 |
| ENSMUSG00000055409 | Nell1 | protein_coding | 0.19729132 | 5.860489754 | 0.5624847 | 0.4639446 | 0.75962137 |
| ENSMUSG00000038296 | Galnt18 | protein_coding | 0.314758416 | 4.65128594 | 0.5624506 | 0.4639579 | 0.75962137 |
| ENSMUSG00000025466 | Fuom | protein_coding | 0.201220891 | 7.227118524 | 0.5624488 | 0.4639586 | 0.75962137 |
| ENSMUSG00000037784 | Dzip1l | protein_coding | -0.662579071 | 2.5623487 | 0.5623901 | 0.4639815 | 0.75962137 |
| ENSMUSG00000087006 | Gm13889 | protein_coding | 0.391382545 | 3.586865845 | 0.5608252 | 0.4645936 | 0.760353512 |
| ENSMUSG00000031594 | Fgl1 | protein_coding | -0.635606153 | 2.721858819 | 0.5607184 | 0.4646354 | 0.760353512 |
| ENSMUSG00000026728 | Vim | protein_coding | -0.386391202 | 4.265749196 | 0.5607044 | 0.4646408 | 0.760353512 |
| ENSMUSG00000037815 | Ctnna1 | protein_coding | -0.243377771 | 5.880425681 | 0.5602239 | 0.4648291 | 0.760533581 |
| ENSMUSG00000059486 | Kbtbd2 | protein_coding | -0.24480639 | 5.815664921 | 0.5600625 | 0.4648923 | 0.760533581 |
| ENSMUSG00000045071 | E130308A19Rik | protein_coding | 0.425117204 | 3.44162894 | 0.5595419 | 0.4650965 | 0.760684068 |
| ENSMUSG00000025451 | Paip1 | protein_coding | -0.348050224 | 8.314837663 | 0.5633062 | 0.4653215 | 0.760684068 |
| ENSMUSG00000037977 | 6430571L13Rik | protein_coding | 0.482795623 | 3.498634964 | 0.5589591 | 0.4653251 | 0.760684068 |
| ENSMUSG00000038618 | Rassf7 | protein_coding | 0.491223783 | 3.373185711 | 0.5587629 | 0.4654022 | 0.760684068 |
| ENSMUSG00000033106 | Slc7a6os | protein_coding | 0.461770253 | 3.552645987 | 0.5587437 | 0.4654097 | 0.760684068 |
| ENSMUSG00000099689 | Zfp383 | protein_coding | -0.431229419 | 3.063736148 | 0.5581505 | 0.4656428 | 0.760684068 |
| ENSMUSG00000051671 | Coa6 | protein_coding | 0.202935332 | 6.509257988 | 0.5579667 | 0.465715 | 0.760684068 |
| ENSMUSG00000022350 | Washc5 | protein_coding | 0.283736692 | 4.438826356 | 0.5579325 | 0.4657284 | 0.760684068 |
| ENSMUSG00000047907 | Tshz2 | protein_coding | -0.291343437 | 4.067112296 | 0.557435 | 0.4659241 | 0.760684068 |
| ENSMUSG00000037166 | Ppp1r14a | protein_coding | -0.42705394 | 3.985494082 | 0.557043 | 0.4660957 | 0.760684068 |
| ENSMUSG00000028100 | Nudt17 | protein_coding | 0.393857648 | 4.793880109 | 0.5571054 | 0.4661371 | 0.760684068 |
| ENSMUSG00000038503 | Mesd | protein_coding | 0.194888109 | 6.570156497 | 0.5567939 | 0.4661764 | 0.760684068 |
| ENSMUSG00000020952 | Scfd1 | protein_coding | -0.18827468 | 6.245587856 | 0.5566995 | 0.4662136 | 0.760684068 |
| ENSMUSG00000047036 | Zfp445 | protein_coding | -0.295693693 | 4.849598076 | 0.556668 | 0.466226 | 0.760684068 |
| ENSMUSG00000020704 | Asic2 | protein_coding | 0.199128802 | 5.867010755 | 0.5565825 | 0.4662597 | 0.760684068 |
| ENSMUSG00000057265 | Bbof1 | protein_coding | -0.725599898 | 2.443925944 | 0.5563743 | 0.4663417 | 0.760684068 |
| ENSMUSG00000021365 | Nedd9 | protein_coding | -0.667380018 | 2.474451511 | 0.5561528 | 0.466429 | 0.760684068 |
| ENSMUSG00000038916 | Soga3 | protein_coding | 0.282740075 | 5.687258894 | 0.5561169 | 0.4664431 | 0.760684068 |
| ENSMUSG00000031709 | Tbc1d9 | protein_coding | -0.263903846 | 4.883991709 | 0.5559981 | 0.4664899 | 0.760684068 |
| ENSMUSG00000043257 | Pigv | protein_coding | -0.580343878 | 2.814847498 | 0.5559711 | 0.4665006 | 0.760684068 |
| ENSMUSG00000002346 | Slc25a42 | protein_coding | 0.392641336 | 4.156457264 | 0.5559133 | 0.4665234 | 0.760684068 |
| ENSMUSG00000024566 | Atp9b | protein_coding | -0.341214365 | 3.993432612 | 0.5558695 | 0.4665406 | 0.760684068 |
| ENSMUSG00000025145 | Lrrc45 | protein_coding | -0.471139171 | 3.403219882 | 0.5547674 | 0.4669755 | 0.761204466 |
| ENSMUSG00000027875 | Hmgcs2 | protein_coding | 0.303901291 | 4.074244339 | 0.5547019 | 0.4670014 | 0.761204466 |
| ENSMUSG00000022707 | Gbe1 | protein_coding | -0.417605552 | 3.4421343 | 0.5545117 | 0.4670765 | 0.761211496 |
| ENSMUSG00000026116 | Tmem131 | protein_coding | -0.336499675 | 3.967843613 | 0.5540866 | 0.4672445 | 0.761369889 |
| ENSMUSG00000034768 | Asb16 | protein_coding | -0.479440204 | 2.949665346 | 0.5537144 | 0.4673916 | 0.761494261 |
| ENSMUSG00000010097 | Nxf1 | protein_coding | 0.158373856 | 7.710549011 | 0.5531712 | 0.4676066 | 0.761646881 |
| ENSMUSG00000021102 | Glrx5 | protein_coding | 0.327104112 | 5.001211171 | 0.5529532 | 0.4676928 | 0.761646881 |
| ENSMUSG00000020115 | Tbk1 | protein_coding | 0.299253534 | 4.763770872 | 0.5529407 | 0.4676978 | 0.761646881 |
| ENSMUSG00000045757 | Zfp764 | protein_coding | -0.984227455 | 2.131660146 | 0.5525997 | 0.4678501 | 0.761779599 |
| ENSMUSG00000021546 | Hnrnpk | protein_coding | -0.200364596 | 8.311756513 | 0.5521128 | 0.4680257 | 0.761950165 |
| ENSMUSG00000024026 | Glo1 | protein_coding | -0.165094539 | 7.490100632 | 0.5519255 | 0.4681 | 0.761955726 |
| ENSMUSG00000001473 | Tubb6 | protein_coding | -0.575782004 | 2.699897975 | 0.5516524 | 0.4682083 | 0.762016624 |
| ENSMUSG00000024091 | Vapa | protein_coding | -0.15789085 | 8.549084186 | 0.5510449 | 0.4684494 | 0.762257061 |
| ENSMUSG00000001415 | Smg5 | protein_coding | 0.271017114 | 4.749983484 | 0.5508306 | 0.4685345 | 0.762257061 |
| ENSMUSG00000029386 | Tctn2 | processed_transcript | -0.27253471 | 4.322253723 | 0.5506793 | 0.4685945 | 0.762257061 |
| ENSMUSG00000029478 | Ncor2 | protein_coding | -0.383221896 | 3.755960441 | 0.5505659 | 0.4686396 | 0.762257061 |
| ENSMUSG00000037463 | Fbxo27 | protein_coding | -0.574005973 | 2.454209972 | 0.5500824 | 0.4688317 | 0.762454258 |
| ENSMUSG00000045193 | Cirbp | protein_coding | 0.173395153 | 9.040230803 | 0.54923 | 0.4691708 | 0.762569542 |
| ENSMUSG00000019890 | Nts | protein_coding | 0.475760133 | 7.780597208 | 0.5628738 | 0.4691895 | 0.762569542 |
| ENSMUSG00000033735 | Spr | protein_coding | 0.369709015 | 5.109772652 | 0.5493166 | 0.469199 | 0.762569542 |
| ENSMUSG00000074797 | Itpa | protein_coding | 0.182123897 | 6.312542748 | 0.5489291 | 0.4692906 | 0.762569542 |
| ENSMUSG00000087184 | Gm11650 | processed_transcript | 0.619717603 | 2.571897185 | 0.5489266 | 0.4692915 | 0.762569542 |
| ENSMUSG00000024187 | Fam234a | protein_coding | -0.397798451 | 4.779985029 | 0.5490671 | 0.4693281 | 0.762569542 |
| ENSMUSG00000027597 | Ahcy | protein_coding | 0.351202386 | 3.452462373 | 0.5484533 | 0.4694801 | 0.762701215 |
| ENSMUSG00000030757 | Zkscan2 | protein_coding | -0.604863486 | 2.533081149 | 0.5479168 | 0.4696939 | 0.76293334 |
| ENSMUSG00000029499 | Pxmp2 | protein_coding | -0.381672536 | 3.719381021 | 0.5476565 | 0.4697977 | 0.762986693 |
| ENSMUSG00000026281 | Dtymk | protein_coding | 0.233542093 | 6.249767161 | 0.5473772 | 0.4699092 | 0.763023608 |
| ENSMUSG00000021892 | Sh3bp5 | protein_coding | -0.293233505 | 4.685489665 | 0.5471191 | 0.4700122 | 0.763023608 |
| ENSMUSG00000010517 | Faf1 | protein_coding | 0.294337931 | 5.091484325 | 0.5470661 | 0.4700333 | 0.763023608 |
| ENSMUSG00000071054 | Safb | protein_coding | 0.29313219 | 4.510427583 | 0.5457505 | 0.4705591 | 0.763666174 |
| ENSMUSG00000058638 | Zfp110 | protein_coding | -0.441130662 | 3.539110823 | 0.5456901 | 0.4705832 | 0.763666174 |
| ENSMUSG00000040250 | Ints13 | protein_coding | 0.365561266 | 3.385165044 | 0.5453262 | 0.4707288 | 0.763666174 |
| ENSMUSG00000029169 | Dhx15 | protein_coding | -0.247757357 | 6.032586644 | 0.5451273 | 0.4708084 | 0.763666174 |
| ENSMUSG00000049281 | Scn3b | protein_coding | 0.183152965 | 6.319144643 | 0.5449768 | 0.4708687 | 0.763666174 |
| ENSMUSG00000037210 | Fam193a | protein_coding | 0.32066529 | 4.910194379 | 0.5448766 | 0.4709088 | 0.763666174 |

| ensembl_gene_id | external_gene_name | gene_biotype | log2 (fold change) | log2 (counts per million) | F-value | p-value | False discovery rate |
|---|---|---|---|---|---|---|---|
| ENSMUSG00000068263 | Efcc1 | protein_coding | -0.359625094 | 3.664385717 | 0.544833 | 0.4709263 | 0.763666174 |
| ENSMUSG00000052105 | Mtcl1 | protein_coding | -0.421712383 | 3.993371631 | 0.5436625 | 0.4713955 | 0.764251346 |
| ENSMUSG00000068740 | Celsr2 | protein_coding | -0.268062298 | 5.933846653 | 0.5435207 | 0.4714524 | 0.764251346 |
| ENSMUSG00000025532 | Crcp | protein_coding | 0.17810325 | 7.105217199 | 0.5434011 | 0.4715004 | 0.764251346 |
| ENSMUSG00000022661 | Cd200 | protein_coding | 0.199149459 | 8.070795326 | 0.5430862 | 0.4716268 | 0.764341042 |
| ENSMUSG00000057522 | Spop | protein_coding | -0.205185629 | 8.272857424 | 0.5427989 | 0.4717422 | 0.764412824 |
| ENSMUSG00000045174 | Amer3 | protein_coding | -0.402290955 | 4.274527975 | 0.5410616 | 0.472458 | 0.765296341 |
| ENSMUSG00000020407 | Upp1 | protein_coding | 0.581065814 | 2.819865986 | 0.5409977 | 0.4724666 | 0.765296341 |
| ENSMUSG00000096199 | Ptrhd1 | protein_coding | 0.247928439 | 5.25331376 | 0.5405956 | 0.4726286 | 0.765296341 |
| ENSMUSG00000038740 | Mvb12b | protein_coding | 0.29513658 | 4.934710172 | 0.540499 | 0.4726675 | 0.765296341 |
| ENSMUSG00000037339 | Fam53a | protein_coding | 0.533990277 | 3.00363387 | 0.5403401 | 0.4727315 | 0.765296341 |
| ENSMUSG00000063760 | Rnf217 | protein_coding | -0.386195266 | 3.511575161 | 0.5400462 | 0.4728501 | 0.765296341 |
| ENSMUSG00000091994 | E130317F20Rik | bidirectional_promote | 0.608254198 | 2.538111552 | 0.539947 | 0.4728901 | 0.765296341 |
| ENSMUSG00000060424 | Pantr1 | lincRNA | -0.385026322 | 7.07974581 | 0.5468567 | 0.4730478 | 0.765296341 |
| ENSMUSG00000032939 | Nup93 | protein_coding | 0.285050057 | 4.417205493 | 0.5395066 | 0.4730677 | 0.765296341 |
| ENSMUSG00000033007 | Asic4 | protein_coding | 0.874494418 | 2.312963814 | 0.5395061 | 0.473085 | 0.765296341 |
| ENSMUSG00000043683 | Fem1a | protein_coding | -0.292346152 | 4.919768379 | 0.5393461 | 0.4731325 | 0.765296341 |
| ENSMUSG00000040373 | Cacng5 | protein_coding | -0.324677197 | 4.872499501 | 0.539324 | 0.4731415 | 0.765296341 |
| ENSMUSG00000045671 | Spred2 | protein_coding | -0.355244841 | 4.025440524 | 0.5390986 | 0.4732324 | 0.765328369 |
| ENSMUSG00000063849 | Ppcdc | protein_coding | 0.237183339 | 5.543887898 | 0.5388379 | 0.4733378 | 0.765365268 |
| ENSMUSG00000003680 | Taf6l | protein_coding | 0.381435389 | 4.353004743 | 0.5386897 | 0.4733976 | 0.765365268 |
| ENSMUSG00000001674 | Ddx18 | protein_coding | 0.28492594 | 4.228747452 | 0.5381583 | 0.4736124 | 0.765597454 |
| ENSMUSG00000052397 | Ezr | protein_coding | -0.331298897 | 4.466363293 | 0.5376508 | 0.4738177 | 0.765748992 |
| ENSMUSG00000075284 | Wipf1 | protein_coding | -0.428293944 | 3.142378486 | 0.5374566 | 0.4738963 | 0.765748992 |
| ENSMUSG00000028099 | Polr3c | protein_coding | 0.30812358 | 5.569447924 | 0.5374049 | 0.4739343 | 0.765748992 |
| ENSMUSG00000020818 | Mfsd11 | protein_coding | 0.23716548 | 5.251139658 | 0.5372227 | 0.473991 | 0.765748992 |
| ENSMUSG00000033808 | Tmem87a | protein_coding | -0.602016578 | 2.727547027 | 0.5365579 | 0.4742603 | 0.766068951 |
| ENSMUSG00000037190 | Cyb561d2 | protein_coding | 0.205886048 | 6.107873697 | 0.5359305 | 0.4745147 | 0.766364758 |
| ENSMUSG00000035297 | Cops4 | protein_coding | 0.16249894 | 8.429446949 | 0.5353673 | 0.4747433 | 0.7666188 |
| ENSMUSG00000046861 | Hectd3 | protein_coding | -0.34399498 | 4.242507124 | 0.5350951 | 0.4748538 | 0.766651097 |
| ENSMUSG00000074607 | Tox2 | protein_coding | 0.37207 | 5.997810365 | 0.5383582 | 0.474916 | 0.766651097 |
| ENSMUSG00000095620 | 2010005H15Rik | protein_coding | 0.449782668 | 3.44310427 | 0.5347432 | 0.4749968 | 0.766651097 |
| ENSMUSG00000021076 | Actr10 | protein_coding | 0.176249667 | 8.860257403 | 0.5345835 | 0.4750617 | 0.766651097 |
| ENSMUSG00000022358 | Fbxo32 | protein_coding | -0.447821774 | 3.737421218 | 0.5341551 | 0.4752359 | 0.766651097 |
| ENSMUSG00000033763 | Mtss1l | protein_coding | -0.22127129 | 5.913357951 | 0.5340849 | 0.4752644 | 0.766651097 |
| ENSMUSG00000035840 | Lysmd3 | protein_coding | -0.348042489 | 3.98831712 | 0.5340551 | 0.4752766 | 0.766651097 |
| ENSMUSG00000021684 | Pde8b | protein_coding | 0.281188325 | 5.163452221 | 0.5339147 | 0.4753337 | 0.766651097 |
| ENSMUSG00000014959 | Gorasp2 | protein_coding | 0.175642521 | 6.525024979 | 0.5331759 | 0.4756344 | 0.76702112 |
| ENSMUSG00000028016 | Ints12 | protein_coding | -0.256117946 | 4.737708314 | 0.5327305 | 0.4758159 | 0.767138412 |
| ENSMUSG00000024539 | Ptpn2 | protein_coding | -0.288576301 | 4.713151037 | 0.5324505 | 0.47593 | 0.767138412 |
| ENSMUSG00000052087 | Rgs14 | protein_coding | -0.271982844 | 5.188153035 | 0.5323001 | 0.4759914 | 0.767138412 |
| ENSMUSG00000067199 | Frat1 | protein_coding | 0.271347167 | 7.740331066 | 0.5339123 | 0.4759925 | 0.767138412 |
| ENSMUSG00000031551 | Ido1 | protein_coding | -0.37683558 | 3.973522036 | 0.531983 | 0.4761207 | 0.767183254 |
| ENSMUSG00000037010 | Apln | protein_coding | 0.429815447 | 3.299783963 | 0.5318425 | 0.476178 | 0.767183254 |
| ENSMUSG00000073705 | Apitd1 | protein_coding | 0.294865927 | 4.583008262 | 0.5314723 | 0.4763291 | 0.767183254 |
| ENSMUSG00000034263 | Ints14 | protein_coding | 0.291649897 | 4.562746553 | 0.5314273 | 0.4763475 | 0.767183254 |
| ENSMUSG00000074918 | Inafm2 | protein_coding | -0.216486081 | 7.812709367 | 0.5312415 | 0.4764234 | 0.767183254 |
| ENSMUSG00000043441 | Gpr149 | protein_coding | 0.414360463 | 3.185985951 | 0.5311803 | 0.4764484 | 0.767183254 |
| ENSMUSG00000023147 | Wrb | protein_coding | 0.161173839 | 8.827525129 | 0.5307321 | 0.4766315 | 0.767363218 |
| ENSMUSG00000020611 | Gna13 | protein_coding | 0.247130069 | 5.400920887 | 0.5302155 | 0.4768428 | 0.767572359 |
| ENSMUSG00000031749 | St3gal2 | protein_coding | -0.247239984 | 5.079419444 | 0.529857 | 0.4769895 | 0.767572359 |
| ENSMUSG00000041203 | Trir | protein_coding | 0.296469602 | 4.336467588 | 0.5298149 | 0.4770067 | 0.767572359 |
| ENSMUSG00000031540 | Kat6a | protein_coding | -0.330047768 | 4.057204075 | 0.5296016 | 0.477094 | 0.767572359 |
| ENSMUSG00000028610 | Dmrtb1 | protein_coding | -0.680897545 | 3.322500982 | 0.5319265 | 0.4771724 | 0.767572359 |
| ENSMUSG00000032355 | Mlip | protein_coding | -0.325882626 | 4.229628769 | 0.5290505 | 0.4773197 | 0.767572359 |
| ENSMUSG00000028284 | Map3k7 | protein_coding | -0.332942183 | 3.835597982 | 0.5289884 | 0.4773452 | 0.767572359 |
| ENSMUSG00000038312 | Edem2 | protein_coding | -0.44844795 | 4.458289532 | 0.5304208 | 0.4774142 | 0.767572359 |
| ENSMUSG00000049439 | Cyp20a1 | protein_coding | 0.32856312 | 4.164389823 | 0.5286435 | 0.4774866 | 0.767572359 |
| ENSMUSG00000028654 | Mycl | protein_coding | 0.514173414 | 2.9758431 | 0.5285898 | 0.4775086 | 0.767572359 |
| ENSMUSG00000022864 | D16Ertd472e | protein_coding | 0.707151716 | 2.559572326 | 0.5284969 | 0.4775466 | 0.767572359 |
| ENSMUSG00000034801 | Sos2 | protein_coding | 0.257997238 | 4.382323573 | 0.5281489 | 0.4776894 | 0.767687046 |
| ENSMUSG00000038696 | Mapkap1 | protein_coding | -0.268846442 | 5.030753625 | 0.5277978 | 0.4778335 | 0.767770517 |
| ENSMUSG00000015542 | Nat9 | protein_coding | 0.237606882 | 5.966383651 | 0.5275839 | 0.4779213 | 0.767770517 |
| ENSMUSG00000034826 | Nup54 | protein_coding | 0.434411502 | 3.960240692 | 0.5275416 | 0.4779555 | 0.767770517 |
| ENSMUSG00000024897 | Apba1 | protein_coding | -0.247008367 | 5.352296882 | 0.5270181 | 0.4781537 | 0.767900024 |
| ENSMUSG00000031796 | Cfap20 | protein_coding | 0.267974567 | 6.061019757 | 0.5269377 | 0.4782036 | 0.767900024 |
| ENSMUSG00000096188 | Cmtm4 | protein_coding | 0.342421897 | 4.718665735 | 0.5267829 | 0.4782504 | 0.767900024 |
| ENSMUSG00000016552 | Foxred2 | protein_coding | 0.521237602 | 2.864188343 | 0.5262624 | 0.4784645 | 0.768129088 |
| ENSMUSG00000030616 | Sytl2 | protein_coding | -0.261074703 | 4.494897183 | 0.5254196 | 0.4788114 | 0.768497222 |
| ENSMUSG00000041119 | Pde9a | protein_coding | 0.272423297 | 5.060572639 | 0.5253582 | 0.4788367 | 0.768497222 |
| ENSMUSG00000037300 | Ttc13 | protein_coding | -0.487492306 | 3.753155692 | 0.5248581 | 0.4791039 | 0.768757143 |
| ENSMUSG00000006423 | C330007P06Rik | protein_coding | -0.303815569 | 4.447876819 | 0.5244607 | 0.4792067 | 0.768757143 |
| ENSMUSG00000034663 | Bmp2k | protein_coding | -0.54531022 | 2.757014937 | 0.5244451 | 0.4792132 | 0.768757143 |
| ENSMUSG00000037685 | Atp8a1 | protein_coding | 0.383188402 | 3.502410305 | 0.5240806 | 0.4793635 | 0.768883676 |
| ENSMUSG00000019868 | Vta1 | protein_coding | 0.172543787 | 6.910586962 | 0.5235339 | 0.4795893 | 0.769131023 |
| ENSMUSG00000085316 | D330050G23Rik | lincRNA | -0.72230195 | 2.556195154 | 0.5224643 | 0.4800315 | 0.769539562 |
| ENSMUSG00000023992 | Trem2 | protein_coding | -0.354408853 | 4.016970492 | 0.5224634 | 0.4800318 | 0.769539562 |
| ENSMUSG00000045435 | Tmem60 | protein_coding | -0.213888794 | 7.072074073 | 0.522209 | 0.4801371 | 0.769539562 |

| ensembl_gene_id | external_gene_name | gene_biotype | log2 (fold change) | log2 (counts per million) | F-value | p-value | False discovery rate |
|---|---|---|---|---|---|---|---|
| ENSMUSG00000061482 | Hist1h4d | protein_coding | -0.310640889 | 3.765522318 | 0.5221235 | 0.4801725 | 0.769539562 |
| ENSMUSG00000070284 | Gmppb | protein_coding | 0.312654361 | 4.677332472 | 0.5220526 | 0.4802018 | 0.769539562 |
| ENSMUSG00000001983 | Taco1 | protein_coding | 0.381042141 | 3.448332275 | 0.5218438 | 0.4802883 | 0.769563386 |
| ENSMUSG00000058070 | Eml1 | protein_coding | -0.423849743 | 4.21443811 | 0.5214224 | 0.4804796 | 0.769672665 |
| ENSMUSG00000038646 | Fam103a1 | protein_coding | -0.17376733 | 6.473152165 | 0.5212086 | 0.4805515 | 0.769672665 |
| ENSMUSG00000021242 | Npc2 | protein_coding | -0.172575186 | 7.42587383 | 0.5211609 | 0.4805712 | 0.769672665 |
| ENSMUSG00000025759 | Mfsd8 | protein_coding | 0.712956531 | 2.851632995 | 0.520441 | 0.4808865 | 0.769950197 |
| ENSMUSG00000004054 | Map3k11 | protein_coding | -0.573476878 | 2.518704053 | 0.5203978 | 0.4808877 | 0.769950197 |
| ENSMUSG00000001100 | Poldip2 | protein_coding | -0.19898225 | 5.884132122 | 0.5197679 | 0.4811493 | 0.770254265 |
| ENSMUSG00000034278 | Dnajc17 | protein_coding | 0.288311772 | 4.320435729 | 0.5186008 | 0.4816344 | 0.77084741 |
| ENSMUSG00000023033 | Scn8a | protein_coding | -0.237971316 | 5.789346112 | 0.5185318 | 0.4816631 | 0.77084741 |
| ENSMUSG00000022779 | Top3b | protein_coding | -0.283393501 | 4.148305104 | 0.5181596 | 0.4818181 | 0.770905947 |
| ENSMUSG00000046722 | Cdc42se1 | protein_coding | -0.232957033 | 5.214550562 | 0.5179738 | 0.4818954 | 0.770905947 |
| ENSMUSG00000038886 | Man2a2 | protein_coding | -0.380823646 | 3.766803742 | 0.5179273 | 0.4819148 | 0.770905947 |
| ENSMUSG00000011096 | Akt1s1 | protein_coding | -0.522285823 | 2.861234585 | 0.5175373 | 0.4820773 | 0.771009349 |
| ENSMUSG00000037735 | 2810032G03Rik | lincRNA | -0.5153552 | 2.874450947 | 0.5173014 | 0.4821756 | 0.771009349 |
| ENSMUSG00000034274 | Thoc5 | protein_coding | 0.258365404 | 5.329991792 | 0.517256 | 0.4821945 | 0.771009349 |
| ENSMUSG00000004791 | Pgf | protein_coding | 0.306481858 | 4.251531182 | 0.5170179 | 0.4822938 | 0.771053471 |
| ENSMUSG00000046519 | Golph3l | protein_coding | -0.326100255 | 4.820095839 | 0.5157266 | 0.4828329 | 0.771743761 |
| ENSMUSG00000029647 | Pan3 | protein_coding | -0.283466278 | 4.615411374 | 0.5156113 | 0.4828811 | 0.771743761 |
| ENSMUSG00000028837 | Psmb2 | protein_coding | 0.162382248 | 8.876893701 | 0.5154469 | 0.4829498 | 0.771743761 |
| ENSMUSG00000046449 | C77370 | protein_coding | -0.302030278 | 4.756877016 | 0.5152964 | 0.4830127 | 0.771743761 |
| ENSMUSG00000030714 | Sgf29 | protein_coding | -0.258549761 | 5.825382955 | 0.5148453 | 0.4832014 | 0.771924867 |
| ENSMUSG00000029330 | Cds1 | protein_coding | -0.281950386 | 4.559815468 | 0.5146822 | 0.4832696 | 0.771924867 |
| ENSMUSG00000029414 | Kntc1 | protein_coding | 0.684956749 | 2.226839101 | 0.5144223 | 0.4833784 | 0.771984016 |
| ENSMUSG00000015217 | Hmgb3 | protein_coding | -0.322018617 | 3.751086296 | 0.5139093 | 0.4835933 | 0.772212436 |
| ENSMUSG00000024063 | Lbh | protein_coding | -0.312934656 | 4.762518515 | 0.5136301 | 0.4837103 | 0.772248182 |
| ENSMUSG00000041763 | Tpp2 | protein_coding | -0.241681661 | 5.151239924 | 0.5134623 | 0.4837806 | 0.772248182 |
| ENSMUSG00000022559 | Fbxl6 | protein_coding | -0.451579068 | 3.018726712 | 0.5131566 | 0.4839088 | 0.772248182 |
| ENSMUSG00000029050 | Ski | protein_coding | -0.290503831 | 5.481192407 | 0.5130649 | 0.4839473 | 0.772248182 |
| ENSMUSG00000024908 | Ppp6r3 | protein_coding | -0.304048053 | 4.469791417 | 0.5129995 | 0.4839748 | 0.772248182 |
| ENSMUSG00000018740 | Slc25a35 | protein_coding | -0.633460846 | 2.782725625 | 0.5127106 | 0.484096 | 0.772284091 |
| ENSMUSG00000032469 | Dbr1 | protein_coding | 0.464444021 | 3.535123546 | 0.5126036 | 0.4841409 | 0.772284091 |
| ENSMUSG00000026925 | Inpp5e | protein_coding | -0.365749672 | 3.412562807 | 0.5118974 | 0.4844375 | 0.772552799 |
| ENSMUSG00000034936 | Arl4d | protein_coding | -0.480588292 | 4.098473613 | 0.5132795 | 0.4845005 | 0.772552799 |
| ENSMUSG00000029343 | Crybb1 | protein_coding | 0.782531401 | 2.065168389 | 0.5114139 | 0.4846408 | 0.772552799 |
| ENSMUSG00000025083 | Afap1l2 | protein_coding | -0.718038594 | 2.378908864 | 0.5113706 | 0.484659 | 0.772552799 |
| ENSMUSG00000004044 | Cavin1 | protein_coding | -0.23516865 | 5.39230093 | 0.5113478 | 0.4846686 | 0.772552799 |
| ENSMUSG00000033161 | Atp1a1 | protein_coding | -0.311125131 | 4.929642025 | 0.5107805 | 0.4849073 | 0.772588739 |
| ENSMUSG00000029730 | Mcm7 | protein_coding | -0.418869615 | 3.548071418 | 0.510749 | 0.4849205 | 0.772588739 |
| ENSMUSG00000029171 | Pgm1 | protein_coding | -0.557855924 | 2.853407313 | 0.5107043 | 0.4849394 | 0.772588739 |
| ENSMUSG00000030254 | Rad18 | protein_coding | -0.416976185 | 3.375191328 | 0.5106113 | 0.4849785 | 0.772588739 |
| ENSMUSG00000007039 | Ddah2 | protein_coding | -0.262819085 | 4.694998761 | 0.5101188 | 0.485186 | 0.77280473 |
| ENSMUSG00000039156 | Stim2 | protein_coding | -0.293698006 | 4.815811894 | 0.5096685 | 0.4853758 | 0.772879077 |
| ENSMUSG00000060152 | Pop5 | protein_coding | 0.2627014 | 5.033105139 | 0.509667 | 0.4853764 | 0.772879077 |
| ENSMUSG00000038738 | Shank1 | protein_coding | -0.29109652 | 5.505999891 | 0.5087369 | 0.4857854 | 0.773144902 |
| ENSMUSG00000068184 | Ndufaf2 | protein_coding | 0.194554357 | 7.34723088 | 0.5086579 | 0.4858022 | 0.773144902 |
| ENSMUSG00000022765 | Snap29 | protein_coding | -0.315215583 | 4.01884437 | 0.5085752 | 0.4858371 | 0.773144902 |
| ENSMUSG00000094786 | Gm14403 | protein_coding | -0.395323756 | 3.592279995 | 0.5085641 | 0.4858418 | 0.773144902 |
| ENSMUSG00000021643 | Serf1 | protein_coding | 0.301531072 | 4.613546209 | 0.5084196 | 0.4859028 | 0.773144902 |
| ENSMUSG00000031516 | Dctn6 | protein_coding | 0.172999611 | 8.032111938 | 0.5079391 | 0.4861059 | 0.773353616 |
| ENSMUSG00000037771 | Slc32a1 | protein_coding | 0.44165205 | 4.616713118 | 0.5091448 | 0.4862369 | 0.773447489 |
| ENSMUSG00000066510 | Ankdd1a | protein_coding | -0.495374639 | 3.148513677 | 0.5071326 | 0.4864471 | 0.773550779 |
| ENSMUSG00000030469 | Zfp719 | protein_coding | -0.360607511 | 3.742661636 | 0.5070459 | 0.4864837 | 0.773550779 |
| ENSMUSG00000039771 | Polr2j | protein_coding | 0.192930331 | 7.22821898 | 0.5069659 | 0.4865176 | 0.773550779 |
| ENSMUSG00000028868 | Wasf2 | protein_coding | -0.269050484 | 4.571624132 | 0.506278 | 0.486809 | 0.773883014 |
| ENSMUSG00000027281 | Slx4ip | protein_coding | -0.360409266 | 3.290612586 | 0.5061329 | 0.4868705 | 0.773883014 |
| ENSMUSG00000038619 | Ensa | protein_coding | 0.169426269 | 9.758406998 | 0.5055307 | 0.4871259 | 0.774070609 |
| ENSMUSG00000042579 | 4632404H12Rik | lincRNA | 0.331289608 | 3.727082463 | 0.5055151 | 0.4871325 | 0.774070609 |
| ENSMUSG00000021895 | Arhgef3 | protein_coding | -0.239107228 | 5.025681311 | 0.5052835 | 0.4872308 | 0.774078195 |
| ENSMUSG00000110841 | AC153368.1 | transcribed_processe | 0.293097639 | 4.00430989 | 0.5051646 | 0.4872813 | 0.774078195 |
| ENSMUSG00000002748 | Baz1b | protein_coding | -0.243288748 | 5.4255037 | 0.5048183 | 0.4874283 | 0.774144447 |
| ENSMUSG00000038170 | Pde4dip | protein_coding | -0.193084286 | 6.043936981 | 0.5044432 | 0.4875877 | 0.774144447 |
| ENSMUSG00000002741 | Ykt6 | protein_coding | 0.26681031 | 5.749168837 | 0.5043826 | 0.4876134 | 0.774144447 |
| ENSMUSG00000020175 | Rab36 | protein_coding | 0.222358226 | 6.417357746 | 0.504163 | 0.4877068 | 0.774144447 |
| ENSMUSG00000000278 | Scpep1 | protein_coding | 0.404790265 | 3.180799759 | 0.5041082 | 0.4877301 | 0.774144447 |
| ENSMUSG00000035067 | Xkr6 | protein_coding | -0.61179626 | 2.614266877 | 0.5040497 | 0.4877549 | 0.774144447 |
| ENSMUSG00000053641 | Dennd4a | protein_coding | -0.299221975 | 4.413775591 | 0.503203 | 0.4881152 | 0.774601962 |
| ENSMUSG00000020211 | Sf3a2 | protein_coding | -0.333496473 | 3.820183956 | 0.5026774 | 0.4883391 | 0.774814076 |
| ENSMUSG00000031628 | Casp3 | protein_coding | -0.50221415 | 3.475953786 | 0.5024338 | 0.4884593 | 0.774814076 |
| ENSMUSG00000024006 | Stk38 | protein_coding | -0.348408429 | 3.52532164 | 0.5023486 | 0.4884792 | 0.774814076 |
| ENSMUSG00000028134 | Ptbp2 | protein_coding | -0.211931658 | 6.239387828 | 0.5022128 | 0.4885371 | 0.774814076 |
| ENSMUSG00000024563 | Smad2 | protein_coding | -0.288425578 | 4.776615974 | 0.5011719 | 0.4889813 | 0.77540419 |
| ENSMUSG00000028469 | Npr2 | protein_coding | 0.42880693 | 3.538515871 | 0.5008601 | 0.4891145 | 0.77550108 |
| ENSMUSG00000020329 | Polrmt | protein_coding | -0.5050683 | 3.309973205 | 0.5002279 | 0.4893847 | 0.77569527 |
| ENSMUSG00000056598 | Drc3 | protein_coding | 0.355524014 | 3.995195366 | 0.5001684 | 0.4894102 | 0.77569527 |
| ENSMUSG00000039987 | Phtf2 | protein_coding | 0.464117701 | 2.961224913 | 0.5000674 | 0.4894534 | 0.77569527 |
| ENSMUSG00000010660 | Plcd1 | protein_coding | 0.632253048 | 3.05950199 | 0.499457 | 0.489774 | 0.776030086 |

| ensembl_gene_id | external_gene_name | gene_biotype | log2 (fold change) | log2 (counts per million) | F-value | p-value | False discovery rate |
|---|---|---|---|---|---|---|---|
| ENSMUSG00000040848 | Sft2d2 | protein_coding | -0.377985797 | 3.875666158 | 0.4992366 | 0.489809 | 0.776030086 |
| ENSMUSG00000057880 | Abat | protein_coding | -0.179999101 | 8.308179823 | 0.498682 | 0.4900467 | 0.776113586 |
| ENSMUSG00000051695 | Pcbp1 | protein_coding | 0.158714611 | 6.761842645 | 0.498664 | 0.4900544 | 0.776113586 |
| ENSMUSG00000025809 | Itgb1 | protein_coding | 0.314474175 | 4.437759961 | 0.4985465 | 0.4901048 | 0.776113586 |
| ENSMUSG00000031847 | 1700030J22Rik | protein_coding | -0.3527862 | 3.621338539 | 0.49844 | 0.4901504 | 0.776113586 |
| ENSMUSG00000057530 | Ece1 | protein_coding | -0.233283096 | 5.311178642 | 0.4980508 | 0.4903174 | 0.77626367 |
| ENSMUSG00000017754 | Pltp | protein_coding | -0.315468825 | 4.931605448 | 0.4978047 | 0.490423 | 0.77631659 |
| ENSMUSG00000031576 | Kcnu1 | protein_coding | 0.655976304 | 2.930505954 | 0.4976622 | 0.4905005 | 0.776324946 |
| ENSMUSG00000022967 | Ifnar1 | protein_coding | 0.331050431 | 3.699766087 | 0.4973837 | 0.4906038 | 0.776374174 |
| ENSMUSG00000019338 | Zfp687 | protein_coding | -0.316843672 | 3.918077177 | 0.4968594 | 0.4908291 | 0.77657888 |
| ENSMUSG00000021540 | Smad5 | protein_coding | 0.251896934 | 4.475325274 | 0.4967466 | 0.4908776 | 0.77657888 |
| ENSMUSG00000035397 | Klf16 | protein_coding | 0.250974664 | 5.288354694 | 0.4965612 | 0.4909573 | 0.77658997 |
| ENSMUSG00000062328 | Rpl17 | protein_coding | 0.383485373 | 3.274014848 | 0.4963944 | 0.491029 | 0.77658997 |
| ENSMUSG00000033278 | Ptprm | protein_coding | -0.318843601 | 4.481869238 | 0.4960116 | 0.4911938 | 0.776714949 |
| ENSMUSG00000019861 | Gopc | protein_coding | -0.239676025 | 4.726471528 | 0.4958751 | 0.4912525 | 0.776714949 |
| ENSMUSG00000028057 | Rit1 | protein_coding | -0.185761425 | 6.135760638 | 0.495506 | 0.4914115 | 0.776852058 |
| ENSMUSG00000013997 | Nit1 | protein_coding | 0.208047538 | 5.735274349 | 0.4949038 | 0.491671 | 0.777148065 |
| ENSMUSG00000025648 | Pfkfb4 | protein_coding | 0.446477261 | 3.353675628 | 0.4946921 | 0.4917623 | 0.777178094 |
| ENSMUSG00000054455 | Vapb | protein_coding | -0.236302377 | 5.438720093 | 0.4943652 | 0.4919033 | 0.777250014 |
| ENSMUSG00000044700 | Tmem201 | protein_coding | -0.289789361 | 4.577270721 | 0.4942516 | 0.4919523 | 0.777250014 |
| ENSMUSG00000085071 | Gm14066 | processed_transcript | -0.526825046 | 3.106832346 | 0.4937025 | 0.4921894 | 0.777462494 |
| ENSMUSG00000055296 | Tmem245 | protein_coding | -0.321839624 | 4.413912717 | 0.4936052 | 0.4922314 | 0.777462494 |
| ENSMUSG00000023015 | Racgap1 | protein_coding | -0.340038519 | 3.922597179 | 0.4928716 | 0.4925485 | 0.777776063 |
| ENSMUSG00000036766 | Dner | protein_coding | -0.220997935 | 6.807641202 | 0.4927491 | 0.4926178 | 0.777776063 |
| ENSMUSG00000028883 | Sema3a | protein_coding | -0.440052512 | 3.199347871 | 0.4923875 | 0.492758 | 0.777776063 |
| ENSMUSG00000003549 | Ercc1 | protein_coding | 0.179341239 | 6.386600643 | 0.4922897 | 0.4928003 | 0.777776063 |
| ENSMUSG00000064326 | Siva1 | protein_coding | 0.245191272 | 5.736593877 | 0.4920094 | 0.4929217 | 0.777776063 |
| ENSMUSG00000026827 | Gpd2 | protein_coding | 0.174987187 | 6.542160794 | 0.4919826 | 0.4929333 | 0.777776063 |
| ENSMUSG00000050822 | Slc29a4 | protein_coding | 0.386424435 | 3.279897872 | 0.4919757 | 0.4929363 | 0.777776063 |
| ENSMUSG00000023262 | Acy1 | protein_coding | 0.296768907 | 4.934477265 | 0.4911585 | 0.4932905 | 0.778143767 |
| ENSMUSG00000021901 | Bap1 | protein_coding | 0.160642476 | 6.542753954 | 0.491104 | 0.4933141 | 0.778143767 |
| ENSMUSG00000040424 | Hipk4 | protein_coding | 0.284447793 | 4.448475181 | 0.4904638 | 0.4935918 | 0.778467704 |
| ENSMUSG00000032498 | Mlh1 | protein_coding | -0.487507397 | 2.910645201 | 0.4900016 | 0.4937926 | 0.778608953 |
| ENSMUSG00000022054 | Nefm | protein_coding | 0.333326014 | 7.018960095 | 0.4935572 | 0.4938262 | 0.778608953 |
| ENSMUSG00000026229 | Psmd1 | protein_coding | 0.185953718 | 7.239032462 | 0.4894465 | 0.4940338 | 0.778710841 |
| ENSMUSG00000001240 | Ramp2 | protein_coding | 0.560141193 | 2.70273353 | 0.489313 | 0.4940918 | 0.778710841 |
| ENSMUSG00000073411 | H2-D1 | protein_coding | 0.473047567 | 4.646071214 | 0.4915402 | 0.4941081 | 0.778710841 |
| ENSMUSG00000027387 | Zc3h8 | protein_coding | 0.321424255 | 3.806664377 | 0.488861 | 0.4942885 | 0.778880984 |
| ENSMUSG00000039191 | Rbpj | protein_coding | -0.383790601 | 3.246021187 | 0.4880685 | 0.4946335 | 0.779217533 |
| ENSMUSG00000059495 | Arhgef12 | protein_coding | -0.222351074 | 5.470293133 | 0.4880377 | 0.494647 | 0.779217533 |
| ENSMUSG00000005609 | Ctr9 | protein_coding | 0.245081923 | 5.359630466 | 0.487638 | 0.4948212 | 0.779238903 |
| ENSMUSG00000017453 | Pipox | protein_coding | -0.576525452 | 2.536586131 | 0.4876221 | 0.4948281 | 0.779238903 |
| ENSMUSG00000072437 | Nanos1 | protein_coding | -0.280999559 | 4.553238287 | 0.4875078 | 0.4948779 | 0.779238903 |
| ENSMUSG00000026309 | Ilkap | protein_coding | 0.250016273 | 5.086444162 | 0.4869639 | 0.4951152 | 0.779498366 |
| ENSMUSG00000042115 | Klhdc8a | protein_coding | 0.272564683 | 5.107572488 | 0.486689 | 0.4952352 | 0.779573183 |
| ENSMUSG00000034035 | Ccdc17 | protein_coding | -0.57131158 | 2.517724767 | 0.486083 | 0.4954999 | 0.779672952 |
| ENSMUSG00000084799 | Ino80dos | lincRNA | -0.4528658 | 2.857561626 | 0.4858355 | 0.4956081 | 0.779672952 |
| ENSMUSG00000007783 | Cpt1c | protein_coding | 0.202444483 | 6.709707017 | 0.4858176 | 0.495616 | 0.779672952 |
| ENSMUSG00000033917 | Gde1 | protein_coding | -0.264945881 | 5.794515996 | 0.4858536 | 0.4956163 | 0.779672952 |
| ENSMUSG00000064068 | Mtx1 | protein_coding | 0.186627645 | 5.985579184 | 0.4857143 | 0.4956611 | 0.779672952 |
| ENSMUSG00000029722 | Agfg2 | protein_coding | -0.250858468 | 5.019407381 | 0.4852423 | 0.4958676 | 0.779783194 |
| ENSMUSG00000002006 | Pdzd4 | protein_coding | -0.328544759 | 4.355802485 | 0.4852225 | 0.4958763 | 0.779783194 |
| ENSMUSG00000020589 | Fam49a | protein_coding | -0.432427653 | 3.25155899 | 0.4850546 | 0.4959497 | 0.779784698 |
| ENSMUSG00000022494 | Shisa9 | protein_coding | -0.262597691 | 4.665970214 | 0.484556 | 0.4961681 | 0.780013953 |
| ENSMUSG00000026812 | Tsc1 | protein_coding | -0.34948272 | 3.868228359 | 0.4839775 | 0.4964216 | 0.780298462 |
| ENSMUSG00000038807 | Rap1gap2 | protein_coding | -0.259164096 | 5.432078849 | 0.4832601 | 0.4967364 | 0.780565683 |
| ENSMUSG00000069895 | Atxn1l | protein_coding | -0.48626032 | 3.812984742 | 0.4838026 | 0.4968014 | 0.780565683 |
| ENSMUSG00000027076 | Timm10 | protein_coding | 0.192586023 | 7.283647256 | 0.4830666 | 0.4968213 | 0.780565683 |
| ENSMUSG00000020250 | Txnrd1 | protein_coding | 0.216695615 | 5.659791275 | 0.4829284 | 0.496882 | 0.780565683 |
| ENSMUSG00000033124 | Atg9a | protein_coding | 0.588051621 | 3.263364428 | 0.4832141 | 0.4970814 | 0.780764896 |
| ENSMUSG00000032624 | Eml4 | protein_coding | -0.31739149 | 3.902866519 | 0.4820529 | 0.4972667 | 0.780880713 |
| ENSMUSG00000040688 | Tbl3 | protein_coding | -0.268946882 | 5.372885284 | 0.4819764 | 0.4973004 | 0.780880713 |
| ENSMUSG00000041777 | Cir1 | protein_coding | -0.40751357 | 2.987626848 | 0.4812101 | 0.4976376 | 0.781196052 |
| ENSMUSG00000055670 | Zzef1 | protein_coding | 0.358631567 | 4.076059421 | 0.481112 | 0.4976808 | 0.781196052 |
| ENSMUSG00000021054 | Sgpp1 | protein_coding | -0.261917439 | 4.598487404 | 0.4810102 | 0.4977257 | 0.781196052 |
| ENSMUSG00000031703 | Itfg1 | protein_coding | -0.229494689 | 6.88123795 | 0.4808302 | 0.4978209 | 0.781196052 |
| ENSMUSG00000041264 | Uspl1 | protein_coding | 0.230309959 | 5.374578541 | 0.4804328 | 0.4979801 | 0.781196052 |
| ENSMUSG00000020900 | Myh10 | protein_coding | 0.219333181 | 6.443934694 | 0.480332 | 0.4980245 | 0.781196052 |
| ENSMUSG00000020218 | Wif1 | protein_coding | -0.528235925 | 3.057694366 | 0.4802931 | 0.4980417 | 0.781196052 |
| ENSMUSG00000020801 | Med31 | protein_coding | 0.224481421 | 6.786437743 | 0.480237 | 0.4980824 | 0.781196052 |
| ENSMUSG00000059187 | Fam19a1 | protein_coding | -0.306425597 | 4.408622389 | 0.4799271 | 0.4982031 | 0.78127146 |
| ENSMUSG00000020577 | Tspan13 | protein_coding | -0.170465646 | 7.317139806 | 0.4797378 | 0.4982867 | 0.781288525 |
| ENSMUSG00000026437 | Cdk18 | protein_coding | -0.313892496 | 5.932591511 | 0.480741 | 0.4984636 | 0.781452035 |
| ENSMUSG00000031198 | Fundc2 | protein_coding | 0.170001589 | 7.531706821 | 0.4789708 | 0.4986254 | 0.781591665 |
| ENSMUSG00000005699 | Pard6a | protein_coding | 0.185976316 | 6.116825136 | 0.4774947 | 0.4992784 | 0.782501206 |
| ENSMUSG00000017837 | Nkiras2 | protein_coding | 0.303689637 | 4.918589916 | 0.4768252 | 0.4995751 | 0.782817868 |
| ENSMUSG00000025040 | Fundc1 | protein_coding | -0.1736808 | 7.658821387 | 0.47648 | 0.4997282 | 0.782817868 |
| ENSMUSG00000022010 | Tsc22d1 | protein_coding | -0.159492585 | 9.219451156 | 0.4764485 | 0.4997421 | 0.782817868 |

| ensembl_gene_id | external_gene_name | gene_biotype | log2 (fold change) | log2 (counts per million) | F-value | p-value | False discovery rate |
|---|---|---|---|---|---|---|---|
| ENSMUSG00000050541 | Adra1b | protein_coding | 0.63085517 | 2.48256062 | 0.4762819 | 0.4998161 | 0.782817868 |
| ENSMUSG00000020496 | Rnf187 | protein_coding | 0.17642094 | 8.020059388 | 0.4761456 | 0.4998766 | 0.782817868 |
| ENSMUSG00000027822 | Slc33a1 | protein_coding | -0.28496656 | 4.249301208 | 0.4759582 | 0.4999598 | 0.782817868 |
| ENSMUSG00000027775 | Mfsd1 | protein_coding | 0.383104164 | 3.452838777 | 0.4758899 | 0.4999901 | 0.782817868 |
| ENSMUSG00000032058 | Ppp2r1b | protein_coding | 0.281267097 | 4.65675267 | 0.4753151 | 0.5002454 | 0.783031437 |
| ENSMUSG00000025764 | Jade1 | protein_coding | -0.196244841 | 5.582317761 | 0.475255 | 0.5002721 | 0.783031437 |
| ENSMUSG00000029179 | Zcchc4 | protein_coding | -0.42774331 | 3.609632238 | 0.4747369 | 0.5005025 | 0.783278009 |
| ENSMUSG00000019761 | Krt10 | protein_coding | 0.240081302 | 5.047705005 | 0.4745564 | 0.5005828 | 0.783289687 |
| ENSMUSG00000032420 | Nt5e | protein_coding | 0.565713036 | 2.641088691 | 0.4741058 | 0.5007833 | 0.783408767 |
| ENSMUSG00000031774 | Fam192a | protein_coding | 0.23703761 | 6.846178738 | 0.4740406 | 0.50087 | 0.783408767 |
| ENSMUSG00000051344 | Plekhm3 | protein_coding | -0.274045929 | 4.396973582 | 0.4738945 | 0.5008775 | 0.783408767 |
| ENSMUSG00000003778 | Brd8 | protein_coding | -0.21319417 | 6.983895678 | 0.4734106 | 0.5011089 | 0.783551062 |
| ENSMUSG00000008393 | Carhsp1 | protein_coding | -0.181645837 | 6.747320037 | 0.4733611 | 0.5011151 | 0.783551062 |
| ENSMUSG00000021010 | Npas3 | protein_coding | -0.62330024 | 2.326806981 | 0.4731998 | 0.501187 | 0.783551062 |
| ENSMUSG00000038042 | Ptpdc1 | protein_coding | 0.33293295 | 4.040815844 | 0.4728858 | 0.5013271 | 0.783656101 |
| ENSMUSG00000021721 | Htr1a | protein_coding | -0.324028848 | 4.1537979 | 0.4722629 | 0.5016051 | 0.783976713 |
| ENSMUSG00000070780 | Rbm47 | protein_coding | -0.775747225 | 2.402039901 | 0.4718391 | 0.5017944 | 0.784127424 |
| ENSMUSG00000029291 | Rufy3 | protein_coding | -0.169686385 | 8.129277051 | 0.4717074 | 0.5018533 | 0.784127424 |
| ENSMUSG00000024011 | Pi16 | protein_coding | -0.607985628 | 3.090579411 | 0.4715447 | 0.5020017 | 0.784127424 |
| ENSMUSG00000056515 | Rab31 | protein_coding | -0.176415487 | 6.519935439 | 0.4712662 | 0.5020506 | 0.784127424 |
| ENSMUSG00000060716 | Plekhh1 | protein_coding | 0.470691669 | 3.439004716 | 0.4712666 | 0.5020662 | 0.784127424 |
| ENSMUSG00000027400 | Pdyn | protein_coding | -0.265568282 | 6.739999228 | 0.4718157 | 0.5024176 | 0.78430021 |
| ENSMUSG00000023286 | Ube2j2 | protein_coding | 0.270880622 | 4.369163291 | 0.4702873 | 0.5024887 | 0.78430021 |
| ENSMUSG00000015342 | Xk | protein_coding | -0.733682634 | 3.146942023 | 0.4730507 | 0.5025026 | 0.78430021 |
| ENSMUSG00000079414 | Gm11110 | protein_coding | 0.642741675 | 2.612340391 | 0.4702299 | 0.5025144 | 0.78430021 |
| ENSMUSG00000023055 | Calcoco1 | protein_coding | -0.224373471 | 6.086909112 | 0.4701695 | 0.5025415 | 0.78430021 |
| ENSMUSG00000031378 | Abcd1 | protein_coding | 0.548995079 | 2.850826298 | 0.4698005 | 0.5027068 | 0.784414728 |
| ENSMUSG00000021186 | Fbln5 | protein_coding | 0.681947774 | 2.269903079 | 0.4696802 | 0.5027607 | 0.784414728 |
| ENSMUSG00000079481 | Nhsl2 | protein_coding | -0.248868011 | 5.12842791 | 0.4692987 | 0.5029319 | 0.784567868 |
| ENSMUSG00000031864 | Ints10 | protein_coding | 0.274328311 | 5.075568629 | 0.468405 | 0.5033331 | 0.785037456 |
| ENSMUSG00000020708 | Psmc5 | protein_coding | 0.162267279 | 9.994707594 | 0.4683031 | 0.5033789 | 0.785037456 |
| ENSMUSG00000038459 | Abhd17c | protein_coding | -0.300654158 | 4.349748203 | 0.4680885 | 0.5034754 | 0.78507404 |
| ENSMUSG00000022867 | Usp25 | protein_coding | -0.321467396 | 4.176363426 | 0.4677636 | 0.5036214 | 0.785187957 |
| ENSMUSG00000058056 | Palld | protein_coding | 0.443545505 | 3.232377554 | 0.4670713 | 0.503933 | 0.785559748 |
| ENSMUSG00000041577 | Prelp | protein_coding | -0.674008411 | 2.359007947 | 0.466102 | 0.5043697 | 0.786126617 |
| ENSMUSG00000004360 | 9330159F19Rik | protein_coding | -0.349291058 | 4.99647521 | 0.4658878 | 0.5045505 | 0.786265614 |
| ENSMUSG00000026088 | Mitd1 | protein_coding | -0.304892973 | 4.400588202 | 0.4655801 | 0.5046051 | 0.786265614 |
| ENSMUSG00000025170 | Rab40b | protein_coding | -0.324592642 | 4.990216302 | 0.4651164 | 0.5048301 | 0.786502258 |
| ENSMUSG00000038212 | Mfsd14b | protein_coding | -0.298580185 | 4.628872732 | 0.4643519 | 0.50516 | 0.786876748 |
| ENSMUSG00000034621 | Gpatch8 | protein_coding | -0.224508814 | 5.727575147 | 0.4641303 | 0.5052602 | 0.786876748 |
| ENSMUSG00000024170 | Telo2 | protein_coding | 0.321489983 | 4.060941039 | 0.4637908 | 0.5054138 | 0.786876748 |
| ENSMUSG00000024140 | Epas1 | protein_coding | -0.342702976 | 4.294059683 | 0.4636308 | 0.5054862 | 0.786876748 |
| ENSMUSG00000006395 | Hyi | polymorphic_pseudog | -0.437883318 | 2.861671572 | 0.4633461 | 0.5056151 | 0.786876748 |
| ENSMUSG00000039065 | Fam173b | protein_coding | -0.24877216 | 4.623880927 | 0.4632761 | 0.5056468 | 0.786876748 |
| ENSMUSG00000049164 | Zfp518a | protein_coding | -0.430712915 | 2.82800712 | 0.4632714 | 0.5056489 | 0.786876748 |
| ENSMUSG00000041235 | Chd7 | protein_coding | -0.34154013 | 3.645943808 | 0.4630882 | 0.505732 | 0.786876748 |
| ENSMUSG00000041459 | Tardbp | protein_coding | 0.183829219 | 6.137299951 | 0.4628971 | 0.5058186 | 0.786876748 |
| ENSMUSG00000007812 | Zfp655 | protein_coding | -0.217945909 | 5.639643847 | 0.4626478 | 0.5059315 | 0.786876748 |
| ENSMUSG00000024456 | Diaph1 | protein_coding | -0.321394936 | 4.603002717 | 0.4626173 | 0.5059454 | 0.786876748 |
| ENSMUSG00000037260 | Hgsnat | protein_coding | 0.285676702 | 4.476158733 | 0.4626101 | 0.5059487 | 0.786876748 |
| ENSMUSG00000020307 | Cdc34 | protein_coding | 0.288788077 | 4.500428073 | 0.4622558 | 0.5061093 | 0.787012825 |
| ENSMUSG00000046668 | Cxxc5 | protein_coding | -0.212924954 | 5.562815778 | 0.4620124 | 0.5062198 | 0.787070787 |
| ENSMUSG00000021090 | Lrrc9 | protein_coding | -0.598397516 | 2.667987494 | 0.4618245 | 0.5063051 | 0.787089622 |
| ENSMUSG00000018239 | Zcchc10 | protein_coding | -0.213661292 | 5.560305535 | 0.4613185 | 0.5065349 | 0.787333064 |
| ENSMUSG00000053799 | Exoc6 | protein_coding | -0.276506988 | 4.011580773 | 0.4610946 | 0.5066367 | 0.787377414 |
| ENSMUSG00000046157 | Tmem229b | protein_coding | 0.366413599 | 3.999142315 | 0.4608145 | 0.506764 | 0.787461478 |
| ENSMUSG00000020647 | Ncoa1 | protein_coding | -0.28187208 | 6.32571021 | 0.4615952 | 0.5070143 | 0.787678565 |
| ENSMUSG00000026289 | Atg16l1 | protein_coding | 0.327937067 | 4.945084035 | 0.4600311 | 0.5071361 | 0.787678565 |
| ENSMUSG00000005566 | Trim28 | protein_coding | 0.165061521 | 6.502968316 | 0.4599414 | 0.5071613 | 0.787678565 |
| ENSMUSG00000034165 | Ccnd3 | protein_coding | 0.316811979 | 3.629713756 | 0.4598383 | 0.5072083 | 0.787678565 |
| ENSMUSG00000045482 | Trrap | protein_coding | -0.33528951 | 4.645502751 | 0.4597371 | 0.50727 | 0.787678565 |
| ENSMUSG00000025156 | Gps1 | protein_coding | 0.192153636 | 5.477462297 | 0.4590954 | 0.5075468 | 0.787994662 |
| ENSMUSG00000024889 | Rce1 | protein_coding | -0.272775201 | 4.453823746 | 0.4585684 | 0.5077873 | 0.788199416 |
| ENSMUSG00000027175 | Tcp11l1 | protein_coding | -0.293972866 | 3.964658979 | 0.4582956 | 0.5079118 | 0.788199416 |
| ENSMUSG00000039671 | Zmynd8 | protein_coding | 0.164609266 | 6.581414069 | 0.4581197 | 0.5079921 | 0.788199416 |
| ENSMUSG00000028274 | Rngtt | protein_coding | 0.27793429 | 5.010765698 | 0.4579036 | 0.5080908 | 0.788199416 |
| ENSMUSG00000008090 | Fgfrl1 | protein_coding | -0.652093158 | 2.626197143 | 0.4578712 | 0.5081056 | 0.788199416 |
| ENSMUSG00000034825 | Nrip3 | protein_coding | -0.261450092 | 7.636221506 | 0.4595893 | 0.5081185 | 0.788199416 |
| ENSMUSG00000052949 | Rnf157 | protein_coding | -0.157250709 | 6.718553286 | 0.4572667 | 0.5083819 | 0.788494315 |
| ENSMUSG00000036644 | Tbc1d9b | protein_coding | 0.245935533 | 5.50299736 | 0.4564254 | 0.508767 | 0.788977718 |
| ENSMUSG00000039234 | Sec24d | protein_coding | 0.415105452 | 3.237169065 | 0.4559992 | 0.5089622 | 0.789135348 |
| ENSMUSG00000022313 | Utp23 | protein_coding | 0.30572398 | 4.466096438 | 0.4558831 | 0.5090154 | 0.789135348 |
| ENSMUSG00000021116 | Eif2s1 | protein_coding | -0.152277608 | 6.639158588 | 0.4556626 | 0.5091165 | 0.78914668 |
| ENSMUSG00000053436 | Mapk14 | protein_coding | -0.246852684 | 5.388833924 | 0.4555471 | 0.5091695 | 0.78914668 |
| ENSMUSG00000031538 | Plat | protein_coding | 0.330142029 | 4.335392688 | 0.455312 | 0.5092773 | 0.789200052 |
| ENSMUSG00000029462 | Vps29 | protein_coding | 0.152501142 | 9.414165902 | 0.4549035 | 0.5094648 | 0.789376825 |
| ENSMUSG00000024212 | Mllt1 | protein_coding | -0.324901922 | 4.066042031 | 0.4545618 | 0.5096217 | 0.789399123 |
| ENSMUSG00000003644 | Rps6ka1 | protein_coding | 0.264652558 | 4.109013313 | 0.4545011 | 0.5096496 | 0.789399123 |

| ensembl_gene_id | external_gene_name | gene_biotype | log2 (fold change) | log2 (counts per million) | F-value | p-value | False discovery rate |
|---|---|---|---|---|---|---|---|
| ENSMUSG00000056413 | Adap1 | protein_coding | -0.214683436 | 6.48871669 | 0.4543926 | 0.5096994 | 0.789399123 |
| ENSMUSG00000052133 | Sema5b | protein_coding | -0.474835007 | 2.685225902 | 0.4541315 | 0.5098194 | 0.789426724 |
| ENSMUSG00000108532 | Gm32647 | lincRNA | -0.653831751 | 2.491066607 | 0.4540207 | 0.5098704 | 0.789426724 |
| ENSMUSG00000005534 | Insr | protein_coding | 0.351455179 | 3.876891473 | 0.4538747 | 0.5099375 | 0.789426724 |
| ENSMUSG00000033020 | Polr2f | protein_coding | 0.188204378 | 7.719445482 | 0.4535656 | 0.5100797 | 0.789533157 |
| ENSMUSG00000048271 | Rbm33 | protein_coding | -0.218808344 | 5.378286856 | 0.4527709 | 0.5104455 | 0.789985732 |
| ENSMUSG00000031446 | Cul4a | protein_coding | 0.325105985 | 4.612522061 | 0.452575 | 0.5105358 | 0.790011714 |
| ENSMUSG00000029777 | Gars | protein_coding | 0.181306667 | 6.473768453 | 0.4523744 | 0.5106283 | 0.790041112 |
| ENSMUSG00000021668 | Polk | protein_coding | -0.400790841 | 3.087683292 | 0.4521391 | 0.5107368 | 0.790095285 |
| ENSMUSG00000044499 | Hs3st5 | protein_coding | -0.371548004 | 3.457253235 | 0.4519432 | 0.5108271 | 0.790121341 |
| ENSMUSG00000019854 | Reps1 | protein_coding | 0.398691197 | 4.639470301 | 0.4524132 | 0.5112085 | 0.79059752 |
| ENSMUSG00000022905 | Kpna1 | protein_coding | -0.176308027 | 6.03260121 | 0.4508899 | 0.5113134 | 0.790639394 |
| ENSMUSG00000048349 | Pou4f1 | protein_coding | -1.864141196 | 1.427937485 | 0.4545917 | 0.5113826 | 0.790639394 |
| ENSMUSG00000027425 | Kat14 | protein_coding | 0.26190921 | 4.035801704 | 0.4505355 | 0.5114772 | 0.79067196 |
| ENSMUSG00000030534 | Vps33b | protein_coding | -0.411441259 | 3.958415376 | 0.4501141 | 0.5116875 | 0.790883337 |
| ENSMUSG00000033940 | Brk1 | protein_coding | 0.146179806 | 9.750060747 | 0.4499167 | 0.5117635 | 0.790887101 |
| ENSMUSG00000062070 | Pgk1 | protein_coding | 0.203804173 | 5.410924104 | 0.4496676 | 0.5118787 | 0.790914442 |
| ENSMUSG00000026470 | Stx6 | protein_coding | 0.231074993 | 5.734930447 | 0.4495607 | 0.5119283 | 0.790914442 |
| ENSMUSG00000021929 | Kpna3 | protein_coding | 0.176003729 | 6.483431306 | 0.4492573 | 0.5120688 | 0.790973467 |
| ENSMUSG00000000282 | Mnt | protein_coding | -0.257072795 | 4.597142169 | 0.4491606 | 0.5121136 | 0.790973467 |
| ENSMUSG00000000959 | Oxa1l | protein_coding | 0.191488729 | 6.370403043 | 0.4489771 | 0.5121986 | 0.790991204 |
| ENSMUSG00000030423 | Pop4 | protein_coding | 0.199652874 | 6.037266014 | 0.4487979 | 0.5122817 | 0.791005883 |
| ENSMUSG00000027359 | Slc27a2 | protein_coding | -0.56378562 | 2.826299836 | 0.448358 | 0.5124857 | 0.791153809 |
| ENSMUSG00000025232 | Hexa | protein_coding | 0.221316103 | 5.996006021 | 0.4482377 | 0.5125416 | 0.791153809 |
| ENSMUSG00000031310 | Zmym3 | protein_coding | 0.164984525 | 6.639066481 | 0.4480606 | 0.5126238 | 0.791153809 |
| ENSMUSG00000021900 | Btd | protein_coding | -0.300399076 | 4.185095894 | 0.447931 | 0.5126839 | 0.791153809 |
| ENSMUSG00000085971 | Gm15411 | processed_transcript | 0.629121746 | 2.348571445 | 0.4477987 | 0.5127454 | 0.791153809 |
| ENSMUSG00000029875 | Ccdc184 | protein_coding | 0.26077807 | 6.12034788 | 0.4474936 | 0.5129429 | 0.791344981 |
| ENSMUSG00000039474 | Wfs1 | protein_coding | 0.199120412 | 6.511998723 | 0.4466446 | 0.5132818 | 0.791644409 |
| ENSMUSG00000021417 | Eci2 | protein_coding | -0.23988521 | 5.131709028 | 0.4465255 | 0.5133373 | 0.791644409 |
| ENSMUSG00000031660 | Brd7 | protein_coding | -0.228873671 | 5.766305864 | 0.4464814 | 0.5133578 | 0.791644409 |
| ENSMUSG00000039166 | Akap7 | protein_coding | 0.187180273 | 6.00025257 | 0.4461887 | 0.5134941 | 0.791740981 |
| ENSMUSG00000071454 | Dtnb | protein_coding | 0.224174924 | 4.995225655 | 0.445786 | 0.5136816 | 0.791833125 |
| ENSMUSG00000015599 | Ttbk1 | protein_coding | 0.401135752 | 3.213615904 | 0.4454981 | 0.5138158 | 0.791833125 |
| ENSMUSG00000038007 | Acer2 | protein_coding | -0.277933669 | 4.297017974 | 0.4454776 | 0.5138254 | 0.791833125 |
| ENSMUSG00000022019 | Tdrd3 | protein_coding | -0.24454016 | 4.73038923 | 0.4454283 | 0.5138484 | 0.791833125 |
| ENSMUSG00000028988 | Ctnnbip1 | protein_coding | 0.385282267 | 3.844294527 | 0.4451039 | 0.5139996 | 0.791952731 |
| ENSMUSG00000021460 | Auh | protein_coding | 0.234666615 | 5.498863594 | 0.44471 | 0.5141834 | 0.791995025 |
| ENSMUSG00000036854 | Hspb6 | protein_coding | 0.257983624 | 5.283787309 | 0.4443676 | 0.5143433 | 0.791995025 |
| ENSMUSG00000048234 | Rnf149 | protein_coding | -0.469206304 | 3.818176469 | 0.4450864 | 0.5144343 | 0.791995025 |
| ENSMUSG00000034807 | Colgalt1 | protein_coding | -0.414473907 | 3.792657305 | 0.4440255 | 0.5145184 | 0.791995025 |
| ENSMUSG00000001999 | Blvra | protein_coding | 0.151350237 | 6.923639702 | 0.4439572 | 0.514535 | 0.791995025 |
| ENSMUSG00000029201 | Ugdh | protein_coding | 0.264056382 | 4.528827159 | 0.4438079 | 0.5146048 | 0.791995025 |
| ENSMUSG00000028643 | Svbp | protein_coding | 0.210329509 | 5.704019942 | 0.4436409 | 0.5146829 | 0.791995025 |
| ENSMUSG00000033416 | Gucd1 | protein_coding | 0.503546805 | 3.277746035 | 0.44367 | 0.5146846 | 0.791995025 |
| ENSMUSG00000037890 | Wdr19 | protein_coding | -0.39965638 | 3.403430134 | 0.4435282 | 0.5147356 | 0.791995025 |
| ENSMUSG00000039633 | Lonrf1 | protein_coding | 0.404654358 | 3.347394235 | 0.4434683 | 0.5147636 | 0.791995025 |
| ENSMUSG00000026096 | Osgepl1 | protein_coding | -0.271853174 | 4.626322671 | 0.4431583 | 0.5149087 | 0.79204072 |
| ENSMUSG00000020280 | Pus10 | protein_coding | -0.334750347 | 3.556230994 | 0.4430899 | 0.5149406 | 0.79204072 |
| ENSMUSG00000028801 | Stpg1 | protein_coding | -0.428058047 | 3.122918865 | 0.4419238 | 0.5154869 | 0.792767585 |
| ENSMUSG00000041057 | Wdr43 | protein_coding | 0.210678285 | 5.112090577 | 0.4409644 | 0.5159372 | 0.793336758 |
| ENSMUSG00000025024 | Smndc1 | protein_coding | -0.259563335 | 4.371692579 | 0.4408208 | 0.5160046 | 0.793336758 |
| ENSMUSG00000027804 | Ppid | protein_coding | 0.147749978 | 6.992843414 | 0.4404845 | 0.5161626 | 0.793466308 |
| ENSMUSG00000040365 | Trim41 | protein_coding | -0.352814229 | 3.930646536 | 0.4391093 | 0.5168098 | 0.794347542 |
| ENSMUSG00000017715 | Pgs1 | protein_coding | 0.267047354 | 4.039491949 | 0.438316 | 0.5171837 | 0.794808708 |
| ENSMUSG00000054008 | Ndst1 | protein_coding | -0.366971687 | 4.169782026 | 0.4380459 | 0.5173263 | 0.794914198 |
| ENSMUSG00000028583 | Pdpn | protein_coding | -0.351745843 | 4.183131412 | 0.4371571 | 0.5177309 | 0.795422281 |
| ENSMUSG00000068921 | Dap3 | protein_coding | 0.157137165 | 7.196531851 | 0.4368145 | 0.5178929 | 0.795557466 |
| ENSMUSG00000025626 | Phf6 | protein_coding | -0.238066432 | 4.926960192 | 0.4362814 | 0.5181451 | 0.795730363 |
| ENSMUSG00000036112 | Metap2 | protein_coding | 0.173196592 | 7.489610262 | 0.4360839 | 0.5182386 | 0.795730363 |
| ENSMUSG00000054423 | Cadps | protein_coding | -0.15316016 | 7.090859496 | 0.4360317 | 0.5182633 | 0.795730363 |
| ENSMUSG00000021585 | Cast | protein_coding | -0.413009729 | 3.004113261 | 0.435831 | 0.5183583 | 0.795730363 |
| ENSMUSG00000002326 | Gmpr2 | protein_coding | 0.240433367 | 5.289476233 | 0.4357354 | 0.5184036 | 0.795730363 |
| ENSMUSG00000053907 | Mat2a | protein_coding | -0.152953358 | 7.41689777 | 0.4356387 | 0.5184494 | 0.795730363 |
| ENSMUSG00000027472 | Pdrg1 | protein_coding | -0.156813704 | 7.819654886 | 0.4348851 | 0.5188068 | 0.796130298 |
| ENSMUSG00000027018 | Hat1 | protein_coding | 0.204959426 | 5.333454055 | 0.4347769 | 0.5188581 | 0.796130298 |
| ENSMUSG00000073207 | Ccdc160 | protein_coding | 0.428885635 | 3.255506659 | 0.4344775 | 0.5190002 | 0.796234661 |
| ENSMUSG00000015850 | Adamtsl4 | protein_coding | -0.605796378 | 2.53228929 | 0.4343064 | 0.5190814 | 0.796245747 |
| ENSMUSG00000056749 | Nfil3 | protein_coding | -0.39461337 | 3.514816785 | 0.4336448 | 0.5193958 | 0.79661428 |
| ENSMUSG00000036167 | Pphln1 | protein_coding | -0.228612999 | 5.291290637 | 0.4330608 | 0.5196735 | 0.796820948 |
| ENSMUSG00000036099 | Vezt | protein_coding | -0.265133377 | 5.092441276 | 0.4330499 | 0.5196787 | 0.796820948 |
| ENSMUSG00000028992 | Nmnat1 | protein_coding | -0.331092968 | 3.606100699 | 0.4328792 | 0.51976 | 0.796831902 |
| ENSMUSG00000028494 | Plin2 | protein_coding | -0.353551262 | 3.848423259 | 0.4321869 | 0.5200897 | 0.79705934 |
| ENSMUSG00000024985 | Tcf7l2 | protein_coding | -0.735364531 | 8.165222098 | 0.4529119 | 0.5201894 | 0.79705934 |
| ENSMUSG00000052310 | Slc39a1 | protein_coding | 0.380665019 | 4.916541823 | 0.4331852 | 0.5201976 | 0.79705934 |
| ENSMUSG00000032300 | 1700017B05Rik | protein_coding | -0.550154495 | 3.158139168 | 0.4318369 | 0.5202716 | 0.79705934 |
| ENSMUSG00000028010 | Gar1 | protein_coding | 0.619726348 | 2.31814993 | 0.4316262 | 0.520357 | 0.79705934 |
| ENSMUSG00000047495 | Dlgap2 | protein_coding | -0.273327645 | 4.648677871 | 0.4315824 | 0.5203779 | 0.79705934 |

| ensembl_gene_id | external_gene_name | gene_biotype | log2 (fold change) | log2 (counts per million) | F-value | p-value | False discovery rate |
|---|---|---|---|---|---|---|---|
| ENSMUSG00000020374 | Rasgef1c | protein_coding | -0.394065962 | 3.537564496 | 0.4314791 | 0.5204272 | 0.79705934 |
| ENSMUSG00000025287 | Acot9 | protein_coding | -0.215641488 | 5.547245871 | 0.4312386 | 0.520542 | 0.797113859 |
| ENSMUSG00000025370 | Cdh9 | protein_coding | -0.61275891 | 2.458546631 | 0.4310939 | 0.5206111 | 0.797113859 |
| ENSMUSG00000040387 | Klhl32 | protein_coding | -0.263240037 | 4.658007617 | 0.4306264 | 0.5208343 | 0.797261793 |
| ENSMUSG00000097102 | 2310069G16Rik | lincRNA | 0.469725684 | 3.187040995 | 0.4305812 | 0.520856 | 0.797261793 |
| ENSMUSG00000038702 | Dsel | protein_coding | -0.422708 | 3.507317858 | 0.4303814 | 0.5209514 | 0.797294409 |
| ENSMUSG00000033102 | Cdc14b | protein_coding | 0.343828878 | 3.484622043 | 0.4299504 | 0.5211575 | 0.797496302 |
| ENSMUSG00000007950 | Abhd8 | protein_coding | 0.188713046 | 6.008048681 | 0.4294525 | 0.5213958 | 0.797747429 |
| ENSMUSG00000041959 | S100a10 | protein_coding | 0.175553377 | 6.685286142 | 0.4289255 | 0.5216482 | 0.797908111 |
| ENSMUSG00000004892 | Bcan | protein_coding | -0.330580931 | 5.297750323 | 0.4296647 | 0.5216959 | 0.797908111 |
| ENSMUSG00000033335 | Dnm2 | protein_coding | -0.436353715 | 4.189013686 | 0.4299054 | 0.5217597 | 0.797908111 |
| ENSMUSG00000055762 | Eef1d | protein_coding | 0.163679674 | 6.868658105 | 0.4284941 | 0.521855 | 0.797908111 |
| ENSMUSG00000032012 | Nectin1 | protein_coding | 0.347814534 | 4.533618981 | 0.4284295 | 0.5219009 | 0.797908111 |
| ENSMUSG00000001016 | Ilf2 | protein_coding | 0.154006203 | 7.841223411 | 0.4283043 | 0.521946 | 0.797908111 |
| ENSMUSG00000039427 | Alg1 | protein_coding | 0.413810806 | 3.42903892 | 0.428038 | 0.5220738 | 0.797937465 |
| ENSMUSG00000036180 | Gatad2a | protein_coding | -0.217803596 | 4.89311502 | 0.4279549 | 0.5221136 | 0.797937465 |
| ENSMUSG00000042198 | Chchd7 | protein_coding | 0.228075807 | 5.275850413 | 0.4272312 | 0.5224612 | 0.798355179 |
| ENSMUSG00000028609 | Magoh | protein_coding | -0.192720382 | 6.642206271 | 0.4267874 | 0.5226746 | 0.798567723 |
| ENSMUSG00000026787 | Gad2 | protein_coding | 0.169847944 | 8.99710077 | 0.4261918 | 0.5229611 | 0.798892056 |
| ENSMUSG00000003418 | St8sia6 | protein_coding | -0.540041728 | 2.527975686 | 0.4256011 | 0.5232456 | 0.799213107 |
| ENSMUSG00000036599 | Chst12 | protein_coding | 0.201999102 | 5.686100521 | 0.4252559 | 0.523412 | 0.799353728 |
| ENSMUSG00000028563 | Tm2d1 | protein_coding | -0.308231891 | 4.406926536 | 0.4250088 | 0.5235312 | 0.799422184 |
| ENSMUSG00000034653 | Ythdc2 | protein_coding | -0.358142767 | 3.328496467 | 0.4244641 | 0.523794 | 0.799709957 |
| ENSMUSG00000056832 | Ttc26 | protein_coding | -0.495824993 | 2.656063282 | 0.4239644 | 0.5240354 | 0.799847576 |
| ENSMUSG00000016831 | Tox4 | protein_coding | 0.207458361 | 6.124848092 | 0.4239423 | 0.5240461 | 0.799847576 |
| ENSMUSG00000024154 | Gtf2a1l | protein_coding | -0.562370458 | 2.657533945 | 0.4238156 | 0.5241073 | 0.799847576 |
| ENSMUSG00000037971 | 1110032A03Rik | protein_coding | -0.141162421 | 7.593220878 | 0.4234832 | 0.524268 | 0.799944506 |
| ENSMUSG00000042507 | Elmsan1 | protein_coding | 0.456916506 | 2.772001058 | 0.4231875 | 0.524411 | 0.799944506 |
| ENSMUSG00000026806 | Ddx31 | protein_coding | 0.517213091 | 2.812605396 | 0.42281 | 0.5245938 | 0.799944506 |
| ENSMUSG00000016757 | Ttll12 | protein_coding | 0.399847986 | 3.701757573 | 0.422688 | 0.5246528 | 0.799944506 |
| ENSMUSG00000044715 | Gskip | protein_coding | -0.146529477 | 7.505913195 | 0.422566 | 0.5247119 | 0.799944506 |
| ENSMUSG00000059713 | Rcan3 | protein_coding | -0.19081662 | 5.330877705 | 0.4223703 | 0.5248067 | 0.799944506 |
| ENSMUSG00000038780 | Smurf1 | protein_coding | 0.27525358 | 3.930851603 | 0.422364 | 0.5248098 | 0.799944506 |
| ENSMUSG00000073468 | Sft2d1 | protein_coding | 0.256060219 | 4.717820799 | 0.4223612 | 0.5248111 | 0.799944506 |
| ENSMUSG00000034842 | Art3 | protein_coding | -0.731491104 | 2.4869802 | 0.4223315 | 0.5248404 | 0.799944506 |
| ENSMUSG00000017132 | Cyth1 | protein_coding | -0.194864154 | 5.502105999 | 0.4220654 | 0.5249545 | 0.800005031 |
| ENSMUSG00000027184 | Caprin1 | protein_coding | -0.176916934 | 6.502732555 | 0.4216446 | 0.5251585 | 0.800161343 |
| ENSMUSG00000037434 | Slc30a1 | protein_coding | 0.371083487 | 3.463613346 | 0.421547 | 0.5252059 | 0.800161343 |
| ENSMUSG00000030990 | Pgap2 | protein_coding | -0.235113837 | 4.838311353 | 0.4210029 | 0.52547 | 0.800450375 |
| ENSMUSG00000025130 | P4hb | protein_coding | 0.183988367 | 6.470404982 | 0.4206728 | 0.5256304 | 0.800474633 |
| ENSMUSG00000045160 | Bola3 | protein_coding | -0.273434961 | 4.126265745 | 0.4205365 | 0.5256966 | 0.800474633 |
| ENSMUSG00000023018 | Smarcd1 | protein_coding | 0.183911006 | 6.309771219 | 0.4205104 | 0.5257093 | 0.800474633 |
| ENSMUSG00000046879 | Irgm1 | protein_coding | 0.386999651 | 4.267996404 | 0.4203001 | 0.5258653 | 0.800511805 |
| ENSMUSG00000029769 | Ccdc136 | protein_coding | -0.251850728 | 6.416344774 | 0.420769 | 0.5259612 | 0.800511805 |
| ENSMUSG00000006024 | Napa | protein_coding | 0.165317649 | 9.942773001 | 0.419718 | 0.5260947 | 0.800511805 |
| ENSMUSG00000040452 | Cdh12 | protein_coding | -0.46050143 | 3.24618239 | 0.4196286 | 0.5261382 | 0.800511805 |
| ENSMUSG00000042462 | Dctpp1 | protein_coding | 0.238678584 | 5.573848654 | 0.4194496 | 0.5262254 | 0.800511805 |
| ENSMUSG00000022096 | Hr | protein_coding | -0.48311114 | 2.578028652 | 0.4193901 | 0.5262543 | 0.800511805 |
| ENSMUSG00000047466 | 8030462N17Rik | protein_coding | -0.381900323 | 3.532953595 | 0.4193891 | 0.5262548 | 0.800511805 |
| ENSMUSG00000040957 | Cables1 | protein_coding | 0.363709043 | 3.644113282 | 0.4189869 | 0.5264508 | 0.800696619 |
| ENSMUSG00000031360 | Ctps2 | protein_coding | 0.271363384 | 5.768637426 | 0.4184662 | 0.5267582 | 0.800711509 |
| ENSMUSG00000030986 | Dhx32 | protein_coding | 0.257458516 | 4.597175037 | 0.418331 | 0.5267706 | 0.800711509 |
| ENSMUSG00000022790 | Igsf11 | protein_coding | 0.393813191 | 3.397509605 | 0.4181604 | 0.5268538 | 0.800711509 |
| ENSMUSG00000027963 | Extl2 | protein_coding | 0.161851604 | 6.745595982 | 0.4178087 | 0.5270255 | 0.800711509 |
| ENSMUSG00000019461 | Plscr3 | protein_coding | 0.26565727 | 4.880976774 | 0.4176974 | 0.5270799 | 0.800711509 |
| ENSMUSG00000071424 | Grid2 | protein_coding | -0.256416681 | 4.632350797 | 0.4176645 | 0.5270959 | 0.800711509 |
| ENSMUSG00000060288 | Ppih | protein_coding | 0.376243474 | 3.363588497 | 0.4176436 | 0.5271061 | 0.800711509 |
| ENSMUSG00000032180 | Tmed1 | protein_coding | 0.19886764 | 5.381310942 | 0.4176223 | 0.5271166 | 0.800711509 |
| ENSMUSG00000024665 | Fads2 | protein_coding | 0.297844293 | 4.030559259 | 0.4175932 | 0.5271307 | 0.800711509 |
| ENSMUSG00000000766 | Oprm1 | protein_coding | -0.241400556 | 5.499168234 | 0.4172564 | 0.5272953 | 0.800735682 |
| ENSMUSG00000031585 | Gtf2e2 | protein_coding | -0.174532636 | 5.868931169 | 0.4172559 | 0.5272956 | 0.800735682 |
| ENSMUSG00000107585 | 3300002P13Rik | lincRNA | 0.486116514 | 2.696054502 | 0.4169447 | 0.5274477 | 0.800853573 |
| ENSMUSG00000040731 | Eif4h | protein_coding | 0.15596512 | 8.360380443 | 0.4165764 | 0.5276279 | 0.801014031 |
| ENSMUSG00000039410 | Prdm16 | protein_coding | 0.2545846 | 5.45472691 | 0.4162607 | 0.5277824 | 0.801135534 |
| ENSMUSG00000033128 | Gga1 | protein_coding | 0.254808434 | 4.297826087 | 0.4157518 | 0.5280317 | 0.801400794 |
| ENSMUSG00000019027 | Dnah1 | protein_coding | 0.285402284 | 5.188644309 | 0.4155021 | 0.5281688 | 0.801408679 |
| ENSMUSG00000045438 | Cox19 | protein_coding | 0.165345537 | 7.156107596 | 0.4152582 | 0.5282736 | 0.801408679 |
| ENSMUSG00000025384 | Faap100 | protein_coding | 0.363805204 | 4.090612436 | 0.4150936 | 0.5283691 | 0.801408679 |
| ENSMUSG00000015363 | Trabd | protein_coding | -0.287808021 | 4.444487945 | 0.4150191 | 0.528391 | 0.801408679 |
| ENSMUSG00000019718 | L3hypdh | protein_coding | 0.3588373 | 3.288056262 | 0.4149005 | 0.5284492 | 0.801408679 |
| ENSMUSG00000042694 | Stn1 | protein_coding | 0.23614046 | 4.440789371 | 0.4148294 | 0.528484 | 0.801408679 |
| ENSMUSG00000054428 | Atpif1 | protein_coding | 0.192920432 | 9.284073866 | 0.4142302 | 0.5287783 | 0.801493834 |
| ENSMUSG00000070509 | Rgma | protein_coding | 0.366177426 | 3.427260483 | 0.4142151 | 0.5287857 | 0.801493834 |
| ENSMUSG00000019877 | Serinc1 | protein_coding | 0.139051716 | 9.253247427 | 0.4140335 | 0.528875 | 0.801493834 |
| ENSMUSG00000042797 | Aqp11 | protein_coding | 0.307924779 | 3.620436067 | 0.4139917 | 0.5288955 | 0.801493834 |
| ENSMUSG00000037989 | Wnk2 | protein_coding | 0.250537122 | 4.822597408 | 0.413604 | 0.5290861 | 0.801493834 |
| ENSMUSG00000039221 | Rpl22l1 | protein_coding | 0.20326639 | 6.996212736 | 0.4136178 | 0.529094 | 0.801493834 |
| ENSMUSG00000032997 | Chpf | protein_coding | 0.250877759 | 5.077606247 | 0.4132593 | 0.5292557 | 0.801493834 |

| ensembl_gene_id | external_gene_name | gene_biotype | log2 (fold change) | log2 (counts per million) | F-value | p-value | False discovery rate |
|---|---|---|---|---|---|---|---|
| ENSMUSG00000006058 | Snf8 | protein_coding | 0.243657923 | 5.101828562 | 0.4130941 | 0.529337 | 0.801493834 |
| ENSMUSG00000026426 | Arl8a | protein_coding | 0.163580515 | 6.998271907 | 0.412953 | 0.5294065 | 0.801493834 |
| ENSMUSG00000050052 | Tdrp | protein_coding | 0.275260971 | 4.529503358 | 0.4128949 | 0.5294351 | 0.801493834 |
| ENSMUSG00000033900 | Map9 | protein_coding | -0.188463829 | 6.439453988 | 0.4125369 | 0.5296114 | 0.801493834 |
| ENSMUSG00000057176 | Ccdc189 | protein_coding | -0.233840196 | 4.692951345 | 0.4123369 | 0.52971 | 0.801493834 |
| ENSMUSG00000030322 | Mbd4 | protein_coding | 0.39581767 | 3.104928882 | 0.4123321 | 0.5297124 | 0.801493834 |
| ENSMUSG00000113216 | AL590503.1 | lincRNA | -0.335678284 | 4.160542779 | 0.412255 | 0.5297504 | 0.801493834 |
| ENSMUSG00000032536 | Trak1 | protein_coding | 0.188523779 | 5.859126942 | 0.4119644 | 0.5298937 | 0.801493834 |
| ENSMUSG00000018707 | Dync1h1 | protein_coding | -0.244625933 | 5.922824251 | 0.4119917 | 0.5298949 | 0.801493834 |
| ENSMUSG00000001127 | Araf | protein_coding | 0.144489087 | 9.716486823 | 0.411908 | 0.5299215 | 0.801493834 |
| ENSMUSG00000103034 | Gm8797 | protein_coding | -0.37326408 | 2.947511634 | 0.4116545 | 0.5300466 | 0.801493834 |
| ENSMUSG00000031387 | Renbp | protein_coding | -0.404565762 | 3.052426427 | 0.4115527 | 0.5300968 | 0.801493834 |
| ENSMUSG00000030103 | Bhlhe40 | protein_coding | -0.214143187 | 6.305069919 | 0.4115134 | 0.5301162 | 0.801493834 |
| ENSMUSG00000051159 | Cited1 | protein_coding | 0.304315261 | 5.499660301 | 0.4120171 | 0.5302406 | 0.801493834 |
| ENSMUSG00000016349 | Eef1a2 | protein_coding | -0.243292975 | 5.491949823 | 0.4112105 | 0.5302658 | 0.801493834 |
| ENSMUSG00000025591 | Tma16 | protein_coding | -0.244666186 | 4.834137173 | 0.4111749 | 0.5302834 | 0.801493834 |
| ENSMUSG00000031231 | Cox7b | protein_coding | 0.190055553 | 9.469297054 | 0.4110824 | 0.5303291 | 0.801493834 |
| ENSMUSG00000094870 | Zfp131 | protein_coding | 0.270973884 | 4.197405735 | 0.4107984 | 0.5304694 | 0.801593227 |
| ENSMUSG00000025825 | Iscu | protein_coding | 0.227842073 | 4.795017122 | 0.4104141 | 0.5306594 | 0.801609112 |
| ENSMUSG00000038902 | Pogz | protein_coding | -0.252737017 | 4.852381412 | 0.4103797 | 0.5306764 | 0.801609112 |
| ENSMUSG00000026596 | Abl2 | protein_coding | -0.253976471 | 4.873128831 | 0.4103248 | 0.5307035 | 0.801609112 |
| ENSMUSG00000033355 | Rtp4 | protein_coding | -0.405169021 | 3.126391149 | 0.4098678 | 0.5309297 | 0.801802931 |
| ENSMUSG00000022193 | Psmb5 | protein_coding | 0.147682616 | 6.843764382 | 0.4097642 | 0.530981 | 0.801802931 |
| ENSMUSG00000039824 | Myl6b | protein_coding | 0.409737142 | 2.879614197 | 0.4095967 | 0.531064 | 0.801815604 |
| ENSMUSG00000029279 | Brdt | protein_coding | 0.344198492 | 3.734904963 | 0.4092466 | 0.5312374 | 0.801964874 |
| ENSMUSG00000030884 | Uqcrc2 | protein_coding | 0.150936282 | 8.887543791 | 0.4089806 | 0.5313693 | 0.802051334 |
| ENSMUSG00000037872 | Ackr1 | protein_coding | 0.261759172 | 4.243581478 | 0.4084587 | 0.5316281 | 0.802247222 |
| ENSMUSG00000028309 | Rnf20 | protein_coding | -0.205802642 | 5.622701641 | 0.4084182 | 0.5316483 | 0.802247222 |
| ENSMUSG00000031010 | Usp9x | protein_coding | -0.26051265 | 4.380907038 | 0.4076285 | 0.5320404 | 0.802651438 |
| ENSMUSG00000063446 | Plppr1 | protein_coding | 0.218550298 | 4.933772752 | 0.4075782 | 0.5320654 | 0.802651438 |
| ENSMUSG00000021916 | Glt8d1 | protein_coding | -0.32935289 | 4.0443837 | 0.4067938 | 0.5324556 | 0.803127318 |
| ENSMUSG00000078794 | Dact3 | protein_coding | -0.266088576 | 4.872785388 | 0.4066112 | 0.5325464 | 0.803148838 |
| ENSMUSG00000040189 | Ccdc114 | protein_coding | -1.040084384 | 2.03408562 | 0.4075969 | 0.5326192 | 0.803148838 |
| ENSMUSG00000048799 | Cep120 | protein_coding | 0.236968112 | 4.952780151 | 0.4062518 | 0.5327254 | 0.803196347 |
| ENSMUSG00000039852 | Rere | protein_coding | -0.192870754 | 6.268421773 | 0.405889 | 0.5329062 | 0.803228544 |
| ENSMUSG00000061838 | Suclg2 | protein_coding | -0.417974446 | 3.038395466 | 0.4057915 | 0.5329548 | 0.803228544 |
| ENSMUSG00000024270 | Slc39a6 | protein_coding | 0.260077397 | 5.133146725 | 0.4056381 | 0.5330313 | 0.803228544 |
| ENSMUSG00000032087 | Dscaml1 | protein_coding | 0.444864928 | 3.56028053 | 0.4056386 | 0.5330456 | 0.803228544 |
| ENSMUSG00000027342 | Pcna | protein_coding | -0.200817437 | 5.227734481 | 0.405347 | 0.5331765 | 0.803313329 |
| ENSMUSG00000026176 | Ctdsp1 | protein_coding | -0.371328748 | 3.061700666 | 0.4049284 | 0.5333855 | 0.803462873 |
| ENSMUSG00000069631 | Strada | protein_coding | -0.288331341 | 4.689361972 | 0.4048488 | 0.5334252 | 0.803462873 |
| ENSMUSG00000044340 | Phlpp1 | protein_coding | -0.211968078 | 5.578806587 | 0.4043974 | 0.5336507 | 0.803646395 |
| ENSMUSG00000059734 | Ndufs8 | protein_coding | 0.154246303 | 9.673630641 | 0.4041788 | 0.53376 | 0.803646395 |
| ENSMUSG00000014426 | Map3k4 | protein_coding | 0.283199626 | 4.223032953 | 0.4037669 | 0.533966 | 0.803646395 |
| ENSMUSG00000069206 | Zfp874a | protein_coding | 0.325781049 | 3.615797756 | 0.4035397 | 0.5340797 | 0.803646395 |
| ENSMUSG00000037426 | Depdc5 | protein_coding | -0.266060558 | 4.586188718 | 0.4033696 | 0.5341649 | 0.803646395 |
| ENSMUSG00000078676 | Casc3 | protein_coding | -0.240248254 | 4.850136266 | 0.4033229 | 0.5341883 | 0.803646395 |
| ENSMUSG00000029147 | Ppm1g | protein_coding | 0.212988021 | 5.239946908 | 0.4032784 | 0.5342106 | 0.803646395 |
| ENSMUSG00000034345 | Gtf2h5 | protein_coding | 0.144950103 | 8.783717357 | 0.4031572 | 0.5342713 | 0.803646395 |
| ENSMUSG00000040147 | Maob | protein_coding | -0.304448931 | 6.05615431 | 0.4048225 | 0.5343173 | 0.803646395 |
| ENSMUSG00000063281 | Zfp35 | protein_coding | 0.237647525 | 5.15766405 | 0.4029745 | 0.5343628 | 0.803646395 |
| ENSMUSG00000022995 | Enah | protein_coding | -0.159095496 | 6.376213725 | 0.402911 | 0.5343946 | 0.803646395 |
| ENSMUSG00000023110 | Prmt5 | protein_coding | -0.15698912 | 6.795089092 | 0.4028126 | 0.5344439 | 0.803646395 |
| ENSMUSG00000031059 | Ndufb11 | protein_coding | 0.178854422 | 9.367674835 | 0.4023641 | 0.5346688 | 0.803660859 |
| ENSMUSG00000022816 | Fstl1 | protein_coding | 0.223826627 | 5.619074361 | 0.4023589 | 0.5346714 | 0.803660859 |
| ENSMUSG00000027498 | Cstf1 | protein_coding | 0.209509441 | 5.412004816 | 0.4023048 | 0.5346985 | 0.803660859 |
| ENSMUSG00000051355 | Commd1 | protein_coding | 0.289885831 | 3.530138769 | 0.4018588 | 0.5349224 | 0.803660859 |
| ENSMUSG00000078161 | Erich3 | protein_coding | -0.321469211 | 4.031065691 | 0.4016918 | 0.5350062 | 0.803660859 |
| ENSMUSG00000031701 | Dnaja2 | protein_coding | 0.167343807 | 7.840731032 | 0.4016595 | 0.5350224 | 0.803660859 |
| ENSMUSG00000024663 | Rab3il1 | protein_coding | -0.375221315 | 3.605714295 | 0.4016158 | 0.5350444 | 0.803660859 |
| ENSMUSG00000028439 | Fam219a | protein_coding | 0.464933502 | 2.820741868 | 0.4016017 | 0.5350514 | 0.803660859 |
| ENSMUSG00000031785 | Adgrg1 | protein_coding | 0.177253217 | 6.230589893 | 0.4011119 | 0.5352975 | 0.80369268 |
| ENSMUSG00000014402 | Tsg101 | protein_coding | 0.145270262 | 6.732788674 | 0.4010933 | 0.5353069 | 0.80369268 |
| ENSMUSG00000055943 | Emc7 | protein_coding | 0.141693077 | 7.536572216 | 0.401037 | 0.5353352 | 0.80369268 |
| ENSMUSG00000028293 | Slc35a1 | protein_coding | 0.216840619 | 5.190766832 | 0.4007286 | 0.5354903 | 0.80369268 |
| ENSMUSG00000001785 | Pwp1 | protein_coding | -0.186741769 | 5.304626079 | 0.4007112 | 0.535499 | 0.80369268 |
| ENSMUSG00000032216 | Nedd4 | protein_coding | 0.176597043 | 8.22302611 | 0.4006675 | 0.5355211 | 0.80369268 |
| ENSMUSG00000030168 | Adipor2 | protein_coding | -0.34149961 | 4.827109208 | 0.4008641 | 0.5356663 | 0.803693911 |
| ENSMUSG00000061981 | Flot2 | protein_coding | 0.177172387 | 6.29793369 | 0.4003688 | 0.5356714 | 0.803693911 |
| ENSMUSG00000098923 | Tmem185b | protein_coding | -0.264403936 | 4.597714848 | 0.400169 | 0.5357719 | 0.803732658 |
| ENSMUSG00000021477 | Ctsl | protein_coding | 0.155608096 | 7.359115582 | 0.3997087 | 0.5360038 | 0.803853835 |
| ENSMUSG00000023074 | Mospd1 | protein_coding | 0.200020335 | 5.229285499 | 0.399598 | 0.5360596 | 0.803853835 |
| ENSMUSG00000057551 | Zfp317 | protein_coding | -0.275000876 | 4.910413976 | 0.3994329 | 0.5361429 | 0.803853835 |
| ENSMUSG00000030898 | Cckbr | protein_coding | 0.449316573 | 3.172540333 | 0.399215 | 0.5362527 | 0.803853835 |
| ENSMUSG00000058966 | Fam57b | protein_coding | 0.365283752 | 4.642708723 | 0.4000365 | 0.5362834 | 0.803853835 |
| ENSMUSG00000022629 | Kif21a | protein_coding | -0.173119223 | 7.104242049 | 0.3991188 | 0.5363013 | 0.803853835 |
| ENSMUSG00000031681 | Smad1 | protein_coding | -0.237141875 | 4.910375733 | 0.3986717 | 0.5365269 | 0.804079952 |
| ENSMUSG00000019876 | Pkib | protein_coding | 0.214907395 | 4.949440417 | 0.3984371 | 0.5366453 | 0.804145401 |

| ensembl_gene_id | external_gene_name | gene_biotype | log2 (fold change) | log2 (counts per million) | F-value | p-value | False discovery rate |
|---|---|---|---|---|---|---|---|
| ENSMUSG00000032281 | Acsbg1 | protein_coding | -0.167381457 | 7.206270198 | 0.3980096 | 0.5368613 | 0.804356961 |
| ENSMUSG00000023007 | Prpf40b | protein_coding | 0.2676606 | 5.256867665 | 0.3975965 | 0.5370702 | 0.804454807 |
| ENSMUSG00000043633 | Fam221b | protein_coding | 0.391625145 | 3.063791925 | 0.3975846 | 0.5370763 | 0.804454807 |
| ENSMUSG00000037331 | Larp1 | protein_coding | -0.220384078 | 5.011395459 | 0.3970438 | 0.5373499 | 0.804752631 |
| ENSMUSG00000024294 | Mib1 | protein_coding | -0.194663945 | 6.045810957 | 0.3968864 | 0.5374296 | 0.804759896 |
| ENSMUSG00000000976 | Heatr6 | protein_coding | 0.251067811 | 4.608350955 | 0.3966903 | 0.537529 | 0.804796592 |
| ENSMUSG00000007610 | Gtpbp3 | protein_coding | -0.417600188 | 2.835083554 | 0.3955435 | 0.5381106 | 0.805433318 |
| ENSMUSG00000029684 | Wasl | protein_coding | -0.171575788 | 6.066731873 | 0.3953481 | 0.5382098 | 0.805433318 |
| ENSMUSG00000053310 | Nrgn | protein_coding | -0.18446498 | 7.189110252 | 0.3951246 | 0.5383233 | 0.805433318 |
| ENSMUSG00000031314 | Taf1 | protein_coding | -0.240599734 | 5.39495505 | 0.395108 | 0.5383318 | 0.805433318 |
| ENSMUSG00000021262 | Evl | protein_coding | 0.354175251 | 3.886538547 | 0.3950818 | 0.5383451 | 0.805433318 |
| ENSMUSG00000028999 | Rint1 | protein_coding | -0.25841062 | 3.966373281 | 0.3949665 | 0.5384037 | 0.805433318 |
| ENSMUSG00000043192 | Gm1840 | processed_pseudoge | 0.707210531 | 2.559594542 | 0.3944426 | 0.5386843 | 0.805741055 |
| ENSMUSG00000002808 | Epdr1 | protein_coding | -0.17645612 | 6.191063094 | 0.3940934 | 0.5388477 | 0.805873409 |
| ENSMUSG00000027340 | Slc23a2 | protein_coding | 0.235291181 | 5.141563334 | 0.3937743 | 0.5390102 | 0.806004286 |
| ENSMUSG00000021226 | Acot2 | protein_coding | 0.274534285 | 5.08055963 | 0.3929761 | 0.539417 | 0.806391264 |
| ENSMUSG00000029191 | Rfc1 | protein_coding | -0.199366603 | 5.548123575 | 0.3928556 | 0.5394785 | 0.806391264 |
| ENSMUSG00000031029 | Eif3f | protein_coding | 0.189488142 | 6.454889722 | 0.392628 | 0.5395946 | 0.806391264 |
| ENSMUSG00000038175 | Mylip | protein_coding | -0.388409151 | 3.224945135 | 0.3926181 | 0.5395996 | 0.806391264 |
| ENSMUSG00000041961 | Znrf3 | protein_coding | -0.253779228 | 5.062820556 | 0.3924602 | 0.5396803 | 0.806391264 |
| ENSMUSG00000038115 | Ano2 | protein_coding | -0.520744467 | 3.114696155 | 0.3923055 | 0.5397734 | 0.806391264 |
| ENSMUSG00000028127 | Abcd3 | protein_coding | 0.220582256 | 5.816527553 | 0.392059 | 0.5398851 | 0.806391264 |
| ENSMUSG00000043432 | Leng9 | protein_coding | 0.25810392 | 4.373271908 | 0.3917821 | 0.5400266 | 0.806391264 |
| ENSMUSG00000031591 | Asah1 | protein_coding | -0.172515006 | 6.231860948 | 0.3917455 | 0.5400453 | 0.806391264 |
| ENSMUSG00000028199 | Cryz | protein_coding | 0.255350081 | 5.182555888 | 0.3916767 | 0.5400805 | 0.806391264 |
| ENSMUSG00000049792 | Bag5 | protein_coding | -0.156627504 | 6.379971003 | 0.3916504 | 0.5400939 | 0.806391264 |
| ENSMUSG00000000301 | Pemt | protein_coding | 0.351398451 | 3.320569159 | 0.3913277 | 0.540259 | 0.806525714 |
| ENSMUSG00000044317 | Gpr4 | protein_coding | -0.611622207 | 2.413926807 | 0.3908622 | 0.5404972 | 0.806759794 |
| ENSMUSG00000038467 | Chmp4b | protein_coding | -0.226799513 | 5.568378355 | 0.3907281 | 0.5405658 | 0.806759794 |
| ENSMUSG00000003068 | Stk11 | protein_coding | -0.22711695 | 5.19273725 | 0.3905462 | 0.540659 | 0.806786811 |
| ENSMUSG00000030424 | Zfp939 | transcribed_unproces | -0.362756894 | 3.067086335 | 0.3900881 | 0.5408937 | 0.806960133 |
| ENSMUSG00000035342 | Lzts2 | protein_coding | 0.481306103 | 3.030487529 | 0.3900267 | 0.5409252 | 0.806960133 |
| ENSMUSG00000046062 | Ppp1r15b | protein_coding | -0.235995338 | 4.111451231 | 0.3896781 | 0.541104 | 0.807080369 |
| ENSMUSG00000014164 | Klhl3 | protein_coding | -0.241560133 | 6.45429583 | 0.3902818 | 0.5411559 | 0.807080369 |
| ENSMUSG00000037563 | Rps16 | protein_coding | 0.414224565 | 2.816220096 | 0.3888677 | 0.5415201 | 0.80741686 |
| ENSMUSG00000024070 | Prkd3 | protein_coding | -0.324741406 | 3.494450604 | 0.3887754 | 0.5415675 | 0.80741686 |
| ENSMUSG00000052707 | Tnrc6a | protein_coding | -0.194083841 | 5.374586533 | 0.388699 | 0.5416068 | 0.80741686 |
| ENSMUSG00000022237 | Ankrd33b | protein_coding | -0.503260173 | 3.099320655 | 0.3885772 | 0.5416835 | 0.807419297 |
| ENSMUSG00000071866 | Ppia | protein_coding | 0.141404538 | 7.158662897 | 0.388296 | 0.541814 | 0.80750184 |
| ENSMUSG00000059248 | 9-Sep | protein_coding | 0.350048065 | 5.15528298 | 0.3895036 | 0.5418992 | 0.807516876 |
| ENSMUSG00000087150 | BC064078 | transcribed_unproces | -0.411616022 | 3.367491776 | 0.387716 | 0.5421124 | 0.80772276 |
| ENSMUSG00000026269 | Rnpepl1 | protein_coding | -0.322793405 | 3.453164609 | 0.3874593 | 0.5422446 | 0.807807737 |
| ENSMUSG00000029657 | Hsph1 | protein_coding | 0.152443687 | 7.818311086 | 0.3866968 | 0.5426377 | 0.808281343 |
| ENSMUSG00000026820 | Ptges2 | protein_coding | -0.200375223 | 5.29739169 | 0.3865284 | 0.5427245 | 0.80829875 |
| ENSMUSG00000030583 | Sipa1l3 | protein_coding | -0.40857452 | 3.080284127 | 0.3859734 | 0.543011 | 0.808509003 |
| ENSMUSG00000008429 | Herpud2 | protein_coding | -0.304147967 | 4.95469928 | 0.3858488 | 0.5430895 | 0.808509003 |
| ENSMUSG00000039176 | Polg | protein_coding | 0.598711588 | 2.443151353 | 0.3858181 | 0.5430913 | 0.808509003 |
| ENSMUSG00000074093 | Svip | protein_coding | -0.196598123 | 6.117589266 | 0.3855773 | 0.5432157 | 0.808582302 |
| ENSMUSG00000020069 | Hnrnph3 | protein_coding | 0.271802524 | 4.443107926 | 0.38535 | 0.5433332 | 0.808645237 |
| ENSMUSG00000026571 | Dcaf6 | protein_coding | 0.230153204 | 4.34606051 | 0.3843606 | 0.5438453 | 0.809253033 |
| ENSMUSG00000028528 | Dnajc6 | protein_coding | -0.147015853 | 6.82525502 | 0.3842702 | 0.5438921 | 0.809253033 |
| ENSMUSG00000025894 | Aasdhppt | protein_coding | 0.172551866 | 6.465648914 | 0.3834675 | 0.5443083 | 0.809760229 |
| ENSMUSG00000069208 | Zfp825 | protein_coding | -0.299246634 | 3.554510274 | 0.3832452 | 0.5444236 | 0.809819808 |
| ENSMUSG00000000056 | Narf | protein_coding | -0.196837202 | 5.785371926 | 0.3827592 | 0.544676 | 0.81002675 |
| ENSMUSG00000002064 | Sdf2 | protein_coding | 0.162385435 | 7.649452526 | 0.3826872 | 0.5447134 | 0.81002675 |
| ENSMUSG00000026080 | Chst10 | protein_coding | 0.188136318 | 5.767554448 | 0.3823106 | 0.5449092 | 0.810114189 |
| ENSMUSG00000004268 | Emg1 | protein_coding | -0.137877307 | 7.510535678 | 0.3822842 | 0.5449229 | 0.810114189 |
| ENSMUSG00000071103 | 1700029J07Rik | protein_coding | -0.755095466 | 2.093148936 | 0.3817221 | 0.5452153 | 0.810426873 |
| ENSMUSG00000031870 | Pgr | protein_coding | -0.234478358 | 4.161279224 | 0.3815902 | 0.545284 | 0.810426873 |
| ENSMUSG00000025878 | Uimc1 | protein_coding | -0.241926629 | 4.440030561 | 0.3812808 | 0.5454451 | 0.810554294 |
| ENSMUSG00000039563 | 2210406O10Rik | lincRNA | -0.347652775 | 3.185292419 | 0.3809584 | 0.5456131 | 0.810691938 |
| ENSMUSG00000050332 | Amer1 | protein_coding | 0.429271744 | 3.496551886 | 0.3806857 | 0.5457692 | 0.810721755 |
| ENSMUSG00000029464 | Gpn3 | protein_coding | 0.228458284 | 5.759942848 | 0.3806307 | 0.5457839 | 0.810721755 |
| ENSMUSG00000024580 | Grpel2 | protein_coding | 0.199140844 | 5.73925752 | 0.3802227 | 0.5459968 | 0.810788021 |
| ENSMUSG00000045613 | Chrm2 | protein_coding | -0.259679424 | 4.4486178 | 0.3801148 | 0.5460531 | 0.810788021 |
| ENSMUSG00000060376 | Bckdha | protein_coding | 0.371203277 | 3.830387795 | 0.3800985 | 0.5460756 | 0.810788021 |
| ENSMUSG00000021750 | Fam107a | protein_coding | -0.189287635 | 6.542335667 | 0.3799673 | 0.5461302 | 0.810788021 |
| ENSMUSG00000031207 | Msn | protein_coding | -0.46692904 | 2.915599636 | 0.3794725 | 0.5463886 | 0.810893221 |
| ENSMUSG00000035944 | Ttc38 | protein_coding | -0.343685259 | 3.406208913 | 0.3794189 | 0.5464167 | 0.810893221 |
| ENSMUSG00000028797 | Tmem234 | protein_coding | 0.132099332 | 7.596368689 | 0.3792632 | 0.546498 | 0.810893221 |
| ENSMUSG00000020198 | Ap3d1 | protein_coding | 0.182872157 | 6.565677307 | 0.3792544 | 0.5465027 | 0.810893221 |
| ENSMUSG00000028295 | Smim8 | protein_coding | -0.182533268 | 6.499060213 | 0.379052 | 0.5466085 | 0.810918073 |
| ENSMUSG00000021932 | Rnaseh2b | protein_coding | 0.204235958 | 5.32804282 | 0.3787209 | 0.5467817 | 0.810918073 |
| ENSMUSG00000057230 | Aak1 | protein_coding | -0.150247898 | 6.234859702 | 0.3785898 | 0.5468503 | 0.810918073 |
| ENSMUSG00000097219 | Gm26551 | lincRNA | -0.316427227 | 3.711355246 | 0.3785325 | 0.5468804 | 0.810918073 |
| ENSMUSG00000035171 | 1110059E24Rik | protein_coding | -0.153963081 | 7.010336727 | 0.3785017 | 0.5468965 | 0.810918073 |
| ENSMUSG00000001794 | Capns1 | protein_coding | 0.140503676 | 7.23377314 | 0.3781432 | 0.5470843 | 0.811084681 |
| ENSMUSG00000061911 | Myt1l | protein_coding | -0.137142905 | 7.934148902 | 0.3772048 | 0.5475763 | 0.811514863 |

| ensembl_gene_id | external_gene_name | gene_biotype | log2 (fold change) | log2 (counts per million) | F-value | p-value | False discovery rate |
|---|---|---|---|---|---|---|---|
| ENSMUSG00000033970 | Rfc3 | protein_coding | -0.203682141 | 4.936916151 | 0.3771864 | 0.547586 | 0.811514863 |
| ENSMUSG00000024260 | Sap130 | protein_coding | -0.325159144 | 3.683428337 | 0.3771581 | 0.5476008 | 0.811514863 |
| ENSMUSG00000060402 | Chst8 | protein_coding | 0.42754996 | 3.486511991 | 0.3768564 | 0.5477732 | 0.811658373 |
| ENSMUSG00000068373 | D430041D05Rik | protein_coding | -0.242211213 | 4.776379239 | 0.3766311 | 0.5478776 | 0.811680568 |
| ENSMUSG00000052302 | Tbc1d30 | protein_coding | -0.192523885 | 5.205387389 | 0.376391 | 0.5480038 | 0.811680568 |
| ENSMUSG00000026558 | Uck2 | protein_coding | 0.311494176 | 3.764361791 | 0.3763704 | 0.5480146 | 0.811680568 |
| ENSMUSG00000021973 | Micu2 | protein_coding | -0.272710591 | 4.362806374 | 0.3760735 | 0.5481707 | 0.811781648 |
| ENSMUSG00000000581 | C1d | protein_coding | -0.162874914 | 9.177309877 | 0.3759535 | 0.5482338 | 0.811781648 |
| ENSMUSG00000032307 | Ube2q2 | protein_coding | -0.191809767 | 5.230685422 | 0.3756724 | 0.5483818 | 0.811888894 |
| ENSMUSG00000006050 | Sra1 | protein_coding | 0.153057241 | 7.857434219 | 0.3753595 | 0.5485465 | 0.812012957 |
| ENSMUSG00000032392 | Parp16 | protein_coding | 0.503574182 | 2.656371158 | 0.3752264 | 0.5486166 | 0.812012957 |
| ENSMUSG00000030536 | Iqgap1 | protein_coding | -0.392984693 | 3.601172273 | 0.3744811 | 0.5490095 | 0.812482615 |
| ENSMUSG00000057421 | Las1l | protein_coding | 0.182954582 | 5.820323439 | 0.3739185 | 0.5493063 | 0.812740688 |
| ENSMUSG00000055322 | Tns1 | protein_coding | -0.406320743 | 3.784379738 | 0.3739023 | 0.5493652 | 0.812740688 |
| ENSMUSG00000055737 | Ghr | protein_coding | -0.354360982 | 2.985612569 | 0.3737212 | 0.5494106 | 0.812740688 |
| ENSMUSG00000002781 | Tmem143 | protein_coding | 0.300742557 | 3.594656534 | 0.3731076 | 0.5497349 | 0.813029412 |
| ENSMUSG00000023079 | Gtf2ird1 | protein_coding | -0.279822687 | 4.057280364 | 0.3728701 | 0.5498605 | 0.813029412 |
| ENSMUSG00000021555 | Naa35 | protein_coding | 0.149962781 | 6.4186992 | 0.3728031 | 0.549896 | 0.813029412 |
| ENSMUSG00000090069 | E430024P14Rik | antisense_RNA | -0.512703401 | 3.360192038 | 0.3737312 | 0.5499082 | 0.813029412 |
| ENSMUSG00000022416 | Cacna1i | protein_coding | -0.444784355 | 2.987274636 | 0.3725822 | 0.5500129 | 0.813043249 |
| ENSMUSG00000055067 | Smyd3 | protein_coding | 0.272068209 | 6.563992212 | 0.3745419 | 0.5500688 | 0.813043249 |
| ENSMUSG00000074170 | Plekhf1 | protein_coding | -0.363918076 | 3.88026458 | 0.3721885 | 0.5502351 | 0.813177351 |
| ENSMUSG00000003031 | Cdkn1b | protein_coding | -0.265881875 | 5.148480046 | 0.3719997 | 0.5503214 | 0.81319304 |
| ENSMUSG00000037058 | Paip2 | protein_coding | 0.128859011 | 7.838311285 | 0.3717706 | 0.5504428 | 0.813205056 |
| ENSMUSG00000030545 | Pex11a | protein_coding | -0.317326627 | 4.157662437 | 0.371699 | 0.5504808 | 0.813205056 |
| ENSMUSG00000056579 | Tug1 | processed_transcript | 0.25589199 | 6.521281845 | 0.3726339 | 0.5506913 | 0.813295838 |
| ENSMUSG00000071757 | Zhx2 | protein_coding | 0.37298279 | 3.222623385 | 0.371298 | 0.5506935 | 0.813295838 |
| ENSMUSG00000050558 | Prokr2 | protein_coding | -0.534874451 | 2.716448841 | 0.3710512 | 0.5508245 | 0.813377572 |
| ENSMUSG00000001767 | Crnkl1 | protein_coding | -0.258093587 | 4.060720349 | 0.370703 | 0.5510094 | 0.813538888 |
| ENSMUSG00000087269 | D330023K18Rik | processed_transcript | 0.192831903 | 5.379798181 | 0.3701458 | 0.5513055 | 0.813864371 |
| ENSMUSG00000022306 | Zfpm2 | protein_coding | 0.544550627 | 2.911789258 | 0.3696037 | 0.5516077 | 0.81401973 |
| ENSMUSG00000028163 | Nfkb1 | protein_coding | 0.358723051 | 3.062743816 | 0.3693994 | 0.5517027 | 0.81401973 |
| ENSMUSG00000092094 | Zfp804b | protein_coding | -0.410534162 | 2.697701864 | 0.3692359 | 0.5517898 | 0.81401973 |
| ENSMUSG00000028664 | Ephb2 | protein_coding | -0.413923265 | 2.859946521 | 0.36919 | 0.5518142 | 0.81401973 |
| ENSMUSG00000035184 | Fam124a | protein_coding | -0.294926267 | 4.021235127 | 0.3691297 | 0.5518464 | 0.81401973 |
| ENSMUSG00000007777 | 0610009B22Rik | protein_coding | -0.169931563 | 7.420469813 | 0.369035 | 0.5518968 | 0.81401973 |
| ENSMUSG00000021427 | Ssr1 | protein_coding | 0.154324478 | 8.64397154 | 0.3688231 | 0.5520097 | 0.81401973 |
| ENSMUSG00000035142 | Nubpl | protein_coding | 0.352641932 | 3.289785091 | 0.3686918 | 0.5520797 | 0.81401973 |
| ENSMUSG00000048388 | Fam171b | protein_coding | -0.147104187 | 7.116365488 | 0.3686687 | 0.5520921 | 0.81401973 |
| ENSMUSG00000054728 | Phactr1 | protein_coding | -0.136776834 | 8.711495992 | 0.3683686 | 0.5522521 | 0.814144131 |
| ENSMUSG00000058886 | Deaf1 | protein_coding | 0.261057896 | 4.746996997 | 0.3678354 | 0.5525368 | 0.814452096 |
| ENSMUSG00000032262 | Elovl4 | protein_coding | 0.170997503 | 5.962057517 | 0.3675869 | 0.5526695 | 0.814536067 |
| ENSMUSG00000019143 | Hars2 | protein_coding | 0.254829886 | 4.390408431 | 0.3665841 | 0.5532058 | 0.815127646 |
| ENSMUSG00000024990 | Rbp4 | protein_coding | -0.44777378 | 2.89401527 | 0.3664868 | 0.5532579 | 0.815127646 |
| ENSMUSG00000025199 | Chuk | protein_coding | -0.46557618 | 3.326487007 | 0.3664369 | 0.5532983 | 0.815127646 |
| ENSMUSG00000049858 | Suox | protein_coding | 0.372186247 | 3.829617284 | 0.3661914 | 0.5534298 | 0.815209668 |
| ENSMUSG00000036510 | Cdh8 | protein_coding | -0.299134409 | 4.200600969 | 0.3658078 | 0.5536217 | 0.815273476 |
| ENSMUSG00000037896 | Rcor1 | protein_coding | -0.304469344 | 4.042405478 | 0.3657485 | 0.5536535 | 0.815273476 |
| ENSMUSG00000091476 | Gm16432 | protein_coding | -0.470420865 | 2.639064719 | 0.3656607 | 0.5537006 | 0.815273476 |
| ENSMUSG00000027016 | Zfp385b | protein_coding | 0.201208572 | 6.379825828 | 0.3654871 | 0.5537937 | 0.81529896 |
| ENSMUSG00000015968 | Cacna1d | protein_coding | 0.289357765 | 3.628391845 | 0.3652666 | 0.553912 | 0.81536147 |
| ENSMUSG00000023353 | Agap3 | protein_coding | 0.171830719 | 5.872220697 | 0.3650076 | 0.554051 | 0.815454496 |
| ENSMUSG00000031154 | Otud5 | protein_coding | -0.231002131 | 5.270545097 | 0.3644835 | 0.5543326 | 0.81565588 |
| ENSMUSG00000028443 | Nudt2 | protein_coding | -0.151559296 | 6.671103307 | 0.3644705 | 0.5543395 | 0.81565588 |
| ENSMUSG00000007850 | Hnrnph1 | protein_coding | -0.148176987 | 6.886565233 | 0.3642317 | 0.5544679 | 0.815690939 |
| ENSMUSG00000036291 | Ap5m1 | protein_coding | -0.303216985 | 3.98614786 | 0.364144 | 0.5545151 | 0.815690939 |
| ENSMUSG00000036391 | Sec24a | protein_coding | -0.295538848 | 3.856373162 | 0.3638789 | 0.5546577 | 0.815789148 |
| ENSMUSG00000029326 | Enoph1 | protein_coding | 0.140240659 | 7.226127742 | 0.3636732 | 0.5547684 | 0.815840382 |
| ENSMUSG00000006705 | Pknox1 | protein_coding | 0.261529445 | 4.007590986 | 0.3634361 | 0.554896 | 0.815916464 |
| ENSMUSG00000107476 | Zfp862-ps | transcribed_unitary_p | -0.422680645 | 2.780247633 | 0.362999 | 0.5551316 | 0.816077903 |
| ENSMUSG00000069072 | Slc7a14 | protein_coding | 0.166995766 | 6.089734951 | 0.3629506 | 0.5551576 | 0.816077903 |
| ENSMUSG00000022375 | Lrrc6 | protein_coding | -0.415735274 | 3.852469351 | 0.3633254 | 0.5553024 | 0.816156879 |
| ENSMUSG00000047897 | Ripply2 | protein_coding | 0.388270791 | 2.911430618 | 0.3625695 | 0.5553631 | 0.816156879 |
| ENSMUSG00000020135 | Apc2 | protein_coding | -0.298173591 | 4.679888566 | 0.3621721 | 0.5555912 | 0.816380408 |
| ENSMUSG00000026254 | Eif4e2 | protein_coding | 0.161861772 | 6.461110761 | 0.3617949 | 0.5557813 | 0.816548143 |
| ENSMUSG00000026933 | Camsap1 | protein_coding | -0.206311933 | 5.571069479 | 0.3600311 | 0.5567357 | 0.817744187 |
| ENSMUSG00000031939 | Taf1d | protein_coding | -0.285400929 | 4.656538511 | 0.3599791 | 0.5567638 | 0.817744187 |
| ENSMUSG00000034111 | Tmed8 | protein_coding | -0.239065681 | 4.609132762 | 0.3598691 | 0.5568235 | 0.817744187 |
| ENSMUSG00000035206 | Sppl2b | protein_coding | 0.290276767 | 3.960929296 | 0.3591146 | 0.5572329 | 0.818167268 |
| ENSMUSG00000026478 | Lamc1 | protein_coding | 0.379060991 | 3.023260391 | 0.3590577 | 0.5572637 | 0.818167268 |
| ENSMUSG00000022263 | Trio | protein_coding | -0.201031553 | 5.447010341 | 0.3587596 | 0.5574257 | 0.818293268 |
| ENSMUSG00000037907 | Ankrd13b | protein_coding | -0.391460296 | 3.239422366 | 0.3583579 | 0.557644 | 0.81850207 |
| ENSMUSG00000029594 | Rbm19 | protein_coding | 0.371541144 | 3.338040783 | 0.3580795 | 0.5577955 | 0.81860077 |
| ENSMUSG00000022419 | Deptor | protein_coding | -0.351196104 | 4.525746661 | 0.358457 | 0.5578635 | 0.81860077 |
| ENSMUSG00000037624 | Kcnk2 | protein_coding | 0.219988383 | 5.383507972 | 0.3574494 | 0.5581384 | 0.818818282 |
| ENSMUSG00000057335 | Cep170 | protein_coding | -0.197158779 | 5.24209487 | 0.3573598 | 0.5581873 | 0.818818282 |
| ENSMUSG00000042211 | Fbxo38 | protein_coding | 0.1938646 | 5.164586381 | 0.3572627 | 0.5582402 | 0.818818282 |
| ENSMUSG00000028032 | Papss1 | protein_coding | 0.139813928 | 6.706910355 | 0.3570045 | 0.5583809 | 0.818912959 |

| ensembl_gene_id | external_gene_name | gene_biotype | log2 (fold change) | log2 (counts per million) | F-value | p-value | False discovery rate |
|---|---|---|---|---|---|---|---|
| ENSMUSG00000001419 | Mef2d | protein_coding | 0.170586333 | 5.675214612 | 0.3564967 | 0.5586579 | 0.819207421 |
| ENSMUSG00000037447 | Arid5a | protein_coding | -0.325348768 | 4.065854634 | 0.3562042 | 0.5588175 | 0.819329839 |
| ENSMUSG00000034121 | Mks1 | protein_coding | 0.939715766 | 2.16448445 | 0.3563096 | 0.559314 | 0.819945937 |
| ENSMUSG00000003810 | Mast2 | protein_coding | -0.186031199 | 5.41546014 | 0.354201 | 0.5599134 | 0.820632643 |
| ENSMUSG00000021798 | Ldb3 | protein_coding | -0.281933379 | 4.315403515 | 0.3540543 | 0.5599939 | 0.820632643 |
| ENSMUSG00000056692 | D17Wsu92e | protein_coding | -0.145137384 | 6.607723939 | 0.3540087 | 0.5600188 | 0.820632643 |
| ENSMUSG00000051864 | Tbc1d22a | protein_coding | 0.251372847 | 4.965717648 | 0.3538831 | 0.5600877 | 0.820632643 |
| ENSMUSG00000030032 | Wdr54 | protein_coding | 0.182488816 | 5.463490421 | 0.3528154 | 0.5606739 | 0.821286261 |
| ENSMUSG00000027551 | Zfp64 | protein_coding | 0.253039746 | 5.124007071 | 0.3527924 | 0.5606865 | 0.821286261 |
| ENSMUSG00000006931 | P3h4 | protein_coding | -0.277400935 | 3.879283003 | 0.3523522 | 0.5609286 | 0.821452938 |
| ENSMUSG00000025159 | Mms19 | protein_coding | 0.267054814 | 4.342493088 | 0.3523076 | 0.5609531 | 0.821452938 |
| ENSMUSG00000021290 | 2010107E04Rik | protein_coding | 0.187676454 | 9.27045432 | 0.3520951 | 0.5610834 | 0.821519499 |
| ENSMUSG00000038784 | Cnot4 | protein_coding | -0.181072579 | 6.426545098 | 0.3518048 | 0.5612299 | 0.821519499 |
| ENSMUSG00000030096 | Slc6a6 | protein_coding | 0.187359617 | 5.537140466 | 0.3517233 | 0.5612747 | 0.821519499 |
| ENSMUSG00000029581 | Fscn1 | protein_coding | -0.174448588 | 5.563897605 | 0.351665 | 0.5613069 | 0.821519499 |
| ENSMUSG00000028671 | Gale | protein_coding | 0.257647842 | 4.816075524 | 0.3515131 | 0.5613905 | 0.821519499 |
| ENSMUSG00000030306 | Tmtc1 | protein_coding | -0.226986152 | 4.54557722 | 0.3510897 | 0.5616239 | 0.821519499 |
| ENSMUSG00000031447 | Lamp1 | protein_coding | 0.307328906 | 3.552951563 | 0.3508972 | 0.5617301 | 0.821519499 |
| ENSMUSG00000059981 | Taok2 | protein_coding | -0.262101259 | 4.662937901 | 0.3507782 | 0.5617957 | 0.821519499 |
| ENSMUSG00000092083 | Kcnb2 | protein_coding | -0.312321394 | 3.41738996 | 0.350775 | 0.5617975 | 0.821519499 |
| ENSMUSG00000030614 | Tmem126b | protein_coding | 0.148246809 | 7.484926532 | 0.3507494 | 0.5618116 | 0.821519499 |
| ENSMUSG00000087075 | A230065H16Rik | protein_coding | -0.173194062 | 6.340546695 | 0.3506999 | 0.561839 | 0.821519499 |
| ENSMUSG00000008384 | Sertad1 | protein_coding | -0.226240091 | 4.178920212 | 0.3503163 | 0.5620507 | 0.821717428 |
| ENSMUSG00000072582 | Ptrh2 | protein_coding | -0.166262301 | 6.181312669 | 0.3496935 | 0.5623949 | 0.822060697 |
| ENSMUSG00000029486 | Mrpl1 | protein_coding | -0.194137264 | 5.919402217 | 0.3496147 | 0.5624384 | 0.822060697 |
| ENSMUSG00000039953 | Clstn1 | protein_coding | 0.142593785 | 7.959736909 | 0.3492643 | 0.5626323 | 0.822232244 |
| ENSMUSG00000004233 | Wars2 | protein_coding | -0.390710888 | 3.088499615 | 0.3489915 | 0.5627833 | 0.822341161 |
| ENSMUSG00000038569 | Rad9b | protein_coding | 0.64378615 | 2.652886334 | 0.348785 | 0.562911 | 0.822343475 |
| ENSMUSG00000022607 | Ptk2 | protein_coding | -0.184570568 | 5.103287679 | 0.3485406 | 0.5630331 | 0.822343475 |
| ENSMUSG00000020372 | Rack1 | protein_coding | 0.145822795 | 8.901453793 | 0.348525 | 0.5630417 | 0.822343475 |
| ENSMUSG00000017801 | Mlx | protein_coding | 0.189852604 | 6.238427675 | 0.3482822 | 0.5631763 | 0.822343475 |
| ENSMUSG00000017210 | Med24 | protein_coding | 0.240450881 | 5.606035835 | 0.3481798 | 0.5632463 | 0.822343475 |
| ENSMUSG00000020075 | Ddx21 | protein_coding | -0.260904265 | 3.940671259 | 0.3479467 | 0.5633624 | 0.822343475 |
| ENSMUSG00000025571 | Tnrc6c | protein_coding | -0.17900875 | 5.659112533 | 0.3479098 | 0.5633829 | 0.822343475 |
| ENSMUSG00000024776 | Stambpl1 | protein_coding | 0.210731576 | 4.764129208 | 0.3478849 | 0.5633967 | 0.822343475 |
| ENSMUSG00000031556 | Tm2d2 | protein_coding | 0.15325381 | 7.927169006 | 0.3468456 | 0.563974 | 0.823059263 |
| ENSMUSG00000046550 | Spin2c | protein_coding | 0.221404866 | 5.370066525 | 0.3467266 | 0.5640401 | 0.823059263 |
| ENSMUSG00000061898 | Rbak | protein_coding | -0.514654489 | 2.679502963 | 0.3462897 | 0.5642832 | 0.823205331 |
| ENSMUSG00000060981 | Hist1h4h | protein_coding | -0.550200246 | 2.313796193 | 0.3461545 | 0.5643585 | 0.823205331 |
| ENSMUSG00000017639 | Rab11fip4 | protein_coding | -0.197227465 | 7.063124303 | 0.3462203 | 0.5643699 | 0.823205331 |
| ENSMUSG00000004846 | Plod3 | protein_coding | -0.342314599 | 3.690383343 | 0.3459285 | 0.5644843 | 0.823260546 |
| ENSMUSG00000031214 | Ophn1 | protein_coding | -0.230133248 | 4.328522202 | 0.3457063 | 0.5646082 | 0.823293301 |
| ENSMUSG00000022940 | Pigp | protein_coding | -0.200864648 | 5.775707716 | 0.3451875 | 0.5648974 | 0.823293301 |
| ENSMUSG00000029413 | Naaa | protein_coding | 0.297430962 | 4.870785773 | 0.3452051 | 0.5649008 | 0.823293301 |
| ENSMUSG00000006576 | Slc4a3 | protein_coding | 0.170861457 | 5.747084477 | 0.3450533 | 0.5649723 | 0.823293301 |
| ENSMUSG00000034723 | Tmx4 | protein_coding | -0.140327991 | 7.659285269 | 0.3449986 | 0.5650028 | 0.823293301 |
| ENSMUSG00000097571 | Jpx | lincRNA | -0.456500013 | 2.721412026 | 0.3449791 | 0.5650137 | 0.823293301 |
| ENSMUSG00000070371 | Prss36 | protein_coding | 0.308535399 | 4.542791938 | 0.3449507 | 0.5650427 | 0.823293301 |
| ENSMUSG00000019710 | Mrpl24 | protein_coding | 0.141349247 | 6.704732725 | 0.3445374 | 0.5652603 | 0.8234361 |
| ENSMUSG00000040867 | Begain | protein_coding | -0.218020165 | 4.989020161 | 0.3444772 | 0.5652939 | 0.8234361 |
| ENSMUSG00000001627 | Ifrd1 | protein_coding | -0.375238664 | 3.542411069 | 0.3443045 | 0.5653904 | 0.823465131 |
| ENSMUSG00000071646 | Mta2 | protein_coding | 0.252307243 | 4.029628737 | 0.3439261 | 0.5656019 | 0.823661611 |
| ENSMUSG00000029119 | Man2b2 | protein_coding | -0.340146474 | 3.495648565 | 0.3426649 | 0.5663081 | 0.824412442 |
| ENSMUSG00000029176 | Anapc4 | protein_coding | 0.164835393 | 6.037861913 | 0.3426163 | 0.5663353 | 0.824412442 |
| ENSMUSG00000040234 | Tm7sf3 | protein_coding | -0.233804628 | 4.603531011 | 0.34257 | 0.5663613 | 0.824412442 |
| ENSMUSG00000036528 | Ppfibp2 | protein_coding | 0.361464531 | 3.651401463 | 0.34241 | 0.566451 | 0.824412442 |
| ENSMUSG00000079427 | Mthfsl | protein_coding | -0.249073331 | 3.962724281 | 0.3423055 | 0.5665097 | 0.824412442 |
| ENSMUSG00000067148 | Polr1c | protein_coding | 0.133633116 | 6.988972938 | 0.3421179 | 0.5666149 | 0.824412442 |
| ENSMUSG00000006304 | Arpc2 | protein_coding | -0.132248071 | 7.838878671 | 0.342048 | 0.5666542 | 0.824412442 |
| ENSMUSG00000021550 | 2210016F16Rik | protein_coding | 0.314263401 | 3.754677248 | 0.3417407 | 0.5668267 | 0.824551908 |
| ENSMUSG00000039908 | Slc26a11 | protein_coding | 0.359779331 | 3.558136159 | 0.3409182 | 0.567289 | 0.825112798 |
| ENSMUSG00000032295 | Man2c1 | protein_coding | -0.247373778 | 5.041759755 | 0.3403117 | 0.5676304 | 0.825497709 |
| ENSMUSG00000023905 | Tnfrsf12a | protein_coding | 0.290262997 | 4.519644428 | 0.3400324 | 0.5677878 | 0.825614896 |
| ENSMUSG00000054920 | Klhl5 | protein_coding | -0.217510134 | 4.89914094 | 0.3398024 | 0.5679175 | 0.82562413 |
| ENSMUSG00000025986 | Slc39a10 | protein_coding | -0.160597094 | 5.661117132 | 0.3396424 | 0.5680076 | 0.82562413 |
| ENSMUSG00000074165 | Zfp788 | protein_coding | -0.252603966 | 4.507406213 | 0.3396126 | 0.5680245 | 0.82562413 |
| ENSMUSG00000048170 | Mcmbp | protein_coding | -0.198012838 | 5.869126509 | 0.33917 | 0.5682742 | 0.825875514 |
| ENSMUSG00000026987 | Baz2b | protein_coding | -0.197506631 | 4.862876181 | 0.3385615 | 0.568618 | 0.826263376 |
| ENSMUSG00000031728 | Zfp821 | protein_coding | 0.257785967 | 4.6341549 | 0.3383146 | 0.5687575 | 0.826354491 |
| ENSMUSG00000030750 | Nsmce1 | protein_coding | -0.151774222 | 6.353034097 | 0.3376993 | 0.5691057 | 0.826663582 |
| ENSMUSG00000097062 | Gm17586 | lincRNA | 0.311911942 | 3.253532886 | 0.3376669 | 0.569124 | 0.826663582 |
| ENSMUSG00000067365 | Tmem128 | protein_coding | 0.165702812 | 6.726846509 | 0.337127 | 0.5694299 | 0.826996172 |
| ENSMUSG00000051185 | Fam174a | protein_coding | -0.205410626 | 5.226008723 | 0.33689 | 0.5695643 | 0.827062839 |
| ENSMUSG00000066900 | Suds3 | protein_coding | 0.172833741 | 5.491229556 | 0.3367748 | 0.5696296 | 0.827062839 |
| ENSMUSG00000020528 | Prpsap2 | protein_coding | -0.213591517 | 5.112066053 | 0.336327 | 0.5698837 | 0.827089928 |
| ENSMUSG00000027937 | Jtb | protein_coding | 0.218631552 | 5.714458474 | 0.3363091 | 0.5698939 | 0.827089928 |
| ENSMUSG00000113679 | AC159633.2 | processed_transcript | -0.296789462 | 3.571613855 | 0.3362352 | 0.5699359 | 0.827089928 |
| ENSMUSG00000036983 | Tfb1m | protein_coding | -0.283566665 | 3.668714948 | 0.3361998 | 0.569956 | 0.827089928 |

| ensembl_gene_id | external_gene_name | gene_biotype | log2 (fold change) | log2 (counts per million) | F-value | p-value | False discovery rate |
|---|---|---|---|---|---|---|---|
| ENSMUSG00000059659 | Gm10069 | antisense_RNA | -0.375623743 | 3.082248651 | 0.3359047 | 0.5701236 | 0.827171297 |
| ENSMUSG00000034308 | Sdr42e1 | protein_coding | -0.412163397 | 3.166429921 | 0.3358303 | 0.5701659 | 0.827171297 |
| ENSMUSG00000024319 | Vps52 | protein_coding | -0.204442675 | 5.761971041 | 0.3356563 | 0.5702648 | 0.827203156 |
| ENSMUSG00000042642 | Flad1 | protein_coding | 0.178339314 | 6.006308765 | 0.3353447 | 0.570442 | 0.827249375 |
| ENSMUSG00000029175 | Slc35f6 | protein_coding | -0.211560447 | 5.03880609 | 0.3353297 | 0.5704505 | 0.827249375 |
| ENSMUSG00000061313 | Ddhd2 | protein_coding | 0.26854255 | 7.095675097 | 0.3371369 | 0.5706873 | 0.827450201 |
| ENSMUSG00000021891 | Mettl6 | protein_coding | 0.199517014 | 5.307868571 | 0.3348161 | 0.5707429 | 0.827450201 |
| ENSMUSG00000026374 | Tsn | protein_coding | -0.141900291 | 8.515474492 | 0.334482 | 0.5709332 | 0.827614523 |
| ENSMUSG00000078974 | Sec61g | protein_coding | 0.207734229 | 5.158550074 | 0.3336252 | 0.5714219 | 0.82809622 |
| ENSMUSG00000044646 | Zbtb7c | protein_coding | 0.433190649 | 2.786383775 | 0.3334871 | 0.5715007 | 0.82809622 |
| ENSMUSG00000018849 | Wwc1 | protein_coding | -0.205404243 | 5.078103555 | 0.3333978 | 0.5715517 | 0.82809622 |
| ENSMUSG00000095567 | Noc2l | protein_coding | 0.234869755 | 4.321235495 | 0.3333251 | 0.5715932 | 0.82809622 |
| ENSMUSG00000071691 | Gm960 | protein_coding | -0.60595191 | 2.546245427 | 0.3332218 | 0.5716651 | 0.82809622 |
| ENSMUSG00000030451 | Herc2 | protein_coding | -0.219881155 | 5.246005761 | 0.33309 | 0.5717275 | 0.82809622 |
| ENSMUSG00000068099 | 1500009C09Rik | protein_coding | 0.160916316 | 5.961928734 | 0.3327432 | 0.5719258 | 0.828118625 |
| ENSMUSG00000046152 | Fut10 | protein_coding | 0.389621835 | 2.851395179 | 0.3325613 | 0.5720298 | 0.828118625 |
| ENSMUSG00000033799 | Fam208b | protein_coding | -0.565701796 | 2.470023968 | 0.3325476 | 0.5720377 | 0.828118625 |
| ENSMUSG00000041729 | Coro2b | protein_coding | 0.177944795 | 5.548722249 | 0.3325241 | 0.5720511 | 0.828118625 |
| ENSMUSG00000020097 | Sgpl1 | protein_coding | 0.373303499 | 3.309907319 | 0.3317716 | 0.5724819 | 0.828521412 |
| ENSMUSG00000023067 | Cdkn1a | protein_coding | -0.402286874 | 4.338982356 | 0.333071 | 0.5725235 | 0.828521412 |
| ENSMUSG00000056679 | Gpr173 | protein_coding | -0.423860168 | 2.789059851 | 0.3315765 | 0.5725937 | 0.828521412 |
| ENSMUSG00000017686 | Rhot1 | protein_coding | -0.199180656 | 5.268118741 | 0.3313964 | 0.572697 | 0.828521412 |
| ENSMUSG00000050272 | Dscam | protein_coding | -0.221751742 | 4.946494286 | 0.3313658 | 0.5727146 | 0.828521412 |
| ENSMUSG00000026278 | Bok | protein_coding | -0.189719361 | 5.664162849 | 0.3311593 | 0.572833 | 0.828581311 |
| ENSMUSG00000040746 | Rnf167 | protein_coding | 0.139525856 | 7.602093668 | 0.3308543 | 0.573008 | 0.828610941 |
| ENSMUSG00000007338 | Mrpl49 | protein_coding | 0.13234762 | 7.222371625 | 0.3307841 | 0.5730483 | 0.828610941 |
| ENSMUSG00000061533 | Cep128 | protein_coding | -0.422314626 | 3.424717076 | 0.3307432 | 0.5730847 | 0.828610941 |
| ENSMUSG00000039670 | Oxld1 | protein_coding | 0.479193018 | 2.546771813 | 0.3304371 | 0.5732476 | 0.828735113 |
| ENSMUSG00000074345 | Tnfaip8l3 | protein_coding | 0.377815502 | 3.454021263 | 0.3300701 | 0.5734586 | 0.828866103 |
| ENSMUSG00000040482 | Dxo | protein_coding | -0.243218061 | 4.449983873 | 0.3300112 | 0.5734924 | 0.828866103 |
| ENSMUSG00000002477 | Snrpd1 | protein_coding | 0.166190126 | 7.826227714 | 0.3297646 | 0.5736343 | 0.828902241 |
| ENSMUSG00000027994 | Mcub | protein_coding | -0.402164016 | 3.54038633 | 0.3297221 | 0.5736716 | 0.828902241 |
| ENSMUSG00000023935 | Spats1 | protein_coding | -0.409294603 | 2.793312523 | 0.3295038 | 0.5737843 | 0.828909224 |
| ENSMUSG00000035189 | Ano4 | protein_coding | -0.23747939 | 4.021598688 | 0.3294235 | 0.5738306 | 0.828909224 |
| ENSMUSG00000093803 | Ppp2r3d | protein_coding | 0.288936333 | 3.402143341 | 0.3287775 | 0.5742027 | 0.829272195 |
| ENSMUSG00000038502 | Ptov1 | protein_coding | 0.174940406 | 6.60639287 | 0.3282598 | 0.5745014 | 0.829272195 |
| ENSMUSG00000036371 | Serbp1 | protein_coding | -0.129963842 | 7.425135801 | 0.3281872 | 0.5745433 | 0.829272195 |
| ENSMUSG00000029392 | Rilpl1 | protein_coding | 0.249431613 | 4.592945239 | 0.3280244 | 0.5746372 | 0.829272195 |
| ENSMUSG00000019055 | Plod1 | protein_coding | 0.289773161 | 3.98051833 | 0.3279637 | 0.5746723 | 0.829272195 |
| ENSMUSG00000025795 | Rassf3 | protein_coding | -0.388006864 | 2.979774299 | 0.3279402 | 0.5746859 | 0.829272195 |
| ENSMUSG00000054766 | Set | protein_coding | -0.164653071 | 8.405427607 | 0.3278071 | 0.5747627 | 0.829272195 |
| ENSMUSG00000056851 | Pcbp2 | protein_coding | -0.132119376 | 6.502847413 | 0.3277502 | 0.5747956 | 0.829272195 |
| ENSMUSG00000024083 | Pja2 | protein_coding | 0.160596985 | 9.105464165 | 0.3276981 | 0.5748257 | 0.829272195 |
| ENSMUSG00000039046 | Usp6nl | protein_coding | -0.264325808 | 3.871457784 | 0.3276507 | 0.5748531 | 0.829272195 |
| ENSMUSG00000042425 | Frmpd3 | protein_coding | -0.443000049 | 2.928074851 | 0.3274389 | 0.5749755 | 0.829337569 |
| ENSMUSG00000035572 | Dcaf10 | protein_coding | 0.305173991 | 3.641918906 | 0.3270003 | 0.5752292 | 0.829518478 |
| ENSMUSG00000073563 | Csnk1g3 | protein_coding | -0.191478189 | 5.131053326 | 0.3269552 | 0.5752552 | 0.829518478 |
| ENSMUSG00000060739 | Nsa2 | protein_coding | 0.270230111 | 3.599785677 | 0.3261966 | 0.5756945 | 0.82978286 |
| ENSMUSG00000025781 | Atp5c1 | protein_coding | 0.129475391 | 10.3395293 | 0.3258769 | 0.5758799 | 0.82978286 |
| ENSMUSG00000035781 | R3hdm4 | protein_coding | 0.160691297 | 7.261553499 | 0.3258728 | 0.5758822 | 0.82978286 |
| ENSMUSG00000007411 | Mark3 | protein_coding | -0.264233231 | 4.971848746 | 0.325834 | 0.5759175 | 0.82978286 |
| ENSMUSG00000047394 | Odf3b | protein_coding | 0.344177855 | 3.508796897 | 0.3257265 | 0.5759671 | 0.82978286 |
| ENSMUSG00000030697 | Ppp4c | protein_coding | -0.168568084 | 5.388606369 | 0.3257125 | 0.5759752 | 0.82978286 |
| ENSMUSG00000035735 | Dagla | protein_coding | -0.317967693 | 3.67213159 | 0.3255832 | 0.5760502 | 0.82978286 |
| ENSMUSG00000039579 | Grin3a | protein_coding | -0.167218566 | 5.533032447 | 0.3255734 | 0.5760559 | 0.82978286 |
| ENSMUSG00000024816 | Frmd8 | protein_coding | -0.254707021 | 5.575737944 | 0.3253461 | 0.5762475 | 0.829947652 |
| ENSMUSG00000015759 | Cnih1 | protein_coding | -0.135722889 | 7.541103114 | 0.3248863 | 0.5764549 | 0.830039501 |
| ENSMUSG00000022505 | Emp2 | protein_coding | -0.403096358 | 3.382255567 | 0.3248898 | 0.5764656 | 0.830039501 |
| ENSMUSG00000039480 | Nt5dc1 | protein_coding | -0.67276552 | 1.833752759 | 0.3245015 | 0.5766786 | 0.830100569 |
| ENSMUSG00000048163 | Selplg | protein_coding | -0.299019794 | 4.155605552 | 0.3244842 | 0.5766887 | 0.830100569 |
| ENSMUSG00000075227 | Znhit2 | protein_coding | 0.30595523 | 3.470282843 | 0.3243966 | 0.5767396 | 0.830100569 |
| ENSMUSG00000027742 | Cog6 | protein_coding | -0.206786027 | 5.242379059 | 0.3241673 | 0.576873 | 0.830181417 |
| ENSMUSG00000022089 | Bin3 | protein_coding | -0.263565971 | 3.915490179 | 0.3239127 | 0.5770212 | 0.830283579 |
| ENSMUSG00000032412 | Atp1b3 | protein_coding | 0.192815547 | 6.144788157 | 0.3235777 | 0.5772164 | 0.830288382 |
| ENSMUSG00000021983 | Atp8a2 | protein_coding | -0.248693412 | 4.255534243 | 0.3234755 | 0.5772759 | 0.830288382 |
| ENSMUSG00000026103 | Gls | protein_coding | -0.191871508 | 6.054018853 | 0.3233167 | 0.5773685 | 0.830288382 |
| ENSMUSG00000029993 | Nfu1 | protein_coding | 0.128078187 | 8.380902641 | 0.3232118 | 0.5774297 | 0.830288382 |
| ENSMUSG00000020752 | Recql5 | protein_coding | 0.310424804 | 3.326451752 | 0.3231331 | 0.5774756 | 0.830288382 |
| ENSMUSG00000027195 | Hsd17b12 | protein_coding | 0.125384324 | 7.438420096 | 0.3229358 | 0.5775906 | 0.830288382 |
| ENSMUSG00000084349 | Rpl3-ps1 | processed_pseudoge | 0.358455537 | 3.374979959 | 0.3229128 | 0.5776041 | 0.830288382 |
| ENSMUSG00000061759 | Armt1 | protein_coding | 0.216278288 | 4.666926094 | 0.3228473 | 0.5776423 | 0.830288382 |
| ENSMUSG00000056999 | Ide | protein_coding | -0.202943093 | 4.665093056 | 0.3224234 | 0.5778898 | 0.830533123 |
| ENSMUSG00000032279 | Idh3a | protein_coding | 0.151174903 | 9.358748694 | 0.3218355 | 0.5782333 | 0.830721683 |
| ENSMUSG00000020882 | Cacnb1 | protein_coding | -0.161069737 | 5.735647018 | 0.3216027 | 0.5783695 | 0.830721683 |
| ENSMUSG00000029128 | Rab28 | protein_coding | 0.168555713 | 6.833190454 | 0.321597 | 0.5783729 | 0.830721683 |
| ENSMUSG00000109398 | Gm3854 | protein_coding | 0.425195789 | 3.106398389 | 0.3215319 | 0.578411 | 0.830721683 |
| ENSMUSG00000043929 | Klhl15 | protein_coding | -0.289765489 | 4.742716628 | 0.3211797 | 0.5786297 | 0.830721683 |
| ENSMUSG00000073616 | Cops9 | protein_coding | -0.186580373 | 8.663054637 | 0.3209903 | 0.5787407 | 0.830721683 |

| ensembl_gene_id | external_gene_name | gene_biotype | log2 (fold change) | log2 (counts per million) | F-value | p-value | False discovery rate |
|---|---|---|---|---|---|---|---|
| ENSMUSG00000023336 | Wfdc1 | protein_coding | 0.601498098 | 2.318903445 | 0.3209452 | 0.5787545 | 0.830721683 |
| ENSMUSG00000020601 | Trib2 | protein_coding | -0.181327572 | 5.577241253 | 0.3208944 | 0.5787842 | 0.830721683 |
| ENSMUSG00000046441 | Cmtr2 | protein_coding | 0.424510262 | 3.056756815 | 0.3207045 | 0.5788955 | 0.830721683 |
| ENSMUSG00000032763 | Ilvbl | protein_coding | 0.256099546 | 5.377961018 | 0.3204977 | 0.5790294 | 0.830721683 |
| ENSMUSG00000019837 | Gtf3c6 | protein_coding | -0.151846037 | 6.844510486 | 0.3204664 | 0.5790351 | 0.830721683 |
| ENSMUSG00000036989 | Trim3 | protein_coding | 0.158396453 | 6.15959805 | 0.3203909 | 0.5790794 | 0.830721683 |
| ENSMUSG00000035629 | Rubcn | protein_coding | 0.216579809 | 5.172054063 | 0.3203494 | 0.5791037 | 0.830721683 |
| ENSMUSG00000023031 | Cela1 | protein_coding | -0.352368767 | 3.943138773 | 0.3200718 | 0.5792792 | 0.830721683 |
| ENSMUSG00000028832 | Stmn1 | protein_coding | -0.133717415 | 7.60688748 | 0.3199918 | 0.5793136 | 0.830721683 |
| ENSMUSG00000021975 | Ints9 | protein_coding | 0.241836069 | 4.656545547 | 0.3198842 | 0.5793768 | 0.830721683 |
| ENSMUSG00000028948 | Nol9 | protein_coding | -0.355792105 | 3.071242924 | 0.3198554 | 0.5793936 | 0.830721683 |
| ENSMUSG00000020166 | Cnot2 | protein_coding | 0.185800408 | 5.701456193 | 0.319731 | 0.5794667 | 0.830721683 |
| ENSMUSG00000052488 | Cherp | protein_coding | 0.401565438 | 4.63832913 | 0.3211519 | 0.5797126 | 0.830721683 |
| ENSMUSG00000024558 | Mapk4 | protein_coding | -0.209677808 | 4.869485368 | 0.3192935 | 0.5797238 | 0.830721683 |
| ENSMUSG00000056216 | Cebpg | protein_coding | 0.207266867 | 4.363496584 | 0.3192703 | 0.5797374 | 0.830721683 |
| ENSMUSG00000037418 | Best1 | protein_coding | 0.514637976 | 2.896601473 | 0.3192824 | 0.5797429 | 0.830721683 |
| ENSMUSG00000081552 | Gm12328 | processed_pseudoge | 0.398706312 | 2.681845599 | 0.3191675 | 0.5797978 | 0.830721683 |
| ENSMUSG00000024462 | Gabbr1 | protein_coding | -0.157674949 | 7.898759309 | 0.3189316 | 0.5799366 | 0.830809793 |
| ENSMUSG00000009418 | Nav1 | protein_coding | -0.179411924 | 5.623536203 | 0.3185159 | 0.5801813 | 0.83104959 |
| ENSMUSG00000027860 | Vangl1 | protein_coding | -0.513398993 | 3.138916828 | 0.3188663 | 0.5802966 | 0.831104046 |
| ENSMUSG00000046058 | Eid2 | protein_coding | -0.234802897 | 4.640219936 | 0.3180984 | 0.5804272 | 0.831180473 |
| ENSMUSG00000053510 | Nrd1 | protein_coding | -0.166086604 | 5.395380543 | 0.317858 | 0.5805689 | 0.831195598 |
| ENSMUSG00000022769 | Sdf2l1 | protein_coding | 0.421119235 | 2.504649534 | 0.3178182 | 0.5805924 | 0.831195598 |
| ENSMUSG00000022525 | Hrasls | protein_coding | 0.175749763 | 6.23717462 | 0.317432 | 0.5808202 | 0.831272974 |
| ENSMUSG00000021771 | Vdac2 | protein_coding | 0.135361661 | 9.130915207 | 0.3173578 | 0.5808641 | 0.831272974 |
| ENSMUSG00000002731 | Prkra | protein_coding | 0.156002406 | 6.252492785 | 0.3172704 | 0.5809156 | 0.831272974 |
| ENSMUSG00000020525 | Ppm1d | protein_coding | 0.333128367 | 3.356789693 | 0.3172026 | 0.5809557 | 0.831272974 |
| ENSMUSG00000042804 | Gpr153 | protein_coding | 0.285463875 | 3.871506597 | 0.3169162 | 0.5811248 | 0.83140438 |
| ENSMUSG00000086943 | 4732414G09Rik | lincRNA | 0.620176534 | 2.822618559 | 0.3166751 | 0.5817539 | 0.831502663 |
| ENSMUSG00000021134 | Srsf5 | protein_coding | 0.136558438 | 7.73424352 | 0.31565 | 0.5818739 | 0.831502663 |
| ENSMUSG00000037933 | Bicd2 | protein_coding | 0.221791022 | 4.4609733 | 0.3154189 | 0.5820108 | 0.831502663 |
| ENSMUSG00000059895 | Ptp4a3 | protein_coding | 0.182348731 | 5.311808711 | 0.3149415 | 0.5822939 | 0.831502663 |
| ENSMUSG00000045294 | Insig1 | protein_coding | -0.195460708 | 5.877428147 | 0.3148967 | 0.5823205 | 0.831502663 |
| ENSMUSG00000039473 | Ubn1 | protein_coding | -0.241794276 | 4.132008918 | 0.3148948 | 0.5823216 | 0.831502663 |
| ENSMUSG00000039286 | Fndc3b | protein_coding | -0.277130625 | 3.988063727 | 0.3146761 | 0.5824514 | 0.831502663 |
| ENSMUSG00000097310 | A930038B10Rik | lincRNA | 0.56082573 | 2.324610215 | 0.3145712 | 0.5825137 | 0.831502663 |
| ENSMUSG00000040586 | Ofd1 | protein_coding | -0.451936126 | 2.848026959 | 0.3145662 | 0.5825167 | 0.831502663 |
| ENSMUSG00000020903 | Stx8 | protein_coding | -0.128875319 | 6.872106506 | 0.3143898 | 0.5826214 | 0.831502663 |
| ENSMUSG00000038267 | Slc22a23 | protein_coding | 0.210617328 | 4.568708152 | 0.3143839 | 0.5826249 | 0.831502663 |
| ENSMUSG00000036390 | Gadd45a | protein_coding | -0.368783455 | 2.947059194 | 0.3143576 | 0.5826405 | 0.831502663 |
| ENSMUSG00000057778 | Cyb5d2 | protein_coding | 0.25163861 | 4.542652472 | 0.3142849 | 0.5826837 | 0.831502663 |
| ENSMUSG00000020629 | Adi1 | protein_coding | 0.169987389 | 6.30891027 | 0.3141889 | 0.5827408 | 0.831502663 |
| ENSMUSG00000011257 | Pabpc4 | protein_coding | -0.31549017 | 3.198286458 | 0.314121 | 0.5827811 | 0.831502663 |
| ENSMUSG00000020886 | Dlg4 | protein_coding | 0.149590833 | 5.726195246 | 0.314117 | 0.5827835 | 0.831502663 |
| ENSMUSG00000039154 | Shd | protein_coding | 0.183601541 | 6.023462431 | 0.3140414 | 0.5828284 | 0.831502663 |
| ENSMUSG00000070880 | Gad1 | protein_coding | 0.137858754 | 8.867156089 | 0.3139392 | 0.5828892 | 0.831502663 |
| ENSMUSG00000041133 | Smc1a | protein_coding | 0.180007859 | 6.176882888 | 0.3138984 | 0.5829134 | 0.831502663 |
| ENSMUSG00000025935 | Tram1 | protein_coding | -0.209928031 | 5.064605052 | 0.3138044 | 0.5829693 | 0.831502663 |
| ENSMUSG00000034442 | Trmt5 | protein_coding | 0.307762466 | 4.143886005 | 0.3137851 | 0.5829933 | 0.831502663 |
| ENSMUSG00000003346 | Abhd17a | protein_coding | 0.243804013 | 4.933499331 | 0.31369 | 0.5830374 | 0.831502663 |
| ENSMUSG00000026791 | Slc2a8 | protein_coding | -0.469511487 | 2.732372097 | 0.3136651 | 0.5830522 | 0.831502663 |
| ENSMUSG00000031497 | Tnfsf13b | protein_coding | -0.393508212 | 3.010661282 | 0.3135515 | 0.5831198 | 0.831502663 |
| ENSMUSG00000074890 | Lcmt2 | protein_coding | 0.39799284 | 3.090681703 | 0.3134161 | 0.5832003 | 0.831502663 |
| ENSMUSG00000024642 | Tle4 | protein_coding | -0.202621043 | 5.585379741 | 0.3134022 | 0.5832086 | 0.831502663 |
| ENSMUSG00000030801 | Kat8 | protein_coding | 0.196151947 | 4.772854295 | 0.31328 | 0.5832814 | 0.831502663 |
| ENSMUSG00000047417 | Rexo1 | protein_coding | 0.255901781 | 4.115535995 | 0.3130352 | 0.5834271 | 0.831600174 |
| ENSMUSG00000071719 | Tmem28 | protein_coding | 0.263179827 | 3.903333358 | 0.3127736 | 0.583583 | 0.831641804 |
| ENSMUSG00000058594 | Fbxo18 | protein_coding | -0.164272403 | 5.709384351 | 0.3127266 | 0.583611 | 0.831641804 |
| ENSMUSG00000042487 | Leo1 | protein_coding | -0.215764956 | 6.18371895 | 0.3124725 | 0.5838078 | 0.831757957 |
| ENSMUSG00000074461 | Gm10699 | TEC | -0.152530168 | 5.997095982 | 0.3123305 | 0.5838472 | 0.831757957 |
| ENSMUSG00000047824 | Pygo2 | protein_coding | 0.228609645 | 4.81998272 | 0.3121415 | 0.58396 | 0.831808395 |
| ENSMUSG00000071659 | Hnrnpul2 | protein_coding | -0.189800437 | 5.875006162 | 0.3116491 | 0.584254 | 0.832111128 |
| ENSMUSG00000051439 | Cd14 | protein_coding | 0.408414507 | 2.931237514 | 0.311155 | 0.5845493 | 0.832111128 |
| ENSMUSG00000031320 | Rps4x | protein_coding | 0.169230081 | 6.337535257 | 0.311153 | 0.5845505 | 0.832111128 |
| ENSMUSG00000029516 | Cit | protein_coding | 0.261561764 | 8.525516425 | 0.3130313 | 0.5846396 | 0.832111128 |
| ENSMUSG00000020828 | Pld2 | protein_coding | -0.292975055 | 3.411782976 | 0.3109987 | 0.5846428 | 0.832111128 |
| ENSMUSG00000027509 | Rae1 | protein_coding | 0.210564409 | 4.86638759 | 0.3109817 | 0.584653 | 0.832111128 |
| ENSMUSG00000025158 | Rfng | protein_coding | -0.240728654 | 4.553115981 | 0.3108794 | 0.5847142 | 0.832111128 |
| ENSMUSG00000051223 | Bzw1 | protein_coding | -0.136492648 | 7.448089155 | 0.3105999 | 0.5848815 | 0.832239083 |
| ENSMUSG00000028901 | Gmeb1 | protein_coding | -0.278035298 | 4.083689922 | 0.3100764 | 0.5851951 | 0.832546892 |
| ENSMUSG00000026644 | Acbd7 | protein_coding | 0.324967288 | 3.201097795 | 0.3099589 | 0.5852656 | 0.832546892 |
| ENSMUSG00000020598 | Nrcam | protein_coding | -0.159646594 | 5.843984081 | 0.3096684 | 0.5854398 | 0.832546892 |
| ENSMUSG00000032076 | Cadm1 | protein_coding | -0.134651619 | 7.275656087 | 0.3093053 | 0.5856577 | 0.832546892 |
| ENSMUSG00000091636 | Akain1 | protein_coding | -0.588092216 | 2.482868168 | 0.3092073 | 0.5857165 | 0.832546892 |
| ENSMUSG00000024480 | Ap3s1 | protein_coding | -0.292631409 | 3.781050752 | 0.3091007 | 0.5857806 | 0.832546892 |
| ENSMUSG00000020657 | Dnajc27 | protein_coding | 0.130765633 | 6.523434476 | 0.3089807 | 0.5858527 | 0.832546892 |
| ENSMUSG00000050132 | Sarm1 | protein_coding | -0.341383005 | 3.056241751 | 0.3088285 | 0.5859442 | 0.832546892 |
| ENSMUSG00000005510 | Ndufs3 | protein_coding | 0.117153051 | 7.416447895 | 0.3085919 | 0.5860864 | 0.832546892 |

| ensembl_gene_id | external_gene_name | gene_biotype | log2 (fold change) | log2 (counts per million) | F-value | p-value | False discovery rate |
|---|---|---|---|---|---|---|---|
| ENSMUSG00000027489 | Necab3 | protein_coding | 0.346157257 | 5.198566932 | 0.3102414 | 0.5861542 | 0.832546892 |
| ENSMUSG00000041046 | Ramp3 | protein_coding | -0.267121651 | 6.002881069 | 0.3096266 | 0.5862177 | 0.832546892 |
| ENSMUSG00000022037 | Clu | protein_coding | 0.158860875 | 9.99337249 | 0.3083199 | 0.58625 | 0.832546892 |
| ENSMUSG00000110277 | Gm45871 | protein_coding | -0.348573516 | 3.264618075 | 0.3082316 | 0.5863031 | 0.832546892 |
| ENSMUSG00000035653 | Lrfn5 | protein_coding | -0.206032772 | 5.506144873 | 0.3081887 | 0.586329 | 0.832546892 |
| ENSMUSG00000053046 | Brsk2 | protein_coding | 0.186956518 | 5.330846141 | 0.3081842 | 0.5863317 | 0.832546892 |
| ENSMUSG00000046997 | Spsb4 | protein_coding | -0.481189853 | 2.451580769 | 0.308176 | 0.5863366 | 0.832546892 |
| ENSMUSG00000038984 | Tspyl5 | protein_coding | -0.179470861 | 6.739903537 | 0.3077019 | 0.5866344 | 0.832699466 |
| ENSMUSG00000040415 | Dtx3 | protein_coding | -0.115192996 | 7.562950715 | 0.3076788 | 0.586636 | 0.832699466 |
| ENSMUSG00000030747 | Dgat2 | protein_coding | -0.183702703 | 5.811963899 | 0.3076118 | 0.5866764 | 0.832699466 |
| ENSMUSG00000037235 | Mxd4 | protein_coding | -0.251629837 | 4.331217165 | 0.307261 | 0.5868878 | 0.832788402 |
| ENSMUSG00000044906 | 4930503L19Rik | protein_coding | -0.493779552 | 2.687705148 | 0.3072509 | 0.5868939 | 0.832788402 |
| ENSMUSG00000027305 | Ndufaf1 | protein_coding | -0.144516819 | 5.910435324 | 0.3068401 | 0.5871418 | 0.832986187 |
| ENSMUSG00000020262 | Adarb1 | protein_coding | -0.221426794 | 5.996686249 | 0.3068373 | 0.5871883 | 0.832986187 |
| ENSMUSG00000000759 | Tubgcp3 | protein_coding | 0.214779466 | 4.827960238 | 0.3061568 | 0.5875545 | 0.833216578 |
| ENSMUSG00000040385 | Ppp1ca | protein_coding | 0.12464174 | 9.192560825 | 0.3061546 | 0.5875558 | 0.833216578 |
| ENSMUSG00000018363 | Smurf2 | protein_coding | -0.235948408 | 4.560869623 | 0.3061095 | 0.5875831 | 0.833216578 |
| ENSMUSG00000004127 | Trmt10a | protein_coding | -0.230478363 | 4.116830788 | 0.3057247 | 0.5878158 | 0.833346278 |
| ENSMUSG00000085208 | Brip1os | antisense_RNA | 0.425469876 | 2.826125638 | 0.305681 | 0.5878423 | 0.833346278 |
| ENSMUSG00000028899 | Taf12 | protein_coding | -0.175877687 | 6.018048433 | 0.3051715 | 0.5881507 | 0.833346278 |
| ENSMUSG00000028218 | Fam92a | protein_coding | -0.155665459 | 5.899350725 | 0.3050593 | 0.5882187 | 0.833346278 |
| ENSMUSG00000026662 | Sephs1 | protein_coding | 0.261891524 | 4.295783792 | 0.30499 | 0.5882607 | 0.833346278 |
| ENSMUSG00000022136 | Dnajc3 | protein_coding | -0.23796868 | 4.487794101 | 0.3049854 | 0.5882635 | 0.833346278 |
| ENSMUSG00000059772 | Slx1b | protein_coding | -0.228088198 | 4.103154165 | 0.3049362 | 0.5882933 | 0.833346278 |
| ENSMUSG00000031657 | Heatr3 | protein_coding | -0.25050143 | 4.264892897 | 0.3049341 | 0.5882946 | 0.833346278 |
| ENSMUSG00000048371 | Pdp2 | protein_coding | -0.290425378 | 5.615301896 | 0.3057938 | 0.5885214 | 0.83355776 |
| ENSMUSG00000029718 | Pcolce | protein_coding | -0.598273365 | 3.082834057 | 0.3052278 | 0.5886233 | 0.833592286 |
| ENSMUSG00000035104 | Eva1a | protein_coding | 0.39993735 | 3.812427907 | 0.3048922 | 0.5887671 | 0.833681348 |
| ENSMUSG00000016256 | Ctsz | protein_coding | -0.184682498 | 6.410510985 | 0.3040532 | 0.5888412 | 0.833681348 |
| ENSMUSG00000026775 | Yme1l1 | protein_coding | -0.178068304 | 6.837194338 | 0.3039065 | 0.5889303 | 0.833697691 |
| ENSMUSG00000037622 | Wdtc1 | protein_coding | 0.210414092 | 4.902716224 | 0.3037453 | 0.589016 | 0.833709245 |
| ENSMUSG00000062075 | Lmnb2 | protein_coding | 0.347827781 | 3.295389858 | 0.3035642 | 0.5891261 | 0.833755258 |
| ENSMUSG00000035247 | Hectd1 | protein_coding | 0.26391027 | 4.236072465 | 0.303377 | 0.5892399 | 0.833758454 |
| ENSMUSG00000074971 | Fibin | protein_coding | 0.403642171 | 2.923401288 | 0.3032975 | 0.5892882 | 0.833758454 |
| ENSMUSG00000030498 | Gas2 | protein_coding | 0.215542713 | 4.081728623 | 0.303178 | 0.5893609 | 0.833758454 |
| ENSMUSG00000024044 | Epb41l3 | protein_coding | -0.160627662 | 7.654101313 | 0.3025922 | 0.5897175 | 0.834153203 |
| ENSMUSG00000018378 | Cuedc1 | protein_coding | 0.258496103 | 3.943362675 | 0.3022042 | 0.589954 | 0.834258505 |
| ENSMUSG00000063239 | Grm4 | protein_coding | -0.21631342 | 4.528026052 | 0.301948 | 0.5901102 | 0.834258505 |
| ENSMUSG00000003541 | Ier3 | protein_coding | 0.184933525 | 4.830210852 | 0.3019265 | 0.5901234 | 0.834258505 |
| ENSMUSG00000033577 | Myo6 | protein_coding | -0.189320054 | 5.153348015 | 0.3018393 | 0.5901766 | 0.834258505 |
| ENSMUSG00000027457 | Snph | protein_coding | -0.159647263 | 6.359652237 | 0.3018019 | 0.5901994 | 0.834258505 |
| ENSMUSG00000016257 | Prelid3b | protein_coding | 0.130950426 | 7.019083206 | 0.3016438 | 0.5902959 | 0.834258505 |
| ENSMUSG00000027395 | Polr1b | protein_coding | 0.457589811 | 2.59214841 | 0.3015796 | 0.5903351 | 0.834258505 |
| ENSMUSG00000057177 | Gsk3a | protein_coding | -0.184414306 | 4.855785757 | 0.3014122 | 0.5904373 | 0.834293338 |
| ENSMUSG00000025262 | Fam120c | protein_coding | -0.423159022 | 3.31235211 | 0.3011068 | 0.5906361 | 0.834441684 |
| ENSMUSG00000017428 | Psmd11 | protein_coding | -0.127686124 | 7.521194612 | 0.3009863 | 0.5906975 | 0.834441684 |
| ENSMUSG00000024831 | Ighmbp2 | protein_coding | 0.412646457 | 3.065338584 | 0.3007913 | 0.5908167 | 0.834460435 |
| ENSMUSG00000021785 | Ngly1 | protein_coding | 0.235610427 | 4.401940673 | 0.3007108 | 0.590866 | 0.834460435 |
| ENSMUSG00000006307 | Kmt2b | protein_coding | -0.409886497 | 3.089273277 | 0.2995921 | 0.5915511 | 0.835261048 |
| ENSMUSG00000005299 | Letm1 | protein_coding | 0.176227849 | 5.787352324 | 0.2995198 | 0.5915954 | 0.835261048 |
| ENSMUSG00000028467 | Gba2 | protein_coding | 0.167290106 | 5.388187898 | 0.2994049 | 0.5916659 | 0.835261048 |
| ENSMUSG00000028550 | Atg4c | protein_coding | -0.184385603 | 5.355859429 | 0.2992235 | 0.5917771 | 0.835308435 |
| ENSMUSG00000028603 | Scp2 | protein_coding | 0.368752263 | 3.075464304 | 0.2985238 | 0.5922068 | 0.835805255 |
| ENSMUSG00000034789 | Rab24 | protein_coding | 0.265715239 | 4.431594642 | 0.2982933 | 0.5923486 | 0.835895537 |
| ENSMUSG00000022892 | App | protein_coding | -0.137279767 | 9.150916813 | 0.2980234 | 0.5925146 | 0.836020148 |
| ENSMUSG00000023830 | Igf2r | protein_coding | -0.308879704 | 3.655759802 | 0.2978729 | 0.5926072 | 0.836041128 |
| ENSMUSG00000063894 | Zkscan8 | protein_coding | 0.311214927 | 3.195151195 | 0.2976665 | 0.5927343 | 0.836110672 |
| ENSMUSG00000000339 | Rtca | protein_coding | 0.285863554 | 4.546336052 | 0.2975202 | 0.5928365 | 0.836145162 |
| ENSMUSG00000032477 | Cdc25a | protein_coding | -0.426178916 | 3.205353042 | 0.2970508 | 0.5931257 | 0.83642141 |
| ENSMUSG00000027800 | Tm4sf1 | protein_coding | 0.236074977 | 4.283675541 | 0.2969304 | 0.5931879 | 0.83642141 |
| ENSMUSG00000028879 | Stx12 | protein_coding | 0.159260167 | 7.947623175 | 0.2968016 | 0.5932673 | 0.836423701 |
| ENSMUSG00000038299 | Wdr36 | protein_coding | 0.24757283 | 4.820468584 | 0.2966617 | 0.5933536 | 0.836435731 |
| ENSMUSG00000015357 | Clpx | protein_coding | 0.182195532 | 5.129757506 | 0.2964512 | 0.5934836 | 0.83650928 |
| ENSMUSG00000040446 | Rprd1a | protein_coding | 0.268929881 | 3.754154012 | 0.296262 | 0.5936004 | 0.836564266 |
| ENSMUSG00000023307 | 5-Mar | protein_coding | 0.195206276 | 5.014800071 | 0.2960985 | 0.5937015 | 0.836597025 |
| ENSMUSG00000022828 | Gtf2e1 | protein_coding | 0.260977261 | 3.880361525 | 0.29563 | 0.593991 | 0.836691462 |
| ENSMUSG00000023043 | Krt18 | protein_coding | -0.434000723 | 3.282090587 | 0.2955679 | 0.5940416 | 0.836691462 |
| ENSMUSG00000028608 | 0610037L13Rik | protein_coding | 0.132204007 | 6.864864031 | 0.2954466 | 0.5941046 | 0.836691462 |
| ENSMUSG00000030291 | Med21 | protein_coding | -0.143634332 | 7.272504161 | 0.2953281 | 0.5941779 | 0.836691462 |
| ENSMUSG00000020090 | Npffr1 | protein_coding | 0.343884752 | 3.041290467 | 0.2951506 | 0.5942878 | 0.836691462 |
| ENSMUSG00000073771 | Btbd19 | protein_coding | -0.336488822 | 3.129784981 | 0.2950407 | 0.5943558 | 0.836691462 |
| ENSMUSG00000038970 | Lmtk2 | protein_coding | -0.237775417 | 4.285647152 | 0.2950038 | 0.5943786 | 0.836691462 |
| ENSMUSG00000001313 | Rnd2 | protein_coding | -0.151726774 | 7.485209106 | 0.2949839 | 0.594391 | 0.836691462 |
| ENSMUSG00000025350 | Rdh5 | protein_coding | -0.458009578 | 3.235796726 | 0.2931985 | 0.5955428 | 0.838139123 |
| ENSMUSG00000022523 | Fgf12 | protein_coding | -0.127197418 | 7.554931054 | 0.293076 | 0.5955753 | 0.838139123 |
| ENSMUSG00000029578 | Wipi2 | protein_coding | -0.132229871 | 6.48268592 | 0.2927817 | 0.5957584 | 0.838287093 |
| ENSMUSG00000073478 | D730003I15Rik | lincRNA | -0.269573875 | 4.288158583 | 0.2923562 | 0.5960234 | 0.838399138 |
| ENSMUSG00000015709 | Arnt2 | protein_coding | -0.150059564 | 6.141255029 | 0.2922588 | 0.5960841 | 0.838399138 |

| ensembl_gene_id | external_gene_name | gene_biotype | log2 (fold change) | log2 (counts per million) | F-value | p-value | False discovery rate |
|---|---|---|---|---|---|---|---|
| ENSMUSG00000028696 | Ipp | protein_coding | 0.332668393 | 3.869324549 | 0.2920655 | 0.5962166 | 0.838399138 |
| ENSMUSG00000078941 | Ak6 | protein_coding | 0.180577599 | 5.52430676 | 0.2919419 | 0.5962817 | 0.838399138 |
| ENSMUSG00000061288 | Taok3 | protein_coding | 0.149428048 | 7.069856673 | 0.2918832 | 0.5963183 | 0.838399138 |
| ENSMUSG00000029780 | Nt5c3 | protein_coding | -0.193788962 | 5.587108197 | 0.2918502 | 0.5963388 | 0.838399138 |
| ENSMUSG00000019370 | Calm3 | protein_coding | 0.123857765 | 9.637650955 | 0.2917781 | 0.5963838 | 0.838399138 |
| ENSMUSG00000023809 | Rps6ka2 | protein_coding | 0.252116283 | 4.225923846 | 0.2914955 | 0.5965602 | 0.838509687 |
| ENSMUSG00000046201 | Scaf8 | protein_coding | -0.234582431 | 4.444221024 | 0.2912736 | 0.5966987 | 0.838509687 |
| ENSMUSG00000060860 | Ube2s | protein_coding | -0.40587742 | 2.896766933 | 0.2911487 | 0.5967768 | 0.838509687 |
| ENSMUSG00000021692 | Dimt1 | protein_coding | -0.228447597 | 3.968309027 | 0.2910953 | 0.5968101 | 0.838509687 |
| ENSMUSG00000021629 | Slc30a5 | protein_coding | -0.233597159 | 4.547295748 | 0.2909422 | 0.5969057 | 0.838509687 |
| ENSMUSG00000074088 | Snrnp40 | protein_coding | 0.21392893 | 5.159838166 | 0.2908245 | 0.5969793 | 0.838509687 |
| ENSMUSG00000028744 | Pqlc2 | protein_coding | 0.24299356 | 4.249229138 | 0.2907782 | 0.5970083 | 0.838509687 |
| ENSMUSG00000020460 | Rps27a | protein_coding | 0.251281109 | 3.584484307 | 0.2903672 | 0.5972654 | 0.838738539 |
| ENSMUSG00000052852 | Reep1 | protein_coding | -0.155185081 | 6.004427529 | 0.290149 | 0.597402 | 0.838738539 |
| ENSMUSG00000100975 | Gm28875 | lincRNA | -0.428936049 | 3.551389565 | 0.2908584 | 0.5974052 | 0.838738539 |
| ENSMUSG00000072964 | Bhlhb9 | protein_coding | -0.163743195 | 5.937865122 | 0.2897828 | 0.5976314 | 0.8389465 |
| ENSMUSG00000020328 | Nudcd2 | protein_coding | -0.163493966 | 5.547238243 | 0.2894959 | 0.5978112 | 0.839089361 |
| ENSMUSG00000038252 | Ncapd2 | protein_coding | -0.584519835 | 2.557168031 | 0.2890004 | 0.598134 | 0.839266647 |
| ENSMUSG00000020648 | Dus4l | protein_coding | -0.424745389 | 2.939779055 | 0.2889511 | 0.598153 | 0.839266647 |
| ENSMUSG00000103103 | 4833445I07Rik | lincRNA | 0.377261358 | 2.77715913 | 0.2889215 | 0.5981716 | 0.839266647 |
| ENSMUSG00000020941 | Map3k14 | protein_coding | 0.592970619 | 2.231274153 | 0.2884417 | 0.598473 | 0.839579921 |
| ENSMUSG00000027465 | Tbc1d20 | protein_coding | -0.19204358 | 5.170926343 | 0.2880138 | 0.5987421 | 0.839847858 |
| ENSMUSG00000092203 | 1110038B12Rik | processed_transcript | 0.174564134 | 5.013220904 | 0.2878054 | 0.5988732 | 0.839922229 |
| ENSMUSG00000019872 | Smpdl3a | protein_coding | -0.224189371 | 4.455349923 | 0.2875454 | 0.599037 | 0.839955342 |
| ENSMUSG00000028555 | Ttc39a | protein_coding | -0.474069981 | 2.707469355 | 0.2873006 | 0.5991912 | 0.839955342 |
| ENSMUSG00000020108 | Ddit4 | protein_coding | 0.343961573 | 3.351009333 | 0.2872961 | 0.599194 | 0.839955342 |
| ENSMUSG00000026018 | Ica1l | protein_coding | -0.199145365 | 4.956794094 | 0.2872719 | 0.5992093 | 0.839955342 |
| ENSMUSG00000024201 | Kdm4b | protein_coding | -0.338055502 | 4.745665976 | 0.2880633 | 0.5994331 | 0.840159532 |
| ENSMUSG00000105843 | Gm19439 | TEC | -0.505700103 | 4.624204269 | 0.29124 | 0.5996774 | 0.840392489 |
| ENSMUSG00000085664 | Atxn7l1os2 | antisense_RNA | 0.391490788 | 2.93795205 | 0.2863369 | 0.5997991 | 0.840453472 |
| ENSMUSG00000022744 | Cldnd1 | protein_coding | -0.122511943 | 7.721319446 | 0.2855422 | 0.6003014 | 0.840866497 |
| ENSMUSG00000003161 | Sri | protein_coding | -0.119332606 | 7.586269207 | 0.2855012 | 0.6003274 | 0.840866497 |
| ENSMUSG00000024137 | E4f1 | protein_coding | 0.343408931 | 3.098904098 | 0.2854995 | 0.6003285 | 0.840866497 |
| ENSMUSG00000039414 | Heatr5b | protein_coding | 0.282935318 | 3.54909801 | 0.2850949 | 0.6005846 | 0.841115651 |
| ENSMUSG00000054099 | Slc25a40 | protein_coding | -0.300411315 | 3.363760727 | 0.28496 | 0.60067 | 0.841125764 |
| ENSMUSG00000028108 | Ecm1 | protein_coding | 0.436422857 | 2.629739521 | 0.2845389 | 0.600937 | 0.841389985 |
| ENSMUSG00000019782 | Rwdd1 | protein_coding | 0.236223779 | 4.027359334 | 0.2841033 | 0.6012133 | 0.841667304 |
| ENSMUSG00000018425 | Dhx40 | protein_coding | -0.193743731 | 4.628552036 | 0.2838265 | 0.6013891 | 0.841711253 |
| ENSMUSG00000021747 | 4930452B06Rik | protein_coding | 0.228448558 | 3.836289135 | 0.2837382 | 0.6014452 | 0.841711253 |
| ENSMUSG00000078716 | Tmem8b | protein_coding | 0.253213104 | 4.758628436 | 0.2835851 | 0.6015425 | 0.841711253 |
| ENSMUSG00000031166 | Wdr13 | protein_coding | -0.134774701 | 5.967335504 | 0.2835609 | 0.6015578 | 0.841711253 |
| ENSMUSG00000019920 | Lims1 | protein_coding | 0.221983924 | 4.735409188 | 0.2833505 | 0.6016916 | 0.841788907 |
| ENSMUSG00000031700 | Gpt2 | protein_coding | -0.222673151 | 4.450509533 | 0.2831109 | 0.6018439 | 0.841892531 |
| ENSMUSG00000026664 | Phyh | protein_coding | -0.110236348 | 8.053546593 | 0.2828143 | 0.6020327 | 0.842026059 |
| ENSMUSG00000007880 | Arid1a | protein_coding | -0.197601474 | 6.022103419 | 0.2827334 | 0.602096 | 0.842026059 |
| ENSMUSG00000000340 | Dbt | protein_coding | -0.352267074 | 3.982134135 | 0.2827714 | 0.6022833 | 0.842178515 |
| ENSMUSG00000029538 | Srsf9 | protein_coding | -0.201016212 | 5.304814752 | 0.281981 | 0.6025637 | 0.842399696 |
| ENSMUSG00000029869 | Ephb6 | protein_coding | -0.280938906 | 4.24987142 | 0.2818651 | 0.6026377 | 0.842399696 |
| ENSMUSG00000085389 | 1700003M07Rik | lincRNA | -0.290116009 | 3.252357839 | 0.2818042 | 0.6026765 | 0.842399696 |
| ENSMUSG00000051346 | Spryd4 | protein_coding | 0.187820608 | 4.964114323 | 0.2812017 | 0.6030612 | 0.842827841 |
| ENSMUSG00000000532 | Acvr1b | protein_coding | 0.233920379 | 4.535971057 | 0.2804041 | 0.6035714 | 0.843431223 |
| ENSMUSG00000040537 | Adam22 | protein_coding | 0.144560026 | 6.220758536 | 0.2799338 | 0.6038726 | 0.843568579 |
| ENSMUSG00000034109 | Golim4 | protein_coding | -0.540909411 | 2.425925254 | 0.279881 | 0.6039064 | 0.843568579 |
| ENSMUSG00000020630 | Rnaseh1 | protein_coding | -0.231378147 | 4.392954136 | 0.2797811 | 0.6039705 | 0.843568579 |
| ENSMUSG00000028882 | Ppp1r8 | protein_coding | 0.164399782 | 5.707958089 | 0.2797608 | 0.6039835 | 0.843568579 |
| ENSMUSG00000054793 | Cadm4 | protein_coding | -0.297408249 | 4.443614982 | 0.2796245 | 0.6041134 | 0.843581996 |
| ENSMUSG00000025076 | Casp7 | protein_coding | -0.523035766 | 2.317533746 | 0.2794962 | 0.6041532 | 0.843581996 |
| ENSMUSG00000036352 | Ubac1 | protein_coding | 0.169301584 | 5.215568652 | 0.2793418 | 0.6042523 | 0.843581996 |
| ENSMUSG00000026049 | Tex30 | protein_coding | -0.214993439 | 4.010276521 | 0.2792566 | 0.6043069 | 0.843581996 |
| ENSMUSG00000016495 | Plgrkt | protein_coding | -0.187071675 | 5.452036591 | 0.2789237 | 0.6045207 | 0.843700159 |
| ENSMUSG00000034574 | Daam1 | protein_coding | -0.218209466 | 4.86905192 | 0.2788804 | 0.6045485 | 0.843700159 |
| ENSMUSG00000036155 | Mgat5 | protein_coding | -0.277808531 | 3.778046629 | 0.2784195 | 0.6048447 | 0.844004072 |
| ENSMUSG00000028158 | Mttp | protein_coding | -0.2841171 | 3.913784523 | 0.2780454 | 0.6050854 | 0.844197749 |
| ENSMUSG00000070476 | Fam217b | protein_coding | 0.143700323 | 5.854551818 | 0.2778647 | 0.6052018 | 0.844197749 |
| ENSMUSG00000020866 | Cacna1g | protein_coding | -0.363340939 | 3.129771186 | 0.2777657 | 0.6052655 | 0.844197749 |
| ENSMUSG00000027488 | Snta1 | protein_coding | 0.215961623 | 4.297580333 | 0.2773491 | 0.605534 | 0.844197749 |
| ENSMUSG00000024132 | Eci1 | protein_coding | 0.244311432 | 5.003862725 | 0.2771887 | 0.6056491 | 0.844197749 |
| ENSMUSG00000019261 | Map1s | protein_coding | 0.315022184 | 4.671903465 | 0.2777509 | 0.6057264 | 0.844197749 |
| ENSMUSG00000045576 | St7l | protein_coding | 0.229736899 | 4.455683809 | 0.2770501 | 0.6057268 | 0.844197749 |
| ENSMUSG00000026620 | Mark1 | protein_coding | 0.211785264 | 4.913049454 | 0.2770437 | 0.6057309 | 0.844197749 |
| ENSMUSG00000038982 | Bloc1s5 | protein_coding | 0.159636122 | 5.813263215 | 0.2770119 | 0.6057515 | 0.844197749 |
| ENSMUSG00000084883 | Ccdc85c | protein_coding | 0.277236182 | 5.217603897 | 0.2775481 | 0.6058318 | 0.844197749 |
| ENSMUSG00000039367 | Sec24c | protein_coding | -0.558970633 | 2.482691938 | 0.2767469 | 0.6059225 | 0.844197749 |
| ENSMUSG00000021696 | Elovl7 | protein_coding | -0.329167041 | 3.896187709 | 0.2767024 | 0.6059629 | 0.844197749 |
| ENSMUSG00000016319 | Slc25a5 | protein_coding | -0.111786833 | 9.055108358 | 0.276361 | 0.6061718 | 0.844197749 |
| ENSMUSG00000017376 | Nlk | protein_coding | -0.178066266 | 5.430241726 | 0.2762447 | 0.6062469 | 0.844197749 |
| ENSMUSG00000045871 | Slitrk6 | protein_coding | -0.521666165 | 2.80035259 | 0.2762366 | 0.6062637 | 0.844197749 |
| ENSMUSG00000074909 | Ranbp6 | protein_coding | -0.21598457 | 5.08870337 | 0.2761174 | 0.6063292 | 0.844197749 |

| ensembl_gene_id | external_gene_name | gene_biotype | log2 (fold change) | log2 (counts per million) | F-value | p-value | False discovery rate |
|---|---|---|---|---|---|---|---|
| ENSMUSG00000044791 | Setd2 | protein_coding | -0.240450594 | 4.536648777 | 0.2760613 | 0.6063655 | 0.844197749 |
| ENSMUSG00000033966 | Cdkl4 | protein_coding | 0.257964807 | 4.235279325 | 0.2759142 | 0.6064606 | 0.844197749 |
| ENSMUSG00000003134 | Tbc1d8 | protein_coding | 0.276659367 | 3.701743693 | 0.275792 | 0.6065397 | 0.844197749 |
| ENSMUSG00000029600 | Rita1 | protein_coding | 0.160445954 | 5.501720603 | 0.2757704 | 0.6065537 | 0.844197749 |
| ENSMUSG00000021285 | Ppp1r13b | protein_coding | 0.302800905 | 3.25431394 | 0.2753609 | 0.6068188 | 0.844339948 |
| ENSMUSG00000048661 | Lemd3 | protein_coding | 0.233747476 | 3.832823905 | 0.2753039 | 0.6068557 | 0.844339948 |
| ENSMUSG00000026750 | Psmb7 | protein_coding | 0.113379903 | 8.167252807 | 0.2752487 | 0.6068914 | 0.844339948 |
| ENSMUSG00000041375 | Ccdc9 | protein_coding | -0.316149078 | 3.067042876 | 0.2749777 | 0.6070671 | 0.844467149 |
| ENSMUSG00000040666 | Sh3bgr | protein_coding | -0.587062417 | 2.521521413 | 0.2746249 | 0.6073074 | 0.844467149 |
| ENSMUSG00000113170 | AC132344.1 | lincRNA | 0.231406665 | 3.751730224 | 0.2745676 | 0.6073331 | 0.844467149 |
| ENSMUSG00000002489 | Tiam1 | protein_coding | -0.182232902 | 5.244388358 | 0.2745291 | 0.6073581 | 0.844467149 |
| ENSMUSG00000072949 | Acot1 | protein_coding | -0.362948951 | 2.791619419 | 0.2745022 | 0.6073755 | 0.844467149 |
| ENSMUSG00000005873 | Reep5 | protein_coding | 0.120934091 | 8.580468128 | 0.2743748 | 0.6074582 | 0.844472946 |
| ENSMUSG00000086477 | Gm15506 | processed_transcript | -0.347977226 | 2.935751709 | 0.2739877 | 0.6077096 | 0.844516293 |
| ENSMUSG00000032109 | Nlrx1 | protein_coding | 0.612622936 | 2.261595834 | 0.2739184 | 0.6077546 | 0.844516293 |
| ENSMUSG00000002372 | Ranbp3 | protein_coding | 0.139479535 | 6.514298845 | 0.2736489 | 0.6079299 | 0.844516293 |
| ENSMUSG00000025486 | Sirt3 | protein_coding | 0.13004943 | 7.720722773 | 0.2735883 | 0.6079693 | 0.844516293 |
| ENSMUSG00000001105 | Ift20 | protein_coding | -0.11257938 | 7.772655175 | 0.2735408 | 0.6080001 | 0.844516293 |
| ENSMUSG00000022748 | Cmss1 | protein_coding | -0.196161222 | 5.107140504 | 0.2735346 | 0.6080042 | 0.844516293 |
| ENSMUSG00000028271 | Gtf2b | protein_coding | -0.13451958 | 6.314934803 | 0.2734808 | 0.6080392 | 0.844516293 |
| ENSMUSG00000045648 | Vwc2l | protein_coding | 0.221931851 | 3.978575379 | 0.2733462 | 0.6081268 | 0.84451683 |
| ENSMUSG00000049657 | Zbtb5 | protein_coding | -0.357410762 | 3.629371052 | 0.2731926 | 0.6082383 | 0.84451683 |
| ENSMUSG00000027397 | Slc20a1 | protein_coding | -0.351681721 | 2.862421355 | 0.2731182 | 0.6082752 | 0.84451683 |
| ENSMUSG00000020869 | Lrrc59 | protein_coding | 0.170520236 | 5.356798914 | 0.2727358 | 0.6085243 | 0.84467805 |
| ENSMUSG00000014353 | Tmem87b | protein_coding | -0.214883126 | 5.005499003 | 0.2722037 | 0.6088712 | 0.84467805 |
| ENSMUSG00000056476 | Med12l | protein_coding | -0.291993407 | 3.301636444 | 0.2721858 | 0.608883 | 0.84467805 |
| ENSMUSG00000031631 | Cfap97 | protein_coding | -0.17320116 | 5.248263559 | 0.2721487 | 0.6089072 | 0.84467805 |
| ENSMUSG00000040865 | Ino80d | protein_coding | 0.247657762 | 3.558900076 | 0.2720685 | 0.6089595 | 0.84467805 |
| ENSMUSG00000031012 | Cask | protein_coding | 0.207096579 | 7.302078276 | 0.273026 | 0.6090355 | 0.84467805 |
| ENSMUSG00000055371 | Stam2 | protein_coding | -0.20822158 | 4.296966598 | 0.2719398 | 0.6090435 | 0.84467805 |
| ENSMUSG00000027810 | Eif2a | protein_coding | -0.121465408 | 6.400073667 | 0.2718729 | 0.6090872 | 0.84467805 |
| ENSMUSG00000074886 | Grk6 | protein_coding | 0.205058638 | 4.896833007 | 0.271856 | 0.6090983 | 0.84467805 |
| ENSMUSG00000043463 | Rab9b | protein_coding | -0.177881022 | 5.675280788 | 0.2712853 | 0.6094712 | 0.84468167 |
| ENSMUSG00000029219 | Slc10a4 | protein_coding | -0.26839657 | 3.807945829 | 0.2711592 | 0.6095537 | 0.84468167 |
| ENSMUSG00000058230 | Arhgap35 | protein_coding | 0.168526802 | 6.117530316 | 0.2711021 | 0.609591 | 0.84468167 |
| ENSMUSG00000042109 | Csdc2 | protein_coding | 0.136697846 | 8.009879572 | 0.2709439 | 0.6096945 | 0.84468167 |
| ENSMUSG00000050447 | Lypd6 | protein_coding | -0.228024485 | 4.566478356 | 0.2709384 | 0.6096982 | 0.84468167 |
| ENSMUSG00000029860 | Zyx | protein_coding | -0.219462235 | 4.171575472 | 0.2709097 | 0.6097169 | 0.84468167 |
| ENSMUSG00000052253 | Zfp622 | protein_coding | 0.212414506 | 5.262507283 | 0.2708151 | 0.6097788 | 0.84468167 |
| ENSMUSG00000032349 | Elovl5 | protein_coding | 0.147647252 | 5.954478647 | 0.2707121 | 0.6098463 | 0.84468167 |
| ENSMUSG00000021400 | Wrnip1 | protein_coding | 0.181367446 | 5.408688002 | 0.2706044 | 0.6099168 | 0.84468167 |
| ENSMUSG00000040502 | 9-Mar | protein_coding | -0.319933042 | 3.218846446 | 0.2703876 | 0.6100589 | 0.84468167 |
| ENSMUSG00000008318 | Relt | protein_coding | -0.358794256 | 3.405330242 | 0.2703592 | 0.6100775 | 0.84468167 |
| ENSMUSG00000058756 | Thra | protein_coding | 0.139519165 | 6.284864811 | 0.2702481 | 0.6101503 | 0.84468167 |
| ENSMUSG00000042682 | Selenok | protein_coding | -0.143691415 | 9.326716745 | 0.2702262 | 0.6101646 | 0.84468167 |
| ENSMUSG00000031290 | Lrch2 | protein_coding | -0.222688192 | 3.809746072 | 0.2701714 | 0.6102006 | 0.84468167 |
| ENSMUSG00000030691 | Fchsd2 | protein_coding | 0.348896176 | 3.146760616 | 0.2695867 | 0.6105842 | 0.84503958 |
| ENSMUSG00000015478 | Rnf5 | protein_coding | 0.120129985 | 6.738153462 | 0.2695047 | 0.6106381 | 0.84503958 |
| ENSMUSG00000030493 | Faap24 | protein_coding | 0.238897464 | 4.212610145 | 0.2694181 | 0.6106949 | 0.84503958 |
| ENSMUSG00000022365 | Derl1 | protein_coding | 0.156360188 | 5.782097291 | 0.2692348 | 0.6108154 | 0.845097514 |
| ENSMUSG00000024617 | Camk2a | protein_coding | -0.165535998 | 7.991297627 | 0.268955 | 0.6110107 | 0.845208323 |
| ENSMUSG00000049421 | Zfp260 | protein_coding | -0.180093244 | 5.497328134 | 0.2687434 | 0.6111385 | 0.845208323 |
| ENSMUSG00000028573 | Fggy | protein_coding | 0.189746536 | 5.243828792 | 0.268695 | 0.6111703 | 0.845208323 |
| ENSMUSG00000028044 | Cks1b | protein_coding | -0.283679088 | 3.204712317 | 0.2686349 | 0.6112099 | 0.845208323 |
| ENSMUSG00000087148 | 1700030C14Rik | antisense_RNA | -0.491439524 | 2.385949082 | 0.2682286 | 0.6114773 | 0.84533123 |
| ENSMUSG00000050312 | Nsun3 | protein_coding | -0.259361416 | 3.963659659 | 0.2682096 | 0.6114899 | 0.84533123 |
| ENSMUSG00000042446 | Zmym4 | protein_coding | -0.421623062 | 2.574492042 | 0.2681417 | 0.6115346 | 0.84533123 |
| ENSMUSG00000040242 | Fgfr1op2 | protein_coding | -0.129180263 | 6.371582917 | 0.2679131 | 0.6116853 | 0.845371036 |
| ENSMUSG00000024758 | Rtn3 | protein_coding | 0.137463694 | 8.667423783 | 0.2678281 | 0.6117413 | 0.845371036 |
| ENSMUSG00000033739 | Fkbpl | protein_coding | 0.18763395 | 5.200513763 | 0.2677402 | 0.6117993 | 0.845371036 |
| ENSMUSG00000027203 | Dut | protein_coding | 0.20434493 | 4.373505902 | 0.2673406 | 0.6120629 | 0.845583524 |
| ENSMUSG00000014547 | Wdfy2 | protein_coding | -0.341161873 | 3.165943596 | 0.2672688 | 0.6121103 | 0.845583524 |
| ENSMUSG00000046679 | C87436 | protein_coding | 0.217924857 | 4.824337619 | 0.2669811 | 0.6123003 | 0.845737391 |
| ENSMUSG00000032373 | Car12 | protein_coding | 0.444510256 | 3.89923021 | 0.2681221 | 0.6124477 | 0.84583229 |
| ENSMUSG00000027475 | Kif3b | protein_coding | 0.187109027 | 4.750649199 | 0.2666352 | 0.6125289 | 0.84583582 |
| ENSMUSG00000021140 | Pcnx | protein_coding | 0.304335518 | 3.71304744 | 0.2662358 | 0.6127931 | 0.846091997 |
| ENSMUSG00000015882 | Lcorl | protein_coding | -0.324625221 | 3.036228734 | 0.2659906 | 0.6129555 | 0.846108187 |
| ENSMUSG00000059975 | Zfp74 | protein_coding | -0.251851922 | 3.668341166 | 0.265979 | 0.6129632 | 0.846108187 |
| ENSMUSG00000022023 | Wbp4 | protein_coding | -0.11531365 | 6.75804316 | 0.2657101 | 0.6131413 | 0.846108187 |
| ENSMUSG00000031953 | Tmem170 | protein_coding | 0.250907833 | 4.277935296 | 0.2656523 | 0.6131796 | 0.846108187 |
| ENSMUSG00000039163 | Cmc1 | protein_coding | 0.219753111 | 4.134331646 | 0.2656241 | 0.6131983 | 0.846108187 |
| ENSMUSG00000039323 | Igfbp2 | protein_coding | -0.37910231 | 3.213648607 | 0.2652966 | 0.6134155 | 0.846283255 |
| ENSMUSG00000034706 | Dnaic2 | protein_coding | -0.245149544 | 3.603919497 | 0.2651091 | 0.6135399 | 0.846283255 |
| ENSMUSG00000021639 | Gtf2h2 | protein_coding | -0.14902299 | 5.981032414 | 0.2649997 | 0.6136125 | 0.846283255 |
| ENSMUSG00000037709 | Fam13a | protein_coding | 0.240982439 | 3.768903823 | 0.2649063 | 0.6136745 | 0.846283255 |
| ENSMUSG00000097554 | Gm26825 | lincRNA | 0.354581988 | 2.700900117 | 0.2648398 | 0.6137187 | 0.846283255 |
| ENSMUSG00000027999 | Pla2g12a | protein_coding | 0.185474504 | 4.434526774 | 0.2646584 | 0.6138391 | 0.84633644 |
| ENSMUSG00000019809 | Pex3 | protein_coding | -0.204789603 | 5.017499214 | 0.2645448 | 0.6139147 | 0.84633644 |

| ensembl_gene_id | external_gene_name | gene_biotype | log2 (fold change) | log2 (counts per million) | F-value | p-value | False discovery rate |
|---|---|---|---|---|---|---|---|
| ENSMUSG00000021203 | Otub2 | protein_coding | 0.41530135 | 3.546582868 | 0.264961 | 0.6140774 | 0.846385697 |
| ENSMUSG00000017466 | Timp2 | protein_coding | 0.118656597 | 8.165904532 | 0.264242 | 0.6141159 | 0.846385697 |
| ENSMUSG00000059602 | Syn3 | protein_coding | 0.292818746 | 3.615109331 | 0.2641359 | 0.6141865 | 0.846385697 |
| ENSMUSG00000030854 | Ptpn5 | protein_coding | 0.12462288 | 8.552007529 | 0.2639839 | 0.6142876 | 0.84641654 |
| ENSMUSG00000027651 | Rprd1b | protein_coding | -0.232294462 | 4.07536025 | 0.2637406 | 0.6144496 | 0.846521092 |
| ENSMUSG00000031391 | L1cam | protein_coding | 0.150502735 | 5.602604218 | 0.2636334 | 0.6145209 | 0.846521092 |
| ENSMUSG00000053293 | Pom121 | protein_coding | 0.279123073 | 4.121406808 | 0.2633027 | 0.6147413 | 0.846678914 |
| ENSMUSG00000041354 | Rgl2 | protein_coding | 0.497938229 | 2.553597632 | 0.2629704 | 0.6149629 | 0.846678914 |
| ENSMUSG00000010095 | Slc3a2 | protein_coding | 0.203267932 | 5.734786904 | 0.2629751 | 0.614971 | 0.846678914 |
| ENSMUSG00000029387 | Gtf2h3 | protein_coding | 0.175420791 | 5.419838996 | 0.2629528 | 0.6149746 | 0.846678914 |
| ENSMUSG00000024535 | Snx24 | protein_coding | -0.240528117 | 4.484463318 | 0.262871 | 0.6150292 | 0.846678914 |
| ENSMUSG00000022895 | Ets2 | protein_coding | 0.226268993 | 5.444192148 | 0.2624735 | 0.6153059 | 0.846821035 |
| ENSMUSG00000026388 | 3110009E18Rik | protein_coding | -0.219953251 | 4.377404602 | 0.2623943 | 0.6153474 | 0.846821035 |
| ENSMUSG00000020396 | Nefh | protein_coding | -0.175612588 | 5.388966316 | 0.2623625 | 0.6153687 | 0.846821035 |
| ENSMUSG00000111088 | AC122428.3 | TEC | -0.377552895 | 3.354417989 | 0.2620367 | 0.6155977 | 0.847027826 |
| ENSMUSG00000031918 | Mtmr2 | protein_coding | -0.185652418 | 5.617265635 | 0.261341 | 0.616052 | 0.847380527 |
| ENSMUSG00000028496 | Mllt3 | protein_coding | -0.165579843 | 5.335618792 | 0.2612828 | 0.616091 | 0.847380527 |
| ENSMUSG00000028869 | Gnl2 | protein_coding | 0.257356041 | 5.825993876 | 0.2622823 | 0.6161066 | 0.847380527 |
| ENSMUSG00000020128 | Vps54 | protein_coding | -0.336766835 | 3.201127812 | 0.2611659 | 0.6161693 | 0.847380527 |
| ENSMUSG00000038422 | Hdhd3 | protein_coding | 0.189883915 | 5.09188441 | 0.2606922 | 0.6164869 | 0.847628412 |
| ENSMUSG00000003863 | Ppfia3 | protein_coding | 0.18543544 | 4.668904969 | 0.2605456 | 0.6165852 | 0.847628412 |
| ENSMUSG00000053819 | Camk2d | protein_coding | -0.122478447 | 7.235924366 | 0.2605444 | 0.616586 | 0.847628412 |
| ENSMUSG00000021495 | Fam193b | protein_coding | -0.362987082 | 3.323254284 | 0.2600154 | 0.6169477 | 0.848017267 |
| ENSMUSG00000021366 | Hivep1 | protein_coding | -0.465223224 | 2.754575673 | 0.2598429 | 0.6170571 | 0.848043852 |
| ENSMUSG00000033596 | Rfwd3 | protein_coding | -0.310708081 | 3.150216625 | 0.2597422 | 0.6171248 | 0.848043852 |
| ENSMUSG00000030180 | Kdm5a | protein_coding | -0.177313889 | 5.70130828 | 0.2595845 | 0.6172309 | 0.848081218 |
| ENSMUSG00000024537 | Psmg2 | protein_coding | 0.130160655 | 6.137538241 | 0.259188 | 0.6174977 | 0.848234528 |
| ENSMUSG00000037465 | Klf10 | protein_coding | 0.23595577 | 3.896625607 | 0.2591843 | 0.6175002 | 0.848234528 |
| ENSMUSG00000027248 | Pdia3 | protein_coding | 0.153087432 | 6.219888738 | 0.2587277 | 0.6178078 | 0.848548657 |
| ENSMUSG00000030397 | Mark4 | protein_coding | -0.220541151 | 4.220877947 | 0.2585566 | 0.6179232 | 0.848598683 |
| ENSMUSG00000027525 | Phactr3 | protein_coding | -0.125405613 | 6.714599074 | 0.2582904 | 0.6181027 | 0.848669973 |
| ENSMUSG00000036782 | Klhl13 | protein_coding | 0.134105982 | 7.218380801 | 0.2582456 | 0.6181329 | 0.848669973 |
| ENSMUSG00000025872 | Thoc3 | protein_coding | 0.185271704 | 5.108572263 | 0.2579998 | 0.6182989 | 0.848789434 |
| ENSMUSG00000032497 | Lrrfip2 | protein_coding | -0.186896958 | 4.629103 | 0.2576211 | 0.6185547 | 0.848860986 |
| ENSMUSG00000018882 | Mrpl45 | protein_coding | -0.122942447 | 7.34872276 | 0.2574909 | 0.6186427 | 0.848860986 |
| ENSMUSG00000027315 | Spint1 | protein_coding | -0.467269034 | 2.528566975 | 0.257484 | 0.6186473 | 0.848860986 |
| ENSMUSG00000016833 | Mrps18c | protein_coding | -0.169006244 | 7.215217141 | 0.2573963 | 0.6187472 | 0.848860986 |
| ENSMUSG00000055725 | Paqr3 | protein_coding | -0.369773035 | 5.331199923 | 0.2599527 | 0.6188936 | 0.848860986 |
| ENSMUSG00000028420 | Tmem38b | protein_coding | 0.34351392 | 3.341440637 | 0.2570932 | 0.6189117 | 0.848860986 |
| ENSMUSG00000062054 | Iah1 | protein_coding | 0.162900566 | 5.522841567 | 0.2569016 | 0.6190414 | 0.848860986 |
| ENSMUSG00000036356 | Csgalnact1 | protein_coding | 0.328779693 | 3.439311521 | 0.2568456 | 0.6190793 | 0.848860986 |
| ENSMUSG00000026848 | Tor1b | protein_coding | -0.214311501 | 4.511419025 | 0.2566402 | 0.6192185 | 0.848860986 |
| ENSMUSG00000031517 | Gpm6a | protein_coding | -0.113481915 | 9.910279319 | 0.2565598 | 0.6192729 | 0.848860986 |
| ENSMUSG00000026575 | Nme7 | protein_coding | -0.216224436 | 4.734404409 | 0.2564879 | 0.6193217 | 0.848860986 |
| ENSMUSG00000056121 | Fez2 | protein_coding | -0.18143038 | 4.617112887 | 0.2564419 | 0.6193528 | 0.848860986 |
| ENSMUSG00000049339 | Retreg2 | protein_coding | 0.13424654 | 6.910979518 | 0.2564059 | 0.6193772 | 0.848860986 |
| ENSMUSG00000021000 | Ctage5 | protein_coding | -0.246225109 | 4.251313208 | 0.2561262 | 0.6195669 | 0.848897435 |
| ENSMUSG00000040681 | Hmgn1 | protein_coding | 0.190304679 | 4.808654613 | 0.2559515 | 0.6196854 | 0.848897435 |
| ENSMUSG00000032507 | Fbxl2 | protein_coding | 0.197317499 | 5.618176864 | 0.2558123 | 0.6197799 | 0.848897435 |
| ENSMUSG00000027652 | Ralgapb | protein_coding | -0.173726131 | 4.935392349 | 0.2555596 | 0.6199515 | 0.848897435 |
| ENSMUSG00000073684 | Faap20 | protein_coding | 0.145948501 | 5.802245919 | 0.2555241 | 0.6199756 | 0.848897435 |
| ENSMUSG00000003948 | Mmd | protein_coding | -0.223030828 | 6.024027836 | 0.2561032 | 0.6200133 | 0.848897435 |
| ENSMUSG00000033565 | Rbfox2 | protein_coding | -0.120831981 | 6.285508904 | 0.2554073 | 0.620055 | 0.848897435 |
| ENSMUSG00000020181 | Nav3 | protein_coding | -0.173765808 | 5.39169619 | 0.2553981 | 0.6200613 | 0.848897435 |
| ENSMUSG00000044676 | Zfp612 | protein_coding | 0.30634519 | 3.313462669 | 0.25532 | 0.6201143 | 0.848897435 |
| ENSMUSG00000060477 | Irak2 | protein_coding | 0.263577756 | 3.333828828 | 0.2551583 | 0.6202243 | 0.848939841 |
| ENSMUSG00000029203 | Ube2k | protein_coding | 0.125860835 | 7.384558187 | 0.2546825 | 0.620548 | 0.849045782 |
| ENSMUSG00000054934 | Kcnmb4 | protein_coding | -0.430187304 | 2.524912605 | 0.254677 | 0.6205517 | 0.849045782 |
| ENSMUSG00000028121 | Bcar3 | protein_coding | -0.255877787 | 4.360199005 | 0.2546326 | 0.620582 | 0.849045782 |
| ENSMUSG00000086841 | 2410006H16Rik | processed_transcript | 0.137484135 | 7.585014337 | 0.2545804 | 0.6206175 | 0.849045782 |
| ENSMUSG00000032199 | Polr2m | protein_coding | -0.122116005 | 6.88665138 | 0.2544542 | 0.6207035 | 0.849055369 |
| ENSMUSG00000018481 | Appbp2 | protein_coding | 0.118121311 | 6.66161444 | 0.2538067 | 0.6211449 | 0.849551084 |
| ENSMUSG00000026227 | 2810459M11Rik | protein_coding | -0.514031418 | 2.526631939 | 0.2536001 | 0.6212859 | 0.849635863 |
| ENSMUSG00000009907 | Vps4b | protein_coding | -0.157534047 | 5.311644651 | 0.2534488 | 0.6213892 | 0.849669069 |
| ENSMUSG00000035078 | Mtmr9 | protein_coding | -0.158933892 | 5.870127254 | 0.2530054 | 0.6216922 | 0.849768409 |
| ENSMUSG00000086272 | BC039966 | antisense_RNA | 0.315445353 | 3.834084902 | 0.2530159 | 0.621696 | 0.849768409 |
| ENSMUSG00000064065 | Ipcef1 | protein_coding | 0.353930231 | 2.627359366 | 0.2529955 | 0.6216989 | 0.849768409 |
| ENSMUSG00000049313 | Sorl1 | protein_coding | 0.139830508 | 6.405108232 | 0.2525189 | 0.6220249 | 0.849817388 |
| ENSMUSG00000060373 | Hnrnpc | protein_coding | -0.110761086 | 7.866966741 | 0.2524454 | 0.6220753 | 0.849817388 |
| ENSMUSG00000041115 | Iqsec2 | protein_coding | 0.212510867 | 4.473498805 | 0.252438 | 0.6220803 | 0.849817388 |
| ENSMUSG00000067567 | Hdac8 | protein_coding | 0.381444604 | 3.922088003 | 0.2531963 | 0.6222324 | 0.849817388 |
| ENSMUSG00000025352 | Gdf11 | protein_coding | 0.218656995 | 4.577487827 | 0.2521445 | 0.6222814 | 0.849817388 |
| ENSMUSG00000032657 | Fam189b | protein_coding | -0.237412121 | 3.935314827 | 0.2521304 | 0.622291 | 0.849817388 |
| ENSMUSG00000020184 | Mdm2 | protein_coding | -0.185033529 | 4.96848928 | 0.252064 | 0.6223365 | 0.849817388 |
| ENSMUSG00000093460 | Six3os1 | processed_transcript | -0.297140779 | 3.60449471 | 0.2519353 | 0.6224247 | 0.849817388 |
| ENSMUSG00000036040 | Adamtsl2 | protein_coding | -0.268563058 | 3.44761555 | 0.2519042 | 0.622446 | 0.849817388 |
| ENSMUSG00000031302 | Nlgn3 | protein_coding | -0.126504008 | 6.648636821 | 0.2516693 | 0.6226071 | 0.849929376 |
| ENSMUSG00000092329 | Gm20388 | protein_coding | 0.10824848 | 6.876614423 | 0.2512928 | 0.6228655 | 0.850174198 |

| ensembl_gene_id | external_gene_name | gene_biotype | log2 (fold change) | log2 (counts per million) | F-value | p-value | False discovery rate |
|---|---|---|---|---|---|---|---|
| ENSMUSG00000031950 | Gabarapl2 | protein_coding | 0.120341267 | 6.722212429 | 0.2511602 | 0.6229565 | 0.850190548 |
| ENSMUSG00000030188 | Magohb | protein_coding | -0.187117064 | 5.418424642 | 0.2508062 | 0.6231998 | 0.850410416 |
| ENSMUSG00000025198 | Erlin1 | protein_coding | -0.26427451 | 3.890880688 | 0.2506956 | 0.6232758 | 0.850410416 |
| ENSMUSG00000031843 | Mphosph6 | protein_coding | -0.15412426 | 6.021009943 | 0.2505424 | 0.6233811 | 0.850446255 |
| ENSMUSG00000032078 | Zpr1 | protein_coding | 0.239717263 | 4.798499595 | 0.2503716 | 0.6235097 | 0.850513673 |
| ENSMUSG00000022186 | Oxct1 | protein_coding | 0.113988694 | 8.154560547 | 0.2499233 | 0.6238074 | 0.850811872 |
| ENSMUSG00000040040 | Ift88 | protein_coding | 0.232439953 | 4.567561074 | 0.2497597 | 0.6239201 | 0.850857662 |
| ENSMUSG00000089832 | Shkbp1 | protein_coding | -0.503906189 | 2.76160648 | 0.2489327 | 0.6245017 | 0.851454551 |
| ENSMUSG00000025909 | Sntg1 | protein_coding | 0.192813553 | 4.783253492 | 0.2488959 | 0.6245161 | 0.851454551 |
| ENSMUSG00000039356 | Exosc2 | protein_coding | 0.245416817 | 3.985303725 | 0.2485772 | 0.6247363 | 0.851646779 |
| ENSMUSG00000002812 | Flii | protein_coding | 0.225956398 | 5.044575425 | 0.2483787 | 0.6248845 | 0.851740784 |
| ENSMUSG00000030754 | Copb1 | protein_coding | -0.170449492 | 5.99782051 | 0.2480492 | 0.6251016 | 0.851879481 |
| ENSMUSG00000024515 | Smad4 | protein_coding | -0.184436114 | 4.955008203 | 0.2479869 | 0.6251447 | 0.851879481 |
| ENSMUSG00000038065 | Mturn | protein_coding | 0.184824419 | 6.059530408 | 0.2478298 | 0.6252645 | 0.851934701 |
| ENSMUSG00000071648 | Rom1 | protein_coding | 0.298140881 | 3.346982988 | 0.2475988 | 0.6254135 | 0.85202988 |
| ENSMUSG00000020432 | Tcn2 | protein_coding | -0.276769725 | 4.297761331 | 0.2473169 | 0.6256199 | 0.852202986 |
| ENSMUSG00000023249 | Parp3 | protein_coding | -0.352667404 | 2.909312316 | 0.2468652 | 0.6259225 | 0.85228099 |
| ENSMUSG00000038072 | Galnt11 | protein_coding | 0.267320142 | 3.718150019 | 0.2468215 | 0.6259528 | 0.85228099 |
| ENSMUSG00000084808 | 9430091E24Rik | processed_transcript | -0.257312498 | 3.72286899 | 0.2467997 | 0.6259679 | 0.85228099 |
| ENSMUSG00000009731 | Kcnd1 | protein_coding | 0.564958084 | 2.365475782 | 0.2467776 | 0.6259942 | 0.85228099 |
| ENSMUSG00000039632 | Ccdc151 | protein_coding | 0.492792814 | 2.469709925 | 0.2462291 | 0.6263644 | 0.852677159 |
| ENSMUSG00000037935 | Smarce1 | protein_coding | 0.170853829 | 5.526289286 | 0.245944 | 0.6265628 | 0.852798204 |
| ENSMUSG00000090291 | Lrrc10b | protein_coding | -0.258640113 | 4.112851668 | 0.2458733 | 0.626612 | 0.852798204 |
| ENSMUSG00000037957 | Wdr20 | protein_coding | -0.22002911 | 4.071077667 | 0.2454853 | 0.6268822 | 0.852861224 |
| ENSMUSG00000035351 | Nup37 | protein_coding | 0.226096109 | 4.096007473 | 0.2453875 | 0.6269504 | 0.852861224 |
| ENSMUSG00000022678 | Nde1 | protein_coding | -0.21668657 | 5.028134558 | 0.2451177 | 0.6271385 | 0.852861224 |
| ENSMUSG00000026087 | Mrpl30 | protein_coding | 0.127338144 | 8.558953754 | 0.2450963 | 0.6271534 | 0.852861224 |
| ENSMUSG00000051768 | Xrcc1 | protein_coding | 0.202661123 | 4.455173051 | 0.2450813 | 0.6271639 | 0.852861224 |
| ENSMUSG00000037902 | Sirpa | protein_coding | -0.127043364 | 6.931019803 | 0.2450346 | 0.6271964 | 0.852861224 |
| ENSMUSG00000039449 | Prpf18 | protein_coding | -0.172330266 | 5.607846361 | 0.2450102 | 0.6272135 | 0.852861224 |
| ENSMUSG00000026556 | Vangl2 | protein_coding | -0.234912085 | 3.631927552 | 0.2448145 | 0.62735 | 0.852897855 |
| ENSMUSG00000054312 | Mrps21 | protein_coding | 0.152395623 | 7.97486002 | 0.2447598 | 0.6273991 | 0.852897855 |
| ENSMUSG00000052656 | Rnf103 | protein_coding | 0.299831714 | 4.638591862 | 0.2451503 | 0.6275359 | 0.85289998 |
| ENSMUSG00000052557 | Gan | protein_coding | 0.275363762 | 3.380614522 | 0.2445148 | 0.6275593 | 0.85289998 |
| ENSMUSG00000028252 | Ccnc | protein_coding | 0.143578375 | 7.047618896 | 0.2440055 | 0.6279153 | 0.853275981 |
| ENSMUSG00000018411 | Mapt | protein_coding | 0.155313226 | 8.897666719 | 0.2430425 | 0.6286004 | 0.8540272 |
| ENSMUSG00000066647 | Gm5113 | protein_coding | 0.168029116 | 4.778625466 | 0.2429893 | 0.6286269 | 0.8540272 |
| ENSMUSG00000060992 | Copz1 | protein_coding | 0.116442032 | 8.602941946 | 0.2423469 | 0.6290778 | 0.854531754 |
| ENSMUSG00000001521 | Tulp3 | protein_coding | 0.293540559 | 3.3495638 | 0.2417553 | 0.6294937 | 0.854988738 |
| ENSMUSG00000025742 | Prps2 | protein_coding | 0.192269831 | 4.834751372 | 0.2415927 | 0.6296081 | 0.855036117 |
| ENSMUSG00000030726 | Pold3 | protein_coding | 0.205463584 | 5.231761122 | 0.2414425 | 0.6297138 | 0.855071722 |
| ENSMUSG00000087461 | C230014O12Rik | lincRNA | 0.36390378 | 2.858043497 | 0.241135 | 0.6299304 | 0.855184432 |
| ENSMUSG00000027286 | Lrrc57 | protein_coding | -0.15546644 | 5.643362814 | 0.2409545 | 0.6300576 | 0.855184432 |
| ENSMUSG00000025558 | Dock9 | protein_coding | -0.242393978 | 4.382703992 | 0.2409343 | 0.6300719 | 0.855184432 |
| ENSMUSG00000053553 | 3110082I17Rik | protein_coding | 0.194270582 | 4.48468091 | 0.2408732 | 0.630115 | 0.855184432 |
| ENSMUSG00000030256 | Bhlhe41 | protein_coding | 0.348203181 | 7.981567153 | 0.2462817 | 0.63021 | 0.855205473 |
| ENSMUSG00000040649 | Rimklb | protein_coding | -0.151087113 | 6.130343826 | 0.2405349 | 0.6303535 | 0.855292327 |
| ENSMUSG00000036636 | Clcn7 | protein_coding | 0.216626283 | 4.562534078 | 0.2402237 | 0.6305733 | 0.855482551 |
| ENSMUSG00000060962 | Dmkn | protein_coding | -0.292482922 | 4.473782362 | 0.2400634 | 0.6309598 | 0.85558267 |
| ENSMUSG00000028136 | Snx27 | protein_coding | 0.137002272 | 6.71847641 | 0.2396352 | 0.6309893 | 0.85558267 |
| ENSMUSG00000005981 | Trap1 | protein_coding | 0.16387918 | 6.09934087 | 0.2396238 | 0.6309973 | 0.85558267 |
| ENSMUSG00000033545 | Znrf1 | protein_coding | -0.161973581 | 5.488763128 | 0.239567 | 0.6310375 | 0.85558267 |
| ENSMUSG00000030329 | Pianp | protein_coding | 0.147604796 | 6.795804784 | 0.2395565 | 0.6310449 | 0.85558267 |
| ENSMUSG00000036435 | Exoc1 | protein_coding | -0.182320555 | 5.289982686 | 0.2393048 | 0.6312231 | 0.855716343 |
| ENSMUSG00000041747 | Utp15 | protein_coding | -0.19746048 | 4.508978226 | 0.2391446 | 0.6313365 | 0.855718714 |
| ENSMUSG00000057506 | Bloc1s2 | protein_coding | -0.157857346 | 5.078684327 | 0.2390131 | 0.6314297 | 0.855718714 |
| ENSMUSG00000008690 | Ncaph2 | protein_coding | 0.153172977 | 7.207697295 | 0.2389803 | 0.6314636 | 0.855718714 |
| ENSMUSG00000078813 | Leng1 | protein_coding | -0.377087212 | 2.817330814 | 0.2387939 | 0.6315851 | 0.855775473 |
| ENSMUSG00000016940 | Kctd2 | protein_coding | 0.180497148 | 5.36580741 | 0.2384594 | 0.6318223 | 0.85584062 |
| ENSMUSG00000048920 | Fkrp | protein_coding | -0.179573416 | 4.62235271 | 0.2383743 | 0.6318827 | 0.85584062 |
| ENSMUSG00000029196 | Tada2b | protein_coding | 0.250752791 | 3.571371922 | 0.2381816 | 0.6320195 | 0.85584062 |
| ENSMUSG00000029207 | Apbb2 | protein_coding | 0.126780553 | 6.354913136 | 0.2381248 | 0.6320598 | 0.85584062 |
| ENSMUSG00000029310 | Nudt9 | protein_coding | 0.160647269 | 5.61468914 | 0.2380909 | 0.6320839 | 0.85584062 |
| ENSMUSG00000054910 | 4931415C17Rik | processed_transcript | -0.442808635 | 2.736321017 | 0.237923 | 0.6322031 | 0.85584062 |
| ENSMUSG00000049971 | Glt1d1 | protein_coding | -0.311956068 | 3.186405473 | 0.2379041 | 0.6322166 | 0.85584062 |
| ENSMUSG00000038403 | Hfe2 | protein_coding | -0.367767411 | 2.767876911 | 0.2378291 | 0.6322699 | 0.85584062 |
| ENSMUSG00000017858 | Ift52 | protein_coding | 0.147739271 | 6.539735167 | 0.2374461 | 0.6325423 | 0.855928031 |
| ENSMUSG00000020538 | Srebf1 | protein_coding | -0.251967937 | 4.174964171 | 0.2372877 | 0.6326549 | 0.855928031 |
| ENSMUSG00000026495 | Efcab2 | protein_coding | -0.172829339 | 5.377614684 | 0.2372597 | 0.6326749 | 0.855928031 |
| ENSMUSG00000023952 | Gtpbp2 | protein_coding | -0.21824272 | 4.165417954 | 0.2371674 | 0.6327406 | 0.855928031 |
| ENSMUSG00000031137 | Fgf13 | protein_coding | 0.161698364 | 6.550759424 | 0.2371084 | 0.6327932 | 0.855928031 |
| ENSMUSG00000022205 | Sub1 | protein_coding | -0.127031785 | 10.23803381 | 0.2370671 | 0.632812 | 0.855928031 |
| ENSMUSG00000021375 | Kif13a | protein_coding | 0.256257539 | 3.691015578 | 0.2369425 | 0.6329008 | 0.855940419 |
| ENSMUSG00000060314 | Zfp941 | protein_coding | 0.137306584 | 6.292525636 | 0.2366501 | 0.6331092 | 0.856055828 |
| ENSMUSG00000049922 | Slc35c1 | protein_coding | -0.235479127 | 4.063966979 | 0.2365993 | 0.6331454 | 0.856055828 |
| ENSMUSG00000050840 | Cdh20 | protein_coding | 0.441973589 | 2.865321039 | 0.2364264 | 0.6332793 | 0.856129328 |
| ENSMUSG00000066640 | Fbxl18 | protein_coding | 0.226494125 | 4.108800012 | 0.2362009 | 0.6334296 | 0.856164107 |
| ENSMUSG00000038366 | Lasp1 | protein_coding | 0.178408139 | 5.692131249 | 0.2361002 | 0.6335015 | 0.856164107 |

| ensembl_gene_id | external_gene_name | gene_biotype | log2 (fold change) | log2 (counts per million) | F-value | p-value | False discovery rate |
|---|---|---|---|---|---|---|---|
| ENSMUSG00000041488 | Stx3 | protein_coding | -0.218851962 | 3.858314056 | 0.2360408 | 0.6335439 | 0.856164107 |
| ENSMUSG00000022203 | Efs | protein_coding | 0.303141343 | 2.928711855 | 0.2356634 | 0.6338135 | 0.856346941 |
| ENSMUSG00000061904 | Slc25a3 | protein_coding | -0.122892559 | 8.245816559 | 0.2356285 | 0.6338385 | 0.856346941 |
| ENSMUSG00000021259 | Cyp46a1 | protein_coding | 0.186373944 | 5.252938892 | 0.2354363 | 0.633976 | 0.856425052 |
| ENSMUSG00000024983 | Vti1a | protein_coding | 0.195533431 | 5.623948769 | 0.235268 | 0.6341069 | 0.856494318 |
| ENSMUSG00000020037 | Rfx4 | protein_coding | -0.205436859 | 3.874908467 | 0.2348347 | 0.6344066 | 0.856791488 |
| ENSMUSG00000041889 | Shisa4 | protein_coding | 0.115753669 | 6.555733141 | 0.2345752 | 0.6345925 | 0.856896361 |
| ENSMUSG00000058740 | Kcnt1 | protein_coding | 0.235169909 | 3.781215717 | 0.2345039 | 0.6346436 | 0.856896361 |
| ENSMUSG00000022340 | Sybu | protein_coding | -0.112853683 | 6.519935658 | 0.2337124 | 0.6352117 | 0.857468401 |
| ENSMUSG00000028655 | Mfsd2a | protein_coding | 0.664396086 | 2.405620596 | 0.2342617 | 0.6352268 | 0.857468401 |
| ENSMUSG00000040028 | Elavl1 | protein_coding | -0.147522368 | 5.450016364 | 0.2334939 | 0.6353688 | 0.857552439 |
| ENSMUSG00000019899 | Lama2 | protein_coding | -0.35487689 | 2.772551931 | 0.2330674 | 0.6356756 | 0.8577688 |
| ENSMUSG00000019960 | Dusp6 | protein_coding | -0.178217201 | 5.189858179 | 0.2329765 | 0.635741 | 0.8577688 |
| ENSMUSG00000021125 | Arg2 | protein_coding | -0.197981633 | 3.883070506 | 0.2329385 | 0.6357684 | 0.8577688 |
| ENSMUSG00000029480 | Dhx37 | protein_coding | 0.258708102 | 3.670939962 | 0.2325109 | 0.6360764 | 0.858076738 |
| ENSMUSG00000010086 | Rnf112 | protein_coding | -0.228857818 | 4.615824824 | 0.232255 | 0.636261 | 0.858218021 |
| ENSMUSG00000052364 | B630019K06Rik | protein_coding | -0.164034954 | 5.775078088 | 0.2319488 | 0.6364819 | 0.858408357 |
| ENSMUSG00000031511 | Arhgef7 | protein_coding | 0.182683406 | 5.673300661 | 0.2318323 | 0.636566 | 0.858414148 |
| ENSMUSG00000015672 | Mrpl32 | protein_coding | -0.141555922 | 6.614622053 | 0.2315674 | 0.6367573 | 0.858531755 |
| ENSMUSG00000032875 | Arhgef17 | protein_coding | 0.122729548 | 6.310172175 | 0.2313896 | 0.6368859 | 0.858531755 |
| ENSMUSG00000032594 | Ip6k1 | protein_coding | 0.118409849 | 7.034763964 | 0.2312887 | 0.6369588 | 0.858531755 |
| ENSMUSG00000046947 | Adck2 | protein_coding | -0.380642448 | 3.16324353 | 0.2312719 | 0.6369815 | 0.858531755 |
| ENSMUSG00000051652 | Lrrc3 | protein_coding | 0.456280379 | 3.470693892 | 0.2324008 | 0.6370524 | 0.858531755 |
| ENSMUSG00000038456 | Dennd2a | protein_coding | -0.257465846 | 3.903286235 | 0.2310201 | 0.6371531 | 0.85855986 |
| ENSMUSG00000022387 | Brd1 | protein_coding | -0.186209969 | 4.943559841 | 0.2307044 | 0.6373817 | 0.85871276 |
| ENSMUSG00000021666 | Gfm2 | protein_coding | 0.246489049 | 3.871473383 | 0.2305068 | 0.6375249 | 0.85871276 |
| ENSMUSG00000061559 | Wdr61 | protein_coding | -0.108861435 | 7.66152338 | 0.2304591 | 0.6375595 | 0.85871276 |
| ENSMUSG00000022437 | Samm50 | protein_coding | 0.141876769 | 6.467781763 | 0.2303252 | 0.6376566 | 0.85871276 |
| ENSMUSG00000028114 | Mettl14 | protein_coding | 0.197903598 | 4.797595066 | 0.2302948 | 0.6376786 | 0.85871276 |
| ENSMUSG00000054733 | Msra | protein_coding | 0.152261657 | 5.175527977 | 0.2300568 | 0.6378512 | 0.85871276 |
| ENSMUSG00000023034 | Nr4a1 | protein_coding | -0.144954 | 5.870254993 | 0.2300145 | 0.6378819 | 0.85871276 |
| ENSMUSG00000044991 | 1110034G24Rik | protein_coding | 0.17234471 | 5.267753978 | 0.2298027 | 0.6380356 | 0.85871276 |
| ENSMUSG00000025289 | Prdx4 | protein_coding | -0.157664727 | 5.927358822 | 0.229735 | 0.6380848 | 0.85871276 |
| ENSMUSG00000054309 | Cpsf3 | protein_coding | 0.162347178 | 6.587729237 | 0.2295967 | 0.6381957 | 0.85871276 |
| ENSMUSG00000028078 | Dclk2 | protein_coding | 0.15743462 | 4.942542086 | 0.2295592 | 0.6382125 | 0.85871276 |
| ENSMUSG00000053470 | Kdm3a | protein_coding | -0.209971147 | 4.227019001 | 0.2294432 | 0.6382968 | 0.85871276 |
| ENSMUSG00000025175 | Fn3k | protein_coding | 0.135479727 | 6.706386204 | 0.2293199 | 0.6383864 | 0.85871276 |
| ENSMUSG00000031730 | Dhodh | protein_coding | 0.138142436 | 6.121563963 | 0.2292914 | 0.6384071 | 0.85871276 |
| ENSMUSG00000074259 | Gramd2 | protein_coding | -0.481263997 | 2.559464492 | 0.2292126 | 0.6384645 | 0.85871276 |
| ENSMUSG00000045287 | Rtn4rl1 | protein_coding | -0.214137309 | 3.935299572 | 0.2290349 | 0.6385937 | 0.858779135 |
| ENSMUSG00000040990 | Sh3kbp1 | protein_coding | -0.143468459 | 5.563578914 | 0.2287332 | 0.6388133 | 0.858967071 |
| ENSMUSG00000045761 | Togaram2 | protein_coding | -0.469960932 | 2.528370165 | 0.228311 | 0.6391209 | 0.859117097 |
| ENSMUSG00000010392 | Gosr1 | protein_coding | -0.198517494 | 5.264274629 | 0.2281539 | 0.6392355 | 0.859117097 |
| ENSMUSG00000030701 | Plekhb1 | protein_coding | 0.131152999 | 9.101479327 | 0.2281412 | 0.6392447 | 0.859117097 |
| ENSMUSG00000052031 | Tagap1 | protein_coding | 0.179719256 | 5.618157086 | 0.2280829 | 0.6392873 | 0.859117097 |
| ENSMUSG00000041245 | Wnk3 | protein_coding | -0.315005689 | 3.176383851 | 0.228032 | 0.6393244 | 0.859117097 |
| ENSMUSG00000035805 | Mlc1 | protein_coding | -0.24290619 | 6.046922441 | 0.2290714 | 0.6394565 | 0.859187255 |
| ENSMUSG00000046613 | Vwa5b2 | protein_coding | 0.341700464 | 2.903223785 | 0.2274014 | 0.6397849 | 0.859521104 |
| ENSMUSG00000020656 | Grhl1 | protein_coding | 0.335977389 | 3.637464104 | 0.2273236 | 0.6398974 | 0.859564855 |
| ENSMUSG00000030683 | Sez6l2 | protein_coding | 0.114192373 | 7.401941601 | 0.2261541 | 0.640698 | 0.860391194 |
| ENSMUSG00000047473 | Zfp30 | protein_coding | 0.17250258 | 4.606516946 | 0.2259603 | 0.6408402 | 0.860391194 |
| ENSMUSG00000073102 | Drc1 | protein_coding | 0.17255263 | 4.882209668 | 0.2259386 | 0.6408561 | 0.860391194 |
| ENSMUSG00000029518 | Rab35 | protein_coding | 0.149903016 | 5.597696146 | 0.2258015 | 0.6409567 | 0.860391194 |
| ENSMUSG00000016262 | Sertad4 | protein_coding | -0.216199146 | 4.413357749 | 0.2257833 | 0.64097 | 0.860391194 |
| ENSMUSG00000056167 | Cnot10 | protein_coding | 0.182387995 | 5.128757025 | 0.2256408 | 0.6410747 | 0.860391194 |
| ENSMUSG00000071265 | 1700086L19Rik | transcribed_unitary_p | 0.141830636 | 5.636011337 | 0.225595 | 0.6411083 | 0.860391194 |
| ENSMUSG00000034066 | Farp2 | protein_coding | 0.314570933 | 2.859174941 | 0.2255346 | 0.6411527 | 0.860391194 |
| ENSMUSG00000031932 | Gpr83 | protein_coding | 0.184247957 | 5.762851993 | 0.2252062 | 0.6414044 | 0.860472256 |
| ENSMUSG00000036721 | Zscan12 | protein_coding | -0.26113056 | 3.671863815 | 0.2250305 | 0.6415232 | 0.860472256 |
| ENSMUSG00000033732 | Sf3b3 | protein_coding | 0.159791468 | 5.440594887 | 0.2250262 | 0.6415264 | 0.860472256 |
| ENSMUSG00000068856 | Sf3b4 | protein_coding | -0.205129135 | 4.936819582 | 0.2247475 | 0.6417315 | 0.860472256 |
| ENSMUSG00000032264 | Zw10 | protein_coding | -0.240100625 | 4.109129274 | 0.2247204 | 0.6417514 | 0.860472256 |
| ENSMUSG00000085436 | Zfp335os | antisense_RNA | -0.252770279 | 3.328951338 | 0.2246245 | 0.641822 | 0.860472256 |
| ENSMUSG00000019433 | Gipc1 | protein_coding | 0.156695927 | 5.693327097 | 0.2245624 | 0.6418678 | 0.860472256 |
| ENSMUSG00000092060 | Bend4 | protein_coding | 0.265014928 | 4.913103167 | 0.2250306 | 0.6419233 | 0.860472256 |
| ENSMUSG00000086607 | 4930511M06Rik | processed_transcript | -0.302768637 | 3.102546833 | 0.2244735 | 0.6419333 | 0.860472256 |
| ENSMUSG00000016528 | Mapkapk2 | protein_coding | 0.416858007 | 2.809409699 | 0.224107 | 0.6422034 | 0.86056342 |
| ENSMUSG00000036006 | Ripor2 | protein_coding | -0.176139559 | 5.540538924 | 0.2239472 | 0.6423212 | 0.86056342 |
| ENSMUSG00000028582 | Cc2d1b | protein_coding | -0.290913039 | 3.403675715 | 0.2239457 | 0.6423223 | 0.86056342 |
| ENSMUSG00000026113 | Inpp4a | protein_coding | 0.202426371 | 9.06552636 | 0.2250107 | 0.6423524 | 0.86056342 |
| ENSMUSG00000030042 | Pole4 | protein_coding | 0.145280816 | 5.649741215 | 0.2237805 | 0.6424442 | 0.86056342 |
| ENSMUSG00000001150 | Mcm3ap | protein_coding | 0.214933367 | 3.73704569 | 0.2236354 | 0.6425514 | 0.86056342 |
| ENSMUSG00000027099 | Mtx2 | protein_coding | -0.179520488 | 5.076598187 | 0.2236217 | 0.6425615 | 0.86056342 |
| ENSMUSG00000071573 | Rnls | protein_coding | 0.264674533 | 3.221765086 | 0.2234741 | 0.6426705 | 0.860597371 |
| ENSMUSG00000060032 | H2afj | protein_coding | 0.318085322 | 2.916554437 | 0.2233707 | 0.6427469 | 0.860597371 |
| ENSMUSG00000032437 | Stt3b | protein_coding | -0.192704814 | 5.189159621 | 0.223184 | 0.6428849 | 0.860675 |
| ENSMUSG00000027108 | Ola1 | protein_coding | 0.109653641 | 7.789073664 | 0.2224338 | 0.6434401 | 0.861302971 |
| ENSMUSG00000008129 | 4930432K21Rik | protein_coding | -0.399922594 | 2.80066687 | 0.2223339 | 0.6435142 | 0.861302971 |

| ensembl_gene_id | external_gene_name | gene_biotype | log2 (fold change) | log2 (counts per million) | F-value | p-value | False discovery rate |
|---|---|---|---|---|---|---|---|
| ENSMUSG00000036707 | Cab39 | protein_coding | 0.140983643 | 6.547629035 | 0.221989 | 0.6437699 | 0.861537951 |
| ENSMUSG00000054027 | Nt5dc3 | protein_coding | -0.205068078 | 4.951660942 | 0.2218202 | 0.6438951 | 0.861598361 |
| ENSMUSG00000021224 | Numb | protein_coding | 0.222145453 | 4.281775495 | 0.2215753 | 0.6440769 | 0.861734339 |
| ENSMUSG00000026051 | 1500015O10Rik | protein_coding | 0.412115215 | 3.884914148 | 0.2216828 | 0.6443407 | 0.861929509 |
| ENSMUSG00000078515 | Ddi2 | protein_coding | 0.168433301 | 5.409686099 | 0.2211537 | 0.6443902 | 0.861929509 |
| ENSMUSG00000046562 | Unc119b | protein_coding | 0.31053766 | 3.136478763 | 0.2209128 | 0.6445694 | 0.861929509 |
| ENSMUSG00000022843 | Clcn2 | protein_coding | 0.429258527 | 2.632605279 | 0.2209123 | 0.6445697 | 0.861929509 |
| ENSMUSG00000073427 | Gm4924 | protein_coding | -0.249720977 | 3.329425983 | 0.2206884 | 0.6447363 | 0.861929509 |
| ENSMUSG00000028894 | Inpp5b | protein_coding | -0.211564208 | 4.148575402 | 0.2204259 | 0.6449319 | 0.861929509 |
| ENSMUSG00000045427 | Hnrnph2 | protein_coding | -0.150674615 | 5.821643913 | 0.2202229 | 0.6450832 | 0.861929509 |
| ENSMUSG00000045031 | Cetn4 | protein_coding | 0.219514435 | 3.838305497 | 0.2201921 | 0.6451061 | 0.861929509 |
| ENSMUSG00000026153 | Fam135a | protein_coding | -0.353554578 | 3.403982412 | 0.2201962 | 0.6451133 | 0.861929509 |
| ENSMUSG00000001569 | Nom1 | protein_coding | 0.183319158 | 4.537747946 | 0.2200757 | 0.6451929 | 0.861929509 |
| ENSMUSG00000069255 | Dusp22 | protein_coding | 0.170334143 | 5.260943692 | 0.2200474 | 0.645214 | 0.861929509 |
| ENSMUSG00000027500 | Stmn2 | protein_coding | 0.10703293 | 10.26118656 | 0.2200434 | 0.645217 | 0.861929509 |
| ENSMUSG00000037617 | Spag1 | protein_coding | -0.427391536 | 2.575763389 | 0.2199793 | 0.6452648 | 0.861929509 |
| ENSMUSG00000030657 | Xylt1 | protein_coding | -0.334824616 | 3.242131493 | 0.2198461 | 0.6453642 | 0.861948044 |
| ENSMUSG00000034312 | Iqsec1 | protein_coding | -0.1683267 | 5.851191908 | 0.2197017 | 0.645472 | 0.861948044 |
| ENSMUSG00000021508 | Cxcl14 | protein_coding | -0.262939337 | 4.651294608 | 0.2198838 | 0.6455192 | 0.861948044 |
| ENSMUSG00000023473 | Celsr3 | protein_coding | 0.386810724 | 2.799257655 | 0.2193908 | 0.6457042 | 0.862088046 |
| ENSMUSG00000026032 | Ndufb3 | protein_coding | 0.148052649 | 8.401086337 | 0.2192132 | 0.645847 | 0.862171711 |
| ENSMUSG00000050663 | Trhde | protein_coding | -0.429131644 | 2.690121764 | 0.2189347 | 0.6460451 | 0.862288918 |
| ENSMUSG00000042408 | Zmym6 | protein_coding | -0.321922269 | 2.98492359 | 0.2188678 | 0.6460952 | 0.862288918 |
| ENSMUSG00000020846 | Rflnb | protein_coding | -0.254795093 | 3.672663658 | 0.218627 | 0.6462755 | 0.862401354 |
| ENSMUSG00000004113 | Cacna1b | protein_coding | 0.20049546 | 4.851315997 | 0.218541 | 0.6463399 | 0.862401354 |
| ENSMUSG00000030753 | Thap12 | protein_coding | -0.167285481 | 4.885944996 | 0.2184002 | 0.6464453 | 0.862435084 |
| ENSMUSG00000031883 | Car7 | protein_coding | 0.238354895 | 3.743672053 | 0.2182753 | 0.6465389 | 0.862445843 |
| ENSMUSG00000019794 | Katna1 | protein_coding | -0.208695273 | 3.812878857 | 0.2181754 | 0.6466138 | 0.862445843 |
| ENSMUSG00000048756 | Foxo3 | protein_coding | 0.181548042 | 5.828438081 | 0.2180073 | 0.64675 | 0.862520487 |
| ENSMUSG00000021241 | Isca2 | protein_coding | -0.097469096 | 8.298252088 | 0.2178135 | 0.6468853 | 0.862569146 |
| ENSMUSG00000028413 | B4galt1 | protein_coding | 0.30648467 | 2.837557655 | 0.2176149 | 0.6470344 | 0.862569146 |
| ENSMUSG00000028910 | Mecr | protein_coding | -0.175464863 | 5.23956045 | 0.2175722 | 0.6470665 | 0.862569146 |
| ENSMUSG00000002102 | Psmc3 | protein_coding | -0.102259454 | 8.549169289 | 0.2175177 | 0.6471073 | 0.862569146 |
| ENSMUSG00000039765 | Cc2d2a | protein_coding | -0.204047768 | 3.858945597 | 0.2172474 | 0.6473105 | 0.86273297 |
| ENSMUSG00000028479 | Gne | protein_coding | 0.174734728 | 4.514314055 | 0.2170998 | 0.6474215 | 0.862773991 |
| ENSMUSG00000071324 | Armc2 | protein_coding | 0.376293001 | 2.970125851 | 0.2168436 | 0.6476142 | 0.862849527 |
| ENSMUSG00000029027 | Dffb | protein_coding | -0.441717399 | 2.44101323 | 0.2167497 | 0.6476849 | 0.862849527 |
| ENSMUSG00000021217 | Tshz3 | protein_coding | -0.280101368 | 2.952394091 | 0.2167045 | 0.6477189 | 0.862849527 |
| ENSMUSG00000002930 | Ppp1r17 | protein_coding | -0.37752123 | 3.752463583 | 0.217336 | 0.6478587 | 0.862918701 |
| ENSMUSG00000003526 | Prodh | protein_coding | -0.261929388 | 4.696585418 | 0.216685 | 0.6479914 | 0.862918701 |
| ENSMUSG00000067367 | Lyar | protein_coding | 0.282898688 | 3.281575278 | 0.2163159 | 0.6480116 | 0.862918701 |
| ENSMUSG00000034930 | Rtkn | protein_coding | -0.18221847 | 4.703298896 | 0.2161803 | 0.6481139 | 0.862948045 |
| ENSMUSG00000027030 | Stk39 | protein_coding | -0.145784059 | 5.963510753 | 0.215695 | 0.64848 | 0.863328625 |
| ENSMUSG00000092274 | Neat1 | lincRNA | -0.27094185 | 3.937045109 | 0.2154476 | 0.6486769 | 0.863483852 |
| ENSMUSG00000038026 | Kcnj9 | protein_coding | 0.288433159 | 4.986539953 | 0.2161767 | 0.6488755 | 0.863641329 |
| ENSMUSG00000033707 | Lrrc24 | protein_coding | 0.227985504 | 4.087936264 | 0.2150521 | 0.6489658 | 0.863654613 |
| ENSMUSG00000037851 | Iars | protein_coding | 0.19251913 | 4.834015032 | 0.2148528 | 0.6491166 | 0.863748403 |
| ENSMUSG00000047649 | Cd3eap | protein_coding | 0.133860185 | 6.083496836 | 0.2146418 | 0.6492764 | 0.863854093 |
| ENSMUSG00000056919 | Cep162 | protein_coding | 0.292789708 | 3.024399606 | 0.2142437 | 0.649578 | 0.864033281 |
| ENSMUSG00000054611 | Kdm2a | protein_coding | -0.161063654 | 4.948489974 | 0.2142372 | 0.6495829 | 0.864033281 |
| ENSMUSG00000062797 | Hikeshi | protein_coding | 0.100229126 | 7.780947549 | 0.2141056 | 0.6496827 | 0.864033281 |
| ENSMUSG00000050721 | Plekho2 | protein_coding | -0.20090135 | 4.310670604 | 0.2139633 | 0.6497907 | 0.864033281 |
| ENSMUSG00000031161 | Hdac6 | protein_coding | 0.203382767 | 4.879867729 | 0.2139341 | 0.6498128 | 0.864033281 |
| ENSMUSG00000030316 | Tamm41 | protein_coding | 0.251199067 | 3.63002157 | 0.2132814 | 0.6503085 | 0.86458553 |
| ENSMUSG00000022394 | L3mbtl2 | protein_coding | 0.244345045 | 4.542254689 | 0.2124545 | 0.6509479 | 0.865073711 |
| ENSMUSG00000055720 | Ubl7 | protein_coding | -0.115936968 | 7.95282427 | 0.2124243 | 0.6509609 | 0.865073711 |
| ENSMUSG00000020519 | Sap30l | protein_coding | -0.314117364 | 2.941717624 | 0.2123562 | 0.6510129 | 0.865073711 |
| ENSMUSG00000022911 | Arl13b | protein_coding | 0.246841727 | 3.263438864 | 0.2121821 | 0.6511456 | 0.865073711 |
| ENSMUSG00000097503 | 3110045C21Rik | antisense_RNA | 0.394665564 | 2.590778942 | 0.2121238 | 0.65119 | 0.865073711 |
| ENSMUSG00000020375 | Rufy1 | protein_coding | -0.163376593 | 4.966513053 | 0.2120095 | 0.6512773 | 0.865073711 |
| ENSMUSG00000025995 | Wdr75 | protein_coding | 0.132217864 | 5.615716546 | 0.2120092 | 0.6512775 | 0.865073711 |
| ENSMUSG00000004383 | Large1 | protein_coding | 0.147669073 | 5.054282434 | 0.2118652 | 0.6513874 | 0.865073711 |
| ENSMUSG00000027201 | Myef2 | protein_coding | -0.347957371 | 2.690352553 | 0.211849 | 0.6513998 | 0.865073711 |
| ENSMUSG00000037098 | Rab11fip3 | protein_coding | -0.190007298 | 4.664841523 | 0.2107889 | 0.6522103 | 0.866043079 |
| ENSMUSG00000098234 | Snhg6 | lincRNA | 0.235453085 | 5.926672747 | 0.2114361 | 0.6525015 | 0.866234104 |
| ENSMUSG00000005907 | Pex1 | protein_coding | 0.319850726 | 3.524648802 | 0.2102516 | 0.6526319 | 0.866234104 |
| ENSMUSG00000037570 | Mcrs1 | protein_coding | -0.159775584 | 4.744328331 | 0.2101706 | 0.6526841 | 0.866234104 |
| ENSMUSG00000022425 | Enpp2 | protein_coding | -0.222595359 | 8.708527452 | 0.2115558 | 0.6527674 | 0.866234104 |
| ENSMUSG00000006412 | Pfdn2 | protein_coding | 0.118948781 | 8.392453779 | 0.2100324 | 0.6527901 | 0.866234104 |
| ENSMUSG00000066551 | Hmgb1 | protein_coding | 0.105117929 | 6.525799609 | 0.2098934 | 0.6528968 | 0.866234104 |
| ENSMUSG00000015879 | Fam184b | protein_coding | -0.265683982 | 3.132677739 | 0.2096462 | 0.6530866 | 0.866234104 |
| ENSMUSG00000035620 | Ric8b | protein_coding | -0.150423139 | 5.915712258 | 0.2096034 | 0.6531195 | 0.866234104 |
| ENSMUSG00000057315 | Arhgap24 | protein_coding | 0.283247158 | 4.943632688 | 0.2103442 | 0.6531546 | 0.866234104 |
| ENSMUSG00000075232 | Amd1 | protein_coding | 0.185064339 | 3.991844883 | 0.2095512 | 0.6531597 | 0.866234104 |
| ENSMUSG00000024378 | Stard4 | protein_coding | -0.24024686 | 4.103663219 | 0.209362 | 0.6533051 | 0.866320109 |
| ENSMUSG00000034675 | Dbn1 | protein_coding | 0.134319084 | 6.113673448 | 0.2091861 | 0.6534404 | 0.866392665 |
| ENSMUSG00000021363 | Mak | protein_coding | 0.547878662 | 2.430673105 | 0.2088443 | 0.6537133 | 0.866643702 |
| ENSMUSG00000057637 | Prdm2 | protein_coding | -0.165120406 | 5.321888145 | 0.2087307 | 0.6537909 | 0.866643702 |

| ensembl_gene_id | external_gene_name | gene_biotype | log2 (fold change) | log2 (counts per million) | F-value | p-value | False discovery rate |
|---|---|---|---|---|---|---|---|
| ENSMUSG00000027882 | Stxbp3 | protein_coding | -0.182943665 | 4.760748714 | 0.2083654 | 0.6540725 | 0.866780711 |
| ENSMUSG00000014243 | Zswim7 | protein_coding | 0.143237786 | 5.390457261 | 0.208363 | 0.6540743 | 0.866780711 |
| ENSMUSG00000035762 | Tmem161b | protein_coding | -0.169642837 | 4.604566558 | 0.2082829 | 0.6541361 | 0.866780711 |
| ENSMUSG00000070462 | Tlnrd1 | protein_coding | -0.227283752 | 4.426052836 | 0.2079659 | 0.6543807 | 0.866998006 |
| ENSMUSG00000026360 | Rgs2 | protein_coding | 0.156538857 | 7.852127661 | 0.2079746 | 0.6545606 | 0.867026502 |
| ENSMUSG00000061589 | Dot1l | protein_coding | -0.248174585 | 3.989008239 | 0.2077293 | 0.6545635 | 0.867026502 |
| ENSMUSG00000022558 | Mroh1 | protein_coding | -0.18296887 | 4.884030128 | 0.2076136 | 0.6546528 | 0.867038049 |
| ENSMUSG00000036678 | Aaas | protein_coding | 0.170755517 | 5.155665353 | 0.2073641 | 0.6548458 | 0.86709617 |
| ENSMUSG00000028015 | Ctso | protein_coding | -0.217864654 | 4.782366102 | 0.207361 | 0.654858 | 0.86709617 |
| ENSMUSG00000033065 | Pfkm | protein_coding | 0.14237782 | 8.237464808 | 0.2070584 | 0.6550921 | 0.867266592 |
| ENSMUSG00000075273 | Ttc30b | protein_coding | -0.175468043 | 4.086097015 | 0.2068878 | 0.6552144 | 0.867266592 |
| ENSMUSG00000023075 | Akirin1 | protein_coding | 0.21257052 | 4.188818032 | 0.2068125 | 0.6552727 | 0.867266592 |
| ENSMUSG00000032239 | Rp9 | protein_coding | 0.165977378 | 4.929225882 | 0.20672 | 0.6553444 | 0.867266592 |
| ENSMUSG00000031505 | Naxd | protein_coding | 0.115900407 | 6.902994987 | 0.2066612 | 0.65539 | 0.867266592 |
| ENSMUSG00000032051 | Fdx1 | protein_coding | -0.238526656 | 3.43317196 | 0.2063051 | 0.6556661 | 0.867515375 |
| ENSMUSG00000026134 | Prim2 | protein_coding | 0.175619925 | 4.413397606 | 0.2061997 | 0.6557478 | 0.867515375 |
| ENSMUSG00000053329 | D10Jhu81e | protein_coding | 0.158918175 | 5.636115395 | 0.2061067 | 0.65582 | 0.867515375 |
| ENSMUSG00000037750 | Fam222b | protein_coding | -0.155109299 | 5.159409597 | 0.2058735 | 0.6560011 | 0.867648234 |
| ENSMUSG00000032116 | Stt3a | protein_coding | -0.134635675 | 5.841603953 | 0.2055003 | 0.6562912 | 0.867822921 |
| ENSMUSG00000097915 | A330009N23Rik | lincRNA | -0.268442915 | 4.085278012 | 0.2055085 | 0.6562946 | 0.867822921 |
| ENSMUSG00000046169 | Adamts6 | protein_coding | 0.26582607 | 3.154787281 | 0.2053826 | 0.6563827 | 0.867827064 |
| ENSMUSG00000051853 | Arf3 | protein_coding | -0.113526761 | 9.245091916 | 0.2052314 | 0.6565004 | 0.867827064 |
| ENSMUSG00000029189 | Sel1l3 | protein_coding | 0.332496758 | 2.90428747 | 0.2051807 | 0.6565399 | 0.867827064 |
| ENSMUSG00000041548 | Hspb8 | protein_coding | -0.180060331 | 4.908712143 | 0.2049451 | 0.6567233 | 0.867927109 |
| ENSMUSG00000036438 | Calm2 | protein_coding | -0.12722201 | 12.2062153 | 0.2048762 | 0.656777 | 0.867927109 |
| ENSMUSG00000049517 | Rps23 | protein_coding | 0.264621049 | 3.084731653 | 0.2046882 | 0.6569235 | 0.86801403 |
| ENSMUSG00000097412 | 1810014B01Rik | transcribed_unproces | -0.341326665 | 2.91370316 | 0.2044797 | 0.6570861 | 0.868122204 |
| ENSMUSG00000078784 | 1810022K09Rik | protein_coding | 0.181968663 | 4.566680193 | 0.2043343 | 0.6571995 | 0.868165389 |
| ENSMUSG00000001507 | Itga3 | protein_coding | 0.223000308 | 3.834342419 | 0.2039103 | 0.6575306 | 0.868379947 |
| ENSMUSG00000034656 | Cacna1a | protein_coding | -0.294532046 | 3.40050763 | 0.2038797 | 0.6575545 | 0.868379947 |
| ENSMUSG00000014075 | Tctex1d2 | protein_coding | -0.111634116 | 6.552979031 | 0.2036964 | 0.6576977 | 0.868379947 |
| ENSMUSG00000039000 | Ube3c | protein_coding | 0.146851709 | 5.459230692 | 0.2036964 | 0.6576977 | 0.868379947 |
| ENSMUSG00000022280 | Rnf19a | protein_coding | -0.195532596 | 4.079640863 | 0.2036095 | 0.6577657 | 0.868379947 |
| ENSMUSG00000028466 | Creb3 | protein_coding | -0.108895394 | 7.342910945 | 0.2034707 | 0.6578743 | 0.868416662 |
| ENSMUSG00000022860 | Chodl | protein_coding | -0.307396624 | 3.315417337 | 0.203228 | 0.6580642 | 0.868560704 |
| ENSMUSG00000046032 | Snx12 | protein_coding | 0.103306081 | 6.618575788 | 0.203043 | 0.6582091 | 0.868610062 |
| ENSMUSG00000055024 | Ep300 | protein_coding | -0.245592669 | 3.593335121 | 0.202974 | 0.6582631 | 0.868610062 |
| ENSMUSG00000035126 | Wdr78 | protein_coding | -0.390785215 | 2.817560419 | 0.2026056 | 0.6585519 | 0.868884505 |
| ENSMUSG00000091478 | Gm10039 | processed_pseudoge | 0.264141246 | 2.898952936 | 0.2022786 | 0.6588086 | 0.869116499 |
| ENSMUSG00000111977 | AC132265.1 | bidirectional_promote | 0.303719221 | 3.175579365 | 0.202147 | 0.6589119 | 0.869122314 |
| ENSMUSG00000098649 | Gm27199 | antisense_RNA | 0.3050867 | 2.751579585 | 0.2020672 | 0.6589746 | 0.869122314 |
| ENSMUSG00000038910 | Plcl2 | protein_coding | -0.186587258 | 5.087735322 | 0.2018634 | 0.6591348 | 0.869150705 |
| ENSMUSG00000070737 | Tmem35b | protein_coding | -0.276563365 | 2.970141968 | 0.2018341 | 0.6591578 | 0.869150705 |
| ENSMUSG00000005672 | Kit | protein_coding | 0.293396405 | 3.267229625 | 0.2013475 | 0.6595407 | 0.869233317 |
| ENSMUSG00000038368 | Focad | protein_coding | -0.311314339 | 2.994303433 | 0.2012228 | 0.6596389 | 0.869233317 |
| ENSMUSG00000042104 | Uggt2 | protein_coding | -0.229551085 | 4.359631651 | 0.2011229 | 0.6597176 | 0.869233317 |
| ENSMUSG00000038827 | Fam206a | protein_coding | -0.162439502 | 4.517711697 | 0.201109 | 0.6597285 | 0.869233317 |
| ENSMUSG00000009995 | Taz | protein_coding | 0.174175683 | 4.485728644 | 0.2010753 | 0.659755 | 0.869233317 |
| ENSMUSG00000023571 | C1qtnf12 | protein_coding | -0.345934036 | 3.022558791 | 0.2010634 | 0.6597644 | 0.869233317 |
| ENSMUSG00000035704 | Alg8 | protein_coding | 0.271713206 | 3.612258277 | 0.2010356 | 0.6597863 | 0.869233317 |
| ENSMUSG00000025509 | Pnpla2 | protein_coding | 0.226976792 | 5.14181246 | 0.2008554 | 0.6601114 | 0.869554995 |
| ENSMUSG00000085795 | Zfp703 | protein_coding | -0.313784545 | 2.951697461 | 0.200093 | 0.6605303 | 0.869900722 |
| ENSMUSG00000045312 | Lhfpl2 | protein_coding | -0.455500787 | 2.521567076 | 0.1997113 | 0.660832 | 0.869900722 |
| ENSMUSG00000026514 | Cnih3 | protein_coding | -0.178868943 | 4.499758929 | 0.1995114 | 0.6609903 | 0.869900722 |
| ENSMUSG00000061451 | Tmem151a | protein_coding | 0.14295377 | 5.821230767 | 0.1994173 | 0.6610648 | 0.869900722 |
| ENSMUSG00000025008 | Tctn3 | protein_coding | 0.377622887 | 3.740737521 | 0.2003603 | 0.6610819 | 0.869900722 |
| ENSMUSG00000025413 | Ttc4 | protein_coding | 0.153034006 | 5.924255249 | 0.1993951 | 0.6610824 | 0.869900722 |
| ENSMUSG00000087380 | 2210408F21Rik | processed_transcript | -0.139712786 | 4.930598638 | 0.199341 | 0.6611252 | 0.869900722 |
| ENSMUSG00000025812 | Pard3 | protein_coding | -0.1670885 | 4.74976956 | 0.1993141 | 0.6611465 | 0.869900722 |
| ENSMUSG00000066880 | Zfp617 | protein_coding | -0.180334528 | 3.954046888 | 0.199267 | 0.6611838 | 0.869900722 |
| ENSMUSG00000027012 | Dync1i2 | protein_coding | -0.104257938 | 7.194547716 | 0.1991739 | 0.6612576 | 0.869900722 |
| ENSMUSG00000055334 | Snupn | protein_coding | 0.147003149 | 5.270604123 | 0.1991662 | 0.6612637 | 0.869900722 |
| ENSMUSG00000028796 | Phc2 | protein_coding | 0.144250484 | 5.428602898 | 0.1990113 | 0.6613866 | 0.869955914 |
| ENSMUSG00000044164 | Rnf182 | protein_coding | -0.298160677 | 3.699817746 | 0.1989128 | 0.6614742 | 0.869962659 |
| ENSMUSG00000047221 | Fam185a | protein_coding | -0.295155584 | 3.551582404 | 0.1988129 | 0.6615535 | 0.869962659 |
| ENSMUSG00000049092 | Gpr137c | protein_coding | -0.330489678 | 2.982284169 | 0.1985365 | 0.6617632 | 0.870132069 |
| ENSMUSG00000037325 | Bbs7 | protein_coding | 0.176132694 | 4.537572092 | 0.1982778 | 0.6619687 | 0.870295865 |
| ENSMUSG00000062580 | Timm17a | protein_coding | 0.094806008 | 9.022382621 | 0.1979918 | 0.6621961 | 0.870401253 |
| ENSMUSG00000036598 | Ccdc113 | protein_coding | -0.17086135 | 4.42532233 | 0.1979734 | 0.6622108 | 0.870401253 |
| ENSMUSG00000025227 | Mfsd13a | protein_coding | 0.166138005 | 4.583608346 | 0.1978368 | 0.6623195 | 0.870437755 |
| ENSMUSG00000067370 | B3galt4 | protein_coding | 0.325274862 | 3.63878684 | 0.1977065 | 0.662668 | 0.870776896 |
| ENSMUSG00000021905 | Dph3 | protein_coding | -0.12934923 | 6.193024074 | 0.1973094 | 0.6627395 | 0.870776896 |
| ENSMUSG00000031148 | Gpkow | protein_coding | -0.188746083 | 5.156053859 | 0.1970175 | 0.6629722 | 0.870976234 |
| ENSMUSG00000087128 | Gm12655 | processed_transcript | 0.369896567 | 2.679446956 | 0.1966836 | 0.6632387 | 0.870994394 |
| ENSMUSG00000031109 | Enox2 | protein_coding | 0.172706401 | 4.325506399 | 0.1966741 | 0.6632463 | 0.870994394 |
| ENSMUSG00000035051 | Dhx57 | protein_coding | -0.190668439 | 4.112709344 | 0.1966643 | 0.6632542 | 0.870994394 |
| ENSMUSG00000025237 | Parp6 | protein_coding | 0.122368452 | 5.991046716 | 0.1965943 | 0.6633101 | 0.870994394 |
| ENSMUSG00000022749 | Tbc1d23 | protein_coding | -0.223369062 | 3.55848293 | 0.1964462 | 0.6634284 | 0.87102774 |

| ensembl_gene_id | external_gene_name | gene_biotype | log2 (fold change) | log2 (counts per million) | F-value | p-value | False discovery rate |
|---|---|---|---|---|---|---|---|
| ENSMUSG00000027257 | Pacsin3 | protein_coding | -0.209855472 | 4.267205859 | 0.1962771 | 0.6635636 | 0.87102774 |
| ENSMUSG00000050323 | Ndufaf6 | protein_coding | -0.339102927 | 2.951008596 | 0.1962585 | 0.6635785 | 0.87102774 |
| ENSMUSG00000001227 | Sema6b | protein_coding | 0.278432363 | 3.384376543 | 0.1958385 | 0.6639145 | 0.871206455 |
| ENSMUSG00000049265 | Kcnk3 | protein_coding | 0.169395224 | 4.463893453 | 0.1957261 | 0.6640045 | 0.871206455 |
| ENSMUSG00000079020 | Slc45a4 | protein_coding | 0.246755259 | 3.704989403 | 0.1957035 | 0.6640226 | 0.871206455 |
| ENSMUSG00000043872 | Zmym1 | protein_coding | 0.255298294 | 3.116851037 | 0.1956834 | 0.6640387 | 0.871206455 |
| ENSMUSG00000022108 | Itm2b | protein_coding | -0.101457313 | 8.615978133 | 0.1952039 | 0.6644231 | 0.87144079 |
| ENSMUSG00000022048 | Dpysl2 | protein_coding | 0.189018137 | 11.92064053 | 0.1956391 | 0.664444 | 0.87144079 |
| ENSMUSG00000024958 | Gpr137 | protein_coding | 0.142721253 | 6.39624649 | 0.1949642 | 0.6646155 | 0.87144079 |
| ENSMUSG00000025510 | Cd151 | protein_coding | 0.273249026 | 3.943163194 | 0.1949649 | 0.6646245 | 0.87144079 |
| ENSMUSG00000075701 | Selenos | protein_coding | -0.136948931 | 7.051738636 | 0.1948095 | 0.6647492 | 0.87144079 |
| ENSMUSG00000020821 | Kif1c | protein_coding | 0.285713347 | 2.868441457 | 0.1947732 | 0.6647689 | 0.87144079 |
| ENSMUSG00000086155 | 9430041J12Rik | antisense_RNA | 0.434100256 | 2.106164436 | 0.1947536 | 0.6647846 | 0.87144079 |
| ENSMUSG00000025873 | Faf2 | protein_coding | 0.136020204 | 5.906230298 | 0.1944907 | 0.6649959 | 0.871484378 |
| ENSMUSG00000029621 | Arpc1a | protein_coding | 0.09291688 | 9.187904977 | 0.1944903 | 0.6649962 | 0.871484378 |
| ENSMUSG00000054604 | Cggbp1 | protein_coding | -0.126965842 | 5.712912326 | 0.1943803 | 0.6650846 | 0.871484378 |
| ENSMUSG00000003421 | Nosip | protein_coding | 0.102101766 | 7.377059727 | 0.1942642 | 0.6651781 | 0.871484378 |
| ENSMUSG00000031772 | Cntnap4 | protein_coding | 0.254246271 | 3.277471789 | 0.1942038 | 0.6652267 | 0.871484378 |
| ENSMUSG00000097156 | Gm3764 | lincRNA | -0.165593497 | 5.037532287 | 0.1940182 | 0.665376 | 0.871484378 |
| ENSMUSG00000024222 | Fkbp5 | protein_coding | -0.207291686 | 4.536240414 | 0.1939322 | 0.6654453 | 0.871484378 |
| ENSMUSG00000004032 | Gstm5 | protein_coding | -0.124354667 | 9.655987292 | 0.1939062 | 0.6654662 | 0.871484378 |
| ENSMUSG00000036949 | Slc39a12 | protein_coding | 0.210150036 | 4.149159605 | 0.1935218 | 0.665776 | 0.87178392 |
| ENSMUSG00000101930 | Gm5441 | antisense_RNA | 0.114432419 | 6.242236178 | 0.1932158 | 0.666023 | 0.87200107 |
| ENSMUSG00000020807 | 4933427D14Rik | protein_coding | 0.218604977 | 3.610852911 | 0.193027 | 0.6661753 | 0.872043732 |
| ENSMUSG00000029623 | Pdap1 | protein_coding | 0.10390661 | 6.886533944 | 0.1929745 | 0.6662177 | 0.872043732 |
| ENSMUSG00000023845 | Lnpep | protein_coding | -0.447673566 | 2.367203165 | 0.19279 | 0.6663668 | 0.872132702 |
| ENSMUSG00000034617 | Mtrr | protein_coding | 0.381754821 | 3.45573141 | 0.1932849 | 0.666452 | 0.872137995 |
| ENSMUSG00000041949 | Tango6 | protein_coding | 0.198930272 | 3.55573917 | 0.192049 | 0.6669665 | 0.872705091 |
| ENSMUSG00000021662 | Arhgef28 | protein_coding | -0.231109998 | 4.535902081 | 0.191941 | 0.6670634 | 0.872725669 |
| ENSMUSG00000010592 | Dazl | protein_coding | -0.590173196 | 2.350461149 | 0.1918831 | 0.6672922 | 0.872858865 |
| ENSMUSG00000039716 | Dock3 | protein_coding | 0.203935051 | 3.952430827 | 0.1916036 | 0.6673275 | 0.872858865 |
| ENSMUSG00000026163 | Sphkap | protein_coding | 0.121076219 | 5.666542243 | 0.1915029 | 0.6674092 | 0.872859569 |
| ENSMUSG00000022353 | Mtss1 | protein_coding | -0.137556808 | 5.152988467 | 0.1912817 | 0.6675888 | 0.872960677 |
| ENSMUSG00000020400 | Tnip1 | protein_coding | -0.178908014 | 5.062295397 | 0.1911335 | 0.6677092 | 0.872960677 |
| ENSMUSG00000029426 | Scarb2 | protein_coding | 0.161175141 | 5.566327779 | 0.1911078 | 0.6677301 | 0.872960677 |
| ENSMUSG00000020732 | Rab37 | protein_coding | -0.451670061 | 4.52075295 | 0.1943298 | 0.6679359 | 0.873123538 |
| ENSMUSG00000029120 | Ppp2r2c | protein_coding | -0.099105552 | 7.304229255 | 0.1906635 | 0.6680913 | 0.873157629 |
| ENSMUSG00000039737 | Prkrip1 | protein_coding | 0.119590959 | 6.35533797 | 0.1906229 | 0.6681243 | 0.873157629 |
| ENSMUSG00000031827 | Cotl1 | protein_coding | 0.114979972 | 5.986020748 | 0.190323 | 0.6683685 | 0.87325946 |
| ENSMUSG00000036779 | Papd5 | protein_coding | -0.189930991 | 4.076825432 | 0.1903162 | 0.668374 | 0.87325946 |
| ENSMUSG00000050989 | Selenon | protein_coding | -0.293957099 | 3.32866713 | 0.190228 | 0.6684459 | 0.87325946 |
| ENSMUSG00000007944 | Ttc9b | protein_coding | 0.202982057 | 4.256573623 | 0.1898888 | 0.6687224 | 0.873514552 |
| ENSMUSG00000018209 | Stk4 | protein_coding | -0.166492076 | 4.73692849 | 0.189417 | 0.6691075 | 0.873911426 |
| ENSMUSG00000019437 | Tlcd1 | protein_coding | 0.204587407 | 4.300287122 | 0.1891336 | 0.6693391 | 0.874107735 |
| ENSMUSG00000058153 | Sez6l | protein_coding | -0.119844789 | 5.904705611 | 0.1888694 | 0.6695552 | 0.874283782 |
| ENSMUSG00000041632 | Mrps27 | protein_coding | 0.171704742 | 5.571094879 | 0.1887238 | 0.6696744 | 0.874333214 |
| ENSMUSG00000022681 | Ntan1 | protein_coding | 0.109439288 | 6.963301012 | 0.1883445 | 0.6699851 | 0.874592726 |
| ENSMUSG00000027655 | Dhx35 | protein_coding | 0.229042584 | 4.10466349 | 0.188233 | 0.6700764 | 0.874592726 |
| ENSMUSG00000061374 | Fiz1 | protein_coding | 0.23343781 | 3.292743738 | 0.1881834 | 0.6701171 | 0.874592726 |
| ENSMUSG00000028757 | Ddost | protein_coding | 0.173552159 | 5.732806411 | 0.1878476 | 0.670402 | 0.87485839 |
| ENSMUSG00000036323 | Srp72 | protein_coding | 0.098458442 | 7.571869453 | 0.1876614 | 0.6705456 | 0.87488137 |
| ENSMUSG00000031555 | Adam9 | protein_coding | -0.209979566 | 3.504916784 | 0.1875124 | 0.670668 | 0.87488137 |
| ENSMUSG00000066621 | Tecpr1 | protein_coding | -0.161756566 | 5.740775339 | 0.187471 | 0.670702 | 0.87488137 |
| ENSMUSG00000043019 | Edem3 | protein_coding | -0.250246474 | 3.210635039 | 0.1873394 | 0.6708102 | 0.87488137 |
| ENSMUSG00000002342 | Tmem161a | protein_coding | 0.193242002 | 4.850548156 | 0.1872155 | 0.6709121 | 0.87488137 |
| ENSMUSG00000029550 | Sppl3 | protein_coding | 0.169332576 | 4.9790143 | 0.1871313 | 0.6709814 | 0.87488137 |
| ENSMUSG00000022718 | Dgcr8 | protein_coding | -0.30785465 | 2.741823488 | 0.1871218 | 0.6709892 | 0.87488137 |
| ENSMUSG00000043153 | Ispd | protein_coding | -0.199340692 | 4.227464938 | 0.1870083 | 0.6710826 | 0.874897112 |
| ENSMUSG00000026520 | Pycr2 | protein_coding | 0.21044942 | 5.07548035 | 0.1868895 | 0.6711897 | 0.874922611 |
| ENSMUSG00000046380 | Jrk | protein_coding | 0.279188154 | 3.524425754 | 0.1866942 | 0.6713413 | 0.874922611 |
| ENSMUSG00000041939 | Mvk | protein_coding | 0.159772483 | 5.485362946 | 0.1866881 | 0.6713463 | 0.874922611 |
| ENSMUSG00000023345 | Poc1a | protein_coding | -0.211603462 | 4.073827309 | 0.1864218 | 0.6715659 | 0.874926729 |
| ENSMUSG00000079434 | Neu2 | protein_coding | -0.293644796 | 3.211015147 | 0.1864101 | 0.6715755 | 0.874926729 |
| ENSMUSG00000034486 | Gbx2 | protein_coding | -0.408181698 | 3.344052447 | 0.1870741 | 0.6715935 | 0.874926729 |
| ENSMUSG00000053925 | Gm9929 | processed_transcript | 0.296728778 | 2.965741851 | 0.1857318 | 0.6721355 | 0.875364508 |
| ENSMUSG00000053950 | Adnp2 | protein_coding | 0.243474547 | 3.565987606 | 0.1857194 | 0.6721457 | 0.875364508 |
| ENSMUSG00000037643 | Prkci | protein_coding | 0.232296953 | 4.427018228 | 0.1856117 | 0.672244 | 0.875364508 |
| ENSMUSG00000043456 | Zfp536 | protein_coding | 0.2580014 | 3.772374222 | 0.1855346 | 0.6722985 | 0.875364508 |
| ENSMUSG00000061979 | Rcc1l | protein_coding | -0.365392782 | 3.091089863 | 0.1854998 | 0.6723366 | 0.875364508 |
| ENSMUSG00000027011 | Ube2e3 | protein_coding | -0.094727181 | 6.868104404 | 0.1852129 | 0.6725647 | 0.875555523 |
| ENSMUSG00000097195 | Snhg5 | processed_transcript | -0.184956126 | 4.857032958 | 0.184806 | 0.6729018 | 0.87585848 |
| ENSMUSG00000032908 | Sgpp2 | protein_coding | 0.599595525 | 2.288440636 | 0.18502 | 0.6729604 | 0.87585848 |
| ENSMUSG00000073702 | Rpl31 | protein_coding | -0.196088814 | 3.938345888 | 0.184547 | 0.6731166 | 0.875955827 |
| ENSMUSG00000027534 | Snx16 | protein_coding | -0.291812756 | 3.755280104 | 0.1844681 | 0.6732156 | 0.875978678 |
| ENSMUSG00000021694 | Ercc8 | protein_coding | -0.221296307 | 4.122049179 | 0.1841385 | 0.6734558 | 0.876062907 |
| ENSMUSG00000078578 | Ube2d3 | protein_coding | -0.093947476 | 7.32672066 | 0.1840074 | 0.6735647 | 0.876062907 |
| ENSMUSG00000060260 | Pwwp2b | protein_coding | 0.217125326 | 4.46380528 | 0.1839498 | 0.6736126 | 0.876062907 |
| ENSMUSG00000000184 | Ccnd2 | protein_coding | 0.160193014 | 5.165602179 | 0.1838871 | 0.6736647 | 0.876062907 |

| ensembl_gene_id | external_gene_name | gene_biotype | log2 (fold change) | log2 (counts per million) | F-value | p-value | False discovery rate |
|---|---|---|---|---|---|---|---|
| ENSMUSG00000032251 | Irak1bp1 | protein_coding | -0.160870933 | 4.895445632 | 0.1838486 | 0.6736967 | 0.876062907 |
| ENSMUSG00000021384 | Susd3 | protein_coding | -0.341079107 | 2.919735398 | 0.1837615 | 0.6737692 | 0.876062907 |
| ENSMUSG00000034194 | R3hcc1 | protein_coding | 0.142135722 | 5.675439554 | 0.1829468 | 0.6744477 | 0.876775558 |
| ENSMUSG00000075590 | Nrbp2 | protein_coding | 0.220649781 | 4.648332518 | 0.1828189 | 0.6745636 | 0.876775558 |
| ENSMUSG00000024991 | Eif3a | protein_coding | -0.150350524 | 6.49308973 | 0.1828015 | 0.6745782 | 0.876775558 |
| ENSMUSG00000031683 | Lsm6 | protein_coding | -0.117083503 | 7.894789187 | 0.1826364 | 0.6747067 | 0.876775558 |
| ENSMUSG00000021103 | Mnat1 | protein_coding | -0.112544302 | 6.011421216 | 0.1826145 | 0.674725 | 0.876775558 |
| ENSMUSG00000042770 | Hebp1 | protein_coding | 0.18783724 | 5.318491374 | 0.1823407 | 0.6749629 | 0.876978694 |
| ENSMUSG00000067873 | Htatsf1 | protein_coding | -0.128652683 | 5.837514164 | 0.1821266 | 0.6751327 | 0.87699173 |
| ENSMUSG00000024909 | Efemp2 | protein_coding | -0.35837144 | 4.049244831 | 0.1831913 | 0.675136 | 0.87699173 |
| ENSMUSG00000033364 | Usp37 | protein_coding | 0.218637438 | 5.195301526 | 0.1820762 | 0.6754086 | 0.87723986 |
| ENSMUSG00000066036 | Ubr4 | protein_coding | 0.158830146 | 5.281390209 | 0.1814193 | 0.6757248 | 0.87744307 |
| ENSMUSG00000031754 | Nudt21 | protein_coding | -0.176622621 | 3.847981517 | 0.1814152 | 0.6757283 | 0.87744307 |
| ENSMUSG00000022827 | Rabl3 | protein_coding | 0.196380711 | 4.725457774 | 0.1812902 | 0.6758331 | 0.877455813 |
| ENSMUSG00000021061 | Sptb | protein_coding | -0.258121623 | 3.921841883 | 0.1810308 | 0.6760599 | 0.877455813 |
| ENSMUSG00000022889 | Mrpl39 | protein_coding | 0.126200952 | 6.207912801 | 0.1810036 | 0.6760735 | 0.877455813 |
| ENSMUSG00000025260 | Hsd17b10 | protein_coding | 0.096410813 | 7.110727815 | 0.1808568 | 0.6761968 | 0.877455813 |
| ENSMUSG00000001143 | Lman2l | protein_coding | 0.1421645 | 5.726341757 | 0.1808178 | 0.6762295 | 0.877455813 |
| ENSMUSG00000050608 | Minos1 | protein_coding | 0.117777093 | 9.318764556 | 0.18074 | 0.6762948 | 0.877455813 |
| ENSMUSG00000092607 | Scnm1 | protein_coding | -0.1037124 | 7.329508744 | 0.1806665 | 0.6763566 | 0.877455813 |
| ENSMUSG00000066438 | Plekhd1 | protein_coding | 0.372580117 | 2.44947377 | 0.1806257 | 0.6763909 | 0.877455813 |
| ENSMUSG00000044813 | Shb | protein_coding | 0.327598339 | 3.022880809 | 0.1805054 | 0.676492 | 0.87748107 |
| ENSMUSG00000028107 | Tars2 | protein_coding | 0.136354422 | 5.71337882 | 0.1804014 | 0.6765794 | 0.877488683 |
| ENSMUSG00000038848 | Ythdf1 | protein_coding | -0.173533702 | 5.274372104 | 0.180283 | 0.6766791 | 0.877512075 |
| ENSMUSG00000021595 | Nsun2 | protein_coding | -0.15198172 | 5.686086394 | 0.1795426 | 0.6773027 | 0.878165539 |
| ENSMUSG00000026127 | Imp4 | protein_coding | -0.151606663 | 4.426090176 | 0.1794425 | 0.6773872 | 0.878165539 |
| ENSMUSG00000041592 | Sdk2 | protein_coding | 0.287919639 | 3.169684741 | 0.1793941 | 0.677428 | 0.878165539 |
| ENSMUSG00000021731 | Mrps30 | protein_coding | -0.208004 | 4.137049958 | 0.1792523 | 0.6775477 | 0.878214839 |
| ENSMUSG00000093738 | AI606473 | processed_transcript | -0.276492556 | 3.792607044 | 0.1790404 | 0.6777357 | 0.878225931 |
| ENSMUSG00000028982 | Slc25a33 | protein_coding | 0.210465799 | 4.497904763 | 0.1790125 | 0.6777502 | 0.878225931 |
| ENSMUSG00000021243 | Fcf1 | protein_coding | -0.161519622 | 5.818224989 | 0.1789628 | 0.6778013 | 0.878225931 |
| ENSMUSG00000052299 | Ltn1 | protein_coding | -0.183605612 | 4.44658231 | 0.1787171 | 0.6779999 | 0.878377424 |
| ENSMUSG00000005698 | Ctcf | protein_coding | -0.126826329 | 6.059742147 | 0.178595 | 0.6781032 | 0.878405416 |
| ENSMUSG00000021051 | Ppp2r5e | protein_coding | 0.117928065 | 5.915104441 | 0.1781795 | 0.6784549 | 0.878675059 |
| ENSMUSG00000029101 | Rgs12 | protein_coding | -0.190387745 | 3.786857235 | 0.1781369 | 0.678491 | 0.878675059 |
| ENSMUSG00000071451 | Psmg4 | protein_coding | 0.14787128 | 5.430961972 | 0.1780104 | 0.6785982 | 0.878675059 |
| ENSMUSG00000034108 | Ccs | protein_coding | -0.112368597 | 6.176152604 | 0.1779084 | 0.6786847 | 0.878675059 |
| ENSMUSG00000062661 | Ncs1 | protein_coding | 0.122793744 | 10.21342779 | 0.1778669 | 0.6787199 | 0.878675059 |
| ENSMUSG00000028771 | Ptpn12 | protein_coding | -0.22738315 | 4.087179303 | 0.1774184 | 0.6791005 | 0.878923951 |
| ENSMUSG00000025917 | Cops5 | protein_coding | -0.105176941 | 7.789224821 | 0.1773974 | 0.6791184 | 0.878923951 |
| ENSMUSG00000059361 | Nrsn2 | protein_coding | 0.237860907 | 3.125861112 | 0.1773515 | 0.6791574 | 0.878923951 |
| ENSMUSG00000002279 | Lmf1 | protein_coding | 0.197765139 | 4.875749006 | 0.1771334 | 0.6793517 | 0.87896418 |
| ENSMUSG00000038542 | Pcid2 | protein_coding | 0.266530118 | 3.686813196 | 0.1771332 | 0.6793519 | 0.87896418 |
| ENSMUSG00000005045 | Chd5 | protein_coding | -0.127007083 | 6.187593633 | 0.1766448 | 0.6797585 | 0.879384371 |
| ENSMUSG00000054263 | Lifr | protein_coding | -0.227521151 | 3.791130107 | 0.1764984 | 0.6798832 | 0.879439875 |
| ENSMUSG00000050910 | Cdr2l | protein_coding | 0.173757127 | 5.033813653 | 0.176375 | 0.6799883 | 0.879470056 |
| ENSMUSG00000020741 | Cluh | protein_coding | 0.136280125 | 5.834445549 | 0.1762687 | 0.6800789 | 0.879481529 |
| ENSMUSG00000020160 | Meis1 | protein_coding | -0.360048497 | 3.539785769 | 0.1767864 | 0.6801853 | 0.879513295 |
| ENSMUSG00000030307 | Slc6a11 | protein_coding | -0.151698906 | 6.060838409 | 0.1758353 | 0.6804577 | 0.879759764 |
| ENSMUSG00000040669 | Phc1 | protein_coding | 0.205698937 | 4.314819874 | 0.1756121 | 0.6806394 | 0.879791031 |
| ENSMUSG00000031642 | Sh3rf1 | protein_coding | -0.13677723 | 4.948869836 | 0.1756049 | 0.6806455 | 0.879791031 |
| ENSMUSG00000087260 | Lamtor5 | protein_coding | -0.10873019 | 7.5296565 | 0.1751682 | 0.681019 | 0.880113967 |
| ENSMUSG00000024902 | Mrpl11 | protein_coding | 0.094614869 | 7.861113835 | 0.1751214 | 0.6810591 | 0.880113967 |
| ENSMUSG00000020516 | Rps6kb1 | protein_coding | -0.158990594 | 5.715167585 | 0.1748103 | 0.6813255 | 0.880273573 |
| ENSMUSG00000052160 | Pld4 | protein_coding | -0.294564362 | 3.550459364 | 0.1746671 | 0.6814572 | 0.880273573 |
| ENSMUSG00000028521 | Slc35d1 | protein_coding | -0.236671641 | 3.083161886 | 0.1745383 | 0.6815587 | 0.880273573 |
| ENSMUSG00000022469 | Rapgef3 | protein_coding | -0.212295379 | 3.899572658 | 0.1744717 | 0.6816158 | 0.880273573 |
| ENSMUSG00000059995 | Atxn7l3 | protein_coding | 0.13975203 | 5.42908172 | 0.1740561 | 0.6819725 | 0.880273573 |
| ENSMUSG00000041477 | Dcp1b | protein_coding | 0.187315259 | 3.722479341 | 0.1740532 | 0.681975 | 0.880273573 |
| ENSMUSG00000028672 | Hmgcl | protein_coding | 0.10501992 | 7.215860882 | 0.1740262 | 0.6819982 | 0.880273573 |
| ENSMUSG00000021852 | Slc35f4 | protein_coding | -0.212979409 | 3.69587323 | 0.1739902 | 0.6820291 | 0.880273573 |
| ENSMUSG00000029650 | Slc46a3 | protein_coding | 0.292274643 | 2.794712814 | 0.1739855 | 0.6820332 | 0.880273573 |
| ENSMUSG00000034037 | Fgd5 | protein_coding | -0.250228834 | 3.056742567 | 0.1739222 | 0.6820876 | 0.880273573 |
| ENSMUSG00000014245 | Pigl | protein_coding | 0.298006691 | 2.891857036 | 0.1738827 | 0.6821215 | 0.880273573 |
| ENSMUSG00000037936 | Scarb1 | protein_coding | -0.18520733 | 4.02383914 | 0.1738322 | 0.6821649 | 0.880273573 |
| ENSMUSG00000025911 | Adhfe1 | protein_coding | -0.198701279 | 3.611597344 | 0.1737246 | 0.6822575 | 0.880287307 |
| ENSMUSG00000051606 | 2010001K21Rik | processed_transcript | 0.502144222 | 2.361829615 | 0.1733666 | 0.6825745 | 0.880494065 |
| ENSMUSG00000050029 | Rap2c | protein_coding | -0.180074778 | 4.081772515 | 0.1732914 | 0.6826303 | 0.880494065 |
| ENSMUSG00000026209 | Dnpep | protein_coding | 0.113793323 | 6.566438219 | 0.1731995 | 0.6827095 | 0.880494065 |
| ENSMUSG00000097486 | Gm17733 | bidirectional_promote | 0.547667276 | 2.426052377 | 0.173535 | 0.6827452 | 0.880494065 |
| ENSMUSG00000019699 | Akt3 | protein_coding | -0.144433248 | 5.479204504 | 0.1729087 | 0.6829601 | 0.880665573 |
| ENSMUSG00000062296 | Trank1 | protein_coding | 0.202835046 | 4.826616183 | 0.1727905 | 0.683071 | 0.880702934 |
| ENSMUSG00000049690 | Nckap5 | protein_coding | 0.435915883 | 2.575471611 | 0.1726743 | 0.6831713 | 0.880726587 |
| ENSMUSG00000022426 | Josd1 | protein_coding | 0.167113578 | 5.076976247 | 0.1724564 | 0.6833504 | 0.880756132 |
| ENSMUSG00000024900 | Cpt1a | protein_coding | -0.430636998 | 2.519854994 | 0.1724042 | 0.6833956 | 0.880756132 |
| ENSMUSG00000028096 | Gpr89 | protein_coding | -0.150513576 | 4.975555012 | 0.1722521 | 0.683527 | 0.880756132 |
| ENSMUSG00000025224 | Gbf1 | protein_coding | 0.144689986 | 5.734216641 | 0.1721315 | 0.6836313 | 0.880756132 |
| ENSMUSG00000040374 | Pex2 | protein_coding | -0.103443768 | 6.584902452 | 0.1720698 | 0.6836846 | 0.880756132 |

| ensembl_gene_id | external_gene_name | gene_biotype | log2 (fold change) | log2 (counts per million) | F-value | p-value | False discovery rate |
|---|---|---|---|---|---|---|---|
| ENSMUSG00000020639 | Pfn4 | protein_coding | 0.194444187 | 4.264502937 | 0.1720001 | 0.6837449 | 0.880756132 |
| ENSMUSG00000027176 | Cstf3 | protein_coding | 0.112044429 | 6.278474817 | 0.171974 | 0.6837675 | 0.880756132 |
| ENSMUSG00000003813 | Rad23a | protein_coding | 0.100554295 | 8.108862216 | 0.1715805 | 0.6841082 | 0.880918718 |
| ENSMUSG00000084843 | B230312C02Rik | lincRNA | -0.253532087 | 3.027621948 | 0.1715737 | 0.6841141 | 0.880918718 |
| ENSMUSG00000097527 | 1700112J16Rik | lincRNA | 0.307203246 | 2.740116355 | 0.1715443 | 0.6841395 | 0.880918718 |
| ENSMUSG00000029636 | Wasf3 | protein_coding | -0.125792816 | 5.556638143 | 0.1713651 | 0.6842949 | 0.881013236 |
| ENSMUSG00000034981 | Parm1 | protein_coding | -0.162815229 | 6.183718437 | 0.1713377 | 0.6844251 | 0.881075455 |
| ENSMUSG00000001366 | Fbxo9 | protein_coding | 0.094230038 | 6.406147436 | 0.1708694 | 0.684725 | 0.881227167 |
| ENSMUSG00000024258 | Polr2d | protein_coding | -0.094758598 | 7.113924232 | 0.1707766 | 0.6848056 | 0.881227167 |
| ENSMUSG00000020664 | Dld | protein_coding | -0.128182946 | 7.390086463 | 0.1707361 | 0.6848496 | 0.881227167 |
| ENSMUSG00000036273 | Lrrk2 | protein_coding | -0.287272777 | 2.917905266 | 0.1707016 | 0.6848708 | 0.881227167 |
| ENSMUSG00000009535 | Rnmt | protein_coding | 0.117209801 | 6.789872447 | 0.1705296 | 0.6850202 | 0.88125437 |
| ENSMUSG00000058761 | Rnf169 | protein_coding | -0.125437757 | 5.562580829 | 0.1704887 | 0.6850558 | 0.88125437 |
| ENSMUSG00000021483 | Cdk20 | protein_coding | 0.156795186 | 4.943556972 | 0.1702541 | 0.6852599 | 0.881271257 |
| ENSMUSG00000038510 | Rpf2 | protein_coding | -0.108428557 | 6.323682725 | 0.1700097 | 0.6854727 | 0.881271257 |
| ENSMUSG00000066152 | Slc31a2 | protein_coding | 0.189197909 | 3.622977023 | 0.169971 | 0.6855064 | 0.881271257 |
| ENSMUSG00000033316 | Galnt9 | protein_coding | -0.184865482 | 4.225352478 | 0.1698993 | 0.6855689 | 0.881271257 |
| ENSMUSG00000021248 | Tmed10 | protein_coding | -0.101963392 | 6.826569887 | 0.1697891 | 0.6856649 | 0.881271257 |
| ENSMUSG00000026767 | Mindy3 | protein_coding | -0.111065468 | 6.240347076 | 0.1696036 | 0.6858267 | 0.881271257 |
| ENSMUSG00000000881 | Dlg3 | protein_coding | 0.137193076 | 5.349493532 | 0.1694371 | 0.6859719 | 0.881271257 |
| ENSMUSG00000025184 | R3hcc1l | protein_coding | 0.20791376 | 3.570688041 | 0.1693837 | 0.6860185 | 0.881271257 |
| ENSMUSG00000021357 | Exoc2 | protein_coding | -0.191197227 | 4.732963836 | 0.1693785 | 0.6860231 | 0.881271257 |
| ENSMUSG00000031393 | Mecp2 | protein_coding | -0.128006134 | 5.871023054 | 0.1693354 | 0.6860607 | 0.881271257 |
| ENSMUSG00000020190 | Mknk2 | protein_coding | 0.216659348 | 3.544430512 | 0.1692786 | 0.6861103 | 0.881271257 |
| ENSMUSG00000040725 | Hnrnpul1 | protein_coding | -0.167156303 | 5.585243096 | 0.1692733 | 0.6861238 | 0.881271257 |
| ENSMUSG00000017548 | Suz12 | protein_coding | -0.158467768 | 4.468041776 | 0.1692511 | 0.6861344 | 0.881271257 |
| ENSMUSG00000043262 | Uevld | protein_coding | -0.317368678 | 2.707248116 | 0.16914 | 0.6862314 | 0.881285156 |
| ENSMUSG00000062014 | Gmfb | protein_coding | 0.122923913 | 7.448109602 | 0.1690569 | 0.6863091 | 0.881285156 |
| ENSMUSG00000036822 | Topors | protein_coding | 0.250576935 | 4.026247427 | 0.168768 | 0.6865654 | 0.881396257 |
| ENSMUSG00000045259 | Klhdc9 | protein_coding | -0.174858157 | 4.511997835 | 0.1687341 | 0.6865862 | 0.881396257 |
| ENSMUSG00000100417 | Gm28578 | lincRNA | -0.398758483 | 2.375551385 | 0.1686375 | 0.6866708 | 0.881396257 |
| ENSMUSG00000029576 | Radil | protein_coding | -0.241285749 | 3.178132561 | 0.1683615 | 0.6869125 | 0.881396257 |
| ENSMUSG00000022420 | Dnal4 | protein_coding | -0.120588226 | 6.838531098 | 0.1682691 | 0.6869934 | 0.881396257 |
| ENSMUSG00000003934 | Efnb3 | protein_coding | -0.104193091 | 7.187374307 | 0.168257 | 0.687004 | 0.881396257 |
| ENSMUSG00000044600 | Smim7 | protein_coding | -0.086407651 | 8.103929156 | 0.1681539 | 0.6870944 | 0.881396257 |
| ENSMUSG00000030218 | Mgp | protein_coding | 0.420074076 | 2.11718742 | 0.1681093 | 0.6871335 | 0.881396257 |
| ENSMUSG00000040006 | Ginm1 | protein_coding | -0.147771963 | 5.331043321 | 0.1680539 | 0.6871821 | 0.881396257 |
| ENSMUSG00000061411 | Nol4l | protein_coding | 0.16575597 | 4.668333996 | 0.167969 | 0.6872565 | 0.881396257 |
| ENSMUSG00000045211 | Nudt18 | protein_coding | 0.136508546 | 5.721261591 | 0.1678664 | 0.6873466 | 0.881396257 |
| ENSMUSG00000076617 | Ighm | IG_C_gene | 0.215930423 | 5.804644818 | 0.1686194 | 0.6873792 | 0.881396257 |
| ENSMUSG00000031760 | Mt3 | protein_coding | 0.124248245 | 10.72742009 | 0.1674115 | 0.6877461 | 0.881607558 |
| ENSMUSG00000031987 | Egln1 | protein_coding | -0.134193553 | 5.548782701 | 0.1673111 | 0.6878344 | 0.881607558 |
| ENSMUSG00000031295 | Phka2 | protein_coding | -0.300351486 | 3.009510972 | 0.1673022 | 0.6878422 | 0.881607558 |
| ENSMUSG00000007646 | Rad51c | protein_coding | -0.281708521 | 3.204790489 | 0.1672683 | 0.687872 | 0.881607558 |
| ENSMUSG00000037280 | Galnt6 | protein_coding | 0.203544785 | 3.473773619 | 0.1671397 | 0.6879852 | 0.881647533 |
| ENSMUSG00000045917 | Tmem268 | protein_coding | 0.18502614 | 4.102658642 | 0.1670231 | 0.6880878 | 0.881673969 |
| ENSMUSG00000031858 | Mau2 | protein_coding | -0.160489015 | 5.019563387 | 0.1666233 | 0.6884399 | 0.881887674 |
| ENSMUSG00000075467 | Dnlz | protein_coding | 0.200971689 | 4.734856396 | 0.1665797 | 0.688487 | 0.881887674 |
| ENSMUSG00000008200 | Fnbp4 | protein_coding | -0.18085782 | 3.868926687 | 0.1665544 | 0.6885006 | 0.881887674 |
| ENSMUSG00000018449 | Rpain | protein_coding | -0.124028708 | 5.882815107 | 0.1664547 | 0.6885885 | 0.881895229 |
| ENSMUSG00000034902 | Pip5k1c | protein_coding | 0.117362849 | 6.460003544 | 0.1662144 | 0.6888005 | 0.882025483 |
| ENSMUSG00000054675 | Tmem119 | protein_coding | 0.288138509 | 3.114579512 | 0.1661536 | 0.6888543 | 0.882025483 |
| ENSMUSG00000055204 | Ankrd17 | protein_coding | 0.109052709 | 6.665545523 | 0.165964 | 0.6890217 | 0.882134763 |
| ENSMUSG00000064302 | Clasp1 | protein_coding | 0.148145813 | 5.440027815 | 0.1652774 | 0.689629 | 0.882807213 |
| ENSMUSG00000038990 | Cables2 | protein_coding | 0.220979811 | 3.377384987 | 0.1650736 | 0.6898096 | 0.882888725 |
| ENSMUSG00000005481 | Ddx39 | protein_coding | 0.157478151 | 5.050695147 | 0.1650202 | 0.6898569 | 0.882888725 |
| ENSMUSG00000040048 | Ndufb10 | protein_coding | 0.100240337 | 10.18834066 | 0.1649049 | 0.6899592 | 0.882914535 |
| ENSMUSG00000029641 | Rasl11a | protein_coding | 0.276662506 | 3.948291261 | 0.1650641 | 0.690109 | 0.88300126 |
| ENSMUSG00000064105 | Cnnm2 | protein_coding | -0.254535068 | 3.6964863 | 0.1644766 | 0.6903479 | 0.883066189 |
| ENSMUSG00000020171 | Yeats4 | protein_coding | -0.184128819 | 4.308757763 | 0.1643655 | 0.6904379 | 0.883066189 |
| ENSMUSG00000057561 | Eif1a | protein_coding | 0.103362363 | 7.456750366 | 0.1643078 | 0.6904892 | 0.883066189 |
| ENSMUSG00000054808 | Actn4 | protein_coding | -0.177842714 | 4.526153361 | 0.16428 | 0.6905139 | 0.883066189 |
| ENSMUSG00000039936 | Pik3cd | protein_coding | 0.336960779 | 2.828825237 | 0.1642164 | 0.6905704 | 0.883066189 |
| ENSMUSG00000031527 | Eri1 | protein_coding | -0.352656763 | 2.766883389 | 0.1639719 | 0.6907879 | 0.883239286 |
| ENSMUSG00000074892 | B3galt5 | protein_coding | 0.14860886 | 4.952348581 | 0.1637952 | 0.6909452 | 0.883335337 |
| ENSMUSG00000037262 | Kin | protein_coding | -0.158358883 | 4.991138795 | 0.1636546 | 0.6910704 | 0.883390358 |
| ENSMUSG00000056429 | Tgoln1 | protein_coding | -0.095264118 | 6.553580373 | 0.1635474 | 0.6911659 | 0.88340747 |
| ENSMUSG00000089687 | Rab42 | protein_coding | -0.286867521 | 2.882036493 | 0.1633108 | 0.6913768 | 0.883572015 |
| ENSMUSG00000034575 | Papd7 | protein_coding | -0.151312598 | 4.694228269 | 0.1632133 | 0.6914637 | 0.883578081 |
| ENSMUSG00000046841 | Ckap4 | protein_coding | -0.167169251 | 4.205987618 | 0.1630491 | 0.6916103 | 0.883587159 |
| ENSMUSG00000032235 | Ice2 | protein_coding | -0.278151547 | 2.950215542 | 0.1628312 | 0.6918049 | 0.883587159 |
| ENSMUSG00000034544 | Rsrc1 | protein_coding | -0.112453232 | 6.519231588 | 0.1627869 | 0.6918445 | 0.883587159 |
| ENSMUSG00000015776 | Med22 | protein_coding | 0.100736667 | 6.323748204 | 0.1627514 | 0.6918763 | 0.883587159 |
| ENSMUSG00000025790 | Slco3a1 | protein_coding | 0.172831251 | 4.733153981 | 0.1627453 | 0.6918817 | 0.883587159 |
| ENSMUSG00000045790 | Ccdc149 | protein_coding | -0.161100487 | 4.636312051 | 0.162464 | 0.6921332 | 0.883609704 |
| ENSMUSG00000032531 | Amotl2 | protein_coding | 0.305468206 | 3.205997303 | 0.1624212 | 0.6921801 | 0.883609704 |
| ENSMUSG00000080727 | C920021L13Rik | antisense_RNA | -0.296659484 | 2.533672648 | 0.1623777 | 0.6922104 | 0.883609704 |
| ENSMUSG00000047434 | Xxylt1 | protein_coding | 0.201481309 | 3.392330409 | 0.162358 | 0.692228 | 0.883609704 |

| ensembl_gene_id | external_gene_name | gene_biotype | log2 (fold change) | log2 (counts per million) | F-value | p-value | False discovery rate |
|---|---|---|---|---|---|---|---|
| ENSMUSG00000026946 | Nmi | protein_coding | -0.230203819 | 3.25338944 | 0.1622533 | 0.6923218 | 0.883624439 |
| ENSMUSG00000032549 | Rab6b | protein_coding | 0.092058808 | 8.76489705 | 0.1619956 | 0.6925525 | 0.883738582 |
| ENSMUSG00000079557 | 2-Mar | protein_coding | -0.138680402 | 5.597889616 | 0.1619699 | 0.6925756 | 0.883738582 |
| ENSMUSG00000033126 | Ybey | protein_coding | 0.194305712 | 4.195006508 | 0.1614803 | 0.6930148 | 0.884194099 |
| ENSMUSG00000029651 | Mtus2 | protein_coding | 0.118608949 | 5.4084515 | 0.161338 | 0.6931425 | 0.884242251 |
| ENSMUSG00000040738 | Ints8 | protein_coding | -0.21236686 | 3.724447073 | 0.1612047 | 0.6932623 | 0.884242251 |
| ENSMUSG00000050846 | Zfp623 | protein_coding | 0.163498686 | 4.333014828 | 0.1610969 | 0.6933592 | 0.884242251 |
| ENSMUSG00000002329 | Mdp1 | protein_coding | 0.087403068 | 7.037462489 | 0.1609243 | 0.6935144 | 0.884242251 |
| ENSMUSG00000062937 | Mtap | protein_coding | 0.149614198 | 5.135717786 | 0.1608789 | 0.6935553 | 0.884242251 |
| ENSMUSG00000027433 | Xrn2 | protein_coding | 0.118250507 | 5.935419471 | 0.1608415 | 0.6935889 | 0.884242251 |
| ENSMUSG00000016018 | Skiv2l2 | protein_coding | 0.138570767 | 5.557120922 | 0.1607979 | 0.6936281 | 0.884242251 |
| ENSMUSG00000047832 | Cdca4 | protein_coding | 0.247281024 | 3.904284938 | 0.1603532 | 0.6940372 | 0.884658862 |
| ENSMUSG00000028668 | Eloa | protein_coding | -0.244862334 | 3.051228497 | 0.160103 | 0.6942543 | 0.884830693 |
| ENSMUSG00000032470 | Mras | protein_coding | -0.100542965 | 6.058891247 | 0.1598762 | 0.6944591 | 0.88496593 |
| ENSMUSG00000031562 | Dctd | protein_coding | -0.206925954 | 4.074161882 | 0.1597958 | 0.6945317 | 0.88496593 |
| ENSMUSG00000007867 | Ift43 | protein_coding | 0.122656304 | 5.788973458 | 0.1597121 | 0.6946073 | 0.88496593 |
| ENSMUSG00000085408 | C530005A16Rik | processed_transcript | -0.235690243 | 3.387807311 | 0.1594319 | 0.6948606 | 0.885031737 |
| ENSMUSG00000049680 | Urgcp | protein_coding | 0.156325036 | 4.460458606 | 0.1594243 | 0.6948675 | 0.885031737 |
| ENSMUSG00000032870 | Smap2 | protein_coding | 0.083168362 | 7.112393624 | 0.1593058 | 0.6949747 | 0.885031737 |
| ENSMUSG00000044018 | Mrpl50 | protein_coding | -0.100644264 | 7.294997489 | 0.15919 | 0.6950795 | 0.885031737 |
| ENSMUSG00000025854 | Fam20c | protein_coding | 0.212355583 | 4.759858499 | 0.1590256 | 0.695269 | 0.885031737 |
| ENSMUSG00000038250 | Usp38 | protein_coding | -0.278218792 | 4.033244723 | 0.1593162 | 0.6952949 | 0.885031737 |
| ENSMUSG00000033554 | Dph5 | protein_coding | -0.121837737 | 5.479971465 | 0.1589077 | 0.6953352 | 0.885031737 |
| ENSMUSG00000035933 | Cog5 | protein_coding | 0.245966255 | 3.09656944 | 0.1589049 | 0.6953377 | 0.885031737 |
| ENSMUSG00000043923 | Ccdc84 | protein_coding | -0.422054052 | 2.271907717 | 0.1588365 | 0.6953997 | 0.885031737 |
| ENSMUSG00000056458 | Mok | protein_coding | 0.194611622 | 4.404013579 | 0.1586126 | 0.6956028 | 0.88518538 |
| ENSMUSG00000015165 | Hnrnpl | protein_coding | -0.105376588 | 6.560718103 | 0.1584481 | 0.6957521 | 0.885270643 |
| ENSMUSG00000001847 | Rac1 | protein_coding | -0.099187277 | 6.536388108 | 0.158089 | 0.6960782 | 0.88553695 |
| ENSMUSG00000070372 | Capza1 | protein_coding | 0.187894932 | 3.747598936 | 0.1580264 | 0.6961352 | 0.88553695 |
| ENSMUSG00000015671 | Psma2 | protein_coding | 0.102806095 | 8.831763041 | 0.1579458 | 0.6962084 | 0.88553695 |
| ENSMUSG00000045106 | Ccdc73 | protein_coding | -0.191023876 | 3.773213786 | 0.1575763 | 0.6965447 | 0.885587748 |
| ENSMUSG00000046814 | Gchfr | protein_coding | 0.212051775 | 3.452972226 | 0.1574674 | 0.6966439 | 0.885587748 |
| ENSMUSG00000037270 | 4932438A13Rik | protein_coding | -0.186549454 | 4.134086467 | 0.1573001 | 0.6967964 | 0.885587748 |
| ENSMUSG00000015087 | Rabl6 | protein_coding | -0.09909632 | 6.476375965 | 0.1572921 | 0.6968037 | 0.885587748 |
| ENSMUSG00000055723 | Rras2 | protein_coding | -0.440304648 | 2.084447914 | 0.1571013 | 0.6969777 | 0.885587748 |
| ENSMUSG00000034269 | Setd5 | protein_coding | 0.156244704 | 4.314935118 | 0.1570639 | 0.6970118 | 0.885587748 |
| ENSMUSG00000031608 | Galnt7 | protein_coding | -0.252694644 | 3.425388949 | 0.1570473 | 0.697027 | 0.885587748 |
| ENSMUSG00000048832 | Vps37c | protein_coding | 0.182295294 | 4.872612485 | 0.1570019 | 0.6970684 | 0.885587748 |
| ENSMUSG00000040560 | Wdr7 | protein_coding | -0.11011252 | 6.03494408 | 0.1569546 | 0.6971115 | 0.885587748 |
| ENSMUSG00000037419 | Endod1 | protein_coding | -0.108005004 | 6.359519041 | 0.1568319 | 0.6972236 | 0.885587748 |
| ENSMUSG00000029185 | Fam114a1 | protein_coding | -0.485337443 | 2.987001015 | 0.1576196 | 0.6972649 | 0.885587748 |
| ENSMUSG00000055692 | Tmem191c | protein_coding | 0.131958246 | 5.673885743 | 0.1565565 | 0.6974752 | 0.885587748 |
| ENSMUSG00000026811 | St6galnac6 | protein_coding | 0.113793189 | 5.915697612 | 0.1565091 | 0.6975185 | 0.885587748 |
| ENSMUSG00000019777 | Hdac2 | protein_coding | -0.159188751 | 5.824625068 | 0.1565001 | 0.6975352 | 0.885587748 |
| ENSMUSG00000032816 | Igdcc4 | protein_coding | 0.249475904 | 3.276347996 | 0.1564596 | 0.6975638 | 0.885587748 |
| ENSMUSG00000022370 | Mrpl13 | protein_coding | 0.096279023 | 7.425329315 | 0.1562214 | 0.6977817 | 0.885587748 |
| ENSMUSG00000021494 | Ddx41 | protein_coding | -0.129726253 | 5.333514977 | 0.1561711 | 0.6978277 | 0.885587748 |
| ENSMUSG00000034748 | Sirt6 | protein_coding | -0.192262023 | 4.765305217 | 0.1559207 | 0.6980655 | 0.885587748 |
| ENSMUSG00000079614 | Seh1l | protein_coding | 0.112807156 | 6.686796075 | 0.1556366 | 0.6983175 | 0.885587748 |
| ENSMUSG00000079179 | Rab10os | antisense_RNA | 0.13470902 | 6.313609855 | 0.1556147 | 0.6983461 | 0.885587748 |
| ENSMUSG00000112343 | AC154734.1 | processed_transcript | -0.408817216 | 2.490998379 | 0.1555666 | 0.6983817 | 0.885587748 |
| ENSMUSG00000027288 | Zfp106 | protein_coding | -0.121880312 | 5.455991824 | 0.1555024 | 0.6984407 | 0.885587748 |
| ENSMUSG00000044934 | Zfp367 | protein_coding | -0.159482482 | 4.7212702 | 0.1554749 | 0.6984659 | 0.885587748 |
| ENSMUSG00000047731 | Wbp1l | protein_coding | -0.209997481 | 4.181229716 | 0.1554573 | 0.6984821 | 0.885587748 |
| ENSMUSG00000015354 | Pcolce2 | protein_coding | -0.297239508 | 2.687559785 | 0.1554169 | 0.6985191 | 0.885587748 |
| ENSMUSG00000040490 | Lrfn2 | protein_coding | -0.241334906 | 2.951923989 | 0.1553467 | 0.6985836 | 0.885587748 |
| ENSMUSG00000051359 | Ncald | protein_coding | 0.108813081 | 8.885252362 | 0.1553309 | 0.6985981 | 0.885587748 |
| ENSMUSG00000024622 | Hmgxb3 | protein_coding | 0.242808855 | 3.1985186 | 0.1552365 | 0.6986848 | 0.885587748 |
| ENSMUSG00000038244 | Mical2 | protein_coding | 0.240765374 | 3.840897518 | 0.1551975 | 0.698729 | 0.885587748 |
| ENSMUSG00000052033 | Pfdn4 | protein_coding | -0.121821961 | 7.429777655 | 0.1551765 | 0.6987484 | 0.885587748 |
| ENSMUSG00000042688 | Mapk6 | protein_coding | -0.169461418 | 5.407855678 | 0.1551187 | 0.6988015 | 0.885587748 |
| ENSMUSG00000086401 | Gm15559 | antisense_RNA | 0.32494343 | 3.268404032 | 0.1548418 | 0.6990926 | 0.885711858 |
| ENSMUSG00000036879 | Phkb | protein_coding | 0.152176621 | 4.607449824 | 0.154711 | 0.699168 | 0.885711858 |
| ENSMUSG00000043510 | Hscb | protein_coding | 0.159104947 | 4.868453037 | 0.1545809 | 0.6992878 | 0.885711858 |
| ENSMUSG00000014856 | Tmem208 | protein_coding | 0.104703977 | 6.816027486 | 0.1544308 | 0.6994261 | 0.885711858 |
| ENSMUSG00000029776 | Hibadh | protein_coding | -0.11846637 | 6.256325144 | 0.1544185 | 0.6994375 | 0.885711858 |
| ENSMUSG00000022763 | Aifm3 | protein_coding | 0.144551903 | 5.137190422 | 0.1543718 | 0.6994805 | 0.885711858 |
| ENSMUSG00000022673 | Mcm4 | protein_coding | -0.221683739 | 3.356208428 | 0.1542484 | 0.6995943 | 0.885711858 |
| ENSMUSG00000025089 | Gfra1 | protein_coding | 0.118961674 | 5.701463054 | 0.1541625 | 0.6996735 | 0.885711858 |
| ENSMUSG00000035024 | Ncapd3 | protein_coding | -0.206148759 | 3.666744051 | 0.1541264 | 0.6997068 | 0.885711858 |
| ENSMUSG00000000915 | Hip1r | protein_coding | -0.186587938 | 4.0628248 | 0.1539569 | 0.6998632 | 0.885711858 |
| ENSMUSG00000026027 | Stradb | protein_coding | -0.159291749 | 4.655099218 | 0.1538881 | 0.6999267 | 0.885711858 |
| ENSMUSG00000026608 | Kctd3 | protein_coding | 0.172163385 | 4.791770273 | 0.1538524 | 0.6999596 | 0.885711858 |
| ENSMUSG00000039523 | Cep104 | protein_coding | 0.199766067 | 4.015936478 | 0.153841 | 0.6999702 | 0.885711858 |
| ENSMUSG00000029397 | Rchy1 | protein_coding | -0.103311402 | 6.608216687 | 0.1537507 | 0.7000535 | 0.885713107 |
| ENSMUSG00000060038 | Dhps | protein_coding | 0.152671817 | 5.142539167 | 0.1535256 | 0.7002616 | 0.885732358 |
| ENSMUSG00000086877 | A230072C01Rik | processed_transcript | -0.147038603 | 4.659431682 | 0.1534819 | 0.700302 | 0.885732358 |
| ENSMUSG00000027998 | Plrg1 | protein_coding | 0.123942614 | 6.257843665 | 0.1534669 | 0.7003159 | 0.885732358 |

| ensembl_gene_id | external_gene_name | gene_biotype | log2 (fold change) | log2 (counts per million) | F-value | p-value | False discovery rate |
|---|---|---|---|---|---|---|---|
| ENSMUSG00000018541 | Cwc25 | protein_coding | 0.143454627 | 4.840670748 | 0.1533257 | 0.7004465 | 0.885793441 |
| ENSMUSG00000017307 | Acot8 | protein_coding | 0.15358204 | 5.065814765 | 0.153189 | 0.7005731 | 0.885849283 |
| ENSMUSG00000035443 | Thyn1 | protein_coding | -0.087796735 | 7.727956212 | 0.1530772 | 0.7006765 | 0.885875899 |
| ENSMUSG00000020697 | Lig3 | protein_coding | 0.298929579 | 2.791081646 | 0.1529513 | 0.7007932 | 0.885919212 |
| ENSMUSG00000059796 | Eif4a1 | protein_coding | 0.096874198 | 6.153011089 | 0.1528026 | 0.700931 | 0.885989328 |
| ENSMUSG00000005687 | Bcas2 | protein_coding | 0.095243595 | 8.263035608 | 0.1526118 | 0.7011079 | 0.886102835 |
| ENSMUSG00000029192 | Tbc1d14 | protein_coding | 0.133781699 | 5.501214275 | 0.1525003 | 0.7012115 | 0.886102835 |
| ENSMUSG00000026766 | Mmadhc | protein_coding | 0.102321175 | 6.389745668 | 0.1524393 | 0.701268 | 0.886102835 |
| ENSMUSG00000036561 | Ppp6r2 | protein_coding | -0.178083879 | 4.045764098 | 0.1521809 | 0.7015081 | 0.886302037 |
| ENSMUSG00000021079 | Timm9 | protein_coding | 0.110924839 | 7.126191626 | 0.1519418 | 0.7017304 | 0.886478784 |
| ENSMUSG00000033961 | Zfp446 | protein_coding | -0.213581342 | 3.355200966 | 0.1516751 | 0.7019786 | 0.886581798 |
| ENSMUSG00000032582 | Rbm6 | protein_coding | -0.155382048 | 4.699735557 | 0.1516154 | 0.7020342 | 0.886581798 |
| ENSMUSG00000027296 | Itpka | protein_coding | -0.209110958 | 5.082096118 | 0.1517809 | 0.7021814 | 0.886581798 |
| ENSMUSG00000070699 | Sars2 | protein_coding | -0.198023413 | 4.124511531 | 0.1511612 | 0.7024575 | 0.886581798 |
| ENSMUSG00000035476 | Tab3 | protein_coding | -0.287798248 | 3.025920837 | 0.151072 | 0.7025408 | 0.886581798 |
| ENSMUSG00000085139 | A730046J19Rik | protein_coding | 0.386710292 | 2.279239719 | 0.1509672 | 0.7026386 | 0.886581798 |
| ENSMUSG00000061536 | Sec22c | protein_coding | -0.137555467 | 4.617431643 | 0.1509614 | 0.702644 | 0.886581798 |
| ENSMUSG00000015341 | Golga7 | protein_coding | -0.108829975 | 5.737273515 | 0.1509613 | 0.7026441 | 0.886581798 |
| ENSMUSG00000004931 | Apba3 | protein_coding | -0.286199471 | 2.61079719 | 0.150945 | 0.7026593 | 0.886581798 |
| ENSMUSG00000021884 | Hacl1 | protein_coding | 0.208547085 | 3.644691839 | 0.1509353 | 0.7026684 | 0.886581798 |
| ENSMUSG00000039753 | Fbxl5 | protein_coding | -0.09314808 | 6.427888956 | 0.1508228 | 0.7027735 | 0.886581798 |
| ENSMUSG00000041598 | Cdc42ep4 | protein_coding | 0.160173896 | 4.979978276 | 0.1506204 | 0.7029626 | 0.886581798 |
| ENSMUSG00000002910 | Arrdc2 | protein_coding | 0.297585756 | 2.682520466 | 0.1505995 | 0.7029821 | 0.886581798 |
| ENSMUSG00000054945 | Gm9958 | protein_coding | -0.556865 | 1.942477758 | 0.1505504 | 0.7030281 | 0.886581798 |
| ENSMUSG00000028437 | Ubap1 | protein_coding | 0.127254214 | 5.993346689 | 0.1504977 | 0.7030774 | 0.886581798 |
| ENSMUSG00000017314 | Mpp2 | protein_coding | 0.110388265 | 6.632723407 | 0.1504402 | 0.7031312 | 0.886581798 |
| ENSMUSG00000033819 | Ppp1r16a | protein_coding | 0.141061541 | 4.734216239 | 0.1502814 | 0.7032797 | 0.886665142 |
| ENSMUSG00000009739 | Pou6f1 | protein_coding | -0.145284194 | 4.692505443 | 0.1501554 | 0.7033977 | 0.886709908 |
| ENSMUSG00000079242 | C730034F03Rik | protein_coding | -0.290044275 | 3.111918437 | 0.1499638 | 0.7035772 | 0.886812517 |
| ENSMUSG00000014074 | Rnf168 | protein_coding | 0.155245883 | 4.749605064 | 0.1498926 | 0.703644 | 0.886812517 |
| ENSMUSG00000034032 | Rpap1 | protein_coding | -0.212766191 | 3.352133866 | 0.1494236 | 0.7040841 | 0.886983545 |
| ENSMUSG00000024640 | Psat1 | protein_coding | -0.08162798 | 7.37906699 | 0.1493729 | 0.7041317 | 0.886983545 |
| ENSMUSG00000041168 | Lonp1 | protein_coding | 0.125854784 | 4.990670459 | 0.1493605 | 0.7041434 | 0.886983545 |
| ENSMUSG00000070394 | Tmem256 | protein_coding | -0.123351846 | 8.328333018 | 0.149352 | 0.7041596 | 0.886983545 |
| ENSMUSG00000060212 | Pcnx2 | protein_coding | -0.209253216 | 3.502275679 | 0.1492264 | 0.7042694 | 0.886983545 |
| ENSMUSG00000044221 | Grsf1 | protein_coding | 0.099720429 | 6.971257597 | 0.1491974 | 0.7042967 | 0.886983545 |
| ENSMUSG00000043621 | Ubxn10 | protein_coding | 0.245165856 | 3.129834677 | 0.1490997 | 0.7043885 | 0.886983545 |
| ENSMUSG00000064215 | Ifi27 | protein_coding | 0.101045426 | 6.399140843 | 0.1490228 | 0.7044608 | 0.886983545 |
| ENSMUSG00000004562 | Arhgef40 | protein_coding | -0.180159076 | 5.044574257 | 0.148951 | 0.7045366 | 0.886983545 |
| ENSMUSG00000087651 | 1500009L16Rik | protein_coding | 0.097583116 | 6.322882621 | 0.14887 | 0.7046046 | 0.886983545 |
| ENSMUSG00000104861 | 3110039M20Rik | lincRNA | 0.23981655 | 3.578127539 | 0.1487209 | 0.704745 | 0.88705641 |
| ENSMUSG00000030638 | Sh3gl3 | protein_coding | -0.15507496 | 5.200236941 | 0.1485108 | 0.704943 | 0.887096165 |
| ENSMUSG00000032518 | Rpsa | protein_coding | 0.179220098 | 3.711441235 | 0.1484478 | 0.7050023 | 0.887096165 |
| ENSMUSG00000031021 | Tmem9b | protein_coding | -0.111362271 | 5.948648697 | 0.1484247 | 0.7050241 | 0.887096165 |
| ENSMUSG00000026213 | Stk11ip | protein_coding | 0.24110115 | 3.316187816 | 0.1481467 | 0.7052864 | 0.887322422 |
| ENSMUSG00000026841 | Fibcd1 | protein_coding | -0.202535796 | 3.830676593 | 0.1480258 | 0.7054005 | 0.887362134 |
| ENSMUSG00000026663 | Atf6 | protein_coding | -0.158016406 | 4.068017822 | 0.1478962 | 0.7055229 | 0.887401253 |
| ENSMUSG00000030584 | Dpf1 | protein_coding | -0.152833684 | 4.907281229 | 0.1478182 | 0.7055966 | 0.887401253 |
| ENSMUSG00000097048 | 1600020E01Rik | antisense_RNA | -0.201399626 | 4.654607397 | 0.1476561 | 0.705758 | 0.887500414 |
| ENSMUSG00000070000 | Fcho1 | protein_coding | 0.199584882 | 3.832681954 | 0.1474077 | 0.7059849 | 0.887681957 |
| ENSMUSG00000025875 | Tspan17 | protein_coding | 0.255003672 | 3.463116363 | 0.1472311 | 0.7061521 | 0.887724462 |
| ENSMUSG00000097638 | Carlr | lincRNA | 0.444250416 | 2.306244157 | 0.1471247 | 0.706253 | 0.887724462 |
| ENSMUSG00000031402 | Mpp1 | protein_coding | 0.122999068 | 6.297692361 | 0.1471105 | 0.7062664 | 0.887724462 |
| ENSMUSG00000022043 | Trim35 | protein_coding | -0.092834987 | 6.707283872 | 0.1467533 | 0.7066052 | 0.888046446 |
| ENSMUSG00000002900 | Lamb1 | protein_coding | 0.235821567 | 3.348845359 | 0.1465485 | 0.7067996 | 0.88818705 |
| ENSMUSG00000058656 | Samd12 | protein_coding | -0.192838056 | 4.076291208 | 0.146442 | 0.7069008 | 0.888199591 |
| ENSMUSG00000037197 | Rbm17 | protein_coding | 0.13379594 | 4.756584771 | 0.1463641 | 0.7069748 | 0.888199591 |
| ENSMUSG00000024878 | Cbwd1 | protein_coding | 0.13347959 | 4.8534708 | 0.1460881 | 0.7072373 | 0.888425574 |
| ENSMUSG00000079197 | Psme2 | protein_coding | 0.097518454 | 6.052629701 | 0.1458506 | 0.7074635 | 0.888605835 |
| ENSMUSG00000067242 | Lgi1 | protein_coding | -0.156579128 | 4.060192259 | 0.1455805 | 0.7077208 | 0.888825295 |
| ENSMUSG00000021536 | Adcy2 | protein_coding | -0.142638936 | 5.192257329 | 0.1453752 | 0.7079167 | 0.888958932 |
| ENSMUSG00000037148 | Arhgap10 | protein_coding | -0.363452324 | 2.855312343 | 0.1453041 | 0.7079926 | 0.888958932 |
| ENSMUSG00000025962 | Fastkd2 | protein_coding | 0.142136418 | 5.107570736 | 0.1451995 | 0.7080843 | 0.888970325 |
| ENSMUSG00000048755 | Mcat | protein_coding | 0.285446357 | 3.814426052 | 0.1454466 | 0.7081818 | 0.888988921 |
| ENSMUSG00000061458 | Nol10 | protein_coding | 0.167085545 | 4.249977819 | 0.1448111 | 0.7084555 | 0.889228692 |
| ENSMUSG00000038205 | Prkab2 | protein_coding | -0.336313425 | 2.740696418 | 0.1446339 | 0.708625 | 0.889337641 |
| ENSMUSG00000047671 | Tctex1d4 | protein_coding | -0.242535922 | 3.345862456 | 0.1444057 | 0.7088435 | 0.889508038 |
| ENSMUSG00000020032 | Nuak1 | protein_coding | 0.151909257 | 4.065982924 | 0.1441328 | 0.7091051 | 0.889567128 |
| ENSMUSG00000027985 | Lef1 | protein_coding | -0.478947423 | 3.205573211 | 0.1453047 | 0.7092554 | 0.889567128 |
| ENSMUSG00000030678 | Maz | protein_coding | 0.169363279 | 4.644659329 | 0.1439568 | 0.7092739 | 0.889567128 |
| ENSMUSG00000024217 | Snrpc | protein_coding | 0.144403813 | 5.049865964 | 0.1439446 | 0.7092856 | 0.889567128 |
| ENSMUSG00000022404 | Slc25a17 | protein_coding | -0.093757605 | 7.268080084 | 0.1439252 | 0.7093042 | 0.889567128 |
| ENSMUSG00000033871 | Ppargc1b | protein_coding | -0.212792767 | 3.937598235 | 0.1435254 | 0.7096883 | 0.889934337 |
| ENSMUSG00000100618 | Gm29595 | lincRNA | -0.293516522 | 2.591720433 | 0.143427 | 0.7097829 | 0.889934337 |
| ENSMUSG00000002289 | Angptl4 | protein_coding | 0.296429786 | 2.480448562 | 0.1432611 | 0.7099425 | 0.889934337 |
| ENSMUSG00000021044 | Adck1 | protein_coding | 0.14709508 | 5.485596723 | 0.1432079 | 0.7099937 | 0.889934337 |
| ENSMUSG00000088595 | Gm24601 | rRNA | -0.16487363 | 4.51822652 | 0.1431901 | 0.7100109 | 0.889934337 |
| ENSMUSG00000081051 | Gm15427 | processed_pseudoge | 0.239165813 | 2.95974261 | 0.1427878 | 0.7103984 | 0.890289583 |

| ensembl_gene_id | external_gene_name | gene_biotype | log2 (fold change) | log2 (counts per million) | F-value | p-value | False discovery rate |
|---|---|---|---|---|---|---|---|
| ENSMUSG00000035062 | Zc4h2 | protein_coding | -0.123838601 | 5.682403937 | 0.1427241 | 0.7104599 | 0.890289583 |
| ENSMUSG00000019054 | Fis1 | protein_coding | 0.09445087 | 10.28494368 | 0.1425735 | 0.7106051 | 0.890354837 |
| ENSMUSG00000026890 | Lhx6 | protein_coding | -0.136730048 | 6.003669517 | 0.1425067 | 0.7106775 | 0.890354837 |
| ENSMUSG00000019883 | Echdc1 | protein_coding | -0.187853945 | 3.896634565 | 0.1423967 | 0.7107758 | 0.890374231 |
| ENSMUSG00000024145 | Pigf | protein_coding | -0.166179933 | 5.052367098 | 0.1421205 | 0.7110426 | 0.890604699 |
| ENSMUSG00000036620 | Mgat4b | protein_coding | -0.307776859 | 2.571468355 | 0.1416593 | 0.711489 | 0.890956671 |
| ENSMUSG00000058620 | Adra2b | protein_coding | 0.219318518 | 3.442978142 | 0.1416412 | 0.7115064 | 0.890956671 |
| ENSMUSG00000024085 | Man2a1 | protein_coding | 0.421431108 | 2.203001725 | 0.1414546 | 0.7116873 | 0.890956671 |
| ENSMUSG00000005357 | Slc1a6 | protein_coding | 0.171963918 | 4.15953709 | 0.1414259 | 0.7117151 | 0.890956671 |
| ENSMUSG00000027273 | Snap25 | protein_coding | 0.121244696 | 12.17103416 | 0.1414024 | 0.7117379 | 0.890956671 |
| ENSMUSG00000034620 | Tmem5 | protein_coding | 0.146140444 | 4.559196807 | 0.1413111 | 0.7118265 | 0.890963799 |
| ENSMUSG00000079508 | Apoo | protein_coding | -0.140595498 | 4.398444904 | 0.1411226 | 0.7120093 | 0.891088899 |
| ENSMUSG00000034295 | Fhod3 | protein_coding | -0.224827016 | 2.940591565 | 0.1409777 | 0.71215 | 0.891161269 |
| ENSMUSG00000027439 | Gzf1 | protein_coding | 0.116017481 | 5.316569817 | 0.1407069 | 0.7124132 | 0.891386854 |
| ENSMUSG00000054408 | Spcs3 | protein_coding | -0.107003892 | 5.892872514 | 0.1406211 | 0.7124966 | 0.891387541 |
| ENSMUSG00000030105 | Arl8b | protein_coding | 0.095711965 | 8.049807547 | 0.1404208 | 0.7126916 | 0.89145989 |
| ENSMUSG00000043542 | Zc2hc1a | protein_coding | 0.087908026 | 7.167439517 | 0.1403897 | 0.7127218 | 0.89145989 |
| ENSMUSG00000055109 | Gm15155 | protein_coding | -0.338004114 | 2.188059548 | 0.1403062 | 0.7128031 | 0.89145989 |
| ENSMUSG00000068749 | Psma5 | protein_coding | 0.084242738 | 6.83374403 | 0.1399354 | 0.7131645 | 0.891745753 |
| ENSMUSG00000021981 | Cab39l | protein_coding | 0.102055532 | 6.511037761 | 0.1399015 | 0.7131976 | 0.891745753 |
| ENSMUSG00000040209 | Zfp704 | protein_coding | -0.274539191 | 2.930957915 | 0.1396723 | 0.7134213 | 0.891921784 |
| ENSMUSG00000048264 | Dip2c | protein_coding | -0.131095021 | 4.975060271 | 0.1395101 | 0.7135797 | 0.89196485 |
| ENSMUSG00000050565 | Tor1aip2 | protein_coding | -0.0949047 | 6.52170787 | 0.1394499 | 0.7136385 | 0.89196485 |
| ENSMUSG00000035764 | Fbxo45 | protein_coding | -0.142557945 | 4.888335855 | 0.1393816 | 0.7137053 | 0.89196485 |
| ENSMUSG00000016637 | Ift27 | protein_coding | -0.095262173 | 7.178958989 | 0.1392975 | 0.7137876 | 0.89196485 |
| ENSMUSG00000042155 | Klhl23 | protein_coding | 0.132222829 | 5.279083359 | 0.1392019 | 0.713881 | 0.891977143 |
| ENSMUSG00000017929 | B4galt5 | protein_coding | 0.136506591 | 4.77261447 | 0.1391178 | 0.7139633 | 0.891977143 |
| ENSMUSG00000005501 | Usp40 | protein_coding | 0.244349229 | 2.889453536 | 0.1389976 | 0.714081 | 0.892020535 |
| ENSMUSG00000036672 | Cenpt | protein_coding | 0.134763443 | 5.315053013 | 0.138773 | 0.7143011 | 0.892141188 |
| ENSMUSG00000026576 | Atp1b1 | protein_coding | 0.108928015 | 8.629832626 | 0.1387297 | 0.7143435 | 0.892141188 |
| ENSMUSG00000051034 | Zfp11 | protein_coding | 0.237559889 | 3.349475125 | 0.1385759 | 0.7144943 | 0.892151935 |
| ENSMUSG00000051246 | Msantd1 | protein_coding | 0.24386361 | 3.377604844 | 0.1384754 | 0.7145929 | 0.892151935 |
| ENSMUSG00000045519 | Zfp560 | protein_coding | 0.1209612 | 5.7476614 | 0.1383906 | 0.7146761 | 0.892151935 |
| ENSMUSG00000036218 | Pdzrn4 | protein_coding | 0.17086459 | 3.917302898 | 0.1383826 | 0.714684 | 0.892151935 |
| ENSMUSG00000070498 | Tmem132b | protein_coding | -0.137382489 | 4.839907472 | 0.1382124 | 0.7148511 | 0.892230239 |
| ENSMUSG00000028358 | Zfp618 | protein_coding | -0.321952921 | 2.889924072 | 0.138099 | 0.7149625 | 0.892230239 |
| ENSMUSG00000037104 | Socs5 | protein_coding | -0.106748229 | 5.970340127 | 0.1380653 | 0.7149956 | 0.892230239 |
| ENSMUSG00000040054 | Baz2a | protein_coding | 0.172409547 | 4.379592687 | 0.1378017 | 0.7152549 | 0.892394227 |
| ENSMUSG00000038975 | Rabggtb | protein_coding | 0.077267417 | 7.224132248 | 0.1377629 | 0.715293 | 0.892394227 |
| ENSMUSG00000021685 | Otp | protein_coding | -0.291690357 | 2.728168457 | 0.1374992 | 0.7155527 | 0.892562877 |
| ENSMUSG00000025217 | Btrc | protein_coding | 0.135795385 | 6.648808445 | 0.1376088 | 0.7155942 | 0.892562877 |
| ENSMUSG00000055835 | Zfp1 | protein_coding | -0.165570961 | 4.897413214 | 0.1372694 | 0.7157792 | 0.892690069 |
| ENSMUSG00000052712 | BC004004 | protein_coding | 0.082155274 | 6.97331961 | 0.136904 | 0.7161399 | 0.893036356 |
| ENSMUSG00000074576 | Mocs3 | protein_coding | 0.36409318 | 2.224110288 | 0.1367376 | 0.7163043 | 0.893044405 |
| ENSMUSG00000028976 | Slc2a5 | protein_coding | 0.457593464 | 2.785864423 | 0.1371179 | 0.7164042 | 0.893044405 |
| ENSMUSG00000028194 | Ddah1 | protein_coding | -0.107076746 | 7.196137723 | 0.1366001 | 0.7164402 | 0.893044405 |
| ENSMUSG00000025578 | Cbx8 | protein_coding | 0.226893099 | 3.14278901 | 0.1364009 | 0.7166373 | 0.893044405 |
| ENSMUSG00000060126 | Tpt1 | protein_coding | 0.184734689 | 3.675308604 | 0.1363988 | 0.7166394 | 0.893044405 |
| ENSMUSG00000000355 | Mcts1 | protein_coding | -0.090074407 | 7.797102568 | 0.1363935 | 0.7166447 | 0.893044405 |
| ENSMUSG00000030279 | C2cd5 | protein_coding | 0.140980775 | 4.359382349 | 0.1360945 | 0.7169408 | 0.893134996 |
| ENSMUSG00000038135 | Crygn | protein_coding | -0.25588995 | 3.045605241 | 0.13609 | 0.7169452 | 0.893134996 |
| ENSMUSG00000029575 | Mmab | protein_coding | 0.133207323 | 5.092512116 | 0.1360685 | 0.7169665 | 0.893134996 |
| ENSMUSG00000048148 | Nwd1 | protein_coding | 0.204484657 | 3.57062 | 0.1355887 | 0.7174426 | 0.893552348 |
| ENSMUSG00000063450 | Syne2 | protein_coding | -0.273970461 | 3.309267407 | 0.1355712 | 0.7174678 | 0.893552348 |
| ENSMUSG00000038398 | Upf3a | protein_coding | -0.155324932 | 4.345430702 | 0.135229 | 0.7178001 | 0.893862726 |
| ENSMUSG00000085218 | BB218582 | processed_transcript | -0.248369019 | 2.970129781 | 0.1349531 | 0.7180747 | 0.893916783 |
| ENSMUSG00000028917 | Plekhm2 | protein_coding | -0.219422084 | 3.915621451 | 0.1349602 | 0.7180754 | 0.893916783 |
| ENSMUSG00000050947 | Amigo1 | protein_coding | -0.145223312 | 4.700277835 | 0.1349348 | 0.7180929 | 0.893916783 |
| ENSMUSG00000097055 | Gm4419 | lincRNA | -0.196413551 | 3.336072077 | 0.1346829 | 0.7183439 | 0.894073016 |
| ENSMUSG00000038274 | Fau | protein_coding | 0.149896754 | 4.187082892 | 0.134642 | 0.7183847 | 0.894073016 |
| ENSMUSG00000032265 | Fam46a | protein_coding | -0.170826567 | 3.586635144 | 0.1345217 | 0.7185047 | 0.894118809 |
| ENSMUSG00000044502 | Bod1 | protein_coding | -0.139276237 | 5.512528293 | 0.1342231 | 0.7188028 | 0.894375158 |
| ENSMUSG00000040710 | St8sia4 | protein_coding | -0.315873942 | 2.980059613 | 0.1341319 | 0.7189016 | 0.894375158 |
| ENSMUSG00000042790 | Rnf214 | protein_coding | -0.112152708 | 5.510956496 | 0.1339814 | 0.7190444 | 0.894375158 |
| ENSMUSG00000041571 | Selenow | protein_coding | 0.111645397 | 12.37679339 | 0.1339215 | 0.7191043 | 0.894375158 |
| ENSMUSG00000028568 | Btf3l4 | protein_coding | -0.129113268 | 4.661104131 | 0.1338737 | 0.7191521 | 0.894375158 |
| ENSMUSG00000020895 | Tmem107 | protein_coding | -0.116438893 | 5.722713389 | 0.1337477 | 0.7192782 | 0.894375158 |
| ENSMUSG00000040699 | Limd2 | protein_coding | -0.149260756 | 5.42800098 | 0.133629 | 0.719397 | 0.894375158 |
| ENSMUSG00000015474 | Ppt2 | protein_coding | -0.217441146 | 2.910200772 | 0.133602 | 0.719424 | 0.894375158 |
| ENSMUSG00000090546 | Cdr1 | protein_coding | -0.239539632 | 4.856864564 | 0.1341289 | 0.7195249 | 0.894375158 |
| ENSMUSG00000024096 | Ralbp1 | protein_coding | 0.104446765 | 5.912335662 | 0.1333941 | 0.7196324 | 0.894375158 |
| ENSMUSG00000018537 | Pcgf2 | protein_coding | 0.208685269 | 3.225708111 | 0.1331801 | 0.719847 | 0.894375158 |
| ENSMUSG00000058064 | Gm10036 | protein_coding | -0.479177728 | 2.964323997 | 0.1339044 | 0.7198712 | 0.894375158 |
| ENSMUSG00000002733 | Plekha3 | protein_coding | -0.139931733 | 4.711575466 | 0.1331554 | 0.7198717 | 0.894375158 |
| ENSMUSG00000027737 | Slc7a11 | protein_coding | 0.239909161 | 3.218321286 | 0.1331521 | 0.7198751 | 0.894375158 |
| ENSMUSG00000050199 | Lgr4 | protein_coding | 0.135847324 | 4.807990148 | 0.1329872 | 0.7200406 | 0.89447743 |
| ENSMUSG00000024824 | Rad9a | protein_coding | 0.158873823 | 4.462150438 | 0.1328353 | 0.7201932 | 0.894492363 |
| ENSMUSG00000028646 | Rragc | protein_coding | -0.111755205 | 5.912414531 | 0.1328097 | 0.720219 | 0.894492363 |

| ensembl_gene_id | external_gene_name | gene_biotype | log2 (fold change) | log2 (counts per million) | F-value | p-value | False discovery rate |
|---|---|---|---|---|---|---|---|
| ENSMUSG00000021192 | Golga5 | protein_coding | 0.174103321 | 3.68689339 | 0.1324817 | 0.7205488 | 0.894795573 |
| ENSMUSG00000029752 | Asns | protein_coding | 0.147789973 | 8.118845226 | 0.1328991 | 0.7206296 | 0.894795573 |
| ENSMUSG00000054715 | Zscan22 | protein_coding | -0.19496351 | 4.282864331 | 0.1321727 | 0.7208677 | 0.894845597 |
| ENSMUSG00000097383 | 1500026H17Rik | bidirectional_promote | 0.153364014 | 3.728712492 | 0.1321237 | 0.7209094 | 0.894845597 |
| ENSMUSG00000027245 | Hypk | protein_coding | -0.154423041 | 4.01193018 | 0.1321137 | 0.7209195 | 0.894845597 |
| ENSMUSG00000037373 | Ctbp1 | protein_coding | 0.096045514 | 6.682706994 | 0.1317891 | 0.7212469 | 0.895084529 |
| ENSMUSG00000020898 | Ctc1 | protein_coding | 0.157941828 | 4.549825919 | 0.1317578 | 0.7212785 | 0.895084529 |
| ENSMUSG00000054237 | Fra10ac1 | protein_coding | -0.114293794 | 5.396569907 | 0.1315739 | 0.7214642 | 0.895211728 |
| ENSMUSG00000025925 | Terf1 | protein_coding | -0.180600774 | 3.790794773 | 0.1314497 | 0.7215898 | 0.895264184 |
| ENSMUSG00000035566 | Pcdh17 | protein_coding | -0.079382983 | 6.755909387 | 0.1313497 | 0.7216908 | 0.895281433 |
| ENSMUSG00000058546 | Rpl23a | protein_coding | -0.199195041 | 3.615805118 | 0.1312713 | 0.7217702 | 0.895281433 |
| ENSMUSG00000002486 | Tchp | protein_coding | 0.237948758 | 2.932409377 | 0.1310433 | 0.7220009 | 0.895321805 |
| ENSMUSG00000022092 | Ppp3cc | protein_coding | -0.243406019 | 3.041517414 | 0.1308845 | 0.7221618 | 0.895321805 |
| ENSMUSG00000024969 | Mark2 | protein_coding | 0.110636115 | 5.771430586 | 0.1308762 | 0.7221702 | 0.895321805 |
| ENSMUSG00000013921 | Clip3 | protein_coding | 0.100879594 | 8.694531453 | 0.1307863 | 0.7222613 | 0.895321805 |
| ENSMUSG00000059439 | Bcas3 | protein_coding | 0.172027168 | 4.684346565 | 0.1307465 | 0.7223017 | 0.895321805 |
| ENSMUSG00000069565 | Dazap1 | protein_coding | -0.20476749 | 3.087351188 | 0.1307459 | 0.7223023 | 0.895321805 |
| ENSMUSG00000020288 | Ahsa2 | protein_coding | 0.196863562 | 3.798943059 | 0.1302884 | 0.7227667 | 0.895733262 |
| ENSMUSG00000028191 | Bcl10 | protein_coding | 0.12276393 | 4.911238581 | 0.1302547 | 0.7228008 | 0.895733262 |
| ENSMUSG00000049122 | Frmd3 | protein_coding | -0.260142437 | 3.188285896 | 0.1301599 | 0.7228973 | 0.895749506 |
| ENSMUSG00000099034 | 2810039B14Rik | transcribed_unitary_p | -0.202752077 | 3.229645535 | 0.1295859 | 0.7234815 | 0.896272233 |
| ENSMUSG00000025580 | Eif4a3 | protein_coding | -0.088724573 | 6.742675217 | 0.1295072 | 0.7235617 | 0.896272233 |
| ENSMUSG00000019826 | Zbtb24 | protein_coding | 0.159616863 | 3.807397015 | 0.1294193 | 0.7236514 | 0.896272233 |
| ENSMUSG00000006369 | Fbln1 | protein_coding | 0.253166453 | 3.155322925 | 0.1293403 | 0.7237319 | 0.896272233 |
| ENSMUSG00000085492 | Trmt61b | processed_transcript | 0.124164049 | 5.168343951 | 0.12926 | 0.7238139 | 0.896272233 |
| ENSMUSG00000018820 | Zfyve27 | protein_coding | -0.119459001 | 5.296735211 | 0.1290579 | 0.7240202 | 0.896272233 |
| ENSMUSG00000003269 | Cyth2 | protein_coding | -0.140214233 | 5.170669015 | 0.1290573 | 0.7240208 | 0.896272233 |
| ENSMUSG00000018548 | Trim37 | protein_coding | -0.08317778 | 6.692543958 | 0.1289904 | 0.7240892 | 0.896272233 |
| ENSMUSG00000034412 | Tbc1d10a | protein_coding | -0.222548533 | 3.277671923 | 0.1288484 | 0.7242343 | 0.896272233 |
| ENSMUSG00000032096 | Arcn1 | protein_coding | 0.104668574 | 5.576845699 | 0.1288418 | 0.724241 | 0.896272233 |
| ENSMUSG00000042121 | Ssh1 | protein_coding | -0.132851865 | 4.392451141 | 0.1288264 | 0.7242568 | 0.896272233 |
| ENSMUSG00000025138 | Sirt7 | protein_coding | 0.249044691 | 3.093681982 | 0.1287653 | 0.7243193 | 0.896272233 |
| ENSMUSG00000075595 | Zfp652 | protein_coding | 0.139884247 | 4.281857141 | 0.1286624 | 0.7244245 | 0.896299277 |
| ENSMUSG00000042032 | Mat2b | protein_coding | 0.083011934 | 7.830334238 | 0.1284657 | 0.724626 | 0.89637283 |
| ENSMUSG00000006699 | Cdc42 | protein_coding | -0.076696242 | 9.861276353 | 0.1282807 | 0.7248155 | 0.89637283 |
| ENSMUSG00000004980 | Hnrnpa2b1 | protein_coding | -0.078465908 | 8.948773507 | 0.1282774 | 0.7248189 | 0.89637283 |
| ENSMUSG00000038760 | Trhr | protein_coding | 0.385359371 | 2.638988981 | 0.1281512 | 0.7249558 | 0.89637283 |
| ENSMUSG00000029763 | Exoc4 | protein_coding | 0.111847319 | 5.708463691 | 0.1280609 | 0.7250408 | 0.89637283 |
| ENSMUSG00000040396 | Abhd13 | protein_coding | -0.171928923 | 3.6968648 | 0.128049 | 0.7250531 | 0.89637283 |
| ENSMUSG00000029229 | Chic2 | protein_coding | 0.192280453 | 3.422371939 | 0.128035 | 0.7250675 | 0.89637283 |
| ENSMUSG00000058267 | Mrps14 | protein_coding | 0.083261984 | 8.757600693 | 0.1279074 | 0.7251984 | 0.896431646 |
| ENSMUSG00000001436 | Slc19a1 | protein_coding | -0.213412286 | 4.7001377 | 0.1278621 | 0.725537 | 0.896530751 |
| ENSMUSG00000079003 | Samd1 | protein_coding | 0.249501883 | 3.283322186 | 0.1274996 | 0.7256175 | 0.896530751 |
| ENSMUSG00000020063 | Sirt1 | protein_coding | -0.22998445 | 2.913816698 | 0.1274399 | 0.7256789 | 0.896530751 |
| ENSMUSG00000029439 | Sfswap | protein_coding | -0.203647625 | 3.823565253 | 0.1274346 | 0.7256844 | 0.896530751 |
| ENSMUSG00000017692 | Rhbdl3 | protein_coding | 0.208871791 | 3.989578344 | 0.1273104 | 0.7258197 | 0.896530751 |
| ENSMUSG00000058174 | Gm5148 | protein_coding | 0.147589321 | 4.361021659 | 0.1271074 | 0.7260213 | 0.896530751 |
| ENSMUSG00000021371 | Mcur1 | protein_coding | -0.22285278 | 3.262011593 | 0.1270419 | 0.7260888 | 0.896530751 |
| ENSMUSG00000032648 | Pygm | protein_coding | -0.166159731 | 4.634923413 | 0.1269831 | 0.7261493 | 0.896530751 |
| ENSMUSG00000005610 | Eif4g2 | protein_coding | -0.104609426 | 8.797451722 | 0.1269686 | 0.7261643 | 0.896530751 |
| ENSMUSG00000020216 | Jsrp1 | protein_coding | 0.228092126 | 3.231822087 | 0.126962 | 0.7261711 | 0.896530751 |
| ENSMUSG00000068966 | Zbtb34 | protein_coding | 0.243188487 | 2.946997983 | 0.1268687 | 0.7262673 | 0.896530751 |
| ENSMUSG00000097413 | A830052D11Rik | lincRNA | -0.286134858 | 2.599959074 | 0.1268573 | 0.7262791 | 0.896530751 |
| ENSMUSG00000073415 | Gm10501 | antisense_RNA | -0.249193497 | 2.689804278 | 0.1267033 | 0.726438 | 0.896623986 |
| ENSMUSG00000021687 | Scamp1 | protein_coding | 0.079675689 | 7.292855386 | 0.1265506 | 0.7265957 | 0.896653426 |
| ENSMUSG00000021115 | Vrk1 | protein_coding | -0.112454002 | 5.201208222 | 0.1265187 | 0.7266287 | 0.896653426 |
| ENSMUSG00000025372 | Baiap2 | protein_coding | -0.098415942 | 7.550732892 | 0.1261042 | 0.7270572 | 0.897079336 |
| ENSMUSG00000022973 | Synj1 | protein_coding | -0.102953186 | 7.54218916 | 0.125776 | 0.7273972 | 0.897395877 |
| ENSMUSG00000041570 | Camsap2 | protein_coding | -0.196657114 | 3.322878241 | 0.1252703 | 0.7279219 | 0.897940158 |
| ENSMUSG00000020027 | Socs2 | protein_coding | 0.195884604 | 3.119462797 | 0.1248937 | 0.7283136 | 0.898195828 |
| ENSMUSG00000015488 | Cacfd1 | protein_coding | 0.156335341 | 4.85552178 | 0.124838 | 0.7283715 | 0.898195828 |
| ENSMUSG00000035045 | Zc3h12b | protein_coding | -0.138113024 | 4.20635853 | 0.1248301 | 0.7283798 | 0.898195828 |
| ENSMUSG00000032562 | Gnai2 | protein_coding | 0.088364798 | 7.278666785 | 0.1246753 | 0.728541 | 0.898291594 |
| ENSMUSG00000054277 | Arfgap3 | protein_coding | -0.130066112 | 5.491942616 | 0.1244949 | 0.728729 | 0.898326095 |
| ENSMUSG00000097393 | D030068K23Rik | lincRNA | 0.147087256 | 5.219533529 | 0.124427 | 0.7287997 | 0.898326095 |
| ENSMUSG00000042328 | Hps4 | protein_coding | -0.167557675 | 3.690043828 | 0.1243755 | 0.7288535 | 0.898326095 |
| ENSMUSG00000060601 | Nr1h2 | protein_coding | -0.105292561 | 5.922656333 | 0.1243052 | 0.7289268 | 0.898326095 |
| ENSMUSG00000053334 | Ficd | protein_coding | -0.214246744 | 3.187597239 | 0.1241923 | 0.7290446 | 0.898326095 |
| ENSMUSG00000022935 | Grik1 | protein_coding | -0.20198668 | 4.004442477 | 0.124105 | 0.7291432 | 0.898326095 |
| ENSMUSG00000018509 | Cenpv | protein_coding | 0.146713593 | 4.041840797 | 0.1240878 | 0.7291537 | 0.898326095 |
| ENSMUSG00000026220 | Slc16a14 | protein_coding | -0.362163693 | 3.227649234 | 0.1243636 | 0.7294513 | 0.898507904 |
| ENSMUSG00000039501 | Znfx1 | protein_coding | -0.172495759 | 3.843822906 | 0.1237532 | 0.7295035 | 0.898507904 |
| ENSMUSG00000036686 | Cc2d1a | protein_coding | -0.163142643 | 4.848330308 | 0.1237069 | 0.729552 | 0.898507904 |
| ENSMUSG00000024921 | Smarca2 | protein_coding | -0.099990696 | 6.0614837 | 0.1235002 | 0.7297683 | 0.898671444 |
| ENSMUSG00000021007 | Spata7 | protein_coding | 0.127663557 | 5.025590544 | 0.1234183 | 0.7298541 | 0.898674122 |
| ENSMUSG00000051518 | Rps19bp1 | protein_coding | 0.158520359 | 4.065044223 | 0.123233 | 0.7300483 | 0.898810315 |
| ENSMUSG00000028256 | Odf2l | protein_coding | 0.241013824 | 3.174813505 | 0.1229277 | 0.7303686 | 0.899068376 |
| ENSMUSG00000047909 | Ankrd16 | protein_coding | 0.303848686 | 2.79214358 | 0.1228499 | 0.7304504 | 0.899068376 |

| ensembl_gene_id | external_gene_name | gene_biotype | log2 (fold change) | log2 (counts per million) | F-value | p-value | False discovery rate |
|---|---|---|---|---|---|---|---|
| ENSMUSG00000027357 | Crls1 | protein_coding | 0.116435922 | 4.835741882 | 0.1227944 | 0.7305087 | 0.899068376 |
| ENSMUSG00000041936 | Agrn | protein_coding | 0.156549374 | 4.319190457 | 0.1224793 | 0.73084 | 0.899373199 |
| ENSMUSG00000044328 | Trp53i13 | protein_coding | 0.167612699 | 3.714643279 | 0.1219148 | 0.7314347 | 0.900001999 |
| ENSMUSG00000031068 | Glrx3 | protein_coding | -0.084027964 | 6.39204857 | 0.1218282 | 0.7315261 | 0.900011482 |
| ENSMUSG00000042354 | Gnl3 | protein_coding | -0.12028993 | 5.427918584 | 0.1217165 | 0.731644 | 0.900053586 |
| ENSMUSG00000010453 | Kansl3 | protein_coding | -0.133391794 | 5.673572314 | 0.1215515 | 0.7318183 | 0.900164996 |
| ENSMUSG00000061665 | Cd2ap | protein_coding | 0.300307294 | 2.505229017 | 0.1211778 | 0.7322135 | 0.900512515 |
| ENSMUSG00000060121 | Gemin2 | protein_coding | -0.123329208 | 4.848682691 | 0.1211261 | 0.7322683 | 0.900512515 |
| ENSMUSG00000041684 | Bivm | protein_coding | -0.138155487 | 4.219318617 | 0.1210106 | 0.7323907 | 0.900559981 |
| ENSMUSG00000032482 | Cspg5 | protein_coding | 0.104056528 | 7.199515376 | 0.1208811 | 0.7325279 | 0.900572822 |
| ENSMUSG00000034707 | Gns | protein_coding | 0.12706515 | 5.097607033 | 0.12084 | 0.7325714 | 0.900572822 |
| ENSMUSG00000021963 | Sap18 | protein_coding | 0.078623623 | 6.985810543 | 0.1207637 | 0.7326524 | 0.900572822 |
| ENSMUSG00000021832 | Psmc6 | protein_coding | -0.081995471 | 7.89661194 | 0.1206675 | 0.7327544 | 0.900595268 |
| ENSMUSG00000044783 | Hjurp | protein_coding | 0.128434813 | 4.757925192 | 0.1204748 | 0.732959 | 0.900642073 |
| ENSMUSG00000084910 | C630043F03Rik | processed_transcript | -0.173134784 | 3.895247764 | 0.1204738 | 0.73296 | 0.900642073 |
| ENSMUSG00000037712 | Fermt2 | protein_coding | -0.116511113 | 5.541511576 | 0.1202527 | 0.733195 | 0.900827859 |
| ENSMUSG00000025134 | Alyref | protein_coding | -0.144759707 | 4.548844003 | 0.1197732 | 0.7337053 | 0.901351897 |
| ENSMUSG00000029470 | P2rx4 | protein_coding | 0.228079998 | 3.457031536 | 0.119341 | 0.7341663 | 0.901647271 |
| ENSMUSG00000057134 | Ado | protein_coding | -0.111438641 | 8.137691572 | 0.1193288 | 0.7341866 | 0.901647271 |
| ENSMUSG00000027341 | Tmem230 | protein_coding | 0.137528608 | 4.33229292 | 0.1191944 | 0.7343229 | 0.901647271 |
| ENSMUSG00000030795 | Fus | protein_coding | -0.118794339 | 5.029174441 | 0.1191236 | 0.7343986 | 0.901647271 |
| ENSMUSG00000001376 | Vps50 | protein_coding | -0.136034451 | 5.090107022 | 0.1190835 | 0.7344414 | 0.901647271 |
| ENSMUSG00000024777 | Ppp2r5b | protein_coding | 0.09840972 | 5.645437289 | 0.1190766 | 0.7344488 | 0.901647271 |
| ENSMUSG00000042178 | Armc5 | protein_coding | -0.247685955 | 3.018035505 | 0.1187193 | 0.7348311 | 0.901792002 |
| ENSMUSG00000054469 | Lclat1 | protein_coding | 0.204921104 | 3.717781153 | 0.1186316 | 0.734925 | 0.901792002 |
| ENSMUSG00000030983 | Bccip | protein_coding | 0.076085389 | 7.669501024 | 0.1185563 | 0.7350058 | 0.901792002 |
| ENSMUSG00000042604 | Kcna4 | protein_coding | -0.189966766 | 3.988612793 | 0.1184806 | 0.7350869 | 0.901792002 |
| ENSMUSG00000056770 | Setd3 | protein_coding | 0.085403199 | 6.51482753 | 0.1184673 | 0.7351012 | 0.901792002 |
| ENSMUSG00000032458 | Copb2 | protein_coding | 0.106059804 | 6.600300104 | 0.1184083 | 0.7351644 | 0.901792002 |
| ENSMUSG00000036430 | Tbcc | protein_coding | -0.197938117 | 3.39072153 | 0.1182889 | 0.7352925 | 0.901792002 |
| ENSMUSG00000030101 | Sumf1 | protein_coding | 0.202895263 | 4.039992714 | 0.1182794 | 0.7353099 | 0.901792002 |
| ENSMUSG00000044352 | Sowaha | protein_coding | -0.17426968 | 5.361425796 | 0.1182919 | 0.7355521 | 0.901792002 |
| ENSMUSG00000028233 | Tgs1 | protein_coding | -0.100634821 | 5.482856638 | 0.1180034 | 0.7355991 | 0.901792002 |
| ENSMUSG00000112964 | CT009718.1 | lincRNA | -0.210259329 | 3.165644942 | 0.1179609 | 0.7356447 | 0.901792002 |
| ENSMUSG00000057367 | Birc2 | protein_coding | 0.137207482 | 5.390641583 | 0.117924 | 0.7356844 | 0.901792002 |
| ENSMUSG00000032786 | Alas1 | protein_coding | 0.139273223 | 6.36808781 | 0.1181509 | 0.7357203 | 0.901792002 |
| ENSMUSG00000026718 | Stam | protein_coding | 0.101713501 | 5.8683243 | 0.1177489 | 0.7358727 | 0.901792002 |
| ENSMUSG00000029465 | Arpc3 | protein_coding | -0.101425009 | 8.650120317 | 0.1177367 | 0.7358858 | 0.901792002 |
| ENSMUSG00000027849 | Syt6 | protein_coding | -0.194919807 | 4.318855024 | 0.1177098 | 0.735922 | 0.901792002 |
| ENSMUSG00000026723 | Trdmt1 | protein_coding | -0.210572483 | 3.159473923 | 0.1176376 | 0.7359924 | 0.901792002 |
| ENSMUSG00000024507 | Hsd17b4 | protein_coding | 0.104364706 | 5.464083413 | 0.1174221 | 0.7362245 | 0.901973624 |
| ENSMUSG00000015829 | Tnr | protein_coding | -0.168887922 | 3.979884262 | 0.1172617 | 0.7363974 | 0.901991011 |
| ENSMUSG00000020513 | Tubd1 | protein_coding | -0.294475963 | 2.857777701 | 0.1172533 | 0.7364065 | 0.901991011 |
| ENSMUSG00000038406 | Scaf1 | protein_coding | 0.144497771 | 4.706982245 | 0.1170497 | 0.7366262 | 0.90210403 |
| ENSMUSG00000022800 | Fyttd1 | protein_coding | -0.093976747 | 6.968082929 | 0.1170123 | 0.7366666 | 0.90210403 |
| ENSMUSG00000005803 | Sqor | protein_coding | -0.175693228 | 3.767947818 | 0.1169275 | 0.7367581 | 0.902113405 |
| ENSMUSG00000032527 | Pccb | protein_coding | -0.120932965 | 5.911814377 | 0.1166784 | 0.7370273 | 0.902340267 |
| ENSMUSG00000038658 | Ric1 | protein_coding | -0.182467545 | 4.409328162 | 0.1165775 | 0.7371436 | 0.902379912 |
| ENSMUSG00000026199 | Ankzf1 | protein_coding | 0.168388997 | 3.976778825 | 0.1162233 | 0.7375199 | 0.90273781 |
| ENSMUSG00000060427 | Zfp868 | protein_coding | 0.176215003 | 4.786684045 | 0.1160468 | 0.7377185 | 0.902875556 |
| ENSMUSG00000055491 | Pprc1 | protein_coding | -0.215305015 | 2.946576924 | 0.115843 | 0.7379325 | 0.902875556 |
| ENSMUSG00000031355 | Arhgap6 | protein_coding | 0.133677132 | 4.359166471 | 0.1157986 | 0.7379807 | 0.902875556 |
| ENSMUSG00000028629 | Exo5 | protein_coding | 0.238751048 | 4.006934492 | 0.115992 | 0.7380476 | 0.902875556 |
| ENSMUSG00000024789 | Jak2 | protein_coding | 0.160172419 | 4.004079368 | 0.1156808 | 0.7381086 | 0.902875556 |
| ENSMUSG00000031652 | N4bp1 | protein_coding | -0.106469449 | 5.833230899 | 0.1156554 | 0.7381362 | 0.902875556 |
| ENSMUSG00000023170 | Gps2 | protein_coding | 0.120734531 | 4.89222568 | 0.1154357 | 0.7383751 | 0.902903426 |
| ENSMUSG00000039033 | Tasp1 | protein_coding | -0.119906096 | 5.575886204 | 0.1153367 | 0.7384829 | 0.902903426 |
| ENSMUSG00000039542 | Ncam1 | protein_coding | -0.092003284 | 8.750398292 | 0.1152148 | 0.7386156 | 0.902903426 |
| ENSMUSG00000025856 | Pdgfa | protein_coding | 0.112284623 | 5.019665444 | 0.1152043 | 0.738627 | 0.902903426 |
| ENSMUSG00000030347 | D6Wsu163e | protein_coding | 0.106209978 | 5.091092312 | 0.1151842 | 0.7386489 | 0.902903426 |
| ENSMUSG00000026333 | Gin1 | protein_coding | -0.164734638 | 4.444129451 | 0.1151714 | 0.7386628 | 0.902903426 |
| ENSMUSG00000041193 | Pla2g5 | protein_coding | -0.2607729 | 2.690346538 | 0.1150759 | 0.7387669 | 0.902927997 |
| ENSMUSG00000043091 | Tuba1c | protein_coding | -0.568140826 | 1.936353087 | 0.1151176 | 0.7389971 | 0.903065801 |
| ENSMUSG00000026491 | Ahctf1 | protein_coding | 0.19705558 | 3.264716847 | 0.1148185 | 0.7390476 | 0.903065801 |
| ENSMUSG00000032462 | Pik3cb | protein_coding | -0.243083077 | 3.052079524 | 0.1147142 | 0.7391615 | 0.903068745 |
| ENSMUSG00000025950 | Idh1 | protein_coding | 0.082471191 | 6.523686016 | 0.1146522 | 0.7392291 | 0.903068745 |
| ENSMUSG00000028962 | Slc4a2 | protein_coding | 0.187815725 | 3.489299007 | 0.1145855 | 0.739302 | 0.903068745 |
| ENSMUSG00000024613 | Tcof1 | protein_coding | 0.143646533 | 4.44965161 | 0.1144809 | 0.7394163 | 0.903105788 |
| ENSMUSG00000011179 | Odc1 | protein_coding | -0.150461406 | 4.258614454 | 0.1142544 | 0.739664 | 0.903305718 |
| ENSMUSG00000021840 | Mapk1ip1l | protein_coding | -0.145886001 | 3.850109201 | 0.114108 | 0.7398242 | 0.903331937 |
| ENSMUSG00000020894 | Vamp2 | protein_coding | 0.080563181 | 9.219983842 | 0.1140813 | 0.7398535 | 0.903331937 |
| ENSMUSG00000021978 | Extl3 | protein_coding | 0.12648291 | 5.404115986 | 0.1135853 | 0.7403972 | 0.903705214 |
| ENSMUSG00000045639 | Zfp629 | protein_coding | -0.142143917 | 4.085104144 | 0.1135072 | 0.740483 | 0.903705214 |
| ENSMUSG00000033373 | Fntb | protein_coding | 0.110739384 | 5.852446831 | 0.1134945 | 0.740497 | 0.903705214 |
| ENSMUSG00000030894 | Tpp1 | protein_coding | -0.12321297 | 6.126023615 | 0.1134274 | 0.7405777 | 0.903705214 |
| ENSMUSG00000021590 | Spata9 | protein_coding | 0.307152387 | 2.518650741 | 0.1133113 | 0.7406982 | 0.903705214 |
| ENSMUSG00000079598 | Clec2l | protein_coding | -0.130226779 | 4.576287823 | 0.1132743 | 0.7407389 | 0.903705214 |
| ENSMUSG00000079055 | Slc8a3 | protein_coding | 0.148525668 | 4.086510186 | 0.1132665 | 0.7407475 | 0.903705214 |

| ensembl_gene_id | external_gene_name | gene_biotype | log2 (fold change) | log2 (counts per million) | F-value | p-value | False discovery rate |
|---|---|---|---|---|---|---|---|
| ENSMUSG00000070056 | Mfhas1 | protein_coding | 0.122811348 | 4.650240112 | 0.1129481 | 0.7410978 | 0.903919232 |
| ENSMUSG00000053519 | Kcnip1 | protein_coding | 0.092578197 | 6.198908436 | 0.1128468 | 0.7412094 | 0.903919232 |
| ENSMUSG00000030922 | Lyrm1 | protein_coding | -0.092283373 | 5.692454423 | 0.112827 | 0.7412312 | 0.903919232 |
| ENSMUSG00000063801 | Ap3s2 | protein_coding | -0.082638176 | 6.881998965 | 0.1128017 | 0.7412592 | 0.903919232 |
| ENSMUSG00000036565 | Ttyh3 | protein_coding | -0.15871968 | 4.395065341 | 0.1125949 | 0.7414871 | 0.9040046 |
| ENSMUSG00000053390 | Zfp952 | protein_coding | 0.157080154 | 4.308508244 | 0.112412 | 0.741689 | 0.9040046 |
| ENSMUSG00000034557 | Zfyve9 | protein_coding | 0.108439256 | 5.23574843 | 0.1123386 | 0.74177 | 0.9040046 |
| ENSMUSG00000070939 | Tgfbrap1 | protein_coding | 0.139721493 | 4.076641298 | 0.112307 | 0.741805 | 0.9040046 |
| ENSMUSG00000023044 | Csad | protein_coding | 0.167733053 | 4.287033396 | 0.112248 | 0.7418701 | 0.9040046 |
| ENSMUSG00000037526 | Atg14 | protein_coding | 0.151517561 | 4.759605011 | 0.1122173 | 0.7419041 | 0.9040046 |
| ENSMUSG00000005069 | Pex5 | protein_coding | 0.138053525 | 4.408127076 | 0.1120271 | 0.7421144 | 0.9040046 |
| ENSMUSG00000024501 | Dpysl3 | protein_coding | -0.11585849 | 6.205575626 | 0.1119629 | 0.7421924 | 0.9040046 |
| ENSMUSG00000084403 | Rps15a-ps8 | processed_pseudoge | -0.576339176 | 1.126234636 | 0.1119556 | 0.7421935 | 0.9040046 |
| ENSMUSG00000030057 | Cnbp | protein_coding | -0.075891666 | 10.42446665 | 0.1118744 | 0.7422834 | 0.9040046 |
| ENSMUSG00000057113 | Npm1 | protein_coding | -0.077162097 | 8.151725743 | 0.1118704 | 0.7422879 | 0.9040046 |
| ENSMUSG00000025279 | Dnase1l3 | protein_coding | -0.441111834 | 1.981800295 | 0.1118251 | 0.742338 | 0.9040046 |
| ENSMUSG00000049866 | Arl4c | protein_coding | 0.110465233 | 5.539892239 | 0.1116288 | 0.7425555 | 0.904167057 |
| ENSMUSG00000028788 | Ptp4a2 | protein_coding | -0.093685468 | 8.424714195 | 0.111465 | 0.7427371 | 0.904285733 |
| ENSMUSG00000003184 | Irf3 | protein_coding | -0.205461232 | 3.253010263 | 0.111383 | 0.7428281 | 0.904294161 |
| ENSMUSG00000045464 | 2810002D19Rik | processed_transcript | 0.189214326 | 4.38772 | 0.1112704 | 0.7429687 | 0.90432681 |
| ENSMUSG00000063714 | Sp3os | antisense_RNA | -0.175564044 | 3.530542364 | 0.1109056 | 0.7433585 | 0.90432681 |
| ENSMUSG00000089889 | 0610040B10Rik | antisense_RNA | -0.165780931 | 3.573180574 | 0.1108375 | 0.7434344 | 0.90432681 |
| ENSMUSG00000013622 | Atraid | protein_coding | -0.089848073 | 7.682198409 | 0.1108036 | 0.7434721 | 0.90432681 |
| ENSMUSG00000097574 | C920006O11Rik | lincRNA | -0.155954977 | 3.671111301 | 0.1106111 | 0.7436864 | 0.90432681 |
| ENSMUSG00000039199 | Zdhhc1 | protein_coding | 0.240952776 | 2.754886922 | 0.1105812 | 0.7437197 | 0.90432681 |
| ENSMUSG00000019817 | Plagl1 | protein_coding | -0.156057053 | 3.547039266 | 0.1105757 | 0.7437259 | 0.90432681 |
| ENSMUSG00000027207 | Galk2 | protein_coding | 0.108853745 | 6.042863571 | 0.1104746 | 0.7438386 | 0.90432681 |
| ENSMUSG00000033282 | Rpgrip1l | protein_coding | 0.158790255 | 4.09526138 | 0.1104259 | 0.7438929 | 0.90432681 |
| ENSMUSG00000085741 | 5430405H02Rik | processed_transcript | -0.182378371 | 3.345413118 | 0.1104 | 0.7439218 | 0.90432681 |
| ENSMUSG00000002778 | Kdelr1 | protein_coding | -0.131979195 | 5.216955976 | 0.1103508 | 0.7439766 | 0.90432681 |
| ENSMUSG00000006526 | Tmem110 | protein_coding | 0.223318008 | 3.326175884 | 0.1103497 | 0.7439779 | 0.90432681 |
| ENSMUSG00000041272 | Tox | protein_coding | 0.241726609 | 6.058983612 | 0.1122 | 0.7439971 | 0.90432681 |
| ENSMUSG00000040820 | Hlcs | protein_coding | 0.166229582 | 3.496479684 | 0.1103009 | 0.7440323 | 0.90432681 |
| ENSMUSG00000035325 | Sec31a | protein_coding | 0.100729112 | 6.04844774 | 0.1100947 | 0.7442625 | 0.904413694 |
| ENSMUSG00000021474 | Sfxn1 | protein_coding | 0.072826391 | 7.44795253 | 0.1100215 | 0.7443443 | 0.904413694 |
| ENSMUSG00000033326 | Kdm4a | protein_coding | 0.273432434 | 2.537096055 | 0.110011 | 0.7443561 | 0.904413694 |
| ENSMUSG00000017286 | Glod4 | protein_coding | 0.072746692 | 7.522430869 | 0.1096638 | 0.7447444 | 0.904783279 |
| ENSMUSG00000024975 | Pdcd4 | protein_coding | 0.144187429 | 7.436718025 | 0.1099985 | 0.7449619 | 0.904868434 |
| ENSMUSG00000030888 | Rrp8 | protein_coding | -0.15435499 | 4.165826031 | 0.1094188 | 0.7450189 | 0.904868434 |
| ENSMUSG00000029233 | Srd5a3 | protein_coding | -0.204331007 | 3.477859204 | 0.1093759 | 0.745067 | 0.904868434 |
| ENSMUSG00000030611 | Mrps11 | protein_coding | -0.078198465 | 7.327358573 | 0.1091943 | 0.7452707 | 0.905013662 |
| ENSMUSG00000029131 | Dnajb6 | protein_coding | 0.134965068 | 4.287201928 | 0.1091155 | 0.7453591 | 0.905018829 |
| ENSMUSG00000029106 | Add1 | protein_coding | 0.075937504 | 7.534351231 | 0.1089917 | 0.7454981 | 0.905085385 |
| ENSMUSG00000021018 | Polr2h | protein_coding | -0.087683927 | 6.12571714 | 0.1088188 | 0.7456925 | 0.905164818 |
| ENSMUSG00000034729 | Mrps10 | protein_coding | 0.086993873 | 6.322368147 | 0.1087837 | 0.7457319 | 0.905164818 |
| ENSMUSG00000033615 | Cplx1 | protein_coding | 0.089250948 | 9.61942052 | 0.1085062 | 0.7460443 | 0.905228769 |
| ENSMUSG00000030088 | Aldh1l1 | protein_coding | 0.181922176 | 4.31170691 | 0.1084995 | 0.7460586 | 0.905228769 |
| ENSMUSG00000030816 | Rnf40 | protein_coding | -0.119538139 | 4.378118553 | 0.10849 | 0.7460624 | 0.905228769 |
| ENSMUSG00000028800 | Hdac1 | protein_coding | -0.14806113 | 3.739194439 | 0.1084378 | 0.7461213 | 0.905228769 |
| ENSMUSG00000032405 | Pias1 | protein_coding | -0.096157945 | 7.093369764 | 0.1081361 | 0.7464615 | 0.905539285 |
| ENSMUSG00000009079 | Ewsr1 | protein_coding | 0.097387555 | 5.932260034 | 0.1078606 | 0.7467725 | 0.905814454 |
| ENSMUSG00000024253 | Dync2li1 | protein_coding | 0.093196246 | 5.827767068 | 0.1074903 | 0.7471913 | 0.906095215 |
| ENSMUSG00000026991 | Pkp4 | protein_coding | -0.072198109 | 6.884406364 | 0.1074148 | 0.7472769 | 0.906095215 |
| ENSMUSG00000032890 | Rims3 | protein_coding | 0.145428202 | 7.353189369 | 0.1079357 | 0.747338 | 0.906095215 |
| ENSMUSG00000039634 | Zfp189 | protein_coding | -0.198257521 | 3.583307246 | 0.1073581 | 0.7473411 | 0.906095215 |
| ENSMUSG00000039703 | Nploc4 | protein_coding | -0.113719976 | 4.976776944 | 0.107193 | 0.7475282 | 0.906219902 |
| ENSMUSG00000024524 | Gnal | protein_coding | -0.087829884 | 7.129695087 | 0.1069098 | 0.7478494 | 0.906507161 |
| ENSMUSG00000060671 | Atp8b2 | protein_coding | -0.17941643 | 3.471157215 | 0.1066935 | 0.7480952 | 0.90670285 |
| ENSMUSG00000026353 | Ubxn4 | protein_coding | -0.089985281 | 5.908342931 | 0.1065617 | 0.7482451 | 0.906782352 |
| ENSMUSG00000010721 | Lmbr1 | protein_coding | 0.110825426 | 4.76906295 | 0.1060989 | 0.7487722 | 0.907318853 |
| ENSMUSG00000022114 | Spry2 | protein_coding | -0.202173142 | 6.703076342 | 0.1073713 | 0.7490932 | 0.907605591 |
| ENSMUSG00000021557 | Agtpbp1 | protein_coding | -0.09266234 | 6.05202309 | 0.1056701 | 0.7492617 | 0.907698038 |
| ENSMUSG00000071341 | Egr4 | protein_coding | -0.409123166 | 1.664414579 | 0.1055807 | 0.7493639 | 0.907698038 |
| ENSMUSG00000041762 | Gpr155 | protein_coding | -0.173573449 | 4.032145918 | 0.1055292 | 0.7494228 | 0.907698038 |
| ENSMUSG00000079330 | Lemd1 | protein_coding | -0.187713056 | 3.620997287 | 0.1052355 | 0.749759 | 0.907997543 |
| ENSMUSG00000097466 | D430036J16Rik | antisense_RNA | 0.200852932 | 3.323470962 | 0.1050713 | 0.7499472 | 0.907997543 |
| ENSMUSG00000022305 | Lrp12 | protein_coding | -0.16341533 | 4.229442977 | 0.1050034 | 0.7500251 | 0.907997543 |
| ENSMUSG00000046341 | Gm11223 | processed_pseudoge | -0.172189464 | 3.20080586 | 0.1049724 | 0.7500606 | 0.907997543 |
| ENSMUSG00000033419 | Snap91 | protein_coding | 0.079973943 | 7.467942265 | 0.1049448 | 0.7500923 | 0.907997543 |
| ENSMUSG00000042428 | Mgat3 | protein_coding | 0.100637423 | 5.993891387 | 0.1048402 | 0.7502123 | 0.908023708 |
| ENSMUSG00000023932 | Cdc5l | protein_coding | 0.100171747 | 5.338712987 | 0.1047417 | 0.7503255 | 0.908023708 |
| ENSMUSG00000041769 | Ppp2r2d | protein_coding | -0.080344669 | 6.312592525 | 0.1046306 | 0.750453 | 0.908023708 |
| ENSMUSG00000044968 | Napepld | protein_coding | -0.221359916 | 3.709934025 | 0.1046134 | 0.7504973 | 0.908023708 |
| ENSMUSG00000026831 | 1700007K13Rik | protein_coding | -0.184646754 | 3.121678407 | 0.1045583 | 0.7505361 | 0.908023708 |
| ENSMUSG00000019803 | Nr2e1 | protein_coding | -0.301260381 | 2.88878223 | 0.1043529 | 0.7507791 | 0.908215512 |
| ENSMUSG00000039257 | Vstm2b | protein_coding | 0.107361154 | 5.263091871 | 0.1042528 | 0.7508876 | 0.908244604 |
| ENSMUSG00000026519 | Tmem63a | protein_coding | 0.15171892 | 4.576866555 | 0.1040653 | 0.7511037 | 0.908403739 |
| ENSMUSG00000087598 | Zfp111 | protein_coding | -0.209221483 | 2.732039677 | 0.1039688 | 0.7512149 | 0.908436102 |

| ensembl_gene_id | external_gene_name | gene_biotype | log2 (fold change) | log2 (counts per million) | F-value | p-value | False discovery rate |
|---|---|---|---|---|---|---|---|
| ENSMUSG00000038630 | Zkscan16 | protein_coding | 0.213302962 | 2.952419971 | 0.10378 | 0.7514328 | 0.90859742 |
| ENSMUSG00000026730 | Pter | protein_coding | -0.200714102 | 4.465772053 | 0.1039177 | 0.7515347 | 0.908618446 |
| ENSMUSG00000001829 | Clpb | protein_coding | 0.094274249 | 5.379357391 | 0.1034605 | 0.7518019 | 0.908839308 |
| ENSMUSG00000061474 | Mrps36 | protein_coding | -0.121428278 | 4.525805202 | 0.1033668 | 0.7519103 | 0.908868152 |
| ENSMUSG00000022401 | Xpnpep3 | protein_coding | -0.177207496 | 3.690002345 | 0.10308 | 0.7522424 | 0.909167391 |
| ENSMUSG00000019891 | Dcbld1 | protein_coding | 0.288414227 | 2.688547111 | 0.1029073 | 0.7524426 | 0.909295518 |
| ENSMUSG00000066571 | 4931406P16Rik | protein_coding | 0.142996099 | 3.750215517 | 0.1028427 | 0.7525175 | 0.909295518 |
| ENSMUSG00000053754 | Chd8 | protein_coding | -0.218722368 | 3.055711784 | 0.1023742 | 0.7530617 | 0.909850855 |
| ENSMUSG00000040570 | Rundc3b | protein_coding | -0.121288967 | 5.015460729 | 0.1018566 | 0.7536646 | 0.910408298 |
| ENSMUSG00000028843 | Sh3bgrl3 | protein_coding | 0.077135698 | 8.961784637 | 0.1016001 | 0.753964 | 0.910408298 |
| ENSMUSG00000015697 | Setdb1 | protein_coding | -0.145784266 | 3.872200855 | 0.1015888 | 0.7539773 | 0.910408298 |
| ENSMUSG00000049882 | Vcpkmt | protein_coding | 0.205233186 | 2.982139181 | 0.1015833 | 0.7539837 | 0.910408298 |
| ENSMUSG00000060579 | Fhit | protein_coding | 0.106957172 | 5.12763996 | 0.1015681 | 0.7540015 | 0.910408298 |
| ENSMUSG00000028416 | Bag1 | protein_coding | 0.086398 | 6.204112354 | 0.1015427 | 0.7540311 | 0.910408298 |
| ENSMUSG00000032504 | Pdcd6ip | protein_coding | -0.118508389 | 5.101560594 | 0.1014105 | 0.7541856 | 0.910492663 |
| ENSMUSG00000022861 | Dgkg | protein_coding | 0.083609899 | 6.621568105 | 0.1012447 | 0.7543796 | 0.910624602 |
| ENSMUSG00000026894 | Morn5 | protein_coding | 0.203771266 | 3.188588865 | 0.1009564 | 0.7547173 | 0.910929962 |
| ENSMUSG00000021819 | Zswim8 | protein_coding | -0.126212577 | 4.343796839 | 0.1008582 | 0.7548324 | 0.910966664 |
| ENSMUSG00000037747 | Phyhipl | protein_coding | -0.083953092 | 7.142047207 | 0.1004724 | 0.7552853 | 0.911410931 |
| ENSMUSG00000022812 | Gsk3b | protein_coding | -0.139702713 | 9.531558881 | 0.1007551 | 0.7555178 | 0.911527491 |
| ENSMUSG00000068742 | Cry2 | protein_coding | -0.098659903 | 5.848385492 | 0.1001066 | 0.7557158 | 0.911527491 |
| ENSMUSG00000020182 | Ddc | protein_coding | 0.100175787 | 5.347214123 | 0.1000773 | 0.7557502 | 0.911527491 |
| ENSMUSG00000071661 | Zbtb3 | protein_coding | 0.201403415 | 3.124408689 | 0.1000436 | 0.7557899 | 0.911527491 |
| ENSMUSG00000043300 | B3galnt1 | protein_coding | 0.091509385 | 6.205817635 | 0.0999723 | 0.7558739 | 0.911527491 |
| ENSMUSG00000002233 | Rhoc | protein_coding | 0.166744285 | 4.492495405 | 0.0998975 | 0.7559686 | 0.911527491 |
| ENSMUSG00000032968 | Inha | protein_coding | -0.283349548 | 2.437307666 | 0.0998863 | 0.7559753 | 0.911527491 |
| ENSMUSG00000043572 | Pars2 | protein_coding | 0.129731361 | 3.973158608 | 0.0996972 | 0.7561984 | 0.911649953 |
| ENSMUSG00000007817 | Zmiz1 | protein_coding | 0.103782111 | 5.827892092 | 0.0996565 | 0.7562464 | 0.911649953 |
| ENSMUSG00000020261 | Slc36a1 | protein_coding | 0.221061083 | 3.689174556 | 0.0995131 | 0.756481 | 0.911732116 |
| ENSMUSG00000055629 | B4galnt4 | protein_coding | 0.102510321 | 5.690109937 | 0.0993539 | 0.7566039 | 0.911732116 |
| ENSMUSG00000041757 | Plekha6 | protein_coding | 0.125252844 | 4.590886825 | 0.0991688 | 0.756823 | 0.911732116 |
| ENSMUSG00000026484 | Rnf2 | protein_coding | -0.108332908 | 5.037563545 | 0.0991644 | 0.7568282 | 0.911732116 |
| ENSMUSG00000034848 | Ttc21b | protein_coding | 0.25282978 | 2.747969089 | 0.0991356 | 0.7568623 | 0.911732116 |
| ENSMUSG00000022820 | Ndufb4 | protein_coding | 0.102072061 | 5.958522135 | 0.0990159 | 0.7570041 | 0.911732116 |
| ENSMUSG00000026111 | Unc50 | protein_coding | -0.099479789 | 5.589462597 | 0.0989678 | 0.7570611 | 0.911732116 |
| ENSMUSG00000024486 | Hbegf | protein_coding | -0.130347709 | 4.366426274 | 0.0989663 | 0.7570628 | 0.911732116 |
| ENSMUSG00000095845 | Gm5741 | protein_coding | -1.094687654 | 2.704893232 | 0.1025506 | 0.757112 | 0.911732116 |
| ENSMUSG00000035131 | Brinp3 | protein_coding | 0.111923178 | 5.172183855 | 0.0988137 | 0.7572437 | 0.911732116 |
| ENSMUSG00000037536 | Fbxo34 | protein_coding | 0.14837691 | 3.509491537 | 0.0988108 | 0.7572472 | 0.911732116 |
| ENSMUSG00000003545 | Fosb | protein_coding | 0.198493142 | 3.305642416 | 0.0983525 | 0.7577916 | 0.912037094 |
| ENSMUSG00000029726 | Mepce | protein_coding | -0.102286954 | 5.396212388 | 0.0983123 | 0.7578394 | 0.912037094 |
| ENSMUSG00000001158 | Snrnp27 | protein_coding | -0.088807434 | 8.072100003 | 0.0982784 | 0.7578797 | 0.912037094 |
| ENSMUSG00000066760 | Psg16 | protein_coding | 0.319948946 | 2.924307113 | 0.0984129 | 0.757895 | 0.912037094 |
| ENSMUSG00000038152 | 5033430I15Rik | lincRNA | 0.220555284 | 2.755846196 | 0.0981537 | 0.758028 | 0.912037094 |
| ENSMUSG00000054280 | Prr14l | protein_coding | -0.200645428 | 3.099167122 | 0.0981103 | 0.7580798 | 0.912037094 |
| ENSMUSG00000039428 | Tmem135 | protein_coding | 0.108212627 | 5.142708133 | 0.0980523 | 0.7581488 | 0.912037094 |
| ENSMUSG00000028709 | Mob3c | protein_coding | 0.369422843 | 2.685665053 | 0.0982217 | 0.7582001 | 0.912037094 |
| ENSMUSG00000045691 | Thtpa | protein_coding | 0.147888123 | 5.335795227 | 0.0979166 | 0.7583455 | 0.912037094 |
| ENSMUSG00000057895 | Zfp105 | protein_coding | -0.234737396 | 2.794361976 | 0.0978471 | 0.7583935 | 0.912037094 |
| ENSMUSG00000028710 | Atpaf1 | protein_coding | -0.177663289 | 3.757950926 | 0.0977872 | 0.7584649 | 0.912037094 |
| ENSMUSG00000027079 | Clp1 | protein_coding | 0.157038011 | 4.119925071 | 0.0977247 | 0.7585395 | 0.912037094 |
| ENSMUSG00000057572 | Zbtb8os | protein_coding | -0.123946226 | 4.87184687 | 0.0976713 | 0.7586032 | 0.912037094 |
| ENSMUSG00000030539 | Sema4b | protein_coding | -0.208939914 | 2.720905013 | 0.0975843 | 0.758707 | 0.912060021 |
| ENSMUSG00000031711 | Zfp330 | protein_coding | -0.080627361 | 5.953053569 | 0.0972999 | 0.759047 | 0.91218473 |
| ENSMUSG00000061080 | Lsamp | protein_coding | -0.064955859 | 7.72007222 | 0.0972642 | 0.7590897 | 0.91218473 |
| ENSMUSG00000045410 | Akr1e1 | protein_coding | -0.078330711 | 6.841923166 | 0.0971962 | 0.7591712 | 0.91218473 |
| ENSMUSG00000019518 | Ap4m1 | protein_coding | -0.209402961 | 2.850539078 | 0.097099 | 0.7592875 | 0.91218473 |
| ENSMUSG00000035293 | G2e3 | protein_coding | -0.264021154 | 2.867473345 | 0.0970622 | 0.7593316 | 0.91218473 |
| ENSMUSG00000036242 | 3632451O06Rik | protein_coding | -0.115843573 | 4.742168553 | 0.0970525 | 0.7593433 | 0.91218473 |
| ENSMUSG00000061111 | Mcrip1 | protein_coding | -0.100888591 | 7.096390871 | 0.0969093 | 0.7595213 | 0.91218473 |
| ENSMUSG00000020766 | Galk1 | protein_coding | 0.147295816 | 4.268817448 | 0.0968657 | 0.7595672 | 0.91218473 |
| ENSMUSG00000030942 | Thumpd1 | protein_coding | -0.124638107 | 5.519415912 | 0.096781 | 0.7596688 | 0.91218473 |
| ENSMUSG00000045404 | Kcnk13 | protein_coding | -0.225146123 | 3.372932918 | 0.0967424 | 0.7597216 | 0.91218473 |
| ENSMUSG00000071860 | 2900055J20Rik | protein_coding | -0.154569645 | 3.835069793 | 0.0967136 | 0.7597496 | 0.91218473 |
| ENSMUSG00000020485 | Supt4a | protein_coding | 0.073513275 | 6.644403082 | 0.0966477 | 0.7598288 | 0.91218473 |
| ENSMUSG00000046637 | Ttc34 | protein_coding | -0.307305735 | 2.504701623 | 0.0963892 | 0.7601393 | 0.912455674 |
| ENSMUSG00000039230 | Tbcd | protein_coding | 0.115980511 | 4.638417579 | 0.0961499 | 0.7604274 | 0.912484573 |
| ENSMUSG00000032356 | Rasgrf1 | protein_coding | 0.092016612 | 8.150316656 | 0.0961363 | 0.7604437 | 0.912484573 |
| ENSMUSG00000054226 | Tprkb | protein_coding | -0.067581342 | 7.424712086 | 0.095964 | 0.7606512 | 0.912484573 |
| ENSMUSG00000027574 | Nkain4 | protein_coding | -0.149843507 | 6.610725985 | 0.0963972 | 0.7607569 | 0.912484573 |
| ENSMUSG00000025956 | Mettl21a | protein_coding | 0.118008009 | 4.838935925 | 0.0958499 | 0.7607889 | 0.912484573 |
| ENSMUSG00000087674 | 4930447M23Rik | lincRNA | 0.177557389 | 3.176993293 | 0.0958483 | 0.7607908 | 0.912484573 |
| ENSMUSG00000004264 | Phb2 | protein_coding | 0.158830741 | 3.591811995 | 0.0958182 | 0.7608272 | 0.912484573 |
| ENSMUSG00000027522 | Stx16 | protein_coding | 0.136964039 | 5.559585882 | 0.0958251 | 0.7608422 | 0.912484573 |
| ENSMUSG00000025404 | R3hdm2 | protein_coding | -0.090266735 | 5.633320548 | 0.0956062 | 0.761083 | 0.91257683 |
| ENSMUSG00000001901 | Kcnh6 | protein_coding | 0.231923599 | 2.875576545 | 0.0955617 | 0.7611368 | 0.91257683 |
| ENSMUSG00000062981 | Mrpl42 | protein_coding | 0.091424281 | 8.561114125 | 0.0955311 | 0.7611738 | 0.91257683 |
| ENSMUSG00000090523 | Gypc | protein_coding | 0.37086656 | 1.789287632 | 0.095389 | 0.7613456 | 0.912681079 |

| ensembl_gene_id | external_gene_name | gene_biotype | log2 (fold change) | log2 (counts per million) | F-value | p-value | False discovery rate |
|---|---|---|---|---|---|---|---|
| ENSMUSG00000015112 | Slc25a13 | protein_coding | -0.192495431 | 3.432724529 | 0.0952617 | 0.7614996 | 0.912763975 |
| ENSMUSG00000029402 | Snrnp35 | protein_coding | -0.111579825 | 5.151248136 | 0.0951829 | 0.761595 | 0.912776563 |
| ENSMUSG00000026594 | Ralgps2 | protein_coding | -0.120172272 | 4.878117885 | 0.095042 | 0.7617657 | 0.912879335 |
| ENSMUSG00000018574 | Acadvl | protein_coding | 0.122733852 | 5.42607439 | 0.0948296 | 0.7620233 | 0.913010558 |
| ENSMUSG00000056666 | Retsat | protein_coding | -0.166557846 | 3.650956387 | 0.094744 | 0.7621271 | 0.913010558 |
| ENSMUSG00000016918 | Sulf1 | protein_coding | 0.213151756 | 3.857128097 | 0.094817 | 0.7621389 | 0.913010558 |
| ENSMUSG00000028133 | Rwdd3 | protein_coding | -0.110929775 | 4.843181953 | 0.0946718 | 0.7622148 | 0.913010558 |
| ENSMUSG00000027134 | Lpcat4 | protein_coding | -0.113955287 | 4.840404694 | 0.0944077 | 0.7625358 | 0.913279314 |
| ENSMUSG00000073481 | 2-Mar | protein_coding | -0.144595353 | 4.460534505 | 0.0943475 | 0.762609 | 0.913279314 |
| ENSMUSG00000028693 | Nasp | protein_coding | -0.130840935 | 4.348576499 | 0.0942745 | 0.7626978 | 0.913283935 |
| ENSMUSG00000003762 | Coq8b | protein_coding | -0.220454189 | 2.662124604 | 0.094122 | 0.7628836 | 0.913404717 |
| ENSMUSG00000036197 | Gxylt1 | protein_coding | -0.166187957 | 3.20701106 | 0.0940504 | 0.7629708 | 0.913407386 |
| ENSMUSG00000039623 | Ap5z1 | protein_coding | -0.196346921 | 3.859692997 | 0.093854 | 0.7632167 | 0.913600034 |
| ENSMUSG00000028890 | Mtf1 | protein_coding | -0.105810735 | 4.908268748 | 0.0937359 | 0.7633546 | 0.913663355 |
| ENSMUSG00000029028 | Lrrc47 | protein_coding | 0.22974917 | 2.925906026 | 0.0934801 | 0.7636671 | 0.913935752 |
| ENSMUSG00000000983 | Wfdc18 | protein_coding | -0.174225867 | 3.370804042 | 0.0932985 | 0.7638893 | 0.914077344 |
| ENSMUSG00000024211 | Grm8 | protein_coding | -0.202775201 | 4.150109232 | 0.0934457 | 0.7639555 | 0.914077344 |
| ENSMUSG00000034282 | Evpl | protein_coding | -0.164065268 | 3.819500099 | 0.0927918 | 0.7645106 | 0.914211327 |
| ENSMUSG00000068479 | Mfap1a | protein_coding | -0.122638458 | 4.009950903 | 0.0927708 | 0.7645364 | 0.914211327 |
| ENSMUSG00000030643 | Rab30 | protein_coding | 0.121342818 | 5.044029253 | 0.0927543 | 0.7645566 | 0.914211327 |
| ENSMUSG00000023236 | Scg5 | protein_coding | -0.077424282 | 9.418817556 | 0.0925642 | 0.7647903 | 0.914211327 |
| ENSMUSG00000051351 | Zfp46 | protein_coding | 0.163176856 | 3.520899645 | 0.092537 | 0.7648238 | 0.914211327 |
| ENSMUSG00000045896 | Paip2b | protein_coding | 0.089634387 | 5.696696941 | 0.0923831 | 0.7650131 | 0.914211327 |
| ENSMUSG00000023977 | Ubr2 | protein_coding | 0.154286371 | 3.830373238 | 0.0923329 | 0.7650749 | 0.914211327 |
| ENSMUSG00000055041 | Commd5 | protein_coding | 0.084339978 | 6.244817319 | 0.0922983 | 0.7651174 | 0.914211327 |
| ENSMUSG00000029625 | Cpsf4 | protein_coding | -0.211234359 | 3.032354332 | 0.0922339 | 0.7651968 | 0.914211327 |
| ENSMUSG00000021418 | Rpp40 | protein_coding | 0.143666684 | 3.772606096 | 0.0921709 | 0.7652745 | 0.914211327 |
| ENSMUSG00000022160 | Mettl3 | protein_coding | 0.161040434 | 3.93605508 | 0.0920869 | 0.765378 | 0.914211327 |
| ENSMUSG00000056014 | A430033K04Rik | protein_coding | -0.138367581 | 4.071842789 | 0.0919061 | 0.765601 | 0.914211327 |
| ENSMUSG00000038563 | Efl1 | protein_coding | -0.106427829 | 4.51451738 | 0.091839 | 0.7656839 | 0.914211327 |
| ENSMUSG00000020087 | Tysnd1 | protein_coding | -0.304196516 | 2.574259294 | 0.0916343 | 0.7659368 | 0.914211327 |
| ENSMUSG00000037509 | Arhgef4 | protein_coding | 0.088937464 | 6.287495244 | 0.0915535 | 0.7660368 | 0.914211327 |
| ENSMUSG00000038718 | Pbx3 | protein_coding | 0.118570401 | 5.39424119 | 0.0914448 | 0.7661713 | 0.914211327 |
| ENSMUSG00000050505 | Pcdh20 | protein_coding | 0.145748367 | 3.923139704 | 0.0914102 | 0.7662141 | 0.914211327 |
| ENSMUSG00000022807 | Osbpl11 | protein_coding | -0.15380459 | 3.436729191 | 0.0913427 | 0.7662977 | 0.914211327 |
| ENSMUSG00000048001 | Hes5 | protein_coding | 0.256673829 | 2.560221355 | 0.0912418 | 0.7664227 | 0.914211327 |
| ENSMUSG00000035274 | Tpbg | protein_coding | 0.217909833 | 3.203808165 | 0.0912304 | 0.7664368 | 0.914211327 |
| ENSMUSG00000027365 | Trpm7 | protein_coding | -0.136789868 | 3.924431 | 0.0912157 | 0.766455 | 0.914211327 |
| ENSMUSG00000024846 | Cst6 | protein_coding | -0.122735801 | 4.547371648 | 0.091174 | 0.7665068 | 0.914211327 |
| ENSMUSG00000032059 | Alg9 | protein_coding | 0.13909491 | 4.222902375 | 0.0911726 | 0.7665085 | 0.914211327 |
| ENSMUSG00000042050 | Wdr60 | protein_coding | -0.154057643 | 3.910344139 | 0.0911277 | 0.7665642 | 0.914211327 |
| ENSMUSG00000035469 | Rcbtb1 | protein_coding | -0.119169613 | 5.131448079 | 0.0909797 | 0.7667478 | 0.914211327 |
| ENSMUSG00000040856 | Dlk1 | protein_coding | -0.076295197 | 6.572223561 | 0.0908986 | 0.7668485 | 0.914211327 |
| ENSMUSG00000023118 | Sympk | protein_coding | -0.133248418 | 4.058834445 | 0.0908357 | 0.7669266 | 0.914211327 |
| ENSMUSG00000031697 | Orc6 | protein_coding | -0.099650632 | 6.86565501 | 0.0908006 | 0.7669765 | 0.914211327 |
| ENSMUSG00000054676 | 1600014C10Rik | protein_coding | 0.12133396 | 4.517361593 | 0.0907541 | 0.7670281 | 0.914211327 |
| ENSMUSG00000028621 | Cyb5rl | protein_coding | 0.148777973 | 4.490891648 | 0.0907086 | 0.7670846 | 0.914211327 |
| ENSMUSG00000038628 | Polr3k | protein_coding | 0.066845554 | 7.76036592 | 0.0907027 | 0.7670919 | 0.914211327 |
| ENSMUSG00000033365 | Ipo13 | protein_coding | -0.109255164 | 5.310807978 | 0.0906178 | 0.7671975 | 0.914211327 |
| ENSMUSG00000030720 | Cln3 | protein_coding | 0.159701271 | 4.129359323 | 0.0905806 | 0.7672438 | 0.914211327 |
| ENSMUSG00000026000 | Lancl1 | protein_coding | 0.071921649 | 6.946182531 | 0.0905615 | 0.7672676 | 0.914211327 |
| ENSMUSG00000039233 | Tbce | protein_coding | 0.124380192 | 5.077393966 | 0.0903985 | 0.7674705 | 0.914211327 |
| ENSMUSG00000026305 | Lrrfip1 | protein_coding | 0.125340442 | 4.908103017 | 0.0903668 | 0.76751 | 0.914211327 |
| ENSMUSG00000029416 | Slc15a4 | protein_coding | 0.125239438 | 4.24000187 | 0.0902986 | 0.767595 | 0.914211327 |
| ENSMUSG00000030849 | Fgfr2 | protein_coding | 0.073141865 | 6.727220766 | 0.0902485 | 0.7676574 | 0.914211327 |
| ENSMUSG00000047238 | Mageh1 | protein_coding | -0.077830729 | 6.191350289 | 0.0902434 | 0.7676638 | 0.914211327 |
| ENSMUSG00000042423 | Fbrs | protein_coding | -0.17123015 | 3.257937636 | 0.0902285 | 0.7676824 | 0.914211327 |
| ENSMUSG00000037519 | Ppfia1 | protein_coding | 0.150526246 | 3.816122203 | 0.0901219 | 0.7678153 | 0.914211327 |
| ENSMUSG00000062202 | Btbd9 | protein_coding | 0.11032906 | 5.808401651 | 0.0900617 | 0.7678904 | 0.914211327 |
| ENSMUSG00000087200 | Lrp8os3 | bidirectional_promote | -0.154619278 | 3.219258825 | 0.0900518 | 0.7679029 | 0.914211327 |
| ENSMUSG00000038879 | Nipal2 | protein_coding | 0.271194117 | 3.33169737 | 0.0902754 | 0.7679194 | 0.914211327 |
| ENSMUSG00000108644 | Gm44920 | lincRNA | -0.271863564 | 2.522787418 | 0.0900239 | 0.7679376 | 0.914211327 |
| ENSMUSG00000042426 | Dhx29 | protein_coding | 0.122117059 | 3.953861934 | 0.0899913 | 0.7679783 | 0.914211327 |
| ENSMUSG00000027006 | Dnajc10 | protein_coding | -0.078906073 | 5.867635521 | 0.0898224 | 0.7681894 | 0.914361344 |
| ENSMUSG00000039463 | Slc9a8 | protein_coding | 0.197829989 | 4.368778682 | 0.0896357 | 0.7687328 | 0.914467066 |
| ENSMUSG00000039384 | Dusp10 | protein_coding | 0.185962274 | 4.084571816 | 0.0893855 | 0.7687695 | 0.914467066 |
| ENSMUSG00000055116 | Arntl | protein_coding | -0.139198312 | 3.893684507 | 0.0893033 | 0.7688393 | 0.914467066 |
| ENSMUSG00000048281 | Dleu7 | protein_coding | 0.137102884 | 3.777676594 | 0.0892313 | 0.7689296 | 0.914467066 |
| ENSMUSG00000018995 | Nars2 | protein_coding | -0.150643677 | 3.687330034 | 0.0891948 | 0.7689754 | 0.914467066 |
| ENSMUSG00000029098 | Acox3 | protein_coding | -0.154215231 | 3.777063605 | 0.0891114 | 0.7690801 | 0.914467066 |
| ENSMUSG00000003355 | Fkbp11 | protein_coding | 0.182534026 | 3.242435879 | 0.0890732 | 0.769128 | 0.914467066 |
| ENSMUSG00000060681 | Slc9a6 | protein_coding | 0.087064801 | 5.87618612 | 0.0890063 | 0.7692121 | 0.914467066 |
| ENSMUSG00000030811 | Fbxl19 | protein_coding | 0.340396641 | 2.807103069 | 0.089171 | 0.7692455 | 0.914467066 |
| ENSMUSG00000041225 | Arhgap12 | protein_coding | -0.08985401 | 5.38148747 | 0.0889489 | 0.7692842 | 0.914467066 |
| ENSMUSG00000025495 | Ptdss2 | protein_coding | -0.117777929 | 5.097319969 | 0.0889376 | 0.7692984 | 0.914467066 |
| ENSMUSG00000024826 | Dpf2 | protein_coding | 0.085267885 | 6.602826591 | 0.0888345 | 0.769428 | 0.914467066 |
| ENSMUSG00000023106 | Denr | protein_coding | -0.069630893 | 7.38599362 | 0.0887539 | 0.7695294 | 0.914467066 |
| ENSMUSG00000028618 | Tmem59 | protein_coding | 0.067959268 | 9.205034219 | 0.0887451 | 0.7695405 | 0.914467066 |

| ensembl_gene_id | external_gene_name | gene_biotype | log2 (fold change) | log2 (counts per million) | F-value | p-value | False discovery rate |
|---|---|---|---|---|---|---|---|
| ENSMUSG00000071856 | Mcc | protein_coding | -0.352142394 | 2.394947163 | 0.0887071 | 0.7695945 | 0.914467066 |
| ENSMUSG00000039652 | Cpeb3 | protein_coding | -0.113025124 | 5.090136494 | 0.0886669 | 0.7696389 | 0.914467066 |
| ENSMUSG00000061132 | Blnk | protein_coding | -0.442592256 | 2.08573582 | 0.0886391 | 0.7697907 | 0.914546384 |
| ENSMUSG00000021185 | 9030617O03Rik | protein_coding | -0.190514388 | 4.068951454 | 0.088436 | 0.770047 | 0.914749773 |
| ENSMUSG00000060397 | Zfp128 | protein_coding | -0.203726232 | 3.274873164 | 0.088178 | 0.7702553 | 0.914896185 |
| ENSMUSG00000025102 | 3110040N11Rik | protein_coding | 0.081192664 | 6.076094014 | 0.0880994 | 0.7703545 | 0.914913006 |
| ENSMUSG00000022978 | Mis18a | protein_coding | 0.130106088 | 3.910452885 | 0.087518 | 0.7710904 | 0.91561812 |
| ENSMUSG00000020868 | Xylt2 | protein_coding | -0.139934817 | 3.705169652 | 0.0874958 | 0.7711185 | 0.91561812 |
| ENSMUSG00000086405 | 9330198N18Rik | lincRNA | 0.197523316 | 2.958387547 | 0.0872337 | 0.7714512 | 0.915893893 |
| ENSMUSG00000057278 | Snrpg | protein_coding | -0.123565055 | 5.204688223 | 0.0871787 | 0.7715211 | 0.915893893 |
| ENSMUSG00000052040 | Klf13 | protein_coding | -0.093988899 | 8.319600223 | 0.0871062 | 0.7716193 | 0.915909327 |
| ENSMUSG00000001962 | Fam50a | protein_coding | -0.166010949 | 4.370050922 | 0.0868886 | 0.7718961 | 0.9161028 |
| ENSMUSG00000054648 | Zfp869 | protein_coding | 0.165310188 | 3.365947779 | 0.0868394 | 0.7719527 | 0.9161028 |
| ENSMUSG00000040029 | Ipo8 | protein_coding | 0.097364517 | 5.01949478 | 0.0867495 | 0.7720672 | 0.916137578 |
| ENSMUSG00000036932 | Aifm1 | protein_coding | -0.108741327 | 5.357849465 | 0.0866262 | 0.7722243 | 0.916222846 |
| ENSMUSG00000055401 | Fbxo6 | protein_coding | -0.091466448 | 5.581647577 | 0.0864839 | 0.7724058 | 0.916337118 |
| ENSMUSG00000073856 | Iqck | protein_coding | -0.154050774 | 3.595928393 | 0.0861783 | 0.7727962 | 0.916352404 |
| ENSMUSG00000097080 | 1700086O06Rik | lincRNA | 0.1591485 | 3.712583447 | 0.0861155 | 0.7728765 | 0.916352404 |
| ENSMUSG00000020183 | Cpm | protein_coding | 0.157156328 | 3.680920737 | 0.0861063 | 0.7728883 | 0.916352404 |
| ENSMUSG00000036606 | Plxnb2 | protein_coding | 0.170150246 | 3.938736052 | 0.0860822 | 0.7729252 | 0.916352404 |
| ENSMUSG00000046567 | 4930430F08Rik | protein_coding | 0.115257447 | 4.367057172 | 0.0860238 | 0.7729939 | 0.916352404 |
| ENSMUSG00000032314 | Etfa | protein_coding | -0.065945828 | 6.758427469 | 0.0859784 | 0.773052 | 0.916352404 |
| ENSMUSG00000043384 | Gprasp1 | protein_coding | 0.203891527 | 3.13095495 | 0.0858459 | 0.7732217 | 0.916352404 |
| ENSMUSG00000031668 | Eif2ak3 | protein_coding | 0.221775613 | 3.672571185 | 0.0860258 | 0.7732269 | 0.916352404 |
| ENSMUSG00000038759 | Nup205 | protein_coding | 0.177187704 | 3.095527149 | 0.0857678 | 0.7733218 | 0.916352404 |
| ENSMUSG00000041460 | Cacna2d4 | protein_coding | -0.250294631 | 2.942424916 | 0.0856885 | 0.7734235 | 0.916352404 |
| ENSMUSG00000070544 | Top1 | protein_coding | 0.088802324 | 6.256643158 | 0.0856815 | 0.7734324 | 0.916352404 |
| ENSMUSG00000031767 | Nudt7 | protein_coding | 0.105205171 | 4.349246878 | 0.0855838 | 0.7735578 | 0.916352404 |
| ENSMUSG00000021136 | Smoc1 | protein_coding | -0.178358017 | 3.176641471 | 0.0855711 | 0.7735741 | 0.916352404 |
| ENSMUSG00000020453 | Patz1 | protein_coding | 0.17418824 | 3.463945877 | 0.0855418 | 0.7736117 | 0.916352404 |
| ENSMUSG00000029022 | Miip | protein_coding | -0.150499975 | 3.589943655 | 0.0853165 | 0.7739012 | 0.916594237 |
| ENSMUSG00000039252 | Lgi2 | protein_coding | -0.107020786 | 4.613793116 | 0.0850209 | 0.7742816 | 0.91694386 |
| ENSMUSG00000031791 | Tmem38a | protein_coding | 0.065869265 | 7.106816414 | 0.0849169 | 0.7744156 | 0.917001514 |
| ENSMUSG00000033955 | Tnks1bp1 | protein_coding | 0.266714059 | 2.644913488 | 0.084514 | 0.7749356 | 0.917516251 |
| ENSMUSG00000039069 | Mtg2 | protein_coding | 0.123521822 | 4.653762963 | 0.0842927 | 0.7752218 | 0.917754007 |
| ENSMUSG00000027487 | Cdk5rap1 | protein_coding | 0.132025467 | 4.290621699 | 0.0841789 | 0.7753691 | 0.917827402 |
| ENSMUSG00000011254 | Thg1l | protein_coding | 0.109713163 | 5.010979857 | 0.0839749 | 0.7756335 | 0.918039316 |
| ENSMUSG00000071655 | Ubxn1 | protein_coding | 0.067107418 | 6.632327151 | 0.0837989 | 0.7758619 | 0.918208533 |
| ENSMUSG00000074064 | Mlycd | protein_coding | -0.138169198 | 4.413333649 | 0.083586 | 0.7761384 | 0.918345226 |
| ENSMUSG00000048603 | Gm9828 | bidirectional_promote | 0.174488572 | 3.293833343 | 0.0835785 | 0.7761482 | 0.918345226 |
| ENSMUSG00000063410 | Stk24 | protein_coding | -0.122558027 | 3.879405367 | 0.0834239 | 0.7763493 | 0.918482086 |
| ENSMUSG00000097350 | 4732491K20Rik | lincRNA | -0.106193953 | 4.549704638 | 0.0833534 | 0.776441 | 0.918489592 |
| ENSMUSG00000049511 | Htr1b | protein_coding | -0.172917529 | 3.790007403 | 0.0832853 | 0.7765298 | 0.91849355 |
| ENSMUSG00000021235 | Coq6 | protein_coding | 0.118601711 | 5.401147564 | 0.0831374 | 0.7767226 | 0.918620527 |
| ENSMUSG00000064127 | Med14 | protein_coding | 0.12940705 | 4.923458654 | 0.0829058 | 0.7770248 | 0.918664846 |
| ENSMUSG00000020783 | Ncbp3 | protein_coding | -0.127697113 | 4.194204079 | 0.0828833 | 0.7770541 | 0.918664846 |
| ENSMUSG00000002578 | Ikzf4 | protein_coding | -0.12390784 | 4.382670822 | 0.0828545 | 0.7770917 | 0.918664846 |
| ENSMUSG00000030083 | Abtb1 | protein_coding | 0.118050636 | 6.035089861 | 0.0829331 | 0.7771018 | 0.918664846 |
| ENSMUSG00000040289 | Hey1 | protein_coding | 0.111850206 | 4.576717294 | 0.0826963 | 0.7772985 | 0.918780243 |
| ENSMUSG00000018387 | Shroom1 | protein_coding | -0.19640188 | 2.888356445 | 0.0826414 | 0.7773703 | 0.918780243 |
| ENSMUSG00000035212 | Leprot | protein_coding | 0.106111519 | 5.555849346 | 0.0825591 | 0.777478 | 0.918806563 |
| ENSMUSG00000014361 | Mertk | protein_coding | -0.250393149 | 2.70795975 | 0.0823613 | 0.7777371 | 0.918959679 |
| ENSMUSG00000061718 | Ppp1r1b | protein_coding | -0.216836976 | 2.587541766 | 0.0823297 | 0.7777785 | 0.918959679 |
| ENSMUSG00000032602 | Slc25a20 | protein_coding | 0.105071512 | 4.862579175 | 0.0822505 | 0.7778824 | 0.918981454 |
| ENSMUSG00000044912 | Syt16 | protein_coding | -0.110473926 | 4.870033124 | 0.0821199 | 0.7780537 | 0.91908288 |
| ENSMUSG00000024241 | Sos1 | protein_coding | 0.158313112 | 3.380003633 | 0.0819958 | 0.7782167 | 0.919140387 |
| ENSMUSG00000032185 | Carm1 | protein_coding | 0.127537412 | 4.119040385 | 0.0819527 | 0.7782733 | 0.919140387 |
| ENSMUSG00000068874 | Selenbp1 | protein_coding | -0.142912704 | 3.323188015 | 0.0817774 | 0.7785038 | 0.919311642 |
| ENSMUSG00000025880 | Smad7 | protein_coding | 0.162713613 | 3.443489442 | 0.0817016 | 0.7786035 | 0.919328367 |
| ENSMUSG00000055305 | Zfp93 | protein_coding | 0.217031975 | 2.681189644 | 0.0815438 | 0.7788114 | 0.919472855 |
| ENSMUSG00000031093 | Dock11 | protein_coding | -0.284642868 | 2.307063063 | 0.081263 | 0.7791818 | 0.919809166 |
| ENSMUSG00000021890 | Eaf1 | protein_coding | -0.183017884 | 3.042197161 | 0.0811926 | 0.7792746 | 0.91981781 |
| ENSMUSG00000006288 | Ttc5 | protein_coding | -0.070224521 | 6.543103202 | 0.0809869 | 0.7795466 | 0.919866864 |
| ENSMUSG00000042197 | Zfp451 | protein_coding | -0.086230165 | 5.442188885 | 0.0809685 | 0.7795709 | 0.919866864 |
| ENSMUSG00000010066 | Cacna2d2 | protein_coding | -0.131654467 | 4.557096645 | 0.0809534 | 0.7795909 | 0.919866864 |
| ENSMUSG00000007570 | Fance | protein_coding | -0.240726286 | 2.876004176 | 0.0809024 | 0.7796584 | 0.919866864 |
| ENSMUSG00000006215 | Zbtb17 | protein_coding | 0.164272903 | 4.625695513 | 0.0807728 | 0.7799876 | 0.920154357 |
| ENSMUSG00000063087 | Gm10125 | processed_transcript | -0.228831478 | 2.386144775 | 0.0805426 | 0.7801351 | 0.920227332 |
| ENSMUSG00000037455 | Slc18b1 | protein_coding | 0.172303714 | 3.00630673 | 0.0804032 | 0.7803201 | 0.920344634 |
| ENSMUSG00000002504 | Slc9a3r2 | protein_coding | 0.100960388 | 4.740477789 | 0.0802954 | 0.7804633 | 0.920412601 |
| ENSMUSG00000032743 | D430042O09Rik | protein_coding | -0.22169784 | 2.810748478 | 0.0801249 | 0.78069 | 0.920535523 |
| ENSMUSG00000029716 | Tfr2 | protein_coding | 0.1373073 | 4.00157809 | 0.0799237 | 0.7809578 | 0.920535523 |
| ENSMUSG00000074754 | Smim26 | protein_coding | 0.075185874 | 6.791577845 | 0.0799231 | 0.7809586 | 0.920535523 |
| ENSMUSG00000025465 | Echs1 | protein_coding | -0.060406619 | 7.770651461 | 0.0798512 | 0.7810544 | 0.920535523 |
| ENSMUSG00000025993 | Slc40a1 | protein_coding | -0.26292246 | 2.315840869 | 0.0798001 | 0.7811225 | 0.920535523 |
| ENSMUSG00000039130 | Zc3hc1 | protein_coding | 0.083200834 | 6.442612538 | 0.0797392 | 0.7812037 | 0.920535523 |
| ENSMUSG00000039531 | Zufsp | protein_coding | 0.124828465 | 3.712014587 | 0.0797004 | 0.7812555 | 0.920535523 |
| ENSMUSG00000035941 | Ibtk | protein_coding | 0.149247218 | 3.603984258 | 0.0796913 | 0.7812676 | 0.920535523 |

| ensembl_gene_id | external_gene_name | gene_biotype | log2 (fold change) | log2 (counts per million) | F-value | p-value | False discovery rate |
|---|---|---|---|---|---|---|---|
| ENSMUSG00000032724 | Abtb2 | protein_coding | 0.262686698 | 2.470819993 | 0.0796105 | 0.7813754 | 0.920535523 |
| ENSMUSG00000029458 | Brap | protein_coding | -0.098843843 | 4.55708519 | 0.0795636 | 0.7814381 | 0.920535523 |
| ENSMUSG00000021264 | Yy1 | protein_coding | -0.128021503 | 4.865081392 | 0.0795103 | 0.7815092 | 0.920535523 |
| ENSMUSG00000097119 | B230354K17Rik | lincRNA | 0.165334744 | 3.440706363 | 0.0792538 | 0.7818522 | 0.920838679 |
| ENSMUSG00000062257 | Opcml | protein_coding | 0.158738774 | 3.674555153 | 0.0791645 | 0.7819717 | 0.92087855 |
| ENSMUSG00000027650 | Tti1 | protein_coding | 0.123040926 | 4.096851908 | 0.0789251 | 0.7822925 | 0.921027977 |
| ENSMUSG00000006715 | Gmnn | protein_coding | -0.153555197 | 4.321437448 | 0.0789201 | 0.7823051 | 0.921027977 |
| ENSMUSG00000030615 | Tmem126a | protein_coding | -0.084754796 | 6.654889712 | 0.078812 | 0.7824444 | 0.921027977 |
| ENSMUSG00000037149 | Ddx1 | protein_coding | 0.066146859 | 7.647847429 | 0.0787867 | 0.7824783 | 0.921027977 |
| ENSMUSG00000002984 | Tomm40 | protein_coding | 0.121171912 | 4.313692104 | 0.0787505 | 0.7825269 | 0.921027977 |
| ENSMUSG00000003438 | Timm50 | protein_coding | 0.1062589 | 4.759467136 | 0.0786545 | 0.7826558 | 0.921078947 |
| ENSMUSG00000021831 | Ero1l | protein_coding | 0.215181938 | 3.134148566 | 0.0785258 | 0.7828289 | 0.921170169 |
| ENSMUSG00000066043 | Phactr4 | protein_coding | -0.197073082 | 2.921263514 | 0.0784343 | 0.782952 | 0.921170169 |
| ENSMUSG00000097353 | A430046D13Rik | lincRNA | -0.125640716 | 3.66636268 | 0.0783929 | 0.7830077 | 0.921170169 |
| ENSMUSG00000002825 | Qtrt1 | protein_coding | 0.094405676 | 5.198670867 | 0.0783421 | 0.783076 | 0.921170169 |
| ENSMUSG00000047139 | Cd24a | protein_coding | -0.133084287 | 4.013841585 | 0.078194 | 0.7832756 | 0.92126521 |
| ENSMUSG00000084215 | Gm11814 | processed_pseudoge | 0.193024117 | 3.054759638 | 0.0781215 | 0.7833733 | 0.92126521 |
| ENSMUSG00000038705 | Gmeb2 | protein_coding | 0.15710685 | 3.403363273 | 0.0780007 | 0.7835364 | 0.92126521 |
| ENSMUSG00000022464 | Slc38a4 | protein_coding | 0.229556176 | 2.634927549 | 0.0778397 | 0.7837538 | 0.92126521 |
| ENSMUSG00000030166 | Rad52 | protein_coding | 0.104070258 | 4.579699803 | 0.0778357 | 0.7837593 | 0.92126521 |
| ENSMUSG00000017843 | Ppp2r5c | protein_coding | 0.070541989 | 7.854451144 | 0.0777896 | 0.7838215 | 0.92126521 |
| ENSMUSG00000075271 | Ttc30a1 | protein_coding | -0.216587072 | 2.943196988 | 0.077753 | 0.783871 | 0.92126521 |
| ENSMUSG00000063334 | Krr1 | protein_coding | 0.111001848 | 4.937537707 | 0.0777174 | 0.7839192 | 0.92126521 |
| ENSMUSG00000050619 | Zscan29 | protein_coding | -0.227756385 | 2.68258433 | 0.0777109 | 0.7839279 | 0.92126521 |
| ENSMUSG00000032456 | Nmnat3 | protein_coding | -0.207371153 | 2.809270267 | 0.0775973 | 0.7840816 | 0.921323833 |
| ENSMUSG00000045374 | Wdr81 | protein_coding | -0.218550662 | 2.921892044 | 0.0775474 | 0.7841491 | 0.921323833 |
| ENSMUSG00000045822 | Zswim3 | protein_coding | -0.151036934 | 3.408235743 | 0.07725 | 0.7845523 | 0.921663046 |
| ENSMUSG00000074748 | Atxn7l3b | protein_coding | -0.060241375 | 7.701684118 | 0.077208 | 0.7846093 | 0.921663046 |
| ENSMUSG00000069135 | Fgfr1op | protein_coding | -0.145287707 | 4.019923772 | 0.0770728 | 0.7847929 | 0.921699113 |
| ENSMUSG00000020902 | Ntn1 | protein_coding | -0.275444923 | 2.661444907 | 0.0769663 | 0.784943 | 0.921699113 |
| ENSMUSG00000042632 | Pla2g6 | protein_coding | -0.09472154 | 5.294236621 | 0.0769349 | 0.7849803 | 0.921699113 |
| ENSMUSG00000055150 | Zfp78 | protein_coding | 0.146764645 | 3.449240101 | 0.0769154 | 0.7850069 | 0.921699113 |
| ENSMUSG00000030598 | Fbxo17 | protein_coding | 0.143657446 | 4.047245717 | 0.0768701 | 0.7850686 | 0.921699113 |
| ENSMUSG00000024410 | 3110002H16Rik | protein_coding | 0.096397533 | 5.443924977 | 0.0767919 | 0.7851749 | 0.921723298 |
| ENSMUSG00000055254 | Ntrk2 | protein_coding | -0.073315668 | 7.519889772 | 0.0764648 | 0.7856208 | 0.921965578 |
| ENSMUSG00000059005 | Hnrnpa3 | protein_coding | -0.109436112 | 3.884910127 | 0.0764208 | 0.7856807 | 0.921965578 |
| ENSMUSG00000031167 | Rbm3 | protein_coding | 0.107989771 | 4.087314132 | 0.0763537 | 0.7857724 | 0.921965578 |
| ENSMUSG00000090247 | Bloc1s1 | protein_coding | -0.095175777 | 5.050698492 | 0.076275 | 0.7858799 | 0.921965578 |
| ENSMUSG00000024008 | Cpne5 | protein_coding | 0.091156991 | 7.202802253 | 0.076255 | 0.7859129 | 0.921965578 |
| ENSMUSG00000067194 | Eif1ax | protein_coding | 0.065511869 | 7.481456596 | 0.0762424 | 0.7859244 | 0.921965578 |
| ENSMUSG00000022013 | Dnajc15 | protein_coding | 0.069458325 | 7.804727818 | 0.0761669 | 0.7860276 | 0.921965578 |
| ENSMUSG00000018921 | Pelp1 | protein_coding | 0.128268003 | 3.524670135 | 0.0761379 | 0.7860672 | 0.921965578 |
| ENSMUSG00000032841 | Prr5l | protein_coding | -0.119702861 | 5.449710752 | 0.0759152 | 0.7863778 | 0.922229294 |
| ENSMUSG00000052962 | Mrpl35 | protein_coding | 0.058224946 | 7.399186012 | 0.0757278 | 0.786629 | 0.922423255 |
| ENSMUSG00000021326 | Trim27 | protein_coding | -0.142906182 | 3.769012894 | 0.0756198 | 0.7867771 | 0.922462313 |
| ENSMUSG00000024997 | Prdx3 | protein_coding | 0.05872031 | 8.346244959 | 0.0755596 | 0.7868598 | 0.922462313 |
| ENSMUSG00000023805 | Synj2 | protein_coding | -0.108594856 | 4.969783912 | 0.075516 | 0.7869196 | 0.922462313 |
| ENSMUSG00000025221 | Kcnip2 | protein_coding | -0.075308572 | 6.323218804 | 0.075279 | 0.7872455 | 0.922621418 |
| ENSMUSG00000025436 | Atp23 | protein_coding | 0.17258273 | 3.030248704 | 0.0752416 | 0.7872969 | 0.922621418 |
| ENSMUSG00000052214 | Opa3 | protein_coding | -0.068233599 | 6.439299332 | 0.0751701 | 0.7873954 | 0.922621418 |
| ENSMUSG00000000605 | Clcn4 | protein_coding | 0.065081465 | 7.351839235 | 0.0751678 | 0.7873986 | 0.922621418 |
| ENSMUSG00000034098 | Fstl5 | protein_coding | 0.095063868 | 5.253125928 | 0.0750689 | 0.7875348 | 0.922680522 |
| ENSMUSG00000040549 | Ckap5 | protein_coding | 0.068324605 | 6.407334351 | 0.0749567 | 0.7876895 | 0.922761212 |
| ENSMUSG00000038351 | Sgsm2 | protein_coding | 0.104426736 | 4.882327647 | 0.0747543 | 0.7879688 | 0.92287354 |
| ENSMUSG00000053550 | Shisa7 | protein_coding | 0.119866489 | 4.438449316 | 0.074736 | 0.787994 | 0.92287354 |
| ENSMUSG00000031834 | Pik3r2 | protein_coding | 0.101701988 | 4.690326057 | 0.0747007 | 0.7880429 | 0.92287354 |
| ENSMUSG00000041608 | Entpd3 | protein_coding | -0.168922695 | 3.918803749 | 0.0745621 | 0.78824 | 0.923003895 |
| ENSMUSG00000032259 | Drd2 | protein_coding | -0.121373634 | 5.144395593 | 0.0744292 | 0.7884239 | 0.923118734 |
| ENSMUSG00000032869 | Psmf1 | protein_coding | 0.088582 | 6.785877124 | 0.0743095 | 0.7885897 | 0.923212318 |
| ENSMUSG00000051113 | Fam71e1 | protein_coding | 0.215334673 | 2.910561462 | 0.0741928 | 0.7887459 | 0.923294623 |
| ENSMUSG00000034858 | Fam214a | protein_coding | 0.116875478 | 4.271265925 | 0.0739959 | 0.7890191 | 0.923513928 |
| ENSMUSG00000003131 | Pafah1b2 | protein_coding | -0.066591972 | 7.544336209 | 0.0738127 | 0.7892738 | 0.923711447 |
| ENSMUSG00000053768 | Chchd3 | protein_coding | -0.057861111 | 6.995113663 | 0.0735845 | 0.7895913 | 0.923977269 |
| ENSMUSG00000021541 | Trpc7 | protein_coding | -0.097731996 | 5.12309614 | 0.0734754 | 0.7897434 | 0.923977269 |
| ENSMUSG00000072235 | Tuba1a | protein_coding | 0.080044903 | 9.916995326 | 0.0734644 | 0.7897587 | 0.923977269 |
| ENSMUSG00000061607 | Mdc1 | protein_coding | -0.309019788 | 2.20541745 | 0.0733344 | 0.78994 | 0.924088879 |
| ENSMUSG00000015126 | Tsr3 | protein_coding | 0.104819981 | 4.581790649 | 0.0732483 | 0.7900603 | 0.924129023 |
| ENSMUSG00000038028 | Tigar | protein_coding | -0.115490417 | 4.351674564 | 0.0729258 | 0.7905112 | 0.924500074 |
| ENSMUSG00000037089 | Slc35b2 | protein_coding | 0.108441638 | 5.014499288 | 0.0728799 | 0.7905754 | 0.924500074 |
| ENSMUSG00000024754 | Tmem2 | protein_coding | 0.184719481 | 3.101771873 | 0.072837 | 0.7906354 | 0.924500074 |
| ENSMUSG00000046909 | Tefm | protein_coding | -0.093128907 | 5.180163379 | 0.0726335 | 0.7909207 | 0.924733094 |
| ENSMUSG00000025980 | Hspd1 | protein_coding | 0.067442883 | 7.043086391 | 0.0725448 | 0.7910452 | 0.924778124 |
| ENSMUSG00000001305 | Rrp15 | protein_coding | -0.112764335 | 4.420875696 | 0.072327 | 0.7913512 | 0.925035245 |
| ENSMUSG00000022151 | Ttc33 | protein_coding | 0.095121622 | 4.688553336 | 0.0720331 | 0.7917648 | 0.925393535 |
| ENSMUSG00000040479 | Dgkz | protein_coding | -0.087759741 | 6.007074155 | 0.071987 | 0.7918298 | 0.925393535 |
| ENSMUSG00000047694 | Yipf6 | protein_coding | -0.134771103 | 4.256792494 | 0.0715029 | 0.7925133 | 0.926091692 |
| ENSMUSG00000027878 | Notch2 | protein_coding | 0.179646002 | 3.242617903 | 0.0714345 | 0.7926102 | 0.926104239 |
| ENSMUSG00000022664 | Slc35a5 | protein_coding | -0.08584079 | 5.115606589 | 0.0711239 | 0.7930503 | 0.926455102 |

| ensembl_gene_id | external_gene_name | gene_biotype | log2 (fold change) | log2 (counts per million) | F-value | p-value | False discovery rate |
|---|---|---|---|---|---|---|---|
| ENSMUSG00000026640 | Plxna2 | protein_coding | -0.106515088 | 4.949512398 | 0.071095 | 0.7930913 | 0.926455102 |
| ENSMUSG00000040918 | Slc19a2 | protein_coding | -0.167536361 | 3.250004622 | 0.0710152 | 0.7932046 | 0.926455102 |
| ENSMUSG00000031239 | Itm2a | protein_coding | 0.089987529 | 5.464426864 | 0.0709796 | 0.7932551 | 0.926455102 |
| ENSMUSG00000024045 | Akap8 | protein_coding | -0.097052351 | 4.802518512 | 0.070736 | 0.7936015 | 0.926758997 |
| ENSMUSG00000036061 | Smug1 | protein_coding | 0.14899529 | 3.632797409 | 0.0706322 | 0.7937491 | 0.926830746 |
| ENSMUSG00000030138 | Bms1 | protein_coding | 0.147879284 | 3.655076753 | 0.0704589 | 0.7939962 | 0.926941715 |
| ENSMUSG00000030835 | Nomo1 | protein_coding | 0.073013519 | 6.283509334 | 0.0704446 | 0.7940166 | 0.926941715 |
| ENSMUSG00000062309 | Rpp25 | protein_coding | 0.10957534 | 4.69093654 | 0.0702969 | 0.7942273 | 0.927087027 |
| ENSMUSG00000011158 | Brf1 | protein_coding | -0.128701441 | 3.943897732 | 0.0700879 | 0.7945259 | 0.927335008 |
| ENSMUSG00000000552 | Zfp385a | protein_coding | 0.12440563 | 4.803536637 | 0.0699186 | 0.7947681 | 0.927499927 |
| ENSMUSG00000034201 | Gas2l1 | protein_coding | -0.171654283 | 3.767728582 | 0.0698724 | 0.7948397 | 0.927499927 |
| ENSMUSG00000028137 | Celf3 | protein_coding | 0.10680024 | 5.27909203 | 0.0696011 | 0.7952233 | 0.927846765 |
| ENSMUSG00000103428 | C130093G08Rik | TEC | -0.18559922 | 4.64571464 | 0.0698445 | 0.7953382 | 0.927880193 |
| ENSMUSG00000011114 | Tbrg1 | protein_coding | 0.072696537 | 7.01306031 | 0.0693165 | 0.7956322 | 0.927920834 |
| ENSMUSG00000044337 | Ackr3 | protein_coding | -0.270518925 | 2.357359568 | 0.0692937 | 0.795665 | 0.927920834 |
| ENSMUSG00000014503 | Pkd2l2 | protein_coding | -0.076097323 | 5.546689647 | 0.0692519 | 0.7957251 | 0.927920834 |
| ENSMUSG00000020754 | Gm20754 | lincRNA | -0.273535919 | 2.397947304 | 0.0692374 | 0.795746 | 0.927920834 |
| ENSMUSG00000020268 | Lyrm7 | protein_coding | 0.072528902 | 5.825074949 | 0.0691968 | 0.7958045 | 0.927920834 |
| ENSMUSG00000047976 | Kcna1 | protein_coding | 0.114478976 | 4.522790484 | 0.0689127 | 0.7962139 | 0.927969372 |
| ENSMUSG00000027612 | Mmp24 | protein_coding | -0.169572116 | 3.06383252 | 0.0689027 | 0.7962284 | 0.927969372 |
| ENSMUSG00000091002 | Tcerg1l | protein_coding | -0.125563526 | 4.286745588 | 0.0688993 | 0.7962333 | 0.927969372 |
| ENSMUSG00000022179 | 4931414P19Rik | protein_coding | -0.216426702 | 2.714418146 | 0.0688096 | 0.7963628 | 0.927969372 |
| ENSMUSG00000056938 | Acbd4 | protein_coding | 0.111163875 | 5.565065564 | 0.0687996 | 0.7963826 | 0.927969372 |
| ENSMUSG00000024286 | Ccny | protein_coding | 0.121895683 | 5.07855017 | 0.0687798 | 0.7964112 | 0.927969372 |
| ENSMUSG00000045471 | Hcrt | protein_coding | -0.161819205 | 10.01304624 | 0.069717 | 0.7964502 | 0.927969372 |
| ENSMUSG00000039717 | Ralyl | protein_coding | -0.061184546 | 7.948291506 | 0.0686836 | 0.7965448 | 0.927979043 |
| ENSMUSG00000035439 | Haus8 | protein_coding | 0.088122793 | 5.555559497 | 0.068558 | 0.7967265 | 0.928090131 |
| ENSMUSG00000074698 | Csnk2a1 | protein_coding | 0.081221009 | 5.274515442 | 0.0684711 | 0.7968523 | 0.928136187 |
| ENSMUSG00000050103 | Agmo | protein_coding | -0.202849301 | 2.957131236 | 0.0679745 | 0.7975728 | 0.928687555 |
| ENSMUSG00000016409 | Nkap | protein_coding | 0.087445381 | 5.150353516 | 0.0679003 | 0.7976806 | 0.928687555 |
| ENSMUSG00000012429 | Mplkip | protein_coding | -0.205429923 | 2.538806212 | 0.0678665 | 0.7977298 | 0.928687555 |
| ENSMUSG00000032547 | Ryk | protein_coding | -0.151514906 | 3.831005932 | 0.067859 | 0.7977407 | 0.928687555 |
| ENSMUSG00000054256 | Msi1 | protein_coding | -0.13889545 | 4.022097204 | 0.0678474 | 0.7977576 | 0.928687555 |
| ENSMUSG00000032342 | Mto1 | protein_coding | 0.128245823 | 3.61128198 | 0.0676477 | 0.7980484 | 0.928698732 |
| ENSMUSG00000027942 | 4933434E20Rik | protein_coding | -0.0568985 | 8.603960631 | 0.0676141 | 0.7980974 | 0.928698732 |
| ENSMUSG00000052428 | Tmco1 | protein_coding | 0.067078502 | 7.076710616 | 0.0675898 | 0.7981329 | 0.928698732 |
| ENSMUSG00000037868 | Egr2 | protein_coding | 0.210849731 | 2.595432528 | 0.0674939 | 0.7982726 | 0.928698732 |
| ENSMUSG00000029994 | Anxa4 | protein_coding | -0.194303814 | 2.78056413 | 0.067446 | 0.7983426 | 0.928698732 |
| ENSMUSG00000036046 | 5031439G07Rik | protein_coding | 0.087922235 | 5.875616793 | 0.0674078 | 0.7983984 | 0.928698732 |
| ENSMUSG00000026077 | Npas2 | protein_coding | 0.119756121 | 3.989062889 | 0.0673978 | 0.7984129 | 0.928698732 |
| ENSMUSG00000041231 | Ublcp1 | protein_coding | 0.075436858 | 5.847100931 | 0.0673669 | 0.7984581 | 0.928698732 |
| ENSMUSG00000036977 | Anapc10 | protein_coding | -0.094438101 | 5.966676204 | 0.067157 | 0.7987702 | 0.928949916 |
| ENSMUSG00000042558 | Adprhl2 | protein_coding | 0.135787479 | 3.937292687 | 0.0670404 | 0.7989356 | 0.928949916 |
| ENSMUSG00000028152 | Tspan5 | protein_coding | 0.067183563 | 6.044116231 | 0.06702 | 0.7989655 | 0.928949916 |
| ENSMUSG00000021431 | Snrnp48 | protein_coding | -0.110378683 | 4.203297408 | 0.0668907 | 0.7991549 | 0.928949916 |
| ENSMUSG00000074825 | Itpripl1 | protein_coding | -0.165895408 | 3.798788421 | 0.0668889 | 0.7991629 | 0.928949916 |
| ENSMUSG00000026765 | Lypd6b | protein_coding | 0.110397084 | 4.229487079 | 0.0668652 | 0.7991924 | 0.928949916 |
| ENSMUSG00000024491 | Rbm27 | protein_coding | -0.089434804 | 5.31306942 | 0.0666274 | 0.7995414 | 0.928970454 |
| ENSMUSG00000005262 | Ufd1 | protein_coding | 0.077643071 | 7.572505418 | 0.0666268 | 0.7995476 | 0.928970454 |
| ENSMUSG00000039450 | Dcxr | protein_coding | -0.126638377 | 4.156763777 | 0.0666081 | 0.7995698 | 0.928970454 |
| ENSMUSG00000041298 | Katnal1 | protein_coding | -0.077661917 | 6.237262926 | 0.0665931 | 0.7995919 | 0.928970454 |
| ENSMUSG00000021725 | Parp8 | protein_coding | -0.139602337 | 3.485293454 | 0.0664288 | 0.7998335 | 0.928970454 |
| ENSMUSG00000034566 | Atp5h | protein_coding | 0.069760314 | 8.487546857 | 0.0664007 | 0.7998748 | 0.928970454 |
| ENSMUSG00000014177 | Tvp23b | protein_coding | -0.06703828 | 6.339747396 | 0.0663841 | 0.7998993 | 0.928970454 |
| ENSMUSG00000015247 | Nipsnap3b | protein_coding | -0.072268607 | 6.272602591 | 0.0663354 | 0.7999709 | 0.928970454 |
| ENSMUSG00000005362 | Crbn | protein_coding | -0.083908583 | 6.155560112 | 0.0662758 | 0.8000587 | 0.928970454 |
| ENSMUSG00000045854 | Lyrm2 | protein_coding | 0.083388612 | 6.16688225 | 0.0662655 | 0.800074 | 0.928970454 |
| ENSMUSG00000045078 | Rnf216 | protein_coding | 0.101202842 | 5.031410019 | 0.066102 | 0.800315 | 0.92915 |
| ENSMUSG00000039043 | Arpin | protein_coding | -0.101448957 | 5.12965249 | 0.0658048 | 0.800754 | 0.929559339 |
| ENSMUSG00000024190 | Dusp1 | protein_coding | 0.109796175 | 4.473585659 | 0.0656247 | 0.8010206 | 0.929768422 |
| ENSMUSG00000027905 | Ddx20 | protein_coding | -0.174791316 | 3.076885073 | 0.0654914 | 0.8012182 | 0.929839717 |
| ENSMUSG00000056204 | Pgpep1 | protein_coding | -0.091489346 | 5.433848232 | 0.0654666 | 0.801255 | 0.929839717 |
| ENSMUSG00000069539 | Scyl2 | protein_coding | -0.178579655 | 2.866869818 | 0.0651761 | 0.8016864 | 0.930091627 |
| ENSMUSG00000008226 | Scrn3 | protein_coding | -0.106397868 | 5.023483344 | 0.0651687 | 0.8016973 | 0.930091627 |
| ENSMUSG00000030122 | Ptms | protein_coding | -0.08620423 | 6.578648641 | 0.065123 | 0.8017706 | 0.930091627 |
| ENSMUSG00000030089 | Slc41a3 | protein_coding | 0.16079841 | 3.496801846 | 0.0650495 | 0.8018747 | 0.930091627 |
| ENSMUSG00000039253 | Fn3krp | protein_coding | 0.226169429 | 3.490451767 | 0.0652438 | 0.8019045 | 0.930091627 |
| ENSMUSG00000032601 | Prkar2a | protein_coding | 0.067172601 | 5.871923199 | 0.064924 | 0.8020616 | 0.93017348 |
| ENSMUSG00000042535 | Gtpbp1 | protein_coding | -0.133896352 | 3.944895231 | 0.0645419 | 0.8026317 | 0.930734287 |
| ENSMUSG00000022414 | Tab1 | protein_coding | -0.120019661 | 5.193363112 | 0.0644047 | 0.802842 | 0.930877757 |
| ENSMUSG00000011148 | Adssl1 | protein_coding | 0.089156175 | 4.719641668 | 0.0641456 | 0.803225 | 0.931131381 |
| ENSMUSG00000018774 | Cd68 | protein_coding | -0.106835114 | 5.002781027 | 0.0640627 | 0.8033493 | 0.931131381 |
| ENSMUSG00000019998 | Stx7 | protein_coding | 0.06204761 | 7.730290443 | 0.0639337 | 0.803543 | 0.931131381 |
| ENSMUSG00000108348 | Gm42372 | protein_coding | 0.06995385 | 6.623810946 | 0.0639321 | 0.8035453 | 0.931131381 |
| ENSMUSG00000046707 | Csnk2a2 | protein_coding | 0.082386546 | 5.413038128 | 0.0638324 | 0.8036951 | 0.931131381 |
| ENSMUSG00000042364 | Snx18 | protein_coding | 0.141245597 | 3.823215141 | 0.0637075 | 0.803883 | 0.931131381 |
| ENSMUSG00000034928 | Rnf44 | protein_coding | -0.082408894 | 5.358573775 | 0.063617 | 0.8040193 | 0.931131381 |
| ENSMUSG00000025172 | Ankrd2 | protein_coding | -0.177962629 | 2.945274011 | 0.0635288 | 0.8041521 | 0.931131381 |

| ensembl_gene_id | external_gene_name | gene_biotype | log2 (fold change) | log2 (counts per million) | F-value | p-value | False discovery rate |
|---|---|---|---|---|---|---|---|
| ENSMUSG00000032301 | Psma4 | protein_coding | -0.065920273 | 7.577675327 | 0.0635251 | 0.8041577 | 0.931131381 |
| ENSMUSG00000024601 | Isoc1 | protein_coding | 0.098803991 | 5.166928495 | 0.0635173 | 0.8041694 | 0.931131381 |
| ENSMUSG00000045932 | Ifit2 | protein_coding | 0.145882017 | 3.092937271 | 0.0634978 | 0.8041988 | 0.931131381 |
| ENSMUSG00000035495 | Tstd2 | protein_coding | -0.135814225 | 3.222864669 | 0.0634416 | 0.8042837 | 0.931131381 |
| ENSMUSG00000029603 | Dtx1 | protein_coding | -0.087175177 | 5.30201011 | 0.0633537 | 0.8044163 | 0.931131381 |
| ENSMUSG00000035048 | Anapc13 | protein_coding | -0.082839503 | 7.441151497 | 0.063291 | 0.8045296 | 0.931131381 |
| ENSMUSG00000031379 | Pir | protein_coding | -0.149225671 | 3.804556121 | 0.063246 | 0.8045788 | 0.931131381 |
| ENSMUSG00000059552 | Trp53 | protein_coding | 0.083046217 | 5.633688644 | 0.0631508 | 0.8047228 | 0.931131381 |
| ENSMUSG00000026942 | Traf2 | protein_coding | -0.105222138 | 4.716629366 | 0.0630599 | 0.8048603 | 0.931131381 |
| ENSMUSG00000001052 | Sec24b | protein_coding | -0.084958199 | 4.881880454 | 0.063031 | 0.804904 | 0.931131381 |
| ENSMUSG00000038535 | Zfp280d | protein_coding | -0.159416777 | 3.146602103 | 0.0630234 | 0.8049154 | 0.931131381 |
| ENSMUSG00000039860 | Srrm3 | protein_coding | -0.159056478 | 2.993255815 | 0.0629146 | 0.8050802 | 0.931131381 |
| ENSMUSG00000078235 | Fam43b | protein_coding | -0.16161646 | 2.797049473 | 0.062906 | 0.8050932 | 0.931131381 |
| ENSMUSG00000085793 | Lin52 | protein_coding | -0.075559628 | 5.72416178 | 0.0627641 | 0.8053084 | 0.931131381 |
| ENSMUSG00000034321 | Exosc1 | protein_coding | -0.075979112 | 6.37568515 | 0.0627399 | 0.8053451 | 0.931131381 |
| ENSMUSG00000001666 | Ddt | protein_coding | -0.080348755 | 5.151267093 | 0.0627066 | 0.8053956 | 0.931131381 |
| ENSMUSG00000038949 | Cnst | protein_coding | 0.122714022 | 4.301464429 | 0.062695 | 0.8054132 | 0.931131381 |
| ENSMUSG00000033400 | Agl | protein_coding | 0.113124478 | 3.815284469 | 0.0626898 | 0.8054211 | 0.931131381 |
| ENSMUSG00000097068 | Gm26760 | lincRNA | 0.192760216 | 2.564848767 | 0.0625457 | 0.8056401 | 0.931131381 |
| ENSMUSG00000043733 | Ptpn11 | protein_coding | -0.09856805 | 5.782791783 | 0.0624702 | 0.8057599 | 0.931131381 |
| ENSMUSG00000002679 | Med6 | protein_coding | -0.084513899 | 5.727165433 | 0.0623757 | 0.8058985 | 0.931131381 |
| ENSMUSG00000001280 | Sp1 | protein_coding | 0.122121682 | 3.948194956 | 0.0623615 | 0.8059202 | 0.931131381 |
| ENSMUSG00000026154 | Sdhaf4 | protein_coding | 0.070242246 | 8.414364103 | 0.0623306 | 0.8059672 | 0.931131381 |
| ENSMUSG00000022244 | Amacr | protein_coding | -0.125185679 | 4.10322344 | 0.0623227 | 0.8059792 | 0.931131381 |
| ENSMUSG00000047409 | Ctdspl | protein_coding | -0.088246753 | 4.700609006 | 0.0623088 | 0.8060004 | 0.931131381 |
| ENSMUSG00000090946 | Ccdc71l | protein_coding | 0.214458148 | 2.466316339 | 0.0623058 | 0.8060049 | 0.931131381 |
| ENSMUSG00000027667 | Zfp639 | protein_coding | -0.082903986 | 5.600760563 | 0.0622406 | 0.8061042 | 0.931146032 |
| ENSMUSG00000035878 | Hykk | protein_coding | 0.301468686 | 2.693860234 | 0.0622024 | 0.8063609 | 0.93134252 |
| ENSMUSG00000030082 | Sec61a1 | protein_coding | 0.084162234 | 4.915062763 | 0.0618795 | 0.8066553 | 0.931582508 |
| ENSMUSG00000044017 | Adgrd1 | protein_coding | 0.145451796 | 2.95835921 | 0.0617627 | 0.806834 | 0.931688802 |
| ENSMUSG00000074555 | Gm10714 | antisense_RNA | -0.210074296 | 2.85445708 | 0.0615653 | 0.8071362 | 0.931937699 |
| ENSMUSG00000012422 | Tmem167 | protein_coding | 0.066682199 | 7.382104096 | 0.061166 | 0.8077493 | 0.932457738 |
| ENSMUSG00000038290 | Smg6 | protein_coding | 0.095861505 | 4.756265061 | 0.061159 | 0.8077601 | 0.932457738 |
| ENSMUSG00000000538 | Tom1l2 | protein_coding | -0.076237611 | 7.247091143 | 0.0609699 | 0.8080562 | 0.932527404 |
| ENSMUSG00000020604 | Arsg | protein_coding | -0.129870422 | 4.808082423 | 0.0609611 | 0.808083 | 0.932527404 |
| ENSMUSG00000028914 | Casp9 | protein_coding | 0.235673188 | 2.50036652 | 0.0609209 | 0.8081266 | 0.932527404 |
| ENSMUSG00000037664 | Cdkn1c | protein_coding | 0.243646663 | 2.328990306 | 0.0608802 | 0.8081894 | 0.932527404 |
| ENSMUSG00000060206 | Zfp462 | protein_coding | 0.107300212 | 4.066285577 | 0.0607742 | 0.8083529 | 0.932527404 |
| ENSMUSG00000051403 | Ppp1r37 | protein_coding | 0.091351609 | 4.556805796 | 0.0607524 | 0.8083865 | 0.932527404 |
| ENSMUSG00000044573 | Acp1 | protein_coding | 0.061340088 | 6.066362044 | 0.0607258 | 0.8084275 | 0.932527404 |
| ENSMUSG00000059851 | Kmt5c | protein_coding | -0.171456821 | 3.019809196 | 0.0606042 | 0.8086154 | 0.932631287 |
| ENSMUSG00000058587 | Tmod3 | protein_coding | 0.062291514 | 6.482394661 | 0.0605484 | 0.8087017 | 0.932631287 |
| ENSMUSG00000027692 | Tnik | protein_coding | -0.106619455 | 5.67519799 | 0.0605111 | 0.8087777 | 0.932631287 |
| ENSMUSG00000025477 | Inpp5a | protein_coding | -0.104483089 | 4.403235227 | 0.0603546 | 0.8090015 | 0.932767097 |
| ENSMUSG00000021767 | Kat6b | protein_coding | -0.105006944 | 4.30740422 | 0.060242 | 0.809176 | 0.932767097 |
| ENSMUSG00000073609 | D2hgdh | protein_coding | 0.164838078 | 3.018181515 | 0.0601347 | 0.8093424 | 0.932767097 |
| ENSMUSG00000041718 | Alg13 | protein_coding | -0.235971309 | 2.759858993 | 0.0601032 | 0.8093964 | 0.932767097 |
| ENSMUSG00000000441 | Raf1 | protein_coding | -0.183582061 | 2.610805292 | 0.0600874 | 0.8094159 | 0.932767097 |
| ENSMUSG00000048486 | Fitm2 | protein_coding | -0.104924188 | 4.319730697 | 0.0600601 | 0.8094583 | 0.932767097 |
| ENSMUSG00000015335 | Zdhhc12 | protein_coding | -0.117641086 | 4.070934075 | 0.0599967 | 0.8095568 | 0.932767097 |
| ENSMUSG00000013701 | Timm23 | protein_coding | 0.139087386 | 3.06188704 | 0.0599362 | 0.8096507 | 0.932767097 |
| ENSMUSG00000028703 | Lrrc41 | protein_coding | 0.130973302 | 3.908571667 | 0.0599199 | 0.8096762 | 0.932767097 |
| ENSMUSG00000020812 | 1810032O08Rik | protein_coding | 0.167023584 | 2.780781175 | 0.0597018 | 0.8100155 | 0.932976735 |
| ENSMUSG00000046417 | Lrrc75a | protein_coding | 0.228031102 | 2.392586218 | 0.0596529 | 0.8100917 | 0.932976735 |
| ENSMUSG00000018666 | Cbx1 | protein_coding | 0.060834326 | 6.193500757 | 0.0596358 | 0.8101185 | 0.932976735 |
| ENSMUSG00000026637 | Traf5 | protein_coding | 0.148156095 | 2.904134733 | 0.0595227 | 0.8102948 | 0.933045672 |
| ENSMUSG00000033308 | Dpyd | protein_coding | -0.174039126 | 2.844302619 | 0.0594861 | 0.8103519 | 0.933045672 |
| ENSMUSG00000073725 | Lmbrd1 | protein_coding | -0.061318812 | 6.696811724 | 0.0594264 | 0.8104451 | 0.933053151 |
| ENSMUSG00000079018 | Ly6c1 | protein_coding | -0.244848501 | 2.631200017 | 0.0592509 | 0.8107195 | 0.933075749 |
| ENSMUSG00000025551 | Fgf14 | protein_coding | -0.096618763 | 5.814159515 | 0.059238 | 0.8107446 | 0.933075749 |
| ENSMUSG00000025898 | Cwf19l2 | protein_coding | -0.107906066 | 4.25217616 | 0.0591577 | 0.8108652 | 0.933075749 |
| ENSMUSG00000027944 | Hax1 | protein_coding | 0.070879603 | 5.327219095 | 0.0591435 | 0.8108876 | 0.933075749 |
| ENSMUSG00000061306 | Slc38a10 | protein_coding | -0.107208145 | 5.193890745 | 0.059132 | 0.8109105 | 0.933075749 |
| ENSMUSG00000024949 | Sf1 | protein_coding | -0.067006723 | 5.964313519 | 0.059081 | 0.8109854 | 0.933075749 |
| ENSMUSG00000023017 | Asic1 | protein_coding | 0.080498568 | 5.062771338 | 0.0589083 | 0.8112562 | 0.933222498 |
| ENSMUSG00000030752 | Kdm8 | protein_coding | 0.203756507 | 2.7555095 | 0.0588889 | 0.8112865 | 0.933222498 |
| ENSMUSG00000087479 | Gm16835 | processed_transcript | -0.139762131 | 3.380178744 | 0.0587666 | 0.8114786 | 0.933343614 |
| ENSMUSG00000032020 | Ubash3b | protein_coding | -0.113583261 | 6.093082325 | 0.0587218 | 0.8118505 | 0.93351498 |
| ENSMUSG00000027531 | Impa1 | protein_coding | -0.051204022 | 6.892489951 | 0.0585145 | 0.811875 | 0.93351498 |
| ENSMUSG00000020660 | Pomc | protein_coding | -0.116519587 | 4.148240725 | 0.0585063 | 0.811888 | 0.93351498 |
| ENSMUSG00000038342 | Mlxip | protein_coding | 0.19955756 | 2.475320782 | 0.0584467 | 0.8119819 | 0.933523095 |
| ENSMUSG00000050164 | Mchr1 | protein_coding | 0.117863598 | 4.577819094 | 0.0582781 | 0.8122477 | 0.933607116 |
| ENSMUSG00000029096 | Htra3 | protein_coding | -0.124917849 | 3.500425552 | 0.0582502 | 0.8122918 | 0.933607116 |
| ENSMUSG00000009073 | Nf2 | protein_coding | 0.090773962 | 4.799165403 | 0.0582352 | 0.8123155 | 0.933607116 |
| ENSMUSG00000106106 | CT010467.1 | rRNA | -0.067836789 | 10.43028008 | 0.0581405 | 0.812465 | 0.933679191 |
| ENSMUSG00000028830 | AU040320 | protein_coding | 0.079205295 | 5.325684901 | 0.0580448 | 0.8126163 | 0.933709791 |
| ENSMUSG00000020070 | Rufy2 | protein_coding | -0.089716365 | 5.062097477 | 0.0579612 | 0.8127485 | 0.933709791 |
| ENSMUSG00000020173 | Cobl | protein_coding | -0.091876679 | 4.979195501 | 0.0579589 | 0.8127521 | 0.933709791 |

| ensembl_gene_id | external_gene_name | gene_biotype | log2 (fold change) | log2 (counts per million) | F-value | p-value | False discovery rate |
|---|---|---|---|---|---|---|---|
| ENSMUSG00000054679 | Srsf12 | protein_coding | -0.108520955 | 5.326645041 | 0.0578899 | 0.8128663 | 0.933741212 |
| ENSMUSG00000015932 | Dstn | protein_coding | -0.057496987 | 8.557411813 | 0.0578274 | 0.8129604 | 0.933749516 |
| ENSMUSG00000034327 | Kctd9 | protein_coding | -0.111796821 | 3.86237387 | 0.0576565 | 0.8132313 | 0.933765055 |
| ENSMUSG00000031174 | Rpgr | protein_coding | -0.089198535 | 4.631533362 | 0.0575673 | 0.813373 | 0.933765055 |
| ENSMUSG00000029697 | Fezf1 | protein_coding | -0.305901176 | 3.055115251 | 0.0578197 | 0.8133925 | 0.933765055 |
| ENSMUSG00000041216 | Clvs1 | protein_coding | 0.108924145 | 4.45779578 | 0.0574246 | 0.8135999 | 0.933765055 |
| ENSMUSG00000091811 | Inafm1 | protein_coding | 0.111360186 | 4.222060814 | 0.0574237 | 0.8136012 | 0.933765055 |
| ENSMUSG00000037905 | Bri3bp | protein_coding | 0.069885257 | 5.625456159 | 0.0573796 | 0.8136713 | 0.933765055 |
| ENSMUSG00000026627 | Tmem206 | protein_coding | 0.116856655 | 4.50691482 | 0.0573586 | 0.8137048 | 0.933765055 |
| ENSMUSG00000051527 | Usp29 | protein_coding | 0.082546882 | 5.338877038 | 0.0573585 | 0.813705 | 0.933765055 |
| ENSMUSG00000020477 | Mrps24 | protein_coding | -0.105301327 | 4.17725496 | 0.0572936 | 0.8138082 | 0.933765055 |
| ENSMUSG00000010048 | Ifrd2 | protein_coding | -0.16083634 | 3.343851751 | 0.0572722 | 0.8138423 | 0.933765055 |
| ENSMUSG00000027272 | Ubr1 | protein_coding | 0.102929756 | 3.851006777 | 0.0571244 | 0.8140779 | 0.93393574 |
| ENSMUSG00000063568 | Jazf1 | protein_coding | -0.085611784 | 4.730185123 | 0.0569459 | 0.8143628 | 0.934162919 |
| ENSMUSG00000053024 | Cntn2 | protein_coding | -0.081694487 | 5.471520605 | 0.0568428 | 0.8145275 | 0.934181932 |
| ENSMUSG00000035967 | Ints6l | protein_coding | 0.180285772 | 2.801393177 | 0.0567677 | 0.8146476 | 0.934181932 |
| ENSMUSG00000021520 | Uqcrb | protein_coding | 0.076159194 | 8.483374154 | 0.0567386 | 0.8146991 | 0.934181932 |
| ENSMUSG00000060743 | H3f3a | protein_coding | -0.098481526 | 3.826424919 | 0.0567043 | 0.8147492 | 0.934181932 |
| ENSMUSG00000089911 | Mfsd14a | protein_coding | -0.11200371 | 4.328370809 | 0.0565772 | 0.8149528 | 0.934181932 |
| ENSMUSG00000057691 | Zfp746 | protein_coding | -0.113210303 | 3.760092009 | 0.0565439 | 0.8150062 | 0.934181932 |
| ENSMUSG00000021133 | Susd6 | protein_coding | -0.134476283 | 3.394380395 | 0.0565174 | 0.8150487 | 0.934181932 |
| ENSMUSG00000043987 | Cep164 | protein_coding | -0.172532434 | 2.793293955 | 0.056487 | 0.8150975 | 0.934181932 |
| ENSMUSG00000032932 | Hspa13 | protein_coding | -0.07815278 | 5.451743719 | 0.0563335 | 0.8153439 | 0.934181932 |
| ENSMUSG00000037852 | Cpe | protein_coding | 0.07105629 | 10.16661404 | 0.0563282 | 0.8153524 | 0.934181932 |
| ENSMUSG00000037134 | Prmt9 | protein_coding | 0.124907401 | 3.579216557 | 0.05629 | 0.8154137 | 0.934181932 |
| ENSMUSG00000037242 | Clic4 | protein_coding | 0.092908258 | 5.37808778 | 0.0562796 | 0.8154305 | 0.934181932 |
| ENSMUSG00000021792 | Fam213a | protein_coding | -0.063947942 | 7.844558865 | 0.0562309 | 0.8155088 | 0.934181932 |
| ENSMUSG00000037278 | Tmem97 | protein_coding | 0.087705019 | 4.261964516 | 0.0561422 | 0.8156515 | 0.934196948 |
| ENSMUSG00000026457 | Adipor1 | protein_coding | -0.076501083 | 5.534447329 | 0.0561148 | 0.8156956 | 0.934196948 |
| ENSMUSG00000038143 | Stox2 | protein_coding | -0.076215358 | 6.042121672 | 0.0560235 | 0.8158427 | 0.934243424 |
| ENSMUSG00000040282 | BC052040 | protein_coding | -0.098223934 | 4.305137647 | 0.0559326 | 0.8159892 | 0.934243424 |
| ENSMUSG00000022550 | Adck5 | protein_coding | 0.127015466 | 4.420545386 | 0.0558871 | 0.8160674 | 0.934243424 |
| ENSMUSG00000029436 | Mmp17 | protein_coding | 0.085661008 | 5.137459336 | 0.055874 | 0.8160837 | 0.934243424 |
| ENSMUSG00000066607 | 6030419C18Rik | protein_coding | 0.173050093 | 2.612149884 | 0.0557837 | 0.8162295 | 0.93425665 |
| ENSMUSG00000031024 | St5 | protein_coding | -0.14333128 | 3.050117708 | 0.0557592 | 0.8162691 | 0.93425665 |
| ENSMUSG00000023064 | Sncg | protein_coding | 0.075839749 | 7.292200882 | 0.0556561 | 0.8164405 | 0.93427493 |
| ENSMUSG00000001034 | Mapk7 | protein_coding | 0.201474113 | 2.897732483 | 0.055591 | 0.8165409 | 0.93427493 |
| ENSMUSG00000017760 | Ctsa | protein_coding | 0.053019873 | 6.797332357 | 0.0555459 | 0.8166139 | 0.93427493 |
| ENSMUSG00000056296 | Synpr | protein_coding | 0.076045697 | 7.160050266 | 0.0555374 | 0.8166326 | 0.93427493 |
| ENSMUSG00000024487 | Yipf5 | protein_coding | -0.068192527 | 5.843376927 | 0.0553774 | 0.8168868 | 0.934415519 |
| ENSMUSG00000035413 | Tmem98 | protein_coding | -0.076842024 | 5.736134457 | 0.0553493 | 0.8169325 | 0.934415519 |
| ENSMUSG00000015846 | Rxra | protein_coding | 0.080024853 | 4.58347294 | 0.0552634 | 0.8170718 | 0.934415519 |
| ENSMUSG00000066595 | Mfsd7b | protein_coding | 0.14771708 | 3.027479416 | 0.0552307 | 0.817125 | 0.934415519 |
| ENSMUSG00000042625 | Safb2 | protein_coding | -0.130550678 | 3.514037681 | 0.055176 | 0.8172137 | 0.934415519 |
| ENSMUSG00000038515 | Grtp1 | protein_coding | -0.150210681 | 3.133808473 | 0.0551372 | 0.8172768 | 0.934415519 |
| ENSMUSG00000024873 | Cnih2 | protein_coding | 0.097848109 | 5.668024471 | 0.0549426 | 0.8175981 | 0.934683392 |
| ENSMUSG00000027180 | Fbxo3 | protein_coding | 0.060231115 | 7.1115003 | 0.0545975 | 0.818156 | 0.935221806 |
| ENSMUSG00000021453 | Gadd45g | protein_coding | -0.092510345 | 5.83083445 | 0.0544945 | 0.8183291 | 0.935286224 |
| ENSMUSG00000038025 | Phf2 | protein_coding | 0.158315934 | 3.106562759 | 0.0544363 | 0.8184195 | 0.935286224 |
| ENSMUSG00000032177 | Pde4a | protein_coding | -0.056427237 | 6.706429982 | 0.0544035 | 0.8184733 | 0.935286224 |
| ENSMUSG00000029452 | Tmem116 | protein_coding | 0.133639225 | 3.387420825 | 0.054118 | 0.8189412 | 0.935310185 |
| ENSMUSG00000029371 | Cxcl5 | protein_coding | -0.312947678 | 2.469770503 | 0.0542233 | 0.8189531 | 0.935310185 |
| ENSMUSG00000035770 | Dync1li2 | protein_coding | -0.063515698 | 6.71919187 | 0.0541105 | 0.8189534 | 0.935310185 |
| ENSMUSG00000004099 | Dnmt1 | protein_coding | 0.091188009 | 4.668233799 | 0.0541027 | 0.8189662 | 0.935310185 |
| ENSMUSG00000035158 | Mitf | protein_coding | 0.207597606 | 2.723060343 | 0.054095 | 0.8189788 | 0.935310185 |
| ENSMUSG00000074923 | Pak6 | protein_coding | -0.087722834 | 4.58149921 | 0.0540723 | 0.8190162 | 0.935310185 |
| ENSMUSG00000032434 | Cmtm6 | protein_coding | 0.193096919 | 2.968209498 | 0.0538116 | 0.8194447 | 0.935572791 |
| ENSMUSG00000029433 | Diablo | protein_coding | -0.075266152 | 4.82074795 | 0.0537955 | 0.8194713 | 0.935572791 |
| ENSMUSG00000060279 | Ap2a1 | protein_coding | 0.20740583 | 4.119981076 | 0.0541544 | 0.8195071 | 0.935572791 |
| ENSMUSG00000068391 | Chrac1 | protein_coding | -0.061312026 | 6.829943269 | 0.0536724 | 0.819674 | 0.935663929 |
| ENSMUSG00000001138 | Cnnm3 | protein_coding | -0.111374899 | 4.023338172 | 0.0532611 | 0.8203533 | 0.936339981 |
| ENSMUSG00000025403 | Shmt2 | protein_coding | 0.120242889 | 4.19618175 | 0.053178 | 0.8204909 | 0.936397684 |
| ENSMUSG00000027455 | Nsfl1c | protein_coding | 0.055037178 | 7.593696853 | 0.0530724 | 0.820666 | 0.936427684 |
| ENSMUSG00000024097 | Srsf7 | protein_coding | 0.059111225 | 8.150589319 | 0.0529764 | 0.8208252 | 0.936427684 |
| ENSMUSG00000033610 | Pank1 | protein_coding | 0.11085945 | 3.737443995 | 0.0529573 | 0.8208568 | 0.936427684 |
| ENSMUSG00000032375 | Aph1b | protein_coding | -0.094713229 | 4.121172696 | 0.0529521 | 0.8208656 | 0.936427684 |
| ENSMUSG00000037649 | H2-DMa | protein_coding | -0.112951895 | 4.709291474 | 0.0528658 | 0.8210136 | 0.93649721 |
| ENSMUSG00000048706 | Lurap1l | protein_coding | -0.094121984 | 4.247336239 | 0.052589 | 0.8214694 | 0.936692333 |
| ENSMUSG00000020928 | Higd1b | protein_coding | -0.168369781 | 2.886518888 | 0.0525047 | 0.8216098 | 0.936692333 |
| ENSMUSG00000039840 | Epg5 | protein_coding | 0.171126164 | 3.388539015 | 0.0525008 | 0.821621 | 0.936692333 |
| ENSMUSG00000029173 | Sepsecs | protein_coding | 0.176607063 | 3.513726781 | 0.0525285 | 0.8216896 | 0.936692333 |
| ENSMUSG00000048701 | Ccdc6 | protein_coding | -0.05738393 | 6.963461909 | 0.052447 | 0.8217061 | 0.936692333 |
| ENSMUSG00000039844 | Rapgef1 | protein_coding | -0.09493154 | 4.591278604 | 0.0524463 | 0.8217073 | 0.936692333 |
| ENSMUSG00000041794 | Myrip | protein_coding | -0.122820267 | 3.511552876 | 0.0523596 | 0.8218521 | 0.936758069 |
| ENSMUSG00000022855 | Senp2 | protein_coding | 0.064695611 | 6.140582818 | 0.0521093 | 0.8222705 | 0.937135596 |
| ENSMUSG00000027495 | Fam210b | protein_coding | -0.090108723 | 4.945300352 | 0.0520008 | 0.8224522 | 0.937243411 |
| ENSMUSG00000020358 | Hnrnpab | protein_coding | 0.169654622 | 2.762757059 | 0.0519384 | 0.8225569 | 0.937263324 |
| ENSMUSG00000050737 | Ptges | protein_coding | -0.1654223 | 2.854792656 | 0.0518018 | 0.8227862 | 0.937425263 |

| ensembl_gene_id | external_gene_name | gene_biotype | log2 (fold change) | log2 (counts per million) | F-value | p-value | False discovery rate |
|---|---|---|---|---|---|---|---|
| ENSMUSG00000036887 | C1qa | protein_coding | -0.144409905 | 3.806809501 | 0.0516908 | 0.8229774 | 0.93743774 |
| ENSMUSG00000035161 | Ints6 | protein_coding | -0.08752073 | 4.27250757 | 0.0516361 | 0.8230647 | 0.93743774 |
| ENSMUSG00000000486 | 1-Sep | protein_coding | -0.150313728 | 2.941187693 | 0.0515196 | 0.8232607 | 0.93743774 |
| ENSMUSG00000015501 | Hivep2 | protein_coding | -0.191403622 | 2.894831611 | 0.0515106 | 0.823276 | 0.93743774 |
| ENSMUSG00000075705 | Msrb1 | protein_coding | 0.079167176 | 4.944493655 | 0.0515089 | 0.8232787 | 0.93743774 |
| ENSMUSG00000039375 | Wdr17 | protein_coding | 0.100505376 | 4.796422682 | 0.0514843 | 0.8233202 | 0.93743774 |
| ENSMUSG00000033684 | Qsox1 | protein_coding | 0.09361796 | 5.093274121 | 0.0514285 | 0.8234143 | 0.93744564 |
| ENSMUSG00000063480 | Snu13 | protein_coding | 0.071732373 | 5.062151622 | 0.0512662 | 0.8236881 | 0.937658022 |
| ENSMUSG00000038876 | Rnf146 | protein_coding | 0.079356523 | 4.722159032 | 0.0507869 | 0.8244995 | 0.938414413 |
| ENSMUSG00000099964 | Gm29514 | antisense_RNA | -0.098085498 | 3.670833723 | 0.0507387 | 0.8245813 | 0.938414413 |
| ENSMUSG00000054405 | Dnajc8 | protein_coding | 0.047088242 | 8.553718443 | 0.0506397 | 0.8247494 | 0.938414413 |
| ENSMUSG00000072946 | Ptgr2 | protein_coding | -0.095118338 | 5.058398107 | 0.050628 | 0.8247695 | 0.938414413 |
| ENSMUSG00000025157 | Zdhhc16 | protein_coding | 0.102982718 | 3.534691485 | 0.0506166 | 0.8247889 | 0.938414413 |
| ENSMUSG00000027890 | Gstm4 | protein_coding | 0.07618398 | 4.802565336 | 0.0505039 | 0.8249805 | 0.938533143 |
| ENSMUSG00000058486 | Wdr91 | protein_coding | 0.117429861 | 4.671450756 | 0.0502528 | 0.8254129 | 0.938915729 |
| ENSMUSG00000040428 | Plekha4 | protein_coding | 0.162145914 | 2.958626033 | 0.0502042 | 0.8254914 | 0.938915729 |
| ENSMUSG00000020827 | Mink1 | protein_coding | -0.077324773 | 4.869979965 | 0.0500988 | 0.8256714 | 0.939021122 |
| ENSMUSG00000037653 | Kctd8 | protein_coding | -0.11781226 | 3.702296862 | 0.0499503 | 0.8259255 | 0.939176289 |
| ENSMUSG00000063972 | Nr6a1 | protein_coding | -0.146496526 | 3.21914049 | 0.0498985 | 0.8260141 | 0.939176289 |
| ENSMUSG00000029193 | Cckar | protein_coding | 0.161772423 | 3.892365771 | 0.0499326 | 0.8261326 | 0.939176289 |
| ENSMUSG00000068015 | Lrch1 | protein_coding | -0.158888936 | 3.917521528 | 0.0498617 | 0.8262539 | 0.939176289 |
| ENSMUSG00000034613 | Ppm1h | protein_coding | 0.102632999 | 7.551319464 | 0.0499351 | 0.8263942 | 0.939176289 |
| ENSMUSG00000020457 | Drg1 | protein_coding | -0.052064208 | 8.483408787 | 0.0496553 | 0.8264313 | 0.939176289 |
| ENSMUSG00000028184 | Adgrl2 | protein_coding | -0.084259239 | 4.233915542 | 0.0496184 | 0.8264946 | 0.939176289 |
| ENSMUSG00000026544 | Dusp23 | protein_coding | 0.119080262 | 3.664859156 | 0.0496056 | 0.8265167 | 0.939176289 |
| ENSMUSG00000030374 | Strn4 | protein_coding | -0.064714408 | 6.040028159 | 0.0495607 | 0.8265939 | 0.939176289 |
| ENSMUSG00000042348 | Arl15 | protein_coding | -0.123830026 | 3.510940519 | 0.0494916 | 0.8267126 | 0.939211913 |
| ENSMUSG00000025220 | Mgea5 | protein_coding | 0.094779504 | 4.184251364 | 0.049367 | 0.8269271 | 0.939356336 |
| ENSMUSG00000004897 | Hdgf | protein_coding | -0.049721894 | 7.34081843 | 0.0491882 | 0.8272355 | 0.939457835 |
| ENSMUSG00000004936 | Map2k1 | protein_coding | 0.059688275 | 7.707290744 | 0.049164 | 0.8272772 | 0.939457835 |
| ENSMUSG00000038069 | Cdkn2aip | protein_coding | -0.127523141 | 3.533200707 | 0.0491418 | 0.8273156 | 0.939457835 |
| ENSMUSG00000079017 | Ifi27l2a | protein_coding | -0.197216916 | 2.489332967 | 0.0490809 | 0.8274207 | 0.939457835 |
| ENSMUSG00000074794 | Arrdc3 | protein_coding | -0.06798295 | 5.529296633 | 0.0490621 | 0.8274533 | 0.939457835 |
| ENSMUSG00000029247 | Paics | protein_coding | -0.050758261 | 7.582638901 | 0.0489183 | 0.8277019 | 0.939640885 |
| ENSMUSG00000028172 | Tacr3 | protein_coding | 0.148440557 | 2.731953969 | 0.0487937 | 0.8279177 | 0.939671179 |
| ENSMUSG00000021947 | Cryl1 | protein_coding | -0.071731577 | 5.011775904 | 0.0487886 | 0.8279266 | 0.939671179 |
| ENSMUSG00000096225 | Lhx8 | protein_coding | 0.101207074 | 4.460769682 | 0.0487257 | 0.8280356 | 0.939671179 |
| ENSMUSG00000036622 | Atp13a2 | protein_coding | -0.091411819 | 5.209432498 | 0.0487012 | 0.8280781 | 0.939671179 |
| ENSMUSG00000025212 | Sfxn3 | protein_coding | 0.069200058 | 7.358371596 | 0.048626 | 0.8282131 | 0.939705152 |
| ENSMUSG00000070570 | Slc17a7 | protein_coding | -0.222198055 | 2.764082517 | 0.0485113 | 0.8284125 | 0.939705152 |
| ENSMUSG00000032479 | Map4 | protein_coding | 0.065633167 | 8.692960752 | 0.0484387 | 0.8285341 | 0.939705152 |
| ENSMUSG00000087178 | A230056P14Rik | processed_transcript | -0.16108392 | 3.420152517 | 0.0483949 | 0.8286149 | 0.939705152 |
| ENSMUSG00000029202 | Pds5a | protein_coding | -0.113263936 | 3.485613207 | 0.0483917 | 0.828616 | 0.939705152 |
| ENSMUSG00000021215 | Net1 | protein_coding | -0.165854917 | 2.936039139 | 0.0483823 | 0.8286324 | 0.939705152 |
| ENSMUSG00000095105 | Edaradd | protein_coding | 0.346205724 | 1.826958411 | 0.0481222 | 0.8290903 | 0.940075202 |
| ENSMUSG00000046182 | Gsg1l | protein_coding | -0.134059013 | 3.759179788 | 0.0480974 | 0.8291336 | 0.940075202 |
| ENSMUSG00000059263 | Usp47 | protein_coding | 0.068835097 | 5.505801422 | 0.0480395 | 0.8292303 | 0.940085769 |
| ENSMUSG00000030711 | Sult1a1 | protein_coding | -0.160757231 | 3.215347347 | 0.0478859 | 0.8294989 | 0.940291127 |
| ENSMUSG00000043323 | Fbrsl1 | protein_coding | 0.122459091 | 3.680549112 | 0.047623 | 0.8299597 | 0.940703407 |
| ENSMUSG00000004895 | Prcc | protein_coding | 0.100179423 | 3.576970569 | 0.0475787 | 0.8300376 | 0.940703407 |
| ENSMUSG00000039126 | Prune2 | protein_coding | -0.118988787 | 8.868720627 | 0.0478567 | 0.8304384 | 0.940946877 |
| ENSMUSG00000039105 | Atp6v1g1 | protein_coding | -0.04734131 | 8.419649537 | 0.0473349 | 0.8304662 | 0.940946877 |
| ENSMUSG00000001056 | Nhp2 | protein_coding | -0.057805889 | 7.596107777 | 0.0472638 | 0.8305916 | 0.940946877 |
| ENSMUSG00000033430 | Terf2ip | protein_coding | -0.072376361 | 5.577778088 | 0.0472576 | 0.8306024 | 0.940946877 |
| ENSMUSG00000045598 | Zfp553 | protein_coding | 0.104231685 | 3.911544166 | 0.0471134 | 0.8308568 | 0.941135875 |
| ENSMUSG00000022451 | Twf1 | protein_coding | -0.171607882 | 2.529935583 | 0.0469486 | 0.831148 | 0.941278043 |
| ENSMUSG00000038828 | Tmem214 | protein_coding | 0.088449645 | 4.820998701 | 0.0469057 | 0.8312239 | 0.941278043 |
| ENSMUSG00000036957 | Lrfn3 | protein_coding | -0.209752829 | 2.342207843 | 0.0468939 | 0.8312449 | 0.941278043 |
| ENSMUSG00000024163 | Mapk8ip3 | protein_coding | 0.048772317 | 7.311188667 | 0.0467657 | 0.8314718 | 0.941435884 |
| ENSMUSG00000027710 | Acad9 | protein_coding | 0.17890292 | 3.779285076 | 0.0468633 | 0.8316043 | 0.941461131 |
| ENSMUSG00000021962 | Dcp1a | protein_coding | -0.105865134 | 3.482967852 | 0.0465639 | 0.8318299 | 0.941461131 |
| ENSMUSG00000033717 | Adra2a | protein_coding | 0.133059694 | 3.305469818 | 0.0465338 | 0.8318834 | 0.941461131 |
| ENSMUSG00000025785 | Exosc7 | protein_coding | 0.053297123 | 7.102316384 | 0.0465204 | 0.8319071 | 0.941461131 |
| ENSMUSG00000030739 | Myh14 | protein_coding | -0.116051301 | 3.342815442 | 0.0464825 | 0.8319747 | 0.941461131 |
| ENSMUSG00000022508 | Bcl6 | protein_coding | 0.148165373 | 3.048045473 | 0.0464573 | 0.8320194 | 0.941461131 |
| ENSMUSG00000001518 | Itfg2 | protein_coding | 0.138140832 | 2.944837538 | 0.0463167 | 0.8322696 | 0.941474231 |
| ENSMUSG00000006134 | Crkl | protein_coding | 0.08994991 | 3.826480559 | 0.0462868 | 0.8323229 | 0.941474231 |
| ENSMUSG00000050627 | Gpd1l | protein_coding | 0.08196868 | 5.678833813 | 0.0462357 | 0.8324139 | 0.941474231 |
| ENSMUSG00000028614 | Ndc1 | protein_coding | 0.168787064 | 3.068110825 | 0.0462223 | 0.8324379 | 0.941474231 |
| ENSMUSG00000020681 | Ace | protein_coding | 0.145646821 | 3.3613673 | 0.046205 | 0.8324688 | 0.941474231 |
| ENSMUSG00000041112 | Elmo1 | protein_coding | 0.053686526 | 8.636083867 | 0.0459436 | 0.8329356 | 0.941810187 |
| ENSMUSG00000055761 | Nkain3 | protein_coding | 0.126231877 | 3.530744899 | 0.045915 | 0.8329868 | 0.941810187 |
| ENSMUSG00000042712 | Tceal9 | protein_coding | -0.058221657 | 9.303714622 | 0.0458485 | 0.8331059 | 0.941810187 |
| ENSMUSG00000037353 | Letmd1 | protein_coding | 0.074736801 | 5.638732501 | 0.0458183 | 0.8331599 | 0.941810187 |
| ENSMUSG00000025019 | Lcor | protein_coding | 0.086449213 | 4.180898007 | 0.0457715 | 0.8332437 | 0.941810187 |
| ENSMUSG00000044033 | Ccdc141 | protein_coding | -0.151237248 | 2.953889257 | 0.0457172 | 0.8333411 | 0.941810187 |
| ENSMUSG00000020893 | Per1 | protein_coding | -0.100198264 | 4.241100739 | 0.0456707 | 0.8334245 | 0.941810187 |
| ENSMUSG00000030688 | Stard10 | protein_coding | -0.083410489 | 4.304015813 | 0.0456455 | 0.8334698 | 0.941810187 |

| ensembl_gene_id | external_gene_name | gene_biotype | log2 (fold change) | log2 (counts per million) | F-value | p-value | False discovery rate |
|---|---|---|---|---|---|---|---|
| ENSMUSG00000024030 | Abcg1 | protein_coding | 0.061766437 | 7.836224138 | 0.0455015 | 0.8337285 | 0.941810187 |
| ENSMUSG00000070643 | Sox13 | protein_coding | -0.197322725 | 2.646483976 | 0.0454381 | 0.8338425 | 0.941810187 |
| ENSMUSG00000020988 | L2hgdh | protein_coding | -0.101418751 | 3.620809624 | 0.0453381 | 0.8340225 | 0.941810187 |
| ENSMUSG00000041268 | Dmxl2 | protein_coding | 0.069682926 | 5.70629224 | 0.0450426 | 0.8345558 | 0.941810187 |
| ENSMUSG00000043639 | Rbm20 | protein_coding | 0.177944761 | 2.429036329 | 0.0450161 | 0.8346036 | 0.941810187 |
| ENSMUSG00000043311 | D17H6S53E | protein_coding | 0.128491471 | 5.399066154 | 0.045216 | 0.834612 | 0.941810187 |
| ENSMUSG00000032555 | Topbp1 | protein_coding | 0.21858254 | 2.651248365 | 0.0449869 | 0.8346607 | 0.941810187 |
| ENSMUSG00000022456 | 3-Sep | protein_coding | -0.051568415 | 8.375689342 | 0.0449765 | 0.8346752 | 0.941810187 |
| ENSMUSG00000038122 | Tbc1d32 | protein_coding | 0.158898223 | 2.883341405 | 0.0449582 | 0.8347084 | 0.941810187 |
| ENSMUSG00000042705 | Commd10 | protein_coding | 0.050083261 | 6.755597794 | 0.0448901 | 0.8348316 | 0.941810187 |
| ENSMUSG00000017999 | Ddx27 | protein_coding | 0.091067733 | 4.486232458 | 0.0448814 | 0.8348474 | 0.941810187 |
| ENSMUSG00000008035 | Mid1ip1 | protein_coding | -0.07440416 | 6.306994888 | 0.0448361 | 0.8349337 | 0.941810187 |
| ENSMUSG00000036501 | Fam13b | protein_coding | 0.098288112 | 3.975044105 | 0.0448242 | 0.834951 | 0.941810187 |
| ENSMUSG00000042506 | Usp22 | protein_coding | -0.068435839 | 8.288347525 | 0.044822 | 0.8349594 | 0.941810187 |
| ENSMUSG00000001323 | Srr | protein_coding | -0.072402872 | 4.668064243 | 0.0448137 | 0.8349701 | 0.941810187 |
| ENSMUSG00000018470 | Kcnab3 | protein_coding | -0.100966618 | 3.594478776 | 0.0448088 | 0.834979 | 0.941810187 |
| ENSMUSG00000023391 | Dlx2 | protein_coding | -0.158092988 | 2.893843547 | 0.0447987 | 0.8349972 | 0.941810187 |
| ENSMUSG00000046364 | Rpl27a | protein_coding | 0.060440412 | 6.757894458 | 0.0447734 | 0.8350431 | 0.941810187 |
| ENSMUSG00000017802 | Retreg3 | protein_coding | -0.095946438 | 3.661015455 | 0.0446406 | 0.8352841 | 0.941831859 |
| ENSMUSG00000018001 | Cyth3 | protein_coding | 0.101733184 | 3.947395335 | 0.0446123 | 0.8353354 | 0.941831859 |
| ENSMUSG00000034800 | Zfp661 | protein_coding | -0.178365973 | 2.819067178 | 0.044592 | 0.8353724 | 0.941831859 |
| ENSMUSG00000029070 | Mxra8 | protein_coding | 0.298323261 | 3.850094941 | 0.045428 | 0.8354127 | 0.941831859 |
| ENSMUSG00000039131 | Gipc2 | protein_coding | 0.106368659 | 3.787851571 | 0.0444857 | 0.8355656 | 0.941871939 |
| ENSMUSG00000052026 | Slc6a7 | protein_coding | 0.111380049 | 3.676002892 | 0.0444539 | 0.8356234 | 0.941871939 |
| ENSMUSG00000049148 | Plcxd3 | protein_coding | 0.081982306 | 5.864056823 | 0.0443979 | 0.8357297 | 0.941893043 |
| ENSMUSG00000036009 | Mettl25 | protein_coding | 0.131822068 | 2.995099745 | 0.0442096 | 0.8360687 | 0.942176369 |
| ENSMUSG00000027324 | Rpusd2 | protein_coding | 0.149021575 | 2.66853841 | 0.0440431 | 0.836373 | 0.94242045 |
| ENSMUSG00000001707 | Eef1e1 | protein_coding | 0.05350265 | 7.812687643 | 0.0438887 | 0.8366556 | 0.94264016 |
| ENSMUSG00000024592 | C330018D20Rik | protein_coding | 0.088555925 | 4.791125602 | 0.0437821 | 0.836851 | 0.942761476 |
| ENSMUSG00000020877 | Scrn2 | protein_coding | 0.134137857 | 3.29208002 | 0.0436976 | 0.8370061 | 0.942837453 |
| ENSMUSG00000032026 | Rexo2 | protein_coding | 0.062884251 | 5.948169437 | 0.0434293 | 0.8374997 | 0.943180191 |
| ENSMUSG00000038240 | Pdss2 | protein_coding | 0.075091456 | 4.928223451 | 0.0434285 | 0.8375012 | 0.943180191 |
| ENSMUSG00000051586 | Mical3 | protein_coding | -0.108032156 | 4.129778632 | 0.0433159 | 0.8377088 | 0.943180191 |
| ENSMUSG00000033948 | Zswim5 | protein_coding | -0.143387546 | 3.294930318 | 0.0432675 | 0.8377979 | 0.943180191 |
| ENSMUSG00000020661 | Dnmt3a | protein_coding | 0.057063485 | 5.956895013 | 0.0432355 | 0.837857 | 0.943180191 |
| ENSMUSG00000025630 | Hprt | protein_coding | 0.046449078 | 9.881867869 | 0.0432022 | 0.8379186 | 0.943180191 |
| ENSMUSG00000084898 | Gm12371 | lincRNA | 0.123085902 | 3.240525916 | 0.0431957 | 0.8379306 | 0.943180191 |
| ENSMUSG00000037894 | H2afz | protein_coding | 0.081583622 | 3.996993766 | 0.0431275 | 0.8380567 | 0.943180191 |
| ENSMUSG00000019894 | Slc6a15 | protein_coding | 0.065873348 | 5.604783337 | 0.0430371 | 0.8382239 | 0.943180191 |
| ENSMUSG00000030822 | Prr14 | protein_coding | 0.083223152 | 5.12615451 | 0.0430334 | 0.8382307 | 0.943180191 |
| ENSMUSG00000084289 | Gm6977 | processed_pseudoge | 0.210647929 | 3.235061878 | 0.0430976 | 0.8383227 | 0.943180191 |
| ENSMUSG00000028051 | Hcn3 | protein_coding | 0.113431063 | 3.320938554 | 0.0429383 | 0.838407 | 0.943180191 |
| ENSMUSG00000074925 | Ptar1 | protein_coding | 0.269180684 | 2.381256175 | 0.0429577 | 0.8384506 | 0.943180191 |
| ENSMUSG00000031467 | Agpat5 | protein_coding | 0.071376124 | 5.353533336 | 0.04281 | 0.8386448 | 0.943299983 |
| ENSMUSG00000027259 | Adal | protein_coding | -0.085168928 | 4.295735333 | 0.0425905 | 0.8390529 | 0.943653461 |
| ENSMUSG00000091475 | 2810468N07Rik | processed_transcript | -0.053076655 | 6.296432369 | 0.0425466 | 0.8391346 | 0.943653461 |
| ENSMUSG00000085881 | Gm15912 | processed_transcript | 0.109387565 | 3.367241813 | 0.04245 | 0.8393146 | 0.943757214 |
| ENSMUSG00000038722 | Bud31 | protein_coding | 0.058817143 | 6.785825901 | 0.0423706 | 0.8394626 | 0.943790103 |
| ENSMUSG00000001774 | Chordc1 | protein_coding | -0.047758751 | 6.496847039 | 0.0423402 | 0.8395194 | 0.943790103 |
| ENSMUSG00000046798 | Cldn12 | protein_coding | -0.080078912 | 4.887008155 | 0.0422766 | 0.8396382 | 0.943824953 |
| ENSMUSG00000039128 | Cdc123 | protein_coding | -0.050656957 | 8.75466243 | 0.0421906 | 0.8397991 | 0.943905914 |
| ENSMUSG00000033377 | Palmd | protein_coding | -0.137560139 | 3.561440039 | 0.0421137 | 0.8399471 | 0.943905914 |
| ENSMUSG00000036835 | Psenen | protein_coding | 0.067220054 | 4.977596897 | 0.0420899 | 0.8399876 | 0.943905914 |
| ENSMUSG00000032537 | Ephb1 | protein_coding | -0.175281761 | 2.574719015 | 0.0420505 | 0.8400613 | 0.943905914 |
| ENSMUSG00000028698 | Pik3r3 | protein_coding | -0.104821077 | 3.624051346 | 0.0419828 | 0.8401881 | 0.943949686 |
| ENSMUSG00000030518 | Fam189a1 | protein_coding | -0.124952994 | 4.028151011 | 0.0418258 | 0.840487 | 0.944186891 |
| ENSMUSG00000023272 | Creld2 | protein_coding | -0.164295453 | 3.29677908 | 0.0417575 | 0.8406264 | 0.944244873 |
| ENSMUSG00000064264 | Zfp428 | protein_coding | 0.109112839 | 3.237660318 | 0.0416969 | 0.8407253 | 0.944257219 |
| ENSMUSG00000017615 | Tnfaip1 | protein_coding | 0.066575242 | 6.625073356 | 0.0416525 | 0.8408131 | 0.944257219 |
| ENSMUSG00000029174 | Tbc1d1 | protein_coding | -0.128739847 | 3.007960314 | 0.0415308 | 0.8410382 | 0.94437177 |
| ENSMUSG00000033047 | Eif3l | protein_coding | 0.049690345 | 7.38292893 | 0.0413591 | 0.8413623 | 0.94437177 |
| ENSMUSG00000063445 | Nmral1 | protein_coding | -0.054791246 | 6.160852941 | 0.0413125 | 0.8414504 | 0.94437177 |
| ENSMUSG00000044285 | Gm1821 | processed_pseudoge | 0.23978755 | 5.447086433 | 0.0425999 | 0.8414762 | 0.94437177 |
| ENSMUSG00000003809 | Gcdh | protein_coding | 0.091724146 | 4.636542732 | 0.0412774 | 0.8415169 | 0.94437177 |
| ENSMUSG00000013787 | Ehmt2 | protein_coding | -0.069743342 | 5.583196149 | 0.0412766 | 0.8415183 | 0.94437177 |
| ENSMUSG00000002221 | Paxip1 | protein_coding | -0.086011297 | 4.563424626 | 0.0412411 | 0.8415856 | 0.94437177 |
| ENSMUSG00000027765 | P2ry1 | protein_coding | -0.125414494 | 3.143338903 | 0.0412241 | 0.8416177 | 0.94437177 |
| ENSMUSG00000033720 | Sfxn5 | protein_coding | -0.079435428 | 8.774874846 | 0.0412453 | 0.8417835 | 0.944372948 |
| ENSMUSG00000062127 | Cttnbp2nl | protein_coding | -0.094627954 | 3.730858449 | 0.0411308 | 0.8417944 | 0.944372948 |
| ENSMUSG00000061086 | Myl4 | protein_coding | -0.13376863 | 2.909886367 | 0.0410263 | 0.8419926 | 0.944496773 |
| ENSMUSG00000020910 | Adprm | protein_coding | -0.064354032 | 5.80374054 | 0.0409419 | 0.842153 | 0.944509073 |
| ENSMUSG00000000869 | Il4 | protein_coding | -0.086791437 | 4.629032743 | 0.040898 | 0.8422364 | 0.944509073 |
| ENSMUSG00000006169 | Clint1 | protein_coding | 0.072027544 | 4.742445546 | 0.0408818 | 0.842267 | 0.944509073 |
| ENSMUSG00000017057 | Il13ra1 | protein_coding | -0.119846027 | 3.208087871 | 0.040823 | 0.842379 | 0.944536169 |
| ENSMUSG00000041625 | Ggact | protein_coding | 0.068507052 | 5.141992694 | 0.0407166 | 0.8425816 | 0.944664741 |
| ENSMUSG00000038482 | Tfdp1 | protein_coding | -0.083450302 | 4.656460446 | 0.0405167 | 0.8429629 | 0.944949019 |
| ENSMUSG00000021645 | Smn1 | protein_coding | -0.062294135 | 5.4629213 | 0.0404916 | 0.8430109 | 0.944949019 |
| ENSMUSG00000028878 | Fam76a | protein_coding | -0.071576029 | 5.551510911 | 0.0402447 | 0.8434835 | 0.945267016 |

| ensembl_gene_id | external_gene_name | gene_biotype | log2 (fold change) | log2 (counts per million) | F-value | p-value | False discovery rate |
|---|---|---|---|---|---|---|---|
| ENSMUSG00000041130 | Zfp598 | protein_coding | -0.139858281 | 3.23157067 | 0.0402287 | 0.8435141 | 0.945267016 |
| ENSMUSG00000056763 | Cspp1 | protein_coding | 0.067445823 | 4.77568364 | 0.0401316 | 0.8437004 | 0.945267016 |
| ENSMUSG00000034168 | Irf2bpl | protein_coding | -0.07735552 | 4.404039221 | 0.0400979 | 0.8437653 | 0.945267016 |
| ENSMUSG00000022265 | Ank | protein_coding | -0.056009086 | 6.578249939 | 0.0400788 | 0.8438019 | 0.945267016 |
| ENSMUSG00000059040 | Eno1b | protein_coding | -0.300796172 | 2.21628682 | 0.0401294 | 0.8438625 | 0.945267016 |
| ENSMUSG00000016664 | Pacsin2 | protein_coding | -0.07786679 | 4.843475171 | 0.0400226 | 0.8439099 | 0.945267016 |
| ENSMUSG00000031442 | Mcf2l | protein_coding | -0.076751482 | 4.428412457 | 0.0399669 | 0.844017 | 0.945288467 |
| ENSMUSG00000053332 | Gas5 | processed_transcript | 0.078995675 | 10.45470529 | 0.0399743 | 0.8441602 | 0.945350396 |
| ENSMUSG00000070822 | Zscan18 | protein_coding | -0.113236246 | 4.896560747 | 0.0398256 | 0.8444464 | 0.945572432 |
| ENSMUSG00000075012 | Fjx1 | protein_coding | -0.142056064 | 3.278786675 | 0.0393703 | 0.8451693 | 0.94604506 |
| ENSMUSG00000062488 | Ifit3b | protein_coding | -0.152226962 | 2.6873316 | 0.039323 | 0.8452611 | 0.94604506 |
| ENSMUSG00000051354 | Samd3 | protein_coding | -0.159036618 | 3.681003651 | 0.0393919 | 0.8452837 | 0.94604506 |
| ENSMUSG00000037196 | Pacrg | protein_coding | -0.051077054 | 6.015750497 | 0.0392731 | 0.8453581 | 0.94604506 |
| ENSMUSG00000003038 | Hmgn2 | protein_coding | 0.0608884 | 5.561308851 | 0.0392687 | 0.8453664 | 0.94604506 |
| ENSMUSG00000021327 | Zkscan3 | protein_coding | 0.094439189 | 3.967106536 | 0.0392533 | 0.8453964 | 0.94604506 |
| ENSMUSG00000026421 | Csrp1 | protein_coding | 0.048460989 | 8.291073707 | 0.0391401 | 0.8456165 | 0.946192963 |
| ENSMUSG00000046532 | Ar | protein_coding | 0.064341538 | 5.82960088 | 0.0390303 | 0.8458302 | 0.946309409 |
| ENSMUSG00000026655 | Fam107b | protein_coding | 0.192561653 | 3.145436871 | 0.0390652 | 0.8459179 | 0.946309409 |
| ENSMUSG00000020319 | Wdpcp | protein_coding | 0.106247349 | 3.361835588 | 0.0389511 | 0.8459846 | 0.946309409 |
| ENSMUSG00000040943 | Tet2 | protein_coding | 0.076197519 | 5.009119942 | 0.0388174 | 0.8462455 | 0.946439684 |
| ENSMUSG00000024534 | Sncaip | protein_coding | -0.178991199 | 2.571521674 | 0.0388012 | 0.8462771 | 0.946439684 |
| ENSMUSG00000057716 | Tmem178b | protein_coding | -0.070247703 | 5.542624092 | 0.0387514 | 0.8463745 | 0.946450194 |
| ENSMUSG00000002455 | Prpf6 | protein_coding | 0.069031551 | 6.141371447 | 0.0386381 | 0.8466002 | 0.946479797 |
| ENSMUSG00000024812 | Tjp2 | protein_coding | -0.10093025 | 3.754612464 | 0.0386228 | 0.8466261 | 0.946479797 |
| ENSMUSG00000054863 | Fam19a5 | protein_coding | 0.055178809 | 6.30314532 | 0.0386029 | 0.846665 | 0.946479797 |
| ENSMUSG00000031786 | Drc7 | protein_coding | 0.183686676 | 3.185635984 | 0.038592 | 0.8468317 | 0.946567705 |
| ENSMUSG00000030647 | Ndufc2 | protein_coding | 0.056067052 | 9.492654567 | 0.038308 | 0.8472439 | 0.946848802 |
| ENSMUSG00000028700 | Pomgnt1 | protein_coding | -0.076434644 | 5.107725066 | 0.0383002 | 0.8472593 | 0.946848802 |
| ENSMUSG00000068747 | Sort1 | protein_coding | 0.071922669 | 5.841445731 | 0.0382276 | 0.8474022 | 0.946910097 |
| ENSMUSG00000028126 | Pip5k1a | protein_coding | -0.062431025 | 5.848862134 | 0.0381302 | 0.8475941 | 0.947015834 |
| ENSMUSG00000036452 | Arhgap26 | protein_coding | 0.065876626 | 5.249432187 | 0.0379908 | 0.8478693 | 0.947015834 |
| ENSMUSG00000024579 | Pcyox1l | protein_coding | -0.088043656 | 3.862839148 | 0.0379795 | 0.8478916 | 0.947015834 |
| ENSMUSG00000023828 | Slc22a3 | protein_coding | -0.160369434 | 2.622116416 | 0.0379484 | 0.847953 | 0.947015834 |
| ENSMUSG00000030417 | Pdcd5 | protein_coding | 0.094154891 | 4.30447361 | 0.0379113 | 0.8480264 | 0.947015834 |
| ENSMUSG00000029705 | Cux1 | protein_coding | -0.058034654 | 6.038084386 | 0.0379013 | 0.8480462 | 0.947015834 |
| ENSMUSG00000045969 | Ing1 | protein_coding | -0.056111198 | 5.778528828 | 0.0378536 | 0.8481405 | 0.947015834 |
| ENSMUSG00000043461 | Sptssb | protein_coding | 0.147588171 | 2.800323848 | 0.0378029 | 0.8482409 | 0.947015834 |
| ENSMUSG00000022109 | Med4 | protein_coding | -0.052471341 | 6.378644572 | 0.0377688 | 0.8483086 | 0.947015834 |
| ENSMUSG00000057766 | Ankrd29 | protein_coding | 0.06191494 | 5.195263539 | 0.037734 | 0.8483775 | 0.947015834 |
| ENSMUSG00000024583 | Txnl1 | protein_coding | -0.046777634 | 7.176357436 | 0.0376757 | 0.848493 | 0.947046445 |
| ENSMUSG00000021752 | Kctd6 | protein_coding | -0.076934565 | 4.238287717 | 0.0375763 | 0.8486905 | 0.947150176 |
| ENSMUSG00000022992 | Kansl2 | protein_coding | 0.137062005 | 2.796807331 | 0.0375402 | 0.8487621 | 0.947150176 |
| ENSMUSG00000050854 | Tmem125 | protein_coding | -0.122160005 | 3.590188888 | 0.0374896 | 0.8488627 | 0.947164124 |
| ENSMUSG00000032120 | C2cd2l | protein_coding | 0.055226862 | 6.011683688 | 0.037376 | 0.8490889 | 0.947318257 |
| ENSMUSG00000021871 | Pnp | protein_coding | 0.132575071 | 2.807099839 | 0.0373027 | 0.849235 | 0.947340284 |
| ENSMUSG00000005973 | Rcn1 | protein_coding | -0.072818601 | 5.132738455 | 0.0371392 | 0.8495614 | 0.947340284 |
| ENSMUSG00000020038 | Cry1 | protein_coding | -0.137159353 | 3.987714875 | 0.0372072 | 0.8495774 | 0.947340284 |
| ENSMUSG00000049532 | Sall2 | protein_coding | -0.057725004 | 5.735768004 | 0.0370924 | 0.8496551 | 0.947340284 |
| ENSMUSG00000031578 | Mak16 | protein_coding | 0.073326292 | 4.675581515 | 0.0370616 | 0.8497166 | 0.947340284 |
| ENSMUSG00000020610 | Amz2 | protein_coding | 0.053162154 | 6.09536867 | 0.0370539 | 0.8497321 | 0.947340284 |
| ENSMUSG00000018909 | Arrb1 | protein_coding | -0.049306021 | 6.224803928 | 0.0370297 | 0.8497805 | 0.947340284 |
| ENSMUSG00000026893 | Gca | protein_coding | -0.08969729 | 4.339121022 | 0.0369847 | 0.8498706 | 0.947340284 |
| ENSMUSG00000095432 | Zfp748 | protein_coding | -0.097422822 | 3.508578526 | 0.0369048 | 0.8500307 | 0.947340284 |
| ENSMUSG00000038615 | Nfe2l1 | protein_coding | -0.043362625 | 6.872197934 | 0.0368853 | 0.85007 | 0.947340284 |
| ENSMUSG00000004085 | Map3k20 | protein_coding | -0.150231672 | 3.11867091 | 0.0368814 | 0.8500778 | 0.947340284 |
| ENSMUSG00000004270 | Lpcat3 | protein_coding | 0.100379635 | 4.037494218 | 0.0366842 | 0.8504739 | 0.94768354 |
| ENSMUSG00000021532 | Fastkd3 | protein_coding | 0.072132951 | 5.034490308 | 0.0364983 | 0.8508485 | 0.947945488 |
| ENSMUSG00000060657 | Marf1 | protein_coding | 0.134452787 | 7.253309898 | 0.0371261 | 0.8509855 | 0.947945488 |
| ENSMUSG00000042608 | Stk40 | protein_coding | -0.112043591 | 3.739329331 | 0.0364016 | 0.8510435 | 0.947945488 |
| ENSMUSG00000028329 | Xpa | protein_coding | 0.093294968 | 3.471350559 | 0.0363927 | 0.8510616 | 0.947945488 |
| ENSMUSG00000029703 | Lrwd1 | protein_coding | -0.116522738 | 3.433185979 | 0.0361106 | 0.8516326 | 0.948483203 |
| ENSMUSG00000020376 | Rnf130 | protein_coding | -0.097100948 | 3.970070322 | 0.0360349 | 0.8517861 | 0.94855593 |
| ENSMUSG00000039183 | Nubp2 | protein_coding | 0.045035494 | 6.605297368 | 0.0359481 | 0.8519625 | 0.948654104 |
| ENSMUSG00000036580 | Spg20 | protein_coding | 0.065852476 | 5.385816295 | 0.0356841 | 0.8525004 | 0.949089111 |
| ENSMUSG00000020709 | Adap2 | protein_coding | -0.154863404 | 2.499203079 | 0.0356697 | 0.8525297 | 0.949089111 |
| ENSMUSG00000058975 | Kcnc1 | protein_coding | 0.063743706 | 5.32414464 | 0.0353695 | 0.8531438 | 0.949674456 |
| ENSMUSG00000032261 | Sh3bgrl2 | protein_coding | 0.091271206 | 5.201580126 | 0.0352936 | 0.8533134 | 0.949683884 |
| ENSMUSG00000074194 | Zfp791 | protein_coding | 0.103999635 | 3.226863141 | 0.0352793 | 0.8533289 | 0.949683884 |
| ENSMUSG00000061650 | Med9 | protein_coding | -0.086375232 | 4.35018074 | 0.0350896 | 0.8537188 | 0.949861936 |
| ENSMUSG00000054836 | Elp6 | protein_coding | -0.055821971 | 5.750769458 | 0.0350353 | 0.8538306 | 0.949861936 |
| ENSMUSG00000022983 | Scaf4 | protein_coding | -0.150620986 | 3.146398055 | 0.0350251 | 0.8538555 | 0.949861936 |
| ENSMUSG00000035557 | Krt17 | protein_coding | 0.132586073 | 2.761113525 | 0.0349751 | 0.8539547 | 0.949861936 |
| ENSMUSG00000029049 | Morn1 | protein_coding | 0.116606271 | 3.800028213 | 0.034918 | 0.8540763 | 0.949861936 |
| ENSMUSG00000018648 | Dusp14 | protein_coding | -0.051386355 | 6.258316206 | 0.0348846 | 0.8541415 | 0.949861936 |
| ENSMUSG00000019577 | Pdk4 | protein_coding | -0.165646769 | 2.415017495 | 0.034866 | 0.8541799 | 0.949861936 |
| ENSMUSG00000019927 | Ube2d1 | protein_coding | 0.049017953 | 6.169708397 | 0.0348584 | 0.8541956 | 0.949861936 |
| ENSMUSG00000021610 | Clptm1l | protein_coding | 0.072853442 | 5.774737366 | 0.0347302 | 0.8544645 | 0.94994963 |
| ENSMUSG00000087197 | Gm13780 | lincRNA | -0.107835459 | 3.154261479 | 0.0346958 | 0.854532 | 0.94994963 |

| ensembl_gene_id | external_gene_name | gene_biotype | log2 (fold change) | log2 (counts per million) | F-value | p-value | False discovery rate |
|---|---|---|---|---|---|---|---|
| ENSMUSG00000020099 | Unc5b | protein_coding | -0.08755914 | 4.237964072 | 0.0346903 | 0.8545434 | 0.94994963 |
| ENSMUSG00000034353 | Ramp1 | protein_coding | -0.079003682 | 6.466903309 | 0.0347327 | 0.8546439 | 0.94994963 |
| ENSMUSG00000039458 | Mtmr12 | protein_coding | -0.097558809 | 3.255889402 | 0.0346069 | 0.8547161 | 0.94994963 |
| ENSMUSG00000025240 | Sacm1l | protein_coding | 0.137275109 | 3.773463814 | 0.0345491 | 0.854982 | 0.950146959 |
| ENSMUSG00000021234 | Fam161b | protein_coding | -0.112386414 | 3.909700963 | 0.0343009 | 0.8553558 | 0.950461028 |
| ENSMUSG00000023938 | Aars2 | protein_coding | -0.155442816 | 2.543370175 | 0.0342129 | 0.8555355 | 0.950461028 |
| ENSMUSG00000034342 | Cbl | protein_coding | 0.071912118 | 4.555585624 | 0.0341872 | 0.8555891 | 0.950461028 |
| ENSMUSG00000068663 | Clec16a | protein_coding | -0.111869486 | 4.878212848 | 0.0342656 | 0.8556536 | 0.950461028 |
| ENSMUSG00000025027 | Xpnpep1 | protein_coding | 0.058722433 | 6.097822581 | 0.0341309 | 0.8557066 | 0.950461028 |
| ENSMUSG00000027188 | Pamr1 | protein_coding | -0.138013607 | 2.718552927 | 0.0340375 | 0.8559018 | 0.950579666 |
| ENSMUSG00000045095 | Magi1 | protein_coding | 0.052335118 | 6.208144879 | 0.033834 | 0.8563282 | 0.95095502 |
| ENSMUSG00000001173 | Ocrl | protein_coding | -0.086507832 | 5.090427923 | 0.0334539 | 0.8571316 | 0.951701984 |
| ENSMUSG00000030243 | Recql | protein_coding | -0.098397451 | 3.821508345 | 0.0332917 | 0.8574707 | 0.951701984 |
| ENSMUSG00000020386 | Sar1b | protein_coding | -0.050695966 | 7.789736804 | 0.0331591 | 0.8577515 | 0.951701984 |
| ENSMUSG00000022772 | Senp5 | protein_coding | 0.090766074 | 3.340804341 | 0.0331376 | 0.8577971 | 0.951701984 |
| ENSMUSG00000010067 | Rassf1 | protein_coding | -0.071401846 | 4.843780091 | 0.0331188 | 0.8578368 | 0.951701984 |
| ENSMUSG00000026914 | Psmd14 | protein_coding | 0.04512743 | 8.344813924 | 0.0330543 | 0.8579738 | 0.951701984 |
| ENSMUSG00000024963 | Dnajc4 | protein_coding | 0.056303469 | 6.0058377 | 0.0330258 | 0.8580343 | 0.951701984 |
| ENSMUSG00000040606 | Kazn | protein_coding | 0.058465065 | 5.968817437 | 0.032924 | 0.8582508 | 0.951701984 |
| ENSMUSG00000049892 | Rasd1 | protein_coding | 0.068477721 | 6.061518301 | 0.0328976 | 0.8583106 | 0.951701984 |
| ENSMUSG00000036545 | Adamts2 | protein_coding | -0.220645007 | 2.421073913 | 0.0328712 | 0.8583766 | 0.951701984 |
| ENSMUSG00000029311 | Hsd17b11 | protein_coding | -0.123534466 | 3.207451278 | 0.0328623 | 0.858382 | 0.951701984 |
| ENSMUSG00000032475 | Nck1 | protein_coding | -0.059233792 | 5.326819621 | 0.0328504 | 0.8584075 | 0.951701984 |
| ENSMUSG00000018974 | Sart3 | protein_coding | -0.079702916 | 4.178009768 | 0.032757 | 0.8586065 | 0.951701984 |
| ENSMUSG00000003848 | Nob1 | protein_coding | 0.056774821 | 6.283997351 | 0.0327364 | 0.8586505 | 0.951701984 |
| ENSMUSG00000056091 | St3gal5 | protein_coding | -0.068019102 | 4.89099431 | 0.032693 | 0.858743 | 0.951701984 |
| ENSMUSG00000031958 | Ldhd | protein_coding | 0.112085754 | 4.17706719 | 0.0326681 | 0.8588656 | 0.951701984 |
| ENSMUSG00000022761 | Lztr1 | protein_coding | -0.049192304 | 6.556894168 | 0.032583 | 0.8589783 | 0.951701984 |
| ENSMUSG00000030872 | Gga2 | protein_coding | 0.129284766 | 3.592282085 | 0.0325359 | 0.8590966 | 0.951701984 |
| ENSMUSG00000024856 | Cdk2ap2 | protein_coding | 0.053994018 | 5.554639604 | 0.0324937 | 0.8591693 | 0.951701984 |
| ENSMUSG00000003849 | Nqo1 | protein_coding | -0.104390306 | 3.758894883 | 0.0324612 | 0.859239 | 0.951701984 |
| ENSMUSG00000027573 | Gid8 | protein_coding | -0.060662167 | 5.507733195 | 0.0324115 | 0.8593457 | 0.951701984 |
| ENSMUSG00000021500 | Ddx46 | protein_coding | 0.056640375 | 6.111591387 | 0.0323553 | 0.8594663 | 0.951701984 |
| ENSMUSG00000100691 | 2010320M18Rik | lincRNA | -0.050553588 | 5.567277269 | 0.0323503 | 0.8594771 | 0.951701984 |
| ENSMUSG00000021701 | Plk2 | protein_coding | 0.043853434 | 7.13169582 | 0.0323053 | 0.8595736 | 0.951701984 |
| ENSMUSG00000048355 | Arxes1 | protein_coding | -0.075692888 | 4.788125196 | 0.0322819 | 0.8596239 | 0.951701984 |
| ENSMUSG00000020333 | Acsl6 | protein_coding | 0.064355015 | 5.723656483 | 0.0322446 | 0.859704 | 0.951701984 |
| ENSMUSG00000038582 | Pptc7 | protein_coding | -0.078914981 | 4.48550398 | 0.0322328 | 0.8597295 | 0.951701984 |
| ENSMUSG00000073293 | Nudt10 | protein_coding | 0.076406559 | 5.579233833 | 0.032218 | 0.859765 | 0.951701984 |
| ENSMUSG00000020922 | Lsm12 | protein_coding | -0.059218809 | 6.049970308 | 0.0322157 | 0.8597662 | 0.951701984 |
| ENSMUSG00000019254 | Ppp1r12c | protein_coding | -0.071567206 | 4.829504984 | 0.0321908 | 0.85982 | 0.951701984 |
| ENSMUSG00000021911 | Parg | protein_coding | -0.074685146 | 4.457585054 | 0.032176 | 0.8598518 | 0.951701984 |
| ENSMUSG00000024712 | Rfk | protein_coding | 0.048204421 | 6.438470408 | 0.0321605 | 0.8598851 | 0.951701984 |
| ENSMUSG00000089809 | Rasgef1b | protein_coding | -0.06822783 | 5.397372802 | 0.0321436 | 0.8599216 | 0.951701984 |
| ENSMUSG00000053093 | Myh7 | protein_coding | -0.116367645 | 2.921770434 | 0.0320221 | 0.8601836 | 0.951894023 |
| ENSMUSG00000058881 | Zfp516 | protein_coding | 0.158330825 | 2.882451343 | 0.0319679 | 0.8603041 | 0.951903835 |
| ENSMUSG00000025969 | Nrp2 | protein_coding | 0.082166189 | 4.422069096 | 0.031936 | 0.8603695 | 0.951903835 |
| ENSMUSG00000032598 | Nckipsd | protein_coding | 0.055585806 | 5.747788628 | 0.0318466 | 0.8605629 | 0.951918786 |
| ENSMUSG00000049526 | Tmem202 | protein_coding | -0.192464522 | 2.35832237 | 0.0318246 | 0.8606105 | 0.951918786 |
| ENSMUSG00000034908 | Sidt2 | protein_coding | 0.075493508 | 4.407146496 | 0.031807 | 0.8606486 | 0.951918786 |
| ENSMUSG00000009030 | Pdcl | protein_coding | 0.062779236 | 5.005014653 | 0.0316828 | 0.860918 | 0.952118787 |
| ENSMUSG00000050234 | Gja4 | protein_coding | 0.158274313 | 2.662100191 | 0.031627 | 0.8610392 | 0.952154941 |
| ENSMUSG00000062098 | Btbd3 | protein_coding | -0.047983672 | 6.552591929 | 0.0315328 | 0.8612439 | 0.952247719 |
| ENSMUSG00000032118 | Fez1 | protein_coding | 0.049515357 | 9.195683626 | 0.0315069 | 0.8613002 | 0.952247719 |
| ENSMUSG00000022677 | Fopnl | protein_coding | -0.041731314 | 8.132747093 | 0.0313957 | 0.8615424 | 0.952417593 |
| ENSMUSG00000026799 | Med27 | protein_coding | -0.049577774 | 5.612580468 | 0.0313319 | 0.8616818 | 0.952473722 |
| ENSMUSG00000029712 | Actl6b | protein_coding | -0.05090719 | 6.161214538 | 0.0311905 | 0.8619907 | 0.952717247 |
| ENSMUSG00000038612 | Mcl1 | protein_coding | -0.081585079 | 4.875636283 | 0.0311181 | 0.8621494 | 0.952794669 |
| ENSMUSG00000021519 | Mterf3 | protein_coding | -0.052489474 | 5.873411583 | 0.0309936 | 0.8624224 | 0.952998484 |
| ENSMUSG00000025885 | Myo5b | protein_coding | -0.074765853 | 4.374766362 | 0.0308946 | 0.8626399 | 0.953062014 |
| ENSMUSG00000038622 | Med30 | protein_coding | 0.093474252 | 3.868644546 | 0.0308867 | 0.8626572 | 0.953062014 |
| ENSMUSG00000026197 | Zfand2b | protein_coding | -0.052750292 | 5.663000998 | 0.0307964 | 0.8628559 | 0.953113277 |
| ENSMUSG00000021373 | Cap2 | protein_coding | -0.046025217 | 7.068830187 | 0.0307614 | 0.8629331 | 0.953113277 |
| ENSMUSG00000022546 | Gpt | protein_coding | -0.089682968 | 4.057947977 | 0.0307449 | 0.8629695 | 0.953113277 |
| ENSMUSG00000028789 | Azin2 | protein_coding | 0.068345464 | 4.471869449 | 0.0305124 | 0.8634829 | 0.95345427 |
| ENSMUSG00000043962 | Thrap3 | protein_coding | -0.081508751 | 5.265451221 | 0.0304706 | 0.8635791 | 0.95345427 |
| ENSMUSG00000032086 | Bace1 | protein_coding | -0.056204876 | 5.419567206 | 0.0304547 | 0.8636108 | 0.95345427 |
| ENSMUSG00000015476 | Prrt1 | protein_coding | 0.055367187 | 5.824490711 | 0.0304084 | 0.8637133 | 0.95345427 |
| ENSMUSG00000028617 | Lrrc42 | protein_coding | 0.0606611 | 4.923167692 | 0.0303745 | 0.8637886 | 0.95345427 |
| ENSMUSG00000048807 | Slc35e4 | protein_coding | -0.077867839 | 4.387235814 | 0.0303014 | 0.8639508 | 0.95345427 |
| ENSMUSG00000030662 | Ipo5 | protein_coding | 0.049989241 | 6.205584337 | 0.0302196 | 0.8641325 | 0.95345427 |
| ENSMUSG00000045333 | Zfp423 | protein_coding | -0.078796958 | 4.549716161 | 0.0302166 | 0.8641392 | 0.95345427 |
| ENSMUSG00000026924 | Sec16a | protein_coding | -0.074872717 | 3.965881154 | 0.0301971 | 0.8641827 | 0.95345427 |
| ENSMUSG00000041890 | Git2 | protein_coding | 0.087762345 | 4.123003064 | 0.0301505 | 0.8642862 | 0.95345427 |
| ENSMUSG00000024039 | Cbs | protein_coding | -0.08035228 | 4.681258939 | 0.0301428 | 0.8643035 | 0.95345427 |
| ENSMUSG00000033287 | Kctd17 | protein_coding | -0.05814571 | 5.488237808 | 0.0300879 | 0.8644257 | 0.95345427 |
| ENSMUSG00000031158 | Timm17b | protein_coding | 0.049063254 | 6.777739209 | 0.0300856 | 0.8644309 | 0.95345427 |
| ENSMUSG00000037525 | Bcdin3d | protein_coding | -0.099183041 | 4.013761634 | 0.0298588 | 0.864941 | 0.953773089 |

| ensembl_gene_id | external_gene_name | gene_biotype | log2 (fold change) | log2 (counts per million) | F-value | p-value | False discovery rate |
|---|---|---|---|---|---|---|---|
| ENSMUSG00000017897 | Eya2 | protein_coding | -0.09696092 | 3.05916692 | 0.0298179 | 0.8650291 | 0.953773089 |
| ENSMUSG00000070687 | Htr1d | protein_coding | -0.204315509 | 2.709892867 | 0.0298737 | 0.8650397 | 0.953773089 |
| ENSMUSG00000067825 | Pex26 | protein_coding | -0.096972182 | 3.494029996 | 0.0297867 | 0.8650989 | 0.953773089 |
| ENSMUSG00000030610 | Det1 | protein_coding | -0.153937875 | 2.628626449 | 0.0297204 | 0.8652476 | 0.953773089 |
| ENSMUSG00000048814 | Lonrf2 | protein_coding | -0.061129219 | 6.857009959 | 0.0297463 | 0.8652522 | 0.953773089 |
| ENSMUSG00000042751 | Nmnat2 | protein_coding | -0.067590704 | 7.270128218 | 0.0295667 | 0.8658041 | 0.954146433 |
| ENSMUSG00000029687 | Ezh2 | protein_coding | -0.136139175 | 3.123296772 | 0.0294491 | 0.8658612 | 0.954146433 |
| ENSMUSG00000051989 | Smim11 | protein_coding | -0.066516994 | 5.259884456 | 0.0294018 | 0.8659643 | 0.954146433 |
| ENSMUSG00000048503 | Tmem136 | protein_coding | -0.081383915 | 3.52924235 | 0.0293909 | 0.8659888 | 0.954146433 |
| ENSMUSG00000029064 | Gnb1 | protein_coding | 0.067370113 | 12.19785351 | 0.0293834 | 0.8660556 | 0.954146433 |
| ENSMUSG00000063787 | Chchd1 | protein_coding | 0.045809277 | 8.251797352 | 0.0293314 | 0.8661233 | 0.954146433 |
| ENSMUSG00000020607 | Fam84a | protein_coding | -0.042133026 | 7.303704423 | 0.0292206 | 0.8663737 | 0.954288418 |
| ENSMUSG00000027671 | Actl6a | protein_coding | -0.070942218 | 4.085812605 | 0.0291637 | 0.8665026 | 0.954288418 |
| ENSMUSG00000000751 | Rpa1 | protein_coding | -0.061930864 | 6.402588931 | 0.0290865 | 0.8666809 | 0.954288418 |
| ENSMUSG00000046111 | Cep295 | protein_coding | -0.110493751 | 2.984215369 | 0.0290222 | 0.8668235 | 0.954288418 |
| ENSMUSG00000021720 | Rnf180 | protein_coding | -0.064289907 | 4.500126184 | 0.0289231 | 0.8670487 | 0.954288418 |
| ENSMUSG00000008348 | Ubc | protein_coding | 0.049110746 | 8.069522431 | 0.0289107 | 0.867077 | 0.954288418 |
| ENSMUSG00000041528 | Rnf123 | protein_coding | 0.181452859 | 2.638792401 | 0.0288858 | 0.8671369 | 0.954288418 |
| ENSMUSG00000040811 | Eml2 | protein_coding | 0.058428207 | 5.45460871 | 0.028798 | 0.8673337 | 0.954288418 |
| ENSMUSG00000003226 | Ranbp2 | protein_coding | 0.083259242 | 3.727733926 | 0.0287763 | 0.8673832 | 0.954288418 |
| ENSMUSG00000039137 | Whrn | protein_coding | 0.079574382 | 4.322699121 | 0.028771 | 0.8673951 | 0.954288418 |
| ENSMUSG00000084885 | 3010001F23Rik | processed_transcript | -0.111483184 | 2.823728381 | 0.028771 | 0.8673952 | 0.954288418 |
| ENSMUSG00000031007 | Atp6ap2 | protein_coding | 0.038083658 | 8.573246147 | 0.0287399 | 0.8674661 | 0.954288418 |
| ENSMUSG00000097039 | Pvt1 | lincRNA | -0.062750858 | 4.786443159 | 0.0287304 | 0.8674879 | 0.954288418 |
| ENSMUSG00000042496 | Prdm10 | protein_coding | -0.101804584 | 3.93350366 | 0.0286323 | 0.8677153 | 0.954288418 |
| ENSMUSG00000000223 | Drp2 | protein_coding | -0.074285154 | 4.106157728 | 0.0286162 | 0.8677487 | 0.954288418 |
| ENSMUSG00000039747 | Orai2 | protein_coding | 0.076031788 | 4.35907682 | 0.0285957 | 0.8677958 | 0.954288418 |
| ENSMUSG00000040420 | Cdh18 | protein_coding | 0.081027846 | 3.766926113 | 0.0285877 | 0.8678141 | 0.954288418 |
| ENSMUSG00000019856 | Fam184a | protein_coding | 0.085916781 | 3.47433877 | 0.028492 | 0.8680333 | 0.954288418 |
| ENSMUSG00000047013 | Fbxo41 | protein_coding | 0.069502633 | 5.231849758 | 0.0284838 | 0.868052 | 0.954288418 |
| ENSMUSG00000103649 | Gm37768 | antisense_RNA | 0.125732517 | 2.671521566 | 0.0284821 | 0.8680559 | 0.954288418 |
| ENSMUSG00000024352 | Spata24 | protein_coding | -0.070927425 | 4.421206474 | 0.0284079 | 0.8682262 | 0.954288418 |
| ENSMUSG00000097213 | Gm26745 | lincRNA | -0.118726123 | 2.703866839 | 0.0283674 | 0.8683193 | 0.954288418 |
| ENSMUSG00000024081 | Cebpz | protein_coding | -0.058949005 | 5.211813581 | 0.0283403 | 0.8683816 | 0.954288418 |
| ENSMUSG00000032680 | 6820408C15Rik | protein_coding | -0.091233968 | 4.215739749 | 0.0283222 | 0.8684265 | 0.954288418 |
| ENSMUSG00000031925 | Maml2 | protein_coding | -0.076768883 | 4.363464793 | 0.0283015 | 0.8684708 | 0.954288418 |
| ENSMUSG00000074781 | Ube2n | protein_coding | -0.044950504 | 6.876333105 | 0.0282561 | 0.8685751 | 0.954305559 |
| ENSMUSG00000055782 | Abcd2 | protein_coding | -0.100363433 | 3.920591442 | 0.0281439 | 0.8688372 | 0.95439994 |
| ENSMUSG00000040720 | Virma | protein_coding | 0.070672993 | 4.250629407 | 0.0281418 | 0.8688386 | 0.95439994 |
| ENSMUSG00000031371 | Haus7 | protein_coding | -0.045286189 | 5.661302932 | 0.0279895 | 0.8691905 | 0.954595445 |
| ENSMUSG00000033159 | Cnppd1 | protein_coding | -0.055586219 | 5.282652657 | 0.027988 | 0.8691941 | 0.954595445 |
| ENSMUSG00000041153 | Osgin2 | protein_coding | -0.102854489 | 3.325612548 | 0.0278802 | 0.8694438 | 0.954772182 |
| ENSMUSG00000033417 | Cacul1 | protein_coding | 0.059943211 | 5.058586534 | 0.0278107 | 0.8696048 | 0.954851489 |
| ENSMUSG00000002265 | Peg3 | protein_coding | -0.046253564 | 8.475969773 | 0.0277132 | 0.8698315 | 0.95493113 |
| ENSMUSG00000040390 | Map3k10 | protein_coding | -0.140402087 | 3.157233329 | 0.0276893 | 0.8698903 | 0.95493113 |
| ENSMUSG00000052512 | Nav2 | protein_coding | -0.065524413 | 5.38433187 | 0.0276649 | 0.8699438 | 0.95493113 |
| ENSMUSG00000032324 | Tspan3 | protein_coding | 0.052365132 | 7.434399191 | 0.0275407 | 0.8702367 | 0.955044169 |
| ENSMUSG00000063919 | Srrm4 | protein_coding | -0.072369685 | 4.491326494 | 0.0275156 | 0.8702919 | 0.955044169 |
| ENSMUSG00000024436 | Mrps18b | protein_coding | -0.062419209 | 4.956158129 | 0.0275065 | 0.8703132 | 0.955044169 |
| ENSMUSG00000026585 | Kifap3 | protein_coding | 0.041034929 | 7.567748367 | 0.0274239 | 0.870506 | 0.955158274 |
| ENSMUSG00000021275 | Tecpr2 | protein_coding | -0.069908266 | 4.203637937 | 0.0272704 | 0.8708656 | 0.955341126 |
| ENSMUSG00000019726 | Lyst | protein_coding | 0.078912468 | 3.96989673 | 0.0271896 | 0.8710552 | 0.955341126 |
| ENSMUSG00000067925 | Cxx1a | protein_coding | 0.072209428 | 3.974048044 | 0.0271779 | 0.8710827 | 0.955341126 |
| ENSMUSG00000086308 | G630016G05Rik | lincRNA | -0.117100952 | 2.940889393 | 0.0271642 | 0.8711147 | 0.955341126 |
| ENSMUSG00000026204 | Ptprn | protein_coding | 0.044585119 | 7.339596512 | 0.0271409 | 0.8711696 | 0.955341126 |
| ENSMUSG00000062929 | Cfl2 | protein_coding | -0.037071556 | 7.9738112 | 0.0271255 | 0.8712057 | 0.955341126 |
| ENSMUSG00000052593 | Adam17 | protein_coding | 0.165295409 | 2.509099436 | 0.0269774 | 0.8715546 | 0.955553639 |
| ENSMUSG00000020929 | Eftud2 | protein_coding | -0.05913012 | 5.697167108 | 0.026931 | 0.8716639 | 0.955553639 |
| ENSMUSG00000028978 | Nos3 | protein_coding | 0.098322844 | 3.220516984 | 0.0269085 | 0.8717171 | 0.955553639 |
| ENSMUSG00000042699 | Dhx9 | protein_coding | 0.068781961 | 4.608989933 | 0.0268924 | 0.871755 | 0.955553639 |
| ENSMUSG00000034751 | Mast4 | protein_coding | -0.086548924 | 3.656852043 | 0.0268342 | 0.8718927 | 0.955607147 |
| ENSMUSG00000022257 | Laptm4b | protein_coding | 0.047382293 | 6.147958706 | 0.0267757 | 0.872031 | 0.955661336 |
| ENSMUSG00000030077 | Chl1 | protein_coding | 0.049125202 | 6.308066585 | 0.0266983 | 0.8722144 | 0.955764959 |
| ENSMUSG00000050074 | Spink8 | protein_coding | -0.113855787 | 3.449041662 | 0.0264483 | 0.8728117 | 0.956174204 |
| ENSMUSG00000050244 | Heatr1 | protein_coding | -0.113435533 | 2.870058153 | 0.0264343 | 0.8728419 | 0.956174204 |
| ENSMUSG00000031963 | Bmper | protein_coding | -0.114144249 | 2.732481153 | 0.0264108 | 0.8728978 | 0.956174204 |
| ENSMUSG00000025521 | Tmem192 | protein_coding | -0.080743328 | 3.808437796 | 0.0263916 | 0.8729436 | 0.956174204 |
| ENSMUSG00000028849 | Map7d1 | protein_coding | -0.041876607 | 6.582567633 | 0.0262657 | 0.8732441 | 0.956343763 |
| ENSMUSG00000022197 | Pdzd2 | protein_coding | -0.091017125 | 3.694721318 | 0.0262523 | 0.8732762 | 0.956343763 |
| ENSMUSG00000070426 | Rnf121 | protein_coding | 0.043904066 | 5.735341833 | 0.0261723 | 0.8734677 | 0.956456042 |
| ENSMUSG00000035151 | Elmod2 | protein_coding | -0.142727562 | 2.45865367 | 0.0261283 | 0.873573 | 0.956473967 |
| ENSMUSG00000005417 | Mprip | protein_coding | 0.055310221 | 5.350901569 | 0.026029 | 0.8738114 | 0.956637528 |
| ENSMUSG00000027427 | Polr3f | protein_coding | -0.067797505 | 4.529543924 | 0.0259804 | 0.8739282 | 0.956667982 |
| ENSMUSG00000001229 | Dpp9 | protein_coding | 0.063974097 | 5.095200661 | 0.0257439 | 0.8744981 | 0.957194439 |
| ENSMUSG00000022672 | Prkdc | protein_coding | -0.167387354 | 2.64344309 | 0.0256937 | 0.8746227 | 0.957233355 |
| ENSMUSG00000020659 | Cbll1 | protein_coding | 0.067053041 | 4.67701495 | 0.0255544 | 0.8749567 | 0.957398485 |
| ENSMUSG00000022378 | Fam49b | protein_coding | -0.038311864 | 6.70146057 | 0.0255374 | 0.874998 | 0.957398485 |
| ENSMUSG00000041895 | Wipi1 | protein_coding | -0.108635581 | 4.434203153 | 0.0255703 | 0.8751142 | 0.957398485 |

| ensembl_gene_id | external_gene_name | gene_biotype | log2 (fold change) | log2 (counts per million) | F-value | p-value | False discovery rate |
|---|---|---|---|---|---|---|---|
| ENSMUSG00000031107 | Rbmx2 | protein_coding | -0.058766891 | 4.255340661 | 0.0253975 | 0.8753378 | 0.957398485 |
| ENSMUSG00000045994 | B3gat1 | protein_coding | 0.061143276 | 4.885530652 | 0.0253648 | 0.8754172 | 0.957398485 |
| ENSMUSG00000032897 | Nfyc | protein_coding | -0.065191775 | 4.118222849 | 0.0253392 | 0.8754797 | 0.957398485 |
| ENSMUSG00000041124 | Msantd4 | protein_coding | 0.044771009 | 6.79558732 | 0.02531 | 0.8755507 | 0.957398485 |
| ENSMUSG00000036372 | Tmem258 | protein_coding | 0.057661825 | 7.304822578 | 0.0253442 | 0.875592 | 0.957398485 |
| ENSMUSG00000020427 | Igfbp3 | protein_coding | 0.119561255 | 2.704675004 | 0.0252154 | 0.8757813 | 0.957398485 |
| ENSMUSG00000042613 | Pbxip1 | protein_coding | -0.075159799 | 3.96877125 | 0.0252089 | 0.8757972 | 0.957398485 |
| ENSMUSG00000022610 | Mapk12 | protein_coding | 0.1583392 | 2.362859331 | 0.0251721 | 0.8758872 | 0.957398485 |
| ENSMUSG00000031839 | Hsbp1 | protein_coding | -0.040080475 | 10.62593126 | 0.0251701 | 0.8758921 | 0.957398485 |
| ENSMUSG00000085042 | Abhd11os | transcribed_unitary_p | -0.060695413 | 5.216477272 | 0.0251429 | 0.8759584 | 0.957398485 |
| ENSMUSG00000066113 | Adamtsl1 | protein_coding | -0.152451313 | 3.038575232 | 0.0251285 | 0.8760753 | 0.957398485 |
| ENSMUSG00000021737 | Psmd6 | protein_coding | -0.037893609 | 8.008013468 | 0.025056 | 0.8761712 | 0.957398485 |
| ENSMUSG00000025050 | Pcgf6 | protein_coding | -0.133739954 | 2.536865775 | 0.025045 | 0.8761981 | 0.957398485 |
| ENSMUSG00000050299 | Gm9843 | protein_coding | -0.058839379 | 4.56489152 | 0.0249414 | 0.8764521 | 0.957571112 |
| ENSMUSG00000058297 | Spock2 | protein_coding | 0.036353907 | 7.721449421 | 0.024908 | 0.8765342 | 0.957571112 |
| ENSMUSG00000030298 | Sec13 | protein_coding | 0.03757067 | 7.532545021 | 0.0248581 | 0.876657 | 0.957607904 |
| ENSMUSG00000026311 | Asb1 | protein_coding | 0.065983716 | 4.529768471 | 0.0247611 | 0.8768955 | 0.957771236 |
| ENSMUSG00000069581 | Tspear | protein_coding | 0.127123779 | 2.714406044 | 0.0246939 | 0.8770613 | 0.957833734 |
| ENSMUSG00000038473 | Nos1ap | protein_coding | 0.049766248 | 5.935121701 | 0.0246361 | 0.8772041 | 0.957833734 |
| ENSMUSG00000024180 | Tmem8 | protein_coding | 0.105601129 | 2.937878498 | 0.0246297 | 0.87722 | 0.957833734 |
| ENSMUSG00000057054 | Inca1 | protein_coding | 0.071400936 | 3.848387541 | 0.0244184 | 0.8777432 | 0.958307655 |
| ENSMUSG00000029521 | Chek2 | protein_coding | -0.093571321 | 3.049746519 | 0.0243516 | 0.877909 | 0.958391411 |
| ENSMUSG00000019689 | Fmc1 | protein_coding | 0.041045769 | 7.439051326 | 0.0242946 | 0.8780509 | 0.958414635 |
| ENSMUSG00000039849 | Pcif1 | protein_coding | -0.041846423 | 6.315142522 | 0.0242714 | 0.8781085 | 0.958414635 |
| ENSMUSG00000044496 | 2510039O18Rik | protein_coding | 0.105197797 | 3.529021307 | 0.0242126 | 0.8782582 | 0.958448748 |
| ENSMUSG00000028656 | Cap1 | protein_coding | -0.047825193 | 6.506580688 | 0.0241873 | 0.878318 | 0.958448748 |
| ENSMUSG00000034163 | Zfc3h1 | protein_coding | -0.119626226 | 2.682463866 | 0.0241169 | 0.8784939 | 0.958455922 |
| ENSMUSG00000027132 | Katnbl1 | protein_coding | 0.083089945 | 3.374344132 | 0.0241133 | 0.8785029 | 0.958455922 |
| ENSMUSG00000042213 | Zfand4 | protein_coding | 0.128303763 | 2.652720383 | 0.0240389 | 0.8786887 | 0.958545355 |
| ENSMUSG00000030663 | 1110004F10Rik | protein_coding | 0.034704875 | 8.6347426 | 0.0240092 | 0.8787631 | 0.958545355 |
| ENSMUSG00000021974 | Fgf9 | protein_coding | -0.055134997 | 5.384579814 | 0.0239317 | 0.8789574 | 0.958566674 |
| ENSMUSG00000066037 | Hnrnpr | protein_coding | 0.039622514 | 6.498009891 | 0.023877 | 0.8790945 | 0.958566674 |
| ENSMUSG00000041930 | Fam222a | protein_coding | -0.155008018 | 2.58983844 | 0.0238761 | 0.8790967 | 0.958566674 |
| ENSMUSG00000022040 | Ephx2 | protein_coding | -0.049865855 | 4.957748281 | 0.023835 | 0.8792002 | 0.958566674 |
| ENSMUSG00000002108 | Nr1h3 | protein_coding | 0.144621784 | 2.680416916 | 0.0238201 | 0.8792374 | 0.958566674 |
| ENSMUSG00000027168 | Pax6 | protein_coding | 0.074258885 | 3.913721954 | 0.0237883 | 0.8793176 | 0.958566674 |
| ENSMUSG00000086555 | Gm13446 | processed_transcript | 0.123002103 | 3.558892642 | 0.0237543 | 0.8795236 | 0.95869411 |
| ENSMUSG00000028782 | Adgrb2 | protein_coding | -0.095807597 | 3.140051897 | 0.0236422 | 0.8796856 | 0.958773487 |
| ENSMUSG00000064325 | Hhip | protein_coding | -0.112392129 | 2.593913966 | 0.0235866 | 0.8798258 | 0.958829088 |
| ENSMUSG00000017144 | Rnd3 | protein_coding | 0.059579291 | 5.151207108 | 0.0235329 | 0.8799616 | 0.95887996 |
| ENSMUSG00000032449 | Slc25a36 | protein_coding | -0.046049642 | 5.766421646 | 0.0234813 | 0.8800923 | 0.958925118 |
| ENSMUSG00000022571 | Pycrl | protein_coding | 0.038605762 | 6.757808045 | 0.0234209 | 0.8802452 | 0.958978251 |
| ENSMUSG00000063362 | Alg11 | protein_coding | -0.04538281 | 6.593971449 | 0.0233917 | 0.8803194 | 0.958978251 |
| ENSMUSG00000035621 | Midn | protein_coding | -0.089060371 | 2.965152088 | 0.0233444 | 0.8804395 | 0.959011945 |
| ENSMUSG00000020614 | Fam20a | protein_coding | 0.145013139 | 2.756788209 | 0.0232652 | 0.8806437 | 0.959025575 |
| ENSMUSG00000020412 | Ascc2 | protein_coding | 0.081386404 | 3.282315563 | 0.0232293 | 0.880732 | 0.959025575 |
| ENSMUSG00000032855 | Pkd1 | protein_coding | 0.116625603 | 2.826235212 | 0.0231575 | 0.880915 | 0.959025575 |
| ENSMUSG00000004110 | Cacna1e | protein_coding | -0.066008093 | 4.923582832 | 0.0231366 | 0.8809684 | 0.959025575 |
| ENSMUSG00000078970 | Wdr92 | protein_coding | 0.072960029 | 4.645180636 | 0.0231266 | 0.8809939 | 0.959025575 |
| ENSMUSG00000024588 | Fech | protein_coding | 0.065228447 | 5.324261847 | 0.0231234 | 0.8810052 | 0.959025575 |
| ENSMUSG00000018796 | Acsl1 | protein_coding | 0.063440206 | 4.888774052 | 0.0230652 | 0.8811508 | 0.959025575 |
| ENSMUSG00000060519 | Tor3a | protein_coding | 0.130561712 | 2.469766756 | 0.0230444 | 0.8812039 | 0.959025575 |
| ENSMUSG00000028902 | Sf3a3 | protein_coding | 0.039037386 | 6.182383047 | 0.0230087 | 0.8812951 | 0.959025575 |
| ENSMUSG00000057133 | Chd6 | protein_coding | -0.047459798 | 5.686323388 | 0.0229897 | 0.8813439 | 0.959025575 |
| ENSMUSG00000094344 | Gm11942 | processed_pseudoge | 0.290492794 | 1.972589933 | 0.0229946 | 0.8814761 | 0.959072428 |
| ENSMUSG00000031438 | Rnf128 | protein_coding | 0.068731298 | 4.235137812 | 0.02283 | 0.8817534 | 0.95912665 |
| ENSMUSG00000020925 | Ccdc43 | protein_coding | 0.045067649 | 5.857816366 | 0.0228283 | 0.8817576 | 0.95912665 |
| ENSMUSG00000022241 | Tars | protein_coding | 0.055999137 | 5.555735571 | 0.0228143 | 0.8817936 | 0.95912665 |
| ENSMUSG00000020076 | Ddx50 | protein_coding | 0.04473768 | 5.727135582 | 0.0226931 | 0.8821055 | 0.959368867 |
| ENSMUSG00000110156 | Gm42067 | antisense_RNA | 0.085339389 | 3.177236938 | 0.0226019 | 0.8823408 | 0.959527803 |
| ENSMUSG00000024892 | Pcx | protein_coding | -0.048052629 | 5.608768198 | 0.0225284 | 0.8825307 | 0.959637261 |
| ENSMUSG00000027643 | Ghrh | protein_coding | 0.093658983 | 3.367773726 | 0.0223843 | 0.882904 | 0.959887075 |
| ENSMUSG00000046020 | Pofut1 | protein_coding | 0.111183537 | 2.626413497 | 0.0223468 | 0.8830015 | 0.959887075 |
| ENSMUSG00000055817 | Mta3 | protein_coding | 0.063279915 | 4.98990718 | 0.0223364 | 0.8830283 | 0.959887075 |
| ENSMUSG00000021178 | Psmc1 | protein_coding | 0.033506295 | 9.082673382 | 0.022275 | 0.8831881 | 0.959963761 |
| ENSMUSG00000071076 | Jund | protein_coding | -0.055974684 | 4.43190559 | 0.0221985 | 0.8833872 | 0.95999215 |
| ENSMUSG00000038141 | Tmem181a | protein_coding | 0.072006489 | 3.611116157 | 0.0221656 | 0.8834729 | 0.95999215 |
| ENSMUSG00000074024 | 4632427E13Rik | bidirectional_promote | 0.200043303 | 2.528839498 | 0.0221907 | 0.8835572 | 0.95999215 |
| ENSMUSG00000034893 | Cog3 | protein_coding | -0.141448799 | 3.031055226 | 0.0221512 | 0.8835713 | 0.95999215 |
| ENSMUSG00000023791 | Pigx | protein_coding | -0.082992646 | 4.053541785 | 0.0220549 | 0.8837651 | 0.960026262 |
| ENSMUSG00000026288 | Inpp5d | protein_coding | -0.198285633 | 2.104741917 | 0.0220169 | 0.8838643 | 0.960026262 |
| ENSMUSG00000063808 | Gpatch1 | protein_coding | -0.049051209 | 5.479376929 | 0.0220134 | 0.8838705 | 0.960026262 |
| ENSMUSG00000032415 | Ube2cbp | protein_coding | -0.075053288 | 3.39040437 | 0.0219736 | 0.8839748 | 0.960042508 |
| ENSMUSG00000070934 | Rraga | protein_coding | -0.050855661 | 6.785571419 | 0.0219407 | 0.8840641 | 0.960042543 |
| ENSMUSG00000036845 | Lin37 | protein_coding | -0.035273166 | 6.935440291 | 0.0218495 | 0.8843005 | 0.960086341 |
| ENSMUSG00000034211 | Mrps17 | protein_coding | 0.035124576 | 7.895572155 | 0.0218336 | 0.8843421 | 0.960086341 |
| ENSMUSG00000023235 | Ccl25 | protein_coding | -0.060328394 | 4.87212232 | 0.0218168 | 0.8843862 | 0.960086341 |
| ENSMUSG00000002908 | Kcnn1 | protein_coding | -0.140213356 | 2.593229118 | 0.0217478 | 0.884568 | 0.960086341 |

| ensembl_gene_id | external_gene_name | gene_biotype | log2 (fold change) | log2 (counts per million) | F-value | p-value | False discovery rate |
|---|---|---|---|---|---|---|---|
| ENSMUSG00000001440 | Kpnb1 | protein_coding | -0.038550645 | 6.058698675 | 0.0217393 | 0.8845902 | 0.960086341 |
| ENSMUSG00000019796 | Lrp11 | protein_coding | -0.052993654 | 5.493221051 | 0.0216983 | 0.8846984 | 0.960086341 |
| ENSMUSG00000020305 | Asb3 | protein_coding | -0.051981983 | 5.64929895 | 0.0216865 | 0.8847294 | 0.960086341 |
| ENSMUSG00000024970 | AI846148 | protein_coding | -0.064239432 | 4.005249017 | 0.0215191 | 0.8851718 | 0.960208015 |
| ENSMUSG00000028991 | Mtor | protein_coding | 0.058115899 | 4.714338801 | 0.0215086 | 0.8851996 | 0.960208015 |
| ENSMUSG00000069094 | Pde7a | protein_coding | 0.089427227 | 3.208383452 | 0.0214896 | 0.88525 | 0.960208015 |
| ENSMUSG00000035910 | Dcdc2a | protein_coding | -0.068302906 | 4.129149183 | 0.0214738 | 0.8852919 | 0.960208015 |
| ENSMUSG00000026155 | Smap1 | protein_coding | 0.076918856 | 3.407709961 | 0.0214623 | 0.8853222 | 0.960208015 |
| ENSMUSG00000086191 | Zfp652os | antisense_RNA | 0.094244479 | 2.934209577 | 0.0214416 | 0.8853773 | 0.960208015 |
| ENSMUSG00000052684 | Jun | protein_coding | 0.078688394 | 3.651238088 | 0.0213759 | 0.8855516 | 0.960300186 |
| ENSMUSG00000031353 | Rbbp7 | protein_coding | -0.038469738 | 7.023521773 | 0.021249 | 0.8858894 | 0.960540167 |
| ENSMUSG00000057469 | E2f6 | protein_coding | -0.0657363 | 4.490085816 | 0.0212256 | 0.8859516 | 0.960540167 |
| ENSMUSG00000031323 | Dmrtc1a | protein_coding | -0.07832379 | 3.489503687 | 0.0211386 | 0.886184 | 0.960580134 |
| ENSMUSG00000018446 | C1qbp | protein_coding | 0.053404211 | 5.541922467 | 0.0211014 | 0.8862834 | 0.960580134 |
| ENSMUSG00000031385 | Plxnb3 | protein_coding | -0.117142097 | 2.818135004 | 0.0210808 | 0.8863385 | 0.960580134 |
| ENSMUSG00000034612 | Chst11 | protein_coding | 0.060569263 | 3.991969294 | 0.0210781 | 0.8863458 | 0.960580134 |
| ENSMUSG00000029622 | Arpc1b | protein_coding | 0.058335243 | 4.752222559 | 0.021038 | 0.8864531 | 0.960599649 |
| ENSMUSG00000038280 | Ostm1 | protein_coding | 0.056206845 | 4.833108994 | 0.0209364 | 0.8867256 | 0.960762913 |
| ENSMUSG00000020491 | 2810021J22Rik | protein_coding | -0.076447037 | 3.813919539 | 0.0209076 | 0.8868028 | 0.960762913 |
| ENSMUSG00000056131 | Pgm3 | protein_coding | -0.082747826 | 4.301884487 | 0.0208641 | 0.8869229 | 0.960762913 |
| ENSMUSG00000042650 | Alkbh5 | protein_coding | 0.06375661 | 6.623995135 | 0.0209042 | 0.887042 | 0.960762913 |
| ENSMUSG00000063439 | B9d2 | protein_coding | -0.034029681 | 7.162228884 | 0.0207817 | 0.8871417 | 0.960762913 |
| ENSMUSG00000028318 | Polr1e | protein_coding | -0.063829789 | 4.451103162 | 0.0207603 | 0.8871993 | 0.960762913 |
| ENSMUSG00000043102 | Qrfp | protein_coding | 0.240734572 | 2.17994674 | 0.0207828 | 0.8872514 | 0.960762913 |
| ENSMUSG00000024122 | Pdpk1 | protein_coding | -0.038295435 | 6.553150578 | 0.0207162 | 0.8873186 | 0.960762913 |
| ENSMUSG00000027361 | Gabpb1 | protein_coding | -0.062488557 | 4.01135822 | 0.0205501 | 0.8877678 | 0.960997116 |
| ENSMUSG00000035585 | Tsen34 | protein_coding | -0.065484977 | 4.467188683 | 0.0205253 | 0.8878351 | 0.960997116 |
| ENSMUSG00000050730 | Arhgap42 | protein_coding | 0.171797079 | 2.211125599 | 0.0205214 | 0.8878455 | 0.960997116 |
| ENSMUSG00000049932 | H2afx | protein_coding | 0.112277296 | 2.595392298 | 0.0204985 | 0.8879079 | 0.960997116 |
| ENSMUSG00000059970 | Hspa2 | protein_coding | -0.061510544 | 5.644125734 | 0.0204659 | 0.8879992 | 0.960997116 |
| ENSMUSG00000018395 | Kif3a | protein_coding | -0.044097404 | 7.687993734 | 0.0203725 | 0.8882533 | 0.960997116 |
| ENSMUSG00000026172 | Bcs1l | protein_coding | 0.042340845 | 6.012627498 | 0.0203688 | 0.8882603 | 0.960997116 |
| ENSMUSG00000036980 | Taf6 | protein_coding | 0.064735465 | 5.728070284 | 0.0203715 | 0.8883334 | 0.960997116 |
| ENSMUSG00000056856 | Jakmip3 | protein_coding | -0.060221934 | 4.059510386 | 0.0203 | 0.8884479 | 0.960997116 |
| ENSMUSG00000025202 | Scd3 | protein_coding | -0.06460794 | 4.335583189 | 0.0202117 | 0.8886889 | 0.960997116 |
| ENSMUSG00000021928 | Ebpl | protein_coding | -0.052704714 | 4.421422157 | 0.0201847 | 0.8887628 | 0.960997116 |
| ENSMUSG00000050373 | Snx21 | protein_coding | 0.060630967 | 4.355608397 | 0.0201317 | 0.8889079 | 0.960997116 |
| ENSMUSG00000032280 | Tle3 | protein_coding | -0.098461783 | 3.152596638 | 0.0201019 | 0.8889895 | 0.960997116 |
| ENSMUSG00000037791 | Phf12 | protein_coding | 0.066766821 | 4.091898863 | 0.0200811 | 0.8890467 | 0.960997116 |
| ENSMUSG00000030275 | Etnk1 | protein_coding | 0.134914274 | 2.811776747 | 0.0200742 | 0.8890681 | 0.960997116 |
| ENSMUSG00000052137 | Rbm12b2 | protein_coding | -0.093949841 | 3.454937519 | 0.0200691 | 0.8890795 | 0.960997116 |
| ENSMUSG00000112390 | CT025654.1 | processed_transcript | 0.098785105 | 3.127733802 | 0.0200611 | 0.8891017 | 0.960997116 |
| ENSMUSG00000063430 | Wscd2 | protein_coding | -0.073714235 | 3.351754842 | 0.0200458 | 0.8891435 | 0.960997116 |
| ENSMUSG00000020720 | Psmd12 | protein_coding | 0.03305995 | 8.112982504 | 0.0199706 | 0.8893503 | 0.961124046 |
| ENSMUSG00000083844 | Ube2d-ps | transcribed_unproces | 0.040763667 | 5.765932777 | 0.0198058 | 0.8898045 | 0.961341154 |
| ENSMUSG00000037737 | Actrt3 | protein_coding | 0.152836106 | 2.315384612 | 0.0197973 | 0.889828 | 0.961341154 |
| ENSMUSG00000027751 | Supt20 | protein_coding | -0.070778949 | 4.469264427 | 0.0197932 | 0.8898392 | 0.961341154 |
| ENSMUSG00000091625 | Lsm5 | protein_coding | -0.089339702 | 3.039674698 | 0.0197379 | 0.8899924 | 0.961341154 |
| ENSMUSG00000073062 | Zxdb | protein_coding | -0.084751693 | 3.790263018 | 0.0197368 | 0.8899983 | 0.961341154 |
| ENSMUSG00000084416 | Rpl10a-ps1 | processed_pseudoge | 0.083184847 | 2.886219556 | 0.0196806 | 0.8901509 | 0.96140942 |
| ENSMUSG00000059447 | Hadhb | protein_coding | 0.059970475 | 3.844665674 | 0.0196232 | 0.8903102 | 0.961484944 |
| ENSMUSG00000020023 | Tmcc3 | protein_coding | -0.152773458 | 2.391541803 | 0.0194884 | 0.8906851 | 0.961744912 |
| ENSMUSG00000059336 | Slc14a1 | protein_coding | -0.11924262 | 2.821442669 | 0.0194246 | 0.8908628 | 0.961744912 |
| ENSMUSG00000035049 | Rrp12 | protein_coding | 0.057252091 | 4.313849598 | 0.0194211 | 0.8908726 | 0.961744912 |
| ENSMUSG00000018923 | Med11 | protein_coding | -0.048221508 | 5.137953717 | 0.0194082 | 0.8909087 | 0.961744912 |
| ENSMUSG00000010663 | Fads1 | protein_coding | 0.04151191 | 5.969591678 | 0.0192928 | 0.8912311 | 0.961958381 |
| ENSMUSG00000057835 | Zfp119a | protein_coding | -0.10308903 | 3.001432612 | 0.0192564 | 0.8913331 | 0.961958381 |
| ENSMUSG00000032062 | 2310030G06Rik | protein_coding | -0.081881958 | 3.466498892 | 0.0192165 | 0.8914451 | 0.961958381 |
| ENSMUSG00000042492 | Tbc1d10b | protein_coding | -0.043290559 | 5.264162678 | 0.0192096 | 0.8914643 | 0.961958381 |
| ENSMUSG00000024927 | Rela | protein_coding | 0.054735281 | 5.16523938 | 0.0190952 | 0.8917857 | 0.962208725 |
| ENSMUSG00000049422 | Chchd10 | protein_coding | -0.052926954 | 5.888721733 | 0.0190415 | 0.89194 | 0.962278613 |
| ENSMUSG00000032127 | Vps11 | protein_coding | 0.053583916 | 6.488605054 | 0.0190401 | 0.8920373 | 0.962287081 |
| ENSMUSG00000065954 | Tacc1 | protein_coding | 0.038841095 | 6.421678638 | 0.0189472 | 0.8922032 | 0.962369431 |
| ENSMUSG00000073700 | Klhl21 | protein_coding | 0.078008556 | 3.31837598 | 0.0188651 | 0.8924356 | 0.962391001 |
| ENSMUSG00000033790 | Tubgcp5 | protein_coding | 0.048706513 | 4.515937438 | 0.0188559 | 0.8924615 | 0.962391001 |
| ENSMUSG00000028990 | Lzic | protein_coding | 0.064202697 | 4.126036374 | 0.0187715 | 0.8927008 | 0.962391001 |
| ENSMUSG00000029127 | Zbtb49 | protein_coding | 0.188131712 | 2.185977151 | 0.0187613 | 0.8927399 | 0.962391001 |
| ENSMUSG00000058440 | Nrf1 | protein_coding | -0.04747169 | 4.588800568 | 0.0187213 | 0.8928434 | 0.962391001 |
| ENSMUSG00000027091 | Zc3h15 | protein_coding | 0.042245403 | 6.253885872 | 0.0187074 | 0.892883 | 0.962391001 |
| ENSMUSG00000027940 | Tpm3 | protein_coding | -0.032263607 | 6.75232842 | 0.0186975 | 0.8929112 | 0.962391001 |
| ENSMUSG00000005374 | Tbl2 | protein_coding | -0.072180692 | 3.460875466 | 0.0186877 | 0.8929392 | 0.962391001 |
| ENSMUSG00000022300 | Dcaf13 | protein_coding | 0.048415216 | 6.142582573 | 0.0186053 | 0.8931764 | 0.962550212 |
| ENSMUSG00000031482 | Slc25a15 | protein_coding | 0.077367596 | 3.020939249 | 0.0185187 | 0.893421 | 0.962653846 |
| ENSMUSG00000003411 | Rab3b | protein_coding | -0.032029659 | 8.485015231 | 0.0184835 | 0.8935217 | 0.962653846 |
| ENSMUSG00000054074 | Skida1 | protein_coding | 0.109524565 | 2.673199539 | 0.0184486 | 0.8936218 | 0.962653846 |
| ENSMUSG00000067150 | Xpo5 | protein_coding | 0.0682305 | 3.970514468 | 0.0184455 | 0.8936307 | 0.962653846 |
| ENSMUSG00000066175 | 2510046G10Rik | antisense_RNA | -0.160571705 | 2.061809455 | 0.0183982 | 0.8937661 | 0.962703321 |
| ENSMUSG00000026135 | Zfp142 | protein_coding | -0.13052235 | 2.531789091 | 0.0183593 | 0.8938779 | 0.962727222 |

| ensembl_gene_id | external_gene_name | gene_biotype | log2 (fold change) | log2 (counts per million) | F-value | p-value | False discovery rate |
|---|---|---|---|---|---|---|---|
| ENSMUSG00000032525 | Nktr | protein_coding | 0.075259691 | 4.031667074 | 0.0182528 | 0.8941841 | 0.96279302 |
| ENSMUSG00000020124 | Usp15 | protein_coding | -0.03715514 | 5.88853095 | 0.0182428 | 0.8942129 | 0.96279302 |
| ENSMUSG00000095041 | AC149090.1 | protein_coding | 0.062459399 | 4.889587051 | 0.0182068 | 0.8943165 | 0.96279302 |
| ENSMUSG00000030008 | Pradc1 | protein_coding | -0.087840555 | 3.403218735 | 0.0181912 | 0.8943617 | 0.96279302 |
| ENSMUSG00000055319 | Sec23ip | protein_coding | -0.049617065 | 5.61426494 | 0.0181825 | 0.8943866 | 0.96279302 |
| ENSMUSG00000020677 | Ddx52 | protein_coding | 0.039800728 | 5.736859524 | 0.0180215 | 0.8948523 | 0.962960851 |
| ENSMUSG00000028680 | Plk3 | protein_coding | 0.082782424 | 3.88673489 | 0.0180201 | 0.8948591 | 0.962960851 |
| ENSMUSG00000053070 | 9230110C19Rik | protein_coding | -0.060273563 | 4.620341163 | 0.0179902 | 0.8949431 | 0.962960851 |
| ENSMUSG00000009070 | Rsph14 | protein_coding | -0.084115151 | 3.214988781 | 0.0179773 | 0.8949806 | 0.962960851 |
| ENSMUSG00000038520 | Tbc1d17 | protein_coding | -0.040869024 | 6.289337849 | 0.0179739 | 0.8949903 | 0.962960851 |
| ENSMUSG00000022471 | Xrcc6 | protein_coding | -0.080852825 | 3.169702503 | 0.0179282 | 0.8951231 | 0.963007395 |
| ENSMUSG00000024955 | Esrra | protein_coding | -0.121383383 | 2.598698475 | 0.017858 | 0.8953275 | 0.963059407 |
| ENSMUSG00000057455 | Rit2 | protein_coding | 0.035010484 | 9.683594199 | 0.01785 | 0.8953506 | 0.963059407 |
| ENSMUSG00000061360 | Phf5a | protein_coding | 0.038927072 | 6.590204573 | 0.0177815 | 0.8955503 | 0.963160382 |
| ENSMUSG00000034324 | Tmem132c | protein_coding | -0.112766647 | 2.717999183 | 0.0177564 | 0.8956236 | 0.963160382 |
| ENSMUSG00000028373 | Astn2 | protein_coding | 0.049119545 | 5.382908985 | 0.0175911 | 0.8961076 | 0.963584496 |
| ENSMUSG00000087403 | Kantr | protein_coding | -0.044825926 | 6.665898781 | 0.0174196 | 0.8966147 | 0.963964318 |
| ENSMUSG00000024130 | Abca3 | protein_coding | 0.093150656 | 2.720833177 | 0.0173875 | 0.896707 | 0.963964318 |
| ENSMUSG00000010175 | Prox1 | protein_coding | 0.070380063 | 4.413979625 | 0.0173763 | 0.8967427 | 0.963964318 |
| ENSMUSG00000036632 | Alg5 | protein_coding | -0.055314657 | 5.020217067 | 0.0173494 | 0.8968194 | 0.963964318 |
| ENSMUSG00000031487 | Brf2 | protein_coding | -0.046539811 | 5.21612389 | 0.0169962 | 0.8978687 | 0.96497026 |
| ENSMUSG00000021111 | Papola | protein_coding | -0.038796082 | 6.108188155 | 0.0169503 | 0.898006 | 0.96497026 |
| ENSMUSG00000000088 | Cox5a | protein_coding | 0.046890217 | 4.60852763 | 0.0169441 | 0.8980245 | 0.96497026 |
| ENSMUSG00000042073 | Abhd14b | protein_coding | -0.069806913 | 3.539141639 | 0.0169111 | 0.8981234 | 0.964980098 |
| ENSMUSG00000019738 | Polr2i | protein_coding | 0.03917677 | 6.638654352 | 0.0168726 | 0.8982385 | 0.965007353 |
| ENSMUSG00000020102 | Slc16a7 | protein_coding | 0.076896094 | 3.21491113 | 0.0167741 | 0.8985342 | 0.965062434 |
| ENSMUSG00000055760 | Gemin6 | protein_coding | -0.063749461 | 4.42354317 | 0.0167563 | 0.8985877 | 0.965062434 |
| ENSMUSG00000034601 | 2700049A03Rik | protein_coding | 0.112282069 | 2.973556882 | 0.0167566 | 0.8985895 | 0.965062434 |
| ENSMUSG00000028597 | Gpx7 | protein_coding | 0.067414214 | 3.864937553 | 0.0167069 | 0.8987364 | 0.965062434 |
| ENSMUSG00000030779 | Rbbp6 | protein_coding | -0.037212611 | 5.982878442 | 0.0166597 | 0.8988788 | 0.965062434 |
| ENSMUSG00000031955 | Bcar1 | protein_coding | -0.056740852 | 4.683054308 | 0.0166507 | 0.8989059 | 0.965062434 |
| ENSMUSG00000042851 | Zc3h6 | protein_coding | 0.070814992 | 3.161167287 | 0.0166467 | 0.898918 | 0.965062434 |
| ENSMUSG00000058351 | Smim4 | protein_coding | 0.053315248 | 5.621326812 | 0.0166019 | 0.8990559 | 0.965114111 |
| ENSMUSG00000093606 | B130034C11Rik | processed_transcript | -0.108447203 | 2.838125638 | 0.0165636 | 0.8991693 | 0.965139504 |
| ENSMUSG00000043079 | Synpo | protein_coding | -0.044674303 | 5.277590938 | 0.0164703 | 0.8994519 | 0.965346504 |
| ENSMUSG00000029918 | Mrps33 | protein_coding | 0.038035251 | 6.964220476 | 0.0163469 | 0.8998272 | 0.965560291 |
| ENSMUSG00000048930 | Tada3 | protein_coding | 0.042439557 | 4.935587301 | 0.0163457 | 0.8998307 | 0.965560291 |
| ENSMUSG00000032267 | Usp28 | protein_coding | 0.120928248 | 2.665603186 | 0.016269 | 0.9000647 | 0.965714969 |
| ENSMUSG00000004880 | Lbr | protein_coding | 0.067183914 | 3.517130989 | 0.0161858 | 0.9003192 | 0.965891727 |
| ENSMUSG00000050954 | Zfp169 | protein_coding | -0.102112481 | 2.607587525 | 0.0161456 | 0.9004424 | 0.965927468 |
| ENSMUSG00000062169 | Cnih4 | protein_coding | 0.038688759 | 5.796278403 | 0.0160941 | 0.9006003 | 0.966000507 |
| ENSMUSG00000073236 | 2500004C02Rik | processed_transcript | -0.048911758 | 4.325522771 | 0.016032 | 0.9007911 | 0.966108759 |
| ENSMUSG00000026932 | Nacc2 | protein_coding | 0.042533354 | 6.10164232 | 0.0159064 | 0.9011785 | 0.966344396 |
| ENSMUSG00000072623 | Zfp9 | protein_coding | 0.07309492 | 4.1590308 | 0.0159033 | 0.9011905 | 0.966344396 |
| ENSMUSG00000006932 | Ctnnb1 | protein_coding | 0.077744472 | 3.323877475 | 0.0156896 | 0.9018504 | 0.966955543 |
| ENSMUSG00000034022 | Cpsf1 | protein_coding | -0.0525154 | 4.900114075 | 0.0155544 | 0.9022719 | 0.967055237 |
| ENSMUSG00000024423 | Impact | protein_coding | -0.042681108 | 10.99090957 | 0.0155526 | 0.9022799 | 0.967055237 |
| ENSMUSG00000037610 | Kcnmb2 | protein_coding | -0.10536932 | 2.971614265 | 0.0155443 | 0.9023059 | 0.967055237 |
| ENSMUSG00000002833 | Hdgfl2 | protein_coding | 0.04717468 | 5.880915318 | 0.0155186 | 0.9023862 | 0.967055237 |
| ENSMUSG00000072494 | Ppp1r3e | protein_coding | -0.071910679 | 3.247803737 | 0.0154874 | 0.9024813 | 0.967055237 |
| ENSMUSG00000031072 | Oraov1 | protein_coding | -0.040055058 | 5.316259961 | 0.0154869 | 0.902483 | 0.967055237 |
| ENSMUSG00000021144 | Mta1 | protein_coding | -0.052922379 | 5.067801162 | 0.0154257 | 0.9026749 | 0.967164503 |
| ENSMUSG00000029093 | Sorcs2 | protein_coding | 0.071469717 | 3.639065966 | 0.0153816 | 0.9028131 | 0.967194874 |
| ENSMUSG00000051238 | Swsap1 | protein_coding | -0.081904325 | 2.90923937 | 0.0153534 | 0.902902 | 0.967194874 |
| ENSMUSG00000040850 | Psme4 | protein_coding | -0.049205323 | 4.122706037 | 0.0153308 | 0.9029731 | 0.967194874 |
| ENSMUSG00000002052 | Supt6 | protein_coding | 0.036098552 | 6.11979242 | 0.0152385 | 0.9032639 | 0.96737687 |
| ENSMUSG00000034190 | Chmp7 | protein_coding | 0.039722787 | 6.459667764 | 0.0151432 | 0.9035652 | 0.96737687 |
| ENSMUSG00000026427 | Eif2d | protein_coding | 0.041801719 | 5.489170157 | 0.0151064 | 0.9036819 | 0.96737687 |
| ENSMUSG00000076928 | Trac | TR_C_gene | -0.138738502 | 2.465869828 | 0.015061 | 0.9038285 | 0.96737687 |
| ENSMUSG00000049134 | Nrap | protein_coding | 0.239084573 | 1.738939819 | 0.0150867 | 0.9038462 | 0.96737687 |
| ENSMUSG00000076432 | Ywhaq | protein_coding | -0.02585264 | 8.08258164 | 0.0150331 | 0.9039147 | 0.96737687 |
| ENSMUSG00000028926 | Cdk14 | protein_coding | 0.044745209 | 7.358194696 | 0.0150532 | 0.9039462 | 0.96737687 |
| ENSMUSG00000031879 | Fam96b | protein_coding | -0.034336207 | 6.989638524 | 0.0150218 | 0.9039505 | 0.96737687 |
| ENSMUSG00000032220 | Myo1e | protein_coding | -0.089609967 | 2.891451718 | 0.0150141 | 0.9039749 | 0.96737687 |
| ENSMUSG00000029401 | Rilpl2 | protein_coding | 0.057418101 | 4.513481641 | 0.0149661 | 0.9041277 | 0.96737687 |
| ENSMUSG00000043531 | Sorcs1 | protein_coding | 0.057199421 | 3.709531103 | 0.0149436 | 0.9041995 | 0.96737687 |
| ENSMUSG00000053870 | Fpgt | protein_coding | 0.073060911 | 3.953645991 | 0.0149372 | 0.9042225 | 0.96737687 |
| ENSMUSG00000029110 | Rnf4 | protein_coding | 0.034017187 | 6.827518734 | 0.0146301 | 0.9052045 | 0.968331102 |
| ENSMUSG00000020312 | Shc2 | protein_coding | 0.066174585 | 3.328318316 | 0.0145869 | 0.9053439 | 0.968342546 |
| ENSMUSG00000049470 | Aff4 | protein_coding | 0.033186462 | 6.631452802 | 0.014571 | 0.9053953 | 0.968342546 |
| ENSMUSG00000021948 | Prkcd | protein_coding | -0.067526813 | 5.889248203 | 0.0145763 | 0.9056133 | 0.96835025 |
| ENSMUSG00000006673 | Qrich1 | protein_coding | 0.033094728 | 5.913338784 | 0.0145002 | 0.9056241 | 0.96835025 |
| ENSMUSG00000067071 | Hes6 | protein_coding | -0.065285273 | 3.233856932 | 0.0144852 | 0.9056727 | 0.96835025 |
| ENSMUSG00000048490 | Nrip1 | protein_coding | -0.049185476 | 4.388132433 | 0.0143874 | 0.90599 | 0.968456395 |
| ENSMUSG00000056158 | Car10 | protein_coding | -0.049433432 | 6.29530553 | 0.0144088 | 0.906014 | 0.968456395 |
| ENSMUSG00000059540 | Tcea2 | protein_coding | 0.05886205 | 4.56843756 | 0.0143714 | 0.9060422 | 0.968456395 |
| ENSMUSG00000023755 | Rhebl1 | protein_coding | -0.072151906 | 4.079893955 | 0.0143197 | 0.9062127 | 0.96846797 |
| ENSMUSG00000028187 | Rpf1 | protein_coding | 0.04705114 | 4.458672323 | 0.0142918 | 0.9063013 | 0.96846797 |

| ensembl_gene_id | external_gene_name | gene_biotype | log2 (fold change) | log2 (counts per million) | F-value | p-value | False discovery rate |
|---|---|---|---|---|---|---|---|
| ENSMUSG00000037060 | Cavin3 | protein_coding | -0.04062342 | 5.077329017 | 0.0142851 | 0.9063232 | 0.96846797 |
| ENSMUSG00000048120 | Entpd1 | protein_coding | -0.062004045 | 3.685527013 | 0.0142004 | 0.9065999 | 0.96866117 |
| ENSMUSG00000095930 | Nim1k | protein_coding | 0.038736825 | 5.445212786 | 0.0141747 | 0.9066842 | 0.96866117 |
| ENSMUSG00000035232 | Pdk3 | protein_coding | 0.085087228 | 4.387157413 | 0.0141396 | 0.9069877 | 0.968889165 |
| ENSMUSG00000021764 | Ndufs4 | protein_coding | 0.037885062 | 8.83922535 | 0.0138338 | 0.9078097 | 0.969656157 |
| ENSMUSG00000056820 | Tsnax | protein_coding | -0.033449561 | 6.500801362 | 0.0137925 | 0.9079447 | 0.969656157 |
| ENSMUSG00000021113 | Snapc1 | protein_coding | 0.034581602 | 5.663202216 | 0.013783 | 0.9079762 | 0.969656157 |
| ENSMUSG00000039683 | Sdk1 | protein_coding | 0.111061605 | 2.433209778 | 0.0137539 | 0.9080729 | 0.969663193 |
| ENSMUSG00000018821 | Avpi1 | protein_coding | -0.030027288 | 7.044228334 | 0.0136401 | 0.9084521 | 0.969971756 |
| ENSMUSG00000023980 | Taf8 | protein_coding | -0.039149588 | 5.220399206 | 0.0136064 | 0.9085646 | 0.969995521 |
| ENSMUSG00000038534 | Osbpl7 | protein_coding | 0.043963044 | 4.381384105 | 0.0135361 | 0.9088002 | 0.970150768 |
| ENSMUSG00000003345 | Csnk1g2 | protein_coding | 0.048779959 | 5.019597493 | 0.0134665 | 0.9090338 | 0.970180856 |
| ENSMUSG00000031751 | Amfr | protein_coding | -0.040269229 | 6.103461629 | 0.0134573 | 0.9090648 | 0.970180856 |
| ENSMUSG00000037151 | Lrrc20 | protein_coding | 0.04809395 | 4.799613182 | 0.0134471 | 0.9090991 | 0.970180856 |
| ENSMUSG00000073755 | 5730409E04Rik | protein_coding | -0.035965464 | 5.339607032 | 0.0133508 | 0.9094236 | 0.970430906 |
| ENSMUSG00000049717 | Lig4 | protein_coding | 0.082340599 | 3.23115119 | 0.0132756 | 0.9096779 | 0.970543163 |
| ENSMUSG00000000631 | Myo18a | protein_coding | -0.040133812 | 5.756674525 | 0.0132588 | 0.9097345 | 0.970543163 |
| ENSMUSG00000032526 | Ss18l2 | protein_coding | 0.026097867 | 8.073288735 | 0.0132334 | 0.9098208 | 0.970543163 |
| ENSMUSG00000020646 | Mboat2 | protein_coding | 0.043604982 | 4.425745491 | 0.0131913 | 0.9099636 | 0.970543163 |
| ENSMUSG00000038855 | Itpkb | protein_coding | -0.053350361 | 4.182160553 | 0.0131855 | 0.9099832 | 0.970543163 |
| ENSMUSG00000017421 | Zfp207 | protein_coding | -0.029365116 | 7.026281644 | 0.0131586 | 0.9100749 | 0.970543163 |
| ENSMUSG00000062376 | Borcs7 | protein_coding | -0.025875693 | 7.115717169 | 0.013126 | 0.9101858 | 0.970543163 |
| ENSMUSG00000050812 | AI314180 | protein_coding | 0.037907749 | 5.528441864 | 0.0131069 | 0.9102509 | 0.970543163 |
| ENSMUSG00000030036 | Mogs | protein_coding | 0.073506491 | 3.324228333 | 0.0130544 | 0.91043 | 0.970556221 |
| ENSMUSG00000002055 | Spag5 | protein_coding | -0.125833457 | 2.194848976 | 0.0130504 | 0.9104437 | 0.970556221 |
| ENSMUSG00000003731 | Kpna6 | protein_coding | 0.041306902 | 4.824788122 | 0.0129575 | 0.9107613 | 0.970677753 |
| ENSMUSG00000026610 | Esrrg | protein_coding | 0.047710502 | 4.177014255 | 0.0129401 | 0.9108211 | 0.970677753 |
| ENSMUSG00000040723 | Rcsd1 | protein_coding | -0.09778738 | 2.563251632 | 0.0129379 | 0.9108285 | 0.970677753 |
| ENSMUSG00000113429 | CT030140.1 | lincRNA | 0.073752524 | 2.950139412 | 0.0128716 | 0.9110564 | 0.970679045 |
| ENSMUSG00000066892 | Fbxl12 | protein_coding | -0.060366213 | 3.706845045 | 0.0128605 | 0.9110945 | 0.970679045 |
| ENSMUSG00000039809 | Gabbr2 | protein_coding | 0.037844409 | 5.834550087 | 0.0128385 | 0.9111702 | 0.970679045 |
| ENSMUSG00000020721 | Helz | protein_coding | 0.033793517 | 5.362374502 | 0.0128137 | 0.9112556 | 0.970679045 |
| ENSMUSG00000004591 | Pkn2 | protein_coding | 0.061371295 | 3.453411419 | 0.012791 | 0.9113341 | 0.970679045 |
| ENSMUSG00000086851 | A430108G06Rik | processed_transcript | -0.156958737 | 2.553515066 | 0.0128202 | 0.9113713 | 0.970679045 |
| ENSMUSG00000020955 | Ap4s1 | protein_coding | -0.035418966 | 5.522627673 | 0.012739 | 0.9115137 | 0.970682659 |
| ENSMUSG00000100164 | 2610306M01Rik | bidirectional_promote | -0.036717575 | 5.269928062 | 0.0126335 | 0.9118792 | 0.970682659 |
| ENSMUSG00000059939 | 9430015G10Rik | protein_coding | 0.052985466 | 3.558765264 | 0.0126279 | 0.9118987 | 0.970682659 |
| ENSMUSG00000020092 | Pald1 | protein_coding | 0.111679164 | 2.379014668 | 0.0126223 | 0.9119182 | 0.970682659 |
| ENSMUSG00000031565 | Fgfr1 | protein_coding | -0.036027109 | 5.45452829 | 0.0126211 | 0.9119224 | 0.970682659 |
| ENSMUSG00000042857 | Gm9776 | bidirectional_promote | -0.083523728 | 2.515001981 | 0.0126202 | 0.9119254 | 0.970682659 |
| ENSMUSG00000004665 | Cnn2 | protein_coding | 0.094293059 | 2.957617468 | 0.0125604 | 0.9121334 | 0.970682659 |
| ENSMUSG00000061689 | Dlgap4 | protein_coding | 0.029685248 | 6.687834921 | 0.0125588 | 0.912139 | 0.970682659 |
| ENSMUSG00000022131 | Gpr180 | protein_coding | 0.059316669 | 3.381276287 | 0.0125007 | 0.9123416 | 0.970682659 |
| ENSMUSG00000020458 | Rtn4 | protein_coding | 0.029310833 | 8.419615773 | 0.0125001 | 0.9123435 | 0.970682659 |
| ENSMUSG00000004317 | Clcn5 | protein_coding | 0.050034968 | 4.298924838 | 0.0124932 | 0.9123677 | 0.970682659 |
| ENSMUSG00000063382 | Bcl9l | protein_coding | 0.05939215 | 4.306735591 | 0.0124404 | 0.9125546 | 0.970785467 |
| ENSMUSG00000022897 | Dyrk1a | protein_coding | -0.03287184 | 6.246967347 | 0.0123536 | 0.9128564 | 0.970934008 |
| ENSMUSG00000024654 | Asrgl1 | protein_coding | -0.025464778 | 7.648444656 | 0.0123427 | 0.9128947 | 0.970934008 |
| ENSMUSG00000047617 | Paxx | protein_coding | -0.025592039 | 7.046361588 | 0.0123227 | 0.9129651 | 0.970934008 |
| ENSMUSG00000030733 | Sh2b1 | protein_coding | -0.037615773 | 4.85364938 | 0.0121626 | 0.91353 | 0.971438682 |
| ENSMUSG00000047712 | Ust | protein_coding | -0.071150571 | 3.075890735 | 0.0121347 | 0.9136285 | 0.971447351 |
| ENSMUSG00000038022 | Mindy4 | protein_coding | -0.068305008 | 3.076901349 | 0.0119158 | 0.9144079 | 0.972179995 |
| ENSMUSG00000053985 | Zfp14 | protein_coding | -0.048902294 | 3.830076174 | 0.0117704 | 0.9149297 | 0.972638574 |
| ENSMUSG00000047879 | Usp14 | protein_coding | 0.03049237 | 7.679212796 | 0.011704 | 0.9151687 | 0.972717407 |
| ENSMUSG00000024875 | Yif1a | protein_coding | 0.028961737 | 6.495635627 | 0.0116747 | 0.9152746 | 0.972717407 |
| ENSMUSG00000012640 | Zfp715 | protein_coding | -0.055421868 | 3.863644095 | 0.0116745 | 0.9152752 | 0.972717407 |
| ENSMUSG00000065922 | n-R5-8s1 | rRNA | -0.04658846 | 5.306719367 | 0.0116073 | 0.9155206 | 0.972816759 |
| ENSMUSG00000022462 | Slc38a2 | protein_coding | 0.035598876 | 5.304103256 | 0.0115771 | 0.915628 | 0.972816759 |
| ENSMUSG00000008859 | Rala | protein_coding | 0.034081506 | 5.393500259 | 0.0115621 | 0.9156824 | 0.972816759 |
| ENSMUSG00000040302 | Rbm48 | protein_coding | -0.041531311 | 4.46789051 | 0.0114928 | 0.9159346 | 0.972816759 |
| ENSMUSG00000032285 | Dnaja4 | protein_coding | 0.076642624 | 2.787858811 | 0.0114886 | 0.9159499 | 0.972816759 |
| ENSMUSG00000033170 | Card10 | protein_coding | -0.155358629 | 2.435956805 | 0.0114834 | 0.9160994 | 0.972816759 |
| ENSMUSG00000062031 | Pgghg | protein_coding | -0.128617276 | 2.076154502 | 0.0114374 | 0.9161367 | 0.972816759 |
| ENSMUSG00000033916 | Chmp2a | protein_coding | -0.022852756 | 9.030813832 | 0.0114058 | 0.9162521 | 0.972816759 |
| ENSMUSG00000040213 | Kyat3 | protein_coding | 0.065497873 | 3.134046576 | 0.011375 | 0.9163648 | 0.972816759 |
| ENSMUSG00000108608 | Gm6916 | transcribed_processe | -0.251159778 | 2.572950252 | 0.0115644 | 0.916445 | 0.972816759 |
| ENSMUSG00000090122 | Kcne1l | protein_coding | -0.106835661 | 2.519501753 | 0.0113175 | 0.9165757 | 0.972816759 |
| ENSMUSG00000073643 | Wdfy1 | protein_coding | -0.10017071 | 4.899426387 | 0.0114683 | 0.9166592 | 0.972816759 |
| ENSMUSG00000019822 | Smpd2 | protein_coding | 0.0524746 | 4.065077862 | 0.0112909 | 0.9166735 | 0.972816759 |
| ENSMUSG00000097311 | Gm26871 | processed_transcript | 0.097219717 | 2.800488916 | 0.0112384 | 0.9168665 | 0.972816759 |
| ENSMUSG00000039372 | 4-Mar | protein_coding | 0.056292388 | 3.590618025 | 0.0112316 | 0.9168916 | 0.972816759 |
| ENSMUSG00000031429 | Psmd10 | protein_coding | -0.026351679 | 6.94919868 | 0.011182 | 0.9170745 | 0.972816759 |
| ENSMUSG00000032803 | Cdv3 | protein_coding | 0.031580959 | 5.753626047 | 0.0111809 | 0.9170786 | 0.972816759 |
| ENSMUSG00000029512 | Ulk1 | protein_coding | 0.037458958 | 4.964379479 | 0.0111694 | 0.917121 | 0.972816759 |
| ENSMUSG00000024430 | Cabyr | protein_coding | 0.091212039 | 2.451006749 | 0.0111625 | 0.9171466 | 0.972816759 |
| ENSMUSG00000040557 | Mettl27 | protein_coding | -0.051226953 | 4.134411456 | 0.0111182 | 0.9173105 | 0.972816759 |
| ENSMUSG00000018678 | Sp2 | protein_coding | -0.109103292 | 2.484319972 | 0.0111068 | 0.9173529 | 0.972816759 |
| ENSMUSG00000033429 | Mcee | protein_coding | -0.029589577 | 6.651822384 | 0.0110403 | 0.9175997 | 0.972816759 |

| ensembl_gene_id | external_gene_name | gene_biotype | log2 (fold change) | log2 (counts per million) | F-value | p-value | False discovery rate |
|---|---|---|---|---|---|---|---|
| ENSMUSG00000032423 | Syncrip | protein_coding | 0.054339517 | 3.30915469 | 0.0110322 | 0.9176296 | 0.972816759 |
| ENSMUSG00000029778 | Adcyap1r1 | protein_coding | -0.0295348 | 6.178430116 | 0.0110247 | 0.9176576 | 0.972816759 |
| ENSMUSG00000112121 | AC163903.1 | lincRNA | 0.082892845 | 2.695447453 | 0.0110235 | 0.9176621 | 0.972816759 |
| ENSMUSG00000032308 | Ulk3 | protein_coding | -0.081344885 | 3.059243722 | 0.0109924 | 0.9177779 | 0.972816759 |
| ENSMUSG00000037788 | Vopp1 | protein_coding | -0.044695637 | 4.910781996 | 0.0109834 | 0.9178114 | 0.972816759 |
| ENSMUSG00000020883 | Fbxl20 | protein_coding | -0.048172756 | 3.978598993 | 0.0109343 | 0.9179946 | 0.972915069 |
| ENSMUSG00000031386 | Hcfc1 | protein_coding | -0.046489608 | 5.197956462 | 0.0108369 | 0.9183615 | 0.972941021 |
| ENSMUSG00000030806 | Stx1b | protein_coding | -0.029112478 | 8.032936804 | 0.0108348 | 0.918367 | 0.972941021 |
| ENSMUSG00000008301 | Phax | protein_coding | 0.021698892 | 7.659439916 | 0.01083 | 0.918385 | 0.972941021 |
| ENSMUSG00000027198 | Ext2 | protein_coding | 0.054182307 | 3.945682192 | 0.0107972 | 0.9185082 | 0.972941021 |
| ENSMUSG00000054484 | Tmem62 | protein_coding | -0.047841046 | 4.323852073 | 0.0107745 | 0.9185939 | 0.972941021 |
| ENSMUSG00000020546 | Stxbp4 | protein_coding | -0.05094415 | 4.780325131 | 0.0107706 | 0.9186103 | 0.972941021 |
| ENSMUSG00000050783 | Htr1f | protein_coding | 0.106773532 | 2.268847071 | 0.0107469 | 0.9186975 | 0.972941021 |
| ENSMUSG00000030257 | Srgap3 | protein_coding | -0.030831838 | 6.169413465 | 0.0107349 | 0.918743 | 0.972941021 |
| ENSMUSG00000022354 | Ndufb9 | protein_coding | 0.031275039 | 10.17307244 | 0.0106945 | 0.9188954 | 0.973006635 |
| ENSMUSG00000028331 | Trmo | protein_coding | 0.108857476 | 3.10188917 | 0.0106373 | 0.9192372 | 0.973191061 |
| ENSMUSG00000026632 | Tatdn3 | protein_coding | -0.041725199 | 4.434896797 | 0.0105762 | 0.9193436 | 0.973191061 |
| ENSMUSG00000028907 | Utp11 | protein_coding | -0.026201382 | 6.560606227 | 0.0105748 | 0.9193488 | 0.973191061 |
| ENSMUSG00000086938 | 4930481A15Rik | antisense_RNA | 0.042827 | 3.896316853 | 0.010553 | 0.9194316 | 0.973191061 |
| ENSMUSG00000020308 | Tpgs1 | protein_coding | 0.068610429 | 4.572860729 | 0.0105022 | 0.9197501 | 0.973325083 |
| ENSMUSG00000020056 | Washc3 | protein_coding | 0.029445381 | 6.423032868 | 0.0104406 | 0.9198603 | 0.973325083 |
| ENSMUSG00000034518 | Hmgxb4 | protein_coding | -0.042727086 | 3.850290875 | 0.0104123 | 0.9199685 | 0.973325083 |
| ENSMUSG00000022848 | Dirc2 | protein_coding | -0.038296766 | 5.275796203 | 0.0104105 | 0.9199754 | 0.973325083 |
| ENSMUSG00000103183 | Gm37090 | TEC | -0.124282956 | 2.365240019 | 0.010397 | 0.920029 | 0.973325083 |
| ENSMUSG00000025086 | Trub1 | protein_coding | 0.036176144 | 5.321156264 | 0.0103607 | 0.9201666 | 0.973325083 |
| ENSMUSG00000039601 | Rcan2 | protein_coding | -0.033579573 | 8.189909291 | 0.0103373 | 0.9202614 | 0.973325083 |
| ENSMUSG00000054034 | Tceal5 | protein_coding | 0.025534559 | 6.457638063 | 0.0103305 | 0.9202824 | 0.973325083 |
| ENSMUSG00000038888 | Ctu1 | protein_coding | -0.087009334 | 3.035940077 | 0.0102702 | 0.9205166 | 0.973352544 |
| ENSMUSG00000069237 | Fam8a1 | protein_coding | -0.034439338 | 5.511208279 | 0.0102658 | 0.9205314 | 0.973352544 |
| ENSMUSG00000036873 | 2410004B18Rik | protein_coding | 0.050414564 | 4.247626815 | 0.0102315 | 0.9206638 | 0.973352544 |
| ENSMUSG00000053774 | Ubxn7 | protein_coding | 0.066478959 | 3.209179744 | 0.0102298 | 0.9206704 | 0.973352544 |
| ENSMUSG00000037553 | Zdhhc18 | protein_coding | -0.048803575 | 4.081426624 | 0.0101724 | 0.9208925 | 0.973372237 |
| ENSMUSG00000004056 | Akt2 | protein_coding | -0.03533782 | 4.845874673 | 0.0101702 | 0.9209011 | 0.973372237 |
| ENSMUSG00000070565 | Rasal2 | protein_coding | -0.039529763 | 5.085976539 | 0.0101549 | 0.9209606 | 0.973372237 |
| ENSMUSG00000015652 | Steap1 | protein_coding | 0.186141304 | 2.66212385 | 0.0101876 | 0.921105 | 0.973411883 |
| ENSMUSG00000036968 | Cnpy4 | protein_coding | -0.037833351 | 5.693023427 | 0.0100791 | 0.921255 | 0.973411883 |
| ENSMUSG00000040466 | Blvrb | protein_coding | 0.030200303 | 5.167728801 | 0.0100753 | 0.9212697 | 0.973411883 |
| ENSMUSG00000104864 | Gm43655 | processed_pseudoge | 0.062738406 | 2.811837455 | 0.0099701 | 0.9216802 | 0.973640473 |
| ENSMUSG00000047030 | Spata2 | protein_coding | 0.047663287 | 4.479292543 | 0.0099374 | 0.9218084 | 0.973640473 |
| ENSMUSG00000040760 | Appl1 | protein_coding | -0.029736764 | 5.821678524 | 0.0099181 | 0.9218839 | 0.973640473 |
| ENSMUSG00000019732 | Calr3 | protein_coding | -0.100307591 | 3.324885192 | 0.0098798 | 0.9222046 | 0.973640473 |
| ENSMUSG00000049804 | Armcx4 | protein_coding | -0.055121877 | 4.416621524 | 0.0098059 | 0.9223275 | 0.973640473 |
| ENSMUSG00000001175 | Calm1 | protein_coding | -0.024688093 | 12.85395451 | 0.0098002 | 0.922348 | 0.973640473 |
| ENSMUSG00000026094 | Stk17b | protein_coding | -0.13533943 | 2.255065074 | 0.0097931 | 0.9223869 | 0.973640473 |
| ENSMUSG00000020679 | Hnf1b | protein_coding | -0.055499086 | 3.596697642 | 0.0097877 | 0.9223976 | 0.973640473 |
| ENSMUSG00000084159 | Gm12696 | processed_pseudoge | 0.062039937 | 2.892748836 | 0.0097716 | 0.9224612 | 0.973640473 |
| ENSMUSG00000038525 | Armc10 | protein_coding | -0.059550107 | 3.275490753 | 0.009769 | 0.9224713 | 0.973640473 |
| ENSMUSG00000031748 | Gnao1 | protein_coding | -0.024243784 | 8.517778331 | 0.0097663 | 0.922482 | 0.973640473 |
| ENSMUSG00000023089 | Ndufa5 | protein_coding | 0.034554154 | 10.15552811 | 0.0097062 | 0.9227274 | 0.97374181 |
| ENSMUSG00000031398 | Plxna3 | protein_coding | 0.071987526 | 2.688513231 | 0.0096964 | 0.9227591 | 0.97374181 |
| ENSMUSG00000050213 | Snip1 | protein_coding | 0.050362699 | 3.608189537 | 0.0096056 | 0.9231205 | 0.973815939 |
| ENSMUSG00000032478 | Nme6 | protein_coding | 0.031605339 | 5.070254869 | 0.0095526 | 0.9233319 | 0.973815939 |
| ENSMUSG00000000876 | Pxmp4 | protein_coding | -0.031170151 | 5.15063373 | 0.0095384 | 0.9233888 | 0.973815939 |
| ENSMUSG00000021254 | Gpatch2l | protein_coding | 0.063578045 | 3.772626543 | 0.0095293 | 0.9234274 | 0.973815939 |
| ENSMUSG00000017176 | Nt5c3b | protein_coding | -0.023629566 | 6.755125826 | 0.0095075 | 0.9235126 | 0.973815939 |
| ENSMUSG00000078485 | Plekhn1 | protein_coding | 0.095471144 | 3.134278056 | 0.0095278 | 0.9235203 | 0.973815939 |
| ENSMUSG00000023092 | Fhl1 | protein_coding | 0.026122901 | 7.233668299 | 0.0095034 | 0.9235293 | 0.973815939 |
| ENSMUSG00000071660 | Ttc9c | protein_coding | -0.027366103 | 5.814119013 | 0.0094972 | 0.9235539 | 0.973815939 |
| ENSMUSG00000007594 | Hapln4 | protein_coding | 0.036847071 | 5.143696213 | 0.0094679 | 0.9236715 | 0.973844475 |
| ENSMUSG00000024830 | Rps6kb2 | protein_coding | -0.088047597 | 3.145732886 | 0.0094268 | 0.9238901 | 0.973885782 |
| ENSMUSG00000021428 | Riok1 | protein_coding | -0.036037061 | 4.878698262 | 0.0094131 | 0.9238918 | 0.973885782 |
| ENSMUSG00000021466 | Ptch1 | protein_coding | 0.040738386 | 4.973170299 | 0.009359 | 0.9241102 | 0.973926769 |
| ENSMUSG00000042331 | Specc1 | protein_coding | 0.040807026 | 5.358759164 | 0.0093591 | 0.9241119 | 0.973926769 |
| ENSMUSG00000021179 | Nrde2 | protein_coding | 0.051077654 | 4.271587522 | 0.0092737 | 0.9244557 | 0.974105687 |
| ENSMUSG00000056427 | Slit3 | protein_coding | -0.06612591 | 2.882208452 | 0.0092674 | 0.9244815 | 0.974105687 |
| ENSMUSG00000046791 | Riox1 | protein_coding | 0.067714691 | 2.938538894 | 0.0092497 | 0.9245534 | 0.974105687 |
| ENSMUSG00000020755 | Sap30bp | protein_coding | -0.021002945 | 6.926630139 | 0.0091854 | 0.924815 | 0.974194496 |
| ENSMUSG00000032637 | Atxn2l | protein_coding | 0.056701712 | 3.237501483 | 0.009148 | 0.924968 | 0.974194496 |
| ENSMUSG00000046480 | Scn4b | protein_coding | -0.043564849 | 5.135063588 | 0.0091453 | 0.924981 | 0.974194496 |
| ENSMUSG00000021770 | Samd8 | protein_coding | -0.033769195 | 5.083330362 | 0.0091055 | 0.9251419 | 0.974194496 |
| ENSMUSG00000044155 | Lsm8 | protein_coding | -0.03218826 | 6.069492638 | 0.0090937 | 0.9251902 | 0.974194496 |
| ENSMUSG00000066406 | Akap13 | protein_coding | -0.061773795 | 3.010877153 | 0.0090805 | 0.9252444 | 0.974194496 |
| ENSMUSG00000067424 | Zfp563 | protein_coding | 0.043534769 | 3.802605472 | 0.0090732 | 0.9252741 | 0.974194496 |
| ENSMUSG00000058420 | Syt17 | protein_coding | 0.038472063 | 5.326430569 | 0.0090323 | 0.9254425 | 0.974194496 |
| ENSMUSG00000018196 | Glrx2 | protein_coding | -0.033328803 | 5.315577468 | 0.0090297 | 0.9254531 | 0.974194496 |
| ENSMUSG00000063600 | Egfem1 | protein_coding | -0.047454038 | 3.503946583 | 0.0089791 | 0.9256617 | 0.974261516 |
| ENSMUSG00000023965 | Fbxl17 | protein_coding | 0.02979639 | 5.385805921 | 0.0089703 | 0.9256979 | 0.974261516 |
| ENSMUSG00000026650 | Meig1 | protein_coding | 0.038570561 | 4.534744567 | 0.0087644 | 0.9265529 | 0.974971197 |

| ensembl_gene_id | external_gene_name | gene_biotype | log2 (fold change) | log2 (counts per million) | F-value | p-value | False discovery rate |
|---|---|---|---|---|---|---|---|
| ENSMUSG00000009687 | Fxyd5 | protein_coding | -0.080444708 | 2.35456016 | 0.0087287 | 0.9267021 | 0.974971197 |
| ENSMUSG00000031848 | Lsm4 | protein_coding | -0.067391248 | 4.547718722 | 0.0087565 | 0.9267338 | 0.974971197 |
| ENSMUSG00000062822 | 4833420G17Rik | protein_coding | 0.033004432 | 5.393469938 | 0.0087139 | 0.926764 | 0.974971197 |
| ENSMUSG00000040618 | Pck2 | protein_coding | 0.033719792 | 5.48517705 | 0.0086694 | 0.926951 | 0.974971197 |
| ENSMUSG00000030087 | Klf15 | protein_coding | -0.053245843 | 3.61185396 | 0.0086646 | 0.926971 | 0.974971197 |
| ENSMUSG00000037287 | Tbcel | protein_coding | -0.036818506 | 4.496771683 | 0.0086432 | 0.9270608 | 0.974971197 |
| ENSMUSG00000071708 | Sms | protein_coding | 0.025243654 | 6.018823401 | 0.0086149 | 0.92718 | 0.974971197 |
| ENSMUSG00000032615 | Nt5m | protein_coding | -0.026199456 | 6.081307107 | 0.0085925 | 0.9272745 | 0.974971197 |
| ENSMUSG00000049800 | Sertad2 | protein_coding | 0.040626032 | 3.710481192 | 0.0085915 | 0.9272789 | 0.974971197 |
| ENSMUSG00000024502 | Jakmip2 | protein_coding | -0.024857456 | 6.198559828 | 0.0085158 | 0.9275989 | 0.975212299 |
| ENSMUSG00000024844 | Banf1 | protein_coding | -0.025589738 | 7.963875365 | 0.0084686 | 0.9277991 | 0.975327409 |
| ENSMUSG00000001802 | Lrp3 | protein_coding | -0.043056768 | 4.149814786 | 0.0084401 | 0.9279205 | 0.975359626 |
| ENSMUSG00000035245 | Eogt | protein_coding | 0.063907935 | 2.911415084 | 0.0083174 | 0.9284449 | 0.975815445 |
| ENSMUSG00000097729 | 2310015A10Rik | processed_transcript | 0.058552426 | 3.141519068 | 0.0082955 | 0.9285388 | 0.975818781 |
| ENSMUSG00000036698 | Ago2 | protein_coding | 0.029206932 | 5.261314727 | 0.0082061 | 0.9289236 | 0.976127727 |
| ENSMUSG00000027894 | Slc6a17 | protein_coding | -0.025885831 | 6.121089919 | 0.0081575 | 0.9291337 | 0.97625311 |
| ENSMUSG00000026918 | Brd3 | protein_coding | 0.022988286 | 6.624802106 | 0.0081191 | 0.9293005 | 0.976280402 |
| ENSMUSG00000051790 | Nlgn2 | protein_coding | 0.024955441 | 6.043379295 | 0.0081097 | 0.9293412 | 0.976280402 |
| ENSMUSG00000070282 | 3000002C10Rik | transcribed_processe | 0.095215526 | 2.154410955 | 0.0080733 | 0.9294994 | 0.976351217 |
| ENSMUSG00000003518 | Dusp3 | protein_coding | 0.030467551 | 8.317145625 | 0.0080532 | 0.9295969 | 0.976358215 |
| ENSMUSG00000104784 | Gm42984 | antisense_RNA | -0.072811256 | 2.66238835 | 0.0079841 | 0.9298888 | 0.976569424 |
| ENSMUSG00000079658 | Eloc | protein_coding | -0.021181888 | 8.764212717 | 0.0079264 | 0.9301422 | 0.976703366 |
| ENSMUSG00000052533 | Nup188 | protein_coding | 0.076638863 | 2.716644853 | 0.0079137 | 0.930198 | 0.976703366 |
| ENSMUSG00000054079 | Utp18 | protein_coding | 0.035468511 | 4.102257036 | 0.0078496 | 0.9304805 | 0.976904552 |
| ENSMUSG00000036885 | Arhgef26 | protein_coding | 0.053937597 | 3.349895326 | 0.0077703 | 0.9308313 | 0.977177492 |
| ENSMUSG00000066149 | Cdc26 | protein_coding | -0.027229871 | 5.822246247 | 0.0077293 | 0.9310135 | 0.97722965 |
| ENSMUSG00000000120 | Ngfr | protein_coding | 0.062725916 | 2.71391218 | 0.0077183 | 0.9310628 | 0.97722965 |
| ENSMUSG00000002266 | Zim1 | protein_coding | 0.048889172 | 3.127123016 | 0.0076735 | 0.9312626 | 0.977284491 |
| ENSMUSG00000031537 | Ikbkb | protein_coding | 0.032852071 | 4.460225865 | 0.0076629 | 0.9313096 | 0.977284491 |
| ENSMUSG00000050556 | Kcnb1 | protein_coding | -0.035311936 | 5.253583299 | 0.0076369 | 0.931426 | 0.977284491 |
| ENSMUSG00000042429 | Adora1 | protein_coding | -0.032555151 | 5.825502439 | 0.0076235 | 0.9314863 | 0.977284491 |
| ENSMUSG00000030271 | Ogg1 | protein_coding | -0.037447872 | 3.96687806 | 0.0075615 | 0.9317645 | 0.977284491 |
| ENSMUSG00000039599 | Fam149b | protein_coding | -0.050398384 | 3.162414155 | 0.0075529 | 0.9318031 | 0.977284491 |
| ENSMUSG00000022789 | Dnm1l | protein_coding | -0.021266282 | 7.672574536 | 0.0075483 | 0.931824 | 0.977284491 |
| ENSMUSG00000025189 | Cnnm1 | protein_coding | 0.023683696 | 5.847748916 | 0.0075313 | 0.9319004 | 0.977284491 |
| ENSMUSG00000050973 | Gdpgp1 | protein_coding | -0.045436674 | 4.282659624 | 0.0075111 | 0.9319919 | 0.977284491 |
| ENSMUSG00000040447 | Spns2 | protein_coding | 0.052568099 | 4.353384316 | 0.0075138 | 0.9320238 | 0.977284491 |
| ENSMUSG00000019929 | Dcn | protein_coding | -0.049959815 | 3.742444667 | 0.0074578 | 0.9322329 | 0.977371047 |
| ENSMUSG00000052605 | Isoc2b | protein_coding | 0.038516991 | 4.50164993 | 0.0074273 | 0.9323711 | 0.977371047 |
| ENSMUSG00000026670 | Uap1 | protein_coding | -0.031033546 | 5.067554371 | 0.0074256 | 0.9323791 | 0.977371047 |
| ENSMUSG00000032679 | Cd59a | protein_coding | -0.03176214 | 5.657192249 | 0.0073562 | 0.932695 | 0.977579721 |
| ENSMUSG00000015149 | Sirt2 | protein_coding | 0.01916695 | 8.34260309 | 0.0073419 | 0.93276 | 0.977579721 |
| ENSMUSG00000048058 | Ldlrad3 | protein_coding | -0.057735479 | 3.027808093 | 0.0072381 | 0.9332358 | 0.977965652 |
| ENSMUSG00000020151 | Ptprr | protein_coding | 0.028886404 | 4.876245521 | 0.0071804 | 0.9335019 | 0.977965652 |
| ENSMUSG00000055866 | Per2 | protein_coding | -0.06293948 | 3.069343328 | 0.0071574 | 0.9336081 | 0.977965652 |
| ENSMUSG00000005442 | Cic | protein_coding | -0.038563505 | 3.706908666 | 0.007135 | 0.9337121 | 0.977965652 |
| ENSMUSG00000019731 | Slc35e1 | protein_coding | -0.04384683 | 4.019405831 | 0.0071327 | 0.9337225 | 0.977965652 |
| ENSMUSG00000028484 | Psip1 | protein_coding | -0.030819156 | 4.976152638 | 0.0071292 | 0.9337387 | 0.977965652 |
| ENSMUSG00000029545 | Acads | protein_coding | -0.050037071 | 3.275808124 | 0.0071169 | 0.9337956 | 0.977965652 |
| ENSMUSG00000020949 | Fkbp3 | protein_coding | -0.019828291 | 9.463429662 | 0.0070929 | 0.9339072 | 0.977965652 |
| ENSMUSG00000026565 | Pou2f1 | protein_coding | -0.056122528 | 3.145914919 | 0.0070844 | 0.9339467 | 0.977965652 |
| ENSMUSG00000035234 | Abraxas1 | protein_coding | 0.057736651 | 2.869221752 | 0.0070197 | 0.9342483 | 0.978095139 |
| ENSMUSG00000030751 | Psma1 | protein_coding | 0.019738537 | 8.507514574 | 0.007007 | 0.9343076 | 0.978095139 |
| ENSMUSG00000033392 | Clasp2 | protein_coding | 0.028549126 | 7.111265289 | 0.0070013 | 0.9343433 | 0.978095139 |
| ENSMUSG00000026988 | Wdsub1 | protein_coding | 0.042273742 | 3.950590657 | 0.0069532 | 0.9345597 | 0.978095748 |
| ENSMUSG00000030515 | Tarsl2 | protein_coding | 0.052937544 | 2.91404841 | 0.0069458 | 0.9345944 | 0.978095748 |
| ENSMUSG00000021807 | 2700060E02Rik | protein_coding | 0.021137369 | 6.484399322 | 0.0068755 | 0.9349257 | 0.978095748 |
| ENSMUSG00000062519 | Zfp398 | protein_coding | -0.04511289 | 3.625922666 | 0.006873 | 0.9349376 | 0.978095748 |
| ENSMUSG00000036097 | Slf2 | protein_coding | -0.028752179 | 5.110730074 | 0.0068638 | 0.9349809 | 0.978095748 |
| ENSMUSG00000034088 | Hdlbp | protein_coding | -0.023538245 | 6.210549682 | 0.0068441 | 0.9350738 | 0.978095748 |
| ENSMUSG00000035215 | Lsm7 | protein_coding | -0.039211536 | 3.812510282 | 0.0068434 | 0.9350773 | 0.978095748 |
| ENSMUSG00000027429 | Sec23b | protein_coding | -0.03159049 | 5.954301935 | 0.0068373 | 0.9351076 | 0.978095748 |
| ENSMUSG00000022421 | Nptxr | protein_coding | -0.030140413 | 6.482046768 | 0.0068267 | 0.9351625 | 0.978095748 |
| ENSMUSG00000028949 | Smarcd3 | protein_coding | 0.039299015 | 3.796072841 | 0.0067973 | 0.9352957 | 0.978139921 |
| ENSMUSG00000038145 | Snrk | protein_coding | -0.035534658 | 4.663585786 | 0.0067765 | 0.9353944 | 0.978148046 |
| ENSMUSG00000025786 | Zdhhc3 | protein_coding | 0.035216881 | 5.349899634 | 0.006751 | 0.9355178 | 0.978181874 |
| ENSMUSG00000017781 | Pitpna | protein_coding | 0.018407171 | 7.228051855 | 0.0066524 | 0.9359875 | 0.978505274 |
| ENSMUSG00000003824 | Syce2 | protein_coding | -0.055391996 | 3.052079266 | 0.0066479 | 0.936009 | 0.978505274 |
| ENSMUSG00000024787 | Snx15 | protein_coding | -0.019572027 | 6.245294448 | 0.0065837 | 0.936318 | 0.97873311 |
| ENSMUSG00000020776 | Fbf1 | protein_coding | -0.039486396 | 3.362885389 | 0.0064434 | 0.9369985 | 0.979349257 |
| ENSMUSG00000029596 | Sdsl | protein_coding | 0.062683714 | 2.705739573 | 0.006357 | 0.9374216 | 0.979689644 |
| ENSMUSG00000078584 | AU022252 | protein_coding | 0.028996188 | 4.762003024 | 0.0063246 | 0.937581 | 0.979689644 |
| ENSMUSG00000027932 | Slc27a3 | protein_coding | -0.071348253 | 2.496547743 | 0.0063121 | 0.9376424 | 0.979689644 |
| ENSMUSG00000021708 | Rasgrf2 | protein_coding | -0.024761735 | 6.114797076 | 0.0062877 | 0.9377631 | 0.979689644 |
| ENSMUSG00000032119 | Hinfp | protein_coding | -0.056988955 | 3.263631182 | 0.0062628 | 0.9378861 | 0.979689644 |
| ENSMUSG00000022412 | Mief1 | protein_coding | 0.034152958 | 5.00375469 | 0.0062427 | 0.9379854 | 0.979689644 |
| ENSMUSG00000047260 | Emc6 | protein_coding | 0.034646324 | 4.534435682 | 0.0062357 | 0.93802 | 0.979689644 |
| ENSMUSG00000029270 | Fam69a | protein_coding | -0.024569477 | 5.543227714 | 0.0062205 | 0.9380957 | 0.979689644 |

| ensembl_gene_id | external_gene_name | gene_biotype | log2 (fold change) | log2 (counts per million) | F-value | p-value | False discovery rate |
|---|---|---|---|---|---|---|---|
| ENSMUSG00000026578 | Ccdc181 | protein_coding | -0.023566785 | 5.90773494 | 0.0062107 | 0.9381442 | 0.979689644 |
| ENSMUSG00000026021 | Sumo1 | protein_coding | -0.018347921 | 8.260689262 | 0.0060944 | 0.9387248 | 0.980158884 |
| ENSMUSG00000032898 | Fbxo21 | protein_coding | 0.023033607 | 6.329678203 | 0.0060684 | 0.9388554 | 0.980158884 |
| ENSMUSG00000028577 | Plaa | protein_coding | 0.029056252 | 4.639243415 | 0.0060661 | 0.938867 | 0.980158884 |
| ENSMUSG00000021024 | Psma6 | protein_coding | 0.018623749 | 9.087785455 | 0.0060199 | 0.9390998 | 0.980269887 |
| ENSMUSG00000055675 | Kbtbd11 | protein_coding | 0.037254045 | 5.211612869 | 0.00601 | 0.9391556 | 0.980269887 |
| ENSMUSG00000028541 | B4galt2 | protein_coding | -0.038640788 | 3.494569312 | 0.0059409 | 0.9395 | 0.980342831 |
| ENSMUSG00000020475 | Pgam2 | protein_coding | -0.03048383 | 4.342649937 | 0.0059383 | 0.9395129 | 0.980342831 |
| ENSMUSG00000057388 | Mrpl18 | protein_coding | 0.017388945 | 8.058049206 | 0.005916 | 0.9396267 | 0.980342831 |
| ENSMUSG00000026458 | Ppfia4 | protein_coding | 0.027244205 | 5.510509538 | 0.0059134 | 0.9396397 | 0.980342831 |
| ENSMUSG00000032244 | Fem1b | protein_coding | 0.020083506 | 6.056165787 | 0.0059052 | 0.9396814 | 0.980342831 |
| ENSMUSG00000036781 | Rps27l | protein_coding | -0.024360855 | 6.179979603 | 0.0058705 | 0.9398589 | 0.980432745 |
| ENSMUSG00000043421 | Hilpda | protein_coding | 0.028039545 | 4.813175792 | 0.0058527 | 0.9399499 | 0.980432745 |
| ENSMUSG00000021919 | Chat | protein_coding | -0.051584375 | 3.470664016 | 0.0058125 | 0.9401576 | 0.980554336 |
| ENSMUSG00000034158 | Lrrc58 | protein_coding | -0.027012065 | 5.653376863 | 0.0057675 | 0.9403872 | 0.98069868 |
| ENSMUSG00000034157 | Cipc | protein_coding | -0.021814455 | 6.034007373 | 0.0057361 | 0.9405496 | 0.9807729 |
| ENSMUSG00000026956 | Uap1l1 | protein_coding | 0.044705892 | 3.614291088 | 0.0057167 | 0.9406501 | 0.980782607 |
| ENSMUSG00000073424 | Cyp4f15 | protein_coding | 0.094132317 | 2.390839071 | 0.0056487 | 0.9410094 | 0.980849774 |
| ENSMUSG00000000861 | Bcl11a | protein_coding | -0.035843241 | 5.098905753 | 0.0056476 | 0.9410106 | 0.980849774 |
| ENSMUSG00000061887 | Ssbp3 | protein_coding | 0.022920728 | 8.011994118 | 0.0056459 | 0.9410196 | 0.980849774 |
| ENSMUSG00000041530 | Ago1 | protein_coding | -0.03123415 | 4.381906915 | 0.0056341 | 0.9410794 | 0.980849774 |
| ENSMUSG00000038437 | Mllt6 | protein_coding | -0.043157423 | 3.657007758 | 0.0056155 | 0.9411767 | 0.980856098 |
| ENSMUSG00000038102 | Trappc11 | protein_coding | 0.026943874 | 4.883764086 | 0.0055362 | 0.9415925 | 0.981109138 |
| ENSMUSG00000057069 | Ero1lb | protein_coding | -0.074857149 | 2.659979662 | 0.0054954 | 0.9418096 | 0.981109138 |
| ENSMUSG00000044674 | Fzd1 | protein_coding | 0.074479012 | 2.389092387 | 0.0054732 | 0.9419254 | 0.981109138 |
| ENSMUSG00000021038 | Vipas39 | protein_coding | -0.023072722 | 6.141613416 | 0.0054647 | 0.9419705 | 0.981109138 |
| ENSMUSG00000109975 | Gm19196 | transcribed_unproces | 0.031490347 | 3.871161809 | 0.0054543 | 0.9420253 | 0.981109138 |
| ENSMUSG00000056076 | Eif3b | protein_coding | -0.021547122 | 5.88964188 | 0.0054524 | 0.9420354 | 0.981109138 |
| ENSMUSG00000017491 | Rarb | protein_coding | 0.03685246 | 3.526248906 | 0.005419 | 0.9422128 | 0.981109138 |
| ENSMUSG00000042873 | Lhfpl4 | protein_coding | -0.019609182 | 6.044644546 | 0.0053853 | 0.9423925 | 0.981109138 |
| ENSMUSG00000042064 | Myo3b | protein_coding | 0.062296123 | 2.505294159 | 0.005385 | 0.9423941 | 0.981109138 |
| ENSMUSG00000026807 | Ak8 | protein_coding | 0.059562467 | 3.527731632 | 0.0053811 | 0.9424718 | 0.981109138 |
| ENSMUSG00000055240 | Zfp101 | protein_coding | 0.04971277 | 3.021592044 | 0.0053665 | 0.942493 | 0.981109138 |
| ENSMUSG00000079036 | Alkbh1 | protein_coding | 0.028748672 | 4.201292992 | 0.0053329 | 0.9426729 | 0.981109138 |
| ENSMUSG00000028405 | Aco1 | protein_coding | 0.037663287 | 4.148423769 | 0.0052952 | 0.9428756 | 0.981109138 |
| ENSMUSG00000017221 | Psmd3 | protein_coding | 0.024190338 | 5.503592171 | 0.005293 | 0.9428873 | 0.981109138 |
| ENSMUSG00000029168 | Dpysl5 | protein_coding | -0.031605975 | 4.853573106 | 0.0052927 | 0.9428889 | 0.981109138 |
| ENSMUSG00000030270 | Cpne9 | protein_coding | 0.061641075 | 5.478852177 | 0.005348 | 0.943025 | 0.981109138 |
| ENSMUSG00000076441 | Ass1 | protein_coding | -0.028053708 | 6.249848399 | 0.0052518 | 0.9431243 | 0.981109138 |
| ENSMUSG00000024516 | Sec11c | protein_coding | 0.022501151 | 7.690142537 | 0.0052434 | 0.9431568 | 0.981109138 |
| ENSMUSG00000056306 | Sertm1 | protein_coding | 0.043298043 | 3.222958381 | 0.0052311 | 0.9432219 | 0.981109138 |
| ENSMUSG00000054770 | Kctd18 | protein_coding | 0.043041814 | 3.477737419 | 0.0052135 | 0.9433172 | 0.981109138 |
| ENSMUSG00000042709 | Atpaf2 | protein_coding | 0.023960537 | 5.953940537 | 0.0051998 | 0.9433918 | 0.981109138 |
| ENSMUSG00000024037 | Wdr4 | protein_coding | 0.02536547 | 4.752591358 | 0.0051933 | 0.9434268 | 0.981109138 |
| ENSMUSG00000042684 | Npl | protein_coding | 0.05090299 | 3.115437471 | 0.0051556 | 0.9436321 | 0.981226972 |
| ENSMUSG00000004508 | Gab2 | protein_coding | 0.031825531 | 3.938016817 | 0.0051352 | 0.9437438 | 0.981226972 |
| ENSMUSG00000028398 | Tmem261 | protein_coding | -0.018546017 | 6.682360992 | 0.005095 | 0.9439639 | 0.981226972 |
| ENSMUSG00000008140 | Emc10 | protein_coding | 0.023831349 | 5.708759003 | 0.0050823 | 0.9440338 | 0.981226972 |
| ENSMUSG00000015189 | Casd1 | protein_coding | 0.023155948 | 5.565796682 | 0.0050705 | 0.9440986 | 0.981226972 |
| ENSMUSG00000001089 | Luzp1 | protein_coding | 0.025618871 | 5.210286883 | 0.0050622 | 0.9441442 | 0.981226972 |
| ENSMUSG00000002625 | Akap8l | protein_coding | 0.022037987 | 5.114808277 | 0.0050559 | 0.9441789 | 0.981226972 |
| ENSMUSG00000038425 | Poli | protein_coding | -0.039968513 | 3.520332659 | 0.0050353 | 0.9442929 | 0.981250661 |
| ENSMUSG00000006378 | Gcat | protein_coding | -0.027593111 | 5.257505579 | 0.0050135 | 0.9444131 | 0.981280754 |
| ENSMUSG00000027419 | Pcsk2 | protein_coding | -0.01808415 | 6.944093578 | 0.0049232 | 0.944915 | 0.981707299 |
| ENSMUSG00000048029 | Eno4 | protein_coding | 0.044120584 | 3.208913066 | 0.0049008 | 0.9450406 | 0.981743002 |
| ENSMUSG00000042404 | Dennd4b | protein_coding | -0.027706545 | 4.202061161 | 0.0048681 | 0.9452238 | 0.981838427 |
| ENSMUSG00000038822 | Hace1 | protein_coding | -0.030928056 | 4.195022349 | 0.0048321 | 0.9454261 | 0.981953727 |
| ENSMUSG00000045625 | Pigz | protein_coding | 0.02616011 | 4.635725031 | 0.0047932 | 0.9456459 | 0.982087145 |
| ENSMUSG00000027506 | Tpd52 | protein_coding | 0.021894506 | 6.305528105 | 0.0047747 | 0.9457506 | 0.982101043 |
| ENSMUSG00000013076 | Amotl1 | protein_coding | 0.039894375 | 7.001349671 | 0.0047424 | 0.9462858 | 0.982561963 |
| ENSMUSG00000024666 | Tmem138 | protein_coding | 0.026428909 | 5.474530119 | 0.0046625 | 0.9463911 | 0.982569301 |
| ENSMUSG00000033478 | Fam160b1 | protein_coding | -0.033033854 | 3.699283433 | 0.0046468 | 0.9464814 | 0.982569301 |
| ENSMUSG00000036712 | Cyld | protein_coding | -0.017091468 | 6.611239859 | 0.0046246 | 0.9466091 | 0.982569301 |
| ENSMUSG00000045176 | Borcs6 | protein_coding | -0.034761461 | 3.728893799 | 0.004616 | 0.9466584 | 0.982569301 |
| ENSMUSG00000022684 | Bfar | protein_coding | 0.020337708 | 5.791579699 | 0.004582 | 0.9468551 | 0.982678558 |
| ENSMUSG00000024302 | Dtna | protein_coding | 0.01723036 | 6.816373177 | 0.0045598 | 0.9469839 | 0.982717793 |
| ENSMUSG00000027349 | Fam98b | protein_coding | 0.017375048 | 6.291079979 | 0.0045147 | 0.9472461 | 0.982845741 |
| ENSMUSG00000026960 | Arl6ip6 | protein_coding | -0.031585875 | 3.486995503 | 0.0045071 | 0.9472904 | 0.982845741 |
| ENSMUSG00000042605 | Atxn2 | protein_coding | -0.019464038 | 5.778144979 | 0.0044626 | 0.9475512 | 0.982950713 |
| ENSMUSG00000049950 | Rpp38 | protein_coding | -0.020400845 | 5.107521382 | 0.00445 | 0.9476246 | 0.982950713 |
| ENSMUSG00000070366 | Plpp4 | protein_coding | 0.033771036 | 4.061904239 | 0.0044431 | 0.9476658 | 0.982950713 |
| ENSMUSG00000003154 | Foxj2 | protein_coding | -0.032291932 | 3.932937719 | 0.0043788 | 0.948045 | 0.982991392 |
| ENSMUSG00000041712 | Ubr7 | protein_coding | -0.026779286 | 4.846411082 | 0.0043685 | 0.9481061 | 0.982991392 |
| ENSMUSG00000000563 | Atp5f1 | protein_coding | 0.014579006 | 9.728584423 | 0.0043628 | 0.9481399 | 0.982991392 |
| ENSMUSG00000019863 | Qrsl1 | protein_coding | 0.033901194 | 4.740587059 | 0.0043575 | 0.9481727 | 0.982991392 |
| ENSMUSG00000022283 | Pabpc1 | protein_coding | -0.021948714 | 5.356366693 | 0.0043466 | 0.9482358 | 0.982991392 |
| ENSMUSG00000068706 | Gm10250 | protein_coding | -0.034322689 | 3.267852304 | 0.0043349 | 0.9483057 | 0.982991392 |
| ENSMUSG00000055065 | Ddx17 | protein_coding | 0.023762106 | 5.506969951 | 0.0043283 | 0.9483449 | 0.982991392 |

| ensembl_gene_id | external_gene_name | gene_biotype | log2 (fold change) | log2 (counts per million) | F-value | p-value | False discovery rate |
|---|---|---|---|---|---|---|---|
| ENSMUSG00000051048 | P4ha3 | protein_coding | -0.060783908 | 2.603268759 | 0.0042655 | 0.9487207 | 0.983286121 |
| ENSMUSG00000041544 | Disp3 | protein_coding | 0.04314939 | 3.256826416 | 0.0042358 | 0.9488991 | 0.98337624 |
| ENSMUSG00000029701 | Rbm28 | protein_coding | 0.023678627 | 5.289942024 | 0.0041548 | 0.9493893 | 0.983405971 |
| ENSMUSG00000034532 | Fbxo16 | protein_coding | 0.022942621 | 5.05428201 | 0.0041432 | 0.9494599 | 0.983405971 |
| ENSMUSG00000058444 | Map2k5 | protein_coding | 0.02605781 | 5.263199779 | 0.0041415 | 0.9494703 | 0.983405971 |
| ENSMUSG00000035064 | Eef2k | protein_coding | -0.029016763 | 3.847565914 | 0.0041327 | 0.9495241 | 0.983405971 |
| ENSMUSG00000001053 | N4bp3 | protein_coding | 0.054954987 | 2.452162379 | 0.0041269 | 0.9495591 | 0.983405971 |
| ENSMUSG00000097924 | A730020E08Rik | bidirectional_promote | -0.048124148 | 2.845040836 | 0.0041257 | 0.9495668 | 0.983405971 |
| ENSMUSG00000001663 | Gstt1 | protein_coding | -0.021433431 | 5.76642151 | 0.0041255 | 0.949568 | 0.983405971 |
| ENSMUSG00000033055 | Ankrd54 | protein_coding | 0.032091708 | 3.488553445 | 0.0040849 | 0.949816 | 0.983568166 |
| ENSMUSG00000002308 | Cd320 | protein_coding | 0.025865399 | 4.901515219 | 0.0040674 | 0.9499236 | 0.983584843 |
| ENSMUSG00000091512 | Lamtor3 | protein_coding | 0.01642845 | 6.806980112 | 0.0039874 | 0.9504181 | 0.983944016 |
| ENSMUSG00000006575 | Rundc3a | protein_coding | 0.018318311 | 6.008340285 | 0.0039635 | 0.9505664 | 0.983944016 |
| ENSMUSG00000073639 | Rab18 | protein_coding | 0.016111724 | 8.258163609 | 0.0039611 | 0.9505813 | 0.983944016 |
| ENSMUSG00000018398 | 8-Sep | protein_coding | -0.013915717 | 6.969690706 | 0.0039523 | 0.9506365 | 0.983944016 |
| ENSMUSG00000039178 | Tbc1d19 | protein_coding | 0.018188263 | 5.520886412 | 0.0038848 | 0.9510593 | 0.984286828 |
| ENSMUSG00000022562 | Oplah | protein_coding | 0.049314268 | 2.775960112 | 0.0038686 | 0.9511613 | 0.984297698 |
| ENSMUSG00000015852 | Fcrls | protein_coding | 0.058625924 | 3.534348842 | 0.0038428 | 0.9514218 | 0.984457996 |
| ENSMUSG00000035226 | Rims4 | protein_coding | -0.023739349 | 5.01318547 | 0.0038127 | 0.9515149 | 0.984457996 |
| ENSMUSG00000032666 | 1700025G04Rik | protein_coding | -0.019767103 | 5.294701351 | 0.0038003 | 0.9515937 | 0.984457996 |
| ENSMUSG00000034813 | Grip1 | protein_coding | 0.022463541 | 4.547246663 | 0.0037863 | 0.9516824 | 0.984457996 |
| ENSMUSG00000019297 | Nop9 | protein_coding | 0.048708982 | 2.89386203 | 0.0037522 | 0.9519001 | 0.984458688 |
| ENSMUSG00000041977 | Arhgef11 | protein_coding | 0.039075745 | 3.297365252 | 0.0037463 | 0.951938 | 0.984458688 |
| ENSMUSG00000032913 | Lrig2 | protein_coding | -0.025801323 | 3.960089999 | 0.0037394 | 0.9519826 | 0.984458688 |
| ENSMUSG00000039307 | Hexdc | protein_coding | -0.029874149 | 3.88887667 | 0.003729 | 0.9520493 | 0.984458688 |
| ENSMUSG00000026959 | Grin1 | protein_coding | 0.048335364 | 2.545413255 | 0.0036805 | 0.9523619 | 0.984628571 |
| ENSMUSG00000035713 | Usp35 | protein_coding | 0.035626564 | 3.237843402 | 0.0036626 | 0.9524776 | 0.984628571 |
| ENSMUSG00000050144 | Slc25a44 | protein_coding | -0.015598845 | 7.042172236 | 0.0036609 | 0.9524883 | 0.984628571 |
| ENSMUSG00000019659 | Ccdc12 | protein_coding | -0.016499307 | 5.902746514 | 0.0036292 | 0.9526943 | 0.984746905 |
| ENSMUSG00000037458 | Azin1 | protein_coding | -0.014993008 | 6.855944283 | 0.0035924 | 0.9529342 | 0.984833422 |
| ENSMUSG00000021252 | Erg28 | protein_coding | -0.014907298 | 7.380017523 | 0.0035661 | 0.9531069 | 0.984833422 |
| ENSMUSG00000037940 | Inpp4b | protein_coding | 0.02973671 | 4.371399073 | 0.0035639 | 0.9531211 | 0.984833422 |
| ENSMUSG00000000805 | Car4 | protein_coding | -0.030811373 | 3.783159165 | 0.0035604 | 0.9531444 | 0.984833422 |
| ENSMUSG00000086968 | 4933431E20Rik | antisense_RNA | -0.030735814 | 4.024833679 | 0.0035402 | 0.9532773 | 0.984876166 |
| ENSMUSG00000028826 | Tmem57 | protein_coding | 0.021840291 | 5.475240805 | 0.0034731 | 0.9537214 | 0.985240311 |
| ENSMUSG00000047844 | Bex4 | protein_coding | 0.018550367 | 5.16928169 | 0.003439 | 0.9539494 | 0.985301608 |
| ENSMUSG00000027274 | Mkks | protein_coding | 0.016912589 | 7.441326678 | 0.0034124 | 0.9541273 | 0.985301608 |
| ENSMUSG00000024790 | Sac3d1 | protein_coding | -0.027736266 | 3.673260541 | 0.0033975 | 0.9542277 | 0.985301608 |
| ENSMUSG00000039218 | Srrm2 | protein_coding | 0.017962789 | 6.124599516 | 0.003368 | 0.9544266 | 0.985301608 |
| ENSMUSG00000096141 | Dnah7a | protein_coding | 0.025176702 | 3.695228633 | 0.003356 | 0.9545078 | 0.985301608 |
| ENSMUSG00000052837 | Junb | protein_coding | 0.044824269 | 2.756355389 | 0.0033465 | 0.9545719 | 0.985301608 |
| ENSMUSG00000014980 | Tsen15 | protein_coding | -0.019931723 | 4.869317829 | 0.0033425 | 0.9545992 | 0.985301608 |
| ENSMUSG00000032040 | Dcps | protein_coding | 0.022433704 | 4.857427172 | 0.0033381 | 0.9546291 | 0.985301608 |
| ENSMUSG00000040785 | Ttc3 | protein_coding | -0.014031862 | 9.858794681 | 0.003322 | 0.9547384 | 0.985301608 |
| ENSMUSG00000030677 | Kif22 | protein_coding | 0.028678035 | 3.465537622 | 0.0033208 | 0.9547463 | 0.985301608 |
| ENSMUSG00000063235 | Ptpmt1 | processed_transcript | 0.025381116 | 3.817350706 | 0.0033027 | 0.95487 | 0.985301608 |
| ENSMUSG00000009905 | Kdsr | protein_coding | -0.018551006 | 5.742732626 | 0.0032813 | 0.9550158 | 0.985301608 |
| ENSMUSG00000020402 | Vdac1 | protein_coding | 0.013352326 | 9.386725854 | 0.0032777 | 0.9550408 | 0.985301608 |
| ENSMUSG00000051373 | Plpp7 | protein_coding | 0.021054196 | 4.706038046 | 0.0032744 | 0.9550636 | 0.985301608 |
| ENSMUSG00000071398 | 2410004P03Rik | protein_coding | -0.033804519 | 3.660535499 | 0.0031837 | 0.9556895 | 0.985784899 |
| ENSMUSG00000051721 | Wdcp | protein_coding | 0.030430587 | 4.719304903 | 0.0031725 | 0.9557735 | 0.985784899 |
| ENSMUSG00000035203 | Epn1 | protein_coding | 0.01678056 | 6.099293271 | 0.0031668 | 0.9558071 | 0.985784899 |
| ENSMUSG00000091537 | Tma7 | protein_coding | 0.012892999 | 7.239049652 | 0.0031315 | 0.9560537 | 0.985855739 |
| ENSMUSG00000047945 | Marcksl1 | protein_coding | 0.022629673 | 4.9562803 | 0.0031307 | 0.9560591 | 0.985855739 |
| ENSMUSG00000049878 | Rlf | protein_coding | 0.027629058 | 3.754233877 | 0.0030673 | 0.956506 | 0.986052941 |
| ENSMUSG00000040270 | Bach2 | protein_coding | 0.033340552 | 3.30761333 | 0.003065 | 0.9565221 | 0.986052941 |
| ENSMUSG00000031513 | Leprotl1 | protein_coding | -0.01888687 | 6.383759213 | 0.0030647 | 0.9565255 | 0.986052941 |
| ENSMUSG00000057132 | Rpgrip1 | protein_coding | 0.033146047 | 3.430311073 | 0.0030087 | 0.9569231 | 0.98634039 |
| ENSMUSG00000027954 | Efna1 | protein_coding | -0.040215234 | 3.056885066 | 0.0029969 | 0.9570077 | 0.98634039 |
| ENSMUSG00000041396 | Mettl18 | protein_coding | -0.021093571 | 4.638651767 | 0.0029812 | 0.9571204 | 0.98634039 |
| ENSMUSG00000028211 | Trp53inp1 | protein_coding | -0.030154554 | 3.28170641 | 0.0029741 | 0.9571712 | 0.98634039 |
| ENSMUSG00000029767 | Calu | protein_coding | -0.01874261 | 5.515481613 | 0.0029386 | 0.9574271 | 0.986509502 |
| ENSMUSG00000036815 | Dpp10 | protein_coding | -0.016326697 | 6.009901135 | 0.0028983 | 0.9577201 | 0.986716863 |
| ENSMUSG00000090698 | Apold1 | protein_coding | -0.038897628 | 3.3684765 | 0.0027988 | 0.958452 | 0.987140615 |
| ENSMUSG00000033029 | 1700088E04Rik | protein_coding | -0.02524814 | 3.823155226 | 0.0027977 | 0.9584592 | 0.987140615 |
| ENSMUSG00000031661 | Nkd1 | protein_coding | -0.05291941 | 2.463176117 | 0.0027919 | 0.9585023 | 0.987140615 |
| ENSMUSG00000013275 | Slc41a1 | protein_coding | 0.020971946 | 4.441277428 | 0.0027804 | 0.9585878 | 0.987140615 |
| ENSMUSG00000024002 | Brd4 | protein_coding | -0.014533894 | 6.56187823 | 0.0027801 | 0.9585904 | 0.987140615 |
| ENSMUSG00000026972 | Arrdc1 | protein_coding | -0.037666829 | 3.728158626 | 0.0027706 | 0.9586881 | 0.987146697 |
| ENSMUSG00000053646 | Plxnb1 | protein_coding | 0.049117721 | 3.351708276 | 0.0027323 | 0.9590107 | 0.987268154 |
| ENSMUSG00000018459 | Slc13a3 | protein_coding | 0.049770724 | 3.046341568 | 0.0027209 | 0.9590621 | 0.987268154 |
| ENSMUSG00000032370 | Lactb | protein_coding | -0.03445901 | 2.970459917 | 0.0027145 | 0.9590815 | 0.987268154 |
| ENSMUSG00000033751 | Gadd45gip1 | protein_coding | -0.040763437 | 3.098671897 | 0.0026863 | 0.9592941 | 0.987285997 |
| ENSMUSG00000046010 | Zfp830 | protein_coding | 0.020253878 | 4.747410748 | 0.002673 | 0.9593946 | 0.987285997 |
| ENSMUSG00000051319 | 1500011K16Rik | protein_coding | 0.012528516 | 7.289265781 | 0.0026703 | 0.9594153 | 0.987285997 |
| ENSMUSG00000018697 | Aatf | protein_coding | 0.030850355 | 3.967412359 | 0.0026638 | 0.9594661 | 0.987285997 |
| ENSMUSG00000020566 | Atp6v1c2 | protein_coding | 0.027636972 | 3.341533126 | 0.0026346 | 0.9596875 | 0.987419367 |
| ENSMUSG00000089665 | Fcor | protein_coding | 0.020947028 | 4.81290421 | 0.0026004 | 0.9599497 | 0.987502851 |

| ensembl_gene_id | external_gene_name | gene_biotype | log2 (fold change) | log2 (counts per million) | F-value | p-value | False discovery rate |
|---|---|---|---|---|---|---|---|
| ENSMUSG00000029009 | Mthfr | protein_coding | 0.039625306 | 3.157893418 | 0.0025978 | 0.9599704 | 0.987502851 |
| ENSMUSG00000060950 | Trmt61a | protein_coding | -0.027270972 | 3.441033552 | 0.0025812 | 0.9600979 | 0.987502851 |
| ENSMUSG00000036986 | Pml | protein_coding | 0.042151688 | 2.724934985 | 0.0025616 | 0.9602493 | 0.987502851 |
| ENSMUSG00000031568 | Rwdd4a | protein_coding | -0.018399445 | 4.71929133 | 0.0025556 | 0.9602959 | 0.987502851 |
| ENSMUSG00000006800 | Sulf2 | protein_coding | 0.0191147 | 5.070675358 | 0.0025525 | 0.9603197 | 0.987502851 |
| ENSMUSG00000028282 | Casp8ap2 | protein_coding | -0.023947765 | 3.979270051 | 0.0025261 | 0.9605252 | 0.987619791 |
| ENSMUSG00000039656 | Rxrb | protein_coding | -0.017381396 | 5.001536529 | 0.0025023 | 0.9607117 | 0.987628616 |
| ENSMUSG00000025817 | Nudt5 | protein_coding | -0.01575194 | 5.502467842 | 0.0024965 | 0.9607575 | 0.987628616 |
| ENSMUSG00000026188 | Tmem169 | protein_coding | 0.022086051 | 3.928872815 | 0.0024877 | 0.9608263 | 0.987628616 |
| ENSMUSG00000053453 | Thoc7 | protein_coding | 0.015458838 | 7.306993074 | 0.0024783 | 0.9609012 | 0.987628616 |
| ENSMUSG00000028247 | Coq3 | protein_coding | 0.015148118 | 6.189198552 | 0.0024399 | 0.9612041 | 0.987845488 |
| ENSMUSG00000037428 | Vgf | protein_coding | 0.025606634 | 4.032652325 | 0.0024227 | 0.9613413 | 0.987892125 |
| ENSMUSG00000041000 | Trim62 | protein_coding | -0.0236045 | 3.93206568 | 0.0023836 | 0.9616541 | 0.988008778 |
| ENSMUSG00000022443 | Myh9 | protein_coding | -0.02516475 | 3.503120734 | 0.0023794 | 0.9616881 | 0.988008778 |
| ENSMUSG00000031935 | Med17 | protein_coding | -0.027666052 | 3.988971524 | 0.0023692 | 0.96177 | 0.988008778 |
| ENSMUSG00000062949 | Atp11c | protein_coding | -0.036717539 | 2.800095287 | 0.0023627 | 0.9618224 | 0.988008778 |
| ENSMUSG00000039686 | Zer1 | protein_coding | 0.02724351 | 5.816096672 | 0.0023505 | 0.9620152 | 0.988029466 |
| ENSMUSG00000003657 | Calb2 | protein_coding | 0.018373877 | 10.09150668 | 0.0023418 | 0.9620263 | 0.988029466 |
| ENSMUSG00000039787 | Cercam | protein_coding | 0.055799335 | 3.018022681 | 0.0023194 | 0.9622556 | 0.98812941 |
| ENSMUSG00000018401 | Mtmr4 | protein_coding | -0.01736806 | 5.241823159 | 0.0023016 | 0.9623187 | 0.98812941 |
| ENSMUSG00000030246 | Ldhb | protein_coding | -0.01100779 | 10.4757629 | 0.0022918 | 0.9623993 | 0.98812941 |
| ENSMUSG00000030286 | Emc3 | protein_coding | -0.014077275 | 6.178071852 | 0.0022803 | 0.9624937 | 0.988131988 |
| ENSMUSG00000047155 | Cyp4x1 | protein_coding | -0.023466526 | 3.539105573 | 0.002255 | 0.9627018 | 0.988251321 |
| ENSMUSG00000018733 | Pex12 | protein_coding | -0.021597883 | 4.360905925 | 0.0021972 | 0.9631824 | 0.988650275 |
| ENSMUSG00000026688 | Mgst3 | protein_coding | 0.010428665 | 8.852648888 | 0.0021748 | 0.9633705 | 0.98874105 |
| ENSMUSG00000052920 | Prkg1 | protein_coding | 0.028139432 | 3.423851385 | 0.0021648 | 0.9634547 | 0.98874105 |
| ENSMUSG00000044748 | Defb1 | protein_coding | -0.058777422 | 3.007935018 | 0.0021573 | 0.9636086 | 0.988804548 |
| ENSMUSG00000045257 | Morn2 | protein_coding | 0.012685718 | 5.872086206 | 0.0021094 | 0.9639253 | 0.988860473 |
| ENSMUSG00000087497 | 2810001G20Rik | antisense_RNA | -0.01712271 | 4.972236344 | 0.0020986 | 0.9640178 | 0.988860473 |
| ENSMUSG00000043059 | Zfp513 | protein_coding | -0.045742945 | 2.252911747 | 0.0020909 | 0.9640837 | 0.988860473 |
| ENSMUSG00000053279 | Aldh1a1 | protein_coding | 0.021485755 | 4.881994317 | 0.0020878 | 0.9641113 | 0.988860473 |
| ENSMUSG00000020196 | Cabin1 | protein_coding | -0.020791644 | 5.004642735 | 0.0020864 | 0.9641229 | 0.988860473 |
| ENSMUSG00000075296 | Aldh3b2 | protein_coding | -0.016285882 | 5.345022307 | 0.0020549 | 0.9643942 | 0.988977152 |
| ENSMUSG00000057156 | Homez | protein_coding | -0.021911031 | 3.484538364 | 0.0020518 | 0.9644206 | 0.988977152 |
| ENSMUSG00000038085 | Cnbd2 | protein_coding | -0.024970161 | 3.309772347 | 0.0020297 | 0.9646125 | 0.989002835 |
| ENSMUSG00000062906 | Hdac10 | protein_coding | -0.037509016 | 3.334555254 | 0.0020305 | 0.9646296 | 0.989002835 |
| ENSMUSG00000036299 | BC031181 | protein_coding | 0.010388132 | 9.008096586 | 0.002016 | 0.9647326 | 0.989014122 |
| ENSMUSG00000030761 | Myo7a | protein_coding | 0.039487196 | 2.559450715 | 0.001994 | 0.9649252 | 0.989086572 |
| ENSMUSG00000028249 | Sdcbp | protein_coding | 0.010537186 | 8.821410057 | 0.0019777 | 0.9650686 | 0.989086572 |
| ENSMUSG00000041278 | Ttc1 | protein_coding | 0.009300919 | 7.870327927 | 0.0019696 | 0.9651402 | 0.989086572 |
| ENSMUSG00000029547 | Ints1 | protein_coding | -0.02504257 | 4.190595333 | 0.0019577 | 0.9652462 | 0.989086572 |
| ENSMUSG00000022836 | Mylk | protein_coding | 0.037030434 | 2.678548687 | 0.0019557 | 0.9652631 | 0.989086572 |
| ENSMUSG00000004319 | Clcn3 | protein_coding | 0.009971719 | 7.199467733 | 0.0019102 | 0.9656698 | 0.989287823 |
| ENSMUSG00000020034 | Tcp11l2 | protein_coding | -0.021559433 | 5.714036698 | 0.0019117 | 0.9656996 | 0.989287823 |
| ENSMUSG00000031352 | Hccs | protein_coding | 0.02059683 | 4.604801972 | 0.0019029 | 0.9657355 | 0.989287823 |
| ENSMUSG00000032046 | Abhd12 | protein_coding | -0.010686193 | 6.967606994 | 0.0018661 | 0.9660678 | 0.989407663 |
| ENSMUSG00000029911 | Ssbp1 | protein_coding | -0.012480126 | 6.047762266 | 0.0018605 | 0.9661188 | 0.989407663 |
| ENSMUSG00000066838 | Zfp772 | protein_coding | -0.027140764 | 3.219859574 | 0.0018541 | 0.9661774 | 0.989407663 |
| ENSMUSG00000030815 | Phkg2 | protein_coding | -0.013895028 | 5.202607546 | 0.0018475 | 0.9662371 | 0.989407663 |
| ENSMUSG00000021039 | Snw1 | protein_coding | -0.010475689 | 6.894242313 | 0.0018393 | 0.9663126 | 0.989407663 |
| ENSMUSG00000020057 | Dram1 | protein_coding | 0.01704196 | 4.04375824 | 0.0017608 | 0.9670386 | 0.989573452 |
| ENSMUSG00000027332 | Ivd | protein_coding | 0.014630653 | 6.87274443 | 0.0017609 | 0.9670418 | 0.989573452 |
| ENSMUSG00000031431 | Tsc22d3 | protein_coding | -0.010285418 | 8.258900288 | 0.0017539 | 0.9671033 | 0.989573452 |
| ENSMUSG00000026566 | Mpzl1 | protein_coding | -0.028127162 | 3.369692735 | 0.0017353 | 0.967278 | 0.989573452 |
| ENSMUSG00000030560 | Ctsc | protein_coding | 0.042996269 | 2.630697253 | 0.0017294 | 0.9673345 | 0.989573452 |
| ENSMUSG00000022526 | Zfp251 | protein_coding | 0.016670473 | 4.270325311 | 0.0017286 | 0.9673409 | 0.989573452 |
| ENSMUSG00000051726 | Kcnf1 | protein_coding | 0.020745854 | 4.390670137 | 0.0017285 | 0.9673422 | 0.989573452 |
| ENSMUSG00000029504 | Ddx51 | protein_coding | 0.051978536 | 2.463090803 | 0.0017302 | 0.9673433 | 0.989573452 |
| ENSMUSG00000040276 | Pacsin1 | protein_coding | 0.010469924 | 7.277581248 | 0.0017264 | 0.967362 | 0.989573452 |
| ENSMUSG00000032633 | Flcn | protein_coding | -0.018224635 | 5.481049435 | 0.001723 | 0.9673948 | 0.989573452 |
| ENSMUSG00000046985 | Tapt1 | protein_coding | 0.019110125 | 3.982898694 | 0.0016573 | 0.9680211 | 0.989958001 |
| ENSMUSG00000032579 | Hemk1 | protein_coding | -0.016625605 | 4.990889154 | 0.0016465 | 0.968126 | 0.989958001 |
| ENSMUSG00000016477 | E2f3 | protein_coding | 0.021058271 | 3.467306497 | 0.0016459 | 0.9681311 | 0.989958001 |
| ENSMUSG00000075703 | Selenoi | protein_coding | 0.019163364 | 5.201008099 | 0.0016429 | 0.9681611 | 0.989958001 |
| ENSMUSG00000052997 | Uba2 | protein_coding | 0.009825794 | 6.448238628 | 0.0016356 | 0.968231 | 0.989958001 |
| ENSMUSG00000034300 | Fam53c | protein_coding | -0.015405731 | 5.380149082 | 0.001617 | 0.9684122 | 0.989964118 |
| ENSMUSG00000029403 | Cdkl2 | protein_coding | -0.013680168 | 5.228739217 | 0.0016161 | 0.9684211 | 0.989964118 |
| ENSMUSG00000052395 | Rft1 | protein_coding | -0.015924959 | 4.127595458 | 0.0015976 | 0.9686026 | 0.990055479 |
| ENSMUSG00000025743 | Sdc3 | protein_coding | -0.022379477 | 3.225559943 | 0.001576 | 0.9688157 | 0.990062314 |
| ENSMUSG00000045348 | Nyap1 | protein_coding | -0.025361358 | 3.3711116 | 0.0015744 | 0.9688311 | 0.990062314 |
| ENSMUSG00000031776 | Arl2bp | protein_coding | -0.009860712 | 6.444304096 | 0.0015689 | 0.9688855 | 0.990062314 |
| ENSMUSG00000051154 | Commd3 | protein_coding | -0.009016846 | 8.214268243 | 0.0015273 | 0.9693009 | 0.990283292 |
| ENSMUSG00000032410 | Xrn1 | protein_coding | 0.01868596 | 4.021214016 | 0.0015215 | 0.969359 | 0.990283292 |
| ENSMUSG00000050830 | Vwc2 | protein_coding | -0.024268234 | 3.167405937 | 0.0015118 | 0.9694562 | 0.990283292 |
| ENSMUSG00000041506 | Rrp9 | protein_coding | -0.013956286 | 4.809833241 | 0.0015069 | 0.9695059 | 0.990283292 |
| ENSMUSG00000019951 | Uhrf1bp1l | protein_coding | 0.012196468 | 5.283490123 | 0.0014992 | 0.9695838 | 0.990283292 |
| ENSMUSG00000045201 | Lrrc3b | protein_coding | 0.01372411 | 4.919093809 | 0.0014776 | 0.969804 | 0.990283292 |
| ENSMUSG00000023963 | Cyp39a1 | protein_coding | -0.048257857 | 2.887792642 | 0.0014816 | 0.9698288 | 0.990283292 |

| ensembl_gene_id | external_gene_name | gene_biotype | log2 (fold change) | log2 (counts per million) | F-value | p-value | False discovery rate |
|---|---|---|---|---|---|---|---|
| ENSMUSG00000025425 | St8sia5 | protein_coding | -0.031210473 | 2.602638481 | 0.0014742 | 0.9698385 | 0.990283292 |
| ENSMUSG00000022783 | Spag6l | protein_coding | 0.019179997 | 3.789404451 | 0.0014494 | 0.9700932 | 0.990449336 |
| ENSMUSG00000042662 | Dusp15 | protein_coding | 0.023952243 | 2.943083448 | 0.0014169 | 0.9704301 | 0.990529575 |
| ENSMUSG00000028165 | Cisd2 | protein_coding | -0.010355367 | 6.175455156 | 0.0014156 | 0.9704436 | 0.990529575 |
| ENSMUSG00000033854 | Kcnk10 | protein_coding | 0.023717163 | 3.331571941 | 0.0014152 | 0.9704482 | 0.990529575 |
| ENSMUSG00000029672 | Fam3c | protein_coding | -0.011867244 | 5.096168148 | 0.0013892 | 0.9707201 | 0.990636206 |
| ENSMUSG00000059669 | Taf1b | protein_coding | 0.015256479 | 4.718518786 | 0.0013826 | 0.9707902 | 0.990636206 |
| ENSMUSG00000032468 | Armc8 | protein_coding | 0.011109664 | 6.763155801 | 0.0013784 | 0.9708343 | 0.990636206 |
| ENSMUSG00000040111 | Gramd1b | protein_coding | -0.011784861 | 5.352504123 | 0.0013702 | 0.9709211 | 0.990636206 |
| ENSMUSG00000094152 | Slc6a16 | protein_coding | -0.02048897 | 3.775682246 | 0.0013482 | 0.9711553 | 0.990754714 |
| ENSMUSG00000003559 | As3mt | protein_coding | -0.012508818 | 5.860821575 | 0.0013378 | 0.9712675 | 0.990754714 |
| ENSMUSG00000058301 | Upf1 | protein_coding | 0.014627847 | 4.106842552 | 0.0013225 | 0.9714322 | 0.990754714 |
| ENSMUSG00000003992 | Ssbp2 | protein_coding | 0.008074437 | 7.282676456 | 0.0013216 | 0.9714413 | 0.990754714 |
| ENSMUSG00000020150 | Gamt | protein_coding | -0.018744423 | 3.771287056 | 0.001307 | 0.9715999 | 0.990754714 |
| ENSMUSG00000049811 | Fam161a | protein_coding | -0.022591816 | 2.959912832 | 0.0012804 | 0.9718898 | 0.990754714 |
| ENSMUSG00000040459 | Arglu1 | protein_coding | 0.010426498 | 7.490944732 | 0.0012797 | 0.9718978 | 0.990754714 |
| ENSMUSG00000032252 | Glce | protein_coding | -0.014345371 | 4.339791961 | 0.0012711 | 0.9719919 | 0.990754714 |
| ENSMUSG00000050931 | Sgms2 | protein_coding | 0.039801346 | 2.269271934 | 0.0012687 | 0.9720185 | 0.990754714 |
| ENSMUSG00000028391 | Wdr31 | protein_coding | -0.016682029 | 5.189647739 | 0.0012661 | 0.972048 | 0.990754714 |
| ENSMUSG00000021270 | Hsp90aa1 | protein_coding | 0.008416168 | 7.648787366 | 0.001258 | 0.9721373 | 0.990754714 |
| ENSMUSG00000032834 | Pwp2 | protein_coding | 0.0185562 | 4.081071624 | 0.0012533 | 0.9721887 | 0.990754714 |
| ENSMUSG00000020621 | Rdh14 | protein_coding | -0.012438465 | 5.326370321 | 0.0012491 | 0.9722351 | 0.990754714 |
| ENSMUSG00000003273 | Car11 | protein_coding | -0.013691113 | 5.660026022 | 0.0012265 | 0.9724875 | 0.990839061 |
| ENSMUSG00000060904 | Arl1 | protein_coding | -0.008104114 | 6.881057215 | 0.001216 | 0.9726055 | 0.990839061 |
| ENSMUSG00000021245 | Mlh3 | protein_coding | -0.018971464 | 3.920570449 | 0.0012114 | 0.9726577 | 0.990839061 |
| ENSMUSG00000026675 | Hsd17b7 | protein_coding | -0.019478637 | 4.474511903 | 0.0012015 | 0.9727727 | 0.990839061 |
| ENSMUSG00000026853 | Crat | protein_coding | -0.014026188 | 4.648621722 | 0.0011839 | 0.9729701 | 0.990839061 |
| ENSMUSG00000017778 | Cox7c | protein_coding | -0.016145824 | 3.971414894 | 0.0011821 | 0.97299 | 0.990839061 |
| ENSMUSG00000036315 | Znrd1 | protein_coding | -0.011309821 | 6.192882707 | 0.0011723 | 0.9731019 | 0.990839061 |
| ENSMUSG00000025085 | Ablim1 | protein_coding | -0.012368111 | 5.591578019 | 0.0011706 | 0.9731217 | 0.990839061 |
| ENSMUSG00000022324 | Matn2 | protein_coding | -0.021564524 | 3.743421757 | 0.0011684 | 0.9731472 | 0.990839061 |
| ENSMUSG00000027984 | Hadh | protein_coding | 0.01051263 | 5.869148006 | 0.0011545 | 0.9733071 | 0.990869838 |
| ENSMUSG00000073155 | 1810058I24Rik | lincRNA | -0.007716869 | 6.7691686 | 0.0011498 | 0.9733617 | 0.990869838 |
| ENSMUSG00000020849 | Ywhae | protein_coding | -0.008748957 | 11.16070672 | 0.0011396 | 0.9734797 | 0.990885912 |
| ENSMUSG00000030823 | 9130019O22Rik | protein_coding | -0.041950832 | 1.986000247 | 0.0011154 | 0.9737631 | 0.990885912 |
| ENSMUSG00000038967 | Pdk2 | protein_coding | 0.011493786 | 6.877811759 | 0.0011143 | 0.9737782 | 0.990885912 |
| ENSMUSG00000028524 | Sgip1 | protein_coding | 0.011964095 | 4.884116224 | 0.0011095 | 0.9738322 | 0.990885912 |
| ENSMUSG00000097316 | Gm10516 | antisense_RNA | -0.030673098 | 2.657396966 | 0.001109 | 0.9738382 | 0.990885912 |
| ENSMUSG00000039308 | Ndst2 | protein_coding | 0.032021812 | 2.597608436 | 0.0010929 | 0.9740282 | 0.99094312 |
| ENSMUSG00000061322 | Dnaic1 | protein_coding | -0.016967771 | 3.965382689 | 0.0010883 | 0.9740832 | 0.99094312 |
| ENSMUSG00000031451 | Gas6 | protein_coding | -0.023440363 | 2.816422772 | 0.001081 | 0.9741709 | 0.99094312 |
| ENSMUSG00000045009 | Prrt3 | protein_coding | -0.025168209 | 2.80428381 | 0.0010614 | 0.9744052 | 0.991078334 |
| ENSMUSG00000041775 | Mapk1ip1 | protein_coding | 0.011094102 | 5.904104184 | 0.0010546 | 0.9744882 | 0.991078334 |
| ENSMUSG00000028634 | Hivep3 | protein_coding | 0.015094447 | 4.62086736 | 0.0010467 | 0.9745839 | 0.991081972 |
| ENSMUSG00000021754 | Map3k1 | protein_coding | -0.01558955 | 3.794129042 | 0.0010286 | 0.9748038 | 0.991211878 |
| ENSMUSG00000036513 | Commd2 | protein_coding | 0.008752305 | 6.116908395 | 0.0010208 | 0.9748999 | 0.991215794 |
| ENSMUSG00000021218 | Gdi2 | protein_coding | -0.008528984 | 8.659815693 | 0.0010004 | 0.9751523 | 0.991378743 |
| ENSMUSG00000030287 | Itpr2 | protein_coding | -0.02470183 | 2.660606631 | 0.0009802 | 0.9754032 | 0.991540046 |
| ENSMUSG00000035834 | Polr3g | protein_coding | -0.036686853 | 1.979969554 | 0.000954 | 0.975734 | 0.991773759 |
| ENSMUSG00000041837 | Pdcd7 | protein_coding | -0.014867033 | 4.364299716 | 0.0009475 | 0.9758176 | 0.991773759 |
| ENSMUSG00000032733 | Snx33 | protein_coding | -0.019841575 | 3.184480107 | 0.0009373 | 0.9759473 | 0.99181182 |
| ENSMUSG00000031924 | Cyb5b | protein_coding | -0.007072241 | 6.754261924 | 0.000911 | 0.9762879 | 0.992030448 |
| ENSMUSG00000044864 | Ankrd50 | protein_coding | 0.012844325 | 4.326761059 | 0.0009063 | 0.9763491 | 0.992030448 |
| ENSMUSG00000025575 | Cant1 | protein_coding | -0.013748984 | 4.199755079 | 0.0008922 | 0.9765331 | 0.992030448 |
| ENSMUSG00000027422 | Rrbp1 | protein_coding | 0.013443832 | 3.931150346 | 0.000887 | 0.9766022 | 0.992030448 |
| ENSMUSG00000092368 | A930015D03Rik | antisense_RNA | -0.020426337 | 2.822723527 | 0.000879 | 0.9767071 | 0.992030448 |
| ENSMUSG00000043207 | Zmpste24 | protein_coding | 0.011870492 | 4.388046085 | 0.000868 | 0.9768543 | 0.992030448 |
| ENSMUSG00000039210 | Gpatch2 | protein_coding | -0.026744704 | 2.641426006 | 0.0008603 | 0.9769569 | 0.992030448 |
| ENSMUSG00000037275 | Gemin5 | protein_coding | 0.016105421 | 3.51674595 | 0.0008501 | 0.977093 | 0.992030448 |
| ENSMUSG00000063320 | 1190007I07Rik | protein_coding | -0.010723569 | 4.499521849 | 0.000847 | 0.9771348 | 0.992030448 |
| ENSMUSG00000015484 | Fam163a | protein_coding | -0.020326508 | 3.346538733 | 0.0008425 | 0.9771958 | 0.992030448 |
| ENSMUSG00000032382 | Snx1 | protein_coding | -0.007139743 | 7.24547317 | 0.0008333 | 0.9773209 | 0.992030448 |
| ENSMUSG00000057554 | Lgals8 | protein_coding | -0.011234534 | 5.631530436 | 0.0008325 | 0.9773324 | 0.992030448 |
| ENSMUSG00000043857 | Mgat5b | protein_coding | -0.012336732 | 4.474945001 | 0.000825 | 0.9774346 | 0.992030448 |
| ENSMUSG00000069020 | Urm1 | protein_coding | -0.006657123 | 6.34336436 | 0.0008236 | 0.977454 | 0.992030448 |
| ENSMUSG00000014195 | Dnajc7 | protein_coding | -0.007157852 | 6.780053273 | 0.0007947 | 0.9778525 | 0.992252406 |
| ENSMUSG00000027996 | Sfrp2 | protein_coding | -0.017386819 | 2.939593684 | 0.0007902 | 0.9779155 | 0.992252406 |
| ENSMUSG00000021036 | Sptlc2 | protein_coding | -0.015018738 | 3.764679548 | 0.0007877 | 0.9779495 | 0.992252406 |
| ENSMUSG00000022022 | Mtrf1 | protein_coding | 0.012964813 | 4.828859959 | 0.0007775 | 0.9780938 | 0.992305186 |
| ENSMUSG00000025007 | Aldh18a1 | protein_coding | -0.017311561 | 4.630399413 | 0.0007616 | 0.9783467 | 0.992397792 |
| ENSMUSG00000062352 | Itgb1bp1 | protein_coding | 0.008415705 | 6.483289988 | 0.000758 | 0.9783697 | 0.992397792 |
| ENSMUSG00000039254 | Pomt1 | protein_coding | -0.018970558 | 2.894868136 | 0.0007374 | 0.9786652 | 0.992603899 |
| ENSMUSG00000043065 | Spice1 | protein_coding | -0.015641637 | 3.476077722 | 0.0007287 | 0.9787911 | 0.992619749 |
| ENSMUSG00000027669 | Gnb4 | protein_coding | -0.007254876 | 6.130793214 | 0.0007236 | 0.9788654 | 0.992619749 |
| ENSMUSG00000031226 | Pbdc1 | protein_coding | -0.006472591 | 6.387891772 | 0.0007025 | 0.979176 | 0.99275594 |
| ENSMUSG00000006599 | Gtf2h1 | protein_coding | -0.011234387 | 4.52194204 | 0.000702 | 0.9791844 | 0.99275594 |
| ENSMUSG00000024817 | Uhrf2 | protein_coding | -0.009753665 | 4.927797077 | 0.0006915 | 0.9793401 | 0.99280513 |
| ENSMUSG00000031065 | Cdk16 | protein_coding | -0.008284127 | 6.001277039 | 0.0006863 | 0.9794175 | 0.99280513 |

| ensembl_gene_id | external_gene_name | gene_biotype | log2 (fold change) | log2 (counts per million) | F-value | p-value | False discovery rate |
|---|---|---|---|---|---|---|---|
| ENSMUSG00000032985 | 5730522E02Rik | protein_coding | -0.018950491 | 3.229163008 | 0.000667 | 0.9797098 | 0.993007791 |
| ENSMUSG00000025314 | Ptprj | protein_coding | -0.018498084 | 2.738237893 | 0.0006416 | 0.9800989 | 0.993308548 |
| ENSMUSG00000078307 | AI593442 | protein_coding | -0.009547324 | 8.096051681 | 0.0006341 | 0.9802363 | 0.993342695 |
| ENSMUSG00000030245 | Golt1b | protein_coding | -0.00681603 | 7.497290907 | 0.0006276 | 0.9803174 | 0.993342695 |
| ENSMUSG00000033157 | Abhd10 | protein_coding | 0.007436579 | 5.680328417 | 0.0006216 | 0.9804116 | 0.993344546 |
| ENSMUSG00000063236 | 1110038F14Rik | protein_coding | 0.007743066 | 5.627800157 | 0.000575 | 0.9811597 | 0.994008874 |
| ENSMUSG00000057409 | Zfp53 | protein_coding | 0.021648408 | 2.409120313 | 0.0005485 | 0.9815997 | 0.99427527 |
| ENSMUSG00000020100 | Slc29a3 | protein_coding | 0.015298481 | 3.130165832 | 0.000548 | 0.9816076 | 0.99427527 |
| ENSMUSG00000040543 | Pitpnm3 | protein_coding | 0.011571645 | 3.960726786 | 0.000528 | 0.981946 | 0.994513592 |
| ENSMUSG00000024413 | Npc1 | protein_coding | 0.010128671 | 4.208555159 | 0.0005232 | 0.9820278 | 0.994513592 |
| ENSMUSG00000050967 | Creg2 | protein_coding | 0.009267407 | 4.896421896 | 0.0005085 | 0.9822831 | 0.994574665 |
| ENSMUSG00000042826 | Fgf11 | protein_coding | 0.01262315 | 3.445099074 | 0.0005059 | 0.9823284 | 0.994574665 |
| ENSMUSG00000062729 | Ppox | protein_coding | 0.011758622 | 4.797051836 | 0.0005039 | 0.9823656 | 0.994574665 |
| ENSMUSG00000061808 | Ttr | protein_coding | -0.026000041 | 11.33229296 | 0.0005146 | 0.9824648 | 0.994581447 |
| ENSMUSG00000021615 | Xrcc4 | protein_coding | -0.009907861 | 4.192460649 | 0.0004796 | 0.9827937 | 0.994820719 |
| ENSMUSG00000049904 | Tmem17 | protein_coding | 0.008924493 | 3.952169071 | 0.0004643 | 0.9830712 | 0.994928119 |
| ENSMUSG00000039715 | Wdr34 | protein_coding | -0.008061251 | 4.733743123 | 0.0004635 | 0.9830848 | 0.994928119 |
| ENSMUSG00000083012 | Fam220a | protein_coding | 0.007582024 | 4.321405995 | 0.0004308 | 0.9836927 | 0.995125678 |
| ENSMUSG00000000148 | Brat1 | protein_coding | 0.011811174 | 4.062798521 | 0.0004266 | 0.9837732 | 0.995125678 |
| ENSMUSG00000048410 | Zfp407 | protein_coding | -0.011301419 | 3.295447265 | 0.0004115 | 0.9840625 | 0.995125678 |
| ENSMUSG00000042505 | Sdhaf3 | protein_coding | -0.005412514 | 6.69178615 | 0.0004055 | 0.9841783 | 0.995125678 |
| ENSMUSG00000053333 | Dis3l2 | protein_coding | -0.014400683 | 2.901809079 | 0.0004045 | 0.9841981 | 0.995125678 |
| ENSMUSG00000030621 | Me3 | protein_coding | 0.010056968 | 4.22441266 | 0.0004027 | 0.9842334 | 0.995125678 |
| ENSMUSG00000038416 | Cdc16 | protein_coding | -0.00733914 | 5.149809649 | 0.0004017 | 0.9842529 | 0.995125678 |
| ENSMUSG00000002043 | Trappc6a | protein_coding | 0.007020132 | 4.975157349 | 0.0004005 | 0.9842767 | 0.995125678 |
| ENSMUSG00000048109 | Rbm15 | protein_coding | -0.01133273 | 5.00797031 | 0.0003962 | 0.9843755 | 0.995125678 |
| ENSMUSG00000035473 | Galm | protein_coding | -0.021868863 | 2.245934502 | 0.0003925 | 0.9844333 | 0.995125678 |
| ENSMUSG00000019763 | Rmnd1 | protein_coding | 0.008094845 | 3.99486324 | 0.0003924 | 0.9844357 | 0.995125678 |
| ENSMUSG00000003847 | Nfat5 | protein_coding | -0.00836814 | 4.026495328 | 0.0003859 | 0.9845659 | 0.995125678 |
| ENSMUSG00000021303 | Gng4 | protein_coding | 0.00484001 | 6.380812427 | 0.0003778 | 0.9847279 | 0.995125678 |
| ENSMUSG00000025277 | Abhd6 | protein_coding | 0.005604895 | 6.217601907 | 0.0003743 | 0.9848 | 0.995125678 |
| ENSMUSG00000021576 | Pdcd6 | protein_coding | 0.004494232 | 7.915904575 | 0.0003737 | 0.9848122 | 0.995125678 |
| ENSMUSG00000064145 | Arih2 | protein_coding | -0.007280082 | 4.752227447 | 0.0003702 | 0.9848827 | 0.995125678 |
| ENSMUSG00000019814 | Ltv1 | protein_coding | -0.007918378 | 4.246042227 | 0.0003679 | 0.9849291 | 0.995125678 |
| ENSMUSG00000042589 | Cux2 | protein_coding | 0.011278362 | 5.710128165 | 0.0003689 | 0.9849458 | 0.995125678 |
| ENSMUSG00000113771 | AC160108.1 | antisense_RNA | -0.012796031 | 3.105544482 | 0.0003548 | 0.9851994 | 0.995205725 |
| ENSMUSG00000030098 | Grip2 | protein_coding | -0.013722036 | 3.227023866 | 0.0003504 | 0.9852933 | 0.995205725 |
| ENSMUSG00000048988 | Elfn1 | protein_coding | 0.013681869 | 3.066622727 | 0.0003483 | 0.9853357 | 0.995205725 |
| ENSMUSG00000108414 | Snhg1 | processed_transcript | 0.007120534 | 4.599979832 | 0.0003455 | 0.9853953 | 0.995205725 |
| ENSMUSG00000027540 | Ptpn1 | protein_coding | -0.007449747 | 4.79176329 | 0.0003281 | 0.9857688 | 0.995406106 |
| ENSMUSG00000000134 | Tfe3 | protein_coding | 0.007136176 | 4.709438247 | 0.000326 | 0.9858129 | 0.995406106 |
| ENSMUSG00000026672 | Optn | protein_coding | 0.009687019 | 3.886804943 | 0.0003234 | 0.9858714 | 0.995406106 |
| ENSMUSG00000028039 | Efna3 | protein_coding | 0.008091516 | 4.154770139 | 0.0003067 | 0.9862408 | 0.995658456 |
| ENSMUSG00000062270 | Morf4l1 | protein_coding | -0.004308735 | 6.593492639 | 0.0002944 | 0.9865189 | 0.995658456 |
| ENSMUSG00000034509 | Mad2l1bp | protein_coding | -0.00568273 | 5.073675527 | 0.0002941 | 0.9865257 | 0.995658456 |
| ENSMUSG00000063870 | Chd4 | protein_coding | 0.004487966 | 6.028503614 | 0.0002918 | 0.9865775 | 0.995658456 |
| ENSMUSG00000031988 | Vps26b | protein_coding | -0.005788894 | 5.71486949 | 0.0002898 | 0.9866252 | 0.995658456 |
| ENSMUSG00000038215 | Cep44 | protein_coding | -0.012159422 | 3.199055763 | 0.0002875 | 0.9866769 | 0.995658456 |
| ENSMUSG00000038271 | Iffo1 | protein_coding | 0.007973319 | 4.008015608 | 0.0002737 | 0.9870008 | 0.99576026 |
| ENSMUSG00000025207 | Sema4g | protein_coding | -0.01216945 | 3.063198317 | 0.0002729 | 0.9870206 | 0.99576026 |
| ENSMUSG00000020048 | Hsp90b1 | protein_coding | -0.003614574 | 7.071487555 | 0.0002714 | 0.9870556 | 0.99576026 |
| ENSMUSG00000042167 | Papd4 | protein_coding | -0.009230972 | 3.182694362 | 0.0002566 | 0.9874138 | 0.996004999 |
| ENSMUSG00000029836 | Cbx3 | protein_coding | 0.007258919 | 3.558430742 | 0.0002538 | 0.9874834 | 0.996004999 |
| ENSMUSG00000025060 | Slk | protein_coding | -0.007881144 | 4.682045416 | 0.0002409 | 0.9878044 | 0.996164242 |
| ENSMUSG00000055044 | Pdlim1 | protein_coding | -0.012527446 | 2.578696262 | 0.00024 | 0.9878266 | 0.996164242 |
| ENSMUSG00000041297 | Cdk13 | protein_coding | -0.00810794 | 4.689720122 | 0.0002309 | 0.9880624 | 0.996268023 |
| ENSMUSG00000047658 | Gal3st3 | protein_coding | -0.010856602 | 3.135643891 | 0.0002288 | 0.9881148 | 0.996268023 |
| ENSMUSG00000052446 | Zfp961 | protein_coding | -0.006528855 | 4.061898998 | 0.0002229 | 0.9882695 | 0.996320526 |
| ENSMUSG00000026142 | Rhbdd1 | protein_coding | -0.015646142 | 2.721468663 | 0.0002181 | 0.9883976 | 0.996320526 |
| ENSMUSG00000038187 | Btbd10 | protein_coding | 0.004631495 | 5.697058457 | 0.0002163 | 0.9884448 | 0.996320526 |
| ENSMUSG00000018899 | Irf1 | protein_coding | 0.011627671 | 2.824912173 | 0.0002126 | 0.9885426 | 0.99632571 |
| ENSMUSG00000026516 | Nvl | protein_coding | 0.005468905 | 5.279776214 | 0.0002085 | 0.9886551 | 0.996333863 |
| ENSMUSG00000025858 | Get4 | protein_coding | -0.006328386 | 3.690348332 | 0.0002055 | 0.988736 | 0.996333863 |
| ENSMUSG00000024381 | Bin1 | protein_coding | -0.004232108 | 6.261509006 | 0.0002005 | 0.9888742 | 0.996379719 |
| ENSMUSG00000041261 | Car8 | protein_coding | 0.006369978 | 4.349591531 | 0.0001789 | 0.9894906 | 0.996732245 |
| ENSMUSG00000021413 | Prpf4b | protein_coding | -0.004516323 | 5.784699678 | 0.0001787 | 0.989497 | 0.996732245 |
| ENSMUSG00000033768 | Nrxn2 | protein_coding | -0.005118282 | 4.840511298 | 0.0001785 | 0.9895022 | 0.996732245 |
| ENSMUSG00000041020 | Map7d2 | protein_coding | -0.002764746 | 7.664329673 | 0.0001748 | 0.9896119 | 0.996749406 |
| ENSMUSG00000041258 | Zfp236 | protein_coding | -0.005207323 | 3.922609002 | 0.0001679 | 0.9898186 | 0.996864233 |
| ENSMUSG00000066637 | Ttc32 | protein_coding | 0.010304952 | 2.762092343 | 0.000157 | 0.9901554 | 0.996977453 |
| ENSMUSG00000001420 | Tmem79 | protein_coding | 0.009202602 | 2.657697046 | 0.0001508 | 0.9903507 | 0.996977453 |
| ENSMUSG00000051401 | Kctd16 | protein_coding | 0.00843272 | 2.779442668 | 0.0001473 | 0.9904622 | 0.996977453 |
| ENSMUSG00000027508 | Pag1 | protein_coding | 0.007814641 | 3.019349105 | 0.0001471 | 0.9904698 | 0.996977453 |
| ENSMUSG00000041879 | Ipo9 | protein_coding | -0.003560525 | 6.119711116 | 0.0001456 | 0.9905186 | 0.996977453 |
| ENSMUSG00000014859 | E2f4 | protein_coding | -0.006025785 | 3.896729469 | 0.0001454 | 0.9905245 | 0.996977453 |
| ENSMUSG00000044134 | Fam109a | protein_coding | 0.013854993 | 2.104216316 | 0.0001371 | 0.9907989 | 0.996977453 |
| ENSMUSG00000034839 | Larp6 | protein_coding | 0.005391132 | 4.866273999 | 0.000137 | 0.9908033 | 0.996977453 |
| ENSMUSG00000035437 | Rabgap1 | protein_coding | 0.007546751 | 3.37899824 | 0.0001323 | 0.9909636 | 0.996977453 |

| ensembl_gene_id | external_gene_name | gene_biotype | log2 (fold change) | log2 (counts per million) | F-value | p-value | False discovery rate |
|---|---|---|---|---|---|---|---|
| ENSMUSG00000029221 | Slc30a9 | protein_coding | 0.00279957 | 6.804376336 | 0.0001278 | 0.9911166 | 0.996977453 |
| ENSMUSG00000017679 | Ttpal | protein_coding | -0.00399261 | 4.909761282 | 0.0001267 | 0.9911562 | 0.996977453 |
| ENSMUSG00000051067 | Lingo3 | protein_coding | -0.007421512 | 3.708246974 | 0.0001257 | 0.9911892 | 0.996977453 |
| ENSMUSG00000004207 | Psap | protein_coding | -0.002839786 | 9.011552708 | 0.0001233 | 0.9912754 | 0.996977453 |
| ENSMUSG00000052459 | Atp6v1a | protein_coding | -0.003054632 | 8.941268298 | 0.0001231 | 0.991281 | 0.996977453 |
| ENSMUSG00000027977 | Ndst3 | protein_coding | -0.008347509 | 2.657612592 | 0.0001195 | 0.991411 | 0.996977453 |
| ENSMUSG00000026283 | Ing5 | protein_coding | -0.003913929 | 4.677544366 | 0.0001162 | 0.9915318 | 0.996977453 |
| ENSMUSG00000031659 | Adcy7 | protein_coding | 0.006486342 | 3.286394113 | 0.0001151 | 0.9915717 | 0.996977453 |
| ENSMUSG00000027253 | Lrp4 | protein_coding | 0.006342784 | 4.078801821 | 0.0001143 | 0.9915999 | 0.996977453 |
| ENSMUSG00000079083 | Jrkl | protein_coding | 0.006804455 | 2.992706332 | 0.0001074 | 0.9918577 | 0.997143432 |
| ENSMUSG00000025026 | Add3 | protein_coding | -0.002900042 | 6.071688875 | 0.0001022 | 0.9920574 | 0.99722686 |
| ENSMUSG00000002602 | Axl | protein_coding | 0.00704384 | 3.171210673 | 0.0001004 | 0.9921262 | 0.99722686 |
| ENSMUSG00000097518 | Gm26694 | lincRNA | 0.004206585 | 4.020025057 | 9.18E-05 | 0.9924697 | 0.9974789 |
| ENSMUSG00000024172 | St6gal2 | protein_coding | -0.00787182 | 2.881335926 | 8.75E-05 | 0.9926495 | 0.997519505 |
| ENSMUSG00000035547 | Capn5 | protein_coding | 0.004080663 | 5.012830379 | 8.53E-05 | 0.9927411 | 0.997519505 |
| ENSMUSG00000097121 | D130020L05Rik | antisense_RNA | -0.004390726 | 3.731361309 | 8.42E-05 | 0.9927884 | 0.997519505 |
| ENSMUSG00000046688 | Tifa | protein_coding | -0.007942206 | 2.772496425 | 8.19E-05 | 0.9928873 | 0.99752562 |
| ENSMUSG00000020802 | Ube2o | protein_coding | -0.003137516 | 4.962172103 | 7.74E-05 | 0.9930853 | 0.997631345 |
| ENSMUSG00000039914 | Coq10a | protein_coding | 0.003380771 | 5.652001736 | 7.36E-05 | 0.9932593 | 0.997713009 |
| ENSMUSG00000097287 | D130017N08Rik | lincRNA | -0.003990637 | 3.860943306 | 6.71E-05 | 0.9935619 | 0.997923658 |
| ENSMUSG00000015461 | Atf6b | protein_coding | 0.004308964 | 4.014220969 | 6.32E-05 | 0.9937547 | 0.997964589 |
| ENSMUSG00000027339 | Rassf2 | protein_coding | -0.003009124 | 4.579458605 | 6.10E-05 | 0.9938628 | 0.997964589 |
| ENSMUSG00000024743 | Syt7 | protein_coding | -0.002463106 | 5.813303596 | 6.06E-05 | 0.993881 | 0.997964589 |
| ENSMUSG00000022808 | Snx4 | protein_coding | 0.002553485 | 5.947934191 | 5.59E-05 | 0.9941259 | 0.998117297 |
| ENSMUSG00000025762 | Larp1b | protein_coding | 0.004611961 | 3.278689258 | 5.14E-05 | 0.9943652 | 0.998192283 |
| ENSMUSG00000034055 | Phka1 | protein_coding | 0.005451577 | 2.741177134 | 4.90E-05 | 0.9945025 | 0.998192283 |
| ENSMUSG00000045658 | Pid1 | protein_coding | 0.002757757 | 4.859347571 | 4.78E-05 | 0.9945684 | 0.998192283 |
| ENSMUSG00000013663 | Pten | protein_coding | -0.002405701 | 5.377594495 | 4.77E-05 | 0.9945719 | 0.998192283 |
| ENSMUSG00000022106 | Rcbtb2 | protein_coding | -0.004428755 | 3.564056249 | 4.61E-05 | 0.9946648 | 0.998192302 |
| ENSMUSG00000032397 | Tipin | protein_coding | -0.002048482 | 6.106471395 | 4.29E-05 | 0.9948541 | 0.998289111 |
| ENSMUSG00000097908 | 4933404O12Rik | lincRNA | 0.003920845 | 3.217324992 | 4.03E-05 | 0.9950145 | 0.998321297 |
| ENSMUSG00000032580 | Rbm5 | protein_coding | 0.001539236 | 6.59973986 | 3.86E-05 | 0.9951157 | 0.998321297 |
| ENSMUSG00000029059 | Fam213b | protein_coding | 0.001886774 | 9.193708674 | 3.79E-05 | 0.9951647 | 0.998321297 |
| ENSMUSG00000068923 | Syt11 | protein_coding | 0.001509531 | 9.840075036 | 3.52E-05 | 0.9953376 | 0.998346973 |
| ENSMUSG00000002799 | Jag2 | protein_coding | 0.005217872 | 2.957578976 | 3.38E-05 | 0.9954303 | 0.998346973 |
| ENSMUSG00000049323 | Smcr8 | protein_coding | 0.002959609 | 3.611316607 | 3.16E-05 | 0.9955824 | 0.998346973 |
| ENSMUSG00000054814 | Usp46 | protein_coding | -0.001713864 | 6.444932736 | 2.85E-05 | 0.9958074 | 0.998346973 |
| ENSMUSG00000016619 | Nup50 | protein_coding | 0.002094314 | 4.860087363 | 2.80E-05 | 0.9958421 | 0.998346973 |
| ENSMUSG00000037470 | Uggt1 | protein_coding | -0.002440228 | 4.094917282 | 2.79E-05 | 0.9958533 | 0.998346973 |
| ENSMUSG00000004631 | Sgce | protein_coding | 0.00142977 | 5.822565911 | 2.70E-05 | 0.9959199 | 0.998346973 |
| ENSMUSG00000032126 | Hmbs | protein_coding | 0.002837731 | 3.707314394 | 2.68E-05 | 0.995933 | 0.998346973 |
| ENSMUSG00000035392 | Dennd1a | protein_coding | 0.001822969 | 4.580423137 | 2.31E-05 | 0.9962207 | 0.998542248 |
| ENSMUSG00000027951 | Adar | protein_coding | -0.003930039 | 3.419891742 | 1.99E-05 | 0.9965018 | 0.998588628 |
| ENSMUSG00000006127 | Inpp5k | protein_coding | 0.001714746 | 5.897336812 | 1.95E-05 | 0.9965313 | 0.998588628 |
| ENSMUSG00000036432 | Siah2 | protein_coding | -0.002636765 | 3.735798607 | 1.93E-05 | 0.9965456 | 0.998588628 |
| ENSMUSG00000026878 | Rab14 | protein_coding | -0.001005043 | 7.850148033 | 1.74E-05 | 0.9967236 | 0.998673958 |
| ENSMUSG00000031875 | Cmtm3 | protein_coding | -0.002243394 | 3.327982326 | 1.59E-05 | 0.996868 | 0.998725587 |
| ENSMUSG00000062591 | Tubb4a | protein_coding | -0.00137731 | 11.07819344 | 1.40E-05 | 0.9970556 | 0.998820427 |
| ENSMUSG00000040151 | Hs2st1 | protein_coding | -0.001471768 | 4.851491335 | 1.17E-05 | 0.9973175 | 0.998989745 |
| ENSMUSG00000030770 | Parva | protein_coding | 0.000982024 | 5.503169889 | 1.00E-05 | 0.9975123 | 0.999007389 |
| ENSMUSG00000029156 | Sgcb | protein_coding | -0.00116475 | 5.265246916 | 9.41E-06 | 0.9975896 | 0.999007389 |
| ENSMUSG00000036278 | Macrod1 | protein_coding | 0.001155385 | 5.257612405 | 9.22E-06 | 0.9976138 | 0.999007389 |
| ENSMUSG00000047423 | AI837181 | protein_coding | 0.000828783 | 6.38112456 | 8.40E-06 | 0.9977231 | 0.999023833 |
| ENSMUSG00000030276 | Ttll3 | protein_coding | 0.002229738 | 2.692870825 | 6.30E-06 | 0.9980277 | 0.999171407 |
| ENSMUSG00000074182 | Znhit6 | protein_coding | -0.000853049 | 4.740682252 | 5.85E-06 | 0.9980993 | 0.999171407 |
| ENSMUSG00000073633 | Fbxo36 | protein_coding | -0.000987557 | 4.715544166 | 5.55E-06 | 0.9981493 | 0.999171407 |
| ENSMUSG00000037408 | Cnnm4 | protein_coding | -0.000660335 | 5.394827322 | 2.25E-06 | 0.9988218 | 0.999751575 |
| ENSMUSG00000078700 | D030028A08Rik | processed_transcript | -0.000976954 | 2.384144999 | 1.28E-06 | 0.999111 | 0.999816075 |
| ENSMUSG00000021573 | Tppp | protein_coding | -0.000273949 | 7.734587106 | 1.27E-06 | 0.9991137 | 0.999816075 |
| ENSMUSG00000006651 | Aplp1 | protein_coding | 0.000298473 | 9.291884173 | 8.51E-07 | 0.9992752 | 0.999816075 |
| ENSMUSG00000003135 | Cnot11 | protein_coding | 0.000510136 | 2.972213039 | 7.75E-07 | 0.9993081 | 0.999816075 |
| ENSMUSG00000020628 | Trappc12 | protein_coding | -0.000189873 | 6.182638544 | 5.41E-07 | 0.999422 | 0.999816075 |
| ENSMUSG00000026317 | Cln8 | protein_coding | -0.000246588 | 5.354004225 | 3.88E-07 | 0.9995109 | 0.999816075 |
| ENSMUSG00000036450 | Hif1an | protein_coding | 0.000276425 | 4.210286783 | 3.47E-07 | 0.9995371 | 0.999816075 |
| ENSMUSG00000035329 | Fbxo33 | protein_coding | 4.06E-05 | 5.444145011 | 1.38E-08 | 0.9999075 | 1 |
| ENSMUSG00000020086 | H2afy2 | protein_coding | -5.62E-05 | 2.575478223 | 5.81E-09 | 0.9999401 | 1 |
| ENSMUSG00000047230 | Cldn2 | protein_coding | 0 | 1.917464903 | 0 | 1 | 1 |
