## Supplemental Table 4 for "Extended amygdala-parabrachial circuits alter threat assessment to regulate feeding"

**TABLE S4**.

| ensembl_gene_id | external_gene_name | gene_biotype | log2 (fold change) | log2 (counts per million) | F-value | p-value | False discovery rate |
|---|---|---|---|---|---|---|---|
| ENSMUSG00000025400 | Tac2 | protein_coding | 4.879840808 | 9.178170504 | 348.95327 | 1.94E-12 | 2.09E-08 |
| ENSMUSG00000024907 | Gal | protein_coding | 2.444546833 | 9.118265128 | 96.315371 | 2.99E-08 | 0.000160937 |
| ENSMUSG00000045731 | Pnoc | protein_coding | 1.57405972 | 6.687693393 | 63.304696 | 5.23E-07 | 0.001873781 |
| ENSMUSG00000047228 | A2ml1 | protein_coding | 1.920154275 | 5.711089496 | 50.521742 | 2.23E-06 | 0.00598548 |
| ENSMUSG00000036578 | Fxyd7 | protein_coding | 1.223069467 | 7.396395722 | 34.394991 | 2.22E-05 | 0.047683583 |
| ENSMUSG00000023150 | Ivns1abp | protein_coding | -1.301431686 | 6.256843816 | 31.728431 | 3.48E-05 | 0.057291697 |
| ENSMUSG00000017707 | Serinc3 | protein_coding | 1.504452705 | 6.07841394 | 31.339129 | 3.73E-05 | 0.057291697 |
| ENSMUSG00000018169 | Mfng | protein_coding | 2.545626059 | 3.752551007 | 29.007311 | 5.68E-05 | 0.076348187 |
| ENSMUSG00000036357 | Gpr101 | protein_coding | 1.799184931 | 4.869678402 | 26.649761 | 8.90E-05 | 0.100841847 |
| ENSMUSG00000045471 | Hcrt | protein_coding | -3.903325682 | 10.01304624 | 34.344446 | 9.38E-05 | 0.100841847 |
| ENSMUSG00000030069 | Prok2 | protein_coding | 2.229458862 | 3.528654604 | 24.595815 | 0.0001344 | 0.126484386 |
| ENSMUSG00000026051 | 1500015O10Rik | protein_coding | -3.915756089 | 3.884914148 | 25.142408 | 0.0001412 | 0.126484386 |
| ENSMUSG00000021091 | Serpina3n | protein_coding | 1.556189386 | 4.794840131 | 23.782644 | 0.0001593 | 0.131736935 |
| ENSMUSG00000000792 | Slc5a5 | protein_coding | 1.220776234 | 6.950427196 | 21.904812 | 0.0002389 | 0.1651074 |
| ENSMUSG00000027270 | Lamp5 | protein_coding | 1.394953881 | 7.01568764 | 21.77817 | 0.0002468 | 0.1651074 |
| ENSMUSG00000008036 | Ap2s1 | protein_coding | 0.938778326 | 8.101066345 | 21.699239 | 0.0002501 | 0.1651074 |
| ENSMUSG00000022425 | Enpp2 | protein_coding | -2.355272958 | 8.708527452 | 23.771409 | 0.000261 | 0.1651074 |
| ENSMUSG00000000214 | Th | protein_coding | 1.750385631 | 5.627137338 | 20.935764 | 0.0002982 | 0.178137404 |
| ENSMUSG00000027620 | Rbm39 | protein_coding | -1.029206578 | 7.1349074 | 20.05974 | 0.0003635 | 0.198798622 |
| ENSMUSG00000030020 | Prickle2 | protein_coding | -1.281447643 | 5.767254028 | 19.986464 | 0.0003698 | 0.198798622 |
| ENSMUSG00000025647 | Shisa5 | protein_coding | 1.090242887 | 7.52527952 | 19.726715 | 0.000393 | 0.201244807 |
| ENSMUSG00000019772 | Vip | protein_coding | 2.338581991 | 3.329568435 | 18.937751 | 0.0004744 | 0.231892042 |
| ENSMUSG00000042258 | Isl1 | protein_coding | 0.998923175 | 6.31340539 | 18.688708 | 0.000504 | 0.235612613 |
| ENSMUSG00000036181 | Hist1h1c | protein_coding | -2.078548761 | 3.712771943 | 18.397107 | 0.0005412 | 0.242476221 |
| ENSMUSG00000001827 | Folr1 | protein_coding | -10.53932424 | 3.469133099 | 21.763453 | 0.0005832 | 0.250865929 |
| ENSMUSG00000030844 | Rgs10 | protein_coding | 0.892156798 | 8.342266456 | 17.393856 | 0.0006951 | 0.274277656 |
| ENSMUSG00000037754 | Ppp1r16b | protein_coding | -1.222678337 | 8.07945623 | 17.383994 | 0.0006991 | 0.274277656 |
| ENSMUSG00000055435 | Maf | protein_coding | -2.46558562 | 3.232228002 | 17.286923 | 0.0007142 | 0.274277656 |
| ENSMUSG00000039672 | Kcne2 | protein_coding | -10.26330514 | 3.019242124 | 19.126619 | 0.000748 | 0.277347076 |
| ENSMUSG00000017978 | Cadps2 | protein_coding | 1.360489068 | 5.975778509 | 16.667299 | 0.00084 | 0.296392265 |
| ENSMUSG00000068220 | Lgals1 | protein_coding | -1.399655199 | 5.289753324 | 16.498195 | 0.0008751 | 0.296392265 |
| ENSMUSG00000046593 | Tmem215 | protein_coding | 1.463265856 | 4.79170462 | 16.467899 | 0.000882 | 0.296392265 |
| ENSMUSG00000004748 | Mtfp1 | protein_coding | 0.913195671 | 7.955389561 | 16.040101 | 0.0009871 | 0.302717024 |
| ENSMUSG00000032172 | Olfm2 | protein_coding | 0.89858603 | 6.814013233 | 15.967399 | 0.0010063 | 0.302717024 |
| ENSMUSG00000028266 | Lmo4 | protein_coding | -1.003885542 | 6.567840984 | 15.952321 | 0.0010104 | 0.302717024 |
| ENSMUSG00000086496 | Gm14204 | processed_transcript | 1.449942871 | 5.415769256 | 15.940799 | 0.0010135 | 0.302717024 |
| ENSMUSG00000024597 | Slc12a2 | protein_coding | -1.428272812 | 5.596955192 | 15.664185 | 0.0010912 | 0.317121616 |
| ENSMUSG00000040118 | Cacna2d1 | protein_coding | -1.076270127 | 5.728301998 | 15.180352 | 0.0012438 | 0.330482782 |
| ENSMUSG00000039202 | Abhd2 | protein_coding | -1.609045989 | 4.758151013 | 15.147828 | 0.0012549 | 0.330482782 |
| ENSMUSG00000061808 | Ttr | protein_coding | -6.935439664 | 11.33229296 | 23.571142 | 0.0013174 | 0.330482782 |
| ENSMUSG00000027495 | Fam210b | protein_coding | -1.435417827 | 4.945300352 | 14.868677 | 0.0013549 | 0.330482782 |
| ENSMUSG00000085336 | Gm11732 | processed_transcript | 2.478230578 | 3.231971655 | 14.629431 | 0.0014477 | 0.330482782 |
| ENSMUSG00000040407 | Akap9 | protein_coding | -1.107335305 | 5.75019992 | 14.584298 | 0.0014661 | 0.330482782 |
| ENSMUSG00000033623 | Pcgf3 | protein_coding | -1.71803018 | 4.33184919 | 14.532628 | 0.0014873 | 0.330482782 |
| ENSMUSG00000032417 | Rwdd2a | protein_coding | 0.826120723 | 7.882959122 | 14.500959 | 0.0015006 | 0.330482782 |
| ENSMUSG00000025869 | Nop16 | protein_coding | 0.791620156 | 7.109687438 | 14.43475 | 0.0015286 | 0.330482782 |
| ENSMUSG00000029334 | Prkg2 | protein_coding | -2.618252975 | 3.431655454 | 14.394398 | 0.001546 | 0.330482782 |
| ENSMUSG00000036880 | Acaa2 | protein_coding | -1.056587768 | 5.883959439 | 14.24841 | 0.0016108 | 0.330482782 |
| ENSMUSG00000024256 | Adcyap1 | protein_coding | -3.014845234 | 3.763489644 | 14.576921 | 0.0016234 | 0.330482782 |
| ENSMUSG00000049796 | Crh | protein_coding | 2.340862791 | 2.83028429 | 14.215585 | 0.0016258 | 0.330482782 |
| ENSMUSG00000045777 | Ifitm10 | protein_coding | 1.002060835 | 5.831106192 | 14.134865 | 0.0016634 | 0.330482782 |
| ENSMUSG00000033730 | Egr3 | protein_coding | -1.631518778 | 4.995065314 | 14.058438 | 0.0016998 | 0.330482782 |
| ENSMUSG00000024610 | Cd74 | protein_coding | -1.935757678 | 4.200338899 | 14.057204 | 0.0017004 | 0.330482782 |
| ENSMUSG00000032253 | Phip | protein_coding | -1.418720672 | 4.797293326 | 14.033397 | 0.001712 | 0.330482782 |
| ENSMUSG00000025862 | Stag2 | protein_coding | -1.344000776 | 4.708151346 | 14.018817 | 0.0017191 | 0.330482782 |
| ENSMUSG00000033307 | Mif | protein_coding | 0.779485902 | 7.599309128 | 14.014663 | 0.0017211 | 0.330482782 |
| ENSMUSG00000031309 | Rps6ka3 | protein_coding | -1.949326986 | 3.483811245 | 13.916526 | 0.0017699 | 0.332297267 |
| ENSMUSG00000079593 | Gm14597 | protein_coding | 0.731333167 | 8.072834169 | 13.872441 | 0.0017924 | 0.332297267 |
| ENSMUSG00000041199 | Rpusd1 | protein_coding | 0.798218104 | 6.885391051 | 13.660402 | 0.0019049 | 0.337196768 |
| ENSMUSG00000007041 | Clic1 | protein_coding | -1.559669386 | 4.372558021 | 13.660204 | 0.001905 | 0.337196768 |
| ENSMUSG00000030306 | Tmtc1 | protein_coding | -1.329005702 | 4.54557722 | 13.645867 | 0.0019129 | 0.337196768 |
| ENSMUSG00000071644 | Eef1g | protein_coding | 0.718733216 | 7.444402866 | 13.510343 | 0.0019893 | 0.341449097 |
| ENSMUSG00000053414 | Hunk | protein_coding | 1.262519722 | 4.479379812 | 13.490993 | 0.0020005 | 0.341449097 |
| ENSMUSG00000033060 | Lmo7 | protein_coding | -2.015903747 | 3.790753184 | 13.423383 | 0.0020402 | 0.342778955 |
| ENSMUSG00000001988 | Npas1 | protein_coding | 2.355703273 | 3.054201747 | 13.061445 | 0.0022682 | 0.371305306 |
| ENSMUSG00000021587 | Pcsk1 | protein_coding | -1.336374321 | 4.154065003 | 13.045373 | 0.002279 | 0.371305306 |
| ENSMUSG00000033068 | Entpd6 | protein_coding | 0.796259553 | 6.758340762 | 12.931441 | 0.0023571 | 0.378292652 |
| ENSMUSG00000062753 | AI413582 | protein_coding | 0.792314537 | 7.381961825 | 12.861432 | 0.0024065 | 0.378721631 |
| ENSMUSG00000021848 | Otx2 | protein_coding | -2.06301536 | 3.240621142 | 12.748176 | 0.0024891 | 0.378721631 |
| ENSMUSG00000003271 | Sult2b1 | protein_coding | 1.74808873 | 3.973833882 | 12.722194 | 0.0025084 | 0.378721631 |
| ENSMUSG00000026247 | Ecel1 | protein_coding | 1.300062188 | 5.931074143 | 12.729433 | 0.0025233 | 0.378721631 |
| ENSMUSG00000013539 | Tango2 | protein_coding | 0.774812899 | 8.030168215 | 12.65717 | 0.0025577 | 0.378721631 |
| ENSMUSG00000005881 | Ergic3 | protein_coding | 0.673043881 | 8.70639423 | 12.639773 | 0.0025711 | 0.378721631 |
| ENSMUSG00000027301 | Oxt | protein_coding | -3.760603686 | 4.513115714 | 14.313108 | 0.0026205 | 0.380779943 |
| ENSMUSG00000022658 | Tagln3 | protein_coding | 0.754978738 | 10.66124834 | 12.427351 | 0.0027408 | 0.39076423 |
| ENSMUSG00000029404 | Arl6ip4 | protein_coding | 0.673532475 | 7.407770941 | 12.401994 | 0.0027618 | 0.39076423 |
| ENSMUSG00000048108 | Tmem72 | protein_coding | -4.736049439 | 2.046472739 | 12.321516 | 0.00283 | 0.392389444 |
| ENSMUSG00000021701 | Plk2 | protein_coding | -0.796218765 | 7.13169582 | 12.302574 | 0.0028463 | 0.392389444 |

| ensembl_gene_id | external_gene_name | gene_biotype | log2 (fold change) | log2 (counts per million) | F-value | p-value | False discovery rate |
|---|---|---|---|---|---|---|---|
| ENSMUSG00000025264 | Tsr2 | protein_coding | 0.792252745 | 7.826594417 | 12.136436 | 0.0029941 | 0.403697086 |
| ENSMUSG00000042078 | Svop | protein_coding | 0.717402285 | 7.24270854 | 12.12623 | 0.0030034 | 0.403697086 |
| ENSMUSG00000085251 | Gm12326 | lincRNA | -2.861337621 | 3.29364466 | 12.050351 | 0.0030804 | 0.408231246 |
| ENSMUSG00000021032 | Ngb | protein_coding | 0.839028513 | 8.359207565 | 11.913578 | 0.0032061 | 0.408231246 |
| ENSMUSG00000026915 | Strbp | protein_coding | -0.767348754 | 6.935576728 | 11.87249 | 0.003247 | 0.408231246 |
| ENSMUSG00000038679 | Trps1 | protein_coding | -1.314423499 | 4.421046374 | 11.82068 | 0.0032994 | 0.408231246 |
| ENSMUSG00000071657 | Bscl2 | protein_coding | 0.72442124 | 7.871977081 | 11.811316 | 0.003309 | 0.408231246 |
| ENSMUSG00000059355 | Wdr83os | protein_coding | 0.74971828 | 6.405259769 | 11.782916 | 0.0033382 | 0.408231246 |
| ENSMUSG00000027347 | Rasgrp1 | protein_coding | -0.843999852 | 6.741384254 | 11.777322 | 0.003344 | 0.408231246 |
| ENSMUSG00000061702 | Tmem91 | protein_coding | 0.760080222 | 7.338949389 | 11.772934 | 0.0033486 | 0.408231246 |
| ENSMUSG00000025792 | Slc25a10 | protein_coding | 1.003564865 | 5.270183151 | 11.647209 | 0.0034821 | 0.408231246 |
| ENSMUSG00000023034 | Nr4a1 | protein_coding | -0.960072103 | 5.870254993 | 11.583546 | 0.0035519 | 0.408231246 |
| ENSMUSG00000052372 | Il1rapl1 | protein_coding | 1.108749537 | 7.527975867 | 11.801472 | 0.0035692 | 0.408231246 |
| ENSMUSG00000096054 | Syne1 | protein_coding | -0.912098671 | 6.143398477 | 11.525639 | 0.0036169 | 0.408231246 |
| ENSMUSG00000024411 | Aqp4 | protein_coding | -1.232102629 | 5.041781743 | 11.493107 | 0.003654 | 0.408231246 |
| ENSMUSG00000057068 | Fam47e | protein_coding | -2.549697505 | 3.11634576 | 11.434621 | 0.0037217 | 0.408231246 |
| ENSMUSG00000032081 | Apoc3 | protein_coding | 1.953019126 | 3.179063449 | 11.405432 | 0.003756 | 0.408231246 |
| ENSMUSG00000008958 | Vps72 | protein_coding | 0.768833738 | 6.102991075 | 11.373034 | 0.0037946 | 0.408231246 |
| ENSMUSG00000061751 | Kalrn | protein_coding | -0.980183566 | 5.524769988 | 11.372774 | 0.0037949 | 0.408231246 |
| ENSMUSG00000030067 | Foxp1 | protein_coding | -0.751710273 | 7.041363291 | 11.365485 | 0.0038036 | 0.408231246 |
| ENSMUSG00000047230 | Cldn2 | protein_coding | -9.050323404 | 1.917464903 | 12.143365 | 0.0038195 | 0.408231246 |
| ENSMUSG00000025666 | Tmem47 | protein_coding | -0.890735322 | 6.171143608 | 11.343061 | 0.0038306 | 0.408231246 |
| ENSMUSG00000037646 | Vps13b | protein_coding | -2.091146357 | 3.46712772 | 11.339927 | 0.0038344 | 0.408231246 |
| ENSMUSG00000047656 | Trpt1 | protein_coding | 0.976075899 | 5.582264213 | 11.280584 | 0.003907 | 0.411882126 |
| ENSMUSG00000030172 | Erc1 | protein_coding | -1.465878415 | 4.023491567 | 11.198299 | 0.0040102 | 0.418662134 |
| ENSMUSG00000022949 | Clic6 | protein_coding | -2.629440937 | 4.046555903 | 11.531702 | 0.0040535 | 0.419108506 |
| ENSMUSG00000039031 | Arhgap18 | protein_coding | -1.841116647 | 3.136624267 | 11.057886 | 0.0041936 | 0.429469035 |
| ENSMUSG00000021478 | Drd1 | protein_coding | -1.325931654 | 5.486676655 | 10.867211 | 0.0044662 | 0.444069718 |
| ENSMUSG00000034998 | Foxn2 | protein_coding | -1.276273162 | 4.48501583 | 10.851807 | 0.0044803 | 0.444069718 |
| ENSMUSG00000039046 | Usp6nl | protein_coding | -1.42296093 | 3.871457784 | 10.791091 | 0.0045689 | 0.444069718 |
| ENSMUSG00000073411 | H2-D1 | protein_coding | -2.131534577 | 4.646071214 | 11.128548 | 0.0045803 | 0.444069718 |
| ENSMUSG00000007721 | Ccdc124 | protein_coding | 0.690986338 | 8.225386948 | 10.721261 | 0.0046733 | 0.444069718 |
| ENSMUSG00000046402 | Rbp1 | protein_coding | -1.34942175 | 5.686358435 | 10.740192 | 0.0046749 | 0.444069718 |
| ENSMUSG00000026090 | 2010300C02Rik | protein_coding | -0.993985824 | 5.642131526 | 10.711857 | 0.0046875 | 0.444069718 |
| ENSMUSG00000045948 | Mrps12 | protein_coding | 0.622113108 | 7.801019874 | 10.639544 | 0.0047989 | 0.444069718 |
| ENSMUSG00000032246 | Calml4 | protein_coding | -2.079037418 | 3.87623688 | 10.6153 | 0.0048454 | 0.444069718 |
| ENSMUSG00000024725 | Ostf1 | protein_coding | -1.879303131 | 3.297632081 | 10.597414 | 0.0048651 | 0.444069718 |
| ENSMUSG00000044167 | Foxo1 | protein_coding | -1.042666708 | 5.322789056 | 10.595188 | 0.0048687 | 0.444069718 |
| ENSMUSG00000075702 | Selenom | protein_coding | 0.727632504 | 9.621468166 | 10.594631 | 0.0048695 | 0.444069718 |
| ENSMUSG00000085609 | 1700016P03Rik | processed_transcript | -1.84290419 | 3.263727687 | 10.592397 | 0.0048731 | 0.444069718 |
| ENSMUSG00000027602 | Map1lc3a | protein_coding | 0.661485971 | 8.415411366 | 10.492976 | 0.0050339 | 0.45486621 |
| ENSMUSG00000027327 | 1700037H04Rik | protein_coding | 0.70624756 | 9.438842398 | 10.458587 | 0.0050909 | 0.456183685 |
| ENSMUSG00000022560 | Slc52a2 | protein_coding | 1.363387143 | 4.264855553 | 10.380497 | 0.005223 | 0.460774543 |
| ENSMUSG00000031647 | Mfap3l | protein_coding | -1.430608046 | 3.545629981 | 10.333579 | 0.0053043 | 0.460774543 |
| ENSMUSG00000058260 | Serpina9 | protein_coding | -1.5387619 | 4.249092998 | 10.302628 | 0.0053587 | 0.460774543 |
| ENSMUSG00000070934 | Rraga | protein_coding | -1.032002227 | 6.785571419 | 10.285761 | 0.0053978 | 0.460774543 |
| ENSMUSG00000072694 | 1500011B03Rik | protein_coding | 0.603210579 | 8.042027602 | 10.269037 | 0.0054185 | 0.460774543 |
| ENSMUSG00000073433 | Arhgdig | protein_coding | 0.944003022 | 5.627176082 | 10.267967 | 0.0054204 | 0.460774543 |
| ENSMUSG00000074872 | Ctxn2 | protein_coding | 0.777298799 | 8.340177171 | 10.25468 | 0.0054443 | 0.460774543 |
| ENSMUSG00000020644 | Id2 | protein_coding | -0.718275943 | 8.614376753 | 10.199009 | 0.0055455 | 0.460774543 |
| ENSMUSG00000011958 | Bnip2 | protein_coding | -1.559917881 | 3.770114211 | 10.161551 | 0.0056149 | 0.460774543 |
| ENSMUSG00000073792 | Alg6 | protein_coding | -1.452424015 | 3.70962646 | 10.122726 | 0.0056878 | 0.460774543 |
| ENSMUSG00000003299 | Mrpl4 | protein_coding | 0.6842492 | 7.4879385 | 10.091198 | 0.0057478 | 0.460774543 |
| ENSMUSG00000032503 | Arpp21 | protein_coding | -1.162039092 | 10.18890769 | 10.247127 | 0.0057969 | 0.460774543 |
| ENSMUSG00000005447 | Pafah1b3 | protein_coding | 0.769049016 | 7.042839988 | 10.063675 | 0.0058007 | 0.460774543 |
| ENSMUSG00000050121 | Opalin | protein_coding | -0.71542619 | 7.410040088 | 10.060175 | 0.0058075 | 0.460774543 |
| ENSMUSG00000028433 | Ubap2 | protein_coding | -1.34600133 | 4.129500924 | 10.023389 | 0.0058792 | 0.460774543 |
| ENSMUSG00000021665 | Hexb | protein_coding | 0.913738763 | 8.13720421 | 9.991156 | 0.0059526 | 0.460774543 |
| ENSMUSG00000024847 | Aip | protein_coding | 0.680777343 | 8.863937582 | 9.9807318 | 0.0059637 | 0.460774543 |
| ENSMUSG00000028670 | Lypla2 | protein_coding | 0.732674092 | 6.34628073 | 9.9277989 | 0.0060704 | 0.460774543 |
| ENSMUSG00000001520 | Nrip2 | protein_coding | 0.972925173 | 6.737210411 | 9.9135997 | 0.0061092 | 0.460774543 |
| ENSMUSG00000052572 | Dlg2 | protein_coding | -0.790947084 | 6.437732956 | 9.9037703 | 0.0061195 | 0.460774543 |
| ENSMUSG00000042745 | Id1 | protein_coding | -1.416265846 | 3.894868094 | 9.8965173 | 0.0061345 | 0.460774543 |
| ENSMUSG00000039345 | Mettl22 | protein_coding | 0.749151017 | 6.464124396 | 9.8548146 | 0.0062211 | 0.460774543 |
| ENSMUSG00000026830 | Ermn | protein_coding | -0.63574149 | 7.384948842 | 9.8346449 | 0.0062635 | 0.460774543 |
| ENSMUSG00000097393 | D030068K23Rik | lincRNA | 1.182456618 | 5.219533529 | 9.8281813 | 0.0062771 | 0.460774543 |
| ENSMUSG00000038664 | Herc1 | protein_coding | -0.985088276 | 5.117133329 | 9.8044856 | 0.0063275 | 0.460774543 |
| ENSMUSG00000002458 | Rgs19 | protein_coding | 0.814899054 | 5.987328493 | 9.8012821 | 0.0063343 | 0.460774543 |
| ENSMUSG00000051504 | Siglech | protein_coding | -1.634035134 | 3.656645677 | 9.7893861 | 0.0063598 | 0.460774543 |
| ENSMUSG00000033423 | Eri3 | protein_coding | 0.602216858 | 8.52696204 | 9.7754435 | 0.0063898 | 0.460774543 |
| ENSMUSG00000034160 | Ogt | protein_coding | -1.091088644 | 5.72437829 | 9.7369181 | 0.0064736 | 0.460774543 |
| ENSMUSG00000031837 | Necab2 | protein_coding | -0.888030902 | 6.648110719 | 9.7236905 | 0.0065026 | 0.460774543 |
| ENSMUSG00000016386 | Mpped2 | protein_coding | -1.018981111 | 5.370563768 | 9.7217253 | 0.0065069 | 0.460774543 |
| ENSMUSG00000039145 | Camk1d | protein_coding | 0.786232764 | 7.706375568 | 9.6842479 | 0.0065901 | 0.460774543 |
| ENSMUSG00000022811 | Zfp148 | protein_coding | -0.728117847 | 6.010483517 | 9.6823504 | 0.0065943 | 0.460774543 |
| ENSMUSG00000023026 | Dip2b | protein_coding | -1.019644662 | 5.396342755 | 9.6802615 | 0.006599 | 0.460774543 |
| ENSMUSG00000006930 | Hap1 | protein_coding | 0.715319927 | 7.80132675 | 9.6152395 | 0.0067464 | 0.464032045 |
| ENSMUSG00000051007 | Gatd1 | protein_coding | 0.760083822 | 7.261766072 | 9.5482412 | 0.0069022 | 0.464032045 |

| ensembl_gene_id | external_gene_name | gene_biotype | log2 (fold change) | log2 (counts per million) | F-value | p-value | False discovery rate |
|---|---|---|---|---|---|---|---|
| ENSMUSG00000003872 | Lin7b | protein_coding | 0.720683003 | 6.138974701 | 9.5476087 | 0.0069037 | 0.464032045 |
| ENSMUSG00000022454 | Nell2 | protein_coding | 0.66388131 | 7.578508581 | 9.537871 | 0.0069267 | 0.464032045 |
| ENSMUSG00000015714 | Cers2 | protein_coding | -0.795043357 | 6.226918209 | 9.5095958 | 0.0069939 | 0.464032045 |
| ENSMUSG00000022587 | Ly6e | protein_coding | 0.744636615 | 7.940055716 | 9.5079992 | 0.0069977 | 0.464032045 |
| ENSMUSG00000031342 | Gpm6b | protein_coding | -0.76894964 | 8.856646969 | 9.4773279 | 0.0070715 | 0.464032045 |
| ENSMUSG00000031848 | Lsm4 | protein_coding | -2.149265657 | 4.547718722 | 9.81871 | 0.007111 | 0.464032045 |
| ENSMUSG00000032366 | Tpm1 | protein_coding | -0.626827027 | 7.779207097 | 9.4479816 | 0.007143 | 0.464032045 |
| ENSMUSG00000033294 | Noc4l | protein_coding | 1.140498717 | 4.910377051 | 9.4450372 | 0.0071502 | 0.464032045 |
| ENSMUSG00000070780 | Rbm47 | protein_coding | -3.415270564 | 2.402039901 | 9.4431281 | 0.0071549 | 0.464032045 |
| ENSMUSG00000033706 | Smyd5 | protein_coding | 0.724625185 | 6.38222208 | 9.4244814 | 0.0072008 | 0.464032045 |
| ENSMUSG00000074746 | Pdzd8 | protein_coding | -0.832905614 | 5.42275172 | 9.4071502 | 0.0072438 | 0.464032045 |
| ENSMUSG00000029718 | Pcolce | protein_coding | -3.309396391 | 3.082834057 | 9.587247 | 0.0072498 | 0.464032045 |
| ENSMUSG00000028082 | Sh3d19 | protein_coding | -1.661952398 | 3.122541729 | 9.3534221 | 0.0073789 | 0.467105857 |
| ENSMUSG00000031255 | Sytl4 | protein_coding | 1.138003224 | 4.055596508 | 9.3011409 | 0.0075131 | 0.467105857 |
| ENSMUSG00000032231 | Anxa2 | protein_coding | -1.232569196 | 4.781141088 | 9.2992409 | 0.007518 | 0.467105857 |
| ENSMUSG00000020263 | Appl2 | protein_coding | -1.022514053 | 4.59630411 | 9.2902152 | 0.0075415 | 0.467105857 |
| ENSMUSG00000061046 | Haghl | protein_coding | 0.645491022 | 7.77156325 | 9.2857063 | 0.0075532 | 0.467105857 |
| ENSMUSG00000037190 | Cyb561d2 | protein_coding | 0.773460249 | 6.107873697 | 9.2836953 | 0.0075585 | 0.467105857 |
| ENSMUSG00000041126 | H2afv | protein_coding | -0.835164385 | 5.493399484 | 9.2545287 | 0.0076351 | 0.467690289 |
| ENSMUSG00000039347 | Atp6v0e2 | protein_coding | 0.686674003 | 10.37809293 | 9.2324835 | 0.0076936 | 0.467690289 |
| ENSMUSG00000024098 | Twsg1 | protein_coding | -1.564876183 | 3.465440633 | 9.2251112 | 0.0077132 | 0.467690289 |
| ENSMUSG00000023964 | Calcr | protein_coding | 1.556527717 | 3.543107807 | 9.1771802 | 0.0078425 | 0.467690289 |
| ENSMUSG00000061762 | Tac1 | protein_coding | -1.233190537 | 8.511015986 | 9.5024704 | 0.0078769 | 0.467690289 |
| ENSMUSG00000010607 | Pigyl | protein_coding | 0.655815608 | 7.440696244 | 9.1531231 | 0.0079083 | 0.467690289 |
| ENSMUSG00000047261 | Gap43 | protein_coding | 0.661092465 | 11.00325107 | 9.150167 | 0.0079165 | 0.467690289 |
| ENSMUSG00000050953 | Gja1 | protein_coding | -0.878983579 | 7.008087782 | 9.1334512 | 0.0079741 | 0.467690289 |
| ENSMUSG00000037400 | Atp11b | protein_coding | -2.290788123 | 3.520111788 | 9.1445121 | 0.0079936 | 0.467690289 |
| ENSMUSG00000003363 | Pld3 | protein_coding | 0.684196955 | 9.931445965 | 9.095564 | 0.0080683 | 0.467690289 |
| ENSMUSG00000026411 | Tmem9 | protein_coding | 0.621541141 | 7.866960137 | 9.0383081 | 0.0082312 | 0.467690289 |
| ENSMUSG00000019132 | BC005537 | protein_coding | -0.719285369 | 6.782680237 | 9.0225336 | 0.0082767 | 0.467690289 |
| ENSMUSG00000041241 | Mul1 | protein_coding | 0.624030549 | 7.04934122 | 9.0158207 | 0.0082962 | 0.467690289 |
| ENSMUSG00000034175 | Rhbdd3 | protein_coding | 1.017834399 | 4.91049185 | 9.0136141 | 0.0083026 | 0.467690289 |
| ENSMUSG00000003949 | Hlf | protein_coding | -0.648166989 | 6.827037216 | 8.9924923 | 0.0083642 | 0.467690289 |
| ENSMUSG00000032376 | Usp3 | protein_coding | -1.103108378 | 4.465359218 | 8.987375 | 0.0083792 | 0.467690289 |
| ENSMUSG00000028622 | Mrpl37 | protein_coding | 0.759882631 | 6.302739605 | 8.9848581 | 0.0083866 | 0.467690289 |
| ENSMUSG00000022108 | Itm2b | protein_coding | -0.639207603 | 8.615978133 | 8.9837892 | 0.0083897 | 0.467690289 |
| ENSMUSG00000041966 | Dcaf17 | protein_coding | -1.369910717 | 3.947191032 | 8.9708828 | 0.0084278 | 0.467690289 |
| ENSMUSG00000000708 | Kat2b | protein_coding | -1.37588919 | 3.592394854 | 8.9603633 | 0.0084589 | 0.467690289 |
| ENSMUSG00000035202 | Lars2 | protein_coding | 0.804780075 | 9.543560959 | 8.9528188 | 0.0084813 | 0.467690289 |
| ENSMUSG00000024500 | Ppp2r2b | protein_coding | 0.600577739 | 7.88228679 | 8.9087182 | 0.0086137 | 0.472255649 |
| ENSMUSG00000048546 | Tob2 | protein_coding | -0.954588906 | 4.887642111 | 8.896128 | 0.0086519 | 0.472255649 |
| ENSMUSG00000110332 | Gm45396 | protein_coding | -1.298967359 | 4.327348264 | 8.8515055 | 0.008789 | 0.473302318 |
| ENSMUSG00000052997 | Uba2 | protein_coding | -0.668693695 | 6.448238628 | 8.8371487 | 0.0088336 | 0.473302318 |
| ENSMUSG00000008575 | Nfib | protein_coding | -1.104017137 | 5.048399068 | 8.7597445 | 0.0090786 | 0.473302318 |
| ENSMUSG00000005360 | Slc1a3 | protein_coding | -0.804173796 | 7.808350894 | 8.7405379 | 0.0091406 | 0.473302318 |
| ENSMUSG00000021951 | Eef1akmt1 | protein_coding | 0.567601506 | 7.540472218 | 8.7335373 | 0.0091633 | 0.473302318 |
| ENSMUSG00000023004 | Tuba1b | protein_coding | 0.639343773 | 7.738139621 | 8.7292395 | 0.0091773 | 0.473302318 |
| ENSMUSG00000021876 | Rnase4 | protein_coding | -1.271486555 | 4.138297218 | 8.7137223 | 0.009228 | 0.473302318 |
| ENSMUSG00000003970 | Rpl8 | protein_coding | 0.672482335 | 10.42760603 | 8.7082743 | 0.0092458 | 0.473302318 |
| ENSMUSG00000037573 | Tob1 | protein_coding | -0.874470492 | 5.356894332 | 8.7039449 | 0.0092601 | 0.473302318 |
| ENSMUSG00000038518 | Jarid2 | protein_coding | 0.834199409 | 8.066763333 | 8.6412908 | 0.0094814 | 0.473302318 |
| ENSMUSG00000017119 | Nbr1 | protein_coding | -0.8056864 | 6.019132822 | 8.6346566 | 0.0094911 | 0.473302318 |
| ENSMUSG00000024194 | Cuta | protein_coding | 0.55828197 | 7.473278638 | 8.6181448 | 0.0095471 | 0.473302318 |
| ENSMUSG00000025868 | Higd2a | protein_coding | 0.577119687 | 8.450003104 | 8.6114098 | 0.0095701 | 0.473302318 |
| ENSMUSG00000028399 | Ptprd | protein_coding | -0.780213176 | 6.425603442 | 8.6004832 | 0.0096075 | 0.473302318 |
| ENSMUSG00000021486 | Prelid1 | protein_coding | 0.66642824 | 7.383620694 | 8.5940003 | 0.0096297 | 0.473302318 |
| ENSMUSG00000036533 | Cdc42ep3 | protein_coding | -1.194872091 | 4.414159543 | 8.5894719 | 0.0096453 | 0.473302318 |
| ENSMUSG00000026335 | Pam | protein_coding | -0.820608601 | 6.194626399 | 8.5851839 | 0.0096601 | 0.473302318 |
| ENSMUSG00000085438 | 1700020I14Rik | lincRNA | -0.635683622 | 6.815821602 | 8.5776982 | 0.009686 | 0.473302318 |
| ENSMUSG00000028176 | Lrrc7 | protein_coding | -1.360990941 | 4.433184604 | 8.570227 | 0.0097119 | 0.473302318 |
| ENSMUSG00000020823 | Sec14l1 | protein_coding | -0.924848822 | 5.454961313 | 8.5592645 | 0.00975 | 0.473302318 |
| ENSMUSG00000030342 | Cd9 | protein_coding | -0.861972714 | 6.841465186 | 8.533894 | 0.0098519 | 0.473302318 |
| ENSMUSG00000030994 | D7Ertd443e | protein_coding | -1.250857344 | 4.277472516 | 8.5300475 | 0.0098525 | 0.473302318 |
| ENSMUSG00000045302 | Preb | protein_coding | 0.80558758 | 5.968168153 | 8.5233982 | 0.009876 | 0.473302318 |
| ENSMUSG00000043388 | Tmem130 | protein_coding | 0.719554324 | 9.615115595 | 8.5223214 | 0.0098798 | 0.473302318 |
| ENSMUSG00000031229 | Atrx | protein_coding | -0.860051425 | 6.221157175 | 8.521427 | 0.0098829 | 0.473302318 |
| ENSMUSG00000086316 | 2210013O21Rik | protein_coding | 0.544796059 | 7.655400986 | 8.5155861 | 0.0099036 | 0.473302318 |
| ENSMUSG00000033063 | Cntnap3 | protein_coding | -2.327743743 | 2.680127865 | 8.5056992 | 0.0099388 | 0.473302318 |
| ENSMUSG00000039007 | Cpq | protein_coding | -1.205901044 | 4.352774455 | 8.5000209 | 0.0099591 | 0.473302318 |
| ENSMUSG00000044734 | Serpinb1a | protein_coding | -0.869049043 | 5.844945195 | 8.4902038 | 0.0099942 | 0.473302318 |
| ENSMUSG00000043419 | Chd3os | protein_coding | 0.568581189 | 8.809065314 | 8.4675046 | 0.010076 | 0.473302318 |
| ENSMUSG00000023939 | Mrpl14 | protein_coding | 0.631490801 | 7.523152473 | 8.4669651 | 0.0100779 | 0.473302318 |
| ENSMUSG00000037625 | Cldn11 | protein_coding | -0.599987799 | 7.407840448 | 8.4570838 | 0.0101138 | 0.473302318 |
| ENSMUSG00000020253 | Ppm1m | protein_coding | 1.17403457 | 4.064014177 | 8.4500363 | 0.0101394 | 0.473302318 |
| ENSMUSG00000044117 | 2900011O08Rik | protein_coding | 0.714098323 | 9.481265429 | 8.442301 | 0.0101677 | 0.473302318 |
| ENSMUSG00000032702 | Kank1 | protein_coding | -1.397932233 | 3.697602007 | 8.4208565 | 0.0102464 | 0.473730231 |
| ENSMUSG00000079523 | Tmsb10 | protein_coding | 0.667531964 | 8.16261568 | 8.4143988 | 0.0102703 | 0.473730231 |
| ENSMUSG00000016179 | Camk1g | protein_coding | 0.820659921 | 6.34527043 | 8.4039469 | 0.010309 | 0.473730231 |

| ensembl_gene_id | external_gene_name | gene_biotype | log2 (fold change) | log2 (counts per million) | F-value | p-value | False discovery rate |
|---|---|---|---|---|---|---|---|
| ENSMUSG00000022594 | Lynx1 | protein_coding | 0.69400925 | 7.52563258 | 8.3879445 | 0.0103687 | 0.474443276 |
| ENSMUSG00000028470 | Hint2 | protein_coding | -1.40306662 | 4.938380281 | 8.3610416 | 0.0105186 | 0.479071945 |
| ENSMUSG00000004667 | Polr2e | protein_coding | 0.594312609 | 8.368268103 | 8.3047616 | 0.0106851 | 0.479071945 |
| ENSMUSG00000023927 | Satb1 | protein_coding | -1.201904701 | 3.755547552 | 8.3004823 | 0.0107016 | 0.479071945 |
| ENSMUSG00000030763 | Lcmt1 | protein_coding | 0.588105094 | 7.912261747 | 8.2866056 | 0.0107556 | 0.479071945 |
| ENSMUSG00000036594 | H2-Aa | protein_coding | -3.754502351 | 2.156284574 | 8.2490449 | 0.0109031 | 0.479071945 |
| ENSMUSG00000097625 | Gm26561 | lincRNA | 0.809728748 | 6.104517801 | 8.248514 | 0.0109052 | 0.479071945 |
| ENSMUSG00000044550 | Tceal3 | protein_coding | 0.544787228 | 7.206967085 | 8.226583 | 0.0109925 | 0.479071945 |
| ENSMUSG00000024007 | Ppil1 | protein_coding | 0.65651644 | 6.774868429 | 8.2258771 | 0.0109953 | 0.479071945 |
| ENSMUSG00000044349 | Snhg11 | protein_coding | -1.008153935 | 7.403342186 | 8.3583401 | 0.011006 | 0.479071945 |
| ENSMUSG00000029614 | Rpl6 | protein_coding | 0.59333383 | 6.90047633 | 8.2189829 | 0.0110229 | 0.479071945 |
| ENSMUSG00000019768 | Esr1 | protein_coding | 1.778693253 | 3.539813594 | 8.1979278 | 0.0111215 | 0.479071945 |
| ENSMUSG00000030102 | Itpr1 | protein_coding | -1.100345097 | 6.325948694 | 8.3179955 | 0.0111315 | 0.479071945 |
| ENSMUSG00000078681 | Tm2d3 | protein_coding | 0.631421823 | 8.32003089 | 8.1830818 | 0.0111679 | 0.479071945 |
| ENSMUSG00000034210 | Efcab14 | protein_coding | -0.74523994 | 5.928916924 | 8.182362 | 0.0111708 | 0.479071945 |
| ENSMUSG00000031104 | Rab33a | protein_coding | 0.668484165 | 6.802206864 | 8.1644476 | 0.011244 | 0.479071945 |
| ENSMUSG00000032959 | Pebp1 | protein_coding | 0.560330083 | 7.635538589 | 8.1563757 | 0.0112771 | 0.479071945 |
| ENSMUSG00000026344 | Lypd1 | protein_coding | 1.763195734 | 2.678574672 | 8.1502199 | 0.0113025 | 0.479071945 |
| ENSMUSG00000025323 | Sp4 | protein_coding | -1.173652232 | 3.837975303 | 8.1490312 | 0.0113074 | 0.479071945 |
| ENSMUSG00000017664 | Slc35c2 | protein_coding | 0.707561657 | 6.684281518 | 8.1468716 | 0.0113163 | 0.479071945 |
| ENSMUSG00000030870 | Ubfd1 | protein_coding | 0.650911214 | 6.262234854 | 8.1323124 | 0.0113766 | 0.479389982 |
| ENSMUSG00000057594 | Arl16 | protein_coding | 0.623647896 | 6.821007018 | 8.1235773 | 0.011413 | 0.479389982 |
| ENSMUSG00000020571 | Pdia6 | protein_coding | 0.556105557 | 7.807010753 | 8.104325 | 0.0114936 | 0.480121056 |
| ENSMUSG00000025492 | Ifitm3 | protein_coding | -1.248406608 | 4.155131299 | 8.0884558 | 0.0115605 | 0.480121056 |
| ENSMUSG00000024944 | Arl2 | protein_coding | 0.572223356 | 9.161043313 | 8.0853594 | 0.0115736 | 0.480121056 |
| ENSMUSG00000018322 | Tomm34 | protein_coding | 0.661074388 | 7.916772627 | 8.0577101 | 0.0116915 | 0.480121056 |
| ENSMUSG00000014551 | Mrps25 | protein_coding | 0.576613201 | 7.710675343 | 8.0078189 | 0.0119076 | 0.480121056 |
| ENSMUSG00000038555 | Reep2 | protein_coding | 0.700398221 | 7.814522904 | 8.0068794 | 0.0119117 | 0.480121056 |
| ENSMUSG00000098973 | Mir6236 | miRNA | 1.20165042 | 8.725991134 | 8.3293609 | 0.0119362 | 0.480121056 |
| ENSMUSG00000020940 | 1700023F06Rik | protein_coding | 0.965008825 | 4.456806218 | 7.9516969 | 0.0121561 | 0.480121056 |
| ENSMUSG00000039745 | Htatip2 | protein_coding | 1.166693589 | 4.621718336 | 7.9504655 | 0.0121616 | 0.480121056 |
| ENSMUSG00000038587 | Akap12 | protein_coding | -1.500960169 | 3.967296444 | 7.9499643 | 0.0121639 | 0.480121056 |
| ENSMUSG00000027309 | 4930402H24Rik | protein_coding | -0.830979332 | 5.915773539 | 7.9493479 | 0.0121666 | 0.480121056 |
| ENSMUSG00000048583 | Igf2 | protein_coding | -2.054840851 | 4.068313374 | 8.1334486 | 0.0122032 | 0.480121056 |
| ENSMUSG00000028857 | Tmem222 | protein_coding | 0.703541159 | 8.220295672 | 7.9035811 | 0.0123739 | 0.480121056 |
| ENSMUSG00000057098 | Ebf1 | protein_coding | 1.089182713 | 3.995808102 | 7.9032803 | 0.0123753 | 0.480121056 |
| ENSMUSG00000034472 | Rasd2 | protein_coding | -0.758770863 | 6.691949808 | 7.8975708 | 0.0124014 | 0.480121056 |
| ENSMUSG00000017485 | Top2b | protein_coding | -0.950718643 | 5.196539979 | 7.8965468 | 0.0124061 | 0.480121056 |
| ENSMUSG00000005089 | Slc1a2 | protein_coding | -1.107302761 | 7.751621344 | 8.1549742 | 0.0124092 | 0.480121056 |
| ENSMUSG00000028868 | Wasf2 | protein_coding | -0.987791579 | 4.571624132 | 7.8773974 | 0.0124943 | 0.480121056 |
| ENSMUSG00000084403 | Rps15a-ps8 | processed_pseudoge | 5.080983167 | 1.126234636 | 7.8677228 | 0.0125391 | 0.480121056 |
| ENSMUSG00000057193 | Slc44a2 | protein_coding | -1.453140168 | 3.950299999 | 7.8659634 | 0.0125473 | 0.480121056 |
| ENSMUSG00000025059 | Gk | protein_coding | -1.093691011 | 4.091340859 | 7.8477188 | 0.0126323 | 0.480121056 |
| ENSMUSG00000062078 | Qk | protein_coding | -0.678605215 | 7.886346701 | 7.8443727 | 0.012648 | 0.480121056 |
| ENSMUSG00000068922 | Msto1 | protein_coding | 0.867380442 | 4.987511977 | 7.8437457 | 0.0126509 | 0.480121056 |
| ENSMUSG00000024387 | Csnk2b | protein_coding | 0.537815911 | 8.273179454 | 7.8428791 | 0.012655 | 0.480121056 |
| ENSMUSG00000039298 | Cdk5rap2 | protein_coding | -1.704777739 | 2.982512472 | 7.8375034 | 0.0126803 | 0.480121056 |
| ENSMUSG00000040289 | Hey1 | protein_coding | -1.010281175 | 4.576717294 | 7.835778 | 0.0126884 | 0.480121056 |
| ENSMUSG00000046178 | Nxph1 | protein_coding | -0.793157026 | 5.835213888 | 7.835479 | 0.0126898 | 0.480121056 |
| ENSMUSG00000070733 | Fryl | protein_coding | -1.213411594 | 3.96075965 | 7.8304258 | 0.0127136 | 0.480121056 |
| ENSMUSG00000029697 | Fezf1 | protein_coding | -3.592533394 | 3.055115251 | 8.1042297 | 0.0127252 | 0.480121056 |
| ENSMUSG00000000276 | Dgke | protein_coding | -1.081187436 | 4.35600267 | 7.7773165 | 0.0129668 | 0.487526059 |
| ENSMUSG00000031292 | Cdkl5 | protein_coding | -0.841056468 | 5.96809459 | 7.7056297 | 0.0133179 | 0.492457994 |
| ENSMUSG00000034757 | Tmub2 | protein_coding | 0.635274695 | 6.644310887 | 7.6930192 | 0.0133807 | 0.492457994 |
| ENSMUSG00000049744 | Arhgap15 | protein_coding | -2.52442338 | 2.51061778 | 7.6929464 | 0.0133811 | 0.492457994 |
| ENSMUSG00000031232 | Magt1 | protein_coding | -1.457045802 | 3.95752326 | 7.6800268 | 0.0134458 | 0.492457994 |
| ENSMUSG00000024446 | Rpp21 | protein_coding | 0.535864604 | 7.644382785 | 7.6791374 | 0.0134503 | 0.492457994 |
| ENSMUSG00000027698 | Nceh1 | protein_coding | 0.531265953 | 7.061547712 | 7.6779834 | 0.0134561 | 0.492457994 |
| ENSMUSG00000027893 | Ahcyl1 | protein_coding | -0.61357476 | 7.54395055 | 7.6690498 | 0.0135011 | 0.492457994 |
| ENSMUSG00000032288 | Imp3 | protein_coding | 1.051494545 | 4.824134145 | 7.6666519 | 0.0135133 | 0.492457994 |
| ENSMUSG00000031820 | Babam1 | protein_coding | 0.657086995 | 7.129763984 | 7.6450488 | 0.013623 | 0.492457994 |
| ENSMUSG00000023277 | Twf2 | protein_coding | 0.792672695 | 5.785012577 | 7.6446928 | 0.0136248 | 0.492457994 |
| ENSMUSG00000026787 | Gad2 | protein_coding | 0.64893122 | 8.99710077 | 7.6329387 | 0.0136849 | 0.492457994 |
| ENSMUSG00000017188 | Coa3 | protein_coding | 0.58525522 | 9.831433548 | 7.6302026 | 0.0136989 | 0.492457994 |
| ENSMUSG00000053565 | Eif3k | protein_coding | 0.617853931 | 9.322931709 | 7.5688859 | 0.0140179 | 0.492457994 |
| ENSMUSG00000103103 | 4833445I07Rik | lincRNA | 1.754278678 | 2.77715913 | 7.5463856 | 0.0141371 | 0.492457994 |
| ENSMUSG00000018040 | Rrp7a | protein_coding | 0.705709215 | 6.211056727 | 7.517856 | 0.0142898 | 0.492457994 |
| ENSMUSG00000020829 | Slc46a1 | protein_coding | 0.781079879 | 5.153076633 | 7.5051157 | 0.0143586 | 0.492457994 |
| ENSMUSG00000041355 | Ssr2 | protein_coding | 0.579257898 | 7.580759846 | 7.4929015 | 0.014425 | 0.492457994 |
| ENSMUSG00000030235 | Slco1c1 | protein_coding | -1.335420156 | 3.677615317 | 7.4927009 | 0.0144261 | 0.492457994 |
| ENSMUSG00000026201 | Stk16 | protein_coding | 0.820152719 | 6.45245225 | 7.4847896 | 0.0144852 | 0.492457994 |
| ENSMUSG00000025208 | Mrpl43 | protein_coding | 0.591361507 | 7.39468562 | 7.4790633 | 0.0145005 | 0.492457994 |
| ENSMUSG00000018906 | P4ha2 | protein_coding | 1.296168963 | 4.267600162 | 7.4743894 | 0.0145262 | 0.492457994 |
| ENSMUSG00000048251 | Bcl11b | protein_coding | -0.760286594 | 6.005800764 | 7.4739006 | 0.0145289 | 0.492457994 |
| ENSMUSG00000028393 | Alad | protein_coding | 0.583161181 | 6.709993692 | 7.4658945 | 0.0145729 | 0.492457994 |
| ENSMUSG00000036398 | Ppp1r11 | protein_coding | 0.547407743 | 9.340274285 | 7.4195268 | 0.0148309 | 0.492457994 |
| ENSMUSG00000041762 | Gpr155 | protein_coding | -1.358033332 | 4.032145918 | 7.4166723 | 0.0148469 | 0.492457994 |
| ENSMUSG00000026781 | Acbd5 | protein_coding | -0.868914487 | 5.32265234 | 7.414789 | 0.0148575 | 0.492457994 |

| ensembl_gene_id | external_gene_name | gene_biotype | log2 (fold change) | log2 (counts per million) | F-value | p-value | False discovery rate |
|---|---|---|---|---|---|---|---|
| ENSMUSG00000031708 | Tecr | protein_coding | 0.60624264 | 10.67126226 | 7.4032813 | 0.0149225 | 0.492457994 |
| ENSMUSG00000041380 | Htr2c | protein_coding | -0.790743375 | 5.492600064 | 7.4023039 | 0.014928 | 0.492457994 |
| ENSMUSG00000026970 | Rbms1 | protein_coding | -1.034101244 | 4.38989333 | 7.4006186 | 0.0149376 | 0.492457994 |
| ENSMUSG00000068580 | Zfyve19 | protein_coding | 0.720127615 | 6.224034757 | 7.3959955 | 0.0149638 | 0.492457994 |
| ENSMUSG00000022564 | Grina | protein_coding | 0.587710596 | 8.836490859 | 7.3875661 | 0.0150118 | 0.492457994 |
| ENSMUSG00000038332 | Sesn1 | protein_coding | -0.837783743 | 5.305426125 | 7.3747673 | 0.0150849 | 0.492457994 |
| ENSMUSG00000020042 | Btbd11 | protein_coding | 0.926933905 | 5.056746215 | 7.3681082 | 0.0151231 | 0.492457994 |
| ENSMUSG00000015749 | Anp32e | protein_coding | -0.579275333 | 6.679126619 | 7.3680805 | 0.0151233 | 0.492457994 |
| ENSMUSG00000018012 | Rac3 | protein_coding | 1.023447713 | 4.290277025 | 7.3649632 | 0.0151412 | 0.492457994 |
| ENSMUSG00000021379 | Id4 | protein_coding | -0.588373677 | 6.834048099 | 7.3648902 | 0.0151416 | 0.492457994 |
| ENSMUSG00000049658 | Bdp1 | protein_coding | -1.27472818 | 3.753330249 | 7.3513381 | 0.0152198 | 0.492457994 |
| ENSMUSG00000050377 | Il31ra | protein_coding | 1.049767607 | 5.067837887 | 7.3389862 | 0.0152915 | 0.492457994 |
| ENSMUSG00000021846 | Peli2 | protein_coding | -1.142661027 | 3.828429506 | 7.335092 | 0.0153142 | 0.492457994 |
| ENSMUSG00000068523 | Gng5 | protein_coding | -1.04380589 | 4.80179362 | 7.3330036 | 0.0153264 | 0.492457994 |
| ENSMUSG00000063594 | Gng8 | protein_coding | -1.14276947 | 4.820213282 | 7.3307353 | 0.0153396 | 0.492457994 |
| ENSMUSG00000024841 | Eif1ad | protein_coding | 0.53050179 | 7.031500112 | 7.3231477 | 0.015384 | 0.492457994 |
| ENSMUSG00000027210 | Meis2 | protein_coding | -0.724404814 | 7.258384013 | 7.3220323 | 0.0153906 | 0.492457994 |
| ENSMUSG00000029695 | Aass | protein_coding | -3.655057905 | 1.638383169 | 7.3070629 | 0.0154786 | 0.492457994 |
| ENSMUSG00000018800 | Abca5 | protein_coding | -1.115326957 | 3.999379252 | 7.3035464 | 0.0154994 | 0.492457994 |
| ENSMUSG00000042453 | Reln | protein_coding | -1.334818849 | 4.225070867 | 7.3006643 | 0.0155164 | 0.492457994 |
| ENSMUSG00000025949 | Pikfyve | protein_coding | -1.08903816 | 4.167803261 | 7.2939218 | 0.0155564 | 0.492457994 |
| ENSMUSG00000020422 | Tns3 | protein_coding | -1.505689542 | 3.221768076 | 7.285658 | 0.0156055 | 0.492457994 |
| ENSMUSG00000049892 | Rasd1 | protein_coding | -0.948964748 | 6.061518301 | 7.2781643 | 0.0156668 | 0.492457994 |
| ENSMUSG00000027613 | Eif6 | protein_coding | 0.571423856 | 6.846657421 | 7.2663251 | 0.0157211 | 0.492457994 |
| ENSMUSG00000023011 | Faim2 | protein_coding | 0.729855962 | 9.330157212 | 7.2616541 | 0.0157492 | 0.492457994 |
| ENSMUSG00000019194 | Scn1b | protein_coding | 0.798304318 | 5.104356915 | 7.259182 | 0.0157641 | 0.492457994 |
| ENSMUSG00000059456 | Ptk2b | protein_coding | -0.684113151 | 7.305700514 | 7.2561049 | 0.0157827 | 0.492457994 |
| ENSMUSG00000040420 | Cdh18 | protein_coding | 1.172417123 | 3.766926113 | 7.2535521 | 0.0157981 | 0.492457994 |
| ENSMUSG00000013997 | Nit1 | protein_coding | 0.718918607 | 5.735274349 | 7.2478265 | 0.0158327 | 0.492457994 |
| ENSMUSG00000020892 | Aloxe3 | protein_coding | 1.401638361 | 4.186535771 | 7.2446716 | 0.0158685 | 0.492457994 |
| ENSMUSG00000032826 | Ank2 | protein_coding | -0.714521531 | 6.614685541 | 7.2369182 | 0.0158989 | 0.492457994 |
| ENSMUSG00000071553 | Cpa2 | protein_coding | 0.940648488 | 4.811288757 | 7.2354358 | 0.0159079 | 0.492457994 |
| ENSMUSG00000029999 | Tgfa | protein_coding | -0.900360492 | 4.891510745 | 7.2344673 | 0.0159138 | 0.492457994 |
| ENSMUSG00000000600 | Krit1 | protein_coding | -1.018479261 | 4.727434989 | 7.2318777 | 0.0159296 | 0.492457994 |
| ENSMUSG00000022139 | Mbnl2 | protein_coding | -0.643355538 | 7.573170751 | 7.2232056 | 0.0159826 | 0.492457994 |
| ENSMUSG00000020153 | Ndufs7 | protein_coding | 0.573239333 | 9.745717511 | 7.2203561 | 0.016 | 0.492457994 |
| ENSMUSG00000037795 | N4bp2 | protein_coding | -2.005307181 | 2.840029752 | 7.2178873 | 0.0160152 | 0.492457994 |
| ENSMUSG00000021591 | Glrx | protein_coding | 0.675644127 | 6.682961607 | 7.2143142 | 0.0160371 | 0.492457994 |
| ENSMUSG00000022119 | Rbm26 | protein_coding | -0.990650108 | 4.824486385 | 7.207323 | 0.0160801 | 0.492457994 |
| ENSMUSG00000038481 | Cdk19 | protein_coding | -0.941591799 | 4.85149738 | 7.1700427 | 0.0163118 | 0.492457994 |
| ENSMUSG00000072501 | Phf20l1 | protein_coding | -1.058102853 | 4.447356078 | 7.1655506 | 0.01634 | 0.492457994 |
| ENSMUSG00000047259 | Mc4r | protein_coding | 1.363661382 | 3.845417213 | 7.1626313 | 0.0163583 | 0.492457994 |
| ENSMUSG00000031796 | Cfap20 | protein_coding | -0.917288092 | 6.061019757 | 7.1636987 | 0.0163686 | 0.492457994 |
| ENSMUSG00000055538 | Zcchc24 | protein_coding | -0.985642987 | 5.011976266 | 7.1532258 | 0.0164175 | 0.492457994 |
| ENSMUSG00000055717 | Slain1 | protein_coding | -0.817570643 | 5.534441336 | 7.1468794 | 0.0164577 | 0.492457994 |
| ENSMUSG00000026869 | Psmd5 | protein_coding | -1.107827196 | 6.091297663 | 7.2510899 | 0.0164648 | 0.492457994 |
| ENSMUSG00000038671 | Arfrp1 | protein_coding | 0.592177198 | 6.881633397 | 7.1439484 | 0.0164762 | 0.492457994 |
| ENSMUSG00000038500 | Prr3 | protein_coding | 0.80685055 | 5.502780232 | 7.1422447 | 0.016487 | 0.492457994 |
| ENSMUSG00000032839 | Trpc1 | protein_coding | -1.504029102 | 3.585666934 | 7.1157886 | 0.0166558 | 0.493316464 |
| ENSMUSG00000069020 | Urm1 | protein_coding | 0.559263906 | 6.34336436 | 7.0947657 | 0.0167913 | 0.493316464 |
| ENSMUSG00000035964 | Tmem59l | protein_coding | 0.594987326 | 9.53239765 | 7.0857828 | 0.0168496 | 0.493316464 |
| ENSMUSG00000019990 | Pde7b | protein_coding | -1.078909004 | 4.13932871 | 7.0826361 | 0.01687 | 0.493316464 |
| ENSMUSG00000092341 | Malat1 | lincRNA | -0.761140553 | 10.80327234 | 7.0816328 | 0.0168766 | 0.493316464 |
| ENSMUSG00000020803 | Txndc17 | protein_coding | 0.513699218 | 7.832674069 | 7.0746655 | 0.016922 | 0.493316464 |
| ENSMUSG00000008658 | Rbfox1 | protein_coding | 0.542634642 | 8.260538718 | 7.0377378 | 0.0171652 | 0.493316464 |
| ENSMUSG00000035754 | Wdr18 | protein_coding | 0.612554251 | 8.189957898 | 7.0313505 | 0.0172077 | 0.493316464 |
| ENSMUSG00000028958 | Tmub1 | protein_coding | 0.608820972 | 6.454854854 | 7.0270309 | 0.0172365 | 0.493316464 |
| ENSMUSG00000039662 | Icmt | protein_coding | -0.974542821 | 4.319494949 | 7.0073267 | 0.0173685 | 0.493316464 |
| ENSMUSG00000060961 | Slc4a4 | protein_coding | -1.060224234 | 5.376427232 | 6.9928875 | 0.0174836 | 0.493316464 |
| ENSMUSG00000028456 | Unc13b | protein_coding | -0.758310383 | 5.591983742 | 6.9832539 | 0.0175314 | 0.493316464 |
| ENSMUSG00000035954 | Dock4 | protein_coding | -1.126493491 | 4.222675166 | 6.9790928 | 0.0175598 | 0.493316464 |
| ENSMUSG00000029123 | Stk32b | protein_coding | 1.470494218 | 4.482213564 | 7.076193 | 0.0175859 | 0.493316464 |
| ENSMUSG00000098557 | Kctd12 | protein_coding | -1.004081248 | 4.537306075 | 6.9701002 | 0.0176212 | 0.493316464 |
| ENSMUSG00000026810 | Dpm2 | protein_coding | 0.49442863 | 7.574163744 | 6.9616736 | 0.0176789 | 0.493316464 |
| ENSMUSG00000037541 | Shank2 | protein_coding | -1.077049666 | 4.28895981 | 6.9535503 | 0.0177348 | 0.493316464 |
| ENSMUSG00000023904 | Hcfc1r1 | protein_coding | 0.510108707 | 8.646237457 | 6.9501137 | 0.0177585 | 0.493316464 |
| ENSMUSG00000046312 | AI464131 | protein_coding | -2.088631507 | 2.567405261 | 6.948838 | 0.0177673 | 0.493316464 |
| ENSMUSG00000000308 | Ckmt1 | protein_coding | 0.786923831 | 8.95899828 | 6.9513772 | 0.0177675 | 0.493316464 |
| ENSMUSG00000036777 | Anln | protein_coding | -0.899663215 | 4.705413853 | 6.9478957 | 0.0177738 | 0.493316464 |
| ENSMUSG00000086370 | Ftx | lincRNA | -1.578754532 | 3.516053268 | 6.9422434 | 0.0178129 | 0.493316464 |
| ENSMUSG00000029840 | Mtpn | protein_coding | -0.659118758 | 7.639171487 | 6.9402351 | 0.0178269 | 0.493316464 |
| ENSMUSG00000020564 | Atxn7l1 | protein_coding | 0.742412615 | 6.226863511 | 6.9337588 | 0.0178718 | 0.493316464 |
| ENSMUSG00000048895 | Cdk5r1 | protein_coding | -0.600293976 | 6.889457101 | 6.8825414 | 0.0182321 | 0.493316464 |
| ENSMUSG00000030516 | Tjp1 | protein_coding | -1.160981608 | 4.111698494 | 6.8770796 | 0.018271 | 0.493316464 |
| ENSMUSG00000022217 | Emc9 | protein_coding | 0.528087135 | 6.944502775 | 6.8538251 | 0.0184376 | 0.493316464 |
| ENSMUSG00000000168 | Dlat | protein_coding | -0.657355942 | 6.283778534 | 6.850871 | 0.0184589 | 0.493316464 |
| ENSMUSG00000037999 | Arap2 | protein_coding | -0.954968176 | 4.659843276 | 6.8476956 | 0.0184819 | 0.493316464 |
| ENSMUSG00000037712 | Fermt2 | protein_coding | -0.815269139 | 5.541511576 | 6.843499 | 0.0185122 | 0.493316464 |

| ensembl_gene_id | external_gene_name | gene_biotype | log2 (fold change) | log2 (counts per million) | F-value | p-value | False discovery rate |
|---|---|---|---|---|---|---|---|
| ENSMUSG00000002985 | Apoe | protein_coding | -1.511188371 | 7.517200894 | 7.5233331 | 0.0185269 | 0.493316464 |
| ENSMUSG00000017943 | Gdap1l1 | protein_coding | 0.634810739 | 7.587768862 | 6.8398925 | 0.0185384 | 0.493316464 |
| ENSMUSG00000022112 | Gpc5 | protein_coding | -1.114421574 | 4.158869144 | 6.8302564 | 0.0186084 | 0.493316464 |
| ENSMUSG00000044285 | Gm1821 | processed_pseudoge | 3.21379226 | 5.447086433 | 8.5291393 | 0.0186217 | 0.493316464 |
| ENSMUSG00000071014 | Ndufb6 | protein_coding | 0.5494343 | 8.649696776 | 6.8274014 | 0.0186292 | 0.493316464 |
| ENSMUSG00000028755 | Cda | protein_coding | 0.998342172 | 4.950900767 | 6.8233633 | 0.0186586 | 0.493316464 |
| ENSMUSG00000007021 | Syngr3 | protein_coding | 0.593729069 | 9.000692261 | 6.8091171 | 0.018763 | 0.493316464 |
| ENSMUSG00000010064 | Slc38a3 | protein_coding | -1.158965798 | 4.503629925 | 6.8027325 | 0.01881 | 0.493316464 |
| ENSMUSG00000048562 | Sp8 | protein_coding | -1.100553536 | 5.007978714 | 6.7974753 | 0.0188488 | 0.493316464 |
| ENSMUSG00000046532 | Ar | protein_coding | 0.767481646 | 5.82960088 | 6.7930516 | 0.0188816 | 0.493316464 |
| ENSMUSG00000024875 | Yif1a | protein_coding | 0.631772361 | 6.495635627 | 6.7927499 | 0.0188838 | 0.493316464 |
| ENSMUSG00000001864 | Aif1l | protein_coding | -0.881385757 | 5.280336379 | 6.7907855 | 0.0188983 | 0.493316464 |
| ENSMUSG00000024999 | Noc3l | protein_coding | -1.842955555 | 2.803657768 | 6.7858317 | 0.0189351 | 0.493316464 |
| ENSMUSG00000028573 | Fggy | protein_coding | 0.86123317 | 5.243828792 | 6.7843993 | 0.0189457 | 0.493316464 |
| ENSMUSG00000020744 | Slc25a19 | protein_coding | 0.726013119 | 5.895115586 | 6.7755295 | 0.0190118 | 0.493316464 |
| ENSMUSG00000002660 | Clpp | protein_coding | 0.518915981 | 8.997684785 | 6.7716908 | 0.0190405 | 0.493316464 |
| ENSMUSG00000052889 | Prkcb | protein_coding | -0.734479477 | 8.464317642 | 6.7636835 | 0.0191188 | 0.493316464 |
| ENSMUSG00000098097 | 6530403H02Rik | lincRNA | -1.874117471 | 2.770981704 | 6.7556092 | 0.0191611 | 0.493316464 |
| ENSMUSG00000000339 | Rtca | protein_coding | -1.26601599 | 4.546336052 | 6.740162 | 0.0192962 | 0.493316464 |
| ENSMUSG00000027944 | Hax1 | protein_coding | 0.683879845 | 5.327219095 | 6.7311684 | 0.0193462 | 0.493316464 |
| ENSMUSG00000048279 | Sacs | protein_coding | -1.166865505 | 4.283864982 | 6.7290746 | 0.0193621 | 0.493316464 |
| ENSMUSG00000046756 | Mrps7 | protein_coding | 0.577404656 | 6.693885339 | 6.7273083 | 0.0193756 | 0.493316464 |
| ENSMUSG00000033228 | Scaf11 | protein_coding | -0.851654963 | 5.105056989 | 6.7269238 | 0.0193785 | 0.493316464 |
| ENSMUSG00000028289 | Epha7 | protein_coding | -0.811843856 | 5.22198334 | 6.7248596 | 0.0193943 | 0.493316464 |
| ENSMUSG00000025889 | Snca | protein_coding | 0.526567458 | 9.380044398 | 6.7158914 | 0.0194629 | 0.493316464 |
| ENSMUSG00000005150 | Wdr83 | protein_coding | 0.68760795 | 6.399766298 | 6.7138968 | 0.0194782 | 0.493316464 |
| ENSMUSG00000035561 | Aldh1b1 | protein_coding | 0.938872803 | 4.460342344 | 6.7078019 | 0.019525 | 0.493316464 |
| ENSMUSG00000005483 | Dnajb1 | protein_coding | -0.625816531 | 6.685269881 | 6.7061111 | 0.019538 | 0.493316464 |
| ENSMUSG00000043391 | 2510009E07Rik | protein_coding | -0.785083132 | 5.119740428 | 6.7013288 | 0.0195749 | 0.493316464 |
| ENSMUSG00000049076 | Acap2 | protein_coding | -0.997476437 | 4.383379589 | 6.7005942 | 0.0195805 | 0.493316464 |
| ENSMUSG00000052581 | Lrrtm4 | protein_coding | -1.054985386 | 5.281600629 | 6.7021038 | 0.0195874 | 0.493316464 |
| ENSMUSG00000022223 | Sdr39u1 | protein_coding | 0.522749753 | 8.470249167 | 6.691492 | 0.0196509 | 0.493316464 |
| ENSMUSG00000032373 | Car12 | protein_coding | -2.084594251 | 3.89923021 | 6.8618504 | 0.0198184 | 0.493316464 |
| ENSMUSG00000030093 | Wnt7a | protein_coding | -1.838467483 | 2.849999121 | 6.6641044 | 0.0198645 | 0.493316464 |
| ENSMUSG00000029635 | Cdk8 | protein_coding | 0.690687322 | 6.496972303 | 6.6604874 | 0.0198929 | 0.493316464 |
| ENSMUSG00000056708 | Ier5 | protein_coding | -1.723708097 | 3.078491672 | 6.655388 | 0.019933 | 0.493316464 |
| ENSMUSG00000094002 | Gm9866 | lincRNA | 0.558578482 | 6.629773502 | 6.6423606 | 0.020036 | 0.493316464 |
| ENSMUSG00000028494 | Plin2 | protein_coding | -1.301620788 | 3.848423259 | 6.6413374 | 0.0200441 | 0.493316464 |
| ENSMUSG00000023456 | Tpi1 | protein_coding | 0.507350073 | 9.877387181 | 6.6335085 | 0.0201062 | 0.493316464 |
| ENSMUSG00000044573 | Acp1 | protein_coding | 0.580088248 | 6.066362044 | 6.6334577 | 0.0201066 | 0.493316464 |
| ENSMUSG00000036437 | Npy1r | protein_coding | 1.00176473 | 4.079519667 | 6.6249857 | 0.0201742 | 0.493316464 |
| ENSMUSG00000029073 | Cptp | protein_coding | 0.889346387 | 6.527474622 | 6.6856317 | 0.0202126 | 0.493316464 |
| ENSMUSG00000021188 | Trip11 | protein_coding | -0.980293025 | 4.587727055 | 6.6144923 | 0.0202582 | 0.493316464 |
| ENSMUSG00000025241 | Fyco1 | protein_coding | -1.626513185 | 3.763028103 | 6.6115541 | 0.0203006 | 0.493316464 |
| ENSMUSG00000048482 | Bdnf | protein_coding | -1.175237871 | 3.991353696 | 6.6013881 | 0.0203636 | 0.493316464 |
| ENSMUSG00000044308 | Ubr3 | protein_coding | -0.902334028 | 6.311988584 | 6.6065437 | 0.0203907 | 0.493316464 |
| ENSMUSG00000018076 | Med13l | protein_coding | -0.871368798 | 5.03335934 | 6.598002 | 0.020391 | 0.493316464 |
| ENSMUSG00000036503 | Rnf13 | protein_coding | -0.561735908 | 7.427291732 | 6.5948333 | 0.0204166 | 0.493316464 |
| ENSMUSG00000020308 | Tpgs1 | protein_coding | -1.636960886 | 4.572860729 | 6.7315939 | 0.0204736 | 0.493316464 |
| ENSMUSG00000015806 | Qdpr | protein_coding | -0.581307697 | 6.779004221 | 6.5870765 | 0.0204796 | 0.493316464 |
| ENSMUSG00000021843 | Ktn1 | protein_coding | -0.805557324 | 5.683007978 | 6.5837123 | 0.0205069 | 0.493316464 |
| ENSMUSG00000049044 | Rapgef4 | protein_coding | -0.866471915 | 6.176587746 | 6.5708587 | 0.0206308 | 0.493316464 |
| ENSMUSG00000057719 | Sh3rf2 | protein_coding | -1.553539219 | 3.397565118 | 6.5638188 | 0.0206695 | 0.493316464 |
| ENSMUSG00000026895 | Ndufa8 | protein_coding | 0.526493552 | 9.504452864 | 6.5585153 | 0.0207131 | 0.493316464 |
| ENSMUSG00000031529 | Tnks | protein_coding | -0.763286784 | 5.229642601 | 6.5575865 | 0.0207208 | 0.493316464 |
| ENSMUSG00000009907 | Vps4b | protein_coding | -0.742649243 | 5.311644651 | 6.5561958 | 0.0207322 | 0.493316464 |
| ENSMUSG00000020778 | Ten1 | protein_coding | 0.525569662 | 7.058388038 | 6.5477987 | 0.0208015 | 0.493316464 |
| ENSMUSG00000024403 | Atp6v1g2 | protein_coding | 0.560209301 | 10.46978886 | 6.5358409 | 0.0209007 | 0.493316464 |
| ENSMUSG00000020178 | Adora2a | protein_coding | -1.305778654 | 4.517630999 | 6.52927 | 0.0209746 | 0.493316464 |
| ENSMUSG00000024248 | Cox7a2l | protein_coding | 0.516767831 | 8.40287394 | 6.5150203 | 0.0210747 | 0.493316464 |
| ENSMUSG00000045007 | Tubg2 | protein_coding | 0.739867133 | 6.830641441 | 6.5114097 | 0.0211242 | 0.493316464 |
| ENSMUSG00000036854 | Hspb6 | protein_coding | 0.891928391 | 5.283787309 | 6.5079937 | 0.0211338 | 0.493316464 |
| ENSMUSG00000035992 | Fnip1 | protein_coding | -0.906259145 | 4.657855697 | 6.4991074 | 0.0212088 | 0.493316464 |
| ENSMUSG00000042157 | Sprr2i | protein_coding | -4.588752725 | 1.905783563 | 6.5072543 | 0.0212431 | 0.493316464 |
| ENSMUSG00000038126 | Mphosph9 | protein_coding | -1.208636748 | 3.540894495 | 6.4942361 | 0.02125 | 0.493316464 |
| ENSMUSG00000009281 | Rarres2 | protein_coding | -1.251304417 | 4.253690316 | 6.4917897 | 0.0212708 | 0.493316464 |
| ENSMUSG00000111471 | AC122463.1 | antisense_RNA | -0.804753341 | 5.102127644 | 6.4908667 | 0.0212786 | 0.493316464 |
| ENSMUSG00000025439 | Clns1a | protein_coding | 0.51318753 | 7.542459201 | 6.4835971 | 0.0213404 | 0.493316464 |
| ENSMUSG00000037736 | Limch1 | protein_coding | -0.709785176 | 5.94337844 | 6.4716545 | 0.0214424 | 0.493316464 |
| ENSMUSG00000074782 | 4833422C13Rik | antisense_RNA | -1.134129303 | 4.016640669 | 6.4708508 | 0.0214493 | 0.493316464 |
| ENSMUSG00000039361 | Picalm | protein_coding | -0.66964252 | 6.279522814 | 6.4632912 | 0.0215142 | 0.493316464 |
| ENSMUSG00000035027 | Map2k2 | protein_coding | 0.685369762 | 5.336854328 | 6.456101 | 0.0215761 | 0.493316464 |
| ENSMUSG00000038538 | Ubn2 | protein_coding | -0.808116858 | 5.661004336 | 6.4552947 | 0.021583 | 0.493316464 |
| ENSMUSG00000074863 | Platr25 | protein_coding | -2.066008871 | 2.442684069 | 6.4522193 | 0.0216096 | 0.493316464 |
| ENSMUSG00000053841 | Txlna | protein_coding | -0.992636452 | 4.507375285 | 6.4465007 | 0.0216591 | 0.493316464 |
| ENSMUSG00000031996 | Aplp2 | protein_coding | -0.553367973 | 7.990537434 | 6.4421634 | 0.0216967 | 0.493316464 |
| ENSMUSG00000097248 | Gm2694 | lincRNA | -1.157987332 | 4.397667393 | 6.4359713 | 0.0217505 | 0.493316464 |
| ENSMUSG00000032399 | Rpl4 | protein_coding | 0.561195862 | 10.60178081 | 6.435731 | 0.0217526 | 0.493316464 |

| ensembl_gene_id | external_gene_name | gene_biotype | log2 (fold change) | log2 (counts per million) | F-value | p-value | False discovery rate |
|---|---|---|---|---|---|---|---|
| ENSMUSG00000058400 | Qrfpr | protein_coding | 1.571009318 | 2.91734529 | 6.427805 | 0.0218217 | 0.493316464 |
| ENSMUSG00000065990 | Aurkaip1 | protein_coding | 0.544166683 | 8.863702639 | 6.4188445 | 0.0219002 | 0.493316464 |
| ENSMUSG00000027400 | Pdyn | protein_coding | -0.92069799 | 6.739999228 | 6.5022611 | 0.0219318 | 0.493316464 |
| ENSMUSG00000050334 | C130071C03Rik | lincRNA | -0.929214699 | 4.538321769 | 6.4149812 | 0.0219341 | 0.493316464 |
| ENSMUSG00000001285 | Myg1 | protein_coding | 0.598229751 | 7.240407338 | 6.4102037 | 0.0219761 | 0.493316464 |
| ENSMUSG00000026034 | Clk1 | protein_coding | -1.048539408 | 5.304603204 | 6.4096382 | 0.0220007 | 0.493316464 |
| ENSMUSG00000026872 | Zeb2 | protein_coding | -0.953703159 | 4.886389772 | 6.4056022 | 0.0220167 | 0.493316464 |
| ENSMUSG00000002012 | Pnck | protein_coding | 0.679972578 | 9.46024771 | 6.3907581 | 0.0221482 | 0.493316464 |
| ENSMUSG00000029755 | Dlx5 | protein_coding | 0.722288365 | 5.722980699 | 6.3810407 | 0.0222347 | 0.493316464 |
| ENSMUSG00000032827 | Ppp1r9a | protein_coding | -0.748337019 | 7.15037439 | 6.3811602 | 0.0222533 | 0.493316464 |
| ENSMUSG00000087211 | Lhx1os | lincRNA | -0.924697505 | 4.070579514 | 6.3725344 | 0.0223108 | 0.493316464 |
| ENSMUSG00000030237 | Slco1a4 | protein_coding | -2.415416853 | 2.316407609 | 6.370211 | 0.0223316 | 0.493316464 |
| ENSMUSG00000051354 | Samd3 | protein_coding | -1.937153103 | 3.681003651 | 6.4557252 | 0.0223354 | 0.493316464 |
| ENSMUSG00000027858 | Tspan2 | protein_coding | -0.646574222 | 6.491047904 | 6.3635366 | 0.0223916 | 0.493316464 |
| ENSMUSG00000025137 | Pcyt2 | protein_coding | 0.64718217 | 7.272872055 | 6.3615376 | 0.0224096 | 0.493316464 |
| ENSMUSG00000086290 | Snhg12 | lincRNA | 0.688224624 | 5.972992914 | 6.3604042 | 0.0224198 | 0.493316464 |
| ENSMUSG00000037742 | Eef1a1 | protein_coding | 0.561825113 | 10.30163358 | 6.3475581 | 0.0225359 | 0.493316464 |
| ENSMUSG00000007872 | Id3 | protein_coding | -0.583111577 | 7.392787143 | 6.3460588 | 0.0225495 | 0.493316464 |
| ENSMUSG00000004071 | Cdip1 | protein_coding | 0.658865328 | 10.04724437 | 6.3453478 | 0.022556 | 0.493316464 |
| ENSMUSG00000031613 | Hpgd | protein_coding | -1.507259896 | 3.347126738 | 6.3434277 | 0.0225734 | 0.493316464 |
| ENSMUSG00000067377 | Tspan6 | protein_coding | 0.59364322 | 6.552946552 | 6.3432904 | 0.0225746 | 0.493316464 |
| ENSMUSG00000018415 | Gid4 | protein_coding | -0.808513018 | 5.948252585 | 6.3360188 | 0.0226408 | 0.493316464 |
| ENSMUSG00000036225 | Kctd1 | protein_coding | -1.077296211 | 5.301943962 | 6.3371575 | 0.0226503 | 0.493316464 |
| ENSMUSG00000063659 | Zbtb18 | protein_coding | -1.072482257 | 3.932208791 | 6.3309846 | 0.0226867 | 0.493316464 |
| ENSMUSG00000028048 | Gba | protein_coding | 0.928597557 | 4.871983274 | 6.3268383 | 0.0227247 | 0.493316464 |
| ENSMUSG00000057411 | Fam173a | protein_coding | 0.564317351 | 8.093885779 | 6.3172252 | 0.0228128 | 0.493316464 |
| ENSMUSG00000021108 | Prkch | protein_coding | -1.171400646 | 4.471156913 | 6.3029552 | 0.0229445 | 0.493316464 |
| ENSMUSG00000035637 | Grhpr | protein_coding | 0.590283804 | 7.164007776 | 6.2975805 | 0.0229943 | 0.493316464 |
| ENSMUSG00000067931 | Zfp948 | protein_coding | -1.258216147 | 3.557058881 | 6.2969667 | 0.0229999 | 0.493316464 |
| ENSMUSG00000006095 | Tbcb | protein_coding | 0.49912212 | 9.009886997 | 6.2943695 | 0.0230241 | 0.493316464 |
| ENSMUSG00000020599 | Rgs9 | protein_coding | -0.665016157 | 6.232323517 | 6.2851307 | 0.0231101 | 0.493316464 |
| ENSMUSG00000069516 | Lyz2 | protein_coding | -1.673820342 | 3.313845195 | 6.2839056 | 0.0231215 | 0.493316464 |
| ENSMUSG00000022124 | Fbxl3 | protein_coding | -0.802246201 | 5.126446197 | 6.283898 | 0.0231216 | 0.493316464 |
| ENSMUSG00000029152 | Ociad1 | protein_coding | 0.568827985 | 8.870748976 | 6.2819967 | 0.0231393 | 0.493316464 |
| ENSMUSG00000039179 | Tekt5 | protein_coding | 0.821637229 | 5.769587978 | 6.2819503 | 0.0231398 | 0.493316464 |
| ENSMUSG00000042215 | Bag2 | protein_coding | 0.891531369 | 4.929207901 | 6.2752084 | 0.0232029 | 0.493316464 |
| ENSMUSG00000046314 | Stxbp6 | protein_coding | -0.663654177 | 5.801587031 | 6.2679278 | 0.0232712 | 0.493316464 |
| ENSMUSG00000042807 | Hecw2 | protein_coding | -1.074060991 | 4.125532092 | 6.2663629 | 0.023286 | 0.493316464 |
| ENSMUSG00000084898 | Gm12371 | lincRNA | 1.346603197 | 3.240525916 | 6.2646646 | 0.023302 | 0.493316464 |
| ENSMUSG00000104597 | B230334C09Rik | antisense_RNA | -1.269197994 | 3.76481505 | 6.2642807 | 0.0233056 | 0.493316464 |
| ENSMUSG00000043372 | Hexim2 | protein_coding | 0.890496292 | 5.644263685 | 6.2601025 | 0.0233651 | 0.493604309 |
| ENSMUSG00000032220 | Myo1e | protein_coding | -1.727522408 | 2.891451718 | 6.2502989 | 0.0234377 | 0.494168184 |
| ENSMUSG00000049521 | Cdc42ep1 | protein_coding | -1.490782853 | 3.740462329 | 6.2391598 | 0.0235638 | 0.494492338 |
| ENSMUSG00000028278 | Rragd | protein_coding | -0.662327492 | 6.228608853 | 6.2345952 | 0.0235872 | 0.494492338 |
| ENSMUSG00000027419 | Pcsk2 | protein_coding | -0.595210712 | 6.944093578 | 6.2341886 | 0.0235911 | 0.494492338 |
| ENSMUSG00000084349 | Rpl3-ps1 | processed_pseudoge | 1.425761678 | 3.374979959 | 6.2163737 | 0.0237619 | 0.497105528 |
| ENSMUSG00000031285 | Dcx | protein_coding | -1.304759419 | 3.778773421 | 6.2042487 | 0.0238791 | 0.498585838 |
| ENSMUSG00000031297 | Slc7a3 | protein_coding | 1.036114613 | 4.918918982 | 6.1893299 | 0.0240445 | 0.498893676 |
| ENSMUSG00000006732 | Mettl1 | protein_coding | 0.643205064 | 6.02174802 | 6.1810733 | 0.0241048 | 0.498893676 |
| ENSMUSG00000020133 | 2310011J03Rik | protein_coding | 0.533478786 | 6.388075915 | 6.1671778 | 0.0242413 | 0.498893676 |
| ENSMUSG00000025203 | Scd2 | protein_coding | -0.623095148 | 8.816656589 | 6.1665219 | 0.0242478 | 0.498893676 |
| ENSMUSG00000033379 | Atp6v0b | protein_coding | 0.588791051 | 8.426530837 | 6.1613732 | 0.0242986 | 0.498893676 |
| ENSMUSG00000062647 | Rpl7a | protein_coding | 0.557271937 | 6.067649454 | 6.1598373 | 0.0243138 | 0.498893676 |
| ENSMUSG00000024227 | Pdzph1 | protein_coding | -1.280196237 | 4.119564187 | 6.1556893 | 0.0243548 | 0.498893676 |
| ENSMUSG00000034154 | Ino80 | protein_coding | -1.273587887 | 3.498932042 | 6.1524256 | 0.0243872 | 0.498893676 |
| ENSMUSG00000003974 | Grm3 | protein_coding | -0.90906998 | 4.983939823 | 6.1495063 | 0.0244162 | 0.498893676 |
| ENSMUSG00000028639 | Ybx1 | protein_coding | -0.930954164 | 5.303736514 | 6.1475431 | 0.0244357 | 0.498893676 |
| ENSMUSG00000002059 | Rab34 | protein_coding | -1.253653696 | 3.398121639 | 6.1450698 | 0.0244603 | 0.498893676 |
| ENSMUSG00000032171 | Pin1 | protein_coding | 0.521422387 | 8.909003625 | 6.1446487 | 0.0244645 | 0.498893676 |
| ENSMUSG00000030983 | Bccip | protein_coding | 0.495418875 | 7.669501024 | 6.1380842 | 0.02453 | 0.498893676 |
| ENSMUSG00000030201 | Lrp6 | protein_coding | -1.177080297 | 3.641438725 | 6.1367472 | 0.0245434 | 0.498893676 |
| ENSMUSG00000028106 | Rprd2 | protein_coding | -0.85851182 | 4.85400875 | 6.1176886 | 0.0247347 | 0.501411371 |
| ENSMUSG00000031157 | Pqbp1 | protein_coding | 0.556179753 | 7.908240987 | 6.1151377 | 0.0247605 | 0.501411371 |
| ENSMUSG00000025579 | Gaa | protein_coding | 0.54856438 | 9.350906561 | 6.1099315 | 0.0248131 | 0.501532806 |
| ENSMUSG00000060601 | Nr1h2 | protein_coding | 0.667737371 | 5.922656333 | 6.0981349 | 0.0249329 | 0.501544911 |
| ENSMUSG00000025508 | Rplp2 | protein_coding | 0.601396162 | 8.84790649 | 6.0968687 | 0.0249458 | 0.501544911 |
| ENSMUSG00000025024 | Smndc1 | protein_coding | -0.894641706 | 4.371692579 | 6.0930931 | 0.0249842 | 0.501544911 |
| ENSMUSG00000014353 | Tmem87b | protein_coding | -0.947251357 | 5.005499003 | 6.0877469 | 0.0250389 | 0.501544911 |
| ENSMUSG00000031626 | Sorbs2 | protein_coding | -0.87827049 | 4.848276821 | 6.0869581 | 0.0250469 | 0.501544911 |
| ENSMUSG00000056854 | Pou3f4 | protein_coding | -1.10134208 | 4.056695907 | 6.0810667 | 0.0251073 | 0.501819209 |
| ENSMUSG00000026202 | Tuba4a | protein_coding | 0.715211836 | 9.099061636 | 6.0707277 | 0.0252344 | 0.502785359 |
| ENSMUSG00000036972 | Zic4 | protein_coding | -1.061438455 | 5.174582636 | 6.0693018 | 0.0252491 | 0.502785359 |
| ENSMUSG00000026179 | Pnkd | protein_coding | 0.53959247 | 8.05180221 | 6.0484211 | 0.0254448 | 0.50305077 |
| ENSMUSG00000034432 | Cops8 | protein_coding | 0.53823706 | 8.788019941 | 6.0430392 | 0.025501 | 0.50305077 |
| ENSMUSG00000030839 | Sergef | protein_coding | 0.575809058 | 6.531287465 | 6.0388647 | 0.0255446 | 0.50305077 |
| ENSMUSG00000049643 | 2310022A10Rik | protein_coding | 0.669572617 | 5.683966829 | 6.0359626 | 0.025575 | 0.50305077 |
| ENSMUSG00000046160 | Olig1 | protein_coding | -0.856155407 | 5.77983526 | 6.0348657 | 0.0255865 | 0.50305077 |
| ENSMUSG00000054850 | Smim10l2a | protein_coding | 0.876081014 | 4.392290456 | 6.0348268 | 0.0255869 | 0.50305077 |

| ensembl_gene_id | external_gene_name | gene_biotype | log2 (fold change) | log2 (counts per million) | F-value | p-value | False discovery rate |
|---|---|---|---|---|---|---|---|
| ENSMUSG00000024346 | Pfdn1 | protein_coding | 0.47038713 | 9.040204816 | 6.028197 | 0.0256566 | 0.50305077 |
| ENSMUSG00000030043 | Tacr1 | protein_coding | 1.280966087 | 3.236117114 | 6.0206524 | 0.0257361 | 0.50305077 |
| ENSMUSG00000022150 | Dab2 | protein_coding | -1.48990788 | 3.048865381 | 6.0101288 | 0.0258474 | 0.50305077 |
| ENSMUSG00000017801 | Mlx | protein_coding | 0.711912712 | 6.238427675 | 5.9997672 | 0.0259576 | 0.50305077 |
| ENSMUSG00000005886 | Ncoa2 | protein_coding | -0.966162642 | 4.780771823 | 5.9991572 | 0.0259641 | 0.50305077 |
| ENSMUSG00000020284 | 1810043G02Rik | protein_coding | 0.622531886 | 6.869623216 | 5.9916961 | 0.0260438 | 0.50305077 |
| ENSMUSG00000059970 | Hspa2 | protein_coding | -0.982538175 | 5.644125734 | 5.9925351 | 0.0260559 | 0.50305077 |
| ENSMUSG00000055553 | Kxd1 | protein_coding | 0.516194525 | 7.517410548 | 5.9875336 | 0.0260884 | 0.50305077 |
| ENSMUSG00000024810 | Il33 | protein_coding | -0.632975854 | 8.05266783 | 5.9805193 | 0.0261637 | 0.50305077 |
| ENSMUSG00000063889 | Crem | protein_coding | -0.785292776 | 5.032857539 | 5.9775516 | 0.0261957 | 0.50305077 |
| ENSMUSG00000031902 | Nfatc3 | protein_coding | -1.34767382 | 3.247578072 | 5.976501 | 0.026207 | 0.50305077 |
| ENSMUSG00000114304 | AC165092.1 | lincRNA | 0.90441312 | 7.087818151 | 6.09517 | 0.0262536 | 0.50305077 |
| ENSMUSG00000006315 | Tmem147 | protein_coding | 0.555855101 | 7.795517713 | 5.9704314 | 0.0262725 | 0.50305077 |
| ENSMUSG00000113029 | AC122190.1 | antisense_RNA | -1.46614976 | 3.301354643 | 5.9646756 | 0.0263348 | 0.50305077 |
| ENSMUSG00000050538 | B230217C12Rik | protein_coding | 0.494166814 | 7.527299184 | 5.9575954 | 0.0264117 | 0.50305077 |
| ENSMUSG00000018585 | Atox1 | protein_coding | 0.547591113 | 8.537547706 | 5.9479935 | 0.0265163 | 0.50305077 |
| ENSMUSG00000043102 | Qrfp | protein_coding | -4.112315625 | 2.17994674 | 6.0237162 | 0.0265207 | 0.50305077 |
| ENSMUSG00000004561 | Mettl17 | protein_coding | 0.894254481 | 4.410173875 | 5.9471932 | 0.0265251 | 0.50305077 |
| ENSMUSG00000001156 | Mxd1 | protein_coding | -1.016277345 | 3.812498202 | 5.9470948 | 0.0265261 | 0.50305077 |
| ENSMUSG00000036333 | Kidins220 | protein_coding | -0.701385431 | 6.645574306 | 5.9462664 | 0.0265352 | 0.50305077 |
| ENSMUSG00000109324 | Prmt1 | protein_coding | 0.536191921 | 7.313579922 | 5.941349 | 0.026589 | 0.50305077 |
| ENSMUSG00000028049 | Scamp3 | protein_coding | 0.568879005 | 6.211898574 | 5.9402326 | 0.0266013 | 0.50305077 |
| ENSMUSG00000034285 | Nipsnap1 | protein_coding | 0.479409925 | 8.165715942 | 5.9317107 | 0.0266949 | 0.50305077 |
| ENSMUSG00000052748 | Swt1 | protein_coding | -1.048179245 | 4.030191547 | 5.9315309 | 0.0266969 | 0.50305077 |
| ENSMUSG00000020882 | Cacnb1 | protein_coding | 0.625994691 | 5.735647018 | 5.9269786 | 0.026747 | 0.50305077 |
| ENSMUSG00000014956 | Ppp1cb | protein_coding | -0.480504601 | 8.33464785 | 5.9194342 | 0.0268304 | 0.50305077 |
| ENSMUSG00000018417 | Myo1b | protein_coding | -0.984004396 | 3.79058272 | 5.9194261 | 0.0268305 | 0.50305077 |
| ENSMUSG00000018677 | Slc25a39 | protein_coding | 0.532918844 | 6.66780126 | 5.9173893 | 0.0268531 | 0.50305077 |
| ENSMUSG00000025810 | Nrp1 | protein_coding | -1.034631946 | 4.129597708 | 5.9031321 | 0.0270116 | 0.504411966 |
| ENSMUSG00000031700 | Gpt2 | protein_coding | -0.944109762 | 4.450509533 | 5.9001418 | 0.027045 | 0.504411966 |
| ENSMUSG00000053297 | AI854703 | protein_coding | 0.770047713 | 5.33239503 | 5.8954719 | 0.0270973 | 0.504411966 |
| ENSMUSG00000038279 | Nop2 | protein_coding | 0.920982744 | 5.250246065 | 5.8940339 | 0.0271134 | 0.504411966 |
| ENSMUSG00000033061 | Resp18 | protein_coding | 0.556853791 | 9.264313916 | 5.8847879 | 0.0272172 | 0.504965403 |
| ENSMUSG00000034714 | Ttyh2 | protein_coding | -0.802231278 | 7.227593125 | 5.9320121 | 0.027237 | 0.504965403 |
| ENSMUSG00000029563 | Foxp2 | protein_coding | -0.854369135 | 5.536918771 | 5.8772496 | 0.0273022 | 0.505302625 |
| ENSMUSG00000025981 | Coq10b | protein_coding | -0.825914413 | 5.300111345 | 5.8657927 | 0.027432 | 0.506589832 |
| ENSMUSG00000049225 | Pdp1 | protein_coding | -0.625224298 | 7.125775586 | 5.8592986 | 0.0275059 | 0.506589832 |
| ENSMUSG00000034891 | Sncb | protein_coding | 0.65671794 | 9.225122231 | 5.8579437 | 0.0275213 | 0.506589832 |
| ENSMUSG00000034845 | Plvap | protein_coding | 0.941757269 | 3.980863032 | 5.8480551 | 0.0276343 | 0.506589832 |
| ENSMUSG00000043284 | Tmem11 | protein_coding | 0.566070639 | 6.622788953 | 5.8402508 | 0.0277239 | 0.506589832 |
| ENSMUSG00000067942 | Zfp160 | protein_coding | -1.151783044 | 3.655992398 | 5.8390148 | 0.0277381 | 0.506589832 |
| ENSMUSG00000020464 | Pnpt1 | protein_coding | -1.314484885 | 3.13808061 | 5.8343733 | 0.0277916 | 0.506589832 |
| ENSMUSG00000026966 | Ssna1 | protein_coding | 0.520242534 | 7.620011426 | 5.830749 | 0.0278334 | 0.506589832 |
| ENSMUSG00000021540 | Smad5 | protein_coding | -0.794373547 | 4.475325274 | 5.8219499 | 0.0279352 | 0.506589832 |
| ENSMUSG00000023064 | Sncg | protein_coding | -0.72102054 | 7.292200882 | 5.8224974 | 0.0279506 | 0.506589832 |
| ENSMUSG00000058488 | Kl | protein_coding | -3.735466084 | 3.713946412 | 6.4425416 | 0.027981 | 0.506589832 |
| ENSMUSG00000007892 | Rplp1 | protein_coding | 0.673509043 | 9.342253724 | 5.8102093 | 0.0280935 | 0.506589832 |
| ENSMUSG00000110027 | C030029H02Rik | lincRNA | -0.922661349 | 4.059750618 | 5.8072573 | 0.0281062 | 0.506589832 |
| ENSMUSG00000038173 | Enpp6 | protein_coding | -1.757340278 | 3.093479553 | 5.805725 | 0.0281241 | 0.506589832 |
| ENSMUSG00000036391 | Sec24a | protein_coding | -1.097527807 | 3.856373162 | 5.8030901 | 0.0281549 | 0.506589832 |
| ENSMUSG00000026721 | Rabgap1l | protein_coding | -0.606377411 | 6.152632248 | 5.7999548 | 0.0281917 | 0.506589832 |
| ENSMUSG00000053004 | Hrh1 | protein_coding | 0.863982771 | 4.110665507 | 5.7958176 | 0.0282402 | 0.506589832 |
| ENSMUSG00000041870 | Ankrd13a | protein_coding | -1.051363703 | 3.997599059 | 5.7953778 | 0.0282454 | 0.506589832 |
| ENSMUSG00000032411 | Tfdp2 | protein_coding | -0.984497648 | 4.105395577 | 5.793545 | 0.0282669 | 0.506589832 |
| ENSMUSG00000028741 | Mrto4 | protein_coding | 0.604445284 | 6.218993079 | 5.7813139 | 0.0284111 | 0.507965605 |
| ENSMUSG00000059474 | Mbtd1 | protein_coding | -1.120653296 | 3.747940993 | 5.7790292 | 0.0284381 | 0.507965605 |
| ENSMUSG00000042388 | Dlgap3 | protein_coding | -1.407222039 | 3.139058895 | 5.7634854 | 0.0286228 | 0.509546784 |
| ENSMUSG00000007338 | Mrpl49 | protein_coding | 0.499651513 | 7.222371625 | 5.760297 | 0.0286609 | 0.509546784 |
| ENSMUSG00000085282 | Gm15663 | processed_transcript | 0.970266448 | 3.665509334 | 5.7596332 | 0.0286688 | 0.509546784 |
| ENSMUSG00000050732 | Vamp8 | protein_coding | -1.19805243 | 5.692453053 | 5.8525073 | 0.0289061 | 0.51061723 |
| ENSMUSG00000022548 | Apod | protein_coding | -0.634278124 | 8.762868782 | 5.7384611 | 0.0289231 | 0.51061723 |
| ENSMUSG00000031807 | Pgls | protein_coding | 0.896205013 | 4.35469058 | 5.735473 | 0.0289592 | 0.51061723 |
| ENSMUSG00000031548 | Sfrp1 | protein_coding | -2.094449413 | 2.630183538 | 5.7349298 | 0.0289657 | 0.51061723 |
| ENSMUSG00000074212 | Dnajb14 | protein_coding | -0.659257814 | 5.796248168 | 5.7296698 | 0.0290294 | 0.51061723 |
| ENSMUSG00000021703 | Serinc5 | protein_coding | -1.058480102 | 4.266735391 | 5.7217442 | 0.0291257 | 0.51061723 |
| ENSMUSG00000038695 | Josd2 | protein_coding | 0.558801125 | 6.239917499 | 5.7215153 | 0.0291285 | 0.51061723 |
| ENSMUSG00000054418 | 2900041M22Rik | processed_transcript | 1.539665589 | 2.730016073 | 5.7131345 | 0.0292307 | 0.51061723 |
| ENSMUSG00000002032 | Tmem25 | protein_coding | 0.722070327 | 6.284978924 | 5.7119678 | 0.0292449 | 0.51061723 |
| ENSMUSG00000021719 | Rgs7bp | protein_coding | -0.496891089 | 7.127353161 | 5.7012558 | 0.0293762 | 0.51061723 |
| ENSMUSG00000024558 | Mapk4 | protein_coding | -0.820728177 | 4.869485368 | 5.6998428 | 0.0293936 | 0.51061723 |
| ENSMUSG00000026058 | Khdrbs2 | protein_coding | -0.784106851 | 4.980876755 | 5.6983388 | 0.0294121 | 0.51061723 |
| ENSMUSG00000030432 | Rpl28 | protein_coding | 0.547764224 | 6.389847888 | 5.6965179 | 0.0294345 | 0.51061723 |
| ENSMUSG00000040929 | Rfx3 | protein_coding | -0.990131979 | 4.279747821 | 5.6945411 | 0.0294588 | 0.51061723 |
| ENSMUSG00000025350 | Rdh5 | protein_coding | -1.921676233 | 3.235796726 | 5.699399 | 0.0294798 | 0.51061723 |
| ENSMUSG00000039018 | Mtg1 | protein_coding | 0.634013724 | 6.316391935 | 5.6921108 | 0.0294888 | 0.51061723 |
| ENSMUSG00000046447 | Camk2n1 | protein_coding | -0.576643517 | 8.814693818 | 5.6841412 | 0.0295874 | 0.510617314 |
| ENSMUSG00000021493 | Pdlim7 | protein_coding | 0.503001431 | 6.876904623 | 5.6733293 | 0.0297217 | 0.510617314 |
| ENSMUSG00000020346 | Mgat1 | protein_coding | -0.760772238 | 5.171762214 | 5.6716413 | 0.0297427 | 0.510617314 |

| ensembl_gene_id | external_gene_name | gene_biotype | log2 (fold change) | log2 (counts per million) | F-value | p-value | False discovery rate |
|---|---|---|---|---|---|---|---|
| ENSMUSG00000024065 | Ehd3 | protein_coding | 0.69335344 | 6.826757013 | 5.6713923 | 0.0297681 | 0.510617314 |
| ENSMUSG00000029211 | Gabra4 | protein_coding | -0.789343848 | 6.717935929 | 5.6771775 | 0.0297933 | 0.510617314 |
| ENSMUSG00000036111 | Lmo1 | protein_coding | 0.753816309 | 4.430360891 | 5.6627071 | 0.0298543 | 0.510617314 |
| ENSMUSG00000026679 | Enkur | protein_coding | -1.078483182 | 4.017114693 | 5.6518257 | 0.0299909 | 0.510617314 |
| ENSMUSG00000032840 | 2410131K14Rik | protein_coding | 0.676068512 | 5.490908084 | 5.6429594 | 0.0301027 | 0.510617314 |
| ENSMUSG00000020572 | Nampt | protein_coding | -0.709183179 | 4.97681015 | 5.6406546 | 0.0301318 | 0.510617314 |
| ENSMUSG00000036949 | Slc39a12 | protein_coding | -1.048123829 | 4.149159605 | 5.6396863 | 0.0301441 | 0.510617314 |
| ENSMUSG00000053470 | Kdm3a | protein_coding | -0.966514749 | 4.227019001 | 5.6340105 | 0.030216 | 0.510617314 |
| ENSMUSG00000023913 | Pla2g7 | protein_coding | -0.609853531 | 7.018557148 | 5.632785 | 0.0302316 | 0.510617314 |
| ENSMUSG00000028333 | Anp32b | protein_coding | -0.796936394 | 5.070327518 | 5.6325922 | 0.030234 | 0.510617314 |
| ENSMUSG00000001039 | B9d1 | protein_coding | 0.651105035 | 5.906474746 | 5.6272806 | 0.0303016 | 0.510617314 |
| ENSMUSG00000021756 | Il6st | protein_coding | -0.994738628 | 4.07293837 | 5.6250991 | 0.0303294 | 0.510617314 |
| ENSMUSG00000027581 | Stmn3 | protein_coding | 0.524229569 | 11.21256015 | 5.62171 | 0.0303726 | 0.510617314 |
| ENSMUSG00000035620 | Ric8b | protein_coding | -0.722044653 | 5.915712258 | 5.621122 | 0.0303801 | 0.510617314 |
| ENSMUSG00000028974 | Dffa | protein_coding | 0.698728658 | 5.691760046 | 5.6202099 | 0.0303918 | 0.510617314 |
| ENSMUSG00000032854 | Ugt8a | protein_coding | -0.648437707 | 5.687384174 | 5.6185371 | 0.0304132 | 0.510617314 |
| ENSMUSG00000037652 | Phc3 | protein_coding | -0.909154712 | 4.092420459 | 5.6165542 | 0.0304385 | 0.510617314 |
| ENSMUSG00000010057 | Nprl2 | protein_coding | 0.679739301 | 6.55218835 | 5.5899056 | 0.0307819 | 0.512637521 |
| ENSMUSG00000020810 | Cygb | protein_coding | 0.507093533 | 7.364913844 | 5.5880783 | 0.0308056 | 0.512637521 |
| ENSMUSG00000021607 | Mrpl36 | protein_coding | 0.447686638 | 7.686975598 | 5.5878806 | 0.0308081 | 0.512637521 |
| ENSMUSG00000039103 | Nexn | protein_coding | -1.210253504 | 4.231479967 | 5.5827479 | 0.0308748 | 0.512637521 |
| ENSMUSG00000032463 | Faim | protein_coding | -0.82951801 | 4.670042742 | 5.5821916 | 0.0308821 | 0.512637521 |
| ENSMUSG00000030161 | Gabarapl1 | protein_coding | 0.541381561 | 10.32629476 | 5.5820896 | 0.0308834 | 0.512637521 |
| ENSMUSG00000050530 | Fam171a1 | protein_coding | -0.674947332 | 5.133659877 | 5.5813766 | 0.0308927 | 0.512637521 |
| ENSMUSG00000047045 | Tmem164 | protein_coding | -1.136297861 | 3.627570521 | 5.5733318 | 0.0309976 | 0.513063307 |
| ENSMUSG00000059288 | Cdyl | protein_coding | -0.976690303 | 4.048326968 | 5.5702476 | 0.031038 | 0.513063307 |
| ENSMUSG00000016918 | Sulf1 | protein_coding | -1.525479509 | 3.857128097 | 5.5935677 | 0.0310892 | 0.513063307 |
| ENSMUSG00000039244 | E130309D02Rik | protein_coding | 0.962050092 | 3.943347011 | 5.5624406 | 0.0311404 | 0.513063307 |
| ENSMUSG00000097736 | 9530059O14Rik | lincRNA | -2.66921773 | 2.537022903 | 5.5605021 | 0.0311886 | 0.513063307 |
| ENSMUSG00000050002 | Idnk | protein_coding | 0.556902233 | 7.140977413 | 5.5572205 | 0.0312091 | 0.513063307 |
| ENSMUSG00000027109 | Sp3 | protein_coding | -0.823324875 | 4.960542373 | 5.5539377 | 0.0312523 | 0.513063307 |
| ENSMUSG00000004270 | Lpcat3 | protein_coding | -1.140714627 | 4.037494218 | 5.5473099 | 0.0313399 | 0.51318193 |
| ENSMUSG00000073423 | Zfp414 | protein_coding | 0.612858172 | 6.22489882 | 5.546171 | 0.031355 | 0.51318193 |
| ENSMUSG00000040138 | Ndp | protein_coding | -0.908682697 | 4.483514932 | 5.5421031 | 0.031409 | 0.513283374 |
| ENSMUSG00000021338 | Carmil1 | protein_coding | -1.174426422 | 3.396781773 | 5.5236231 | 0.0316553 | 0.51463533 |
| ENSMUSG00000053931 | Cnn3 | protein_coding | -0.987455481 | 4.961678493 | 5.5235761 | 0.0316559 | 0.51463533 |
| ENSMUSG00000039419 | Cntnap2 | protein_coding | 0.502583044 | 7.646214872 | 5.521985 | 0.0316773 | 0.51463533 |
| ENSMUSG00000038418 | Egr1 | protein_coding | -0.677229824 | 6.746067033 | 5.5203134 | 0.0316997 | 0.51463533 |
| ENSMUSG00000029135 | Fosl2 | protein_coding | -1.495135752 | 3.461338492 | 5.5179799 | 0.031731 | 0.51463533 |
| ENSMUSG00000067274 | Rplp0 | protein_coding | 0.468384577 | 8.351305797 | 5.5125505 | 0.031804 | 0.515042491 |
| ENSMUSG00000026674 | Ddr2 | protein_coding | -1.482961728 | 2.860090175 | 5.498559 | 0.031993 | 0.515918487 |
| ENSMUSG00000022537 | Tmem44 | protein_coding | -1.2173088 | 3.575541263 | 5.4971082 | 0.0320127 | 0.515918487 |
| ENSMUSG00000040250 | Ints13 | protein_coding | -1.068665821 | 3.385165044 | 5.4933574 | 0.0320636 | 0.515918487 |
| ENSMUSG00000024502 | Jakmip2 | protein_coding | 0.571160301 | 6.198559828 | 5.485217 | 0.0321744 | 0.515918487 |
| ENSMUSG00000040591 | 1110051M20Rik | protein_coding | 0.584185534 | 8.374392058 | 5.4817746 | 0.0322215 | 0.515918487 |
| ENSMUSG00000033174 | Mgll | protein_coding | -0.684785174 | 6.842980407 | 5.4823949 | 0.032236 | 0.515918487 |
| ENSMUSG00000029088 | Kcnip4 | protein_coding | -0.647216872 | 5.534930803 | 5.4769262 | 0.0322878 | 0.515918487 |
| ENSMUSG00000078429 | Ctdsp2 | protein_coding | -0.960596897 | 4.81346619 | 5.4759265 | 0.0323015 | 0.515918487 |
| ENSMUSG00000028367 | Txn1 | protein_coding | 0.498502137 | 8.31226565 | 5.4748792 | 0.0323159 | 0.515918487 |
| ENSMUSG00000032802 | Srxn1 | protein_coding | 0.601130865 | 6.536059391 | 5.4687852 | 0.0323995 | 0.515918487 |
| ENSMUSG00000087674 | 4930447M23Rik | lincRNA | 1.21399471 | 3.176993293 | 5.4679827 | 0.0324106 | 0.515918487 |
| ENSMUSG00000018648 | Dusp14 | protein_coding | -0.594006903 | 6.258316206 | 5.4650435 | 0.0324511 | 0.515918487 |
| ENSMUSG00000024006 | Stk38 | protein_coding | -1.069031401 | 3.52532164 | 5.4569386 | 0.032563 | 0.515918487 |
| ENSMUSG00000032599 | Ip6k2 | protein_coding | 0.518391148 | 6.790141189 | 5.4525492 | 0.0326237 | 0.515918487 |
| ENSMUSG00000033295 | Ptprf | protein_coding | 0.607278247 | 5.670309118 | 5.4464192 | 0.0327089 | 0.515918487 |
| ENSMUSG00000031851 | Ntpcr | protein_coding | 0.531633706 | 6.280122565 | 5.4445357 | 0.0327351 | 0.515918487 |
| ENSMUSG00000016252 | Atp5e | protein_coding | 0.56164113 | 9.763074852 | 5.4423913 | 0.0327649 | 0.515918487 |
| ENSMUSG00000003813 | Rad23a | protein_coding | 0.512145492 | 8.108862216 | 5.4399142 | 0.0327995 | 0.515918487 |
| ENSMUSG00000032320 | Rcn2 | protein_coding | -0.785271447 | 6.618201067 | 5.4443832 | 0.0328615 | 0.515918487 |
| ENSMUSG00000022272 | Myo10 | protein_coding | -0.83238945 | 4.921313803 | 5.4350237 | 0.0328678 | 0.515918487 |
| ENSMUSG00000049090 | Zadh2 | protein_coding | -1.006459746 | 3.667105702 | 5.4337219 | 0.032886 | 0.515918487 |
| ENSMUSG00000018451 | 6330403K07Rik | lincRNA | 0.510951607 | 10.97421339 | 5.429252 | 0.0329486 | 0.515918487 |
| ENSMUSG00000015224 | Cyp2j9 | protein_coding | -0.945602615 | 4.778190594 | 5.4268328 | 0.0329825 | 0.515918487 |
| ENSMUSG00000021534 | 1700001L19Rik | protein_coding | 0.63712356 | 6.731267775 | 5.4240926 | 0.033021 | 0.515918487 |
| ENSMUSG00000025531 | Chm | protein_coding | -0.892157908 | 4.489383728 | 5.4209592 | 0.0330651 | 0.515918487 |
| ENSMUSG00000042506 | Usp22 | protein_coding | 0.679603059 | 8.288347525 | 5.4179658 | 0.0331306 | 0.515918487 |
| ENSMUSG00000009394 | Syn2 | protein_coding | 0.491114738 | 8.464177171 | 5.4117451 | 0.0331951 | 0.515918487 |
| ENSMUSG00000026600 | Soat1 | protein_coding | -0.938452259 | 4.214065536 | 5.4112958 | 0.0332015 | 0.515918487 |
| ENSMUSG00000038539 | Atf5 | protein_coding | -1.814216499 | 2.495895301 | 5.4008926 | 0.033349 | 0.516731759 |
| ENSMUSG00000052423 | B4galt3 | protein_coding | 0.680356989 | 5.498150261 | 5.3967848 | 0.0334075 | 0.516731759 |
| ENSMUSG00000039701 | Usp53 | protein_coding | -1.291941447 | 3.229360718 | 5.3888575 | 0.0335207 | 0.516731759 |
| ENSMUSG00000006289 | Osgep | protein_coding | -0.979449267 | 4.89572541 | 5.3880717 | 0.0335319 | 0.516731759 |
| ENSMUSG00000025658 | Cnksr2 | protein_coding | -0.881436777 | 5.375949277 | 5.3878826 | 0.0335346 | 0.516731759 |
| ENSMUSG00000020623 | Map2k6 | protein_coding | -1.073927076 | 4.021855182 | 5.3873565 | 0.0335422 | 0.516731759 |
| ENSMUSG00000025132 | Arhgdia | protein_coding | 0.536191111 | 9.882980384 | 5.3737913 | 0.033737 | 0.518989578 |
| ENSMUSG00000067847 | Romo1 | protein_coding | 0.529662025 | 8.20766826 | 5.3626949 | 0.0338973 | 0.520711177 |
| ENSMUSG00000040565 | Btaf1 | protein_coding | -1.222415654 | 3.801692078 | 5.3553022 | 0.0340046 | 0.520947638 |
| ENSMUSG00000045103 | Dmd | protein_coding | -0.744606195 | 4.828921698 | 5.3549599 | 0.0340096 | 0.520947638 |

| ensembl_gene_id | external_gene_name | gene_biotype | log2 (fold change) | log2 (counts per million) | F-value | p-value | False discovery rate |
|---|---|---|---|---|---|---|---|
| ENSMUSG00000043572 | Pars2 | protein_coding | 0.860198615 | 3.973158608 | 5.3466693 | 0.0341304 | 0.521419663 |
| ENSMUSG00000032452 | Clstn2 | protein_coding | -0.871618485 | 4.601326026 | 5.3461921 | 0.0341374 | 0.521419663 |
| ENSMUSG00000022500 | Litaf | protein_coding | -0.543641034 | 6.764656062 | 5.3400693 | 0.034227 | 0.521966705 |
| ENSMUSG00000030660 | Pik3c2a | protein_coding | -1.697406551 | 2.710355144 | 5.329266 | 0.0343857 | 0.521966705 |
| ENSMUSG00000027668 | Mfn1 | protein_coding | -1.021934155 | 4.57648401 | 5.3285483 | 0.0343963 | 0.521966705 |
| ENSMUSG00000019984 | Med23 | protein_coding | -0.968155425 | 3.592601219 | 5.3285438 | 0.0343963 | 0.521966705 |
| ENSMUSG00000070802 | Pnmal2 | protein_coding | 0.572403266 | 8.621589177 | 5.327214 | 0.0344159 | 0.521966705 |
| ENSMUSG00000017667 | Zfp334 | protein_coding | -0.779901368 | 4.632684367 | 5.3166489 | 0.0345721 | 0.523049121 |
| ENSMUSG00000002379 | Ndufa11 | protein_coding | 0.522192159 | 9.817108866 | 5.3100164 | 0.0346706 | 0.523049121 |
| ENSMUSG00000027133 | Nop10 | protein_coding | 0.628085656 | 9.048247012 | 5.3092953 | 0.0346813 | 0.523049121 |
| ENSMUSG00000057130 | Txnl4a | protein_coding | 0.591383017 | 6.757728205 | 5.3092578 | 0.0346819 | 0.523049121 |
| ENSMUSG00000046798 | Cldn12 | protein_coding | -0.831541219 | 4.887008155 | 5.3015015 | 0.0347975 | 0.523228333 |
| ENSMUSG00000023800 | Tiam2 | protein_coding | -1.021497298 | 4.999777228 | 5.3003381 | 0.0348386 | 0.523228333 |
| ENSMUSG00000036966 | Spryd3 | protein_coding | 0.627793479 | 8.680516596 | 5.2954377 | 0.0348882 | 0.523228333 |
| ENSMUSG00000063849 | Ppcdc | protein_coding | 0.671383817 | 5.543887898 | 5.2928095 | 0.0349275 | 0.523228333 |
| ENSMUSG00000035354 | Uvrag | protein_coding | -1.097752948 | 3.360669855 | 5.2921768 | 0.034937 | 0.523228333 |
| ENSMUSG00000008690 | Ncaph2 | protein_coding | 0.651064894 | 7.207697295 | 5.2892454 | 0.0350048 | 0.523514527 |
| ENSMUSG00000057841 | Rpl32 | protein_coding | 0.615460883 | 9.869372076 | 5.2828857 | 0.0350767 | 0.523643587 |
| ENSMUSG00000033910 | Gucy1a3 | protein_coding | -0.466293746 | 6.664581123 | 5.2806239 | 0.0351109 | 0.523643587 |
| ENSMUSG00000042182 | Bend6 | protein_coding | 0.502780596 | 7.256237673 | 5.2753519 | 0.0351905 | 0.524011771 |
| ENSMUSG00000031872 | Bean1 | protein_coding | 1.55631821 | 2.534409631 | 5.2725445 | 0.035233 | 0.524011771 |
| ENSMUSG00000017561 | Crlf3 | protein_coding | -1.314405692 | 3.450944053 | 5.2547261 | 0.0355041 | 0.527314893 |
| ENSMUSG00000020737 | Jpt1 | protein_coding | 0.474374236 | 8.50463391 | 5.2483792 | 0.0356013 | 0.527882998 |
| ENSMUSG00000027248 | Pdia3 | protein_coding | -0.636403523 | 6.219888738 | 5.2451743 | 0.0356505 | 0.527882998 |
| ENSMUSG00000005625 | Psmd4 | protein_coding | 0.479166422 | 8.643510323 | 5.2417645 | 0.0357029 | 0.527882998 |
| ENSMUSG00000022570 | Tsta3 | protein_coding | 0.470807133 | 6.72415395 | 5.2394334 | 0.0357388 | 0.527882998 |
| ENSMUSG00000048218 | Amigo2 | protein_coding | 0.838332547 | 4.458766099 | 5.2313235 | 0.0358639 | 0.528561347 |
| ENSMUSG00000026255 | Efhd1 | protein_coding | -1.051636125 | 4.00969525 | 5.2300894 | 0.035883 | 0.528561347 |
| ENSMUSG00000001017 | Chtop | protein_coding | -0.550105345 | 5.922652074 | 5.2167735 | 0.0360897 | 0.529465532 |
| ENSMUSG00000016024 | Lbp | protein_coding | -2.226898201 | 2.970601539 | 5.245591 | 0.0360974 | 0.529465532 |
| ENSMUSG00000079435 | Rpl36a | protein_coding | 0.766549585 | 5.304824514 | 5.2153865 | 0.0361113 | 0.529465532 |
| ENSMUSG00000096472 | Cdkn2d | protein_coding | 0.446165335 | 8.239729479 | 5.2134611 | 0.0361413 | 0.529465532 |
| ENSMUSG00000024565 | Sall3 | protein_coding | -1.789254181 | 2.707476908 | 5.2068944 | 0.0362439 | 0.529671383 |
| ENSMUSG00000039197 | Adk | protein_coding | -0.464024054 | 7.492440639 | 5.2029177 | 0.0363062 | 0.529671383 |
| ENSMUSG00000040624 | Plekhg1 | protein_coding | -1.148058958 | 3.843939393 | 5.1956081 | 0.0364211 | 0.529671383 |
| ENSMUSG00000038486 | Sv2a | protein_coding | 0.462972106 | 8.389974562 | 5.1930481 | 0.0364614 | 0.529671383 |
| ENSMUSG00000074825 | Itpripl1 | protein_coding | -1.362845573 | 3.798788421 | 5.1898251 | 0.0365364 | 0.529671383 |
| ENSMUSG00000041112 | Elmo1 | protein_coding | 0.515913966 | 8.636083867 | 5.1853055 | 0.0365837 | 0.529671383 |
| ENSMUSG00000034353 | Ramp1 | protein_coding | -0.904871593 | 6.466903309 | 5.2548974 | 0.0366994 | 0.529671383 |
| ENSMUSG00000021606 | Ndufs6 | protein_coding | 0.538497443 | 8.767019294 | 5.17478 | 0.0367506 | 0.529671383 |
| ENSMUSG00000033863 | Klf9 | protein_coding | -0.50809192 | 6.174847207 | 5.1708748 | 0.0368128 | 0.529671383 |
| ENSMUSG00000032553 | Srprb | protein_coding | 0.609192016 | 5.847270819 | 5.1676502 | 0.0368642 | 0.529671383 |
| ENSMUSG00000038170 | Pde4dip | protein_coding | -0.570643603 | 6.043936981 | 5.1671539 | 0.0368721 | 0.529671383 |
| ENSMUSG00000019579 | Mydgf | protein_coding | 0.498209433 | 7.3175133 | 5.1578442 | 0.0370211 | 0.529671383 |
| ENSMUSG00000030707 | Coro1a | protein_coding | 0.500042426 | 7.390618051 | 5.1547536 | 0.0370707 | 0.529671383 |
| ENSMUSG00000069045 | Ddx3y | protein_coding | -0.615924072 | 6.104745614 | 5.154602 | 0.0370732 | 0.529671383 |
| ENSMUSG00000015305 | Sash1 | protein_coding | -1.032662899 | 4.177191885 | 5.1461696 | 0.0372089 | 0.529671383 |
| ENSMUSG00000051323 | Pcdh19 | protein_coding | -0.692755432 | 5.411491613 | 5.1447059 | 0.0372326 | 0.529671383 |
| ENSMUSG00000021268 | Meg3 | lincRNA | -1.293909706 | 7.508684122 | 5.5530173 | 0.0372525 | 0.529671383 |
| ENSMUSG00000023868 | Pde10a | protein_coding | -0.780951561 | 6.422342543 | 5.1474752 | 0.0372763 | 0.529671383 |
| ENSMUSG00000023025 | Larp4 | protein_coding | -0.741128069 | 4.523088923 | 5.1385207 | 0.0373326 | 0.529671383 |
| ENSMUSG00000002489 | Tiam1 | protein_coding | -0.729762461 | 5.244388358 | 5.1380324 | 0.0373405 | 0.529671383 |
| ENSMUSG00000038530 | Rgs4 | protein_coding | -0.487723795 | 8.482805222 | 5.1363847 | 0.0373672 | 0.529671383 |
| ENSMUSG00000084289 | Gm6977 | processed_pseudoge | -2.171993407 | 3.235061878 | 5.2104284 | 0.0373951 | 0.529671383 |
| ENSMUSG00000060012 | Kif13b | protein_coding | -1.088412282 | 3.707502773 | 5.1257932 | 0.0375394 | 0.529671383 |
| ENSMUSG00000058317 | Ube2e2 | protein_coding | 0.503297531 | 6.828385677 | 5.1237941 | 0.037572 | 0.529671383 |
| ENSMUSG00000022956 | Atp5o | protein_coding | 0.45200804 | 8.53452819 | 5.1204601 | 0.0376264 | 0.529671383 |
| ENSMUSG00000070306 | Ccdc153 | protein_coding | -0.82354413 | 5.855935236 | 5.1209129 | 0.0376434 | 0.529671383 |
| ENSMUSG00000021676 | Iqgap2 | protein_coding | -1.51963411 | 3.054315862 | 5.114492 | 0.0377241 | 0.529671383 |
| ENSMUSG00000027430 | Dtd1 | protein_coding | 0.438345909 | 8.941783108 | 5.1117595 | 0.0377689 | 0.529671383 |
| ENSMUSG00000057132 | Rpgrip1 | protein_coding | 1.240750344 | 3.430311073 | 5.1111223 | 0.0377794 | 0.529671383 |
| ENSMUSG00000063457 | Rps15 | protein_coding | 0.605771 | 8.929819648 | 5.1102006 | 0.0377945 | 0.529671383 |
| ENSMUSG00000022673 | Mcm4 | protein_coding | -1.193560914 | 3.356208428 | 5.1094686 | 0.0378065 | 0.529671383 |
| ENSMUSG00000032766 | Gng11 | protein_coding | -0.619252558 | 6.068835467 | 5.1086187 | 0.0378205 | 0.529671383 |
| ENSMUSG00000074129 | Rpl13a | protein_coding | 0.650230534 | 5.367134901 | 5.1054504 | 0.0378726 | 0.529671383 |
| ENSMUSG00000023068 | Nus1 | protein_coding | -0.614650202 | 5.801789935 | 5.1052885 | 0.0378753 | 0.529671383 |
| ENSMUSG00000078713 | Tomm5 | protein_coding | 0.519446332 | 6.628975089 | 5.1042617 | 0.0378922 | 0.529671383 |
| ENSMUSG00000073676 | Hspe1 | protein_coding | 0.522328234 | 7.452301563 | 5.0970302 | 0.0380116 | 0.529671383 |
| ENSMUSG00000022021 | Diaph3 | protein_coding | 0.891371226 | 4.311244021 | 5.0938991 | 0.0380634 | 0.529671383 |
| ENSMUSG00000019872 | Smpdl3a | protein_coding | -0.874260044 | 4.455349923 | 5.0928748 | 0.0380803 | 0.529671383 |
| ENSMUSG00000037351 | Actr1b | protein_coding | 0.600001751 | 8.367555443 | 5.0897211 | 0.0381326 | 0.529671383 |
| ENSMUSG00000041329 | Atp1b2 | protein_coding | -0.676195582 | 7.359503043 | 5.0837456 | 0.0382566 | 0.529671383 |
| ENSMUSG00000028937 | Acot7 | protein_coding | 0.631678279 | 10.6617693 | 5.0793906 | 0.0383045 | 0.529671383 |
| ENSMUSG00000044288 | Cnr1 | protein_coding | 0.585130412 | 6.287251988 | 5.0733883 | 0.0384048 | 0.529671383 |
| ENSMUSG00000056553 | Ptprn2 | protein_coding | -1.115511994 | 3.91967776 | 5.0717046 | 0.038433 | 0.529671383 |
| ENSMUSG00000041736 | Tspo | protein_coding | -1.158193821 | 3.492472395 | 5.0694579 | 0.0384707 | 0.529671383 |
| ENSMUSG00000036752 | Tubb4b | protein_coding | 0.573917339 | 7.668649757 | 5.0689532 | 0.0384791 | 0.529671383 |
| ENSMUSG00000029617 | Ccz1 | protein_coding | -0.93771205 | 4.257759713 | 5.0622199 | 0.0385923 | 0.529671383 |

| ensembl_gene_id | external_gene_name | gene_biotype | log2 (fold change) | log2 (counts per million) | F-value | p-value | False discovery rate |
|---|---|---|---|---|---|---|---|
| ENSMUSG00000025135 | Anapc11 | protein_coding | 0.44012055 | 9.282356923 | 5.0617971 | 0.0385994 | 0.529671383 |
| ENSMUSG00000047250 | Ptgs1 | protein_coding | 0.974713824 | 3.772499142 | 5.0616405 | 0.038602 | 0.529671383 |
| ENSMUSG00000024033 | Rsph1 | protein_coding | -0.620238766 | 5.553946355 | 5.060862 | 0.0386151 | 0.529671383 |
| ENSMUSG00000053801 | Grwd1 | protein_coding | 0.644033838 | 6.023791231 | 5.0588228 | 0.0386495 | 0.529671383 |
| ENSMUSG00000032384 | Csnk1g1 | protein_coding | -0.742002295 | 5.506981906 | 5.0576805 | 0.0386687 | 0.529671383 |
| ENSMUSG00000029838 | Ptn | protein_coding | -0.68123544 | 8.78621346 | 5.0507816 | 0.03881 | 0.529671383 |
| ENSMUSG00000027829 | Ccnl1 | protein_coding | -1.079806896 | 3.801533746 | 5.0488685 | 0.0388177 | 0.529671383 |
| ENSMUSG00000007038 | Neu1 | protein_coding | 0.703112424 | 5.20445601 | 5.0483013 | 0.0388273 | 0.529671383 |
| ENSMUSG00000026857 | Ntmt1 | protein_coding | 0.483218443 | 7.068004118 | 5.0461033 | 0.0388646 | 0.529671383 |
| ENSMUSG00000028444 | Cntfr | protein_coding | -1.144733602 | 3.91942776 | 5.0392996 | 0.0389802 | 0.529744875 |
| ENSMUSG00000025931 | Paqr8 | protein_coding | -0.935939291 | 4.716172339 | 5.0338794 | 0.0390726 | 0.529744875 |
| ENSMUSG00000031365 | Zfp275 | protein_coding | -0.889914534 | 4.555549452 | 5.0336423 | 0.0390766 | 0.529744875 |
| ENSMUSG00000068859 | Sp9 | protein_coding | -0.959978016 | 3.855387841 | 5.0321452 | 0.0391022 | 0.529744875 |
| ENSMUSG00000003382 | Etv3 | protein_coding | -0.890801034 | 3.984042676 | 5.0313215 | 0.0391163 | 0.529744875 |
| ENSMUSG00000041120 | Nbl1 | protein_coding | -1.046813729 | 3.95341608 | 5.0247525 | 0.0392288 | 0.529928489 |
| ENSMUSG00000002031 | Ift46 | protein_coding | 0.578867775 | 7.052698323 | 5.0235789 | 0.0392489 | 0.529928489 |
| ENSMUSG00000018199 | Trove2 | protein_coding | -0.612404112 | 5.844787989 | 5.0131172 | 0.0394289 | 0.529928489 |
| ENSMUSG00000036769 | Wdr44 | protein_coding | -1.206873934 | 4.062501324 | 5.0125343 | 0.0394638 | 0.529928489 |
| ENSMUSG00000046432 | Bex3 | protein_coding | 0.509833295 | 9.760962919 | 5.0062536 | 0.0395475 | 0.529928489 |
| ENSMUSG00000029001 | Fbxo44 | protein_coding | 0.424584543 | 7.966833578 | 5.003441 | 0.0395962 | 0.529928489 |
| ENSMUSG00000039911 | Spsb1 | protein_coding | -1.398250658 | 2.936041677 | 4.9975089 | 0.0396992 | 0.529928489 |
| ENSMUSG00000033099 | Nol12 | protein_coding | 0.436790257 | 6.686652263 | 4.9965465 | 0.0397159 | 0.529928489 |
| ENSMUSG00000106775 | C130093G08Rik | TEC | -1.502261659 | 4.64571464 | 5.1193803 | 0.0397433 | 0.529928489 |
| ENSMUSG00000022671 | Mzt2 | protein_coding | 0.506312402 | 6.378649776 | 4.9925703 | 0.0397852 | 0.529928489 |
| ENSMUSG00000020315 | Sptbn1 | protein_coding | -0.6335642 | 7.420372831 | 4.9937672 | 0.0397892 | 0.529928489 |
| ENSMUSG00000029642 | Polr1d | protein_coding | 0.509719964 | 6.929829932 | 4.9908663 | 0.0398149 | 0.529928489 |
| ENSMUSG00000024176 | Sox8 | protein_coding | -0.885796247 | 4.254497487 | 4.9876872 | 0.0398704 | 0.529928489 |
| ENSMUSG00000070319 | Eif3g | protein_coding | 0.438916342 | 7.844693706 | 4.9839168 | 0.0399363 | 0.529928489 |
| ENSMUSG00000108778 | Gm20083 | lincRNA | -1.920036465 | 2.465986827 | 4.9838825 | 0.0399369 | 0.529928489 |
| ENSMUSG00000025209 | Twnk | protein_coding | -1.373301969 | 3.574506826 | 4.9768361 | 0.0400605 | 0.529928489 |
| ENSMUSG00000037605 | Adgrl3 | protein_coding | -0.900206408 | 5.248406716 | 4.968731 | 0.0402032 | 0.529928489 |
| ENSMUSG00000027677 | Ttc14 | protein_coding | -1.283083236 | 3.715954324 | 4.9683254 | 0.0402104 | 0.529928489 |
| ENSMUSG00000037031 | Tspan15 | protein_coding | -1.009961021 | 4.183674783 | 4.9606718 | 0.0403457 | 0.529928489 |
| ENSMUSG00000059412 | Fxyd2 | protein_coding | 0.77154871 | 5.13876124 | 4.9559546 | 0.0404294 | 0.529928489 |
| ENSMUSG00000067722 | BC003965 | protein_coding | 0.495710244 | 6.375988644 | 4.9548247 | 0.0404494 | 0.529928489 |
| ENSMUSG00000003528 | Slc25a1 | protein_coding | 0.883545758 | 5.133553603 | 4.9559829 | 0.0404539 | 0.529928489 |
| ENSMUSG00000020774 | Aspa | protein_coding | -0.758524973 | 5.993678956 | 4.9544561 | 0.0404811 | 0.529928489 |
| ENSMUSG00000022489 | Pde1b | protein_coding | -0.527165039 | 7.601957761 | 4.9510829 | 0.040516 | 0.529928489 |
| ENSMUSG00000032299 | Commd4 | protein_coding | 0.514244165 | 7.677940495 | 4.9489101 | 0.0405547 | 0.529928489 |
| ENSMUSG00000020134 | Peli1 | protein_coding | -0.787641922 | 5.027350715 | 4.933278 | 0.0408344 | 0.529928489 |
| ENSMUSG00000009681 | Bcr | protein_coding | -0.54180566 | 6.228285443 | 4.9280303 | 0.0409288 | 0.529928489 |
| ENSMUSG00000030842 | Lamtor1 | protein_coding | 0.44897903 | 7.343612553 | 4.926735 | 0.0409521 | 0.529928489 |
| ENSMUSG00000059323 | Tonsl | protein_coding | 0.45255471 | 9.574008816 | 4.9247308 | 0.0409882 | 0.529928489 |
| ENSMUSG00000050875 | A730017C20Rik | protein_coding | 0.569991544 | 8.826962204 | 4.9238501 | 0.0410041 | 0.529928489 |
| ENSMUSG00000029559 | 2210016L21Rik | protein_coding | 0.488293635 | 8.426601944 | 4.9224802 | 0.0410289 | 0.529928489 |
| ENSMUSG00000021377 | Dek | protein_coding | -0.702472161 | 5.116669153 | 4.9217317 | 0.0410424 | 0.529928489 |
| ENSMUSG00000012848 | Rps5 | protein_coding | 0.570792521 | 10.12689909 | 4.9120986 | 0.0412169 | 0.529928489 |
| ENSMUSG00000029162 | Khk | protein_coding | 0.621953502 | 5.107388417 | 4.9074757 | 0.0413009 | 0.529928489 |
| ENSMUSG00000029616 | Erp29 | protein_coding | 0.455716837 | 7.669565449 | 4.905385 | 0.041339 | 0.529928489 |
| ENSMUSG00000033632 | AW554918 | protein_coding | -0.885085566 | 3.894266693 | 4.905164 | 0.041343 | 0.529928489 |
| ENSMUSG00000022617 | Chkb | protein_coding | 0.846707643 | 3.886913244 | 4.9030222 | 0.0413821 | 0.529928489 |
| ENSMUSG00000034024 | Cct2 | protein_coding | 0.469935278 | 8.002043955 | 4.9014388 | 0.041411 | 0.529928489 |
| ENSMUSG00000055839 | Elob | protein_coding | 0.506529381 | 9.024725631 | 4.8981753 | 0.0414706 | 0.529928489 |
| ENSMUSG00000014226 | Cacybp | protein_coding | -0.552283597 | 7.549578527 | 4.896909 | 0.0414938 | 0.529928489 |
| ENSMUSG00000037979 | Ccdc92 | protein_coding | 0.575785406 | 6.192813252 | 4.8944889 | 0.0415381 | 0.529928489 |
| ENSMUSG00000042246 | Tmc7 | protein_coding | -1.62989619 | 3.010522308 | 4.8916369 | 0.0415904 | 0.529928489 |
| ENSMUSG00000031901 | Dus2 | protein_coding | 0.716750069 | 5.148643558 | 4.8890512 | 0.0416378 | 0.529928489 |
| ENSMUSG00000027366 | Sppl2a | protein_coding | -0.763326053 | 4.958975193 | 4.8866875 | 0.0416813 | 0.529928489 |
| ENSMUSG00000042750 | Bex2 | protein_coding | 0.476098922 | 10.3553781 | 4.8866346 | 0.0416823 | 0.529928489 |
| ENSMUSG00000041308 | Sntb2 | protein_coding | -1.863009681 | 2.552772567 | 4.8794242 | 0.0418152 | 0.529928489 |
| ENSMUSG00000021303 | Gng4 | protein_coding | 0.498053279 | 6.380812427 | 4.8792887 | 0.0418177 | 0.529928489 |
| ENSMUSG00000028798 | Eif3i | protein_coding | 0.451777702 | 9.126720684 | 4.8748261 | 0.0419001 | 0.529928489 |
| ENSMUSG00000026817 | Ak1 | protein_coding | 0.53505429 | 9.070943259 | 4.8725354 | 0.0419426 | 0.529928489 |
| ENSMUSG00000071064 | Zfp827 | protein_coding | -0.842920946 | 4.348245981 | 4.8711487 | 0.0419683 | 0.529928489 |
| ENSMUSG00000035242 | Oaz1 | protein_coding | 0.455025651 | 8.286674294 | 4.8699727 | 0.0419901 | 0.529928489 |
| ENSMUSG00000033208 | S100b | protein_coding | -0.826992488 | 6.003589543 | 4.8744656 | 0.0419991 | 0.529928489 |
| ENSMUSG00000000567 | Sox9 | protein_coding | -0.682623494 | 5.735015818 | 4.8686381 | 0.0420148 | 0.529928489 |
| ENSMUSG00000025468 | Caly | protein_coding | 0.483839737 | 9.560052941 | 4.8683526 | 0.0420202 | 0.529928489 |
| ENSMUSG00000013150 | Gfod2 | protein_coding | 0.473712035 | 6.447896045 | 4.867611 | 0.0420339 | 0.529928489 |
| ENSMUSG00000030697 | Ppp4c | protein_coding | -0.601606869 | 5.388606369 | 4.8659062 | 0.0420656 | 0.529928489 |
| ENSMUSG00000040746 | Rnf167 | protein_coding | 0.483992303 | 7.602093668 | 4.8633457 | 0.0421132 | 0.529928489 |
| ENSMUSG00000006740 | Kif5b | protein_coding | -0.599018678 | 6.115875941 | 4.8632833 | 0.0421144 | 0.529928489 |
| ENSMUSG00000030086 | Chchd6 | protein_coding | 0.610199608 | 7.819536396 | 4.8642696 | 0.0421215 | 0.529928489 |
| ENSMUSG00000044894 | Uqcrq | protein_coding | 0.539804639 | 10.39844636 | 4.8610733 | 0.0421556 | 0.529928489 |
| ENSMUSG00000028648 | Ndufs5 | protein_coding | 0.484710397 | 7.458033273 | 4.8592572 | 0.0421894 | 0.529928489 |
| ENSMUSG00000062210 | Tnfaip8 | protein_coding | -1.063140371 | 4.226963676 | 4.8569248 | 0.042233 | 0.529928489 |
| ENSMUSG00000029414 | Kntc1 | protein_coding | 1.9029653 | 2.226839101 | 4.8568369 | 0.0422346 | 0.529928489 |
| ENSMUSG00000020003 | Pex7 | protein_coding | -0.83713283 | 4.586048451 | 4.8507301 | 0.0423488 | 0.530435329 |

| ensembl_gene_id | external_gene_name | gene_biotype | log2 (fold change) | log2 (counts per million) | F-value | p-value | False discovery rate |
|---|---|---|---|---|---|---|---|
| ENSMUSG00000037266 | Rsrp1 | protein_coding | -1.787355167 | 5.794475185 | 5.2974265 | 0.0423906 | 0.530435329 |
| ENSMUSG00000000776 | Polr3d | protein_coding | -1.572146087 | 3.890417899 | 4.9032324 | 0.0424974 | 0.530435329 |
| ENSMUSG00000033014 | Trim33 | protein_coding | -0.73691768 | 4.776449862 | 4.8406371 | 0.0425384 | 0.530435329 |
| ENSMUSG00000022342 | Kcnv1 | protein_coding | -1.300298981 | 3.770552748 | 4.8396802 | 0.042582 | 0.530435329 |
| ENSMUSG00000057329 | Bcl2 | protein_coding | -1.137123757 | 3.884976949 | 4.8367193 | 0.0426123 | 0.530435329 |
| ENSMUSG00000000305 | Cdh4 | protein_coding | 0.625691198 | 5.535970234 | 4.8362952 | 0.0426203 | 0.530435329 |
| ENSMUSG00000047507 | Baiap3 | protein_coding | 0.785639145 | 6.667296313 | 4.8819778 | 0.0427537 | 0.53059314 |
| ENSMUSG00000029535 | Triap1 | protein_coding | 0.541254893 | 6.337030239 | 4.825349 | 0.0428275 | 0.53059314 |
| ENSMUSG00000042210 | Abhd14a | protein_coding | 0.685093402 | 6.311576668 | 4.8252635 | 0.0428547 | 0.53059314 |
| ENSMUSG00000040794 | C1qtnf4 | protein_coding | -1.363979314 | 3.310333493 | 4.8226852 | 0.0428781 | 0.53059314 |
| ENSMUSG00000037463 | Fbxo27 | protein_coding | -1.618194513 | 2.454209972 | 4.8208431 | 0.0429131 | 0.53059314 |
| ENSMUSG00000043004 | Gng2 | protein_coding | 0.536385063 | 7.764402545 | 4.8200065 | 0.042929 | 0.53059314 |
| ENSMUSG00000027406 | Idh3b | protein_coding | 0.477440609 | 9.161263292 | 4.8084785 | 0.0431491 | 0.532536426 |
| ENSMUSG00000031130 | Brs3 | protein_coding | 2.252748204 | 3.223265056 | 4.9392187 | 0.0431853 | 0.532536426 |
| ENSMUSG00000030204 | Ddx47 | protein_coding | 0.519727034 | 6.951213676 | 4.7985192 | 0.0433403 | 0.533835272 |
| ENSMUSG00000027287 | Snap23 | protein_coding | -1.028678726 | 4.287965476 | 4.7891448 | 0.0435211 | 0.534112568 |
| ENSMUSG00000020098 | Pcbd1 | protein_coding | 0.462157157 | 7.694197222 | 4.7890589 | 0.0435228 | 0.534112568 |
| ENSMUSG00000039759 | Thap3 | protein_coding | 0.553247605 | 6.023403703 | 4.7881825 | 0.0435398 | 0.534112568 |
| ENSMUSG00000085980 | Gm12408 | lincRNA | 1.39411544 | 2.757608371 | 4.785273 | 0.0435961 | 0.534112568 |
| ENSMUSG00000060992 | Copz1 | protein_coding | 0.468094496 | 8.602941946 | 4.782709 | 0.0436458 | 0.534112568 |
| ENSMUSG00000055862 | Izumo4 | protein_coding | 0.523795379 | 6.276003752 | 4.781291 | 0.0436733 | 0.534112568 |
| ENSMUSG00000025239 | Limd1 | protein_coding | -0.939507475 | 4.056360226 | 4.7793771 | 0.0437105 | 0.534112568 |
| ENSMUSG00000034021 | Pds5b | protein_coding | -0.540462695 | 6.033313339 | 4.7721289 | 0.0438516 | 0.534435448 |
| ENSMUSG00000104939 | Gm33651 | lincRNA | -1.381401436 | 3.545577896 | 4.7710325 | 0.0438988 | 0.534435448 |
| ENSMUSG00000030386 | Zfp606 | protein_coding | -1.117984767 | 3.567408348 | 4.7689559 | 0.0439136 | 0.534435448 |
| ENSMUSG00000071547 | Nt5dc2 | protein_coding | 0.984425556 | 3.340278492 | 4.7675827 | 0.0439404 | 0.534435448 |
| ENSMUSG00000021071 | Trim9 | protein_coding | -0.878538369 | 5.423972199 | 4.7666015 | 0.0439854 | 0.534435448 |
| ENSMUSG00000049760 | 2410015M20Rik | protein_coding | 0.510722066 | 8.724607971 | 4.7571961 | 0.0441441 | 0.53457889 |
| ENSMUSG00000053907 | Mat2a | protein_coding | -0.46626043 | 7.41689777 | 4.7556652 | 0.0441742 | 0.53457889 |
| ENSMUSG00000035566 | Pcdh17 | protein_coding | 0.432727341 | 6.755909387 | 4.7513279 | 0.0442596 | 0.53457889 |
| ENSMUSG00000025915 | Sgk3 | protein_coding | -0.711344069 | 5.458691514 | 4.7507455 | 0.0442711 | 0.53457889 |
| ENSMUSG00000049583 | Grm5 | protein_coding | -0.67326066 | 7.011997702 | 4.7513345 | 0.0442854 | 0.53457889 |
| ENSMUSG00000005354 | Txn2 | protein_coding | 0.469168621 | 9.312066701 | 4.7473259 | 0.0443386 | 0.53457889 |
| ENSMUSG00000058076 | Sdhc | protein_coding | 0.457354586 | 9.022960936 | 4.7469905 | 0.0443452 | 0.53457889 |
| ENSMUSG00000002107 | Celf2 | protein_coding | -0.492031601 | 7.824730066 | 4.7424847 | 0.0444344 | 0.535053755 |
| ENSMUSG00000037628 | Cdkn3 | protein_coding | 1.489586577 | 2.570795995 | 4.7359485 | 0.0445641 | 0.535296577 |
| ENSMUSG00000026851 | BC005624 | protein_coding | 0.455890006 | 7.926438961 | 4.7289941 | 0.0447026 | 0.535296577 |
| ENSMUSG00000028772 | Zcchc17 | protein_coding | 0.413917639 | 8.941094121 | 4.72708 | 0.0447408 | 0.535296577 |
| ENSMUSG00000001036 | Epn2 | protein_coding | -0.531374218 | 6.300248836 | 4.7251443 | 0.0447794 | 0.535296577 |
| ENSMUSG00000054079 | Utp18 | protein_coding | -0.801840062 | 4.102257036 | 4.7248842 | 0.0447846 | 0.535296577 |
| ENSMUSG00000041079 | Rwdd2b | protein_coding | -0.823693373 | 4.046549818 | 4.7240635 | 0.044801 | 0.535296577 |
| ENSMUSG00000034254 | Agpat1 | protein_coding | 0.592186184 | 5.640121654 | 4.7239641 | 0.044803 | 0.535296577 |
| ENSMUSG00000022443 | Myh9 | protein_coding | -1.038319555 | 3.503120734 | 4.7173509 | 0.0449355 | 0.535387792 |
| ENSMUSG00000025283 | Sat1 | protein_coding | -0.549368134 | 6.838112687 | 4.7165095 | 0.0449524 | 0.535387792 |
| ENSMUSG00000024963 | Dnajc4 | protein_coding | 0.608637262 | 6.0058377 | 4.7137687 | 0.0450075 | 0.535387792 |
| ENSMUSG00000051232 | Tmem199 | protein_coding | 0.45679235 | 7.900208875 | 4.7136518 | 0.0450098 | 0.535387792 |
| ENSMUSG00000064247 | Plcxd1 | protein_coding | 0.692216264 | 4.633945197 | 4.7071419 | 0.0451409 | 0.53635404 |
| ENSMUSG00000074591 | Ankub1 | protein_coding | -1.886378517 | 2.561682603 | 4.7012878 | 0.0452592 | 0.536860232 |
| ENSMUSG00000001524 | Gtf2h4 | protein_coding | 0.687658307 | 5.099909183 | 4.6988161 | 0.0453093 | 0.536860232 |
| ENSMUSG00000027660 | Skil | protein_coding | -0.517751377 | 6.513864919 | 4.6953095 | 0.0453804 | 0.536860232 |
| ENSMUSG00000005506 | Celf1 | protein_coding | -0.478240202 | 6.52656855 | 4.6947448 | 0.0453918 | 0.536860232 |
| ENSMUSG00000009470 | Tnpo1 | protein_coding | -0.85552955 | 4.766316474 | 4.6918155 | 0.0454514 | 0.536860232 |
| ENSMUSG00000020267 | Hint1 | protein_coding | 0.482394682 | 10.43714047 | 4.6898118 | 0.0454921 | 0.536860232 |
| ENSMUSG00000039959 | Hip1 | protein_coding | -0.738942838 | 4.775215907 | 4.6878042 | 0.045533 | 0.536860232 |
| ENSMUSG00000037364 | Srrt | protein_coding | 0.604717355 | 5.570223714 | 4.6762645 | 0.0457689 | 0.538575156 |
| ENSMUSG00000032413 | Rasa2 | protein_coding | -0.840251539 | 4.582664916 | 4.6757883 | 0.0457786 | 0.538575156 |
| ENSMUSG00000071855 | Ccdc112 | protein_coding | -0.981606845 | 4.039742645 | 4.6654874 | 0.0459904 | 0.540042322 |
| ENSMUSG00000039716 | Dock3 | protein_coding | -0.927130384 | 3.952430827 | 4.6606464 | 0.0460903 | 0.540042322 |
| ENSMUSG00000041660 | Bbox1 | protein_coding | -1.521963015 | 3.012470954 | 4.6554118 | 0.0461987 | 0.540042322 |
| ENSMUSG00000004285 | Atp6v1f | protein_coding | 0.464668101 | 9.236162641 | 4.6518451 | 0.0462727 | 0.540042322 |
| ENSMUSG00000048458 | Fam212b | protein_coding | -0.895310518 | 5.087369993 | 4.6506589 | 0.0462973 | 0.540042322 |
| ENSMUSG00000025151 | Maged1 | protein_coding | 0.672655628 | 10.0312578 | 4.6514768 | 0.0463066 | 0.540042322 |
| ENSMUSG00000031913 | Vps4a | protein_coding | -1.04601011 | 5.448352667 | 4.6987317 | 0.0463237 | 0.540042322 |
| ENSMUSG00000034480 | Diaph2 | protein_coding | -1.011529897 | 4.128041007 | 4.6467934 | 0.0463777 | 0.540042322 |
| ENSMUSG00000005871 | Apc | protein_coding | -0.72973705 | 5.403525281 | 4.6400434 | 0.0465184 | 0.540042322 |
| ENSMUSG00000020526 | Znhit3 | protein_coding | 0.452222199 | 7.147062682 | 4.6394984 | 0.0465298 | 0.540042322 |
| ENSMUSG00000034259 | Exosc4 | protein_coding | 0.537453038 | 6.135253347 | 4.6379922 | 0.0465613 | 0.540042322 |
| ENSMUSG00000016534 | Lamp2 | protein_coding | -0.594310871 | 6.080716892 | 4.6348785 | 0.0466265 | 0.540042322 |
| ENSMUSG00000024829 | Mrpl21 | protein_coding | 0.43716678 | 7.109861112 | 4.63071 | 0.0467139 | 0.540042322 |
| ENSMUSG00000045534 | Kcna5 | protein_coding | -1.041652607 | 3.796053901 | 4.6296614 | 0.0467359 | 0.540042322 |
| ENSMUSG00000033799 | Fam208b | protein_coding | -2.01957138 | 2.470023968 | 4.6233845 | 0.046868 | 0.540042322 |
| ENSMUSG00000031714 | Gab1 | protein_coding | -0.809484156 | 4.408208257 | 4.6222011 | 0.0468929 | 0.540042322 |
| ENSMUSG00000031709 | Tbc1d9 | protein_coding | -0.703347325 | 4.883991709 | 4.6169989 | 0.0470027 | 0.540042322 |
| ENSMUSG00000020230 | Prmt2 | protein_coding | 0.659527127 | 9.243598961 | 4.6187259 | 0.0470621 | 0.540042322 |
| ENSMUSG00000031937 | Vstm5 | protein_coding | 0.849662503 | 4.592154197 | 4.6138906 | 0.0470685 | 0.540042322 |
| ENSMUSG00000024900 | Cpt1a | protein_coding | -2.145222062 | 2.519854994 | 4.6125495 | 0.0470969 | 0.540042322 |
| ENSMUSG00000037493 | Cib2 | protein_coding | 0.633866525 | 5.405799481 | 4.6089573 | 0.0471731 | 0.540042322 |
| ENSMUSG00000049184 | Purg | protein_coding | -0.733431102 | 4.934903238 | 4.6088403 | 0.0471756 | 0.540042322 |

| ensembl_gene_id | external_gene_name | gene_biotype | log2 (fold change) | log2 (counts per million) | F-value | p-value | False discovery rate |
|---|---|---|---|---|---|---|---|
| ENSMUSG00000100826 | Snhg14 | processed_transcript | -0.59445792 | 6.081352785 | 4.6072726 | 0.0472089 | 0.540042322 |
| ENSMUSG00000002820 | Atg4d | protein_coding | -1.185395865 | 3.81406085 | 4.6008472 | 0.0473457 | 0.540042322 |
| ENSMUSG00000030337 | Vamp1 | protein_coding | 0.640026774 | 7.451851913 | 4.5974065 | 0.0475153 | 0.540042322 |
| ENSMUSG00000059070 | Rpl18 | protein_coding | 0.569012234 | 6.174859405 | 4.5891781 | 0.0475952 | 0.540042322 |
| ENSMUSG00000042380 | Smim12 | protein_coding | 0.48972208 | 7.240841027 | 4.5874763 | 0.0476318 | 0.540042322 |
| ENSMUSG00000020354 | Sgcd | protein_coding | -1.636698796 | 2.665280259 | 4.5843191 | 0.0476996 | 0.540042322 |
| ENSMUSG00000039168 | Dap | protein_coding | -0.83240513 | 4.225448907 | 4.5837748 | 0.0477113 | 0.540042322 |
| ENSMUSG00000032083 | Apoa1 | protein_coding | 1.138491678 | 3.085571856 | 4.5803807 | 0.0477844 | 0.540042322 |
| ENSMUSG00000044252 | Osbpl1a | protein_coding | -0.478912894 | 6.867745508 | 4.5794862 | 0.0478037 | 0.540042322 |
| ENSMUSG00000097326 | A330048O09Rik | lincRNA | -1.372613481 | 3.431106651 | 4.5791311 | 0.0478113 | 0.540042322 |
| ENSMUSG00000068696 | Gpr88 | protein_coding | -1.36804419 | 6.335451933 | 4.9093233 | 0.047832 | 0.540042322 |
| ENSMUSG00000114309 | AC163038.1 | bidirectional_promote | 1.43562027 | 3.07934206 | 4.5748805 | 0.0479031 | 0.540042322 |
| ENSMUSG00000025739 | Gng13 | protein_coding | 0.774305884 | 4.249985549 | 4.5743023 | 0.0479156 | 0.540042322 |
| ENSMUSG00000030275 | Etnk1 | protein_coding | -1.921548713 | 2.811776747 | 4.5751092 | 0.0479216 | 0.540042322 |
| ENSMUSG00000006728 | Cdk4 | protein_coding | 0.459672686 | 7.036813166 | 4.5693817 | 0.0480221 | 0.540042322 |
| ENSMUSG00000054013 | Tmem179 | protein_coding | 0.532787925 | 7.989735247 | 4.5687609 | 0.0480356 | 0.540042322 |
| ENSMUSG00000057229 | Atp5sl | protein_coding | 0.533199153 | 6.897571732 | 4.5678004 | 0.0480564 | 0.540042322 |
| ENSMUSG00000062908 | Acadm | protein_coding | -0.580330893 | 5.884754575 | 4.5664142 | 0.0480865 | 0.540042322 |
| ENSMUSG00000020189 | Osbpl8 | protein_coding | -0.676208636 | 5.708584989 | 4.5662833 | 0.0480893 | 0.540042322 |
| ENSMUSG00000002015 | Bcap31 | protein_coding | 0.461363644 | 7.856845416 | 4.565775 | 0.0481004 | 0.540042322 |
| ENSMUSG00000032641 | Gpr19 | protein_coding | 0.534748006 | 5.796899798 | 4.562925 | 0.0481623 | 0.540042322 |
| ENSMUSG00000006941 | Eif1b | protein_coding | 0.442550925 | 8.260969803 | 4.5626866 | 0.0481675 | 0.540042322 |
| ENSMUSG00000024858 | Grk2 | protein_coding | 0.551273345 | 6.36618441 | 4.5598947 | 0.0482283 | 0.540042322 |
| ENSMUSG00000078348 | Sf3b5 | protein_coding | 0.466111036 | 8.019502921 | 4.5595192 | 0.0482364 | 0.540042322 |
| ENSMUSG00000043411 | Usp48 | protein_coding | -0.755862993 | 4.625503793 | 4.5553867 | 0.0483266 | 0.540042322 |
| ENSMUSG00000018166 | Erbb3 | protein_coding | -1.991339824 | 2.661932717 | 4.5544181 | 0.0483477 | 0.540042322 |
| ENSMUSG00000025724 | Sec11a | protein_coding | 0.540687322 | 6.941212833 | 4.549229 | 0.0484613 | 0.540042322 |
| ENSMUSG00000032667 | Pon2 | protein_coding | -0.733784024 | 5.017661309 | 4.546873 | 0.0485129 | 0.540042322 |
| ENSMUSG00000039740 | Alg2 | protein_coding | 0.520725224 | 7.151120417 | 4.5449559 | 0.048555 | 0.540042322 |
| ENSMUSG00000028033 | Kcnq5 | protein_coding | -0.793253249 | 4.809074714 | 4.5431732 | 0.0485941 | 0.540042322 |
| ENSMUSG00000006262 | Mob1b | protein_coding | -0.8109891 | 3.822096656 | 4.5421777 | 0.048616 | 0.540042322 |
| ENSMUSG00000026860 | Sh3glb2 | protein_coding | 0.523300179 | 8.027200636 | 4.5407471 | 0.0486475 | 0.540042322 |
| ENSMUSG00000022635 | Zcrb1 | protein_coding | -0.697641114 | 5.522701848 | 4.5368548 | 0.0487332 | 0.540042322 |
| ENSMUSG00000032051 | Fdx1 | protein_coding | -1.038795034 | 3.43317196 | 4.5341003 | 0.048794 | 0.540042322 |
| ENSMUSG00000011589 | Fsd1 | protein_coding | 0.601460098 | 7.226805919 | 4.5331487 | 0.0488417 | 0.540042322 |
| ENSMUSG00000026728 | Vim | protein_coding | -1.021978451 | 4.265749196 | 4.5310326 | 0.0488618 | 0.540042322 |
| ENSMUSG00000021287 | Xrcc3 | protein_coding | 1.393822991 | 2.800430518 | 4.5308199 | 0.0488665 | 0.540042322 |
| ENSMUSG00000025102 | 3110040N11Rik | protein_coding | 0.52620249 | 6.076094014 | 4.5188964 | 0.049131 | 0.542005849 |
| ENSMUSG00000028692 | Akr1a1 | protein_coding | 0.47614752 | 10.57220494 | 4.5154381 | 0.0492081 | 0.542005849 |
| ENSMUSG00000028906 | Epb41 | protein_coding | -0.702831591 | 4.773790182 | 4.5140079 | 0.04924 | 0.542005849 |
| ENSMUSG00000030329 | Pianp | protein_coding | 0.579099323 | 6.795804784 | 4.5137486 | 0.0492458 | 0.542005849 |
| ENSMUSG00000052369 | Tmem106c | protein_coding | 0.424462865 | 7.361881729 | 4.5062948 | 0.0494124 | 0.542161186 |
| ENSMUSG00000000416 | Cttnbp2 | protein_coding | -0.81572541 | 5.046223057 | 4.5032338 | 0.0494811 | 0.542161186 |
| ENSMUSG00000019945 | 1700040L02Rik | protein_coding | -1.028938074 | 3.569373878 | 4.5004273 | 0.0495441 | 0.542161186 |
| ENSMUSG00000026678 | Rgs5 | protein_coding | -1.014985273 | 4.246764995 | 4.5003553 | 0.0495457 | 0.542161186 |
| ENSMUSG00000038845 | Phb | protein_coding | 0.603103846 | 6.715993829 | 4.501372 | 0.0495497 | 0.542161186 |
| ENSMUSG00000011831 | Evi5 | protein_coding | -0.76969092 | 4.949072978 | 4.4993589 | 0.0495681 | 0.542161186 |
| ENSMUSG00000011884 | Gltp | protein_coding | -0.744749941 | 6.231484029 | 4.4983101 | 0.0496186 | 0.542161186 |
| ENSMUSG00000036067 | Slc2a6 | protein_coding | 0.949500186 | 4.741101066 | 4.4962943 | 0.0496639 | 0.542161186 |
| ENSMUSG00000046480 | Scn4b | protein_coding | -0.897234963 | 5.135063588 | 4.4940888 | 0.0497137 | 0.542161186 |
| ENSMUSG00000061904 | Slc25a3 | protein_coding | -0.495689649 | 8.245816559 | 4.4869871 | 0.0498473 | 0.542813147 |
| ENSMUSG00000030189 | Ybx3 | protein_coding | -0.948650658 | 3.853006636 | 4.4857886 | 0.0498744 | 0.542813147 |
| ENSMUSG00000019890 | Nts | protein_coding | 1.258557157 | 7.780597208 | 4.8717817 | 0.0499758 | 0.542966182 |
| ENSMUSG00000001445 | Mrpl10 | protein_coding | 0.538579731 | 6.93874724 | 4.4807158 | 0.0499894 | 0.542966182 |
| ENSMUSG00000030878 | Cdr2 | protein_coding | 0.998554296 | 3.563215937 | 4.475232 | 0.0501142 | 0.543472995 |
| ENSMUSG00000026788 | Zbtb43 | protein_coding | -0.758426775 | 4.339041362 | 4.4710364 | 0.0502098 | 0.543472995 |
| ENSMUSG00000025364 | Pa2g4 | protein_coding | 0.503453532 | 7.354736714 | 4.4703127 | 0.0502264 | 0.543472995 |
| ENSMUSG00000001380 | Hars | protein_coding | 0.493057689 | 8.238366538 | 4.4697907 | 0.0502383 | 0.543472995 |
| ENSMUSG00000028059 | Arhgef2 | protein_coding | -0.830834263 | 4.502508929 | 4.4620837 | 0.0504147 | 0.54483313 |
| ENSMUSG00000007987 | Ift22 | protein_coding | 0.42022011 | 8.388343311 | 4.4540193 | 0.0506 | 0.545181739 |
| ENSMUSG00000027575 | Arfgap1 | protein_coding | 0.509592922 | 6.255567342 | 4.4522595 | 0.0506406 | 0.545181739 |
| ENSMUSG00000035274 | Tpbg | protein_coding | -1.420717318 | 3.203808165 | 4.4488646 | 0.0507189 | 0.545181739 |
| ENSMUSG00000044881 | Coa4 | protein_coding | 0.732566958 | 4.985126148 | 4.4480911 | 0.0507368 | 0.545181739 |
| ENSMUSG00000023961 | Enpp4 | protein_coding | -0.721822816 | 5.192161877 | 4.4466026 | 0.0507712 | 0.545181739 |
| ENSMUSG00000030838 | Ush1c | protein_coding | 2.467709354 | 2.411824723 | 4.4633099 | 0.0508963 | 0.545181739 |
| ENSMUSG00000059213 | Ddn | protein_coding | -1.07905553 | 5.670490936 | 4.5336018 | 0.0509204 | 0.545181739 |
| ENSMUSG00000007603 | Dus3l | protein_coding | 0.560988392 | 6.250032188 | 4.4396911 | 0.0509313 | 0.545181739 |
| ENSMUSG00000023367 | Tmem176a | protein_coding | -0.794976717 | 5.30402105 | 4.4387881 | 0.0509523 | 0.545181739 |
| ENSMUSG00000054612 | Mgmt | protein_coding | 1.134906743 | 3.160805417 | 4.4361577 | 0.0510134 | 0.545181739 |
| ENSMUSG00000015980 | Lrrc27 | protein_coding | 0.917367654 | 4.809992088 | 4.4369636 | 0.0510217 | 0.545181739 |
| ENSMUSG00000020794 | Ube2g1 | protein_coding | -0.576824584 | 5.654516639 | 4.434354 | 0.0510553 | 0.545181739 |
| ENSMUSG00000036820 | Amdhd2 | protein_coding | 0.728351843 | 5.454167321 | 4.4302071 | 0.051152 | 0.54567159 |
| ENSMUSG00000091985 | Gm17354 | lincRNA | -1.610832094 | 2.537460446 | 4.4258732 | 0.0512532 | 0.54571851 |
| ENSMUSG00000031762 | Mt2 | protein_coding | -1.034975179 | 5.28709843 | 4.4692674 | 0.0512962 | 0.54571851 |
| ENSMUSG00000078958 | Atp6ap1l | protein_coding | -1.129309814 | 3.491809677 | 4.4205742 | 0.0513772 | 0.54571851 |
| ENSMUSG00000000127 | Fer | protein_coding | -0.819347906 | 4.309187938 | 4.4202422 | 0.051385 | 0.54571851 |
| ENSMUSG00000015095 | Fbxw5 | protein_coding | 0.490480564 | 7.437499802 | 4.419172 | 0.0514101 | 0.54571851 |
| ENSMUSG00000031983 | 2310022B05Rik | protein_coding | -0.777503459 | 4.985790845 | 4.4162635 | 0.0514784 | 0.545904534 |

| ensembl_gene_id | external_gene_name | gene_biotype | log2 (fold change) | log2 (counts per million) | F-value | p-value | False discovery rate |
|---|---|---|---|---|---|---|---|
| ENSMUSG00000050558 | Prokr2 | protein_coding | -1.765472473 | 2.716448841 | 4.4099897 | 0.0516261 | 0.54601582 |
| ENSMUSG00000024974 | Smc3 | protein_coding | -0.648186382 | 5.852107548 | 4.4079326 | 0.0516746 | 0.54601582 |
| ENSMUSG00000036916 | Zfp280c | protein_coding | -1.280295267 | 3.532355404 | 4.4044063 | 0.0517579 | 0.54601582 |
| ENSMUSG00000058589 | Anks1b | protein_coding | -0.405675682 | 7.130867999 | 4.4020674 | 0.0518132 | 0.54601582 |
| ENSMUSG00000036298 | Slc2a13 | protein_coding | -0.799442731 | 5.510649288 | 4.4028575 | 0.0518217 | 0.54601582 |
| ENSMUSG00000002808 | Epdr1 | protein_coding | -0.54413599 | 6.191063094 | 4.3991529 | 0.0518823 | 0.54601582 |
| ENSMUSG00000040813 | Tex264 | protein_coding | 0.455535775 | 6.528082836 | 4.3961895 | 0.0519526 | 0.54601582 |
| ENSMUSG00000001729 | Akt1 | protein_coding | 0.4424476 | 6.455986 | 4.3944164 | 0.0519948 | 0.54601582 |
| ENSMUSG00000030905 | Crym | protein_coding | -0.925416066 | 7.711231282 | 4.5070022 | 0.0520105 | 0.54601582 |
| ENSMUSG00000058638 | Zfp110 | protein_coding | -1.170475974 | 3.539110823 | 4.3866924 | 0.0521788 | 0.54601582 |
| ENSMUSG00000029223 | Uchl1 | protein_coding | 0.440732674 | 10.33781782 | 4.3791125 | 0.0523601 | 0.54601582 |
| ENSMUSG00000018865 | Sult4a1 | protein_coding | 0.52809111 | 9.330066523 | 4.3768024 | 0.0524155 | 0.54601582 |
| ENSMUSG00000025268 | Maged2 | protein_coding | 0.634384144 | 7.775685718 | 4.3820443 | 0.052436 | 0.54601582 |
| ENSMUSG00000026649 | Cfap126 | protein_coding | -0.903960342 | 4.686892357 | 4.3750195 | 0.0524583 | 0.54601582 |
| ENSMUSG00000056870 | Gulp1 | protein_coding | -1.251615947 | 3.06054886 | 4.3714643 | 0.0525437 | 0.54601582 |
| ENSMUSG00000042505 | Sdhaf3 | protein_coding | 0.508781264 | 6.69178615 | 4.3701795 | 0.0525747 | 0.54601582 |
| ENSMUSG00000043831 | Lysmd4 | protein_coding | 0.452117573 | 6.736433411 | 4.3577108 | 0.0528759 | 0.54601582 |
| ENSMUSG00000046364 | Rpl27a | protein_coding | 0.53935754 | 6.757894458 | 4.3556623 | 0.0529256 | 0.54601582 |
| ENSMUSG00000010095 | Slc3a2 | protein_coding | -0.76549148 | 5.734786904 | 4.3539229 | 0.0529952 | 0.54601582 |
| ENSMUSG00000026383 | Epb41l5 | protein_coding | -0.975410554 | 3.350534585 | 4.3522341 | 0.0530089 | 0.54601582 |
| ENSMUSG00000022884 | Eif4a2 | protein_coding | 0.506131269 | 10.53247066 | 4.3498534 | 0.0530668 | 0.54601582 |
| ENSMUSG00000069601 | Ank3 | protein_coding | -0.483660132 | 6.505575988 | 4.3485954 | 0.0530974 | 0.54601582 |
| ENSMUSG00000029528 | Pxn | protein_coding | -1.726714045 | 2.558738636 | 4.3443107 | 0.0532019 | 0.54601582 |
| ENSMUSG00000049299 | Trappc1 | protein_coding | 0.47993732 | 7.355933209 | 4.3427624 | 0.0532398 | 0.54601582 |
| ENSMUSG00000058715 | Fcer1g | protein_coding | -1.060923184 | 4.652242436 | 4.3452722 | 0.053278 | 0.54601582 |
| ENSMUSG00000029344 | Tpst2 | protein_coding | -1.048847148 | 3.349081953 | 4.3410992 | 0.0532804 | 0.54601582 |
| ENSMUSG00000024924 | Vldlr | protein_coding | -0.810181771 | 5.759781989 | 4.3403687 | 0.0533257 | 0.54601582 |
| ENSMUSG00000013089 | Etv5 | protein_coding | -0.831878807 | 4.202614693 | 4.3326047 | 0.0534886 | 0.54601582 |
| ENSMUSG00000001025 | S100a6 | protein_coding | -0.817229974 | 5.360026536 | 4.328219 | 0.0535965 | 0.54601582 |
| ENSMUSG00000079426 | Arpc4 | protein_coding | 0.465887395 | 8.860484356 | 4.3279166 | 0.053604 | 0.54601582 |
| ENSMUSG00000005823 | Gpr108 | protein_coding | 0.651490105 | 5.589183926 | 4.3272727 | 0.0536198 | 0.54601582 |
| ENSMUSG00000028565 | Nfia | protein_coding | -0.597492554 | 6.998809161 | 4.3266745 | 0.053662 | 0.54601582 |
| ENSMUSG00000022076 | Klhl1 | protein_coding | 0.90289345 | 4.332187391 | 4.3231667 | 0.0537211 | 0.54601582 |
| ENSMUSG00000030095 | Tmem43 | protein_coding | 0.574409585 | 5.636489023 | 4.3201545 | 0.0537956 | 0.54601582 |
| ENSMUSG00000032560 | Dnajc13 | protein_coding | -0.936097921 | 4.102390063 | 4.3169067 | 0.053876 | 0.54601582 |
| ENSMUSG00000073725 | Lmbrd1 | protein_coding | -0.481671694 | 6.696811724 | 4.3165873 | 0.0538839 | 0.54601582 |
| ENSMUSG00000038417 | Fig4 | protein_coding | -0.791521026 | 4.988663656 | 4.3159055 | 0.0539008 | 0.54601582 |
| ENSMUSG00000056459 | Zbtb25 | protein_coding | -1.101085753 | 3.542563016 | 4.311325 | 0.0540145 | 0.54601582 |
| ENSMUSG00000019303 | Psmc3ip | protein_coding | -0.926880395 | 3.827689081 | 4.3104086 | 0.0540372 | 0.54601582 |
| ENSMUSG00000060002 | Chpt1 | protein_coding | -0.89392781 | 3.839606672 | 4.3094876 | 0.0540601 | 0.54601582 |
| ENSMUSG00000028969 | Cdk5 | protein_coding | 0.43052597 | 8.546438567 | 4.3066419 | 0.054131 | 0.54601582 |
| ENSMUSG00000024309 | Pfdn6 | protein_coding | 0.398058579 | 8.166682519 | 4.306532 | 0.0541337 | 0.54601582 |
| ENSMUSG00000034667 | Xpot | protein_coding | -0.733461676 | 4.929697942 | 4.3053386 | 0.0541635 | 0.54601582 |
| ENSMUSG00000028081 | Rps3a1 | protein_coding | 0.567629534 | 7.926400516 | 4.3062716 | 0.0541677 | 0.54601582 |
| ENSMUSG00000020083 | 2010107G23Rik | protein_coding | 0.47391168 | 6.60916479 | 4.3047629 | 0.0541778 | 0.54601582 |
| ENSMUSG00000021374 | Nup153 | protein_coding | -1.033494209 | 3.758688117 | 4.3033397 | 0.0542133 | 0.54601582 |
| ENSMUSG00000100147 | 1700047M11Rik | lincRNA | -0.836181399 | 4.785041665 | 4.3032423 | 0.0542157 | 0.54601582 |
| ENSMUSG00000029103 | Lrpap1 | protein_coding | 0.413560911 | 9.146954222 | 4.3029224 | 0.0542237 | 0.54601582 |
| ENSMUSG00000037071 | Scd1 | protein_coding | -0.719336426 | 5.692765077 | 4.302532 | 0.0542335 | 0.54601582 |
| ENSMUSG00000058672 | Tubb2a | protein_coding | 0.547373949 | 9.439810375 | 4.3019671 | 0.0542476 | 0.54601582 |
| ENSMUSG00000037902 | Sirpa | protein_coding | -0.490902322 | 6.931019803 | 4.3002129 | 0.0542914 | 0.54601582 |
| ENSMUSG00000037876 | Jmjd1c | protein_coding | -0.822875626 | 4.82454241 | 4.2983364 | 0.0543384 | 0.54601582 |
| ENSMUSG00000029198 | Grpel1 | protein_coding | 0.434493535 | 8.453275361 | 4.295881 | 0.0543998 | 0.54601582 |
| ENSMUSG00000027820 | Mme | protein_coding | -1.481420773 | 2.654254081 | 4.2952622 | 0.0544153 | 0.54601582 |
| ENSMUSG00000040659 | Efhd2 | protein_coding | 0.515047712 | 6.788462381 | 4.2951593 | 0.0544179 | 0.54601582 |
| ENSMUSG00000020681 | Ace | protein_coding | -1.304561718 | 3.3613673 | 4.2948837 | 0.0544248 | 0.54601582 |
| ENSMUSG00000055320 | Tead1 | protein_coding | -0.879713163 | 3.646694706 | 4.2939646 | 0.0544479 | 0.54601582 |
| ENSMUSG00000022528 | Hes1 | protein_coding | -1.258536482 | 2.960252542 | 4.2906351 | 0.0545315 | 0.54601582 |
| ENSMUSG00000060301 | 2610008E11Rik | protein_coding | -0.956312171 | 3.531408418 | 4.2890559 | 0.0545712 | 0.54601582 |
| ENSMUSG00000028412 | Slc44a1 | protein_coding | -0.679833153 | 5.617019208 | 4.2872282 | 0.0546172 | 0.54601582 |
| ENSMUSG00000005378 | Wbscr22 | protein_coding | 0.482745163 | 6.300554617 | 4.2851498 | 0.0546695 | 0.54601582 |
| ENSMUSG00000063428 | Ddo | protein_coding | -0.769515745 | 4.108612222 | 4.28454 | 0.0546849 | 0.54601582 |
| ENSMUSG00000067787 | Blcap | protein_coding | 0.452282927 | 8.873666099 | 4.2828808 | 0.0547267 | 0.54601582 |
| ENSMUSG00000039114 | Nrn1 | protein_coding | -1.048575242 | 6.156109319 | 4.3892323 | 0.0547734 | 0.54601582 |
| ENSMUSG00000026587 | Astn1 | protein_coding | -0.551116962 | 6.196100042 | 4.2802441 | 0.0547933 | 0.54601582 |
| ENSMUSG00000027375 | Mal | protein_coding | -0.513392448 | 7.874490494 | 4.2783876 | 0.0548402 | 0.54601582 |
| ENSMUSG00000110834 | CAAA01180111.2 | lincRNA | 0.741113368 | 4.268034083 | 4.2746789 | 0.0549341 | 0.546129442 |
| ENSMUSG00000039621 | Prex1 | protein_coding | -1.112840445 | 4.392303219 | 4.2761489 | 0.0549974 | 0.546129442 |
| ENSMUSG00000067851 | Arfgef1 | protein_coding | -0.702001022 | 5.310221381 | 4.2719238 | 0.055004 | 0.546129442 |
| ENSMUSG00000007458 | M6pr | protein_coding | 0.485992763 | 7.138541407 | 4.2621162 | 0.0552536 | 0.548101478 |
| ENSMUSG00000015852 | Fcrls | protein_coding | -1.85463014 | 3.534348842 | 4.3444276 | 0.0553687 | 0.548603659 |
| ENSMUSG00000063297 | Luzp2 | protein_coding | -0.654644333 | 6.133680057 | 4.2561428 | 0.0554063 | 0.548603659 |
| ENSMUSG00000075706 | Gpx4 | protein_coding | 0.450115629 | 8.737888227 | 4.2479071 | 0.0556176 | 0.54942072 |
| ENSMUSG00000030323 | Ift122 | protein_coding | -0.749971825 | 4.801334896 | 4.2475122 | 0.0556277 | 0.54942072 |
| ENSMUSG00000021069 | Pygl | protein_coding | -1.820466958 | 2.657915326 | 4.2469538 | 0.0556421 | 0.54942072 |
| ENSMUSG00000010045 | Tmem115 | protein_coding | 0.583765811 | 5.388791014 | 4.2414538 | 0.0557838 | 0.550314543 |
| ENSMUSG00000015090 | Ptgds | protein_coding | -0.683826405 | 9.368961098 | 4.2577508 | 0.055889 | 0.550671528 |
| ENSMUSG00000002900 | Lamb1 | protein_coding | -1.175572977 | 3.348845359 | 4.2342644 | 0.0559696 | 0.550671528 |

| ensembl_gene_id | external_gene_name | gene_biotype | log2 (fold change) | log2 (counts per million) | F-value | p-value | False discovery rate |
|---|---|---|---|---|---|---|---|
| ENSMUSG00000005892 | Trh | protein_coding | -0.745392196 | 7.312990643 | 4.2893763 | 0.0559783 | 0.550671528 |
| ENSMUSG00000022216 | Psme1 | protein_coding | 0.463371716 | 7.421260387 | 4.2286924 | 0.0561142 | 0.550671528 |
| ENSMUSG00000029328 | Hnrnpdl | protein_coding | -0.443544589 | 6.69906303 | 4.2275362 | 0.0561442 | 0.550671528 |
| ENSMUSG00000027405 | Nop56 | protein_coding | 0.508743251 | 7.276843481 | 4.2260261 | 0.0561835 | 0.550671528 |
| ENSMUSG00000087651 | 1500009L16Rik | protein_coding | 0.470669699 | 6.322882621 | 4.2249532 | 0.0562114 | 0.550671528 |
| ENSMUSG00000077450 | Rab11b | protein_coding | 0.501762649 | 7.827942109 | 4.2239362 | 0.0562379 | 0.550671528 |
| ENSMUSG00000026826 | Nr4a2 | protein_coding | -1.438946458 | 2.621287073 | 4.2214166 | 0.0563036 | 0.550671528 |
| ENSMUSG00000019923 | Zwint | protein_coding | 0.536348748 | 12.04740912 | 4.2186207 | 0.0563765 | 0.550671528 |
| ENSMUSG00000060803 | Gstp1 | protein_coding | 0.414779492 | 8.774563866 | 4.2183627 | 0.0563833 | 0.550671528 |
| ENSMUSG00000022475 | Hdac7 | protein_coding | 0.962099162 | 4.279906756 | 4.2136764 | 0.0565059 | 0.550896968 |
| ENSMUSG00000026469 | Xpr1 | protein_coding | -0.511965918 | 6.068042782 | 4.2135632 | 0.0565088 | 0.550896968 |
| ENSMUSG00000053025 | Sv2b | protein_coding | -0.665261664 | 5.410793827 | 4.2097643 | 0.0566084 | 0.550961883 |
| ENSMUSG00000035198 | Tubg1 | protein_coding | 0.653501951 | 6.697199905 | 4.2139206 | 0.0566297 | 0.550961883 |
| ENSMUSG00000026836 | Acvr1 | protein_coding | -0.953293246 | 3.561656619 | 4.2074505 | 0.0566692 | 0.550961883 |
| ENSMUSG00000041540 | Sox5 | protein_coding | -0.861315779 | 4.527138898 | 4.1996256 | 0.0568753 | 0.552271263 |
| ENSMUSG00000057406 | Nsd2 | protein_coding | -0.587431761 | 5.332635908 | 4.198439 | 0.0569066 | 0.552271263 |
| ENSMUSG00000032606 | Nicn1 | protein_coding | 0.465501755 | 8.336555782 | 4.1890223 | 0.0571559 | 0.55237738 |
| ENSMUSG00000041828 | Abca8a | protein_coding | 1.193549507 | 3.14266088 | 4.1880148 | 0.0571826 | 0.55237738 |
| ENSMUSG00000004451 | Ralb | protein_coding | 0.632291862 | 4.742937865 | 4.1827761 | 0.057322 | 0.55237738 |
| ENSMUSG00000037664 | Cdkn1c | protein_coding | -1.911382581 | 2.328990306 | 4.181929 | 0.0573445 | 0.55237738 |
| ENSMUSG00000038369 | Ncoa6 | protein_coding | -0.916546856 | 4.480358588 | 4.177552 | 0.0574613 | 0.55237738 |
| ENSMUSG00000025381 | Cnpy2 | protein_coding | 0.427386224 | 7.272330767 | 4.1751912 | 0.0575244 | 0.55237738 |
| ENSMUSG00000023186 | Vwa5a | protein_coding | -1.285212249 | 2.970044968 | 4.1748725 | 0.0575329 | 0.55237738 |
| ENSMUSG00000006403 | Adamts4 | protein_coding | -1.234157941 | 3.873262184 | 4.1747372 | 0.0575645 | 0.55237738 |
| ENSMUSG00000067430 | Zfp763 | protein_coding | -0.860605328 | 4.197765692 | 4.1728184 | 0.0575879 | 0.55237738 |
| ENSMUSG00000049154 | Fam183b | protein_coding | -0.512653623 | 5.985182853 | 4.1721602 | 0.0576055 | 0.55237738 |
| ENSMUSG00000027312 | Atrn | protein_coding | -0.833367429 | 5.29521595 | 4.1718128 | 0.0576428 | 0.55237738 |
| ENSMUSG00000070283 | Ndufaf3 | protein_coding | 0.446152321 | 8.101443415 | 4.1668645 | 0.0577476 | 0.55237738 |
| ENSMUSG00000024059 | Clip4 | protein_coding | -0.570875095 | 5.871477717 | 4.1663294 | 0.0577619 | 0.55237738 |
| ENSMUSG00000025855 | Prkar1b | protein_coding | 0.661464204 | 10.12034065 | 4.1718695 | 0.0577635 | 0.55237738 |
| ENSMUSG00000008153 | Clstn3 | protein_coding | 0.482592555 | 8.246138266 | 4.165648 | 0.0577803 | 0.55237738 |
| ENSMUSG00000044465 | Fam160a2 | protein_coding | -0.741995802 | 4.499321294 | 4.1643378 | 0.0578155 | 0.55237738 |
| ENSMUSG00000031210 | Gpr165 | protein_coding | 1.233857871 | 3.673757385 | 4.1648139 | 0.0578307 | 0.55237738 |
| ENSMUSG00000036751 | Cox6b1 | protein_coding | 0.508242728 | 10.25496019 | 4.1633461 | 0.0578422 | 0.55237738 |
| ENSMUSG00000027673 | Ndufb5 | protein_coding | 0.442285599 | 9.868636205 | 4.160943 | 0.0579069 | 0.552504764 |
| ENSMUSG00000029772 | Ahcyl2 | protein_coding | -0.555173485 | 5.592556027 | 4.156755 | 0.0580199 | 0.553092101 |
| ENSMUSG00000070866 | Zfp804a | protein_coding | -0.905627684 | 4.426000685 | 4.1517158 | 0.0581562 | 0.553900393 |
| ENSMUSG00000044629 | Cnrip1 | protein_coding | 0.402938869 | 7.54233379 | 4.1459893 | 0.0583115 | 0.553974417 |
| ENSMUSG00000059291 | Rpl11 | protein_coding | 0.679929857 | 4.430452932 | 4.1426621 | 0.058402 | 0.553974417 |
| ENSMUSG00000096956 | Snhg18 | lincRNA | -1.644930592 | 2.863060401 | 4.1403674 | 0.0584645 | 0.553974417 |
| ENSMUSG00000040548 | Tex2 | protein_coding | 0.479442761 | 6.582425221 | 4.1351317 | 0.0586074 | 0.553974417 |
| ENSMUSG00000031409 | Tceal6 | protein_coding | 0.444156107 | 6.238640912 | 4.1327787 | 0.0586718 | 0.553974417 |
| ENSMUSG00000034488 | Edil3 | protein_coding | -0.446863749 | 6.477288724 | 4.1323345 | 0.0586839 | 0.553974417 |
| ENSMUSG00000040025 | Ythdf2 | protein_coding | -0.492754184 | 6.242873669 | 4.13231 | 0.0586846 | 0.553974417 |
| ENSMUSG00000026028 | Trak2 | protein_coding | -0.767720812 | 5.226720733 | 4.1275395 | 0.0588153 | 0.553974417 |
| ENSMUSG00000040462 | Os9 | protein_coding | 0.420635836 | 6.832922169 | 4.1274533 | 0.0588177 | 0.553974417 |
| ENSMUSG00000034708 | Grn | protein_coding | 0.528612397 | 5.539982496 | 4.1253151 | 0.0588764 | 0.553974417 |
| ENSMUSG00000032640 | Chsy1 | protein_coding | -0.864092076 | 3.73724095 | 4.1219865 | 0.058968 | 0.553974417 |
| ENSMUSG00000047409 | Ctdspl | protein_coding | -0.662596647 | 4.700609006 | 4.1215034 | 0.0589813 | 0.553974417 |
| ENSMUSG00000039323 | Igfbp2 | protein_coding | -1.399726399 | 3.213648607 | 4.1212101 | 0.0589893 | 0.553974417 |
| ENSMUSG00000097392 | D930016D06Rik | lincRNA | -1.627615482 | 2.721452666 | 4.1200167 | 0.0590222 | 0.553974417 |
| ENSMUSG00000020137 | Thap2 | protein_coding | -1.024374817 | 3.538827538 | 4.119691 | 0.0590312 | 0.553974417 |
| ENSMUSG00000031344 | Gabrq | protein_coding | 0.913226746 | 4.170744822 | 4.1134457 | 0.0592036 | 0.553974417 |
| ENSMUSG00000019947 | Arid5b | protein_coding | -0.682059194 | 5.545585943 | 4.1116523 | 0.0592532 | 0.553974417 |
| ENSMUSG00000035493 | Tgfbi | protein_coding | -1.940937199 | 2.223095746 | 4.1094176 | 0.0593151 | 0.553974417 |
| ENSMUSG00000028521 | Slc35d1 | protein_coding | -1.068021083 | 3.083161886 | 4.109375 | 0.0593163 | 0.553974417 |
| ENSMUSG00000007617 | Homer1 | protein_coding | -0.651408186 | 5.929049542 | 4.1091395 | 0.0593228 | 0.553974417 |
| ENSMUSG00000042228 | Lyn | protein_coding | -1.990826351 | 2.375049205 | 4.1060325 | 0.059409 | 0.553974417 |
| ENSMUSG00000040459 | Arglu1 | protein_coding | -0.545956138 | 7.490944732 | 4.1058654 | 0.0594137 | 0.553974417 |
| ENSMUSG00000030127 | Cops7a | protein_coding | 0.418784288 | 8.366976052 | 4.1024473 | 0.0595087 | 0.553974417 |
| ENSMUSG00000021895 | Arhgef3 | protein_coding | -0.629273697 | 5.025681311 | 4.102143 | 0.0595172 | 0.553974417 |
| ENSMUSG00000027199 | Gatm | protein_coding | -0.528383188 | 8.833057624 | 4.1004674 | 0.0595638 | 0.553974417 |
| ENSMUSG00000053062 | Jam2 | protein_coding | -0.806904906 | 3.995873277 | 4.0999283 | 0.0595788 | 0.553974417 |
| ENSMUSG00000015396 | Cd83 | protein_coding | 0.567680833 | 6.530667424 | 4.0966008 | 0.0596716 | 0.553974417 |
| ENSMUSG00000037486 | Asxl2 | protein_coding | -0.817876432 | 4.385081825 | 4.0942674 | 0.0597368 | 0.553974417 |
| ENSMUSG00000034647 | Ankrd12 | protein_coding | -0.681693903 | 6.328542907 | 4.094358 | 0.0597625 | 0.553974417 |
| ENSMUSG00000021466 | Ptch1 | protein_coding | -0.78721532 | 4.973170299 | 4.0925269 | 0.0597855 | 0.553974417 |
| ENSMUSG00000017734 | Dbndd2 | protein_coding | -0.671841814 | 5.176144182 | 4.0905174 | 0.0598417 | 0.553974417 |
| ENSMUSG00000026904 | Slc4a10 | protein_coding | -0.658019266 | 5.330997393 | 4.0892413 | 0.0598774 | 0.553974417 |
| ENSMUSG00000020015 | Cdk17 | protein_coding | -0.617018672 | 6.730462063 | 4.0889828 | 0.059913 | 0.553974417 |
| ENSMUSG00000028945 | Rheb | protein_coding | -0.448958398 | 6.803813429 | 4.0867389 | 0.0599476 | 0.553974417 |
| ENSMUSG00000026171 | Rnf25 | protein_coding | 0.548441002 | 6.474817037 | 4.0835861 | 0.0600362 | 0.553974417 |
| ENSMUSG00000024120 | Lrpprc | protein_coding | -0.53775089 | 5.719920287 | 4.0824899 | 0.060067 | 0.553974417 |
| ENSMUSG00000078671 | Chd2 | protein_coding | -0.692459059 | 4.680624463 | 4.0822425 | 0.060074 | 0.553974417 |
| ENSMUSG00000068205 | Macrod2 | protein_coding | -0.690765006 | 5.345425192 | 4.0788689 | 0.0601689 | 0.553974417 |
| ENSMUSG00000047423 | AI837181 | protein_coding | 0.522991708 | 6.38112456 | 4.0775036 | 0.0602074 | 0.553974417 |
| ENSMUSG00000031783 | Polr2c | protein_coding | 0.485962762 | 5.726039436 | 4.0757891 | 0.0602558 | 0.553974417 |
| ENSMUSG00000030016 | Zfp638 | protein_coding | -0.727463366 | 5.660084374 | 4.0746424 | 0.0602882 | 0.553974417 |

| ensembl_gene_id | external_gene_name | gene_biotype | log2 (fold change) | log2 (counts per million) | F-value | p-value | False discovery rate |
|---|---|---|---|---|---|---|---|
| ENSMUSG00000014554 | Dguok | protein_coding | 0.544441261 | 5.686858152 | 4.0690135 | 0.0604474 | 0.553974417 |
| ENSMUSG00000067148 | Polr1c | protein_coding | 0.417250763 | 6.988972938 | 4.0680791 | 0.0604739 | 0.553974417 |
| ENSMUSG00000024012 | Mtch1 | protein_coding | 0.431270307 | 9.233100313 | 4.067587 | 0.0604879 | 0.553974417 |
| ENSMUSG00000044080 | S100a1 | protein_coding | -0.804307918 | 8.199146881 | 4.1533892 | 0.060488 | 0.553974417 |
| ENSMUSG00000031833 | Mast3 | protein_coding | -0.68576499 | 5.532815314 | 4.0659684 | 0.0605338 | 0.553974417 |
| ENSMUSG00000062372 | Otof | protein_coding | 1.182448937 | 3.280649575 | 4.060785 | 0.0606811 | 0.554850466 |
| ENSMUSG00000032679 | Cd59a | protein_coding | 0.674825191 | 5.657192249 | 4.0537629 | 0.0608814 | 0.55541407 |
| ENSMUSG00000026743 | Mllt10 | protein_coding | -0.704539455 | 4.706873075 | 4.053193 | 0.0608977 | 0.55541407 |
| ENSMUSG00000041264 | Uspl1 | protein_coding | 0.604355138 | 5.374578541 | 4.0528 | 0.0609089 | 0.55541407 |
| ENSMUSG00000076498 | Trbc2 | TR_C_gene | -1.452270202 | 2.604877757 | 4.0513849 | 0.0609494 | 0.55541407 |
| ENSMUSG00000009640 | Dmap1 | protein_coding | 0.5318977 | 5.696855416 | 4.046186 | 0.0610983 | 0.556187846 |
| ENSMUSG00000047459 | Dynlrb1 | protein_coding | 0.488455864 | 10.28289572 | 4.0448131 | 0.0611377 | 0.556187846 |
| ENSMUSG00000019831 | Wasf1 | protein_coding | -0.451154925 | 7.054750933 | 4.0380937 | 0.061331 | 0.557017869 |
| ENSMUSG00000027346 | Gpcpd1 | protein_coding | -0.643492043 | 4.817931451 | 4.0379351 | 0.0613356 | 0.557017869 |
| ENSMUSG00000009894 | Snap47 | protein_coding | 0.593146803 | 10.21676749 | 4.037231 | 0.0613844 | 0.557017869 |
| ENSMUSG00000045438 | Cox19 | protein_coding | 0.466197374 | 7.156107596 | 4.0330587 | 0.0614763 | 0.557376802 |
| ENSMUSG00000001750 | Tcirg1 | protein_coding | 1.273260103 | 2.990143412 | 4.0312031 | 0.06153 | 0.557376802 |
| ENSMUSG00000028843 | Sh3bgrl3 | protein_coding | 0.439764944 | 8.961784637 | 4.0294948 | 0.0615794 | 0.557376802 |
| ENSMUSG00000052430 | Bmpr1b | protein_coding | -1.221163577 | 3.043873098 | 4.0238811 | 0.0617422 | 0.557729618 |
| ENSMUSG00000030942 | Thumpd1 | protein_coding | -0.745491571 | 5.519415912 | 4.0215852 | 0.061809 | 0.557729618 |
| ENSMUSG00000003039 | Fam32a | protein_coding | 0.393364814 | 8.117999032 | 4.0210158 | 0.0618255 | 0.557729618 |
| ENSMUSG00000071042 | Rasgrp3 | protein_coding | -1.251493668 | 3.738488421 | 4.0205944 | 0.0618663 | 0.557729618 |
| ENSMUSG00000014837 | 4931428F04Rik | protein_coding | 0.928325791 | 5.338054235 | 4.0554637 | 0.0619356 | 0.557729618 |
| ENSMUSG00000029234 | Tmem165 | protein_coding | -1.26793746 | 3.647534826 | 4.0160034 | 0.0620001 | 0.557729618 |
| ENSMUSG00000029919 | Hpgds | protein_coding | -1.209960179 | 3.063518813 | 4.0135616 | 0.0620428 | 0.557729618 |
| ENSMUSG00000032368 | Zic1 | protein_coding | -0.543715482 | 6.105026732 | 4.0129096 | 0.0620619 | 0.557729618 |
| ENSMUSG00000027087 | Itgav | protein_coding | -1.18085165 | 5.549576828 | 4.1086974 | 0.0620852 | 0.557729618 |
| ENSMUSG00000028480 | Glipr2 | protein_coding | -1.603144236 | 2.854064298 | 4.0004051 | 0.0624285 | 0.55856111 |
| ENSMUSG00000056602 | Fry | protein_coding | -0.588945115 | 5.901134414 | 4.0002515 | 0.0624331 | 0.55856111 |
| ENSMUSG00000042426 | Dhx29 | protein_coding | -0.749422573 | 3.953861934 | 3.9987367 | 0.0624776 | 0.55856111 |
| ENSMUSG00000097080 | 1700086O06Rik | lincRNA | 0.982633732 | 3.712583447 | 3.9969125 | 0.0625314 | 0.55856111 |
| ENSMUSG00000002343 | Armc6 | protein_coding | 0.641387267 | 6.423998086 | 3.9938969 | 0.062649 | 0.55856111 |
| ENSMUSG00000028121 | Bcar3 | protein_coding | -0.942850124 | 4.360199005 | 3.9927041 | 0.0626556 | 0.55856111 |
| ENSMUSG00000056895 | Hist3h2ba | protein_coding | 0.924447369 | 3.746305997 | 3.9921427 | 0.0626722 | 0.55856111 |
| ENSMUSG00000045967 | Gpr158 | protein_coding | -0.636764495 | 5.215833255 | 3.9896844 | 0.0627449 | 0.55856111 |
| ENSMUSG00000024425 | Ndfip1 | protein_coding | -0.434498971 | 6.807682116 | 3.9874193 | 0.062812 | 0.55856111 |
| ENSMUSG00000042700 | Sipa1l1 | protein_coding | -0.643171869 | 6.147735034 | 3.9865895 | 0.0628366 | 0.55856111 |
| ENSMUSG00000019943 | Atp2b1 | protein_coding | -0.520939257 | 8.203782665 | 3.9861017 | 0.0628511 | 0.55856111 |
| ENSMUSG00000040640 | Erc2 | protein_coding | -0.55548112 | 6.308424434 | 3.9845985 | 0.0628957 | 0.55856111 |
| ENSMUSG00000059273 | Zc3h4 | protein_coding | -0.683320616 | 4.300804903 | 3.9832075 | 0.062937 | 0.55856111 |
| ENSMUSG00000024290 | Rock1 | protein_coding | -1.041805365 | 3.749513411 | 3.9825419 | 0.0629567 | 0.55856111 |
| ENSMUSG00000025470 | Zfp511 | protein_coding | 0.67685257 | 4.322987393 | 3.9818047 | 0.0629786 | 0.55856111 |
| ENSMUSG00000058355 | Abce1 | protein_coding | -0.546108314 | 5.983356726 | 3.9769444 | 0.0631233 | 0.55856111 |
| ENSMUSG00000055137 | Sugct | protein_coding | -1.194744603 | 2.948896426 | 3.9768788 | 0.0631253 | 0.55856111 |
| ENSMUSG00000029484 | Anxa3 | protein_coding | -1.068410921 | 3.495999761 | 3.9760137 | 0.0631511 | 0.55856111 |
| ENSMUSG00000101621 | Gm28905 | lincRNA | -1.823336716 | 2.29363569 | 3.973197 | 0.0632351 | 0.55856111 |
| ENSMUSG00000025857 | Dnaaf5 | protein_coding | -4.151869084 | 1.77534841 | 4.0097677 | 0.0632625 | 0.55856111 |
| ENSMUSG00000038576 | Susd4 | protein_coding | 0.480583201 | 7.183129061 | 3.9700191 | 0.0633301 | 0.55856111 |
| ENSMUSG00000033020 | Polr2f | protein_coding | 0.503418268 | 7.719445482 | 3.9690351 | 0.0633596 | 0.55856111 |
| ENSMUSG00000003037 | Rab8a | protein_coding | 0.424781945 | 7.495020344 | 3.9686035 | 0.0633725 | 0.55856111 |
| ENSMUSG00000020364 | Zfp354a | protein_coding | -1.177876632 | 3.351531826 | 3.9600917 | 0.063628 | 0.56035361 |
| ENSMUSG00000066705 | Fxyd6 | protein_coding | 0.43473687 | 9.078928226 | 3.9564782 | 0.0637368 | 0.560819201 |
| ENSMUSG00000032757 | Bet1 | protein_coding | -0.428478703 | 6.576747972 | 3.9530125 | 0.0638414 | 0.560819201 |
| ENSMUSG00000058254 | Tspan7 | protein_coding | 0.421031326 | 11.05145679 | 3.9502498 | 0.0639249 | 0.560819201 |
| ENSMUSG00000021932 | Rnaseh2b | protein_coding | 0.596092557 | 5.32804282 | 3.949852 | 0.0639369 | 0.560819201 |
| ENSMUSG00000019920 | Lims1 | protein_coding | -0.763170811 | 4.735409188 | 3.9496968 | 0.0639416 | 0.560819201 |
| ENSMUSG00000022961 | Son | protein_coding | -0.526487445 | 7.369054877 | 3.9476945 | 0.0640023 | 0.560893399 |
| ENSMUSG00000029780 | Nt5c3 | protein_coding | -0.658749902 | 5.587108197 | 3.9382673 | 0.0642886 | 0.562570913 |
| ENSMUSG00000013822 | Elof1 | protein_coding | 0.398880136 | 8.163650204 | 3.9379479 | 0.0642983 | 0.562570913 |
| ENSMUSG00000021098 | 4930447C04Rik | protein_coding | -1.401460872 | 2.70016333 | 3.9307208 | 0.0645189 | 0.564041713 |
| ENSMUSG00000015575 | Atp6v0e | protein_coding | -0.616785081 | 7.084785121 | 3.9301353 | 0.0646416 | 0.564655587 |
| ENSMUSG00000028004 | Npy2r | protein_coding | 1.221360433 | 4.064201765 | 3.9585157 | 0.0647907 | 0.565048908 |
| ENSMUSG00000020799 | Tekt1 | protein_coding | -0.826895625 | 4.468957633 | 3.920506 | 0.0648321 | 0.565048908 |
| ENSMUSG00000037815 | Ctnna1 | protein_coding | -0.594896475 | 5.880425681 | 3.917719 | 0.0649179 | 0.565048908 |
| ENSMUSG00000036617 | Etl4 | protein_coding | -0.880754041 | 4.367446043 | 3.9167881 | 0.0649466 | 0.565048908 |
| ENSMUSG00000027007 | Ssfa2 | protein_coding | -1.413989989 | 3.402114785 | 3.9149183 | 0.0650331 | 0.565048908 |
| ENSMUSG00000078137 | Ankrd63 | protein_coding | -0.879334779 | 4.836188436 | 3.9127729 | 0.0650993 | 0.565048908 |
| ENSMUSG00000033578 | Tmem35a | protein_coding | 0.449771903 | 6.739431291 | 3.911428 | 0.065112 | 0.565048908 |
| ENSMUSG00000005566 | Trim28 | protein_coding | 0.435452984 | 6.502968316 | 3.9089364 | 0.0651891 | 0.565048908 |
| ENSMUSG00000036402 | Gng12 | protein_coding | -0.741653614 | 4.976671488 | 3.9083043 | 0.0652086 | 0.565048908 |
| ENSMUSG00000010609 | Psen2 | protein_coding | 0.695154265 | 4.709192614 | 3.9071154 | 0.0652454 | 0.565048908 |
| ENSMUSG00000038056 | Kmt2c | protein_coding | -0.699285974 | 4.777930851 | 3.9049736 | 0.0653118 | 0.565048908 |
| ENSMUSG00000040183 | Ankrd6 | protein_coding | 0.638074434 | 5.466469459 | 3.9048013 | 0.0653172 | 0.565048908 |
| ENSMUSG00000017491 | Rarb | protein_coding | -0.909743876 | 3.526248906 | 3.9022186 | 0.0653974 | 0.565196663 |
| ENSMUSG00000022964 | Tmem50b | protein_coding | 0.514539602 | 7.304724141 | 3.9008664 | 0.0654394 | 0.565196663 |
| ENSMUSG00000025967 | Eef1b2 | protein_coding | 0.500207352 | 9.00386979 | 3.8991237 | 0.0654936 | 0.565210897 |
| ENSMUSG00000008668 | Rps18 | protein_coding | 0.524768432 | 7.167909004 | 3.894722 | 0.0656308 | 0.565940306 |
| ENSMUSG00000057378 | Ryr3 | protein_coding | -1.18412775 | 3.667007566 | 3.8914274 | 0.0657626 | 0.566622775 |

| ensembl_gene_id | external_gene_name | gene_biotype | log2 (fold change) | log2 (counts per million) | F-value | p-value | False discovery rate |
|---|---|---|---|---|---|---|---|
| ENSMUSG00000027739 | Rab33b | protein_coding | -0.58083637 | 5.633893349 | 3.8845581 | 0.0659487 | 0.567106148 |
| ENSMUSG00000020776 | Fbf1 | protein_coding | -0.896498396 | 3.362885389 | 3.8821547 | 0.0660242 | 0.567106148 |
| ENSMUSG00000040260 | Daam2 | protein_coding | -0.769855547 | 4.34735342 | 3.8815901 | 0.0660419 | 0.567106148 |
| ENSMUSG00000027953 | Slc50a1 | protein_coding | 0.459402226 | 5.91594276 | 3.8798613 | 0.0660963 | 0.567106148 |
| ENSMUSG00000029066 | Mrpl20 | protein_coding | 0.448304327 | 8.825360776 | 3.8776391 | 0.0661662 | 0.567106148 |
| ENSMUSG00000030177 | Ccdc77 | protein_coding | -1.731961039 | 2.536813618 | 3.8751334 | 0.0662452 | 0.567106148 |
| ENSMUSG00000064043 | Trerf1 | processed_transcript | -0.707278828 | 4.211828547 | 3.8742315 | 0.0662737 | 0.567106148 |
| ENSMUSG00000021690 | Jmy | protein_coding | -0.827563395 | 4.531544115 | 3.8733899 | 0.0663002 | 0.567106148 |
| ENSMUSG00000053141 | Ptprt | protein_coding | -0.823967125 | 4.442839227 | 3.8731275 | 0.0663085 | 0.567106148 |
| ENSMUSG00000064115 | Cadm2 | protein_coding | -0.659084242 | 5.531232018 | 3.8714964 | 0.06636 | 0.567106148 |
| ENSMUSG00000006675 | P4htm | protein_coding | 0.504331157 | 6.530913841 | 3.8659894 | 0.0665344 | 0.567106148 |
| ENSMUSG00000019996 | Map7 | protein_coding | -0.988825394 | 3.207849493 | 3.8643389 | 0.0665867 | 0.567106148 |
| ENSMUSG00000027562 | Car2 | protein_coding | -0.464189818 | 10.41162244 | 3.8615828 | 0.0666742 | 0.567106148 |
| ENSMUSG00000029169 | Dhx15 | protein_coding | -0.609215341 | 6.032586644 | 3.8610865 | 0.06669 | 0.567106148 |
| ENSMUSG00000020577 | Tspan13 | protein_coding | -0.445870574 | 7.317139806 | 3.8607055 | 0.0667021 | 0.567106148 |
| ENSMUSG00000040771 | Oard1 | protein_coding | -0.475466559 | 6.457537831 | 3.8602453 | 0.0667167 | 0.567106148 |
| ENSMUSG00000039838 | Slc45a1 | protein_coding | 0.64218897 | 6.048189818 | 3.85773 | 0.0668259 | 0.567106148 |
| ENSMUSG00000024427 | Spry4 | protein_coding | -0.74946345 | 4.300934509 | 3.8562673 | 0.0668434 | 0.567106148 |
| ENSMUSG00000031523 | Dlc1 | protein_coding | -1.12364951 | 3.484914263 | 3.8562086 | 0.0668453 | 0.567106148 |
| ENSMUSG00000000085 | Scmh1 | protein_coding | -0.6544877 | 5.071774487 | 3.8553235 | 0.0668735 | 0.567106148 |
| ENSMUSG00000035441 | Myo1d | protein_coding | -0.724749933 | 4.208263697 | 3.8507599 | 0.0670192 | 0.567678828 |
| ENSMUSG00000026694 | Mettl13 | protein_coding | -1.013627468 | 3.717727346 | 3.8499033 | 0.0670466 | 0.567678828 |
| ENSMUSG00000027882 | Stxbp3 | protein_coding | -0.725578515 | 4.760748714 | 3.8438843 | 0.0672395 | 0.568428135 |
| ENSMUSG00000030782 | Tgfb1i1 | protein_coding | 0.79143048 | 4.468546827 | 3.8438417 | 0.0672408 | 0.568428135 |
| ENSMUSG00000034467 | Dynlrb2 | protein_coding | -0.951361893 | 4.88891651 | 3.844049 | 0.06734 | 0.568559677 |
| ENSMUSG00000021125 | Arg2 | protein_coding | -0.742619219 | 3.883070506 | 3.8374091 | 0.0674477 | 0.568559677 |
| ENSMUSG00000010803 | Gabra1 | protein_coding | -0.424324654 | 7.603223781 | 3.8370873 | 0.067458 | 0.568559677 |
| ENSMUSG00000029182 | 1700001C02Rik | protein_coding | -1.05210877 | 3.55603683 | 3.8350986 | 0.0675221 | 0.568559677 |
| ENSMUSG00000028886 | Eya3 | protein_coding | 1.203671115 | 3.124177612 | 3.8338764 | 0.0675616 | 0.568559677 |
| ENSMUSG00000031730 | Dhodh | protein_coding | 0.511106438 | 6.121563963 | 3.8332647 | 0.0675813 | 0.568559677 |
| ENSMUSG00000006050 | Sra1 | protein_coding | 0.442577681 | 7.857434219 | 3.8313277 | 0.0676439 | 0.568559677 |
| ENSMUSG00000038059 | Smim3 | protein_coding | -0.925752259 | 3.914729862 | 3.8288009 | 0.0677256 | 0.568559677 |
| ENSMUSG00000029823 | Luc7l2 | protein_coding | -0.58626987 | 5.357817522 | 3.8285956 | 0.0677323 | 0.568559677 |
| ENSMUSG00000038526 | Car14 | protein_coding | -0.895231305 | 4.815138323 | 3.826672 | 0.0678237 | 0.568756786 |
| ENSMUSG00000036053 | Fmnl2 | protein_coding | -0.840866852 | 4.242882389 | 3.8246068 | 0.0678615 | 0.568756786 |
| ENSMUSG00000045294 | Insig1 | protein_coding | -0.629229862 | 5.877428147 | 3.8187724 | 0.0680511 | 0.569406621 |
| ENSMUSG00000022016 | Akap11 | protein_coding | -0.711866894 | 6.148210347 | 3.8218647 | 0.0680903 | 0.569406621 |
| ENSMUSG00000022311 | Csmd3 | protein_coding | -0.907757535 | 4.866877705 | 3.8160729 | 0.0681683 | 0.569406621 |
| ENSMUSG00000041710 | Trpc5 | protein_coding | 1.114994451 | 3.258501206 | 3.8125534 | 0.0682539 | 0.569406621 |
| ENSMUSG00000025170 | Rab40b | protein_coding | -0.86352214 | 4.990216302 | 3.8122789 | 0.0682921 | 0.569406621 |
| ENSMUSG00000027313 | Chac1 | protein_coding | 0.750452922 | 4.619900276 | 3.8101781 | 0.0683315 | 0.569406621 |
| ENSMUSG00000040620 | Dhx33 | protein_coding | -1.257914591 | 2.79761533 | 3.8084879 | 0.0683868 | 0.569406621 |
| ENSMUSG00000016554 | Eif3d | protein_coding | 0.422925595 | 6.399336327 | 3.8070508 | 0.0684339 | 0.569406621 |
| ENSMUSG00000060098 | Prmt7 | protein_coding | 0.936012213 | 5.276261344 | 3.8405018 | 0.0684781 | 0.569406621 |
| ENSMUSG00000027222 | Pex16 | protein_coding | 0.519293254 | 6.157758827 | 3.8055203 | 0.0684841 | 0.569406621 |
| ENSMUSG00000041193 | Pla2g5 | protein_coding | -1.407177352 | 2.690346538 | 3.8043783 | 0.0685215 | 0.569406621 |
| ENSMUSG00000018102 | Hist1h2bc | protein_coding | -0.812056102 | 5.968762913 | 3.8298579 | 0.0688227 | 0.569869549 |
| ENSMUSG00000062901 | Klhl24 | protein_coding | -0.655560439 | 5.141138767 | 3.7942158 | 0.068856 | 0.569869549 |
| ENSMUSG00000000409 | Lck | protein_coding | 1.072885105 | 3.777898361 | 3.794746 | 0.0688677 | 0.569869549 |
| ENSMUSG00000057375 | Yipf1 | protein_coding | 0.453436699 | 6.431579522 | 3.7930475 | 0.0688945 | 0.569869549 |
| ENSMUSG00000033938 | Ndufb7 | protein_coding | 0.449120655 | 9.54847961 | 3.7930197 | 0.0688955 | 0.569869549 |
| ENSMUSG00000054199 | Gon4l | protein_coding | -1.191376547 | 4.01507341 | 3.8026833 | 0.0689289 | 0.569869549 |
| ENSMUSG00000032050 | Rdx | protein_coding | -0.66792982 | 5.48007083 | 3.789189 | 0.0690221 | 0.569869549 |
| ENSMUSG00000022136 | Dnajc3 | protein_coding | -0.775833976 | 4.487794101 | 3.7888899 | 0.069032 | 0.569869549 |
| ENSMUSG00000004934 | Pias4 | protein_coding | -2.227451253 | 2.134810029 | 3.7861328 | 0.0691233 | 0.569869549 |
| ENSMUSG00000038690 | Atp5j2 | protein_coding | 0.53015113 | 9.890778011 | 3.7851325 | 0.0691565 | 0.569869549 |
| ENSMUSG00000044716 | Dok7 | protein_coding | 1.07760195 | 3.466412865 | 3.7836498 | 0.0692057 | 0.569869549 |
| ENSMUSG00000015879 | Fam184b | protein_coding | -1.053173063 | 3.132677739 | 3.7819465 | 0.0692623 | 0.569869549 |
| ENSMUSG00000033965 | Slc16a2 | protein_coding | -1.215108586 | 3.931101434 | 3.7900465 | 0.0693459 | 0.569869549 |
| ENSMUSG00000033382 | Trappc8 | protein_coding | -0.947603509 | 3.871704066 | 3.77912 | 0.0693563 | 0.569869549 |
| ENSMUSG00000029673 | Auts2 | protein_coding | -0.420937753 | 6.258686383 | 3.7786419 | 0.0693722 | 0.569869549 |
| ENSMUSG00000005899 | Smpd4 | protein_coding | -1.041324583 | 3.595173308 | 3.7755311 | 0.0694759 | 0.570285398 |
| ENSMUSG00000041992 | Rapgef5 | protein_coding | -0.609749681 | 5.380962526 | 3.7727803 | 0.0695677 | 0.570394624 |
| ENSMUSG00000028341 | Nr4a3 | protein_coding | -1.369147361 | 3.210663218 | 3.7687418 | 0.0697027 | 0.570394624 |
| ENSMUSG00000039983 | Ccdc32 | protein_coding | 0.412462886 | 7.137188915 | 3.7681361 | 0.069723 | 0.570394624 |
| ENSMUSG00000030979 | Uros | protein_coding | 0.445054369 | 6.304832327 | 3.7679862 | 0.069728 | 0.570394624 |
| ENSMUSG00000030881 | Arfip2 | protein_coding | 0.454109453 | 6.476021251 | 3.7671991 | 0.0697544 | 0.570394624 |
| ENSMUSG00000002205 | Vrk3 | protein_coding | 0.671968816 | 5.766803364 | 3.7581966 | 0.0700862 | 0.57233256 |
| ENSMUSG00000041740 | Rnf10 | protein_coding | 0.528926824 | 5.532524674 | 3.756886 | 0.070101 | 0.57233256 |
| ENSMUSG00000070498 | Tmem132b | protein_coding | -0.660553565 | 4.839907472 | 3.7554003 | 0.0701511 | 0.57233256 |
| ENSMUSG00000031066 | Usp11 | protein_coding | 0.623632246 | 7.700349473 | 3.7819722 | 0.0702458 | 0.57267084 |
| ENSMUSG00000006418 | Rnf114 | protein_coding | 0.444092974 | 7.368095382 | 3.7474843 | 0.0704187 | 0.573218503 |
| ENSMUSG00000038400 | Pmepa1 | protein_coding | -0.731186992 | 4.561801788 | 3.7474578 | 0.0704196 | 0.573218503 |
| ENSMUSG00000039620 | 6430573F11Rik | protein_coding | -0.920405581 | 3.508994447 | 3.7444452 | 0.0705217 | 0.573615896 |
| ENSMUSG00000031627 | Irf2 | protein_coding | -0.568633066 | 4.944026901 | 3.7427395 | 0.0705797 | 0.573628465 |
| ENSMUSG00000027878 | Notch2 | protein_coding | -1.204678052 | 3.242617903 | 3.7399056 | 0.070676 | 0.573628465 |
| ENSMUSG00000024132 | Eci1 | protein_coding | -0.827532381 | 5.003862725 | 3.7403328 | 0.0706909 | 0.573628465 |
| ENSMUSG00000001768 | Rin2 | protein_coding | -0.782776676 | 4.377031292 | 3.7370985 | 0.0707716 | 0.573628465 |

| ensembl_gene_id | external_gene_name | gene_biotype | log2 (fold change) | log2 (counts per million) | F-value | p-value | False discovery rate |
|---|---|---|---|---|---|---|---|
| ENSMUSG00000043498 | 9330132A10Rik | pseudogene | -0.917156707 | 3.451029808 | 3.7355211 | 0.0708254 | 0.573628465 |
| ENSMUSG00000015652 | Steap1 | protein_coding | -3.414525019 | 2.66212385 | 3.8535395 | 0.0708434 | 0.573628465 |
| ENSMUSG00000022604 | Cep97 | protein_coding | -0.846059417 | 3.975601437 | 3.7330359 | 0.0709102 | 0.573737889 |
| ENSMUSG00000002732 | Fkbp7 | protein_coding | -1.341194684 | 2.801679118 | 3.7292047 | 0.0710412 | 0.574046548 |
| ENSMUSG00000051344 | Plekhm3 | protein_coding | -0.709398919 | 4.396973582 | 3.7287988 | 0.0710551 | 0.574046548 |
| ENSMUSG00000061589 | Dot1l | protein_coding | -0.975977355 | 3.989008239 | 3.7259132 | 0.071154 | 0.574321468 |
| ENSMUSG00000015839 | Nfe2l2 | protein_coding | -0.910920668 | 4.716166321 | 3.7255494 | 0.071196 | 0.574321468 |
| ENSMUSG00000020821 | Kif1c | protein_coding | -1.151807986 | 2.868441457 | 3.7203099 | 0.0713465 | 0.575104316 |
| ENSMUSG00000041708 | Mpped1 | protein_coding | 0.546779599 | 5.241067827 | 3.7170699 | 0.0714581 | 0.575326366 |
| ENSMUSG00000029627 | Zkscan14 | protein_coding | 0.616133745 | 4.75515435 | 3.7164036 | 0.0714811 | 0.575326366 |
| ENSMUSG00000004849 | Ap1s1 | protein_coding | 0.413360262 | 9.61928205 | 3.7085644 | 0.071752 | 0.575508393 |
| ENSMUSG00000021198 | Unc79 | protein_coding | -0.958620822 | 3.788694174 | 3.7077413 | 0.0717805 | 0.575508393 |
| ENSMUSG00000030058 | Copg1 | protein_coding | 0.52394848 | 7.934758162 | 3.7084982 | 0.0717838 | 0.575508393 |
| ENSMUSG00000031171 | Ftsj1 | protein_coding | 0.570999678 | 5.746934887 | 3.706553 | 0.0718217 | 0.575508393 |
| ENSMUSG00000002580 | Mien1 | protein_coding | 0.479503924 | 7.03482004 | 3.7059391 | 0.071843 | 0.575508393 |
| ENSMUSG00000027423 | Snx5 | protein_coding | -0.498031487 | 6.114143889 | 3.7058273 | 0.0718469 | 0.575508393 |
| ENSMUSG00000078439 | Smim24 | protein_coding | 1.289940299 | 2.738258183 | 3.7035346 | 0.0719265 | 0.575508393 |
| ENSMUSG00000027384 | Ndufaf5 | protein_coding | 0.420321207 | 7.220808295 | 3.7033795 | 0.0719319 | 0.575508393 |
| ENSMUSG00000022090 | Pdlim2 | protein_coding | -0.496072026 | 6.69297983 | 3.6990442 | 0.0720826 | 0.576045465 |
| ENSMUSG00000021288 | Klc1 | protein_coding | -0.494003594 | 7.094391431 | 3.6973397 | 0.072142 | 0.576045465 |
| ENSMUSG00000035024 | Ncapd3 | protein_coding | -0.93630549 | 3.666744051 | 3.6950453 | 0.0722221 | 0.576045465 |
| ENSMUSG00000047371 | Zfp768 | protein_coding | -1.762465602 | 2.477287658 | 3.6944505 | 0.0722428 | 0.576045465 |
| ENSMUSG00000036879 | Phkb | protein_coding | -0.683593421 | 4.607449824 | 3.6934209 | 0.0722788 | 0.576045465 |
| ENSMUSG00000024526 | Cidea | protein_coding | -0.954203226 | 3.764927347 | 3.6922301 | 0.0723204 | 0.576045465 |
| ENSMUSG00000045092 | S1pr1 | protein_coding | -0.682892872 | 5.733062734 | 3.6881575 | 0.072463 | 0.576482475 |
| ENSMUSG00000032009 | Sesn3 | protein_coding | -0.569164953 | 6.287416602 | 3.6868496 | 0.0725088 | 0.576482475 |
| ENSMUSG00000002602 | Axl | protein_coding | -1.26106362 | 3.171210673 | 3.6829121 | 0.072647 | 0.576482475 |
| ENSMUSG00000061288 | Taok3 | protein_coding | 0.480330296 | 7.069856673 | 3.6825226 | 0.0726607 | 0.576482475 |
| ENSMUSG00000042045 | Sln | protein_coding | -1.085653578 | 6.315763863 | 3.8294684 | 0.0726994 | 0.576482475 |
| ENSMUSG00000032500 | Dclk3 | protein_coding | -0.927486231 | 4.730840745 | 3.6817041 | 0.0727191 | 0.576482475 |
| ENSMUSG00000029290 | Zfp326 | protein_coding | -0.755950688 | 4.101239666 | 3.679157 | 0.0727792 | 0.576482475 |
| ENSMUSG00000045287 | Rtn4rl1 | protein_coding | -0.791937339 | 3.935299572 | 3.6779446 | 0.0728219 | 0.576482475 |
| ENSMUSG00000007207 | Stx1a | protein_coding | 0.677400426 | 7.031155404 | 3.7160106 | 0.0729678 | 0.576482475 |
| ENSMUSG00000028522 | Mier1 | protein_coding | -0.552751125 | 5.329494853 | 3.6713907 | 0.0730533 | 0.576482475 |
| ENSMUSG00000063142 | Kcnma1 | protein_coding | -0.457104778 | 6.222946109 | 3.6702242 | 0.0730946 | 0.576482475 |
| ENSMUSG00000060675 | Pla2g16 | protein_coding | -0.665889805 | 6.167451262 | 3.6707346 | 0.0731061 | 0.576482475 |
| ENSMUSG00000020219 | Timm13 | protein_coding | 0.413604583 | 8.92112637 | 3.669353 | 0.0731254 | 0.576482475 |
| ENSMUSG00000031149 | Praf2 | protein_coding | 0.490868066 | 6.38582939 | 3.6679711 | 0.0731744 | 0.576482475 |
| ENSMUSG00000026959 | Grin1 | protein_coding | -1.430166562 | 2.545413255 | 3.6677584 | 0.0731819 | 0.576482475 |
| ENSMUSG00000028517 | Plpp3 | protein_coding | -0.702948995 | 6.071146533 | 3.6691156 | 0.0732416 | 0.576482475 |
| ENSMUSG00000028773 | Fabp3 | protein_coding | 0.43329851 | 6.611636062 | 3.6636644 | 0.0733272 | 0.576482475 |
| ENSMUSG00000033352 | Map2k4 | protein_coding | 0.416797113 | 7.402422423 | 3.6631478 | 0.0733456 | 0.576482475 |
| ENSMUSG00000042401 | Crtac1 | protein_coding | -0.922994963 | 4.769151047 | 3.6626235 | 0.0733939 | 0.576482475 |
| ENSMUSG00000003031 | Cdkn1b | protein_coding | -0.772465966 | 5.148480046 | 3.6592266 | 0.0734851 | 0.576538996 |
| ENSMUSG00000025189 | Cnnm1 | protein_coding | 0.472620567 | 5.847748916 | 3.6585752 | 0.0735083 | 0.576538996 |
| ENSMUSG00000034880 | Mrpl34 | protein_coding | 0.456353864 | 7.009545127 | 3.6547607 | 0.0736444 | 0.576818065 |
| ENSMUSG00000035696 | Rnf38 | protein_coding | -0.827788279 | 4.031856508 | 3.6541696 | 0.0736655 | 0.576818065 |
| ENSMUSG00000030747 | Dgat2 | protein_coding | -0.5842102 | 5.811963899 | 3.6516632 | 0.0737551 | 0.576818065 |
| ENSMUSG00000034059 | Ypel4 | protein_coding | 0.456085414 | 6.631333858 | 3.6507211 | 0.0737888 | 0.576818065 |
| ENSMUSG00000061118 | Dnajc30 | protein_coding | 0.556012082 | 5.483751363 | 3.6485362 | 0.0738671 | 0.576818065 |
| ENSMUSG00000024914 | Drap1 | protein_coding | 0.412155725 | 6.442584975 | 3.6485312 | 0.0738673 | 0.576818065 |
| ENSMUSG00000060935 | Tmem263 | protein_coding | -0.447365485 | 7.272686587 | 3.6436935 | 0.0740409 | 0.576818065 |
| ENSMUSG00000036570 | Fxyd1 | protein_coding | -0.886436325 | 6.27820526 | 3.715124 | 0.0740473 | 0.576818065 |
| ENSMUSG00000020919 | Stat5b | protein_coding | -0.772134334 | 4.271669641 | 3.6434122 | 0.074051 | 0.576818065 |
| ENSMUSG00000027371 | Fahd2a | protein_coding | 0.47029077 | 6.322687499 | 3.6424563 | 0.0740854 | 0.576818065 |
| ENSMUSG00000001525 | Tubb5 | protein_coding | 0.539820295 | 10.28704004 | 3.6411059 | 0.074134 | 0.576818065 |
| ENSMUSG00000024785 | Rcl1 | protein_coding | -0.859226498 | 4.641329716 | 3.6376532 | 0.0742584 | 0.577012474 |
| ENSMUSG00000022231 | Sema5a | protein_coding | -0.793199955 | 4.641101121 | 3.6374339 | 0.0742663 | 0.577012474 |
| ENSMUSG00000034525 | Ice1 | protein_coding | -0.774593714 | 4.221830255 | 3.630646 | 0.0745116 | 0.578500388 |
| ENSMUSG00000029003 | Mad2l2 | protein_coding | 0.617628451 | 5.318783669 | 3.6281026 | 0.0746037 | 0.57879801 |
| ENSMUSG00000032740 | Ccdc88a | protein_coding | -0.606720654 | 5.344034453 | 3.6254811 | 0.0746989 | 0.579056638 |
| ENSMUSG00000031715 | Smarca5 | protein_coding | -0.62052686 | 5.237627904 | 3.6231964 | 0.0747819 | 0.579056638 |
| ENSMUSG00000050965 | Prkca | protein_coding | -0.555401041 | 5.234489138 | 3.6217944 | 0.0748329 | 0.579056638 |
| ENSMUSG00000023094 | Msrb2 | protein_coding | 0.476690009 | 5.95553684 | 3.6194471 | 0.0749184 | 0.579056638 |
| ENSMUSG00000036634 | Mag | protein_coding | -0.418029831 | 7.307725008 | 3.6161703 | 0.0750379 | 0.579056638 |
| ENSMUSG00000084786 | Ubl5 | protein_coding | 0.5030248 | 9.224991029 | 3.6157039 | 0.0750549 | 0.579056638 |
| ENSMUSG00000034484 | Snx2 | protein_coding | 0.577419929 | 7.439300765 | 3.6177465 | 0.0751401 | 0.579056638 |
| ENSMUSG00000028690 | Mmachc | protein_coding | 0.703992349 | 4.731167092 | 3.6118392 | 0.0751962 | 0.579056638 |
| ENSMUSG00000002010 | Idh3g | protein_coding | 0.390088256 | 8.415720531 | 3.6112276 | 0.0752186 | 0.579056638 |
| ENSMUSG00000034958 | Atcay | protein_coding | 0.640961861 | 6.108660041 | 3.6107821 | 0.0752648 | 0.579056638 |
| ENSMUSG00000029757 | Dync1i1 | protein_coding | 0.511382437 | 7.483071199 | 3.6096273 | 0.0752772 | 0.579056638 |
| ENSMUSG00000045624 | Esf1 | protein_coding | -0.640229119 | 5.188180045 | 3.6094623 | 0.0752833 | 0.579056638 |
| ENSMUSG00000034910 | Pygo1 | protein_coding | -0.693441144 | 4.763878088 | 3.6050846 | 0.0754439 | 0.579465998 |
| ENSMUSG00000021037 | Ahsa1 | protein_coding | 0.396984107 | 7.846547904 | 3.6040994 | 0.0754801 | 0.579465998 |
| ENSMUSG00000035686 | Thrsp | protein_coding | -0.687988003 | 6.131810158 | 3.6042135 | 0.0755844 | 0.579465998 |
| ENSMUSG00000016262 | Sertad4 | protein_coding | -0.796408692 | 4.413357749 | 3.6011543 | 0.0755885 | 0.579465998 |
| ENSMUSG00000026663 | Atf6 | protein_coding | -0.718137179 | 4.068017822 | 3.5994519 | 0.0756512 | 0.579465998 |
| ENSMUSG00000036699 | Zcchc12 | protein_coding | 0.799181656 | 9.901316203 | 3.6696704 | 0.0757837 | 0.579465998 |

| ensembl_gene_id | external_gene_name | gene_biotype | log2 (fold change) | log2 (counts per million) | F-value | p-value | False discovery rate |
|---|---|---|---|---|---|---|---|
| ENSMUSG00000021368 | Tbc1d7 | protein_coding | 0.445030054 | 6.389363671 | 3.5955085 | 0.0757967 | 0.579465998 |
| ENSMUSG00000040415 | Dtx3 | protein_coding | 0.357071773 | 7.562950715 | 3.595449 | 0.0757989 | 0.579465998 |
| ENSMUSG00000032534 | Cep63 | protein_coding | 0.665218215 | 4.898120913 | 3.5936031 | 0.0758672 | 0.579465998 |
| ENSMUSG00000031786 | Drc7 | protein_coding | -1.670149439 | 3.185635984 | 3.6229572 | 0.0758754 | 0.579465998 |
| ENSMUSG00000030588 | Yif1b | protein_coding | 0.438802849 | 6.243423051 | 3.5895362 | 0.0760178 | 0.579927087 |
| ENSMUSG00000059498 | Fcgr3 | protein_coding | -1.106269731 | 4.39498967 | 3.6160343 | 0.0760436 | 0.579927087 |
| ENSMUSG00000038095 | Sbno1 | protein_coding | -0.522514001 | 6.455760382 | 3.5863582 | 0.0761357 | 0.579948939 |
| ENSMUSG00000031399 | Fam3a | protein_coding | 0.578488746 | 5.110490453 | 3.5847622 | 0.076195 | 0.579948939 |
| ENSMUSG00000042719 | Naa25 | protein_coding | 0.942248339 | 3.837378938 | 3.5844044 | 0.0762083 | 0.579948939 |
| ENSMUSG00000034744 | Nagk | protein_coding | 0.435063086 | 7.326640144 | 3.5808574 | 0.0763403 | 0.580436887 |
| ENSMUSG00000031781 | Ciapin1 | protein_coding | 0.413233646 | 7.512321031 | 3.5788692 | 0.0764144 | 0.580436887 |
| ENSMUSG00000038451 | Spsb2 | protein_coding | -0.788795202 | 3.76742242 | 3.5751054 | 0.076555 | 0.580436887 |
| ENSMUSG00000056201 | Cfl1 | protein_coding | 0.39590234 | 10.34545569 | 3.5709584 | 0.0767102 | 0.580436887 |
| ENSMUSG00000040952 | Rps19 | protein_coding | 0.523611062 | 8.554367268 | 3.5716042 | 0.0767159 | 0.580436887 |
| ENSMUSG00000069769 | Msi2 | protein_coding | -0.551478503 | 5.347387036 | 3.5706911 | 0.0767202 | 0.580436887 |
| ENSMUSG00000045962 | Wnk1 | protein_coding | -0.582744677 | 5.624088413 | 3.570532 | 0.0767262 | 0.580436887 |
| ENSMUSG00000036611 | Eepd1 | protein_coding | -0.986585505 | 3.164771061 | 3.5690624 | 0.0767813 | 0.580436887 |
| ENSMUSG00000020863 | Luc7l3 | protein_coding | -0.52223369 | 6.400903113 | 3.5682093 | 0.0768133 | 0.580436887 |
| ENSMUSG00000052456 | Asna1 | protein_coding | 0.380289982 | 8.054059734 | 3.5648297 | 0.0769402 | 0.580436887 |
| ENSMUSG00000049148 | Plcxd3 | protein_coding | 0.664721241 | 5.864056823 | 3.5652465 | 0.0769545 | 0.580436887 |
| ENSMUSG00000018770 | Atp5g3 | protein_coding | 0.409733354 | 10.45908229 | 3.5627961 | 0.0770167 | 0.580436887 |
| ENSMUSG00000063887 | Nlgn1 | protein_coding | -0.523435668 | 6.333251997 | 3.5621026 | 0.0770429 | 0.580436887 |
| ENSMUSG00000043441 | Gpr149 | protein_coding | 0.966380534 | 3.185985951 | 3.5599753 | 0.077123 | 0.580436887 |
| ENSMUSG00000091537 | Tma7 | protein_coding | 0.393872166 | 7.239049652 | 3.5585882 | 0.0771753 | 0.580436887 |
| ENSMUSG00000014633 | Cmc2 | protein_coding | 0.499871551 | 5.842276473 | 3.5551762 | 0.0773042 | 0.580436887 |
| ENSMUSG00000020537 | Drg2 | protein_coding | 0.519931953 | 7.094896345 | 3.5548356 | 0.0773471 | 0.580436887 |
| ENSMUSG00000020598 | Nrcam | protein_coding | -0.498359595 | 5.843984081 | 3.5521966 | 0.077417 | 0.580436887 |
| ENSMUSG00000058799 | Nap1l1 | protein_coding | -0.403895121 | 6.744221483 | 3.5520958 | 0.0774208 | 0.580436887 |
| ENSMUSG00000028959 | Fastk | protein_coding | 0.47143573 | 6.605509675 | 3.5504937 | 0.0774815 | 0.580436887 |
| ENSMUSG00000028572 | Hook1 | protein_coding | -0.867287739 | 3.667438773 | 3.5472893 | 0.0776031 | 0.580436887 |
| ENSMUSG00000002459 | Rgs20 | protein_coding | -0.536722095 | 6.075147803 | 3.5444054 | 0.0777128 | 0.580436887 |
| ENSMUSG00000039630 | Hnrnpu | protein_coding | -0.367372855 | 7.5492458 | 3.5441443 | 0.0777227 | 0.580436887 |
| ENSMUSG00000020956 | Dtd2 | protein_coding | -1.08925245 | 5.50627464 | 3.6217076 | 0.0777328 | 0.580436887 |
| ENSMUSG00000062110 | Scfd2 | protein_coding | 0.854161673 | 4.006111283 | 3.5435244 | 0.0777463 | 0.580436887 |
| ENSMUSG00000024589 | Nedd4l | protein_coding | -0.531770911 | 6.077579682 | 3.5434376 | 0.0777496 | 0.580436887 |
| ENSMUSG00000039735 | Fnbp1l | protein_coding | -0.618969275 | 4.974653609 | 3.5422612 | 0.0777944 | 0.580436887 |
| ENSMUSG00000030310 | Slc6a1 | protein_coding | -0.523248017 | 7.111988007 | 3.5421111 | 0.0778001 | 0.580436887 |
| ENSMUSG00000021994 | Wnt5a | protein_coding | -0.988720751 | 3.22810331 | 3.5388476 | 0.0779246 | 0.580436887 |
| ENSMUSG00000015243 | Abca1 | protein_coding | -1.018920575 | 3.073190224 | 3.5381272 | 0.0779521 | 0.580436887 |
| ENSMUSG00000036810 | Cnep1r1 | protein_coding | -0.60078632 | 5.715660577 | 3.5356325 | 0.0780474 | 0.580436887 |
| ENSMUSG00000068039 | Tcp1 | protein_coding | 0.449698213 | 8.356202542 | 3.5355747 | 0.0780496 | 0.580436887 |
| ENSMUSG00000032621 | Srek1 | protein_coding | -0.60426534 | 5.54780392 | 3.5346886 | 0.0780835 | 0.580436887 |
| ENSMUSG00000025204 | Ndufb8 | protein_coding | 0.447280948 | 9.711001295 | 3.534056 | 0.0781077 | 0.580436887 |
| ENSMUSG00000028223 | Decr1 | protein_coding | -0.646904923 | 5.156132739 | 3.53089 | 0.078229 | 0.580651233 |
| ENSMUSG00000023147 | Wrb | protein_coding | 0.376577104 | 8.827525129 | 3.5304154 | 0.0782472 | 0.580651233 |
| ENSMUSG00000061981 | Flot2 | protein_coding | 0.476003498 | 6.29793369 | 3.52894 | 0.0783038 | 0.580651233 |
| ENSMUSG00000025147 | Mob2 | protein_coding | 0.396335945 | 6.615878038 | 3.5270993 | 0.0783745 | 0.580651233 |
| ENSMUSG00000001891 | Ugp2 | protein_coding | -0.428530664 | 6.183373685 | 3.5244232 | 0.0784774 | 0.580651233 |
| ENSMUSG00000019804 | Snx3 | protein_coding | 0.378626951 | 8.357148891 | 3.5238179 | 0.0785007 | 0.580651233 |
| ENSMUSG00000039246 | Lyplal1 | protein_coding | 0.647862394 | 4.955391354 | 3.5234594 | 0.0785145 | 0.580651233 |
| ENSMUSG00000019790 | Stxbp5 | protein_coding | -0.594607273 | 5.111077448 | 3.5218637 | 0.078576 | 0.580706506 |
| ENSMUSG00000042978 | Sbk1 | protein_coding | -0.864316678 | 4.205062604 | 3.5184823 | 0.0787065 | 0.581128241 |
| ENSMUSG00000025583 | Rptor | protein_coding | 0.578819714 | 4.777241995 | 3.5160331 | 0.0788011 | 0.581128241 |
| ENSMUSG00000025351 | Cd63 | protein_coding | -0.646400687 | 6.778323389 | 3.5373216 | 0.0788134 | 0.581128241 |
| ENSMUSG00000038383 | Pigu | protein_coding | 0.627007998 | 5.496662148 | 3.5136433 | 0.0788936 | 0.581128241 |
| ENSMUSG00000049792 | Bag5 | protein_coding | -0.431724767 | 6.379971003 | 3.5133939 | 0.0789033 | 0.581128241 |
| ENSMUSG00000061207 | Stk19 | protein_coding | 0.436837482 | 6.162909754 | 3.5116781 | 0.0789698 | 0.58121987 |
| ENSMUSG00000023087 | Noct | protein_coding | -0.619275214 | 4.931298851 | 3.5003271 | 0.0794114 | 0.583959881 |
| ENSMUSG00000028174 | Rpe65 | protein_coding | -3.049700567 | 2.05384736 | 3.5150063 | 0.0794507 | 0.583959881 |
| ENSMUSG00000073139 | Tmem185a | protein_coding | 0.485642776 | 5.789414709 | 3.4966733 | 0.0795541 | 0.58432075 |
| ENSMUSG00000096199 | Ptrhd1 | protein_coding | 0.569147584 | 5.25331376 | 3.4919077 | 0.0797408 | 0.585291917 |
| ENSMUSG00000031166 | Wdr13 | protein_coding | -0.43479625 | 5.967335504 | 3.48658 | 0.0799501 | 0.586427737 |
| ENSMUSG00000057738 | Sptan1 | protein_coding | -0.537002199 | 7.14596439 | 3.4815484 | 0.0801785 | 0.586701613 |
| ENSMUSG00000031583 | Wrn | protein_coding | -1.188652318 | 3.024138021 | 3.4789769 | 0.0802499 | 0.586701613 |
| ENSMUSG00000034898 | Filip1 | protein_coding | -0.792695557 | 4.020894749 | 3.4772701 | 0.0803173 | 0.586701613 |
| ENSMUSG00000039546 | Ajap1 | protein_coding | -0.499332236 | 5.796713387 | 3.4750289 | 0.080406 | 0.586701613 |
| ENSMUSG00000004394 | Tmed4 | protein_coding | 0.387923037 | 8.130152195 | 3.4747926 | 0.0804154 | 0.586701613 |
| ENSMUSG00000079657 | Rab26 | protein_coding | 0.797728895 | 5.200088954 | 3.4748987 | 0.0804884 | 0.586701613 |
| ENSMUSG00000063802 | Hspbp1 | protein_coding | 0.455995578 | 7.8338441 | 3.4718258 | 0.080533 | 0.586701613 |
| ENSMUSG00000028964 | Park7 | protein_coding | 0.373415462 | 10.18779602 | 3.4660473 | 0.0807627 | 0.586701613 |
| ENSMUSG00000029166 | Mapre3 | protein_coding | 0.408630904 | 6.801726034 | 3.4656623 | 0.080778 | 0.586701613 |
| ENSMUSG00000036745 | Ttll7 | protein_coding | -0.409333404 | 7.030682103 | 3.4651182 | 0.0807997 | 0.586701613 |
| ENSMUSG00000028793 | Rnf19b | protein_coding | 0.623845601 | 4.540440979 | 3.4588348 | 0.0810505 | 0.586701613 |
| ENSMUSG00000014547 | Wdfy2 | protein_coding | -1.143317991 | 3.165943596 | 3.4584939 | 0.0810641 | 0.586701613 |
| ENSMUSG00000028389 | Zfp37 | protein_coding | -0.68118318 | 4.774334343 | 3.4564429 | 0.0811462 | 0.586701613 |
| ENSMUSG00000042444 | Mindy2 | protein_coding | -0.690743869 | 5.504832714 | 3.4545621 | 0.0812215 | 0.586701613 |
| ENSMUSG00000061731 | Ext1 | protein_coding | -0.718055707 | 4.01353138 | 3.4545392 | 0.0812224 | 0.586701613 |
| ENSMUSG00000028753 | Vwa5b1 | protein_coding | -0.900964709 | 3.62853526 | 3.4540094 | 0.0812437 | 0.586701613 |

| ensembl_gene_id | external_gene_name | gene_biotype | log2 (fold change) | log2 (counts per million) | F-value | p-value | False discovery rate |
|---|---|---|---|---|---|---|---|
| ENSMUSG00000066456 | Hmgn3 | protein_coding | -0.515835642 | 7.04535961 | 3.4533697 | 0.0812693 | 0.586701613 |
| ENSMUSG00000027962 | Vcam1 | protein_coding | -1.083501036 | 3.937435322 | 3.4539763 | 0.0812752 | 0.586701613 |
| ENSMUSG00000002617 | Zfp40 | protein_coding | -1.066311588 | 3.713377652 | 3.4526603 | 0.0812978 | 0.586701613 |
| ENSMUSG00000025144 | Cenpx | protein_coding | 0.486472482 | 6.391339461 | 3.4524769 | 0.0813052 | 0.586701613 |
| ENSMUSG00000026361 | Cdc73 | protein_coding | -0.631316568 | 4.876864289 | 3.4504712 | 0.0813857 | 0.586701613 |
| ENSMUSG00000017776 | Crk | protein_coding | -0.537950427 | 6.282211737 | 3.4489371 | 0.0814474 | 0.586701613 |
| ENSMUSG00000010136 | Pifo | protein_coding | -0.683124428 | 4.485323866 | 3.448355 | 0.0814708 | 0.586701613 |
| ENSMUSG00000086657 | Stamos | processed_transcript | -1.479947712 | 2.476351875 | 3.4468781 | 0.0815302 | 0.586701613 |
| ENSMUSG00000064215 | Ifi27 | protein_coding | -0.446578448 | 6.399140843 | 3.4463192 | 0.0815527 | 0.586701613 |
| ENSMUSG00000035649 | Zcchc7 | protein_coding | -0.655438708 | 4.421059595 | 3.4453253 | 0.0815928 | 0.586701613 |
| ENSMUSG00000046062 | Ppp1r15b | protein_coding | -0.64719922 | 4.111451231 | 3.4452468 | 0.0815959 | 0.586701613 |
| ENSMUSG00000041040 | Fam117b | protein_coding | -1.42045804 | 2.756767461 | 3.4450344 | 0.0816045 | 0.586701613 |
| ENSMUSG00000026520 | Pycr2 | protein_coding | -0.836389116 | 5.07548035 | 3.4445963 | 0.0816524 | 0.586701613 |
| ENSMUSG00000052698 | Tln2 | protein_coding | -0.577802864 | 5.70148989 | 3.442715 | 0.081698 | 0.586701613 |
| ENSMUSG00000024397 | Aif1 | protein_coding | -1.046693795 | 4.614596879 | 3.4716204 | 0.0817122 | 0.586701613 |
| ENSMUSG00000021647 | Cartpt | protein_coding | 0.632111364 | 6.853129418 | 3.4704203 | 0.08176 | 0.586701613 |
| ENSMUSG00000021660 | Btf3 | protein_coding | 0.469280865 | 6.743615765 | 3.4404885 | 0.0817879 | 0.586701613 |
| ENSMUSG00000006611 | Hfe | protein_coding | -1.339012234 | 2.95804863 | 3.438838 | 0.0818547 | 0.586739446 |
| ENSMUSG00000060771 | Tsga10 | protein_coding | -1.158354807 | 3.247839513 | 3.4365723 | 0.0819464 | 0.586739446 |
| ENSMUSG00000033735 | Spr | protein_coding | -0.853553583 | 5.109772652 | 3.4390316 | 0.0819569 | 0.586739446 |
| ENSMUSG00000027952 | Pmvk | protein_coding | 0.382749299 | 8.011103667 | 3.4263151 | 0.0823631 | 0.58873252 |
| ENSMUSG00000055491 | Pprc1 | protein_coding | -1.090894303 | 2.946576924 | 3.425638 | 0.0823907 | 0.58873252 |
| ENSMUSG00000021709 | Erbin | protein_coding | -0.683451145 | 4.967908066 | 3.4242904 | 0.0824456 | 0.58873252 |
| ENSMUSG00000059363 | Fxn | protein_coding | -1.193518751 | 2.881427468 | 3.4240781 | 0.0824543 | 0.58873252 |
| ENSMUSG00000016344 | Ppdpf | protein_coding | 0.549724673 | 5.202265819 | 3.4188425 | 0.0826683 | 0.589868443 |
| ENSMUSG00000037679 | Inf2 | protein_coding | -0.689987011 | 4.847573096 | 3.4167878 | 0.0827524 | 0.589920991 |
| ENSMUSG00000015943 | Bola1 | protein_coding | 0.397829781 | 6.70976838 | 3.415984 | 0.0827853 | 0.589920991 |
| ENSMUSG00000043164 | Tmem212 | protein_coding | -1.087963154 | 3.784289306 | 3.4135433 | 0.0829158 | 0.590459441 |
| ENSMUSG00000041506 | Rrp9 | protein_coding | 0.601803199 | 4.809833241 | 3.4112352 | 0.0829803 | 0.590463092 |
| ENSMUSG00000021194 | Chga | protein_coding | 0.41554051 | 7.63615483 | 3.4094742 | 0.0830527 | 0.590463092 |
| ENSMUSG00000020152 | Actr2 | protein_coding | -0.411501052 | 9.058469489 | 3.4087856 | 0.0830811 | 0.590463092 |
| ENSMUSG00000033728 | Lrrc14 | protein_coding | -0.731381362 | 4.191297207 | 3.4054182 | 0.0832198 | 0.591058654 |
| ENSMUSG00000030655 | Smg1 | protein_coding | -0.857748961 | 3.940697427 | 3.4006449 | 0.083417 | 0.591235625 |
| ENSMUSG00000027669 | Gnb4 | protein_coding | -0.457491086 | 6.130793214 | 3.400528 | 0.0834218 | 0.591235625 |
| ENSMUSG00000038628 | Polr3k | protein_coding | 0.370894356 | 7.76036592 | 3.3996544 | 0.083458 | 0.591235625 |
| ENSMUSG00000006299 | Aamp | protein_coding | 0.495774335 | 9.100446211 | 3.3982335 | 0.0835168 | 0.591235625 |
| ENSMUSG00000051811 | Cox6b2 | protein_coding | -0.748570776 | 4.008247307 | 3.3981649 | 0.0835197 | 0.591235625 |
| ENSMUSG00000035199 | Arl6ip5 | protein_coding | 0.358022617 | 8.05023803 | 3.3948559 | 0.0836569 | 0.591817307 |
| ENSMUSG00000060036 | Rpl3 | protein_coding | 0.485251835 | 7.69626689 | 3.3934172 | 0.0837166 | 0.591850536 |
| ENSMUSG00000026484 | Rnf2 | protein_coding | -0.583721789 | 5.037563545 | 3.3878171 | 0.0839496 | 0.592018423 |
| ENSMUSG00000063015 | Ccni | protein_coding | -0.520211735 | 6.356034087 | 3.3870103 | 0.0839833 | 0.592018423 |
| ENSMUSG00000027778 | Ift80 | protein_coding | -0.878523903 | 3.804707773 | 3.3867331 | 0.0839948 | 0.592018423 |
| ENSMUSG00000022337 | Emc2 | protein_coding | -0.39884201 | 7.531188737 | 3.3840096 | 0.0841085 | 0.592018423 |
| ENSMUSG00000024621 | Csf1r | protein_coding | -0.761296717 | 4.449327018 | 3.3834146 | 0.0841333 | 0.592018423 |
| ENSMUSG00000034532 | Fbxo16 | protein_coding | 0.593651104 | 5.05428201 | 3.3820267 | 0.0841914 | 0.592018423 |
| ENSMUSG00000059554 | Ccdc28a | protein_coding | 0.44872855 | 6.973733931 | 3.3815005 | 0.0842134 | 0.592018423 |
| ENSMUSG00000037773 | Pced1a | protein_coding | 0.985129722 | 3.918021502 | 3.3813103 | 0.0842517 | 0.592018423 |
| ENSMUSG00000062209 | Erbb4 | protein_coding | -0.736565272 | 4.623836947 | 3.3795797 | 0.0842938 | 0.592018423 |
| ENSMUSG00000009291 | Pttg1ip | protein_coding | -0.482417114 | 6.171831776 | 3.3787879 | 0.0843269 | 0.592018423 |
| ENSMUSG00000027163 | Commd9 | protein_coding | 0.479302557 | 7.208490425 | 3.3780924 | 0.0843561 | 0.592018423 |
| ENSMUSG00000021650 | Ptcd2 | protein_coding | 0.471718798 | 6.435827119 | 3.3756551 | 0.0844583 | 0.592018423 |
| ENSMUSG00000036980 | Taf6 | protein_coding | 0.755939696 | 5.728070284 | 3.3955414 | 0.0844765 | 0.592018423 |
| ENSMUSG00000021474 | Sfxn1 | protein_coding | 0.365522299 | 7.44795253 | 3.3743972 | 0.0845111 | 0.592018423 |
| ENSMUSG00000000682 | Cd52 | protein_coding | -1.601570714 | 2.638470569 | 3.3719083 | 0.0846158 | 0.592041099 |
| ENSMUSG00000028066 | Pmf1 | protein_coding | 0.630182052 | 4.495058868 | 3.3704482 | 0.0846772 | 0.592041099 |
| ENSMUSG00000019917 | 10-Sep | protein_coding | -1.096156861 | 3.195238907 | 3.3703928 | 0.0846796 | 0.592041099 |
| ENSMUSG00000032469 | Dbr1 | protein_coding | -1.100095734 | 3.535123546 | 3.3676502 | 0.0847951 | 0.592463922 |
| ENSMUSG00000020212 | Mdm1 | protein_coding | -0.90864404 | 3.198376845 | 3.363087 | 0.0849878 | 0.593082612 |
| ENSMUSG00000035725 | Prkx | protein_coding | -1.156187845 | 3.582011779 | 3.3631648 | 0.085015 | 0.593082612 |
| ENSMUSG00000038000 | Acd | protein_coding | 0.540915829 | 6.959643338 | 3.3616141 | 0.0850806 | 0.593082612 |
| ENSMUSG00000038910 | Plcl2 | protein_coding | -0.704210423 | 5.087735322 | 3.3568107 | 0.0852537 | 0.593082612 |
| ENSMUSG00000057110 | Cntrl | protein_coding | -1.803889777 | 2.464555657 | 3.3564759 | 0.0852679 | 0.593082612 |
| ENSMUSG00000024208 | Uqcc2 | protein_coding | 0.428981017 | 9.520094615 | 3.3558214 | 0.0852957 | 0.593082612 |
| ENSMUSG00000022255 | Mtdh | protein_coding | -0.725909038 | 4.957280925 | 3.3552303 | 0.0853208 | 0.593082612 |
| ENSMUSG00000035228 | Ccdc106 | protein_coding | 0.634697425 | 5.471957661 | 3.3551344 | 0.0853249 | 0.593082612 |
| ENSMUSG00000028334 | Nans | protein_coding | 0.852937103 | 3.834587374 | 3.3523203 | 0.0854446 | 0.59353068 |
| ENSMUSG00000026764 | Kif5c | protein_coding | -0.485836468 | 7.28772482 | 3.3458615 | 0.08572 | 0.593673586 |
| ENSMUSG00000023572 | Ccndbp1 | protein_coding | 0.398072037 | 7.556442304 | 3.3453954 | 0.0857399 | 0.593673586 |
| ENSMUSG00000031787 | Katnb1 | protein_coding | 0.664115108 | 6.125970279 | 3.3575905 | 0.0857966 | 0.593673586 |
| ENSMUSG00000022885 | St6gal1 | protein_coding | -1.029374704 | 3.565684672 | 3.3436816 | 0.0858131 | 0.593673586 |
| ENSMUSG00000039607 | Rbms3 | protein_coding | -0.657783819 | 5.626298058 | 3.343174 | 0.0858349 | 0.593673586 |
| ENSMUSG00000067629 | Syngap1 | protein_coding | -0.840013066 | 3.426343774 | 3.3430537 | 0.08584 | 0.593673586 |
| ENSMUSG00000029141 | Slc4a1ap | protein_coding | -0.628665513 | 4.239304345 | 3.3427825 | 0.0858516 | 0.593673586 |
| ENSMUSG00000039164 | Naif1 | protein_coding | 1.000934335 | 3.153482618 | 3.34147 | 0.0859078 | 0.593680352 |
| ENSMUSG00000020604 | Arsg | protein_coding | -0.893768748 | 4.808082423 | 3.3421586 | 0.0859892 | 0.593861525 |
| ENSMUSG00000108621 | Gm37494 | lincRNA | -1.154082328 | 2.913498847 | 3.3374106 | 0.0860819 | 0.594119665 |
| ENSMUSG00000055681 | Cope | protein_coding | 0.355987313 | 9.279655957 | 3.3346203 | 0.0862018 | 0.594565525 |
| ENSMUSG00000040021 | Lats1 | protein_coding | -1.009522424 | 3.305346806 | 3.3327041 | 0.0862842 | 0.594752701 |

| ensembl_gene_id | external_gene_name | gene_biotype | log2 (fold change) | log2 (counts per million) | F-value | p-value | False discovery rate |
|---|---|---|---|---|---|---|---|
| ENSMUSG00000029126 | Nsg1 | protein_coding | 0.390706213 | 9.756698452 | 3.3295095 | 0.0864219 | 0.595238699 |
| ENSMUSG00000060512 | 0610040J01Rik | protein_coding | -2.199629675 | 2.003837371 | 3.3276928 | 0.0865003 | 0.595238699 |
| ENSMUSG00000030064 | Frmd4b | protein_coding | -0.985988503 | 3.473205466 | 3.327218 | 0.0865208 | 0.595238699 |
| ENSMUSG00000021327 | Zkscan3 | protein_coding | -0.799648171 | 3.967106536 | 3.3259219 | 0.0865768 | 0.59524314 |
| ENSMUSG00000066026 | Dhrs3 | protein_coding | -0.591691235 | 5.994403766 | 3.3225782 | 0.0867215 | 0.595479228 |
| ENSMUSG00000042737 | Dpm3 | protein_coding | 0.488105551 | 6.893419205 | 3.3203425 | 0.0868184 | 0.595479228 |
| ENSMUSG00000031605 | Klhl2 | protein_coding | -0.485844255 | 6.092562795 | 3.3192508 | 0.0868657 | 0.595479228 |
| ENSMUSG00000022428 | Cby1 | protein_coding | 0.403666459 | 7.321213859 | 3.3189824 | 0.0868774 | 0.595479228 |
| ENSMUSG00000028863 | Meaf6 | protein_coding | 0.421007873 | 7.874483094 | 3.318737 | 0.086888 | 0.595479228 |
| ENSMUSG00000032316 | Clk3 | protein_coding | 0.56075388 | 5.255660421 | 3.3146307 | 0.0870665 | 0.596322245 |
| ENSMUSG00000025169 | Ogfod3 | protein_coding | -0.817960156 | 4.034516583 | 3.3114096 | 0.0872068 | 0.596902967 |
| ENSMUSG00000031577 | Tti2 | protein_coding | 0.843182902 | 3.608366245 | 3.308394 | 0.0873384 | 0.597423388 |
| ENSMUSG00000019179 | Mdh2 | protein_coding | 0.446467755 | 10.78112278 | 3.3047132 | 0.0874993 | 0.597751103 |
| ENSMUSG00000024576 | Csnk1a1 | protein_coding | -0.492465567 | 7.22639045 | 3.3046317 | 0.0875029 | 0.597751103 |
| ENSMUSG00000039239 | Tgfb2 | protein_coding | -0.789882603 | 4.117506558 | 3.3034858 | 0.0875531 | 0.597751103 |
| ENSMUSG00000001473 | Tubb6 | protein_coding | -1.326261476 | 2.699897975 | 3.3021633 | 0.087611 | 0.597767248 |
| ENSMUSG00000026834 | Acvr1c | protein_coding | -0.755124958 | 3.894645852 | 3.2995049 | 0.0877276 | 0.598183437 |
| ENSMUSG00000024359 | Hspa9 | protein_coding | -0.459527277 | 6.974329792 | 3.29559 | 0.0878997 | 0.598675095 |
| ENSMUSG00000040370 | Etfrf1 | protein_coding | -0.452878718 | 6.488541711 | 3.2948843 | 0.0879308 | 0.598675095 |
| ENSMUSG00000027365 | Trpm7 | protein_coding | -0.757723391 | 3.924431 | 3.2940672 | 0.0879668 | 0.598675095 |
| ENSMUSG00000022285 | Ywhaz | protein_coding | -0.409358387 | 10.25803832 | 3.2908886 | 0.0881069 | 0.599249605 |
| ENSMUSG00000029822 | Osbpl3 | protein_coding | -1.263139787 | 2.998039118 | 3.2843604 | 0.0883956 | 0.600357018 |
| ENSMUSG00000017009 | Sdc4 | protein_coding | -0.575270777 | 7.306298795 | 3.2918985 | 0.0884307 | 0.600357018 |
| ENSMUSG00000031537 | Ikbkb | protein_coding | -0.625333091 | 4.460225865 | 3.2822073 | 0.088491 | 0.600357018 |
| ENSMUSG00000096914 | Galntl6 | protein_coding | -0.571287 | 5.198842689 | 3.2821612 | 0.0884931 | 0.600357018 |
| ENSMUSG00000006356 | Crip2 | protein_coding | 0.392660141 | 8.839673363 | 3.2759128 | 0.0887708 | 0.60186127 |
| ENSMUSG00000058571 | Gpc6 | protein_coding | -0.970491786 | 3.164254474 | 3.2718275 | 0.0889529 | 0.601893469 |
| ENSMUSG00000029467 | Atp2a2 | protein_coding | -0.469305942 | 7.904779447 | 3.2714572 | 0.0889694 | 0.601893469 |
| ENSMUSG00000030357 | Fkbp4 | protein_coding | 0.386459287 | 7.238306529 | 3.2707874 | 0.0889993 | 0.601893469 |
| ENSMUSG00000063047 | Zfp780b | protein_coding | -0.582868088 | 4.690286311 | 3.2695967 | 0.0890525 | 0.601893469 |
| ENSMUSG00000063320 | 1190007I07Rik | protein_coding | 0.603594455 | 4.499521849 | 3.2695326 | 0.0890554 | 0.601893469 |
| ENSMUSG00000030350 | Prmt8 | protein_coding | -0.464157344 | 6.015413048 | 3.2674431 | 0.0891488 | 0.602146568 |
| ENSMUSG00000023019 | Gpd1 | protein_coding | -0.847772908 | 3.716180756 | 3.2649113 | 0.0892622 | 0.602533948 |
| ENSMUSG00000039480 | Nt5dc1 | protein_coding | -2.06486124 | 1.833752759 | 3.2618401 | 0.0894 | 0.602948436 |
| ENSMUSG00000057816 | 1700007G11Rik | protein_coding | -0.791817825 | 3.541309499 | 3.2609958 | 0.0894379 | 0.602948436 |
| ENSMUSG00000034245 | Hdac11 | protein_coding | 0.430966342 | 8.240556123 | 3.2597199 | 0.0894952 | 0.602948436 |
| ENSMUSG00000039640 | Mrpl12 | protein_coding | 0.474556452 | 6.935011068 | 3.2585488 | 0.0895479 | 0.602948436 |
| ENSMUSG00000038658 | Ric1 | protein_coding | -0.893170043 | 4.409328162 | 3.2560118 | 0.0896928 | 0.603016116 |
| ENSMUSG00000109946 | D2Bwg1423e | protein_coding | 0.420668837 | 6.58515184 | 3.2532356 | 0.0897873 | 0.603016116 |
| ENSMUSG00000061740 | Cyp2d22 | protein_coding | -1.005118691 | 3.458521513 | 3.2516035 | 0.089861 | 0.603016116 |
| ENSMUSG00000033671 | Cep350 | protein_coding | -1.266095229 | 2.879776363 | 3.2492227 | 0.0899686 | 0.603016116 |
| ENSMUSG00000022602 | Arc | protein_coding | -0.70955233 | 5.510500588 | 3.2478245 | 0.0900626 | 0.603016116 |
| ENSMUSG00000040729 | Cep126 | protein_coding | -1.488532817 | 3.213043619 | 3.2537767 | 0.0900676 | 0.603016116 |
| ENSMUSG00000020719 | Ddx5 | protein_coding | -0.39952282 | 8.230626734 | 3.2464661 | 0.0900934 | 0.603016116 |
| ENSMUSG00000028137 | Celf3 | protein_coding | 0.659889369 | 5.27909203 | 3.246062 | 0.0901118 | 0.603016116 |
| ENSMUSG00000094344 | Gm11942 | processed_pseudoge | -3.327722659 | 1.972589933 | 3.2779271 | 0.090134 | 0.603016116 |
| ENSMUSG00000020570 | Sypl | protein_coding | -0.464760251 | 6.263794843 | 3.2452795 | 0.0901472 | 0.603016116 |
| ENSMUSG00000044229 | Nxpe4 | protein_coding | -1.22991569 | 3.301324228 | 3.244671 | 0.0901748 | 0.603016116 |
| ENSMUSG00000026447 | Pik3c2b | protein_coding | -0.774949082 | 4.035664441 | 3.2415688 | 0.0903157 | 0.603450286 |
| ENSMUSG00000030609 | Aen | protein_coding | 0.818990303 | 4.770893489 | 3.239978 | 0.0904188 | 0.603450286 |
| ENSMUSG00000057863 | Rpl36 | protein_coding | 0.475632304 | 5.553229542 | 3.2383462 | 0.0904623 | 0.603450286 |
| ENSMUSG00000076439 | Mog | protein_coding | -0.426604306 | 7.849745662 | 3.2383033 | 0.0904642 | 0.603450286 |
| ENSMUSG00000051557 | Pusl1 | protein_coding | 0.861293966 | 4.982828726 | 3.2522562 | 0.0906153 | 0.604083108 |
| ENSMUSG00000041881 | Ndufa7 | protein_coding | 0.467716758 | 9.535408849 | 3.2331154 | 0.0907008 | 0.604267464 |
| ENSMUSG00000030714 | Sgf29 | protein_coding | 0.586775031 | 5.825382955 | 3.2314233 | 0.0907781 | 0.604267464 |
| ENSMUSG00000023911 | Flywch2 | protein_coding | 0.474117803 | 5.777949659 | 3.230628 | 0.0908145 | 0.604267464 |
| ENSMUSG00000026131 | Dst | protein_coding | -0.666789015 | 5.014893336 | 3.2294657 | 0.0908677 | 0.604267464 |
| ENSMUSG00000021451 | Sema4d | protein_coding | -0.749790351 | 3.612911937 | 3.2274566 | 0.0909597 | 0.604505549 |
| ENSMUSG00000040048 | Ndufb10 | protein_coding | 0.401579081 | 10.18834066 | 3.226125 | 0.0910208 | 0.604537676 |
| ENSMUSG00000022523 | Fgf12 | protein_coding | 0.382334483 | 7.554931054 | 3.2221312 | 0.0912042 | 0.605061726 |
| ENSMUSG00000027834 | Serpini1 | protein_coding | -0.424188607 | 6.329136714 | 3.2219564 | 0.0912122 | 0.605061726 |
| ENSMUSG00000001783 | Rtcb | protein_coding | 0.503233143 | 6.885349732 | 3.2185492 | 0.0913998 | 0.605484806 |
| ENSMUSG00000063531 | Sema3e | protein_coding | -1.166587951 | 2.783433438 | 3.2146498 | 0.0915489 | 0.605484806 |
| ENSMUSG00000026203 | Dnajb2 | protein_coding | 0.37814369 | 6.674401916 | 3.2145616 | 0.091553 | 0.605484806 |
| ENSMUSG00000028184 | Adgrl2 | protein_coding | -0.624103348 | 4.233915542 | 3.2144689 | 0.0915573 | 0.605484806 |
| ENSMUSG00000060988 | Galnt13 | protein_coding | -0.60812859 | 5.405734543 | 3.2144632 | 0.0915575 | 0.605484806 |
| ENSMUSG00000028847 | Trappc3 | protein_coding | 0.34761698 | 7.886045481 | 3.2123501 | 0.0916552 | 0.605758107 |
| ENSMUSG00000029359 | Tesc | protein_coding | -0.808663789 | 4.607347057 | 3.2087605 | 0.0918214 | 0.606425834 |
| ENSMUSG00000027534 | Snx16 | protein_coding | -1.132028474 | 3.755280104 | 3.2090514 | 0.0919199 | 0.606425834 |
| ENSMUSG00000030284 | Creld1 | protein_coding | 0.483978193 | 6.220781924 | 3.2065173 | 0.0919254 | 0.606425834 |
| ENSMUSG00000022841 | Ap2m1 | protein_coding | 0.464509408 | 7.793836002 | 3.2046621 | 0.0920116 | 0.606621978 |
| ENSMUSG00000040877 | Wdr25 | protein_coding | 1.13777761 | 3.857686571 | 3.2262235 | 0.0921431 | 0.606748111 |
| ENSMUSG00000044788 | Fads6 | protein_coding | -0.814840067 | 4.033293867 | 3.2000764 | 0.0922249 | 0.606748111 |
| ENSMUSG00000035521 | Gnptg | protein_coding | 0.374363597 | 7.368568313 | 3.199197 | 0.0922659 | 0.606748111 |
| ENSMUSG00000024357 | Sil1 | protein_coding | 0.466846644 | 6.185084574 | 3.198851 | 0.092282 | 0.606748111 |
| ENSMUSG00000059278 | Naa38 | protein_coding | 0.469713501 | 7.56424352 | 3.1978758 | 0.0923275 | 0.606748111 |
| ENSMUSG00000002319 | Ipo4 | protein_coding | 0.540967339 | 6.218564489 | 3.196079 | 0.0924114 | 0.606748111 |
| ENSMUSG00000020736 | Nt5c | protein_coding | 0.346647198 | 7.34385362 | 3.1957726 | 0.0924257 | 0.606748111 |

| ensembl_gene_id | external_gene_name | gene_biotype | log2 (fold change) | log2 (counts per million) | F-value | p-value | False discovery rate |
|---|---|---|---|---|---|---|---|
| ENSMUSG00000062006 | Rpl34 | protein_coding | 0.506715011 | 7.859558926 | 3.1938018 | 0.0925486 | 0.607184543 |
| ENSMUSG00000047613 | A430005L14Rik | protein_coding | 0.347501609 | 7.307185625 | 3.1912494 | 0.0926372 | 0.607357782 |
| ENSMUSG00000026750 | Psmb7 | protein_coding | 0.349852152 | 8.167252807 | 3.190166 | 0.092688 | 0.607357782 |
| ENSMUSG00000056820 | Tsnax | protein_coding | -0.468051117 | 6.500801362 | 3.1834838 | 0.0930018 | 0.608998095 |
| ENSMUSG00000039853 | Trim14 | protein_coding | -1.972009402 | 2.136635728 | 3.1824258 | 0.0930516 | 0.608998095 |
| ENSMUSG00000026219 | Trip12 | protein_coding | -0.436363493 | 6.114001535 | 3.1806978 | 0.093133 | 0.609160104 |
| ENSMUSG00000031750 | Il34 | protein_coding | 0.535605853 | 5.729122828 | 3.1768164 | 0.0933162 | 0.609857702 |
| ENSMUSG00000022471 | Xrcc6 | protein_coding | -1.000865073 | 3.169702503 | 3.1757895 | 0.0933647 | 0.609857702 |
| ENSMUSG00000024798 | Htr7 | protein_coding | 0.742347865 | 4.349140398 | 3.1743735 | 0.0934317 | 0.609857702 |
| ENSMUSG00000022174 | Dad1 | protein_coding | 0.395145379 | 9.050245605 | 3.1736369 | 0.0934665 | 0.609857702 |
| ENSMUSG00000052137 | Rbm12b2 | protein_coding | -1.093406654 | 3.454937519 | 3.1704226 | 0.0936188 | 0.61016903 |
| ENSMUSG00000029817 | Tra2a | protein_coding | -0.615730866 | 4.997497304 | 3.1702348 | 0.0936277 | 0.61016903 |
| ENSMUSG00000029093 | Sorcs2 | protein_coding | -0.951062567 | 3.639065966 | 3.1672767 | 0.0937682 | 0.610714093 |
| ENSMUSG00000030541 | Idh2 | protein_coding | -0.592505408 | 4.56323447 | 3.164594 | 0.0938957 | 0.610816868 |
| ENSMUSG00000023043 | Krt18 | protein_coding | -1.332001709 | 3.282090587 | 3.1651967 | 0.0938979 | 0.610816868 |
| ENSMUSG00000041035 | Gm17018 | protein_coding | 0.367738963 | 8.814747378 | 3.1633627 | 0.0939543 | 0.610816868 |
| ENSMUSG00000027339 | Rassf2 | protein_coding | -0.630421596 | 4.579458605 | 3.1603458 | 0.0940982 | 0.611046934 |
| ENSMUSG00000024579 | Pcyox1l | protein_coding | -0.741246115 | 3.862839148 | 3.1593206 | 0.0941471 | 0.611046934 |
| ENSMUSG00000038642 | Ctss | protein_coding | -0.763597405 | 6.462792108 | 3.2107827 | 0.0941602 | 0.611046934 |
| ENSMUSG00000019494 | Cops6 | protein_coding | 0.345914267 | 9.482982842 | 3.1577764 | 0.0942209 | 0.611071727 |
| ENSMUSG00000058833 | 2810428I15Rik | protein_coding | 0.438221176 | 7.558352681 | 3.1550079 | 0.0943533 | 0.611295418 |
| ENSMUSG00000031584 | Gsr | protein_coding | -0.53846492 | 5.228194358 | 3.1546784 | 0.094369 | 0.611295418 |
| ENSMUSG00000108456 | 4732496C06Rik | lincRNA | -0.99148075 | 3.32660542 | 3.1531093 | 0.0944442 | 0.611316717 |
| ENSMUSG00000043733 | Ptpn11 | protein_coding | -0.647068182 | 5.782791783 | 3.1528816 | 0.094486 | 0.611316717 |
| ENSMUSG00000031333 | Abcb7 | protein_coding | -0.714392299 | 4.066281437 | 3.1463279 | 0.0947699 | 0.611364633 |
| ENSMUSG00000022197 | Pdzd2 | protein_coding | -0.920277941 | 3.694721318 | 3.1453616 | 0.0948164 | 0.611364633 |
| ENSMUSG00000035835 | Plppr3 | protein_coding | 0.55171946 | 5.406949151 | 3.14392 | 0.0948858 | 0.611364633 |
| ENSMUSG00000027166 | Dnajc24 | protein_coding | -0.69340233 | 5.076899597 | 3.143244 | 0.0949184 | 0.611364633 |
| ENSMUSG00000024844 | Banf1 | protein_coding | 0.44640373 | 7.963875365 | 3.1430118 | 0.0949296 | 0.611364633 |
| ENSMUSG00000002475 | Abhd3 | protein_coding | -0.620069005 | 5.700202732 | 3.1429571 | 0.0949322 | 0.611364633 |
| ENSMUSG00000028960 | Ube4b | protein_coding | -0.539793807 | 5.598998595 | 3.142802 | 0.0949397 | 0.611364633 |
| ENSMUSG00000023992 | Trem2 | protein_coding | -0.803683521 | 4.016970492 | 3.1424035 | 0.0949589 | 0.611364633 |
| ENSMUSG00000026596 | Abl2 | protein_coding | -0.64873162 | 4.873128831 | 3.1414456 | 0.0950051 | 0.611364633 |
| ENSMUSG00000022387 | Brd1 | protein_coding | -0.633949167 | 4.943559841 | 3.1388558 | 0.0951302 | 0.611564027 |
| ENSMUSG00000048175 | Asb8 | protein_coding | 0.44467143 | 7.244738122 | 3.1372833 | 0.0952063 | 0.611564027 |
| ENSMUSG00000003585 | Sec14l2 | protein_coding | -0.716379603 | 3.81467034 | 3.1372735 | 0.0952067 | 0.611564027 |
| ENSMUSG00000000594 | Gm2a | protein_coding | -0.414842254 | 7.111728144 | 3.1348195 | 0.0953256 | 0.611961737 |
| ENSMUSG00000097383 | 1500026H17Rik | bidirectional_promote | 0.675179269 | 3.728712492 | 3.1332717 | 0.0954006 | 0.612078007 |
| ENSMUSG00000039563 | 2210406O10Rik | lincRNA | -0.925136641 | 3.185292419 | 3.1301626 | 0.0955515 | 0.612680824 |
| ENSMUSG00000029174 | Tbc1d1 | protein_coding | -1.038776572 | 3.007960314 | 3.1265603 | 0.0957268 | 0.613373873 |
| ENSMUSG00000029554 | Mad1l1 | protein_coding | 0.505852189 | 5.278598499 | 3.1255719 | 0.0957749 | 0.613373873 |
| ENSMUSG00000034659 | Tmem109 | protein_coding | 0.508758891 | 6.521911539 | 3.123607 | 0.0958707 | 0.613373873 |
| ENSMUSG00000022619 | Mapk8ip2 | protein_coding | -0.507793328 | 6.320424847 | 3.1230415 | 0.0958983 | 0.613373873 |
| ENSMUSG00000029524 | Sirt4 | protein_coding | -0.737553284 | 3.985969658 | 3.1211045 | 0.0959929 | 0.613373873 |
| ENSMUSG00000018405 | Mrm1 | protein_coding | 0.945228304 | 3.018384625 | 3.1191558 | 0.0960882 | 0.613373873 |
| ENSMUSG00000031617 | Tmem184c | protein_coding | -0.717857735 | 4.827262625 | 3.1187106 | 0.09611 | 0.613373873 |
| ENSMUSG00000037787 | Apopt1 | protein_coding | 0.360318647 | 6.798427294 | 3.1185878 | 0.096116 | 0.613373873 |
| ENSMUSG00000066568 | Lsm14a | protein_coding | -0.479603045 | 5.43454583 | 3.1130143 | 0.0963892 | 0.614752439 |
| ENSMUSG00000041096 | Tspyl2 | protein_coding | 0.428141199 | 6.912719448 | 3.1093451 | 0.0965695 | 0.615537636 |
| ENSMUSG00000021945 | Zmym2 | protein_coding | -0.546997803 | 5.437061708 | 3.1068846 | 0.0966907 | 0.615944853 |
| ENSMUSG00000055447 | Cd47 | protein_coding | -0.397321029 | 7.192665503 | 3.1040687 | 0.0968296 | 0.616464452 |
| ENSMUSG00000036850 | Mrpl41 | protein_coding | 0.347096384 | 8.315883781 | 3.1016305 | 0.09695 | 0.616834325 |
| ENSMUSG00000029598 | Plbd2 | protein_coding | 0.602729479 | 5.096875469 | 3.0990764 | 0.0970764 | 0.616834325 |
| ENSMUSG00000028779 | Pef1 | protein_coding | 0.39547939 | 7.551714064 | 3.0982847 | 0.0971156 | 0.616834325 |
| ENSMUSG00000104861 | 3110039M20Rik | lincRNA | 0.989351601 | 3.578127539 | 3.0982541 | 0.0971171 | 0.616834325 |
| ENSMUSG00000026159 | Agfg1 | protein_coding | -0.537072396 | 5.152506144 | 3.0964121 | 0.0972084 | 0.616884175 |
| ENSMUSG00000057101 | Zfp180 | protein_coding | -0.552199013 | 5.044179791 | 3.0957815 | 0.0972397 | 0.616884175 |
| ENSMUSG00000023336 | Wfdc1 | protein_coding | 1.689415861 | 2.318903445 | 3.0930832 | 0.0973737 | 0.616991542 |
| ENSMUSG00000029686 | Cul1 | protein_coding | -0.43620131 | 6.502790169 | 3.0928573 | 0.0973849 | 0.616991542 |
| ENSMUSG00000024486 | Hbegf | protein_coding | 0.660798372 | 4.366426274 | 3.0919754 | 0.0974288 | 0.616991542 |
| ENSMUSG00000028729 | Ebna1bp2 | protein_coding | 0.426763848 | 6.966038955 | 3.0905082 | 0.0975018 | 0.617090412 |
| ENSMUSG00000048240 | Gng7 | protein_coding | -0.489138357 | 8.62343048 | 3.0890459 | 0.0975746 | 0.617188019 |
| ENSMUSG00000027881 | Prpf38b | protein_coding | -0.883458266 | 3.818091983 | 3.0858757 | 0.0977327 | 0.617824539 |
| ENSMUSG00000005681 | Apoa2 | protein_coding | 0.820652088 | 3.382292466 | 3.0814209 | 0.0979553 | 0.618868298 |
| ENSMUSG00000019808 | Adat2 | protein_coding | 0.999041947 | 2.972440111 | 3.0762401 | 0.0982151 | 0.619556581 |
| ENSMUSG00000055866 | Per2 | protein_coding | -1.215772869 | 3.069343328 | 3.0762374 | 0.0982152 | 0.619556581 |
| ENSMUSG00000027804 | Ppid | protein_coding | 0.353697197 | 6.992843414 | 3.0743938 | 0.0983078 | 0.619556581 |
| ENSMUSG00000035151 | Elmod2 | protein_coding | -1.46180899 | 2.45865367 | 3.0738938 | 0.0983329 | 0.619556581 |
| ENSMUSG00000026235 | Epha4 | protein_coding | -0.587024289 | 6.211637622 | 3.0738631 | 0.0983656 | 0.619556581 |
| ENSMUSG00000029430 | Ran | protein_coding | 0.390596023 | 8.036845059 | 3.0723623 | 0.09841 | 0.619556581 |
| ENSMUSG00000021685 | Otp | protein_coding | -1.297021935 | 2.728168457 | 3.07114 | 0.0984715 | 0.619581252 |
| ENSMUSG00000024218 | Taf11 | protein_coding | 0.368861623 | 6.62879639 | 3.0694047 | 0.098559 | 0.619719909 |
| ENSMUSG00000028842 | Ago3 | protein_coding | -0.841404352 | 4.372173985 | 3.0676307 | 0.0986485 | 0.619719909 |
| ENSMUSG00000054945 | Gm9958 | protein_coding | -2.43198435 | 1.942477758 | 3.0672741 | 0.0986665 | 0.619719909 |
| ENSMUSG00000039611 | Tmem246 | protein_coding | 0.457997482 | 6.775406665 | 3.0640816 | 0.0988278 | 0.620370852 |
| ENSMUSG00000085891 | Gm14634 | processed_transcript | -1.533704856 | 2.374153954 | 3.061182 | 0.0989746 | 0.620465652 |
| ENSMUSG00000010376 | Nedd8 | protein_coding | 0.402925396 | 9.396075068 | 3.0605989 | 0.0990041 | 0.620465652 |
| ENSMUSG00000003662 | Ciao1 | protein_coding | 0.453707356 | 6.247825886 | 3.0603316 | 0.0990177 | 0.620465652 |

| ensembl_gene_id | external_gene_name | gene_biotype | log2 (fold change) | log2 (counts per million) | F-value | p-value | False discovery rate |
|---|---|---|---|---|---|---|---|
| ENSMUSG00000046027 | Stard5 | protein_coding | -0.667984422 | 4.094386568 | 3.0592272 | 0.0990737 | 0.620465652 |
| ENSMUSG00000039001 | Rps21 | protein_coding | 0.509347273 | 8.753982477 | 3.0564303 | 0.0992468 | 0.621080796 |
| ENSMUSG00000032333 | Stoml1 | protein_coding | 0.506400373 | 6.128324996 | 3.0550195 | 0.0992874 | 0.621080796 |
| ENSMUSG00000021939 | Ctsb | protein_coding | 0.37783771 | 10.06288877 | 3.0509565 | 0.0994944 | 0.621269154 |
| ENSMUSG00000026740 | Dnajc1 | protein_coding | -1.030955318 | 4.213419847 | 3.0633367 | 0.0994953 | 0.621269154 |
| ENSMUSG00000027546 | Atp9a | protein_coding | 0.377356123 | 7.260064044 | 3.0505609 | 0.0995145 | 0.621269154 |
| ENSMUSG00000009216 | Fam163b | protein_coding | 0.68341358 | 4.936179858 | 3.0498918 | 0.0995487 | 0.621269154 |
| ENSMUSG00000045659 | Plekha7 | protein_coding | -1.072506557 | 3.350783315 | 3.047335 | 0.0996792 | 0.62147499 |
| ENSMUSG00000052848 | C130026L21Rik | lincRNA | 0.602446027 | 4.471491748 | 3.0432957 | 0.0998859 | 0.62147499 |
| ENSMUSG00000039952 | Dag1 | protein_coding | -0.598324429 | 4.91240035 | 3.0376246 | 0.1001769 | 0.62147499 |
| ENSMUSG00000020953 | Coch | protein_coding | -0.820559183 | 3.620251044 | 3.0367201 | 0.1002234 | 0.62147499 |
| ENSMUSG00000071662 | Polr2g | protein_coding | 0.416091877 | 8.576597452 | 3.0328762 | 0.1004214 | 0.62147499 |
| ENSMUSG00000027999 | Pla2g12a | protein_coding | 0.567843418 | 4.434526774 | 3.0324558 | 0.100443 | 0.62147499 |
| ENSMUSG00000024695 | Zfp91 | protein_coding | -0.46037096 | 5.647609796 | 3.0321369 | 0.1004595 | 0.62147499 |
| ENSMUSG00000019796 | Lrp11 | protein_coding | -0.578017986 | 5.493221051 | 3.030885 | 0.1005241 | 0.62147499 |
| ENSMUSG00000052852 | Reep1 | protein_coding | -0.46183072 | 6.004427529 | 3.0291758 | 0.1006124 | 0.62147499 |
| ENSMUSG00000021536 | Adcy2 | protein_coding | -0.601007671 | 5.192257329 | 3.0261382 | 0.1007695 | 0.62147499 |
| ENSMUSG00000054256 | Msi1 | protein_coding | -0.857469635 | 4.022097204 | 3.0245488 | 0.1008518 | 0.62147499 |
| ENSMUSG00000025278 | Flnb | protein_coding | -1.367719054 | 3.310572923 | 3.0271846 | 0.1008822 | 0.62147499 |
| ENSMUSG00000038235 | F11r | protein_coding | -0.959507407 | 3.493792847 | 3.0230555 | 0.1009292 | 0.62147499 |
| ENSMUSG00000038156 | Spon1 | protein_coding | -0.500804186 | 5.425957612 | 3.0222089 | 0.1009731 | 0.62147499 |
| ENSMUSG00000018287 | Spag7 | protein_coding | 0.329533996 | 8.922908867 | 3.0206528 | 0.1010539 | 0.62147499 |
| ENSMUSG00000063787 | Chchd1 | protein_coding | 0.420997734 | 8.251797352 | 3.0204491 | 0.1010645 | 0.62147499 |
| ENSMUSG00000019590 | Cyb561 | protein_coding | 0.422901069 | 7.175003432 | 3.019009 | 0.1011394 | 0.62147499 |
| ENSMUSG00000010663 | Fads1 | protein_coding | -0.477659133 | 5.969591678 | 3.0189885 | 0.1011404 | 0.62147499 |
| ENSMUSG00000024414 | Mrpl27 | protein_coding | 0.379104501 | 7.467434165 | 3.0176884 | 0.101208 | 0.62147499 |
| ENSMUSG00000046922 | Gpr6 | protein_coding | -1.025313672 | 3.465183134 | 3.0176801 | 0.1012085 | 0.62147499 |
| ENSMUSG00000039715 | Wdr34 | protein_coding | -0.599188788 | 4.733743123 | 3.0165045 | 0.1012697 | 0.62147499 |
| ENSMUSG00000037965 | Zc3h7a | protein_coding | 0.525500406 | 6.426642557 | 3.0166027 | 0.1012957 | 0.62147499 |
| ENSMUSG00000027935 | Rab13 | protein_coding | 0.560575426 | 4.98848082 | 3.0145232 | 0.1013729 | 0.62147499 |
| ENSMUSG00000004892 | Bcan | protein_coding | -0.815269222 | 5.297750323 | 3.0291237 | 0.1014501 | 0.62147499 |
| ENSMUSG00000028809 | Srrm1 | protein_coding | -0.376525438 | 7.215659472 | 3.0127028 | 0.1014679 | 0.62147499 |
| ENSMUSG00000038121 | Fam210a | protein_coding | -0.561682578 | 5.42499516 | 3.0125303 | 0.1014769 | 0.62147499 |
| ENSMUSG00000033066 | Gas7 | protein_coding | -0.568324454 | 6.236110823 | 3.0108342 | 0.1015966 | 0.62147499 |
| ENSMUSG00000013698 | Pea15a | protein_coding | 0.450222846 | 10.3333805 | 3.0063554 | 0.1017999 | 0.62147499 |
| ENSMUSG00000041235 | Chd7 | protein_coding | -0.804575419 | 3.645943808 | 3.0060819 | 0.1018142 | 0.62147499 |
| ENSMUSG00000018339 | Gpx3 | protein_coding | 0.451323921 | 9.533675749 | 3.0056211 | 0.1018383 | 0.62147499 |
| ENSMUSG00000044667 | Plppr4 | protein_coding | -0.412095219 | 6.517486683 | 3.005288 | 0.1018558 | 0.62147499 |
| ENSMUSG00000025422 | Agap2 | protein_coding | -0.925122154 | 3.675226007 | 3.0037107 | 0.1019386 | 0.62147499 |
| ENSMUSG00000034928 | Rnf44 | protein_coding | -0.521712815 | 5.358573775 | 3.0036166 | 0.1019435 | 0.62147499 |
| ENSMUSG00000022013 | Dnajc15 | protein_coding | 0.395126525 | 7.804727818 | 3.0034457 | 0.1019525 | 0.62147499 |
| ENSMUSG00000000632 | Sez6 | protein_coding | -0.638492768 | 6.559710408 | 3.0261428 | 0.1019762 | 0.62147499 |
| ENSMUSG00000059742 | Kcnh7 | protein_coding | -0.711208465 | 3.864897987 | 3.0024329 | 0.1020056 | 0.62147499 |
| ENSMUSG00000005583 | Mef2c | protein_coding | -0.505259472 | 6.856169045 | 3.0015651 | 0.1020824 | 0.62147499 |
| ENSMUSG00000032348 | Gsta4 | protein_coding | 0.372304226 | 7.262439524 | 2.9990566 | 0.1021832 | 0.62147499 |
| ENSMUSG00000042410 | Agps | protein_coding | -0.653362335 | 4.159991566 | 2.998461 | 0.1022145 | 0.62147499 |
| ENSMUSG00000040936 | Ulk4 | protein_coding | -0.798439214 | 3.71758564 | 2.9975903 | 0.1022604 | 0.62147499 |
| ENSMUSG00000019998 | Stx7 | protein_coding | 0.384957627 | 7.730290443 | 2.9970993 | 0.1022863 | 0.62147499 |
| ENSMUSG00000047454 | Gphn | protein_coding | 0.392888138 | 7.188718565 | 2.9963917 | 0.1023236 | 0.62147499 |
| ENSMUSG00000035139 | Secisbp2 | protein_coding | -0.645634756 | 4.315297583 | 2.9944989 | 0.1024234 | 0.62147499 |
| ENSMUSG00000020918 | Kat2a | protein_coding | 0.492829106 | 5.123732653 | 2.9943675 | 0.1024304 | 0.62147499 |
| ENSMUSG00000007989 | Fzd3 | protein_coding | -0.468316156 | 5.708642128 | 2.9938794 | 0.1024561 | 0.62147499 |
| ENSMUSG00000019889 | Ptprk | protein_coding | -1.018557948 | 3.102611883 | 2.9937827 | 0.1024612 | 0.62147499 |
| ENSMUSG00000073565 | Prr16 | protein_coding | -1.628526237 | 2.509511292 | 2.9932606 | 0.1024888 | 0.62147499 |
| ENSMUSG00000059920 | 4930453N24Rik | protein_coding | -0.633312291 | 4.780918003 | 2.9885814 | 0.1027363 | 0.62147499 |
| ENSMUSG00000022314 | Rad21 | protein_coding | -0.497489732 | 5.920039703 | 2.9879684 | 0.1027688 | 0.62147499 |
| ENSMUSG00000031982 | Arv1 | protein_coding | 0.853169426 | 3.505064802 | 2.9869166 | 0.1028246 | 0.62147499 |
| ENSMUSG00000084808 | 9430091E24Rik | processed_transcript | -0.828975516 | 3.72286899 | 2.9843861 | 0.1029589 | 0.62147499 |
| ENSMUSG00000021982 | Cdadc1 | protein_coding | 0.413774795 | 6.603513022 | 2.9838744 | 0.1029861 | 0.62147499 |
| ENSMUSG00000061838 | Suclg2 | protein_coding | -1.056408968 | 3.038395466 | 2.9823805 | 0.1030655 | 0.62147499 |
| ENSMUSG00000032437 | Stt3b | protein_coding | -0.651370628 | 5.189159621 | 2.9820655 | 0.1030822 | 0.62147499 |
| ENSMUSG00000049555 | Tmie | protein_coding | 0.707796602 | 4.130788794 | 2.9818708 | 0.1030926 | 0.62147499 |
| ENSMUSG00000042650 | Alkbh5 | protein_coding | -0.710785101 | 6.623995135 | 3.0287332 | 0.1030928 | 0.62147499 |
| ENSMUSG00000021057 | Akap5 | protein_coding | -0.435854888 | 6.766825998 | 2.9817791 | 0.1030975 | 0.62147499 |
| ENSMUSG00000028173 | Wls | protein_coding | -0.62014485 | 4.860362235 | 2.9817114 | 0.1031011 | 0.62147499 |
| ENSMUSG00000019428 | Fkbp8 | protein_coding | 0.349361989 | 8.438425054 | 2.9809222 | 0.1031431 | 0.62147499 |
| ENSMUSG00000025933 | Tmem14a | protein_coding | 0.402949336 | 6.769581659 | 2.9808203 | 0.1031485 | 0.62147499 |
| ENSMUSG00000008683 | Rps15a | protein_coding | 0.476196069 | 7.305842225 | 2.9805499 | 0.1031941 | 0.62147499 |
| ENSMUSG00000031214 | Ophn1 | protein_coding | -0.622605986 | 4.328522202 | 2.9793958 | 0.1032244 | 0.62147499 |
| ENSMUSG00000026841 | Fibcd1 | protein_coding | -0.843142351 | 3.830676593 | 2.9790502 | 0.1032428 | 0.62147499 |
| ENSMUSG00000054667 | Irs4 | protein_coding | 0.707376025 | 4.720075632 | 2.9780896 | 0.103294 | 0.62147499 |
| ENSMUSG00000035704 | Alg8 | protein_coding | 0.943368273 | 3.612258277 | 2.9754861 | 0.1034329 | 0.62147499 |
| ENSMUSG00000040351 | Ankib1 | protein_coding | -0.667257703 | 4.844242296 | 2.9743927 | 0.1034913 | 0.62147499 |
| ENSMUSG00000022421 | Nptxr | protein_coding | -0.581459987 | 6.482046768 | 2.9762025 | 0.1035081 | 0.62147499 |
| ENSMUSG00000029428 | Stx2 | protein_coding | 1.125115911 | 2.756617378 | 2.9736492 | 0.1035311 | 0.62147499 |
| ENSMUSG00000022099 | Dmtn | protein_coding | 0.383374451 | 7.798985178 | 2.97293 | 0.1035695 | 0.62147499 |
| ENSMUSG00000033004 | Mycbp2 | protein_coding | -0.556668935 | 6.269808914 | 2.9702387 | 0.1037448 | 0.621999575 |
| ENSMUSG00000022793 | B4galt4 | protein_coding | -0.98106187 | 3.48912952 | 2.9687307 | 0.1037945 | 0.621999575 |

| ensembl_gene_id | external_gene_name | gene_biotype | log2 (fold change) | log2 (counts per million) | F-value | p-value | False discovery rate |
|---|---|---|---|---|---|---|---|
| ENSMUSG00000102215 | Gm2464 | lincRNA | 0.625640329 | 4.146585718 | 2.9680589 | 0.1038305 | 0.621999575 |
| ENSMUSG00000034560 | Washc4 | protein_coding | -0.807536083 | 4.050259752 | 2.9629812 | 0.1041033 | 0.622578049 |
| ENSMUSG00000025656 | Arhgef9 | protein_coding | -0.37184351 | 7.063057312 | 2.9590398 | 0.1043157 | 0.622578049 |
| ENSMUSG00000020248 | Nfyb | protein_coding | -0.53163302 | 5.796719178 | 2.9587635 | 0.1043306 | 0.622578049 |
| ENSMUSG00000022462 | Slc38a2 | protein_coding | -0.52382156 | 5.304103256 | 2.9579439 | 0.1043749 | 0.622578049 |
| ENSMUSG00000024812 | Tjp2 | protein_coding | -0.816062394 | 3.754612464 | 2.9578678 | 0.104379 | 0.622578049 |
| ENSMUSG00000047671 | Tctex1d4 | protein_coding | -1.015723674 | 3.345862456 | 2.9571358 | 0.1044185 | 0.622578049 |
| ENSMUSG00000003849 | Nqo1 | protein_coding | -0.920574187 | 3.758894883 | 2.9568945 | 0.1044316 | 0.622578049 |
| ENSMUSG00000085042 | Abhd11os | transcribed_unitary_p | -0.607232404 | 5.216477272 | 2.9568723 | 0.1044328 | 0.622578049 |
| ENSMUSG00000049252 | Lrp1b | protein_coding | -0.73662886 | 5.20458909 | 2.9564565 | 0.1044865 | 0.622578049 |
| ENSMUSG00000036158 | Prickle1 | protein_coding | -0.628050767 | 4.668239792 | 2.9555164 | 0.104506 | 0.622578049 |
| ENSMUSG00000038742 | Angptl6 | protein_coding | 0.780305069 | 4.581476581 | 2.9541244 | 0.1046126 | 0.622867692 |
| ENSMUSG00000025172 | Ankrd2 | protein_coding | -1.135360128 | 2.945274011 | 2.9521224 | 0.1046897 | 0.622982158 |
| ENSMUSG00000102252 | Snrpn | protein_coding | 0.518691978 | 5.158850636 | 2.9506499 | 0.1047696 | 0.623112345 |
| ENSMUSG00000022623 | Shank3 | protein_coding | -1.088835375 | 2.872524293 | 2.9485535 | 0.1048833 | 0.623248184 |
| ENSMUSG00000062661 | Ncs1 | protein_coding | 0.452329944 | 10.21342779 | 2.946597 | 0.1049896 | 0.623248184 |
| ENSMUSG00000022751 | Nit2 | protein_coding | 0.457653855 | 5.828133495 | 2.9458797 | 0.1050286 | 0.623248184 |
| ENSMUSG00000046982 | Tshz1 | protein_coding | -0.622098504 | 4.906798199 | 2.9450071 | 0.1050761 | 0.623248184 |
| ENSMUSG00000074282 | Zfp94 | protein_coding | -0.584135303 | 4.398316985 | 2.9444528 | 0.1051063 | 0.623248184 |
| ENSMUSG00000020328 | Nudcd2 | protein_coding | 0.472702993 | 5.547238243 | 2.9436847 | 0.1051481 | 0.623248184 |
| ENSMUSG00000004018 | Fancl | protein_coding | 0.596341489 | 4.841292438 | 2.9417929 | 0.1052512 | 0.623248184 |
| ENSMUSG00000051469 | Zfp24 | protein_coding | -0.521473023 | 5.445761587 | 2.9417037 | 0.1052561 | 0.623248184 |
| ENSMUSG00000061032 | Rrp1 | protein_coding | 0.428825194 | 8.565164267 | 2.9405429 | 0.1053194 | 0.623279926 |
| ENSMUSG00000046822 | Slc39a3 | protein_coding | -0.766429624 | 4.066614776 | 2.9392978 | 0.1053874 | 0.623339149 |
| ENSMUSG00000008373 | Prpf31 | protein_coding | 0.366349581 | 6.59998186 | 2.9373496 | 0.1054939 | 0.623625848 |
| ENSMUSG00000032714 | Syde1 | protein_coding | -1.815461358 | 2.675543045 | 2.9371461 | 0.1056186 | 0.623957021 |
| ENSMUSG00000031095 | Cul4b | protein_coding | -0.849856591 | 3.859403634 | 2.9342058 | 0.1056659 | 0.623957021 |
| ENSMUSG00000031425 | Plp1 | protein_coding | -0.429364521 | 10.61790917 | 2.9306257 | 0.1058623 | 0.624573717 |
| ENSMUSG00000020886 | Dlg4 | protein_coding | 0.413583853 | 5.726195246 | 2.9301843 | 0.1058865 | 0.624573717 |
| ENSMUSG00000025512 | Chid1 | protein_coding | 0.406049488 | 6.30292777 | 2.9276518 | 0.1060257 | 0.624813329 |
| ENSMUSG00000007039 | Ddah2 | protein_coding | -0.580176136 | 4.694998761 | 2.9273314 | 0.1060434 | 0.624813329 |
| ENSMUSG00000054519 | Zfp867 | protein_coding | -0.781972903 | 3.704421995 | 2.9257549 | 0.1061302 | 0.624905233 |
| ENSMUSG00000022312 | Eif3h | protein_coding | 0.356197488 | 8.916982619 | 2.9239194 | 0.1062313 | 0.624905233 |
| ENSMUSG00000087549 | Gm13427 | antisense_RNA | -0.846447204 | 6.941726025 | 3.0229976 | 0.1062333 | 0.624905233 |
| ENSMUSG00000031327 | Chic1 | protein_coding | -0.501950228 | 5.439736471 | 2.9211629 | 0.1063834 | 0.624985435 |
| ENSMUSG00000000838 | Fmr1 | protein_coding | -0.612387246 | 4.646406992 | 2.9208164 | 0.1064026 | 0.624985435 |
| ENSMUSG00000063632 | Sox11 | protein_coding | -0.465299153 | 6.041335213 | 2.9204771 | 0.1064213 | 0.624985435 |
| ENSMUSG00000028995 | Fam126a | protein_coding | -0.993111781 | 3.874948119 | 2.9191893 | 0.1065238 | 0.625097656 |
| ENSMUSG00000049659 | Aftph | protein_coding | -0.487186029 | 5.669652592 | 2.9174007 | 0.1065915 | 0.625097656 |
| ENSMUSG00000026748 | Plxdc2 | protein_coding | -0.697721913 | 4.74135173 | 2.916806 | 0.1066244 | 0.625097656 |
| ENSMUSG00000020362 | Cnot6 | protein_coding | -0.593919339 | 5.369454094 | 2.9159295 | 0.1066729 | 0.625097656 |
| ENSMUSG00000074994 | Qser1 | protein_coding | -1.4157758 | 2.57466188 | 2.91199 | 0.1068915 | 0.626037313 |
| ENSMUSG00000056185 | Snx32 | protein_coding | 0.376529701 | 7.503683569 | 2.9099743 | 0.1070036 | 0.626352336 |
| ENSMUSG00000039771 | Polr2j | protein_coding | 0.418178188 | 7.22821898 | 2.9078712 | 0.1071206 | 0.626488543 |
| ENSMUSG00000032232 | Cgnl1 | protein_coding | -2.052252987 | 1.933576761 | 2.9068447 | 0.1071778 | 0.626488543 |
| ENSMUSG00000022244 | Amacr | protein_coding | -0.791616747 | 4.10322344 | 2.9064174 | 0.1072016 | 0.626488543 |
| ENSMUSG00000027298 | Tyro3 | protein_coding | -0.565064453 | 5.381614131 | 2.9043606 | 0.1073163 | 0.626818316 |
| ENSMUSG00000026031 | Cflar | protein_coding | -0.8670552 | 3.42663038 | 2.9013578 | 0.1074841 | 0.627094241 |
| ENSMUSG00000055078 | Gabra5 | protein_coding | 0.938427759 | 3.08861533 | 2.8999557 | 0.1075625 | 0.627094241 |
| ENSMUSG00000062580 | Timm17a | protein_coding | 0.328917789 | 9.022382621 | 2.8983659 | 0.1076515 | 0.627094241 |
| ENSMUSG00000041084 | Ostc | protein_coding | 0.339162772 | 7.486781233 | 2.8974982 | 0.1077002 | 0.627094241 |
| ENSMUSG00000025885 | Myo5b | protein_coding | -0.666261732 | 4.374766362 | 2.8968156 | 0.1077384 | 0.627094241 |
| ENSMUSG00000036822 | Topors | protein_coding | -0.957476122 | 4.026247427 | 2.8970409 | 0.1077571 | 0.627094241 |
| ENSMUSG00000009876 | Cox4i2 | protein_coding | -1.272482868 | 2.777996292 | 2.8962207 | 0.1077718 | 0.627094241 |
| ENSMUSG00000010797 | Wnt2 | protein_coding | -1.084233997 | 3.163097649 | 2.8938438 | 0.1079052 | 0.627199012 |
| ENSMUSG00000015214 | Mtmr1 | protein_coding | -0.679903104 | 4.185210537 | 2.8934673 | 0.1079264 | 0.627199012 |
| ENSMUSG00000022391 | Rangap1 | protein_coding | 0.38898293 | 7.905234947 | 2.8913421 | 0.1080459 | 0.627199012 |
| ENSMUSG00000019230 | Lhx9 | protein_coding | -1.50953822 | 4.271214065 | 2.9877429 | 0.108103 | 0.627199012 |
| ENSMUSG00000028411 | Aptx | protein_coding | 0.369655848 | 6.507321975 | 2.8889736 | 0.1081793 | 0.627199012 |
| ENSMUSG00000018858 | Mrpl58 | protein_coding | 0.344319193 | 7.702655499 | 2.8886735 | 0.1081962 | 0.627199012 |
| ENSMUSG00000024908 | Ppp6r3 | protein_coding | -0.665722498 | 4.469791417 | 2.8876855 | 0.1082519 | 0.627199012 |
| ENSMUSG00000006241 | Ccdc159 | protein_coding | 0.81396389 | 3.402527741 | 2.8866928 | 0.1083079 | 0.627199012 |
| ENSMUSG00000028607 | Cpt2 | protein_coding | -0.776907757 | 3.913188687 | 2.886571 | 0.1083148 | 0.627199012 |
| ENSMUSG00000022544 | Eef2kmt | protein_coding | -0.836809067 | 3.838473285 | 2.8841306 | 0.1084526 | 0.627436938 |
| ENSMUSG00000023020 | Cox14 | protein_coding | 0.443556581 | 8.847201036 | 2.8837775 | 0.1084725 | 0.627436938 |
| ENSMUSG00000035268 | Pkig | protein_coding | -0.387529408 | 7.141990585 | 2.8821237 | 0.1085661 | 0.62757384 |
| ENSMUSG00000054871 | Tmem158 | protein_coding | -0.83834623 | 4.795902813 | 2.8826541 | 0.10865 | 0.62757384 |
| ENSMUSG00000055319 | Sec23ip | protein_coding | -0.576175341 | 5.61426494 | 2.8798692 | 0.1086938 | 0.62757384 |
| ENSMUSG00000023106 | Denr | protein_coding | 0.359522714 | 7.38599362 | 2.8781373 | 0.108792 | 0.62757384 |
| ENSMUSG00000024181 | Mrpl28 | protein_coding | 0.34870471 | 8.842048939 | 2.8759223 | 0.1089177 | 0.62757384 |
| ENSMUSG00000034203 | Chchd4 | protein_coding | 0.346153147 | 7.239234446 | 2.874613 | 0.1089921 | 0.62757384 |
| ENSMUSG00000034255 | Arhgap27 | protein_coding | 0.547652659 | 5.421061186 | 2.8743724 | 0.1090058 | 0.62757384 |
| ENSMUSG00000022114 | Spry2 | protein_coding | -1.002782546 | 6.703076342 | 3.0209193 | 0.1090882 | 0.62757384 |
| ENSMUSG00000033184 | Tmed7 | protein_coding | -0.643076894 | 5.589289033 | 2.8717498 | 0.1091864 | 0.62757384 |
| ENSMUSG00000042292 | Mkl1 | protein_coding | -0.593780465 | 4.91846856 | 2.8708675 | 0.1092054 | 0.62757384 |
| ENSMUSG00000020562 | Efcab10 | protein_coding | 0.483716243 | 5.368158711 | 2.8699243 | 0.1092591 | 0.62757384 |
| ENSMUSG00000024667 | Tmem216 | protein_coding | 0.502131592 | 5.050720056 | 2.8690121 | 0.1093112 | 0.62757384 |
| ENSMUSG00000074060 | Fbxw15 | protein_coding | -0.915250722 | 3.577313877 | 2.868765 | 0.1093253 | 0.62757384 |

| ensembl_gene_id | external_gene_name | gene_biotype | log2 (fold change) | log2 (counts per million) | F-value | p-value | False discovery rate |
|---|---|---|---|---|---|---|---|
| ENSMUSG00000024082 | Ndufaf7 | protein_coding | 0.362804059 | 6.749561846 | 2.8682443 | 0.109355 | 0.62757384 |
| ENSMUSG00000059834 | Sclt1 | protein_coding | -1.038842828 | 3.183665427 | 2.8679523 | 0.1093717 | 0.62757384 |
| ENSMUSG00000033335 | Dnm2 | protein_coding | -1.055558867 | 4.189013686 | 2.886992 | 0.1095051 | 0.627575522 |
| ENSMUSG00000045100 | Slc25a26 | protein_coding | 0.723996088 | 4.702150218 | 2.8655739 | 0.1095075 | 0.627575522 |
| ENSMUSG00000003344 | Btbd2 | protein_coding | 0.418499222 | 6.043958302 | 2.8648836 | 0.109547 | 0.627575522 |
| ENSMUSG00000033960 | Jcad | protein_coding | -0.894532332 | 4.348085359 | 2.8636555 | 0.1096487 | 0.627823262 |
| ENSMUSG00000030316 | Tamm41 | protein_coding | -0.848267328 | 3.63002157 | 2.860461 | 0.1098004 | 0.628357355 |
| ENSMUSG00000063229 | Ldha | protein_coding | 0.352928053 | 8.55512679 | 2.8573447 | 0.1099793 | 0.629046622 |
| ENSMUSG00000020471 | Pold2 | protein_coding | 0.553179627 | 5.39565547 | 2.8535365 | 0.1101985 | 0.629516046 |
| ENSMUSG00000039221 | Rpl22l1 | protein_coding | 0.482760702 | 6.996212736 | 2.85377 | 0.1102164 | 0.629516046 |
| ENSMUSG00000046598 | Bdh1 | protein_coding | 0.503993592 | 5.806172135 | 2.8516915 | 0.1103048 | 0.629516046 |
| ENSMUSG00000024766 | Lipo3 | protein_coding | -0.923747296 | 3.2692085 | 2.8503079 | 0.1103847 | 0.629516046 |
| ENSMUSG00000040269 | Mrps28 | protein_coding | 0.361119211 | 6.852699931 | 2.8485404 | 0.1104868 | 0.629516046 |
| ENSMUSG00000030689 | Ino80e | protein_coding | 0.367492558 | 6.461476471 | 2.8478635 | 0.1105259 | 0.629516046 |
| ENSMUSG00000027797 | Dclk1 | protein_coding | -0.378487698 | 7.263284849 | 2.8473328 | 0.1105566 | 0.629516046 |
| ENSMUSG00000024197 | Plin3 | protein_coding | -0.596782945 | 5.200013871 | 2.8471715 | 0.1105659 | 0.629516046 |
| ENSMUSG00000031422 | Morf4l2 | protein_coding | 0.411413833 | 8.396678086 | 2.8444632 | 0.1107227 | 0.629516046 |
| ENSMUSG00000050244 | Heatr1 | protein_coding | 1.067892886 | 2.870058153 | 2.8432786 | 0.1107914 | 0.629516046 |
| ENSMUSG00000019978 | Epb41l2 | protein_coding | -0.627350894 | 4.435299091 | 2.8428955 | 0.1108136 | 0.629516046 |
| ENSMUSG00000031557 | Plekha2 | protein_coding | -1.4236667 | 2.553370924 | 2.8428626 | 0.1108155 | 0.629516046 |
| ENSMUSG00000058690 | Ccser2 | protein_coding | -0.472317776 | 5.493256827 | 2.8414443 | 0.1108978 | 0.629516046 |
| ENSMUSG00000038668 | Lpar1 | protein_coding | -0.604937223 | 5.332223296 | 2.8368112 | 0.1111672 | 0.629516046 |
| ENSMUSG00000040532 | Abhd11 | protein_coding | 0.396668555 | 6.511618318 | 2.8361418 | 0.1112061 | 0.629516046 |
| ENSMUSG00000047496 | Rnf152 | protein_coding | -0.845843529 | 3.959589499 | 2.8356643 | 0.111234 | 0.629516046 |
| ENSMUSG00000064293 | Cntn4 | protein_coding | -0.641861608 | 4.833247032 | 2.8347516 | 0.1112871 | 0.629516046 |
| ENSMUSG00000063439 | B9d2 | protein_coding | 0.360361369 | 7.162228884 | 2.8341368 | 0.111323 | 0.629516046 |
| ENSMUSG00000003623 | Crot | protein_coding | -0.627306612 | 4.772818687 | 2.833091 | 0.111384 | 0.629516046 |
| ENSMUSG00000031105 | Slc25a14 | protein_coding | 0.361228816 | 7.199902593 | 2.8330799 | 0.1113847 | 0.629516046 |
| ENSMUSG00000000078 | Klf6 | protein_coding | -0.51244145 | 5.934176 | 2.8323617 | 0.1114266 | 0.629516046 |
| ENSMUSG00000022791 | Tnk2 | protein_coding | -0.592576696 | 5.128254215 | 2.8309209 | 0.1115107 | 0.629516046 |
| ENSMUSG00000022442 | Ttll1 | protein_coding | 0.531326305 | 7.433576324 | 2.8445422 | 0.1115184 | 0.629516046 |
| ENSMUSG00000037214 | Thap1 | protein_coding | -0.667063104 | 3.795572481 | 2.8306903 | 0.1115242 | 0.629516046 |
| ENSMUSG00000021629 | Slc30a5 | protein_coding | -0.672516032 | 4.547295748 | 2.8306772 | 0.111525 | 0.629516046 |
| ENSMUSG00000037058 | Paip2 | protein_coding | 0.322185255 | 7.838311285 | 2.8269741 | 0.1117417 | 0.630003614 |
| ENSMUSG00000037270 | 4932438A13Rik | protein_coding | -0.730086244 | 4.134086467 | 2.82613 | 0.1117911 | 0.630003614 |
| ENSMUSG00000024767 | Otub1 | protein_coding | 0.422964552 | 10.83118066 | 2.8255887 | 0.1118229 | 0.630003614 |
| ENSMUSG00000018565 | Elp5 | protein_coding | 0.34519723 | 8.18573126 | 2.825199 | 0.1118457 | 0.630003614 |
| ENSMUSG00000002871 | Tpra1 | protein_coding | 0.516909262 | 5.906543138 | 2.822935 | 0.1119786 | 0.63034967 |
| ENSMUSG00000090523 | Gypc | protein_coding | -1.915796332 | 1.789287632 | 2.821736 | 0.112049 | 0.63034967 |
| ENSMUSG00000053192 | Mllt11 | protein_coding | 0.515703544 | 6.47669126 | 2.8216921 | 0.112083 | 0.63034967 |
| ENSMUSG00000013833 | Med16 | protein_coding | -0.643582282 | 3.914232501 | 2.8186721 | 0.1122293 | 0.63071189 |
| ENSMUSG00000020132 | Rab21 | protein_coding | -0.364653826 | 6.693586283 | 2.8176393 | 0.1122901 | 0.63071189 |
| ENSMUSG00000028745 | Capzb | protein_coding | 0.358924884 | 8.992086046 | 2.816199 | 0.112375 | 0.63071189 |
| ENSMUSG00000090112 | Shprh | protein_coding | -0.728192137 | 4.266955647 | 2.8160806 | 0.112382 | 0.63071189 |
| ENSMUSG00000033488 | BC026585 | protein_coding | -0.800864845 | 4.166418027 | 2.8150262 | 0.1124442 | 0.630731845 |
| ENSMUSG00000029759 | Pon3 | protein_coding | -2.062169367 | 2.371750793 | 2.8133352 | 0.1125755 | 0.631138943 |
| ENSMUSG00000055334 | Snupn | protein_coding | 0.499392895 | 5.270604123 | 2.8091243 | 0.1127933 | 0.631779973 |
| ENSMUSG00000049336 | Tenm2 | protein_coding | -0.596820727 | 5.487232192 | 2.8083267 | 0.1128405 | 0.631779973 |
| ENSMUSG00000038174 | Fam126b | protein_coding | -0.410563811 | 6.424129411 | 2.8078238 | 0.1128703 | 0.631779973 |
| ENSMUSG00000030591 | Psmd8 | protein_coding | 0.357208284 | 8.434043169 | 2.8054901 | 0.1130088 | 0.631779973 |
| ENSMUSG00000022865 | Cxadr | protein_coding | -1.012247254 | 2.899589865 | 2.8044846 | 0.1130685 | 0.631779973 |
| ENSMUSG00000049119 | Fam110b | protein_coding | 0.389599304 | 6.340096032 | 2.80319 | 0.1131455 | 0.631779973 |
| ENSMUSG00000024883 | Rin1 | protein_coding | -0.968771845 | 3.607593129 | 2.802787 | 0.1131695 | 0.631779973 |
| ENSMUSG00000028684 | Urod | protein_coding | 0.44293365 | 7.131386531 | 2.8019204 | 0.113221 | 0.631779973 |
| ENSMUSG00000015755 | Tab2 | protein_coding | -0.651224009 | 5.182467895 | 2.8010616 | 0.1132722 | 0.631779973 |
| ENSMUSG00000044857 | Lemd2 | protein_coding | -1.040059246 | 3.499210004 | 2.8009736 | 0.1132774 | 0.631779973 |
| ENSMUSG00000002147 | Stat6 | protein_coding | -1.218944547 | 2.770354408 | 2.7989164 | 0.1134 | 0.632113279 |
| ENSMUSG00000042229 | Rabif | protein_coding | 0.376126725 | 6.905860524 | 2.7979986 | 0.1134547 | 0.632113279 |
| ENSMUSG00000028064 | Sema4a | protein_coding | 0.750520946 | 4.49220998 | 2.7963365 | 0.1135539 | 0.632338444 |
| ENSMUSG00000030105 | Arl8b | protein_coding | 0.386804026 | 8.049807547 | 2.7946744 | 0.1136533 | 0.632525663 |
| ENSMUSG00000058454 | Dhcr7 | protein_coding | 0.448823468 | 5.621171495 | 2.7938059 | 0.1137052 | 0.632525663 |
| ENSMUSG00000025066 | Sfr1 | protein_coding | 0.321689 | 8.415339589 | 2.7916882 | 0.113832 | 0.632689609 |
| ENSMUSG00000006373 | Pgrmc1 | protein_coding | 0.358499622 | 9.279987835 | 2.791348 | 0.1138524 | 0.632689609 |
| ENSMUSG00000030126 | Tmcc1 | protein_coding | -0.557446729 | 5.601854664 | 2.7852996 | 0.1142154 | 0.634379416 |
| ENSMUSG00000036022 | Fam122b | protein_coding | -0.905690337 | 3.233161775 | 2.7824055 | 0.1143897 | 0.634692926 |
| ENSMUSG00000022797 | Tfrc | protein_coding | -0.626142019 | 4.404060591 | 2.7820501 | 0.1144111 | 0.634692926 |
| ENSMUSG00000022577 | Ly6h | protein_coding | 0.3814852 | 6.972127586 | 2.7809929 | 0.1144748 | 0.634692926 |
| ENSMUSG00000045128 | Rpl18a | protein_coding | 0.443244 | 8.363054074 | 2.780443 | 0.114508 | 0.634692926 |
| ENSMUSG00000058056 | Palld | protein_coding | 0.974744678 | 3.232377554 | 2.7774263 | 0.1146901 | 0.635375153 |
| ENSMUSG00000031168 | Ebp | protein_coding | 0.445224919 | 5.573726174 | 2.7729823 | 0.1149591 | 0.636311429 |
| ENSMUSG00000057409 | Zfp53 | protein_coding | 1.39458837 | 2.409120313 | 2.7724405 | 0.114992 | 0.636311429 |
| ENSMUSG00000020639 | Pfn4 | protein_coding | 0.705554496 | 4.264502937 | 2.7713961 | 0.1150553 | 0.636311429 |
| ENSMUSG00000038365 | Fbxo25 | protein_coding | 0.44881713 | 6.579606532 | 2.7702324 | 0.115126 | 0.636311429 |
| ENSMUSG00000006576 | Slc4a3 | protein_coding | 0.437880554 | 5.747084477 | 2.769493 | 0.1151709 | 0.636311429 |
| ENSMUSG00000042541 | Sem1 | protein_coding | 0.462632697 | 9.098111375 | 2.7665298 | 0.1153825 | 0.636311429 |
| ENSMUSG00000030168 | Adipor2 | protein_coding | -0.834195857 | 4.827109208 | 2.7736689 | 0.1154507 | 0.636311429 |
| ENSMUSG00000026456 | Cyb5r1 | protein_coding | 0.545437342 | 5.561723233 | 2.7645146 | 0.1154738 | 0.636311429 |
| ENSMUSG00000036572 | Upf3b | protein_coding | -0.487776957 | 5.476546506 | 2.7635928 | 0.11553 | 0.636311429 |

| ensembl_gene_id | external_gene_name | gene_biotype | log2 (fold change) | log2 (counts per million) | F-value | p-value | False discovery rate |
|---|---|---|---|---|---|---|---|
| ENSMUSG00000033577 | Myo6 | protein_coding | -0.527580241 | 5.153348015 | 2.7632517 | 0.1155508 | 0.636311429 |
| ENSMUSG00000040681 | Hmgn1 | protein_coding | -0.573824608 | 4.808654613 | 2.762084 | 0.115622 | 0.636311429 |
| ENSMUSG00000036006 | Ripor2 | protein_coding | -0.570643274 | 5.540538924 | 2.7610184 | 0.1156871 | 0.636311429 |
| ENSMUSG00000024937 | Ehbp1l1 | protein_coding | 0.468506403 | 5.694630554 | 2.7602754 | 0.1157325 | 0.636311429 |
| ENSMUSG00000038495 | Otud7b | protein_coding | -0.467097979 | 5.380455749 | 2.7602424 | 0.1157345 | 0.636311429 |
| ENSMUSG00000006304 | Arpc2 | protein_coding | -0.345499475 | 7.838878671 | 2.7585372 | 0.1158388 | 0.636311429 |
| ENSMUSG00000015757 | Ppil4 | protein_coding | -0.436936551 | 5.927004705 | 2.7583885 | 0.1158479 | 0.636311429 |
| ENSMUSG00000020758 | Itgb4 | protein_coding | -0.824632547 | 3.371077176 | 2.7581061 | 0.1158651 | 0.636311429 |
| ENSMUSG00000000934 | Top1mt | protein_coding | 0.565841554 | 5.3546275 | 2.7567493 | 0.1159482 | 0.636442503 |
| ENSMUSG00000019158 | Tmem160 | protein_coding | 0.704124168 | 4.02644066 | 2.7518344 | 0.1162497 | 0.637628968 |
| ENSMUSG00000055436 | Srsf11 | protein_coding | -0.488237365 | 5.796884674 | 2.7512929 | 0.1162829 | 0.637628968 |
| ENSMUSG00000071176 | Arhgef10 | protein_coding | -0.644159153 | 4.175205598 | 2.7461012 | 0.1166025 | 0.639055499 |
| ENSMUSG00000042747 | Krtcap2 | protein_coding | 0.387903373 | 7.698861476 | 2.739305 | 0.1170225 | 0.640840441 |
| ENSMUSG00000037938 | Chchd5 | protein_coding | 0.362730934 | 6.292169243 | 2.7388045 | 0.1170535 | 0.640840441 |
| ENSMUSG00000041328 | Pcf11 | protein_coding | -0.808177123 | 3.966971098 | 2.7365091 | 0.1171958 | 0.640840441 |
| ENSMUSG00000005674 | Tomm40l | protein_coding | 0.553021848 | 5.84229215 | 2.7363131 | 0.1172079 | 0.640840441 |
| ENSMUSG00000022829 | Stxbp5l | protein_coding | -0.767230283 | 3.902236103 | 2.7356649 | 0.1172481 | 0.640840441 |
| ENSMUSG00000022840 | Adcy5 | protein_coding | -0.53157812 | 6.869010957 | 2.7374858 | 0.1172858 | 0.640840441 |
| ENSMUSG00000021519 | Mterf3 | protein_coding | 0.446887268 | 5.873411583 | 2.7323499 | 0.1174541 | 0.640869435 |
| ENSMUSG00000024958 | Gpr137 | protein_coding | 0.483433459 | 6.39624649 | 2.7320458 | 0.117473 | 0.640869435 |
| ENSMUSG00000003235 | Eif2b5 | protein_coding | 0.533274824 | 5.373775988 | 2.7312038 | 0.1175254 | 0.640869435 |
| ENSMUSG00000028096 | Gpr89 | protein_coding | 0.54305612 | 4.975555012 | 2.7294721 | 0.1176333 | 0.640869435 |
| ENSMUSG00000014077 | Chp1 | protein_coding | 0.408388387 | 7.876562314 | 2.7291331 | 0.1176544 | 0.640869435 |
| ENSMUSG00000037386 | Rims2 | protein_coding | -0.383212375 | 6.12669985 | 2.726665 | 0.1178084 | 0.640869435 |
| ENSMUSG00000003119 | Cdk12 | protein_coding | -0.70358946 | 4.176047042 | 2.7236969 | 0.1179938 | 0.640869435 |
| ENSMUSG00000048758 | Rpl29 | protein_coding | 0.443601553 | 5.486489102 | 2.7232721 | 0.1180204 | 0.640869435 |
| ENSMUSG00000038371 | Sbf2 | protein_coding | -0.600290777 | 4.293764089 | 2.7207701 | 0.1181771 | 0.640869435 |
| ENSMUSG00000031198 | Fundc2 | protein_coding | 0.366856491 | 7.531706821 | 2.7187824 | 0.1183017 | 0.640869435 |
| ENSMUSG00000024038 | Ndufv3 | protein_coding | 0.383257057 | 9.184277111 | 2.7183528 | 0.1183287 | 0.640869435 |
| ENSMUSG00000048304 | Slitrk3 | protein_coding | -0.698275374 | 4.373265654 | 2.7179308 | 0.1183551 | 0.640869435 |
| ENSMUSG00000025477 | Inpp5a | protein_coding | -0.646039258 | 4.403235227 | 2.7173091 | 0.1183942 | 0.640869435 |
| ENSMUSG00000044364 | Tmem74b | protein_coding | 0.751505927 | 3.394795497 | 2.7171903 | 0.1184016 | 0.640869435 |
| ENSMUSG00000016253 | Nelfcd | protein_coding | 0.514564428 | 6.811902554 | 2.7175766 | 0.1184918 | 0.640869435 |
| ENSMUSG00000020921 | Tmem101 | protein_coding | 0.464045394 | 5.629054628 | 2.7157457 | 0.1184924 | 0.640869435 |
| ENSMUSG00000023827 | Agpat4 | protein_coding | 0.393697418 | 7.118018798 | 2.7151087 | 0.1185325 | 0.640869435 |
| ENSMUSG00000033940 | Brk1 | protein_coding | 0.325523602 | 9.750060747 | 2.7138806 | 0.1186097 | 0.640869435 |
| ENSMUSG00000031358 | Msl3 | protein_coding | 0.600239225 | 4.63740218 | 2.7134386 | 0.1186376 | 0.640869435 |
| ENSMUSG00000035953 | Tmem55b | protein_coding | 0.511195855 | 5.558917797 | 2.7125114 | 0.1186959 | 0.640869435 |
| ENSMUSG00000041483 | Zfp281 | protein_coding | -0.634864981 | 4.817946654 | 2.7124085 | 0.1187024 | 0.640869435 |
| ENSMUSG00000071337 | Tia1 | protein_coding | -0.784252642 | 3.8261691 | 2.7112974 | 0.1187724 | 0.640869435 |
| ENSMUSG00000028107 | Tars2 | protein_coding | 0.478394716 | 5.71337882 | 2.7106642 | 0.1188124 | 0.640869435 |
| ENSMUSG00000053580 | Tanc2 | protein_coding | -0.465908374 | 5.970584882 | 2.7100563 | 0.1188507 | 0.640869435 |
| ENSMUSG00000040123 | Zmym5 | protein_coding | -0.493001504 | 5.017706106 | 2.7094999 | 0.1188858 | 0.640869435 |
| ENSMUSG00000042323 | Pbrm1 | protein_coding | -0.490189667 | 5.235378555 | 2.7093319 | 0.1188964 | 0.640869435 |
| ENSMUSG00000027560 | Dok5 | protein_coding | 0.477814518 | 5.700823367 | 2.7092713 | 0.1189003 | 0.640869435 |
| ENSMUSG00000097785 | B230217O12Rik | lincRNA | 0.565722178 | 5.528692352 | 2.7074806 | 0.1190134 | 0.641032543 |
| ENSMUSG00000033054 | Npat | protein_coding | -0.629345918 | 4.384894328 | 2.7064089 | 0.1190811 | 0.641032543 |
| ENSMUSG00000027905 | Ddx20 | protein_coding | -1.044833627 | 3.076885073 | 2.7055035 | 0.1191384 | 0.641032543 |
| ENSMUSG00000026879 | Gsn | protein_coding | -0.380268887 | 6.200328088 | 2.7050201 | 0.119169 | 0.641032543 |
| ENSMUSG00000068263 | Efcc1 | protein_coding | 0.728317728 | 3.664385717 | 2.7036813 | 0.1192538 | 0.641167829 |
| ENSMUSG00000020646 | Mboat2 | protein_coding | -0.573105469 | 4.425745491 | 2.7021647 | 0.1193499 | 0.641363964 |
| ENSMUSG00000048109 | Rbm15 | protein_coding | -0.869517951 | 5.00797031 | 2.7190128 | 0.1195068 | 0.641886333 |
| ENSMUSG00000022571 | Pycrl | protein_coding | 0.375331407 | 6.757808045 | 2.6938201 | 0.1198805 | 0.64251008 |
| ENSMUSG00000061859 | Patj | protein_coding | 0.895838966 | 5.584489827 | 2.7419285 | 0.1199125 | 0.64251008 |
| ENSMUSG00000032637 | Atxn2l | protein_coding | -0.899027078 | 3.237501483 | 2.6926656 | 0.1199541 | 0.64251008 |
| ENSMUSG00000024966 | Stip1 | protein_coding | 0.450369781 | 8.728754636 | 2.692622 | 0.1199569 | 0.64251008 |
| ENSMUSG00000030045 | Mrpl19 | protein_coding | 0.331811186 | 7.065750676 | 2.6915396 | 0.1200259 | 0.64251008 |
| ENSMUSG00000049755 | Zfp672 | protein_coding | 0.545889738 | 5.688238535 | 2.6915288 | 0.1200266 | 0.64251008 |
| ENSMUSG00000081534 | Slc48a1 | protein_coding | -0.460971968 | 7.330296116 | 2.6906183 | 0.1201163 | 0.64251008 |
| ENSMUSG00000044408 | Sptssa | protein_coding | -0.395500863 | 6.727464877 | 2.6894225 | 0.1201612 | 0.64251008 |
| ENSMUSG00000024811 | Tnks2 | protein_coding | -0.636730612 | 5.379517139 | 2.6878511 | 0.1202917 | 0.64251008 |
| ENSMUSG00000045257 | Morn2 | protein_coding | 0.410256588 | 5.872086206 | 2.6866443 | 0.120339 | 0.64251008 |
| ENSMUSG00000033949 | Trim36 | protein_coding | -0.8633335 | 3.119246863 | 2.6863556 | 0.1203575 | 0.64251008 |
| ENSMUSG00000026755 | Arpc5l | protein_coding | 0.317344488 | 8.091165685 | 2.6855951 | 0.1204062 | 0.64251008 |
| ENSMUSG00000079562 | Maea | protein_coding | 0.41092832 | 7.573083674 | 2.6817263 | 0.1206545 | 0.64251008 |
| ENSMUSG00000043131 | Mob1a | protein_coding | -0.841770005 | 3.664848471 | 2.6795451 | 0.1207947 | 0.64251008 |
| ENSMUSG00000022240 | Ctnnd2 | protein_coding | -0.344140746 | 7.273596759 | 2.6768012 | 0.1209714 | 0.64251008 |
| ENSMUSG00000062151 | Unc13c | protein_coding | -0.571936322 | 6.355882941 | 2.6791595 | 0.1209883 | 0.64251008 |
| ENSMUSG00000030032 | Wdr54 | protein_coding | 0.45465754 | 5.463490421 | 2.6755471 | 0.1210522 | 0.64251008 |
| ENSMUSG00000022526 | Zfp251 | protein_coding | 0.594017372 | 4.270325311 | 2.6748917 | 0.1210945 | 0.64251008 |
| ENSMUSG00000051111 | Sv2c | protein_coding | 0.594340438 | 4.970558256 | 2.6746791 | 0.1211083 | 0.64251008 |
| ENSMUSG00000030231 | Plekha5 | protein_coding | -0.475461801 | 5.603126478 | 2.671819 | 0.121293 | 0.64251008 |
| ENSMUSG00000008384 | Sertad1 | protein_coding | -0.574666249 | 4.178920212 | 2.6710622 | 0.121342 | 0.64251008 |
| ENSMUSG00000026098 | Pms1 | protein_coding | -1.35122137 | 2.571576675 | 2.670538 | 0.1213759 | 0.64251008 |
| ENSMUSG00000022742 | Cpox | protein_coding | -0.571207743 | 5.632461472 | 2.6696216 | 0.1214352 | 0.64251008 |
| ENSMUSG00000022018 | Rgcc | protein_coding | -1.163928886 | 3.124808967 | 2.669178 | 0.1214639 | 0.64251008 |
| ENSMUSG00000020283 | Pex13 | protein_coding | -0.64989317 | 4.641362991 | 2.6691429 | 0.1214662 | 0.64251008 |
| ENSMUSG00000054894 | Atp5s | protein_coding | 0.446358295 | 5.555725959 | 2.6690793 | 0.1214703 | 0.64251008 |

| ensembl_gene_id | external_gene_name | gene_biotype | log2 (fold change) | log2 (counts per million) | F-value | p-value | False discovery rate |
|---|---|---|---|---|---|---|---|
| ENSMUSG00000018669 | Cdk5rap3 | protein_coding | 0.464592941 | 6.585328663 | 2.6684828 | 0.121509 | 0.64251008 |
| ENSMUSG00000024191 | Bnip1 | protein_coding | 0.472098918 | 5.806986693 | 2.6682457 | 0.1215243 | 0.64251008 |
| ENSMUSG00000018412 | Kansl1 | protein_coding | -0.5656871 | 5.409290668 | 2.6673377 | 0.1215832 | 0.64251008 |
| ENSMUSG00000025813 | Homer2 | protein_coding | -0.664935032 | 4.084011976 | 2.6660111 | 0.1216693 | 0.64251008 |
| ENSMUSG00000109536 | 9330162G02Rik | TEC | -0.782829956 | 3.428943043 | 2.6652205 | 0.1217206 | 0.64251008 |
| ENSMUSG00000036748 | Cuedc2 | protein_coding | 0.321552936 | 8.642863803 | 2.6648571 | 0.1217442 | 0.64251008 |
| ENSMUSG00000026854 | Usp20 | protein_coding | 0.478614435 | 5.97601616 | 2.6644167 | 0.1217728 | 0.64251008 |
| ENSMUSG00000053475 | Tnfaip6 | protein_coding | -0.561411327 | 5.348703418 | 2.664148 | 0.1217902 | 0.64251008 |
| ENSMUSG00000026547 | Tagln2 | protein_coding | -0.66552794 | 4.606795362 | 2.6623722 | 0.1219057 | 0.64251008 |
| ENSMUSG00000024431 | Nr3c1 | protein_coding | -0.366960632 | 6.420682602 | 2.6613277 | 0.1219737 | 0.64251008 |
| ENSMUSG00000004880 | Lbr | protein_coding | -0.794306591 | 3.517130989 | 2.6612808 | 0.1219767 | 0.64251008 |
| ENSMUSG00000038806 | Sde2 | protein_coding | 0.368064014 | 6.14126393 | 2.6606756 | 0.1220161 | 0.64251008 |
| ENSMUSG00000002661 | Alkbh7 | protein_coding | 0.370953479 | 6.488680113 | 2.6594876 | 0.1220935 | 0.64251008 |
| ENSMUSG00000039842 | Mcph1 | protein_coding | 0.62867216 | 4.052234881 | 2.6593897 | 0.1220999 | 0.64251008 |
| ENSMUSG00000026701 | Prdx6 | protein_coding | -0.377075041 | 9.748615925 | 2.6593271 | 0.122104 | 0.64251008 |
| ENSMUSG00000069520 | Tmem19 | protein_coding | 0.446437141 | 5.323552314 | 2.6588899 | 0.1221325 | 0.64251008 |
| ENSMUSG00000030047 | Arhgap25 | protein_coding | -1.630910238 | 2.611281715 | 2.6566764 | 0.1223084 | 0.643058983 |
| ENSMUSG00000023330 | Dtwd1 | protein_coding | 0.427932682 | 5.508982348 | 2.6554589 | 0.1223564 | 0.643058983 |
| ENSMUSG00000035900 | Gramd4 | protein_coding | -0.808543934 | 3.297468782 | 2.6519988 | 0.1225828 | 0.643933878 |
| ENSMUSG00000042207 | Kdm5b | protein_coding | -0.44824803 | 5.486204669 | 2.6475304 | 0.1228758 | 0.644206098 |
| ENSMUSG00000027076 | Timm10 | protein_coding | 0.408020332 | 7.283647256 | 2.6463979 | 0.1229502 | 0.644206098 |
| ENSMUSG00000054474 | Thnsl2 | protein_coding | 0.935401013 | 3.354334824 | 2.646105 | 0.1229695 | 0.644206098 |
| ENSMUSG00000004393 | Ddx56 | protein_coding | 0.434533534 | 7.161439461 | 2.6450383 | 0.1230396 | 0.644206098 |
| ENSMUSG00000030917 | Tmem159 | protein_coding | 0.447188268 | 5.45839797 | 2.6444994 | 0.1230751 | 0.644206098 |
| ENSMUSG00000047945 | Marcksl1 | protein_coding | 0.595228418 | 4.9562803 | 2.6437694 | 0.1231232 | 0.644206098 |
| ENSMUSG00000047721 | Bola2 | protein_coding | 0.4805869 | 7.920219984 | 2.6437172 | 0.1231581 | 0.644206098 |
| ENSMUSG00000041607 | Mbp | protein_coding | -0.513909346 | 11.80476767 | 2.6422793 | 0.1232528 | 0.644206098 |
| ENSMUSG00000062098 | Btbd3 | protein_coding | -0.403865266 | 6.552591929 | 2.6400124 | 0.1233708 | 0.644206098 |
| ENSMUSG00000025794 | Rpl14 | protein_coding | 0.372128281 | 9.060645753 | 2.6399464 | 0.1233752 | 0.644206098 |
| ENSMUSG00000071234 | Syndig1l | protein_coding | -0.658275627 | 5.554383131 | 2.6388784 | 0.1234772 | 0.644206098 |
| ENSMUSG00000059534 | Uqcr10 | protein_coding | 0.431004016 | 10.21776316 | 2.6379764 | 0.1235053 | 0.644206098 |
| ENSMUSG00000032913 | Lrig2 | protein_coding | -0.629943982 | 3.960089999 | 2.6377042 | 0.1235233 | 0.644206098 |
| ENSMUSG00000033618 | Map3k13 | protein_coding | -0.94558222 | 2.837090241 | 2.6369961 | 0.1235701 | 0.644206098 |
| ENSMUSG00000041132 | N4bp2l1 | protein_coding | -0.536211132 | 5.07898896 | 2.6369858 | 0.1235708 | 0.644206098 |
| ENSMUSG00000042396 | Rbm7 | protein_coding | -0.458319993 | 5.8744885 | 2.6353662 | 0.1236779 | 0.644206098 |
| ENSMUSG00000009090 | Ap1b1 | protein_coding | 0.450459792 | 7.029784393 | 2.635497 | 0.1237008 | 0.644206098 |
| ENSMUSG00000087128 | Gm12655 | processed_transcript | 1.224303902 | 2.679446956 | 2.6346145 | 0.1237277 | 0.644206098 |
| ENSMUSG00000019970 | Sgk1 | protein_coding | -0.584692671 | 7.726937866 | 2.6579447 | 0.123857 | 0.644206098 |
| ENSMUSG00000034157 | Cipc | protein_coding | -0.429803474 | 6.034007373 | 2.6322066 | 0.1238873 | 0.644206098 |
| ENSMUSG00000004268 | Emg1 | protein_coding | 0.328187294 | 7.510535678 | 2.6321249 | 0.1238927 | 0.644206098 |
| ENSMUSG00000034724 | Cnot6l | protein_coding | -0.527496349 | 5.221231872 | 2.6293843 | 0.1240747 | 0.644545655 |
| ENSMUSG00000036896 | C1qc | protein_coding | -1.099038853 | 4.251901366 | 2.65839 | 0.1240779 | 0.644545655 |
| ENSMUSG00000004843 | Chmp2b | protein_coding | -0.522567493 | 5.617063188 | 2.6266945 | 0.1242536 | 0.645146597 |
| ENSMUSG00000005897 | Nr2c1 | protein_coding | -1.133955157 | 2.943919074 | 2.6252501 | 0.1243498 | 0.645192735 |
| ENSMUSG00000033389 | Arhgap44 | protein_coding | 0.539625249 | 8.415104961 | 2.6416881 | 0.124481 | 0.645192735 |
| ENSMUSG00000025290 | Rps24 | protein_coding | 0.46919513 | 8.061290589 | 2.6228585 | 0.1245407 | 0.645192735 |
| ENSMUSG00000028556 | Dock7 | protein_coding | -0.765494404 | 3.640016014 | 2.6220552 | 0.1245629 | 0.645192735 |
| ENSMUSG00000025898 | Cwf19l2 | protein_coding | -0.661306038 | 4.25217616 | 2.621434 | 0.1246044 | 0.645192735 |
| ENSMUSG00000030682 | Cdipt | protein_coding | 0.397815248 | 6.957219419 | 2.6211629 | 0.1246225 | 0.645192735 |
| ENSMUSG00000028292 | Rars2 | protein_coding | -0.479015632 | 5.453790549 | 2.6188807 | 0.124775 | 0.645365606 |
| ENSMUSG00000039768 | Dnajc11 | protein_coding | 0.552747615 | 6.270257955 | 2.6202317 | 0.1247992 | 0.645365606 |
| ENSMUSG00000026782 | Abi2 | protein_coding | -0.371275907 | 7.19286247 | 2.6179713 | 0.1248359 | 0.645365606 |
| ENSMUSG00000105867 | Gm42517 | protein_coding | -1.009584738 | 2.859212297 | 2.6142026 | 0.1250885 | 0.645926115 |
| ENSMUSG00000039706 | Ldb2 | protein_coding | -0.418928169 | 6.280817788 | 2.6140876 | 0.1250962 | 0.645926115 |
| ENSMUSG00000020163 | Uqcr11 | protein_coding | 0.450905349 | 9.283705867 | 2.613666 | 0.1251245 | 0.645926115 |
| ENSMUSG00000020708 | Psmc5 | protein_coding | 0.347312733 | 9.994707594 | 2.6116561 | 0.1252596 | 0.646312835 |
| ENSMUSG00000059921 | Unc5c | protein_coding | -0.772333572 | 4.057897879 | 2.6101873 | 0.1253583 | 0.646512301 |
| ENSMUSG00000022427 | Tomm22 | protein_coding | 0.319641371 | 7.662045527 | 2.6083115 | 0.1254846 | 0.646853375 |
| ENSMUSG00000033596 | Rfwd3 | protein_coding | -0.913832398 | 3.150216625 | 2.6063059 | 0.1256198 | 0.6470515 |
| ENSMUSG00000068299 | Nat8f4 | protein_coding | -0.832920037 | 3.405434141 | 2.604866 | 0.125717 | 0.6470515 |
| ENSMUSG00000027806 | Tsc22d2 | protein_coding | -0.599425587 | 4.615468649 | 2.60432 | 0.1257539 | 0.6470515 |
| ENSMUSG00000019080 | Mfsd3 | protein_coding | 0.710748407 | 4.22252676 | 2.6041737 | 0.1257638 | 0.6470515 |
| ENSMUSG00000058246 | Gm10037 | protein_coding | 0.596672275 | 4.268719226 | 2.6031563 | 0.1258325 | 0.647084725 |
| ENSMUSG00000015776 | Med22 | protein_coding | 0.365004306 | 6.323748204 | 2.6022975 | 0.1258906 | 0.647084725 |
| ENSMUSG00000030465 | Psd3 | protein_coding | -0.391247019 | 6.568826009 | 2.5986708 | 0.1261361 | 0.648037197 |
| ENSMUSG00000046201 | Scaf8 | protein_coding | -0.646164848 | 4.444221024 | 2.5967865 | 0.1262639 | 0.648164531 |
| ENSMUSG00000031990 | Jam3 | protein_coding | -0.535937558 | 4.968066662 | 2.5965283 | 0.1262815 | 0.648164531 |
| ENSMUSG00000047675 | Rps8 | protein_coding | 0.463142934 | 6.217113011 | 2.594048 | 0.12645 | 0.648305246 |
| ENSMUSG00000034875 | Nudt19 | protein_coding | 0.318102794 | 6.684341927 | 2.5934704 | 0.1264893 | 0.648305246 |
| ENSMUSG00000034158 | Lrrc58 | protein_coding | -0.527937333 | 5.653376863 | 2.5934631 | 0.1264898 | 0.648305246 |
| ENSMUSG00000029084 | Cd38 | protein_coding | -0.932872199 | 3.063188531 | 2.5912911 | 0.1266376 | 0.648753797 |
| ENSMUSG00000026568 | Mpc2 | protein_coding | -0.737445949 | 6.934030928 | 2.6406859 | 0.1269394 | 0.649523226 |
| ENSMUSG00000020459 | Mtif2 | protein_coding | 0.500452718 | 5.176760627 | 2.5865592 | 0.1269604 | 0.649523226 |
| ENSMUSG00000030002 | Dusp11 | protein_coding | -0.667083132 | 3.916512805 | 2.5864335 | 0.126969 | 0.649523226 |
| ENSMUSG00000064210 | Ano6 | protein_coding | -0.81330119 | 3.670365067 | 2.5848536 | 0.127077 | 0.649766677 |
| ENSMUSG00000071341 | Egr4 | protein_coding | -1.937299286 | 1.664414579 | 2.5819927 | 0.1272729 | 0.649907984 |
| ENSMUSG00000039629 | Strip2 | protein_coding | -0.773218192 | 4.260741772 | 2.5818017 | 0.127286 | 0.649907984 |
| ENSMUSG00000071646 | Mta2 | protein_coding | -0.634900542 | 4.029628737 | 2.5816705 | 0.127295 | 0.649907984 |

| ensembl_gene_id | external_gene_name | gene_biotype | log2 (fold change) | log2 (counts per million) | F-value | p-value | False discovery rate |
|---|---|---|---|---|---|---|---|
| ENSMUSG00000042712 | Tceal9 | protein_coding | 0.395714751 | 9.303714622 | 2.5803603 | 0.1273848 | 0.649907984 |
| ENSMUSG00000026434 | Nucks1 | protein_coding | -0.375984504 | 7.003684231 | 2.5793159 | 0.1274565 | 0.649907984 |
| ENSMUSG00000025078 | Nhlrc2 | protein_coding | -1.07981624 | 2.748052881 | 2.5791587 | 0.1274673 | 0.649907984 |
| ENSMUSG00000028669 | Pithd1 | protein_coding | 0.364222754 | 6.480969073 | 2.5768554 | 0.1276256 | 0.650406481 |
| ENSMUSG00000032570 | Atp2c1 | protein_coding | -0.536690899 | 5.012945453 | 2.5726301 | 0.1279165 | 0.651060522 |
| ENSMUSG00000029309 | Sparcl1 | protein_coding | -0.414503746 | 9.275472997 | 2.5723902 | 0.127933 | 0.651060522 |
| ENSMUSG00000040028 | Elavl1 | protein_coding | -0.450264061 | 5.450016364 | 2.5707309 | 0.1280475 | 0.651060522 |
| ENSMUSG00000054808 | Actn4 | protein_coding | -0.648173345 | 4.526153361 | 2.5681386 | 0.1282266 | 0.651060522 |
| ENSMUSG00000045867 | Cradd | protein_coding | 0.491380399 | 4.529806171 | 2.568033 | 0.1282339 | 0.651060522 |
| ENSMUSG00000024792 | Zfpl1 | protein_coding | -0.95824228 | 3.797924973 | 2.5680078 | 0.1282672 | 0.651060522 |
| ENSMUSG00000047888 | Tnrc6b | protein_coding | -0.516634034 | 4.947256467 | 2.5670776 | 0.1283 | 0.651060522 |
| ENSMUSG00000008763 | Man1a2 | protein_coding | -0.562821471 | 5.403876338 | 2.5664689 | 0.1283422 | 0.651060522 |
| ENSMUSG00000037872 | Ackr1 | protein_coding | 0.592191842 | 4.243581478 | 2.5655449 | 0.1284062 | 0.651060522 |
| ENSMUSG00000035051 | Dhx57 | protein_coding | 0.624448777 | 4.112709344 | 2.5652657 | 0.1284255 | 0.651060522 |
| ENSMUSG00000024245 | Tmem178 | protein_coding | -1.176869498 | 3.659488846 | 2.5814846 | 0.1285287 | 0.651060522 |
| ENSMUSG00000021500 | Ddx46 | protein_coding | -0.46411451 | 6.111591387 | 2.563554 | 0.1285442 | 0.651060522 |
| ENSMUSG00000030876 | Mettl9 | protein_coding | -0.592949633 | 5.042005265 | 2.5635174 | 0.1285467 | 0.651060522 |
| ENSMUSG00000039578 | Ccser1 | protein_coding | -0.615657183 | 4.043679451 | 2.5627266 | 0.1286016 | 0.651060522 |
| ENSMUSG00000029993 | Nfu1 | protein_coding | 0.326918876 | 8.380902641 | 2.5614231 | 0.1286921 | 0.651212094 |
| ENSMUSG00000027589 | Pcmtd2 | protein_coding | -0.482583719 | 5.765213064 | 2.5575572 | 0.1289609 | 0.651767013 |
| ENSMUSG00000034211 | Mrps17 | protein_coding | 0.344659583 | 7.895572155 | 2.5573879 | 0.1289727 | 0.651767013 |
| ENSMUSG00000015812 | Gnrh1 | protein_coding | -0.952671856 | 5.32651247 | 2.6005524 | 0.1290654 | 0.651767013 |
| ENSMUSG00000029146 | Snx17 | protein_coding | 0.409987636 | 8.228083589 | 2.5557774 | 0.1290849 | 0.651767013 |
| ENSMUSG00000024668 | Sdhaf2 | protein_coding | 0.330274661 | 6.812997948 | 2.5554924 | 0.1291048 | 0.651767013 |
| ENSMUSG00000037174 | Elf2 | protein_coding | -0.699384001 | 3.947263548 | 2.5527306 | 0.1292975 | 0.651802194 |
| ENSMUSG00000047193 | Dync2h1 | protein_coding | -0.555565386 | 5.172442651 | 2.5496571 | 0.1295124 | 0.651802194 |
| ENSMUSG00000026452 | Syt2 | protein_coding | 0.581869479 | 4.272954533 | 2.549144 | 0.1295484 | 0.651802194 |
| ENSMUSG00000020372 | Rack1 | protein_coding | 0.357336691 | 8.901453793 | 2.5487159 | 0.1295783 | 0.651802194 |
| ENSMUSG00000007783 | Cpt1c | protein_coding | 0.419648765 | 6.709707017 | 2.5479513 | 0.1296319 | 0.651802194 |
| ENSMUSG00000037119 | Fam91a1 | protein_coding | -0.577160543 | 4.981655417 | 2.5469745 | 0.1297004 | 0.651802194 |
| ENSMUSG00000039917 | Rhbdd2 | protein_coding | -0.996134346 | 6.527210063 | 2.6813789 | 0.1297509 | 0.651802194 |
| ENSMUSG00000027215 | Cd82 | protein_coding | -0.471894614 | 6.399149265 | 2.5457601 | 0.1297856 | 0.651802194 |
| ENSMUSG00000023919 | Cenpq | protein_coding | -1.031891264 | 3.48608627 | 2.5449096 | 0.1298768 | 0.651802194 |
| ENSMUSG00000037204 | Atg101 | protein_coding | 0.3285826 | 6.702083432 | 2.5443929 | 0.1298815 | 0.651802194 |
| ENSMUSG00000032306 | Mpi | protein_coding | 0.3780038 | 7.010687497 | 2.5437781 | 0.1299247 | 0.651802194 |
| ENSMUSG00000036934 | 4921524J17Rik | protein_coding | -0.488025987 | 5.390113529 | 2.5431003 | 0.1299724 | 0.651802194 |
| ENSMUSG00000061607 | Mdc1 | protein_coding | -1.734267411 | 2.20541745 | 2.5430463 | 0.1299762 | 0.651802194 |
| ENSMUSG00000006010 | BC003331 | protein_coding | -0.447768903 | 5.896520046 | 2.542089 | 0.1300435 | 0.651802194 |
| ENSMUSG00000025986 | Slc39a10 | protein_coding | -0.403827016 | 5.661117132 | 2.5419154 | 0.1300557 | 0.651802194 |
| ENSMUSG00000033237 | Arid2 | protein_coding | -0.58904414 | 4.539216334 | 2.540859 | 0.13013 | 0.651802194 |
| ENSMUSG00000057335 | Cep170 | protein_coding | -0.483888391 | 5.24209487 | 2.5392546 | 0.130243 | 0.651802194 |
| ENSMUSG00000042298 | Ttc19 | protein_coding | -0.502082482 | 5.377593297 | 2.5380981 | 0.1303246 | 0.651802194 |
| ENSMUSG00000020432 | Tcn2 | protein_coding | -0.820417671 | 4.297761331 | 2.5381905 | 0.1303496 | 0.651802194 |
| ENSMUSG00000032735 | Ablim3 | protein_coding | 0.564901937 | 4.937772889 | 2.5375795 | 0.1303612 | 0.651802194 |
| ENSMUSG00000041935 | AW549877 | protein_coding | -0.478495064 | 5.234525089 | 2.5370114 | 0.1304012 | 0.651802194 |
| ENSMUSG00000028256 | Odf2l | protein_coding | 0.991143261 | 3.174813505 | 2.5356585 | 0.1304968 | 0.651802194 |
| ENSMUSG00000026641 | Usf1 | protein_coding | 0.378583425 | 6.31063339 | 2.5347788 | 0.1305589 | 0.651802194 |
| ENSMUSG00000043432 | Leng9 | protein_coding | 0.593372207 | 4.373271908 | 2.5340497 | 0.1306105 | 0.651802194 |
| ENSMUSG00000041453 | Rpl21 | protein_coding | 0.452697438 | 5.190588427 | 2.532076 | 0.1307501 | 0.651802194 |
| ENSMUSG00000029776 | Hibadh | protein_coding | -0.441503517 | 6.256325144 | 2.5307771 | 0.1308421 | 0.651802194 |
| ENSMUSG00000026172 | Bcs1l | protein_coding | 0.42742862 | 6.012627498 | 2.5289073 | 0.1309747 | 0.651802194 |
| ENSMUSG00000017288 | Vps53 | protein_coding | 0.504839627 | 6.081862636 | 2.528563 | 0.1309992 | 0.651802194 |
| ENSMUSG00000071037 | Camkmt | protein_coding | -0.685214996 | 4.431726802 | 2.5277943 | 0.1310537 | 0.651802194 |
| ENSMUSG00000020514 | Mrpl22 | protein_coding | 0.38850261 | 6.224653797 | 2.5277841 | 0.1310545 | 0.651802194 |
| ENSMUSG00000036534 | Slc38a7 | protein_coding | 0.924934382 | 3.731879295 | 2.5281486 | 0.1310601 | 0.651802194 |
| ENSMUSG00000054855 | Rnd1 | protein_coding | 0.57036751 | 5.994944056 | 2.5286085 | 0.1311105 | 0.651802194 |
| ENSMUSG00000021027 | Ralgapa1 | protein_coding | -0.49498898 | 5.251444168 | 2.5269728 | 0.1311121 | 0.651802194 |
| ENSMUSG00000024959 | Bad | protein_coding | 0.333383477 | 7.611176989 | 2.5238908 | 0.1313313 | 0.652439618 |
| ENSMUSG00000022708 | Zbtb20 | protein_coding | -0.532133848 | 6.260583793 | 2.523733 | 0.131374 | 0.652439618 |
| ENSMUSG00000019849 | Prep | protein_coding | -0.624504612 | 3.773300172 | 2.5224998 | 0.1314303 | 0.652439618 |
| ENSMUSG00000020521 | Rnft1 | protein_coding | -0.639899401 | 4.037475334 | 2.5217607 | 0.131483 | 0.652439618 |
| ENSMUSG00000039294 | BC017643 | protein_coding | 0.882551836 | 4.386000687 | 2.5370108 | 0.1315478 | 0.652460184 |
| ENSMUSG00000097929 | Tunar | protein_coding | 0.541494139 | 4.680065021 | 2.5161529 | 0.1318835 | 0.653566736 |
| ENSMUSG00000036198 | Arhgap36 | protein_coding | 0.511784317 | 5.900011865 | 2.5160274 | 0.1318925 | 0.653566736 |
| ENSMUSG00000020704 | Asic2 | protein_coding | 0.382945362 | 5.867010755 | 2.5123443 | 0.1321564 | 0.654572667 |
| ENSMUSG00000025584 | Pde8a | protein_coding | -0.756719149 | 4.010309423 | 2.5108882 | 0.1322608 | 0.654788619 |
| ENSMUSG00000032517 | Mobp | protein_coding | -0.348333298 | 7.82379215 | 2.5093478 | 0.1323715 | 0.655034881 |
| ENSMUSG00000059708 | Akap17b | protein_coding | -0.646351375 | 3.972768792 | 2.5049892 | 0.1326852 | 0.656211542 |
| ENSMUSG00000040943 | Tet2 | protein_coding | -0.563231424 | 5.009119942 | 2.5038194 | 0.1327696 | 0.656211542 |
| ENSMUSG00000037936 | Scarb1 | protein_coding | -0.647560849 | 4.02383914 | 2.5035034 | 0.1327924 | 0.656211542 |
| ENSMUSG00000041801 | Phlda3 | protein_coding | 0.478133399 | 5.543786398 | 2.50231 | 0.1328785 | 0.656335575 |
| ENSMUSG00000002332 | Dhrs1 | protein_coding | 0.346551841 | 7.310267359 | 2.5012273 | 0.1329567 | 0.656420343 |
| ENSMUSG00000092416 | Zfp141 | protein_coding | -0.911832492 | 3.008401772 | 2.4999581 | 0.1330485 | 0.65657187 |
| ENSMUSG00000023723 | Mrps23 | protein_coding | 0.345009465 | 6.937090636 | 2.4985587 | 0.1331497 | 0.656770106 |
| ENSMUSG00000012422 | Tmem167 | protein_coding | 0.385907931 | 7.382104096 | 2.4954806 | 0.1333728 | 0.657287194 |
| ENSMUSG00000010097 | Nxf1 | protein_coding | 0.305003945 | 7.710549011 | 2.4954249 | 0.1333768 | 0.657287194 |
| ENSMUSG00000021420 | Fars2 | protein_coding | 0.417193785 | 5.774157552 | 2.4922218 | 0.1336094 | 0.658131738 |
| ENSMUSG00000027782 | Kpna4 | protein_coding | -0.423557272 | 5.463499398 | 2.4913192 | 0.133675 | 0.658146974 |

| ensembl_gene_id | external_gene_name | gene_biotype | log2 (fold change) | log2 (counts per million) | F-value | p-value | False discovery rate |
|---|---|---|---|---|---|---|---|
| ENSMUSG00000040339 | Fam102b | protein_coding | -0.426204039 | 5.639958171 | 2.4904962 | 0.1337349 | 0.658146974 |
| ENSMUSG00000022323 | Rida | protein_coding | -0.460374402 | 7.338296693 | 2.4895582 | 0.1338347 | 0.658336734 |
| ENSMUSG00000097023 | Mir9-3hg | lincRNA | -0.755987913 | 3.721432233 | 2.4836002 | 0.1342379 | 0.659821662 |
| ENSMUSG00000030788 | Rnf141 | protein_coding | -0.483105909 | 5.494603547 | 2.482443 | 0.1343225 | 0.659821662 |
| ENSMUSG00000038344 | Txlng | protein_coding | -0.793023515 | 3.688826837 | 2.4822669 | 0.1343354 | 0.659821662 |
| ENSMUSG00000035093 | Secisbp2l | protein_coding | -0.455026878 | 6.285429562 | 2.4803593 | 0.134475 | 0.659821662 |
| ENSMUSG00000006527 | Sfmbt1 | protein_coding | -0.56188133 | 4.659079906 | 2.4798428 | 0.1345129 | 0.659821662 |
| ENSMUSG00000052981 | Ube2ql1 | protein_coding | 0.381939228 | 6.85934638 | 2.479619 | 0.1345293 | 0.659821662 |
| ENSMUSG00000033751 | Gadd45gip1 | protein_coding | -1.156471126 | 3.098671897 | 2.4791175 | 0.1345661 | 0.659821662 |
| ENSMUSG00000025860 | Xiap | protein_coding | -0.39308272 | 6.136989412 | 2.4752372 | 0.1348509 | 0.660272716 |
| ENSMUSG00000024990 | Rbp4 | protein_coding | -1.082636185 | 2.89401527 | 2.4734494 | 0.1349824 | 0.660272716 |
| ENSMUSG00000024976 | Shoc2 | protein_coding | -0.47624989 | 8.430827714 | 2.473105 | 0.1350393 | 0.660272716 |
| ENSMUSG00000005371 | Fbxo11 | protein_coding | -0.376443493 | 6.474362649 | 2.4725198 | 0.1350509 | 0.660272716 |
| ENSMUSG00000016664 | Pacsin2 | protein_coding | -0.563551368 | 4.843475171 | 2.4723477 | 0.1350636 | 0.660272716 |
| ENSMUSG00000007944 | Ttc9b | protein_coding | 0.661667875 | 4.256573623 | 2.4717644 | 0.1351065 | 0.660272716 |
| ENSMUSG00000020491 | 2810021J22Rik | protein_coding | 0.755108309 | 3.813919539 | 2.4708069 | 0.1351771 | 0.660272716 |
| ENSMUSG00000001138 | Cnnm3 | protein_coding | -0.698248527 | 4.023338172 | 2.4698454 | 0.135248 | 0.660272716 |
| ENSMUSG00000054162 | Spock3 | protein_coding | -0.341536581 | 7.329591639 | 2.4697376 | 0.135256 | 0.660272716 |
| ENSMUSG00000020790 | Ankfy1 | protein_coding | -0.654245333 | 4.178651392 | 2.4690992 | 0.1353031 | 0.660272716 |
| ENSMUSG00000031207 | Msn | protein_coding | -1.11596682 | 2.915599636 | 2.4686871 | 0.1353335 | 0.660272716 |
| ENSMUSG00000034793 | G6pc3 | protein_coding | 0.47962518 | 6.389852792 | 2.4675868 | 0.1354462 | 0.660523131 |
| ENSMUSG00000020056 | Washc3 | protein_coding | 0.410065129 | 6.423032868 | 2.4661189 | 0.1355233 | 0.660599156 |
| ENSMUSG00000031883 | Car7 | protein_coding | 0.723615084 | 3.743672053 | 2.4650698 | 0.1356009 | 0.660638547 |
| ENSMUSG00000021972 | Hmbox1 | protein_coding | -0.7197762 | 3.662169594 | 2.4639284 | 0.1356854 | 0.660638547 |
| ENSMUSG00000035164 | Zc3h12c | protein_coding | -0.852101845 | 3.641309505 | 2.4635197 | 0.1357157 | 0.660638547 |
| ENSMUSG00000087679 | Tmem250-ps | lincRNA | 0.434332381 | 9.365723529 | 2.4618593 | 0.1358387 | 0.660881703 |
| ENSMUSG00000105265 | Sox2ot | processed_transcript | -0.365904579 | 6.116986292 | 2.4611881 | 0.1358885 | 0.660881703 |
| ENSMUSG00000042831 | Alkbh6 | protein_coding | 0.529627113 | 7.43084631 | 2.4766775 | 0.1359653 | 0.660955895 |
| ENSMUSG00000047414 | Flrt2 | protein_coding | -0.62270185 | 4.531359265 | 2.4582277 | 0.1361084 | 0.661122833 |
| ENSMUSG00000048142 | Nat8l | protein_coding | 0.353342725 | 6.965309063 | 2.4580374 | 0.1361226 | 0.661122833 |
| ENSMUSG00000040855 | Reps2 | protein_coding | -0.461815657 | 6.145058229 | 2.4560018 | 0.136274 | 0.661559627 |
| ENSMUSG00000060703 | Cd302 | protein_coding | -0.616214865 | 5.575286645 | 2.4554169 | 0.1363491 | 0.661625217 |
| ENSMUSG00000023175 | Bsg | protein_coding | -0.579247776 | 8.302447301 | 2.4771896 | 0.1366196 | 0.662543073 |
| ENSMUSG00000037355 | Uvssa | protein_coding | -1.118968948 | 2.934807809 | 2.4508074 | 0.1366614 | 0.662543073 |
| ENSMUSG00000021033 | Gstz1 | protein_coding | 0.336863926 | 7.07085356 | 2.4477479 | 0.1368903 | 0.662923799 |
| ENSMUSG00000111947 | AC131120.1 | lincRNA | -1.414050026 | 2.78943609 | 2.4480795 | 0.1368969 | 0.662923799 |
| ENSMUSG00000063019 | Manbal | protein_coding | 0.335280343 | 7.243091372 | 2.4465933 | 0.1369767 | 0.662923799 |
| ENSMUSG00000021282 | Eif5 | protein_coding | -0.328730127 | 6.971456499 | 2.446462 | 0.1369866 | 0.662923799 |
| ENSMUSG00000021559 | Dapk1 | protein_coding | -0.468893715 | 5.818781129 | 2.441059 | 0.1373921 | 0.663608378 |
| ENSMUSG00000074736 | Syndig1 | protein_coding | 0.407745634 | 6.409774563 | 2.4389383 | 0.1375517 | 0.663608378 |
| ENSMUSG00000034765 | Dusp5 | protein_coding | -1.293895296 | 2.497909975 | 2.4381437 | 0.1376116 | 0.663608378 |
| ENSMUSG00000045078 | Rnf216 | protein_coding | -0.565802791 | 5.031410019 | 2.4372781 | 0.1376768 | 0.663608378 |
| ENSMUSG00000047361 | Gm973 | protein_coding | -1.472335265 | 2.290108878 | 2.4370713 | 0.1376924 | 0.663608378 |
| ENSMUSG00000028322 | Exosc3 | protein_coding | -0.852894623 | 3.460895537 | 2.4365331 | 0.137733 | 0.663608378 |
| ENSMUSG00000085069 | Gm13111 | antisense_RNA | -1.247074249 | 2.502866299 | 2.4361023 | 0.1377655 | 0.663608378 |
| ENSMUSG00000057497 | Fam136a | protein_coding | 0.347894259 | 6.854044648 | 2.4358579 | 0.137784 | 0.663608378 |
| ENSMUSG00000034912 | Mdga2 | protein_coding | -0.646876129 | 5.501504469 | 2.4354648 | 0.1378451 | 0.663608378 |
| ENSMUSG00000030530 | Furin | protein_coding | -0.960416154 | 2.998263109 | 2.4349991 | 0.1378488 | 0.663608378 |
| ENSMUSG00000072692 | Rpl37rt | transcribed_processe | 0.581899292 | 5.42589792 | 2.4353529 | 0.1378535 | 0.663608378 |
| ENSMUSG00000051855 | Mest | protein_coding | -0.878479114 | 3.702198442 | 2.4347365 | 0.1378686 | 0.663608378 |
| ENSMUSG00000058325 | Dock1 | protein_coding | -0.672452365 | 4.34361608 | 2.4332967 | 0.1379774 | 0.663798656 |
| ENSMUSG00000030209 | Grin2b | protein_coding | -0.43415209 | 6.206190231 | 2.4325797 | 0.1380316 | 0.663798656 |
| ENSMUSG00000027282 | Mtch2 | protein_coding | 0.294104783 | 7.647824628 | 2.4309192 | 0.1381572 | 0.664105892 |
| ENSMUSG00000046699 | Slitrk4 | protein_coding | -0.572506433 | 4.671661976 | 2.4294637 | 0.1382675 | 0.664338872 |
| ENSMUSG00000025332 | Kdm5c | protein_coding | -0.612359489 | 4.416504247 | 2.4285627 | 0.1383358 | 0.664370198 |
| ENSMUSG00000020923 | Ubtf | protein_coding | -0.424635434 | 6.328263927 | 2.4271115 | 0.1384459 | 0.664602132 |
| ENSMUSG00000008429 | Herpud2 | protein_coding | -0.705435644 | 4.95469928 | 2.426163 | 0.1385494 | 0.664757554 |
| ENSMUSG00000042302 | Ehbp1 | protein_coding | -0.713172053 | 4.174940949 | 2.4244348 | 0.1386492 | 0.664757554 |
| ENSMUSG00000021313 | Ryr2 | protein_coding | -0.876170023 | 4.29272841 | 2.4292264 | 0.1386637 | 0.664757554 |
| ENSMUSG00000113429 | CT030140.1 | lincRNA | 0.916371966 | 2.950139412 | 2.4229324 | 0.1387636 | 0.664939603 |
| ENSMUSG00000026385 | Dbi | protein_coding | -0.56015702 | 9.643997402 | 2.4369425 | 0.1389215 | 0.665202438 |
| ENSMUSG00000021555 | Naa35 | protein_coding | -0.350557767 | 6.4186992 | 2.4201549 | 0.1389752 | 0.665202438 |
| ENSMUSG00000031954 | Cfdp1 | protein_coding | 0.299389797 | 8.68150032 | 2.4197769 | 0.139004 | 0.665202438 |
| ENSMUSG00000040037 | Negr1 | protein_coding | -0.46188691 | 6.345821179 | 2.4160454 | 0.139289 | 0.666269686 |
| ENSMUSG00000074733 | Zfp950 | protein_coding | -0.494347576 | 5.169887919 | 2.4131699 | 0.1395091 | 0.667025756 |
| ENSMUSG00000001120 | Pcbp3 | protein_coding | -0.749915024 | 6.028750481 | 2.4436188 | 0.1398824 | 0.668513764 |
| ENSMUSG00000025538 | Sumf2 | protein_coding | 0.656425516 | 5.000445832 | 2.4072644 | 0.1399938 | 0.66874892 |
| ENSMUSG00000037419 | Endod1 | protein_coding | -0.388758583 | 6.359519041 | 2.4038419 | 0.140226 | 0.669326092 |
| ENSMUSG00000049985 | Ankrd55 | protein_coding | 0.828328734 | 3.601224438 | 2.4033486 | 0.140264 | 0.669326092 |
| ENSMUSG00000030059 | Tmf1 | protein_coding | -0.692668962 | 6.469920041 | 2.4373168 | 0.1403562 | 0.669326092 |
| ENSMUSG00000073155 | 1810058I24Rik | lincRNA | 0.319771014 | 6.7691686 | 2.3995384 | 0.1405582 | 0.669326092 |
| ENSMUSG00000079641 | Rpl39 | protein_coding | 0.512065302 | 6.23679866 | 2.3996842 | 0.1405784 | 0.669326092 |
| ENSMUSG00000029371 | Cxcl5 | protein_coding | -2.005342005 | 2.469770503 | 2.4151182 | 0.1405787 | 0.669326092 |
| ENSMUSG00000022180 | Slc7a8 | protein_coding | -0.508872678 | 4.673576006 | 2.3987196 | 0.1406215 | 0.669326092 |
| ENSMUSG00000042043 | Tbca | protein_coding | 0.430747867 | 7.893308059 | 2.399095 | 0.1406239 | 0.669326092 |
| ENSMUSG00000074896 | Ifit3 | protein_coding | -0.892847189 | 3.128248239 | 2.3974133 | 0.1407226 | 0.669326092 |
| ENSMUSG00000022947 | Cbr3 | protein_coding | -0.917174061 | 3.36984053 | 2.3972261 | 0.1407371 | 0.669326092 |
| ENSMUSG00000025373 | Rnf41 | protein_coding | 0.369605173 | 7.783121997 | 2.3961018 | 0.1408242 | 0.669444219 |

| ensembl_gene_id | external_gene_name | gene_biotype | log2 (fold change) | log2 (counts per million) | F-value | p-value | False discovery rate |
|---|---|---|---|---|---|---|---|
| ENSMUSG00000038295 | Atg9b | protein_coding | 0.627577489 | 4.819288226 | 2.3952548 | 0.1408899 | 0.66946037 |
| ENSMUSG00000001700 | Gramd3 | protein_coding | -0.834827056 | 3.769375656 | 2.3930639 | 0.1410599 | 0.669972128 |
| ENSMUSG00000006673 | Qrich1 | protein_coding | -0.390378568 | 5.913338784 | 2.3912296 | 0.1412024 | 0.670099415 |
| ENSMUSG00000092310 | Gm20509 | protein_coding | -0.81987411 | 3.420100164 | 2.3908905 | 0.1412288 | 0.670099415 |
| ENSMUSG00000040907 | Atp1a3 | protein_coding | 0.322688107 | 9.92749935 | 2.3899351 | 0.1413031 | 0.670099415 |
| ENSMUSG00000029038 | Ssu72 | protein_coding | 0.302405523 | 7.920760484 | 2.3888752 | 0.1413856 | 0.670099415 |
| ENSMUSG00000022621 | Rabl2 | protein_coding | 0.439023761 | 6.451840936 | 2.3887127 | 0.1413983 | 0.670099415 |
| ENSMUSG00000031760 | Mt3 | protein_coding | 0.424805701 | 10.72742009 | 2.3874386 | 0.1414975 | 0.67018141 |
| ENSMUSG00000038570 | Saxo2 | protein_coding | 0.473354972 | 4.880712069 | 2.3868911 | 0.1415402 | 0.67018141 |
| ENSMUSG00000037896 | Rcor1 | protein_coding | -0.716432503 | 4.042405478 | 2.3836763 | 0.1417912 | 0.670910244 |
| ENSMUSG00000021696 | Elovl7 | protein_coding | -0.895037935 | 3.896187709 | 2.3836588 | 0.141824 | 0.670910244 |
| ENSMUSG00000032883 | Acsl3 | protein_coding | -0.482451412 | 5.964220527 | 2.3825231 | 0.1418813 | 0.670910244 |
| ENSMUSG00000033499 | Larp4b | protein_coding | -0.382482965 | 6.176354794 | 2.3781285 | 0.1422255 | 0.671181327 |
| ENSMUSG00000029722 | Agfg2 | protein_coding | -0.511275875 | 5.019407381 | 2.37774 | 0.142256 | 0.671181327 |
| ENSMUSG00000035086 | Becn1 | protein_coding | 0.707861511 | 3.339431092 | 2.3772992 | 0.1422906 | 0.671181327 |
| ENSMUSG00000044709 | Gemin7 | protein_coding | 0.415123023 | 5.674839794 | 2.3754897 | 0.1424327 | 0.671181327 |
| ENSMUSG00000004945 | Tmem242 | protein_coding | 0.330441809 | 6.377803356 | 2.3751748 | 0.1424574 | 0.671181327 |
| ENSMUSG00000039154 | Shd | protein_coding | 0.456732134 | 6.023462431 | 2.3733415 | 0.1426016 | 0.671181327 |
| ENSMUSG00000001082 | Mfsd10 | protein_coding | 0.88941722 | 3.679602123 | 2.372416 | 0.1427059 | 0.671181327 |
| ENSMUSG00000029361 | Nos1 | protein_coding | 0.56119043 | 6.17370665 | 2.3800118 | 0.1427179 | 0.671181327 |
| ENSMUSG00000050908 | Tvp23a | protein_coding | 0.649175541 | 3.957495416 | 2.3713235 | 0.1427605 | 0.671181327 |
| ENSMUSG00000055150 | Zfp78 | protein_coding | 0.737248369 | 3.449240101 | 2.370478 | 0.1428272 | 0.671181327 |
| ENSMUSG00000019966 | Kitl | protein_coding | -0.72579505 | 4.71695056 | 2.3705572 | 0.1428523 | 0.671181327 |
| ENSMUSG00000029022 | Miip | protein_coding | -0.729515035 | 3.589943655 | 2.3701269 | 0.1428548 | 0.671181327 |
| ENSMUSG00000036192 | Rorb | protein_coding | -0.996622006 | 5.784684724 | 2.4503599 | 0.1428894 | 0.671181327 |
| ENSMUSG00000040520 | Manea | protein_coding | -0.639030665 | 4.031956797 | 2.368251 | 0.1430029 | 0.671181327 |
| ENSMUSG00000069895 | Atxn1l | protein_coding | -0.998193567 | 3.812984742 | 2.3757283 | 0.1430076 | 0.671181327 |
| ENSMUSG00000022769 | Sdf2l1 | protein_coding | -1.061148084 | 2.504649534 | 2.3673461 | 0.1430744 | 0.671181327 |
| ENSMUSG00000044352 | Sowaha | protein_coding | -0.723279986 | 5.361425796 | 2.3812234 | 0.1431266 | 0.671181327 |
| ENSMUSG00000053111 | Fank1 | protein_coding | 0.680881468 | 4.313828583 | 2.3666572 | 0.1431288 | 0.671181327 |
| ENSMUSG00000045790 | Ccdc149 | protein_coding | -0.566118761 | 4.636312051 | 2.3662535 | 0.1431607 | 0.671181327 |
| ENSMUSG00000023806 | Rsph3b | protein_coding | 0.874553916 | 2.821497084 | 2.365712 | 0.1432035 | 0.671181327 |
| ENSMUSG00000050213 | Snip1 | protein_coding | -0.727474892 | 3.608189537 | 2.3646135 | 0.1432904 | 0.671181327 |
| ENSMUSG00000047767 | Atg16l2 | protein_coding | 0.772893228 | 4.074407515 | 2.3645685 | 0.1433254 | 0.671181327 |
| ENSMUSG00000003410 | Elavl3 | protein_coding | 0.330540558 | 6.897769516 | 2.3635029 | 0.1433784 | 0.671181327 |
| ENSMUSG00000059857 | Ntng1 | protein_coding | -0.691548961 | 5.235176209 | 2.3646207 | 0.1434367 | 0.671181327 |
| ENSMUSG00000041733 | Coq5 | protein_coding | 0.357132032 | 6.582183234 | 2.3616924 | 0.1435219 | 0.671184123 |
| ENSMUSG00000064246 | Chil1 | protein_coding | -0.730577632 | 4.459438099 | 2.360968 | 0.1436108 | 0.671184123 |
| ENSMUSG00000042962 | Gm5436 | processed_pseudoge | 0.628172516 | 3.503938644 | 2.3590666 | 0.1437303 | 0.671184123 |
| ENSMUSG00000054640 | Slc8a1 | protein_coding | -0.894080981 | 3.27902357 | 2.3584103 | 0.1437825 | 0.671184123 |
| ENSMUSG00000028328 | Tmod1 | protein_coding | -0.55776798 | 4.726566109 | 2.358121 | 0.1438055 | 0.671184123 |
| ENSMUSG00000052712 | BC004004 | protein_coding | 0.309189835 | 6.97331961 | 2.3580418 | 0.1438118 | 0.671184123 |
| ENSMUSG00000001105 | Ift20 | protein_coding | 0.299895747 | 7.772655175 | 2.3570147 | 0.1438935 | 0.671274031 |
| ENSMUSG00000039539 | Sgcz | protein_coding | -0.763624888 | 4.028245037 | 2.3557677 | 0.1439927 | 0.671445776 |
| ENSMUSG00000071649 | B3gat3 | protein_coding | 0.353240975 | 6.366993141 | 2.3539438 | 0.1441381 | 0.671832105 |
| ENSMUSG00000015478 | Rnf5 | protein_coding | 0.321759005 | 6.738153462 | 2.3524725 | 0.1442554 | 0.671871847 |
| ENSMUSG00000008167 | Fbxw9 | protein_coding | 0.691316728 | 4.700894323 | 2.3526639 | 0.1442716 | 0.671871847 |
| ENSMUSG00000036990 | Otud4 | protein_coding | -0.693388416 | 4.057387711 | 2.3478692 | 0.1446234 | 0.672005359 |
| ENSMUSG00000032112 | Trappc4 | protein_coding | 0.294254353 | 8.58449716 | 2.3477501 | 0.1446329 | 0.672005359 |
| ENSMUSG00000034903 | Cobll1 | protein_coding | -1.574366083 | 2.680774634 | 2.3478948 | 0.1446527 | 0.672005359 |
| ENSMUSG00000051048 | P4ha3 | protein_coding | 1.293876772 | 2.603268759 | 2.3470982 | 0.1446851 | 0.672005359 |
| ENSMUSG00000032712 | 2810474O19Rik | protein_coding | -0.637554017 | 3.9110502 | 2.3456396 | 0.144802 | 0.672005359 |
| ENSMUSG00000036087 | Slain2 | protein_coding | -1.146435939 | 3.484801061 | 2.354701 | 0.1448786 | 0.672005359 |
| ENSMUSG00000044646 | Zbtb7c | protein_coding | 1.035171278 | 2.786383775 | 2.3442623 | 0.1449125 | 0.672005359 |
| ENSMUSG00000021431 | Snrnp48 | protein_coding | -0.601173746 | 4.203297408 | 2.3438339 | 0.1449469 | 0.672005359 |
| ENSMUSG00000024851 | Pitpnm1 | protein_coding | 0.430695653 | 5.789043041 | 2.342078 | 0.1450879 | 0.672005359 |
| ENSMUSG00000044134 | Fam109a | protein_coding | -1.71940792 | 2.104216316 | 2.3403776 | 0.1452247 | 0.672005359 |
| ENSMUSG00000029863 | Casp2 | protein_coding | -0.947346957 | 2.862350715 | 2.3401582 | 0.1452423 | 0.672005359 |
| ENSMUSG00000037010 | Apln | protein_coding | -0.826925283 | 3.299783963 | 2.3397532 | 0.1452749 | 0.672005359 |
| ENSMUSG00000043881 | Kbtbd7 | protein_coding | -0.589454892 | 4.593530467 | 2.3383271 | 0.1453898 | 0.672005359 |
| ENSMUSG00000041571 | Selenow | protein_coding | 0.422333475 | 12.37679339 | 2.3375621 | 0.1454514 | 0.672005359 |
| ENSMUSG00000024926 | Kat5 | protein_coding | 0.439127842 | 6.447044186 | 2.337449 | 0.1454605 | 0.672005359 |
| ENSMUSG00000027438 | Napb | protein_coding | 0.388046223 | 8.827453936 | 2.3367914 | 0.1455136 | 0.672005359 |
| ENSMUSG00000075284 | Wipf1 | protein_coding | -0.82730421 | 3.142378486 | 2.3366863 | 0.145522 | 0.672005359 |
| ENSMUSG00000006333 | Rps9 | protein_coding | 0.350706448 | 9.87165058 | 2.3365503 | 0.145533 | 0.672005359 |
| ENSMUSG00000039515 | Ptpa | protein_coding | 0.374308676 | 8.410019744 | 2.3359835 | 0.1455788 | 0.672005359 |
| ENSMUSG00000041278 | Ttc1 | protein_coding | 0.290591549 | 7.870327927 | 2.3345852 | 0.1456917 | 0.672005359 |
| ENSMUSG00000021759 | Plpp1 | protein_coding | 0.486632593 | 4.696786493 | 2.33442 | 0.145705 | 0.672005359 |
| ENSMUSG00000032872 | Cyb5r4 | protein_coding | -0.681170575 | 3.598388793 | 2.3340617 | 0.145734 | 0.672005359 |
| ENSMUSG00000022474 | Pmm1 | protein_coding | 0.295097216 | 8.903386654 | 2.3340165 | 0.1457376 | 0.672005359 |
| ENSMUSG00000052707 | Tnrc6a | protein_coding | -0.43734492 | 5.374586533 | 2.3316122 | 0.1459321 | 0.672495568 |
| ENSMUSG00000025372 | Baiap2 | protein_coding | -0.389462817 | 7.550732892 | 2.3304165 | 0.1460289 | 0.672495568 |
| ENSMUSG00000061298 | Agbl4 | protein_coding | 0.602129889 | 4.51388219 | 2.3303837 | 0.1460315 | 0.672495568 |
| ENSMUSG00000001506 | Col1a1 | protein_coding | -1.818049863 | 2.015746299 | 2.32888 | 0.1461534 | 0.672559492 |
| ENSMUSG00000042705 | Commd10 | protein_coding | 0.327113066 | 6.755597794 | 2.3286692 | 0.1461705 | 0.672559492 |
| ENSMUSG00000074698 | Csnk2a1 | protein_coding | -0.434834906 | 5.274515442 | 2.3267687 | 0.1463248 | 0.672981222 |
| ENSMUSG00000039278 | Pcsk1n | protein_coding | 0.474507579 | 5.94998412 | 2.3233891 | 0.1465995 | 0.673799559 |
| ENSMUSG00000020873 | Slc35b1 | protein_coding | 0.31487464 | 7.164479832 | 2.3223197 | 0.1466866 | 0.673799559 |

| ensembl_gene_id | external_gene_name | gene_biotype | log2 (fold change) | log2 (counts per million) | F-value | p-value | False discovery rate |
|---|---|---|---|---|---|---|---|
| ENSMUSG00000061603 | Akap6 | protein_coding | -0.419475866 | 5.891526183 | 2.322121 | 0.1467028 | 0.673799559 |
| ENSMUSG00000069495 | Epc2 | protein_coding | -0.604876905 | 4.693634474 | 2.3215008 | 0.1467533 | 0.673799559 |
| ENSMUSG00000093483 | AA465934 | processed_transcript | -0.815373887 | 3.409305018 | 2.3200837 | 0.1468689 | 0.674042309 |
| ENSMUSG00000068747 | Sort1 | protein_coding | -0.515238218 | 5.841445731 | 2.3160921 | 0.147195 | 0.675062662 |
| ENSMUSG00000026705 | Klhl20 | protein_coding | -0.581995856 | 3.967137319 | 2.3158255 | 0.1472168 | 0.675062662 |
| ENSMUSG00000059495 | Arhgef12 | protein_coding | -0.445568393 | 5.470293133 | 2.3140477 | 0.1473623 | 0.675407932 |
| ENSMUSG00000034403 | Pja1 | protein_coding | 0.463896535 | 8.745547709 | 2.3142266 | 0.1474617 | 0.675407932 |
| ENSMUSG00000017344 | Vtn | protein_coding | -0.589641253 | 4.202792313 | 2.3126056 | 0.1474805 | 0.675407932 |
| ENSMUSG00000028005 | Gucy1b3 | protein_coding | -0.352639651 | 7.007594023 | 2.3118381 | 0.1475435 | 0.675408588 |
| ENSMUSG00000031939 | Taf1d | protein_coding | 0.656653747 | 4.656538511 | 2.3100024 | 0.1476941 | 0.675810687 |
| ENSMUSG00000029819 | Npy | protein_coding | -0.740243088 | 6.897415766 | 2.3691763 | 0.1478677 | 0.676138313 |
| ENSMUSG00000024661 | Fth1 | protein_coding | -1.817597526 | 12.1728887 | 2.5733691 | 0.1478975 | 0.676138313 |
| ENSMUSG00000031948 | Kars | protein_coding | 0.45761676 | 7.945595081 | 2.3082228 | 0.1479544 | 0.676138313 |
| ENSMUSG00000022762 | Ncam2 | protein_coding | -0.730248975 | 4.609970633 | 2.3046966 | 0.1481621 | 0.676755024 |
| ENSMUSG00000044600 | Smim7 | protein_coding | 0.290262721 | 8.103929156 | 2.3036727 | 0.1482152 | 0.676755024 |
| ENSMUSG00000067604 | Nms | protein_coding | -1.093022565 | 2.993198315 | 2.3013089 | 0.1484104 | 0.677358502 |
| ENSMUSG00000019810 | Fuca2 | protein_coding | 0.415606034 | 6.321853766 | 2.2989163 | 0.1486082 | 0.677515546 |
| ENSMUSG00000065922 | n-R5-8s1 | rRNA | 0.593848114 | 5.306719367 | 2.2983109 | 0.1486897 | 0.677515546 |
| ENSMUSG00000065954 | Tacc1 | protein_coding | -0.393024856 | 6.421678638 | 2.297514 | 0.1487243 | 0.677515546 |
| ENSMUSG00000001786 | Fbxo7 | protein_coding | -0.477761936 | 5.935291912 | 2.2962899 | 0.1488258 | 0.677515546 |
| ENSMUSG00000030770 | Parva | protein_coding | -0.432034172 | 5.503169889 | 2.2951633 | 0.1489192 | 0.677515546 |
| ENSMUSG00000030301 | Ccdc91 | protein_coding | 0.410690123 | 5.880553558 | 2.2936717 | 0.1490431 | 0.677515546 |
| ENSMUSG00000036202 | Rif1 | protein_coding | -0.647807616 | 4.189042573 | 2.2934996 | 0.1490574 | 0.677515546 |
| ENSMUSG00000022463 | Srebf2 | protein_coding | -0.388827682 | 5.999905884 | 2.2924222 | 0.1491469 | 0.677515546 |
| ENSMUSG00000005621 | Zfp592 | protein_coding | -0.800300591 | 3.739771541 | 2.2922542 | 0.1491609 | 0.677515546 |
| ENSMUSG00000056515 | Rab31 | protein_coding | -0.357509152 | 6.519935439 | 2.2915437 | 0.14922 | 0.677515546 |
| ENSMUSG00000059119 | Nap1l4 | protein_coding | 0.444806492 | 8.400587276 | 2.2909108 | 0.149304 | 0.677515546 |
| ENSMUSG00000051285 | Pcmtd1 | protein_coding | -0.475864905 | 5.094840284 | 2.2904115 | 0.1493142 | 0.677515546 |
| ENSMUSG00000034413 | Neurl1b | protein_coding | -0.967749645 | 3.442703507 | 2.2893209 | 0.149405 | 0.677515546 |
| ENSMUSG00000037253 | Mex3c | protein_coding | -0.788923454 | 3.938099964 | 2.2889115 | 0.1494391 | 0.677515546 |
| ENSMUSG00000026977 | 7-Mar | protein_coding | -0.434154606 | 5.295410902 | 2.2888998 | 0.1494401 | 0.677515546 |
| ENSMUSG00000010047 | Hyal2 | protein_coding | 0.586743695 | 4.857066359 | 2.2887463 | 0.1494529 | 0.677515546 |
| ENSMUSG00000028100 | Nudt17 | protein_coding | 0.71992149 | 4.793880109 | 2.2891883 | 0.149566 | 0.677533269 |
| ENSMUSG00000030880 | Polr3e | protein_coding | -0.787474243 | 4.903822005 | 2.299872 | 0.1496696 | 0.677533269 |
| ENSMUSG00000015501 | Hivep2 | protein_coding | -1.190798822 | 2.894831611 | 2.2855527 | 0.1497193 | 0.677533269 |
| ENSMUSG00000025499 | Hras | protein_coding | 0.417042799 | 6.458936108 | 2.2855288 | 0.1497213 | 0.677533269 |
| ENSMUSG00000021608 | Lpcat1 | protein_coding | 0.585364458 | 4.229477468 | 2.2849242 | 0.1497718 | 0.677533269 |
| ENSMUSG00000028683 | Eif2b3 | protein_coding | 0.388328927 | 6.287355809 | 2.2828262 | 0.1499473 | 0.677686156 |
| ENSMUSG00000046442 | Ppm1e | protein_coding | -0.441458305 | 5.906311378 | 2.2827874 | 0.1499505 | 0.677686156 |
| ENSMUSG00000002797 | Ggct | protein_coding | 0.461255602 | 6.464061391 | 2.2822184 | 0.1500295 | 0.677686156 |
| ENSMUSG00000019039 | Dalrd3 | protein_coding | 0.530524368 | 5.753914959 | 2.2818809 | 0.1500577 | 0.677686156 |
| ENSMUSG00000049800 | Sertad2 | protein_coding | -0.606429764 | 3.710481192 | 2.2794609 | 0.1502292 | 0.678175805 |
| ENSMUSG00000084830 | Gm14539 | processed_pseudoge | 0.616558237 | 3.93243946 | 2.2781596 | 0.1503384 | 0.678383931 |
| ENSMUSG00000000399 | Ndufa9 | protein_coding | 0.305188376 | 9.01370572 | 2.2766181 | 0.1504679 | 0.678683341 |
| ENSMUSG00000019936 | Epyc | protein_coding | -1.061767269 | 2.734746908 | 2.2747962 | 0.1506211 | 0.678892619 |
| ENSMUSG00000026185 | Igfbp5 | protein_coding | -0.547641276 | 5.404350889 | 2.2737181 | 0.1507118 | 0.678892619 |
| ENSMUSG00000032402 | Smad3 | protein_coding | -0.632750641 | 4.130400917 | 2.2725065 | 0.1508139 | 0.678892619 |
| ENSMUSG00000035632 | Cnot3 | protein_coding | 0.623004531 | 3.568040834 | 2.272121 | 0.1508464 | 0.678892619 |
| ENSMUSG00000025571 | Tnrc6c | protein_coding | -0.420679812 | 5.659112533 | 2.2718434 | 0.1508698 | 0.678892619 |
| ENSMUSG00000055639 | Dach1 | protein_coding | -0.643393822 | 3.902452405 | 2.2710303 | 0.1509384 | 0.678892619 |
| ENSMUSG00000034842 | Art3 | protein_coding | -1.60481133 | 2.4869802 | 2.2709789 | 0.150974 | 0.678892619 |
| ENSMUSG00000078307 | AI593442 | protein_coding | -0.53010696 | 8.096051681 | 2.2850523 | 0.1510518 | 0.678892619 |
| ENSMUSG00000029610 | Aimp2 | protein_coding | 0.338498864 | 6.109981577 | 2.2693066 | 0.1510839 | 0.678892619 |
| ENSMUSG00000020841 | Cpd | protein_coding | -0.583457045 | 4.600038782 | 2.2679278 | 0.1512004 | 0.678892619 |
| ENSMUSG00000048720 | Tbc1d12 | protein_coding | -0.799081267 | 3.270310205 | 2.2676632 | 0.1512227 | 0.678892619 |
| ENSMUSG00000022389 | Tef | protein_coding | -0.289357764 | 7.112904169 | 2.2668548 | 0.1512911 | 0.678892619 |
| ENSMUSG00000025025 | Mxi1 | protein_coding | -0.477355013 | 5.435271417 | 2.2654098 | 0.1514134 | 0.678892619 |
| ENSMUSG00000025362 | Rps26 | protein_coding | 0.448914568 | 7.665758863 | 2.2658853 | 0.1514869 | 0.678892619 |
| ENSMUSG00000028876 | Epha10 | protein_coding | 0.758462831 | 3.683833959 | 2.2643956 | 0.1514993 | 0.678892619 |
| ENSMUSG00000074657 | Kif5a | protein_coding | -0.430114326 | 8.897045906 | 2.2641968 | 0.1515475 | 0.678892619 |
| ENSMUSG00000008206 | Cers4 | protein_coding | 0.430839673 | 5.832784365 | 2.2627882 | 0.1516356 | 0.678892619 |
| ENSMUSG00000074738 | Fndc10 | protein_coding | 0.588886139 | 4.143632959 | 2.2626105 | 0.1516507 | 0.678892619 |
| ENSMUSG00000048416 | Mlf1 | protein_coding | -0.702344878 | 3.661680414 | 2.2617397 | 0.1517246 | 0.67894084 |
| ENSMUSG00000001962 | Fam50a | protein_coding | -0.78378354 | 4.370050922 | 2.2575885 | 0.1521088 | 0.680377065 |
| ENSMUSG00000025789 | St8sia2 | protein_coding | -1.163826562 | 2.608282858 | 2.2546269 | 0.15233 | 0.681030716 |
| ENSMUSG00000014294 | Ndufa2 | protein_coding | 0.408069246 | 9.642070955 | 2.2540213 | 0.1523817 | 0.681030716 |
| ENSMUSG00000024831 | Ighmbp2 | protein_coding | 1.016130808 | 3.065338584 | 2.2514876 | 0.1525981 | 0.681714702 |
| ENSMUSG00000039166 | Akap7 | protein_coding | 0.380389317 | 6.00025257 | 2.2492409 | 0.1527903 | 0.682135546 |
| ENSMUSG00000028414 | Fktn | protein_coding | -0.44449722 | 4.980741172 | 2.2489042 | 0.1528192 | 0.682135546 |
| ENSMUSG00000022450 | Ndufa6 | protein_coding | 0.370895238 | 10.01225996 | 2.2474102 | 0.1529472 | 0.682423747 |
| ENSMUSG00000039660 | Spout1 | protein_coding | 0.430410566 | 5.892875434 | 2.2450013 | 0.1531539 | 0.682595847 |
| ENSMUSG00000025588 | Nat1 | protein_coding | -1.386340129 | 2.428411009 | 2.244277 | 0.1532161 | 0.682595847 |
| ENSMUSG00000061778 | Mospd2 | protein_coding | -0.683673285 | 3.429067216 | 2.2424333 | 0.1533747 | 0.682595847 |
| ENSMUSG00000020926 | Adam11 | protein_coding | 0.486562385 | 4.884671681 | 2.2414837 | 0.1534564 | 0.682595847 |
| ENSMUSG00000052305 | Hbb-bs | protein_coding | -0.568850505 | 5.942453252 | 2.2427725 | 0.1534591 | 0.682595847 |
| ENSMUSG00000020713 | Gh | protein_coding | -1.286793421 | 3.159073707 | 2.2490084 | 0.1536078 | 0.682595847 |
| ENSMUSG00000009555 | Cdk9 | protein_coding | 0.574892117 | 4.755560864 | 2.2393815 | 0.1536375 | 0.682595847 |
| ENSMUSG00000097924 | A730020E08Rik | bidirectional_promote | 1.018452699 | 2.845040836 | 2.2387311 | 0.1536936 | 0.682595847 |

| ensembl_gene_id | external_gene_name | gene_biotype | log2 (fold change) | log2 (counts per million) | F-value | p-value | False discovery rate |
|---|---|---|---|---|---|---|---|
| ENSMUSG00000022210 | Dhrs4 | protein_coding | -0.591520361 | 4.737723596 | 2.2383372 | 0.1537276 | 0.682595847 |
| ENSMUSG00000032575 | Manf | protein_coding | -0.407681014 | 6.02686512 | 2.2383118 | 0.1537298 | 0.682595847 |
| ENSMUSG00000006360 | Crip1 | protein_coding | -0.813548771 | 4.025108628 | 2.2360422 | 0.153957 | 0.682595847 |
| ENSMUSG00000094483 | Purb | protein_coding | -0.313289223 | 8.418996873 | 2.2346424 | 0.1540468 | 0.682595847 |
| ENSMUSG00000008393 | Carhsp1 | protein_coding | -0.362567869 | 6.747320037 | 2.232867 | 0.1542004 | 0.682595847 |
| ENSMUSG00000020684 | Rasl10b | protein_coding | -0.658396446 | 5.728973139 | 2.246272 | 0.1542562 | 0.682595847 |
| ENSMUSG00000028383 | Hsdl2 | protein_coding | -0.398447693 | 5.550690746 | 2.231669 | 0.1543042 | 0.682595847 |
| ENSMUSG00000097789 | Gm2115 | protein_coding | -0.77031583 | 3.277335339 | 2.2313968 | 0.1543278 | 0.682595847 |
| ENSMUSG00000046079 | Lrrc8d | protein_coding | -0.516726224 | 4.44451498 | 2.2306357 | 0.1543938 | 0.682595847 |
| ENSMUSG00000014873 | Surf2 | protein_coding | 0.387446463 | 6.407623706 | 2.2303215 | 0.1544211 | 0.682595847 |
| ENSMUSG00000029193 | Cckar | protein_coding | -1.006067205 | 3.892365771 | 2.2438182 | 0.1544668 | 0.682595847 |
| ENSMUSG00000028581 | Laptm5 | protein_coding | -0.516074058 | 5.352613588 | 2.2288428 | 0.1545494 | 0.682595847 |
| ENSMUSG00000052151 | Plpp2 | protein_coding | 0.523765612 | 4.438587461 | 2.2285244 | 0.1545771 | 0.682595847 |
| ENSMUSG00000028232 | Tmem68 | protein_coding | -0.400351826 | 5.896256195 | 2.2283842 | 0.1545893 | 0.682595847 |
| ENSMUSG00000047935 | Gm5607 | sense_overlapping | -0.500524682 | 4.563505048 | 2.2264525 | 0.1547572 | 0.682595847 |
| ENSMUSG00000000120 | Ngfr | protein_coding | -0.990192422 | 2.71391218 | 2.2260783 | 0.1547897 | 0.682595847 |
| ENSMUSG00000020802 | Ube2o | protein_coding | 0.482014223 | 4.962172103 | 2.2257688 | 0.1548167 | 0.682595847 |
| ENSMUSG00000032330 | Cox7a2 | protein_coding | 0.409561219 | 10.09163773 | 2.2254359 | 0.1548456 | 0.682595847 |
| ENSMUSG00000000916 | Nsun5 | protein_coding | 0.759115354 | 3.846835917 | 2.2238697 | 0.1549821 | 0.682595847 |
| ENSMUSG00000042607 | Asb4 | protein_coding | 0.732810725 | 4.45846655 | 2.2223263 | 0.1551479 | 0.682595847 |
| ENSMUSG00000059316 | Slc27a4 | protein_coding | 0.468819964 | 5.374491072 | 2.2211042 | 0.1552233 | 0.682595847 |
| ENSMUSG00000034121 | Mks1 | protein_coding | 2.130429774 | 2.16448445 | 2.2353217 | 0.1552757 | 0.682595847 |
| ENSMUSG00000032579 | Hemk1 | protein_coding | -0.561951723 | 4.990889154 | 2.2200049 | 0.1553193 | 0.682595847 |
| ENSMUSG00000024073 | Birc6 | protein_coding | -0.47528368 | 5.592319642 | 2.2197866 | 0.1553384 | 0.682595847 |
| ENSMUSG00000024644 | Cndp2 | protein_coding | 0.506244058 | 6.333924575 | 2.2208493 | 0.155395 | 0.682595847 |
| ENSMUSG00000028060 | 2810403A07Rik | protein_coding | -0.574466196 | 4.220048193 | 2.2179638 | 0.1554978 | 0.682595847 |
| ENSMUSG00000024074 | Crim1 | protein_coding | -0.420419956 | 5.537589851 | 2.2177087 | 0.1555201 | 0.682595847 |
| ENSMUSG00000022521 | Crebbp | protein_coding | -0.405128742 | 7.026471536 | 2.2172509 | 0.1555602 | 0.682595847 |
| ENSMUSG00000041353 | Tmem29 | protein_coding | 0.334965866 | 6.878532586 | 2.2171681 | 0.1555674 | 0.682595847 |
| ENSMUSG00000012117 | Dhdds | protein_coding | 0.388264242 | 6.15705973 | 2.2171069 | 0.1555728 | 0.682595847 |
| ENSMUSG00000049327 | Kmt5a | protein_coding | -0.57722949 | 4.106663803 | 2.2148202 | 0.1557732 | 0.682595847 |
| ENSMUSG00000061013 | Mkx | protein_coding | -1.258444683 | 2.553565789 | 2.2137563 | 0.1558665 | 0.682595847 |
| ENSMUSG00000002997 | Prkar2b | protein_coding | -0.431204987 | 6.045390933 | 2.2130998 | 0.1559241 | 0.682595847 |
| ENSMUSG00000016257 | Prelid3b | protein_coding | 0.321546302 | 7.019083206 | 2.2125827 | 0.1559695 | 0.682595847 |
| ENSMUSG00000029458 | Brap | protein_coding | -0.478855734 | 4.55708519 | 2.211958 | 0.1560244 | 0.682595847 |
| ENSMUSG00000001630 | Stk38l | protein_coding | -0.455504485 | 5.132107652 | 2.2105364 | 0.1561494 | 0.682595847 |
| ENSMUSG00000042532 | Golga7b | protein_coding | 0.401159943 | 7.965838073 | 2.2103509 | 0.1561657 | 0.682595847 |
| ENSMUSG00000087461 | C230014O12Rik | lincRNA | 0.99310721 | 2.858043497 | 2.2100801 | 0.1561895 | 0.682595847 |
| ENSMUSG00000063856 | Gpx1 | protein_coding | 0.303071863 | 9.114042541 | 2.2068936 | 0.1564702 | 0.682595847 |
| ENSMUSG00000042109 | Csdc2 | protein_coding | 0.353558189 | 8.009879572 | 2.2066107 | 0.1564951 | 0.682595847 |
| ENSMUSG00000035640 | Cbarp | protein_coding | 0.385398083 | 6.86161817 | 2.205888 | 0.1565589 | 0.682595847 |
| ENSMUSG00000105843 | Gm19439 | TEC | -1.332000144 | 4.624204269 | 2.2962953 | 0.1565629 | 0.682595847 |
| ENSMUSG00000042548 | Asxl1 | protein_coding | -0.662229567 | 3.71268328 | 2.2046653 | 0.1566668 | 0.682595847 |
| ENSMUSG00000022400 | Rbx1 | protein_coding | 0.3056702 | 9.326508806 | 2.2037535 | 0.1567474 | 0.682595847 |
| ENSMUSG00000026457 | Adipor1 | protein_coding | -0.440691504 | 5.534447329 | 2.2034963 | 0.1567701 | 0.682595847 |
| ENSMUSG00000042097 | Zfp239 | protein_coding | 0.377613285 | 6.28642213 | 2.2011439 | 0.1569782 | 0.682595847 |
| ENSMUSG00000044150 | A830080D01Rik | protein_coding | -0.941326487 | 3.460706837 | 2.1993081 | 0.1571408 | 0.682595847 |
| ENSMUSG00000018340 | Anxa6 | protein_coding | 0.400452768 | 6.697405451 | 2.1990342 | 0.1571651 | 0.682595847 |
| ENSMUSG00000030657 | Xylt1 | protein_coding | -0.985973586 | 3.242131493 | 2.1989167 | 0.1571755 | 0.682595847 |
| ENSMUSG00000022858 | Tra2b | protein_coding | -0.376796011 | 6.079785217 | 2.1984608 | 0.1572159 | 0.682595847 |
| ENSMUSG00000032846 | Zswim6 | processed_transcript | -0.929609107 | 3.842680476 | 2.199972 | 0.1572492 | 0.682595847 |
| ENSMUSG00000055567 | Unc80 | protein_coding | -0.388463415 | 6.175450892 | 2.1978422 | 0.1572708 | 0.682595847 |
| ENSMUSG00000020691 | Mettl2 | protein_coding | 0.677111619 | 4.728498721 | 2.1976499 | 0.1573191 | 0.682595847 |
| ENSMUSG00000046311 | Zfp62 | protein_coding | -0.706478893 | 3.823587933 | 2.1956508 | 0.1574654 | 0.682595847 |
| ENSMUSG00000034729 | Mrps10 | protein_coding | 0.354196159 | 6.322368147 | 2.194545 | 0.1575637 | 0.682595847 |
| ENSMUSG00000050071 | Bex1 | protein_coding | 0.317824767 | 8.358075561 | 2.1935453 | 0.1576527 | 0.682595847 |
| ENSMUSG00000053641 | Dennd4a | protein_coding | -0.575138654 | 4.413775591 | 2.1933302 | 0.1576718 | 0.682595847 |
| ENSMUSG00000037926 | Ssh2 | protein_coding | -0.557060585 | 4.286228006 | 2.1926772 | 0.1577299 | 0.682595847 |
| ENSMUSG00000036948 | BC037034 | protein_coding | 0.665786277 | 3.583442074 | 2.1925114 | 0.1577447 | 0.682595847 |
| ENSMUSG00000037703 | Lzts3 | protein_coding | -0.468430988 | 5.886241849 | 2.1918803 | 0.1578009 | 0.682595847 |
| ENSMUSG00000044628 | Rnf208 | protein_coding | 0.388293947 | 6.932068985 | 2.1917014 | 0.1578169 | 0.682595847 |
| ENSMUSG00000069184 | Zfp72 | protein_coding | -0.880200775 | 3.149835557 | 2.1915145 | 0.1578335 | 0.682595847 |
| ENSMUSG00000039850 | Endov | protein_coding | 0.499423407 | 4.222574245 | 2.1907861 | 0.1578985 | 0.682595847 |
| ENSMUSG00000014769 | Psmb1 | protein_coding | 0.299604316 | 10.02840023 | 2.1904481 | 0.1579286 | 0.682595847 |
| ENSMUSG00000037972 | Snn | protein_coding | -0.436740093 | 5.734400238 | 2.1903949 | 0.1579333 | 0.682595847 |
| ENSMUSG00000052273 | Dnah3 | protein_coding | -1.149358931 | 2.341484185 | 2.1902759 | 0.157944 | 0.682595847 |
| ENSMUSG00000038816 | Ctnnal1 | protein_coding | -0.664652661 | 3.863867728 | 2.189647 | 0.1580001 | 0.682595847 |
| ENSMUSG00000027189 | Trim44 | protein_coding | -0.358764632 | 6.742039693 | 2.1881732 | 0.1581317 | 0.682595847 |
| ENSMUSG00000032604 | Qars | protein_coding | 0.427613265 | 5.644562786 | 2.1880734 | 0.1581406 | 0.682595847 |
| ENSMUSG00000027243 | Harbi1 | protein_coding | 0.719216522 | 4.173488341 | 2.1875019 | 0.1581916 | 0.682595847 |
| ENSMUSG00000046671 | Mtfr1l | protein_coding | 0.313089472 | 7.657374727 | 2.1862446 | 0.1583041 | 0.682595847 |
| ENSMUSG00000024642 | Tle4 | protein_coding | -0.492840725 | 5.585379741 | 2.1858638 | 0.1583381 | 0.682595847 |
| ENSMUSG00000037032 | Apbb1 | protein_coding | 0.399266818 | 9.16330702 | 2.1857121 | 0.1583517 | 0.682595847 |
| ENSMUSG00000031879 | Fam96b | protein_coding | 0.375424174 | 6.989638524 | 2.1853762 | 0.1583818 | 0.682595847 |
| ENSMUSG00000058159 | T2 | processed_transcript | 1.192363592 | 2.287451488 | 2.184299 | 0.1584782 | 0.682595847 |
| ENSMUSG00000002068 | Ccne1 | protein_coding | 0.725666769 | 3.513192971 | 2.1840747 | 0.1584983 | 0.682595847 |
| ENSMUSG00000040181 | Fmo1 | protein_coding | -0.860792962 | 3.594445406 | 2.1820317 | 0.1586815 | 0.682595847 |
| ENSMUSG00000105879 | Gm6204 | processed_pseudoge | 0.619917029 | 3.852311768 | 2.1814505 | 0.1587336 | 0.682595847 |

| ensembl_gene_id | external_gene_name | gene_biotype | log2 (fold change) | log2 (counts per million) | F-value | p-value | False discovery rate |
|---|---|---|---|---|---|---|---|
| ENSMUSG00000021687 | Scamp1 | protein_coding | -0.303577465 | 7.292855386 | 2.1801862 | 0.1588471 | 0.682595847 |
| ENSMUSG00000060240 | Cend1 | protein_coding | 0.393308498 | 9.860547497 | 2.1797184 | 0.1588892 | 0.682595847 |
| ENSMUSG00000030612 | Mrpl46 | protein_coding | 0.291071178 | 7.79607444 | 2.1796188 | 0.1588981 | 0.682595847 |
| ENSMUSG00000038683 | Pak1ip1 | protein_coding | 0.376379151 | 7.238185548 | 2.1784256 | 0.1590054 | 0.682595847 |
| ENSMUSG00000028413 | B4galt1 | protein_coding | -0.892402185 | 2.837557655 | 2.177068 | 0.1591275 | 0.682595847 |
| ENSMUSG00000028649 | Macf1 | protein_coding | -0.630703833 | 5.171982175 | 2.1770405 | 0.1591612 | 0.682595847 |
| ENSMUSG00000020964 | Sel1l | protein_coding | -0.372163863 | 5.974239127 | 2.1763708 | 0.1591903 | 0.682595847 |
| ENSMUSG00000036452 | Arhgap26 | protein_coding | 0.451710932 | 5.249432187 | 2.1756242 | 0.1592576 | 0.682595847 |
| ENSMUSG00000042523 | Dnal1 | protein_coding | -0.481519601 | 5.155331092 | 2.1746548 | 0.159345 | 0.682595847 |
| ENSMUSG00000029804 | Herc3 | protein_coding | 0.403205013 | 5.997854704 | 2.1737892 | 0.1594231 | 0.682595847 |
| ENSMUSG00000038014 | Fam120a | protein_coding | -0.502981083 | 5.576354021 | 2.1734463 | 0.159454 | 0.682595847 |
| ENSMUSG00000032352 | Lrrc1 | protein_coding | -0.769681357 | 3.422985212 | 2.1725483 | 0.1595351 | 0.682595847 |
| ENSMUSG00000086822 | 5330413P13Rik | lincRNA | -1.411572468 | 2.127317112 | 2.1725461 | 0.1595353 | 0.682595847 |
| ENSMUSG00000021962 | Dcp1a | protein_coding | -0.666891981 | 3.482967852 | 2.1722954 | 0.1595579 | 0.682595847 |
| ENSMUSG00000048076 | Arf1 | protein_coding | 0.333564914 | 9.201643865 | 2.1714796 | 0.1596316 | 0.682595847 |
| ENSMUSG00000049624 | Slc17a5 | protein_coding | 0.570123878 | 4.188864397 | 2.1703309 | 0.1597355 | 0.682595847 |
| ENSMUSG00000039860 | Srrm3 | protein_coding | -0.863865707 | 2.993255815 | 2.1687908 | 0.1598749 | 0.682595847 |
| ENSMUSG00000024393 | Prrc2a | protein_coding | 0.509552829 | 4.643013439 | 2.1683376 | 0.1599159 | 0.682595847 |
| ENSMUSG00000061524 | Zic2 | protein_coding | -1.146503588 | 2.582367582 | 2.168093 | 0.1599381 | 0.682595847 |
| ENSMUSG00000017721 | Pigt | protein_coding | 0.613311311 | 4.034348605 | 2.1667356 | 0.1600611 | 0.682595847 |
| ENSMUSG00000062252 | Lhfpl5 | protein_coding | 0.363464233 | 6.03676927 | 2.1662557 | 0.1601046 | 0.682595847 |
| ENSMUSG00000039068 | Zzz3 | protein_coding | -0.551716853 | 4.790596783 | 2.1661115 | 0.1601177 | 0.682595847 |
| ENSMUSG00000095325 | Zfp870 | protein_coding | -0.749184366 | 3.837029677 | 2.1659201 | 0.1601351 | 0.682595847 |
| ENSMUSG00000092329 | Gm20388 | protein_coding | 0.288129348 | 6.876614423 | 2.1640642 | 0.1603036 | 0.682595847 |
| ENSMUSG00000035898 | Uba6 | protein_coding | -0.711026749 | 4.259359949 | 2.1640586 | 0.1603041 | 0.682595847 |
| ENSMUSG00000064373 | Selenop | protein_coding | -0.440844996 | 7.836042949 | 2.1642484 | 0.160318 | 0.682595847 |
| ENSMUSG00000072214 | 5-Sep | protein_coding | 0.354841916 | 7.012611073 | 2.1637178 | 0.160335 | 0.682595847 |
| ENSMUSG00000026211 | Obsl1 | protein_coding | 0.908788368 | 2.832879547 | 2.1629281 | 0.1604068 | 0.682595847 |
| ENSMUSG00000022419 | Deptor | protein_coding | -0.800451437 | 4.525746661 | 2.169828 | 0.1604118 | 0.682595847 |
| ENSMUSG00000017802 | Retreg3 | protein_coding | -0.615275282 | 3.661015455 | 2.1627189 | 0.1604258 | 0.682595847 |
| ENSMUSG00000020064 | Herc4 | protein_coding | -0.732515837 | 3.692384802 | 2.1620783 | 0.1604841 | 0.682595847 |
| ENSMUSG00000062963 | Ufc1 | protein_coding | 0.311165641 | 8.435340463 | 2.1613011 | 0.1605548 | 0.682595847 |
| ENSMUSG00000001507 | Itga3 | protein_coding | 0.655797062 | 3.834342419 | 2.1598491 | 0.1606871 | 0.682595847 |
| ENSMUSG00000096943 | Gm26721 | lincRNA | -1.048479279 | 2.809165358 | 2.1596301 | 0.160707 | 0.682595847 |
| ENSMUSG00000001143 | Lman2l | protein_coding | 0.444885684 | 5.726341757 | 2.1595414 | 0.1607151 | 0.682595847 |
| ENSMUSG00000022386 | Trmu | protein_coding | 0.528052043 | 5.064305744 | 2.1587086 | 0.160791 | 0.682595847 |
| ENSMUSG00000069662 | Marcks | protein_coding | -0.296285439 | 7.706302332 | 2.1586785 | 0.1607938 | 0.682595847 |
| ENSMUSG00000028528 | Dnajc6 | protein_coding | -0.319958834 | 6.82525502 | 2.1570692 | 0.1609406 | 0.682760509 |
| ENSMUSG00000028034 | Fubp1 | protein_coding | 0.374721184 | 6.719120052 | 2.155786 | 0.1610578 | 0.682760509 |
| ENSMUSG00000003380 | Rabac1 | protein_coding | 0.419999874 | 8.737440246 | 2.1551815 | 0.1611442 | 0.682760509 |
| ENSMUSG00000037126 | Psd | protein_coding | -0.512059662 | 5.103726207 | 2.1523011 | 0.1613766 | 0.682760509 |
| ENSMUSG00000044098 | Rsbn1 | protein_coding | -0.952708922 | 2.860981529 | 2.1521894 | 0.1613868 | 0.682760509 |
| ENSMUSG00000046324 | Ermp1 | protein_coding | -0.713919002 | 4.23663807 | 2.1511926 | 0.1614782 | 0.682760509 |
| ENSMUSG00000037735 | 2810032G03Rik | lincRNA | -0.979513854 | 2.874450947 | 2.1506082 | 0.1615318 | 0.682760509 |
| ENSMUSG00000029111 | Nelfa | protein_coding | 0.562830789 | 5.752998423 | 2.1511693 | 0.1615934 | 0.682760509 |
| ENSMUSG00000024515 | Smad4 | protein_coding | -0.499803971 | 4.955008203 | 2.1496966 | 0.1616154 | 0.682760509 |
| ENSMUSG00000024072 | Yipf4 | protein_coding | -0.499304014 | 5.55137118 | 2.1486526 | 0.1617112 | 0.682760509 |
| ENSMUSG00000023999 | Kif6 | protein_coding | -1.321308886 | 2.428414314 | 2.1484682 | 0.1617282 | 0.682760509 |
| ENSMUSG00000008140 | Emc10 | protein_coding | -0.450061853 | 5.708759003 | 2.1476809 | 0.1618005 | 0.682760509 |
| ENSMUSG00000030652 | Coq7 | protein_coding | 0.279934525 | 7.84327492 | 2.1467503 | 0.1618861 | 0.682760509 |
| ENSMUSG00000057234 | Mettl15 | protein_coding | -0.683513891 | 3.394642558 | 2.1464499 | 0.1619137 | 0.682760509 |
| ENSMUSG00000001424 | Snd1 | protein_coding | 0.403813551 | 5.71235447 | 2.1463172 | 0.1619259 | 0.682760509 |
| ENSMUSG00000027122 | Arl14ep | protein_coding | -0.42745201 | 6.619910862 | 2.1451455 | 0.1620337 | 0.682760509 |
| ENSMUSG00000027247 | Arhgap1 | protein_coding | 0.475281893 | 6.71735667 | 2.1497806 | 0.1620553 | 0.682760509 |
| ENSMUSG00000068856 | Sf3b4 | protein_coding | 0.575126693 | 4.936819582 | 2.142464 | 0.1622808 | 0.682760509 |
| ENSMUSG00000042742 | Bmt2 | protein_coding | -0.559816346 | 4.507488992 | 2.1421083 | 0.1623136 | 0.682760509 |
| ENSMUSG00000023571 | C1qtnf12 | protein_coding | -1.05149492 | 3.022558791 | 2.1420949 | 0.1623148 | 0.682760509 |
| ENSMUSG00000005054 | Cstb | protein_coding | 0.346166163 | 8.031710396 | 2.1419221 | 0.1623308 | 0.682760509 |
| ENSMUSG00000074923 | Pak6 | protein_coding | -0.506744968 | 4.58149921 | 2.1411575 | 0.1624014 | 0.682760509 |
| ENSMUSG00000002345 | Borcs8 | protein_coding | 0.35285197 | 7.178621538 | 2.1408052 | 0.1624339 | 0.682760509 |
| ENSMUSG00000004035 | Gstm7 | protein_coding | -0.298070811 | 7.183323043 | 2.1407989 | 0.1624345 | 0.682760509 |
| ENSMUSG00000061315 | Naca | protein_coding | 0.361391498 | 6.815709084 | 2.1399222 | 0.1625154 | 0.682760509 |
| ENSMUSG00000005986 | Ankrd13d | protein_coding | 0.532857973 | 5.30003387 | 2.1395182 | 0.1625528 | 0.682760509 |
| ENSMUSG00000024298 | Zfp871 | protein_coding | -0.551713094 | 4.790642478 | 2.1390789 | 0.1625934 | 0.682760509 |
| ENSMUSG00000037062 | Sh3glb1 | protein_coding | -0.433745456 | 5.881245522 | 2.1379497 | 0.1626978 | 0.682760509 |
| ENSMUSG00000032407 | U2surp | protein_coding | -0.341877352 | 6.560169412 | 2.1379473 | 0.1626981 | 0.682760509 |
| ENSMUSG00000003814 | Calr | protein_coding | 0.311877827 | 8.158670102 | 2.1374331 | 0.1627456 | 0.682760509 |
| ENSMUSG00000022696 | Sidt1 | protein_coding | -0.853727907 | 3.069529831 | 2.1363419 | 0.1628467 | 0.682760509 |
| ENSMUSG00000009293 | Ube2g2 | protein_coding | 0.369234048 | 6.617847669 | 2.1361508 | 0.1628644 | 0.682760509 |
| ENSMUSG00000026843 | Fubp3 | protein_coding | -0.517195152 | 4.951118832 | 2.1351263 | 0.1629593 | 0.682892293 |
| ENSMUSG00000024378 | Stard4 | protein_coding | -0.70856782 | 4.103663219 | 2.1308132 | 0.1633598 | 0.684303855 |
| ENSMUSG00000044148 | 1810030O07Rik | protein_coding | 0.408435711 | 5.633291775 | 2.1269691 | 0.1637178 | 0.684711119 |
| ENSMUSG00000022779 | Top3b | protein_coding | -0.528467802 | 4.148305104 | 2.1265658 | 0.1637554 | 0.684711119 |
| ENSMUSG00000030619 | Eed | protein_coding | -0.651543043 | 4.505695003 | 2.1260638 | 0.1638022 | 0.684711119 |
| ENSMUSG00000037172 | E330009J07Rik | protein_coding | -0.549863229 | 4.419615746 | 2.1256991 | 0.1638362 | 0.684711119 |
| ENSMUSG00000027933 | Ints3 | protein_coding | 0.502168045 | 4.900079861 | 2.1256862 | 0.1638374 | 0.684711119 |
| ENSMUSG00000060924 | Csmd1 | protein_coding | -0.740572051 | 4.672874283 | 2.1269014 | 0.1638692 | 0.684711119 |
| ENSMUSG00000049571 | Cfap46 | protein_coding | 0.901685305 | 3.510504677 | 2.1253189 | 0.1639028 | 0.684711119 |

| ensembl_gene_id | external_gene_name | gene_biotype | log2 (fold change) | log2 (counts per million) | F-value | p-value | False discovery rate |
|---|---|---|---|---|---|---|---|
| ENSMUSG00000078640 | Gm11627 | protein_coding | -1.005790068 | 3.102746571 | 2.123728 | 0.1640203 | 0.684936136 |
| ENSMUSG00000017764 | Zswim1 | protein_coding | 0.512829792 | 5.908942062 | 2.121854 | 0.1642266 | 0.685057784 |
| ENSMUSG00000027624 | Epb41l1 | protein_coding | 0.350492536 | 6.805994717 | 2.1214283 | 0.1642354 | 0.685057784 |
| ENSMUSG00000012405 | Rpl15 | protein_coding | 0.315552363 | 6.940425499 | 2.1193026 | 0.1644345 | 0.685057784 |
| ENSMUSG00000037286 | Stag1 | protein_coding | -0.469715747 | 5.854711725 | 2.1184188 | 0.1645174 | 0.685057784 |
| ENSMUSG00000036478 | Btg1 | protein_coding | -0.544977393 | 5.003005241 | 2.1183189 | 0.1645268 | 0.685057784 |
| ENSMUSG00000049339 | Retreg2 | protein_coding | 0.349554767 | 6.910979518 | 2.1182612 | 0.1645322 | 0.685057784 |
| ENSMUSG00000061718 | Ppp1r1b | protein_coding | -1.021407392 | 2.587541766 | 2.1182591 | 0.1645324 | 0.685057784 |
| ENSMUSG00000070730 | Rmdn3 | protein_coding | 0.573913724 | 4.036268146 | 2.1179745 | 0.1645591 | 0.685057784 |
| ENSMUSG00000041645 | Ddx24 | protein_coding | 0.326905929 | 8.638056338 | 2.1165841 | 0.1646897 | 0.685335883 |
| ENSMUSG00000034839 | Larp6 | protein_coding | -0.617749228 | 4.866273999 | 2.1133603 | 0.1649928 | 0.68590569 |
| ENSMUSG00000089809 | Rasgef1b | protein_coding | -0.509756573 | 5.397372802 | 2.113255 | 0.1650027 | 0.68590569 |
| ENSMUSG00000031158 | Timm17b | protein_coding | 0.372662442 | 6.777739209 | 2.1125073 | 0.1650731 | 0.68590569 |
| ENSMUSG00000027601 | Mtfr1 | protein_coding | -0.710655471 | 3.960358547 | 2.112416 | 0.1650817 | 0.68590569 |
| ENSMUSG00000044986 | Tst | protein_coding | -0.507192775 | 6.522073545 | 2.1164443 | 0.1653314 | 0.686112282 |
| ENSMUSG00000037487 | Ubr5 | protein_coding | -0.440589502 | 5.318454565 | 2.1085426 | 0.1654471 | 0.686112282 |
| ENSMUSG00000021820 | Camk2g | protein_coding | 0.280091438 | 7.244062361 | 2.1083487 | 0.1654655 | 0.686112282 |
| ENSMUSG00000034751 | Mast4 | protein_coding | -0.706699647 | 3.656852043 | 2.1075683 | 0.1655392 | 0.686112282 |
| ENSMUSG00000032422 | Snx14 | protein_coding | -0.600802347 | 4.587299046 | 2.1071492 | 0.1655788 | 0.686112282 |
| ENSMUSG00000068748 | Ptprz1 | protein_coding | -0.710260159 | 5.973749057 | 2.1353422 | 0.1655862 | 0.686112282 |
| ENSMUSG00000034278 | Dnajc17 | protein_coding | 0.524537108 | 4.320435729 | 2.1058572 | 0.1657011 | 0.686112282 |
| ENSMUSG00000039483 | Asb6 | protein_coding | 0.411172044 | 6.025324549 | 2.1057971 | 0.1657067 | 0.686112282 |
| ENSMUSG00000028053 | Ash1l | protein_coding | -0.407870354 | 6.306372161 | 2.1053746 | 0.1657467 | 0.686112282 |
| ENSMUSG00000021803 | Cdhr1 | protein_coding | -1.093238297 | 2.655449623 | 2.105134 | 0.1657695 | 0.686112282 |
| ENSMUSG00000032383 | Ppib | protein_coding | 0.353491362 | 6.18592516 | 2.1023107 | 0.1660371 | 0.686665159 |
| ENSMUSG00000028069 | Gpatch4 | protein_coding | 0.337558684 | 6.731263987 | 2.1021727 | 0.1660502 | 0.686665159 |
| ENSMUSG00000054057 | A930004D18Rik | protein_coding | 0.619382618 | 3.860349917 | 2.1012868 | 0.1661343 | 0.686665159 |
| ENSMUSG00000051890 | Klhdc1 | protein_coding | -0.52573454 | 4.471963416 | 2.100423 | 0.1662164 | 0.686665159 |
| ENSMUSG00000029613 | Eif2ak1 | protein_coding | -0.503624758 | 4.672484427 | 2.0996004 | 0.1662945 | 0.686665159 |
| ENSMUSG00000038145 | Snrk | protein_coding | -0.575360221 | 4.663585786 | 2.0995844 | 0.1662961 | 0.686665159 |
| ENSMUSG00000100691 | 2010320M18Rik | lincRNA | 0.369324709 | 5.567277269 | 2.099016 | 0.1663501 | 0.686665159 |
| ENSMUSG00000033411 | Ctdspl2 | protein_coding | -0.595583899 | 4.635409623 | 2.097426 | 0.1665014 | 0.687025952 |
| ENSMUSG00000107283 | Mpv17 | protein_coding | 0.321122191 | 6.948560434 | 2.0965358 | 0.1665862 | 0.687112131 |
| ENSMUSG00000024269 | Tpgs2 | protein_coding | 0.315129287 | 6.612250471 | 2.0936081 | 0.1668654 | 0.687459435 |
| ENSMUSG00000034349 | Smc4 | protein_coding | -1.22272758 | 2.374126917 | 2.0916041 | 0.1670569 | 0.687459435 |
| ENSMUSG00000026727 | Rsu1 | protein_coding | -0.476015635 | 5.540845219 | 2.0903409 | 0.1671777 | 0.687459435 |
| ENSMUSG00000022095 | Fam160b2 | protein_coding | -0.609816197 | 3.768285915 | 2.090206 | 0.1671906 | 0.687459435 |
| ENSMUSG00000031112 | Stk26 | protein_coding | -1.284426183 | 2.454219891 | 2.0895238 | 0.1672559 | 0.687459435 |
| ENSMUSG00000026558 | Uck2 | protein_coding | -0.672259458 | 3.764361791 | 2.0886891 | 0.1673358 | 0.687459435 |
| ENSMUSG00000048486 | Fitm2 | protein_coding | -0.568835742 | 4.319730697 | 2.0886787 | 0.1673368 | 0.687459435 |
| ENSMUSG00000020876 | Snx11 | protein_coding | 0.670175674 | 4.596379397 | 2.0887728 | 0.1673588 | 0.687459435 |
| ENSMUSG00000041014 | Nrg3 | protein_coding | -0.545955903 | 4.681865863 | 2.0883723 | 0.1673662 | 0.687459435 |
| ENSMUSG00000024885 | Aldh3b1 | protein_coding | -0.70453511 | 3.921266022 | 2.0876054 | 0.1674397 | 0.687459435 |
| ENSMUSG00000038319 | Kcnh2 | protein_coding | 0.861640915 | 2.905448428 | 2.0863016 | 0.1675648 | 0.687459435 |
| ENSMUSG00000068264 | Ap5s1 | protein_coding | 0.374668578 | 6.289212354 | 2.0859143 | 0.1676019 | 0.687459435 |
| ENSMUSG00000038370 | Pcp4l1 | protein_coding | -0.451937954 | 8.590326869 | 2.0868058 | 0.167637 | 0.687459435 |
| ENSMUSG00000002804 | Nudt14 | protein_coding | 0.380614319 | 5.569466343 | 2.0850028 | 0.1676895 | 0.687459435 |
| ENSMUSG00000052387 | Trpm3 | protein_coding | -0.904340472 | 4.38765766 | 2.0995799 | 0.1678812 | 0.687459435 |
| ENSMUSG00000022905 | Kpna1 | protein_coding | -0.34795236 | 6.03260121 | 2.0819671 | 0.1679814 | 0.687459435 |
| ENSMUSG00000061601 | Pclo | protein_coding | -0.521046617 | 5.385969975 | 2.0804883 | 0.1681238 | 0.687459435 |
| ENSMUSG00000012609 | Ttll5 | protein_coding | 0.567419563 | 5.153133943 | 2.0799426 | 0.1681764 | 0.687459435 |
| ENSMUSG00000034640 | Tiparp | protein_coding | -0.675937275 | 4.451759757 | 2.0797377 | 0.1681961 | 0.687459435 |
| ENSMUSG00000027162 | Lin7c | protein_coding | -0.317405251 | 6.417872567 | 2.079556 | 0.1682137 | 0.687459435 |
| ENSMUSG00000059820 | AU019823 | protein_coding | -0.766838623 | 3.350332117 | 2.0791242 | 0.1682553 | 0.687459435 |
| ENSMUSG00000021991 | Cacna2d3 | protein_coding | -0.571408297 | 5.559323087 | 2.0790555 | 0.1682929 | 0.687459435 |
| ENSMUSG00000051579 | Tceal8 | protein_coding | -0.329684736 | 7.465920246 | 2.0783899 | 0.1683262 | 0.687459435 |
| ENSMUSG00000022752 | Tomm70a | protein_coding | -0.318271793 | 7.013920706 | 2.0782925 | 0.1683355 | 0.687459435 |
| ENSMUSG00000033096 | Apmap | protein_coding | 0.358584198 | 5.88880109 | 2.07807 | 0.168357 | 0.687459435 |
| ENSMUSG00000078695 | Cisd3 | protein_coding | 0.302671015 | 6.448157169 | 2.0776417 | 0.1683984 | 0.687459435 |
| ENSMUSG00000087006 | Gm13889 | protein_coding | -0.686694923 | 3.586865845 | 2.0771186 | 0.1684489 | 0.687459435 |
| ENSMUSG00000050394 | Armcx6 | protein_coding | 0.778461907 | 3.473449672 | 2.0767049 | 0.1684889 | 0.687459435 |
| ENSMUSG00000019235 | Rps6kl1 | protein_coding | 0.43756437 | 5.743658113 | 2.0763369 | 0.1685244 | 0.687459435 |
| ENSMUSG00000019528 | Gyg | protein_coding | 0.373309421 | 6.934157067 | 2.0753129 | 0.1686234 | 0.687602445 |
| ENSMUSG00000032353 | Tmed3 | protein_coding | 0.303482523 | 7.049319492 | 2.0708119 | 0.1690594 | 0.689119122 |
| ENSMUSG00000003033 | Ap1m1 | protein_coding | 0.42801967 | 7.141286704 | 2.0695716 | 0.1692108 | 0.689474525 |
| ENSMUSG00000087380 | 2210408F21Rik | processed_transcript | 0.408078021 | 4.930598638 | 2.066807 | 0.1694486 | 0.689877799 |
| ENSMUSG00000015981 | Stk32c | protein_coding | -0.886089747 | 3.19143195 | 2.0665876 | 0.1694699 | 0.689877799 |
| ENSMUSG00000032478 | Nme6 | protein_coding | 0.421265576 | 5.070254869 | 2.065701 | 0.1695563 | 0.689877799 |
| ENSMUSG00000021124 | Vti1b | protein_coding | -0.691574075 | 8.008059592 | 2.1208991 | 0.1695664 | 0.689877799 |
| ENSMUSG00000007564 | Ppp2r1a | protein_coding | 0.361763476 | 9.864951749 | 2.0648084 | 0.1696432 | 0.689929427 |
| ENSMUSG00000073888 | Ccl27a | protein_coding | 0.376811828 | 6.095853882 | 2.0641093 | 0.1697113 | 0.689945598 |
| ENSMUSG00000097486 | Gm17733 | bidirectional_promote | 1.713756396 | 2.426052377 | 2.0737588 | 0.1698988 | 0.690346871 |
| ENSMUSG00000066175 | 2510046G10Rik | antisense_RNA | 1.548449989 | 2.061809455 | 2.0617818 | 0.1699385 | 0.690346871 |
| ENSMUSG00000026527 | Rgs7 | protein_coding | -0.444505996 | 5.555931476 | 2.0594158 | 0.1701697 | 0.691025212 |
| ENSMUSG00000067279 | Ppp1r3c | protein_coding | -0.587231416 | 4.881953259 | 2.0587528 | 0.1702346 | 0.691027687 |
| ENSMUSG00000020840 | Blmh | protein_coding | 0.322764546 | 7.288699791 | 2.0578551 | 0.1703225 | 0.691109437 |
| ENSMUSG00000042726 | Trafd1 | protein_coding | -0.586494626 | 4.927957212 | 2.0568805 | 0.1704179 | 0.691109437 |
| ENSMUSG00000032375 | Aph1b | protein_coding | -0.542330027 | 4.121172696 | 2.0565785 | 0.1704475 | 0.691109437 |

| ensembl_gene_id | external_gene_name | gene_biotype | log2 (fold change) | log2 (counts per million) | F-value | p-value | False discovery rate |
|---|---|---|---|---|---|---|---|
| ENSMUSG00000030538 | Cib1 | protein_coding | -0.570784758 | 4.759552767 | 2.0557969 | 0.1705242 | 0.691155363 |
| ENSMUSG00000073415 | Gm10501 | antisense_RNA | 0.909303787 | 2.689804278 | 2.0551522 | 0.1705874 | 0.691155363 |
| ENSMUSG00000020427 | Igfbp3 | protein_coding | -1.000626341 | 2.704675004 | 2.0538949 | 0.1707108 | 0.691183652 |
| ENSMUSG00000024360 | Etf1 | protein_coding | -0.370986844 | 6.051335038 | 2.0526728 | 0.1708309 | 0.691183652 |
| ENSMUSG00000032131 | Abcg4 | protein_coding | 0.451896866 | 5.149193701 | 2.0514179 | 0.1709543 | 0.691183652 |
| ENSMUSG00000017400 | Stac2 | protein_coding | -0.694447664 | 4.566754954 | 2.0516455 | 0.1709628 | 0.691183652 |
| ENSMUSG00000045733 | Sprn | protein_coding | -0.595228923 | 4.718273447 | 2.0511988 | 0.1709758 | 0.691183652 |
| ENSMUSG00000020156 | Mum1 | protein_coding | 0.513514726 | 5.315632701 | 2.0511472 | 0.1709809 | 0.691183652 |
| ENSMUSG00000001289 | Pfdn5 | protein_coding | 0.395255914 | 9.841359291 | 2.050503 | 0.1710443 | 0.691183652 |
| ENSMUSG00000024953 | Prdx5 | protein_coding | 0.293440321 | 10.25842496 | 2.0492228 | 0.1711704 | 0.691396704 |
| ENSMUSG00000037369 | Kdm6a | protein_coding | -0.68061709 | 3.870583074 | 2.0486624 | 0.1712257 | 0.691396704 |
| ENSMUSG00000001517 | Foxm1 | protein_coding | 0.922518121 | 2.814224342 | 2.0476579 | 0.1713247 | 0.691519876 |
| ENSMUSG00000030978 | Rrm1 | protein_coding | -0.712398989 | 4.168672564 | 2.0464515 | 0.1714438 | 0.691519876 |
| ENSMUSG00000028998 | Tomm7 | protein_coding | 0.396666795 | 8.606368385 | 2.0467101 | 0.1714491 | 0.691519876 |
| ENSMUSG00000062234 | Gak | protein_coding | -0.409664653 | 5.490160043 | 2.0456767 | 0.1715203 | 0.691547616 |
| ENSMUSG00000040433 | Zbtb38 | protein_coding | -0.490752727 | 7.245201503 | 2.0569439 | 0.171608 | 0.691641824 |
| ENSMUSG00000028402 | Mpdz | protein_coding | -0.666526579 | 3.322529361 | 2.0427978 | 0.171805 | 0.69202973 |
| ENSMUSG00000098108 | Gm27008 | antisense_RNA | 0.878215688 | 3.144980135 | 2.0425154 | 0.1718329 | 0.69202973 |
| ENSMUSG00000044340 | Phlpp1 | protein_coding | -0.437881908 | 5.578806587 | 2.0394849 | 0.1721333 | 0.692798364 |
| ENSMUSG00000069917 | Hba-a2 | protein_coding | -0.755441024 | 3.810027504 | 2.0386495 | 0.1722162 | 0.692798364 |
| ENSMUSG00000063317 | Usp31 | protein_coding | -0.464547658 | 5.091182228 | 2.0381505 | 0.1722658 | 0.692798364 |
| ENSMUSG00000055022 | Cntn1 | protein_coding | -0.29225937 | 7.891555497 | 2.0379922 | 0.1722815 | 0.692798364 |
| ENSMUSG00000020903 | Stx8 | protein_coding | 0.297776474 | 6.872106506 | 2.0373376 | 0.1723465 | 0.692800822 |
| ENSMUSG00000018001 | Cyth3 | protein_coding | -0.630724638 | 3.947395335 | 2.034515 | 0.1726273 | 0.693670225 |
| ENSMUSG00000002372 | Ranbp3 | protein_coding | 0.344608688 | 6.514298845 | 2.0335723 | 0.1727212 | 0.693788309 |
| ENSMUSG00000035278 | Plekhj1 | protein_coding | 0.322901712 | 6.151296234 | 2.0314296 | 0.1729349 | 0.694387208 |
| ENSMUSG00000031592 | Pcm1 | protein_coding | -0.496017266 | 5.501152417 | 2.0299057 | 0.1730871 | 0.694615452 |
| ENSMUSG00000031864 | Ints10 | protein_coding | -0.524189268 | 5.075568629 | 2.0288243 | 0.1731952 | 0.694615452 |
| ENSMUSG00000004056 | Akt2 | protein_coding | -0.457684894 | 4.845874673 | 2.0283332 | 0.1732443 | 0.694615452 |
| ENSMUSG00000038459 | Abhd17c | protein_coding | -0.576631661 | 4.349748203 | 2.0282745 | 0.1732501 | 0.694615452 |
| ENSMUSG00000025175 | Fn3k | protein_coding | 0.364776273 | 6.706386204 | 2.0260006 | 0.1734778 | 0.69507951 |
| ENSMUSG00000029446 | Psph | protein_coding | -0.438568057 | 5.792015047 | 2.0254386 | 0.1735341 | 0.69507951 |
| ENSMUSG00000049493 | Pls1 | protein_coding | -0.638034334 | 4.987409284 | 2.0238876 | 0.1737204 | 0.69507951 |
| ENSMUSG00000027533 | Fabp5 | protein_coding | 0.305866458 | 6.901610174 | 2.0233899 | 0.1737396 | 0.69507951 |
| ENSMUSG00000029810 | Tmem176b | protein_coding | -0.339601248 | 6.570724312 | 2.0230018 | 0.1737785 | 0.69507951 |
| ENSMUSG00000031310 | Zmym3 | protein_coding | 0.317697303 | 6.639066481 | 2.0229153 | 0.1737872 | 0.69507951 |
| ENSMUSG00000021290 | 2010107E04Rik | protein_coding | 0.40723408 | 9.27045432 | 2.0224473 | 0.173865 | 0.69507951 |
| ENSMUSG00000039901 | 9130011E15Rik | protein_coding | 0.676018453 | 3.936812485 | 2.0211595 | 0.1739636 | 0.69507951 |
| ENSMUSG00000025010 | Ccnj | protein_coding | 0.871528678 | 3.045179781 | 2.020572 | 0.1740227 | 0.69507951 |
| ENSMUSG00000017831 | Rab5a | protein_coding | -0.593078731 | 5.016698875 | 2.0201274 | 0.1740674 | 0.69507951 |
| ENSMUSG00000021114 | Atp6v1d | protein_coding | 0.286590573 | 9.903445503 | 2.0200333 | 0.1740769 | 0.69507951 |
| ENSMUSG00000041351 | Rap1gap | protein_coding | -0.453281393 | 6.109828099 | 2.0184257 | 0.1742388 | 0.695127056 |
| ENSMUSG00000021226 | Acot2 | protein_coding | -0.571164377 | 5.08055963 | 2.0173577 | 0.1743464 | 0.695127056 |
| ENSMUSG00000022270 | Retreg1 | protein_coding | -0.377496482 | 5.980789228 | 2.0171933 | 0.174363 | 0.695127056 |
| ENSMUSG00000020190 | Mknk2 | protein_coding | -0.685350476 | 3.544430512 | 2.0167253 | 0.1744102 | 0.695127056 |
| ENSMUSG00000033373 | Fntb | protein_coding | 0.422863062 | 5.852446831 | 2.0167066 | 0.1744121 | 0.695127056 |
| ENSMUSG00000002103 | Acp2 | protein_coding | 0.740734629 | 4.573478707 | 2.0236223 | 0.1745014 | 0.695144957 |
| ENSMUSG00000031969 | Acad8 | protein_coding | -0.450021551 | 5.460269292 | 2.0153808 | 0.1745458 | 0.695144957 |
| ENSMUSG00000021252 | Erg28 | protein_coding | 0.321377885 | 7.380017523 | 2.0144186 | 0.174643 | 0.69527449 |
| ENSMUSG00000029647 | Pan3 | protein_coding | -0.515364834 | 4.615411374 | 2.0133999 | 0.174746 | 0.695426891 |
| ENSMUSG00000021361 | Tmem14c | protein_coding | 0.354419558 | 6.955817858 | 2.0114499 | 0.1749433 | 0.695954467 |
| ENSMUSG00000034663 | Bmp2k | protein_coding | -0.996398978 | 2.757014937 | 2.0106923 | 0.17502 | 0.696002157 |
| ENSMUSG00000005804 | Bloc1s6 | protein_coding | 0.531482002 | 6.412023216 | 2.0213023 | 0.1751418 | 0.696229052 |
| ENSMUSG00000034371 | Tkfc | protein_coding | 0.594470981 | 3.857554488 | 2.0084659 | 0.1752457 | 0.69638471 |
| ENSMUSG00000028156 | Eif4e | protein_coding | 0.299811576 | 7.869576667 | 2.0042524 | 0.1756739 | 0.69763015 |
| ENSMUSG00000024287 | Thoc1 | protein_coding | -0.359196562 | 5.986577429 | 2.0039069 | 0.175709 | 0.69763015 |
| ENSMUSG00000022390 | Zc3h7b | protein_coding | -0.516035665 | 5.210253924 | 2.0033901 | 0.1757617 | 0.69763015 |
| ENSMUSG00000021870 | Slmap | protein_coding | -0.516786005 | 5.716524717 | 2.0029725 | 0.1758349 | 0.69763015 |
| ENSMUSG00000038648 | Creb3l2 | protein_coding | -1.447640795 | 2.243356346 | 2.002194 | 0.1758835 | 0.69763015 |
| ENSMUSG00000024039 | Cbs | protein_coding | -0.602764953 | 4.681258939 | 2.0001944 | 0.1760875 | 0.698181604 |
| ENSMUSG00000033510 | Otud7a | protein_coding | -1.032263194 | 2.772559865 | 1.9985948 | 0.1762508 | 0.698333021 |
| ENSMUSG00000097587 | 4930578M01Rik | lincRNA | -0.805347287 | 3.936053073 | 1.9980838 | 0.1763338 | 0.698333021 |
| ENSMUSG00000003868 | Ruvbl2 | protein_coding | 0.42582782 | 6.01352007 | 1.9974772 | 0.1763651 | 0.698333021 |
| ENSMUSG00000026349 | Ccnt2 | protein_coding | -0.517293359 | 4.920341983 | 1.997194 | 0.1763941 | 0.698333021 |
| ENSMUSG00000031290 | Lrch2 | protein_coding | -0.555752932 | 3.809746072 | 1.9950388 | 0.1766147 | 0.698333021 |
| ENSMUSG00000030695 | Aldoa | protein_coding | 0.338418398 | 11.9914057 | 1.9946447 | 0.1766551 | 0.698333021 |
| ENSMUSG00000061578 | Ksr2 | protein_coding | -0.527828058 | 4.148114964 | 1.9930547 | 0.1768181 | 0.698333021 |
| ENSMUSG00000023988 | Bysl | protein_coding | 0.460840356 | 5.257597428 | 1.9929816 | 0.1768256 | 0.698333021 |
| ENSMUSG00000100975 | Gm28875 | lincRNA | -1.045434656 | 3.551389565 | 2.0023868 | 0.1770125 | 0.698333021 |
| ENSMUSG00000036098 | Myrf | protein_coding | -0.544325003 | 4.412695229 | 1.990988 | 0.1770303 | 0.698333021 |
| ENSMUSG00000021012 | Zc3h14 | protein_coding | -0.549020175 | 4.454841109 | 1.9894182 | 0.1771917 | 0.698333021 |
| ENSMUSG00000037443 | Cep85 | protein_coding | -0.664220263 | 3.550220314 | 1.9886073 | 0.1772751 | 0.698333021 |
| ENSMUSG00000028519 | Dab1 | protein_coding | 0.327723604 | 5.940746159 | 1.9876467 | 0.177374 | 0.698333021 |
| ENSMUSG00000041684 | Bivm | protein_coding | -0.514773535 | 4.219318617 | 1.9872494 | 0.177415 | 0.698333021 |
| ENSMUSG00000052656 | Rnf103 | protein_coding | -0.786192436 | 4.638591862 | 1.9982576 | 0.1774752 | 0.698333021 |
| ENSMUSG00000028222 | Calb1 | protein_coding | -0.380401467 | 7.475597524 | 1.9865681 | 0.1774852 | 0.698333021 |
| ENSMUSG00000047036 | Zfp445 | protein_coding | -0.513742948 | 4.849598076 | 1.985572 | 0.1775879 | 0.698333021 |
| ENSMUSG00000029640 | Usp12 | protein_coding | -0.844131276 | 3.012577258 | 1.9852934 | 0.1776166 | 0.698333021 |

| ensembl_gene_id | external_gene_name | gene_biotype | log2 (fold change) | log2 (counts per million) | F-value | p-value | False discovery rate |
|---|---|---|---|---|---|---|---|
| ENSMUSG00000031381 | Piga | protein_coding | -0.710153326 | 3.304020973 | 1.9845689 | 0.1776914 | 0.698333021 |
| ENSMUSG00000052681 | Rap1b | protein_coding | -0.439082882 | 5.520195589 | 1.9844378 | 0.177705 | 0.698333021 |
| ENSMUSG00000054836 | Elp6 | protein_coding | 0.380944401 | 5.750769458 | 1.983917 | 0.1777587 | 0.698333021 |
| ENSMUSG00000027805 | Pfn2 | protein_coding | 0.340193758 | 10.00691878 | 1.9837098 | 0.1777801 | 0.698333021 |
| ENSMUSG00000053279 | Aldh1a1 | protein_coding | -0.609954003 | 4.881994317 | 1.9821321 | 0.1779739 | 0.698333021 |
| ENSMUSG00000031931 | Ankrd49 | protein_coding | -0.471026042 | 4.469545221 | 1.9817575 | 0.1779819 | 0.698333021 |
| ENSMUSG00000040713 | Creg1 | protein_coding | -0.61878026 | 4.165173706 | 1.9815718 | 0.1780011 | 0.698333021 |
| ENSMUSG00000042772 | Smg7 | protein_coding | -0.312427328 | 6.953742699 | 1.9810016 | 0.1780601 | 0.698333021 |
| ENSMUSG00000046204 | Pnma2 | protein_coding | 0.336767884 | 7.583794278 | 1.9809027 | 0.1780704 | 0.698333021 |
| ENSMUSG00000053317 | Sec61b | protein_coding | 0.313304248 | 6.311453429 | 1.9805417 | 0.1781078 | 0.698333021 |
| ENSMUSG00000038453 | Srcin1 | protein_coding | -0.388320455 | 5.644276852 | 1.9797844 | 0.1781862 | 0.698333021 |
| ENSMUSG00000040472 | Rabggta | protein_coding | 0.368220292 | 6.178422067 | 1.9785436 | 0.1783148 | 0.698333021 |
| ENSMUSG00000049097 | Ankrd34a | protein_coding | 0.427579198 | 5.665853992 | 1.977862 | 0.1783855 | 0.698333021 |
| ENSMUSG00000036613 | Tssc1 | protein_coding | 0.398977983 | 6.756816031 | 1.9779219 | 0.1784099 | 0.698333021 |
| ENSMUSG00000051034 | Zfp11 | protein_coding | 0.811020698 | 3.349475125 | 1.9775902 | 0.1784137 | 0.698333021 |
| ENSMUSG00000057541 | Pus7 | protein_coding | -0.959147882 | 2.835257955 | 1.9775804 | 0.1784147 | 0.698333021 |
| ENSMUSG00000036062 | Phf24 | protein_coding | -0.346631354 | 6.335359416 | 1.9771783 | 0.1784564 | 0.698333021 |
| ENSMUSG00000037236 | Matr3 | protein_coding | -0.309623634 | 8.047291825 | 1.9766647 | 0.1785097 | 0.698333021 |
| ENSMUSG00000045636 | Mtus1 | protein_coding | -0.495184578 | 5.259558151 | 1.9761884 | 0.1785592 | 0.698333021 |
| ENSMUSG00000030313 | Dennd5b | protein_coding | -0.406413251 | 4.922465207 | 1.9755176 | 0.1786289 | 0.698333021 |
| ENSMUSG00000036641 | Ccdc148 | protein_coding | -0.482310613 | 5.152694454 | 1.9747444 | 0.1787092 | 0.698333021 |
| ENSMUSG00000024395 | Lims2 | protein_coding | -0.474064268 | 5.325717595 | 1.9732181 | 0.178868 | 0.698333021 |
| ENSMUSG00000050428 | Fbxo46 | protein_coding | -0.466694035 | 4.897242929 | 1.9730943 | 0.1788809 | 0.698333021 |
| ENSMUSG00000096370 | Gm21992 | protein_coding | -0.868550545 | 3.226450449 | 1.9723045 | 0.1789631 | 0.698333021 |
| ENSMUSG00000059586 | Nsmce2 | protein_coding | 0.372577003 | 6.325690648 | 1.9722016 | 0.1789739 | 0.698333021 |
| ENSMUSG00000037251 | Pomk | protein_coding | 0.411063649 | 5.944052646 | 1.9719877 | 0.1789961 | 0.698333021 |
| ENSMUSG00000052917 | Senp7 | protein_coding | -0.400500541 | 5.549636345 | 1.971266 | 0.1790714 | 0.698333021 |
| ENSMUSG00000062797 | Hikeshi | protein_coding | 0.27595269 | 7.780947549 | 1.9708661 | 0.179113 | 0.698333021 |
| ENSMUSG00000032060 | Cryab | protein_coding | -0.337932335 | 9.198418857 | 1.967414 | 0.1794734 | 0.699484484 |
| ENSMUSG00000087269 | D330023K18Rik | processed_transcript | 0.402147661 | 5.379798181 | 1.9659599 | 0.1796255 | 0.699823538 |
| ENSMUSG00000022679 | Mpv17l | protein_coding | -0.43574745 | 5.775120625 | 1.9629226 | 0.1799437 | 0.700809178 |
| ENSMUSG00000070284 | Gmppb | protein_coding | 0.546880454 | 4.677332472 | 1.9600039 | 0.1802501 | 0.701133849 |
| ENSMUSG00000030216 | Wbp11 | protein_coding | 0.326490416 | 6.936768788 | 1.959651 | 0.1802872 | 0.701133849 |
| ENSMUSG00000030322 | Mbd4 | protein_coding | 0.776876537 | 3.104928882 | 1.959581 | 0.1802945 | 0.701133849 |
| ENSMUSG00000002395 | Use1 | protein_coding | 0.313940583 | 8.642709291 | 1.9592943 | 0.1803247 | 0.701133849 |
| ENSMUSG00000024067 | Dpy30 | protein_coding | 0.323947075 | 8.230608169 | 1.9586087 | 0.1803968 | 0.701133849 |
| ENSMUSG00000042901 | Aida | protein_coding | 0.392717928 | 5.122552065 | 1.9584045 | 0.1804182 | 0.701133849 |
| ENSMUSG00000004642 | Slbp | protein_coding | 0.396487706 | 6.246675971 | 1.9559353 | 0.1806783 | 0.701555207 |
| ENSMUSG00000020381 | Mrnip | protein_coding | 0.440837041 | 5.218172705 | 1.9551861 | 0.1807573 | 0.701555207 |
| ENSMUSG00000046876 | Atxn1 | protein_coding | -0.58629518 | 6.992813897 | 1.9813667 | 0.1807749 | 0.701555207 |
| ENSMUSG00000055373 | Fut9 | protein_coding | -0.45412403 | 5.562331203 | 1.9548164 | 0.1807963 | 0.701555207 |
| ENSMUSG00000020114 | Cand1 | protein_coding | -0.351330028 | 6.489287104 | 1.9539341 | 0.1808894 | 0.701555207 |
| ENSMUSG00000080268 | Brms1 | protein_coding | 0.324237884 | 6.284589523 | 1.9519939 | 0.1810943 | 0.701555207 |
| ENSMUSG00000042312 | S100a13 | protein_coding | -0.571574691 | 6.416420582 | 1.9689107 | 0.181176 | 0.701555207 |
| ENSMUSG00000035285 | Nat14 | protein_coding | 0.397163838 | 6.095206322 | 1.949643 | 0.181343 | 0.701555207 |
| ENSMUSG00000030007 | Cct7 | protein_coding | 0.367259537 | 9.448790663 | 1.9492511 | 0.1813845 | 0.701555207 |
| ENSMUSG00000029632 | Ndufa4 | protein_coding | 0.384723206 | 11.03033526 | 1.9488105 | 0.1814312 | 0.701555207 |
| ENSMUSG00000068739 | Sars | protein_coding | 0.356485996 | 8.568066418 | 1.9487918 | 0.1814332 | 0.701555207 |
| ENSMUSG00000033721 | Vav3 | protein_coding | -0.786627742 | 3.443028302 | 1.9473205 | 0.1815892 | 0.701555207 |
| ENSMUSG00000059552 | Trp53 | protein_coding | 0.417754115 | 5.633688644 | 1.9471399 | 0.1816083 | 0.701555207 |
| ENSMUSG00000020475 | Pgam2 | protein_coding | -0.506544313 | 4.342649937 | 1.9462506 | 0.1817027 | 0.701555207 |
| ENSMUSG00000042129 | Rassf4 | protein_coding | -0.676386721 | 3.448326088 | 1.9457931 | 0.1817513 | 0.701555207 |
| ENSMUSG00000040867 | Begain | protein_coding | 0.470017532 | 4.989020161 | 1.9456293 | 0.1817687 | 0.701555207 |
| ENSMUSG00000060843 | Ctnna3 | protein_coding | -0.808620971 | 2.978928468 | 1.9455655 | 0.1817754 | 0.701555207 |
| ENSMUSG00000087574 | C030037D09Rik | lincRNA | -0.746420935 | 3.542319725 | 1.9451182 | 0.181823 | 0.701555207 |
| ENSMUSG00000045087 | S1pr5 | protein_coding | -1.020339191 | 2.742858132 | 1.944937 | 0.1818422 | 0.701555207 |
| ENSMUSG00000026478 | Lamc1 | protein_coding | -0.809576361 | 3.023260391 | 1.9430496 | 0.1820429 | 0.701555207 |
| ENSMUSG00000035171 | 1110059E24Rik | protein_coding | 0.31652301 | 7.010336727 | 1.9419509 | 0.1821598 | 0.701555207 |
| ENSMUSG00000013593 | Ndufs2 | protein_coding | 0.309851336 | 8.539366537 | 1.9396286 | 0.1824073 | 0.701555207 |
| ENSMUSG00000001383 | Zmat2 | protein_coding | 0.308354744 | 9.115223408 | 1.9395885 | 0.1824116 | 0.701555207 |
| ENSMUSG00000041268 | Dmxl2 | protein_coding | -0.419876308 | 5.70629224 | 1.9389888 | 0.1824756 | 0.701555207 |
| ENSMUSG00000021497 | Txndc15 | protein_coding | 0.347266058 | 6.261456886 | 1.938555 | 0.1825219 | 0.701555207 |
| ENSMUSG00000002190 | Clgn | protein_coding | -1.170654368 | 2.905563367 | 1.9378046 | 0.1826326 | 0.701555207 |
| ENSMUSG00000039130 | Zc3hc1 | protein_coding | 0.371601202 | 6.442612538 | 1.9367419 | 0.1827156 | 0.701555207 |
| ENSMUSG00000069844 | Sco1 | protein_coding | -1.071719987 | 3.304374672 | 1.9400925 | 0.1827249 | 0.701555207 |
| ENSMUSG00000071856 | Mcc | protein_coding | -1.557789675 | 2.394947163 | 1.9367267 | 0.1827477 | 0.701555207 |
| ENSMUSG00000002844 | Adprh | protein_coding | 0.325638677 | 7.485039426 | 1.9362501 | 0.1827682 | 0.701555207 |
| ENSMUSG00000042616 | Oscp1 | protein_coding | 0.358274812 | 6.296014459 | 1.9358164 | 0.1828145 | 0.701555207 |
| ENSMUSG00000109156 | Gm45194 | protein_coding | -0.655395671 | 4.031249565 | 1.9341135 | 0.1829968 | 0.701555207 |
| ENSMUSG00000029017 | Pmpcb | protein_coding | 0.297656929 | 7.100038551 | 1.9337445 | 0.1830363 | 0.701555207 |
| ENSMUSG00000040824 | Snrpd2 | protein_coding | 0.367728389 | 7.163819088 | 1.9332744 | 0.1830867 | 0.701555207 |
| ENSMUSG00000029864 | Gstk1 | protein_coding | -0.816272567 | 4.460523854 | 1.9438871 | 0.1831371 | 0.701555207 |
| ENSMUSG00000027459 | Fam110a | protein_coding | 0.602456946 | 5.031921134 | 1.9330186 | 0.1831446 | 0.701555207 |
| ENSMUSG00000026248 | Mrpl44 | protein_coding | 0.302185529 | 7.060938854 | 1.9319387 | 0.1832299 | 0.701555207 |
| ENSMUSG00000027639 | Samhd1 | protein_coding | -0.701642704 | 3.30342135 | 1.9318608 | 0.1832383 | 0.701555207 |
| ENSMUSG00000106981 | Gm45495 | lincRNA | -0.931280654 | 3.014933517 | 1.9314886 | 0.1832782 | 0.701555207 |
| ENSMUSG00000023966 | Rsph9 | protein_coding | 0.298328023 | 6.664529236 | 1.9309254 | 0.1833387 | 0.701555207 |
| ENSMUSG00000069171 | Nr2f1 | protein_coding | 0.468002111 | 4.585679755 | 1.9302264 | 0.1834137 | 0.701555207 |

| ensembl_gene_id | external_gene_name | gene_biotype | log2 (fold change) | log2 (counts per million) | F-value | p-value | False discovery rate |
|---|---|---|---|---|---|---|---|
| ENSMUSG00000026077 | Npas2 | protein_coding | -0.587012952 | 3.989062889 | 1.9297996 | 0.1834596 | 0.701555207 |
| ENSMUSG00000029625 | Cpsf4 | protein_coding | -0.897555664 | 3.032354332 | 1.9295431 | 0.1834871 | 0.701555207 |
| ENSMUSG00000031570 | Plpp5 | protein_coding | 0.485808926 | 4.85331234 | 1.9283166 | 0.183619 | 0.701555207 |
| ENSMUSG00000013523 | Bcas1 | protein_coding | -0.615322188 | 5.245615683 | 1.9292242 | 0.1836323 | 0.701555207 |
| ENSMUSG00000037640 | Zfp60 | protein_coding | -0.520275595 | 4.904706894 | 1.9280425 | 0.1836485 | 0.701555207 |
| ENSMUSG00000020205 | Phlda1 | protein_coding | -0.611442678 | 4.042854638 | 1.927868 | 0.1836672 | 0.701555207 |
| ENSMUSG00000045414 | 1190002N15Rik | protein_coding | -0.619665429 | 3.857464086 | 1.9274565 | 0.1837115 | 0.701555207 |
| ENSMUSG00000039943 | Plcb4 | protein_coding | -0.380677268 | 6.39625355 | 1.9273446 | 0.1837236 | 0.701555207 |
| ENSMUSG00000027808 | Serp1 | protein_coding | -0.551570875 | 4.652541343 | 1.9256955 | 0.1839012 | 0.701691831 |
| ENSMUSG00000030335 | Mrpl51 | protein_coding | 0.285155076 | 6.91307832 | 1.9256532 | 0.1839057 | 0.701691831 |
| ENSMUSG00000019471 | Cdc37 | protein_coding | 0.266258866 | 7.155075556 | 1.9249876 | 0.1839775 | 0.701691831 |
| ENSMUSG00000055493 | Epm2a | protein_coding | -0.555791559 | 4.133105188 | 1.9245898 | 0.1840204 | 0.701691831 |
| ENSMUSG00000027890 | Gstm4 | protein_coding | 0.425983047 | 4.802565336 | 1.9233929 | 0.1841495 | 0.70193538 |
| ENSMUSG00000027584 | Oprl1 | protein_coding | 0.388259377 | 6.424172654 | 1.9221546 | 0.1842832 | 0.702190185 |
| ENSMUSG00000030868 | Dctn5 | protein_coding | 0.346819685 | 5.84964316 | 1.9209742 | 0.1844108 | 0.702190185 |
| ENSMUSG00000000374 | Trappc10 | protein_coding | -0.489545812 | 4.455311871 | 1.9209609 | 0.1844123 | 0.702190185 |
| ENSMUSG00000000340 | Dbt | protein_coding | -0.852229778 | 3.982134135 | 1.9249236 | 0.1845629 | 0.702514896 |
| ENSMUSG00000039016 | Timm8b | protein_coding | 0.413020213 | 9.334659322 | 1.9185865 | 0.1846997 | 0.702787006 |
| ENSMUSG00000075376 | Rc3h2 | protein_coding | -0.585094998 | 5.503473446 | 1.9182043 | 0.184853 | 0.703023133 |
| ENSMUSG00000031681 | Smad1 | protein_coding | -0.478358323 | 4.910375733 | 1.9165269 | 0.1848925 | 0.703023133 |
| ENSMUSG00000005299 | Letm1 | protein_coding | -0.408638816 | 5.787352324 | 1.9145164 | 0.1851108 | 0.703049472 |
| ENSMUSG00000000787 | Ddx3x | protein_coding | -0.327125434 | 6.582524005 | 1.9137736 | 0.1851916 | 0.703049472 |
| ENSMUSG00000058174 | Gm5148 | protein_coding | 0.518456521 | 4.361021659 | 1.9137551 | 0.1851936 | 0.703049472 |
| ENSMUSG00000027452 | Acss1 | protein_coding | -0.953739795 | 3.118755559 | 1.9116693 | 0.1854205 | 0.703049472 |
| ENSMUSG00000015357 | Clpx | protein_coding | -0.4240612 | 5.129757506 | 1.9115731 | 0.185431 | 0.703049472 |
| ENSMUSG00000051671 | Coa6 | protein_coding | 0.340140267 | 6.509257988 | 1.9111998 | 0.1854716 | 0.703049472 |
| ENSMUSG00000028710 | Atpaf1 | protein_coding | -0.723269331 | 3.757950926 | 1.9111236 | 0.1854799 | 0.703049472 |
| ENSMUSG00000021957 | Tkt | protein_coding | 0.381838403 | 8.534920959 | 1.9113676 | 0.1854838 | 0.703049472 |
| ENSMUSG00000034135 | Sik3 | protein_coding | -0.306915728 | 6.39715125 | 1.9104277 | 0.1855557 | 0.703049472 |
| ENSMUSG00000087056 | Gm14004 | processed_transcript | 0.529994015 | 5.122867176 | 1.9098556 | 0.1856181 | 0.703049472 |
| ENSMUSG00000001911 | Nfix | protein_coding | -0.445925846 | 5.602617967 | 1.9098269 | 0.1856212 | 0.703049472 |
| ENSMUSG00000070436 | Serpinh1 | protein_coding | -0.866182583 | 3.617527688 | 1.9094729 | 0.1856902 | 0.703049472 |
| ENSMUSG00000023017 | Asic1 | protein_coding | 0.414940438 | 5.062771338 | 1.9084754 | 0.1857687 | 0.703049472 |
| ENSMUSG00000028778 | Hcrtr1 | protein_coding | 1.976690417 | 3.40245049 | 2.0122507 | 0.1858148 | 0.703049472 |
| ENSMUSG00000055421 | Pcdh9 | protein_coding | -0.328790458 | 6.98252191 | 1.9071182 | 0.1859168 | 0.703118054 |
| ENSMUSG00000022201 | Zfr | protein_coding | -0.289890649 | 8.293012069 | 1.9066893 | 0.1859637 | 0.703118054 |
| ENSMUSG00000036502 | Tmem255a | protein_coding | 0.371417226 | 6.151645705 | 1.9056415 | 0.1860783 | 0.703169734 |
| ENSMUSG00000032396 | Dis3l | protein_coding | 0.500629844 | 4.497877359 | 1.9043376 | 0.1862209 | 0.703169734 |
| ENSMUSG00000039904 | Gpr37 | protein_coding | -0.422613778 | 6.394693251 | 1.9040172 | 0.1862737 | 0.703169734 |
| ENSMUSG00000029202 | Pds5a | protein_coding | -0.652514604 | 3.485613207 | 1.9035062 | 0.186312 | 0.703169734 |
| ENSMUSG00000031367 | Ap1s2 | protein_coding | 0.405385024 | 6.820562455 | 1.9032744 | 0.1863678 | 0.703169734 |
| ENSMUSG00000022199 | Slc22a17 | protein_coding | 0.320813922 | 8.546739703 | 1.9027242 | 0.1863976 | 0.703169734 |
| ENSMUSG00000058013 | 11-Sep | protein_coding | -0.36385983 | 6.660549514 | 1.9023823 | 0.1864351 | 0.703169734 |
| ENSMUSG00000024870 | Rab1b | protein_coding | 0.313940795 | 8.0398978 | 1.9006112 | 0.1866294 | 0.703291304 |
| ENSMUSG00000004937 | Sgta | protein_coding | 0.458980854 | 8.72233234 | 1.9111452 | 0.1866565 | 0.703291304 |
| ENSMUSG00000020289 | Nprl3 | protein_coding | 0.415112803 | 5.770974429 | 1.8991282 | 0.1867923 | 0.703291304 |
| ENSMUSG00000058835 | Abi1 | protein_coding | -0.309689121 | 6.933388157 | 1.8984486 | 0.186867 | 0.703291304 |
| ENSMUSG00000029304 | Spp1 | protein_coding | -0.708720114 | 4.246992768 | 1.8985582 | 0.1868853 | 0.703291304 |
| ENSMUSG00000044042 | Fmn1 | protein_coding | -0.680201884 | 3.170494221 | 1.8970393 | 0.187022 | 0.703291304 |
| ENSMUSG00000015804 | Med28 | protein_coding | 0.382066642 | 8.268673354 | 1.8970778 | 0.1870482 | 0.703291304 |
| ENSMUSG00000026283 | Ing5 | protein_coding | -0.459497405 | 4.677544366 | 1.8966119 | 0.1870691 | 0.703291304 |
| ENSMUSG00000038462 | Uqcrfs1 | protein_coding | -0.450216155 | 6.788603422 | 1.8992694 | 0.1870782 | 0.703291304 |
| ENSMUSG00000037514 | Pank2 | protein_coding | -0.582083304 | 4.32589004 | 1.8957501 | 0.187164 | 0.703291304 |
| ENSMUSG00000041360 | Pum3 | protein_coding | 0.386662641 | 5.666340357 | 1.8951462 | 0.1872305 | 0.703291304 |
| ENSMUSG00000020496 | Rnf187 | protein_coding | 0.318965522 | 8.020059388 | 1.8942194 | 0.1873327 | 0.703291304 |
| ENSMUSG00000063800 | Prpf38a | protein_coding | -0.426510526 | 6.011208155 | 1.8930006 | 0.1874672 | 0.703291304 |
| ENSMUSG00000037916 | Ndufv1 | protein_coding | 0.322997552 | 9.217131399 | 1.8912839 | 0.1876568 | 0.703291304 |
| ENSMUSG00000058709 | Egln2 | protein_coding | 0.345488971 | 6.016477242 | 1.8902581 | 0.1877702 | 0.703291304 |
| ENSMUSG00000054611 | Kdm2a | protein_coding | -0.439525819 | 4.948489974 | 1.8901889 | 0.1877779 | 0.703291304 |
| ENSMUSG00000002413 | Braf | protein_coding | -0.380998231 | 6.161610104 | 1.8900248 | 0.1877961 | 0.703291304 |
| ENSMUSG00000038831 | Ralgps1 | protein_coding | -0.619269004 | 4.464031335 | 1.8899914 | 0.1877997 | 0.703291304 |
| ENSMUSG00000010608 | Rbm25 | protein_coding | -0.307432753 | 6.623945702 | 1.8899776 | 0.1878013 | 0.703291304 |
| ENSMUSG00000058444 | Map2k5 | protein_coding | 0.504244976 | 5.263199779 | 1.8897325 | 0.1878284 | 0.703291304 |
| ENSMUSG00000020022 | Ndufa12 | protein_coding | 0.281522574 | 8.523937266 | 1.8888867 | 0.187922 | 0.703291304 |
| ENSMUSG00000038866 | Zcchc2 | protein_coding | -0.778992841 | 2.898668744 | 1.8873863 | 0.1880882 | 0.703291304 |
| ENSMUSG00000032589 | Bsn | protein_coding | -0.604940233 | 4.550189453 | 1.8873348 | 0.188094 | 0.703291304 |
| ENSMUSG00000025555 | Farp1 | protein_coding | -0.555008654 | 4.541455002 | 1.8869581 | 0.1881357 | 0.703291304 |
| ENSMUSG00000027603 | Ggt7 | protein_coding | 0.36057753 | 7.073577587 | 1.8869326 | 0.1881385 | 0.703291304 |
| ENSMUSG00000052609 | Plekhg3 | protein_coding | -0.702429584 | 3.484572256 | 1.8866481 | 0.1881701 | 0.703291304 |
| ENSMUSG00000048897 | Zfp710 | protein_coding | -0.508378968 | 4.031453304 | 1.8855603 | 0.1882908 | 0.703291304 |
| ENSMUSG00000096847 | Tmem151b | protein_coding | 0.394532929 | 6.425331316 | 1.8847132 | 0.1883849 | 0.703291304 |
| ENSMUSG00000026856 | Dolpp1 | protein_coding | 0.471431619 | 4.741127617 | 1.8841042 | 0.1884525 | 0.703291304 |
| ENSMUSG00000029655 | N4bp2l2 | protein_coding | -0.537077743 | 5.050342002 | 1.8840937 | 0.1884537 | 0.703291304 |
| ENSMUSG00000043153 | Ispd | protein_coding | 0.574806049 | 4.227464938 | 1.8837229 | 0.1884949 | 0.703291304 |
| ENSMUSG00000026657 | Frmd4a | protein_coding | -0.462295365 | 4.795458317 | 1.8828077 | 0.1885966 | 0.703426907 |
| ENSMUSG00000033669 | Zfp7 | protein_coding | 0.7189492 | 3.10848337 | 1.8812504 | 0.18877 | 0.703829213 |
| ENSMUSG00000022684 | Bfar | protein_coding | 0.37350411 | 5.791579699 | 1.8796843 | 0.1889445 | 0.70409847 |
| ENSMUSG00000010406 | Mrpl52 | protein_coding | 0.338329383 | 7.395968681 | 1.8785001 | 0.1890765 | 0.70409847 |

| ensembl_gene_id | external_gene_name | gene_biotype | log2 (fold change) | log2 (counts per million) | F-value | p-value | False discovery rate |
|---|---|---|---|---|---|---|---|
| ENSMUSG00000021772 | Nkiras1 | protein_coding | 0.267001716 | 8.258030439 | 1.8779953 | 0.1891329 | 0.70409847 |
| ENSMUSG00000039693 | Msantd3 | protein_coding | -0.619183555 | 3.699884254 | 1.8776418 | 0.1891723 | 0.70409847 |
| ENSMUSG00000033377 | Palmd | protein_coding | -0.844792481 | 3.561440039 | 1.8777125 | 0.1891948 | 0.70409847 |
| ENSMUSG00000024142 | Mlst8 | protein_coding | 0.617055619 | 4.379486391 | 1.8766282 | 0.1892856 | 0.70409847 |
| ENSMUSG00000063882 | Uqcrh | protein_coding | 0.399695942 | 10.25289796 | 1.8767654 | 0.1893005 | 0.70409847 |
| ENSMUSG00000002014 | Ssr4 | protein_coding | 0.368913912 | 7.614212691 | 1.8756776 | 0.1893918 | 0.704194374 |
| ENSMUSG00000031633 | Slc25a4 | protein_coding | 0.29916443 | 11.73553787 | 1.8733689 | 0.1896502 | 0.704799469 |
| ENSMUSG00000019647 | Sema6a | protein_coding | -0.536379219 | 4.673341871 | 1.872093 | 0.1897932 | 0.704799469 |
| ENSMUSG00000043461 | Sptssb | protein_coding | 0.939466187 | 2.800323848 | 1.8720841 | 0.1897942 | 0.704799469 |
| ENSMUSG00000036860 | Mrpl55 | protein_coding | 0.293290017 | 7.83164215 | 1.8699233 | 0.1900366 | 0.704799469 |
| ENSMUSG00000063698 | Sfxn4 | protein_coding | -0.387481291 | 5.734147057 | 1.8697959 | 0.1900509 | 0.704799469 |
| ENSMUSG00000042460 | C1galt1 | protein_coding | -1.115610527 | 2.674108706 | 1.8689382 | 0.1901473 | 0.704799469 |
| ENSMUSG00000020954 | Strn3 | protein_coding | -0.453184807 | 5.765313245 | 1.8678045 | 0.1902747 | 0.704799469 |
| ENSMUSG00000054008 | Ndst1 | protein_coding | -0.700845879 | 4.169782026 | 1.8679428 | 0.1902895 | 0.704799469 |
| ENSMUSG00000031600 | Vps37a | protein_coding | -0.390495262 | 5.384052491 | 1.8673725 | 0.1903233 | 0.704799469 |
| ENSMUSG00000018919 | Tm4sf5 | protein_coding | 0.733855853 | 3.347728474 | 1.8673478 | 0.1903261 | 0.704799469 |
| ENSMUSG00000066406 | Akap13 | protein_coding | -0.816482841 | 3.010877153 | 1.8670177 | 0.1903633 | 0.704799469 |
| ENSMUSG00000067288 | Rps28 | protein_coding | 0.533048955 | 4.775434699 | 1.8667939 | 0.1903884 | 0.704799469 |
| ENSMUSG00000026339 | Ccdc93 | protein_coding | -0.869966974 | 2.93749674 | 1.8659829 | 0.1904798 | 0.704799469 |
| ENSMUSG00000048249 | Crebrf | protein_coding | -0.315712061 | 6.30893955 | 1.8658052 | 0.1904998 | 0.704799469 |
| ENSMUSG00000055254 | Ntrk2 | protein_coding | -0.332865829 | 7.519889772 | 1.8652542 | 0.1905618 | 0.704799469 |
| ENSMUSG00000042744 | Hectd4 | protein_coding | -0.355957443 | 6.342138632 | 1.8647119 | 0.190623 | 0.704799469 |
| ENSMUSG00000032475 | Nck1 | protein_coding | -0.409782685 | 5.326819621 | 1.8634543 | 0.1907648 | 0.704799469 |
| ENSMUSG00000002881 | Nab1 | protein_coding | -0.585205575 | 4.198576992 | 1.8627781 | 0.1908411 | 0.704799469 |
| ENSMUSG00000019975 | Ikbip | protein_coding | -0.65950655 | 3.845814942 | 1.862075 | 0.1909205 | 0.704799469 |
| ENSMUSG00000036782 | Klhl13 | protein_coding | 0.326392523 | 7.218380801 | 1.8619735 | 0.190932 | 0.704799469 |
| ENSMUSG00000041556 | Fbxo2 | protein_coding | 0.488989464 | 4.92481782 | 1.8616566 | 0.1909678 | 0.704799469 |
| ENSMUSG00000012126 | Ubxn11 | protein_coding | 0.330502705 | 5.614019991 | 1.8611204 | 0.1910284 | 0.704799469 |
| ENSMUSG00000024382 | Ercc3 | protein_coding | 0.490605865 | 4.83417328 | 1.8601172 | 0.1911418 | 0.704799469 |
| ENSMUSG00000036887 | C1qa | protein_coding | -0.79906496 | 3.806809501 | 1.8579837 | 0.1914136 | 0.704799469 |
| ENSMUSG00000053825 | Ppfia2 | protein_coding | -0.33538654 | 6.398884823 | 1.8573167 | 0.1914589 | 0.704799469 |
| ENSMUSG00000013921 | Clip3 | protein_coding | 0.344604789 | 8.694531453 | 1.8570396 | 0.1914904 | 0.704799469 |
| ENSMUSG00000028560 | Usp1 | protein_coding | -0.811011242 | 3.026843183 | 1.8569844 | 0.1914966 | 0.704799469 |
| ENSMUSG00000009863 | Sdhb | protein_coding | 0.301435372 | 10.22007054 | 1.8565721 | 0.1915434 | 0.704799469 |
| ENSMUSG00000021996 | Esd | protein_coding | 0.279906573 | 8.145446247 | 1.8556684 | 0.1916459 | 0.704799469 |
| ENSMUSG00000034602 | Mon2 | protein_coding | -0.522876401 | 4.907468043 | 1.8555859 | 0.1916553 | 0.704799469 |
| ENSMUSG00000049323 | Smcr8 | protein_coding | -0.659788306 | 3.611316607 | 1.8550245 | 0.191719 | 0.704799469 |
| ENSMUSG00000033685 | Ucp2 | protein_coding | -0.389066272 | 6.573361326 | 1.8549276 | 0.19173 | 0.704799469 |
| ENSMUSG00000022760 | Thap7 | protein_coding | 0.361971845 | 6.147698245 | 1.8542064 | 0.1918119 | 0.704799469 |
| ENSMUSG00000023861 | Mpc1 | protein_coding | -0.306336791 | 6.224462493 | 1.8541458 | 0.1918188 | 0.704799469 |
| ENSMUSG00000030603 | Psmc4 | protein_coding | 0.27716986 | 8.493986337 | 1.8537123 | 0.1918681 | 0.704799469 |
| ENSMUSG00000024420 | Zfp521 | protein_coding | 0.565723501 | 4.269042558 | 1.853307 | 0.1919141 | 0.704799469 |
| ENSMUSG00000060371 | Caln1 | protein_coding | -0.434788461 | 6.668395795 | 1.8528026 | 0.1920017 | 0.704799621 |
| ENSMUSG00000034336 | Ina | protein_coding | 0.284568788 | 7.552055652 | 1.8518212 | 0.1920832 | 0.704799621 |
| ENSMUSG00000022663 | Atg3 | protein_coding | -0.421944507 | 5.160404018 | 1.8515782 | 0.1921108 | 0.704799621 |
| ENSMUSG00000018334 | Ksr1 | protein_coding | -0.746501645 | 3.4061664 | 1.8496934 | 0.1923255 | 0.705054506 |
| ENSMUSG00000048307 | Ankrd46 | protein_coding | 0.35972201 | 8.673423792 | 1.8492772 | 0.192373 | 0.705054506 |
| ENSMUSG00000087200 | Lrp8os3 | bidirectional_promote | 0.635035188 | 3.219258825 | 1.8492421 | 0.192377 | 0.705054506 |
| ENSMUSG00000004961 | Syt5 | protein_coding | 0.267828189 | 8.28337684 | 1.8486183 | 0.1924482 | 0.705074973 |
| ENSMUSG00000054752 | Fsd1l | protein_coding | -0.610763207 | 3.702558382 | 1.8479546 | 0.1925239 | 0.705112217 |
| ENSMUSG00000023156 | Rpp14 | protein_coding | 0.292562578 | 6.519071113 | 1.8469728 | 0.192636 | 0.705132713 |
| ENSMUSG00000026809 | Spaca9 | protein_coding | -1.028263787 | 2.80355454 | 1.8465153 | 0.1926883 | 0.705132713 |
| ENSMUSG00000038888 | Ctu1 | protein_coding | -1.081680808 | 3.035940077 | 1.8458328 | 0.1927965 | 0.705132713 |
| ENSMUSG00000085436 | Zfp335os | antisense_RNA | 0.658093874 | 3.328951338 | 1.8441033 | 0.1929642 | 0.705132713 |
| ENSMUSG00000070953 | Rabepk | protein_coding | 0.29738048 | 6.597109698 | 1.8436936 | 0.1930111 | 0.705132713 |
| ENSMUSG00000051695 | Pcbp1 | protein_coding | -0.279559493 | 6.761842645 | 1.843582 | 0.1930239 | 0.705132713 |
| ENSMUSG00000039257 | Vstm2b | protein_coding | 0.408847782 | 5.263091871 | 1.8434326 | 0.193041 | 0.705132713 |
| ENSMUSG00000044447 | Dock5 | protein_coding | -1.142134418 | 2.442328391 | 1.8431917 | 0.1930686 | 0.705132713 |
| ENSMUSG00000032370 | Lactb | protein_coding | -0.831364861 | 2.970459917 | 1.8425417 | 0.193143 | 0.705132713 |
| ENSMUSG00000029462 | Vps29 | protein_coding | 0.278340926 | 9.414165902 | 1.8405274 | 0.1933741 | 0.705132713 |
| ENSMUSG00000063334 | Krr1 | protein_coding | 0.489122037 | 4.937537707 | 1.8394099 | 0.1935024 | 0.705132713 |
| ENSMUSG00000063049 | Ing2 | protein_coding | -0.600167961 | 4.181450923 | 1.8392851 | 0.1935167 | 0.705132713 |
| ENSMUSG00000032294 | Pkm | protein_coding | 0.419543672 | 10.22861395 | 1.8391435 | 0.1935632 | 0.705132713 |
| ENSMUSG00000070705 | Eid2b | protein_coding | 0.625286459 | 3.510431073 | 1.8382813 | 0.1936321 | 0.705132713 |
| ENSMUSG00000020273 | Papolg | protein_coding | -0.558637144 | 4.466994118 | 1.8382567 | 0.1936349 | 0.705132713 |
| ENSMUSG00000026987 | Baz2b | protein_coding | -0.422755084 | 4.862876181 | 1.8380709 | 0.1936563 | 0.705132713 |
| ENSMUSG00000059824 | Dbp | protein_coding | -0.596582848 | 3.487219614 | 1.83693 | 0.1937876 | 0.705132713 |
| ENSMUSG00000019362 | D8Ertd738e | protein_coding | 0.272193657 | 8.80868539 | 1.8368342 | 0.1937986 | 0.705132713 |
| ENSMUSG00000053477 | Tcf4 | protein_coding | -0.319722185 | 6.748185216 | 1.8354245 | 0.1939609 | 0.705132713 |
| ENSMUSG00000036327 | Qsox2 | protein_coding | -0.995852723 | 2.993632608 | 1.8349466 | 0.194016 | 0.705132713 |
| ENSMUSG00000021079 | Timm9 | protein_coding | 0.349362516 | 7.126191626 | 1.8349133 | 0.1940198 | 0.705132713 |
| ENSMUSG00000024170 | Telo2 | protein_coding | 0.57738295 | 4.060941039 | 1.8345853 | 0.1940577 | 0.705132713 |
| ENSMUSG00000062867 | Impdh2 | protein_coding | 0.555756802 | 4.062557865 | 1.8343617 | 0.1940834 | 0.705132713 |
| ENSMUSG00000048058 | Ldlrad3 | protein_coding | -0.850867698 | 3.027808093 | 1.8341895 | 0.1941033 | 0.705132713 |
| ENSMUSG00000053553 | 3110082I17Rik | protein_coding | 0.48517495 | 4.48468091 | 1.8335842 | 0.1941731 | 0.705148128 |
| ENSMUSG00000001763 | Tspan33 | protein_coding | 0.720253081 | 3.839518618 | 1.8327672 | 0.1942976 | 0.705250321 |
| ENSMUSG00000056211 | R3hdm1 | protein_coding | -0.283015193 | 6.656809729 | 1.8321008 | 0.1943444 | 0.705250321 |
| ENSMUSG00000071723 | Gspt2 | protein_coding | 0.393570371 | 5.012475864 | 1.8316365 | 0.194398 | 0.705250321 |

| ensembl_gene_id | external_gene_name | gene_biotype | log2 (fold change) | log2 (counts per million) | F-value | p-value | False discovery rate |
|---|---|---|---|---|---|---|---|
| ENSMUSG00000022507 | 1810013L24Rik | protein_coding | -0.295581133 | 6.407993132 | 1.8254385 | 0.1951159 | 0.707615889 |
| ENSMUSG00000022358 | Fbxo32 | protein_coding | -0.764888935 | 3.737421218 | 1.8237536 | 0.1953116 | 0.708086897 |
| ENSMUSG00000028822 | Tmem50a | protein_coding | 0.272389697 | 8.468053722 | 1.8188465 | 0.1958831 | 0.709878476 |
| ENSMUSG00000047712 | Ust | protein_coding | -0.802803903 | 3.075890735 | 1.8176419 | 0.1960237 | 0.709878476 |
| ENSMUSG00000024234 | Mtpap | protein_coding | -0.436793119 | 5.238940671 | 1.8170577 | 0.1960919 | 0.709878476 |
| ENSMUSG00000027889 | Ampd2 | protein_coding | 0.429013178 | 6.126254103 | 1.8170176 | 0.1961267 | 0.709878476 |
| ENSMUSG00000051107 | Gm15440 | protein_coding | -0.543390483 | 3.974827409 | 1.8166814 | 0.1961359 | 0.709878476 |
| ENSMUSG00000026393 | Nek7 | protein_coding | -0.452934681 | 6.044481512 | 1.8158271 | 0.1962658 | 0.710109754 |
| ENSMUSG00000029600 | Rita1 | protein_coding | 0.37271874 | 5.501720603 | 1.8149311 | 0.1963405 | 0.710141218 |
| ENSMUSG00000035236 | Scai | protein_coding | -0.528820695 | 4.68379217 | 1.8133387 | 0.196527 | 0.710462703 |
| ENSMUSG00000029231 | Pdgfra | protein_coding | -0.859701749 | 3.623955412 | 1.8105942 | 0.1968789 | 0.710462703 |
| ENSMUSG00000069835 | Sat2 | protein_coding | 0.373228315 | 5.743220845 | 1.8100516 | 0.1969126 | 0.710462703 |
| ENSMUSG00000053950 | Adnp2 | protein_coding | -0.697185155 | 3.565987606 | 1.8099322 | 0.1969266 | 0.710462703 |
| ENSMUSG00000029920 | Smarcad1 | protein_coding | -0.634753454 | 3.78108318 | 1.8096412 | 0.1969608 | 0.710462703 |
| ENSMUSG00000031134 | Rbmx | protein_coding | -0.395580251 | 6.32331531 | 1.8086455 | 0.1970778 | 0.710462703 |
| ENSMUSG00000061479 | Snrpa | protein_coding | 0.417421463 | 6.147516646 | 1.8075376 | 0.1972081 | 0.710462703 |
| ENSMUSG00000022248 | Rad1 | protein_coding | -0.41768326 | 4.790968335 | 1.8074353 | 0.1972201 | 0.710462703 |
| ENSMUSG00000024566 | Atp9b | protein_coding | -0.566272318 | 3.993432612 | 1.8071944 | 0.1972485 | 0.710462703 |
| ENSMUSG00000015709 | Arnt2 | protein_coding | -0.342567281 | 6.141255029 | 1.8065941 | 0.1973192 | 0.710462703 |
| ENSMUSG00000054364 | Rhob | protein_coding | -0.293917082 | 7.058097079 | 1.8065617 | 0.197323 | 0.710462703 |
| ENSMUSG00000026473 | Glul | protein_coding | -0.304323396 | 8.303558326 | 1.8059411 | 0.1973961 | 0.710462703 |
| ENSMUSG00000041836 | Ptpre | protein_coding | -0.521278629 | 4.376135082 | 1.8056632 | 0.1974288 | 0.710462703 |
| ENSMUSG00000028527 | Ak4 | protein_coding | 0.455138723 | 4.634569626 | 1.8046319 | 0.1975504 | 0.710462703 |
| ENSMUSG00000005262 | Ufd1 | protein_coding | 0.366280816 | 7.572505418 | 1.804498 | 0.1975962 | 0.710462703 |
| ENSMUSG00000054843 | Atrnl1 | protein_coding | -0.470623567 | 4.743823263 | 1.8036699 | 0.1976639 | 0.710462703 |
| ENSMUSG00000014355 | Anapc1 | protein_coding | -0.579040011 | 4.217643733 | 1.8033174 | 0.1977055 | 0.710462703 |
| ENSMUSG00000024608 | Rps14 | protein_coding | 0.455693605 | 10.3060541 | 1.8087469 | 0.1978318 | 0.710462703 |
| ENSMUSG00000064138 | Fam172a | protein_coding | -0.713466334 | 3.102864087 | 1.8020312 | 0.1978574 | 0.710462703 |
| ENSMUSG00000002980 | Bcam | protein_coding | 1.099334711 | 3.430448586 | 1.8172869 | 0.1978904 | 0.710462703 |
| ENSMUSG00000019139 | Isyna1 | protein_coding | -0.581028589 | 5.3840446 | 1.8022286 | 0.1979949 | 0.710462703 |
| ENSMUSG00000059839 | Zfp874b | protein_coding | -0.82421203 | 3.056650539 | 1.8006073 | 0.1980257 | 0.710462703 |
| ENSMUSG00000046230 | Vps13a | protein_coding | -0.602616118 | 3.501451692 | 1.7999829 | 0.1980996 | 0.710462703 |
| ENSMUSG00000040738 | Ints8 | protein_coding | -0.653027183 | 3.724447073 | 1.7997857 | 0.1981229 | 0.710462703 |
| ENSMUSG00000032481 | Smarcc1 | protein_coding | 0.386654123 | 5.557874425 | 1.7993931 | 0.1981694 | 0.710462703 |
| ENSMUSG00000017837 | Nkiras2 | protein_coding | 0.532568518 | 4.918589916 | 1.799322 | 0.1981778 | 0.710462703 |
| ENSMUSG00000024135 | Srbd1 | protein_coding | -0.857930474 | 3.426965269 | 1.7984119 | 0.1983156 | 0.710462703 |
| ENSMUSG00000024617 | Camk2a | protein_coding | -0.394324278 | 7.991297627 | 1.7983154 | 0.198327 | 0.710462703 |
| ENSMUSG00000053510 | Nrd1 | protein_coding | -0.362839589 | 5.395380543 | 1.7979064 | 0.1983455 | 0.710462703 |
| ENSMUSG00000078117 | Gm16485 | protein_coding | 1.112592612 | 2.420546414 | 1.7959177 | 0.1985814 | 0.711002195 |
| ENSMUSG00000097431 | Gm26782 | lincRNA | 0.386033836 | 5.107348508 | 1.7955221 | 0.1986283 | 0.711002195 |
| ENSMUSG00000033790 | Tubgcp5 | protein_coding | -0.435512463 | 4.515937438 | 1.7943346 | 0.1987694 | 0.711050657 |
| ENSMUSG00000052560 | Cpne8 | protein_coding | -0.427787101 | 5.446482968 | 1.7937371 | 0.1988404 | 0.711050657 |
| ENSMUSG00000025421 | Hdhd2 | protein_coding | 0.270183203 | 8.431281938 | 1.7937142 | 0.1988432 | 0.711050657 |
| ENSMUSG00000021275 | Tecpr2 | protein_coding | -0.52006172 | 4.203637937 | 1.7931064 | 0.1989154 | 0.711050657 |
| ENSMUSG00000041841 | Rpl37 | protein_coding | 0.458616938 | 8.0548986 | 1.8012376 | 0.1991873 | 0.711050657 |
| ENSMUSG00000020828 | Pld2 | protein_coding | -0.648034268 | 3.411782976 | 1.7907363 | 0.1991976 | 0.711050657 |
| ENSMUSG00000023046 | Igfbp6 | protein_coding | -0.61735837 | 4.68971003 | 1.7900916 | 0.1993044 | 0.711050657 |
| ENSMUSG00000090841 | Myl6 | protein_coding | 0.293260799 | 7.729844937 | 1.7894685 | 0.1993488 | 0.711050657 |
| ENSMUSG00000070923 | Klhl9 | protein_coding | -0.306503846 | 8.232878104 | 1.7892764 | 0.1993717 | 0.711050657 |
| ENSMUSG00000032504 | Pdcd6ip | protein_coding | -0.457532078 | 5.101560594 | 1.7877787 | 0.1995505 | 0.711050657 |
| ENSMUSG00000027852 | Nras | protein_coding | -0.270327 | 7.041033276 | 1.7877181 | 0.1995577 | 0.711050657 |
| ENSMUSG00000005802 | Slc30a4 | protein_coding | -0.900037955 | 3.385005898 | 1.7867182 | 0.1997072 | 0.711050657 |
| ENSMUSG00000028441 | 1110017D15Rik | protein_coding | -0.385934142 | 5.39075987 | 1.786238 | 0.1997346 | 0.711050657 |
| ENSMUSG00000022451 | Twf1 | protein_coding | -0.986667397 | 2.529935583 | 1.7859211 | 0.1997725 | 0.711050657 |
| ENSMUSG00000032328 | Tmem30a | protein_coding | -0.270745057 | 7.184056146 | 1.7855424 | 0.1998178 | 0.711050657 |
| ENSMUSG00000024131 | Slc3a1 | protein_coding | 0.301378002 | 7.649142147 | 1.7850847 | 0.1998726 | 0.711050657 |
| ENSMUSG00000024424 | Ttc39c | protein_coding | 0.601620335 | 3.912295957 | 1.7850548 | 0.1998762 | 0.711050657 |
| ENSMUSG00000024483 | Ankhd1 | protein_coding | -0.434365449 | 4.962412996 | 1.7846203 | 0.1999282 | 0.711050657 |
| ENSMUSG00000022516 | Nudt16l1 | protein_coding | 0.287194981 | 7.037987779 | 1.7823554 | 0.2001996 | 0.711050657 |
| ENSMUSG00000013160 | Atp6v0d1 | protein_coding | 0.25570762 | 8.515038157 | 1.7817393 | 0.2002735 | 0.711050657 |
| ENSMUSG00000003316 | Glg1 | protein_coding | 0.274139049 | 6.681295146 | 1.7808214 | 0.2003836 | 0.711050657 |
| ENSMUSG00000029121 | Crmp1 | protein_coding | 0.579470142 | 4.814668844 | 1.7806295 | 0.2004366 | 0.711050657 |
| ENSMUSG00000004565 | Pnpla6 | protein_coding | 0.464139309 | 4.932264495 | 1.7793332 | 0.2005624 | 0.711050657 |
| ENSMUSG00000024764 | Naa40 | protein_coding | -0.660988229 | 3.508491131 | 1.777572 | 0.2007743 | 0.711050657 |
| ENSMUSG00000030898 | Cckbr | protein_coding | 0.853535893 | 3.172540333 | 1.7773806 | 0.2007973 | 0.711050657 |
| ENSMUSG00000028180 | Zranb2 | protein_coding | -0.289084138 | 7.576783338 | 1.7773272 | 0.2008038 | 0.711050657 |
| ENSMUSG00000072640 | Lyrm9 | protein_coding | 0.39837323 | 6.788462245 | 1.7775584 | 0.2008058 | 0.711050657 |
| ENSMUSG00000037990 | Sh3rf3 | protein_coding | -0.573214375 | 3.771848816 | 1.7771374 | 0.2008266 | 0.711050657 |
| ENSMUSG00000033943 | Mga | protein_coding | -0.479526134 | 5.35196869 | 1.7770085 | 0.2008421 | 0.711050657 |
| ENSMUSG00000037536 | Fbxo34 | protein_coding | -0.576472037 | 3.509491537 | 1.776924 | 0.2008523 | 0.711050657 |
| ENSMUSG00000053617 | Sh3pxd2a | protein_coding | -0.962941249 | 3.092813162 | 1.7766083 | 0.2008903 | 0.711050657 |
| ENSMUSG00000055214 | Pld5 | protein_coding | -0.406792454 | 5.240425973 | 1.7756456 | 0.2010063 | 0.711050657 |
| ENSMUSG00000038517 | Tbkbp1 | protein_coding | -0.873320508 | 3.123857561 | 1.7753682 | 0.2010398 | 0.711050657 |
| ENSMUSG00000021680 | Crhbp | protein_coding | -0.505757831 | 4.435789994 | 1.7753124 | 0.2010465 | 0.711050657 |
| ENSMUSG00000003865 | Gys1 | protein_coding | -0.596065418 | 4.209137295 | 1.7751678 | 0.2010639 | 0.711050657 |
| ENSMUSG00000022339 | Ebag9 | protein_coding | -0.455104036 | 4.976404661 | 1.7750791 | 0.2010746 | 0.711050657 |
| ENSMUSG00000039765 | Cc2d2a | protein_coding | -0.536474515 | 3.858945597 | 1.7749638 | 0.2010885 | 0.711050657 |
| ENSMUSG00000041773 | Enc1 | protein_coding | 0.366940022 | 8.466463369 | 1.7742409 | 0.2012056 | 0.711230846 |

| ensembl_gene_id | external_gene_name | gene_biotype | log2 (fold change) | log2 (counts per million) | F-value | p-value | False discovery rate |
|---|---|---|---|---|---|---|---|
| ENSMUSG00000030536 | Iqgap1 | protein_coding | -0.790215702 | 3.601172273 | 1.773071 | 0.201317 | 0.711271634 |
| ENSMUSG00000050945 | Zfp438 | protein_coding | -0.958176166 | 3.362616456 | 1.7731633 | 0.2014145 | 0.711271634 |
| ENSMUSG00000026087 | Mrpl30 | protein_coding | 0.310457165 | 8.558953754 | 1.7714136 | 0.2015173 | 0.711271634 |
| ENSMUSG00000049960 | Mrps16 | protein_coding | 0.282250117 | 6.888654785 | 1.770621 | 0.2016132 | 0.711271634 |
| ENSMUSG00000015790 | Surf1 | protein_coding | 0.287146722 | 8.21324202 | 1.7702307 | 0.2016604 | 0.711271634 |
| ENSMUSG00000090061 | Nwd2 | protein_coding | -1.147443911 | 2.857913007 | 1.7703742 | 0.2016729 | 0.711271634 |
| ENSMUSG00000030286 | Emc3 | protein_coding | -0.360209435 | 6.178071852 | 1.7697126 | 0.2017231 | 0.711271634 |
| ENSMUSG00000101493 | 2810405F17Rik | lincRNA | -1.006747526 | 2.5205655 | 1.7688628 | 0.2018261 | 0.711271634 |
| ENSMUSG00000052299 | Ltn1 | protein_coding | -0.531620249 | 4.44658231 | 1.7686102 | 0.2018567 | 0.711271634 |
| ENSMUSG00000069744 | Psmb3 | protein_coding | 0.307598137 | 7.452217863 | 1.7682241 | 0.2019035 | 0.711271634 |
| ENSMUSG00000027875 | Hmgcs2 | protein_coding | -0.494668899 | 4.074244339 | 1.7676731 | 0.2019703 | 0.711271634 |
| ENSMUSG00000039910 | Cited2 | protein_coding | -0.451286691 | 4.336699034 | 1.766897 | 0.2020644 | 0.711271634 |
| ENSMUSG00000015222 | Map2 | protein_coding | -0.271953752 | 8.291809547 | 1.7666013 | 0.2021003 | 0.711271634 |
| ENSMUSG00000025475 | Adgra1 | protein_coding | 0.390549658 | 6.688221718 | 1.7664937 | 0.2021432 | 0.711271634 |
| ENSMUSG00000030213 | Atf7ip | protein_coding | -0.464637711 | 4.988359292 | 1.7655528 | 0.2022276 | 0.711335822 |
| ENSMUSG00000047146 | Tet1 | protein_coding | -0.605370194 | 3.973923499 | 1.7635616 | 0.2024697 | 0.711944337 |
| ENSMUSG00000042363 | Lgalsl | protein_coding | -0.30707006 | 6.228389488 | 1.762555 | 0.2025922 | 0.711944337 |
| ENSMUSG00000024064 | Galnt14 | protein_coding | 0.522683688 | 3.847379419 | 1.7618575 | 0.2026771 | 0.711944337 |
| ENSMUSG00000018567 | Gabarap | protein_coding | 0.290381852 | 9.783309314 | 1.7615727 | 0.2027118 | 0.711944337 |
| ENSMUSG00000025609 | Mkln1 | protein_coding | -0.362684323 | 5.956628802 | 1.7610116 | 0.2027802 | 0.711944337 |
| ENSMUSG00000068921 | Dap3 | protein_coding | 0.286139029 | 7.196531851 | 1.7607373 | 0.2028137 | 0.711944337 |
| ENSMUSG00000073940 | Hbb-bt | protein_coding | -0.730950604 | 3.790202896 | 1.7603237 | 0.2028641 | 0.711944337 |
| ENSMUSG00000025326 | Ube3a | protein_coding | -0.363663115 | 5.993583477 | 1.7591525 | 0.203007 | 0.712213391 |
| ENSMUSG00000014402 | Tsg101 | protein_coding | -0.278691952 | 6.732788674 | 1.7585935 | 0.2030752 | 0.712220456 |
| ENSMUSG00000002428 | Hltf | protein_coding | -0.473470406 | 4.522616814 | 1.7558682 | 0.2034084 | 0.713140335 |
| ENSMUSG00000050315 | Synpo2 | protein_coding | -0.835937686 | 4.186734964 | 1.7648416 | 0.2034702 | 0.713140335 |
| ENSMUSG00000006024 | Napa | protein_coding | 0.306528875 | 9.942773001 | 1.7542605 | 0.2036053 | 0.713381318 |
| ENSMUSG00000027176 | Cstf3 | protein_coding | 0.324258392 | 6.278474817 | 1.7530161 | 0.2037578 | 0.713475268 |
| ENSMUSG00000022142 | Nup155 | protein_coding | -0.76237581 | 2.915053981 | 1.7528291 | 0.2037807 | 0.713475268 |
| ENSMUSG00000056493 | Foxk1 | protein_coding | -0.641305485 | 3.329235493 | 1.7524183 | 0.2038311 | 0.713475268 |
| ENSMUSG00000038473 | Nos1ap | protein_coding | 0.380450589 | 5.935121701 | 1.7515763 | 0.2039345 | 0.713604712 |
| ENSMUSG00000025532 | Crcp | protein_coding | 0.289877824 | 7.105217199 | 1.750846 | 0.2040242 | 0.71368632 |
| ENSMUSG00000026356 | Dars | protein_coding | -0.411658692 | 6.447264805 | 1.7505409 | 0.2040914 | 0.71368942 |
| ENSMUSG00000050608 | Minos1 | protein_coding | 0.332212432 | 9.318764556 | 1.7493574 | 0.2042071 | 0.713861881 |
| ENSMUSG00000020333 | Acsl6 | protein_coding | -0.435592254 | 5.723656483 | 1.7487761 | 0.2042786 | 0.713879787 |
| ENSMUSG00000113216 | AL590503.1 | lincRNA | -0.636902984 | 4.160542779 | 1.7479271 | 0.2043831 | 0.71401291 |
| ENSMUSG00000025757 | Hspa4l | protein_coding | -0.363572017 | 6.93759472 | 1.7426605 | 0.2050328 | 0.715859965 |
| ENSMUSG00000027466 | Rbck1 | protein_coding | 0.358391895 | 6.158664113 | 1.7416878 | 0.2051531 | 0.715859965 |
| ENSMUSG00000004446 | Bid | protein_coding | 0.457020853 | 5.242118399 | 1.740819 | 0.2052606 | 0.715859965 |
| ENSMUSG00000032580 | Rbm5 | protein_coding | -0.299609212 | 6.59973986 | 1.7397105 | 0.2053979 | 0.715859965 |
| ENSMUSG00000039217 | Il18 | protein_coding | -0.530959693 | 5.980826172 | 1.7479774 | 0.2054599 | 0.715859965 |
| ENSMUSG00000018841 | Rad51d | protein_coding | 0.424778485 | 5.734623597 | 1.7378734 | 0.2056257 | 0.715859965 |
| ENSMUSG00000057134 | Ado | protein_coding | -0.391863502 | 8.137691572 | 1.7378146 | 0.2056627 | 0.715859965 |
| ENSMUSG00000042111 | Ccdc115 | protein_coding | 0.384422959 | 5.416773609 | 1.7373768 | 0.2056873 | 0.715859965 |
| ENSMUSG00000020171 | Yeats4 | protein_coding | -0.550830815 | 4.308757763 | 1.7370206 | 0.2057315 | 0.715859965 |
| ENSMUSG00000058600 | Rpl30 | protein_coding | 0.396192833 | 6.950298277 | 1.7372116 | 0.2057375 | 0.715859965 |
| ENSMUSG00000029076 | Sdf4 | protein_coding | 0.265604097 | 8.512245284 | 1.7361892 | 0.2058348 | 0.715859965 |
| ENSMUSG00000032174 | Icam5 | protein_coding | -1.296471579 | 2.171267815 | 1.7360008 | 0.2058582 | 0.715859965 |
| ENSMUSG00000034614 | Pik3ip1 | protein_coding | -0.574575128 | 4.147793106 | 1.7357351 | 0.2058912 | 0.715859965 |
| ENSMUSG00000042105 | Inpp5f | protein_coding | 0.359583378 | 8.659097926 | 1.7356398 | 0.2059031 | 0.715859965 |
| ENSMUSG00000032936 | Camkv | protein_coding | -0.31302787 | 8.008859896 | 1.7355805 | 0.2059104 | 0.715859965 |
| ENSMUSG00000051256 | Jagn1 | protein_coding | 0.303356227 | 6.482956711 | 1.7344156 | 0.2060553 | 0.715874336 |
| ENSMUSG00000027193 | Api5 | protein_coding | -0.350632602 | 5.813265692 | 1.7337464 | 0.2061385 | 0.715874336 |
| ENSMUSG00000034994 | Eef2 | protein_coding | 0.34292094 | 10.00423559 | 1.7335135 | 0.2061675 | 0.715874336 |
| ENSMUSG00000079017 | Ifi27l2a | protein_coding | -1.091888045 | 2.489332967 | 1.7324721 | 0.2062972 | 0.715874336 |
| ENSMUSG00000040928 | S100pbp | protein_coding | -0.401658165 | 5.027817843 | 1.732309 | 0.2063176 | 0.715874336 |
| ENSMUSG00000109321 | A330076H08Rik | processed_transcript | -0.584168377 | 3.772797869 | 1.7320749 | 0.2063467 | 0.715874336 |
| ENSMUSG00000032300 | 1700017B05Rik | protein_coding | -1.027994428 | 3.158139168 | 1.7320414 | 0.2063806 | 0.715874336 |
| ENSMUSG00000059939 | 9430015G10Rik | protein_coding | 0.561720769 | 3.558765264 | 1.7312079 | 0.2064548 | 0.715900858 |
| ENSMUSG00000025486 | Sirt3 | protein_coding | 0.296590955 | 7.720722773 | 1.729482 | 0.2066702 | 0.716416644 |
| ENSMUSG00000046722 | Cdc42se1 | protein_coding | -0.391041799 | 5.214550562 | 1.7284379 | 0.2068006 | 0.716637783 |
| ENSMUSG00000036591 | Arhgap21 | protein_coding | -0.568680686 | 4.502984032 | 1.7274192 | 0.206928 | 0.716846472 |
| ENSMUSG00000025478 | Dpysl4 | protein_coding | 0.477731363 | 5.052584115 | 1.7265029 | 0.2070426 | 0.716846472 |
| ENSMUSG00000034445 | Cyb561a3 | protein_coding | 0.977386998 | 2.512051655 | 1.7261986 | 0.2070807 | 0.716846472 |
| ENSMUSG00000027439 | Gzf1 | protein_coding | -0.372456864 | 5.316569817 | 1.7258249 | 0.2071275 | 0.716846472 |
| ENSMUSG00000034295 | Fhod3 | protein_coding | -0.727815382 | 2.940591565 | 1.7252181 | 0.2072035 | 0.716878265 |
| ENSMUSG00000095620 | 2010005H15Rik | protein_coding | 0.727308375 | 3.44310427 | 1.7246872 | 0.20727 | 0.716878265 |
| ENSMUSG00000041477 | Dcp1b | protein_coding | 0.53249537 | 3.722479341 | 1.7230882 | 0.2074706 | 0.717140374 |
| ENSMUSG00000071415 | Rpl23 | protein_coding | 0.371671993 | 7.520585782 | 1.722516 | 0.207572 | 0.717140374 |
| ENSMUSG00000039830 | Olig2 | protein_coding | -0.707180454 | 3.316571828 | 1.7216119 | 0.2076559 | 0.717140374 |
| ENSMUSG00000043535 | Setx | protein_coding | -0.503208086 | 4.54475546 | 1.7214673 | 0.2076741 | 0.717140374 |
| ENSMUSG00000019710 | Mrpl24 | protein_coding | 0.286497582 | 6.704732725 | 1.7214261 | 0.2076793 | 0.717140374 |
| ENSMUSG00000028347 | Tmeff1 | protein_coding | -0.371979142 | 5.739402037 | 1.7204653 | 0.2078 | 0.717327054 |
| ENSMUSG00000026610 | Esrrg | protein_coding | -0.50420947 | 4.177014255 | 1.719444 | 0.2079285 | 0.717450218 |
| ENSMUSG00000010453 | Kansl3 | protein_coding | -0.461874622 | 5.673572314 | 1.719121 | 0.2079692 | 0.717450218 |
| ENSMUSG00000048154 | Kmt2d | protein_coding | -0.514047899 | 6.139164003 | 1.7261727 | 0.2081693 | 0.717811544 |
| ENSMUSG00000037349 | Nudt22 | protein_coding | 0.41857887 | 4.836511979 | 1.7162567 | 0.2083301 | 0.717811544 |
| ENSMUSG00000005161 | Prdx2 | protein_coding | 0.255831084 | 9.48512235 | 1.715862 | 0.2083799 | 0.717811544 |

| ensembl_gene_id | external_gene_name | gene_biotype | log2 (fold change) | log2 (counts per million) | F-value | p-value | False discovery rate |
|---|---|---|---|---|---|---|---|
| ENSMUSG00000022488 | Nckap1l | protein_coding | -1.677093823 | 2.372215698 | 1.7203723 | 0.2084122 | 0.717811544 |
| ENSMUSG00000086841 | 2410006H16Rik | processed_transcript | 0.323484125 | 7.585014337 | 1.7153322 | 0.2084468 | 0.717811544 |
| ENSMUSG00000031770 | Herpud1 | protein_coding | 0.315618254 | 6.77839581 | 1.7147676 | 0.2085181 | 0.717811544 |
| ENSMUSG00000035248 | Zcchc6 | protein_coding | -0.711371303 | 3.076259494 | 1.7141337 | 0.2085981 | 0.717811544 |
| ENSMUSG00000087165 | 2010001A14Rik | processed_transcript | -0.529992754 | 4.217475595 | 1.7140562 | 0.2086079 | 0.717811544 |
| ENSMUSG00000041057 | Wdr43 | protein_coding | -0.380280652 | 5.112090577 | 1.7131651 | 0.2087206 | 0.717922146 |
| ENSMUSG00000032870 | Smap2 | protein_coding | 0.247515885 | 7.112393624 | 1.7125244 | 0.2088016 | 0.717922146 |
| ENSMUSG00000030125 | Lrrc23 | protein_coding | 0.6026858 | 3.572157851 | 1.7116618 | 0.2089108 | 0.717922146 |
| ENSMUSG00000032212 | Sltm | protein_coding | -0.374834218 | 5.581230775 | 1.7116525 | 0.208912 | 0.717922146 |
| ENSMUSG00000026672 | Optn | protein_coding | -0.648493328 | 3.886804943 | 1.7101966 | 0.2090964 | 0.717922146 |
| ENSMUSG00000040006 | Ginm1 | protein_coding | -0.433624514 | 5.331043321 | 1.7101261 | 0.2091053 | 0.717922146 |
| ENSMUSG00000019054 | Fis1 | protein_coding | 0.296756998 | 10.28494368 | 1.7097918 | 0.2091477 | 0.717922146 |
| ENSMUSG00000028419 | Chmp5 | protein_coding | 0.26416022 | 8.829841351 | 1.7095828 | 0.2091742 | 0.717922146 |
| ENSMUSG00000024867 | Pip5k1b | protein_coding | -0.410146888 | 5.233252831 | 1.708224 | 0.2093466 | 0.717959589 |
| ENSMUSG00000069729 | Arid1b | protein_coding | -0.390112157 | 5.325468977 | 1.7080502 | 0.2093686 | 0.717959589 |
| ENSMUSG00000021774 | Ube2e1 | protein_coding | 0.352324695 | 5.5198214 | 1.7079073 | 0.2093868 | 0.717959589 |
| ENSMUSG00000041703 | Zic5 | protein_coding | -0.794323311 | 3.725062182 | 1.706719 | 0.2095673 | 0.717959589 |
| ENSMUSG00000068036 | Afdn | protein_coding | -0.467768381 | 5.070946904 | 1.7048508 | 0.2097753 | 0.717959589 |
| ENSMUSG00000028780 | Sema3c | protein_coding | -1.154768564 | 2.825147627 | 1.704702 | 0.2098238 | 0.717959589 |
| ENSMUSG00000021094 | Dhrs7 | protein_coding | -0.418245798 | 6.063722617 | 1.7035603 | 0.2099396 | 0.717959589 |
| ENSMUSG00000022750 | Klhl22 | protein_coding | 0.315129304 | 6.821613233 | 1.7033541 | 0.2099659 | 0.717959589 |
| ENSMUSG00000027883 | Gpsm2 | protein_coding | -0.707583785 | 2.962965676 | 1.7032676 | 0.2099769 | 0.717959589 |
| ENSMUSG00000049957 | Ccdc137 | protein_coding | 0.404032583 | 4.829238784 | 1.7031548 | 0.2099913 | 0.717959589 |
| ENSMUSG00000041771 | Slc24a4 | protein_coding | -1.024721455 | 2.504585725 | 1.7027838 | 0.2100386 | 0.717959589 |
| ENSMUSG00000024122 | Pdpk1 | protein_coding | -0.318607749 | 6.553150578 | 1.7025483 | 0.2100686 | 0.717959589 |
| ENSMUSG00000000253 | Gmpr | protein_coding | 0.2600883 | 7.122825237 | 1.7023592 | 0.2100927 | 0.717959589 |
| ENSMUSG00000111375 | Btbd8 | protein_coding | -0.486399501 | 5.097936744 | 1.7016738 | 0.2101801 | 0.717959589 |
| ENSMUSG00000027315 | Spint1 | protein_coding | -1.12587129 | 2.528566975 | 1.701623 | 0.2101866 | 0.717959589 |
| ENSMUSG00000025558 | Dock9 | protein_coding | -0.592979868 | 4.382703992 | 1.7000034 | 0.2103934 | 0.718042411 |
| ENSMUSG00000051166 | Eml5 | protein_coding | -0.556542565 | 3.710090778 | 1.6994475 | 0.2104644 | 0.718042411 |
| ENSMUSG00000022364 | Tbc1d31 | protein_coding | -1.148128574 | 2.316712019 | 1.6992087 | 0.2104949 | 0.718042411 |
| ENSMUSG00000022332 | Khdrbs3 | protein_coding | 0.325672547 | 6.130064683 | 1.698946 | 0.2105285 | 0.718042411 |
| ENSMUSG00000029778 | Adcyap1r1 | protein_coding | -0.33653275 | 6.178430116 | 1.6988191 | 0.2105448 | 0.718042411 |
| ENSMUSG00000023170 | Gps2 | protein_coding | -0.423983413 | 4.89222568 | 1.6964262 | 0.210851 | 0.718719272 |
| ENSMUSG00000019952 | Poc1b | protein_coding | -1.085322515 | 2.66438283 | 1.696224 | 0.2108769 | 0.718719272 |
| ENSMUSG00000028851 | Nudc | protein_coding | -0.551904276 | 5.935962443 | 1.7041572 | 0.2110085 | 0.718779484 |
| ENSMUSG00000047238 | Mageh1 | protein_coding | 0.306185458 | 6.191350289 | 1.6950432 | 0.2110283 | 0.718779484 |
| ENSMUSG00000048001 | Hes5 | protein_coding | -1.020412427 | 2.560221355 | 1.6937296 | 0.2111968 | 0.718926096 |
| ENSMUSG00000031371 | Haus7 | protein_coding | 0.319673042 | 5.661302932 | 1.6936656 | 0.211205 | 0.718926096 |
| ENSMUSG00000079055 | Slc8a3 | protein_coding | 0.519262449 | 4.086510186 | 1.6927223 | 0.2113262 | 0.71892825 |
| ENSMUSG00000022913 | Psmg1 | protein_coding | 0.426368108 | 4.951692239 | 1.6924879 | 0.2113563 | 0.71892825 |
| ENSMUSG00000060487 | Samd5 | protein_coding | -0.475744283 | 4.481120482 | 1.6919288 | 0.2114281 | 0.71892825 |
| ENSMUSG00000028518 | Prkaa2 | protein_coding | -0.487477748 | 4.330244399 | 1.691128 | 0.2115311 | 0.71892825 |
| ENSMUSG00000058153 | Sez6l | protein_coding | 0.325398917 | 5.904705611 | 1.6910593 | 0.2115399 | 0.71892825 |
| ENSMUSG00000093674 | Rpl41 | protein_coding | 0.400183298 | 9.105067067 | 1.6907912 | 0.2116816 | 0.719182435 |
| ENSMUSG00000025212 | Sfxn3 | protein_coding | 0.369478128 | 7.358371596 | 1.6874003 | 0.2120408 | 0.719832731 |
| ENSMUSG00000061119 | Prcp | protein_coding | -0.889299924 | 3.268052999 | 1.6870668 | 0.2120838 | 0.719832731 |
| ENSMUSG00000033739 | Fkbpl | protein_coding | 0.426073165 | 5.200513763 | 1.6858293 | 0.2122141 | 0.719832731 |
| ENSMUSG00000061136 | Prpf40a | protein_coding | -0.427074142 | 5.872851531 | 1.6856287 | 0.21224 | 0.719832731 |
| ENSMUSG00000037152 | Ndufc1 | protein_coding | 0.334984219 | 9.412866579 | 1.6853696 | 0.2122735 | 0.719832731 |
| ENSMUSG00000034473 | Sec22a | protein_coding | 0.304887328 | 6.196040873 | 1.6853602 | 0.2122747 | 0.719832731 |
| ENSMUSG00000034595 | Ppp1r18 | protein_coding | -1.0287211 | 2.624881968 | 1.684594 | 0.2123737 | 0.71994147 |
| ENSMUSG00000027770 | Dhx36 | protein_coding | -0.393751268 | 6.075141452 | 1.6833005 | 0.212541 | 0.720281582 |
| ENSMUSG00000059187 | Fam19a1 | protein_coding | -0.528704575 | 4.408622389 | 1.6827231 | 0.2126158 | 0.720307886 |
| ENSMUSG00000005779 | Psmb4 | protein_coding | 0.289861787 | 7.730041608 | 1.6821315 | 0.2126924 | 0.720340456 |
| ENSMUSG00000037855 | Zfp365 | protein_coding | -0.272644176 | 7.027593811 | 1.6806963 | 0.2128784 | 0.720743424 |
| ENSMUSG00000108434 | Gm45768 | lincRNA | -0.928967248 | 3.28471313 | 1.6790234 | 0.2131249 | 0.721326466 |
| ENSMUSG00000035735 | Dagla | protein_coding | -0.665626709 | 3.67213159 | 1.6774014 | 0.2133062 | 0.721326466 |
| ENSMUSG00000041073 | Nacad | protein_coding | 0.452070628 | 4.855918819 | 1.6767959 | 0.2133849 | 0.721326466 |
| ENSMUSG00000029148 | Nrbp1 | protein_coding | 0.39795105 | 6.827885153 | 1.6765605 | 0.213445 | 0.721326466 |
| ENSMUSG00000030447 | Cyfip1 | protein_coding | -0.41989561 | 5.532259004 | 1.6758081 | 0.2135135 | 0.721326466 |
| ENSMUSG00000022176 | Rem2 | protein_coding | -1.309506052 | 2.818293959 | 1.6802103 | 0.2135378 | 0.721326466 |
| ENSMUSG00000025995 | Wdr75 | protein_coding | 0.336764924 | 5.615716546 | 1.6753484 | 0.2135733 | 0.721326466 |
| ENSMUSG00000034681 | Rnps1 | protein_coding | 0.349194158 | 6.11300136 | 1.6751601 | 0.2135978 | 0.721326466 |
| ENSMUSG00000083282 | Ctsf | protein_coding | -0.489014691 | 5.385650902 | 1.6747265 | 0.2136543 | 0.721326466 |
| ENSMUSG00000020431 | Adcy1 | protein_coding | -0.599324266 | 4.762574829 | 1.6742553 | 0.2137451 | 0.721402402 |
| ENSMUSG00000022261 | Sdc2 | protein_coding | -0.345436959 | 6.526697747 | 1.673415 | 0.2138253 | 0.721402402 |
| ENSMUSG00000015656 | Hspa8 | protein_coding | 0.439935092 | 8.910845924 | 1.6817826 | 0.2138781 | 0.721402402 |
| ENSMUSG00000036023 | Parp2 | protein_coding | -0.347443377 | 5.474415451 | 1.6723927 | 0.2139586 | 0.721429424 |
| ENSMUSG00000042660 | Wdr55 | protein_coding | 0.413407959 | 4.827518776 | 1.6719207 | 0.2140203 | 0.721429424 |
| ENSMUSG00000031370 | Zrsr2 | protein_coding | 0.303454678 | 6.645237934 | 1.6702813 | 0.2142344 | 0.721864764 |
| ENSMUSG00000001300 | Efnb2 | protein_coding | -0.610655773 | 4.524014235 | 1.6701295 | 0.2142837 | 0.721864764 |
| ENSMUSG00000046550 | Spin2c | protein_coding | 0.439217206 | 5.370066525 | 1.6679811 | 0.2145354 | 0.72248413 |
| ENSMUSG00000038515 | Grtp1 | protein_coding | -0.764177638 | 3.133808473 | 1.6674736 | 0.2146019 | 0.72248413 |
| ENSMUSG00000028878 | Fam76a | protein_coding | -0.423416746 | 5.551510911 | 1.6664723 | 0.2147331 | 0.722699685 |
| ENSMUSG00000034216 | Vps18 | protein_coding | 0.602948665 | 5.153911227 | 1.6716036 | 0.2148496 | 0.722865267 |
| ENSMUSG00000029209 | Gnpda2 | protein_coding | -0.340006432 | 5.92283896 | 1.6638036 | 0.2150834 | 0.723036794 |
| ENSMUSG00000020225 | Tmbim4 | protein_coding | 0.296354084 | 6.932846328 | 1.6636447 | 0.2151043 | 0.723036794 |

| ensembl_gene_id | external_gene_name | gene_biotype | log2 (fold change) | log2 (counts per million) | F-value | p-value | False discovery rate |
|---|---|---|---|---|---|---|---|
| ENSMUSG00000004044 | Cavin1 | protein_coding | -0.389910272 | 5.39230093 | 1.6634756 | 0.2151265 | 0.723036794 |
| ENSMUSG00000022770 | Dlg1 | protein_coding | -0.386352704 | 5.375225563 | 1.6629925 | 0.2151901 | 0.723036794 |
| ENSMUSG00000021366 | Hivep1 | protein_coding | -1.097177987 | 2.754575673 | 1.6624331 | 0.2152636 | 0.723036794 |
| ENSMUSG00000002944 | Cd36 | protein_coding | -1.365199154 | 2.164736826 | 1.6619201 | 0.2153311 | 0.723036794 |
| ENSMUSG00000053063 | Clec12a | protein_coding | -1.451240882 | 2.502526598 | 1.6624258 | 0.2153712 | 0.723036794 |
| ENSMUSG00000031723 | Txnl4b | protein_coding | 0.39959205 | 6.709429716 | 1.6608876 | 0.2154964 | 0.72323109 |
| ENSMUSG00000079018 | Ly6c1 | protein_coding | -1.214136721 | 2.631200017 | 1.6591615 | 0.2156945 | 0.723460074 |
| ENSMUSG00000017144 | Rnd3 | protein_coding | -0.459799416 | 5.151207108 | 1.6591263 | 0.2156992 | 0.723460074 |
| ENSMUSG00000027253 | Lrp4 | protein_coding | -0.703781096 | 4.078801821 | 1.6586458 | 0.2157919 | 0.723545303 |
| ENSMUSG00000002222 | Rmnd5a | protein_coding | -0.390078326 | 5.427513093 | 1.6576939 | 0.2158882 | 0.723640665 |
| ENSMUSG00000034818 | Celf5 | protein_coding | -0.312392154 | 6.515981532 | 1.6571886 | 0.2159549 | 0.723640665 |
| ENSMUSG00000027695 | Pld1 | protein_coding | -1.123754346 | 2.534102462 | 1.6553316 | 0.2162003 | 0.724109004 |
| ENSMUSG00000036672 | Cenpt | protein_coding | 0.421289623 | 5.315053013 | 1.6547059 | 0.2162831 | 0.724109004 |
| ENSMUSG00000022672 | Prkdc | protein_coding | -1.255921247 | 2.64344309 | 1.6545378 | 0.2163347 | 0.724109004 |
| ENSMUSG00000073856 | Iqck | protein_coding | 0.613816114 | 3.595928393 | 1.6540947 | 0.216364 | 0.724109004 |
| ENSMUSG00000032583 | Mon1a | protein_coding | 0.494105655 | 4.200117861 | 1.653041 | 0.2165036 | 0.724345781 |
| ENSMUSG00000016637 | Ift27 | protein_coding | 0.297731546 | 7.178958989 | 1.6519058 | 0.2166541 | 0.724345781 |
| ENSMUSG00000031349 | Nsdhl | protein_coding | 0.547281562 | 5.791566562 | 1.6601238 | 0.2167037 | 0.724345781 |
| ENSMUSG00000052144 | Ppp4r2 | protein_coding | -0.406704502 | 4.914873638 | 1.6514196 | 0.2167186 | 0.724345781 |
| ENSMUSG00000024570 | Rbfa | protein_coding | 0.612070707 | 3.664244014 | 1.6510202 | 0.2167716 | 0.724345781 |
| ENSMUSG00000074794 | Arrdc3 | protein_coding | -0.361812095 | 5.529296633 | 1.6490414 | 0.2170344 | 0.724789759 |
| ENSMUSG00000034848 | Ttc21b | protein_coding | -0.954586376 | 2.747969089 | 1.6487025 | 0.2170795 | 0.724789759 |
| ENSMUSG00000028466 | Creb3 | protein_coding | 0.28135165 | 7.342910945 | 1.6479699 | 0.2171769 | 0.724789759 |
| ENSMUSG00000038406 | Scaf1 | protein_coding | 0.490652353 | 4.706982245 | 1.647741 | 0.2172073 | 0.724789759 |
| ENSMUSG00000054452 | Aes | protein_coding | 0.282809175 | 7.901939294 | 1.6471559 | 0.2172852 | 0.724789759 |
| ENSMUSG00000021791 | Dydc2 | protein_coding | -0.811378818 | 2.879453552 | 1.6465048 | 0.2173719 | 0.724789759 |
| ENSMUSG00000046768 | Rhoj | protein_coding | 0.937504881 | 2.680213226 | 1.6462318 | 0.2174083 | 0.724789759 |
| ENSMUSG00000028437 | Ubap1 | protein_coding | 0.38112377 | 5.993346689 | 1.6456677 | 0.2174834 | 0.724789759 |
| ENSMUSG00000039316 | Rftn1 | protein_coding | -0.443820857 | 4.447628496 | 1.6449199 | 0.2175831 | 0.724789759 |
| ENSMUSG00000075266 | Cenpw | protein_coding | -0.876874977 | 3.134142398 | 1.6445228 | 0.2176361 | 0.724789759 |
| ENSMUSG00000021639 | Gtf2h2 | protein_coding | 0.336887945 | 5.981032414 | 1.6444495 | 0.2176459 | 0.724789759 |
| ENSMUSG00000028910 | Mecr | protein_coding | 0.437413251 | 5.23956045 | 1.6439098 | 0.2177179 | 0.724805128 |
| ENSMUSG00000019087 | Atp6ap1 | protein_coding | 0.260525053 | 9.103616349 | 1.642521 | 0.2179034 | 0.724836429 |
| ENSMUSG00000084883 | Ccdc85c | protein_coding | -0.620933909 | 5.217603897 | 1.6496483 | 0.2180247 | 0.724836429 |
| ENSMUSG00000040466 | Blvrb | protein_coding | -0.353248417 | 5.167728801 | 1.6414286 | 0.2180494 | 0.724836429 |
| ENSMUSG00000035772 | Mrps2 | protein_coding | 0.360615664 | 5.785814546 | 1.6404508 | 0.2181802 | 0.724836429 |
| ENSMUSG00000061576 | Dpp6 | protein_coding | 0.297209402 | 7.247168666 | 1.6395313 | 0.2183033 | 0.724836429 |
| ENSMUSG00000086859 | Snhg20 | lincRNA | 0.607188413 | 3.822661677 | 1.6393587 | 0.2183264 | 0.724836429 |
| ENSMUSG00000024891 | Slc29a2 | protein_coding | 0.48107056 | 4.347253447 | 1.6393024 | 0.2183339 | 0.724836429 |
| ENSMUSG00000059974 | Ntm | protein_coding | -0.27768628 | 7.518673752 | 1.6391534 | 0.2183539 | 0.724836429 |
| ENSMUSG00000078451 | Ppil6 | protein_coding | -0.828022384 | 2.86356817 | 1.63888 | 0.2183905 | 0.724836429 |
| ENSMUSG00000003352 | Cacnb3 | protein_coding | 0.34232786 | 5.509101422 | 1.6387989 | 0.2184014 | 0.724836429 |
| ENSMUSG00000028837 | Psmb2 | protein_coding | 0.262593223 | 8.876893701 | 1.6366028 | 0.2186959 | 0.725417456 |
| ENSMUSG00000034462 | Pkd2 | protein_coding | -0.823530848 | 2.742760216 | 1.6364877 | 0.2187114 | 0.725417456 |
| ENSMUSG00000039456 | Morc3 | protein_coding | -0.621198387 | 4.556758285 | 1.6344022 | 0.2190208 | 0.726036375 |
| ENSMUSG00000109088 | Gm44593 | lincRNA | -0.867419454 | 2.679908493 | 1.6339748 | 0.219049 | 0.726036375 |
| ENSMUSG00000009927 | Rps25 | protein_coding | 0.339088263 | 6.467116773 | 1.6335921 | 0.2191005 | 0.726036375 |
| ENSMUSG00000022620 | Arsa | protein_coding | 0.504209169 | 4.693227459 | 1.6325995 | 0.2192341 | 0.72625531 |
| ENSMUSG00000096257 | Ccer2 | protein_coding | 0.956093847 | 2.987224756 | 1.6308444 | 0.2194998 | 0.726574054 |
| ENSMUSG00000009418 | Nav1 | protein_coding | -0.37274371 | 5.623536203 | 1.6305994 | 0.2195037 | 0.726574054 |
| ENSMUSG00000010911 | Apip | protein_coding | 0.334411836 | 5.986840214 | 1.6299063 | 0.2195972 | 0.726574054 |
| ENSMUSG00000031862 | Atp13a1 | protein_coding | -0.73440608 | 3.360370246 | 1.6295611 | 0.2196438 | 0.726574054 |
| ENSMUSG00000021832 | Psmc6 | protein_coding | 0.273564949 | 7.89661194 | 1.62938 | 0.2196682 | 0.726574054 |
| ENSMUSG00000020246 | Hcfc2 | protein_coding | -0.480268065 | 4.458769391 | 1.6274493 | 0.2199291 | 0.726727643 |
| ENSMUSG00000027374 | Mrps5 | protein_coding | 0.333404289 | 5.855082703 | 1.6271625 | 0.2199678 | 0.726727643 |
| ENSMUSG00000038354 | Ankrd35 | protein_coding | -1.10818095 | 2.39288361 | 1.6263096 | 0.2200832 | 0.726727643 |
| ENSMUSG00000108112 | Gm45193 | processed_transcript | -0.64513627 | 3.974443416 | 1.6249942 | 0.2202613 | 0.726727643 |
| ENSMUSG00000033126 | Ybey | protein_coding | 0.557099013 | 4.195006508 | 1.6242076 | 0.2203679 | 0.726727643 |
| ENSMUSG00000020656 | Grhl1 | protein_coding | 0.812236957 | 3.637464104 | 1.6252786 | 0.220379 | 0.726727643 |
| ENSMUSG00000056629 | Fkbp2 | protein_coding | 0.304238341 | 9.108188008 | 1.6238053 | 0.2204225 | 0.726727643 |
| ENSMUSG00000023828 | Slc22a3 | protein_coding | -0.977770974 | 2.622116416 | 1.6236393 | 0.220445 | 0.726727643 |
| ENSMUSG00000025034 | Trim8 | protein_coding | -0.374445967 | 5.425123894 | 1.6230907 | 0.2205194 | 0.726727643 |
| ENSMUSG00000040945 | Rcc2 | protein_coding | 0.311283582 | 5.898254329 | 1.622385 | 0.2206152 | 0.726727643 |
| ENSMUSG00000021061 | Sptb | protein_coding | -0.71155087 | 3.921841883 | 1.6216595 | 0.2207428 | 0.726727643 |
| ENSMUSG00000017868 | Sgk2 | protein_coding | -0.927474238 | 3.413828535 | 1.6225594 | 0.2207476 | 0.726727643 |
| ENSMUSG00000021911 | Parg | protein_coding | -0.486462265 | 4.457585054 | 1.62129 | 0.2207638 | 0.726727643 |
| ENSMUSG00000020679 | Hnf1b | protein_coding | 0.647716367 | 3.596697642 | 1.6208453 | 0.2208243 | 0.726727643 |
| ENSMUSG00000027525 | Phactr3 | protein_coding | -0.288140779 | 6.714599074 | 1.6208202 | 0.2208277 | 0.726727643 |
| ENSMUSG00000002028 | Kmt2a | protein_coding | -0.476672623 | 5.263140623 | 1.6205959 | 0.2208582 | 0.726727643 |
| ENSMUSG00000025907 | Rb1cc1 | protein_coding | -0.379169696 | 5.252218328 | 1.6202374 | 0.2209069 | 0.726727643 |
| ENSMUSG00000030744 | Rps3 | protein_coding | 0.323018806 | 10.97915624 | 1.619182 | 0.2210505 | 0.726727643 |
| ENSMUSG00000026112 | Coa5 | protein_coding | 0.283525396 | 6.58948572 | 1.6187204 | 0.2211133 | 0.726727643 |
| ENSMUSG00000020300 | Cpeb4 | protein_coding | -0.386524105 | 5.245710737 | 1.6187074 | 0.2211151 | 0.726727643 |
| ENSMUSG00000031953 | Tmem170 | protein_coding | -0.565183143 | 4.277935296 | 1.6185693 | 0.2211339 | 0.726727643 |
| ENSMUSG00000003469 | Phyhip | protein_coding | 0.282108064 | 8.650892577 | 1.6173586 | 0.2212988 | 0.726944068 |
| ENSMUSG00000020962 | Gtf2a1 | protein_coding | -0.35108932 | 5.943670029 | 1.6170129 | 0.2213459 | 0.726944068 |
| ENSMUSG00000035864 | Syt1 | protein_coding | -0.285579121 | 9.368887695 | 1.6165817 | 0.2214047 | 0.726944068 |
| ENSMUSG00000057649 | Brd9 | protein_coding | 0.364480832 | 5.955178553 | 1.6156555 | 0.2215311 | 0.726944068 |

| ensembl_gene_id | external_gene_name | gene_biotype | log2 (fold change) | log2 (counts per million) | F-value | p-value | False discovery rate |
|---|---|---|---|---|---|---|---|
| ENSMUSG00000050234 | Gja4 | protein_coding | 1.022520189 | 2.662100191 | 1.6152852 | 0.2215816 | 0.726944068 |
| ENSMUSG00000060212 | Pcnx2 | protein_coding | -0.634494777 | 3.502275679 | 1.6143257 | 0.2217127 | 0.726944068 |
| ENSMUSG00000043913 | Ccdc60 | protein_coding | -1.153498343 | 2.315341886 | 1.6138867 | 0.2217726 | 0.726944068 |
| ENSMUSG00000021917 | Spcs1 | protein_coding | 0.344294222 | 8.833726053 | 1.6132789 | 0.2218557 | 0.726944068 |
| ENSMUSG00000031918 | Mtmr2 | protein_coding | -0.424094418 | 5.617265635 | 1.6129189 | 0.221905 | 0.726944068 |
| ENSMUSG00000057388 | Mrpl18 | protein_coding | 0.260648398 | 8.058049206 | 1.6125826 | 0.221951 | 0.726944068 |
| ENSMUSG00000021222 | Dcaf4 | protein_coding | 0.365149834 | 6.168689986 | 1.6120714 | 0.2220209 | 0.726944068 |
| ENSMUSG00000020611 | Gna13 | protein_coding | -0.394540859 | 5.400920887 | 1.6118966 | 0.2220448 | 0.726944068 |
| ENSMUSG00000059436 | Max | protein_coding | -0.540680886 | 5.952105046 | 1.6199128 | 0.2220786 | 0.726944068 |
| ENSMUSG00000020687 | Cdc27 | protein_coding | -0.372304668 | 5.098841015 | 1.6089321 | 0.2224511 | 0.727614687 |
| ENSMUSG00000021716 | Srek1ip1 | protein_coding | -0.332681382 | 6.026478648 | 1.6087938 | 0.2224701 | 0.727614687 |
| ENSMUSG00000026504 | Sdccag8 | protein_coding | 0.609194671 | 4.75536557 | 1.6094439 | 0.2224865 | 0.727614687 |
| ENSMUSG00000020990 | Cdkl1 | protein_coding | 0.485440568 | 4.588056942 | 1.6079958 | 0.2225796 | 0.727697929 |
| ENSMUSG00000044952 | Kctd21 | protein_coding | 0.553409328 | 4.021924645 | 1.6060945 | 0.2228409 | 0.727704387 |
| ENSMUSG00000022973 | Synj1 | protein_coding | -0.338205938 | 7.54218916 | 1.6054381 | 0.2229311 | 0.727704387 |
| ENSMUSG00000023452 | Pisd | protein_coding | 0.325431865 | 5.562063979 | 1.6049944 | 0.2229922 | 0.727704387 |
| ENSMUSG00000037669 | Ldah | protein_coding | -0.286294742 | 6.544297754 | 1.6049124 | 0.2230035 | 0.727704387 |
| ENSMUSG00000020130 | Tbc1d15 | protein_coding | -0.367426188 | 5.682578696 | 1.6040793 | 0.2231182 | 0.727704387 |
| ENSMUSG00000007036 | Abhd16a | protein_coding | 0.35695862 | 5.766196043 | 1.6040709 | 0.2231194 | 0.727704387 |
| ENSMUSG00000034768 | Asb16 | protein_coding | 0.740865698 | 2.949665346 | 1.603852 | 0.2231495 | 0.727704387 |
| ENSMUSG00000020716 | Nf1 | protein_coding | -0.435362168 | 5.142647013 | 1.6036788 | 0.2231734 | 0.727704387 |
| ENSMUSG00000027712 | Anxa5 | protein_coding | -0.33985511 | 7.370112413 | 1.6021552 | 0.2233834 | 0.727704387 |
| ENSMUSG00000036943 | Rab8b | protein_coding | -0.480486383 | 4.885630103 | 1.6019457 | 0.2234123 | 0.727704387 |
| ENSMUSG00000056966 | Gjc3 | protein_coding | -0.384988576 | 5.573142038 | 1.600778 | 0.2235735 | 0.727704387 |
| ENSMUSG00000108348 | Gm42372 | protein_coding | 0.317422338 | 6.623810946 | 1.600014 | 0.223679 | 0.727704387 |
| ENSMUSG00000004100 | Ppan | protein_coding | 0.633678326 | 3.711390121 | 1.5996054 | 0.2237355 | 0.727704387 |
| ENSMUSG00000073838 | Tufm | protein_coding | 0.516109049 | 3.580335509 | 1.5994881 | 0.2237517 | 0.727704387 |
| ENSMUSG00000047115 | Fam221a | protein_coding | 1.131216466 | 2.264758872 | 1.5994169 | 0.2237615 | 0.727704387 |
| ENSMUSG00000032540 | Abhd5 | protein_coding | -0.575389261 | 3.705300625 | 1.5989844 | 0.2238213 | 0.727704387 |
| ENSMUSG00000001313 | Rnd2 | protein_coding | 0.320445073 | 7.485209106 | 1.5986429 | 0.2238686 | 0.727704387 |
| ENSMUSG00000019775 | Rgs17 | protein_coding | 0.294563844 | 7.274097657 | 1.5968723 | 0.2241137 | 0.727704387 |
| ENSMUSG00000113902 | AC121965.1 | bidirectional_promote | 0.372461134 | 9.115190095 | 1.5964132 | 0.2242063 | 0.727704387 |
| ENSMUSG00000026275 | Ppp1r7 | protein_coding | -0.272204571 | 7.815914867 | 1.5959714 | 0.2242385 | 0.727704387 |
| ENSMUSG00000030423 | Pop4 | protein_coding | 0.340807311 | 6.037266014 | 1.5953665 | 0.2243224 | 0.727704387 |
| ENSMUSG00000052397 | Ezr | protein_coding | -0.525526536 | 4.466363293 | 1.5947054 | 0.2244141 | 0.727704387 |
| ENSMUSG00000027220 | Syt13 | protein_coding | 0.293391617 | 6.202347776 | 1.5941546 | 0.2244906 | 0.727704387 |
| ENSMUSG00000005268 | Prlr | protein_coding | 0.799086897 | 3.490090336 | 1.5932539 | 0.2246446 | 0.727704387 |
| ENSMUSG00000019432 | Ddx39b | protein_coding | 0.287256177 | 7.952588544 | 1.5929622 | 0.2246562 | 0.727704387 |
| ENSMUSG00000046562 | Unc119b | protein_coding | -0.766401727 | 3.136478763 | 1.5913422 | 0.2248814 | 0.727704387 |
| ENSMUSG00000046947 | Adck2 | protein_coding | -0.927212892 | 3.16324353 | 1.5913253 | 0.2249127 | 0.727704387 |
| ENSMUSG00000049100 | Pcdh10 | protein_coding | -0.567203725 | 5.010605996 | 1.5903139 | 0.2250535 | 0.727704387 |
| ENSMUSG00000085622 | 3110056K07Rik | processed_transcript | -0.619395451 | 4.492442256 | 1.5895446 | 0.2251606 | 0.727704387 |
| ENSMUSG00000036218 | Pdzrn4 | protein_coding | 0.523595436 | 3.917302898 | 1.5883312 | 0.2253009 | 0.727704387 |
| ENSMUSG00000027002 | Nckap1 | protein_coding | -0.294958389 | 7.916296501 | 1.5873782 | 0.2254338 | 0.727704387 |
| ENSMUSG00000037049 | Smpd1 | protein_coding | 0.30919509 | 6.9351457 | 1.5872828 | 0.2254471 | 0.727704387 |
| ENSMUSG00000044763 | Trmt10c | protein_coding | -0.505437433 | 4.636811629 | 1.5872587 | 0.2254505 | 0.727704387 |
| ENSMUSG00000005534 | Insr | protein_coding | 0.591513704 | 3.876891473 | 1.5864537 | 0.2255629 | 0.727704387 |
| ENSMUSG00000028871 | Rspo1 | protein_coding | -0.901910199 | 3.104288689 | 1.5850404 | 0.2257604 | 0.727704387 |
| ENSMUSG00000025231 | Sufu | protein_coding | -0.791279244 | 3.201262282 | 1.5848874 | 0.2257818 | 0.727704387 |
| ENSMUSG00000030094 | Xpc | protein_coding | -0.60436936 | 3.523634083 | 1.5847482 | 0.2258013 | 0.727704387 |
| ENSMUSG00000021775 | Nr1d2 | protein_coding | -0.359512668 | 5.451792979 | 1.584588 | 0.2258237 | 0.727704387 |
| ENSMUSG00000024430 | Cabyr | protein_coding | -1.011040191 | 2.451006749 | 1.5844996 | 0.2258361 | 0.727704387 |
| ENSMUSG00000008976 | Gabpa | protein_coding | -0.723862574 | 2.888751968 | 1.5844719 | 0.2258399 | 0.727704387 |
| ENSMUSG00000029050 | Ski | protein_coding | -0.469907065 | 5.481192407 | 1.5843902 | 0.2258514 | 0.727704387 |
| ENSMUSG00000050144 | Slc25a44 | protein_coding | 0.294431563 | 7.042172236 | 1.5843418 | 0.2258581 | 0.727704387 |
| ENSMUSG00000033819 | Ppp1r16a | protein_coding | 0.414770431 | 4.734216239 | 1.5843141 | 0.225862 | 0.727704387 |
| ENSMUSG00000058966 | Fam57b | protein_coding | 0.65679265 | 4.642708723 | 1.5905551 | 0.2258861 | 0.727704387 |
| ENSMUSG00000033981 | Gria2 | protein_coding | -0.317944677 | 8.100468188 | 1.5834962 | 0.2259765 | 0.727704387 |
| ENSMUSG00000021365 | Nedd9 | protein_coding | -1.053058055 | 2.474451511 | 1.5829337 | 0.2260553 | 0.727704387 |
| ENSMUSG00000018572 | Phf23 | protein_coding | 0.344148107 | 5.843592551 | 1.5821697 | 0.2261623 | 0.727704387 |
| ENSMUSG00000087370 | Tmem170b | protein_coding | -0.457209154 | 5.335219299 | 1.5818312 | 0.2262098 | 0.727704387 |
| ENSMUSG00000033278 | Ptprm | protein_coding | -0.524482588 | 4.481869238 | 1.5816987 | 0.2262284 | 0.727704387 |
| ENSMUSG00000028744 | Pqlc2 | protein_coding | 0.512371872 | 4.249229138 | 1.5816569 | 0.2262342 | 0.727704387 |
| ENSMUSG00000039100 | 6-Mar | protein_coding | -0.342579588 | 6.267039497 | 1.5813602 | 0.2262758 | 0.727704387 |
| ENSMUSG00000006057 | Atp5g1 | protein_coding | 0.297844263 | 5.899637889 | 1.5809937 | 0.2263272 | 0.727704387 |
| ENSMUSG00000041695 | Kcnj2 | protein_coding | -1.485942078 | 2.236459618 | 1.5806445 | 0.2264051 | 0.727704387 |
| ENSMUSG00000022151 | Ttc33 | protein_coding | 0.403627971 | 4.688553336 | 1.5800021 | 0.2264664 | 0.727704387 |
| ENSMUSG00000026347 | Tmem163 | protein_coding | -0.580334293 | 4.287206901 | 1.5792072 | 0.226578 | 0.727704387 |
| ENSMUSG00000029062 | Cdk11b | protein_coding | -0.380225743 | 4.978969342 | 1.5787725 | 0.2266391 | 0.727704387 |
| ENSMUSG00000032519 | Slc25a38 | protein_coding | -0.427864014 | 4.960370605 | 1.576967 | 0.226893 | 0.727704387 |
| ENSMUSG00000002221 | Paxip1 | protein_coding | -0.488396249 | 4.563424626 | 1.5762414 | 0.2269952 | 0.727704387 |
| ENSMUSG00000027475 | Kif3b | protein_coding | -0.416000066 | 4.750649199 | 1.5760526 | 0.2270218 | 0.727704387 |
| ENSMUSG00000048106 | 4632415L05Rik | lincRNA | -0.644724401 | 3.111521231 | 1.575477 | 0.2271029 | 0.727704387 |
| ENSMUSG00000003660 | Snrnp200 | protein_coding | -0.407511604 | 5.519479476 | 1.5750858 | 0.227158 | 0.727704387 |
| ENSMUSG00000029363 | Rfc5 | protein_coding | -0.897253791 | 2.78036276 | 1.574735 | 0.2272075 | 0.727704387 |
| ENSMUSG00000033075 | Senp1 | protein_coding | -0.788615429 | 3.370551733 | 1.5741652 | 0.2272878 | 0.727704387 |
| ENSMUSG00000042557 | Sin3a | protein_coding | -0.81023203 | 2.824291115 | 1.5738395 | 0.2273338 | 0.727704387 |
| ENSMUSG00000020547 | Bzw2 | protein_coding | 0.304203666 | 6.436693677 | 1.5735135 | 0.2273798 | 0.727704387 |

| ensembl_gene_id | external_gene_name | gene_biotype | log2 (fold change) | log2 (counts per million) | F-value | p-value | False discovery rate |
|---|---|---|---|---|---|---|---|
| ENSMUSG00000040029 | Ipo8 | protein_coding | -0.379964857 | 5.01949478 | 1.5734569 | 0.2273878 | 0.727704387 |
| ENSMUSG00000056204 | Pgpep1 | protein_coding | -0.41211752 | 5.433848232 | 1.5733332 | 0.2274052 | 0.727704387 |
| ENSMUSG00000068917 | Clk2 | protein_coding | -0.868933191 | 3.160121422 | 1.5731939 | 0.2274249 | 0.727704387 |
| ENSMUSG00000038608 | Dock10 | protein_coding | -0.550954066 | 4.563913217 | 1.5727888 | 0.2274821 | 0.727704387 |
| ENSMUSG00000051435 | Fhad1 | protein_coding | -0.356985308 | 5.190793183 | 1.5726725 | 0.2274985 | 0.727704387 |
| ENSMUSG00000022051 | Bnip3l | protein_coding | -0.247010097 | 7.068522962 | 1.5726504 | 0.2275016 | 0.727704387 |
| ENSMUSG00000074305 | Peak1 | protein_coding | -0.706508341 | 3.140702015 | 1.5724393 | 0.2275314 | 0.727704387 |
| ENSMUSG00000059486 | Kbtbd2 | protein_coding | -0.376932636 | 5.815664921 | 1.5719205 | 0.2276047 | 0.727704387 |
| ENSMUSG00000060147 | Serpinb6a | protein_coding | -0.366678671 | 6.124552557 | 1.5716858 | 0.2276379 | 0.727704387 |
| ENSMUSG00000029056 | Pank4 | protein_coding | 0.413516912 | 4.982483717 | 1.5715496 | 0.2276572 | 0.727704387 |
| ENSMUSG00000086697 | 4933427G23Rik | transcribed_processe | -0.949799565 | 2.445828553 | 1.5707769 | 0.2277664 | 0.727766895 |
| ENSMUSG00000046096 | BC030336 | protein_coding | -0.359546971 | 5.853572325 | 1.5704541 | 0.2278121 | 0.727766895 |
| ENSMUSG00000024853 | Sf3b2 | protein_coding | 0.293248606 | 9.008018132 | 1.5678644 | 0.2281789 | 0.728675837 |
| ENSMUSG00000025909 | Sntg1 | protein_coding | 0.437647712 | 4.783253492 | 1.5671508 | 0.2282801 | 0.728675837 |
| ENSMUSG00000022711 | Pmm2 | protein_coding | -0.387295719 | 5.048380884 | 1.5670109 | 0.2282999 | 0.728675837 |
| ENSMUSG00000045409 | Trim39 | protein_coding | -0.630027602 | 3.885929486 | 1.5648275 | 0.2286099 | 0.729292908 |
| ENSMUSG00000022111 | Uchl3 | protein_coding | 0.320687815 | 6.159675216 | 1.5646942 | 0.2286289 | 0.729292908 |
| ENSMUSG00000053838 | Nudcd3 | protein_coding | 0.360461489 | 7.133423902 | 1.5632553 | 0.2288623 | 0.72942087 |
| ENSMUSG00000064068 | Mtx1 | protein_coding | 0.303405636 | 5.985579184 | 1.5629499 | 0.228877 | 0.72942087 |
| ENSMUSG00000090125 | Pou3f1 | protein_coding | -0.559189255 | 3.977710097 | 1.5628179 | 0.2288958 | 0.72942087 |
| ENSMUSG00000036882 | Arhgap33 | protein_coding | -0.410373777 | 5.61869211 | 1.5623389 | 0.228964 | 0.72942087 |
| ENSMUSG00000026883 | Dab2ip | protein_coding | -0.493792457 | 4.455905101 | 1.5620286 | 0.2290082 | 0.72942087 |
| ENSMUSG00000037843 | Vstm2l | protein_coding | -0.904709923 | 2.609806328 | 1.5613011 | 0.2291118 | 0.729534938 |
| ENSMUSG00000031665 | Sall1 | protein_coding | -0.759165214 | 3.208844785 | 1.5603376 | 0.2292492 | 0.729756257 |
| ENSMUSG00000025035 | Arl3 | protein_coding | 0.273003398 | 9.134159162 | 1.5594951 | 0.2293694 | 0.729899168 |
| ENSMUSG00000028187 | Rpf1 | protein_coding | 0.445060252 | 4.458672323 | 1.5590718 | 0.2294299 | 0.729899168 |
| ENSMUSG00000055629 | B4galnt4 | protein_coding | 0.367641641 | 5.690109937 | 1.5571799 | 0.2297002 | 0.730501627 |
| ENSMUSG00000034675 | Dbn1 | protein_coding | -0.33555152 | 6.113673448 | 1.5556696 | 0.2299163 | 0.730501627 |
| ENSMUSG00000041236 | Vps41 | protein_coding | 0.326481476 | 7.767959497 | 1.5556533 | 0.2299186 | 0.730501627 |
| ENSMUSG00000029581 | Fscn1 | protein_coding | -0.336604068 | 5.563897605 | 1.5554764 | 0.2299439 | 0.730501627 |
| ENSMUSG00000018042 | Cyb5r3 | protein_coding | 0.319473753 | 7.649355006 | 1.555372 | 0.2299589 | 0.730501627 |
| ENSMUSG00000005533 | Igf1r | protein_coding | -0.511331815 | 4.398673351 | 1.5541386 | 0.2301356 | 0.730752454 |
| ENSMUSG00000034275 | Igsf9b | protein_coding | -0.552084941 | 4.741212726 | 1.5537512 | 0.2302199 | 0.730752454 |
| ENSMUSG00000001089 | Luzp1 | protein_coding | -0.411934369 | 5.210286883 | 1.5523083 | 0.2303982 | 0.730752454 |
| ENSMUSG00000001260 | Gabrg1 | protein_coding | 0.315643976 | 6.075152125 | 1.5520561 | 0.2304344 | 0.730752454 |
| ENSMUSG00000023944 | Hsp90ab1 | protein_coding | 0.306759632 | 11.18772475 | 1.5505924 | 0.2306447 | 0.730752454 |
| ENSMUSG00000005716 | Pvalb | protein_coding | -0.50399484 | 7.802006038 | 1.5676905 | 0.2306569 | 0.730752454 |
| ENSMUSG00000045374 | Wdr81 | protein_coding | -0.905567326 | 2.921892044 | 1.5498218 | 0.2307555 | 0.730752454 |
| ENSMUSG00000114362 | AC124433.1 | antisense_RNA | -1.281374998 | 2.15005337 | 1.5489791 | 0.2308768 | 0.730752454 |
| ENSMUSG00000041544 | Disp3 | protein_coding | 0.746703754 | 3.256826416 | 1.5480199 | 0.2310149 | 0.730752454 |
| ENSMUSG00000055660 | Mettl4 | protein_coding | -0.888447897 | 2.489487307 | 1.5475382 | 0.2310843 | 0.730752454 |
| ENSMUSG00000029283 | Cdc7 | protein_coding | 0.924142487 | 2.676501028 | 1.5473768 | 0.2311076 | 0.730752454 |
| ENSMUSG00000039218 | Srrm2 | protein_coding | -0.35325993 | 6.124599516 | 1.5461882 | 0.231279 | 0.730752454 |
| ENSMUSG00000001150 | Mcm3ap | protein_coding | -0.516995927 | 3.73704569 | 1.5460513 | 0.2312987 | 0.730752454 |
| ENSMUSG00000056962 | Jmjd6 | protein_coding | 0.353516993 | 5.364287745 | 1.5456002 | 0.2313638 | 0.730752454 |
| ENSMUSG00000031320 | Rps4x | protein_coding | 0.341597465 | 6.337535257 | 1.5454921 | 0.2313794 | 0.730752454 |
| ENSMUSG00000071252 | 2210408I21Rik | protein_coding | -0.424958945 | 6.947680985 | 1.5527098 | 0.231385 | 0.730752454 |
| ENSMUSG00000022040 | Ephx2 | protein_coding | -0.36813598 | 4.957748281 | 1.5451008 | 0.2314359 | 0.730752454 |
| ENSMUSG00000039753 | Fbxl5 | protein_coding | -0.273298759 | 6.427888956 | 1.5449485 | 0.2314579 | 0.730752454 |
| ENSMUSG00000040759 | Cmtm5 | protein_coding | -0.378521209 | 7.823720568 | 1.5453551 | 0.2315034 | 0.730752454 |
| ENSMUSG00000026153 | Fam135a | protein_coding | -0.867379508 | 3.403982412 | 1.5445173 | 0.2315488 | 0.730752454 |
| ENSMUSG00000021830 | Txndc16 | protein_coding | -0.367311647 | 5.510028045 | 1.5440513 | 0.2315875 | 0.730752454 |
| ENSMUSG00000048878 | Hexim1 | protein_coding | -0.440053721 | 4.585483857 | 1.5435685 | 0.2316573 | 0.730752454 |
| ENSMUSG00000005357 | Slc1a6 | protein_coding | 0.513941382 | 4.15953709 | 1.5433199 | 0.2316932 | 0.730752454 |
| ENSMUSG00000021264 | Yy1 | protein_coding | -0.518496073 | 4.865081392 | 1.5431557 | 0.231717 | 0.730752454 |
| ENSMUSG00000000247 | Lhx2 | protein_coding | -0.942767445 | 3.10998722 | 1.5432169 | 0.2317368 | 0.730752454 |
| ENSMUSG00000051319 | 1500011K16Rik | protein_coding | 0.273282341 | 7.289265781 | 1.5422439 | 0.2318489 | 0.730891634 |
| ENSMUSG00000037996 | Slc24a2 | protein_coding | -0.379232226 | 6.09735924 | 1.5409744 | 0.2320328 | 0.731256798 |
| ENSMUSG00000026259 | Ngef | protein_coding | -0.500825615 | 6.559742315 | 1.5528693 | 0.2321746 | 0.731489526 |
| ENSMUSG00000027011 | Ube2e3 | protein_coding | -0.250216882 | 6.868104404 | 1.5376078 | 0.2325212 | 0.732305464 |
| ENSMUSG00000069670 | Nkain2 | protein_coding | -0.52543066 | 5.140302414 | 1.5374289 | 0.2325758 | 0.732305464 |
| ENSMUSG00000032036 | Kirrel3 | protein_coding | -0.397149304 | 5.479960075 | 1.5368045 | 0.2326379 | 0.732305464 |
| ENSMUSG00000023967 | Mrps18a | protein_coding | 0.342548369 | 5.128822715 | 1.5353327 | 0.232852 | 0.732701581 |
| ENSMUSG00000024325 | Ring1 | protein_coding | 0.81132581 | 2.785337148 | 1.5350027 | 0.2329 | 0.732701581 |
| ENSMUSG00000026280 | Atg4b | protein_coding | 0.315046738 | 6.247778244 | 1.5329558 | 0.2331983 | 0.733425273 |
| ENSMUSG00000030654 | Arl6ip1 | protein_coding | 0.250052772 | 10.05420097 | 1.5320743 | 0.2333269 | 0.73361385 |
| ENSMUSG00000024902 | Mrpl11 | protein_coding | 0.253988474 | 7.861113835 | 1.5314492 | 0.2334181 | 0.73361385 |
| ENSMUSG00000041046 | Ramp3 | protein_coding | 0.540403556 | 6.002881069 | 1.5427914 | 0.2335084 | 0.73361385 |
| ENSMUSG00000031325 | 4930519F16Rik | transcribed_unitary_p | -0.919726565 | 2.588531123 | 1.5306482 | 0.2335351 | 0.73361385 |
| ENSMUSG00000044708 | Kcnj10 | protein_coding | -0.494697207 | 4.889760016 | 1.5301571 | 0.2336069 | 0.73361385 |
| ENSMUSG00000043633 | Fam221b | protein_coding | 0.691693194 | 3.063791925 | 1.5294942 | 0.2337038 | 0.73361385 |
| ENSMUSG00000001018 | Snapin | protein_coding | -0.434936991 | 6.521484819 | 1.5342706 | 0.2337358 | 0.73361385 |
| ENSMUSG00000032456 | Nmnat3 | protein_coding | 0.834806837 | 2.809270267 | 1.5285988 | 0.2338348 | 0.733710327 |
| ENSMUSG00000013495 | Tmem175 | protein_coding | 0.43213517 | 5.834361965 | 1.5279402 | 0.2339597 | 0.733851347 |
| ENSMUSG00000031774 | Fam192a | protein_coding | 0.385240521 | 6.846178738 | 1.5279942 | 0.2340271 | 0.733851347 |
| ENSMUSG00000014496 | Ankrd28 | protein_coding | -0.605893805 | 3.551917331 | 1.5268373 | 0.2340927 | 0.733851347 |
| ENSMUSG00000027942 | 4933434E20Rik | protein_coding | 0.245530026 | 8.603960631 | 1.5264282 | 0.2341527 | 0.733851347 |
| ENSMUSG00000068184 | Ndufaf2 | protein_coding | 0.305215364 | 7.34723088 | 1.5236913 | 0.2345544 | 0.734466399 |

| ensembl_gene_id | external_gene_name | gene_biotype | log2 (fold change) | log2 (counts per million) | F-value | p-value | False discovery rate |
|---|---|---|---|---|---|---|---|
| ENSMUSG00000024824 | Rad9a | protein_coding | 0.486674779 | 4.462150438 | 1.5233247 | 0.2346082 | 0.734466399 |
| ENSMUSG00000021266 | Wars | protein_coding | 0.51804935 | 7.014786767 | 1.5423216 | 0.2346139 | 0.734466399 |
| ENSMUSG00000028992 | Nmnat1 | protein_coding | -0.572102305 | 3.606100699 | 1.5229532 | 0.2346629 | 0.734466399 |
| ENSMUSG00000064145 | Arih2 | protein_coding | -0.428539455 | 4.752227447 | 1.5220348 | 0.2347979 | 0.734466399 |
| ENSMUSG00000031023 | Akip1 | protein_coding | -0.56314772 | 3.610570956 | 1.5219727 | 0.2348071 | 0.734466399 |
| ENSMUSG00000039349 | C130074G19Rik | protein_coding | -0.822098431 | 3.535374711 | 1.522005 | 0.2348308 | 0.734466399 |
| ENSMUSG00000029536 | Gatc | protein_coding | 0.292109696 | 5.99262506 | 1.5207646 | 0.2349849 | 0.734466399 |
| ENSMUSG00000068457 | Uty | protein_coding | -0.612712228 | 3.355291775 | 1.5207536 | 0.2349865 | 0.734466399 |
| ENSMUSG00000050552 | Lamtor4 | protein_coding | 0.302517811 | 7.544119523 | 1.520445 | 0.235032 | 0.734466399 |
| ENSMUSG00000004508 | Gab2 | protein_coding | -0.500898136 | 3.938016817 | 1.5196124 | 0.2351547 | 0.734636297 |
| ENSMUSG00000026639 | Lamb3 | protein_coding | -0.866855697 | 3.199071581 | 1.5186687 | 0.2352938 | 0.73485759 |
| ENSMUSG00000021373 | Cap2 | protein_coding | -0.296710819 | 7.068830187 | 1.5174711 | 0.2354706 | 0.735041074 |
| ENSMUSG00000005813 | Metap1 | protein_coding | 0.365516384 | 5.827359175 | 1.5169943 | 0.235541 | 0.735041074 |
| ENSMUSG00000061232 | H2-K1 | protein_coding | -0.761037963 | 3.521797605 | 1.5170746 | 0.2355577 | 0.735041074 |
| ENSMUSG00000041205 | Map6d1 | protein_coding | -0.484625976 | 4.165930521 | 1.5154927 | 0.235763 | 0.735468397 |
| ENSMUSG00000086363 | A330102I10Rik | lincRNA | 0.448514355 | 4.731109168 | 1.5145597 | 0.2359011 | 0.735685632 |
| ENSMUSG00000063077 | Kif1b | protein_coding | -0.26346421 | 7.914263442 | 1.5135377 | 0.2360524 | 0.735744255 |
| ENSMUSG00000048376 | F2r | protein_coding | -0.907360147 | 2.762143809 | 1.5128963 | 0.2361475 | 0.735744255 |
| ENSMUSG00000034781 | Gna11 | protein_coding | 0.315049287 | 8.147263377 | 1.5126116 | 0.2361897 | 0.735744255 |
| ENSMUSG00000036435 | Exoc1 | protein_coding | -0.42137213 | 5.289982686 | 1.5125854 | 0.2361935 | 0.735744255 |
| ENSMUSG00000038046 | Mrm3 | protein_coding | 0.332117425 | 5.840858269 | 1.5106395 | 0.2364823 | 0.736251553 |
| ENSMUSG00000021990 | Spata13 | protein_coding | -0.561235337 | 4.088362386 | 1.5098221 | 0.2366037 | 0.736251553 |
| ENSMUSG00000026245 | Farsb | protein_coding | 0.292820926 | 7.306000075 | 1.5097574 | 0.2366133 | 0.736251553 |
| ENSMUSG00000022416 | Cacna1i | protein_coding | -0.830832164 | 2.987274636 | 1.5096432 | 0.2366303 | 0.736251553 |
| ENSMUSG00000029405 | G3bp2 | protein_coding | 0.359396351 | 7.275277119 | 1.5087109 | 0.2367973 | 0.736351419 |
| ENSMUSG00000025505 | Tmem80 | protein_coding | 0.360811619 | 6.001325798 | 1.5078005 | 0.2369043 | 0.736351419 |
| ENSMUSG00000085084 | 4930570G19Rik | lincRNA | -0.461120856 | 4.099744963 | 1.5072468 | 0.2369868 | 0.736351419 |
| ENSMUSG00000020235 | Fzr1 | protein_coding | 0.304725711 | 6.145964474 | 1.5069396 | 0.2370325 | 0.736351419 |
| ENSMUSG00000020182 | Ddc | protein_coding | 0.352222071 | 5.347214123 | 1.5068409 | 0.2370472 | 0.736351419 |
| ENSMUSG00000030166 | Rad52 | protein_coding | 0.414783367 | 4.579699803 | 1.505686 | 0.2372194 | 0.736351419 |
| ENSMUSG00000038347 | Tcte2 | protein_coding | -0.47434121 | 4.131523896 | 1.5051654 | 0.237297 | 0.736351419 |
| ENSMUSG00000074649 | BC029722 | protein_coding | -0.549299527 | 4.083220011 | 1.5050776 | 0.2373101 | 0.736351419 |
| ENSMUSG00000018501 | Ncor1 | protein_coding | -0.320532657 | 5.848514274 | 1.5042526 | 0.2374332 | 0.736351419 |
| ENSMUSG00000035383 | Pmch | protein_coding | 1.171287271 | 6.988678681 | 1.6336256 | 0.2375447 | 0.736351419 |
| ENSMUSG00000051518 | Rps19bp1 | protein_coding | -0.506795489 | 4.065044223 | 1.5031247 | 0.2376016 | 0.736351419 |
| ENSMUSG00000053226 | Dand5 | protein_coding | 0.412959601 | 4.903094634 | 1.5029236 | 0.2376317 | 0.736351419 |
| ENSMUSG00000021180 | Rps6ka5 | protein_coding | -0.67859797 | 4.180148764 | 1.5031297 | 0.2376735 | 0.736351419 |
| ENSMUSG00000025921 | Rdh10 | protein_coding | -0.868821559 | 2.745578129 | 1.5025893 | 0.2376817 | 0.736351419 |
| ENSMUSG00000020700 | Map3k3 | protein_coding | -0.710124151 | 3.35893511 | 1.5025365 | 0.2376896 | 0.736351419 |
| ENSMUSG00000019820 | Utrn | protein_coding | -0.405814977 | 5.03436791 | 1.5017071 | 0.2378136 | 0.736395884 |
| ENSMUSG00000020374 | Rasgef1c | protein_coding | -0.67797039 | 3.537564496 | 1.5015247 | 0.2378409 | 0.736395884 |
| ENSMUSG00000040888 | Gfer | protein_coding | 0.395476656 | 4.47667064 | 1.4994492 | 0.2381516 | 0.737107044 |
| ENSMUSG00000046668 | Cxxc5 | protein_coding | -0.352192443 | 5.562815778 | 1.4990755 | 0.2382077 | 0.737107044 |
| ENSMUSG00000052373 | Mpp3 | protein_coding | 0.446952176 | 5.007328505 | 1.4983679 | 0.2383138 | 0.737223226 |
| ENSMUSG00000031878 | Nae1 | protein_coding | -0.3163829 | 6.584111225 | 1.4971195 | 0.2385011 | 0.737590614 |
| ENSMUSG00000030731 | Syt3 | protein_coding | 0.397106235 | 4.880763602 | 1.4952223 | 0.2387862 | 0.737920838 |
| ENSMUSG00000054302 | Eapp | protein_coding | -0.275216288 | 7.104930646 | 1.4948097 | 0.2388483 | 0.737920838 |
| ENSMUSG00000056222 | Spock1 | protein_coding | 0.257588017 | 6.523959085 | 1.4947224 | 0.2388614 | 0.737920838 |
| ENSMUSG00000034593 | Myo5a | protein_coding | -0.344315969 | 7.076444363 | 1.4945028 | 0.2389228 | 0.737920838 |
| ENSMUSG00000020485 | Supt4a | protein_coding | 0.262249814 | 6.644403082 | 1.4938484 | 0.2389929 | 0.737920838 |
| ENSMUSG00000052087 | Rgs14 | protein_coding | -0.418508974 | 5.188153035 | 1.493671 | 0.2390196 | 0.737920838 |
| ENSMUSG00000035439 | Haus8 | protein_coding | 0.372590277 | 5.555559497 | 1.4921487 | 0.239249 | 0.738343143 |
| ENSMUSG00000073236 | 2500004C02Rik | processed_transcript | -0.432177989 | 4.325522771 | 1.4918519 | 0.2392938 | 0.738343143 |
| ENSMUSG00000026074 | Map4k4 | protein_coding | -0.470636045 | 4.142938693 | 1.4902529 | 0.2395351 | 0.738875634 |
| ENSMUSG00000026634 | Angel2 | protein_coding | -0.344573254 | 5.422101788 | 1.4895777 | 0.239637 | 0.738877016 |
| ENSMUSG00000090291 | Lrrc10b | protein_coding | -0.585210254 | 4.112851668 | 1.4893402 | 0.2396729 | 0.738877016 |
| ENSMUSG00000029471 | Camkk2 | protein_coding | -0.459205755 | 4.662695683 | 1.4886855 | 0.2397719 | 0.738945616 |
| ENSMUSG00000085071 | Gm14066 | processed_transcript | -0.84724614 | 3.106832346 | 1.4882838 | 0.2398326 | 0.738945616 |
| ENSMUSG00000042707 | Dnali1 | protein_coding | -0.77196288 | 3.274985517 | 1.4876506 | 0.2399284 | 0.739029031 |
| ENSMUSG00000022280 | Rnf19a | protein_coding | -0.484974581 | 4.079640863 | 1.4861494 | 0.2401557 | 0.739435936 |
| ENSMUSG00000056268 | Dennd1b | protein_coding | -0.926630144 | 2.790479501 | 1.4854938 | 0.2402551 | 0.739435936 |
| ENSMUSG00000112121 | AC163903.1 | lincRNA | 0.870338319 | 2.695447453 | 1.484762 | 0.2403661 | 0.739435936 |
| ENSMUSG00000037461 | Ints7 | protein_coding | 0.403429387 | 5.262373825 | 1.4837624 | 0.2405177 | 0.739435936 |
| ENSMUSG00000028797 | Tmem234 | protein_coding | 0.237181323 | 7.596368689 | 1.4837423 | 0.2405208 | 0.739435936 |
| ENSMUSG00000022477 | Aco2 | protein_coding | 0.292756777 | 8.493533715 | 1.4836457 | 0.2405355 | 0.739435936 |
| ENSMUSG00000070639 | Lrrc8b | protein_coding | -0.462087124 | 4.292064938 | 1.4833159 | 0.2405855 | 0.739435936 |
| ENSMUSG00000031438 | Rnf128 | protein_coding | 0.501540055 | 4.235137812 | 1.482586 | 0.2406964 | 0.739435936 |
| ENSMUSG00000032902 | Slc16a1 | protein_coding | -0.44155882 | 4.132903036 | 1.4820419 | 0.2407791 | 0.739435936 |
| ENSMUSG00000072769 | Gm10419 | lincRNA | -0.295348166 | 6.107039761 | 1.4819013 | 0.2408005 | 0.739435936 |
| ENSMUSG00000034777 | Vax2 | protein_coding | -0.926942662 | 2.562757672 | 1.4814978 | 0.2408618 | 0.739435936 |
| ENSMUSG00000030717 | Nupr1 | protein_coding | -0.670428498 | 5.056776348 | 1.4922178 | 0.2409573 | 0.739435936 |
| ENSMUSG00000027823 | Gmps | protein_coding | -0.383295097 | 7.305183761 | 1.4829001 | 0.2409884 | 0.739435936 |
| ENSMUSG00000029110 | Rnf4 | protein_coding | 0.310472446 | 6.827518734 | 1.4804371 | 0.2410232 | 0.739435936 |
| ENSMUSG00000038805 | Six3 | protein_coding | -0.451924789 | 5.097093561 | 1.4796276 | 0.2411465 | 0.739603061 |
| ENSMUSG00000051224 | Tceanc | protein_coding | -0.946922295 | 2.64279464 | 1.4777446 | 0.2414336 | 0.740272317 |
| ENSMUSG00000078812 | Eif5a | protein_coding | 0.283433056 | 9.469711168 | 1.4770599 | 0.2415381 | 0.740376795 |
| ENSMUSG00000022664 | Slc35a5 | protein_coding | -0.358808309 | 5.115606589 | 1.4763844 | 0.2416412 | 0.740376795 |
| ENSMUSG00000000184 | Ccnd2 | protein_coding | -0.415971017 | 5.165602179 | 1.4758019 | 0.2417302 | 0.740376795 |

| ensembl_gene_id | external_gene_name | gene_biotype | log2 (fold change) | log2 (counts per million) | F-value | p-value | False discovery rate |
|---|---|---|---|---|---|---|---|
| ENSMUSG00000046138 | 9930021J03Rik | protein_coding | -0.457722762 | 5.848909932 | 1.4759023 | 0.241743 | 0.740376795 |
| ENSMUSG00000026229 | Psmd1 | protein_coding | 0.292701925 | 7.239032462 | 1.4748837 | 0.2418705 | 0.74047699 |
| ENSMUSG00000059003 | Grin2a | protein_coding | -0.573110046 | 4.417501533 | 1.4745956 | 0.2419146 | 0.74047699 |
| ENSMUSG00000038446 | Cdc40 | protein_coding | -0.500844208 | 5.089583073 | 1.4736602 | 0.2420859 | 0.74047699 |
| ENSMUSG00000009013 | Dynll1 | protein_coding | 0.340484212 | 10.1079042 | 1.4732016 | 0.2421279 | 0.74047699 |
| ENSMUSG00000060538 | Tmem219 | protein_coding | 0.429083367 | 4.342771791 | 1.4730166 | 0.2421563 | 0.74047699 |
| ENSMUSG00000024513 | Mbd2 | protein_coding | -0.550443356 | 3.827740814 | 1.4723978 | 0.2422511 | 0.74047699 |
| ENSMUSG00000031618 | Nr3c2 | protein_coding | -0.557174501 | 4.117893082 | 1.4716163 | 0.2423709 | 0.74047699 |
| ENSMUSG00000085971 | Gm15411 | processed_transcript | 1.024866806 | 2.348571445 | 1.4715029 | 0.2423883 | 0.74047699 |
| ENSMUSG00000031765 | Mt1 | protein_coding | -0.528499203 | 10.16083644 | 1.488126 | 0.2423955 | 0.74047699 |
| ENSMUSG00000038352 | Arl5c | protein_coding | -0.779765194 | 4.748157757 | 1.4859455 | 0.2425953 | 0.740776452 |
| ENSMUSG00000063018 | 2010204K13Rik | processed_transcript | 0.300060743 | 7.865832667 | 1.4690763 | 0.2427608 | 0.740776452 |
| ENSMUSG00000016018 | Skiv2l2 | protein_coding | -0.384189492 | 5.557120922 | 1.469058 | 0.2427636 | 0.740776452 |
| ENSMUSG00000018449 | Rpain | protein_coding | 0.33438398 | 5.882815107 | 1.4687749 | 0.2428071 | 0.740776452 |
| ENSMUSG00000025403 | Shmt2 | protein_coding | 0.572588136 | 4.19618175 | 1.4685738 | 0.242838 | 0.740776452 |
| ENSMUSG00000097287 | D130017N08Rik | lincRNA | -0.541679823 | 3.860943306 | 1.46795 | 0.2429339 | 0.74085885 |
| ENSMUSG00000027500 | Stmn2 | protein_coding | 0.250867474 | 10.26118656 | 1.4661785 | 0.2432066 | 0.741177587 |
| ENSMUSG00000026842 | Abl1 | protein_coding | -0.62773109 | 4.118710143 | 1.4660241 | 0.2432585 | 0.741177587 |
| ENSMUSG00000028577 | Plaa | protein_coding | -0.413512663 | 4.639243415 | 1.4652715 | 0.2433463 | 0.741177587 |
| ENSMUSG00000066441 | Rdh11 | protein_coding | -0.421277078 | 4.662928458 | 1.4649802 | 0.2433912 | 0.741177587 |
| ENSMUSG00000090553 | Snrpe | protein_coding | 0.34264416 | 7.173520655 | 1.4645736 | 0.2434821 | 0.741177587 |
| ENSMUSG00000046854 | Pip5kl1 | protein_coding | 1.578027752 | 2.460729268 | 1.4748246 | 0.2435911 | 0.741177587 |
| ENSMUSG00000015542 | Nat9 | protein_coding | 0.358139106 | 5.966383651 | 1.4631011 | 0.2436812 | 0.741177587 |
| ENSMUSG00000041429 | Nthl1 | protein_coding | 0.675744792 | 3.525973992 | 1.4629437 | 0.2437055 | 0.741177587 |
| ENSMUSG00000038696 | Mapkap1 | protein_coding | -0.411229575 | 5.030753625 | 1.4622405 | 0.2438141 | 0.741177587 |
| ENSMUSG00000059518 | Znhit1 | protein_coding | 0.265237636 | 7.934591468 | 1.4620494 | 0.2438437 | 0.741177587 |
| ENSMUSG00000021242 | Npc2 | protein_coding | -0.265188922 | 7.42587383 | 1.4614613 | 0.2439346 | 0.741177587 |
| ENSMUSG00000035798 | Zdhhc17 | protein_coding | -0.342379856 | 5.272496121 | 1.4593137 | 0.244267 | 0.741177587 |
| ENSMUSG00000027799 | Nbea | protein_coding | -0.30177996 | 6.13149796 | 1.4584393 | 0.2444025 | 0.741177587 |
| ENSMUSG00000035629 | Rubcn | protein_coding | -0.423241297 | 5.172054063 | 1.457479 | 0.2445514 | 0.741177587 |
| ENSMUSG00000012483 | Rpa3 | protein_coding | -0.524703917 | 4.794647199 | 1.4571678 | 0.2445997 | 0.741177587 |
| ENSMUSG00000004631 | Sgce | protein_coding | -0.304562052 | 5.822565911 | 1.4567591 | 0.2446631 | 0.741177587 |
| ENSMUSG00000066189 | Cacng3 | protein_coding | 0.250131884 | 7.436770443 | 1.4566875 | 0.2446742 | 0.741177587 |
| ENSMUSG00000062627 | Mysm1 | protein_coding | -0.36571821 | 5.109075998 | 1.4561972 | 0.2447503 | 0.741177587 |
| ENSMUSG00000085793 | Lin52 | protein_coding | -0.333959092 | 5.72416178 | 1.4557132 | 0.2448255 | 0.741177587 |
| ENSMUSG00000022790 | Igsf11 | protein_coding | -0.666731834 | 3.397509605 | 1.4556125 | 0.2448412 | 0.741177587 |
| ENSMUSG00000035849 | Krt222 | protein_coding | -0.346702432 | 5.881823538 | 1.455612 | 0.2448413 | 0.741177587 |
| ENSMUSG00000026482 | Rgl1 | protein_coding | 0.305465673 | 5.569146892 | 1.4555339 | 0.2448534 | 0.741177587 |
| ENSMUSG00000041303 | Gtf3c3 | protein_coding | -0.371504704 | 5.121092947 | 1.4554621 | 0.2448645 | 0.741177587 |
| ENSMUSG00000036206 | Sh3bp4 | protein_coding | -0.653316816 | 3.751942726 | 1.4552126 | 0.2449033 | 0.741177587 |
| ENSMUSG00000040022 | Rab11fip2 | protein_coding | -0.400223427 | 4.496235975 | 1.4544168 | 0.2450271 | 0.741177587 |
| ENSMUSG00000038774 | Ascc3 | protein_coding | -0.703526387 | 3.242007027 | 1.4539415 | 0.245101 | 0.741177587 |
| ENSMUSG00000015711 | Prune1 | protein_coding | 0.492629365 | 5.363997029 | 1.4540042 | 0.2451193 | 0.741177587 |
| ENSMUSG00000005575 | Ube2m | protein_coding | 0.309325995 | 6.758331894 | 1.4526657 | 0.2452996 | 0.741177587 |
| ENSMUSG00000022031 | Elp3 | protein_coding | 0.383928918 | 6.756834004 | 1.4535125 | 0.2453182 | 0.741177587 |
| ENSMUSG00000061533 | Cep128 | protein_coding | -0.819615809 | 3.424717076 | 1.4517677 | 0.2454676 | 0.741177587 |
| ENSMUSG00000073987 | Ggh | protein_coding | -0.413392547 | 5.054664238 | 1.4515799 | 0.2454688 | 0.741177587 |
| ENSMUSG00000037348 | Paqr7 | protein_coding | 0.330979051 | 6.590218466 | 1.451238 | 0.2455221 | 0.741177587 |
| ENSMUSG00000001542 | Ell2 | protein_coding | -0.61642268 | 3.6254633 | 1.4512148 | 0.2455258 | 0.741177587 |
| ENSMUSG00000031144 | Syp | protein_coding | 0.4233966 | 6.858672657 | 1.4575229 | 0.2456323 | 0.741177587 |
| ENSMUSG00000045482 | Trrap | protein_coding | -0.548308079 | 4.645502751 | 1.4504176 | 0.2456781 | 0.741177587 |
| ENSMUSG00000022054 | Nefm | protein_coding | -0.529616959 | 7.018960095 | 1.4678202 | 0.2456824 | 0.741177587 |
| ENSMUSG00000037737 | Actrt3 | protein_coding | 1.185146505 | 2.315384612 | 1.4501069 | 0.2456986 | 0.741177587 |
| ENSMUSG00000049764 | Zfp280b | protein_coding | -0.597251528 | 3.189917158 | 1.4499348 | 0.2457255 | 0.741177587 |
| ENSMUSG00000088595 | Gm24601 | rRNA | 0.476392958 | 4.51822652 | 1.4497713 | 0.245751 | 0.741177587 |
| ENSMUSG00000020224 | Llph | protein_coding | 0.390294767 | 4.444461609 | 1.4489836 | 0.245874 | 0.741177587 |
| ENSMUSG00000035824 | Tk2 | protein_coding | 0.409262844 | 5.365178531 | 1.4486942 | 0.2459193 | 0.741177587 |
| ENSMUSG00000032215 | Rsl24d1 | protein_coding | 0.269004677 | 6.40011498 | 1.4478003 | 0.246059 | 0.741177587 |
| ENSMUSG00000037138 | Aff3 | protein_coding | -0.469337247 | 4.817062394 | 1.4477916 | 0.2460604 | 0.741177587 |
| ENSMUSG00000037278 | Tmem97 | protein_coding | 0.4033853 | 4.261964516 | 1.4477222 | 0.2460712 | 0.741177587 |
| ENSMUSG00000057614 | Gnai1 | protein_coding | -0.255232016 | 6.97412425 | 1.4445276 | 0.2465716 | 0.741450416 |
| ENSMUSG00000020451 | Limk2 | protein_coding | 0.378452493 | 6.679281051 | 1.4450617 | 0.2465896 | 0.741450416 |
| ENSMUSG00000112964 | CT009718.1 | lincRNA | 0.667190754 | 3.165644942 | 1.4442673 | 0.2466124 | 0.741450416 |
| ENSMUSG00000026003 | Acadl | protein_coding | -0.456534236 | 5.020147576 | 1.4437799 | 0.2466888 | 0.741450416 |
| ENSMUSG00000052942 | Glis3 | protein_coding | -0.626663544 | 3.356678143 | 1.4436575 | 0.2467081 | 0.741450416 |
| ENSMUSG00000039652 | Cpeb3 | protein_coding | -0.418793016 | 5.090136494 | 1.4431239 | 0.2467918 | 0.741450416 |
| ENSMUSG00000021794 | Glud1 | protein_coding | -0.332193624 | 6.788155854 | 1.4428002 | 0.2468426 | 0.741450416 |
| ENSMUSG00000014075 | Tctex1d2 | protein_coding | 0.269736141 | 6.552979031 | 1.4425213 | 0.2468865 | 0.741450416 |
| ENSMUSG00000059689 | Zfp637 | protein_coding | 0.237547986 | 7.198845951 | 1.442449 | 0.2468978 | 0.741450416 |
| ENSMUSG00000017314 | Mpp2 | protein_coding | 0.309946424 | 6.632723407 | 1.4418962 | 0.2469847 | 0.741450416 |
| ENSMUSG00000029924 | Slc37a3 | protein_coding | 0.447095434 | 5.162331966 | 1.4418911 | 0.2469855 | 0.741450416 |
| ENSMUSG00000048379 | Socs4 | protein_coding | -0.902181667 | 2.690753874 | 1.4418671 | 0.2469892 | 0.741450416 |
| ENSMUSG00000035790 | Cep19 | protein_coding | 0.243764816 | 7.53305366 | 1.4399667 | 0.2472882 | 0.74214058 |
| ENSMUSG00000056486 | Chn1 | protein_coding | 0.241280399 | 7.21260601 | 1.4392863 | 0.2473953 | 0.742254972 |
| ENSMUSG00000064181 | Rab3ip | protein_coding | 0.411677683 | 4.458143797 | 1.4384341 | 0.2475296 | 0.742341329 |
| ENSMUSG00000020811 | Wscd1 | protein_coding | -0.346023024 | 5.535521252 | 1.43817 | 0.2475712 | 0.742341329 |
| ENSMUSG00000025420 | Katnal2 | protein_coding | -0.602653168 | 3.421721063 | 1.4376113 | 0.2476593 | 0.742341329 |
| ENSMUSG00000045594 | Glb1 | protein_coding | 0.567322297 | 5.061458579 | 1.4413278 | 0.2477872 | 0.742341329 |

| ensembl_gene_id | external_gene_name | gene_biotype | log2 (fold change) | log2 (counts per million) | F-value | p-value | False discovery rate |
|---|---|---|---|---|---|---|---|
| ENSMUSG00000003929 | Zfp81 | protein_coding | -0.676157307 | 3.661235405 | 1.4365921 | 0.2478202 | 0.742341329 |
| ENSMUSG00000097055 | Gm4419 | lincRNA | 0.581934085 | 3.336072077 | 1.4364771 | 0.2478383 | 0.742341329 |
| ENSMUSG00000024781 | Lipa | protein_coding | 0.369514331 | 5.126524875 | 1.4355213 | 0.2479893 | 0.742551349 |
| ENSMUSG00000050856 | Atp5k | protein_coding | 0.365892282 | 9.580473233 | 1.4352415 | 0.2480614 | 0.742551349 |
| ENSMUSG00000053898 | Ech1 | protein_coding | -0.24657303 | 7.728086844 | 1.4335047 | 0.2483082 | 0.742551349 |
| ENSMUSG00000058656 | Samd12 | protein_coding | -0.554508261 | 4.076291208 | 1.4329448 | 0.2483968 | 0.742551349 |
| ENSMUSG00000031155 | Pim2 | protein_coding | -0.591431825 | 5.437067109 | 1.4417558 | 0.248437 | 0.742551349 |
| ENSMUSG00000029729 | Zkscan1 | protein_coding | -0.35058184 | 5.761704343 | 1.4326328 | 0.2484462 | 0.742551349 |
| ENSMUSG00000074345 | Tnfaip8l3 | protein_coding | 0.709321709 | 3.454021263 | 1.4321548 | 0.2485219 | 0.742551349 |
| ENSMUSG00000066235 | Pomgnt2 | protein_coding | 0.405084125 | 5.394538667 | 1.431888 | 0.2485642 | 0.742551349 |
| ENSMUSG00000042225 | Ammecr1 | protein_coding | -0.60823032 | 3.249757824 | 1.4310295 | 0.2487003 | 0.742551349 |
| ENSMUSG00000037098 | Rab11fip3 | protein_coding | -0.454441496 | 4.664841523 | 1.4309675 | 0.2487102 | 0.742551349 |
| ENSMUSG00000030605 | Mfge8 | protein_coding | -0.387762594 | 7.272475873 | 1.4354635 | 0.2487105 | 0.742551349 |
| ENSMUSG00000102805 | Gm37240 | protein_coding | -0.655419297 | 3.117351977 | 1.4303761 | 0.248804 | 0.742551349 |
| ENSMUSG00000036002 | Fam214b | protein_coding | 0.451170631 | 4.200672327 | 1.4302757 | 0.2488199 | 0.742551349 |
| ENSMUSG00000031139 | Mcf2 | protein_coding | 0.715168031 | 3.053798652 | 1.4299275 | 0.2488752 | 0.742551349 |
| ENSMUSG00000044934 | Zfp367 | protein_coding | 0.439004792 | 4.7212702 | 1.4290636 | 0.2490124 | 0.742754595 |
| ENSMUSG00000021007 | Spata7 | protein_coding | -0.397881485 | 5.025590544 | 1.4283674 | 0.249123 | 0.742832038 |
| ENSMUSG00000013465 | Nelfb | protein_coding | 0.283749328 | 5.997050314 | 1.4277006 | 0.249229 | 0.742832038 |
| ENSMUSG00000032359 | Ctsh | protein_coding | -0.655724541 | 3.698743639 | 1.4275832 | 0.2492477 | 0.742832038 |
| ENSMUSG00000025777 | Gdap1 | protein_coding | -0.268811893 | 6.892165658 | 1.4268317 | 0.2493673 | 0.742832038 |
| ENSMUSG00000024302 | Dtna | protein_coding | -0.279537117 | 6.816373177 | 1.4267282 | 0.2493838 | 0.742832038 |
| ENSMUSG00000042632 | Pla2g6 | protein_coding | -0.374381659 | 5.294236621 | 1.4254693 | 0.2495843 | 0.743223101 |
| ENSMUSG00000006599 | Gtf2h1 | protein_coding | -0.464596047 | 4.52194204 | 1.4248785 | 0.2496784 | 0.743223101 |
| ENSMUSG00000004535 | Tax1bp1 | protein_coding | -0.23279162 | 7.530073695 | 1.4232021 | 0.2499459 | 0.743223101 |
| ENSMUSG00000022437 | Samm50 | protein_coding | 0.319698677 | 6.467781763 | 1.4231284 | 0.2499576 | 0.743223101 |
| ENSMUSG00000024154 | Gtf2a1l | protein_coding | -0.960634802 | 2.657533945 | 1.4227616 | 0.2500162 | 0.743223101 |
| ENSMUSG00000068762 | Gstm6 | protein_coding | 0.298900364 | 6.187682757 | 1.4226302 | 0.2500372 | 0.743223101 |
| ENSMUSG00000003153 | Slc2a3 | protein_coding | 0.249615167 | 6.362208726 | 1.422333 | 0.2500847 | 0.743223101 |
| ENSMUSG00000032135 | Mcam | protein_coding | -1.072678495 | 2.90400085 | 1.4212975 | 0.250278 | 0.743223101 |
| ENSMUSG00000004040 | Stat3 | protein_coding | -0.572813433 | 4.838964581 | 1.4215444 | 0.2503118 | 0.743223101 |
| ENSMUSG00000063273 | Naa15 | protein_coding | -0.424013189 | 5.168552921 | 1.4207345 | 0.2503402 | 0.743223101 |
| ENSMUSG00000029422 | Rsrc2 | protein_coding | -0.278619698 | 7.29315906 | 1.420472 | 0.2503822 | 0.743223101 |
| ENSMUSG00000022043 | Trim35 | protein_coding | -0.264780157 | 6.707283872 | 1.420173 | 0.2504301 | 0.743223101 |
| ENSMUSG00000035007 | Rundc1 | protein_coding | -0.564945643 | 4.464754984 | 1.4199188 | 0.2504986 | 0.743223101 |
| ENSMUSG00000097195 | Snhg5 | processed_transcript | -0.470775572 | 4.857032958 | 1.4197021 | 0.2505054 | 0.743223101 |
| ENSMUSG00000050668 | Gpatch11 | protein_coding | 0.272046575 | 6.136313262 | 1.4194125 | 0.2505518 | 0.743223101 |
| ENSMUSG00000034951 | Cog7 | protein_coding | 0.395353993 | 5.961421117 | 1.4182257 | 0.2507698 | 0.74354127 |
| ENSMUSG00000046240 | Hepacam | protein_coding | -0.571024603 | 4.693450718 | 1.4175447 | 0.250879 | 0.74354127 |
| ENSMUSG00000067365 | Tmem128 | protein_coding | 0.307932094 | 6.726846509 | 1.4170463 | 0.2509312 | 0.74354127 |
| ENSMUSG00000050188 | Lsm10 | protein_coding | 0.346140592 | 5.930104451 | 1.4164027 | 0.2510345 | 0.74354127 |
| ENSMUSG00000020142 | Slc1a4 | protein_coding | -0.436468085 | 4.882143168 | 1.4153385 | 0.2512054 | 0.74354127 |
| ENSMUSG00000035376 | Hacd2 | protein_coding | -0.544573557 | 3.720250957 | 1.4149549 | 0.2512671 | 0.74354127 |
| ENSMUSG00000073680 | Tmem88b | protein_coding | -0.427287898 | 5.823505136 | 1.4148636 | 0.2513095 | 0.74354127 |
| ENSMUSG00000030759 | Far1 | protein_coding | -0.346893301 | 5.452490102 | 1.4146727 | 0.2513124 | 0.74354127 |
| ENSMUSG00000032586 | Traip | protein_coding | 0.405691575 | 4.521045935 | 1.4145992 | 0.2513243 | 0.74354127 |
| ENSMUSG00000007891 | Ctsd | protein_coding | 0.305332112 | 6.348899188 | 1.4144357 | 0.2513506 | 0.74354127 |
| ENSMUSG00000003099 | Ppp5c | protein_coding | 0.330914477 | 7.03799125 | 1.4136463 | 0.2515053 | 0.743694855 |
| ENSMUSG00000028790 | Khdrbs1 | protein_coding | -0.367346231 | 5.447666043 | 1.4129635 | 0.2515875 | 0.743694855 |
| ENSMUSG00000029068 | Ccnl2 | protein_coding | -0.497476335 | 3.8551514 | 1.4121416 | 0.2517199 | 0.743694855 |
| ENSMUSG00000052139 | Babam2 | protein_coding | 0.291997978 | 6.603073595 | 1.4120838 | 0.2517292 | 0.743694855 |
| ENSMUSG00000052102 | Gnpda1 | protein_coding | 0.40870314 | 5.444722279 | 1.4117648 | 0.2517806 | 0.743694855 |
| ENSMUSG00000074030 | Exoc8 | protein_coding | 0.393800833 | 4.648191444 | 1.4112193 | 0.2518685 | 0.743694855 |
| ENSMUSG00000020077 | Srgn | protein_coding | -0.926226362 | 2.586494069 | 1.4108705 | 0.2519248 | 0.743694855 |
| ENSMUSG00000022120 | Rnf219 | protein_coding | -0.512593511 | 4.369410088 | 1.4103438 | 0.2520098 | 0.743694855 |
| ENSMUSG00000056367 | Actr3b | protein_coding | -0.302966404 | 6.426638227 | 1.4100612 | 0.2520554 | 0.743694855 |
| ENSMUSG00000030061 | Uba3 | protein_coding | -0.274896419 | 6.785816306 | 1.4097625 | 0.2521036 | 0.743694855 |
| ENSMUSG00000038544 | Inip | protein_coding | -0.356386624 | 5.42137329 | 1.4093932 | 0.2521633 | 0.743694855 |
| ENSMUSG00000027030 | Stk39 | protein_coding | -0.341914435 | 5.963510753 | 1.4087465 | 0.2522677 | 0.743744538 |
| ENSMUSG00000062591 | Tubb4a | protein_coding | 0.395189207 | 11.07819344 | 1.4088111 | 0.2523581 | 0.743744538 |
| ENSMUSG00000042010 | Acacb | protein_coding | 0.639217472 | 4.116900543 | 1.4067184 | 0.2526235 | 0.743744538 |
| ENSMUSG00000051950 | B3glct | protein_coding | -0.751361074 | 3.011145977 | 1.4062133 | 0.2526775 | 0.743744538 |
| ENSMUSG00000074758 | Gm5535 | lincRNA | -0.815669062 | 2.770397321 | 1.4061181 | 0.2526929 | 0.743744538 |
| ENSMUSG00000002266 | Zim1 | protein_coding | 0.598835485 | 3.127123016 | 1.4059016 | 0.252728 | 0.743744538 |
| ENSMUSG00000038760 | Trhr | protein_coding | 1.150272181 | 2.638988981 | 1.4057896 | 0.2527739 | 0.743744538 |
| ENSMUSG00000070366 | Plpp4 | protein_coding | 0.544257574 | 4.061904239 | 1.4053866 | 0.2528115 | 0.743744538 |
| ENSMUSG00000027359 | Slc27a2 | protein_coding | -0.928891839 | 2.826299836 | 1.4052379 | 0.2528356 | 0.743744538 |
| ENSMUSG00000074971 | Fibin | protein_coding | 0.781951648 | 2.923401288 | 1.404951 | 0.2528821 | 0.743744538 |
| ENSMUSG00000078974 | Sec61g | protein_coding | 0.385845704 | 5.158550074 | 1.404588 | 0.2529409 | 0.743744538 |
| ENSMUSG00000014361 | Mertk | protein_coding | -0.959949792 | 2.70795975 | 1.4021483 | 0.253337 | 0.743982392 |
| ENSMUSG00000044122 | Proca1 | protein_coding | -0.511627148 | 3.87143142 | 1.4018105 | 0.2533919 | 0.743982392 |
| ENSMUSG00000028619 | Tceanc2 | protein_coding | 0.545643384 | 3.57439876 | 1.4014653 | 0.253448 | 0.743982392 |
| ENSMUSG00000113701 | AC121583.1 | transcribed_processe | -0.423089671 | 4.360972706 | 1.4013355 | 0.2534691 | 0.743982392 |
| ENSMUSG00000032109 | Nlrx1 | protein_coding | 1.24512896 | 2.261595834 | 1.4011087 | 0.253506 | 0.743982392 |
| ENSMUSG00000006641 | Slc5a6 | protein_coding | 0.443482846 | 4.429235679 | 1.4006706 | 0.2535772 | 0.743982392 |
| ENSMUSG00000072915 | Gm12258 | protein_coding | -0.577162123 | 3.798291442 | 1.4005307 | 0.2536 | 0.743982392 |
| ENSMUSG00000038128 | Camk4 | protein_coding | -0.383647875 | 6.331167359 | 1.4004219 | 0.2536454 | 0.743982392 |
| ENSMUSG00000018427 | Ypel2 | protein_coding | -0.258711799 | 6.565486051 | 1.3999271 | 0.2536983 | 0.743982392 |

| ensembl_gene_id | external_gene_name | gene_biotype | log2 (fold change) | log2 (counts per million) | F-value | p-value | False discovery rate |
|---|---|---|---|---|---|---|---|
| ENSMUSG00000020048 | Hsp90b1 | protein_coding | -0.237900518 | 7.071487555 | 1.3995789 | 0.253755 | 0.743982392 |
| ENSMUSG00000024826 | Dpf2 | protein_coding | 0.307017263 | 6.602826591 | 1.3994074 | 0.2537829 | 0.743982392 |
| ENSMUSG00000028381 | Ugcg | protein_coding | -0.351208323 | 5.413771416 | 1.3976304 | 0.2540725 | 0.744177548 |
| ENSMUSG00000032946 | Rasgrp2 | protein_coding | 0.343752454 | 5.225065832 | 1.3973981 | 0.2541104 | 0.744177548 |
| ENSMUSG00000054720 | Lrrc8c | protein_coding | -0.480486933 | 3.937073174 | 1.3971303 | 0.2541541 | 0.744177548 |
| ENSMUSG00000001436 | Slc19a1 | protein_coding | 0.641960023 | 4.7001377 | 1.4028255 | 0.2542993 | 0.744177548 |
| ENSMUSG00000040818 | Dennd6a | protein_coding | 0.498418777 | 3.729116788 | 1.3961227 | 0.2543186 | 0.744177548 |
| ENSMUSG00000048100 | Taf13 | protein_coding | -0.400377488 | 5.28099885 | 1.3956801 | 0.254391 | 0.744177548 |
| ENSMUSG00000025370 | Cdh9 | protein_coding | -1.029458756 | 2.458546631 | 1.3955671 | 0.2544094 | 0.744177548 |
| ENSMUSG00000097537 | 2610020C07Rik | lincRNA | -0.408319081 | 4.705334267 | 1.3948893 | 0.2545202 | 0.744177548 |
| ENSMUSG00000023272 | Creld2 | protein_coding | -0.880576859 | 3.29677908 | 1.3952405 | 0.2545632 | 0.744177548 |
| ENSMUSG00000033701 | Acbd6 | protein_coding | 0.353529843 | 5.386860559 | 1.3943548 | 0.2546076 | 0.744177548 |
| ENSMUSG00000048603 | Gm9828 | bidirectional_promote | 0.644971009 | 3.293833343 | 1.3940504 | 0.2546574 | 0.744177548 |
| ENSMUSG00000024369 | Nelfe | protein_coding | 0.305470995 | 6.009581249 | 1.3934677 | 0.2547528 | 0.744177548 |
| ENSMUSG00000022193 | Psmb5 | protein_coding | 0.247071527 | 6.843764382 | 1.3933055 | 0.2547793 | 0.744177548 |
| ENSMUSG00000052544 | St6galnac3 | protein_coding | -0.541836661 | 3.529795754 | 1.3930242 | 0.2548254 | 0.744177548 |
| ENSMUSG00000027203 | Dut | protein_coding | -0.425869704 | 4.373505902 | 1.3924563 | 0.2549184 | 0.744177548 |
| ENSMUSG00000008999 | Bmp7 | protein_coding | -0.726917796 | 3.158227919 | 1.3912954 | 0.2551087 | 0.744177548 |
| ENSMUSG00000021504 | B4galt7 | protein_coding | 0.432663252 | 4.868777202 | 1.3902963 | 0.2552726 | 0.744177548 |
| ENSMUSG00000019470 | Xab2 | protein_coding | 0.491879883 | 5.777824972 | 1.3967924 | 0.2552799 | 0.744177548 |
| ENSMUSG00000019874 | Fabp7 | protein_coding | -0.543956661 | 6.935996241 | 1.4106663 | 0.2553056 | 0.744177548 |
| ENSMUSG00000004558 | Ndrg2 | protein_coding | -0.297892479 | 9.746559123 | 1.390012 | 0.2553193 | 0.744177548 |
| ENSMUSG00000058997 | Vwa8 | protein_coding | -0.704564203 | 4.08039628 | 1.3934108 | 0.2553215 | 0.744177548 |
| ENSMUSG00000037784 | Dzip1l | protein_coding | -0.970641158 | 2.5623487 | 1.3890865 | 0.2554713 | 0.744177548 |
| ENSMUSG00000067370 | B3galt4 | protein_coding | 0.779359174 | 3.63878684 | 1.3931194 | 0.2555139 | 0.744177548 |
| ENSMUSG00000028149 | Rap1gds1 | protein_coding | 0.247535042 | 7.472948559 | 1.3885361 | 0.2555618 | 0.744177548 |
| ENSMUSG00000026799 | Med27 | protein_coding | 0.299498636 | 5.612580468 | 1.3884276 | 0.2555796 | 0.744177548 |
| ENSMUSG00000024350 | Dnajc18 | protein_coding | 0.316899633 | 7.567937018 | 1.3853401 | 0.2560879 | 0.745284021 |
| ENSMUSG00000006442 | Srm | protein_coding | 0.384189382 | 5.229121767 | 1.3850534 | 0.2561351 | 0.745284021 |
| ENSMUSG00000045107 | Saysd1 | protein_coding | 0.35568788 | 5.068478322 | 1.3847723 | 0.2561815 | 0.745284021 |
| ENSMUSG00000031775 | Pllp | protein_coding | -0.415705528 | 4.431402865 | 1.3842769 | 0.2562632 | 0.745284021 |
| ENSMUSG00000021235 | Coq6 | protein_coding | 0.438414488 | 5.401147564 | 1.3840165 | 0.2563062 | 0.745284021 |
| ENSMUSG00000034799 | Unc13a | protein_coding | -0.513528748 | 3.774578642 | 1.3823876 | 0.2565752 | 0.745818858 |
| ENSMUSG00000031231 | Cox7b | protein_coding | 0.315930379 | 9.469297054 | 1.3818314 | 0.2566671 | 0.745818858 |
| ENSMUSG00000010392 | Gosr1 | protein_coding | -0.449628294 | 5.264274629 | 1.3816433 | 0.2566982 | 0.745818858 |
| ENSMUSG00000034341 | Wbp2 | protein_coding | 0.347092169 | 10.2406158 | 1.3813468 | 0.2567748 | 0.7458398 |
| ENSMUSG00000032238 | Rora | protein_coding | -0.404842161 | 6.226476772 | 1.3796374 | 0.2570577 | 0.746390075 |
| ENSMUSG00000063524 | Eno1 | protein_coding | 0.315565169 | 8.641938171 | 1.3788641 | 0.2571583 | 0.746390075 |
| ENSMUSG00000025651 | Uqcrc1 | protein_coding | 0.382628636 | 9.629316734 | 1.3819195 | 0.2571724 | 0.746390075 |
| ENSMUSG00000020108 | Ddit4 | protein_coding | -0.691432757 | 3.351009333 | 1.3774895 | 0.2573863 | 0.746390148 |
| ENSMUSG00000029054 | Gabrd | protein_coding | -0.44518891 | 5.524120706 | 1.3766378 | 0.2575551 | 0.746390148 |
| ENSMUSG00000076432 | Ywhaq | protein_coding | -0.226921029 | 8.08258164 | 1.3764707 | 0.2575554 | 0.746390148 |
| ENSMUSG00000050229 | Pigm | protein_coding | -0.715559379 | 2.779106092 | 1.3761523 | 0.2576083 | 0.746390148 |
| ENSMUSG00000041623 | D11Wsu47e | protein_coding | 0.573375545 | 3.499927593 | 1.375817 | 0.257664 | 0.746390148 |
| ENSMUSG00000032492 | Pth1r | protein_coding | 0.638570273 | 3.535233835 | 1.3751013 | 0.257783 | 0.746390148 |
| ENSMUSG00000050855 | Zfp940 | protein_coding | 0.452292711 | 4.580419847 | 1.3748193 | 0.2578299 | 0.746390148 |
| ENSMUSG00000028228 | Cpne3 | protein_coding | -0.737985991 | 3.502049258 | 1.3747754 | 0.2578646 | 0.746390148 |
| ENSMUSG00000039968 | Rsbn1l | protein_coding | -0.596412599 | 3.902598808 | 1.3743694 | 0.2579047 | 0.746390148 |
| ENSMUSG00000078435 | AU041133 | protein_coding | -0.785943635 | 2.924900475 | 1.3740525 | 0.2579574 | 0.746390148 |
| ENSMUSG00000038115 | Ano2 | protein_coding | -0.905046734 | 3.114696155 | 1.3740215 | 0.25799 | 0.746390148 |
| ENSMUSG00000029470 | P2rx4 | protein_coding | -0.711266102 | 3.457031536 | 1.3730346 | 0.2581269 | 0.746390148 |
| ENSMUSG00000026333 | Gin1 | protein_coding | -0.522176511 | 4.444129451 | 1.3728193 | 0.2581628 | 0.746390148 |
| ENSMUSG00000047617 | Paxx | protein_coding | 0.24526342 | 7.046361588 | 1.3725699 | 0.2582043 | 0.746390148 |
| ENSMUSG00000035382 | Pcsk7 | protein_coding | 0.65162981 | 3.056818934 | 1.372285 | 0.2582518 | 0.746390148 |
| ENSMUSG00000031227 | Magee1 | protein_coding | 0.236364234 | 6.897244192 | 1.3719996 | 0.2582994 | 0.746390148 |
| ENSMUSG00000001052 | Sec24b | protein_coding | -0.363453488 | 4.881880454 | 1.3716812 | 0.2583525 | 0.746390148 |
| ENSMUSG00000024593 | Megf10 | protein_coding | -0.853445807 | 2.874228502 | 1.3710694 | 0.2584545 | 0.746475204 |
| ENSMUSG00000006717 | Acot13 | protein_coding | 0.266245537 | 8.428009377 | 1.3706107 | 0.2585311 | 0.746475204 |
| ENSMUSG00000031604 | Msmo1 | protein_coding | -0.243297844 | 7.175170499 | 1.3702567 | 0.2585902 | 0.746475204 |
| ENSMUSG00000052833 | Sae1 | protein_coding | 0.331861376 | 7.901327901 | 1.3696472 | 0.2587194 | 0.746525456 |
| ENSMUSG00000000627 | Sema4f | protein_coding | 0.490619804 | 4.304261615 | 1.3687356 | 0.2588443 | 0.746525456 |
| ENSMUSG00000045589 | Frrs1l | protein_coding | -0.261213383 | 6.858846421 | 1.3678333 | 0.2589952 | 0.746525456 |
| ENSMUSG00000001833 | 7-Sep | protein_coding | -0.275374838 | 9.081041745 | 1.3676002 | 0.2590342 | 0.746525456 |
| ENSMUSG00000020423 | Btg2 | protein_coding | -0.691752274 | 5.95360029 | 1.395502 | 0.2591193 | 0.746525456 |
| ENSMUSG00000054021 | Sirt5 | protein_coding | 0.394292063 | 5.581060247 | 1.3669504 | 0.259143 | 0.746525456 |
| ENSMUSG00000028530 | Jak1 | protein_coding | -0.260202281 | 6.645700066 | 1.3666199 | 0.2591983 | 0.746525456 |
| ENSMUSG00000025221 | Kcnip2 | protein_coding | -0.294253515 | 6.323218804 | 1.366547 | 0.2592105 | 0.746525456 |
| ENSMUSG00000004110 | Cacna1e | protein_coding | -0.466295144 | 4.923582832 | 1.3664165 | 0.2592324 | 0.746525456 |
| ENSMUSG00000039942 | Ptger4 | protein_coding | -0.514152136 | 4.093155842 | 1.3658293 | 0.2593308 | 0.746608876 |
| ENSMUSG00000089945 | Pakap | protein_coding | -0.692894871 | 3.012438572 | 1.3652287 | 0.2594315 | 0.746698796 |
| ENSMUSG00000036432 | Siah2 | protein_coding | -0.642709312 | 3.735798607 | 1.364787 | 0.2595056 | 0.746712151 |
| ENSMUSG00000022019 | Tdrd3 | protein_coding | -0.392759045 | 4.73038923 | 1.3639131 | 0.2596522 | 0.746934279 |
| ENSMUSG00000027787 | Nmd3 | protein_coding | -0.311369341 | 5.983076922 | 1.3629011 | 0.2598222 | 0.747142167 |
| ENSMUSG00000030098 | Grip2 | protein_coding | 0.774656907 | 3.227023866 | 1.3617551 | 0.2600148 | 0.747142167 |
| ENSMUSG00000006335 | Tfpt | protein_coding | 0.299367321 | 5.883506186 | 1.3616451 | 0.2600333 | 0.747142167 |
| ENSMUSG00000061411 | Nol4l | protein_coding | -0.432353574 | 4.668333996 | 1.3611248 | 0.2601209 | 0.747142167 |
| ENSMUSG00000020380 | Rad50 | protein_coding | -0.496597952 | 3.776395721 | 1.3607267 | 0.2601879 | 0.747142167 |
| ENSMUSG00000040549 | Ckap5 | protein_coding | -0.266629159 | 6.407334351 | 1.3606217 | 0.2602055 | 0.747142167 |

| ensembl_gene_id | external_gene_name | gene_biotype | log2 (fold change) | log2 (counts per million) | F-value | p-value | False discovery rate |
|---|---|---|---|---|---|---|---|
| ENSMUSG00000022674 | Ube2v2 | protein_coding | -0.385795316 | 4.830762363 | 1.3605901 | 0.2602109 | 0.747142167 |
| ENSMUSG00000054459 | Vsnl1 | protein_coding | 0.286751687 | 10.53233629 | 1.3595947 | 0.2603785 | 0.747231891 |
| ENSMUSG00000021400 | Wrnip1 | protein_coding | -0.372276398 | 5.408688002 | 1.3595794 | 0.2603811 | 0.747231891 |
| ENSMUSG00000022130 | Tgds | protein_coding | -0.956254599 | 2.404967611 | 1.3577302 | 0.2606929 | 0.747244127 |
| ENSMUSG00000041775 | Mapk1ip1 | protein_coding | 0.360969118 | 5.904104184 | 1.3574358 | 0.2607426 | 0.747244127 |
| ENSMUSG00000028062 | Lamtor2 | protein_coding | 0.287457619 | 9.130326099 | 1.3574149 | 0.2607461 | 0.747244127 |
| ENSMUSG00000067424 | Zfp563 | protein_coding | -0.48718606 | 3.802605472 | 1.3571361 | 0.2607932 | 0.747244127 |
| ENSMUSG00000031546 | Gins4 | protein_coding | 0.337882171 | 4.980697551 | 1.3570903 | 0.2608009 | 0.747244127 |
| ENSMUSG00000035770 | Dync1li2 | protein_coding | -0.291936248 | 6.71919187 | 1.3570823 | 0.2608023 | 0.747244127 |
| ENSMUSG00000035158 | Mitf | protein_coding | -0.96105275 | 2.723060343 | 1.3562685 | 0.2609398 | 0.747305456 |
| ENSMUSG00000028132 | Tmem56 | protein_coding | -0.528522147 | 4.357039653 | 1.3559277 | 0.2609974 | 0.747305456 |
| ENSMUSG00000030223 | Ptpro | protein_coding | 0.356131938 | 4.842292511 | 1.355095 | 0.2611382 | 0.747305456 |
| ENSMUSG00000033697 | Arhgap39 | protein_coding | -0.855952258 | 2.727258943 | 1.3547628 | 0.2611944 | 0.747305456 |
| ENSMUSG00000025060 | Slk | protein_coding | -0.542036779 | 4.682045416 | 1.35477 | 0.2612205 | 0.747305456 |
| ENSMUSG00000048787 | Dcun1d3 | protein_coding | -0.530636398 | 3.568965369 | 1.3544263 | 0.2612513 | 0.747305456 |
| ENSMUSG00000018698 | Lhx1 | protein_coding | -0.86839626 | 2.559117788 | 1.3540784 | 0.2613102 | 0.747305456 |
| ENSMUSG00000027330 | Cdc25b | protein_coding | -0.7298362 | 3.054587964 | 1.3485524 | 0.262248 | 0.749787891 |
| ENSMUSG00000001985 | Grik3 | protein_coding | 0.416809369 | 4.438517667 | 1.3479963 | 0.2623426 | 0.749859034 |
| ENSMUSG00000020181 | Nav3 | protein_coding | -0.366301642 | 5.39169619 | 1.3473974 | 0.2624445 | 0.74994418 |
| ENSMUSG00000031671 | Setd6 | protein_coding | 0.763750269 | 3.194264437 | 1.3470021 | 0.2625118 | 0.74994418 |
| ENSMUSG00000027651 | Rprd1b | protein_coding | -0.481566634 | 4.07536025 | 1.3455712 | 0.2627557 | 0.750271909 |
| ENSMUSG00000046997 | Spsb4 | protein_coding | -0.935542233 | 2.451580769 | 1.3451018 | 0.2628358 | 0.750271909 |
| ENSMUSG00000050310 | Rictor | protein_coding | -0.411800981 | 4.044007343 | 1.3447059 | 0.2629034 | 0.750271909 |
| ENSMUSG00000032507 | Fbxl2 | protein_coding | 0.409332878 | 5.618176864 | 1.3444527 | 0.2629466 | 0.750271909 |
| ENSMUSG00000078453 | Abracl | protein_coding | 0.457399427 | 4.923847524 | 1.3442836 | 0.2629754 | 0.750271909 |
| ENSMUSG00000028583 | Pdpn | protein_coding | -0.567369109 | 4.183131412 | 1.3434503 | 0.2631178 | 0.75047883 |
| ENSMUSG00000043259 | Fam13c | protein_coding | 0.304403673 | 5.533974813 | 1.3429392 | 0.2632051 | 0.750528904 |
| ENSMUSG00000025262 | Fam120c | protein_coding | -0.829056971 | 3.31235211 | 1.3419916 | 0.2633944 | 0.750869423 |
| ENSMUSG00000024037 | Wdr4 | protein_coding | -0.373601992 | 4.752591358 | 1.3409342 | 0.2635481 | 0.751108623 |
| ENSMUSG00000001100 | Poldip2 | protein_coding | -0.293061174 | 5.884132122 | 1.340017 | 0.2637052 | 0.751357239 |
| ENSMUSG00000028261 | Ndufaf4 | protein_coding | -0.282507735 | 6.343887909 | 1.3394348 | 0.263805 | 0.751442465 |
| ENSMUSG00000041638 | Gcn1l1 | protein_coding | 0.418527528 | 4.07384481 | 1.338578 | 0.263952 | 0.751661933 |
| ENSMUSG00000046994 | Mars2 | protein_coding | -0.724128592 | 3.103361942 | 1.3375346 | 0.2641311 | 0.751866217 |
| ENSMUSG00000025159 | Mms19 | protein_coding | -0.474905701 | 4.342493088 | 1.3373455 | 0.2641635 | 0.751866217 |
| ENSMUSG00000036036 | Zfp57 | protein_coding | 1.288633636 | 2.74899328 | 1.3464307 | 0.2642627 | 0.75194298 |
| ENSMUSG00000066877 | Nck2 | protein_coding | -0.503948969 | 3.486586342 | 1.3363746 | 0.2643304 | 0.75194298 |
| ENSMUSG00000023991 | Foxp4 | protein_coding | 0.598563724 | 3.535904737 | 1.3347226 | 0.2646146 | 0.752139982 |
| ENSMUSG00000032228 | Tcf12 | protein_coding | -0.425039563 | 4.905642339 | 1.3343432 | 0.2646799 | 0.752139982 |
| ENSMUSG00000026858 | Miga2 | protein_coding | 0.355432999 | 5.634868065 | 1.3342301 | 0.2646993 | 0.752139982 |
| ENSMUSG00000026626 | Ppp2r5a | protein_coding | -0.496116337 | 4.44567116 | 1.3340803 | 0.2647251 | 0.752139982 |
| ENSMUSG00000002416 | Ndufb2 | protein_coding | 0.269428968 | 6.208844997 | 1.3339398 | 0.2647494 | 0.752139982 |
| ENSMUSG00000041313 | Slc7a1 | protein_coding | -0.486112253 | 3.810194806 | 1.333325 | 0.2648553 | 0.752242163 |
| ENSMUSG00000037720 | Tmem33 | protein_coding | -0.340181753 | 6.126478516 | 1.3316242 | 0.2651486 | 0.752789895 |
| ENSMUSG00000004698 | Hdac9 | protein_coding | -0.383133652 | 5.084762076 | 1.3309909 | 0.265258 | 0.752789895 |
| ENSMUSG00000008301 | Phax | protein_coding | 0.21854001 | 7.659439916 | 1.3309899 | 0.2652582 | 0.752789895 |
| ENSMUSG00000035851 | Ythdc1 | protein_coding | -0.368962902 | 5.114196526 | 1.330232 | 0.2653891 | 0.752851303 |
| ENSMUSG00000026377 | Nifk | protein_coding | 0.295946818 | 5.969500023 | 1.3300542 | 0.2654198 | 0.752851303 |
| ENSMUSG00000022849 | Hspbap1 | protein_coding | -0.780557591 | 3.208849033 | 1.3295037 | 0.265515 | 0.752922682 |
| ENSMUSG00000021494 | Ddx41 | protein_coding | 0.343412726 | 5.333514977 | 1.3287835 | 0.2656396 | 0.753077326 |
| ENSMUSG00000025958 | Creb1 | protein_coding | -0.405922811 | 5.110658551 | 1.3278594 | 0.2657995 | 0.753332211 |
| ENSMUSG00000067870 | Rpl31-ps8 | processed_pseudoge | 0.562601353 | 3.545993671 | 1.3257826 | 0.2661595 | 0.754153691 |
| ENSMUSG00000062115 | Rai1 | protein_coding | 0.400698899 | 5.042439665 | 1.3252297 | 0.2662555 | 0.754226795 |
| ENSMUSG00000040896 | Kcnd3 | protein_coding | -0.527781916 | 4.172259452 | 1.3245208 | 0.2663785 | 0.754376701 |
| ENSMUSG00000040734 | Ppp1r13l | protein_coding | -0.885887677 | 2.762359993 | 1.3229569 | 0.2666503 | 0.754935984 |
| ENSMUSG00000041958 | Pigs | protein_coding | 0.372316694 | 5.845747439 | 1.3225769 | 0.2667164 | 0.754935984 |
| ENSMUSG00000037148 | Arhgap10 | protein_coding | -1.02043184 | 2.855312343 | 1.3220863 | 0.2668288 | 0.755055404 |
| ENSMUSG00000017677 | Wsb1 | protein_coding | -0.510726053 | 4.565200691 | 1.3213267 | 0.266934 | 0.755154382 |
| ENSMUSG00000040564 | Apoc1 | protein_coding | -0.654488355 | 3.600949766 | 1.3192403 | 0.2672977 | 0.755625736 |
| ENSMUSG00000028706 | Nsun4 | protein_coding | 0.494400764 | 4.674578843 | 1.3185869 | 0.2674118 | 0.755625736 |
| ENSMUSG00000041720 | Pi4ka | protein_coding | 0.467648494 | 4.881560216 | 1.3181847 | 0.267482 | 0.755625736 |
| ENSMUSG00000033306 | Lpp | protein_coding | -0.545116004 | 3.491464405 | 1.3177971 | 0.2675497 | 0.755625736 |
| ENSMUSG00000039058 | Ak5 | protein_coding | 0.338209129 | 5.474999154 | 1.3177273 | 0.2675619 | 0.755625736 |
| ENSMUSG00000062397 | Zfp706 | protein_coding | -0.260399764 | 8.195450523 | 1.3175547 | 0.267592 | 0.755625736 |
| ENSMUSG00000041774 | Ydjc | protein_coding | 0.654273213 | 3.276588869 | 1.3175517 | 0.2675926 | 0.755625736 |
| ENSMUSG00000044933 | Sstr3 | protein_coding | 1.062997801 | 1.991282608 | 1.3232432 | 0.2676648 | 0.755631327 |
| ENSMUSG00000022370 | Mrpl13 | protein_coding | 0.253633715 | 7.425329315 | 1.3162575 | 0.2678188 | 0.755635609 |
| ENSMUSG00000054312 | Mrps21 | protein_coding | 0.320379638 | 7.97486002 | 1.3163246 | 0.2678341 | 0.755635609 |
| ENSMUSG00000040385 | Ppp1ca | protein_coding | 0.234645373 | 9.192560825 | 1.315589 | 0.2679358 | 0.755635609 |
| ENSMUSG00000029034 | Ints11 | protein_coding | -0.696030819 | 3.329845632 | 1.3152137 | 0.2680015 | 0.755635609 |
| ENSMUSG00000038970 | Lmtk2 | protein_coding | -0.461259534 | 4.285647152 | 1.3151212 | 0.2680177 | 0.755635609 |
| ENSMUSG00000001924 | Uba1 | protein_coding | 0.251931075 | 7.770458192 | 1.3133851 | 0.2683219 | 0.75623028 |
| ENSMUSG00000018923 | Med11 | protein_coding | 0.35982096 | 5.137953717 | 1.312885 | 0.2684096 | 0.75623028 |
| ENSMUSG00000042213 | Zfand4 | protein_coding | 0.857499791 | 2.652720383 | 1.3125555 | 0.2684675 | 0.75623028 |
| ENSMUSG00000031644 | Nek1 | protein_coding | -0.670646166 | 3.354982581 | 1.3121843 | 0.2685326 | 0.75623028 |
| ENSMUSG00000078515 | Ddi2 | protein_coding | -0.376035274 | 5.409686099 | 1.3116879 | 0.2686198 | 0.75623028 |
| ENSMUSG00000035310 | Lin54 | protein_coding | -0.694458309 | 2.820126201 | 1.3115125 | 0.2686506 | 0.75623028 |
| ENSMUSG00000038181 | Chpf2 | protein_coding | 1.147335712 | 2.365429595 | 1.3097016 | 0.2689689 | 0.756928244 |
| ENSMUSG00000097310 | A930038B10Rik | lincRNA | 1.028997987 | 2.324610215 | 1.3082695 | 0.269221 | 0.756962818 |

| ensembl_gene_id | external_gene_name | gene_biotype | log2 (fold change) | log2 (counts per million) | F-value | p-value | False discovery rate |
|---|---|---|---|---|---|---|---|
| ENSMUSG00000036905 | C1qb | protein_coding | -0.57448935 | 6.327932409 | 1.3265343 | 0.2692314 | 0.756962818 |
| ENSMUSG00000034274 | Thoc5 | protein_coding | 0.371725608 | 5.329991792 | 1.3075814 | 0.2693423 | 0.756962818 |
| ENSMUSG00000033916 | Chmp2a | protein_coding | 0.222310968 | 9.030813832 | 1.3075146 | 0.2693541 | 0.756962818 |
| ENSMUSG00000026797 | Stxbp1 | protein_coding | 0.370221431 | 9.183460742 | 1.310548 | 0.2693661 | 0.756962818 |
| ENSMUSG00000015850 | Adamtsl4 | protein_coding | -0.977498703 | 2.53228929 | 1.3068966 | 0.269463 | 0.756962818 |
| ENSMUSG00000022324 | Matn2 | protein_coding | -0.664476449 | 3.743421757 | 1.3069871 | 0.269474 | 0.756962818 |
| ENSMUSG00000025234 | Arih1 | protein_coding | -0.402793851 | 5.035524536 | 1.3053193 | 0.2697414 | 0.757095329 |
| ENSMUSG00000070462 | Tlnrd1 | protein_coding | -0.523422236 | 4.426052836 | 1.3050911 | 0.2697817 | 0.757095329 |
| ENSMUSG00000022185 | Acin1 | protein_coding | -0.455252957 | 3.949173567 | 1.3049639 | 0.2698042 | 0.757095329 |
| ENSMUSG00000020032 | Nuak1 | protein_coding | -0.417408851 | 4.065982924 | 1.3048551 | 0.2698234 | 0.757095329 |
| ENSMUSG00000006435 | Neurl1a | protein_coding | -0.370105631 | 5.92054398 | 1.3045732 | 0.2698732 | 0.757095329 |
| ENSMUSG00000013646 | Sh3bp5l | protein_coding | 0.320113748 | 5.210169796 | 1.3004563 | 0.270602 | 0.758917946 |
| ENSMUSG00000023307 | 5-Mar | protein_coding | 0.369890865 | 5.014800071 | 1.2997232 | 0.2707321 | 0.758917946 |
| ENSMUSG00000020946 | Gosr2 | protein_coding | 0.292619657 | 6.890727503 | 1.2997091 | 0.2707346 | 0.758917946 |
| ENSMUSG00000028673 | Fuca1 | protein_coding | 0.244971584 | 6.853897675 | 1.2978383 | 0.2710669 | 0.759476101 |
| ENSMUSG00000020801 | Med31 | protein_coding | 0.334190589 | 6.786437743 | 1.2979438 | 0.271075 | 0.759476101 |
| ENSMUSG00000045608 | Dbx2 | protein_coding | -0.525069249 | 3.945594001 | 1.2970562 | 0.271206 | 0.759645165 |
| ENSMUSG00000039270 | Megf9 | protein_coding | -0.357088215 | 5.528848811 | 1.2954768 | 0.2714871 | 0.759939418 |
| ENSMUSG00000027752 | Exosc8 | protein_coding | 0.294295697 | 5.884115606 | 1.2953663 | 0.2715068 | 0.759939418 |
| ENSMUSG00000021541 | Trpc7 | protein_coding | 0.372187518 | 5.12309614 | 1.2952752 | 0.271523 | 0.759939418 |
| ENSMUSG00000003847 | Nfat5 | protein_coding | -0.442811075 | 4.026495328 | 1.2940712 | 0.2717377 | 0.760342196 |
| ENSMUSG00000048772 | Tmem53 | protein_coding | 0.438726389 | 4.221569174 | 1.2932323 | 0.2718873 | 0.760563085 |
| ENSMUSG00000033769 | Exoc6b | protein_coding | 0.345362917 | 5.277434501 | 1.2922279 | 0.2720667 | 0.760752395 |
| ENSMUSG00000027175 | Tcp11l1 | protein_coding | 0.448558299 | 3.964658979 | 1.292061 | 0.2720965 | 0.760752395 |
| ENSMUSG00000030539 | Sema4b | protein_coding | -0.702537158 | 2.720905013 | 1.289923 | 0.2724789 | 0.761622043 |
| ENSMUSG00000022208 | Jph4 | protein_coding | -0.284960583 | 6.716425145 | 1.2894901 | 0.2725564 | 0.761622043 |
| ENSMUSG00000038291 | Snx25 | protein_coding | -0.464116993 | 4.481618643 | 1.2891346 | 0.27262 | 0.761622043 |
| ENSMUSG00000028782 | Adgrb2 | protein_coding | -0.652240984 | 3.140051897 | 1.2880744 | 0.27281 | 0.76166148 |
| ENSMUSG00000097545 | Mir124a-1hg | lincRNA | -0.60652849 | 3.619828558 | 1.2880702 | 0.2728108 | 0.76166148 |
| ENSMUSG00000031847 | 1700030J22Rik | protein_coding | 0.51576949 | 3.621338539 | 1.2878703 | 0.2728466 | 0.76166148 |
| ENSMUSG00000008348 | Ubc | protein_coding | 0.297327771 | 8.069522431 | 1.2871813 | 0.2729702 | 0.761808693 |
| ENSMUSG00000026942 | Traf2 | protein_coding | 0.431189525 | 4.716629366 | 1.2866888 | 0.2730586 | 0.761857639 |
| ENSMUSG00000030298 | Sec13 | protein_coding | 0.245373667 | 7.532545021 | 1.2859751 | 0.2731868 | 0.761912208 |
| ENSMUSG00000083844 | Ube2d-ps | transcribed_unproces | 0.297946 | 5.765932777 | 1.2855429 | 0.2732644 | 0.761912208 |
| ENSMUSG00000027333 | Smox | protein_coding | -0.376998764 | 4.904103028 | 1.2853965 | 0.2732907 | 0.761912208 |
| ENSMUSG00000015697 | Setdb1 | protein_coding | -0.475675565 | 3.872200855 | 1.2844924 | 0.2734533 | 0.762057126 |
| ENSMUSG00000048776 | Pthlh | protein_coding | -1.032947174 | 2.503974901 | 1.2840696 | 0.2735294 | 0.762057126 |
| ENSMUSG00000041417 | Pik3r1 | protein_coding | -0.365949136 | 5.653165169 | 1.2839252 | 0.2735553 | 0.762057126 |
| ENSMUSG00000018379 | Srsf1 | protein_coding | -0.29690432 | 5.664583714 | 1.2827633 | 0.2737645 | 0.762243148 |
| ENSMUSG00000052613 | Pcdh15 | protein_coding | -0.506424768 | 4.051659865 | 1.2819037 | 0.2739195 | 0.762243148 |
| ENSMUSG00000044835 | Ankrd45 | protein_coding | -0.349954213 | 5.516952363 | 1.2816769 | 0.2739603 | 0.762243148 |
| ENSMUSG00000055675 | Kbtbd11 | protein_coding | -0.500017265 | 5.211612869 | 1.2821502 | 0.2739721 | 0.762243148 |
| ENSMUSG00000047658 | Gal3st3 | protein_coding | 0.735715007 | 3.135643891 | 1.2814266 | 0.2740055 | 0.762243148 |
| ENSMUSG00000021700 | Rab3c | protein_coding | -0.392301106 | 8.36536048 | 1.2865859 | 0.2741132 | 0.762243148 |
| ENSMUSG00000028398 | Tmem261 | protein_coding | 0.2668771 | 6.682360992 | 1.280141 | 0.2742375 | 0.762243148 |
| ENSMUSG00000052146 | Rps10 | protein_coding | 0.339091627 | 5.612174742 | 1.2798539 | 0.2742893 | 0.762243148 |
| ENSMUSG00000040856 | Dlk1 | protein_coding | -0.262452704 | 6.572223561 | 1.2796793 | 0.2743208 | 0.762243148 |
| ENSMUSG00000058135 | Gstm1 | protein_coding | -0.282312787 | 8.942497461 | 1.2796232 | 0.274331 | 0.762243148 |
| ENSMUSG00000031078 | Cttn | protein_coding | 0.356014797 | 6.768797952 | 1.2790069 | 0.2745393 | 0.76244347 |
| ENSMUSG00000013275 | Slc41a1 | protein_coding | -0.411712209 | 4.441277428 | 1.2784396 | 0.2745449 | 0.76244347 |
| ENSMUSG00000048807 | Slc35e4 | protein_coding | -0.464118136 | 4.387235814 | 1.2776505 | 0.2746876 | 0.762505581 |
| ENSMUSG00000029455 | Aldh2 | protein_coding | -0.35587616 | 5.05775773 | 1.2763572 | 0.2749218 | 0.762505581 |
| ENSMUSG00000068391 | Chrac1 | protein_coding | 0.271416552 | 6.829943269 | 1.2760039 | 0.2749858 | 0.762505581 |
| ENSMUSG00000018425 | Dhx40 | protein_coding | -0.376850144 | 4.628552036 | 1.2759824 | 0.2749897 | 0.762505581 |
| ENSMUSG00000027864 | Ptgfrn | protein_coding | -0.548395739 | 4.140524915 | 1.2759318 | 0.2749989 | 0.762505581 |
| ENSMUSG00000041375 | Ccdc9 | protein_coding | -0.627109223 | 3.067042876 | 1.2755486 | 0.2750683 | 0.762505581 |
| ENSMUSG00000022235 | Cmbl | protein_coding | 0.341036847 | 5.518609554 | 1.2747358 | 0.2752157 | 0.762505581 |
| ENSMUSG00000016308 | Ube2a | protein_coding | 0.237367011 | 7.517151785 | 1.2743691 | 0.2752823 | 0.762505581 |
| ENSMUSG00000074622 | Mafb | protein_coding | -0.955214358 | 2.802937455 | 1.2740944 | 0.2753321 | 0.762505581 |
| ENSMUSG00000034345 | Gtf2h5 | protein_coding | 0.233966536 | 8.783717357 | 1.2738873 | 0.2753697 | 0.762505581 |
| ENSMUSG00000009828 | Ick | protein_coding | -0.634104609 | 3.469201804 | 1.2737502 | 0.2753946 | 0.762505581 |
| ENSMUSG00000074194 | Zfp791 | protein_coding | -0.573286829 | 3.226863141 | 1.2736204 | 0.2754182 | 0.762505581 |
| ENSMUSG00000024054 | Smchd1 | protein_coding | -0.588883833 | 3.379495894 | 1.2731501 | 0.2755036 | 0.762545772 |
| ENSMUSG00000026924 | Sec16a | protein_coding | -0.445338405 | 3.965881154 | 1.2719702 | 0.2757181 | 0.762739798 |
| ENSMUSG00000038429 | Usp5 | protein_coding | 0.409011012 | 8.407859596 | 1.280668 | 0.2757625 | 0.762739798 |
| ENSMUSG00000056310 | Tyw1 | protein_coding | 0.416620039 | 4.485944164 | 1.271594 | 0.2757865 | 0.762739798 |
| ENSMUSG00000026127 | Imp4 | protein_coding | -0.370014073 | 4.426090176 | 1.2709437 | 0.2759049 | 0.762870881 |
| ENSMUSG00000026820 | Ptges2 | protein_coding | 0.32972095 | 5.29739169 | 1.2702365 | 0.2760336 | 0.763030733 |
| ENSMUSG00000020368 | Canx | protein_coding | -0.221159927 | 9.236388523 | 1.2692057 | 0.2762214 | 0.763262619 |
| ENSMUSG00000043702 | Pde12 | protein_coding | -0.929848256 | 2.91466877 | 1.2688683 | 0.2763096 | 0.763262619 |
| ENSMUSG00000033740 | St18 | protein_coding | -0.449391316 | 3.954821333 | 1.2681653 | 0.2764112 | 0.763262619 |
| ENSMUSG00000024535 | Snx24 | protein_coding | -0.48593669 | 4.484463318 | 1.2679178 | 0.2764564 | 0.763262619 |
| ENSMUSG00000021010 | Npas3 | protein_coding | -0.94951201 | 2.326806981 | 1.26783 | 0.2764724 | 0.763262619 |
| ENSMUSG00000064367 | mt-Nd5 | protein_coding | 0.380499766 | 9.814911841 | 1.2711659 | 0.2766117 | 0.763451237 |
| ENSMUSG00000027544 | Nfatc2 | protein_coding | 0.496754966 | 4.198646993 | 1.2660296 | 0.2768013 | 0.763600743 |
| ENSMUSG00000038872 | Zfhx3 | protein_coding | 0.300326597 | 6.113214942 | 1.2653492 | 0.2769257 | 0.763600743 |
| ENSMUSG00000048701 | Ccdc6 | protein_coding | 0.255934899 | 6.963461909 | 1.2648575 | 0.2770157 | 0.763600743 |
| ENSMUSG00000052921 | Arhgef15 | protein_coding | 0.310549138 | 6.334668996 | 1.2641525 | 0.2771448 | 0.763600743 |

| ensembl_gene_id | external_gene_name | gene_biotype | log2 (fold change) | log2 (counts per million) | F-value | p-value | False discovery rate |
|---|---|---|---|---|---|---|---|
| ENSMUSG00000069206 | Zfp874a | protein_coding | 0.520054743 | 3.615797756 | 1.2640181 | 0.2771694 | 0.763600743 |
| ENSMUSG00000038172 | Ttc39b | protein_coding | -0.433067143 | 4.595200301 | 1.2637402 | 0.2772203 | 0.763600743 |
| ENSMUSG00000061898 | Rbak | protein_coding | -0.91463732 | 2.679502963 | 1.2635944 | 0.277247 | 0.763600743 |
| ENSMUSG00000020238 | Ncln | protein_coding | 0.427615985 | 4.257319797 | 1.2630209 | 0.2773521 | 0.763600743 |
| ENSMUSG00000030532 | Hddc3 | protein_coding | -0.501204522 | 5.144441123 | 1.2633482 | 0.27736 | 0.763600743 |
| ENSMUSG00000020486 | 4-Sep | protein_coding | -0.263690686 | 8.938024311 | 1.2628904 | 0.277376 | 0.763600743 |
| ENSMUSG00000070280 | Slc22a14 | protein_coding | -0.75718972 | 2.627265896 | 1.2622805 | 0.2774879 | 0.763667445 |
| ENSMUSG00000024099 | Ndufv2 | protein_coding | 0.249958684 | 10.3424644 | 1.261453 | 0.2776397 | 0.763667445 |
| ENSMUSG00000031176 | Dynlt3 | protein_coding | 0.241271879 | 8.581354738 | 1.2613733 | 0.2776544 | 0.763667445 |
| ENSMUSG00000079317 | Trappc2 | protein_coding | 0.242084671 | 6.535422888 | 1.2609598 | 0.2777303 | 0.763667445 |
| ENSMUSG00000030287 | Itpr2 | protein_coding | 0.801247714 | 2.660606631 | 1.2604287 | 0.2778278 | 0.763667445 |
| ENSMUSG00000042439 | Zfp532 | protein_coding | -0.367251574 | 4.906140566 | 1.2590133 | 0.2780881 | 0.763667445 |
| ENSMUSG00000009035 | Tmem184b | protein_coding | -0.493673839 | 4.128339592 | 1.2587148 | 0.278143 | 0.763667445 |
| ENSMUSG00000052794 | 1700030K09Rik | protein_coding | 0.471626508 | 4.025646093 | 1.2585057 | 0.2781815 | 0.763667445 |
| ENSMUSG00000075318 | Scn2a | protein_coding | -0.234589711 | 6.959248327 | 1.2581305 | 0.2782505 | 0.763667445 |
| ENSMUSG00000036499 | Eea1 | protein_coding | -0.442495735 | 4.669997963 | 1.2576334 | 0.2783421 | 0.763667445 |
| ENSMUSG00000026312 | Cdh7 | protein_coding | -0.417815476 | 4.282803202 | 1.2574778 | 0.2783707 | 0.763667445 |
| ENSMUSG00000022008 | Gpalpp1 | protein_coding | -0.390500516 | 4.981242341 | 1.2574266 | 0.2783802 | 0.763667445 |
| ENSMUSG00000027763 | Mbnl1 | protein_coding | -0.292173829 | 6.597685114 | 1.2572983 | 0.2784038 | 0.763667445 |
| ENSMUSG00000069763 | Tmem100 | protein_coding | -0.450977693 | 5.158979621 | 1.2566403 | 0.2785251 | 0.763667445 |
| ENSMUSG00000021482 | Aaed1 | protein_coding | -0.453674095 | 4.010025099 | 1.2566082 | 0.278531 | 0.763667445 |
| ENSMUSG00000015291 | Gdi1 | protein_coding | 0.375686333 | 10.42006291 | 1.2589671 | 0.2785972 | 0.763667445 |
| ENSMUSG00000041323 | Ak7 | protein_coding | -0.843241249 | 2.577050978 | 1.2561342 | 0.2786184 | 0.763667445 |
| ENSMUSG00000023266 | Frs3 | protein_coding | 0.427261403 | 4.120808735 | 1.2554826 | 0.2787386 | 0.763667445 |
| ENSMUSG00000021276 | Cinp | protein_coding | 0.240848037 | 6.88534505 | 1.2554229 | 0.2787496 | 0.763667445 |
| ENSMUSG00000002748 | Baz1b | protein_coding | -0.352201307 | 5.4255037 | 1.2548377 | 0.2788577 | 0.763768821 |
| ENSMUSG00000020608 | Smc6 | protein_coding | -0.31411564 | 5.399207557 | 1.2523423 | 0.279319 | 0.764776078 |
| ENSMUSG00000003444 | Med29 | protein_coding | 0.42776726 | 3.908901613 | 1.2519822 | 0.2793856 | 0.764776078 |
| ENSMUSG00000025533 | Asl | protein_coding | 0.250125414 | 6.434689736 | 1.2505399 | 0.2796528 | 0.764776078 |
| ENSMUSG00000091625 | Lsm5 | protein_coding | 0.64431801 | 3.039674698 | 1.2500729 | 0.2797394 | 0.764776078 |
| ENSMUSG00000019146 | Cacng2 | protein_coding | 0.3473817 | 5.129436191 | 1.249516 | 0.2798427 | 0.764776078 |
| ENSMUSG00000090733 | Rps27 | protein_coding | 0.413858532 | 4.099858358 | 1.2491394 | 0.2799125 | 0.764776078 |
| ENSMUSG00000018401 | Mtmr4 | protein_coding | -0.371249876 | 5.241823159 | 1.2490519 | 0.2799288 | 0.764776078 |
| ENSMUSG00000054986 | Sec14l3 | protein_coding | -0.903941056 | 2.385090534 | 1.248799 | 0.2799757 | 0.764776078 |
| ENSMUSG00000041654 | Slc39a11 | protein_coding | 0.522632896 | 4.16824922 | 1.2486658 | 0.2800005 | 0.764776078 |
| ENSMUSG00000000827 | Tpd52l2 | protein_coding | 0.241207055 | 7.251869599 | 1.2470724 | 0.2802965 | 0.764776078 |
| ENSMUSG00000015961 | Adss | protein_coding | -0.290341142 | 5.890868046 | 1.2468489 | 0.2803381 | 0.764776078 |
| ENSMUSG00000052738 | Suclg1 | protein_coding | 0.221507685 | 9.129198051 | 1.2468069 | 0.2803459 | 0.764776078 |
| ENSMUSG00000102869 | 2900097C17Rik | lincRNA | 0.301347308 | 7.970443406 | 1.2467682 | 0.2803531 | 0.764776078 |
| ENSMUSG00000033009 | Ogfod1 | protein_coding | 0.274557397 | 6.537881971 | 1.2466864 | 0.2803683 | 0.764776078 |
| ENSMUSG00000031393 | Mecp2 | protein_coding | -0.318467487 | 5.871023054 | 1.2463685 | 0.2804275 | 0.764776078 |
| ENSMUSG00000047388 | Atmin | protein_coding | 0.301344649 | 5.551702336 | 1.2462834 | 0.2804433 | 0.764776078 |
| ENSMUSG00000031508 | Ankrd10 | protein_coding | -0.49389997 | 4.519871206 | 1.2461215 | 0.2804734 | 0.764776078 |
| ENSMUSG00000031590 | Frg1 | protein_coding | 0.244833012 | 7.061392723 | 1.2459485 | 0.2805056 | 0.764776078 |
| ENSMUSG00000030042 | Pole4 | protein_coding | 0.310687464 | 5.649741215 | 1.245342 | 0.2806185 | 0.764889949 |
| ENSMUSG00000015568 | Lpl | protein_coding | -0.63388336 | 3.603553369 | 1.2440774 | 0.2808541 | 0.765303076 |
| ENSMUSG00000031772 | Cntnap4 | protein_coding | 0.581318622 | 3.277471789 | 1.2437649 | 0.2809124 | 0.765303076 |
| ENSMUSG00000079215 | Zfp664 | protein_coding | -0.362171382 | 4.983712089 | 1.2432185 | 0.2810143 | 0.765321137 |
| ENSMUSG00000029390 | Tmed2 | protein_coding | -0.587501556 | 3.063734541 | 1.2426156 | 0.2811269 | 0.765321137 |
| ENSMUSG00000086003 | B230206L02Rik | processed_transcript | -0.448197065 | 4.057111651 | 1.242585 | 0.2811326 | 0.765321137 |
| ENSMUSG00000069539 | Scyl2 | protein_coding | 0.706335693 | 2.866869818 | 1.2421162 | 0.2812201 | 0.765365694 |
| ENSMUSG00000003549 | Ercc1 | protein_coding | 0.258221808 | 6.386600643 | 1.2415106 | 0.2813332 | 0.765412567 |
| ENSMUSG00000034111 | Tmed8 | protein_coding | 0.403157351 | 4.609132762 | 1.241262 | 0.2813797 | 0.765412567 |
| ENSMUSG00000004356 | Utp20 | protein_coding | 0.928968608 | 2.419253035 | 1.2406168 | 0.2815003 | 0.765547066 |
| ENSMUSG00000046519 | Golph3l | protein_coding | -0.465376272 | 4.820095839 | 1.2397585 | 0.2816609 | 0.765650594 |
| ENSMUSG00000024539 | Ptpn2 | protein_coding | -0.404343555 | 4.713151037 | 1.2396522 | 0.2816808 | 0.765650594 |
| ENSMUSG00000052105 | Mtcl1 | protein_coding | -0.585129359 | 3.993371631 | 1.238721 | 0.2818552 | 0.765775804 |
| ENSMUSG00000026526 | Fh1 | protein_coding | 0.239769719 | 7.555256967 | 1.2386456 | 0.2818693 | 0.765775804 |
| ENSMUSG00000022771 | Ppil2 | protein_coding | 0.298454709 | 6.472391962 | 1.2371551 | 0.2821487 | 0.765859854 |
| ENSMUSG00000042133 | Ppig | protein_coding | -0.297187663 | 6.474703558 | 1.2368593 | 0.2822042 | 0.765859854 |
| ENSMUSG00000028838 | Extl1 | protein_coding | 0.379241399 | 4.424538236 | 1.2367924 | 0.2822168 | 0.765859854 |
| ENSMUSG00000036580 | Spg20 | protein_coding | -0.355424208 | 5.385816295 | 1.2365988 | 0.2822531 | 0.765859854 |
| ENSMUSG00000057788 | Ddx49 | protein_coding | 0.432873848 | 5.217780257 | 1.2365467 | 0.2822629 | 0.765859854 |
| ENSMUSG00000032263 | Bckdhb | protein_coding | 0.445471434 | 3.918022141 | 1.2358641 | 0.2823911 | 0.765859854 |
| ENSMUSG00000022295 | Atp6v1c1 | protein_coding | 0.246297976 | 9.301728821 | 1.2346469 | 0.2826198 | 0.765859854 |
| ENSMUSG00000048728 | Zfp454 | protein_coding | 0.543826242 | 3.464650488 | 1.2346181 | 0.2826252 | 0.765859854 |
| ENSMUSG00000049536 | Tceal1 | protein_coding | 0.214460923 | 7.576970249 | 1.2344039 | 0.2826655 | 0.765859854 |
| ENSMUSG00000024491 | Rbm27 | protein_coding | -0.353106626 | 5.31306942 | 1.2343314 | 0.2826791 | 0.765859854 |
| ENSMUSG00000057219 | Armc7 | protein_coding | 0.417411517 | 4.448733052 | 1.234307 | 0.2826837 | 0.765859854 |
| ENSMUSG00000056050 | Mia3 | protein_coding | -0.323608348 | 5.465371818 | 1.2335857 | 0.2828194 | 0.766004754 |
| ENSMUSG00000026867 | Gapvd1 | protein_coding | -0.314644352 | 5.793790567 | 1.2324543 | 0.2830324 | 0.766004754 |
| ENSMUSG00000040321 | Zfp770 | protein_coding | -0.618246355 | 3.689128903 | 1.2324331 | 0.2830364 | 0.766004754 |
| ENSMUSG00000024231 | Cul2 | protein_coding | -0.368664006 | 5.685802949 | 1.2322827 | 0.2830648 | 0.766004754 |
| ENSMUSG00000059923 | Grb2 | protein_coding | 0.218558253 | 6.953409347 | 1.2321146 | 0.2830965 | 0.766004754 |
| ENSMUSG00000028445 | Enho | protein_coding | -0.359098975 | 6.246349103 | 1.2318923 | 0.2831646 | 0.766004754 |
| ENSMUSG00000034664 | Itga2b | protein_coding | 0.4503835 | 4.189217898 | 1.2310738 | 0.2832927 | 0.766158484 |
| ENSMUSG00000055128 | Cgrrf1 | protein_coding | 0.286900338 | 6.209589568 | 1.2306336 | 0.2833757 | 0.766190389 |
| ENSMUSG00000041840 | Haus1 | protein_coding | -0.270146152 | 5.787045799 | 1.2292792 | 0.2836314 | 0.766689029 |

| ensembl_gene_id | external_gene_name | gene_biotype | log2 (fold change) | log2 (counts per million) | F-value | p-value | False discovery rate |
|---|---|---|---|---|---|---|---|
| ENSMUSG00000018634 | Crhr1 | protein_coding | 0.770055508 | 2.681756768 | 1.228464 | 0.2837855 | 0.766790445 |
| ENSMUSG00000049134 | Nrap | protein_coding | -2.062832159 | 1.738939819 | 1.2338204 | 0.2838578 | 0.766790445 |
| ENSMUSG00000025793 | Hgs | protein_coding | 0.30846768 | 5.683935347 | 1.2274572 | 0.283976 | 0.766790445 |
| ENSMUSG00000057315 | Arhgap24 | protein_coding | -0.63182915 | 4.943632688 | 1.2357793 | 0.284003 | 0.766790445 |
| ENSMUSG00000029192 | Tbc1d14 | protein_coding | -0.347952962 | 5.501214275 | 1.2266356 | 0.2841315 | 0.766790445 |
| ENSMUSG00000036427 | Gpi1 | protein_coding | 0.326916592 | 9.013078224 | 1.2266101 | 0.2841625 | 0.766790445 |
| ENSMUSG00000026687 | Aldh9a1 | protein_coding | 0.306867457 | 6.127759362 | 1.2264422 | 0.2841681 | 0.766790445 |
| ENSMUSG00000028243 | Ubxn2b | protein_coding | -0.330936848 | 5.753515982 | 1.225466 | 0.2843531 | 0.767097109 |
| ENSMUSG00000071645 | Tut1 | protein_coding | 0.379395858 | 5.181577807 | 1.2242055 | 0.2845922 | 0.767549525 |
| ENSMUSG00000033720 | Sfxn5 | protein_coding | -0.399965246 | 8.774874846 | 1.2301085 | 0.2847826 | 0.767810372 |
| ENSMUSG00000028923 | Necap2 | protein_coding | -0.339353164 | 5.508739971 | 1.2226419 | 0.2848892 | 0.767810372 |
| ENSMUSG00000107872 | Gm44511 | protein_coding | -0.672238943 | 3.305188604 | 1.2225685 | 0.2849031 | 0.767810372 |
| ENSMUSG00000033416 | Gucd1 | protein_coding | -0.763341545 | 3.277746035 | 1.2217959 | 0.2850762 | 0.767863889 |
| ENSMUSG00000051022 | Hs3st1 | protein_coding | -0.419020963 | 4.575954066 | 1.2216363 | 0.2850804 | 0.767863889 |
| ENSMUSG00000097585 | E230029C05Rik | processed_transcript | -0.553446484 | 3.481865834 | 1.2213377 | 0.2851372 | 0.767863889 |
| ENSMUSG00000068099 | 1500009C09Rik | protein_coding | 0.279272159 | 5.961928734 | 1.2197051 | 0.2854481 | 0.768508608 |
| ENSMUSG00000059146 | Ntrk3 | protein_coding | -0.283272113 | 6.111669655 | 1.2191522 | 0.2855535 | 0.768599909 |
| ENSMUSG00000054484 | Tmem62 | protein_coding | -0.466803789 | 4.323852073 | 1.2187066 | 0.2856385 | 0.768636257 |
| ENSMUSG00000031703 | Itfg1 | protein_coding | -0.33540016 | 6.88123795 | 1.2172982 | 0.2859334 | 0.768968631 |
| ENSMUSG00000075020 | Mir670hg | lincRNA | -0.57848114 | 4.707947878 | 1.2204221 | 0.2859785 | 0.768968631 |
| ENSMUSG00000000486 | 1-Sep | protein_coding | 0.663555148 | 2.941187693 | 1.2161436 | 0.2861279 | 0.768968631 |
| ENSMUSG00000109336 | Samd4b | protein_coding | -0.409293381 | 5.297529667 | 1.2158866 | 0.286177 | 0.768968631 |
| ENSMUSG00000028211 | Trp53inp1 | protein_coding | -0.560691141 | 3.28170641 | 1.2158275 | 0.2861883 | 0.768968631 |
| ENSMUSG00000021750 | Fam107a | protein_coding | -0.3107989 | 6.542335667 | 1.2158133 | 0.2861911 | 0.768968631 |
| ENSMUSG00000031561 | Tenm3 | protein_coding | -0.58654364 | 3.863586045 | 1.2144496 | 0.286452 | 0.769242065 |
| ENSMUSG00000029171 | Pgm1 | protein_coding | -0.796777116 | 2.853407313 | 1.2134707 | 0.2866395 | 0.769242065 |
| ENSMUSG00000062991 | Nrg1 | protein_coding | -0.330247226 | 5.108745726 | 1.2134622 | 0.2866412 | 0.769242065 |
| ENSMUSG00000059714 | Flot1 | protein_coding | 0.29075315 | 6.609385333 | 1.2131303 | 0.2867048 | 0.769242065 |
| ENSMUSG00000020740 | Gga3 | protein_coding | 0.320614539 | 5.906229905 | 1.2129228 | 0.2867446 | 0.769242065 |
| ENSMUSG00000040373 | Cacng5 | protein_coding | -0.447969909 | 4.872499501 | 1.2127296 | 0.2867816 | 0.769242065 |
| ENSMUSG00000038250 | Usp38 | protein_coding | 0.697894854 | 4.033244723 | 1.2177343 | 0.2868362 | 0.769242065 |
| ENSMUSG00000026048 | Ercc5 | protein_coding | -0.574327807 | 3.90961275 | 1.212132 | 0.2868962 | 0.769242065 |
| ENSMUSG00000029512 | Ulk1 | protein_coding | -0.357543079 | 4.964379479 | 1.2117861 | 0.2869626 | 0.769242065 |
| ENSMUSG00000026209 | Dnpep | protein_coding | 0.272992413 | 6.566438219 | 1.2114789 | 0.2870216 | 0.769242065 |
| ENSMUSG00000025665 | Rps6ka6 | protein_coding | 0.625210313 | 3.323199076 | 1.2111761 | 0.2870797 | 0.769242065 |
| ENSMUSG00000112390 | CT025654.1 | processed_transcript | 0.693909289 | 3.127733802 | 1.2089565 | 0.2875064 | 0.770193484 |
| ENSMUSG00000048537 | Phldb1 | protein_coding | -0.371255821 | 4.846870434 | 1.2066215 | 0.2879562 | 0.770828174 |
| ENSMUSG00000062012 | Zfp13 | protein_coding | 0.346955423 | 5.172760166 | 1.2064695 | 0.2879855 | 0.770828174 |
| ENSMUSG00000016921 | Srsf6 | protein_coding | -0.42077664 | 5.095380755 | 1.2064023 | 0.2879985 | 0.770828174 |
| ENSMUSG00000036989 | Trim3 | protein_coding | 0.278635733 | 6.15959805 | 1.2062385 | 0.2880301 | 0.770828174 |
| ENSMUSG00000096225 | Lhx8 | protein_coding | 0.455786409 | 4.460769682 | 1.2052094 | 0.2882287 | 0.771081751 |
| ENSMUSG00000003226 | Ranbp2 | protein_coding | -0.494049618 | 3.727733926 | 1.2047019 | 0.2883267 | 0.771081751 |
| ENSMUSG00000060519 | Tor3a | protein_coding | -0.875224216 | 2.469766756 | 1.2045329 | 0.2883594 | 0.771081751 |
| ENSMUSG00000005442 | Cic | protein_coding | -0.458932084 | 3.706908666 | 1.2042622 | 0.2884117 | 0.771081751 |
| ENSMUSG00000003199 | Mpnd | protein_coding | 0.323926751 | 5.730802558 | 1.2026621 | 0.2887211 | 0.771717222 |
| ENSMUSG00000068798 | Rap1a | protein_coding | -0.344522443 | 4.606959822 | 1.2022317 | 0.2888044 | 0.771748054 |
| ENSMUSG00000024150 | Mcfd2 | protein_coding | 0.23341911 | 7.672831725 | 1.2010046 | 0.2890422 | 0.772017994 |
| ENSMUSG00000017969 | Ptgis | protein_coding | 0.844814837 | 2.848786097 | 1.200969 | 0.2890491 | 0.772017994 |
| ENSMUSG00000043257 | Pigv | protein_coding | -0.79011509 | 2.814847498 | 1.2002633 | 0.2891859 | 0.772191685 |
| ENSMUSG00000034801 | Sos2 | protein_coding | 0.351454799 | 4.382323573 | 1.1988369 | 0.2894627 | 0.772558978 |
| ENSMUSG00000050390 | C77080 | protein_coding | -0.471587655 | 4.336900118 | 1.1983289 | 0.2895614 | 0.772558978 |
| ENSMUSG00000051510 | Mafg | protein_coding | 0.29581507 | 5.839539025 | 1.1982736 | 0.2895722 | 0.772558978 |
| ENSMUSG00000018707 | Dync1h1 | protein_coding | -0.383214088 | 5.922824251 | 1.1982081 | 0.2896108 | 0.772558978 |
| ENSMUSG00000097277 | 2900076A07Rik | lincRNA | -0.461690452 | 3.852739501 | 1.1973445 | 0.2897528 | 0.77274593 |
| ENSMUSG00000032224 | Fam81a | protein_coding | -0.349711139 | 8.374444457 | 1.1983195 | 0.2900534 | 0.773230312 |
| ENSMUSG00000005846 | Rsl1d1 | protein_coding | 0.261670419 | 7.142042132 | 1.1955149 | 0.2901089 | 0.773230312 |
| ENSMUSG00000048222 | Mfap1b | protein_coding | -0.348761769 | 4.988416576 | 1.1950539 | 0.2901987 | 0.773230312 |
| ENSMUSG00000012640 | Zfp715 | protein_coding | -0.513774783 | 3.863644095 | 1.1948603 | 0.2902364 | 0.773230312 |
| ENSMUSG00000030802 | Bckdk | protein_coding | 0.336473819 | 6.812872469 | 1.19446 | 0.2903404 | 0.773230312 |
| ENSMUSG00000030688 | Stard10 | protein_coding | -0.390784768 | 4.304015813 | 1.1941965 | 0.2903659 | 0.773230312 |
| ENSMUSG00000029090 | Adgra3 | protein_coding | 1.165511601 | 2.282222908 | 1.1936017 | 0.2905078 | 0.773416794 |
| ENSMUSG00000021375 | Kif13a | protein_coding | -0.52472024 | 3.691015578 | 1.1927095 | 0.2906561 | 0.773619992 |
| ENSMUSG00000029767 | Calu | protein_coding | -0.34636863 | 5.515481613 | 1.1921064 | 0.2907739 | 0.7736888 |
| ENSMUSG00000025157 | Zdhhc16 | protein_coding | 0.452683388 | 3.534691485 | 1.1918406 | 0.2908258 | 0.7736888 |
| ENSMUSG00000031429 | Psmd10 | protein_coding | 0.247008802 | 6.94919868 | 1.1909333 | 0.2910032 | 0.773969268 |
| ENSMUSG00000037617 | Spag1 | protein_coding | -0.925766365 | 2.575763389 | 1.1904661 | 0.2910946 | 0.774020955 |
| ENSMUSG00000109284 | B230311B06Rik | lincRNA | -0.593821754 | 3.309155098 | 1.1894847 | 0.2912868 | 0.774264824 |
| ENSMUSG00000015126 | Tsr3 | protein_coding | -0.386313553 | 4.581790649 | 1.1888497 | 0.2914112 | 0.774264824 |
| ENSMUSG00000006705 | Pknox1 | protein_coding | -0.431034215 | 4.007590986 | 1.1888367 | 0.2914138 | 0.774264824 |
| ENSMUSG00000059895 | Ptp4a3 | protein_coding | 0.320938821 | 5.311808711 | 1.1884524 | 0.2914891 | 0.774264824 |
| ENSMUSG00000068329 | Htra2 | protein_coding | 0.400949645 | 4.739776655 | 1.1879895 | 0.2915799 | 0.774264824 |
| ENSMUSG00000041685 | Fcho2 | protein_coding | -0.464896736 | 3.901869086 | 1.1877932 | 0.2916184 | 0.774264824 |
| ENSMUSG00000068373 | D430041D05Rik | protein_coding | -0.394611898 | 4.776379239 | 1.1862803 | 0.2919154 | 0.774862061 |
| ENSMUSG00000033436 | Armcx2 | protein_coding | 0.283373488 | 6.195995287 | 1.1857877 | 0.2920122 | 0.774927683 |
| ENSMUSG00000038437 | Mllt6 | protein_coding | -0.574263922 | 3.657007758 | 1.1842948 | 0.2923058 | 0.775515481 |
| ENSMUSG00000006586 | Runx1t1 | protein_coding | -0.381605311 | 6.322733554 | 1.1841258 | 0.2924774 | 0.775779407 |
| ENSMUSG00000063870 | Chd4 | protein_coding | -0.261665372 | 6.028503614 | 1.1823024 | 0.2926983 | 0.776172003 |
| ENSMUSG00000025269 | Apex2 | protein_coding | 0.592719997 | 3.326395185 | 1.1819397 | 0.2927698 | 0.776172003 |

| ensembl_gene_id | external_gene_name | gene_biotype | log2 (fold change) | log2 (counts per million) | F-value | p-value | False discovery rate |
|---|---|---|---|---|---|---|---|
| ENSMUSG00000044988 | Ucn3 | protein_coding | -0.688039491 | 3.261059349 | 1.1805052 | 0.2930529 | 0.776287825 |
| ENSMUSG00000037971 | 1110032A03Rik | protein_coding | 0.214131498 | 7.593220878 | 1.1800239 | 0.293148 | 0.776287825 |
| ENSMUSG00000024158 | Hagh | protein_coding | 0.232005492 | 8.304436477 | 1.1799519 | 0.2931622 | 0.776287825 |
| ENSMUSG00000005893 | Nr2c2 | protein_coding | -0.398791509 | 4.941019602 | 1.1799479 | 0.293163 | 0.776287825 |
| ENSMUSG00000022456 | 3-Sep | protein_coding | 0.239902171 | 8.375689342 | 1.1798507 | 0.2931822 | 0.776287825 |
| ENSMUSG00000025939 | Ube2w | protein_coding | 0.225750002 | 8.205577493 | 1.1782297 | 0.2935028 | 0.776287825 |
| ENSMUSG00000033809 | Alg3 | protein_coding | 0.378815445 | 4.917568556 | 1.1779946 | 0.2935493 | 0.776287825 |
| ENSMUSG00000031558 | Slit2 | protein_coding | -0.575505668 | 3.494476546 | 1.1778572 | 0.2935765 | 0.776287825 |
| ENSMUSG00000034936 | Arl4d | protein_coding | -0.672724378 | 4.098473613 | 1.1828483 | 0.29359 | 0.776287825 |
| ENSMUSG00000068101 | Cenpm | protein_coding | 0.833244686 | 2.727712796 | 1.1777046 | 0.2936067 | 0.776287825 |
| ENSMUSG00000004360 | 9330159F19Rik | protein_coding | -0.511836487 | 4.99647521 | 1.1783978 | 0.2936076 | 0.776287825 |
| ENSMUSG00000038457 | Tmem255b | protein_coding | -1.239366599 | 2.342129433 | 1.1767864 | 0.2938144 | 0.776643387 |
| ENSMUSG00000022132 | Cldn10 | protein_coding | -0.594738877 | 5.794280525 | 1.1887926 | 0.2939284 | 0.776643387 |
| ENSMUSG00000001964 | Emd | protein_coding | -0.388615635 | 5.350165237 | 1.1759282 | 0.2939588 | 0.776643387 |
| ENSMUSG00000026073 | Il1r2 | protein_coding | 1.138951075 | 2.054757729 | 1.1753305 | 0.2940773 | 0.776691887 |
| ENSMUSG00000051427 | Ccdc157 | protein_coding | -0.649847218 | 2.943598465 | 1.1748596 | 0.2941708 | 0.776691887 |
| ENSMUSG00000031539 | Ap3m2 | protein_coding | 0.314285537 | 7.416457161 | 1.1747552 | 0.2942173 | 0.776691887 |
| ENSMUSG00000063145 | Bbs5 | protein_coding | -0.393615354 | 4.886398385 | 1.1739926 | 0.294343 | 0.776691887 |
| ENSMUSG00000014195 | Dnajc7 | protein_coding | 0.249832889 | 6.780053273 | 1.1739064 | 0.2943601 | 0.776691887 |
| ENSMUSG00000030268 | Bcat1 | protein_coding | 0.331228185 | 7.583348572 | 1.1741941 | 0.2945014 | 0.776691887 |
| ENSMUSG00000019961 | Tmpo | protein_coding | -0.358808517 | 4.458552505 | 1.1726548 | 0.2946089 | 0.776691887 |
| ENSMUSG00000000355 | Mcts1 | protein_coding | 0.239891452 | 7.797102568 | 1.1726498 | 0.2946099 | 0.776691887 |
| ENSMUSG00000087260 | Lamtor5 | protein_coding | 0.255438868 | 7.5296565 | 1.1724154 | 0.2946566 | 0.776691887 |
| ENSMUSG00000009575 | Cbx5 | protein_coding | 0.215246301 | 8.581273462 | 1.1715223 | 0.2948343 | 0.776691887 |
| ENSMUSG00000000282 | Mnt | protein_coding | 0.376877765 | 4.597142169 | 1.171295 | 0.2948796 | 0.776691887 |
| ENSMUSG00000027479 | Mapre1 | protein_coding | -0.230673297 | 7.318382081 | 1.1693374 | 0.2952699 | 0.776691887 |
| ENSMUSG00000021573 | Tppp | protein_coding | -0.240901141 | 7.734587106 | 1.1691293 | 0.2953114 | 0.776691887 |
| ENSMUSG00000036097 | Slf2 | protein_coding | -0.343893722 | 5.110730074 | 1.1688242 | 0.2953723 | 0.776691887 |
| ENSMUSG00000026207 | Speg | protein_coding | -0.570014825 | 3.661828598 | 1.168614 | 0.2954143 | 0.776691887 |
| ENSMUSG00000056856 | Jakmip3 | protein_coding | 0.414423404 | 4.059510386 | 1.1675941 | 0.295618 | 0.776691887 |
| ENSMUSG00000053137 | Mapk11 | protein_coding | 0.490898837 | 3.555196215 | 1.1672252 | 0.2956917 | 0.776691887 |
| ENSMUSG00000027737 | Slc7a11 | protein_coding | -0.649738854 | 3.218321286 | 1.1671563 | 0.2957055 | 0.776691887 |
| ENSMUSG00000026617 | Bpnt1 | protein_coding | 0.242116431 | 6.995437354 | 1.1671394 | 0.2957089 | 0.776691887 |
| ENSMUSG00000030720 | Cln3 | protein_coding | 0.519020132 | 4.129359323 | 1.1664693 | 0.2958429 | 0.776691887 |
| ENSMUSG00000032324 | Tspan3 | protein_coding | -0.312965358 | 7.434399191 | 1.1665866 | 0.2958451 | 0.776691887 |
| ENSMUSG00000024889 | Rce1 | protein_coding | -0.39938585 | 4.453823746 | 1.1660992 | 0.295917 | 0.776691887 |
| ENSMUSG00000031545 | Gpat4 | protein_coding | 0.319661276 | 6.547037953 | 1.1660702 | 0.2959484 | 0.776691887 |
| ENSMUSG00000056608 | Chd9 | protein_coding | -0.35961276 | 5.372321809 | 1.1658753 | 0.2959618 | 0.776691887 |
| ENSMUSG00000039523 | Cep104 | protein_coding | 0.496715711 | 4.015936478 | 1.1656785 | 0.2960012 | 0.776691887 |
| ENSMUSG00000064037 | Gpn1 | protein_coding | 0.230852875 | 6.995836777 | 1.1654128 | 0.2960544 | 0.776691887 |
| ENSMUSG00000029649 | Pomp | protein_coding | 0.244635949 | 9.62776888 | 1.1650798 | 0.2961211 | 0.776691887 |
| ENSMUSG00000034858 | Fam214a | protein_coding | -0.42382286 | 4.271265925 | 1.1649582 | 0.2961455 | 0.776691887 |
| ENSMUSG00000043091 | Tuba1c | protein_coding | 1.646590685 | 1.936353087 | 1.1697902 | 0.296174 | 0.776691887 |
| ENSMUSG00000029130 | Rnf32 | protein_coding | -0.374528697 | 4.393035802 | 1.1640436 | 0.2963288 | 0.776691887 |
| ENSMUSG00000033400 | Agl | protein_coding | -0.446321139 | 3.815284469 | 1.163938 | 0.2963499 | 0.776691887 |
| ENSMUSG00000032096 | Arcn1 | protein_coding | -0.287749753 | 5.576845699 | 1.1637243 | 0.2963928 | 0.776691887 |
| ENSMUSG00000024477 | Pggt1b | protein_coding | -0.526954227 | 3.324418447 | 1.1635278 | 0.2964322 | 0.776691887 |
| ENSMUSG00000014426 | Map3k4 | protein_coding | -0.439133344 | 4.223032953 | 1.1634252 | 0.2964528 | 0.776691887 |
| ENSMUSG00000027506 | Tpd52 | protein_coding | 0.31015713 | 6.305528105 | 1.1631322 | 0.2965116 | 0.776691887 |
| ENSMUSG00000005514 | Por | protein_coding | 0.32293946 | 6.626504381 | 1.1629318 | 0.2965774 | 0.776691887 |
| ENSMUSG00000005103 | Wdr1 | protein_coding | 0.35505904 | 7.835162807 | 1.1665396 | 0.296822 | 0.776904749 |
| ENSMUSG00000042492 | Tbc1d10b | protein_coding | 0.305430336 | 5.264162678 | 1.1613681 | 0.2968659 | 0.776904749 |
| ENSMUSG00000031154 | Otud5 | protein_coding | -0.378698869 | 5.270545097 | 1.1611471 | 0.2969103 | 0.776904749 |
| ENSMUSG00000020186 | Csrp2 | protein_coding | 0.351765532 | 4.560132812 | 1.1609613 | 0.2969477 | 0.776904749 |
| ENSMUSG00000015942 | Gtf2ird2 | protein_coding | 0.502983896 | 3.549050209 | 1.1599504 | 0.2971511 | 0.777242446 |
| ENSMUSG00000014856 | Tmem208 | protein_coding | 0.260264827 | 6.816027486 | 1.1593738 | 0.2972672 | 0.777242446 |
| ENSMUSG00000014767 | Tbp | protein_coding | 0.322865159 | 5.374297906 | 1.158045 | 0.2975349 | 0.777242446 |
| ENSMUSG00000026890 | Lhx6 | protein_coding | 0.353654855 | 6.003669517 | 1.1579492 | 0.2975798 | 0.777242446 |
| ENSMUSG00000029501 | Ankle2 | protein_coding | -0.400063876 | 4.398547778 | 1.1574734 | 0.2976502 | 0.777242446 |
| ENSMUSG00000048118 | Arid4a | protein_coding | -0.303380644 | 6.112381513 | 1.1570329 | 0.2977391 | 0.777242446 |
| ENSMUSG00000029571 | Tmem106b | protein_coding | -0.313593598 | 5.82573429 | 1.1568812 | 0.2977697 | 0.777242446 |
| ENSMUSG00000030137 | Tuba8 | protein_coding | 0.727216696 | 3.06575667 | 1.1566737 | 0.2978116 | 0.777242446 |
| ENSMUSG00000022607 | Ptk2 | protein_coding | -0.308342687 | 5.103287679 | 1.1565977 | 0.2978269 | 0.777242446 |
| ENSMUSG00000045441 | Gprin3 | protein_coding | -0.423991284 | 4.460587543 | 1.1564416 | 0.2978584 | 0.777242446 |
| ENSMUSG00000067995 | Gtf2f2 | protein_coding | -0.302671445 | 5.655540052 | 1.1562968 | 0.2978877 | 0.777242446 |
| ENSMUSG00000007827 | Ankrd26 | protein_coding | -0.478089139 | 4.709306544 | 1.1561437 | 0.2979441 | 0.777242446 |
| ENSMUSG00000028416 | Bag1 | protein_coding | 0.264358189 | 6.204112354 | 1.1551222 | 0.298125 | 0.777413253 |
| ENSMUSG00000040464 | Gtpbp10 | protein_coding | -0.758140089 | 2.572768242 | 1.1549607 | 0.2981577 | 0.777413253 |
| ENSMUSG00000022765 | Snap29 | protein_coding | -0.436253344 | 4.01884437 | 1.1546163 | 0.2982273 | 0.777413253 |
| ENSMUSG00000000751 | Rpa1 | protein_coding | -0.358566864 | 6.402588931 | 1.1542311 | 0.2983308 | 0.777413253 |
| ENSMUSG00000033256 | Shf | protein_coding | -0.737099832 | 2.677663578 | 1.1539058 | 0.2983711 | 0.777413253 |
| ENSMUSG00000041261 | Car8 | protein_coding | 0.463717899 | 4.349591531 | 1.1531121 | 0.2985318 | 0.777425378 |
| ENSMUSG00000056476 | Med12l | protein_coding | -0.553155938 | 3.301636444 | 1.1524172 | 0.2986726 | 0.777425378 |
| ENSMUSG00000041997 | Tlk1 | protein_coding | -0.297099405 | 5.412870422 | 1.151696 | 0.2988189 | 0.777425378 |
| ENSMUSG00000030826 | Bcat2 | protein_coding | 0.44933111 | 4.651427735 | 1.1512055 | 0.2989184 | 0.777425378 |
| ENSMUSG00000041891 | Lman1 | protein_coding | -0.536562784 | 3.955768525 | 1.1511217 | 0.2989354 | 0.777425378 |
| ENSMUSG00000018761 | Mpdu1 | protein_coding | 0.252737291 | 6.826506948 | 1.150845 | 0.2989915 | 0.777425378 |
| ENSMUSG00000039697 | Ncoa7 | protein_coding | -0.264819354 | 5.966730322 | 1.1508073 | 0.2989992 | 0.777425378 |

| ensembl_gene_id | external_gene_name | gene_biotype | log2 (fold change) | log2 (counts per million) | F-value | p-value | False discovery rate |
|---|---|---|---|---|---|---|---|
| ENSMUSG00000025261 | Huwe1 | protein_coding | -0.269083065 | 5.898105929 | 1.1506437 | 0.2990324 | 0.777425378 |
| ENSMUSG00000020696 | Rffl | protein_coding | -0.653609545 | 3.31766545 | 1.1503337 | 0.2990953 | 0.777425378 |
| ENSMUSG00000045691 | Thtpa | protein_coding | 0.459114449 | 5.335795227 | 1.1509899 | 0.2990987 | 0.777425378 |
| ENSMUSG00000042590 | Ipo11 | protein_coding | -0.543306207 | 3.43643476 | 1.1494785 | 0.2992691 | 0.777548255 |
| ENSMUSG00000000301 | Pemt | protein_coding | 0.543043715 | 3.320569159 | 1.1493724 | 0.2992906 | 0.777548255 |
| ENSMUSG00000051177 | Plcb1 | protein_coding | -0.54980394 | 3.797051818 | 1.1486832 | 0.2994308 | 0.777724436 |
| ENSMUSG00000024640 | Psat1 | protein_coding | -0.207366024 | 7.37906699 | 1.1479953 | 0.2995707 | 0.777900036 |
| ENSMUSG00000070047 | Fat1 | protein_coding | -0.602760726 | 3.598370895 | 1.1470699 | 0.2997592 | 0.777995655 |
| ENSMUSG00000026384 | Ptpn4 | protein_coding | -0.293542746 | 5.596980002 | 1.146974 | 0.2997787 | 0.777995655 |
| ENSMUSG00000021824 | Ap3m1 | protein_coding | -0.336534178 | 4.904119096 | 1.1467486 | 0.2998246 | 0.777995655 |
| ENSMUSG00000019082 | Slc25a22 | protein_coding | 0.348534888 | 7.807801147 | 1.1501375 | 0.300062 | 0.778423821 |
| ENSMUSG00000000568 | Hnrnpd | protein_coding | -0.268628731 | 6.889537929 | 1.1449898 | 0.3001833 | 0.778550566 |
| ENSMUSG00000036390 | Gadd45a | protein_coding | -0.649085712 | 2.947059194 | 1.1432582 | 0.300537 | 0.778884947 |
| ENSMUSG00000043496 | Tril | protein_coding | -0.702567533 | 4.010766033 | 1.1477082 | 0.3006151 | 0.778884947 |
| ENSMUSG00000038803 | Ost4 | protein_coding | 0.280564592 | 6.636590519 | 1.1426469 | 0.300662 | 0.778884947 |
| ENSMUSG00000045664 | Cdc42ep2 | protein_coding | -0.373419084 | 5.455680769 | 1.1422548 | 0.3007422 | 0.778884947 |
| ENSMUSG00000031673 | Cdh11 | protein_coding | -0.383712909 | 4.760155225 | 1.1416791 | 0.30086 | 0.778884947 |
| ENSMUSG00000025781 | Atp5c1 | protein_coding | 0.220125807 | 10.3395293 | 1.1413437 | 0.3009287 | 0.778884947 |
| ENSMUSG00000006395 | Hyi | polymorphic_pseudog | -0.63412633 | 2.861671572 | 1.141057 | 0.3009874 | 0.778884947 |
| ENSMUSG00000031917 | Nip7 | protein_coding | 0.274847701 | 6.922530447 | 1.1408732 | 0.3010251 | 0.778884947 |
| ENSMUSG00000035891 | Cerk | protein_coding | -0.402986969 | 4.706977048 | 1.1403712 | 0.3011279 | 0.778884947 |
| ENSMUSG00000035517 | Tdrd7 | protein_coding | -0.490571997 | 4.36072328 | 1.1398505 | 0.3012347 | 0.778884947 |
| ENSMUSG00000041577 | Prelp | protein_coding | -0.981735271 | 2.359007947 | 1.1396212 | 0.3012817 | 0.778884947 |
| ENSMUSG00000040055 | Gjb6 | protein_coding | -0.418277814 | 5.585771799 | 1.1395357 | 0.3013246 | 0.778884947 |
| ENSMUSG00000022774 | Ncbp2 | protein_coding | 0.208587732 | 7.268332187 | 1.1387593 | 0.3014586 | 0.778884947 |
| ENSMUSG00000016541 | Atxn10 | protein_coding | 0.276600217 | 9.478745114 | 1.138626 | 0.301486 | 0.778884947 |
| ENSMUSG00000001844 | Zdhhc4 | protein_coding | 0.305943047 | 5.724670165 | 1.1385621 | 0.3014991 | 0.778884947 |
| ENSMUSG00000001707 | Eef1e1 | protein_coding | 0.247385501 | 7.812687643 | 1.1381619 | 0.3015813 | 0.778884947 |
| ENSMUSG00000052684 | Jun | protein_coding | -0.526912599 | 3.651238088 | 1.1378443 | 0.3016465 | 0.778884947 |
| ENSMUSG00000033955 | Tnks1bp1 | protein_coding | -0.901122598 | 2.644913488 | 1.137706 | 0.3016749 | 0.778884947 |
| ENSMUSG00000033526 | Ppip5k1 | protein_coding | 0.370513392 | 4.9857567 | 1.1376402 | 0.3016884 | 0.778884947 |
| ENSMUSG00000075316 | Scn9a | protein_coding | -0.459396111 | 3.5078237 | 1.1369107 | 0.3018384 | 0.778896008 |
| ENSMUSG00000034321 | Exosc1 | protein_coding | 0.29358661 | 6.37568515 | 1.136609 | 0.3019004 | 0.778896008 |
| ENSMUSG00000019659 | Ccdc12 | protein_coding | 0.265000145 | 5.902746514 | 1.1365624 | 0.30191 | 0.778896008 |
| ENSMUSG00000005469 | Prkaca | protein_coding | 0.268269426 | 7.918285294 | 1.1352247 | 0.3021854 | 0.779142815 |
| ENSMUSG00000024212 | Mllt1 | protein_coding | -0.471608311 | 4.066042031 | 1.1352062 | 0.3021892 | 0.779142815 |
| ENSMUSG00000027177 | Hipk3 | protein_coding | -0.355903381 | 5.20658307 | 1.1350417 | 0.3022231 | 0.779142815 |
| ENSMUSG00000031672 | Got2 | protein_coding | 0.326756779 | 7.03732811 | 1.1343339 | 0.302461 | 0.779370529 |
| ENSMUSG00000029131 | Dnajb6 | protein_coding | -0.397540304 | 4.287201928 | 1.1336479 | 0.3025104 | 0.779370529 |
| ENSMUSG00000040396 | Abhd13 | protein_coding | -0.469426151 | 3.6968648 | 1.1335585 | 0.3025288 | 0.779370529 |
| ENSMUSG00000029179 | Zcchc4 | protein_coding | -0.608062264 | 3.609632238 | 1.1328828 | 0.3026683 | 0.779404708 |
| ENSMUSG00000028431 | Ikbkap | protein_coding | -0.31801436 | 5.093809927 | 1.1327917 | 0.3026871 | 0.779404708 |
| ENSMUSG00000064325 | Hhip | protein_coding | -0.718445675 | 2.593913966 | 1.1316962 | 0.3029133 | 0.779699912 |
| ENSMUSG00000042032 | Mat2b | protein_coding | -0.225869126 | 7.830334238 | 1.1315346 | 0.3029467 | 0.779699912 |
| ENSMUSG00000021785 | Ngly1 | protein_coding | -0.417480249 | 4.401940673 | 1.1305146 | 0.3031576 | 0.780053339 |
| ENSMUSG00000028771 | Ptpn12 | protein_coding | -0.527291812 | 4.087179303 | 1.1301581 | 0.3032314 | 0.780053339 |
| ENSMUSG00000024911 | Fibp | protein_coding | 0.244053166 | 6.886098027 | 1.1298186 | 0.3033017 | 0.780053339 |
| ENSMUSG00000061482 | Hist1h4d | protein_coding | -0.419277713 | 3.765522318 | 1.1290548 | 0.3034598 | 0.780273469 |
| ENSMUSG00000033216 | Eefsec | protein_coding | 0.445866628 | 4.607461669 | 1.1285084 | 0.3035731 | 0.780378022 |
| ENSMUSG00000049130 | C5ar1 | protein_coding | -1.200679397 | 2.071131458 | 1.127748 | 0.3037307 | 0.780534228 |
| ENSMUSG00000039568 | Ubald1 | protein_coding | 0.249957885 | 6.558367213 | 1.1275152 | 0.303779 | 0.780534228 |
| ENSMUSG00000046962 | Zbtb21 | protein_coding | -0.320513549 | 5.212693497 | 1.1269505 | 0.3038962 | 0.780645957 |
| ENSMUSG00000053870 | Fpgt | protein_coding | -0.580673619 | 3.953645991 | 1.1265006 | 0.3040148 | 0.780645957 |
| ENSMUSG00000025156 | Gps1 | protein_coding | 0.272747331 | 5.477462297 | 1.1262566 | 0.3040403 | 0.780645957 |
| ENSMUSG00000027822 | Slc33a1 | protein_coding | -0.402024985 | 4.249301208 | 1.1250194 | 0.3042974 | 0.780829215 |
| ENSMUSG00000031133 | Arhgef6 | protein_coding | -0.76249882 | 2.854956478 | 1.1246661 | 0.3043709 | 0.780829215 |
| ENSMUSG00000043251 | Exoc3l | protein_coding | 0.973741965 | 2.066138384 | 1.1245457 | 0.3043959 | 0.780829215 |
| ENSMUSG00000028514 | Usp24 | protein_coding | -0.680216355 | 3.687224854 | 1.1244899 | 0.3044992 | 0.780829215 |
| ENSMUSG00000074093 | Svip | protein_coding | -0.308090493 | 6.117589266 | 1.1239745 | 0.3045148 | 0.780829215 |
| ENSMUSG00000019907 | Ppp1r12a | protein_coding | -0.287320033 | 6.455738273 | 1.1233218 | 0.3046507 | 0.780829215 |
| ENSMUSG00000029249 | Rest | protein_coding | -1.016841268 | 2.127344686 | 1.1229976 | 0.3047182 | 0.780829215 |
| ENSMUSG00000019210 | Atp6v1e1 | protein_coding | 0.213144716 | 10.1101088 | 1.1227897 | 0.3047616 | 0.780829215 |
| ENSMUSG00000025722 | Wdr73 | protein_coding | 0.296873659 | 5.634725503 | 1.1227722 | 0.3047652 | 0.780829215 |
| ENSMUSG00000003131 | Pafah1b2 | protein_coding | -0.238154262 | 7.544336209 | 1.1220243 | 0.3049211 | 0.781042553 |
| ENSMUSG00000041598 | Cdc42ep4 | protein_coding | -0.400280542 | 4.979978276 | 1.120331 | 0.3052745 | 0.781761532 |
| ENSMUSG00000046711 | Hmga1 | protein_coding | 0.443287376 | 3.853199304 | 1.1192061 | 0.3055096 | 0.781972934 |
| ENSMUSG00000020541 | Tom1l1 | protein_coding | -0.316026089 | 5.201902742 | 1.1184711 | 0.3056633 | 0.781972934 |
| ENSMUSG00000027710 | Acad9 | protein_coding | -0.806308278 | 3.779285076 | 1.1264736 | 0.3057403 | 0.781972934 |
| ENSMUSG00000036299 | BC031181 | protein_coding | 0.222267194 | 9.008096586 | 1.1180696 | 0.3057473 | 0.781972934 |
| ENSMUSG00000025790 | Slco3a1 | protein_coding | 0.409544829 | 4.733153981 | 1.1177651 | 0.3058111 | 0.781972934 |
| ENSMUSG00000055805 | Fmnl1 | protein_coding | -0.427620318 | 4.428708604 | 1.117399 | 0.3058878 | 0.781972934 |
| ENSMUSG00000067818 | Myl9 | protein_coding | -0.397730956 | 4.61961519 | 1.1164041 | 0.3060962 | 0.781972934 |
| ENSMUSG00000004788 | Eif2b2 | protein_coding | 0.248954666 | 6.234374159 | 1.1163832 | 0.3061006 | 0.781972934 |
| ENSMUSG00000029469 | Ift81 | protein_coding | -0.338108666 | 5.347854405 | 1.1162471 | 0.3061291 | 0.781972934 |
| ENSMUSG00000085829 | Gm4285 | antisense_RNA | 0.934934822 | 2.39625023 | 1.1159838 | 0.3061844 | 0.781972934 |
| ENSMUSG00000097908 | 4933404O12Rik | lincRNA | 0.591484629 | 3.217324992 | 1.1158253 | 0.3062176 | 0.781972934 |
| ENSMUSG00000034247 | Plekhm1 | protein_coding | -0.463188334 | 3.720036447 | 1.1157676 | 0.3062297 | 0.781972934 |
| ENSMUSG00000090399 | Gm38399 | lincRNA | -0.664509486 | 2.784710787 | 1.1150733 | 0.3063754 | 0.782033887 |

| ensembl_gene_id | external_gene_name | gene_biotype | log2 (fold change) | log2 (counts per million) | F-value | p-value | False discovery rate |
|---|---|---|---|---|---|---|---|
| ENSMUSG00000029861 | Fam131b | protein_coding | -0.53718014 | 3.633777531 | 1.1147471 | 0.3064439 | 0.782033887 |
| ENSMUSG00000043183 | Simc1 | protein_coding | 0.529994058 | 3.341186191 | 1.1134483 | 0.3067168 | 0.782033887 |
| ENSMUSG00000029199 | Lias | protein_coding | 0.240343415 | 6.878861536 | 1.113392 | 0.3067286 | 0.782033887 |
| ENSMUSG00000042787 | Exog | protein_coding | 0.419541148 | 5.026639843 | 1.1132773 | 0.3067527 | 0.782033887 |
| ENSMUSG00000036550 | Cnot1 | protein_coding | -0.386306947 | 4.584060115 | 1.1132265 | 0.3067634 | 0.782033887 |
| ENSMUSG00000039683 | Sdk1 | protein_coding | -0.92635988 | 2.433209778 | 1.1129604 | 0.3068193 | 0.782033887 |
| ENSMUSG00000032038 | St3gal4 | protein_coding | 0.367182785 | 4.551804892 | 1.1126949 | 0.3068752 | 0.782033887 |
| ENSMUSG00000084416 | Rpl10a-ps1 | processed_pseudoge | 0.565943003 | 2.886219556 | 1.1125384 | 0.3069081 | 0.782033887 |
| ENSMUSG00000039219 | Arid4b | protein_coding | -0.300623834 | 5.786670601 | 1.1119853 | 0.3070245 | 0.78210485 |
| ENSMUSG00000055541 | Lair1 | protein_coding | -0.615431852 | 3.076907307 | 1.111715 | 0.3070814 | 0.78210485 |
| ENSMUSG00000028101 | Pias3 | protein_coding | 0.472596915 | 5.432729769 | 1.1144716 | 0.3074205 | 0.782592156 |
| ENSMUSG00000045237 | 1110012L19Rik | protein_coding | 0.256135992 | 5.830513754 | 1.1099082 | 0.3074622 | 0.782592156 |
| ENSMUSG00000061286 | Exosc5 | protein_coding | 0.268297674 | 6.021888694 | 1.1097713 | 0.3074911 | 0.782592156 |
| ENSMUSG00000061111 | Mcrip1 | protein_coding | -0.313538617 | 7.096390871 | 1.1090197 | 0.3076748 | 0.782790591 |
| ENSMUSG00000040688 | Tbl3 | protein_coding | 0.370428187 | 5.372885284 | 1.1081544 | 0.3078325 | 0.782790591 |
| ENSMUSG00000041658 | Rragb | protein_coding | -0.893535815 | 5.215019542 | 1.144184 | 0.3078409 | 0.782790591 |
| ENSMUSG00000015968 | Cacna1d | protein_coding | 0.45472673 | 3.628391845 | 1.1078414 | 0.3078986 | 0.782790591 |
| ENSMUSG00000053819 | Camk2d | protein_coding | -0.231492972 | 7.235924366 | 1.107507 | 0.3079693 | 0.782790591 |
| ENSMUSG00000015937 | H2afy | protein_coding | 0.480203897 | 4.298275487 | 1.1071152 | 0.3080522 | 0.782790591 |
| ENSMUSG00000015247 | Nipsnap3b | protein_coding | 0.268086397 | 6.272602591 | 1.1069899 | 0.3080787 | 0.782790591 |
| ENSMUSG00000000171 | Sdhd | protein_coding | 0.220559319 | 9.071525417 | 1.1061984 | 0.3082461 | 0.783022127 |
| ENSMUSG00000057093 | Zfp607b | protein_coding | 0.525009881 | 3.5890363 | 1.1057151 | 0.3083485 | 0.783022127 |
| ENSMUSG00000099696 | 2900052N01Rik | lincRNA | -0.294098763 | 5.699290823 | 1.1055273 | 0.3083882 | 0.783022127 |
| ENSMUSG00000097868 | A330069E16Rik | lincRNA | -0.526523713 | 4.657656107 | 1.1051066 | 0.3085683 | 0.783294354 |
| ENSMUSG00000049971 | Glt1d1 | protein_coding | -0.619142922 | 3.186405473 | 1.1040622 | 0.3086988 | 0.78344066 |
| ENSMUSG00000037852 | Cpe | protein_coding | -0.289070009 | 10.16661404 | 1.1027721 | 0.3089726 | 0.783742687 |
| ENSMUSG00000042390 | Gatad2b | protein_coding | -0.290811277 | 5.723837878 | 1.1024498 | 0.309041 | 0.783742687 |
| ENSMUSG00000029213 | Commd8 | protein_coding | 0.240651632 | 7.193414177 | 1.1023086 | 0.3090711 | 0.783742687 |
| ENSMUSG00000044730 | 9930104L06Rik | processed_transcript | -0.598806795 | 3.161767278 | 1.1014031 | 0.3092635 | 0.783742687 |
| ENSMUSG00000032309 | Fbxo22 | protein_coding | 0.427303159 | 4.51369685 | 1.1012845 | 0.3092887 | 0.783742687 |
| ENSMUSG00000000088 | Cox5a | protein_coding | -0.345680088 | 4.60852763 | 1.1003582 | 0.3094858 | 0.783742687 |
| ENSMUSG00000064372 | mt-Tp | Mt_tRNA | -0.479741474 | 4.066601396 | 1.1000627 | 0.3095487 | 0.783742687 |
| ENSMUSG00000041229 | Phf8 | protein_coding | -0.396255514 | 4.269503119 | 1.0999947 | 0.3095632 | 0.783742687 |
| ENSMUSG00000097503 | 3110045C21Rik | antisense_RNA | 0.811438978 | 2.590778942 | 1.0999581 | 0.309571 | 0.783742687 |
| ENSMUSG00000069565 | Dazap1 | protein_coding | -0.547186893 | 3.087351188 | 1.0995975 | 0.3096478 | 0.783742687 |
| ENSMUSG00000097062 | Gm17586 | lincRNA | 0.50832972 | 3.253532886 | 1.0985701 | 0.3098667 | 0.783742687 |
| ENSMUSG00000043192 | Gm1840 | processed_pseudoge | -1.086119431 | 2.559594542 | 1.0985092 | 0.3099047 | 0.783742687 |
| ENSMUSG00000020783 | Ncbp3 | protein_coding | -0.426712353 | 4.194204079 | 1.097382 | 0.3101202 | 0.783742687 |
| ENSMUSG00000022464 | Slc38a4 | protein_coding | 0.777384742 | 2.634927549 | 1.0967986 | 0.3102448 | 0.783742687 |
| ENSMUSG00000049265 | Kcnk3 | protein_coding | 0.363116511 | 4.463893453 | 1.0967382 | 0.3102577 | 0.783742687 |
| ENSMUSG00000037653 | Kctd8 | protein_coding | -0.506187444 | 3.702296862 | 1.0967341 | 0.3102586 | 0.783742687 |
| ENSMUSG00000037325 | Bbs7 | protein_coding | 0.375147507 | 4.537572092 | 1.0966856 | 0.3102689 | 0.783742687 |
| ENSMUSG00000029551 | Psmg3 | protein_coding | 0.642320362 | 2.713556447 | 1.0965881 | 0.3102897 | 0.783742687 |
| ENSMUSG00000005949 | Ctns | protein_coding | -0.587424938 | 3.660435599 | 1.0965808 | 0.3102913 | 0.783742687 |
| ENSMUSG00000079197 | Psme2 | protein_coding | 0.24257514 | 6.052629701 | 1.0962815 | 0.3103553 | 0.783742687 |
| ENSMUSG00000031575 | Ash2l | protein_coding | -0.384297503 | 4.821138281 | 1.09612 | 0.3103898 | 0.783742687 |
| ENSMUSG00000104864 | Gm43655 | processed_pseudoge | 0.595209924 | 2.811837455 | 1.0959013 | 0.3104365 | 0.783742687 |
| ENSMUSG00000001552 | Jup | protein_coding | -0.369874828 | 4.976714421 | 1.0956316 | 0.3104942 | 0.783742687 |
| ENSMUSG00000004994 | Ccdc130 | protein_coding | 0.319112689 | 5.078397919 | 1.0949131 | 0.3106478 | 0.783906763 |
| ENSMUSG00000033021 | Gmppa | protein_coding | 0.352837346 | 5.686105646 | 1.0932749 | 0.3109986 | 0.783906763 |
| ENSMUSG00000037458 | Azin1 | protein_coding | -0.23969392 | 6.855944283 | 1.0930429 | 0.3110484 | 0.783906763 |
| ENSMUSG00000017132 | Cyth1 | protein_coding | -0.287408624 | 5.502105999 | 1.0928853 | 0.3110821 | 0.783906763 |
| ENSMUSG00000036941 | Elac1 | protein_coding | -0.380686968 | 4.921002979 | 1.0928205 | 0.311096 | 0.783906763 |
| ENSMUSG00000057113 | Npm1 | protein_coding | 0.219112594 | 8.151725743 | 1.0926488 | 0.3111328 | 0.783906763 |
| ENSMUSG00000024735 | Prpf19 | protein_coding | 0.28469259 | 5.857345862 | 1.0925453 | 0.3111551 | 0.783906763 |
| ENSMUSG00000027671 | Actl6a | protein_coding | -0.397613383 | 4.085812605 | 1.091757 | 0.3113242 | 0.783906763 |
| ENSMUSG00000020720 | Psmd12 | protein_coding | 0.222057801 | 8.112982504 | 1.0916138 | 0.3113549 | 0.783906763 |
| ENSMUSG00000013155 | Enkd1 | protein_coding | 0.473159592 | 3.709672084 | 1.0913764 | 0.3114059 | 0.783906763 |
| ENSMUSG00000044014 | Npy5r | protein_coding | 0.778532308 | 2.983356318 | 1.0912292 | 0.3114375 | 0.783906763 |
| ENSMUSG00000043962 | Thrap3 | protein_coding | -0.448979509 | 5.265451221 | 1.0911608 | 0.311477 | 0.783906763 |
| ENSMUSG00000021133 | Susd6 | protein_coding | -0.543585799 | 3.394380395 | 1.0908778 | 0.3115129 | 0.783906763 |
| ENSMUSG00000027774 | Gfm1 | protein_coding | -0.368641113 | 4.742264326 | 1.0905665 | 0.3115798 | 0.783906763 |
| ENSMUSG00000001173 | Ocrl | protein_coding | -0.453967853 | 5.090427923 | 1.0901502 | 0.3116941 | 0.783962976 |
| ENSMUSG00000025428 | Atp5a1 | protein_coding | 0.288997331 | 10.5517582 | 1.0897842 | 0.3117479 | 0.783962976 |
| ENSMUSG00000111977 | AC132265.1 | bidirectional_promote | 0.635296981 | 3.175579365 | 1.0884031 | 0.3120451 | 0.784526768 |
| ENSMUSG00000062866 | Phactr2 | protein_coding | 0.317815126 | 5.45711007 | 1.087187 | 0.3123071 | 0.784922769 |
| ENSMUSG00000040918 | Slc19a2 | protein_coding | 0.594137671 | 3.250004622 | 1.0869943 | 0.3123486 | 0.784922769 |
| ENSMUSG00000036955 | Kif1bp | protein_coding | -0.276758095 | 6.236492011 | 1.0864646 | 0.3124629 | 0.784950288 |
| ENSMUSG00000053768 | Chchd3 | protein_coding | 0.201993144 | 6.995113663 | 1.0857108 | 0.3126255 | 0.784950288 |
| ENSMUSG00000020859 | Spag9 | protein_coding | -0.252879357 | 6.745140716 | 1.0856359 | 0.3126417 | 0.784950288 |
| ENSMUSG00000006411 | Nectin4 | protein_coding | 0.865263901 | 3.299685051 | 1.0914583 | 0.3126601 | 0.784950288 |
| ENSMUSG00000045980 | Tmem104 | protein_coding | 0.549043737 | 3.024620185 | 1.0848597 | 0.3128093 | 0.784950288 |
| ENSMUSG00000016487 | Ppfibp1 | protein_coding | -0.597496001 | 3.142430266 | 1.0847914 | 0.312824 | 0.784950288 |
| ENSMUSG00000025809 | Itgb1 | protein_coding | -0.422709379 | 4.437759961 | 1.0843397 | 0.3129216 | 0.784950288 |
| ENSMUSG00000030443 | Zfp583 | protein_coding | 0.466136614 | 3.713609321 | 1.0842384 | 0.3129435 | 0.784950288 |
| ENSMUSG00000031986 | Sprtn | protein_coding | -0.901496969 | 2.526641802 | 1.0833095 | 0.3131444 | 0.78496808 |
| ENSMUSG00000023286 | Ube2j2 | protein_coding | 0.371172542 | 4.369163291 | 1.0822543 | 0.3133728 | 0.78496808 |
| ENSMUSG00000020376 | Rnf130 | protein_coding | -0.487608397 | 3.970070322 | 1.0822267 | 0.3133788 | 0.78496808 |

| ensembl_gene_id | external_gene_name | gene_biotype | log2 (fold change) | log2 (counts per million) | F-value | p-value | False discovery rate |
|---|---|---|---|---|---|---|---|
| ENSMUSG00000021285 | Ppp1r13b | protein_coding | -0.549610347 | 3.25431394 | 1.0819856 | 0.313431 | 0.78496808 |
| ENSMUSG00000085925 | Rtl1 | protein_coding | -0.475918279 | 4.657096858 | 1.0820636 | 0.3134389 | 0.78496808 |
| ENSMUSG00000028035 | Dnajb4 | protein_coding | -0.26387094 | 6.584486295 | 1.0815494 | 0.3135255 | 0.78496808 |
| ENSMUSG00000066829 | Zfp810 | protein_coding | -0.666478411 | 3.844884453 | 1.0837767 | 0.3135456 | 0.78496808 |
| ENSMUSG00000022012 | Enox1 | protein_coding | -0.464147681 | 4.283157381 | 1.0813466 | 0.3135695 | 0.78496808 |
| ENSMUSG00000026319 | 2310035C23Rik | protein_coding | -0.359005527 | 5.275347878 | 1.0811707 | 0.3136076 | 0.78496808 |
| ENSMUSG00000049680 | Urgcp | protein_coding | 0.368511488 | 4.460458606 | 1.0800721 | 0.3138459 | 0.785221953 |
| ENSMUSG00000084215 | Gm11814 | processed_pseudoge | -0.659990314 | 3.054759638 | 1.0795433 | 0.3139607 | 0.785221953 |
| ENSMUSG00000039967 | Zfp292 | protein_coding | -0.427083354 | 4.74236974 | 1.0789441 | 0.3140909 | 0.785221953 |
| ENSMUSG00000005078 | Jkamp | protein_coding | 0.212360348 | 7.001393794 | 1.0783307 | 0.3142242 | 0.785221953 |
| ENSMUSG00000024101 | Washc1 | protein_coding | 0.317920677 | 5.043493892 | 1.0783188 | 0.3142267 | 0.785221953 |
| ENSMUSG00000018286 | Psmb6 | protein_coding | 0.231266735 | 9.771318944 | 1.0781652 | 0.3142602 | 0.785221953 |
| ENSMUSG00000074754 | Smim26 | protein_coding | 0.250616038 | 6.791577845 | 1.0780083 | 0.3142943 | 0.785221953 |
| ENSMUSG00000029875 | Ccdc184 | protein_coding | 0.366343176 | 6.12034788 | 1.0780116 | 0.3143834 | 0.785221953 |
| ENSMUSG00000040767 | Snrnp25 | protein_coding | 0.239314174 | 6.89303129 | 1.077593 | 0.3143846 | 0.785221953 |
| ENSMUSG00000040652 | Oaz2 | protein_coding | 0.248668259 | 6.202725126 | 1.0773416 | 0.3144393 | 0.785221953 |
| ENSMUSG00000035934 | Pknox2 | protein_coding | 0.323407052 | 5.223197322 | 1.0763777 | 0.3146491 | 0.785369174 |
| ENSMUSG00000064065 | Ipcef1 | protein_coding | 0.657784359 | 2.627359366 | 1.0763399 | 0.3146574 | 0.785369174 |
| ENSMUSG00000086649 | Gm15286 | processed_transcript | 0.651025625 | 2.672795255 | 1.0760645 | 0.3147174 | 0.785369174 |
| ENSMUSG00000015094 | Npdc1 | protein_coding | 0.313533349 | 6.540215449 | 1.0757612 | 0.3148081 | 0.785413422 |
| ENSMUSG00000000805 | Car4 | protein_coding | 0.485786269 | 3.783159165 | 1.0747898 | 0.3149952 | 0.785518431 |
| ENSMUSG00000019969 | Psen1 | protein_coding | 0.29515262 | 5.820382091 | 1.0747848 | 0.3149963 | 0.785518431 |
| ENSMUSG00000042079 | Hnrnpf | protein_coding | -0.571064964 | 2.968149836 | 1.0736541 | 0.3152431 | 0.785951563 |
| ENSMUSG00000033933 | Vhl | protein_coding | -0.366644956 | 5.045703462 | 1.0723406 | 0.3155301 | 0.786316535 |
| ENSMUSG00000036264 | Fstl4 | protein_coding | -0.650645844 | 3.592194445 | 1.0724276 | 0.3155357 | 0.786316535 |
| ENSMUSG00000060938 | Rpl26 | protein_coding | 0.297686725 | 8.23068949 | 1.071413 | 0.3157577 | 0.786432193 |
| ENSMUSG00000006464 | Bbs1 | protein_coding | -0.624901332 | 3.40991514 | 1.0711504 | 0.3157905 | 0.786432193 |
| ENSMUSG00000035805 | Mlc1 | protein_coding | -0.48608134 | 6.046922441 | 1.0807228 | 0.3158136 | 0.786432193 |
| ENSMUSG00000006782 | Cnp | protein_coding | -0.322500556 | 9.811024305 | 1.0702447 | 0.3160135 | 0.786432193 |
| ENSMUSG00000110723 | AC153912.1 | protein_coding | -0.734243138 | 2.861603887 | 1.0696569 | 0.3161176 | 0.786432193 |
| ENSMUSG00000067071 | Hes6 | protein_coding | -0.515552895 | 3.233856932 | 1.0695846 | 0.3161335 | 0.786432193 |
| ENSMUSG00000027598 | Itch | protein_coding | -0.344478485 | 4.714672349 | 1.069496 | 0.3161529 | 0.786432193 |
| ENSMUSG00000048997 | Atxn7l2 | protein_coding | -0.626919103 | 2.92928557 | 1.0689483 | 0.316273 | 0.786432193 |
| ENSMUSG00000027132 | Katnbl1 | protein_coding | 0.500967636 | 3.374344132 | 1.0687894 | 0.3163079 | 0.786432193 |
| ENSMUSG00000025511 | Tspan4 | protein_coding | 0.434506127 | 4.134808784 | 1.0687638 | 0.3163135 | 0.786432193 |
| ENSMUSG00000018372 | Cep95 | protein_coding | 0.39629582 | 3.87349461 | 1.065707 | 0.3169852 | 0.787809712 |
| ENSMUSG00000026914 | Psmd14 | protein_coding | 0.232738041 | 8.344813924 | 1.065572 | 0.3170149 | 0.787809712 |
| ENSMUSG00000045215 | Asxl3 | protein_coding | -0.547749153 | 3.130231723 | 1.0652427 | 0.3170874 | 0.787809712 |
| ENSMUSG00000021177 | Tdp1 | protein_coding | 0.393611025 | 4.693445551 | 1.0647479 | 0.3171963 | 0.787811671 |
| ENSMUSG00000022089 | Bin3 | protein_coding | -0.439292842 | 3.915490179 | 1.0645737 | 0.3172347 | 0.787811671 |
| ENSMUSG00000026709 | Dars2 | protein_coding | 0.407612673 | 4.442531 | 1.0633328 | 0.3175082 | 0.787909544 |
| ENSMUSG00000029478 | Ncor2 | protein_coding | -0.489234282 | 3.755960441 | 1.0630487 | 0.3175709 | 0.787909544 |
| ENSMUSG00000025213 | Kazald1 | protein_coding | -0.684786507 | 2.824612093 | 1.0630338 | 0.3175742 | 0.787909544 |
| ENSMUSG00000006342 | Susd2 | protein_coding | 0.51116647 | 4.02023654 | 1.0626189 | 0.3176657 | 0.787909544 |
| ENSMUSG00000047766 | Lrrc49 | protein_coding | 0.27692441 | 6.572946789 | 1.0624838 | 0.3176956 | 0.787909544 |
| ENSMUSG00000024137 | E4f1 | protein_coding | -0.606776994 | 3.098904098 | 1.0624014 | 0.3177137 | 0.787909544 |
| ENSMUSG00000006191 | Cdkal1 | protein_coding | -0.380824777 | 4.641648167 | 1.0614046 | 0.3179339 | 0.78820428 |
| ENSMUSG00000022505 | Emp2 | protein_coding | -0.672053338 | 3.382255567 | 1.0613107 | 0.3179792 | 0.78820428 |
| ENSMUSG00000028676 | Srsf10 | protein_coding | -0.281383638 | 5.481555705 | 1.0605698 | 0.3181184 | 0.788276049 |
| ENSMUSG00000019055 | Plod1 | protein_coding | -0.474887248 | 3.98051833 | 1.0601404 | 0.3182134 | 0.788276049 |
| ENSMUSG00000041959 | S100a10 | protein_coding | -0.25258377 | 6.685286142 | 1.0600057 | 0.3182432 | 0.788276049 |
| ENSMUSG00000001280 | Sp1 | protein_coding | -0.460396325 | 3.948194956 | 1.059661 | 0.3183195 | 0.788276049 |
| ENSMUSG00000019837 | Gtf3c6 | protein_coding | -0.253113068 | 6.844510486 | 1.0594118 | 0.3183747 | 0.788276049 |
| ENSMUSG00000048264 | Dip2c | protein_coding | -0.330966235 | 4.975060271 | 1.0581057 | 0.318664 | 0.788439781 |
| ENSMUSG00000070372 | Capza1 | protein_coding | -0.444557923 | 3.747598936 | 1.0580638 | 0.3186733 | 0.788439781 |
| ENSMUSG00000039328 | Rnf122 | protein_coding | -0.414479757 | 4.062670197 | 1.0571706 | 0.3188714 | 0.788439781 |
| ENSMUSG00000036093 | Arl5a | protein_coding | 0.364559461 | 4.645275477 | 1.0563882 | 0.3190451 | 0.788439781 |
| ENSMUSG00000019971 | Cep290 | protein_coding | -0.565169252 | 3.458790192 | 1.056183 | 0.3190906 | 0.788439781 |
| ENSMUSG00000036371 | Serbp1 | protein_coding | -0.213583594 | 7.425135801 | 1.0551738 | 0.3193149 | 0.788439781 |
| ENSMUSG00000040035 | Disp2 | protein_coding | 0.378784471 | 7.018399155 | 1.0615678 | 0.3193877 | 0.788439781 |
| ENSMUSG00000006005 | Tpr | protein_coding | -0.332638764 | 5.924069098 | 1.0546402 | 0.3194335 | 0.788439781 |
| ENSMUSG00000102674 | 8030442B05Rik | lincRNA | -0.612187466 | 3.225545928 | 1.0544583 | 0.319474 | 0.788439781 |
| ENSMUSG00000000420 | Galnt1 | protein_coding | -0.40074567 | 4.118701884 | 1.0542533 | 0.3195196 | 0.788439781 |
| ENSMUSG00000020462 | Cfap36 | protein_coding | 0.219668511 | 7.174457757 | 1.0535691 | 0.3196719 | 0.788439781 |
| ENSMUSG00000052133 | Sema5b | protein_coding | -0.668154921 | 2.685225902 | 1.0535016 | 0.3196869 | 0.788439781 |
| ENSMUSG00000048371 | Pdp2 | protein_coding | -0.498535806 | 5.615301896 | 1.0600779 | 0.3197161 | 0.788439781 |
| ENSMUSG00000095567 | Noc2l | protein_coding | -0.38123637 | 4.321235495 | 1.0533305 | 0.319725 | 0.788439781 |
| ENSMUSG00000028293 | Slc35a1 | protein_coding | 0.318207319 | 5.190766832 | 1.0524617 | 0.3199186 | 0.788439781 |
| ENSMUSG00000053253 | Ndfip2 | protein_coding | -0.27130599 | 5.975936153 | 1.0524475 | 0.3199218 | 0.788439781 |
| ENSMUSG00000009112 | Bcl2l13 | protein_coding | 0.447077365 | 4.002064157 | 1.0519317 | 0.3200368 | 0.788439781 |
| ENSMUSG00000019577 | Pdk4 | protein_coding | -0.843898641 | 2.415017495 | 1.0516147 | 0.3201075 | 0.788439781 |
| ENSMUSG00000050132 | Sarm1 | protein_coding | -0.58048711 | 3.056241751 | 1.0511442 | 0.3202125 | 0.788439781 |
| ENSMUSG00000025094 | Slc18a2 | protein_coding | -0.655557653 | 3.023974236 | 1.0509064 | 0.3202655 | 0.788439781 |
| ENSMUSG00000020982 | Nemf | protein_coding | -0.242192392 | 6.868900378 | 1.0507826 | 0.3202932 | 0.788439781 |
| ENSMUSG00000020847 | Rph3al | protein_coding | 0.564792401 | 3.492750588 | 1.0499482 | 0.3204795 | 0.788439781 |
| ENSMUSG00000061306 | Slc38a10 | protein_coding | 0.409804158 | 5.193890745 | 1.0492885 | 0.3206514 | 0.788439781 |
| ENSMUSG00000002504 | Slc9a3r2 | protein_coding | 0.330710688 | 4.740477789 | 1.0490606 | 0.320678 | 0.788439781 |
| ENSMUSG00000028986 | Klhl7 | protein_coding | -0.269725622 | 6.381126515 | 1.0489502 | 0.3207026 | 0.788439781 |

| ensembl_gene_id | external_gene_name | gene_biotype | log2 (fold change) | log2 (counts per million) | F-value | p-value | False discovery rate |
|---|---|---|---|---|---|---|---|
| ENSMUSG00000029385 | Ccng2 | protein_coding | -0.421374215 | 4.500731803 | 1.0487116 | 0.320756 | 0.788439781 |
| ENSMUSG00000061080 | Lsamp | protein_coding | 0.193871122 | 7.72007222 | 1.0485324 | 0.3207961 | 0.788439781 |
| ENSMUSG00000071659 | Hnrnpul2 | protein_coding | -0.319361413 | 5.875006162 | 1.0484782 | 0.3208082 | 0.788439781 |
| ENSMUSG00000035189 | Ano4 | protein_coding | -0.388454355 | 4.021598688 | 1.0480586 | 0.3209022 | 0.788439781 |
| ENSMUSG00000028128 | F3 | protein_coding | -0.419974805 | 5.787861744 | 1.050324 | 0.3210184 | 0.788439781 |
| ENSMUSG00000042606 | Hirip3 | protein_coding | 0.398759807 | 4.380832311 | 1.0473658 | 0.3210573 | 0.788439781 |
| ENSMUSG00000059248 | 9-Sep | protein_coding | 0.519302956 | 5.15528298 | 1.0528072 | 0.32106 | 0.788439781 |
| ENSMUSG00000041220 | Elovl6 | protein_coding | -0.317416412 | 5.424769524 | 1.0470077 | 0.3211375 | 0.788439781 |
| ENSMUSG00000004933 | Matk | protein_coding | 0.257725835 | 7.258529824 | 1.0467819 | 0.3211881 | 0.788439781 |
| ENSMUSG00000074001 | Klhl40 | protein_coding | -1.042105544 | 2.161219232 | 1.0467572 | 0.3211937 | 0.788439781 |
| ENSMUSG00000025436 | Atp23 | protein_coding | 0.581899784 | 3.030248704 | 1.0463666 | 0.3212812 | 0.788439781 |
| ENSMUSG00000020184 | Mdm2 | protein_coding | -0.345913743 | 4.96848928 | 1.0458221 | 0.3214034 | 0.788439781 |
| ENSMUSG00000021715 | Cwc27 | protein_coding | -0.516415881 | 3.686163553 | 1.0457443 | 0.3214208 | 0.788439781 |
| ENSMUSG00000067199 | Frat1 | protein_coding | 0.344529397 | 7.740331066 | 1.0493818 | 0.3215518 | 0.788439781 |
| ENSMUSG00000020770 | Unk | protein_coding | -0.768946197 | 3.006113874 | 1.0450755 | 0.3215709 | 0.788439781 |
| ENSMUSG00000035885 | Cox8a | protein_coding | 0.402580798 | 9.631013762 | 1.0533178 | 0.3216557 | 0.788439781 |
| ENSMUSG00000090150 | Acad11 | protein_coding | -0.56034416 | 2.967308539 | 1.0446407 | 0.3216686 | 0.788439781 |
| ENSMUSG00000029253 | Cenpc1 | protein_coding | -0.408049803 | 4.863560757 | 1.0444283 | 0.3217162 | 0.788439781 |
| ENSMUSG00000028483 | Snapc3 | protein_coding | 0.281561788 | 5.714900084 | 1.044415 | 0.3217193 | 0.788439781 |
| ENSMUSG00000040842 | Szrd1 | protein_coding | -0.33377016 | 5.368235244 | 1.0435244 | 0.3219194 | 0.788439781 |
| ENSMUSG00000000959 | Oxa1l | protein_coding | 0.264620551 | 6.370403043 | 1.0432125 | 0.3219896 | 0.788439781 |
| ENSMUSG00000113722 | AC140477.3 | lincRNA | 0.335215052 | 4.754180436 | 1.0431559 | 0.3220023 | 0.788439781 |
| ENSMUSG00000043398 | Gpr135 | protein_coding | 0.569351914 | 3.060723677 | 1.0430484 | 0.3220265 | 0.788439781 |
| ENSMUSG00000024533 | Spire1 | protein_coding | -0.321634259 | 5.694342266 | 1.0430166 | 0.3220336 | 0.788439781 |
| ENSMUSG00000002320 | Tm9sf1 | protein_coding | 0.530096244 | 3.637929839 | 1.0412269 | 0.3224366 | 0.788592686 |
| ENSMUSG00000056073 | Grik2 | protein_coding | -0.330213673 | 5.420068935 | 1.0411265 | 0.3224592 | 0.788592686 |
| ENSMUSG00000034390 | Cmip | protein_coding | 0.309915277 | 7.457598778 | 1.0415909 | 0.322471 | 0.788592686 |
| ENSMUSG00000048000 | Gigyf2 | protein_coding | -0.2405151 | 6.073082081 | 1.0410198 | 0.3224833 | 0.788592686 |
| ENSMUSG00000085885 | Gm11906 | processed_transcript | -0.37719663 | 4.577046964 | 1.0409383 | 0.3225016 | 0.788592686 |
| ENSMUSG00000028653 | Trit1 | protein_coding | 0.382381663 | 4.177133886 | 1.0402788 | 0.3226503 | 0.788592686 |
| ENSMUSG00000027751 | Supt20 | protein_coding | -0.471059685 | 4.469264427 | 1.0401063 | 0.3226893 | 0.788592686 |
| ENSMUSG00000038489 | Polr2l | protein_coding | 0.308350068 | 4.985064175 | 1.0397067 | 0.3227794 | 0.788592686 |
| ENSMUSG00000021000 | Ctage5 | protein_coding | -0.455515687 | 4.251313208 | 1.039545 | 0.3228159 | 0.788592686 |
| ENSMUSG00000043154 | Ppp2r3a | protein_coding | -0.373276477 | 4.875235619 | 1.0392015 | 0.3228935 | 0.788592686 |
| ENSMUSG00000039801 | 2410089E03Rik | protein_coding | -0.580116212 | 3.251676142 | 1.0386533 | 0.3230173 | 0.788592686 |
| ENSMUSG00000019891 | Dcbld1 | protein_coding | -0.846562202 | 2.688547111 | 1.0385497 | 0.3230407 | 0.788592686 |
| ENSMUSG00000000902 | Smarcb1 | protein_coding | 0.261207585 | 7.733330558 | 1.0378148 | 0.3232068 | 0.788592686 |
| ENSMUSG00000031513 | Leprotl1 | protein_coding | -0.319010123 | 6.383759213 | 1.0378944 | 0.3232131 | 0.788592686 |
| ENSMUSG00000021782 | Dlg5 | protein_coding | -0.48400907 | 3.431185895 | 1.0377848 | 0.3232136 | 0.788592686 |
| ENSMUSG00000032434 | Cmtm6 | protein_coding | -0.780854067 | 2.968209498 | 1.0371043 | 0.3233675 | 0.788592686 |
| ENSMUSG00000049670 | Morn4 | protein_coding | 0.198086504 | 7.601645334 | 1.0370955 | 0.3233695 | 0.788592686 |
| ENSMUSG00000039671 | Zmynd8 | protein_coding | -0.226524023 | 6.581414069 | 1.0367549 | 0.3234466 | 0.788592686 |
| ENSMUSG00000030393 | Zik1 | protein_coding | -0.529216132 | 3.691285119 | 1.0363951 | 0.3235281 | 0.788592686 |
| ENSMUSG00000020014 | Cfap54 | protein_coding | -0.726363482 | 2.646951021 | 1.0360155 | 0.323614 | 0.788592686 |
| ENSMUSG00000048310 | Pskh1 | protein_coding | -0.673640817 | 2.831746232 | 1.0359177 | 0.3236362 | 0.788592686 |
| ENSMUSG00000071398 | 2410004P03Rik | protein_coding | -0.5595937 | 3.660535499 | 1.0350941 | 0.3238228 | 0.788617249 |
| ENSMUSG00000022890 | Atp5j | protein_coding | 0.246412177 | 10.19202003 | 1.0350148 | 0.3238408 | 0.788617249 |
| ENSMUSG00000038068 | Rnf144b | protein_coding | -0.501851782 | 3.326195331 | 1.0340078 | 0.3240692 | 0.788617249 |
| ENSMUSG00000061130 | Ppm1b | protein_coding | -0.308339396 | 5.23265105 | 1.0339573 | 0.3240807 | 0.788617249 |
| ENSMUSG00000021508 | Cxcl14 | protein_coding | -0.524555572 | 4.651294608 | 1.0356972 | 0.3241219 | 0.788617249 |
| ENSMUSG00000028139 | Riiad1 | protein_coding | -0.34232144 | 5.2233371 | 1.0335076 | 0.3241828 | 0.788617249 |
| ENSMUSG00000030327 | Necap1 | protein_coding | 0.286450619 | 9.043497889 | 1.0334606 | 0.3242177 | 0.788617249 |
| ENSMUSG00000039183 | Nubp2 | protein_coding | 0.219255771 | 6.605297368 | 1.0332867 | 0.3242329 | 0.788617249 |
| ENSMUSG00000074457 | S100a16 | protein_coding | -0.271957253 | 8.358842546 | 1.0327346 | 0.3243584 | 0.788657029 |
| ENSMUSG00000062232 | Rapgef2 | protein_coding | -0.330758631 | 5.343580898 | 1.0318239 | 0.3245654 | 0.788657029 |
| ENSMUSG00000006931 | P3h4 | protein_coding | -0.435865941 | 3.879283003 | 1.0316741 | 0.3245995 | 0.788657029 |
| ENSMUSG00000037111 | Setd7 | protein_coding | -0.324817475 | 4.984668212 | 1.0316654 | 0.3246015 | 0.788657029 |
| ENSMUSG00000029426 | Scarb2 | protein_coding | -0.342853391 | 5.566327779 | 1.0316014 | 0.324616 | 0.788657029 |
| ENSMUSG00000029145 | Eif2b4 | protein_coding | 0.291184784 | 5.925867278 | 1.0307379 | 0.3248125 | 0.788665548 |
| ENSMUSG00000024943 | Smc5 | protein_coding | -0.489567935 | 4.320390629 | 1.0305514 | 0.324855 | 0.788665548 |
| ENSMUSG00000048170 | Mcmbp | protein_coding | -0.316727048 | 5.869126509 | 1.0304108 | 0.324887 | 0.788665548 |
| ENSMUSG00000029723 | Tsc22d4 | protein_coding | -0.397652589 | 7.49352734 | 1.0394264 | 0.3249129 | 0.788665548 |
| ENSMUSG00000001794 | Capns1 | protein_coding | 0.21053269 | 7.23377314 | 1.0298992 | 0.3250036 | 0.788707634 |
| ENSMUSG00000004568 | Arhgef18 | protein_coding | -0.571388318 | 3.849706534 | 1.0289802 | 0.3252373 | 0.789096682 |
| ENSMUSG00000063568 | Jazf1 | protein_coding | 0.330132176 | 4.730185123 | 1.0281036 | 0.3254131 | 0.789334431 |
| ENSMUSG00000024472 | Dcp2 | protein_coding | 0.3770026 | 4.493941162 | 1.0278015 | 0.3254821 | 0.789334431 |
| ENSMUSG00000020701 | Tmem132e | protein_coding | -0.550971593 | 3.462792717 | 1.0253241 | 0.3260485 | 0.790509086 |
| ENSMUSG00000046111 | Cep295 | protein_coding | -0.604246241 | 2.984215369 | 1.0250402 | 0.3261135 | 0.790509086 |
| ENSMUSG00000020091 | Eif4ebp2 | protein_coding | 0.329355073 | 4.73625418 | 1.0245373 | 0.3262287 | 0.79061009 |
| ENSMUSG00000068966 | Zbtb34 | protein_coding | 0.623379034 | 2.946997983 | 1.0234684 | 0.3264737 | 0.79101002 |
| ENSMUSG00000051041 | Olfml1 | protein_coding | -0.370429418 | 5.013972872 | 1.0231756 | 0.3265408 | 0.79101002 |
| ENSMUSG00000027716 | Trpc3 | protein_coding | -0.752970637 | 2.94328663 | 1.0223969 | 0.3267195 | 0.79126465 |
| ENSMUSG00000029053 | Prkcz | protein_coding | 0.22148873 | 7.089822294 | 1.021612 | 0.3268998 | 0.791462666 |
| ENSMUSG00000025993 | Slc40a1 | protein_coding | -0.875993398 | 2.315840869 | 1.0209388 | 0.3270545 | 0.791462666 |
| ENSMUSG00000038827 | Fam206a | protein_coding | 0.332564481 | 4.517711697 | 1.0207821 | 0.3270905 | 0.791462666 |
| ENSMUSG00000061518 | Cox5b | protein_coding | 0.283188638 | 8.013800626 | 1.020517 | 0.3271756 | 0.791462666 |
| ENSMUSG00000034758 | Tle6 | protein_coding | 1.186174235 | 2.579938696 | 1.0244139 | 0.3274594 | 0.791462666 |
| ENSMUSG00000032531 | Amotl2 | protein_coding | -0.703321359 | 3.205997303 | 1.0185255 | 0.327634 | 0.791462666 |

| ensembl_gene_id | external_gene_name | gene_biotype | log2 (fold change) | log2 (counts per million) | F-value | p-value | False discovery rate |
|---|---|---|---|---|---|---|---|
| ENSMUSG00000034064 | Poglut1 | protein_coding | 0.28748242 | 6.366769192 | 1.0183918 | 0.3276408 | 0.791462666 |
| ENSMUSG00000070570 | Slc17a7 | protein_coding | -0.942854222 | 2.764082517 | 1.0181484 | 0.327721 | 0.791462666 |
| ENSMUSG00000036712 | Cyld | protein_coding | -0.232187403 | 6.611239859 | 1.0178128 | 0.3277743 | 0.791462666 |
| ENSMUSG00000027678 | Ncoa3 | protein_coding | -0.420910634 | 4.86053662 | 1.0178039 | 0.3277763 | 0.791462666 |
| ENSMUSG00000014164 | Klhl3 | protein_coding | -0.359008486 | 6.45429583 | 1.0201648 | 0.3278466 | 0.791462666 |
| ENSMUSG00000042670 | Immp1l | protein_coding | -0.298083797 | 5.770079126 | 1.017386 | 0.3278727 | 0.791462666 |
| ENSMUSG00000026827 | Gpd2 | protein_coding | 0.228300044 | 6.542160794 | 1.0172817 | 0.3278968 | 0.791462666 |
| ENSMUSG00000007812 | Zfp655 | protein_coding | 0.293603976 | 5.639643847 | 1.0169192 | 0.3279804 | 0.791462666 |
| ENSMUSG00000039128 | Cdc123 | protein_coding | 0.225927264 | 8.75466243 | 1.0167314 | 0.3280238 | 0.791462666 |
| ENSMUSG00000028145 | Them4 | protein_coding | 0.257246385 | 6.586980391 | 1.0161316 | 0.3281623 | 0.791462666 |
| ENSMUSG00000025104 | Hdgfl3 | protein_coding | -0.277796943 | 6.422532832 | 1.0159906 | 0.3281949 | 0.791462666 |
| ENSMUSG00000004798 | Ulk2 | protein_coding | -0.350284082 | 5.441127595 | 1.0157167 | 0.3282582 | 0.791462666 |
| ENSMUSG00000032412 | Atp1b3 | protein_coding | -0.312619771 | 6.144788157 | 1.015415 | 0.3283279 | 0.791462666 |
| ENSMUSG00000005699 | Pard6a | protein_coding | 0.245885792 | 6.116825136 | 1.0152246 | 0.3283719 | 0.791462666 |
| ENSMUSG00000059447 | Hadhb | protein_coding | -0.394709949 | 3.844665674 | 1.0152188 | 0.3283733 | 0.791462666 |
| ENSMUSG00000003464 | Pex19 | protein_coding | 0.256242829 | 7.878205664 | 1.0146844 | 0.3284968 | 0.791462666 |
| ENSMUSG00000030203 | Dusp16 | protein_coding | -0.5649337 | 3.462242681 | 1.0142993 | 0.3285859 | 0.791462666 |
| ENSMUSG00000029165 | Agbl5 | protein_coding | 0.590577124 | 2.977751785 | 1.0140448 | 0.3286448 | 0.791462666 |
| ENSMUSG00000020124 | Usp15 | protein_coding | -0.253727348 | 5.88853095 | 1.0137765 | 0.328707 | 0.791462666 |
| ENSMUSG00000021022 | Ppp2r3c | protein_coding | 0.333204333 | 4.722584434 | 1.0137418 | 0.328715 | 0.791462666 |
| ENSMUSG00000032040 | Dcps | protein_coding | 0.354391056 | 4.857427172 | 1.0130383 | 0.328878 | 0.791638292 |
| ENSMUSG00000021549 | Rasa1 | protein_coding | -0.443526026 | 3.628780971 | 1.0125603 | 0.3289887 | 0.791638292 |
| ENSMUSG00000072623 | Zfp9 | protein_coding | -0.53456885 | 4.1590308 | 1.0125773 | 0.3290088 | 0.791638292 |
| ENSMUSG00000038582 | Pptc7 | protein_coding | -0.405085522 | 4.48550398 | 1.0119097 | 0.3291396 | 0.791775881 |
| ENSMUSG00000054675 | Tmem119 | protein_coding | -0.650800692 | 3.114579512 | 1.0109958 | 0.3293517 | 0.791897546 |
| ENSMUSG00000053119 | Chmp3 | protein_coding | 0.220438127 | 7.855505258 | 1.0109034 | 0.3293731 | 0.791897546 |
| ENSMUSG00000030629 | Zfand6 | protein_coding | -0.225118638 | 7.01366206 | 1.0107399 | 0.3294111 | 0.791897546 |
| ENSMUSG00000066735 | Vkorc1l1 | protein_coding | -0.346411283 | 4.973310593 | 1.0103816 | 0.3294943 | 0.791920557 |
| ENSMUSG00000047907 | Tshz2 | protein_coding | 0.356562848 | 4.067112296 | 1.0098341 | 0.3296216 | 0.791973303 |
| ENSMUSG00000020027 | Socs2 | protein_coding | -0.508650626 | 3.119462797 | 1.0088101 | 0.3298597 | 0.791973303 |
| ENSMUSG00000037818 | Abhd18 | protein_coding | -0.549924225 | 3.23669657 | 1.0080698 | 0.330032 | 0.791973303 |
| ENSMUSG00000063430 | Wscd2 | protein_coding | 0.47391647 | 3.351754842 | 1.0079986 | 0.3300486 | 0.791973303 |
| ENSMUSG00000020029 | Nudt4 | protein_coding | 0.241782048 | 6.206252499 | 1.0078963 | 0.3300724 | 0.791973303 |
| ENSMUSG00000041650 | Pcca | protein_coding | -0.422729659 | 4.350311959 | 1.0078539 | 0.3300823 | 0.791973303 |
| ENSMUSG00000045211 | Nudt18 | protein_coding | 0.303182502 | 5.721261591 | 1.007352 | 0.3301992 | 0.791973303 |
| ENSMUSG00000050854 | Tmem125 | protein_coding | -0.581160307 | 3.590188888 | 1.0066445 | 0.3303642 | 0.791973303 |
| ENSMUSG00000039976 | Tbc1d16 | protein_coding | -0.367984047 | 4.131890637 | 1.006357 | 0.3304312 | 0.791973303 |
| ENSMUSG00000007594 | Hapln4 | protein_coding | 0.344189549 | 5.143696213 | 1.0053054 | 0.3306766 | 0.791973303 |
| ENSMUSG00000030604 | Zfp626 | protein_coding | -0.713317094 | 2.699631647 | 1.0050971 | 0.3307253 | 0.791973303 |
| ENSMUSG00000020792 | Exoc7 | protein_coding | 0.273075973 | 7.062563813 | 1.0048634 | 0.3307798 | 0.791973303 |
| ENSMUSG00000015766 | Eps8 | protein_coding | -0.436723823 | 4.559607813 | 1.0048252 | 0.3307888 | 0.791973303 |
| ENSMUSG00000028990 | Lzic | protein_coding | -0.429093837 | 4.126036374 | 1.0038742 | 0.3310111 | 0.791973303 |
| ENSMUSG00000057963 | Itpk1 | protein_coding | 0.280282313 | 5.643085714 | 1.0034245 | 0.3311162 | 0.791973303 |
| ENSMUSG00000060904 | Arl1 | protein_coding | 0.211485949 | 6.881057215 | 1.0029228 | 0.3312336 | 0.791973303 |
| ENSMUSG00000014602 | Kif1a | protein_coding | 0.225335465 | 8.857309886 | 1.0027815 | 0.3312667 | 0.791973303 |
| ENSMUSG00000030718 | Ppme1 | protein_coding | 0.317809945 | 7.3473376 | 1.0045465 | 0.3314634 | 0.791973303 |
| ENSMUSG00000001248 | Gramd1a | protein_coding | 0.430460411 | 4.273829548 | 1.001921 | 0.3314682 | 0.791973303 |
| ENSMUSG00000027568 | Ntsr1 | protein_coding | 0.707995377 | 2.674081521 | 1.0013833 | 0.3315942 | 0.791973303 |
| ENSMUSG00000028557 | Rnf11 | protein_coding | 0.19933693 | 7.771785049 | 1.0010796 | 0.3316654 | 0.791973303 |
| ENSMUSG00000036678 | Aaas | protein_coding | 0.339881025 | 5.155665353 | 1.0010691 | 0.3316679 | 0.791973303 |
| ENSMUSG00000030471 | Zdhhc13 | protein_coding | -0.493671719 | 3.432563223 | 1.0009948 | 0.3316853 | 0.791973303 |
| ENSMUSG00000030649 | Anapc15 | protein_coding | 0.450295756 | 3.334285731 | 1.0004005 | 0.3318247 | 0.791973303 |
| ENSMUSG00000025173 | Wdr45b | protein_coding | 0.22461068 | 6.486144189 | 1.0001466 | 0.3318843 | 0.791973303 |
| ENSMUSG00000034083 | Ccdc174 | protein_coding | -0.245058841 | 5.939229856 | 0.9999518 | 0.33193 | 0.791973303 |
| ENSMUSG00000031576 | Kcnu1 | protein_coding | 0.833846727 | 2.930505954 | 1.0000081 | 0.3319407 | 0.791973303 |
| ENSMUSG00000036885 | Arhgef26 | protein_coding | -0.560714262 | 3.349895326 | 0.9997556 | 0.3319761 | 0.791973303 |
| ENSMUSG00000031024 | St5 | protein_coding | -0.558347753 | 3.050117708 | 0.9994734 | 0.3320423 | 0.791973303 |
| ENSMUSG00000069814 | Ccdc92b | protein_coding | -0.400847703 | 4.020722003 | 0.9989829 | 0.3321576 | 0.791973303 |
| ENSMUSG00000017390 | Aldoc | protein_coding | 0.22493002 | 10.98811899 | 0.998666 | 0.3322321 | 0.791973303 |
| ENSMUSG00000054226 | Tprkb | protein_coding | -0.199719968 | 7.424712086 | 0.998629 | 0.3322407 | 0.791973303 |
| ENSMUSG00000040359 | Ufl1 | protein_coding | -0.549635819 | 2.978506641 | 0.9983049 | 0.332317 | 0.791973303 |
| ENSMUSG00000047635 | 2810006K23Rik | protein_coding | 0.321890436 | 5.227095628 | 0.9980738 | 0.3323713 | 0.791973303 |
| ENSMUSG00000026510 | Trp53bp2 | protein_coding | -0.382292146 | 4.712471166 | 0.997856 | 0.3324225 | 0.791973303 |
| ENSMUSG00000039844 | Rapgef1 | protein_coding | -0.379440481 | 4.591278604 | 0.9978128 | 0.3324327 | 0.791973303 |
| ENSMUSG00000060962 | Dmkn | protein_coding | -0.549231523 | 4.473782362 | 0.9999953 | 0.332528 | 0.791973303 |
| ENSMUSG00000020777 | Acox1 | protein_coding | -0.261256694 | 6.792100537 | 0.9967937 | 0.3326725 | 0.791973303 |
| ENSMUSG00000004317 | Clcn5 | protein_coding | -0.409434372 | 4.298924838 | 0.9964207 | 0.3327604 | 0.791973303 |
| ENSMUSG00000031754 | Nudt21 | protein_coding | -0.379115074 | 3.847981517 | 0.9949859 | 0.3330987 | 0.791973303 |
| ENSMUSG00000042211 | Fbxo38 | protein_coding | -0.295736536 | 5.164586381 | 0.9949091 | 0.3331168 | 0.791973303 |
| ENSMUSG00000038619 | Ensa | protein_coding | 0.215824957 | 9.758406998 | 0.9941916 | 0.3332861 | 0.791973303 |
| ENSMUSG00000037519 | Ppfia1 | protein_coding | -0.456227263 | 3.816122203 | 0.99407 | 0.3333148 | 0.791973303 |
| ENSMUSG00000032011 | Thy1 | protein_coding | 0.274810419 | 7.286844666 | 0.9940287 | 0.3333484 | 0.791973303 |
| ENSMUSG00000030595 | Nfkbib | protein_coding | 0.333610582 | 5.707432319 | 0.9937237 | 0.3333966 | 0.791973303 |
| ENSMUSG00000025950 | Idh1 | protein_coding | -0.222178689 | 6.523686016 | 0.9932921 | 0.3334986 | 0.791973303 |
| ENSMUSG00000047379 | B4gat1 | protein_coding | 0.284554134 | 7.438646218 | 0.9930738 | 0.333574 | 0.791973303 |
| ENSMUSG00000021592 | Arsk | protein_coding | 0.632190915 | 3.16525007 | 0.99295 | 0.3335795 | 0.791973303 |
| ENSMUSG00000048581 | E130311K13Rik | protein_coding | -0.817216203 | 2.635295738 | 0.9928373 | 0.3336061 | 0.791973303 |
| ENSMUSG00000028234 | Rps20 | protein_coding | 0.279771752 | 7.008257084 | 0.9925356 | 0.3337012 | 0.791973303 |

| ensembl_gene_id | external_gene_name | gene_biotype | log2 (fold change) | log2 (counts per million) | F-value | p-value | False discovery rate |
|---|---|---|---|---|---|---|---|
| ENSMUSG00000051335 | Gfod1 | protein_coding | -0.413388316 | 3.964177096 | 0.9921409 | 0.3337708 | 0.791973303 |
| ENSMUSG00000001416 | Cct3 | protein_coding | 0.227810501 | 7.875360436 | 0.9921009 | 0.3337803 | 0.791973303 |
| ENSMUSG00000000711 | Rab5b | protein_coding | 0.359423775 | 6.431023966 | 0.9959782 | 0.3337867 | 0.791973303 |
| ENSMUSG00000048965 | Mrgpre | protein_coding | 0.399408965 | 4.43760468 | 0.9919557 | 0.3338146 | 0.791973303 |
| ENSMUSG00000025487 | Psmd13 | protein_coding | 0.209082899 | 7.360820135 | 0.9917744 | 0.3338575 | 0.791973303 |
| ENSMUSG00000022498 | Txndc11 | protein_coding | -0.592730479 | 2.905277887 | 0.9916252 | 0.3338929 | 0.791973303 |
| ENSMUSG00000021040 | Slirp | protein_coding | 0.365259245 | 4.494074905 | 0.9913186 | 0.3339655 | 0.791973303 |
| ENSMUSG00000029659 | Medag | protein_coding | -0.994658876 | 2.621532017 | 0.991327 | 0.3339872 | 0.791973303 |
| ENSMUSG00000021149 | Gtpbp4 | protein_coding | 0.240799024 | 6.593417034 | 0.9911156 | 0.3340135 | 0.791973303 |
| ENSMUSG00000039070 | Cpa4 | protein_coding | -0.98485424 | 2.618529772 | 0.9911157 | 0.3340372 | 0.791973303 |
| ENSMUSG00000019861 | Gopc | protein_coding | -0.310562165 | 4.726471528 | 0.9909242 | 0.3340589 | 0.791973303 |
| ENSMUSG00000027649 | Ctnnbl1 | protein_coding | 0.284429031 | 6.228980183 | 0.9906956 | 0.334113 | 0.791973303 |
| ENSMUSG00000049517 | Rps23 | protein_coding | 0.525333527 | 3.084731653 | 0.9902091 | 0.3342283 | 0.791973303 |
| ENSMUSG00000052406 | Rexo4 | protein_coding | 0.354335397 | 5.485134177 | 0.9900372 | 0.3342691 | 0.791973303 |
| ENSMUSG00000031010 | Usp9x | protein_coding | -0.372541008 | 4.380907038 | 0.9898915 | 0.3343036 | 0.791973303 |
| ENSMUSG00000026035 | Ppil3 | protein_coding | 0.28390514 | 6.186644103 | 0.9892502 | 0.3344557 | 0.792131506 |
| ENSMUSG00000023977 | Ubr2 | protein_coding | -0.460868429 | 3.830373238 | 0.9889889 | 0.3345177 | 0.792131506 |
| ENSMUSG00000041164 | Zmiz2 | protein_coding | 0.246687768 | 6.669697262 | 0.9885102 | 0.3346314 | 0.792226164 |
| ENSMUSG00000021635 | Rad17 | protein_coding | -0.356325918 | 4.620835006 | 0.9881099 | 0.3347265 | 0.792276823 |
| ENSMUSG00000038902 | Pogz | protein_coding | -0.359823732 | 4.852381412 | 0.9869654 | 0.3349985 | 0.792746249 |
| ENSMUSG00000020591 | Ntsr2 | protein_coding | -0.50926056 | 5.596747364 | 0.9938375 | 0.3352441 | 0.793152801 |
| ENSMUSG00000028013 | Ppa2 | protein_coding | 0.253997419 | 6.441135138 | 0.9852178 | 0.3354145 | 0.793256899 |
| ENSMUSG00000022680 | Pdxdc1 | protein_coding | 0.316940669 | 5.315056667 | 0.9851292 | 0.3354356 | 0.793256899 |
| ENSMUSG00000032305 | Fam219b | protein_coding | -0.344506515 | 5.791319871 | 0.984222 | 0.3356519 | 0.793593757 |
| ENSMUSG00000021730 | Hcn1 | protein_coding | -0.334932804 | 4.651421315 | 0.9830429 | 0.3359332 | 0.793780136 |
| ENSMUSG00000058793 | Cds2 | protein_coding | -0.216517143 | 7.027360207 | 0.9829908 | 0.3359457 | 0.793780136 |
| ENSMUSG00000034118 | Tpst1 | protein_coding | -0.405340409 | 4.475285247 | 0.9829637 | 0.3359521 | 0.793780136 |
| ENSMUSG00000022407 | Adsl | protein_coding | 0.282953606 | 6.061447949 | 0.9825006 | 0.3360627 | 0.793867004 |
| ENSMUSG00000062949 | Atp11c | protein_coding | -0.6877818 | 2.800095287 | 0.9814254 | 0.3363197 | 0.793968378 |
| ENSMUSG00000034636 | Zyg11b | protein_coding | -0.233538783 | 5.867173384 | 0.9811555 | 0.3363843 | 0.793968378 |
| ENSMUSG00000051934 | Spats2 | protein_coding | -0.322374796 | 4.648541293 | 0.9804058 | 0.3365637 | 0.793968378 |
| ENSMUSG00000038976 | Ppp1r9b | protein_coding | -0.297217422 | 7.021831273 | 0.980069 | 0.336668 | 0.793968378 |
| ENSMUSG00000028743 | Akr7a5 | protein_coding | 0.404265213 | 3.887303776 | 0.9798483 | 0.3366972 | 0.793968378 |
| ENSMUSG00000031028 | Tub | protein_coding | 0.266663374 | 5.704199397 | 0.9798037 | 0.3367079 | 0.793968378 |
| ENSMUSG00000025026 | Add3 | protein_coding | -0.260053406 | 6.071688875 | 0.9793258 | 0.3368224 | 0.793968378 |
| ENSMUSG00000003809 | Gcdh | protein_coding | 0.404535095 | 4.636542732 | 0.9788993 | 0.3369247 | 0.793968378 |
| ENSMUSG00000028309 | Rnf20 | protein_coding | -0.292254967 | 5.622701641 | 0.9788832 | 0.3369285 | 0.793968378 |
| ENSMUSG00000062691 | Cebpzos | protein_coding | 0.313952193 | 4.80714867 | 0.9786247 | 0.3369905 | 0.793968378 |
| ENSMUSG00000040424 | Hipk4 | protein_coding | 0.362971079 | 4.448475181 | 0.9785721 | 0.3370031 | 0.793968378 |
| ENSMUSG00000070544 | Top1 | protein_coding | -0.274617045 | 6.256643158 | 0.9781964 | 0.3370933 | 0.793968378 |
| ENSMUSG00000057691 | Zfp746 | protein_coding | 0.427206564 | 3.760092009 | 0.9778014 | 0.3371881 | 0.793968378 |
| ENSMUSG00000035674 | Ndufa3 | protein_coding | 0.305009999 | 10.02601084 | 0.9770131 | 0.3374009 | 0.793968378 |
| ENSMUSG00000022892 | App | protein_coding | -0.228067261 | 9.150916813 | 0.9767882 | 0.3374314 | 0.793968378 |
| ENSMUSG00000025395 | Prim1 | protein_coding | -0.433173302 | 4.083517732 | 0.9766017 | 0.3374762 | 0.793968378 |
| ENSMUSG00000030134 | Rasgef1a | protein_coding | -0.283530731 | 7.583743779 | 0.9765062 | 0.3375227 | 0.793968378 |
| ENSMUSG00000010505 | Myt1 | protein_coding | -0.711486671 | 2.486515823 | 0.9761814 | 0.3375772 | 0.793968378 |
| ENSMUSG00000025138 | Sirt7 | protein_coding | -0.629326001 | 3.093681982 | 0.9756821 | 0.3376973 | 0.793968378 |
| ENSMUSG00000020988 | L2hgdh | protein_coding | -0.431002149 | 3.620809624 | 0.9748112 | 0.3379069 | 0.793968378 |
| ENSMUSG00000068328 | Aup1 | protein_coding | 0.425555872 | 6.236428025 | 0.9825001 | 0.3379266 | 0.793968378 |
| ENSMUSG00000038248 | Sobp | protein_coding | -0.360111763 | 4.816461611 | 0.9746923 | 0.3379355 | 0.793968378 |
| ENSMUSG00000011877 | Git1 | protein_coding | -0.218547367 | 6.426263458 | 0.9746582 | 0.3379437 | 0.793968378 |
| ENSMUSG00000030615 | Tmem126a | protein_coding | 0.270567145 | 6.654889712 | 0.9746001 | 0.3379577 | 0.793968378 |
| ENSMUSG00000025871 | 4833439L19Rik | protein_coding | 0.274986898 | 8.268381697 | 0.9744315 | 0.3380219 | 0.793968378 |
| ENSMUSG00000043460 | Elfn2 | protein_coding | 0.537467134 | 3.91564847 | 0.9741103 | 0.3380993 | 0.793968378 |
| ENSMUSG00000022052 | Ppp2r2a | protein_coding | -0.288914837 | 6.583689077 | 0.9739377 | 0.3381173 | 0.793968378 |
| ENSMUSG00000026027 | Stradb | protein_coding | -0.367722221 | 4.655099218 | 0.9737063 | 0.3381731 | 0.793968378 |
| ENSMUSG00000005936 | Kctd20 | protein_coding | -0.322923533 | 5.242159555 | 0.9730209 | 0.3383383 | 0.794004798 |
| ENSMUSG00000037949 | Ano10 | protein_coding | -0.445993264 | 4.049037285 | 0.9724266 | 0.3384817 | 0.794004798 |
| ENSMUSG00000097102 | 2310069G16Rik | lincRNA | -0.643858478 | 3.187040995 | 0.9721289 | 0.3385536 | 0.794004798 |
| ENSMUSG00000024862 | Klc2 | protein_coding | 0.398353324 | 5.55964312 | 0.972207 | 0.3386251 | 0.794004798 |
| ENSMUSG00000028104 | Polr3gl | protein_coding | 0.420398556 | 3.652093066 | 0.9717895 | 0.3386356 | 0.794004798 |
| ENSMUSG00000070643 | Sox13 | protein_coding | -0.84760556 | 2.646483976 | 0.9715738 | 0.3386877 | 0.794004798 |
| ENSMUSG00000060579 | Fhit | protein_coding | 0.299913542 | 5.12763996 | 0.9715 | 0.3387055 | 0.794004798 |
| ENSMUSG00000045658 | Pid1 | protein_coding | 0.356321423 | 4.859347571 | 0.9703345 | 0.3389872 | 0.794476637 |
| ENSMUSG00000020532 | Acaca | protein_coding | -0.432135223 | 3.55394166 | 0.9700562 | 0.3390545 | 0.794476637 |
| ENSMUSG00000110841 | AC153368.1 | transcribed_processe | 0.366710622 | 4.00430989 | 0.9683996 | 0.3394556 | 0.795243251 |
| ENSMUSG00000025495 | Ptdss2 | protein_coding | -0.356526543 | 5.097319969 | 0.9678953 | 0.3395779 | 0.795356351 |
| ENSMUSG00000026437 | Cdk18 | protein_coding | 0.404784682 | 5.932591511 | 0.9711591 | 0.3396985 | 0.795465513 |
| ENSMUSG00000031093 | Dock11 | protein_coding | -0.912455754 | 2.307063063 | 0.966146 | 0.3400024 | 0.795478783 |
| ENSMUSG00000006021 | Kptn | protein_coding | 0.394067912 | 5.224893488 | 0.9657714 | 0.3401169 | 0.795478783 |
| ENSMUSG00000022160 | Mettl3 | protein_coding | -0.476641582 | 3.93605508 | 0.9654943 | 0.3401608 | 0.795478783 |
| ENSMUSG00000024750 | Zfand5 | protein_coding | -0.251107656 | 6.018019954 | 0.9637419 | 0.3405871 | 0.795478783 |
| ENSMUSG00000032523 | Hhatl | protein_coding | -0.875728329 | 3.133468773 | 0.9672535 | 0.3406423 | 0.795478783 |
| ENSMUSG00000024045 | Akap8 | protein_coding | -0.328244275 | 4.802518512 | 0.962448 | 0.3409023 | 0.795478783 |
| ENSMUSG00000051154 | Commd3 | protein_coding | 0.205697369 | 8.214268243 | 0.9624062 | 0.3409125 | 0.795478783 |
| ENSMUSG00000051705 | Senp8 | protein_coding | -0.387242627 | 4.582202443 | 0.9623286 | 0.3409314 | 0.795478783 |
| ENSMUSG00000027742 | Cog6 | protein_coding | -0.326831778 | 5.242379059 | 0.9622995 | 0.3409386 | 0.795478783 |
| ENSMUSG00000022401 | Xpnpep3 | protein_coding | -0.49760711 | 3.690002345 | 0.9622972 | 0.3409391 | 0.795478783 |

| ensembl_gene_id | external_gene_name | gene_biotype | log2 (fold change) | log2 (counts per million) | F-value | p-value | False discovery rate |
|---|---|---|---|---|---|---|---|
| ENSMUSG00000039725 | Trp53rka | protein_coding | -0.672213476 | 3.216718572 | 0.9620896 | 0.3409897 | 0.795478783 |
| ENSMUSG00000028496 | Mllt3 | protein_coding | -0.291264359 | 5.335618792 | 0.9618878 | 0.341039 | 0.795478783 |
| ENSMUSG00000051062 | Fbll1 | protein_coding | 0.323561109 | 5.381324325 | 0.9618754 | 0.341042 | 0.795478783 |
| ENSMUSG00000070697 | Utp3 | protein_coding | 0.216973355 | 6.400541001 | 0.9612768 | 0.3411881 | 0.795478783 |
| ENSMUSG00000058486 | Wdr91 | protein_coding | 0.46486917 | 4.671450756 | 0.9607556 | 0.3413387 | 0.795478783 |
| ENSMUSG00000018507 | Trpv2 | protein_coding | 0.509994222 | 4.695499816 | 0.9641398 | 0.3413423 | 0.795478783 |
| ENSMUSG00000015092 | Edf1 | protein_coding | 0.225562241 | 7.716842567 | 0.9604772 | 0.3413833 | 0.795478783 |
| ENSMUSG00000034687 | Fras1 | protein_coding | -1.098767396 | 2.484782701 | 0.9607829 | 0.3413936 | 0.795478783 |
| ENSMUSG00000020375 | Rufy1 | protein_coding | -0.318796355 | 4.966513053 | 0.9603232 | 0.341421 | 0.795478783 |
| ENSMUSG00000022204 | Ngdn | protein_coding | -0.235235534 | 6.862385799 | 0.9602615 | 0.341436 | 0.795478783 |
| ENSMUSG00000037475 | Thoc2 | protein_coding | -0.399011283 | 4.708718785 | 0.9601008 | 0.3414753 | 0.795478783 |
| ENSMUSG00000021745 | Ptprg | protein_coding | -0.462878958 | 4.195560502 | 0.960075 | 0.3414816 | 0.795478783 |
| ENSMUSG00000040848 | Sft2d2 | protein_coding | -0.480972447 | 3.875666158 | 0.9598761 | 0.3415302 | 0.795478783 |
| ENSMUSG00000039512 | Uhrf1bp1 | protein_coding | -0.519509336 | 3.174498448 | 0.9597841 | 0.3415527 | 0.795478783 |
| ENSMUSG00000021140 | Pcnx | protein_coding | -0.526984647 | 3.71304744 | 0.9597806 | 0.3415536 | 0.795478783 |
| ENSMUSG00000036054 | Sugp2 | protein_coding | -0.291840597 | 4.989714878 | 0.9587659 | 0.3418018 | 0.795765235 |
| ENSMUSG00000029238 | Clock | protein_coding | -0.339692564 | 5.392085765 | 0.9585462 | 0.3418555 | 0.795765235 |
| ENSMUSG00000021645 | Smn1 | protein_coding | 0.275132362 | 5.4629213 | 0.9581363 | 0.3419559 | 0.795765235 |
| ENSMUSG00000036833 | Pnpla7 | protein_coding | -0.764218297 | 2.789872672 | 0.9574682 | 0.3421196 | 0.795765235 |
| ENSMUSG00000035297 | Cops4 | protein_coding | 0.197434349 | 8.429446949 | 0.9574281 | 0.3421294 | 0.795765235 |
| ENSMUSG00000028078 | Dclk2 | protein_coding | -0.29379794 | 4.942542086 | 0.9572034 | 0.3421845 | 0.795765235 |
| ENSMUSG00000095930 | Nim1k | protein_coding | -0.291583011 | 5.445212786 | 0.9570444 | 0.3422235 | 0.795765235 |
| ENSMUSG00000034616 | Ssh3 | protein_coding | 0.566572238 | 3.420697223 | 0.9561498 | 0.3424429 | 0.795765235 |
| ENSMUSG00000032525 | Nktr | protein_coding | -0.498121827 | 4.031667074 | 0.9561147 | 0.3424515 | 0.795765235 |
| ENSMUSG00000066415 | Msl2 | protein_coding | -0.448624917 | 4.015005728 | 0.956091 | 0.3424573 | 0.795765235 |
| ENSMUSG00000042688 | Mapk6 | protein_coding | -0.38654845 | 5.407855678 | 0.9554247 | 0.3426442 | 0.795765235 |
| ENSMUSG00000066621 | Tecpr1 | protein_coding | 0.331421405 | 5.740775339 | 0.9550904 | 0.342703 | 0.795765235 |
| ENSMUSG00000064337 | mt-Rnr1 | Mt_rRNA | -0.50576765 | 10.86487464 | 0.9714832 | 0.3427077 | 0.795765235 |
| ENSMUSG00000032527 | Pccb | protein_coding | -0.31760311 | 5.911814377 | 0.9546381 | 0.3428141 | 0.795765235 |
| ENSMUSG00000019899 | Lama2 | protein_coding | -0.662586128 | 2.772551931 | 0.9543015 | 0.3428969 | 0.795765235 |
| ENSMUSG00000070858 | Gm1673 | protein_coding | 0.468323532 | 3.401330592 | 0.9540484 | 0.3429591 | 0.795765235 |
| ENSMUSG00000054309 | Cpsf3 | protein_coding | 0.300004614 | 6.587729237 | 0.9535197 | 0.3431124 | 0.795765235 |
| ENSMUSG00000054619 | Mettl7a1 | protein_coding | -0.521869595 | 3.794749842 | 0.9534186 | 0.343114 | 0.795765235 |
| ENSMUSG00000093738 | AI606473 | processed_transcript | 0.579297067 | 3.792607044 | 0.9533279 | 0.3431596 | 0.795765235 |
| ENSMUSG00000086191 | Zfp652os | antisense_RNA | -0.576098569 | 2.934209577 | 0.9532251 | 0.3431616 | 0.795765235 |
| ENSMUSG00000039717 | Ralyl | protein_coding | 0.207123757 | 7.948291506 | 0.9529448 | 0.3432306 | 0.795765235 |
| ENSMUSG00000053390 | Zfp952 | protein_coding | -0.418421025 | 4.308508244 | 0.9523647 | 0.3433735 | 0.795924793 |
| ENSMUSG00000038861 | Pi4kb | protein_coding | 0.468918605 | 3.527739879 | 0.9510195 | 0.3437051 | 0.796365283 |
| ENSMUSG00000038150 | Ormdl3 | protein_coding | 0.24443426 | 6.097584382 | 0.9507068 | 0.3437823 | 0.796365283 |
| ENSMUSG00000008859 | Rala | protein_coding | -0.282913276 | 5.393500259 | 0.9503926 | 0.3438598 | 0.796365283 |
| ENSMUSG00000051768 | Xrcc1 | protein_coding | 0.361044011 | 4.455173051 | 0.950206 | 0.3439059 | 0.796365283 |
| ENSMUSG00000062352 | Itgb1bp1 | protein_coding | 0.270371647 | 6.483289988 | 0.9500928 | 0.3439338 | 0.796365283 |
| ENSMUSG00000064339 | mt-Rnr2 | Mt_rRNA | -0.459019772 | 13.83599611 | 0.9570788 | 0.3440546 | 0.79647353 |
| ENSMUSG00000031153 | Gripap1 | protein_coding | 0.211001034 | 7.663267864 | 0.9481432 | 0.3444157 | 0.796839529 |
| ENSMUSG00000028252 | Ccnc | protein_coding | 0.256691253 | 7.047618896 | 0.947443 | 0.344589 | 0.796839529 |
| ENSMUSG00000022807 | Osbpl11 | protein_coding | -0.454402711 | 3.436729191 | 0.9471644 | 0.344658 | 0.796839529 |
| ENSMUSG00000056383 | AI987944 | protein_coding | -0.682791797 | 3.222489646 | 0.9471927 | 0.3446742 | 0.796839529 |
| ENSMUSG00000045031 | Cetn4 | protein_coding | -0.415686617 | 3.838305497 | 0.946742 | 0.3447626 | 0.796839529 |
| ENSMUSG00000039233 | Tbce | protein_coding | -0.368979062 | 5.077393966 | 0.9467348 | 0.3447644 | 0.796839529 |
| ENSMUSG00000002910 | Arrdc2 | protein_coding | -0.685980975 | 2.682520466 | 0.9466757 | 0.344779 | 0.796839529 |
| ENSMUSG00000020812 | 1810032O08Rik | protein_coding | 0.600788548 | 2.780781175 | 0.9464389 | 0.3448377 | 0.796839529 |
| ENSMUSG00000020850 | Prpf8 | protein_coding | -0.237462988 | 6.328230262 | 0.9459229 | 0.3449657 | 0.796839529 |
| ENSMUSG00000079467 | Gm14966 | antisense_RNA | -0.95183302 | 1.889569052 | 0.9458006 | 0.344996 | 0.796839529 |
| ENSMUSG00000041986 | Elmod1 | protein_coding | -0.385560686 | 6.195305318 | 0.9491639 | 0.3450636 | 0.796839529 |
| ENSMUSG00000062794 | Zfp599 | protein_coding | -0.457010803 | 13.5488801 | 0.9526113 | 0.3451481 | 0.796839529 |
| ENSMUSG00000048388 | Fam171b | protein_coding | -0.215746348 | 7.116365488 | 0.9447568 | 0.345255 | 0.796839529 |
| ENSMUSG00000079242 | C730034F03Rik | protein_coding | -0.670534432 | 3.111918437 | 0.9447258 | 0.3452627 | 0.796839529 |
| ENSMUSG00000044950 | Pwwp2a | protein_coding | -0.383786978 | 4.588338942 | 0.9444777 | 0.3453243 | 0.796839529 |
| ENSMUSG00000050931 | Sgms2 | protein_coding | -1.007627003 | 2.269271934 | 0.9440239 | 0.345437 | 0.796928612 |
| ENSMUSG00000008540 | Mgst1 | protein_coding | -0.608259267 | 5.354484766 | 0.9541769 | 0.3456831 | 0.797201842 |
| ENSMUSG00000030823 | 9130019O22Rik | protein_coding | -1.142804362 | 1.986000247 | 0.9429511 | 0.3457037 | 0.797201842 |
| ENSMUSG00000097156 | Gm3764 | lincRNA | -0.334514585 | 5.037532287 | 0.9422543 | 0.3458771 | 0.797308565 |
| ENSMUSG00000023932 | Cdc5l | protein_coding | -0.274876442 | 5.338712987 | 0.9420437 | 0.3459295 | 0.797308565 |
| ENSMUSG00000041921 | Metap1d | protein_coding | 0.444169664 | 3.832582105 | 0.9418714 | 0.3459725 | 0.797308565 |
| ENSMUSG00000017740 | Slc12a5 | protein_coding | -0.390200777 | 6.468301671 | 0.9467687 | 0.3461648 | 0.797439979 |
| ENSMUSG00000070695 | Cntnap5a | protein_coding | -0.369307781 | 4.667547053 | 0.9409748 | 0.3461958 | 0.797439979 |
| ENSMUSG00000039488 | Cntn5 | protein_coding | -0.584525712 | 3.316897843 | 0.9406792 | 0.3462695 | 0.797439979 |
| ENSMUSG00000096916 | Zfp850 | protein_coding | 0.914118792 | 2.376784484 | 0.9400509 | 0.3464262 | 0.797439979 |
| ENSMUSG00000063296 | Tmem117 | protein_coding | -0.648384236 | 3.037931534 | 0.9398965 | 0.3464647 | 0.797439979 |
| ENSMUSG00000071708 | Sms | protein_coding | 0.239367621 | 6.018823401 | 0.9396585 | 0.3465241 | 0.797439979 |
| ENSMUSG00000020087 | Tysnd1 | protein_coding | -0.906816711 | 2.574259294 | 0.939442 | 0.3465782 | 0.797439979 |
| ENSMUSG00000064030 | Pym1 | protein_coding | 0.325810522 | 4.753457262 | 0.9392634 | 0.3466228 | 0.797439979 |
| ENSMUSG00000021178 | Psmc1 | protein_coding | 0.197677917 | 9.082673382 | 0.938396 | 0.3468394 | 0.797554952 |
| ENSMUSG00000044772 | Sntn | protein_coding | -0.542503779 | 3.33696982 | 0.9380054 | 0.3469371 | 0.797554952 |
| ENSMUSG00000029826 | Zc3hav1 | protein_coding | -0.480090411 | 3.925240295 | 0.9377939 | 0.34699 | 0.797554952 |
| ENSMUSG00000024273 | 2700062C07Rik | protein_coding | -0.276521729 | 5.358026639 | 0.9376692 | 0.3470211 | 0.797554952 |
| ENSMUSG00000033707 | Lrrc24 | protein_coding | 0.431027947 | 4.087936264 | 0.9371772 | 0.3471442 | 0.797554952 |
| ENSMUSG00000079184 | Mphosph8 | protein_coding | -0.216140079 | 6.514185979 | 0.9370947 | 0.3471649 | 0.797554952 |

| ensembl_gene_id | external_gene_name | gene_biotype | log2 (fold change) | log2 (counts per million) | F-value | p-value | False discovery rate |
|---|---|---|---|---|---|---|---|
| ENSMUSG00000003534 | Ddr1 | protein_coding | -0.352403386 | 4.699261605 | 0.9369699 | 0.3471961 | 0.797554952 |
| ENSMUSG00000064351 | mt-Co1 | protein_coding | -0.403949011 | 11.50948277 | 0.9424202 | 0.3472661 | 0.797554952 |
| ENSMUSG00000024537 | Psmg2 | protein_coding | 0.2244243 | 6.137538241 | 0.9359198 | 0.3474591 | 0.797655113 |
| ENSMUSG00000024294 | Mib1 | protein_coding | -0.274014711 | 6.045810957 | 0.9356153 | 0.3475354 | 0.797655113 |
| ENSMUSG00000057615 | Ldoc1 | protein_coding | 0.439423318 | 3.467957453 | 0.9355825 | 0.3475436 | 0.797655113 |
| ENSMUSG00000014606 | Slc25a11 | protein_coding | 0.203843783 | 8.391474826 | 0.9350652 | 0.3476733 | 0.797655113 |
| ENSMUSG00000043300 | B3galnt1 | protein_coding | 0.253946053 | 6.205817635 | 0.9344858 | 0.3478186 | 0.797655113 |
| ENSMUSG00000042507 | Elmsan1 | protein_coding | -0.620638526 | 2.772001058 | 0.9344635 | 0.3478242 | 0.797655113 |
| ENSMUSG00000022179 | 4931414P19Rik | protein_coding | -0.73792369 | 2.714418146 | 0.9343502 | 0.3478527 | 0.797655113 |
| ENSMUSG00000022843 | Clcn2 | protein_coding | -0.81097969 | 2.632605279 | 0.9339907 | 0.3479429 | 0.797655113 |
| ENSMUSG00000047789 | Slc38a9 | protein_coding | -0.376056492 | 3.844259702 | 0.9338536 | 0.3479773 | 0.797655113 |
| ENSMUSG00000029466 | Anapc7 | protein_coding | 0.28330848 | 5.210385149 | 0.9324599 | 0.3483275 | 0.798287676 |
| ENSMUSG00000046607 | Hrk | protein_coding | -0.351229985 | 4.154284163 | 0.930217 | 0.3488922 | 0.799411388 |
| ENSMUSG00000034973 | Dopey1 | protein_coding | -0.550340741 | 3.857142208 | 0.9298913 | 0.3489973 | 0.799451649 |
| ENSMUSG00000041377 | Ninj2 | protein_coding | -0.394970916 | 4.6352863 | 0.9295575 | 0.3490584 | 0.799451649 |
| ENSMUSG00000026509 | Capn2 | protein_coding | -0.352288924 | 4.854152845 | 0.9289076 | 0.3492224 | 0.799606352 |
| ENSMUSG00000020580 | Rock2 | protein_coding | -0.232565818 | 6.259630581 | 0.9272867 | 0.3496319 | 0.799606352 |
| ENSMUSG00000012076 | Brms1l | protein_coding | -0.261431319 | 5.794557637 | 0.9271821 | 0.3496583 | 0.799606352 |
| ENSMUSG00000011154 | Cfap161 | protein_coding | 0.372330433 | 4.412085731 | 0.9269923 | 0.3497063 | 0.799606352 |
| ENSMUSG00000016481 | Cr1l | protein_coding | -0.412205325 | 3.992040996 | 0.9264709 | 0.3498382 | 0.799606352 |
| ENSMUSG00000019232 | Etnppl | protein_coding | -0.273929516 | 5.211151314 | 0.9261424 | 0.3499213 | 0.799606352 |
| ENSMUSG00000043301 | Kcnj6 | protein_coding | -0.599259144 | 2.906168264 | 0.92591 | 0.3499801 | 0.799606352 |
| ENSMUSG00000056267 | Cep70 | protein_coding | -0.354939876 | 4.42457313 | 0.9256928 | 0.3500352 | 0.799606352 |
| ENSMUSG00000075705 | Msrb1 | protein_coding | 0.304390013 | 4.944493655 | 0.9255902 | 0.3500611 | 0.799606352 |
| ENSMUSG00000005069 | Pex5 | protein_coding | -0.36268825 | 4.408127076 | 0.9255043 | 0.3500829 | 0.799606352 |
| ENSMUSG00000034300 | Fam53c | protein_coding | -0.337807493 | 5.380149082 | 0.9250247 | 0.3502044 | 0.799606352 |
| ENSMUSG00000009687 | Fxyd5 | protein_coding | 0.751654681 | 2.35456016 | 0.9250076 | 0.3502087 | 0.799606352 |
| ENSMUSG00000035919 | Bbs9 | protein_coding | 0.34481008 | 4.403336251 | 0.9249247 | 0.3502298 | 0.799606352 |
| ENSMUSG00000023067 | Cdkn1a | protein_coding | -0.619116552 | 4.338982356 | 0.9302763 | 0.3502736 | 0.799606352 |
| ENSMUSG00000028108 | Ecm1 | protein_coding | 0.708052412 | 2.629739521 | 0.9244327 | 0.3503545 | 0.799606352 |
| ENSMUSG00000048747 | E130114P18Rik | protein_coding | -0.479571393 | 4.640997814 | 0.9246345 | 0.350362 | 0.799606352 |
| ENSMUSG00000079339 | Ifit1bl1 | protein_coding | 1.276296184 | 2.273117401 | 0.9278078 | 0.3503901 | 0.799606352 |
| ENSMUSG00000062376 | Borcs7 | protein_coding | 0.197309738 | 7.115717169 | 0.9239232 | 0.3504837 | 0.799650121 |
| ENSMUSG00000032181 | Scg3 | protein_coding | -0.258008498 | 7.058210045 | 0.9235492 | 0.3505786 | 0.799696983 |
| ENSMUSG00000028282 | Casp8ap2 | protein_coding | 0.414800532 | 3.979270051 | 0.9221503 | 0.3509339 | 0.800038066 |
| ENSMUSG00000001767 | Crnkl1 | protein_coding | -0.373327005 | 4.060720349 | 0.9221316 | 0.3509386 | 0.800038066 |
| ENSMUSG00000100417 | Gm28578 | lincRNA | -0.864449125 | 2.375551385 | 0.9220817 | 0.3509513 | 0.800038066 |
| ENSMUSG00000020709 | Adap2 | protein_coding | -0.729695209 | 2.499203079 | 0.92175 | 0.3510357 | 0.800060727 |
| ENSMUSG00000054091 | 1810037I17Rik | protein_coding | -0.257171922 | 7.637371474 | 0.9210645 | 0.35121 | 0.800288521 |
| ENSMUSG00000038084 | Opa1 | protein_coding | -0.253565143 | 5.952056589 | 0.9206408 | 0.3513179 | 0.800364638 |
| ENSMUSG00000021254 | Gpatch2l | protein_coding | -0.57338171 | 3.772626543 | 0.9198455 | 0.3515433 | 0.80063072 |
| ENSMUSG00000043531 | Sorcs1 | protein_coding | -0.40978921 | 3.709531103 | 0.9190767 | 0.3517164 | 0.80063072 |
| ENSMUSG00000056427 | Slit3 | protein_coding | -0.605910575 | 2.882208452 | 0.9190577 | 0.3517212 | 0.80063072 |
| ENSMUSG00000022848 | Dirc2 | protein_coding | -0.329612836 | 5.275796203 | 0.9190135 | 0.3517325 | 0.80063072 |
| ENSMUSG00000035126 | Wdr78 | protein_coding | -0.770163047 | 2.817560419 | 0.9184389 | 0.3518791 | 0.800685462 |
| ENSMUSG00000036599 | Chst12 | protein_coding | 0.269101945 | 5.686100521 | 0.9182036 | 0.3519391 | 0.800685462 |
| ENSMUSG00000047881 | Rell1 | protein_coding | -0.877275226 | 2.650813724 | 0.9179221 | 0.3520109 | 0.800685462 |
| ENSMUSG00000059182 | Skap2 | protein_coding | -0.414499381 | 4.404641125 | 0.917752 | 0.3520544 | 0.800685462 |
| ENSMUSG00000056596 | Trnp1 | protein_coding | -0.321676151 | 5.774710519 | 0.9173579 | 0.352155 | 0.800745005 |
| ENSMUSG00000001127 | Araf | protein_coding | 0.195927304 | 9.716486823 | 0.9170281 | 0.3522393 | 0.800767296 |
| ENSMUSG00000007777 | 0610009B22Rik | protein_coding | 0.243232321 | 7.420469813 | 0.9162676 | 0.3524337 | 0.80103987 |
| ENSMUSG00000068250 | Amn1 | protein_coding | 0.276246715 | 5.656343951 | 0.9155858 | 0.3526081 | 0.801266986 |
| ENSMUSG00000029675 | Eln | protein_coding | -1.037926836 | 3.211596894 | 0.9218069 | 0.3532409 | 0.802535231 |
| ENSMUSG00000033732 | Sf3b3 | protein_coding | 0.291521033 | 5.440594887 | 0.9121608 | 0.3534862 | 0.80274105 |
| ENSMUSG00000000148 | Brat1 | protein_coding | 0.495827173 | 4.062798521 | 0.9120722 | 0.3535318 | 0.80274105 |
| ENSMUSG00000045251 | Zfp688 | protein_coding | 0.308729993 | 4.787095876 | 0.9112038 | 0.3537322 | 0.80274105 |
| ENSMUSG00000018651 | Tada2a | protein_coding | 0.372065741 | 5.035906239 | 0.9102646 | 0.3539738 | 0.80274105 |
| ENSMUSG00000055200 | Sertad3 | protein_coding | -0.589022329 | 3.194498465 | 0.9102044 | 0.3539892 | 0.80274105 |
| ENSMUSG00000005687 | Bcas2 | protein_coding | 0.211315137 | 8.263035608 | 0.9101136 | 0.3540126 | 0.80274105 |
| ENSMUSG00000056999 | Ide | protein_coding | -0.31227536 | 4.665093056 | 0.9091864 | 0.3542514 | 0.80274105 |
| ENSMUSG00000074918 | Inafm2 | protein_coding | -0.259790144 | 7.812709367 | 0.908686 | 0.3543804 | 0.80274105 |
| ENSMUSG00000026784 | Pdss1 | protein_coding | -0.498484991 | 3.362814986 | 0.9084917 | 0.3544304 | 0.80274105 |
| ENSMUSG00000038141 | Tmem181a | protein_coding | -0.4221043 | 3.611116157 | 0.908246 | 0.3544938 | 0.80274105 |
| ENSMUSG00000030806 | Stx1b | protein_coding | 0.242072019 | 8.032936804 | 0.9080865 | 0.354535 | 0.80274105 |
| ENSMUSG00000056167 | Cnot10 | protein_coding | 0.331199345 | 5.128757025 | 0.9077273 | 0.3546276 | 0.80274105 |
| ENSMUSG00000042699 | Dhx9 | protein_coding | -0.365632173 | 4.608989933 | 0.9076242 | 0.3546542 | 0.80274105 |
| ENSMUSG00000020843 | Timm22 | protein_coding | 0.239575189 | 7.608335653 | 0.9074423 | 0.3547012 | 0.80274105 |
| ENSMUSG00000028420 | Tmem38b | protein_coding | 0.581747396 | 3.341440637 | 0.9073164 | 0.3547337 | 0.80274105 |
| ENSMUSG00000027227 | Sord | protein_coding | 0.353511121 | 5.580487141 | 0.9070196 | 0.354833 | 0.80274105 |
| ENSMUSG00000054733 | Msra | protein_coding | 0.273990595 | 5.175527977 | 0.9066248 | 0.3549123 | 0.80274105 |
| ENSMUSG00000039501 | Znfx1 | protein_coding | -0.428081065 | 3.843822906 | 0.9063503 | 0.3549832 | 0.80274105 |
| ENSMUSG00000042613 | Pbxip1 | protein_coding | -0.41232842 | 3.96877125 | 0.9063466 | 0.3549842 | 0.80274105 |
| ENSMUSG00000018474 | Chd3 | protein_coding | -0.215123687 | 6.673162646 | 0.9060248 | 0.3550673 | 0.80274105 |
| ENSMUSG00000079610 | Ankrd39 | protein_coding | 0.400507321 | 3.616515507 | 0.9060133 | 0.3550703 | 0.80274105 |
| ENSMUSG00000039552 | Rsph4a | protein_coding | 0.758282993 | 2.769764578 | 0.9059332 | 0.355091 | 0.80274105 |
| ENSMUSG00000025702 | 8-Mar | protein_coding | -0.437466124 | 4.250494278 | 0.9055945 | 0.3551786 | 0.80274105 |
| ENSMUSG00000024955 | Esrra | protein_coding | -0.800810585 | 2.598698475 | 0.90551 | 0.3552005 | 0.80274105 |
| ENSMUSG00000054693 | Adam10 | protein_coding | -0.236387196 | 5.802093242 | 0.9050726 | 0.3553136 | 0.80274105 |

| ensembl_gene_id | external_gene_name | gene_biotype | log2 (fold change) | log2 (counts per million) | F-value | p-value | False discovery rate |
|---|---|---|---|---|---|---|---|
| ENSMUSG00000023030 | Slc11a2 | protein_coding | 0.420496795 | 4.586060056 | 0.9047869 | 0.3553876 | 0.80274105 |
| ENSMUSG00000027455 | Nsfl1c | protein_coding | 0.206463822 | 7.593696853 | 0.9046119 | 0.3554329 | 0.80274105 |
| ENSMUSG00000020634 | Ubxn2a | protein_coding | -0.269007131 | 5.43505583 | 0.9043757 | 0.355494 | 0.80274105 |
| ENSMUSG00000041879 | Ipo9 | protein_coding | 0.254804578 | 6.119711116 | 0.9043665 | 0.3554964 | 0.80274105 |
| ENSMUSG00000027489 | Necab3 | protein_coding | 0.535925125 | 5.198566932 | 0.9105005 | 0.3558517 | 0.80324867 |
| ENSMUSG00000043670 | Diras1 | protein_coding | 0.410701257 | 5.99051786 | 0.9079037 | 0.3559613 | 0.80324867 |
| ENSMUSG00000024530 | Prelid3a | protein_coding | 0.230002393 | 6.054081993 | 0.9023773 | 0.3560121 | 0.80324867 |
| ENSMUSG00000043463 | Rab9b | protein_coding | -0.297485235 | 5.675280788 | 0.9015494 | 0.356227 | 0.80324867 |
| ENSMUSG00000032078 | Zpr1 | protein_coding | 0.411096634 | 4.798499595 | 0.9013771 | 0.3562944 | 0.80324867 |
| ENSMUSG00000020610 | Amz2 | protein_coding | 0.238021176 | 6.09536867 | 0.901116 | 0.3563396 | 0.80324867 |
| ENSMUSG00000040760 | Appl1 | protein_coding | -0.2596333 | 5.821678524 | 0.9008297 | 0.356414 | 0.80324867 |
| ENSMUSG00000002109 | Ddb2 | protein_coding | -0.672316445 | 2.829398467 | 0.9007517 | 0.3564343 | 0.80324867 |
| ENSMUSG00000050705 | 2310061I04Rik | protein_coding | 0.259415517 | 5.638533226 | 0.9007141 | 0.3564441 | 0.80324867 |
| ENSMUSG00000021546 | Hnrnpk | protein_coding | -0.234761654 | 8.311756513 | 0.9003295 | 0.3565441 | 0.80324867 |
| ENSMUSG00000053289 | Ddx10 | protein_coding | -0.35676717 | 5.026413245 | 0.9001313 | 0.3565956 | 0.80324867 |
| ENSMUSG00000037624 | Kcnk2 | protein_coding | 0.315811509 | 5.383507972 | 0.8991726 | 0.3568451 | 0.80324867 |
| ENSMUSG00000035152 | Ap2b1 | protein_coding | -0.244166265 | 6.570604096 | 0.8990507 | 0.3568769 | 0.80324867 |
| ENSMUSG00000004460 | Dnajb11 | protein_coding | 0.316959046 | 4.93697652 | 0.8989026 | 0.3569154 | 0.80324867 |
| ENSMUSG00000015829 | Tnr | protein_coding | -0.428518764 | 3.979884262 | 0.8988945 | 0.3569176 | 0.80324867 |
| ENSMUSG00000046807 | Lrrc75b | protein_coding | 0.381713342 | 4.251037385 | 0.8988069 | 0.3569404 | 0.80324867 |
| ENSMUSG00000031107 | Rbmx2 | protein_coding | -0.319865897 | 4.255340661 | 0.8986121 | 0.3569911 | 0.80324867 |
| ENSMUSG00000041594 | Tmtc4 | protein_coding | 0.792014613 | 2.540256851 | 0.8976275 | 0.3572478 | 0.803398904 |
| ENSMUSG00000028032 | Papss1 | protein_coding | -0.202720369 | 6.706910355 | 0.8975187 | 0.3572762 | 0.803398904 |
| ENSMUSG00000029029 | Wrap73 | protein_coding | 0.257144337 | 5.753226431 | 0.89729 | 0.3573358 | 0.803398904 |
| ENSMUSG00000079283 | 2310009B15Rik | protein_coding | 0.351109526 | 4.54184601 | 0.8972098 | 0.3573567 | 0.803398904 |
| ENSMUSG00000019977 | Hbs1l | protein_coding | 0.304915385 | 5.924672831 | 0.896888 | 0.3574407 | 0.803419745 |
| ENSMUSG00000003541 | Ier3 | protein_coding | 0.28867412 | 4.830210852 | 0.8962235 | 0.3576142 | 0.803641765 |
| ENSMUSG00000035615 | Frmpd1 | protein_coding | 0.953154751 | 2.221045393 | 0.8956526 | 0.3577634 | 0.803753447 |
| ENSMUSG00000033569 | Adgrb3 | protein_coding | -0.332663022 | 5.961160046 | 0.8955475 | 0.3578134 | 0.803753447 |
| ENSMUSG00000026094 | Stk17b | protein_coding | -1.205139462 | 2.255065074 | 0.8952823 | 0.3579811 | 0.803866734 |
| ENSMUSG00000027447 | Cst3 | protein_coding | -0.418865975 | 11.15737554 | 0.9015927 | 0.3581447 | 0.803866734 |
| ENSMUSG00000028134 | Ptbp2 | protein_coding | -0.259213524 | 6.239387828 | 0.8940836 | 0.3581738 | 0.803866734 |
| ENSMUSG00000067578 | Cbln4 | protein_coding | -0.460710919 | 4.364327424 | 0.8938351 | 0.3582614 | 0.803866734 |
| ENSMUSG00000044528 | Tram1l1 | protein_coding | 0.193292859 | 6.654881795 | 0.8937065 | 0.3582726 | 0.803866734 |
| ENSMUSG00000098649 | Gm27199 | antisense_RNA | -0.588909435 | 2.751579585 | 0.8931657 | 0.3584142 | 0.803866734 |
| ENSMUSG00000097416 | Gm26670 | antisense_RNA | -0.756069032 | 2.352260028 | 0.8930044 | 0.3584565 | 0.803866734 |
| ENSMUSG00000019802 | Sec63 | protein_coding | -0.416849026 | 4.528129988 | 0.8929541 | 0.3584697 | 0.803866734 |
| ENSMUSG00000052302 | Tbc1d30 | protein_coding | -0.271602792 | 5.205387389 | 0.8923788 | 0.3586205 | 0.803866734 |
| ENSMUSG00000028567 | Txndc12 | protein_coding | 0.232570847 | 6.104744527 | 0.8921048 | 0.3586924 | 0.803866734 |
| ENSMUSG00000053310 | Nrgn | protein_coding | 0.251640999 | 7.189110252 | 0.8917176 | 0.358794 | 0.803866734 |
| ENSMUSG00000059811 | Atl2 | protein_coding | -0.301754505 | 4.999606337 | 0.8916326 | 0.3588163 | 0.803866734 |
| ENSMUSG00000060216 | Arrb2 | protein_coding | 0.383034398 | 4.17512589 | 0.8914608 | 0.3588614 | 0.803866734 |
| ENSMUSG00000025423 | Pias2 | protein_coding | -0.271098307 | 5.407554787 | 0.891274 | 0.3589105 | 0.803866734 |
| ENSMUSG00000003234 | Abcf3 | protein_coding | 0.28343296 | 7.064597713 | 0.8900233 | 0.3592616 | 0.804485561 |
| ENSMUSG00000000278 | Scpep1 | protein_coding | 0.485049828 | 3.180799759 | 0.889544 | 0.3593651 | 0.804549937 |
| ENSMUSG00000035829 | Ppp1r26 | protein_coding | 0.421757524 | 3.6414229 | 0.8890474 | 0.3594958 | 0.80467494 |
| ENSMUSG00000064127 | Med14 | protein_coding | 0.383413242 | 4.923458654 | 0.8878347 | 0.3598152 | 0.80522225 |
| ENSMUSG00000001666 | Ddt | protein_coding | 0.274399571 | 5.151267093 | 0.8871948 | 0.3599839 | 0.805282899 |
| ENSMUSG00000024180 | Tmem8 | protein_coding | 0.573338433 | 2.937878498 | 0.8869547 | 0.3600472 | 0.805282899 |
| ENSMUSG00000001829 | Clpb | protein_coding | 0.250397304 | 5.379357391 | 0.8868274 | 0.3600808 | 0.805282899 |
| ENSMUSG00000022263 | Trio | protein_coding | -0.289640531 | 5.447010341 | 0.8864419 | 0.3601826 | 0.805282899 |
| ENSMUSG00000026546 | Cfap45 | protein_coding | 0.463309185 | 3.53867362 | 0.8863123 | 0.3602168 | 0.805282899 |
| ENSMUSG00000028464 | Tpm2 | protein_coding | -0.422484462 | 4.452468545 | 0.8857637 | 0.3603616 | 0.805328446 |
| ENSMUSG00000048022 | Tmem229a | protein_coding | -0.413572923 | 4.868359132 | 0.8857528 | 0.3603869 | 0.805328446 |
| ENSMUSG00000026163 | Sphkap | protein_coding | -0.237866073 | 5.666542243 | 0.8850401 | 0.3605528 | 0.805531729 |
| ENSMUSG00000029569 | Tmem168 | protein_coding | -0.631239329 | 3.063842105 | 0.8843825 | 0.3607267 | 0.805752805 |
| ENSMUSG00000060475 | Wtap | protein_coding | -0.514107693 | 5.476082691 | 0.8910014 | 0.3608993 | 0.805874724 |
| ENSMUSG00000000125 | Wnt3 | protein_coding | -0.522603824 | 4.082509671 | 0.8836945 | 0.3609312 | 0.805874724 |
| ENSMUSG00000037455 | Slc18b1 | protein_coding | -0.523511945 | 3.00630673 | 0.8830558 | 0.3610779 | 0.805938827 |
| ENSMUSG00000036315 | Znrd1 | protein_coding | 0.281638712 | 6.192882707 | 0.8829354 | 0.3611098 | 0.805938827 |
| ENSMUSG00000030218 | Mgp | protein_coding | -0.883745688 | 2.11718742 | 0.8823325 | 0.3612696 | 0.80611581 |
| ENSMUSG00000051483 | Cbr1 | protein_coding | 0.23722669 | 7.257001137 | 0.8812304 | 0.3615619 | 0.80611581 |
| ENSMUSG00000020747 | Tmem94 | protein_coding | -0.31307358 | 4.770877502 | 0.8812214 | 0.3615643 | 0.80611581 |
| ENSMUSG00000014245 | Pigl | protein_coding | -0.61455093 | 2.891857036 | 0.8802479 | 0.3618228 | 0.80611581 |
| ENSMUSG00000037710 | Cisd1 | protein_coding | 0.283366597 | 8.734266486 | 0.8798683 | 0.361946 | 0.80611581 |
| ENSMUSG00000022668 | Gtpbp8 | protein_coding | 0.321952891 | 4.736968523 | 0.8797811 | 0.3619468 | 0.80611581 |
| ENSMUSG00000030533 | Unc45a | protein_coding | 0.353275937 | 4.486014819 | 0.8796566 | 0.3619799 | 0.80611581 |
| ENSMUSG00000041362 | Shtn1 | protein_coding | -0.287578776 | 5.496170292 | 0.8791262 | 0.362121 | 0.80611581 |
| ENSMUSG00000040482 | Dxo | protein_coding | 0.360475163 | 4.449983873 | 0.879061 | 0.3621383 | 0.80611581 |
| ENSMUSG00000020463 | Ppp4r3b | protein_coding | -0.290280613 | 5.015230073 | 0.8787922 | 0.3622098 | 0.80611581 |
| ENSMUSG00000047213 | Ythdf3 | protein_coding | -0.357962943 | 4.409674026 | 0.8784993 | 0.3622878 | 0.80611581 |
| ENSMUSG00000001227 | Sema6b | protein_coding | 0.531954589 | 3.384376543 | 0.8783418 | 0.3623297 | 0.80611581 |
| ENSMUSG00000002833 | Hdgfl2 | protein_coding | 0.321974366 | 5.880915318 | 0.8781553 | 0.3624017 | 0.80611581 |
| ENSMUSG00000036528 | Ppfibp2 | protein_coding | -0.529315977 | 3.651401463 | 0.8778409 | 0.3624631 | 0.80611581 |
| ENSMUSG00000079330 | Lemd1 | protein_coding | -0.498491145 | 3.620997287 | 0.877807 | 0.3624722 | 0.80611581 |
| ENSMUSG00000028909 | Ptpru | protein_coding | 0.549593031 | 3.507827806 | 0.8778025 | 0.3624734 | 0.80611581 |
| ENSMUSG00000034007 | Scaper | protein_coding | -0.267766773 | 5.731327711 | 0.8777244 | 0.3624942 | 0.80611581 |
| ENSMUSG00000038132 | Rbm24 | protein_coding | -0.368510664 | 4.232803002 | 0.8775581 | 0.3625385 | 0.80611581 |

| ensembl_gene_id | external_gene_name | gene_biotype | log2 (fold change) | log2 (counts per million) | F-value | p-value | False discovery rate |
|---|---|---|---|---|---|---|---|
| ENSMUSG00000033615 | Cplx1 | protein_coding | 0.230434269 | 9.61942052 | 0.8766159 | 0.3627896 | 0.806188608 |
| ENSMUSG00000109836 | Gm45884 | TEC | -0.417304707 | 3.596234696 | 0.8765589 | 0.3628049 | 0.806188608 |
| ENSMUSG00000023249 | Parp3 | protein_coding | -0.613113642 | 2.909312316 | 0.8764829 | 0.3628251 | 0.806188608 |
| ENSMUSG00000026821 | Ralgds | protein_coding | -0.297091166 | 5.58292015 | 0.8763106 | 0.3628711 | 0.806188608 |
| ENSMUSG00000020396 | Nefh | protein_coding | -0.294122326 | 5.388966316 | 0.8750795 | 0.3631998 | 0.806687591 |
| ENSMUSG00000025227 | Mfsd13a | protein_coding | -0.318783754 | 4.583608346 | 0.8746532 | 0.3633137 | 0.806687591 |
| ENSMUSG00000046182 | Gsg1l | protein_coding | -0.52429156 | 3.759179788 | 0.8747103 | 0.3633207 | 0.806687591 |
| ENSMUSG00000017493 | Igfbp4 | protein_coding | -0.523197725 | 4.478549762 | 0.8763239 | 0.3636403 | 0.807230462 |
| ENSMUSG00000024772 | Ehd1 | protein_coding | -0.295826488 | 4.516743433 | 0.8719336 | 0.3640417 | 0.807954652 |
| ENSMUSG00000027006 | Dnajc10 | protein_coding | -0.224995094 | 5.867635521 | 0.8714186 | 0.3641798 | 0.80807033 |
| ENSMUSG00000061322 | Dnaic1 | protein_coding | -0.438978724 | 3.965382689 | 0.871085 | 0.3642693 | 0.80807033 |
| ENSMUSG00000029725 | Ppp1r35 | protein_coding | 0.620325521 | 2.750171303 | 0.8703342 | 0.3644708 | 0.80807033 |
| ENSMUSG00000079477 | Rab7 | protein_coding | 0.284253011 | 6.059395973 | 0.8698915 | 0.3645897 | 0.80807033 |
| ENSMUSG00000006736 | Tspan31 | protein_coding | 0.205706341 | 6.733065879 | 0.8695233 | 0.3646887 | 0.80807033 |
| ENSMUSG00000032077 | Bud13 | protein_coding | 0.533205923 | 4.458012439 | 0.872606 | 0.3647239 | 0.80807033 |
| ENSMUSG00000026199 | Ankzf1 | protein_coding | 0.416800218 | 3.976778825 | 0.8690769 | 0.3648087 | 0.80807033 |
| ENSMUSG00000024370 | Cdc23 | protein_coding | 0.375368771 | 5.487269264 | 0.8690842 | 0.3648289 | 0.80807033 |
| ENSMUSG00000031622 | Sin3b | protein_coding | 0.193144282 | 6.505434759 | 0.8689372 | 0.3648463 | 0.80807033 |
| ENSMUSG00000028479 | Gne | protein_coding | 0.316629336 | 4.514314055 | 0.8689212 | 0.3648505 | 0.80807033 |
| ENSMUSG00000036009 | Mettl25 | protein_coding | 0.528598635 | 2.995099745 | 0.8682631 | 0.3650276 | 0.80807033 |
| ENSMUSG00000006998 | Psmd2 | protein_coding | 0.257121831 | 7.72167652 | 0.8682823 | 0.3650447 | 0.80807033 |
| ENSMUSG00000026878 | Rab14 | protein_coding | -0.205813082 | 7.850148033 | 0.8677945 | 0.3651538 | 0.80807033 |
| ENSMUSG00000027424 | Mgme1 | protein_coding | 0.357146543 | 3.799947461 | 0.8676252 | 0.3651994 | 0.80807033 |
| ENSMUSG00000028948 | Nol9 | protein_coding | -0.539796612 | 3.071242924 | 0.8675447 | 0.365221 | 0.80807033 |
| ENSMUSG00000030519 | Apba2 | protein_coding | 0.318332809 | 5.802000429 | 0.8657942 | 0.3657151 | 0.808829117 |
| ENSMUSG00000030222 | Rerg | protein_coding | -0.265565125 | 5.623906843 | 0.8654171 | 0.3657947 | 0.808829117 |
| ENSMUSG00000010110 | Stx5a | protein_coding | 0.222695481 | 6.442202614 | 0.8652591 | 0.3658374 | 0.808829117 |
| ENSMUSG00000027184 | Caprin1 | protein_coding | -0.232162952 | 6.502732555 | 0.8651575 | 0.3658649 | 0.808829117 |
| ENSMUSG00000097574 | C920006O11Rik | lincRNA | 0.395842608 | 3.671111301 | 0.8646056 | 0.3660139 | 0.808831087 |
| ENSMUSG00000031360 | Ctps2 | protein_coding | 0.35303984 | 5.768637426 | 0.8648954 | 0.3660162 | 0.808831087 |
| ENSMUSG00000079083 | Jrkl | protein_coding | 0.553586457 | 2.992706332 | 0.863607 | 0.3662839 | 0.80925635 |
| ENSMUSG00000062944 | 9130023H24Rik | protein_coding | -0.479263942 | 3.946583191 | 0.8631172 | 0.3664164 | 0.809346463 |
| ENSMUSG00000020530 | Ggnbp2 | protein_coding | -0.20170072 | 7.034951486 | 0.8629 | 0.3664752 | 0.809346463 |
| ENSMUSG00000032532 | Cck | protein_coding | -0.562094136 | 3.604041295 | 0.862462 | 0.3666159 | 0.809490992 |
| ENSMUSG00000030000 | Add2 | protein_coding | 0.219454705 | 7.219291826 | 0.8616572 | 0.3668119 | 0.809757368 |
| ENSMUSG00000037470 | Uggt1 | protein_coding | -0.392987673 | 4.094917282 | 0.8604308 | 0.3671445 | 0.810244248 |
| ENSMUSG00000030583 | Sipa1l3 | protein_coding | -0.561614781 | 3.080284127 | 0.8602019 | 0.3672067 | 0.810244248 |
| ENSMUSG00000045284 | Dcaf12l1 | protein_coding | -0.235213393 | 6.420948291 | 0.8600111 | 0.3672585 | 0.810244248 |
| ENSMUSG00000053166 | Cdh22 | protein_coding | -0.669691465 | 2.911345034 | 0.8597157 | 0.3673387 | 0.810254967 |
| ENSMUSG00000028932 | Psmc2 | protein_coding | 0.202331274 | 8.499905704 | 0.8593308 | 0.3674433 | 0.810277913 |
| ENSMUSG00000026885 | Ttll11 | protein_coding | 0.359896844 | 4.24246534 | 0.8588124 | 0.3675842 | 0.810277913 |
| ENSMUSG00000030264 | Thumpd3 | protein_coding | -0.316492921 | 4.766676556 | 0.8586195 | 0.3676366 | 0.810277913 |
| ENSMUSG00000026000 | Lancl1 | protein_coding | -0.202654209 | 6.946182531 | 0.857998 | 0.3678057 | 0.810277913 |
| ENSMUSG00000051730 | Mettl5 | protein_coding | -0.220737819 | 6.12775311 | 0.8574169 | 0.3679639 | 0.810277913 |
| ENSMUSG00000021520 | Uqcrb | protein_coding | 0.268565427 | 8.483374154 | 0.8564779 | 0.3682418 | 0.810277913 |
| ENSMUSG00000049751 | Rpl36al | protein_coding | 0.482260894 | 3.012442042 | 0.8563111 | 0.3682652 | 0.810277913 |
| ENSMUSG00000002731 | Prkra | protein_coding | 0.232462238 | 6.252492785 | 0.8557687 | 0.3684132 | 0.810277913 |
| ENSMUSG00000041438 | Utp4 | protein_coding | 0.339635642 | 5.843235692 | 0.8557951 | 0.368428 | 0.810277913 |
| ENSMUSG00000023960 | Enpp5 | protein_coding | -0.175805194 | 7.834927433 | 0.8555257 | 0.3684795 | 0.810277913 |
| ENSMUSG00000043059 | Zfp513 | protein_coding | -0.857530233 | 2.252911747 | 0.8551008 | 0.3685954 | 0.810277913 |
| ENSMUSG00000027618 | Nfs1 | protein_coding | 0.325838927 | 4.773280756 | 0.8545711 | 0.3687401 | 0.810277913 |
| ENSMUSG00000111966 | AL589661.1 | processed_pseudoge | 0.483567646 | 3.01471568 | 0.8544514 | 0.3687728 | 0.810277913 |
| ENSMUSG00000087184 | Gm11650 | processed_transcript | 0.693674308 | 2.571897185 | 0.8543932 | 0.3687887 | 0.810277913 |
| ENSMUSG00000024561 | Mbd1 | protein_coding | 0.315466256 | 5.181178733 | 0.8537699 | 0.368959 | 0.810277913 |
| ENSMUSG00000027577 | Chrna4 | protein_coding | 0.57302663 | 4.640889132 | 0.8605566 | 0.3689934 | 0.810277913 |
| ENSMUSG00000003154 | Foxj2 | protein_coding | -0.41209148 | 3.932937719 | 0.8535798 | 0.369011 | 0.810277913 |
| ENSMUSG00000042608 | Stk40 | protein_coding | -0.497671805 | 3.739329331 | 0.8535238 | 0.3690263 | 0.810277913 |
| ENSMUSG00000029463 | Fam216a | protein_coding | 0.391436838 | 4.12607671 | 0.8533683 | 0.3690689 | 0.810277913 |
| ENSMUSG00000032185 | Carm1 | protein_coding | 0.372484393 | 4.119040385 | 0.8532307 | 0.3691065 | 0.810277913 |
| ENSMUSG00000026188 | Tmem169 | protein_coding | 0.371046305 | 3.928872815 | 0.8530884 | 0.3691454 | 0.810277913 |
| ENSMUSG00000027879 | Sec22b | protein_coding | -0.20277902 | 7.907287196 | 0.8529201 | 0.3691915 | 0.810277913 |
| ENSMUSG00000040697 | Dnajc16 | protein_coding | -0.460495047 | 4.041502549 | 0.8527987 | 0.3692247 | 0.810277913 |
| ENSMUSG00000038065 | Mturn | protein_coding | 0.310707235 | 6.059530408 | 0.8523404 | 0.3693722 | 0.810277913 |
| ENSMUSG00000071267 | Zfp942 | protein_coding | 0.471530067 | 3.31002476 | 0.8519967 | 0.3694443 | 0.810277913 |
| ENSMUSG00000025648 | Pfkfb4 | protein_coding | 0.528076131 | 3.353675628 | 0.8519388 | 0.3694602 | 0.810277913 |
| ENSMUSG00000030101 | Sumf1 | protein_coding | -0.498453849 | 4.039992714 | 0.8518393 | 0.3695094 | 0.810277913 |
| ENSMUSG00000020910 | Adprm | protein_coding | 0.266613774 | 5.80374054 | 0.8517168 | 0.369521 | 0.810277913 |
| ENSMUSG00000020668 | Kif3c | protein_coding | 0.769564291 | 2.57057705 | 0.851668 | 0.3695344 | 0.810277913 |
| ENSMUSG00000001418 | Glmp | protein_coding | 0.22835505 | 6.461224664 | 0.8513679 | 0.3696166 | 0.810293039 |
| ENSMUSG00000036112 | Metap2 | protein_coding | 0.219609157 | 7.489610262 | 0.850767 | 0.3697814 | 0.810403705 |
| ENSMUSG00000040054 | Baz2a | protein_coding | -0.391154444 | 4.379592687 | 0.8502722 | 0.3699171 | 0.810403705 |
| ENSMUSG00000053929 | Cyhr1 | protein_coding | 0.193186459 | 6.914207001 | 0.8500041 | 0.3699907 | 0.810403705 |
| ENSMUSG00000026305 | Lrrfip1 | protein_coding | -0.351563846 | 4.908103017 | 0.849855 | 0.3700317 | 0.810403705 |
| ENSMUSG00000049775 | Tmsb4x | protein_coding | 0.285945935 | 11.71089182 | 0.8498904 | 0.3700439 | 0.810403705 |
| ENSMUSG00000028099 | Polr3c | protein_coding | 0.350467775 | 5.569447924 | 0.849585 | 0.3701278 | 0.810422309 |
| ENSMUSG00000024845 | Tmem134 | processed_transcript | 0.415166922 | 3.994797671 | 0.848343 | 0.3704472 | 0.810811523 |
| ENSMUSG00000055720 | Ubl7 | protein_coding | 0.210552128 | 7.95282427 | 0.8482358 | 0.3704766 | 0.810811523 |
| ENSMUSG00000044864 | Ankrd50 | protein_coding | 0.356082466 | 4.326761059 | 0.8477741 | 0.3706037 | 0.810811523 |

| ensembl_gene_id | external_gene_name | gene_biotype | log2 (fold change) | log2 (counts per million) | F-value | p-value | False discovery rate |
|---|---|---|---|---|---|---|---|
| ENSMUSG00000079020 | Slc45a4 | protein_coding | 0.46377194 | 3.704989403 | 0.8473693 | 0.3707151 | 0.810811523 |
| ENSMUSG00000020268 | Lyrm7 | protein_coding | 0.230422238 | 5.825074949 | 0.8473311 | 0.3707256 | 0.810811523 |
| ENSMUSG00000031072 | Oraov1 | protein_coding | 0.26886991 | 5.316259961 | 0.8469874 | 0.3708203 | 0.810811523 |
| ENSMUSG00000022234 | Cct5 | protein_coding | 0.219573317 | 9.145480461 | 0.8469399 | 0.3708334 | 0.810811523 |
| ENSMUSG00000093436 | Gm20646 | sense_overlapping | -0.405358876 | 3.643241201 | 0.8464677 | 0.3709635 | 0.810930636 |
| ENSMUSG00000064349 | mt-Tc | Mt_tRNA | -0.562522897 | 2.982255113 | 0.8456861 | 0.3711789 | 0.810930636 |
| ENSMUSG00000030341 | Tnfrsf1a | protein_coding | -0.701420508 | 2.560831926 | 0.8454502 | 0.371244 | 0.810930636 |
| ENSMUSG00000071679 | Zcchc16 | protein_coding | 0.615964842 | 2.834051564 | 0.8452912 | 0.3712879 | 0.810930636 |
| ENSMUSG00000050272 | Dscam | protein_coding | 0.32148588 | 4.946494286 | 0.8441336 | 0.3716075 | 0.810930636 |
| ENSMUSG00000055737 | Ghr | protein_coding | -0.489367996 | 2.985612569 | 0.843927 | 0.3716646 | 0.810930636 |
| ENSMUSG00000069631 | Strada | protein_coding | -0.382256366 | 4.689361972 | 0.8437022 | 0.3717268 | 0.810930636 |
| ENSMUSG00000024998 | Plce1 | protein_coding | -0.52514407 | 2.957736556 | 0.8436942 | 0.371729 | 0.810930636 |
| ENSMUSG00000026615 | Eprs | protein_coding | -0.253400061 | 6.566048291 | 0.8430722 | 0.371901 | 0.810930636 |
| ENSMUSG00000028803 | Nipal3 | protein_coding | 0.291412145 | 5.249474232 | 0.8429475 | 0.3719355 | 0.810930636 |
| ENSMUSG00000031226 | Pbdc1 | protein_coding | 0.203675543 | 6.387891772 | 0.8428956 | 0.3719498 | 0.810930636 |
| ENSMUSG00000009549 | Srp14 | protein_coding | 0.209471044 | 9.519047073 | 0.8424559 | 0.3720715 | 0.810930636 |
| ENSMUSG00000054428 | Atpif1 | protein_coding | 0.249671337 | 9.284073866 | 0.8421218 | 0.372164 | 0.810930636 |
| ENSMUSG00000071866 | Ppia | protein_coding | 0.189118597 | 7.158662897 | 0.8420364 | 0.3721877 | 0.810930636 |
| ENSMUSG00000017858 | Ift52 | protein_coding | 0.252382951 | 6.539735167 | 0.8417438 | 0.3722687 | 0.810930636 |
| ENSMUSG00000053702 | Nebl | protein_coding | -0.291682276 | 4.805923661 | 0.8416325 | 0.3722995 | 0.810930636 |
| ENSMUSG00000021796 | Bmpr1a | protein_coding | -0.356424123 | 4.532161124 | 0.8414506 | 0.3723499 | 0.810930636 |
| ENSMUSG00000039579 | Grin3a | protein_coding | 0.244227233 | 5.533032447 | 0.8409798 | 0.3724804 | 0.810930636 |
| ENSMUSG00000029602 | Rasal1 | protein_coding | 0.341155568 | 5.027993756 | 0.8408834 | 0.3725072 | 0.810930636 |
| ENSMUSG00000021413 | Prpf4b | protein_coding | -0.283919089 | 5.784699678 | 0.8407432 | 0.372546 | 0.810930636 |
| ENSMUSG00000024283 | Wac | protein_coding | -0.247069649 | 5.65565543 | 0.8405429 | 0.3726016 | 0.810930636 |
| ENSMUSG00000021072 | Tmx1 | protein_coding | 0.215936982 | 7.163245087 | 0.8405413 | 0.372602 | 0.810930636 |
| ENSMUSG00000060143 | Gm10076 | protein_coding | 0.499442164 | 3.08257954 | 0.8404045 | 0.37264 | 0.810930636 |
| ENSMUSG00000033392 | Clasp2 | protein_coding | -0.287056098 | 7.111265289 | 0.8406181 | 0.3726978 | 0.810930636 |
| ENSMUSG00000097675 | 1700101I11Rik | processed_transcript | 0.548846869 | 2.873337155 | 0.839837 | 0.3727975 | 0.810983489 |
| ENSMUSG00000060548 | Tnfrsf19 | protein_coding | -0.499230156 | 3.548605899 | 0.8394542 | 0.3729038 | 0.811050628 |
| ENSMUSG00000022525 | Hrasls | protein_coding | 0.259029267 | 6.23717462 | 0.8388848 | 0.373062 | 0.811230588 |
| ENSMUSG00000021468 | Sptlc1 | protein_coding | -0.306655052 | 4.391982916 | 0.8382657 | 0.373234 | 0.811282419 |
| ENSMUSG00000056771 | Gm10010 | lincRNA | 0.903762043 | 2.306700587 | 0.8382562 | 0.3732367 | 0.811282419 |
| ENSMUSG00000053716 | Dusp7 | protein_coding | -0.362335459 | 4.516290361 | 0.8378156 | 0.3733593 | 0.811301457 |
| ENSMUSG00000026864 | Hspa5 | protein_coding | 0.179767364 | 8.346771058 | 0.8376823 | 0.3733963 | 0.811301457 |
| ENSMUSG00000027360 | Hdc | protein_coding | -0.499203488 | 3.202417784 | 0.836895 | 0.3736155 | 0.811613675 |
| ENSMUSG00000030849 | Fgfr2 | protein_coding | 0.202165027 | 6.727220766 | 0.8358659 | 0.3739023 | 0.812055994 |
| ENSMUSG00000060572 | Mfap2 | protein_coding | 0.386618259 | 4.044965662 | 0.8356225 | 0.3739702 | 0.812055994 |
| ENSMUSG00000038766 | Gabpb2 | protein_coding | -0.666763111 | 2.501157165 | 0.8347949 | 0.3742011 | 0.812353378 |
| ENSMUSG00000021423 | Ly86 | protein_coding | -0.403974862 | 4.544423758 | 0.834406 | 0.3743097 | 0.812353378 |
| ENSMUSG00000028671 | Gale | protein_coding | 0.358919724 | 4.816075524 | 0.8335187 | 0.3745576 | 0.812353378 |
| ENSMUSG00000020090 | Npffr1 | protein_coding | 0.521257441 | 3.041290467 | 0.8333205 | 0.374613 | 0.812353378 |
| ENSMUSG00000047246 | Hist1h2be | protein_coding | -0.598592514 | 2.832654143 | 0.832759 | 0.37477 | 0.812353378 |
| ENSMUSG00000001366 | Fbxo9 | protein_coding | 0.188895832 | 6.406147436 | 0.8324708 | 0.3748507 | 0.812353378 |
| ENSMUSG00000048578 | Mlec | protein_coding | 0.253399362 | 5.514728006 | 0.832253 | 0.3749117 | 0.812353378 |
| ENSMUSG00000024413 | Npc1 | protein_coding | -0.369490505 | 4.208555159 | 0.8321363 | 0.3749443 | 0.812353378 |
| ENSMUSG00000029376 | Mthfd2l | protein_coding | -0.331238296 | 4.601838328 | 0.8320369 | 0.3749721 | 0.812353378 |
| ENSMUSG00000020290 | Xpo1 | protein_coding | -0.304597371 | 5.05689535 | 0.8318479 | 0.3750251 | 0.812353378 |
| ENSMUSG00000022698 | Naa50 | protein_coding | -0.219094032 | 5.952970426 | 0.8318155 | 0.3750341 | 0.812353378 |
| ENSMUSG00000032966 | Fkbp1a | protein_coding | -0.200760782 | 9.154511293 | 0.8314181 | 0.3751455 | 0.812353378 |
| ENSMUSG00000025534 | Gusb | protein_coding | 0.264872884 | 5.25349744 | 0.8314095 | 0.3751479 | 0.812353378 |
| ENSMUSG00000038244 | Mical2 | protein_coding | -0.509279525 | 3.840897518 | 0.8310011 | 0.375284 | 0.812353378 |
| ENSMUSG00000022025 | Cnmd | protein_coding | -1.056115466 | 2.401022234 | 0.8306651 | 0.375412 | 0.812353378 |
| ENSMUSG00000025083 | Afap1l2 | protein_coding | -0.849220709 | 2.378908864 | 0.8301422 | 0.3755032 | 0.812353378 |
| ENSMUSG00000027273 | Snap25 | protein_coding | 0.266655745 | 12.17103416 | 0.8300637 | 0.3755253 | 0.812353378 |
| ENSMUSG00000042816 | Gpr151 | protein_coding | -1.672952396 | 2.063609126 | 0.8363953 | 0.3756099 | 0.812353378 |
| ENSMUSG00000033773 | Rpap2 | protein_coding | 0.425723776 | 4.518197972 | 0.8294776 | 0.3757115 | 0.812353378 |
| ENSMUSG00000059173 | Pde1a | protein_coding | 0.239617225 | 6.814877576 | 0.8292706 | 0.3757479 | 0.812353378 |
| ENSMUSG00000056004 | 9330182L06Rik | protein_coding | -0.300759575 | 5.064102773 | 0.8285051 | 0.375963 | 0.812353378 |
| ENSMUSG00000035776 | Cd99l2 | protein_coding | 0.181598093 | 7.23010765 | 0.8284713 | 0.3759725 | 0.812353378 |
| ENSMUSG00000047155 | Cyp4x1 | protein_coding | 0.408498847 | 3.539105573 | 0.8282687 | 0.3760295 | 0.812353378 |
| ENSMUSG00000040972 | Igsf21 | protein_coding | -0.527468158 | 4.150300322 | 0.829646 | 0.3760521 | 0.812353378 |
| ENSMUSG00000054708 | Ankrd24 | protein_coding | 0.410049554 | 3.893537642 | 0.8281397 | 0.3760657 | 0.812353378 |
| ENSMUSG00000023495 | Pcbp4 | protein_coding | 0.320887435 | 5.32965233 | 0.8281198 | 0.3760713 | 0.812353378 |
| ENSMUSG00000001687 | Ubl3 | protein_coding | -0.190220414 | 8.338305244 | 0.8267967 | 0.3764437 | 0.812994388 |
| ENSMUSG00000036339 | Tmem260 | protein_coding | -0.479045483 | 3.371221441 | 0.8264617 | 0.3765381 | 0.813034888 |
| ENSMUSG00000029672 | Fam3c | protein_coding | -0.264694032 | 5.096168148 | 0.8257142 | 0.3767487 | 0.813219509 |
| ENSMUSG00000021051 | Ppp2r5e | protein_coding | 0.230358995 | 5.915104441 | 0.8256218 | 0.3767748 | 0.813219509 |
| ENSMUSG00000031402 | Mpp1 | protein_coding | 0.264347382 | 6.297692361 | 0.8253316 | 0.3768567 | 0.813232929 |
| ENSMUSG00000033308 | Dpyd | protein_coding | -0.59870039 | 2.844302619 | 0.8250257 | 0.3769429 | 0.813255927 |
| ENSMUSG00000026496 | Parp1 | protein_coding | -0.266486636 | 5.257757209 | 0.8246004 | 0.377063 | 0.813273251 |
| ENSMUSG00000038215 | Cep44 | protein_coding | -0.598052796 | 3.199055763 | 0.8244614 | 0.3771022 | 0.813273251 |
| ENSMUSG00000022641 | Bbx | protein_coding | -0.308148365 | 5.438121776 | 0.8239409 | 0.3772492 | 0.813421937 |
| ENSMUSG00000031555 | Adam9 | protein_coding | -0.403282145 | 3.504916784 | 0.8236818 | 0.3773225 | 0.813421937 |
| ENSMUSG00000097729 | 2310015A10Rik | processed_transcript | -0.533950319 | 3.141519068 | 0.823244 | 0.3774462 | 0.81350805 |
| ENSMUSG00000035671 | Zswim4 | protein_coding | -0.360883708 | 3.825529101 | 0.8227988 | 0.3775722 | 0.81350805 |
| ENSMUSG00000022064 | Pibf1 | protein_coding | -0.51542809 | 3.199530729 | 0.8227378 | 0.3775894 | 0.81350805 |
| ENSMUSG00000040560 | Wdr7 | protein_coding | -0.230766365 | 6.03494408 | 0.8220131 | 0.3777945 | 0.813768614 |

| ensembl_gene_id | external_gene_name | gene_biotype | log2 (fold change) | log2 (counts per million) | F-value | p-value | False discovery rate |
|---|---|---|---|---|---|---|---|
| ENSMUSG00000025404 | R3hdm2 | protein_coding | -0.242318369 | 5.633320548 | 0.8217759 | 0.3778617 | 0.813768614 |
| ENSMUSG00000045136 | Tubb2b | protein_coding | 0.251120118 | 5.423477762 | 0.8205887 | 0.3781981 | 0.814330093 |
| ENSMUSG00000042625 | Safb2 | protein_coding | -0.462142784 | 3.514037681 | 0.8197758 | 0.3784288 | 0.814663549 |
| ENSMUSG00000019795 | Pcmt1 | protein_coding | 0.195154339 | 8.737572817 | 0.8189921 | 0.3786513 | 0.814762907 |
| ENSMUSG00000019689 | Fmc1 | protein_coding | 0.216450619 | 7.439051326 | 0.8187269 | 0.3787266 | 0.814762907 |
| ENSMUSG00000086607 | 4930511M06Rik | processed_transcript | -0.531492884 | 3.102546833 | 0.8186827 | 0.3787392 | 0.814762907 |
| ENSMUSG00000036376 | Abt1 | protein_coding | 0.261666775 | 6.069733086 | 0.8185462 | 0.378778 | 0.814762907 |
| ENSMUSG00000029769 | Ccdc136 | protein_coding | 0.319241272 | 6.416344774 | 0.8202144 | 0.3788538 | 0.814763067 |
| ENSMUSG00000033813 | Tcea1 | protein_coding | -0.313879333 | 4.076756085 | 0.8168841 | 0.3792509 | 0.815325991 |
| ENSMUSG00000021900 | Btd | protein_coding | -0.3717202 | 4.185095894 | 0.8168266 | 0.3792672 | 0.815325991 |
| ENSMUSG00000031758 | Cdyl2 | protein_coding | -0.547446471 | 4.238801782 | 0.8191534 | 0.3794397 | 0.815533804 |
| ENSMUSG00000050069 | Grem2 | protein_coding | 0.451937797 | 3.546617738 | 0.8152998 | 0.3797024 | 0.815770229 |
| ENSMUSG00000062995 | Ica1 | protein_coding | 0.291573649 | 6.464474724 | 0.8149908 | 0.379812 | 0.815770229 |
| ENSMUSG00000031295 | Phka2 | protein_coding | -0.611549174 | 3.009510972 | 0.8149097 | 0.3798137 | 0.815770229 |
| ENSMUSG00000037275 | Gemin5 | protein_coding | -0.45637167 | 3.51674595 | 0.8147713 | 0.3798532 | 0.815770229 |
| ENSMUSG00000031792 | Usb1 | protein_coding | 0.338144348 | 4.800074597 | 0.8144258 | 0.3799518 | 0.815819104 |
| ENSMUSG00000073542 | Cep76 | protein_coding | 0.548654731 | 2.759413771 | 0.8135317 | 0.3802073 | 0.815941055 |
| ENSMUSG00000046449 | C77370 | protein_coding | -0.347785397 | 4.756877016 | 0.8132126 | 0.3802985 | 0.815941055 |
| ENSMUSG00000052253 | Zfp622 | protein_coding | 0.333034342 | 5.262507283 | 0.8131797 | 0.3803079 | 0.815941055 |
| ENSMUSG00000047989 | Ino80c | protein_coding | -0.25101294 | 6.052578601 | 0.8131649 | 0.3803122 | 0.815941055 |
| ENSMUSG00000024454 | Hdac3 | protein_coding | 0.204019302 | 7.052944055 | 0.8114535 | 0.380802 | 0.816664964 |
| ENSMUSG00000070880 | Gad1 | protein_coding | 0.201338514 | 8.867156089 | 0.8111205 | 0.3808974 | 0.816664964 |
| ENSMUSG00000052428 | Tmco1 | protein_coding | 0.211044373 | 7.076710616 | 0.8110831 | 0.3809081 | 0.816664964 |
| ENSMUSG00000029047 | Pex10 | protein_coding | 0.275322037 | 4.967391868 | 0.8109253 | 0.3809534 | 0.816664964 |
| ENSMUSG00000020448 | Rnf185 | protein_coding | 0.322153229 | 5.802407097 | 0.8105223 | 0.3810903 | 0.816740504 |
| ENSMUSG00000037108 | Zcwpw1 | protein_coding | 0.40463915 | 3.452655585 | 0.8102727 | 0.3811405 | 0.816740504 |
| ENSMUSG00000019852 | Arfgef3 | protein_coding | -0.260558242 | 5.896840829 | 0.8092661 | 0.3814294 | 0.817092247 |
| ENSMUSG00000042446 | Zmym4 | protein_coding | -0.676709395 | 2.574492042 | 0.808939 | 0.3815234 | 0.817092247 |
| ENSMUSG00000026958 | Dpp7 | protein_coding | 0.322001256 | 4.544340655 | 0.8086692 | 0.3816009 | 0.817092247 |
| ENSMUSG00000043940 | Wdfy3 | protein_coding | -0.317696566 | 5.725792591 | 0.8080841 | 0.3817691 | 0.817092247 |
| ENSMUSG00000003545 | Fosb | protein_coding | -0.518812322 | 3.305642416 | 0.8077095 | 0.3818769 | 0.817092247 |
| ENSMUSG00000019948 | Actr6 | protein_coding | -0.217265725 | 6.397790488 | 0.807683 | 0.3818845 | 0.817092247 |
| ENSMUSG00000004931 | Apba3 | protein_coding | -0.611435528 | 2.61079719 | 0.8073955 | 0.3819672 | 0.817092247 |
| ENSMUSG00000025485 | Ric8a | protein_coding | 0.278416926 | 5.790532906 | 0.8064242 | 0.3822469 | 0.817092247 |
| ENSMUSG00000050332 | Amer1 | protein_coding | 0.564338112 | 3.496551886 | 0.8064839 | 0.3822511 | 0.817092247 |
| ENSMUSG00000087174 | 5530601H04Rik | processed_transcript | 0.263102699 | 5.187209242 | 0.8061554 | 0.3823244 | 0.817092247 |
| ENSMUSG00000068284 | Usf3 | protein_coding | -0.475469794 | 3.504776773 | 0.8057736 | 0.3824345 | 0.817092247 |
| ENSMUSG00000040007 | Bahd1 | protein_coding | -0.564259644 | 3.270660705 | 0.805576 | 0.3824914 | 0.817092247 |
| ENSMUSG00000041570 | Camsap2 | protein_coding | -0.45782566 | 3.322878241 | 0.8049873 | 0.3826613 | 0.817092247 |
| ENSMUSG00000021713 | Ppwd1 | protein_coding | -0.892882203 | 2.292183636 | 0.8046578 | 0.3827564 | 0.817092247 |
| ENSMUSG00000032598 | Nckipsd | protein_coding | 0.253577962 | 5.747788628 | 0.8044431 | 0.3828184 | 0.817092247 |
| ENSMUSG00000074797 | Itpa | protein_coding | 0.200032402 | 6.312542748 | 0.8042603 | 0.3828712 | 0.817092247 |
| ENSMUSG00000050473 | Slc35d3 | protein_coding | -0.595003861 | 3.052097934 | 0.8041029 | 0.3829166 | 0.817092247 |
| ENSMUSG00000078607 | 1810010H24Rik | protein_coding | -0.937959491 | 2.574770644 | 0.8030519 | 0.3832417 | 0.817092247 |
| ENSMUSG00000027552 | E2f5 | protein_coding | -0.494631958 | 2.977907243 | 0.8027138 | 0.3833182 | 0.817092247 |
| ENSMUSG00000031109 | Enox2 | protein_coding | 0.316109266 | 4.325506399 | 0.8026287 | 0.3833428 | 0.817092247 |
| ENSMUSG00000038957 | Edc3 | protein_coding | 0.306409407 | 5.546286261 | 0.8024534 | 0.3833936 | 0.817092247 |
| ENSMUSG00000029781 | Fkbp9 | protein_coding | -0.571994456 | 3.42077307 | 0.802458 | 0.3834135 | 0.817092247 |
| ENSMUSG00000028273 | Pdlim5 | protein_coding | -0.438030824 | 4.080537185 | 0.8019457 | 0.3835405 | 0.817092247 |
| ENSMUSG00000004902 | Slc25a18 | protein_coding | -0.258997761 | 6.351347819 | 0.8018436 | 0.3835701 | 0.817092247 |
| ENSMUSG00000029250 | Polr2b | protein_coding | -0.261228641 | 5.613263363 | 0.801611 | 0.3836375 | 0.817092247 |
| ENSMUSG00000028070 | Naxe | protein_coding | 0.245635639 | 7.422686091 | 0.8011397 | 0.3837953 | 0.817092247 |
| ENSMUSG00000030884 | Uqcrc2 | protein_coding | 0.191953185 | 8.887543791 | 0.8009588 | 0.3838265 | 0.817092247 |
| ENSMUSG00000026273 | Mterf4 | protein_coding | 0.25164949 | 5.478050371 | 0.8008813 | 0.383849 | 0.817092247 |
| ENSMUSG00000036513 | Commd2 | protein_coding | 0.22257263 | 6.116908395 | 0.8007291 | 0.3838931 | 0.817092247 |
| ENSMUSG00000028455 | Stoml2 | protein_coding | 0.241086065 | 7.123481909 | 0.8004938 | 0.3839613 | 0.817092247 |
| ENSMUSG00000038612 | Mcl1 | protein_coding | -0.379318367 | 4.875636283 | 0.8001574 | 0.3840589 | 0.817092247 |
| ENSMUSG00000036422 | Pcdh8 | protein_coding | -0.407510035 | 4.455041384 | 0.8000987 | 0.3840759 | 0.817092247 |
| ENSMUSG00000038776 | Ephx1 | protein_coding | 0.348657163 | 4.720512754 | 0.7999527 | 0.3841183 | 0.817092247 |
| ENSMUSG00000026603 | Smyd2 | protein_coding | 0.401151318 | 3.360309474 | 0.799622 | 0.3842143 | 0.817092247 |
| ENSMUSG00000063358 | Mapk1 | protein_coding | -0.232555592 | 8.06731653 | 0.7995961 | 0.3842218 | 0.817092247 |
| ENSMUSG00000053080 | 2700081O15Rik | protein_coding | -0.598849875 | 2.671389807 | 0.7995386 | 0.3842385 | 0.817092247 |
| ENSMUSG00000023010 | Tmbim6 | protein_coding | 0.206234081 | 9.694523127 | 0.7994909 | 0.3842524 | 0.817092247 |
| ENSMUSG00000104674 | Gm42756 | antisense_RNA | -0.429158379 | 3.81264865 | 0.7994024 | 0.3842781 | 0.817092247 |
| ENSMUSG00000042743 | Sgtb | protein_coding | -0.288144546 | 5.508618151 | 0.7991044 | 0.3843646 | 0.817092247 |
| ENSMUSG00000020585 | Laptm4a | protein_coding | -0.219980131 | 8.703104482 | 0.7982992 | 0.3845987 | 0.817092247 |
| ENSMUSG00000031516 | Dctn6 | protein_coding | 0.196933472 | 8.032111938 | 0.7978222 | 0.3847374 | 0.817092247 |
| ENSMUSG00000026463 | Atp2b4 | protein_coding | -0.235964329 | 6.025280533 | 0.7977471 | 0.3847593 | 0.817092247 |
| ENSMUSG00000023830 | Igf2r | protein_coding | -0.463974244 | 3.655759802 | 0.7971704 | 0.3849271 | 0.817092247 |
| ENSMUSG00000004267 | Eno2 | protein_coding | 0.252951772 | 10.23750331 | 0.7971323 | 0.3849382 | 0.817092247 |
| ENSMUSG00000025979 | Mob4 | protein_coding | -0.193431465 | 7.801513554 | 0.7969351 | 0.3849956 | 0.817092247 |
| ENSMUSG00000097121 | D130020L05Rik | antisense_RNA | 0.386915796 | 3.731361309 | 0.7967959 | 0.3850362 | 0.817092247 |
| ENSMUSG00000003355 | Fkbp11 | protein_coding | 0.493668723 | 3.242435879 | 0.7967791 | 0.3850411 | 0.817092247 |
| ENSMUSG00000101892 | 9130401M01Rik | protein_coding | 0.287540126 | 4.625813844 | 0.7966558 | 0.385077 | 0.817092247 |
| ENSMUSG00000026739 | Bmi1 | protein_coding | -0.347592421 | 4.297099157 | 0.7958921 | 0.3852995 | 0.817092247 |
| ENSMUSG00000026888 | Grb14 | protein_coding | -0.28709459 | 6.202078848 | 0.7959031 | 0.3853176 | 0.817092247 |
| ENSMUSG00000005682 | Pan2 | protein_coding | -0.759041584 | 2.311549658 | 0.7957218 | 0.3853492 | 0.817092247 |
| ENSMUSG00000069089 | Cdk7 | protein_coding | -0.270228649 | 5.207456483 | 0.795691 | 0.3853582 | 0.817092247 |

| ensembl_gene_id | external_gene_name | gene_biotype | log2 (fold change) | log2 (counts per million) | F-value | p-value | False discovery rate |
|---|---|---|---|---|---|---|---|
| ENSMUSG00000107002 | 0610012G03Rik | protein_coding | 0.343841076 | 4.20281554 | 0.7954277 | 0.385435 | 0.817092247 |
| ENSMUSG00000095105 | Edaradd | protein_coding | -1.317274696 | 1.826958411 | 0.7954012 | 0.3854639 | 0.817092247 |
| ENSMUSG00000008892 | Vdac3 | protein_coding | 0.175785521 | 7.591641338 | 0.795194 | 0.3855032 | 0.817092247 |
| ENSMUSG00000032184 | Lysmd2 | protein_coding | 0.199379476 | 6.451402441 | 0.7943464 | 0.3857506 | 0.817092247 |
| ENSMUSG00000045210 | Vcpip1 | protein_coding | 0.243015717 | 5.468057633 | 0.7943265 | 0.3857564 | 0.817092247 |
| ENSMUSG00000027291 | Vps39 | protein_coding | 0.351954422 | 5.778991377 | 0.795191 | 0.3859045 | 0.817092247 |
| ENSMUSG00000075334 | Rprm | protein_coding | 0.380239206 | 5.63172732 | 0.7960589 | 0.3860226 | 0.817092247 |
| ENSMUSG00000030435 | U2af2 | protein_coding | -0.419675356 | 3.440949511 | 0.7931026 | 0.3861142 | 0.817092247 |
| ENSMUSG00000035227 | Spcs2 | protein_coding | 0.205337547 | 8.275560973 | 0.7928114 | 0.3861993 | 0.817092247 |
| ENSMUSG00000039367 | Sec24c | protein_coding | -0.878894171 | 2.482691938 | 0.7927113 | 0.3862287 | 0.817092247 |
| ENSMUSG00000044627 | Swi5 | protein_coding | 0.223789812 | 9.426210742 | 0.7926259 | 0.3862536 | 0.817092247 |
| ENSMUSG00000030534 | Vps33b | protein_coding | -0.50318184 | 3.958415376 | 0.7924601 | 0.3863233 | 0.817092247 |
| ENSMUSG00000020109 | Dnajb12 | protein_coding | 0.244873378 | 5.728433243 | 0.7923266 | 0.3863412 | 0.817092247 |
| ENSMUSG00000058070 | Eml1 | protein_coding | -0.481329646 | 4.21443811 | 0.7922395 | 0.3863879 | 0.817092247 |
| ENSMUSG00000067336 | Bmpr2 | protein_coding | -0.24835437 | 5.509089548 | 0.7918571 | 0.3864787 | 0.817092247 |
| ENSMUSG00000019726 | Lyst | protein_coding | -0.389127912 | 3.96989673 | 0.7914858 | 0.3865875 | 0.817092247 |
| ENSMUSG00000029009 | Mthfr | protein_coding | -0.635703574 | 3.157893418 | 0.7909689 | 0.3867602 | 0.817092247 |
| ENSMUSG00000017176 | Nt5c3b | protein_coding | 0.195842471 | 6.755125826 | 0.7907785 | 0.3867948 | 0.817092247 |
| ENSMUSG00000037771 | Slc32a1 | protein_coding | -0.501587668 | 4.616713118 | 0.7931502 | 0.3869203 | 0.817092247 |
| ENSMUSG00000053684 | BC048403 | protein_coding | 0.588498085 | 2.762959654 | 0.7903077 | 0.386933 | 0.817092247 |
| ENSMUSG00000036639 | Nudt1 | protein_coding | 0.310762669 | 4.102397497 | 0.7901228 | 0.3869872 | 0.817092247 |
| ENSMUSG00000001034 | Mapk7 | protein_coding | 0.685872579 | 2.897732483 | 0.7900021 | 0.3870226 | 0.817092247 |
| ENSMUSG00000029434 | Vps33a | protein_coding | -0.31889582 | 5.761476132 | 0.7899532 | 0.3870581 | 0.817092247 |
| ENSMUSG00000043542 | Zc2hc1a | protein_coding | 0.189412975 | 7.167439517 | 0.7896233 | 0.3871339 | 0.817092247 |
| ENSMUSG00000027167 | Elp4 | protein_coding | -0.336581225 | 3.932947935 | 0.7895282 | 0.3871618 | 0.817092247 |
| ENSMUSG00000000804 | Usp32 | protein_coding | -0.235477538 | 5.593076024 | 0.7892903 | 0.3872317 | 0.817092247 |
| ENSMUSG00000038070 | Cntln | protein_coding | -0.462603261 | 3.697882556 | 0.788713 | 0.3874013 | 0.817183134 |
| ENSMUSG00000040003 | Magi2 | protein_coding | -0.260766952 | 5.226439022 | 0.7884066 | 0.3874914 | 0.817183134 |
| ENSMUSG00000027080 | Med19 | protein_coding | 0.236339792 | 6.069591869 | 0.7870253 | 0.3878979 | 0.817183134 |
| ENSMUSG00000050846 | Zfp623 | protein_coding | 0.327479887 | 4.333014828 | 0.786812 | 0.3879607 | 0.817183134 |
| ENSMUSG00000034825 | Nrip3 | protein_coding | -0.315158524 | 7.636221506 | 0.7904113 | 0.3879832 | 0.817183134 |
| ENSMUSG00000022075 | Rhobtb2 | protein_coding | -0.278918719 | 5.279647874 | 0.7863567 | 0.3880949 | 0.817183134 |
| ENSMUSG00000022972 | 1110004E09Rik | protein_coding | -0.386355833 | 6.017678376 | 0.7905188 | 0.3881537 | 0.817183134 |
| ENSMUSG00000000869 | Il4 | protein_coding | -0.348598066 | 4.629032743 | 0.7860362 | 0.3881894 | 0.817183134 |
| ENSMUSG00000079442 | St6galnac4 | protein_coding | 0.390680544 | 4.861569362 | 0.786011 | 0.3882179 | 0.817183134 |
| ENSMUSG00000078816 | Prkcg | protein_coding | -0.323063237 | 7.171257509 | 0.7902476 | 0.3882331 | 0.817183134 |
| ENSMUSG00000015290 | Ubl4a | protein_coding | 0.208486757 | 7.399745294 | 0.7858673 | 0.3882392 | 0.817183134 |
| ENSMUSG00000037499 | Nenf | protein_coding | 0.249401789 | 5.674159859 | 0.7858297 | 0.3882503 | 0.817183134 |
| ENSMUSG00000021048 | Mthfd1 | protein_coding | -0.346637868 | 4.127595466 | 0.7856258 | 0.3883104 | 0.817183134 |
| ENSMUSG00000053985 | Zfp14 | protein_coding | -0.365934474 | 3.830076174 | 0.7851589 | 0.3884482 | 0.817183134 |
| ENSMUSG00000063260 | Syt10 | protein_coding | -0.375788799 | 3.929621671 | 0.7848164 | 0.3885493 | 0.817183134 |
| ENSMUSG00000032497 | Lrrfip2 | protein_coding | -0.298813613 | 4.629103 | 0.7846747 | 0.3885911 | 0.817183134 |
| ENSMUSG00000037012 | Hk1 | protein_coding | 0.373710543 | 7.295175808 | 0.7911597 | 0.3887326 | 0.817183134 |
| ENSMUSG00000041037 | Irgq | protein_coding | 0.307012887 | 4.537448549 | 0.7841643 | 0.3887419 | 0.817183134 |
| ENSMUSG00000028811 | Yars | protein_coding | 0.254933539 | 7.068007096 | 0.7841119 | 0.3887784 | 0.817183134 |
| ENSMUSG00000000296 | Tpd52l1 | protein_coding | 0.175220759 | 7.58623538 | 0.783929 | 0.3888114 | 0.817183134 |
| ENSMUSG00000042428 | Mgat3 | protein_coding | 0.249594797 | 5.993891387 | 0.7835752 | 0.3889161 | 0.817183134 |
| ENSMUSG00000059734 | Ndufs8 | protein_coding | 0.195156573 | 9.673630641 | 0.7834523 | 0.3889524 | 0.817183134 |
| ENSMUSG00000001123 | Lgals9 | protein_coding | -0.480085638 | 3.411263067 | 0.7832148 | 0.3890226 | 0.817183134 |
| ENSMUSG00000032754 | Slc8b1 | protein_coding | -0.617333617 | 2.605505296 | 0.7822944 | 0.389295 | 0.817595599 |
| ENSMUSG00000039849 | Pcif1 | protein_coding | 0.215807819 | 6.315142522 | 0.7818289 | 0.3894329 | 0.817725516 |
| ENSMUSG00000014905 | Dnajb9 | protein_coding | -0.314520536 | 5.484625982 | 0.7807508 | 0.3897526 | 0.818091203 |
| ENSMUSG00000020638 | Cmpk2 | protein_coding | 0.256631171 | 5.485329651 | 0.7804833 | 0.3898319 | 0.818091203 |
| ENSMUSG00000021754 | Map3k1 | protein_coding | -0.393373162 | 3.794129042 | 0.7803167 | 0.3898814 | 0.818091203 |
| ENSMUSG00000020458 | Rtn4 | protein_coding | -0.212339761 | 8.419615773 | 0.7802154 | 0.3899114 | 0.818091203 |
| ENSMUSG00000019188 | H13 | protein_coding | 0.374930773 | 5.09278234 | 0.7799887 | 0.3899997 | 0.818116799 |
| ENSMUSG00000023460 | Rab12 | protein_coding | -0.240175741 | 5.846956722 | 0.7791713 | 0.3902215 | 0.818422453 |
| ENSMUSG00000028541 | B4galt2 | protein_coding | 0.40131978 | 3.494569312 | 0.7785937 | 0.3903932 | 0.818595527 |
| ENSMUSG00000016150 | Tenm1 | protein_coding | -0.359093336 | 4.381687609 | 0.7781953 | 0.3905117 | 0.818595527 |
| ENSMUSG00000028789 | Azin2 | protein_coding | 0.312872566 | 4.471869449 | 0.7781258 | 0.3905324 | 0.818595527 |
| ENSMUSG00000054509 | Parp4 | protein_coding | -0.693918746 | 2.787169199 | 0.7768801 | 0.3909033 | 0.819213291 |
| ENSMUSG00000024949 | Sf1 | protein_coding | -0.22236996 | 5.964313519 | 0.7764566 | 0.3910296 | 0.819318165 |
| ENSMUSG00000040270 | Bach2 | protein_coding | -0.486549439 | 3.30761333 | 0.775924 | 0.3911884 | 0.819491224 |
| ENSMUSG00000024436 | Mrps18b | protein_coding | 0.300740977 | 4.956158129 | 0.7745972 | 0.3915845 | 0.819788188 |
| ENSMUSG00000031839 | Hsbp1 | protein_coding | 0.20209411 | 10.62593126 | 0.7745594 | 0.3915958 | 0.819788188 |
| ENSMUSG00000003992 | Ssbp2 | protein_coding | 0.177678607 | 7.282676456 | 0.7744482 | 0.391629 | 0.819788188 |
| ENSMUSG00000016552 | Foxred2 | protein_coding | 0.568835583 | 2.864188343 | 0.7744278 | 0.3916351 | 0.819788188 |
| ENSMUSG00000004637 | Wwox | protein_coding | 0.287323029 | 6.345004564 | 0.7739582 | 0.3917963 | 0.81988147 |
| ENSMUSG00000029518 | Rab35 | protein_coding | 0.251513376 | 5.597696146 | 0.7735167 | 0.3919074 | 0.81988147 |
| ENSMUSG00000020474 | Polm | protein_coding | 0.564539886 | 3.367098616 | 0.7731216 | 0.3920466 | 0.81988147 |
| ENSMUSG00000017453 | Pipox | protein_coding | 0.657138699 | 2.536586131 | 0.7729382 | 0.3920806 | 0.81988147 |
| ENSMUSG00000037085 | Trmt12 | protein_coding | 0.285538494 | 5.502149929 | 0.772769 | 0.3921312 | 0.81988147 |
| ENSMUSG00000038116 | Phf20 | protein_coding | 0.195861187 | 6.675524015 | 0.7727493 | 0.3921371 | 0.81988147 |
| ENSMUSG00000063434 | Sorcs3 | protein_coding | 0.279038055 | 5.116461354 | 0.7714993 | 0.3925116 | 0.820353764 |
| ENSMUSG00000039253 | Fn3krp | protein_coding | -0.717481015 | 3.490451767 | 0.7757314 | 0.3925156 | 0.820353764 |
| ENSMUSG00000025607 | Copg2 | protein_coding | -0.248208675 | 7.752464664 | 0.7708298 | 0.3927333 | 0.820548966 |
| ENSMUSG00000016427 | Ndufa1 | protein_coding | 0.255246206 | 9.064829798 | 0.7704964 | 0.3928333 | 0.820548966 |
| ENSMUSG00000028111 | Ctsk | protein_coding | -0.827672207 | 3.124977368 | 0.7739784 | 0.3928379 | 0.820548966 |

| ensembl_gene_id | external_gene_name | gene_biotype | log2 (fold change) | log2 (counts per million) | F-value | p-value | False discovery rate |
|---|---|---|---|---|---|---|---|
| ENSMUSG00000031568 | Rwdd4a | protein_coding | -0.292452116 | 4.71929133 | 0.7700195 | 0.3929557 | 0.820572354 |
| ENSMUSG00000026986 | Hnmt | protein_coding | -0.190766065 | 6.539923914 | 0.7696412 | 0.3930693 | 0.820572354 |
| ENSMUSG00000029657 | Hsph1 | protein_coding | 0.195340451 | 7.818311086 | 0.7693972 | 0.3931426 | 0.820572354 |
| ENSMUSG00000023007 | Prpf40b | protein_coding | -0.34047772 | 5.256867665 | 0.7693582 | 0.3931544 | 0.820572354 |
| ENSMUSG00000033763 | Mtss1l | protein_coding | -0.243249652 | 5.913357951 | 0.7687391 | 0.3933405 | 0.820735555 |
| ENSMUSG00000003746 | Man1a | protein_coding | -0.319993055 | 4.613703272 | 0.7684461 | 0.3934287 | 0.820735555 |
| ENSMUSG00000028430 | Nol6 | protein_coding | 0.25167104 | 6.081971905 | 0.768294 | 0.3934744 | 0.820735555 |
| ENSMUSG00000052926 | Rnaseh2a | protein_coding | 0.239872349 | 5.930473243 | 0.7680832 | 0.3935379 | 0.820735555 |
| ENSMUSG00000062981 | Mrpl42 | protein_coding | 0.235288577 | 8.561114125 | 0.7672298 | 0.3937949 | 0.821112356 |
| ENSMUSG00000068551 | Zfp467 | protein_coding | -0.350631821 | 4.430571319 | 0.7669425 | 0.3938815 | 0.821133654 |
| ENSMUSG00000003411 | Rab3b | protein_coding | -0.189077728 | 8.485015231 | 0.766548 | 0.3940004 | 0.821222338 |
| ENSMUSG00000067925 | Cxx1a | protein_coding | -0.350345971 | 3.974048044 | 0.7655489 | 0.3943018 | 0.821448197 |
| ENSMUSG00000027018 | Hat1 | protein_coding | -0.248236171 | 5.333454055 | 0.7650951 | 0.3944389 | 0.821448197 |
| ENSMUSG00000039661 | Dusp26 | protein_coding | 0.195008101 | 7.594853447 | 0.7648994 | 0.394498 | 0.821448197 |
| ENSMUSG00000035596 | Mboat7 | protein_coding | 0.21561044 | 6.622442919 | 0.764631 | 0.3945791 | 0.821448197 |
| ENSMUSG00000004151 | Etv1 | protein_coding | -0.266989623 | 5.695036509 | 0.7641618 | 0.3947209 | 0.821448197 |
| ENSMUSG00000041343 | Ankrd42 | protein_coding | -0.492224853 | 3.169587784 | 0.763933 | 0.3947901 | 0.821448197 |
| ENSMUSG00000006169 | Clint1 | protein_coding | 0.282276282 | 4.742445546 | 0.7637605 | 0.3948423 | 0.821448197 |
| ENSMUSG00000036845 | Lin37 | protein_coding | 0.189487277 | 6.935440291 | 0.7634312 | 0.3949419 | 0.821448197 |
| ENSMUSG00000020069 | Hnrnph3 | protein_coding | -0.349082724 | 4.443107926 | 0.763331 | 0.3949723 | 0.821448197 |
| ENSMUSG00000028080 | Lrba | protein_coding | -0.476585758 | 3.827978285 | 0.763027 | 0.3950643 | 0.821448197 |
| ENSMUSG00000011427 | Zfp790 | protein_coding | -0.47534306 | 3.195718129 | 0.7629517 | 0.3950871 | 0.821448197 |
| ENSMUSG00000047694 | Yipf6 | protein_coding | -0.403896116 | 4.256792494 | 0.7624399 | 0.3952421 | 0.821448197 |
| ENSMUSG00000043415 | Otud1 | protein_coding | -0.524598536 | 3.314810434 | 0.7624245 | 0.3952468 | 0.821448197 |
| ENSMUSG00000028657 | Ppt1 | protein_coding | 0.191171993 | 7.447321557 | 0.7622381 | 0.3953033 | 0.821448197 |
| ENSMUSG00000027411 | Vps16 | protein_coding | 0.332648454 | 5.552159739 | 0.7620553 | 0.3953794 | 0.821448197 |
| ENSMUSG00000021661 | Ankra2 | protein_coding | 0.247698052 | 5.411818967 | 0.7618742 | 0.3954136 | 0.821448197 |
| ENSMUSG00000029802 | Abcg2 | protein_coding | 0.548562083 | 2.876853871 | 0.7614888 | 0.3955304 | 0.821448197 |
| ENSMUSG00000036775 | Decr2 | protein_coding | 0.194549912 | 6.638058043 | 0.7612759 | 0.395595 | 0.821448197 |
| ENSMUSG00000036686 | Cc2d1a | protein_coding | 0.367616304 | 4.848330308 | 0.7609676 | 0.3956886 | 0.821448197 |
| ENSMUSG00000061650 | Med9 | protein_coding | 0.364608979 | 4.35018074 | 0.7599384 | 0.3960012 | 0.821448197 |
| ENSMUSG00000056579 | Tug1 | processed_transcript | -0.335702881 | 6.521281845 | 0.7633914 | 0.3960146 | 0.821448197 |
| ENSMUSG00000035910 | Dcdc2a | protein_coding | -0.371642499 | 4.129149183 | 0.7596896 | 0.3960768 | 0.821448197 |
| ENSMUSG00000023092 | Fhl1 | protein_coding | 0.212177025 | 7.233668299 | 0.7595289 | 0.3961256 | 0.821448197 |
| ENSMUSG00000001986 | Gria3 | protein_coding | 0.214983808 | 5.891999901 | 0.7595149 | 0.3961299 | 0.821448197 |
| ENSMUSG00000019843 | Fyn | protein_coding | 0.300073135 | 4.647893775 | 0.7594832 | 0.3961395 | 0.821448197 |
| ENSMUSG00000047603 | Zfp235 | protein_coding | 0.287156513 | 4.396018757 | 0.7593202 | 0.3961891 | 0.821448197 |
| ENSMUSG00000022350 | Washc5 | protein_coding | 0.299179999 | 4.438826356 | 0.7590748 | 0.3962637 | 0.821448197 |
| ENSMUSG00000022681 | Ntan1 | protein_coding | 0.199526398 | 6.963301012 | 0.7587336 | 0.3963675 | 0.821448197 |
| ENSMUSG00000030854 | Ptpn5 | protein_coding | 0.191989701 | 8.552007529 | 0.758685 | 0.3963823 | 0.821448197 |
| ENSMUSG00000056537 | Rlim | protein_coding | -0.305436992 | 5.508096725 | 0.7586251 | 0.3964005 | 0.821448197 |
| ENSMUSG00000030614 | Tmem126b | protein_coding | 0.197905217 | 7.484926532 | 0.7576534 | 0.3966964 | 0.821902973 |
| ENSMUSG00000025580 | Eif4a3 | protein_coding | 0.194962232 | 6.742675217 | 0.7565718 | 0.3970262 | 0.822427716 |
| ENSMUSG00000031782 | Coq9 | protein_coding | 0.204892881 | 6.965553373 | 0.7561074 | 0.3971679 | 0.822562842 |
| ENSMUSG00000052364 | B630019K06Rik | protein_coding | 0.268767939 | 5.775078088 | 0.7551456 | 0.3974616 | 0.82285613 |
| ENSMUSG00000030512 | Snrpa1 | protein_coding | -0.183517944 | 6.516635699 | 0.7548814 | 0.3975424 | 0.82285613 |
| ENSMUSG00000053550 | Shisa7 | protein_coding | 0.344809993 | 4.438449316 | 0.7548011 | 0.3975669 | 0.82285613 |
| ENSMUSG00000089872 | Rps6kc1 | protein_coding | 0.275192626 | 4.90180226 | 0.7545901 | 0.3976314 | 0.82285613 |
| ENSMUSG00000021102 | Glrx5 | protein_coding | 0.344946616 | 5.001211171 | 0.7541526 | 0.3977652 | 0.82285613 |
| ENSMUSG00000025630 | Hprt | protein_coding | 0.176474338 | 9.881867869 | 0.7541414 | 0.3977686 | 0.82285613 |
| ENSMUSG00000030302 | Atp2b2 | protein_coding | -0.206408138 | 8.150041567 | 0.7535768 | 0.3979414 | 0.823055234 |
| ENSMUSG00000018509 | Cenpv | protein_coding | 0.327113182 | 4.041840797 | 0.7530003 | 0.398118 | 0.823157985 |
| ENSMUSG00000020151 | Ptprr | protein_coding | 0.268314476 | 4.876245521 | 0.7529147 | 0.3981442 | 0.823157985 |
| ENSMUSG00000043518 | Rai2 | protein_coding | -0.391915415 | 4.267342959 | 0.7526559 | 0.3982235 | 0.823163696 |
| ENSMUSG00000038112 | AW551984 | protein_coding | 0.29570697 | 6.598811039 | 0.7540314 | 0.3983276 | 0.823179841 |
| ENSMUSG00000032193 | Ldlr | protein_coding | 0.548275072 | 3.684855885 | 0.7527968 | 0.3983844 | 0.823179841 |
| ENSMUSG00000036442 | Thap11 | protein_coding | 0.338214069 | 4.678581292 | 0.7515372 | 0.3985666 | 0.823258102 |
| ENSMUSG00000001518 | Itfg2 | protein_coding | 0.503814427 | 2.944837538 | 0.75148 | 0.3985841 | 0.823258102 |
| ENSMUSG00000024140 | Epas1 | protein_coding | -0.400287936 | 4.294059683 | 0.7512589 | 0.398652 | 0.823258102 |
| ENSMUSG00000063646 | Jakmip1 | protein_coding | -0.207322982 | 5.848733938 | 0.7503306 | 0.3989371 | 0.823563185 |
| ENSMUSG00000030616 | Sytl2 | protein_coding | -0.285882632 | 4.494897183 | 0.7502792 | 0.3989529 | 0.823563185 |
| ENSMUSG00000048490 | Nrip1 | protein_coding | -0.324898633 | 4.388132433 | 0.7499157 | 0.3990646 | 0.823635691 |
| ENSMUSG00000025190 | Got1 | protein_coding | 0.250239039 | 9.60610963 | 0.7492903 | 0.3992775 | 0.823693088 |
| ENSMUSG00000002279 | Lmf1 | protein_coding | 0.368177242 | 4.875749006 | 0.7491253 | 0.3993283 | 0.823693088 |
| ENSMUSG00000022637 | Cblb | protein_coding | -0.631091877 | 3.005102669 | 0.7485251 | 0.3994925 | 0.823693088 |
| ENSMUSG00000026754 | Golga1 | protein_coding | -0.261282947 | 4.866080909 | 0.7484551 | 0.3995141 | 0.823693088 |
| ENSMUSG00000032648 | Pygm | protein_coding | -0.369984631 | 4.634923413 | 0.7482127 | 0.3995888 | 0.823693088 |
| ENSMUSG00000002578 | Ikzf4 | protein_coding | -0.341014612 | 4.382670822 | 0.7480306 | 0.3996449 | 0.823693088 |
| ENSMUSG00000000056 | Narf | protein_coding | -0.252117691 | 5.785371926 | 0.7477992 | 0.3997162 | 0.823693088 |
| ENSMUSG00000031934 | Panx1 | protein_coding | -0.421918979 | 3.856228081 | 0.7476212 | 0.399771 | 0.823693088 |
| ENSMUSG00000038205 | Prkab2 | protein_coding | -0.707209636 | 2.740696418 | 0.7475861 | 0.3997818 | 0.823693088 |
| ENSMUSG00000026887 | Mrrf | protein_coding | 0.262279143 | 5.462209478 | 0.7467267 | 0.400047 | 0.824071332 |
| ENSMUSG00000022545 | Ercc4 | protein_coding | 0.531478812 | 4.237128675 | 0.7498163 | 0.4001187 | 0.824071332 |
| ENSMUSG00000025178 | Pi4k2a | protein_coding | 0.254405026 | 6.30671777 | 0.7461245 | 0.4002329 | 0.824109402 |
| ENSMUSG00000062116 | Zfp954 | protein_coding | 0.333852211 | 5.134182017 | 0.7458418 | 0.4003202 | 0.824109402 |
| ENSMUSG00000014776 | Nol3 | protein_coding | 0.371879111 | 5.167329487 | 0.7457562 | 0.4003671 | 0.824109402 |
| ENSMUSG00000052031 | Tagap1 | protein_coding | 0.29439957 | 5.618157086 | 0.744805 | 0.4006407 | 0.824391821 |
| ENSMUSG00000097040 | 2610316D01Rik | processed_transcript | 0.401383554 | 4.550037084 | 0.7446542 | 0.4007078 | 0.824391821 |

| ensembl_gene_id | external_gene_name | gene_biotype | log2 (fold change) | log2 (counts per million) | F-value | p-value | False discovery rate |
|---|---|---|---|---|---|---|---|
| ENSMUSG00000035585 | Tsen34 | protein_coding | -0.36113838 | 4.467188683 | 0.7445023 | 0.4007343 | 0.824391821 |
| ENSMUSG00000005505 | Kbtbd4 | protein_coding | 0.422090701 | 3.955570353 | 0.7442161 | 0.4008229 | 0.824416322 |
| ENSMUSG00000078235 | Fam43b | protein_coding | 0.50373903 | 2.797049473 | 0.7433987 | 0.401076 | 0.824639795 |
| ENSMUSG00000029474 | Rnf34 | protein_coding | 0.227140582 | 6.829315571 | 0.7432424 | 0.4011244 | 0.824639795 |
| ENSMUSG00000041354 | Rgl2 | protein_coding | 0.753417818 | 2.553597632 | 0.7430436 | 0.4011861 | 0.824639795 |
| ENSMUSG00000031776 | Arl2bp | protein_coding | 0.194916551 | 6.444304096 | 0.7426651 | 0.4013034 | 0.824639795 |
| ENSMUSG00000030304 | Ergic2 | protein_coding | 0.196581941 | 6.167755877 | 0.7426277 | 0.401315 | 0.824639795 |
| ENSMUSG00000019864 | Rtn4ip1 | protein_coding | 0.323573706 | 4.815927102 | 0.7421493 | 0.4014634 | 0.824787139 |
| ENSMUSG00000071648 | Rom1 | protein_coding | -0.471456473 | 3.346982988 | 0.7417786 | 0.4015784 | 0.824865858 |
| ENSMUSG00000028546 | Elavl4 | protein_coding | 0.239624093 | 7.37055083 | 0.7415383 | 0.4016734 | 0.824903448 |
| ENSMUSG00000034037 | Fgd5 | protein_coding | 0.469132944 | 3.056742567 | 0.7409706 | 0.4018294 | 0.825066117 |
| ENSMUSG00000025220 | Mgea5 | protein_coding | -0.335306724 | 4.184251364 | 0.7404335 | 0.4019963 | 0.825251285 |
| ENSMUSG00000015023 | Ddx19a | protein_coding | 0.494410984 | 2.818227446 | 0.7401534 | 0.4020834 | 0.825271819 |
| ENSMUSG00000004127 | Trmt10a | protein_coding | 0.326307568 | 4.116830788 | 0.7399078 | 0.4021598 | 0.825271819 |
| ENSMUSG00000082536 | Gm13456 | processed_pseudoge | 0.66003327 | 2.286606122 | 0.7387503 | 0.4025202 | 0.825656194 |
| ENSMUSG00000032114 | Slc37a4 | protein_coding | -0.362242956 | 4.078771897 | 0.7386943 | 0.4025376 | 0.825656194 |
| ENSMUSG00000032297 | Celf6 | protein_coding | 0.305266875 | 5.126596465 | 0.7385663 | 0.4025775 | 0.825656194 |
| ENSMUSG00000030986 | Dhx32 | protein_coding | -0.311550927 | 4.597175037 | 0.7377143 | 0.4028431 | 0.826028499 |
| ENSMUSG00000045519 | Zfp560 | protein_coding | 0.253244005 | 5.7476614 | 0.7370071 | 0.4030637 | 0.826028499 |
| ENSMUSG00000054728 | Phactr1 | protein_coding | -0.177250035 | 8.711495992 | 0.7367247 | 0.4031519 | 0.826028499 |
| ENSMUSG00000030079 | Ruvbl1 | protein_coding | 0.274932758 | 5.052224155 | 0.7364067 | 0.4032512 | 0.826028499 |
| ENSMUSG00000005125 | Ndrg1 | protein_coding | -0.266007764 | 5.389788109 | 0.7362071 | 0.4033136 | 0.826028499 |
| ENSMUSG00000022565 | Plec | protein_coding | -0.293078571 | 4.769189975 | 0.7360634 | 0.4033585 | 0.826028499 |
| ENSMUSG00000041147 | Brca2 | protein_coding | -0.485083989 | 3.318215813 | 0.7354893 | 0.403538 | 0.826028499 |
| ENSMUSG00000032564 | Cpne4 | protein_coding | -0.283493873 | 5.771340579 | 0.7354829 | 0.40354 | 0.826028499 |
| ENSMUSG00000052593 | Adam17 | protein_coding | -0.796294837 | 2.509099436 | 0.7350298 | 0.4036817 | 0.826028499 |
| ENSMUSG00000032890 | Rims3 | protein_coding | 0.345399653 | 7.353189369 | 0.7411411 | 0.4037065 | 0.826028499 |
| ENSMUSG00000024219 | Anks1 | protein_coding | -0.430609926 | 3.484281808 | 0.7347648 | 0.4037647 | 0.826028499 |
| ENSMUSG00000022721 | Trmt2a | protein_coding | 0.340728248 | 5.02992924 | 0.7340374 | 0.4039925 | 0.826028499 |
| ENSMUSG00000021259 | Cyp46a1 | protein_coding | 0.297860381 | 5.252938892 | 0.7338679 | 0.4040456 | 0.826028499 |
| ENSMUSG00000032217 | Rnf111 | protein_coding | -0.308617673 | 4.463264588 | 0.7337259 | 0.40409 | 0.826028499 |
| ENSMUSG00000038351 | Sgsm2 | protein_coding | -0.298979517 | 4.882327647 | 0.7335866 | 0.4041337 | 0.826028499 |
| ENSMUSG00000021681 | Aggf1 | protein_coding | -0.292770338 | 4.869183155 | 0.7331805 | 0.404261 | 0.826028499 |
| ENSMUSG00000023032 | Slc4a8 | protein_coding | 0.292783068 | 4.621661121 | 0.7331503 | 0.4042705 | 0.826028499 |
| ENSMUSG00000027884 | Clcc1 | protein_coding | 0.285117713 | 4.867255731 | 0.7328588 | 0.4043619 | 0.826028499 |
| ENSMUSG00000064329 | Scn1a | protein_coding | -0.227090184 | 6.450607601 | 0.732692 | 0.4044143 | 0.826028499 |
| ENSMUSG00000034163 | Zfc3h1 | protein_coding | -0.606393052 | 2.682463866 | 0.7326499 | 0.4044275 | 0.826028499 |
| ENSMUSG00000041625 | Ggact | protein_coding | 0.263689461 | 5.141992694 | 0.7326074 | 0.4044408 | 0.826028499 |
| ENSMUSG00000027748 | Trpc4 | protein_coding | 0.307904441 | 4.408237184 | 0.7325813 | 0.404449 | 0.826028499 |
| ENSMUSG00000023104 | Rfc2 | protein_coding | 0.19323042 | 6.384480116 | 0.7313285 | 0.4048424 | 0.826508427 |
| ENSMUSG00000025155 | Dus1l | protein_coding | 0.364832483 | 4.475230336 | 0.7312219 | 0.404876 | 0.826508427 |
| ENSMUSG00000025544 | Tm9sf2 | protein_coding | -0.353037783 | 5.955620366 | 0.7333622 | 0.4049901 | 0.826508427 |
| ENSMUSG00000058748 | Zfp958 | protein_coding | 0.43184232 | 3.542261307 | 0.7308036 | 0.4050075 | 0.826508427 |
| ENSMUSG00000073639 | Rab18 | protein_coding | 0.198861991 | 8.258163609 | 0.730379 | 0.405141 | 0.826508427 |
| ENSMUSG00000063480 | Snu13 | protein_coding | 0.245671975 | 5.062151622 | 0.7303659 | 0.4051452 | 0.826508427 |
| ENSMUSG00000018848 | Rars | protein_coding | 0.219358164 | 6.479045316 | 0.7298817 | 0.4052975 | 0.826539196 |
| ENSMUSG00000031659 | Adcy7 | protein_coding | 0.468518645 | 3.286394113 | 0.7296021 | 0.4053856 | 0.826539196 |
| ENSMUSG00000098702 | 1500015A07Rik | antisense_RNA | 0.458634964 | 3.304849174 | 0.7293512 | 0.4054646 | 0.826539196 |
| ENSMUSG00000034993 | Vat1 | protein_coding | 0.226951958 | 6.200072375 | 0.7289478 | 0.4055917 | 0.826539196 |
| ENSMUSG00000005510 | Ndufs3 | protein_coding | 0.163640868 | 7.416447895 | 0.7287807 | 0.4056444 | 0.826539196 |
| ENSMUSG00000022614 | Lmf2 | protein_coding | 0.447784902 | 3.527637532 | 0.7286481 | 0.4056862 | 0.826539196 |
| ENSMUSG00000007815 | Rhoa | protein_coding | -0.264891969 | 5.819720118 | 0.7283808 | 0.4057705 | 0.826539196 |
| ENSMUSG00000020128 | Vps54 | protein_coding | 0.5110743 | 3.201127812 | 0.728366 | 0.4057752 | 0.826539196 |
| ENSMUSG00000022191 | Drosha | protein_coding | -0.2261184 | 6.414153116 | 0.7277912 | 0.4059565 | 0.826682493 |
| ENSMUSG00000025162 | Csnk1d | protein_coding | 0.221376397 | 6.384389758 | 0.7274751 | 0.4060563 | 0.826682493 |
| ENSMUSG00000049321 | Zfp2 | protein_coding | 0.348402175 | 4.650543512 | 0.7269378 | 0.406226 | 0.826682493 |
| ENSMUSG00000037416 | Dmxl1 | protein_coding | -0.414399498 | 3.915605645 | 0.7268916 | 0.4062406 | 0.826682493 |
| ENSMUSG00000008129 | 4930432K21Rik | protein_coding | 0.65597691 | 2.80066687 | 0.726717 | 0.4062958 | 0.826682493 |
| ENSMUSG00000037260 | Hgsnat | protein_coding | -0.326382337 | 4.476158733 | 0.7266821 | 0.4063068 | 0.826682493 |
| ENSMUSG00000055407 | Map6 | protein_coding | -0.293297971 | 5.469052074 | 0.7257831 | 0.4065911 | 0.826851102 |
| ENSMUSG00000047719 | Ubiad1 | protein_coding | -0.440119223 | 3.207248485 | 0.7255194 | 0.4066745 | 0.826851102 |
| ENSMUSG00000014074 | Rnf168 | protein_coding | -0.312150826 | 4.749605064 | 0.7254078 | 0.4067098 | 0.826851102 |
| ENSMUSG00000021003 | Galc | protein_coding | -0.495068301 | 3.282337328 | 0.725333 | 0.4067335 | 0.826851102 |
| ENSMUSG00000097207 | 6030443J06Rik | lincRNA | -0.462109027 | 3.104366737 | 0.7248914 | 0.4068733 | 0.826851102 |
| ENSMUSG00000048285 | Frmd6 | protein_coding | -0.662427423 | 2.624950927 | 0.7247345 | 0.406923 | 0.826851102 |
| ENSMUSG00000044066 | Cep68 | protein_coding | -0.403399852 | 3.705952981 | 0.724719 | 0.4069279 | 0.826851102 |
| ENSMUSG00000068479 | Mfap1a | protein_coding | -0.313503116 | 4.009950903 | 0.7241267 | 0.4071156 | 0.827076244 |
| ENSMUSG00000039234 | Sec24d | protein_coding | 0.471761859 | 3.237169065 | 0.7235323 | 0.4073041 | 0.827190736 |
| ENSMUSG00000091478 | Gm10039 | processed_pseudoge | 0.451244569 | 2.898952936 | 0.723262 | 0.4073899 | 0.827190736 |
| ENSMUSG00000041396 | Mettl18 | protein_coding | -0.300400449 | 4.638651767 | 0.7230436 | 0.4074592 | 0.827190736 |
| ENSMUSG00000030138 | Bms1 | protein_coding | -0.432751445 | 3.655076753 | 0.722979 | 0.4074797 | 0.827190736 |
| ENSMUSG00000025982 | Sf3b1 | protein_coding | -0.174166119 | 7.046822769 | 0.7220999 | 0.4077589 | 0.827320028 |
| ENSMUSG00000030584 | Dpf1 | protein_coding | -0.309379151 | 4.907281229 | 0.7219067 | 0.4078203 | 0.827320028 |
| ENSMUSG00000022555 | Dgat1 | protein_coding | 0.460322391 | 3.529617965 | 0.721883 | 0.4078279 | 0.827320028 |
| ENSMUSG00000047342 | Zfp286 | protein_coding | 0.277030215 | 5.013948161 | 0.7218097 | 0.4078512 | 0.827320028 |
| ENSMUSG00000027160 | Ccdc34 | protein_coding | -0.507944352 | 3.2779481 | 0.7204684 | 0.4082779 | 0.827962099 |
| ENSMUSG00000027628 | Aar2 | protein_coding | 0.216526992 | 6.484043331 | 0.7201087 | 0.4083924 | 0.827962099 |
| ENSMUSG00000020669 | Sh3yl1 | protein_coding | 0.4071014 | 4.384485981 | 0.7200169 | 0.4084417 | 0.827962099 |

| ensembl_gene_id | external_gene_name | gene_biotype | log2 (fold change) | log2 (counts per million) | F-value | p-value | False discovery rate |
|---|---|---|---|---|---|---|---|
| ENSMUSG00000089917 | Uckl1 | protein_coding | 0.295599368 | 5.065466382 | 0.7198474 | 0.4084757 | 0.827962099 |
| ENSMUSG00000064254 | Ethe1 | protein_coding | 0.33047909 | 4.879331073 | 0.7196001 | 0.4085545 | 0.827965776 |
| ENSMUSG00000021590 | Spata9 | protein_coding | 0.698702364 | 2.518650741 | 0.7190229 | 0.4087385 | 0.828117328 |
| ENSMUSG00000009731 | Kcnd1 | protein_coding | 0.867231942 | 2.365475782 | 0.718664 | 0.4088731 | 0.828117328 |
| ENSMUSG00000093460 | Six3os1 | processed_transcript | -0.460892861 | 3.60449471 | 0.7185853 | 0.4088781 | 0.828117328 |
| ENSMUSG00000097881 | Celrr | lincRNA | -0.747436676 | 2.872962828 | 0.7180252 | 0.409077 | 0.828117328 |
| ENSMUSG00000028608 | 0610037L13Rik | protein_coding | 0.187125679 | 6.864864031 | 0.7176625 | 0.4091728 | 0.828117328 |
| ENSMUSG00000022881 | Rfc4 | protein_coding | -0.530945628 | 4.263347673 | 0.7206832 | 0.4092031 | 0.828117328 |
| ENSMUSG00000031634 | Ufsp2 | protein_coding | 0.169327698 | 7.256195566 | 0.7175379 | 0.4092126 | 0.828117328 |
| ENSMUSG00000034926 | Dhcr24 | protein_coding | 0.49266142 | 3.241174384 | 0.7174353 | 0.4092454 | 0.828117328 |
| ENSMUSG00000042675 | Ypel3 | protein_coding | 0.235706386 | 6.043903776 | 0.7168026 | 0.4094476 | 0.828188936 |
| ENSMUSG00000001773 | Folh1 | protein_coding | -0.584266682 | 2.654719932 | 0.7162855 | 0.4096131 | 0.828188936 |
| ENSMUSG00000058975 | Kcnc1 | protein_coding | 0.260226726 | 5.32414464 | 0.7156587 | 0.4098137 | 0.828188936 |
| ENSMUSG00000040524 | Zfp609 | protein_coding | -0.389174132 | 3.534331021 | 0.715562 | 0.4098447 | 0.828188936 |
| ENSMUSG00000026662 | Sephs1 | protein_coding | -0.365873337 | 4.295783792 | 0.7155405 | 0.4098516 | 0.828188936 |
| ENSMUSG00000036622 | Atp13a2 | protein_coding | 0.317952451 | 5.209432498 | 0.7151033 | 0.4099916 | 0.828188936 |
| ENSMUSG00000028896 | Rcc1 | protein_coding | 0.473605897 | 3.877977304 | 0.7151422 | 0.4099992 | 0.828188936 |
| ENSMUSG00000029027 | Dffb | protein_coding | -0.743603957 | 2.44101323 | 0.7150732 | 0.4100013 | 0.828188936 |
| ENSMUSG00000037730 | Mynn | protein_coding | -0.325553282 | 4.006506073 | 0.7149721 | 0.4100337 | 0.828188936 |
| ENSMUSG00000019817 | Plagl1 | protein_coding | 0.360445696 | 3.547039266 | 0.7149183 | 0.410051 | 0.828188936 |
| ENSMUSG00000023952 | Gtpbp2 | protein_coding | 0.343963915 | 4.165417954 | 0.7143992 | 0.4102174 | 0.828369482 |
| ENSMUSG00000039713 | Plekhg5 | protein_coding | 0.323545683 | 4.772621883 | 0.7137588 | 0.4104229 | 0.82862879 |
| ENSMUSG00000017999 | Ddx27 | protein_coding | -0.332085786 | 4.486232458 | 0.7130897 | 0.4106377 | 0.828756724 |
| ENSMUSG00000086405 | 9330198N18Rik | lincRNA | 0.509878297 | 2.958387547 | 0.7130133 | 0.4106622 | 0.828756724 |
| ENSMUSG00000049176 | Frmpd4 | protein_coding | -0.27332784 | 5.270504808 | 0.7123778 | 0.4108665 | 0.828756724 |
| ENSMUSG00000018411 | Mapt | protein_coding | 0.241307229 | 8.897666719 | 0.7116913 | 0.4111073 | 0.828756724 |
| ENSMUSG00000025959 | Klf7 | protein_coding | -0.241032732 | 5.180760094 | 0.7112742 | 0.4112216 | 0.828756724 |
| ENSMUSG00000046434 | Hnrnpa1 | protein_coding | 0.317481137 | 3.84562228 | 0.7112692 | 0.4112232 | 0.828756724 |
| ENSMUSG00000042272 | Sestd1 | protein_coding | -0.432225365 | 3.482998893 | 0.7110833 | 0.4112831 | 0.828756724 |
| ENSMUSG00000011179 | Odc1 | protein_coding | -0.343788444 | 4.258614454 | 0.7110639 | 0.4112893 | 0.828756724 |
| ENSMUSG00000023031 | Cela1 | protein_coding | -0.482644966 | 3.943138773 | 0.7110134 | 0.4113255 | 0.828756724 |
| ENSMUSG00000095362 | Gm14325 | protein_coding | -0.270332028 | 5.72133313 | 0.7100685 | 0.41161 | 0.828756724 |
| ENSMUSG00000033318 | Gstt2 | protein_coding | 0.404425516 | 3.994900866 | 0.7100569 | 0.4116138 | 0.828756724 |
| ENSMUSG00000068686 | Cd59b | protein_coding | 0.524446506 | 2.836200096 | 0.7098332 | 0.4116859 | 0.828756724 |
| ENSMUSG00000028434 | Epb41l4b | protein_coding | -0.276000636 | 4.797368241 | 0.7097243 | 0.4117211 | 0.828756724 |
| ENSMUSG00000014846 | Tppp3 | protein_coding | 0.219457158 | 8.447401721 | 0.7093238 | 0.4118503 | 0.828756724 |
| ENSMUSG00000042757 | Tmem108 | protein_coding | -0.43587208 | 3.452302513 | 0.7093195 | 0.4118516 | 0.828756724 |
| ENSMUSG00000030638 | Sh3gl3 | protein_coding | -0.310387164 | 5.200236941 | 0.7090736 | 0.411931 | 0.828756724 |
| ENSMUSG00000039195 | 1110008P14Rik | protein_coding | 0.353607306 | 4.218693784 | 0.7089918 | 0.4119574 | 0.828756724 |
| ENSMUSG00000028820 | Sfpq | protein_coding | -0.227788997 | 5.63432158 | 0.7089712 | 0.4119641 | 0.828756724 |
| ENSMUSG00000042462 | Dctpp1 | protein_coding | 0.281048626 | 5.573848654 | 0.7088346 | 0.4120082 | 0.828756724 |
| ENSMUSG00000000581 | C1d | protein_coding | 0.203233696 | 9.177309877 | 0.7087742 | 0.4120277 | 0.828756724 |
| ENSMUSG00000026632 | Tatdn3 | protein_coding | -0.312296988 | 4.434896797 | 0.7084466 | 0.4121335 | 0.828814521 |
| ENSMUSG00000052310 | Slc39a1 | protein_coding | -0.44507243 | 4.916541823 | 0.7105192 | 0.4122373 | 0.828868333 |
| ENSMUSG00000018217 | Pmp22 | protein_coding | -0.28867457 | 5.840557124 | 0.7073888 | 0.4124754 | 0.829161603 |
| ENSMUSG00000051098 | Mblac2 | protein_coding | 0.301486381 | 5.543081478 | 0.7071973 | 0.4125374 | 0.829161603 |
| ENSMUSG00000020661 | Dnmt3a | protein_coding | -0.211068869 | 5.956895013 | 0.7069341 | 0.4126226 | 0.829177784 |
| ENSMUSG00000025371 | Chmp6 | protein_coding | 0.282894237 | 5.972846728 | 0.7059159 | 0.4129722 | 0.829321175 |
| ENSMUSG00000026798 | Coq4 | protein_coding | 0.210365452 | 5.82078392 | 0.7058406 | 0.4129767 | 0.829321175 |
| ENSMUSG00000039633 | Lonrf1 | protein_coding | -0.464277745 | 3.347394235 | 0.7056279 | 0.4130456 | 0.829321175 |
| ENSMUSG00000021732 | Fgf10 | protein_coding | -0.677949926 | 2.872667115 | 0.7054539 | 0.4131021 | 0.829321175 |
| ENSMUSG00000030990 | Pgap2 | protein_coding | -0.278708336 | 4.838311353 | 0.7052427 | 0.4131705 | 0.829321175 |
| ENSMUSG00000048978 | Nrsn1 | protein_coding | -0.174451743 | 9.176900978 | 0.705081 | 0.413223 | 0.829321175 |
| ENSMUSG00000062296 | Trank1 | protein_coding | -0.374653301 | 4.826616183 | 0.7050683 | 0.413247 | 0.829321175 |
| ENSMUSG00000051451 | Crebzf | protein_coding | -0.61773477 | 2.589739195 | 0.7043306 | 0.4134665 | 0.829321175 |
| ENSMUSG00000026155 | Smap1 | protein_coding | -0.403200105 | 3.407709961 | 0.7042431 | 0.4134949 | 0.829321175 |
| ENSMUSG00000004626 | Stxbp2 | protein_coding | -0.350061175 | 4.167826793 | 0.7038001 | 0.4136388 | 0.829321175 |
| ENSMUSG00000010277 | 2610507B11Rik | protein_coding | 0.294810597 | 5.370121584 | 0.7036951 | 0.4136729 | 0.829321175 |
| ENSMUSG00000060166 | Zdhhc8 | protein_coding | 0.369301939 | 3.884171771 | 0.7036331 | 0.4136931 | 0.829321175 |
| ENSMUSG00000041697 | Cox6a1 | protein_coding | 0.281023396 | 9.755176941 | 0.7045416 | 0.4139042 | 0.829321175 |
| ENSMUSG00000026849 | Tor1a | protein_coding | 0.289637499 | 5.716089504 | 0.7029173 | 0.4139258 | 0.829321175 |
| ENSMUSG00000025235 | Bbs4 | protein_coding | 0.274481349 | 6.460608854 | 0.7031263 | 0.4139299 | 0.829321175 |
| ENSMUSG00000028693 | Nasp | protein_coding | 0.324388138 | 4.348576499 | 0.7026929 | 0.4139988 | 0.829321175 |
| ENSMUSG00000035476 | Tab3 | protein_coding | 0.56392335 | 3.025920837 | 0.7024538 | 0.4140766 | 0.829321175 |
| ENSMUSG00000006800 | Sulf2 | protein_coding | -0.290107307 | 5.070675358 | 0.701936 | 0.4142451 | 0.829321175 |
| ENSMUSG00000063087 | Gm10125 | processed_transcript | -0.622430436 | 2.386144775 | 0.7019104 | 0.4142535 | 0.829321175 |
| ENSMUSG00000018481 | Appbp2 | protein_coding | -0.179589992 | 6.66161444 | 0.701857 | 0.4142709 | 0.829321175 |
| ENSMUSG00000040249 | Lrp1 | protein_coding | -0.26277718 | 5.675659049 | 0.701726 | 0.4143135 | 0.829321175 |
| ENSMUSG00000035392 | Dennd1a | protein_coding | -0.290159983 | 4.580423137 | 0.7011558 | 0.4144993 | 0.8295099 |
| ENSMUSG00000029178 | Klf3 | protein_coding | -0.41355851 | 3.876269742 | 0.7009632 | 0.4145621 | 0.8295099 |
| ENSMUSG00000033565 | Rbfox2 | protein_coding | 0.181975339 | 6.285508904 | 0.7007148 | 0.4146431 | 0.829517634 |
| ENSMUSG00000042579 | 4632404H12Rik | lincRNA | 0.351846852 | 3.727082463 | 0.7003357 | 0.4147668 | 0.829610672 |
| ENSMUSG00000031447 | Lamp1 | protein_coding | -0.395363442 | 3.552951563 | 0.6998122 | 0.4149376 | 0.829798036 |
| ENSMUSG00000047824 | Pygo2 | protein_coding | 0.309418177 | 4.81998272 | 0.6990214 | 0.4151959 | 0.830160187 |
| ENSMUSG00000029014 | Dnajc2 | protein_coding | -0.285072068 | 4.619513144 | 0.6987666 | 0.4152792 | 0.830172373 |
| ENSMUSG00000072437 | Nanos1 | protein_coding | -0.307988452 | 4.553238287 | 0.6978828 | 0.4155682 | 0.830415645 |
| ENSMUSG00000093930 | Hmgcs1 | protein_coding | -0.223007194 | 7.080878806 | 0.6977715 | 0.4156047 | 0.830415645 |
| ENSMUSG00000097166 | 9330179D12Rik | antisense_RNA | -0.458670983 | 3.657326077 | 0.6976421 | 0.415647 | 0.830415645 |

| ensembl_gene_id | external_gene_name | gene_biotype | log2 (fold change) | log2 (counts per million) | F-value | p-value | False discovery rate |
|---|---|---|---|---|---|---|---|
| ENSMUSG00000075590 | Nrbp2 | protein_coding | -0.394131294 | 4.648332518 | 0.69726 | 0.4157919 | 0.830415645 |
| ENSMUSG00000034893 | Cog3 | protein_coding | -0.734392645 | 3.031055226 | 0.698418 | 0.4157934 | 0.830415645 |
| ENSMUSG00000075595 | Zfp652 | protein_coding | -0.29706871 | 4.281857141 | 0.6967534 | 0.4159381 | 0.830415645 |
| ENSMUSG00000024498 | Tcerg1 | protein_coding | -0.275233406 | 5.115507489 | 0.6967431 | 0.4159415 | 0.830415645 |
| ENSMUSG00000008035 | Mid1ip1 | protein_coding | -0.268836795 | 6.306994888 | 0.6960004 | 0.4162047 | 0.830786904 |
| ENSMUSG00000036873 | 2410004B18Rik | protein_coding | -0.379419671 | 4.247626815 | 0.695666 | 0.4162947 | 0.830795042 |
| ENSMUSG00000040234 | Tm7sf3 | protein_coding | -0.305180616 | 4.603531011 | 0.6954569 | 0.4163633 | 0.830795042 |
| ENSMUSG00000012429 | Mplkip | protein_coding | 0.596792296 | 2.538806212 | 0.694716 | 0.4166066 | 0.830989319 |
| ENSMUSG00000052798 | Nup107 | protein_coding | 0.291052409 | 4.677629792 | 0.6944036 | 0.4167092 | 0.830989319 |
| ENSMUSG00000020832 | Eral1 | protein_coding | 0.291178615 | 5.595521897 | 0.6943746 | 0.4167188 | 0.830989319 |
| ENSMUSG00000049717 | Lig4 | protein_coding | 0.538936857 | 3.23115119 | 0.6940106 | 0.4168384 | 0.830989319 |
| ENSMUSG00000029207 | Apbb2 | protein_coding | -0.197820894 | 6.354913136 | 0.6937167 | 0.4169351 | 0.830989319 |
| ENSMUSG00000038650 | Rnh1 | protein_coding | 0.217575119 | 7.089744045 | 0.6935182 | 0.4170004 | 0.830989319 |
| ENSMUSG00000034126 | Pomt2 | protein_coding | 0.717174834 | 2.751272529 | 0.6935743 | 0.4170016 | 0.830989319 |
| ENSMUSG00000028525 | Pde4b | protein_coding | -0.203744469 | 7.408735093 | 0.6929179 | 0.417198 | 0.831226547 |
| ENSMUSG00000026812 | Tsc1 | protein_coding | -0.38304198 | 3.868228359 | 0.6921026 | 0.4174665 | 0.831607562 |
| ENSMUSG00000044337 | Ackr3 | protein_coding | -0.7919222 | 2.357359568 | 0.6916092 | 0.4176292 | 0.831777505 |
| ENSMUSG00000018160 | Med1 | protein_coding | -0.250158848 | 6.664700456 | 0.6907016 | 0.4179483 | 0.832113408 |
| ENSMUSG00000032267 | Usp28 | protein_coding | -0.723503217 | 2.665603186 | 0.690629 | 0.4179526 | 0.832113408 |
| ENSMUSG00000037960 | Card19 | protein_coding | -0.339980728 | 4.101817204 | 0.6897659 | 0.4182377 | 0.832342936 |
| ENSMUSG00000028953 | Abcf2 | protein_coding | 0.336801442 | 6.907046021 | 0.6948304 | 0.4182475 | 0.832342936 |
| ENSMUSG00000029787 | Avl9 | protein_coding | -0.298314677 | 4.490577068 | 0.6895772 | 0.4183001 | 0.832342936 |
| ENSMUSG00000056342 | Usp34 | protein_coding | -0.225903825 | 5.862036858 | 0.6882007 | 0.4187556 | 0.833095117 |
| ENSMUSG00000051246 | Msantd1 | protein_coding | 0.490702168 | 3.377604844 | 0.6877579 | 0.4189023 | 0.833232785 |
| ENSMUSG00000026109 | Tmeff2 | protein_coding | 0.200598232 | 7.305166309 | 0.6864242 | 0.4193446 | 0.833828596 |
| ENSMUSG00000016831 | Tox4 | protein_coding | 0.239373759 | 6.124848092 | 0.6861675 | 0.4194298 | 0.833828596 |
| ENSMUSG00000007411 | Mark3 | protein_coding | -0.351783862 | 4.971848746 | 0.6862123 | 0.4194345 | 0.833828596 |
| ENSMUSG00000032459 | Mrps22 | protein_coding | 0.287885748 | 5.211931843 | 0.6856389 | 0.4196053 | 0.833853927 |
| ENSMUSG00000043683 | Fem1a | protein_coding | -0.302077895 | 4.919768379 | 0.6855345 | 0.41964 | 0.833853927 |
| ENSMUSG00000045994 | B3gat1 | protein_coding | -0.290509522 | 4.885530652 | 0.6854146 | 0.4196799 | 0.833853927 |
| ENSMUSG00000023274 | Cd4 | protein_coding | -0.305579836 | 4.681910182 | 0.6844953 | 0.4199855 | 0.833952642 |
| ENSMUSG00000001016 | Ilf2 | protein_coding | 0.176921706 | 7.841223411 | 0.6844629 | 0.4199963 | 0.833952642 |
| ENSMUSG00000046750 | Selenov | protein_coding | 0.286362433 | 4.165583799 | 0.6843172 | 0.4200448 | 0.833952642 |
| ENSMUSG00000037933 | Bicd2 | protein_coding | -0.297556474 | 4.4609733 | 0.6840446 | 0.4201355 | 0.833952642 |
| ENSMUSG00000074575 | Kcng1 | protein_coding | -0.435045958 | 3.102649966 | 0.683904 | 0.4201823 | 0.833952642 |
| ENSMUSG00000040720 | Virma | protein_coding | -0.318459485 | 4.250629407 | 0.6837896 | 0.4202204 | 0.833952642 |
| ENSMUSG00000004771 | Rab11a | protein_coding | -0.167226522 | 6.915055833 | 0.6834339 | 0.4203389 | 0.833952642 |
| ENSMUSG00000005501 | Usp40 | protein_coding | -0.496018928 | 2.889453536 | 0.6832702 | 0.4203934 | 0.833952642 |
| ENSMUSG00000038025 | Phf2 | protein_coding | -0.513056446 | 3.106562759 | 0.6831678 | 0.4204275 | 0.833952642 |
| ENSMUSG00000023826 | Park2 | protein_coding | -0.297408243 | 4.804067349 | 0.6828118 | 0.4205462 | 0.834034184 |
| ENSMUSG00000030772 | Dkk3 | protein_coding | -0.316638456 | 5.609323161 | 0.6823134 | 0.4207319 | 0.834061651 |
| ENSMUSG00000025140 | Pycr1 | protein_coding | -0.459236251 | 3.771223146 | 0.6817742 | 0.4208923 | 0.834061651 |
| ENSMUSG00000029452 | Tmem116 | protein_coding | -0.43288782 | 3.387420825 | 0.6816886 | 0.4209209 | 0.834061651 |
| ENSMUSG00000046862 | Pramef8 | protein_coding | 0.357742295 | 4.561603272 | 0.6816882 | 0.4209211 | 0.834061651 |
| ENSMUSG00000005732 | Ranbp1 | protein_coding | 0.1900493 | 7.170496367 | 0.6815358 | 0.4209719 | 0.834061651 |
| ENSMUSG00000035614 | Togaram1 | protein_coding | -0.323399126 | 4.576164842 | 0.681283 | 0.4210564 | 0.834061651 |
| ENSMUSG00000026790 | Odf2 | protein_coding | 0.274955064 | 5.846598924 | 0.6807196 | 0.4212446 | 0.834061651 |
| ENSMUSG00000028677 | Rnf220 | protein_coding | 0.176288984 | 6.594261474 | 0.6806336 | 0.4212734 | 0.834061651 |
| ENSMUSG00000026404 | Ddx59 | protein_coding | -0.470743889 | 2.920960976 | 0.6805717 | 0.4212941 | 0.834061651 |
| ENSMUSG00000056014 | A430033K04Rik | protein_coding | -0.344436912 | 4.071842789 | 0.6804471 | 0.4213357 | 0.834061651 |
| ENSMUSG00000052928 | Ctif | protein_coding | 0.278292428 | 4.956431126 | 0.6799302 | 0.4215086 | 0.834250375 |
| ENSMUSG00000038594 | Cep85l | protein_coding | -0.62580176 | 3.084781103 | 0.679491 | 0.4216751 | 0.834426213 |
| ENSMUSG00000032097 | Ddx6 | protein_coding | -0.181637364 | 6.223176284 | 0.6787976 | 0.4218878 | 0.834607236 |
| ENSMUSG00000020755 | Sap30bp | protein_coding | 0.16416269 | 6.926630139 | 0.6786963 | 0.4219218 | 0.834607236 |
| ENSMUSG00000024878 | Cbwd1 | protein_coding | 0.260519412 | 4.8534708 | 0.6769606 | 0.422504 | 0.835605253 |
| ENSMUSG00000036469 | 1-Mar | protein_coding | 0.240557223 | 4.910146209 | 0.6764094 | 0.4226892 | 0.835676271 |
| ENSMUSG00000040412 | 5330417C22Rik | protein_coding | -0.319612762 | 4.990527782 | 0.6761687 | 0.4227701 | 0.835676271 |
| ENSMUSG00000063354 | Slc39a4 | protein_coding | -1.915819962 | 1.36567172 | 0.6833907 | 0.4228789 | 0.835676271 |
| ENSMUSG00000026796 | Fam129b | protein_coding | 0.461162572 | 2.90440795 | 0.6755733 | 0.4229703 | 0.835676271 |
| ENSMUSG00000024260 | Sap130 | protein_coding | -0.399186104 | 3.683428337 | 0.6754706 | 0.4230049 | 0.835676271 |
| ENSMUSG00000023990 | Tfeb | protein_coding | -0.814022732 | 2.436470241 | 0.6753284 | 0.4230527 | 0.835676271 |
| ENSMUSG00000036510 | Cdh8 | protein_coding | -0.372841706 | 4.200600969 | 0.6752356 | 0.4230839 | 0.835676271 |
| ENSMUSG00000071073 | Lrrc73 | protein_coding | 0.345752386 | 3.816430592 | 0.6746889 | 0.423268 | 0.835886277 |
| ENSMUSG00000048351 | Coa7 | protein_coding | 0.211913894 | 5.885505597 | 0.6741341 | 0.4234549 | 0.836078302 |
| ENSMUSG00000020476 | Dbnl | protein_coding | 0.179899953 | 6.914025836 | 0.673935 | 0.423522 | 0.836078302 |
| ENSMUSG00000039782 | Cpeb2 | protein_coding | -0.278845396 | 4.842610896 | 0.6734411 | 0.4236886 | 0.836078302 |
| ENSMUSG00000057561 | Eif1a | protein_coding | 0.190147276 | 7.456750366 | 0.6734254 | 0.4236939 | 0.836078302 |
| ENSMUSG00000030493 | Faap24 | protein_coding | 0.34106851 | 4.212610145 | 0.6732471 | 0.423754 | 0.836078302 |
| ENSMUSG00000063065 | Mapk3 | protein_coding | 0.163576282 | 7.28755616 | 0.6729779 | 0.4238448 | 0.836104117 |
| ENSMUSG00000023079 | Gtf2ird1 | protein_coding | -0.344566835 | 4.057280364 | 0.6726587 | 0.4239526 | 0.836145096 |
| ENSMUSG00000043668 | Tox3 | protein_coding | -0.424026869 | 3.709354237 | 0.6723874 | 0.4240442 | 0.836145096 |
| ENSMUSG00000025408 | Ddit3 | protein_coding | -0.214881032 | 6.742839347 | 0.6722256 | 0.4240989 | 0.836145096 |
| ENSMUSG00000037553 | Zdhhc18 | protein_coding | -0.362349852 | 4.081426624 | 0.6713978 | 0.4243787 | 0.836543312 |
| ENSMUSG00000031392 | Irak1 | protein_coding | -0.246079311 | 5.165498465 | 0.6711388 | 0.4244663 | 0.836562649 |
| ENSMUSG00000058093 | Zfp729b | protein_coding | -0.670184691 | 2.732909415 | 0.6706474 | 0.4246325 | 0.836736994 |
| ENSMUSG00000038764 | Ptpn3 | protein_coding | -0.524683446 | 4.327253601 | 0.6733896 | 0.4248811 | 0.836835287 |
| ENSMUSG00000021721 | Htr1a | protein_coding | -0.354094195 | 4.1537979 | 0.6697831 | 0.4249253 | 0.836835287 |
| ENSMUSG00000036120 | Rfxank | protein_coding | 0.271858215 | 4.926668222 | 0.6697001 | 0.4249534 | 0.836835287 |

| ensembl_gene_id | external_gene_name | gene_biotype | log2 (fold change) | log2 (counts per million) | F-value | p-value | False discovery rate |
|---|---|---|---|---|---|---|---|
| ENSMUSG00000047804 | Akap10 | protein_coding | -0.592727877 | 2.627325968 | 0.6694584 | 0.4250353 | 0.836835287 |
| ENSMUSG00000029432 | Nipsnap2 | protein_coding | -0.242913284 | 8.179459623 | 0.6694084 | 0.4250715 | 0.836835287 |
| ENSMUSG00000034908 | Sidt2 | protein_coding | -0.315451886 | 4.407146496 | 0.6674822 | 0.4257061 | 0.837885471 |
| ENSMUSG00000053334 | Ficd | protein_coding | -0.456484154 | 3.187597239 | 0.6671795 | 0.425809 | 0.837885471 |
| ENSMUSG00000025228 | Actr1a | protein_coding | 0.277905314 | 7.759377898 | 0.6691423 | 0.4258898 | 0.837885471 |
| ENSMUSG00000034518 | Hmgxb4 | protein_coding | -0.312569452 | 3.850290875 | 0.6668093 | 0.4259348 | 0.837885471 |
| ENSMUSG00000035621 | Midn | protein_coding | -0.436599556 | 2.965152088 | 0.6665085 | 0.4260372 | 0.837885471 |
| ENSMUSG00000060681 | Slc9a6 | protein_coding | 0.216201 | 5.87618612 | 0.6660033 | 0.4262091 | 0.837885471 |
| ENSMUSG00000055943 | Emc7 | protein_coding | 0.165907288 | 7.536572216 | 0.6659538 | 0.426226 | 0.837885471 |
| ENSMUSG00000024270 | Slc39a6 | protein_coding | -0.304347508 | 5.133146725 | 0.6659468 | 0.4262284 | 0.837885471 |
| ENSMUSG00000029096 | Htra3 | protein_coding | 0.383162492 | 3.500425552 | 0.6654833 | 0.4263862 | 0.838042557 |
| ENSMUSG00000000384 | Tbrg4 | protein_coding | 0.315702978 | 4.916363317 | 0.664887 | 0.4265895 | 0.838288822 |
| ENSMUSG00000046034 | Otulin | protein_coding | 0.407400194 | 3.192789776 | 0.6646045 | 0.4266858 | 0.838324948 |
| ENSMUSG00000028382 | Ptbp3 | protein_coding | -0.356923069 | 3.931620566 | 0.6638456 | 0.4269447 | 0.838577283 |
| ENSMUSG00000020728 | Cep112 | protein_coding | 0.482671603 | 2.978589964 | 0.663771 | 0.4269702 | 0.838577283 |
| ENSMUSG00000025050 | Pcgf6 | protein_coding | -0.634593862 | 2.536865775 | 0.6628353 | 0.4272899 | 0.838978227 |
| ENSMUSG00000037366 | Pafah2 | protein_coding | -0.764777643 | 2.283650416 | 0.6627168 | 0.4273304 | 0.838978227 |
| ENSMUSG00000024827 | Gldc | protein_coding | -0.690499648 | 2.589045251 | 0.6616168 | 0.4277067 | 0.839486888 |
| ENSMUSG00000022940 | Pigp | protein_coding | -0.254808947 | 5.775707716 | 0.6615032 | 0.4277456 | 0.839486888 |
| ENSMUSG00000022861 | Dgkg | protein_coding | -0.195418698 | 6.621568105 | 0.6612024 | 0.4278487 | 0.839531123 |
| ENSMUSG00000037857 | Nufip2 | protein_coding | -0.253815186 | 6.648928208 | 0.6610375 | 0.4279243 | 0.839531123 |
| ENSMUSG00000003421 | Nosip | protein_coding | 0.171096123 | 7.377059727 | 0.6604653 | 0.4281013 | 0.839582297 |
| ENSMUSG00000045767 | B230219D22Rik | protein_coding | -0.222140789 | 5.908005349 | 0.6602442 | 0.4281771 | 0.839582297 |
| ENSMUSG00000031963 | Bmper | protein_coding | -0.525360454 | 2.732481153 | 0.6600886 | 0.4282304 | 0.839582297 |
| ENSMUSG00000028869 | Gnl2 | protein_coding | 0.370582256 | 5.825993876 | 0.6633991 | 0.4282627 | 0.839582297 |
| ENSMUSG00000045404 | Kcnk13 | protein_coding | -0.540163324 | 3.372932918 | 0.6589424 | 0.428643 | 0.84007741 |
| ENSMUSG00000025597 | Klhl4 | protein_coding | -0.328978033 | 3.970025512 | 0.658726 | 0.4286983 | 0.84007741 |
| ENSMUSG00000022469 | Rapgef3 | protein_coding | -0.378296423 | 3.899572658 | 0.6575832 | 0.4290912 | 0.84007741 |
| ENSMUSG00000046818 | Ddit4l | protein_coding | -0.322006915 | 4.730396204 | 0.6575362 | 0.4291074 | 0.84007741 |
| ENSMUSG00000022338 | Eny2 | protein_coding | -0.180705745 | 7.138504758 | 0.6572885 | 0.4291927 | 0.84007741 |
| ENSMUSG00000028397 | Kdm4c | protein_coding | -0.326130916 | 4.255247997 | 0.6571336 | 0.429246 | 0.84007741 |
| ENSMUSG00000048874 | Phf3 | protein_coding | -0.254295322 | 5.110300952 | 0.6570822 | 0.4292637 | 0.84007741 |
| ENSMUSG00000014039 | Prdm15 | protein_coding | -0.671661347 | 2.37350164 | 0.6570764 | 0.4292657 | 0.84007741 |
| ENSMUSG00000028098 | Rnf115 | protein_coding | 0.211060382 | 5.421290025 | 0.6569808 | 0.4292986 | 0.84007741 |
| ENSMUSG00000026281 | Dtymk | protein_coding | 0.23199745 | 6.249767161 | 0.6569405 | 0.4293125 | 0.84007741 |
| ENSMUSG00000023353 | Agap3 | protein_coding | 0.209051993 | 5.872220697 | 0.6567601 | 0.4293746 | 0.84007741 |
| ENSMUSG00000021190 | Lgmn | protein_coding | 0.2084665 | 7.624437304 | 0.6562228 | 0.4295598 | 0.840209379 |
| ENSMUSG00000034789 | Rab24 | protein_coding | 0.356100949 | 4.431594642 | 0.6560928 | 0.4296046 | 0.840209379 |
| ENSMUSG00000037940 | Inpp4b | protein_coding | -0.369510689 | 4.371399073 | 0.6558843 | 0.4296765 | 0.840209379 |
| ENSMUSG00000022899 | Slc15a2 | protein_coding | -0.460014083 | 3.892640066 | 0.6553341 | 0.4298854 | 0.840219641 |
| ENSMUSG00000028248 | Pnisr | protein_coding | -0.235691582 | 5.828569493 | 0.6551241 | 0.4299388 | 0.840219641 |
| ENSMUSG00000003360 | Ddx23 | protein_coding | -0.367269068 | 3.709420481 | 0.6550331 | 0.4299702 | 0.840219641 |
| ENSMUSG00000031375 | Bgn | protein_coding | -0.508422161 | 3.133467997 | 0.6549633 | 0.4299943 | 0.840219641 |
| ENSMUSG00000085396 | Firre | processed_transcript | -0.414087537 | 3.688050705 | 0.6545025 | 0.4301534 | 0.840313976 |
| ENSMUSG00000016181 | Diexf | protein_coding | -0.594235319 | 3.022252251 | 0.6541675 | 0.4302692 | 0.840313976 |
| ENSMUSG00000041977 | Arhgef11 | protein_coding | 0.468020473 | 3.297365252 | 0.654022 | 0.4303195 | 0.840313976 |
| ENSMUSG00000071528 | Usmg5 | protein_coding | 0.230580116 | 7.503552794 | 0.6539739 | 0.4303552 | 0.840313976 |
| ENSMUSG00000024479 | Mal2 | protein_coding | -0.227988617 | 5.853344285 | 0.6535943 | 0.4304674 | 0.840380475 |
| ENSMUSG00000027296 | Itpka | protein_coding | -0.398645218 | 5.082096118 | 0.655263 | 0.4305846 | 0.840429723 |
| ENSMUSG00000070394 | Tmem256 | protein_coding | 0.234299083 | 8.328333018 | 0.6531246 | 0.4306489 | 0.840429723 |
| ENSMUSG00000028889 | Yrdc | protein_coding | -0.363700376 | 4.415054992 | 0.6514856 | 0.4311977 | 0.840852418 |
| ENSMUSG00000044676 | Zfp612 | protein_coding | 0.441951276 | 3.313462669 | 0.6509955 | 0.4313677 | 0.840852418 |
| ENSMUSG00000070576 | Mn1 | protein_coding | 0.307515357 | 8.317535647 | 0.6551076 | 0.4314477 | 0.840852418 |
| ENSMUSG00000045205 | Dpy19l4 | protein_coding | -0.339884381 | 3.949632797 | 0.6506834 | 0.431476 | 0.840852418 |
| ENSMUSG00000040280 | Ndufa4l2 | protein_coding | -0.641448332 | 2.881207013 | 0.6501067 | 0.4316763 | 0.840852418 |
| ENSMUSG00000024193 | Phf1 | protein_coding | 0.334525557 | 4.191752132 | 0.65008 | 0.4316856 | 0.840852418 |
| ENSMUSG00000030075 | Cntn3 | protein_coding | -0.407811027 | 3.320332865 | 0.6498338 | 0.4317711 | 0.840852418 |
| ENSMUSG00000024527 | Afg3l2 | protein_coding | 0.264951792 | 5.199336318 | 0.6497633 | 0.4317957 | 0.840852418 |
| ENSMUSG00000031403 | Dkc1 | protein_coding | -0.217375932 | 5.864226478 | 0.6495764 | 0.4318606 | 0.840852418 |
| ENSMUSG00000039637 | Coro7 | protein_coding | -0.259109869 | 4.868359133 | 0.649563 | 0.4318653 | 0.840852418 |
| ENSMUSG00000040883 | Tmem205 | protein_coding | 0.231313163 | 6.500854608 | 0.6495444 | 0.4318717 | 0.840852418 |
| ENSMUSG00000022415 | Syngr1 | protein_coding | 0.15422357 | 7.596962092 | 0.6494398 | 0.4319081 | 0.840852418 |
| ENSMUSG00000050312 | Nsun3 | protein_coding | -0.370465129 | 3.963659659 | 0.6493468 | 0.4319404 | 0.840852418 |
| ENSMUSG00000022960 | Donson | protein_coding | -0.945228096 | 2.715215233 | 0.6519112 | 0.4320255 | 0.840852418 |
| ENSMUSG00000022237 | Ankrd33b | protein_coding | -0.598039904 | 3.099320655 | 0.6491195 | 0.4320385 | 0.840852418 |
| ENSMUSG00000084957 | Bbip1 | protein_coding | 0.177422995 | 7.193835222 | 0.6487464 | 0.4321493 | 0.840854027 |
| ENSMUSG00000048078 | Tenm4 | protein_coding | -0.354374484 | 4.19453274 | 0.6486132 | 0.4321957 | 0.840854027 |
| ENSMUSG00000024085 | Man2a1 | protein_coding | -0.828998396 | 2.203001725 | 0.6480736 | 0.4323836 | 0.840953197 |
| ENSMUSG00000024190 | Dusp1 | protein_coding | -0.315366647 | 4.473585659 | 0.6480176 | 0.4324031 | 0.840953197 |
| ENSMUSG00000019738 | Polr2i | protein_coding | 0.220426205 | 6.638654352 | 0.6476122 | 0.4325443 | 0.841070271 |
| ENSMUSG00000017002 | Slpi | protein_coding | -0.566731057 | 2.919325958 | 0.6470705 | 0.4327332 | 0.841070271 |
| ENSMUSG00000026455 | Klhl12 | protein_coding | -0.364065289 | 4.628527613 | 0.6470214 | 0.4327692 | 0.841070271 |
| ENSMUSG00000022131 | Gpr180 | protein_coding | -0.390482464 | 3.381276287 | 0.6469474 | 0.4327761 | 0.841070271 |
| ENSMUSG00000020388 | Pdlim4 | protein_coding | -0.429619878 | 3.714481648 | 0.6464503 | 0.4329496 | 0.84121919 |
| ENSMUSG00000029464 | Gpn3 | protein_coding | 0.26968537 | 5.759942848 | 0.6458917 | 0.4331446 | 0.84121919 |
| ENSMUSG00000051727 | Kctd14 | protein_coding | -0.767512438 | 2.14395777 | 0.6456172 | 0.4332405 | 0.84121919 |
| ENSMUSG00000021134 | Srsf5 | protein_coding | 0.177431172 | 7.73424352 | 0.6451917 | 0.4333892 | 0.84121919 |
| ENSMUSG00000092607 | Scnm1 | protein_coding | 0.17819457 | 7.329508744 | 0.6451405 | 0.4334071 | 0.84121919 |

| ensembl_gene_id | external_gene_name | gene_biotype | log2 (fold change) | log2 (counts per million) | F-value | p-value | False discovery rate |
|---|---|---|---|---|---|---|---|
| ENSMUSG00000020135 | Apc2 | protein_coding | -0.365370378 | 4.679888566 | 0.6450702 | 0.4334506 | 0.84121919 |
| ENSMUSG00000022476 | Polr3h | protein_coding | 0.211692631 | 6.335432108 | 0.6448226 | 0.4335183 | 0.84121919 |
| ENSMUSG00000031533 | Mrps31 | protein_coding | 0.258628669 | 6.396049181 | 0.6446879 | 0.4335843 | 0.84121919 |
| ENSMUSG00000024242 | Map4k3 | protein_coding | -0.289223039 | 5.042346066 | 0.6445603 | 0.4336101 | 0.84121919 |
| ENSMUSG00000030276 | Ttll3 | protein_coding | 0.64588744 | 2.692870825 | 0.6443946 | 0.4336681 | 0.84121919 |
| ENSMUSG00000025218 | Poll | protein_coding | 0.279952205 | 6.066550268 | 0.6441038 | 0.4337887 | 0.84121919 |
| ENSMUSG00000028136 | Snx27 | protein_coding | -0.205467042 | 6.71847641 | 0.6440419 | 0.4337915 | 0.84121919 |
| ENSMUSG00000037110 | Ralgapa2 | protein_coding | -0.642042936 | 2.806873281 | 0.6435479 | 0.4339645 | 0.841402956 |
| ENSMUSG00000043602 | Zfp3 | protein_coding | 0.340149709 | 3.811429342 | 0.6432035 | 0.4340852 | 0.841485195 |
| ENSMUSG00000034110 | Kctd7 | protein_coding | -0.320776596 | 4.071132289 | 0.6427017 | 0.4342612 | 0.841618846 |
| ENSMUSG00000010175 | Prox1 | protein_coding | -0.391394853 | 4.413979625 | 0.6424652 | 0.434363 | 0.841618846 |
| ENSMUSG00000026173 | Plcd4 | protein_coding | 0.390401242 | 3.898973072 | 0.6423374 | 0.434389 | 0.841618846 |
| ENSMUSG00000028936 | Rpl22 | protein_coding | 0.203594607 | 7.214161766 | 0.6420245 | 0.4344988 | 0.841680002 |
| ENSMUSG00000006711 | D130043K22Rik | protein_coding | -0.484299801 | 2.919162373 | 0.6416893 | 0.4346165 | 0.84175629 |
| ENSMUSG00000037257 | Aagab | protein_coding | 0.24981872 | 6.616531165 | 0.6412023 | 0.4348064 | 0.841972526 |
| ENSMUSG00000028439 | Fam219a | protein_coding | -0.535554232 | 2.820741868 | 0.6405359 | 0.4350219 | 0.842224712 |
| ENSMUSG00000024780 | Cdc37l1 | protein_coding | -0.179931078 | 6.340182071 | 0.6400483 | 0.4351934 | 0.842224712 |
| ENSMUSG00000034981 | Parm1 | protein_coding | 0.285945442 | 6.183718437 | 0.6406757 | 0.435199 | 0.842224712 |
| ENSMUSG00000035246 | Pcyt1b | protein_coding | -0.295690328 | 4.864259464 | 0.6398876 | 0.43525 | 0.842224712 |
| ENSMUSG00000025969 | Nrp2 | protein_coding | 0.333106649 | 4.422069096 | 0.6394444 | 0.435406 | 0.842259416 |
| ENSMUSG00000020949 | Fkbp3 | protein_coding | 0.17121579 | 9.463429662 | 0.6393918 | 0.4354245 | 0.842259416 |
| ENSMUSG00000025466 | Fuom | protein_coding | 0.19461169 | 7.227118524 | 0.6386141 | 0.4356986 | 0.842355896 |
| ENSMUSG00000026917 | Wdr5 | protein_coding | 0.377796547 | 3.894247691 | 0.6385795 | 0.4357108 | 0.842355896 |
| ENSMUSG00000021013 | Ttc8 | protein_coding | -0.237264814 | 5.003376252 | 0.6385388 | 0.4357252 | 0.842355896 |
| ENSMUSG00000039000 | Ube3c | protein_coding | -0.237658476 | 5.459230692 | 0.6383613 | 0.4357878 | 0.842355896 |
| ENSMUSG00000058756 | Thra | protein_coding | 0.194529987 | 6.284864811 | 0.6379479 | 0.4359336 | 0.842486389 |
| ENSMUSG00000061613 | U2af1 | protein_coding | 0.281984717 | 4.06321943 | 0.637268 | 0.4361736 | 0.842798773 |
| ENSMUSG00000045690 | Wdr89 | protein_coding | 0.311595124 | 4.393169587 | 0.6365251 | 0.4364362 | 0.843154573 |
| ENSMUSG00000004789 | Dlst | protein_coding | 0.207846909 | 6.92417559 | 0.6362353 | 0.4365387 | 0.84317389 |
| ENSMUSG00000032087 | Dscaml1 | protein_coding | 0.501660393 | 3.56028053 | 0.6355866 | 0.4367869 | 0.84317389 |
| ENSMUSG00000026753 | Ppp6c | protein_coding | -0.175726288 | 6.718598413 | 0.6354386 | 0.4368206 | 0.84317389 |
| ENSMUSG00000038324 | Trpc4ap | protein_coding | 0.175879463 | 6.560717276 | 0.6353989 | 0.4368346 | 0.84317389 |
| ENSMUSG00000036606 | Plxnb2 | protein_coding | 0.417975356 | 3.938736052 | 0.6354416 | 0.4368383 | 0.84317389 |
| ENSMUSG00000024921 | Smarca2 | protein_coding | 0.206081619 | 6.0614837 | 0.6349744 | 0.436985 | 0.843181629 |
| ENSMUSG00000029245 | Epha5 | protein_coding | -0.279203313 | 5.230722747 | 0.6349345 | 0.4369991 | 0.843181629 |
| ENSMUSG00000031497 | Tnfsf13b | protein_coding | 0.509437323 | 3.010661282 | 0.6346981 | 0.4370829 | 0.843191983 |
| ENSMUSG00000028291 | Akirin2 | protein_coding | -0.249847915 | 5.350126835 | 0.6343237 | 0.4372156 | 0.843296684 |
| ENSMUSG00000020218 | Wif1 | protein_coding | -0.558624068 | 3.057694366 | 0.6339778 | 0.4373382 | 0.843382015 |
| ENSMUSG00000063052 | Lrrc40 | protein_coding | -0.274419232 | 5.179749987 | 0.6329548 | 0.4377014 | 0.843832207 |
| ENSMUSG00000022313 | Utp23 | protein_coding | 0.325664135 | 4.466096438 | 0.632878 | 0.4377286 | 0.843832207 |
| ENSMUSG00000003527 | Dgcr14 | protein_coding | 0.245626395 | 5.291420031 | 0.6319137 | 0.4380714 | 0.84399759 |
| ENSMUSG00000031534 | Smim19 | protein_coding | -0.288482435 | 4.746452526 | 0.6317341 | 0.4381353 | 0.84399759 |
| ENSMUSG00000024957 | Kcnk4 | protein_coding | 0.651227276 | 2.800689289 | 0.6317223 | 0.4381395 | 0.84399759 |
| ENSMUSG00000031378 | Abcd1 | protein_coding | 0.5718129 | 2.850826298 | 0.6314896 | 0.4382223 | 0.84399759 |
| ENSMUSG00000040621 | Gemin8 | protein_coding | -0.316659283 | 3.708095508 | 0.6313064 | 0.4382875 | 0.84399759 |
| ENSMUSG00000037196 | Pacrg | protein_coding | 0.186097603 | 6.015750497 | 0.6311289 | 0.4383507 | 0.84399759 |
| ENSMUSG00000045009 | Prrt3 | protein_coding | -0.562569599 | 2.80428381 | 0.6310607 | 0.438375 | 0.84399759 |
| ENSMUSG00000039835 | Nhsl1 | protein_coding | -0.443323399 | 3.785744422 | 0.6308972 | 0.4384518 | 0.84399759 |
| ENSMUSG00000028550 | Atg4c | protein_coding | -0.245100743 | 5.355859429 | 0.6302631 | 0.4386591 | 0.84399759 |
| ENSMUSG00000038077 | Kcna6 | protein_coding | 0.253090212 | 5.795123717 | 0.6300458 | 0.4387366 | 0.84399759 |
| ENSMUSG00000017747 | Ghdc | protein_coding | -0.747344666 | 2.546931946 | 0.6299947 | 0.4387548 | 0.84399759 |
| ENSMUSG00000059851 | Kmt5c | protein_coding | -0.508632223 | 3.019809196 | 0.6299685 | 0.4387641 | 0.84399759 |
| ENSMUSG00000025893 | Kbtbd3 | protein_coding | -0.253896482 | 5.278127537 | 0.6297703 | 0.4388348 | 0.84399759 |
| ENSMUSG00000059142 | Zfp945 | protein_coding | -0.285599165 | 4.555713559 | 0.6295298 | 0.4389206 | 0.844011678 |
| ENSMUSG00000059602 | Syn3 | protein_coding | 0.408484618 | 3.615109331 | 0.6281092 | 0.4394279 | 0.844306062 |
| ENSMUSG00000038975 | Rabggtb | protein_coding | 0.149989672 | 7.224132248 | 0.6279662 | 0.439479 | 0.844306062 |
| ENSMUSG00000026103 | Gls | protein_coding | -0.245075397 | 6.054018853 | 0.627961 | 0.4394809 | 0.844306062 |
| ENSMUSG00000002006 | Pdzd4 | protein_coding | -0.342660854 | 4.355802485 | 0.6279589 | 0.4394816 | 0.844306062 |
| ENSMUSG00000028785 | Hpca | protein_coding | 0.194379787 | 9.923674582 | 0.6278197 | 0.4395314 | 0.844306062 |
| ENSMUSG00000026566 | Mpzl1 | protein_coding | -0.490148615 | 3.369692735 | 0.6277822 | 0.4395448 | 0.844306062 |
| ENSMUSG00000027351 | Spred1 | protein_coding | -0.190910366 | 6.353192015 | 0.6274729 | 0.4396554 | 0.844367731 |
| ENSMUSG00000027361 | Gabpb1 | protein_coding | -0.315584456 | 4.01135822 | 0.6270616 | 0.4398026 | 0.84439779 |
| ENSMUSG00000028436 | Dcaf12 | protein_coding | -0.265075787 | 4.935003927 | 0.6269903 | 0.4398281 | 0.84439779 |
| ENSMUSG00000003423 | Pih1d1 | protein_coding | 0.259530188 | 5.332552352 | 0.626645 | 0.4399517 | 0.844399156 |
| ENSMUSG00000056763 | Cspp1 | protein_coding | 0.241743259 | 4.77568364 | 0.6265497 | 0.4399859 | 0.844399156 |
| ENSMUSG00000030774 | Pak1 | protein_coding | 0.196420447 | 8.523614792 | 0.6262679 | 0.4400868 | 0.844442165 |
| ENSMUSG00000035443 | Thyn1 | protein_coding | 0.161479401 | 7.727956212 | 0.6259313 | 0.4402075 | 0.844496108 |
| ENSMUSG00000024647 | Cbln2 | protein_coding | 0.276380632 | 4.755811278 | 0.625336 | 0.440421 | 0.844496108 |
| ENSMUSG00000032280 | Tle3 | protein_coding | -0.503860254 | 3.152596638 | 0.624942 | 0.4405623 | 0.844496108 |
| ENSMUSG00000048355 | Arxes1 | protein_coding | -0.304907777 | 4.788125196 | 0.6248542 | 0.4405938 | 0.844496108 |
| ENSMUSG00000062444 | Ap3b2 | protein_coding | 0.222888638 | 6.66180657 | 0.6244657 | 0.4407334 | 0.844496108 |
| ENSMUSG00000032705 | Exd2 | protein_coding | 0.309001728 | 4.414390876 | 0.6244397 | 0.4407427 | 0.844496108 |
| ENSMUSG00000064326 | Siva1 | protein_coding | 0.250126092 | 5.736593877 | 0.6243119 | 0.4407886 | 0.844496108 |
| ENSMUSG00000021248 | Tmed10 | protein_coding | -0.178855694 | 6.826569887 | 0.6237733 | 0.4409821 | 0.844496108 |
| ENSMUSG00000036282 | Naa30 | protein_coding | -0.339640255 | 4.880531943 | 0.6235617 | 0.4410767 | 0.844496108 |
| ENSMUSG00000055531 | Cpsf6 | protein_coding | -0.234899944 | 5.774154191 | 0.6233493 | 0.4411346 | 0.844496108 |
| ENSMUSG00000020038 | Cry1 | protein_coding | -0.515795379 | 3.987714875 | 0.6251834 | 0.4411952 | 0.844496108 |
| ENSMUSG00000006390 | Elovl1 | protein_coding | -0.270390476 | 4.990622228 | 0.6226323 | 0.4413926 | 0.844496108 |

| ensembl_gene_id | external_gene_name | gene_biotype | log2 (fold change) | log2 (counts per million) | F-value | p-value | False discovery rate |
|---|---|---|---|---|---|---|---|
| ENSMUSG00000037523 | Mavs | protein_coding | 0.360012513 | 4.380389987 | 0.6224455 | 0.4414599 | 0.844496108 |
| ENSMUSG00000002341 | Ncan | protein_coding | -0.18141717 | 7.218162338 | 0.6223021 | 0.4415115 | 0.844496108 |
| ENSMUSG00000028051 | Hcn3 | protein_coding | -0.395379214 | 3.320938554 | 0.6221943 | 0.4415503 | 0.844496108 |
| ENSMUSG00000101585 | 1600010M07Rik | antisense_RNA | -0.551537812 | 2.830697685 | 0.6215211 | 0.4417929 | 0.844496108 |
| ENSMUSG00000032002 | Dcun1d5 | protein_coding | 0.195647253 | 5.856712684 | 0.6212676 | 0.4418843 | 0.844496108 |
| ENSMUSG00000029708 | Gcc1 | protein_coding | -0.327735958 | 3.788800149 | 0.6211203 | 0.4419375 | 0.844496108 |
| ENSMUSG00000060716 | Plekhh1 | protein_coding | -0.492495393 | 3.439004716 | 0.6209382 | 0.4420217 | 0.844496108 |
| ENSMUSG00000017754 | Pltp | protein_coding | -0.323073381 | 4.931605448 | 0.6206589 | 0.442104 | 0.844496108 |
| ENSMUSG00000038271 | Iffo1 | protein_coding | -0.346929175 | 4.008015608 | 0.6203514 | 0.442215 | 0.844496108 |
| ENSMUSG00000040997 | Abhd4 | protein_coding | -0.183094413 | 6.797734357 | 0.6202913 | 0.4422367 | 0.844496108 |
| ENSMUSG00000021720 | Rnf180 | protein_coding | 0.270404993 | 4.500126184 | 0.619904 | 0.4423766 | 0.844496108 |
| ENSMUSG00000012114 | Med15 | protein_coding | 0.251908115 | 5.038707628 | 0.6192945 | 0.4425969 | 0.844496108 |
| ENSMUSG00000050148 | Ubqln2 | protein_coding | 0.167184087 | 6.796164998 | 0.6189613 | 0.4427174 | 0.844496108 |
| ENSMUSG00000040747 | Cd53 | protein_coding | -0.387223477 | 4.078831472 | 0.6187555 | 0.4427919 | 0.844496108 |
| ENSMUSG00000037966 | Ninj1 | protein_coding | 0.273021973 | 4.936713085 | 0.6185471 | 0.4428673 | 0.844496108 |
| ENSMUSG00000026197 | Zfand2b | protein_coding | 0.214613388 | 5.663000998 | 0.6176825 | 0.4431804 | 0.844496108 |
| ENSMUSG00000021814 | Anxa7 | protein_coding | 0.209516529 | 6.427187044 | 0.6175289 | 0.4432361 | 0.844496108 |
| ENSMUSG00000099964 | Gm29514 | antisense_RNA | -0.313046901 | 3.670833723 | 0.6174725 | 0.4432565 | 0.844496108 |
| ENSMUSG00000040463 | Mybbp1a | protein_coding | 0.266531792 | 4.832715963 | 0.6172114 | 0.4433512 | 0.844496108 |
| ENSMUSG00000031059 | Ndufb11 | protein_coding | 0.201235803 | 9.367674835 | 0.6171039 | 0.4433901 | 0.844496108 |
| ENSMUSG00000035168 | Tanc1 | protein_coding | -0.343582592 | 3.707261857 | 0.6170657 | 0.443404 | 0.844496108 |
| ENSMUSG00000048755 | Mcat | protein_coding | 0.531048514 | 3.814426052 | 0.6190392 | 0.4434502 | 0.844496108 |
| ENSMUSG00000047714 | Ppp1r2 | protein_coding | -0.256614558 | 5.611116276 | 0.6168698 | 0.443475 | 0.844496108 |
| ENSMUSG00000039754 | Alkbh4 | protein_coding | 0.37879445 | 4.552657248 | 0.6166966 | 0.4435563 | 0.844496108 |
| ENSMUSG00000037847 | Nmrk1 | protein_coding | -0.416758659 | 3.704456929 | 0.6165404 | 0.4435946 | 0.844496108 |
| ENSMUSG00000050914 | Ankrd37 | protein_coding | -0.248602481 | 4.687636827 | 0.6164327 | 0.4436337 | 0.844496108 |
| ENSMUSG00000027886 | 1700013F07Rik | protein_coding | -0.864847709 | 2.351329396 | 0.6161877 | 0.443741 | 0.844496108 |
| ENSMUSG00000045854 | Lyrm2 | protein_coding | 0.230744474 | 6.16688225 | 0.6159487 | 0.4438094 | 0.844496108 |
| ENSMUSG00000042761 | Mrap2 | protein_coding | 0.245743083 | 5.658619907 | 0.6156053 | 0.4439341 | 0.844496108 |
| ENSMUSG00000028643 | Svbp | protein_coding | 0.224566319 | 5.704019942 | 0.6154176 | 0.4440024 | 0.844496108 |
| ENSMUSG00000035840 | Lysmd3 | protein_coding | -0.342171826 | 3.98831712 | 0.6153735 | 0.4440184 | 0.844496108 |
| ENSMUSG00000039206 | Daglb | protein_coding | 0.248338666 | 4.974757951 | 0.6153699 | 0.4440197 | 0.844496108 |
| ENSMUSG00000032458 | Copb2 | protein_coding | -0.221244875 | 6.600300104 | 0.6149628 | 0.4441677 | 0.844496108 |
| ENSMUSG00000033543 | Gtf2a2 | protein_coding | 0.177724236 | 7.381250823 | 0.6147693 | 0.4442381 | 0.844496108 |
| ENSMUSG00000023832 | Acat2 | protein_coding | 0.209617855 | 6.774164154 | 0.6147522 | 0.4442443 | 0.844496108 |
| ENSMUSG00000021068 | Nin | protein_coding | -0.420754068 | 3.182937395 | 0.6147516 | 0.4442445 | 0.844496108 |
| ENSMUSG00000024985 | Tcf7l2 | protein_coding | -0.818600452 | 8.165222098 | 0.6484934 | 0.4442572 | 0.844496108 |
| ENSMUSG00000032615 | Nt5m | protein_coding | 0.201346611 | 6.081307107 | 0.6145951 | 0.4443015 | 0.844496108 |
| ENSMUSG00000048799 | Cep120 | protein_coding | -0.265998174 | 4.952780151 | 0.6145842 | 0.4443054 | 0.844496108 |
| ENSMUSG00000058655 | Eif4b | protein_coding | 0.243990753 | 7.164427178 | 0.6151242 | 0.4443224 | 0.844496108 |
| ENSMUSG00000021120 | Pigh | protein_coding | 0.48429576 | 3.330616155 | 0.6145271 | 0.4443262 | 0.844496108 |
| ENSMUSG00000032925 | Itgbl1 | protein_coding | -0.358807873 | 3.819230925 | 0.6141643 | 0.4444583 | 0.844496108 |
| ENSMUSG00000018846 | Pank3 | protein_coding | -0.262312911 | 4.737048058 | 0.6140344 | 0.4445055 | 0.844496108 |
| ENSMUSG00000028978 | Nos3 | protein_coding | 0.425356725 | 3.220516984 | 0.614014 | 0.444513 | 0.844496108 |
| ENSMUSG00000018209 | Stk4 | protein_coding | -0.27449784 | 4.73692849 | 0.6137963 | 0.4445922 | 0.844497511 |
| ENSMUSG00000046814 | Gchfr | protein_coding | -0.382212908 | 3.452972226 | 0.6130187 | 0.4448756 | 0.844847013 |
| ENSMUSG00000025551 | Fgf14 | protein_coding | -0.285016164 | 5.814159515 | 0.6129107 | 0.4449334 | 0.844847013 |
| ENSMUSG00000054099 | Slc25a40 | protein_coding | -0.40415924 | 3.363760727 | 0.6121818 | 0.445181 | 0.845076248 |
| ENSMUSG00000003848 | Nob1 | protein_coding | 0.222893083 | 6.283997351 | 0.6120987 | 0.4452113 | 0.845076248 |
| ENSMUSG00000039531 | Zufsp | protein_coding | 0.31323796 | 3.712014587 | 0.6116255 | 0.4453841 | 0.845236285 |
| ENSMUSG00000038256 | Bcl9 | protein_coding | 0.343538358 | 4.243007085 | 0.6112606 | 0.4455174 | 0.845236285 |
| ENSMUSG00000021597 | Slf1 | protein_coding | -0.312225603 | 4.400162418 | 0.610897 | 0.4456503 | 0.845236285 |
| ENSMUSG00000020257 | Wdr82 | protein_coding | -0.195167631 | 6.41093458 | 0.6108011 | 0.4456854 | 0.845236285 |
| ENSMUSG00000018548 | Trim37 | protein_coding | -0.165619198 | 6.692543958 | 0.6107761 | 0.4456945 | 0.845236285 |
| ENSMUSG00000027395 | Polr1b | protein_coding | -0.595503925 | 2.59214841 | 0.6103724 | 0.4458422 | 0.845236285 |
| ENSMUSG00000035397 | Klf16 | protein_coding | -0.253853371 | 5.288354694 | 0.6103625 | 0.4458458 | 0.845236285 |
| ENSMUSG00000035834 | Polr3g | protein_coding | -0.860295228 | 1.979969554 | 0.6099591 | 0.4459935 | 0.845237978 |
| ENSMUSG00000046330 | Rpl37a | protein_coding | 0.283800799 | 7.908050191 | 0.6129822 | 0.4460039 | 0.845237978 |
| ENSMUSG00000047897 | Ripply2 | protein_coding | 0.453959339 | 2.911430618 | 0.608732 | 0.4464431 | 0.845921135 |
| ENSMUSG00000031284 | Pak3 | protein_coding | 0.161685801 | 7.575495308 | 0.6083839 | 0.4465708 | 0.846013995 |
| ENSMUSG00000046873 | Mbtps2 | protein_coding | -0.356171841 | 3.778127387 | 0.6081478 | 0.4466574 | 0.846029044 |
| ENSMUSG00000039963 | Ccdc40 | protein_coding | 0.407135452 | 3.778706287 | 0.6070668 | 0.4470544 | 0.846631901 |
| ENSMUSG00000034317 | Trim59 | protein_coding | -0.442058191 | 4.038572998 | 0.6068255 | 0.4471614 | 0.84668535 |
| ENSMUSG00000032199 | Polr2m | protein_coding | 0.171397648 | 6.88665138 | 0.6063709 | 0.4473103 | 0.846818178 |
| ENSMUSG00000041837 | Pdcd7 | protein_coding | 0.340775073 | 4.364299716 | 0.6059299 | 0.4474725 | 0.846850052 |
| ENSMUSG00000037697 | Ddhd1 | protein_coding | -0.619381338 | 2.963253179 | 0.6059467 | 0.4474846 | 0.846850052 |
| ENSMUSG00000021993 | Mipep | protein_coding | 0.286471139 | 4.298738987 | 0.6052256 | 0.4477319 | 0.847168891 |
| ENSMUSG00000018909 | Arrb1 | protein_coding | -0.182295771 | 6.224803928 | 0.6047291 | 0.4479149 | 0.847273098 |
| ENSMUSG00000091636 | Akain1 | protein_coding | -0.76279757 | 2.482868168 | 0.6045614 | 0.4479767 | 0.847273098 |
| ENSMUSG00000018167 | Stard3 | protein_coding | 0.287795652 | 5.476916645 | 0.6044844 | 0.4480233 | 0.847273098 |
| ENSMUSG00000020895 | Tmem107 | protein_coding | 0.224827291 | 5.722713389 | 0.6038807 | 0.4482278 | 0.847377898 |
| ENSMUSG00000029101 | Rgs12 | protein_coding | -0.320840083 | 3.786857235 | 0.6038576 | 0.4482364 | 0.847377898 |
| ENSMUSG00000049657 | Zbtb5 | protein_coding | -0.487873919 | 3.629371052 | 0.6034151 | 0.4484179 | 0.847539265 |
| ENSMUSG00000057342 | Sphk2 | protein_coding | 0.289532214 | 4.929433609 | 0.603117 | 0.4485098 | 0.847539265 |
| ENSMUSG00000040481 | Bptf | protein_coding | -0.214214963 | 6.101804355 | 0.6029862 | 0.4485582 | 0.847539265 |
| ENSMUSG00000025321 | Itgb8 | protein_coding | -0.439474722 | 3.993392886 | 0.6024367 | 0.4487795 | 0.847689771 |
| ENSMUSG00000062542 | Syt9 | protein_coding | 0.337483618 | 4.246402326 | 0.6023442 | 0.4487955 | 0.847689771 |
| ENSMUSG00000098923 | Tmem185b | protein_coding | -0.297275954 | 4.597714848 | 0.6017 | 0.4490339 | 0.847866819 |

| ensembl_gene_id | external_gene_name | gene_biotype | log2 (fold change) | log2 (counts per million) | F-value | p-value | False discovery rate |
|---|---|---|---|---|---|---|---|
| ENSMUSG00000020175 | Rab36 | protein_coding | 0.220298234 | 6.417357746 | 0.6014567 | 0.4491239 | 0.847866819 |
| ENSMUSG00000078716 | Tmem8b | protein_coding | 0.333185629 | 4.758628436 | 0.6013191 | 0.4491749 | 0.847866819 |
| ENSMUSG00000030598 | Fbxo17 | protein_coding | -0.367018382 | 4.047245717 | 0.6012388 | 0.4492046 | 0.847866819 |
| ENSMUSG00000073678 | Pgap1 | protein_coding | -0.27064934 | 4.58660552 | 0.6006242 | 0.4494324 | 0.848147809 |
| ENSMUSG00000025089 | Gfra1 | protein_coding | 0.213024171 | 5.701463054 | 0.6000004 | 0.4496637 | 0.848435466 |
| ENSMUSG00000048782 | Insc | protein_coding | -0.664653603 | 2.653834911 | 0.5997462 | 0.449758 | 0.848441099 |
| ENSMUSG00000039656 | Rxrb | protein_coding | 0.244200528 | 5.001536529 | 0.5994994 | 0.4498496 | 0.848441099 |
| ENSMUSG00000022707 | Gbe1 | protein_coding | -0.398546708 | 3.4421343 | 0.5993547 | 0.4499034 | 0.848441099 |
| ENSMUSG00000025217 | Btrc | protein_coding | 0.257156764 | 6.648808445 | 0.5996799 | 0.4501281 | 0.848512475 |
| ENSMUSG00000047205 | Dusp18 | protein_coding | 0.291762218 | 4.869226442 | 0.598029 | 0.4503961 | 0.848512475 |
| ENSMUSG00000070644 | Etnk2 | protein_coding | 0.554226089 | 2.743692198 | 0.5974993 | 0.4505932 | 0.848512475 |
| ENSMUSG00000031517 | Gpm6a | protein_coding | -0.158663108 | 9.910279319 | 0.5974764 | 0.4506017 | 0.848512475 |
| ENSMUSG00000000915 | Hip1r | protein_coding | -0.336510738 | 4.0628248 | 0.5973764 | 0.450639 | 0.848512475 |
| ENSMUSG00000029265 | Dr1 | protein_coding | -0.331926424 | 4.416622029 | 0.5973309 | 0.4506559 | 0.848512475 |
| ENSMUSG00000029716 | Tfr2 | protein_coding | 0.340136419 | 4.00157809 | 0.5971886 | 0.4507089 | 0.848512475 |
| ENSMUSG00000030401 | Rtn2 | protein_coding | 0.262301071 | 5.087100593 | 0.5968156 | 0.4508478 | 0.848512475 |
| ENSMUSG00000071550 | Cfap44 | protein_coding | -0.695905235 | 2.155988903 | 0.5966268 | 0.4509182 | 0.848512475 |
| ENSMUSG00000040451 | Sgms1 | protein_coding | -0.315615321 | 4.122359206 | 0.5962259 | 0.4510677 | 0.848512475 |
| ENSMUSG00000034587 | 8430429K09Rik | processed_transcript | 0.327439282 | 4.422276254 | 0.5957523 | 0.4512443 | 0.848512475 |
| ENSMUSG00000011254 | Thg1l | protein_coding | 0.265124118 | 5.010979857 | 0.5956503 | 0.4512824 | 0.848512475 |
| ENSMUSG00000021752 | Kctd6 | protein_coding | -0.280215495 | 4.238287717 | 0.5954578 | 0.4513542 | 0.848512475 |
| ENSMUSG00000004031 | Brinp2 | protein_coding | 0.255100452 | 4.603007488 | 0.5952342 | 0.4514377 | 0.848512475 |
| ENSMUSG00000031431 | Tsc22d3 | protein_coding | -0.173592824 | 8.258900288 | 0.5952305 | 0.4514391 | 0.848512475 |
| ENSMUSG00000020457 | Drg1 | protein_coding | 0.163953784 | 8.483408787 | 0.5951873 | 0.4514552 | 0.848512475 |
| ENSMUSG00000083674 | Zfp133-ps | transcribed_unitary_p | 0.477665944 | 3.335927274 | 0.5950976 | 0.4514887 | 0.848512475 |
| ENSMUSG00000049550 | Clip1 | protein_coding | 0.195234202 | 6.116884207 | 0.5948997 | 0.4515626 | 0.848512475 |
| ENSMUSG00000029538 | Srsf9 | protein_coding | -0.267365197 | 5.304814752 | 0.5945325 | 0.4516999 | 0.848512475 |
| ENSMUSG00000054387 | Mdm4 | protein_coding | -0.215685895 | 5.099620466 | 0.5944575 | 0.4517279 | 0.848512475 |
| ENSMUSG00000047446 | Arl4a | protein_coding | -0.288007796 | 6.549602201 | 0.5960248 | 0.451843 | 0.848512475 |
| ENSMUSG00000033170 | Card10 | protein_coding | 1.012992311 | 2.435956805 | 0.5970483 | 0.4518597 | 0.848512475 |
| ENSMUSG00000030376 | Slc8a2 | protein_coding | -0.530468026 | 2.93078696 | 0.5939626 | 0.4519129 | 0.848512475 |
| ENSMUSG00000031521 | Aga | protein_coding | -0.337932959 | 4.549416809 | 0.5938563 | 0.4519527 | 0.848512475 |
| ENSMUSG00000028672 | Hmgcl | protein_coding | 0.176260965 | 7.215860882 | 0.5938046 | 0.451972 | 0.848512475 |
| ENSMUSG00000041078 | Grid1 | protein_coding | 0.219643168 | 5.258684022 | 0.5937489 | 0.4519929 | 0.848512475 |
| ENSMUSG00000078350 | Smim1 | protein_coding | 0.22860029 | 5.456210545 | 0.5932222 | 0.45219 | 0.848734395 |
| ENSMUSG00000028218 | Fam92a | protein_coding | 0.197226907 | 5.899350725 | 0.5923061 | 0.4525333 | 0.849153653 |
| ENSMUSG00000020519 | Sap30l | protein_coding | -0.482273622 | 2.941717624 | 0.5922045 | 0.4525713 | 0.849153653 |
| ENSMUSG00000051910 | Sox6 | protein_coding | -0.41934754 | 3.748186388 | 0.5918955 | 0.4526872 | 0.849222869 |
| ENSMUSG00000027465 | Tbc1d20 | protein_coding | 0.249945097 | 5.170926343 | 0.5914441 | 0.4528566 | 0.849392432 |
| ENSMUSG00000053754 | Chd8 | protein_coding | -0.482650028 | 3.055711784 | 0.5911367 | 0.452972 | 0.849426493 |
| ENSMUSG00000025812 | Pard3 | protein_coding | -0.263191443 | 4.74976956 | 0.5907572 | 0.4531145 | 0.849426493 |
| ENSMUSG00000032115 | Hyou1 | protein_coding | 0.381485537 | 4.211311351 | 0.5906875 | 0.4531587 | 0.849426493 |
| ENSMUSG00000025962 | Fastkd2 | protein_coding | 0.259868674 | 5.107570736 | 0.5903965 | 0.45325 | 0.849426493 |
| ENSMUSG00000058388 | Phtf1 | protein_coding | -0.346670602 | 4.01175604 | 0.590245 | 0.453307 | 0.849426493 |
| ENSMUSG00000104283 | Gm37459 | antisense_RNA | -0.299617244 | 4.559865584 | 0.5899803 | 0.4534065 | 0.849426493 |
| ENSMUSG00000040731 | Eif4h | protein_coding | 0.168668911 | 8.360380443 | 0.5897015 | 0.4535114 | 0.849426493 |
| ENSMUSG00000070348 | Ccnd1 | protein_coding | -0.686005626 | 2.568116953 | 0.5896874 | 0.4535167 | 0.849426493 |
| ENSMUSG00000043557 | Mdga1 | protein_coding | -0.463590579 | 3.316663069 | 0.5894152 | 0.4536191 | 0.849426493 |
| ENSMUSG00000021024 | Psma6 | protein_coding | 0.167583191 | 9.087785455 | 0.5892941 | 0.4536647 | 0.849426493 |
| ENSMUSG00000027012 | Dync1i2 | protein_coding | -0.164071591 | 7.194547716 | 0.5887786 | 0.4538588 | 0.849642001 |
| ENSMUSG00000054196 | Cthrc1 | protein_coding | 0.202830259 | 5.38972456 | 0.5880597 | 0.4541298 | 0.849705446 |
| ENSMUSG00000025384 | Faap100 | protein_coding | -0.393685824 | 4.090612436 | 0.5872898 | 0.4544381 | 0.849705446 |
| ENSMUSG00000024941 | Scyl1 | protein_coding | 0.224411167 | 5.49164501 | 0.5871984 | 0.4544547 | 0.849705446 |
| ENSMUSG00000026088 | Mitd1 | protein_coding | -0.313837608 | 4.400588202 | 0.5869781 | 0.4545379 | 0.849705446 |
| ENSMUSG00000062604 | Srpk2 | protein_coding | -0.181771938 | 7.969830073 | 0.5867187 | 0.4546359 | 0.849705446 |
| ENSMUSG00000022820 | Ndufb4 | protein_coding | 0.225428242 | 5.958522135 | 0.5866302 | 0.4546693 | 0.849705446 |
| ENSMUSG00000036545 | Adamts2 | protein_coding | -0.867982131 | 2.421073913 | 0.5866322 | 0.4547336 | 0.849705446 |
| ENSMUSG00000042520 | Ubap2l | protein_coding | -0.351098337 | 4.45753949 | 0.5864455 | 0.4547391 | 0.849705446 |
| ENSMUSG00000072235 | Tuba1a | protein_coding | 0.20548813 | 9.916995326 | 0.5861833 | 0.4548381 | 0.849705446 |
| ENSMUSG00000053007 | Creb5 | protein_coding | -0.428534017 | 3.695559275 | 0.5861078 | 0.4548667 | 0.849705446 |
| ENSMUSG00000035401 | Emsy | protein_coding | -0.285827096 | 4.647654301 | 0.5859582 | 0.4549233 | 0.849705446 |
| ENSMUSG00000035226 | Rims4 | protein_coding | 0.266912723 | 5.01318547 | 0.5855452 | 0.4550795 | 0.849705446 |
| ENSMUSG00000032763 | Ilvbl | protein_coding | 0.313499759 | 5.377961018 | 0.5855126 | 0.4551097 | 0.849705446 |
| ENSMUSG00000080316 | Spaca6 | protein_coding | -0.345308365 | 3.488998129 | 0.5854307 | 0.4551228 | 0.849705446 |
| ENSMUSG00000041939 | Mvk | protein_coding | 0.256627646 | 5.485362946 | 0.5853889 | 0.4551386 | 0.849705446 |
| ENSMUSG00000078784 | 1810022K09Rik | protein_coding | 0.27897017 | 4.566680193 | 0.5853403 | 0.455157 | 0.849705446 |
| ENSMUSG00000037994 | Slc9b2 | protein_coding | 0.519065964 | 2.693653155 | 0.5849942 | 0.455288 | 0.849802531 |
| ENSMUSG00000032625 | Thsd7a | protein_coding | -0.430679124 | 4.466602786 | 0.586454 | 0.4553876 | 0.849834158 |
| ENSMUSG00000052551 | Adarb2 | protein_coding | -0.281163379 | 4.072963191 | 0.5845321 | 0.4554631 | 0.849834158 |
| ENSMUSG00000022639 | Dubr | lincRNA | -0.422629996 | 3.784583327 | 0.5838872 | 0.4557075 | 0.849930493 |
| ENSMUSG00000005656 | Snx6 | protein_coding | -0.176561602 | 7.146636293 | 0.5838461 | 0.4557231 | 0.849930493 |
| ENSMUSG00000032386 | Trip4 | protein_coding | 0.234094423 | 4.950811401 | 0.5836715 | 0.4557893 | 0.849930493 |
| ENSMUSG00000023707 | Ogfod2 | protein_coding | 0.245949448 | 5.971557086 | 0.5832738 | 0.4559402 | 0.849930493 |
| ENSMUSG00000062760 | 1810041L15Rik | protein_coding | -0.293119774 | 5.549646128 | 0.5831801 | 0.4559936 | 0.849930493 |
| ENSMUSG00000087687 | Pet100 | protein_coding | 0.229750568 | 6.789267763 | 0.5829389 | 0.4560852 | 0.849930493 |
| ENSMUSG00000029201 | Ugdh | protein_coding | -0.275269497 | 4.528827159 | 0.582762 | 0.4561345 | 0.849930493 |
| ENSMUSG00000048503 | Tmem136 | protein_coding | -0.331668678 | 3.52924235 | 0.582596 | 0.4561976 | 0.849930493 |
| ENSMUSG00000024997 | Prdx3 | protein_coding | 0.148248938 | 8.346244959 | 0.5820379 | 0.4564097 | 0.849930493 |

| ensembl_gene_id | external_gene_name | gene_biotype | log2 (fold change) | log2 (counts per million) | F-value | p-value | False discovery rate |
|---|---|---|---|---|---|---|---|
| ENSMUSG00000016349 | Eef1a2 | protein_coding | 0.262864307 | 5.491949823 | 0.5819388 | 0.4564473 | 0.849930493 |
| ENSMUSG00000039108 | Lsm14b | protein_coding | -0.296220451 | 4.354056581 | 0.58188 | 0.4564697 | 0.849930493 |
| ENSMUSG00000033845 | Mrpl15 | protein_coding | 0.15045826 | 7.412366499 | 0.581516 | 0.4566082 | 0.849930493 |
| ENSMUSG00000063146 | Clip2 | protein_coding | 0.368664891 | 3.68875427 | 0.5814861 | 0.4566195 | 0.849930493 |
| ENSMUSG00000018574 | Acadvl | protein_coding | 0.275715195 | 5.42607439 | 0.5814815 | 0.4566213 | 0.849930493 |
| ENSMUSG00000033964 | Zbtb41 | protein_coding | -0.407669881 | 3.457689117 | 0.581227 | 0.4567181 | 0.849963662 |
| ENSMUSG00000024261 | Syt4 | protein_coding | -0.170897639 | 8.374893494 | 0.5806022 | 0.456956 | 0.850246591 |
| ENSMUSG00000062257 | Opcml | protein_coding | -0.391492696 | 3.674555153 | 0.5804125 | 0.4570283 | 0.850246591 |
| ENSMUSG00000030982 | 9030624J02Rik | protein_coding | 0.295254938 | 6.165526476 | 0.5819075 | 0.4571502 | 0.850326193 |
| ENSMUSG00000031129 | Slc9a9 | protein_coding | -0.328023542 | 3.493578143 | 0.5796393 | 0.457323 | 0.850346488 |
| ENSMUSG00000020893 | Per1 | protein_coding | -0.326803218 | 4.241100739 | 0.5795798 | 0.4573458 | 0.850346488 |
| ENSMUSG00000046417 | Lrrc75a | protein_coding | -0.651979728 | 2.392586218 | 0.579442 | 0.4573983 | 0.850346488 |
| ENSMUSG00000049489 | Fam58b | protein_coding | 0.254985371 | 4.903408993 | 0.5787491 | 0.4576628 | 0.850648556 |
| ENSMUSG00000072704 | Smim10l1 | protein_coding | -0.226390215 | 8.020245595 | 0.5785253 | 0.4577661 | 0.850648556 |
| ENSMUSG00000025578 | Cbx8 | protein_coding | 0.422823702 | 3.14278901 | 0.5783949 | 0.4577981 | 0.850648556 |
| ENSMUSG00000030551 | Nr2f2 | protein_coding | 0.287152658 | 6.558364138 | 0.5795371 | 0.4582721 | 0.850987604 |
| ENSMUSG00000028029 | Aimp1 | protein_coding | 0.191313104 | 7.773119981 | 0.5764213 | 0.4585532 | 0.850987604 |
| ENSMUSG00000046329 | Slc25a23 | protein_coding | -0.308323097 | 3.69950055 | 0.5762407 | 0.4586224 | 0.850987604 |
| ENSMUSG00000044147 | Arf6 | protein_coding | 0.22203122 | 5.076528552 | 0.5760892 | 0.4586805 | 0.850987604 |
| ENSMUSG00000062393 | Dgkk | protein_coding | 0.296389751 | 4.040301523 | 0.5760803 | 0.4586839 | 0.850987604 |
| ENSMUSG00000029227 | Fip1l1 | protein_coding | 0.363565072 | 4.152033898 | 0.5759087 | 0.4587497 | 0.850987604 |
| ENSMUSG00000042055 | Wdr11 | protein_coding | -0.257806673 | 4.757641252 | 0.575819 | 0.4587841 | 0.850987604 |
| ENSMUSG00000068206 | Pick1 | protein_coding | 0.279650208 | 4.937320041 | 0.5758077 | 0.4587884 | 0.850987604 |
| ENSMUSG00000027630 | Tbl1xr1 | protein_coding | -0.233796514 | 5.024175438 | 0.5757642 | 0.4588051 | 0.850987604 |
| ENSMUSG00000045180 | Shroom2 | protein_coding | 0.211219671 | 6.313908682 | 0.5756866 | 0.4588348 | 0.850987604 |
| ENSMUSG00000045665 | Mfsd5 | protein_coding | 0.241945169 | 4.838395001 | 0.5753294 | 0.4589719 | 0.850987604 |
| ENSMUSG00000058587 | Tmod3 | protein_coding | 0.17446574 | 6.482394661 | 0.5752464 | 0.4590037 | 0.850987604 |
| ENSMUSG00000022781 | Pak2 | protein_coding | -0.252348377 | 5.053092836 | 0.5751287 | 0.4590489 | 0.850987604 |
| ENSMUSG00000033326 | Kdm4a | protein_coding | -0.573733193 | 2.537096055 | 0.5750253 | 0.4590885 | 0.850987604 |
| ENSMUSG00000022951 | Rcan1 | protein_coding | 0.199349547 | 5.367254783 | 0.5744685 | 0.4593024 | 0.851119675 |
| ENSMUSG00000034165 | Ccnd3 | protein_coding | 0.319458712 | 3.629713756 | 0.5737954 | 0.4595611 | 0.851119675 |
| ENSMUSG00000024084 | Qpct | protein_coding | -0.181064967 | 6.503661628 | 0.5736722 | 0.4596085 | 0.851119675 |
| ENSMUSG00000034813 | Grip1 | protein_coding | 0.250792976 | 4.547246663 | 0.573444 | 0.4596962 | 0.851119675 |
| ENSMUSG00000019845 | Tube1 | protein_coding | -0.66845626 | 2.32141918 | 0.5733949 | 0.4597151 | 0.851119675 |
| ENSMUSG00000027305 | Ndufaf1 | protein_coding | 0.179625821 | 5.910435324 | 0.5732117 | 0.4597856 | 0.851119675 |
| ENSMUSG00000005672 | Kit | protein_coding | 0.446752057 | 3.267229625 | 0.5731386 | 0.4598137 | 0.851119675 |
| ENSMUSG00000005803 | Sqor | protein_coding | -0.356799768 | 3.767947818 | 0.5730484 | 0.4598484 | 0.851119675 |
| ENSMUSG00000019505 | Ubb | protein_coding | 0.1751756 | 10.89795493 | 0.5728075 | 0.4599412 | 0.851119675 |
| ENSMUSG00000037722 | Gnpnat1 | protein_coding | 0.269080554 | 4.307557478 | 0.5727289 | 0.4599714 | 0.851119675 |
| ENSMUSG00000046985 | Tapt1 | protein_coding | -0.324505536 | 3.982898694 | 0.5725757 | 0.4600305 | 0.851119675 |
| ENSMUSG00000020922 | Lsm12 | protein_coding | 0.226646516 | 6.049970308 | 0.5717461 | 0.4603502 | 0.851505131 |
| ENSMUSG00000028221 | Tmem55a | protein_coding | -0.156572261 | 7.032470232 | 0.5716242 | 0.4603972 | 0.851505131 |
| ENSMUSG00000054405 | Dnajc8 | protein_coding | 0.143866114 | 8.553718443 | 0.5710786 | 0.4606077 | 0.851588957 |
| ENSMUSG00000069135 | Fgfr1op | protein_coding | -0.36245464 | 4.019923772 | 0.5707117 | 0.4607493 | 0.851588957 |
| ENSMUSG00000032536 | Trak1 | protein_coding | -0.202552523 | 5.859126942 | 0.5704698 | 0.4608427 | 0.851588957 |
| ENSMUSG00000030509 | Asb7 | protein_coding | -0.278564204 | 4.074378218 | 0.5698478 | 0.4610831 | 0.851588957 |
| ENSMUSG00000020385 | Clk4 | protein_coding | -0.239557944 | 5.68838557 | 0.5698219 | 0.4610931 | 0.851588957 |
| ENSMUSG00000045410 | Akr1e1 | protein_coding | -0.173533102 | 6.841923166 | 0.5698158 | 0.4610955 | 0.851588957 |
| ENSMUSG00000038497 | Tmco3 | protein_coding | 0.27552201 | 5.312085953 | 0.5696701 | 0.4611519 | 0.851588957 |
| ENSMUSG00000050786 | Ccdc126 | protein_coding | 0.340414032 | 3.822217161 | 0.56959 | 0.4611828 | 0.851588957 |
| ENSMUSG00000047368 | Abhd17b | protein_coding | -0.25636284 | 5.259547045 | 0.5695659 | 0.4611922 | 0.851588957 |
| ENSMUSG00000018774 | Cd68 | protein_coding | 0.288998695 | 5.002781027 | 0.569312 | 0.4612904 | 0.851588957 |
| ENSMUSG00000045969 | Ing1 | protein_coding | -0.1990498 | 5.778528828 | 0.5689931 | 0.4614138 | 0.851588957 |
| ENSMUSG00000024258 | Polr2d | protein_coding | 0.157258834 | 7.113924232 | 0.5685567 | 0.4615828 | 0.851588957 |
| ENSMUSG00000014504 | Srp19 | protein_coding | 0.189355198 | 8.272780773 | 0.5684783 | 0.4616131 | 0.851588957 |
| ENSMUSG00000036620 | Mgat4b | protein_coding | -0.567000142 | 2.571468355 | 0.5679784 | 0.4618068 | 0.851588957 |
| ENSMUSG00000027433 | Xrn2 | protein_coding | -0.20312529 | 5.935419471 | 0.5679449 | 0.4618198 | 0.851588957 |
| ENSMUSG00000020407 | Upp1 | protein_coding | 0.533323005 | 2.819865986 | 0.567795 | 0.461878 | 0.851588957 |
| ENSMUSG00000032869 | Psmf1 | protein_coding | 0.222361338 | 6.785877124 | 0.5678238 | 0.4618844 | 0.851588957 |
| ENSMUSG00000003420 | Fcgrt | protein_coding | -0.453884245 | 3.467224619 | 0.567448 | 0.4620125 | 0.851588957 |
| ENSMUSG00000091474 | 2610021A01Rik | protein_coding | 0.375238364 | 3.469323336 | 0.5671473 | 0.4621292 | 0.851588957 |
| ENSMUSG00000097638 | Carlr | lincRNA | 0.786360343 | 2.306244157 | 0.5668934 | 0.4622278 | 0.851588957 |
| ENSMUSG00000037533 | Rapgef6 | protein_coding | -0.258848461 | 5.157853183 | 0.5667118 | 0.4622983 | 0.851588957 |
| ENSMUSG00000039449 | Prpf18 | protein_coding | -0.240056185 | 5.607846361 | 0.5666827 | 0.4623095 | 0.851588957 |
| ENSMUSG00000031137 | Fgf13 | protein_coding | 0.226848654 | 6.550759424 | 0.5663117 | 0.4624712 | 0.851588957 |
| ENSMUSG00000021747 | 4930452B06Rik | protein_coding | 0.29170205 | 3.836289135 | 0.5661636 | 0.4625112 | 0.851588957 |
| ENSMUSG00000020467 | Efemp1 | protein_coding | -0.430401015 | 3.055751695 | 0.5658461 | 0.4626346 | 0.851588957 |
| ENSMUSG00000024683 | Mrpl16 | protein_coding | 0.180297286 | 6.373796248 | 0.5656476 | 0.4627118 | 0.851588957 |
| ENSMUSG00000019803 | Nr2e1 | protein_coding | -0.646024104 | 2.88878223 | 0.5656442 | 0.4627307 | 0.851588957 |
| ENSMUSG00000018442 | Derl2 | protein_coding | -0.215927804 | 5.580025005 | 0.5654748 | 0.462779 | 0.851588957 |
| ENSMUSG00000024901 | Peli3 | protein_coding | 0.303701893 | 3.903887518 | 0.5652523 | 0.4628656 | 0.851588957 |
| ENSMUSG00000032932 | Hspa13 | protein_coding | -0.226623463 | 5.451743719 | 0.565027 | 0.4629532 | 0.851588957 |
| ENSMUSG00000024165 | Hn1l | protein_coding | 0.313002327 | 4.286297872 | 0.5649803 | 0.4629714 | 0.851588957 |
| ENSMUSG00000100594 | 2810414N06Rik | lincRNA | -0.419699682 | 3.632830618 | 0.5649666 | 0.4629768 | 0.851588957 |
| ENSMUSG00000074671 | Tspyl3 | protein_coding | 0.256805261 | 5.022841253 | 0.5647212 | 0.4630723 | 0.851619068 |
| ENSMUSG00000031949 | Adat1 | protein_coding | 0.356569494 | 3.422585029 | 0.5644086 | 0.463194 | 0.851697263 |
| ENSMUSG00000020444 | Guk1 | protein_coding | -0.263158322 | 7.484912509 | 0.5660265 | 0.4634395 | 0.85186017 |
| ENSMUSG00000001419 | Mef2d | protein_coding | 0.194624148 | 5.675214612 | 0.5637748 | 0.4634411 | 0.85186017 |

| ensembl_gene_id | external_gene_name | gene_biotype | log2 (fold change) | log2 (counts per million) | F-value | p-value | False discovery rate |
|---|---|---|---|---|---|---|---|
| ENSMUSG00000035437 | Rabgap1 | protein_coding | -0.451532127 | 3.37899824 | 0.5633055 | 0.4636241 | 0.852050977 |
| ENSMUSG00000038388 | Mpp6 | protein_coding | -0.26749095 | 4.397720151 | 0.5629631 | 0.4637578 | 0.852150918 |
| ENSMUSG00000054910 | 4931415C17Rik | processed_transcript | -0.627439269 | 2.736321017 | 0.5622502 | 0.4640362 | 0.852509102 |
| ENSMUSG00000019370 | Calm3 | protein_coding | 0.156351686 | 9.637650955 | 0.5620581 | 0.4641113 | 0.852509102 |
| ENSMUSG00000046152 | Fut10 | protein_coding | 0.456618381 | 2.851395179 | 0.5617197 | 0.4642435 | 0.852606447 |
| ENSMUSG00000036186 | Fam69b | protein_coding | 0.28930759 | 4.7655652 | 0.5614936 | 0.464332 | 0.852623217 |
| ENSMUSG00000024925 | Rnaseh2c | protein_coding | 0.447804602 | 4.817628596 | 0.5650174 | 0.4644515 | 0.852633387 |
| ENSMUSG00000022817 | Itgb5 | protein_coding | -0.491099214 | 2.879385228 | 0.5610742 | 0.4644961 | 0.852633387 |
| ENSMUSG00000032388 | Spg21 | protein_coding | -0.238760908 | 5.239548365 | 0.5607381 | 0.4646277 | 0.852729366 |
| ENSMUSG00000005312 | Ubqln1 | protein_coding | 0.151269619 | 7.206045527 | 0.5603583 | 0.4647764 | 0.852773643 |
| ENSMUSG00000000861 | Bcl11a | protein_coding | 0.323777207 | 5.098905753 | 0.560123 | 0.4648861 | 0.852773643 |
| ENSMUSG00000021712 | Trim23 | protein_coding | -0.294258864 | 5.5530116 | 0.5599703 | 0.4649459 | 0.852773643 |
| ENSMUSG00000063450 | Syne2 | protein_coding | -0.512095207 | 3.309267407 | 0.5599114 | 0.464969 | 0.852773643 |
| ENSMUSG00000032235 | Ice2 | protein_coding | 0.468802131 | 2.950215542 | 0.5595167 | 0.4651063 | 0.852880032 |
| ENSMUSG00000022759 | Lrrc74b | protein_coding | -0.425481787 | 3.16521939 | 0.5588253 | 0.4653777 | 0.853232049 |
| ENSMUSG00000033365 | Ipo13 | protein_coding | -0.24826271 | 5.310807978 | 0.5583439 | 0.4655668 | 0.853433257 |
| ENSMUSG00000042064 | Myo3b | protein_coding | -0.579643501 | 2.505294159 | 0.5578321 | 0.4657679 | 0.85351951 |
| ENSMUSG00000026578 | Ccdc181 | protein_coding | 0.202942067 | 5.90773494 | 0.557657 | 0.4658368 | 0.85351951 |
| ENSMUSG00000009356 | Lpo | protein_coding | -1.001753526 | 2.031690728 | 0.5576627 | 0.4658519 | 0.85351951 |
| ENSMUSG00000034424 | Gcsh | protein_coding | 0.158545869 | 7.694820109 | 0.556899 | 0.466135 | 0.853813157 |
| ENSMUSG00000050074 | Spink8 | protein_coding | -0.478178513 | 3.449041662 | 0.5568518 | 0.466171 | 0.853813157 |
| ENSMUSG00000026224 | 4933407L21Rik | protein_coding | 0.283169572 | 4.27700674 | 0.5565216 | 0.4662836 | 0.853859548 |
| ENSMUSG00000002658 | Gtf2f1 | protein_coding | 0.19987931 | 6.596767364 | 0.5563231 | 0.4663619 | 0.853859548 |
| ENSMUSG00000028063 | Lmna | protein_coding | -0.227002175 | 5.095959939 | 0.5560387 | 0.4664739 | 0.853859548 |
| ENSMUSG00000032322 | Pstpip1 | protein_coding | 0.657372353 | 2.260251507 | 0.5559371 | 0.466514 | 0.853859548 |
| ENSMUSG00000073664 | Nbeal1 | protein_coding | -0.375740563 | 3.613402452 | 0.5551512 | 0.466824 | 0.854203293 |
| ENSMUSG00000001441 | Npepps | protein_coding | -0.158446417 | 6.845432145 | 0.5549499 | 0.4669034 | 0.854203293 |
| ENSMUSG00000048271 | Rbm33 | protein_coding | -0.221722967 | 5.378286856 | 0.5547645 | 0.4669767 | 0.854203293 |
| ENSMUSG00000026544 | Dusp23 | protein_coding | 0.36091631 | 3.664859156 | 0.5546559 | 0.4670195 | 0.854203293 |
| ENSMUSG00000032733 | Snx33 | protein_coding | -0.442701913 | 3.184480107 | 0.5543535 | 0.467139 | 0.85427649 |
| ENSMUSG00000025076 | Casp7 | protein_coding | 0.669266439 | 2.317533746 | 0.553919 | 0.4673108 | 0.854445254 |
| ENSMUSG00000039682 | Lap3 | protein_coding | -0.183004106 | 6.696552077 | 0.5534053 | 0.4675139 | 0.854457996 |
| ENSMUSG00000086155 | 9430041J12Rik | antisense_RNA | 0.657895711 | 2.106164436 | 0.5531723 | 0.4676061 | 0.854457996 |
| ENSMUSG00000091002 | Tcerg1l | protein_coding | -0.325327351 | 4.286745588 | 0.5522426 | 0.4679743 | 0.854457996 |
| ENSMUSG00000028999 | Rint1 | protein_coding | -0.279773487 | 3.966373281 | 0.5522317 | 0.4679786 | 0.854457996 |
| ENSMUSG00000022203 | Efs | protein_coding | 0.418497365 | 2.928711855 | 0.5512276 | 0.4683768 | 0.854457996 |
| ENSMUSG00000037608 | Bclaf1 | protein_coding | -0.186815372 | 7.408676833 | 0.5511016 | 0.4684268 | 0.854457996 |
| ENSMUSG00000047832 | Cdca4 | protein_coding | 0.414410581 | 3.904284938 | 0.5509271 | 0.4685134 | 0.854457996 |
| ENSMUSG00000038160 | Atg5 | protein_coding | 0.162594643 | 6.381312863 | 0.5506624 | 0.4686013 | 0.854457996 |
| ENSMUSG00000029228 | Lnx1 | protein_coding | 0.360770391 | 4.27486884 | 0.5506676 | 0.4686165 | 0.854457996 |
| ENSMUSG00000026766 | Mmadhc | protein_coding | 0.176633011 | 6.389745668 | 0.55031 | 0.4687412 | 0.854457996 |
| ENSMUSG00000035559 | Mpv17l2 | protein_coding | 0.455321086 | 3.124717197 | 0.550175 | 0.4687949 | 0.854457996 |
| ENSMUSG00000027800 | Tm4sf1 | protein_coding | -0.292318074 | 4.283675541 | 0.5500046 | 0.4688626 | 0.854457996 |
| ENSMUSG00000005667 | Mthfd2 | protein_coding | 0.226982638 | 5.701857152 | 0.5497578 | 0.4689608 | 0.854457996 |
| ENSMUSG00000042389 | Tsen2 | protein_coding | 0.222807863 | 5.267071006 | 0.5496845 | 0.4689899 | 0.854457996 |
| ENSMUSG00000040490 | Lrfn2 | protein_coding | -0.416698565 | 2.951923989 | 0.5494513 | 0.4690827 | 0.854457996 |
| ENSMUSG00000064368 | mt-Nd6 | protein_coding | 0.270932481 | 8.052755857 | 0.5520285 | 0.4691099 | 0.854457996 |
| ENSMUSG00000069678 | Pcgf1 | protein_coding | 0.209698244 | 5.263945238 | 0.5486835 | 0.4693883 | 0.854457996 |
| ENSMUSG00000026585 | Kifap3 | protein_coding | -0.16801959 | 7.567748367 | 0.5486737 | 0.4693923 | 0.854457996 |
| ENSMUSG00000043445 | Pgp | protein_coding | 0.264425711 | 4.355717368 | 0.5485904 | 0.4694254 | 0.854457996 |
| ENSMUSG00000024511 | Rab27b | protein_coding | 0.241139229 | 6.756997328 | 0.5490908 | 0.4695188 | 0.854457996 |
| ENSMUSG00000024383 | Map3k2 | protein_coding | -0.331331864 | 3.76212729 | 0.548081 | 0.4696284 | 0.854457996 |
| ENSMUSG00000086938 | 4930481A15Rik | antisense_RNA | 0.279952515 | 3.896316853 | 0.5478786 | 0.4697091 | 0.854457996 |
| ENSMUSG00000021038 | Vipas39 | protein_coding | 0.209911679 | 6.141613416 | 0.5478335 | 0.4697272 | 0.854457996 |
| ENSMUSG00000005087 | Cd44 | protein_coding | -0.569003513 | 2.696449333 | 0.5472503 | 0.4699598 | 0.854457996 |
| ENSMUSG00000031778 | Cx3cl1 | protein_coding | -0.169134005 | 7.131790328 | 0.5469822 | 0.4700668 | 0.854457996 |
| ENSMUSG00000109121 | Gm44644 | lincRNA | -0.434651064 | 3.242296823 | 0.546906 | 0.4700973 | 0.854457996 |
| ENSMUSG00000051331 | Cacna1c | protein_coding | -0.388452485 | 3.874655236 | 0.5467024 | 0.4701786 | 0.854457996 |
| ENSMUSG00000023938 | Aars2 | protein_coding | -0.572732617 | 2.543370175 | 0.5466458 | 0.4702012 | 0.854457996 |
| ENSMUSG00000022247 | Brix1 | protein_coding | 0.162917281 | 6.667334184 | 0.5465787 | 0.470228 | 0.854457996 |
| ENSMUSG00000051586 | Mical3 | protein_coding | -0.351186813 | 4.129778632 | 0.5463302 | 0.4703273 | 0.854457996 |
| ENSMUSG00000035681 | Kcnc2 | protein_coding | -0.267115574 | 5.261467745 | 0.5460437 | 0.4704418 | 0.854457996 |
| ENSMUSG00000021890 | Eaf1 | protein_coding | 0.430803206 | 3.042197161 | 0.5459914 | 0.4704627 | 0.854457996 |
| ENSMUSG00000034574 | Daam1 | protein_coding | -0.279706611 | 4.86905192 | 0.5456157 | 0.470613 | 0.854457996 |
| ENSMUSG00000053286 | Trmt1l | protein_coding | -0.351733758 | 3.963026734 | 0.5454794 | 0.4706675 | 0.854457996 |
| ENSMUSG00000030410 | Dmwd | protein_coding | 0.200650477 | 6.561215865 | 0.544806 | 0.4709371 | 0.854457996 |
| ENSMUSG00000026667 | Uhmk1 | protein_coding | 0.168871942 | 6.742017928 | 0.5443423 | 0.4711229 | 0.854457996 |
| ENSMUSG00000037331 | Larp1 | protein_coding | -0.236366062 | 5.011395459 | 0.5443266 | 0.4711292 | 0.854457996 |
| ENSMUSG00000018476 | Kdm6b | protein_coding | -0.401464204 | 5.4998987 | 0.5477615 | 0.4712242 | 0.854457996 |
| ENSMUSG00000026110 | Mgat4a | protein_coding | -0.2855218 | 4.364640138 | 0.543743 | 0.4713632 | 0.854457996 |
| ENSMUSG00000011752 | Pgam1 | protein_coding | 0.287223571 | 3.725204375 | 0.5436138 | 0.471415 | 0.854457996 |
| ENSMUSG00000039427 | Alg1 | protein_coding | -0.424822263 | 3.42903892 | 0.5435277 | 0.4714496 | 0.854457996 |
| ENSMUSG00000042535 | Gtpbp1 | protein_coding | -0.355805837 | 3.944895231 | 0.5434923 | 0.4714638 | 0.854457996 |
| ENSMUSG00000002546 | Golga2 | protein_coding | -0.249499589 | 5.61340806 | 0.5433699 | 0.4715129 | 0.854457996 |
| ENSMUSG00000022656 | Nectin3 | protein_coding | -0.494159284 | 3.045823265 | 0.5431145 | 0.4716154 | 0.854457996 |
| ENSMUSG00000068735 | Trp53i11 | protein_coding | -0.220224625 | 5.634389852 | 0.5431075 | 0.4716182 | 0.854457996 |
| ENSMUSG00000022265 | Ank | protein_coding | -0.188634011 | 6.578249939 | 0.5430283 | 0.47165 | 0.854457996 |
| ENSMUSG00000026360 | Rgs2 | protein_coding | 0.229750697 | 7.852127661 | 0.5437284 | 0.4716932 | 0.854457996 |

| ensembl_gene_id | external_gene_name | gene_biotype | log2 (fold change) | log2 (counts per million) | F-value | p-value | False discovery rate |
|---|---|---|---|---|---|---|---|
| ENSMUSG00000040536 | Necab1 | protein_coding | -0.309169461 | 5.67572211 | 0.5439306 | 0.4717245 | 0.854457996 |
| ENSMUSG00000026254 | Eif4e2 | protein_coding | 0.180011918 | 6.461110761 | 0.5428257 | 0.4717314 | 0.854457996 |
| ENSMUSG00000030760 | Acer3 | protein_coding | -0.223087014 | 4.766273208 | 0.5427332 | 0.4717685 | 0.854457996 |
| ENSMUSG00000025937 | Lactb2 | protein_coding | -0.362999651 | 3.577857389 | 0.5427302 | 0.4717698 | 0.854457996 |
| ENSMUSG00000018796 | Acsl1 | protein_coding | 0.279195209 | 4.888774052 | 0.5424429 | 0.4718852 | 0.854457996 |
| ENSMUSG00000028385 | Snx30 | protein_coding | -0.229864985 | 4.822640067 | 0.5422891 | 0.471947 | 0.854457996 |
| ENSMUSG00000038290 | Smg6 | protein_coding | -0.260805106 | 4.756265061 | 0.5422233 | 0.4719734 | 0.854457996 |
| ENSMUSG00000021738 | Atxn7 | protein_coding | -0.367612052 | 3.312715562 | 0.5420824 | 0.4720301 | 0.854457996 |
| ENSMUSG00000108461 | AV356131 | bidirectional_promote | 0.559214518 | 2.681893611 | 0.5420641 | 0.4720375 | 0.854457996 |
| ENSMUSG00000079108 | Srp54b | protein_coding | -0.376525276 | 3.34042388 | 0.5420351 | 0.4720491 | 0.854457996 |
| ENSMUSG00000071424 | Grid2 | protein_coding | -0.267106071 | 4.632350797 | 0.5417226 | 0.4721748 | 0.854457996 |
| ENSMUSG00000038011 | Dnah10 | protein_coding | 0.508029799 | 2.859504059 | 0.5417027 | 0.4721828 | 0.854457996 |
| ENSMUSG00000041889 | Shisa4 | protein_coding | 0.159784944 | 6.555733141 | 0.54155 | 0.4722443 | 0.854457996 |
| ENSMUSG00000020705 | Ddx42 | protein_coding | -0.193897432 | 5.811682995 | 0.5414912 | 0.4722679 | 0.854457996 |
| ENSMUSG00000040387 | Klhl32 | protein_coding | 0.268459724 | 4.658007617 | 0.5414393 | 0.4722888 | 0.854457996 |
| ENSMUSG00000029388 | Eif2b1 | protein_coding | 0.152336784 | 6.75653842 | 0.5412336 | 0.4723716 | 0.854457996 |
| ENSMUSG00000070476 | Fam217b | protein_coding | 0.18202421 | 5.854551818 | 0.5410276 | 0.4724546 | 0.854457996 |
| ENSMUSG00000042604 | Kcna4 | protein_coding | -0.372067948 | 3.988612793 | 0.5409575 | 0.4724828 | 0.854457996 |
| ENSMUSG00000016528 | Mapkapk2 | protein_coding | -0.592544647 | 2.809409699 | 0.5404923 | 0.4726702 | 0.854469962 |
| ENSMUSG00000019518 | Ap4m1 | protein_coding | -0.453927375 | 2.850539078 | 0.5403692 | 0.4727198 | 0.854469962 |
| ENSMUSG00000030498 | Gas2 | protein_coding | -0.261676504 | 4.081728623 | 0.5403494 | 0.4727278 | 0.854469962 |
| ENSMUSG00000029439 | Sfswap | protein_coding | -0.384181795 | 3.823565253 | 0.5399846 | 0.4728749 | 0.854592234 |
| ENSMUSG00000053799 | Exoc6 | protein_coding | -0.274033281 | 4.011580773 | 0.5396708 | 0.4730015 | 0.854677361 |
| ENSMUSG00000052336 | Cx3cr1 | protein_coding | -0.341408549 | 4.189887564 | 0.5389594 | 0.4732887 | 0.855052611 |
| ENSMUSG00000024384 | Iws1 | protein_coding | -0.32811719 | 3.589956295 | 0.5384807 | 0.4734821 | 0.855165613 |
| ENSMUSG00000021892 | Sh3bp5 | protein_coding | -0.266538028 | 4.685489665 | 0.5384109 | 0.4735103 | 0.855165613 |
| ENSMUSG00000039059 | Hrh3 | protein_coding | -0.321171189 | 3.775898326 | 0.5379253 | 0.4737066 | 0.855341818 |
| ENSMUSG00000004264 | Phb2 | protein_coding | -0.343815237 | 3.591811995 | 0.537688 | 0.4738027 | 0.855341818 |
| ENSMUSG00000007656 | Arpp19 | protein_coding | -0.208720243 | 9.503377922 | 0.5375436 | 0.4738611 | 0.855341818 |
| ENSMUSG00000000766 | Oprm1 | protein_coding | 0.249009053 | 5.499168234 | 0.536841 | 0.4741456 | 0.855341818 |
| ENSMUSG00000027596 | a | protein_coding | -0.478670959 | 2.687726089 | 0.5365943 | 0.4742456 | 0.855341818 |
| ENSMUSG00000027997 | Casp6 | protein_coding | -0.485754385 | 2.845442278 | 0.5363704 | 0.4743363 | 0.855341818 |
| ENSMUSG00000073478 | D730003I15Rik | lincRNA | -0.334273944 | 4.288158583 | 0.5360767 | 0.4744554 | 0.855341818 |
| ENSMUSG00000031660 | Brd7 | protein_coding | -0.229704567 | 5.766305864 | 0.5358908 | 0.4745308 | 0.855341818 |
| ENSMUSG00000033808 | Tmem87a | protein_coding | -0.554008405 | 2.727547027 | 0.5358122 | 0.4745627 | 0.855341818 |
| ENSMUSG00000003273 | Car11 | protein_coding | -0.261761349 | 5.660026022 | 0.5353213 | 0.474762 | 0.855341818 |
| ENSMUSG00000031485 | Prosc | protein_coding | 0.184397548 | 6.440097532 | 0.5352165 | 0.4748045 | 0.855341818 |
| ENSMUSG00000020864 | Ankrd40 | protein_coding | -0.205533466 | 5.976955726 | 0.5350693 | 0.4748643 | 0.855341818 |
| ENSMUSG00000022946 | Dopey2 | protein_coding | 0.293817827 | 4.038381557 | 0.534995 | 0.4748945 | 0.855341818 |
| ENSMUSG00000038982 | Bloc1s5 | protein_coding | 0.201304801 | 5.813263215 | 0.5348399 | 0.4749575 | 0.855341818 |
| ENSMUSG00000041468 | Gpr12 | protein_coding | -0.278840688 | 4.7505083 | 0.5348108 | 0.4749693 | 0.855341818 |
| ENSMUSG00000034324 | Tmem132c | protein_coding | -0.569193705 | 2.717999183 | 0.5342871 | 0.4751822 | 0.855341818 |
| ENSMUSG00000048930 | Tada3 | protein_coding | 0.220293966 | 4.935587301 | 0.5342154 | 0.4752114 | 0.855341818 |
| ENSMUSG00000064225 | Paqr9 | protein_coding | 0.316540271 | 4.614825467 | 0.5342098 | 0.4752136 | 0.855341818 |
| ENSMUSG00000032571 | Pik3r4 | protein_coding | 0.32876954 | 4.210568491 | 0.534115 | 0.4752522 | 0.855341818 |
| ENSMUSG00000079038 | D130040H23Rik | protein_coding | -0.60462974 | 2.546525431 | 0.5340716 | 0.4752698 | 0.855341818 |
| ENSMUSG00000031608 | Galnt7 | protein_coding | -0.427953521 | 3.425388949 | 0.5338586 | 0.4753565 | 0.855341818 |
| ENSMUSG00000046093 | Hpcal4 | protein_coding | -0.160941523 | 8.117955572 | 0.5338371 | 0.4753652 | 0.855341818 |
| ENSMUSG00000022867 | Usp25 | protein_coding | -0.314129647 | 4.176363426 | 0.5336543 | 0.4754397 | 0.855341818 |
| ENSMUSG00000030545 | Pex11a | protein_coding | -0.348607747 | 4.157662437 | 0.5332826 | 0.475591 | 0.855341818 |
| ENSMUSG00000026824 | Kcnj3 | protein_coding | 0.240841325 | 4.63725668 | 0.5328317 | 0.4757747 | 0.855341818 |
| ENSMUSG00000049907 | Rasl11b | protein_coding | 0.266598974 | 4.109289556 | 0.5327593 | 0.4758041 | 0.855341818 |
| ENSMUSG00000042156 | Dzip1 | protein_coding | -0.248851263 | 5.309869513 | 0.5323791 | 0.4759592 | 0.855341818 |
| ENSMUSG00000045348 | Nyap1 | protein_coding | -0.427187522 | 3.3711116 | 0.5323128 | 0.4759862 | 0.855341818 |
| ENSMUSG00000056296 | Synpr | protein_coding | 0.213776719 | 7.160050266 | 0.5322837 | 0.476015 | 0.855341818 |
| ENSMUSG00000040102 | Klhl42 | protein_coding | 0.328325883 | 4.246095242 | 0.5321498 | 0.4760527 | 0.855341818 |
| ENSMUSG00000025872 | Thoc3 | protein_coding | 0.241135127 | 5.108572263 | 0.5319471 | 0.4761353 | 0.855341818 |
| ENSMUSG00000017417 | Plxdc1 | protein_coding | 0.300684284 | 3.993092198 | 0.5317721 | 0.4762068 | 0.855341818 |
| ENSMUSG00000022354 | Ndufb9 | protein_coding | 0.200418871 | 10.17307244 | 0.531584 | 0.4762835 | 0.855341818 |
| ENSMUSG00000048485 | Zbtb8b | protein_coding | 0.460213504 | 2.654409053 | 0.5312049 | 0.4764384 | 0.855341818 |
| ENSMUSG00000035960 | Apex1 | protein_coding | 0.205515523 | 5.558135195 | 0.5311587 | 0.4764572 | 0.855341818 |
| ENSMUSG00000024036 | Slc37a1 | protein_coding | -0.460220485 | 2.945747937 | 0.5311239 | 0.4764714 | 0.855341818 |
| ENSMUSG00000028300 | 3110043O21Rik | protein_coding | 0.24195187 | 5.315404608 | 0.5306166 | 0.4766787 | 0.855571102 |
| ENSMUSG00000055653 | Gpc3 | protein_coding | 0.258675678 | 4.439939438 | 0.5302058 | 0.4768468 | 0.855729843 |
| ENSMUSG00000059810 | Rgs3 | protein_coding | -0.371385141 | 3.626404954 | 0.5299325 | 0.4769586 | 0.85578768 |
| ENSMUSG00000021619 | Atg10 | protein_coding | 0.174800495 | 5.97382329 | 0.5297361 | 0.477039 | 0.855789113 |
| ENSMUSG00000030282 | Cmas | protein_coding | 0.14877982 | 6.74470459 | 0.5293674 | 0.4771899 | 0.855917114 |
| ENSMUSG00000039372 | 4-Mar | protein_coding | 0.350193071 | 3.590618025 | 0.5288374 | 0.4774071 | 0.856112285 |
| ENSMUSG00000027316 | Gfra4 | protein_coding | 0.226187128 | 5.032923443 | 0.5287132 | 0.477458 | 0.856112285 |
| ENSMUSG00000002963 | Pnkp | protein_coding | 0.223580979 | 4.95504095 | 0.5278218 | 0.4778236 | 0.856517035 |
| ENSMUSG00000022965 | Ifngr2 | protein_coding | 0.214905748 | 5.18798113 | 0.5277746 | 0.477843 | 0.856517035 |
| ENSMUSG00000031980 | Agt | protein_coding | -0.205846568 | 6.828260512 | 0.5275292 | 0.4779437 | 0.856554856 |
| ENSMUSG00000020658 | Efr3b | protein_coding | -0.192620884 | 5.894749229 | 0.5269458 | 0.4781834 | 0.856830867 |
| ENSMUSG00000024712 | Rfk | protein_coding | 0.177239189 | 6.438470408 | 0.5266194 | 0.4783176 | 0.856830867 |
| ENSMUSG00000022517 | Mgrn1 | protein_coding | 0.350516004 | 6.009870723 | 0.5303572 | 0.478385 | 0.856830867 |
| ENSMUSG00000030257 | Srgap3 | protein_coding | -0.197506736 | 6.169413465 | 0.526379 | 0.4784165 | 0.856830867 |
| ENSMUSG00000019856 | Fam184a | protein_coding | -0.337319953 | 3.47433877 | 0.5261106 | 0.4785269 | 0.856885948 |
| ENSMUSG00000034343 | Ube2f | protein_coding | 0.170045241 | 6.296843582 | 0.5258315 | 0.4786418 | 0.856948947 |

| ensembl_gene_id | external_gene_name | gene_biotype | log2 (fold change) | log2 (counts per million) | F-value | p-value | False discovery rate |
|---|---|---|---|---|---|---|---|
| ENSMUSG00000022009 | Nufip1 | protein_coding | 0.293247433 | 3.825513868 | 0.5254018 | 0.4788188 | 0.856992897 |
| ENSMUSG00000057133 | Chd6 | protein_coding | -0.207537972 | 5.686323388 | 0.5250754 | 0.4789532 | 0.856992897 |
| ENSMUSG00000039737 | Prkrip1 | protein_coding | 0.180161834 | 6.35533797 | 0.5246924 | 0.4791112 | 0.856992897 |
| ENSMUSG00000024817 | Uhrf2 | protein_coding | -0.245265025 | 4.927797077 | 0.5245011 | 0.47919 | 0.856992897 |
| ENSMUSG00000029650 | Slc46a3 | protein_coding | 0.458189553 | 2.794712814 | 0.5243081 | 0.4792697 | 0.856992897 |
| ENSMUSG00000025410 | Dctn2 | protein_coding | 0.192748206 | 8.007037595 | 0.5242091 | 0.4793105 | 0.856992897 |
| ENSMUSG00000021986 | Amer2 | protein_coding | -0.265423969 | 5.07212423 | 0.5241072 | 0.4793526 | 0.856992897 |
| ENSMUSG00000048332 | Lhfp | protein_coding | -0.438964237 | 3.587935089 | 0.5241334 | 0.4793586 | 0.856992897 |
| ENSMUSG00000037134 | Prmt9 | protein_coding | -0.347909685 | 3.579216557 | 0.5240319 | 0.4793836 | 0.856992897 |
| ENSMUSG00000072825 | Cep170b | protein_coding | 0.194490196 | 6.618849116 | 0.5236185 | 0.4795543 | 0.857155531 |
| ENSMUSG00000058402 | Zfp420 | protein_coding | -0.289687609 | 3.993449678 | 0.523207 | 0.4797243 | 0.857316895 |
| ENSMUSG00000062590 | Armc9 | protein_coding | 0.21595736 | 5.167580818 | 0.5227086 | 0.4799304 | 0.857542667 |
| ENSMUSG00000002781 | Tmem143 | protein_coding | 0.321589921 | 3.594656534 | 0.5224344 | 0.4800438 | 0.857553497 |
| ENSMUSG00000031148 | Gpkow | protein_coding | -0.281844413 | 5.156053859 | 0.5221822 | 0.4801482 | 0.857553497 |
| ENSMUSG00000011658 | Fuz | protein_coding | 0.509708551 | 2.572785091 | 0.5218809 | 0.4802729 | 0.857553497 |
| ENSMUSG00000029406 | Pitpnm2 | protein_coding | -0.242769646 | 5.065491646 | 0.5214945 | 0.480433 | 0.857553497 |
| ENSMUSG00000032349 | Elovl5 | protein_coding | -0.187167215 | 5.954478647 | 0.5214766 | 0.4804404 | 0.857553497 |
| ENSMUSG00000026655 | Fam107b | protein_coding | -0.645697805 | 3.145436871 | 0.5230448 | 0.4804416 | 0.857553497 |
| ENSMUSG00000071172 | Srsf3 | protein_coding | -0.160195516 | 7.133198226 | 0.5211977 | 0.480556 | 0.857553497 |
| ENSMUSG00000074220 | Zfp382 | protein_coding | 0.386130195 | 3.186077791 | 0.521153 | 0.4805745 | 0.857553497 |
| ENSMUSG00000021039 | Snw1 | protein_coding | -0.161232675 | 6.894242313 | 0.5204585 | 0.4808625 | 0.857808165 |
| ENSMUSG00000044768 | D1Ertd622e | protein_coding | -0.273417479 | 4.60672709 | 0.520233 | 0.4809561 | 0.857808165 |
| ENSMUSG00000021518 | Ptdss1 | protein_coding | 0.35018428 | 4.259465503 | 0.5202723 | 0.4809565 | 0.857808165 |
| ENSMUSG00000030754 | Copb1 | protein_coding | -0.225960609 | 5.99782051 | 0.5196808 | 0.4811854 | 0.85792905 |
| ENSMUSG00000038836 | Agbl3 | protein_coding | -0.21911751 | 5.249206097 | 0.519514 | 0.4812547 | 0.85792905 |
| ENSMUSG00000027285 | Haus2 | protein_coding | 0.188698038 | 6.045529529 | 0.5194925 | 0.4812636 | 0.85792905 |
| ENSMUSG00000053094 | Tmem248 | protein_coding | 0.251888618 | 4.729966442 | 0.519077 | 0.4814364 | 0.858094691 |
| ENSMUSG00000001323 | Srr | protein_coding | 0.223734289 | 4.668064243 | 0.5186123 | 0.4816296 | 0.858173 |
| ENSMUSG00000097466 | D430036J16Rik | antisense_RNA | -0.407623425 | 3.323470962 | 0.5185308 | 0.4816636 | 0.858173 |
| ENSMUSG00000037235 | Mxd4 | protein_coding | -0.299138011 | 4.331217165 | 0.5183625 | 0.4817336 | 0.858173 |
| ENSMUSG00000030579 | Tyrobp | protein_coding | -0.353061423 | 5.019160979 | 0.5192285 | 0.4817995 | 0.858173 |
| ENSMUSG00000038552 | Fndc4 | protein_coding | -0.248205805 | 5.636499316 | 0.5178142 | 0.4819619 | 0.858200611 |
| ENSMUSG00000031376 | Atp2b3 | protein_coding | 0.340786246 | 3.234894051 | 0.5177045 | 0.4820076 | 0.858200611 |
| ENSMUSG00000058420 | Syt17 | protein_coding | -0.266720032 | 5.326430569 | 0.5172785 | 0.4821851 | 0.858200611 |
| ENSMUSG00000046753 | Ccdc66 | protein_coding | -0.415625394 | 3.093185447 | 0.5171446 | 0.482241 | 0.858200611 |
| ENSMUSG00000026730 | Pter | protein_coding | -0.411985552 | 4.465772053 | 0.5185393 | 0.482307 | 0.858200611 |
| ENSMUSG00000057778 | Cyb5d2 | protein_coding | 0.291803579 | 4.542652472 | 0.5169736 | 0.4823123 | 0.858200611 |
| ENSMUSG00000041798 | Gck | protein_coding | 0.29444105 | 5.389062002 | 0.5162875 | 0.4826152 | 0.858200611 |
| ENSMUSG00000009079 | Ewsr1 | protein_coding | 0.193422252 | 5.932260034 | 0.5159847 | 0.4827251 | 0.858200611 |
| ENSMUSG00000109006 | B230209E15Rik | lincRNA | 0.196154535 | 6.39717013 | 0.5159676 | 0.4827322 | 0.858200611 |
| ENSMUSG00000020743 | Mif4gd | protein_coding | -0.223898324 | 5.246410867 | 0.5158686 | 0.4827736 | 0.858200611 |
| ENSMUSG00000020037 | Rfx4 | protein_coding | -0.278401367 | 3.874908467 | 0.5157123 | 0.4828389 | 0.858200611 |
| ENSMUSG00000062075 | Lmnb2 | protein_coding | -0.413151649 | 3.295389858 | 0.5156939 | 0.4828466 | 0.858200611 |
| ENSMUSG00000025374 | Nabp2 | protein_coding | 0.236985552 | 5.12100011 | 0.5154903 | 0.4829316 | 0.858200611 |
| ENSMUSG00000056673 | Kdm5d | protein_coding | -0.41512997 | 2.893253497 | 0.5154885 | 0.4829324 | 0.858200611 |
| ENSMUSG00000026078 | Pdcl3 | protein_coding | -0.154895066 | 7.318520387 | 0.5151473 | 0.483075 | 0.858232701 |
| ENSMUSG00000056260 | Lrif1 | protein_coding | 0.218436051 | 5.03457639 | 0.5150636 | 0.4831101 | 0.858232701 |
| ENSMUSG00000047648 | Fbxo30 | protein_coding | -0.473375715 | 2.810722436 | 0.5147096 | 0.4832582 | 0.858288933 |
| ENSMUSG00000025240 | Sacm1l | protein_coding | -0.48595398 | 3.773463814 | 0.5161177 | 0.4833147 | 0.858288933 |
| ENSMUSG00000041153 | Osgin2 | protein_coding | -0.404540699 | 3.325612548 | 0.5141853 | 0.4834777 | 0.858288933 |
| ENSMUSG00000026123 | Plekhb2 | protein_coding | 0.291496996 | 9.104807935 | 0.5175378 | 0.4834986 | 0.858288933 |
| ENSMUSG00000073705 | Apitd1 | protein_coding | 0.262404255 | 4.583008262 | 0.5137975 | 0.4836401 | 0.858288933 |
| ENSMUSG00000020319 | Wdpcp | protein_coding | -0.352884195 | 3.361835588 | 0.5136828 | 0.4836882 | 0.858288933 |
| ENSMUSG00000020024 | Cep83 | protein_coding | -0.305371731 | 4.004862667 | 0.5136536 | 0.4837004 | 0.858288933 |
| ENSMUSG00000027165 | B230118H07Rik | protein_coding | 0.16615607 | 7.438964117 | 0.5124003 | 0.4842263 | 0.858814154 |
| ENSMUSG00000025451 | Paip1 | protein_coding | -0.306104918 | 8.314837663 | 0.5161976 | 0.4842512 | 0.858814154 |
| ENSMUSG00000061758 | Akr1b10 | protein_coding | -0.205992846 | 5.577998856 | 0.5123045 | 0.4842665 | 0.858814154 |
| ENSMUSG00000021193 | Pitrm1 | protein_coding | 0.301310957 | 4.532985356 | 0.5114784 | 0.4846137 | 0.858814154 |
| ENSMUSG00000034105 | Tldc1 | protein_coding | 0.324122108 | 3.480613273 | 0.5114694 | 0.4846174 | 0.858814154 |
| ENSMUSG00000075478 | Slitrk1 | protein_coding | 0.267015516 | 4.949897252 | 0.5113659 | 0.484661 | 0.858814154 |
| ENSMUSG00000069919 | Hba-a1 | protein_coding | -0.363651821 | 4.437089775 | 0.5111328 | 0.4847756 | 0.858814154 |
| ENSMUSG00000036893 | Ehmt1 | protein_coding | 0.212243168 | 5.202402411 | 0.5108283 | 0.4848872 | 0.858814154 |
| ENSMUSG00000001983 | Taco1 | protein_coding | 0.340078445 | 3.448332275 | 0.5107706 | 0.4849114 | 0.858814154 |
| ENSMUSG00000023192 | Grm2 | protein_coding | -0.522899152 | 3.081456582 | 0.5104707 | 0.4850377 | 0.858814154 |
| ENSMUSG00000037747 | Phyhipl | protein_coding | -0.173211262 | 7.142047207 | 0.5104706 | 0.4850378 | 0.858814154 |
| ENSMUSG00000002326 | Gmpr2 | protein_coding | 0.235881159 | 5.289476233 | 0.5104519 | 0.4850456 | 0.858814154 |
| ENSMUSG00000028403 | Zdhhc21 | protein_coding | -0.2784224 | 4.489614356 | 0.510451 | 0.485046 | 0.858814154 |
| ENSMUSG00000041530 | Ago1 | protein_coding | 0.269694299 | 4.381906915 | 0.5100213 | 0.4852271 | 0.858814154 |
| ENSMUSG00000020955 | Ap4s1 | protein_coding | -0.204857482 | 5.522627673 | 0.509948 | 0.485258 | 0.858814154 |
| ENSMUSG00000029651 | Mtus2 | protein_coding | -0.192458774 | 5.4084515 | 0.5094208 | 0.4854802 | 0.858814154 |
| ENSMUSG00000037868 | Egr2 | protein_coding | -0.530887798 | 2.595432528 | 0.50913 | 0.4856029 | 0.858814154 |
| ENSMUSG00000021983 | Atp8a2 | protein_coding | -0.285770502 | 4.255534243 | 0.5090011 | 0.4856573 | 0.858814154 |
| ENSMUSG00000079614 | Seh1l | protein_coding | 0.185310119 | 6.686796075 | 0.5088562 | 0.4857185 | 0.858814154 |
| ENSMUSG00000034243 | Golgb1 | protein_coding | -0.223521031 | 5.276950784 | 0.5088072 | 0.4857392 | 0.858814154 |
| ENSMUSG00000057069 | Ero1lb | protein_coding | -0.665092308 | 2.659979662 | 0.5088082 | 0.4857553 | 0.858814154 |
| ENSMUSG00000040506 | Ambra1 | protein_coding | -0.262089576 | 5.066546137 | 0.5086251 | 0.485816 | 0.858814154 |
| ENSMUSG00000059439 | Bcas3 | protein_coding | -0.310224707 | 4.684346565 | 0.5085841 | 0.4858334 | 0.858814154 |
| ENSMUSG00000044912 | Syt16 | protein_coding | 0.249454574 | 4.870033124 | 0.5078675 | 0.4861362 | 0.859208184 |

| ensembl_gene_id | external_gene_name | gene_biotype | log2 (fold change) | log2 (counts per million) | F-value | p-value | False discovery rate |
|---|---|---|---|---|---|---|---|
| ENSMUSG00000023110 | Prmt5 | protein_coding | 0.160318407 | 6.795089092 | 0.5075407 | 0.4862744 | 0.859311199 |
| ENSMUSG00000019868 | Vta1 | protein_coding | 0.154239751 | 6.910586962 | 0.5068852 | 0.4865518 | 0.859513476 |
| ENSMUSG00000028158 | Mttp | protein_coding | -0.351326747 | 3.913784523 | 0.5066488 | 0.4866519 | 0.859513476 |
| ENSMUSG00000014349 | Ube2z | protein_coding | -0.176495226 | 6.026995833 | 0.5064536 | 0.4867346 | 0.859513476 |
| ENSMUSG00000027255 | Arfgap2 | protein_coding | 0.301052821 | 4.615170239 | 0.5064043 | 0.4867555 | 0.859513476 |
| ENSMUSG00000009647 | Mcu | protein_coding | -0.182568858 | 6.260621807 | 0.5063263 | 0.4867885 | 0.859513476 |
| ENSMUSG00000006307 | Kmt2b | protein_coding | -0.489636992 | 3.089273277 | 0.5057018 | 0.4870533 | 0.859815242 |
| ENSMUSG00000026134 | Prim2 | protein_coding | -0.250841347 | 4.413397606 | 0.5055461 | 0.4871194 | 0.859815242 |
| ENSMUSG00000049946 | BC030500 | protein_coding | 0.264556867 | 5.154877365 | 0.5052142 | 0.4872602 | 0.859888566 |
| ENSMUSG00000032462 | Pik3cb | protein_coding | -0.468665977 | 3.052079524 | 0.5050508 | 0.4873296 | 0.859888566 |
| ENSMUSG00000037070 | Rbmxl1 | protein_coding | 0.256837762 | 4.031584788 | 0.5048831 | 0.4874008 | 0.859888566 |
| ENSMUSG00000030077 | Chl1 | protein_coding | 0.193954776 | 6.308066585 | 0.5043853 | 0.4876123 | 0.860046748 |
| ENSMUSG00000068114 | Ccdc134 | protein_coding | -0.425480967 | 3.203838353 | 0.5039964 | 0.4877776 | 0.860046748 |
| ENSMUSG00000019763 | Rmnd1 | protein_coding | 0.263338422 | 3.99486324 | 0.5039116 | 0.4878137 | 0.860046748 |
| ENSMUSG00000023942 | Slc29a1 | protein_coding | 0.268499181 | 4.982933342 | 0.5037276 | 0.4878919 | 0.860046748 |
| ENSMUSG00000033557 | Fam20b | protein_coding | 0.227133134 | 5.503172762 | 0.503365 | 0.4880463 | 0.860046748 |
| ENSMUSG00000039275 | Foxk2 | protein_coding | -0.24118697 | 5.098792257 | 0.5032909 | 0.4880778 | 0.860046748 |
| ENSMUSG00000021973 | Micu2 | protein_coding | -0.288707599 | 4.362806374 | 0.5032731 | 0.4880854 | 0.860046748 |
| ENSMUSG00000031337 | Mtm1 | protein_coding | -0.594273702 | 2.849705972 | 0.5031675 | 0.4881303 | 0.860046748 |
| ENSMUSG00000097412 | 1810014B01Rik | transcribed_unproces | 0.485729674 | 2.91370316 | 0.5029634 | 0.4882172 | 0.860058991 |
| ENSMUSG00000027540 | Ptpn1 | protein_coding | -0.266507996 | 4.79176329 | 0.5025123 | 0.4884094 | 0.860067336 |
| ENSMUSG00000060261 | Gtf2i | protein_coding | -0.19195349 | 6.664395869 | 0.5024627 | 0.4884306 | 0.860067336 |
| ENSMUSG00000016346 | Kcnq2 | protein_coding | 0.175705124 | 5.984644413 | 0.5022336 | 0.4885283 | 0.860067336 |
| ENSMUSG00000032673 | Prorsd1 | protein_coding | -0.247188321 | 4.986987756 | 0.5022015 | 0.4885419 | 0.860067336 |
| ENSMUSG00000029777 | Gars | protein_coding | -0.174397677 | 6.473768453 | 0.5017187 | 0.4887479 | 0.860173626 |
| ENSMUSG00000054252 | Fgfr3 | protein_coding | -0.274953911 | 4.760512676 | 0.5016706 | 0.4887684 | 0.860173626 |
| ENSMUSG00000021250 | Fos | protein_coding | -0.234608643 | 6.540849949 | 0.5016769 | 0.4888423 | 0.860173626 |
| ENSMUSG00000026187 | Xrcc5 | protein_coding | 0.211545782 | 5.202809155 | 0.5000896 | 0.4894439 | 0.860879516 |
| ENSMUSG00000081552 | Gm12328 | processed_pseudoge | 0.44989595 | 2.681845599 | 0.5000046 | 0.4894803 | 0.860879516 |
| ENSMUSG00000042734 | Ttc9 | protein_coding | -0.235704001 | 5.301513792 | 0.4999967 | 0.4894836 | 0.860879516 |
| ENSMUSG00000015971 | Actr8 | protein_coding | 0.184825998 | 6.486960432 | 0.4991796 | 0.4898335 | 0.861353906 |
| ENSMUSG00000025354 | Dnajc14 | protein_coding | 0.269394097 | 4.999085832 | 0.4985632 | 0.4900976 | 0.861677488 |
| ENSMUSG00000027667 | Zfp639 | protein_coding | -0.214612956 | 5.600760563 | 0.497816 | 0.4904182 | 0.86198746 |
| ENSMUSG00000020640 | Itsn2 | protein_coding | -0.296844894 | 3.894198218 | 0.4977786 | 0.4904342 | 0.86198746 |
| ENSMUSG00000027257 | Pacsin3 | protein_coding | -0.305907914 | 4.267205859 | 0.4971198 | 0.4907171 | 0.862197124 |
| ENSMUSG00000039804 | Ncoa5 | protein_coding | -0.276702424 | 4.500782273 | 0.4968325 | 0.4908406 | 0.862197124 |
| ENSMUSG00000032261 | Sh3bgrl2 | protein_coding | -0.313656717 | 5.201580126 | 0.4968178 | 0.4909062 | 0.862197124 |
| ENSMUSG00000026313 | Hdac4 | protein_coding | -0.213905227 | 5.850926802 | 0.4966172 | 0.4909332 | 0.862197124 |
| ENSMUSG00000040174 | Alkbh3 | protein_coding | 0.42595663 | 2.89348082 | 0.4965679 | 0.4909544 | 0.862197124 |
| ENSMUSG00000057531 | Dtnbp1 | protein_coding | 0.176332985 | 6.867280888 | 0.4959921 | 0.4912022 | 0.862386733 |
| ENSMUSG00000050556 | Kcnb1 | protein_coding | -0.260311768 | 5.253583299 | 0.4959263 | 0.4912305 | 0.862386733 |
| ENSMUSG00000036169 | Sostdc1 | protein_coding | -0.419918728 | 5.072898366 | 0.4989987 | 0.491303 | 0.862386733 |
| ENSMUSG00000035828 | Pim3 | protein_coding | -0.24484251 | 5.045713111 | 0.4953937 | 0.4914598 | 0.862417728 |
| ENSMUSG00000030611 | Mrps11 | protein_coding | 0.151502188 | 7.327358573 | 0.4953445 | 0.4914811 | 0.862417728 |
| ENSMUSG00000033610 | Pank1 | protein_coding | -0.309385207 | 3.737443995 | 0.4950481 | 0.4916088 | 0.862434478 |
| ENSMUSG00000021215 | Net1 | protein_coding | -0.487714641 | 2.936039139 | 0.4949501 | 0.491651 | 0.862434478 |
| ENSMUSG00000021217 | Tshz3 | protein_coding | -0.38780688 | 2.952394091 | 0.4934607 | 0.4922939 | 0.863396967 |
| ENSMUSG00000071573 | Rnls | protein_coding | 0.355591377 | 3.221765086 | 0.4932458 | 0.4923867 | 0.863396967 |
| ENSMUSG00000028804 | Csmd2 | protein_coding | 0.419379414 | 3.077264071 | 0.4930148 | 0.4924866 | 0.863396967 |
| ENSMUSG00000033444 | Specc1l | protein_coding | -0.239566638 | 4.588919484 | 0.4929356 | 0.4925209 | 0.863396967 |
| ENSMUSG00000030091 | Nup210 | protein_coding | -0.529525791 | 2.789625532 | 0.4919038 | 0.4929674 | 0.86355612 |
| ENSMUSG00000039086 | Ss18l1 | protein_coding | 0.228307625 | 4.984078631 | 0.4918592 | 0.4929868 | 0.86355612 |
| ENSMUSG00000051234 | Rnf7 | protein_coding | 0.136118195 | 7.855072363 | 0.4918429 | 0.4929938 | 0.86355612 |
| ENSMUSG00000042642 | Flad1 | protein_coding | 0.196037081 | 6.006308765 | 0.4917213 | 0.4930465 | 0.86355612 |
| ENSMUSG00000021585 | Cast | protein_coding | -0.40304854 | 3.004113261 | 0.4915339 | 0.4931277 | 0.86355612 |
| ENSMUSG00000034035 | Ccdc17 | protein_coding | -0.529569425 | 2.517724767 | 0.4914318 | 0.4931719 | 0.86355612 |
| ENSMUSG00000050605 | Zfp61 | protein_coding | -0.297129511 | 4.759084114 | 0.4913547 | 0.4932054 | 0.86355612 |
| ENSMUSG00000037907 | Ankrd13b | protein_coding | -0.420890116 | 3.239422366 | 0.4912083 | 0.4932689 | 0.86355612 |
| ENSMUSG00000032042 | Srpr | protein_coding | 0.182051271 | 5.738268041 | 0.4908733 | 0.4934141 | 0.86355612 |
| ENSMUSG00000021621 | Zcchc9 | protein_coding | 0.275255085 | 4.912392051 | 0.4905616 | 0.4935494 | 0.86355612 |
| ENSMUSG00000003378 | Grik5 | protein_coding | -0.417357166 | 3.244811165 | 0.490305 | 0.4936608 | 0.86355612 |
| ENSMUSG00000057895 | Zfp105 | protein_coding | -0.482743201 | 2.794361976 | 0.4902005 | 0.4937062 | 0.86355612 |
| ENSMUSG00000041961 | Znrf3 | protein_coding | -0.25984197 | 5.062820556 | 0.4901878 | 0.4937117 | 0.86355612 |
| ENSMUSG00000044748 | Defb1 | protein_coding | -0.822937557 | 3.007935018 | 0.493767 | 0.493736 | 0.86355612 |
| ENSMUSG00000031950 | Gabarapl2 | protein_coding | 0.152681051 | 6.722212429 | 0.4895405 | 0.4939929 | 0.863782968 |
| ENSMUSG00000027422 | Rrbp1 | protein_coding | -0.288115506 | 3.931150346 | 0.4894636 | 0.4940263 | 0.863782968 |
| ENSMUSG00000022533 | Atp13a3 | protein_coding | -0.294900288 | 4.077991839 | 0.4892418 | 0.4941228 | 0.863811174 |
| ENSMUSG00000026696 | Vamp4 | protein_coding | 0.175043085 | 7.902477043 | 0.4887657 | 0.49433 | 0.863812777 |
| ENSMUSG00000097145 | 9230114K14Rik | processed_transcript | -0.514769782 | 2.639944073 | 0.4883165 | 0.4945255 | 0.863812777 |
| ENSMUSG00000021483 | Cdk20 | protein_coding | 0.240894301 | 4.943556972 | 0.4882846 | 0.4945394 | 0.863812777 |
| ENSMUSG00000078485 | Plekhn1 | protein_coding | 0.619349412 | 3.134278056 | 0.489649 | 0.4946823 | 0.863812777 |
| ENSMUSG00000021476 | Habp4 | protein_coding | 0.172773007 | 6.94883417 | 0.4876117 | 0.4948326 | 0.863812777 |
| ENSMUSG00000052763 | Zfp212 | protein_coding | 0.346130021 | 3.652756293 | 0.4875902 | 0.494842 | 0.863812777 |
| ENSMUSG00000028691 | Prdx1 | protein_coding | -0.162504436 | 9.35749687 | 0.4874641 | 0.494897 | 0.863812777 |
| ENSMUSG00000074578 | Zfas1 | processed_transcript | 0.207902765 | 7.938346674 | 0.4873992 | 0.4949414 | 0.863812777 |
| ENSMUSG00000021273 | Fdft1 | protein_coding | 0.172572338 | 7.190815928 | 0.486895 | 0.4951453 | 0.863812777 |
| ENSMUSG00000040612 | Ildr2 | protein_coding | -0.240129026 | 5.880273793 | 0.4867608 | 0.49522 | 0.863812777 |
| ENSMUSG00000010755 | Cars | protein_coding | 0.3015927 | 5.871903777 | 0.4885264 | 0.4952384 | 0.863812777 |

| ensembl_gene_id | external_gene_name | gene_biotype | log2 (fold change) | log2 (counts per million) | F-value | p-value | False discovery rate |
|---|---|---|---|---|---|---|---|
| ENSMUSG00000041341 | Atg2b | protein_coding | 0.305518064 | 3.948557634 | 0.4865388 | 0.4953008 | 0.863812777 |
| ENSMUSG00000073207 | Ccdc160 | protein_coding | 0.409662141 | 3.255506659 | 0.4864346 | 0.4953463 | 0.863812777 |
| ENSMUSG00000054263 | Lifr | protein_coding | -0.346044461 | 3.791130107 | 0.4864014 | 0.4953608 | 0.863812777 |
| ENSMUSG00000031585 | Gtf2e2 | protein_coding | 0.171373182 | 5.868931169 | 0.4862281 | 0.4954365 | 0.863812777 |
| ENSMUSG00000029279 | Brdt | protein_coding | -0.341139812 | 3.734904963 | 0.4861935 | 0.4954517 | 0.863812777 |
| ENSMUSG00000034551 | Hdx | protein_coding | -0.770574924 | 2.241436869 | 0.4853181 | 0.4958344 | 0.863812777 |
| ENSMUSG00000027650 | Tti1 | protein_coding | 0.276434154 | 4.096851908 | 0.4849859 | 0.4959798 | 0.863812777 |
| ENSMUSG00000031610 | Scrg1 | protein_coding | -0.5057669 | 5.703641688 | 0.4956388 | 0.4960368 | 0.863812777 |
| ENSMUSG00000111080 | AC153899.1 | lincRNA | 0.194984096 | 5.911816474 | 0.4847251 | 0.496094 | 0.863812777 |
| ENSMUSG00000031753 | Cog4 | protein_coding | 0.33391354 | 4.126550012 | 0.4845846 | 0.4961556 | 0.863812777 |
| ENSMUSG00000061436 | Hipk2 | protein_coding | -0.365894846 | 4.56739696 | 0.4850781 | 0.496243 | 0.863812777 |
| ENSMUSG00000030030 | 1700003E16Rik | protein_coding | 0.218830847 | 5.293670112 | 0.4842386 | 0.4963072 | 0.863812777 |
| ENSMUSG00000038485 | Socs7 | protein_coding | -0.222998424 | 5.370109791 | 0.4842073 | 0.4963209 | 0.863812777 |
| ENSMUSG00000041670 | Rims1 | protein_coding | -0.17285877 | 6.013708503 | 0.48415 | 0.496346 | 0.863812777 |
| ENSMUSG00000038085 | Cnbd2 | protein_coding | -0.351986265 | 3.309772347 | 0.4839492 | 0.496434 | 0.863812777 |
| ENSMUSG00000037339 | Fam53a | protein_coding | 0.453695811 | 3.00363387 | 0.483868 | 0.4964696 | 0.863812777 |
| ENSMUSG00000034566 | Atp5h | protein_coding | 0.171203054 | 8.487546857 | 0.4837713 | 0.496512 | 0.863812777 |
| ENSMUSG00000030621 | Me3 | protein_coding | -0.31837396 | 4.22441266 | 0.4837593 | 0.4965173 | 0.863812777 |
| ENSMUSG00000020647 | Ncoa1 | protein_coding | -0.264868435 | 6.32571021 | 0.485143 | 0.4965337 | 0.863812777 |
| ENSMUSG00000032425 | Zfp949 | protein_coding | 0.244690958 | 5.079971543 | 0.4831681 | 0.4967767 | 0.864095795 |
| ENSMUSG00000026944 | Abca2 | protein_coding | -0.211207826 | 6.357412888 | 0.4827458 | 0.4969622 | 0.864106209 |
| ENSMUSG00000015653 | Steap2 | protein_coding | -0.31091167 | 3.443828162 | 0.4827184 | 0.4969742 | 0.864106209 |
| ENSMUSG00000037526 | Atg14 | protein_coding | -0.286986741 | 4.759605011 | 0.4825756 | 0.4970369 | 0.864106209 |
| ENSMUSG00000024376 | Epb41l4a | protein_coding | -0.369736849 | 3.386500114 | 0.482365 | 0.4971295 | 0.864106209 |
| ENSMUSG00000030521 | Mphosph10 | protein_coding | 0.23098119 | 4.697835945 | 0.4821347 | 0.4972308 | 0.864106209 |
| ENSMUSG00000097875 | Gm26704 | lincRNA | 0.340866522 | 3.844733875 | 0.4820571 | 0.4972649 | 0.864106209 |
| ENSMUSG00000041141 | Pnmal1 | protein_coding | 0.197603984 | 7.956656141 | 0.4817696 | 0.4974073 | 0.864214117 |
| ENSMUSG00000034168 | Irf2bpl | protein_coding | -0.244802927 | 4.404039221 | 0.4808496 | 0.4977964 | 0.864750317 |
| ENSMUSG00000098659 | 1110015O18Rik | lincRNA | 0.492208042 | 2.736696433 | 0.4801867 | 0.4980886 | 0.864901521 |
| ENSMUSG00000050379 | 6-Sep | protein_coding | 0.229018945 | 6.389171914 | 0.4802151 | 0.4981169 | 0.864901521 |
| ENSMUSG00000036598 | Ccdc113 | protein_coding | 0.241749812 | 4.42532233 | 0.4801049 | 0.4981247 | 0.864901521 |
| ENSMUSG00000069049 | Eif2s3y | protein_coding | -0.260099044 | 5.038632738 | 0.4796998 | 0.4983034 | 0.864973031 |
| ENSMUSG00000034252 | Senp6 | protein_coding | -0.238771281 | 5.136205662 | 0.4795841 | 0.4983545 | 0.864973031 |
| ENSMUSG00000024614 | Tmx3 | protein_coding | -0.194173602 | 5.685585436 | 0.4790952 | 0.4985704 | 0.864973031 |
| ENSMUSG00000038274 | Fau | protein_coding | 0.256160322 | 4.187082892 | 0.4787303 | 0.4987317 | 0.864973031 |
| ENSMUSG00000031840 | Rab3a | protein_coding | -0.250591745 | 9.649739403 | 0.480136 | 0.4987693 | 0.864973031 |
| ENSMUSG00000002058 | Unc119 | protein_coding | 0.26548633 | 4.628577171 | 0.4785461 | 0.4988131 | 0.864973031 |
| ENSMUSG00000014748 | Tex261 | protein_coding | -0.23282131 | 5.957894525 | 0.4784603 | 0.498867 | 0.864973031 |
| ENSMUSG00000079427 | Mthfsl | protein_coding | 0.267620078 | 3.962724281 | 0.4781746 | 0.4989775 | 0.864973031 |
| ENSMUSG00000060032 | H2afj | protein_coding | 0.420149567 | 2.916554437 | 0.4778421 | 0.4991246 | 0.864973031 |
| ENSMUSG00000001802 | Lrp3 | protein_coding | 0.29395568 | 4.149814786 | 0.4778203 | 0.4991342 | 0.864973031 |
| ENSMUSG00000047123 | Ticam1 | protein_coding | -0.556613942 | 2.958278918 | 0.4778346 | 0.4991438 | 0.864973031 |
| ENSMUSG00000031355 | Arhgap6 | protein_coding | 0.245934884 | 4.359166471 | 0.4777163 | 0.4991803 | 0.864973031 |
| ENSMUSG00000020697 | Lig3 | protein_coding | -0.483306926 | 2.791081646 | 0.4776455 | 0.4992116 | 0.864973031 |
| ENSMUSG00000021947 | Cryl1 | protein_coding | -0.205162924 | 5.011775904 | 0.4772212 | 0.4993996 | 0.865159304 |
| ENSMUSG00000036026 | Tmem63b | protein_coding | 0.219274243 | 5.229959865 | 0.4767961 | 0.499588 | 0.865229147 |
| ENSMUSG00000038740 | Mvb12b | protein_coding | -0.252473727 | 4.934710172 | 0.4767671 | 0.4996008 | 0.865229147 |
| ENSMUSG00000058594 | Fbxo18 | protein_coding | -0.185510051 | 5.709384351 | 0.4764235 | 0.4997533 | 0.865353742 |
| ENSMUSG00000042487 | Leo1 | protein_coding | 0.242060509 | 6.18371895 | 0.4762281 | 0.4998977 | 0.865464544 |
| ENSMUSG00000002325 | Irf9 | protein_coding | -0.354327277 | 3.572885205 | 0.474937 | 0.5004135 | 0.865970861 |
| ENSMUSG00000022677 | Fopnl | protein_coding | 0.147666072 | 8.132747093 | 0.4749246 | 0.500419 | 0.865970861 |
| ENSMUSG00000005873 | Reep5 | protein_coding | 0.144726033 | 8.580468128 | 0.4748959 | 0.5004318 | 0.865970861 |
| ENSMUSG00000035372 | 1810055G02Rik | protein_coding | 0.268441302 | 4.514499838 | 0.4744985 | 0.5006086 | 0.866126268 |
| ENSMUSG00000030225 | Dera | protein_coding | -0.37862255 | 3.302145592 | 0.4743319 | 0.5006827 | 0.866126268 |
| ENSMUSG00000098188 | Sowahc | protein_coding | -0.382993581 | 3.065199733 | 0.4739192 | 0.5008664 | 0.86613974 |
| ENSMUSG00000021391 | Cenpp | protein_coding | -0.337101058 | 3.797325587 | 0.4737274 | 0.5009519 | 0.86613974 |
| ENSMUSG00000070565 | Rasal2 | protein_coding | -0.246736252 | 5.085976539 | 0.4737067 | 0.5009611 | 0.86613974 |
| ENSMUSG00000032816 | Igdcc4 | protein_coding | -0.395164324 | 3.276347996 | 0.4733184 | 0.5011342 | 0.86613974 |
| ENSMUSG00000034430 | Zxdc | protein_coding | 0.287371562 | 3.944649992 | 0.4731317 | 0.5012174 | 0.86613974 |
| ENSMUSG00000063796 | Slc22a8 | protein_coding | -0.668605585 | 2.651915332 | 0.4728669 | 0.5013513 | 0.86613974 |
| ENSMUSG00000019302 | Atp6v0a1 | protein_coding | 0.165702577 | 7.714074091 | 0.472768 | 0.5013797 | 0.86613974 |
| ENSMUSG00000074890 | Lcmt2 | protein_coding | 0.440486671 | 3.090681703 | 0.472629 | 0.5014417 | 0.86613974 |
| ENSMUSG00000042050 | Wdr60 | protein_coding | -0.320721266 | 3.910344139 | 0.4724816 | 0.5015074 | 0.86613974 |
| ENSMUSG00000030610 | Det1 | protein_coding | -0.566273162 | 2.628626449 | 0.4723298 | 0.5015752 | 0.86613974 |
| ENSMUSG00000060981 | Hist1h4h | protein_coding | -0.591901547 | 2.313796193 | 0.472327 | 0.5015765 | 0.86613974 |
| ENSMUSG00000053398 | Phgdh | protein_coding | -0.21403592 | 5.139658463 | 0.4719214 | 0.5017577 | 0.866232212 |
| ENSMUSG00000028093 | Acp6 | protein_coding | 0.356602838 | 3.547751919 | 0.4718464 | 0.5017912 | 0.866232212 |
| ENSMUSG00000033124 | Atg9a | protein_coding | 0.522359793 | 3.263364428 | 0.4723417 | 0.5018906 | 0.866264725 |
| ENSMUSG00000024604 | Rbm22 | protein_coding | -0.160693511 | 5.880171391 | 0.4710426 | 0.5021505 | 0.86645418 |
| ENSMUSG00000028167 | Bdh2 | protein_coding | -0.322162745 | 3.684436894 | 0.4709534 | 0.5021905 | 0.86645418 |
| ENSMUSG00000030701 | Plekhb1 | protein_coding | -0.172620351 | 9.101479327 | 0.4708381 | 0.5022421 | 0.86645418 |
| ENSMUSG00000046785 | Epm2aip1 | protein_coding | -0.164149584 | 6.769392348 | 0.4705499 | 0.5023711 | 0.866537741 |
| ENSMUSG00000032398 | Snapc5 | protein_coding | -0.184900931 | 6.535597671 | 0.4702499 | 0.5025054 | 0.86663046 |
| ENSMUSG00000004567 | Mcoln1 | protein_coding | 0.261742788 | 4.725137671 | 0.4698851 | 0.5026689 | 0.866773342 |
| ENSMUSG00000097006 | 9530082P21Rik | lincRNA | 0.203285549 | 8.267374426 | 0.4693988 | 0.5029027 | 0.867016323 |
| ENSMUSG00000039263 | Npepl1 | protein_coding | -0.885480845 | 2.256624466 | 0.4695226 | 0.5031308 | 0.867016323 |
| ENSMUSG00000014867 | Surf4 | protein_coding | 0.210740158 | 5.453202944 | 0.4687746 | 0.5031671 | 0.867016323 |
| ENSMUSG00000072582 | Ptrh2 | protein_coding | 0.175051081 | 6.181312669 | 0.4686684 | 0.5032148 | 0.867016323 |

| ensembl_gene_id | external_gene_name | gene_biotype | log2 (fold change) | log2 (counts per million) | F-value | p-value | False discovery rate |
|---|---|---|---|---|---|---|---|
| ENSMUSG00000026608 | Kctd3 | protein_coding | -0.274174234 | 4.791770273 | 0.4685167 | 0.5032829 | 0.867016323 |
| ENSMUSG00000026771 | Spopl | protein_coding | -0.351174735 | 3.182406445 | 0.468493 | 0.5032936 | 0.867016323 |
| ENSMUSG00000056832 | Ttc26 | protein_coding | -0.479600054 | 2.656063282 | 0.4681793 | 0.5034345 | 0.86712018 |
| ENSMUSG00000094910 | D430019H16Rik | protein_coding | -0.265018425 | 5.818413957 | 0.4680385 | 0.5035732 | 0.867220122 |
| ENSMUSG00000075703 | Selenoi | protein_coding | -0.295899868 | 5.201008099 | 0.4673163 | 0.5038384 | 0.86749408 |
| ENSMUSG00000057530 | Ece1 | protein_coding | -0.206744749 | 5.311178642 | 0.4670486 | 0.5039432 | 0.86749408 |
| ENSMUSG00000026458 | Ppfia4 | protein_coding | -0.221434145 | 5.510509538 | 0.4667535 | 0.5040761 | 0.86749408 |
| ENSMUSG00000033249 | Hsf4 | protein_coding | 0.446622422 | 2.823140125 | 0.4667205 | 0.5040909 | 0.86749408 |
| ENSMUSG00000026064 | Ptp4a1 | protein_coding | 0.259878928 | 4.085936593 | 0.4664391 | 0.5042177 | 0.86749408 |
| ENSMUSG00000076617 | Ighm | IG_C_gene | -0.330138642 | 5.804644818 | 0.4687402 | 0.5044251 | 0.86749408 |
| ENSMUSG00000024480 | Ap3s1 | protein_coding | -0.32882861 | 3.781050752 | 0.465646 | 0.5045754 | 0.86749408 |
| ENSMUSG00000047842 | Diras2 | protein_coding | -0.234380533 | 5.535112795 | 0.4651974 | 0.5047779 | 0.86749408 |
| ENSMUSG00000022718 | Dgcr8 | protein_coding | -0.446305535 | 2.741823488 | 0.4651528 | 0.504798 | 0.86749408 |
| ENSMUSG00000032024 | Clmp | protein_coding | 0.357940091 | 3.363255867 | 0.4650791 | 0.5048313 | 0.86749408 |
| ENSMUSG00000066798 | Zbtb6 | protein_coding | -0.328965607 | 3.804663878 | 0.4649221 | 0.5049023 | 0.86749408 |
| ENSMUSG00000079045 | Prox1os | antisense_RNA | -0.399075118 | 4.162944071 | 0.465564 | 0.5049304 | 0.86749408 |
| ENSMUSG00000059669 | Taf1b | protein_coding | 0.253925283 | 4.718518786 | 0.4648576 | 0.5049314 | 0.86749408 |
| ENSMUSG00000040147 | Maob | protein_coding | -0.300148399 | 6.05615431 | 0.4668908 | 0.5049686 | 0.86749408 |
| ENSMUSG00000090698 | Apold1 | protein_coding | -0.458932685 | 3.3684765 | 0.4646478 | 0.5050419 | 0.86749408 |
| ENSMUSG00000021767 | Kat6b | protein_coding | 0.264843599 | 4.30740422 | 0.4644008 | 0.5051378 | 0.86749408 |
| ENSMUSG00000025980 | Hspd1 | protein_coding | 0.155032695 | 7.043086391 | 0.4637402 | 0.5054367 | 0.86749408 |
| ENSMUSG00000045106 | Ccdc73 | protein_coding | -0.299477183 | 3.773213786 | 0.4634566 | 0.5055651 | 0.86749408 |
| ENSMUSG00000020413 | Hus1 | protein_coding | -0.404320623 | 3.090170719 | 0.4634167 | 0.5055831 | 0.86749408 |
| ENSMUSG00000036372 | Tmem258 | protein_coding | 0.224206753 | 7.304822578 | 0.4646821 | 0.5056181 | 0.86749408 |
| ENSMUSG00000050812 | AI314180 | protein_coding | -0.205977399 | 5.528441864 | 0.4632664 | 0.5056512 | 0.86749408 |
| ENSMUSG00000018541 | Cwc25 | protein_coding | -0.227679229 | 4.840670748 | 0.4630819 | 0.5057348 | 0.86749408 |
| ENSMUSG00000028578 | Caap1 | protein_coding | -0.591901118 | 2.471576403 | 0.463037 | 0.5057552 | 0.86749408 |
| ENSMUSG00000024069 | Slc30a6 | protein_coding | -0.287833457 | 4.55533128 | 0.462927 | 0.505805 | 0.86749408 |
| ENSMUSG00000026554 | Dcaf8 | protein_coding | 0.19828152 | 7.101564655 | 0.4628166 | 0.5058707 | 0.86749408 |
| ENSMUSG00000078676 | Casc3 | protein_coding | -0.235523631 | 4.850136266 | 0.4627249 | 0.5058966 | 0.86749408 |
| ENSMUSG00000034187 | Nsf | protein_coding | 0.245521826 | 9.831542962 | 0.4641197 | 0.5059105 | 0.86749408 |
| ENSMUSG00000035351 | Nup37 | protein_coding | 0.281190367 | 4.096007473 | 0.4624845 | 0.5060056 | 0.867518874 |
| ENSMUSG00000040128 | Pnrc1 | protein_coding | -0.183418926 | 5.564301215 | 0.4614684 | 0.5064668 | 0.867994944 |
| ENSMUSG00000032652 | Crebl2 | protein_coding | -0.234351958 | 7.062361033 | 0.4625276 | 0.5065918 | 0.867994944 |
| ENSMUSG00000035325 | Sec31a | protein_coding | 0.187200824 | 6.04844774 | 0.4611883 | 0.5065941 | 0.867994944 |
| ENSMUSG00000097518 | Gm26694 | lincRNA | 0.270325798 | 4.020025057 | 0.4611617 | 0.5066062 | 0.867994944 |
| ENSMUSG00000015176 | Nolc1 | protein_coding | 0.157004767 | 6.376195778 | 0.4606874 | 0.5068218 | 0.868046704 |
| ENSMUSG00000021495 | Fam193b | protein_coding | -0.443926308 | 3.323254284 | 0.4606171 | 0.5068629 | 0.868046704 |
| ENSMUSG00000017778 | Cox7c | protein_coding | 0.289043642 | 3.971414894 | 0.4605626 | 0.5068786 | 0.868046704 |
| ENSMUSG00000057469 | E2f6 | protein_coding | -0.279735094 | 4.490085816 | 0.4596963 | 0.507273 | 0.868509715 |
| ENSMUSG00000008435 | Rdh13 | protein_coding | 0.182106837 | 6.406501572 | 0.4592007 | 0.5074988 | 0.868509715 |
| ENSMUSG00000033885 | Pxk | protein_coding | 0.241187452 | 5.072323438 | 0.4591681 | 0.5075137 | 0.868509715 |
| ENSMUSG00000042851 | Zc3h6 | protein_coding | -0.339277374 | 3.161167287 | 0.4591106 | 0.5075399 | 0.868509715 |
| ENSMUSG00000020085 | Aifm2 | protein_coding | 0.339579785 | 4.33042888 | 0.4589709 | 0.5076192 | 0.868509715 |
| ENSMUSG00000023923 | Tbc1d5 | protein_coding | -0.250099226 | 4.134017351 | 0.4589053 | 0.5076335 | 0.868509715 |
| ENSMUSG00000034800 | Zfp661 | protein_coding | -0.527276958 | 2.819067178 | 0.4585954 | 0.5077749 | 0.868613376 |
| ENSMUSG00000097353 | A430046D13Rik | lincRNA | 0.275956291 | 3.66636268 | 0.4583006 | 0.5079095 | 0.86870539 |
| ENSMUSG00000039199 | Zdhhc1 | protein_coding | 0.442651702 | 2.754886922 | 0.4574689 | 0.5082895 | 0.869083804 |
| ENSMUSG00000025766 | D3Ertd751e | protein_coding | 0.276764023 | 4.628732869 | 0.4573712 | 0.5083342 | 0.869083804 |
| ENSMUSG00000026664 | Phyh | protein_coding | -0.128267513 | 8.053546593 | 0.4572199 | 0.5084034 | 0.869083804 |
| ENSMUSG00000026341 | Actr3 | protein_coding | -0.155693282 | 6.223189003 | 0.4571091 | 0.508454 | 0.869083804 |
| ENSMUSG00000029402 | Snrnp35 | protein_coding | -0.223766365 | 5.151248136 | 0.4567529 | 0.508617 | 0.869224267 |
| ENSMUSG00000044715 | Gskip | protein_coding | 0.138569802 | 7.505913195 | 0.4563438 | 0.5088044 | 0.869319663 |
| ENSMUSG00000024588 | Fech | protein_coding | -0.264850104 | 5.324261847 | 0.4559743 | 0.5089892 | 0.869319663 |
| ENSMUSG00000024797 | Vps51 | protein_coding | 0.252216016 | 4.582024347 | 0.4559263 | 0.5089957 | 0.869319663 |
| ENSMUSG00000040596 | Pogk | protein_coding | -0.245345285 | 4.689284965 | 0.4557762 | 0.5090644 | 0.869319663 |
| ENSMUSG00000037709 | Fam13a | protein_coding | 0.286147928 | 3.768903823 | 0.4557003 | 0.5090992 | 0.869319663 |
| ENSMUSG00000000881 | Dlg3 | protein_coding | -0.205353158 | 5.349493532 | 0.4553926 | 0.5092404 | 0.869319663 |
| ENSMUSG00000006715 | Gmnn | protein_coding | -0.337739461 | 4.321437448 | 0.4550159 | 0.5094287 | 0.869319663 |
| ENSMUSG00000059183 | Mtfmt | protein_coding | 0.269980746 | 4.232497729 | 0.4547844 | 0.5095195 | 0.869319663 |
| ENSMUSG00000032178 | Ilf3 | protein_coding | 0.26493085 | 7.480326421 | 0.4575068 | 0.5095963 | 0.869319663 |
| ENSMUSG00000020412 | Ascc2 | protein_coding | -0.329161797 | 3.282315563 | 0.4545311 | 0.5096358 | 0.869319663 |
| ENSMUSG00000037224 | Zfyve28 | protein_coding | -0.263797089 | 3.891816337 | 0.4543473 | 0.5097203 | 0.869319663 |
| ENSMUSG00000035547 | Capn5 | protein_coding | -0.27261511 | 5.012830379 | 0.454276 | 0.509753 | 0.869319663 |
| ENSMUSG00000055991 | Zkscan5 | protein_coding | 0.31742714 | 3.547191066 | 0.4542352 | 0.5097718 | 0.869319663 |
| ENSMUSG00000034429 | Zfp707 | protein_coding | 0.304488839 | 4.951956453 | 0.4541972 | 0.5098047 | 0.869319663 |
| ENSMUSG00000042515 | Mum1l1 | protein_coding | -0.413999305 | 3.670943387 | 0.4537598 | 0.5100058 | 0.869524717 |
| ENSMUSG00000033909 | Usp36 | protein_coding | -0.251521373 | 4.481393143 | 0.4530189 | 0.5103313 | 0.869941784 |
| ENSMUSG00000021488 | Nsd1 | protein_coding | -0.175294516 | 6.332371186 | 0.4523011 | 0.5106621 | 0.870033837 |
| ENSMUSG00000027187 | Cat | protein_coding | -0.225499487 | 5.842333269 | 0.4521612 | 0.5107266 | 0.870033837 |
| ENSMUSG00000022783 | Spag6l | protein_coding | 0.307034045 | 3.789404451 | 0.4520109 | 0.5107958 | 0.870033837 |
| ENSMUSG00000022636 | Alcam | protein_coding | -0.185310984 | 7.059554592 | 0.4519607 | 0.510819 | 0.870033837 |
| ENSMUSG00000031925 | Maml2 | protein_coding | 0.278482775 | 4.363464793 | 0.4516898 | 0.510944 | 0.870033837 |
| ENSMUSG00000027932 | Slc27a3 | protein_coding | 0.547321779 | 2.496547743 | 0.4515658 | 0.5110012 | 0.870033837 |
| ENSMUSG00000030518 | Fam189a1 | protein_coding | -0.375858168 | 4.028151011 | 0.4515761 | 0.5110119 | 0.870033837 |
| ENSMUSG00000038718 | Pbx3 | protein_coding | 0.238784872 | 5.39424119 | 0.4513691 | 0.511092 | 0.870033837 |
| ENSMUSG00000032336 | Nptn | protein_coding | -0.143639207 | 8.112727407 | 0.4511225 | 0.5112059 | 0.870033837 |
| ENSMUSG00000033658 | Ddx19b | protein_coding | 0.372953645 | 2.951968457 | 0.4507073 | 0.5113978 | 0.870033837 |

| ensembl_gene_id | external_gene_name | gene_biotype | log2 (fold change) | log2 (counts per million) | F-value | p-value | False discovery rate |
|---|---|---|---|---|---|---|---|
| ENSMUSG00000060181 | Slc35e3 | protein_coding | 0.32588331 | 3.131692399 | 0.4505534 | 0.5114689 | 0.870033837 |
| ENSMUSG00000038305 | Spats2l | protein_coding | -0.281338945 | 4.865048663 | 0.4504365 | 0.511523 | 0.870033837 |
| ENSMUSG00000036466 | Megf11 | protein_coding | -0.239237325 | 4.842826329 | 0.4504275 | 0.5115271 | 0.870033837 |
| ENSMUSG00000029576 | Radil | protein_coding | 0.358927993 | 3.178132561 | 0.4503858 | 0.5115464 | 0.870033837 |
| ENSMUSG00000025192 | Entpd7 | protein_coding | 0.351939662 | 3.783401664 | 0.4502039 | 0.5116306 | 0.870033837 |
| ENSMUSG00000060314 | Zfp941 | protein_coding | 0.172000859 | 6.292525636 | 0.4500972 | 0.5116799 | 0.870033837 |
| ENSMUSG00000034570 | Inpp5j | protein_coding | -0.38751741 | 3.112371007 | 0.449607 | 0.5119068 | 0.870282075 |
| ENSMUSG00000051615 | Rap2a | protein_coding | 0.194085564 | 5.783770997 | 0.4493508 | 0.5120255 | 0.870300374 |
| ENSMUSG00000033768 | Nrxn2 | protein_coding | 0.232983597 | 4.840511298 | 0.4492342 | 0.5120795 | 0.870300374 |
| ENSMUSG00000018377 | Vezf1 | protein_coding | -0.209271735 | 5.292392694 | 0.4490309 | 0.5121737 | 0.87032295 |
| ENSMUSG00000022574 | Naprt | protein_coding | 0.425315743 | 3.163063662 | 0.4485688 | 0.512388 | 0.870549511 |
| ENSMUSG00000007836 | Hnrnpa0 | protein_coding | -0.167330884 | 7.269052776 | 0.4482898 | 0.5125174 | 0.870631848 |
| ENSMUSG00000049421 | Zfp260 | protein_coding | 0.211453276 | 5.497328134 | 0.4480347 | 0.5126358 | 0.87069542 |
| ENSMUSG00000018470 | Kcnab3 | protein_coding | -0.291595351 | 3.594478776 | 0.447237 | 0.5130063 | 0.871062577 |
| ENSMUSG00000003923 | Tfam | protein_coding | 0.18840334 | 5.862559057 | 0.4472079 | 0.5130199 | 0.871062577 |
| ENSMUSG00000061759 | Armt1 | protein_coding | 0.230520014 | 4.666926094 | 0.4470464 | 0.513095 | 0.871062577 |
| ENSMUSG00000036834 | Plch1 | protein_coding | 0.428699778 | 3.273452275 | 0.4466077 | 0.513299 | 0.871139592 |
| ENSMUSG00000071753 | C230004F18Rik | processed_transcript | 0.245564092 | 4.836514358 | 0.4465371 | 0.5133319 | 0.871139592 |
| ENSMUSG00000021097 | Clmn | protein_coding | -0.226785993 | 5.483942147 | 0.4464264 | 0.5133834 | 0.871139592 |
| ENSMUSG00000042790 | Rnf214 | protein_coding | -0.186951217 | 5.510956496 | 0.4445742 | 0.5142468 | 0.872467052 |
| ENSMUSG00000004562 | Arhgef40 | protein_coding | -0.284707301 | 5.044574257 | 0.4437028 | 0.5146692 | 0.872886051 |
| ENSMUSG00000083307 | AA414768 | processed_pseudoge | -0.389835609 | 2.909304346 | 0.4435674 | 0.5147173 | 0.872886051 |
| ENSMUSG00000027324 | Rpusd2 | protein_coding | -0.433832072 | 2.66853841 | 0.4435245 | 0.5147373 | 0.872886051 |
| ENSMUSG00000029545 | Acads | protein_coding | 0.358237862 | 3.275808124 | 0.4430206 | 0.5149731 | 0.872900438 |
| ENSMUSG00000036352 | Ubac1 | protein_coding | 0.193437624 | 5.215568652 | 0.4429873 | 0.5149887 | 0.872900438 |
| ENSMUSG00000031628 | Casp3 | protein_coding | 0.42910099 | 3.475953786 | 0.4430185 | 0.5149893 | 0.872900438 |
| ENSMUSG00000047747 | Rnf150 | protein_coding | -0.307230172 | 4.084373128 | 0.4427619 | 0.5150942 | 0.872940612 |
| ENSMUSG00000035378 | Shq1 | protein_coding | -0.423008924 | 2.995015407 | 0.4424959 | 0.5152188 | 0.873006751 |
| ENSMUSG00000025402 | Nab2 | protein_coding | -0.346892747 | 3.529868613 | 0.4423319 | 0.5152956 | 0.873006751 |
| ENSMUSG00000029521 | Chek2 | protein_coding | 0.36167311 | 3.049746519 | 0.4416187 | 0.51563 | 0.873249255 |
| ENSMUSG00000028461 | Ccdc107 | protein_coding | 0.383553739 | 5.916558428 | 0.4471606 | 0.515653 | 0.873249255 |
| ENSMUSG00000044792 | Isca1 | protein_coding | 0.153705241 | 6.796535888 | 0.4415071 | 0.5156824 | 0.873249255 |
| ENSMUSG00000039375 | Wdr17 | protein_coding | -0.268596788 | 4.796422682 | 0.4408807 | 0.5159765 | 0.873486527 |
| ENSMUSG00000022897 | Dyrk1a | protein_coding | -0.179538699 | 6.246967347 | 0.4405756 | 0.5161198 | 0.873486527 |
| ENSMUSG00000046909 | Tefm | protein_coding | 0.208513481 | 5.180163379 | 0.4405579 | 0.5161282 | 0.873486527 |
| ENSMUSG00000032479 | Map4 | protein_coding | -0.181384068 | 8.692960752 | 0.440459 | 0.5161746 | 0.873486527 |
| ENSMUSG00000107585 | 3300002P13Rik | lincRNA | 0.449359375 | 2.696054502 | 0.440344 | 0.5162287 | 0.873486527 |
| ENSMUSG00000063919 | Srrm4 | protein_coding | 0.263151338 | 4.491326494 | 0.4401602 | 0.5163151 | 0.873495296 |
| ENSMUSG00000046215 | Rprml | protein_coding | 0.378616143 | 3.693100388 | 0.4398379 | 0.5164819 | 0.873553073 |
| ENSMUSG00000034639 | Setmar | protein_coding | 0.436691865 | 2.899282814 | 0.4396788 | 0.5165416 | 0.873553073 |
| ENSMUSG00000028651 | Ppie | protein_coding | 0.215959944 | 4.584048901 | 0.4394272 | 0.51666 | 0.873553073 |
| ENSMUSG00000073609 | D2hgdh | protein_coding | -0.407357392 | 3.018181515 | 0.4392525 | 0.5167423 | 0.873553073 |
| ENSMUSG00000038503 | Mesd | protein_coding | 0.157204083 | 6.570156497 | 0.4392054 | 0.5167645 | 0.873553073 |
| ENSMUSG00000032624 | Eml4 | protein_coding | -0.277434608 | 3.902866519 | 0.4390522 | 0.5168367 | 0.873553073 |
| ENSMUSG00000086555 | Gm13446 | processed_transcript | 0.480016257 | 3.558892642 | 0.4400871 | 0.5169381 | 0.873587174 |
| ENSMUSG00000002733 | Plekha3 | protein_coding | -0.232158896 | 4.711575466 | 0.438565 | 0.5170663 | 0.87365296 |
| ENSMUSG00000026637 | Traf5 | protein_coding | 0.364065257 | 2.904134733 | 0.4384097 | 0.5171395 | 0.87365296 |
| ENSMUSG00000001901 | Kcnh6 | protein_coding | 0.449252335 | 2.875576545 | 0.4379999 | 0.5173329 | 0.873744661 |
| ENSMUSG00000038257 | Glra3 | protein_coding | -0.341344806 | 3.159077533 | 0.4379503 | 0.5173563 | 0.873744661 |
| ENSMUSG00000024172 | St6gal2 | protein_coding | -0.510464599 | 2.881335926 | 0.4377355 | 0.5174577 | 0.873778635 |
| ENSMUSG00000032965 | Ift57 | protein_coding | -0.41114822 | 3.069249186 | 0.4371455 | 0.5177364 | 0.87399732 |
| ENSMUSG00000040189 | Ccdc114 | protein_coding | -1.001989838 | 2.03408562 | 0.4383327 | 0.5177631 | 0.87399732 |
| ENSMUSG00000019960 | Dusp6 | protein_coding | -0.223324333 | 5.189858179 | 0.4367516 | 0.5179226 | 0.87399732 |
| ENSMUSG00000024222 | Fkbp5 | protein_coding | -0.284609487 | 4.536240414 | 0.4365929 | 0.5179977 | 0.87399732 |
| ENSMUSG00000016619 | Nup50 | protein_coding | 0.237323837 | 4.860087363 | 0.4364343 | 0.5180727 | 0.87399732 |
| ENSMUSG00000024487 | Yipf5 | protein_coding | -0.175042189 | 5.843376927 | 0.4362417 | 0.5181639 | 0.87399732 |
| ENSMUSG00000020100 | Slc29a3 | protein_coding | -0.395317591 | 3.130165832 | 0.4360359 | 0.5182613 | 0.87399732 |
| ENSMUSG00000020741 | Cluh | protein_coding | 0.194572949 | 5.834445549 | 0.4359363 | 0.5183085 | 0.87399732 |
| ENSMUSG00000019715 | Gle1 | protein_coding | 0.232749923 | 6.066302344 | 0.4357668 | 0.5184038 | 0.87399732 |
| ENSMUSG00000014959 | Gorasp2 | protein_coding | 0.144195383 | 6.525024979 | 0.435682 | 0.5184289 | 0.87399732 |
| ENSMUSG00000050587 | Lrrc4c | protein_coding | -0.136702326 | 6.84101994 | 0.4355716 | 0.5184813 | 0.87399732 |
| ENSMUSG00000018661 | Cog1 | protein_coding | 0.308143718 | 3.643888327 | 0.4353015 | 0.5186093 | 0.874076109 |
| ENSMUSG00000031865 | Dctn1 | protein_coding | 0.187838097 | 6.87130974 | 0.4351228 | 0.5187091 | 0.874107294 |
| ENSMUSG00000021811 | Dnajc9 | protein_coding | 0.208231488 | 6.019361797 | 0.4346387 | 0.5189237 | 0.874289805 |
| ENSMUSG00000032259 | Drd2 | protein_coding | -0.268335786 | 5.144395593 | 0.4342039 | 0.5191452 | 0.874289805 |
| ENSMUSG00000026707 | Nsun6 | protein_coding | -0.369762901 | 3.048525799 | 0.4337184 | 0.5193608 | 0.874289805 |
| ENSMUSG00000031935 | Med17 | protein_coding | -0.341724988 | 3.988971524 | 0.4335647 | 0.5194339 | 0.874289805 |
| ENSMUSG00000055041 | Commd5 | protein_coding | 0.16609772 | 6.244817319 | 0.4335412 | 0.519445 | 0.874289805 |
| ENSMUSG00000020912 | Krt12 | protein_coding | -0.508580788 | 2.457496946 | 0.4334861 | 0.5194712 | 0.874289805 |
| ENSMUSG00000035495 | Tstd2 | protein_coding | -0.325076502 | 3.222864669 | 0.4330655 | 0.5196713 | 0.874289805 |
| ENSMUSG00000015837 | Sqstm1 | protein_coding | 0.151388731 | 8.745918331 | 0.4329828 | 0.5197106 | 0.874289805 |
| ENSMUSG00000036214 | Znrd1as | protein_coding | -0.316363498 | 3.733411621 | 0.4328308 | 0.519783 | 0.874289805 |
| ENSMUSG00000048483 | Zdhhc22 | protein_coding | 0.340047786 | 7.562365395 | 0.4409225 | 0.5198331 | 0.874289805 |
| ENSMUSG00000007659 | Bcl2l1 | protein_coding | 0.144826519 | 8.134873366 | 0.4324768 | 0.5199516 | 0.874289805 |
| ENSMUSG00000078652 | Psme3 | protein_coding | -0.150684384 | 6.240512895 | 0.4323677 | 0.5200035 | 0.874289805 |
| ENSMUSG00000030272 | Camk1 | protein_coding | 0.273999002 | 3.713426668 | 0.4320949 | 0.5201335 | 0.874289805 |
| ENSMUSG00000027086 | Fastkd1 | protein_coding | -0.246841069 | 4.208332391 | 0.4320504 | 0.5201548 | 0.874289805 |
| ENSMUSG00000066510 | Ankdd1a | protein_coding | 0.415450194 | 3.148513677 | 0.4320142 | 0.520172 | 0.874289805 |

| ensembl_gene_id | external_gene_name | gene_biotype | log2 (fold change) | log2 (counts per million) | F-value | p-value | False discovery rate |
|---|---|---|---|---|---|---|---|
| ENSMUSG00000005610 | Eif4g2 | protein_coding | -0.176814787 | 8.797451722 | 0.4318551 | 0.5202479 | 0.874289805 |
| ENSMUSG00000029094 | Afap1 | protein_coding | 0.231981259 | 4.558819411 | 0.4318068 | 0.5202709 | 0.874289805 |
| ENSMUSG00000097316 | Gm10516 | antisense_RNA | 0.548659168 | 2.657396966 | 0.4317288 | 0.5203081 | 0.874289805 |
| ENSMUSG00000026688 | Mgst3 | protein_coding | 0.133637669 | 8.852648888 | 0.4311924 | 0.520564 | 0.874289805 |
| ENSMUSG00000029670 | Ing3 | protein_coding | -0.202574758 | 5.85921492 | 0.4309795 | 0.5206657 | 0.874289805 |
| ENSMUSG00000027200 | Sema6d | protein_coding | -0.221955439 | 4.817091646 | 0.430967 | 0.5206716 | 0.874289805 |
| ENSMUSG00000025876 | Unc5a | protein_coding | 0.320868315 | 4.321028332 | 0.4306901 | 0.5208189 | 0.874289805 |
| ENSMUSG00000085385 | Snhg17 | processed_transcript | -0.558818434 | 2.346316515 | 0.4304658 | 0.5209111 | 0.874289805 |
| ENSMUSG00000020522 | Mfap3 | protein_coding | 0.241077357 | 4.986215384 | 0.4304215 | 0.5209323 | 0.874289805 |
| ENSMUSG00000003644 | Rps6ka1 | protein_coding | 0.232967716 | 4.109013313 | 0.4303809 | 0.5209517 | 0.874289805 |
| ENSMUSG00000026991 | Pkp4 | protein_coding | -0.132120495 | 6.884406364 | 0.4302331 | 0.5210223 | 0.874289805 |
| ENSMUSG00000017760 | Ctsa | protein_coding | 0.134144534 | 6.797332357 | 0.4300146 | 0.5211268 | 0.874289805 |
| ENSMUSG00000039704 | Lmbrd2 | protein_coding | -0.552863825 | 2.339368941 | 0.4299171 | 0.5211734 | 0.874289805 |
| ENSMUSG00000002477 | Snrpd1 | protein_coding | 0.172426248 | 7.826227714 | 0.4299133 | 0.5211753 | 0.874289805 |
| ENSMUSG00000028756 | Pink1 | protein_coding | 0.158670098 | 7.55903131 | 0.4296042 | 0.5213232 | 0.874401493 |
| ENSMUSG00000026490 | Cdc42bpa | protein_coding | -0.183367407 | 6.237104324 | 0.4292435 | 0.5214959 | 0.874446869 |
| ENSMUSG00000045095 | Magi1 | protein_coding | 0.169372985 | 6.208144879 | 0.429208 | 0.5215129 | 0.874446869 |
| ENSMUSG00000028618 | Tmem59 | protein_coding | 0.13588588 | 9.205034219 | 0.4284631 | 0.5218698 | 0.874909009 |
| ENSMUSG00000036275 | 9530068E07Rik | protein_coding | 0.17377385 | 6.025936184 | 0.4281506 | 0.5220197 | 0.875023867 |
| ENSMUSG00000033161 | Atp1a1 | protein_coding | -0.260777735 | 4.929642025 | 0.427872 | 0.5221535 | 0.875111628 |
| ENSMUSG00000018750 | Zbtb4 | protein_coding | -0.146998902 | 6.130943418 | 0.4276084 | 0.52228 | 0.875187321 |
| ENSMUSG00000045996 | Polr2k | protein_coding | 0.248011806 | 4.235161033 | 0.4272123 | 0.5224703 | 0.875369766 |
| ENSMUSG00000018899 | Irf1 | protein_coding | 0.472253444 | 2.824912173 | 0.4269566 | 0.5225932 | 0.875439238 |
| ENSMUSG00000026342 | Slc35f5 | protein_coding | -0.457862223 | 3.268655608 | 0.426543 | 0.522807 | 0.87566107 |
| ENSMUSG00000051113 | Fam71e1 | protein_coding | 0.466840313 | 2.910561462 | 0.4258748 | 0.5231138 | 0.876038357 |
| ENSMUSG00000034620 | Tmem5 | protein_coding | -0.231117016 | 4.559196807 | 0.4251536 | 0.5234614 | 0.876041288 |
| ENSMUSG00000022601 | Zbtb11 | protein_coding | 0.307358457 | 3.69462832 | 0.4251079 | 0.5234834 | 0.876041288 |
| ENSMUSG00000015671 | Psma2 | protein_coding | 0.153356426 | 8.831763041 | 0.4248751 | 0.5235957 | 0.876041288 |
| ENSMUSG00000036632 | Alg5 | protein_coding | -0.250467946 | 5.020217067 | 0.4246854 | 0.5236872 | 0.876041288 |
| ENSMUSG00000011960 | Ccnt1 | protein_coding | 0.200934838 | 5.296576263 | 0.4246658 | 0.5236967 | 0.876041288 |
| ENSMUSG00000087075 | A230065H16Rik | protein_coding | -0.174342586 | 6.340546695 | 0.4245481 | 0.5237535 | 0.876041288 |
| ENSMUSG00000045193 | Cirbp | protein_coding | 0.138638976 | 9.040230803 | 0.4243479 | 0.5238501 | 0.876041288 |
| ENSMUSG00000026220 | Slc16a14 | protein_coding | -0.619298727 | 3.227649234 | 0.4267773 | 0.5238564 | 0.876041288 |
| ENSMUSG00000026353 | Ubxn4 | protein_coding | -0.16413721 | 5.908342931 | 0.4240289 | 0.5240042 | 0.876041288 |
| ENSMUSG00000025645 | Ccdc51 | protein_coding | -0.323733655 | 3.507253039 | 0.4239692 | 0.5240331 | 0.876041288 |
| ENSMUSG00000021357 | Exoc2 | protein_coding | -0.277085927 | 4.732963836 | 0.4239571 | 0.5240389 | 0.876041288 |
| ENSMUSG00000024560 | Cxxc1 | protein_coding | -0.344370137 | 4.553512164 | 0.4243616 | 0.5241465 | 0.876041288 |
| ENSMUSG00000038665 | Dgki | protein_coding | -0.313418307 | 3.621750182 | 0.4235846 | 0.524219 | 0.876041288 |
| ENSMUSG00000027655 | Dhx35 | protein_coding | 0.310816179 | 4.10466349 | 0.4235079 | 0.5242561 | 0.876041288 |
| ENSMUSG00000040389 | Wdr47 | protein_coding | -0.172798666 | 6.156199779 | 0.4227864 | 0.5246052 | 0.876287458 |
| ENSMUSG00000067028 | Cntnap5b | protein_coding | -0.410548599 | 2.91609367 | 0.4227265 | 0.5246342 | 0.876287458 |
| ENSMUSG00000030922 | Lyrm1 | protein_coding | 0.162419131 | 5.692454423 | 0.42259 | 0.5247003 | 0.876287458 |
| ENSMUSG00000070371 | Prss36 | protein_coding | 0.308797293 | 4.542791938 | 0.4224638 | 0.5247763 | 0.876287458 |
| ENSMUSG00000096935 | 1700113A16Rik | antisense_RNA | 0.53961416 | 2.436714362 | 0.4223617 | 0.5248109 | 0.876287458 |
| ENSMUSG00000021711 | Trappc13 | protein_coding | -0.137925102 | 7.177096875 | 0.4215945 | 0.5251829 | 0.876552088 |
| ENSMUSG00000021240 | Abcd4 | protein_coding | -0.457224337 | 2.905909648 | 0.4214834 | 0.5252367 | 0.876552088 |
| ENSMUSG00000038736 | Nudcd1 | protein_coding | -0.277565977 | 4.382402137 | 0.4213755 | 0.5252891 | 0.876552088 |
| ENSMUSG00000071719 | Tmem28 | protein_coding | 0.276177405 | 3.903333358 | 0.4211021 | 0.5254218 | 0.876552088 |
| ENSMUSG00000071253 | Slc25a16 | protein_coding | -0.232056123 | 4.729946211 | 0.4210655 | 0.5254396 | 0.876552088 |
| ENSMUSG00000027180 | Fbxo3 | protein_coding | -0.152981353 | 7.1115003 | 0.42096 | 0.5254908 | 0.876552088 |
| ENSMUSG00000029505 | Ep400 | protein_coding | -0.279898021 | 4.822313806 | 0.4208667 | 0.525551 | 0.876552088 |
| ENSMUSG00000032291 | Crabp1 | protein_coding | -0.288300238 | 4.617059342 | 0.4206911 | 0.5256215 | 0.876552088 |
| ENSMUSG00000033417 | Cacul1 | protein_coding | -0.212931026 | 5.058586534 | 0.4204669 | 0.5257304 | 0.876597848 |
| ENSMUSG00000024097 | Srsf7 | protein_coding | -0.152415601 | 8.150589319 | 0.4202671 | 0.5258276 | 0.876623855 |
| ENSMUSG00000021241 | Isca2 | protein_coding | 0.123177466 | 8.298252088 | 0.4198633 | 0.526024 | 0.876815342 |
| ENSMUSG00000039599 | Fam149b | protein_coding | 0.341188895 | 3.162414155 | 0.4196285 | 0.5261383 | 0.876869968 |
| ENSMUSG00000027010 | Slc25a12 | protein_coding | 0.217750813 | 7.805494318 | 0.4199583 | 0.5265506 | 0.877421215 |
| ENSMUSG00000024952 | Rps6ka4 | protein_coding | 0.25419482 | 4.084557071 | 0.4184621 | 0.5267066 | 0.877439026 |
| ENSMUSG00000039536 | Stau1 | protein_coding | -0.203822046 | 5.958777813 | 0.4184254 | 0.5267245 | 0.877439026 |
| ENSMUSG00000051359 | Ncald | protein_coding | -0.163451028 | 8.885252362 | 0.4176986 | 0.5270793 | 0.877486341 |
| ENSMUSG00000030291 | Med21 | protein_coding | 0.155385841 | 7.272504161 | 0.4176608 | 0.5270978 | 0.877486341 |
| ENSMUSG00000004364 | Cul3 | protein_coding | -0.148881346 | 6.451955058 | 0.4174525 | 0.5271995 | 0.877486341 |
| ENSMUSG00000008318 | Relt | protein_coding | -0.409382955 | 3.405330242 | 0.4174488 | 0.5272013 | 0.877486341 |
| ENSMUSG00000050891 | Tatdn1 | protein_coding | 0.181469107 | 5.328396989 | 0.417417 | 0.5272168 | 0.877486341 |
| ENSMUSG00000074247 | Dda1 | protein_coding | -0.215813139 | 5.191936078 | 0.4173644 | 0.5272426 | 0.877486341 |
| ENSMUSG00000040712 | Camta2 | protein_coding | 0.252309845 | 6.010244327 | 0.4181607 | 0.5273903 | 0.877487296 |
| ENSMUSG00000029502 | Golga3 | protein_coding | 0.210602642 | 5.084133763 | 0.4170293 | 0.5274063 | 0.877487296 |
| ENSMUSG00000028848 | Gpn2 | protein_coding | -0.418755416 | 2.981742726 | 0.4166191 | 0.527607 | 0.877653218 |
| ENSMUSG00000049807 | Arhgap23 | protein_coding | -0.270162472 | 4.26667607 | 0.4164917 | 0.5276693 | 0.877653218 |
| ENSMUSG00000036167 | Pphln1 | protein_coding | -0.205096739 | 5.291290637 | 0.4156235 | 0.5280945 | 0.877838704 |
| ENSMUSG00000025576 | Rbfox3 | protein_coding | 0.142654507 | 6.697180347 | 0.4155282 | 0.5281413 | 0.877838704 |
| ENSMUSG00000036555 | Iqce | protein_coding | -0.289019398 | 4.140155216 | 0.4154706 | 0.5281695 | 0.877838704 |
| ENSMUSG00000007610 | Gtpbp3 | protein_coding | -0.39308859 | 2.835083554 | 0.4154361 | 0.5281864 | 0.877838704 |
| ENSMUSG00000073733 | Rsg1 | protein_coding | 0.553136698 | 2.950123744 | 0.4155743 | 0.528189 | 0.877838704 |
| ENSMUSG00000051412 | Vamp7 | protein_coding | -0.169958754 | 6.712283676 | 0.4150337 | 0.5283838 | 0.877947627 |
| ENSMUSG00000021432 | Slc35b3 | protein_coding | 0.378962973 | 3.149178668 | 0.414917 | 0.528441 | 0.877947627 |
| ENSMUSG00000020260 | Pofut2 | protein_coding | 0.241149281 | 5.643181593 | 0.4148278 | 0.5284995 | 0.877947627 |
| ENSMUSG00000038535 | Zfp280d | protein_coding | 0.371851818 | 3.146602103 | 0.4145501 | 0.5286211 | 0.878014084 |

| ensembl_gene_id | external_gene_name | gene_biotype | log2 (fold change) | log2 (counts per million) | F-value | p-value | False discovery rate |
|---|---|---|---|---|---|---|---|
| ENSMUSG00000005225 | Plekha8 | protein_coding | -0.342905952 | 3.064504677 | 0.414117 | 0.5288339 | 0.878063977 |
| ENSMUSG00000024548 | Setbp1 | protein_coding | -0.350602885 | 3.466152354 | 0.4140168 | 0.5288831 | 0.878063977 |
| ENSMUSG00000003068 | Stk11 | protein_coding | 0.212494677 | 5.19273725 | 0.4139903 | 0.5288962 | 0.878063977 |
| ENSMUSG00000020288 | Ahsa2 | protein_coding | 0.317344521 | 3.798943059 | 0.4131753 | 0.5292971 | 0.878511766 |
| ENSMUSG00000005417 | Mprip | protein_coding | -0.201329734 | 5.350901569 | 0.4131098 | 0.5293293 | 0.878511766 |
| ENSMUSG00000022788 | Fgd4 | protein_coding | -0.252273315 | 3.77936611 | 0.4127498 | 0.5295065 | 0.87852783 |
| ENSMUSG00000030704 | Rab6a | protein_coding | -0.131395815 | 8.306047026 | 0.4124222 | 0.529668 | 0.87852783 |
| ENSMUSG00000002728 | Naa20 | protein_coding | 0.1376333 | 7.108867017 | 0.4123154 | 0.5297206 | 0.87852783 |
| ENSMUSG00000031574 | Star | protein_coding | -0.406286921 | 2.702912723 | 0.4123093 | 0.5297236 | 0.87852783 |
| ENSMUSG00000001376 | Vps50 | protein_coding | -0.231570401 | 5.090107022 | 0.4122609 | 0.5297475 | 0.87852783 |
| ENSMUSG00000018809 | Smyd4 | protein_coding | 0.275964957 | 4.364534529 | 0.411968 | 0.5298919 | 0.878581013 |
| ENSMUSG00000028410 | Dnaja1 | protein_coding | -0.128753838 | 7.796491924 | 0.4118115 | 0.5299691 | 0.878581013 |
| ENSMUSG00000020436 | Gabrg2 | protein_coding | 0.250413622 | 3.993808384 | 0.411699 | 0.5300246 | 0.878581013 |
| ENSMUSG00000050627 | Gpd1l | protein_coding | -0.223779123 | 5.678833813 | 0.4115242 | 0.5301109 | 0.878588554 |
| ENSMUSG00000030207 | Fam234b | protein_coding | 0.194109728 | 6.781779912 | 0.4105881 | 0.530588 | 0.878943676 |
| ENSMUSG00000003955 | Fam162a | protein_coding | 0.192418877 | 6.252638151 | 0.4103498 | 0.5306912 | 0.878943676 |
| ENSMUSG00000050373 | Snx21 | protein_coding | -0.249963091 | 4.355608397 | 0.4101326 | 0.5307986 | 0.878943676 |
| ENSMUSG00000063884 | Ptcd3 | protein_coding | -0.206558763 | 5.521540319 | 0.4100915 | 0.530819 | 0.878943676 |
| ENSMUSG00000006932 | Ctnnb1 | protein_coding | -0.363817231 | 3.323877475 | 0.4100704 | 0.5308294 | 0.878943676 |
| ENSMUSG00000026004 | Kansl1l | protein_coding | -0.384623066 | 3.449601336 | 0.4096821 | 0.5310216 | 0.878943676 |
| ENSMUSG00000022305 | Lrp12 | protein_coding | -0.295247199 | 4.229442977 | 0.4095847 | 0.5310699 | 0.878943676 |
| ENSMUSG00000037343 | Taf2 | protein_coding | 0.271433167 | 3.868903352 | 0.4095651 | 0.5310796 | 0.878943676 |
| ENSMUSG00000039395 | Mreg | protein_coding | 0.467458049 | 2.458589269 | 0.4094549 | 0.5311342 | 0.878943676 |
| ENSMUSG00000033233 | Trim45 | protein_coding | 0.409291325 | 3.97313716 | 0.4105573 | 0.531154 | 0.878943676 |
| ENSMUSG00000035203 | Epn1 | protein_coding | 0.173286855 | 6.099293271 | 0.409273 | 0.5312243 | 0.878943676 |
| ENSMUSG00000054408 | Spcs3 | protein_coding | -0.166829648 | 5.892872514 | 0.4085932 | 0.5315614 | 0.879259453 |
| ENSMUSG00000040990 | Sh3kbp1 | protein_coding | -0.175361044 | 5.563578914 | 0.4084984 | 0.5316084 | 0.879259453 |
| ENSMUSG00000047126 | Cltc | protein_coding | -0.131096527 | 8.0806273 | 0.4083936 | 0.5316605 | 0.879259453 |
| ENSMUSG00000044813 | Shb | protein_coding | -0.449322464 | 3.022880809 | 0.4081089 | 0.5318018 | 0.879357881 |
| ENSMUSG00000029189 | Sel1l3 | protein_coding | -0.427943154 | 2.90428747 | 0.4079189 | 0.5318962 | 0.879363171 |
| ENSMUSG00000029629 | Phf14 | protein_coding | -0.234499971 | 4.886092772 | 0.4077732 | 0.5319685 | 0.879363171 |
| ENSMUSG00000060126 | Tpt1 | protein_coding | -0.290564336 | 3.675308604 | 0.4063332 | 0.5326849 | 0.880196444 |
| ENSMUSG00000040423 | Rc3h1 | protein_coding | -0.199313611 | 4.843479795 | 0.4062122 | 0.5327452 | 0.880196444 |
| ENSMUSG00000112639 | AC134329.1 | lincRNA | -0.379661142 | 2.834979682 | 0.4060756 | 0.5328132 | 0.880196444 |
| ENSMUSG00000038267 | Slc22a23 | protein_coding | -0.21797538 | 4.568708152 | 0.4058316 | 0.5329348 | 0.880196444 |
| ENSMUSG00000035078 | Mtmr9 | protein_coding | -0.184137648 | 5.870127254 | 0.4057327 | 0.5329842 | 0.880196444 |
| ENSMUSG00000030706 | Mrpl48 | protein_coding | 0.134342744 | 7.445085293 | 0.4055655 | 0.5330676 | 0.880196444 |
| ENSMUSG00000020524 | Gria1 | protein_coding | 0.134141874 | 8.10925162 | 0.4051741 | 0.5332628 | 0.880196444 |
| ENSMUSG00000031221 | Igbp1 | protein_coding | 0.203569956 | 6.132569825 | 0.4049447 | 0.5333918 | 0.880196444 |
| ENSMUSG00000032119 | Hinfp | protein_coding | 0.415050091 | 3.263631182 | 0.4047868 | 0.5334562 | 0.880196444 |
| ENSMUSG00000112280 | AC153962.3 | processed_transcript | -0.272033684 | 3.952618323 | 0.4046272 | 0.5335359 | 0.880196444 |
| ENSMUSG00000048330 | Ric3 | protein_coding | 0.221790066 | 5.306662517 | 0.4044161 | 0.5336414 | 0.880196444 |
| ENSMUSG00000058881 | Zfp516 | protein_coding | -0.516894448 | 2.882451343 | 0.404365 | 0.5336814 | 0.880196444 |
| ENSMUSG00000036957 | Lrfn3 | protein_coding | -0.56749104 | 2.342207843 | 0.4041035 | 0.5337977 | 0.880196444 |
| ENSMUSG00000022307 | Oxr1 | protein_coding | -0.198191124 | 7.482523217 | 0.4043883 | 0.5338231 | 0.880196444 |
| ENSMUSG00000032786 | Alas1 | protein_coding | 0.234093455 | 6.36808781 | 0.405138 | 0.5338419 | 0.880196444 |
| ENSMUSG00000072946 | Ptgr2 | protein_coding | 0.244120812 | 5.058398107 | 0.403704 | 0.5339975 | 0.880196444 |
| ENSMUSG00000003824 | Syce2 | protein_coding | 0.391223194 | 3.052079266 | 0.403387 | 0.5341562 | 0.880196444 |
| ENSMUSG00000058897 | Col25a1 | protein_coding | 0.226829157 | 4.848781409 | 0.403309 | 0.5341952 | 0.880196444 |
| ENSMUSG00000022212 | Cpne6 | protein_coding | 0.227011994 | 7.179004098 | 0.4049494 | 0.5342541 | 0.880196444 |
| ENSMUSG00000043991 | Pura | protein_coding | -0.152749235 | 6.957728223 | 0.4030562 | 0.5343219 | 0.880196444 |
| ENSMUSG00000057858 | Fam204a | protein_coding | -0.199100286 | 5.679382187 | 0.4029148 | 0.5343927 | 0.880196444 |
| ENSMUSG00000052752 | Traf7 | protein_coding | 0.191326675 | 4.874099156 | 0.4028604 | 0.5344199 | 0.880196444 |
| ENSMUSG00000040710 | St8sia4 | protein_coding | -0.503154825 | 2.980059613 | 0.4028646 | 0.5344323 | 0.880196444 |
| ENSMUSG00000022855 | Senp2 | protein_coding | -0.164271402 | 6.140582818 | 0.4025587 | 0.5345712 | 0.880196444 |
| ENSMUSG00000029761 | Cald1 | protein_coding | -0.355451821 | 4.789121677 | 0.4038537 | 0.5346432 | 0.880196444 |
| ENSMUSG00000039148 | Sart1 | protein_coding | 0.297036769 | 4.561525235 | 0.402415 | 0.5346577 | 0.880196444 |
| ENSMUSG00000030630 | Fah | protein_coding | 0.189121791 | 5.376407871 | 0.4023364 | 0.5346827 | 0.880196444 |
| ENSMUSG00000096210 | H1f0 | protein_coding | -0.157880135 | 5.834766493 | 0.4017196 | 0.5349922 | 0.880533035 |
| ENSMUSG00000033917 | Gde1 | protein_coding | -0.22060061 | 5.794515996 | 0.4016314 | 0.5350509 | 0.880533035 |
| ENSMUSG00000028127 | Abcd3 | protein_coding | -0.20364954 | 5.816527553 | 0.4012885 | 0.5352088 | 0.880658021 |
| ENSMUSG00000004113 | Cacna1b | protein_coding | 0.24589906 | 4.851315997 | 0.4009452 | 0.5353814 | 0.880763715 |
| ENSMUSG00000019828 | Grm1 | protein_coding | -0.184248766 | 5.464318844 | 0.4007748 | 0.5354671 | 0.880763715 |
| ENSMUSG00000028295 | Smim8 | protein_coding | 0.170673638 | 6.499060213 | 0.4005962 | 0.5355569 | 0.880763715 |
| ENSMUSG00000097451 | Rian | processed_transcript | -0.175928949 | 5.684275856 | 0.4004415 | 0.5356348 | 0.880763715 |
| ENSMUSG00000037563 | Rps16 | protein_coding | 0.379274349 | 2.816220096 | 0.4003465 | 0.5356826 | 0.880763715 |
| ENSMUSG00000028138 | Adh5 | protein_coding | 0.129529814 | 7.589360747 | 0.3996736 | 0.5360215 | 0.881186228 |
| ENSMUSG00000027642 | Rpn2 | protein_coding | 0.118437782 | 7.653253879 | 0.3992092 | 0.5362557 | 0.881353838 |
| ENSMUSG00000001054 | Rmnd5b | protein_coding | 0.182352679 | 5.63977067 | 0.3990378 | 0.5363421 | 0.881353838 |
| ENSMUSG00000078941 | Ak6 | protein_coding | 0.191420222 | 5.52430676 | 0.3986349 | 0.5365454 | 0.881353838 |
| ENSMUSG00000027207 | Galk2 | protein_coding | 0.187751776 | 6.042863571 | 0.3984283 | 0.5366498 | 0.881353838 |
| ENSMUSG00000044647 | Csrnp3 | protein_coding | -0.153840026 | 6.295928373 | 0.3983546 | 0.536687 | 0.881353838 |
| ENSMUSG00000030757 | Zkscan2 | protein_coding | -0.474677285 | 2.533081149 | 0.3982559 | 0.5367369 | 0.881353838 |
| ENSMUSG00000020917 | Acly | protein_coding | 0.176067291 | 7.114004496 | 0.3982108 | 0.536774 | 0.881353838 |
| ENSMUSG00000033157 | Abhd10 | protein_coding | 0.171088624 | 5.680328417 | 0.3980337 | 0.5368492 | 0.881353838 |
| ENSMUSG00000018974 | Sart3 | protein_coding | -0.253885185 | 4.178009768 | 0.3976703 | 0.5370329 | 0.881353838 |
| ENSMUSG00000052033 | Pfdn4 | protein_coding | 0.177330653 | 7.429777655 | 0.3976959 | 0.5370343 | 0.881353838 |
| ENSMUSG00000049300 | Prmt6 | protein_coding | 0.280238398 | 3.436988072 | 0.3975903 | 0.5370734 | 0.881353838 |

| ensembl_gene_id | external_gene_name | gene_biotype | log2 (fold change) | log2 (counts per million) | F-value | p-value | False discovery rate |
|---|---|---|---|---|---|---|---|
| ENSMUSG00000093904 | Tomm20 | protein_coding | -0.144992618 | 6.853735272 | 0.3975238 | 0.537107 | 0.881353838 |
| ENSMUSG00000037958 | Nsrp1 | protein_coding | 0.176589255 | 6.112309632 | 0.3972238 | 0.5372588 | 0.881468421 |
| ENSMUSG00000033579 | Fa2h | protein_coding | 0.219291469 | 4.900477609 | 0.3967909 | 0.537478 | 0.881693564 |
| ENSMUSG00000059991 | Nptx2 | protein_coding | 0.343123696 | 3.168592128 | 0.3963056 | 0.537724 | 0.881834487 |
| ENSMUSG00000044783 | Hjurp | protein_coding | 0.211317681 | 4.757925192 | 0.3960898 | 0.5378334 | 0.881834487 |
| ENSMUSG00000030421 | Uri1 | protein_coding | -0.198070359 | 4.919120289 | 0.3960694 | 0.5378438 | 0.881834487 |
| ENSMUSG00000029345 | Tfip11 | protein_coding | 0.304711076 | 4.892480515 | 0.3965459 | 0.537892 | 0.881834487 |
| ENSMUSG00000035234 | Abraxas1 | protein_coding | 0.393190754 | 2.869221752 | 0.395763 | 0.5379992 | 0.881875782 |
| ENSMUSG00000038949 | Cnst | protein_coding | 0.279212502 | 4.301464429 | 0.3948046 | 0.538486 | 0.882539166 |
| ENSMUSG00000021025 | Nfkbia | protein_coding | 0.20882235 | 5.816459171 | 0.3940938 | 0.5388475 | 0.882997156 |
| ENSMUSG00000007613 | Tgfbr1 | protein_coding | -0.551670551 | 3.187294785 | 0.3949536 | 0.5389631 | 0.883052041 |
| ENSMUSG00000034171 | Faah | protein_coding | -0.246974616 | 4.293621047 | 0.3934993 | 0.5391503 | 0.88322408 |
| ENSMUSG00000030683 | Sez6l2 | protein_coding | 0.136834735 | 7.401941601 | 0.3928318 | 0.5394906 | 0.883463634 |
| ENSMUSG00000048756 | Foxo3 | protein_coding | -0.222555709 | 5.828438081 | 0.3926683 | 0.5395882 | 0.883463634 |
| ENSMUSG00000046897 | Zfp740 | protein_coding | -0.198761855 | 4.845189861 | 0.3926301 | 0.5395935 | 0.883463634 |
| ENSMUSG00000026708 | Cenpl | protein_coding | 0.472914225 | 2.71994695 | 0.3925285 | 0.5396454 | 0.883463634 |
| ENSMUSG00000038784 | Cnot4 | protein_coding | -0.174969182 | 6.426545098 | 0.3924072 | 0.5397073 | 0.883463634 |
| ENSMUSG00000021958 | Pinx1 | protein_coding | -0.216031521 | 4.210913605 | 0.3915864 | 0.5401266 | 0.883718067 |
| ENSMUSG00000001229 | Dpp9 | protein_coding | 0.226454862 | 5.095200661 | 0.3915075 | 0.540167 | 0.883718067 |
| ENSMUSG00000052331 | Ankrd44 | protein_coding | -0.521110349 | 2.457030023 | 0.390328 | 0.5407708 | 0.883718067 |
| ENSMUSG00000049882 | Vcpkmt | protein_coding | -0.367897317 | 2.982139181 | 0.3899535 | 0.5409628 | 0.883718067 |
| ENSMUSG00000028766 | Alpl | protein_coding | -0.621298286 | 2.546182813 | 0.3897267 | 0.5410932 | 0.883718067 |
| ENSMUSG00000021666 | Gfm2 | protein_coding | 0.289878488 | 3.871473383 | 0.3895463 | 0.5411717 | 0.883718067 |
| ENSMUSG00000028890 | Mtf1 | protein_coding | 0.196013765 | 4.908268748 | 0.3894497 | 0.5412212 | 0.883718067 |
| ENSMUSG00000049038 | Mterf2 | protein_coding | -0.376604397 | 3.173659394 | 0.389253 | 0.5413222 | 0.883718067 |
| ENSMUSG00000025184 | R3hcc1l | protein_coding | -0.287370626 | 3.570688041 | 0.3892301 | 0.5413339 | 0.883718067 |
| ENSMUSG00000001056 | Nhp2 | protein_coding | 0.150838048 | 7.596107777 | 0.3889447 | 0.5414805 | 0.883718067 |
| ENSMUSG00000021271 | Zfp839 | protein_coding | -0.239630206 | 3.894737059 | 0.3889196 | 0.5414934 | 0.883718067 |
| ENSMUSG00000030243 | Recql | protein_coding | 0.305545711 | 3.821508345 | 0.3888886 | 0.5415093 | 0.883718067 |
| ENSMUSG00000086308 | G630016G05Rik | lincRNA | 0.401589302 | 2.940889393 | 0.3887823 | 0.5415639 | 0.883718067 |
| ENSMUSG00000029389 | Ddx55 | protein_coding | 0.1879682 | 5.187898048 | 0.3887759 | 0.5415673 | 0.883718067 |
| ENSMUSG00000002984 | Tomm40 | protein_coding | 0.243811122 | 4.313692104 | 0.3886621 | 0.5416257 | 0.883718067 |
| ENSMUSG00000054051 | Ercc6 | protein_coding | -0.498133701 | 2.562444111 | 0.3886321 | 0.5416412 | 0.883718067 |
| ENSMUSG00000001627 | Ifrd1 | protein_coding | -0.36540764 | 3.542411069 | 0.3885682 | 0.541674 | 0.883718067 |
| ENSMUSG00000001569 | Nom1 | protein_coding | -0.222064984 | 4.537747946 | 0.3885079 | 0.541705 | 0.883718067 |
| ENSMUSG00000036693 | Nop14 | protein_coding | 0.218049577 | 5.11560451 | 0.388469 | 0.541725 | 0.883718067 |
| ENSMUSG00000020075 | Ddx21 | protein_coding | -0.252123093 | 3.940671259 | 0.388165 | 0.5418813 | 0.883718067 |
| ENSMUSG00000057367 | Birc2 | protein_coding | -0.227360159 | 5.390641583 | 0.3879267 | 0.542004 | 0.883718067 |
| ENSMUSG00000032127 | Vps11 | protein_coding | 0.219951535 | 6.488605054 | 0.3888066 | 0.5420413 | 0.883718067 |
| ENSMUSG00000009905 | Kdsr | protein_coding | -0.184409643 | 5.742732626 | 0.3877208 | 0.54211 | 0.883718067 |
| ENSMUSG00000047495 | Dlgap2 | protein_coding | -0.236940978 | 4.648677871 | 0.3876968 | 0.5421223 | 0.883718067 |
| ENSMUSG00000028358 | Zfp618 | protein_coding | -0.496950689 | 2.889924072 | 0.3874589 | 0.5422448 | 0.883718067 |
| ENSMUSG00000067369 | Trmt2b | protein_coding | -0.273029886 | 4.47209768 | 0.3873729 | 0.5422891 | 0.883718067 |
| ENSMUSG00000030036 | Mogs | protein_coding | 0.362771077 | 3.324228333 | 0.3873651 | 0.5422932 | 0.883718067 |
| ENSMUSG00000021831 | Ero1l | protein_coding | -0.436538857 | 3.134148566 | 0.3871179 | 0.5424206 | 0.883718067 |
| ENSMUSG00000055109 | Gm15155 | protein_coding | 0.509358713 | 2.188059548 | 0.3870633 | 0.5424487 | 0.883718067 |
| ENSMUSG00000035575 | Utp6 | protein_coding | -0.227060037 | 4.648040619 | 0.3869054 | 0.5425301 | 0.883718067 |
| ENSMUSG00000036932 | Aifm1 | protein_coding | -0.210336279 | 5.357849465 | 0.3868905 | 0.5425378 | 0.883718067 |
| ENSMUSG00000048410 | Zfp407 | protein_coding | -0.316341396 | 3.295447265 | 0.3867301 | 0.5426205 | 0.883718067 |
| ENSMUSG00000097911 | Gm26691 | antisense_RNA | -0.408284273 | 2.904787901 | 0.3863094 | 0.5428376 | 0.883718067 |
| ENSMUSG00000031224 | Magee2 | protein_coding | 0.241944448 | 5.034566242 | 0.3860368 | 0.5429783 | 0.883718067 |
| ENSMUSG00000009739 | Pou6f1 | protein_coding | 0.211596072 | 4.692505443 | 0.3857776 | 0.5431122 | 0.883718067 |
| ENSMUSG00000015112 | Slc25a13 | protein_coding | -0.354391798 | 3.432724529 | 0.3856273 | 0.5431899 | 0.883718067 |
| ENSMUSG00000056153 | Socs6 | protein_coding | -0.236140718 | 4.079099879 | 0.3853668 | 0.5433245 | 0.883718067 |
| ENSMUSG00000039286 | Fndc3b | protein_coding | -0.280520355 | 3.988063727 | 0.3853575 | 0.5433293 | 0.883718067 |
| ENSMUSG00000050890 | Pdik1l | protein_coding | -0.245678954 | 4.28095881 | 0.3853295 | 0.5433438 | 0.883718067 |
| ENSMUSG00000034248 | Slc25a37 | protein_coding | -0.428446292 | 2.954380726 | 0.3849233 | 0.5435539 | 0.883718067 |
| ENSMUSG00000039194 | Rlbp1 | protein_coding | -0.334874969 | 4.133638272 | 0.3849229 | 0.5435682 | 0.883718067 |
| ENSMUSG00000028354 | Fmn2 | protein_coding | 0.24684896 | 4.408030175 | 0.3847838 | 0.5436261 | 0.883718067 |
| ENSMUSG00000030505 | Prmt3 | protein_coding | -0.296204385 | 3.326721743 | 0.384728 | 0.543655 | 0.883718067 |
| ENSMUSG00000029221 | Slc30a9 | protein_coding | 0.139659464 | 6.804376336 | 0.3846049 | 0.5437187 | 0.883718067 |
| ENSMUSG00000085795 | Zfp703 | protein_coding | -0.399842313 | 2.951697461 | 0.3845273 | 0.5437589 | 0.883718067 |
| ENSMUSG00000033991 | Ttc37 | protein_coding | 0.338724737 | 3.353452265 | 0.3844826 | 0.5437821 | 0.883718067 |
| ENSMUSG00000021340 | Gpld1 | protein_coding | 0.292627351 | 3.527496448 | 0.3843839 | 0.5438332 | 0.883718067 |
| ENSMUSG00000026036 | Nif3l1 | protein_coding | 0.218394933 | 6.00136296 | 0.384336 | 0.543872 | 0.883718067 |
| ENSMUSG00000070282 | 3000002C10Rik | transcribed_processe | 0.594102323 | 2.154410955 | 0.3842748 | 0.5438897 | 0.883718067 |
| ENSMUSG00000021596 | Mctp1 | protein_coding | -0.248934042 | 6.10910159 | 0.3850719 | 0.5440219 | 0.883771037 |
| ENSMUSG00000086943 | 4732414G09Rik | lincRNA | 0.612727327 | 2.822618559 | 0.3848318 | 0.544146 | 0.883771037 |
| ENSMUSG00000029128 | Rab28 | protein_coding | -0.168238689 | 6.833190454 | 0.3835747 | 0.5442526 | 0.883771037 |
| ENSMUSG00000046613 | Vwa5b2 | protein_coding | 0.40107337 | 2.903223785 | 0.3835315 | 0.544275 | 0.883771037 |
| ENSMUSG00000021748 | Pdhb | protein_coding | 0.119058516 | 7.858428937 | 0.3832134 | 0.5444401 | 0.883771037 |
| ENSMUSG00000026278 | Bok | protein_coding | -0.186736233 | 5.664162849 | 0.3829238 | 0.5445905 | 0.883771037 |
| ENSMUSG00000028696 | Ipp | protein_coding | -0.346954767 | 3.869324549 | 0.382729 | 0.5447057 | 0.883771037 |
| ENSMUSG00000024846 | Cst6 | protein_coding | -0.229841426 | 4.547371648 | 0.3826155 | 0.5447507 | 0.883771037 |
| ENSMUSG00000049422 | Chchd10 | protein_coding | -0.217014582 | 5.888721733 | 0.3825156 | 0.5448166 | 0.883771037 |
| ENSMUSG00000023791 | Pigx | protein_coding | 0.313469386 | 4.053541785 | 0.3822848 | 0.5449366 | 0.883771037 |
| ENSMUSG00000057156 | Homez | protein_coding | -0.27268535 | 3.484538364 | 0.3821834 | 0.5449753 | 0.883771037 |
| ENSMUSG00000044005 | Gls2 | protein_coding | 0.284828093 | 3.5291459 | 0.3821609 | 0.544987 | 0.883771037 |

| ensembl_gene_id | external_gene_name | gene_biotype | log2 (fold change) | log2 (counts per million) | F-value | p-value | False discovery rate |
|---|---|---|---|---|---|---|---|
| ENSMUSG00000032307 | Ube2q2 | protein_coding | -0.177003421 | 5.230685422 | 0.3821537 | 0.5449908 | 0.883771037 |
| ENSMUSG00000063931 | Pepd | protein_coding | 0.186398356 | 6.764992624 | 0.381845 | 0.5451653 | 0.883851197 |
| ENSMUSG00000022092 | Ppp3cc | protein_coding | -0.381152127 | 3.041517414 | 0.3816237 | 0.5452665 | 0.883851197 |
| ENSMUSG00000031538 | Plat | protein_coding | -0.275056775 | 4.335392688 | 0.3815848 | 0.5452868 | 0.883851197 |
| ENSMUSG00000071369 | Map3k5 | protein_coding | -0.225616035 | 4.299181997 | 0.3810823 | 0.5455485 | 0.884012986 |
| ENSMUSG00000024392 | Bag6 | protein_coding | 0.196307671 | 5.578006025 | 0.3810775 | 0.545551 | 0.884012986 |
| ENSMUSG00000024580 | Grpel2 | protein_coding | 0.180744877 | 5.73925752 | 0.3806648 | 0.5457662 | 0.884228356 |
| ENSMUSG00000004980 | Hnrnpa2b1 | protein_coding | -0.123513162 | 8.948773507 | 0.3794029 | 0.546425 | 0.884775589 |
| ENSMUSG00000017897 | Eya2 | protein_coding | -0.316606057 | 3.05916692 | 0.3794016 | 0.5464257 | 0.884775589 |
| ENSMUSG00000075324 | Fign | protein_coding | -0.343224422 | 4.297662235 | 0.3798209 | 0.5464686 | 0.884775589 |
| ENSMUSG00000027895 | Kcnc4 | protein_coding | 0.190777418 | 5.077709349 | 0.3791437 | 0.5465605 | 0.884775589 |
| ENSMUSG00000024666 | Tmem138 | protein_coding | 0.216414628 | 5.474530119 | 0.3790387 | 0.5466155 | 0.884775589 |
| ENSMUSG00000035557 | Krt17 | protein_coding | 0.395126673 | 2.761113525 | 0.3790134 | 0.5466287 | 0.884775589 |
| ENSMUSG00000066357 | Wdr6 | protein_coding | 0.358764441 | 7.794394425 | 0.387302 | 0.5466799 | 0.884775589 |
| ENSMUSG00000050751 | Pgbd5 | protein_coding | -0.143489008 | 6.575469217 | 0.3782845 | 0.5470103 | 0.88505491 |
| ENSMUSG00000048988 | Elfn1 | protein_coding | 0.408704222 | 3.066622727 | 0.3782714 | 0.5470171 | 0.88505491 |
| ENSMUSG00000020818 | Mfsd11 | protein_coding | 0.180413931 | 5.251139658 | 0.3779109 | 0.547206 | 0.885149225 |
| ENSMUSG00000078877 | Gm14295 | protein_coding | -0.275138435 | 3.695255168 | 0.377774 | 0.5472777 | 0.885149225 |
| ENSMUSG00000057176 | Ccdc189 | protein_coding | -0.204663214 | 4.692951345 | 0.3775744 | 0.5473824 | 0.885149225 |
| ENSMUSG00000042348 | Arl15 | protein_coding | -0.312219789 | 3.510940519 | 0.377532 | 0.5474047 | 0.885149225 |
| ENSMUSG00000097119 | B230354K17Rik | lincRNA | 0.326647201 | 3.440706363 | 0.3772224 | 0.5475671 | 0.885278797 |
| ENSMUSG00000097536 | 2610037D02Rik | lincRNA | -0.497706854 | 2.550039944 | 0.377047 | 0.5476592 | 0.885294519 |
| ENSMUSG00000091512 | Lamtor3 | protein_coding | 0.14523323 | 6.806980112 | 0.3768715 | 0.5477513 | 0.885304397 |
| ENSMUSG00000044317 | Gpr4 | protein_coding | -0.553068188 | 2.413926807 | 0.3766745 | 0.5478548 | 0.885304397 |
| ENSMUSG00000085067 | Gm15631 | lincRNA | -0.24715597 | 5.229903911 | 0.3764781 | 0.5479719 | 0.885304397 |
| ENSMUSG00000046658 | Zfp316 | protein_coding | -0.425259198 | 2.786277545 | 0.3762634 | 0.5480708 | 0.885304397 |
| ENSMUSG00000021234 | Fam161b | protein_coding | -0.340825933 | 3.909700963 | 0.3762408 | 0.5480966 | 0.885304397 |
| ENSMUSG00000020079 | Supv3l1 | protein_coding | 0.428910053 | 3.067613605 | 0.3760952 | 0.5481593 | 0.885304397 |
| ENSMUSG00000032123 | Dpagt1 | protein_coding | 0.385184711 | 3.034197488 | 0.3755854 | 0.5484276 | 0.8854593 |
| ENSMUSG00000049411 | Tmem241 | protein_coding | 0.327542786 | 4.27090545 | 0.3755924 | 0.5484902 | 0.8854593 |
| ENSMUSG00000040297 | Suco | protein_coding | -0.214395366 | 4.478071577 | 0.3752944 | 0.5485808 | 0.8854593 |
| ENSMUSG00000051065 | Mb21d2 | protein_coding | 0.202473504 | 4.829982531 | 0.3752872 | 0.5485846 | 0.8854593 |
| ENSMUSG00000042303 | Sgsm3 | protein_coding | 0.19813167 | 6.572978962 | 0.3749347 | 0.5488206 | 0.885514771 |
| ENSMUSG00000046709 | Mapk10 | protein_coding | 0.154228763 | 8.463617021 | 0.3748113 | 0.5488353 | 0.885514771 |
| ENSMUSG00000021186 | Fbln5 | protein_coding | -0.553453256 | 2.269903079 | 0.3747531 | 0.548866 | 0.885514771 |
| ENSMUSG00000034220 | Gpc1 | protein_coding | 0.236215441 | 4.569326509 | 0.3743368 | 0.5490856 | 0.88563622 |
| ENSMUSG00000024241 | Sos1 | protein_coding | 0.306582579 | 3.380003633 | 0.3741671 | 0.5491751 | 0.88563622 |
| ENSMUSG00000037509 | Arhgef4 | protein_coding | 0.163286259 | 6.287495244 | 0.3741421 | 0.5491883 | 0.88563622 |
| ENSMUSG00000061371 | Zfp873 | protein_coding | -0.270179158 | 4.0858188 | 0.3737317 | 0.549405 | 0.885720126 |
| ENSMUSG00000020088 | Sar1a | protein_coding | 0.12997248 | 7.259950516 | 0.3736186 | 0.5494648 | 0.885720126 |
| ENSMUSG00000023945 | Slc5a7 | protein_coding | -0.319309589 | 3.343747888 | 0.3735519 | 0.5495 | 0.885720126 |
| ENSMUSG00000022300 | Dcaf13 | protein_coding | 0.196975267 | 6.142582573 | 0.3734459 | 0.5495699 | 0.885720126 |
| ENSMUSG00000005045 | Chd5 | protein_coding | 0.167899473 | 6.187593633 | 0.3730598 | 0.5497601 | 0.885738462 |
| ENSMUSG00000053436 | Mapk14 | protein_coding | -0.204406386 | 5.388833924 | 0.3730248 | 0.5497787 | 0.885738462 |
| ENSMUSG00000060534 | Dcc | protein_coding | -0.370844911 | 3.846926997 | 0.3730257 | 0.5498283 | 0.885738462 |
| ENSMUSG00000039585 | Myo9a | protein_coding | 0.402041828 | 2.871054901 | 0.3727528 | 0.5499226 | 0.885757564 |
| ENSMUSG00000038212 | Mfsd14b | protein_coding | -0.244513551 | 4.628872732 | 0.3721617 | 0.5502356 | 0.885799905 |
| ENSMUSG00000041415 | Dicer1 | protein_coding | -0.257253519 | 4.0657958 | 0.3720962 | 0.5502702 | 0.885799905 |
| ENSMUSG00000052374 | Actn2 | protein_coding | -0.479332534 | 3.558390301 | 0.3735934 | 0.5503171 | 0.885799905 |
| ENSMUSG00000002307 | Daxx | protein_coding | 0.229588089 | 5.234720564 | 0.3719776 | 0.5503331 | 0.885799905 |
| ENSMUSG00000021185 | 9030617O03Rik | protein_coding | -0.358569008 | 4.068951454 | 0.3724196 | 0.5503608 | 0.885799905 |
| ENSMUSG00000032526 | Ss18l2 | protein_coding | 0.125877687 | 8.073288735 | 0.3717579 | 0.5504496 | 0.885810265 |
| ENSMUSG00000038007 | Acer2 | protein_coding | 0.230899856 | 4.297017974 | 0.3714732 | 0.5506005 | 0.885920578 |
| ENSMUSG00000022159 | Rab2b | protein_coding | 0.178704718 | 4.849298695 | 0.3712807 | 0.5507027 | 0.885952369 |
| ENSMUSG00000070509 | Rgma | protein_coding | 0.312737747 | 3.427260483 | 0.3710282 | 0.5508367 | 0.88603229 |
| ENSMUSG00000043671 | Dpy19l3 | protein_coding | -0.272242793 | 4.19884492 | 0.3707711 | 0.5509732 | 0.88603229 |
| ENSMUSG00000000266 | Mid2 | protein_coding | -0.288750099 | 4.204205419 | 0.3704253 | 0.5511569 | 0.88603229 |
| ENSMUSG00000066637 | Ttc32 | protein_coding | -0.458115179 | 2.762092343 | 0.3701242 | 0.551317 | 0.88603229 |
| ENSMUSG00000026819 | Slc25a25 | protein_coding | 0.158502251 | 5.904882908 | 0.3700459 | 0.5513587 | 0.88603229 |
| ENSMUSG00000023912 | Slc25a27 | protein_coding | 0.182190093 | 4.990672139 | 0.3699059 | 0.5514331 | 0.88603229 |
| ENSMUSG00000055762 | Eef1d | protein_coding | 0.138183572 | 6.868658105 | 0.3696915 | 0.5515472 | 0.88603229 |
| ENSMUSG00000015165 | Hnrnpl | protein_coding | -0.14716729 | 6.560718103 | 0.3696834 | 0.5515515 | 0.88603229 |
| ENSMUSG00000020635 | Fkbp1b | protein_coding | -0.242105375 | 4.312844413 | 0.3694738 | 0.5516631 | 0.88603229 |
| ENSMUSG00000057554 | Lgals8 | protein_coding | 0.21507352 | 5.631530436 | 0.3692387 | 0.5517883 | 0.88603229 |
| ENSMUSG00000028920 | Fbxo42 | protein_coding | -0.262660567 | 3.880096418 | 0.3691662 | 0.5518269 | 0.88603229 |
| ENSMUSG00000032279 | Idh3a | protein_coding | 0.147266978 | 9.358748694 | 0.369127 | 0.5518478 | 0.88603229 |
| ENSMUSG00000017404 | Rpl19 | protein_coding | 0.267721324 | 3.484945015 | 0.3690526 | 0.5518874 | 0.88603229 |
| ENSMUSG00000021219 | Rgs6 | protein_coding | -0.274279657 | 4.009210378 | 0.3690179 | 0.5519059 | 0.88603229 |
| ENSMUSG00000029310 | Nudt9 | protein_coding | 0.181363651 | 5.61468914 | 0.3687625 | 0.552042 | 0.886118537 |
| ENSMUSG00000023089 | Ndufa5 | protein_coding | 0.193498328 | 10.15552811 | 0.3683193 | 0.5523283 | 0.886249529 |
| ENSMUSG00000055322 | Tns1 | protein_coding | -0.370366729 | 3.784379738 | 0.3682323 | 0.5523747 | 0.886249529 |
| ENSMUSG00000028563 | Tm2d1 | protein_coding | 0.260849046 | 4.406926536 | 0.3679854 | 0.5524567 | 0.886249529 |
| ENSMUSG00000036095 | Dgkb | protein_coding | -0.16366212 | 7.206414384 | 0.36795 | 0.5524756 | 0.886249529 |
| ENSMUSG00000087197 | Gm13780 | lincRNA | -0.321054824 | 3.154261479 | 0.3678373 | 0.5525357 | 0.886249529 |
| ENSMUSG00000021209 | Ppp4r4 | protein_coding | -0.235114218 | 4.206087628 | 0.367519 | 0.5527058 | 0.88639004 |
| ENSMUSG00000028962 | Slc4a2 | protein_coding | -0.305671341 | 3.489299007 | 0.3671014 | 0.552929 | 0.886504676 |
| ENSMUSG00000002055 | Spag5 | protein_coding | -0.611624475 | 2.194848976 | 0.3670769 | 0.5529421 | 0.886504676 |
| ENSMUSG00000056215 | Lrguk | protein_coding | -0.520182407 | 2.904019546 | 0.3665123 | 0.5532579 | 0.886878749 |

| ensembl_gene_id | external_gene_name | gene_biotype | log2 (fold change) | log2 (counts per million) | F-value | p-value | False discovery rate |
|---|---|---|---|---|---|---|---|
| ENSMUSG00000074682 | Zcchc3 | protein_coding | -0.300444161 | 3.928975272 | 0.3663155 | 0.5533497 | 0.886893591 |
| ENSMUSG00000021981 | Cab39l | protein_coding | -0.150635337 | 6.511037761 | 0.3653222 | 0.5538822 | 0.887614729 |
| ENSMUSG00000087618 | Gm14114 | lincRNA | -0.443270846 | 2.454754012 | 0.3650482 | 0.5540293 | 0.88771816 |
| ENSMUSG00000027254 | Map1a | protein_coding | 0.176526009 | 7.385341799 | 0.3648419 | 0.5541537 | 0.887785218 |
| ENSMUSG00000006378 | Gcat | protein_coding | 0.213692533 | 5.257505579 | 0.3643758 | 0.5543905 | 0.888032302 |
| ENSMUSG00000038936 | Sccpdh | protein_coding | 0.144157795 | 7.117289642 | 0.36396 | 0.554614 | 0.888165421 |
| ENSMUSG00000025337 | Sbds | protein_coding | 0.117543075 | 8.185856683 | 0.3636772 | 0.5547662 | 0.888165421 |
| ENSMUSG00000059336 | Slc14a1 | protein_coding | -0.474466548 | 2.821442669 | 0.3634413 | 0.5548932 | 0.888165421 |
| ENSMUSG00000059493 | Nhs | protein_coding | -0.294166493 | 3.366856046 | 0.3633039 | 0.5549673 | 0.888165421 |
| ENSMUSG00000021277 | Traf3 | protein_coding | 0.283137199 | 3.856128044 | 0.3632524 | 0.554995 | 0.888165421 |
| ENSMUSG00000040725 | Hnrnpul1 | protein_coding | -0.224188131 | 5.585243096 | 0.3632244 | 0.5550237 | 0.888165421 |
| ENSMUSG00000017679 | Ttpal | protein_coding | -0.195326818 | 4.909761282 | 0.363147 | 0.5550518 | 0.888165421 |
| ENSMUSG00000038991 | Txndc5 | protein_coding | 0.228230085 | 5.050696154 | 0.3629791 | 0.5551423 | 0.888178089 |
| ENSMUSG00000027427 | Polr3f | protein_coding | -0.231542841 | 4.529543924 | 0.3625903 | 0.5553519 | 0.888341389 |
| ENSMUSG00000041444 | Arhgap32 | protein_coding | -0.18140448 | 4.925505771 | 0.3624742 | 0.5554146 | 0.888341389 |
| ENSMUSG00000020671 | Rab10 | protein_coding | -0.356460134 | 2.972997929 | 0.3623303 | 0.5554922 | 0.888341389 |
| ENSMUSG00000040479 | Dgkz | protein_coding | -0.180012853 | 6.007074155 | 0.3620151 | 0.5556623 | 0.88836486 |
| ENSMUSG00000019066 | Rab3d | protein_coding | -0.636844626 | 2.520365655 | 0.3609789 | 0.556236 | 0.88836486 |
| ENSMUSG00000032666 | 1700025G04Rik | protein_coding | 0.175110666 | 5.294701351 | 0.3609516 | 0.5562372 | 0.88836486 |
| ENSMUSG00000033059 | Pygb | protein_coding | 0.194975367 | 6.971582932 | 0.3615669 | 0.5562502 | 0.88836486 |
| ENSMUSG00000059975 | Zfp74 | protein_coding | 0.267184889 | 3.668341166 | 0.3608789 | 0.5562765 | 0.88836486 |
| ENSMUSG00000021910 | Nisch | protein_coding | 0.124555322 | 7.4462223 | 0.3608653 | 0.5562839 | 0.88836486 |
| ENSMUSG00000015733 | Capza2 | protein_coding | -0.125328994 | 9.262856819 | 0.3608457 | 0.5562945 | 0.88836486 |
| ENSMUSG00000055024 | Ep300 | protein_coding | -0.299338908 | 3.593335121 | 0.3607481 | 0.5563473 | 0.88836486 |
| ENSMUSG00000037376 | Trmt6 | protein_coding | -0.193109329 | 5.655510675 | 0.3606562 | 0.5563971 | 0.88836486 |
| ENSMUSG00000038909 | Kat7 | protein_coding | -0.187888317 | 6.342927976 | 0.3606502 | 0.5564139 | 0.88836486 |
| ENSMUSG00000076928 | Trac | TR_C_gene | 0.615104666 | 2.465869828 | 0.3604782 | 0.556507 | 0.88836486 |
| ENSMUSG00000025086 | Trub1 | protein_coding | 0.193798619 | 5.321156264 | 0.3603267 | 0.5565755 | 0.88836486 |
| ENSMUSG00000031266 | Gla | protein_coding | 0.312290823 | 3.426680206 | 0.3603168 | 0.5565809 | 0.88836486 |
| ENSMUSG00000019558 | Slc6a8 | protein_coding | -0.185494412 | 5.170336647 | 0.3601399 | 0.5566767 | 0.888385943 |
| ENSMUSG00000031889 | D230025D16Rik | protein_coding | 0.228608385 | 4.334105351 | 0.3599168 | 0.5567976 | 0.888447046 |
| ENSMUSG00000024805 | Pcgf5 | protein_coding | -0.216228777 | 5.309325547 | 0.3589578 | 0.557318 | 0.889010109 |
| ENSMUSG00000045071 | E130308A19Rik | protein_coding | -0.308226917 | 3.44162894 | 0.3589338 | 0.557331 | 0.889010109 |
| ENSMUSG00000018589 | Glra2 | protein_coding | 0.216796254 | 5.258017311 | 0.3586101 | 0.5575069 | 0.889010109 |
| ENSMUSG00000043241 | Upf2 | protein_coding | -0.23055543 | 5.165862436 | 0.3585906 | 0.5575175 | 0.889010109 |
| ENSMUSG00000061374 | Fiz1 | protein_coding | 0.291409686 | 3.292743738 | 0.3583828 | 0.5576305 | 0.889010109 |
| ENSMUSG00000032340 | Neo1 | protein_coding | -0.229934419 | 4.574868656 | 0.3577275 | 0.557987 | 0.889010109 |
| ENSMUSG00000087598 | Zfp111 | protein_coding | 0.352261274 | 2.732039677 | 0.357569 | 0.5580733 | 0.889010109 |
| ENSMUSG00000028467 | Gba2 | protein_coding | 0.165894449 | 5.388187898 | 0.3574411 | 0.558143 | 0.889010109 |
| ENSMUSG00000038884 | A230050P20Rik | protein_coding | 0.319534511 | 4.417019797 | 0.3578032 | 0.5582016 | 0.889010109 |
| ENSMUSG00000060260 | Pwwp2b | protein_coding | -0.275962483 | 4.46380528 | 0.3572609 | 0.5582412 | 0.889010109 |
| ENSMUSG00000096146 | Kcnj11 | protein_coding | 0.272782033 | 3.954955941 | 0.3572148 | 0.5582663 | 0.889010109 |
| ENSMUSG00000049122 | Frmd3 | protein_coding | 0.391619301 | 3.188285896 | 0.3571972 | 0.5582759 | 0.889010109 |
| ENSMUSG00000033319 | Fem1c | protein_coding | -0.249825654 | 3.986097007 | 0.3571512 | 0.5583009 | 0.889010109 |
| ENSMUSG00000055932 | Fto | protein_coding | 0.249582358 | 8.114774257 | 0.3594696 | 0.5583509 | 0.889010109 |
| ENSMUSG00000003402 | Prkcsh | protein_coding | 0.247007869 | 5.289178934 | 0.3569152 | 0.5584431 | 0.889010109 |
| ENSMUSG00000022404 | Slc25a17 | protein_coding | -0.135047918 | 7.268080084 | 0.3567565 | 0.5585161 | 0.889010109 |
| ENSMUSG00000022466 | Rpap3 | protein_coding | 0.166174514 | 5.866990366 | 0.3566835 | 0.558556 | 0.889010109 |
| ENSMUSG00000038677 | Scube3 | protein_coding | -0.455580809 | 2.59670893 | 0.3563306 | 0.5587485 | 0.889184952 |
| ENSMUSG00000020846 | Rflnb | protein_coding | -0.297676247 | 3.672663658 | 0.3558644 | 0.5590031 | 0.889254384 |
| ENSMUSG00000030669 | Calca | protein_coding | -0.644408502 | 3.089485496 | 0.3573278 | 0.5593526 | 0.889254384 |
| ENSMUSG00000048481 | Mypop | protein_coding | 0.229727743 | 3.951406015 | 0.3550363 | 0.559456 | 0.889254384 |
| ENSMUSG00000038403 | Hfe2 | protein_coding | 0.408060395 | 2.767876911 | 0.3548782 | 0.5595425 | 0.889254384 |
| ENSMUSG00000042834 | Nrep | protein_coding | -0.15765283 | 6.433058837 | 0.3547746 | 0.5595992 | 0.889254384 |
| ENSMUSG00000026577 | Blzf1 | protein_coding | -0.193882578 | 5.141870769 | 0.3547719 | 0.5596007 | 0.889254384 |
| ENSMUSG00000029621 | Arpc1a | protein_coding | 0.114224104 | 9.187904977 | 0.3546816 | 0.5596501 | 0.889254384 |
| ENSMUSG00000050271 | Prag1 | protein_coding | -0.298944956 | 3.463287593 | 0.3546805 | 0.5596507 | 0.889254384 |
| ENSMUSG00000000346 | Dazap2 | protein_coding | -0.129836021 | 8.570230436 | 0.3546481 | 0.5596685 | 0.889254384 |
| ENSMUSG00000027016 | Zfp385b | protein_coding | 0.179856097 | 6.379825828 | 0.3545299 | 0.5597332 | 0.889254384 |
| ENSMUSG00000024855 | Pacs1 | protein_coding | 0.184022607 | 5.570134098 | 0.3543668 | 0.5598226 | 0.889254384 |
| ENSMUSG00000025130 | P4hb | protein_coding | -0.154215186 | 6.470404982 | 0.3542338 | 0.5598954 | 0.889254384 |
| ENSMUSG00000019578 | Ubxn6 | protein_coding | -0.14405158 | 6.444794577 | 0.3541604 | 0.5599357 | 0.889254384 |
| ENSMUSG00000024109 | Nrxn1 | protein_coding | -0.150655005 | 8.156305757 | 0.354129 | 0.5599529 | 0.889254384 |
| ENSMUSG00000018347 | Zkscan6 | protein_coding | 0.476922096 | 3.488809905 | 0.3554651 | 0.560051 | 0.889254384 |
| ENSMUSG00000053093 | Myh7 | protein_coding | -0.35484888 | 2.921770434 | 0.3537392 | 0.5601666 | 0.889254384 |
| ENSMUSG00000020231 | Dip2a | protein_coding | -0.234676905 | 4.17885028 | 0.3535807 | 0.5602536 | 0.889254384 |
| ENSMUSG00000020894 | Vamp2 | protein_coding | 0.129065235 | 9.219983842 | 0.3535313 | 0.5602807 | 0.889254384 |
| ENSMUSG00000023072 | Cep89 | protein_coding | -0.284082832 | 3.758035032 | 0.3532199 | 0.5604517 | 0.889394482 |
| ENSMUSG00000040502 | 9-Mar | protein_coding | -0.334748898 | 3.218846446 | 0.3528899 | 0.5606329 | 0.889550831 |
| ENSMUSG00000040018 | Cox15 | protein_coding | 0.304306766 | 3.750012357 | 0.3526109 | 0.5607863 | 0.889662959 |
| ENSMUSG00000006276 | Eps15l1 | protein_coding | 0.17633567 | 5.558917699 | 0.3522939 | 0.5609607 | 0.88977544 |
| ENSMUSG00000027965 | Olfm3 | protein_coding | 0.220513842 | 4.733912521 | 0.3519407 | 0.561155 | 0.88977544 |
| ENSMUSG00000013236 | Ptprs | protein_coding | -0.151403277 | 7.138867334 | 0.3518353 | 0.5612131 | 0.88977544 |
| ENSMUSG00000030613 | Ccdc90b | protein_coding | -0.182642461 | 5.437788353 | 0.3515033 | 0.561396 | 0.88977544 |
| ENSMUSG00000000441 | Raf1 | protein_coding | -0.407373268 | 2.610805292 | 0.3514805 | 0.5614085 | 0.88977544 |
| ENSMUSG00000020070 | Rufy2 | protein_coding | 0.200850613 | 5.062097477 | 0.351336 | 0.5614881 | 0.88977544 |
| ENSMUSG00000026439 | Rbbp5 | protein_coding | -0.19067787 | 5.231769556 | 0.3512908 | 0.561513 | 0.88977544 |
| ENSMUSG00000052214 | Opa3 | protein_coding | -0.134792046 | 6.439299332 | 0.3511669 | 0.5615814 | 0.88977544 |

| ensembl_gene_id | external_gene_name | gene_biotype | log2 (fold change) | log2 (counts per million) | F-value | p-value | False discovery rate |
|---|---|---|---|---|---|---|---|
| ENSMUSG00000047466 | 8030462N17Rik | protein_coding | 0.317452662 | 3.532953595 | 0.3510269 | 0.5616586 | 0.88977544 |
| ENSMUSG00000068617 | Efcab1 | protein_coding | 0.167111404 | 5.490461706 | 0.35091 | 0.561723 | 0.88977544 |
| ENSMUSG00000053329 | D10Jhu81e | protein_coding | -0.189321158 | 5.636115395 | 0.3508295 | 0.5617675 | 0.88977544 |
| ENSMUSG00000042364 | Snx18 | protein_coding | -0.30172932 | 3.823215141 | 0.350655 | 0.5618637 | 0.889796852 |
| ENSMUSG00000060568 | Fam78b | protein_coding | -0.307143714 | 3.485171774 | 0.3504522 | 0.5619757 | 0.889843073 |
| ENSMUSG00000026491 | Ahctf1 | protein_coding | -0.313823768 | 3.264716847 | 0.3498936 | 0.5622843 | 0.890200613 |
| ENSMUSG00000059995 | Atxn7l3 | protein_coding | 0.179733745 | 5.42908172 | 0.349635 | 0.5624272 | 0.890295894 |
| ENSMUSG00000063550 | Nup98 | protein_coding | -0.171321098 | 5.737977979 | 0.3494651 | 0.5625212 | 0.89031355 |
| ENSMUSG00000038252 | Ncapd2 | protein_coding | -0.591568812 | 2.557168031 | 0.3490425 | 0.5627684 | 0.890573668 |
| ENSMUSG00000033676 | Gabrb3 | protein_coding | -0.137609074 | 6.4470507 | 0.348699 | 0.5629453 | 0.890615018 |
| ENSMUSG00000058569 | Tmed9 | protein_coding | -0.205662606 | 8.118184108 | 0.3496623 | 0.5629601 | 0.890615018 |
| ENSMUSG00000021902 | Phf7 | protein_coding | -0.643339534 | 2.481842382 | 0.3478465 | 0.5634519 | 0.89103709 |
| ENSMUSG00000024501 | Dpysl3 | protein_coding | 0.185672318 | 6.205575626 | 0.3477137 | 0.563505 | 0.89103709 |
| ENSMUSG00000015747 | Vps45 | protein_coding | 0.143085889 | 6.472370461 | 0.3476484 | 0.563528 | 0.89103709 |
| ENSMUSG00000021552 | Gkap1 | protein_coding | -0.313619454 | 3.370369014 | 0.347419 | 0.5636553 | 0.89103709 |
| ENSMUSG00000009073 | Nf2 | protein_coding | -0.2023132 | 4.799165403 | 0.3471861 | 0.5637847 | 0.89103709 |
| ENSMUSG00000006392 | Med8 | protein_coding | -0.146584346 | 6.431049831 | 0.3471542 | 0.5638024 | 0.89103709 |
| ENSMUSG00000085881 | Gm15912 | processed_transcript | 0.283433691 | 3.367241813 | 0.347146 | 0.563807 | 0.89103709 |
| ENSMUSG00000051978 | Erich1 | protein_coding | 0.236836371 | 4.202474229 | 0.3466294 | 0.5640942 | 0.891272147 |
| ENSMUSG00000052337 | Immt | protein_coding | 0.16174807 | 7.219270007 | 0.3466042 | 0.5641215 | 0.891272147 |
| ENSMUSG00000045427 | Hnrnph2 | protein_coding | -0.172881664 | 5.821643913 | 0.3463138 | 0.5642698 | 0.891375536 |
| ENSMUSG00000041895 | Wipi1 | protein_coding | -0.366917684 | 4.434203153 | 0.3475489 | 0.5643879 | 0.891431086 |
| ENSMUSG00000007670 | Khsrp | protein_coding | -0.175872756 | 5.071241666 | 0.3451167 | 0.5649369 | 0.892071934 |
| ENSMUSG00000026791 | Slc2a8 | protein_coding | -0.452672455 | 2.732372097 | 0.3450761 | 0.5649595 | 0.892071934 |
| ENSMUSG00000040276 | Pacsin1 | protein_coding | -0.135339985 | 7.277581248 | 0.3449093 | 0.5650527 | 0.89208799 |
| ENSMUSG00000046185 | Zfp84 | protein_coding | -0.348872577 | 3.379679398 | 0.3447596 | 0.5651362 | 0.892088901 |
| ENSMUSG00000034522 | Zfp395 | protein_coding | 0.257887368 | 3.499835818 | 0.3444476 | 0.5653105 | 0.892232991 |
| ENSMUSG00000024664 | Fads3 | protein_coding | 0.32774997 | 4.01031925 | 0.3441494 | 0.565525 | 0.892391321 |
| ENSMUSG00000042066 | Tmcc2 | protein_coding | -0.173425515 | 6.075979298 | 0.3439711 | 0.5655768 | 0.892391321 |
| ENSMUSG00000039519 | Cyp7b1 | protein_coding | -0.329218369 | 3.486738706 | 0.3436483 | 0.5657573 | 0.892545242 |
| ENSMUSG00000021245 | Mlh3 | protein_coding | -0.292172747 | 3.920570449 | 0.3434624 | 0.5658614 | 0.892578469 |
| ENSMUSG00000021576 | Pdcd6 | protein_coding | -0.124587313 | 7.915904575 | 0.3431398 | 0.566042 | 0.892674509 |
| ENSMUSG00000025207 | Sema4g | protein_coding | 0.391316516 | 3.063198317 | 0.3427469 | 0.5662622 | 0.892674509 |
| ENSMUSG00000025899 | Alkbh8 | protein_coding | -0.168925638 | 5.523790829 | 0.3427158 | 0.5662796 | 0.892674509 |
| ENSMUSG00000019894 | Slc6a15 | protein_coding | -0.169868982 | 5.604783337 | 0.3426134 | 0.566337 | 0.892674509 |
| ENSMUSG00000079481 | Nhsl2 | protein_coding | -0.194535152 | 5.12842791 | 0.3426127 | 0.5663373 | 0.892674509 |
| ENSMUSG00000022376 | Adcy8 | protein_coding | 0.334926446 | 2.998032649 | 0.3424184 | 0.5664463 | 0.892688441 |
| ENSMUSG00000027971 | Ndst4 | protein_coding | -0.285935786 | 3.514078181 | 0.3423009 | 0.5665122 | 0.892688441 |
| ENSMUSG00000039648 | Kyat1 | protein_coding | 0.276881718 | 4.543496185 | 0.341868 | 0.5667683 | 0.892765717 |
| ENSMUSG00000036057 | Ptpn23 | protein_coding | 0.273445408 | 3.875969448 | 0.341724 | 0.5668361 | 0.892765717 |
| ENSMUSG00000033439 | Trmt13 | protein_coding | -0.383412635 | 2.783158122 | 0.3416443 | 0.5668809 | 0.892765717 |
| ENSMUSG00000020628 | Trappc12 | protein_coding | 0.137146453 | 6.182638544 | 0.341622 | 0.5668934 | 0.892765717 |
| ENSMUSG00000032293 | Ireb2 | protein_coding | -0.213613913 | 4.662883172 | 0.3411564 | 0.5671551 | 0.892784953 |
| ENSMUSG00000005102 | Eif2ak4 | protein_coding | -0.288499591 | 3.634315011 | 0.3411189 | 0.5671762 | 0.892784953 |
| ENSMUSG00000052676 | Zmat1 | protein_coding | 0.241198545 | 4.490497069 | 0.3407412 | 0.5673886 | 0.892784953 |
| ENSMUSG00000026767 | Mindy3 | protein_coding | 0.143241789 | 6.240347076 | 0.3407357 | 0.5673917 | 0.892784953 |
| ENSMUSG00000004187 | Kifc2 | protein_coding | -0.155654553 | 6.489650591 | 0.3407107 | 0.5674058 | 0.892784953 |
| ENSMUSG00000055202 | Zfp811 | protein_coding | -0.237467268 | 4.399401755 | 0.3406169 | 0.5674586 | 0.892784953 |
| ENSMUSG00000056305 | Usp39 | protein_coding | 0.20055764 | 4.208385166 | 0.3403663 | 0.5675997 | 0.892784953 |
| ENSMUSG00000016940 | Kctd2 | protein_coding | 0.195509542 | 5.36580741 | 0.3401969 | 0.5676951 | 0.892784953 |
| ENSMUSG00000051159 | Cited1 | protein_coding | 0.25056277 | 5.499660301 | 0.3405213 | 0.5678464 | 0.892784953 |
| ENSMUSG00000029348 | Asphd2 | protein_coding | -0.171672765 | 5.911058479 | 0.3398293 | 0.5679023 | 0.892784953 |
| ENSMUSG00000040653 | Ppp1r14c | protein_coding | 0.28098108 | 3.440860724 | 0.3396184 | 0.5680212 | 0.892784953 |
| ENSMUSG00000006127 | Inpp5k | protein_coding | 0.20564913 | 5.897336812 | 0.3395413 | 0.5680777 | 0.892784953 |
| ENSMUSG00000020948 | Klhl28 | protein_coding | 0.17769048 | 4.843608937 | 0.3392942 | 0.5682041 | 0.892784953 |
| ENSMUSG00000006423 | C330007P06Rik | protein_coding | -0.223621036 | 4.447876819 | 0.3391858 | 0.5682653 | 0.892784953 |
| ENSMUSG00000004207 | Psap | protein_coding | -0.136316752 | 9.011552708 | 0.3389292 | 0.5684102 | 0.892784953 |
| ENSMUSG00000055305 | Zfp93 | protein_coding | 0.400574638 | 2.681189644 | 0.3388159 | 0.5684742 | 0.892784953 |
| ENSMUSG00000035765 | Dym | protein_coding | 0.156391029 | 5.538047024 | 0.3384465 | 0.5686829 | 0.892784953 |
| ENSMUSG00000039686 | Zer1 | protein_coding | -0.300216544 | 5.816096672 | 0.3404 | 0.5688508 | 0.892784953 |
| ENSMUSG00000001674 | Ddx18 | protein_coding | 0.20426228 | 4.228747452 | 0.3380878 | 0.5688858 | 0.892784953 |
| ENSMUSG00000025920 | Stau2 | protein_coding | -0.116369315 | 7.064248822 | 0.3379558 | 0.5689605 | 0.892784953 |
| ENSMUSG00000051951 | Xkr4 | protein_coding | -0.347602459 | 3.610802665 | 0.337885 | 0.5690136 | 0.892784953 |
| ENSMUSG00000031661 | Nkd1 | protein_coding | 0.52681102 | 2.463176117 | 0.3378277 | 0.569033 | 0.892784953 |
| ENSMUSG00000033282 | Rpgrip1l | protein_coding | 0.251329382 | 4.09526138 | 0.3378203 | 0.5690372 | 0.892784953 |
| ENSMUSG00000028405 | Aco1 | protein_coding | -0.275055917 | 4.148423769 | 0.3377551 | 0.5690741 | 0.892784953 |
| ENSMUSG00000110156 | Gm42067 | antisense_RNA | 0.299002667 | 3.177236938 | 0.3377458 | 0.5690794 | 0.892784953 |
| ENSMUSG00000041949 | Tango6 | protein_coding | -0.239949202 | 3.55573917 | 0.3376403 | 0.5691391 | 0.892784953 |
| ENSMUSG00000018189 | Uchl5 | protein_coding | 0.122911148 | 6.939610943 | 0.3376258 | 0.5691473 | 0.892784953 |
| ENSMUSG00000019927 | Ube2d1 | protein_coding | 0.138580498 | 6.169708397 | 0.3372423 | 0.5693646 | 0.892982653 |
| ENSMUSG00000033684 | Qsox1 | protein_coding | 0.217459509 | 5.093274121 | 0.337014 | 0.5694939 | 0.892982653 |
| ENSMUSG00000003731 | Kpna6 | protein_coding | -0.192267945 | 4.824788122 | 0.3369637 | 0.5695225 | 0.892982653 |
| ENSMUSG00000066392 | Nrxn3 | protein_coding | 0.134940976 | 7.907882488 | 0.336429 | 0.5698258 | 0.893328062 |
| ENSMUSG00000033323 | Ctdp1 | protein_coding | -0.552066673 | 2.510626562 | 0.3362356 | 0.5699357 | 0.893369994 |
| ENSMUSG00000055897 | Ppp4r1l-ps | transcribed_unproces | -0.286061915 | 3.477519279 | 0.3360531 | 0.5700393 | 0.893402221 |
| ENSMUSG00000026213 | Stk11ip | protein_coding | 0.328548243 | 3.316187816 | 0.335773 | 0.5701984 | 0.893437924 |
| ENSMUSG00000020484 | Xbp1 | protein_coding | -0.1861201 | 5.883448601 | 0.3355291 | 0.5703371 | 0.893437924 |
| ENSMUSG00000036835 | Psenen | protein_coding | 0.172294715 | 4.977596897 | 0.3353021 | 0.5704662 | 0.893437924 |

| ensembl_gene_id | external_gene_name | gene_biotype | log2 (fold change) | log2 (counts per million) | F-value | p-value | False discovery rate |
|---|---|---|---|---|---|---|---|
| ENSMUSG00000054455 | Vapb | protein_coding | 0.176910118 | 5.438720093 | 0.3351882 | 0.570531 | 0.893437924 |
| ENSMUSG00000034586 | Hid1 | protein_coding | 0.182815418 | 6.134864464 | 0.3351382 | 0.5705595 | 0.893437924 |
| ENSMUSG00000063873 | Slc24a3 | protein_coding | -0.179478632 | 6.203920954 | 0.3351362 | 0.5705606 | 0.893437924 |
| ENSMUSG00000058441 | Panx2 | protein_coding | -0.244338648 | 4.069168426 | 0.3347855 | 0.5707603 | 0.893592686 |
| ENSMUSG00000056679 | Gpr173 | protein_coding | -0.391057375 | 2.789059851 | 0.3346604 | 0.5708315 | 0.893592686 |
| ENSMUSG00000038002 | Cramp1l | protein_coding | -0.293185376 | 3.327462723 | 0.3345249 | 0.5709087 | 0.893592686 |
| ENSMUSG00000021728 | Emb | protein_coding | 0.179601112 | 5.713795227 | 0.3342421 | 0.5710699 | 0.893714873 |
| ENSMUSG00000029167 | Ppargc1a | protein_coding | -0.174774079 | 5.318008391 | 0.3337933 | 0.5713259 | 0.893985347 |
| ENSMUSG00000032998 | Foxj3 | protein_coding | -0.182991515 | 5.043641327 | 0.3336425 | 0.571412 | 0.893989942 |
| ENSMUSG00000027263 | Tubgcp4 | protein_coding | -0.325981341 | 3.016042096 | 0.3334967 | 0.5714952 | 0.893990068 |
| ENSMUSG00000049305 | Ccdc71 | protein_coding | 0.195210131 | 5.392133224 | 0.3332908 | 0.5716128 | 0.894043952 |
| ENSMUSG00000000552 | Zfp385a | protein_coding | 0.246497169 | 4.803536637 | 0.3331218 | 0.5717093 | 0.894064927 |
| ENSMUSG00000024909 | Efemp2 | protein_coding | -0.445117233 | 4.049244831 | 0.334522 | 0.5721677 | 0.894467496 |
| ENSMUSG00000053040 | Aph1c | protein_coding | 0.284811763 | 3.38231935 | 0.3322985 | 0.5721802 | 0.894467496 |
| ENSMUSG00000055723 | Rras2 | protein_coding | -0.590427043 | 2.084447914 | 0.332046 | 0.5723248 | 0.894467496 |
| ENSMUSG00000094707 | A830019P07Rik | lincRNA | -0.51639348 | 2.449636593 | 0.3319012 | 0.5724077 | 0.894467496 |
| ENSMUSG00000001420 | Tmem79 | protein_coding | 0.391048542 | 2.657697046 | 0.3317391 | 0.5725005 | 0.894467496 |
| ENSMUSG00000022508 | Bcl6 | protein_coding | -0.361476886 | 3.048045473 | 0.331626 | 0.5725654 | 0.894467496 |
| ENSMUSG00000020334 | Slc22a4 | protein_coding | 0.336024839 | 3.121545902 | 0.3315835 | 0.5725898 | 0.894467496 |
| ENSMUSG00000021930 | Spryd7 | protein_coding | -0.120222097 | 6.699836475 | 0.3315094 | 0.5726322 | 0.894467496 |
| ENSMUSG00000073154 | 9330158H04Rik | lincRNA | -0.36860368 | 2.924406404 | 0.3312329 | 0.5727908 | 0.894538506 |
| ENSMUSG00000044906 | 4930503L19Rik | protein_coding | 0.466549794 | 2.687705148 | 0.33114 | 0.5728441 | 0.894538506 |
| ENSMUSG00000027985 | Lef1 | protein_coding | -0.671988816 | 3.205573211 | 0.3343663 | 0.5730366 | 0.894626718 |
| ENSMUSG00000062906 | Hdac10 | protein_coding | 0.434944438 | 3.334555254 | 0.3313541 | 0.5730669 | 0.894626718 |
| ENSMUSG00000030232 | Aebp2 | protein_coding | -0.181425287 | 5.211686 | 0.3303373 | 0.573305 | 0.894776291 |
| ENSMUSG00000007029 | Vars | protein_coding | -0.3249569 | 3.074595686 | 0.3302952 | 0.5733292 | 0.894776291 |
| ENSMUSG00000053347 | Zfp943 | protein_coding | -0.344850987 | 4.009394861 | 0.330393 | 0.5735161 | 0.894782721 |
| ENSMUSG00000020513 | Tubd1 | protein_coding | -0.454191557 | 2.857777701 | 0.3298003 | 0.5736137 | 0.894782721 |
| ENSMUSG00000028719 | Cmpk1 | protein_coding | -0.139111195 | 7.30232154 | 0.3295445 | 0.5737609 | 0.894782721 |
| ENSMUSG00000031551 | Ido1 | protein_coding | -0.271477162 | 3.973522036 | 0.3294088 | 0.5738391 | 0.894782721 |
| ENSMUSG00000022433 | Csnk1e | protein_coding | -0.203352153 | 5.460540342 | 0.3292954 | 0.5739043 | 0.894782721 |
| ENSMUSG00000029420 | Rimbp2 | protein_coding | -0.269996471 | 3.871527703 | 0.3292586 | 0.5739255 | 0.894782721 |
| ENSMUSG00000025287 | Acot9 | protein_coding | -0.17236426 | 5.547245871 | 0.3289008 | 0.5741316 | 0.894782721 |
| ENSMUSG00000028407 | Toporsos | protein_coding | 0.262424315 | 3.603594937 | 0.3288831 | 0.5741418 | 0.894782721 |
| ENSMUSG00000025314 | Ptprj | protein_coding | -0.383694853 | 2.738237893 | 0.3288528 | 0.5741594 | 0.894782721 |
| ENSMUSG00000015932 | Dstn | protein_coding | -0.125485657 | 8.557411813 | 0.3288422 | 0.5741654 | 0.894782721 |
| ENSMUSG00000037239 | Spred3 | protein_coding | 0.327333379 | 2.938059179 | 0.3285183 | 0.5743522 | 0.894930981 |
| ENSMUSG00000062190 | Lancl2 | protein_coding | 0.450381332 | 3.077568906 | 0.3284109 | 0.574427 | 0.894930981 |
| ENSMUSG00000015668 | Pdzd11 | protein_coding | 0.115296215 | 7.534398526 | 0.3280051 | 0.5746484 | 0.895146171 |
| ENSMUSG00000022661 | Cd200 | protein_coding | 0.14066292 | 8.070795326 | 0.3277151 | 0.5748159 | 0.895277413 |
| ENSMUSG00000021779 | Thrb | protein_coding | -0.214093619 | 5.798525101 | 0.3274854 | 0.5749614 | 0.895313909 |
| ENSMUSG00000043987 | Cep164 | protein_coding | 0.376768977 | 2.793293955 | 0.3273631 | 0.5750193 | 0.895313909 |
| ENSMUSG00000039678 | Tbc1d13 | protein_coding | -0.270111349 | 3.440285826 | 0.3271811 | 0.5751245 | 0.895313909 |
| ENSMUSG00000026932 | Nacc2 | protein_coding | 0.175232987 | 6.10164232 | 0.3270081 | 0.5752246 | 0.895313909 |
| ENSMUSG00000020594 | Pum2 | protein_coding | -0.125322991 | 7.253718798 | 0.3269545 | 0.5752556 | 0.895313909 |
| ENSMUSG00000057265 | Bbof1 | protein_coding | -0.51244461 | 2.443925944 | 0.326646 | 0.5754342 | 0.895462182 |
| ENSMUSG00000049164 | Zfp518a | protein_coding | -0.331621412 | 2.82800712 | 0.3264315 | 0.5755584 | 0.895525955 |
| ENSMUSG00000015377 | Dennd6b | protein_coding | 0.282988236 | 3.564111178 | 0.3262045 | 0.5756899 | 0.895600331 |
| ENSMUSG00000034709 | Ppp1r21 | protein_coding | -0.161815028 | 5.805066777 | 0.3260447 | 0.5757825 | 0.895600331 |
| ENSMUSG00000030199 | Etv6 | protein_coding | -0.4233957 | 3.028764161 | 0.325804 | 0.5759221 | 0.895600331 |
| ENSMUSG00000021036 | Sptlc2 | protein_coding | 0.277330848 | 3.764679548 | 0.3257743 | 0.5759394 | 0.895600331 |
| ENSMUSG00000030500 | Slc17a6 | protein_coding | -0.599558053 | 6.462160859 | 0.3399587 | 0.5761459 | 0.895740061 |
| ENSMUSG00000032968 | Inha | protein_coding | -0.470095537 | 2.437307666 | 0.3251737 | 0.576288 | 0.895740061 |
| ENSMUSG00000026466 | Tor1aip1 | protein_coding | -0.239446529 | 3.687747395 | 0.3250802 | 0.5763423 | 0.895740061 |
| ENSMUSG00000027833 | Shox2 | protein_coding | -0.805144176 | 4.859036932 | 0.3395647 | 0.5763625 | 0.895740061 |
| ENSMUSG00000015484 | Fam163a | protein_coding | 0.362954128 | 3.346538733 | 0.3248521 | 0.5764748 | 0.895785178 |
| ENSMUSG00000030647 | Ndufc2 | protein_coding | 0.148436191 | 9.492654567 | 0.3243973 | 0.5767392 | 0.896066556 |
| ENSMUSG00000026443 | Lrrn2 | protein_coding | 0.150975365 | 5.853491411 | 0.3240769 | 0.5769257 | 0.896078086 |
| ENSMUSG00000034949 | Zfr2 | protein_coding | 0.258975193 | 3.528318463 | 0.3239446 | 0.5770027 | 0.896078086 |
| ENSMUSG00000065126 | Snord104 | snoRNA | 0.322783854 | 3.124400437 | 0.3237956 | 0.5770895 | 0.896078086 |
| ENSMUSG00000000983 | Wfdc18 | protein_coding | -0.296739771 | 3.370804042 | 0.3237047 | 0.5771424 | 0.896078086 |
| ENSMUSG00000042121 | Ssh1 | protein_coding | -0.192353345 | 4.392451141 | 0.3236688 | 0.5771633 | 0.896078086 |
| ENSMUSG00000043207 | Zmpste24 | protein_coding | 0.20790461 | 4.388046085 | 0.3231298 | 0.5774775 | 0.896436488 |
| ENSMUSG00000027956 | Tmem144 | protein_coding | -0.278011041 | 3.501868317 | 0.3229016 | 0.5776106 | 0.896490105 |
| ENSMUSG00000033355 | Rtp4 | protein_coding | 0.327208443 | 3.126391149 | 0.3226631 | 0.5777498 | 0.896490105 |
| ENSMUSG00000044328 | Trp53i13 | protein_coding | 0.2470735 | 3.714643279 | 0.3226419 | 0.5777622 | 0.896490105 |
| ENSMUSG00000024188 | Luc7l | protein_coding | -0.128603587 | 6.515031364 | 0.3224959 | 0.5778474 | 0.896493046 |
| ENSMUSG00000000759 | Tubgcp3 | protein_coding | 0.199774819 | 4.827960238 | 0.3222068 | 0.5780163 | 0.896527324 |
| ENSMUSG00000021807 | 2700060E02Rik | protein_coding | -0.132222611 | 6.484399322 | 0.3221726 | 0.5780363 | 0.896527324 |
| ENSMUSG00000060743 | H3f3a | protein_coding | -0.214137327 | 3.826424919 | 0.3216503 | 0.5783417 | 0.896701543 |
| ENSMUSG00000018733 | Pex12 | protein_coding | 0.23761159 | 4.360905925 | 0.3215906 | 0.5783766 | 0.896701543 |
| ENSMUSG00000036270 | Edc4 | protein_coding | -0.21845697 | 4.747502669 | 0.3214743 | 0.5784446 | 0.896701543 |
| ENSMUSG00000022453 | Naga | protein_coding | 0.196549342 | 5.324071958 | 0.3211231 | 0.5786502 | 0.896701543 |
| ENSMUSG00000021699 | Pde4d | protein_coding | -0.154597933 | 5.435247316 | 0.3210547 | 0.5786903 | 0.896701543 |
| ENSMUSG00000015882 | Lcorl | protein_coding | 0.324100658 | 3.036228734 | 0.3208075 | 0.5788351 | 0.896701543 |
| ENSMUSG00000034648 | Lrrn1 | protein_coding | -0.242040373 | 4.623744272 | 0.3207875 | 0.5788468 | 0.896701543 |
| ENSMUSG00000029534 | St7 | protein_coding | 0.192790326 | 4.278496417 | 0.3207793 | 0.5788516 | 0.896701543 |
| ENSMUSG00000045761 | Togaram2 | protein_coding | -0.513235231 | 2.528370165 | 0.3206951 | 0.578901 | 0.896701543 |

| ensembl_gene_id | external_gene_name | gene_biotype | log2 (fold change) | log2 (counts per million) | F-value | p-value | False discovery rate |
|---|---|---|---|---|---|---|---|
| ENSMUSG00000027429 | Sec23b | protein_coding | -0.198016748 | 5.954301935 | 0.3204088 | 0.5790815 | 0.896701543 |
| ENSMUSG00000028549 | Itgb3bp | protein_coding | 0.19969194 | 4.194755483 | 0.3203612 | 0.5790968 | 0.896701543 |
| ENSMUSG00000035769 | Xylb | protein_coding | 0.31110071 | 3.277310768 | 0.3199665 | 0.5793284 | 0.896701543 |
| ENSMUSG00000021928 | Ebpl | protein_coding | -0.191638103 | 4.421422157 | 0.3198062 | 0.5794225 | 0.896701543 |
| ENSMUSG00000044365 | Cxxc4 | protein_coding | -0.181771696 | 5.379609679 | 0.3197545 | 0.5794529 | 0.896701543 |
| ENSMUSG00000097068 | Gm26760 | lincRNA | -0.39829919 | 2.564848767 | 0.3196443 | 0.5795176 | 0.896701543 |
| ENSMUSG00000073002 | Vamp5 | protein_coding | -0.351874288 | 3.080028733 | 0.3196022 | 0.5795424 | 0.896701543 |
| ENSMUSG00000042208 | 0610010F05Rik | protein_coding | -0.166622701 | 4.933790896 | 0.3195616 | 0.5795662 | 0.896701543 |
| ENSMUSG00000020928 | Higd1b | protein_coding | 0.37712433 | 2.886518888 | 0.3193028 | 0.5797183 | 0.896716916 |
| ENSMUSG00000027194 | Ttc17 | protein_coding | -0.22040502 | 3.704452578 | 0.319161 | 0.5798017 | 0.896716916 |
| ENSMUSG00000024410 | 3110002H16Rik | protein_coding | 0.178469208 | 5.443924977 | 0.3191191 | 0.5798263 | 0.896716916 |
| ENSMUSG00000019295 | Tmem129 | protein_coding | 0.183002739 | 5.344032066 | 0.3188306 | 0.5799961 | 0.896850393 |
| ENSMUSG00000008226 | Scrn3 | protein_coding | -0.215187766 | 5.023483344 | 0.3186763 | 0.5800868 | 0.896861777 |
| ENSMUSG00000024927 | Rela | protein_coding | -0.204230538 | 5.16523938 | 0.3183285 | 0.5802916 | 0.896979917 |
| ENSMUSG00000026150 | Mff | protein_coding | -0.13856797 | 7.330525342 | 0.3182633 | 0.5803301 | 0.896979917 |
| ENSMUSG00000020359 | Phykpl | protein_coding | 0.25121022 | 3.634499591 | 0.3179898 | 0.5804912 | 0.89710006 |
| ENSMUSG00000054893 | Zfp667 | protein_coding | -0.178736658 | 5.479913346 | 0.3176246 | 0.5807066 | 0.897224202 |
| ENSMUSG00000063694 | Cycs | protein_coding | 0.166422105 | 5.154449995 | 0.3174355 | 0.5808182 | 0.897224202 |
| ENSMUSG00000040618 | Pck2 | protein_coding | 0.185426167 | 5.48517705 | 0.3173126 | 0.5808907 | 0.897224202 |
| ENSMUSG00000056917 | Sipa1 | protein_coding | 0.262236034 | 4.245049254 | 0.3172879 | 0.5809053 | 0.897224202 |
| ENSMUSG00000061474 | Mrps36 | protein_coding | 0.193313122 | 4.525805202 | 0.3171039 | 0.5810139 | 0.89726308 |
| ENSMUSG00000002486 | Tchp | protein_coding | 0.334743697 | 2.932409377 | 0.3168748 | 0.5811493 | 0.897343223 |
| ENSMUSG00000018378 | Cuedc1 | protein_coding | 0.238997538 | 3.943362675 | 0.3161908 | 0.5815537 | 0.897590392 |
| ENSMUSG00000051413 | Plagl2 | protein_coding | -0.306472868 | 2.963117602 | 0.3160083 | 0.5816617 | 0.897590392 |
| ENSMUSG00000027894 | Slc6a17 | protein_coding | 0.146411798 | 6.121089919 | 0.3157046 | 0.5818416 | 0.897590392 |
| ENSMUSG00000071655 | Ubxn1 | protein_coding | -0.11895282 | 6.632327151 | 0.3156379 | 0.5818811 | 0.897590392 |
| ENSMUSG00000026427 | Eif2d | protein_coding | -0.174565884 | 5.489170157 | 0.3156142 | 0.5818951 | 0.897590392 |
| ENSMUSG00000032295 | Man2c1 | protein_coding | 0.216680746 | 5.041759755 | 0.3152523 | 0.5821096 | 0.897590392 |
| ENSMUSG00000039987 | Phtf2 | protein_coding | 0.331611518 | 2.961224913 | 0.3151894 | 0.5821469 | 0.897590392 |
| ENSMUSG00000009207 | Lnpk | protein_coding | -0.163457818 | 5.532369626 | 0.3151271 | 0.5821838 | 0.897590392 |
| ENSMUSG00000026894 | Morn5 | protein_coding | -0.328539919 | 3.188588865 | 0.3151095 | 0.5821942 | 0.897590392 |
| ENSMUSG00000034673 | Pbx2 | protein_coding | -0.204146821 | 4.721255609 | 0.3150199 | 0.5822474 | 0.897590392 |
| ENSMUSG00000021771 | Vdac2 | protein_coding | 0.122716712 | 9.130915207 | 0.3149274 | 0.5823023 | 0.897590392 |
| ENSMUSG00000035958 | Tdp2 | protein_coding | -0.235702018 | 3.702396804 | 0.3147409 | 0.5824129 | 0.897590392 |
| ENSMUSG00000028882 | Ppp1r8 | protein_coding | 0.158226378 | 5.707958089 | 0.3144998 | 0.5825561 | 0.897590392 |
| ENSMUSG00000024854 | Pold4 | protein_coding | -0.163010699 | 5.691467525 | 0.3141874 | 0.5827416 | 0.897590392 |
| ENSMUSG00000039156 | Stim2 | protein_coding | 0.209590249 | 4.815811894 | 0.3141493 | 0.5827643 | 0.897590392 |
| ENSMUSG00000013736 | Trnt1 | protein_coding | 0.177367848 | 6.412082725 | 0.3141353 | 0.5827851 | 0.897590392 |
| ENSMUSG00000040797 | Iqsec3 | protein_coding | -0.206217374 | 5.413933185 | 0.3139487 | 0.5828835 | 0.897590392 |
| ENSMUSG00000042429 | Adora1 | protein_coding | -0.191115656 | 5.825502439 | 0.3138782 | 0.5829254 | 0.897590392 |
| ENSMUSG00000078622 | Ccdc47 | protein_coding | 0.183203546 | 5.700669994 | 0.3137059 | 0.5830279 | 0.897590392 |
| ENSMUSG00000020775 | Mrpl38 | protein_coding | 0.193213821 | 6.000364874 | 0.3136028 | 0.5831018 | 0.897590392 |
| ENSMUSG00000070056 | Mfhas1 | protein_coding | -0.186515749 | 4.650240112 | 0.313061 | 0.5834117 | 0.897590392 |
| ENSMUSG00000036537 | Rnf113a1 | protein_coding | 0.188900751 | 5.094406699 | 0.3130537 | 0.5834161 | 0.897590392 |
| ENSMUSG00000026519 | Tmem63a | protein_coding | -0.23975761 | 4.576866555 | 0.3129116 | 0.5835008 | 0.897590392 |
| ENSMUSG00000042425 | Frmpd3 | protein_coding | -0.397306207 | 2.928074851 | 0.3128572 | 0.5835332 | 0.897590392 |
| ENSMUSG00000016757 | Ttll12 | protein_coding | -0.31128555 | 3.701757573 | 0.3126607 | 0.5836503 | 0.897590392 |
| ENSMUSG00000073563 | Csnk1g3 | protein_coding | -0.171197805 | 5.131053326 | 0.312607 | 0.5836823 | 0.897590392 |
| ENSMUSG00000025964 | Adam23 | protein_coding | 0.141182742 | 6.261421837 | 0.312485 | 0.5837551 | 0.897590392 |
| ENSMUSG00000029701 | Rbm28 | protein_coding | -0.187519648 | 5.289942024 | 0.3121867 | 0.583933 | 0.897590392 |
| ENSMUSG00000036099 | Vezt | protein_coding | -0.205996174 | 5.092441276 | 0.3120782 | 0.5839977 | 0.897590392 |
| ENSMUSG00000021773 | Comtd1 | protein_coding | 0.328052093 | 3.079362413 | 0.3120382 | 0.5840216 | 0.897590392 |
| ENSMUSG00000025867 | Cplx2 | protein_coding | -0.123385502 | 8.226536737 | 0.3119712 | 0.5840616 | 0.897590392 |
| ENSMUSG00000026553 | Copa | protein_coding | 0.164603321 | 6.898510448 | 0.3116652 | 0.5842568 | 0.897590392 |
| ENSMUSG00000033460 | Armcx1 | protein_coding | 0.148918808 | 6.736641341 | 0.3116436 | 0.5842573 | 0.897590392 |
| ENSMUSG00000045917 | Tmem268 | protein_coding | -0.230560241 | 4.102658642 | 0.3115452 | 0.5843161 | 0.897590392 |
| ENSMUSG00000026227 | 2810459M11Rik | protein_coding | -0.52501849 | 2.526631939 | 0.3114361 | 0.5843813 | 0.897590392 |
| ENSMUSG00000025875 | Tspan17 | protein_coding | 0.335306679 | 3.463116363 | 0.3114154 | 0.5843937 | 0.897590392 |
| ENSMUSG00000026933 | Camsap1 | protein_coding | -0.175515412 | 5.571069479 | 0.3113414 | 0.5844379 | 0.897590392 |
| ENSMUSG00000058267 | Mrps14 | protein_coding | 0.118212853 | 8.757600693 | 0.3112097 | 0.5845166 | 0.897590392 |
| ENSMUSG00000034832 | Tet3 | protein_coding | -0.201727114 | 4.60386671 | 0.3109464 | 0.5846741 | 0.897590392 |
| ENSMUSG00000006412 | Pfdn2 | protein_coding | 0.131642288 | 8.392453779 | 0.3108399 | 0.5847378 | 0.897590392 |
| ENSMUSG00000024947 | Men1 | protein_coding | -0.177802694 | 5.013387401 | 0.3107812 | 0.584773 | 0.897590392 |
| ENSMUSG00000030096 | Slc6a6 | protein_coding | -0.160594522 | 5.537140466 | 0.3106143 | 0.5848729 | 0.897590392 |
| ENSMUSG00000023852 | Chd1 | protein_coding | -0.11524421 | 6.615970631 | 0.3104611 | 0.5849646 | 0.897590392 |
| ENSMUSG00000021477 | Ctsl | protein_coding | -0.125300344 | 7.359115582 | 0.3103674 | 0.5850207 | 0.897590392 |
| ENSMUSG00000044224 | Dnajc21 | protein_coding | 0.204058591 | 4.343740295 | 0.3102924 | 0.5850657 | 0.897590392 |
| ENSMUSG00000027679 | Dnajc19 | protein_coding | 0.138827468 | 6.113242534 | 0.3098061 | 0.5853572 | 0.897909586 |
| ENSMUSG00000032202 | Rab27a | protein_coding | 0.278019132 | 3.891648485 | 0.3095672 | 0.5855005 | 0.897912404 |
| ENSMUSG00000020785 | Camkk1 | protein_coding | 0.179361308 | 5.651688003 | 0.3093886 | 0.5856077 | 0.897912404 |
| ENSMUSG00000074064 | Mlycd | protein_coding | 0.241530319 | 4.413333649 | 0.3093855 | 0.5856096 | 0.897912404 |
| ENSMUSG00000035783 | Acta2 | protein_coding | -0.418600242 | 3.195979537 | 0.3085734 | 0.5861099 | 0.89855142 |
| ENSMUSG00000013076 | Amotl1 | protein_coding | 0.294433079 | 7.001349671 | 0.313642 | 0.5862505 | 0.898638849 |
| ENSMUSG00000035247 | Hectd1 | protein_coding | -0.242167192 | 4.236072465 | 0.3079727 | 0.586459 | 0.898724608 |
| ENSMUSG00000025237 | Parp6 | protein_coding | -0.139778628 | 5.991046716 | 0.3079484 | 0.5864736 | 0.898724608 |
| ENSMUSG00000027078 | Ube2l6 | protein_coding | 0.220710408 | 4.029431324 | 0.3077379 | 0.5866004 | 0.898790829 |
| ENSMUSG00000020415 | Pttg1 | protein_coding | -0.105178864 | 7.486446678 | 0.3074152 | 0.5867948 | 0.898838637 |
| ENSMUSG00000049940 | Pgrmc2 | protein_coding | -0.167791201 | 5.786279103 | 0.3071916 | 0.5869297 | 0.898838637 |

| ensembl_gene_id | external_gene_name | gene_biotype | log2 (fold change) | log2 (counts per million) | F-value | p-value | False discovery rate |
|---|---|---|---|---|---|---|---|
| ENSMUSG00000103183 | Gm37090 | TEC | 0.612428256 | 2.365240019 | 0.3071736 | 0.5869529 | 0.898838637 |
| ENSMUSG00000036061 | Smug1 | protein_coding | 0.281740278 | 3.632797409 | 0.3071315 | 0.586966 | 0.898838637 |
| ENSMUSG00000031189 | Aff2 | protein_coding | -0.364223773 | 2.77269957 | 0.3063515 | 0.5874368 | 0.899353047 |
| ENSMUSG00000027502 | Rtfdc1 | protein_coding | 0.129138045 | 7.356181442 | 0.3061569 | 0.5875544 | 0.899353047 |
| ENSMUSG00000035764 | Fbxo45 | protein_coding | 0.192021141 | 4.888335855 | 0.3060727 | 0.5876054 | 0.899353047 |
| ENSMUSG00000022787 | Wdr53 | protein_coding | 0.182017654 | 4.781929122 | 0.3060213 | 0.5876364 | 0.899353047 |
| ENSMUSG00000024091 | Vapa | protein_coding | -0.107573438 | 8.549084186 | 0.3057005 | 0.5878305 | 0.899518705 |
| ENSMUSG00000027304 | Rtf1 | protein_coding | -0.184884542 | 5.315831213 | 0.3053469 | 0.5880445 | 0.899518705 |
| ENSMUSG00000051351 | Zfp46 | protein_coding | 0.268653229 | 3.520899645 | 0.3051624 | 0.5881562 | 0.899518705 |
| ENSMUSG00000032126 | Hmbs | protein_coding | -0.276876504 | 3.707314394 | 0.3050744 | 0.5882096 | 0.899518705 |
| ENSMUSG00000034889 | Cactin | protein_coding | 0.448215155 | 2.749299879 | 0.3050668 | 0.5882142 | 0.899518705 |
| ENSMUSG00000057715 | A830018L16Rik | protein_coding | -0.19668622 | 4.937030067 | 0.3050055 | 0.5882513 | 0.899518705 |
| ENSMUSG00000042035 | Igsf3 | protein_coding | 0.173568451 | 5.47660172 | 0.3047896 | 0.5883822 | 0.899518705 |
| ENSMUSG00000028161 | Ppp3ca | protein_coding | -0.111922437 | 9.571254196 | 0.3047373 | 0.5884139 | 0.899518705 |
| ENSMUSG00000062101 | Zfp119b | protein_coding | -0.490410831 | 2.779244344 | 0.3045747 | 0.5885248 | 0.899560291 |
| ENSMUSG00000010307 | Tmem86a | protein_coding | -0.234298724 | 4.325601805 | 0.3043182 | 0.5886681 | 0.899651543 |
| ENSMUSG00000002625 | Akap8l | protein_coding | 0.155306792 | 5.114808277 | 0.3040797 | 0.5888129 | 0.899744942 |
| ENSMUSG00000027519 | Rab22a | protein_coding | -0.144516204 | 5.427502453 | 0.3037924 | 0.5889874 | 0.899833026 |
| ENSMUSG00000044636 | Csrnp2 | protein_coding | -0.243331636 | 4.041868163 | 0.3036019 | 0.5891031 | 0.899833026 |
| ENSMUSG00000044149 | Nkrf | protein_coding | 0.214717639 | 4.948426562 | 0.303359 | 0.5892508 | 0.899833026 |
| ENSMUSG00000074607 | Tox2 | protein_coding | -0.255476732 | 5.997810365 | 0.3048332 | 0.589353 | 0.899833026 |
| ENSMUSG00000063535 | Zfp773 | protein_coding | -0.555202024 | 2.375556669 | 0.3031738 | 0.5893635 | 0.899833026 |
| ENSMUSG00000045038 | Prkce | protein_coding | -0.124587512 | 7.599534377 | 0.3031587 | 0.5893726 | 0.899833026 |
| ENSMUSG00000054423 | Cadps | protein_coding | 0.116193839 | 7.090859496 | 0.3028689 | 0.589549 | 0.899974566 |
| ENSMUSG00000026425 | Srgap2 | protein_coding | -0.158971121 | 5.390911162 | 0.3026148 | 0.5897038 | 0.900025962 |
| ENSMUSG00000038860 | Garnl3 | protein_coding | 0.20113861 | 5.181698089 | 0.3023838 | 0.5898445 | 0.900025962 |
| ENSMUSG00000026021 | Sumo1 | protein_coding | 0.11763718 | 8.260689262 | 0.3023269 | 0.5898792 | 0.900025962 |
| ENSMUSG00000061393 | Acvr2b | protein_coding | -0.40188241 | 2.642841186 | 0.3022641 | 0.5899175 | 0.900025962 |
| ENSMUSG00000053453 | Thoc7 | protein_coding | 0.155193316 | 7.306993074 | 0.3020163 | 0.5900808 | 0.900147374 |
| ENSMUSG00000051790 | Nlgn2 | protein_coding | 0.138348958 | 6.043379295 | 0.3015889 | 0.5903294 | 0.900398811 |
| ENSMUSG00000044155 | Lsm8 | protein_coding | 0.168420041 | 6.069492638 | 0.3012274 | 0.5905501 | 0.90060781 |
| ENSMUSG00000023809 | Rps6ka2 | protein_coding | 0.231671348 | 4.225923846 | 0.3004746 | 0.5910105 | 0.901084916 |
| ENSMUSG00000059659 | Gm10069 | antisense_RNA | 0.32287286 | 3.082248651 | 0.3004417 | 0.5910306 | 0.901084916 |
| ENSMUSG00000025133 | Ints4 | protein_coding | -0.207059822 | 4.854161641 | 0.3001845 | 0.5911881 | 0.90119727 |
| ENSMUSG00000061887 | Ssbp3 | protein_coding | 0.151851228 | 8.011994118 | 0.2994788 | 0.5916327 | 0.901405016 |
| ENSMUSG00000085389 | 1700003M07Rik | lincRNA | 0.271877972 | 3.252357839 | 0.2990771 | 0.591867 | 0.901405016 |
| ENSMUSG00000074088 | Snrnp40 | protein_coding | 0.196797803 | 5.159838166 | 0.2990534 | 0.5918816 | 0.901405016 |
| ENSMUSG00000027634 | Ndrg3 | protein_coding | 0.167807088 | 9.406259515 | 0.2989708 | 0.5919444 | 0.901405016 |
| ENSMUSG00000007880 | Arid1a | protein_coding | -0.18583389 | 6.022103419 | 0.2989661 | 0.5919473 | 0.901405016 |
| ENSMUSG00000026765 | Lypd6b | protein_coding | 0.211586864 | 4.229487079 | 0.298928 | 0.5919585 | 0.901405016 |
| ENSMUSG00000036790 | Slitrk2 | protein_coding | -0.248611359 | 3.373395421 | 0.2989091 | 0.5919702 | 0.901405016 |
| ENSMUSG00000033454 | Zbtb1 | protein_coding | -0.393313927 | 2.644817207 | 0.2988687 | 0.591995 | 0.901405016 |
| ENSMUSG00000050029 | Rap2c | protein_coding | -0.215830648 | 4.081772515 | 0.2985757 | 0.592175 | 0.90155143 |
| ENSMUSG00000072875 | Gpr27 | protein_coding | -0.242850053 | 4.238632282 | 0.2982565 | 0.5923712 | 0.901690498 |
| ENSMUSG00000106106 | CT010467.1 | rRNA | 0.139799903 | 10.43028008 | 0.2981543 | 0.5924341 | 0.901690498 |
| ENSMUSG00000024962 | Vegfb | protein_coding | 0.175560857 | 5.333725805 | 0.2978096 | 0.5926461 | 0.901765044 |
| ENSMUSG00000062961 | Ccdc177 | protein_coding | -0.251087439 | 3.767734098 | 0.2977798 | 0.5926645 | 0.901765044 |
| ENSMUSG00000055371 | Stam2 | protein_coding | -0.199119347 | 4.296966598 | 0.2976659 | 0.5927346 | 0.901765044 |
| ENSMUSG00000014850 | Msh3 | protein_coding | -0.243692001 | 3.962347534 | 0.2972252 | 0.5930061 | 0.902050516 |
| ENSMUSG00000024381 | Bin1 | protein_coding | -0.148853331 | 6.261509006 | 0.296899 | 0.5932072 | 0.902228755 |
| ENSMUSG00000038954 | Supt3 | protein_coding | 0.19882417 | 4.465072397 | 0.2966913 | 0.5933354 | 0.902296049 |
| ENSMUSG00000020280 | Pus10 | protein_coding | 0.24899735 | 3.556230994 | 0.2963674 | 0.5935353 | 0.902424526 |
| ENSMUSG00000028443 | Nudt2 | protein_coding | 0.124360378 | 6.671103307 | 0.2962562 | 0.593604 | 0.902424526 |
| ENSMUSG00000027981 | Rnpc3 | protein_coding | -0.30853464 | 2.974047067 | 0.2961002 | 0.5937004 | 0.902424526 |
| ENSMUSG00000034543 | Morc2a | protein_coding | 0.213835785 | 3.987447383 | 0.2960109 | 0.5937556 | 0.902424526 |
| ENSMUSG00000040430 | Pitpnc1 | protein_coding | 0.14321685 | 7.311017842 | 0.2958225 | 0.593872 | 0.902473988 |
| ENSMUSG00000022538 | Lsg1 | protein_coding | -0.151004872 | 5.240216218 | 0.2956504 | 0.5939784 | 0.902508126 |
| ENSMUSG00000021076 | Actr10 | protein_coding | 0.119167144 | 8.860257403 | 0.2951546 | 0.5942853 | 0.902554595 |
| ENSMUSG00000086968 | 4933431E20Rik | antisense_RNA | -0.25637935 | 4.024833679 | 0.295054 | 0.5943475 | 0.902554595 |
| ENSMUSG00000035232 | Pdk3 | protein_coding | -0.356122379 | 4.387157413 | 0.2965707 | 0.5943637 | 0.902554595 |
| ENSMUSG00000075271 | Ttc30a1 | protein_coding | -0.387033419 | 2.943196988 | 0.2948222 | 0.5944912 | 0.902554595 |
| ENSMUSG00000054942 | Miga1 | protein_coding | -0.156681536 | 5.453316848 | 0.2947879 | 0.5945124 | 0.902554595 |
| ENSMUSG00000043421 | Hilpda | protein_coding | 0.180755425 | 4.813175792 | 0.2945899 | 0.5946352 | 0.902554595 |
| ENSMUSG00000101930 | Gm5441 | antisense_RNA | 0.128332015 | 6.242236178 | 0.2943221 | 0.5948012 | 0.902554595 |
| ENSMUSG00000021103 | Mnat1 | protein_coding | 0.129988806 | 6.011421216 | 0.2942908 | 0.5948206 | 0.902554595 |
| ENSMUSG00000029603 | Dtx1 | protein_coding | -0.171726325 | 5.30201011 | 0.294157 | 0.5949037 | 0.902554595 |
| ENSMUSG00000062127 | Cttnbp2nl | protein_coding | -0.231108472 | 3.730858449 | 0.2941219 | 0.5949254 | 0.902554595 |
| ENSMUSG00000029413 | Naaa | protein_coding | 0.248508164 | 4.870785773 | 0.2941302 | 0.5949323 | 0.902554595 |
| ENSMUSG00000057522 | Spop | protein_coding | -0.138073023 | 8.272857424 | 0.2933681 | 0.5953937 | 0.903074198 |
| ENSMUSG00000021453 | Gadd45g | protein_coding | -0.196359476 | 5.83083445 | 0.2933084 | 0.5954428 | 0.903074198 |
| ENSMUSG00000020657 | Dnajc27 | protein_coding | -0.116209812 | 6.523434476 | 0.2929565 | 0.5956496 | 0.903089156 |
| ENSMUSG00000031631 | Cfap97 | protein_coding | -0.164311456 | 5.248263559 | 0.2928908 | 0.5956905 | 0.903089156 |
| ENSMUSG00000018593 | Sparc | protein_coding | -0.21912007 | 9.204871844 | 0.2942821 | 0.5958523 | 0.903089156 |
| ENSMUSG00000048794 | Cfap100 | protein_coding | 0.442464972 | 2.308252603 | 0.2924554 | 0.5959616 | 0.903089156 |
| ENSMUSG00000022375 | Lrrc6 | protein_coding | 0.339355467 | 3.852469351 | 0.2928749 | 0.5960066 | 0.903089156 |
| ENSMUSG00000064345 | mt-Nd2 | protein_coding | 0.195460942 | 10.31322046 | 0.2929538 | 0.5961166 | 0.903089156 |
| ENSMUSG00000024077 | Strn | protein_coding | -0.134258293 | 6.419242611 | 0.2921887 | 0.5961277 | 0.903089156 |
| ENSMUSG00000057322 | Rpl38 | protein_coding | 0.202319479 | 4.578530249 | 0.2921709 | 0.5961389 | 0.903089156 |

| ensembl_gene_id | external_gene_name | gene_biotype | log2 (fold change) | log2 (counts per million) | F-value | p-value | False discovery rate |
|---|---|---|---|---|---|---|---|
| ENSMUSG00000020216 | Jsrp1 | protein_coding | -0.314629174 | 3.231822087 | 0.2920592 | 0.5962085 | 0.903089156 |
| ENSMUSG00000047417 | Rexo1 | protein_coding | 0.223040044 | 4.115535995 | 0.2916549 | 0.5964606 | 0.90334384 |
| ENSMUSG00000022827 | Rabl3 | protein_coding | 0.225627461 | 4.725457774 | 0.2914368 | 0.5965968 | 0.903344333 |
| ENSMUSG00000024646 | Cyb5a | protein_coding | -0.136392585 | 8.28430428 | 0.2912905 | 0.5966882 | 0.903344333 |
| ENSMUSG00000033918 | Parl | protein_coding | 0.151249775 | 6.333580154 | 0.2912197 | 0.5967324 | 0.903344333 |
| ENSMUSG00000032418 | Me1 | protein_coding | -0.198567096 | 4.417696933 | 0.2909524 | 0.5968994 | 0.903344333 |
| ENSMUSG00000022377 | Asap1 | protein_coding | -0.182862985 | 5.303138934 | 0.2906807 | 0.5970692 | 0.903344333 |
| ENSMUSG00000098112 | Bin2 | protein_coding | -0.566103087 | 2.31569101 | 0.2906704 | 0.5970757 | 0.903344333 |
| ENSMUSG00000057054 | Inca1 | protein_coding | 0.223382766 | 3.848387541 | 0.2904983 | 0.5971833 | 0.903344333 |
| ENSMUSG00000093803 | Ppp2r3d | protein_coding | 0.245470445 | 3.402143341 | 0.290288 | 0.597315 | 0.903344333 |
| ENSMUSG00000027014 | Cwc22 | protein_coding | -0.284754937 | 3.187849467 | 0.2902519 | 0.5973375 | 0.903344333 |
| ENSMUSG00000032018 | Sc5d | protein_coding | 0.119610363 | 6.268769391 | 0.2901987 | 0.5973709 | 0.903344333 |
| ENSMUSG00000035901 | Dennd5a | protein_coding | -0.124257906 | 6.827601737 | 0.290176 | 0.5973851 | 0.903344333 |
| ENSMUSG00000024429 | Gnl1 | protein_coding | 0.123623224 | 6.405451743 | 0.2900213 | 0.597482 | 0.903363821 |
| ENSMUSG00000026926 | Pmpca | protein_coding | -0.183186782 | 6.246171186 | 0.2896539 | 0.5977241 | 0.903575304 |
| ENSMUSG00000029547 | Ints1 | protein_coding | 0.276291233 | 4.190595333 | 0.2895489 | 0.5977899 | 0.903575304 |
| ENSMUSG00000035277 | Arx | protein_coding | -0.298741197 | 3.12653984 | 0.2892995 | 0.5979344 | 0.903666678 |
| ENSMUSG00000021376 | Tpmt | protein_coding | -0.208542481 | 3.970889341 | 0.2889667 | 0.5981432 | 0.903855228 |
| ENSMUSG00000037072 | Selenof | protein_coding | -0.12920841 | 8.841278563 | 0.2878669 | 0.5988345 | 0.904445931 |
| ENSMUSG00000030057 | Cnbp | protein_coding | 0.110847313 | 10.42446665 | 0.2878527 | 0.5988435 | 0.904445931 |
| ENSMUSG00000056755 | Grm7 | protein_coding | -0.185968172 | 5.387431684 | 0.2878453 | 0.5988481 | 0.904445931 |
| ENSMUSG00000024068 | Spast | protein_coding | -0.227970613 | 4.623684832 | 0.2878097 | 0.5988706 | 0.904445931 |
| ENSMUSG00000022546 | Gpt | protein_coding | 0.249214974 | 4.057947977 | 0.287503 | 0.5990637 | 0.904610551 |
| ENSMUSG00000033880 | Lgals3bp | protein_coding | 0.297772635 | 4.716344849 | 0.2879664 | 0.5993647 | 0.904881818 |
| ENSMUSG00000026249 | Serpine2 | protein_coding | 0.161835335 | 7.864507227 | 0.2869301 | 0.5994679 | 0.904881818 |
| ENSMUSG00000041215 | Yeats2 | protein_coding | -0.297398905 | 5.218110279 | 0.2884286 | 0.5994958 | 0.904881818 |
| ENSMUSG00000026181 | Ppm1f | protein_coding | 0.178276301 | 5.832316627 | 0.2866822 | 0.5995811 | 0.904883596 |
| ENSMUSG00000083695 | Rnf138rt1 | protein_coding | 0.315653691 | 3.793234133 | 0.2863821 | 0.5997824 | 0.904989824 |
| ENSMUSG00000028796 | Phc2 | protein_coding | -0.157853322 | 5.428602898 | 0.2861304 | 0.5999295 | 0.904989824 |
| ENSMUSG00000034755 | Pcdh11x | protein_coding | 0.171492417 | 5.168562025 | 0.2859223 | 0.6000611 | 0.904989824 |
| ENSMUSG00000061360 | Phf5a | protein_coding | 0.141920451 | 6.590204573 | 0.2858767 | 0.6000899 | 0.904989824 |
| ENSMUSG00000029821 | Dfna5 | protein_coding | -0.181878541 | 7.575944082 | 0.2865009 | 0.6001549 | 0.904989824 |
| ENSMUSG00000031556 | Tm2d2 | protein_coding | 0.126522611 | 7.927169006 | 0.2857277 | 0.6001841 | 0.904989824 |
| ENSMUSG00000047153 | Khnyn | protein_coding | 0.42255773 | 2.890848492 | 0.2856571 | 0.6002406 | 0.904989824 |
| ENSMUSG00000045259 | Klhdc9 | protein_coding | 0.206800131 | 4.511997835 | 0.2853473 | 0.6004248 | 0.905140578 |
| ENSMUSG00000021371 | Mcur1 | protein_coding | -0.305608048 | 3.262011593 | 0.2847141 | 0.6008259 | 0.905496153 |
| ENSMUSG00000059361 | Nrsn2 | protein_coding | 0.272796846 | 3.125861112 | 0.2847062 | 0.6008309 | 0.905496153 |
| ENSMUSG00000071757 | Zhx2 | protein_coding | -0.296643834 | 3.222623385 | 0.2844195 | 0.6010127 | 0.905496153 |
| ENSMUSG00000024970 | AI846148 | protein_coding | -0.213360202 | 4.005249017 | 0.284258 | 0.6011151 | 0.905496153 |
| ENSMUSG00000039936 | Pik3cd | protein_coding | 0.400024743 | 2.828825237 | 0.2841109 | 0.6012085 | 0.905496153 |
| ENSMUSG00000029311 | Hsd17b11 | protein_coding | -0.332208218 | 3.207451278 | 0.2840967 | 0.6012175 | 0.905496153 |
| ENSMUSG00000034263 | Ints14 | protein_coding | 0.192967397 | 4.562746553 | 0.2838687 | 0.6013622 | 0.905496153 |
| ENSMUSG00000029343 | Crybb1 | protein_coding | -0.524733924 | 2.065168389 | 0.2832929 | 0.6017282 | 0.905496153 |
| ENSMUSG00000039410 | Prdm16 | protein_coding | 0.19022939 | 5.45472691 | 0.2830822 | 0.6018622 | 0.905496153 |
| ENSMUSG00000050730 | Arhgap42 | protein_coding | -0.585597748 | 2.211125599 | 0.2829888 | 0.6019216 | 0.905496153 |
| ENSMUSG00000068742 | Cry2 | protein_coding | -0.151575255 | 5.848385492 | 0.2826905 | 0.6021116 | 0.905496153 |
| ENSMUSG00000025223 | Ldb1 | protein_coding | 0.167500177 | 5.148441697 | 0.2826715 | 0.6021237 | 0.905496153 |
| ENSMUSG00000040964 | Arhgef10l | protein_coding | -0.239214487 | 3.819511373 | 0.282391 | 0.6023023 | 0.905496153 |
| ENSMUSG00000029366 | Dck | protein_coding | -0.197276605 | 4.599533717 | 0.282346 | 0.602331 | 0.905496153 |
| ENSMUSG00000029151 | Slc30a3 | protein_coding | -0.606616491 | 2.706361497 | 0.2832345 | 0.6023525 | 0.905496153 |
| ENSMUSG00000020814 | Mxra7 | protein_coding | -0.227808814 | 3.90545563 | 0.2822907 | 0.6023662 | 0.905496153 |
| ENSMUSG00000045576 | St7l | protein_coding | -0.211305042 | 4.455683809 | 0.2822771 | 0.6023749 | 0.905496153 |
| ENSMUSG00000026102 | Inpp1 | protein_coding | 0.143467854 | 5.806926255 | 0.2821367 | 0.6024644 | 0.905496153 |
| ENSMUSG00000048029 | Eno4 | protein_coding | -0.306044054 | 3.208913066 | 0.282078 | 0.6025019 | 0.905496153 |
| ENSMUSG00000028698 | Pik3r3 | protein_coding | -0.248545905 | 3.624051346 | 0.2819586 | 0.602578 | 0.905496153 |
| ENSMUSG00000000295 | Hddc2 | protein_coding | 0.133299389 | 7.101080233 | 0.2818503 | 0.6026471 | 0.905496153 |
| ENSMUSG00000039230 | Tbcd | protein_coding | -0.181110879 | 4.638417579 | 0.2817186 | 0.6027312 | 0.905496153 |
| ENSMUSG00000060373 | Hnrnpc | protein_coding | 0.10651132 | 7.866966741 | 0.2815775 | 0.6028212 | 0.905496153 |
| ENSMUSG00000047976 | Kcna1 | protein_coding | -0.21082796 | 4.522790484 | 0.2815713 | 0.6028252 | 0.905496153 |
| ENSMUSG00000029064 | Gnb1 | protein_coding | 0.189660984 | 12.19785351 | 0.2817497 | 0.6028803 | 0.905496153 |
| ENSMUSG00000022763 | Aifm3 | protein_coding | -0.178228472 | 5.137190422 | 0.2813739 | 0.6029512 | 0.905496153 |
| ENSMUSG00000027288 | Zfp106 | protein_coding | -0.149811013 | 5.455991824 | 0.2812034 | 0.6030601 | 0.905496153 |
| ENSMUSG00000024182 | Axin1 | protein_coding | -0.196645054 | 4.05334865 | 0.2811222 | 0.603112 | 0.905496153 |
| ENSMUSG00000037418 | Best1 | protein_coding | -0.440540197 | 2.896601473 | 0.2810428 | 0.6031745 | 0.905496153 |
| ENSMUSG00000027993 | Trim2 | protein_coding | -0.110046836 | 7.573465912 | 0.2810051 | 0.6031869 | 0.905496153 |
| ENSMUSG00000039463 | Slc9a8 | protein_coding | 0.317552937 | 4.368778682 | 0.2815341 | 0.603434 | 0.905505734 |
| ENSMUSG00000056938 | Acbd4 | protein_coding | 0.203834781 | 5.565065564 | 0.2806175 | 0.6034465 | 0.905505734 |
| ENSMUSG00000039533 | Mmd2 | protein_coding | -0.157260848 | 7.968709712 | 0.2805322 | 0.6035011 | 0.905505734 |
| ENSMUSG00000041975 | Mettl8 | protein_coding | 0.28016664 | 4.448576166 | 0.2807761 | 0.6035547 | 0.905505734 |
| ENSMUSG00000035478 | Mbd3 | protein_coding | -0.195664888 | 5.413148314 | 0.2802359 | 0.6036791 | 0.905505734 |
| ENSMUSG00000085139 | A730046J19Rik | protein_coding | 0.475407606 | 2.279239719 | 0.2801424 | 0.6037389 | 0.905505734 |
| ENSMUSG00000034931 | Dhx8 | protein_coding | -0.2009587 | 4.399326705 | 0.280074 | 0.6037828 | 0.905505734 |
| ENSMUSG00000057637 | Prdm2 | protein_coding | -0.174865415 | 5.321888145 | 0.2798425 | 0.6039311 | 0.905547674 |
| ENSMUSG00000022515 | Anks3 | protein_coding | -0.239626768 | 4.971730314 | 0.2796858 | 0.6040433 | 0.905547674 |
| ENSMUSG00000108532 | Gm32647 | lincRNA | -0.471583899 | 2.491066607 | 0.2796362 | 0.6040634 | 0.905547674 |
| ENSMUSG00000057982 | Zfp809 | protein_coding | -0.230723105 | 4.043580567 | 0.2794674 | 0.6041716 | 0.905583729 |
| ENSMUSG00000063511 | Snrnp70 | protein_coding | -0.176254296 | 4.871768543 | 0.2792854 | 0.6042884 | 0.905609493 |
| ENSMUSG00000032551 | 1110059G10Rik | protein_coding | -0.153461316 | 5.850311521 | 0.2791782 | 0.6043573 | 0.905609493 |

| ensembl_gene_id | external_gene_name | gene_biotype | log2 (fold change) | log2 (counts per million) | F-value | p-value | False discovery rate |
|---|---|---|---|---|---|---|---|
| ENSMUSG00000046836 | Brox | protein_coding | 0.176049183 | 4.588628176 | 0.2789791 | 0.6044851 | 0.905674801 |
| ENSMUSG00000034442 | Trmt5 | protein_coding | 0.261510893 | 4.143886005 | 0.2782266 | 0.6049805 | 0.906197111 |
| ENSMUSG00000038602 | Slc35f1 | protein_coding | 0.165022549 | 5.437663363 | 0.2781612 | 0.6050109 | 0.906197111 |
| ENSMUSG00000048216 | Gpr85 | protein_coding | 0.14823218 | 6.060611206 | 0.2780437 | 0.6050865 | 0.906197111 |
| ENSMUSG00000034940 | Synrg | protein_coding | -0.135024988 | 6.096106445 | 0.2778705 | 0.605198 | 0.906237928 |
| ENSMUSG00000029575 | Mmab | protein_coding | 0.172783404 | 5.092512116 | 0.2777219 | 0.6052937 | 0.906254991 |
| ENSMUSG00000074259 | Gramd2 | protein_coding | -0.487293856 | 2.559464492 | 0.277207 | 0.6056256 | 0.906463971 |
| ENSMUSG00000024011 | Pi16 | protein_coding | 0.423770979 | 3.090579411 | 0.2772263 | 0.6056689 | 0.906463971 |
| ENSMUSG00000069581 | Tspear | protein_coding | -0.389532541 | 2.714406044 | 0.2769065 | 0.6058195 | 0.906463971 |
| ENSMUSG00000045064 | Zc2hc1c | protein_coding | 0.331049183 | 3.334567697 | 0.2768639 | 0.605847 | 0.906463971 |
| ENSMUSG00000037296 | Lsm1 | protein_coding | 0.166979853 | 6.027523324 | 0.2767383 | 0.6059281 | 0.906463971 |
| ENSMUSG00000022967 | Ifnar1 | protein_coding | -0.224022482 | 3.699766087 | 0.2766192 | 0.6060049 | 0.906463971 |
| ENSMUSG00000034379 | Wdr5b | protein_coding | 0.282256205 | 3.381513871 | 0.27659 | 0.6060238 | 0.906463971 |
| ENSMUSG00000029472 | Anapc5 | protein_coding | 0.136193137 | 7.048916681 | 0.2764602 | 0.6061077 | 0.906463971 |
| ENSMUSG00000062519 | Zfp398 | protein_coding | -0.261259128 | 3.625922666 | 0.2762531 | 0.6062415 | 0.906537985 |
| ENSMUSG00000034269 | Setd5 | protein_coding | -0.188644351 | 4.314935118 | 0.2758363 | 0.606511 | 0.906797561 |
| ENSMUSG00000031922 | Cep57 | protein_coding | -0.149812552 | 5.132438851 | 0.2755401 | 0.6067027 | 0.906797561 |
| ENSMUSG00000003435 | Supt5 | protein_coding | 0.205815626 | 5.680561552 | 0.2755143 | 0.6067616 | 0.906797561 |
| ENSMUSG00000024319 | Vps52 | protein_coding | 0.168450406 | 5.761971041 | 0.2754475 | 0.6067627 | 0.906797561 |
| ENSMUSG00000025762 | Larp1b | protein_coding | -0.308535898 | 3.278689258 | 0.2752842 | 0.6068684 | 0.906797561 |
| ENSMUSG00000069094 | Pde7a | protein_coding | 0.290228653 | 3.208383452 | 0.275203 | 0.606921 | 0.906797561 |
| ENSMUSG00000066152 | Slc31a2 | protein_coding | 0.218045885 | 3.622977023 | 0.2750456 | 0.6070231 | 0.906823989 |
| ENSMUSG00000026279 | Thap4 | protein_coding | 0.160153653 | 4.762737754 | 0.2745995 | 0.6073124 | 0.907130141 |
| ENSMUSG00000051343 | Rab11fip5 | protein_coding | 0.165140907 | 5.378459488 | 0.2742739 | 0.6075237 | 0.907242487 |
| ENSMUSG00000049470 | Aff4 | protein_coding | -0.131536361 | 6.631452802 | 0.2742237 | 0.6075563 | 0.907242487 |
| ENSMUSG00000036977 | Anapc10 | protein_coding | 0.17330882 | 5.966676204 | 0.27374 | 0.6078822 | 0.907472879 |
| ENSMUSG00000031343 | Gabra3 | protein_coding | 0.165168336 | 5.587913627 | 0.2736149 | 0.607952 | 0.907472879 |
| ENSMUSG00000039205 | Ciz1 | protein_coding | 0.216582024 | 4.429933006 | 0.2735049 | 0.6080235 | 0.907472879 |
| ENSMUSG00000037428 | Vgf | protein_coding | -0.248129397 | 4.032652325 | 0.2734301 | 0.6080722 | 0.907472879 |
| ENSMUSG00000035847 | Ids | protein_coding | -0.128458614 | 6.718583316 | 0.2733373 | 0.6081326 | 0.907472879 |
| ENSMUSG00000033055 | Ankrd54 | protein_coding | 0.238138062 | 3.488553445 | 0.2731153 | 0.6082771 | 0.907562538 |
| ENSMUSG00000032044 | Rpusd4 | protein_coding | 0.199154593 | 4.453725906 | 0.272963 | 0.6083762 | 0.907584569 |
| ENSMUSG00000048047 | Zbtb33 | protein_coding | -0.360295299 | 2.590932394 | 0.2727149 | 0.6085379 | 0.907646352 |
| ENSMUSG00000008604 | Ubqln4 | protein_coding | -0.143411238 | 5.605717211 | 0.2726404 | 0.6085865 | 0.907646352 |
| ENSMUSG00000020361 | Hspa4 | protein_coding | -0.154418077 | 5.412967386 | 0.2723851 | 0.608753 | 0.907768763 |
| ENSMUSG00000036902 | Neto2 | protein_coding | -0.212782509 | 4.898348325 | 0.2721846 | 0.6088837 | 0.90783786 |
| ENSMUSG00000026775 | Yme1l1 | protein_coding | -0.154064099 | 6.837194338 | 0.271713 | 0.6092032 | 0.9081882 |
| ENSMUSG00000029561 | Oasl2 | protein_coding | 0.706327874 | 3.05310636 | 0.275799 | 0.6093184 | 0.908234152 |
| ENSMUSG00000032470 | Mras | protein_coding | -0.119613828 | 6.058891247 | 0.2711447 | 0.6095632 | 0.908320312 |
| ENSMUSG00000001774 | Chordc1 | protein_coding | -0.110412357 | 6.496847039 | 0.271106 | 0.6095885 | 0.908320312 |
| ENSMUSG00000022394 | L3mbtl2 | protein_coding | 0.249995114 | 4.542254689 | 0.2710606 | 0.6096297 | 0.908320312 |
| ENSMUSG00000003873 | Bax | protein_coding | 0.121111002 | 6.257187138 | 0.2708651 | 0.6097461 | 0.908367976 |
| ENSMUSG00000027665 | Pik3ca | protein_coding | -0.149548465 | 5.507951303 | 0.2702336 | 0.6101598 | 0.908858321 |
| ENSMUSG00000047554 | Tmem41b | protein_coding | -0.184879296 | 4.389520327 | 0.2699038 | 0.6103761 | 0.908998408 |
| ENSMUSG00000084843 | B230312C02Rik | lincRNA | -0.291155002 | 3.027621948 | 0.2697066 | 0.6105055 | 0.908998408 |
| ENSMUSG00000008090 | Fgfrl1 | protein_coding | 0.453349199 | 2.626197143 | 0.2697037 | 0.6105074 | 0.908998408 |
| ENSMUSG00000019189 | Rnf145 | protein_coding | 0.176679936 | 5.142568878 | 0.269005 | 0.6109664 | 0.909416715 |
| ENSMUSG00000026627 | Tmem206 | protein_coding | -0.231029401 | 4.50691482 | 0.2683344 | 0.6114077 | 0.909416715 |
| ENSMUSG00000024456 | Diaph1 | protein_coding | -0.223857059 | 4.603002717 | 0.2682236 | 0.6114806 | 0.909416715 |
| ENSMUSG00000032030 | Cul5 | protein_coding | -0.153022257 | 5.90373286 | 0.2681806 | 0.611509 | 0.909416715 |
| ENSMUSG00000054426 | A930005H10Rik | processed_transcript | 0.169727876 | 4.744549992 | 0.2681295 | 0.6115426 | 0.909416715 |
| ENSMUSG00000008489 | Elavl2 | protein_coding | -0.130717701 | 6.592545263 | 0.2681139 | 0.6115529 | 0.909416715 |
| ENSMUSG00000069072 | Slc7a14 | protein_coding | -0.130949978 | 6.089734951 | 0.2680233 | 0.6116126 | 0.909416715 |
| ENSMUSG00000074519 | Zfp971 | protein_coding | 0.263853001 | 3.113243149 | 0.2679906 | 0.6116342 | 0.909416715 |
| ENSMUSG00000022257 | Laptm4b | protein_coding | -0.13686765 | 6.147958706 | 0.2679433 | 0.6116653 | 0.909416715 |
| ENSMUSG00000074892 | B3galt5 | protein_coding | 0.172545387 | 4.952348581 | 0.2677493 | 0.6117932 | 0.909416715 |
| ENSMUSG00000056019 | Zfp709 | protein_coding | -0.348940382 | 2.776760095 | 0.2675583 | 0.6119192 | 0.909416715 |
| ENSMUSG00000020121 | Srgap1 | protein_coding | -0.245192432 | 4.487032466 | 0.2675503 | 0.6119359 | 0.909416715 |
| ENSMUSG00000006731 | B4galnt1 | protein_coding | -0.239164809 | 3.284752277 | 0.2675206 | 0.6119441 | 0.909416715 |
| ENSMUSG00000026975 | Dph7 | protein_coding | 0.12534235 | 6.119075733 | 0.2672387 | 0.6121302 | 0.909416715 |
| ENSMUSG00000024613 | Tcof1 | protein_coding | -0.19980913 | 4.44965161 | 0.2672267 | 0.6121381 | 0.909416715 |
| ENSMUSG00000032116 | Stt3a | protein_coding | -0.140351068 | 5.841603953 | 0.2672215 | 0.6121416 | 0.909416715 |
| ENSMUSG00000004846 | Plod3 | protein_coding | -0.275489378 | 3.690383343 | 0.2669538 | 0.6123184 | 0.909538022 |
| ENSMUSG00000038708 | Golga4 | protein_coding | -0.148554567 | 5.144764638 | 0.2668418 | 0.6123924 | 0.909538022 |
| ENSMUSG00000033653 | Vps8 | protein_coding | 0.207278286 | 4.796670176 | 0.2665087 | 0.6126126 | 0.909605785 |
| ENSMUSG00000020340 | Cyfip2 | protein_coding | 0.125005374 | 8.355776164 | 0.2663135 | 0.6127417 | 0.909605785 |
| ENSMUSG00000071691 | Gm960 | protein_coding | -0.498784347 | 2.546245427 | 0.2663194 | 0.6127492 | 0.909605785 |
| ENSMUSG00000037287 | Tbcel | protein_coding | -0.186576295 | 4.496771683 | 0.2662612 | 0.6127764 | 0.909605785 |
| ENSMUSG00000022292 | Rrm2b | protein_coding | -0.179608314 | 4.879641011 | 0.2660048 | 0.6129461 | 0.909608715 |
| ENSMUSG00000056458 | Mok | protein_coding | 0.228431446 | 4.404013579 | 0.2658094 | 0.6130755 | 0.909608715 |
| ENSMUSG00000003354 | Ccdc65 | protein_coding | 0.218278175 | 4.403472492 | 0.2657768 | 0.6130971 | 0.909608715 |
| ENSMUSG00000033540 | Idua | protein_coding | -0.210606861 | 4.047153992 | 0.265702 | 0.6131467 | 0.909608715 |
| ENSMUSG00000032985 | 5730522E02Rik | protein_coding | -0.345509735 | 3.229163008 | 0.2655996 | 0.6132146 | 0.909608715 |
| ENSMUSG00000003062 | Stard3nl | protein_coding | 0.108429287 | 7.038505656 | 0.2651496 | 0.613513 | 0.909608715 |
| ENSMUSG00000017548 | Suz12 | protein_coding | -0.181114604 | 4.468041776 | 0.2651481 | 0.613514 | 0.909608715 |
| ENSMUSG00000033904 | Ccp110 | protein_coding | -0.144874163 | 5.276146718 | 0.2649378 | 0.6136536 | 0.909608715 |
| ENSMUSG00000055044 | Pdlim1 | protein_coding | -0.381103227 | 2.578696262 | 0.2649362 | 0.6136546 | 0.909608715 |
| ENSMUSG00000090546 | Cdr1 | protein_coding | -0.30947496 | 4.856864564 | 0.2662568 | 0.6137019 | 0.909608715 |

| ensembl_gene_id | external_gene_name | gene_biotype | log2 (fold change) | log2 (counts per million) | F-value | p-value | False discovery rate |
|---|---|---|---|---|---|---|---|
| ENSMUSG00000040390 | Map3k10 | protein_coding | -0.397592191 | 3.157233329 | 0.2648716 | 0.6137088 | 0.909608715 |
| ENSMUSG00000042293 | Gm5617 | protein_coding | 0.243057427 | 4.872114521 | 0.2644094 | 0.6140588 | 0.910001982 |
| ENSMUSG00000070287 | Slc35g2 | protein_coding | -0.164831435 | 5.014595835 | 0.2638958 | 0.6143463 | 0.910080307 |
| ENSMUSG00000070939 | Tgfbrap1 | protein_coding | -0.19485498 | 4.076641298 | 0.2637849 | 0.6144201 | 0.910080307 |
| ENSMUSG00000020561 | Twistnb | protein_coding | 0.23656133 | 3.707681291 | 0.2637694 | 0.6144304 | 0.910080307 |
| ENSMUSG00000032997 | Chpf | protein_coding | 0.181520506 | 5.077606247 | 0.2637301 | 0.6144566 | 0.910080307 |
| ENSMUSG00000021905 | Dph3 | protein_coding | -0.136677254 | 6.193024074 | 0.2635028 | 0.6146079 | 0.910080307 |
| ENSMUSG00000029147 | Ppm1g | protein_coding | 0.156240552 | 5.239946908 | 0.2634855 | 0.6146195 | 0.910080307 |
| ENSMUSG00000021548 | Ccnh | protein_coding | -0.110850479 | 7.14178214 | 0.2633127 | 0.6147346 | 0.910125478 |
| ENSMUSG00000050192 | Eif5a2 | protein_coding | 0.146867987 | 5.390785133 | 0.2628909 | 0.615016 | 0.910416654 |
| ENSMUSG00000030254 | Rad18 | protein_coding | -0.273994127 | 3.375191328 | 0.2625376 | 0.6152517 | 0.910526282 |
| ENSMUSG00000030753 | Thap12 | protein_coding | -0.16761765 | 4.885944996 | 0.2625262 | 0.6152594 | 0.910526282 |
| ENSMUSG00000020817 | Rabep1 | protein_coding | -0.123704752 | 6.820638269 | 0.2623609 | 0.6153697 | 0.91056432 |
| ENSMUSG00000028552 | Eps15 | protein_coding | -0.119027517 | 6.687543463 | 0.2616453 | 0.6158483 | 0.910986378 |
| ENSMUSG00000014313 | Cox6c | protein_coding | 0.142215614 | 10.12988312 | 0.2614026 | 0.6160108 | 0.910986378 |
| ENSMUSG00000037685 | Atp8a1 | protein_coding | -0.245829746 | 3.502410305 | 0.2612046 | 0.6161433 | 0.910986378 |
| ENSMUSG00000039065 | Fam173b | protein_coding | -0.17078555 | 4.623880927 | 0.2610711 | 0.6162328 | 0.910986378 |
| ENSMUSG00000031482 | Slc25a15 | protein_coding | 0.263276464 | 3.020939249 | 0.2609886 | 0.6162881 | 0.910986378 |
| ENSMUSG00000058192 | Zfp846 | protein_coding | -0.235582681 | 4.151514707 | 0.2609164 | 0.6163365 | 0.910986378 |
| ENSMUSG00000031835 | Mbtps1 | protein_coding | 0.151711852 | 5.428851872 | 0.2608089 | 0.6164086 | 0.910986378 |
| ENSMUSG00000078700 | D030028A08Rik | processed_transcript | 0.401479208 | 2.384144999 | 0.2605376 | 0.6165906 | 0.910986378 |
| ENSMUSG00000037750 | Fam222b | protein_coding | -0.159532823 | 5.159409597 | 0.2603965 | 0.6166853 | 0.910986378 |
| ENSMUSG00000073293 | Nudt10 | protein_coding | -0.198573538 | 5.579233833 | 0.2603984 | 0.6166952 | 0.910986378 |
| ENSMUSG00000024899 | Papss2 | protein_coding | 0.249601465 | 3.466976227 | 0.2600904 | 0.6168909 | 0.910986378 |
| ENSMUSG00000022822 | Abcc5 | protein_coding | 0.118050786 | 6.360321778 | 0.2600642 | 0.6169084 | 0.910986378 |
| ENSMUSG00000005362 | Crbn | protein_coding | 0.151217419 | 6.155560112 | 0.2599505 | 0.6169849 | 0.910986378 |
| ENSMUSG00000045932 | Ifit2 | protein_coding | 0.267497224 | 3.092937271 | 0.2599325 | 0.6169969 | 0.910986378 |
| ENSMUSG00000028865 | Cd164l2 | protein_coding | 0.194739317 | 3.901543846 | 0.259884 | 0.6170295 | 0.910986378 |
| ENSMUSG00000016256 | Ctsz | protein_coding | -0.156152984 | 6.410510985 | 0.2598529 | 0.6170616 | 0.910986378 |
| ENSMUSG00000041000 | Trim62 | protein_coding | 0.223647721 | 3.93206568 | 0.2597863 | 0.6170952 | 0.910986378 |
| ENSMUSG00000032680 | 6820408C15Rik | protein_coding | -0.252212544 | 4.215739749 | 0.2590699 | 0.6175884 | 0.91149413 |
| ENSMUSG00000021969 | Zdhhc20 | protein_coding | -0.127437746 | 6.100182519 | 0.2590232 | 0.6176087 | 0.91149413 |
| ENSMUSG00000031239 | Itm2a | protein_coding | 0.156022314 | 5.464426864 | 0.2588084 | 0.6177534 | 0.911553689 |
| ENSMUSG00000032897 | Nfyc | protein_coding | -0.19009515 | 4.118222849 | 0.2584914 | 0.6179671 | 0.911553689 |
| ENSMUSG00000024151 | Msh2 | protein_coding | -0.219834341 | 5.128752653 | 0.25833 | 0.6180872 | 0.911553689 |
| ENSMUSG00000032841 | Prr5l | protein_coding | -0.201998288 | 5.449710752 | 0.2582037 | 0.6181724 | 0.911553689 |
| ENSMUSG00000043510 | Hscb | protein_coding | 0.186497953 | 4.868453037 | 0.2581702 | 0.6181838 | 0.911553689 |
| ENSMUSG00000045871 | Slitrk6 | protein_coding | -0.463851938 | 2.80035259 | 0.2574995 | 0.6186481 | 0.911553689 |
| ENSMUSG00000000948 | Gm38393 | protein_coding | 0.175580165 | 4.792417404 | 0.2573993 | 0.6187046 | 0.911553689 |
| ENSMUSG00000062031 | Pgghg | protein_coding | -0.56147479 | 2.076154502 | 0.2573864 | 0.6187134 | 0.911553689 |
| ENSMUSG00000028944 | Prkag2 | protein_coding | 0.131903451 | 5.982183124 | 0.2573547 | 0.6187348 | 0.911553689 |
| ENSMUSG00000027746 | Ufm1 | protein_coding | 0.127074361 | 6.165844384 | 0.2573418 | 0.6187435 | 0.911553689 |
| ENSMUSG00000020952 | Scfd1 | protein_coding | 0.116492212 | 6.245587856 | 0.2572456 | 0.6188086 | 0.911553689 |
| ENSMUSG00000078591 | Hs3st4 | protein_coding | 0.151409987 | 5.317790841 | 0.2568567 | 0.6190718 | 0.911553689 |
| ENSMUSG00000068151 | A230006K03Rik | lincRNA | 0.179901698 | 4.432775736 | 0.256714 | 0.6191684 | 0.911553689 |
| ENSMUSG00000044477 | Zfand3 | protein_coding | -0.178080386 | 4.897481481 | 0.2566807 | 0.619191 | 0.911553689 |
| ENSMUSG00000031988 | Vps26b | protein_coding | 0.156609091 | 5.71486949 | 0.2565819 | 0.6192579 | 0.911553689 |
| ENSMUSG00000051817 | Sox12 | protein_coding | 0.231644988 | 3.284725338 | 0.2564079 | 0.6193758 | 0.911553689 |
| ENSMUSG00000035642 | Aamdc | protein_coding | 0.120250919 | 7.16647457 | 0.2563451 | 0.6194185 | 0.911553689 |
| ENSMUSG00000029427 | Zcchc8 | protein_coding | -0.232806613 | 3.828438522 | 0.2563183 | 0.6194366 | 0.911553689 |
| ENSMUSG00000063808 | Gpatch1 | protein_coding | 0.152146214 | 5.479376929 | 0.2561536 | 0.6195483 | 0.911553689 |
| ENSMUSG00000100599 | 1700120C14Rik | processed_transcript | -0.374008454 | 2.57138685 | 0.2561487 | 0.6195517 | 0.911553689 |
| ENSMUSG00000057506 | Bloc1s2 | protein_coding | 0.148630286 | 5.078684327 | 0.2560501 | 0.6196185 | 0.911553689 |
| ENSMUSG00000049511 | Htr1b | protein_coding | -0.277090176 | 3.790007403 | 0.2559928 | 0.6196574 | 0.911553689 |
| ENSMUSG00000026277 | Stk25 | protein_coding | -0.118369522 | 7.586585753 | 0.2559815 | 0.6196651 | 0.911553689 |
| ENSMUSG00000021147 | Wdr37 | protein_coding | -0.133156707 | 6.485592369 | 0.2556904 | 0.6198627 | 0.911553689 |
| ENSMUSG00000024442 | 0610009O20Rik | protein_coding | 0.127317438 | 5.770186713 | 0.255613 | 0.6199153 | 0.911553689 |
| ENSMUSG00000086477 | Gm15506 | processed_transcript | -0.308251066 | 2.935751709 | 0.2554592 | 0.6200198 | 0.911553689 |
| ENSMUSG00000047363 | Cstad | protein_coding | 0.232311408 | 3.822675004 | 0.2554289 | 0.6200403 | 0.911553689 |
| ENSMUSG00000031916 | Cog8 | protein_coding | -0.425751467 | 2.432140051 | 0.2552743 | 0.6201454 | 0.911553689 |
| ENSMUSG00000042770 | Hebp1 | protein_coding | 0.201446365 | 5.318491374 | 0.255267 | 0.6201615 | 0.911553689 |
| ENSMUSG00000028954 | Nub1 | protein_coding | 0.130804123 | 6.897385071 | 0.2551051 | 0.6202604 | 0.911553689 |
| ENSMUSG00000025971 | Maip1 | protein_coding | 0.15481218 | 4.989493481 | 0.2549135 | 0.6203908 | 0.911553689 |
| ENSMUSG00000070737 | Tmem35b | protein_coding | 0.282417605 | 2.970141968 | 0.2548663 | 0.6204229 | 0.911553689 |
| ENSMUSG00000004791 | Pgf | protein_coding | 0.194491773 | 4.251531182 | 0.2547243 | 0.6205195 | 0.911553689 |
| ENSMUSG00000001999 | Blvra | protein_coding | 0.104239728 | 6.923639702 | 0.2546931 | 0.6205408 | 0.911553689 |
| ENSMUSG00000024800 | Rpp30 | protein_coding | 0.120998771 | 6.361405398 | 0.2545825 | 0.6206161 | 0.911553689 |
| ENSMUSG00000055817 | Mta3 | protein_coding | -0.195038345 | 4.98990718 | 0.2540474 | 0.6209807 | 0.911590968 |
| ENSMUSG00000060450 | Rnf14 | protein_coding | 0.147056971 | 8.835925022 | 0.2533446 | 0.6214714 | 0.911590968 |
| ENSMUSG00000032435 | Dync1li1 | protein_coding | -0.176277961 | 7.157449133 | 0.2538426 | 0.6215993 | 0.911590968 |
| ENSMUSG00000024856 | Cdk2ap2 | protein_coding | -0.137576086 | 5.554639604 | 0.2531003 | 0.6216273 | 0.911590968 |
| ENSMUSG00000034290 | Nek9 | protein_coding | -0.247056991 | 3.854974297 | 0.2530889 | 0.6216351 | 0.911590968 |
| ENSMUSG00000026565 | Pou2f1 | protein_coding | -0.306918797 | 3.145914919 | 0.25303 | 0.6216753 | 0.911590968 |
| ENSMUSG00000020549 | Elac2 | protein_coding | 0.236107641 | 3.621125108 | 0.2529853 | 0.6217059 | 0.911590968 |
| ENSMUSG00000028532 | Cachd1 | protein_coding | 0.229806934 | 3.756836825 | 0.2529834 | 0.6217072 | 0.911590968 |
| ENSMUSG00000014601 | Strip1 | protein_coding | 0.241607535 | 5.067694756 | 0.2533352 | 0.6218053 | 0.911590968 |
| ENSMUSG00000026644 | Acbd7 | protein_coding | -0.266825506 | 3.201097795 | 0.2525294 | 0.6220177 | 0.911590968 |
| ENSMUSG00000071604 | Fam189a2 | protein_coding | 0.38347131 | 2.759313938 | 0.2524673 | 0.6220603 | 0.911590968 |

| ensembl_gene_id | external_gene_name | gene_biotype | log2 (fold change) | log2 (counts per million) | F-value | p-value | False discovery rate |
|---|---|---|---|---|---|---|---|
| ENSMUSG00000003161 | Sri | protein_coding | 0.102186489 | 7.586269207 | 0.2524664 | 0.6220609 | 0.911590968 |
| ENSMUSG00000081225 | Cyp2j12 | protein_coding | -0.159748066 | 4.902637011 | 0.252259 | 0.6222029 | 0.911590968 |
| ENSMUSG00000029063 | Nadk | protein_coding | -0.168002216 | 4.830288005 | 0.2518913 | 0.6224548 | 0.911590968 |
| ENSMUSG00000021258 | Ccnk | protein_coding | -0.288652774 | 3.107589317 | 0.2518758 | 0.6224655 | 0.911590968 |
| ENSMUSG00000024317 | Rnf138 | protein_coding | -0.179307893 | 4.513559622 | 0.2518569 | 0.6224785 | 0.911590968 |
| ENSMUSG00000022346 | Myc | protein_coding | 0.32635464 | 3.014558506 | 0.2516583 | 0.6226146 | 0.911590968 |
| ENSMUSG00000028758 | Kif17 | protein_coding | 0.176294626 | 4.78534031 | 0.2515171 | 0.6227115 | 0.911590968 |
| ENSMUSG00000073468 | Sft2d1 | protein_coding | 0.178892199 | 4.717820799 | 0.2513909 | 0.6227981 | 0.911590968 |
| ENSMUSG00000038398 | Upf3a | protein_coding | -0.193410419 | 4.345430702 | 0.2513636 | 0.6228169 | 0.911590968 |
| ENSMUSG00000029416 | Slc15a4 | protein_coding | -0.19008699 | 4.24000187 | 0.2512147 | 0.6229191 | 0.911590968 |
| ENSMUSG00000038425 | Poli | protein_coding | -0.257803063 | 3.520332659 | 0.2510223 | 0.6230512 | 0.911590968 |
| ENSMUSG00000032601 | Prkar2a | protein_coding | 0.120064572 | 5.871923199 | 0.2509896 | 0.6230737 | 0.911590968 |
| ENSMUSG00000030122 | Ptms | protein_coding | 0.153862555 | 6.578648641 | 0.250983 | 0.6230893 | 0.911590968 |
| ENSMUSG00000081051 | Gm15427 | processed_pseudoge | 0.28684169 | 2.95974261 | 0.2507807 | 0.6232173 | 0.911590968 |
| ENSMUSG00000038206 | Fbxo8 | protein_coding | -0.181242196 | 4.913619376 | 0.2507634 | 0.6232292 | 0.911590968 |
| ENSMUSG00000019791 | Hint3 | protein_coding | 0.196006426 | 4.098868876 | 0.2503929 | 0.623484 | 0.911590968 |
| ENSMUSG00000072980 | Oip5 | protein_coding | -0.254140818 | 3.608398567 | 0.2503234 | 0.6235319 | 0.911590968 |
| ENSMUSG00000027498 | Cstf1 | protein_coding | -0.150685458 | 5.412004816 | 0.2502924 | 0.6235532 | 0.911590968 |
| ENSMUSG00000047606 | Ankrd34c | protein_coding | -0.262503707 | 3.019823095 | 0.2502691 | 0.6235693 | 0.911590968 |
| ENSMUSG00000020515 | Cnot8 | protein_coding | 0.133934697 | 6.864369852 | 0.2502043 | 0.6236138 | 0.911590968 |
| ENSMUSG00000024892 | Pcx | protein_coding | -0.146283728 | 5.608768198 | 0.2500284 | 0.623735 | 0.911590968 |
| ENSMUSG00000073643 | Wdfy1 | protein_coding | 0.428892117 | 4.899426387 | 0.2546125 | 0.623871 | 0.911590968 |
| ENSMUSG00000000884 | Gnb1l | protein_coding | 0.216142779 | 5.353469977 | 0.2499155 | 0.6239442 | 0.911590968 |
| ENSMUSG00000044674 | Fzd1 | protein_coding | -0.461381009 | 2.389092387 | 0.2496315 | 0.6240085 | 0.911590968 |
| ENSMUSG00000013622 | Atraid | protein_coding | 0.122685523 | 7.682198409 | 0.2494353 | 0.6241438 | 0.911590968 |
| ENSMUSG00000022982 | Sod1 | protein_coding | 0.13289437 | 10.04575214 | 0.2493171 | 0.6242253 | 0.911590968 |
| ENSMUSG00000038042 | Ptpdc1 | protein_coding | 0.218560154 | 4.040815844 | 0.2492751 | 0.6242543 | 0.911590968 |
| ENSMUSG00000005374 | Tbl2 | protein_coding | 0.239278035 | 3.460875466 | 0.2489977 | 0.6244458 | 0.911590968 |
| ENSMUSG00000024238 | Zeb1 | protein_coding | -0.121401898 | 6.277915847 | 0.2489081 | 0.6245077 | 0.911590968 |
| ENSMUSG00000032498 | Mlh1 | protein_coding | 0.316082516 | 2.910645201 | 0.2486442 | 0.62469 | 0.911590968 |
| ENSMUSG00000040365 | Trim41 | protein_coding | 0.241287751 | 3.930646536 | 0.2484306 | 0.6248377 | 0.911590968 |
| ENSMUSG00000071302 | 2610044O15Rik8 | protein_coding | 0.164408273 | 4.952829646 | 0.2483966 | 0.6248612 | 0.911590968 |
| ENSMUSG00000029992 | Gfpt1 | protein_coding | -0.182074401 | 5.066774403 | 0.2481713 | 0.6250171 | 0.911590968 |
| ENSMUSG00000003934 | Efnb3 | protein_coding | -0.11567081 | 7.187374307 | 0.2481487 | 0.6250327 | 0.911590968 |
| ENSMUSG00000022403 | St13 | protein_coding | -0.127885265 | 9.582884206 | 0.2481002 | 0.6250663 | 0.911590968 |
| ENSMUSG00000029833 | Trim24 | protein_coding | -0.198838025 | 3.973895543 | 0.2480145 | 0.6251256 | 0.911590968 |
| ENSMUSG00000001755 | Coasy | protein_coding | 0.177339177 | 5.350843705 | 0.2479585 | 0.6251644 | 0.911590968 |
| ENSMUSG00000047649 | Cd3eap | protein_coding | 0.130694449 | 6.083496836 | 0.2478614 | 0.6252316 | 0.911590968 |
| ENSMUSG00000022214 | Dcaf11 | protein_coding | 0.169426444 | 5.684362016 | 0.2478509 | 0.6252389 | 0.911590968 |
| ENSMUSG00000050711 | Scg2 | protein_coding | -0.107935583 | 9.620289236 | 0.2477664 | 0.6252975 | 0.911590968 |
| ENSMUSG00000026946 | Nmi | protein_coding | -0.26019912 | 3.25338944 | 0.2477609 | 0.6253012 | 0.911590968 |
| ENSMUSG00000050321 | Neto1 | protein_coding | -0.206835368 | 4.702651805 | 0.2477279 | 0.6253241 | 0.911590968 |
| ENSMUSG00000029726 | Mepce | protein_coding | 0.14765834 | 5.396212388 | 0.2477187 | 0.6253305 | 0.911590968 |
| ENSMUSG00000005198 | Polr2a | protein_coding | -0.234194412 | 4.727635432 | 0.2476611 | 0.6253813 | 0.911590968 |
| ENSMUSG00000027108 | Ola1 | protein_coding | 0.105288012 | 7.789073664 | 0.2476086 | 0.6254068 | 0.911590968 |
| ENSMUSG00000032121 | Tmem218 | protein_coding | -0.146478376 | 5.524120132 | 0.2475121 | 0.6254737 | 0.911590968 |
| ENSMUSG00000070738 | Dgkd | protein_coding | 0.206403051 | 4.513298388 | 0.2472385 | 0.6256633 | 0.911596851 |
| ENSMUSG00000024104 | Washc2 | protein_coding | -0.188575136 | 5.126617546 | 0.2471484 | 0.6257259 | 0.911596851 |
| ENSMUSG00000033047 | Eif3l | protein_coding | 0.110533609 | 7.38292893 | 0.2470772 | 0.6257753 | 0.911596851 |
| ENSMUSG00000029161 | Cgref1 | protein_coding | 0.166323493 | 5.114376841 | 0.2470174 | 0.6258168 | 0.911596851 |
| ENSMUSG00000032281 | Acsbg1 | protein_coding | -0.120545552 | 7.206270198 | 0.2468632 | 0.6259238 | 0.911629241 |
| ENSMUSG00000010914 | Pdhx | protein_coding | 0.121768134 | 6.463653855 | 0.2465618 | 0.6261332 | 0.911810731 |
| ENSMUSG00000021385 | Ippk | protein_coding | -0.215570169 | 4.175136875 | 0.2462165 | 0.6263732 | 0.911848823 |
| ENSMUSG00000025878 | Uimc1 | protein_coding | -0.177729785 | 4.440030561 | 0.2461484 | 0.6264206 | 0.911848823 |
| ENSMUSG00000029330 | Cds1 | protein_coding | -0.178217498 | 4.559815468 | 0.2460136 | 0.6265144 | 0.911848823 |
| ENSMUSG00000027566 | Psma7 | protein_coding | 0.11206392 | 9.496865509 | 0.2459144 | 0.6265834 | 0.911848823 |
| ENSMUSG00000047921 | Trappc9 | protein_coding | 0.156423255 | 6.268041293 | 0.2459107 | 0.6265968 | 0.911848823 |
| ENSMUSG00000022438 | Parvb | protein_coding | -0.210369556 | 3.865977338 | 0.2457286 | 0.6267128 | 0.911848823 |
| ENSMUSG00000002102 | Psmc3 | protein_coding | 0.098969291 | 8.549169289 | 0.2456708 | 0.626753 | 0.911848823 |
| ENSMUSG00000028610 | Dmrtb1 | protein_coding | -0.426508633 | 3.322500982 | 0.246256 | 0.6270059 | 0.912011753 |
| ENSMUSG00000040236 | Trappc5 | protein_coding | 0.115667358 | 6.273591413 | 0.2451348 | 0.6271266 | 0.912011753 |
| ENSMUSG00000025375 | Aatk | protein_coding | -0.156805155 | 5.730584382 | 0.2449592 | 0.627249 | 0.912011753 |
| ENSMUSG00000020590 | Snx13 | protein_coding | -0.16805615 | 4.746669632 | 0.2448457 | 0.6273283 | 0.912011753 |
| ENSMUSG00000022948 | Setd4 | protein_coding | 0.392199088 | 3.210991503 | 0.2451593 | 0.627386 | 0.912011753 |
| ENSMUSG00000042185 | Nfrkb | protein_coding | -0.269294001 | 3.61727441 | 0.2445635 | 0.6275252 | 0.912011753 |
| ENSMUSG00000025279 | Dnase1l3 | protein_coding | -0.600713046 | 1.981800295 | 0.2445216 | 0.6275545 | 0.912011753 |
| ENSMUSG00000042063 | Zfp386 | protein_coding | 0.172477715 | 5.182603598 | 0.244427 | 0.6276206 | 0.912011753 |
| ENSMUSG00000070822 | Zscan18 | protein_coding | -0.257088775 | 4.896560747 | 0.2447398 | 0.6278219 | 0.912011753 |
| ENSMUSG00000031731 | Ap1g1 | protein_coding | -0.283175454 | 3.130153161 | 0.2440863 | 0.6278587 | 0.912011753 |
| ENSMUSG00000034850 | Tmem127 | protein_coding | 0.168232494 | 6.47245845 | 0.2439907 | 0.6281299 | 0.912011753 |
| ENSMUSG00000085208 | Brip1os | antisense_RNA | -0.345911368 | 2.826125638 | 0.2436803 | 0.6281428 | 0.912011753 |
| ENSMUSG00000061666 | Gdpd1 | protein_coding | 0.104344777 | 6.787748303 | 0.2436539 | 0.6281613 | 0.912011753 |
| ENSMUSG00000005142 | Man2b1 | protein_coding | 0.175080532 | 4.325568572 | 0.2432088 | 0.6284731 | 0.912011753 |
| ENSMUSG00000027692 | Tnik | protein_coding | -0.195488053 | 5.67519799 | 0.2431667 | 0.6285418 | 0.912011753 |
| ENSMUSG00000067873 | Htatsf1 | protein_coding | -0.13580574 | 5.837514164 | 0.2430609 | 0.6285767 | 0.912011753 |
| ENSMUSG00000039384 | Dusp10 | protein_coding | 0.27767392 | 4.084571816 | 0.2430507 | 0.6286418 | 0.912011753 |
| ENSMUSG00000053581 | Zfand2a | protein_coding | 0.164613163 | 6.114160358 | 0.2427224 | 0.6288249 | 0.912011753 |
| ENSMUSG00000019699 | Akt3 | protein_coding | -0.156342382 | 5.479204504 | 0.2425913 | 0.6289062 | 0.912011753 |

| ensembl_gene_id | external_gene_name | gene_biotype | log2 (fold change) | log2 (counts per million) | F-value | p-value | False discovery rate |
|---|---|---|---|---|---|---|---|
| ENSMUSG00000029687 | Ezh2 | protein_coding | 0.354257471 | 3.123296772 | 0.2424269 | 0.6290324 | 0.912011753 |
| ENSMUSG00000094626 | Cecr6 | protein_coding | 0.343335233 | 3.036081441 | 0.2419059 | 0.6293877 | 0.912011753 |
| ENSMUSG00000024603 | Dctn4 | protein_coding | -0.168008719 | 7.585918399 | 0.2424366 | 0.6294323 | 0.912011753 |
| ENSMUSG00000019878 | Hsf2 | protein_coding | -0.21837464 | 5.061571402 | 0.2416782 | 0.6295587 | 0.912011753 |
| ENSMUSG00000059878 | Zfp422 | protein_coding | 0.159460776 | 5.255728432 | 0.2416403 | 0.6295746 | 0.912011753 |
| ENSMUSG00000029071 | Dvl1 | protein_coding | -0.181573068 | 4.1831954 | 0.2414405 | 0.6297152 | 0.912011753 |
| ENSMUSG00000083012 | Fam220a | protein_coding | 0.16285388 | 4.321405995 | 0.2411154 | 0.6299442 | 0.912011753 |
| ENSMUSG00000028224 | Nbn | protein_coding | -0.264263262 | 3.964025293 | 0.2410713 | 0.629986 | 0.912011753 |
| ENSMUSG00000027865 | Gdap2 | protein_coding | -0.194301994 | 5.846192724 | 0.2413452 | 0.6300001 | 0.912011753 |
| ENSMUSG00000022558 | Mroh1 | protein_coding | 0.179165572 | 4.884030128 | 0.2407143 | 0.630227 | 0.912011753 |
| ENSMUSG00000074749 | Kiz | protein_coding | -0.215133029 | 4.178520559 | 0.2406687 | 0.6302592 | 0.912011753 |
| ENSMUSG00000036040 | Adamtsl2 | protein_coding | 0.23889227 | 3.44761555 | 0.2405032 | 0.6303759 | 0.912011753 |
| ENSMUSG00000020614 | Fam20a | protein_coding | -0.426298698 | 2.756788209 | 0.2404881 | 0.6303973 | 0.912011753 |
| ENSMUSG00000041923 | Nol4 | protein_coding | -0.123243264 | 6.112688642 | 0.2402747 | 0.6305373 | 0.912011753 |
| ENSMUSG00000027809 | Etfdh | protein_coding | -0.169841315 | 4.954823637 | 0.2401681 | 0.6306126 | 0.912011753 |
| ENSMUSG00000015217 | Hmgb3 | protein_coding | -0.201221234 | 3.751086296 | 0.2399159 | 0.6307908 | 0.912011753 |
| ENSMUSG00000073481 | 2-Mar | protein_coding | -0.210654148 | 4.460534505 | 0.2398074 | 0.6308675 | 0.912011753 |
| ENSMUSG00000074793 | Hspa12b | protein_coding | -0.309963871 | 2.950965648 | 0.2397081 | 0.6309377 | 0.912011753 |
| ENSMUSG00000024271 | Elp2 | protein_coding | 0.136730911 | 6.922050333 | 0.2396981 | 0.6309548 | 0.912011753 |
| ENSMUSG00000020430 | Pes1 | protein_coding | 0.178473755 | 5.057633856 | 0.2395685 | 0.6310365 | 0.912011753 |
| ENSMUSG00000066442 | Mthfs | protein_coding | 0.28876788 | 2.857450493 | 0.2393966 | 0.6311581 | 0.912011753 |
| ENSMUSG00000017615 | Tnfaip1 | protein_coding | 0.145062464 | 6.625073356 | 0.2391853 | 0.6313184 | 0.912011753 |
| ENSMUSG00000028524 | Sgip1 | protein_coding | -0.160365942 | 4.884116224 | 0.2390806 | 0.6313819 | 0.912011753 |
| ENSMUSG00000002233 | Rhoc | protein_coding | -0.234910927 | 4.492495405 | 0.2387518 | 0.6316256 | 0.912011753 |
| ENSMUSG00000021978 | Extl3 | protein_coding | 0.166471653 | 5.404115986 | 0.2386855 | 0.6316619 | 0.912011753 |
| ENSMUSG00000028538 | St3gal3 | protein_coding | -0.257410827 | 3.241666298 | 0.238604 | 0.6317197 | 0.912011753 |
| ENSMUSG00000025016 | Tm9sf3 | protein_coding | -0.14274149 | 6.079951102 | 0.2385821 | 0.6317353 | 0.912011753 |
| ENSMUSG00000026576 | Atp1b1 | protein_coding | -0.130678052 | 8.629832626 | 0.2384388 | 0.6318369 | 0.912011753 |
| ENSMUSG00000041028 | Ghitm | protein_coding | -0.109604907 | 9.508313207 | 0.2382799 | 0.6319497 | 0.912011753 |
| ENSMUSG00000028826 | Tmem57 | protein_coding | -0.165264533 | 5.475240805 | 0.2382069 | 0.6320015 | 0.912011753 |
| ENSMUSG00000029153 | Ociad2 | protein_coding | -0.092580257 | 7.775326845 | 0.238134 | 0.6320533 | 0.912011753 |
| ENSMUSG00000022551 | Cyc1 | protein_coding | 0.111574347 | 6.715909011 | 0.2380331 | 0.6321249 | 0.912011753 |
| ENSMUSG00000073424 | Cyp4f15 | protein_coding | -0.562354627 | 2.390839071 | 0.2379854 | 0.6322004 | 0.912011753 |
| ENSMUSG00000085492 | Trmt61b | processed_transcript | 0.15288075 | 5.168343951 | 0.2379006 | 0.6322191 | 0.912011753 |
| ENSMUSG00000028114 | Mettl14 | protein_coding | -0.183271771 | 4.797595066 | 0.2378005 | 0.6322902 | 0.912011753 |
| ENSMUSG00000063804 | Lin28b | protein_coding | -0.270623239 | 3.228588473 | 0.2376527 | 0.6323953 | 0.912011753 |
| ENSMUSG00000006920 | Ezh1 | protein_coding | 0.11229377 | 6.198793368 | 0.23764 | 0.6324043 | 0.912011753 |
| ENSMUSG00000085156 | Snhg15 | processed_transcript | 0.2607373 | 3.023525146 | 0.2375435 | 0.632473 | 0.912011753 |
| ENSMUSG00000064036 | Mro | protein_coding | -0.136767525 | 5.294121439 | 0.2375029 | 0.6325018 | 0.912011753 |
| ENSMUSG00000024906 | Mus81 | protein_coding | 0.185410672 | 4.973403595 | 0.2373804 | 0.632589 | 0.912011753 |
| ENSMUSG00000035509 | Fbxl21 | protein_coding | -0.206798771 | 3.861763145 | 0.2372841 | 0.6326576 | 0.912011753 |
| ENSMUSG00000036019 | Tmtc2 | protein_coding | 0.234541737 | 3.387662185 | 0.2371632 | 0.6327436 | 0.912011753 |
| ENSMUSG00000037887 | Dusp8 | protein_coding | 0.133706206 | 8.228678214 | 0.2371603 | 0.6327563 | 0.912011753 |
| ENSMUSG00000037395 | Rcor3 | protein_coding | -0.236136567 | 3.439315693 | 0.2370071 | 0.6328547 | 0.912011753 |
| ENSMUSG00000021686 | Ap3b1 | protein_coding | -0.175570974 | 4.754380161 | 0.2367911 | 0.6330086 | 0.912011753 |
| ENSMUSG00000034000 | Neu4 | protein_coding | 0.19558102 | 4.023571202 | 0.2367415 | 0.633044 | 0.912011753 |
| ENSMUSG00000050619 | Zscan29 | protein_coding | 0.36120235 | 2.68258433 | 0.236684 | 0.633085 | 0.912011753 |
| ENSMUSG00000042423 | Fbrs | protein_coding | -0.253490581 | 3.257937636 | 0.2364639 | 0.633242 | 0.912011753 |
| ENSMUSG00000054863 | Fam19a5 | protein_coding | -0.124688113 | 6.30314532 | 0.2364529 | 0.6332498 | 0.912011753 |
| ENSMUSG00000038152 | 5033430I15Rik | lincRNA | 0.30968607 | 2.755846196 | 0.2364515 | 0.6332508 | 0.912011753 |
| ENSMUSG00000040865 | Ino80d | protein_coding | -0.209939285 | 3.558900076 | 0.2364437 | 0.6332563 | 0.912011753 |
| ENSMUSG00000038506 | Dcun1d2 | protein_coding | 0.159710963 | 5.279136708 | 0.236443 | 0.6332569 | 0.912011753 |
| ENSMUSG00000022895 | Ets2 | protein_coding | -0.195709565 | 5.444192148 | 0.2364043 | 0.6332951 | 0.912011753 |
| ENSMUSG00000010362 | Rdm1 | protein_coding | -0.186428787 | 5.33457854 | 0.2363672 | 0.633311 | 0.912011753 |
| ENSMUSG00000026288 | Inpp5d | protein_coding | -0.599078573 | 2.104741917 | 0.2362972 | 0.6333715 | 0.912011753 |
| ENSMUSG00000078794 | Dact3 | protein_coding | -0.185465481 | 4.872785388 | 0.2362679 | 0.6333818 | 0.912011753 |
| ENSMUSG00000030256 | Bhlhe41 | protein_coding | -0.316419814 | 7.981567153 | 0.2416422 | 0.6334136 | 0.912011753 |
| ENSMUSG00000063765 | Chadl | protein_coding | 0.219444681 | 3.341909579 | 0.2359969 | 0.6335753 | 0.912011753 |
| ENSMUSG00000031503 | Col4a2 | protein_coding | 0.386884861 | 2.444763669 | 0.2359425 | 0.6336141 | 0.912011753 |
| ENSMUSG00000038366 | Lasp1 | protein_coding | -0.162759745 | 5.692131249 | 0.2359167 | 0.6336325 | 0.912011753 |
| ENSMUSG00000018697 | Aatf | protein_coding | -0.264888548 | 3.967412359 | 0.2359069 | 0.6336501 | 0.912011753 |
| ENSMUSG00000041319 | Thoc6 | protein_coding | 0.248154394 | 4.222495636 | 0.2353682 | 0.6340352 | 0.912380476 |
| ENSMUSG00000026489 | Coq8a | protein_coding | -0.158052106 | 5.611937598 | 0.2352964 | 0.634076 | 0.912380476 |
| ENSMUSG00000039913 | Pak7 | protein_coding | -0.159963618 | 4.788729336 | 0.234927 | 0.6343404 | 0.912577206 |
| ENSMUSG00000020456 | Ogdh | protein_coding | 0.143743683 | 8.134128311 | 0.2348831 | 0.6343825 | 0.912577206 |
| ENSMUSG00000055065 | Ddx17 | protein_coding | -0.159498184 | 5.506969951 | 0.2339804 | 0.6350192 | 0.912743409 |
| ENSMUSG00000055761 | Nkain3 | protein_coding | 0.257985707 | 3.530744899 | 0.2339611 | 0.6350331 | 0.912743409 |
| ENSMUSG00000002210 | Smg9 | protein_coding | 0.186687655 | 4.2138044 | 0.2338871 | 0.6350862 | 0.912743409 |
| ENSMUSG00000037465 | Klf10 | protein_coding | -0.204028043 | 3.896625607 | 0.2338182 | 0.6351357 | 0.912743409 |
| ENSMUSG00000001909 | Trmt1 | protein_coding | 0.182167586 | 5.004510961 | 0.2337396 | 0.6351922 | 0.912743409 |
| ENSMUSG00000024816 | Frmd8 | protein_coding | 0.196292416 | 5.575737944 | 0.233758 | 0.6352283 | 0.912743409 |
| ENSMUSG00000021481 | Zfp346 | protein_coding | -0.188902548 | 4.227970794 | 0.2336167 | 0.6352805 | 0.912743409 |
| ENSMUSG00000037013 | Ss18 | protein_coding | -0.216539053 | 4.372798749 | 0.2335407 | 0.6353351 | 0.912743409 |
| ENSMUSG00000044367 | Slc16a13 | protein_coding | 0.567506796 | 2.677996523 | 0.2343244 | 0.6354141 | 0.912743409 |
| ENSMUSG00000028759 | Hp1bp3 | protein_coding | -0.119318248 | 7.591971683 | 0.233263 | 0.6355348 | 0.912743409 |
| ENSMUSG00000034152 | Exoc3 | protein_coding | 0.109940717 | 7.219471058 | 0.2331888 | 0.6355882 | 0.912743409 |
| ENSMUSG00000058006 | Mdn1 | protein_coding | -0.323241348 | 3.343406399 | 0.2331624 | 0.6356178 | 0.912743409 |
| ENSMUSG00000021392 | Nol8 | protein_coding | -0.170117493 | 4.738627855 | 0.2330933 | 0.6356569 | 0.912743409 |

| ensembl_gene_id | external_gene_name | gene_biotype | log2 (fold change) | log2 (counts per million) | F-value | p-value | False discovery rate |
|---|---|---|---|---|---|---|---|
| ENSMUSG00000059277 | R74862 | processed_transcript | -0.325004477 | 2.99845578 | 0.2330524 | 0.6356864 | 0.912743409 |
| ENSMUSG00000024896 | Minpp1 | protein_coding | -0.182423828 | 4.77867413 | 0.2327217 | 0.6359245 | 0.912963425 |
| ENSMUSG00000055963 | Triqk | protein_coding | 0.116818511 | 5.890380288 | 0.2322703 | 0.6362499 | 0.913148975 |
| ENSMUSG00000030733 | Sh2b1 | protein_coding | -0.150121654 | 4.85364938 | 0.2321536 | 0.6363341 | 0.913148975 |
| ENSMUSG00000026563 | Tada1 | protein_coding | 0.340089178 | 2.606274316 | 0.2320784 | 0.6363884 | 0.913148975 |
| ENSMUSG00000041702 | Btbd7 | protein_coding | -0.229562755 | 3.842871206 | 0.2320714 | 0.6363934 | 0.913148975 |
| ENSMUSG00000054823 | Nsd3 | protein_coding | 0.132345086 | 6.613822115 | 0.2316556 | 0.6366936 | 0.913457845 |
| ENSMUSG00000041781 | Cpsf2 | protein_coding | -0.129078147 | 5.601195972 | 0.2311419 | 0.637065 | 0.913868769 |
| ENSMUSG00000029070 | Mxra8 | protein_coding | -0.624812735 | 3.850094941 | 0.2360977 | 0.6372933 | 0.914052105 |
| ENSMUSG00000050299 | Gm9843 | protein_coding | 0.162588996 | 4.56489152 | 0.2306055 | 0.6374534 | 0.914052105 |
| ENSMUSG00000017291 | Taok1 | protein_coding | -0.190376515 | 3.751936543 | 0.2305578 | 0.6374879 | 0.914052105 |
| ENSMUSG00000087148 | 1700030C14Rik | antisense_RNA | 0.413451086 | 2.385949082 | 0.2304893 | 0.6375375 | 0.914052105 |
| ENSMUSG00000039110 | Mycbpap | protein_coding | -0.318950224 | 2.801243388 | 0.2303785 | 0.6376179 | 0.914052105 |
| ENSMUSG00000052516 | Robo2 | protein_coding | 0.184144159 | 4.285642096 | 0.230048 | 0.6378576 | 0.914273929 |
| ENSMUSG00000009995 | Taz | protein_coding | 0.168753184 | 4.485728644 | 0.2297421 | 0.6380797 | 0.914360443 |
| ENSMUSG00000057880 | Abat | protein_coding | -0.111758139 | 8.308179823 | 0.2297305 | 0.638088 | 0.914360443 |
| ENSMUSG00000097867 | Lppos | lincRNA | -0.321695816 | 2.724646276 | 0.2295603 | 0.6382117 | 0.91441577 |
| ENSMUSG00000023873 | 1700010I14Rik | protein_coding | 0.380929004 | 2.972098165 | 0.2291704 | 0.6385056 | 0.914629812 |
| ENSMUSG00000004069 | Dnaja3 | protein_coding | 0.126255927 | 6.564819385 | 0.2291208 | 0.6385312 | 0.914629812 |
| ENSMUSG00000035933 | Cog5 | protein_coding | -0.268274841 | 3.09656944 | 0.2286908 | 0.6388442 | 0.91495627 |
| ENSMUSG00000022836 | Mylk | protein_coding | -0.365875863 | 2.678548687 | 0.2284586 | 0.6390134 | 0.915076697 |
| ENSMUSG00000034377 | Tulp4 | protein_coding | -0.127506756 | 5.511357516 | 0.2282683 | 0.6391521 | 0.915153447 |
| ENSMUSG00000027520 | Zdbf2 | protein_coding | -0.200923428 | 4.992381259 | 0.228062 | 0.6393129 | 0.915192897 |
| ENSMUSG00000066571 | 4931406P16Rik | protein_coding | -0.19385192 | 3.750215517 | 0.2277906 | 0.6395006 | 0.915192897 |
| ENSMUSG00000022241 | Tars | protein_coding | -0.161729966 | 5.555735571 | 0.2277727 | 0.6395137 | 0.915192897 |
| ENSMUSG00000018932 | Map2k3 | protein_coding | -0.280746292 | 2.856853344 | 0.2276608 | 0.6395954 | 0.915192897 |
| ENSMUSG00000028007 | Snx7 | protein_coding | -0.208522789 | 3.934943451 | 0.2275176 | 0.6397 | 0.915192897 |
| ENSMUSG00000027968 | Larp7 | protein_coding | -0.152037447 | 5.413909771 | 0.2272158 | 0.6399206 | 0.915192897 |
| ENSMUSG00000039908 | Slc26a11 | protein_coding | 0.265562464 | 3.558136159 | 0.227006 | 0.640074 | 0.915192897 |
| ENSMUSG00000023259 | Slc26a6 | protein_coding | 0.420891465 | 2.561602246 | 0.2268638 | 0.6401781 | 0.915192897 |
| ENSMUSG00000021279 | Cdc42bpb | protein_coding | -0.241292923 | 3.217432205 | 0.2268512 | 0.6401873 | 0.915192897 |
| ENSMUSG00000029156 | Sgcb | protein_coding | -0.165310534 | 5.265246916 | 0.2268487 | 0.6401891 | 0.915192897 |
| ENSMUSG00000022468 | Endou | protein_coding | -0.577477293 | 3.049065569 | 0.2281105 | 0.6405199 | 0.915192897 |
| ENSMUSG00000091475 | 2810468N07Rik | processed_transcript | -0.111815084 | 6.296432369 | 0.226329 | 0.6405697 | 0.915192897 |
| ENSMUSG00000013419 | Zfp651 | protein_coding | 0.19950951 | 4.179712255 | 0.2261978 | 0.640666 | 0.915192897 |
| ENSMUSG00000029176 | Anapc4 | protein_coding | -0.122200937 | 6.037861913 | 0.2260894 | 0.6407455 | 0.915192897 |
| ENSMUSG00000032657 | Fam189b | protein_coding | 0.204673437 | 3.935314827 | 0.2259427 | 0.640853 | 0.915192897 |
| ENSMUSG00000030560 | Ctsc | protein_coding | -0.450566368 | 2.630697253 | 0.2259261 | 0.6408756 | 0.915192897 |
| ENSMUSG00000031398 | Plxna3 | protein_coding | 0.314999835 | 2.688513231 | 0.2259015 | 0.6408833 | 0.915192897 |
| ENSMUSG00000033713 | Foxn3 | protein_coding | -0.14478764 | 6.87413152 | 0.2257759 | 0.6409859 | 0.915192897 |
| ENSMUSG00000037643 | Prkci | protein_coding | -0.232936981 | 4.427018228 | 0.2257282 | 0.6410209 | 0.915192897 |
| ENSMUSG00000020677 | Ddx52 | protein_coding | 0.128048615 | 5.736859524 | 0.2256673 | 0.6410552 | 0.915192897 |
| ENSMUSG00000033793 | Atp6v1h | protein_coding | 0.117998649 | 8.77313999 | 0.2256256 | 0.6410858 | 0.915192897 |
| ENSMUSG00000026889 | Rbm18 | protein_coding | 0.105135939 | 6.793040632 | 0.2256053 | 0.6411007 | 0.915192897 |
| ENSMUSG00000030407 | Qpctl | protein_coding | 0.155535861 | 4.971526964 | 0.2255332 | 0.6411537 | 0.915192897 |
| ENSMUSG00000025437 | Usp33 | protein_coding | -0.293173124 | 3.952541438 | 0.225766 | 0.6412466 | 0.915192897 |
| ENSMUSG00000026269 | Rnpepl1 | protein_coding | -0.22478893 | 3.453164609 | 0.2251428 | 0.6414407 | 0.915192897 |
| ENSMUSG00000075296 | Aldh3b2 | protein_coding | 0.154917021 | 5.345022307 | 0.2250464 | 0.6415115 | 0.915192897 |
| ENSMUSG00000047514 | Tspyl1 | protein_coding | -0.118087368 | 8.789173962 | 0.2250206 | 0.6415305 | 0.915192897 |
| ENSMUSG00000066306 | Numa1 | protein_coding | 0.168503891 | 4.956374096 | 0.2249195 | 0.6416049 | 0.915192897 |
| ENSMUSG00000042406 | Atf4 | protein_coding | -0.106751003 | 7.274473083 | 0.2247117 | 0.6417579 | 0.915192897 |
| ENSMUSG00000002496 | Tsc2 | protein_coding | -0.186503365 | 4.776192968 | 0.2246465 | 0.6418058 | 0.915192897 |
| ENSMUSG00000024259 | Slc25a46 | protein_coding | -0.137130027 | 5.508648155 | 0.2245225 | 0.6418971 | 0.915192897 |
| ENSMUSG00000029836 | Cbx3 | protein_coding | 0.195961172 | 3.558430742 | 0.224103 | 0.6422063 | 0.915192897 |
| ENSMUSG00000038412 | Higd1a | protein_coding | -0.102525266 | 8.510514161 | 0.2240715 | 0.6422295 | 0.915192897 |
| ENSMUSG00000017929 | B4galt5 | protein_coding | 0.157218392 | 4.77261447 | 0.2239747 | 0.642301 | 0.915192897 |
| ENSMUSG00000030711 | Sult1a1 | protein_coding | -0.318444511 | 3.215347347 | 0.2239715 | 0.6423034 | 0.915192897 |
| ENSMUSG00000031065 | Cdk16 | protein_coding | -0.136683971 | 6.001277039 | 0.223938 | 0.642328 | 0.915192897 |
| ENSMUSG00000052446 | Zfp961 | protein_coding | 0.188025296 | 4.061898998 | 0.2237552 | 0.6424629 | 0.915192897 |
| ENSMUSG00000032392 | Parp16 | protein_coding | 0.350138042 | 2.656371158 | 0.2237299 | 0.6424816 | 0.915192897 |
| ENSMUSG00000054976 | Nyap2 | protein_coding | 0.173167638 | 4.313598731 | 0.2236981 | 0.6425051 | 0.915192897 |
| ENSMUSG00000028312 | Smc2 | protein_coding | 0.343426418 | 2.554614109 | 0.2234017 | 0.642724 | 0.915192897 |
| ENSMUSG00000030087 | Klf15 | protein_coding | -0.247082834 | 3.61185396 | 0.223399 | 0.642726 | 0.915192897 |
| ENSMUSG00000038780 | Smurf1 | protein_coding | 0.181175625 | 3.930851603 | 0.2233607 | 0.6427543 | 0.915192897 |
| ENSMUSG00000032285 | Dnaja4 | protein_coding | -0.30859248 | 2.787858811 | 0.2227644 | 0.6431953 | 0.915699618 |
| ENSMUSG00000092083 | Kcnb2 | protein_coding | 0.226294147 | 3.41738996 | 0.222207 | 0.6436082 | 0.91612775 |
| ENSMUSG00000031562 | Dctd | protein_coding | -0.222826468 | 4.074161882 | 0.2220967 | 0.64369 | 0.91612775 |
| ENSMUSG00000026111 | Unc50 | protein_coding | -0.136034744 | 5.589462597 | 0.221799 | 0.6439109 | 0.91612775 |
| ENSMUSG00000041907 | Gpr45 | protein_coding | -0.243450788 | 3.999234521 | 0.2217109 | 0.6439763 | 0.91612775 |
| ENSMUSG00000002608 | Ccdc97 | protein_coding | 0.166654745 | 5.891737021 | 0.2216692 | 0.6440175 | 0.91612775 |
| ENSMUSG00000024169 | Ift140 | protein_coding | -0.27537633 | 2.989982275 | 0.221625 | 0.64404 | 0.91612775 |
| ENSMUSG00000027286 | Lrrc57 | protein_coding | -0.136304745 | 5.643362814 | 0.2215544 | 0.6440924 | 0.91612775 |
| ENSMUSG00000060950 | Trmt61a | protein_coding | 0.229211669 | 3.441033552 | 0.2213401 | 0.6442516 | 0.916233014 |
| ENSMUSG00000087497 | 2810001G20Rik | antisense_RNA | 0.159615494 | 4.972236344 | 0.2208922 | 0.6445847 | 0.916585442 |
| ENSMUSG00000048040 | Arxes2 | protein_coding | 0.116865302 | 6.371574271 | 0.2206736 | 0.6447474 | 0.916695548 |
| ENSMUSG00000066196 | Spag8 | protein_coding | -0.267904472 | 3.189612317 | 0.2203113 | 0.6450173 | 0.916958094 |
| ENSMUSG00000026018 | Ica1l | protein_coding | -0.159323575 | 4.956794094 | 0.2200699 | 0.6451972 | 0.91708874 |
| ENSMUSG00000075486 | Commd6 | protein_coding | -0.126271718 | 7.794029343 | 0.2199592 | 0.6452798 | 0.91708874 |

| ensembl_gene_id | external_gene_name | gene_biotype | log2 (fold change) | log2 (counts per million) | F-value | p-value | False discovery rate |
|---|---|---|---|---|---|---|---|
| ENSMUSG00000032086 | Bace1 | protein_coding | -0.137975685 | 5.419567206 | 0.2198175 | 0.6453855 | 0.917117815 |
| ENSMUSG00000030314 | Atg7 | protein_coding | -0.210449993 | 6.424224491 | 0.2206753 | 0.6457369 | 0.917495929 |
| ENSMUSG00000057729 | Prtn3 | protein_coding | 0.486475988 | 2.570975663 | 0.2192086 | 0.6458949 | 0.917599086 |
| ENSMUSG00000008333 | Snrpb2 | protein_coding | -0.128395683 | 6.805207652 | 0.2188333 | 0.6461211 | 0.917799175 |
| ENSMUSG00000031458 | Coprs | protein_coding | 0.151752986 | 6.282476966 | 0.2185367 | 0.6463532 | 0.917912122 |
| ENSMUSG00000025892 | Gria4 | protein_coding | -0.14791867 | 5.401965913 | 0.218472 | 0.6463916 | 0.917912122 |
| ENSMUSG00000022253 | Nadk2 | protein_coding | -0.167306724 | 4.544090733 | 0.2182911 | 0.6465271 | 0.917912122 |
| ENSMUSG00000027367 | Stard7 | protein_coding | 0.10075324 | 6.49919492 | 0.2182441 | 0.6465623 | 0.917912122 |
| ENSMUSG00000043716 | Rpl7 | protein_coding | 0.122035247 | 8.566012322 | 0.2181573 | 0.6466274 | 0.917912122 |
| ENSMUSG00000090223 | Pcp4 | protein_coding | -0.151120864 | 8.449282519 | 0.2183376 | 0.6467512 | 0.917966675 |
| ENSMUSG00000020482 | Ccdc117 | protein_coding | -0.267494939 | 3.228345873 | 0.2175047 | 0.6471171 | 0.918228437 |
| ENSMUSG00000098332 | 2310009A05Rik | processed_transcript | 0.155348514 | 5.400427499 | 0.2174692 | 0.6471438 | 0.918228437 |
| ENSMUSG00000035184 | Fam124a | protein_coding | -0.207171329 | 4.021235127 | 0.217359 | 0.6472266 | 0.918228437 |
| ENSMUSG00000031751 | Amfr | protein_coding | -0.147824872 | 6.103461629 | 0.2172054 | 0.6473421 | 0.918228437 |
| ENSMUSG00000009030 | Pdcl | protein_coding | -0.150016128 | 5.005014653 | 0.2170788 | 0.6474373 | 0.918228437 |
| ENSMUSG00000005882 | Uqcc1 | protein_coding | 0.117386889 | 7.093360736 | 0.2170646 | 0.647448 | 0.918228437 |
| ENSMUSG00000039356 | Exosc2 | protein_coding | 0.207573803 | 3.985303725 | 0.216688 | 0.6477313 | 0.918509151 |
| ENSMUSG00000033629 | Hacd3 | protein_coding | 0.095876862 | 7.534286954 | 0.2162062 | 0.6480943 | 0.918839287 |
| ENSMUSG00000022761 | Lztr1 | protein_coding | -0.115819042 | 6.556894168 | 0.2161522 | 0.648135 | 0.918839287 |
| ENSMUSG00000066151 | Fkbp15 | protein_coding | -0.155758313 | 4.607863867 | 0.2157527 | 0.6484364 | 0.919145351 |
| ENSMUSG00000059263 | Usp47 | protein_coding | -0.132996937 | 5.505801422 | 0.2153382 | 0.6487495 | 0.919198308 |
| ENSMUSG00000034731 | Dgkh | protein_coding | -0.245046449 | 3.239338009 | 0.2152725 | 0.6487992 | 0.919198308 |
| ENSMUSG00000031641 | Cbr4 | protein_coding | -0.149620942 | 5.240956239 | 0.215254 | 0.6488132 | 0.919198308 |
| ENSMUSG00000026853 | Crat | protein_coding | -0.172563921 | 4.648621722 | 0.2149794 | 0.6490208 | 0.919198308 |
| ENSMUSG00000073910 | Mob3b | protein_coding | 0.256507091 | 3.951102461 | 0.2149548 | 0.6490495 | 0.919198308 |
| ENSMUSG00000021196 | Pfkp | protein_coding | 0.136984413 | 7.676349815 | 0.214949 | 0.6490539 | 0.919198308 |
| ENSMUSG00000032264 | Zw10 | protein_coding | -0.214785256 | 4.109129274 | 0.2149116 | 0.6490721 | 0.919198308 |
| ENSMUSG00000056131 | Pgm3 | protein_coding | -0.242969842 | 4.301884487 | 0.2147768 | 0.6491842 | 0.919235897 |
| ENSMUSG00000031642 | Sh3rf1 | protein_coding | -0.13813265 | 4.948869836 | 0.2145791 | 0.6493238 | 0.919312602 |
| ENSMUSG00000047003 | Zfp41 | protein_coding | -0.42138839 | 2.686236001 | 0.2143192 | 0.6495308 | 0.919333746 |
| ENSMUSG00000018363 | Smurf2 | protein_coding | -0.180401045 | 4.560869623 | 0.2142171 | 0.6495982 | 0.919333746 |
| ENSMUSG00000027104 | Atf2 | protein_coding | -0.118420453 | 7.127111128 | 0.214133 | 0.6496619 | 0.919333746 |
| ENSMUSG00000039105 | Atp6v1g1 | protein_coding | 0.091678562 | 8.419649537 | 0.2139801 | 0.649778 | 0.919333746 |
| ENSMUSG00000037151 | Lrrc20 | protein_coding | -0.175066434 | 4.799613182 | 0.2139585 | 0.6497943 | 0.919333746 |
| ENSMUSG00000005683 | Cs | protein_coding | 0.137382778 | 8.087842053 | 0.2138005 | 0.6499242 | 0.919333746 |
| ENSMUSG00000024404 | Riok3 | protein_coding | -0.184515284 | 4.928221499 | 0.2137702 | 0.6499372 | 0.919333746 |
| ENSMUSG00000023027 | Atf1 | protein_coding | -0.356132623 | 2.517143718 | 0.2136544 | 0.6500251 | 0.919337108 |
| ENSMUSG00000021171 | Esyt2 | protein_coding | -0.176140439 | 4.758010707 | 0.2134318 | 0.6501943 | 0.919347216 |
| ENSMUSG00000032356 | Rasgrf1 | protein_coding | -0.125432034 | 8.150316656 | 0.21342 | 0.6502033 | 0.919347216 |
| ENSMUSG00000028039 | Efna3 | protein_coding | 0.193553084 | 4.154770139 | 0.2130394 | 0.6504926 | 0.919376683 |
| ENSMUSG00000036086 | Zranb3 | protein_coding | 0.363606267 | 2.837932009 | 0.2129013 | 0.6505977 | 0.919376683 |
| ENSMUSG00000020520 | Galnt10 | protein_coding | -0.223435114 | 3.456535794 | 0.2128482 | 0.6506381 | 0.919376683 |
| ENSMUSG00000021192 | Golga5 | protein_coding | -0.200833353 | 3.68689339 | 0.2127256 | 0.6507314 | 0.919376683 |
| ENSMUSG00000022722 | Arl6 | protein_coding | 0.111295775 | 7.395387216 | 0.2127162 | 0.6507386 | 0.919376683 |
| ENSMUSG00000038323 | 1700066M21Rik | protein_coding | -0.299180271 | 3.207639769 | 0.2126912 | 0.6507576 | 0.919376683 |
| ENSMUSG00000061911 | Myt1l | protein_coding | 0.093761802 | 7.934148902 | 0.2126059 | 0.6508226 | 0.919376683 |
| ENSMUSG00000038143 | Stox2 | protein_coding | -0.135564305 | 6.042121672 | 0.2123547 | 0.651014 | 0.9194048 |
| ENSMUSG00000018740 | Slc25a35 | protein_coding | -0.374487825 | 2.782725625 | 0.21217 | 0.6511548 | 0.9194048 |
| ENSMUSG00000022550 | Adck5 | protein_coding | 0.224638598 | 4.420545386 | 0.2120449 | 0.6512602 | 0.9194048 |
| ENSMUSG00000033220 | Rac2 | protein_coding | 0.34697544 | 2.77756677 | 0.2118421 | 0.651405 | 0.9194048 |
| ENSMUSG00000053483 | Usp21 | protein_coding | 0.15721903 | 5.18299884 | 0.2118006 | 0.6514367 | 0.9194048 |
| ENSMUSG00000073418 | C4b | protein_coding | -0.460986237 | 2.333341342 | 0.2117253 | 0.6514942 | 0.9194048 |
| ENSMUSG00000026389 | Steap3 | protein_coding | -0.429739073 | 2.068060088 | 0.2121412 | 0.6515698 | 0.9194048 |
| ENSMUSG00000020869 | Lrrc59 | protein_coding | 0.136340384 | 5.356798914 | 0.2115814 | 0.6516041 | 0.9194048 |
| ENSMUSG00000033159 | Cnppd1 | protein_coding | -0.139560919 | 5.282652657 | 0.2115711 | 0.651612 | 0.9194048 |
| ENSMUSG00000003134 | Tbc1d8 | protein_coding | -0.219660658 | 3.701743693 | 0.2108811 | 0.6521396 | 0.919786678 |
| ENSMUSG00000092203 | 1110038B12Rik | processed_transcript | 0.135589707 | 5.013220904 | 0.210831 | 0.652178 | 0.919786678 |
| ENSMUSG00000026388 | 3110009E18Rik | protein_coding | -0.18004319 | 4.377404602 | 0.210738 | 0.6522492 | 0.919786678 |
| ENSMUSG00000035293 | G2e3 | protein_coding | -0.35727695 | 2.867473345 | 0.2107163 | 0.6522658 | 0.919786678 |
| ENSMUSG00000021694 | Ercc8 | protein_coding | 0.215330124 | 4.122049179 | 0.2106582 | 0.6523104 | 0.919786678 |
| ENSMUSG00000021262 | Evl | protein_coding | 0.233667923 | 3.886538547 | 0.2104202 | 0.6524927 | 0.919902981 |
| ENSMUSG00000002324 | Rec8 | protein_coding | -0.29135762 | 3.110103132 | 0.2102639 | 0.6526125 | 0.919902981 |
| ENSMUSG00000025272 | Tro | protein_coding | -0.128340208 | 7.082353565 | 0.2100428 | 0.6527921 | 0.919902981 |
| ENSMUSG00000033316 | Galnt9 | protein_coding | 0.186879392 | 4.225352478 | 0.2097549 | 0.6530031 | 0.919902981 |
| ENSMUSG00000066129 | Kndc1 | protein_coding | 0.171808725 | 4.97263177 | 0.2097366 | 0.6530172 | 0.919902981 |
| ENSMUSG00000071477 | Zfp777 | protein_coding | 0.218241839 | 3.500699086 | 0.2096108 | 0.6531139 | 0.919902981 |
| ENSMUSG00000026037 | Orc2 | protein_coding | -0.412606748 | 2.489259787 | 0.2095506 | 0.6531601 | 0.919902981 |
| ENSMUSG00000034902 | Pip5k1c | protein_coding | 0.119727367 | 6.460003544 | 0.2095335 | 0.6531733 | 0.919902981 |
| ENSMUSG00000027605 | Acss2 | protein_coding | 0.228192869 | 3.965329002 | 0.2094009 | 0.6532752 | 0.919902981 |
| ENSMUSG00000031732 | Phlpp2 | protein_coding | -0.185768988 | 4.01577232 | 0.2093733 | 0.6532964 | 0.919902981 |
| ENSMUSG00000036707 | Cab39 | protein_coding | 0.124393756 | 6.547629035 | 0.2093246 | 0.6533339 | 0.919902981 |
| ENSMUSG00000026192 | Atic | protein_coding | 0.16006224 | 5.524916594 | 0.2090394 | 0.6535532 | 0.920006516 |
| ENSMUSG00000039307 | Hexdc | protein_coding | 0.20290461 | 3.88887667 | 0.2090066 | 0.6535785 | 0.920006516 |
| ENSMUSG00000031696 | Vps35 | protein_coding | -0.093983559 | 7.691568384 | 0.2082563 | 0.6541566 | 0.92050824 |
| ENSMUSG00000038644 | Pold1 | protein_coding | 0.37098033 | 2.400839149 | 0.2079901 | 0.654362 | 0.92050824 |
| ENSMUSG00000028453 | Fancg | protein_coding | 0.184920377 | 3.980522971 | 0.2079853 | 0.6543658 | 0.92050824 |
| ENSMUSG00000038844 | Kif16b | protein_coding | -0.263263052 | 3.107738631 | 0.2078086 | 0.6545022 | 0.92050824 |
| ENSMUSG00000027456 | Sdcbp2 | protein_coding | -0.337062297 | 3.053528154 | 0.207772 | 0.6545305 | 0.92050824 |

| ensembl_gene_id | external_gene_name | gene_biotype | log2 (fold change) | log2 (counts per million) | F-value | p-value | False discovery rate |
|---|---|---|---|---|---|---|---|
| ENSMUSG00000036819 | Jmjd4 | protein_coding | 0.177718756 | 4.267717668 | 0.2077692 | 0.6545326 | 0.92050824 |
| ENSMUSG00000003233 | Dvl3 | protein_coding | 0.216436649 | 4.062868915 | 0.2076613 | 0.654616 | 0.92050824 |
| ENSMUSG00000049690 | Nckap5 | protein_coding | -0.434716753 | 2.575471611 | 0.2076435 | 0.6546396 | 0.92050824 |
| ENSMUSG00000031813 | Mvb12a | protein_coding | -0.17903416 | 4.721394405 | 0.2073609 | 0.6548483 | 0.92050824 |
| ENSMUSG00000019906 | Lin7a | protein_coding | 0.128134883 | 6.307667229 | 0.2072455 | 0.6549375 | 0.92050824 |
| ENSMUSG00000027206 | Cops2 | protein_coding | 0.113732119 | 7.228814751 | 0.2072259 | 0.6549527 | 0.92050824 |
| ENSMUSG00000038422 | Hdhd3 | protein_coding | 0.153593972 | 5.09188441 | 0.2072136 | 0.6549622 | 0.92050824 |
| ENSMUSG00000085218 | BB218582 | processed_transcript | -0.281849442 | 2.970129781 | 0.2070304 | 0.6551039 | 0.920587132 |
| ENSMUSG00000066647 | Gm5113 | protein_coding | -0.141199148 | 4.778625466 | 0.2068516 | 0.6552424 | 0.920636748 |
| ENSMUSG00000003680 | Taf6l | protein_coding | -0.214319772 | 4.353004743 | 0.2066246 | 0.6554183 | 0.920636748 |
| ENSMUSG00000032249 | Anp32a | protein_coding | -0.103759876 | 6.482187072 | 0.2065095 | 0.6555075 | 0.920636748 |
| ENSMUSG00000047085 | Lrrc4b | protein_coding | 0.159271015 | 4.831499957 | 0.2063799 | 0.6556081 | 0.920636748 |
| ENSMUSG00000024975 | Pdcd4 | protein_coding | -0.181157621 | 7.436718025 | 0.207394 | 0.6556633 | 0.920636748 |
| ENSMUSG00000023021 | Cers5 | protein_coding | -0.151137606 | 5.106722379 | 0.2062091 | 0.6557405 | 0.920636748 |
| ENSMUSG00000050447 | Lypd6 | protein_coding | 0.180848705 | 4.566478356 | 0.2061202 | 0.6558095 | 0.920636748 |
| ENSMUSG00000044018 | Mrpl50 | protein_coding | 0.104266706 | 7.294997489 | 0.2061013 | 0.6558242 | 0.920636748 |
| ENSMUSG00000021090 | Lrrc9 | protein_coding | -0.366572614 | 2.667987494 | 0.205487 | 0.6563016 | 0.920842539 |
| ENSMUSG00000006906 | Stambp | protein_coding | 0.10978283 | 6.173512635 | 0.2054195 | 0.656354 | 0.920842539 |
| ENSMUSG00000054400 | Cklf | protein_coding | 0.328068816 | 2.540702746 | 0.2053893 | 0.6563775 | 0.920842539 |
| ENSMUSG00000024474 | Ik | protein_coding | 0.114995169 | 7.511914059 | 0.2052917 | 0.6564535 | 0.920842539 |
| ENSMUSG00000042763 | Maneal | protein_coding | 0.187522728 | 4.41117024 | 0.2052509 | 0.6564852 | 0.920842539 |
| ENSMUSG00000020660 | Pomc | protein_coding | 0.198477186 | 4.148240725 | 0.2051015 | 0.6566015 | 0.920842539 |
| ENSMUSG00000061665 | Cd2ap | protein_coding | -0.354953745 | 2.505229017 | 0.2046727 | 0.6569356 | 0.920842539 |
| ENSMUSG00000020827 | Mink1 | protein_coding | -0.14262194 | 4.869979965 | 0.2046484 | 0.6569545 | 0.920842539 |
| ENSMUSG00000000753 | Serpinf1 | protein_coding | 0.211717065 | 4.379507595 | 0.2044458 | 0.6571223 | 0.920842539 |
| ENSMUSG00000024030 | Abcg1 | protein_coding | -0.119751815 | 7.836224138 | 0.2044317 | 0.6571235 | 0.920842539 |
| ENSMUSG00000053046 | Brsk2 | protein_coding | 0.138220611 | 5.330846141 | 0.2044185 | 0.6571338 | 0.920842539 |
| ENSMUSG00000055670 | Zzef1 | protein_coding | 0.210644654 | 4.076059421 | 0.2043159 | 0.6572138 | 0.920842539 |
| ENSMUSG00000040785 | Ttc3 | protein_coding | -0.100680818 | 9.858794681 | 0.2043085 | 0.6572196 | 0.920842539 |
| ENSMUSG00000040666 | Sh3bgr | protein_coding | 0.459211167 | 2.521521413 | 0.2043182 | 0.6572219 | 0.920842539 |
| ENSMUSG00000010721 | Lmbr1 | protein_coding | 0.139591887 | 4.76906295 | 0.2040576 | 0.6574155 | 0.920842539 |
| ENSMUSG00000000223 | Drp2 | protein_coding | 0.180420422 | 4.106157728 | 0.2040572 | 0.6574158 | 0.920842539 |
| ENSMUSG00000021326 | Trim27 | protein_coding | -0.214273488 | 3.769012894 | 0.2040434 | 0.6574266 | 0.920842539 |
| ENSMUSG00000092274 | Neat1 | lincRNA | -0.240945338 | 3.937045109 | 0.2036908 | 0.6577119 | 0.920908066 |
| ENSMUSG00000030822 | Prr14 | protein_coding | -0.165225181 | 5.12615451 | 0.2036693 | 0.6577189 | 0.920908066 |
| ENSMUSG00000055409 | Nell1 | protein_coding | 0.107831905 | 5.860489754 | 0.2035837 | 0.6577859 | 0.920908066 |
| ENSMUSG00000006273 | Atp6v1b2 | protein_coding | 0.108326896 | 9.505910288 | 0.2035019 | 0.6578499 | 0.920908066 |
| ENSMUSG00000060038 | Dhps | protein_coding | -0.160468422 | 5.142539167 | 0.2034358 | 0.6579016 | 0.920908066 |
| ENSMUSG00000097039 | Pvt1 | lincRNA | 0.151828819 | 4.786443159 | 0.2033196 | 0.6579925 | 0.920915477 |
| ENSMUSG00000051721 | Wdcp | protein_coding | -0.222077097 | 4.719304903 | 0.2023367 | 0.6588151 | 0.921861382 |
| ENSMUSG00000031755 | Bbs2 | protein_coding | 0.194708416 | 4.505617287 | 0.2021173 | 0.6589352 | 0.921861382 |
| ENSMUSG00000032064 | Dixdc1 | protein_coding | -0.102359068 | 6.123348435 | 0.2020199 | 0.6590117 | 0.921861382 |
| ENSMUSG00000021065 | Fut8 | protein_coding | 0.150377957 | 4.914717648 | 0.2019844 | 0.6590397 | 0.921861382 |
| ENSMUSG00000040843 | Tiprl | protein_coding | 0.105560203 | 7.344423892 | 0.2019114 | 0.659097 | 0.921861382 |
| ENSMUSG00000018666 | Cbx1 | protein_coding | -0.102119636 | 6.193500757 | 0.2017078 | 0.6592571 | 0.921920447 |
| ENSMUSG00000004032 | Gstm5 | protein_coding | -0.116013369 | 9.655987292 | 0.2015146 | 0.6594091 | 0.921920447 |
| ENSMUSG00000041488 | Stx3 | protein_coding | -0.184699077 | 3.858314056 | 0.2013867 | 0.6595098 | 0.921920447 |
| ENSMUSG00000029910 | Mad2l1 | protein_coding | 0.187035122 | 4.158878799 | 0.201368 | 0.6595245 | 0.921920447 |
| ENSMUSG00000074886 | Grk6 | protein_coding | 0.160089045 | 4.896833007 | 0.2011212 | 0.6597189 | 0.921920447 |
| ENSMUSG00000079056 | Kcnip3 | protein_coding | -0.09970935 | 6.305152198 | 0.2010436 | 0.65978 | 0.921920447 |
| ENSMUSG00000029703 | Lrwd1 | protein_coding | -0.25133524 | 3.433185979 | 0.2009509 | 0.6598531 | 0.921920447 |
| ENSMUSG00000001942 | Siae | protein_coding | 0.16286267 | 4.628530044 | 0.2008636 | 0.6599219 | 0.921920447 |
| ENSMUSG00000021179 | Nrde2 | protein_coding | -0.216673279 | 4.271587522 | 0.2007044 | 0.6600474 | 0.921920447 |
| ENSMUSG00000043384 | Gprasp1 | protein_coding | 0.282261423 | 3.13095495 | 0.2006736 | 0.6600718 | 0.921920447 |
| ENSMUSG00000029287 | Tgfbr3 | protein_coding | -0.400514439 | 2.925349882 | 0.2007048 | 0.6600824 | 0.921920447 |
| ENSMUSG00000039954 | Stk32a | protein_coding | -0.343222518 | 3.783346815 | 0.2008736 | 0.660678 | 0.922332084 |
| ENSMUSG00000024423 | Impact | protein_coding | 0.139259458 | 10.99090957 | 0.1999048 | 0.6606887 | 0.922332084 |
| ENSMUSG00000031748 | Gnao1 | protein_coding | 0.099880863 | 8.517778331 | 0.1998883 | 0.6606921 | 0.922332084 |
| ENSMUSG00000038127 | Ccdc50 | protein_coding | -0.117262497 | 5.518560666 | 0.1995377 | 0.6609695 | 0.922332084 |
| ENSMUSG00000038705 | Gmeb2 | protein_coding | -0.228744603 | 3.403363273 | 0.1994591 | 0.6610317 | 0.922332084 |
| ENSMUSG00000097527 | 1700112J16Rik | lincRNA | -0.302110917 | 2.740116355 | 0.1994574 | 0.661033 | 0.922332084 |
| ENSMUSG00000024991 | Eif3a | protein_coding | -0.143593093 | 6.49308973 | 0.1993652 | 0.6611157 | 0.922332084 |
| ENSMUSG00000037492 | Zmat4 | protein_coding | -0.170933093 | 5.848493678 | 0.1993599 | 0.661154 | 0.922332084 |
| ENSMUSG00000042857 | Gm9776 | bidirectional_promote | -0.303510538 | 2.515001981 | 0.1992718 | 0.66118 | 0.922332084 |
| ENSMUSG00000027849 | Syt6 | protein_coding | -0.23179825 | 4.318855024 | 0.1992092 | 0.6612393 | 0.922332084 |
| ENSMUSG00000031027 | Stk33 | protein_coding | -0.255232922 | 3.030573786 | 0.1990343 | 0.6613683 | 0.922332084 |
| ENSMUSG00000035845 | Alg12 | protein_coding | 0.220812422 | 3.806612406 | 0.1988999 | 0.6614749 | 0.922332084 |
| ENSMUSG00000023236 | Scg5 | protein_coding | 0.103349032 | 9.418817556 | 0.1988781 | 0.6614921 | 0.922332084 |
| ENSMUSG00000040648 | Ppip5k2 | protein_coding | 0.158915777 | 4.174445386 | 0.1986068 | 0.6617074 | 0.922512623 |
| ENSMUSG00000078670 | Fam174b | protein_coding | 0.278279109 | 3.171188857 | 0.1979247 | 0.6622495 | 0.922586811 |
| ENSMUSG00000022744 | Cldnd1 | protein_coding | 0.092847463 | 7.721319446 | 0.1977384 | 0.6623978 | 0.922586811 |
| ENSMUSG00000021221 | Dpf3 | protein_coding | -0.240351305 | 3.478082958 | 0.1976906 | 0.6624358 | 0.922586811 |
| ENSMUSG00000102437 | Gm38048 | sense_overlapping | -0.276528463 | 3.008105649 | 0.1975886 | 0.662517 | 0.922586811 |
| ENSMUSG00000024790 | Sac3d1 | protein_coding | 0.192280753 | 3.673260541 | 0.197552 | 0.6625462 | 0.922586811 |
| ENSMUSG00000041143 | Tmco4 | protein_coding | 0.351510574 | 2.414615072 | 0.1974452 | 0.6626313 | 0.922586811 |
| ENSMUSG00000071103 | 1700029J07Rik | protein_coding | -0.499886509 | 2.093148936 | 0.1972468 | 0.6627894 | 0.922586811 |
| ENSMUSG00000004936 | Map2k1 | protein_coding | -0.109162759 | 7.707290744 | 0.1967767 | 0.6631644 | 0.922586811 |
| ENSMUSG00000050947 | Amigo1 | protein_coding | 0.159376927 | 4.700277835 | 0.1966664 | 0.6632525 | 0.922586811 |

| ensembl_gene_id | external_gene_name | gene_biotype | log2 (fold change) | log2 (counts per million) | F-value | p-value | False discovery rate |
|---|---|---|---|---|---|---|---|
| ENSMUSG00000032312 | Csk | protein_coding | 0.181116693 | 4.611753996 | 0.1966593 | 0.6632582 | 0.922586811 |
| ENSMUSG00000024217 | Snrpc | protein_coding | -0.153905234 | 5.049865964 | 0.1966421 | 0.6632719 | 0.922586811 |
| ENSMUSG00000055302 | Mrfap1 | protein_coding | 0.091944056 | 7.938451238 | 0.1965692 | 0.6633301 | 0.922586811 |
| ENSMUSG00000026096 | Osgepl1 | protein_coding | -0.165548406 | 4.626322671 | 0.1964282 | 0.6634428 | 0.922586811 |
| ENSMUSG00000042694 | Stn1 | protein_coding | 0.147198598 | 4.440789371 | 0.1963699 | 0.6634894 | 0.922586811 |
| ENSMUSG00000039254 | Pomt1 | protein_coding | 0.28102983 | 2.894868136 | 0.1963232 | 0.6635267 | 0.922586811 |
| ENSMUSG00000036986 | Pml | protein_coding | 0.334810826 | 2.724934985 | 0.1962851 | 0.6635572 | 0.922586811 |
| ENSMUSG00000037306 | Man1c1 | protein_coding | 0.246697393 | 3.687824388 | 0.1962447 | 0.6635895 | 0.922586811 |
| ENSMUSG00000055835 | Zfp1 | protein_coding | -0.180928838 | 4.897413214 | 0.1960865 | 0.663716 | 0.922586811 |
| ENSMUSG00000029098 | Acox3 | protein_coding | -0.209149628 | 3.777063605 | 0.196081 | 0.6637205 | 0.922586811 |
| ENSMUSG00000032180 | Tmed1 | protein_coding | 0.123639489 | 5.381310942 | 0.1959474 | 0.6638274 | 0.922586811 |
| ENSMUSG00000079235 | Ccdc13 | protein_coding | 0.218473858 | 3.461573038 | 0.1958959 | 0.6638685 | 0.922586811 |
| ENSMUSG00000003038 | Hmgn2 | protein_coding | 0.123610454 | 5.561308851 | 0.1958514 | 0.6639042 | 0.922586811 |
| ENSMUSG00000030663 | 1110004F10Rik | protein_coding | 0.090263004 | 8.6347426 | 0.1957883 | 0.6639547 | 0.922586811 |
| ENSMUSG00000028163 | Nfkb1 | protein_coding | 0.235786109 | 3.062743816 | 0.1956289 | 0.6640824 | 0.922586811 |
| ENSMUSG00000027652 | Ralgapb | protein_coding | 0.138289999 | 4.935392349 | 0.1956273 | 0.6640836 | 0.922586811 |
| ENSMUSG00000000326 | Comt | protein_coding | 0.094378486 | 7.220056283 | 0.1954984 | 0.664187 | 0.922586811 |
| ENSMUSG00000018931 | Natd1 | protein_coding | -0.312649646 | 2.723686703 | 0.1953704 | 0.6642896 | 0.922586811 |
| ENSMUSG00000020627 | Klhl29 | protein_coding | -0.20640898 | 4.320473232 | 0.1952037 | 0.6644232 | 0.922586811 |
| ENSMUSG00000039634 | Zfp189 | protein_coding | 0.242730808 | 3.583307246 | 0.1951948 | 0.6644304 | 0.922586811 |
| ENSMUSG00000057363 | Uxs1 | protein_coding | -0.212398489 | 3.824973209 | 0.1951659 | 0.6644536 | 0.922586811 |
| ENSMUSG00000030835 | Nomo1 | protein_coding | 0.110490949 | 6.283509334 | 0.1951232 | 0.6644879 | 0.922586811 |
| ENSMUSG00000037824 | Tspan14 | protein_coding | -0.16020937 | 4.513875636 | 0.1950346 | 0.664559 | 0.922586811 |
| ENSMUSG00000021470 | Ercc6l2 | protein_coding | -0.219428798 | 3.592632967 | 0.1945467 | 0.6649509 | 0.922586811 |
| ENSMUSG00000051726 | Kcnf1 | protein_coding | -0.200704783 | 4.390670137 | 0.1944158 | 0.6650561 | 0.922586811 |
| ENSMUSG00000057716 | Tmem178b | protein_coding | 0.143050763 | 5.542624092 | 0.1942125 | 0.6652197 | 0.922586811 |
| ENSMUSG00000006058 | Snf8 | protein_coding | 0.151318044 | 5.101828562 | 0.1940326 | 0.6653645 | 0.922586811 |
| ENSMUSG00000003452 | Bicd1 | protein_coding | -0.121765238 | 5.506908073 | 0.1939915 | 0.6653976 | 0.922586811 |
| ENSMUSG00000024457 | Trim26 | protein_coding | 0.161942231 | 4.668947304 | 0.1939674 | 0.665417 | 0.922586811 |
| ENSMUSG00000073131 | Vma21 | protein_coding | -0.100945539 | 6.259907015 | 0.1939329 | 0.6654447 | 0.922586811 |
| ENSMUSG00000054723 | Vmac | protein_coding | -0.219396277 | 3.522435072 | 0.1938526 | 0.6655094 | 0.922586811 |
| ENSMUSG00000037610 | Kcnmb2 | protein_coding | 0.338241703 | 2.971614265 | 0.1938328 | 0.6655348 | 0.922586811 |
| ENSMUSG00000025085 | Ablim1 | protein_coding | 0.144653554 | 5.591578019 | 0.1937235 | 0.6656134 | 0.922586811 |
| ENSMUSG00000031278 | Acsl4 | protein_coding | -0.124653467 | 6.224603325 | 0.1935177 | 0.6657793 | 0.922586811 |
| ENSMUSG00000066640 | Fbxl18 | protein_coding | -0.186708376 | 4.108800012 | 0.1935089 | 0.6657865 | 0.922586811 |
| ENSMUSG00000078546 | Zfp995 | protein_coding | -0.273174757 | 2.667089842 | 0.1934657 | 0.6658213 | 0.922586811 |
| ENSMUSG00000003559 | As3mt | protein_coding | -0.137508808 | 5.860821575 | 0.1934538 | 0.6658309 | 0.922586811 |
| ENSMUSG00000024594 | Prrc1 | protein_coding | -0.443855075 | 2.087124265 | 0.1934255 | 0.6658537 | 0.922586811 |
| ENSMUSG00000111088 | AC122428.3 | TEC | -0.296835624 | 3.354417989 | 0.1932709 | 0.6659879 | 0.922586811 |
| ENSMUSG00000027293 | Ehd4 | protein_coding | -0.204688684 | 3.606209092 | 0.1931953 | 0.6660395 | 0.922586811 |
| ENSMUSG00000040675 | Mthfd1l | protein_coding | 0.225059161 | 3.427437564 | 0.1931816 | 0.6660505 | 0.922586811 |
| ENSMUSG00000094152 | Slc6a16 | protein_coding | 0.222736373 | 3.775682246 | 0.1927379 | 0.666409 | 0.922607478 |
| ENSMUSG00000047844 | Bex4 | protein_coding | 0.126252289 | 5.16928169 | 0.1927041 | 0.6664363 | 0.922607478 |
| ENSMUSG00000112343 | AC154734.1 | processed_transcript | 0.412645639 | 2.490998379 | 0.1925716 | 0.6665434 | 0.922607478 |
| ENSMUSG00000064284 | Cdpf1 | protein_coding | 0.120545222 | 6.128265536 | 0.1925642 | 0.6665494 | 0.922607478 |
| ENSMUSG00000034653 | Ythdc2 | protein_coding | -0.220596484 | 3.328496467 | 0.1923759 | 0.6667017 | 0.922607478 |
| ENSMUSG00000020495 | Smg8 | protein_coding | 0.193398544 | 3.617206367 | 0.1922988 | 0.6667641 | 0.922607478 |
| ENSMUSG00000086272 | BC039966 | antisense_RNA | 0.248582372 | 3.834084902 | 0.1922462 | 0.6668162 | 0.922607478 |
| ENSMUSG00000000538 | Tom1l2 | protein_coding | -0.123815279 | 7.247091143 | 0.1922254 | 0.6668331 | 0.922607478 |
| ENSMUSG00000020877 | Scrn2 | protein_coding | -0.256540571 | 3.29208002 | 0.192208 | 0.6668377 | 0.922607478 |
| ENSMUSG00000070426 | Rnf121 | protein_coding | -0.108555185 | 5.735341833 | 0.1920821 | 0.6669396 | 0.92262988 |
| ENSMUSG00000007837 | Prrg2 | protein_coding | -0.224773476 | 3.673805446 | 0.191593 | 0.6673361 | 0.923048017 |
| ENSMUSG00000090386 | Mir99ahg | processed_transcript | -0.199902937 | 4.000903403 | 0.1914975 | 0.6674136 | 0.923048017 |
| ENSMUSG00000045034 | Ankrd34b | protein_coding | -0.20534231 | 3.798458628 | 0.1913502 | 0.6675332 | 0.923079034 |
| ENSMUSG00000021460 | Auh | protein_coding | 0.139527262 | 5.498863594 | 0.1911124 | 0.6677263 | 0.923079034 |
| ENSMUSG00000050967 | Creg2 | protein_coding | 0.163206921 | 4.896421896 | 0.1910379 | 0.6677869 | 0.923079034 |
| ENSMUSG00000029229 | Chic2 | protein_coding | 0.212587693 | 3.422371939 | 0.1910061 | 0.6678127 | 0.923079034 |
| ENSMUSG00000038510 | Rpf2 | protein_coding | -0.104996613 | 6.323682725 | 0.1909415 | 0.6678652 | 0.923079034 |
| ENSMUSG00000028654 | Mycl | protein_coding | -0.280216826 | 2.9758431 | 0.1905911 | 0.6681502 | 0.923120302 |
| ENSMUSG00000042709 | Atpaf2 | protein_coding | 0.131956876 | 5.953940537 | 0.1905688 | 0.6681684 | 0.923120302 |
| ENSMUSG00000057421 | Las1l | protein_coding | -0.119108823 | 5.820323439 | 0.1905115 | 0.668215 | 0.923120302 |
| ENSMUSG00000032750 | Gab3 | protein_coding | 0.215032085 | 3.593346028 | 0.1904827 | 0.6682385 | 0.923120302 |
| ENSMUSG00000021610 | Clptm1l | protein_coding | -0.155707766 | 5.774737366 | 0.1903152 | 0.6683842 | 0.923146009 |
| ENSMUSG00000042198 | Chchd7 | protein_coding | -0.138646788 | 5.275850413 | 0.190249 | 0.6684288 | 0.923146009 |
| ENSMUSG00000063160 | Numbl | protein_coding | -0.213987726 | 3.21841183 | 0.1900651 | 0.6685787 | 0.923156287 |
| ENSMUSG00000020848 | Doc2b | protein_coding | 0.191372799 | 3.522851538 | 0.1900292 | 0.6686079 | 0.923156287 |
| ENSMUSG00000021770 | Samd8 | protein_coding | -0.140531934 | 5.083330362 | 0.1893132 | 0.6691923 | 0.923844467 |
| ENSMUSG00000021891 | Mettl6 | protein_coding | -0.136741983 | 5.307868571 | 0.1891997 | 0.669285 | 0.923853919 |
| ENSMUSG00000038848 | Ythdf1 | protein_coding | -0.162317003 | 5.274372104 | 0.1888195 | 0.6695961 | 0.923927727 |
| ENSMUSG00000062169 | Cnih4 | protein_coding | 0.120450068 | 5.796278403 | 0.1888057 | 0.6696073 | 0.923927727 |
| ENSMUSG00000027245 | Hypk | protein_coding | 0.168000383 | 4.01193018 | 0.1887687 | 0.6696376 | 0.923927727 |
| ENSMUSG00000019841 | Rev3l | protein_coding | -0.201018747 | 4.223288506 | 0.1885033 | 0.669855 | 0.923927727 |
| ENSMUSG00000035215 | Lsm7 | protein_coding | -0.187716145 | 3.812510282 | 0.188406 | 0.6699347 | 0.923927727 |
| ENSMUSG00000020198 | Ap3d1 | protein_coding | 0.11707846 | 6.565677307 | 0.188321 | 0.6700043 | 0.923927727 |
| ENSMUSG00000027994 | Mcub | protein_coding | 0.277229666 | 3.54038633 | 0.1882286 | 0.6700894 | 0.923927727 |
| ENSMUSG00000007739 | Cct4 | protein_coding | 0.09603323 | 7.595742519 | 0.1882058 | 0.6700988 | 0.923927727 |
| ENSMUSG00000025956 | Mettl21a | protein_coding | 0.150213132 | 4.838935925 | 0.1881899 | 0.6701118 | 0.923927727 |
| ENSMUSG00000031379 | Pir | protein_coding | -0.235206786 | 3.804556121 | 0.1878802 | 0.6703659 | 0.924122674 |

| ensembl_gene_id | external_gene_name | gene_biotype | log2 (fold change) | log2 (counts per million) | F-value | p-value | False discovery rate |
|---|---|---|---|---|---|---|---|
| ENSMUSG00000036285 | Noa1 | protein_coding | -0.223985432 | 3.49091992 | 0.1878081 | 0.6704251 | 0.924122674 |
| ENSMUSG00000038722 | Bud31 | protein_coding | 0.112590606 | 6.785825901 | 0.18767 | 0.6705385 | 0.924160533 |
| ENSMUSG00000020455 | Trim11 | protein_coding | 0.217332277 | 3.555612013 | 0.1873925 | 0.6707666 | 0.92432651 |
| ENSMUSG00000025139 | Tollip | protein_coding | -0.116993121 | 7.020217951 | 0.1873143 | 0.6708308 | 0.92432651 |
| ENSMUSG00000019822 | Smpd2 | protein_coding | 0.1939356 | 4.065077862 | 0.1870494 | 0.6710488 | 0.924458581 |
| ENSMUSG00000021243 | Fcf1 | protein_coding | 0.150064934 | 5.818224989 | 0.1866892 | 0.6713547 | 0.924458581 |
| ENSMUSG00000063235 | Ptpmt1 | processed_transcript | 0.173319699 | 3.817350706 | 0.1866634 | 0.6713667 | 0.924458581 |
| ENSMUSG00000002845 | Tmem39a | protein_coding | 0.268891649 | 3.255610213 | 0.1865822 | 0.6714336 | 0.924458581 |
| ENSMUSG00000070687 | Htr1d | protein_coding | -0.469564011 | 2.709892867 | 0.1870129 | 0.6714393 | 0.924458581 |
| ENSMUSG00000033948 | Zswim5 | protein_coding | -0.272187274 | 3.294930318 | 0.1865619 | 0.6714503 | 0.924458581 |
| ENSMUSG00000032898 | Fbxo21 | protein_coding | -0.116599401 | 6.329678203 | 0.1864671 | 0.6715285 | 0.924458581 |
| ENSMUSG00000078813 | Leng1 | protein_coding | 0.30228029 | 2.817330814 | 0.1859562 | 0.6719501 | 0.924842558 |
| ENSMUSG00000026516 | Nvl | protein_coding | -0.149195263 | 5.279776214 | 0.1858907 | 0.6720042 | 0.924842558 |
| ENSMUSG00000035505 | Cox18 | protein_coding | 0.194427527 | 3.559834666 | 0.1858166 | 0.6720654 | 0.924842558 |
| ENSMUSG00000037706 | Cd81 | protein_coding | 0.097253328 | 9.865788007 | 0.185442 | 0.6723751 | 0.924930576 |
| ENSMUSG00000078566 | Bnip3 | protein_coding | -0.134435244 | 5.546691871 | 0.1851809 | 0.6725912 | 0.924930576 |
| ENSMUSG00000039262 | Prrc2b | protein_coding | -0.129581023 | 5.773422834 | 0.1850096 | 0.6727331 | 0.924930576 |
| ENSMUSG00000032192 | Gnb5 | protein_coding | -0.087390828 | 7.735530839 | 0.1848095 | 0.6728989 | 0.924930576 |
| ENSMUSG00000026289 | Atg16l1 | protein_coding | -0.188915712 | 4.945084035 | 0.1848196 | 0.6728998 | 0.924930576 |
| ENSMUSG00000047731 | Wbp1l | protein_coding | 0.208203444 | 4.181229716 | 0.1847505 | 0.6729478 | 0.924930576 |
| ENSMUSG00000020453 | Patz1 | protein_coding | 0.232004938 | 3.463945877 | 0.1846987 | 0.6729908 | 0.924930576 |
| ENSMUSG00000028967 | Errfi1 | protein_coding | -0.179220795 | 4.431978901 | 0.1846696 | 0.6730149 | 0.924930576 |
| ENSMUSG00000049281 | Scn3b | protein_coding | -0.097219404 | 6.319144643 | 0.1845918 | 0.6730794 | 0.924930576 |
| ENSMUSG00000044566 | Cage1 | protein_coding | 0.208200733 | 3.494822054 | 0.1845226 | 0.6731369 | 0.924930576 |
| ENSMUSG00000000563 | Atp5f1 | protein_coding | 0.086368726 | 9.728584423 | 0.184486 | 0.6731672 | 0.924930576 |
| ENSMUSG00000050063 | Klk6 | protein_coding | -0.254812309 | 3.952359432 | 0.1845435 | 0.673169 | 0.924930576 |
| ENSMUSG00000049106 | Dcaf5 | protein_coding | -0.113969269 | 5.500459631 | 0.184334 | 0.6732934 | 0.924930576 |
| ENSMUSG00000020642 | Rnf144a | protein_coding | -0.215208982 | 3.501786462 | 0.1842856 | 0.6733336 | 0.924930576 |
| ENSMUSG00000030352 | Tspan9 | protein_coding | -0.227704051 | 3.740494061 | 0.1837726 | 0.6737599 | 0.925369167 |
| ENSMUSG00000029397 | Rchy1 | protein_coding | 0.102796787 | 6.608216687 | 0.1835918 | 0.6739104 | 0.925369167 |
| ENSMUSG00000063063 | Ctnna2 | protein_coding | -0.100590419 | 6.555486821 | 0.1834855 | 0.6739989 | 0.925369167 |
| ENSMUSG00000018459 | Slc13a3 | protein_coding | 0.371111551 | 3.046341568 | 0.1836086 | 0.6741509 | 0.925369167 |
| ENSMUSG00000038286 | Bphl | protein_coding | 0.087383356 | 6.518229383 | 0.183286 | 0.674165 | 0.925369167 |
| ENSMUSG00000060860 | Ube2s | protein_coding | 0.292453269 | 2.896766933 | 0.1831843 | 0.6742497 | 0.925369167 |
| ENSMUSG00000068740 | Celsr2 | protein_coding | 0.141595429 | 5.933846653 | 0.1831776 | 0.6742553 | 0.925369167 |
| ENSMUSG00000029775 | Klhdc10 | protein_coding | 0.11546166 | 5.376062898 | 0.1829798 | 0.6744202 | 0.925371802 |
| ENSMUSG00000021589 | Rhobtb3 | protein_coding | 0.193597508 | 3.888652789 | 0.1829688 | 0.6744294 | 0.925371802 |
| ENSMUSG00000067567 | Hdac8 | protein_coding | 0.293345016 | 3.922088003 | 0.1832506 | 0.6747531 | 0.925697854 |
| ENSMUSG00000021286 | Zfyve21 | protein_coding | -0.146399248 | 5.29721669 | 0.1822275 | 0.6750483 | 0.925839173 |
| ENSMUSG00000031357 | Syap1 | protein_coding | -0.199432146 | 3.793191762 | 0.1821583 | 0.6751062 | 0.925839173 |
| ENSMUSG00000001467 | Cyp51 | protein_coding | 0.119800926 | 5.708776918 | 0.1821485 | 0.6751144 | 0.925839173 |
| ENSMUSG00000028015 | Ctso | protein_coding | -0.1863906 | 4.782366102 | 0.1817652 | 0.6754442 | 0.926173409 |
| ENSMUSG00000023965 | Fbxl17 | protein_coding | -0.122133707 | 5.385805921 | 0.1810583 | 0.6760276 | 0.926855181 |
| ENSMUSG00000044968 | Napepld | protein_coding | -0.266667154 | 3.709934025 | 0.1807449 | 0.6763239 | 0.927113508 |
| ENSMUSG00000004591 | Pkn2 | protein_coding | -0.210191698 | 3.453411419 | 0.1806285 | 0.6763885 | 0.927113508 |
| ENSMUSG00000032076 | Cadm1 | protein_coding | -0.093996504 | 7.275656087 | 0.1804828 | 0.676511 | 0.927163267 |
| ENSMUSG00000022561 | Gpaa1 | protein_coding | 0.145866366 | 5.135706021 | 0.1801918 | 0.6767558 | 0.927302181 |
| ENSMUSG00000054277 | Arfgap3 | protein_coding | 0.142399041 | 5.491942616 | 0.1801573 | 0.6767849 | 0.927302181 |
| ENSMUSG00000038013 | Wipf2 | protein_coding | 0.099096514 | 6.463170301 | 0.1799571 | 0.6769534 | 0.927414922 |
| ENSMUSG00000073427 | Gm4924 | protein_coding | 0.205094291 | 3.329425983 | 0.1798183 | 0.6770703 | 0.927456895 |
| ENSMUSG00000003575 | Crtc1 | protein_coding | 0.106962425 | 5.899694798 | 0.1796136 | 0.6772428 | 0.927575066 |
| ENSMUSG00000059981 | Taok2 | protein_coding | -0.171337806 | 4.662937901 | 0.1792849 | 0.6775201 | 0.927637564 |
| ENSMUSG00000048814 | Lonrf2 | protein_coding | 0.13656609 | 6.857009959 | 0.1793723 | 0.6776097 | 0.927637564 |
| ENSMUSG00000021692 | Dimt1 | protein_coding | 0.163012426 | 3.968309027 | 0.1789687 | 0.6777872 | 0.927637564 |
| ENSMUSG00000027244 | Atg13 | protein_coding | 0.121712913 | 6.63133618 | 0.1788457 | 0.6778911 | 0.927637564 |
| ENSMUSG00000024982 | Zdhhc6 | protein_coding | 0.128749759 | 4.825078853 | 0.1787769 | 0.6779493 | 0.927637564 |
| ENSMUSG00000021670 | Hmgcr | protein_coding | -0.110329963 | 5.731228708 | 0.1786843 | 0.6780276 | 0.927637564 |
| ENSMUSG00000039428 | Tmem135 | protein_coding | -0.133196091 | 5.142708133 | 0.1786533 | 0.6780538 | 0.927637564 |
| ENSMUSG00000028664 | Ephb2 | protein_coding | -0.263718321 | 2.859946521 | 0.178641 | 0.6780643 | 0.927637564 |
| ENSMUSG00000038525 | Armc10 | protein_coding | 0.231058972 | 3.275490753 | 0.1786217 | 0.6780805 | 0.927637564 |
| ENSMUSG00000074218 | Cox7a1 | protein_coding | 0.169182787 | 3.850723237 | 0.1785241 | 0.6781631 | 0.927637564 |
| ENSMUSG00000014932 | Yes1 | protein_coding | -0.343637796 | 2.407524458 | 0.1783868 | 0.6782794 | 0.927637564 |
| ENSMUSG00000091811 | Inafm1 | protein_coding | -0.178576368 | 4.222060814 | 0.1783251 | 0.6783316 | 0.927637564 |
| ENSMUSG00000040649 | Rimklb | protein_coding | -0.11888995 | 6.130343826 | 0.1782326 | 0.6784099 | 0.927637564 |
| ENSMUSG00000023118 | Sympk | protein_coding | -0.170102674 | 4.058834445 | 0.1779321 | 0.6786646 | 0.927768031 |
| ENSMUSG00000035984 | Nme5 | protein_coding | -0.111213733 | 6.20005046 | 0.1776642 | 0.6788919 | 0.927768031 |
| ENSMUSG00000009292 | Trpm2 | protein_coding | 0.230679386 | 3.85111753 | 0.1776614 | 0.6789033 | 0.927768031 |
| ENSMUSG00000024581 | Napg | protein_coding | 0.096592576 | 7.799076739 | 0.1776393 | 0.678913 | 0.927768031 |
| ENSMUSG00000024146 | Cript | protein_coding | 0.09687945 | 8.259973186 | 0.1776113 | 0.6789367 | 0.927768031 |
| ENSMUSG00000071669 | Snx29 | protein_coding | -0.22189118 | 3.290606917 | 0.1773543 | 0.679155 | 0.927948345 |
| ENSMUSG00000032177 | Pde4a | protein_coding | -0.093000838 | 6.706429982 | 0.1771023 | 0.6793692 | 0.928112662 |
| ENSMUSG00000108350 | Gm44950 | TEC | -0.178863262 | 4.493757093 | 0.1770098 | 0.6794479 | 0.928112662 |
| ENSMUSG00000063446 | Plppr1 | protein_coding | -0.131169029 | 4.933772752 | 0.176845 | 0.679588 | 0.928113459 |
| ENSMUSG00000021646 | Mccc2 | protein_coding | 0.159053654 | 5.453756839 | 0.1768092 | 0.6796276 | 0.928113459 |
| ENSMUSG00000030815 | Phkg2 | protein_coding | 0.123588749 | 5.202607546 | 0.176663 | 0.679743 | 0.928113459 |
| ENSMUSG00000060427 | Zfp868 | protein_coding | -0.198476054 | 4.786684045 | 0.176586 | 0.6798176 | 0.928113459 |
| ENSMUSG00000040521 | Tsfm | protein_coding | 0.107593117 | 6.541211415 | 0.1763762 | 0.6799873 | 0.928113459 |
| ENSMUSG00000021270 | Hsp90aa1 | protein_coding | -0.091067688 | 7.648787366 | 0.1763334 | 0.6800238 | 0.928113459 |

| ensembl_gene_id | external_gene_name | gene_biotype | log2 (fold change) | log2 (counts per million) | F-value | p-value | False discovery rate |
|---|---|---|---|---|---|---|---|
| ENSMUSG00000031511 | Arhgef7 | protein_coding | -0.145337456 | 5.673300661 | 0.1762551 | 0.6800906 | 0.928113459 |
| ENSMUSG00000028351 | Brinp1 | protein_coding | 0.101351681 | 6.494578386 | 0.175943 | 0.6803568 | 0.928113459 |
| ENSMUSG00000019433 | Gipc1 | protein_coding | 0.125923613 | 5.693327097 | 0.1756398 | 0.6806157 | 0.928113459 |
| ENSMUSG00000078584 | AU022252 | protein_coding | -0.139355578 | 4.762003024 | 0.1756393 | 0.6806162 | 0.928113459 |
| ENSMUSG00000030662 | Ipo5 | protein_coding | -0.109992095 | 6.205584337 | 0.1755523 | 0.6806905 | 0.928113459 |
| ENSMUSG00000028898 | Trnau1ap | protein_coding | 0.178184352 | 4.140814043 | 0.1755448 | 0.6806969 | 0.928113459 |
| ENSMUSG00000051029 | Serpinb1b | protein_coding | -0.148778883 | 4.661446212 | 0.1754653 | 0.6807649 | 0.928113459 |
| ENSMUSG00000026956 | Uap1l1 | protein_coding | 0.224286516 | 3.614291088 | 0.1752689 | 0.6809328 | 0.928113459 |
| ENSMUSG00000025854 | Fam20c | protein_coding | 0.201956209 | 4.759858499 | 0.1753074 | 0.6809428 | 0.928113459 |
| ENSMUSG00000034342 | Cbl | protein_coding | -0.148227593 | 4.555585624 | 0.1752529 | 0.6809465 | 0.928113459 |
| ENSMUSG00000037692 | Ahdc1 | protein_coding | -0.20785534 | 3.881753752 | 0.1750645 | 0.6811078 | 0.928113459 |
| ENSMUSG00000064264 | Zfp428 | protein_coding | -0.203846332 | 3.237660318 | 0.1750485 | 0.6811214 | 0.928113459 |
| ENSMUSG00000031302 | Nlgn3 | protein_coding | -0.096351533 | 6.648636821 | 0.1748562 | 0.6812862 | 0.928113459 |
| ENSMUSG00000044700 | Tmem201 | protein_coding | 0.156750669 | 4.577270721 | 0.1748185 | 0.6813185 | 0.928113459 |
| ENSMUSG00000019888 | Mgat4c | protein_coding | -0.172275731 | 4.655284636 | 0.1747605 | 0.6813681 | 0.928113459 |
| ENSMUSG00000037104 | Socs5 | protein_coding | -0.109699355 | 5.970340127 | 0.1747291 | 0.681395 | 0.928113459 |
| ENSMUSG00000062488 | Ifit3b | protein_coding | -0.294229871 | 2.6873316 | 0.174588 | 0.681516 | 0.928113459 |
| ENSMUSG00000031652 | N4bp1 | protein_coding | 0.118957925 | 5.833230899 | 0.1745324 | 0.6815637 | 0.928113459 |
| ENSMUSG00000097290 | 1300002E11Rik | lincRNA | -0.151012332 | 4.73278305 | 0.1744828 | 0.6816062 | 0.928113459 |
| ENSMUSG00000032186 | Tmod2 | protein_coding | -0.088053715 | 8.400191226 | 0.1743156 | 0.6817497 | 0.928185787 |
| ENSMUSG00000066687 | Zbtb16 | protein_coding | -0.304723289 | 2.852946376 | 0.1741444 | 0.6818967 | 0.928185787 |
| ENSMUSG00000059866 | Tnip2 | protein_coding | -0.275075882 | 2.952295257 | 0.1741192 | 0.6819183 | 0.928185787 |
| ENSMUSG00000062270 | Morf4l1 | protein_coding | 0.095339138 | 6.593492639 | 0.173997 | 0.6820233 | 0.928211219 |
| ENSMUSG00000047248 | C2cd3 | protein_coding | -0.43670286 | 2.391018639 | 0.1737918 | 0.6822086 | 0.928260995 |
| ENSMUSG00000029168 | Dpysl5 | protein_coding | 0.164699861 | 4.853573106 | 0.1736981 | 0.6822803 | 0.928260995 |
| ENSMUSG00000038372 | Gmds | protein_coding | 0.172340248 | 3.71384034 | 0.17364 | 0.6823303 | 0.928260995 |
| ENSMUSG00000030259 | Rassf8 | protein_coding | -0.192554136 | 3.946340098 | 0.1735529 | 0.6824052 | 0.928260995 |
| ENSMUSG00000024587 | Nars | protein_coding | 0.105552153 | 8.572052115 | 0.1734398 | 0.6825025 | 0.928275905 |
| ENSMUSG00000034930 | Rtkn | protein_coding | 0.148457579 | 4.703298896 | 0.1731344 | 0.6827655 | 0.928354868 |
| ENSMUSG00000073176 | Zfp449 | protein_coding | -0.181347945 | 3.378219399 | 0.1730839 | 0.682809 | 0.928354868 |
| ENSMUSG00000036564 | Ndrg4 | protein_coding | -0.1423744 | 11.22154694 | 0.1731272 | 0.6828196 | 0.928354868 |
| ENSMUSG00000020023 | Tmcc3 | protein_coding | 0.4118698 | 2.391541803 | 0.1727668 | 0.6830825 | 0.928559956 |
| ENSMUSG00000057894 | Zfp329 | protein_coding | -0.141864435 | 5.127169057 | 0.1726966 | 0.6831431 | 0.928559956 |
| ENSMUSG00000020120 | Plek | protein_coding | -0.197140451 | 3.509583045 | 0.1725887 | 0.6832362 | 0.92856917 |
| ENSMUSG00000039431 | Mtmr7 | protein_coding | 0.113251851 | 6.083458304 | 0.1722835 | 0.6834999 | 0.92881006 |
| ENSMUSG00000018882 | Mrpl45 | protein_coding | 0.091516689 | 7.34872276 | 0.1720317 | 0.6837176 | 0.928968951 |
| ENSMUSG00000038260 | Trpm4 | protein_coding | 0.151814218 | 4.247393398 | 0.1718713 | 0.6838564 | 0.928968951 |
| ENSMUSG00000020739 | Nup85 | protein_coding | -0.143860006 | 5.327087493 | 0.1716078 | 0.6840845 | 0.928968951 |
| ENSMUSG00000030739 | Myh14 | protein_coding | -0.20343725 | 3.342815442 | 0.1714939 | 0.6841832 | 0.928968951 |
| ENSMUSG00000025940 | Tmem70 | protein_coding | -0.140181817 | 4.937732559 | 0.1714839 | 0.6841919 | 0.928968951 |
| ENSMUSG00000022100 | Xpo7 | protein_coding | -0.12221147 | 5.314992387 | 0.1714711 | 0.6842029 | 0.928968951 |
| ENSMUSG00000014668 | Chfr | protein_coding | -0.418506583 | 2.161270994 | 0.1712945 | 0.6843561 | 0.928968951 |
| ENSMUSG00000024163 | Mapk8ip3 | protein_coding | 0.084964548 | 7.311188667 | 0.1712618 | 0.6843844 | 0.928968951 |
| ENSMUSG00000005981 | Trap1 | protein_coding | 0.125895257 | 6.09934087 | 0.1712504 | 0.6843943 | 0.928968951 |
| ENSMUSG00000067150 | Xpo5 | protein_coding | 0.188683413 | 3.970514468 | 0.1709982 | 0.6846131 | 0.928989092 |
| ENSMUSG00000030082 | Sec61a1 | protein_coding | 0.127120558 | 4.915062763 | 0.1709777 | 0.684631 | 0.928989092 |
| ENSMUSG00000055917 | Zfp277 | protein_coding | 0.168959867 | 5.231259473 | 0.1706964 | 0.6848841 | 0.928989092 |
| ENSMUSG00000039982 | Dtx4 | protein_coding | 0.161453738 | 4.140135314 | 0.1705428 | 0.6850088 | 0.928989092 |
| ENSMUSG00000039714 | Cplx3 | protein_coding | 0.321449903 | 3.717022402 | 0.1713214 | 0.6850314 | 0.928989092 |
| ENSMUSG00000060090 | Rp2 | protein_coding | -0.303463303 | 2.647798057 | 0.1704898 | 0.6850549 | 0.928989092 |
| ENSMUSG00000033022 | Cdo1 | protein_coding | -0.123415269 | 5.595285487 | 0.1703733 | 0.6851562 | 0.928989092 |
| ENSMUSG00000091896 | Ube2d2a | protein_coding | -0.113156588 | 5.745542378 | 0.1703112 | 0.6852102 | 0.928989092 |
| ENSMUSG00000017428 | Psmd11 | protein_coding | 0.087466296 | 7.521194612 | 0.1702799 | 0.6852375 | 0.928989092 |
| ENSMUSG00000050079 | Rspry1 | protein_coding | -0.140123335 | 4.873429617 | 0.170239 | 0.6852731 | 0.928989092 |
| ENSMUSG00000078619 | Smarcd2 | protein_coding | -0.383535138 | 2.691362152 | 0.1698392 | 0.6856301 | 0.929310386 |
| ENSMUSG00000038828 | Tmem214 | protein_coding | -0.15354918 | 4.820998701 | 0.1697026 | 0.6857404 | 0.929310386 |
| ENSMUSG00000003184 | Irf3 | protein_coding | -0.231939943 | 3.253010263 | 0.1696011 | 0.6858289 | 0.929310386 |
| ENSMUSG00000004896 | Rrnad1 | protein_coding | 0.18203063 | 3.883037571 | 0.1695702 | 0.6858558 | 0.929310386 |
| ENSMUSG00000045268 | Zfp691 | protein_coding | -0.32929692 | 2.370564015 | 0.169281 | 0.6861082 | 0.929463611 |
| ENSMUSG00000066042 | Med18 | protein_coding | 0.140707401 | 4.473760807 | 0.1691374 | 0.6862336 | 0.929463611 |
| ENSMUSG00000001305 | Rrp15 | protein_coding | 0.156804219 | 4.420875696 | 0.1691303 | 0.6862398 | 0.929463611 |
| ENSMUSG00000020170 | Frs2 | protein_coding | -0.152562338 | 4.04767324 | 0.1690448 | 0.6863146 | 0.929463611 |
| ENSMUSG00000056076 | Eif3b | protein_coding | -0.109431203 | 5.88964188 | 0.168759 | 0.6865645 | 0.929684944 |
| ENSMUSG00000022842 | Ece2 | protein_coding | 0.134253721 | 5.364220344 | 0.1686032 | 0.6867008 | 0.929752478 |
| ENSMUSG00000095432 | Zfp748 | protein_coding | 0.188976744 | 3.508578526 | 0.1682248 | 0.6870323 | 0.929928718 |
| ENSMUSG00000034610 | Zcchc11 | protein_coding | -0.176777351 | 4.181040372 | 0.168136 | 0.6871101 | 0.929928718 |
| ENSMUSG00000024639 | Gnaq | protein_coding | 0.119560915 | 8.272937565 | 0.1680859 | 0.6871628 | 0.929928718 |
| ENSMUSG00000021704 | Mtx3 | protein_coding | 0.150193231 | 4.285084892 | 0.1680598 | 0.6871769 | 0.929928718 |
| ENSMUSG00000090247 | Bloc1s1 | protein_coding | -0.128935399 | 5.050698492 | 0.167928 | 0.6872925 | 0.929968058 |
| ENSMUSG00000020180 | Snrpd3 | protein_coding | 0.097239347 | 8.641579732 | 0.1675079 | 0.6876614 | 0.929989157 |
| ENSMUSG00000034145 | Tmem63c | protein_coding | 0.20946283 | 3.439357991 | 0.1674632 | 0.6877007 | 0.929989157 |
| ENSMUSG00000032120 | C2cd2l | protein_coding | -0.106590621 | 6.011683688 | 0.1672427 | 0.6878945 | 0.929989157 |
| ENSMUSG00000054874 | Pcnx3 | protein_coding | 0.323866938 | 2.79137171 | 0.1672318 | 0.6879041 | 0.929989157 |
| ENSMUSG00000024293 | Esco1 | protein_coding | -0.1389433 | 4.417822002 | 0.1671745 | 0.6879545 | 0.929989157 |
| ENSMUSG00000026309 | Ilkap | protein_coding | -0.133366409 | 5.086444162 | 0.1671214 | 0.6880013 | 0.929989157 |
| ENSMUSG00000034707 | Gns | protein_coding | -0.136273992 | 5.097607033 | 0.167037 | 0.6880755 | 0.929989157 |
| ENSMUSG00000022962 | Gart | protein_coding | -0.173107084 | 4.633510336 | 0.1669805 | 0.6881252 | 0.929989157 |
| ENSMUSG00000022678 | Nde1 | protein_coding | -0.16349844 | 5.028134558 | 0.1669565 | 0.6881464 | 0.929989157 |

| ensembl_gene_id | external_gene_name | gene_biotype | log2 (fold change) | log2 (counts per million) | F-value | p-value | False discovery rate |
|---|---|---|---|---|---|---|---|
| ENSMUSG00000063785 | Utp14a | protein_coding | -0.244694288 | 3.571688047 | 0.1668791 | 0.6882232 | 0.929989157 |
| ENSMUSG00000032565 | Nudt16 | protein_coding | -0.151292723 | 4.08789586 | 0.1668281 | 0.6882594 | 0.929989157 |
| ENSMUSG00000031958 | Ldhd | protein_coding | 0.229884816 | 4.17706719 | 0.1667933 | 0.688455 | 0.930136464 |
| ENSMUSG00000034201 | Gas2l1 | protein_coding | 0.240807663 | 3.767728582 | 0.1663522 | 0.6886877 | 0.930333971 |
| ENSMUSG00000039208 | Metrnl | protein_coding | -0.292473989 | 2.675614047 | 0.1662215 | 0.6887943 | 0.930361138 |
| ENSMUSG00000000976 | Heatr6 | protein_coding | -0.147589481 | 4.608350955 | 0.1655239 | 0.6894108 | 0.930998698 |
| ENSMUSG00000043872 | Zmym1 | protein_coding | 0.21242696 | 3.116851037 | 0.1652225 | 0.6896777 | 0.930998698 |
| ENSMUSG00000041426 | Hibch | protein_coding | 0.130026325 | 5.204312644 | 0.1650301 | 0.6898482 | 0.930998698 |
| ENSMUSG00000042216 | Sgsm1 | protein_coding | 0.124947802 | 6.090930906 | 0.164938 | 0.6899298 | 0.930998698 |
| ENSMUSG00000031903 | Pla2g15 | protein_coding | 0.14253967 | 4.10737651 | 0.1648652 | 0.6899943 | 0.930998698 |
| ENSMUSG00000004895 | Prcc | protein_coding | -0.16982293 | 3.576970569 | 0.1647582 | 0.6900893 | 0.930998698 |
| ENSMUSG00000024524 | Gnal | protein_coding | -0.099639263 | 7.129695087 | 0.164737 | 0.690108 | 0.930998698 |
| ENSMUSG00000047139 | Cd24a | protein_coding | 0.175725684 | 4.013841585 | 0.1647105 | 0.6901316 | 0.930998698 |
| ENSMUSG00000034098 | Fstl5 | protein_coding | 0.127945648 | 5.253125928 | 0.1646715 | 0.6901662 | 0.930998698 |
| ENSMUSG00000038990 | Cables2 | protein_coding | 0.200150589 | 3.377384987 | 0.164623 | 0.6902092 | 0.930998698 |
| ENSMUSG00000060733 | Ipmk | protein_coding | -0.157163215 | 3.940777536 | 0.1646124 | 0.6902187 | 0.930998698 |
| ENSMUSG00000039458 | Mtmr12 | protein_coding | -0.194088482 | 3.255889402 | 0.1644023 | 0.6904053 | 0.931003134 |
| ENSMUSG00000038546 | Ranbp9 | protein_coding | -0.184808986 | 3.615199335 | 0.1641777 | 0.6906049 | 0.931003134 |
| ENSMUSG00000024286 | Ccny | protein_coding | -0.171565259 | 5.07855017 | 0.1641836 | 0.6906083 | 0.931003134 |
| ENSMUSG00000021301 | Hecw1 | protein_coding | -0.130904238 | 5.212150038 | 0.1640649 | 0.6907052 | 0.931003134 |
| ENSMUSG00000031245 | Hmgn5 | protein_coding | 0.205365587 | 3.447118563 | 0.1639628 | 0.690796 | 0.931003134 |
| ENSMUSG00000041890 | Git2 | protein_coding | -0.186274361 | 4.123003064 | 0.1639538 | 0.690804 | 0.931003134 |
| ENSMUSG00000078427 | Sarnp | protein_coding | -0.104630247 | 6.551563415 | 0.1639268 | 0.690828 | 0.931003134 |
| ENSMUSG00000031387 | Renbp | protein_coding | -0.233649627 | 3.052426427 | 0.1637344 | 0.6909993 | 0.931050776 |
| ENSMUSG00000052395 | Rft1 | protein_coding | -0.146899229 | 4.127595458 | 0.1636667 | 0.6910596 | 0.931050776 |
| ENSMUSG00000029386 | Tctn2 | processed_transcript | -0.135702247 | 4.322253723 | 0.1635953 | 0.6911232 | 0.931050776 |
| ENSMUSG00000061244 | Exoc5 | protein_coding | -0.136769569 | 4.714899718 | 0.163438 | 0.6912634 | 0.931122998 |
| ENSMUSG00000032537 | Ephb1 | protein_coding | 0.313584224 | 2.574719015 | 0.1633254 | 0.6913638 | 0.931141588 |
| ENSMUSG00000039988 | Ankrd13c | protein_coding | -0.105235222 | 5.982011039 | 0.1631774 | 0.6914958 | 0.931202762 |
| ENSMUSG00000006215 | Zbtb17 | protein_coding | 0.211373802 | 4.625695513 | 0.1629916 | 0.6918927 | 0.931620566 |
| ENSMUSG00000019761 | Krt10 | protein_coding | 0.127394979 | 5.047705005 | 0.1625727 | 0.692036 | 0.931690193 |
| ENSMUSG00000015846 | Rxra | protein_coding | 0.124562979 | 4.58347294 | 0.1623763 | 0.6922117 | 0.931690193 |
| ENSMUSG00000031111 | Igsf1 | protein_coding | 0.124443224 | 5.180803012 | 0.1621817 | 0.6923858 | 0.931690193 |
| ENSMUSG00000023074 | Mospd1 | protein_coding | 0.115632996 | 5.229285499 | 0.1621629 | 0.6924027 | 0.931690193 |
| ENSMUSG00000027387 | Zc3h8 | protein_coding | 0.167344481 | 3.806664377 | 0.1620842 | 0.6924731 | 0.931690193 |
| ENSMUSG00000022110 | Sucla2 | protein_coding | -0.096217112 | 8.265718248 | 0.1619299 | 0.6926114 | 0.931690193 |
| ENSMUSG00000074771 | Ankef1 | protein_coding | 0.298181798 | 2.771985152 | 0.1618999 | 0.6926383 | 0.931690193 |
| ENSMUSG00000026260 | Ndufa10 | protein_coding | -0.129272536 | 8.476402603 | 0.1620738 | 0.6926565 | 0.931690193 |
| ENSMUSG00000031887 | Tradd | protein_coding | 0.277125096 | 2.985530773 | 0.1617592 | 0.6927645 | 0.931690193 |
| ENSMUSG00000003200 | Sh3gl1 | protein_coding | -0.1833257 | 4.02389838 | 0.1616268 | 0.6928832 | 0.931690193 |
| ENSMUSG00000071266 | Zfp946 | protein_coding | 0.268590367 | 2.83477977 | 0.161611 | 0.6928975 | 0.931690193 |
| ENSMUSG00000022141 | Nipbl | protein_coding | -0.206400958 | 4.416390134 | 0.1613385 | 0.6931507 | 0.931750895 |
| ENSMUSG00000020366 | Mapk9 | protein_coding | 0.088573626 | 6.710022446 | 0.1613091 | 0.6931685 | 0.931750895 |
| ENSMUSG00000033530 | Ttc7b | protein_coding | 0.200184018 | 4.516190074 | 0.1613058 | 0.6932026 | 0.931750895 |
| ENSMUSG00000031385 | Plxnb3 | protein_coding | 0.293924521 | 2.818135004 | 0.1608379 | 0.6935922 | 0.932116345 |
| ENSMUSG00000050796 | B3galt6 | protein_coding | 0.196503328 | 3.461346429 | 0.160776 | 0.6936478 | 0.932116345 |
| ENSMUSG00000053574 | 4930563E22Rik | processed_transcript | -0.210640022 | 3.28713829 | 0.1606766 | 0.6937374 | 0.932120172 |
| ENSMUSG00000060739 | Nsa2 | protein_coding | 0.171721681 | 3.599785677 | 0.1605524 | 0.6938492 | 0.93215401 |
| ENSMUSG00000023235 | Ccl25 | protein_coding | 0.148610842 | 4.87212232 | 0.1602109 | 0.694157 | 0.932407961 |
| ENSMUSG00000032940 | Rbm11 | protein_coding | 0.155299199 | 4.260309132 | 0.1601073 | 0.6942505 | 0.932407961 |
| ENSMUSG00000025978 | Rftn2 | protein_coding | -0.13349851 | 4.74961476 | 0.1599828 | 0.6943629 | 0.932407961 |
| ENSMUSG00000022353 | Mtss1 | protein_coding | -0.114865226 | 5.152988467 | 0.1598239 | 0.6945063 | 0.932407961 |
| ENSMUSG00000042797 | Aqp11 | protein_coding | 0.173034467 | 3.620436067 | 0.1594666 | 0.6948293 | 0.932407961 |
| ENSMUSG00000041729 | Coro2b | protein_coding | 0.111918625 | 5.548722249 | 0.1594645 | 0.6948311 | 0.932407961 |
| ENSMUSG00000025613 | Cct8 | protein_coding | 0.112337914 | 8.639490942 | 0.1594233 | 0.6948769 | 0.932407961 |
| ENSMUSG00000028426 | Rad23b | protein_coding | -0.110792894 | 5.85898496 | 0.1593781 | 0.6949092 | 0.932407961 |
| ENSMUSG00000033083 | Tbc1d4 | protein_coding | -0.211285267 | 3.271941174 | 0.1593028 | 0.6949774 | 0.932407961 |
| ENSMUSG00000064105 | Cnnm2 | protein_coding | 0.227729681 | 3.6964863 | 0.1592597 | 0.6950249 | 0.932407961 |
| ENSMUSG00000044164 | Rnf182 | protein_coding | -0.244017336 | 3.699817746 | 0.1591077 | 0.6951625 | 0.932407961 |
| ENSMUSG00000033871 | Ppargc1b | protein_coding | 0.203605073 | 3.937598235 | 0.1589135 | 0.6953299 | 0.932407961 |
| ENSMUSG00000021577 | Sdha | protein_coding | 0.122890938 | 9.626183273 | 0.158922 | 0.6953307 | 0.932407961 |
| ENSMUSG00000054604 | Cggbp1 | protein_coding | -0.104745878 | 5.712912326 | 0.1586574 | 0.6955621 | 0.932407961 |
| ENSMUSG00000034453 | Polr3b | protein_coding | -0.184273207 | 3.459292314 | 0.1585724 | 0.6956392 | 0.932407961 |
| ENSMUSG00000029007 | Agtrap | protein_coding | -0.162076123 | 4.356498875 | 0.1585591 | 0.6956513 | 0.932407961 |
| ENSMUSG00000052155 | Acvr2a | protein_coding | -0.173739301 | 3.817366856 | 0.1583207 | 0.6958677 | 0.932407961 |
| ENSMUSG00000027239 | Mdk | protein_coding | -0.242826388 | 4.884046734 | 0.1591131 | 0.6958739 | 0.932407961 |
| ENSMUSG00000079003 | Samd1 | protein_coding | -0.252506145 | 3.283322186 | 0.1583059 | 0.6958812 | 0.932407961 |
| ENSMUSG00000037373 | Ctbp1 | protein_coding | -0.096029232 | 6.682706994 | 0.1582028 | 0.6959748 | 0.932407961 |
| ENSMUSG00000030869 | Ndufab1 | protein_coding | 0.145595439 | 4.057246265 | 0.1579996 | 0.6961596 | 0.932407961 |
| ENSMUSG00000057969 | Sema3b | protein_coding | 0.47584278 | 2.483645804 | 0.1583098 | 0.6962061 | 0.932407961 |
| ENSMUSG00000003208 | Ccdc94 | protein_coding | -0.216399823 | 3.964551078 | 0.1579299 | 0.6962314 | 0.932407961 |
| ENSMUSG00000042751 | Nmnat2 | protein_coding | -0.143162361 | 7.270128218 | 0.158401 | 0.6963098 | 0.932407961 |
| ENSMUSG00000021731 | Mrps30 | protein_coding | -0.178101257 | 4.137049958 | 0.1577816 | 0.6963578 | 0.932407961 |
| ENSMUSG00000047786 | Lix1 | protein_coding | -0.139407893 | 5.070151099 | 0.1576928 | 0.6964386 | 0.932407961 |
| ENSMUSG00000072872 | Rybp | protein_coding | 0.278664156 | 2.674842252 | 0.1576652 | 0.6964638 | 0.932407961 |
| ENSMUSG00000027067 | Ssrp1 | protein_coding | -0.097227519 | 6.496944194 | 0.1575507 | 0.696568 | 0.932407961 |
| ENSMUSG00000020941 | Map3k14 | protein_coding | -0.397094678 | 2.231274153 | 0.157431 | 0.6966771 | 0.932407961 |
| ENSMUSG00000029592 | Usp30 | protein_coding | -0.170810652 | 4.847948825 | 0.1574265 | 0.6966897 | 0.932407961 |

| ensembl_gene_id | external_gene_name | gene_biotype | log2 (fold change) | log2 (counts per million) | F-value | p-value | False discovery rate |
|---|---|---|---|---|---|---|---|
| ENSMUSG00000018849 | Wwc1 | protein_coding | -0.128919395 | 5.078103555 | 0.157377 | 0.6967263 | 0.932407961 |
| ENSMUSG00000017412 | Cacnb4 | protein_coding | 0.104614533 | 5.601837244 | 0.1572624 | 0.6968307 | 0.932431664 |
| ENSMUSG00000022529 | Zfp263 | protein_coding | -0.152027911 | 3.891581902 | 0.1571403 | 0.6969421 | 0.932464603 |
| ENSMUSG00000013707 | Tnfaip8l2 | protein_coding | -0.343391712 | 3.013897389 | 0.1570229 | 0.6971008 | 0.932560983 |
| ENSMUSG00000020149 | Rab1a | protein_coding | -0.094069114 | 8.128892063 | 0.1564309 | 0.69759 | 0.93304071 |
| ENSMUSG00000028832 | Stmn1 | protein_coding | 0.085148874 | 7.60688748 | 0.156384 | 0.697633 | 0.93304071 |
| ENSMUSG00000023206 | Il15ra | protein_coding | 0.213578478 | 3.391109557 | 0.1561949 | 0.697806 | 0.933043618 |
| ENSMUSG00000030029 | Lrig1 | protein_coding | -0.209653851 | 3.127578595 | 0.1561508 | 0.6978463 | 0.933043618 |
| ENSMUSG00000038467 | Chmp4b | protein_coding | -0.130998597 | 5.568378355 | 0.1560972 | 0.6978955 | 0.933043618 |
| ENSMUSG00000022000 | Zc3h13 | protein_coding | -0.124235601 | 5.432840269 | 0.155822 | 0.6981475 | 0.933264556 |
| ENSMUSG00000000131 | Xpo6 | protein_coding | 0.184837835 | 5.281730071 | 0.1559614 | 0.6983456 | 0.933413295 |
| ENSMUSG00000026080 | Chst10 | protein_coding | -0.109236558 | 5.767554448 | 0.1552764 | 0.6986481 | 0.933623579 |
| ENSMUSG00000015994 | Fnta | protein_coding | 0.125653242 | 5.94889738 | 0.1552455 | 0.6986765 | 0.933623579 |
| ENSMUSG00000025289 | Prdx4 | protein_coding | -0.118368402 | 5.927358822 | 0.1550485 | 0.6988576 | 0.933749435 |
| ENSMUSG00000061455 | Stx17 | protein_coding | -0.142915988 | 4.505653527 | 0.1548798 | 0.6990127 | 0.933840708 |
| ENSMUSG00000030750 | Nsmce1 | protein_coding | 0.09351743 | 6.353034097 | 0.1546483 | 0.6992258 | 0.934007176 |
| ENSMUSG00000028597 | Gpx7 | protein_coding | 0.186056422 | 3.864937553 | 0.1543993 | 0.6994551 | 0.934007176 |
| ENSMUSG00000075700 | Selenot | protein_coding | 0.121914489 | 5.747603591 | 0.1543458 | 0.6995045 | 0.934007176 |
| ENSMUSG00000045598 | Zfp553 | protein_coding | 0.171309532 | 3.911544166 | 0.1543319 | 0.6995173 | 0.934007176 |
| ENSMUSG00000037022 | Mmaa | protein_coding | -0.138689226 | 4.865514758 | 0.1542729 | 0.6995716 | 0.934007176 |
| ENSMUSG00000029705 | Cux1 | protein_coding | 0.106462093 | 6.038084386 | 0.1540389 | 0.6997875 | 0.934171184 |
| ENSMUSG00000032034 | Kcnj5 | protein_coding | 0.305940546 | 2.643768098 | 0.1539514 | 0.6998682 | 0.934171184 |
| ENSMUSG00000035530 | Eif1 | protein_coding | 0.104185143 | 9.399045328 | 0.153619 | 0.7001753 | 0.934342861 |
| ENSMUSG00000050103 | Agmo | protein_coding | -0.279340356 | 2.957131236 | 0.1534677 | 0.7003151 | 0.934342861 |
| ENSMUSG00000030226 | Lmo3 | protein_coding | -0.088876014 | 7.101001143 | 0.1534421 | 0.7003389 | 0.934342861 |
| ENSMUSG00000038072 | Galnt11 | protein_coding | -0.191590368 | 3.718150019 | 0.1533171 | 0.7004545 | 0.934342861 |
| ENSMUSG00000052837 | Junb | protein_coding | -0.276753402 | 2.756355389 | 0.1532731 | 0.7004951 | 0.934342861 |
| ENSMUSG00000027810 | Eif2a | protein_coding | 0.083033625 | 6.400073667 | 0.1531939 | 0.7005685 | 0.934342861 |
| ENSMUSG00000020849 | Ywhae | protein_coding | 0.09236723 | 11.16070672 | 0.1530623 | 0.7006903 | 0.934342861 |
| ENSMUSG00000015522 | Arnt | protein_coding | -0.188633782 | 3.379198166 | 0.1528928 | 0.7008474 | 0.934342861 |
| ENSMUSG00000036792 | Mbd5 | protein_coding | -0.124864984 | 4.79709081 | 0.1528395 | 0.7008968 | 0.934342861 |
| ENSMUSG00000010048 | Ifrd2 | protein_coding | -0.240288904 | 3.343851751 | 0.152833 | 0.7009028 | 0.934342861 |
| ENSMUSG00000089911 | Mfsd14a | protein_coding | -0.167925936 | 4.328370809 | 0.1526985 | 0.7010275 | 0.934342861 |
| ENSMUSG00000038811 | Gngt2 | protein_coding | -0.270211198 | 2.841497445 | 0.1526142 | 0.7011057 | 0.934342861 |
| ENSMUSG00000045983 | Eif4g1 | protein_coding | -0.12158398 | 5.20967708 | 0.1524286 | 0.701278 | 0.934342861 |
| ENSMUSG00000087403 | Kantr | protein_coding | -0.121182157 | 6.665898781 | 0.1523567 | 0.7013531 | 0.934342861 |
| ENSMUSG00000059897 | Zfp930 | protein_coding | 0.197106163 | 3.991080406 | 0.1523415 | 0.7013589 | 0.934342861 |
| ENSMUSG00000012535 | Tnpo3 | protein_coding | 0.158013244 | 5.407096462 | 0.15232 | 0.7013871 | 0.934342861 |
| ENSMUSG00000015745 | Plekho1 | protein_coding | -0.233552424 | 3.719621721 | 0.1521543 | 0.7015411 | 0.934432284 |
| ENSMUSG00000058706 | 0610030E20Rik | protein_coding | -0.257651225 | 2.85645559 | 0.1517959 | 0.7018661 | 0.934749362 |
| ENSMUSG00000039461 | Tcta | protein_coding | 0.103205192 | 6.791598511 | 0.1515003 | 0.7021414 | 0.934991064 |
| ENSMUSG00000004085 | Map3k20 | protein_coding | -0.278114558 | 3.11867091 | 0.1510178 | 0.7025914 | 0.934991064 |
| ENSMUSG00000037426 | Depdc5 | protein_coding | 0.148334389 | 4.586188718 | 0.1510158 | 0.7025932 | 0.934991064 |
| ENSMUSG00000029918 | Mrps33 | protein_coding | 0.105165345 | 6.964220476 | 0.150965 | 0.7026406 | 0.934991064 |
| ENSMUSG00000050565 | Tor1aip2 | protein_coding | -0.090211093 | 6.52170787 | 0.1509556 | 0.7026495 | 0.934991064 |
| ENSMUSG00000031683 | Lsm6 | protein_coding | -0.097293036 | 7.894789187 | 0.1508594 | 0.7027393 | 0.934991064 |
| ENSMUSG00000060477 | Irak2 | protein_coding | 0.183335347 | 3.333828828 | 0.150798 | 0.7027967 | 0.934991064 |
| ENSMUSG00000025142 | Aspscr1 | protein_coding | 0.203730053 | 3.519149347 | 0.150688 | 0.7028994 | 0.934991064 |
| ENSMUSG00000090946 | Ccdc71l | protein_coding | -0.304230937 | 2.466316339 | 0.1505951 | 0.7029863 | 0.934991064 |
| ENSMUSG00000018537 | Pcgf2 | protein_coding | -0.201623864 | 3.225708111 | 0.150539 | 0.7030387 | 0.934991064 |
| ENSMUSG00000032308 | Ulk3 | protein_coding | 0.27285936 | 3.059243722 | 0.1504435 | 0.7031281 | 0.934991064 |
| ENSMUSG00000045973 | Slc25a51 | protein_coding | -0.101936613 | 6.216469553 | 0.1504266 | 0.7031439 | 0.934991064 |
| ENSMUSG00000066149 | Cdc26 | protein_coding | 0.109249239 | 5.822246247 | 0.1503288 | 0.7032354 | 0.934991064 |
| ENSMUSG00000021707 | Dhfr | protein_coding | 0.213291533 | 3.269350618 | 0.1502973 | 0.7032649 | 0.934991064 |
| ENSMUSG00000049354 | Dcaf7 | protein_coding | -0.089662389 | 6.43481068 | 0.1500499 | 0.7034966 | 0.93518343 |
| ENSMUSG00000031985 | Gnpat | protein_coding | -0.127311928 | 5.414486197 | 0.1498498 | 0.7036841 | 0.935317093 |
| ENSMUSG00000015289 | Lage3 | protein_coding | -0.119523085 | 5.787581657 | 0.1494633 | 0.7040468 | 0.935579439 |
| ENSMUSG00000042502 | Cd2bp2 | protein_coding | 0.108924844 | 6.020395609 | 0.149454 | 0.7040555 | 0.935579439 |
| ENSMUSG00000026176 | Ctdsp1 | protein_coding | -0.206126878 | 3.061700666 | 0.1492884 | 0.7042111 | 0.935670534 |
| ENSMUSG00000029048 | Rer1 | protein_coding | 0.095175897 | 6.511122146 | 0.1490495 | 0.7044357 | 0.935811393 |
| ENSMUSG00000103034 | Gm8797 | protein_coding | 0.204307061 | 2.947511634 | 0.148914 | 0.7045632 | 0.935811393 |
| ENSMUSG00000014813 | Stc1 | protein_coding | 0.187859614 | 3.672506606 | 0.1488341 | 0.7046384 | 0.935811393 |
| ENSMUSG00000075229 | Ccdc58 | protein_coding | -0.134416036 | 4.814919982 | 0.1486743 | 0.7047889 | 0.935811393 |
| ENSMUSG00000021852 | Slc35f4 | protein_coding | 0.179001346 | 3.69587323 | 0.1485882 | 0.70487 | 0.935811393 |
| ENSMUSG00000006235 | Epor | protein_coding | -0.224291848 | 2.937804514 | 0.1485351 | 0.70492 | 0.935811393 |
| ENSMUSG00000025821 | Zfp282 | protein_coding | 0.2339006 | 3.051024875 | 0.1484419 | 0.7050079 | 0.935811393 |
| ENSMUSG00000034544 | Rsrc1 | protein_coding | -0.098122032 | 6.519231588 | 0.1484362 | 0.7050133 | 0.935811393 |
| ENSMUSG00000018821 | Avpi1 | protein_coding | -0.090385785 | 7.044228334 | 0.1480821 | 0.7053474 | 0.935928564 |
| ENSMUSG00000074748 | Atxn7l3b | protein_coding | 0.075953141 | 7.701684118 | 0.1479334 | 0.7054877 | 0.935928564 |
| ENSMUSG00000011306 | Sugp1 | protein_coding | 0.114528027 | 6.264808811 | 0.1479213 | 0.7054993 | 0.935928564 |
| ENSMUSG00000020538 | Srebf1 | protein_coding | -0.181761134 | 4.174964171 | 0.1478894 | 0.7055294 | 0.935928564 |
| ENSMUSG00000042249 | Grk3 | protein_coding | -0.176817717 | 4.939557764 | 0.1478059 | 0.7056165 | 0.935928564 |
| ENSMUSG00000057551 | Zfp317 | protein_coding | -0.15296455 | 4.910413976 | 0.1477894 | 0.7056238 | 0.935928564 |
| ENSMUSG00000051067 | Lingo3 | protein_coding | -0.231672818 | 3.708246974 | 0.1475858 | 0.7058245 | 0.936079323 |
| ENSMUSG00000038633 | Degs1 | protein_coding | 0.075163162 | 7.198477339 | 0.1473719 | 0.7060188 | 0.93612645 |
| ENSMUSG00000066438 | Plekhd1 | protein_coding | -0.306999587 | 2.44947377 | 0.1472645 | 0.7061205 | 0.93612645 |
| ENSMUSG00000026049 | Tex30 | protein_coding | -0.142562629 | 4.010276521 | 0.1472637 | 0.7061212 | 0.93612645 |
| ENSMUSG00000031026 | Trim66 | protein_coding | -0.172575264 | 3.792633113 | 0.1470124 | 0.7063594 | 0.936134215 |

| ensembl_gene_id | external_gene_name | gene_biotype | log2 (fold change) | log2 (counts per million) | F-value | p-value | False discovery rate |
|---|---|---|---|---|---|---|---|
| ENSMUSG00000020115 | Tbk1 | protein_coding | 0.139832422 | 4.763770872 | 0.1469995 | 0.7063717 | 0.936134215 |
| ENSMUSG00000022814 | Umps | protein_coding | 0.135555939 | 5.473291853 | 0.1469104 | 0.7064562 | 0.936134215 |
| ENSMUSG00000037905 | Bri3bp | protein_coding | -0.101994572 | 5.625456159 | 0.1468536 | 0.70651 | 0.936134215 |
| ENSMUSG00000055900 | Tmem69 | protein_coding | -0.173903405 | 3.362338281 | 0.1465847 | 0.7067652 | 0.936134215 |
| ENSMUSG00000020576 | Nbas | protein_coding | -0.168768073 | 4.351054025 | 0.1465602 | 0.7067885 | 0.936134215 |
| ENSMUSG00000029440 | Psmd9 | protein_coding | 0.094176313 | 6.265659844 | 0.1462232 | 0.7071088 | 0.936134215 |
| ENSMUSG00000039601 | Rcan2 | protein_coding | -0.115565778 | 8.189909291 | 0.1462403 | 0.7071133 | 0.936134215 |
| ENSMUSG00000028555 | Ttc39a | protein_coding | -0.310203341 | 2.707469355 | 0.1461978 | 0.7071329 | 0.936134215 |
| ENSMUSG00000061273 | Mmgt1 | protein_coding | -0.135153497 | 4.400901462 | 0.1460031 | 0.7073182 | 0.936134215 |
| ENSMUSG00000033966 | Cdkl4 | protein_coding | -0.170592577 | 4.235279325 | 0.1458708 | 0.7074442 | 0.936134215 |
| ENSMUSG00000049950 | Rpp38 | protein_coding | 0.106167735 | 5.107521382 | 0.1457411 | 0.7075678 | 0.936134215 |
| ENSMUSG00000025413 | Ttc4 | protein_coding | -0.119300255 | 5.924255249 | 0.1455986 | 0.7077036 | 0.936134215 |
| ENSMUSG00000044471 | Lncpint | lincRNA | -0.372254081 | 2.492313321 | 0.1455793 | 0.7077219 | 0.936134215 |
| ENSMUSG00000063239 | Grm4 | protein_coding | 0.13662211 | 4.528026052 | 0.1455297 | 0.7077693 | 0.936134215 |
| ENSMUSG00000039887 | Alg14 | protein_coding | -0.099827383 | 5.90807092 | 0.1455145 | 0.7077837 | 0.936134215 |
| ENSMUSG00000079084 | Ccdc82 | protein_coding | -0.158089933 | 4.377176218 | 0.1453721 | 0.7079195 | 0.936134215 |
| ENSMUSG00000058440 | Nrf1 | protein_coding | 0.12024327 | 4.588800568 | 0.1452519 | 0.7080343 | 0.936134215 |
| ENSMUSG00000043061 | Tmem18 | protein_coding | -0.131334006 | 4.516317477 | 0.1451484 | 0.7081332 | 0.936134215 |
| ENSMUSG00000025202 | Scd3 | protein_coding | 0.157464172 | 4.335583189 | 0.145116 | 0.7081641 | 0.936134215 |
| ENSMUSG00000030515 | Tarsl2 | protein_coding | -0.220741664 | 2.91404841 | 0.1450388 | 0.7082378 | 0.936134215 |
| ENSMUSG00000039670 | Oxld1 | protein_coding | -0.288530388 | 2.546771813 | 0.1449317 | 0.7083402 | 0.936134215 |
| ENSMUSG00000055725 | Paqr3 | protein_coding | -0.254291121 | 5.331199923 | 0.1463568 | 0.7083649 | 0.936134215 |
| ENSMUSG00000035131 | Brinp3 | protein_coding | 0.123182206 | 5.172183855 | 0.1448958 | 0.7083745 | 0.936134215 |
| ENSMUSG00000037656 | Slc20a2 | protein_coding | -0.145140445 | 4.918344931 | 0.1448112 | 0.7084554 | 0.936134215 |
| ENSMUSG00000052713 | Zfp608 | protein_coding | -0.159162123 | 4.004855985 | 0.1448061 | 0.7084603 | 0.936134215 |
| ENSMUSG00000048234 | Rnf149 | protein_coding | -0.245347778 | 3.818176469 | 0.1450236 | 0.7084777 | 0.936134215 |
| ENSMUSG00000039542 | Ncam1 | protein_coding | 0.093895509 | 8.750398292 | 0.1446492 | 0.7086104 | 0.936194517 |
| ENSMUSG00000040452 | Cdh12 | protein_coding | -0.247706776 | 3.24618239 | 0.1444387 | 0.708812 | 0.936256802 |
| ENSMUSG00000020745 | Pafah1b1 | protein_coding | -0.083316122 | 7.459927703 | 0.1443094 | 0.7089358 | 0.936256802 |
| ENSMUSG00000024095 | Hnrnpll | protein_coding | -0.143433911 | 5.037191559 | 0.1442658 | 0.7089776 | 0.936256802 |
| ENSMUSG00000021224 | Numb | protein_coding | 0.162474045 | 4.281775495 | 0.1442364 | 0.7090058 | 0.936256802 |
| ENSMUSG00000104784 | Gm42984 | antisense_RNA | 0.280517991 | 2.66238835 | 0.1440905 | 0.7091457 | 0.936295342 |
| ENSMUSG00000028988 | Ctnnbip1 | protein_coding | 0.197770625 | 3.844294527 | 0.1438888 | 0.7093392 | 0.936295342 |
| ENSMUSG00000046500 | Fam19a4 | protein_coding | -0.319787125 | 2.344135323 | 0.1438694 | 0.7093578 | 0.936295342 |
| ENSMUSG00000027303 | Ptpra | protein_coding | 0.129393813 | 4.260738267 | 0.1438171 | 0.709408 | 0.936295342 |
| ENSMUSG00000028373 | Astn2 | protein_coding | 0.127618868 | 5.382908985 | 0.1436756 | 0.7095439 | 0.936295342 |
| ENSMUSG00000052914 | Cyp2j6 | protein_coding | -0.198195679 | 4.464166553 | 0.1437064 | 0.7095574 | 0.936295342 |
| ENSMUSG00000020150 | Gamt | protein_coding | -0.178685219 | 3.771287056 | 0.1433709 | 0.7098368 | 0.936530027 |
| ENSMUSG00000038879 | Nipal2 | protein_coding | -0.311771944 | 3.33169737 | 0.1436911 | 0.7099094 | 0.936530027 |
| ENSMUSG00000039852 | Rere | protein_coding | -0.104552152 | 6.268421773 | 0.1429171 | 0.7102737 | 0.936588664 |
| ENSMUSG00000028165 | Cisd2 | protein_coding | -0.094986039 | 6.175455156 | 0.1428415 | 0.7103466 | 0.936588664 |
| ENSMUSG00000031790 | Mmp15 | protein_coding | -0.161778489 | 3.531035954 | 0.1427695 | 0.710416 | 0.936588664 |
| ENSMUSG00000063275 | Hacd1 | protein_coding | -0.289518235 | 2.469023942 | 0.142698 | 0.710485 | 0.936588664 |
| ENSMUSG00000058301 | Upf1 | protein_coding | -0.138398721 | 4.106842552 | 0.1425579 | 0.7106202 | 0.936588664 |
| ENSMUSG00000052512 | Nav2 | protein_coding | -0.135841685 | 5.38433187 | 0.1425396 | 0.7106378 | 0.936588664 |
| ENSMUSG00000051864 | Tbc1d22a | protein_coding | 0.144635473 | 4.965717648 | 0.1425374 | 0.7106399 | 0.936588664 |
| ENSMUSG00000018196 | Glrx2 | protein_coding | 0.120350498 | 5.315577468 | 0.1423486 | 0.7108223 | 0.936588664 |
| ENSMUSG00000031921 | Terf2 | protein_coding | 0.128591449 | 4.788759732 | 0.1422886 | 0.7108802 | 0.936588664 |
| ENSMUSG00000051674 | Dcun1d4 | protein_coding | -0.100322216 | 6.531076334 | 0.142282 | 0.7108865 | 0.936588664 |
| ENSMUSG00000076441 | Ass1 | protein_coding | -0.133505172 | 6.249848399 | 0.1423187 | 0.7109304 | 0.936588664 |
| ENSMUSG00000037740 | Mrps26 | protein_coding | 0.1162846 | 4.850606707 | 0.1421656 | 0.7109991 | 0.936588664 |
| ENSMUSG00000013701 | Timm23 | protein_coding | 0.194166372 | 3.06188704 | 0.1418446 | 0.7113096 | 0.936868724 |
| ENSMUSG00000040040 | Ift88 | protein_coding | 0.158646202 | 4.567561074 | 0.1417656 | 0.711386 | 0.936868724 |
| ENSMUSG00000053963 | Stum | protein_coding | 0.125485241 | 4.746367882 | 0.1413887 | 0.7117512 | 0.937102894 |
| ENSMUSG00000029480 | Dhx37 | protein_coding | 0.182709275 | 3.670939962 | 0.1413607 | 0.7117783 | 0.937102894 |
| ENSMUSG00000036959 | Bcorl1 | protein_coding | -0.171664199 | 3.776822993 | 0.1413124 | 0.7118252 | 0.937102894 |
| ENSMUSG00000053091 | Lins1 | protein_coding | -0.21425798 | 3.081821021 | 0.1408908 | 0.7122344 | 0.937526795 |
| ENSMUSG00000025817 | Nudt5 | protein_coding | -0.107958392 | 5.502467842 | 0.1406729 | 0.7124462 | 0.937690853 |
| ENSMUSG00000029415 | Sdad1 | protein_coding | 0.171977055 | 4.236613644 | 0.1405257 | 0.7125894 | 0.937764543 |
| ENSMUSG00000021484 | Lman2 | protein_coding | 0.126634693 | 5.333730486 | 0.1403604 | 0.7127503 | 0.937861466 |
| ENSMUSG00000027573 | Gid8 | protein_coding | 0.114806811 | 5.507733195 | 0.1402209 | 0.7128862 | 0.937862931 |
| ENSMUSG00000056900 | Usp13 | protein_coding | -0.159501023 | 4.009411994 | 0.1400932 | 0.7130107 | 0.937862931 |
| ENSMUSG00000033065 | Pfkm | protein_coding | 0.106549274 | 8.237464808 | 0.1399816 | 0.7131274 | 0.937862931 |
| ENSMUSG00000041642 | Kif21b | protein_coding | 0.183117441 | 3.805694586 | 0.1398899 | 0.7132089 | 0.937862931 |
| ENSMUSG00000028961 | Pgd | protein_coding | 0.124306048 | 5.789461953 | 0.1398461 | 0.7132516 | 0.937862931 |
| ENSMUSG00000039474 | Wfs1 | protein_coding | -0.101523824 | 6.511998723 | 0.1396973 | 0.7133968 | 0.937862931 |
| ENSMUSG00000042042 | Csgalnact2 | protein_coding | -0.270022276 | 2.890965698 | 0.1396577 | 0.7134355 | 0.937862931 |
| ENSMUSG00000025742 | Prps2 | protein_coding | -0.133065946 | 4.834751372 | 0.1395486 | 0.7135421 | 0.937862931 |
| ENSMUSG00000039953 | Clstn1 | protein_coding | 0.082010894 | 7.959736909 | 0.139405 | 0.7136824 | 0.937862931 |
| ENSMUSG00000052812 | Atad2b | protein_coding | -0.221949341 | 3.200018374 | 0.1392855 | 0.7137992 | 0.937862931 |
| ENSMUSG00000030527 | Crtc3 | protein_coding | 0.244411998 | 3.106435 | 0.139212 | 0.7138711 | 0.937862931 |
| ENSMUSG00000016382 | Pls3 | protein_coding | -0.091295577 | 6.281365613 | 0.1391796 | 0.7139029 | 0.937862931 |
| ENSMUSG00000030424 | Zfp939 | transcribed_unproces | -0.198169571 | 3.067086335 | 0.1391042 | 0.7139766 | 0.937862931 |
| ENSMUSG00000003458 | Ncstn | protein_coding | 0.119304328 | 5.951255116 | 0.1390283 | 0.7140509 | 0.937862931 |
| ENSMUSG00000071650 | Ganab | protein_coding | 0.091663561 | 5.829963717 | 0.1389588 | 0.714119 | 0.937862931 |
| ENSMUSG00000021311 | Mtr | protein_coding | -0.226110239 | 3.017225109 | 0.1387575 | 0.7143162 | 0.937862931 |
| ENSMUSG00000030678 | Maz | protein_coding | 0.150887801 | 4.644659329 | 0.13873 | 0.7143431 | 0.937862931 |
| ENSMUSG00000085913 | Gm15601 | antisense_RNA | -0.261668867 | 2.707428456 | 0.138722 | 0.714351 | 0.937862931 |

| ensembl_gene_id | external_gene_name | gene_biotype | log2 (fold change) | log2 (counts per million) | F-value | p-value | False discovery rate |
|---|---|---|---|---|---|---|---|
| ENSMUSG00000011832 | Evi5l | protein_coding | -0.110412547 | 4.973127487 | 0.1386633 | 0.7144086 | 0.937862931 |
| ENSMUSG00000071860 | 2900055J20Rik | protein_coding | 0.168225525 | 3.835069793 | 0.1384628 | 0.7146052 | 0.938006591 |
| ENSMUSG00000022426 | Josd1 | protein_coding | 0.135652405 | 5.076976247 | 0.1380624 | 0.7149985 | 0.938030488 |
| ENSMUSG00000020766 | Galk1 | protein_coding | 0.159509586 | 4.268817448 | 0.137991 | 0.7150686 | 0.938030488 |
| ENSMUSG00000038615 | Nfe2l1 | protein_coding | -0.076540489 | 6.872197934 | 0.1377921 | 0.7152643 | 0.938030488 |
| ENSMUSG00000057914 | Cacnb2 | protein_coding | -0.156124033 | 4.629845211 | 0.137637 | 0.715417 | 0.938030488 |
| ENSMUSG00000023284 | Zfp605 | protein_coding | -0.195214915 | 3.455143663 | 0.1376369 | 0.7154171 | 0.938030488 |
| ENSMUSG00000025626 | Phf6 | protein_coding | -0.122086112 | 4.926960192 | 0.1373888 | 0.7156615 | 0.938030488 |
| ENSMUSG00000047044 | D030056L22Rik | protein_coding | 0.155671564 | 3.731754147 | 0.1373305 | 0.715719 | 0.938030488 |
| ENSMUSG00000040822 | 1700123O20Rik | protein_coding | 0.08741601 | 6.609926099 | 0.1368572 | 0.7161861 | 0.938030488 |
| ENSMUSG00000042496 | Prdm10 | protein_coding | -0.203425068 | 3.93350366 | 0.1367055 | 0.7163438 | 0.938030488 |
| ENSMUSG00000010592 | Dazl | protein_coding | 0.454727414 | 2.350461149 | 0.1367804 | 0.7164205 | 0.938030488 |
| ENSMUSG00000041020 | Map7d2 | protein_coding | 0.070400896 | 7.664329673 | 0.1365855 | 0.7164547 | 0.938030488 |
| ENSMUSG00000029641 | Rasl11a | protein_coding | 0.227973994 | 3.948291261 | 0.1368456 | 0.71646 | 0.938030488 |
| ENSMUSG00000026749 | Nek6 | protein_coding | -0.151384758 | 4.331659509 | 0.136379 | 0.716659 | 0.938030488 |
| ENSMUSG00000062184 | Hs6st2 | protein_coding | -0.108471355 | 5.469406566 | 0.1362253 | 0.7168112 | 0.938030488 |
| ENSMUSG00000029599 | Ddx54 | protein_coding | -0.206834574 | 3.394293668 | 0.1361502 | 0.7168855 | 0.938030488 |
| ENSMUSG00000037001 | Zfp39 | protein_coding | -0.169042933 | 3.344581473 | 0.1359857 | 0.7170486 | 0.938030488 |
| ENSMUSG00000067860 | Zic3 | protein_coding | -0.14320611 | 3.910919561 | 0.1358948 | 0.7171388 | 0.938030488 |
| ENSMUSG00000016946 | Kctd5 | protein_coding | -0.308575855 | 3.874375361 | 0.1366403 | 0.7171586 | 0.938030488 |
| ENSMUSG00000029916 | Agk | protein_coding | 0.112138099 | 4.77577093 | 0.1357303 | 0.7173021 | 0.938030488 |
| ENSMUSG00000086429 | Gt(ROSA)26Sor | antisense_RNA | -0.150549099 | 5.287847624 | 0.1356406 | 0.7173989 | 0.938030488 |
| ENSMUSG00000042155 | Klhl23 | protein_coding | -0.118914094 | 5.279083359 | 0.1354236 | 0.7176066 | 0.938030488 |
| ENSMUSG00000001334 | Fndc5 | protein_coding | 0.128886721 | 5.763481412 | 0.1353843 | 0.7176534 | 0.938030488 |
| ENSMUSG00000033781 | Asb13 | protein_coding | 0.098906349 | 5.697185403 | 0.1353157 | 0.7177139 | 0.938030488 |
| ENSMUSG00000025134 | Alyref | protein_coding | -0.140401028 | 4.548844003 | 0.1352625 | 0.7177668 | 0.938030488 |
| ENSMUSG00000075704 | Txnrd2 | protein_coding | 0.177851964 | 3.669811175 | 0.1352451 | 0.7177841 | 0.938030488 |
| ENSMUSG00000040447 | Spns2 | protein_coding | 0.202452028 | 4.353384316 | 0.1354441 | 0.7177846 | 0.938030488 |
| ENSMUSG00000041297 | Cdk13 | protein_coding | -0.178971274 | 4.689720122 | 0.1351818 | 0.7178753 | 0.938030488 |
| ENSMUSG00000024070 | Prkd3 | protein_coding | 0.174228297 | 3.494450604 | 0.135081 | 0.7179474 | 0.938030488 |
| ENSMUSG00000047242 | Taf9b | protein_coding | -0.101145107 | 6.562037021 | 0.1350492 | 0.717979 | 0.938030488 |
| ENSMUSG00000028821 | Syf2 | protein_coding | -0.086705882 | 7.347891965 | 0.1349301 | 0.7180976 | 0.938030488 |
| ENSMUSG00000061983 | Rps12 | protein_coding | 0.19241546 | 3.176514267 | 0.1348773 | 0.7181502 | 0.938030488 |
| ENSMUSG00000058761 | Rnf169 | protein_coding | 0.101480813 | 5.562580829 | 0.1347266 | 0.7183004 | 0.938030488 |
| ENSMUSG00000057177 | Gsk3a | protein_coding | 0.112197025 | 4.855785757 | 0.1346726 | 0.7183542 | 0.938030488 |
| ENSMUSG00000026773 | Pfkfb3 | protein_coding | 0.205806895 | 3.827403228 | 0.1345194 | 0.7185148 | 0.938030488 |
| ENSMUSG00000066148 | Prpf4 | protein_coding | 0.128376929 | 4.626979966 | 0.1344955 | 0.7185309 | 0.938030488 |
| ENSMUSG00000021938 | Pspc1 | protein_coding | -0.149302316 | 4.338091377 | 0.1343613 | 0.7186648 | 0.938030488 |
| ENSMUSG00000018008 | Cyth4 | protein_coding | 0.160027357 | 3.659424417 | 0.13432 | 0.7187061 | 0.938030488 |
| ENSMUSG00000013662 | Atad1 | protein_coding | -0.091904635 | 6.835617778 | 0.134264 | 0.7187619 | 0.938030488 |
| ENSMUSG00000020305 | Asb3 | protein_coding | 0.117634281 | 5.64929895 | 0.1341964 | 0.7188295 | 0.938030488 |
| ENSMUSG00000079316 | Rab9 | protein_coding | 0.093318755 | 6.864754682 | 0.1340882 | 0.7189376 | 0.938030488 |
| ENSMUSG00000071379 | Hpcal1 | protein_coding | 0.140537543 | 7.646691765 | 0.1346545 | 0.7190332 | 0.938030488 |
| ENSMUSG00000028478 | Clta | protein_coding | 0.09605144 | 6.887830154 | 0.1339415 | 0.7190843 | 0.938030488 |
| ENSMUSG00000018965 | Ywhah | protein_coding | 0.074629907 | 8.259081373 | 0.1339165 | 0.7191093 | 0.938030488 |
| ENSMUSG00000056124 | B4galt6 | protein_coding | -0.086150583 | 6.352546689 | 0.133871 | 0.7191548 | 0.938030488 |
| ENSMUSG00000038335 | Tsr1 | protein_coding | -0.122604605 | 5.146566259 | 0.1338305 | 0.7191953 | 0.938030488 |
| ENSMUSG00000024220 | Zfp523 | protein_coding | -0.157111326 | 4.664263773 | 0.1337597 | 0.7192662 | 0.938030488 |
| ENSMUSG00000031767 | Nudt7 | protein_coding | -0.11978051 | 4.349246878 | 0.1337478 | 0.719278 | 0.938030488 |
| ENSMUSG00000041679 | Lrrc29 | protein_coding | 0.235341229 | 2.848384261 | 0.1336544 | 0.7193716 | 0.938030488 |
| ENSMUSG00000028249 | Sdcbp | protein_coding | -0.079140566 | 8.821410057 | 0.1334649 | 0.7195614 | 0.938030488 |
| ENSMUSG00000044442 | N6amt1 | protein_coding | -0.07831823 | 6.41823056 | 0.1331947 | 0.7198324 | 0.938030488 |
| ENSMUSG00000038876 | Rnf146 | protein_coding | 0.116757328 | 4.722159032 | 0.1331376 | 0.7198897 | 0.938030488 |
| ENSMUSG00000050910 | Cdr2l | protein_coding | 0.136972411 | 5.033813653 | 0.1330429 | 0.7199847 | 0.938030488 |
| ENSMUSG00000021047 | Nova1 | protein_coding | 0.092950895 | 6.313334427 | 0.1330396 | 0.7199881 | 0.938030488 |
| ENSMUSG00000043262 | Uevld | protein_coding | -0.258190251 | 2.707248116 | 0.1330344 | 0.7199933 | 0.938030488 |
| ENSMUSG00000024807 | Syvn1 | protein_coding | 0.135484326 | 4.645465583 | 0.1329543 | 0.7200737 | 0.938030488 |
| ENSMUSG00000020220 | Vps13d | protein_coding | 0.163851399 | 4.030778721 | 0.1328648 | 0.7201636 | 0.938030488 |
| ENSMUSG00000052727 | Map1b | protein_coding | -0.178931662 | 9.563344979 | 0.1341126 | 0.7202313 | 0.938030488 |
| ENSMUSG00000021136 | Smoc1 | protein_coding | -0.203215406 | 3.176641471 | 0.1327326 | 0.7202964 | 0.938030488 |
| ENSMUSG00000022789 | Dnm1l | protein_coding | 0.081205233 | 7.672574536 | 0.1326614 | 0.7203681 | 0.938030488 |
| ENSMUSG00000004347 | Pde1c | protein_coding | -0.130856541 | 5.082863341 | 0.1324876 | 0.7205429 | 0.938030488 |
| ENSMUSG00000041225 | Arhgap12 | protein_coding | -0.100146791 | 5.38148747 | 0.1324835 | 0.7205471 | 0.938030488 |
| ENSMUSG00000039795 | Zfand1 | protein_coding | -0.10583675 | 5.231060953 | 0.1324564 | 0.7205743 | 0.938030488 |
| ENSMUSG00000034108 | Ccs | protein_coding | -0.088530095 | 6.176152604 | 0.1324465 | 0.7205843 | 0.938030488 |
| ENSMUSG00000039220 | Ppp1r10 | protein_coding | -0.138253094 | 5.239057029 | 0.1323911 | 0.72064 | 0.938030488 |
| ENSMUSG00000032867 | Fbxw8 | protein_coding | -0.260432177 | 2.483064752 | 0.1322153 | 0.7208171 | 0.938030488 |
| ENSMUSG00000092035 | Peg10 | protein_coding | 0.124024067 | 4.61487077 | 0.1321532 | 0.7208797 | 0.938030488 |
| ENSMUSG00000028883 | Sema3a | protein_coding | -0.208489067 | 3.199347871 | 0.1321412 | 0.7208918 | 0.938030488 |
| ENSMUSG00000046567 | 4930430F08Rik | protein_coding | 0.12959225 | 4.367057172 | 0.1321007 | 0.7209325 | 0.938030488 |
| ENSMUSG00000003437 | Paf1 | protein_coding | -0.165161236 | 5.326865428 | 0.1322657 | 0.7209621 | 0.938030488 |
| ENSMUSG00000074182 | Znhit6 | protein_coding | 0.116450128 | 4.740682252 | 0.1320653 | 0.7209682 | 0.938030488 |
| ENSMUSG00000044071 | Fam19a2 | protein_coding | -0.142604767 | 4.660714796 | 0.1318667 | 0.7211685 | 0.938030488 |
| ENSMUSG00000031442 | Mcf2l | protein_coding | -0.127193139 | 4.428412457 | 0.1317689 | 0.7212673 | 0.938030488 |
| ENSMUSG00000050721 | Plekho2 | protein_coding | -0.144089725 | 4.310670604 | 0.1317551 | 0.7212813 | 0.938030488 |
| ENSMUSG00000026116 | Tmem131 | protein_coding | -0.149771454 | 3.967843613 | 0.1316164 | 0.7214213 | 0.938030488 |
| ENSMUSG00000024758 | Rtn3 | protein_coding | -0.088067026 | 8.667423783 | 0.1315658 | 0.7214724 | 0.938030488 |
| ENSMUSG00000028931 | Kcnab2 | protein_coding | 0.080664686 | 7.385451449 | 0.1314319 | 0.7216077 | 0.938030488 |

| ensembl_gene_id | external_gene_name | gene_biotype | log2 (fold change) | log2 (counts per million) | F-value | p-value | False discovery rate |
|---|---|---|---|---|---|---|---|
| ENSMUSG00000063316 | Rpl27 | protein_coding | 0.150597437 | 3.778532114 | 0.131388 | 0.7216521 | 0.938030488 |
| ENSMUSG00000062421 | Arf2 | protein_coding | 0.10341122 | 6.704694964 | 0.1312947 | 0.7217465 | 0.938030488 |
| ENSMUSG00000007646 | Rad51c | protein_coding | 0.227024346 | 3.204790489 | 0.1311723 | 0.7218703 | 0.938030488 |
| ENSMUSG00000046572 | Zfp518b | protein_coding | -0.302264229 | 2.427074338 | 0.131158 | 0.7218848 | 0.938030488 |
| ENSMUSG00000052584 | Serp2 | protein_coding | 0.08089103 | 6.425509516 | 0.1311429 | 0.7219 | 0.938030488 |
| ENSMUSG00000042569 | Dhrs7b | protein_coding | 0.107827726 | 5.481559796 | 0.1310698 | 0.7219741 | 0.938030488 |
| ENSMUSG00000029004 | Kmt2e | protein_coding | 0.105413177 | 6.199021232 | 0.1310231 | 0.7220214 | 0.938030488 |
| ENSMUSG00000061689 | Dlgap4 | protein_coding | -0.087517966 | 6.687834921 | 0.1309786 | 0.7220665 | 0.938030488 |
| ENSMUSG00000031353 | Rbbp7 | protein_coding | 0.086939621 | 7.023521773 | 0.13095 | 0.7220954 | 0.938030488 |
| ENSMUSG00000016319 | Slc25a5 | protein_coding | -0.070315676 | 9.055108358 | 0.1308612 | 0.7221854 | 0.938030488 |
| ENSMUSG00000043448 | Gjc2 | protein_coding | -0.325242402 | 2.313014584 | 0.1308342 | 0.7222128 | 0.938030488 |
| ENSMUSG00000041133 | Smc1a | protein_coding | -0.105921188 | 6.176882888 | 0.1306903 | 0.7223586 | 0.938051581 |
| ENSMUSG00000042167 | Papd4 | protein_coding | -0.190068477 | 3.182694362 | 0.1306461 | 0.7224035 | 0.938051581 |
| ENSMUSG00000032549 | Rab6b | protein_coding | 0.075280945 | 8.76489705 | 0.1305399 | 0.7225113 | 0.938078189 |
| ENSMUSG00000073771 | Btbd19 | protein_coding | -0.204336427 | 3.129784981 | 0.1301768 | 0.7228801 | 0.938443734 |
| ENSMUSG00000075410 | Prcd | protein_coding | -0.273329046 | 3.463079104 | 0.1301144 | 0.7232383 | 0.938795474 |
| ENSMUSG00000068706 | Gm10250 | protein_coding | 0.170582968 | 3.267852304 | 0.1297074 | 0.7233577 | 0.938837048 |
| ENSMUSG00000025591 | Tma16 | protein_coding | -0.125486686 | 4.834137173 | 0.1295294 | 0.7235391 | 0.938959232 |
| ENSMUSG00000026418 | Tnni1 | protein_coding | -0.110051869 | 4.722029144 | 0.1293573 | 0.7237146 | 0.939073589 |
| ENSMUSG00000053519 | Kcnip1 | protein_coding | 0.090108682 | 6.198908436 | 0.1291866 | 0.7238887 | 0.93914049 |
| ENSMUSG00000007415 | Gatad1 | protein_coding | 0.092702985 | 7.549900113 | 0.1291356 | 0.7239408 | 0.93914049 |
| ENSMUSG00000060073 | Psma3 | protein_coding | 0.091325988 | 6.12581061 | 0.1288482 | 0.7242345 | 0.939408189 |
| ENSMUSG00000032547 | Ryk | protein_coding | -0.190534794 | 3.831005932 | 0.1286602 | 0.7244268 | 0.93944686 |
| ENSMUSG00000029482 | Aacs | protein_coding | -0.14611665 | 4.888742531 | 0.1286482 | 0.7244391 | 0.93944686 |
| ENSMUSG00000006476 | Nsmf | protein_coding | -0.132702119 | 6.355059657 | 0.1285889 | 0.7247927 | 0.939792067 |
| ENSMUSG00000003581 | Rnf215 | protein_coding | 0.154283751 | 3.454155927 | 0.1279733 | 0.7251308 | 0.940064937 |
| ENSMUSG00000049091 | Sephs2 | protein_coding | -0.156338521 | 5.725464198 | 0.1282124 | 0.725178 | 0.940064937 |
| ENSMUSG00000058407 | Txndc9 | protein_coding | -0.076155672 | 7.320790892 | 0.1277578 | 0.7253521 | 0.940177332 |
| ENSMUSG00000029291 | Rufy3 | protein_coding | 0.080365378 | 8.129277051 | 0.1274976 | 0.7256196 | 0.940245705 |
| ENSMUSG00000022812 | Gsk3b | protein_coding | -0.144251776 | 9.531558881 | 0.1280894 | 0.7256547 | 0.940245705 |
| ENSMUSG00000061461 | Smim20 | protein_coding | -0.072825126 | 7.151173044 | 0.1274513 | 0.7256672 | 0.940245705 |
| ENSMUSG00000051401 | Kctd16 | protein_coding | -0.226182281 | 2.779442668 | 0.1272009 | 0.7259249 | 0.94042124 |
| ENSMUSG00000015748 | Prpf3 | protein_coding | 0.149666149 | 4.036123094 | 0.1269703 | 0.7261626 | 0.94042124 |
| ENSMUSG00000048796 | Cyb561d1 | protein_coding | 0.182215846 | 3.55124083 | 0.1268667 | 0.7262694 | 0.94042124 |
| ENSMUSG00000031451 | Gas6 | protein_coding | 0.230673384 | 2.816422772 | 0.1267919 | 0.7263465 | 0.94042124 |
| ENSMUSG00000047797 | Gjb1 | protein_coding | -0.148526253 | 6.266397011 | 0.127148 | 0.7264357 | 0.94042124 |
| ENSMUSG00000027963 | Extl2 | protein_coding | 0.081061851 | 6.745595982 | 0.126661 | 0.7264817 | 0.94042124 |
| ENSMUSG00000064302 | Clasp1 | protein_coding | 0.117765073 | 5.440027815 | 0.1266338 | 0.7265097 | 0.94042124 |
| ENSMUSG00000024787 | Snx15 | protein_coding | 0.078111465 | 6.245294448 | 0.1265812 | 0.7265641 | 0.94042124 |
| ENSMUSG00000022634 | Yaf2 | protein_coding | -0.08121565 | 6.893968802 | 0.1265415 | 0.7266051 | 0.94042124 |
| ENSMUSG00000024493 | Lars | protein_coding | -0.111201072 | 5.57129113 | 0.1264717 | 0.7266772 | 0.94042124 |
| ENSMUSG00000032743 | D430042O09Rik | protein_coding | 0.253030692 | 2.810748478 | 0.1260883 | 0.7270737 | 0.940599818 |
| ENSMUSG00000032477 | Cdc25a | protein_coding | -0.254612315 | 3.205353042 | 0.1260749 | 0.7270951 | 0.940599818 |
| ENSMUSG00000005204 | Senp3 | protein_coding | -0.118032401 | 5.074071555 | 0.1259782 | 0.7271877 | 0.940599818 |
| ENSMUSG00000048960 | Prex2 | protein_coding | -0.326278873 | 3.109215943 | 0.1261839 | 0.7272929 | 0.940599818 |
| ENSMUSG00000035595 | 1600002K03Rik | protein_coding | 0.16367532 | 3.89745917 | 0.1258086 | 0.7273634 | 0.940599818 |
| ENSMUSG00000022747 | St3gal6 | protein_coding | -0.172748851 | 3.685599646 | 0.1257764 | 0.7273968 | 0.940599818 |
| ENSMUSG00000029233 | Srd5a3 | protein_coding | 0.199104669 | 3.477859204 | 0.1256839 | 0.7274927 | 0.940599818 |
| ENSMUSG00000027397 | Slc20a1 | protein_coding | -0.21857646 | 2.862421355 | 0.1256624 | 0.727515 | 0.940599818 |
| ENSMUSG00000050164 | Mchr1 | protein_coding | 0.156849406 | 4.577819094 | 0.1255412 | 0.7276407 | 0.94062629 |
| ENSMUSG00000058298 | Mcm9 | protein_coding | -0.220674255 | 2.917679339 | 0.125474 | 0.7277104 | 0.94062629 |
| ENSMUSG00000000605 | Clcn4 | protein_coding | 0.076414835 | 7.351839235 | 0.1249648 | 0.7282396 | 0.941197149 |
| ENSMUSG00000021384 | Susd3 | protein_coding | -0.257295659 | 2.919735398 | 0.1247694 | 0.7284429 | 0.941257549 |
| ENSMUSG00000044991 | 1110034G24Rik | protein_coding | 0.115277316 | 5.267753978 | 0.1246947 | 0.7285207 | 0.941257549 |
| ENSMUSG00000033595 | Lgi3 | protein_coding | 0.096366938 | 5.970040407 | 0.1246004 | 0.7286189 | 0.941257549 |
| ENSMUSG00000078786 | BC024978 | protein_coding | 0.147220858 | 4.375174023 | 0.1244149 | 0.7288124 | 0.941257549 |
| ENSMUSG00000027722 | Spata5 | protein_coding | 0.168943189 | 3.663025969 | 0.1243889 | 0.7288394 | 0.941257549 |
| ENSMUSG00000046844 | Vat1l | protein_coding | 0.08964398 | 7.407664796 | 0.1243483 | 0.7288819 | 0.941257549 |
| ENSMUSG00000025246 | Tbl1x | protein_coding | -0.093983345 | 6.155743411 | 0.1243318 | 0.7288991 | 0.941257549 |
| ENSMUSG00000022044 | Stmn4 | protein_coding | -0.094517848 | 8.71982359 | 0.1240843 | 0.7291574 | 0.941368869 |
| ENSMUSG00000028345 | Tex10 | protein_coding | -0.147607857 | 4.033069419 | 0.1240815 | 0.7291604 | 0.941368869 |
| ENSMUSG00000021112 | Mpp5 | protein_coding | -0.090036473 | 5.843263303 | 0.1238366 | 0.7294163 | 0.94157136 |
| ENSMUSG00000037017 | Zscan21 | protein_coding | 0.140990766 | 4.961300475 | 0.1236737 | 0.7295867 | 0.94157136 |
| ENSMUSG00000031453 | Rasa3 | protein_coding | 0.115336573 | 4.952652906 | 0.1236132 | 0.72965 | 0.94157136 |
| ENSMUSG00000021936 | Mapk8 | protein_coding | -0.104449363 | 5.316795201 | 0.1235632 | 0.7297023 | 0.94157136 |
| ENSMUSG00000025503 | Taldo1 | protein_coding | 0.069323131 | 8.388558491 | 0.1235129 | 0.729755 | 0.94157136 |
| ENSMUSG00000033214 | Slitrk5 | protein_coding | -0.115913978 | 4.334733499 | 0.1232735 | 0.7300058 | 0.941781959 |
| ENSMUSG00000040428 | Plekha4 | protein_coding | -0.230801005 | 2.958626033 | 0.1230892 | 0.7301992 | 0.941918362 |
| ENSMUSG00000022340 | Sybu | protein_coding | -0.074771229 | 6.519935658 | 0.1230031 | 0.7302896 | 0.94192199 |
| ENSMUSG00000011837 | Snapc2 | protein_coding | 0.115238557 | 5.29531142 | 0.1226968 | 0.7306113 | 0.942223953 |
| ENSMUSG00000024782 | Ak3 | protein_coding | -0.209829387 | 2.964821122 | 0.1226046 | 0.7307082 | 0.942235909 |
| ENSMUSG00000005413 | Hmox1 | protein_coding | 0.168421916 | 3.43502437 | 0.1223628 | 0.7309626 | 0.942272846 |
| ENSMUSG00000038916 | Soga3 | protein_coding | -0.120791769 | 5.687258894 | 0.1223545 | 0.7309714 | 0.942272846 |
| ENSMUSG00000054679 | Srsf12 | protein_coding | -0.144258868 | 5.326645041 | 0.1222496 | 0.7310891 | 0.942272846 |
| ENSMUSG00000038886 | Man2a2 | protein_coding | -0.169328539 | 3.766803742 | 0.1221361 | 0.7312014 | 0.942272846 |
| ENSMUSG00000037622 | Wdtc1 | protein_coding | 0.12096831 | 4.902716224 | 0.121928 | 0.7314208 | 0.942272846 |
| ENSMUSG00000039178 | Tbc1d19 | protein_coding | 0.092688598 | 5.520886412 | 0.1219051 | 0.731445 | 0.942272846 |
| ENSMUSG00000096974 | Gm26881 | antisense_RNA | -0.197240059 | 2.940711554 | 0.1218075 | 0.731548 | 0.942272846 |

| ensembl_gene_id | external_gene_name | gene_biotype | log2 (fold change) | log2 (counts per million) | F-value | p-value | False discovery rate |
|---|---|---|---|---|---|---|---|
| ENSMUSG00000060657 | Marf1 | protein_coding | -0.225178915 | 7.253309898 | 0.124152 | 0.7317357 | 0.942272846 |
| ENSMUSG00000055692 | Tmem191c | protein_coding | -0.106066891 | 5.673885743 | 0.1215277 | 0.7318435 | 0.942272846 |
| ENSMUSG00000028484 | Psip1 | protein_coding | 0.115648826 | 4.976152638 | 0.1215007 | 0.731872 | 0.942272846 |
| ENSMUSG00000046791 | Riox1 | protein_coding | 0.222535921 | 2.938538894 | 0.1214969 | 0.7318761 | 0.942272846 |
| ENSMUSG00000094786 | Gm14403 | protein_coding | -0.176788139 | 3.592279995 | 0.1214557 | 0.7319196 | 0.942272846 |
| ENSMUSG00000070699 | Sars2 | protein_coding | 0.161556327 | 4.124511531 | 0.1213831 | 0.7319964 | 0.942272846 |
| ENSMUSG00000040606 | Kazn | protein_coding | 0.102012342 | 5.968817437 | 0.1213313 | 0.7320511 | 0.942272846 |
| ENSMUSG00000041817 | Fam169a | protein_coding | -0.118355576 | 5.870143142 | 0.1212796 | 0.7321059 | 0.942272846 |
| ENSMUSG00000026925 | Inpp5e | protein_coding | 0.16212024 | 3.412562807 | 0.1212484 | 0.7321389 | 0.942272846 |
| ENSMUSG00000022559 | Fbxl6 | protein_coding | -0.200738313 | 3.018726712 | 0.1211589 | 0.7322336 | 0.942281918 |
| ENSMUSG00000031843 | Mphosph6 | protein_coding | -0.097821094 | 6.021009943 | 0.1210142 | 0.7323868 | 0.942346664 |
| ENSMUSG00000041168 | Lonp1 | protein_coding | -0.103212285 | 4.990670459 | 0.1209459 | 0.7324592 | 0.942346664 |
| ENSMUSG00000017286 | Glod4 | protein_coding | 0.069501944 | 7.522430869 | 0.1206866 | 0.7327342 | 0.942510871 |
| ENSMUSG00000018995 | Nars2 | protein_coding | -0.15979546 | 3.687330034 | 0.1206409 | 0.7327826 | 0.942510871 |
| ENSMUSG00000022037 | Clu | protein_coding | 0.090471699 | 9.99337249 | 0.1205777 | 0.7328498 | 0.942510871 |
| ENSMUSG00000028982 | Slc25a33 | protein_coding | 0.156326046 | 4.497904763 | 0.1204024 | 0.7330359 | 0.942587613 |
| ENSMUSG00000031479 | Vps36 | protein_coding | -0.105511594 | 5.319831092 | 0.1203421 | 0.7331 | 0.942587613 |
| ENSMUSG00000021901 | Bap1 | protein_coding | -0.072486354 | 6.542753954 | 0.120235 | 0.7332138 | 0.942587613 |
| ENSMUSG00000038233 | Fam198a | protein_coding | 0.338763429 | 2.170744089 | 0.1201114 | 0.7333452 | 0.942587613 |
| ENSMUSG00000066036 | Ubr4 | protein_coding | -0.11787089 | 5.281390209 | 0.1200778 | 0.733381 | 0.942587613 |
| ENSMUSG00000038533 | Cbfa2t2 | protein_coding | 0.139908342 | 6.28692202 | 0.1204433 | 0.7334354 | 0.942587613 |
| ENSMUSG00000036766 | Dner | protein_coding | -0.099550682 | 6.807641202 | 0.1197494 | 0.733738 | 0.942863853 |
| ENSMUSG00000042589 | Cux2 | protein_coding | -0.186067649 | 5.710128165 | 0.1201829 | 0.7339759 | 0.943056836 |
| ENSMUSG00000040111 | Gramd1b | protein_coding | -0.100298984 | 5.352504123 | 0.1192029 | 0.7343138 | 0.943290515 |
| ENSMUSG00000021066 | Atl1 | protein_coding | 0.103150659 | 6.57713241 | 0.1191318 | 0.7343971 | 0.943290515 |
| ENSMUSG00000020092 | Pald1 | protein_coding | 0.311335094 | 2.379014668 | 0.1191027 | 0.7344209 | 0.943290515 |
| ENSMUSG00000041592 | Sdk2 | protein_coding | 0.212041127 | 3.169684741 | 0.118933 | 0.7346024 | 0.943410856 |
| ENSMUSG00000034706 | Dnaic2 | protein_coding | -0.149737391 | 3.603919497 | 0.1186452 | 0.7349105 | 0.943686557 |
| ENSMUSG00000020111 | Micu1 | protein_coding | 0.079402164 | 7.00800492 | 0.1184903 | 0.7350765 | 0.943686557 |
| ENSMUSG00000041548 | Hspb8 | protein_coding | -0.12494804 | 4.908712143 | 0.1181181 | 0.7354758 | 0.943686557 |
| ENSMUSG00000001415 | Smg5 | protein_coding | 0.113685351 | 4.749983484 | 0.1180448 | 0.7355546 | 0.943686557 |
| ENSMUSG00000029246 | Ppat | protein_coding | 0.186902676 | 4.337523247 | 0.1181628 | 0.7355648 | 0.943686557 |
| ENSMUSG00000024201 | Kdm4b | protein_coding | -0.198115123 | 4.745665976 | 0.1183553 | 0.7356599 | 0.943686557 |
| ENSMUSG00000032397 | Tipin | protein_coding | 0.097671021 | 6.106471395 | 0.1178808 | 0.7357308 | 0.943686557 |
| ENSMUSG00000019732 | Calr3 | protein_coding | 0.315555884 | 3.324885192 | 0.1184534 | 0.7357322 | 0.943686557 |
| ENSMUSG00000060279 | Ap2a1 | protein_coding | 0.278233442 | 4.119981076 | 0.1187397 | 0.7357605 | 0.943686557 |
| ENSMUSG00000058620 | Adra2b | protein_coding | 0.181337864 | 3.442978142 | 0.1178523 | 0.7357615 | 0.943686557 |
| ENSMUSG00000003808 | Farsa | protein_coding | 0.121070127 | 5.889888517 | 0.1177152 | 0.7359161 | 0.943686557 |
| ENSMUSG00000023169 | Slc38a1 | protein_coding | -0.080837405 | 7.064916432 | 0.1176386 | 0.7359914 | 0.943686557 |
| ENSMUSG00000079179 | Rab10os | antisense_RNA | 0.10648319 | 6.313609855 | 0.1175539 | 0.7360897 | 0.943686557 |
| ENSMUSG00000026960 | Arl6ip6 | protein_coding | -0.146876447 | 3.486995503 | 0.1174097 | 0.7362378 | 0.943686557 |
| ENSMUSG00000015672 | Mrpl32 | protein_coding | 0.091760132 | 6.614622053 | 0.1173982 | 0.7362503 | 0.943686557 |
| ENSMUSG00000001270 | Ckb | protein_coding | -0.097237797 | 10.89649594 | 0.117386 | 0.7362634 | 0.943686557 |
| ENSMUSG00000027860 | Vangl1 | protein_coding | -0.285871727 | 3.138916828 | 0.1174752 | 0.7363508 | 0.943686557 |
| ENSMUSG00000004054 | Map3k11 | protein_coding | -0.249139459 | 2.518704053 | 0.1172624 | 0.7363967 | 0.943686557 |
| ENSMUSG00000097571 | Jpx | lincRNA | -0.24331473 | 2.721412026 | 0.1169719 | 0.7367101 | 0.943865038 |
| ENSMUSG00000030451 | Herc2 | protein_coding | -0.118977959 | 5.246005761 | 0.1169706 | 0.7367115 | 0.943865038 |
| ENSMUSG00000027854 | Sike1 | protein_coding | -0.16304245 | 4.124325647 | 0.1167231 | 0.7369789 | 0.943865561 |
| ENSMUSG00000028089 | Chd1l | protein_coding | 0.204860827 | 3.109815379 | 0.1166955 | 0.7370088 | 0.943865561 |
| ENSMUSG00000026620 | Mark1 | protein_coding | -0.125020386 | 4.913049454 | 0.116564 | 0.737151 | 0.943865561 |
| ENSMUSG00000033429 | Mcee | protein_coding | 0.087476718 | 6.651822384 | 0.1164582 | 0.7372655 | 0.943865561 |
| ENSMUSG00000091264 | Smim13 | protein_coding | -0.074473209 | 6.418624663 | 0.1164217 | 0.737305 | 0.943865561 |
| ENSMUSG00000022864 | D16Ertd472e | protein_coding | 0.297996216 | 2.559572326 | 0.1164152 | 0.7373121 | 0.943865561 |
| ENSMUSG00000025474 | Tubgcp2 | protein_coding | 0.120234273 | 5.413787032 | 0.1162893 | 0.7374484 | 0.943865561 |
| ENSMUSG00000006651 | Aplp1 | protein_coding | -0.100913178 | 9.291884173 | 0.1162282 | 0.7375218 | 0.943865561 |
| ENSMUSG00000060397 | Zfp128 | protein_coding | 0.212224753 | 3.274873164 | 0.1159504 | 0.7378159 | 0.943865561 |
| ENSMUSG00000020312 | Shc2 | protein_coding | 0.169408086 | 3.328318316 | 0.1159177 | 0.7378514 | 0.943865561 |
| ENSMUSG00000020834 | Dhrs13 | protein_coding | 0.265751495 | 2.554185785 | 0.1159155 | 0.7378538 | 0.943865561 |
| ENSMUSG00000032842 | Abcc10 | protein_coding | 0.281449674 | 2.920291954 | 0.1158209 | 0.7379637 | 0.943865561 |
| ENSMUSG00000025791 | Pgm2 | protein_coding | 0.102786527 | 7.022991 | 0.1155989 | 0.7382048 | 0.943865561 |
| ENSMUSG00000041203 | Trir | protein_coding | -0.125714488 | 4.336467588 | 0.1155684 | 0.7382308 | 0.943865561 |
| ENSMUSG00000019297 | Nop9 | protein_coding | -0.246429523 | 2.89386203 | 0.115543 | 0.7382584 | 0.943865561 |
| ENSMUSG00000005034 | Prkacb | protein_coding | 0.082358071 | 8.785418905 | 0.1155249 | 0.7382781 | 0.943865561 |
| ENSMUSG00000092060 | Bend4 | protein_coding | 0.172207262 | 4.913103167 | 0.1157298 | 0.7383319 | 0.943865561 |
| ENSMUSG00000034361 | Cpne2 | protein_coding | 0.108157045 | 5.225440251 | 0.1154565 | 0.7383525 | 0.943865561 |
| ENSMUSG00000038894 | Irs2 | protein_coding | 0.112219947 | 5.25886737 | 0.1153804 | 0.7384353 | 0.943865561 |
| ENSMUSG00000028907 | Utp11 | protein_coding | -0.07898863 | 6.560606227 | 0.1153508 | 0.7384675 | 0.943865561 |
| ENSMUSG00000085316 | D330050G23Rik | lincRNA | -0.310184177 | 2.556195154 | 0.115174 | 0.73866 | 0.943999375 |
| ENSMUSG00000022494 | Shisa9 | protein_coding | -0.11673364 | 4.665970214 | 0.1148799 | 0.7389805 | 0.94429682 |
| ENSMUSG00000017466 | Timp2 | protein_coding | 0.07116205 | 8.165904532 | 0.1145808 | 0.7393071 | 0.94460188 |
| ENSMUSG00000003721 | Insig2 | protein_coding | -0.070907053 | 6.989022052 | 0.1144812 | 0.7394159 | 0.944623041 |
| ENSMUSG00000074461 | Gm10699 | TEC | -0.084302628 | 5.997095982 | 0.1144049 | 0.7394994 | 0.944623041 |
| ENSMUSG00000032449 | Slc25a36 | protein_coding | 0.092471399 | 5.766421646 | 0.1142633 | 0.7396543 | 0.944649937 |
| ENSMUSG00000047013 | Fbxo41 | protein_coding | 0.126418512 | 5.231849758 | 0.114225 | 0.7396961 | 0.944649937 |
| ENSMUSG00000026825 | Dnm1 | protein_coding | 0.085756799 | 6.683177301 | 0.1140892 | 0.7398448 | 0.944727604 |
| ENSMUSG00000035011 | Zbtb7a | protein_coding | -0.117538112 | 5.645804851 | 0.1140078 | 0.7399339 | 0.944729166 |
| ENSMUSG00000044795 | Cyb5d1 | protein_coding | 0.230327964 | 3.322605512 | 0.1138537 | 0.7401099 | 0.944780671 |
| ENSMUSG00000022103 | Gfra2 | protein_coding | 0.121411702 | 5.329226926 | 0.1138107 | 0.7401499 | 0.944780671 |

| ensembl_gene_id | external_gene_name | gene_biotype | log2 (fold change) | log2 (counts per million) | F-value | p-value | False discovery rate |
|---|---|---|---|---|---|---|---|
| ENSMUSG00000004633 | Chn2 | protein_coding | -0.27323995 | 2.262162866 | 0.1131892 | 0.7408325 | 0.945520282 |
| ENSMUSG00000014232 | Cluap1 | protein_coding | -0.102775187 | 4.900014006 | 0.1128878 | 0.7411643 | 0.945520282 |
| ENSMUSG00000026831 | 1700007K13Rik | protein_coding | 0.174657435 | 3.121678407 | 0.1128798 | 0.7411731 | 0.945520282 |
| ENSMUSG00000029578 | Wipi2 | protein_coding | 0.074753677 | 6.48268592 | 0.1128407 | 0.7412162 | 0.945520282 |
| ENSMUSG00000056116 | H2-T22 | protein_coding | 0.148275501 | 4.006138147 | 0.1128345 | 0.741223 | 0.945520282 |
| ENSMUSG00000030103 | Bhlhe40 | protein_coding | -0.102459164 | 6.305069919 | 0.1127248 | 0.7413439 | 0.945520282 |
| ENSMUSG00000020034 | Tcp11l2 | protein_coding | 0.150853939 | 5.714036698 | 0.1129918 | 0.7414008 | 0.945520282 |
| ENSMUSG00000050963 | Kcns2 | protein_coding | 0.245461588 | 2.687444094 | 0.1125906 | 0.7414919 | 0.945520282 |
| ENSMUSG00000037851 | Iars | protein_coding | -0.126811064 | 4.834015032 | 0.11244 | 0.7416581 | 0.945520282 |
| ENSMUSG00000034501 | Pcnx4 | protein_coding | -0.119773853 | 4.806627932 | 0.1123107 | 0.7418009 | 0.945520282 |
| ENSMUSG00000037234 | Hook3 | protein_coding | -0.14183666 | 4.477801318 | 0.1122542 | 0.7418633 | 0.945520282 |
| ENSMUSG00000033364 | Usp37 | protein_coding | -0.156577085 | 5.195301526 | 0.112393 | 0.7418891 | 0.945520282 |
| ENSMUSG00000058704 | Memo1 | protein_coding | 0.129059542 | 4.05571093 | 0.1122214 | 0.7418996 | 0.945520282 |
| ENSMUSG00000027984 | Hadh | protein_coding | 0.094227141 | 5.869148006 | 0.1120538 | 0.7420849 | 0.945520282 |
| ENSMUSG00000040043 | Rbms2 | protein_coding | -0.155428536 | 3.612148986 | 0.1120329 | 0.742108 | 0.945520282 |
| ENSMUSG00000030872 | Gga2 | protein_coding | -0.218714542 | 3.592282085 | 0.1120308 | 0.7421439 | 0.945520282 |
| ENSMUSG00000038055 | Dexi | protein_coding | -0.137262163 | 4.185808353 | 0.111797 | 0.7423691 | 0.945520282 |
| ENSMUSG00000002799 | Jag2 | protein_coding | 0.272147288 | 2.957578976 | 0.1117858 | 0.7423885 | 0.945520282 |
| ENSMUSG00000028582 | Cc2d1b | protein_coding | -0.188131017 | 3.403675715 | 0.1117691 | 0.7424001 | 0.945520282 |
| ENSMUSG00000030659 | Nucb2 | protein_coding | -0.094509788 | 5.219319898 | 0.1115486 | 0.7426444 | 0.945570807 |
| ENSMUSG00000001521 | Tulp3 | protein_coding | -0.180742911 | 3.3495638 | 0.1115124 | 0.7426846 | 0.945570807 |
| ENSMUSG00000027996 | Sfrp2 | protein_coding | -0.188412733 | 2.939593684 | 0.1112741 | 0.742949 | 0.945570807 |
| ENSMUSG00000001240 | Ramp2 | protein_coding | 0.240787698 | 2.70273353 | 0.1112661 | 0.7429578 | 0.945570807 |
| ENSMUSG00000037957 | Wdr20 | protein_coding | 0.134851012 | 4.071077667 | 0.1112126 | 0.7430173 | 0.945570807 |
| ENSMUSG00000003518 | Dusp3 | protein_coding | -0.103513884 | 8.317145625 | 0.1111293 | 0.7431472 | 0.945570807 |
| ENSMUSG00000027341 | Tmem230 | protein_coding | -0.120818969 | 4.33229292 | 0.1110532 | 0.7431944 | 0.945570807 |
| ENSMUSG00000061313 | Ddhd2 | protein_coding | 0.140354336 | 7.095675097 | 0.1116467 | 0.7432145 | 0.945570807 |
| ENSMUSG00000024081 | Cebpz | protein_coding | -0.106519138 | 5.211813581 | 0.1110168 | 0.7432349 | 0.945570807 |
| ENSMUSG00000018433 | Nol11 | protein_coding | -0.112724908 | 5.266250352 | 0.1108942 | 0.7433713 | 0.945570807 |
| ENSMUSG00000052557 | Gan | protein_coding | -0.168580654 | 3.380614522 | 0.1108621 | 0.743407 | 0.945570807 |
| ENSMUSG00000024777 | Ppp2r5b | protein_coding | 0.086287178 | 5.645437289 | 0.1107421 | 0.7435406 | 0.945628815 |
| ENSMUSG00000031808 | Slc27a1 | protein_coding | -0.125134233 | 5.635908775 | 0.1106331 | 0.7436689 | 0.945680157 |
| ENSMUSG00000035413 | Tmem98 | protein_coding | -0.099086759 | 5.736134457 | 0.1104595 | 0.7438554 | 0.945732677 |
| ENSMUSG00000024983 | Vti1a | protein_coding | 0.121694251 | 5.623948769 | 0.1104175 | 0.7439092 | 0.945732677 |
| ENSMUSG00000028830 | AU040320 | protein_coding | 0.09929699 | 5.325684901 | 0.1103532 | 0.743974 | 0.945732677 |
| ENSMUSG00000060671 | Atp8b2 | protein_coding | -0.166417626 | 3.471157215 | 0.1101137 | 0.7442413 | 0.945960663 |
| ENSMUSG00000003072 | Atp5d | protein_coding | -0.168362587 | 8.518672717 | 0.1113644 | 0.7443711 | 0.946013705 |
| ENSMUSG00000032468 | Armc8 | protein_coding | -0.090446236 | 6.763155801 | 0.1095454 | 0.7448771 | 0.94613757 |
| ENSMUSG00000028792 | Ak2 | protein_coding | 0.101237887 | 5.347829628 | 0.1094052 | 0.7450342 | 0.94613757 |
| ENSMUSG00000031347 | Cetn2 | protein_coding | -0.084404866 | 7.865143906 | 0.109239 | 0.7452205 | 0.94613757 |
| ENSMUSG00000004366 | Sst | protein_coding | -0.101916252 | 8.91773104 | 0.1091317 | 0.745366 | 0.94613757 |
| ENSMUSG00000024654 | Asrgl1 | protein_coding | -0.069150974 | 7.648444656 | 0.1090561 | 0.7454259 | 0.94613757 |
| ENSMUSG00000039096 | Rsad1 | protein_coding | -0.211095525 | 3.080677049 | 0.1090066 | 0.7454815 | 0.94613757 |
| ENSMUSG00000047777 | Phf13 | protein_coding | 0.149641734 | 3.441621594 | 0.1089561 | 0.7455382 | 0.94613757 |
| ENSMUSG00000038280 | Ostm1 | protein_coding | 0.116407206 | 4.833108994 | 0.1088566 | 0.74565 | 0.94613757 |
| ENSMUSG00000056919 | Cep162 | protein_coding | -0.189773605 | 3.024399606 | 0.1087685 | 0.745749 | 0.94613757 |
| ENSMUSG00000064370 | mt-Cytb | protein_coding | -0.1330594 | 12.19260763 | 0.1089448 | 0.7458951 | 0.94613757 |
| ENSMUSG00000032342 | Mto1 | protein_coding | -0.147787136 | 3.61128198 | 0.1085297 | 0.7460178 | 0.94613757 |
| ENSMUSG00000038374 | Rbm8a | protein_coding | 0.069612175 | 7.190096563 | 0.108313 | 0.7462619 | 0.94613757 |
| ENSMUSG00000036368 | Rmdn2 | protein_coding | -0.250030019 | 2.618125722 | 0.1082346 | 0.7463504 | 0.94613757 |
| ENSMUSG00000032251 | Irak1bp1 | protein_coding | -0.112568136 | 4.895445632 | 0.1080104 | 0.7466033 | 0.94613757 |
| ENSMUSG00000032420 | Nt5e | protein_coding | 0.243260123 | 2.641088691 | 0.1079617 | 0.7466583 | 0.94613757 |
| ENSMUSG00000047428 | Dlk2 | protein_coding | 0.190417038 | 3.196595907 | 0.1079469 | 0.746675 | 0.94613757 |
| ENSMUSG00000028708 | Mknk1 | protein_coding | 0.120913195 | 4.546987891 | 0.1079284 | 0.7466959 | 0.94613757 |
| ENSMUSG00000026965 | Anapc2 | protein_coding | -0.082638244 | 6.07619777 | 0.1078996 | 0.7467284 | 0.94613757 |
| ENSMUSG00000024400 | Wdr33 | protein_coding | 0.078123267 | 6.93051735 | 0.1077249 | 0.7469259 | 0.94613757 |
| ENSMUSG00000086075 | Gm15728 | lincRNA | 0.187736542 | 3.171824099 | 0.1076328 | 0.7470301 | 0.94613757 |
| ENSMUSG00000023845 | Lnpep | protein_coding | 0.303667567 | 2.367203165 | 0.1076148 | 0.7470504 | 0.94613757 |
| ENSMUSG00000086629 | 2810403D21Rik | processed_transcript | -0.175430957 | 3.35964998 | 0.1076103 | 0.7470555 | 0.94613757 |
| ENSMUSG00000086877 | A230072C01Rik | processed_transcript | -0.112394028 | 4.659431682 | 0.1075607 | 0.7471117 | 0.94613757 |
| ENSMUSG00000036561 | Ppp6r2 | protein_coding | -0.136431199 | 4.045764098 | 0.1073932 | 0.7473013 | 0.94613757 |
| ENSMUSG00000037791 | Phf12 | protein_coding | -0.140308625 | 4.091898863 | 0.1071819 | 0.7475407 | 0.94613757 |
| ENSMUSG00000030469 | Zfp719 | protein_coding | 0.151103352 | 3.742661636 | 0.107125 | 0.7476053 | 0.94613757 |
| ENSMUSG00000026154 | Sdhaf4 | protein_coding | 0.083852935 | 8.414364103 | 0.1070834 | 0.7476525 | 0.94613757 |
| ENSMUSG00000034723 | Tmx4 | protein_coding | -0.071427815 | 7.659285269 | 0.1070729 | 0.7476644 | 0.94613757 |
| ENSMUSG00000025153 | Fasn | protein_coding | -0.105179219 | 6.044390897 | 0.1070459 | 0.747695 | 0.94613757 |
| ENSMUSG00000035245 | Eogt | protein_coding | -0.209372342 | 2.911415084 | 0.1070284 | 0.7477149 | 0.94613757 |
| ENSMUSG00000014980 | Tsen15 | protein_coding | 0.102614217 | 4.869317829 | 0.1070262 | 0.7477173 | 0.94613757 |
| ENSMUSG00000055053 | Nfic | protein_coding | 0.115398991 | 5.752961924 | 0.1070254 | 0.7477251 | 0.94613757 |
| ENSMUSG00000019794 | Katna1 | protein_coding | -0.133299862 | 3.812878857 | 0.1068573 | 0.7479091 | 0.94613757 |
| ENSMUSG00000022800 | Fyttd1 | protein_coding | 0.081796565 | 6.968082929 | 0.1068132 | 0.7479592 | 0.94613757 |
| ENSMUSG00000026223 | Itm2c | protein_coding | 0.081226802 | 9.311222768 | 0.1067763 | 0.7480011 | 0.94613757 |
| ENSMUSG00000022091 | Sorbs3 | protein_coding | 0.107794223 | 4.802148404 | 0.1067537 | 0.7480269 | 0.94613757 |
| ENSMUSG00000021156 | Zmynd11 | protein_coding | -0.12751968 | 4.511812128 | 0.1066898 | 0.7480994 | 0.94613757 |
| ENSMUSG00000108644 | Gm44920 | lincRNA | -0.270852972 | 2.522787418 | 0.1066459 | 0.7481494 | 0.94613757 |
| ENSMUSG00000064202 | 4430402I18Rik | protein_coding | -0.206038234 | 3.057452635 | 0.106643 | 0.7481527 | 0.94613757 |
| ENSMUSG00000074829 | 2010315B03Rik | protein_coding | 0.133970496 | 4.00988942 | 0.1065142 | 0.7482992 | 0.94613757 |
| ENSMUSG00000005609 | Ctr9 | protein_coding | 0.103962146 | 5.359630466 | 0.1064246 | 0.7484012 | 0.94613757 |

| ensembl_gene_id | external_gene_name | gene_biotype | log2 (fold change) | log2 (counts per million) | F-value | p-value | False discovery rate |
|---|---|---|---|---|---|---|---|
| ENSMUSG00000016495 | Plgrkt | protein_coding | 0.105094325 | 5.452036591 | 0.1062719 | 0.748575 | 0.94613757 |
| ENSMUSG00000004221 | Ikbkg | protein_coding | -0.126911016 | 5.073525441 | 0.1061844 | 0.7486747 | 0.94613757 |
| ENSMUSG00000022992 | Kansl2 | protein_coding | 0.208796801 | 2.796807331 | 0.1060748 | 0.7487997 | 0.94613757 |
| ENSMUSG00000055026 | Gabrg3 | protein_coding | -0.10717809 | 4.684126774 | 0.1060663 | 0.7488094 | 0.94613757 |
| ENSMUSG00000029624 | Ptcd1 | protein_coding | 0.154122118 | 3.945842654 | 0.1059783 | 0.7489098 | 0.94613757 |
| ENSMUSG00000031924 | Cyb5b | protein_coding | -0.069645375 | 6.754261924 | 0.1059581 | 0.7489328 | 0.94613757 |
| ENSMUSG00000034075 | Zdhhc5 | protein_coding | -0.151258231 | 3.375278264 | 0.1058888 | 0.749012 | 0.94613757 |
| ENSMUSG00000073700 | Klhl21 | protein_coding | 0.167869276 | 3.31837598 | 0.1058774 | 0.7490249 | 0.94613757 |
| ENSMUSG00000031299 | Pdha1 | protein_coding | 0.135904829 | 7.285462606 | 0.1064538 | 0.7490296 | 0.94613757 |
| ENSMUSG00000046573 | Lyrm4 | protein_coding | 0.092349803 | 7.735868849 | 0.1058356 | 0.7490795 | 0.94613757 |
| ENSMUSG00000031068 | Glrx3 | protein_coding | -0.071452325 | 6.39204857 | 0.1057148 | 0.7492107 | 0.94613757 |
| ENSMUSG00000047030 | Spata2 | protein_coding | 0.141176141 | 4.479292543 | 0.1057137 | 0.7492119 | 0.94613757 |
| ENSMUSG00000031594 | Fgl1 | protein_coding | -0.252754617 | 2.721858819 | 0.1057067 | 0.7492199 | 0.94613757 |
| ENSMUSG00000035206 | Sppl2b | protein_coding | -0.143005289 | 3.960929296 | 0.1055913 | 0.7493518 | 0.946193041 |
| ENSMUSG00000034591 | Slc41a2 | protein_coding | -0.127085013 | 3.769568938 | 0.1053108 | 0.7496727 | 0.946229557 |
| ENSMUSG00000048120 | Entpd1 | protein_coding | -0.153911682 | 3.685527013 | 0.1050927 | 0.7499227 | 0.946229557 |
| ENSMUSG00000029587 | Zfp12 | protein_coding | -0.26639208 | 3.779491193 | 0.1055885 | 0.7500137 | 0.946229557 |
| ENSMUSG00000061143 | Maml3 | protein_coding | -0.182775791 | 3.033189784 | 0.104651 | 0.7504296 | 0.946229557 |
| ENSMUSG00000029994 | Anxa4 | protein_coding | -0.221603911 | 2.78056413 | 0.1045885 | 0.7505014 | 0.946229557 |
| ENSMUSG00000022436 | Sh3bp1 | protein_coding | 0.072029708 | 7.128496643 | 0.1045298 | 0.750569 | 0.946229557 |
| ENSMUSG00000031691 | Tnpo2 | protein_coding | -0.19815251 | 3.370183233 | 0.1045217 | 0.7505782 | 0.946229557 |
| ENSMUSG00000028646 | Rragc | protein_coding | -0.090507484 | 5.912414531 | 0.1045043 | 0.7505982 | 0.946229557 |
| ENSMUSG00000003438 | Timm50 | protein_coding | 0.111275977 | 4.759467136 | 0.1044648 | 0.7506437 | 0.946229557 |
| ENSMUSG00000031834 | Pik3r2 | protein_coding | -0.109538974 | 4.690326057 | 0.1044327 | 0.7506806 | 0.946229557 |
| ENSMUSG00000049232 | Tigd2 | protein_coding | -0.157657166 | 4.020976715 | 0.1043762 | 0.7507457 | 0.946229557 |
| ENSMUSG00000063904 | 4930533O14Rik | protein_coding | 0.135228142 | 5.120789091 | 0.104232 | 0.7509183 | 0.946229557 |
| ENSMUSG00000024268 | Celf4 | protein_coding | 0.094801407 | 9.063432229 | 0.1041942 | 0.7509619 | 0.946229557 |
| ENSMUSG00000052459 | Atp6v1a | protein_coding | -0.081238448 | 8.941268298 | 0.104168 | 0.7509853 | 0.946229557 |
| ENSMUSG00000079036 | Alkbh1 | protein_coding | -0.115694306 | 4.201292992 | 0.104121 | 0.7510395 | 0.946229557 |
| ENSMUSG00000033931 | Rbm34 | protein_coding | -0.11194727 | 4.615637206 | 0.1039421 | 0.7512457 | 0.946229557 |
| ENSMUSG00000040363 | Bcor | protein_coding | -0.101462088 | 4.932756 | 0.1038275 | 0.7513779 | 0.946229557 |
| ENSMUSG00000031242 | 2610002M06Rik | protein_coding | -0.083773399 | 5.652742903 | 0.103793 | 0.7514178 | 0.946229557 |
| ENSMUSG00000038967 | Pdk2 | protein_coding | -0.10136479 | 6.877811759 | 0.1037759 | 0.751462 | 0.946229557 |
| ENSMUSG00000029403 | Cdkl2 | protein_coding | 0.099728155 | 5.228739217 | 0.1037541 | 0.7514626 | 0.946229557 |
| ENSMUSG00000031667 | Aktip | protein_coding | 0.076605704 | 8.165522375 | 0.1037541 | 0.7514626 | 0.946229557 |
| ENSMUSG00000028020 | Glrb | protein_coding | -0.067707775 | 6.764037551 | 0.1037381 | 0.7514812 | 0.946229557 |
| ENSMUSG00000037434 | Slc30a1 | protein_coding | -0.166665662 | 3.463613346 | 0.1036984 | 0.751527 | 0.946229557 |
| ENSMUSG00000000876 | Pxmp4 | protein_coding | 0.093498053 | 5.15063373 | 0.1036724 | 0.7515571 | 0.946229557 |
| ENSMUSG00000021111 | Papola | protein_coding | 0.087335749 | 6.108188155 | 0.1036519 | 0.7515806 | 0.946229557 |
| ENSMUSG00000043587 | Pxylp1 | protein_coding | -0.144204855 | 3.931386906 | 0.1035047 | 0.7517508 | 0.946238156 |
| ENSMUSG00000003500 | Impdh1 | protein_coding | -0.159835547 | 4.04516045 | 0.1034875 | 0.7517707 | 0.946238156 |
| ENSMUSG00000084910 | C630043F03Rik | processed_transcript | -0.146486677 | 3.895247764 | 0.103354 | 0.7519251 | 0.946238156 |
| ENSMUSG00000003762 | Coq8b | protein_coding | -0.211454628 | 2.662124604 | 0.1033416 | 0.7519395 | 0.946238156 |
| ENSMUSG00000045757 | Zfp764 | protein_coding | -0.389430684 | 2.131660146 | 0.1032192 | 0.7520878 | 0.946314119 |
| ENSMUSG00000037217 | Syn1 | protein_coding | 0.083580083 | 6.267628301 | 0.1029719 | 0.7523676 | 0.946504363 |
| ENSMUSG00000033006 | Sox10 | protein_coding | -0.113012776 | 5.169216814 | 0.1028996 | 0.7524515 | 0.946504363 |
| ENSMUSG00000032788 | Pdxk | protein_coding | 0.126610282 | 7.679197173 | 0.103349 | 0.7525031 | 0.946504363 |
| ENSMUSG00000032878 | Ccdc85a | protein_coding | -0.093683391 | 5.289002769 | 0.1026307 | 0.7527637 | 0.946532643 |
| ENSMUSG00000043323 | Fbrsl1 | protein_coding | 0.162850698 | 3.680549112 | 0.1025946 | 0.7528055 | 0.946532643 |
| ENSMUSG00000113679 | AC159633.2 | processed_transcript | -0.149588267 | 3.571613855 | 0.1024522 | 0.7529711 | 0.946532643 |
| ENSMUSG00000041552 | Ptchd1 | protein_coding | -0.158200747 | 3.196184494 | 0.1023196 | 0.7531252 | 0.946532643 |
| ENSMUSG00000054280 | Prr14l | protein_coding | 0.186153126 | 3.099167122 | 0.1023146 | 0.753131 | 0.946532643 |
| ENSMUSG00000047221 | Fam185a | protein_coding | -0.193482516 | 3.551582404 | 0.1022726 | 0.7531867 | 0.946532643 |
| ENSMUSG00000064343 | mt-Tq | Mt_tRNA | -0.23876055 | 2.419131349 | 0.1022503 | 0.753206 | 0.946532643 |
| ENSMUSG00000073889 | Il11ra1 | protein_coding | 0.125410179 | 3.906025596 | 0.1022298 | 0.7532298 | 0.946532643 |
| ENSMUSG00000046032 | Snx12 | protein_coding | 0.066593294 | 6.618575788 | 0.1018555 | 0.753666 | 0.94689509 |
| ENSMUSG00000028030 | Tbck | protein_coding | -0.128652843 | 6.406016081 | 0.1021787 | 0.7536943 | 0.94689509 |
| ENSMUSG00000109776 | Gm30400 | processed_transcript | 0.267261956 | 2.474284492 | 0.1016688 | 0.7538838 | 0.946976113 |
| ENSMUSG00000107476 | Zfp862-ps | transcribed_unitary_p | 0.203269063 | 2.780247633 | 0.1015126 | 0.7540663 | 0.946976113 |
| ENSMUSG00000029020 | Mfn2 | protein_coding | 0.105353775 | 7.370764413 | 0.101528 | 0.7543059 | 0.946976113 |
| ENSMUSG00000019806 | Aig1 | protein_coding | -0.102528496 | 6.224464513 | 0.1012887 | 0.7543347 | 0.946976113 |
| ENSMUSG00000021877 | Arf4 | protein_coding | -0.079780798 | 8.805973161 | 0.1012597 | 0.754362 | 0.946976113 |
| ENSMUSG00000041439 | Mfsd6 | protein_coding | 0.089469067 | 5.493051545 | 0.1012447 | 0.7543796 | 0.946976113 |
| ENSMUSG00000027954 | Efna1 | protein_coding | -0.213199004 | 3.056885066 | 0.1011886 | 0.7544452 | 0.946976113 |
| ENSMUSG00000078970 | Wdr92 | protein_coding | -0.139145172 | 4.645180636 | 0.1011731 | 0.7544633 | 0.946976113 |
| ENSMUSG00000022335 | Zfat | protein_coding | 0.281983922 | 2.241048006 | 0.1010221 | 0.7546402 | 0.947087572 |
| ENSMUSG00000056692 | D17Wsu92e | protein_coding | -0.070746046 | 6.607723939 | 0.1008931 | 0.7547915 | 0.947118474 |
| ENSMUSG00000090093 | Gm14399 | protein_coding | 0.216853952 | 2.701887579 | 0.1008509 | 0.754841 | 0.947118474 |
| ENSMUSG00000035653 | Lrfn5 | protein_coding | -0.107546571 | 5.506144873 | 0.1005716 | 0.7551688 | 0.947232838 |
| ENSMUSG00000028044 | Cks1b | protein_coding | -0.158610621 | 3.204712317 | 0.1005194 | 0.7552302 | 0.947232838 |
| ENSMUSG00000022200 | Golph3 | protein_coding | -0.100824414 | 5.640855745 | 0.1004879 | 0.7552671 | 0.947232838 |
| ENSMUSG00000044221 | Grsf1 | protein_coding | 0.07449297 | 6.971257597 | 0.1004731 | 0.7552845 | 0.947232838 |
| ENSMUSG00000038079 | Tmem237 | protein_coding | 0.096198038 | 6.25793534 | 0.0999266 | 0.7559278 | 0.947542638 |
| ENSMUSG00000074358 | Ccdc61 | protein_coding | 0.145215683 | 3.424313251 | 0.0998617 | 0.7560043 | 0.947542638 |
| ENSMUSG00000090641 | Zfp712 | protein_coding | 0.244120418 | 2.588525085 | 0.0998382 | 0.7560321 | 0.947542638 |
| ENSMUSG00000042046 | Dstyk | protein_coding | -0.095696754 | 5.169482576 | 0.0998071 | 0.7560687 | 0.947542638 |
| ENSMUSG00000062054 | Iah1 | protein_coding | 0.09219041 | 5.522841567 | 0.0996587 | 0.7562438 | 0.947542638 |
| ENSMUSG00000021427 | Ssr1 | protein_coding | 0.073053952 | 8.64397154 | 0.099623 | 0.756286 | 0.947542638 |

| ensembl_gene_id | external_gene_name | gene_biotype | log2 (fold change) | log2 (counts per million) | F-value | p-value | False discovery rate |
|---|---|---|---|---|---|---|---|
| ENSMUSG00000027940 | Tpm3 | protein_coding | 0.067787313 | 6.75232842 | 0.0995094 | 0.7564202 | 0.947542638 |
| ENSMUSG00000034826 | Nup54 | protein_coding | 0.170199712 | 3.960240692 | 0.0995 | 0.7564378 | 0.947542638 |
| ENSMUSG00000003604 | Aven | protein_coding | -0.147174731 | 4.143550156 | 0.0994458 | 0.7564953 | 0.947542638 |
| ENSMUSG00000034807 | Colgalt1 | protein_coding | -0.179262934 | 3.792657305 | 0.0994403 | 0.7565085 | 0.947542638 |
| ENSMUSG00000039157 | Fam102a | protein_coding | 0.074249249 | 6.538388626 | 0.0994272 | 0.7565173 | 0.947542638 |
| ENSMUSG00000030738 | Eif3c | protein_coding | -0.076550061 | 7.789052658 | 0.0993444 | 0.7566152 | 0.947542638 |
| ENSMUSG00000029735 | Tpk1 | protein_coding | -0.305410699 | 2.61527718 | 0.0992485 | 0.7567352 | 0.947542638 |
| ENSMUSG00000021028 | Mbip | protein_coding | -0.116228161 | 4.987960437 | 0.0992176 | 0.7567652 | 0.947542638 |
| ENSMUSG00000020435 | Osbp2 | protein_coding | -0.16105882 | 3.854302169 | 0.0989502 | 0.7570819 | 0.947629793 |
| ENSMUSG00000024528 | Srfbp1 | protein_coding | 0.11893896 | 4.73445239 | 0.098865 | 0.7571829 | 0.947629793 |
| ENSMUSG00000020902 | Ntn1 | protein_coding | -0.285411791 | 2.661444907 | 0.0988655 | 0.7571885 | 0.947629793 |
| ENSMUSG00000017692 | Rhbdl3 | protein_coding | 0.16673856 | 3.989578344 | 0.0988142 | 0.7572498 | 0.947629793 |
| ENSMUSG00000040599 | Mis12 | protein_coding | -0.116785164 | 4.846130486 | 0.098735 | 0.7573371 | 0.947629793 |
| ENSMUSG00000038028 | Tigar | protein_coding | 0.122179669 | 4.351674564 | 0.0986707 | 0.7574135 | 0.947629793 |
| ENSMUSG00000011148 | Adssl1 | protein_coding | 0.100448092 | 4.719641668 | 0.0986385 | 0.7574517 | 0.947629793 |
| ENSMUSG00000034820 | Cpsf7 | protein_coding | 0.288448369 | 2.343303687 | 0.098456 | 0.7576686 | 0.947667835 |
| ENSMUSG00000042385 | Gzmk | protein_coding | -0.311826729 | 2.587285795 | 0.0984437 | 0.7576897 | 0.947667835 |
| ENSMUSG00000049881 | 2810025M15Rik | transcribed_processe | -0.084878063 | 5.804134644 | 0.0981726 | 0.7580056 | 0.947667835 |
| ENSMUSG00000031432 | Prps1 | protein_coding | 0.062736015 | 7.358218619 | 0.0980617 | 0.7581377 | 0.947667835 |
| ENSMUSG00000032252 | Glce | protein_coding | 0.11445869 | 4.339791961 | 0.0980485 | 0.7581534 | 0.947667835 |
| ENSMUSG00000039680 | Mrps6 | protein_coding | -0.097499017 | 6.241458897 | 0.0979228 | 0.7583032 | 0.947667835 |
| ENSMUSG00000029060 | Mib2 | protein_coding | 0.129232991 | 4.132679786 | 0.0978833 | 0.7583503 | 0.947667835 |
| ENSMUSG00000067194 | Eif1ax | protein_coding | -0.067782067 | 7.481456596 | 0.0978517 | 0.758388 | 0.947667835 |
| ENSMUSG00000031256 | Cstf2 | protein_coding | -0.136184749 | 4.311827684 | 0.0978509 | 0.7583889 | 0.947667835 |
| ENSMUSG00000029108 | Pcdh7 | protein_coding | -0.090403166 | 5.97182435 | 0.0978032 | 0.7584458 | 0.947667835 |
| ENSMUSG00000019986 | Ahi1 | protein_coding | 0.084620994 | 8.7779824 | 0.0977984 | 0.7584515 | 0.947667835 |
| ENSMUSG00000000142 | Axin2 | protein_coding | -0.160597491 | 3.539760292 | 0.0977121 | 0.7585545 | 0.947686404 |
| ENSMUSG00000023075 | Akirin1 | protein_coding | -0.132692616 | 4.188818032 | 0.0976215 | 0.7586627 | 0.947694525 |
| ENSMUSG00000096910 | Zfp955b | protein_coding | 0.215482147 | 2.486901328 | 0.0974114 | 0.7589137 | 0.947694525 |
| ENSMUSG00000027201 | Myef2 | protein_coding | -0.21576436 | 2.690352553 | 0.0973643 | 0.75897 | 0.947694525 |
| ENSMUSG00000004947 | Dtx2 | protein_coding | -0.221183173 | 2.572193851 | 0.097357 | 0.7589788 | 0.947694525 |
| ENSMUSG00000028700 | Pomgnt1 | protein_coding | -0.111189006 | 5.107725066 | 0.0973352 | 0.7590048 | 0.947694525 |
| ENSMUSG00000027342 | Pcna | protein_coding | -0.089790908 | 5.227734481 | 0.0971176 | 0.7592652 | 0.947694525 |
| ENSMUSG00000038646 | Fam103a1 | protein_coding | 0.068296799 | 6.473152165 | 0.0970216 | 0.7593803 | 0.947694525 |
| ENSMUSG00000079658 | Eloc | protein_coding | 0.067529605 | 8.764212717 | 0.0970112 | 0.7593927 | 0.947694525 |
| ENSMUSG00000097213 | Gm26745 | lincRNA | 0.199348909 | 2.703866839 | 0.0969844 | 0.7594249 | 0.947694525 |
| ENSMUSG00000026234 | Ncl | protein_coding | -0.069350707 | 8.209735329 | 0.0969698 | 0.7594424 | 0.947694525 |
| ENSMUSG00000021929 | Kpna3 | protein_coding | 0.074254696 | 6.483431306 | 0.0966872 | 0.7597814 | 0.947843607 |
| ENSMUSG00000027079 | Clp1 | protein_coding | -0.142097891 | 4.119925071 | 0.0965945 | 0.7598926 | 0.947843607 |
| ENSMUSG00000028914 | Casp9 | protein_coding | -0.270923209 | 2.50036652 | 0.0965327 | 0.7599669 | 0.947843607 |
| ENSMUSG00000042359 | Osbpl6 | protein_coding | -0.111661475 | 5.156949153 | 0.0964372 | 0.7600816 | 0.947843607 |
| ENSMUSG00000046793 | Gpr61 | protein_coding | -0.123544295 | 4.355467775 | 0.0964293 | 0.7600911 | 0.947843607 |
| ENSMUSG00000074579 | Lekr1 | protein_coding | -0.17241928 | 3.043518446 | 0.0964258 | 0.7600954 | 0.947843607 |
| ENSMUSG00000028709 | Mob3c | protein_coding | 0.33076722 | 2.685665053 | 0.0965233 | 0.7602288 | 0.947843607 |
| ENSMUSG00000022889 | Mrpl39 | protein_coding | 0.083715626 | 6.207912801 | 0.0962683 | 0.7602848 | 0.947843607 |
| ENSMUSG00000024251 | Thada | protein_coding | 0.151555288 | 3.259667911 | 0.0962098 | 0.7603552 | 0.947843607 |
| ENSMUSG00000028010 | Gar1 | protein_coding | -0.264427667 | 2.31814993 | 0.0959911 | 0.7606187 | 0.947988035 |
| ENSMUSG00000040356 | Skiv2l | protein_coding | 0.105212619 | 4.674149618 | 0.0959673 | 0.7606473 | 0.947988035 |
| ENSMUSG00000032908 | Sgpp2 | protein_coding | -0.392943737 | 2.288440636 | 0.0959586 | 0.7608223 | 0.948096198 |
| ENSMUSG00000030816 | Rnf40 | protein_coding | 0.102092426 | 4.378118553 | 0.0956261 | 0.761059 | 0.948281302 |
| ENSMUSG00000045875 | Adra1a | protein_coding | -0.155123584 | 3.405582033 | 0.0955459 | 0.7611559 | 0.948292102 |
| ENSMUSG00000052488 | Cherp | protein_coding | 0.198374992 | 4.63832913 | 0.0957287 | 0.7614842 | 0.94836546 |
| ENSMUSG00000033430 | Terf2ip | protein_coding | 0.093502681 | 5.577778088 | 0.0952649 | 0.7614957 | 0.94836546 |
| ENSMUSG00000019897 | Ccdc59 | protein_coding | 0.06629429 | 7.036182763 | 0.0952014 | 0.7615726 | 0.94836546 |
| ENSMUSG00000014243 | Zswim7 | protein_coding | 0.087938949 | 5.390457261 | 0.0951055 | 0.7616888 | 0.94836546 |
| ENSMUSG00000036890 | Gtdc1 | protein_coding | -0.124703769 | 4.693488783 | 0.0950726 | 0.7617286 | 0.94836546 |
| ENSMUSG00000045896 | Paip2b | protein_coding | -0.082892122 | 5.696696941 | 0.09499 | 0.7618288 | 0.94836546 |
| ENSMUSG00000005615 | Pcyt1a | protein_coding | -0.116789282 | 5.212254866 | 0.0949128 | 0.7619223 | 0.94836546 |
| ENSMUSG00000039047 | Pigk | protein_coding | 0.080000483 | 6.552210073 | 0.0947953 | 0.7620648 | 0.94836546 |
| ENSMUSG00000026670 | Uap1 | protein_coding | -0.101115669 | 5.067554371 | 0.0947783 | 0.7620855 | 0.94836546 |
| ENSMUSG00000038569 | Rad9b | protein_coding | -0.304101118 | 2.652886334 | 0.094696 | 0.7621918 | 0.94836546 |
| ENSMUSG00000033545 | Znrf1 | protein_coding | -0.092911219 | 5.488763128 | 0.0945929 | 0.7623106 | 0.94836546 |
| ENSMUSG00000039159 | Ube2h | protein_coding | 0.087290344 | 6.566999314 | 0.0945532 | 0.7623589 | 0.94836546 |
| ENSMUSG00000021236 | Entpd5 | protein_coding | 0.114099879 | 4.612089594 | 0.0945141 | 0.7624064 | 0.94836546 |
| ENSMUSG00000100164 | 2610306M01Rik | bidirectional_promote | 0.09138035 | 5.269928062 | 0.0944787 | 0.7624495 | 0.94836546 |
| ENSMUSG00000020889 | Nr1d1 | protein_coding | 0.10292778 | 6.574850309 | 0.0943735 | 0.7626872 | 0.948480136 |
| ENSMUSG00000024347 | Psd2 | protein_coding | -0.152096241 | 4.892576603 | 0.0943918 | 0.7627181 | 0.948480136 |
| ENSMUSG00000061477 | Rps7 | protein_coding | 0.09242496 | 5.938776465 | 0.0941739 | 0.7628203 | 0.948497582 |
| ENSMUSG00000021963 | Sap18 | protein_coding | -0.063297146 | 6.985810543 | 0.0939542 | 0.7630881 | 0.948718463 |
| ENSMUSG00000032939 | Nup93 | protein_coding | -0.107985426 | 4.417205493 | 0.0938019 | 0.763274 | 0.948718463 |
| ENSMUSG00000029752 | Asns | protein_coding | 0.113192517 | 8.118845226 | 0.0941338 | 0.7632842 | 0.948718463 |
| ENSMUSG00000041912 | Tdrkh | protein_coding | 0.079666758 | 5.926393016 | 0.0937389 | 0.7633509 | 0.948718463 |
| ENSMUSG00000055540 | Epha6 | protein_coding | -0.155579984 | 3.574033654 | 0.093647 | 0.7634632 | 0.948748354 |
| ENSMUSG00000019261 | Map1s | protein_coding | -0.166473574 | 4.671903465 | 0.0936673 | 0.763685 | 0.948755137 |
| ENSMUSG00000002329 | Mdp1 | protein_coding | 0.060599308 | 7.037462489 | 0.0933005 | 0.7638869 | 0.948755137 |
| ENSMUSG00000024008 | Cpne5 | protein_coding | 0.091664461 | 7.202802253 | 0.0931726 | 0.7640499 | 0.948755137 |
| ENSMUSG00000039210 | Gpatch2 | protein_coding | -0.254752068 | 2.641426006 | 0.093162 | 0.7640565 | 0.948755137 |
| ENSMUSG00000022883 | Robo1 | protein_coding | -0.120090487 | 3.967815851 | 0.0931501 | 0.7640711 | 0.948755137 |

| ensembl_gene_id | external_gene_name | gene_biotype | log2 (fold change) | log2 (counts per million) | F-value | p-value | False discovery rate |
|---|---|---|---|---|---|---|---|
| ENSMUSG00000061024 | Rrs1 | protein_coding | -0.147556289 | 3.855579368 | 0.0931452 | 0.7640771 | 0.948755137 |
| ENSMUSG00000020255 | D10Wsu102e | protein_coding | -0.110899978 | 5.333206271 | 0.0931212 | 0.7641065 | 0.948755137 |
| ENSMUSG00000023328 | Ache | protein_coding | 0.072691899 | 6.236611995 | 0.0930657 | 0.7641745 | 0.948755137 |
| ENSMUSG00000030729 | Pgm2l1 | protein_coding | -0.079647156 | 6.782313586 | 0.0928294 | 0.7644645 | 0.949005607 |
| ENSMUSG00000073702 | Rpl31 | protein_coding | -0.126804632 | 3.938345888 | 0.0927254 | 0.7645921 | 0.949038408 |
| ENSMUSG00000022629 | Kif21a | protein_coding | -0.076189998 | 7.104242049 | 0.0926229 | 0.7647181 | 0.949038408 |
| ENSMUSG00000025825 | Iscu | protein_coding | 0.098169524 | 4.795017122 | 0.0925923 | 0.7647557 | 0.949038408 |
| ENSMUSG00000028576 | Ift74 | protein_coding | 0.08007344 | 5.909639603 | 0.0924219 | 0.7649653 | 0.949189027 |
| ENSMUSG00000029059 | Fam213b | protein_coding | 0.084841483 | 9.193708674 | 0.0922657 | 0.7651576 | 0.949291496 |
| ENSMUSG00000031932 | Gpr83 | protein_coding | 0.10706309 | 5.762851993 | 0.0922166 | 0.7652245 | 0.949291496 |
| ENSMUSG00000018293 | Pfn1 | protein_coding | 0.075777094 | 6.820414673 | 0.0921054 | 0.7653552 | 0.949344096 |
| ENSMUSG00000028133 | Rwdd3 | protein_coding | 0.099421809 | 4.843181953 | 0.0918283 | 0.7656971 | 0.949576299 |
| ENSMUSG00000024835 | Coro1b | protein_coding | -0.066582145 | 6.337349066 | 0.0917717 | 0.7657671 | 0.949576299 |
| ENSMUSG00000049439 | Cyp20a1 | protein_coding | 0.123840268 | 4.164389823 | 0.0917349 | 0.7658125 | 0.949576299 |
| ENSMUSG00000039738 | Slx4 | protein_coding | 0.189340368 | 2.92523349 | 0.0914385 | 0.766179 | 0.949576299 |
| ENSMUSG00000023473 | Celsr3 | protein_coding | 0.225469405 | 2.799257655 | 0.0913899 | 0.7662392 | 0.949576299 |
| ENSMUSG00000037594 | BC022687 | protein_coding | 0.162995369 | 3.430828508 | 0.0913437 | 0.7662964 | 0.949576299 |
| ENSMUSG00000055733 | Nap1l3 | protein_coding | -0.093296415 | 5.768470532 | 0.0913408 | 0.7663 | 0.949576299 |
| ENSMUSG00000038039 | Gcc2 | protein_coding | -0.080822057 | 5.948137453 | 0.0913335 | 0.7663091 | 0.949576299 |
| ENSMUSG00000038822 | Hace1 | protein_coding | -0.122502932 | 4.195022349 | 0.0913019 | 0.7663483 | 0.949576299 |
| ENSMUSG00000034923 | Ly6g6f | protein_coding | 0.180601113 | 3.087477309 | 0.0910829 | 0.7666198 | 0.949576299 |
| ENSMUSG00000110185 | Igip | protein_coding | -0.107240251 | 5.364580679 | 0.0909219 | 0.7668196 | 0.949576299 |
| ENSMUSG00000059713 | Rcan3 | protein_coding | -0.080766085 | 5.330877705 | 0.0907183 | 0.7670725 | 0.949576299 |
| ENSMUSG00000029676 | Pot1a | protein_coding | -0.133848285 | 4.502546478 | 0.0906863 | 0.7671123 | 0.949576299 |
| ENSMUSG00000025027 | Xpnpep1 | protein_coding | -0.087358383 | 6.097822581 | 0.0906528 | 0.767154 | 0.949576299 |
| ENSMUSG00000015002 | Efr3a | protein_coding | -0.077701489 | 5.889607461 | 0.090628 | 0.7671848 | 0.949576299 |
| ENSMUSG00000046688 | Tifa | protein_coding | -0.2409852 | 2.772496425 | 0.090581 | 0.7672433 | 0.949576299 |
| ENSMUSG00000011158 | Brf1 | protein_coding | -0.133515311 | 3.943897732 | 0.0905666 | 0.7672612 | 0.949576299 |
| ENSMUSG00000029684 | Wasl | protein_coding | -0.074938111 | 6.066731873 | 0.0904501 | 0.7674063 | 0.949576299 |
| ENSMUSG00000021906 | Oxnad1 | protein_coding | -0.108892778 | 4.883129261 | 0.0903615 | 0.7675166 | 0.949576299 |
| ENSMUSG00000062929 | Cfl2 | protein_coding | 0.061641768 | 7.9738112 | 0.0903061 | 0.7675857 | 0.949576299 |
| ENSMUSG00000056306 | Sertm1 | protein_coding | 0.163131515 | 3.222958381 | 0.0902567 | 0.7676472 | 0.949576299 |
| ENSMUSG00000039747 | Orai2 | protein_coding | -0.12300492 | 4.35907682 | 0.0902504 | 0.7676551 | 0.949576299 |
| ENSMUSG00000028636 | Ppcs | protein_coding | 0.163586421 | 4.117834814 | 0.0901981 | 0.7677429 | 0.949576299 |
| ENSMUSG00000029438 | Bcl7a | protein_coding | -0.10520993 | 5.314470997 | 0.0899279 | 0.7680576 | 0.949576299 |
| ENSMUSG00000045312 | Lhfpl2 | protein_coding | -0.280095303 | 2.521567076 | 0.0898859 | 0.76811 | 0.949576299 |
| ENSMUSG00000034192 | Lsm3 | protein_coding | 0.1040344 | 6.577785955 | 0.0899551 | 0.7682138 | 0.949576299 |
| ENSMUSG00000022336 | Eif3e | protein_coding | 0.075113228 | 7.651434213 | 0.0897288 | 0.7683064 | 0.949576299 |
| ENSMUSG00000027487 | Cdk5rap1 | protein_coding | 0.123547188 | 4.290621699 | 0.0895752 | 0.7684986 | 0.949576299 |
| ENSMUSG00000046879 | Irgm1 | protein_coding | -0.161692988 | 4.267996404 | 0.0895384 | 0.7685673 | 0.949576299 |
| ENSMUSG00000041608 | Entpd3 | protein_coding | -0.169230818 | 3.918803749 | 0.0894931 | 0.7686076 | 0.949576299 |
| ENSMUSG00000020650 | Bcap29 | protein_coding | 0.076874679 | 6.324469534 | 0.0894597 | 0.7686432 | 0.949576299 |
| ENSMUSG00000060121 | Gemin2 | protein_coding | 0.096462245 | 4.848682691 | 0.0894512 | 0.7686539 | 0.949576299 |
| ENSMUSG00000046380 | Jrk | protein_coding | -0.175829616 | 3.524425754 | 0.0893975 | 0.7687213 | 0.949576299 |
| ENSMUSG00000039050 | Osbpl2 | protein_coding | -0.083241533 | 5.793113036 | 0.08935 | 0.7687807 | 0.949576299 |
| ENSMUSG00000024665 | Fads2 | protein_coding | 0.124664342 | 4.030559259 | 0.0892914 | 0.7688542 | 0.949576299 |
| ENSMUSG00000019027 | Dnah1 | protein_coding | 0.119957106 | 5.188644309 | 0.089165 | 0.769019 | 0.949576299 |
| ENSMUSG00000004768 | Rab23 | protein_coding | -0.139603542 | 4.504457909 | 0.08899 | 0.7692388 | 0.949576299 |
| ENSMUSG00000032555 | Topbp1 | protein_coding | -0.280028369 | 2.651248365 | 0.0889628 | 0.7692729 | 0.949576299 |
| ENSMUSG00000038390 | Gpr162 | protein_coding | 0.120441622 | 4.424986241 | 0.0889395 | 0.769296 | 0.949576299 |
| ENSMUSG00000096141 | Dnah7a | protein_coding | -0.118024993 | 3.695228633 | 0.0888562 | 0.7694007 | 0.949576299 |
| ENSMUSG00000021514 | Zfp369 | protein_coding | -0.163835095 | 3.534012811 | 0.0887578 | 0.7695245 | 0.949576299 |
| ENSMUSG00000028568 | Btf3l4 | protein_coding | -0.095923223 | 4.661104131 | 0.0887465 | 0.7695388 | 0.949576299 |
| ENSMUSG00000027508 | Pag1 | protein_coding | 0.174153921 | 3.019349105 | 0.0887127 | 0.7695813 | 0.949576299 |
| ENSMUSG00000075033 | Nxpe3 | protein_coding | -0.161132369 | 3.370713129 | 0.0886137 | 0.7697058 | 0.949576299 |
| ENSMUSG00000022109 | Med4 | protein_coding | 0.073140704 | 6.378644572 | 0.088477 | 0.7698781 | 0.949576299 |
| ENSMUSG00000032602 | Slc25a20 | protein_coding | 0.098928769 | 4.862579175 | 0.0883819 | 0.769998 | 0.949576299 |
| ENSMUSG00000025575 | Cant1 | protein_coding | 0.124305779 | 4.199755079 | 0.0883248 | 0.77007 | 0.949576299 |
| ENSMUSG00000027272 | Ubr1 | protein_coding | -0.116453347 | 3.851006777 | 0.0882603 | 0.7701515 | 0.949576299 |
| ENSMUSG00000033186 | Mzt1 | protein_coding | -0.063714182 | 7.298392549 | 0.0882469 | 0.7701684 | 0.949576299 |
| ENSMUSG00000008855 | Hdac5 | protein_coding | -0.100935374 | 5.34606645 | 0.0882357 | 0.7701825 | 0.949576299 |
| ENSMUSG00000058741 | Prr19 | protein_coding | 0.297073305 | 2.448788106 | 0.0882149 | 0.7702149 | 0.949576299 |
| ENSMUSG00000022106 | Rcbtb2 | protein_coding | -0.176945993 | 3.564056249 | 0.0881828 | 0.7702554 | 0.949576299 |
| ENSMUSG00000051502 | Ufsp1 | protein_coding | 0.125506846 | 3.98069418 | 0.0880399 | 0.7704298 | 0.949576299 |
| ENSMUSG00000050830 | Vwc2 | protein_coding | -0.168998039 | 3.167405937 | 0.088012 | 0.770465 | 0.949576299 |
| ENSMUSG00000020891 | Alox8 | protein_coding | 0.241013651 | 2.964018178 | 0.0879554 | 0.7705428 | 0.949576299 |
| ENSMUSG00000039057 | Myo16 | protein_coding | -0.216252785 | 2.929518244 | 0.0879248 | 0.7705753 | 0.949576299 |
| ENSMUSG00000028454 | Pigo | protein_coding | 0.181554864 | 3.200219405 | 0.0878968 | 0.7706107 | 0.949576299 |
| ENSMUSG00000039676 | Capsl | protein_coding | -0.154254915 | 4.066368369 | 0.0878594 | 0.7706642 | 0.949576299 |
| ENSMUSG00000004865 | Srpk1 | protein_coding | -0.078642449 | 6.884929363 | 0.0876852 | 0.7708785 | 0.949581927 |
| ENSMUSG00000043639 | Rbm20 | protein_coding | -0.226413557 | 2.429036329 | 0.0876794 | 0.7708859 | 0.949581927 |
| ENSMUSG00000021314 | Amph | protein_coding | 0.102726174 | 7.151884838 | 0.0878388 | 0.7709375 | 0.949581927 |
| ENSMUSG00000027936 | Crtc2 | protein_coding | -0.183815097 | 3.167573966 | 0.087572 | 0.771022 | 0.949581927 |
| ENSMUSG00000037060 | Cavin3 | protein_coding | 0.09148231 | 5.077329017 | 0.0874584 | 0.771166 | 0.949623418 |
| ENSMUSG00000022816 | Fstl1 | protein_coding | 0.09466504 | 5.619074361 | 0.0873455 | 0.7713092 | 0.949623418 |
| ENSMUSG00000054237 | Fra10ac1 | protein_coding | -0.08500807 | 5.396569907 | 0.0872779 | 0.7713951 | 0.949623418 |
| ENSMUSG00000029730 | Mcm7 | protein_coding | 0.157513896 | 3.548071418 | 0.0871092 | 0.7716094 | 0.949623418 |
| ENSMUSG00000020173 | Cobl | protein_coding | 0.102489457 | 4.979195501 | 0.0871074 | 0.7716117 | 0.949623418 |

| ensembl_gene_id | external_gene_name | gene_biotype | log2 (fold change) | log2 (counts per million) | F-value | p-value | False discovery rate |
|---|---|---|---|---|---|---|---|
| ENSMUSG00000021018 | Polr2h | protein_coding | -0.071612768 | 6.12571714 | 0.087093 | 0.77163 | 0.949623418 |
| ENSMUSG00000026640 | Plxna2 | protein_coding | -0.107486179 | 4.949512398 | 0.0869042 | 0.7718702 | 0.949623418 |
| ENSMUSG00000039809 | Gabbr2 | protein_coding | -0.089780912 | 5.834550087 | 0.0867982 | 0.7720051 | 0.949623418 |
| ENSMUSG00000035967 | Ints6l | protein_coding | -0.203171391 | 2.801393177 | 0.0867606 | 0.772053 | 0.949623418 |
| ENSMUSG00000023018 | Smarcd1 | protein_coding | 0.075796958 | 6.309771219 | 0.0865239 | 0.7723548 | 0.949623418 |
| ENSMUSG00000051373 | Plpp7 | protein_coding | 0.098337229 | 4.706038046 | 0.0864012 | 0.7725114 | 0.949623418 |
| ENSMUSG00000022378 | Fam49b | protein_coding | 0.064150369 | 6.70146057 | 0.0863649 | 0.7725578 | 0.949623418 |
| ENSMUSG00000032562 | Gnai2 | protein_coding | 0.066945285 | 7.278666785 | 0.0863574 | 0.7725673 | 0.949623418 |
| ENSMUSG00000019877 | Serinc1 | protein_coding | -0.057986073 | 9.253247427 | 0.0862541 | 0.7726993 | 0.949623418 |
| ENSMUSG00000028394 | Pole3 | protein_coding | 0.06653626 | 6.041170929 | 0.0862085 | 0.7727576 | 0.949623418 |
| ENSMUSG00000045827 | Serpinb9 | protein_coding | 0.135080844 | 3.913009374 | 0.0861438 | 0.7728403 | 0.949623418 |
| ENSMUSG00000038456 | Dennd2a | protein_coding | -0.143378626 | 3.903286235 | 0.0860684 | 0.7729368 | 0.949623418 |
| ENSMUSG00000032482 | Cspg5 | protein_coding | 0.079901121 | 7.199515376 | 0.0860683 | 0.7729369 | 0.949623418 |
| ENSMUSG00000050199 | Lgr4 | protein_coding | 0.099213903 | 4.807990148 | 0.0860602 | 0.7729473 | 0.949623418 |
| ENSMUSG00000026806 | Ddx31 | protein_coding | 0.210276964 | 2.812605396 | 0.0860058 | 0.773017 | 0.949623418 |
| ENSMUSG00000026893 | Gca | protein_coding | -0.124782153 | 4.339121022 | 0.0859253 | 0.77312 | 0.949623418 |
| ENSMUSG00000066892 | Fbxl12 | protein_coding | -0.142146526 | 3.706845045 | 0.085899 | 0.7731536 | 0.949623418 |
| ENSMUSG00000042328 | Hps4 | protein_coding | -0.12709225 | 3.690043828 | 0.0858215 | 0.7732529 | 0.949623418 |
| ENSMUSG00000026880 | Stom | protein_coding | 0.211340687 | 2.608445582 | 0.0857298 | 0.7733706 | 0.949623418 |
| ENSMUSG00000000794 | Kcnn3 | protein_coding | -0.163427031 | 3.752232933 | 0.0856293 | 0.7735054 | 0.949623418 |
| ENSMUSG00000090110 | Cmc4 | protein_coding | -0.119334881 | 3.930517205 | 0.0855963 | 0.7735418 | 0.949623418 |
| ENSMUSG00000040446 | Rprd1a | protein_coding | -0.131026048 | 3.754154012 | 0.085256 | 0.7739789 | 0.949623418 |
| ENSMUSG00000032590 | Apeh | protein_coding | 0.113157871 | 4.439069016 | 0.0852033 | 0.7740468 | 0.949623418 |
| ENSMUSG00000025743 | Sdc3 | protein_coding | 0.149312786 | 3.225559943 | 0.0851134 | 0.7741624 | 0.949623418 |
| ENSMUSG00000016200 | Syt14 | protein_coding | -0.155828177 | 3.44593663 | 0.0851019 | 0.7741772 | 0.949623418 |
| ENSMUSG00000028603 | Scp2 | protein_coding | 0.177837032 | 3.075464304 | 0.0850661 | 0.7742233 | 0.949623418 |
| ENSMUSG00000024112 | Cacna1h | protein_coding | -0.253746263 | 2.669602741 | 0.0850603 | 0.7742308 | 0.949623418 |
| ENSMUSG00000046707 | Csnk2a2 | protein_coding | 0.086482634 | 5.413038128 | 0.0850595 | 0.7742318 | 0.949623418 |
| ENSMUSG00000048834 | Vstm2a | protein_coding | -0.066334861 | 6.941873829 | 0.0849738 | 0.7743423 | 0.949623418 |
| ENSMUSG00000028540 | Dph2 | protein_coding | 0.121416241 | 4.539993622 | 0.0849521 | 0.7743702 | 0.949623418 |
| ENSMUSG00000043885 | Slc36a4 | protein_coding | 0.097860267 | 5.17974341 | 0.084934 | 0.7743935 | 0.949623418 |
| ENSMUSG00000055204 | Ankrd17 | protein_coding | -0.071144044 | 6.665545523 | 0.0849089 | 0.7744259 | 0.949623418 |
| ENSMUSG00000048163 | Selplg | protein_coding | -0.139588061 | 4.155605552 | 0.0848387 | 0.7745165 | 0.949623418 |
| ENSMUSG00000023055 | Calcoco1 | protein_coding | -0.087063855 | 6.086909112 | 0.084796 | 0.7745715 | 0.949623418 |
| ENSMUSG00000030726 | Pold3 | protein_coding | 0.110559344 | 5.231761122 | 0.084783 | 0.7745882 | 0.949623418 |
| ENSMUSG00000007908 | Hmgcll1 | protein_coding | -0.124808734 | 4.293571825 | 0.0846104 | 0.7748111 | 0.949788391 |
| ENSMUSG00000024026 | Glo1 | protein_coding | -0.059007986 | 7.490100632 | 0.0844999 | 0.7749538 | 0.949819563 |
| ENSMUSG00000041815 | Poldip3 | protein_coding | -0.063232896 | 8.061233683 | 0.0844267 | 0.7750485 | 0.949819563 |
| ENSMUSG00000074576 | Mocs3 | protein_coding | 0.258308278 | 2.224110288 | 0.0843856 | 0.7751015 | 0.949819563 |
| ENSMUSG00000034656 | Cacna1a | protein_coding | -0.172929155 | 3.40050763 | 0.0842538 | 0.7752722 | 0.949920421 |
| ENSMUSG00000028331 | Trmo | protein_coding | -0.279298651 | 3.10188917 | 0.0839046 | 0.776089 | 0.950757844 |
| ENSMUSG00000038375 | Trp53inp2 | protein_coding | -0.087070068 | 5.434666145 | 0.0835906 | 0.7761325 | 0.950757844 |
| ENSMUSG00000020994 | Pnn | protein_coding | 0.085576969 | 5.979140039 | 0.0833975 | 0.7763836 | 0.950852402 |
| ENSMUSG00000029095 | Ablim2 | protein_coding | -0.070657334 | 6.51450988 | 0.0833953 | 0.7763865 | 0.950852402 |
| ENSMUSG00000024335 | Brd2 | protein_coding | -0.064852511 | 6.404587949 | 0.0831156 | 0.7767509 | 0.951142647 |
| ENSMUSG00000073147 | 5031425E22Rik | lincRNA | 0.156769529 | 3.38855646 | 0.0830777 | 0.7768004 | 0.951142647 |
| ENSMUSG00000029191 | Rfc1 | protein_coding | 0.083388376 | 5.548123575 | 0.0828647 | 0.7770785 | 0.951374779 |
| ENSMUSG00000036840 | Siah1a | protein_coding | 0.128813239 | 3.974477674 | 0.0827916 | 0.7771739 | 0.951383356 |
| ENSMUSG00000066880 | Zfp617 | protein_coding | 0.105752956 | 3.954046888 | 0.0826778 | 0.7773227 | 0.951457129 |
| ENSMUSG00000044847 | Lsm11 | protein_coding | -0.177037171 | 3.172878441 | 0.0823267 | 0.7777824 | 0.951855221 |
| ENSMUSG00000059890 | Ube4a | protein_coding | -0.122574825 | 3.81874335 | 0.0822089 | 0.777937 | 0.951855221 |
| ENSMUSG00000032911 | Cspg4 | protein_coding | 0.162280289 | 3.823510684 | 0.0821581 | 0.7780095 | 0.951855221 |
| ENSMUSG00000032609 | Klhdc8b | protein_coding | 0.093713183 | 5.787174411 | 0.0820033 | 0.7782068 | 0.951855221 |
| ENSMUSG00000024240 | Epc1 | protein_coding | -0.099090383 | 4.111177096 | 0.0818704 | 0.7783815 | 0.951855221 |
| ENSMUSG00000001157 | Gmcl1 | protein_coding | -0.134713026 | 4.452459458 | 0.0817952 | 0.7784862 | 0.951855221 |
| ENSMUSG00000072494 | Ppp1r3e | protein_coding | -0.1507702 | 3.247803737 | 0.081778 | 0.778503 | 0.951855221 |
| ENSMUSG00000030315 | Vgll4 | protein_coding | 0.088525581 | 4.932348267 | 0.0817154 | 0.7785853 | 0.951855221 |
| ENSMUSG00000009734 | Pou6f2 | protein_coding | -0.281010834 | 2.355691237 | 0.0816833 | 0.7786276 | 0.951855221 |
| ENSMUSG00000036273 | Lrrk2 | protein_coding | 0.180756586 | 2.917905266 | 0.08164 | 0.7786847 | 0.951855221 |
| ENSMUSG00000027523 | Gnas | protein_coding | -0.127293151 | 3.948568539 | 0.0816026 | 0.7787339 | 0.951855221 |
| ENSMUSG00000017307 | Acot8 | protein_coding | 0.101804235 | 5.065814765 | 0.0815599 | 0.7787901 | 0.951855221 |
| ENSMUSG00000035399 | Oser1 | protein_coding | 0.108464888 | 3.785850373 | 0.0815534 | 0.7787987 | 0.951855221 |
| ENSMUSG00000036501 | Fam13b | protein_coding | 0.120209458 | 3.975044105 | 0.081393 | 0.7790101 | 0.951926101 |
| ENSMUSG00000028140 | Mrpl9 | protein_coding | 0.066719634 | 6.042274355 | 0.0813751 | 0.7790337 | 0.951926101 |
| ENSMUSG00000037149 | Ddx1 | protein_coding | -0.06133814 | 7.647847429 | 0.0812215 | 0.7792365 | 0.951933227 |
| ENSMUSG00000021244 | Ylpm1 | protein_coding | -0.083459333 | 5.454123486 | 0.0811766 | 0.7792958 | 0.951933227 |
| ENSMUSG00000038534 | Osbpl7 | protein_coding | 0.09782591 | 4.381384105 | 0.0811695 | 0.7793051 | 0.951933227 |
| ENSMUSG00000024601 | Isoc1 | protein_coding | 0.101436103 | 5.166928495 | 0.0810787 | 0.7794252 | 0.95197178 |
| ENSMUSG00000044475 | Ascc1 | protein_coding | 0.061657427 | 6.756643452 | 0.0809105 | 0.7796476 | 0.952005655 |
| ENSMUSG00000031156 | Slc35a2 | protein_coding | 0.082457764 | 5.421110488 | 0.0807869 | 0.7798113 | 0.952005655 |
| ENSMUSG00000059326 | Csf2ra | protein_coding | -0.189468006 | 2.661422324 | 0.080784 | 0.7798151 | 0.952005655 |
| ENSMUSG00000027457 | Snph | protein_coding | 0.075189565 | 6.359652237 | 0.0807329 | 0.7798828 | 0.952005655 |
| ENSMUSG00000021265 | Slc25a29 | protein_coding | 0.147301515 | 3.224824225 | 0.0806443 | 0.7800001 | 0.952005655 |
| ENSMUSG00000033209 | Ttc28 | protein_coding | 0.18529249 | 2.80421923 | 0.0805523 | 0.7801221 | 0.952005655 |
| ENSMUSG00000109865 | Hspa14 | protein_coding | -0.0800551 | 5.770359006 | 0.0805468 | 0.7801295 | 0.952005655 |
| ENSMUSG00000021918 | Nek4 | protein_coding | 0.116980893 | 4.061334243 | 0.0805229 | 0.7801612 | 0.952005655 |
| ENSMUSG00000021669 | Col4a3bp | protein_coding | 0.090665033 | 4.648174305 | 0.0802435 | 0.7805322 | 0.952262481 |
| ENSMUSG00000040225 | Prrc2c | protein_coding | -0.074519375 | 6.235597442 | 0.080231 | 0.7805488 | 0.952262481 |

| ensembl_gene_id | external_gene_name | gene_biotype | log2 (fold change) | log2 (counts per million) | F-value | p-value | False discovery rate |
|---|---|---|---|---|---|---|---|
| ENSMUSG00000036644 | Tbc1d9b | protein_coding | 0.093430243 | 5.50299736 | 0.0799349 | 0.7809428 | 0.952581195 |
| ENSMUSG00000030083 | Abtb1 | protein_coding | 0.105435501 | 6.035089861 | 0.0799845 | 0.7809872 | 0.952581195 |
| ENSMUSG00000041609 | Bicdl1 | protein_coding | 0.127279309 | 4.482857563 | 0.0796855 | 0.7812754 | 0.952727204 |
| ENSMUSG00000051355 | Commd1 | protein_coding | 0.11681776 | 3.530138769 | 0.0796789 | 0.7812842 | 0.952727204 |
| ENSMUSG00000075289 | Carns1 | protein_coding | 0.149447844 | 3.162253939 | 0.0794487 | 0.7815916 | 0.952994029 |
| ENSMUSG00000016477 | E2f3 | protein_coding | -0.133296783 | 3.467306497 | 0.0793434 | 0.7817324 | 0.953057647 |
| ENSMUSG00000026723 | Trdmt1 | protein_coding | -0.157728243 | 3.159473923 | 0.0790815 | 0.7820829 | 0.953376892 |
| ENSMUSG00000029401 | Rilpl2 | protein_coding | 0.119710861 | 4.513481641 | 0.0788732 | 0.7823622 | 0.953609195 |
| ENSMUSG00000040151 | Hs2st1 | protein_coding | 0.109851234 | 4.851491335 | 0.0786597 | 0.7826489 | 0.953721581 |
| ENSMUSG00000032875 | Arhgef17 | protein_coding | 0.06507003 | 6.310172175 | 0.0786184 | 0.7827044 | 0.953721581 |
| ENSMUSG00000027811 | 4930579G24Rik | protein_coding | -0.222755076 | 2.607655136 | 0.0786064 | 0.7827204 | 0.953721581 |
| ENSMUSG00000021916 | Glt8d1 | protein_coding | -0.132210019 | 4.0443837 | 0.0784822 | 0.7828875 | 0.95374075 |
| ENSMUSG00000019809 | Pex3 | protein_coding | 0.101404872 | 5.017499214 | 0.078178 | 0.7832972 | 0.95374075 |
| ENSMUSG00000031565 | Fgfr1 | protein_coding | -0.081805301 | 5.45452829 | 0.0781666 | 0.7833126 | 0.95374075 |
| ENSMUSG00000049112 | Oxtr | protein_coding | 0.194320502 | 2.82457283 | 0.0780956 | 0.7834083 | 0.95374075 |
| ENSMUSG00000028423 | Nfx1 | protein_coding | -0.079236225 | 6.440801969 | 0.0780105 | 0.7835232 | 0.95374075 |
| ENSMUSG00000087178 | A230056P14Rik | processed_transcript | 0.186069528 | 3.420152517 | 0.0779596 | 0.7835976 | 0.95374075 |
| ENSMUSG00000052906 | Ubxn8 | protein_coding | -0.073876486 | 6.051718057 | 0.0777852 | 0.7838275 | 0.95374075 |
| ENSMUSG00000028016 | Ints12 | protein_coding | 0.089145059 | 4.737708314 | 0.0777599 | 0.7838616 | 0.95374075 |
| ENSMUSG00000074505 | Fat3 | protein_coding | 0.129294215 | 4.060654349 | 0.0776932 | 0.7839518 | 0.95374075 |
| ENSMUSG00000045176 | Borcs6 | protein_coding | 0.12955517 | 3.728893799 | 0.0775055 | 0.7842059 | 0.95374075 |
| ENSMUSG00000026317 | Cln8 | protein_coding | -0.100179853 | 5.354004225 | 0.0774774 | 0.784244 | 0.95374075 |
| ENSMUSG00000038122 | Tbc1d32 | protein_coding | 0.188845576 | 2.883341405 | 0.0773307 | 0.7844429 | 0.95374075 |
| ENSMUSG00000062729 | Ppox | protein_coding | 0.132501886 | 4.797051836 | 0.0773383 | 0.7844535 | 0.95374075 |
| ENSMUSG00000044562 | Rasip1 | protein_coding | 0.198949495 | 3.036528528 | 0.0773204 | 0.7844568 | 0.95374075 |
| ENSMUSG00000032855 | Pkd1 | protein_coding | -0.193810753 | 2.826235212 | 0.0772265 | 0.7845842 | 0.95374075 |
| ENSMUSG00000041769 | Ppp2r2d | protein_coding | -0.063000926 | 6.312592525 | 0.0772093 | 0.7846075 | 0.95374075 |
| ENSMUSG00000086119 | Gm2415 | antisense_RNA | -0.194209485 | 2.633322965 | 0.0771987 | 0.784622 | 0.95374075 |
| ENSMUSG00000028629 | Exo5 | protein_coding | 0.176133364 | 4.006934492 | 0.0773372 | 0.7846567 | 0.95374075 |
| ENSMUSG00000111336 | AC160338.2 | processed_pseudoge | 0.189670288 | 2.495703663 | 0.0770636 | 0.7848055 | 0.95374075 |
| ENSMUSG00000029392 | Rilpl1 | protein_coding | -0.109857113 | 4.592945239 | 0.0770428 | 0.7848337 | 0.95374075 |
| ENSMUSG00000029499 | Pxmp2 | protein_coding | 0.130338071 | 3.719381021 | 0.0770245 | 0.7848586 | 0.95374075 |
| ENSMUSG00000001576 | Ergic1 | protein_coding | 0.08450005 | 5.837601065 | 0.0768746 | 0.7850623 | 0.95374075 |
| ENSMUSG00000024055 | Cyp4f13 | protein_coding | -0.215211587 | 2.949855005 | 0.0767728 | 0.7852009 | 0.95374075 |
| ENSMUSG00000056121 | Fez2 | protein_coding | -0.090568462 | 4.617112887 | 0.0767134 | 0.7852818 | 0.95374075 |
| ENSMUSG00000049516 | Spty2d1 | protein_coding | -0.117926337 | 3.873395935 | 0.0766816 | 0.7853251 | 0.95374075 |
| ENSMUSG00000049878 | Rlf | protein_coding | -0.125600328 | 3.754233877 | 0.0765702 | 0.785477 | 0.95374075 |
| ENSMUSG00000028893 | Sesn2 | protein_coding | -0.160237152 | 3.422479289 | 0.0765547 | 0.785498 | 0.95374075 |
| ENSMUSG00000033900 | Map9 | protein_coding | -0.07401591 | 6.439453988 | 0.0762481 | 0.7859166 | 0.95374075 |
| ENSMUSG00000035032 | Nek11 | protein_coding | -0.215084285 | 2.447111599 | 0.0761323 | 0.7860749 | 0.95374075 |
| ENSMUSG00000027951 | Adar | protein_coding | -0.222875124 | 3.419891742 | 0.0763437 | 0.7860779 | 0.95374075 |
| ENSMUSG00000024969 | Mark2 | protein_coding | 0.076663688 | 5.771430586 | 0.0759893 | 0.7862706 | 0.95374075 |
| ENSMUSG00000025856 | Pdgfa | protein_coding | 0.082861879 | 5.019665444 | 0.0759575 | 0.7863141 | 0.95374075 |
| ENSMUSG00000040543 | Pitpnm3 | protein_coding | -0.12629338 | 3.960726786 | 0.075816 | 0.7865079 | 0.95374075 |
| ENSMUSG00000063253 | Scoc | protein_coding | 0.060913478 | 7.516493739 | 0.0757824 | 0.7865541 | 0.95374075 |
| ENSMUSG00000019853 | Hebp2 | protein_coding | 0.106272826 | 3.909298826 | 0.0757224 | 0.7866364 | 0.95374075 |
| ENSMUSG00000025586 | Cpeb1 | protein_coding | 0.119626051 | 5.172911502 | 0.0756631 | 0.7867321 | 0.95374075 |
| ENSMUSG00000089759 | 3632454L22Rik | processed_transcript | -0.195280284 | 2.614671891 | 0.0755454 | 0.7868792 | 0.95374075 |
| ENSMUSG00000091476 | Gm16432 | protein_coding | -0.195860626 | 2.639064719 | 0.075535 | 0.7868935 | 0.95374075 |
| ENSMUSG00000022035 | Ccdc25 | protein_coding | 0.064478261 | 6.514769902 | 0.0753292 | 0.7871764 | 0.95374075 |
| ENSMUSG00000004099 | Dnmt1 | protein_coding | -0.098008541 | 4.668233799 | 0.0753101 | 0.7872027 | 0.95374075 |
| ENSMUSG00000052298 | Cdc42se2 | protein_coding | 0.076421988 | 7.224808403 | 0.0752276 | 0.7873218 | 0.95374075 |
| ENSMUSG00000050821 | Fam131a | protein_coding | 0.057034082 | 6.95714607 | 0.0752102 | 0.7873401 | 0.95374075 |
| ENSMUSG00000050989 | Selenon | protein_coding | -0.168946579 | 3.32866713 | 0.0751955 | 0.7873604 | 0.95374075 |
| ENSMUSG00000032724 | Abtb2 | protein_coding | -0.232234993 | 2.470819993 | 0.0751614 | 0.7874074 | 0.95374075 |
| ENSMUSG00000027221 | Chst1 | protein_coding | 0.066612604 | 6.988503255 | 0.075061 | 0.7875457 | 0.95374075 |
| ENSMUSG00000036907 | C1ql2 | protein_coding | 0.240792855 | 2.369910729 | 0.0750355 | 0.7875808 | 0.95374075 |
| ENSMUSG00000003863 | Ppfia3 | protein_coding | 0.090276528 | 4.668904969 | 0.0750211 | 0.7876007 | 0.95374075 |
| ENSMUSG00000025521 | Tmem192 | protein_coding | 0.12377079 | 3.808437796 | 0.0750155 | 0.7876084 | 0.95374075 |
| ENSMUSG00000037935 | Smarce1 | protein_coding | 0.085695825 | 5.526289286 | 0.0749608 | 0.7876838 | 0.95374075 |
| ENSMUSG00000040782 | Rfwd2 | protein_coding | -0.166113821 | 3.500210466 | 0.0748523 | 0.7878334 | 0.95374075 |
| ENSMUSG00000040188 | Scamp2 | protein_coding | 0.082942679 | 5.15352161 | 0.0748152 | 0.7878847 | 0.95374075 |
| ENSMUSG00000027663 | Zmat3 | protein_coding | -0.069568395 | 5.696992462 | 0.0748022 | 0.7879026 | 0.95374075 |
| ENSMUSG00000031314 | Taf1 | protein_coding | -0.095539404 | 5.39495505 | 0.0747216 | 0.788014 | 0.95374075 |
| ENSMUSG00000037788 | Vopp1 | protein_coding | -0.106398031 | 4.910781996 | 0.0747175 | 0.7880196 | 0.95374075 |
| ENSMUSG00000003279 | Dlgap1 | protein_coding | -0.066952126 | 6.176423712 | 0.0747085 | 0.7880321 | 0.95374075 |
| ENSMUSG00000025764 | Jade1 | protein_coding | -0.070997188 | 5.582317761 | 0.0746459 | 0.7881186 | 0.95374075 |
| ENSMUSG00000021537 | Cetn3 | protein_coding | -0.076450913 | 7.85686863 | 0.0746371 | 0.7881364 | 0.95374075 |
| ENSMUSG00000025369 | Smarcc2 | protein_coding | 0.0671417 | 6.432757822 | 0.0745358 | 0.7882707 | 0.95374075 |
| ENSMUSG00000027835 | Pdcd10 | protein_coding | -0.070330279 | 6.234613368 | 0.0744558 | 0.7883815 | 0.95374075 |
| ENSMUSG00000028469 | Npr2 | protein_coding | -0.149612152 | 3.538515871 | 0.074349 | 0.7885294 | 0.95374075 |
| ENSMUSG00000030089 | Slc41a3 | protein_coding | 0.156085305 | 3.496801846 | 0.0743293 | 0.7885567 | 0.95374075 |
| ENSMUSG00000022420 | Dnal4 | protein_coding | -0.073038937 | 6.838531098 | 0.0740462 | 0.7889493 | 0.95374075 |
| ENSMUSG00000066551 | Hmgb1 | protein_coding | 0.056785356 | 6.525799609 | 0.0739636 | 0.7890641 | 0.95374075 |
| ENSMUSG00000020621 | Rdh14 | protein_coding | -0.087127823 | 5.326370321 | 0.0736758 | 0.7894642 | 0.95374075 |
| ENSMUSG00000060679 | Mrps9 | protein_coding | 0.065293895 | 6.640901099 | 0.0736624 | 0.7894829 | 0.95374075 |
| ENSMUSG00000025616 | Usp16 | protein_coding | -0.091857778 | 4.813608305 | 0.0736432 | 0.7895096 | 0.95374075 |
| ENSMUSG00000040483 | Xaf1 | protein_coding | -0.107580359 | 4.266145799 | 0.0736331 | 0.7895237 | 0.95374075 |

| ensembl_gene_id | external_gene_name | gene_biotype | log2 (fold change) | log2 (counts per million) | F-value | p-value | False discovery rate |
|---|---|---|---|---|---|---|---|
| ENSMUSG00000049932 | H2afx | protein_coding | -0.194172948 | 2.595392298 | 0.0736186 | 0.7895439 | 0.95374075 |
| ENSMUSG00000047473 | Zfp30 | protein_coding | -0.08952271 | 4.606516946 | 0.0735032 | 0.7897047 | 0.95374075 |
| ENSMUSG00000027708 | Dcun1d1 | protein_coding | -0.081454411 | 5.159536717 | 0.0734364 | 0.7897978 | 0.95374075 |
| ENSMUSG00000025019 | Lcor | protein_coding | 0.099394108 | 4.180898007 | 0.0734283 | 0.7898091 | 0.95374075 |
| ENSMUSG00000033102 | Cdc14b | protein_coding | -0.128536926 | 3.484622043 | 0.073321 | 0.7899588 | 0.95374075 |
| ENSMUSG00000062040 | Zfp27 | protein_coding | -0.117745318 | 4.00792207 | 0.0733057 | 0.7899801 | 0.95374075 |
| ENSMUSG00000036940 | Kdm1a | protein_coding | -0.13622429 | 3.272368553 | 0.0731485 | 0.7901997 | 0.95374075 |
| ENSMUSG00000019832 | Rab32 | protein_coding | -0.352672198 | 1.917970587 | 0.0731342 | 0.7902252 | 0.95374075 |
| ENSMUSG00000043252 | Tmem64 | protein_coding | -0.112320003 | 4.869118991 | 0.0730933 | 0.7902769 | 0.95374075 |
| ENSMUSG00000109398 | Gm3854 | protein_coding | 0.182955978 | 3.106398389 | 0.0730875 | 0.7902849 | 0.95374075 |
| ENSMUSG00000040112 | Mrps35 | protein_coding | 0.110907112 | 4.382661176 | 0.0730862 | 0.7902867 | 0.95374075 |
| ENSMUSG00000024422 | Dhx16 | protein_coding | 0.172624228 | 2.864319705 | 0.0730561 | 0.7903288 | 0.95374075 |
| ENSMUSG00000024754 | Tmem2 | protein_coding | -0.168242607 | 3.101771873 | 0.0729289 | 0.7905068 | 0.95374075 |
| ENSMUSG00000015536 | Mocs2 | protein_coding | -0.054900771 | 7.941274613 | 0.072904 | 0.7905416 | 0.95374075 |
| ENSMUSG00000031657 | Heatr3 | protein_coding | -0.111802102 | 4.264892897 | 0.0728257 | 0.7906513 | 0.95374075 |
| ENSMUSG00000049556 | Lingo1 | protein_coding | 0.086698716 | 5.534299387 | 0.072807 | 0.7906776 | 0.95374075 |
| ENSMUSG00000038026 | Kcnj9 | protein_coding | -0.152824923 | 4.986539953 | 0.0730806 | 0.7907137 | 0.95374075 |
| ENSMUSG00000036104 | Rab3gap1 | protein_coding | -0.078578444 | 6.15508334 | 0.0727772 | 0.7907193 | 0.95374075 |
| ENSMUSG00000090626 | Tex9 | protein_coding | 0.12843141 | 3.860997671 | 0.0727501 | 0.7907572 | 0.95374075 |
| ENSMUSG00000027706 | Sec62 | protein_coding | -0.05608578 | 7.909474174 | 0.0727427 | 0.7907676 | 0.95374075 |
| ENSMUSG00000058886 | Deaf1 | protein_coding | -0.105430072 | 4.746996997 | 0.0726614 | 0.7908816 | 0.95374075 |
| ENSMUSG00000047181 | Samd14 | protein_coding | 0.083624571 | 5.095027515 | 0.0726522 | 0.7908945 | 0.95374075 |
| ENSMUSG00000029922 | Mkrn1 | protein_coding | -0.0599381 | 6.384668255 | 0.0726485 | 0.7908996 | 0.95374075 |
| ENSMUSG00000035069 | Oma1 | protein_coding | -0.131609712 | 3.725733586 | 0.0726405 | 0.7909109 | 0.95374075 |
| ENSMUSG00000030092 | Cntn6 | protein_coding | -0.115160907 | 4.10731142 | 0.0726224 | 0.7909363 | 0.95374075 |
| ENSMUSG00000041460 | Cacna2d4 | protein_coding | -0.211332232 | 2.942424916 | 0.0725878 | 0.7909848 | 0.95374075 |
| ENSMUSG00000043140 | Tmem186 | protein_coding | 0.106320515 | 5.17310996 | 0.0724027 | 0.7912447 | 0.953846064 |
| ENSMUSG00000020057 | Dram1 | protein_coding | -0.099543983 | 4.04375824 | 0.0723811 | 0.7912751 | 0.953846064 |
| ENSMUSG00000066043 | Phactr4 | protein_coding | -0.172966239 | 2.921263514 | 0.0723361 | 0.7913383 | 0.953846064 |
| ENSMUSG00000078862 | Gm14326 | protein_coding | -0.107739388 | 4.483002058 | 0.0722669 | 0.7914356 | 0.953856449 |
| ENSMUSG00000092536 | Gm20501 | bidirectional_promote | 0.119076199 | 3.668349907 | 0.0721246 | 0.7916359 | 0.953861189 |
| ENSMUSG00000004665 | Cnn2 | protein_coding | -0.206440452 | 2.957617468 | 0.072122 | 0.7916396 | 0.953861189 |
| ENSMUSG00000024462 | Gabbr1 | protein_coding | 0.068267745 | 7.898759309 | 0.0720275 | 0.7917727 | 0.953861189 |
| ENSMUSG00000058297 | Spock2 | protein_coding | 0.05629145 | 7.721449421 | 0.0719444 | 0.7918899 | 0.953861189 |
| ENSMUSG00000030980 | Knop1 | protein_coding | 0.074169685 | 5.433076153 | 0.0718464 | 0.792028 | 0.953861189 |
| ENSMUSG00000022935 | Grik1 | protein_coding | -0.140497284 | 4.004442477 | 0.0718303 | 0.7920564 | 0.953861189 |
| ENSMUSG00000051223 | Bzw1 | protein_coding | -0.059945578 | 7.448089155 | 0.0717767 | 0.7921264 | 0.953861189 |
| ENSMUSG00000027977 | Ndst3 | protein_coding | -0.186949594 | 2.657612592 | 0.0717606 | 0.7921492 | 0.953861189 |
| ENSMUSG00000053134 | Supt7l | protein_coding | 0.091635679 | 4.77275833 | 0.0715544 | 0.7924406 | 0.954105204 |
| ENSMUSG00000070002 | Ell | protein_coding | -0.089924399 | 5.35422909 | 0.0714692 | 0.7925611 | 0.954143483 |
| ENSMUSG00000002812 | Flii | protein_coding | 0.10981464 | 5.044575425 | 0.071193 | 0.7929578 | 0.954514219 |
| ENSMUSG00000028613 | Lrp8 | protein_coding | -0.127564592 | 4.297031855 | 0.0710745 | 0.7931204 | 0.954603093 |
| ENSMUSG00000043909 | Trp53bp1 | protein_coding | 0.103242695 | 4.852830253 | 0.0709936 | 0.7932353 | 0.954634466 |
| ENSMUSG00000029364 | Wsb2 | protein_coding | 0.076150655 | 5.49505858 | 0.0708891 | 0.7933838 | 0.954706319 |
| ENSMUSG00000050323 | Ndufaf6 | protein_coding | -0.186101001 | 2.951008596 | 0.0707668 | 0.7935577 | 0.954743491 |
| ENSMUSG00000038618 | Rassf7 | protein_coding | 0.157588166 | 3.373185711 | 0.0706863 | 0.7936722 | 0.954743491 |
| ENSMUSG00000020536 | Llgl1 | protein_coding | -0.093922806 | 4.550861631 | 0.0706301 | 0.7937522 | 0.954743491 |
| ENSMUSG00000025377 | Tepsin | protein_coding | -0.168070017 | 2.691341674 | 0.0705102 | 0.793923 | 0.954743491 |
| ENSMUSG00000062014 | Gmfb | protein_coding | 0.072240782 | 7.448109602 | 0.0704468 | 0.7940165 | 0.954743491 |
| ENSMUSG00000041216 | Clvs1 | protein_coding | 0.109610531 | 4.45779578 | 0.0704142 | 0.7940599 | 0.954743491 |
| ENSMUSG00000033007 | Asic4 | protein_coding | -0.283349321 | 2.312963814 | 0.0703758 | 0.7941202 | 0.954743491 |
| ENSMUSG00000058318 | Phf21a | protein_coding | -0.091391123 | 4.990714443 | 0.0703686 | 0.794125 | 0.954743491 |
| ENSMUSG00000036781 | Rps27l | protein_coding | 0.076607532 | 6.179979603 | 0.0701003 | 0.7945082 | 0.955097395 |
| ENSMUSG00000030417 | Pdcd5 | protein_coding | -0.116370825 | 4.30447361 | 0.0699407 | 0.7947365 | 0.955179644 |
| ENSMUSG00000048970 | C1galt1c1 | protein_coding | -0.084901263 | 5.775163264 | 0.0699046 | 0.7947882 | 0.955179644 |
| ENSMUSG00000004383 | Large1 | protein_coding | -0.077208406 | 5.054282434 | 0.0698663 | 0.7948431 | 0.955179644 |
| ENSMUSG00000027188 | Pamr1 | protein_coding | 0.17975343 | 2.718552927 | 0.0697477 | 0.795013 | 0.955213548 |
| ENSMUSG00000030088 | Aldh1l1 | protein_coding | 0.13232352 | 4.31170691 | 0.0697264 | 0.7950489 | 0.955213548 |
| ENSMUSG00000030380 | Mzf1 | protein_coding | -0.229975104 | 2.223890863 | 0.0695985 | 0.795227 | 0.955240991 |
| ENSMUSG00000032216 | Nedd4 | protein_coding | 0.067021169 | 8.22302611 | 0.0695829 | 0.7952495 | 0.955240991 |
| ENSMUSG00000055745 | Ldoc1l | protein_coding | 0.078356062 | 5.336780888 | 0.0695001 | 0.7953683 | 0.955277068 |
| ENSMUSG00000028614 | Ndc1 | protein_coding | 0.187446292 | 3.068110825 | 0.0693011 | 0.7956544 | 0.955346499 |
| ENSMUSG00000035342 | Lzts2 | protein_coding | -0.183232698 | 3.030487529 | 0.0692313 | 0.7957548 | 0.955346499 |
| ENSMUSG00000056091 | St3gal5 | protein_coding | -0.090308329 | 4.89099431 | 0.0691983 | 0.7958023 | 0.955346499 |
| ENSMUSG00000004233 | Wars2 | protein_coding | -0.158987324 | 3.088499615 | 0.0691608 | 0.7958563 | 0.955346499 |
| ENSMUSG00000027164 | Traf6 | protein_coding | -0.184962249 | 2.683158561 | 0.0691202 | 0.7959148 | 0.955346499 |
| ENSMUSG00000042115 | Klhdc8a | protein_coding | -0.093434203 | 5.107572488 | 0.0690894 | 0.7959592 | 0.955346499 |
| ENSMUSG00000015476 | Prrt1 | protein_coding | 0.075756578 | 5.824490711 | 0.0688135 | 0.7963572 | 0.95565887 |
| ENSMUSG00000071076 | Jund | protein_coding | 0.089623366 | 4.43190559 | 0.0687858 | 0.7963972 | 0.95565887 |
| ENSMUSG00000028854 | Slc9a1 | protein_coding | -0.099294024 | 4.565030444 | 0.0686818 | 0.7965475 | 0.955732567 |
| ENSMUSG00000031487 | Brf2 | protein_coding | 0.085074646 | 5.21612389 | 0.0685833 | 0.79669 | 0.955796855 |
| ENSMUSG00000022983 | Scaf4 | protein_coding | 0.191249566 | 3.146398055 | 0.0684389 | 0.7969043 | 0.955878155 |
| ENSMUSG00000025232 | Hexa | protein_coding | 0.078577355 | 5.996006021 | 0.0684136 | 0.7969355 | 0.955878155 |
| ENSMUSG00000031701 | Dnaja2 | protein_coding | 0.062859831 | 7.840731032 | 0.0683514 | 0.7970257 | 0.955879708 |
| ENSMUSG00000034156 | Tspoap1 | protein_coding | -0.115127182 | 4.054410639 | 0.0681675 | 0.7972924 | 0.955897923 |
| ENSMUSG00000054934 | Kcnmb4 | protein_coding | -0.203255061 | 2.524912605 | 0.0680578 | 0.7974516 | 0.955897923 |
| ENSMUSG00000021987 | Mtmr6 | protein_coding | -0.072224226 | 6.883620978 | 0.0680513 | 0.7974612 | 0.955897923 |
| ENSMUSG00000028894 | Inpp5b | protein_coding | 0.10704749 | 4.148575402 | 0.067973 | 0.7975749 | 0.955897923 |

| ensembl_gene_id | external_gene_name | gene_biotype | log2 (fold change) | log2 (counts per million) | F-value | p-value | False discovery rate |
|---|---|---|---|---|---|---|---|
| ENSMUSG00000025525 | Apool | protein_coding | -0.116797972 | 4.103545178 | 0.0678959 | 0.7976871 | 0.955897923 |
| ENSMUSG00000031446 | Cul4a | protein_coding | -0.114349798 | 4.612522061 | 0.0678881 | 0.7976983 | 0.955897923 |
| ENSMUSG00000059423 | Zfp933 | protein_coding | -0.163956338 | 3.282108402 | 0.0678868 | 0.7977003 | 0.955897923 |
| ENSMUSG00000066613 | Zfp932 | protein_coding | -0.109505155 | 3.590341987 | 0.0678512 | 0.7977521 | 0.955897923 |
| ENSMUSG00000054074 | Skida1 | protein_coding | 0.190115454 | 2.673199539 | 0.0676763 | 0.7980068 | 0.956091231 |
| ENSMUSG00000028899 | Taf12 | protein_coding | -0.075607405 | 6.018048433 | 0.0676183 | 0.7980912 | 0.956091231 |
| ENSMUSG00000112117 | RMST_1 | lincRNA | -0.119337135 | 3.455362308 | 0.0675249 | 0.7982274 | 0.956147873 |
| ENSMUSG00000063801 | Ap3s2 | protein_coding | 0.05819726 | 6.881998965 | 0.067397 | 0.7984141 | 0.956169837 |
| ENSMUSG00000040550 | Otud6b | protein_coding | -0.081028057 | 5.995032259 | 0.0673905 | 0.7984236 | 0.956169837 |
| ENSMUSG00000018820 | Zfyve27 | protein_coding | -0.078719016 | 5.296735211 | 0.0672006 | 0.7987011 | 0.956395691 |
| ENSMUSG00000026241 | Nppc | protein_coding | -0.144006919 | 3.262840325 | 0.0670845 | 0.798871 | 0.956492564 |
| ENSMUSG00000047547 | Cltb | protein_coding | -0.069294449 | 7.68489647 | 0.0669915 | 0.7990072 | 0.956549174 |
| ENSMUSG00000039703 | Nploc4 | protein_coding | -0.081933583 | 4.976776944 | 0.0668481 | 0.7992174 | 0.956607487 |
| ENSMUSG00000039414 | Heatr5b | protein_coding | -0.124215845 | 3.54909801 | 0.0668369 | 0.7992339 | 0.956607487 |
| ENSMUSG00000037259 | Dzank1 | protein_coding | -0.112321556 | 4.315431019 | 0.0667059 | 0.7994262 | 0.956731181 |
| ENSMUSG00000026974 | Zmynd19 | protein_coding | 0.10062754 | 3.884006846 | 0.0666125 | 0.7995633 | 0.956740523 |
| ENSMUSG00000032262 | Elovl4 | protein_coding | 0.066131321 | 5.962057517 | 0.0664735 | 0.7997676 | 0.956740523 |
| ENSMUSG00000027827 | Kcnab1 | protein_coding | -0.05615094 | 6.617753753 | 0.0664561 | 0.7997932 | 0.956740523 |
| ENSMUSG00000019718 | L3hypdh | protein_coding | 0.129826089 | 3.288056262 | 0.0662562 | 0.8000876 | 0.956740523 |
| ENSMUSG00000037674 | Rfx7 | protein_coding | -0.1095749 | 3.692786048 | 0.0662453 | 0.8001037 | 0.956740523 |
| ENSMUSG00000072653 | Zfp783 | transcribed_unproces | 0.195292078 | 2.885958419 | 0.0661955 | 0.8001771 | 0.956740523 |
| ENSMUSG00000048833 | Slc39a9 | protein_coding | 0.07785918 | 4.910425204 | 0.0661593 | 0.8002304 | 0.956740523 |
| ENSMUSG00000028191 | Bcl10 | protein_coding | -0.079683598 | 4.911238581 | 0.0661449 | 0.8002517 | 0.956740523 |
| ENSMUSG00000038807 | Rap1gap2 | protein_coding | 0.08731962 | 5.432078849 | 0.0661413 | 0.8002571 | 0.956740523 |
| ENSMUSG00000020307 | Cdc34 | protein_coding | -0.099198863 | 4.500428073 | 0.0660961 | 0.8003237 | 0.956740523 |
| ENSMUSG00000034282 | Evpl | protein_coding | -0.126324649 | 3.819500099 | 0.0660308 | 0.8004201 | 0.956749411 |
| ENSMUSG00000026571 | Dcaf6 | protein_coding | -0.086432939 | 4.34606051 | 0.0657849 | 0.8007835 | 0.957077412 |
| ENSMUSG00000033983 | Coil | protein_coding | -0.091899991 | 4.528481232 | 0.065684 | 0.8009329 | 0.95714709 |
| ENSMUSG00000020400 | Tnip1 | protein_coding | 0.095438344 | 5.062295397 | 0.0655766 | 0.8010919 | 0.95714709 |
| ENSMUSG00000021302 | Ggps1 | protein_coding | 0.054977958 | 6.849266201 | 0.0654241 | 0.8013181 | 0.95714709 |
| ENSMUSG00000020929 | Eftud2 | protein_coding | 0.083881231 | 5.697167108 | 0.065387 | 0.8013732 | 0.95714709 |
| ENSMUSG00000035878 | Hykk | protein_coding | 0.279937865 | 2.693860234 | 0.0654913 | 0.8014222 | 0.95714709 |
| ENSMUSG00000042404 | Dennd4b | protein_coding | -0.09240299 | 4.202061161 | 0.065266 | 0.8015528 | 0.95714709 |
| ENSMUSG00000079109 | Pms2 | protein_coding | 0.206657936 | 2.451916495 | 0.0652127 | 0.801632 | 0.95714709 |
| ENSMUSG00000034780 | B3galt1 | protein_coding | -0.076008903 | 5.905181875 | 0.0651829 | 0.8016763 | 0.95714709 |
| ENSMUSG00000040563 | Plppr2 | protein_coding | 0.091817476 | 5.355403126 | 0.0651473 | 0.8017292 | 0.95714709 |
| ENSMUSG00000109975 | Gm19196 | transcribed_unproces | 0.098717627 | 3.871161809 | 0.065034 | 0.8018978 | 0.95714709 |
| ENSMUSG00000054676 | 1600014C10Rik | protein_coding | -0.093476421 | 4.517361593 | 0.0650083 | 0.801936 | 0.95714709 |
| ENSMUSG00000064289 | Tank | protein_coding | -0.073737128 | 5.599823498 | 0.064968 | 0.8019959 | 0.95714709 |
| ENSMUSG00000026424 | Gpr37l1 | protein_coding | -0.073613857 | 6.583081309 | 0.064966 | 0.801999 | 0.95714709 |
| ENSMUSG00000035597 | Prpf39 | protein_coding | -0.102170318 | 4.118613969 | 0.0648186 | 0.8022186 | 0.957302973 |
| ENSMUSG00000024663 | Rab3il1 | protein_coding | 0.137190293 | 3.605714295 | 0.0647537 | 0.8023154 | 0.957312247 |
| ENSMUSG00000047067 | Dusp28 | protein_coding | 0.097094311 | 3.811464744 | 0.0646788 | 0.8024272 | 0.957335418 |
| ENSMUSG00000037166 | Ppp1r14a | protein_coding | -0.133081733 | 3.985494082 | 0.0646249 | 0.8025129 | 0.957335418 |
| ENSMUSG00000001440 | Kpnb1 | protein_coding | -0.060618677 | 6.058698675 | 0.0645313 | 0.8026474 | 0.957389684 |
| ENSMUSG00000066595 | Mfsd7b | protein_coding | 0.144513377 | 3.027479416 | 0.0644212 | 0.8028122 | 0.957480014 |
| ENSMUSG00000016559 | H3f3b | protein_coding | 0.055403987 | 9.584166665 | 0.0643423 | 0.8029302 | 0.957514538 |
| ENSMUSG00000026918 | Brd3 | protein_coding | -0.058995816 | 6.624802106 | 0.0642226 | 0.8031094 | 0.957622056 |
| ENSMUSG00000015120 | Ube2i | protein_coding | 0.117755095 | 3.638052954 | 0.0641058 | 0.8032846 | 0.957652906 |
| ENSMUSG00000036779 | Papd5 | protein_coding | 0.100196294 | 4.076825432 | 0.0639385 | 0.8035358 | 0.957652906 |
| ENSMUSG00000053333 | Dis3l2 | protein_coding | -0.16556726 | 2.901809079 | 0.0639311 | 0.8035468 | 0.957652906 |
| ENSMUSG00000037415 | Ranbp10 | protein_coding | 0.154141728 | 3.407233385 | 0.0637927 | 0.8037548 | 0.957652906 |
| ENSMUSG00000002342 | Tmem161a | protein_coding | -0.10263283 | 4.850548156 | 0.0637237 | 0.8038586 | 0.957652906 |
| ENSMUSG00000022359 | Wdyhv1 | protein_coding | 0.063905374 | 5.77310825 | 0.0637235 | 0.803859 | 0.957652906 |
| ENSMUSG00000039840 | Epg5 | protein_coding | -0.171530127 | 3.388539015 | 0.063705 | 0.803892 | 0.957652906 |
| ENSMUSG00000031467 | Agpat5 | protein_coding | 0.079120167 | 5.353533336 | 0.0636669 | 0.8039442 | 0.957652906 |
| ENSMUSG00000030224 | Strap | protein_coding | 0.05586335 | 6.523884665 | 0.0636563 | 0.8039601 | 0.957652906 |
| ENSMUSG00000004096 | Cwc15 | protein_coding | 0.058796836 | 8.159173265 | 0.0636101 | 0.8040297 | 0.957652906 |
| ENSMUSG00000027499 | Pkia | protein_coding | -0.056350882 | 7.19892695 | 0.0635535 | 0.804115 | 0.957652906 |
| ENSMUSG00000031532 | Saraf | protein_coding | -0.067108402 | 7.232649969 | 0.0633055 | 0.804489 | 0.957949565 |
| ENSMUSG00000029033 | Acap3 | protein_coding | -0.082926251 | 4.663657039 | 0.0632703 | 0.8045422 | 0.957949565 |
| ENSMUSG00000056394 | Lig1 | protein_coding | -0.097942274 | 4.074865149 | 0.0632015 | 0.8046462 | 0.957967236 |
| ENSMUSG00000029634 | Rnf6 | protein_coding | 0.078414357 | 7.05288655 | 0.0630857 | 0.8048489 | 0.958015149 |
| ENSMUSG00000041119 | Pde9a | protein_coding | 0.085627304 | 5.060572639 | 0.0630283 | 0.804908 | 0.958015149 |
| ENSMUSG00000022682 | Rrn3 | protein_coding | 0.100228025 | 7.475947597 | 0.0631294 | 0.8052186 | 0.958015149 |
| ENSMUSG00000036278 | Macrod1 | protein_coding | 0.086738434 | 5.257612405 | 0.0628045 | 0.8052472 | 0.958015149 |
| ENSMUSG00000019373 | Cops3 | protein_coding | 0.048680157 | 7.154969852 | 0.062729 | 0.8053617 | 0.958015149 |
| ENSMUSG00000037363 | Letm2 | protein_coding | -0.128778565 | 3.508788132 | 0.0626863 | 0.8054265 | 0.958015149 |
| ENSMUSG00000099689 | Zfp383 | protein_coding | -0.132022374 | 3.063736148 | 0.0626777 | 0.8054395 | 0.958015149 |
| ENSMUSG00000039033 | Tasp1 | protein_coding | 0.080399919 | 5.575886204 | 0.062581 | 0.8055864 | 0.958015149 |
| ENSMUSG00000110080 | Gm6145 | processed_transcript | 0.21765081 | 3.08788904 | 0.062699 | 0.8055942 | 0.958015149 |
| ENSMUSG00000008305 | Tle1 | protein_coding | -0.088916366 | 4.754083591 | 0.0625747 | 0.805596 | 0.958015149 |
| ENSMUSG00000028973 | Abcb8 | protein_coding | 0.063968115 | 6.129467242 | 0.0625283 | 0.8056664 | 0.958015149 |
| ENSMUSG00000048911 | Rnf24 | protein_coding | -0.15714683 | 3.282949124 | 0.0623402 | 0.8059526 | 0.958249428 |
| ENSMUSG00000055489 | Ano5 | protein_coding | -0.157573265 | 4.58816733 | 0.0625074 | 0.8061031 | 0.958322513 |
| ENSMUSG00000034449 | Dhrs11 | protein_coding | 0.134210608 | 3.101245341 | 0.0620096 | 0.8064566 | 0.958393206 |
| ENSMUSG00000103649 | Gm37768 | antisense_RNA | -0.1692126 | 2.671521566 | 0.0619722 | 0.8065137 | 0.958393206 |
| ENSMUSG00000019876 | Pkib | protein_coding | -0.07712642 | 4.949440417 | 0.0619291 | 0.8065795 | 0.958393206 |

| ensembl_gene_id | external_gene_name | gene_biotype | log2 (fold change) | log2 (counts per million) | F-value | p-value | False discovery rate |
|---|---|---|---|---|---|---|---|
| ENSMUSG00000026014 | Raph1 | protein_coding | -0.079620327 | 4.442371162 | 0.0618726 | 0.8066658 | 0.958393206 |
| ENSMUSG00000023905 | Tnfrsf12a | protein_coding | 0.112011687 | 4.519644428 | 0.0617243 | 0.8068927 | 0.958393206 |
| ENSMUSG00000071072 | Ptges3 | protein_coding | -0.050208803 | 8.537608677 | 0.0616981 | 0.8069328 | 0.958393206 |
| ENSMUSG00000100618 | Gm29595 | lincRNA | -0.175942352 | 2.591720433 | 0.0616922 | 0.8069419 | 0.958393206 |
| ENSMUSG00000045464 | 2810002D19Rik | processed_transcript | -0.128332277 | 4.38772 | 0.0616952 | 0.8069486 | 0.958393206 |
| ENSMUSG00000057835 | Zfp119a | protein_coding | 0.167801084 | 3.001432612 | 0.0616772 | 0.8069648 | 0.958393206 |
| ENSMUSG00000013663 | Pten | protein_coding | 0.078597035 | 5.377594495 | 0.0615712 | 0.8071271 | 0.958480166 |
| ENSMUSG00000057455 | Rit2 | protein_coding | 0.05923411 | 9.683594199 | 0.0614925 | 0.8072479 | 0.958496561 |
| ENSMUSG00000046574 | Prr12 | protein_coding | 0.113731327 | 4.780336741 | 0.0612528 | 0.8076208 | 0.958496561 |
| ENSMUSG00000049866 | Arl4c | protein_coding | 0.074414301 | 5.539892239 | 0.0612442 | 0.807629 | 0.958496561 |
| ENSMUSG00000035572 | Dcaf10 | protein_coding | 0.118879848 | 3.641918906 | 0.0607859 | 0.8083348 | 0.958496561 |
| ENSMUSG00000028680 | Plk3 | protein_coding | -0.138417651 | 3.88673489 | 0.0607833 | 0.8083438 | 0.958496561 |
| ENSMUSG00000021948 | Prkcd | protein_coding | -0.126348063 | 5.889248203 | 0.061079 | 0.808375 | 0.958496561 |
| ENSMUSG00000026017 | Carf | protein_coding | -0.111211782 | 4.410506606 | 0.0606835 | 0.8084928 | 0.958496561 |
| ENSMUSG00000015461 | Atf6b | protein_coding | -0.121380515 | 4.014220969 | 0.0604687 | 0.808825 | 0.958496561 |
| ENSMUSG00000000197 | Nalcn | protein_coding | 0.087082524 | 5.587749691 | 0.0604405 | 0.8088686 | 0.958496561 |
| ENSMUSG00000002996 | Hbp1 | protein_coding | 0.106950626 | 3.632420087 | 0.0603801 | 0.808962 | 0.958496561 |
| ENSMUSG00000028233 | Tgs1 | protein_coding | -0.065670253 | 5.482856638 | 0.0603712 | 0.8089758 | 0.958496561 |
| ENSMUSG00000037361 | Sf3b6 | protein_coding | 0.058929334 | 8.375932533 | 0.0603309 | 0.8090382 | 0.958496561 |
| ENSMUSG00000091994 | E130317F20Rik | bidirectional_promote | 0.18265378 | 2.538111552 | 0.0602732 | 0.8091276 | 0.958496561 |
| ENSMUSG00000050663 | Trhde | protein_coding | -0.205915739 | 2.690121764 | 0.0602615 | 0.8091458 | 0.958496561 |
| ENSMUSG00000040302 | Rbm48 | protein_coding | -0.086664132 | 4.46789051 | 0.0601677 | 0.8092913 | 0.958496561 |
| ENSMUSG00000042662 | Dusp15 | protein_coding | 0.141590286 | 2.943083448 | 0.0600365 | 0.8094949 | 0.958496561 |
| ENSMUSG00000057230 | Aak1 | protein_coding | -0.054584066 | 6.234859702 | 0.059969 | 0.8095998 | 0.958496561 |
| ENSMUSG00000046679 | C87436 | protein_coding | -0.093965813 | 4.824337619 | 0.0599178 | 0.8096794 | 0.958496561 |
| ENSMUSG00000022536 | Glyr1 | protein_coding | 0.064124724 | 7.648161917 | 0.0598703 | 0.8097533 | 0.958496561 |
| ENSMUSG00000057604 | Lmcd1 | protein_coding | -0.143535023 | 3.760950453 | 0.0598275 | 0.8098248 | 0.958496561 |
| ENSMUSG00000041528 | Rnf123 | protein_coding | -0.238396237 | 2.638792401 | 0.0598055 | 0.809859 | 0.958496561 |
| ENSMUSG00000021809 | Nudt13 | protein_coding | 0.105044313 | 4.202712886 | 0.0597819 | 0.8098908 | 0.958496561 |
| ENSMUSG00000024083 | Pja2 | protein_coding | -0.062656178 | 9.105464165 | 0.0597525 | 0.8099366 | 0.958496561 |
| ENSMUSG00000029447 | Cct6a | processed_transcript | 0.05231169 | 8.111927685 | 0.0597303 | 0.8099712 | 0.958496561 |
| ENSMUSG00000006676 | Usp19 | protein_coding | 0.093554109 | 3.875609456 | 0.0597212 | 0.8099853 | 0.958496561 |
| ENSMUSG00000030801 | Kat8 | protein_coding | -0.077917083 | 4.772854295 | 0.0597141 | 0.8099965 | 0.958496561 |
| ENSMUSG00000038502 | Ptov1 | protein_coding | 0.067825236 | 6.60639287 | 0.0596924 | 0.8100302 | 0.958496561 |
| ENSMUSG00000020099 | Unc5b | protein_coding | -0.104721664 | 4.237964072 | 0.0596478 | 0.8100997 | 0.958496561 |
| ENSMUSG00000021840 | Mapk1ip1l | protein_coding | 0.096000134 | 3.850109201 | 0.0596308 | 0.8101262 | 0.958496561 |
| ENSMUSG00000039478 | Micu3 | protein_coding | 0.088436318 | 8.724196797 | 0.0597847 | 0.810191 | 0.958496561 |
| ENSMUSG00000043456 | Zfp536 | protein_coding | -0.132777274 | 3.772374222 | 0.05958 | 0.8102054 | 0.958496561 |
| ENSMUSG00000003308 | Keap1 | protein_coding | 0.121377991 | 4.751555809 | 0.0596078 | 0.8102571 | 0.958496561 |
| ENSMUSG00000044770 | Scml4 | protein_coding | 0.129271992 | 3.261107086 | 0.0595239 | 0.8102929 | 0.958496561 |
| ENSMUSG00000035601 | Trmt10b | protein_coding | -0.060725344 | 5.834775679 | 0.0594937 | 0.8103401 | 0.958496561 |
| ENSMUSG00000027168 | Pax6 | protein_coding | -0.106853837 | 3.913721954 | 0.0594922 | 0.8103424 | 0.958496561 |
| ENSMUSG00000025858 | Get4 | protein_coding | 0.09787082 | 3.690348332 | 0.0594265 | 0.8104449 | 0.958496561 |
| ENSMUSG00000027775 | Mfsd1 | protein_coding | 0.122040034 | 3.452838777 | 0.0593814 | 0.8105154 | 0.958496561 |
| ENSMUSG00000041346 | Wrap53 | protein_coding | -0.076826776 | 4.948486831 | 0.0593733 | 0.8105282 | 0.958496561 |
| ENSMUSG00000028164 | Manba | protein_coding | -0.165964738 | 2.762128959 | 0.0592441 | 0.81073 | 0.95854962 |
| ENSMUSG00000066324 | Impad1 | protein_coding | -0.049881937 | 6.810191745 | 0.059188 | 0.8108179 | 0.95854962 |
| ENSMUSG00000040722 | Scamp5 | protein_coding | 0.050167292 | 7.379337979 | 0.0590744 | 0.8109957 | 0.95854962 |
| ENSMUSG00000063760 | Rnf217 | protein_coding | 0.116326358 | 3.511575161 | 0.0590671 | 0.8110071 | 0.95854962 |
| ENSMUSG00000024352 | Spata24 | protein_coding | -0.093181909 | 4.421206474 | 0.0590597 | 0.8110187 | 0.95854962 |
| ENSMUSG00000020102 | Slc16a7 | protein_coding | -0.131141322 | 3.21491113 | 0.0588149 | 0.8114028 | 0.958768239 |
| ENSMUSG00000006471 | Ndor1 | protein_coding | 0.129171056 | 3.286308488 | 0.0587379 | 0.8115237 | 0.958768239 |
| ENSMUSG00000018143 | Mafk | protein_coding | -0.147355265 | 3.372681478 | 0.0586928 | 0.8115946 | 0.958768239 |
| ENSMUSG00000024993 | Fam45a | protein_coding | 0.060080042 | 6.807848676 | 0.0585492 | 0.8118204 | 0.958768239 |
| ENSMUSG00000052160 | Pld4 | protein_coding | 0.155229815 | 3.550459364 | 0.0585463 | 0.8118301 | 0.958768239 |
| ENSMUSG00000079434 | Neu2 | protein_coding | -0.150249318 | 3.211015147 | 0.0585374 | 0.811839 | 0.958768239 |
| ENSMUSG00000031938 | 4931406C07Rik | protein_coding | -0.124705176 | 3.821503406 | 0.0584508 | 0.8119754 | 0.958768239 |
| ENSMUSG00000033883 | D3Ertd254e | protein_coding | -0.093471427 | 4.246837968 | 0.058437 | 0.8119971 | 0.958768239 |
| ENSMUSG00000026972 | Arrdc1 | protein_coding | 0.157279919 | 3.728158626 | 0.0585115 | 0.8120062 | 0.958768239 |
| ENSMUSG00000020648 | Dus4l | protein_coding | -0.174103771 | 2.939779055 | 0.058312 | 0.8121942 | 0.958786782 |
| ENSMUSG00000075012 | Fjx1 | protein_coding | 0.157032874 | 3.278786675 | 0.0583028 | 0.8122087 | 0.958786782 |
| ENSMUSG00000027531 | Impa1 | protein_coding | -0.046636769 | 6.892489951 | 0.0582517 | 0.8122894 | 0.958786782 |
| ENSMUSG00000087590 | Epb41l4aos | lincRNA | 0.112629534 | 3.339230179 | 0.0581396 | 0.8124663 | 0.9588904 |
| ENSMUSG00000050088 | 1600012H06Rik | protein_coding | 0.251199404 | 2.783557498 | 0.0579756 | 0.8129168 | 0.95903818 |
| ENSMUSG00000036529 | Sbf1 | protein_coding | 0.074548096 | 5.353329883 | 0.0578256 | 0.8129631 | 0.95903818 |
| ENSMUSG00000035529 | Prdm4 | protein_coding | 0.120306438 | 3.940585413 | 0.0578176 | 0.8129759 | 0.95903818 |
| ENSMUSG00000050840 | Cdh20 | protein_coding | -0.198472275 | 2.865321039 | 0.0577864 | 0.8130303 | 0.95903818 |
| ENSMUSG00000073755 | 5730409E04Rik | protein_coding | 0.068066338 | 5.339607032 | 0.0576972 | 0.8131667 | 0.95903818 |
| ENSMUSG00000035863 | Palm | protein_coding | -0.06994337 | 5.592000077 | 0.0576749 | 0.8132021 | 0.95903818 |
| ENSMUSG00000062203 | Gspt1 | protein_coding | -0.147732603 | 2.725209974 | 0.0576403 | 0.8132572 | 0.95903818 |
| ENSMUSG00000090015 | Gm15446 | protein_coding | -0.16861134 | 3.140716799 | 0.0576101 | 0.813305 | 0.95903818 |
| ENSMUSG00000034748 | Sirt6 | protein_coding | -0.106620137 | 4.765305217 | 0.0574348 | 0.8135884 | 0.959046083 |
| ENSMUSG00000019437 | Tlcd1 | protein_coding | 0.102161849 | 4.300287122 | 0.0574204 | 0.8136065 | 0.959046083 |
| ENSMUSG00000039473 | Ubn1 | protein_coding | 0.093939296 | 4.132008918 | 0.0573738 | 0.8136806 | 0.959046083 |
| ENSMUSG00000063362 | Alg11 | protein_coding | -0.064889285 | 6.593971449 | 0.0573368 | 0.8137395 | 0.959046083 |
| ENSMUSG00000070814 | 6330408A02Rik | protein_coding | -0.146564788 | 3.603768717 | 0.057292 | 0.8138157 | 0.959046083 |
| ENSMUSG00000037300 | Ttc13 | protein_coding | -0.147129973 | 3.753155692 | 0.0572805 | 0.8138469 | 0.959046083 |
| ENSMUSG00000060376 | Bckdha | protein_coding | -0.130353819 | 3.830387795 | 0.0570971 | 0.8141263 | 0.959246938 |

| ensembl_gene_id | external_gene_name | gene_biotype | log2 (fold change) | log2 (counts per million) | F-value | p-value | False discovery rate |
|---|---|---|---|---|---|---|---|
| ENSMUSG00000042804 | Gpr153 | protein_coding | -0.109861922 | 3.871506597 | 0.0569981 | 0.8142795 | 0.959246938 |
| ENSMUSG00000025266 | Gnl3l | protein_coding | 0.098992073 | 8.742299325 | 0.0572744 | 0.8143144 | 0.959246938 |
| ENSMUSG00000069208 | Zfp825 | protein_coding | -0.105314307 | 3.554510274 | 0.0568707 | 0.8144829 | 0.959246938 |
| ENSMUSG00000049526 | Tmem202 | protein_coding | -0.236047043 | 2.35832237 | 0.0568676 | 0.8144879 | 0.959246938 |
| ENSMUSG00000033272 | Slc35a4 | protein_coding | 0.066078065 | 6.143286942 | 0.0568272 | 0.8145526 | 0.959246938 |
| ENSMUSG00000078572 | Ndufaf8 | protein_coding | 0.068571275 | 6.474336998 | 0.0567084 | 0.8147426 | 0.959286056 |
| ENSMUSG00000018736 | Ndel1 | protein_coding | 0.057310267 | 6.757699419 | 0.0565498 | 0.8149968 | 0.959286056 |
| ENSMUSG00000025158 | Rfng | protein_coding | -0.093713045 | 4.553115981 | 0.0565314 | 0.8150262 | 0.959286056 |
| ENSMUSG00000027574 | Nkain4 | protein_coding | -0.105046772 | 6.610725985 | 0.0567985 | 0.815072 | 0.959286056 |
| ENSMUSG00000068115 | Ninl | protein_coding | -0.250055985 | 2.241747413 | 0.0564771 | 0.8151133 | 0.959286056 |
| ENSMUSG00000064179 | Tnnt1 | protein_coding | -0.16601155 | 4.643140755 | 0.0567677 | 0.8151211 | 0.959286056 |
| ENSMUSG00000097915 | A330009N23Rik | lincRNA | 0.128141975 | 4.085278012 | 0.0563988 | 0.8152439 | 0.959325636 |
| ENSMUSG00000037997 | Parp11 | protein_coding | -0.089517 | 4.771992288 | 0.056239 | 0.8154957 | 0.959421128 |
| ENSMUSG00000058392 | Rrp1b | protein_coding | 0.080047809 | 4.924290776 | 0.0562342 | 0.8155035 | 0.959421128 |
| ENSMUSG00000036815 | Dpp10 | protein_coding | -0.065526722 | 6.009901135 | 0.056107 | 0.8157081 | 0.959556909 |
| ENSMUSG00000015342 | Xk | protein_coding | -0.231104333 | 3.146942023 | 0.0561747 | 0.8159819 | 0.95973082 |
| ENSMUSG00000031352 | Hccs | protein_coding | 0.101368211 | 4.604801972 | 0.0558253 | 0.8161622 | 0.95973082 |
| ENSMUSG00000035064 | Eef2k | protein_coding | -0.097122871 | 3.847565914 | 0.0558053 | 0.8161946 | 0.95973082 |
| ENSMUSG00000022757 | Tfg | protein_coding | -0.073702152 | 6.460969884 | 0.0557584 | 0.8162752 | 0.95973082 |
| ENSMUSG00000029203 | Ube2k | protein_coding | 0.053608089 | 7.384558187 | 0.0557386 | 0.8163022 | 0.95973082 |
| ENSMUSG00000045822 | Zswim3 | protein_coding | 0.116634295 | 3.408235743 | 0.0556762 | 0.8164032 | 0.959744591 |
| ENSMUSG00000033487 | Fndc3a | protein_coding | -0.065007003 | 5.590292676 | 0.0555932 | 0.8165373 | 0.959797321 |
| ENSMUSG00000035142 | Nubpl | protein_coding | 0.123786143 | 3.289785091 | 0.0554801 | 0.8167205 | 0.959885249 |
| ENSMUSG00000029911 | Ssbp1 | protein_coding | 0.062006054 | 6.047762266 | 0.0554368 | 0.8167907 | 0.959885249 |
| ENSMUSG00000020387 | Jade2 | protein_coding | -0.064929488 | 5.322138839 | 0.0550986 | 0.8173394 | 0.96027419 |
| ENSMUSG00000041777 | Cir1 | protein_coding | -0.126021324 | 2.987626848 | 0.0550979 | 0.8173406 | 0.96027419 |
| ENSMUSG00000054793 | Cadm4 | protein_coding | -0.120395604 | 4.443614982 | 0.0550389 | 0.8174541 | 0.96027419 |
| ENSMUSG00000049401 | Ogfr | protein_coding | -0.099633409 | 3.834392804 | 0.0550129 | 0.8174788 | 0.96027419 |
| ENSMUSG00000040859 | Bsdc1 | protein_coding | 0.057483636 | 5.986133482 | 0.0549193 | 0.8176312 | 0.960348234 |
| ENSMUSG00000006818 | Sod2 | protein_coding | 0.052390152 | 7.601455699 | 0.0548172 | 0.8177976 | 0.960388631 |
| ENSMUSG00000037007 | Zfp113 | protein_coding | -0.126396475 | 3.324830141 | 0.0546892 | 0.8180064 | 0.960388631 |
| ENSMUSG00000028933 | Xrcc2 | protein_coding | 0.110838192 | 3.25736211 | 0.0546131 | 0.8181306 | 0.960388631 |
| ENSMUSG00000036242 | 3632451O06Rik | protein_coding | 0.079107938 | 4.742168553 | 0.0545797 | 0.8181851 | 0.960388631 |
| ENSMUSG00000049313 | Sorl1 | protein_coding | -0.059247587 | 6.405108232 | 0.0545466 | 0.8182392 | 0.960388631 |
| ENSMUSG00000030245 | Golt1b | protein_coding | 0.057901728 | 7.497290907 | 0.0545417 | 0.8182472 | 0.960388631 |
| ENSMUSG00000039285 | Azi2 | protein_coding | 0.071790688 | 5.542731768 | 0.0545151 | 0.8182908 | 0.960388631 |
| ENSMUSG00000045333 | Zfp423 | protein_coding | -0.096460085 | 4.549716161 | 0.054385 | 0.8185035 | 0.96046306 |
| ENSMUSG00000025816 | Sec61a2 | protein_coding | -0.081849151 | 4.587601177 | 0.0543356 | 0.8185844 | 0.96046306 |
| ENSMUSG00000020183 | Cpm | protein_coding | 0.113089928 | 3.680920737 | 0.0542724 | 0.818688 | 0.96046306 |
| ENSMUSG00000019814 | Ltv1 | protein_coding | 0.087417799 | 4.246042227 | 0.0542372 | 0.8187455 | 0.96046306 |
| ENSMUSG00000041130 | Zfp598 | protein_coding | 0.14747655 | 3.23157067 | 0.0541793 | 0.8188405 | 0.96046306 |
| ENSMUSG00000039137 | Whrn | protein_coding | -0.099297709 | 4.322699121 | 0.0541491 | 0.8188901 | 0.96046306 |
| ENSMUSG00000068663 | Clec16a | protein_coding | 0.127899023 | 4.878212848 | 0.0541383 | 0.8191968 | 0.960685116 |
| ENSMUSG00000039067 | Psmd7 | protein_coding | 0.045507945 | 8.601291436 | 0.053925 | 0.8192581 | 0.960685116 |
| ENSMUSG00000033906 | Zdhhc15 | protein_coding | -0.202710314 | 2.51327674 | 0.0538143 | 0.8194403 | 0.960793924 |
| ENSMUSG00000024622 | Hmgxb3 | protein_coding | -0.129751709 | 3.1985186 | 0.0537317 | 0.8195764 | 0.960848736 |
| ENSMUSG00000030110 | Ret | protein_coding | -0.107901058 | 3.503712827 | 0.0536686 | 0.8196803 | 0.9608658 |
| ENSMUSG00000054027 | Nt5dc3 | protein_coding | -0.091988567 | 4.951660942 | 0.0535281 | 0.819912 | 0.961032724 |
| ENSMUSG00000059772 | Slx1b | protein_coding | -0.087119384 | 4.103154165 | 0.0534039 | 0.8201171 | 0.961066495 |
| ENSMUSG00000017210 | Med24 | protein_coding | 0.085522753 | 5.606035835 | 0.0534053 | 0.8201196 | 0.961066495 |
| ENSMUSG00000020523 | Fam114a2 | protein_coding | 0.060992365 | 6.023009542 | 0.0532037 | 0.8204483 | 0.961219803 |
| ENSMUSG00000045201 | Lrrc3b | protein_coding | 0.074927694 | 4.919093809 | 0.053199 | 0.8204561 | 0.961219803 |
| ENSMUSG00000053332 | Gas5 | processed_transcript | -0.083395786 | 10.45470529 | 0.0532717 | 0.8205186 | 0.961219803 |
| ENSMUSG00000024304 | Cdh2 | protein_coding | -0.102108753 | 3.838995901 | 0.0530871 | 0.8206416 | 0.961230225 |
| ENSMUSG00000108414 | Snhg1 | processed_transcript | -0.080315784 | 4.599979832 | 0.0530003 | 0.8207855 | 0.961230225 |
| ENSMUSG00000028833 | Ncdn | protein_coding | -0.055402194 | 8.830321483 | 0.0529942 | 0.8207957 | 0.961230225 |
| ENSMUSG00000027404 | Snrpb | protein_coding | 0.056946712 | 7.354725503 | 0.0529212 | 0.8209168 | 0.961267334 |
| ENSMUSG00000052534 | Pbx1 | protein_coding | 0.052385056 | 7.052924281 | 0.0528495 | 0.8210359 | 0.961269782 |
| ENSMUSG00000018326 | Ywhab | protein_coding | 0.045219954 | 9.025884273 | 0.0527912 | 0.8211329 | 0.961269782 |
| ENSMUSG00000006763 | Saal1 | protein_coding | 0.086100265 | 4.029950953 | 0.0527586 | 0.821187 | 0.961269782 |
| ENSMUSG00000003657 | Calb2 | protein_coding | -0.079728996 | 10.09150668 | 0.0527243 | 0.821415 | 0.961410354 |
| ENSMUSG00000001855 | Nup214 | protein_coding | -0.075725845 | 4.491393077 | 0.0525049 | 0.8216096 | 0.961410354 |
| ENSMUSG00000025873 | Faf2 | protein_coding | -0.06437578 | 5.906230298 | 0.0523896 | 0.8218019 | 0.961410354 |
| ENSMUSG00000042684 | Npl | protein_coding | 0.147181183 | 3.115437471 | 0.0522561 | 0.822025 | 0.961410354 |
| ENSMUSG00000097048 | 1600020E01Rik | antisense_RNA | -0.109302505 | 4.654607397 | 0.0522462 | 0.8220462 | 0.961410354 |
| ENSMUSG00000031749 | St3gal2 | protein_coding | 0.070693444 | 5.079419444 | 0.05222 | 0.8220854 | 0.961410354 |
| ENSMUSG00000003418 | St8sia6 | protein_coding | 0.171852482 | 2.527975686 | 0.0521638 | 0.8221794 | 0.961410354 |
| ENSMUSG00000050052 | Tdrp | protein_coding | 0.088662088 | 4.529503358 | 0.0520982 | 0.8222891 | 0.961410354 |
| ENSMUSG00000027643 | Ghrh | protein_coding | 0.129796169 | 3.367773726 | 0.052094 | 0.8222961 | 0.961410354 |
| ENSMUSG00000035473 | Galm | protein_coding | 0.229101096 | 2.245934502 | 0.0520836 | 0.8223136 | 0.961410354 |
| ENSMUSG00000048915 | Efna5 | protein_coding | 0.097496195 | 6.530370452 | 0.0523123 | 0.8223296 | 0.961410354 |
| ENSMUSG00000038482 | Tfdp1 | protein_coding | 0.086061237 | 4.656460446 | 0.0520439 | 0.82238 | 0.961410354 |
| ENSMUSG00000043289 | Mei4 | protein_coding | 0.198262448 | 2.37639446 | 0.0519335 | 0.8225651 | 0.961440813 |
| ENSMUSG00000066037 | Hnrnpr | protein_coding | -0.053296919 | 6.498009891 | 0.0518891 | 0.8226397 | 0.961440813 |
| ENSMUSG00000014786 | Slc9a5 | protein_coding | -0.115561658 | 3.415460126 | 0.0518571 | 0.8226933 | 0.961440813 |
| ENSMUSG00000090122 | Kcne1l | protein_coding | -0.209041248 | 2.519501753 | 0.0516599 | 0.8230246 | 0.961440813 |
| ENSMUSG00000029270 | Fam69a | protein_coding | 0.064451354 | 5.543227714 | 0.0516287 | 0.8230772 | 0.961440813 |
| ENSMUSG00000040374 | Pex2 | protein_coding | 0.051626618 | 6.584902452 | 0.0516183 | 0.8230946 | 0.961440813 |

| ensembl_gene_id | external_gene_name | gene_biotype | log2 (fold change) | log2 (counts per million) | F-value | p-value | False discovery rate |
|---|---|---|---|---|---|---|---|
| ENSMUSG00000030805 | Stx4a | protein_coding | -0.061772037 | 6.131729887 | 0.0516051 | 0.8231168 | 0.961440813 |
| ENSMUSG00000040265 | Dnm3 | protein_coding | 0.071732366 | 5.663832161 | 0.0515918 | 0.8231393 | 0.961440813 |
| ENSMUSG00000041298 | Katnal1 | protein_coding | 0.062222895 | 6.237262926 | 0.0515207 | 0.823259 | 0.961440813 |
| ENSMUSG00000002289 | Angptl4 | protein_coding | 0.160930592 | 2.480448562 | 0.0514962 | 0.8233002 | 0.961440813 |
| ENSMUSG00000007570 | Fance | protein_coding | -0.175116318 | 2.876004176 | 0.0512951 | 0.8236394 | 0.961732496 |
| ENSMUSG00000029504 | Ddx51 | protein_coding | -0.258705538 | 2.463090803 | 0.0512095 | 0.8238776 | 0.961906134 |
| ENSMUSG00000031785 | Adgrg1 | protein_coding | -0.057629013 | 6.230589893 | 0.0510256 | 0.8240949 | 0.961923642 |
| ENSMUSG00000042073 | Abhd14b | protein_coding | -0.110596437 | 3.539141639 | 0.0510229 | 0.8240995 | 0.961923642 |
| ENSMUSG00000042354 | Gnl3 | protein_coding | -0.071030191 | 5.427918584 | 0.0509294 | 0.8242578 | 0.961923642 |
| ENSMUSG00000037353 | Letmd1 | protein_coding | -0.07182165 | 5.638732501 | 0.0508986 | 0.82431 | 0.961923642 |
| ENSMUSG00000001910 | Nacc1 | protein_coding | 0.080014258 | 4.97235212 | 0.050881 | 0.8243398 | 0.961923642 |
| ENSMUSG00000034617 | Mtrr | protein_coding | 0.177221708 | 3.45573141 | 0.0509584 | 0.8244446 | 0.961941538 |
| ENSMUSG00000055760 | Gemin6 | protein_coding | 0.100750928 | 4.42354317 | 0.0506025 | 0.8248127 | 0.962181142 |
| ENSMUSG00000069255 | Dusp22 | protein_coding | 0.074223862 | 5.260943692 | 0.050593 | 0.824829 | 0.962181142 |
| ENSMUSG00000005907 | Pex1 | protein_coding | -0.142309406 | 3.524648802 | 0.0505325 | 0.8249364 | 0.962202075 |
| ENSMUSG00000103428 | Gm20754 | lincRNA | -0.212354836 | 2.397947304 | 0.0501023 | 0.8256655 | 0.962784846 |
| ENSMUSG00000033106 | Slc7a6os | protein_coding | -0.125107973 | 3.552645987 | 0.0500275 | 0.8257933 | 0.962784846 |
| ENSMUSG00000064061 | Dzip3 | protein_coding | -0.059224222 | 5.567912203 | 0.050026 | 0.825796 | 0.962784846 |
| ENSMUSG00000010554 | Mettl16 | protein_coding | 0.054532527 | 6.035521682 | 0.049989 | 0.8258593 | 0.962784846 |
| ENSMUSG00000020386 | Sar1b | protein_coding | 0.056709246 | 7.789736804 | 0.0499638 | 0.8259024 | 0.962784846 |
| ENSMUSG00000041617 | Ccdc74a | protein_coding | -0.114332443 | 3.585730457 | 0.0498799 | 0.8260461 | 0.962784846 |
| ENSMUSG00000001175 | Calm1 | protein_coding | 0.050780089 | 12.85395451 | 0.0498701 | 0.8260628 | 0.962784846 |
| ENSMUSG00000027599 | Armc1 | protein_coding | 0.054673278 | 7.614774014 | 0.0497511 | 0.8262669 | 0.962918341 |
| ENSMUSG00000068823 | Csde1 | protein_coding | -0.064734741 | 8.718473416 | 0.0496931 | 0.826371 | 0.96293532 |
| ENSMUSG00000040820 | Hlcs | protein_coding | 0.100878084 | 3.496479684 | 0.0494893 | 0.8267166 | 0.963233669 |
| ENSMUSG00000023033 | Scn8a | protein_coding | -0.066972743 | 5.789346112 | 0.049273 | 0.8270891 | 0.963390691 |
| ENSMUSG00000001995 | Sipa1l2 | protein_coding | 0.112539736 | 3.881671861 | 0.0491601 | 0.827284 | 0.963390691 |
| ENSMUSG00000041423 | Paqr6 | protein_coding | -0.095607296 | 3.762613481 | 0.0491502 | 0.8273011 | 0.963390691 |
| ENSMUSG00000020674 | Pxdn | protein_coding | -0.218166412 | 2.519421216 | 0.0490973 | 0.827397 | 0.963390691 |
| ENSMUSG00000022365 | Derl1 | protein_coding | -0.06086407 | 5.782097291 | 0.0490935 | 0.8273989 | 0.963390691 |
| ENSMUSG00000013787 | Ehmt2 | protein_coding | 0.06916511 | 5.583196149 | 0.0490018 | 0.8275575 | 0.963390691 |
| ENSMUSG00000028559 | Osbpl9 | protein_coding | -0.055108334 | 5.932342339 | 0.0489654 | 0.8276204 | 0.963390691 |
| ENSMUSG00000066838 | Zfp772 | protein_coding | 0.126687777 | 3.219859574 | 0.0488922 | 0.8277471 | 0.963390691 |
| ENSMUSG00000019929 | Dcn | protein_coding | 0.116248457 | 3.742444667 | 0.0488519 | 0.8278169 | 0.963390691 |
| ENSMUSG00000027509 | Rae1 | protein_coding | 0.075708903 | 4.86638759 | 0.0488194 | 0.8278732 | 0.963390691 |
| ENSMUSG00000028343 | Erp44 | protein_coding | -0.061422073 | 5.414849164 | 0.0487066 | 0.8280687 | 0.963390691 |
| ENSMUSG00000020460 | Rps27a | protein_coding | 0.093216137 | 3.584484307 | 0.0486719 | 0.8281289 | 0.963390691 |
| ENSMUSG00000048495 | Tyw5 | protein_coding | -0.072284612 | 4.875694732 | 0.0486302 | 0.8282014 | 0.963390691 |
| ENSMUSG00000025427 | Rnf165 | protein_coding | -0.094583999 | 4.695807413 | 0.0486033 | 0.8282482 | 0.963390691 |
| ENSMUSG00000022911 | Arl13b | protein_coding | 0.106942331 | 3.263438864 | 0.0485646 | 0.8283154 | 0.963390691 |
| ENSMUSG00000023044 | Csad | protein_coding | -0.100349619 | 4.287033396 | 0.0485078 | 0.8284141 | 0.963390691 |
| ENSMUSG00000073633 | Fbxo36 | protein_coding | 0.083874481 | 4.715544166 | 0.0483867 | 0.8286246 | 0.963390691 |
| ENSMUSG00000032369 | Plscr1 | protein_coding | -0.187512689 | 2.352033512 | 0.0483844 | 0.8286287 | 0.963390691 |
| ENSMUSG00000084799 | Ino80dos | lincRNA | -0.13052512 | 2.857561626 | 0.0483423 | 0.8287019 | 0.963390691 |
| ENSMUSG00000004462 | Tbccd1 | protein_coding | -0.097750812 | 4.312858335 | 0.0483318 | 0.8287204 | 0.963390691 |
| ENSMUSG00000021578 | Ccdc127 | protein_coding | 0.051628691 | 6.335168064 | 0.0483216 | 0.8287381 | 0.963390691 |
| ENSMUSG00000021871 | Pnp | protein_coding | -0.137314254 | 2.807099839 | 0.048217 | 0.8289204 | 0.963390691 |
| ENSMUSG00000024211 | Grm8 | protein_coding | -0.133174192 | 4.150109232 | 0.0482995 | 0.8289503 | 0.963390691 |
| ENSMUSG00000032518 | Rpsa | protein_coding | 0.092518428 | 3.711441235 | 0.0481612 | 0.8290178 | 0.963390691 |
| ENSMUSG00000035941 | Ibtk | protein_coding | -0.105495385 | 3.603984258 | 0.0481191 | 0.8290912 | 0.963390691 |
| ENSMUSG00000005370 | Msh6 | protein_coding | -0.212667765 | 2.915605067 | 0.0481514 | 0.8292083 | 0.963422677 |
| ENSMUSG00000024732 | Ccdc86 | protein_coding | 0.088456903 | 4.064614931 | 0.0479982 | 0.8293025 | 0.963428018 |
| ENSMUSG00000036908 | Unc93b1 | protein_coding | -0.183319424 | 3.136844393 | 0.0480064 | 0.8294015 | 0.963438987 |
| ENSMUSG00000050783 | Htr1f | protein_coding | -0.204020289 | 2.268847071 | 0.0476274 | 0.829952 | 0.963974257 |
| ENSMUSG00000073616 | Cops9 | protein_coding | 0.065339437 | 8.663054637 | 0.0474196 | 0.8303217 | 0.964299463 |
| ENSMUSG00000022685 | Parn | protein_coding | 0.077087197 | 4.910161498 | 0.0471592 | 0.8307759 | 0.964441009 |
| ENSMUSG00000038102 | Trappc11 | protein_coding | -0.071640517 | 4.883764086 | 0.0471358 | 0.8308172 | 0.964441009 |
| ENSMUSG00000002679 | Med6 | protein_coding | -0.067026611 | 5.727165433 | 0.0471275 | 0.8308319 | 0.964441009 |
| ENSMUSG00000041560 | Nop53 | protein_coding | 0.073874687 | 6.640298729 | 0.0471569 | 0.830893 | 0.964441009 |
| ENSMUSG00000020723 | Cacng4 | protein_coding | -0.073500692 | 5.752116786 | 0.0470863 | 0.8309046 | 0.964441009 |
| ENSMUSG00000003348 | Mob3a | protein_coding | -0.173636322 | 2.797709961 | 0.0470002 | 0.8310567 | 0.964441009 |
| ENSMUSG00000028271 | Gtf2b | protein_coding | -0.050866269 | 6.314934803 | 0.0469332 | 0.8311754 | 0.964441009 |
| ENSMUSG00000022704 | Qtrt2 | protein_coding | -0.111310576 | 3.32029933 | 0.0468792 | 0.8312708 | 0.964441009 |
| ENSMUSG00000000194 | Gpr107 | protein_coding | -0.085373708 | 4.487791172 | 0.0468568 | 0.8313105 | 0.964441009 |
| ENSMUSG00000030035 | Wbp1 | protein_coding | 0.096372614 | 4.942696081 | 0.0468424 | 0.8313404 | 0.964441009 |
| ENSMUSG00000029790 | Cep41 | protein_coding | -0.139024752 | 3.099118787 | 0.046705 | 0.8315796 | 0.964614348 |
| ENSMUSG00000058291 | Zfp68 | protein_coding | -0.062600043 | 5.270613456 | 0.0466367 | 0.8317006 | 0.964650696 |
| ENSMUSG00000066900 | Suds3 | protein_coding | -0.058417432 | 5.491229556 | 0.0464314 | 0.8320655 | 0.964969824 |
| ENSMUSG00000021710 | Nln | protein_coding | -0.084163369 | 4.26161093 | 0.0463677 | 0.8321788 | 0.964997202 |
| ENSMUSG00000027198 | Ext2 | protein_coding | -0.101811074 | 3.945682192 | 0.0461067 | 0.8326442 | 0.965432693 |
| ENSMUSG00000051391 | Ywhag | protein_coding | 0.059417255 | 9.147746212 | 0.0458085 | 0.8331774 | 0.965667857 |
| ENSMUSG00000073102 | Drc1 | protein_coding | -0.070718436 | 4.882209668 | 0.0457698 | 0.8332469 | 0.965667857 |
| ENSMUSG00000030847 | Bag3 | protein_coding | 0.139092241 | 2.816494219 | 0.0456903 | 0.8333895 | 0.965667857 |
| ENSMUSG00000034621 | Gpatch8 | protein_coding | -0.064306725 | 5.727575147 | 0.0456546 | 0.8334534 | 0.965667857 |
| ENSMUSG00000027612 | Mmp24 | protein_coding | 0.125464834 | 3.06383252 | 0.0456348 | 0.8334891 | 0.965667857 |
| ENSMUSG00000027646 | Src | protein_coding | -0.097690612 | 4.171514877 | 0.0456317 | 0.8334946 | 0.965667857 |
| ENSMUSG00000020262 | Adarb1 | protein_coding | 0.077775049 | 5.996686249 | 0.045611 | 0.8335476 | 0.965667857 |
| ENSMUSG00000066760 | Psg16 | protein_coding | -0.198671798 | 2.924307113 | 0.0456573 | 0.8335654 | 0.965667857 |

| ensembl_gene_id | external_gene_name | gene_biotype | log2 (fold change) | log2 (counts per million) | F-value | p-value | False discovery rate |
|---|---|---|---|---|---|---|---|
| ENSMUSG00000051185 | Fam174a | protein_coding | 0.068750529 | 5.226008723 | 0.0454909 | 0.8337475 | 0.965739443 |
| ENSMUSG00000030727 | Rabep2 | protein_coding | -0.114613451 | 3.876506452 | 0.0454419 | 0.8338397 | 0.965739443 |
| ENSMUSG00000039958 | Etfbkmt | protein_coding | -0.094234245 | 4.189351122 | 0.045408 | 0.8338967 | 0.965739443 |
| ENSMUSG00000019143 | Hars2 | protein_coding | -0.081300859 | 4.390408431 | 0.0451771 | 0.8343128 | 0.966056199 |
| ENSMUSG00000035666 | Gtf3c4 | protein_coding | 0.100412484 | 3.779221143 | 0.0450478 | 0.8345463 | 0.966056199 |
| ENSMUSG00000085882 | 2610507I01Rik | processed_transcript | -0.116215188 | 3.233352781 | 0.0450413 | 0.8345581 | 0.966056199 |
| ENSMUSG00000036975 | Tmem177 | protein_coding | 0.09804247 | 3.831928398 | 0.045 | 0.8346328 | 0.966056199 |
| ENSMUSG00000095041 | AC149090.1 | protein_coding | -0.089486428 | 4.889587051 | 0.0449884 | 0.8346537 | 0.966056199 |
| ENSMUSG00000089832 | Shkbp1 | protein_coding | 0.195002508 | 2.76160648 | 0.0449596 | 0.8347101 | 0.966056199 |
| ENSMUSG00000031540 | Kat6a | protein_coding | -0.087669643 | 4.057204075 | 0.0448115 | 0.834974 | 0.966056199 |
| ENSMUSG00000084885 | 3010001F23Rik | processed_transcript | 0.126504655 | 2.823728381 | 0.0448048 | 0.8349861 | 0.966056199 |
| ENSMUSG00000045174 | Amer3 | protein_coding | -0.105805652 | 4.274527975 | 0.044804 | 0.834992 | 0.966056199 |
| ENSMUSG00000029433 | Diablo | protein_coding | 0.062480686 | 4.82074795 | 0.0447593 | 0.8350686 | 0.966056199 |
| ENSMUSG00000095139 | Pou3f2 | protein_coding | -0.069439238 | 4.942613641 | 0.0446 | 0.8353578 | 0.966195827 |
| ENSMUSG00000052949 | Rnf157 | protein_coding | 0.044739576 | 6.718553286 | 0.0445713 | 0.83541 | 0.966195827 |
| ENSMUSG00000032059 | Alg9 | protein_coding | 0.088301694 | 4.222902375 | 0.0445444 | 0.8354588 | 0.966195827 |
| ENSMUSG00000063600 | Egfem1 | protein_coding | 0.095996157 | 3.503946583 | 0.0444363 | 0.8356555 | 0.966319308 |
| ENSMUSG00000046287 | Pnma3 | protein_coding | 0.098970107 | 4.538704125 | 0.0443684 | 0.8357834 | 0.966363298 |
| ENSMUSG00000034867 | Ankrd27 | protein_coding | 0.09390823 | 4.345784258 | 0.0442658 | 0.8359662 | 0.966470709 |
| ENSMUSG00000075273 | Ttc30b | protein_coding | -0.07393726 | 4.086097015 | 0.0441149 | 0.8362416 | 0.966685265 |
| ENSMUSG00000002908 | Kcnn1 | protein_coding | -0.182434911 | 2.593229118 | 0.0440491 | 0.836362 | 0.966720491 |
| ENSMUSG00000007817 | Zmiz1 | protein_coding | 0.062691281 | 5.827892092 | 0.0439459 | 0.8365508 | 0.966797816 |
| ENSMUSG00000057236 | Rbbp4 | protein_coding | 0.058443829 | 7.282346161 | 0.0438579 | 0.8367163 | 0.966797816 |
| ENSMUSG00000028656 | Cap1 | protein_coding | 0.058621116 | 6.506580688 | 0.0437983 | 0.8368213 | 0.966797816 |
| ENSMUSG00000062822 | 4833420G17Rik | protein_coding | -0.067471035 | 5.393469938 | 0.0437883 | 0.8368396 | 0.966797816 |
| ENSMUSG00000033597 | Caskin1 | protein_coding | -0.079780613 | 5.013224174 | 0.0437393 | 0.8369295 | 0.966797816 |
| ENSMUSG00000015085 | Entpd2 | protein_coding | -0.125157314 | 2.870821829 | 0.0437182 | 0.8369684 | 0.966797816 |
| ENSMUSG00000031311 | Nono | protein_coding | 0.072000618 | 6.397071582 | 0.0436788 | 0.8371106 | 0.966830899 |
| ENSMUSG00000027714 | Exosc9 | protein_coding | 0.048503804 | 6.244827269 | 0.0436047 | 0.8371768 | 0.966830899 |
| ENSMUSG00000021708 | Rasgrf2 | protein_coding | -0.059415386 | 6.114797076 | 0.0434551 | 0.8374521 | 0.966958314 |
| ENSMUSG00000097059 | Fam120aos | lincRNA | 0.132708623 | 2.730748456 | 0.0434272 | 0.8375036 | 0.966958314 |
| ENSMUSG00000038301 | Snx10 | protein_coding | -0.053073486 | 8.472617737 | 0.0433982 | 0.8375569 | 0.966958314 |
| ENSMUSG00000005148 | Klf5 | protein_coding | -0.104164911 | 3.456679139 | 0.0433244 | 0.837693 | 0.967011582 |
| ENSMUSG00000030888 | Rrp8 | protein_coding | -0.088451679 | 4.165826031 | 0.043223 | 0.8378803 | 0.967105422 |
| ENSMUSG00000089665 | Fcor | protein_coding | 0.077553819 | 4.81290421 | 0.0431098 | 0.8380894 | 0.967105422 |
| ENSMUSG00000041420 | Meis3 | protein_coding | 0.075562219 | 4.46361733 | 0.043021 | 0.8382537 | 0.967105422 |
| ENSMUSG00000020277 | Pfkl | protein_coding | 0.070355924 | 7.5166223 | 0.0430641 | 0.8383375 | 0.967105422 |
| ENSMUSG00000044231 | Nhlrc1 | protein_coding | -0.100234026 | 3.692155523 | 0.0429394 | 0.8384049 | 0.967105422 |
| ENSMUSG00000038178 | Slc43a2 | protein_coding | -0.079405923 | 4.133100289 | 0.0429098 | 0.8384598 | 0.967105422 |
| ENSMUSG00000022710 | Usp7 | protein_coding | -0.054444024 | 5.755904599 | 0.0428964 | 0.8384846 | 0.967105422 |
| ENSMUSG00000057182 | Scn3a | protein_coding | -0.058121294 | 5.406071106 | 0.0428785 | 0.8385177 | 0.967105422 |
| ENSMUSG00000031841 | Cdh13 | protein_coding | 0.040279567 | 7.328179184 | 0.0428429 | 0.8385837 | 0.967105422 |
| ENSMUSG00000085181 | Gm12709 | antisense_RNA | -0.068048961 | 5.139537362 | 0.0427459 | 0.8387639 | 0.967209422 |
| ENSMUSG00000045252 | Zfp574 | protein_coding | -0.088273079 | 4.009958688 | 0.0425264 | 0.8391722 | 0.967258478 |
| ENSMUSG00000061979 | Rcc1l | protein_coding | -0.15973695 | 3.091089863 | 0.0424464 | 0.8393254 | 0.967258478 |
| ENSMUSG00000045639 | Zfp629 | protein_coding | -0.07916226 | 4.085104144 | 0.042433 | 0.8393462 | 0.967258478 |
| ENSMUSG00000087479 | Gm16835 | processed_transcript | -0.108377557 | 3.380178744 | 0.0424225 | 0.8393658 | 0.967258478 |
| ENSMUSG00000020566 | Atp6v1c2 | protein_coding | -0.101087555 | 3.341533126 | 0.0423965 | 0.8394145 | 0.967258478 |
| ENSMUSG00000041763 | Tpp2 | protein_coding | -0.063319864 | 5.151239924 | 0.0422695 | 0.8396514 | 0.967258478 |
| ENSMUSG00000017264 | Exosc10 | protein_coding | 0.072561928 | 5.261205355 | 0.0422268 | 0.8397313 | 0.967258478 |
| ENSMUSG00000021023 | 1110008L16Rik | protein_coding | -0.107191876 | 3.348714097 | 0.0422022 | 0.8397772 | 0.967258478 |
| ENSMUSG00000006134 | Crkl | protein_coding | -0.078111712 | 3.826480559 | 0.042106 | 0.8399574 | 0.967258478 |
| ENSMUSG00000061451 | Tmem151a | protein_coding | -0.059826852 | 5.821230767 | 0.0420891 | 0.839989 | 0.967258478 |
| ENSMUSG00000052605 | Isoc2b | protein_coding | 0.083219364 | 4.50164993 | 0.042073 | 0.8400191 | 0.967258478 |
| ENSMUSG00000055240 | Zfp101 | protein_coding | -0.12689052 | 3.021592044 | 0.0420669 | 0.8400305 | 0.967258478 |
| ENSMUSG00000026643 | Nmt2 | protein_coding | 0.059353221 | 8.632761936 | 0.0420544 | 0.8400582 | 0.967258478 |
| ENSMUSG00000049553 | Polr1a | protein_coding | 0.102902666 | 3.578568344 | 0.0420243 | 0.8401103 | 0.967258478 |
| ENSMUSG00000055782 | Abcd2 | protein_coding | -0.111852541 | 3.920591442 | 0.0420023 | 0.8401557 | 0.967258478 |
| ENSMUSG00000091864 | Gm17102 | lincRNA | 0.117574567 | 3.367061646 | 0.0418811 | 0.840379 | 0.967393919 |
| ENSMUSG00000057766 | Ankrd29 | protein_coding | -0.059381943 | 5.195263539 | 0.041819 | 0.8404956 | 0.967393919 |
| ENSMUSG00000071265 | 1700086L19Rik | transcribed_unitary_p | 0.055496792 | 5.636011337 | 0.041757 | 0.8406121 | 0.967393919 |
| ENSMUSG00000038416 | Cdc16 | protein_coding | -0.068137951 | 5.149809649 | 0.0416292 | 0.8408528 | 0.967393919 |
| ENSMUSG00000035067 | Xkr6 | protein_coding | 0.159929075 | 2.614266877 | 0.0415958 | 0.8409157 | 0.967393919 |
| ENSMUSG00000047909 | Ankrd16 | protein_coding | 0.160149478 | 2.79214358 | 0.0415628 | 0.8409778 | 0.967393919 |
| ENSMUSG00000071652 | Ints5 | protein_coding | -0.090256958 | 3.756312583 | 0.0415518 | 0.8409986 | 0.967393919 |
| ENSMUSG00000038368 | Focad | protein_coding | -0.129174622 | 2.994303433 | 0.0415504 | 0.8410012 | 0.967393919 |
| ENSMUSG00000019863 | Qrsl1 | protein_coding | 0.095074291 | 4.740587059 | 0.0414774 | 0.8411431 | 0.967393919 |
| ENSMUSG00000075701 | Selenos | protein_coding | -0.05769629 | 7.051738636 | 0.0414615 | 0.841173 | 0.967393919 |
| ENSMUSG00000025425 | St8sia5 | protein_coding | -0.150904499 | 2.602638481 | 0.0413128 | 0.8414498 | 0.96748394 |
| ENSMUSG00000009630 | Ppp2cb | protein_coding | -0.05195728 | 6.220185394 | 0.0412535 | 0.8415621 | 0.96748394 |
| ENSMUSG00000025145 | Lrrc45 | protein_coding | -0.117418386 | 3.403219882 | 0.0412028 | 0.8416581 | 0.96748394 |
| ENSMUSG00000042529 | Kcnj12 | protein_coding | 0.115273254 | 3.044335872 | 0.0411124 | 0.8418294 | 0.96748394 |
| ENSMUSG00000078908 | Mon1b | protein_coding | 0.168948574 | 2.834909832 | 0.0410976 | 0.8418615 | 0.96748394 |
| ENSMUSG00000022195 | 6030458C11Rik | protein_coding | 0.084518898 | 5.026614836 | 0.0410959 | 0.8418647 | 0.96748394 |
| ENSMUSG00000024544 | Ldlrad4 | protein_coding | 0.105431081 | 4.545320812 | 0.0411262 | 0.8418811 | 0.96748394 |
| ENSMUSG00000005481 | Ddx39 | protein_coding | 0.071369535 | 5.050695147 | 0.0410147 | 0.8420146 | 0.967534009 |
| ENSMUSG00000074781 | Ube2n | protein_coding | 0.049265976 | 6.876333105 | 0.0408762 | 0.8422777 | 0.967537187 |
| ENSMUSG00000043162 | Pyurf | protein_coding | -0.111444886 | 5.055327612 | 0.0409263 | 0.8424488 | 0.967537187 |

| ensembl_gene_id | external_gene_name | gene_biotype | log2 (fold change) | log2 (counts per million) | F-value | p-value | False discovery rate |
|---|---|---|---|---|---|---|---|
| ENSMUSG00000038135 | Crygn | protein_coding | -0.127884135 | 3.045605241 | 0.0407763 | 0.8424679 | 0.967537187 |
| ENSMUSG00000047710 | Champ1 | protein_coding | 0.076407825 | 4.904989309 | 0.0407712 | 0.8424775 | 0.967537187 |
| ENSMUSG00000035781 | R3hdm4 | protein_coding | -0.05179803 | 7.261553499 | 0.0407183 | 0.8425784 | 0.967537187 |
| ENSMUSG00000033361 | Prrg3 | protein_coding | 0.090436889 | 4.063571141 | 0.0405155 | 0.8429653 | 0.967537187 |
| ENSMUSG00000097820 | E530011L22Rik | antisense_RNA | 0.155123946 | 2.661232199 | 0.040449 | 0.8430924 | 0.967537187 |
| ENSMUSG00000061654 | Spry3 | protein_coding | -0.109182568 | 3.475000625 | 0.0403234 | 0.8433327 | 0.967537187 |
| ENSMUSG00000026556 | Vangl2 | protein_coding | -0.08679065 | 3.631927552 | 0.0401722 | 0.8436226 | 0.967537187 |
| ENSMUSG00000029516 | Cit | protein_coding | -0.085790683 | 8.525516425 | 0.040358 | 0.8436627 | 0.967537187 |
| ENSMUSG00000029407 | Uso1 | protein_coding | -0.060037281 | 5.557244643 | 0.0401274 | 0.8437085 | 0.967537187 |
| ENSMUSG00000051403 | Ppp1r37 | protein_coding | -0.067535841 | 4.556805796 | 0.0400636 | 0.843831 | 0.967537187 |
| ENSMUSG00000037606 | Osbpl5 | protein_coding | 0.078375342 | 4.151074454 | 0.0400358 | 0.8438845 | 0.967537187 |
| ENSMUSG00000043311 | D17H6S53E | protein_coding | -0.110477029 | 5.399066154 | 0.0402005 | 0.8439158 | 0.967537187 |
| ENSMUSG00000022010 | Tsc22d1 | protein_coding | 0.042134231 | 9.219451156 | 0.0400052 | 0.8439433 | 0.967537187 |
| ENSMUSG00000019254 | Ppp1r12c | protein_coding | -0.072742633 | 4.829504984 | 0.0399759 | 0.8439997 | 0.967537187 |
| ENSMUSG00000034265 | Zdhhc14 | protein_coding | 0.087951207 | 4.272817134 | 0.0399321 | 0.8440839 | 0.967537187 |
| ENSMUSG00000038187 | Btbd10 | protein_coding | -0.057415734 | 5.697058457 | 0.0399234 | 0.8441008 | 0.967537187 |
| ENSMUSG00000042148 | Cox10 | protein_coding | 0.059774778 | 5.255918177 | 0.0398383 | 0.8442646 | 0.967537187 |
| ENSMUSG00000038022 | Mindy4 | protein_coding | 0.113504578 | 3.076901349 | 0.0398027 | 0.8443333 | 0.967537187 |
| ENSMUSG00000026083 | Eif5b | protein_coding | -0.061858882 | 6.243742382 | 0.0397271 | 0.8444791 | 0.967537187 |
| ENSMUSG00000070044 | Fam149a | protein_coding | -0.075083838 | 4.159628302 | 0.0397238 | 0.8444856 | 0.967537187 |
| ENSMUSG00000025357 | Dgka | protein_coding | -0.165872399 | 2.571706719 | 0.0396998 | 0.8445318 | 0.967537187 |
| ENSMUSG00000021143 | Pacs2 | protein_coding | -0.061285186 | 5.061377253 | 0.0396984 | 0.8445345 | 0.967537187 |
| ENSMUSG00000042992 | Borcs5 | protein_coding | 0.080043621 | 5.352748953 | 0.039672 | 0.8445896 | 0.967537187 |
| ENSMUSG00000027950 | Chrnb2 | protein_coding | -0.075088318 | 4.510799278 | 0.039669 | 0.8445913 | 0.967537187 |
| ENSMUSG00000037977 | 6430571L13Rik | protein_coding | 0.116250901 | 3.498634964 | 0.0396216 | 0.8446829 | 0.967537187 |
| ENSMUSG00000024044 | Epb41l3 | protein_coding | -0.053093638 | 7.654101313 | 0.0396206 | 0.8446848 | 0.967537187 |
| ENSMUSG00000039824 | Myl6b | protein_coding | 0.115022742 | 2.879614197 | 0.0395604 | 0.8448012 | 0.967537187 |
| ENSMUSG00000032265 | Fam46a | protein_coding | 0.084385752 | 3.586635144 | 0.0395366 | 0.8448472 | 0.967537187 |
| ENSMUSG00000030741 | Spns1 | protein_coding | -0.063882362 | 5.835921161 | 0.0395314 | 0.8448574 | 0.967537187 |
| ENSMUSG00000036898 | Zfp157 | protein_coding | -0.073992981 | 4.376131062 | 0.0395111 | 0.8448967 | 0.967537187 |
| ENSMUSG00000058546 | Rpl23a | protein_coding | 0.099304204 | 3.615805118 | 0.0393912 | 0.8451288 | 0.967699931 |
| ENSMUSG00000019173 | Rab5c | protein_coding | 0.068453768 | 5.459684264 | 0.0393113 | 0.8452838 | 0.967774375 |
| ENSMUSG00000045435 | Tmem60 | protein_coding | 0.053414375 | 7.072074073 | 0.0392238 | 0.8454537 | 0.967817827 |
| ENSMUSG00000002265 | Peg3 | protein_coding | 0.050134021 | 8.475969773 | 0.0391769 | 0.8455448 | 0.967817827 |
| ENSMUSG00000024213 | Nudt3 | protein_coding | 0.044127415 | 8.135176112 | 0.0391528 | 0.8455918 | 0.967817827 |
| ENSMUSG00000052419 | 2610001J05Rik | protein_coding | -0.042645007 | 7.768716308 | 0.0390992 | 0.845696 | 0.967833938 |
| ENSMUSG00000028675 | Pnrc2 | protein_coding | -0.062111772 | 6.454867682 | 0.0390551 | 0.8457859 | 0.967833938 |
| ENSMUSG00000022048 | Dpysl2 | protein_coding | -0.077110679 | 11.92064053 | 0.0389592 | 0.8461242 | 0.968044981 |
| ENSMUSG00000028977 | Casz1 | protein_coding | -0.083434621 | 3.652476258 | 0.0388661 | 0.8461504 | 0.968044981 |
| ENSMUSG00000036188 | Ankmy2 | protein_coding | 0.047426537 | 6.939125546 | 0.038671 | 0.8465317 | 0.968301116 |
| ENSMUSG00000037892 | Pcdh18 | protein_coding | 0.105925529 | 3.706613125 | 0.0386595 | 0.8465543 | 0.968301116 |
| ENSMUSG00000024002 | Brd4 | protein_coding | 0.049326801 | 6.56187823 | 0.0386016 | 0.8466676 | 0.96832772 |
| ENSMUSG00000036291 | Ap5m1 | protein_coding | 0.090005232 | 3.98614786 | 0.0385538 | 0.8467613 | 0.968331812 |
| ENSMUSG00000021810 | Ecd | protein_coding | 0.06050325 | 6.148060357 | 0.0384846 | 0.846897 | 0.968366784 |
| ENSMUSG00000057181 | 5730455P16Rik | protein_coding | -0.05163619 | 6.975994754 | 0.0384464 | 0.8469721 | 0.968366784 |
| ENSMUSG00000071661 | Zbtb3 | protein_coding | -0.113447415 | 3.124408689 | 0.0384006 | 0.847062 | 0.968366784 |
| ENSMUSG00000029127 | Zbtb49 | protein_coding | 0.243752334 | 2.185977151 | 0.038281 | 0.8473116 | 0.968549119 |
| ENSMUSG00000027035 | Cers6 | protein_coding | 0.073743734 | 4.366479279 | 0.038162 | 0.8475315 | 0.968622412 |
| ENSMUSG00000029608 | Rph3a | protein_coding | 0.06516701 | 7.193579012 | 0.0381655 | 0.8476784 | 0.968622412 |
| ENSMUSG00000020925 | Ccdc43 | protein_coding | 0.052972073 | 5.857816366 | 0.0380439 | 0.8477643 | 0.968622412 |
| ENSMUSG00000025925 | Terf1 | protein_coding | -0.088609026 | 3.790794773 | 0.0380439 | 0.8477644 | 0.968622412 |
| ENSMUSG00000027099 | Mtx2 | protein_coding | -0.067491548 | 5.076598187 | 0.0379718 | 0.8479068 | 0.968622412 |
| ENSMUSG00000029636 | Wasf3 | protein_coding | 0.053925859 | 5.556638143 | 0.0379558 | 0.8479384 | 0.968622412 |
| ENSMUSG00000006463 | Zdhhc24 | protein_coding | -0.089322976 | 3.984992583 | 0.0379215 | 0.8480063 | 0.968622412 |
| ENSMUSG00000075227 | Znhit2 | protein_coding | 0.09448938 | 3.470282843 | 0.0378553 | 0.8481373 | 0.968669173 |
| ENSMUSG00000038914 | Dido1 | protein_coding | -0.060161739 | 4.996768125 | 0.0376856 | 0.8484734 | 0.968768322 |
| ENSMUSG00000003526 | Prodh | protein_coding | -0.099757163 | 4.696585418 | 0.037638 | 0.8486684 | 0.968768322 |
| ENSMUSG00000052539 | Magi3 | protein_coding | -0.134580333 | 2.938889907 | 0.0375558 | 0.8487312 | 0.968768322 |
| ENSMUSG00000033478 | Fam160b1 | protein_coding | -0.085493905 | 3.699283433 | 0.0375492 | 0.8487442 | 0.968768322 |
| ENSMUSG00000028465 | Tln1 | protein_coding | 0.059342828 | 5.298574854 | 0.0375476 | 0.8487474 | 0.968768322 |
| ENSMUSG00000050017 | Pitpnb | protein_coding | 0.049773193 | 6.832640067 | 0.0375062 | 0.8488297 | 0.968768322 |
| ENSMUSG00000036764 | Dnajc12 | protein_coding | -0.039006911 | 6.958223742 | 0.0374936 | 0.8488548 | 0.968768322 |
| ENSMUSG00000095845 | Gm5741 | protein_coding | 0.610147468 | 2.704893232 | 0.0385628 | 0.8492845 | 0.96906774 |
| ENSMUSG00000000439 | Mkrn2 | protein_coding | -0.055457195 | 5.914693123 | 0.0372715 | 0.8492973 | 0.96906774 |
| ENSMUSG00000039041 | Adrm1 | protein_coding | 0.074202632 | 4.483912201 | 0.037223 | 0.8493941 | 0.969075334 |
| ENSMUSG00000022995 | Enah | protein_coding | 0.043977535 | 6.376213725 | 0.0370712 | 0.8496976 | 0.969318676 |
| ENSMUSG00000029763 | Exoc4 | protein_coding | -0.054716014 | 5.708463691 | 0.0369352 | 0.8499699 | 0.969386995 |
| ENSMUSG00000027357 | Crls1 | protein_coding | -0.058156177 | 4.835741882 | 0.0369336 | 0.849973 | 0.969386995 |
| ENSMUSG00000030846 | Tial1 | protein_coding | 0.051175213 | 6.111302703 | 0.0369063 | 0.8500279 | 0.969386995 |
| ENSMUSG00000033961 | Zfp446 | protein_coding | -0.096155131 | 3.355200966 | 0.0368221 | 0.8501968 | 0.96938916 |
| ENSMUSG00000034327 | Kctd9 | protein_coding | 0.081215437 | 3.86237387 | 0.0367913 | 0.8502586 | 0.96938916 |
| ENSMUSG00000042873 | Lhfpl4 | protein_coding | -0.046734094 | 6.044644546 | 0.0367706 | 0.8503002 | 0.96938916 |
| ENSMUSG00000042810 | Krba1 | protein_coding | 0.078445448 | 3.77363301 | 0.0366998 | 0.8504426 | 0.969396098 |
| ENSMUSG00000004319 | Clcn3 | protein_coding | -0.039875776 | 7.199467733 | 0.0366779 | 0.8504866 | 0.969396098 |
| ENSMUSG00000054770 | Kctd18 | protein_coding | -0.103782664 | 3.477737419 | 0.0364862 | 0.8508727 | 0.969590634 |
| ENSMUSG00000028861 | Mrps15 | protein_coding | -0.053960401 | 6.071185528 | 0.0364356 | 0.8509749 | 0.969590634 |
| ENSMUSG00000064363 | mt-Nd4 | protein_coding | 0.066648773 | 10.34042568 | 0.0365006 | 0.8509794 | 0.969590634 |
| ENSMUSG00000049739 | Zfp646 | protein_coding | -0.09970781 | 3.184393989 | 0.0363936 | 0.8510598 | 0.969590634 |

| ensembl_gene_id | external_gene_name | gene_biotype | log2 (fold change) | log2 (counts per million) | F-value | p-value | False discovery rate |
|---|---|---|---|---|---|---|---|
| ENSMUSG00000032298 | Neil1 | protein_coding | -0.148865293 | 3.125484663 | 0.036347 | 0.8511577 | 0.969590634 |
| ENSMUSG00000025255 | Zfhx4 | protein_coding | -0.066044456 | 4.583105789 | 0.036325 | 0.8511983 | 0.969590634 |
| ENSMUSG00000026189 | Pecr | protein_coding | -0.117008495 | 2.841673926 | 0.0360367 | 0.8517826 | 0.970114744 |
| ENSMUSG00000024897 | Apba1 | protein_coding | 0.058820209 | 5.352296882 | 0.0359883 | 0.8518808 | 0.970114744 |
| ENSMUSG00000074227 | Spint2 | protein_coding | 0.045621745 | 5.881870537 | 0.0358259 | 0.8522111 | 0.970114744 |
| ENSMUSG00000056214 | Pard6g | protein_coding | -0.109818204 | 3.154507109 | 0.0357716 | 0.8523218 | 0.970114744 |
| ENSMUSG00000049728 | Zfp668 | protein_coding | 0.089470432 | 3.787904068 | 0.0357572 | 0.8523513 | 0.970114744 |
| ENSMUSG00000024253 | Dync2li1 | protein_coding | -0.04893864 | 5.827767068 | 0.0356876 | 0.8524932 | 0.970114744 |
| ENSMUSG00000028609 | Magoh | protein_coding | 0.05077171 | 6.642206271 | 0.0356722 | 0.8525246 | 0.970114744 |
| ENSMUSG00000007050 | Lsm2 | protein_coding | 0.099117339 | 3.413094897 | 0.035611 | 0.8526496 | 0.970114744 |
| ENSMUSG00000021130 | Galnt16 | protein_coding | 0.04618881 | 6.390037046 | 0.0355931 | 0.8526861 | 0.970114744 |
| ENSMUSG00000021337 | Scgn | protein_coding | 0.11432701 | 3.085266767 | 0.0355789 | 0.8527151 | 0.970114744 |
| ENSMUSG00000029661 | Col1a2 | protein_coding | -0.15313985 | 3.187938435 | 0.0355658 | 0.8528195 | 0.970114744 |
| ENSMUSG00000074925 | Ptar1 | protein_coding | -0.223616742 | 2.381256175 | 0.0355341 | 0.852879 | 0.970114744 |
| ENSMUSG00000058446 | Znrf2 | protein_coding | -0.085675735 | 3.726275141 | 0.0354632 | 0.8529519 | 0.970114744 |
| ENSMUSG00000092368 | A930015D03Rik | antisense_RNA | -0.118042082 | 2.822723527 | 0.0354229 | 0.8530343 | 0.970114744 |
| ENSMUSG00000025200 | Cwf19l1 | protein_coding | 0.068112788 | 4.413403784 | 0.0354085 | 0.8530639 | 0.970114744 |
| ENSMUSG00000038630 | Zkscan16 | protein_coding | -0.113439473 | 2.952419971 | 0.0353899 | 0.8531019 | 0.970114744 |
| ENSMUSG00000072964 | Bhlhb9 | protein_coding | 0.051995135 | 5.937865122 | 0.0351986 | 0.8534946 | 0.970326684 |
| ENSMUSG00000029028 | Lrrc47 | protein_coding | 0.127689775 | 2.925906026 | 0.0351892 | 0.853514 | 0.970326684 |
| ENSMUSG00000068732 | Tmem167b | protein_coding | -0.069373809 | 5.617912655 | 0.0351537 | 0.8535908 | 0.970326684 |
| ENSMUSG00000031951 | Tmem231 | protein_coding | 0.077000087 | 4.565835644 | 0.0351234 | 0.8536493 | 0.970326684 |
| ENSMUSG00000074582 | Arfgef2 | protein_coding | -0.091820612 | 4.268294228 | 0.0350094 | 0.8538879 | 0.970441474 |
| ENSMUSG00000033717 | Adra2a | protein_coding | 0.104714698 | 3.305469818 | 0.0349712 | 0.8539628 | 0.970441474 |
| ENSMUSG00000022024 | Sugt1 | protein_coding | -0.038060645 | 7.235566152 | 0.034943 | 0.854021 | 0.970441474 |
| ENSMUSG00000002550 | Uck1 | protein_coding | -0.095484464 | 3.433031781 | 0.0348984 | 0.8541129 | 0.970443406 |
| ENSMUSG00000079557 | 2-Mar | protein_coding | 0.058544637 | 5.597889616 | 0.0348211 | 0.8542727 | 0.970495343 |
| ENSMUSG00000029185 | Fam114a1 | protein_coding | 0.208278987 | 2.987001015 | 0.0349539 | 0.8543392 | 0.970495343 |
| ENSMUSG00000026792 | Lrsam1 | protein_coding | -0.092396808 | 4.490933125 | 0.0345346 | 0.8548699 | 0.970737006 |
| ENSMUSG00000019782 | Rwdd1 | protein_coding | 0.074625471 | 4.027359334 | 0.0345125 | 0.854912 | 0.970737006 |
| ENSMUSG00000024276 | Zfp397 | protein_coding | -0.082976274 | 4.089525751 | 0.0344021 | 0.8551414 | 0.970737006 |
| ENSMUSG00000055401 | Fbxo6 | protein_coding | 0.052559631 | 5.581647577 | 0.0344018 | 0.855142 | 0.970737006 |
| ENSMUSG00000030279 | C2cd5 | protein_coding | 0.064338276 | 4.359382349 | 0.0343649 | 0.8552188 | 0.970737006 |
| ENSMUSG00000029465 | Arpc3 | protein_coding | 0.049928424 | 8.650120317 | 0.0343382 | 0.8552744 | 0.970737006 |
| ENSMUSG00000029012 | Orc5 | protein_coding | -0.059946695 | 5.490140977 | 0.0342856 | 0.8553839 | 0.970737006 |
| ENSMUSG00000037989 | Wnk2 | protein_coding | 0.065399615 | 4.822597408 | 0.0342691 | 0.8554184 | 0.970737006 |
| ENSMUSG00000035212 | Leprot | protein_coding | 0.062140045 | 5.555849346 | 0.0342157 | 0.8555298 | 0.970737006 |
| ENSMUSG00000002741 | Ykt6 | protein_coding | -0.063235314 | 5.749168837 | 0.0341956 | 0.8555716 | 0.970737006 |
| ENSMUSG00000026426 | Arl8a | protein_coding | -0.042893533 | 6.998271907 | 0.0341731 | 0.8556187 | 0.970737006 |
| ENSMUSG00000040044 | Orc3 | protein_coding | 0.057882419 | 6.001959922 | 0.0340751 | 0.8558233 | 0.970737006 |
| ENSMUSG00000047022 | Mipol1 | protein_coding | -0.077181768 | 4.213269487 | 0.034041 | 0.8558945 | 0.970737006 |
| ENSMUSG00000064120 | Mocs1 | protein_coding | 0.174361421 | 2.373372985 | 0.0340266 | 0.8559247 | 0.970737006 |
| ENSMUSG00000064341 | mt-Nd1 | protein_coding | -0.067796738 | 12.02550762 | 0.0339672 | 0.8561442 | 0.970737006 |
| ENSMUSG00000029771 | Irf5 | protein_coding | 0.127338079 | 2.881099797 | 0.0339139 | 0.8561606 | 0.970737006 |
| ENSMUSG00000046169 | Adamts6 | protein_coding | 0.097669023 | 3.154787281 | 0.0338737 | 0.856245 | 0.970737006 |
| ENSMUSG00000021096 | Ppm1a | protein_coding | 0.034806688 | 7.718985064 | 0.0338421 | 0.8563112 | 0.970737006 |
| ENSMUSG00000042197 | Zfp451 | protein_coding | -0.050840459 | 5.442188885 | 0.0338213 | 0.8563549 | 0.970737006 |
| ENSMUSG00000028634 | Hivep3 | protein_coding | -0.078030121 | 4.62086736 | 0.0337368 | 0.8565322 | 0.970737006 |
| ENSMUSG00000020009 | Ifngr1 | protein_coding | -0.073737069 | 4.016042888 | 0.0337131 | 0.8565821 | 0.970737006 |
| ENSMUSG00000045625 | Pigz | protein_coding | 0.063005347 | 4.635725031 | 0.0335924 | 0.8568359 | 0.970737006 |
| ENSMUSG00000033128 | Gga1 | protein_coding | -0.065693235 | 4.297826087 | 0.0335771 | 0.8568681 | 0.970737006 |
| ENSMUSG00000038537 | Mc3r | protein_coding | 0.183404406 | 2.441413846 | 0.0335698 | 0.8568873 | 0.970737006 |
| ENSMUSG00000000325 | Arvcf | protein_coding | -0.083154362 | 3.429181469 | 0.0335165 | 0.856996 | 0.970737006 |
| ENSMUSG00000073557 | Ppp1r12b | protein_coding | 0.137113568 | 2.64936118 | 0.0334559 | 0.8571237 | 0.970737006 |
| ENSMUSG00000032062 | 2310030G06Rik | protein_coding | -0.098581878 | 3.466498892 | 0.0334467 | 0.8571431 | 0.970737006 |
| ENSMUSG00000008682 | Rpl10 | protein_coding | -0.099278166 | 3.033878391 | 0.0334085 | 0.8572238 | 0.970737006 |
| ENSMUSG00000024740 | Ddb1 | protein_coding | -0.059195636 | 7.069296325 | 0.0334478 | 0.8572352 | 0.970737006 |
| ENSMUSG00000063972 | Nr6a1 | protein_coding | 0.108921658 | 3.21914049 | 0.0333381 | 0.8573725 | 0.970737006 |
| ENSMUSG00000032633 | Flcn | protein_coding | 0.0729665 | 5.481049435 | 0.0333098 | 0.857436 | 0.970737006 |
| ENSMUSG00000089889 | 0610040B10Rik | antisense_RNA | 0.082626334 | 3.573180574 | 0.0332766 | 0.8575025 | 0.970737006 |
| ENSMUSG00000056258 | Kcnq3 | protein_coding | -0.070712579 | 5.110848953 | 0.0332632 | 0.857531 | 0.970737006 |
| ENSMUSG00000032020 | Ubash3b | protein_coding | 0.077859264 | 6.093082325 | 0.0332655 | 0.8577506 | 0.970746997 |
| ENSMUSG00000019731 | Slc35e1 | protein_coding | -0.086067307 | 4.019405831 | 0.0331124 | 0.8578504 | 0.970746997 |
| ENSMUSG00000026761 | Orc4 | protein_coding | -0.045782136 | 6.233257525 | 0.0330986 | 0.8578797 | 0.970746997 |
| ENSMUSG00000043079 | Synpo | protein_coding | -0.057730725 | 5.277590938 | 0.0330886 | 0.8579009 | 0.970746997 |
| ENSMUSG00000035969 | Rusc2 | protein_coding | -0.068214737 | 4.488377439 | 0.033036 | 0.8580126 | 0.970771245 |
| ENSMUSG00000001847 | Rac1 | protein_coding | -0.041296734 | 6.536388108 | 0.0329153 | 0.8582693 | 0.970782893 |
| ENSMUSG00000041231 | Ublcp1 | protein_coding | 0.047983534 | 5.847100931 | 0.0328985 | 0.858305 | 0.970782893 |
| ENSMUSG00000002129 | Sf3a1 | protein_coding | 0.053729821 | 6.106236639 | 0.0328705 | 0.8583647 | 0.970782893 |
| ENSMUSG00000036615 | Rfxap | protein_coding | 0.06821746 | 4.236576245 | 0.0328218 | 0.8584683 | 0.970782893 |
| ENSMUSG00000037408 | Cnnm4 | protein_coding | 0.072683561 | 5.394827322 | 0.0328024 | 0.8585133 | 0.970782893 |
| ENSMUSG00000022553 | Maf1 | protein_coding | -0.075432453 | 5.636313139 | 0.0328048 | 0.858579 | 0.970782893 |
| ENSMUSG00000042558 | Adprhl2 | protein_coding | -0.086151738 | 3.937292687 | 0.0327343 | 0.8586549 | 0.970782893 |
| ENSMUSG00000002930 | Ppp1r17 | protein_coding | -0.134038851 | 3.752463583 | 0.0327682 | 0.8588053 | 0.970835634 |
| ENSMUSG00000030060 | Hmces | protein_coding | -0.062988504 | 5.224205205 | 0.0325941 | 0.8589545 | 0.970835634 |
| ENSMUSG00000002017 | Fam98a | protein_coding | 0.064482951 | 4.176499394 | 0.032575 | 0.8589953 | 0.970835634 |
| ENSMUSG00000039043 | Arpin | protein_coding | 0.064998687 | 5.12965249 | 0.0325435 | 0.8590627 | 0.970835634 |
| ENSMUSG00000015149 | Sirt2 | protein_coding | -0.036815584 | 8.34260309 | 0.0324804 | 0.8591979 | 0.970856188 |

| ensembl_gene_id | external_gene_name | gene_biotype | log2 (fold change) | log2 (counts per million) | F-value | p-value | False discovery rate |
|---|---|---|---|---|---|---|---|
| ENSMUSG00000060923 | Acyp2 | protein_coding | 0.037259726 | 7.248511188 | 0.0323146 | 0.8595537 | 0.970856188 |
| ENSMUSG00000026238 | Ptma | protein_coding | 0.041639561 | 6.344461929 | 0.0322413 | 0.8597112 | 0.970856188 |
| ENSMUSG00000020477 | Mrps24 | protein_coding | 0.071828938 | 4.17725496 | 0.0322262 | 0.8597436 | 0.970856188 |
| ENSMUSG00000047260 | Emc6 | protein_coding | 0.071593311 | 4.534435682 | 0.0322124 | 0.8597735 | 0.970856188 |
| ENSMUSG00000048706 | Lurap1l | protein_coding | 0.067037729 | 4.247336239 | 0.0322105 | 0.8597774 | 0.970856188 |
| ENSMUSG00000021189 | Atxn3 | protein_coding | -0.08257842 | 3.618118477 | 0.0322074 | 0.8597841 | 0.970856188 |
| ENSMUSG00000072949 | Acot1 | protein_coding | 0.113162717 | 2.791619419 | 0.0321649 | 0.8598756 | 0.970856188 |
| ENSMUSG00000045083 | Lingo2 | protein_coding | -0.05735273 | 4.931962584 | 0.0321566 | 0.8598935 | 0.970856188 |
| ENSMUSG00000023391 | Dlx2 | protein_coding | 0.121329953 | 2.893843547 | 0.0319848 | 0.8602641 | 0.971172725 |
| ENSMUSG00000071074 | Yipf3 | protein_coding | -0.061148913 | 5.260315035 | 0.0319126 | 0.86042 | 0.971240478 |
| ENSMUSG00000031505 | Naxd | protein_coding | 0.041408132 | 6.902994987 | 0.0318291 | 0.8606009 | 0.971240478 |
| ENSMUSG00000032480 | Dhx30 | protein_coding | 0.049744922 | 6.567190402 | 0.0317845 | 0.8606975 | 0.971240478 |
| ENSMUSG00000036257 | Pnpla8 | protein_coding | 0.041974253 | 6.179466432 | 0.0317691 | 0.8607307 | 0.971240478 |
| ENSMUSG00000041794 | Myrip | protein_coding | -0.087060778 | 3.511552876 | 0.0316874 | 0.8609079 | 0.971240478 |
| ENSMUSG00000026032 | Ndufb3 | protein_coding | 0.051268546 | 8.401086337 | 0.031668 | 0.8609537 | 0.971240478 |
| ENSMUSG00000026839 | Upp2 | protein_coding | 0.07395652 | 4.29042745 | 0.0316651 | 0.8609564 | 0.971240478 |
| ENSMUSG00000024799 | Tm7sf2 | protein_coding | -0.083229498 | 4.114726198 | 0.0315709 | 0.8611611 | 0.97136945 |
| ENSMUSG00000027900 | Dram2 | protein_coding | 0.050724107 | 5.679446548 | 0.0314764 | 0.8613667 | 0.971405856 |
| ENSMUSG00000031987 | Egln1 | protein_coding | -0.053048944 | 5.548782701 | 0.0314282 | 0.8614717 | 0.971405856 |
| ENSMUSG00000047996 | Prrg1 | protein_coding | -0.08503216 | 3.350186742 | 0.0314058 | 0.8615206 | 0.971405856 |
| ENSMUSG00000020733 | Slc9a3r1 | protein_coding | -0.066143648 | 4.971427444 | 0.031308 | 0.8617339 | 0.971405856 |
| ENSMUSG00000029765 | Plxna4 | protein_coding | -0.066983274 | 4.416682776 | 0.0312819 | 0.8617909 | 0.971405856 |
| ENSMUSG00000036046 | 5031439G07Rik | protein_coding | 0.054468972 | 5.875616793 | 0.0312651 | 0.8618277 | 0.971405856 |
| ENSMUSG00000048707 | Tprn | protein_coding | 0.071346603 | 4.150384956 | 0.0312269 | 0.8619112 | 0.971405856 |
| ENSMUSG00000028902 | Sf3a3 | protein_coding | 0.041411021 | 6.182383047 | 0.0312247 | 0.861916 | 0.971405856 |
| ENSMUSG00000017781 | Pitpna | protein_coding | 0.036233096 | 7.228051855 | 0.0310492 | 0.8623004 | 0.971649725 |
| ENSMUSG00000031870 | Pgr | protein_coding | 0.060917293 | 4.161279224 | 0.0310434 | 0.8623132 | 0.971649725 |
| ENSMUSG00000023755 | Rhebl1 | protein_coding | 0.096493176 | 4.079893955 | 0.0310013 | 0.862409 | 0.971655855 |
| ENSMUSG00000026374 | Tsn | protein_coding | -0.039387899 | 8.515474492 | 0.0308945 | 0.8626401 | 0.971814502 |
| ENSMUSG00000032355 | Mlip | protein_coding | -0.071808156 | 4.229628769 | 0.0308314 | 0.8627789 | 0.971837035 |
| ENSMUSG00000047583 | Tyw3 | protein_coding | -0.058839191 | 4.365579562 | 0.0307968 | 0.8628551 | 0.971837035 |
| ENSMUSG00000062761 | Zfp512 | protein_coding | 0.04606401 | 5.651797819 | 0.0307392 | 0.8629819 | 0.971837035 |
| ENSMUSG00000042472 | Zfp410 | protein_coding | -0.055899145 | 5.151333987 | 0.0306705 | 0.8631335 | 0.971837035 |
| ENSMUSG00000028799 | Zfp362 | protein_coding | -0.098748153 | 2.955070288 | 0.0306556 | 0.8631664 | 0.971837035 |
| ENSMUSG00000035762 | Tmem161b | protein_coding | 0.059248429 | 4.604566558 | 0.0306004 | 0.8632883 | 0.971837035 |
| ENSMUSG00000045466 | Zfp956 | protein_coding | 0.077470136 | 4.374954297 | 0.0305984 | 0.8632928 | 0.971837035 |
| ENSMUSG00000056899 | Immp2l | protein_coding | 0.073950951 | 4.525311142 | 0.0304836 | 0.8635467 | 0.97193544 |
| ENSMUSG00000028452 | Vcp | protein_coding | -0.121001835 | 2.751292978 | 0.0304541 | 0.8636121 | 0.97193544 |
| ENSMUSG00000049804 | Armcx4 | protein_coding | 0.088334845 | 4.416621524 | 0.0304379 | 0.8636514 | 0.97193544 |
| ENSMUSG00000025537 | Phkg1 | protein_coding | -0.097399267 | 3.493481303 | 0.0303711 | 0.863796 | 0.971996501 |
| ENSMUSG00000017421 | Zfp207 | protein_coding | -0.040647412 | 7.026281644 | 0.0302657 | 0.8640301 | 0.972101334 |
| ENSMUSG00000031527 | Eri1 | protein_coding | 0.137735994 | 2.766883389 | 0.0302259 | 0.8641186 | 0.972101334 |
| ENSMUSG00000020107 | Anapc16 | protein_coding | 0.034619399 | 7.218092901 | 0.0301675 | 0.8642486 | 0.972101334 |
| ENSMUSG00000020196 | Cabin1 | protein_coding | -0.072021792 | 5.004642735 | 0.0301318 | 0.8643315 | 0.972101334 |
| ENSMUSG00000032067 | Pts | protein_coding | -0.047076086 | 7.21680756 | 0.0301259 | 0.8643412 | 0.972101334 |
| ENSMUSG00000025007 | Aldh18a1 | protein_coding | 0.099142269 | 4.630399413 | 0.0301643 | 0.8644355 | 0.9721057 |
| ENSMUSG00000021974 | Fgf9 | protein_coding | 0.056238192 | 5.384579814 | 0.0299968 | 0.8646291 | 0.972213002 |
| ENSMUSG00000050822 | Slc29a4 | protein_coding | -0.086387174 | 3.279897872 | 0.0299598 | 0.8647117 | 0.972213002 |
| ENSMUSG00000036306 | Lzts1 | protein_coding | -0.040645686 | 8.608718317 | 0.0298641 | 0.8649257 | 0.972256097 |
| ENSMUSG00000018599 | Mief2 | protein_coding | 0.063194421 | 4.81715464 | 0.0298346 | 0.8649918 | 0.972256097 |
| ENSMUSG00000024583 | Txnl1 | protein_coding | -0.037993162 | 7.176357436 | 0.0298214 | 0.8650213 | 0.972256097 |
| ENSMUSG00000027582 | Zgpat | protein_coding | 0.144515178 | 2.583425405 | 0.0295859 | 0.8655496 | 0.972727642 |
| ENSMUSG00000031979 | Cog2 | protein_coding | 0.062074436 | 4.659814181 | 0.0295419 | 0.8656487 | 0.972727642 |
| ENSMUSG00000024079 | Eif2ak2 | protein_coding | -0.244408607 | 1.681537871 | 0.0294826 | 0.8657857 | 0.972727642 |
| ENSMUSG00000031697 | Orc6 | protein_coding | 0.051709747 | 6.86565501 | 0.0294559 | 0.8658458 | 0.972727642 |
| ENSMUSG00000020978 | Klhdc2 | protein_coding | 0.039131493 | 6.554160516 | 0.0293622 | 0.8660537 | 0.972727642 |
| ENSMUSG00000025785 | Exosc7 | protein_coding | 0.038559594 | 7.102316384 | 0.0293348 | 0.8661155 | 0.972727642 |
| ENSMUSG00000068882 | Ssb | protein_coding | -0.043033888 | 7.356923846 | 0.0292541 | 0.8662979 | 0.972727642 |
| ENSMUSG00000098176 | Ccdc166 | protein_coding | -0.081292835 | 4.102231282 | 0.0292505 | 0.8663061 | 0.972727642 |
| ENSMUSG00000028630 | Dyrk2 | protein_coding | 0.073015581 | 3.865607839 | 0.0292088 | 0.8664004 | 0.972727642 |
| ENSMUSG00000022772 | Senp5 | protein_coding | 0.077308722 | 3.340804341 | 0.0291627 | 0.8665049 | 0.972727642 |
| ENSMUSG00000034071 | Zfp551 | protein_coding | -0.196748834 | 2.569405987 | 0.0291422 | 0.866685 | 0.972727642 |
| ENSMUSG00000021203 | Otub2 | protein_coding | 0.124340685 | 3.546582868 | 0.0291414 | 0.8666868 | 0.972727642 |
| ENSMUSG00000022556 | Hsf1 | protein_coding | 0.06862373 | 4.899558544 | 0.0290824 | 0.8666868 | 0.972727642 |
| ENSMUSG00000038759 | Nup205 | protein_coding | -0.09384571 | 3.095527149 | 0.0290734 | 0.8667073 | 0.972727642 |
| ENSMUSG00000040774 | Cept1 | protein_coding | 0.039235462 | 6.258797858 | 0.0289911 | 0.866894 | 0.972835644 |
| ENSMUSG00000031668 | Eif2ak3 | protein_coding | -0.116693081 | 3.672571185 | 0.0289454 | 0.8671313 | 0.972856907 |
| ENSMUSG00000053293 | Pom121 | protein_coding | 0.083708359 | 4.121406808 | 0.0288739 | 0.8671606 | 0.972856907 |
| ENSMUSG00000028991 | Mtor | protein_coding | -0.061294322 | 4.714338801 | 0.0288635 | 0.8671844 | 0.972856907 |
| ENSMUSG00000029712 | Actl6b | protein_coding | -0.044617844 | 6.161214538 | 0.0287879 | 0.8673566 | 0.972948613 |
| ENSMUSG00000025935 | Tram1 | protein_coding | 0.057781825 | 5.064605052 | 0.0286689 | 0.8676284 | 0.973044455 |
| ENSMUSG00000032350 | Gclc | protein_coding | -0.05229185 | 5.5127045 | 0.0286394 | 0.8676958 | 0.973044455 |
| ENSMUSG00000028152 | Tspan5 | protein_coding | -0.039978623 | 6.044116231 | 0.0285686 | 0.8678577 | 0.973044455 |
| ENSMUSG00000025006 | Sorbs1 | protein_coding | -0.042194235 | 6.236332475 | 0.0285505 | 0.8678992 | 0.973044455 |
| ENSMUSG00000050737 | Ptges | protein_coding | 0.111528425 | 2.854792656 | 0.0284937 | 0.8680292 | 0.973044455 |
| ENSMUSG00000002308 | Cd320 | protein_coding | -0.062265345 | 4.901515219 | 0.0284654 | 0.8680942 | 0.973044455 |
| ENSMUSG00000018547 | Pip4k2b | protein_coding | 0.068255253 | 4.264914796 | 0.0283482 | 0.8683633 | 0.973044455 |
| ENSMUSG00000041936 | Agrn | protein_coding | 0.068257275 | 4.319190457 | 0.0283166 | 0.8684359 | 0.973044455 |

| ensembl_gene_id | external_gene_name | gene_biotype | log2 (fold change) | log2 (counts per million) | F-value | p-value | False discovery rate |
|---|---|---|---|---|---|---|---|
| ENSMUSG00000018171 | Vmp1 | protein_coding | -0.036062457 | 7.624051019 | 0.0283128 | 0.8684448 | 0.973044455 |
| ENSMUSG00000025795 | Rassf3 | protein_coding | -0.104201613 | 2.979774299 | 0.0282974 | 0.8684801 | 0.973044455 |
| ENSMUSG00000045039 | Megf8 | protein_coding | -0.071899754 | 4.716612134 | 0.028281 | 0.8685179 | 0.973044455 |
| ENSMUSG00000033970 | Rfc3 | protein_coding | -0.050835576 | 4.936916151 | 0.0282033 | 0.8686968 | 0.973044455 |
| ENSMUSG00000025092 | Hspa12a | protein_coding | -0.042272025 | 5.960480157 | 0.0281674 | 0.8687796 | 0.973044455 |
| ENSMUSG00000063236 | 1110038F14Rik | protein_coding | 0.04932745 | 5.627800157 | 0.0281594 | 0.8687981 | 0.973044455 |
| ENSMUSG00000030411 | Nova2 | protein_coding | -0.054797846 | 5.176793 | 0.0281588 | 0.8687994 | 0.973044455 |
| ENSMUSG00000031169 | Porcn | protein_coding | 0.062361175 | 5.106679883 | 0.0279326 | 0.8693223 | 0.97317227 |
| ENSMUSG00000022552 | Sharpin | protein_coding | -0.076557977 | 5.492261759 | 0.0279818 | 0.8693394 | 0.97317227 |
| ENSMUSG00000029869 | Ephb6 | protein_coding | 0.080287461 | 4.24987142 | 0.0277464 | 0.8697543 | 0.97317227 |
| ENSMUSG00000059013 | Sh2d3c | protein_coding | 0.07500627 | 4.324343061 | 0.0277345 | 0.869782 | 0.97317227 |
| ENSMUSG00000044786 | Zfp36 | protein_coding | 0.108812158 | 2.927527003 | 0.0277328 | 0.869786 | 0.97317227 |
| ENSMUSG00000031930 | Wwp2 | protein_coding | -0.063216618 | 5.607010301 | 0.0276967 | 0.8698732 | 0.97317227 |
| ENSMUSG00000100837 | 1700063D05Rik | lincRNA | 0.084479587 | 3.539659676 | 0.0276492 | 0.8699803 | 0.97317227 |
| ENSMUSG00000001158 | Snrnp27 | protein_coding | -0.042979938 | 8.072100003 | 0.0276002 | 0.8700945 | 0.97317227 |
| ENSMUSG00000042572 | Ube2q1 | protein_coding | -0.048806132 | 5.737948593 | 0.0275411 | 0.8702323 | 0.97317227 |
| ENSMUSG00000026442 | Nfasc | protein_coding | -0.039921348 | 5.774130877 | 0.0274754 | 0.8703858 | 0.97317227 |
| ENSMUSG00000038564 | Ift172 | protein_coding | -0.086621715 | 3.500283351 | 0.0274003 | 0.8705613 | 0.97317227 |
| ENSMUSG00000014503 | Pkd2l2 | protein_coding | 0.043600018 | 5.546689647 | 0.0273897 | 0.8705861 | 0.97317227 |
| ENSMUSG00000033712 | Ccar2 | protein_coding | 0.059415614 | 5.48746704 | 0.0273845 | 0.8705982 | 0.97317227 |
| ENSMUSG00000028637 | Ccdc30 | protein_coding | 0.04971058 | 5.308539103 | 0.0272655 | 0.870877 | 0.97317227 |
| ENSMUSG00000020593 | Lpin1 | protein_coding | 0.068309515 | 4.67513294 | 0.0272225 | 0.870978 | 0.97317227 |
| ENSMUSG00000024308 | Tapbp | protein_coding | 0.051923092 | 4.960714711 | 0.027141 | 0.8711692 | 0.97317227 |
| ENSMUSG00000025199 | Chuk | protein_coding | -0.115612483 | 3.326487007 | 0.0271207 | 0.8712204 | 0.97317227 |
| ENSMUSG00000056941 | Commd7 | protein_coding | -0.053197806 | 6.256936883 | 0.0271174 | 0.8712281 | 0.97317227 |
| ENSMUSG00000026020 | Nop58 | protein_coding | -0.041999867 | 6.335977242 | 0.0271019 | 0.8712614 | 0.97317227 |
| ENSMUSG00000028949 | Smarcd3 | protein_coding | -0.071191377 | 3.796072841 | 0.0269393 | 0.8716444 | 0.97317227 |
| ENSMUSG00000045991 | Onecut2 | protein_coding | 0.056898961 | 4.74678289 | 0.0269088 | 0.8717164 | 0.97317227 |
| ENSMUSG00000060152 | Pop5 | protein_coding | 0.054743163 | 5.033105139 | 0.0268914 | 0.8717574 | 0.97317227 |
| ENSMUSG00000003346 | Abhd17a | protein_coding | -0.064840415 | 4.933499331 | 0.0268695 | 0.8718093 | 0.97317227 |
| ENSMUSG00000028917 | Plekhm2 | protein_coding | 0.089158456 | 3.915621451 | 0.0268673 | 0.8718177 | 0.97317227 |
| ENSMUSG00000089687 | Rab42 | protein_coding | 0.105980188 | 2.882036493 | 0.0268518 | 0.8718509 | 0.97317227 |
| ENSMUSG00000020630 | Rnaseh1 | protein_coding | 0.065341552 | 4.392954136 | 0.0268512 | 0.8718523 | 0.97317227 |
| ENSMUSG00000022434 | Fam118a | protein_coding | 0.050086829 | 5.011475363 | 0.0267906 | 0.8719958 | 0.97317227 |
| ENSMUSG00000032594 | Ip6k1 | protein_coding | 0.036653982 | 7.034763964 | 0.0267263 | 0.872148 | 0.97317227 |
| ENSMUSG00000048920 | Fkrp | protein_coding | 0.054783119 | 4.62235271 | 0.0267206 | 0.8721615 | 0.97317227 |
| ENSMUSG00000020160 | Meis1 | protein_coding | 0.127747789 | 3.539785769 | 0.0268022 | 0.8721691 | 0.97317227 |
| ENSMUSG00000032607 | Amt | protein_coding | -0.089811279 | 5.327476593 | 0.0268139 | 0.8722228 | 0.97317227 |
| ENSMUSG00000021595 | Nsun2 | protein_coding | -0.053463451 | 5.686086394 | 0.0266909 | 0.872232 | 0.97317227 |
| ENSMUSG00000039470 | Zdhhc2 | protein_coding | 0.055178623 | 5.22588489 | 0.0266166 | 0.8724082 | 0.97317227 |
| ENSMUSG00000078161 | Erich3 | protein_coding | -0.075550464 | 4.031065691 | 0.0265997 | 0.8724483 | 0.97317227 |
| ENSMUSG00000028602 | Tnfrsf8 | protein_coding | 0.196320898 | 1.854737579 | 0.0265985 | 0.8724513 | 0.97317227 |
| ENSMUSG00000021087 | Rtn1 | protein_coding | 0.033657286 | 10.12214384 | 0.0264675 | 0.8727628 | 0.97317227 |
| ENSMUSG00000044876 | Zfp444 | protein_coding | 0.058272064 | 5.120859974 | 0.0264421 | 0.8728231 | 0.97317227 |
| ENSMUSG00000021919 | Chat | protein_coding | -0.100288486 | 3.470664016 | 0.0264381 | 0.8728361 | 0.97317227 |
| ENSMUSG00000027349 | Fam98b | protein_coding | 0.038271919 | 6.291079979 | 0.0264131 | 0.8728924 | 0.97317227 |
| ENSMUSG00000042682 | Selenok | protein_coding | -0.041010351 | 9.326716745 | 0.0263822 | 0.8729659 | 0.97317227 |
| ENSMUSG00000028655 | Mfsd2a | protein_coding | -0.20119888 | 2.405620596 | 0.0264197 | 0.8730039 | 0.97317227 |
| ENSMUSG00000026811 | St6galnac6 | protein_coding | -0.04253462 | 5.915697612 | 0.0263562 | 0.8730281 | 0.97317227 |
| ENSMUSG00000065979 | Cpped1 | protein_coding | -0.045447191 | 6.910962486 | 0.0261827 | 0.8734429 | 0.97317227 |
| ENSMUSG00000029196 | Tada2b | protein_coding | 0.075292877 | 3.571371922 | 0.0261712 | 0.8734704 | 0.97317227 |
| ENSMUSG00000019124 | Scrn1 | protein_coding | 0.04320191 | 7.737756782 | 0.0260974 | 0.8736471 | 0.97317227 |
| ENSMUSG00000113170 | AC132344.1 | lincRNA | -0.064733992 | 3.751730224 | 0.026097 | 0.8736483 | 0.97317227 |
| ENSMUSG00000031591 | Asah1 | protein_coding | -0.040632322 | 6.231860948 | 0.0260876 | 0.8736707 | 0.97317227 |
| ENSMUSG00000030556 | Lrrc28 | protein_coding | 0.039922578 | 5.877708572 | 0.0260489 | 0.8737636 | 0.97317227 |
| ENSMUSG00000027674 | Pex5l | protein_coding | -0.083423995 | 3.225459184 | 0.0260414 | 0.8737815 | 0.97317227 |
| ENSMUSG00000034509 | Mad2l1bp | protein_coding | 0.048624131 | 5.073675527 | 0.0259753 | 0.8739404 | 0.97317227 |
| ENSMUSG00000041115 | Iqsec2 | protein_coding | -0.061955135 | 4.473498805 | 0.0259613 | 0.873974 | 0.97317227 |
| ENSMUSG00000055850 | Rnf181 | protein_coding | 0.032167178 | 7.940958936 | 0.0259583 | 0.8739814 | 0.97317227 |
| ENSMUSG00000030795 | Fus | protein_coding | -0.050538994 | 5.029174441 | 0.0259251 | 0.8740613 | 0.97317227 |
| ENSMUSG00000026623 | Lpgat1 | protein_coding | 0.036013983 | 6.42459055 | 0.0258985 | 0.8741252 | 0.97317227 |
| ENSMUSG00000050541 | Adra1b | protein_coding | 0.132460452 | 2.48256062 | 0.0258819 | 0.8741651 | 0.97317227 |
| ENSMUSG00000022808 | Snx4 | protein_coding | 0.05003977 | 5.947934191 | 0.0258728 | 0.874187 | 0.97317227 |
| ENSMUSG00000047141 | Zfp654 | protein_coding | -0.07132879 | 3.831966821 | 0.0258693 | 0.8741955 | 0.97317227 |
| ENSMUSG00000003778 | Brd8 | protein_coding | -0.045488656 | 6.983895678 | 0.0258516 | 0.8742414 | 0.97317227 |
| ENSMUSG00000104960 | Snhg8 | processed_transcript | 0.06484333 | 6.492113512 | 0.025941 | 0.8742532 | 0.97317227 |
| ENSMUSG00000046157 | Tmem229b | protein_coding | 0.07822866 | 3.999142315 | 0.0257039 | 0.8745946 | 0.973276209 |
| ENSMUSG00000027350 | Chgb | protein_coding | 0.035601948 | 7.543770136 | 0.0256525 | 0.874719 | 0.973276209 |
| ENSMUSG00000040537 | Adam22 | protein_coding | 0.039798109 | 6.220758536 | 0.0256197 | 0.8747985 | 0.973276209 |
| ENSMUSG00000044533 | Rps2 | protein_coding | 0.062484704 | 4.659442267 | 0.0256194 | 0.8747993 | 0.973276209 |
| ENSMUSG00000029998 | Pcyox1 | protein_coding | 0.039736706 | 6.168019748 | 0.0255893 | 0.8748721 | 0.973276209 |
| ENSMUSG00000048644 | Ctxn1 | protein_coding | -0.030500234 | 7.951408505 | 0.0255821 | 0.8748896 | 0.973276209 |
| ENSMUSG00000028926 | Cdk14 | protein_coding | 0.053089636 | 7.358194696 | 0.0255231 | 0.8751575 | 0.973473555 |
| ENSMUSG00000036356 | Csgalnact1 | protein_coding | -0.093867035 | 3.439311521 | 0.0254028 | 0.8753248 | 0.973501686 |
| ENSMUSG00000024830 | Rps6kb2 | protein_coding | 0.131530275 | 3.145732886 | 0.0254231 | 0.8753639 | 0.973501686 |
| ENSMUSG00000011114 | Tbrg1 | protein_coding | -0.040036265 | 7.01306031 | 0.0252483 | 0.8757011 | 0.973776011 |
| ENSMUSG00000105891 | A230001M10Rik | lincRNA | 0.077747618 | 3.815658261 | 0.0251761 | 0.8758775 | 0.973871451 |
| ENSMUSG00000025352 | Gdf11 | protein_coding | 0.062610951 | 4.577487827 | 0.0250839 | 0.876103 | 0.974015593 |

| ensembl_gene_id | external_gene_name | gene_biotype | log2 (fold change) | log2 (counts per million) | F-value | p-value | False discovery rate |
|---|---|---|---|---|---|---|---|
| ENSMUSG00000034189 | Hsdl1 | protein_coding | 0.042057014 | 6.774531571 | 0.025049 | 0.8761883 | 0.974015593 |
| ENSMUSG00000024145 | Pigf | protein_coding | 0.063346415 | 5.052367098 | 0.024898 | 0.8765588 | 0.974326674 |
| ENSMUSG00000021395 | Spin1 | protein_coding | 0.035891049 | 7.491724962 | 0.0248268 | 0.876734 | 0.974358277 |
| ENSMUSG00000005470 | Asf1b | protein_coding | 0.207540637 | 2.029709115 | 0.0247622 | 0.8769046 | 0.974358277 |
| ENSMUSG00000022749 | Tbc1d23 | protein_coding | -0.072362148 | 3.55848293 | 0.0247455 | 0.8769342 | 0.974358277 |
| ENSMUSG00000022557 | Bop1 | protein_coding | 0.060180765 | 5.784828613 | 0.0247211 | 0.8770058 | 0.974358277 |
| ENSMUSG00000019826 | Zbtb24 | protein_coding | 0.063277521 | 3.807397015 | 0.0247025 | 0.8770402 | 0.974358277 |
| ENSMUSG00000075467 | Dnlz | protein_coding | 0.069995457 | 4.734856396 | 0.0245992 | 0.8772984 | 0.974430955 |
| ENSMUSG00000022329 | Stk3 | protein_coding | -0.07298524 | 4.112228634 | 0.024589 | 0.8773206 | 0.974430955 |
| ENSMUSG00000048832 | Vps37c | protein_coding | -0.065679879 | 4.872612485 | 0.024566 | 0.8773775 | 0.974430955 |
| ENSMUSG00000039886 | Tmem120a | protein_coding | 0.055563553 | 4.318940806 | 0.0245254 | 0.877478 | 0.974441898 |
| ENSMUSG00000034173 | Zbed5 | protein_coding | -0.079677477 | 3.221061799 | 0.0244241 | 0.8777289 | 0.974502174 |
| ENSMUSG00000027134 | Lpcat4 | protein_coding | 0.052701096 | 4.840404694 | 0.0243411 | 0.8779353 | 0.974502174 |
| ENSMUSG00000050505 | Pcdh20 | protein_coding | 0.068239397 | 3.923139704 | 0.0243388 | 0.8779409 | 0.974502174 |
| ENSMUSG00000075415 | Fnbp1 | protein_coding | -0.041745118 | 6.990679437 | 0.0242997 | 0.8780381 | 0.974502174 |
| ENSMUSG00000092094 | Zfp804b | protein_coding | -0.096219265 | 2.697701864 | 0.0242689 | 0.8781148 | 0.974502174 |
| ENSMUSG00000026142 | Rhbdd1 | protein_coding | 0.14999121 | 2.721468663 | 0.024273 | 0.878116 | 0.974502174 |
| ENSMUSG00000038611 | Phrf1 | protein_coding | 0.058850327 | 4.427450614 | 0.0242481 | 0.8781667 | 0.974502174 |
| ENSMUSG00000003345 | Csnk1g2 | protein_coding | 0.059317268 | 5.019597493 | 0.0240911 | 0.8785584 | 0.974836257 |
| ENSMUSG00000027394 | Ttl | protein_coding | -0.049186577 | 4.823554834 | 0.0239539 | 0.8789018 | 0.974836271 |
| ENSMUSG00000032314 | Etfa | protein_coding | -0.031749142 | 6.758427469 | 0.0239345 | 0.8789502 | 0.974836271 |
| ENSMUSG00000036636 | Clcn7 | protein_coding | -0.062172869 | 4.562534078 | 0.0239271 | 0.8789688 | 0.974836271 |
| ENSMUSG00000031885 | Cbfb | protein_coding | -0.058308834 | 4.202384283 | 0.0239146 | 0.8790002 | 0.974836271 |
| ENSMUSG00000024312 | Wdr46 | protein_coding | 0.101246044 | 2.900480382 | 0.02391 | 0.8790117 | 0.974836271 |
| ENSMUSG00000019813 | Cep57l1 | protein_coding | -0.084777551 | 3.313911245 | 0.0237265 | 0.8794731 | 0.975117758 |
| ENSMUSG00000029312 | Klhl8 | protein_coding | -0.054644658 | 4.255544535 | 0.0236026 | 0.8797856 | 0.975117758 |
| ENSMUSG00000024789 | Jak2 | protein_coding | 0.065507135 | 4.004079368 | 0.0235689 | 0.8798706 | 0.975117758 |
| ENSMUSG00000113314 | AC122013.1 | lincRNA | 0.120567888 | 2.498443276 | 0.0235633 | 0.8798847 | 0.975117758 |
| ENSMUSG00000034032 | Rpap1 | protein_coding | 0.076918184 | 3.352133866 | 0.023549 | 0.8799209 | 0.975117758 |
| ENSMUSG00000036052 | Dnajb5 | protein_coding | 0.05515851 | 5.281467032 | 0.0234913 | 0.8800668 | 0.975117758 |
| ENSMUSG00000021115 | Vrk1 | protein_coding | -0.044147729 | 5.201208222 | 0.0234255 | 0.8802335 | 0.975117758 |
| ENSMUSG00000039252 | Lgi2 | protein_coding | 0.051100792 | 4.613793116 | 0.0234049 | 0.8802859 | 0.975117758 |
| ENSMUSG00000010660 | Plcd1 | protein_coding | -0.123529804 | 3.05950199 | 0.0233915 | 0.880331 | 0.975117758 |
| ENSMUSG00000090069 | E430024P14Rik | antisense_RNA | -0.117229461 | 3.360192038 | 0.0234105 | 0.880384 | 0.975117758 |
| ENSMUSG00000063410 | Stk24 | protein_coding | -0.059107734 | 3.879405367 | 0.0233638 | 0.8803902 | 0.975117758 |
| ENSMUSG00000030779 | Rbbp6 | protein_coding | -0.040169983 | 5.982878442 | 0.0233455 | 0.8804366 | 0.975117758 |
| ENSMUSG00000022412 | Mief1 | protein_coding | -0.060216354 | 5.00375469 | 0.023327 | 0.8804835 | 0.975117758 |
| ENSMUSG00000026833 | Olfm1 | protein_coding | 0.033911815 | 7.687292613 | 0.0232988 | 0.8805553 | 0.975117758 |
| ENSMUSG00000030303 | Far2 | protein_coding | 0.086509172 | 4.74885854 | 0.0233049 | 0.8807268 | 0.975117758 |
| ENSMUSG00000008734 | Gprc5b | protein_coding | 0.063355758 | 4.896268762 | 0.0232007 | 0.8808048 | 0.975117758 |
| ENSMUSG00000085664 | Atxn7l1os2 | antisense_RNA | 0.100609486 | 2.93795205 | 0.0231998 | 0.8808071 | 0.975117758 |
| ENSMUSG00000033854 | Kcnk10 | protein_coding | 0.08680651 | 3.331571941 | 0.0229629 | 0.8814125 | 0.975687509 |
| ENSMUSG00000020409 | Slu7 | protein_coding | 0.035761326 | 7.116265078 | 0.0229083 | 0.8815524 | 0.975741959 |
| ENSMUSG00000030374 | Strn4 | protein_coding | 0.039963156 | 6.040028159 | 0.0227933 | 0.8818476 | 0.975764134 |
| ENSMUSG00000021615 | Xrcc4 | protein_coding | -0.062078881 | 4.192460649 | 0.0227751 | 0.8818943 | 0.975764134 |
| ENSMUSG00000040354 | Mars | protein_coding | 0.064875084 | 4.800313522 | 0.0227762 | 0.8818946 | 0.975764134 |
| ENSMUSG00000021668 | Polk | protein_coding | -0.082085719 | 3.087683292 | 0.0227592 | 0.8819354 | 0.975764134 |
| ENSMUSG00000031827 | Cotl1 | protein_coding | -0.036100494 | 5.986020748 | 0.0226076 | 0.882326 | 0.975826789 |
| ENSMUSG00000024507 | Hsd17b4 | protein_coding | -0.041690092 | 5.464083413 | 0.022573 | 0.8824153 | 0.975826789 |
| ENSMUSG00000040723 | Rcsd1 | protein_coding | -0.117838353 | 2.563251632 | 0.0225543 | 0.8824637 | 0.975826789 |
| ENSMUSG00000030811 | Fbxl19 | protein_coding | 0.155019275 | 2.807103069 | 0.0225447 | 0.8826058 | 0.975826789 |
| ENSMUSG00000039530 | Tusc3 | protein_coding | -0.042336676 | 7.370832467 | 0.0224217 | 0.88281 | 0.975826789 |
| ENSMUSG00000036106 | Prr5 | protein_coding | -0.083456596 | 3.567331264 | 0.0224017 | 0.8828589 | 0.975826789 |
| ENSMUSG00000031728 | Zfp821 | protein_coding | 0.060085805 | 4.6341549 | 0.0223445 | 0.8830073 | 0.975826789 |
| ENSMUSG00000034088 | Hdlbp | protein_coding | -0.038792543 | 6.210549682 | 0.0223237 | 0.8830614 | 0.975826789 |
| ENSMUSG00000063894 | Zkscan8 | protein_coding | -0.077250069 | 3.195151195 | 0.0222631 | 0.883219 | 0.975826789 |
| ENSMUSG00000020321 | Mdh1 | protein_coding | 0.03385617 | 11.90593286 | 0.0222058 | 0.8833683 | 0.975826789 |
| ENSMUSG00000062070 | Pgk1 | protein_coding | 0.041153567 | 5.410924104 | 0.0221832 | 0.883427 | 0.975826789 |
| ENSMUSG00000085741 | 5430405H02Rik | processed_transcript | -0.074582954 | 3.345413118 | 0.0221552 | 0.8835 | 0.975826789 |
| ENSMUSG00000049858 | Suox | protein_coding | 0.082801175 | 3.829617284 | 0.022128 | 0.883574 | 0.975826789 |
| ENSMUSG00000027488 | Snta1 | protein_coding | -0.055380327 | 4.297580333 | 0.0221199 | 0.8835922 | 0.975826789 |
| ENSMUSG00000006494 | Pdk1 | protein_coding | -0.070455705 | 4.046963452 | 0.0221099 | 0.8836184 | 0.975826789 |
| ENSMUSG00000009076 | Zmat5 | protein_coding | 0.042316966 | 5.897945453 | 0.0221081 | 0.883623 | 0.975826789 |
| ENSMUSG00000030486 | Zfp108 | protein_coding | -0.112506436 | 2.501826416 | 0.0221021 | 0.8836387 | 0.975826789 |
| ENSMUSG00000007850 | Hnrnph1 | protein_coding | -0.033289526 | 6.886565233 | 0.0220693 | 0.8837243 | 0.975826789 |
| ENSMUSG00000027238 | Frmd5 | protein_coding | 0.054982582 | 4.258166199 | 0.0220446 | 0.883789 | 0.975826789 |
| ENSMUSG00000002524 | Puf60 | protein_coding | 0.032184165 | 8.566875765 | 0.0219691 | 0.8839866 | 0.975826789 |
| ENSMUSG00000032329 | Hmg20a | protein_coding | 0.041451377 | 6.190941691 | 0.0219584 | 0.8840147 | 0.975826789 |
| ENSMUSG00000022351 | Sqle | protein_coding | 0.047899828 | 5.66798087 | 0.0218198 | 0.8843786 | 0.975826789 |
| ENSMUSG00000038240 | Pdss2 | protein_coding | -0.048465954 | 4.928223451 | 0.0217974 | 0.8844374 | 0.975826789 |
| ENSMUSG00000053774 | Ubxn7 | protein_coding | 0.088196472 | 3.209179744 | 0.0217759 | 0.884494 | 0.975826789 |
| ENSMUSG00000033419 | Snap91 | protein_coding | 0.033192565 | 7.467942265 | 0.0217735 | 0.8845003 | 0.975826789 |
| ENSMUSG00000032834 | Pwp2 | protein_coding | -0.070493447 | 4.081071624 | 0.0217592 | 0.8845379 | 0.975826789 |
| ENSMUSG00000024078 | Ttc27 | protein_coding | 0.082832592 | 4.069331207 | 0.0217425 | 0.8845957 | 0.975826789 |
| ENSMUSG00000050973 | Gdpgp1 | protein_coding | -0.070461681 | 4.282659624 | 0.0217279 | 0.8846204 | 0.975826789 |
| ENSMUSG00000063626 | Unc5d | protein_coding | 0.047860437 | 5.112167552 | 0.0217266 | 0.8846238 | 0.975826789 |
| ENSMUSG00000022283 | Pabpc1 | protein_coding | 0.044623365 | 5.356366693 | 0.0216727 | 0.884766 | 0.975883503 |
| ENSMUSG00000021218 | Gdi2 | protein_coding | 0.036122166 | 8.659815693 | 0.0215758 | 0.8850218 | 0.976040014 |

| ensembl_gene_id | external_gene_name | gene_biotype | log2 (fold change) | log2 (counts per million) | F-value | p-value | False discovery rate |
|---|---|---|---|---|---|---|---|
| ENSMUSG00000021819 | Zswim8 | protein_coding | -0.05314442 | 4.343796839 | 0.0215433 | 0.8851076 | 0.976040014 |
| ENSMUSG00000044502 | Bod1 | protein_coding | 0.050794327 | 5.512528293 | 0.021516 | 0.8851802 | 0.976040014 |
| ENSMUSG00000031012 | Cask | protein_coding | 0.052934139 | 7.302078276 | 0.0215287 | 0.8853261 | 0.976100872 |
| ENSMUSG00000028229 | Rmdn1 | protein_coding | -0.055033067 | 4.287708467 | 0.0213644 | 0.8855821 | 0.976283034 |
| ENSMUSG00000029267 | Mtf2 | protein_coding | -0.050762118 | 4.569289192 | 0.021263 | 0.8858519 | 0.976335246 |
| ENSMUSG00000015341 | Golga7 | protein_coding | 0.03719474 | 5.737273515 | 0.0212465 | 0.885896 | 0.976335246 |
| ENSMUSG00000045763 | Basp1 | protein_coding | 0.030468386 | 7.471415446 | 0.0212443 | 0.8859019 | 0.976335246 |
| ENSMUSG00000067367 | Lyar | protein_coding | 0.08019264 | 3.281575278 | 0.0211348 | 0.8861941 | 0.976472976 |
| ENSMUSG00000002280 | Narfl | protein_coding | -0.074978991 | 3.723069689 | 0.0211294 | 0.8862085 | 0.976472976 |
| ENSMUSG00000042104 | Uggt2 | protein_coding | 0.067735898 | 4.359631651 | 0.0210672 | 0.8863749 | 0.976551603 |
| ENSMUSG00000020441 | 2310033P09Rik | protein_coding | 0.063536196 | 3.652092943 | 0.0210349 | 0.8864615 | 0.976551603 |
| ENSMUSG00000021144 | Mta1 | protein_coding | -0.056201615 | 5.067801162 | 0.0209103 | 0.8867957 | 0.976773279 |
| ENSMUSG00000043923 | Ccdc84 | protein_coding | -0.139975636 | 2.271907717 | 0.0208922 | 0.8868444 | 0.976773279 |
| ENSMUSG00000043929 | Klhl15 | protein_coding | -0.067444711 | 4.742716628 | 0.0208193 | 0.8870435 | 0.976811781 |
| ENSMUSG00000027522 | Stx16 | protein_coding | -0.058162899 | 5.559585882 | 0.0208156 | 0.887061 | 0.976811781 |
| ENSMUSG00000031993 | Snx19 | protein_coding | 0.047466468 | 5.034120209 | 0.0207126 | 0.8873281 | 0.976951805 |
| ENSMUSG00000042271 | Nxt2 | protein_coding | 0.041760791 | 6.680657324 | 0.0206982 | 0.8873699 | 0.976951805 |
| ENSMUSG00000040850 | Psme4 | protein_coding | 0.051886187 | 4.122706037 | 0.0205836 | 0.8876771 | 0.976987087 |
| ENSMUSG00000033628 | Pik3c3 | protein_coding | 0.045412091 | 5.531721256 | 0.0205765 | 0.8876962 | 0.976987087 |
| ENSMUSG00000044496 | 2510039O18Rik | protein_coding | -0.088213678 | 3.529021307 | 0.0205727 | 0.8877095 | 0.976987087 |
| ENSMUSG00000020250 | Txnrd1 | protein_coding | -0.040645501 | 5.659791275 | 0.0205194 | 0.8878511 | 0.976987087 |
| ENSMUSG00000029674 | Limk1 | protein_coding | -0.117759353 | 2.789344449 | 0.0204889 | 0.8879338 | 0.976987087 |
| ENSMUSG00000036983 | Tfb1m | protein_coding | 0.063708681 | 3.668714948 | 0.0204526 | 0.8880325 | 0.976987087 |
| ENSMUSG00000021884 | Hacl1 | protein_coding | -0.069644382 | 3.644691839 | 0.0204506 | 0.8880379 | 0.976987087 |
| ENSMUSG00000059540 | Tcea2 | protein_coding | 0.063707163 | 4.56843756 | 0.0203167 | 0.8884024 | 0.977288039 |
| ENSMUSG00000030120 | Mlf2 | protein_coding | 0.0268421 | 7.984126122 | 0.0202681 | 0.8885349 | 0.977333837 |
| ENSMUSG00000097709 | 2810429I04Rik | lincRNA | -0.073954058 | 3.477889688 | 0.0201673 | 0.8888106 | 0.977349138 |
| ENSMUSG00000031536 | Polb | protein_coding | 0.027185228 | 7.048115771 | 0.0201538 | 0.8888475 | 0.977349138 |
| ENSMUSG00000036698 | Ago2 | protein_coding | -0.041713821 | 5.261314727 | 0.0201419 | 0.88888 | 0.977349138 |
| ENSMUSG00000040124 | Gorab | protein_coding | -0.05070639 | 4.695669483 | 0.0201301 | 0.8889124 | 0.977349138 |
| ENSMUSG00000022554 | Hgh1 | protein_coding | -0.094314278 | 2.639369943 | 0.0200697 | 0.889078 | 0.977405443 |
| ENSMUSG00000078789 | Dph1 | protein_coding | 0.075735259 | 3.712056252 | 0.0200452 | 0.8891454 | 0.977405443 |
| ENSMUSG00000028391 | Wdr31 | protein_coding | 0.060136686 | 5.189647739 | 0.0198398 | 0.8897135 | 0.97788527 |
| ENSMUSG00000053411 | Cbx7 | protein_coding | -0.042885065 | 5.132280649 | 0.0198062 | 0.8898035 | 0.97788527 |
| ENSMUSG00000061086 | Myl4 | protein_coding | 0.084554562 | 2.909886367 | 0.0197749 | 0.8898899 | 0.97788527 |
| ENSMUSG00000029049 | Morn1 | protein_coding | -0.079711666 | 3.800028213 | 0.0197558 | 0.8899456 | 0.97788527 |
| ENSMUSG00000062328 | Rpl17 | protein_coding | 0.068762783 | 3.274014848 | 0.0195537 | 0.8905032 | 0.978174655 |
| ENSMUSG00000024127 | Prepl | protein_coding | 0.089373193 | 7.106366955 | 0.0198769 | 0.8906229 | 0.978174655 |
| ENSMUSG00000019461 | Plscr3 | protein_coding | -0.052165454 | 4.880976774 | 0.0194927 | 0.8906731 | 0.978174655 |
| ENSMUSG00000037570 | Mcrs1 | protein_coding | 0.044305284 | 4.744328331 | 0.0194908 | 0.8906784 | 0.978174655 |
| ENSMUSG00000024277 | Mapre2 | protein_coding | 0.03652326 | 8.384898168 | 0.0193774 | 0.8909947 | 0.978174655 |
| ENSMUSG00000030180 | Kdm5a | protein_coding | -0.044200253 | 5.70130828 | 0.0193681 | 0.8910206 | 0.978174655 |
| ENSMUSG00000044348 | Slc25a53 | protein_coding | 0.046258021 | 4.529963874 | 0.0193367 | 0.8911083 | 0.978174655 |
| ENSMUSG00000006288 | Ttc5 | protein_coding | 0.0312709 | 6.543103202 | 0.0193321 | 0.8911212 | 0.978174655 |
| ENSMUSG00000084159 | Gm12696 | processed_pseudoge | -0.079508152 | 2.892748836 | 0.0193192 | 0.8911572 | 0.978174655 |
| ENSMUSG00000062309 | Rpp25 | protein_coding | 0.052095238 | 4.69093654 | 0.0192533 | 0.8913419 | 0.978174655 |
| ENSMUSG00000062202 | Btbd9 | protein_coding | 0.046407178 | 5.808401651 | 0.0192491 | 0.8913537 | 0.978174655 |
| ENSMUSG00000027425 | Kat14 | protein_coding | 0.04907067 | 4.035801704 | 0.0192429 | 0.8913711 | 0.978174655 |
| ENSMUSG00000000378 | Ccm2 | protein_coding | 0.034402108 | 5.708089225 | 0.0191758 | 0.8915591 | 0.978174655 |
| ENSMUSG00000037103 | Dcaf15 | protein_coding | -0.086019923 | 3.497198755 | 0.0191641 | 0.891595 | 0.978174655 |
| ENSMUSG00000020664 | Dld | protein_coding | -0.039193386 | 7.390086463 | 0.0191437 | 0.8916522 | 0.978174655 |
| ENSMUSG00000024873 | Cnih2 | protein_coding | 0.052497035 | 5.668024471 | 0.0191217 | 0.8917142 | 0.978174655 |
| ENSMUSG00000024759 | Atl3 | protein_coding | -0.084075097 | 3.070248104 | 0.019106 | 0.8917554 | 0.978174655 |
| ENSMUSG00000049672 | Zbtb14 | protein_coding | 0.069759857 | 3.195792007 | 0.019035 | 0.8919555 | 0.97829429 |
| ENSMUSG00000029436 | Mmp17 | protein_coding | 0.045332496 | 5.137459336 | 0.0189161 | 0.8922911 | 0.978562643 |
| ENSMUSG00000030270 | Cpne9 | protein_coding | 0.106217416 | 5.478852177 | 0.0191644 | 0.8924158 | 0.978573709 |
| ENSMUSG00000017418 | Arl5b | protein_coding | -0.079954613 | 3.134239237 | 0.01884 | 0.8925065 | 0.978573709 |
| ENSMUSG00000022125 | Cln5 | protein_coding | -0.073523171 | 3.400346536 | 0.0187531 | 0.8927531 | 0.978573709 |
| ENSMUSG00000021706 | Zfyve16 | protein_coding | -0.071231171 | 3.227398287 | 0.0187488 | 0.8927654 | 0.978573709 |
| ENSMUSG00000030269 | Mtmr14 | protein_coding | 0.075915134 | 2.959908207 | 0.0187225 | 0.89284 | 0.978573709 |
| ENSMUSG00000029669 | Tspan12 | protein_coding | -0.042842914 | 6.187878731 | 0.01872 | 0.8928473 | 0.978573709 |
| ENSMUSG00000010066 | Cacna2d2 | protein_coding | -0.057645756 | 4.557096645 | 0.0186861 | 0.8929436 | 0.978579522 |
| ENSMUSG00000034109 | Golim4 | protein_coding | 0.126889769 | 2.425925254 | 0.0185608 | 0.8933007 | 0.978871154 |
| ENSMUSG00000029319 | Coq2 | protein_coding | 0.043228188 | 4.903824083 | 0.0184701 | 0.8935602 | 0.978971709 |
| ENSMUSG00000053646 | Plxnb1 | protein_coding | -0.116153343 | 3.351708276 | 0.0184981 | 0.8936464 | 0.978971709 |
| ENSMUSG00000047539 | Fbxo28 | protein_coding | -0.04466134 | 4.946111766 | 0.0184313 | 0.8936713 | 0.978971709 |
| ENSMUSG00000018446 | C1qbp | protein_coding | -0.045407382 | 5.541922467 | 0.0183727 | 0.8938392 | 0.978971709 |
| ENSMUSG00000031563 | Wwc2 | protein_coding | 0.053610509 | 4.512205655 | 0.0183353 | 0.8939468 | 0.978971709 |
| ENSMUSG00000056069 | Fam105a | protein_coding | -0.071722785 | 3.276289901 | 0.018284 | 0.8940943 | 0.978971709 |
| ENSMUSG00000020211 | Sf3a2 | protein_coding | 0.057959567 | 3.820183956 | 0.0182788 | 0.8941092 | 0.978971709 |
| ENSMUSG00000030265 | Kras | protein_coding | -0.043435881 | 5.529993642 | 0.0182748 | 0.8941208 | 0.978971709 |
| ENSMUSG00000036721 | Zscan12 | protein_coding | 0.067787343 | 3.671863815 | 0.0182365 | 0.8942309 | 0.978992592 |
| ENSMUSG00000030761 | Myo7a | protein_coding | -0.108554459 | 2.559450715 | 0.0181647 | 0.8944381 | 0.97911973 |
| ENSMUSG00000037197 | Rbm17 | protein_coding | 0.042739458 | 4.756584771 | 0.0180953 | 0.8946388 | 0.97917852 |
| ENSMUSG00000030677 | Kif22 | protein_coding | 0.060793919 | 3.465537622 | 0.0180757 | 0.8946953 | 0.97917852 |
| ENSMUSG00000002981 | Clptm1 | protein_coding | -0.035393093 | 5.646695765 | 0.0180308 | 0.8948255 | 0.97917852 |
| ENSMUSG00000080727 | C920021L13Rik | antisense_RNA | 0.090021473 | 2.533672648 | 0.0180202 | 0.8948561 | 0.97917852 |
| ENSMUSG00000059208 | Hnrnpm | protein_coding | 0.037516324 | 6.265211582 | 0.0179678 | 0.8950082 | 0.979245382 |

| ensembl_gene_id | external_gene_name | gene_biotype | log2 (fold change) | log2 (counts per million) | F-value | p-value | False discovery rate |
|---|---|---|---|---|---|---|---|
| ENSMUSG00000028194 | Ddah1 | protein_coding | -0.035356125 | 7.196137723 | 0.0178722 | 0.8952862 | 0.979420218 |
| ENSMUSG00000030062 | Rpn1 | protein_coding | -0.026830364 | 6.965638885 | 0.0178502 | 0.8953502 | 0.979420218 |
| ENSMUSG00000075232 | Amd1 | protein_coding | 0.048925371 | 3.991844883 | 0.0177999 | 0.8954967 | 0.979480806 |
| ENSMUSG00000032047 | Acat1 | protein_coding | 0.02851862 | 8.079841106 | 0.0177494 | 0.8956443 | 0.979542621 |
| ENSMUSG00000015335 | Zdhhc12 | protein_coding | -0.05822744 | 4.070934075 | 0.0177038 | 0.8957774 | 0.979588538 |
| ENSMUSG00000028645 | Slc2a1 | protein_coding | 0.071913092 | 3.207892489 | 0.0176704 | 0.8958751 | 0.979595754 |
| ENSMUSG00000002346 | Slc25a42 | protein_coding | 0.06318066 | 4.156457264 | 0.0176079 | 0.8960585 | 0.979660701 |
| ENSMUSG00000015843 | Rxrg | protein_coding | 0.030183025 | 6.702866231 | 0.017588 | 0.8961167 | 0.979660701 |
| ENSMUSG00000032485 | Scap | protein_coding | 0.050068335 | 4.527274056 | 0.0174745 | 0.8964504 | 0.979705335 |
| ENSMUSG00000028199 | Cryz | protein_coding | -0.049014632 | 5.182555888 | 0.0174399 | 0.8965523 | 0.979705335 |
| ENSMUSG00000032410 | Xrn1 | protein_coding | -0.057456405 | 4.021214016 | 0.0173911 | 0.8966964 | 0.979705335 |
| ENSMUSG00000038175 | Mylip | protein_coding | 0.074362656 | 3.224945135 | 0.0172964 | 0.8969761 | 0.979705335 |
| ENSMUSG00000041632 | Mrps27 | protein_coding | 0.047281891 | 5.571094879 | 0.0172794 | 0.8970265 | 0.979705335 |
| ENSMUSG00000002957 | Ap2a2 | protein_coding | 0.037435259 | 7.263747972 | 0.0172677 | 0.8970639 | 0.979705335 |
| ENSMUSG00000022860 | Chodl | protein_coding | 0.081651545 | 3.315417337 | 0.017247 | 0.8971226 | 0.979705335 |
| ENSMUSG00000028468 | Rgp1 | protein_coding | -0.069071384 | 3.156965295 | 0.0171993 | 0.8972642 | 0.979705335 |
| ENSMUSG00000040651 | Fam208a | protein_coding | 0.068258014 | 3.492858956 | 0.0171604 | 0.8973797 | 0.979705335 |
| ENSMUSG00000060882 | Kcnd2 | protein_coding | -0.029723863 | 6.631503232 | 0.0171567 | 0.8973906 | 0.979705335 |
| ENSMUSG00000026502 | Desi2 | protein_coding | 0.049653438 | 4.263983837 | 0.0171261 | 0.8974817 | 0.979705335 |
| ENSMUSG00000036918 | Ttc7 | protein_coding | -0.114248333 | 2.377443965 | 0.0171076 | 0.8975366 | 0.979705335 |
| ENSMUSG00000058740 | Kcnt1 | protein_coding | 0.057362955 | 3.781215717 | 0.017026 | 0.8977798 | 0.979705335 |
| ENSMUSG00000029016 | Clcn6 | protein_coding | -0.036197894 | 5.548644841 | 0.0170235 | 0.8977874 | 0.979705335 |
| ENSMUSG00000030008 | Pradc1 | protein_coding | -0.077211768 | 3.403218735 | 0.017002 | 0.8978513 | 0.979705335 |
| ENSMUSG00000047412 | Zbtb44 | protein_coding | -0.07005903 | 3.245284772 | 0.0169999 | 0.8978578 | 0.979705335 |
| ENSMUSG00000079111 | Kdelr2 | protein_coding | 0.041097874 | 6.432431385 | 0.016988 | 0.8978958 | 0.979705335 |
| ENSMUSG00000108030 | 9530062K07Rik | bidirectional_promote | -0.114113627 | 3.071294766 | 0.0169814 | 0.897999 | 0.979705335 |
| ENSMUSG00000024187 | Fam234a | protein_coding | -0.063738075 | 4.779985029 | 0.0169107 | 0.8981384 | 0.979705335 |
| ENSMUSG00000100252 | Mir124-2hg | lincRNA | -0.039364626 | 5.424679353 | 0.016863 | 0.8982675 | 0.979705335 |
| ENSMUSG00000032058 | Ppp2r1b | protein_coding | -0.048097184 | 4.65675267 | 0.0168581 | 0.898282 | 0.979705335 |
| ENSMUSG00000028617 | Lrrc42 | protein_coding | 0.041096212 | 4.923167692 | 0.0168453 | 0.8983204 | 0.979705335 |
| ENSMUSG00000053925 | Gm9929 | processed_transcript | 0.080869826 | 2.965741851 | 0.0168107 | 0.8984243 | 0.979705335 |
| ENSMUSG00000003269 | Cyth2 | protein_coding | -0.04609897 | 5.170669015 | 0.0167795 | 0.8985181 | 0.979705335 |
| ENSMUSG00000037221 | Mospd3 | protein_coding | -0.082499821 | 4.650157983 | 0.016825 | 0.8986038 | 0.979705335 |
| ENSMUSG00000025510 | Cd151 | protein_coding | 0.072529905 | 3.943163194 | 0.01675 | 0.8986093 | 0.979705335 |
| ENSMUSG00000025236 | Adpgk | protein_coding | -0.085997339 | 3.119365889 | 0.0167209 | 0.8986944 | 0.979705335 |
| ENSMUSG00000036450 | Hif1an | protein_coding | -0.055205717 | 4.210286783 | 0.0167162 | 0.8987086 | 0.979705335 |
| ENSMUSG00000020589 | Fam49a | protein_coding | 0.073067125 | 3.25155899 | 0.0166736 | 0.898837 | 0.979746011 |
| ENSMUSG00000009535 | Rnmt | protein_coding | 0.033333573 | 6.789872447 | 0.0166335 | 0.8989578 | 0.97977837 |
| ENSMUSG00000020694 | Tlk2 | protein_coding | -0.038381846 | 5.276387066 | 0.0165677 | 0.8991567 | 0.979844462 |
| ENSMUSG00000025193 | Cutc | protein_coding | -0.046069551 | 4.378400117 | 0.0165532 | 0.8992007 | 0.979844462 |
| ENSMUSG00000036646 | Man1b1 | protein_coding | -0.037897544 | 5.165876527 | 0.0163148 | 0.8999251 | 0.980400048 |
| ENSMUSG00000025911 | Adhfe1 | protein_coding | 0.05543283 | 3.611597344 | 0.0162995 | 0.8999718 | 0.980400048 |
| ENSMUSG00000050240 | Hic2 | protein_coding | -0.108744362 | 3.345082412 | 0.0162848 | 0.9001724 | 0.980400048 |
| ENSMUSG00000086851 | A430108G06Rik | processed_transcript | -0.161018508 | 2.553515066 | 0.0162469 | 0.9002879 | 0.980400048 |
| ENSMUSG00000035486 | Plk5 | protein_coding | 0.070117782 | 3.165917379 | 0.016142 | 0.9004533 | 0.980400048 |
| ENSMUSG00000003135 | Cnot11 | protein_coding | -0.067044831 | 2.972213039 | 0.0161217 | 0.9005156 | 0.980400048 |
| ENSMUSG00000040414 | Slc25a28 | protein_coding | 0.056727862 | 4.194077487 | 0.0161155 | 0.9005347 | 0.980400048 |
| ENSMUSG00000023088 | Abcc1 | protein_coding | -0.144691318 | 2.924059921 | 0.0161794 | 0.9005569 | 0.980400048 |
| ENSMUSG00000011096 | Akt1s1 | protein_coding | -0.083976718 | 2.861234585 | 0.0160271 | 0.9008062 | 0.980400048 |
| ENSMUSG00000012296 | Tjap1 | protein_coding | -0.103898435 | 2.745423782 | 0.016014 | 0.9008466 | 0.980400048 |
| ENSMUSG00000025340 | Rabgef1 | protein_coding | 0.036905274 | 5.840807769 | 0.0160045 | 0.9008758 | 0.980400048 |
| ENSMUSG00000032745 | Gpbp1 | protein_coding | -0.034598563 | 6.97775992 | 0.0159848 | 0.9009365 | 0.980400048 |
| ENSMUSG00000018765 | Fxr2 | protein_coding | 0.035016843 | 5.496278609 | 0.0159393 | 0.9010768 | 0.980400048 |
| ENSMUSG00000068290 | Ddrgk1 | protein_coding | 0.030990964 | 7.561920124 | 0.0159009 | 0.9011952 | 0.980400048 |
| ENSMUSG00000068749 | Psma5 | protein_coding | 0.025849896 | 6.83374403 | 0.0158814 | 0.9012557 | 0.980400048 |
| ENSMUSG00000073096 | Lrrc61 | protein_coding | 0.061566795 | 4.166503939 | 0.0158333 | 0.9014046 | 0.980400048 |
| ENSMUSG00000015474 | Ppt2 | protein_coding | -0.068321855 | 2.910200772 | 0.015818 | 0.9014519 | 0.980400048 |
| ENSMUSG00000054715 | Zscan22 | protein_coding | -0.061543803 | 4.282864331 | 0.0158165 | 0.9014589 | 0.980400048 |
| ENSMUSG00000036377 | C530008M17Rik | protein_coding | -0.04583281 | 4.964826786 | 0.0157458 | 0.9016758 | 0.980400048 |
| ENSMUSG00000022186 | Oxct1 | protein_coding | 0.026077373 | 8.154560547 | 0.0157392 | 0.9016962 | 0.980400048 |
| ENSMUSG00000037262 | Kin | protein_coding | -0.044657913 | 4.991138795 | 0.0156382 | 0.9020104 | 0.980400048 |
| ENSMUSG00000028207 | Asph | protein_coding | -0.039354261 | 5.442789133 | 0.015561 | 0.9022511 | 0.980400048 |
| ENSMUSG00000021044 | Adck1 | protein_coding | 0.044063482 | 5.485596723 | 0.0155355 | 0.902331 | 0.980400048 |
| ENSMUSG00000068134 | Zfp120 | protein_coding | 0.052947555 | 4.44365837 | 0.0155226 | 0.9023713 | 0.980400048 |
| ENSMUSG00000027274 | Mkks | protein_coding | 0.032858302 | 7.441326678 | 0.0155074 | 0.9024187 | 0.980400048 |
| ENSMUSG00000040557 | Mettl27 | protein_coding | -0.055013169 | 4.134411456 | 0.0154425 | 0.9026219 | 0.980400048 |
| ENSMUSG00000020986 | Sec23a | protein_coding | -0.063234727 | 4.69112239 | 0.0154525 | 0.9026414 | 0.980400048 |
| ENSMUSG00000074165 | Zfp788 | protein_coding | -0.049047704 | 4.507406213 | 0.0153951 | 0.9027709 | 0.980400048 |
| ENSMUSG00000071533 | Pcnp | protein_coding | 0.02585371 | 8.028674101 | 0.0153904 | 0.9027856 | 0.980400048 |
| ENSMUSG00000024773 | Atg2a | protein_coding | 0.067003678 | 3.524198162 | 0.0153729 | 0.9028405 | 0.980400048 |
| ENSMUSG00000029212 | Gabrb1 | protein_coding | -0.0317062 | 6.368171514 | 0.0153338 | 0.9029637 | 0.980400048 |
| ENSMUSG00000021418 | Rpp40 | protein_coding | 0.053197797 | 3.772606096 | 0.015333 | 0.9029661 | 0.980400048 |
| ENSMUSG00000048905 | 4930539E08Rik | protein_coding | -0.105324371 | 2.498641152 | 0.0153183 | 0.9030124 | 0.980400048 |
| ENSMUSG00000049922 | Slc35c1 | protein_coding | 0.054609877 | 4.063966979 | 0.0152931 | 0.9030918 | 0.980400048 |
| ENSMUSG00000027185 | Nat10 | protein_coding | -0.055276927 | 3.557744579 | 0.015269 | 0.9031678 | 0.980400048 |
| ENSMUSG00000054792 | Klhl18 | protein_coding | 0.043244748 | 4.768380638 | 0.0152612 | 0.9031922 | 0.980400048 |
| ENSMUSG00000051497 | Kcnj16 | protein_coding | -0.065906527 | 3.231320837 | 0.0152498 | 0.9032282 | 0.980400048 |
| ENSMUSG00000060402 | Chst8 | protein_coding | -0.077772687 | 3.486511991 | 0.0152008 | 0.9033854 | 0.980400048 |

| ensembl_gene_id | external_gene_name | gene_biotype | log2 (fold change) | log2 (counts per million) | F-value | p-value | False discovery rate |
|---|---|---|---|---|---|---|---|
| ENSMUSG00000053024 | Cntn2 | protein_coding | -0.038506063 | 5.471520605 | 0.0151728 | 0.9034715 | 0.980400048 |
| ENSMUSG00000024743 | Syt7 | protein_coding | -0.035506622 | 5.813303596 | 0.0151562 | 0.903524 | 0.980400048 |
| ENSMUSG00000029247 | Paics | protein_coding | -0.025780558 | 7.582638901 | 0.015147 | 0.9035531 | 0.980400048 |
| ENSMUSG00000022452 | Smdt1 | protein_coding | 0.029014233 | 8.365848453 | 0.015121 | 0.9036357 | 0.980400048 |
| ENSMUSG00000018428 | Akap1 | protein_coding | -0.064553122 | 3.861081418 | 0.0150566 | 0.9038401 | 0.980400048 |
| ENSMUSG00000079598 | Clec2l | protein_coding | 0.043219935 | 4.576287823 | 0.015048 | 0.9038673 | 0.980400048 |
| ENSMUSG00000028757 | Ddost | protein_coding | -0.044653627 | 5.732806411 | 0.0150033 | 0.9040118 | 0.980400048 |
| ENSMUSG00000014859 | E2f4 | protein_coding | 0.055548559 | 3.896729469 | 0.0149939 | 0.9040394 | 0.980400048 |
| ENSMUSG00000034486 | Gbx2 | protein_coding | 0.10548792 | 3.344052447 | 0.0150203 | 0.9041048 | 0.980400048 |
| ENSMUSG00000019777 | Hdac2 | protein_coding | -0.044914457 | 5.824625068 | 0.0149662 | 0.9041301 | 0.980400048 |
| ENSMUSG00000006699 | Cdc42 | protein_coding | 0.023874615 | 9.861276353 | 0.014937 | 0.9042206 | 0.980400048 |
| ENSMUSG00000029173 | Sepsecs | protein_coding | 0.085207585 | 3.513726781 | 0.0148612 | 0.904525 | 0.980400048 |
| ENSMUSG00000040724 | Kcna2 | protein_coding | -0.041648964 | 5.230897557 | 0.0147798 | 0.9047233 | 0.980400048 |
| ENSMUSG00000039069 | Mtg2 | protein_coding | -0.047022668 | 4.653762963 | 0.0147424 | 0.9048433 | 0.980400048 |
| ENSMUSG00000062646 | Ganc | protein_coding | -0.048924137 | 3.615466265 | 0.0147174 | 0.9049237 | 0.980400048 |
| ENSMUSG00000040097 | Flywch1 | protein_coding | -0.079230059 | 7.261020424 | 0.014992 | 0.9049244 | 0.980400048 |
| ENSMUSG00000020402 | Vdac1 | protein_coding | -0.0257843 | 9.386725854 | 0.0146548 | 0.9051251 | 0.980400048 |
| ENSMUSG00000020721 | Helz | protein_coding | -0.032918037 | 5.362374502 | 0.0146379 | 0.9051794 | 0.980400048 |
| ENSMUSG00000041712 | Ubr7 | protein_coding | -0.044664581 | 4.846411082 | 0.0146257 | 0.9052186 | 0.980400048 |
| ENSMUSG00000017167 | Cntnap1 | protein_coding | -0.044517021 | 4.798864447 | 0.0145902 | 0.9053334 | 0.980400048 |
| ENSMUSG00000035284 | Vps13c | protein_coding | -0.045286498 | 4.709763561 | 0.0145412 | 0.9054914 | 0.980400048 |
| ENSMUSG00000019854 | Reps1 | protein_coding | 0.064616772 | 4.639470301 | 0.0145271 | 0.9056308 | 0.980400048 |
| ENSMUSG00000031711 | Zfp330 | protein_coding | 0.02835719 | 5.953053569 | 0.0144926 | 0.9056486 | 0.980400048 |
| ENSMUSG00000025894 | Aasdhppt | protein_coding | 0.030516188 | 6.465648914 | 0.0144788 | 0.9056934 | 0.980400048 |
| ENSMUSG00000014599 | Csf1 | protein_coding | -0.06813178 | 2.841161253 | 0.0144231 | 0.9058741 | 0.980400048 |
| ENSMUSG00000096995 | 2810029C07Rik | bidirectional_promote | 0.053896684 | 3.791981806 | 0.0143707 | 0.9060443 | 0.980400048 |
| ENSMUSG00000023909 | Paqr4 | protein_coding | -0.049808926 | 4.688858335 | 0.0143529 | 0.9061023 | 0.980400048 |
| ENSMUSG00000034575 | Papd7 | protein_coding | 0.040832707 | 4.694228269 | 0.0143421 | 0.9061375 | 0.980400048 |
| ENSMUSG00000005267 | Zfp287 | protein_coding | -0.059925714 | 3.255660461 | 0.014316 | 0.9062224 | 0.980400048 |
| ENSMUSG00000086382 | Chrna1os | antisense_RNA | -0.077713158 | 2.764194342 | 0.0143062 | 0.9062545 | 0.980400048 |
| ENSMUSG00000001098 | Kctd10 | protein_coding | 0.074491076 | 3.615844524 | 0.01423 | 0.9065056 | 0.980400048 |
| ENSMUSG00000036817 | Sun1 | protein_coding | 0.048958654 | 5.080491014 | 0.0142035 | 0.9065923 | 0.980400048 |
| ENSMUSG00000038622 | Med30 | protein_coding | -0.057570546 | 3.868644546 | 0.0142017 | 0.9065956 | 0.980400048 |
| ENSMUSG00000000134 | Tfe3 | protein_coding | -0.042878507 | 4.709438247 | 0.0141957 | 0.9066152 | 0.980400048 |
| ENSMUSG00000001151 | Pcnt | protein_coding | 0.073750933 | 3.436726609 | 0.0141052 | 0.9069145 | 0.980400048 |
| ENSMUSG00000071324 | Armc2 | protein_coding | 0.086917021 | 2.970125851 | 0.0141013 | 0.9069249 | 0.980400048 |
| ENSMUSG00000056770 | Setd3 | protein_coding | -0.026853761 | 6.51482753 | 0.014095 | 0.9069455 | 0.980400048 |
| ENSMUSG00000040033 | Stat2 | protein_coding | 0.092278037 | 2.603208494 | 0.0140703 | 0.9070265 | 0.980400048 |
| ENSMUSG00000034066 | Farp2 | protein_coding | 0.071139864 | 2.859174941 | 0.0140527 | 0.9070845 | 0.980400048 |
| ENSMUSG00000034354 | Mtmr3 | protein_coding | -0.044310805 | 5.133218822 | 0.0140491 | 0.9070964 | 0.980400048 |
| ENSMUSG00000031858 | Mau2 | protein_coding | -0.042500032 | 5.019563387 | 0.014042 | 0.9071197 | 0.980400048 |
| ENSMUSG00000020868 | Xylt2 | protein_coding | 0.050972456 | 3.705169652 | 0.0140141 | 0.9072116 | 0.980400048 |
| ENSMUSG00000028849 | Map7d1 | protein_coding | -0.027869848 | 6.582567633 | 0.013987 | 0.907301 | 0.980400048 |
| ENSMUSG00000046020 | Pofut1 | protein_coding | 0.079850198 | 2.626413497 | 0.0139663 | 0.9073691 | 0.980400048 |
| ENSMUSG00000030307 | Slc6a11 | protein_coding | -0.039003137 | 6.060838409 | 0.0139666 | 0.9073705 | 0.980400048 |
| ENSMUSG00000025465 | Echs1 | protein_coding | 0.023022962 | 7.770651461 | 0.01395 | 0.9074231 | 0.980400048 |
| ENSMUSG00000007589 | Tinf2 | protein_coding | -0.057163862 | 3.47534983 | 0.0138636 | 0.9077086 | 0.980400048 |
| ENSMUSG00000034194 | R3hcc1 | protein_coding | -0.035631384 | 5.675439554 | 0.0138534 | 0.9077426 | 0.980400048 |
| ENSMUSG00000023050 | Map3k12 | protein_coding | -0.052078676 | 4.076085082 | 0.0138381 | 0.9077933 | 0.980400048 |
| ENSMUSG00000042595 | Fam199x | protein_coding | -0.05775261 | 3.526595474 | 0.0138162 | 0.9078657 | 0.980400048 |
| ENSMUSG00000000631 | Myo18a | protein_coding | 0.037284238 | 5.756674525 | 0.0138061 | 0.9078994 | 0.980400048 |
| ENSMUSG00000029120 | Ppp2r2c | protein_coding | -0.024287148 | 7.304229255 | 0.0137491 | 0.9080887 | 0.980400048 |
| ENSMUSG00000024050 | Wiz | protein_coding | 0.056161733 | 3.622251832 | 0.0137376 | 0.908127 | 0.980400048 |
| ENSMUSG00000031386 | Hcfc1 | protein_coding | -0.047613464 | 5.197956462 | 0.0136548 | 0.9084052 | 0.980400048 |
| ENSMUSG00000028329 | Xpa | protein_coding | 0.051820618 | 3.471350559 | 0.0136187 | 0.9085234 | 0.980400048 |
| ENSMUSG00000038773 | Kdm3b | protein_coding | -0.042902767 | 3.911557916 | 0.0136008 | 0.9085836 | 0.980400048 |
| ENSMUSG00000025260 | Hsd17b10 | protein_coding | 0.024078619 | 7.110727815 | 0.0135964 | 0.9085981 | 0.980400048 |
| ENSMUSG00000033029 | 1700088E04Rik | protein_coding | 0.050581823 | 3.823155226 | 0.0135955 | 0.9086012 | 0.980400048 |
| ENSMUSG00000054414 | Slc30a7 | protein_coding | 0.068081273 | 3.354132197 | 0.0135885 | 0.9086245 | 0.980400048 |
| ENSMUSG00000017843 | Ppp2r5c | protein_coding | -0.026854926 | 7.854451144 | 0.0135347 | 0.9088047 | 0.980400048 |
| ENSMUSG00000097350 | 4732491K20Rik | lincRNA | -0.038942636 | 4.549704638 | 0.0134869 | 0.908965 | 0.980400048 |
| ENSMUSG00000020097 | Sgpl1 | protein_coding | -0.068116492 | 3.309907319 | 0.0134596 | 0.9090571 | 0.980400048 |
| ENSMUSG00000036968 | Cnpy4 | protein_coding | -0.039862673 | 5.693023427 | 0.0134586 | 0.9090602 | 0.980400048 |
| ENSMUSG00000026421 | Csrp1 | protein_coding | -0.025922851 | 8.291073707 | 0.0134404 | 0.9091216 | 0.980400048 |
| ENSMUSG00000032727 | Mier3 | protein_coding | 0.033611849 | 5.296214816 | 0.0134296 | 0.909158 | 0.980400048 |
| ENSMUSG00000039450 | Dcxr | protein_coding | 0.051767382 | 4.156763777 | 0.0134192 | 0.9091927 | 0.980400048 |
| ENSMUSG00000031875 | Cmtm3 | protein_coding | -0.059341871 | 3.327982326 | 0.0133644 | 0.9093776 | 0.980501116 |
| ENSMUSG00000046010 | Zfp830 | protein_coding | 0.04114099 | 4.747410748 | 0.0133301 | 0.9094935 | 0.98052776 |
| ENSMUSG00000038005 | Hpf1 | protein_coding | -0.027448831 | 5.994374637 | 0.0132927 | 0.9096198 | 0.980547408 |
| ENSMUSG00000031374 | Zfp92 | protein_coding | 0.076687131 | 2.641578821 | 0.0132402 | 0.9097975 | 0.980547408 |
| ENSMUSG00000036825 | Ssx2ip | protein_coding | -0.031721287 | 5.533175361 | 0.0132191 | 0.9098693 | 0.980547408 |
| ENSMUSG00000056598 | Drc3 | protein_coding | 0.052246508 | 3.995195366 | 0.013217 | 0.9098765 | 0.980547408 |
| ENSMUSG00000021684 | Pde8b | protein_coding | -0.040053132 | 5.163452221 | 0.0131153 | 0.9102223 | 0.980781408 |
| ENSMUSG00000018239 | Zcchc10 | protein_coding | -0.032853324 | 5.560305535 | 0.0130995 | 0.9102761 | 0.980781408 |
| ENSMUSG00000087620 | 5330434G04Rik | lincRNA | 0.048666636 | 4.666725903 | 0.0130629 | 0.9104007 | 0.980817441 |
| ENSMUSG00000097560 | Gm26904 | lincRNA | 0.067691103 | 3.585983041 | 0.0129846 | 0.9106709 | 0.980908327 |
| ENSMUSG00000038733 | Wdr26 | protein_coding | -0.037448166 | 6.151534961 | 0.0129632 | 0.9107441 | 0.980908327 |
| ENSMUSG00000000560 | Gabra2 | protein_coding | -0.051797507 | 6.153070811 | 0.0130153 | 0.9107587 | 0.980908327 |

| ensembl_gene_id | external_gene_name | gene_biotype | log2 (fold change) | log2 (counts per million) | F-value | p-value | False discovery rate |
|---|---|---|---|---|---|---|---|
| ENSMUSG00000052962 | Mrpl35 | protein_coding | -0.021937621 | 7.399186012 | 0.0129172 | 0.9108996 | 0.980961804 |
| ENSMUSG00000043190 | Rfesd | protein_coding | -0.034230411 | 5.511361375 | 0.012828 | 0.9112064 | 0.981093236 |
| ENSMUSG00000022610 | Mapk12 | protein_coding | 0.102752825 | 2.362859331 | 0.0128202 | 0.9112333 | 0.981093236 |
| ENSMUSG00000033998 | Kcnk1 | protein_coding | -0.060675894 | 3.67952757 | 0.0127697 | 0.9114075 | 0.981093236 |
| ENSMUSG00000029461 | Fam168a | protein_coding | -0.030654187 | 7.503392389 | 0.0127663 | 0.9114193 | 0.981093236 |
| ENSMUSG00000027340 | Slc23a2 | protein_coding | 0.038438459 | 5.141563334 | 0.0127493 | 0.9114779 | 0.981093236 |
| ENSMUSG00000027597 | Ahcy | protein_coding | -0.048388757 | 3.452462373 | 0.0126695 | 0.9117542 | 0.981207739 |
| ENSMUSG00000033285 | Wdr3 | protein_coding | 0.103589013 | 2.434928954 | 0.0126222 | 0.9119184 | 0.981207739 |
| ENSMUSG00000020131 | Pcsk4 | protein_coding | 0.06288077 | 3.616326002 | 0.0126167 | 0.9119374 | 0.981207739 |
| ENSMUSG00000020516 | Rps6kb1 | protein_coding | -0.038937363 | 5.715167585 | 0.0125839 | 0.9120516 | 0.981207739 |
| ENSMUSG00000029486 | Mrpl1 | protein_coding | 0.033540344 | 5.919402217 | 0.0125693 | 0.9121023 | 0.981207739 |
| ENSMUSG00000031391 | L1cam | protein_coding | -0.029913098 | 5.602604218 | 0.0125609 | 0.9121317 | 0.981207739 |
| ENSMUSG00000040383 | Aqr | protein_coding | 0.043194914 | 4.519892556 | 0.0124713 | 0.912444 | 0.981248504 |
| ENSMUSG00000038542 | Pcid2 | protein_coding | 0.063998686 | 3.686813196 | 0.0124332 | 0.9125798 | 0.981248504 |
| ENSMUSG00000055239 | Kcmf1 | protein_coding | -0.034794344 | 5.772593006 | 0.0124009 | 0.9126906 | 0.981248504 |
| ENSMUSG00000063445 | Nmral1 | protein_coding | 0.027337189 | 6.160852941 | 0.0123922 | 0.9127212 | 0.981248504 |
| ENSMUSG00000004897 | Hdgf | protein_coding | 0.022748286 | 7.34081843 | 0.0123843 | 0.9127486 | 0.981248504 |
| ENSMUSG00000042331 | Specc1 | protein_coding | 0.042715097 | 5.358759164 | 0.0123721 | 0.9127939 | 0.981248504 |
| ENSMUSG00000031029 | Eif3f | protein_coding | 0.030556035 | 6.454889722 | 0.0123412 | 0.9129002 | 0.981248504 |
| ENSMUSG00000020883 | Fbxl20 | protein_coding | -0.04657811 | 3.978598993 | 0.0123223 | 0.9129664 | 0.981248504 |
| ENSMUSG00000034312 | Iqsec1 | protein_coding | 0.036323241 | 5.851191908 | 0.0123153 | 0.9129909 | 0.981248504 |
| ENSMUSG00000074170 | Plekhf1 | protein_coding | 0.060224246 | 3.88026458 | 0.0122654 | 0.9131689 | 0.981261862 |
| ENSMUSG00000044030 | Irf2bp1 | protein_coding | 0.03957509 | 4.346356885 | 0.01226 | 0.9131859 | 0.981261862 |
| ENSMUSG00000039354 | Smarcal1 | protein_coding | 0.050389362 | 4.604776588 | 0.0121618 | 0.913535 | 0.981538943 |
| ENSMUSG00000035104 | Eva1a | protein_coding | 0.071840354 | 3.812427907 | 0.0121186 | 0.9137661 | 0.981689159 |
| ENSMUSG00000048349 | Pou4f1 | protein_coding | 0.273257064 | 1.427937485 | 0.0120798 | 0.9140765 | 0.981690165 |
| ENSMUSG00000032423 | Syncrip | protein_coding | -0.051511856 | 3.30915469 | 0.0119978 | 0.914115 | 0.981690165 |
| ENSMUSG00000027983 | Cyp2u1 | protein_coding | -0.044763543 | 4.690115659 | 0.0119738 | 0.9142009 | 0.981690165 |
| ENSMUSG00000020089 | Ppa1 | protein_coding | -0.027650166 | 7.898838711 | 0.0119519 | 0.9142789 | 0.981690165 |
| ENSMUSG00000031197 | Vbp1 | protein_coding | -0.022261521 | 7.867598489 | 0.0119061 | 0.9144426 | 0.981690165 |
| ENSMUSG00000026999 | Nup35 | protein_coding | 0.039221689 | 4.37657614 | 0.0118926 | 0.9144911 | 0.981690165 |
| ENSMUSG00000033350 | Chst2 | protein_coding | 0.022619528 | 6.92829808 | 0.0118828 | 0.9145259 | 0.981690165 |
| ENSMUSG00000026575 | Nme7 | protein_coding | -0.042432798 | 4.734404409 | 0.0118685 | 0.9145774 | 0.981690165 |
| ENSMUSG00000027569 | Mrgbp | protein_coding | -0.064462222 | 3.122264797 | 0.0118653 | 0.9145887 | 0.981690165 |
| ENSMUSG00000021816 | Ppp3cb | protein_coding | 0.024736201 | 7.762420812 | 0.0117875 | 0.9148682 | 0.981728254 |
| ENSMUSG00000097164 | Cep83os | bidirectional_promote | 0.033752103 | 5.223949307 | 0.011781 | 0.9148913 | 0.981728254 |
| ENSMUSG00000021532 | Fastkd3 | protein_coding | -0.037313339 | 5.034490308 | 0.0117792 | 0.9148981 | 0.981728254 |
| ENSMUSG00000022205 | Sub1 | protein_coding | -0.025801213 | 10.23803381 | 0.0117268 | 0.9150865 | 0.981832468 |
| ENSMUSG00000059040 | Eno1b | protein_coding | 0.147780937 | 2.21628682 | 0.0117089 | 0.9152354 | 0.981864701 |
| ENSMUSG00000025509 | Pnpla2 | protein_coding | 0.049617033 | 5.14181246 | 0.0116209 | 0.9155103 | 0.981864701 |
| ENSMUSG00000097311 | Gm26871 | processed_transcript | -0.089869934 | 2.800488916 | 0.0115854 | 0.9155981 | 0.981864701 |
| ENSMUSG00000024516 | Sec11c | protein_coding | -0.030503982 | 7.690142537 | 0.0115672 | 0.9156662 | 0.981864701 |
| ENSMUSG00000023915 | Tnfrsf21 | protein_coding | -0.030814817 | 5.548326239 | 0.011564 | 0.9156755 | 0.981864701 |
| ENSMUSG00000039477 | Tnrc18 | protein_coding | -0.047750031 | 3.305940623 | 0.0115493 | 0.9157291 | 0.981864701 |
| ENSMUSG00000059401 | Mamld1 | protein_coding | -0.04218727 | 4.822807118 | 0.011542 | 0.9157557 | 0.981864701 |
| ENSMUSG00000054648 | Zfp869 | protein_coding | 0.054545564 | 3.365947779 | 0.0114801 | 0.9159807 | 0.982008058 |
| ENSMUSG00000023345 | Poc1a | protein_coding | -0.047829367 | 4.073827309 | 0.0114156 | 0.9162164 | 0.982162793 |
| ENSMUSG00000037503 | Fam168b | protein_coding | -0.024655735 | 6.432525751 | 0.011389 | 0.9163137 | 0.982169176 |
| ENSMUSG00000020086 | H2afy2 | protein_coding | 0.083901938 | 2.575478223 | 0.0113628 | 0.9164095 | 0.982174008 |
| ENSMUSG00000023963 | Cyp39a1 | protein_coding | -0.121946665 | 2.887792642 | 0.0113461 | 0.9166528 | 0.982241827 |
| ENSMUSG00000046341 | Gm11223 | processed_pseudoge | 0.051433592 | 3.20080586 | 0.0112894 | 0.9166787 | 0.982241827 |
| ENSMUSG00000014177 | Tvp23b | protein_coding | -0.02520259 | 6.339747396 | 0.011269 | 0.9167538 | 0.982241827 |
| ENSMUSG00000078578 | Ube2d3 | protein_coding | -0.021177768 | 7.32672066 | 0.0112263 | 0.9169111 | 0.982241827 |
| ENSMUSG00000035179 | Ppp1r32 | protein_coding | 0.058006016 | 3.234266445 | 0.0112213 | 0.9169296 | 0.982241827 |
| ENSMUSG00000032046 | Abhd12 | protein_coding | 0.023750639 | 6.967606994 | 0.0110969 | 0.9173896 | 0.982561089 |
| ENSMUSG00000060424 | Pantr1 | lincRNA | 0.050129152 | 7.07974581 | 0.0111786 | 0.9174486 | 0.982561089 |
| ENSMUSG00000031201 | Brcc3 | protein_coding | 0.025903397 | 7.292192666 | 0.0110289 | 0.9176418 | 0.982561089 |
| ENSMUSG00000035045 | Zc3h12b | protein_coding | -0.03742448 | 4.20635853 | 0.0110213 | 0.9176702 | 0.982561089 |
| ENSMUSG00000020900 | Myh10 | protein_coding | -0.030247263 | 6.443934694 | 0.0110124 | 0.9177032 | 0.982561089 |
| ENSMUSG00000017221 | Psmd3 | protein_coding | 0.031728285 | 5.503592171 | 0.0109929 | 0.9177758 | 0.982561089 |
| ENSMUSG00000022914 | Brwd1 | protein_coding | -0.025510818 | 6.174824678 | 0.0108124 | 0.9184511 | 0.982656896 |
| ENSMUSG00000025224 | Gbf1 | protein_coding | -0.032916664 | 5.734216641 | 0.0107464 | 0.9186994 | 0.982656896 |
| ENSMUSG00000009070 | Rsph14 | protein_coding | -0.059219596 | 3.214988781 | 0.010713 | 0.9188256 | 0.982656896 |
| ENSMUSG00000022724 | Riox2 | protein_coding | 0.056442507 | 3.831922771 | 0.0107122 | 0.9188306 | 0.982656896 |
| ENSMUSG00000030814 | Bcl7c | protein_coding | -0.068990238 | 2.647911309 | 0.010699 | 0.9188785 | 0.982656896 |
| ENSMUSG00000034612 | Chst11 | protein_coding | 0.039199473 | 3.991969294 | 0.0106915 | 0.9189067 | 0.982656896 |
| ENSMUSG00000020297 | Nsg2 | protein_coding | 0.020577409 | 9.103249512 | 0.0106753 | 0.9189677 | 0.982656896 |
| ENSMUSG00000029560 | Snx8 | protein_coding | 0.049385217 | 4.573281168 | 0.0106708 | 0.9189872 | 0.982656896 |
| ENSMUSG00000029125 | Stx18 | protein_coding | -0.030448248 | 5.479894927 | 0.0106654 | 0.9190054 | 0.982656896 |
| ENSMUSG00000054920 | Klhl5 | protein_coding | -0.035151159 | 4.89914094 | 0.0106635 | 0.9190124 | 0.982656896 |
| ENSMUSG00000035726 | Supt16 | protein_coding | -0.033522523 | 5.624381339 | 0.010655 | 0.9190446 | 0.982656896 |
| ENSMUSG00000031959 | Wdr59 | protein_coding | -0.045005585 | 3.81467286 | 0.0106159 | 0.9191929 | 0.982656896 |
| ENSMUSG00000057147 | Dph6 | protein_coding | 0.029600164 | 5.277548766 | 0.0106061 | 0.91923 | 0.982656896 |
| ENSMUSG00000038855 | Itpkb | protein_coding | -0.043539845 | 4.182160553 | 0.0105703 | 0.9193659 | 0.982656896 |
| ENSMUSG00000073062 | Zxdb | protein_coding | -0.056441264 | 3.790263018 | 0.0105587 | 0.9194122 | 0.982656896 |
| ENSMUSG00000018583 | G3bp1 | protein_coding | 0.026251409 | 6.057077749 | 0.0105254 | 0.9195366 | 0.982656896 |
| ENSMUSG00000034042 | Gpbp1l1 | protein_coding | -0.046429881 | 3.696995942 | 0.010524 | 0.919542 | 0.982656896 |
| ENSMUSG00000018395 | Kif3a | protein_coding | -0.02889552 | 7.687993734 | 0.0104942 | 0.9196576 | 0.982656896 |

| ensembl_gene_id | external_gene_name | gene_biotype | log2 (fold change) | log2 (counts per million) | F-value | p-value | False discovery rate |
|---|---|---|---|---|---|---|---|
| ENSMUSG00000010067 | Rassf1 | protein_coding | -0.036634512 | 4.843780091 | 0.0104823 | 0.919701 | 0.982656896 |
| ENSMUSG00000032187 | Smarca4 | protein_coding | -0.033087392 | 5.949081 | 0.0104734 | 0.9197348 | 0.982656896 |
| ENSMUSG00000049904 | Tmem17 | protein_coding | -0.038595558 | 3.952169071 | 0.0104605 | 0.9197844 | 0.982656896 |
| ENSMUSG00000000489 | Pdgfb | protein_coding | -0.053943466 | 3.367482814 | 0.0104227 | 0.9199289 | 0.982693817 |
| ENSMUSG00000059796 | Eif4a1 | protein_coding | -0.022996204 | 6.153011089 | 0.0103747 | 0.9201125 | 0.982693817 |
| ENSMUSG00000022802 | Lmln | protein_coding | -0.082760299 | 2.455071012 | 0.0103611 | 0.9201647 | 0.982693817 |
| ENSMUSG00000057897 | Camk2b | protein_coding | 0.022538667 | 8.455445347 | 0.0103508 | 0.9202044 | 0.982693817 |
| ENSMUSG00000021501 | Caml | protein_coding | 0.026122267 | 6.034386963 | 0.0103322 | 0.9202759 | 0.982693817 |
| ENSMUSG00000027300 | Ubox5 | protein_coding | 0.036574946 | 4.192564187 | 0.0102672 | 0.9205259 | 0.982752511 |
| ENSMUSG00000039191 | Rbpj | protein_coding | 0.050719937 | 3.246021187 | 0.0102612 | 0.9205493 | 0.982752511 |
| ENSMUSG00000042680 | Garem1 | protein_coding | -0.048620167 | 3.877893025 | 0.0102297 | 0.920671 | 0.982752511 |
| ENSMUSG00000099034 | 2810039B14Rik | transcribed_unitary_p | 0.051859123 | 3.229645535 | 0.0102231 | 0.9206964 | 0.982752511 |
| ENSMUSG00000087259 | 2610035D17Rik | lincRNA | -0.029760356 | 5.366582611 | 0.0101823 | 0.9208544 | 0.982766021 |
| ENSMUSG00000034308 | Sdr42e1 | protein_coding | -0.06556202 | 3.166429921 | 0.0101677 | 0.9209107 | 0.982766021 |
| ENSMUSG00000031167 | Rbm3 | protein_coding | 0.035679374 | 4.087314132 | 0.0101075 | 0.9211444 | 0.982766021 |
| ENSMUSG00000026005 | Rpe | protein_coding | -0.044769767 | 4.424036185 | 0.0100802 | 0.9212507 | 0.982766021 |
| ENSMUSG00000039163 | Cmc1 | protein_coding | 0.038799007 | 4.134331646 | 0.0100751 | 0.9212705 | 0.982766021 |
| ENSMUSG00000062778 | Chia1 | protein_coding | -0.110412037 | 2.152560485 | 0.0100674 | 0.9213005 | 0.982766021 |
| ENSMUSG00000068015 | Lrch1 | protein_coding | 0.065089378 | 3.917521528 | 0.010075 | 0.9213489 | 0.982766021 |
| ENSMUSG00000004473 | Clec11a | protein_coding | 0.047087001 | 3.374650093 | 0.0100259 | 0.9214623 | 0.982767659 |
| ENSMUSG00000002043 | Trappc6a | protein_coding | -0.031985562 | 4.975157349 | 0.0100077 | 0.9215332 | 0.982767659 |
| ENSMUSG00000019916 | P4ha1 | protein_coding | -0.039177469 | 4.435432049 | 0.009916 | 0.9218923 | 0.982880383 |
| ENSMUSG00000049092 | Gpr137c | protein_coding | -0.0672416 | 2.982284169 | 0.0098766 | 0.9220473 | 0.982880383 |
| ENSMUSG00000057666 | Gapdh | protein_coding | -0.038266928 | 4.429047748 | 0.0098725 | 0.9220633 | 0.982880383 |
| ENSMUSG00000061755 | Bod1l | protein_coding | 0.035403336 | 5.3902274 | 0.0098724 | 0.9220635 | 0.982880383 |
| ENSMUSG00000029106 | Add1 | protein_coding | -0.020816639 | 7.534351231 | 0.0098411 | 0.9221868 | 0.982880383 |
| ENSMUSG00000007653 | Gabrb2 | protein_coding | 0.049292573 | 4.267264857 | 0.0098398 | 0.9221939 | 0.982880383 |
| ENSMUSG00000051652 | Lrrc3 | protein_coding | 0.084907384 | 3.470693892 | 0.0098609 | 0.9222787 | 0.982880383 |
| ENSMUSG00000039308 | Ndst2 | protein_coding | -0.086909277 | 2.597608436 | 0.0096958 | 0.9227615 | 0.983297457 |
| ENSMUSG00000025917 | Cops5 | protein_coding | 0.022394819 | 7.789224821 | 0.0096693 | 0.9228669 | 0.98331128 |
| ENSMUSG00000035049 | Rrp12 | protein_coding | -0.03666295 | 4.313849598 | 0.0096133 | 0.9230896 | 0.98331128 |
| ENSMUSG00000071359 | Tbpl1 | protein_coding | -0.027119088 | 6.377242744 | 0.0095992 | 0.923146 | 0.98331128 |
| ENSMUSG00000031161 | Hdac6 | protein_coding | 0.0390232 | 4.879867729 | 0.0095408 | 0.9233793 | 0.98331128 |
| ENSMUSG00000055296 | Tmem245 | protein_coding | 0.040756497 | 4.413912717 | 0.0095333 | 0.9234091 | 0.98331128 |
| ENSMUSG00000016526 | Dyrk3 | protein_coding | -0.086839407 | 2.25077061 | 0.0095048 | 0.9235235 | 0.98331128 |
| ENSMUSG00000035713 | Usp35 | protein_coding | 0.052106471 | 3.237843402 | 0.0094812 | 0.9236183 | 0.98331128 |
| ENSMUSG00000063172 | Hspb11 | protein_coding | -0.026076027 | 5.895855498 | 0.0094531 | 0.9237309 | 0.98331128 |
| ENSMUSG00000098087 | Gm17750 | lincRNA | -0.049120286 | 3.3932452 | 0.0094433 | 0.9237706 | 0.98331128 |
| ENSMUSG00000063714 | Sp3os | antisense_RNA | -0.046717837 | 3.530542364 | 0.0094309 | 0.9238205 | 0.98331128 |
| ENSMUSG00000035944 | Ttc38 | protein_coding | 0.049402033 | 3.406208913 | 0.0094265 | 0.923838 | 0.98331128 |
| ENSMUSG00000003810 | Mast2 | protein_coding | 0.027649721 | 5.41546014 | 0.0094181 | 0.9238718 | 0.98331128 |
| ENSMUSG00000028801 | Stpg1 | protein_coding | 0.056659735 | 3.122918865 | 0.0093114 | 0.9243029 | 0.983336695 |
| ENSMUSG00000066113 | Adamtsl1 | protein_coding | -0.084630942 | 3.038575232 | 0.0092964 | 0.924413 | 0.983336695 |
| ENSMUSG00000027195 | Hsd17b12 | protein_coding | -0.019377825 | 7.438420096 | 0.0092723 | 0.9244613 | 0.983336695 |
| ENSMUSG00000015759 | Cnih1 | protein_coding | 0.020901618 | 7.541103114 | 0.0092639 | 0.9244955 | 0.983336695 |
| ENSMUSG00000037525 | Bcdin3d | protein_coding | -0.050187684 | 4.013761634 | 0.009214 | 0.9247006 | 0.983336695 |
| ENSMUSG00000041272 | Tox | protein_coding | -0.063495246 | 6.058983612 | 0.0093331 | 0.9247872 | 0.983336695 |
| ENSMUSG00000110277 | Gm45871 | protein_coding | 0.054848585 | 3.264618075 | 0.0091729 | 0.9248662 | 0.983336695 |
| ENSMUSG00000025786 | Zdhhc3 | protein_coding | -0.037417932 | 5.349899634 | 0.0091727 | 0.9248688 | 0.983336695 |
| ENSMUSG00000024940 | Ltbp3 | protein_coding | 0.064716246 | 2.779765272 | 0.009167 | 0.9248902 | 0.983336695 |
| ENSMUSG00000038074 | Fkbp14 | protein_coding | -0.04825499 | 3.382830478 | 0.0091362 | 0.9250163 | 0.983336695 |
| ENSMUSG00000086040 | Wipf3 | protein_coding | -0.066080084 | 2.898000175 | 0.009127 | 0.9250536 | 0.983336695 |
| ENSMUSG00000034601 | 2700049A03Rik | protein_coding | -0.075540545 | 2.973556882 | 0.0091186 | 0.92509 | 0.983336695 |
| ENSMUSG00000020629 | Adi1 | protein_coding | 0.026284754 | 6.30891027 | 0.0090712 | 0.9252824 | 0.983336695 |
| ENSMUSG00000046637 | Ttc34 | protein_coding | -0.086108779 | 2.504701623 | 0.0090497 | 0.9253708 | 0.983336695 |
| ENSMUSG00000036992 | Nxt1 | protein_coding | -0.03182672 | 4.667078708 | 0.009049 | 0.9253738 | 0.983336695 |
| ENSMUSG00000025277 | Abhd6 | protein_coding | 0.02499925 | 6.217601907 | 0.0089691 | 0.9257028 | 0.983336695 |
| ENSMUSG00000097219 | Gm26551 | lincRNA | 0.044389461 | 3.711355246 | 0.0089561 | 0.9257566 | 0.983336695 |
| ENSMUSG00000016206 | H2-M3 | protein_coding | 0.060135054 | 3.008335465 | 0.0089436 | 0.9258083 | 0.983336695 |
| ENSMUSG00000024534 | Sncaip | protein_coding | 0.077952485 | 2.571521674 | 0.0088911 | 0.9260255 | 0.983336695 |
| ENSMUSG00000022122 | Ednrb | protein_coding | -0.029226726 | 6.1008618 | 0.0088786 | 0.9260776 | 0.983336695 |
| ENSMUSG00000026930 | Gpsm1 | protein_coding | 0.026207484 | 6.004158475 | 0.0088741 | 0.9260959 | 0.983336695 |
| ENSMUSG00000041930 | Fam222a | protein_coding | -0.086218631 | 2.58983844 | 0.0088544 | 0.9261777 | 0.983336695 |
| ENSMUSG00000015087 | Rabl6 | protein_coding | 0.021397896 | 6.476375965 | 0.0088288 | 0.9262842 | 0.983336695 |
| ENSMUSG00000037242 | Clic4 | protein_coding | -0.033495587 | 5.37808778 | 0.0088255 | 0.9262981 | 0.983336695 |
| ENSMUSG00000096433 | Zfp994 | protein_coding | 0.07694433 | 2.366309007 | 0.0088103 | 0.9263614 | 0.983336695 |
| ENSMUSG00000042605 | Atxn2 | protein_coding | 0.02488695 | 5.778144979 | 0.0087967 | 0.9264182 | 0.983336695 |
| ENSMUSG00000069237 | Fam8a1 | protein_coding | -0.029054828 | 5.511208279 | 0.0087916 | 0.9264394 | 0.983336695 |
| ENSMUSG00000055116 | Arntl | protein_coding | -0.039743028 | 3.893684507 | 0.0087716 | 0.9265228 | 0.983336695 |
| ENSMUSG00000048550 | Thnsl1 | protein_coding | -0.031941234 | 5.112092337 | 0.0087534 | 0.926599 | 0.983336695 |
| ENSMUSG00000025198 | Erlin1 | protein_coding | -0.04499592 | 3.890880688 | 0.0087438 | 0.9266391 | 0.983336695 |
| ENSMUSG00000020844 | Nxn | protein_coding | -0.042029615 | 3.816429731 | 0.0086838 | 0.9268902 | 0.983460788 |
| ENSMUSG00000021737 | Psmd6 | protein_coding | 0.020333454 | 8.008013468 | 0.0086722 | 0.926939 | 0.983460788 |
| ENSMUSG00000068876 | Cgn | protein_coding | 0.048522406 | 3.525450988 | 0.0085908 | 0.9272815 | 0.983727099 |
| ENSMUSG00000020358 | Hnrnpab | protein_coding | 0.062360218 | 2.762757059 | 0.0085262 | 0.9275548 | 0.983884166 |
| ENSMUSG00000058351 | Smim4 | protein_coding | 0.034740336 | 5.621326812 | 0.008508 | 0.9276337 | 0.983884166 |
| ENSMUSG00000020706 | Ftsj3 | protein_coding | -0.03930933 | 4.824402061 | 0.0084914 | 0.927704 | 0.983884166 |
| ENSMUSG00000005698 | Ctcf | protein_coding | -0.025180125 | 6.059742147 | 0.0084594 | 0.9278383 | 0.983929502 |

| ensembl_gene_id | external_gene_name | gene_biotype | log2 (fold change) | log2 (counts per million) | F-value | p-value | False discovery rate |
|---|---|---|---|---|---|---|---|
| ENSMUSG00000029219 | Slc10a4 | protein_coding | 0.043115252 | 3.807945829 | 0.0084309 | 0.9279595 | 0.98396103 |
| ENSMUSG00000071654 | Uqcc3 | protein_coding | 0.024213773 | 6.871597521 | 0.0083614 | 0.9282562 | 0.984038771 |
| ENSMUSG00000047394 | Odf3b | protein_coding | 0.049868812 | 3.508796897 | 0.0083464 | 0.9283206 | 0.984038771 |
| ENSMUSG00000041669 | Prima1 | protein_coding | 0.075012803 | 2.634618313 | 0.0083141 | 0.928459 | 0.984038771 |
| ENSMUSG00000024687 | Osbp | protein_coding | -0.039011233 | 4.320719299 | 0.0082997 | 0.9285207 | 0.984038771 |
| ENSMUSG00000079414 | Gm11110 | protein_coding | 0.076801429 | 2.612340391 | 0.008291 | 0.9285581 | 0.984038771 |
| ENSMUSG00000073982 | Rhog | protein_coding | 0.026717662 | 6.224358474 | 0.0082628 | 0.9286792 | 0.984038771 |
| ENSMUSG00000032118 | Fez1 | protein_coding | 0.023127052 | 9.195683626 | 0.0082608 | 0.9286878 | 0.984038771 |
| ENSMUSG00000046841 | Ckap4 | protein_coding | -0.034262792 | 4.205987618 | 0.0082429 | 0.9287649 | 0.984038771 |
| ENSMUSG00000032382 | Snx1 | protein_coding | -0.020429941 | 7.24547317 | 0.0081932 | 0.9289795 | 0.984169149 |
| ENSMUSG00000038299 | Wdr36 | protein_coding | 0.037279877 | 4.820468584 | 0.0081506 | 0.9291638 | 0.984258247 |
| ENSMUSG00000038349 | Plcl1 | protein_coding | -0.029221451 | 4.898905079 | 0.0081119 | 0.9293314 | 0.984258247 |
| ENSMUSG00000020053 | Igf1 | protein_coding | 0.093932378 | 2.51168405 | 0.0081108 | 0.9293382 | 0.984258247 |
| ENSMUSG00000067924 | Cxx1b | protein_coding | -0.026563834 | 4.928086095 | 0.0080609 | 0.9295537 | 0.984389468 |
| ENSMUSG00000015363 | Trabd | protein_coding | -0.036444496 | 4.444487945 | 0.0079965 | 0.9298345 | 0.984589887 |
| ENSMUSG00000006575 | Rundc3a | protein_coding | 0.023560057 | 6.008340285 | 0.0079098 | 0.9302152 | 0.984654155 |
| ENSMUSG00000026848 | Tor1b | protein_coding | 0.034283096 | 4.511419025 | 0.0079055 | 0.9302341 | 0.984654155 |
| ENSMUSG00000029175 | Slc35f6 | protein_coding | -0.029637315 | 5.03880609 | 0.007899 | 0.9302625 | 0.984654155 |
| ENSMUSG00000022828 | Gtf2e1 | protein_coding | 0.038661912 | 3.880361525 | 0.0078879 | 0.9303115 | 0.984654155 |
| ENSMUSG00000109554 | A230057D06Rik | lincRNA | 0.079757744 | 2.69635699 | 0.0078789 | 0.930353 | 0.984654155 |
| ENSMUSG00000015189 | Casd1 | protein_coding | 0.0262 | 5.565796682 | 0.007833 | 0.9305538 | 0.984749443 |
| ENSMUSG00000074415 | 2610203C20Rik | processed_transcript | -0.02601135 | 5.094700376 | 0.0077965 | 0.9307152 | 0.984749443 |
| ENSMUSG00000057103 | Nat8f1 | protein_coding | -0.031607241 | 4.708141597 | 0.0077866 | 0.9307591 | 0.984749443 |
| ENSMUSG00000021428 | Riok1 | protein_coding | 0.029840358 | 4.878698262 | 0.0077753 | 0.9308095 | 0.984749443 |
| ENSMUSG00000021116 | Eif2s1 | protein_coding | -0.018124906 | 6.639158588 | 0.0077546 | 0.930901 | 0.984749443 |
| ENSMUSG00000060288 | Ppih | protein_coding | -0.046325256 | 3.363588497 | 0.0077337 | 0.9309942 | 0.984751203 |
| ENSMUSG00000009733 | Tfcp2 | protein_coding | 0.031946883 | 4.626596057 | 0.0076571 | 0.9313356 | 0.984884994 |
| ENSMUSG00000038020 | Rapgefl1 | protein_coding | 0.031838183 | 5.041302395 | 0.0076509 | 0.9313635 | 0.984884994 |
| ENSMUSG00000097554 | Gm26825 | lincRNA | -0.054512724 | 2.700900117 | 0.0076025 | 0.9315802 | 0.984884994 |
| ENSMUSG00000020899 | Pfas | protein_coding | -0.029599388 | 5.876812559 | 0.0075936 | 0.9316219 | 0.984884994 |
| ENSMUSG00000020607 | Fam84a | protein_coding | -0.019523087 | 7.303704423 | 0.0075365 | 0.9318771 | 0.984884994 |
| ENSMUSG00000035161 | Ints6 | protein_coding | -0.03045068 | 4.27250757 | 0.007535 | 0.931884 | 0.984884994 |
| ENSMUSG00000040209 | Zfp704 | protein_coding | -0.058000668 | 2.930957915 | 0.0075109 | 0.9319926 | 0.984884994 |
| ENSMUSG00000002108 | Nr1h3 | protein_coding | 0.073478143 | 2.680416916 | 0.0075093 | 0.9319998 | 0.984884994 |
| ENSMUSG00000038069 | Cdkn2aip | protein_coding | 0.045204996 | 3.533200707 | 0.0074485 | 0.932275 | 0.984884994 |
| ENSMUSG00000061132 | Blnk | protein_coding | -0.117052347 | 2.08573582 | 0.0074366 | 0.9323616 | 0.984884994 |
| ENSMUSG00000062937 | Mtap | protein_coding | -0.029264815 | 5.135717786 | 0.0074197 | 0.9324058 | 0.984884994 |
| ENSMUSG00000087478 | 4930506C21Rik | processed_transcript | -0.062201439 | 2.649936518 | 0.0074122 | 0.9324397 | 0.984884994 |
| ENSMUSG00000086401 | Gm15559 | antisense_RNA | -0.064464884 | 3.268404032 | 0.0074072 | 0.9324719 | 0.984884994 |
| ENSMUSG00000028760 | Eif4g3 | protein_coding | -0.025411851 | 5.792021111 | 0.007402 | 0.9324862 | 0.984884994 |
| ENSMUSG00000037531 | Mrpl47 | protein_coding | 0.025594981 | 5.937371966 | 0.0074002 | 0.9324946 | 0.984884994 |
| ENSMUSG00000014763 | Fam120b | protein_coding | -0.021992635 | 6.325271253 | 0.0073357 | 0.9327884 | 0.984912639 |
| ENSMUSG00000040687 | Madd | protein_coding | 0.023536045 | 5.786761892 | 0.0073244 | 0.93284 | 0.984912639 |
| ENSMUSG00000035048 | Anapc13 | protein_coding | -0.025704131 | 7.441151497 | 0.0073169 | 0.9328806 | 0.984912639 |
| ENSMUSG00000040282 | BC052040 | protein_coding | 0.032316888 | 4.305137647 | 0.0073104 | 0.932904 | 0.984912639 |
| ENSMUSG00000021671 | Poc5 | protein_coding | -0.030576918 | 4.472090136 | 0.0072746 | 0.9330682 | 0.984912639 |
| ENSMUSG00000052533 | Nup188 | protein_coding | 0.066709988 | 2.716644853 | 0.0072678 | 0.9330996 | 0.984912639 |
| ENSMUSG00000052026 | Slc6a7 | protein_coding | -0.04085891 | 3.676002892 | 0.0072542 | 0.9331619 | 0.984912639 |
| ENSMUSG00000047284 | Neurl4 | protein_coding | -0.033660922 | 5.23873819 | 0.0072109 | 0.9333627 | 0.984927302 |
| ENSMUSG00000005338 | Cadm3 | protein_coding | 0.025680231 | 5.57829691 | 0.0071997 | 0.9334127 | 0.984927302 |
| ENSMUSG00000024096 | Ralbp1 | protein_coding | -0.022084924 | 5.912335662 | 0.0071915 | 0.9334506 | 0.984927302 |
| ENSMUSG00000014550 | Rbsn | protein_coding | 0.026580207 | 5.928368641 | 0.007141 | 0.933684 | 0.985076961 |
| ENSMUSG00000055067 | Smyd3 | protein_coding | 0.033931686 | 6.563992212 | 0.0070426 | 0.934274 | 0.985397787 |
| ENSMUSG00000027676 | Ccdc39 | protein_coding | -0.052139119 | 3.229858903 | 0.0069814 | 0.9344275 | 0.985397787 |
| ENSMUSG00000026471 | Mr1 | protein_coding | -0.117806643 | 2.113181124 | 0.0069555 | 0.9346127 | 0.985397787 |
| ENSMUSG00000029681 | Bcl7b | protein_coding | 0.023016201 | 6.484288191 | 0.0068757 | 0.9349245 | 0.985397787 |
| ENSMUSG00000008822 | Acyp1 | protein_coding | -0.019285082 | 6.363257055 | 0.0068463 | 0.9350636 | 0.985397787 |
| ENSMUSG00000041852 | Tcf20 | protein_coding | 0.027989971 | 5.792420449 | 0.0068428 | 0.93508 | 0.985397787 |
| ENSMUSG00000096188 | Cmtm4 | protein_coding | -0.03533656 | 4.718665735 | 0.0068187 | 0.9351944 | 0.985397787 |
| ENSMUSG00000042305 | Tmem183a | protein_coding | 0.025124478 | 5.834486723 | 0.0068166 | 0.935204 | 0.985397787 |
| ENSMUSG00000029860 | Zyx | protein_coding | 0.03171938 | 4.171575472 | 0.0068094 | 0.9352382 | 0.985397787 |
| ENSMUSG00000002718 | Cse1l | protein_coding | 0.022544098 | 5.626755388 | 0.0067905 | 0.9353279 | 0.985397787 |
| ENSMUSG00000020078 | Vps26a | protein_coding | -0.020871421 | 6.546155547 | 0.0067797 | 0.9353794 | 0.985397787 |
| ENSMUSG00000029326 | Enoph1 | protein_coding | -0.017449439 | 7.226127742 | 0.0067746 | 0.9354035 | 0.985397787 |
| ENSMUSG00000038871 | Bpgm | protein_coding | 0.017577879 | 7.396211372 | 0.0067714 | 0.9354187 | 0.985397787 |
| ENSMUSG00000047866 | Lonp2 | protein_coding | 0.020264996 | 6.782646359 | 0.0067339 | 0.9355977 | 0.985397787 |
| ENSMUSG00000024592 | C330018D20Rik | protein_coding | -0.031540342 | 4.791125602 | 0.0067097 | 0.9357131 | 0.985397787 |
| ENSMUSG00000032012 | Nectin1 | protein_coding | 0.039369373 | 4.533618981 | 0.0067079 | 0.9357233 | 0.985397787 |
| ENSMUSG00000085408 | C530005A16Rik | processed_transcript | -0.044176062 | 3.387807311 | 0.0067049 | 0.9357358 | 0.985397787 |
| ENSMUSG00000020546 | Stxbp4 | protein_coding | 0.036520974 | 4.780325131 | 0.006677 | 0.935871 | 0.985397787 |
| ENSMUSG00000025582 | Nptx1 | protein_coding | -0.030404878 | 4.495206793 | 0.0066712 | 0.9358972 | 0.985397787 |
| ENSMUSG00000017376 | Nlk | protein_coding | 0.025166034 | 5.430241726 | 0.0066467 | 0.9360148 | 0.985397787 |
| ENSMUSG00000021131 | Erh | protein_coding | 0.034623135 | 3.646676036 | 0.0066287 | 0.9361014 | 0.985397787 |
| ENSMUSG00000071054 | Safb | protein_coding | 0.029263567 | 4.510427583 | 0.0066284 | 0.936103 | 0.985397787 |
| ENSMUSG00000033972 | Zfp944 | protein_coding | -0.033505425 | 3.843087249 | 0.0066277 | 0.9361061 | 0.985397787 |
| ENSMUSG00000017715 | Pgs1 | protein_coding | -0.029737311 | 4.039491949 | 0.0066108 | 0.9361875 | 0.985397787 |
| ENSMUSG00000041915 | Ammecr1l | protein_coding | 0.042705464 | 3.297961726 | 0.0065387 | 0.9365357 | 0.985482403 |
| ENSMUSG00000032381 | Fam96a | protein_coding | -0.018820775 | 6.727894162 | 0.0065311 | 0.9365723 | 0.985482403 |

| ensembl_gene_id | external_gene_name | gene_biotype | log2 (fold change) | log2 (counts per million) | F-value | p-value | False discovery rate |
|---|---|---|---|---|---|---|---|
| ENSMUSG00000026204 | Ptprn | protein_coding | 0.019919406 | 7.339596512 | 0.0065224 | 0.9366148 | 0.985482403 |
| ENSMUSG00000066057 | Gm1976 | lincRNA | -0.04870963 | 3.030686164 | 0.0065183 | 0.9366344 | 0.985482403 |
| ENSMUSG00000048429 | Timm29 | protein_coding | 0.026899644 | 5.196733817 | 0.0064437 | 0.9369973 | 0.985767772 |
| ENSMUSG00000108608 | Gm6916 | transcribed_processe | -0.17197509 | 2.572950252 | 0.0065244 | 0.9371825 | 0.985817087 |
| ENSMUSG00000015312 | Gadd45b | protein_coding | 0.037123699 | 3.546688812 | 0.0063966 | 0.9372276 | 0.985817087 |
| ENSMUSG00000031060 | Rbm10 | protein_coding | -0.02999907 | 4.31472965 | 0.0063303 | 0.9375528 | 0.985948385 |
| ENSMUSG00000020335 | Zfp354b | protein_coding | -0.042546394 | 3.46258822 | 0.0063169 | 0.9376188 | 0.985948385 |
| ENSMUSG00000039990 | Edrf1 | protein_coding | -0.048131473 | 3.266259873 | 0.0063091 | 0.9376571 | 0.985948385 |
| ENSMUSG00000030397 | Mark4 | protein_coding | -0.031341578 | 4.220877947 | 0.0062882 | 0.9377603 | 0.985948385 |
| ENSMUSG00000025759 | Mfsd8 | protein_coding | 0.070218337 | 2.851632995 | 0.006264 | 0.9378816 | 0.985948385 |
| ENSMUSG00000108776 | Gm45169 | lincRNA | 0.041941068 | 3.493116215 | 0.0062594 | 0.9379025 | 0.985948385 |
| ENSMUSG00000021643 | Serf1 | protein_coding | 0.030197445 | 4.613546209 | 0.0062015 | 0.9381901 | 0.986154308 |
| ENSMUSG00000028274 | Rngtt | protein_coding | -0.029337319 | 5.010765698 | 0.0061816 | 0.9382891 | 0.986161903 |
| ENSMUSG00000028284 | Map3k7 | protein_coding | 0.032719001 | 3.835597982 | 0.0061391 | 0.938501 | 0.986288241 |
| ENSMUSG00000043065 | Spice1 | protein_coding | -0.04126692 | 3.476077722 | 0.0060995 | 0.9386995 | 0.986333791 |
| ENSMUSG00000025037 | Maoa | protein_coding | -0.02369598 | 5.56564449 | 0.0060938 | 0.9387278 | 0.986333791 |
| ENSMUSG00000024556 | Me2 | protein_coding | 0.030449647 | 4.003990473 | 0.006053 | 0.9389331 | 0.986453121 |
| ENSMUSG00000019792 | Trmt11 | protein_coding | 0.036649545 | 4.368417646 | 0.0059637 | 0.9393856 | 0.986529215 |
| ENSMUSG00000020993 | Trappc6b | protein_coding | -0.02275885 | 5.564164391 | 0.005961 | 0.9393978 | 0.986529215 |
| ENSMUSG00000026778 | Prkcq | protein_coding | -0.042602206 | 3.361686752 | 0.0059572 | 0.9394169 | 0.986529215 |
| ENSMUSG00000028247 | Coq3 | protein_coding | 0.021546072 | 6.189198552 | 0.0059455 | 0.9394764 | 0.986529215 |
| ENSMUSG00000016510 | Mtif3 | protein_coding | 0.02385147 | 5.049693734 | 0.0059364 | 0.9395228 | 0.986529215 |
| ENSMUSG00000034932 | Mrpl54 | protein_coding | 0.033115602 | 6.746762172 | 0.0059551 | 0.9395773 | 0.986529215 |
| ENSMUSG00000025880 | Smad7 | protein_coding | 0.03948164 | 3.443489442 | 0.0058517 | 0.9399546 | 0.986529215 |
| ENSMUSG00000027680 | Fxr1 | protein_coding | -0.027554726 | 5.052254019 | 0.0058506 | 0.9399604 | 0.986529215 |
| ENSMUSG00000042447 | Mios | protein_coding | -0.037646376 | 3.605521355 | 0.005828 | 0.9400761 | 0.986529215 |
| ENSMUSG00000031007 | Atp6ap2 | protein_coding | 0.015631668 | 8.573246147 | 0.0058185 | 0.940125 | 0.986529215 |
| ENSMUSG00000018559 | Ctdnep1 | protein_coding | 0.02353232 | 5.025039211 | 0.0057756 | 0.9403458 | 0.986529215 |
| ENSMUSG00000057440 | Mpp7 | protein_coding | -0.041568034 | 3.335569132 | 0.0057732 | 0.9403582 | 0.986529215 |
| ENSMUSG00000049721 | Gal3st1 | protein_coding | -0.047327313 | 3.6048914 | 0.0057092 | 0.9406905 | 0.986529215 |
| ENSMUSG00000055403 | 4933427D06Rik | lincRNA | 0.051206264 | 3.222625254 | 0.0057072 | 0.9407011 | 0.986529215 |
| ENSMUSG00000033554 | Dph5 | protein_coding | -0.021056408 | 5.479971465 | 0.0057001 | 0.9407364 | 0.986529215 |
| ENSMUSG00000019951 | Uhrf1bp1l | protein_coding | -0.021664231 | 5.283490123 | 0.0056938 | 0.9407688 | 0.986529215 |
| ENSMUSG00000024948 | Map4k2 | protein_coding | -0.041157818 | 4.428703934 | 0.0056808 | 0.9408748 | 0.986529215 |
| ENSMUSG00000051346 | Spryd4 | protein_coding | 0.02423849 | 4.964114323 | 0.0056701 | 0.9408917 | 0.986529215 |
| ENSMUSG00000039787 | Cercam | protein_coding | 0.079452575 | 3.018022681 | 0.0056911 | 0.9409115 | 0.986529215 |
| ENSMUSG00000032405 | Pias1 | protein_coding | 0.01999673 | 7.093369764 | 0.0056223 | 0.941141 | 0.986529215 |
| ENSMUSG00000020166 | Cnot2 | protein_coding | -0.022423163 | 5.701456193 | 0.0056206 | 0.9411501 | 0.986529215 |
| ENSMUSG00000048148 | Nwd1 | protein_coding | -0.037650961 | 3.57062 | 0.0056029 | 0.9412426 | 0.986529215 |
| ENSMUSG00000025393 | Atp5b | protein_coding | -0.020390156 | 10.30612259 | 0.0055966 | 0.9412756 | 0.986529215 |
| ENSMUSG00000030246 | Ldhb | protein_coding | 0.015690196 | 10.4757629 | 0.0055922 | 0.9412988 | 0.986529215 |
| ENSMUSG00000027937 | Jtb | protein_coding | -0.025516945 | 5.714458474 | 0.0055343 | 0.9416025 | 0.986529215 |
| ENSMUSG00000021364 | Elovl2 | protein_coding | -0.070637375 | 2.667073904 | 0.0055298 | 0.9416278 | 0.986529215 |
| ENSMUSG00000041747 | Utp15 | protein_coding | -0.027370637 | 4.508978226 | 0.0055161 | 0.9416985 | 0.986529215 |
| ENSMUSG00000056698 | Elmod3 | protein_coding | -0.021676803 | 5.625959417 | 0.0054861 | 0.9418569 | 0.986529215 |
| ENSMUSG00000110444 | Gm10033 | transcribed_unproces | 0.053503033 | 3.567408666 | 0.0054554 | 0.9420768 | 0.986529215 |
| ENSMUSG00000034107 | Ano7 | protein_coding | -0.063472356 | 2.574093117 | 0.0054418 | 0.9420921 | 0.986529215 |
| ENSMUSG00000044250 | Pced1b | protein_coding | -0.042838135 | 2.992362881 | 0.0054416 | 0.942093 | 0.986529215 |
| ENSMUSG00000054814 | Usp46 | protein_coding | 0.021552494 | 6.444932736 | 0.0054288 | 0.9421608 | 0.986529215 |
| ENSMUSG00000055923 | Aasdh | protein_coding | -0.044990744 | 3.069737384 | 0.0053838 | 0.9424005 | 0.986529215 |
| ENSMUSG00000022414 | Tab1 | protein_coding | -0.031527736 | 5.193363112 | 0.0053376 | 0.9426492 | 0.986529215 |
| ENSMUSG00000020612 | Prkar1a | protein_coding | -0.026221328 | 9.700402142 | 0.0053402 | 0.9426907 | 0.986529215 |
| ENSMUSG00000027765 | P2ry1 | protein_coding | -0.04094565 | 3.143338903 | 0.0053124 | 0.942783 | 0.986529215 |
| ENSMUSG00000066643 | Wdr35 | protein_coding | 0.024266926 | 5.363477915 | 0.0053061 | 0.9428171 | 0.986529215 |
| ENSMUSG00000028172 | Tacr3 | protein_coding | 0.044446264 | 2.731953969 | 0.005304 | 0.9428286 | 0.986529215 |
| ENSMUSG00000032744 | Heyl | protein_coding | 0.0392802 | 3.977557591 | 0.0052537 | 0.943101 | 0.986529215 |
| ENSMUSG00000051238 | Swsap1 | protein_coding | 0.043590159 | 2.90923937 | 0.0052468 | 0.9431371 | 0.986529215 |
| ENSMUSG00000026470 | Stx6 | protein_coding | -0.022666177 | 5.734930447 | 0.0052323 | 0.9432153 | 0.986529215 |
| ENSMUSG00000093606 | B130034C11Rik | processed_transcript | 0.055406491 | 2.838125638 | 0.0052313 | 0.943221 | 0.986529215 |
| ENSMUSG00000036155 | Mgat5 | protein_coding | 0.034659627 | 3.778046629 | 0.0052168 | 0.9432996 | 0.986529215 |
| ENSMUSG00000035314 | Gdpd5 | protein_coding | 0.031635919 | 3.631621812 | 0.0051872 | 0.9434601 | 0.986529215 |
| ENSMUSG00000033486 | Catsper2 | protein_coding | -0.03862719 | 3.225605393 | 0.0051398 | 0.9437184 | 0.986529215 |
| ENSMUSG00000021967 | Mrpl57 | protein_coding | -0.019387197 | 6.480578311 | 0.0051303 | 0.9437704 | 0.986529215 |
| ENSMUSG00000023805 | Synj2 | protein_coding | -0.02563382 | 4.969783912 | 0.0050551 | 0.9441837 | 0.986529215 |
| ENSMUSG00000023935 | Spats1 | protein_coding | 0.046176669 | 2.793312523 | 0.0050514 | 0.9442036 | 0.986529215 |
| ENSMUSG00000014418 | Hps5 | protein_coding | -0.043813342 | 3.081212267 | 0.0050217 | 0.9443676 | 0.986529215 |
| ENSMUSG00000037089 | Slc35b2 | protein_coding | -0.025887748 | 5.014499288 | 0.0050089 | 0.9444388 | 0.986529215 |
| ENSMUSG00000031928 | Mre11a | protein_coding | -0.037755944 | 3.140148877 | 0.0050033 | 0.9444697 | 0.986529215 |
| ENSMUSG00000042628 | Zfyve1 | protein_coding | -0.024638742 | 4.594394296 | 0.0050022 | 0.9444756 | 0.986529215 |
| ENSMUSG00000043621 | Ubxn10 | protein_coding | -0.040769033 | 3.129834677 | 0.004999 | 0.9444933 | 0.986529215 |
| ENSMUSG00000020601 | Trib2 | protein_coding | -0.020645217 | 5.577241253 | 0.0049966 | 0.9445068 | 0.986529215 |
| ENSMUSG00000036667 | Tcaf1 | protein_coding | 0.018372231 | 7.530609626 | 0.0049922 | 0.9445311 | 0.986529215 |
| ENSMUSG00000054034 | Tceal5 | protein_coding | 0.016168286 | 6.457638063 | 0.0049899 | 0.9445442 | 0.986529215 |
| ENSMUSG00000026988 | Wdsub1 | protein_coding | 0.03254441 | 3.950590657 | 0.0049813 | 0.9445918 | 0.986529215 |
| ENSMUSG00000029577 | Ube3b | protein_coding | 0.023230037 | 4.872432996 | 0.0049764 | 0.9446189 | 0.986529215 |
| ENSMUSG00000039318 | Rab3gap2 | protein_coding | -0.032081348 | 4.163688123 | 0.0049759 | 0.9446215 | 0.986529215 |
| ENSMUSG00000074909 | Ranbp6 | protein_coding | 0.02641356 | 5.08870337 | 0.0049675 | 0.9446682 | 0.986529215 |
| ENSMUSG00000021257 | Angel1 | protein_coding | 0.048998339 | 3.090701824 | 0.0049556 | 0.9447343 | 0.986529215 |

| ensembl_gene_id | external_gene_name | gene_biotype | log2 (fold change) | log2 (counts per million) | F-value | p-value | False discovery rate |
|---|---|---|---|---|---|---|---|
| ENSMUSG00000061950 | Ppp4r1 | protein_coding | -0.040730501 | 3.309485181 | 0.0049383 | 0.9448308 | 0.986529215 |
| ENSMUSG00000040586 | Ofd1 | protein_coding | -0.051455164 | 2.848026959 | 0.0048965 | 0.9450644 | 0.986529215 |
| ENSMUSG00000010290 | AI597479 | protein_coding | 0.023929438 | 5.439774411 | 0.0048802 | 0.9451558 | 0.986529215 |
| ENSMUSG00000091705 | H2-Q2 | protein_coding | -0.052743183 | 2.492497233 | 0.0048797 | 0.9451587 | 0.986529215 |
| ENSMUSG00000023015 | Racgap1 | protein_coding | 0.030812276 | 3.922597179 | 0.0048737 | 0.9451921 | 0.986529215 |
| ENSMUSG00000047098 | Rnf31 | protein_coding | -0.032568628 | 4.120447558 | 0.0048647 | 0.945243 | 0.986529215 |
| ENSMUSG00000032244 | Fem1b | protein_coding | 0.016605645 | 6.056165787 | 0.0048645 | 0.9452442 | 0.986529215 |
| ENSMUSG00000001663 | Gstt1 | protein_coding | 0.02116943 | 5.76642151 | 0.004848 | 0.9453367 | 0.986530074 |
| ENSMUSG00000035329 | Fbxo33 | protein_coding | -0.021698516 | 5.444145011 | 0.0047363 | 0.9459691 | 0.986886717 |
| ENSMUSG00000040410 | Fbxl4 | protein_coding | -0.022681868 | 4.835194913 | 0.0047308 | 0.9460003 | 0.986886717 |
| ENSMUSG00000055013 | Agap1 | protein_coding | -0.016882982 | 6.385730368 | 0.0047277 | 0.946018 | 0.986886717 |
| ENSMUSG00000038637 | Lrrc56 | protein_coding | 0.048838158 | 2.744091276 | 0.0047229 | 0.9460456 | 0.986886717 |
| ENSMUSG00000026675 | Hsd17b7 | protein_coding | 0.035004306 | 4.474511903 | 0.0046786 | 0.946304 | 0.986915144 |
| ENSMUSG00000044469 | Tnfaip8l1 | protein_coding | -0.037433224 | 3.351556379 | 0.004654 | 0.9464396 | 0.986915144 |
| ENSMUSG00000030751 | Psma1 | protein_coding | -0.014681202 | 8.507514574 | 0.0046522 | 0.9464499 | 0.986915144 |
| ENSMUSG00000003948 | Mmd | protein_coding | -0.027385308 | 6.024027836 | 0.0046426 | 0.9465579 | 0.986915144 |
| ENSMUSG00000028126 | Pip5k1a | protein_coding | -0.019795504 | 5.848862134 | 0.0046152 | 0.9466629 | 0.986915144 |
| ENSMUSG00000062931 | Zfp938 | protein_coding | 0.015034017 | 6.348800424 | 0.0046113 | 0.9466856 | 0.986915144 |
| ENSMUSG00000040455 | Usp45 | protein_coding | -0.025105925 | 4.613921663 | 0.0045987 | 0.9467583 | 0.986915144 |
| ENSMUSG00000001053 | N4bp3 | protein_coding | -0.052720542 | 2.452162379 | 0.0045903 | 0.9468071 | 0.986915144 |
| ENSMUSG00000036180 | Gatad2a | protein_coding | -0.020468929 | 4.89311502 | 0.0045421 | 0.9470864 | 0.987110571 |
| ENSMUSG00000026803 | Ttf1 | protein_coding | 0.030347371 | 3.734458087 | 0.0044876 | 0.9474046 | 0.987260679 |
| ENSMUSG00000045648 | Vwc2l | protein_coding | -0.025832339 | 3.978575379 | 0.004486 | 0.947414 | 0.987260679 |
| ENSMUSG00000024063 | Lbh | protein_coding | -0.026593332 | 4.762518515 | 0.0044584 | 0.9475758 | 0.987281518 |
| ENSMUSG00000020525 | Ppm1d | protein_coding | 0.035648315 | 3.356789693 | 0.0044383 | 0.9476937 | 0.987281518 |
| ENSMUSG00000040699 | Limd2 | protein_coding | 0.024784248 | 5.42800098 | 0.0044356 | 0.9477095 | 0.987281518 |
| ENSMUSG00000046058 | Eid2 | protein_coding | -0.025128039 | 4.640219936 | 0.0043845 | 0.9480113 | 0.987395887 |
| ENSMUSG00000022023 | Wbp4 | protein_coding | 0.013502545 | 6.75804316 | 0.0043816 | 0.9480286 | 0.987395887 |
| ENSMUSG00000047434 | Xxylt1 | protein_coding | 0.029909733 | 3.392330409 | 0.0043523 | 0.9482021 | 0.987395887 |
| ENSMUSG00000021363 | Mak | protein_coding | 0.071558132 | 2.430673105 | 0.0043525 | 0.9482025 | 0.987395887 |
| ENSMUSG00000026019 | Wdr12 | protein_coding | 0.014949957 | 6.68670352 | 0.0043395 | 0.9482784 | 0.987395887 |
| ENSMUSG00000063281 | Zfp35 | protein_coding | -0.022316303 | 5.15766405 | 0.0043039 | 0.9484903 | 0.987485656 |
| ENSMUSG00000040957 | Cables1 | protein_coding | -0.033251069 | 3.644113282 | 0.0042942 | 0.9485483 | 0.987485656 |
| ENSMUSG00000024130 | Abca3 | protein_coding | -0.041882631 | 2.720833177 | 0.0042378 | 0.9488869 | 0.987635876 |
| ENSMUSG00000026495 | Efcab2 | protein_coding | 0.020952205 | 5.377614684 | 0.0041941 | 0.9491507 | 0.987635876 |
| ENSMUSG00000037890 | Wdr19 | protein_coding | 0.035426796 | 3.403430134 | 0.004189 | 0.9491816 | 0.987635876 |
| ENSMUSG00000027091 | Zc3h15 | protein_coding | 0.018191417 | 6.253885872 | 0.0041816 | 0.9492266 | 0.987635876 |
| ENSMUSG00000049532 | Sall2 | protein_coding | 0.017657927 | 5.735768004 | 0.0041801 | 0.9492354 | 0.987635876 |
| ENSMUSG00000020176 | Grb10 | protein_coding | -0.019624393 | 5.982098793 | 0.0041648 | 0.9493282 | 0.987635876 |
| ENSMUSG00000045671 | Spred2 | protein_coding | 0.028479812 | 4.025440524 | 0.0041637 | 0.9493355 | 0.987635876 |
| ENSMUSG00000004530 | Coro1c | protein_coding | 0.024626184 | 6.0219746 | 0.0041315 | 0.9495639 | 0.987777965 |
| ENSMUSG00000074785 | Plxnc1 | protein_coding | 0.019705674 | 5.013057301 | 0.0041073 | 0.9496789 | 0.987802051 |
| ENSMUSG00000046441 | Cmtr2 | protein_coding | -0.043374676 | 3.056756815 | 0.0040854 | 0.949813 | 0.987845992 |
| ENSMUSG00000022175 | Lrp10 | protein_coding | -0.032182892 | 3.736932533 | 0.0040379 | 0.9501054 | 0.9879743 |
| ENSMUSG00000047879 | Usp14 | protein_coding | 0.016293518 | 7.679212796 | 0.0040183 | 0.9502266 | 0.9879743 |
| ENSMUSG00000041459 | Tardbp | protein_coding | 0.015569499 | 6.137299951 | 0.0040118 | 0.9502668 | 0.9879743 |
| ENSMUSG00000001785 | Pwp1 | protein_coding | -0.017028668 | 5.304626079 | 0.0040058 | 0.9503039 | 0.9879743 |
| ENSMUSG00000045160 | Bola3 | protein_coding | 0.024219249 | 4.126265745 | 0.0039663 | 0.9505489 | 0.988020661 |
| ENSMUSG00000021557 | Agtpbp1 | protein_coding | 0.016348977 | 6.05202309 | 0.0039594 | 0.9505921 | 0.988020661 |
| ENSMUSG00000090266 | Mettl23 | protein_coding | 0.025887726 | 4.816072364 | 0.0039297 | 0.9507777 | 0.988020661 |
| ENSMUSG00000000532 | Acvr1b | protein_coding | 0.02508519 | 4.535971057 | 0.0039247 | 0.9508088 | 0.988020661 |
| ENSMUSG00000037894 | H2afz | protein_coding | 0.022335063 | 3.996993766 | 0.0039127 | 0.9508839 | 0.988020661 |
| ENSMUSG00000027848 | Olfml3 | protein_coding | -0.025341154 | 4.462495399 | 0.0039006 | 0.9509596 | 0.988020661 |
| ENSMUSG00000037752 | Xkr8 | protein_coding | -0.041154127 | 2.798958383 | 0.0038955 | 0.9509917 | 0.988020661 |
| ENSMUSG00000068040 | Tm9sf4 | protein_coding | -0.015611375 | 6.085282906 | 0.0038736 | 0.9511296 | 0.988068496 |
| ENSMUSG00000031099 | Smarca1 | protein_coding | -0.020842061 | 5.353965234 | 0.0038581 | 0.9512275 | 0.988074727 |
| ENSMUSG00000005973 | Rcn1 | protein_coding | -0.021313019 | 5.132738455 | 0.0038281 | 0.951417 | 0.988176135 |
| ENSMUSG00000002983 | Relb | protein_coding | 0.046393275 | 2.645367851 | 0.00379 | 0.9516588 | 0.988331729 |
| ENSMUSG00000029119 | Man2b2 | protein_coding | -0.03255996 | 3.495648565 | 0.0037623 | 0.951836 | 0.988375447 |
| ENSMUSG00000033287 | Kctd17 | protein_coding | 0.018618974 | 5.488237808 | 0.0037192 | 0.9521124 | 0.988375447 |
| ENSMUSG00000032415 | Ube2cbp | protein_coding | 0.028086184 | 3.39040437 | 0.0037156 | 0.9521354 | 0.988375447 |
| ENSMUSG00000029623 | Pdap1 | protein_coding | 0.013127967 | 6.886533944 | 0.0037114 | 0.9521625 | 0.988375447 |
| ENSMUSG00000028884 | Rpa2 | protein_coding | 0.020955635 | 5.220254905 | 0.0036841 | 0.9523386 | 0.988375447 |
| ENSMUSG00000021725 | Parp8 | protein_coding | -0.029898551 | 3.485293454 | 0.0036675 | 0.952446 | 0.988375447 |
| ENSMUSG00000026239 | Pde6d | protein_coding | 0.018530207 | 4.990558208 | 0.0036602 | 0.9524927 | 0.988375447 |
| ENSMUSG00000016409 | Nkap | protein_coding | -0.018491929 | 5.150353516 | 0.0036602 | 0.9524933 | 0.988375447 |
| ENSMUSG00000015599 | Ttbk1 | protein_coding | -0.03274358 | 3.213615904 | 0.0036439 | 0.9525988 | 0.988375447 |
| ENSMUSG00000085558 | 4930412C18Rik | processed_transcript | 0.029830152 | 3.611819385 | 0.0036268 | 0.9527102 | 0.988375447 |
| ENSMUSG00000051306 | Usp42 | protein_coding | 0.024663491 | 4.038454282 | 0.0036265 | 0.9527119 | 0.988375447 |
| ENSMUSG00000018398 | 8-Sep | protein_coding | -0.012099459 | 6.969690706 | 0.0035925 | 0.9529337 | 0.988443334 |
| ENSMUSG00000044017 | Adgrd1 | protein_coding | -0.031782301 | 2.95835921 | 0.0035685 | 0.9530911 | 0.988443334 |
| ENSMUSG00000038034 | Igsf8 | protein_coding | -0.015032281 | 5.971416614 | 0.0035639 | 0.953121 | 0.988443334 |
| ENSMUSG00000056158 | Car10 | protein_coding | -0.022342562 | 6.29530553 | 0.0035345 | 0.9533608 | 0.988443334 |
| ENSMUSG00000027550 | Lrrcc1 | protein_coding | -0.047413996 | 4.236047588 | 0.0035331 | 0.9534736 | 0.988443334 |
| ENSMUSG00000074024 | 4632427E13Rik | bidirectional_promote | 0.07220701 | 2.528839498 | 0.0035129 | 0.9535167 | 0.988443334 |
| ENSMUSG00000067825 | Pex26 | protein_coding | 0.030140926 | 3.494029996 | 0.0034705 | 0.9537391 | 0.988443334 |
| ENSMUSG00000002052 | Supt6 | protein_coding | -0.015632206 | 6.11979242 | 0.0034403 | 0.9539401 | 0.988443334 |
| ENSMUSG00000022978 | Mis18a | protein_coding | -0.023372837 | 3.910452885 | 0.0034283 | 0.9540204 | 0.988443334 |

| ensembl_gene_id | external_gene_name | gene_biotype | log2 (fold change) | log2 (counts per million) | F-value | p-value | False discovery rate |
|---|---|---|---|---|---|---|---|
| ENSMUSG00000026107 | Nabp1 | protein_coding | -0.02890813 | 3.476302538 | 0.0033996 | 0.9542134 | 0.988443334 |
| ENSMUSG00000051736 | Fam229b | protein_coding | 0.016809377 | 5.959750319 | 0.003396 | 0.9542372 | 0.988443334 |
| ENSMUSG00000038984 | Tspyl5 | protein_coding | -0.017203636 | 6.739903537 | 0.0033961 | 0.9542379 | 0.988443334 |
| ENSMUSG00000060735 | Rxfp3 | protein_coding | 0.045732771 | 2.619317142 | 0.0033488 | 0.954556 | 0.988443334 |
| ENSMUSG00000017670 | Elmo2 | protein_coding | -0.016876527 | 6.908676478 | 0.0033436 | 0.9545925 | 0.988443334 |
| ENSMUSG00000035949 | Fbxw2 | protein_coding | 0.023596525 | 5.108968232 | 0.0032783 | 0.9550377 | 0.988443334 |
| ENSMUSG00000034006 | Pqlc1 | protein_coding | 0.020494963 | 5.558147472 | 0.0032629 | 0.9551424 | 0.988443334 |
| ENSMUSG00000053070 | 9230110C19Rik | protein_coding | 0.023345146 | 4.620341163 | 0.0032572 | 0.9551816 | 0.988443334 |
| ENSMUSG00000005580 | Adcy9 | protein_coding | -0.030938623 | 4.012218975 | 0.003252 | 0.9552185 | 0.988443334 |
| ENSMUSG00000022022 | Mtrf1 | protein_coding | 0.024092785 | 4.828859959 | 0.0032387 | 0.9553086 | 0.988443334 |
| ENSMUSG00000040570 | Rundc3b | protein_coding | 0.019683986 | 5.015460729 | 0.0032339 | 0.9553415 | 0.988443334 |
| ENSMUSG00000038296 | Galnt18 | protein_coding | -0.021645912 | 4.65128594 | 0.0032287 | 0.9553776 | 0.988443334 |
| ENSMUSG00000021550 | 2210016F16Rik | protein_coding | 0.027557784 | 3.754677248 | 0.0032104 | 0.9555045 | 0.988443334 |
| ENSMUSG00000020483 | Dynll2 | protein_coding | -0.01810504 | 8.058738353 | 0.0032071 | 0.955556 | 0.988443334 |
| ENSMUSG00000051606 | 2010001K21Rik | processed_transcript | 0.061104996 | 2.361829615 | 0.0031975 | 0.9555945 | 0.988443334 |
| ENSMUSG00000040213 | Kyat3 | protein_coding | 0.03148972 | 3.134046576 | 0.0031814 | 0.9557054 | 0.988443334 |
| ENSMUSG00000026737 | Pip4k2a | protein_coding | -0.014711583 | 5.635831493 | 0.0031721 | 0.9557701 | 0.988443334 |
| ENSMUSG00000011257 | Pabpc4 | protein_coding | -0.028903564 | 3.198286458 | 0.0031657 | 0.9558146 | 0.988443334 |
| ENSMUSG00000028409 | Smu1 | protein_coding | 0.013205825 | 6.920026931 | 0.0031646 | 0.9558226 | 0.988443334 |
| ENSMUSG00000074555 | Gm10714 | antisense_RNA | 0.04334888 | 2.85445708 | 0.0031612 | 0.9558458 | 0.988443334 |
| ENSMUSG00000022306 | Zfpm2 | protein_coding | -0.045158705 | 2.911789258 | 0.0031113 | 0.9561966 | 0.988443334 |
| ENSMUSG00000032905 | Atg12 | protein_coding | -0.01147299 | 6.55226275 | 0.0031049 | 0.9562407 | 0.988443334 |
| ENSMUSG00000000826 | Dnajc5 | protein_coding | 0.012233609 | 8.25432264 | 0.0031035 | 0.9562506 | 0.988443334 |
| ENSMUSG00000004929 | Thop1 | protein_coding | -0.027505241 | 4.018110319 | 0.003081 | 0.9564089 | 0.988443334 |
| ENSMUSG00000030271 | Ogg1 | protein_coding | -0.021761224 | 3.96687806 | 0.0030808 | 0.9564107 | 0.988443334 |
| ENSMUSG00000051853 | Arf3 | protein_coding | -0.012662495 | 9.245091916 | 0.0030638 | 0.9565306 | 0.988443334 |
| ENSMUSG00000028621 | Cyb5rl | protein_coding | 0.024842171 | 4.490891648 | 0.0030591 | 0.9565643 | 0.988443334 |
| ENSMUSG00000026650 | Meig1 | protein_coding | -0.020658754 | 4.534744567 | 0.003041 | 0.9566924 | 0.988443334 |
| ENSMUSG00000015889 | Lta4h | protein_coding | 0.014109746 | 6.221510192 | 0.0030337 | 0.9567449 | 0.988443334 |
| ENSMUSG00000113771 | AC160108.1 | antisense_RNA | -0.03398269 | 3.105544482 | 0.0030187 | 0.9568513 | 0.988443334 |
| ENSMUSG00000008200 | Fnbp4 | protein_coding | 0.022069221 | 3.868926687 | 0.0029893 | 0.9570622 | 0.988443334 |
| ENSMUSG00000026807 | Ak8 | protein_coding | -0.040318245 | 3.527731632 | 0.0029798 | 0.9571725 | 0.988443334 |
| ENSMUSG00000044033 | Ccdc141 | protein_coding | 0.03515272 | 2.953889257 | 0.0029725 | 0.9571827 | 0.988443334 |
| ENSMUSG00000044551 | 9930012K11Rik | protein_coding | 0.046513103 | 2.885556435 | 0.0029646 | 0.9572409 | 0.988443334 |
| ENSMUSG00000045573 | Penk | protein_coding | -0.019545024 | 10.40961684 | 0.0029549 | 0.9573517 | 0.988443334 |
| ENSMUSG00000029550 | Sppl3 | protein_coding | -0.019324587 | 4.9790143 | 0.002949 | 0.957352 | 0.988443334 |
| ENSMUSG00000040136 | Abcc8 | protein_coding | 0.078121486 | 2.080395375 | 0.002947 | 0.9574084 | 0.988443334 |
| ENSMUSG00000010025 | Aldh3a2 | protein_coding | 0.022122932 | 5.385840034 | 0.0029316 | 0.9574792 | 0.988443334 |
| ENSMUSG00000028277 | Ube2j1 | protein_coding | 0.014868298 | 6.01648963 | 0.0029284 | 0.9575012 | 0.988443334 |
| ENSMUSG00000025008 | Tctn3 | protein_coding | 0.041396304 | 3.740737521 | 0.0029378 | 0.9575187 | 0.988443334 |
| ENSMUSG00000000167 | Pih1d2 | protein_coding | -0.023124922 | 3.671753911 | 0.0029185 | 0.9575727 | 0.988443334 |
| ENSMUSG00000041718 | Alg13 | protein_coding | 0.047224174 | 2.759858993 | 0.0029097 | 0.9576382 | 0.988443334 |
| ENSMUSG00000026158 | Ogfrl1 | protein_coding | -0.015465685 | 6.023285108 | 0.0028886 | 0.957791 | 0.988443334 |
| ENSMUSG00000086350 | B230369F24Rik | antisense_RNA | -0.023712921 | 3.605258578 | 0.0028862 | 0.9578083 | 0.988443334 |
| ENSMUSG00000056749 | Nfil3 | protein_coding | 0.029350247 | 3.514816785 | 0.0028781 | 0.9578673 | 0.988443334 |
| ENSMUSG00000071451 | Psmg4 | protein_coding | -0.017108191 | 5.430961972 | 0.0028777 | 0.9578705 | 0.988443334 |
| ENSMUSG00000037280 | Galnt6 | protein_coding | -0.024187758 | 3.473773619 | 0.0028581 | 0.9580136 | 0.988443334 |
| ENSMUSG00000024563 | Smad2 | protein_coding | -0.01984319 | 4.776615974 | 0.0028458 | 0.958104 | 0.988443334 |
| ENSMUSG00000025545 | Clybl | protein_coding | -0.011397714 | 7.084661771 | 0.0028452 | 0.9581089 | 0.988443334 |
| ENSMUSG00000043019 | Edem3 | protein_coding | 0.027847963 | 3.210635039 | 0.0027953 | 0.9584772 | 0.988728453 |
| ENSMUSG00000020752 | Recql5 | protein_coding | -0.026106508 | 3.326451752 | 0.0027815 | 0.9585801 | 0.98873974 |
| ENSMUSG00000071454 | Dtnb | protein_coding | -0.015988258 | 4.995225655 | 0.0027469 | 0.9588383 | 0.988911208 |
| ENSMUSG00000056216 | Cebpg | protein_coding | 0.01722003 | 4.363496584 | 0.0026755 | 0.959376 | 0.989144205 |
| ENSMUSG00000058624 | Gda | protein_coding | 0.01326754 | 6.490384451 | 0.0026742 | 0.9593856 | 0.989144205 |
| ENSMUSG00000028800 | Hdac1 | protein_coding | 0.021129622 | 3.739194439 | 0.0026613 | 0.9594838 | 0.989144205 |
| ENSMUSG00000027428 | Rbbp9 | protein_coding | -0.015290875 | 6.297842946 | 0.0026519 | 0.9595552 | 0.989144205 |
| ENSMUSG00000041258 | Zfp236 | protein_coding | -0.018822643 | 3.922609002 | 0.0026497 | 0.9595725 | 0.989144205 |
| ENSMUSG00000034040 | Wbscr17 | protein_coding | -0.020142483 | 4.346021108 | 0.002641 | 0.9596384 | 0.989144205 |
| ENSMUSG00000032239 | Rp9 | protein_coding | 0.017017869 | 4.929225882 | 0.0026319 | 0.9597081 | 0.989144205 |
| ENSMUSG00000053730 | Tmem39b | protein_coding | -0.030526497 | 3.168198016 | 0.0025727 | 0.9601632 | 0.989233497 |
| ENSMUSG00000031090 | Nadsyn1 | protein_coding | -0.038019526 | 2.574717907 | 0.0025713 | 0.9601738 | 0.989233497 |
| ENSMUSG00000021693 | Kif2a | protein_coding | -0.012660212 | 5.858300644 | 0.0025701 | 0.9601832 | 0.989233497 |
| ENSMUSG00000022792 | Yars2 | protein_coding | 0.020573263 | 3.835039822 | 0.0025686 | 0.960195 | 0.989233497 |
| ENSMUSG00000039826 | Trub2 | protein_coding | -0.023442476 | 5.013049099 | 0.0025616 | 0.9602547 | 0.989233497 |
| ENSMUSG00000038302 | Afg1l | protein_coding | 0.019609322 | 4.72901734 | 0.0025474 | 0.9603592 | 0.989246295 |
| ENSMUSG00000026135 | Zfp142 | protein_coding | -0.043831576 | 2.531789091 | 0.0024946 | 0.9607723 | 0.989345381 |
| ENSMUSG00000022094 | Slc39a14 | protein_coding | 0.032189285 | 3.064125829 | 0.0024899 | 0.960809 | 0.989345381 |
| ENSMUSG00000028788 | Ptp4a2 | protein_coding | -0.012751831 | 8.424714195 | 0.0024784 | 0.9608997 | 0.989345381 |
| ENSMUSG00000038520 | Tbc1d17 | protein_coding | -0.013832875 | 6.289337849 | 0.0024753 | 0.9609243 | 0.989345381 |
| ENSMUSG00000006522 | Itih3 | protein_coding | -0.021480118 | 5.66703038 | 0.0024727 | 0.9609828 | 0.989345381 |
| ENSMUSG00000031955 | Bcar1 | protein_coding | -0.019897818 | 4.683054308 | 0.0024647 | 0.9610074 | 0.989345381 |
| ENSMUSG00000097413 | A830052D11Rik | lincRNA | 0.035732505 | 2.599959074 | 0.0023798 | 0.9616845 | 0.989901997 |
| ENSMUSG00000053181 | A830005F24Rik | protein_coding | -0.032322643 | 3.006694887 | 0.0023672 | 0.9617859 | 0.989901997 |
| ENSMUSG00000054737 | Zfp182 | protein_coding | 0.036353502 | 2.357138798 | 0.0023539 | 0.9618934 | 0.989901997 |
| ENSMUSG00000025905 | Oprk1 | protein_coding | -0.021597629 | 4.134242713 | 0.0023409 | 0.9619989 | 0.989901997 |
| ENSMUSG00000063406 | Tmed5 | protein_coding | 0.017251739 | 5.217522506 | 0.0023397 | 0.9620084 | 0.989901997 |
| ENSMUSG00000020866 | Cacna1g | protein_coding | -0.030272083 | 3.129771186 | 0.0023147 | 0.9622115 | 0.989954552 |
| ENSMUSG00000055707 | Klhl26 | protein_coding | 0.021115049 | 5.529094558 | 0.0023069 | 0.9623108 | 0.989954552 |

| ensembl_gene_id | external_gene_name | gene_biotype | log2 (fold change) | log2 (counts per million) | F-value | p-value | False discovery rate |
|---|---|---|---|---|---|---|---|
| ENSMUSG00000008136 | Fhl2 | protein_coding | 0.036045056 | 3.259917399 | 0.0023007 | 0.9623356 | 0.989954552 |
| ENSMUSG00000014592 | Camta1 | protein_coding | -0.010520886 | 7.884971833 | 0.0022484 | 0.9627568 | 0.990293123 |
| ENSMUSG00000040667 | Nup88 | protein_coding | -0.013912587 | 6.057076029 | 0.0022203 | 0.9629895 | 0.990413718 |
| ENSMUSG00000051339 | 2900026A02Rik | protein_coding | 0.020334948 | 4.055889803 | 0.0021951 | 0.9632005 | 0.990413718 |
| ENSMUSG00000023262 | Acy1 | protein_coding | 0.017989517 | 4.934477265 | 0.002193 | 0.9632177 | 0.990413718 |
| ENSMUSG00000035173 | Ccdc186 | protein_coding | -0.015753655 | 5.177513361 | 0.002181 | 0.9633186 | 0.990413718 |
| ENSMUSG00000087143 | A830082K12Rik | antisense_RNA | -0.020053883 | 3.85780153 | 0.0021791 | 0.9633346 | 0.990413718 |
| ENSMUSG00000039770 | Ypel5 | protein_coding | 0.009232827 | 8.954682556 | 0.0021428 | 0.9636408 | 0.990542662 |
| ENSMUSG00000025782 | Taf3 | protein_coding | -0.015196938 | 4.81405377 | 0.0021325 | 0.9637283 | 0.990542662 |
| ENSMUSG00000031202 | Rab39b | protein_coding | 0.016815516 | 4.459768071 | 0.0021316 | 0.9637364 | 0.990542662 |
| ENSMUSG00000007867 | Ift43 | protein_coding | -0.012784869 | 5.788973458 | 0.002094 | 0.9640571 | 0.990568111 |
| ENSMUSG00000047187 | Rab2a | protein_coding | -0.00914123 | 8.097974959 | 0.002093 | 0.9640659 | 0.990568111 |
| ENSMUSG00000030752 | Kdm8 | protein_coding | -0.034927752 | 2.7555095 | 0.0020926 | 0.9640689 | 0.990568111 |
| ENSMUSG00000040669 | Phc1 | protein_coding | -0.020331934 | 4.314819874 | 0.0020814 | 0.9641649 | 0.990568111 |
| ENSMUSG00000022096 | Hr | protein_coding | -0.030991506 | 2.578028652 | 0.0020728 | 0.9642395 | 0.990568111 |
| ENSMUSG00000028633 | Ctps | protein_coding | 0.013990049 | 5.140252002 | 0.0020642 | 0.9643139 | 0.990568111 |
| ENSMUSG00000032777 | Gtf3c1 | protein_coding | -0.016667729 | 5.043845491 | 0.0020326 | 0.9645872 | 0.990654354 |
| ENSMUSG00000046139 | Patl1 | protein_coding | -0.02076064 | 3.763403618 | 0.0020256 | 0.9646486 | 0.990654354 |
| ENSMUSG00000027784 | Ppm1l | protein_coding | -0.01361209 | 5.153257303 | 0.0020138 | 0.9647516 | 0.990654354 |
| ENSMUSG00000020261 | Slc36a1 | protein_coding | 0.028527739 | 3.689174556 | 0.0020131 | 0.9647663 | 0.990654354 |
| ENSMUSG00000048661 | Lemd3 | protein_coding | 0.018042309 | 3.832823905 | 0.0019968 | 0.9649009 | 0.990697948 |
| ENSMUSG00000079508 | Apoo | protein_coding | 0.015149977 | 4.398444904 | 0.0019746 | 0.9650965 | 0.990754249 |
| ENSMUSG00000032582 | Rbm6 | protein_coding | -0.016147919 | 4.699735557 | 0.0019696 | 0.96514 | 0.990754249 |
| ENSMUSG00000042350 | Arel1 | protein_coding | 0.019173524 | 4.131847776 | 0.0019498 | 0.9653161 | 0.990756154 |
| ENSMUSG00000027332 | Ivd | protein_coding | -0.014008367 | 6.87274443 | 0.0019407 | 0.9654014 | 0.990756154 |
| ENSMUSG00000002846 | Timmdc1 | protein_coding | -0.018602213 | 5.077553258 | 0.0019384 | 0.9654183 | 0.990756154 |
| ENSMUSG00000040435 | Ppp1r15a | protein_coding | -0.018370914 | 4.902967388 | 0.0019211 | 0.9655722 | 0.990819497 |
| ENSMUSG00000028668 | Eloa | protein_coding | 0.024284175 | 3.051228497 | 0.0018927 | 0.9658271 | 0.99095694 |
| ENSMUSG00000032850 | Rnft2 | protein_coding | -0.016374267 | 4.516265606 | 0.0018857 | 0.9658904 | 0.99095694 |
| ENSMUSG00000052040 | Klf13 | protein_coding | -0.012372983 | 8.319600223 | 0.0018121 | 0.9665633 | 0.991547402 |
| ENSMUSG00000057278 | Snrpg | protein_coding | -0.016155549 | 5.204688223 | 0.0017937 | 0.966732 | 0.991547402 |
| ENSMUSG00000071660 | Ttc9c | protein_coding | -0.01084148 | 5.814119013 | 0.0017926 | 0.9667426 | 0.991547402 |
| ENSMUSG00000034292 | Traf3ip1 | protein_coding | 0.021467857 | 3.585416351 | 0.0017765 | 0.9668917 | 0.99160574 |
| ENSMUSG00000039176 | Polg | protein_coding | -0.036526264 | 2.443151353 | 0.0017608 | 0.9670385 | 0.991661731 |
| ENSMUSG00000040272 | Accs | protein_coding | -0.04278287 | 2.430935469 | 0.001733 | 0.9673009 | 0.991695597 |
| ENSMUSG00000026414 | Tnnt2 | protein_coding | 0.019237502 | 3.784445905 | 0.001721 | 0.9674128 | 0.991695597 |
| ENSMUSG00000026494 | Kif26b | protein_coding | 0.02171689 | 3.373505931 | 0.001719 | 0.9674323 | 0.991695597 |
| ENSMUSG00000026614 | Slc30a10 | protein_coding | -0.027232496 | 3.199459821 | 0.0017181 | 0.9674404 | 0.991695597 |
| ENSMUSG00000031860 | Pbx4 | protein_coding | -0.015532442 | 4.024994436 | 0.0016826 | 0.9677786 | 0.991926495 |
| ENSMUSG00000044807 | Zfp354c | protein_coding | 0.015540886 | 4.196527264 | 0.0016707 | 0.9678926 | 0.991926495 |
| ENSMUSG00000072258 | Taf1a | protein_coding | -0.0161382 | 4.182932386 | 0.0016655 | 0.9679424 | 0.991926495 |
| ENSMUSG00000032773 | Chrm1 | protein_coding | 0.015422022 | 5.403171046 | 0.0016172 | 0.9684108 | 0.992201868 |
| ENSMUSG00000046269 | Usp27x | protein_coding | -0.017104131 | 3.983395356 | 0.0016155 | 0.9684274 | 0.992201868 |
| ENSMUSG00000042570 | Mier2 | protein_coding | 0.02352383 | 3.993611786 | 0.0016086 | 0.9685026 | 0.992201868 |
| ENSMUSG00000021662 | Arhgef28 | protein_coding | -0.019246768 | 4.535902081 | 0.0015999 | 0.9685802 | 0.992201868 |
| ENSMUSG00000050954 | Zfp169 | protein_coding | -0.029082585 | 2.607587525 | 0.0015824 | 0.9687515 | 0.992282845 |
| ENSMUSG00000051439 | Cd14 | protein_coding | -0.026187675 | 2.931237514 | 0.0015697 | 0.9688779 | 0.992317789 |
| ENSMUSG00000022321 | Cdh10 | protein_coding | 0.015120884 | 4.203458714 | 0.0015541 | 0.9690322 | 0.992381289 |
| ENSMUSG00000058230 | Arhgap35 | protein_coding | 0.011533998 | 6.117530316 | 0.0015329 | 0.9692439 | 0.99239648 |
| ENSMUSG00000059482 | 2610301B20Rik | protein_coding | -0.008832059 | 6.793202852 | 0.0015293 | 0.9692807 | 0.99239648 |
| ENSMUSG00000032026 | Rexo2 | protein_coding | -0.010733324 | 5.948169437 | 0.0015248 | 0.9693258 | 0.99239648 |
| ENSMUSG00000044499 | Hs3st5 | protein_coding | 0.019628359 | 3.457253235 | 0.0015133 | 0.9694416 | 0.99239648 |
| ENSMUSG00000017057 | Il13ra1 | protein_coding | 0.020970101 | 3.208087871 | 0.0015048 | 0.9695272 | 0.99239648 |
| ENSMUSG00000037447 | Arid5a | protein_coding | 0.019196052 | 4.065854634 | 0.0014891 | 0.9696868 | 0.99239648 |
| ENSMUSG00000002949 | Timm44 | protein_coding | -0.010873593 | 6.385036777 | 0.0014885 | 0.9696931 | 0.99239648 |
| ENSMUSG00000028057 | Rit1 | protein_coding | -0.009238067 | 6.135760638 | 0.001473 | 0.9698507 | 0.992463277 |
| ENSMUSG00000015944 | Gatsl2 | protein_coding | -0.00897079 | 6.851639751 | 0.0014636 | 0.9699467 | 0.992467141 |
| ENSMUSG00000094870 | Zfp131 | protein_coding | 0.014553094 | 4.197405735 | 0.0014451 | 0.970138 | 0.99256836 |
| ENSMUSG00000010936 | Vac14 | protein_coding | 0.013333259 | 4.58232797 | 0.0013969 | 0.9706396 | 0.992987121 |
| ENSMUSG00000026311 | Asb1 | protein_coding | -0.014191521 | 4.529768471 | 0.0013859 | 0.9707558 | 0.993011546 |
| ENSMUSG00000048281 | Dleu7 | protein_coding | 0.015423773 | 3.777676594 | 0.0013701 | 0.9709224 | 0.993087434 |
| ENSMUSG00000043857 | Mgat5b | protein_coding | -0.014420413 | 4.474945001 | 0.0013592 | 0.9710386 | 0.993111821 |
| ENSMUSG00000036197 | Gxylt1 | protein_coding | 0.017983157 | 3.20701106 | 0.001327 | 0.971383 | 0.993242374 |
| ENSMUSG00000030691 | Fchsd2 | protein_coding | -0.022093138 | 3.146760616 | 0.0013114 | 0.9715524 | 0.993242374 |
| ENSMUSG00000030347 | D6Wsu163e | protein_coding | 0.010307337 | 5.091092312 | 0.0013095 | 0.971573 | 0.993242374 |
| ENSMUSG00000074968 | Ano3 | protein_coding | -0.014592328 | 5.341786411 | 0.0013051 | 0.9716211 | 0.993242374 |
| ENSMUSG00000029596 | Sdsl | protein_coding | -0.025751672 | 2.705739573 | 0.0012995 | 0.9716809 | 0.993242374 |
| ENSMUSG00000027378 | Nphp1 | protein_coding | -0.017552589 | 3.735808116 | 0.001289 | 0.9717961 | 0.993242374 |
| ENSMUSG00000041765 | Ubac2 | protein_coding | -0.010631153 | 5.337075299 | 0.0012771 | 0.9719266 | 0.993242374 |
| ENSMUSG00000042408 | Zmym6 | protein_coding | 0.022394471 | 2.98492359 | 0.0012711 | 0.971992 | 0.993242374 |
| ENSMUSG00000021417 | Eci2 | protein_coding | 0.011562942 | 5.131709028 | 0.0012471 | 0.9722574 | 0.993242374 |
| ENSMUSG00000027998 | Plrg1 | protein_coding | -0.010148827 | 6.257843665 | 0.0012402 | 0.9723341 | 0.993242374 |
| ENSMUSG00000031729 | Ist1 | protein_coding | 0.012944332 | 5.561109839 | 0.0012232 | 0.9725253 | 0.993242374 |
| ENSMUSG00000057060 | Slc35f3 | protein_coding | 0.01719846 | 3.816031417 | 0.001195 | 0.9728433 | 0.993242374 |
| ENSMUSG00000039914 | Coq10a | protein_coding | -0.012375315 | 5.652001736 | 0.0011871 | 0.972934 | 0.993242374 |
| ENSMUSG00000024776 | Stambpl1 | protein_coding | -0.011167454 | 4.764129208 | 0.0011833 | 0.9729767 | 0.993242374 |
| ENSMUSG00000018861 | Fdxr | protein_coding | -0.015157668 | 3.581501674 | 0.0011791 | 0.9730244 | 0.993242374 |
| ENSMUSG00000025040 | Fundc1 | protein_coding | -0.007875795 | 7.658821387 | 0.0011765 | 0.9730546 | 0.993242374 |

| ensembl_gene_id | external_gene_name | gene_biotype | log2 (fold change) | log2 (counts per million) | F-value | p-value | False discovery rate |
|---|---|---|---|---|---|---|---|
| ENSMUSG00000006369 | Fbln1 | protein_coding | -0.021865545 | 3.155322925 | 0.0011713 | 0.973114 | 0.993242374 |
| ENSMUSG00000027281 | Slx4ip | protein_coding | -0.015798504 | 3.290612586 | 0.0011681 | 0.9731501 | 0.993242374 |
| ENSMUSG00000043964 | Orai3 | protein_coding | 0.016384273 | 3.812328674 | 0.0011654 | 0.9731818 | 0.993242374 |
| ENSMUSG00000038738 | Shank1 | protein_coding | -0.012681042 | 5.505999891 | 0.0011599 | 0.9732451 | 0.993242374 |
| ENSMUSG00000041408 | Wapl | protein_coding | 0.007379508 | 6.719766122 | 0.0011592 | 0.973253 | 0.993242374 |
| ENSMUSG00000029089 | 5730480H06Rik | protein_coding | -0.018141261 | 3.863149373 | 0.0011565 | 0.9732847 | 0.993242374 |
| ENSMUSG00000055676 | Gm5069 | transcribed_processe | -0.021307834 | 2.771218341 | 0.0011559 | 0.9732907 | 0.993242374 |
| ENSMUSG00000026718 | Stam | protein_coding | -0.009127247 | 5.8683243 | 0.0011428 | 0.973443 | 0.993303497 |
| ENSMUSG00000047084 | Ngrn | protein_coding | 0.009007794 | 6.385211894 | 0.0011198 | 0.9737117 | 0.993483452 |
| ENSMUSG00000066607 | 6030419C18Rik | protein_coding | -0.022081389 | 2.612149884 | 0.0011028 | 0.9739118 | 0.993499856 |
| ENSMUSG00000027472 | Pdrg1 | protein_coding | -0.007205317 | 7.819654886 | 0.0011027 | 0.9739126 | 0.993499856 |
| ENSMUSG00000051527 | Usp29 | protein_coding | -0.010193341 | 5.338877038 | 0.0010534 | 0.9745018 | 0.994006632 |
| ENSMUSG00000054469 | Lclat1 | protein_coding | -0.0173946 | 3.717781153 | 0.0010356 | 0.9747185 | 0.994045528 |
| ENSMUSG00000029622 | Arpc1b | protein_coding | -0.011723027 | 4.752222559 | 0.0010281 | 0.9748105 | 0.994045528 |
| ENSMUSG00000044791 | Setd2 | protein_coding | 0.01337178 | 4.536648777 | 0.0010275 | 0.9748173 | 0.994045528 |
| ENSMUSG00000029036 | Atad3a | protein_coding | 0.015767727 | 3.588426218 | 0.0009717 | 0.9755105 | 0.994610262 |
| ENSMUSG00000056413 | Adap1 | protein_coding | 0.00903466 | 6.48871669 | 0.0009681 | 0.9755561 | 0.994610262 |
| ENSMUSG00000039623 | Ap5z1 | protein_coding | -0.018011374 | 3.859692997 | 0.0009497 | 0.9757905 | 0.994651376 |
| ENSMUSG00000097616 | 1110019D14Rik | lincRNA | 0.017326696 | 3.245475465 | 0.000943 | 0.9758745 | 0.994651376 |
| ENSMUSG00000029580 | Actb | protein_coding | 0.011780048 | 12.74089718 | 0.000938 | 0.9759613 | 0.994651376 |
| ENSMUSG00000023980 | Taf8 | protein_coding | 0.009336915 | 5.220399206 | 0.0009322 | 0.9760139 | 0.994651376 |
| ENSMUSG00000028488 | Sh3gl2 | protein_coding | -0.007157721 | 7.615826739 | 0.0009287 | 0.9760589 | 0.994651376 |
| ENSMUSG00000036438 | Calm2 | protein_coding | 0.007733468 | 12.2062153 | 0.0009091 | 0.9763126 | 0.994815611 |
| ENSMUSG00000018678 | Sp2 | protein_coding | 0.02804565 | 2.484319972 | 0.0008868 | 0.9766049 | 0.995019229 |
| ENSMUSG00000027828 | Ssr3 | protein_coding | -0.005588231 | 7.518890899 | 0.0008781 | 0.9767201 | 0.995042236 |
| ENSMUSG00000028580 | Pum1 | protein_coding | 0.009345394 | 6.789635868 | 0.0008708 | 0.9768192 | 0.995048977 |
| ENSMUSG00000045795 | Whamm | protein_coding | -0.022315319 | 2.505882743 | 0.0008433 | 0.9771855 | 0.995327766 |
| ENSMUSG00000010086 | Rnf112 | protein_coding | -0.012222715 | 4.615824824 | 0.0007982 | 0.9778036 | 0.995863094 |
| ENSMUSG00000017639 | Rab11fip4 | protein_coding | 0.008581767 | 7.063124303 | 0.000788 | 0.9779486 | 0.995916423 |
| ENSMUSG00000002455 | Prpf6 | protein_coding | -0.008926863 | 6.141371447 | 0.0007777 | 0.9780903 | 0.995966429 |
| ENSMUSG00000002778 | Kdelr1 | protein_coding | -0.010004083 | 5.216955976 | 0.000764 | 0.9782838 | 0.996069076 |
| ENSMUSG00000073684 | Faap20 | protein_coding | 0.007172139 | 5.802245919 | 0.0007449 | 0.978558 | 0.996149128 |
| ENSMUSG00000097717 | 1700066J03Rik | lincRNA | 0.018736656 | 3.00396811 | 0.0007373 | 0.9786668 | 0.996149128 |
| ENSMUSG00000048039 | Isg20l2 | protein_coding | 0.012732485 | 3.647626949 | 0.0007366 | 0.9786768 | 0.996149128 |
| ENSMUSG00000087150 | BC064078 | transcribed_unproces | -0.01628524 | 3.367491776 | 0.0007327 | 0.978733 | 0.996149128 |
| ENSMUSG00000026618 | Iars2 | protein_coding | 0.013569092 | 3.856479725 | 0.0007116 | 0.979042 | 0.996239591 |
| ENSMUSG00000070000 | Fcho1 | protein_coding | 0.012567264 | 3.832681954 | 0.0007096 | 0.979072 | 0.996239591 |
| ENSMUSG00000051989 | Smim11 | protein_coding | -0.009399087 | 5.259884456 | 0.000707 | 0.9791104 | 0.996239591 |
| ENSMUSG00000041124 | Msantd4 | protein_coding | 0.006754164 | 6.79558732 | 0.0006932 | 0.9793149 | 0.996239591 |
| ENSMUSG00000039126 | Prune2 | protein_coding | 0.013063698 | 8.868720627 | 0.000693 | 0.9794225 | 0.996239591 |
| ENSMUSG00000020780 | Srp68 | protein_coding | -0.006088078 | 6.480610392 | 0.0006839 | 0.9794531 | 0.996239591 |
| ENSMUSG00000042178 | Armc5 | protein_coding | 0.017069107 | 3.018035505 | 0.000678 | 0.9795427 | 0.996239591 |
| ENSMUSG00000034022 | Cpsf1 | protein_coding | -0.009911951 | 4.900114075 | 0.0006671 | 0.9797072 | 0.996239591 |
| ENSMUSG00000056023 | Gm9989 | processed_transcript | -0.014271877 | 3.638110107 | 0.000667 | 0.9797086 | 0.996239591 |
| ENSMUSG00000026240 | Cops7b | protein_coding | 0.00911482 | 4.814157823 | 0.000664 | 0.9797546 | 0.996239591 |
| ENSMUSG00000032258 | Lca5 | protein_coding | -0.02857915 | 2.502113867 | 0.000659 | 0.979841 | 0.996239591 |
| ENSMUSG00000106379 | Lhfpl3 | protein_coding | 0.006511712 | 6.027799078 | 0.0006212 | 0.9804175 | 0.996553267 |
| ENSMUSG00000026594 | Ralgps2 | protein_coding | -0.008844278 | 4.878117885 | 0.0006193 | 0.9804481 | 0.996553267 |
| ENSMUSG00000030188 | Magohb | protein_coding | 0.008468895 | 5.418424642 | 0.0006184 | 0.9804625 | 0.996553267 |
| ENSMUSG00000036241 | Ube2r2 | protein_coding | 0.005378696 | 6.414309663 | 0.0006147 | 0.9805202 | 0.996553267 |
| ENSMUSG00000018921 | Pelp1 | protein_coding | -0.010398847 | 3.524670135 | 0.0006059 | 0.980661 | 0.996556666 |
| ENSMUSG00000060206 | Zfp462 | protein_coding | -0.009705611 | 4.066285577 | 0.0006022 | 0.9807199 | 0.996556666 |
| ENSMUSG00000039879 | Heca | protein_coding | 0.009734023 | 3.829112711 | 0.0005895 | 0.9809243 | 0.996556666 |
| ENSMUSG00000029594 | Rbm19 | protein_coding | -0.013605247 | 3.338040783 | 0.0005887 | 0.9809375 | 0.996556666 |
| ENSMUSG00000019883 | Echdc1 | protein_coding | -0.010975702 | 3.896634565 | 0.0005836 | 0.9810191 | 0.996556666 |
| ENSMUSG00000062198 | 2700097O09Rik | protein_coding | 0.008974256 | 4.295031524 | 0.000572 | 0.9812091 | 0.996556666 |
| ENSMUSG00000021764 | Ndufs4 | protein_coding | -0.007027289 | 8.83922535 | 0.0005714 | 0.9812192 | 0.996556666 |
| ENSMUSG00000053644 | Aldh7a1 | protein_coding | -0.006202256 | 5.997732818 | 0.0005686 | 0.9812649 | 0.996556666 |
| ENSMUSG00000058064 | Gm10036 | protein_coding | -0.028317351 | 2.964323997 | 0.0005629 | 0.9814061 | 0.996605881 |
| ENSMUSG00000006526 | Tmem110 | protein_coding | -0.014224853 | 3.326175884 | 0.0005435 | 0.9816838 | 0.996793708 |
| ENSMUSG00000030145 | Zfp248 | protein_coding | -0.00940598 | 4.432422608 | 0.0005359 | 0.981812 | 0.996829843 |
| ENSMUSG00000069227 | Gprin1 | protein_coding | 0.009491905 | 3.98846622 | 0.0005187 | 0.9821059 | 0.996976339 |
| ENSMUSG00000021798 | Ldb3 | protein_coding | 0.009794455 | 4.315403515 | 0.0005141 | 0.9821848 | 0.996976339 |
| ENSMUSG00000042826 | Fgf11 | protein_coding | -0.011511782 | 3.445099074 | 0.0005077 | 0.9822966 | 0.996976339 |
| ENSMUSG00000042766 | Trim46 | protein_coding | 0.005899044 | 5.439437472 | 0.000502 | 0.9823966 | 0.996976339 |
| ENSMUSG00000034310 | Tmem132d | protein_coding | 0.01327437 | 2.98634704 | 0.0004999 | 0.9824328 | 0.996976339 |
| ENSMUSG00000020659 | Cbll1 | protein_coding | 0.00846773 | 4.67701495 | 0.0004919 | 0.9825742 | 0.996976339 |
| ENSMUSG00000019818 | Cd164 | protein_coding | 0.005902657 | 6.237024336 | 0.0004902 | 0.9826053 | 0.996976339 |
| ENSMUSG00000052920 | Prkg1 | protein_coding | 0.011963594 | 3.423851385 | 0.0004729 | 0.982914 | 0.997195383 |
| ENSMUSG00000015354 | Pcolce2 | protein_coding | 0.014743942 | 2.687559785 | 0.0004596 | 0.9831567 | 0.997217586 |
| ENSMUSG00000032101 | Ddx25 | protein_coding | 0.005030029 | 7.373943566 | 0.0004574 | 0.9831967 | 0.997217586 |
| ENSMUSG00000020411 | Nipal4 | protein_coding | -0.014934688 | 2.931015967 | 0.0004564 | 0.9832141 | 0.997217586 |
| ENSMUSG00000016833 | Mrps18c | protein_coding | 0.006354529 | 7.215217141 | 0.0004375 | 0.9835673 | 0.99737235 |
| ENSMUSG00000026117 | Zap70 | protein_coding | 0.022762188 | 2.329439338 | 0.0004311 | 0.9836869 | 0.99737235 |
| ENSMUSG00000036894 | Rap2b | protein_coding | -0.006459227 | 6.864916024 | 0.0004303 | 0.983704 | 0.99737235 |
| ENSMUSG00000048004 | Tmem196 | protein_coding | -0.009140335 | 4.160727152 | 0.0004242 | 0.9838181 | 0.99737235 |
| ENSMUSG00000020329 | Polrmt | protein_coding | -0.013443467 | 3.309973205 | 0.0004235 | 0.9838304 | 0.99737235 |
| ENSMUSG00000001211 | Agpat3 | protein_coding | 0.006785895 | 5.813345945 | 0.0003833 | 0.984618 | 0.997775653 |

| ensembl_gene_id | external_gene_name | gene_biotype | log2 (fold change) | log2 (counts per million) | F-value | p-value | False discovery rate |
|---|---|---|---|---|---|---|---|
| ENSMUSG00000061458 | Nol10 | protein_coding | -0.007788766 | 4.249977819 | 0.0003816 | 0.9846515 | 0.997775653 |
| ENSMUSG00000038312 | Edem2 | protein_coding | 0.010899751 | 4.458289532 | 0.0003762 | 0.9847756 | 0.997775653 |
| ENSMUSG00000030643 | Rab30 | protein_coding | -0.00689401 | 5.044029253 | 0.0003615 | 0.9850619 | 0.997775653 |
| ENSMUSG00000022856 | Tmem41a | protein_coding | -0.006536296 | 4.42514257 | 0.0003548 | 0.9852009 | 0.997775653 |
| ENSMUSG00000034412 | Tbc1d10a | protein_coding | -0.010627011 | 3.277671923 | 0.0003544 | 0.9852085 | 0.997775653 |
| ENSMUSG00000020692 | Nle1 | protein_coding | -0.009345554 | 4.406804354 | 0.0003527 | 0.9852438 | 0.997775653 |
| ENSMUSG00000034557 | Zfyve9 | protein_coding | -0.005480937 | 5.23574843 | 0.0003464 | 0.9853776 | 0.997775653 |
| ENSMUSG00000034190 | Chmp7 | protein_coding | -0.00535425 | 6.459667764 | 0.0003311 | 0.9857035 | 0.997775653 |
| ENSMUSG00000025745 | Hadha | protein_coding | 0.005369352 | 5.641655946 | 0.000326 | 0.9858145 | 0.997775653 |
| ENSMUSG00000002968 | Med25 | protein_coding | -0.007002113 | 4.253980917 | 0.0003153 | 0.986048 | 0.997775653 |
| ENSMUSG00000039089 | L3mbtl3 | protein_coding | 0.006243312 | 4.024540819 | 0.0003108 | 0.9861492 | 0.997775653 |
| ENSMUSG00000020326 | Ccng1 | protein_coding | -0.006162303 | 4.812654868 | 0.0003095 | 0.9861768 | 0.997775653 |
| ENSMUSG00000040761 | Spen | protein_coding | 0.00882777 | 4.045662314 | 0.0003017 | 0.9863538 | 0.997775653 |
| ENSMUSG00000067242 | Lgi1 | protein_coding | 0.006491478 | 4.060192259 | 0.0003015 | 0.9863565 | 0.997775653 |
| ENSMUSG00000068874 | Selenbp1 | protein_coding | 0.007869724 | 3.323188015 | 0.0002986 | 0.9864221 | 0.997775653 |
| ENSMUSG00000028703 | Lrrc41 | protein_coding | 0.008386175 | 3.908571667 | 0.0002976 | 0.9864454 | 0.997775653 |
| ENSMUSG00000038563 | Efl1 | protein_coding | -0.005513606 | 4.51451738 | 0.0002968 | 0.9864647 | 0.997775653 |
| ENSMUSG00000038762 | Abcf1 | protein_coding | 0.004550563 | 6.924859348 | 0.0002947 | 0.986511 | 0.997775653 |
| ENSMUSG00000022562 | Oplah | protein_coding | -0.01231026 | 2.775960112 | 0.0002912 | 0.9865919 | 0.997775653 |
| ENSMUSG00000010517 | Faf1 | protein_coding | 0.006091242 | 5.091484325 | 0.0002842 | 0.9867547 | 0.997775653 |
| ENSMUSG00000031021 | Tmem9b | protein_coding | -0.004429424 | 5.948648697 | 0.0002824 | 0.986796 | 0.997775653 |
| ENSMUSG00000025280 | Polr3a | protein_coding | 0.007767756 | 3.709208896 | 0.0002817 | 0.9868134 | 0.997775653 |
| ENSMUSG00000026433 | Rab29 | protein_coding | 0.011439322 | 3.270456569 | 0.0002802 | 0.9868492 | 0.997775653 |
| ENSMUSG00000009406 | Elk1 | protein_coding | -0.006562192 | 4.138443608 | 0.0002776 | 0.9869089 | 0.997775653 |
| ENSMUSG00000034055 | Phka1 | protein_coding | 0.011814558 | 2.741177134 | 0.0002775 | 0.9869117 | 0.997775653 |
| ENSMUSG00000048921 | Zfp689 | protein_coding | -0.007856068 | 3.831255787 | 0.0002771 | 0.9869204 | 0.997775653 |
| ENSMUSG00000036323 | Srp72 | protein_coding | -0.003344944 | 7.571869453 | 0.0002604 | 0.98732 | 0.997775653 |
| ENSMUSG00000032803 | Cdv3 | protein_coding | 0.00429458 | 5.753626047 | 0.0002492 | 0.9875978 | 0.997775653 |
| ENSMUSG00000029878 | Dbpht2 | lincRNA | 0.00639761 | 8.515574096 | 0.0002469 | 0.9876814 | 0.997775653 |
| ENSMUSG00000025429 | Pstpip2 | protein_coding | -0.012155193 | 2.547898882 | 0.0002443 | 0.9877179 | 0.997775653 |
| ENSMUSG00000037210 | Fam193a | protein_coding | 0.006144027 | 4.910194379 | 0.0002432 | 0.9877474 | 0.997775653 |
| ENSMUSG00000046861 | Hectd3 | protein_coding | -0.006665813 | 4.242507124 | 0.0002417 | 0.9877858 | 0.997775653 |
| ENSMUSG00000020074 | Ccar1 | protein_coding | -0.00466002 | 5.314641543 | 0.0002413 | 0.9877951 | 0.997775653 |
| ENSMUSG00000079157 | Fam155a | protein_coding | 0.004091172 | 5.533616211 | 0.0002273 | 0.9881542 | 0.997775653 |
| ENSMUSG00000028879 | Stx12 | protein_coding | 0.003959734 | 7.947623175 | 0.0002207 | 0.9883277 | 0.997775653 |
| ENSMUSG00000037993 | Dhx38 | protein_coding | -0.006800672 | 4.389647886 | 0.0002197 | 0.9883538 | 0.997775653 |
| ENSMUSG00000019785 | Clvs2 | protein_coding | -0.00412422 | 5.983015569 | 0.0002136 | 0.9885172 | 0.997775653 |
| ENSMUSG00000028226 | Mmp16 | protein_coding | -0.006313713 | 4.054769991 | 0.0002116 | 0.9885701 | 0.997775653 |
| ENSMUSG00000021792 | Fam213a | protein_coding | 0.003563181 | 7.844558865 | 0.0002097 | 0.9886206 | 0.997775653 |
| ENSMUSG00000002635 | Pdcd2l | protein_coding | 0.005721843 | 4.720181586 | 0.0002024 | 0.9888228 | 0.997775653 |
| ENSMUSG00000050022 | Amz1 | protein_coding | 0.005100038 | 4.858056728 | 0.000201 | 0.9888594 | 0.997775653 |
| ENSMUSG00000048497 | Mmgt2 | protein_coding | 0.007414153 | 3.594320019 | 0.0002007 | 0.9888698 | 0.997775653 |
| ENSMUSG00000020898 | Ctc1 | protein_coding | 0.005573155 | 4.549825919 | 0.000199 | 0.9889154 | 0.997775653 |
| ENSMUSG00000032301 | Psma4 | protein_coding | 0.003360663 | 7.577675327 | 0.0001984 | 0.988932 | 0.997775653 |
| ENSMUSG00000056666 | Retsat | protein_coding | 0.006890426 | 3.650956387 | 0.0001953 | 0.9890184 | 0.997775653 |
| ENSMUSG00000028976 | Slc2a5 | protein_coding | 0.015634646 | 2.785864423 | 0.0001959 | 0.9890208 | 0.997775653 |
| ENSMUSG00000031578 | Mak16 | protein_coding | 0.004838758 | 4.675581515 | 0.0001949 | 0.989032 | 0.997775653 |
| ENSMUSG00000086171 | Pcsk2os1 | antisense_RNA | -0.010019471 | 2.510343127 | 0.0001917 | 0.9891216 | 0.997775653 |
| ENSMUSG00000025041 | Nt5c2 | protein_coding | -0.004159459 | 5.858135852 | 0.0001867 | 0.9892635 | 0.997775653 |
| ENSMUSG00000022490 | Ppp1r1a | protein_coding | 0.004265311 | 5.34981323 | 0.0001858 | 0.9892899 | 0.997775653 |
| ENSMUSG00000035469 | Rcbtb1 | protein_coding | -0.00490526 | 5.131448079 | 0.0001856 | 0.9892958 | 0.997775653 |
| ENSMUSG00000028952 | Zbtb48 | protein_coding | 0.005504846 | 4.685661917 | 0.000182 | 0.9893998 | 0.997775653 |
| ENSMUSG00000058503 | Fam133b | protein_coding | 0.003395291 | 6.062374776 | 0.0001764 | 0.9895634 | 0.997775653 |
| ENSMUSG00000028901 | Gmeb1 | protein_coding | -0.006012735 | 4.083689922 | 0.0001747 | 0.9896149 | 0.997775653 |
| ENSMUSG00000019338 | Zfp687 | protein_coding | -0.005296044 | 3.918077177 | 0.0001663 | 0.9898676 | 0.997775653 |
| ENSMUSG00000015143 | Actn1 | protein_coding | -0.004096864 | 4.509325244 | 0.0001591 | 0.9900891 | 0.997775653 |
| ENSMUSG00000039632 | Ccdc151 | protein_coding | 0.01132392 | 2.469709925 | 0.0001586 | 0.9901037 | 0.997775653 |
| ENSMUSG00000037649 | H2-DMa | protein_coding | -0.005533329 | 4.709291474 | 0.0001532 | 0.9902748 | 0.997775653 |
| ENSMUSG00000024843 | Chka | protein_coding | 0.004933187 | 4.607991209 | 0.0001515 | 0.9903287 | 0.997775653 |
| ENSMUSG00000057572 | Zbtb8os | protein_coding | 0.004441308 | 4.87184687 | 0.0001511 | 0.990343 | 0.997775653 |
| ENSMUSG00000031483 | Erlin2 | protein_coding | -0.003859618 | 5.141166293 | 0.0001496 | 0.990391 | 0.997775653 |
| ENSMUSG00000032612 | Usp4 | protein_coding | -0.002980451 | 6.156896791 | 0.0001491 | 0.9904055 | 0.997775653 |
| ENSMUSG00000056429 | Tgoln1 | protein_coding | -0.002584834 | 6.553580373 | 0.0001448 | 0.9905447 | 0.997775653 |
| ENSMUSG00000041757 | Plekha6 | protein_coding | 0.004335885 | 4.590886825 | 0.0001437 | 0.990581 | 0.997775653 |
| ENSMUSG00000030987 | Stim1 | protein_coding | 0.003188947 | 5.982933369 | 0.0001407 | 0.9906802 | 0.997775653 |
| ENSMUSG00000036249 | Rbm43 | protein_coding | -0.004286262 | 5.19496541 | 0.0001402 | 0.9906954 | 0.997775653 |
| ENSMUSG00000062373 | Tmem65 | protein_coding | -0.003405677 | 5.907264928 | 0.0001395 | 0.9907212 | 0.997775653 |
| ENSMUSG00000038342 | Mlxip | protein_coding | 0.008762368 | 2.475320782 | 0.0001362 | 0.9908291 | 0.997775653 |
| ENSMUSG00000017631 | Abr | protein_coding | 0.002675338 | 7.151394272 | 0.0001359 | 0.9908399 | 0.997775653 |
| ENSMUSG00000040811 | Eml2 | protein_coding | -0.003652578 | 5.45460871 | 0.0001359 | 0.9908417 | 0.997775653 |
| ENSMUSG00000008475 | Arpc5 | protein_coding | 0.002339373 | 7.254298124 | 0.0001327 | 0.9909482 | 0.997775653 |
| ENSMUSG00000020605 | Hs1bp3 | protein_coding | 0.004939961 | 3.979064474 | 0.0001311 | 0.991002 | 0.997775653 |
| ENSMUSG00000090063 | Dlx6os1 | antisense_RNA | -0.005698947 | 4.559342189 | 0.0001268 | 0.9911519 | 0.997791191 |
| ENSMUSG00000028318 | Polr1e | protein_coding | -0.004482434 | 4.451103162 | 0.0001232 | 0.9912797 | 0.997791191 |
| ENSMUSG00000062526 | Mppe1 | protein_coding | -0.007299423 | 2.980219548 | 0.0001227 | 0.9912958 | 0.997791191 |
| ENSMUSG00000054477 | Kcnn2 | protein_coding | -0.004594006 | 3.612487642 | 0.0001172 | 0.9914941 | 0.997897462 |
| ENSMUSG00000039828 | Wdr70 | protein_coding | -0.002888338 | 5.70048989 | 0.000113 | 0.991649 | 0.99795989 |
| ENSMUSG00000038712 | Mindy1 | protein_coding | 0.004298779 | 4.8086224 | 0.0001029 | 0.9920302 | 0.997982831 |

| ensembl_gene_id | external_gene_name | gene_biotype | log2 (fold change) | log2 (counts per million) | F-value | p-value | False discovery rate |
|---|---|---|---|---|---|---|---|
| ENSMUSG00000063200 | Nol7 | protein_coding | 0.002709485 | 6.24434449 | 9.88E-05 | 0.9921905 | 0.997982831 |
| ENSMUSG00000035062 | Zc4h2 | protein_coding | -0.002954214 | 5.682403937 | 9.77E-05 | 0.9922346 | 0.997982831 |
| ENSMUSG00000020807 | 4933427D14Rik | protein_coding | 0.004438005 | 3.610852911 | 9.66E-05 | 0.9922778 | 0.997982831 |
| ENSMUSG00000060227 | Casc4 | protein_coding | -0.002903829 | 5.343589844 | 9.61E-05 | 0.9922965 | 0.997982831 |
| ENSMUSG00000020271 | Fbxw11 | protein_coding | 0.002632095 | 6.205915144 | 9.61E-05 | 0.9922983 | 0.997982831 |
| ENSMUSG00000029782 | Tmem209 | protein_coding | 0.004225283 | 3.896533446 | 9.55E-05 | 0.9923214 | 0.997982831 |
| ENSMUSG00000017686 | Rhot1 | protein_coding | -0.003032675 | 5.268118741 | 9.24E-05 | 0.9924464 | 0.997990223 |
| ENSMUSG00000021113 | Snapc1 | protein_coding | -0.002556997 | 5.663202216 | 9.08E-05 | 0.9925144 | 0.997990223 |
| ENSMUSG00000054766 | Set | protein_coding | -0.002444737 | 8.405427607 | 8.68E-05 | 0.9926804 | 0.997997386 |
| ENSMUSG00000036565 | Ttyh3 | protein_coding | -0.003992884 | 4.395065341 | 8.61E-05 | 0.9927072 | 0.997997386 |
| ENSMUSG00000015597 | Zfp318 | protein_coding | 0.003222075 | 4.926958998 | 8.29E-05 | 0.9928462 | 0.998007693 |
| ENSMUSG00000027088 | Phospho2 | protein_coding | -0.002494585 | 5.56835317 | 8.16E-05 | 0.992903 | 0.998007693 |
| ENSMUSG00000034235 | Usp54 | protein_coding | -0.007461818 | 2.469394203 | 7.41E-05 | 0.9932362 | 0.998229661 |
| ENSMUSG00000078570 | 1110065P20Rik | protein_coding | -0.00632624 | 2.752640373 | 7.25E-05 | 0.9933095 | 0.998229661 |
| ENSMUSG00000026113 | Inpp4a | protein_coding | 0.003015913 | 9.06552636 | 6.00E-05 | 0.9939239 | 0.998318206 |
| ENSMUSG00000061536 | Sec22c | protein_coding | 0.002482995 | 4.617431643 | 5.92E-05 | 0.9939529 | 0.998318206 |
| ENSMUSG00000044072 | Eml6 | protein_coding | -0.004752475 | 2.775507336 | 5.86E-05 | 0.9939829 | 0.998318206 |
| ENSMUSG00000026511 | Srp9 | protein_coding | 0.001563672 | 7.03976577 | 5.86E-05 | 0.9939832 | 0.998318206 |
| ENSMUSG00000039131 | Gipc2 | protein_coding | 0.003458983 | 3.787851571 | 5.69E-05 | 0.9940705 | 0.998318206 |
| ENSMUSG00000043629 | 1700019D03Rik | protein_coding | 0.003517554 | 3.571831769 | 5.50E-05 | 0.9941743 | 0.998318206 |
| ENSMUSG00000020076 | Ddx50 | protein_coding | 0.001949996 | 5.727135582 | 5.20E-05 | 0.9943336 | 0.998318206 |
| ENSMUSG00000033342 | Plppr5 | protein_coding | -0.002482126 | 4.639042414 | 5.18E-05 | 0.9943442 | 0.998318206 |
| ENSMUSG00000031791 | Tmem38a | protein_coding | 0.001481797 | 7.106816414 | 5.17E-05 | 0.9943498 | 0.998318206 |
| ENSMUSG00000045613 | Chrm2 | protein_coding | 0.002743519 | 4.4486178 | 5.10E-05 | 0.994386 | 0.998318206 |
| ENSMUSG00000015488 | Cacfd1 | protein_coding | 0.002856812 | 4.85552178 | 5.05E-05 | 0.9944189 | 0.998318206 |
| ENSMUSG00000020063 | Sirt1 | protein_coding | -0.004077588 | 2.913816698 | 4.87E-05 | 0.9945178 | 0.998324294 |
| ENSMUSG00000019467 | Arhgef25 | protein_coding | -0.002457153 | 4.862343665 | 4.66E-05 | 0.9946375 | 0.998333432 |
| ENSMUSG00000053460 | Ggcx | protein_coding | 0.004903515 | 3.244690604 | 4.53E-05 | 0.9947126 | 0.998333432 |
| ENSMUSG00000031174 | Rpgr | protein_coding | -0.002220606 | 4.631533362 | 4.29E-05 | 0.9948518 | 0.998350856 |
| ENSMUSG00000031323 | Dmrtc1a | protein_coding | -0.003150631 | 3.489503687 | 4.13E-05 | 0.9949512 | 0.998350856 |
| ENSMUSG00000034226 | Rhov | protein_coding | 0.004841904 | 2.758434946 | 4.04E-05 | 0.9950085 | 0.998350856 |
| ENSMUSG00000002064 | Sdf2 | protein_coding | -0.001467865 | 7.649452526 | 3.76E-05 | 0.9951807 | 0.99838707 |
| ENSMUSG00000018387 | Shroom1 | protein_coding | 0.00378069 | 2.888356445 | 3.68E-05 | 0.9952303 | 0.99838707 |
| ENSMUSG00000042671 | Rgs8 | protein_coding | -0.001767673 | 6.430973914 | 3.19E-05 | 0.99556 | 0.998562103 |
| ENSMUSG00000027551 | Zfp64 | protein_coding | 0.002171659 | 5.124007071 | 3.15E-05 | 0.9955905 | 0.998562103 |
| ENSMUSG00000021054 | Sgpp1 | protein_coding | -0.001695282 | 4.598487404 | 2.43E-05 | 0.9961295 | 0.998978696 |
| ENSMUSG00000061559 | Wdr61 | protein_coding | -0.001002752 | 7.66152338 | 2.35E-05 | 0.9961916 | 0.998978696 |
| ENSMUSG00000025823 | Pdia4 | protein_coding | 0.001153404 | 5.920511761 | 1.89E-05 | 0.9965839 | 0.99912015 |
| ENSMUSG00000029387 | Gtf2h3 | protein_coding | 0.001348298 | 5.419838996 | 1.88E-05 | 0.9965948 | 0.99912015 |
| ENSMUSG00000078861 | Zfp931 | protein_coding | -0.001849927 | 5.300697294 | 1.86E-05 | 0.9966114 | 0.99912015 |
| ENSMUSG00000056851 | Pcbp2 | protein_coding | -0.000854903 | 6.502847413 | 1.65E-05 | 0.9968075 | 0.999213012 |
| ENSMUSG00000031246 | Sh3bgrl | protein_coding | 0.001106031 | 7.149897123 | 1.57E-05 | 0.9968899 | 0.999213012 |
| ENSMUSG00000026514 | Cnih3 | protein_coding | 0.001262649 | 4.499758929 | 1.19E-05 | 0.9972863 | 0.999484066 |
| ENSMUSG00000025968 | Ndufs1 | protein_coding | 0.001203118 | 5.702321585 | 1.05E-05 | 0.9974512 | 0.999484066 |
| ENSMUSG00000036430 | Tbcc | protein_coding | 0.001628938 | 3.39072153 | 9.58E-06 | 0.9975676 | 0.999484066 |
| ENSMUSG00000036160 | Surf6 | protein_coding | 0.001280869 | 4.498404518 | 8.70E-06 | 0.9976818 | 0.999484066 |
| ENSMUSG00000030894 | Tpp1 | protein_coding | -0.000889054 | 6.126023615 | 7.10E-06 | 0.9979059 | 0.999484066 |
| ENSMUSG00000022748 | Cmss1 | protein_coding | -0.000796061 | 5.107140504 | 5.39E-06 | 0.998175 | 0.999484066 |
| ENSMUSG00000068923 | Syt11 | protein_coding | 0.000534318 | 9.840075036 | 5.29E-06 | 0.9981919 | 0.999484066 |
| ENSMUSG00000023921 | Mut | protein_coding | -0.00069476 | 5.10118876 | 4.51E-06 | 0.9983306 | 0.999484066 |
| ENSMUSG00000027259 | Adal | protein_coding | 0.000793413 | 4.295735333 | 4.48E-06 | 0.9983373 | 0.999484066 |
| ENSMUSG00000059005 | Hnrnpa3 | protein_coding | -0.000760784 | 3.884910127 | 4.46E-06 | 0.9983398 | 0.999484066 |
| ENSMUSG00000030255 | Sspn | protein_coding | -0.001560112 | 2.634035781 | 4.43E-06 | 0.9983457 | 0.999484066 |
| ENSMUSG00000021975 | Ints9 | protein_coding | -0.000739452 | 4.656545547 | 3.61E-06 | 0.9985068 | 0.999484066 |
| ENSMUSG00000046727 | Cystm1 | protein_coding | 0.000369268 | 8.856223718 | 3.11E-06 | 0.9986137 | 0.999484066 |
| ENSMUSG00000007950 | Abhd8 | protein_coding | 0.000445296 | 6.008048681 | 2.89E-06 | 0.998665 | 0.999484066 |
| ENSMUSG00000031960 | Aars | protein_coding | 0.000477205 | 7.021415259 | 2.72E-06 | 0.9987048 | 0.999484066 |
| ENSMUSG00000034613 | Ppm1h | protein_coding | 0.000659654 | 7.551319464 | 2.48E-06 | 0.9987652 | 0.999484066 |
| ENSMUSG00000040242 | Fgfr1op2 | protein_coding | 0.000314197 | 6.371582917 | 1.91E-06 | 0.9989148 | 0.999484066 |
| ENSMUSG00000041245 | Wnk3 | protein_coding | 0.000797992 | 3.176383851 | 1.73E-06 | 0.9989669 | 0.999484066 |
| ENSMUSG00000098234 | Snhg6 | lincRNA | -0.000569636 | 5.926672747 | 1.50E-06 | 0.9990385 | 0.999484066 |
| ENSMUSG00000020732 | Rab37 | protein_coding | 0.001064011 | 4.52075295 | 1.30E-06 | 0.9991096 | 0.999484066 |
| ENSMUSG00000049811 | Fam161a | protein_coding | 0.000584402 | 2.959912832 | 1.02E-06 | 0.9992051 | 0.999484066 |
| ENSMUSG00000038702 | Dsel | protein_coding | -0.000551374 | 3.507317858 | 8.89E-07 | 0.9992591 | 0.999484066 |
| ENSMUSG00000020528 | Prpsap2 | protein_coding | 0.000300538 | 5.112066053 | 7.98E-07 | 0.9992982 | 0.999484066 |
| ENSMUSG00000063382 | Bcl9l | protein_coding | 0.000319177 | 4.306735591 | 4.31E-07 | 0.9994844 | 0.999577341 |
| ENSMUSG00000002825 | Qtrt1 | protein_coding | 4.16E-05 | 5.198670867 | 1.85E-08 | 0.999893 | 0.999893035 |
