## Supplemental Table 5 for "Extended amygdala-parabrachial circuits alter threat assessment to regulate feeding"

### TABLE S5

| ensembl_gene_id | external_gene_name | gene_biotype | log2 (fold change) | log2 (counts per million) | F-value | p-value | False discovery rate |
|---|---|---|---|---|---|---|---|
| ENSMUSG00000025400 | Tac2 | protein_coding | 4.325582504 | 9.178170504 | 146.06814 | 1.48E-09 | 1.59E-05 |
| ENSMUSG00000045471 | Hcrt | protein_coding | -6.123358423 | 10.01304624 | 50.089962 | 1.67E-05 | 0.090012175 |
| ENSMUSG00000003872 | Lin7b | protein_coding | 1.701091154 | 6.138974701 | 25.33619 | 0.0001156 | 0.414265157 |
| ENSMUSG00000044933 | Sstr3 | protein_coding | 9.894550688 | 1.991282608 | 24.477447 | 0.0001653 | 0.444481553 |
| ENSMUSG00000036357 | Gpr101 | protein_coding | 2.247204506 | 4.869678402 | 19.622944 | 0.0004028 | 0.866200183 |
| ENSMUSG00000026389 | Steap3 | protein_coding | 8.750722364 | 2.068060088 | 19.324721 | 0.0004978 | 0.89216578 |
| ENSMUSG00000008036 | Ap2s1 | protein_coding | 1.22921003 | 8.101066345 | 17.94313 | 0.0006055 | 0.930110595 |
| ENSMUSG00000030844 | Rgs10 | protein_coding | 1.239537624 | 8.342266456 | 16.191429 | 0.0009484 | 0.999175987 |
| ENSMUSG00000040118 | Cacna2d1 | protein_coding | -1.519624089 | 5.728301998 | 16.113251 | 0.0009682 | 0.999175987 |
| ENSMUSG00000048644 | Ctxn1 | protein_coding | 1.058082906 | 7.951408505 | 15.275494 | 0.001212 | 0.999175987 |
| ENSMUSG00000024907 | Gal | protein_coding | 1.378810037 | 9.118265128 | 15.162462 | 0.0012499 | 0.999175987 |
| ENSMUSG00000022577 | Ly6h | protein_coding | 1.260887743 | 6.972127586 | 14.750013 | 0.0014001 | 0.999175987 |
| ENSMUSG00000024256 | Adcyap1 | protein_coding | -3.994858322 | 3.763489644 | 15.086245 | 0.0014163 | 0.999175987 |
| ENSMUSG00000010045 | Tmem115 | protein_coding | 1.562590031 | 5.388791014 | 14.618319 | 0.0014522 | 0.999175987 |
| ENSMUSG00000042258 | Isl1 | protein_coding | 1.258224574 | 6.31340539 | 14.281663 | 0.0015958 | 0.999175987 |
| ENSMUSG00000061518 | Cox5b | protein_coding | 1.491057602 | 8.013800626 | 13.754644 | 0.0018584 | 0.999175987 |
| ENSMUSG00000045731 | Pnoc | protein_coding | 1.047495314 | 6.687693393 | 13.523636 | 0.0019817 | 0.999175987 |
| ENSMUSG00000018169 | Mfng | protein_coding | 2.477581796 | 3.752551007 | 13.107586 | 0.0022376 | 0.999175987 |
| ENSMUSG00000029635 | Cdk8 | protein_coding | 1.378483852 | 6.496972303 | 12.828808 | 0.00243 | 0.999175987 |
| ENSMUSG00000030069 | Prok2 | protein_coding | 2.267701808 | 3.528654604 | 12.186322 | 0.0029488 | 0.999175987 |
| ENSMUSG00000036578 | Fxyd7 | protein_coding | 1.031286148 | 7.396395722 | 11.803901 | 0.0033166 | 0.999175987 |
| ENSMUSG00000027466 | Rbck1 | protein_coding | 1.324788245 | 6.158664113 | 11.584536 | 0.0035509 | 0.999175987 |
| ENSMUSG00000078117 | Gm16485 | protein_coding | 4.222526581 | 2.420546414 | 11.245472 | 0.0039507 | 0.999175987 |
| ENSMUSG00000053414 | Hunk | protein_coding | 1.671480532 | 4.479379812 | 11.184629 | 0.0040277 | 0.999175987 |
| ENSMUSG00000027270 | Lamp5 | protein_coding | 1.439307316 | 7.01568764 | 11.106525 | 0.0041368 | 0.999175987 |
| ENSMUSG00000085282 | Gm15663 | processed_transcript | 1.953821419 | 3.665509334 | 11.011079 | 0.0042569 | 0.999175987 |
| ENSMUSG00000005892 | Trh | protein_coding | -1.685425278 | 7.312990643 | 11.272779 | 0.0043079 | 0.999175987 |
| ENSMUSG00000039662 | Icmt | protein_coding | -1.688137631 | 4.319494949 | 10.928626 | 0.0043709 | 0.999175987 |
| ENSMUSG00000037628 | Cdkn3 | protein_coding | 3.325195914 | 2.570795995 | 10.49 | 0.0050388 | 0.999175987 |
| ENSMUSG00000041966 | Dcaf17 | protein_coding | -2.000646425 | 3.947191032 | 10.461771 | 0.0050856 | 0.999175987 |
| ENSMUSG00000024526 | Cidea | protein_coding | -2.230842483 | 3.764927347 | 10.382468 | 0.0052197 | 0.999175987 |
| ENSMUSG00000068580 | Zfyve19 | protein_coding | 1.223588414 | 6.224034757 | 10.297036 | 0.0053686 | 0.999175987 |
| ENSMUSG00000045302 | Preb | protein_coding | 1.276090416 | 5.968168153 | 10.27255 | 0.0054122 | 0.999175987 |
| ENSMUSG00000023277 | Twf2 | protein_coding | 1.319009392 | 5.785012577 | 10.193931 | 0.0055549 | 0.999175987 |
| ENSMUSG00000047228 | A2ml1 | protein_coding | 1.229301493 | 5.711089496 | 10.07189 | 0.0057848 | 0.999175987 |
| ENSMUSG00000021876 | Rnase4 | protein_coding | -1.865682216 | 4.138297218 | 10.027463 | 0.0058712 | 0.999175987 |
| ENSMUSG00000024952 | Rps6ka4 | protein_coding | 1.804781656 | 4.084557071 | 10.00446 | 0.0059165 | 0.999175987 |
| ENSMUSG00000049744 | Arhgap15 | protein_coding | -3.763520192 | 2.51061778 | 9.8977599 | 0.0061319 | 0.999175987 |
| ENSMUSG00000074738 | Fndc10 | protein_coding | 1.772129813 | 4.143632959 | 9.6438754 | 0.006681 | 0.999175987 |
| ENSMUSG00000020940 | 1700023F06Rik | protein_coding | 1.542609286 | 4.456806218 | 9.6435913 | 0.0066817 | 0.999175987 |
| ENSMUSG00000028670 | Lypla2 | protein_coding | 1.036096292 | 6.34628073 | 9.624004 | 0.0067263 | 0.999175987 |
| ENSMUSG00000020464 | Pnpt1 | protein_coding | -2.316560463 | 3.13808061 | 9.5973483 | 0.0067876 | 0.999175987 |
| ENSMUSG00000000792 | Slc5a5 | protein_coding | 1.160627909 | 6.950427196 | 9.4851366 | 0.0070527 | 0.999175987 |
| ENSMUSG00000042157 | Sprr2i | protein_coding | -7.130264609 | 1.905783563 | 9.4807736 | 0.0071207 | 0.999175987 |
| ENSMUSG00000008958 | Vps72 | protein_coding | 1.006663939 | 6.102991075 | 9.4543262 | 0.0071275 | 0.999175987 |
| ENSMUSG00000022021 | Diaph3 | protein_coding | 1.7634064 | 4.311244021 | 9.4139369 | 0.0072269 | 0.999175987 |
| ENSMUSG00000057863 | Rpl36 | protein_coding | 1.162494918 | 5.553229542 | 9.3830367 | 0.0073041 | 0.999175987 |
| ENSMUSG00000020133 | 2310011J03Rik | protein_coding | 0.943968433 | 6.388075915 | 9.3802681 | 0.007311 | 0.999175987 |
| ENSMUSG00000035992 | Fnip1 | protein_coding | -1.49839825 | 4.657855697 | 9.3302407 | 0.0074381 | 0.999175987 |
| ENSMUSG00000037999 | Arap2 | protein_coding | -1.534380663 | 4.659843276 | 9.3180856 | 0.0074693 | 0.999175987 |
| ENSMUSG00000039850 | Endov | protein_coding | 1.479317203 | 4.222574245 | 9.1811004 | 0.0078318 | 0.999175987 |
| ENSMUSG00000023064 | Sncg | protein_coding | -1.258072758 | 7.292200882 | 9.1274269 | 0.0079908 | 0.999175987 |
| ENSMUSG00000031309 | Rps6ka3 | protein_coding | -2.079247243 | 3.483811245 | 9.0830219 | 0.0081037 | 0.999175987 |
| ENSMUSG00000025645 | Ccdc51 | protein_coding | -2.113055958 | 3.507253039 | 8.9697401 | 0.0084311 | 0.999175987 |
| ENSMUSG00000025792 | Slc25a10 | protein_coding | 1.253965863 | 5.270183151 | 8.8041684 | 0.0089371 | 0.999175987 |
| ENSMUSG00000031949 | Adat1 | protein_coding | 2.056778804 | 3.422585029 | 8.8018079 | 0.0089445 | 0.999175987 |
| ENSMUSG00000086363 | A330102I10Rik | lincRNA | 1.549654254 | 4.731109168 | 8.6648635 | 0.0093896 | 0.999175987 |
| ENSMUSG00000007721 | Ccdc124 | protein_coding | 0.88241541 | 8.225386948 | 8.5029734 | 0.0099485 | 0.999175987 |
| ENSMUSG00000034254 | Agpat1 | protein_coding | 1.139677853 | 5.640121654 | 8.4712006 | 0.0100626 | 0.999175987 |
| ENSMUSG00000035285 | Nat14 | protein_coding | 1.18426473 | 6.095206322 | 8.4160456 | 0.0102642 | 0.999175987 |
| ENSMUSG00000004748 | Mtfp1 | protein_coding | 0.949894542 | 7.955389561 | 8.3951021 | 0.0103419 | 0.999175987 |
| ENSMUSG00000022617 | Chkb | protein_coding | 1.600971481 | 3.886913244 | 8.3708746 | 0.0104327 | 0.999175987 |
| ENSMUSG00000022089 | Bin3 | protein_coding | -1.743747556 | 3.915490179 | 8.3663295 | 0.0104498 | 0.999175987 |
| ENSMUSG00000003352 | Cacnb3 | protein_coding | 1.107786712 | 5.509101422 | 8.3452739 | 0.0105296 | 0.999175987 |
| ENSMUSG00000023150 | Ivns1abp | protein_coding | -0.903204916 | 6.256843816 | 8.2977729 | 0.0107121 | 0.999175987 |
| ENSMUSG00000035228 | Ccdc106 | protein_coding | 1.43044161 | 5.471957661 | 8.2224097 | 0.0110092 | 0.999175987 |
| ENSMUSG00000037876 | Jmjd1c | protein_coding | -1.580793156 | 4.82454241 | 8.2081136 | 0.0110666 | 0.999175987 |
| ENSMUSG00000042380 | Smim12 | protein_coding | 0.936021143 | 7.240841027 | 8.1641897 | 0.011245 | 0.999175987 |
| ENSMUSG00000000214 | Th | protein_coding | 1.571967476 | 5.627137338 | 8.1402571 | 0.0113577 | 0.999175987 |
| ENSMUSG00000001750 | Tcirg1 | protein_coding | 2.651151497 | 2.990143412 | 8.119409 | 0.0114304 | 0.999175987 |
| ENSMUSG00000019768 | Esr1 | protein_coding | 2.649697797 | 3.539813594 | 8.1018824 | 0.011518 | 0.999175987 |
| ENSMUSG00000049796 | Crh | protein_coding | 2.532635277 | 2.83028429 | 8.0565238 | 0.0116966 | 0.999175987 |
| ENSMUSG00000006611 | Hfe | protein_coding | -2.827222275 | 2.95804863 | 8.0280347 | 0.0118195 | 0.999175987 |
| ENSMUSG00000020042 | Btbd11 | protein_coding | 1.391869436 | 5.056746215 | 7.9037866 | 0.0123729 | 0.999175987 |
| ENSMUSG00000057342 | Sphk2 | protein_coding | 1.495582307 | 4.929433609 | 7.7675647 | 0.013014 | 0.999175987 |
| ENSMUSG00000028673 | Fuca1 | protein_coding | 0.854400962 | 6.853897675 | 7.7299563 | 0.0131975 | 0.999175987 |
| ENSMUSG00000049097 | Ankrd34a | protein_coding | 1.209684041 | 5.665853992 | 7.7090188 | 0.013301 | 0.999175987 |
| ENSMUSG00000056394 | Lig1 | protein_coding | -1.535994282 | 4.074865149 | 7.6867076 | 0.0134123 | 0.999175987 |

| ensembl_gene_id | external_gene_name | gene_biotype | log2 (fold change) | log2 (counts per million) | F-value | p-value | False discovery rate |
|---|---|---|---|---|---|---|---|
| ENSMUSG00000035372 | 1810055G02Rik | protein_coding | 1.551793113 | 4.514499838 | 7.6152925 | 0.0137757 | 0.999175987 |
| ENSMUSG00000027778 | Ift80 | protein_coding | -1.817540588 | 3.804707773 | 7.6098159 | 0.0138041 | 0.999175987 |
| ENSMUSG00000025647 | Shisa5 | protein_coding | 0.974376072 | 7.52527952 | 7.5962525 | 0.0138745 | 0.999175987 |
| ENSMUSG00000027602 | Map1lc3a | protein_coding | 0.803728042 | 8.415411366 | 7.5413942 | 0.0141636 | 0.999175987 |
| ENSMUSG00000035835 | Plppr3 | protein_coding | 1.231704547 | 5.406949151 | 7.5309768 | 0.0142193 | 0.999175987 |
| ENSMUSG00000007036 | Abhd16a | protein_coding | 1.109250845 | 5.766196043 | 7.5251695 | 0.0142505 | 0.999175987 |
| ENSMUSG00000021608 | Lpcat1 | protein_coding | 1.537379772 | 4.229477468 | 7.4942138 | 0.0144178 | 0.999175987 |
| ENSMUSG00000025208 | Mrpl43 | protein_coding | 0.847735631 | 7.39468562 | 7.4816484 | 0.0144864 | 0.999175987 |
| ENSMUSG00000022760 | Thap7 | protein_coding | 1.037918398 | 6.147698245 | 7.4576321 | 0.0146185 | 0.999175987 |
| ENSMUSG00000002944 | Cd36 | protein_coding | -4.090855011 | 2.164736826 | 7.4423453 | 0.0147033 | 0.999175987 |
| ENSMUSG00000043411 | Usp48 | protein_coding | -1.338231037 | 4.625503793 | 7.388982 | 0.0150037 | 0.999175987 |
| ENSMUSG00000022500 | Litaf | protein_coding | -0.888023448 | 6.764656062 | 7.3341412 | 0.0153197 | 0.999175987 |
| ENSMUSG00000023927 | Satb1 | protein_coding | -1.537325347 | 3.755547552 | 7.3132279 | 0.0154423 | 0.999175987 |
| ENSMUSG00000031229 | Atrx | protein_coding | -1.098804126 | 6.221157175 | 7.3096562 | 0.0154633 | 0.999175987 |
| ENSMUSG00000053841 | Txlna | protein_coding | -1.449433462 | 4.507375285 | 7.2393368 | 0.0158842 | 0.999175987 |
| ENSMUSG00000030443 | Zfp583 | protein_coding | 1.741482314 | 3.713609321 | 7.2233558 | 0.0159817 | 0.999175987 |
| ENSMUSG00000001156 | Mxd1 | protein_coding | -1.55189941 | 3.812498202 | 7.2215873 | 0.0159925 | 0.999175987 |
| ENSMUSG00000027088 | Phospho2 | protein_coding | 1.052568848 | 5.56835317 | 7.1651156 | 0.0163427 | 0.999175987 |
| ENSMUSG00000033949 | Trim36 | protein_coding | -1.949155892 | 3.119246863 | 7.1559091 | 0.0164006 | 0.999175987 |
| ENSMUSG00000020238 | Ncln | protein_coding | 1.463690778 | 4.257319797 | 7.1450295 | 0.0164694 | 0.999175987 |
| ENSMUSG00000054894 | Atp5s | protein_coding | 1.042828077 | 5.555725959 | 7.1273649 | 0.0165817 | 0.999175987 |
| ENSMUSG00000027327 | 1700037H04Rik | protein_coding | 0.835674918 | 9.438842398 | 7.1240002 | 0.0166032 | 0.999175987 |
| ENSMUSG00000053574 | 4930563E22Rik | processed_transcript | -1.986826525 | 3.28713829 | 7.1214864 | 0.0166193 | 0.999175987 |
| ENSMUSG00000032583 | Mon1a | protein_coding | 1.475360667 | 4.200117861 | 7.1184814 | 0.0166385 | 0.999175987 |
| ENSMUSG00000021587 | Pcsk1 | protein_coding | -1.343450006 | 4.154065003 | 7.113276 | 0.0166719 | 0.999175987 |
| ENSMUSG00000024500 | Ppp2r2b | protein_coding | 0.768225941 | 7.88228679 | 7.1067786 | 0.0167137 | 0.999175987 |
| ENSMUSG00000034560 | Washc4 | protein_coding | -1.724331853 | 4.050259752 | 7.045725 | 0.0171123 | 0.999175987 |
| ENSMUSG00000007029 | Vars | protein_coding | 2.17395143 | 3.074595686 | 7.01868 | 0.0172923 | 0.999175987 |
| ENSMUSG00000037966 | Ninj1 | protein_coding | 1.322377392 | 4.936713085 | 6.9878705 | 0.0175 | 0.999175987 |
| ENSMUSG00000028098 | Rnf115 | protein_coding | 0.980382595 | 5.421290025 | 6.956899 | 0.0177117 | 0.999175987 |
| ENSMUSG00000021536 | Adcy2 | protein_coding | -1.276206834 | 5.192257329 | 6.9471771 | 0.0177788 | 0.999175987 |
| ENSMUSG00000006941 | Eif1b | protein_coding | 0.779842159 | 8.260969803 | 6.9305338 | 0.0178943 | 0.999175987 |
| ENSMUSG00000023495 | Pcbp4 | protein_coding | 1.324212843 | 5.32965233 | 6.9008725 | 0.0181022 | 0.999175987 |
| ENSMUSG00000039244 | E130309D02Rik | protein_coding | 1.54846486 | 3.943347011 | 6.8960295 | 0.0181364 | 0.999175987 |
| ENSMUSG00000039886 | Tmem120a | protein_coding | 1.342808459 | 4.318940806 | 6.8825327 | 0.0182321 | 0.999175987 |
| ENSMUSG00000038173 | Enpp6 | protein_coding | -2.582381568 | 3.093479553 | 6.8738827 | 0.0182938 | 0.999175987 |
| ENSMUSG00000027884 | Clcc1 | protein_coding | 1.255414663 | 4.867255731 | 6.8727418 | 0.0183019 | 0.999175987 |
| ENSMUSG00000026201 | Stk16 | protein_coding | 1.130316364 | 6.45245225 | 6.8669431 | 0.0183614 | 0.999175987 |
| ENSMUSG00000025862 | Stag2 | protein_coding | -1.268103327 | 4.708151346 | 6.8444367 | 0.0185054 | 0.999175987 |
| ENSMUSG00000039278 | Pcsk1n | protein_coding | 1.167048553 | 5.94998412 | 6.8247886 | 0.0186482 | 0.999175987 |
| ENSMUSG00000001630 | Stk38l | protein_coding | -1.122342907 | 5.132107652 | 6.8243812 | 0.0186512 | 0.999175987 |
| ENSMUSG00000036106 | Prr5 | protein_coding | 2.121416252 | 3.567331264 | 6.8133903 | 0.0187316 | 0.999175987 |
| ENSMUSG00000039759 | Thap3 | protein_coding | 0.944351045 | 6.023403703 | 6.7893553 | 0.0189089 | 0.999175987 |
| ENSMUSG00000030020 | Prickle2 | protein_coding | -1.014703687 | 5.767254028 | 6.7851529 | 0.0189401 | 0.999175987 |
| ENSMUSG00000002968 | Med25 | protein_coding | 1.472391159 | 4.253980917 | 6.7587076 | 0.0191378 | 0.999175987 |
| ENSMUSG00000069045 | Ddx3y | protein_coding | -0.977220974 | 6.104745614 | 6.7252167 | 0.0193915 | 0.999175987 |
| ENSMUSG00000057130 | Txnl4a | protein_coding | 0.955221817 | 6.757728205 | 6.7087713 | 0.0195175 | 0.999175987 |
| ENSMUSG00000079657 | Rab26 | protein_coding | 1.611433387 | 5.200088954 | 6.7055026 | 0.01959 | 0.999175987 |
| ENSMUSG00000043602 | Zfp3 | protein_coding | 1.588363846 | 3.811429342 | 6.6901535 | 0.0196613 | 0.999175987 |
| ENSMUSG00000030086 | Chchd6 | protein_coding | 1.025552459 | 7.819536396 | 6.6756959 | 0.0197924 | 0.999175987 |
| ENSMUSG00000030172 | Erc1 | protein_coding | -1.515480344 | 4.023491567 | 6.642362 | 0.0200359 | 0.999175987 |
| ENSMUSG00000059182 | Skap2 | protein_coding | -1.564182707 | 4.404641125 | 6.6214587 | 0.0202024 | 0.999175987 |
| ENSMUSG00000038235 | F11r | protein_coding | -1.959860291 | 3.493792847 | 6.6090027 | 0.0203023 | 0.999175987 |
| ENSMUSG00000029810 | Tmem176b | protein_coding | -0.861109454 | 6.570724312 | 6.5986286 | 0.0203859 | 0.999175987 |
| ENSMUSG00000041560 | Nop53 | protein_coding | 1.253371687 | 6.640298729 | 6.6561773 | 0.0204054 | 0.999175987 |
| ENSMUSG00000028393 | Alad | protein_coding | 0.782132981 | 6.709993692 | 6.5436446 | 0.0208359 | 0.999175987 |
| ENSMUSG00000063889 | Crem | protein_coding | -1.136144103 | 5.032857539 | 6.5361717 | 0.020898 | 0.999175987 |
| ENSMUSG00000057193 | Slc44a2 | protein_coding | -1.79936839 | 3.950299999 | 6.5323509 | 0.0209298 | 0.999175987 |
| ENSMUSG00000020131 | Pcsk4 | protein_coding | 2.09059872 | 3.616326002 | 6.5202447 | 0.0210309 | 0.999175987 |
| ENSMUSG00000080268 | Brms1 | protein_coding | 0.843196563 | 6.284589523 | 6.4622643 | 0.021523 | 0.999175987 |
| ENSMUSG00000028480 | Glipr2 | protein_coding | -2.761984793 | 2.854064298 | 6.4590509 | 0.0215507 | 0.999175987 |
| ENSMUSG00000035382 | Pcsk7 | protein_coding | 2.05585614 | 3.056818934 | 6.4547408 | 0.0215878 | 0.999175987 |
| ENSMUSG00000028653 | Trit1 | protein_coding | 1.377258569 | 4.177133886 | 6.4545265 | 0.0215897 | 0.999175987 |
| ENSMUSG00000024835 | Coro1b | protein_coding | 0.788372787 | 6.337349066 | 6.3944154 | 0.0221157 | 0.999175987 |
| ENSMUSG00000061119 | Prcp | protein_coding | -2.412919089 | 3.268052999 | 6.3866601 | 0.0222043 | 0.999175987 |
| ENSMUSG00000037979 | Ccdc92 | protein_coding | 0.942731794 | 6.192813252 | 6.3841402 | 0.0222071 | 0.999175987 |
| ENSMUSG00000066441 | Rdh11 | protein_coding | -1.231086727 | 4.662928458 | 6.3610015 | 0.0224144 | 0.999175987 |
| ENSMUSG00000040563 | Plppr2 | protein_coding | 1.296559497 | 5.355403126 | 6.3465161 | 0.0225454 | 0.999175987 |
| ENSMUSG00000032114 | Slc37a4 | protein_coding | -1.493789368 | 4.078771897 | 6.3168806 | 0.022816 | 0.999175987 |
| ENSMUSG00000056895 | Hist3h2ba | protein_coding | 1.674649159 | 3.746305997 | 6.2940051 | 0.0230275 | 0.999175987 |
| ENSMUSG00000020775 | Mrpl38 | protein_coding | 1.242179985 | 6.000364874 | 6.2918631 | 0.0230674 | 0.999175987 |
| ENSMUSG00000041660 | Bbox1 | protein_coding | -2.401069368 | 3.012470954 | 6.2799422 | 0.0231586 | 0.999175987 |
| ENSMUSG00000020422 | Tns3 | protein_coding | -1.890942944 | 3.221768076 | 6.276542 | 0.0231904 | 0.999175987 |
| ENSMUSG00000024841 | Eif1ad | protein_coding | 0.702344043 | 7.031500112 | 6.2702269 | 0.0232496 | 0.999175987 |
| ENSMUSG00000063698 | Sfxn4 | protein_coding | -0.996635477 | 5.734147057 | 6.2634439 | 0.0233135 | 0.999175987 |
| ENSMUSG00000022099 | Dmtn | protein_coding | 0.792878243 | 7.798985178 | 6.2356389 | 0.0235772 | 0.999175987 |
| ENSMUSG00000028959 | Fastk | protein_coding | 0.891211412 | 6.605509675 | 6.2058821 | 0.0238633 | 0.999175987 |
| ENSMUSG00000028082 | Sh3d19 | protein_coding | -1.811617133 | 3.122541729 | 6.1965795 | 0.0239535 | 0.999175987 |

| ensembl_gene_id | external_gene_name | gene_biotype | log2 (fold change) | log2 (counts per million) | F-value | p-value | False discovery rate |
|---|---|---|---|---|---|---|---|
| ENSMUSG00000027309 | 4930402H24Rik | protein_coding | -1.01228401 | 5.915773539 | 6.1800693 | 0.0241146 | 0.999175987 |
| ENSMUSG00000026754 | Golga1 | protein_coding | -1.054704387 | 4.866080909 | 6.1667441 | 0.0242456 | 0.999175987 |
| ENSMUSG00000020459 | Mtif2 | protein_coding | 1.108699681 | 5.176760627 | 6.1468812 | 0.0244423 | 0.999175987 |
| ENSMUSG00000031232 | Magt1 | protein_coding | -1.757268109 | 3.95752326 | 6.1351773 | 0.0245591 | 0.999175987 |
| ENSMUSG00000052372 | Il1rapl1 | protein_coding | 1.15807813 | 7.527975867 | 6.1970595 | 0.0246968 | 0.999175987 |
| ENSMUSG00000007038 | Neu1 | protein_coding | 1.114612875 | 5.20445601 | 6.0858776 | 0.025058 | 0.999175987 |
| ENSMUSG00000004961 | Syt5 | protein_coding | 0.692174221 | 8.28337684 | 6.0802445 | 0.0251157 | 0.999175987 |
| ENSMUSG00000026383 | Epb41l5 | protein_coding | -1.599527004 | 3.350534585 | 6.0793165 | 0.0251253 | 0.999175987 |
| ENSMUSG00000034932 | Mrpl54 | protein_coding | 1.540994603 | 6.746762172 | 6.2708638 | 0.0252177 | 0.999175987 |
| ENSMUSG00000029404 | Arl6ip4 | protein_coding | 0.675171524 | 7.407770941 | 6.0560133 | 0.0253659 | 0.999175987 |
| ENSMUSG00000036181 | Hist1h1c | protein_coding | -1.571596163 | 3.712771943 | 6.0554366 | 0.0253719 | 0.999175987 |
| ENSMUSG00000031529 | Tnks | protein_coding | -1.012004224 | 5.229642601 | 6.0432006 | 0.0254993 | 0.999175987 |
| ENSMUSG00000037646 | Vps13b | protein_coding | -2.014303064 | 3.46712772 | 6.0404359 | 0.0255282 | 0.999175987 |
| ENSMUSG00000038530 | Rgs4 | protein_coding | -0.7388237 | 8.482805222 | 6.0350485 | 0.0255846 | 0.999175987 |
| ENSMUSG00000038587 | Akap12 | protein_coding | -1.755439936 | 3.967296444 | 6.0039616 | 0.0259129 | 0.999175987 |
| ENSMUSG00000054723 | Vmac | protein_coding | -1.718807196 | 3.522435072 | 5.954621 | 0.026444 | 0.999175987 |
| ENSMUSG00000057098 | Ebf1 | protein_coding | 1.363765148 | 3.995808102 | 5.9491555 | 0.0265036 | 0.999175987 |
| ENSMUSG00000040548 | Tex2 | protein_coding | 0.821388705 | 6.582425221 | 5.9311463 | 0.0267011 | 0.999175987 |
| ENSMUSG00000015749 | Anp32e | protein_coding | -0.723217309 | 6.679126619 | 5.9071626 | 0.0269667 | 0.999175987 |
| ENSMUSG00000034445 | Cyb561a3 | protein_coding | 2.664368765 | 2.512051655 | 5.9037477 | 0.0270048 | 0.999175987 |
| ENSMUSG00000026879 | Gsn | protein_coding | -0.78662599 | 6.200328088 | 5.8946301 | 0.0271067 | 0.999175987 |
| ENSMUSG00000071477 | Zfp777 | protein_coding | 1.67104853 | 3.500699086 | 5.883152 | 0.0272356 | 0.999175987 |
| ENSMUSG00000028974 | Dffa | protein_coding | 1.028060715 | 5.691760046 | 5.8801423 | 0.0272696 | 0.999175987 |
| ENSMUSG00000037400 | Atp11b | protein_coding | -2.39446366 | 3.520111788 | 5.8868794 | 0.027309 | 0.999175987 |
| ENSMUSG00000025739 | Gng13 | protein_coding | 1.27167447 | 4.249985549 | 5.8741617 | 0.0273371 | 0.999175987 |
| ENSMUSG00000036834 | Plch1 | protein_coding | 2.250253298 | 3.273452275 | 5.836642 | 0.0277654 | 0.999175987 |
| ENSMUSG00000035561 | Aldh1b1 | protein_coding | 1.266994338 | 4.460342344 | 5.834801 | 0.0277866 | 0.999175987 |
| ENSMUSG00000031570 | Plpp5 | protein_coding | 1.211417091 | 4.85331234 | 5.8318666 | 0.0278205 | 0.999175987 |
| ENSMUSG00000054986 | Sec14l3 | protein_coding | -2.736423098 | 2.385090534 | 5.830085 | 0.027841 | 0.999175987 |
| ENSMUSG00000001524 | Gtf2h4 | protein_coding | 1.100577974 | 5.099909183 | 5.8246251 | 0.0279042 | 0.999175987 |
| ENSMUSG00000035954 | Dock4 | protein_coding | -1.408709873 | 4.222675166 | 5.8225646 | 0.0279281 | 0.999175987 |
| ENSMUSG00000040771 | Oard1 | protein_coding | -0.816490479 | 6.457537831 | 5.8217207 | 0.0279379 | 0.999175987 |
| ENSMUSG00000008658 | Rbfox1 | protein_coding | 0.704342488 | 8.260538718 | 5.7969579 | 0.0282268 | 0.999175987 |
| ENSMUSG00000014767 | Tbp | protein_coding | 1.03400279 | 5.374297906 | 5.7966656 | 0.0282302 | 0.999175987 |
| ENSMUSG00000021451 | Sema4d | protein_coding | -1.392097926 | 3.612911937 | 5.777301 | 0.0284586 | 0.999175987 |
| ENSMUSG00000028857 | Tmem222 | protein_coding | 0.861988079 | 8.220295672 | 5.7649038 | 0.0286059 | 0.999175987 |
| ENSMUSG00000028766 | Alpl | protein_coding | 3.599265811 | 2.546182813 | 5.7603577 | 0.0286821 | 0.999175987 |
| ENSMUSG00000033307 | Mif | protein_coding | 0.716533612 | 7.599309128 | 5.7550212 | 0.028724 | 0.999175987 |
| ENSMUSG00000048728 | Zfp454 | protein_coding | 1.696833695 | 3.464650488 | 5.753296 | 0.0287447 | 0.999175987 |
| ENSMUSG00000020892 | Aloxe3 | protein_coding | 1.819028297 | 4.186535771 | 5.7532997 | 0.0287666 | 0.999175987 |
| ENSMUSG00000044167 | Foxo1 | protein_coding | -1.053811137 | 5.322789056 | 5.7459652 | 0.0288327 | 0.999175987 |
| ENSMUSG00000034636 | Zyg11b | protein_coding | -0.795012162 | 5.867173384 | 5.7365694 | 0.0289459 | 0.999175987 |
| ENSMUSG00000108461 | AV356131 | bidirectional_promote | 2.677619601 | 2.681893611 | 5.7282409 | 0.0290467 | 0.999175987 |
| ENSMUSG00000017667 | Zfp334 | protein_coding | -1.122348491 | 4.632684367 | 5.7257969 | 0.0290764 | 0.999175987 |
| ENSMUSG00000025155 | Dus1l | protein_coding | 1.475643362 | 4.475230336 | 5.7013602 | 0.0293749 | 0.999175987 |
| ENSMUSG00000007041 | Clic1 | protein_coding | -1.351943731 | 4.372558021 | 5.6958652 | 0.0294425 | 0.999175987 |
| ENSMUSG00000050530 | Fam171a1 | protein_coding | -0.947577689 | 5.133659877 | 5.6813471 | 0.029622 | 0.999175987 |
| ENSMUSG00000023939 | Mrpl14 | protein_coding | 0.740728423 | 7.523152473 | 5.6764619 | 0.0296827 | 0.999175987 |
| ENSMUSG00000018012 | Rac3 | protein_coding | 1.308543724 | 4.290277025 | 5.6751653 | 0.0296988 | 0.999175987 |
| ENSMUSG00000035725 | Prkx | protein_coding | -2.064032329 | 3.582011779 | 5.6607576 | 0.0299011 | 0.999175987 |
| ENSMUSG00000041079 | Rwdd2b | protein_coding | -1.248643095 | 4.046549818 | 5.6458691 | 0.030066 | 0.999175987 |
| ENSMUSG00000035027 | Map2k2 | protein_coding | 0.917899785 | 5.336854328 | 5.6448643 | 0.0300786 | 0.999175987 |
| ENSMUSG00000062115 | Rai1 | protein_coding | 1.195510569 | 5.042439665 | 5.6319729 | 0.0302419 | 0.999175987 |
| ENSMUSG00000034175 | Rhbdd3 | protein_coding | 1.157652935 | 4.91049185 | 5.6255051 | 0.0303242 | 0.999175987 |
| ENSMUSG00000035649 | Zcchc7 | protein_coding | -1.162826483 | 4.421059595 | 5.616558 | 0.0304385 | 0.999175987 |
| ENSMUSG00000033706 | Smyd5 | protein_coding | 0.801937727 | 6.38222208 | 5.6127567 | 0.0304872 | 0.999175987 |
| ENSMUSG00000083307 | AA414768 | processed_pseudoge | -1.956913109 | 2.909304346 | 5.6063166 | 0.0305699 | 0.999175987 |
| ENSMUSG00000032827 | Ppp1r9a | protein_coding | -0.97231029 | 7.15037439 | 5.6056931 | 0.0306005 | 0.999175987 |
| ENSMUSG00000018906 | P4ha2 | protein_coding | 1.630523962 | 4.267600162 | 5.5962445 | 0.0306998 | 0.999175987 |
| ENSMUSG00000086496 | Gm14204 | processed_transcript | 1.236675164 | 5.415769256 | 5.5902342 | 0.0307776 | 0.999175987 |
| ENSMUSG00000071646 | Mta2 | protein_coding | -1.313554881 | 4.029628737 | 5.5747456 | 0.0309792 | 0.999175987 |
| ENSMUSG00000039316 | Rftn1 | protein_coding | -1.146169632 | 4.447628496 | 5.5710522 | 0.0310275 | 0.999175987 |
| ENSMUSG00000029062 | Cdk11b | protein_coding | -1.004128836 | 4.978969342 | 5.5688796 | 0.0310559 | 0.999175987 |
| ENSMUSG00000020926 | Adam11 | protein_coding | 1.098616779 | 4.884671681 | 5.564461 | 0.0311138 | 0.999175987 |
| ENSMUSG00000020003 | Pex7 | protein_coding | -1.235589877 | 4.586048451 | 5.5620491 | 0.0311455 | 0.999175987 |
| ENSMUSG00000041360 | Pum3 | protein_coding | 0.948472591 | 5.666340357 | 5.5531214 | 0.0312631 | 0.999175987 |
| ENSMUSG00000078671 | Chd2 | protein_coding | -1.121407815 | 4.680624463 | 5.550871 | 0.0312928 | 0.999175987 |
| ENSMUSG00000009112 | Bcl2l13 | protein_coding | 1.482406538 | 4.002064157 | 5.5299437 | 0.0315708 | 0.999175987 |
| ENSMUSG00000111375 | Btbd8 | protein_coding | -1.229126246 | 5.097936744 | 5.5254978 | 0.0316302 | 0.999175987 |
| ENSMUSG00000059355 | Wdr83os | protein_coding | 0.735499372 | 6.405259769 | 5.5049639 | 0.0319063 | 0.999175987 |
| ENSMUSG00000041975 | Mettl8 | protein_coding | 1.808840652 | 4.448576166 | 5.5267238 | 0.0320466 | 0.999175987 |
| ENSMUSG00000028433 | Ubap2 | protein_coding | -1.357131987 | 4.129500924 | 5.4910394 | 0.0320951 | 0.999175987 |
| ENSMUSG00000026223 | Itm2c | protein_coding | 0.827020861 | 9.311222768 | 5.4874241 | 0.0321443 | 0.999175987 |
| ENSMUSG00000015224 | Cyp2j9 | protein_coding | -1.306266615 | 4.778190594 | 5.4789807 | 0.0322597 | 0.999175987 |
| ENSMUSG00000035851 | Ythdc1 | protein_coding | -1.049959775 | 5.114196526 | 5.469292 | 0.0323926 | 0.999175987 |
| ENSMUSG00000025209 | Twnk | protein_coding | -1.955715745 | 3.574506826 | 5.4653616 | 0.0324467 | 0.999175987 |
| ENSMUSG00000031604 | Msmo1 | protein_coding | -0.683265284 | 7.175170499 | 5.4580764 | 0.0325472 | 0.999175987 |
| ENSMUSG00000031878 | Nae1 | protein_coding | -0.846393664 | 6.584111225 | 5.4446482 | 0.0327335 | 0.999175987 |

| ensembl_gene_id | external_gene_name | gene_biotype | log2 (fold change) | log2 (counts per million) | F-value | p-value | False discovery rate |
|---|---|---|---|---|---|---|---|
| ENSMUSG00000028345 | Tex10 | protein_coding | 1.418815724 | 4.033069419 | 5.4421965 | 0.0327676 | 0.999175987 |
| ENSMUSG00000039197 | Adk | protein_coding | -0.662122054 | 7.492440639 | 5.4224982 | 0.0330434 | 0.999175987 |
| ENSMUSG00000034757 | Tmub2 | protein_coding | 0.763389999 | 6.644310887 | 5.4072072 | 0.0332594 | 0.999175987 |
| ENSMUSG00000041229 | Phf8 | protein_coding | -1.230022 | 4.269503119 | 5.4067539 | 0.0332658 | 0.999175987 |
| ENSMUSG00000021993 | Mipep | protein_coding | 1.237031775 | 4.298738987 | 5.3976704 | 0.0333949 | 0.999175987 |
| ENSMUSG00000074212 | Dnajb14 | protein_coding | -0.887868406 | 5.796248168 | 5.3784239 | 0.0336703 | 0.999175987 |
| ENSMUSG00000051323 | Pcdh19 | protein_coding | -0.984126435 | 5.411491613 | 5.3783148 | 0.0336719 | 0.999175987 |
| ENSMUSG00000029536 | Gatc | protein_coding | 0.783250446 | 5.99262506 | 5.3713756 | 0.0337718 | 0.999175987 |
| ENSMUSG00000002416 | Ndufb2 | protein_coding | 0.771520319 | 6.208844997 | 5.3605885 | 0.0339278 | 0.999175987 |
| ENSMUSG00000054013 | Tmem179 | protein_coding | 0.82666895 | 7.989735247 | 5.3571573 | 0.0339777 | 0.999175987 |
| ENSMUSG00000021478 | Drd1 | protein_coding | -1.264997969 | 5.486676655 | 5.3528519 | 0.0340638 | 0.999175987 |
| ENSMUSG00000026347 | Tmem163 | protein_coding | -1.492914401 | 4.287206901 | 5.3235787 | 0.0344696 | 0.999175987 |
| ENSMUSG00000047205 | Dusp18 | protein_coding | 1.252372479 | 4.869226442 | 5.3140043 | 0.0346113 | 0.999175987 |
| ENSMUSG00000005534 | Insr | protein_coding | 1.554007426 | 3.876891473 | 5.3111074 | 0.0346544 | 0.999175987 |
| ENSMUSG00000038319 | Kcnh2 | protein_coding | 1.990421905 | 2.905448428 | 5.3058424 | 0.0347327 | 0.999175987 |
| ENSMUSG00000038644 | Pold1 | protein_coding | 2.776041877 | 2.400839149 | 5.300969 | 0.0348054 | 0.999175987 |
| ENSMUSG00000085251 | Gm12326 | lincRNA | -2.412157787 | 3.29364466 | 5.2994362 | 0.0348521 | 0.999175987 |
| ENSMUSG00000039911 | Spsb1 | protein_coding | -1.960265741 | 2.936041677 | 5.2832108 | 0.0350718 | 0.999175987 |
| ENSMUSG00000052698 | Tln2 | protein_coding | -0.998223833 | 5.70148989 | 5.2828158 | 0.0350778 | 0.999175987 |
| ENSMUSG00000034173 | Zbed5 | protein_coding | 1.70502504 | 3.221061799 | 5.2719102 | 0.0352426 | 0.999175987 |
| ENSMUSG00000032959 | Pebp1 | protein_coding | 0.644723 | 7.635538589 | 5.2704191 | 0.0352652 | 0.999175987 |
| ENSMUSG00000031503 | Col4a2 | protein_coding | 2.703174143 | 2.444763669 | 5.2652462 | 0.0353438 | 0.999175987 |
| ENSMUSG00000035202 | Lars2 | protein_coding | 0.885683109 | 9.543560959 | 5.2631003 | 0.0353764 | 0.999175987 |
| ENSMUSG00000041720 | Pi4ka | protein_coding | 1.338438684 | 4.881560216 | 5.253464 | 0.0355234 | 0.999175987 |
| ENSMUSG00000046160 | Olig1 | protein_coding | -1.101676215 | 5.77983526 | 5.2499052 | 0.0355779 | 0.999175987 |
| ENSMUSG00000035772 | Mrps2 | protein_coding | 0.924772308 | 5.785814546 | 5.2490857 | 0.0355905 | 0.999175987 |
| ENSMUSG00000028743 | Akr7a5 | protein_coding | 1.354822871 | 3.887303776 | 5.2471546 | 0.0356201 | 0.999175987 |
| ENSMUSG00000032624 | Eml4 | protein_coding | -1.355101172 | 3.902866519 | 5.2281632 | 0.0359128 | 0.999175987 |
| ENSMUSG00000017978 | Cadps2 | protein_coding | 1.101808909 | 5.975778509 | 5.2294747 | 0.0359165 | 0.999175987 |
| ENSMUSG00000021125 | Arg2 | protein_coding | -1.209856515 | 3.883070506 | 5.2221246 | 0.0360065 | 0.999175987 |
| ENSMUSG00000026203 | Dnajb2 | protein_coding | 0.688552918 | 6.674401916 | 5.2201176 | 0.0360377 | 0.999175987 |
| ENSMUSG00000025470 | Zfp511 | protein_coding | 1.11789211 | 4.322987393 | 5.2090988 | 0.0362095 | 0.999175987 |
| ENSMUSG00000031782 | Coq9 | protein_coding | 0.76663368 | 6.965553373 | 5.2024917 | 0.0363129 | 0.999175987 |
| ENSMUSG00000024870 | Rab1b | protein_coding | 0.740530172 | 8.0398978 | 5.2004927 | 0.0363443 | 0.999175987 |
| ENSMUSG00000032172 | Olfm2 | protein_coding | 0.736414377 | 6.814013233 | 5.1945715 | 0.0364374 | 0.999175987 |
| ENSMUSG00000071553 | Cpa2 | protein_coding | 1.146398059 | 4.811288757 | 5.1905899 | 0.0365002 | 0.999175987 |
| ENSMUSG00000029617 | Ccz1 | protein_coding | -1.307920023 | 4.257759713 | 5.1897768 | 0.036513 | 0.999175987 |
| ENSMUSG00000058355 | Abce1 | protein_coding | -0.87077434 | 5.983356726 | 5.1855269 | 0.0365802 | 0.999175987 |
| ENSMUSG00000016344 | Ppdpf | protein_coding | 0.970869367 | 5.202265819 | 5.1826849 | 0.0366252 | 0.999175987 |
| ENSMUSG00000041540 | Sox5 | protein_coding | -1.32011917 | 4.527138898 | 5.1783475 | 0.0366939 | 0.999175987 |
| ENSMUSG00000032288 | Imp3 | protein_coding | 1.245163162 | 4.824134145 | 5.1742603 | 0.0367589 | 0.999175987 |
| ENSMUSG00000058833 | 2810428I15Rik | protein_coding | 0.802206067 | 7.558352681 | 5.1740512 | 0.0367622 | 0.999175987 |
| ENSMUSG00000028104 | Polr3gl | protein_coding | 1.397858761 | 3.652093066 | 5.1657992 | 0.0368938 | 0.999175987 |
| ENSMUSG00000060301 | 2610008E11Rik | protein_coding | -1.435703859 | 3.531408418 | 5.1463242 | 0.0372064 | 0.999175987 |
| ENSMUSG00000029484 | Anxa3 | protein_coding | -1.653653838 | 3.495999761 | 5.1381683 | 0.0373383 | 0.999175987 |
| ENSMUSG00000031903 | Pla2g15 | protein_coding | 1.145757756 | 4.10737651 | 5.1228976 | 0.0375866 | 0.999175987 |
| ENSMUSG00000038664 | Herc1 | protein_coding | -0.978718808 | 5.117133329 | 5.1130693 | 0.0377474 | 0.999175987 |
| ENSMUSG00000063594 | Gng8 | protein_coding | -1.310835403 | 4.820213282 | 5.1130487 | 0.0377477 | 0.999175987 |
| ENSMUSG00000026743 | Mllt10 | protein_coding | -1.095333044 | 4.706873075 | 5.0909437 | 0.0381124 | 0.999175987 |
| ENSMUSG00000033382 | Trappc8 | protein_coding | -1.516942152 | 3.871704066 | 5.087183 | 0.0381748 | 0.999175987 |
| ENSMUSG00000092341 | Malat1 | lincRNA | -0.892566347 | 10.80327234 | 5.0751043 | 0.0383761 | 0.999175987 |
| ENSMUSG00000033068 | Entpd6 | protein_coding | 0.71647791 | 6.758340762 | 5.0734566 | 0.0384037 | 0.999175987 |
| ENSMUSG00000021069 | Pygl | protein_coding | -2.677396554 | 2.657915326 | 5.0666659 | 0.0385175 | 0.999175987 |
| ENSMUSG00000026219 | Trip12 | protein_coding | -0.770479814 | 6.114001535 | 5.0588287 | 0.0386494 | 0.999175987 |
| ENSMUSG00000058454 | Dhcr7 | protein_coding | 0.864640477 | 5.621171495 | 5.0564537 | 0.0386894 | 0.999175987 |
| ENSMUSG00000106981 | Gm45495 | lincRNA | -2.092505027 | 3.014933517 | 5.0357346 | 0.0390409 | 0.999175987 |
| ENSMUSG00000012126 | Ubxn11 | protein_coding | 0.776819983 | 5.614019991 | 5.0278392 | 0.0391759 | 0.999175987 |
| ENSMUSG00000018405 | Mrm1 | protein_coding | 1.739926958 | 3.018384625 | 5.0274896 | 0.0391818 | 0.999175987 |
| ENSMUSG00000034171 | Faah | protein_coding | 1.260588286 | 4.293621047 | 5.0118214 | 0.0394512 | 0.999175987 |
| ENSMUSG00000022112 | Gpc5 | protein_coding | -1.307381438 | 4.158869144 | 5.0082803 | 0.0395124 | 0.999175987 |
| ENSMUSG00000021188 | Trip11 | protein_coding | -1.173246082 | 4.587727055 | 5.0040391 | 0.0395858 | 0.999175987 |
| ENSMUSG00000026755 | Arpc5l | protein_coding | 0.617247161 | 8.091165685 | 5.0026747 | 0.0396095 | 0.999175987 |
| ENSMUSG00000018166 | Erbb3 | protein_coding | -2.773199665 | 2.661932717 | 4.9938565 | 0.0397628 | 0.999175987 |
| ENSMUSG00000038555 | Reep2 | protein_coding | 0.793063036 | 7.814522904 | 4.9912215 | 0.0398087 | 0.999175987 |
| ENSMUSG00000024925 | Rnaseh2c | protein_coding | 1.973017439 | 4.817628596 | 5.1119144 | 0.039868 | 0.999175987 |
| ENSMUSG00000051107 | Gm15440 | protein_coding | -1.257716548 | 3.974827409 | 4.983949 | 0.0399358 | 0.999175987 |
| ENSMUSG00000079435 | Rpl36a | protein_coding | 1.076242021 | 5.304824514 | 4.9799273 | 0.0400063 | 0.999175987 |
| ENSMUSG00000030093 | Wnt7a | protein_coding | -2.120292162 | 2.849999121 | 4.9782949 | 0.0400349 | 0.999175987 |
| ENSMUSG00000029283 | Cdc7 | protein_coding | 2.430026988 | 2.676501028 | 4.9698577 | 0.0401834 | 0.999175987 |
| ENSMUSG00000028282 | Casp8ap2 | protein_coding | 1.371072892 | 3.979270051 | 4.9569893 | 0.040411 | 0.999175987 |
| ENSMUSG00000036169 | Sostdc1 | protein_coding | 1.926466036 | 5.072898366 | 5.078322 | 0.040435 | 0.999175987 |
| ENSMUSG00000057060 | Slc35f3 | protein_coding | 1.580643482 | 3.816031417 | 4.9467451 | 0.0405933 | 0.999175987 |
| ENSMUSG00000068299 | Nat8f4 | protein_coding | -1.589268044 | 3.405434141 | 4.9322234 | 0.0408533 | 0.999175987 |
| ENSMUSG00000046139 | Patl1 | protein_coding | -1.457514393 | 3.763403618 | 4.9306379 | 0.0408818 | 0.999175987 |
| ENSMUSG00000039086 | Ss18l1 | protein_coding | 1.032712229 | 4.984078631 | 4.9067198 | 0.0413147 | 0.999175987 |
| ENSMUSG00000030516 | Tjp1 | protein_coding | -1.335731413 | 4.111698494 | 4.9035477 | 0.0413725 | 0.999175987 |
| ENSMUSG00000029060 | Mib2 | protein_coding | 1.318143609 | 4.132679786 | 4.8784913 | 0.0418324 | 0.999175987 |
| ENSMUSG00000087211 | Lhx1os | lincRNA | -1.117283837 | 4.070579514 | 4.8746144 | 0.0419041 | 0.999175987 |

| ensembl_gene_id | external_gene_name | gene_biotype | log2 (fold change) | log2 (counts per million) | F-value | p-value | False discovery rate |
|---|---|---|---|---|---|---|---|
| ENSMUSG00000006392 | Med8 | protein_coding | -0.774270117 | 6.431049831 | 4.8608711 | 0.0421593 | 0.999175987 |
| ENSMUSG00000029577 | Ube3b | protein_coding | 1.035294787 | 4.872432996 | 4.8559731 | 0.0422507 | 0.999175987 |
| ENSMUSG00000028266 | Lmo4 | protein_coding | -0.759776318 | 6.567840984 | 4.8552555 | 0.0422642 | 0.999175987 |
| ENSMUSG00000024597 | Slc12a2 | protein_coding | -1.07378474 | 5.596955192 | 4.8539695 | 0.0422882 | 0.999175987 |
| ENSMUSG00000000127 | Fer | protein_coding | -1.185685046 | 4.309187938 | 4.8506096 | 0.0423511 | 0.999175987 |
| ENSMUSG00000029209 | Gnpda2 | protein_coding | -0.815125431 | 5.92283896 | 4.8505865 | 0.0423515 | 0.999175987 |
| ENSMUSG00000032826 | Ank2 | protein_coding | -0.810006197 | 6.614685541 | 4.8342276 | 0.0426594 | 0.999175987 |
| ENSMUSG00000015804 | Med28 | protein_coding | 0.871413057 | 8.268673354 | 4.8313456 | 0.0427394 | 0.999175987 |
| ENSMUSG00000032024 | Clmp | protein_coding | 1.660860546 | 3.363255867 | 4.8221956 | 0.0428874 | 0.999175987 |
| ENSMUSG00000010607 | Pigyl | protein_coding | 0.681105839 | 7.440696244 | 4.8164184 | 0.0429974 | 0.999175987 |
| ENSMUSG00000021025 | Nfkbia | protein_coding | 1.046090231 | 5.816459171 | 4.8099192 | 0.0431215 | 0.999175987 |
| ENSMUSG00000026019 | Wdr12 | protein_coding | -0.704427744 | 6.68670352 | 4.8020029 | 0.0432733 | 0.999175987 |
| ENSMUSG00000036764 | Dnajc12 | protein_coding | -0.624425397 | 6.958223742 | 4.8010322 | 0.043292 | 0.999175987 |
| ENSMUSG00000042215 | Bag2 | protein_coding | 1.124569435 | 4.929207901 | 4.7982048 | 0.0433463 | 0.999175987 |
| ENSMUSG00000053510 | Nrd1 | protein_coding | -0.833393287 | 5.395380543 | 4.7981474 | 0.0433475 | 0.999175987 |
| ENSMUSG00000026411 | Tmem9 | protein_coding | 0.647256002 | 7.866960137 | 4.7739991 | 0.0438152 | 0.999175987 |
| ENSMUSG00000074863 | Platr25 | protein_coding | -2.321340322 | 2.442684069 | 4.7552111 | 0.0441831 | 0.999175987 |
| ENSMUSG00000038518 | Jarid2 | protein_coding | 0.888950076 | 8.066763333 | 4.7461495 | 0.0443878 | 0.999175987 |
| ENSMUSG00000074060 | Fbxw15 | protein_coding | -1.6196659 | 3.577313877 | 4.7405534 | 0.0444727 | 0.999175987 |
| ENSMUSG00000032840 | 2410131K14Rik | protein_coding | 0.889479113 | 5.490908084 | 4.7389467 | 0.0445045 | 0.999175987 |
| ENSMUSG00000002981 | Clptm1 | protein_coding | 0.815183107 | 5.646695765 | 4.7385874 | 0.0445117 | 0.999175987 |
| ENSMUSG00000039345 | Mettl22 | protein_coding | 0.745588614 | 6.464124396 | 4.7385124 | 0.0445132 | 0.999175987 |
| ENSMUSG00000033632 | AW554918 | protein_coding | -1.19843246 | 3.894266693 | 4.7382791 | 0.0445178 | 0.999175987 |
| ENSMUSG00000044788 | Fads6 | protein_coding | -1.372406754 | 4.033293867 | 4.7235589 | 0.0448111 | 0.999175987 |
| ENSMUSG00000020130 | Tbc1d15 | protein_coding | -0.884400417 | 5.682578696 | 4.7151375 | 0.04498 | 0.999175987 |
| ENSMUSG00000025993 | Slc40a1 | protein_coding | -2.654555668 | 2.315840869 | 4.7106398 | 0.0450704 | 0.999175987 |
| ENSMUSG00000027459 | Fam110a | protein_coding | 1.357015335 | 5.031921134 | 4.7099285 | 0.0451108 | 0.999175987 |
| ENSMUSG00000035919 | Bbs9 | protein_coding | 1.118195357 | 4.403336251 | 4.702579 | 0.0452331 | 0.999175987 |
| ENSMUSG00000062753 | AI413582 | protein_coding | 0.686943635 | 7.381961825 | 4.6966989 | 0.0453522 | 0.999175987 |
| ENSMUSG00000000125 | Wnt3 | protein_coding | -1.693540733 | 4.082509671 | 4.6889853 | 0.0455351 | 0.999175987 |
| ENSMUSG00000042831 | Alkbh6 | protein_coding | 1.050123459 | 7.43084631 | 4.7328976 | 0.0456325 | 0.999175987 |
| ENSMUSG00000034525 | Ice1 | protein_coding | -1.215824642 | 4.221830255 | 4.6709363 | 0.0458783 | 0.999175987 |
| ENSMUSG00000024194 | Cuta | protein_coding | 0.587280703 | 7.473278638 | 4.6607221 | 0.0460888 | 0.999175987 |
| ENSMUSG00000028648 | Ndufs5 | protein_coding | 0.678533046 | 7.458033273 | 4.6594258 | 0.0461156 | 0.999175987 |
| ENSMUSG00000039620 | 6430573F11Rik | protein_coding | -1.414212883 | 3.508994447 | 4.6362349 | 0.0465981 | 0.999175987 |
| ENSMUSG00000043670 | Diras1 | protein_coding | 1.341272543 | 5.99051786 | 4.7044959 | 0.0467825 | 0.999175987 |
| ENSMUSG00000025609 | Mkln1 | protein_coding | -0.824337251 | 5.956628802 | 4.6271628 | 0.0467884 | 0.999175987 |
| ENSMUSG00000039842 | Mcph1 | protein_coding | 1.199325852 | 4.052234881 | 4.6213752 | 0.0469103 | 0.999175987 |
| ENSMUSG00000034659 | Tmem109 | protein_coding | 0.885944632 | 6.521911539 | 4.6182119 | 0.0469771 | 0.999175987 |
| ENSMUSG00000037071 | Scd1 | protein_coding | -1.035178192 | 5.692765077 | 4.6137361 | 0.0470718 | 0.999175987 |
| ENSMUSG00000028053 | Ash1l | protein_coding | -0.844125415 | 6.306372161 | 4.6031941 | 0.0472957 | 0.999175987 |
| ENSMUSG00000020156 | Mum1 | protein_coding | 1.108061154 | 5.315632701 | 4.5977706 | 0.0474113 | 0.999175987 |
| ENSMUSG00000023919 | Cenpq | protein_coding | -1.901177749 | 3.48608627 | 4.5975075 | 0.0474434 | 0.999175987 |
| ENSMUSG00000030994 | D7Ertd443e | protein_coding | -1.248942345 | 4.277472516 | 4.582741 | 0.0477336 | 0.999175987 |
| ENSMUSG00000025508 | Rplp2 | protein_coding | 0.746329134 | 8.84790649 | 4.581317 | 0.0477642 | 0.999175987 |
| ENSMUSG00000043102 | Qrfp | protein_coding | -4.760725175 | 2.17994674 | 4.6289477 | 0.0477738 | 0.999175987 |
| ENSMUSG00000026796 | Fam129b | protein_coding | 1.738120415 | 2.90440795 | 4.5763421 | 0.0478715 | 0.999175987 |
| ENSMUSG00000003299 | Mrpl4 | protein_coding | 0.65982391 | 7.4879385 | 4.5710429 | 0.0479861 | 0.999175987 |
| ENSMUSG00000029598 | Plbd2 | protein_coding | 1.051473197 | 5.096875469 | 4.5705004 | 0.0479979 | 0.999175987 |
| ENSMUSG00000027811 | 4930579G24Rik | protein_coding | 2.496036836 | 2.607655136 | 4.5655505 | 0.0481052 | 0.999175987 |
| ENSMUSG00000032640 | Chsy1 | protein_coding | -1.254525255 | 3.73724095 | 4.5654048 | 0.0481084 | 0.999175987 |
| ENSMUSG00000029193 | Cckar | protein_coding | -2.021566872 | 3.892365771 | 4.6086793 | 0.0482049 | 0.999175987 |
| ENSMUSG00000028436 | Dcaf12 | protein_coding | -1.006409365 | 4.935003927 | 4.5579803 | 0.04827 | 0.999175987 |
| ENSMUSG00000027301 | Oxt | protein_coding | -2.741410433 | 4.513115714 | 4.8353492 | 0.0482879 | 0.999175987 |
| ENSMUSG00000022538 | Lsg1 | protein_coding | -0.836388585 | 5.240216218 | 4.5462391 | 0.0485268 | 0.999175987 |
| ENSMUSG00000067279 | Ppp1r3c | protein_coding | -1.215892332 | 4.881953259 | 4.5355564 | 0.0487618 | 0.999175987 |
| ENSMUSG00000031458 | Coprs | protein_coding | 0.987338029 | 6.282476966 | 4.5253674 | 0.049014 | 0.999175987 |
| ENSMUSG00000035781 | R3hdm4 | protein_coding | 0.771429246 | 7.261553499 | 4.5087594 | 0.0493573 | 0.999175987 |
| ENSMUSG00000036242 | 3632451O06Rik | protein_coding | -1.023105028 | 4.742168553 | 4.504109 | 0.0494614 | 0.999175987 |
| ENSMUSG00000006288 | Ttc5 | protein_coding | -0.676548403 | 6.543103202 | 4.4978131 | 0.0496029 | 0.999175987 |
| ENSMUSG00000024999 | Noc3l | protein_coding | -2.003797666 | 2.803657768 | 4.4961083 | 0.0496413 | 0.999175987 |
| ENSMUSG00000029182 | 1700001C02Rik | protein_coding | -1.558542151 | 3.55603683 | 4.4938164 | 0.049693 | 0.999175987 |
| ENSMUSG00000015757 | Ppil4 | protein_coding | -0.780665546 | 5.927004705 | 4.4912227 | 0.0497515 | 0.999175987 |
| ENSMUSG00000067430 | Zfp763 | protein_coding | -1.231249784 | 4.197765692 | 4.48783 | 0.0498282 | 0.999175987 |
| ENSMUSG00000071042 | Rasgrp3 | protein_coding | -1.794623053 | 3.738488421 | 4.4890029 | 0.0498285 | 0.999175987 |
| ENSMUSG00000021377 | Dek | protein_coding | -0.925440712 | 5.116669153 | 4.4830697 | 0.049936 | 0.999175987 |
| ENSMUSG00000034667 | Xpot | protein_coding | -1.039289017 | 4.929697942 | 4.4808185 | 0.0499871 | 0.999175987 |
| ENSMUSG00000021774 | Ube2e1 | protein_coding | 0.817149533 | 5.5198214 | 4.4804482 | 0.0499955 | 0.999175987 |
| ENSMUSG00000033294 | Noc4l | protein_coding | 1.128922647 | 4.910377051 | 4.4727667 | 0.0501704 | 0.999175987 |
| ENSMUSG00000044252 | Osbpl1a | protein_coding | -0.660948986 | 6.867745508 | 4.4718613 | 0.050191 | 0.999175987 |
| ENSMUSG00000057738 | Sptan1 | protein_coding | -0.848913631 | 7.14596439 | 4.4674451 | 0.0503188 | 0.999175987 |
| ENSMUSG00000026694 | Mettl13 | protein_coding | -1.50146692 | 3.717727346 | 4.4539718 | 0.0506011 | 0.999175987 |
| ENSMUSG00000013150 | Gfod2 | protein_coding | 0.648083292 | 6.447896045 | 4.4538098 | 0.0506048 | 0.999175987 |
| ENSMUSG00000027519 | Rab22a | protein_coding | -0.780223601 | 5.427502453 | 4.4492524 | 0.05071 | 0.999175987 |
| ENSMUSG00000015092 | Edf1 | protein_coding | 0.691197927 | 7.716842567 | 4.4439607 | 0.0508323 | 0.999175987 |
| ENSMUSG00000024781 | Lipa | protein_coding | 0.931067704 | 5.126524875 | 4.4416373 | 0.0508862 | 0.999175987 |
| ENSMUSG00000028622 | Mrpl37 | protein_coding | 0.766546698 | 6.302739605 | 4.4382735 | 0.0509642 | 0.999175987 |
| ENSMUSG00000034126 | Pomt2 | protein_coding | 2.667623801 | 2.751272529 | 4.4356811 | 0.0510515 | 0.999175987 |

| ensembl_gene_id | external_gene_name | gene_biotype | log2 (fold change) | log2 (counts per million) | F-value | p-value | False discovery rate |
|---|---|---|---|---|---|---|---|
| ENSMUSG00000046269 | Usp27x | protein_coding | 1.277075307 | 3.983395356 | 4.4234922 | 0.0513089 | 0.999175987 |
| ENSMUSG00000036111 | Lmo1 | protein_coding | 0.963740351 | 4.430360891 | 4.4202669 | 0.0513844 | 0.999175987 |
| ENSMUSG00000070733 | Fryl | protein_coding | -1.243213839 | 3.96075965 | 4.4200257 | 0.0513901 | 0.999175987 |
| ENSMUSG00000020114 | Cand1 | protein_coding | -0.741797773 | 6.489287104 | 4.41838 | 0.0514287 | 0.999175987 |
| ENSMUSG00000045777 | Ifitm10 | protein_coding | 0.80634788 | 5.831106192 | 4.41829 | 0.0514308 | 0.999175987 |
| ENSMUSG00000028559 | Osbpl9 | protein_coding | 0.740997449 | 5.932342339 | 4.4024191 | 0.0518049 | 0.999175987 |
| ENSMUSG00000039246 | Lyplal1 | protein_coding | 1.041800193 | 4.955391354 | 4.3972562 | 0.0519273 | 0.999175987 |
| ENSMUSG00000021707 | Dhfr | protein_coding | 1.66307597 | 3.269350618 | 4.3929376 | 0.0520299 | 0.999175987 |
| ENSMUSG00000040653 | Ppp1r14c | protein_coding | 1.446255621 | 3.440860724 | 4.3823245 | 0.0522831 | 0.999175987 |
| ENSMUSG00000076498 | Trbc2 | TR_C_gene | -2.059370701 | 2.604877757 | 4.3739078 | 0.052485 | 0.999175987 |
| ENSMUSG00000026587 | Astn1 | protein_coding | -0.77768038 | 6.196100042 | 4.3723024 | 0.0525236 | 0.999175987 |
| ENSMUSG00000063047 | Zfp780b | protein_coding | -0.937094896 | 4.690286311 | 4.3713659 | 0.0525461 | 0.999175987 |
| ENSMUSG00000021098 | 4930447C04Rik | protein_coding | -1.983406749 | 2.70016333 | 4.3662164 | 0.0526702 | 0.999175987 |
| ENSMUSG00000024797 | Vps51 | protein_coding | 1.118262889 | 4.582024347 | 4.362452 | 0.0527611 | 0.999175987 |
| ENSMUSG00000037773 | Pced1a | protein_coding | 1.633798204 | 3.918021502 | 4.3620566 | 0.052798 | 0.999175987 |
| ENSMUSG00000031902 | Nfatc3 | protein_coding | -1.560190328 | 3.247578072 | 4.3584812 | 0.0528573 | 0.999175987 |
| ENSMUSG00000024120 | Lrpprc | protein_coding | -0.775769911 | 5.719920287 | 4.3576378 | 0.0528777 | 0.999175987 |
| ENSMUSG00000097431 | Gm26782 | lincRNA | 0.859064579 | 5.107348508 | 4.3575053 | 0.0528809 | 0.999175987 |
| ENSMUSG00000073433 | Arhgdig | protein_coding | 0.883728542 | 5.627176082 | 4.3552481 | 0.0529357 | 0.999175987 |
| ENSMUSG00000059288 | Cdyl | protein_coding | -1.181463645 | 4.048326968 | 4.3517354 | 0.053021 | 0.999175987 |
| ENSMUSG00000041199 | Rpusd1 | protein_coding | 0.646595882 | 6.885391051 | 4.3504179 | 0.0530531 | 0.999175987 |
| ENSMUSG00000079523 | Tmsb10 | protein_coding | 0.687364052 | 8.16261568 | 4.3494598 | 0.0530764 | 0.999175987 |
| ENSMUSG00000042078 | Svop | protein_coding | 0.61628015 | 7.24270854 | 4.3483219 | 0.0531041 | 0.999175987 |
| ENSMUSG00000026255 | Efhd1 | protein_coding | -1.312097339 | 4.00969525 | 4.3468487 | 0.05314 | 0.999175987 |
| ENSMUSG00000025813 | Homer2 | protein_coding | -1.177819211 | 4.084011976 | 4.343596 | 0.0532194 | 0.999175987 |
| ENSMUSG00000022619 | Mapk8ip2 | protein_coding | -0.835864672 | 6.320424847 | 4.342358 | 0.0532496 | 0.999175987 |
| ENSMUSG00000036972 | Zic4 | protein_coding | -1.229281566 | 5.174582636 | 4.3359356 | 0.0534343 | 0.999175987 |
| ENSMUSG00000036225 | Kctd1 | protein_coding | -1.220571094 | 5.301943962 | 4.333077 | 0.0535045 | 0.999175987 |
| ENSMUSG00000005682 | Pan2 | protein_coding | -2.488074886 | 2.311549658 | 4.3248659 | 0.0536792 | 0.999175987 |
| ENSMUSG00000030036 | Mogs | protein_coding | -1.752344846 | 3.324228333 | 4.321994 | 0.0537501 | 0.999175987 |
| ENSMUSG00000052584 | Serp2 | protein_coding | 0.659643641 | 6.425509516 | 4.3143803 | 0.0539386 | 0.999175987 |
| ENSMUSG00000061731 | Ext1 | protein_coding | -1.108553152 | 4.01353138 | 4.3082653 | 0.0540906 | 0.999175987 |
| ENSMUSG00000019804 | Snx3 | protein_coding | 0.596809297 | 8.357148891 | 4.2997986 | 0.0543018 | 0.999175987 |
| ENSMUSG00000046311 | Zfp62 | protein_coding | -1.363266896 | 3.823587933 | 4.2932953 | 0.0544647 | 0.999175987 |
| ENSMUSG00000008999 | Bmp7 | protein_coding | -1.786184892 | 3.158227919 | 4.291911 | 0.0544994 | 0.999175987 |
| ENSMUSG00000035900 | Gramd4 | protein_coding | -1.427000533 | 3.297468782 | 4.2847122 | 0.0546805 | 0.999175987 |
| ENSMUSG00000034845 | Plvap | protein_coding | 1.166336148 | 3.980863032 | 4.2812803 | 0.0547671 | 0.999175987 |
| ENSMUSG00000027568 | Ntsr1 | protein_coding | 2.176941193 | 2.674081521 | 4.2747577 | 0.0549321 | 0.999175987 |
| ENSMUSG00000035754 | Wdr18 | protein_coding | 0.682880073 | 8.189957898 | 4.2616836 | 0.0552646 | 0.999175987 |
| ENSMUSG00000031837 | Necab2 | protein_coding | -0.810021774 | 6.648110719 | 4.2573167 | 0.0553762 | 0.999175987 |
| ENSMUSG00000037224 | Zfyve28 | protein_coding | -1.134955798 | 3.891816337 | 4.2458046 | 0.0556716 | 0.999175987 |
| ENSMUSG00000045948 | Mrps12 | protein_coding | 0.562446575 | 7.801019874 | 4.2425725 | 0.0557549 | 0.999175987 |
| ENSMUSG00000022560 | Slc52a2 | protein_coding | 1.257027563 | 4.264855553 | 4.2412816 | 0.0557882 | 0.999175987 |
| ENSMUSG00000020455 | Trim11 | protein_coding | 1.489049934 | 3.555612013 | 4.238501 | 0.05586 | 0.999175987 |
| ENSMUSG00000097625 | Gm26561 | lincRNA | 0.834610475 | 6.104517801 | 4.2330201 | 0.0560019 | 0.999175987 |
| ENSMUSG00000028483 | Snapc3 | protein_coding | 0.808148876 | 5.714900084 | 4.224277 | 0.056229 | 0.999175987 |
| ENSMUSG00000021108 | Prkch | protein_coding | -1.307148368 | 4.471156913 | 4.2239514 | 0.0562375 | 0.999175987 |
| ENSMUSG00000078789 | Dph1 | protein_coding | 1.589759888 | 3.712056252 | 4.2192631 | 0.0563598 | 0.999175987 |
| ENSMUSG00000045665 | Mfsd5 | protein_coding | 0.939501257 | 4.838395001 | 4.2157233 | 0.0564523 | 0.999175987 |
| ENSMUSG00000063884 | Ptcd3 | protein_coding | 0.939162611 | 5.521540319 | 4.2107204 | 0.0565833 | 0.999175987 |
| ENSMUSG00000017119 | Nbr1 | protein_coding | -0.77927314 | 6.019132822 | 4.2103628 | 0.0565927 | 0.999175987 |
| ENSMUSG00000049354 | Dcaf7 | protein_coding | -0.670375021 | 6.43481068 | 4.2035313 | 0.0567723 | 0.999175987 |
| ENSMUSG00000022811 | Zfp148 | protein_coding | -0.664781567 | 6.010483517 | 4.2034718 | 0.0567739 | 0.999175987 |
| ENSMUSG00000045980 | Tmem104 | protein_coding | 1.578628479 | 3.024620185 | 4.2018573 | 0.0568164 | 0.999175987 |
| ENSMUSG00000026708 | Cenpl | protein_coding | 2.250473183 | 2.71994695 | 4.1969019 | 0.0569472 | 0.999175987 |
| ENSMUSG00000058013 | 11-Sep | protein_coding | -0.757816944 | 6.660549514 | 4.1933576 | 0.057041 | 0.999175987 |
| ENSMUSG00000020644 | Id2 | protein_coding | -0.637966142 | 8.614376753 | 4.187642 | 0.0571925 | 0.999175987 |
| ENSMUSG00000075706 | Gpx4 | protein_coding | 0.637737663 | 8.737888227 | 4.1811428 | 0.0573655 | 0.999175987 |
| ENSMUSG00000046607 | Hrk | protein_coding | -1.043120513 | 4.154284163 | 4.1732819 | 0.0575755 | 0.999175987 |
| ENSMUSG00000035885 | Cox8a | protein_coding | 1.157056509 | 9.631013762 | 4.2490564 | 0.0577843 | 0.999175987 |
| ENSMUSG00000027222 | Pex16 | protein_coding | 0.776613359 | 6.157758827 | 4.1619701 | 0.0578792 | 0.999175987 |
| ENSMUSG00000052707 | Tnrc6a | protein_coding | -0.819534503 | 5.374586533 | 4.1602975 | 0.0579243 | 0.999175987 |
| ENSMUSG00000020886 | Dlg4 | protein_coding | 0.702949547 | 5.726195246 | 4.1594174 | 0.057948 | 0.999175987 |
| ENSMUSG00000045268 | Zfp691 | protein_coding | 2.370844132 | 2.370564015 | 4.1586058 | 0.0579699 | 0.999175987 |
| ENSMUSG00000074030 | Exoc8 | protein_coding | 0.966764671 | 4.648191444 | 4.1524979 | 0.058135 | 0.999175987 |
| ENSMUSG00000044807 | Zfp354c | protein_coding | -1.095434272 | 4.196527264 | 4.1507637 | 0.058182 | 0.999175987 |
| ENSMUSG00000024621 | Csf1r | protein_coding | -1.165512036 | 4.449327018 | 4.1459832 | 0.0583117 | 0.999175987 |
| ENSMUSG00000006930 | Hap1 | protein_coding | 0.673095915 | 7.80132675 | 4.142955 | 0.0583941 | 0.999175987 |
| ENSMUSG00000037710 | Cisd1 | protein_coding | 0.875774072 | 8.734266486 | 4.1289459 | 0.0588049 | 0.999175987 |
| ENSMUSG00000027858 | Tspan2 | protein_coding | -0.722054252 | 6.491047904 | 4.1265737 | 0.0588419 | 0.999175987 |
| ENSMUSG00000054051 | Ercc6 | protein_coding | 2.362934549 | 2.562444111 | 4.1178895 | 0.0590809 | 0.999175987 |
| ENSMUSG00000035891 | Cerk | protein_coding | -1.073414199 | 4.706977048 | 4.1175827 | 0.0590893 | 0.999175987 |
| ENSMUSG00000034707 | Gns | protein_coding | -0.959352118 | 5.097607033 | 4.1146194 | 0.0591712 | 0.999175987 |
| ENSMUSG00000025059 | Gk | protein_coding | -1.081039291 | 4.091340859 | 4.1086914 | 0.0593353 | 0.999175987 |
| ENSMUSG00000021680 | Crhbp | protein_coding | -1.076725722 | 4.435789994 | 4.1084886 | 0.0593409 | 0.999175987 |
| ENSMUSG00000020564 | Atxn7l1 | protein_coding | 0.820968236 | 6.226863511 | 4.1081742 | 0.0593496 | 0.999175987 |
| ENSMUSG00000025172 | Ankrd2 | protein_coding | -1.854555781 | 2.945274011 | 4.1034028 | 0.0594821 | 0.999175987 |
| ENSMUSG00000031807 | Pgls | protein_coding | 1.097946596 | 4.35469058 | 4.099653 | 0.0595865 | 0.999175987 |

| ensembl_gene_id | external_gene_name | gene_biotype | log2 (fold change) | log2 (counts per million) | F-value | p-value | False discovery rate |
|---|---|---|---|---|---|---|---|
| ENSMUSG00000028385 | Snx30 | protein_coding | -0.889819727 | 4.822640067 | 4.0873918 | 0.0599293 | 0.999175987 |
| ENSMUSG00000029094 | Afap1 | protein_coding | 1.018859743 | 4.558819411 | 4.0857281 | 0.059976 | 0.999175987 |
| ENSMUSG00000025958 | Creb1 | protein_coding | -0.994791956 | 5.110658551 | 4.0762845 | 0.0602418 | 0.999175987 |
| ENSMUSG00000030978 | Rrm1 | protein_coding | -1.398122302 | 4.168672564 | 4.0748885 | 0.0602812 | 0.999175987 |
| ENSMUSG00000025203 | Scd2 | protein_coding | -0.704647802 | 8.816656589 | 4.0729175 | 0.0603369 | 0.999175987 |
| ENSMUSG00000021338 | Carmil1 | protein_coding | -1.376798817 | 3.396781773 | 4.0626516 | 0.060628 | 0.999175987 |
| ENSMUSG00000031255 | Sytl4 | protein_coding | 1.078955062 | 4.055596508 | 4.0572183 | 0.0607827 | 0.999175987 |
| ENSMUSG00000040407 | Akap9 | protein_coding | -0.798508537 | 5.75019992 | 4.0563316 | 0.060808 | 0.999175987 |
| ENSMUSG00000029535 | Triap1 | protein_coding | 0.710260503 | 6.337030239 | 4.0531306 | 0.0608995 | 0.999175987 |
| ENSMUSG00000043913 | Ccdc60 | protein_coding | -2.508614901 | 2.315341886 | 4.0530708 | 0.0609012 | 0.999175987 |
| ENSMUSG00000004100 | Ppan | protein_coding | 1.46316013 | 3.711390121 | 4.0530564 | 0.0609016 | 0.999175987 |
| ENSMUSG00000027752 | Exosc8 | protein_coding | 0.74273167 | 5.884115606 | 4.0524694 | 0.0609184 | 0.999175987 |
| ENSMUSG00000067847 | Romo1 | protein_coding | 0.658273275 | 8.20766826 | 4.0489948 | 0.0610178 | 0.999175987 |
| ENSMUSG00000053477 | Tcf4 | protein_coding | -0.664704568 | 6.748185216 | 4.0400979 | 0.0612733 | 0.999175987 |
| ENSMUSG00000042579 | 4632404H12Rik | lincRNA | 1.206295758 | 3.727082463 | 4.0312902 | 0.0615275 | 0.999175987 |
| ENSMUSG00000037847 | Nmrk1 | protein_coding | -1.490800781 | 3.704456929 | 4.0234543 | 0.0617546 | 0.999175987 |
| ENSMUSG00000032735 | Ablim3 | protein_coding | 1.021240433 | 4.937772889 | 4.0102504 | 0.0621396 | 0.999175987 |
| ENSMUSG00000090150 | Acad11 | protein_coding | -1.535484812 | 2.967308539 | 4.0082956 | 0.0621969 | 0.999175987 |
| ENSMUSG00000046178 | Nxph1 | protein_coding | -0.782682331 | 5.835213888 | 4.0028771 | 0.0623558 | 0.999175987 |
| ENSMUSG00000040021 | Lats1 | protein_coding | -1.52302297 | 3.305346806 | 4.0020772 | 0.0623794 | 0.999175987 |
| ENSMUSG00000027712 | Anxa5 | protein_coding | -0.75400618 | 7.370112413 | 3.9996841 | 0.0624498 | 0.999175987 |
| ENSMUSG00000003271 | Sult2b1 | protein_coding | 1.41210096 | 3.973833882 | 3.997759 | 0.0625064 | 0.999175987 |
| ENSMUSG00000027238 | Frmd5 | protein_coding | 1.056720355 | 4.258166199 | 3.9970161 | 0.0625283 | 0.999175987 |
| ENSMUSG00000046314 | Stxbp6 | protein_coding | -0.733643017 | 5.801587031 | 3.9968394 | 0.0625336 | 0.999175987 |
| ENSMUSG00000050121 | Opalin | protein_coding | -0.624498924 | 7.410040088 | 3.996643 | 0.0625393 | 0.999175987 |
| ENSMUSG00000063785 | Utp14a | protein_coding | -1.692601994 | 3.571688047 | 3.9959167 | 0.0625894 | 0.999175987 |
| ENSMUSG00000027162 | Lin7c | protein_coding | -0.615739459 | 6.417872567 | 3.9918052 | 0.0626822 | 0.999175987 |
| ENSMUSG00000038776 | Ephx1 | protein_coding | 1.127296746 | 4.720512754 | 3.9904746 | 0.0627215 | 0.999175987 |
| ENSMUSG00000045100 | Slc25a26 | protein_coding | 1.225995783 | 4.702150218 | 3.981909 | 0.0629755 | 0.999175987 |
| ENSMUSG00000032368 | Zic1 | protein_coding | -0.753292803 | 6.105026732 | 3.9786562 | 0.0630723 | 0.999175987 |
| ENSMUSG00000020841 | Cpd | protein_coding | -1.077656235 | 4.600038782 | 3.9781655 | 0.0630869 | 0.999175987 |
| ENSMUSG00000074872 | Ctxn2 | protein_coding | 0.693980656 | 8.340177171 | 3.9759996 | 0.0631515 | 0.999175987 |
| ENSMUSG00000029817 | Tra2a | protein_coding | -0.961959897 | 4.997497304 | 3.9744324 | 0.0631982 | 0.999175987 |
| ENSMUSG00000037108 | Zcwpw1 | protein_coding | 1.283017884 | 3.452655585 | 3.971116 | 0.0632973 | 0.999175987 |
| ENSMUSG00000071369 | Map3k5 | protein_coding | -1.0277385 | 4.299181997 | 3.9708837 | 0.0633043 | 0.999175987 |
| ENSMUSG00000047045 | Tmem164 | protein_coding | -1.313474354 | 3.627570521 | 3.9680927 | 0.0633878 | 0.999175987 |
| ENSMUSG00000020634 | Ubxn2a | protein_coding | -0.788804412 | 5.43505583 | 3.9602498 | 0.0636232 | 0.999175987 |
| ENSMUSG00000024563 | Smad2 | protein_coding | -1.049463482 | 4.776615974 | 3.9550798 | 0.063779 | 0.999175987 |
| ENSMUSG00000027366 | Sppl2a | protein_coding | -0.952059577 | 4.958975193 | 3.9549222 | 0.0637837 | 0.999175987 |
| ENSMUSG00000021665 | Hexb | protein_coding | 0.826119558 | 8.13720421 | 3.9523832 | 0.0638892 | 0.999175987 |
| ENSMUSG00000075702 | Selenom | protein_coding | 0.636439293 | 9.621468166 | 3.9495758 | 0.0639453 | 0.999175987 |
| ENSMUSG00000039145 | Camk1d | protein_coding | 0.720240103 | 7.706375568 | 3.9450505 | 0.0640824 | 0.999175987 |
| ENSMUSG00000085609 | 1700016P03Rik | processed_transcript | -1.488356025 | 3.263727687 | 3.9400569 | 0.0642341 | 0.999175987 |
| ENSMUSG00000038156 | Spon1 | protein_coding | -0.800019766 | 5.425957612 | 3.9387872 | 0.0642727 | 0.999175987 |
| ENSMUSG00000039166 | Akap7 | protein_coding | 0.71740653 | 6.00025257 | 3.9322439 | 0.0644723 | 0.999175987 |
| ENSMUSG00000031937 | Vstm5 | protein_coding | 1.129403386 | 4.592154197 | 3.9319148 | 0.0644824 | 0.999175987 |
| ENSMUSG00000044709 | Gemin7 | protein_coding | 0.763932844 | 5.674839794 | 3.9307196 | 0.0645189 | 0.999175987 |
| ENSMUSG00000027304 | Rtf1 | protein_coding | -0.935539447 | 5.315831213 | 3.9288249 | 0.0645769 | 0.999175987 |
| ENSMUSG00000042185 | Nfrkb | protein_coding | -1.522594349 | 3.61727441 | 3.928105 | 0.0645989 | 0.999175987 |
| ENSMUSG00000050953 | Gja1 | protein_coding | -0.794373984 | 7.008087782 | 3.9243352 | 0.0647433 | 0.999175987 |
| ENSMUSG00000058709 | Egln2 | protein_coding | 0.709897876 | 6.016477242 | 3.9175523 | 0.064923 | 0.999175987 |
| ENSMUSG00000032604 | Qars | protein_coding | -0.820502916 | 5.644562786 | 3.9162065 | 0.0649645 | 0.999175987 |
| ENSMUSG00000020733 | Slc9a3r1 | protein_coding | 1.051202151 | 4.971427444 | 3.915931 | 0.064973 | 0.999175987 |
| ENSMUSG00000021962 | Dcp1a | protein_coding | -1.254570433 | 3.482967852 | 3.9158206 | 0.0649764 | 0.999175987 |
| ENSMUSG00000031375 | Bgn | protein_coding | -1.737974101 | 3.133467997 | 3.9153158 | 0.064992 | 0.999175987 |
| ENSMUSG00000038570 | Saxo2 | protein_coding | 0.868157524 | 4.880712069 | 3.9130871 | 0.0650607 | 0.999175987 |
| ENSMUSG00000038370 | Pcp4l1 | protein_coding | 0.866631425 | 8.590326869 | 3.9101713 | 0.0652635 | 0.999175987 |
| ENSMUSG00000058799 | Nap1l1 | protein_coding | -0.592750101 | 6.744221483 | 3.9059922 | 0.0652803 | 0.999175987 |
| ENSMUSG00000062234 | Gak | protein_coding | -0.793557815 | 5.490160043 | 3.905831 | 0.0652853 | 0.999175987 |
| ENSMUSG00000055373 | Fut9 | protein_coding | -0.890098033 | 5.562331203 | 3.8964984 | 0.0655754 | 0.999175987 |
| ENSMUSG00000026841 | Fibcd1 | protein_coding | -1.342637727 | 3.830676593 | 3.8960334 | 0.0655899 | 0.999175987 |
| ENSMUSG00000019917 | 10-Sep | protein_coding | -1.62395825 | 3.195238907 | 3.8958295 | 0.0655962 | 0.999175987 |
| ENSMUSG00000062012 | Zfp13 | protein_coding | 0.890854793 | 5.172760166 | 3.8952912 | 0.065613 | 0.999175987 |
| ENSMUSG00000078429 | Ctdsp2 | protein_coding | -1.11364684 | 4.81346619 | 3.8948951 | 0.0656254 | 0.999175987 |
| ENSMUSG00000024424 | Ttc39c | protein_coding | 1.279768512 | 3.912295957 | 3.8893671 | 0.0657981 | 0.999175987 |
| ENSMUSG00000097248 | Gm2694 | lincRNA | -1.230044947 | 4.397667393 | 3.8761857 | 0.066212 | 0.999175987 |
| ENSMUSG00000039640 | Mrpl12 | protein_coding | 0.741059109 | 6.935011068 | 3.8752153 | 0.0662426 | 0.999175987 |
| ENSMUSG00000018585 | Atox1 | protein_coding | 0.631893255 | 8.537547706 | 3.8735113 | 0.0662964 | 0.999175987 |
| ENSMUSG00000024807 | Syvn1 | protein_coding | 1.044185487 | 4.645465583 | 3.8664472 | 0.0665198 | 0.999175987 |
| ENSMUSG00000019975 | Ikbip | protein_coding | -1.320457 | 3.845814942 | 3.8643567 | 0.0665861 | 0.999175987 |
| ENSMUSG00000024234 | Mtpap | protein_coding | -0.888651204 | 5.238940671 | 3.8522032 | 0.0669731 | 0.999175987 |
| ENSMUSG00000056854 | Pou3f4 | protein_coding | -1.201592585 | 4.056695907 | 3.8483055 | 0.0670977 | 0.999175987 |
| ENSMUSG00000021039 | Snw1 | protein_coding | -0.618454777 | 6.894242313 | 3.8423176 | 0.0672898 | 0.999175987 |
| ENSMUSG00000031865 | Dctn1 | protein_coding | 0.795282699 | 6.87130974 | 3.84311 | 0.0672934 | 0.999175987 |
| ENSMUSG00000002147 | Stat6 | protein_coding | -1.959796252 | 2.770354408 | 3.8410647 | 0.06733 | 0.999175987 |
| ENSMUSG00000030126 | Tmcc1 | protein_coding | -0.911625171 | 5.601854664 | 3.83801 | 0.0674283 | 0.999175987 |
| ENSMUSG00000006527 | Sfmbt1 | protein_coding | -0.975603722 | 4.659079906 | 3.8331119 | 0.0675862 | 0.999175987 |
| ENSMUSG00000046822 | Slc39a3 | protein_coding | -1.211191675 | 4.066614776 | 3.831236 | 0.0676468 | 0.999175987 |

| ensembl_gene_id | external_gene_name | gene_biotype | log2 (fold change) | log2 (counts per million) | F-value | p-value | False discovery rate |
|---|---|---|---|---|---|---|---|
| ENSMUSG00000039168 | Dap | protein_coding | -1.051029109 | 4.225448907 | 3.8255244 | 0.0678318 | 0.999175987 |
| ENSMUSG00000042515 | Mum1l1 | protein_coding | -1.684313748 | 3.670943387 | 3.8221335 | 0.067971 | 0.999175987 |
| ENSMUSG00000039103 | Nexn | protein_coding | -1.352740353 | 4.231479967 | 3.8132474 | 0.0682312 | 0.999175987 |
| ENSMUSG00000042323 | Pbrm1 | protein_coding | -0.811382559 | 5.235378555 | 3.8020226 | 0.0685989 | 0.999175987 |
| ENSMUSG00000044730 | 9930104L06Rik | processed_transcript | -1.552901625 | 3.161767278 | 3.8002033 | 0.0686587 | 0.999175987 |
| ENSMUSG00000085336 | Gm11732 | processed_transcript | 1.817305743 | 3.231971655 | 3.8000339 | 0.0686643 | 0.999175987 |
| ENSMUSG00000026641 | Usf1 | protein_coding | 0.662821963 | 6.31063339 | 3.7970736 | 0.0687617 | 0.999175987 |
| ENSMUSG00000030213 | Atf7ip | protein_coding | -0.952496186 | 4.988359292 | 3.7924737 | 0.0689135 | 0.999175987 |
| ENSMUSG00000000776 | Polr3d | protein_coding | -1.885553459 | 3.890417899 | 3.82675 | 0.0690807 | 0.999175987 |
| ENSMUSG00000036907 | C1ql2 | protein_coding | 2.581669949 | 2.369910729 | 3.7867445 | 0.0691031 | 0.999175987 |
| ENSMUSG00000024589 | Nedd4l | protein_coding | -0.767159462 | 6.077579682 | 3.7843362 | 0.0691829 | 0.999175987 |
| ENSMUSG00000058317 | Ube2e2 | protein_coding | 0.617907309 | 6.828385677 | 3.7823077 | 0.0692503 | 0.999175987 |
| ENSMUSG00000028261 | Ndufaf4 | protein_coding | -0.666204027 | 6.343887909 | 3.7806079 | 0.0693068 | 0.999175987 |
| ENSMUSG00000000708 | Kat2b | protein_coding | -1.206090518 | 3.592394854 | 3.773023 | 0.0695596 | 0.999175987 |
| ENSMUSG00000034891 | Sncb | protein_coding | 0.754251223 | 9.225122231 | 3.7650177 | 0.0698275 | 0.999175987 |
| ENSMUSG00000035632 | Cnot3 | protein_coding | 1.156483092 | 3.568040834 | 3.7634553 | 0.06988 | 0.999175987 |
| ENSMUSG00000036437 | Npy1r | protein_coding | 1.09625544 | 4.079519667 | 3.7618976 | 0.0699323 | 0.999175987 |
| ENSMUSG00000027287 | Snap23 | protein_coding | -1.241926468 | 4.287965476 | 3.7557604 | 0.0701389 | 0.999175987 |
| ENSMUSG00000054843 | Atrnl1 | protein_coding | -0.947415011 | 4.743823263 | 3.7524426 | 0.0702509 | 0.999175987 |
| ENSMUSG00000036880 | Acaa2 | protein_coding | -0.744012576 | 5.883959439 | 3.7508766 | 0.0703038 | 0.999175987 |
| ENSMUSG00000029729 | Zkscan1 | protein_coding | -0.796166862 | 5.761704343 | 3.750438 | 0.0703187 | 0.999175987 |
| ENSMUSG00000031577 | Tti2 | protein_coding | 1.288916224 | 3.608366245 | 3.7479736 | 0.0704021 | 0.999175987 |
| ENSMUSG00000044005 | Gls2 | protein_coding | -1.292106474 | 3.5291459 | 3.7468764 | 0.0704393 | 0.999175987 |
| ENSMUSG00000022008 | Gpalpp1 | protein_coding | -0.945880732 | 4.981242341 | 3.7446381 | 0.0705152 | 0.999175987 |
| ENSMUSG00000104674 | Gm42756 | antisense_RNA | -1.294587135 | 3.81264865 | 3.7430981 | 0.0705675 | 0.999175987 |
| ENSMUSG00000016181 | Diexf | protein_coding | -1.995166415 | 3.022252251 | 3.7430816 | 0.070568 | 0.999175987 |
| ENSMUSG00000026787 | Gad2 | protein_coding | 0.649772312 | 8.99710077 | 3.7356755 | 0.0708201 | 0.999175987 |
| ENSMUSG00000061298 | Agbl4 | protein_coding | 1.097910742 | 4.51388219 | 3.734467 | 0.0708614 | 0.999175987 |
| ENSMUSG00000021277 | Traf3 | protein_coding | 1.315080841 | 3.856128044 | 3.7342356 | 0.0708693 | 0.999175987 |
| ENSMUSG00000105265 | Sox2ot | processed_transcript | -0.629267435 | 6.116986292 | 3.7262283 | 0.0711432 | 0.999175987 |
| ENSMUSG00000025869 | Nop16 | protein_coding | 0.576709013 | 7.109687438 | 3.7209474 | 0.0713246 | 0.999175987 |
| ENSMUSG00000031996 | Aplp2 | protein_coding | -0.586274572 | 7.990537434 | 3.7198015 | 0.071364 | 0.999175987 |
| ENSMUSG00000060487 | Samd5 | protein_coding | -0.979841534 | 4.481120482 | 3.7196039 | 0.0713708 | 0.999175987 |
| ENSMUSG00000003949 | Hlf | protein_coding | -0.578450691 | 6.827037216 | 3.7193569 | 0.0713793 | 0.999175987 |
| ENSMUSG00000047250 | Ptgs1 | protein_coding | 1.209408823 | 3.772499142 | 3.7184037 | 0.0714121 | 0.999175987 |
| ENSMUSG00000027080 | Med19 | protein_coding | 0.732396865 | 6.069591869 | 3.7181398 | 0.0714212 | 0.999175987 |
| ENSMUSG00000092536 | Gm20501 | bidirectional_promote | 1.220049302 | 3.668349907 | 3.7174918 | 0.0714436 | 0.999175987 |
| ENSMUSG00000030689 | Ino80e | protein_coding | 0.598260818 | 6.461476471 | 3.712736 | 0.0716077 | 0.999175987 |
| ENSMUSG00000027953 | Slc50a1 | protein_coding | 0.641488416 | 5.91594276 | 3.7108648 | 0.0716724 | 0.999175987 |
| ENSMUSG00000030083 | Abtb1 | protein_coding | -1.023992868 | 6.035089861 | 3.7148282 | 0.0720933 | 0.999175987 |
| ENSMUSG00000004535 | Tax1bp1 | protein_coding | -0.527787895 | 7.530073695 | 3.6981024 | 0.0721155 | 0.999175987 |
| ENSMUSG00000026678 | Rgs5 | protein_coding | -1.257010288 | 4.246764995 | 3.6977701 | 0.072127 | 0.999175987 |
| ENSMUSG00000032452 | Clstn2 | protein_coding | -1.004693284 | 4.601326026 | 3.6968872 | 0.0721578 | 0.999175987 |
| ENSMUSG00000041444 | Arhgap32 | protein_coding | 0.821705493 | 4.925505771 | 3.6919583 | 0.0723299 | 0.999175987 |
| ENSMUSG00000059003 | Grin2a | protein_coding | -1.262815588 | 4.417501533 | 3.6915946 | 0.0723426 | 0.999175987 |
| ENSMUSG00000023707 | Ogfod2 | protein_coding | 0.885943845 | 5.971557086 | 3.6891461 | 0.0724283 | 0.999175987 |
| ENSMUSG00000031292 | Cdkl5 | protein_coding | -0.801849878 | 5.96809459 | 3.6852593 | 0.0725646 | 0.999175987 |
| ENSMUSG00000023964 | Calcr | protein_coding | 1.453895971 | 3.543107807 | 3.6841077 | 0.072605 | 0.999175987 |
| ENSMUSG00000039672 | Kcne2 | protein_coding | -5.959761029 | 3.019242124 | 3.8113162 | 0.0727103 | 0.999175987 |
| ENSMUSG00000031714 | Gab1 | protein_coding | -0.998606606 | 4.408208257 | 3.6777457 | 0.0728289 | 0.999175987 |
| ENSMUSG00000028630 | Dyrk2 | protein_coding | 1.172224742 | 3.865607839 | 3.6747974 | 0.0729329 | 0.999175987 |
| ENSMUSG00000058246 | Gm10037 | protein_coding | 1.015646489 | 4.268719226 | 3.6715826 | 0.0730465 | 0.999175987 |
| ENSMUSG00000004473 | Clec11a | protein_coding | 1.288658896 | 3.374650093 | 3.6703435 | 0.0730903 | 0.999175987 |
| ENSMUSG00000027523 | Gnas | protein_coding | 1.216808827 | 3.948568539 | 3.6701273 | 0.073098 | 0.999175987 |
| ENSMUSG00000027935 | Rab13 | protein_coding | 0.887273445 | 4.98848082 | 3.670006 | 0.0731023 | 0.999175987 |
| ENSMUSG00000052852 | Reep1 | protein_coding | -0.71162715 | 6.004427529 | 3.6647527 | 0.0732886 | 0.999175987 |
| ENSMUSG00000049521 | Cdc42ep1 | protein_coding | -1.544798548 | 3.740462329 | 3.6644667 | 0.0733284 | 0.999175987 |
| ENSMUSG00000028886 | Eya3 | protein_coding | 1.705296366 | 3.124177612 | 3.6625508 | 0.0733668 | 0.999175987 |
| ENSMUSG00000042447 | Mios | protein_coding | 1.364447889 | 3.605521355 | 3.6614414 | 0.0734063 | 0.999175987 |
| ENSMUSG00000037787 | Apopt1 | protein_coding | 0.556745139 | 6.798427294 | 3.6542451 | 0.0736628 | 0.999175987 |
| ENSMUSG00000053111 | Fank1 | protein_coding | 1.21993792 | 4.313828583 | 3.6527089 | 0.0737177 | 0.999175987 |
| ENSMUSG00000004035 | Gstm7 | protein_coding | -0.546083806 | 7.183323043 | 3.6494493 | 0.0738344 | 0.999175987 |
| ENSMUSG00000036275 | 9530068E07Rik | protein_coding | 0.722693072 | 6.025936184 | 3.6444166 | 0.0740149 | 0.999175987 |
| ENSMUSG00000059183 | Mtfmt | protein_coding | 1.098506343 | 4.232497729 | 3.6428258 | 0.0740721 | 0.999175987 |
| ENSMUSG00000037331 | Larp1 | protein_coding | -0.862389572 | 5.011395459 | 3.6417134 | 0.0741121 | 0.999175987 |
| ENSMUSG00000014633 | Cmc2 | protein_coding | 0.723573441 | 5.842276473 | 3.6412397 | 0.0741292 | 0.999175987 |
| ENSMUSG00000046593 | Tmem215 | protein_coding | 0.987926606 | 4.79170462 | 3.632827 | 0.0744327 | 0.999175987 |
| ENSMUSG00000024059 | Clip4 | protein_coding | -0.743684062 | 5.871477717 | 3.6315896 | 0.0744774 | 0.999175987 |
| ENSMUSG00000038126 | Mphosph9 | protein_coding | -1.231779875 | 3.540894495 | 3.61787 | 0.0749759 | 0.999175987 |
| ENSMUSG00000026211 | Obsl1 | protein_coding | 1.702568771 | 2.832879547 | 3.6088369 | 0.0753062 | 0.999175987 |
| ENSMUSG00000035354 | Uvrag | protein_coding | -1.244546973 | 3.360669855 | 3.6081097 | 0.0753329 | 0.999175987 |
| ENSMUSG00000020248 | Nfyb | protein_coding | -0.818292354 | 5.796719178 | 3.6051005 | 0.0754433 | 0.999175987 |
| ENSMUSG00000003226 | Ranbp2 | protein_coding | -1.204294971 | 3.727733926 | 3.6000963 | 0.0756275 | 0.999175987 |
| ENSMUSG00000061758 | Akr1b10 | protein_coding | -0.769093352 | 5.577998856 | 3.5999218 | 0.0756339 | 0.999175987 |
| ENSMUSG00000044768 | D1Ertd622e | protein_coding | -1.009316885 | 4.60672709 | 3.5987996 | 0.0756753 | 0.999175987 |
| ENSMUSG00000021500 | Ddx46 | protein_coding | -0.770984552 | 6.111591387 | 3.5968487 | 0.0757473 | 0.999175987 |
| ENSMUSG00000027347 | Rasgrp1 | protein_coding | -0.642632376 | 6.741384254 | 3.5928609 | 0.0758946 | 0.999175987 |
| ENSMUSG00000022779 | Top3b | protein_coding | -0.962056548 | 4.148305104 | 3.5905389 | 0.0759806 | 0.999175987 |

| ensembl_gene_id | external_gene_name | gene_biotype | log2 (fold change) | log2 (counts per million) | F-value | p-value | False discovery rate |
|---|---|---|---|---|---|---|---|
| ENSMUSG00000043183 | Simc1 | protein_coding | 1.365521741 | 3.341186191 | 3.5846474 | 0.0761993 | 0.999175987 |
| ENSMUSG00000018677 | Slc25a39 | protein_coding | 0.593513056 | 6.66780126 | 3.582502 | 0.0762791 | 0.999175987 |
| ENSMUSG00000000902 | Smarcb1 | protein_coding | 0.691323873 | 7.733330558 | 3.5778983 | 0.0764507 | 0.999175987 |
| ENSMUSG00000003970 | Rpl8 | protein_coding | 0.617028903 | 10.42760603 | 3.5765947 | 0.0764993 | 0.999175987 |
| ENSMUSG00000070953 | Rabepk | protein_coding | 0.590711734 | 6.597109698 | 3.5761689 | 0.0765152 | 0.999175987 |
| ENSMUSG00000024845 | Tmem134 | processed_transcript | 1.220462543 | 3.994797671 | 3.5757686 | 0.0765302 | 0.999175987 |
| ENSMUSG00000036820 | Amdhd2 | protein_coding | 0.93931766 | 5.454167321 | 3.5756861 | 0.0765333 | 0.999175987 |
| ENSMUSG00000034998 | Foxn2 | protein_coding | -0.989068685 | 4.48501583 | 3.5745126 | 0.0765771 | 0.999175987 |
| ENSMUSG00000019194 | Scn1b | protein_coding | 0.803540214 | 5.104356915 | 3.5742886 | 0.0765855 | 0.999175987 |
| ENSMUSG00000074736 | Syndig1 | protein_coding | 0.704480465 | 6.409774563 | 3.5700652 | 0.0767437 | 0.999175987 |
| ENSMUSG00000089945 | Pakap | protein_coding | -1.553015001 | 3.012438572 | 3.5631631 | 0.0770029 | 0.999175987 |
| ENSMUSG00000040124 | Gorab | protein_coding | 0.959912435 | 4.695669483 | 3.5615879 | 0.0770622 | 0.999175987 |
| ENSMUSG00000019158 | Tmem160 | protein_coding | 1.148160666 | 4.02644066 | 3.5567364 | 0.0772453 | 0.999175987 |
| ENSMUSG00000028292 | Rars2 | protein_coding | -0.779515368 | 5.453790549 | 3.553611 | 0.0773634 | 0.999175987 |
| ENSMUSG00000025094 | Slc18a2 | protein_coding | -1.675466414 | 3.023974236 | 3.5482994 | 0.0775648 | 0.999175987 |
| ENSMUSG00000023087 | Noct | protein_coding | -0.865494401 | 4.931298851 | 3.5430327 | 0.077765 | 0.999175987 |
| ENSMUSG00000016386 | Mpped2 | protein_coding | -0.843383901 | 5.370563768 | 3.5423218 | 0.0777921 | 0.999175987 |
| ENSMUSG00000044786 | Zfp36 | protein_coding | 1.772551681 | 2.927527003 | 3.5420778 | 0.0778014 | 0.999175987 |
| ENSMUSG00000045733 | Sprn | protein_coding | -1.089237309 | 4.718273447 | 3.5365803 | 0.0780112 | 0.999175987 |
| ENSMUSG00000022594 | Lynx1 | protein_coding | 0.645490497 | 7.52563258 | 3.5306916 | 0.0782366 | 0.999175987 |
| ENSMUSG00000020219 | Timm13 | protein_coding | 0.578200678 | 8.92112637 | 3.5215463 | 0.0785883 | 0.999175987 |
| ENSMUSG00000026679 | Enkur | protein_coding | -1.156634597 | 4.017114693 | 3.5210461 | 0.0786075 | 0.999175987 |
| ENSMUSG00000017664 | Slc35c2 | protein_coding | 0.666177005 | 6.684281518 | 3.5108363 | 0.0790024 | 0.999175987 |
| ENSMUSG00000086405 | 9330198N18Rik | lincRNA | 1.622184909 | 2.958387547 | 3.5107395 | 0.0790062 | 0.999175987 |
| ENSMUSG00000028690 | Mmachc | protein_coding | 0.998274394 | 4.731167092 | 3.5098194 | 0.0790419 | 0.999175987 |
| ENSMUSG00000070858 | Gm1673 | protein_coding | 1.292492144 | 3.401330592 | 3.5094527 | 0.0790561 | 0.999175987 |
| ENSMUSG00000001988 | Npas1 | protein_coding | 1.751934706 | 3.054201747 | 3.5086137 | 0.0790887 | 0.999175987 |
| ENSMUSG00000053907 | Mat2a | protein_coding | -0.559615337 | 7.41689777 | 3.501722 | 0.079357 | 0.999175987 |
| ENSMUSG00000067604 | Nms | protein_coding | -1.849787794 | 2.993198315 | 3.5013234 | 0.0793725 | 0.999175987 |
| ENSMUSG00000021759 | Plpp1 | protein_coding | 0.858684695 | 4.696786493 | 3.4982342 | 0.0794931 | 0.999175987 |
| ENSMUSG00000019132 | BC005537 | protein_coding | -0.619894855 | 6.782680237 | 3.4946465 | 0.0796335 | 0.999175987 |
| ENSMUSG00000041697 | Cox6a1 | protein_coding | 0.894742899 | 9.755176941 | 3.5134953 | 0.0796671 | 0.999175987 |
| ENSMUSG00000047507 | Baiap3 | protein_coding | 0.958180893 | 6.667296313 | 3.5212513 | 0.0797656 | 0.999175987 |
| ENSMUSG00000026547 | Tagln2 | protein_coding | -1.058107407 | 4.606795362 | 3.4880248 | 0.0798932 | 0.999175987 |
| ENSMUSG00000033623 | Pcgf3 | protein_coding | -1.125691166 | 4.33184919 | 3.4871815 | 0.0799264 | 0.999175987 |
| ENSMUSG00000032050 | Rdx | protein_coding | -0.891605316 | 5.48007083 | 3.484354 | 0.0800377 | 0.999175987 |
| ENSMUSG00000024069 | Slc30a6 | protein_coding | -1.111068769 | 4.55533128 | 3.483801 | 0.0800595 | 0.999175987 |
| ENSMUSG00000041328 | Pcf11 | protein_coding | -1.260963637 | 3.966971098 | 3.4822451 | 0.0801208 | 0.999175987 |
| ENSMUSG00000060166 | Zdhhc8 | protein_coding | 1.179538872 | 3.884171771 | 3.4811889 | 0.0801625 | 0.999175987 |
| ENSMUSG00000035596 | Mboat7 | protein_coding | 0.656462075 | 6.622442919 | 3.478584 | 0.0802654 | 0.999175987 |
| ENSMUSG00000026915 | Strbp | protein_coding | -0.574133325 | 6.935576728 | 3.478386 | 0.0802732 | 0.999175987 |
| ENSMUSG00000033918 | Parl | protein_coding | 0.744623626 | 6.333580154 | 3.4773326 | 0.0803149 | 0.999175987 |
| ENSMUSG00000007603 | Dus3l | protein_coding | 0.711171995 | 6.250032188 | 3.476744 | 0.0803381 | 0.999175987 |
| ENSMUSG00000032376 | Usp3 | protein_coding | -0.940844988 | 4.465359218 | 3.4744252 | 0.08043 | 0.999175987 |
| ENSMUSG00000040624 | Plekhg1 | protein_coding | -1.276077569 | 3.843939393 | 3.474124 | 0.0804419 | 0.999175987 |
| ENSMUSG00000020530 | Ggnbp2 | protein_coding | -0.569811983 | 7.034951486 | 3.4732823 | 0.0804753 | 0.999175987 |
| ENSMUSG00000031149 | Praf2 | protein_coding | 0.684520903 | 6.38582939 | 3.4731791 | 0.0804793 | 0.999175987 |
| ENSMUSG00000026127 | Imp4 | protein_coding | -0.860888408 | 4.426090176 | 3.4729127 | 0.0804899 | 0.999175987 |
| ENSMUSG00000036036 | Zfp57 | protein_coding | 3.116850765 | 2.74899328 | 3.5143714 | 0.0806948 | 0.999175987 |
| ENSMUSG00000020598 | Nrcam | protein_coding | -0.688391965 | 5.843984081 | 3.4615315 | 0.0809427 | 0.999175987 |
| ENSMUSG00000027582 | Zgpat | protein_coding | 2.299821951 | 2.583425405 | 3.4607899 | 0.0809724 | 0.999175987 |
| ENSMUSG00000047547 | Cltb | protein_coding | 0.704635172 | 7.68489647 | 3.4580118 | 0.0810834 | 0.999175987 |
| ENSMUSG00000024084 | Qpct | protein_coding | -0.626695921 | 6.503661628 | 3.45546 | 0.0811855 | 0.999175987 |
| ENSMUSG00000042790 | Rnf214 | protein_coding | -0.735145796 | 5.510956496 | 3.4488618 | 0.0814504 | 0.999175987 |
| ENSMUSG00000016252 | Atp5e | protein_coding | 0.639263492 | 9.763074852 | 3.4480163 | 0.0814844 | 0.999175987 |
| ENSMUSG00000024208 | Uqcc2 | protein_coding | 0.620710308 | 9.520094615 | 3.4479609 | 0.0814866 | 0.999175987 |
| ENSMUSG00000054057 | A930004D18Rik | protein_coding | 1.142367596 | 3.860349917 | 3.4415676 | 0.0817443 | 0.999175987 |
| ENSMUSG00000058318 | Phf21a | protein_coding | -0.904080253 | 4.990714443 | 3.4413664 | 0.0817525 | 0.999175987 |
| ENSMUSG00000029505 | Ep400 | protein_coding | -1.128213992 | 4.822313806 | 3.4367249 | 0.0819705 | 0.999175987 |
| ENSMUSG00000038774 | Ascc3 | protein_coding | -1.506375705 | 3.242007027 | 3.4318083 | 0.0821396 | 0.999175987 |
| ENSMUSG00000011958 | Bnip2 | protein_coding | -1.213958919 | 3.770114211 | 3.4268382 | 0.0823418 | 0.999175987 |
| ENSMUSG00000035964 | Tmem59l | protein_coding | 0.591661947 | 9.53239765 | 3.4231212 | 0.0824934 | 0.999175987 |
| ENSMUSG00000017485 | Top2b | protein_coding | -0.857966707 | 5.196539979 | 3.4129767 | 0.0829087 | 0.999175987 |
| ENSMUSG00000048497 | Mmgt2 | protein_coding | 1.402489828 | 3.594320019 | 3.4110581 | 0.0829876 | 0.999175987 |
| ENSMUSG00000058756 | Thra | protein_coding | 0.638912412 | 6.284864811 | 3.4048119 | 0.0832448 | 0.999175987 |
| ENSMUSG00000014554 | Dguok | protein_coding | 0.711969676 | 5.686858152 | 3.4034294 | 0.0833019 | 0.999175987 |
| ENSMUSG00000085891 | Gm14634 | processed_transcript | -2.196570886 | 2.374153954 | 3.3993654 | 0.0834699 | 0.999175987 |
| ENSMUSG00000017802 | Retreg3 | protein_coding | -1.080194929 | 3.661015455 | 3.3959874 | 0.0836099 | 0.999175987 |
| ENSMUSG00000034616 | Ssh3 | protein_coding | 1.536101632 | 3.420697223 | 3.3827212 | 0.0841623 | 0.999175987 |
| ENSMUSG00000027639 | Samhd1 | protein_coding | -1.290367299 | 3.30342135 | 3.3812671 | 0.0842231 | 0.999175987 |
| ENSMUSG00000043284 | Tmem11 | protein_coding | 0.616111515 | 6.622788953 | 3.3777605 | 0.08437 | 0.999175987 |
| ENSMUSG00000033819 | Ppp1r16a | protein_coding | 0.863513906 | 4.734216239 | 3.3763081 | 0.0844309 | 0.999175987 |
| ENSMUSG00000024610 | Cd74 | protein_coding | -1.253735294 | 4.200338899 | 3.3749569 | 0.0844876 | 0.999175987 |
| ENSMUSG00000027620 | Rbm39 | protein_coding | -0.578157622 | 7.1349074 | 3.3625678 | 0.0850098 | 0.999175987 |
| ENSMUSG00000025142 | Aspscr1 | protein_coding | 1.392638004 | 3.519149347 | 3.3540395 | 0.0853715 | 0.999175987 |
| ENSMUSG00000026869 | Psmd5 | protein_coding | -1.036737625 | 6.091297663 | 3.3808965 | 0.0854184 | 0.999175987 |
| ENSMUSG00000039578 | Ccser1 | protein_coding | -0.977801066 | 4.043679451 | 3.3525146 | 0.0854363 | 0.999175987 |
| ENSMUSG00000003382 | Etv3 | protein_coding | -1.006531451 | 3.984042676 | 3.3522474 | 0.0854477 | 0.999175987 |

| ensembl_gene_id | external_gene_name | gene_biotype | log2 (fold change) | log2 (counts per million) | F-value | p-value | False discovery rate |
|---|---|---|---|---|---|---|---|
| ENSMUSG00000023026 | Dip2b | protein_coding | -0.822295531 | 5.396342755 | 3.3518577 | 0.0854643 | 0.999175987 |
| ENSMUSG00000008373 | Prpf31 | protein_coding | 0.558180282 | 6.59998186 | 3.3512064 | 0.085492 | 0.999175987 |
| ENSMUSG00000098557 | Kctd12 | protein_coding | -0.951195316 | 4.537306075 | 3.3489439 | 0.0855884 | 0.999175987 |
| ENSMUSG00000032038 | St3gal4 | protein_coding | 0.913921477 | 4.551804892 | 3.3470245 | 0.0856703 | 0.999175987 |
| ENSMUSG00000038369 | Ncoa6 | protein_coding | -1.130164616 | 4.480358588 | 3.34532 | 0.0857431 | 0.999175987 |
| ENSMUSG00000031641 | Cbr4 | protein_coding | -0.833236376 | 5.240956239 | 3.3443468 | 0.0857847 | 0.999175987 |
| ENSMUSG00000049764 | Zfp280b | protein_coding | -1.269497131 | 3.189917158 | 3.3442658 | 0.0857882 | 0.999175987 |
| ENSMUSG00000035521 | Gnptg | protein_coding | 0.546253392 | 7.368568313 | 3.3441298 | 0.085794 | 0.999175987 |
| ENSMUSG00000020448 | Rnf185 | protein_coding | 0.938475311 | 5.802407097 | 3.3444896 | 0.0858091 | 0.999175987 |
| ENSMUSG00000031613 | Hpgd | protein_coding | -1.465114433 | 3.347126738 | 3.3429544 | 0.0858443 | 0.999175987 |
| ENSMUSG00000030189 | Ybx3 | protein_coding | -1.132466229 | 3.853006636 | 3.342249 | 0.0858744 | 0.999175987 |
| ENSMUSG00000025576 | Rbfox3 | protein_coding | 0.575454225 | 6.697180347 | 3.34175 | 0.0858958 | 0.999175987 |
| ENSMUSG00000035530 | Eif1 | protein_coding | 0.689459404 | 9.399045328 | 3.3375392 | 0.0860764 | 0.999175987 |
| ENSMUSG00000102215 | Gm2464 | lincRNA | 0.956100644 | 4.146585718 | 3.3320171 | 0.0863138 | 0.999175987 |
| ENSMUSG00000073910 | Mob3b | protein_coding | 1.45857803 | 3.951102461 | 3.3323261 | 0.086331 | 0.999175987 |
| ENSMUSG00000041220 | Elovl6 | protein_coding | -0.793078086 | 5.424769524 | 3.3291469 | 0.0864375 | 0.999175987 |
| ENSMUSG00000025439 | Clns1a | protein_coding | 0.525753984 | 7.542459201 | 3.3287269 | 0.0864556 | 0.999175987 |
| ENSMUSG00000020186 | Csrp2 | protein_coding | 0.852178508 | 4.560132812 | 3.3269769 | 0.0865312 | 0.999175987 |
| ENSMUSG00000027823 | Gmps | protein_coding | -0.805528886 | 7.305183761 | 3.3313286 | 0.0867107 | 0.999175987 |
| ENSMUSG00000044716 | Dok7 | protein_coding | 1.452884461 | 3.466412865 | 3.3185703 | 0.0868953 | 0.999175987 |
| ENSMUSG00000028399 | Ptprd | protein_coding | -0.669733566 | 6.425603442 | 3.3176841 | 0.0869337 | 0.999175987 |
| ENSMUSG00000024055 | Cyp4f13 | protein_coding | 2.031937385 | 2.949855005 | 3.3165867 | 0.0869814 | 0.999175987 |
| ENSMUSG00000062691 | Cebpzos | protein_coding | 0.827250815 | 4.80714867 | 3.3158229 | 0.0870146 | 0.999175987 |
| ENSMUSG00000022076 | Klhl1 | protein_coding | 1.145662366 | 4.332187391 | 3.3092617 | 0.0873005 | 0.999175987 |
| ENSMUSG00000071550 | Cfap44 | protein_coding | 2.343085362 | 2.155988903 | 3.3074661 | 0.0873789 | 0.999175987 |
| ENSMUSG00000042444 | Mindy2 | protein_coding | -0.935347555 | 5.504832714 | 3.3055583 | 0.0874623 | 0.999175987 |
| ENSMUSG00000008668 | Rps18 | protein_coding | 0.691529108 | 7.167909004 | 3.3042773 | 0.0875184 | 0.999175987 |
| ENSMUSG00000056493 | Foxk1 | protein_coding | -1.227902823 | 3.329235493 | 3.3026009 | 0.0875918 | 0.999175987 |
| ENSMUSG00000039632 | Ccdc151 | protein_coding | -2.374066178 | 2.469709925 | 3.3019495 | 0.0876204 | 0.999175987 |
| ENSMUSG00000034135 | Sik3 | protein_coding | -0.566378167 | 6.39715125 | 3.3004561 | 0.0876859 | 0.999175987 |
| ENSMUSG00000006335 | Tfpt | protein_coding | 0.664272998 | 5.883506186 | 3.2997527 | 0.0877168 | 0.999175987 |
| ENSMUSG00000026803 | Ttf1 | protein_coding | 1.173289568 | 3.734458087 | 3.2978483 | 0.0878004 | 0.999175987 |
| ENSMUSG00000020810 | Cygb | protein_coding | 0.556715201 | 7.364913844 | 3.29587 | 0.0878874 | 0.999175987 |
| ENSMUSG00000034248 | Slc25a37 | protein_coding | 1.798522689 | 2.954380726 | 3.2954498 | 0.0879059 | 0.999175987 |
| ENSMUSG00000021486 | Prelid1 | protein_coding | 0.590797311 | 7.383620694 | 3.2950303 | 0.0879244 | 0.999175987 |
| ENSMUSG00000091985 | Gm17354 | lincRNA | -1.880189141 | 2.537460446 | 3.2940667 | 0.0879668 | 0.999175987 |
| ENSMUSG00000021918 | Nek4 | protein_coding | 1.07236455 | 4.061334243 | 3.2885315 | 0.088211 | 0.999175987 |
| ENSMUSG00000007827 | Ankrd26 | protein_coding | -1.130067799 | 4.709306544 | 3.2861262 | 0.088348 | 0.999175987 |
| ENSMUSG00000026482 | Rgl1 | protein_coding | 0.655078858 | 5.569146892 | 3.284255 | 0.0884002 | 0.999175987 |
| ENSMUSG00000026860 | Sh3glb2 | protein_coding | 0.636411049 | 8.027200636 | 3.2833905 | 0.0884386 | 0.999175987 |
| ENSMUSG00000063434 | Sorcs3 | protein_coding | 0.821149021 | 5.116461354 | 3.2773313 | 0.0887076 | 0.999175987 |
| ENSMUSG00000037669 | Ldah | protein_coding | -0.574631315 | 6.544297754 | 3.2762699 | 0.0887549 | 0.999175987 |
| ENSMUSG00000006356 | Crip2 | protein_coding | 0.560096976 | 8.839673363 | 3.2753055 | 0.0887978 | 0.999175987 |
| ENSMUSG00000025658 | Cnksr2 | protein_coding | -0.947885435 | 5.375949277 | 3.2749133 | 0.0888153 | 0.999175987 |
| ENSMUSG00000048118 | Arid4a | protein_coding | -0.71505218 | 6.112381513 | 3.2744203 | 0.0888372 | 0.999175987 |
| ENSMUSG00000031983 | 2310022B05Rik | protein_coding | -0.92654024 | 4.985790845 | 3.2724629 | 0.0889245 | 0.999175987 |
| ENSMUSG00000031256 | Cstf2 | protein_coding | -1.11267708 | 4.311827684 | 3.2703674 | 0.0890181 | 0.999175987 |
| ENSMUSG00000059498 | Fcgr3 | protein_coding | -1.444489899 | 4.39498967 | 3.2920294 | 0.0890848 | 0.999175987 |
| ENSMUSG00000038170 | Pde4dip | protein_coding | -0.633497877 | 6.043936981 | 3.2688031 | 0.089088 | 0.999175987 |
| ENSMUSG00000036333 | Kidins220 | protein_coding | -0.721625332 | 6.645574306 | 3.2629087 | 0.089352 | 0.999175987 |
| ENSMUSG00000059895 | Ptp4a3 | protein_coding | 0.757499841 | 5.311808711 | 3.2614127 | 0.0894192 | 0.999175987 |
| ENSMUSG00000050377 | Il31ra | protein_coding | 1.013879834 | 5.067837887 | 3.2587872 | 0.0895372 | 0.999175987 |
| ENSMUSG00000030847 | Bag3 | protein_coding | 1.711475887 | 2.816494219 | 3.2575712 | 0.0895919 | 0.999175987 |
| ENSMUSG00000023913 | Pla2g7 | protein_coding | -0.645075161 | 7.018557148 | 3.2559796 | 0.0896636 | 0.999175987 |
| ENSMUSG00000026649 | Cfap126 | protein_coding | -1.072588681 | 4.686892357 | 3.2520908 | 0.089839 | 0.999175987 |
| ENSMUSG00000020137 | Thap2 | protein_coding | -1.245316163 | 3.538827538 | 3.249168 | 0.0899711 | 0.999175987 |
| ENSMUSG00000005897 | Nr2c1 | protein_coding | -1.72179897 | 2.943919074 | 3.2454308 | 0.0901404 | 0.999175987 |
| ENSMUSG00000044308 | Ubr3 | protein_coding | -0.877497841 | 6.311988584 | 3.247486 | 0.0901589 | 0.999175987 |
| ENSMUSG00000022119 | Rbm26 | protein_coding | -0.906553701 | 4.824486385 | 3.2413522 | 0.0903255 | 0.999175987 |
| ENSMUSG00000105867 | Gm42517 | protein_coding | -1.552269959 | 2.859212297 | 3.2405928 | 0.09036 | 0.999175987 |
| ENSMUSG00000085069 | Gm13111 | antisense_RNA | -1.966611236 | 2.502866299 | 3.238404 | 0.0904596 | 0.999175987 |
| ENSMUSG00000003099 | Ppp5c | protein_coding | 0.715490124 | 7.03799125 | 3.2386132 | 0.0904809 | 0.999175987 |
| ENSMUSG00000015396 | Cd83 | protein_coding | 0.724926164 | 6.530667424 | 3.2345104 | 0.0906371 | 0.999175987 |
| ENSMUSG00000026377 | Nifk | protein_coding | 0.660245294 | 5.969500023 | 3.2321181 | 0.0907464 | 0.999175987 |
| ENSMUSG00000001909 | Trmt1 | protein_coding | 0.9673735 | 5.004510961 | 3.2320859 | 0.0907479 | 0.999175987 |
| ENSMUSG00000025423 | Pias2 | protein_coding | -0.723523813 | 5.407554787 | 3.2319609 | 0.0907536 | 0.999175987 |
| ENSMUSG00000040883 | Tmem205 | protein_coding | 0.735173739 | 6.500854608 | 3.230325 | 0.0908284 | 0.999175987 |
| ENSMUSG00000021967 | Mrpl57 | protein_coding | 0.689825768 | 6.480578311 | 3.2275186 | 0.0909569 | 0.999175987 |
| ENSMUSG00000045087 | S1pr5 | protein_coding | -1.81415225 | 2.742858132 | 3.2222987 | 0.0911965 | 0.999175987 |
| ENSMUSG00000028455 | Stoml2 | protein_coding | 0.690189085 | 7.123481909 | 3.2217333 | 0.0912225 | 0.999175987 |
| ENSMUSG00000087590 | Epb41l4aos | lincRNA | 1.200947933 | 3.339230179 | 3.2184764 | 0.0913724 | 0.999175987 |
| ENSMUSG00000059708 | Akap17b | protein_coding | -1.016123041 | 3.972768792 | 3.215061 | 0.0915299 | 0.999175987 |
| ENSMUSG00000062908 | Acadm | protein_coding | -0.677422301 | 5.884754575 | 3.2147317 | 0.0915451 | 0.999175987 |
| ENSMUSG00000003072 | Atp5d | protein_coding | 1.31231533 | 8.518672717 | 3.3646159 | 0.0915827 | 0.999175987 |
| ENSMUSG00000026447 | Pik3c2b | protein_coding | -1.070264663 | 4.035664441 | 3.2094048 | 0.0917915 | 0.999175987 |
| ENSMUSG00000052430 | Bmpr1b | protein_coding | -1.489492847 | 3.043873098 | 3.2088082 | 0.0918192 | 0.999175987 |
| ENSMUSG00000027316 | Gfra4 | protein_coding | 0.793834822 | 5.032923443 | 3.2064463 | 0.0919287 | 0.999175987 |
| ENSMUSG00000062210 | Tnfaip8 | protein_coding | -1.180572672 | 4.226963676 | 3.2062558 | 0.0919376 | 0.999175987 |

| ensembl_gene_id | external_gene_name | gene_biotype | log2 (fold change) | log2 (counts per million) | F-value | p-value | False discovery rate |
|---|---|---|---|---|---|---|---|
| ENSMUSG00000046402 | Rbp1 | protein_coding | -0.9980624 | 5.686358435 | 3.2074008 | 0.0919964 | 0.999175987 |
| ENSMUSG00000028833 | Ncdn | protein_coding | 0.609174804 | 8.830321483 | 3.2044096 | 0.0920233 | 0.999175987 |
| ENSMUSG00000047804 | Akap10 | protein_coding | -1.819732375 | 2.627325968 | 3.2041255 | 0.0920365 | 0.999175987 |
| ENSMUSG00000031358 | Msl3 | protein_coding | 0.935040417 | 4.63740218 | 3.1991195 | 0.0922695 | 0.999175987 |
| ENSMUSG00000022797 | Tfrc | protein_coding | -0.933138072 | 4.404060591 | 3.198985 | 0.0922758 | 0.999175987 |
| ENSMUSG00000034259 | Exosc4 | protein_coding | 0.638940418 | 6.135253347 | 3.1986294 | 0.0922924 | 0.999175987 |
| ENSMUSG00000034902 | Pip5k1c | protein_coding | 0.663934812 | 6.460003544 | 3.196303 | 0.0924009 | 0.999175987 |
| ENSMUSG00000042228 | Lyn | protein_coding | -2.32691773 | 2.375049205 | 3.1928587 | 0.0925619 | 0.999175987 |
| ENSMUSG00000074129 | Rpl13a | protein_coding | 0.739268254 | 5.367134901 | 3.191756 | 0.0926135 | 0.999175987 |
| ENSMUSG00000027400 | Pdyn | protein_coding | -0.894109311 | 6.739999228 | 3.2172138 | 0.0926244 | 0.999175987 |
| ENSMUSG00000026842 | Abl1 | protein_coding | -1.296448719 | 4.118710143 | 3.1915158 | 0.0926556 | 0.999175987 |
| ENSMUSG00000030286 | Emc3 | protein_coding | -0.679549834 | 6.178071852 | 3.1882811 | 0.0927764 | 0.999175987 |
| ENSMUSG00000074227 | Spint2 | protein_coding | 0.612550396 | 5.881870537 | 3.1882312 | 0.0927787 | 0.999175987 |
| ENSMUSG00000041556 | Fbxo2 | protein_coding | 0.916215888 | 4.92481782 | 3.1878988 | 0.0927943 | 0.999175987 |
| ENSMUSG00000022564 | Grina | protein_coding | 0.551945474 | 8.836490859 | 3.183817 | 0.0929861 | 0.999175987 |
| ENSMUSG00000071644 | Eef1g | protein_coding | 0.500047414 | 7.444402866 | 3.1836036 | 0.0929962 | 0.999175987 |
| ENSMUSG00000024940 | Ltbp3 | protein_coding | 1.760063814 | 2.779765272 | 3.1805397 | 0.0931405 | 0.999175987 |
| ENSMUSG00000032171 | Pin1 | protein_coding | 0.535949028 | 8.909003625 | 3.1796427 | 0.0931828 | 0.999175987 |
| ENSMUSG00000002068 | Ccne1 | protein_coding | 1.266898901 | 3.513192971 | 3.1785203 | 0.0932357 | 0.999175987 |
| ENSMUSG00000024220 | Zfp523 | protein_coding | 1.08646483 | 4.664263773 | 3.1773955 | 0.0932888 | 0.999175987 |
| ENSMUSG00000020692 | Nle1 | protein_coding | 1.268944309 | 4.406804354 | 3.1756208 | 0.0934035 | 0.999175987 |
| ENSMUSG00000024810 | Il33 | protein_coding | -0.640395544 | 8.05266783 | 3.1704763 | 0.0936163 | 0.999175987 |
| ENSMUSG00000033282 | Rpgrip1l | protein_coding | 1.096612428 | 4.09526138 | 3.1693964 | 0.0936675 | 0.999175987 |
| ENSMUSG00000041828 | Abca8a | protein_coding | 1.502818817 | 3.14266088 | 3.1692452 | 0.0936747 | 0.999175987 |
| ENSMUSG00000004031 | Brinp2 | protein_coding | 0.842007578 | 4.603007488 | 3.168992 | 0.0936867 | 0.999175987 |
| ENSMUSG00000042401 | Crtac1 | protein_coding | -1.185162814 | 4.769151047 | 3.1681769 | 0.0937563 | 0.999175987 |
| ENSMUSG00000031775 | Pllp | protein_coding | 0.886074194 | 4.431402865 | 3.1627037 | 0.0939857 | 0.999175987 |
| ENSMUSG00000069844 | Sco1 | protein_coding | -1.89008696 | 3.304374672 | 3.1638903 | 0.0943007 | 0.999175987 |
| ENSMUSG00000030231 | Plekha5 | protein_coding | -0.720895635 | 5.603126478 | 3.1553972 | 0.0943346 | 0.999175987 |
| ENSMUSG00000022016 | Akap11 | protein_coding | -0.899749941 | 6.148210347 | 3.1574529 | 0.0943843 | 0.999175987 |
| ENSMUSG00000104597 | B230334C09Rik | antisense_RNA | -1.218255114 | 3.76481505 | 3.1524015 | 0.0944781 | 0.999175987 |
| ENSMUSG00000039001 | Rps21 | protein_coding | 0.739509781 | 8.753982477 | 3.1513468 | 0.0945596 | 0.999175987 |
| ENSMUSG00000029141 | Slc4a1ap | protein_coding | -0.84941651 | 4.239304345 | 3.1490811 | 0.0946375 | 0.999175987 |
| ENSMUSG00000030707 | Coro1a | protein_coding | 0.558562244 | 7.390618051 | 3.1485662 | 0.0946622 | 0.999175987 |
| ENSMUSG00000029093 | Sorcs2 | protein_coding | -1.319871825 | 3.639065966 | 3.1479496 | 0.0946919 | 0.999175987 |
| ENSMUSG00000079593 | Gm14597 | protein_coding | 0.499083302 | 8.072834169 | 3.1470525 | 0.094735 | 0.999175987 |
| ENSMUSG00000043441 | Gpr149 | protein_coding | 1.305460938 | 3.185985951 | 3.1451472 | 0.0948267 | 0.999175987 |
| ENSMUSG00000016179 | Camk1g | protein_coding | 0.720949913 | 6.34527043 | 3.1436238 | 0.0949001 | 0.999175987 |
| ENSMUSG00000042388 | Dlgap3 | protein_coding | -1.40781386 | 3.139058895 | 3.1429651 | 0.0949318 | 0.999175987 |
| ENSMUSG00000025239 | Limd1 | protein_coding | -1.046520074 | 4.056360226 | 3.1394416 | 0.0951019 | 0.999175987 |
| ENSMUSG00000029088 | Kcnip4 | protein_coding | -0.678550925 | 5.534930803 | 3.1382626 | 0.0951589 | 0.999175987 |
| ENSMUSG00000013539 | Tango2 | protein_coding | 0.553232363 | 8.030168215 | 3.1374939 | 0.0951961 | 0.999175987 |
| ENSMUSG00000018865 | Sult4a1 | protein_coding | 0.639256695 | 9.330066523 | 3.1364205 | 0.095248 | 0.999175987 |
| ENSMUSG00000046432 | Bex3 | protein_coding | 0.576570895 | 9.760962919 | 3.1351952 | 0.0953074 | 0.999175987 |
| ENSMUSG00000031023 | Akip1 | protein_coding | -1.130170044 | 3.610570956 | 3.1342926 | 0.0953511 | 0.999175987 |
| ENSMUSG00000047259 | Mc4r | protein_coding | 1.304552059 | 3.845417213 | 3.1341802 | 0.0953566 | 0.999175987 |
| ENSMUSG00000046312 | AI464131 | protein_coding | -1.842265185 | 2.567405261 | 3.1339909 | 0.0953657 | 0.999175987 |
| ENSMUSG00000066148 | Prpf4 | protein_coding | -0.886093687 | 4.626979966 | 3.1328097 | 0.095423 | 0.999175987 |
| ENSMUSG00000098702 | 1500015A07Rik | antisense_RNA | 1.366878977 | 3.304849174 | 3.1313545 | 0.0954936 | 0.999175987 |
| ENSMUSG00000111947 | AC131120.1 | lincRNA | -2.161760068 | 2.78943609 | 3.13104 | 0.0955399 | 0.999175987 |
| ENSMUSG00000060012 | Kif13b | protein_coding | -1.161731216 | 3.707502773 | 3.130199 | 0.0955498 | 0.999175987 |
| ENSMUSG00000022529 | Zfp263 | protein_coding | 0.964791811 | 3.891581902 | 3.1281928 | 0.0956473 | 0.999175987 |
| ENSMUSG00000046598 | Bdh1 | protein_coding | 0.757656864 | 5.806172135 | 3.1239626 | 0.0958534 | 0.999175987 |
| ENSMUSG00000063800 | Prpf38a | protein_coding | -0.766631425 | 6.011208155 | 3.1231037 | 0.0958953 | 0.999175987 |
| ENSMUSG00000019761 | Krt10 | protein_coding | 0.793672698 | 5.047705005 | 3.1226533 | 0.0959173 | 0.999175987 |
| ENSMUSG00000051007 | Gatd1 | protein_coding | 0.622601236 | 7.261766072 | 3.1213407 | 0.0959814 | 0.999175987 |
| ENSMUSG00000057841 | Rpl32 | protein_coding | 0.676935316 | 9.869372076 | 3.1199059 | 0.0960515 | 0.999175987 |
| ENSMUSG00000027748 | Trpc4 | protein_coding | 0.907331698 | 4.408237184 | 3.1163753 | 0.0962243 | 0.999175987 |
| ENSMUSG00000018160 | Med1 | protein_coding | -0.747834611 | 6.664700456 | 3.1151489 | 0.0963154 | 0.999175987 |
| ENSMUSG00000026615 | Eprs | protein_coding | -0.685559008 | 6.566048291 | 3.1139785 | 0.0963418 | 0.999175987 |
| ENSMUSG00000000247 | Lhx2 | protein_coding | -1.854897437 | 3.10998722 | 3.11321 | 0.0964105 | 0.999175987 |
| ENSMUSG00000022203 | Efs | protein_coding | 1.423581926 | 2.928711855 | 3.1115425 | 0.0964615 | 0.999175987 |
| ENSMUSG00000034165 | Ccnd3 | protein_coding | 1.061655793 | 3.629713756 | 3.1086954 | 0.0966015 | 0.999175987 |
| ENSMUSG00000032702 | Kank1 | protein_coding | -1.15077778 | 3.697602007 | 3.1074662 | 0.096662 | 0.999175987 |
| ENSMUSG00000024325 | Ring1 | protein_coding | 1.669054837 | 2.785337148 | 3.1072108 | 0.0966746 | 0.999175987 |
| ENSMUSG00000032534 | Cep63 | protein_coding | 0.889808863 | 4.898120913 | 3.1063758 | 0.0967158 | 0.999175987 |
| ENSMUSG00000063317 | Usp31 | protein_coding | -0.801653012 | 5.091182228 | 3.1057017 | 0.096749 | 0.999175987 |
| ENSMUSG00000058600 | Rpl30 | protein_coding | 0.756517988 | 6.950298277 | 3.1043508 | 0.0968467 | 0.999175987 |
| ENSMUSG00000003411 | Rab3b | protein_coding | -0.535741543 | 8.485015231 | 3.0991378 | 0.0970734 | 0.999175987 |
| ENSMUSG00000035726 | Supt16 | protein_coding | -0.808849255 | 5.624381339 | 3.097349 | 0.097162 | 0.999175987 |
| ENSMUSG00000028412 | Slc44a1 | protein_coding | -0.801725664 | 5.617019208 | 3.0972683 | 0.097166 | 0.999175987 |
| ENSMUSG00000020132 | Rab21 | protein_coding | -0.535339726 | 6.693586283 | 3.0963998 | 0.097209 | 0.999175987 |
| ENSMUSG00000018800 | Abca5 | protein_coding | -0.993563672 | 3.999379252 | 3.0941535 | 0.0973205 | 0.999175987 |
| ENSMUSG00000045409 | Trim39 | protein_coding | -1.238003443 | 3.885929486 | 3.0903796 | 0.0975082 | 0.999175987 |
| ENSMUSG00000032527 | Pccb | protein_coding | -0.803792436 | 5.911814377 | 3.0881106 | 0.0976212 | 0.999175987 |
| ENSMUSG00000030782 | Tgfb1i1 | protein_coding | 1.023397608 | 4.468546827 | 3.0875514 | 0.0976491 | 0.999175987 |
| ENSMUSG00000049299 | Trappc1 | protein_coding | 0.578030901 | 7.355933209 | 3.0847663 | 0.0977881 | 0.999175987 |
| ENSMUSG00000025981 | Coq10b | protein_coding | -0.827171564 | 5.300111345 | 3.083891 | 0.0978318 | 0.999175987 |

| ensembl_gene_id | external_gene_name | gene_biotype | log2 (fold change) | log2 (counts per million) | F-value | p-value | False discovery rate |
|---|---|---|---|---|---|---|---|
| ENSMUSG00000078453 | Abracl | protein_coding | 0.991113709 | 4.923847524 | 3.0830784 | 0.0978724 | 0.999175987 |
| ENSMUSG00000034088 | Hdlbp | protein_coding | -0.645444946 | 6.210549682 | 3.0811665 | 0.0979681 | 0.999175987 |
| ENSMUSG00000029772 | Ahcyl2 | protein_coding | -0.664935309 | 5.592556027 | 3.0798654 | 0.0980332 | 0.999175987 |
| ENSMUSG00000027983 | Cyp2u1 | protein_coding | 1.020482652 | 4.690115659 | 3.0769334 | 0.0981803 | 0.999175987 |
| ENSMUSG00000061046 | Haghl | protein_coding | 0.531731468 | 7.77156325 | 3.075964 | 0.0982289 | 0.999175987 |
| ENSMUSG00000030588 | Yif1b | protein_coding | 0.579702662 | 6.243423051 | 3.0723125 | 0.0984125 | 0.999175987 |
| ENSMUSG00000063457 | Rps15 | protein_coding | 0.672125111 | 8.929819648 | 3.0712276 | 0.0984671 | 0.999175987 |
| ENSMUSG00000026764 | Kif5c | protein_coding | -0.650422029 | 7.28772482 | 3.06904 | 0.0985774 | 0.999175987 |
| ENSMUSG00000020823 | Sec14l1 | protein_coding | -0.763217554 | 5.454961313 | 3.0688327 | 0.0985878 | 0.999175987 |
| ENSMUSG00000020471 | Pold2 | protein_coding | 0.822818978 | 5.39565547 | 3.0671712 | 0.0986717 | 0.999175987 |
| ENSMUSG00000024378 | Stard4 | protein_coding | -1.185671379 | 4.103663219 | 3.0613703 | 0.0989651 | 0.999175987 |
| ENSMUSG00000021555 | Naa35 | protein_coding | -0.553953902 | 6.4186992 | 3.0572312 | 0.099175 | 0.999175987 |
| ENSMUSG00000033965 | Slc16a2 | protein_coding | -1.491646762 | 3.931101434 | 3.0634329 | 0.0992343 | 0.999175987 |
| ENSMUSG00000056763 | Cspp1 | protein_coding | 0.759101652 | 4.77568364 | 3.0545449 | 0.0993116 | 0.999175987 |
| ENSMUSG00000022978 | Mis18a | protein_coding | 0.987460241 | 3.910452885 | 3.0464651 | 0.0997237 | 0.999175987 |
| ENSMUSG00000031755 | Bbs2 | protein_coding | 1.088460063 | 4.505617287 | 3.0462316 | 0.0997356 | 0.999175987 |
| ENSMUSG00000027359 | Slc27a2 | protein_coding | -1.903960298 | 2.826299836 | 3.04389 | 0.0998555 | 0.999175987 |
| ENSMUSG00000064210 | Ano6 | protein_coding | -1.220878319 | 3.670365067 | 3.0430571 | 0.0998981 | 0.999175987 |
| ENSMUSG00000033669 | Zfp7 | protein_coding | 1.327689827 | 3.10848337 | 3.0426385 | 0.0999196 | 0.999175987 |
| ENSMUSG00000043498 | 9330132A10Rik | pseudogene | -1.124720893 | 3.451029808 | 3.0369495 | 0.1002116 | 0.999175987 |
| ENSMUSG00000055435 | Maf | protein_coding | -1.333921869 | 3.232228002 | 3.0346797 | 0.1003284 | 0.999175987 |
| ENSMUSG00000047656 | Trpt1 | protein_coding | 0.728301929 | 5.582264213 | 3.0324628 | 0.1004427 | 0.999175987 |
| ENSMUSG00000047454 | Gphn | protein_coding | 0.564100703 | 7.188718565 | 3.0316719 | 0.1004835 | 0.999175987 |
| ENSMUSG00000049076 | Acap2 | protein_coding | -0.918123822 | 4.383379589 | 3.0314482 | 0.100495 | 0.999175987 |
| ENSMUSG00000024395 | Lims2 | protein_coding | -0.818765149 | 5.325717595 | 3.023847 | 0.1008882 | 0.999175987 |
| ENSMUSG00000031381 | Piga | protein_coding | -1.215681836 | 3.304020973 | 3.0205671 | 0.1010584 | 0.999175987 |
| ENSMUSG00000020623 | Map2k6 | protein_coding | -1.106203398 | 4.021855182 | 3.0199608 | 0.1010899 | 0.999175987 |
| ENSMUSG00000000296 | Tpd52l1 | protein_coding | 0.488622521 | 7.58623538 | 3.0191919 | 0.1011298 | 0.999175987 |
| ENSMUSG00000059412 | Fxyd2 | protein_coding | 0.865415725 | 5.13876124 | 3.019171 | 0.1011309 | 0.999175987 |
| ENSMUSG00000004558 | Ndrg2 | protein_coding | -0.616469993 | 9.746559123 | 3.0161626 | 0.1012875 | 0.999175987 |
| ENSMUSG00000039765 | Cc2d2a | protein_coding | -0.979458575 | 3.858945597 | 3.0091063 | 0.1016558 | 0.999175987 |
| ENSMUSG00000085795 | Zfp703 | protein_coding | -1.579060991 | 2.951697461 | 3.0045799 | 0.1018929 | 0.999175987 |
| ENSMUSG00000031887 | Tradd | protein_coding | 1.72698386 | 2.985530773 | 2.9987663 | 0.1021985 | 0.999175987 |
| ENSMUSG00000006095 | Tbcb | protein_coding | 0.492088291 | 9.009886997 | 2.9986497 | 0.1022046 | 0.999175987 |
| ENSMUSG00000010505 | Myt1 | protein_coding | 1.772617514 | 2.486515823 | 2.9967047 | 0.1023071 | 0.999175987 |
| ENSMUSG00000055538 | Zcchc24 | protein_coding | -0.874511768 | 5.011976266 | 2.995154 | 0.1023888 | 0.999175987 |
| ENSMUSG00000031297 | Slc7a3 | protein_coding | 1.048864223 | 4.918918982 | 2.9943094 | 0.1024646 | 0.999175987 |
| ENSMUSG00000025821 | Zfp282 | protein_coding | 1.51024131 | 3.051024875 | 2.9911243 | 0.1026017 | 0.999175987 |
| ENSMUSG00000022090 | Pdlim2 | protein_coding | -0.622152291 | 6.69297983 | 2.9899276 | 0.1026651 | 0.999175987 |
| ENSMUSG00000022204 | Ngdn | protein_coding | -0.584483134 | 6.862385799 | 2.9898047 | 0.1026716 | 0.999175987 |
| ENSMUSG00000021027 | Ralgapa1 | protein_coding | -0.750391966 | 5.251444168 | 2.9841432 | 0.1029718 | 0.999175987 |
| ENSMUSG00000029676 | Pot1a | protein_coding | 1.092254794 | 4.502546478 | 2.982613 | 0.1030531 | 0.999175987 |
| ENSMUSG00000037119 | Fam91a1 | protein_coding | -0.869298149 | 4.981655417 | 2.9815642 | 0.1031089 | 0.999175987 |
| ENSMUSG00000086308 | G630016G05Rik | lincRNA | 1.58738379 | 2.940889393 | 2.9803145 | 0.1031754 | 0.999175987 |
| ENSMUSG00000032123 | Dpagt1 | protein_coding | 1.580484896 | 3.034197488 | 2.9799963 | 0.1031924 | 0.999175987 |
| ENSMUSG00000047583 | Tyw3 | protein_coding | -0.817197924 | 4.365579562 | 2.979681 | 0.1032092 | 0.999175987 |
| ENSMUSG00000015312 | Gadd45b | protein_coding | 1.146300794 | 3.546688812 | 2.97737 | 0.1033324 | 0.999175987 |
| ENSMUSG00000026857 | Ntmt1 | protein_coding | 0.530648935 | 7.068004118 | 2.9765949 | 0.1033737 | 0.999175987 |
| ENSMUSG00000025889 | Snca | protein_coding | 0.500600548 | 9.380044398 | 2.972916 | 0.1035703 | 0.999175987 |
| ENSMUSG00000026925 | Inpp5e | protein_coding | -1.14359013 | 3.412562807 | 2.9728941 | 0.1035715 | 0.999175987 |
| ENSMUSG00000033237 | Arid2 | protein_coding | -0.886821313 | 4.539216334 | 2.9726716 | 0.1035834 | 0.999175987 |
| ENSMUSG00000034744 | Nagk | protein_coding | 0.566217769 | 7.326640144 | 2.9724185 | 0.1035969 | 0.999175987 |
| ENSMUSG00000045867 | Cradd | protein_coding | 0.757983986 | 4.529806171 | 2.9718468 | 0.1036275 | 0.999175987 |
| ENSMUSG00000068115 | Ninl | protein_coding | 2.622271626 | 2.241747413 | 2.9697572 | 0.1037394 | 0.999175987 |
| ENSMUSG00000026836 | Acvr1 | protein_coding | -1.101547551 | 3.561656619 | 2.9691225 | 0.1037734 | 0.999175987 |
| ENSMUSG00000034614 | Pik3ip1 | protein_coding | -1.04757069 | 4.147793106 | 2.9673761 | 0.1038671 | 0.999175987 |
| ENSMUSG00000005823 | Gpr108 | protein_coding | 0.774633088 | 5.589183926 | 2.9662978 | 0.103925 | 0.999175987 |
| ENSMUSG00000030313 | Dennd5b | protein_coding | -0.696621347 | 4.922465207 | 2.9648791 | 0.1040013 | 0.999175987 |
| ENSMUSG00000085913 | Gm15601 | antisense_RNA | -1.709540652 | 2.707428456 | 2.9625871 | 0.1041246 | 0.999175987 |
| ENSMUSG00000029334 | Prkg2 | protein_coding | -1.526546201 | 3.431655454 | 2.9599864 | 0.1042647 | 0.999175987 |
| ENSMUSG00000057411 | Fam173a | protein_coding | 0.552543734 | 8.093885779 | 2.9597102 | 0.1042796 | 0.999175987 |
| ENSMUSG00000027643 | Ghrh | protein_coding | -1.401465063 | 3.367773726 | 2.9590151 | 0.1043171 | 0.999175987 |
| ENSMUSG00000035696 | Rnf38 | protein_coding | -1.030993387 | 4.031856508 | 2.9575314 | 0.1043972 | 0.999175987 |
| ENSMUSG00000058897 | Col25a1 | protein_coding | 0.87608172 | 4.848781409 | 2.9509575 | 0.1047529 | 0.999175987 |
| ENSMUSG00000014353 | Tmem87b | protein_coding | -0.91097607 | 5.005499003 | 2.9505261 | 0.1047763 | 0.999175987 |
| ENSMUSG00000020260 | Pofut2 | protein_coding | 0.91870486 | 5.643181593 | 2.9463878 | 0.1050323 | 0.999175987 |
| ENSMUSG00000040929 | Rfx3 | protein_coding | -0.971376101 | 4.279747821 | 2.9454609 | 0.1050514 | 0.999175987 |
| ENSMUSG00000034880 | Mrpl34 | protein_coding | 0.585694508 | 7.009545127 | 2.9447348 | 0.1050909 | 0.999175987 |
| ENSMUSG00000048482 | Bdnf | protein_coding | -1.074448121 | 3.991353696 | 2.9427557 | 0.1051987 | 0.999175987 |
| ENSMUSG00000014956 | Ppp1cb | protein_coding | -0.472846198 | 8.33464785 | 2.9413247 | 0.1052768 | 0.999175987 |
| ENSMUSG00000027132 | Katnbl1 | protein_coding | 1.186677297 | 3.374344132 | 2.9409826 | 0.1052954 | 0.999175987 |
| ENSMUSG00000025584 | Pde8a | protein_coding | -1.131244279 | 4.010309423 | 2.9360642 | 0.1055642 | 0.999175987 |
| ENSMUSG00000020744 | Slc25a19 | protein_coding | 0.685335053 | 5.895115586 | 2.9354696 | 0.1055967 | 0.999175987 |
| ENSMUSG00000024098 | Twsg1 | protein_coding | -1.182111028 | 3.465440633 | 2.9341253 | 0.1056703 | 0.999175987 |
| ENSMUSG00000032872 | Cyb5r4 | protein_coding | -1.061742221 | 3.598388793 | 2.9340745 | 0.1056731 | 0.999175987 |
| ENSMUSG00000028081 | Rps3a1 | protein_coding | 0.670202466 | 7.926400516 | 2.9345434 | 0.1056787 | 0.999175987 |
| ENSMUSG00000055707 | Klhl26 | protein_coding | 1.079590431 | 5.529094558 | 2.9540456 | 0.1057228 | 0.999175987 |
| ENSMUSG00000059070 | Rpl18 | protein_coding | 0.651650778 | 6.174859405 | 2.93291 | 0.1057369 | 0.999175987 |

| ensembl_gene_id | external_gene_name | gene_biotype | log2 (fold change) | log2 (counts per million) | F-value | p-value | False discovery rate |
|---|---|---|---|---|---|---|---|
| ENSMUSG00000033060 | Lmo7 | protein_coding | -1.244971454 | 3.790753184 | 2.9324657 | 0.1057613 | 0.999175987 |
| ENSMUSG00000027371 | Fahd2a | protein_coding | 0.601291944 | 6.322687499 | 2.9209159 | 0.1063971 | 0.999175987 |
| ENSMUSG00000063659 | Zbtb18 | protein_coding | -0.995587247 | 3.932208791 | 2.9176757 | 0.1065762 | 0.999175987 |
| ENSMUSG00000027787 | Nmd3 | protein_coding | -0.638548274 | 5.983076922 | 2.9176252 | 0.106579 | 0.999175987 |
| ENSMUSG00000067377 | Tspan6 | protein_coding | 0.576590947 | 6.552946552 | 2.916507 | 0.106641 | 0.999175987 |
| ENSMUSG00000097924 | A730020E08Rik | bidirectional_promote | 1.664987899 | 2.845040836 | 2.9145547 | 0.1067492 | 0.999175987 |
| ENSMUSG00000038871 | Bpgm | protein_coding | -0.516808722 | 7.396211372 | 2.9133575 | 0.1068156 | 0.999175987 |
| ENSMUSG00000030682 | Cdipt | protein_coding | 0.600037167 | 6.957219419 | 2.9131489 | 0.1068272 | 0.999175987 |
| ENSMUSG00000037174 | Elf2 | protein_coding | -1.035505653 | 3.947263548 | 2.9109721 | 0.1069481 | 0.999175987 |
| ENSMUSG00000002732 | Fkbp7 | protein_coding | -1.605503103 | 2.801679118 | 2.9070069 | 0.1071688 | 0.999175987 |
| ENSMUSG00000041329 | Atp1b2 | protein_coding | -0.709974925 | 7.359503043 | 2.9069238 | 0.1072047 | 0.999175987 |
| ENSMUSG00000038181 | Chpf2 | protein_coding | 2.51341763 | 2.365429595 | 2.9050293 | 0.107279 | 0.999175987 |
| ENSMUSG00000038056 | Kmt2c | protein_coding | -0.836505445 | 4.777930851 | 2.8994996 | 0.107588 | 0.999175987 |
| ENSMUSG00000039701 | Usp53 | protein_coding | -1.283554984 | 3.229360718 | 2.8976394 | 0.1076923 | 0.999175987 |
| ENSMUSG00000096914 | Galntl6 | protein_coding | -0.747175171 | 5.198842689 | 2.8973237 | 0.1077099 | 0.999175987 |
| ENSMUSG00000050786 | Ccdc126 | protein_coding | 1.104200057 | 3.822217161 | 2.896899 | 0.1077338 | 0.999175987 |
| ENSMUSG00000019428 | Fkbp8 | protein_coding | 0.490804193 | 8.438425054 | 2.8956965 | 0.1078012 | 0.999175987 |
| ENSMUSG00000049907 | Rasl11b | protein_coding | 0.887382963 | 4.109289556 | 2.8954975 | 0.1078124 | 0.999175987 |
| ENSMUSG00000018322 | Tomm34 | protein_coding | 0.567165499 | 7.916772627 | 2.8927166 | 0.1079686 | 0.999175987 |
| ENSMUSG00000028070 | Naxe | protein_coding | 0.665220366 | 7.422686091 | 2.8924127 | 0.108017 | 0.999175987 |
| ENSMUSG00000046862 | Pramef8 | protein_coding | 1.060183197 | 4.561603272 | 2.8902471 | 0.1081075 | 0.999175987 |
| ENSMUSG00000022561 | Gpaa1 | protein_coding | 0.832772641 | 5.135706021 | 2.8873838 | 0.1082689 | 0.999175987 |
| ENSMUSG00000027663 | Zmat3 | protein_coding | -0.610880899 | 5.696992462 | 2.8856819 | 0.1083649 | 0.999175987 |
| ENSMUSG00000038648 | Creb3l2 | protein_coding | -2.352756169 | 2.243356346 | 2.8815343 | 0.1085995 | 0.999175987 |
| ENSMUSG00000032936 | Camkv | protein_coding | 0.567035392 | 8.008859896 | 2.881103 | 0.1086239 | 0.999175987 |
| ENSMUSG00000031133 | Arhgef6 | protein_coding | -1.687916533 | 2.854956478 | 2.8808436 | 0.1086386 | 0.999175987 |
| ENSMUSG00000027133 | Nop10 | protein_coding | 0.662184475 | 9.048247012 | 2.8800202 | 0.1086852 | 0.999175987 |
| ENSMUSG00000038451 | Spsb2 | protein_coding | -0.979591618 | 3.76742242 | 2.8795619 | 0.1087112 | 0.999175987 |
| ENSMUSG00000052748 | Swt1 | protein_coding | -0.999234896 | 4.030191547 | 2.8795344 | 0.1087127 | 0.999175987 |
| ENSMUSG00000038084 | Opa1 | protein_coding | -0.630963516 | 5.952056589 | 2.8792341 | 0.1087298 | 0.999175987 |
| ENSMUSG00000026966 | Ssna1 | protein_coding | 0.522454295 | 7.620011426 | 2.8777599 | 0.1088134 | 0.999175987 |
| ENSMUSG00000075318 | Scn2a | protein_coding | -0.498534624 | 6.959248327 | 2.874132 | 0.1090195 | 0.999175987 |
| ENSMUSG00000044364 | Tmem74b | protein_coding | 1.113761936 | 3.394795497 | 2.8725544 | 0.1091093 | 0.999175987 |
| ENSMUSG00000067288 | Rps28 | protein_coding | 0.947482627 | 4.775434699 | 2.8696433 | 0.1092752 | 0.999175987 |
| ENSMUSG00000062372 | Otof | protein_coding | 1.440720374 | 3.280649575 | 2.8688264 | 0.1093218 | 0.999175987 |
| ENSMUSG00000051034 | Zfp11 | protein_coding | 1.400376385 | 3.349475125 | 2.867245 | 0.109412 | 0.999175987 |
| ENSMUSG00000047115 | Fam221a | protein_coding | 2.212750272 | 2.264758872 | 2.8657015 | 0.1095003 | 0.999175987 |
| ENSMUSG00000098973 | Mir6236 | miRNA | 1.023936518 | 8.725991134 | 2.9179737 | 0.109597 | 0.999175987 |
| ENSMUSG00000026707 | Nsun6 | protein_coding | -1.335176426 | 3.048525799 | 2.8619991 | 0.1097122 | 0.999175987 |
| ENSMUSG00000069729 | Arid1b | protein_coding | -0.705767661 | 5.325468977 | 2.8614843 | 0.1097417 | 0.999175987 |
| ENSMUSG00000036023 | Parp2 | protein_coding | -0.637071957 | 5.474415451 | 2.8611044 | 0.1097635 | 0.999175987 |
| ENSMUSG00000052572 | Dlg2 | protein_coding | -0.587127272 | 6.437732956 | 2.8600645 | 0.1098231 | 0.999175987 |
| ENSMUSG00000103103 | 4833445I07Rik | lincRNA | 1.546862334 | 2.77715913 | 2.8600384 | 0.1098246 | 0.999175987 |
| ENSMUSG00000028494 | Plin2 | protein_coding | -1.167158288 | 3.848423259 | 2.858331 | 0.1099227 | 0.999175987 |
| ENSMUSG00000041355 | Ssr2 | protein_coding | 0.511692441 | 7.580759846 | 2.8582645 | 0.1099265 | 0.999175987 |
| ENSMUSG00000035849 | Krt222 | protein_coding | -0.681071659 | 5.881823538 | 2.8574603 | 0.1099727 | 0.999175987 |
| ENSMUSG00000024298 | Zfp871 | protein_coding | -0.889810894 | 4.790642478 | 2.854175 | 0.1101617 | 0.999175987 |
| ENSMUSG00000050229 | Pigm | protein_coding | -1.434852395 | 2.779106092 | 2.8540839 | 0.1101669 | 0.999175987 |
| ENSMUSG00000036893 | Ehmt1 | protein_coding | 0.715399217 | 5.202402411 | 2.8528937 | 0.1102355 | 0.999175987 |
| ENSMUSG00000030264 | Thumpd3 | protein_coding | -0.810439077 | 4.766676556 | 2.8520178 | 0.110286 | 0.999175987 |
| ENSMUSG00000051504 | Siglech | protein_coding | -1.168019814 | 3.656645677 | 2.8504508 | 0.1103764 | 0.999175987 |
| ENSMUSG00000037487 | Ubr5 | protein_coding | -0.716530296 | 5.318454565 | 2.84838 | 0.110496 | 0.999175987 |
| ENSMUSG00000026339 | Ccdc93 | protein_coding | -1.484665346 | 2.93749674 | 2.8473805 | 0.1105538 | 0.999175987 |
| ENSMUSG00000025915 | Sgk3 | protein_coding | -0.761150183 | 5.458691514 | 2.8472351 | 0.1105622 | 0.999175987 |
| ENSMUSG00000019984 | Med23 | protein_coding | -0.972053135 | 3.592601219 | 2.8433239 | 0.1107887 | 0.999175987 |
| ENSMUSG00000029054 | Gabrd | protein_coding | 0.899246957 | 5.524120706 | 2.840199 | 0.1110015 | 0.999175987 |
| ENSMUSG00000004929 | Thop1 | protein_coding | 1.194495163 | 4.018110319 | 2.8388073 | 0.111051 | 0.999175987 |
| ENSMUSG00000032422 | Snx14 | protein_coding | -0.973693581 | 4.587299046 | 2.8385493 | 0.111066 | 0.999175987 |
| ENSMUSG00000002748 | Baz1b | protein_coding | -0.744216989 | 5.4255037 | 2.8373408 | 0.1111363 | 0.999175987 |
| ENSMUSG00000004667 | Polr2e | protein_coding | 0.496843925 | 8.368268103 | 2.8370922 | 0.1111508 | 0.999175987 |
| ENSMUSG00000000805 | Car4 | protein_coding | 1.124692742 | 3.783159165 | 2.8367828 | 0.1111688 | 0.999175987 |
| ENSMUSG00000021711 | Trappc13 | protein_coding | -0.504325581 | 7.177096875 | 2.8354892 | 0.1112442 | 0.999175987 |
| ENSMUSG00000009216 | Fam163b | protein_coding | 0.947459545 | 4.936179858 | 2.8352575 | 0.1112577 | 0.999175987 |
| ENSMUSG00000056153 | Socs6 | protein_coding | -0.894942955 | 4.079099879 | 2.8342947 | 0.1113138 | 0.999175987 |
| ENSMUSG00000051950 | B3glct | protein_coding | -1.479385502 | 3.011145977 | 2.8310379 | 0.1115039 | 0.999175987 |
| ENSMUSG00000032231 | Anxa2 | protein_coding | -0.925482608 | 4.781141088 | 2.8305485 | 0.1115325 | 0.999175987 |
| ENSMUSG00000026469 | Xpr1 | protein_coding | -0.584292237 | 6.068042782 | 2.829495 | 0.1115941 | 0.999175987 |
| ENSMUSG00000022668 | Gtpbp8 | protein_coding | 0.824326004 | 4.736968523 | 2.828426 | 0.1116566 | 0.999175987 |
| ENSMUSG00000085438 | 1700020I14Rik | lincRNA | -0.506753182 | 6.815821602 | 2.8245441 | 0.1118841 | 0.999175987 |
| ENSMUSG00000008683 | Rps15a | protein_coding | 0.66286166 | 7.305842225 | 2.8248696 | 0.1118964 | 0.999175987 |
| ENSMUSG00000001855 | Nup214 | protein_coding | -0.785605114 | 4.491393077 | 2.8222767 | 0.1120173 | 0.999175987 |
| ENSMUSG00000048772 | Tmem53 | protein_coding | 0.931672307 | 4.221569174 | 2.8214278 | 0.1120671 | 0.999175987 |
| ENSMUSG00000111471 | AC122463.1 | antisense_RNA | -0.732514923 | 5.102127644 | 2.8196595 | 0.1121712 | 0.999175987 |
| ENSMUSG00000020152 | Actr2 | protein_coding | -0.523418258 | 9.058469489 | 2.8172867 | 0.1123109 | 0.999175987 |
| ENSMUSG00000037815 | Ctnna1 | protein_coding | -0.703422429 | 5.880425681 | 2.8165235 | 0.1123559 | 0.999175987 |
| ENSMUSG00000038517 | Tbkbp1 | protein_coding | -1.516604244 | 3.123857561 | 2.8152624 | 0.1124303 | 0.999175987 |
| ENSMUSG00000000838 | Fmr1 | protein_coding | -0.828842412 | 4.646406992 | 2.814524 | 0.1124739 | 0.999175987 |
| ENSMUSG00000069769 | Msi2 | protein_coding | -0.678690317 | 5.347387036 | 2.8140292 | 0.1125031 | 0.999175987 |

| ensembl_gene_id | external_gene_name | gene_biotype | log2 (fold change) | log2 (counts per million) | F-value | p-value | False discovery rate |
|---|---|---|---|---|---|---|---|
| ENSMUSG00000041429 | Nthl1 | protein_coding | 1.354489455 | 3.525973992 | 2.8122348 | 0.1126092 | 0.999175987 |
| ENSMUSG00000031227 | Magee1 | protein_coding | 0.481795949 | 6.897244192 | 2.8118921 | 0.1126294 | 0.999175987 |
| ENSMUSG00000036186 | Fam69b | protein_coding | 0.925975339 | 4.7655652 | 2.8101072 | 0.112735 | 0.999175987 |
| ENSMUSG00000050069 | Grem2 | protein_coding | 1.20405756 | 3.546617738 | 2.8098235 | 0.1127518 | 0.999175987 |
| ENSMUSG00000022604 | Cep97 | protein_coding | -1.009133831 | 3.975601437 | 2.8091841 | 0.1127897 | 0.999175987 |
| ENSMUSG00000030102 | Itpr1 | protein_coding | -0.877584545 | 6.325948694 | 2.8273596 | 0.1128615 | 0.999175987 |
| ENSMUSG00000028048 | Gba | protein_coding | 0.892085646 | 4.871983274 | 2.8029348 | 0.1131607 | 0.999175987 |
| ENSMUSG00000053644 | Aldh7a1 | protein_coding | -0.615668172 | 5.997732818 | 2.7989738 | 0.1133966 | 0.999175987 |
| ENSMUSG00000051427 | Ccdc157 | protein_coding | -1.398868135 | 2.943598465 | 2.7972021 | 0.1135023 | 0.999175987 |
| ENSMUSG00000029390 | Tmed2 | protein_coding | -1.226169493 | 3.063734541 | 2.7822307 | 0.1144002 | 0.999175987 |
| ENSMUSG00000052889 | Prkcb | protein_coding | -0.652770512 | 8.464317642 | 2.7820783 | 0.1144408 | 0.999175987 |
| ENSMUSG00000023017 | Asic1 | protein_coding | -0.716314118 | 5.062771338 | 2.7800291 | 0.114533 | 0.999175987 |
| ENSMUSG00000036641 | Ccdc148 | protein_coding | -0.796401144 | 5.152694454 | 2.7797582 | 0.1145493 | 0.999175987 |
| ENSMUSG00000006373 | Pgrmc1 | protein_coding | 0.509718198 | 9.279987835 | 2.7766325 | 0.1147381 | 0.999175987 |
| ENSMUSG00000021379 | Id4 | protein_coding | -0.501503932 | 6.834048099 | 2.7714218 | 0.1150538 | 0.999175987 |
| ENSMUSG00000027800 | Tm4sf1 | protein_coding | 0.924561756 | 4.283675541 | 2.7655603 | 0.1154101 | 0.999175987 |
| ENSMUSG00000024446 | Rpp21 | protein_coding | 0.459343222 | 7.644382785 | 2.7620322 | 0.1156252 | 0.999175987 |
| ENSMUSG00000009555 | Cdk9 | protein_coding | 0.916951953 | 4.755560864 | 2.7604487 | 0.1157219 | 0.999175987 |
| ENSMUSG00000007892 | Rplp1 | protein_coding | 0.664644339 | 9.342253724 | 2.7587774 | 0.1158555 | 0.999175987 |
| ENSMUSG00000095325 | Zfp870 | protein_coding | -1.173059052 | 3.837029677 | 2.7528131 | 0.1161896 | 0.999175987 |
| ENSMUSG00000024493 | Lars | protein_coding | -0.732393377 | 5.57129113 | 2.7516256 | 0.1162625 | 0.999175987 |
| ENSMUSG00000037795 | N4bp2 | protein_coding | -1.628751701 | 2.840029752 | 2.749617 | 0.116386 | 0.999175987 |
| ENSMUSG00000032135 | Mcam | protein_coding | -2.056363095 | 2.90400085 | 2.7496626 | 0.1164146 | 0.999175987 |
| ENSMUSG00000034154 | Ino80 | protein_coding | -1.159481661 | 3.498932042 | 2.7483615 | 0.1164632 | 0.999175987 |
| ENSMUSG00000021337 | Scgn | protein_coding | -1.429931538 | 3.085266767 | 2.7482883 | 0.1164678 | 0.999175987 |
| ENSMUSG00000063856 | Gpx1 | protein_coding | 0.481278905 | 9.114042541 | 2.745099 | 0.1166643 | 0.999175987 |
| ENSMUSG00000062760 | 1810041L15Rik | protein_coding | -0.892370944 | 5.549646128 | 2.7418935 | 0.1168938 | 0.999175987 |
| ENSMUSG00000004317 | Clcn5 | protein_coding | -0.956895711 | 4.298924838 | 2.7387713 | 0.1170555 | 0.999175987 |
| ENSMUSG00000020774 | Aspa | protein_coding | -0.779057697 | 5.993678956 | 2.738759 | 0.1170877 | 0.999175987 |
| ENSMUSG00000097080 | 1700086O06Rik | lincRNA | 1.163620184 | 3.712583447 | 2.7346152 | 0.1173133 | 0.999175987 |
| ENSMUSG00000041841 | Rpl37 | protein_coding | 0.810943243 | 8.0548986 | 2.7550383 | 0.1173492 | 0.999175987 |
| ENSMUSG00000073792 | Alg6 | protein_coding | -1.022440974 | 3.70962646 | 2.7323291 | 0.1174554 | 0.999175987 |
| ENSMUSG00000028518 | Prkaa2 | protein_coding | -0.862190615 | 4.330244399 | 2.7309421 | 0.1175417 | 0.999175987 |
| ENSMUSG00000056537 | Rlim | protein_coding | -0.814410142 | 5.508096725 | 2.7292951 | 0.1176443 | 0.999175987 |
| ENSMUSG00000038417 | Fig4 | protein_coding | -0.872275867 | 4.988663656 | 2.7290422 | 0.1176601 | 0.999175987 |
| ENSMUSG00000025724 | Sec11a | protein_coding | 0.597653058 | 6.941212833 | 2.719842 | 0.1182352 | 0.999175987 |
| ENSMUSG00000021811 | Dnajc9 | protein_coding | 0.744274902 | 6.019361797 | 2.717432 | 0.1183865 | 0.999175987 |
| ENSMUSG00000011831 | Evi5 | protein_coding | -0.825985866 | 4.949072978 | 2.7172789 | 0.1183961 | 0.999175987 |
| ENSMUSG00000026687 | Aldh9a1 | protein_coding | 0.653397773 | 6.127759362 | 2.7171821 | 0.1184022 | 0.999175987 |
| ENSMUSG00000109121 | Gm44644 | lincRNA | -1.345843389 | 3.242296823 | 2.7157252 | 0.1184937 | 0.999175987 |
| ENSMUSG00000096943 | Gm26721 | lincRNA | -1.618595286 | 2.809165358 | 2.7113969 | 0.1187662 | 0.999175987 |
| ENSMUSG00000035863 | Palm | protein_coding | 0.679118344 | 5.592000077 | 2.7086641 | 0.1189386 | 0.999175987 |
| ENSMUSG00000069516 | Lyz2 | protein_coding | -1.461789456 | 3.313845195 | 2.7067336 | 0.1190606 | 0.999175987 |
| ENSMUSG00000001517 | Foxm1 | protein_coding | 1.535877358 | 2.814224342 | 2.7050061 | 0.1191699 | 0.999175987 |
| ENSMUSG00000007589 | Tinf2 | protein_coding | 1.142523989 | 3.47534983 | 2.7016303 | 0.1193838 | 0.999175987 |
| ENSMUSG00000031210 | Gpr165 | protein_coding | 1.452562483 | 3.673757385 | 2.698512 | 0.1196133 | 0.999175987 |
| ENSMUSG00000042570 | Mier2 | protein_coding | 1.375112341 | 3.993611786 | 2.6963792 | 0.1200297 | 0.999175987 |
| ENSMUSG00000030125 | Lrrc23 | protein_coding | 1.086140509 | 3.572157851 | 2.6914801 | 0.1200297 | 0.999175987 |
| ENSMUSG00000024411 | Aqp4 | protein_coding | -0.810080604 | 5.041781743 | 2.6899487 | 0.1201276 | 0.999175987 |
| ENSMUSG00000062647 | Rpl7a | protein_coding | 0.527035424 | 6.067649454 | 2.6893797 | 0.1201639 | 0.999175987 |
| ENSMUSG00000039480 | Nt5dc1 | protein_coding | -2.510840642 | 1.833752759 | 2.6884478 | 0.1202235 | 0.999175987 |
| ENSMUSG00000022335 | Zfat | protein_coding | 2.112960027 | 2.241048006 | 2.6850805 | 0.1204392 | 0.999175987 |
| ENSMUSG00000031626 | Sorbs2 | protein_coding | -0.802639365 | 4.848276821 | 2.6817136 | 0.1206553 | 0.999175987 |
| ENSMUSG00000020799 | Tekt1 | protein_coding | -0.946106534 | 4.468957633 | 2.6807896 | 0.1207147 | 0.999175987 |
| ENSMUSG00000031546 | Gins4 | protein_coding | 0.678117801 | 4.980697551 | 2.6799528 | 0.1207685 | 0.999175987 |
| ENSMUSG00000053080 | 2700081O15Rik | protein_coding | -1.53545618 | 2.671389807 | 2.6781054 | 0.1208874 | 0.999175987 |
| ENSMUSG00000072692 | Rpl37rt | transcribed_processe | 0.875106295 | 5.42589792 | 2.6762495 | 0.1210384 | 0.999175987 |
| ENSMUSG00000014776 | Nol3 | protein_coding | 1.014455407 | 5.167329487 | 2.6756997 | 0.1210739 | 0.999175987 |
| ENSMUSG00000046027 | Stard5 | protein_coding | -0.864102389 | 4.094386568 | 2.6726543 | 0.121239 | 0.999175987 |
| ENSMUSG00000020474 | Polm | protein_coding | 1.525057118 | 3.367098616 | 2.672099 | 0.1213064 | 0.999175987 |
| ENSMUSG00000025362 | Rps26 | protein_coding | 0.697033795 | 7.665758863 | 2.6731178 | 0.1213235 | 0.999175987 |
| ENSMUSG00000026958 | Dpp7 | protein_coding | 0.838311679 | 4.544340655 | 2.6713293 | 0.1213247 | 0.999175987 |
| ENSMUSG00000037364 | Srrt | protein_coding | 0.654900002 | 5.570223714 | 2.6709373 | 0.12135 | 0.999175987 |
| ENSMUSG00000036990 | Otud4 | protein_coding | -1.025713081 | 4.057387711 | 2.6702741 | 0.121393 | 0.999175987 |
| ENSMUSG00000032459 | Mrps22 | protein_coding | 0.811955799 | 5.211931843 | 2.6657456 | 0.1216865 | 0.999175987 |
| ENSMUSG00000106106 | CT010467.1 | rRNA | 0.593509131 | 10.43028008 | 2.6656136 | 0.121695 | 0.999175987 |
| ENSMUSG00000038668 | Lpar1 | protein_coding | -0.816426789 | 5.332223296 | 2.6645756 | 0.1217625 | 0.999175987 |
| ENSMUSG00000032503 | Arpp21 | protein_coding | -0.810199959 | 10.18890769 | 2.6789797 | 0.121977 | 0.999175987 |
| ENSMUSG00000006576 | Slc4a3 | protein_coding | 0.612474535 | 5.747084477 | 2.6603426 | 0.1220378 | 0.999175987 |
| ENSMUSG00000034361 | Cpne2 | protein_coding | 0.739573974 | 5.225440251 | 2.6592213 | 0.1221109 | 0.999175987 |
| ENSMUSG00000028174 | Rpe65 | protein_coding | -3.365641527 | 2.05384736 | 2.6670823 | 0.1222379 | 0.999175987 |
| ENSMUSG00000037138 | Aff3 | protein_coding | -0.890661977 | 4.817062394 | 2.6571685 | 0.1222448 | 0.999175987 |
| ENSMUSG00000029098 | Acox3 | protein_coding | -1.089465954 | 3.777063605 | 2.6551735 | 0.1223751 | 0.999175987 |
| ENSMUSG00000034472 | Rasd2 | protein_coding | -0.609054583 | 6.691949808 | 2.6521266 | 0.1225744 | 0.999175987 |
| ENSMUSG00000032396 | Dis3l | protein_coding | 0.849556406 | 4.497877359 | 2.650423 | 0.122686 | 0.999175987 |
| ENSMUSG00000015968 | Cacna1d | protein_coding | 1.005286944 | 3.628391845 | 2.6496416 | 0.1227373 | 0.999175987 |
| ENSMUSG00000010803 | Gabra1 | protein_coding | -0.492636726 | 7.603223781 | 2.6487672 | 0.1227946 | 0.999175987 |
| ENSMUSG00000005899 | Smpd4 | protein_coding | -1.198552717 | 3.595173308 | 2.6485024 | 0.122812 | 0.999175987 |

| ensembl_gene_id | external_gene_name | gene_biotype | log2 (fold change) | log2 (counts per million) | F-value | p-value | False discovery rate |
|---|---|---|---|---|---|---|---|
| ENSMUSG00000079641 | Rpl39 | protein_coding | 0.769062485 | 6.23679866 | 2.6470201 | 0.1229409 | 0.999175987 |
| ENSMUSG00000053137 | Mapk11 | protein_coding | 1.063619275 | 3.555196215 | 2.6430111 | 0.1231731 | 0.999175987 |
| ENSMUSG00000104861 | 3110039M20Rik | lincRNA | 1.310992635 | 3.578127539 | 2.642914 | 0.1231795 | 0.999175987 |
| ENSMUSG00000039693 | Msantd3 | protein_coding | -1.020762035 | 3.699884254 | 2.6401018 | 0.1233649 | 0.999175987 |
| ENSMUSG00000041958 | Pigs | protein_coding | 0.754671881 | 5.845747439 | 2.6388771 | 0.1234458 | 0.999175987 |
| ENSMUSG00000061207 | Stk19 | protein_coding | 0.540414457 | 6.162909754 | 2.6309517 | 0.1239706 | 0.999175987 |
| ENSMUSG00000038497 | Tmco3 | protein_coding | 0.846429231 | 5.312085953 | 2.6305598 | 0.1239966 | 0.999175987 |
| ENSMUSG00000028861 | Mrps15 | protein_coding | 0.650141407 | 6.071185528 | 2.6292045 | 0.1240866 | 0.999175987 |
| ENSMUSG00000023110 | Prmt5 | protein_coding | -0.518198994 | 6.795089092 | 2.6287313 | 0.1241181 | 0.999175987 |
| ENSMUSG00000035769 | Xylb | protein_coding | 1.281788818 | 3.277310768 | 2.6282555 | 0.1241497 | 0.999175987 |
| ENSMUSG00000036052 | Dnajb5 | protein_coding | 0.828426732 | 5.281467032 | 2.6255002 | 0.1243331 | 0.999175987 |
| ENSMUSG00000020163 | Uqcr11 | protein_coding | 0.645200608 | 9.283705867 | 2.6239832 | 0.1244342 | 0.999175987 |
| ENSMUSG00000032839 | Trpc1 | protein_coding | -1.226744476 | 3.585666934 | 2.6198735 | 0.1247086 | 0.999175987 |
| ENSMUSG00000019990 | Pde7b | protein_coding | -0.894141311 | 4.13932871 | 2.6187602 | 0.1247831 | 0.999175987 |
| ENSMUSG00000038628 | Polr3k | protein_coding | 0.463730865 | 7.76036592 | 2.6171258 | 0.1248925 | 0.999175987 |
| ENSMUSG00000050965 | Prkca | protein_coding | -0.657147955 | 5.234489138 | 2.6161719 | 0.1249564 | 0.999175987 |
| ENSMUSG00000031715 | Smarca5 | protein_coding | -0.733083251 | 5.237627904 | 2.6157579 | 0.1249842 | 0.999175987 |
| ENSMUSG00000039031 | Arhgap18 | protein_coding | -1.186993229 | 3.136624267 | 2.6146184 | 0.1250606 | 0.999175987 |
| ENSMUSG00000013155 | Enkd1 | protein_coding | 1.05037409 | 3.709672084 | 2.6145782 | 0.1250633 | 0.999175987 |
| ENSMUSG00000040774 | Cept1 | protein_coding | 0.528906572 | 6.258797858 | 2.6135799 | 0.1251303 | 0.999175987 |
| ENSMUSG00000028906 | Epb41 | protein_coding | -0.740147703 | 4.773790182 | 2.6127772 | 0.1251842 | 0.999175987 |
| ENSMUSG00000002059 | Rab34 | protein_coding | -1.10752005 | 3.398121639 | 2.6100807 | 0.1253655 | 0.999175987 |
| ENSMUSG00000003308 | Keap1 | protein_coding | 1.158302905 | 4.751555809 | 2.6180927 | 0.1254258 | 0.999175987 |
| ENSMUSG00000035765 | Dym | protein_coding | 0.618005226 | 5.538047024 | 2.6079493 | 0.125509 | 0.999175987 |
| ENSMUSG00000027829 | Ccnl1 | protein_coding | -1.063600492 | 3.801533746 | 2.6046872 | 0.1257291 | 0.999175987 |
| ENSMUSG00000031342 | Gpm6b | protein_coding | -0.5574986 | 8.856646969 | 2.6041628 | 0.1257645 | 0.999175987 |
| ENSMUSG00000037463 | Fbxo27 | protein_coding | -1.607084842 | 2.454209972 | 2.6031819 | 0.1258308 | 0.999175987 |
| ENSMUSG00000030030 | 1700003E16Rik | protein_coding | 0.723887928 | 5.293670112 | 2.6029498 | 0.1258465 | 0.999175987 |
| ENSMUSG00000024227 | Pdzph1 | protein_coding | -1.125074605 | 4.119564187 | 2.6008281 | 0.12599 | 0.999175987 |
| ENSMUSG00000031156 | Slc35a2 | protein_coding | 0.66556195 | 5.421110488 | 2.6007739 | 0.1259937 | 0.999175987 |
| ENSMUSG00000006740 | Kif5b | protein_coding | -0.608682915 | 6.115875941 | 2.6000577 | 0.1260422 | 0.999175987 |
| ENSMUSG00000028581 | Laptm5 | protein_coding | -0.775981472 | 5.352613588 | 2.599273 | 0.1260953 | 0.999175987 |
| ENSMUSG00000020109 | Dnajb12 | protein_coding | 0.63213451 | 5.728433243 | 2.5983515 | 0.1261578 | 0.999175987 |
| ENSMUSG00000032589 | Bsn | protein_coding | -0.989120441 | 4.550189453 | 2.5976122 | 0.1262079 | 0.999175987 |
| ENSMUSG00000042210 | Abhd14a | protein_coding | 0.719770079 | 6.311576668 | 2.5977065 | 0.126233 | 0.999175987 |
| ENSMUSG00000021981 | Cab39l | protein_coding | 0.566712291 | 6.511037761 | 2.5969258 | 0.1262545 | 0.999175987 |
| ENSMUSG00000043633 | Fam221b | protein_coding | 1.292944229 | 3.063791925 | 2.5943069 | 0.1264324 | 0.999175987 |
| ENSMUSG00000029201 | Ugdh | protein_coding | 0.818524918 | 4.528827159 | 2.5878685 | 0.126871 | 0.999175987 |
| ENSMUSG00000062078 | Qk | protein_coding | -0.539996472 | 7.886346701 | 2.5864535 | 0.1269676 | 0.999175987 |
| ENSMUSG00000067942 | Zfp160 | protein_coding | -1.039181551 | 3.655992398 | 2.5863893 | 0.126972 | 0.999175987 |
| ENSMUSG00000034839 | Larp6 | protein_coding | -0.955846274 | 4.866273999 | 2.5861403 | 0.1269891 | 0.999175987 |
| ENSMUSG00000029730 | Mcm7 | protein_coding | -1.223210328 | 3.548071418 | 2.5849945 | 0.1270674 | 0.999175987 |
| ENSMUSG00000021647 | Cartpt | protein_coding | 0.784153517 | 6.853129418 | 2.6000302 | 0.127265 | 0.999175987 |
| ENSMUSG00000028093 | Acp6 | protein_coding | 1.210299277 | 3.547751919 | 2.578582 | 0.1275069 | 0.999175987 |
| ENSMUSG00000053475 | Tnfaip6 | protein_coding | -0.769240746 | 5.348703418 | 2.5783861 | 0.1275204 | 0.999175987 |
| ENSMUSG00000028876 | Epha10 | protein_coding | 1.163662198 | 3.683833959 | 2.5778631 | 0.1275563 | 0.999175987 |
| ENSMUSG00000036882 | Arhgap33 | protein_coding | -0.737597834 | 5.61869211 | 2.5750083 | 0.1277526 | 0.999175987 |
| ENSMUSG00000038375 | Trp53inp2 | protein_coding | -0.683151742 | 5.434666145 | 2.5732028 | 0.127877 | 0.999175987 |
| ENSMUSG00000021003 | Galc | protein_coding | 1.326698586 | 3.282337328 | 2.5725242 | 0.1279238 | 0.999175987 |
| ENSMUSG00000068036 | Afdn | protein_coding | -0.805001721 | 5.070946904 | 2.5725193 | 0.1279241 | 0.999175987 |
| ENSMUSG00000079283 | 2310009B15Rik | protein_coding | 0.850277389 | 4.54184601 | 2.5722635 | 0.1279418 | 0.999175987 |
| ENSMUSG00000029344 | Tpst2 | protein_coding | -1.106326105 | 3.349081953 | 2.5709289 | 0.1280339 | 0.999175987 |
| ENSMUSG00000055493 | Epm2a | protein_coding | -0.894537208 | 4.133105188 | 2.5707192 | 0.1280483 | 0.999175987 |
| ENSMUSG00000022961 | Son | protein_coding | -0.592866955 | 7.369054877 | 2.5688105 | 0.1281802 | 0.999175987 |
| ENSMUSG00000053094 | Tmem248 | protein_coding | 0.803402321 | 4.729966442 | 2.56838 | 0.1282099 | 0.999175987 |
| ENSMUSG00000049553 | Polr1a | protein_coding | 1.151135793 | 3.578568344 | 2.5654019 | 0.1284161 | 0.999175987 |
| ENSMUSG00000018167 | Stard3 | protein_coding | 0.846901471 | 5.476916645 | 2.5652005 | 0.1284615 | 0.999175987 |
| ENSMUSG00000020811 | Wscd1 | protein_coding | -0.648306936 | 5.535521252 | 2.5639138 | 0.1285192 | 0.999175987 |
| ENSMUSG00000058159 | T2 | processed_transcript | 1.877967298 | 2.287451488 | 2.5627747 | 0.1285982 | 0.999175987 |
| ENSMUSG00000029755 | Dlx5 | protein_coding | 0.656509453 | 5.722980699 | 2.5627153 | 0.1286023 | 0.999175987 |
| ENSMUSG00000029563 | Foxp2 | protein_coding | -0.775413204 | 5.536918771 | 2.5610252 | 0.1287197 | 0.999175987 |
| ENSMUSG00000063887 | Nlgn1 | protein_coding | -0.61706008 | 6.333251997 | 2.5609069 | 0.1287279 | 0.999175987 |
| ENSMUSG00000028033 | Kcnq5 | protein_coding | -0.821847316 | 4.809074714 | 2.5603463 | 0.1287669 | 0.999175987 |
| ENSMUSG00000024811 | Tnks2 | protein_coding | -0.862584806 | 5.379517139 | 2.5539097 | 0.1292452 | 0.999175987 |
| ENSMUSG00000029012 | Orc5 | protein_coding | -0.731408078 | 5.490140977 | 2.552371 | 0.1293227 | 0.999175987 |
| ENSMUSG00000097023 | Mir9-3hg | lincRNA | -1.058945101 | 3.721432233 | 2.5520038 | 0.1293483 | 0.999175987 |
| ENSMUSG00000042750 | Bex2 | protein_coding | 0.490910753 | 10.3553781 | 2.5480654 | 0.1296239 | 0.999175987 |
| ENSMUSG00000037419 | Endod1 | protein_coding | -0.561526388 | 6.359519041 | 2.5478761 | 0.1296372 | 0.999175987 |
| ENSMUSG00000002341 | Ncan | protein_coding | -0.517265911 | 7.218162338 | 2.5471853 | 0.1296856 | 0.999175987 |
| ENSMUSG00000047606 | Ankrd34c | protein_coding | 1.190276457 | 3.019823095 | 2.5459741 | 0.1297705 | 0.999175987 |
| ENSMUSG00000054752 | Fsd1l | protein_coding | -0.993763429 | 3.702558382 | 2.5456035 | 0.1297965 | 0.999175987 |
| ENSMUSG00000020451 | Limk2 | protein_coding | 0.718730032 | 6.679281051 | 2.5464371 | 0.1298526 | 0.999175987 |
| ENSMUSG00000012114 | Med15 | protein_coding | 0.727594754 | 5.038707628 | 2.5428541 | 0.1299897 | 0.999175987 |
| ENSMUSG00000015214 | Mtmr1 | protein_coding | -0.881822058 | 4.185210537 | 2.5402437 | 0.1301734 | 0.999175987 |
| ENSMUSG00000078784 | 1810022K09Rik | protein_coding | 0.827510132 | 4.566680193 | 2.5399297 | 0.1301955 | 0.999175987 |
| ENSMUSG00000009687 | Fxyd5 | protein_coding | 1.785073332 | 2.35456016 | 2.5394615 | 0.1302285 | 0.999175987 |
| ENSMUSG00000031370 | Zrsr2 | protein_coding | 0.533258402 | 6.645237934 | 2.5372196 | 0.1303866 | 0.999175987 |
| ENSMUSG00000038059 | Smim3 | protein_coding | -1.039723084 | 3.914729862 | 2.5356414 | 0.130498 | 0.999175987 |

| ensembl_gene_id | external_gene_name | gene_biotype | log2 (fold change) | log2 (counts per million) | F-value | p-value | False discovery rate |
|---|---|---|---|---|---|---|---|
| ENSMUSG00000042548 | Asxl1 | protein_coding | -0.987892949 | 3.71268328 | 2.5344454 | 0.1305825 | 0.999175987 |
| ENSMUSG00000034021 | Pds5b | protein_coding | -0.548919194 | 6.033313339 | 2.5332634 | 0.1306661 | 0.999175987 |
| ENSMUSG00000044628 | Rnf208 | protein_coding | 0.597005586 | 6.932068985 | 2.5331057 | 0.1306772 | 0.999175987 |
| ENSMUSG00000025420 | Katnal2 | protein_coding | -1.115036475 | 3.421721063 | 2.5299221 | 0.1309028 | 0.999175987 |
| ENSMUSG00000020029 | Nudt4 | protein_coding | 0.543520973 | 6.206252499 | 2.5284003 | 0.1310107 | 0.999175987 |
| ENSMUSG00000041000 | Trim62 | protein_coding | -0.997107175 | 3.93206568 | 2.5280248 | 0.1310374 | 0.999175987 |
| ENSMUSG00000019590 | Cyb561 | protein_coding | 0.553891889 | 7.175003432 | 2.5260873 | 0.131175 | 0.999175987 |
| ENSMUSG00000043162 | Pyurf | protein_coding | 1.257256564 | 5.055327612 | 2.5530322 | 0.1313156 | 0.999175987 |
| ENSMUSG00000030538 | Cib1 | protein_coding | -0.878582052 | 4.759552767 | 2.5232765 | 0.131375 | 0.999175987 |
| ENSMUSG00000028278 | Rragd | protein_coding | -0.585134005 | 6.228608853 | 2.5229932 | 0.1313952 | 0.999175987 |
| ENSMUSG00000005225 | Plekha8 | protein_coding | -1.186976706 | 3.064504677 | 2.5208723 | 0.1315463 | 0.999175987 |
| ENSMUSG00000005681 | Apoa2 | protein_coding | 1.072763781 | 3.382292466 | 2.5199482 | 0.1316123 | 0.999175987 |
| ENSMUSG00000035969 | Rusc2 | protein_coding | -0.840119837 | 4.488377439 | 2.518128 | 0.1317423 | 0.999175987 |
| ENSMUSG00000055403 | 4933427D06Rik | lincRNA | -1.530577339 | 3.222625254 | 2.516987 | 0.1318553 | 0.999175987 |
| ENSMUSG00000021497 | Txndc15 | protein_coding | 0.564402422 | 6.261456886 | 2.5163162 | 0.1318718 | 0.999175987 |
| ENSMUSG00000087165 | 2010001A14Rik | processed_transcript | -0.894569661 | 4.217475595 | 2.5137324 | 0.1320568 | 0.999175987 |
| ENSMUSG00000026834 | Acvr1c | protein_coding | -0.906458687 | 3.894645852 | 2.5130035 | 0.1321091 | 0.999175987 |
| ENSMUSG00000030465 | Psd3 | protein_coding | -0.537992142 | 6.568826009 | 2.511124 | 0.1322439 | 0.999175987 |
| ENSMUSG00000028519 | Dab1 | protein_coding | 0.524858237 | 5.940746159 | 2.5110791 | 0.1322471 | 0.999175987 |
| ENSMUSG00000053580 | Tanc2 | protein_coding | -0.626683425 | 5.970584882 | 2.508868 | 0.132406 | 0.999175987 |
| ENSMUSG00000021621 | Zcchc9 | protein_coding | 0.891842111 | 4.912392051 | 2.5085123 | 0.1324316 | 0.999175987 |
| ENSMUSG00000053483 | Usp21 | protein_coding | 0.769916655 | 5.18299884 | 2.5060936 | 0.1326056 | 0.999175987 |
| ENSMUSG00000027695 | Pld1 | protein_coding | -1.896760922 | 2.534102462 | 2.5022094 | 0.1328858 | 0.999175987 |
| ENSMUSG00000062542 | Syt9 | protein_coding | 0.9909109 | 4.246402326 | 2.5010113 | 0.1329723 | 0.999175987 |
| ENSMUSG00000006005 | Tpr | protein_coding | -0.718449407 | 5.924069098 | 2.5005346 | 0.1330068 | 0.999175987 |
| ENSMUSG00000022790 | Igsf11 | protein_coding | 1.228298894 | 3.397509605 | 2.500366 | 0.133019 | 0.999175987 |
| ENSMUSG00000039910 | Cited2 | protein_coding | -0.74748765 | 4.336699034 | 2.4988975 | 0.1331252 | 0.999175987 |
| ENSMUSG00000029165 | Agbl5 | protein_coding | 1.337565935 | 2.977751785 | 2.4981978 | 0.1331758 | 0.999175987 |
| ENSMUSG00000029446 | Psph | protein_coding | -0.681161267 | 5.792015047 | 2.4979055 | 0.133197 | 0.999175987 |
| ENSMUSG00000074785 | Plxnc1 | protein_coding | -0.687287224 | 5.013057301 | 2.4974776 | 0.133228 | 0.999175987 |
| ENSMUSG00000052423 | B4galt3 | protein_coding | 0.662382121 | 5.498150261 | 2.4956941 | 0.1333573 | 0.999175987 |
| ENSMUSG00000028434 | Epb41l4b | protein_coding | -0.727133366 | 4.797368241 | 2.4948315 | 0.1334198 | 0.999175987 |
| ENSMUSG00000026344 | Lypd1 | protein_coding | 1.410945119 | 2.678574672 | 2.4946905 | 0.1334301 | 0.999175987 |
| ENSMUSG00000029467 | Atp2a2 | protein_coding | -0.572643437 | 7.904779447 | 2.493967 | 0.1334826 | 0.999175987 |
| ENSMUSG00000025086 | Trub1 | protein_coding | -0.726239704 | 5.321156264 | 2.4930452 | 0.1335495 | 0.999175987 |
| ENSMUSG00000003528 | Slc25a1 | protein_coding | 0.902893216 | 5.133553603 | 2.4908463 | 0.1337409 | 0.999175987 |
| ENSMUSG00000004040 | Stat3 | protein_coding | -1.05911068 | 4.838964581 | 2.4864785 | 0.1341421 | 0.999175987 |
| ENSMUSG00000075334 | Rprm | protein_coding | 0.970098678 | 5.63172732 | 2.4994976 | 0.1342279 | 0.999175987 |
| ENSMUSG00000020601 | Trib2 | protein_coding | -0.651668769 | 5.577241253 | 2.4826294 | 0.1343089 | 0.999175987 |
| ENSMUSG00000021690 | Jmy | protein_coding | -0.915311142 | 4.531544115 | 2.4810204 | 0.1344266 | 0.999175987 |
| ENSMUSG00000062006 | Rpl34 | protein_coding | 0.6382642 | 7.859558926 | 2.4798695 | 0.1345424 | 0.999175987 |
| ENSMUSG00000025533 | Asl | protein_coding | -0.500758943 | 6.434689736 | 2.4779461 | 0.134652 | 0.999175987 |
| ENSMUSG00000015942 | Gtf2ird2 | protein_coding | 1.060044059 | 3.549050209 | 2.4774148 | 0.134691 | 0.999175987 |
| ENSMUSG00000062209 | Erbb4 | protein_coding | -0.871061165 | 4.623836947 | 2.4769552 | 0.1347247 | 0.999175987 |
| ENSMUSG00000057219 | Armc7 | protein_coding | 0.853110417 | 4.448733052 | 2.4763419 | 0.1347698 | 0.999175987 |
| ENSMUSG00000001864 | Aif1l | protein_coding | -0.733390237 | 5.280336379 | 2.4755879 | 0.1348252 | 0.999175987 |
| ENSMUSG00000024422 | Dhx16 | protein_coding | 1.439775247 | 2.864319705 | 2.4748018 | 0.134883 | 0.999175987 |
| ENSMUSG00000020380 | Rad50 | protein_coding | -0.93690941 | 3.776395721 | 2.4733195 | 0.134992 | 0.999175987 |
| ENSMUSG00000038679 | Trps1 | protein_coding | -0.813848023 | 4.421046374 | 2.4723523 | 0.1350632 | 0.999175987 |
| ENSMUSG00000037031 | Tspan15 | protein_coding | -0.974441662 | 4.183674783 | 2.4711819 | 0.1351495 | 0.999175987 |
| ENSMUSG00000039208 | Metrnl | protein_coding | 1.622373958 | 2.675614047 | 2.4707782 | 0.1351792 | 0.999175987 |
| ENSMUSG00000025144 | Cenpx | protein_coding | 0.588134338 | 6.391339461 | 2.4704103 | 0.1352063 | 0.999175987 |
| ENSMUSG00000027679 | Dnajc19 | protein_coding | 0.556939017 | 6.113242534 | 2.468478 | 0.1353489 | 0.999175987 |
| ENSMUSG00000074746 | Pdzd8 | protein_coding | -0.585600338 | 5.42275172 | 2.4660284 | 0.13553 | 0.999175987 |
| ENSMUSG00000005621 | Zfp592 | protein_coding | -1.150293889 | 3.739771541 | 2.4623866 | 0.1357996 | 0.999175987 |
| ENSMUSG00000030533 | Unc45a | protein_coding | 0.845563165 | 4.486014819 | 2.4623053 | 0.1358057 | 0.999175987 |
| ENSMUSG00000068328 | Aup1 | protein_coding | 0.971341514 | 6.236428025 | 2.4903017 | 0.1362331 | 0.999175987 |
| ENSMUSG00000000384 | Tbrg4 | protein_coding | 0.867573461 | 4.916363317 | 2.4547881 | 0.1363644 | 0.999175987 |
| ENSMUSG00000020486 | 4-Sep | protein_coding | -0.516828255 | 8.938024311 | 2.4543362 | 0.1363981 | 0.999175987 |
| ENSMUSG00000029833 | Trim24 | protein_coding | -0.87858788 | 3.973895543 | 2.4537794 | 0.1364396 | 0.999175987 |
| ENSMUSG00000012422 | Tmem167 | protein_coding | 0.545473198 | 7.382104096 | 2.4534314 | 0.1364656 | 0.999175987 |
| ENSMUSG00000001833 | 7-Sep | protein_coding | -0.517918787 | 9.081041745 | 2.4512906 | 0.1366254 | 0.999175987 |
| ENSMUSG00000035547 | Capn5 | protein_coding | -0.893946341 | 5.012830379 | 2.45124 | 0.1366291 | 0.999175987 |
| ENSMUSG00000048776 | Pthlh | protein_coding | -1.971331242 | 2.503974901 | 2.4510974 | 0.1366398 | 0.999175987 |
| ENSMUSG00000019936 | Epyc | protein_coding | -1.494915394 | 2.734746908 | 2.4507877 | 0.1366629 | 0.999175987 |
| ENSMUSG00000094870 | Zfp131 | protein_coding | 0.850657728 | 4.197405735 | 2.4498058 | 0.1367363 | 0.999175987 |
| ENSMUSG00000032580 | Rbm5 | protein_coding | -0.499959816 | 6.59973986 | 2.4486181 | 0.1368251 | 0.999175987 |
| ENSMUSG00000038481 | Cdk19 | protein_coding | -0.754913906 | 4.85149738 | 2.4473371 | 0.136921 | 0.999175987 |
| ENSMUSG00000042293 | Gm5617 | protein_coding | 1.055931584 | 4.872114521 | 2.4493176 | 0.1369235 | 0.999175987 |
| ENSMUSG00000017747 | Ghdc | protein_coding | -2.056353734 | 2.546931946 | 2.4460556 | 0.137017 | 0.999175987 |
| ENSMUSG00000028863 | Meaf6 | protein_coding | 0.516048027 | 7.874483094 | 2.4440394 | 0.1371683 | 0.999175987 |
| ENSMUSG00000033272 | Slc35a4 | protein_coding | 0.616095678 | 6.143286942 | 2.4437447 | 0.1371904 | 0.999175987 |
| ENSMUSG00000021287 | Xrcc3 | protein_coding | 1.482306763 | 2.800430518 | 2.4435712 | 0.1372034 | 0.999175987 |
| ENSMUSG00000048720 | Tbc1d12 | protein_coding | -1.143762863 | 3.270310205 | 2.4433592 | 0.1372193 | 0.999175987 |
| ENSMUSG00000028223 | Decr1 | protein_coding | -0.747601728 | 5.156132739 | 2.4418672 | 0.1373314 | 0.999175987 |
| ENSMUSG00000098188 | Sowahc | protein_coding | -1.218782452 | 3.065199733 | 2.4413116 | 0.1373732 | 0.999175987 |
| ENSMUSG00000036769 | Wdr44 | protein_coding | -1.142698102 | 4.062501324 | 2.4387634 | 0.1375964 | 0.999175987 |
| ENSMUSG00000067722 | BC003965 | protein_coding | 0.497189873 | 6.375988644 | 2.4362547 | 0.137754 | 0.999175987 |

| ensembl_gene_id | external_gene_name | gene_biotype | log2 (fold change) | log2 (counts per million) | F-value | p-value | False discovery rate |
|---|---|---|---|---|---|---|---|
| ENSMUSG00000031028 | Tub | protein_coding | 0.599290586 | 5.704199397 | 2.4348177 | 0.1378625 | 0.999175987 |
| ENSMUSG00000063428 | Ddo | protein_coding | -0.804184402 | 4.108612222 | 2.4347139 | 0.1378703 | 0.999175987 |
| ENSMUSG00000086155 | 9430041J12Rik | antisense_RNA | 1.989731808 | 2.106164436 | 2.4306379 | 0.1381786 | 0.999175987 |
| ENSMUSG00000040640 | Erc2 | protein_coding | -0.603883113 | 6.308424434 | 2.430065 | 0.1382219 | 0.999175987 |
| ENSMUSG00000026207 | Speg | protein_coding | 1.159678792 | 3.661828598 | 2.429972 | 0.138229 | 0.999175987 |
| ENSMUSG00000000148 | Brat1 | protein_coding | 1.152585295 | 4.062798521 | 2.4271898 | 0.1384714 | 0.999175987 |
| ENSMUSG00000010608 | Rbm25 | protein_coding | -0.488000989 | 6.623945702 | 2.4263491 | 0.1385038 | 0.999175987 |
| ENSMUSG00000037296 | Lsm1 | protein_coding | 0.704035475 | 6.027523324 | 2.4232528 | 0.1387392 | 0.999175987 |
| ENSMUSG00000038886 | Man2a2 | protein_coding | -1.065956472 | 3.766803742 | 2.4232049 | 0.1387428 | 0.999175987 |
| ENSMUSG00000025323 | Sp4 | protein_coding | -0.869364865 | 3.837975303 | 2.418266 | 0.1391193 | 0.999175987 |
| ENSMUSG00000036533 | Cdc42ep3 | protein_coding | -0.859381766 | 4.414159543 | 2.4167145 | 0.1392378 | 0.999175987 |
| ENSMUSG00000010025 | Aldh3a2 | protein_coding | 0.906173088 | 5.385840034 | 2.4163575 | 0.1392966 | 0.999175987 |
| ENSMUSG00000061118 | Dnajc30 | protein_coding | 0.648236413 | 5.483751363 | 2.4148603 | 0.1393796 | 0.999175987 |
| ENSMUSG00000030942 | Thumpd1 | protein_coding | -0.801681098 | 5.519415912 | 2.4087655 | 0.139847 | 0.999175987 |
| ENSMUSG00000057329 | Bcl2 | protein_coding | -1.098255268 | 3.884976949 | 2.4080255 | 0.1399039 | 0.999175987 |
| ENSMUSG00000003119 | Cdk12 | protein_coding | -0.913015638 | 4.176047042 | 2.407537 | 0.1399414 | 0.999175987 |
| ENSMUSG00000029838 | Ptn | protein_coding | -0.652509257 | 8.78621346 | 2.4067458 | 0.1400337 | 0.999175987 |
| ENSMUSG00000062110 | Scfd2 | protein_coding | 1.009868965 | 4.006111283 | 2.4063012 | 0.1400365 | 0.999175987 |
| ENSMUSG00000033948 | Zswim5 | protein_coding | -1.386404325 | 3.294930318 | 2.4053638 | 0.1401087 | 0.999175987 |
| ENSMUSG00000044465 | Fam160a2 | protein_coding | -0.779423808 | 4.499321294 | 2.4039187 | 0.14022 | 0.999175987 |
| ENSMUSG00000056870 | Gulp1 | protein_coding | -1.256422482 | 3.06054886 | 2.4031146 | 0.140282 | 0.999175987 |
| ENSMUSG00000005089 | Slc1a2 | protein_coding | -0.827463908 | 7.751621344 | 2.4348395 | 0.1403426 | 0.999175987 |
| ENSMUSG00000015812 | Gnrh1 | protein_coding | -1.271151277 | 5.32651247 | 2.4416196 | 0.1404024 | 0.999175987 |
| ENSMUSG00000069565 | Dazap1 | protein_coding | -1.133824107 | 3.087351188 | 2.4009455 | 0.1404495 | 0.999175987 |
| ENSMUSG00000024191 | Bnip1 | protein_coding | 0.640069446 | 5.806986693 | 2.4005345 | 0.1404812 | 0.999175987 |
| ENSMUSG00000018541 | Cwc25 | protein_coding | -0.732615668 | 4.840670748 | 2.3990419 | 0.1405966 | 0.999175987 |
| ENSMUSG00000016481 | Cr1l | protein_coding | -0.928782898 | 3.992040996 | 2.3983581 | 0.1406495 | 0.999175987 |
| ENSMUSG00000047417 | Rexo1 | protein_coding | 0.911741806 | 4.115535995 | 2.3981331 | 0.1406669 | 0.999175987 |
| ENSMUSG00000028173 | Wls | protein_coding | -0.77341038 | 4.860362235 | 2.3980128 | 0.1406762 | 0.999175987 |
| ENSMUSG00000023050 | Map3k12 | protein_coding | -0.969311171 | 4.076085082 | 2.3965993 | 0.1407857 | 0.999175987 |
| ENSMUSG00000049252 | Lrp1b | protein_coding | -0.915853145 | 5.20458909 | 2.3944891 | 0.1409807 | 0.999175987 |
| ENSMUSG00000028871 | Rspo1 | protein_coding | 1.564236381 | 3.104288689 | 2.3907225 | 0.1412418 | 0.999175987 |
| ENSMUSG00000070280 | Slc22a14 | protein_coding | -1.446798959 | 2.627265896 | 2.3842336 | 0.1417476 | 0.999175987 |
| ENSMUSG00000055670 | Zzef1 | protein_coding | 1.026846659 | 4.076059421 | 2.3834898 | 0.1418058 | 0.999175987 |
| ENSMUSG00000068205 | Macrod2 | protein_coding | -0.73261232 | 5.345425192 | 2.3826026 | 0.1418751 | 0.999175987 |
| ENSMUSG00000101892 | 9130401M01Rik | protein_coding | 0.71110588 | 4.625813844 | 2.3815823 | 0.1419549 | 0.999175987 |
| ENSMUSG00000034075 | Zdhhc5 | protein_coding | -1.010211423 | 3.375278264 | 2.3806623 | 0.1420269 | 0.999175987 |
| ENSMUSG00000018599 | Mief2 | protein_coding | 0.802999258 | 4.81715464 | 2.379319 | 0.1421322 | 0.999175987 |
| ENSMUSG00000055900 | Tmem69 | protein_coding | 0.995389909 | 3.362338281 | 2.3740316 | 0.1425473 | 0.999175987 |
| ENSMUSG00000024958 | Gpr137 | protein_coding | 0.643275163 | 6.39624649 | 2.3735221 | 0.1425874 | 0.999175987 |
| ENSMUSG00000005360 | Slc1a3 | protein_coding | -0.579006366 | 7.808350894 | 2.3730653 | 0.1426233 | 0.999175987 |
| ENSMUSG00000062901 | Klhl24 | protein_coding | -0.720078853 | 5.141138767 | 2.370764 | 0.1428046 | 0.999175987 |
| ENSMUSG00000033597 | Caskin1 | protein_coding | 0.83433167 | 5.013224174 | 2.3704173 | 0.1428319 | 0.999175987 |
| ENSMUSG00000051606 | 2010001K21Rik | processed_transcript | 2.398775532 | 2.361829615 | 2.3704531 | 0.1428605 | 0.999175987 |
| ENSMUSG00000006315 | Tmem147 | protein_coding | 0.501009225 | 7.795517713 | 2.3689615 | 0.1429468 | 0.999175987 |
| ENSMUSG00000004451 | Ralb | protein_coding | 0.682928645 | 4.742937865 | 2.3688922 | 0.1429523 | 0.999175987 |
| ENSMUSG00000033379 | Atp6v0b | protein_coding | 0.522114776 | 8.426530837 | 2.3680419 | 0.1430194 | 0.999175987 |
| ENSMUSG00000009927 | Rps25 | protein_coding | 0.582611358 | 6.467116773 | 2.3667281 | 0.1431232 | 0.999175987 |
| ENSMUSG00000028180 | Zranb2 | protein_coding | -0.468346508 | 7.576783338 | 2.3654414 | 0.1432249 | 0.999175987 |
| ENSMUSG00000034210 | Efcab14 | protein_coding | -0.55443763 | 5.928916924 | 2.3642728 | 0.1433174 | 0.999175987 |
| ENSMUSG00000024425 | Ndfip1 | protein_coding | -0.467902118 | 6.807682116 | 2.3611586 | 0.1435642 | 0.999175987 |
| ENSMUSG00000040856 | Dlk1 | protein_coding | -0.501556474 | 6.572223561 | 2.3592479 | 0.1437159 | 0.999175987 |
| ENSMUSG00000036617 | Etl4 | protein_coding | -0.939687515 | 4.367446043 | 2.3586585 | 0.1437628 | 0.999175987 |
| ENSMUSG00000028555 | Ttc39a | protein_coding | -1.765472541 | 2.707469355 | 2.3586151 | 0.1437662 | 0.999175987 |
| ENSMUSG00000024231 | Cul2 | protein_coding | -0.714551531 | 5.685802949 | 2.3563131 | 0.1439493 | 0.999175987 |
| ENSMUSG00000055660 | Mettl4 | protein_coding | -1.513148984 | 2.489487307 | 2.355796 | 0.1439905 | 0.999175987 |
| ENSMUSG00000063796 | Slc22a8 | protein_coding | 2.164311459 | 2.651915332 | 2.3559496 | 0.1440097 | 0.999175987 |
| ENSMUSG00000035505 | Cox18 | protein_coding | 0.98785097 | 3.559834666 | 2.3553798 | 0.1440236 | 0.999175987 |
| ENSMUSG00000031387 | Renbp | protein_coding | -1.252563693 | 3.052426427 | 2.3551713 | 0.1440402 | 0.999175987 |
| ENSMUSG00000039515 | Ptpa | protein_coding | 0.535917406 | 8.410019744 | 2.3534127 | 0.1441804 | 0.999175987 |
| ENSMUSG00000010277 | 2610507B11Rik | protein_coding | 0.769875955 | 5.370121584 | 2.3530232 | 0.1442115 | 0.999175987 |
| ENSMUSG00000029823 | Luc7l2 | protein_coding | -0.637883708 | 5.357817522 | 2.3522841 | 0.1442705 | 0.999175987 |
| ENSMUSG00000027166 | Dnajc24 | protein_coding | -0.830500504 | 5.076899597 | 2.3519751 | 0.1442951 | 0.999175987 |
| ENSMUSG00000020407 | Upp1 | protein_coding | 1.562298929 | 2.819865986 | 2.3508678 | 0.1443836 | 0.999175987 |
| ENSMUSG00000033411 | Ctdspl2 | protein_coding | -0.876612809 | 4.635409623 | 2.3476364 | 0.144642 | 0.999175987 |
| ENSMUSG00000029999 | Tgfa | protein_coding | -0.706168409 | 4.891510745 | 2.3475089 | 0.1446522 | 0.999175987 |
| ENSMUSG00000039621 | Prex1 | protein_coding | -1.130428996 | 4.392303219 | 2.34757 | 0.1447615 | 0.999175987 |
| ENSMUSG00000010797 | Wnt2 | protein_coding | -1.337364684 | 3.163097649 | 2.345345 | 0.1448256 | 0.999175987 |
| ENSMUSG00000031026 | Trim66 | protein_coding | -0.973436199 | 3.792633113 | 2.3447747 | 0.1448714 | 0.999175987 |
| ENSMUSG00000039967 | Zfp292 | protein_coding | -0.879817085 | 4.74236974 | 2.3446036 | 0.1448851 | 0.999175987 |
| ENSMUSG00000027782 | Kpna4 | protein_coding | -0.575320358 | 5.463499398 | 2.3443264 | 0.1449073 | 0.999175987 |
| ENSMUSG00000041241 | Mul1 | protein_coding | 0.455430259 | 7.04934122 | 2.3442279 | 0.1449153 | 0.999175987 |
| ENSMUSG00000039007 | Cpq | protein_coding | -0.86546781 | 4.352774455 | 2.3432453 | 0.1449941 | 0.999175987 |
| ENSMUSG00000007653 | Gabrb2 | protein_coding | -1.076974884 | 4.267264857 | 2.3425322 | 0.1450828 | 0.999175987 |
| ENSMUSG00000020728 | Cep112 | protein_coding | 1.304855677 | 2.978589964 | 2.3407833 | 0.145192 | 0.999175987 |
| ENSMUSG00000027164 | Traf6 | protein_coding | 1.537324789 | 2.683158561 | 2.3402385 | 0.1452359 | 0.999175987 |
| ENSMUSG00000056966 | Gjc3 | protein_coding | -0.650371495 | 5.573142038 | 2.3386948 | 0.1453601 | 0.999175987 |
| ENSMUSG00000076617 | Ighm | IG_C_gene | -1.047341787 | 5.804644818 | 2.3692302 | 0.1454032 | 0.999175987 |

| ensembl_gene_id | external_gene_name | gene_biotype | log2 (fold change) | log2 (counts per million) | F-value | p-value | False discovery rate |
|---|---|---|---|---|---|---|---|
| ENSMUSG00000084786 | Ubl5 | protein_coding | 0.577635102 | 9.224991029 | 2.3354749 | 0.1456198 | 0.999175987 |
| ENSMUSG00000031864 | Ints10 | protein_coding | -0.789453452 | 5.075568629 | 2.3350877 | 0.1456511 | 0.999175987 |
| ENSMUSG00000029524 | Sirt4 | protein_coding | -0.880980187 | 3.985969658 | 2.3344358 | 0.1457037 | 0.999175987 |
| ENSMUSG00000032293 | Ireb2 | protein_coding | -0.784283426 | 4.662883172 | 2.3343414 | 0.1457114 | 0.999175987 |
| ENSMUSG00000049336 | Tenm2 | protein_coding | -0.754991576 | 5.487232192 | 2.3332161 | 0.1458023 | 0.999175987 |
| ENSMUSG00000086859 | Snhg20 | lincRNA | 1.043232427 | 3.822661677 | 2.3324695 | 0.1458627 | 0.999175987 |
| ENSMUSG00000003469 | Phyhip | protein_coding | 0.482167791 | 8.650892577 | 2.331747 | 0.1459211 | 0.999175987 |
| ENSMUSG00000031207 | Msn | protein_coding | -1.492311882 | 2.915599636 | 2.3270863 | 0.146299 | 0.999175987 |
| ENSMUSG00000022450 | Ndufa6 | protein_coding | 0.538088166 | 10.01225996 | 2.3267768 | 0.1463241 | 0.999175987 |
| ENSMUSG00000064329 | Scn1a | protein_coding | -0.569755853 | 6.450607601 | 2.326198 | 0.1463711 | 0.999175987 |
| ENSMUSG00000029343 | Crybb1 | protein_coding | 2.139607843 | 2.065168389 | 2.3258658 | 0.1463981 | 0.999175987 |
| ENSMUSG00000026527 | Rgs7 | protein_coding | -0.659610362 | 5.555931476 | 2.3243142 | 0.1465243 | 0.999175987 |
| ENSMUSG00000029571 | Tmem106b | protein_coding | -0.622615103 | 5.82573429 | 2.3235972 | 0.1465826 | 0.999175987 |
| ENSMUSG00000100826 | Snhg14 | processed_transcript | -0.583712897 | 6.081352785 | 2.3227637 | 0.1466504 | 0.999175987 |
| ENSMUSG00000033618 | Map3k13 | protein_coding | -1.22241093 | 2.837090241 | 2.3191913 | 0.1469417 | 0.999175987 |
| ENSMUSG00000032712 | 2810474O19Rik | protein_coding | -0.878054414 | 3.9110502 | 2.3179675 | 0.1470416 | 0.999175987 |
| ENSMUSG00000028232 | Tmem68 | protein_coding | -0.570187013 | 5.896256195 | 2.3164351 | 0.1471669 | 0.999175987 |
| ENSMUSG00000024732 | Ccdc86 | protein_coding | 0.875528068 | 4.064614931 | 2.3161389 | 0.1471911 | 0.999175987 |
| ENSMUSG00000038332 | Sesn1 | protein_coding | -0.64953879 | 5.305426125 | 2.315318 | 0.1472583 | 0.999175987 |
| ENSMUSG00000035069 | Oma1 | protein_coding | -1.049040181 | 3.725733586 | 2.3127779 | 0.1474664 | 0.999175987 |
| ENSMUSG00000068220 | Lgals1 | protein_coding | -0.707988542 | 5.289753324 | 2.311381 | 0.147581 | 0.999175987 |
| ENSMUSG00000015176 | Nolc1 | protein_coding | -0.500170853 | 6.376195778 | 2.310361 | 0.1476647 | 0.999175987 |
| ENSMUSG00000028967 | Errfi1 | protein_coding | -0.893530122 | 4.431978901 | 2.3098653 | 0.1477054 | 0.999175987 |
| ENSMUSG00000021288 | Klc1 | protein_coding | 0.547081779 | 7.094391431 | 2.3089685 | 0.1477791 | 0.999175987 |
| ENSMUSG00000074305 | Peak1 | protein_coding | -1.18705803 | 3.140702015 | 2.3075336 | 0.1478971 | 0.999175987 |
| ENSMUSG00000027716 | Trpc3 | protein_coding | -1.577184445 | 2.94328663 | 2.3069283 | 0.1479469 | 0.999175987 |
| ENSMUSG00000041303 | Gtf3c3 | protein_coding | -0.65554167 | 5.121092947 | 2.3063204 | 0.147997 | 0.999175987 |
| ENSMUSG00000032238 | Rora | protein_coding | -0.73118448 | 6.226476772 | 2.3061534 | 0.1480421 | 0.999175987 |
| ENSMUSG00000024387 | Csnk2b | protein_coding | 0.416650365 | 8.273179454 | 2.305741 | 0.1480447 | 0.999175987 |
| ENSMUSG00000042359 | Osbpl6 | protein_coding | -0.771129925 | 5.156949153 | 2.3047324 | 0.1481278 | 0.999175987 |
| ENSMUSG00000021374 | Nup153 | protein_coding | -1.041533632 | 3.758688117 | 2.3038553 | 0.1482001 | 0.999175987 |
| ENSMUSG00000021972 | Hmbox1 | protein_coding | -0.966431584 | 3.662169594 | 2.3036235 | 0.1482193 | 0.999175987 |
| ENSMUSG00000002808 | Epdr1 | protein_coding | -0.549484712 | 6.191063094 | 2.3009514 | 0.1484399 | 0.999175987 |
| ENSMUSG00000026247 | Ecel1 | protein_coding | 0.796918024 | 5.931074143 | 2.3014663 | 0.1485113 | 0.999175987 |
| ENSMUSG00000055401 | Fbxo6 | protein_coding | -0.609424394 | 5.581647577 | 2.298406 | 0.1486505 | 0.999175987 |
| ENSMUSG00000067274 | Rplp0 | protein_coding | 0.43163071 | 8.351305797 | 2.2977526 | 0.1487046 | 0.999175987 |
| ENSMUSG00000028797 | Tmem234 | protein_coding | 0.419619377 | 7.596368689 | 2.2967376 | 0.1487887 | 0.999175987 |
| ENSMUSG00000021703 | Serinc5 | protein_coding | -0.914586123 | 4.266735391 | 2.2919984 | 0.1491821 | 0.999175987 |
| ENSMUSG00000046364 | Rpl27a | protein_coding | 0.558633626 | 6.757894458 | 2.2911659 | 0.1492514 | 0.999175987 |
| ENSMUSG00000027394 | Ttl | protein_coding | 0.682512497 | 4.823554834 | 2.2903753 | 0.1493172 | 0.999175987 |
| ENSMUSG00000039298 | Cdk5rap2 | protein_coding | -1.234655923 | 2.982512472 | 2.2889139 | 0.1494389 | 0.999175987 |
| ENSMUSG00000053565 | Eif3k | protein_coding | 0.485883457 | 9.322931709 | 2.2877845 | 0.1495331 | 0.999175987 |
| ENSMUSG00000031758 | Cdyl2 | protein_coding | -1.283424881 | 4.238801782 | 2.301634 | 0.1495979 | 0.999175987 |
| ENSMUSG00000078439 | Smim24 | protein_coding | 1.468728807 | 2.738258183 | 2.2865789 | 0.1496336 | 0.999175987 |
| ENSMUSG00000035246 | Pcyt1b | protein_coding | -0.78507799 | 4.864259464 | 2.286153 | 0.1496692 | 0.999175987 |
| ENSMUSG00000051359 | Ncald | protein_coding | -0.539053452 | 8.885252362 | 2.2847016 | 0.1497904 | 0.999175987 |
| ENSMUSG00000022673 | Mcm4 | protein_coding | -1.096120017 | 3.356208428 | 2.2826777 | 0.1499597 | 0.999175987 |
| ENSMUSG00000030199 | Etv6 | protein_coding | 1.599976594 | 3.028764161 | 2.2818399 | 0.1500298 | 0.999175987 |
| ENSMUSG00000000276 | Dgke | protein_coding | -0.803160431 | 4.35600267 | 2.2807094 | 0.1501245 | 0.999175987 |
| ENSMUSG00000020015 | Cdk17 | protein_coding | -0.640915374 | 6.730462063 | 2.2801501 | 0.1502027 | 0.999175987 |
| ENSMUSG00000033228 | Scaf11 | protein_coding | -0.684782215 | 5.105056989 | 2.2781543 | 0.1503389 | 0.999175987 |
| ENSMUSG00000039395 | Mreg | protein_coding | 1.591728576 | 2.458589269 | 2.2777204 | 0.1503753 | 0.999175987 |
| ENSMUSG00000024883 | Rin1 | protein_coding | -1.202453268 | 3.607593129 | 2.2760147 | 0.1505186 | 0.999175987 |
| ENSMUSG00000021259 | Cyp46a1 | protein_coding | 0.747041569 | 5.252938892 | 2.2729862 | 0.1507735 | 0.999175987 |
| ENSMUSG00000047261 | Gap43 | protein_coding | 0.471422561 | 11.00325107 | 2.2726792 | 0.1507993 | 0.999175987 |
| ENSMUSG00000025909 | Sntg1 | protein_coding | 0.751112684 | 4.783253492 | 2.2661626 | 0.1513497 | 0.999175987 |
| ENSMUSG00000002718 | Cse1l | protein_coding | 0.58382619 | 5.626755388 | 2.2633485 | 0.1515881 | 0.999175987 |
| ENSMUSG00000029840 | Mtpn | protein_coding | -0.522465478 | 7.639171487 | 2.2632987 | 0.1515923 | 0.999175987 |
| ENSMUSG00000049154 | Fam183b | protein_coding | -0.527310151 | 5.985182853 | 2.262496 | 0.1516604 | 0.999175987 |
| ENSMUSG00000042064 | Myo3b | protein_coding | 1.640781589 | 2.505294159 | 2.2609715 | 0.1517899 | 0.999175987 |
| ENSMUSG00000043131 | Mob1a | protein_coding | -1.069401798 | 3.664848471 | 2.2597535 | 0.1518934 | 0.999175987 |
| ENSMUSG00000031198 | Fundc2 | protein_coding | 0.476654141 | 7.531706821 | 2.2591491 | 0.1519448 | 0.999175987 |
| ENSMUSG00000048142 | Nat8l | protein_coding | 0.483418401 | 6.965309063 | 2.2587163 | 0.1519816 | 0.999175987 |
| ENSMUSG00000020267 | Hint1 | protein_coding | 0.477958971 | 10.43714047 | 2.2581027 | 0.1520338 | 0.999175987 |
| ENSMUSG00000030471 | Zdhhc13 | protein_coding | -1.038697559 | 3.432563223 | 2.25751 | 0.1520842 | 0.999175987 |
| ENSMUSG00000036745 | Ttll7 | protein_coding | -0.462038042 | 7.030682103 | 2.2569338 | 0.1521333 | 0.999175987 |
| ENSMUSG00000058402 | Zfp420 | protein_coding | -0.843076584 | 3.993449678 | 2.2560572 | 0.152208 | 0.999175987 |
| ENSMUSG00000015980 | Lrrc27 | protein_coding | 0.942671726 | 4.809992088 | 2.2555915 | 0.152279 | 0.999175987 |
| ENSMUSG00000068859 | Sp9 | protein_coding | -0.881447591 | 3.855387841 | 2.2539927 | 0.1523841 | 0.999175987 |
| ENSMUSG00000026826 | Nr4a2 | protein_coding | -1.4124972 | 2.621287073 | 2.2536468 | 0.1524136 | 0.999175987 |
| ENSMUSG00000044288 | Cnr1 | protein_coding | 0.557259323 | 6.287251988 | 2.2518767 | 0.1525648 | 0.999175987 |
| ENSMUSG00000033769 | Exoc6b | protein_coding | 0.651176016 | 5.277434501 | 2.2516062 | 0.152588 | 0.999175987 |
| ENSMUSG00000019796 | Lrp11 | protein_coding | -0.696028613 | 5.493221051 | 2.2514571 | 0.1526007 | 0.999175987 |
| ENSMUSG00000023011 | Faim2 | protein_coding | 0.582112918 | 9.330157212 | 2.2511888 | 0.1526236 | 0.999175987 |
| ENSMUSG00000028755 | Cda | protein_coding | 0.825515508 | 4.950900767 | 2.2503415 | 0.1526961 | 0.999175987 |
| ENSMUSG00000041733 | Coq5 | protein_coding | 0.497668713 | 6.582183234 | 2.2493442 | 0.1527815 | 0.999175987 |
| ENSMUSG00000027215 | Cd82 | protein_coding | -0.618494796 | 6.399149265 | 2.2456702 | 0.1530965 | 0.999175987 |
| ENSMUSG00000026675 | Hsd17b7 | protein_coding | -1.093011103 | 4.474511903 | 2.2468254 | 0.1531192 | 0.999175987 |

| ensembl_gene_id | external_gene_name | gene_biotype | log2 (fold change) | log2 (counts per million) | F-value | p-value | False discovery rate |
|---|---|---|---|---|---|---|---|
| ENSMUSG00000002108 | Nr1h3 | protein_coding | 1.81417082 | 2.680416916 | 2.245213 | 0.1531357 | 0.999175987 |
| ENSMUSG00000073676 | Hspe1 | protein_coding | 0.49553071 | 7.452301563 | 2.2448128 | 0.1531701 | 0.999175987 |
| ENSMUSG00000024219 | Anks1 | protein_coding | 1.070769735 | 3.484281808 | 2.2425668 | 0.1533632 | 0.999175987 |
| ENSMUSG00000042675 | Ypel3 | protein_coding | 0.594654885 | 6.043903776 | 2.2422876 | 0.1533872 | 0.999175987 |
| ENSMUSG00000002032 | Tmem25 | protein_coding | 0.649503252 | 6.284978924 | 2.2420044 | 0.1534116 | 0.999175987 |
| ENSMUSG00000037625 | Cldn11 | protein_coding | -0.429610716 | 7.407840448 | 2.2418692 | 0.1534232 | 0.999175987 |
| ENSMUSG00000048550 | Thnsl1 | protein_coding | -0.722894298 | 5.112092337 | 2.2411063 | 0.1534889 | 0.999175987 |
| ENSMUSG00000012296 | Tjap1 | protein_coding | 1.769323355 | 2.745423782 | 2.2404162 | 0.1535483 | 0.999175987 |
| ENSMUSG00000033096 | Apmap | protein_coding | 0.532159823 | 5.88880109 | 2.2393439 | 0.1536407 | 0.999175987 |
| ENSMUSG00000031425 | Plp1 | protein_coding | -0.524944037 | 10.61790917 | 2.2390742 | 0.153664 | 0.999175987 |
| ENSMUSG00000024911 | Fibp | protein_coding | 0.489356859 | 6.886098027 | 2.2388541 | 0.153683 | 0.999175987 |
| ENSMUSG00000059493 | Nhs | protein_coding | -1.023397585 | 3.366856046 | 2.2386509 | 0.1537005 | 0.999175987 |
| ENSMUSG00000033308 | Dpyd | protein_coding | -1.383256765 | 2.844302619 | 2.2381082 | 0.1537473 | 0.999175987 |
| ENSMUSG00000030839 | Sergef | protein_coding | 0.501855921 | 6.531287465 | 2.2379245 | 0.1537632 | 0.999175987 |
| ENSMUSG00000028960 | Ube4b | protein_coding | -0.635085863 | 5.598998595 | 2.2352703 | 0.1539925 | 0.999175987 |
| ENSMUSG00000021244 | Ylpm1 | protein_coding | 0.620615838 | 5.454123486 | 2.2346895 | 0.1540427 | 0.999175987 |
| ENSMUSG00000030759 | Far1 | protein_coding | -0.609690453 | 5.452490102 | 2.233377 | 0.1541563 | 0.999175987 |
| ENSMUSG00000025393 | Atp5b | protein_coding | 0.576395928 | 10.30612259 | 2.2328194 | 0.1542045 | 0.999175987 |
| ENSMUSG00000047767 | Atg16l2 | protein_coding | 1.084231274 | 4.074407515 | 2.2323362 | 0.1542776 | 0.999175987 |
| ENSMUSG00000030660 | Pik3c2a | protein_coding | -1.46860699 | 2.710355144 | 2.2309515 | 0.1543664 | 0.999175987 |
| ENSMUSG00000022761 | Lztr1 | protein_coding | -0.525091952 | 6.556894168 | 2.2263831 | 0.1547632 | 0.999175987 |
| ENSMUSG00000022475 | Hdac7 | protein_coding | 1.008676404 | 4.279906756 | 2.2259866 | 0.1547977 | 0.999175987 |
| ENSMUSG00000085622 | 3110056K07Rik | processed_transcript | -1.018684052 | 4.492442256 | 2.2252578 | 0.1548924 | 0.999175987 |
| ENSMUSG00000084830 | Gm14539 | processed_pseudoge | 0.872836537 | 3.93243946 | 2.2236888 | 0.1549978 | 0.999175987 |
| ENSMUSG00000062526 | Mppe1 | protein_coding | 1.409477004 | 2.980219548 | 2.2222359 | 0.1551245 | 0.999175987 |
| ENSMUSG00000035104 | Eva1a | protein_coding | 1.392249813 | 3.812427907 | 2.2350957 | 0.1551446 | 0.999175987 |
| ENSMUSG00000039361 | Picalm | protein_coding | -0.544722617 | 6.279522814 | 2.2204817 | 0.1552776 | 0.999175987 |
| ENSMUSG00000067870 | Rpl31-ps8 | processed_pseudoge | 1.044974546 | 3.545993671 | 2.2196319 | 0.1553519 | 0.999175987 |
| ENSMUSG00000073154 | 9330158H04Rik | lincRNA | -1.3427104 | 2.924406404 | 2.2194967 | 0.1553637 | 0.999175987 |
| ENSMUSG00000009293 | Ube2g2 | protein_coding | 0.53672762 | 6.617847669 | 2.2192216 | 0.1553878 | 0.999175987 |
| ENSMUSG00000034799 | Unc13a | protein_coding | -0.906928393 | 3.774578642 | 2.2152989 | 0.1557312 | 0.999175987 |
| ENSMUSG00000051285 | Pcmtd1 | protein_coding | -0.650435889 | 5.094840284 | 2.2150628 | 0.1557519 | 0.999175987 |
| ENSMUSG00000030380 | Mzf1 | protein_coding | 1.852168484 | 2.223890863 | 2.2125965 | 0.1559683 | 0.999175987 |
| ENSMUSG00000063334 | Krr1 | protein_coding | 0.765246626 | 4.937537707 | 2.2119987 | 0.1560208 | 0.999175987 |
| ENSMUSG00000040952 | Rps19 | protein_coding | 0.589006955 | 8.554367268 | 2.2117824 | 0.156071 | 0.999175987 |
| ENSMUSG00000032946 | Rasgrp2 | protein_coding | 0.617556108 | 5.225065832 | 2.2111645 | 0.1560941 | 0.999175987 |
| ENSMUSG00000003585 | Sec14l2 | protein_coding | -0.834149287 | 3.81467034 | 2.2095144 | 0.1562393 | 0.999175987 |
| ENSMUSG00000063297 | Luzp2 | protein_coding | -0.653580097 | 6.133680057 | 2.2089371 | 0.1562901 | 0.999175987 |
| ENSMUSG00000002985 | Apoe | protein_coding | -1.123872644 | 7.517200894 | 2.3016146 | 0.1563701 | 0.999175987 |
| ENSMUSG00000025931 | Paqr8 | protein_coding | -0.853802678 | 4.716172339 | 2.2063215 | 0.1565206 | 0.999175987 |
| ENSMUSG00000040506 | Ambra1 | protein_coding | -0.767834662 | 5.066546137 | 2.2058356 | 0.1565635 | 0.999175987 |
| ENSMUSG00000027544 | Nfatc2 | protein_coding | 0.941835987 | 4.198646993 | 2.2043901 | 0.1566911 | 0.999175987 |
| ENSMUSG00000029759 | Pon3 | protein_coding | -2.400819278 | 2.371750793 | 2.2042384 | 0.1567357 | 0.999175987 |
| ENSMUSG00000001036 | Epn2 | protein_coding | -0.505837123 | 6.300248836 | 2.2007627 | 0.1570119 | 0.999175987 |
| ENSMUSG00000110834 | CAAA01180111.2 | lincRNA | 0.763444326 | 4.268034083 | 2.1995817 | 0.1571166 | 0.999175987 |
| ENSMUSG00000046756 | Mrps7 | protein_coding | 0.472244093 | 6.693885339 | 2.1990782 | 0.1571612 | 0.999175987 |
| ENSMUSG00000025223 | Ldb1 | protein_coding | 0.666804362 | 5.148441697 | 2.1989421 | 0.1571733 | 0.999175987 |
| ENSMUSG00000013593 | Ndufs2 | protein_coding | 0.470000843 | 8.539366537 | 2.1988638 | 0.1571802 | 0.999175987 |
| ENSMUSG00000022452 | Smdt1 | protein_coding | 0.495714682 | 8.365848453 | 2.1986764 | 0.1571968 | 0.999175987 |
| ENSMUSG00000021715 | Cwc27 | protein_coding | -1.045060415 | 3.686163553 | 2.1985354 | 0.1572093 | 0.999175987 |
| ENSMUSG00000037415 | Ranbp10 | protein_coding | 1.292934388 | 3.407233385 | 2.197443 | 0.1573062 | 0.999175987 |
| ENSMUSG00000040463 | Mybbp1a | protein_coding | 0.717123675 | 4.832715963 | 2.1963086 | 0.157407 | 0.999175987 |
| ENSMUSG00000017801 | Mlx | protein_coding | 0.616087312 | 6.238427675 | 2.1946473 | 0.1575546 | 0.999175987 |
| ENSMUSG00000031575 | Ash2l | protein_coding | -0.763315846 | 4.821138281 | 2.1939738 | 0.1576145 | 0.999175987 |
| ENSMUSG00000039349 | C130074G19Rik | protein_coding | -1.36225823 | 3.535374711 | 2.1940151 | 0.157642 | 0.999175987 |
| ENSMUSG00000001827 | Folr1 | protein_coding | -5.108927328 | 3.469133099 | 2.2790552 | 0.1576444 | 0.999175987 |
| ENSMUSG00000066036 | Ubr4 | protein_coding | -0.713347835 | 5.281390209 | 2.1923042 | 0.1577632 | 0.999175987 |
| ENSMUSG00000027562 | Car2 | protein_coding | -0.488635028 | 10.41162244 | 2.1917495 | 0.1578126 | 0.999175987 |
| ENSMUSG00000032253 | Phip | protein_coding | -0.752832016 | 4.797293326 | 2.1916374 | 0.1578226 | 0.999175987 |
| ENSMUSG00000047371 | Zfp768 | protein_coding | -1.809505787 | 2.477287658 | 2.1908966 | 0.1578886 | 0.999175987 |
| ENSMUSG00000027122 | Arl14ep | protein_coding | -0.602996342 | 6.619910862 | 2.1904216 | 0.157931 | 0.999175987 |
| ENSMUSG00000096916 | Zfp850 | protein_coding | 2.031445793 | 2.376784484 | 2.1901773 | 0.1579528 | 0.999175987 |
| ENSMUSG00000023307 | 5-Mar | protein_coding | 0.684711146 | 5.014800071 | 2.188841 | 0.158072 | 0.999175987 |
| ENSMUSG00000026740 | Dnajc1 | protein_coding | -1.194036675 | 4.213419847 | 2.1958395 | 0.1580805 | 0.999175987 |
| ENSMUSG00000042328 | Hps4 | protein_coding | -0.908921993 | 3.690043828 | 2.1887074 | 0.1580839 | 0.999175987 |
| ENSMUSG00000031990 | Jam3 | protein_coding | -0.684252961 | 4.968066662 | 2.188426 | 0.1581091 | 0.999175987 |
| ENSMUSG00000027221 | Chst1 | protein_coding | 0.510404388 | 6.988503255 | 2.1883407 | 0.1581167 | 0.999175987 |
| ENSMUSG00000031982 | Arv1 | protein_coding | 1.051366991 | 3.505064802 | 2.1874348 | 0.1581976 | 0.999175987 |
| ENSMUSG00000045589 | Frrs1l | protein_coding | -0.46349997 | 6.858846421 | 2.1871162 | 0.1582261 | 0.999175987 |
| ENSMUSG00000044708 | Kcnj10 | protein_coding | -0.827248622 | 4.889760016 | 2.1858582 | 0.1583386 | 0.999175987 |
| ENSMUSG00000020319 | Wdpcp | protein_coding | -1.028419979 | 3.361835588 | 2.1853637 | 0.1583829 | 0.999175987 |
| ENSMUSG00000021071 | Trim9 | protein_coding | -0.820574366 | 5.423972199 | 2.1841172 | 0.1585257 | 0.999175987 |
| ENSMUSG00000019849 | Prep | protein_coding | -0.80837358 | 3.773300172 | 2.1835215 | 0.1585479 | 0.999175987 |
| ENSMUSG00000043415 | Otud1 | protein_coding | 1.254722219 | 3.314810434 | 2.1828191 | 0.1586108 | 0.999175987 |
| ENSMUSG00000063354 | Slc39a4 | protein_coding | 6.405917175 | 1.36567172 | 2.2277588 | 0.158686 | 0.999175987 |
| ENSMUSG00000043518 | Rai2 | protein_coding | -0.936599016 | 4.267342959 | 2.1815776 | 0.1587222 | 0.999175987 |
| ENSMUSG00000042737 | Dpm3 | protein_coding | 0.565656306 | 6.893419205 | 2.1815443 | 0.1587252 | 0.999175987 |
| ENSMUSG00000051671 | Coa6 | protein_coding | 0.517795099 | 6.509257988 | 2.1811261 | 0.1587627 | 0.999175987 |

| ensembl_gene_id | external_gene_name | gene_biotype | log2 (fold change) | log2 (counts per million) | F-value | p-value | False discovery rate |
|---|---|---|---|---|---|---|---|
| ENSMUSG00000037965 | Zc3h7a | protein_coding | 0.639604791 | 6.426642557 | 2.1796026 | 0.1589307 | 0.999175987 |
| ENSMUSG00000002845 | Tmem39a | protein_coding | 1.329167051 | 3.255610213 | 2.1787219 | 0.1589787 | 0.999175987 |
| ENSMUSG00000029001 | Fbxo44 | protein_coding | 0.399731939 | 7.966833578 | 2.1785577 | 0.1589935 | 0.999175987 |
| ENSMUSG00000001552 | Jup | protein_coding | -0.729274567 | 4.976714421 | 2.1775451 | 0.1590846 | 0.999175987 |
| ENSMUSG00000045827 | Serpinb9 | protein_coding | 0.977945136 | 3.913009374 | 2.176827 | 0.1591492 | 0.999175987 |
| ENSMUSG00000114304 | AC165092.1 | lincRNA | 0.782641351 | 7.087818151 | 2.1960438 | 0.1593569 | 0.999175987 |
| ENSMUSG00000027452 | Acss1 | protein_coding | -1.406534569 | 3.118755559 | 2.1741816 | 0.1593877 | 0.999175987 |
| ENSMUSG00000034110 | Kctd7 | protein_coding | -0.82903425 | 4.071132289 | 2.1737295 | 0.1594284 | 0.999175987 |
| ENSMUSG00000005802 | Slc30a4 | protein_coding | -1.370199541 | 3.385005898 | 2.1738816 | 0.1594459 | 0.999175987 |
| ENSMUSG00000031139 | Mcf2 | protein_coding | 1.271465808 | 3.053798652 | 2.1733166 | 0.1594657 | 0.999175987 |
| ENSMUSG00000032673 | Prorsd1 | protein_coding | -0.722160659 | 4.986987756 | 2.1731832 | 0.1594778 | 0.999175987 |
| ENSMUSG00000016206 | H2-M3 | protein_coding | 1.348351983 | 3.008335465 | 2.1727343 | 0.1595183 | 0.999175987 |
| ENSMUSG00000025326 | Ube3a | protein_coding | -0.565267027 | 5.993583477 | 2.1713192 | 0.1596461 | 0.999175987 |
| ENSMUSG00000036158 | Prickle1 | protein_coding | -0.74964112 | 4.668239792 | 2.1709067 | 0.1596834 | 0.999175987 |
| ENSMUSG00000031760 | Mt3 | protein_coding | 0.577976901 | 10.72742009 | 2.1702982 | 0.1597384 | 0.999175987 |
| ENSMUSG00000046062 | Ppp1r15b | protein_coding | -0.716365534 | 4.111451231 | 2.1700278 | 0.1597629 | 0.999175987 |
| ENSMUSG00000098097 | 6530403H02Rik | lincRNA | -1.405134981 | 2.770981704 | 2.1694263 | 0.1598173 | 0.999175987 |
| ENSMUSG00000022711 | Pmm2 | protein_coding | -0.63793166 | 5.048380884 | 2.1684483 | 0.1599059 | 0.999175987 |
| ENSMUSG00000034777 | Vax2 | protein_coding | -1.528746662 | 2.562757672 | 2.1628826 | 0.1604109 | 0.999175987 |
| ENSMUSG00000025395 | Prim1 | protein_coding | -0.903673096 | 4.083517732 | 2.1614378 | 0.1605424 | 0.999175987 |
| ENSMUSG00000086697 | 4933427G23Rik | transcribed_processe | -1.541840995 | 2.445828553 | 2.1601827 | 0.1606567 | 0.999175987 |
| ENSMUSG00000027668 | Mfn1 | protein_coding | -0.894067631 | 4.57648401 | 2.1601169 | 0.1606627 | 0.999175987 |
| ENSMUSG00000025982 | Sf3b1 | protein_coding | -0.424451832 | 7.046822769 | 2.1594653 | 0.160722 | 0.999175987 |
| ENSMUSG00000037251 | Pomk | protein_coding | 0.615209887 | 5.944052646 | 2.1594254 | 0.1607257 | 0.999175987 |
| ENSMUSG00000040183 | Ankrd6 | protein_coding | 0.678813633 | 5.466469459 | 2.1588479 | 0.1607783 | 0.999175987 |
| ENSMUSG00000025234 | Arih1 | protein_coding | -0.723856048 | 5.035524536 | 2.1575934 | 0.1608928 | 0.999175987 |
| ENSMUSG00000038736 | Nudcd1 | protein_coding | -0.881388867 | 4.382402137 | 2.1555722 | 0.1610773 | 0.999175987 |
| ENSMUSG00000030878 | Cdr2 | protein_coding | 0.999508691 | 3.563215937 | 2.1545996 | 0.1611662 | 0.999175987 |
| ENSMUSG00000033557 | Fam20b | protein_coding | 0.669589892 | 5.503172762 | 2.1544633 | 0.1611787 | 0.999175987 |
| ENSMUSG00000027581 | Stmn3 | protein_coding | 0.462817661 | 11.21256015 | 2.1470346 | 0.1618599 | 0.999175987 |
| ENSMUSG00000051256 | Jagn1 | protein_coding | 0.481495563 | 6.482956711 | 2.146616 | 0.1618984 | 0.999175987 |
| ENSMUSG00000039607 | Rbms3 | protein_coding | -0.728877555 | 5.626298058 | 2.1465826 | 0.1619015 | 0.999175987 |
| ENSMUSG00000075704 | Txnrd2 | protein_coding | 1.01543753 | 3.669811175 | 2.1424969 | 0.1622778 | 0.999175987 |
| ENSMUSG00000020413 | Hus1 | protein_coding | -1.219361295 | 3.090170719 | 2.1423468 | 0.1622916 | 0.999175987 |
| ENSMUSG00000024926 | Kat5 | protein_coding | 0.602194651 | 6.447044186 | 2.1410157 | 0.1624145 | 0.999175987 |
| ENSMUSG00000074758 | Gm5535 | lincRNA | -1.394478441 | 2.770397321 | 2.1393238 | 0.1625708 | 0.999175987 |
| ENSMUSG00000033910 | Gucy1a3 | protein_coding | -0.414768168 | 6.664581123 | 2.1376365 | 0.1627268 | 0.999175987 |
| ENSMUSG00000052374 | Actn2 | protein_coding | 1.638718192 | 3.558390301 | 2.1566248 | 0.1628696 | 0.999175987 |
| ENSMUSG00000026414 | Tnnt2 | protein_coding | 0.971360072 | 3.784445905 | 2.1359071 | 0.162887 | 0.999175987 |
| ENSMUSG00000001910 | Nacc1 | protein_coding | 0.736588801 | 4.97235212 | 2.1359019 | 0.1628874 | 0.999175987 |
| ENSMUSG00000038046 | Mrm3 | protein_coding | 0.564271918 | 5.840858269 | 2.1337775 | 0.1630844 | 0.999175987 |
| ENSMUSG00000003380 | Rabac1 | protein_coding | 0.596315814 | 8.737440246 | 2.1323152 | 0.1632513 | 0.999175987 |
| ENSMUSG00000037972 | Snn | protein_coding | -0.602733839 | 5.734400238 | 2.1316093 | 0.1632858 | 0.999175987 |
| ENSMUSG00000030629 | Zfand6 | protein_coding | -0.459241334 | 7.01366206 | 2.1293321 | 0.1634976 | 0.999175987 |
| ENSMUSG00000016308 | Ube2a | protein_coding | 0.436149981 | 7.517151785 | 2.1261043 | 0.1637984 | 0.999175987 |
| ENSMUSG00000020608 | Smc6 | protein_coding | -0.572505821 | 5.399207557 | 2.1257873 | 0.163828 | 0.999175987 |
| ENSMUSG00000085408 | C530005A16Rik | processed_transcript | -1.115852616 | 3.387807311 | 2.1244807 | 0.16395 | 0.999175987 |
| ENSMUSG00000028572 | Hook1 | protein_coding | -0.924378674 | 3.667438773 | 2.1241869 | 0.1639774 | 0.999175987 |
| ENSMUSG00000038489 | Polr2l | protein_coding | 0.629471343 | 4.985064175 | 2.1236223 | 0.1640302 | 0.999175987 |
| ENSMUSG00000026883 | Dab2ip | protein_coding | -0.804128989 | 4.455905101 | 2.1231747 | 0.164072 | 0.999175987 |
| ENSMUSG00000032854 | Ugt8a | protein_coding | -0.551992395 | 5.687384174 | 2.1228714 | 0.1641004 | 0.999175987 |
| ENSMUSG00000021290 | 2010107E04Rik | protein_coding | 0.595241415 | 9.27045432 | 2.122762 | 0.1641417 | 0.999175987 |
| ENSMUSG00000054414 | Slc30a7 | protein_coding | -1.205089411 | 3.354132197 | 2.1220955 | 0.164173 | 0.999175987 |
| ENSMUSG00000039754 | Alkbh4 | protein_coding | 1.006084826 | 4.552657248 | 2.120595 | 0.1643445 | 0.999175987 |
| ENSMUSG00000059277 | R74862 | processed_transcript | -1.372929073 | 2.99845578 | 2.1186886 | 0.1644921 | 0.999175987 |
| ENSMUSG00000021493 | Pdlim7 | protein_coding | 0.439180913 | 6.876904623 | 2.1178331 | 0.1645724 | 0.999175987 |
| ENSMUSG00000093674 | Rpl41 | protein_coding | 0.639234659 | 9.105067067 | 2.1182283 | 0.1646481 | 0.999175987 |
| ENSMUSG00000053390 | Zfp952 | protein_coding | -0.879615133 | 4.308508244 | 2.1164553 | 0.1647018 | 0.999175987 |
| ENSMUSG00000020521 | Rnft1 | protein_coding | -0.811087072 | 4.037475334 | 2.1163987 | 0.1647071 | 0.999175987 |
| ENSMUSG00000073139 | Tmem185a | protein_coding | 0.540162311 | 5.789414709 | 2.1147009 | 0.1648667 | 0.999175987 |
| ENSMUSG00000024036 | Slc37a1 | protein_coding | -1.288708753 | 2.945747937 | 2.114147 | 0.1649188 | 0.999175987 |
| ENSMUSG00000048416 | Mlf1 | protein_coding | -0.938888371 | 3.661680414 | 2.1130388 | 0.1650231 | 0.999175987 |
| ENSMUSG00000030659 | Nucb2 | protein_coding | 0.582623185 | 5.219319898 | 2.1117771 | 0.1651419 | 0.999175987 |
| ENSMUSG00000040896 | Kcnd3 | protein_coding | -0.928866824 | 4.172259452 | 2.1114533 | 0.1651725 | 0.999175987 |
| ENSMUSG00000022454 | Nell2 | protein_coding | 0.447548659 | 7.578508581 | 2.1109454 | 0.1652204 | 0.999175987 |
| ENSMUSG00000025175 | Fn3k | protein_coding | 0.530606398 | 6.706386204 | 2.1104068 | 0.1652712 | 0.999175987 |
| ENSMUSG00000036615 | Rfxap | protein_coding | 0.779763111 | 4.236576245 | 2.1093123 | 0.1653744 | 0.999175987 |
| ENSMUSG00000027652 | Ralgapb | protein_coding | -0.645250739 | 4.935392349 | 2.1089414 | 0.1654095 | 0.999175987 |
| ENSMUSG00000059534 | Uqcr10 | protein_coding | 0.549894778 | 10.21776316 | 2.1084756 | 0.1654535 | 0.999175987 |
| ENSMUSG00000017776 | Crk | protein_coding | -0.585713592 | 6.282211737 | 2.1061655 | 0.1656719 | 0.999175987 |
| ENSMUSG00000020246 | Hcfc2 | protein_coding | -0.764871741 | 4.458769391 | 2.1061508 | 0.1656733 | 0.999175987 |
| ENSMUSG00000035278 | Plekhj1 | protein_coding | 0.468718305 | 6.151296234 | 2.1052057 | 0.1657627 | 0.999175987 |
| ENSMUSG00000101621 | Gm28905 | lincRNA | -1.746277148 | 2.29363569 | 2.1049724 | 0.1657848 | 0.999175987 |
| ENSMUSG00000003235 | Eif2b5 | protein_coding | 0.67087815 | 5.373775988 | 2.1019076 | 0.1660754 | 0.999175987 |
| ENSMUSG00000032740 | Ccdc88a | protein_coding | -0.638340784 | 5.344034453 | 2.0991483 | 0.1663375 | 0.999175987 |
| ENSMUSG00000065990 | Aurkaip1 | protein_coding | 0.44467899 | 8.863702639 | 2.0986046 | 0.1663892 | 0.999175987 |
| ENSMUSG00000005583 | Mef2c | protein_coding | -0.58962922 | 6.856169045 | 2.09829 | 0.1664502 | 0.999175987 |
| ENSMUSG00000073733 | Rsg1 | protein_coding | 1.804234249 | 2.950123744 | 2.0987832 | 0.1665206 | 0.999175987 |

| ensembl_gene_id | external_gene_name | gene_biotype | log2 (fold change) | log2 (counts per million) | F-value | p-value | False discovery rate |
|---|---|---|---|---|---|---|---|
| ENSMUSG00000042246 | Tmc7 | protein_coding | -1.428056494 | 3.010522308 | 2.0968077 | 0.1665603 | 0.999175987 |
| ENSMUSG00000026895 | Ndufa8 | protein_coding | 0.425296855 | 9.504452864 | 2.0968072 | 0.1665603 | 0.999175987 |
| ENSMUSG00000026109 | Tmeff2 | protein_coding | 0.498630812 | 7.305166309 | 2.0964194 | 0.1665973 | 0.999175987 |
| ENSMUSG00000079426 | Arpc4 | protein_coding | 0.462583808 | 8.860484356 | 2.0935314 | 0.1668727 | 0.999175987 |
| ENSMUSG00000039419 | Cntnap2 | protein_coding | 0.441569151 | 7.646214872 | 2.0899579 | 0.1672143 | 0.999175987 |
| ENSMUSG00000037348 | Paqr7 | protein_coding | 0.566280406 | 6.590218466 | 2.0869888 | 0.1674988 | 0.999175987 |
| ENSMUSG00000017002 | Slpi | protein_coding | -1.619081326 | 2.919325958 | 2.0865881 | 0.1675373 | 0.999175987 |
| ENSMUSG00000055202 | Zfp811 | protein_coding | -0.822563492 | 4.399401755 | 2.0827536 | 0.1679057 | 0.999175987 |
| ENSMUSG00000005779 | Psmb4 | protein_coding | 0.459360906 | 7.730041608 | 2.0811272 | 0.1680622 | 0.999175987 |
| ENSMUSG00000004356 | Utp20 | protein_coding | 1.751193365 | 2.419253035 | 2.0809655 | 0.1680778 | 0.999175987 |
| ENSMUSG00000054474 | Thnsl2 | protein_coding | 1.205700134 | 3.354334824 | 2.0794506 | 0.1682238 | 0.999175987 |
| ENSMUSG00000018931 | Natd1 | protein_coding | 1.475505298 | 2.723686703 | 2.0781265 | 0.1683516 | 0.999175987 |
| ENSMUSG00000108434 | Gm45768 | lincRNA | -1.429576359 | 3.28471313 | 2.0779315 | 0.1684013 | 0.999175987 |
| ENSMUSG00000028060 | 2810403A07Rik | protein_coding | -0.774554376 | 4.220048193 | 2.0772101 | 0.1684401 | 0.999175987 |
| ENSMUSG00000022816 | Fstl1 | protein_coding | 0.655119164 | 5.619074361 | 2.0750111 | 0.1686526 | 0.999175987 |
| ENSMUSG00000044894 | Uqcrq | protein_coding | 0.503870429 | 10.39844636 | 2.0739605 | 0.1687543 | 0.999175987 |
| ENSMUSG00000028936 | Rpl22 | protein_coding | 0.52083735 | 7.214161766 | 2.0737928 | 0.1687705 | 0.999175987 |
| ENSMUSG00000021198 | Unc79 | protein_coding | -0.985118208 | 3.788694174 | 2.0737627 | 0.1687734 | 0.999175987 |
| ENSMUSG00000049401 | Ogfr | protein_coding | -0.862710834 | 3.834392804 | 2.0731199 | 0.1688357 | 0.999175987 |
| ENSMUSG00000023259 | Slc26a6 | protein_coding | 1.840787352 | 2.561602246 | 2.0721144 | 0.1689331 | 0.999175987 |
| ENSMUSG00000048039 | Isg20l2 | protein_coding | 0.964365609 | 3.647626949 | 2.0674407 | 0.1693869 | 0.999175987 |
| ENSMUSG00000031833 | Mast3 | protein_coding | -0.677737682 | 5.532815314 | 2.0664019 | 0.169488 | 0.999175987 |
| ENSMUSG00000097392 | D930016D06Rik | lincRNA | -1.528978249 | 2.721452666 | 2.0663748 | 0.1694907 | 0.999175987 |
| ENSMUSG00000059586 | Nsmce2 | protein_coding | 0.544314213 | 6.325690648 | 2.0658551 | 0.1695412 | 0.999175987 |
| ENSMUSG00000041453 | Rpl21 | protein_coding | 0.583992201 | 5.190588427 | 2.0650893 | 0.1696158 | 0.999175987 |
| ENSMUSG00000031939 | Taf1d | protein_coding | 0.884176798 | 4.656538511 | 2.0643408 | 0.1696888 | 0.999175987 |
| ENSMUSG00000041119 | Pde9a | protein_coding | -0.697797851 | 5.060572639 | 2.0637941 | 0.1697421 | 0.999175987 |
| ENSMUSG00000021038 | Vipas39 | protein_coding | -0.57969415 | 6.141613416 | 2.0633328 | 0.1697871 | 0.999175987 |
| ENSMUSG00000033287 | Kctd17 | protein_coding | 0.620713343 | 5.488237808 | 2.0629711 | 0.1698224 | 0.999175987 |
| ENSMUSG00000026260 | Ndufa10 | protein_coding | 0.652611905 | 8.476402603 | 2.0690545 | 0.1698563 | 0.999175987 |
| ENSMUSG00000050711 | Scg2 | protein_coding | -0.439629563 | 9.620289236 | 2.0626198 | 0.1698566 | 0.999175987 |
| ENSMUSG00000053025 | Sv2b | protein_coding | -0.647408625 | 5.410793827 | 2.0623947 | 0.1698786 | 0.999175987 |
| ENSMUSG00000049643 | 2310022A10Rik | protein_coding | 0.561493614 | 5.683966829 | 2.0617158 | 0.1699449 | 0.999175987 |
| ENSMUSG00000006675 | P4htm | protein_coding | 0.527588333 | 6.530913841 | 2.0612459 | 0.1699908 | 0.999175987 |
| ENSMUSG00000020495 | Smg8 | protein_coding | 0.904142827 | 3.617206367 | 2.0609285 | 0.1700218 | 0.999175987 |
| ENSMUSG00000022438 | Parvb | protein_coding | -0.855095764 | 3.865977338 | 2.0602957 | 0.1700837 | 0.999175987 |
| ENSMUSG00000048997 | Atxn7l2 | protein_coding | -1.213407937 | 2.92928557 | 2.0575399 | 0.1703533 | 0.999175987 |
| ENSMUSG00000024735 | Prpf19 | protein_coding | 0.558577205 | 5.857345862 | 2.0570096 | 0.1704053 | 0.999175987 |
| ENSMUSG00000045237 | 1110012L19Rik | protein_coding | 0.49709593 | 5.830513754 | 2.0561315 | 0.1704914 | 0.999175987 |
| ENSMUSG00000057229 | Atp5sl | protein_coding | 0.511905935 | 6.897571732 | 2.0552504 | 0.1705778 | 0.999175987 |
| ENSMUSG00000061778 | Mospd2 | protein_coding | -0.909416243 | 3.429067216 | 2.0545667 | 0.1706449 | 0.999175987 |
| ENSMUSG00000021606 | Ndufs6 | protein_coding | 0.48474366 | 8.767019294 | 2.0534001 | 0.1707594 | 0.999175987 |
| ENSMUSG00000057554 | Lgals8 | protein_coding | -0.721660086 | 5.631530436 | 2.0500313 | 0.1710908 | 0.999175987 |
| ENSMUSG00000031438 | Rnf128 | protein_coding | 0.841133154 | 4.235137812 | 2.0487232 | 0.1712197 | 0.999175987 |
| ENSMUSG00000034424 | Gcsh | protein_coding | 0.431600958 | 7.694820109 | 2.043803 | 0.1717055 | 0.999175987 |
| ENSMUSG00000029231 | Pdgfra | protein_coding | -1.263241558 | 3.623955412 | 2.044036 | 0.1717133 | 0.999175987 |
| ENSMUSG00000003974 | Grm3 | protein_coding | -0.72289675 | 4.983939823 | 2.0427974 | 0.171805 | 0.999175987 |
| ENSMUSG00000036916 | Zfp280c | protein_coding | -1.177994965 | 3.532355404 | 2.0417106 | 0.1719126 | 0.999175987 |
| ENSMUSG00000020253 | Ppm1m | protein_coding | 0.830227404 | 4.064014177 | 2.0415226 | 0.1719313 | 0.999175987 |
| ENSMUSG00000019970 | Sgk1 | protein_coding | -0.716481546 | 7.726937866 | 2.0576016 | 0.1719769 | 0.999175987 |
| ENSMUSG00000070822 | Zscan18 | protein_coding | -1.051999742 | 4.896560747 | 2.0527513 | 0.172019 | 0.999175987 |
| ENSMUSG00000086290 | Snhg12 | lincRNA | 0.558379029 | 5.972992914 | 2.0403836 | 0.1720442 | 0.999175987 |
| ENSMUSG00000038884 | A230050P20Rik | protein_coding | 1.094091339 | 4.417019797 | 2.0461451 | 0.1720595 | 0.999175987 |
| ENSMUSG00000032842 | Abcc10 | protein_coding | 1.726843833 | 2.920291954 | 2.0398473 | 0.1721282 | 0.999175987 |
| ENSMUSG00000073542 | Cep76 | protein_coding | 1.246580824 | 2.759413771 | 2.0391131 | 0.1721702 | 0.999175987 |
| ENSMUSG00000033964 | Zbtb41 | protein_coding | -1.067383102 | 3.457689117 | 2.0379364 | 0.172287 | 0.999175987 |
| ENSMUSG00000024304 | Cdh2 | protein_coding | -0.893170244 | 3.838995901 | 2.0343189 | 0.1726468 | 0.999175987 |
| ENSMUSG00000031665 | Sall1 | protein_coding | -1.199217772 | 3.208844785 | 2.0328042 | 0.1727978 | 0.999175987 |
| ENSMUSG00000025531 | Chm | protein_coding | -0.751957796 | 4.489383728 | 2.0326406 | 0.1728141 | 0.999175987 |
| ENSMUSG00000029053 | Prkcz | protein_coding | 0.444515793 | 7.089822294 | 2.0298167 | 0.173096 | 0.999175987 |
| ENSMUSG00000042426 | Dhx29 | protein_coding | -0.745436359 | 3.953861934 | 2.0294867 | 0.1731289 | 0.999175987 |
| ENSMUSG00000026634 | Angel2 | protein_coding | -0.563175454 | 5.422101788 | 2.0287956 | 0.173198 | 0.999175987 |
| ENSMUSG00000042541 | Sem1 | protein_coding | 0.565698199 | 9.098111375 | 2.0290323 | 0.1732052 | 0.999175987 |
| ENSMUSG00000042757 | Tmem108 | protein_coding | 1.040413022 | 3.452302513 | 2.0267882 | 0.1733989 | 0.999175987 |
| ENSMUSG00000078348 | Sf3b5 | protein_coding | 0.443465497 | 8.019502921 | 2.0250712 | 0.1735709 | 0.999175987 |
| ENSMUSG00000041921 | Metap1d | protein_coding | 0.931736247 | 3.832582105 | 2.0245074 | 0.1736274 | 0.999175987 |
| ENSMUSG00000057963 | Itpk1 | protein_coding | 0.567459021 | 5.643085714 | 2.0224538 | 0.1738336 | 0.999175987 |
| ENSMUSG00000016262 | Sertad4 | protein_coding | 0.827328384 | 4.413357749 | 2.0221568 | 0.1738634 | 0.999175987 |
| ENSMUSG00000067851 | Arfgef1 | protein_coding | -0.668451517 | 5.310221381 | 2.0221144 | 0.1738677 | 0.999175987 |
| ENSMUSG00000097929 | Tunar | protein_coding | 0.694478161 | 4.680065021 | 2.0210978 | 0.1739698 | 0.999175987 |
| ENSMUSG00000002996 | Hbp1 | protein_coding | 0.88530768 | 3.632420087 | 2.0206986 | 0.17401 | 0.999175987 |
| ENSMUSG00000037339 | Fam53a | protein_coding | 1.330593991 | 3.00363387 | 2.0205776 | 0.1740222 | 0.999175987 |
| ENSMUSG00000063765 | Chadl | protein_coding | 0.919201646 | 3.341909579 | 2.0204982 | 0.1740301 | 0.999175987 |
| ENSMUSG00000026159 | Agfg1 | protein_coding | -0.604168913 | 5.152506144 | 2.0203833 | 0.1740417 | 0.999175987 |
| ENSMUSG00000040794 | C1qtnf4 | protein_coding | -1.197870915 | 3.310333493 | 2.0181736 | 0.1742642 | 0.999175987 |
| ENSMUSG00000061143 | Maml3 | protein_coding | 1.140320182 | 3.033189784 | 2.0177863 | 0.1743032 | 0.999175987 |
| ENSMUSG00000057406 | Nsd2 | protein_coding | -0.567162463 | 5.332635908 | 2.0169597 | 0.1743865 | 0.999175987 |
| ENSMUSG00000030534 | Vps33b | protein_coding | -1.123736633 | 3.958415376 | 2.0125776 | 0.1748599 | 0.999175987 |

| ensembl_gene_id | external_gene_name | gene_biotype | log2 (fold change) | log2 (counts per million) | F-value | p-value | False discovery rate |
|---|---|---|---|---|---|---|---|
| ENSMUSG00000056069 | Fam105a | protein_coding | 1.073920405 | 3.276289901 | 2.0099086 | 0.1750994 | 0.999175987 |
| ENSMUSG00000049550 | Clip1 | protein_coding | 0.51019707 | 6.116884207 | 2.0095523 | 0.1751355 | 0.999175987 |
| ENSMUSG00000110027 | C030029H02Rik | lincRNA | -0.74137863 | 4.059750618 | 2.0095087 | 0.1751399 | 0.999175987 |
| ENSMUSG00000021271 | Zfp839 | protein_coding | -0.766993605 | 3.894737059 | 2.0080979 | 0.175283 | 0.999175987 |
| ENSMUSG00000045414 | 1190002N15Rik | protein_coding | -0.869851285 | 3.857464086 | 2.0071178 | 0.1753825 | 0.999175987 |
| ENSMUSG00000097518 | Gm26694 | lincRNA | 0.801870423 | 4.020025057 | 2.0061174 | 0.1754842 | 0.999175987 |
| ENSMUSG00000055717 | Slain1 | protein_coding | -0.597683719 | 5.534441336 | 2.005432 | 0.1755539 | 0.999175987 |
| ENSMUSG00000021072 | Tmx1 | protein_coding | 0.474616376 | 7.163245087 | 2.0050451 | 0.1755932 | 0.999175987 |
| ENSMUSG00000003575 | Crtc1 | protein_coding | 0.507685523 | 5.899694798 | 2.0045732 | 0.1756412 | 0.999175987 |
| ENSMUSG00000024901 | Peli3 | protein_coding | 0.819511949 | 3.903887518 | 2.0027128 | 0.1758306 | 0.999175987 |
| ENSMUSG00000051502 | Ufsp1 | protein_coding | 0.858017857 | 3.98069418 | 2.0016657 | 0.1759374 | 0.999175987 |
| ENSMUSG00000023175 | Bsg | protein_coding | 0.73057604 | 8.302447301 | 2.0200048 | 0.1759481 | 0.999175987 |
| ENSMUSG00000022671 | Mzt2 | protein_coding | 0.458233008 | 6.378649776 | 2.0012099 | 0.1759838 | 0.999175987 |
| ENSMUSG00000033510 | Otud7a | protein_coding | -1.404088201 | 2.772559865 | 1.9999856 | 0.1761088 | 0.999175987 |
| ENSMUSG00000069814 | Ccdc92b | protein_coding | -0.792940445 | 4.020722003 | 1.9983026 | 0.1762807 | 0.999175987 |
| ENSMUSG00000020346 | Mgat1 | protein_coding | -0.625309349 | 5.171762214 | 1.9978225 | 0.1763298 | 0.999175987 |
| ENSMUSG00000028405 | Aco1 | protein_coding | -0.945952689 | 4.148423769 | 1.9974483 | 0.176368 | 0.999175987 |
| ENSMUSG00000037486 | Asxl2 | protein_coding | -0.787681193 | 4.385081825 | 1.9973346 | 0.1763797 | 0.999175987 |
| ENSMUSG00000040181 | Fmo1 | protein_coding | -1.13251998 | 3.594445406 | 1.9966885 | 0.1764458 | 0.999175987 |
| ENSMUSG00000086822 | 5330413P13Rik | lincRNA | -1.82172755 | 2.127317112 | 1.9966267 | 0.1764521 | 0.999175987 |
| ENSMUSG00000017868 | Sgk2 | protein_coding | -1.416522471 | 3.413828535 | 1.9961631 | 0.1766642 | 0.999175987 |
| ENSMUSG00000039660 | Spout1 | protein_coding | 0.579657803 | 5.892875434 | 1.994318 | 0.1766885 | 0.999175987 |
| ENSMUSG00000020547 | Bzw2 | protein_coding | 0.487853734 | 6.436693677 | 1.993961 | 0.1767251 | 0.999175987 |
| ENSMUSG00000024074 | Crim1 | protein_coding | -0.556748071 | 5.537589851 | 1.99357 | 0.1767652 | 0.999175987 |
| ENSMUSG00000078974 | Sec61g | protein_coding | 0.655267361 | 5.158550074 | 1.9924119 | 0.1768841 | 0.999175987 |
| ENSMUSG00000030255 | Sspn | protein_coding | -1.479455519 | 2.634035781 | 1.991998 | 0.1769266 | 0.999175987 |
| ENSMUSG00000005881 | Ergic3 | protein_coding | 0.382138436 | 8.70639423 | 1.9896946 | 0.1771633 | 0.999175987 |
| ENSMUSG00000034875 | Nudt19 | protein_coding | 0.397125351 | 6.684341927 | 1.9889836 | 0.1772364 | 0.999175987 |
| ENSMUSG00000009907 | Vps4b | protein_coding | -0.567925567 | 5.311644651 | 1.9886792 | 0.1772677 | 0.999175987 |
| ENSMUSG00000042726 | Trafd1 | protein_coding | -0.80275815 | 4.927957212 | 1.9881609 | 0.1773211 | 0.999175987 |
| ENSMUSG00000027014 | Cwc22 | protein_coding | -1.047865559 | 3.187849467 | 1.9870895 | 0.1774314 | 0.999175987 |
| ENSMUSG00000031376 | Atp2b3 | protein_coding | 0.958397491 | 3.234894051 | 1.9864536 | 0.177497 | 0.999175987 |
| ENSMUSG00000043962 | Thrap3 | protein_coding | -0.849983141 | 5.265451221 | 1.9863056 | 0.1775429 | 0.999175987 |
| ENSMUSG00000051497 | Kcnj16 | protein_coding | -1.059178185 | 3.231320837 | 1.9857981 | 0.1775646 | 0.999175987 |
| ENSMUSG00000034118 | Tpst1 | protein_coding | -0.805553451 | 4.475285247 | 1.9854209 | 0.1776035 | 0.999175987 |
| ENSMUSG00000027429 | Sec23b | protein_coding | -0.69555587 | 5.954301935 | 1.9855632 | 0.1776195 | 0.999175987 |
| ENSMUSG00000020083 | 2010107G23Rik | protein_coding | 0.460302536 | 6.60916479 | 1.984772 | 0.1776705 | 0.999175987 |
| ENSMUSG00000021136 | Smoc1 | protein_coding | -1.111952733 | 3.176641471 | 1.9839228 | 0.1777581 | 0.999175987 |
| ENSMUSG00000039523 | Cep104 | protein_coding | 0.925785297 | 4.015936478 | 1.983781 | 0.1777728 | 0.999175987 |
| ENSMUSG00000041132 | N4bp2l1 | protein_coding | -0.648132566 | 5.07898896 | 1.983 | 0.1778535 | 0.999175987 |
| ENSMUSG00000039740 | Alg2 | protein_coding | 0.492231934 | 7.151120417 | 1.9820514 | 0.1779515 | 0.999175987 |
| ENSMUSG00000026020 | Nop58 | protein_coding | -0.507610048 | 6.335977242 | 1.9811601 | 0.1780438 | 0.999175987 |
| ENSMUSG00000109324 | Prmt1 | protein_coding | 0.443100521 | 7.313579922 | 1.9787744 | 0.1782909 | 0.999175987 |
| ENSMUSG00000018736 | Ndel1 | protein_coding | 0.481130615 | 6.757699419 | 1.9779453 | 0.1783768 | 0.999175987 |
| ENSMUSG00000037266 | Rsrp1 | protein_coding | -1.49052106 | 5.794475185 | 2.0715428 | 0.1785006 | 0.999175987 |
| ENSMUSG00000063077 | Kif1b | protein_coding | -0.423096666 | 7.914263442 | 1.9765716 | 0.1785194 | 0.999175987 |
| ENSMUSG00000029655 | N4bp2l2 | protein_coding | -0.766968786 | 5.050342002 | 1.9761713 | 0.178561 | 0.999175987 |
| ENSMUSG00000020076 | Ddx50 | protein_coding | 0.538186428 | 5.727135582 | 1.9760845 | 0.17857 | 0.999175987 |
| ENSMUSG00000022244 | Amacr | protein_coding | -0.907807602 | 4.10322344 | 1.9755966 | 0.1786207 | 0.999175987 |
| ENSMUSG00000000168 | Dlat | protein_coding | -0.49084321 | 6.283778534 | 1.9755775 | 0.1786227 | 0.999175987 |
| ENSMUSG00000029675 | Eln | protein_coding | -2.087418579 | 3.211596894 | 2.0039551 | 0.1786833 | 0.999175987 |
| ENSMUSG00000079083 | Jrkl | protein_coding | 1.193593267 | 2.992706332 | 1.9699551 | 0.1792081 | 0.999175987 |
| ENSMUSG00000038695 | Josd2 | protein_coding | 0.467466057 | 6.239917499 | 1.9686488 | 0.1793444 | 0.999175987 |
| ENSMUSG00000053730 | Tmem39b | protein_coding | 1.215575902 | 3.168198016 | 1.9664074 | 0.1795787 | 0.999175987 |
| ENSMUSG00000049904 | Tmem17 | protein_coding | -0.751005626 | 3.952169071 | 1.9663567 | 0.179584 | 0.999175987 |
| ENSMUSG00000039114 | Nrn1 | protein_coding | -0.969598753 | 6.156109319 | 1.9950488 | 0.1795903 | 0.999175987 |
| ENSMUSG00000040321 | Zfp770 | protein_coding | -1.086102298 | 3.689128903 | 1.9661871 | 0.1796017 | 0.999175987 |
| ENSMUSG00000027360 | Hdc | protein_coding | 1.078848974 | 3.202417784 | 1.9653713 | 0.1796871 | 0.999175987 |
| ENSMUSG00000041477 | Dcp1b | protein_coding | 0.812186212 | 3.722479341 | 1.9633782 | 0.1798959 | 0.999175987 |
| ENSMUSG00000056383 | AI987944 | protein_coding | -1.361514665 | 3.222489646 | 1.9635425 | 0.1799093 | 0.999175987 |
| ENSMUSG00000110332 | Gm45396 | protein_coding | -0.827918142 | 4.327348264 | 1.9625755 | 0.1799801 | 0.999175987 |
| ENSMUSG00000041870 | Ankrd13a | protein_coding | -0.840566825 | 3.997599059 | 1.9608829 | 0.1801577 | 0.999175987 |
| ENSMUSG00000035179 | Ppp1r32 | protein_coding | 1.101848784 | 3.234266445 | 1.9605157 | 0.1801963 | 0.999175987 |
| ENSMUSG00000005447 | Pafah1b3 | protein_coding | 0.487116522 | 7.042839988 | 1.9602276 | 0.1802266 | 0.999175987 |
| ENSMUSG00000026112 | Coa5 | protein_coding | 0.444266226 | 6.58948572 | 1.9597327 | 0.1802786 | 0.999175987 |
| ENSMUSG00000098234 | Snhg6 | lincRNA | 0.929308565 | 5.926672747 | 1.9822191 | 0.1803518 | 0.999175987 |
| ENSMUSG00000043460 | Elfn2 | protein_coding | 1.093794922 | 3.91564847 | 1.9589729 | 0.1803891 | 0.999175987 |
| ENSMUSG00000019907 | Ppp1r12a | protein_coding | -0.532126216 | 6.455738273 | 1.9584797 | 0.1804103 | 0.999175987 |
| ENSMUSG00000060036 | Rpl3 | protein_coding | 0.526920924 | 7.69626689 | 1.9573712 | 0.180527 | 0.999175987 |
| ENSMUSG00000064181 | Rab3ip | protein_coding | 0.688353851 | 4.458143797 | 1.9571532 | 0.18055 | 0.999175987 |
| ENSMUSG00000032500 | Dclk3 | protein_coding | -0.93067441 | 4.730840745 | 1.9566243 | 0.1806363 | 0.999175987 |
| ENSMUSG00000002459 | Rgs20 | protein_coding | -0.556292473 | 6.075147803 | 1.9559014 | 0.1806819 | 0.999175987 |
| ENSMUSG00000024580 | Grpel2 | protein_coding | 0.582428941 | 5.73925752 | 1.9558775 | 0.1806844 | 0.999175987 |
| ENSMUSG00000047989 | Ino80c | protein_coding | -0.547491725 | 6.052578601 | 1.9545514 | 0.1808242 | 0.999175987 |
| ENSMUSG00000028556 | Dock7 | protein_coding | -0.912178099 | 3.640016014 | 1.9534587 | 0.1809396 | 0.999175987 |
| ENSMUSG00000078435 | AU041133 | protein_coding | -1.293453156 | 2.924900475 | 1.9534416 | 0.1809414 | 0.999175987 |
| ENSMUSG00000030306 | Tmtc1 | protein_coding | -0.685083893 | 4.54557722 | 1.9521351 | 0.1810794 | 0.999175987 |
| ENSMUSG00000015599 | Ttbk1 | protein_coding | 1.077989396 | 3.213615904 | 1.9509579 | 0.1812039 | 0.999175987 |

| ensembl_gene_id | external_gene_name | gene_biotype | log2 (fold change) | log2 (counts per million) | F-value | p-value | False discovery rate |
|---|---|---|---|---|---|---|---|
| ENSMUSG00000020289 | Nprl3 | protein_coding | 0.600885169 | 5.770974429 | 1.9488699 | 0.1814249 | 0.999175987 |
| ENSMUSG00000027599 | Armc1 | protein_coding | 0.484809276 | 7.614774014 | 1.946808 | 0.1816435 | 0.999175987 |
| ENSMUSG00000070730 | Rmdn3 | protein_coding | 0.790921126 | 4.036268146 | 1.9465752 | 0.1816682 | 0.999175987 |
| ENSMUSG00000036959 | Bcorl1 | protein_coding | -0.89921507 | 3.776822993 | 1.9465048 | 0.1816757 | 0.999175987 |
| ENSMUSG00000031540 | Kat6a | protein_coding | -0.81740545 | 4.057204075 | 1.945961 | 0.1817334 | 0.999175987 |
| ENSMUSG00000033423 | Eri3 | protein_coding | 0.384023817 | 8.52696204 | 1.944145 | 0.1819264 | 0.999175987 |
| ENSMUSG00000046722 | Cdc42se1 | protein_coding | -0.581727414 | 5.214550562 | 1.9440744 | 0.1819339 | 0.999175987 |
| ENSMUSG00000003363 | Pld3 | protein_coding | 0.452478193 | 9.931445965 | 1.9415704 | 0.1822004 | 0.999175987 |
| ENSMUSG00000044477 | Zfand3 | protein_coding | 0.693755854 | 4.897481481 | 1.9390087 | 0.1824735 | 0.999175987 |
| ENSMUSG00000029152 | Ociad1 | protein_coding | 0.451610359 | 8.870748976 | 1.9367642 | 0.1827132 | 0.999175987 |
| ENSMUSG00000034681 | Rnps1 | protein_coding | 0.537318513 | 6.11300136 | 1.9356919 | 0.1828279 | 0.999175987 |
| ENSMUSG00000027952 | Pmvk | protein_coding | 0.41023839 | 8.011103667 | 1.9355582 | 0.1828422 | 0.999175987 |
| ENSMUSG00000040936 | Ulk4 | protein_coding | -0.887321293 | 3.71758564 | 1.9352739 | 0.1828726 | 0.999175987 |
| ENSMUSG00000019943 | Atp2b1 | protein_coding | -0.506016706 | 8.203782665 | 1.9341226 | 0.1829958 | 0.999175987 |
| ENSMUSG00000001227 | Sema6b | protein_coding | 1.129280444 | 3.384376543 | 1.9325618 | 0.1831631 | 0.999175987 |
| ENSMUSG00000051111 | Sv2c | protein_coding | 0.726073557 | 4.970558256 | 1.931647 | 0.1832612 | 0.999175987 |
| ENSMUSG00000021258 | Ccnk | protein_coding | -1.126615957 | 3.107589317 | 1.931474 | 0.1832798 | 0.999175987 |
| ENSMUSG00000024725 | Ostf1 | protein_coding | -1.068251171 | 3.297632081 | 1.9308935 | 0.1833421 | 0.999175987 |
| ENSMUSG00000045205 | Dpy19l4 | protein_coding | -0.821230097 | 3.949632797 | 1.9303422 | 0.1834013 | 0.999175987 |
| ENSMUSG00000023020 | Cox14 | protein_coding | 0.517507467 | 8.847201036 | 1.9264454 | 0.1838204 | 0.999175987 |
| ENSMUSG00000032411 | Tfdp2 | protein_coding | -0.779312432 | 4.105395577 | 1.9256777 | 0.1839031 | 0.999175987 |
| ENSMUSG00000021803 | Cdhr1 | protein_coding | -1.429169025 | 2.655449623 | 1.9255017 | 0.1839221 | 0.999175987 |
| ENSMUSG00000013833 | Med16 | protein_coding | 0.74935331 | 3.914232501 | 1.9253905 | 0.183934 | 0.999175987 |
| ENSMUSG00000034706 | Dnaic2 | protein_coding | -0.853020099 | 3.603919497 | 1.9240411 | 0.1840796 | 0.999175987 |
| ENSMUSG00000045790 | Ccdc149 | protein_coding | -0.713858206 | 4.636312051 | 1.9237926 | 0.1841064 | 0.999175987 |
| ENSMUSG00000059278 | Naa38 | protein_coding | 0.520353755 | 7.56424352 | 1.9230148 | 0.1841903 | 0.999175987 |
| ENSMUSG00000031144 | Syp | protein_coding | 0.698273985 | 6.858672657 | 1.9323116 | 0.1843717 | 0.999175987 |
| ENSMUSG00000024937 | Ehbp1l1 | protein_coding | 0.559252447 | 5.694630554 | 1.9203053 | 0.1844832 | 0.999175987 |
| ENSMUSG00000020766 | Galk1 | protein_coding | 0.845125109 | 4.268817448 | 1.9198446 | 0.184533 | 0.999175987 |
| ENSMUSG00000018882 | Mrpl45 | protein_coding | -0.433513349 | 7.34872276 | 1.9195876 | 0.1845609 | 0.999175987 |
| ENSMUSG00000031931 | Ankrd49 | protein_coding | 0.646778128 | 4.469545221 | 1.9183979 | 0.1846897 | 0.999175987 |
| ENSMUSG00000068457 | Uty | protein_coding | -0.950809353 | 3.355291775 | 1.9142302 | 0.1851419 | 0.999175987 |
| ENSMUSG00000026098 | Pms1 | protein_coding | -1.540849146 | 2.571576675 | 1.9139935 | 0.1851676 | 0.999175987 |
| ENSMUSG00000086003 | B230206L02Rik | processed_transcript | -0.775106394 | 4.057111651 | 1.912725 | 0.1853056 | 0.999175987 |
| ENSMUSG00000028602 | Tnfrsf8 | protein_coding | 2.535911005 | 1.854737579 | 1.9125178 | 0.1853281 | 0.999175987 |
| ENSMUSG00000051412 | Vamp7 | protein_coding | -0.512857518 | 6.712283676 | 1.9120881 | 0.1853749 | 0.999175987 |
| ENSMUSG00000036022 | Fam122b | protein_coding | -1.032958679 | 3.233161775 | 1.912052 | 0.1853788 | 0.999175987 |
| ENSMUSG00000051041 | Olfml1 | protein_coding | -0.709184663 | 5.013972872 | 1.9118948 | 0.1853959 | 0.999175987 |
| ENSMUSG00000060548 | Tnfrsf19 | protein_coding | -1.054529371 | 3.548605899 | 1.9113674 | 0.1854534 | 0.999175987 |
| ENSMUSG00000021814 | Anxa7 | protein_coding | 0.525072037 | 6.427187044 | 1.9096157 | 0.1856443 | 0.999175987 |
| ENSMUSG00000059974 | Ntm | protein_coding | -0.421185427 | 7.518673752 | 1.9094749 | 0.1856596 | 0.999175987 |
| ENSMUSG00000024135 | Srbd1 | protein_coding | -1.222543719 | 3.426965269 | 1.9095377 | 0.1856832 | 0.999175987 |
| ENSMUSG00000003344 | Btbd2 | protein_coding | 0.487918426 | 6.043958302 | 1.9088396 | 0.1857289 | 0.999175987 |
| ENSMUSG00000030770 | Parva | protein_coding | -0.552531859 | 5.503169889 | 1.9087465 | 0.1857391 | 0.999175987 |
| ENSMUSG00000039347 | Atp6v0e2 | protein_coding | 0.446766054 | 10.37809293 | 1.9077862 | 0.1858439 | 0.999175987 |
| ENSMUSG00000051817 | Sox12 | protein_coding | 0.902430234 | 3.284725338 | 1.907065 | 0.1859227 | 0.999175987 |
| ENSMUSG00000047423 | AI837181 | protein_coding | 0.510250544 | 6.38112456 | 1.9063503 | 0.1860008 | 0.999175987 |
| ENSMUSG00000007021 | Syngr3 | protein_coding | 0.449361773 | 9.000692261 | 1.9063418 | 0.1860017 | 0.999175987 |
| ENSMUSG00000078681 | Tm2d3 | protein_coding | 0.436130998 | 8.32003089 | 1.9062621 | 0.1860104 | 0.999175987 |
| ENSMUSG00000061762 | Tac1 | protein_coding | -0.755140628 | 8.511015986 | 1.9279594 | 0.1860611 | 0.999175987 |
| ENSMUSG00000041837 | Pdcd7 | protein_coding | 0.860121786 | 4.364299716 | 1.9052823 | 0.1861175 | 0.999175987 |
| ENSMUSG00000028273 | Pdlim5 | protein_coding | -0.946516852 | 4.080537185 | 1.9042221 | 0.1862336 | 0.999175987 |
| ENSMUSG00000042453 | Reln | protein_coding | -0.926274952 | 4.225070867 | 1.9039985 | 0.1862581 | 0.999175987 |
| ENSMUSG00000021488 | Nsd1 | protein_coding | -0.506622195 | 6.332371186 | 1.9037254 | 0.186288 | 0.999175987 |
| ENSMUSG00000044164 | Rnf182 | protein_coding | -1.192893503 | 3.699817746 | 1.900181 | 0.186707 | 0.999175987 |
| ENSMUSG00000019927 | Ube2d1 | protein_coding | 0.466570514 | 6.169708397 | 1.8979512 | 0.1869217 | 0.999175987 |
| ENSMUSG00000040022 | Rab11fip2 | protein_coding | -0.638046603 | 4.496235975 | 1.8979074 | 0.1869265 | 0.999175987 |
| ENSMUSG00000026473 | Glul | protein_coding | -0.437741445 | 8.303558326 | 1.8956207 | 0.1871782 | 0.999175987 |
| ENSMUSG00000024764 | Naa40 | protein_coding | -0.946883774 | 3.508491131 | 1.8925502 | 0.1875169 | 0.999175987 |
| ENSMUSG00000034371 | Tkfc | protein_coding | 0.83094252 | 3.857554488 | 1.8914017 | 0.1876438 | 0.999175987 |
| ENSMUSG00000035704 | Alg8 | protein_coding | 1.078915978 | 3.612258277 | 1.8907616 | 0.1877146 | 0.999175987 |
| ENSMUSG00000087184 | Gm11650 | processed_transcript | 1.484270201 | 2.571897185 | 1.8884601 | 0.1879693 | 0.999175987 |
| ENSMUSG00000020919 | Stat5b | protein_coding | -0.770195193 | 4.271669641 | 1.8883866 | 0.1879774 | 0.999175987 |
| ENSMUSG00000045107 | Saysd1 | protein_coding | 0.593293063 | 5.068478322 | 1.8883404 | 0.1879825 | 0.999175987 |
| ENSMUSG00000057110 | Cntrl | protein_coding | -1.798720525 | 2.464555657 | 1.8872804 | 0.1881 | 0.999175987 |
| ENSMUSG00000061702 | Tmem91 | protein_coding | 0.436103097 | 7.338949389 | 1.8871812 | 0.188111 | 0.999175987 |
| ENSMUSG00000032369 | Plscr1 | protein_coding | -1.656408139 | 2.352033512 | 1.8864335 | 0.1881939 | 0.999175987 |
| ENSMUSG00000000253 | Gmpr | protein_coding | 0.389594891 | 7.122825237 | 1.8860611 | 0.1882352 | 0.999175987 |
| ENSMUSG00000014551 | Mrps25 | protein_coding | 0.400179869 | 7.710675343 | 1.8854378 | 0.1883044 | 0.999175987 |
| ENSMUSG00000034949 | Zfr2 | protein_coding | 0.89204328 | 3.528318463 | 1.8849349 | 0.1883602 | 0.999175987 |
| ENSMUSG00000029534 | St7 | protein_coding | 0.666502005 | 4.278496417 | 1.8848015 | 0.1883751 | 0.999175987 |
| ENSMUSG00000010047 | Hyal2 | protein_coding | 0.764447567 | 4.857066359 | 1.8843962 | 0.1884201 | 0.999175987 |
| ENSMUSG00000095567 | Noc2l | protein_coding | -0.719022506 | 4.321235495 | 1.8819416 | 0.188693 | 0.999175987 |
| ENSMUSG00000042298 | Ttc19 | protein_coding | -0.603642702 | 5.377593297 | 1.8811087 | 0.1887857 | 0.999175987 |
| ENSMUSG00000020212 | Mdm1 | protein_coding | -0.937058095 | 3.198376845 | 1.8810119 | 0.1887965 | 0.999175987 |
| ENSMUSG00000063260 | Syt10 | protein_coding | -0.814414291 | 3.929621671 | 1.8796948 | 0.1889433 | 0.999175987 |
| ENSMUSG00000040242 | Fgfr1op2 | protein_coding | 0.440897939 | 6.371582917 | 1.8769747 | 0.1892468 | 0.999175987 |
| ENSMUSG00000005370 | Msh6 | protein_coding | -1.888947571 | 2.915605067 | 1.8863829 | 0.189375 | 0.999175987 |

| ensembl_gene_id | external_gene_name | gene_biotype | log2 (fold change) | log2 (counts per million) | F-value | p-value | False discovery rate |
|---|---|---|---|---|---|---|---|
| ENSMUSG00000041997 | Tlk1 | protein_coding | -0.531774602 | 5.412870422 | 1.8752949 | 0.1894346 | 0.999175987 |
| ENSMUSG00000037742 | Eef1a1 | protein_coding | 0.436313363 | 10.30163358 | 1.8741569 | 0.1895619 | 0.999175987 |
| ENSMUSG00000032193 | Ldlr | protein_coding | 1.248137918 | 3.684855885 | 1.8749233 | 0.1897767 | 0.999175987 |
| ENSMUSG00000062101 | Zfp119b | protein_coding | -1.692931382 | 2.779244344 | 1.871715 | 0.1898658 | 0.999175987 |
| ENSMUSG00000041889 | Shisa4 | protein_coding | 0.421585205 | 6.555733141 | 1.8696992 | 0.1900618 | 0.999175987 |
| ENSMUSG00000018415 | Gid4 | protein_coding | -0.60608618 | 5.948252585 | 1.8690942 | 0.1901297 | 0.999175987 |
| ENSMUSG00000034343 | Ube2f | protein_coding | 0.455999249 | 6.296843582 | 1.8682326 | 0.1902266 | 0.999175987 |
| ENSMUSG00000040620 | Dhx33 | protein_coding | -1.195040241 | 2.79761533 | 1.8673854 | 0.1903219 | 0.999175987 |
| ENSMUSG00000036339 | Tmem260 | protein_coding | 1.015258842 | 3.371221441 | 1.8667849 | 0.1903895 | 0.999175987 |
| ENSMUSG00000021032 | Ngb | protein_coding | 0.475757252 | 8.359207565 | 1.8619772 | 0.1909316 | 0.999175987 |
| ENSMUSG00000092416 | Zfp141 | protein_coding | -1.081359781 | 3.008401772 | 1.8612911 | 0.1910091 | 0.999175987 |
| ENSMUSG00000025370 | Cdh9 | protein_coding | -1.644184056 | 2.458546631 | 1.8611139 | 0.1910291 | 0.999175987 |
| ENSMUSG00000052560 | Cpne8 | protein_coding | -0.608943858 | 5.446482968 | 1.8604277 | 0.1911067 | 0.999175987 |
| ENSMUSG00000059810 | Rgs3 | protein_coding | 0.985638886 | 3.626404954 | 1.859346 | 0.1912291 | 0.999175987 |
| ENSMUSG00000027968 | Larp7 | protein_coding | -0.612856159 | 5.413909771 | 1.8588512 | 0.1912851 | 0.999175987 |
| ENSMUSG00000027698 | Nceh1 | protein_coding | 0.373583747 | 7.061547712 | 1.8575241 | 0.1914354 | 0.999175987 |
| ENSMUSG00000029466 | Anapc7 | protein_coding | 0.571496146 | 5.210385149 | 1.8569986 | 0.191495 | 0.999175987 |
| ENSMUSG00000031557 | Plekha2 | protein_coding | -1.556267613 | 2.553370924 | 1.8542616 | 0.1918056 | 0.999175987 |
| ENSMUSG00000025860 | Xiap | protein_coding | -0.474117813 | 6.136989412 | 1.8542438 | 0.1918077 | 0.999175987 |
| ENSMUSG00000026163 | Sphkap | protein_coding | 0.484918673 | 5.666542243 | 1.8538466 | 0.1918528 | 0.999175987 |
| ENSMUSG00000022548 | Apod | protein_coding | -0.500687876 | 8.762868782 | 1.8529974 | 0.1919494 | 0.999175987 |
| ENSMUSG00000021932 | Rnaseh2b | protein_coding | 0.583462539 | 5.32804282 | 1.8521327 | 0.1920477 | 0.999175987 |
| ENSMUSG00000042505 | Sdhaf3 | protein_coding | 0.472584995 | 6.69178615 | 1.8521285 | 0.1920482 | 0.999175987 |
| ENSMUSG00000005949 | Ctns | protein_coding | -1.066030786 | 3.660435599 | 1.8521079 | 0.1920505 | 0.999175987 |
| ENSMUSG00000050002 | Idnk | protein_coding | 0.459307655 | 7.140977413 | 1.8489203 | 0.1924137 | 0.999175987 |
| ENSMUSG00000055839 | Elob | protein_coding | 0.444400801 | 9.024725631 | 1.847704 | 0.1925525 | 0.999175987 |
| ENSMUSG00000023186 | Vwa5a | protein_coding | -1.166266403 | 2.970044968 | 1.8471454 | 0.1926163 | 0.999175987 |
| ENSMUSG00000009356 | Lpo | protein_coding | -2.519127715 | 2.031690728 | 1.8464554 | 0.1927253 | 0.999175987 |
| ENSMUSG00000040410 | Fbxl4 | protein_coding | 0.63743927 | 4.835194913 | 1.845706 | 0.1927808 | 0.999175987 |
| ENSMUSG00000033361 | Prrg3 | protein_coding | -0.865061258 | 4.063571141 | 1.8456586 | 0.1927862 | 0.999175987 |
| ENSMUSG00000055026 | Gabrg3 | protein_coding | 0.633352327 | 4.684126774 | 1.8455346 | 0.1928004 | 0.999175987 |
| ENSMUSG00000002475 | Abhd3 | protein_coding | -0.661426067 | 5.700202732 | 1.8451496 | 0.1928444 | 0.999175987 |
| ENSMUSG00000026664 | Phyh | protein_coding | -0.363102338 | 8.053546593 | 1.8412338 | 0.193293 | 0.999175987 |
| ENSMUSG00000000751 | Rpa1 | protein_coding | -0.635856928 | 6.402588931 | 1.8401276 | 0.1934501 | 0.999175987 |
| ENSMUSG00000034252 | Senp6 | protein_coding | -0.657185984 | 5.136205662 | 1.8384514 | 0.1936125 | 0.999175987 |
| ENSMUSG00000022340 | Sybu | protein_coding | -0.408494399 | 6.519935658 | 1.8383016 | 0.1936298 | 0.999175987 |
| ENSMUSG00000039202 | Abhd2 | protein_coding | -0.750694067 | 4.758151013 | 1.8377018 | 0.1936988 | 0.999175987 |
| ENSMUSG00000027834 | Serpini1 | protein_coding | -0.448157881 | 6.329136714 | 1.8375111 | 0.1937207 | 0.999175987 |
| ENSMUSG00000025283 | Sat1 | protein_coding | -0.477196791 | 6.838112687 | 1.837232 | 0.1937528 | 0.999175987 |
| ENSMUSG00000037351 | Actr1b | protein_coding | 0.515695144 | 8.367555443 | 1.8369234 | 0.1937883 | 0.999175987 |
| ENSMUSG00000030067 | Foxp1 | protein_coding | -0.417290886 | 7.041363291 | 1.8355279 | 0.193949 | 0.999175987 |
| ENSMUSG00000032399 | Rpl4 | protein_coding | 0.428121653 | 10.60178081 | 1.8337797 | 0.1941506 | 0.999175987 |
| ENSMUSG00000024427 | Spry4 | protein_coding | -0.71528551 | 4.300934509 | 1.8333119 | 0.1942046 | 0.999175987 |
| ENSMUSG00000046574 | Prr12 | protein_coding | 0.903832248 | 4.780336741 | 1.833233 | 0.1942438 | 0.999175987 |
| ENSMUSG00000028256 | Odf2l | protein_coding | 1.207306219 | 3.174813505 | 1.8317911 | 0.1943802 | 0.999175987 |
| ENSMUSG00000071253 | Slc25a16 | protein_coding | -0.680901001 | 4.729946211 | 1.8311642 | 0.1944526 | 0.999175987 |
| ENSMUSG00000034687 | Fras1 | protein_coding | -2.087127936 | 2.484782701 | 1.8315258 | 0.1945203 | 0.999175987 |
| ENSMUSG00000038095 | Sbno1 | protein_coding | -0.520690218 | 6.455760382 | 1.8293963 | 0.1946571 | 0.999175987 |
| ENSMUSG00000021143 | Pacs2 | protein_coding | 0.589579137 | 5.061377253 | 1.8291203 | 0.1946891 | 0.999175987 |
| ENSMUSG00000073423 | Zfp414 | protein_coding | 0.503274669 | 6.22489882 | 1.8275205 | 0.1948744 | 0.999175987 |
| ENSMUSG00000040428 | Plekha4 | protein_coding | 1.257776944 | 2.958626033 | 1.826884 | 0.1949482 | 0.999175987 |
| ENSMUSG00000021091 | Serpina3n | protein_coding | 0.623192627 | 4.794840131 | 1.8265059 | 0.194992 | 0.999175987 |
| ENSMUSG00000067931 | Zfp948 | protein_coding | -0.916089767 | 3.557058881 | 1.8262275 | 0.1950243 | 0.999175987 |
| ENSMUSG00000041891 | Lman1 | protein_coding | 0.956903594 | 3.955768525 | 1.826082 | 0.1950412 | 0.999175987 |
| ENSMUSG00000020770 | Unk | protein_coding | -1.409395738 | 3.006113874 | 1.8225863 | 0.1954474 | 0.999175987 |
| ENSMUSG00000037752 | Xkr8 | protein_coding | -1.297743269 | 2.798958383 | 1.8216944 | 0.1955512 | 0.999175987 |
| ENSMUSG00000038342 | Mlxip | protein_coding | -1.44765975 | 2.475320782 | 1.8213955 | 0.195586 | 0.999175987 |
| ENSMUSG00000048251 | Bcl11b | protein_coding | -0.518378423 | 6.005800764 | 1.8199141 | 0.1957586 | 0.999175987 |
| ENSMUSG00000024065 | Ehd3 | protein_coding | 0.563182873 | 6.826757013 | 1.8184067 | 0.1959645 | 0.999175987 |
| ENSMUSG00000018669 | Cdk5rap3 | protein_coding | 0.548384653 | 6.585328663 | 1.8170465 | 0.1960932 | 0.999175987 |
| ENSMUSG00000024914 | Drap1 | protein_coding | 0.415301541 | 6.442584975 | 1.8166572 | 0.1961387 | 0.999175987 |
| ENSMUSG00000039059 | Hrh3 | protein_coding | 0.835047339 | 3.775898326 | 1.8165729 | 0.1961485 | 0.999175987 |
| ENSMUSG00000040007 | Bahd1 | protein_coding | -1.18258142 | 3.270660705 | 1.8145433 | 0.1963859 | 0.999175987 |
| ENSMUSG00000026965 | Anapc2 | protein_coding | -0.478724779 | 6.07619777 | 1.8145205 | 0.1963886 | 0.999175987 |
| ENSMUSG00000037735 | 2810032G03Rik | lincRNA | -1.242277513 | 2.874450947 | 1.81415 | 0.196432 | 0.999175987 |
| ENSMUSG00000041143 | Tmco4 | protein_coding | 1.534974371 | 2.414615072 | 1.8115732 | 0.196734 | 0.999175987 |
| ENSMUSG00000047747 | Rnf150 | protein_coding | -0.868758621 | 4.084373128 | 1.8112746 | 0.196769 | 0.999175987 |
| ENSMUSG00000025899 | Alkbh8 | protein_coding | -0.546213693 | 5.523790829 | 1.8111301 | 0.1967859 | 0.999175987 |
| ENSMUSG00000022949 | Clic6 | protein_coding | -1.34308766 | 4.046555903 | 1.8259256 | 0.1968868 | 0.999175987 |
| ENSMUSG00000027487 | Cdk5rap1 | protein_coding | -0.792081356 | 4.290621699 | 1.8099245 | 0.1969275 | 0.999175987 |
| ENSMUSG00000020629 | Adi1 | protein_coding | 0.525939101 | 6.30891027 | 1.8092883 | 0.1970022 | 0.999175987 |
| ENSMUSG00000015943 | Bola1 | protein_coding | 0.413019011 | 6.70976838 | 1.8079619 | 0.1971582 | 0.999175987 |
| ENSMUSG00000061536 | Sec22c | protein_coding | -0.614868993 | 4.617431643 | 1.8077575 | 0.1971822 | 0.999175987 |
| ENSMUSG00000021219 | Rgs6 | protein_coding | 0.858589825 | 4.009210378 | 1.8063402 | 0.197349 | 0.999175987 |
| ENSMUSG00000021794 | Glud1 | protein_coding | -0.521493888 | 6.788155854 | 1.804059 | 0.1976179 | 0.999175987 |
| ENSMUSG00000036860 | Mrpl55 | protein_coding | 0.41022989 | 7.83164215 | 1.8035449 | 0.1976786 | 0.999175987 |
| ENSMUSG00000054519 | Zfp867 | protein_coding | -0.846469122 | 3.704421995 | 1.8023624 | 0.1978182 | 0.999175987 |
| ENSMUSG00000024498 | Tcerg1 | protein_coding | -0.621482205 | 5.115507489 | 1.802333 | 0.1978217 | 0.999175987 |

| ensembl_gene_id | external_gene_name | gene_biotype | log2 (fold change) | log2 (counts per million) | F-value | p-value | False discovery rate |
|---|---|---|---|---|---|---|---|
| ENSMUSG00000022885 | St6gal1 | protein_coding | -1.034247318 | 3.565684672 | 1.8005927 | 0.1980274 | 0.999175987 |
| ENSMUSG00000041126 | H2afv | protein_coding | -0.50868017 | 5.493399484 | 1.8003527 | 0.1980558 | 0.999175987 |
| ENSMUSG00000027315 | Spint1 | protein_coding | -1.595179888 | 2.528566975 | 1.799459 | 0.1981616 | 0.999175987 |
| ENSMUSG00000064326 | Siva1 | protein_coding | 0.604545623 | 5.736593877 | 1.7989331 | 0.1982238 | 0.999175987 |
| ENSMUSG00000002617 | Zfp40 | protein_coding | -1.040634584 | 3.713377652 | 1.798368 | 0.1982908 | 0.999175987 |
| ENSMUSG00000017344 | Vtn | protein_coding | -0.722850667 | 4.202792313 | 1.7978306 | 0.1983545 | 0.999175987 |
| ENSMUSG00000060512 | 0610040J01Rik | protein_coding | -2.112789033 | 2.003837371 | 1.7977064 | 0.1983692 | 0.999175987 |
| ENSMUSG00000020235 | Fzr1 | protein_coding | 0.473987112 | 6.145964474 | 1.796886 | 0.1984665 | 0.999175987 |
| ENSMUSG00000055128 | Cgrrf1 | protein_coding | -0.493002287 | 6.209589568 | 1.7964449 | 0.1985188 | 0.999175987 |
| ENSMUSG00000022124 | Fbxl3 | protein_coding | -0.587882986 | 5.126446197 | 1.7962462 | 0.1985424 | 0.999175987 |
| ENSMUSG00000109536 | 9330162G02Rik | TEC | -0.885393138 | 3.428943043 | 1.7953613 | 0.1986474 | 0.999175987 |
| ENSMUSG00000026753 | Ppp6c | protein_coding | -0.416217805 | 6.718598413 | 1.7953601 | 0.1986476 | 0.999175987 |
| ENSMUSG00000026701 | Prdx6 | protein_coding | -0.433738153 | 9.748615925 | 1.7931999 | 0.1989043 | 0.999175987 |
| ENSMUSG00000066595 | Mfsd7b | protein_coding | 1.084745342 | 3.027479416 | 1.7931158 | 0.1989143 | 0.999175987 |
| ENSMUSG00000022802 | Lmln | protein_coding | 1.565277792 | 2.455071012 | 1.7927705 | 0.1989554 | 0.999175987 |
| ENSMUSG00000028792 | Ak2 | protein_coding | 0.583050416 | 5.347829628 | 1.7915062 | 0.1991059 | 0.999175987 |
| ENSMUSG00000068922 | Msto1 | protein_coding | 0.595083806 | 4.987511977 | 1.7914659 | 0.1991107 | 0.999175987 |
| ENSMUSG00000031667 | Aktip | protein_coding | 0.45138609 | 8.165522375 | 1.7905704 | 0.1992174 | 0.999175987 |
| ENSMUSG00000041881 | Ndufa7 | protein_coding | 0.496721879 | 9.535408849 | 1.789204 | 0.1993803 | 0.999175987 |
| ENSMUSG00000041439 | Mfsd6 | protein_coding | 0.53501299 | 5.493051545 | 1.7889075 | 0.1994157 | 0.999175987 |
| ENSMUSG00000014837 | 4931428F04Rik | protein_coding | 0.888157939 | 5.338054235 | 1.7982268 | 0.1994694 | 0.999175987 |
| ENSMUSG00000047153 | Khnyn | protein_coding | 1.536127073 | 2.890848492 | 1.7879329 | 0.199562 | 0.999175987 |
| ENSMUSG00000019039 | Dalrd3 | protein_coding | 0.671793997 | 5.753914959 | 1.7871036 | 0.1996611 | 0.999175987 |
| ENSMUSG00000029781 | Fkbp9 | protein_coding | 1.207887655 | 3.42077307 | 1.787046 | 0.199668 | 0.999175987 |
| ENSMUSG00000031792 | Usb1 | protein_coding | 0.716486446 | 4.800074597 | 1.7865091 | 0.1997022 | 0.999175987 |
| ENSMUSG00000046215 | Rprml | protein_coding | 1.095373219 | 3.693100388 | 1.7857782 | 0.1998195 | 0.999175987 |
| ENSMUSG00000100147 | 1700047M11Rik | lincRNA | -0.738578208 | 4.785041665 | 1.7851855 | 0.1998605 | 0.999175987 |
| ENSMUSG00000001100 | Poldip2 | protein_coding | -0.475448 | 5.884132122 | 1.7850712 | 0.1998742 | 0.999175987 |
| ENSMUSG00000060373 | Hnrnpc | protein_coding | -0.380142039 | 7.866966741 | 1.7831501 | 0.2001043 | 0.999175987 |
| ENSMUSG00000060147 | Serpinb6a | protein_coding | -0.547774659 | 6.124552557 | 1.7830805 | 0.2001126 | 0.999175987 |
| ENSMUSG00000031600 | Vps37a | protein_coding | -0.533015293 | 5.384052491 | 1.7827188 | 0.200156 | 0.999175987 |
| ENSMUSG00000026830 | Ermn | protein_coding | -0.375513433 | 7.384948842 | 1.7827144 | 0.2001565 | 0.999175987 |
| ENSMUSG00000057729 | Prtn3 | protein_coding | 2.040968007 | 2.570975663 | 1.7836474 | 0.2002051 | 0.999175987 |
| ENSMUSG00000024576 | Csnk1a1 | protein_coding | -0.504804404 | 7.22639045 | 1.7822363 | 0.2002139 | 0.999175987 |
| ENSMUSG00000057541 | Pus7 | protein_coding | -1.256652553 | 2.835257955 | 1.780584 | 0.2004122 | 0.999175987 |
| ENSMUSG00000051354 | Samd3 | protein_coding | -1.369034992 | 3.681003651 | 1.7890036 | 0.2005646 | 0.999175987 |
| ENSMUSG00000033256 | Shf | protein_coding | -1.267754533 | 2.677663578 | 1.7789021 | 0.2006143 | 0.999175987 |
| ENSMUSG00000025218 | Poll | protein_coding | 0.662757143 | 6.066550268 | 1.7780399 | 0.2007479 | 0.999175987 |
| ENSMUSG00000028643 | Svbp | protein_coding | 0.542967024 | 5.704019942 | 1.7776186 | 0.2007687 | 0.999175987 |
| ENSMUSG00000064254 | Ethe1 | protein_coding | 0.743885723 | 4.879331073 | 1.7747638 | 0.2011127 | 0.999175987 |
| ENSMUSG00000071014 | Ndufb6 | protein_coding | 0.400275061 | 8.649696776 | 1.7740471 | 0.2011991 | 0.999175987 |
| ENSMUSG00000052456 | Asna1 | protein_coding | 0.382349695 | 8.054059734 | 1.7727524 | 0.2013555 | 0.999175987 |
| ENSMUSG00000040820 | Hlcs | protein_coding | 0.857837334 | 3.496479684 | 1.7724808 | 0.2013883 | 0.999175987 |
| ENSMUSG00000040037 | Negr1 | protein_coding | -0.551608511 | 6.345821179 | 1.7710764 | 0.2015581 | 0.999175987 |
| ENSMUSG00000030604 | Zfp626 | protein_coding | -1.312351304 | 2.699631647 | 1.7709131 | 0.2015778 | 0.999175987 |
| ENSMUSG00000113029 | AC122190.1 | antisense_RNA | -1.077148993 | 3.301354643 | 1.7699538 | 0.2016939 | 0.999175987 |
| ENSMUSG00000029048 | Rer1 | protein_coding | 0.465162925 | 6.511122146 | 1.7672504 | 0.2020216 | 0.999175987 |
| ENSMUSG00000032766 | Gng11 | protein_coding | -0.504436685 | 6.068835467 | 1.7672372 | 0.2020232 | 0.999175987 |
| ENSMUSG00000019841 | Rev3l | protein_coding | -0.862323559 | 4.223288506 | 1.7664134 | 0.2021231 | 0.999175987 |
| ENSMUSG00000055723 | Rras2 | protein_coding | -1.919099059 | 2.084447914 | 1.7657001 | 0.2022097 | 0.999175987 |
| ENSMUSG00000024847 | Aip | protein_coding | 0.409654144 | 8.863937582 | 1.7646381 | 0.2023388 | 0.999175987 |
| ENSMUSG00000071649 | B3gat3 | protein_coding | 0.436456386 | 6.366993141 | 1.7644613 | 0.2023603 | 0.999175987 |
| ENSMUSG00000024924 | Vldlr | protein_coding | -0.713618732 | 5.759781989 | 1.7646716 | 0.2023645 | 0.999175987 |
| ENSMUSG00000047193 | Dync2h1 | protein_coding | -0.64431811 | 5.172442651 | 1.7639327 | 0.2024245 | 0.999175987 |
| ENSMUSG00000021737 | Psmd6 | protein_coding | -0.410547684 | 8.008013468 | 1.7634467 | 0.2024837 | 0.999175987 |
| ENSMUSG00000062627 | Mysm1 | protein_coding | -0.560071673 | 5.109075998 | 1.7619855 | 0.2026615 | 0.999175987 |
| ENSMUSG00000078812 | Eif5a | protein_coding | 0.440584736 | 9.469711168 | 1.7614048 | 0.2027323 | 0.999175987 |
| ENSMUSG00000030498 | Gas2 | protein_coding | 0.66644874 | 4.081728623 | 1.7606043 | 0.2028299 | 0.999175987 |
| ENSMUSG00000000223 | Drp2 | protein_coding | -0.754157095 | 4.106157728 | 1.7586259 | 0.2030713 | 0.999175987 |
| ENSMUSG00000033730 | Egr3 | protein_coding | -0.772009812 | 4.995065314 | 1.7581572 | 0.2031285 | 0.999175987 |
| ENSMUSG00000059456 | Ptk2b | protein_coding | -0.467080589 | 7.305700514 | 1.7575229 | 0.203206 | 0.999175987 |
| ENSMUSG00000037514 | Pank2 | protein_coding | -0.780341658 | 4.32589004 | 1.7568259 | 0.2032912 | 0.999175987 |
| ENSMUSG00000035566 | Pcdh17 | protein_coding | 0.374810333 | 6.755909387 | 1.7555645 | 0.2034456 | 0.999175987 |
| ENSMUSG00000041506 | Rrp9 | protein_coding | 0.615850933 | 4.809833241 | 1.755036 | 0.2035103 | 0.999175987 |
| ENSMUSG00000004665 | Cnn2 | protein_coding | -1.448727961 | 2.957617468 | 1.7546237 | 0.2035608 | 0.999175987 |
| ENSMUSG00000025868 | Higd2a | protein_coding | 0.37227285 | 8.450003104 | 1.7544325 | 0.2035842 | 0.999175987 |
| ENSMUSG00000028383 | Hsdl2 | protein_coding | -0.494212356 | 5.550690746 | 1.7523256 | 0.2038425 | 0.999175987 |
| ENSMUSG00000025876 | Unc5a | protein_coding | 0.927053234 | 4.321028332 | 1.7500213 | 0.2041553 | 0.999175987 |
| ENSMUSG00000024604 | Rbm22 | protein_coding | -0.436523227 | 5.880171391 | 1.7489085 | 0.2042623 | 0.999175987 |
| ENSMUSG00000048410 | Zfp407 | protein_coding | 0.946453677 | 3.295447265 | 1.7477929 | 0.2043997 | 0.999175987 |
| ENSMUSG00000051557 | Pusl1 | protein_coding | 0.910464473 | 4.982828726 | 1.7531236 | 0.2045043 | 0.999175987 |
| ENSMUSG00000039456 | Morc3 | protein_coding | -0.895150076 | 4.556758285 | 1.7469907 | 0.2045282 | 0.999175987 |
| ENSMUSG00000038121 | Fam210a | protein_coding | -0.595783823 | 5.42499516 | 1.7463885 | 0.2045727 | 0.999175987 |
| ENSMUSG00000020400 | Tnip1 | protein_coding | -0.699234064 | 5.062295397 | 1.7460131 | 0.2046189 | 0.999175987 |
| ENSMUSG00000030310 | Slc6a1 | protein_coding | -0.512260148 | 7.111988007 | 1.7453307 | 0.2047031 | 0.999175987 |
| ENSMUSG00000061455 | Stx17 | protein_coding | -0.676932869 | 4.505653527 | 1.7446399 | 0.2047883 | 0.999175987 |
| ENSMUSG00000040272 | Accs | protein_coding | 1.959845319 | 2.430935469 | 1.7441595 | 0.2048774 | 0.999175987 |
| ENSMUSG00000032298 | Neil1 | protein_coding | -1.451273779 | 3.125484663 | 1.7431318 | 0.2050043 | 0.999175987 |

| ensembl_gene_id | external_gene_name | gene_biotype | log2 (fold change) | log2 (counts per million) | F-value | p-value | False discovery rate |
|---|---|---|---|---|---|---|---|
| ENSMUSG00000059316 | Slc27a4 | protein_coding | 0.594973229 | 5.374491072 | 1.7428716 | 0.2050068 | 0.999175987 |
| ENSMUSG00000086841 | 2410006H16Rik | processed_transcript | 0.464357215 | 7.585014337 | 1.7424473 | 0.2050592 | 0.999175987 |
| ENSMUSG00000100594 | 2810414N06Rik | lincRNA | -1.027722147 | 3.632830618 | 1.7402445 | 0.2053318 | 0.999175987 |
| ENSMUSG00000029790 | Cep41 | protein_coding | 1.209007679 | 3.099118787 | 1.7387015 | 0.205523 | 0.999175987 |
| ENSMUSG00000029725 | Ppp1r35 | protein_coding | 1.266093819 | 2.750171303 | 1.7385701 | 0.2055393 | 0.999175987 |
| ENSMUSG00000028780 | Sema3c | protein_coding | -1.600738149 | 2.825147627 | 1.7379977 | 0.20564 | 0.999175987 |
| ENSMUSG00000020964 | Sel1l | protein_coding | -0.466155323 | 5.974239127 | 1.7371082 | 0.2057207 | 0.999175987 |
| ENSMUSG00000042389 | Tsen2 | protein_coding | 0.565406863 | 5.267071006 | 1.736992 | 0.2057351 | 0.999175987 |
| ENSMUSG00000063296 | Tmem117 | protein_coding | -1.229292105 | 3.037931534 | 1.7360762 | 0.2058488 | 0.999175987 |
| ENSMUSG00000042109 | Csdc2 | protein_coding | 0.446724485 | 8.009879572 | 1.7353795 | 0.2059354 | 0.999175987 |
| ENSMUSG00000032081 | Apoc3 | protein_coding | 1.094641374 | 3.179063449 | 1.7340869 | 0.2060962 | 0.999175987 |
| ENSMUSG00000044471 | Lncpint | lincRNA | 1.838789492 | 2.492313321 | 1.7336875 | 0.2061459 | 0.999175987 |
| ENSMUSG00000053553 | 3110082I17Rik | protein_coding | 0.673130607 | 4.48468091 | 1.7336416 | 0.2061516 | 0.999175987 |
| ENSMUSG00000012405 | Rpl15 | protein_coding | 0.406369026 | 6.940425499 | 1.7301062 | 0.2065923 | 0.999175987 |
| ENSMUSG00000024829 | Mrpl21 | protein_coding | 0.381399842 | 7.109861112 | 1.72989 | 0.2066192 | 0.999175987 |
| ENSMUSG00000027775 | Mfsd1 | protein_coding | 0.938680128 | 3.452838777 | 1.729137 | 0.2067133 | 0.999175987 |
| ENSMUSG00000024772 | Ehd1 | protein_coding | -0.584176574 | 4.516743433 | 1.7266844 | 0.2070199 | 0.999175987 |
| ENSMUSG00000022248 | Rad1 | protein_coding | -0.571817772 | 4.790968335 | 1.7250695 | 0.2072221 | 0.999175987 |
| ENSMUSG00000019230 | Lhx9 | protein_coding | -1.569191226 | 4.271214065 | 1.7659708 | 0.2072546 | 0.999175987 |
| ENSMUSG00000022009 | Nufip1 | protein_coding | 0.761210981 | 3.825513868 | 1.7241128 | 0.207342 | 0.999175987 |
| ENSMUSG00000046287 | Pnma3 | protein_coding | -0.875348194 | 4.538704125 | 1.7241531 | 0.2073666 | 0.999175987 |
| ENSMUSG00000021519 | Mterf3 | protein_coding | 0.505751663 | 5.873411583 | 1.7238502 | 0.207375 | 0.999175987 |
| ENSMUSG00000007837 | Prrg2 | protein_coding | -0.9442681 | 3.673805446 | 1.7235238 | 0.2074159 | 0.999175987 |
| ENSMUSG00000001039 | B9d1 | protein_coding | 0.515758191 | 5.906474746 | 1.7230707 | 0.2074728 | 0.999175987 |
| ENSMUSG00000042055 | Wdr11 | protein_coding | 0.633110869 | 4.757641252 | 1.722319 | 0.2075671 | 0.999175987 |
| ENSMUSG00000074749 | Kiz | protein_coding | -0.810235116 | 4.178520559 | 1.7210419 | 0.2077275 | 0.999175987 |
| ENSMUSG00000029130 | Rnf32 | protein_coding | -0.638925799 | 4.393035802 | 1.7200046 | 0.207858 | 0.999175987 |
| ENSMUSG00000026617 | Bpnt1 | protein_coding | 0.418290679 | 6.995437354 | 1.7191583 | 0.2079645 | 0.999175987 |
| ENSMUSG00000030042 | Pole4 | protein_coding | 0.519378319 | 5.649741215 | 1.7169797 | 0.2082389 | 0.999175987 |
| ENSMUSG00000040359 | Ufl1 | protein_coding | -1.002497956 | 2.978506641 | 1.7168389 | 0.2082567 | 0.999175987 |
| ENSMUSG00000056296 | Synpr | protein_coding | 0.545790301 | 7.160050266 | 1.7168134 | 0.2082895 | 0.999175987 |
| ENSMUSG00000038910 | Plcl2 | protein_coding | -0.700269977 | 5.087735322 | 1.7151389 | 0.2084712 | 0.999175987 |
| ENSMUSG00000070923 | Klhl9 | protein_coding | -0.421074859 | 8.232878104 | 1.7145302 | 0.208548 | 0.999175987 |
| ENSMUSG00000032413 | Rasa2 | protein_coding | -0.699138205 | 4.582664916 | 1.7143653 | 0.2085689 | 0.999175987 |
| ENSMUSG00000029569 | Tmem168 | protein_coding | -1.219664709 | 3.063842105 | 1.7136516 | 0.2086591 | 0.999175987 |
| ENSMUSG00000052369 | Tmem106c | protein_coding | 0.373518998 | 7.361881729 | 1.7124998 | 0.2088047 | 0.999175987 |
| ENSMUSG00000024142 | Mlst8 | protein_coding | 0.848569354 | 4.379486391 | 1.7122068 | 0.2088418 | 0.999175987 |
| ENSMUSG00000034160 | Ogt | protein_coding | -0.625453205 | 5.72437829 | 1.7112856 | 0.2089584 | 0.999175987 |
| ENSMUSG00000024457 | Trim26 | protein_coding | 0.687064874 | 4.668947304 | 1.7110808 | 0.2089843 | 0.999175987 |
| ENSMUSG00000033099 | Nol12 | protein_coding | 0.36517448 | 6.686652263 | 1.7105403 | 0.2090528 | 0.999175987 |
| ENSMUSG00000000567 | Sox9 | protein_coding | -0.558770287 | 5.735015818 | 1.7100068 | 0.2091204 | 0.999175987 |
| ENSMUSG00000021843 | Ktn1 | protein_coding | -0.567098737 | 5.683007978 | 1.7096664 | 0.2091636 | 0.999175987 |
| ENSMUSG00000025145 | Lrrc45 | protein_coding | -1.068876304 | 3.403219882 | 1.7070532 | 0.2094953 | 0.999175987 |
| ENSMUSG00000037674 | Rfx7 | protein_coding | 0.782030696 | 3.692786048 | 1.7069382 | 0.2095099 | 0.999175987 |
| ENSMUSG00000023800 | Tiam2 | protein_coding | -0.796466663 | 4.999777228 | 1.7046526 | 0.2098301 | 0.999175987 |
| ENSMUSG00000012848 | Rps5 | protein_coding | 0.480623609 | 10.12689909 | 1.7043269 | 0.209842 | 0.999175987 |
| ENSMUSG00000058420 | Syt17 | protein_coding | -0.682208749 | 5.326430569 | 1.703438 | 0.2099552 | 0.999175987 |
| ENSMUSG00000040356 | Skiv2l | protein_coding | 0.630892408 | 4.674149618 | 1.703273 | 0.2099762 | 0.999175987 |
| ENSMUSG00000071073 | Lrrc73 | protein_coding | 0.787912862 | 3.816430592 | 1.7030968 | 0.2099987 | 0.999175987 |
| ENSMUSG00000007950 | Abhd8 | protein_coding | 0.484253028 | 6.008048681 | 1.7030851 | 0.2100002 | 0.999175987 |
| ENSMUSG00000049184 | Purg | protein_coding | -0.613941845 | 4.934903238 | 1.702954 | 0.2100169 | 0.999175987 |
| ENSMUSG00000023088 | Abcc1 | protein_coding | 2.195931215 | 2.924059921 | 1.7222348 | 0.210103 | 0.999175987 |
| ENSMUSG00000050705 | 2310061I04Rik | protein_coding | 0.509166845 | 5.638533226 | 1.7021268 | 0.2101224 | 0.999175987 |
| ENSMUSG00000040269 | Mrps28 | protein_coding | 0.39784314 | 6.852699931 | 1.7007616 | 0.2102966 | 0.999175987 |
| ENSMUSG00000072640 | Lyrm9 | protein_coding | 0.555611439 | 6.788462245 | 1.6991314 | 0.2105344 | 0.999175987 |
| ENSMUSG00000042043 | Tbca | protein_coding | 0.517339751 | 7.893308059 | 1.6986756 | 0.2105926 | 0.999175987 |
| ENSMUSG00000047824 | Pygo2 | protein_coding | 0.68745444 | 4.81998272 | 1.6980123 | 0.2106479 | 0.999175987 |
| ENSMUSG00000036398 | Ppp1r11 | protein_coding | 0.374171372 | 9.340274285 | 1.6978589 | 0.2106676 | 0.999175987 |
| ENSMUSG00000062797 | Hikeshi | protein_coding | 0.364312495 | 7.780947549 | 1.6971387 | 0.2107598 | 0.999175987 |
| ENSMUSG00000031548 | Sfrp1 | protein_coding | -1.506979726 | 2.630183538 | 1.6966764 | 0.210819 | 0.999175987 |
| ENSMUSG00000058135 | Gstm1 | protein_coding | -0.456579078 | 8.942497461 | 1.6966679 | 0.21082 | 0.999175987 |
| ENSMUSG00000022176 | Rem2 | protein_coding | -1.792824917 | 2.818293959 | 1.7012918 | 0.2108274 | 0.999175987 |
| ENSMUSG00000102437 | Gm38048 | sense_overlapping | 1.154256568 | 3.008105649 | 1.6963414 | 0.2108619 | 0.999175987 |
| ENSMUSG00000030401 | Rtn2 | protein_coding | 0.630118976 | 5.087100593 | 1.6961452 | 0.210887 | 0.999175987 |
| ENSMUSG00000038022 | Mindy4 | protein_coding | -1.056455581 | 3.076901349 | 1.6957074 | 0.2109431 | 0.999175987 |
| ENSMUSG00000023367 | Tmem176a | protein_coding | -0.680052767 | 5.30402105 | 1.6939211 | 0.2111722 | 0.999175987 |
| ENSMUSG00000020220 | Vps13d | protein_coding | 0.836019712 | 4.030778721 | 1.6936547 | 0.2112064 | 0.999175987 |
| ENSMUSG00000043391 | 2510009E07Rik | protein_coding | -0.54530427 | 5.119740428 | 1.693193 | 0.2112657 | 0.999175987 |
| ENSMUSG00000009734 | Pou6f2 | protein_coding | -1.790108401 | 2.355691237 | 1.692702 | 0.2113288 | 0.999175987 |
| ENSMUSG00000058006 | Mdn1 | protein_coding | 1.249032944 | 3.343406399 | 1.6925021 | 0.2113839 | 0.999175987 |
| ENSMUSG00000027189 | Trim44 | protein_coding | -0.441902266 | 6.742039693 | 1.6918519 | 0.211438 | 0.999175987 |
| ENSMUSG00000032300 | 1700017B05Rik | protein_coding | -1.403430964 | 3.158139168 | 1.6918733 | 0.2114648 | 0.999175987 |
| ENSMUSG00000028121 | Bcar3 | protein_coding | -0.847889295 | 4.360199005 | 1.6914413 | 0.2114908 | 0.999175987 |
| ENSMUSG00000026924 | Sec16a | protein_coding | -0.721666373 | 3.965881154 | 1.6908315 | 0.2115692 | 0.999175987 |
| ENSMUSG00000039637 | Coro7 | protein_coding | 0.589654295 | 4.868359133 | 1.689775 | 0.2117052 | 0.999175987 |
| ENSMUSG00000086657 | Stamos | processed_transcript | -1.388057074 | 2.476351875 | 1.6893907 | 0.2117547 | 0.999175987 |
| ENSMUSG00000020063 | Sirt1 | protein_coding | 1.078652768 | 2.913816698 | 1.6893856 | 0.2117554 | 0.999175987 |
| ENSMUSG00000032202 | Rab27a | protein_coding | 0.932606473 | 3.891648485 | 1.6890811 | 0.2117946 | 0.999175987 |

| ensembl_gene_id | external_gene_name | gene_biotype | log2 (fold change) | log2 (counts per million) | F-value | p-value | False discovery rate |
|---|---|---|---|---|---|---|---|
| ENSMUSG00000028051 | Hcn3 | protein_coding | -0.918569618 | 3.320938554 | 1.688901 | 0.2118178 | 0.999175987 |
| ENSMUSG00000021371 | Mcur1 | protein_coding | -1.049963698 | 3.262011593 | 1.6877157 | 0.2119706 | 0.999175987 |
| ENSMUSG00000005575 | Ube2m | protein_coding | 0.474934189 | 6.758331894 | 1.6864673 | 0.2121317 | 0.999175987 |
| ENSMUSG00000114309 | AC163038.1 | bidirectional_promote | 1.252788054 | 3.07934206 | 1.6860243 | 0.2121889 | 0.999175987 |
| ENSMUSG00000050608 | Minos1 | protein_coding | 0.464507742 | 9.318764556 | 1.6855437 | 0.212251 | 0.999175987 |
| ENSMUSG00000038633 | Degs1 | protein_coding | 0.360530185 | 7.198477339 | 1.6854321 | 0.2122654 | 0.999175987 |
| ENSMUSG00000042707 | Dnali1 | protein_coding | -1.133014176 | 3.274985517 | 1.6836443 | 0.2124965 | 0.999175987 |
| ENSMUSG00000017858 | Ift52 | protein_coding | -0.508134931 | 6.539735167 | 1.6817027 | 0.2127479 | 0.999175987 |
| ENSMUSG00000035164 | Zc3h12c | protein_coding | -0.968088085 | 3.641309505 | 1.6816825 | 0.2127505 | 0.999175987 |
| ENSMUSG00000068263 | Efcc1 | protein_coding | 0.816980836 | 3.664385717 | 1.6811222 | 0.2128231 | 0.999175987 |
| ENSMUSG00000030168 | Adipor2 | protein_coding | -0.900882544 | 4.827109208 | 1.684108 | 0.2129361 | 0.999175987 |
| ENSMUSG00000022386 | Trmu | protein_coding | 0.667316737 | 5.064305744 | 1.6785781 | 0.2131533 | 0.999175987 |
| ENSMUSG00000031883 | Car7 | protein_coding | 0.85466201 | 3.743672053 | 1.6784423 | 0.2131709 | 0.999175987 |
| ENSMUSG00000054302 | Eapp | protein_coding | -0.409690155 | 7.104930646 | 1.6774091 | 0.2133052 | 0.999175987 |
| ENSMUSG00000002064 | Sdf2 | protein_coding | -0.438715776 | 7.649452526 | 1.6757109 | 0.2135261 | 0.999175987 |
| ENSMUSG00000039568 | Ubald1 | protein_coding | 0.434236371 | 6.558367213 | 1.6756308 | 0.2135365 | 0.999175987 |
| ENSMUSG00000002379 | Ndufa11 | protein_coding | 0.418972707 | 9.817108866 | 1.6753831 | 0.2135688 | 0.999175987 |
| ENSMUSG00000039070 | Cpa4 | protein_coding | -1.768070579 | 2.618529772 | 1.6730979 | 0.213896 | 0.999175987 |
| ENSMUSG00000038690 | Atp5j2 | protein_coding | 0.503524982 | 9.890778011 | 1.6723927 | 0.2139586 | 0.999175987 |
| ENSMUSG00000035517 | Tdrd7 | protein_coding | 0.837628188 | 4.36072328 | 1.6721348 | 0.2139923 | 0.999175987 |
| ENSMUSG00000062393 | Dgkk | protein_coding | 0.722320949 | 4.040301523 | 1.6720351 | 0.2140053 | 0.999175987 |
| ENSMUSG00000019578 | Ubxn6 | protein_coding | 0.442435457 | 6.444794577 | 1.671614 | 0.2140603 | 0.999175987 |
| ENSMUSG00000028522 | Mier1 | protein_coding | -0.516976075 | 5.329494853 | 1.6714036 | 0.2140878 | 0.999175987 |
| ENSMUSG00000056515 | Rab31 | protein_coding | -0.428315828 | 6.519935439 | 1.6710728 | 0.214131 | 0.999175987 |
| ENSMUSG00000031521 | Aga | protein_coding | -0.795858912 | 4.549416809 | 1.6692986 | 0.214363 | 0.999175987 |
| ENSMUSG00000046982 | Tshz1 | protein_coding | -0.650054417 | 4.906798199 | 1.6691867 | 0.2143776 | 0.999175987 |
| ENSMUSG00000028367 | Txn1 | protein_coding | 0.392999282 | 8.31226565 | 1.6683258 | 0.2144903 | 0.999175987 |
| ENSMUSG00000022364 | Tbc1d31 | protein_coding | -1.561265938 | 2.316712019 | 1.6671569 | 0.2146434 | 0.999175987 |
| ENSMUSG00000029622 | Arpc1b | protein_coding | 0.668527031 | 4.752222559 | 1.6668425 | 0.2146846 | 0.999175987 |
| ENSMUSG00000041078 | Grid1 | protein_coding | 0.524542444 | 5.258684022 | 1.6662518 | 0.2147621 | 0.999175987 |
| ENSMUSG00000022043 | Trim35 | protein_coding | -0.403466031 | 6.707283872 | 1.6661942 | 0.2147696 | 0.999175987 |
| ENSMUSG00000044573 | Acp1 | protein_coding | 0.415161718 | 6.066362044 | 1.666154 | 0.2147749 | 0.999175987 |
| ENSMUSG00000026556 | Vangl2 | protein_coding | -0.791084326 | 3.631927552 | 1.6642377 | 0.2150264 | 0.999175987 |
| ENSMUSG00000050908 | Tvp23a | protein_coding | 0.780355587 | 3.957495416 | 1.6632236 | 0.2151597 | 0.999175987 |
| ENSMUSG00000038526 | Car14 | protein_coding | -0.811569892 | 4.815138323 | 1.6631177 | 0.2152029 | 0.999175987 |
| ENSMUSG00000020444 | Guk1 | protein_coding | 0.637539529 | 7.484912509 | 1.6735511 | 0.2152725 | 0.999175987 |
| ENSMUSG00000003662 | Ciao1 | protein_coding | 0.478013039 | 6.247825886 | 1.6622871 | 0.2152828 | 0.999175987 |
| ENSMUSG00000049760 | 2410015M20Rik | protein_coding | 0.431228925 | 8.724607971 | 1.6617755 | 0.2153501 | 0.999175987 |
| ENSMUSG00000034639 | Setmar | protein_coding | 1.223524897 | 2.899282814 | 1.6589049 | 0.2157283 | 0.999175987 |
| ENSMUSG00000042810 | Krba1 | protein_coding | 0.75191692 | 3.77363301 | 1.6579158 | 0.2158589 | 0.999175987 |
| ENSMUSG00000015714 | Cers2 | protein_coding | -0.458729908 | 6.226918209 | 1.6575998 | 0.2159006 | 0.999175987 |
| ENSMUSG00000033209 | Ttc28 | protein_coding | 1.201301321 | 2.80421923 | 1.6570842 | 0.2159687 | 0.999175987 |
| ENSMUSG00000031154 | Otud5 | protein_coding | -0.635009817 | 5.270545097 | 1.6570062 | 0.215979 | 0.999175987 |
| ENSMUSG00000032998 | Foxj3 | protein_coding | -0.576709673 | 5.043641327 | 1.6562258 | 0.2160821 | 0.999175987 |
| ENSMUSG00000019437 | Tlcd1 | protein_coding | 0.77940491 | 4.300287122 | 1.6560319 | 0.2161077 | 0.999175987 |
| ENSMUSG00000040260 | Daam2 | protein_coding | -0.695736114 | 4.34735342 | 1.6557875 | 0.21614 | 0.999175987 |
| ENSMUSG00000000266 | Mid2 | protein_coding | -0.853608072 | 4.204205419 | 1.6556519 | 0.216158 | 0.999175987 |
| ENSMUSG00000038390 | Gpr162 | protein_coding | 0.741552072 | 4.424986241 | 1.6552329 | 0.2162134 | 0.999175987 |
| ENSMUSG00000031820 | Babam1 | protein_coding | 0.437575888 | 7.129763984 | 1.6546784 | 0.2162868 | 0.999175987 |
| ENSMUSG00000025505 | Tmem80 | protein_coding | 0.539791825 | 6.001325798 | 1.6546627 | 0.2162888 | 0.999175987 |
| ENSMUSG00000030551 | Nr2f2 | protein_coding | 0.694052654 | 6.558364138 | 1.6658063 | 0.2163218 | 0.999175987 |
| ENSMUSG00000002031 | Ift46 | protein_coding | 0.475428731 | 7.052698323 | 1.6540807 | 0.2163659 | 0.999175987 |
| ENSMUSG00000045007 | Tubg2 | protein_coding | 0.535039399 | 6.830641441 | 1.6527082 | 0.216577 | 0.999175987 |
| ENSMUSG00000042302 | Ehbp1 | protein_coding | -0.81081206 | 4.174940949 | 1.6517387 | 0.2166762 | 0.999175987 |
| ENSMUSG00000028753 | Vwa5b1 | protein_coding | -0.861430676 | 3.62853526 | 1.6515825 | 0.216697 | 0.999175987 |
| ENSMUSG00000057156 | Homez | protein_coding | 0.798213734 | 3.484538364 | 1.6513797 | 0.2167239 | 0.999175987 |
| ENSMUSG00000024608 | Rps14 | protein_coding | 0.622681178 | 10.3060541 | 1.6563182 | 0.2168173 | 0.999175987 |
| ENSMUSG00000021676 | Iqgap2 | protein_coding | -1.161379109 | 3.054315862 | 1.6504132 | 0.2168522 | 0.999175987 |
| ENSMUSG00000050552 | Lamtor4 | protein_coding | 0.448657422 | 7.544119523 | 1.6484091 | 0.2171185 | 0.999175987 |
| ENSMUSG00000033809 | Alg3 | protein_coding | 0.643897291 | 4.917568556 | 1.6483088 | 0.2171318 | 0.999175987 |
| ENSMUSG00000045128 | Rpl18a | protein_coding | 0.487089322 | 8.363054074 | 1.6478812 | 0.2171887 | 0.999175987 |
| ENSMUSG00000030203 | Dusp16 | protein_coding | -1.005656442 | 3.462242681 | 1.647725 | 0.2172095 | 0.999175987 |
| ENSMUSG00000022972 | 1110004E09Rik | protein_coding | 0.789691716 | 6.017678376 | 1.6609743 | 0.2172359 | 0.999175987 |
| ENSMUSG00000041632 | Mrps27 | protein_coding | -0.655471135 | 5.571094879 | 1.6459521 | 0.2174456 | 0.999175987 |
| ENSMUSG00000000568 | Hnrnpd | protein_coding | -0.452020599 | 6.889537929 | 1.6457824 | 0.2174682 | 0.999175987 |
| ENSMUSG00000105879 | Gm6204 | processed_pseudoge | 0.771782024 | 3.852311768 | 1.6443741 | 0.2176559 | 0.999175987 |
| ENSMUSG00000053898 | Ech1 | protein_coding | -0.371231553 | 7.728086844 | 1.6439999 | 0.2177059 | 0.999175987 |
| ENSMUSG00000030315 | Vgll4 | protein_coding | 0.566536198 | 4.932348267 | 1.642238 | 0.2179412 | 0.999175987 |
| ENSMUSG00000029778 | Adcyap1r1 | protein_coding | -0.464858605 | 6.178430116 | 1.6419566 | 0.2179788 | 0.999175987 |
| ENSMUSG00000044149 | Nkrf | protein_coding | 0.712744766 | 4.948426562 | 1.6418351 | 0.217995 | 0.999175987 |
| ENSMUSG00000033740 | St18 | protein_coding | -0.714072658 | 3.954821333 | 1.6407509 | 0.21814 | 0.999175987 |
| ENSMUSG00000062590 | Armc9 | protein_coding | 0.545146904 | 5.167580818 | 1.6392401 | 0.2183423 | 0.999175987 |
| ENSMUSG00000024773 | Atg2a | protein_coding | 0.989303056 | 3.524198162 | 1.6389508 | 0.218381 | 0.999175987 |
| ENSMUSG00000025290 | Rps24 | protein_coding | 0.529407702 | 8.061290589 | 1.6374381 | 0.218613 | 0.999175987 |
| ENSMUSG00000028528 | Dnajc6 | protein_coding | -0.391296421 | 6.82525502 | 1.6365704 | 0.2187003 | 0.999175987 |
| ENSMUSG00000028080 | Lrba | protein_coding | -0.970957132 | 3.827978285 | 1.636183 | 0.2187523 | 0.999175987 |
| ENSMUSG00000057103 | Nat8f1 | protein_coding | -0.648292817 | 4.708141597 | 1.6355059 | 0.2188432 | 0.999175987 |
| ENSMUSG00000034675 | Dbn1 | protein_coding | 0.483312445 | 6.113673448 | 1.6350796 | 0.2189005 | 0.999175987 |

| ensembl_gene_id | external_gene_name | gene_biotype | log2 (fold change) | log2 (counts per million) | F-value | p-value | False discovery rate |
|---|---|---|---|---|---|---|---|
| ENSMUSG00000054452 | Aes | protein_coding | 0.401062319 | 7.901939294 | 1.634916 | 0.2189225 | 0.999175987 |
| ENSMUSG00000072694 | 1500011B03Rik | protein_coding | 0.344148713 | 8.042027602 | 1.6344211 | 0.218989 | 0.999175987 |
| ENSMUSG00000049321 | Zfp2 | protein_coding | 0.74642881 | 4.650543512 | 1.6340235 | 0.2190425 | 0.999175987 |
| ENSMUSG00000025262 | Fam120c | protein_coding | -1.270792144 | 3.31235211 | 1.6333882 | 0.2191572 | 0.999175987 |
| ENSMUSG00000038457 | Tmem255b | protein_coding | -1.985931203 | 2.342129433 | 1.6328821 | 0.2192253 | 0.999175987 |
| ENSMUSG00000015709 | Arnt2 | protein_coding | -0.457249938 | 6.141255029 | 1.6326055 | 0.2192333 | 0.999175987 |
| ENSMUSG00000041708 | Mpped1 | protein_coding | 0.519137013 | 5.241067827 | 1.632191 | 0.2192891 | 0.999175987 |
| ENSMUSG00000006728 | Cdk4 | protein_coding | 0.392356201 | 7.036813166 | 1.631979 | 0.2193177 | 0.999175987 |
| ENSMUSG00000041740 | Rnf10 | protein_coding | 0.499650628 | 5.532524674 | 1.6319341 | 0.2193238 | 0.999175987 |
| ENSMUSG00000085664 | Atxn7l1os2 | antisense_RNA | 1.202952357 | 2.93795205 | 1.6318222 | 0.2193388 | 0.999175987 |
| ENSMUSG00000015243 | Abca1 | protein_coding | -0.947705787 | 3.073190224 | 1.6313868 | 0.2193975 | 0.999175987 |
| ENSMUSG00000074578 | Zfas1 | processed_transcript | 0.540778692 | 7.938346674 | 1.6303294 | 0.2195693 | 0.999175987 |
| ENSMUSG00000021670 | Hmgcr | protein_coding | -0.470295807 | 5.731228708 | 1.6300871 | 0.2195728 | 0.999175987 |
| ENSMUSG00000040472 | Rabggta | protein_coding | 0.476826643 | 6.178422067 | 1.6297277 | 0.2196213 | 0.999175987 |
| ENSMUSG00000055214 | Pld5 | protein_coding | -0.544390485 | 5.240425973 | 1.6288796 | 0.2197358 | 0.999175987 |
| ENSMUSG00000031671 | Setd6 | protein_coding | 1.217485636 | 3.194264437 | 1.6287737 | 0.2197501 | 0.999175987 |
| ENSMUSG00000038503 | Mesd | protein_coding | -0.430429179 | 6.570156497 | 1.6286555 | 0.2197661 | 0.999175987 |
| ENSMUSG00000034083 | Ccdc174 | protein_coding | -0.43772923 | 5.939229856 | 1.6284764 | 0.2197902 | 0.999175987 |
| ENSMUSG00000015839 | Nfe2l2 | protein_coding | -0.82688179 | 4.716166321 | 1.627834 | 0.2199062 | 0.999175987 |
| ENSMUSG00000022332 | Khdrbs3 | protein_coding | 0.453839875 | 6.130064683 | 1.6248848 | 0.2202761 | 0.999175987 |
| ENSMUSG00000028382 | Ptbp3 | protein_coding | -0.781626462 | 3.931620566 | 1.6239315 | 0.2204053 | 0.999175987 |
| ENSMUSG00000056820 | Tsnax | protein_coding | -0.467851593 | 6.500801362 | 1.6235036 | 0.2204634 | 0.999175987 |
| ENSMUSG00000029033 | Acap3 | protein_coding | 0.596462942 | 4.663657039 | 1.6230302 | 0.2205276 | 0.999175987 |
| ENSMUSG00000028998 | Tomm7 | protein_coding | 0.503832499 | 8.606368385 | 1.6231679 | 0.220538 | 0.999175987 |
| ENSMUSG00000027971 | Ndst4 | protein_coding | -0.875628807 | 3.514078181 | 1.6223062 | 0.2206258 | 0.999175987 |
| ENSMUSG00000025477 | Inpp5a | protein_coding | -0.696967284 | 4.403235227 | 1.6219069 | 0.2206801 | 0.999175987 |
| ENSMUSG00000003623 | Crot | protein_coding | -0.660869899 | 4.772818687 | 1.6216783 | 0.2207111 | 0.999175987 |
| ENSMUSG00000040651 | Fam208a | protein_coding | -0.939255066 | 3.492858956 | 1.6208177 | 0.220828 | 0.999175987 |
| ENSMUSG00000063802 | Hspbp1 | protein_coding | 0.44476559 | 7.8338441 | 1.6207284 | 0.2208402 | 0.999175987 |
| ENSMUSG00000018199 | Trove2 | protein_coding | -0.48260078 | 5.844787989 | 1.6204822 | 0.2208736 | 0.999175987 |
| ENSMUSG00000022389 | Tef | protein_coding | -0.343526551 | 7.112904169 | 1.6193328 | 0.22103 | 0.999175987 |
| ENSMUSG00000021983 | Atp8a2 | protein_coding | -0.717581904 | 4.255534243 | 1.6185649 | 0.2211345 | 0.999175987 |
| ENSMUSG00000044447 | Dock5 | protein_coding | -1.454956975 | 2.442328391 | 1.6184868 | 0.2211451 | 0.999175987 |
| ENSMUSG00000027330 | Cdc25b | protein_coding | -1.107315384 | 3.054587964 | 1.6180689 | 0.221202 | 0.999175987 |
| ENSMUSG00000073427 | Gm4924 | protein_coding | -0.876072638 | 3.329425983 | 1.6177212 | 0.2212494 | 0.999175987 |
| ENSMUSG00000006676 | Usp19 | protein_coding | 0.696389447 | 3.875609456 | 1.6174916 | 0.2212807 | 0.999175987 |
| ENSMUSG00000031772 | Cntnap4 | protein_coding | -0.952903086 | 3.277471789 | 1.617277 | 0.2213099 | 0.999175987 |
| ENSMUSG00000022469 | Rapgef3 | protein_coding | -0.833937179 | 3.899572658 | 1.6161649 | 0.2214616 | 0.999175987 |
| ENSMUSG00000043964 | Orai3 | protein_coding | 0.875302442 | 3.812328674 | 1.6157897 | 0.2215128 | 0.999175987 |
| ENSMUSG00000002308 | Cd320 | protein_coding | 0.663600655 | 4.901515219 | 1.6156736 | 0.2215286 | 0.999175987 |
| ENSMUSG00000029463 | Fam216a | protein_coding | 0.773702332 | 4.12607671 | 1.6156138 | 0.2215368 | 0.999175987 |
| ENSMUSG00000020700 | Map3k3 | protein_coding | 1.039920776 | 3.35893511 | 1.6140845 | 0.2217456 | 0.999175987 |
| ENSMUSG00000048833 | Slc39a9 | protein_coding | 0.548746098 | 4.910425204 | 1.6131286 | 0.2218763 | 0.999175987 |
| ENSMUSG00000001786 | Fbxo7 | protein_coding | -0.559849494 | 5.935291912 | 1.6128318 | 0.2219169 | 0.999175987 |
| ENSMUSG00000056367 | Actr3b | protein_coding | -0.454433007 | 6.426638227 | 1.6123553 | 0.2219821 | 0.999175987 |
| ENSMUSG00000026374 | Tsn | protein_coding | -0.401941359 | 8.515474492 | 1.6098151 | 0.22233 | 0.999175987 |
| ENSMUSG00000029695 | Aass | protein_coding | -2.105525182 | 1.638383169 | 1.6085199 | 0.2225077 | 0.999175987 |
| ENSMUSG00000024393 | Prrc2a | protein_coding | 0.62870756 | 4.643013439 | 1.6077281 | 0.2226164 | 0.999175987 |
| ENSMUSG00000041992 | Rapgef5 | protein_coding | -0.55378603 | 5.380962526 | 1.6076245 | 0.2226306 | 0.999175987 |
| ENSMUSG00000031327 | Chic1 | protein_coding | -0.517595794 | 5.439736471 | 1.6063698 | 0.222803 | 0.999175987 |
| ENSMUSG00000028389 | Zfp37 | protein_coding | -0.639522896 | 4.774334343 | 1.6051749 | 0.2229674 | 0.999175987 |
| ENSMUSG00000026491 | Ahctf1 | protein_coding | -0.950816863 | 3.264716847 | 1.6038045 | 0.223156 | 0.999175987 |
| ENSMUSG00000036751 | Cox6b1 | protein_coding | 0.450487896 | 10.25496019 | 1.6034532 | 0.2232044 | 0.999175987 |
| ENSMUSG00000040659 | Efhd2 | protein_coding | 0.449927529 | 6.788462381 | 1.603045 | 0.2232607 | 0.999175987 |
| ENSMUSG00000024566 | Atp9b | protein_coding | -0.745157664 | 3.993432612 | 1.602137 | 0.2233859 | 0.999175987 |
| ENSMUSG00000029919 | Hpgds | protein_coding | -1.035042452 | 3.063518813 | 1.601572 | 0.2234639 | 0.999175987 |
| ENSMUSG00000025597 | Klhl4 | protein_coding | -0.718427622 | 3.970025512 | 1.6012765 | 0.2235047 | 0.999175987 |
| ENSMUSG00000039068 | Zzz3 | protein_coding | -0.662016314 | 4.790596783 | 1.6012479 | 0.2235086 | 0.999175987 |
| ENSMUSG00000039239 | Tgfb2 | protein_coding | -0.759985479 | 4.117506558 | 1.6009978 | 0.2235431 | 0.999175987 |
| ENSMUSG00000001518 | Itfg2 | protein_coding | 1.049464491 | 2.944837538 | 1.5994741 | 0.2237536 | 0.999175987 |
| ENSMUSG00000039205 | Ciz1 | protein_coding | 0.747308059 | 4.429933006 | 1.5993233 | 0.2237745 | 0.999175987 |
| ENSMUSG00000061286 | Exosc5 | protein_coding | 0.458456418 | 6.021888694 | 1.5988957 | 0.2238336 | 0.999175987 |
| ENSMUSG00000037376 | Trmt6 | protein_coding | -0.573067196 | 5.655510675 | 1.5987418 | 0.2238549 | 0.999175987 |
| ENSMUSG00000048100 | Taf13 | protein_coding | -0.598514472 | 5.28099885 | 1.5982384 | 0.2239245 | 0.999175987 |
| ENSMUSG00000063882 | Uqcrh | protein_coding | 0.526041352 | 10.25289796 | 1.5979121 | 0.2239987 | 0.999175987 |
| ENSMUSG00000038462 | Uqcrfs1 | protein_coding | -0.578012345 | 6.788603422 | 1.5992692 | 0.2240695 | 0.999175987 |
| ENSMUSG00000030802 | Bckdk | protein_coding | 0.555291172 | 6.812872469 | 1.5971647 | 0.2241022 | 0.999175987 |
| ENSMUSG00000028295 | Smim8 | protein_coding | 0.483334268 | 6.499060213 | 1.5966536 | 0.224144 | 0.999175987 |
| ENSMUSG00000046185 | Zfp84 | protein_coding | -1.055300548 | 3.379679398 | 1.5962193 | 0.2242042 | 0.999175987 |
| ENSMUSG00000048930 | Tada3 | protein_coding | 0.540956883 | 4.935587301 | 1.595808 | 0.2242612 | 0.999175987 |
| ENSMUSG00000029029 | Wrap73 | protein_coding | 0.488164855 | 5.753226431 | 1.5947822 | 0.2244034 | 0.999175987 |
| ENSMUSG00000022552 | Sharpin | protein_coding | 0.822943998 | 5.492261759 | 1.6026986 | 0.2244322 | 0.999175987 |
| ENSMUSG00000029253 | Cenpc1 | protein_coding | -0.70565205 | 4.863560757 | 1.5940078 | 0.2245109 | 0.999175987 |
| ENSMUSG00000032352 | Lrrc1 | protein_coding | -0.911173014 | 3.422985212 | 1.5939575 | 0.2245179 | 0.999175987 |
| ENSMUSG00000053004 | Hrh1 | protein_coding | 0.654197409 | 4.110665507 | 1.5935361 | 0.2245764 | 0.999175987 |
| ENSMUSG00000055137 | Sugct | protein_coding | -1.027722329 | 2.948896426 | 1.5932028 | 0.2246227 | 0.999175987 |
| ENSMUSG00000032480 | Dhx30 | protein_coding | 0.500366182 | 6.567190402 | 1.5923301 | 0.224744 | 0.999175987 |
| ENSMUSG00000020354 | Sgcd | protein_coding | -1.290467986 | 2.665280259 | 1.590387 | 0.2250144 | 0.999175987 |

| ensembl_gene_id | external_gene_name | gene_biotype | log2 (fold change) | log2 (counts per million) | F-value | p-value | False discovery rate |
|---|---|---|---|---|---|---|---|
| ENSMUSG00000007207 | Stx1a | protein_coding | 0.63739989 | 7.031155404 | 1.6008058 | 0.2250146 | 0.999175987 |
| ENSMUSG00000060090 | Rp2 | protein_coding | -1.298744038 | 2.647798057 | 1.5888502 | 0.2252285 | 0.999175987 |
| ENSMUSG00000024668 | Sdhaf2 | protein_coding | 0.370972798 | 6.812997948 | 1.5873496 | 0.2254378 | 0.999175987 |
| ENSMUSG00000004849 | Ap1s1 | protein_coding | 0.385673479 | 9.61928205 | 1.5873467 | 0.2254382 | 0.999175987 |
| ENSMUSG00000068290 | Ddrgk1 | protein_coding | 0.438549723 | 7.561920124 | 1.5865525 | 0.2255491 | 0.999175987 |
| ENSMUSG00000047721 | Bola2 | protein_coding | 0.531757983 | 7.920219984 | 1.5866384 | 0.225566 | 0.999175987 |
| ENSMUSG00000041308 | Sntb2 | protein_coding | -1.413873247 | 2.552772567 | 1.5861145 | 0.2256103 | 0.999175987 |
| ENSMUSG00000026727 | Rsu1 | protein_coding | -0.579214524 | 5.540845219 | 1.58592 | 0.2256375 | 0.999175987 |
| ENSMUSG00000044117 | 2900011O08Rik | protein_coding | 0.443028744 | 9.481265429 | 1.5853393 | 0.2257186 | 0.999175987 |
| ENSMUSG00000022312 | Eif3h | protein_coding | 0.37368863 | 8.916982619 | 1.5852293 | 0.225734 | 0.999175987 |
| ENSMUSG00000020462 | Cfap36 | protein_coding | 0.382184572 | 7.174457757 | 1.5833747 | 0.2259935 | 0.999175987 |
| ENSMUSG00000015377 | Dennd6b | protein_coding | 0.898939146 | 3.564111178 | 1.5830199 | 0.2260432 | 0.999175987 |
| ENSMUSG00000013646 | Sh3bp5l | protein_coding | 0.503470044 | 5.210169796 | 1.5830025 | 0.2260456 | 0.999175987 |
| ENSMUSG00000002580 | Mien1 | protein_coding | 0.44725048 | 7.03482004 | 1.5810258 | 0.2263227 | 0.999175987 |
| ENSMUSG00000072214 | 5-Sep | protein_coding | 0.432798193 | 7.012611073 | 1.5804456 | 0.2264041 | 0.999175987 |
| ENSMUSG00000028111 | Ctsk | protein_coding | -1.648281626 | 3.124977368 | 1.5907309 | 0.2264464 | 0.999175987 |
| ENSMUSG00000067150 | Xpo5 | protein_coding | -0.818135847 | 3.970514468 | 1.5800315 | 0.2264623 | 0.999175987 |
| ENSMUSG00000046380 | Jrk | protein_coding | -1.050788383 | 3.524425754 | 1.5797543 | 0.2265012 | 0.999175987 |
| ENSMUSG00000015981 | Stk32c | protein_coding | -1.060387537 | 3.19143195 | 1.5796493 | 0.2265159 | 0.999175987 |
| ENSMUSG00000006442 | Srm | protein_coding | 0.585618168 | 5.229121767 | 1.5785546 | 0.2266697 | 0.999175987 |
| ENSMUSG00000030763 | Lcmt1 | protein_coding | 0.367011565 | 7.912261747 | 1.5783108 | 0.226704 | 0.999175987 |
| ENSMUSG00000024827 | Gldc | protein_coding | -1.486972181 | 2.589045251 | 1.5781623 | 0.2267249 | 0.999175987 |
| ENSMUSG00000102805 | Gm37240 | protein_coding | -0.955368556 | 3.117351977 | 1.5770538 | 0.2268808 | 0.999175987 |
| ENSMUSG00000029614 | Rpl6 | protein_coding | 0.371906277 | 6.90047633 | 1.5765466 | 0.2269522 | 0.999175987 |
| ENSMUSG00000028809 | Srrm1 | protein_coding | -0.382001676 | 7.215659472 | 1.5752072 | 0.2271409 | 0.999175987 |
| ENSMUSG00000033278 | Ptprm | protein_coding | -0.731401366 | 4.481869238 | 1.5733605 | 0.2274014 | 0.999175987 |
| ENSMUSG00000036206 | Sh3bp4 | protein_coding | -0.946077769 | 3.751942726 | 1.5728434 | 0.2274744 | 0.999175987 |
| ENSMUSG00000018501 | Ncor1 | protein_coding | -0.459191475 | 5.848514274 | 1.5724982 | 0.2275231 | 0.999175987 |
| ENSMUSG00000041417 | Pik3r1 | protein_coding | -0.566402501 | 5.653165169 | 1.5723864 | 0.2275389 | 0.999175987 |
| ENSMUSG00000029101 | Rgs12 | protein_coding | -0.729023681 | 3.786857235 | 1.5716701 | 0.2276401 | 0.999175987 |
| ENSMUSG00000039942 | Ptger4 | protein_coding | -0.766006109 | 4.093155842 | 1.5697475 | 0.2279121 | 0.999175987 |
| ENSMUSG00000068523 | Gng5 | protein_coding | -0.660900398 | 4.80179362 | 1.5695854 | 0.227935 | 0.999175987 |
| ENSMUSG00000027134 | Lpcat4 | protein_coding | -0.600283309 | 4.840404694 | 1.5688056 | 0.2280455 | 0.999175987 |
| ENSMUSG00000029659 | Medag | protein_coding | -1.723659937 | 2.621532017 | 1.5685602 | 0.228109 | 0.999175987 |
| ENSMUSG00000070544 | Top1 | protein_coding | 0.489157147 | 6.256643158 | 1.5679554 | 0.228166 | 0.999175987 |
| ENSMUSG00000037605 | Adgrl3 | protein_coding | -0.693155836 | 5.248406716 | 1.566924 | 0.2283122 | 0.999175987 |
| ENSMUSG00000026799 | Med27 | protein_coding | -0.453166955 | 5.612580468 | 1.5667075 | 0.228343 | 0.999175987 |
| ENSMUSG00000031860 | Pbx4 | protein_coding | 0.678567461 | 4.024994436 | 1.566316 | 0.2283985 | 0.999175987 |
| ENSMUSG00000063145 | Bbs5 | protein_coding | -0.635797264 | 4.886398385 | 1.5656171 | 0.2284978 | 0.999175987 |
| ENSMUSG00000025907 | Rb1cc1 | protein_coding | -0.51921742 | 5.252218328 | 1.5650788 | 0.2285742 | 0.999175987 |
| ENSMUSG00000028064 | Sema4a | protein_coding | 0.807823347 | 4.49220998 | 1.5649512 | 0.2285924 | 0.999175987 |
| ENSMUSG00000024805 | Pcgf5 | protein_coding | -0.632406782 | 5.309325547 | 1.564777 | 0.2286171 | 0.999175987 |
| ENSMUSG00000054008 | Ndst1 | protein_coding | -0.892643294 | 4.169782026 | 1.5646924 | 0.2286579 | 0.999175987 |
| ENSMUSG00000040370 | Etfrf1 | protein_coding | -0.435294009 | 6.488541711 | 1.5640099 | 0.2287262 | 0.999175987 |
| ENSMUSG00000065922 | n-R5-8s1 | rRNA | 0.699182572 | 5.306719367 | 1.5635537 | 0.2288199 | 0.999175987 |
| ENSMUSG00000033938 | Ndufb7 | protein_coding | 0.411481353 | 9.54847961 | 1.5633144 | 0.2288251 | 0.999175987 |
| ENSMUSG00000029594 | Rbm19 | protein_coding | 0.996867602 | 3.338040783 | 1.5626702 | 0.2289168 | 0.999175987 |
| ENSMUSG00000015879 | Fam184b | protein_coding | -0.932288006 | 3.132677739 | 1.5623861 | 0.2289573 | 0.999175987 |
| ENSMUSG00000078591 | Hs3st4 | protein_coding | 0.531468284 | 5.317790841 | 1.5614472 | 0.229091 | 0.999175987 |
| ENSMUSG00000026972 | Arrdc1 | protein_coding | -1.163132627 | 3.728158626 | 1.5663574 | 0.2291269 | 0.999175987 |
| ENSMUSG00000019659 | Ccdc12 | protein_coding | 0.441558015 | 5.902746514 | 1.560859 | 0.2291749 | 0.999175987 |
| ENSMUSG00000037994 | Slc9b2 | protein_coding | 1.220469655 | 2.693653155 | 1.5605593 | 0.2292176 | 0.999175987 |
| ENSMUSG00000054612 | Mgmt | protein_coding | 0.976741394 | 3.160805417 | 1.5603801 | 0.2292431 | 0.999175987 |
| ENSMUSG00000026000 | Lancl1 | protein_coding | -0.384940208 | 6.946182531 | 1.5584037 | 0.2295253 | 0.999175987 |
| ENSMUSG00000026810 | Dpm2 | protein_coding | 0.333978199 | 7.574163744 | 1.557813 | 0.2296097 | 0.999175987 |
| ENSMUSG00000042208 | 0610010F05Rik | protein_coding | -0.515546968 | 4.933790896 | 1.5560022 | 0.2298687 | 0.999175987 |
| ENSMUSG00000113902 | AC121965.1 | bidirectional_promote | 0.524300885 | 9.115190095 | 1.5561373 | 0.2298781 | 0.999175987 |
| ENSMUSG00000045174 | Amer3 | protein_coding | -0.880666697 | 4.274527975 | 1.5536143 | 0.2302395 | 0.999175987 |
| ENSMUSG00000025757 | Hspa4l | protein_coding | -0.480985914 | 6.93759472 | 1.5533635 | 0.2302467 | 0.999175987 |
| ENSMUSG00000049327 | Kmt5a | protein_coding | -0.672184371 | 4.106663803 | 1.5525844 | 0.2303585 | 0.999175987 |
| ENSMUSG00000034928 | Rnf44 | protein_coding | -0.52451699 | 5.358573775 | 1.5525651 | 0.2303613 | 0.999175987 |
| ENSMUSG00000027387 | Zc3h8 | protein_coding | 0.736991941 | 3.806664377 | 1.5521211 | 0.2304251 | 0.999175987 |
| ENSMUSG00000009035 | Tmem184b | protein_coding | -0.766899532 | 4.128339592 | 1.5518951 | 0.2304575 | 0.999175987 |
| ENSMUSG00000029166 | Mapre3 | protein_coding | 0.390257875 | 6.801726034 | 1.5517959 | 0.2304718 | 0.999175987 |
| ENSMUSG00000059326 | Csf2ra | protein_coding | 1.18163768 | 2.661422324 | 1.5510813 | 0.2305744 | 0.999175987 |
| ENSMUSG00000029014 | Dnajc2 | protein_coding | -0.594447349 | 4.619513144 | 1.5506687 | 0.2306337 | 0.999175987 |
| ENSMUSG00000024858 | Grk2 | protein_coding | 0.459559608 | 6.36618441 | 1.5503777 | 0.2306755 | 0.999175987 |
| ENSMUSG00000030075 | Cntn3 | protein_coding | -0.879604821 | 3.320332865 | 1.5501733 | 0.2307049 | 0.999175987 |
| ENSMUSG00000002797 | Ggct | protein_coding | 0.542921786 | 6.464061391 | 1.5492112 | 0.2308721 | 0.999175987 |
| ENSMUSG00000039943 | Plcb4 | protein_coding | -0.477014642 | 6.39625355 | 1.5485536 | 0.230938 | 0.999175987 |
| ENSMUSG00000032096 | Arcn1 | protein_coding | 0.466956404 | 5.576845699 | 1.5484493 | 0.230953 | 0.999175987 |
| ENSMUSG00000031681 | Smad1 | protein_coding | -0.601923995 | 4.910375733 | 1.5482631 | 0.2309799 | 0.999175987 |
| ENSMUSG00000010554 | Mettl16 | protein_coding | -0.43144826 | 6.035521682 | 1.5481825 | 0.2309915 | 0.999175987 |
| ENSMUSG00000042816 | Gpr151 | protein_coding | -3.126920244 | 2.063609126 | 1.5646974 | 0.2310803 | 0.999175987 |
| ENSMUSG00000050783 | Htr1f | protein_coding | 1.65221448 | 2.268847071 | 1.5475376 | 0.2310844 | 0.999175987 |
| ENSMUSG00000022404 | Slc25a17 | protein_coding | -0.396549323 | 7.268080084 | 1.5453139 | 0.2314051 | 0.999175987 |
| ENSMUSG00000092329 | Gm20388 | protein_coding | 0.346353231 | 6.876614423 | 1.5443054 | 0.2315508 | 0.999175987 |
| ENSMUSG00000022136 | Dnajc3 | protein_coding | -0.687965393 | 4.487794101 | 1.5437179 | 0.2316357 | 0.999175987 |

| ensembl_gene_id | external_gene_name | gene_biotype | log2 (fold change) | log2 (counts per million) | F-value | p-value | False discovery rate |
|---|---|---|---|---|---|---|---|
| ENSMUSG00000024063 | Lbh | protein_coding | -0.700365843 | 4.762518515 | 1.5426652 | 0.2317879 | 0.999175987 |
| ENSMUSG00000036503 | Rnf13 | protein_coding | -0.377891214 | 7.427291732 | 1.5426347 | 0.2317924 | 0.999175987 |
| ENSMUSG00000043004 | Gng2 | protein_coding | 0.433736101 | 7.764402545 | 1.5425291 | 0.2318076 | 0.999175987 |
| ENSMUSG00000022793 | B4galt4 | protein_coding | -0.975273911 | 3.48912952 | 1.5419282 | 0.2318946 | 0.999175987 |
| ENSMUSG00000025579 | Gaa | protein_coding | 0.393736887 | 9.350906561 | 1.5412715 | 0.2319897 | 0.999175987 |
| ENSMUSG00000062794 | Zfp599 | protein_coding | -0.817449188 | 13.5488801 | 1.5565224 | 0.232059 | 0.999175987 |
| ENSMUSG00000033075 | Senp1 | protein_coding | -1.076506148 | 3.370551733 | 1.5405951 | 0.2320877 | 0.999175987 |
| ENSMUSG00000026443 | Lrrn2 | protein_coding | 0.468412342 | 5.853491411 | 1.5403531 | 0.2321228 | 0.999175987 |
| ENSMUSG00000030094 | Xpc | protein_coding | -0.830798514 | 3.523634083 | 1.5402005 | 0.2321449 | 0.999175987 |
| ENSMUSG00000109946 | D2Bwg1423e | protein_coding | 0.413713062 | 6.58515184 | 1.5401824 | 0.2321475 | 0.999175987 |
| ENSMUSG00000005699 | Pard6a | protein_coding | 0.430809003 | 6.116825136 | 1.5398311 | 0.2321985 | 0.999175987 |
| ENSMUSG00000051098 | Mblac2 | protein_coding | 0.633908051 | 5.543081478 | 1.5383216 | 0.2324175 | 0.999175987 |
| ENSMUSG00000045624 | Esf1 | protein_coding | -0.579481181 | 5.188180045 | 1.5381906 | 0.2324365 | 0.999175987 |
| ENSMUSG00000095620 | 2010005H15Rik | protein_coding | -0.991162348 | 3.44310427 | 1.5377935 | 0.2324942 | 0.999175987 |
| ENSMUSG00000048970 | C1galt1c1 | protein_coding | -0.561020341 | 5.775163264 | 1.5336442 | 0.2330979 | 0.999175987 |
| ENSMUSG00000002205 | Vrk3 | protein_coding | 0.615451312 | 5.766803364 | 1.5335774 | 0.2331363 | 0.999175987 |
| ENSMUSG00000002372 | Ranbp3 | protein_coding | 0.426218631 | 6.514298845 | 1.5333503 | 0.2331408 | 0.999175987 |
| ENSMUSG00000073176 | Zfp449 | protein_coding | -0.759408077 | 3.378219399 | 1.5327807 | 0.2332238 | 0.999175987 |
| ENSMUSG00000019235 | Rps6kl1 | protein_coding | 0.536967639 | 5.743658113 | 1.5315326 | 0.2334059 | 0.999175987 |
| ENSMUSG00000045917 | Tmem268 | protein_coding | -0.723000326 | 4.102658642 | 1.53054 | 0.2335509 | 0.999175987 |
| ENSMUSG00000061576 | Dpp6 | protein_coding | 0.409560378 | 7.247168666 | 1.529868 | 0.2336491 | 0.999175987 |
| ENSMUSG00000028779 | Pef1 | protein_coding | 0.396346143 | 7.551714064 | 1.5297271 | 0.2336697 | 0.999175987 |
| ENSMUSG00000024287 | Thoc1 | protein_coding | -0.438873083 | 5.986577429 | 1.529722 | 0.2336705 | 0.999175987 |
| ENSMUSG00000023961 | Enpp4 | protein_coding | -0.587563912 | 5.192161877 | 1.5292912 | 0.2337335 | 0.999175987 |
| ENSMUSG00000048040 | Arxes2 | protein_coding | 0.437536055 | 6.371574271 | 1.5290567 | 0.2337678 | 0.999175987 |
| ENSMUSG00000038872 | Zfhx3 | protein_coding | 0.468356131 | 6.113214942 | 1.5287506 | 0.2338125 | 0.999175987 |
| ENSMUSG00000071652 | Ints5 | protein_coding | -0.772978689 | 3.756312583 | 1.5280828 | 0.2339103 | 0.999175987 |
| ENSMUSG00000034587 | 8430429K09Rik | processed_transcript | 0.749778265 | 4.422276254 | 1.5276071 | 0.23398 | 0.999175987 |
| ENSMUSG00000021102 | Glrx5 | protein_coding | 0.700590345 | 5.001211171 | 1.5271245 | 0.2340506 | 0.999175987 |
| ENSMUSG00000036934 | 4921524J17Rik | protein_coding | -0.526923442 | 5.390113529 | 1.5261049 | 0.2342001 | 0.999175987 |
| ENSMUSG00000046034 | Otulin | protein_coding | 0.883052579 | 3.192789776 | 1.5256523 | 0.2342665 | 0.999175987 |
| ENSMUSG00000001768 | Rin2 | protein_coding | -0.691988522 | 4.377031292 | 1.525122 | 0.2343443 | 0.999175987 |
| ENSMUSG00000049680 | Urgcp | protein_coding | 0.623991795 | 4.460458606 | 1.5250784 | 0.2343507 | 0.999175987 |
| ENSMUSG00000021969 | Zdhhc20 | protein_coding | -0.435510082 | 6.100182519 | 1.5244213 | 0.2344472 | 0.999175987 |
| ENSMUSG00000038371 | Sbf2 | protein_coding | -0.623419486 | 4.293764089 | 1.5230274 | 0.234652 | 0.999175987 |
| ENSMUSG00000042520 | Ubap2l | protein_coding | -0.793181295 | 4.45753949 | 1.5200939 | 0.2350837 | 0.999175987 |
| ENSMUSG00000052539 | Magi3 | protein_coding | -1.204716317 | 2.938889907 | 1.5200514 | 0.23509 | 0.999175987 |
| ENSMUSG00000039714 | Cplx3 | protein_coding | -1.367193557 | 3.717022402 | 1.535421 | 0.2350935 | 0.999175987 |
| ENSMUSG00000050605 | Zfp61 | protein_coding | -0.734738658 | 4.759084114 | 1.5197309 | 0.2351372 | 0.999175987 |
| ENSMUSG00000029718 | Pcolce | protein_coding | -1.680918324 | 3.082834057 | 1.5270044 | 0.2352443 | 0.999175987 |
| ENSMUSG00000071528 | Usmg5 | protein_coding | 0.499927635 | 7.503552794 | 1.5182194 | 0.2353886 | 0.999175987 |
| ENSMUSG00000029687 | Ezh2 | protein_coding | 1.26265102 | 3.123296772 | 1.5180318 | 0.2354163 | 0.999175987 |
| ENSMUSG00000104939 | Gm33651 | lincRNA | -1.049387162 | 3.545577896 | 1.5172205 | 0.2355361 | 0.999175987 |
| ENSMUSG00000053226 | Dand5 | protein_coding | 0.593508277 | 4.903094634 | 1.5167769 | 0.2355732 | 0.999175987 |
| ENSMUSG00000034663 | Bmp2k | protein_coding | -1.192939952 | 2.757014937 | 1.5158984 | 0.235703 | 0.999175987 |
| ENSMUSG00000026048 | Ercc5 | protein_coding | -0.893781438 | 3.90961275 | 1.5151929 | 0.2358074 | 0.999175987 |
| ENSMUSG00000059363 | Fxn | protein_coding | -1.085709743 | 2.881427468 | 1.5149862 | 0.2358379 | 0.999175987 |
| ENSMUSG00000058997 | Vwa8 | protein_coding | -1.022677858 | 4.08039628 | 1.5182206 | 0.2359372 | 0.999175987 |
| ENSMUSG00000001313 | Rnd2 | protein_coding | 0.443626043 | 7.485209106 | 1.5129378 | 0.2361413 | 0.999175987 |
| ENSMUSG00000039294 | BC017643 | protein_coding | 0.988000608 | 4.386000687 | 1.5195482 | 0.2362009 | 0.999175987 |
| ENSMUSG00000028069 | Gpatch4 | protein_coding | 0.407981579 | 6.731263987 | 1.5124376 | 0.2362155 | 0.999175987 |
| ENSMUSG00000053470 | Kdm3a | protein_coding | -0.691604948 | 4.227019001 | 1.5122327 | 0.2362458 | 0.999175987 |
| ENSMUSG00000023074 | Mospd1 | protein_coding | 0.50140803 | 5.229285499 | 1.5121163 | 0.2362631 | 0.999175987 |
| ENSMUSG00000038068 | Rnf144b | protein_coding | -0.847183009 | 3.326195331 | 1.5115713 | 0.236344 | 0.999175987 |
| ENSMUSG00000042182 | Bend6 | protein_coding | 0.383742723 | 7.256237673 | 1.5112301 | 0.2363946 | 0.999175987 |
| ENSMUSG00000006471 | Ndor1 | protein_coding | 0.939211664 | 3.286308488 | 1.5107693 | 0.236463 | 0.999175987 |
| ENSMUSG00000049583 | Grm5 | protein_coding | -0.525788036 | 7.011997702 | 1.510039 | 0.2365999 | 0.999175987 |
| ENSMUSG00000030610 | Det1 | protein_coding | -1.421303347 | 2.628626449 | 1.5091437 | 0.2367045 | 0.999175987 |
| ENSMUSG00000027799 | Nbea | protein_coding | -0.429754475 | 6.13149796 | 1.50909 | 0.2367125 | 0.999175987 |
| ENSMUSG00000009733 | Tfcp2 | protein_coding | 0.637996784 | 4.626596057 | 1.5088284 | 0.2367514 | 0.999175987 |
| ENSMUSG00000050855 | Zfp940 | protein_coding | 0.677196885 | 4.580419847 | 1.5087975 | 0.236756 | 0.999175987 |
| ENSMUSG00000047361 | Gm973 | protein_coding | -1.568949002 | 2.290108878 | 1.5071984 | 0.236994 | 0.999175987 |
| ENSMUSG00000034145 | Tmem63c | protein_coding | 0.918156235 | 3.439357991 | 1.506985 | 0.2370258 | 0.999175987 |
| ENSMUSG00000021420 | Fars2 | protein_coding | 0.463007883 | 5.774157552 | 1.5065376 | 0.2370924 | 0.999175987 |
| ENSMUSG00000019818 | Cd164 | protein_coding | 0.463820273 | 6.237024336 | 1.5062628 | 0.2371334 | 0.999175987 |
| ENSMUSG00000069601 | Ank3 | protein_coding | -0.396722976 | 6.505575988 | 1.5060344 | 0.2371674 | 0.999175987 |
| ENSMUSG00000059811 | Atl2 | protein_coding | -0.549422938 | 4.999606337 | 1.5060118 | 0.2371708 | 0.999175987 |
| ENSMUSG00000024798 | Htr7 | protein_coding | 0.732116342 | 4.349140398 | 1.5054195 | 0.2372591 | 0.999175987 |
| ENSMUSG00000041168 | Lonp1 | protein_coding | -0.515262423 | 4.990670459 | 1.5038023 | 0.2375004 | 0.999175987 |
| ENSMUSG00000004568 | Arhgef18 | protein_coding | -0.963501915 | 3.849706534 | 1.5034707 | 0.2375784 | 0.999175987 |
| ENSMUSG00000063362 | Alg11 | protein_coding | -0.469785044 | 6.593971449 | 1.5032574 | 0.2375818 | 0.999175987 |
| ENSMUSG00000025544 | Tm9sf2 | protein_coding | -0.711355391 | 5.955620366 | 1.5105427 | 0.2375983 | 0.999175987 |
| ENSMUSG00000029910 | Mad2l1 | protein_coding | 0.728129238 | 4.158878799 | 1.5027167 | 0.2376626 | 0.999175987 |
| ENSMUSG00000042772 | Smg7 | protein_coding | -0.38195463 | 6.953742699 | 1.5025751 | 0.2376838 | 0.999175987 |
| ENSMUSG00000062981 | Mrpl42 | protein_coding | 0.468216946 | 8.561114125 | 1.5024846 | 0.2376973 | 0.999175987 |
| ENSMUSG00000087679 | Tmem250-ps | lincRNA | 0.484157401 | 9.365723529 | 1.502009 | 0.2377684 | 0.999175987 |
| ENSMUSG00000024683 | Mrpl16 | protein_coding | 0.417704988 | 6.373796248 | 1.5003856 | 0.2380114 | 0.999175987 |
| ENSMUSG00000055322 | Tns1 | protein_coding | -1.05150042 | 3.784379738 | 1.5003279 | 0.2381231 | 0.999175987 |

| ensembl_gene_id | external_gene_name | gene_biotype | log2 (fold change) | log2 (counts per million) | F-value | p-value | False discovery rate |
|---|---|---|---|---|---|---|---|
| ENSMUSG00000005986 | Ankrd13d | protein_coding | 0.639222589 | 5.30003387 | 1.4993529 | 0.2381661 | 0.999175987 |
| ENSMUSG00000022629 | Kif21a | protein_coding | -0.433067449 | 7.104242049 | 1.4991555 | 0.2381957 | 0.999175987 |
| ENSMUSG00000040225 | Prrc2c | protein_coding | -0.45503348 | 6.235597442 | 1.499092 | 0.2382052 | 0.999175987 |
| ENSMUSG00000040188 | Scamp2 | protein_coding | -0.525985555 | 5.15352161 | 1.4988944 | 0.2382348 | 0.999175987 |
| ENSMUSG00000028565 | Nfia | protein_coding | -0.486974477 | 6.998809161 | 1.4989191 | 0.2382595 | 0.999175987 |
| ENSMUSG00000010307 | Tmem86a | protein_coding | 0.734339085 | 4.325601805 | 1.4935742 | 0.2390342 | 0.999175987 |
| ENSMUSG00000030350 | Prmt8 | protein_coding | -0.438205832 | 6.015413048 | 1.4931736 | 0.2390946 | 0.999175987 |
| ENSMUSG00000061136 | Prpf40a | protein_coding | -0.561833825 | 5.872851531 | 1.4931665 | 0.2390956 | 0.999175987 |
| ENSMUSG00000037989 | Wnk2 | protein_coding | 0.61285148 | 4.822597408 | 1.4931097 | 0.2391042 | 0.999175987 |
| ENSMUSG00000025422 | Agap2 | protein_coding | -0.897891574 | 3.675226007 | 1.492643 | 0.2391745 | 0.999175987 |
| ENSMUSG00000026970 | Rbms1 | protein_coding | -0.634683947 | 4.38989333 | 1.4924199 | 0.2392081 | 0.999175987 |
| ENSMUSG00000039987 | Phtf2 | protein_coding | 1.03202447 | 2.961224913 | 1.4916815 | 0.2393195 | 0.999175987 |
| ENSMUSG00000024593 | Megf10 | protein_coding | -1.231355771 | 2.874228502 | 1.4910073 | 0.2394212 | 0.999175987 |
| ENSMUSG00000038416 | Cdc16 | protein_coding | -0.577395324 | 5.149809649 | 1.4907807 | 0.2394554 | 0.999175987 |
| ENSMUSG00000027605 | Acss2 | protein_coding | 0.875833493 | 3.965329002 | 1.4907466 | 0.2394605 | 0.999175987 |
| ENSMUSG00000018698 | Lhx1 | protein_coding | -1.255280182 | 2.559117788 | 1.4902321 | 0.2395382 | 0.999175987 |
| ENSMUSG00000038538 | Ubn2 | protein_coding | -0.534761218 | 5.661004336 | 1.4900363 | 0.2395678 | 0.999175987 |
| ENSMUSG00000050856 | Atp5k | protein_coding | 0.531514724 | 9.580473233 | 1.4901221 | 0.2395831 | 0.999175987 |
| ENSMUSG00000020120 | Plek | protein_coding | -0.813465154 | 3.509583045 | 1.489595 | 0.2396344 | 0.999175987 |
| ENSMUSG00000029309 | Sparcl1 | protein_coding | -0.441125932 | 9.275472997 | 1.4882943 | 0.239831 | 0.999175987 |
| ENSMUSG00000033335 | Dnm2 | protein_coding | -1.045503962 | 4.189013686 | 1.4952451 | 0.2398805 | 0.999175987 |
| ENSMUSG00000039539 | Sgcz | protein_coding | -0.827345723 | 4.028245037 | 1.4870148 | 0.2400247 | 0.999175987 |
| ENSMUSG00000035242 | Oaz1 | protein_coding | 0.358812246 | 8.286674294 | 1.4869199 | 0.240039 | 0.999175987 |
| ENSMUSG00000034220 | Gpc1 | protein_coding | 0.673733845 | 4.569326509 | 1.4865136 | 0.2401006 | 0.999175987 |
| ENSMUSG00000043419 | Chd3os | protein_coding | 0.340429338 | 8.809065314 | 1.486428 | 0.2401135 | 0.999175987 |
| ENSMUSG00000020577 | Tspan13 | protein_coding | -0.386889543 | 7.317139806 | 1.4858194 | 0.2402057 | 0.999175987 |
| ENSMUSG00000000420 | Galnt1 | protein_coding | -0.666512363 | 4.118701884 | 1.4856815 | 0.2402266 | 0.999175987 |
| ENSMUSG00000041763 | Tpp2 | protein_coding | -0.530549613 | 5.151239924 | 1.485218 | 0.2402969 | 0.999175987 |
| ENSMUSG00000029775 | Klhdc10 | protein_coding | 0.468862165 | 5.376062898 | 1.4838968 | 0.2404973 | 0.999175987 |
| ENSMUSG00000009281 | Rarres2 | protein_coding | -0.812332037 | 4.253690316 | 1.4830569 | 0.2406249 | 0.999175987 |
| ENSMUSG00000033318 | Gstt2 | protein_coding | 0.836614256 | 3.994900866 | 1.4828193 | 0.240661 | 0.999175987 |
| ENSMUSG00000002660 | Clpp | protein_coding | 0.346712176 | 8.997684785 | 1.4820711 | 0.2407747 | 0.999175987 |
| ENSMUSG00000017943 | Gdap1l1 | protein_coding | 0.423080018 | 7.587768862 | 1.4818201 | 0.2408128 | 0.999175987 |
| ENSMUSG00000031508 | Ankrd10 | protein_coding | -0.752484935 | 4.519871206 | 1.4816285 | 0.240842 | 0.999175987 |
| ENSMUSG00000031171 | Ftsj1 | protein_coding | 0.516594899 | 5.746934887 | 1.4812122 | 0.2409053 | 0.999175987 |
| ENSMUSG00000020674 | Pxdn | protein_coding | 1.740716878 | 2.519421216 | 1.4806966 | 0.241012 | 0.999175987 |
| ENSMUSG00000022488 | Nckap1l | protein_coding | -2.089386655 | 2.372215698 | 1.4838631 | 0.2410751 | 0.999175987 |
| ENSMUSG00000060098 | Prmt7 | protein_coding | 0.839032195 | 5.276261344 | 1.4868679 | 0.2411425 | 0.999175987 |
| ENSMUSG00000035378 | Shq1 | protein_coding | 1.100736351 | 2.995015407 | 1.4795859 | 0.2411529 | 0.999175987 |
| ENSMUSG00000058400 | Qrfpr | protein_coding | 1.08444391 | 2.91734529 | 1.4784931 | 0.2413194 | 0.999175987 |
| ENSMUSG00000025810 | Nrp1 | protein_coding | -0.71175001 | 4.129597708 | 1.4781739 | 0.2413681 | 0.999175987 |
| ENSMUSG00000024889 | Rce1 | protein_coding | -0.630653793 | 4.453823746 | 1.4773255 | 0.2414975 | 0.999175987 |
| ENSMUSG00000021820 | Camk2g | protein_coding | 0.333841196 | 7.244062361 | 1.4756388 | 0.2417551 | 0.999175987 |
| ENSMUSG00000023904 | Hcfc1r1 | protein_coding | 0.335575621 | 8.646237457 | 1.4751449 | 0.2418306 | 0.999175987 |
| ENSMUSG00000041205 | Map6d1 | protein_coding | -0.667111887 | 4.165930521 | 1.4740222 | 0.2420023 | 0.999175987 |
| ENSMUSG00000020863 | Luc7l3 | protein_coding | -0.466606304 | 6.400903113 | 1.4738511 | 0.2420285 | 0.999175987 |
| ENSMUSG00000003847 | Nfat5 | protein_coding | 0.662672337 | 4.026495328 | 1.4734692 | 0.242087 | 0.999175987 |
| ENSMUSG00000025555 | Farp1 | protein_coding | -0.68376004 | 4.541455002 | 1.4729725 | 0.242163 | 0.999175987 |
| ENSMUSG00000034000 | Neu4 | protein_coding | -0.690149653 | 4.023571202 | 1.4720969 | 0.2422972 | 0.999175987 |
| ENSMUSG00000032407 | U2surp | protein_coding | -0.397566806 | 6.560169412 | 1.4719051 | 0.2423266 | 0.999175987 |
| ENSMUSG00000035572 | Dcaf10 | protein_coding | 0.832389528 | 3.641918906 | 1.4715063 | 0.2423877 | 0.999175987 |
| ENSMUSG00000033128 | Gga1 | protein_coding | 0.615948991 | 4.297826087 | 1.4703904 | 0.2425589 | 0.999175987 |
| ENSMUSG00000021534 | 1700001L19Rik | protein_coding | 0.475209794 | 6.731267775 | 1.4702669 | 0.2425779 | 0.999175987 |
| ENSMUSG00000028013 | Ppa2 | protein_coding | 0.442112233 | 6.441135138 | 1.4701797 | 0.2425913 | 0.999175987 |
| ENSMUSG00000005078 | Jkamp | protein_coding | 0.352762273 | 7.001393794 | 1.4697809 | 0.2426525 | 0.999175987 |
| ENSMUSG00000026179 | Pnkd | protein_coding | 0.380000513 | 8.05180221 | 1.4695161 | 0.2426932 | 0.999175987 |
| ENSMUSG00000050132 | Sarm1 | protein_coding | -0.959626791 | 3.056241751 | 1.4691217 | 0.2427538 | 0.999175987 |
| ENSMUSG00000056832 | Ttc26 | protein_coding | -1.191448204 | 2.656063282 | 1.4688802 | 0.2427909 | 0.999175987 |
| ENSMUSG00000045252 | Zfp574 | protein_coding | 0.737970066 | 4.009958688 | 1.4687353 | 0.2428132 | 0.999175987 |
| ENSMUSG00000031231 | Cox7b | protein_coding | 0.463728798 | 9.469297054 | 1.4679569 | 0.2429328 | 0.999175987 |
| ENSMUSG00000042744 | Hectd4 | protein_coding | -0.44159842 | 6.342138632 | 1.4672961 | 0.2430345 | 0.999175987 |
| ENSMUSG00000030064 | Frmd4b | protein_coding | -0.895985579 | 3.473205466 | 1.4671119 | 0.2430629 | 0.999175987 |
| ENSMUSG00000063146 | Clip2 | protein_coding | 0.838341245 | 3.68875427 | 1.4655464 | 0.243304 | 0.999175987 |
| ENSMUSG00000053825 | Ppfia2 | protein_coding | -0.417628386 | 6.398884823 | 1.4655161 | 0.2433086 | 0.999175987 |
| ENSMUSG00000060002 | Chpt1 | protein_coding | -0.709599745 | 3.839606672 | 1.4653699 | 0.2433312 | 0.999175987 |
| ENSMUSG00000024403 | Atp6v1g2 | protein_coding | 0.378866859 | 10.46978886 | 1.4639814 | 0.2435453 | 0.999175987 |
| ENSMUSG00000038831 | Ralgps1 | protein_coding | -0.756984927 | 4.464031335 | 1.4636862 | 0.2435909 | 0.999175987 |
| ENSMUSG00000031365 | Zfp275 | protein_coding | -0.662920937 | 4.555549452 | 1.4632523 | 0.2436578 | 0.999175987 |
| ENSMUSG00000038077 | Kcna6 | protein_coding | 0.549373926 | 5.795123717 | 1.4631205 | 0.2436782 | 0.999175987 |
| ENSMUSG00000036782 | Klhl13 | protein_coding | 0.411995603 | 7.218380801 | 1.4628273 | 0.2437235 | 0.999175987 |
| ENSMUSG00000019146 | Cacng2 | protein_coding | 0.52731099 | 5.129436191 | 1.4603902 | 0.2441003 | 0.999175987 |
| ENSMUSG00000022427 | Tomm22 | protein_coding | 0.34069329 | 7.662045527 | 1.4600989 | 0.2441454 | 0.999175987 |
| ENSMUSG00000068882 | Ssb | protein_coding | -0.429145899 | 7.356923846 | 1.4594976 | 0.2442385 | 0.999175987 |
| ENSMUSG00000032067 | Pts | protein_coding | -0.463144545 | 7.21680756 | 1.4593286 | 0.2442646 | 0.999175987 |
| ENSMUSG00000041571 | Selenow | protein_coding | 0.475972063 | 12.37679339 | 1.4589623 | 0.2443214 | 0.999175987 |
| ENSMUSG00000029559 | 2210016L21Rik | protein_coding | 0.379657835 | 8.426601944 | 1.4588518 | 0.2443385 | 0.999175987 |
| ENSMUSG00000003581 | Rnf215 | protein_coding | 0.74675729 | 3.454155927 | 1.4573351 | 0.2445737 | 0.999175987 |
| ENSMUSG00000029428 | Stx2 | protein_coding | 1.119367364 | 2.756617378 | 1.455275 | 0.2448936 | 0.999175987 |

| ensembl_gene_id | external_gene_name | gene_biotype | log2 (fold change) | log2 (counts per million) | F-value | p-value | False discovery rate |
|---|---|---|---|---|---|---|---|
| ENSMUSG00000039137 | Whrn | protein_coding | 0.727682557 | 4.322699121 | 1.4549447 | 0.244945 | 0.999175987 |
| ENSMUSG00000028402 | Mpdz | protein_coding | -0.782714654 | 3.322529361 | 1.4540214 | 0.2450886 | 0.999175987 |
| ENSMUSG00000031367 | Ap1s2 | protein_coding | 0.505788243 | 6.820562455 | 1.4536207 | 0.245179 | 0.999175987 |
| ENSMUSG00000020312 | Shc2 | protein_coding | -0.855802128 | 3.328318316 | 1.4534391 | 0.2451792 | 0.999175987 |
| ENSMUSG00000052997 | Uba2 | protein_coding | -0.377153745 | 6.448238628 | 1.453069 | 0.2452368 | 0.999175987 |
| ENSMUSG00000048758 | Rpl29 | protein_coding | 0.462886405 | 5.486489102 | 1.4524083 | 0.2453397 | 0.999175987 |
| ENSMUSG00000025967 | Eef1b2 | protein_coding | 0.435964444 | 9.00386979 | 1.4521097 | 0.2453863 | 0.999175987 |
| ENSMUSG00000036478 | Btg1 | protein_coding | -0.627231242 | 5.003005241 | 1.4514484 | 0.2454893 | 0.999175987 |
| ENSMUSG00000003660 | Snrnp200 | protein_coding | -0.547841772 | 5.519479476 | 1.4502643 | 0.245674 | 0.999175987 |
| ENSMUSG00000021221 | Dpf3 | protein_coding | -0.913890399 | 3.478082958 | 1.4466748 | 0.2462351 | 0.999175987 |
| ENSMUSG00000025656 | Arhgef9 | protein_coding | -0.363882984 | 7.063057312 | 1.4466236 | 0.2462431 | 0.999175987 |
| ENSMUSG00000034910 | Pygo1 | protein_coding | -0.606946204 | 4.763878088 | 1.4457048 | 0.246387 | 0.999175987 |
| ENSMUSG00000032375 | Aph1b | protein_coding | -0.63685014 | 4.121172696 | 1.4455987 | 0.2464037 | 0.999175987 |
| ENSMUSG00000007989 | Fzd3 | protein_coding | -0.453004917 | 5.708642128 | 1.4451296 | 0.2464772 | 0.999175987 |
| ENSMUSG00000023235 | Ccl25 | protein_coding | 0.632655806 | 4.87212232 | 1.4428881 | 0.2468288 | 0.999175987 |
| ENSMUSG00000039715 | Wdr34 | protein_coding | -0.578869386 | 4.733743123 | 1.442518 | 0.246887 | 0.999175987 |
| ENSMUSG00000037720 | Tmem33 | protein_coding | -0.496496786 | 6.126478516 | 1.4416968 | 0.247016 | 0.999175987 |
| ENSMUSG00000038766 | Gabpb2 | protein_coding | -1.212897463 | 2.501157165 | 1.441528 | 0.2470426 | 0.999175987 |
| ENSMUSG00000054823 | Nsd3 | protein_coding | 0.468688458 | 6.613822115 | 1.4405115 | 0.2472024 | 0.999175987 |
| ENSMUSG00000030591 | Psmd8 | protein_coding | 0.364449375 | 8.434043169 | 1.4371784 | 0.2477276 | 0.999175987 |
| ENSMUSG00000050558 | Prokr2 | protein_coding | -1.350136988 | 2.716448841 | 1.4365081 | 0.2478334 | 0.999175987 |
| ENSMUSG00000044014 | Npy5r | protein_coding | 1.284140741 | 2.983356318 | 1.4360899 | 0.2478995 | 0.999175987 |
| ENSMUSG00000039221 | Rpl22l1 | protein_coding | 0.488943213 | 6.996212736 | 1.4358088 | 0.2479718 | 0.999175987 |
| ENSMUSG00000071267 | Zfp942 | protein_coding | 0.859592171 | 3.31002476 | 1.4351913 | 0.2480414 | 0.999175987 |
| ENSMUSG00000026064 | Ptp4a1 | protein_coding | 0.644575805 | 4.085936593 | 1.4348484 | 0.2480956 | 0.999175987 |
| ENSMUSG00000042460 | C1galt1 | protein_coding | -1.325807901 | 2.674108706 | 1.4347163 | 0.2481165 | 0.999175987 |
| ENSMUSG00000024957 | Kcnk4 | protein_coding | 1.41429711 | 2.800689289 | 1.4336879 | 0.2482792 | 0.999175987 |
| ENSMUSG00000020716 | Nf1 | protein_coding | -0.574933982 | 5.142647013 | 1.4331643 | 0.2483621 | 0.999175987 |
| ENSMUSG00000023025 | Larp4 | protein_coding | -0.542059379 | 4.523088923 | 1.4327216 | 0.2484322 | 0.999175987 |
| ENSMUSG00000062098 | Btbd3 | protein_coding | -0.417043825 | 6.552591929 | 1.4326915 | 0.2484369 | 0.999175987 |
| ENSMUSG00000030237 | Slco1a4 | protein_coding | -1.487157183 | 2.316407609 | 1.4324299 | 0.2484784 | 0.999175987 |
| ENSMUSG00000021094 | Dhrs7 | protein_coding | -0.536898612 | 6.063722617 | 1.4322066 | 0.2485137 | 0.999175987 |
| ENSMUSG00000045594 | Glb1 | protein_coding | 0.813995003 | 5.061458579 | 1.4347439 | 0.2488241 | 0.999175987 |
| ENSMUSG00000027346 | Gpcpd1 | protein_coding | -0.531547732 | 4.817931451 | 1.4287016 | 0.2490699 | 0.999175987 |
| ENSMUSG00000079184 | Mphosph8 | protein_coding | -0.373821566 | 6.514185979 | 1.4279235 | 0.2491936 | 0.999175987 |
| ENSMUSG00000024442 | 0610009O20Rik | protein_coding | -0.426419321 | 5.770186713 | 1.4276769 | 0.2492328 | 0.999175987 |
| ENSMUSG00000040138 | Ndp | protein_coding | -0.634208149 | 4.483514932 | 1.4274591 | 0.2492675 | 0.999175987 |
| ENSMUSG00000032235 | Ice2 | protein_coding | 1.06470172 | 2.950215542 | 1.4267612 | 0.2493785 | 0.999175987 |
| ENSMUSG00000001018 | Snapin | protein_coding | 0.58997817 | 6.521484819 | 1.431188 | 0.2493865 | 0.999175987 |
| ENSMUSG00000043190 | Rfesd | protein_coding | -0.510227555 | 5.511361375 | 1.4265284 | 0.2494156 | 0.999175987 |
| ENSMUSG00000027677 | Ttc14 | protein_coding | -0.92008879 | 3.715954324 | 1.4262967 | 0.2494525 | 0.999175987 |
| ENSMUSG00000040813 | Tex264 | protein_coding | 0.370999133 | 6.528082836 | 1.4249162 | 0.2496724 | 0.999175987 |
| ENSMUSG00000030731 | Syt3 | protein_coding | 0.553534677 | 4.880763602 | 1.4247377 | 0.2497009 | 0.999175987 |
| ENSMUSG00000035934 | Pknox2 | protein_coding | 0.531437361 | 5.223197322 | 1.4245889 | 0.2497246 | 0.999175987 |
| ENSMUSG00000058254 | Tspan7 | protein_coding | 0.360599361 | 11.05145679 | 1.4244801 | 0.2497419 | 0.999175987 |
| ENSMUSG00000032366 | Tpm1 | protein_coding | -0.33812831 | 7.779207097 | 1.4241975 | 0.249787 | 0.999175987 |
| ENSMUSG00000022656 | Nectin3 | protein_coding | -1.116645089 | 3.045823265 | 1.4240721 | 0.249807 | 0.999175987 |
| ENSMUSG00000020877 | Scrn2 | protein_coding | -0.989266609 | 3.29208002 | 1.4238279 | 0.249846 | 0.999175987 |
| ENSMUSG00000030842 | Lamtor1 | protein_coding | 0.344372893 | 7.343612553 | 1.4235329 | 0.2498931 | 0.999175987 |
| ENSMUSG00000027210 | Meis2 | protein_coding | -0.441944259 | 7.258384013 | 1.4231498 | 0.2499542 | 0.999175987 |
| ENSMUSG00000029632 | Ndufa4 | protein_coding | 0.46868236 | 11.03033526 | 1.4228441 | 0.250003 | 0.999175987 |
| ENSMUSG00000025666 | Tmem47 | protein_coding | -0.433299657 | 6.171143608 | 1.4227577 | 0.2500168 | 0.999175987 |
| ENSMUSG00000022718 | Dgcr8 | protein_coding | -1.096390967 | 2.741823488 | 1.4222296 | 0.2501012 | 0.999175987 |
| ENSMUSG00000014601 | Strip1 | protein_coding | 0.818696392 | 5.067694756 | 1.427091 | 0.2501797 | 0.999175987 |
| ENSMUSG00000047409 | Ctdspl | protein_coding | -0.541930017 | 4.700609006 | 1.4198834 | 0.2504764 | 0.999175987 |
| ENSMUSG00000096054 | Syne1 | protein_coding | -0.440199667 | 6.143398477 | 1.4191564 | 0.2505928 | 0.999175987 |
| ENSMUSG00000027963 | Extl2 | protein_coding | -0.384992826 | 6.745595982 | 1.4185413 | 0.2506914 | 0.999175987 |
| ENSMUSG00000087549 | Gm13427 | antisense_RNA | -0.807646517 | 6.941726025 | 1.4480112 | 0.2507793 | 0.999175987 |
| ENSMUSG00000063049 | Ing2 | protein_coding | -0.732809266 | 4.181450923 | 1.4162731 | 0.2510553 | 0.999175987 |
| ENSMUSG00000070866 | Zfp804a | protein_coding | -0.724634949 | 4.426000685 | 1.4162669 | 0.2510563 | 0.999175987 |
| ENSMUSG00000026073 | Il1r2 | protein_coding | 1.815934354 | 2.054757729 | 1.4153321 | 0.2512064 | 0.999175987 |
| ENSMUSG00000020014 | Cfap54 | protein_coding | -1.17391259 | 2.646951021 | 1.4152818 | 0.2512145 | 0.999175987 |
| ENSMUSG00000033540 | Idua | protein_coding | -0.681410935 | 4.047153992 | 1.4125592 | 0.2516526 | 0.999175987 |
| ENSMUSG00000022235 | Cmbl | protein_coding | -0.510941672 | 5.518609554 | 1.412285 | 0.2516968 | 0.999175987 |
| ENSMUSG00000050148 | Ubqln2 | protein_coding | 0.357920253 | 6.796164998 | 1.4120853 | 0.2517289 | 0.999175987 |
| ENSMUSG00000035615 | Frmpd1 | protein_coding | 1.734644855 | 2.221045393 | 1.4117091 | 0.2517896 | 0.999175987 |
| ENSMUSG00000044734 | Serpinb1a | protein_coding | -0.488504205 | 5.844945195 | 1.4115112 | 0.2518215 | 0.999175987 |
| ENSMUSG00000006299 | Aamp | protein_coding | 0.456124482 | 9.100446211 | 1.4098253 | 0.2520935 | 0.999175987 |
| ENSMUSG00000032997 | Chpf | protein_coding | 0.596965816 | 5.077606247 | 1.409339 | 0.252172 | 0.999175987 |
| ENSMUSG00000042216 | Sgsm1 | protein_coding | 0.519826276 | 6.090930906 | 1.4090371 | 0.2522208 | 0.999175987 |
| ENSMUSG00000044551 | 9930012K11Rik | protein_coding | 1.459712787 | 2.885556435 | 1.408119 | 0.2523969 | 0.999175987 |
| ENSMUSG00000041420 | Meis3 | protein_coding | 0.61516984 | 4.46361733 | 1.4078286 | 0.2524161 | 0.999175987 |
| ENSMUSG00000033400 | Agl | protein_coding | -0.69020911 | 3.815284469 | 1.4064718 | 0.2526357 | 0.999175987 |
| ENSMUSG00000006262 | Mob1b | protein_coding | -0.619324765 | 3.822096656 | 1.4047362 | 0.2529169 | 0.999175987 |
| ENSMUSG00000110841 | AC153368.1 | transcribed_processe | 0.630097488 | 4.00430989 | 1.4046988 | 0.2529229 | 0.999175987 |
| ENSMUSG00000020124 | Usp15 | protein_coding | -0.420452133 | 5.88853095 | 1.4045407 | 0.2529486 | 0.999175987 |
| ENSMUSG00000022762 | Ncam2 | protein_coding | -0.78555805 | 4.609970633 | 1.4045179 | 0.25298 | 0.999175987 |
| ENSMUSG00000025468 | Caly | protein_coding | 0.37088629 | 9.560052941 | 1.4033963 | 0.2531343 | 0.999175987 |

| ensembl_gene_id | external_gene_name | gene_biotype | log2 (fold change) | log2 (counts per million) | F-value | p-value | False discovery rate |
|---|---|---|---|---|---|---|---|
| ENSMUSG00000029073 | Cptp | protein_coding | 0.586849398 | 6.527474622 | 1.4078434 | 0.25318 | 0.999175987 |
| ENSMUSG00000035236 | Scai | protein_coding | -0.644049756 | 4.68379217 | 1.4015069 | 0.2534412 | 0.999175987 |
| ENSMUSG00000021892 | Sh3bp5 | protein_coding | -0.604739801 | 4.685489665 | 1.3999629 | 0.2536924 | 0.999175987 |
| ENSMUSG00000050967 | Creg2 | protein_coding | 0.626912854 | 4.896421896 | 1.399301 | 0.2538002 | 0.999175987 |
| ENSMUSG00000014243 | Zswim7 | protein_coding | -0.479193371 | 5.390457261 | 1.3987554 | 0.2538891 | 0.999175987 |
| ENSMUSG00000041747 | Utp15 | protein_coding | -0.616794289 | 4.508978226 | 1.3985137 | 0.2539285 | 0.999175987 |
| ENSMUSG00000038803 | Ost4 | protein_coding | 0.441659777 | 6.636590519 | 1.3963637 | 0.2542793 | 0.999175987 |
| ENSMUSG00000074591 | Ankub1 | protein_coding | -1.357876426 | 2.561682603 | 1.3960857 | 0.2543247 | 0.999175987 |
| ENSMUSG00000061578 | Ksr2 | protein_coding | -0.616711654 | 4.148114964 | 1.3950981 | 0.2544861 | 0.999175987 |
| ENSMUSG00000043372 | Hexim2 | protein_coding | 0.605915078 | 5.644263685 | 1.3951944 | 0.254498 | 0.999175987 |
| ENSMUSG00000027378 | Nphp1 | protein_coding | 0.82329853 | 3.735808116 | 1.3947851 | 0.2545372 | 0.999175987 |
| ENSMUSG00000018076 | Med13l | protein_coding | -0.552842243 | 5.03335934 | 1.3938525 | 0.2546898 | 0.999175987 |
| ENSMUSG00000006464 | Bbs1 | protein_coding | -0.994185964 | 3.40991514 | 1.393145 | 0.2548056 | 0.999175987 |
| ENSMUSG00000033249 | Hsf4 | protein_coding | 1.110056688 | 2.823140125 | 1.3931201 | 0.2548097 | 0.999175987 |
| ENSMUSG00000051727 | Kctd14 | protein_coding | -1.567936377 | 2.14395777 | 1.390704 | 0.2552057 | 0.999175987 |
| ENSMUSG00000020821 | Kif1c | protein_coding | 0.972495333 | 2.868441457 | 1.3905088 | 0.2552378 | 0.999175987 |
| ENSMUSG00000070056 | Mfhas1 | protein_coding | -0.555296267 | 4.650240112 | 1.3878663 | 0.2556719 | 0.999175987 |
| ENSMUSG00000059714 | Flot1 | protein_coding | 0.443546153 | 6.609385333 | 1.3878246 | 0.2556788 | 0.999175987 |
| ENSMUSG00000022210 | Dhrs4 | protein_coding | -0.648673949 | 4.737723596 | 1.3867021 | 0.2558635 | 0.999175987 |
| ENSMUSG00000020562 | Efcab10 | protein_coding | 0.481526908 | 5.368158711 | 1.386448 | 0.2559053 | 0.999175987 |
| ENSMUSG00000054715 | Zscan22 | protein_coding | -0.816661471 | 4.282864331 | 1.3864642 | 0.2559302 | 0.999175987 |
| ENSMUSG00000067995 | Gtf2f2 | protein_coding | -0.465305843 | 5.655540052 | 1.3858917 | 0.255997 | 0.999175987 |
| ENSMUSG00000024012 | Mtch1 | protein_coding | 0.359091926 | 9.233100313 | 1.3856983 | 0.2560288 | 0.999175987 |
| ENSMUSG00000043398 | Gpr135 | protein_coding | 0.941621486 | 3.060723677 | 1.3853266 | 0.2560901 | 0.999175987 |
| ENSMUSG00000033208 | S100b | protein_coding | -0.607995057 | 6.003589543 | 1.3858951 | 0.2560966 | 0.999175987 |
| ENSMUSG00000059839 | Zfp874b | protein_coding | -0.998867988 | 3.056650539 | 1.3851816 | 0.256114 | 0.999175987 |
| ENSMUSG00000025665 | Rps6ka6 | protein_coding | 0.959170348 | 3.323199076 | 1.3843801 | 0.2562462 | 0.999175987 |
| ENSMUSG00000095432 | Zfp748 | protein_coding | 0.768754576 | 3.508578526 | 1.3832858 | 0.2564268 | 0.999175987 |
| ENSMUSG00000060961 | Slc4a4 | protein_coding | -0.644521702 | 5.376427232 | 1.3831866 | 0.2564707 | 0.999175987 |
| ENSMUSG00000025289 | Prdx4 | protein_coding | -0.499196264 | 5.927358822 | 1.3827815 | 0.2565101 | 0.999175987 |
| ENSMUSG00000029036 | Atad3a | protein_coding | 0.848495097 | 3.588426218 | 1.3822561 | 0.2565969 | 0.999175987 |
| ENSMUSG00000029864 | Gstk1 | protein_coding | -0.952532887 | 4.460523854 | 1.3886669 | 0.2566085 | 0.999175987 |
| ENSMUSG00000009681 | Bcr | protein_coding | -0.400159033 | 6.228285443 | 1.3817546 | 0.2566798 | 0.999175987 |
| ENSMUSG00000034825 | Nrip3 | protein_coding | -0.586953061 | 7.636221506 | 1.3902547 | 0.2566902 | 0.999175987 |
| ENSMUSG00000026171 | Rnf25 | protein_coding | 0.457438307 | 6.474817037 | 1.3813308 | 0.2567499 | 0.999175987 |
| ENSMUSG00000017837 | Nkiras2 | protein_coding | 0.666798688 | 4.918589916 | 1.3799889 | 0.2569719 | 0.999175987 |
| ENSMUSG00000039016 | Timm8b | protein_coding | 0.499695501 | 9.334659322 | 1.3796154 | 0.2570613 | 0.999175987 |
| ENSMUSG00000020476 | Dbnl | protein_coding | 0.365703026 | 6.914025836 | 1.3792266 | 0.2570982 | 0.999175987 |
| ENSMUSG00000030043 | Tacr1 | protein_coding | 0.88957783 | 3.236117114 | 1.3784709 | 0.2572235 | 0.999175987 |
| ENSMUSG00000023274 | Cd4 | protein_coding | 0.612192575 | 4.681910182 | 1.3780551 | 0.2572924 | 0.999175987 |
| ENSMUSG00000002661 | Alkbh7 | protein_coding | 0.380789842 | 6.488680113 | 1.3778891 | 0.25732 | 0.999175987 |
| ENSMUSG00000022704 | Qtrt2 | protein_coding | 0.863178539 | 3.32029933 | 1.3775414 | 0.2573776 | 0.999175987 |
| ENSMUSG00000007944 | Ttc9b | protein_coding | 0.706484639 | 4.256573623 | 1.3775166 | 0.2573818 | 0.999175987 |
| ENSMUSG00000053963 | Stum | protein_coding | 0.553376948 | 4.746367882 | 1.3772863 | 0.25742 | 0.999175987 |
| ENSMUSG00000060371 | Caln1 | protein_coding | -0.523492135 | 6.668395795 | 1.3765338 | 0.2575724 | 0.999175987 |
| ENSMUSG00000057133 | Chd6 | protein_coding | -0.473678743 | 5.686323388 | 1.3758343 | 0.2576611 | 0.999175987 |
| ENSMUSG00000028066 | Pmf1 | protein_coding | 0.576599088 | 4.495058868 | 1.3746706 | 0.2578546 | 0.999175987 |
| ENSMUSG00000030754 | Copb1 | protein_coding | -0.517600069 | 5.99782051 | 1.3743369 | 0.2579101 | 0.999175987 |
| ENSMUSG00000030376 | Slc8a2 | protein_coding | -1.119613199 | 2.93078696 | 1.3740961 | 0.2579502 | 0.999175987 |
| ENSMUSG00000029427 | Zcchc8 | protein_coding | 0.764704295 | 3.828438522 | 1.3731989 | 0.2580995 | 0.999175987 |
| ENSMUSG00000038372 | Gmds | protein_coding | 0.69015167 | 3.71384034 | 1.3725064 | 0.2582149 | 0.999175987 |
| ENSMUSG00000078451 | Ppil6 | protein_coding | -1.046385143 | 2.86356817 | 1.3720441 | 0.258292 | 0.999175987 |
| ENSMUSG00000096847 | Tmem151b | protein_coding | 0.480258356 | 6.425331316 | 1.3714719 | 0.2583874 | 0.999175987 |
| ENSMUSG00000022751 | Nit2 | protein_coding | 0.446623493 | 5.828133495 | 1.3705801 | 0.2585362 | 0.999175987 |
| ENSMUSG00000037239 | Spred3 | protein_coding | 0.958565489 | 2.938059179 | 1.3704853 | 0.258552 | 0.999175987 |
| ENSMUSG00000022946 | Dopey2 | protein_coding | 0.67298322 | 4.038381557 | 1.3684731 | 0.2588882 | 0.999175987 |
| ENSMUSG00000090733 | Rps27 | protein_coding | 0.619663236 | 4.099858358 | 1.3675655 | 0.25904 | 0.999175987 |
| ENSMUSG00000078652 | Psme3 | protein_coding | -0.377241086 | 6.240512895 | 1.3659198 | 0.2593156 | 0.999175987 |
| ENSMUSG00000027624 | Epb41l1 | protein_coding | -0.400575998 | 6.805994717 | 1.3657232 | 0.2593486 | 0.999175987 |
| ENSMUSG00000047459 | Dynlrb1 | protein_coding | 0.405194131 | 10.28289572 | 1.3655381 | 0.2593796 | 0.999175987 |
| ENSMUSG00000021643 | Serf1 | protein_coding | 0.635720108 | 4.613546209 | 1.3649647 | 0.2594758 | 0.999175987 |
| ENSMUSG00000038535 | Zfp280d | protein_coding | 0.958613713 | 3.146602103 | 1.3641772 | 0.2596079 | 0.999175987 |
| ENSMUSG00000017286 | Glod4 | protein_coding | -0.331257937 | 7.522430869 | 1.3640933 | 0.259622 | 0.999175987 |
| ENSMUSG00000021798 | Ldb3 | protein_coding | -0.714988818 | 4.315403515 | 1.3637591 | 0.2596781 | 0.999175987 |
| ENSMUSG00000041935 | AW549877 | protein_coding | -0.488372853 | 5.234525089 | 1.3636077 | 0.2597035 | 0.999175987 |
| ENSMUSG00000027016 | Zfp385b | protein_coding | 0.50120924 | 6.379825828 | 1.3631571 | 0.2597792 | 0.999175987 |
| ENSMUSG00000036992 | Nxt1 | protein_coding | -0.552247657 | 4.667078708 | 1.3625435 | 0.2598823 | 0.999175987 |
| ENSMUSG00000020128 | Vps54 | protein_coding | 0.99390149 | 3.201127812 | 1.3624478 | 0.2598984 | 0.999175987 |
| ENSMUSG00000021650 | Ptcd2 | protein_coding | 0.427873098 | 6.435827119 | 1.3617459 | 0.2600164 | 0.999175987 |
| ENSMUSG00000040373 | Cacng5 | protein_coding | -0.664571168 | 4.872499501 | 1.3613105 | 0.2600896 | 0.999175987 |
| ENSMUSG00000025789 | St8sia2 | protein_coding | -1.223981768 | 2.608282858 | 1.3600367 | 0.2603041 | 0.999175987 |
| ENSMUSG00000000600 | Krit1 | protein_coding | -0.604534002 | 4.727434989 | 1.3591368 | 0.2604557 | 0.999175987 |
| ENSMUSG00000036748 | Cuedc2 | protein_coding | 0.326975134 | 8.642863803 | 1.3590974 | 0.2604623 | 0.999175987 |
| ENSMUSG00000026078 | Pdcl3 | protein_coding | -0.354114515 | 7.318520387 | 1.3584512 | 0.2605713 | 0.999175987 |
| ENSMUSG00000025170 | Rab40b | protein_coding | -0.712988753 | 4.990216302 | 1.3580208 | 0.2606712 | 0.999175987 |
| ENSMUSG00000026568 | Mpc2 | protein_coding | 0.740869547 | 6.934030928 | 1.3763755 | 0.2607193 | 0.999175987 |
| ENSMUSG00000033487 | Fndc3a | protein_coding | -0.453546576 | 5.590292676 | 1.3575452 | 0.2607242 | 0.999175987 |
| ENSMUSG00000028222 | Calb1 | protein_coding | -0.440097697 | 7.475597524 | 1.3564541 | 0.2609084 | 0.999175987 |

| ensembl_gene_id | external_gene_name | gene_biotype | log2 (fold change) | log2 (counts per million) | F-value | p-value | False discovery rate |
|---|---|---|---|---|---|---|---|
| ENSMUSG00000027613 | Eif6 | protein_coding | 0.352743315 | 6.846657421 | 1.3560993 | 0.2609684 | 0.999175987 |
| ENSMUSG00000021782 | Dlg5 | protein_coding | -0.771714296 | 3.431185895 | 1.3560457 | 0.2609774 | 0.999175987 |
| ENSMUSG00000022614 | Lmf2 | protein_coding | 0.875771299 | 3.527637532 | 1.3559398 | 0.2609953 | 0.999175987 |
| ENSMUSG00000056596 | Trnp1 | protein_coding | -0.547766127 | 5.774710519 | 1.3555746 | 0.2610571 | 0.999175987 |
| ENSMUSG00000020785 | Camkk1 | protein_coding | 0.534842705 | 5.651688003 | 1.3553256 | 0.2610992 | 0.999175987 |
| ENSMUSG00000039164 | Naif1 | protein_coding | 0.91786521 | 3.153482618 | 1.3546604 | 0.2612117 | 0.999175987 |
| ENSMUSG00000032333 | Stoml1 | protein_coding | 0.482604001 | 6.128324996 | 1.3539737 | 0.2613279 | 0.999175987 |
| ENSMUSG00000029090 | Adgra3 | protein_coding | 1.797858137 | 2.282222908 | 1.3537997 | 0.2613847 | 0.999175987 |
| ENSMUSG00000003378 | Grik5 | protein_coding | 0.979472308 | 3.244811165 | 1.3529355 | 0.2615038 | 0.999175987 |
| ENSMUSG00000032519 | Slc25a38 | protein_coding | 0.557783616 | 4.960370605 | 1.3522583 | 0.2616186 | 0.999175987 |
| ENSMUSG00000011832 | Evi5l | protein_coding | -0.486269008 | 4.973127487 | 1.351964 | 0.2616685 | 0.999175987 |
| ENSMUSG00000020658 | Efr3b | protein_coding | 0.436440265 | 5.894749229 | 1.351963 | 0.2616686 | 0.999175987 |
| ENSMUSG00000097040 | 2610316D01Rik | processed_transcript | 0.77399546 | 4.550037084 | 1.3518582 | 0.2617137 | 0.999175987 |
| ENSMUSG00000022025 | Cnmd | protein_coding | -1.851297329 | 2.401022234 | 1.3510986 | 0.2618851 | 0.999175987 |
| ENSMUSG00000026090 | 2010300C02Rik | protein_coding | -0.48432619 | 5.642131526 | 1.3504009 | 0.2619338 | 0.999175987 |
| ENSMUSG00000075289 | Carns1 | protein_coding | 0.882798685 | 3.162253939 | 1.3503426 | 0.2619437 | 0.999175987 |
| ENSMUSG00000047675 | Rps8 | protein_coding | 0.477182688 | 6.217113011 | 1.3499269 | 0.2620143 | 0.999175987 |
| ENSMUSG00000078619 | Smarcd2 | protein_coding | -1.518849874 | 2.691362152 | 1.3498929 | 0.2620473 | 0.999175987 |
| ENSMUSG00000050821 | Fam131a | protein_coding | 0.342436801 | 6.95714607 | 1.3490144 | 0.2621694 | 0.999175987 |
| ENSMUSG00000028322 | Exosc3 | protein_coding | -0.875248716 | 3.460895537 | 1.3486899 | 0.2622246 | 0.999175987 |
| ENSMUSG00000070509 | Rgma | protein_coding | 0.851019693 | 3.427260483 | 1.3484566 | 0.2622643 | 0.999175987 |
| ENSMUSG00000052917 | Senp7 | protein_coding | -0.463292972 | 5.549636345 | 1.3479308 | 0.2623537 | 0.999175987 |
| ENSMUSG00000008129 | 4930432K21Rik | protein_coding | -1.282563601 | 2.80066687 | 1.3479097 | 0.2623573 | 0.999175987 |
| ENSMUSG00000016409 | Nkap | protein_coding | -0.502986308 | 5.150353516 | 1.3476261 | 0.2624056 | 0.999175987 |
| ENSMUSG00000020441 | 2310033P09Rik | protein_coding | 0.730561266 | 3.652092943 | 1.3473535 | 0.262452 | 0.999175987 |
| ENSMUSG00000050334 | C130071C03Rik | lincRNA | -0.583529451 | 4.538321769 | 1.3471093 | 0.2624936 | 0.999175987 |
| ENSMUSG00000038914 | Dido1 | protein_coding | -0.507611642 | 4.996768125 | 1.3468721 | 0.262534 | 0.999175987 |
| ENSMUSG00000052837 | Junb | protein_coding | 1.156954554 | 2.756355389 | 1.3466826 | 0.2625663 | 0.999175987 |
| ENSMUSG00000026843 | Fubp3 | protein_coding | -0.570697279 | 4.951118832 | 1.3455047 | 0.2627671 | 0.999175987 |
| ENSMUSG00000073856 | Iqck | protein_coding | 0.787681341 | 3.595928393 | 1.3451441 | 0.2628286 | 0.999175987 |
| ENSMUSG00000024515 | Smad4 | protein_coding | -0.553011105 | 4.955008203 | 1.344467 | 0.2629441 | 0.999175987 |
| ENSMUSG00000066607 | 6030419C18Rik | protein_coding | 1.094584719 | 2.612149884 | 1.344415 | 0.262953 | 0.999175987 |
| ENSMUSG00000035161 | Ints6 | protein_coding | 0.574665098 | 4.27250757 | 1.3438996 | 0.263041 | 0.999175987 |
| ENSMUSG00000034595 | Ppp1r18 | protein_coding | -1.253195094 | 2.624881968 | 1.3434763 | 0.2631133 | 0.999175987 |
| ENSMUSG00000038642 | Ctss | protein_coding | -0.686041814 | 6.462792108 | 1.355859 | 0.2631624 | 0.999175987 |
| ENSMUSG00000002458 | Rgs19 | protein_coding | 0.432389931 | 5.987328493 | 1.3406366 | 0.2635991 | 0.999175987 |
| ENSMUSG00000018770 | Atp5g3 | protein_coding | 0.358337949 | 10.45908229 | 1.3401697 | 0.2636791 | 0.999175987 |
| ENSMUSG00000048988 | Elfn1 | protein_coding | 1.095913157 | 3.066622727 | 1.339774 | 0.2637469 | 0.999175987 |
| ENSMUSG00000022601 | Zbtb11 | protein_coding | 0.784260114 | 3.69462832 | 1.3393921 | 0.2638123 | 0.999175987 |
| ENSMUSG00000034342 | Cbl | protein_coding | 0.578455504 | 4.555585624 | 1.3381741 | 0.2640213 | 0.999175987 |
| ENSMUSG00000025135 | Anapc11 | protein_coding | 0.322764396 | 9.282356923 | 1.3376636 | 0.2641089 | 0.999175987 |
| ENSMUSG00000021392 | Nol8 | protein_coding | -0.573954759 | 4.738627855 | 1.3371129 | 0.2642035 | 0.999175987 |
| ENSMUSG00000071757 | Zhx2 | protein_coding | 0.907820481 | 3.222623385 | 1.3366692 | 0.2642797 | 0.999175987 |
| ENSMUSG00000042272 | Sestd1 | protein_coding | -0.827289637 | 3.482998893 | 1.336433 | 0.2643203 | 0.999175987 |
| ENSMUSG00000047434 | Xxylt1 | protein_coding | 0.74362569 | 3.392330409 | 1.3362952 | 0.264344 | 0.999175987 |
| ENSMUSG00000026977 | 7-Mar | protein_coding | -0.463768229 | 5.295410902 | 1.3359163 | 0.2644092 | 0.999175987 |
| ENSMUSG00000032036 | Kirrel3 | protein_coding | -0.518267258 | 5.479960075 | 1.3356877 | 0.2644485 | 0.999175987 |
| ENSMUSG00000037824 | Tspan14 | protein_coding | -0.59050633 | 4.513875636 | 1.3353065 | 0.2645141 | 0.999175987 |
| ENSMUSG00000034349 | Smc4 | protein_coding | -1.322398523 | 2.374126917 | 1.3348376 | 0.2645947 | 0.999175987 |
| ENSMUSG00000008489 | Elavl2 | protein_coding | -0.410491423 | 6.592545263 | 1.334117 | 0.2647188 | 0.999175987 |
| ENSMUSG00000048222 | Mfap1b | protein_coding | -0.516500448 | 4.988416576 | 1.3337281 | 0.2647858 | 0.999175987 |
| ENSMUSG00000051469 | Zfp24 | protein_coding | -0.48913763 | 5.445761587 | 1.3332809 | 0.2648629 | 0.999175987 |
| ENSMUSG00000036402 | Gng12 | protein_coding | -0.599408145 | 4.976671488 | 1.3332218 | 0.2648731 | 0.999175987 |
| ENSMUSG00000027722 | Spata5 | protein_coding | 0.794286955 | 3.663025969 | 1.3323402 | 0.2650251 | 0.999175987 |
| ENSMUSG00000022911 | Arl13b | protein_coding | 0.79616382 | 3.263438864 | 1.3321712 | 0.2650543 | 0.999175987 |
| ENSMUSG00000100252 | Mir124-2hg | lincRNA | 0.492379631 | 5.424679353 | 1.3315593 | 0.2651599 | 0.999175987 |
| ENSMUSG00000097350 | 4732491K20Rik | lincRNA | 0.546432227 | 4.549704638 | 1.3312642 | 0.2652108 | 0.999175987 |
| ENSMUSG00000036510 | Cdh8 | protein_coding | -0.735055483 | 4.200600969 | 1.3297068 | 0.2654799 | 0.999175987 |
| ENSMUSG00000030881 | Arfip2 | protein_coding | 0.385514281 | 6.476021251 | 1.3288872 | 0.2656216 | 0.999175987 |
| ENSMUSG00000049775 | Tmsb4x | protein_coding | 0.508902381 | 11.71089182 | 1.3285421 | 0.2657085 | 0.999175987 |
| ENSMUSG00000018919 | Tm4sf5 | protein_coding | 0.889171683 | 3.347728474 | 1.3274774 | 0.2658657 | 0.999175987 |
| ENSMUSG00000000916 | Nsun5 | protein_coding | 0.841046808 | 3.846835917 | 1.3263374 | 0.2660633 | 0.999175987 |
| ENSMUSG00000042742 | Bmt2 | protein_coding | -0.609304193 | 4.507488992 | 1.326235 | 0.266081 | 0.999175987 |
| ENSMUSG00000033554 | Dph5 | protein_coding | -0.454443922 | 5.479971465 | 1.3260961 | 0.2661051 | 0.999175987 |
| ENSMUSG00000029084 | Cd38 | protein_coding | -0.918282125 | 3.063188531 | 1.3252624 | 0.2662498 | 0.999175987 |
| ENSMUSG00000071855 | Ccdc112 | protein_coding | -0.714746459 | 4.039742645 | 1.3246209 | 0.2663611 | 0.999175987 |
| ENSMUSG00000022982 | Sod1 | protein_coding | 0.434669162 | 10.04575214 | 1.3240106 | 0.2664672 | 0.999175987 |
| ENSMUSG00000043621 | Ubxn10 | protein_coding | 0.940032108 | 3.129834677 | 1.3235319 | 0.2665504 | 0.999175987 |
| ENSMUSG00000032925 | Itgbl1 | protein_coding | -0.733647764 | 3.819230925 | 1.3212269 | 0.2669514 | 0.999175987 |
| ENSMUSG00000063550 | Nup98 | protein_coding | 0.472473374 | 5.737977979 | 1.3203476 | 0.2671046 | 0.999175987 |
| ENSMUSG00000085436 | Zfp335os | antisense_RNA | 0.792518738 | 3.328951338 | 1.3197808 | 0.2672034 | 0.999175987 |
| ENSMUSG00000061898 | Rbak | protein_coding | -1.29430654 | 2.679502963 | 1.318926 | 0.2673526 | 0.999175987 |
| ENSMUSG00000060679 | Mrps9 | protein_coding | -0.391081938 | 6.640901099 | 1.3166075 | 0.2677576 | 0.999175987 |
| ENSMUSG00000028164 | Manba | protein_coding | 1.119802512 | 2.762128959 | 1.3165861 | 0.2677614 | 0.999175987 |
| ENSMUSG00000039830 | Olig2 | protein_coding | -0.854833408 | 3.316571828 | 1.3158718 | 0.2678863 | 0.999175987 |
| ENSMUSG00000028698 | Pik3r3 | protein_coding | -0.757421146 | 3.624051346 | 1.3152409 | 0.2679967 | 0.999175987 |
| ENSMUSG00000021303 | Gng4 | protein_coding | 0.368747506 | 6.380812427 | 1.3143666 | 0.2681499 | 0.999175987 |
| ENSMUSG00000022185 | Acin1 | protein_coding | -0.637792279 | 3.949173567 | 1.3143077 | 0.2681602 | 0.999175987 |

| ensembl_gene_id | external_gene_name | gene_biotype | log2 (fold change) | log2 (counts per million) | F-value | p-value | False discovery rate |
|---|---|---|---|---|---|---|---|
| ENSMUSG00000044348 | Slc25a53 | protein_coding | -0.539929977 | 4.529963874 | 1.3142807 | 0.2681649 | 0.999175987 |
| ENSMUSG00000039046 | Usp6nl | protein_coding | -0.675496442 | 3.871457784 | 1.3139126 | 0.2682294 | 0.999175987 |
| ENSMUSG00000021501 | Caml | protein_coding | 0.417579339 | 6.034386963 | 1.3139001 | 0.2682316 | 0.999175987 |
| ENSMUSG00000002012 | Pnck | protein_coding | 0.441247235 | 9.46024771 | 1.313775 | 0.2682536 | 0.999175987 |
| ENSMUSG00000031583 | Wrn | protein_coding | -0.996030871 | 3.024138021 | 1.3135051 | 0.2683009 | 0.999175987 |
| ENSMUSG00000025651 | Uqcrc1 | protein_coding | 0.532575869 | 9.629316734 | 1.3161622 | 0.2683455 | 0.999175987 |
| ENSMUSG00000083012 | Fam220a | protein_coding | 0.539435983 | 4.321405995 | 1.3132477 | 0.268346 | 0.999175987 |
| ENSMUSG00000020064 | Herc4 | protein_coding | -0.790482853 | 3.692384802 | 1.3119835 | 0.2685679 | 0.999175987 |
| ENSMUSG00000049300 | Prmt6 | protein_coding | 0.720179295 | 3.436988072 | 1.3113046 | 0.2686871 | 0.999175987 |
| ENSMUSG00000005674 | Tomm40l | protein_coding | 0.54805868 | 5.84229215 | 1.3109454 | 0.2687502 | 0.999175987 |
| ENSMUSG00000045251 | Zfp688 | protein_coding | 0.527674 | 4.787095876 | 1.3104595 | 0.2688356 | 0.999175987 |
| ENSMUSG00000033701 | Acbd6 | protein_coding | 0.48852794 | 5.386860559 | 1.3092397 | 0.2690502 | 0.999175987 |
| ENSMUSG00000028156 | Eif4e | protein_coding | 0.3447651 | 7.869576667 | 1.3078636 | 0.2692925 | 0.999175987 |
| ENSMUSG00000059975 | Zfp74 | protein_coding | 0.720518896 | 3.668341166 | 1.3078577 | 0.2692936 | 0.999175987 |
| ENSMUSG00000022867 | Usp25 | protein_coding | 0.689123681 | 4.176363426 | 1.306139 | 0.2695967 | 0.999175987 |
| ENSMUSG00000018809 | Smyd4 | protein_coding | -0.703434284 | 4.364534529 | 1.3059957 | 0.269622 | 0.999175987 |
| ENSMUSG00000028931 | Kcnab2 | protein_coding | 0.360627485 | 7.385451449 | 1.3059491 | 0.2696302 | 0.999175987 |
| ENSMUSG00000035277 | Arx | protein_coding | -0.889160722 | 3.12653984 | 1.3047529 | 0.2698414 | 0.999175987 |
| ENSMUSG00000050271 | Prag1 | protein_coding | -0.806175287 | 3.463287593 | 1.3042896 | 0.2699233 | 0.999175987 |
| ENSMUSG00000045690 | Wdr89 | protein_coding | 0.636637172 | 4.393169587 | 1.3040627 | 0.2699634 | 0.999175987 |
| ENSMUSG00000058620 | Adra2b | protein_coding | 0.857580301 | 3.442978142 | 1.3039166 | 0.2699892 | 0.999175987 |
| ENSMUSG00000026187 | Xrcc5 | protein_coding | 0.486051359 | 5.202809155 | 1.303558 | 0.2700527 | 0.999175987 |
| ENSMUSG00000028729 | Ebna1bp2 | protein_coding | 0.395662469 | 6.966038955 | 1.3032564 | 0.270106 | 0.999175987 |
| ENSMUSG00000001415 | Smg5 | protein_coding | 0.536625395 | 4.749983484 | 1.3025872 | 0.2702245 | 0.999175987 |
| ENSMUSG00000036545 | Adamts2 | protein_coding | -1.80502206 | 2.421073913 | 1.3022313 | 0.2703852 | 0.999175987 |
| ENSMUSG00000062604 | Srpk2 | protein_coding | -0.381234005 | 7.969830073 | 1.3011645 | 0.2704765 | 0.999175987 |
| ENSMUSG00000026748 | Plxdc2 | protein_coding | -0.646047077 | 4.74135173 | 1.3004936 | 0.2705954 | 0.999175987 |
| ENSMUSG00000019817 | Plagl1 | protein_coding | 0.690547507 | 3.547039266 | 1.3001312 | 0.2706597 | 0.999175987 |
| ENSMUSG00000044475 | Ascc1 | protein_coding | -0.349965859 | 6.756643452 | 1.3001069 | 0.270664 | 0.999175987 |
| ENSMUSG00000032898 | Fbxo21 | protein_coding | -0.435057978 | 6.329678203 | 1.300008 | 0.2706816 | 0.999175987 |
| ENSMUSG00000071679 | Zcchc16 | protein_coding | -1.089398325 | 2.834051564 | 1.2985618 | 0.2709383 | 0.999175987 |
| ENSMUSG00000059273 | Zc3h4 | protein_coding | -0.539162482 | 4.300804903 | 1.2983759 | 0.2709714 | 0.999175987 |
| ENSMUSG00000022914 | Brwd1 | protein_coding | -0.395349572 | 6.174824678 | 1.2983698 | 0.2709724 | 0.999175987 |
| ENSMUSG00000022433 | Csnk1e | protein_coding | -0.568271633 | 5.460540342 | 1.2981957 | 0.2710034 | 0.999175987 |
| ENSMUSG00000035640 | Cbarp | protein_coding | 0.421541966 | 6.86161817 | 1.2978235 | 0.2710695 | 0.999175987 |
| ENSMUSG00000038486 | Sv2a | protein_coding | 0.330238882 | 8.389974562 | 1.2972152 | 0.2711777 | 0.999175987 |
| ENSMUSG00000017491 | Rarb | protein_coding | 0.727559479 | 3.526248906 | 1.2971907 | 0.2711821 | 0.999175987 |
| ENSMUSG00000029361 | Nos1 | protein_coding | 0.593004307 | 6.17370665 | 1.2997915 | 0.2712669 | 0.999175987 |
| ENSMUSG00000025264 | Tsr2 | protein_coding | 0.371022188 | 7.826594417 | 1.2962157 | 0.2713556 | 0.999175987 |
| ENSMUSG00000045034 | Ankrd34b | protein_coding | -0.752449411 | 3.798458628 | 1.2954458 | 0.2714926 | 0.999175987 |
| ENSMUSG00000033350 | Chst2 | protein_coding | 0.334529401 | 6.92829808 | 1.2953114 | 0.2715166 | 0.999175987 |
| ENSMUSG00000031148 | Gpkow | protein_coding | -0.624390926 | 5.156053859 | 1.2949665 | 0.271578 | 0.999175987 |
| ENSMUSG00000050963 | Kcns2 | protein_coding | 1.196491152 | 2.687444094 | 1.2945532 | 0.2716517 | 0.999175987 |
| ENSMUSG00000025132 | Arhgdia | protein_coding | 0.375720974 | 9.882980384 | 1.2928421 | 0.271957 | 0.999175987 |
| ENSMUSG00000028961 | Pgd | protein_coding | -0.538175766 | 5.789461953 | 1.2925974 | 0.2720007 | 0.999175987 |
| ENSMUSG00000028914 | Casp9 | protein_coding | -1.407820727 | 2.50036652 | 1.2917004 | 0.2721609 | 0.999175987 |
| ENSMUSG00000019579 | Mydgf | protein_coding | 0.356156214 | 7.3175133 | 1.2915448 | 0.2721888 | 0.999175987 |
| ENSMUSG00000003559 | As3mt | protein_coding | -0.501777041 | 5.860821575 | 1.2914688 | 0.2722023 | 0.999175987 |
| ENSMUSG00000039680 | Mrps6 | protein_coding | 0.500800217 | 6.241458897 | 1.2911724 | 0.2722553 | 0.999175987 |
| ENSMUSG00000041346 | Wrap53 | protein_coding | -0.504141216 | 4.948486831 | 1.2888504 | 0.272671 | 0.999175987 |
| ENSMUSG00000017561 | Crlf3 | protein_coding | -0.88330554 | 3.450944053 | 1.2883691 | 0.2727572 | 0.999175987 |
| ENSMUSG00000019689 | Fmc1 | protein_coding | 0.385844447 | 7.439051326 | 1.288329 | 0.2727644 | 0.999175987 |
| ENSMUSG00000033676 | Gabrb3 | protein_coding | 0.372752895 | 6.4470507 | 1.2883056 | 0.2727686 | 0.999175987 |
| ENSMUSG00000030983 | Bccip | protein_coding | 0.323899851 | 7.669501024 | 1.2881376 | 0.2727987 | 0.999175987 |
| ENSMUSG00000026831 | 1700007K13Rik | protein_coding | 0.836045459 | 3.121678407 | 1.2879176 | 0.2728382 | 0.999175987 |
| ENSMUSG00000027601 | Mtfr1 | protein_coding | -0.769153757 | 3.960358547 | 1.287617 | 0.2728921 | 0.999175987 |
| ENSMUSG00000037652 | Phc3 | protein_coding | -0.598482472 | 4.092420459 | 1.2874334 | 0.272925 | 0.999175987 |
| ENSMUSG00000040957 | Cables1 | protein_coding | 0.817974361 | 3.644113282 | 1.2863625 | 0.2731172 | 0.999175987 |
| ENSMUSG00000040280 | Ndufa4l2 | protein_coding | -1.247440094 | 2.881207013 | 1.2861853 | 0.273149 | 0.999175987 |
| ENSMUSG00000014164 | Klhl3 | protein_coding | -0.566463752 | 6.45429583 | 1.2899639 | 0.2731543 | 0.999175987 |
| ENSMUSG00000076439 | Mog | protein_coding | -0.375945597 | 7.849745662 | 1.2857041 | 0.2732355 | 0.999175987 |
| ENSMUSG00000022174 | Dad1 | protein_coding | 0.358612734 | 9.050245605 | 1.2855178 | 0.273269 | 0.999175987 |
| ENSMUSG00000033233 | Trim45 | protein_coding | 1.052243268 | 3.97313716 | 1.2911166 | 0.2733029 | 0.999175987 |
| ENSMUSG00000037461 | Ints7 | protein_coding | 0.536137282 | 5.262373825 | 1.2847219 | 0.273412 | 0.999175987 |
| ENSMUSG00000025227 | Mfsd13a | protein_coding | 0.543662844 | 4.583608346 | 1.284451 | 0.2734607 | 0.999175987 |
| ENSMUSG00000021700 | Rab3c | protein_coding | -0.549904858 | 8.36536048 | 1.2893776 | 0.2736135 | 0.999175987 |
| ENSMUSG00000042851 | Zc3h6 | protein_coding | 0.797994961 | 3.161167287 | 1.2824189 | 0.2738266 | 0.999175987 |
| ENSMUSG00000044229 | Nxpe4 | protein_coding | -1.048653847 | 3.301324228 | 1.2820791 | 0.2738878 | 0.999175987 |
| ENSMUSG00000037750 | Fam222b | protein_coding | -0.498956837 | 5.159409597 | 1.2795214 | 0.2743494 | 0.999175987 |
| ENSMUSG00000023068 | Nus1 | protein_coding | -0.426923737 | 5.801789935 | 1.2794936 | 0.2743544 | 0.999175987 |
| ENSMUSG00000038347 | Tcte2 | protein_coding | -0.609189283 | 4.131523896 | 1.2784694 | 0.2745395 | 0.999175987 |
| ENSMUSG00000048581 | E130311K13Rik | protein_coding | 1.312744897 | 2.635295738 | 1.2777864 | 0.274663 | 0.999175987 |
| ENSMUSG00000031986 | Sprtn | protein_coding | -1.352023309 | 2.526641802 | 1.2777013 | 0.2746784 | 0.999175987 |
| ENSMUSG00000011179 | Odc1 | protein_coding | -0.64840332 | 4.258614454 | 1.277657 | 0.2746864 | 0.999175987 |
| ENSMUSG00000025204 | Ndufb8 | protein_coding | 0.383704621 | 9.711001295 | 1.2774314 | 0.2747273 | 0.999175987 |
| ENSMUSG00000020696 | Rffl | protein_coding | 0.974266476 | 3.31766545 | 1.2769843 | 0.2748082 | 0.999175987 |
| ENSMUSG00000020263 | Appl2 | protein_coding | -0.520030256 | 4.59630411 | 1.2768868 | 0.2748259 | 0.999175987 |
| ENSMUSG00000010592 | Dazl | protein_coding | 1.979405697 | 2.350461149 | 1.2794751 | 0.2748984 | 0.999175987 |

| ensembl_gene_id | external_gene_name | gene_biotype | log2 (fold change) | log2 (counts per million) | F-value | p-value | False discovery rate |
|---|---|---|---|---|---|---|---|
| ENSMUSG00000054364 | Rhob | protein_coding | -0.346715443 | 7.058097079 | 1.2761722 | 0.2749553 | 0.999175987 |
| ENSMUSG00000005803 | Sqor | protein_coding | -0.748472163 | 3.767947818 | 1.2750704 | 0.275155 | 0.999175987 |
| ENSMUSG00000020834 | Dhrs13 | protein_coding | 1.276279968 | 2.554185785 | 1.2742159 | 0.2753101 | 0.999175987 |
| ENSMUSG00000035401 | Emsy | protein_coding | -0.590943473 | 4.647654301 | 1.2739255 | 0.2753628 | 0.999175987 |
| ENSMUSG00000071415 | Rpl23 | protein_coding | 0.455737111 | 7.520585782 | 1.2733262 | 0.2754982 | 0.999175987 |
| ENSMUSG00000033685 | Ucp2 | protein_coding | -0.451423433 | 6.573361326 | 1.272245 | 0.2756681 | 0.999175987 |
| ENSMUSG00000027937 | Jtb | protein_coding | 0.547684431 | 5.714458474 | 1.2716948 | 0.2757682 | 0.999175987 |
| ENSMUSG00000031347 | Cetn2 | protein_coding | -0.406458432 | 7.865143906 | 1.2710005 | 0.2758945 | 0.999175987 |
| ENSMUSG00000070934 | Rraga | protein_coding | -0.498120086 | 6.785571419 | 1.2705194 | 0.2760087 | 0.999175987 |
| ENSMUSG00000020876 | Snx11 | protein_coding | 0.749593266 | 4.596379397 | 1.2700285 | 0.2760981 | 0.999175987 |
| ENSMUSG00000052812 | Atad2b | protein_coding | -0.943789898 | 3.200018374 | 1.2697911 | 0.2761148 | 0.999175987 |
| ENSMUSG00000015335 | Zdhhc12 | protein_coding | -0.698935278 | 4.070934075 | 1.2692059 | 0.2762214 | 0.999175987 |
| ENSMUSG00000038954 | Supt3 | protein_coding | 0.586528832 | 4.465072397 | 1.2681085 | 0.2764216 | 0.999175987 |
| ENSMUSG00000007739 | Cct4 | protein_coding | 0.353806428 | 7.595742519 | 1.267802 | 0.2764775 | 0.999175987 |
| ENSMUSG00000030615 | Tmem126a | protein_coding | 0.438724786 | 6.654889712 | 1.2676428 | 0.2765066 | 0.999175987 |
| ENSMUSG00000036054 | Sugp2 | protein_coding | 0.473561581 | 4.989714878 | 1.2673097 | 0.2765674 | 0.999175987 |
| ENSMUSG00000024472 | Dcp2 | protein_coding | -0.595324039 | 4.493941162 | 1.2673034 | 0.2765685 | 0.999175987 |
| ENSMUSG00000029191 | Rfc1 | protein_coding | -0.462277309 | 5.548123575 | 1.2669404 | 0.2766349 | 0.999175987 |
| ENSMUSG00000017188 | Coa3 | protein_coding | 0.340720699 | 9.831433548 | 1.265934 | 0.2768188 | 0.999175987 |
| ENSMUSG00000022544 | Eef2kmt | protein_coding | -0.766529625 | 3.838473285 | 1.2654481 | 0.2769076 | 0.999175987 |
| ENSMUSG00000048911 | Rnf24 | protein_coding | 1.010552863 | 3.282949124 | 1.2622169 | 0.2774995 | 0.999175987 |
| ENSMUSG00000021477 | Ctsl | protein_coding | -0.356796004 | 7.359115582 | 1.2621709 | 0.277508 | 0.999175987 |
| ENSMUSG00000024397 | Aif1 | protein_coding | -0.864626586 | 4.614596879 | 1.267301 | 0.2775986 | 0.999175987 |
| ENSMUSG00000002504 | Slc9a3r2 | protein_coding | 0.51635016 | 4.740477789 | 1.2614645 | 0.2776376 | 0.999175987 |
| ENSMUSG00000042625 | Safb2 | protein_coding | -0.804861706 | 3.514037681 | 1.2611242 | 0.2777001 | 0.999175987 |
| ENSMUSG00000030619 | Eed | protein_coding | -0.694095133 | 4.505695003 | 1.2597523 | 0.2779521 | 0.999175987 |
| ENSMUSG00000042978 | Sbk1 | protein_coding | -0.712587296 | 4.205062604 | 1.2597108 | 0.2779598 | 0.999175987 |
| ENSMUSG00000003534 | Ddr1 | protein_coding | -0.572991559 | 4.699261605 | 1.2594022 | 0.2780165 | 0.999175987 |
| ENSMUSG00000025010 | Ccnj | protein_coding | 0.990944635 | 3.045179781 | 1.2593487 | 0.2780264 | 0.999175987 |
| ENSMUSG00000042700 | Sipa1l1 | protein_coding | -0.502350048 | 6.147735034 | 1.258946 | 0.2781004 | 0.999175987 |
| ENSMUSG00000037747 | Phyhipl | protein_coding | -0.38343218 | 7.142047207 | 1.2586448 | 0.2781559 | 0.999175987 |
| ENSMUSG00000036599 | Chst12 | protein_coding | -0.448952483 | 5.686100521 | 1.258331 | 0.2782136 | 0.999175987 |
| ENSMUSG00000038206 | Fbxo8 | protein_coding | -0.569703756 | 4.913619376 | 1.257857 | 0.2783009 | 0.999175987 |
| ENSMUSG00000050315 | Synpo2 | protein_coding | -0.97530641 | 4.186734964 | 1.2633016 | 0.2783288 | 0.999175987 |
| ENSMUSG00000047888 | Tnrc6b | protein_coding | -0.504740598 | 4.947256467 | 1.2576208 | 0.2783444 | 0.999175987 |
| ENSMUSG00000033004 | Mycbp2 | protein_coding | -0.504867208 | 6.269808914 | 1.2572468 | 0.2784398 | 0.999175987 |
| ENSMUSG00000019080 | Mfsd3 | protein_coding | 0.710728601 | 4.22252676 | 1.2570168 | 0.2784557 | 0.999175987 |
| ENSMUSG00000010911 | Apip | protein_coding | 0.418547893 | 5.986840214 | 1.2569497 | 0.278468 | 0.999175987 |
| ENSMUSG00000007411 | Mark3 | protein_coding | -0.668935179 | 4.971848746 | 1.2569883 | 0.2784874 | 0.999175987 |
| ENSMUSG00000043535 | Setx | protein_coding | -0.601043091 | 4.54475546 | 1.2568194 | 0.2784921 | 0.999175987 |
| ENSMUSG00000025626 | Phf6 | protein_coding | 0.520165688 | 4.926960192 | 1.2567192 | 0.2785105 | 0.999175987 |
| ENSMUSG00000059291 | Rpl11 | protein_coding | 0.537529463 | 4.430452932 | 1.2565075 | 0.2785496 | 0.999175987 |
| ENSMUSG00000001248 | Gramd1a | protein_coding | 0.694654867 | 4.273829548 | 1.2562016 | 0.278606 | 0.999175987 |
| ENSMUSG00000001763 | Tspan33 | protein_coding | 0.864034462 | 3.839518618 | 1.2551923 | 0.2788187 | 0.999175987 |
| ENSMUSG00000022321 | Cdh10 | protein_coding | -0.605930758 | 4.203458714 | 1.2547282 | 0.2788779 | 0.999175987 |
| ENSMUSG00000007458 | M6pr | protein_coding | 0.377183616 | 7.138541407 | 1.2546048 | 0.2789007 | 0.999175987 |
| ENSMUSG00000053063 | Clec12a | protein_coding | -1.707089778 | 2.502526598 | 1.2548062 | 0.2789597 | 0.999175987 |
| ENSMUSG00000051022 | Hs3st1 | protein_coding | 0.597623557 | 4.575954066 | 1.2533067 | 0.2791406 | 0.999175987 |
| ENSMUSG00000001143 | Lman2l | protein_coding | 0.483305708 | 5.726341757 | 1.2530959 | 0.2791795 | 0.999175987 |
| ENSMUSG00000036442 | Thap11 | protein_coding | 0.624764498 | 4.678581292 | 1.2526067 | 0.27927 | 0.999175987 |
| ENSMUSG00000036825 | Ssx2ip | protein_coding | 0.437695081 | 5.533175361 | 1.2519357 | 0.2793942 | 0.999175987 |
| ENSMUSG00000044080 | S100a1 | protein_coding | -0.61269721 | 8.199146881 | 1.2635766 | 0.2794877 | 0.999175987 |
| ENSMUSG00000049517 | Rps23 | protein_coding | 0.843789992 | 3.084731653 | 1.2492841 | 0.2798857 | 0.999175987 |
| ENSMUSG00000011589 | Fsd1 | protein_coding | 0.451964687 | 7.226805919 | 1.2491901 | 0.2799296 | 0.999175987 |
| ENSMUSG00000014294 | Ndufa2 | protein_coding | 0.433110812 | 9.642070955 | 1.2482502 | 0.2800777 | 0.999175987 |
| ENSMUSG00000022551 | Cyc1 | protein_coding | 0.36236869 | 6.715909011 | 1.2463315 | 0.2804343 | 0.999175987 |
| ENSMUSG00000044667 | Plppr4 | protein_coding | -0.370345056 | 6.517486683 | 1.2461153 | 0.2804746 | 0.999175987 |
| ENSMUSG00000034612 | Chst11 | protein_coding | -0.60157704 | 3.991969294 | 1.246081 | 0.280481 | 0.999175987 |
| ENSMUSG00000038500 | Prr3 | protein_coding | 0.484289235 | 5.502780232 | 1.2459715 | 0.2805013 | 0.999175987 |
| ENSMUSG00000042743 | Sgtb | protein_coding | -0.504914064 | 5.508618151 | 1.2455047 | 0.2805882 | 0.999175987 |
| ENSMUSG00000003316 | Glg1 | protein_coding | 0.326769334 | 6.681295146 | 1.2450982 | 0.2806639 | 0.999175987 |
| ENSMUSG00000052155 | Acvr2a | protein_coding | -0.683638672 | 3.817366856 | 1.2443493 | 0.2808035 | 0.999175987 |
| ENSMUSG00000028289 | Epha7 | protein_coding | -0.481041254 | 5.22198334 | 1.2441948 | 0.2808323 | 0.999175987 |
| ENSMUSG00000028923 | Necap2 | protein_coding | -0.479137589 | 5.508739971 | 1.2432294 | 0.2810123 | 0.999175987 |
| ENSMUSG00000045659 | Plekha7 | protein_coding | -0.937653203 | 3.350783315 | 1.2431623 | 0.2810248 | 0.999175987 |
| ENSMUSG00000019861 | Gopc | protein_coding | -0.488831386 | 4.726471528 | 1.2430441 | 0.2810469 | 0.999175987 |
| ENSMUSG00000026005 | Rpe | protein_coding | -0.699809464 | 4.424036185 | 1.2428278 | 0.2810873 | 0.999175987 |
| ENSMUSG00000027881 | Prpf38b | protein_coding | -0.770839442 | 3.818091983 | 1.2426085 | 0.2811282 | 0.999175987 |
| ENSMUSG00000026817 | Ak1 | protein_coding | 0.385994509 | 9.070943259 | 1.2421097 | 0.2812213 | 0.999175987 |
| ENSMUSG00000052551 | Adarb2 | protein_coding | -0.575159048 | 4.072963191 | 1.2420892 | 0.2812252 | 0.999175987 |
| ENSMUSG00000032485 | Scap | protein_coding | 0.600598121 | 4.527274056 | 1.2420406 | 0.2812342 | 0.999175987 |
| ENSMUSG00000068264 | Ap5s1 | protein_coding | 0.412032451 | 6.289212354 | 1.2409788 | 0.2814327 | 0.999175987 |
| ENSMUSG00000021022 | Ppp2r3c | protein_coding | 0.525709685 | 4.722584434 | 1.2393412 | 0.281739 | 0.999175987 |
| ENSMUSG00000085492 | Trmt61b | processed_transcript | 0.495160123 | 5.168343951 | 1.2370213 | 0.2821738 | 0.999175987 |
| ENSMUSG00000015790 | Surf1 | protein_coding | 0.341553063 | 8.21324202 | 1.2364035 | 0.2822898 | 0.999175987 |
| ENSMUSG00000045967 | Gpr158 | protein_coding | -0.491988331 | 5.215833255 | 1.2358825 | 0.2823876 | 0.999175987 |
| ENSMUSG00000028176 | Lrrc7 | protein_coding | -0.690889229 | 4.433184604 | 1.2356408 | 0.282433 | 0.999175987 |
| ENSMUSG00000010064 | Slc38a3 | protein_coding | -0.674080321 | 4.503629925 | 1.2349464 | 0.2825635 | 0.999175987 |

| ensembl_gene_id | external_gene_name | gene_biotype | log2 (fold change) | log2 (counts per million) | F-value | p-value | False discovery rate |
|---|---|---|---|---|---|---|---|
| ENSMUSG00000024054 | Smchd1 | protein_coding | -0.806454689 | 3.379495894 | 1.2345947 | 0.2826296 | 0.999175987 |
| ENSMUSG00000034480 | Diaph2 | protein_coding | -0.712260017 | 4.128041007 | 1.2342261 | 0.2826989 | 0.999175987 |
| ENSMUSG00000022858 | Tra2b | protein_coding | -0.395205735 | 6.079785217 | 1.2339486 | 0.2827511 | 0.999175987 |
| ENSMUSG00000032330 | Cox7a2 | protein_coding | 0.434867845 | 10.09163773 | 1.233511 | 0.2828335 | 0.999175987 |
| ENSMUSG00000020135 | Apc2 | protein_coding | -0.709280943 | 4.679888566 | 1.2327896 | 0.2829956 | 0.999175987 |
| ENSMUSG00000002307 | Daxx | protein_coding | -0.596129209 | 5.234720564 | 1.232343 | 0.2830534 | 0.999175987 |
| ENSMUSG00000022064 | Pibf1 | protein_coding | -0.877777371 | 3.199530729 | 1.2318095 | 0.283154 | 0.999175987 |
| ENSMUSG00000047363 | Cstad | protein_coding | 0.729676754 | 3.822675004 | 1.2314755 | 0.2832169 | 0.999175987 |
| ENSMUSG00000028944 | Prkag2 | protein_coding | 0.410061298 | 5.982183124 | 1.2312573 | 0.2832581 | 0.999175987 |
| ENSMUSG00000042901 | Aida | protein_coding | 0.442578429 | 5.122552065 | 1.2312327 | 0.2832627 | 0.999175987 |
| ENSMUSG00000033065 | Pfkm | protein_coding | -0.448215559 | 8.237464808 | 1.2303414 | 0.2834571 | 0.999175987 |
| ENSMUSG00000024507 | Hsd17b4 | protein_coding | -0.435854486 | 5.464083413 | 1.2297384 | 0.2835447 | 0.999175987 |
| ENSMUSG00000021635 | Rad17 | protein_coding | -0.556748358 | 4.620835006 | 1.2291392 | 0.2836579 | 0.999175987 |
| ENSMUSG00000034813 | Grip1 | protein_coding | 0.521905451 | 4.547246663 | 1.2288603 | 0.2837106 | 0.999175987 |
| ENSMUSG00000061013 | Mkx | protein_coding | -1.27422911 | 2.553565789 | 1.2287964 | 0.2837227 | 0.999175987 |
| ENSMUSG00000090553 | Snrpe | protein_coding | 0.447050131 | 7.173520655 | 1.2274885 | 0.2839963 | 0.999175987 |
| ENSMUSG00000024548 | Setbp1 | protein_coding | -0.846196291 | 3.466152354 | 1.2269729 | 0.2840676 | 0.999175987 |
| ENSMUSG00000031168 | Ebp | protein_coding | 0.422804905 | 5.573726174 | 1.2267587 | 0.2841082 | 0.999175987 |
| ENSMUSG00000004264 | Phb2 | protein_coding | -0.732776257 | 3.591811995 | 1.2266974 | 0.2841198 | 0.999175987 |
| ENSMUSG00000030374 | Strn4 | protein_coding | 0.414915694 | 6.040028159 | 1.2256853 | 0.2843115 | 0.999175987 |
| ENSMUSG00000044072 | Eml6 | protein_coding | 0.978875266 | 2.775507336 | 1.2252089 | 0.2844019 | 0.999175987 |
| ENSMUSG00000031333 | Abcb7 | protein_coding | -0.616956803 | 4.066281437 | 1.2243648 | 0.284562 | 0.999175987 |
| ENSMUSG00000030269 | Mtmr14 | protein_coding | 0.87542374 | 2.959908207 | 1.2207733 | 0.2852446 | 0.999175987 |
| ENSMUSG00000028461 | Ccdc107 | protein_coding | 0.9160528 | 5.916558428 | 1.2434273 | 0.2854601 | 0.999175987 |
| ENSMUSG00000031202 | Rab39b | protein_coding | -0.566456003 | 4.459768071 | 1.2194296 | 0.2855006 | 0.999175987 |
| ENSMUSG00000035184 | Fam124a | protein_coding | -0.691518329 | 4.021235127 | 1.2191662 | 0.2855508 | 0.999175987 |
| ENSMUSG00000033960 | Jcad | protein_coding | -0.801099343 | 4.348085359 | 1.2192937 | 0.2855527 | 0.999175987 |
| ENSMUSG00000020431 | Adcy1 | protein_coding | -0.709372373 | 4.762574829 | 1.2191181 | 0.2855861 | 0.999175987 |
| ENSMUSG00000028271 | Gtf2b | protein_coding | -0.366367466 | 6.314934803 | 1.2183283 | 0.2857106 | 0.999175987 |
| ENSMUSG00000022742 | Cpox | protein_coding | -0.536199616 | 5.632461472 | 1.2175709 | 0.2858552 | 0.999175987 |
| ENSMUSG00000045664 | Cdc42ep2 | protein_coding | -0.53936281 | 5.455680769 | 1.2169641 | 0.2859711 | 0.999175987 |
| ENSMUSG00000086649 | Gm15286 | processed_transcript | 0.99525765 | 2.672795255 | 1.2167344 | 0.286015 | 0.999175987 |
| ENSMUSG00000056938 | Acbd4 | protein_coding | 0.603207034 | 5.565065564 | 1.2167861 | 0.2860312 | 0.999175987 |
| ENSMUSG00000064247 | Plcxd1 | protein_coding | 0.505305791 | 4.633945197 | 1.2157476 | 0.2862036 | 0.999175987 |
| ENSMUSG00000036513 | Commd2 | protein_coding | 0.389676865 | 6.116908395 | 1.2155053 | 0.28625 | 0.999175987 |
| ENSMUSG00000018209 | Stk4 | protein_coding | -0.54369348 | 4.73692849 | 1.215206 | 0.2863072 | 0.999175987 |
| ENSMUSG00000053985 | Zfp14 | protein_coding | -0.640382427 | 3.830076174 | 1.2148791 | 0.2863698 | 0.999175987 |
| ENSMUSG00000032083 | Apoa1 | protein_coding | 0.843570973 | 3.085571856 | 1.2140773 | 0.2865233 | 0.999175987 |
| ENSMUSG00000025153 | Fasn | protein_coding | 0.501343107 | 6.044390897 | 1.2133548 | 0.2866617 | 0.999175987 |
| ENSMUSG00000022425 | Enpp2 | protein_coding | -0.691427952 | 8.708527452 | 1.2290219 | 0.2866854 | 0.999175987 |
| ENSMUSG00000054850 | Smim10l2a | protein_coding | 0.565184456 | 4.392290456 | 1.2128487 | 0.2867588 | 0.999175987 |
| ENSMUSG00000047368 | Abhd17b | protein_coding | -0.524045874 | 5.259547045 | 1.2120898 | 0.2869043 | 0.999175987 |
| ENSMUSG00000073888 | Ccl27a | protein_coding | 0.41175236 | 6.095853882 | 1.2118707 | 0.2869464 | 0.999175987 |
| ENSMUSG00000022681 | Ntan1 | protein_coding | 0.358241773 | 6.963301012 | 1.2118017 | 0.2869596 | 0.999175987 |
| ENSMUSG00000046711 | Hmga1 | protein_coding | 0.660668594 | 3.853199304 | 1.2110158 | 0.2871105 | 0.999175987 |
| ENSMUSG00000030849 | Fgfr2 | protein_coding | 0.345636681 | 6.727220766 | 1.2103795 | 0.2872328 | 0.999175987 |
| ENSMUSG00000027531 | Impa1 | protein_coding | -0.300378938 | 6.892489951 | 1.2090228 | 0.2874937 | 0.999175987 |
| ENSMUSG00000016757 | Ttll12 | protein_coding | 0.865434403 | 3.701757573 | 1.2088487 | 0.2875272 | 0.999175987 |
| ENSMUSG00000039771 | Polr2j | protein_coding | 0.384558682 | 7.22821898 | 1.2088476 | 0.2875274 | 0.999175987 |
| ENSMUSG00000049130 | C5ar1 | protein_coding | -1.692985667 | 2.071131458 | 1.2086158 | 0.287572 | 0.999175987 |
| ENSMUSG00000029095 | Ablim2 | protein_coding | 0.380402108 | 6.51450988 | 1.2085814 | 0.2875786 | 0.999175987 |
| ENSMUSG00000061374 | Fiz1 | protein_coding | 0.762513069 | 3.292743738 | 1.2084691 | 0.2876002 | 0.999175987 |
| ENSMUSG00000036822 | Topors | protein_coding | -0.860635865 | 4.026247427 | 1.2085273 | 0.2876151 | 0.999175987 |
| ENSMUSG00000018750 | Zbtb4 | protein_coding | -0.347507874 | 6.130943418 | 1.2061615 | 0.288045 | 0.999175987 |
| ENSMUSG00000025007 | Aldh18a1 | protein_coding | -0.894606648 | 4.630399413 | 1.212115 | 0.288233 | 0.999175987 |
| ENSMUSG00000031134 | Rbmx | protein_coding | -0.451466159 | 6.32331531 | 1.2051006 | 0.2882497 | 0.999175987 |
| ENSMUSG00000091994 | E130317F20Rik | bidirectional_promote | 1.167386616 | 2.538111552 | 1.2043442 | 0.2883958 | 0.999175987 |
| ENSMUSG00000069678 | Pcgf1 | protein_coding | 0.442327073 | 5.263945238 | 1.2037638 | 0.288508 | 0.999175987 |
| ENSMUSG00000042213 | Zfand4 | protein_coding | 1.176631812 | 2.652720383 | 1.2037339 | 0.2885138 | 0.999175987 |
| ENSMUSG00000036986 | Pml | protein_coding | -1.187146041 | 2.724934985 | 1.203606 | 0.2885385 | 0.999175987 |
| ENSMUSG00000002107 | Celf2 | protein_coding | -0.345501556 | 7.824730066 | 1.2032251 | 0.2886122 | 0.999175987 |
| ENSMUSG00000028820 | Sfpq | protein_coding | -0.417238735 | 5.63432158 | 1.2024346 | 0.2887652 | 0.999175987 |
| ENSMUSG00000027936 | Crtc2 | protein_coding | 0.968309164 | 3.167573966 | 1.2016159 | 0.2889237 | 0.999175987 |
| ENSMUSG00000021079 | Timm9 | protein_coding | 0.402552768 | 7.126191626 | 1.2006241 | 0.2891159 | 0.999175987 |
| ENSMUSG00000018567 | Gabarap | protein_coding | 0.341138141 | 9.783309314 | 1.2004368 | 0.2891522 | 0.999175987 |
| ENSMUSG00000011837 | Snapc2 | protein_coding | 0.51260418 | 5.29531142 | 1.1994643 | 0.2893409 | 0.999175987 |
| ENSMUSG00000032469 | Dbr1 | protein_coding | -0.910214414 | 3.535123546 | 1.1982068 | 0.2895851 | 0.999175987 |
| ENSMUSG00000041775 | Mapk1ip1 | protein_coding | -0.483600156 | 5.904104184 | 1.1979313 | 0.2896387 | 0.999175987 |
| ENSMUSG00000025905 | Oprk1 | protein_coding | 0.695908908 | 4.134242713 | 1.1974495 | 0.2897323 | 0.999175987 |
| ENSMUSG00000034801 | Sos2 | protein_coding | 0.500633255 | 4.382323573 | 1.1962652 | 0.2899628 | 0.999175987 |
| ENSMUSG00000026878 | Rab14 | protein_coding | -0.340215996 | 7.850148033 | 1.1959028 | 0.2900333 | 0.999175987 |
| ENSMUSG00000029203 | Ube2k | protein_coding | 0.351819626 | 7.384558187 | 1.1957331 | 0.2900664 | 0.999175987 |
| ENSMUSG00000039488 | Cntn5 | protein_coding | -0.914905217 | 3.316897843 | 1.1956799 | 0.2900767 | 0.999175987 |
| ENSMUSG00000019845 | Tube1 | protein_coding | -1.337272636 | 2.32141918 | 1.195516 | 0.2901087 | 0.999175987 |
| ENSMUSG00000030643 | Rab30 | protein_coding | 0.561185182 | 5.044029253 | 1.1950648 | 0.2901966 | 0.999175987 |
| ENSMUSG00000047496 | Rnf152 | protein_coding | -0.758438332 | 3.959589499 | 1.1949395 | 0.290221 | 0.999175987 |
| ENSMUSG00000031221 | Igbp1 | protein_coding | 0.497599601 | 6.132569825 | 1.1946532 | 0.2903027 | 0.999175987 |
| ENSMUSG00000040524 | Zfp609 | protein_coding | -0.70355844 | 3.534331021 | 1.1944483 | 0.2903167 | 0.999175987 |

| ensembl_gene_id | external_gene_name | gene_biotype | log2 (fold change) | log2 (counts per million) | F-value | p-value | False discovery rate |
|---|---|---|---|---|---|---|---|
| ENSMUSG00000058690 | Ccser2 | protein_coding | -0.427655284 | 5.493256827 | 1.1937786 | 0.2904474 | 0.999175987 |
| ENSMUSG00000035901 | Dennd5a | protein_coding | -0.355291648 | 6.827601737 | 1.1931523 | 0.2905696 | 0.999175987 |
| ENSMUSG00000049792 | Bag5 | protein_coding | -0.352204869 | 6.379971003 | 1.1929759 | 0.290604 | 0.999175987 |
| ENSMUSG00000028868 | Wasf2 | protein_coding | -0.529552127 | 4.571624132 | 1.1929031 | 0.2906183 | 0.999175987 |
| ENSMUSG00000021721 | Htr1a | protein_coding | -0.66340115 | 4.1537979 | 1.1924348 | 0.2907097 | 0.999175987 |
| ENSMUSG00000018820 | Zfyve27 | protein_coding | -0.468641199 | 5.296735211 | 1.1921719 | 0.2907611 | 0.999175987 |
| ENSMUSG00000020777 | Acox1 | protein_coding | -0.402334929 | 6.792100537 | 1.1910293 | 0.2909845 | 0.999175987 |
| ENSMUSG00000036026 | Tmem63b | protein_coding | 0.493111029 | 5.229959865 | 1.1909259 | 0.2910047 | 0.999175987 |
| ENSMUSG00000033323 | Ctdp1 | protein_coding | -1.454481257 | 2.510626562 | 1.1906692 | 0.2910549 | 0.999175987 |
| ENSMUSG00000022051 | Bnip3l | protein_coding | -0.301742824 | 7.068522962 | 1.1903742 | 0.2911126 | 0.999175987 |
| ENSMUSG00000033020 | Polr2f | protein_coding | 0.393645696 | 7.719445482 | 1.1902984 | 0.2911275 | 0.999175987 |
| ENSMUSG00000097393 | D030068K23Rik | lincRNA | 0.589436451 | 5.219533529 | 1.1897806 | 0.2912288 | 0.999175987 |
| ENSMUSG00000005268 | Prlr | protein_coding | 0.993992817 | 3.490090336 | 1.1883836 | 0.2915284 | 0.999175987 |
| ENSMUSG00000065126 | Snord104 | snoRNA | 0.886747048 | 3.124400437 | 1.1881088 | 0.2915565 | 0.999175987 |
| ENSMUSG00000045519 | Zfp560 | protein_coding | 0.457260943 | 5.7476614 | 1.1878618 | 0.2916049 | 0.999175987 |
| ENSMUSG00000029822 | Osbpl3 | protein_coding | -1.031090701 | 2.998039118 | 1.1864033 | 0.2918912 | 0.999175987 |
| ENSMUSG00000038248 | Sobp | protein_coding | -0.55369523 | 4.816461611 | 1.1859831 | 0.2919738 | 0.999175987 |
| ENSMUSG00000042249 | Grk3 | protein_coding | -0.704527626 | 4.939557764 | 1.1854617 | 0.2921021 | 0.999175987 |
| ENSMUSG00000019782 | Rwdd1 | protein_coding | 0.620685152 | 4.027359334 | 1.1846613 | 0.2922337 | 0.999175987 |
| ENSMUSG00000021752 | Kctd6 | protein_coding | -0.556859951 | 4.238287717 | 1.1844489 | 0.2922755 | 0.999175987 |
| ENSMUSG00000032458 | Copb2 | protein_coding | -0.43240686 | 6.600300104 | 1.182619 | 0.2926359 | 0.999175987 |
| ENSMUSG00000021917 | Spcs1 | protein_coding | 0.41999531 | 8.833726053 | 1.1825228 | 0.2926548 | 0.999175987 |
| ENSMUSG00000062070 | Pgk1 | protein_coding | -0.426478761 | 5.410924104 | 1.1822236 | 0.2927138 | 0.999175987 |
| ENSMUSG00000022272 | Myo10 | protein_coding | -0.534831869 | 4.921313803 | 1.1813604 | 0.2928841 | 0.999175987 |
| ENSMUSG00000027175 | Tcp11l1 | protein_coding | 0.608783266 | 3.964658979 | 1.1800006 | 0.2931526 | 0.999175987 |
| ENSMUSG00000112121 | AC163903.1 | lincRNA | 1.111183617 | 2.695447453 | 1.1795829 | 0.2932352 | 0.999175987 |
| ENSMUSG00000020941 | Map3k14 | protein_coding | 1.538902875 | 2.231274153 | 1.1793769 | 0.2932759 | 0.999175987 |
| ENSMUSG00000041343 | Ankrd42 | protein_coding | 0.862294676 | 3.169587784 | 1.1793089 | 0.2932893 | 0.999175987 |
| ENSMUSG00000000489 | Pdgfb | protein_coding | 0.817400703 | 3.367482814 | 1.1792774 | 0.2932955 | 0.999175987 |
| ENSMUSG00000030386 | Zfp606 | protein_coding | -0.762492458 | 3.567408348 | 1.1790146 | 0.2933475 | 0.999175987 |
| ENSMUSG00000092274 | Neat1 | lincRNA | 0.813980108 | 3.937045109 | 1.1788048 | 0.2934148 | 0.999175987 |
| ENSMUSG00000049112 | Oxtr | protein_coding | 1.077822501 | 2.82457283 | 1.1764651 | 0.2938523 | 0.999175987 |
| ENSMUSG00000017764 | Zswim1 | protein_coding | 0.547544492 | 5.908942062 | 1.1765394 | 0.2938633 | 0.999175987 |
| ENSMUSG00000025193 | Cutc | protein_coding | -0.546695359 | 4.378400117 | 1.1751253 | 0.2941181 | 0.999175987 |
| ENSMUSG00000051339 | 2900026A02Rik | protein_coding | -0.665344713 | 4.055889803 | 1.1749731 | 0.2941483 | 0.999175987 |
| ENSMUSG00000046169 | Adamts6 | protein_coding | 0.817876754 | 3.154787281 | 1.174465 | 0.2942491 | 0.999175987 |
| ENSMUSG00000068856 | Sf3b4 | protein_coding | 0.606331903 | 4.936819582 | 1.1743282 | 0.2942763 | 0.999175987 |
| ENSMUSG00000073565 | Prr16 | protein_coding | -1.353275062 | 2.509511292 | 1.1739074 | 0.2943599 | 0.999175987 |
| ENSMUSG00000046876 | Atxn1 | protein_coding | -0.631383655 | 6.992813897 | 1.1850973 | 0.2944653 | 0.999175987 |
| ENSMUSG00000040025 | Ythdf2 | protein_coding | -0.366400725 | 6.242873669 | 1.1731391 | 0.2945126 | 0.999175987 |
| ENSMUSG00000026546 | Cfap45 | protein_coding | 0.762218537 | 3.53867362 | 1.1719056 | 0.294758 | 0.999175987 |
| ENSMUSG00000054196 | Cthrc1 | protein_coding | 0.409014261 | 5.38972456 | 1.1719034 | 0.2947585 | 0.999175987 |
| ENSMUSG00000028394 | Pole3 | protein_coding | 0.348049364 | 6.041170929 | 1.1716535 | 0.2948082 | 0.999175987 |
| ENSMUSG00000029076 | Sdf4 | protein_coding | 0.310382867 | 8.512245284 | 1.1709173 | 0.2949549 | 0.999175987 |
| ENSMUSG00000039838 | Slc45a1 | protein_coding | 0.506633415 | 6.048189818 | 1.1709107 | 0.2949819 | 0.999175987 |
| ENSMUSG00000059187 | Fam19a1 | protein_coding | -0.615598009 | 4.408622389 | 1.1688632 | 0.2953645 | 0.999175987 |
| ENSMUSG00000086382 | Chrna1os | antisense_RNA | -0.990009998 | 2.764194342 | 1.1687462 | 0.2953879 | 0.999175987 |
| ENSMUSG00000028741 | Mrto4 | protein_coding | 0.389168252 | 6.218993079 | 1.1684462 | 0.2954478 | 0.999175987 |
| ENSMUSG00000015937 | H2afy | protein_coding | 0.707562321 | 4.298275487 | 1.1676884 | 0.2955992 | 0.999175987 |
| ENSMUSG00000025777 | Gdap1 | protein_coding | -0.341402875 | 6.892165658 | 1.1669485 | 0.2957471 | 0.999175987 |
| ENSMUSG00000022466 | Rpap3 | protein_coding | 0.428441499 | 5.866990366 | 1.1663921 | 0.2958584 | 0.999175987 |
| ENSMUSG00000035798 | Zdhhc17 | protein_coding | -0.42933285 | 5.272496121 | 1.1663043 | 0.2958759 | 0.999175987 |
| ENSMUSG00000098087 | Gm17750 | lincRNA | 0.768344713 | 3.3932452 | 1.1652163 | 0.2960937 | 0.999175987 |
| ENSMUSG00000040606 | Kazn | protein_coding | 0.448439309 | 5.968817437 | 1.1651545 | 0.2961061 | 0.999175987 |
| ENSMUSG00000003410 | Elavl3 | protein_coding | 0.329571121 | 6.897769516 | 1.1651277 | 0.2961115 | 0.999175987 |
| ENSMUSG00000022516 | Nudt16l1 | protein_coding | 0.330475882 | 7.037987779 | 1.1645938 | 0.2962185 | 0.999175987 |
| ENSMUSG00000031409 | Tceal6 | protein_coding | 0.337064225 | 6.238640912 | 1.1641296 | 0.2963115 | 0.999175987 |
| ENSMUSG00000055150 | Zfp78 | protein_coding | 0.738671881 | 3.449240101 | 1.16393 | 0.2963515 | 0.999175987 |
| ENSMUSG00000034484 | Snx2 | protein_coding | 0.468253309 | 7.439300765 | 1.1645259 | 0.2963694 | 0.999175987 |
| ENSMUSG00000026281 | Dtymk | protein_coding | 0.439075678 | 6.249767161 | 1.1630051 | 0.2965371 | 0.999175987 |
| ENSMUSG00000053024 | Cntn2 | protein_coding | -0.477053386 | 5.471520605 | 1.1629052 | 0.2965571 | 0.999175987 |
| ENSMUSG00000032902 | Slc16a1 | protein_coding | -0.547154295 | 4.132903036 | 1.1628411 | 0.29657 | 0.999175987 |
| ENSMUSG00000030177 | Ccdc77 | protein_coding | -1.262181717 | 2.536813618 | 1.1613418 | 0.2968712 | 0.999175987 |
| ENSMUSG00000037300 | Ttc13 | protein_coding | -0.935928918 | 3.753155692 | 1.1610792 | 0.297017 | 0.999175987 |
| ENSMUSG00000024309 | Pfdn6 | protein_coding | 0.294743495 | 8.166682519 | 1.1605679 | 0.2970268 | 0.999175987 |
| ENSMUSG00000025949 | Pikfyve | protein_coding | -0.594033194 | 4.167803261 | 1.1583961 | 0.2974641 | 0.999175987 |
| ENSMUSG00000038301 | Snx10 | protein_coding | -0.387291278 | 8.472617737 | 1.1578859 | 0.297567 | 0.999175987 |
| ENSMUSG00000022883 | Robo1 | protein_coding | 0.601478921 | 3.967815851 | 1.1574064 | 0.2976637 | 0.999175987 |
| ENSMUSG00000027598 | Itch | protein_coding | -0.502493482 | 4.714672349 | 1.1568201 | 0.297782 | 0.999175987 |
| ENSMUSG00000030201 | Lrp6 | protein_coding | -0.69859959 | 3.641438725 | 1.1564259 | 0.2978616 | 0.999175987 |
| ENSMUSG00000029192 | Tbc1d14 | protein_coding | -0.475005059 | 5.501214275 | 1.1563936 | 0.2978681 | 0.999175987 |
| ENSMUSG00000024082 | Ndufaf7 | protein_coding | 0.328756783 | 6.749561846 | 1.1558606 | 0.2979758 | 0.999175987 |
| ENSMUSG00000022748 | Cmss1 | protein_coding | -0.520718962 | 5.107140504 | 1.1557084 | 0.2980066 | 0.999175987 |
| ENSMUSG00000024790 | Sac3d1 | protein_coding | 0.659513752 | 3.673260541 | 1.1551933 | 0.2981107 | 0.999175987 |
| ENSMUSG00000020315 | Sptbn1 | protein_coding | -0.422908103 | 7.420372831 | 1.1550888 | 0.2981573 | 0.999175987 |
| ENSMUSG00000028409 | Smu1 | protein_coding | -0.357504251 | 6.920026931 | 1.1546668 | 0.2982171 | 0.999175987 |
| ENSMUSG00000024959 | Bad | protein_coding | 0.320859231 | 7.611176989 | 1.1524605 | 0.2986638 | 0.999175987 |
| ENSMUSG00000068551 | Zfp467 | protein_coding | -0.602589214 | 4.430571319 | 1.1519132 | 0.2987748 | 0.999175987 |

| ensembl_gene_id | external_gene_name | gene_biotype | log2 (fold change) | log2 (counts per million) | F-value | p-value | False discovery rate |
|---|---|---|---|---|---|---|---|
| ENSMUSG00000025871 | 4833439L19Rik | protein_coding | 0.425613171 | 8.268381697 | 1.1514566 | 0.2988929 | 0.999175987 |
| ENSMUSG00000029528 | Pxn | protein_coding | -1.183220682 | 2.558738636 | 1.1503152 | 0.2990991 | 0.999175987 |
| ENSMUSG00000029330 | Cds1 | protein_coding | -0.543115319 | 4.559815468 | 1.1491935 | 0.299327 | 0.999175987 |
| ENSMUSG00000031709 | Tbc1d9 | protein_coding | -0.487607197 | 4.883991709 | 1.1489807 | 0.2993703 | 0.999175987 |
| ENSMUSG00000061904 | Slc25a3 | protein_coding | -0.35007111 | 8.245816559 | 1.1482775 | 0.2995133 | 0.999175987 |
| ENSMUSG00000024429 | Gnl1 | protein_coding | 0.349620591 | 6.405451743 | 1.1480376 | 0.2995621 | 0.999175987 |
| ENSMUSG00000007670 | Khsrp | protein_coding | -0.451472159 | 5.071241666 | 1.1467866 | 0.2998169 | 0.999175987 |
| ENSMUSG00000029003 | Mad2l2 | protein_coding | 0.496664678 | 5.318783669 | 1.1462832 | 0.2999195 | 0.999175987 |
| ENSMUSG00000040855 | Reps2 | protein_coding | -0.440230882 | 6.145058229 | 1.1460047 | 0.2999762 | 0.999175987 |
| ENSMUSG00000016534 | Lamp2 | protein_coding | -0.411329696 | 6.080716892 | 1.1459579 | 0.2999858 | 0.999175987 |
| ENSMUSG00000055943 | Emc7 | protein_coding | 0.309058161 | 7.536572216 | 1.1459466 | 0.2999881 | 0.999175987 |
| ENSMUSG00000024483 | Ankhd1 | protein_coding | -0.486044937 | 4.962412996 | 1.1453077 | 0.3001184 | 0.999175987 |
| ENSMUSG00000028106 | Rprd2 | protein_coding | -0.510985214 | 4.85400875 | 1.1451849 | 0.3001435 | 0.999175987 |
| ENSMUSG00000022537 | Tmem44 | protein_coding | -0.759350479 | 3.575541263 | 1.1447868 | 0.3002247 | 0.999175987 |
| ENSMUSG00000010453 | Kansl3 | protein_coding | -0.52762572 | 5.673572314 | 1.1447285 | 0.3002366 | 0.999175987 |
| ENSMUSG00000020848 | Doc2b | protein_coding | 0.672526782 | 3.522851538 | 1.1444648 | 0.3002904 | 0.999175987 |
| ENSMUSG00000031767 | Nudt7 | protein_coding | -0.496081945 | 4.349246878 | 1.1439598 | 0.3003936 | 0.999175987 |
| ENSMUSG00000079111 | Kdelr2 | protein_coding | 0.479331018 | 6.432431385 | 1.1438548 | 0.3004405 | 0.999175987 |
| ENSMUSG00000034708 | Grn | protein_coding | 0.39817656 | 5.539982496 | 1.143529 | 0.3004816 | 0.999175987 |
| ENSMUSG00000035958 | Tdp2 | protein_coding | -0.630428936 | 3.702396804 | 1.1434531 | 0.3004971 | 0.999175987 |
| ENSMUSG00000019813 | Cep57l1 | protein_coding | 0.835572473 | 3.313911245 | 1.1419252 | 0.3008096 | 0.999175987 |
| ENSMUSG00000018761 | Mpdu1 | protein_coding | 0.358576815 | 6.826506948 | 1.1418519 | 0.3008246 | 0.999175987 |
| ENSMUSG00000039801 | 2410089E03Rik | protein_coding | 0.856586499 | 3.251676142 | 1.1415552 | 0.3008854 | 0.999175987 |
| ENSMUSG00000027895 | Kcnc4 | protein_coding | 0.46836886 | 5.077709349 | 1.1407996 | 0.3010401 | 0.999175987 |
| ENSMUSG00000087370 | Tmem170b | protein_coding | -0.541958129 | 5.335219299 | 1.1397578 | 0.3012537 | 0.999175987 |
| ENSMUSG00000112117 | RMST_1 | lincRNA | -0.68903542 | 3.455362308 | 1.1394044 | 0.3013262 | 0.999175987 |
| ENSMUSG00000032258 | Lca5 | protein_coding | -1.687072099 | 2.502113867 | 1.1416285 | 0.3013265 | 0.999175987 |
| ENSMUSG00000019806 | Aig1 | protein_coding | -0.485349029 | 6.224464513 | 1.1390361 | 0.3014271 | 0.999175987 |
| ENSMUSG00000020484 | Xbp1 | protein_coding | -0.482843736 | 5.883448601 | 1.1386147 | 0.3014883 | 0.999175987 |
| ENSMUSG00000056698 | Elmod3 | protein_coding | -0.440282922 | 5.625959417 | 1.1382689 | 0.3015593 | 0.999175987 |
| ENSMUSG00000019082 | Slc25a22 | protein_coding | 0.495058324 | 7.807801147 | 1.1419205 | 0.3017344 | 0.999175987 |
| ENSMUSG00000040729 | Cep126 | protein_coding | -1.181798881 | 3.213043619 | 1.138372 | 0.3017896 | 0.999175987 |
| ENSMUSG00000037997 | Parp11 | protein_coding | -0.571499481 | 4.771992288 | 1.1369521 | 0.3018299 | 0.999175987 |
| ENSMUSG00000024785 | Rcl1 | protein_coding | 0.67417498 | 4.641329716 | 1.1365393 | 0.3019148 | 0.999175987 |
| ENSMUSG00000040824 | Snrpd2 | protein_coding | 0.402005068 | 7.163819088 | 1.1364617 | 0.3019308 | 0.999175987 |
| ENSMUSG00000030254 | Rad18 | protein_coding | -0.802137715 | 3.375191328 | 1.136194 | 0.3019858 | 0.999175987 |
| ENSMUSG00000027018 | Hat1 | protein_coding | 0.426079818 | 5.333454055 | 1.1354617 | 0.3021366 | 0.999175987 |
| ENSMUSG00000034121 | Mks1 | protein_coding | 2.187530589 | 2.16448445 | 1.1398946 | 0.302271 | 0.999175987 |
| ENSMUSG00000032802 | Srxn1 | protein_coding | 0.391876126 | 6.536059391 | 1.1345135 | 0.3023319 | 0.999175987 |
| ENSMUSG00000053007 | Creb5 | protein_coding | -0.828565993 | 3.695559275 | 1.1342524 | 0.3023857 | 0.999175987 |
| ENSMUSG00000004980 | Hnrnpa2b1 | protein_coding | -0.301209019 | 8.948773507 | 1.1341079 | 0.3024155 | 0.999175987 |
| ENSMUSG00000022261 | Sdc2 | protein_coding | -0.39862406 | 6.526697747 | 1.1324534 | 0.3027569 | 0.999175987 |
| ENSMUSG00000034007 | Scaper | protein_coding | -0.42661961 | 5.731327711 | 1.132025 | 0.3028454 | 0.999175987 |
| ENSMUSG00000026418 | Tnni1 | protein_coding | -0.456720046 | 4.722029144 | 1.1316026 | 0.3029327 | 0.999175987 |
| ENSMUSG00000023266 | Frs3 | protein_coding | 0.582186335 | 4.120808735 | 1.1315331 | 0.302947 | 0.999175987 |
| ENSMUSG00000024817 | Uhrf2 | protein_coding | 0.507263421 | 4.927797077 | 1.1313421 | 0.3029865 | 0.999175987 |
| ENSMUSG00000045201 | Lrrc3b | protein_coding | 0.489414487 | 4.919093809 | 1.1307402 | 0.303111 | 0.999175987 |
| ENSMUSG00000022285 | Ywhaz | protein_coding | -0.335698529 | 10.25803832 | 1.1305042 | 0.3031598 | 0.999175987 |
| ENSMUSG00000038115 | Ano2 | protein_coding | -1.136669058 | 3.114696155 | 1.1302883 | 0.3032297 | 0.999175987 |
| ENSMUSG00000036905 | C1qb | protein_coding | -0.741393428 | 6.327932409 | 1.1440443 | 0.3033816 | 0.999175987 |
| ENSMUSG00000033306 | Lpp | protein_coding | -0.7034195 | 3.491464405 | 1.1293402 | 0.3034007 | 0.999175987 |
| ENSMUSG00000015961 | Adss | protein_coding | -0.38778281 | 5.890868046 | 1.127605 | 0.3037604 | 0.999175987 |
| ENSMUSG00000040433 | Zbtb38 | protein_coding | -0.508255716 | 7.245201503 | 1.1322973 | 0.3037691 | 0.999175987 |
| ENSMUSG00000006010 | BC003331 | protein_coding | -0.414941703 | 5.896520046 | 1.1272108 | 0.3038422 | 0.999175987 |
| ENSMUSG00000018387 | Shroom1 | protein_coding | -0.938972944 | 2.888356445 | 1.1271798 | 0.3038486 | 0.999175987 |
| ENSMUSG00000035764 | Fbxo45 | protein_coding | 0.522709364 | 4.888335855 | 1.1268855 | 0.3039097 | 0.999175987 |
| ENSMUSG00000000278 | Scpep1 | protein_coding | -0.78345021 | 3.180799759 | 1.1267511 | 0.3039376 | 0.999175987 |
| ENSMUSG00000032418 | Me1 | protein_coding | -0.550530094 | 4.417696933 | 1.1264412 | 0.3040019 | 0.999175987 |
| ENSMUSG00000022437 | Samm50 | protein_coding | 0.404924216 | 6.467781763 | 1.1262314 | 0.3040455 | 0.999175987 |
| ENSMUSG00000038172 | Ttc39b | protein_coding | -0.570482232 | 4.595200301 | 1.1254353 | 0.3042109 | 0.999175987 |
| ENSMUSG00000057440 | Mpp7 | protein_coding | -0.815399519 | 3.335569132 | 1.125292 | 0.3042407 | 0.999175987 |
| ENSMUSG00000039477 | Tnrc18 | protein_coding | 0.672946886 | 3.305940623 | 1.1251047 | 0.3042797 | 0.999175987 |
| ENSMUSG00000006241 | Ccdc159 | protein_coding | 0.731507425 | 3.402527741 | 1.1243082 | 0.3044453 | 0.999175987 |
| ENSMUSG00000032384 | Csnk1g1 | protein_coding | -0.483657571 | 5.506981906 | 1.1230337 | 0.3047107 | 0.999175987 |
| ENSMUSG00000034602 | Mon2 | protein_coding | -0.567521499 | 4.907468043 | 1.1224533 | 0.3048317 | 0.999175987 |
| ENSMUSG00000031591 | Asah1 | protein_coding | -0.376602612 | 6.231860948 | 1.1211053 | 0.3051128 | 0.999175987 |
| ENSMUSG00000025217 | Btrc | protein_coding | -0.501364343 | 6.648808445 | 1.1232841 | 0.3051386 | 0.999175987 |
| ENSMUSG00000042712 | Tceal9 | protein_coding | 0.371703999 | 9.303714622 | 1.1204988 | 0.3052395 | 0.999175987 |
| ENSMUSG00000008475 | Arpc5 | protein_coding | 0.304232977 | 7.254298124 | 1.1199773 | 0.3053484 | 0.999175987 |
| ENSMUSG00000035620 | Ric8b | protein_coding | -0.447387563 | 5.915712258 | 1.119526 | 0.3054427 | 0.999175987 |
| ENSMUSG00000055862 | Izumo4 | protein_coding | 0.362836555 | 6.276003752 | 1.1189915 | 0.3055545 | 0.999175987 |
| ENSMUSG00000043557 | Mdga1 | protein_coding | -0.893503214 | 3.316663069 | 1.1184862 | 0.3056602 | 0.999175987 |
| ENSMUSG00000025025 | Mxi1 | protein_coding | -0.468394864 | 5.435271417 | 1.1183234 | 0.3056942 | 0.999175987 |
| ENSMUSG00000023094 | Msrb2 | protein_coding | 0.378881547 | 5.95553684 | 1.1175118 | 0.3058641 | 0.999175987 |
| ENSMUSG00000037926 | Ssh2 | protein_coding | -0.552978052 | 4.286228006 | 1.1172705 | 0.3059147 | 0.999175987 |
| ENSMUSG00000026600 | Soat1 | protein_coding | -0.593188659 | 4.214065536 | 1.1160664 | 0.306167 | 0.999175987 |
| ENSMUSG00000028525 | Pde4b | protein_coding | -0.36385528 | 7.408735093 | 1.1156868 | 0.3062467 | 0.999175987 |
| ENSMUSG00000031708 | Tecr | protein_coding | 0.336442693 | 10.67126226 | 1.1156773 | 0.3062487 | 0.999175987 |

| ensembl_gene_id | external_gene_name | gene_biotype | log2 (fold change) | log2 (counts per million) | F-value | p-value | False discovery rate |
|---|---|---|---|---|---|---|---|
| ENSMUSG00000028999 | Rint1 | protein_coding | -0.559494069 | 3.966373281 | 1.1155957 | 0.3062658 | 0.999175987 |
| ENSMUSG00000034709 | Ppp1r21 | protein_coding | -0.421610357 | 5.805066777 | 1.1154643 | 0.3062933 | 0.999175987 |
| ENSMUSG00000053334 | Ficd | protein_coding | -0.82785036 | 3.187597239 | 1.1147319 | 0.3064471 | 0.999175987 |
| ENSMUSG00000015090 | Ptgds | protein_coding | -0.485748176 | 9.368961098 | 1.1169937 | 0.3064475 | 0.999175987 |
| ENSMUSG00000060216 | Arrb2 | protein_coding | 0.612990817 | 4.17512589 | 1.1144772 | 0.3065006 | 0.999175987 |
| ENSMUSG00000060735 | Rxfp3 | protein_coding | -1.180596354 | 2.619317142 | 1.1144076 | 0.3065152 | 0.999175987 |
| ENSMUSG00000035139 | Secisbp2 | protein_coding | 0.554014533 | 4.315297583 | 1.1140371 | 0.306593 | 0.999175987 |
| ENSMUSG00000030649 | Anapc15 | protein_coding | 0.681114478 | 3.334285731 | 1.1139797 | 0.3066051 | 0.999175987 |
| ENSMUSG00000078713 | Tomm5 | protein_coding | 0.346697005 | 6.628975089 | 1.1139589 | 0.3066094 | 0.999175987 |
| ENSMUSG00000020571 | Pdia6 | protein_coding | 0.294838057 | 7.807010753 | 1.1136204 | 0.3066806 | 0.999175987 |
| ENSMUSG00000001289 | Pfdn5 | protein_coding | 0.415283164 | 9.841359291 | 1.1134003 | 0.3067268 | 0.999175987 |
| ENSMUSG00000035864 | Syt1 | protein_coding | -0.332672559 | 9.368887695 | 1.1123414 | 0.3069496 | 0.999175987 |
| ENSMUSG00000024180 | Tmem8 | protein_coding | 0.917113351 | 2.937878498 | 1.1121128 | 0.3069977 | 0.999175987 |
| ENSMUSG00000053950 | Adnp2 | protein_coding | -0.765599275 | 3.565987606 | 1.1115686 | 0.3071123 | 0.999175987 |
| ENSMUSG00000016018 | Skiv2l2 | protein_coding | -0.46940933 | 5.557120922 | 1.1111349 | 0.3072036 | 0.999175987 |
| ENSMUSG00000106775 | C130093G08Rik | TEC | -0.953192267 | 4.64571464 | 1.1211668 | 0.3072452 | 0.999175987 |
| ENSMUSG00000030605 | Mfge8 | protein_coding | -0.478124419 | 7.272475873 | 1.1136618 | 0.307312 | 0.999175987 |
| ENSMUSG00000037606 | Osbpl5 | protein_coding | 0.587703905 | 4.151074454 | 1.1103199 | 0.3073754 | 0.999175987 |
| ENSMUSG00000001773 | Folh1 | protein_coding | -1.005839546 | 2.654719932 | 1.1095475 | 0.3075383 | 0.999175987 |
| ENSMUSG00000020107 | Anapc16 | protein_coding | 0.297558468 | 7.218092901 | 1.1094 | 0.3075695 | 0.999175987 |
| ENSMUSG00000021120 | Pigh | protein_coding | 0.934147136 | 3.330616155 | 1.1073371 | 0.3080052 | 0.999175987 |
| ENSMUSG00000022546 | Gpt | protein_coding | 0.69410961 | 4.057947977 | 1.1072633 | 0.3080208 | 0.999175987 |
| ENSMUSG00000004933 | Matk | protein_coding | 0.377172834 | 7.258529824 | 1.1070314 | 0.3080699 | 0.999175987 |
| ENSMUSG00000000766 | Oprm1 | protein_coding | 0.507292759 | 5.499168234 | 1.1065559 | 0.3081705 | 0.999175987 |
| ENSMUSG00000053684 | BC048403 | protein_coding | 0.998644014 | 2.762959654 | 1.1064228 | 0.3081986 | 0.999175987 |
| ENSMUSG00000000827 | Tpd52l2 | protein_coding | 0.323306364 | 7.251869599 | 1.1063605 | 0.3082118 | 0.999175987 |
| ENSMUSG00000079562 | Maea | protein_coding | 0.376756412 | 7.573083674 | 1.1062607 | 0.308233 | 0.999175987 |
| ENSMUSG00000041112 | Elmo1 | protein_coding | 0.340163891 | 8.636083867 | 1.105844 | 0.3083212 | 0.999175987 |
| ENSMUSG00000005813 | Metap1 | protein_coding | 0.446306295 | 5.827359175 | 1.1053883 | 0.3084177 | 0.999175987 |
| ENSMUSG00000093904 | Tomm20 | protein_coding | -0.340838225 | 6.853735272 | 1.1046905 | 0.3085656 | 0.999175987 |
| ENSMUSG00000000374 | Trappc10 | protein_coding | -0.516858932 | 4.455311871 | 1.1038999 | 0.3087332 | 0.999175987 |
| ENSMUSG00000061859 | Patj | protein_coding | 0.822220217 | 5.584489827 | 1.1150102 | 0.3088233 | 0.999175987 |
| ENSMUSG00000034724 | Cnot6l | protein_coding | -0.476087583 | 5.221231872 | 1.1032308 | 0.3088752 | 0.999175987 |
| ENSMUSG00000086477 | Gm15506 | processed_transcript | -0.901493859 | 2.935751709 | 1.1029732 | 0.3089299 | 0.999175987 |
| ENSMUSG00000028107 | Tars2 | protein_coding | 0.435414282 | 5.71337882 | 1.1029161 | 0.308942 | 0.999175987 |
| ENSMUSG00000035910 | Dcdc2a | protein_coding | 0.628130117 | 4.129149183 | 1.1025872 | 0.3090119 | 0.999175987 |
| ENSMUSG00000028621 | Cyb5rl | protein_coding | -0.670185556 | 4.490891648 | 1.1020198 | 0.3091324 | 0.999175987 |
| ENSMUSG00000032526 | Ss18l2 | protein_coding | 0.30735563 | 8.073288735 | 1.1015098 | 0.3092408 | 0.999175987 |
| ENSMUSG00000045071 | E130308A19Rik | protein_coding | 0.763292192 | 3.44162894 | 1.1010186 | 0.3093453 | 0.999175987 |
| ENSMUSG00000053253 | Ndfip2 | protein_coding | -0.389645762 | 5.975936153 | 1.1008668 | 0.3093776 | 0.999175987 |
| ENSMUSG00000096995 | 2810029C07Rik | bidirectional_promote | 0.671392551 | 3.791981806 | 1.1008341 | 0.3093846 | 0.999175987 |
| ENSMUSG00000055109 | Gm15155 | protein_coding | 1.22438622 | 2.188059548 | 1.1004769 | 0.3094605 | 0.999175987 |
| ENSMUSG00000052406 | Rexo4 | protein_coding | 0.53295489 | 5.485134177 | 1.0985239 | 0.3098766 | 0.999175987 |
| ENSMUSG00000036053 | Fmnl2 | protein_coding | -0.62069437 | 4.242882389 | 1.0983132 | 0.3099215 | 0.999175987 |
| ENSMUSG00000078570 | 1110065P20Rik | protein_coding | 1.110687437 | 2.752640373 | 1.0982307 | 0.3099391 | 0.999175987 |
| ENSMUSG00000030016 | Zfp638 | protein_coding | -0.522951044 | 5.660084374 | 1.0975633 | 0.3100815 | 0.999175987 |
| ENSMUSG00000068101 | Cenpm | protein_coding | 1.16430203 | 2.727712796 | 1.0973083 | 0.3101359 | 0.999175987 |
| ENSMUSG00000028804 | Csmd2 | protein_coding | 0.89610825 | 3.077264071 | 1.0969098 | 0.310221 | 0.999175987 |
| ENSMUSG00000026393 | Nek7 | protein_coding | -0.491566696 | 6.044481512 | 1.0967867 | 0.3102723 | 0.999175987 |
| ENSMUSG00000032185 | Carm1 | protein_coding | 0.601184056 | 4.119040385 | 1.0940935 | 0.3108233 | 0.999175987 |
| ENSMUSG00000097585 | E230029C05Rik | processed_transcript | -0.725655656 | 3.481865834 | 1.0940683 | 0.3108287 | 0.999175987 |
| ENSMUSG00000063511 | Snrnp70 | protein_coding | -0.490773653 | 4.871768543 | 1.0940553 | 0.3108315 | 0.999175987 |
| ENSMUSG00000034958 | Atcay | protein_coding | 0.505071754 | 6.108660041 | 1.0938551 | 0.3108992 | 0.999175987 |
| ENSMUSG00000030410 | Dmwd | protein_coding | 0.404171517 | 6.561215865 | 1.0934385 | 0.3109636 | 0.999175987 |
| ENSMUSG00000030246 | Ldhb | protein_coding | -0.310482165 | 10.4757629 | 1.0934168 | 0.3109682 | 0.999175987 |
| ENSMUSG00000044988 | Ucn3 | protein_coding | -0.917620897 | 3.261059349 | 1.0934031 | 0.3109712 | 0.999175987 |
| ENSMUSG00000051439 | Cd14 | protein_coding | 0.983323919 | 2.931237514 | 1.093052 | 0.3110464 | 0.999175987 |
| ENSMUSG00000052581 | Lrrtm4 | protein_coding | -0.58221076 | 5.281600629 | 1.0921993 | 0.3112541 | 0.999175987 |
| ENSMUSG00000036693 | Nop14 | protein_coding | 0.521209891 | 5.11560451 | 1.0918059 | 0.3113137 | 0.999175987 |
| ENSMUSG00000025427 | Rnf165 | protein_coding | -0.632920822 | 4.695807413 | 1.0910099 | 0.3114845 | 0.999175987 |
| ENSMUSG00000027546 | Atp9a | protein_coding | 0.321973101 | 7.260064044 | 1.0906112 | 0.3115702 | 0.999175987 |
| ENSMUSG00000029007 | Agtrap | protein_coding | 0.601130735 | 4.356498875 | 1.0900585 | 0.311689 | 0.999175987 |
| ENSMUSG00000032392 | Parp16 | protein_coding | 1.105706068 | 2.656371158 | 1.0898561 | 0.3117325 | 0.999175987 |
| ENSMUSG00000055407 | Map6 | protein_coding | -0.504426299 | 5.469052074 | 1.088875 | 0.3119435 | 0.999175987 |
| ENSMUSG00000029211 | Gabra4 | protein_coding | -0.477033446 | 6.717935929 | 1.0893496 | 0.3119746 | 0.999175987 |
| ENSMUSG00000026791 | Slc2a8 | protein_coding | -1.128952152 | 2.732372097 | 1.0886894 | 0.3119835 | 0.999175987 |
| ENSMUSG00000048583 | Igf2 | protein_coding | -0.992936321 | 4.068313374 | 1.0946739 | 0.3121501 | 0.999175987 |
| ENSMUSG00000022217 | Emc9 | protein_coding | 0.300605776 | 6.944502775 | 1.087717 | 0.3121929 | 0.999175987 |
| ENSMUSG00000034820 | Cpsf7 | protein_coding | 1.38301811 | 2.343303687 | 1.0874984 | 0.31224 | 0.999175987 |
| ENSMUSG00000025583 | Rptor | protein_coding | -0.462008527 | 4.777241995 | 1.087396 | 0.312262 | 0.999175987 |
| ENSMUSG00000019978 | Epb41l2 | protein_coding | -0.539695771 | 4.435299091 | 1.0873625 | 0.3122692 | 0.999175987 |
| ENSMUSG00000028619 | Tceanc2 | protein_coding | 0.690206379 | 3.57439876 | 1.0860348 | 0.3125556 | 0.999175987 |
| ENSMUSG00000002820 | Atg4d | protein_coding | -0.783758911 | 3.81406085 | 1.0860163 | 0.3125596 | 0.999175987 |
| ENSMUSG00000015994 | Fnta | protein_coding | 0.472549981 | 5.94889738 | 1.0859525 | 0.3125733 | 0.999175987 |
| ENSMUSG00000030744 | Rps3 | protein_coding | 0.376664019 | 10.97915624 | 1.0856671 | 0.3126349 | 0.999175987 |
| ENSMUSG00000029364 | Wsb2 | protein_coding | 0.423525293 | 5.49505858 | 1.085456 | 0.3126805 | 0.999175987 |
| ENSMUSG00000023830 | Igf2r | protein_coding | -0.758947709 | 3.655759802 | 1.0851786 | 0.3127404 | 0.999175987 |
| ENSMUSG00000021796 | Bmpr1a | protein_coding | -0.565619223 | 4.532161124 | 1.0841406 | 0.3129647 | 0.999175987 |

| ensembl_gene_id | external_gene_name | gene_biotype | log2 (fold change) | log2 (counts per million) | F-value | p-value | False discovery rate |
|---|---|---|---|---|---|---|---|
| ENSMUSG00000074415 | 2610203C20Rik | processed_transcript | 0.432546618 | 5.094700376 | 1.0831851 | 0.3131713 | 0.999175987 |
| ENSMUSG00000024026 | Glo1 | protein_coding | -0.298403076 | 7.490100632 | 1.0827052 | 0.3132752 | 0.999175987 |
| ENSMUSG00000039546 | Ajap1 | protein_coding | -0.388931702 | 5.796713387 | 1.0823016 | 0.3133626 | 0.999175987 |
| ENSMUSG00000071708 | Sms | protein_coding | 0.36506935 | 6.018823401 | 1.0820849 | 0.3134095 | 0.999175987 |
| ENSMUSG00000108778 | Gm20083 | lincRNA | -1.173692065 | 2.465986827 | 1.0819893 | 0.3134302 | 0.999175987 |
| ENSMUSG00000048000 | Gigyf2 | protein_coding | -0.344299036 | 6.073082081 | 1.0817139 | 0.3134899 | 0.999175987 |
| ENSMUSG00000030209 | Grin2b | protein_coding | -0.404052218 | 6.206190231 | 1.0807299 | 0.3137032 | 0.999175987 |
| ENSMUSG00000024273 | 2700062C07Rik | protein_coding | -0.41718309 | 5.358026639 | 1.0801529 | 0.3138284 | 0.999175987 |
| ENSMUSG00000024002 | Brd4 | protein_coding | -0.369832401 | 6.56187823 | 1.0799339 | 0.3138759 | 0.999175987 |
| ENSMUSG00000063320 | 1190007I07Rik | protein_coding | 0.494925725 | 4.499521849 | 1.0797907 | 0.313907 | 0.999175987 |
| ENSMUSG00000001520 | Nrip2 | protein_coding | 0.461002043 | 6.737210411 | 1.0796414 | 0.3139642 | 0.999175987 |
| ENSMUSG00000000948 | Gm38393 | protein_coding | 0.508237824 | 4.792417404 | 1.0792392 | 0.3140267 | 0.999175987 |
| ENSMUSG00000045427 | Hnrnph2 | protein_coding | -0.430121141 | 5.821643913 | 1.0783824 | 0.3142129 | 0.999175987 |
| ENSMUSG00000036775 | Decr2 | protein_coding | -0.329322392 | 6.638058043 | 1.0780487 | 0.3142855 | 0.999175987 |
| ENSMUSG00000009828 | Ick | protein_coding | -0.811159377 | 3.469201804 | 1.0779849 | 0.3142994 | 0.999175987 |
| ENSMUSG00000039853 | Trim14 | protein_coding | -1.509770684 | 2.136635728 | 1.0771729 | 0.314476 | 0.999175987 |
| ENSMUSG00000052146 | Rps10 | protein_coding | 0.444168868 | 5.612174742 | 1.0769399 | 0.3145267 | 0.999175987 |
| ENSMUSG00000000078 | Klf6 | protein_coding | -0.440407592 | 5.934176 | 1.0764214 | 0.3146396 | 0.999175987 |
| ENSMUSG00000024851 | Pitpnm1 | protein_coding | 0.416864707 | 5.789043041 | 1.0759261 | 0.3147475 | 0.999175987 |
| ENSMUSG00000005615 | Pcyt1a | protein_coding | -0.554359478 | 5.212254866 | 1.0755788 | 0.3148232 | 0.999175987 |
| ENSMUSG00000035674 | Ndufa3 | protein_coding | 0.455672502 | 10.02601084 | 1.075598 | 0.3148437 | 0.999175987 |
| ENSMUSG00000037890 | Wdr19 | protein_coding | -0.804320379 | 3.403430134 | 1.0751968 | 0.3149065 | 0.999175987 |
| ENSMUSG00000024687 | Osbp | protein_coding | 0.631148808 | 4.320719299 | 1.0745485 | 0.3150479 | 0.999175987 |
| ENSMUSG00000021033 | Gstz1 | protein_coding | 0.317978427 | 7.07085356 | 1.0740133 | 0.3151647 | 0.999175987 |
| ENSMUSG00000026781 | Acbd5 | protein_coding | -0.453524379 | 5.32265234 | 1.0726623 | 0.3154598 | 0.999175987 |
| ENSMUSG00000049751 | Rpl36al | protein_coding | 0.774715248 | 3.012442042 | 1.0725257 | 0.3154896 | 0.999175987 |
| ENSMUSG00000030352 | Tspan9 | protein_coding | -0.77390259 | 3.740494061 | 1.0718167 | 0.3156447 | 0.999175987 |
| ENSMUSG00000030655 | Smg1 | protein_coding | -0.660987302 | 3.940697427 | 1.0714981 | 0.3157144 | 0.999175987 |
| ENSMUSG00000061461 | Smim20 | protein_coding | -0.297899386 | 7.151173044 | 1.0705687 | 0.3159179 | 0.999175987 |
| ENSMUSG00000032212 | Sltm | protein_coding | -0.414532084 | 5.581230775 | 1.0702895 | 0.315979 | 0.999175987 |
| ENSMUSG00000060938 | Rpl26 | protein_coding | 0.423552752 | 8.23068949 | 1.0697759 | 0.3161162 | 0.999175987 |
| ENSMUSG00000025487 | Psmd13 | protein_coding | -0.308791132 | 7.360820135 | 1.0692119 | 0.3162152 | 0.999175987 |
| ENSMUSG00000018593 | Sparc | protein_coding | -0.590553065 | 9.204871844 | 1.0788169 | 0.3162256 | 0.999175987 |
| ENSMUSG00000025374 | Nabp2 | protein_coding | 0.486288495 | 5.12100011 | 1.0684399 | 0.3163846 | 0.999175987 |
| ENSMUSG00000022407 | Adsl | protein_coding | -0.421041068 | 6.061447949 | 1.0678703 | 0.3165096 | 0.999175987 |
| ENSMUSG00000085980 | Gm12408 | lincRNA | 0.952350433 | 2.757608371 | 1.0675785 | 0.3165737 | 0.999175987 |
| ENSMUSG00000031563 | Wwc2 | protein_coding | -0.579069623 | 4.512205655 | 1.0670632 | 0.3166869 | 0.999175987 |
| ENSMUSG00000037012 | Hk1 | protein_coding | 0.622862399 | 7.295175808 | 1.0775462 | 0.3167909 | 0.999175987 |
| ENSMUSG00000026833 | Olfm1 | protein_coding | 0.325037137 | 7.687292613 | 1.0665298 | 0.3168042 | 0.999175987 |
| ENSMUSG00000068206 | Pick1 | protein_coding | 0.543672149 | 4.937320041 | 1.0664703 | 0.3168173 | 0.999175987 |
| ENSMUSG00000042363 | Lgalsl | protein_coding | -0.334929487 | 6.228389488 | 1.0662006 | 0.3168766 | 0.999175987 |
| ENSMUSG00000022565 | Plec | protein_coding | -0.493967789 | 4.769189975 | 1.0660081 | 0.3169189 | 0.999175987 |
| ENSMUSG00000044066 | Cep68 | protein_coding | 0.69569204 | 3.705952981 | 1.0654057 | 0.3170515 | 0.999175987 |
| ENSMUSG00000040652 | Oaz2 | protein_coding | 0.352103513 | 6.202725126 | 1.0644785 | 0.3172557 | 0.999175987 |
| ENSMUSG00000111966 | AL589661.1 | processed_pseudoge | 0.773533693 | 3.01471568 | 1.0640907 | 0.3173411 | 0.999175987 |
| ENSMUSG00000019863 | Qrsl1 | protein_coding | -0.685014784 | 4.740587059 | 1.0639069 | 0.3174062 | 0.999175987 |
| ENSMUSG00000058070 | Eml1 | protein_coding | -0.779817663 | 4.21443811 | 1.0635182 | 0.3174919 | 0.999175987 |
| ENSMUSG00000021708 | Rasgrf2 | protein_coding | -0.415613484 | 6.114797076 | 1.0631282 | 0.3175534 | 0.999175987 |
| ENSMUSG00000032249 | Anp32a | protein_coding | -0.331532251 | 6.482187072 | 1.06282 | 0.3176213 | 0.999175987 |
| ENSMUSG00000030337 | Vamp1 | protein_coding | 0.440813178 | 7.451851913 | 1.0626797 | 0.3177416 | 0.999175987 |
| ENSMUSG00000059834 | Sclt1 | protein_coding | -0.861313134 | 3.183665427 | 1.0617587 | 0.3178557 | 0.999175987 |
| ENSMUSG00000028478 | Clta | protein_coding | 0.383901102 | 6.887830154 | 1.0616504 | 0.3178796 | 0.999175987 |
| ENSMUSG00000037996 | Slc24a2 | protein_coding | -0.439922106 | 6.09735924 | 1.0616411 | 0.3178816 | 0.999175987 |
| ENSMUSG00000076441 | Ass1 | protein_coding | -0.515177317 | 6.249848399 | 1.0625932 | 0.3179149 | 0.999175987 |
| ENSMUSG00000015745 | Plekho1 | protein_coding | -0.866469672 | 3.719621721 | 1.0615457 | 0.3179273 | 0.999175987 |
| ENSMUSG00000022208 | Jph4 | protein_coding | -0.36266652 | 6.716425145 | 1.061128 | 0.317995 | 0.999175987 |
| ENSMUSG00000020153 | Ndufs7 | protein_coding | 0.313961038 | 9.745717511 | 1.0605922 | 0.3181135 | 0.999175987 |
| ENSMUSG00000021756 | Il6st | protein_coding | -0.593387124 | 4.07293837 | 1.0602894 | 0.3181805 | 0.999175987 |
| ENSMUSG00000059824 | Dbp | protein_coding | -0.631224011 | 3.487219614 | 1.0601832 | 0.3182039 | 0.999175987 |
| ENSMUSG00000022370 | Mrpl13 | protein_coding | 0.323656628 | 7.425329315 | 1.0597245 | 0.3183055 | 0.999175987 |
| ENSMUSG00000026872 | Zeb2 | protein_coding | -0.530453973 | 4.886389772 | 1.0593542 | 0.3183874 | 0.999175987 |
| ENSMUSG00000037134 | Prmt9 | protein_coding | -0.698146455 | 3.579216557 | 1.0592572 | 0.3184089 | 0.999175987 |
| ENSMUSG00000024787 | Snx15 | protein_coding | 0.320106414 | 6.245294448 | 1.0588587 | 0.3184971 | 0.999175987 |
| ENSMUSG00000023018 | Smarcd1 | protein_coding | 0.376131948 | 6.309771219 | 1.0587464 | 0.318522 | 0.999175987 |
| ENSMUSG00000078350 | Smim1 | protein_coding | 0.43476118 | 5.456210545 | 1.0578711 | 0.318716 | 0.999175987 |
| ENSMUSG00000028969 | Cdk5 | protein_coding | 0.304381039 | 8.546438567 | 1.0575644 | 0.318784 | 0.999175987 |
| ENSMUSG00000024948 | Map4k2 | protein_coding | 0.800993622 | 4.428703934 | 1.0600687 | 0.3188548 | 0.999175987 |
| ENSMUSG00000024565 | Sall3 | protein_coding | -1.068927964 | 2.707476908 | 1.0569382 | 0.318923 | 0.999175987 |
| ENSMUSG00000001829 | Clpb | protein_coding | 0.388692473 | 5.379357391 | 1.0567145 | 0.3189726 | 0.999175987 |
| ENSMUSG00000020411 | Nipal4 | protein_coding | -1.014675243 | 2.931015967 | 1.0557057 | 0.3191967 | 0.999175987 |
| ENSMUSG00000025995 | Wdr75 | protein_coding | 0.380486412 | 5.615716546 | 1.0542255 | 0.3195258 | 0.999175987 |
| ENSMUSG00000027996 | Sfrp2 | protein_coding | -0.820366118 | 2.939593684 | 1.0541675 | 0.3195387 | 0.999175987 |
| ENSMUSG00000057234 | Mettl15 | protein_coding | -0.661374417 | 3.394642558 | 1.0541049 | 0.3195526 | 0.999175987 |
| ENSMUSG00000035310 | Lin54 | protein_coding | -0.865131261 | 2.820126201 | 1.0539474 | 0.3195877 | 0.999175987 |
| ENSMUSG00000039156 | Stim2 | protein_coding | -0.54524921 | 4.815811894 | 1.052299 | 0.3199549 | 0.999175987 |
| ENSMUSG00000063316 | Rpl27 | protein_coding | 0.606980043 | 3.778532114 | 1.0515429 | 0.3201235 | 0.999175987 |
| ENSMUSG00000025873 | Faf2 | protein_coding | -0.408104681 | 5.906230298 | 1.0513367 | 0.3201695 | 0.999175987 |
| ENSMUSG00000033917 | Gde1 | protein_coding | -0.502541876 | 5.794515996 | 1.0511993 | 0.3202246 | 0.999175987 |

| ensembl_gene_id | external_gene_name | gene_biotype | log2 (fold change) | log2 (counts per million) | F-value | p-value | False discovery rate |
|---|---|---|---|---|---|---|---|
| ENSMUSG00000027506 | Tpd52 | protein_coding | 0.419395292 | 6.305528105 | 1.0509016 | 0.3202666 | 0.999175987 |
| ENSMUSG00000017167 | Cntnap1 | protein_coding | 0.535875721 | 4.798864447 | 1.0507252 | 0.320306 | 0.999175987 |
| ENSMUSG00000056019 | Zfp709 | protein_coding | -0.966199845 | 2.776760095 | 1.0491827 | 0.3206506 | 0.999175987 |
| ENSMUSG00000002635 | Pdcd2l | protein_coding | 0.586973328 | 4.720181586 | 1.0486233 | 0.3207758 | 0.999175987 |
| ENSMUSG00000010406 | Mrpl52 | protein_coding | 0.360295115 | 7.395968681 | 1.0483596 | 0.3208348 | 0.999175987 |
| ENSMUSG00000060538 | Tmem219 | protein_coding | 0.518969351 | 4.342771791 | 1.0480648 | 0.3209008 | 0.999175987 |
| ENSMUSG00000033943 | Mga | protein_coding | -0.512485042 | 5.35196869 | 1.0456847 | 0.3214342 | 0.999175987 |
| ENSMUSG00000027886 | 1700013F07Rik | protein_coding | -1.560707574 | 2.351329396 | 1.0451191 | 0.3215855 | 0.999175987 |
| ENSMUSG00000022587 | Ly6e | protein_coding | 0.353612044 | 7.940055716 | 1.0449267 | 0.3216043 | 0.999175987 |
| ENSMUSG00000040462 | Os9 | protein_coding | 0.302008462 | 6.832922169 | 1.0446826 | 0.3216591 | 0.999175987 |
| ENSMUSG00000022416 | Cacna1i | protein_coding | -0.957497633 | 2.987274636 | 1.0440177 | 0.3218085 | 0.999175987 |
| ENSMUSG00000018733 | Pex12 | protein_coding | -0.609561387 | 4.360905925 | 1.0438089 | 0.3218555 | 0.999175987 |
| ENSMUSG00000030062 | Rpn1 | protein_coding | 0.291162761 | 6.965638885 | 1.0437187 | 0.3218757 | 0.999175987 |
| ENSMUSG00000038552 | Fndc4 | protein_coding | -0.49481648 | 5.636499316 | 1.0437051 | 0.3218788 | 0.999175987 |
| ENSMUSG00000026028 | Trak2 | protein_coding | -0.534321753 | 5.226720733 | 1.0430039 | 0.3220365 | 0.999175987 |
| ENSMUSG00000034430 | Zxdc | protein_coding | 0.611450791 | 3.944649992 | 1.0428999 | 0.3220599 | 0.999175987 |
| ENSMUSG00000032348 | Gsta4 | protein_coding | 0.312944197 | 7.262439524 | 1.042328 | 0.3221886 | 0.999175987 |
| ENSMUSG00000046922 | Gpr6 | protein_coding | -0.828215687 | 3.465183134 | 1.0420299 | 0.3222557 | 0.999175987 |
| ENSMUSG00000002658 | Gtf2f1 | protein_coding | 0.389393349 | 6.596767364 | 1.0415695 | 0.3223594 | 0.999175987 |
| ENSMUSG00000024248 | Cox7a2l | protein_coding | 0.295056882 | 8.40287394 | 1.0411145 | 0.3224619 | 0.999175987 |
| ENSMUSG00000022663 | Atg3 | protein_coding | -0.442196589 | 5.160404018 | 1.0407104 | 0.322553 | 0.999175987 |
| ENSMUSG00000001999 | Blvra | protein_coding | -0.298925698 | 6.923639702 | 1.0396006 | 0.3228034 | 0.999175987 |
| ENSMUSG00000028005 | Gucy1b3 | protein_coding | -0.331561959 | 7.007594023 | 1.0394596 | 0.3228352 | 0.999175987 |
| ENSMUSG00000020661 | Dnmt3a | protein_coding | -0.360815822 | 5.956895013 | 1.0394201 | 0.3228441 | 0.999175987 |
| ENSMUSG00000042129 | Rassf4 | protein_coding | -0.68558218 | 3.448326088 | 1.0390124 | 0.3229362 | 0.999175987 |
| ENSMUSG00000053040 | Aph1c | protein_coding | 0.719373926 | 3.38231935 | 1.0377517 | 0.3232211 | 0.999175987 |
| ENSMUSG00000042097 | Zfp239 | protein_coding | 0.370780037 | 6.28642213 | 1.0373501 | 0.3233119 | 0.999175987 |
| ENSMUSG00000015971 | Actr8 | protein_coding | -0.379047758 | 6.486960432 | 1.0364821 | 0.3235083 | 0.999175987 |
| ENSMUSG00000026798 | Coq4 | protein_coding | 0.362373235 | 5.82078392 | 1.0357249 | 0.3236798 | 0.999175987 |
| ENSMUSG00000029111 | Nelfa | protein_coding | 0.559228127 | 5.752998423 | 1.0358211 | 0.3237462 | 0.999175987 |
| ENSMUSG00000041354 | Rgl2 | protein_coding | 1.276499199 | 2.553597632 | 1.034402 | 0.3239798 | 0.999175987 |
| ENSMUSG00000031851 | Ntpcr | protein_coding | 0.330917734 | 6.280122565 | 1.0340575 | 0.3240579 | 0.999175987 |
| ENSMUSG00000087574 | C030037D09Rik | lincRNA | -0.750631895 | 3.542319725 | 1.0334705 | 0.3241912 | 0.999175987 |
| ENSMUSG00000000631 | Myo18a | protein_coding | 0.456726844 | 5.756674525 | 1.0333749 | 0.3242129 | 0.999175987 |
| ENSMUSG00000046985 | Tapt1 | protein_coding | -0.614851637 | 3.982898694 | 1.0333394 | 0.324221 | 0.999175987 |
| ENSMUSG00000025534 | Gusb | protein_coding | 0.421080154 | 5.25349744 | 1.0330505 | 0.3242866 | 0.999175987 |
| ENSMUSG00000018661 | Cog1 | protein_coding | 0.684023767 | 3.643888327 | 1.0325114 | 0.3244091 | 0.999175987 |
| ENSMUSG00000068923 | Syt11 | protein_coding | -0.333786602 | 9.840075036 | 1.0323335 | 0.3244495 | 0.999175987 |
| ENSMUSG00000020783 | Ncbp3 | protein_coding | -0.580423623 | 4.194204079 | 1.0316387 | 0.3246075 | 0.999175987 |
| ENSMUSG00000020257 | Wdr82 | protein_coding | -0.356857149 | 6.41093458 | 1.031056 | 0.3247401 | 0.999175987 |
| ENSMUSG00000020273 | Papolg | protein_coding | -0.582577807 | 4.466994118 | 1.0307307 | 0.3248142 | 0.999175987 |
| ENSMUSG00000028898 | Trnau1ap | protein_coding | 0.615487224 | 4.140814043 | 1.0302141 | 0.3249318 | 0.999175987 |
| ENSMUSG00000062252 | Lhfpl5 | protein_coding | -0.35850722 | 6.03676927 | 1.029534 | 0.3250868 | 0.999175987 |
| ENSMUSG00000041073 | Nacad | protein_coding | 0.506651189 | 4.855918819 | 1.0294894 | 0.325097 | 0.999175987 |
| ENSMUSG00000021974 | Fgf9 | protein_coding | 0.466300507 | 5.384579814 | 1.0291611 | 0.3251718 | 0.999175987 |
| ENSMUSG00000055681 | Cope | protein_coding | 0.281673199 | 9.279655957 | 1.0286783 | 0.325282 | 0.999175987 |
| ENSMUSG00000048218 | Amigo2 | protein_coding | 0.535518107 | 4.458766099 | 1.0286185 | 0.3252956 | 0.999175987 |
| ENSMUSG00000031700 | Gpt2 | protein_coding | -0.544242595 | 4.450509533 | 1.0285951 | 0.3253009 | 0.999175987 |
| ENSMUSG00000055817 | Mta3 | protein_coding | -0.554082219 | 4.98990718 | 1.0281335 | 0.3254063 | 0.999175987 |
| ENSMUSG00000024414 | Mrpl27 | protein_coding | 0.315281644 | 7.467434165 | 1.0273775 | 0.3255789 | 0.999175987 |
| ENSMUSG00000038291 | Snx25 | protein_coding | -0.57907011 | 4.481618643 | 1.0273775 | 0.3255789 | 0.999175987 |
| ENSMUSG00000071076 | Jund | protein_coding | 0.490537082 | 4.43190559 | 1.0269961 | 0.3256661 | 0.999175987 |
| ENSMUSG00000025764 | Jade1 | protein_coding | 0.371490486 | 5.582317761 | 1.0261645 | 0.3258562 | 0.999175987 |
| ENSMUSG00000053716 | Dusp7 | protein_coding | -0.562049363 | 4.516290361 | 1.0256009 | 0.3259851 | 0.999175987 |
| ENSMUSG00000018909 | Arrb1 | protein_coding | -0.334431799 | 6.224803928 | 1.0240764 | 0.3263343 | 0.999175987 |
| ENSMUSG00000022428 | Cby1 | protein_coding | 0.319948629 | 7.321213859 | 1.0237088 | 0.3264185 | 0.999175987 |
| ENSMUSG00000035559 | Mpv17l2 | protein_coding | 0.890374545 | 3.124717197 | 1.022909 | 0.326602 | 0.999175987 |
| ENSMUSG00000037703 | Lzts3 | protein_coding | -0.4464707 | 5.886241849 | 1.0229073 | 0.3266024 | 0.999175987 |
| ENSMUSG00000096956 | Snhg18 | lincRNA | -1.092074228 | 2.863060401 | 1.0228988 | 0.3266043 | 0.999175987 |
| ENSMUSG00000046442 | Ppm1e | protein_coding | -0.411655461 | 5.906311378 | 1.0219291 | 0.3268269 | 0.999175987 |
| ENSMUSG00000019890 | Nts | protein_coding | 0.841319999 | 7.780597208 | 1.055224 | 0.3268415 | 0.999175987 |
| ENSMUSG00000022463 | Srebf2 | protein_coding | -0.36347488 | 5.999905884 | 1.0213372 | 0.3269629 | 0.999175987 |
| ENSMUSG00000087174 | 5530601H04Rik | processed_transcript | 0.419670762 | 5.187209242 | 1.0210412 | 0.327031 | 0.999175987 |
| ENSMUSG00000072501 | Phf20l1 | protein_coding | -0.544660203 | 4.447356078 | 1.0197738 | 0.3273225 | 0.999175987 |
| ENSMUSG00000011884 | Gltp | protein_coding | -0.490048259 | 6.231484029 | 1.0191635 | 0.3274871 | 0.999175987 |
| ENSMUSG00000001755 | Coasy | protein_coding | 0.511867961 | 5.350843705 | 1.0189504 | 0.3275121 | 0.999175987 |
| ENSMUSG00000063015 | Ccni | protein_coding | -0.397298863 | 6.356034087 | 1.0183102 | 0.3276596 | 0.999175987 |
| ENSMUSG00000055200 | Sertad3 | protein_coding | -0.867122271 | 3.194498465 | 1.0175067 | 0.3278449 | 0.999175987 |
| ENSMUSG00000039356 | Exosc2 | protein_coding | -0.641816986 | 3.985303725 | 1.0163459 | 0.3281128 | 0.999175987 |
| ENSMUSG00000028234 | Rps20 | protein_coding | 0.403326756 | 7.008257084 | 1.0162103 | 0.3281682 | 0.999175987 |
| ENSMUSG00000027439 | Gzf1 | protein_coding | -0.40152845 | 5.316569817 | 1.0156473 | 0.3282742 | 0.999175987 |
| ENSMUSG00000021177 | Tdp1 | protein_coding | 0.548995705 | 4.693445551 | 1.0149241 | 0.3284414 | 0.999175987 |
| ENSMUSG00000026235 | Epha4 | protein_coding | -0.469206973 | 6.211637622 | 1.0149897 | 0.3284503 | 0.999175987 |
| ENSMUSG00000097316 | Gm10516 | antisense_RNA | 1.201243873 | 2.657396966 | 1.0147279 | 0.3284868 | 0.999175987 |
| ENSMUSG00000063281 | Zfp35 | protein_coding | 0.485398398 | 5.15766405 | 1.014569 | 0.3285235 | 0.999175987 |
| ENSMUSG00000011752 | Pgam1 | protein_coding | 0.561335495 | 3.725204375 | 1.0145232 | 0.3285341 | 0.999175987 |
| ENSMUSG00000045103 | Dmd | protein_coding | -0.446594578 | 4.828921698 | 1.0141802 | 0.3286135 | 0.999175987 |
| ENSMUSG00000024420 | Zfp521 | protein_coding | 0.60222028 | 4.269042558 | 1.0140756 | 0.3286377 | 0.999175987 |

| ensembl_gene_id | external_gene_name | gene_biotype | log2 (fold change) | log2 (counts per million) | F-value | p-value | False discovery rate |
|---|---|---|---|---|---|---|---|
| ENSMUSG00000020091 | Eif4ebp2 | protein_coding | 0.465178114 | 4.73625418 | 1.0136392 | 0.3287387 | 0.999175987 |
| ENSMUSG00000029151 | Slc30a3 | protein_coding | 1.629076013 | 2.706361497 | 1.0187325 | 0.3287615 | 0.999175987 |
| ENSMUSG00000020576 | Nbas | protein_coding | -0.625650888 | 4.351054025 | 1.0131743 | 0.3288464 | 0.999175987 |
| ENSMUSG00000036019 | Tmtc2 | protein_coding | 0.693871531 | 3.387662185 | 1.0131401 | 0.3288543 | 0.999175987 |
| ENSMUSG00000001285 | Myg1 | protein_coding | 0.340295321 | 7.240407338 | 1.0131358 | 0.3288553 | 0.999175987 |
| ENSMUSG00000097006 | 9530082P21Rik | lincRNA | 0.424744468 | 8.267374426 | 1.0124848 | 0.3290302 | 0.999175987 |
| ENSMUSG00000075316 | Scn9a | protein_coding | -0.605658633 | 3.5078237 | 1.0105046 | 0.3294658 | 0.999175987 |
| ENSMUSG00000016427 | Ndufa1 | protein_coding | 0.415800962 | 9.064829798 | 1.0103005 | 0.3295372 | 0.999175987 |
| ENSMUSG00000014226 | Cacybp | protein_coding | -0.349372854 | 7.549578527 | 1.0100149 | 0.3295796 | 0.999175987 |
| ENSMUSG00000034664 | Itga2b | protein_coding | 0.586175411 | 4.189217898 | 1.0097021 | 0.3296523 | 0.999175987 |
| ENSMUSG00000058761 | Rnf169 | protein_coding | -0.394097483 | 5.562580829 | 1.0095644 | 0.3296843 | 0.999175987 |
| ENSMUSG00000042229 | Rabif | protein_coding | 0.322070354 | 6.905860524 | 1.0093565 | 0.3297326 | 0.999175987 |
| ENSMUSG00000003437 | Paf1 | protein_coding | 0.64780043 | 5.326865428 | 1.0118139 | 0.3297735 | 0.999175987 |
| ENSMUSG00000031129 | Slc9a9 | protein_coding | -0.605605053 | 3.493578143 | 1.0078577 | 0.3300814 | 0.999175987 |
| ENSMUSG00000006057 | Atp5g1 | protein_coding | 0.3385656 | 5.899637889 | 1.0068787 | 0.3303095 | 0.999175987 |
| ENSMUSG00000044469 | Tnfaip8l1 | protein_coding | -0.777178547 | 3.351556379 | 1.0067066 | 0.3303497 | 0.999175987 |
| ENSMUSG00000016319 | Slc25a5 | protein_coding | -0.275416937 | 9.055108358 | 1.0066162 | 0.3303707 | 0.999175987 |
| ENSMUSG00000005732 | Ranbp1 | protein_coding | 0.328455563 | 7.170496367 | 1.0056249 | 0.330602 | 0.999175987 |
| ENSMUSG00000026709 | Dars2 | protein_coding | 0.567274507 | 4.442531 | 1.0046422 | 0.3308315 | 0.999175987 |
| ENSMUSG00000027167 | Elp4 | protein_coding | -0.531393827 | 3.932947935 | 1.0044068 | 0.3308865 | 0.999175987 |
| ENSMUSG00000022637 | Cblb | protein_coding | -1.016687461 | 3.005102669 | 1.0033156 | 0.3311417 | 0.999175987 |
| ENSMUSG00000055676 | Gm5069 | transcribed_processe | 0.893485437 | 2.771218341 | 1.002815 | 0.3312588 | 0.999175987 |
| ENSMUSG00000060579 | Fhit | protein_coding | 0.43366993 | 5.12763996 | 1.0027964 | 0.3312632 | 0.999175987 |
| ENSMUSG00000038515 | Grtp1 | protein_coding | -0.824316917 | 3.133808473 | 1.0023051 | 0.3313782 | 0.999175987 |
| ENSMUSG00000050234 | Gja4 | protein_coding | 1.154532261 | 2.662100191 | 1.0017841 | 0.3315003 | 0.999175987 |
| ENSMUSG00000033871 | Ppargc1b | protein_coding | 0.72443775 | 3.937598235 | 1.0017516 | 0.3315079 | 0.999175987 |
| ENSMUSG00000025078 | Nhlrc2 | protein_coding | -0.918327481 | 2.748052881 | 1.0014838 | 0.3315707 | 0.999175987 |
| ENSMUSG00000028576 | Ift74 | protein_coding | -0.371898057 | 5.909639603 | 1.0010959 | 0.3316616 | 0.999175987 |
| ENSMUSG00000006411 | Nectin4 | protein_coding | 1.203861118 | 3.299685051 | 1.0056302 | 0.3318238 | 0.999175987 |
| ENSMUSG00000033751 | Gadd45gip1 | protein_coding | -1.010968446 | 3.098671897 | 0.9999172 | 0.3319381 | 0.999175987 |
| ENSMUSG00000006050 | Sra1 | protein_coding | 0.322484459 | 7.857434219 | 0.9996998 | 0.3319892 | 0.999175987 |
| ENSMUSG00000027160 | Ccdc34 | protein_coding | 0.842289369 | 3.2779481 | 0.9994215 | 0.3320546 | 0.999175987 |
| ENSMUSG00000051510 | Mafg | protein_coding | -0.384071879 | 5.839539025 | 0.9992656 | 0.3320912 | 0.999175987 |
| ENSMUSG00000040945 | Rcc2 | protein_coding | 0.347898338 | 5.898254329 | 0.9990981 | 0.3321305 | 0.999175987 |
| ENSMUSG00000050213 | Snip1 | protein_coding | -0.659975428 | 3.608189537 | 0.9990455 | 0.3321429 | 0.999175987 |
| ENSMUSG00000020513 | Tubd1 | protein_coding | -1.110100851 | 2.857777701 | 0.9986745 | 0.3322301 | 0.999175987 |
| ENSMUSG00000024843 | Chka | protein_coding | 0.570246183 | 4.607991209 | 0.9986528 | 0.3322352 | 0.999175987 |
| ENSMUSG00000020758 | Itgb4 | protein_coding | -0.684498267 | 3.371077176 | 0.9981386 | 0.3323561 | 0.999175987 |
| ENSMUSG00000087674 | 4930447M23Rik | lincRNA | 0.742536792 | 3.176993293 | 0.997033 | 0.3326162 | 0.999175987 |
| ENSMUSG00000056131 | Pgm3 | protein_coding | -0.738358458 | 4.301884487 | 0.9967255 | 0.3327124 | 0.999175987 |
| ENSMUSG00000027282 | Mtch2 | protein_coding | 0.268074449 | 7.647824628 | 0.9963491 | 0.3327773 | 0.999175987 |
| ENSMUSG00000041225 | Arhgap12 | protein_coding | -0.387951439 | 5.38148747 | 0.996169 | 0.3328197 | 0.999175987 |
| ENSMUSG00000098176 | Ccdc166 | protein_coding | 0.677743493 | 4.102231282 | 0.9961568 | 0.3328226 | 0.999175987 |
| ENSMUSG00000041483 | Zfp281 | protein_coding | -0.532229749 | 4.817946654 | 0.9958632 | 0.3328918 | 0.999175987 |
| ENSMUSG00000058638 | Zfp110 | protein_coding | -0.762577896 | 3.539110823 | 0.9945785 | 0.3331948 | 0.999175987 |
| ENSMUSG00000093803 | Ppp2r3d | protein_coding | 0.647206275 | 3.402143341 | 0.9945074 | 0.3332116 | 0.999175987 |
| ENSMUSG00000028044 | Cks1b | protein_coding | -0.704375629 | 3.204712317 | 0.9944654 | 0.3332215 | 0.999175987 |
| ENSMUSG00000032553 | Srprb | protein_coding | 0.383306411 | 5.847270819 | 0.9936636 | 0.3334108 | 0.999175987 |
| ENSMUSG00000028414 | Fktn | protein_coding | -0.410941521 | 4.980741172 | 0.9929615 | 0.3335768 | 0.999175987 |
| ENSMUSG00000029673 | Auts2 | protein_coding | -0.301799159 | 6.258686383 | 0.99268 | 0.3336433 | 0.999175987 |
| ENSMUSG00000037149 | Ddx1 | protein_coding | -0.303032673 | 7.647847429 | 0.9924608 | 0.3336951 | 0.999175987 |
| ENSMUSG00000025287 | Acot9 | protein_coding | -0.421930669 | 5.547245871 | 0.9924016 | 0.3337091 | 0.999175987 |
| ENSMUSG00000057497 | Fam136a | protein_coding | 0.316426722 | 6.854044648 | 0.9920366 | 0.3337955 | 0.999175987 |
| ENSMUSG00000005142 | Man2b1 | protein_coding | 0.506282151 | 4.325568572 | 0.9920337 | 0.3337962 | 0.999175987 |
| ENSMUSG00000021087 | Rtn1 | protein_coding | 0.291635321 | 10.12214384 | 0.991212 | 0.3339907 | 0.999175987 |
| ENSMUSG00000045648 | Vwc2l | protein_coding | -0.54539816 | 3.978575379 | 0.991163 | 0.3340023 | 0.999175987 |
| ENSMUSG00000040699 | Limd2 | protein_coding | -0.52495752 | 5.42800098 | 0.9910491 | 0.3340293 | 0.999175987 |
| ENSMUSG00000057101 | Zfp180 | protein_coding | -0.435569946 | 5.044179791 | 0.9907834 | 0.3340922 | 0.999175987 |
| ENSMUSG00000028669 | Pithd1 | protein_coding | 0.322200288 | 6.480969073 | 0.9906965 | 0.3341128 | 0.999175987 |
| ENSMUSG00000026319 | 2310035C23Rik | protein_coding | -0.481142741 | 5.275347878 | 0.9903666 | 0.334191 | 0.999175987 |
| ENSMUSG00000022421 | Nptxr | protein_coding | -0.467948811 | 6.482046768 | 0.9906865 | 0.3342015 | 0.999175987 |
| ENSMUSG00000005413 | Hmox1 | protein_coding | 0.68638344 | 3.43502437 | 0.9902957 | 0.3342078 | 0.999175987 |
| ENSMUSG00000073700 | Klhl21 | protein_coding | -0.731354216 | 3.31837598 | 0.990087 | 0.3342573 | 0.999175987 |
| ENSMUSG00000027603 | Ggt7 | protein_coding | 0.372565095 | 7.073577587 | 0.9899192 | 0.334297 | 0.999175987 |
| ENSMUSG00000051319 | 1500011K16Rik | protein_coding | 0.311318284 | 7.289265781 | 0.9894465 | 0.3344092 | 0.999175987 |
| ENSMUSG00000062661 | Ncs1 | protein_coding | 0.373830814 | 10.21342779 | 0.9886297 | 0.334603 | 0.999175987 |
| ENSMUSG00000032417 | Rwdd2a | protein_coding | 0.309315717 | 7.882959122 | 0.9885292 | 0.3346269 | 0.999175987 |
| ENSMUSG00000063430 | Wscd2 | protein_coding | 0.668390251 | 3.351754842 | 0.9884687 | 0.3346412 | 0.999175987 |
| ENSMUSG00000070695 | Cntnap5a | protein_coding | -0.53046199 | 4.667547053 | 0.9884071 | 0.3346559 | 0.999175987 |
| ENSMUSG00000019528 | Gyg | protein_coding | 0.367486792 | 6.934157067 | 0.9879315 | 0.3347688 | 0.999175987 |
| ENSMUSG00000028444 | Cntfr | protein_coding | -0.685351654 | 3.91942776 | 0.986955 | 0.335001 | 0.999175987 |
| ENSMUSG00000022311 | Csmd3 | protein_coding | -0.633992936 | 4.866877705 | 0.9870012 | 0.3350137 | 0.999175987 |
| ENSMUSG00000078607 | 1810010H24Rik | protein_coding | -1.437044975 | 2.574770644 | 0.986776 | 0.3350672 | 0.999175987 |
| ENSMUSG00000068748 | Ptprz1 | protein_coding | -0.671724622 | 5.973749057 | 0.9951826 | 0.3350792 | 0.999175987 |
| ENSMUSG00000054418 | 2900041M22Rik | processed_transcript | 0.923018509 | 2.730016073 | 0.9855682 | 0.335331 | 0.999175987 |
| ENSMUSG00000036833 | Pnpla7 | protein_coding | -1.070752411 | 2.789872672 | 0.9855681 | 0.3353311 | 0.999175987 |
| ENSMUSG00000022658 | Tagln3 | protein_coding | 0.304363789 | 10.66124834 | 0.9854736 | 0.3353536 | 0.999175987 |
| ENSMUSG00000006920 | Ezh1 | protein_coding | 0.325138996 | 6.198793368 | 0.9854411 | 0.3353613 | 0.999175987 |

| ensembl_gene_id | external_gene_name | gene_biotype | log2 (fold change) | log2 (counts per million) | F-value | p-value | False discovery rate |
|---|---|---|---|---|---|---|---|
| ENSMUSG00000079215 | Zfp664 | protein_coding | -0.451839702 | 4.983712089 | 0.9854288 | 0.3353642 | 0.999175987 |
| ENSMUSG00000026974 | Zmynd19 | protein_coding | 0.551093198 | 3.884006846 | 0.9853943 | 0.3353725 | 0.999175987 |
| ENSMUSG00000022935 | Grik1 | protein_coding | -0.735184641 | 4.004442477 | 0.9852614 | 0.3354278 | 0.999175987 |
| ENSMUSG00000057181 | 5730455P16Rik | protein_coding | -0.368943164 | 6.975994754 | 0.9850016 | 0.335466 | 0.999175987 |
| ENSMUSG00000032437 | Stt3b | protein_coding | -0.520921766 | 5.189159621 | 0.9843703 | 0.3356165 | 0.999175987 |
| ENSMUSG00000048834 | Vstm2a | protein_coding | -0.318509345 | 6.941873829 | 0.9842449 | 0.3356464 | 0.999175987 |
| ENSMUSG00000027770 | Dhx36 | protein_coding | -0.421747735 | 6.075141452 | 0.983541 | 0.3358143 | 0.999175987 |
| ENSMUSG00000053799 | Exoc6 | protein_coding | -0.520111943 | 4.011580773 | 0.9829531 | 0.3359547 | 0.999175987 |
| ENSMUSG00000032733 | Snx33 | protein_coding | -0.828742766 | 3.184480107 | 0.982757 | 0.3360015 | 0.999175987 |
| ENSMUSG00000015522 | Arnt | protein_coding | 0.679717018 | 3.379198166 | 0.9821641 | 0.3361432 | 0.999175987 |
| ENSMUSG00000041718 | Alg13 | protein_coding | 1.229620706 | 2.759858993 | 0.9818323 | 0.3362461 | 0.999175987 |
| ENSMUSG00000018974 | Sart3 | protein_coding | -0.56242372 | 4.178009768 | 0.9816234 | 0.3362724 | 0.999175987 |
| ENSMUSG00000055053 | Nfic | protein_coding | 0.496506342 | 5.752961924 | 0.9815243 | 0.3363197 | 0.999175987 |
| ENSMUSG00000030225 | Dera | protein_coding | -0.760763997 | 3.302145592 | 0.9811034 | 0.3363968 | 0.999175987 |
| ENSMUSG00000025158 | Rfng | protein_coding | -0.551356215 | 4.553115981 | 0.9796698 | 0.33674 | 0.999175987 |
| ENSMUSG00000026388 | 3110009E18Rik | protein_coding | 0.546204672 | 4.377404602 | 0.9792796 | 0.3368335 | 0.999175987 |
| ENSMUSG00000002028 | Kmt2a | protein_coding | -0.516943238 | 5.263140623 | 0.9792763 | 0.3368343 | 0.999175987 |
| ENSMUSG00000028358 | Zfp618 | protein_coding | -1.108157036 | 2.889924072 | 0.9792618 | 0.3368378 | 0.999175987 |
| ENSMUSG00000041771 | Slc24a4 | protein_coding | -1.062828042 | 2.504585725 | 0.9789511 | 0.3369122 | 0.999175987 |
| ENSMUSG00000036777 | Anln | protein_coding | -0.463863156 | 4.705413853 | 0.9776948 | 0.3372136 | 0.999175987 |
| ENSMUSG00000010609 | Psen2 | protein_coding | 0.498631364 | 4.709192614 | 0.9772024 | 0.3373319 | 0.999175987 |
| ENSMUSG00000078640 | Gm11627 | protein_coding | -0.932016237 | 3.102746571 | 0.976883 | 0.3374086 | 0.999175987 |
| ENSMUSG00000063535 | Zfp773 | protein_coding | -1.392421659 | 2.375556669 | 0.976456 | 0.3375112 | 0.999175987 |
| ENSMUSG00000059689 | Zfp637 | protein_coding | 0.277973648 | 7.198845951 | 0.9758618 | 0.3376541 | 0.999175987 |
| ENSMUSG00000036002 | Fam214b | protein_coding | 0.532485512 | 4.200672327 | 0.9757576 | 0.3376792 | 0.999175987 |
| ENSMUSG00000038855 | Itpkb | protein_coding | -0.592629252 | 4.182160553 | 0.9756046 | 0.337716 | 0.999175987 |
| ENSMUSG00000032192 | Gnb5 | protein_coding | -0.28341782 | 7.735530839 | 0.9754332 | 0.3377572 | 0.999175987 |
| ENSMUSG00000015095 | Fbxw5 | protein_coding | 0.329106954 | 7.437499802 | 0.9745333 | 0.3379738 | 0.999175987 |
| ENSMUSG00000028096 | Gpr89 | protein_coding | -0.463488682 | 4.975555012 | 0.9744679 | 0.3379896 | 0.999175987 |
| ENSMUSG00000055116 | Arntl | protein_coding | -0.593583024 | 3.893684507 | 0.9742804 | 0.3380348 | 0.999175987 |
| ENSMUSG00000001211 | Agpat3 | protein_coding | 0.484982537 | 5.813345945 | 0.9739619 | 0.338135 | 0.999175987 |
| ENSMUSG00000033526 | Ppip5k1 | protein_coding | 0.489955501 | 4.9857567 | 0.9735465 | 0.3382116 | 0.999175987 |
| ENSMUSG00000025794 | Rpl14 | protein_coding | 0.322030994 | 9.060645753 | 0.9734985 | 0.3382232 | 0.999175987 |
| ENSMUSG00000026667 | Uhmk1 | protein_coding | 0.320936682 | 6.742017928 | 0.9731959 | 0.3382961 | 0.999175987 |
| ENSMUSG00000039179 | Tekt5 | protein_coding | 0.464078753 | 5.769587978 | 0.9729154 | 0.3383638 | 0.999175987 |
| ENSMUSG00000002109 | Ddb2 | protein_coding | -0.970917906 | 2.829398467 | 0.9725517 | 0.3384516 | 0.999175987 |
| ENSMUSG00000032181 | Scg3 | protein_coding | -0.372162271 | 7.058210045 | 0.9721689 | 0.3385439 | 0.999175987 |
| ENSMUSG00000031105 | Slc25a14 | protein_coding | 0.301565512 | 7.199902593 | 0.971785 | 0.3386366 | 0.999175987 |
| ENSMUSG00000007613 | Tgfbr1 | protein_coding | 1.223262514 | 3.187294785 | 0.9753501 | 0.3386898 | 0.999175987 |
| ENSMUSG00000020709 | Adap2 | protein_coding | -1.043385284 | 2.499203079 | 0.9711992 | 0.3387782 | 0.999175987 |
| ENSMUSG00000041040 | Fam117b | protein_coding | -1.016892033 | 2.756767461 | 0.9710242 | 0.3388204 | 0.999175987 |
| ENSMUSG00000002324 | Rec8 | protein_coding | 0.891899839 | 3.110103132 | 0.9709885 | 0.3388291 | 0.999175987 |
| ENSMUSG00000028527 | Ak4 | protein_coding | 0.478816402 | 4.634569626 | 0.9708115 | 0.3388719 | 0.999175987 |
| ENSMUSG00000039678 | Tbc1d13 | protein_coding | -0.652373181 | 3.440285826 | 0.970539 | 0.3389378 | 0.999175987 |
| ENSMUSG00000034442 | Trmt5 | protein_coding | 0.696544479 | 4.143886005 | 0.970485 | 0.3389743 | 0.999175987 |
| ENSMUSG00000001663 | Gstt1 | protein_coding | -0.424256054 | 5.76642151 | 0.9695536 | 0.3391762 | 0.999175987 |
| ENSMUSG00000053801 | Grwd1 | protein_coding | 0.404429923 | 6.023791231 | 0.9694965 | 0.33919 | 0.999175987 |
| ENSMUSG00000020415 | Pttg1 | protein_coding | -0.263250757 | 7.486446678 | 0.9692451 | 0.3392508 | 0.999175987 |
| ENSMUSG00000056553 | Ptprn2 | protein_coding | -0.664997127 | 3.91967776 | 0.9687823 | 0.3393629 | 0.999175987 |
| ENSMUSG00000036790 | Slitrk2 | protein_coding | 0.633293122 | 3.373395421 | 0.9684179 | 0.3394512 | 0.999175987 |
| ENSMUSG00000073705 | Apitd1 | protein_coding | 0.512899431 | 4.583008262 | 0.9680788 | 0.3395334 | 0.999175987 |
| ENSMUSG00000001767 | Crnkl1 | protein_coding | -0.536877368 | 4.060720349 | 0.9680087 | 0.3395504 | 0.999175987 |
| ENSMUSG00000041710 | Trpc5 | protein_coding | 0.812202795 | 3.258501206 | 0.9679987 | 0.3395528 | 0.999175987 |
| ENSMUSG00000001270 | Ckb | protein_coding | 0.39409059 | 10.89649594 | 0.9676269 | 0.339643 | 0.999175987 |
| ENSMUSG00000020150 | Gamt | protein_coding | 0.654890238 | 3.771287056 | 0.9674198 | 0.3396932 | 0.999175987 |
| ENSMUSG00000031673 | Cdh11 | protein_coding | -0.492213434 | 4.760155225 | 0.9672121 | 0.3397436 | 0.999175987 |
| ENSMUSG00000005354 | Txn2 | protein_coding | 0.302184515 | 9.312066701 | 0.9670714 | 0.3397777 | 0.999175987 |
| ENSMUSG00000015305 | Sash1 | protein_coding | -0.613603383 | 4.177191885 | 0.9669139 | 0.339816 | 0.999175987 |
| ENSMUSG00000078676 | Casc3 | protein_coding | -0.479354084 | 4.850136266 | 0.9668157 | 0.3398398 | 0.999175987 |
| ENSMUSG00000022840 | Adcy5 | protein_coding | -0.440128116 | 6.869010957 | 0.9663454 | 0.3400662 | 0.999175987 |
| ENSMUSG00000028986 | Klhl7 | protein_coding | -0.363570251 | 6.381126515 | 0.9652629 | 0.340217 | 0.999175987 |
| ENSMUSG00000015290 | Ubl4a | protein_coding | 0.328735491 | 7.399745294 | 0.9644714 | 0.3404095 | 0.999175987 |
| ENSMUSG00000022884 | Eif4a2 | protein_coding | 0.3400164 | 10.53247066 | 0.9629373 | 0.3407831 | 0.999175987 |
| ENSMUSG00000029175 | Slc35f6 | protein_coding | -0.462745775 | 5.03880609 | 0.9628917 | 0.3407942 | 0.999175987 |
| ENSMUSG00000039634 | Zfp189 | protein_coding | 0.764942964 | 3.583307246 | 0.9625689 | 0.3408729 | 0.999175987 |
| ENSMUSG00000028228 | Cpne3 | protein_coding | -0.852449016 | 3.502049258 | 0.9618874 | 0.3410624 | 0.999175987 |
| ENSMUSG00000055653 | Gpc3 | protein_coding | 0.498008317 | 4.439939438 | 0.9610793 | 0.3412363 | 0.999175987 |
| ENSMUSG00000027312 | Atrn | protein_coding | -0.553158701 | 5.29521595 | 0.9608287 | 0.3413208 | 0.999175987 |
| ENSMUSG00000063760 | Rnf217 | protein_coding | -0.665485315 | 3.511575161 | 0.9605694 | 0.3413608 | 0.999175987 |
| ENSMUSG00000035614 | Togaram1 | protein_coding | 0.541929859 | 4.576164842 | 0.9603646 | 0.3414108 | 0.999175987 |
| ENSMUSG00000016552 | Foxred2 | protein_coding | -0.911567072 | 2.864188343 | 0.9602206 | 0.341446 | 0.999175987 |
| ENSMUSG00000028059 | Arhgef2 | protein_coding | -0.530052988 | 4.502508929 | 0.9591608 | 0.3417051 | 0.999175987 |
| ENSMUSG00000002963 | Pnkp | protein_coding | 0.42916809 | 4.95504095 | 0.9587007 | 0.3418177 | 0.999175987 |
| ENSMUSG00000031555 | Adam9 | protein_coding | 0.608783288 | 3.504916784 | 0.9571534 | 0.3421967 | 0.999175987 |
| ENSMUSG00000028692 | Akr1a1 | protein_coding | 0.312861739 | 10.57220494 | 0.9570065 | 0.3422327 | 0.999175987 |
| ENSMUSG00000030729 | Pgm2l1 | protein_coding | -0.359813228 | 6.782313586 | 0.9551448 | 0.3426896 | 0.999175987 |
| ENSMUSG00000030880 | Polr3e | protein_coding | -0.702228643 | 4.903822005 | 0.9582828 | 0.3427696 | 0.999175987 |
| ENSMUSG00000013160 | Atp6v0d1 | protein_coding | 0.266210982 | 8.515038157 | 0.9546106 | 0.3428209 | 0.999175987 |

| ensembl_gene_id | external_gene_name | gene_biotype | log2 (fold change) | log2 (counts per million) | F-value | p-value | False discovery rate |
|---|---|---|---|---|---|---|---|
| ENSMUSG00000057335 | Cep170 | protein_coding | -0.414808226 | 5.24209487 | 0.954229 | 0.3429147 | 0.999175987 |
| ENSMUSG00000039474 | Wfs1 | protein_coding | 0.374898038 | 6.511998723 | 0.9540212 | 0.3429658 | 0.999175987 |
| ENSMUSG00000032425 | Zfp949 | protein_coding | 0.490570862 | 5.079971543 | 0.9533588 | 0.3431287 | 0.999175987 |
| ENSMUSG00000029147 | Ppm1g | protein_coding | -0.42283634 | 5.239946908 | 0.9532524 | 0.3431549 | 0.999175987 |
| ENSMUSG00000097560 | Gm26904 | lincRNA | 0.828210946 | 3.585983041 | 0.9532178 | 0.3431867 | 0.999175987 |
| ENSMUSG00000028785 | Hpca | protein_coding | 0.339968342 | 9.923674582 | 0.9515167 | 0.3435825 | 0.999175987 |
| ENSMUSG00000029627 | Zkscan14 | protein_coding | 0.447604712 | 4.75515435 | 0.9514567 | 0.3435973 | 0.999175987 |
| ENSMUSG00000025140 | Pycr1 | protein_coding | -0.754218753 | 3.771223146 | 0.9514051 | 0.34361 | 0.999175987 |
| ENSMUSG00000032727 | Mier3 | protein_coding | -0.398615988 | 5.296214816 | 0.9511706 | 0.3436678 | 0.999175987 |
| ENSMUSG00000024238 | Zeb1 | protein_coding | -0.334020958 | 6.277915847 | 0.9500103 | 0.3439542 | 0.999175987 |
| ENSMUSG00000029123 | Stk32b | protein_coding | 0.776742435 | 4.482213564 | 0.953552 | 0.3439835 | 0.999175987 |
| ENSMUSG00000057378 | Ryr3 | protein_coding | -0.793815633 | 3.667007566 | 0.9497874 | 0.3440325 | 0.999175987 |
| ENSMUSG00000047085 | Lrrc4b | protein_coding | 0.483891638 | 4.831499957 | 0.9495793 | 0.3440606 | 0.999175987 |
| ENSMUSG00000060935 | Tmem263 | protein_coding | -0.318741022 | 7.272686587 | 0.9495287 | 0.3440731 | 0.999175987 |
| ENSMUSG00000016946 | Kctd5 | protein_coding | -1.147162395 | 3.874375361 | 0.9584344 | 0.3441466 | 0.999175987 |
| ENSMUSG00000012609 | Ttll5 | protein_coding | 0.54829447 | 5.153133943 | 0.9489434 | 0.3442178 | 0.999175987 |
| ENSMUSG00000028057 | Rit1 | protein_coding | -0.331646438 | 6.135760638 | 0.9487636 | 0.3442622 | 0.999175987 |
| ENSMUSG00000001157 | Gmcl1 | protein_coding | -0.646199881 | 4.452459458 | 0.9487818 | 0.344281 | 0.999175987 |
| ENSMUSG00000045441 | Gprin3 | protein_coding | -0.534363615 | 4.460587543 | 0.9486585 | 0.3442882 | 0.999175987 |
| ENSMUSG00000039065 | Fam173b | protein_coding | -0.458954152 | 4.623880927 | 0.9486427 | 0.3442921 | 0.999175987 |
| ENSMUSG00000020668 | Kif3c | protein_coding | 1.172848494 | 2.57057705 | 0.9481751 | 0.3444078 | 0.999175987 |
| ENSMUSG00000060143 | Gm10076 | protein_coding | 0.759990482 | 3.08257954 | 0.9477914 | 0.3445027 | 0.999175987 |
| ENSMUSG00000058407 | Txndc9 | protein_coding | -0.292616964 | 7.320790892 | 0.9464874 | 0.3448257 | 0.999175987 |
| ENSMUSG00000031284 | Pak3 | protein_coding | -0.286112442 | 7.575495308 | 0.9464062 | 0.3448458 | 0.999175987 |
| ENSMUSG00000031109 | Enox2 | protein_coding | 0.488652699 | 4.325506399 | 0.9462883 | 0.3448751 | 0.999175987 |
| ENSMUSG00000088595 | Gm24601 | rRNA | 0.547317757 | 4.51822652 | 0.9452779 | 0.3451257 | 0.999175987 |
| ENSMUSG00000089809 | Rasgef1b | protein_coding | -0.476677949 | 5.397372802 | 0.9452362 | 0.345136 | 0.999175987 |
| ENSMUSG00000039262 | Prrc2b | protein_coding | 0.414533425 | 5.773422834 | 0.9451181 | 0.3451653 | 0.999175987 |
| ENSMUSG00000017404 | Rpl19 | protein_coding | 0.613154908 | 3.484945015 | 0.9449161 | 0.3452155 | 0.999175987 |
| ENSMUSG00000026650 | Meig1 | protein_coding | 0.515369595 | 4.534744567 | 0.944566 | 0.3453024 | 0.999175987 |
| ENSMUSG00000110723 | AC153912.1 | protein_coding | -0.953464558 | 2.861603887 | 0.9445598 | 0.3453039 | 0.999175987 |
| ENSMUSG00000033629 | Hacd3 | protein_coding | 0.284775334 | 7.534286954 | 0.9443448 | 0.3453573 | 0.999175987 |
| ENSMUSG00000006731 | B4galnt1 | protein_coding | -0.631071277 | 3.284752277 | 0.9441371 | 0.3454089 | 0.999175987 |
| ENSMUSG00000031112 | Stk26 | protein_coding | -1.17286675 | 2.454219891 | 0.9435003 | 0.3455672 | 0.999175987 |
| ENSMUSG00000037070 | Rbmxl1 | protein_coding | 0.502030733 | 4.031584788 | 0.9433966 | 0.3455929 | 0.999175987 |
| ENSMUSG00000037902 | Sirpa | protein_coding | -0.321186359 | 6.931019803 | 0.9429603 | 0.3457014 | 0.999175987 |
| ENSMUSG00000017418 | Arl5b | protein_coding | -0.798842175 | 3.134239237 | 0.9425311 | 0.3458082 | 0.999175987 |
| ENSMUSG00000074782 | 4833422C13Rik | antisense_RNA | -0.587203647 | 4.016640669 | 0.9423774 | 0.3458465 | 0.999175987 |
| ENSMUSG00000052516 | Robo2 | protein_coding | 0.525239027 | 4.285642096 | 0.9409214 | 0.3462091 | 0.999175987 |
| ENSMUSG00000077450 | Rab11b | protein_coding | 0.338201162 | 7.827942109 | 0.9408926 | 0.3462163 | 0.999175987 |
| ENSMUSG00000048108 | Tmem72 | protein_coding | 1.863960014 | 2.046472739 | 0.9407861 | 0.3462428 | 0.999175987 |
| ENSMUSG00000018293 | Pfn1 | protein_coding | 0.343509574 | 6.820414673 | 0.9406237 | 0.3462834 | 0.999175987 |
| ENSMUSG00000042198 | Chchd7 | protein_coding | 0.435519032 | 5.275850413 | 0.9405778 | 0.3462948 | 0.999175987 |
| ENSMUSG00000058793 | Cds2 | protein_coding | -0.298053242 | 7.027360207 | 0.9405136 | 0.3463108 | 0.999175987 |
| ENSMUSG00000014177 | Tvp23b | protein_coding | -0.325475474 | 6.339747396 | 0.9393042 | 0.3466126 | 0.999175987 |
| ENSMUSG00000030616 | Sytl2 | protein_coding | -0.449598604 | 4.494897183 | 0.9392558 | 0.3466247 | 0.999175987 |
| ENSMUSG00000046798 | Cldn12 | protein_coding | -0.485014714 | 4.887008155 | 0.9392147 | 0.3466349 | 0.999175987 |
| ENSMUSG00000044647 | Csrnp3 | protein_coding | -0.332347633 | 6.295928373 | 0.9389489 | 0.3467013 | 0.999175987 |
| ENSMUSG00000037366 | Pafah2 | protein_coding | -1.286079471 | 2.283650416 | 0.93862 | 0.3467835 | 0.999175987 |
| ENSMUSG00000031584 | Gsr | protein_coding | -0.40951775 | 5.228194358 | 0.9377302 | 0.3470059 | 0.999175987 |
| ENSMUSG00000023004 | Tuba1b | protein_coding | 0.299864693 | 7.738139621 | 0.937639 | 0.3470287 | 0.999175987 |
| ENSMUSG00000050947 | Amigo1 | protein_coding | -0.494487906 | 4.700277835 | 0.9372955 | 0.3471146 | 0.999175987 |
| ENSMUSG00000020396 | Nefh | protein_coding | -0.427659118 | 5.388966316 | 0.9364972 | 0.3473144 | 0.999175987 |
| ENSMUSG00000010110 | Stx5a | protein_coding | 0.329520351 | 6.442202614 | 0.9362948 | 0.3473651 | 0.999175987 |
| ENSMUSG00000027893 | Ahcyl1 | protein_coding | -0.297982699 | 7.54395055 | 0.9360891 | 0.3474166 | 0.999175987 |
| ENSMUSG00000027981 | Rnpc3 | protein_coding | 0.76846657 | 2.974047067 | 0.935521 | 0.347559 | 0.999175987 |
| ENSMUSG00000041548 | Hspb8 | protein_coding | -0.496248824 | 4.908712143 | 0.935039 | 0.3476799 | 0.999175987 |
| ENSMUSG00000034245 | Hdac11 | protein_coding | 0.329301609 | 8.240556123 | 0.934657 | 0.3477757 | 0.999175987 |
| ENSMUSG00000036634 | Mag | protein_coding | -0.297363024 | 7.307725008 | 0.9346087 | 0.3477878 | 0.999175987 |
| ENSMUSG00000113429 | CT030140.1 | lincRNA | 0.813948857 | 2.950139412 | 0.9343223 | 0.3478597 | 0.999175987 |
| ENSMUSG00000038696 | Mapkap1 | protein_coding | -0.460488213 | 5.030753625 | 0.9342761 | 0.3478712 | 0.999175987 |
| ENSMUSG00000050188 | Lsm10 | protein_coding | 0.401033768 | 5.930104451 | 0.9342602 | 0.3478752 | 0.999175987 |
| ENSMUSG00000031485 | Prosc | protein_coding | 0.346894055 | 6.440097532 | 0.9337795 | 0.3479959 | 0.999175987 |
| ENSMUSG00000061559 | Wdr61 | protein_coding | -0.282811753 | 7.66152338 | 0.9335577 | 0.3480516 | 0.999175987 |
| ENSMUSG00000021546 | Hnrnpk | protein_coding | -0.336207193 | 8.311756513 | 0.9335074 | 0.3480643 | 0.999175987 |
| ENSMUSG00000038805 | Six3 | protein_coding | -0.499842065 | 5.097093561 | 0.9334702 | 0.3480736 | 0.999175987 |
| ENSMUSG00000055078 | Gabra5 | protein_coding | 0.767077174 | 3.08861533 | 0.932069 | 0.3484258 | 0.999175987 |
| ENSMUSG00000022471 | Xrcc6 | protein_coding | -0.748349274 | 3.169702503 | 0.9309087 | 0.3487179 | 0.999175987 |
| ENSMUSG00000017679 | Ttpal | protein_coding | -0.44126757 | 4.909761282 | 0.9307993 | 0.3487455 | 0.999175987 |
| ENSMUSG00000009013 | Dynll1 | protein_coding | 0.385482801 | 10.1079042 | 0.9306233 | 0.3487898 | 0.999175987 |
| ENSMUSG00000003808 | Farsa | protein_coding | 0.483803179 | 5.889888517 | 0.9304736 | 0.3488505 | 0.999175987 |
| ENSMUSG00000031467 | Agpat5 | protein_coding | 0.428819105 | 5.353533336 | 0.9301057 | 0.3489202 | 0.999175987 |
| ENSMUSG00000005506 | Celf1 | protein_coding | -0.297268931 | 6.52656855 | 0.9295684 | 0.3490557 | 0.999175987 |
| ENSMUSG00000096146 | Kcnj11 | protein_coding | 0.62985479 | 3.954955941 | 0.9291728 | 0.3491555 | 0.999175987 |
| ENSMUSG00000042770 | Hebp1 | protein_coding | 0.546624444 | 5.318491374 | 0.929176 | 0.3491777 | 0.999175987 |
| ENSMUSG00000087618 | Gm14114 | lincRNA | -0.987683943 | 2.454754012 | 0.9290412 | 0.3491887 | 0.999175987 |
| ENSMUSG00000039089 | L3mbtl3 | protein_coding | 0.484633421 | 4.024540819 | 0.9289687 | 0.349207 | 0.999175987 |
| ENSMUSG00000026037 | Orc2 | protein_coding | 1.236156348 | 2.489259787 | 0.9287416 | 0.3492643 | 0.999175987 |

| ensembl_gene_id | external_gene_name | gene_biotype | log2 (fold change) | log2 (counts per million) | F-value | p-value | False discovery rate |
|---|---|---|---|---|---|---|---|
| ENSMUSG00000040565 | Btaf1 | protein_coding | -0.691495925 | 3.801692078 | 0.9285181 | 0.3493207 | 0.999175987 |
| ENSMUSG00000020280 | Pus10 | protein_coding | 0.624204083 | 3.556230994 | 0.9278794 | 0.3494821 | 0.999175987 |
| ENSMUSG00000021646 | Mccc2 | protein_coding | 0.518936445 | 5.453756839 | 0.9278114 | 0.3495222 | 0.999175987 |
| ENSMUSG00000035199 | Arl6ip5 | protein_coding | 0.266924023 | 8.05023803 | 0.927578 | 0.3495582 | 0.999175987 |
| ENSMUSG00000021365 | Nedd9 | protein_coding | -1.107024805 | 2.474451511 | 0.9269504 | 0.3497169 | 0.999175987 |
| ENSMUSG00000028773 | Fabp3 | protein_coding | 0.310937241 | 6.611636062 | 0.9269282 | 0.3497225 | 0.999175987 |
| ENSMUSG00000043164 | Tmem212 | protein_coding | -0.772117346 | 3.784289306 | 0.926846 | 0.3497663 | 0.999175987 |
| ENSMUSG00000097412 | 1810014B01Rik | transcribed_unproces | 0.937769643 | 2.91370316 | 0.9260162 | 0.3499533 | 0.999175987 |
| ENSMUSG00000025537 | Phkg1 | protein_coding | 0.765963338 | 3.493481303 | 0.9247853 | 0.3502651 | 0.999175987 |
| ENSMUSG00000017740 | Slc12a5 | protein_coding | -0.542016905 | 6.468301671 | 0.9298769 | 0.350374 | 0.999175987 |
| ENSMUSG00000061601 | Pclo | protein_coding | -0.481755231 | 5.385969975 | 0.9242989 | 0.3503884 | 0.999175987 |
| ENSMUSG00000005981 | Trap1 | protein_coding | -0.415576528 | 6.09934087 | 0.9241174 | 0.3504344 | 0.999175987 |
| ENSMUSG00000034923 | Ly6g6f | protein_coding | -0.816140215 | 3.087477309 | 0.9239589 | 0.3504746 | 0.999175987 |
| ENSMUSG00000026571 | Dcaf6 | protein_coding | 0.458536395 | 4.34606051 | 0.9236503 | 0.3505529 | 0.999175987 |
| ENSMUSG00000034413 | Neurl1b | protein_coding | -0.839887394 | 3.442703507 | 0.9228826 | 0.3507478 | 0.999175987 |
| ENSMUSG00000038028 | Tigar | protein_coding | -0.530716377 | 4.351674564 | 0.9223787 | 0.3508759 | 0.999175987 |
| ENSMUSG00000020290 | Xpo1 | protein_coding | -0.449254402 | 5.05689535 | 0.9217174 | 0.3510439 | 0.999175987 |
| ENSMUSG00000020463 | Ppp4r3b | protein_coding | -0.415864125 | 5.015230073 | 0.9205512 | 0.3513407 | 0.999175987 |
| ENSMUSG00000025790 | Slco3a1 | protein_coding | 0.530207389 | 4.733153981 | 0.9204373 | 0.3513697 | 0.999175987 |
| ENSMUSG00000049960 | Mrps16 | protein_coding | 0.289958179 | 6.888654785 | 0.9201901 | 0.3514326 | 0.999175987 |
| ENSMUSG00000062257 | Opcml | protein_coding | 0.69350775 | 3.674555153 | 0.9200934 | 0.3514573 | 0.999175987 |
| ENSMUSG00000032186 | Tmod2 | protein_coding | -0.285455141 | 8.400191226 | 0.9197424 | 0.3515467 | 0.999175987 |
| ENSMUSG00000033488 | BC026585 | protein_coding | -0.633264869 | 4.166418027 | 0.9192149 | 0.3516811 | 0.999175987 |
| ENSMUSG00000026424 | Gpr37l1 | protein_coding | 0.390495416 | 6.583081309 | 0.9188197 | 0.3517819 | 0.999175987 |
| ENSMUSG00000056919 | Cep162 | protein_coding | -0.782216031 | 3.024399606 | 0.9186304 | 0.3518302 | 0.999175987 |
| ENSMUSG00000029027 | Dffb | protein_coding | -1.173108388 | 2.44101323 | 0.9185102 | 0.3518609 | 0.999175987 |
| ENSMUSG00000066175 | 2510046G10Rik | antisense_RNA | 1.477056956 | 2.061809455 | 0.9184471 | 0.351877 | 0.999175987 |
| ENSMUSG00000005469 | Prkaca | protein_coding | 0.34343709 | 7.918285294 | 0.9182906 | 0.3519169 | 0.999175987 |
| ENSMUSG00000020982 | Nemf | protein_coding | -0.31837341 | 6.868900378 | 0.9182786 | 0.3519199 | 0.999175987 |
| ENSMUSG00000032320 | Rcn2 | protein_coding | -0.446260783 | 6.618201067 | 0.9181148 | 0.3520843 | 0.999175987 |
| ENSMUSG00000087479 | Gm16835 | processed_transcript | -0.713383865 | 3.380178744 | 0.9173521 | 0.3521565 | 0.999175987 |
| ENSMUSG00000038412 | Higd1a | protein_coding | -0.292668622 | 8.510514161 | 0.917172 | 0.3522025 | 0.999175987 |
| ENSMUSG00000061273 | Mmgt1 | protein_coding | -0.477262622 | 4.400901462 | 0.9164645 | 0.3523834 | 0.999175987 |
| ENSMUSG00000030854 | Ptpn5 | protein_coding | 0.299669377 | 8.552007529 | 0.9160541 | 0.3524883 | 0.999175987 |
| ENSMUSG00000058704 | Memo1 | protein_coding | 0.5207272 | 4.05571093 | 0.9147367 | 0.3528255 | 0.999175987 |
| ENSMUSG00000054312 | Mrps21 | protein_coding | 0.380323764 | 7.97486002 | 0.9147678 | 0.3528404 | 0.999175987 |
| ENSMUSG00000022490 | Ppp1r1a | protein_coding | 0.424200879 | 5.34981323 | 0.9138784 | 0.3530455 | 0.999175987 |
| ENSMUSG00000069135 | Fgfr1op | protein_coding | -0.644775539 | 4.019923772 | 0.913744 | 0.35308 | 0.999175987 |
| ENSMUSG00000049038 | Mterf2 | protein_coding | -0.807548158 | 3.173659394 | 0.9135972 | 0.3531176 | 0.999175987 |
| ENSMUSG00000001986 | Gria3 | protein_coding | 0.334470423 | 5.891999901 | 0.9132089 | 0.3532172 | 0.999175987 |
| ENSMUSG00000025241 | Fyco1 | protein_coding | -0.812336252 | 3.763028103 | 0.9132321 | 0.353234 | 0.999175987 |
| ENSMUSG00000025255 | Zfhx4 | protein_coding | -0.465720574 | 4.583105789 | 0.9131046 | 0.3532439 | 0.999175987 |
| ENSMUSG00000070283 | Ndufaf3 | protein_coding | 0.297912994 | 8.101443415 | 0.9125614 | 0.3533834 | 0.999175987 |
| ENSMUSG00000030122 | Ptms | protein_coding | 0.416297177 | 6.578648641 | 0.9125871 | 0.3533996 | 0.999175987 |
| ENSMUSG00000022337 | Emc2 | protein_coding | -0.28981727 | 7.531188737 | 0.9124371 | 0.3534153 | 0.999175987 |
| ENSMUSG00000002486 | Tchp | protein_coding | 0.809815833 | 2.932409377 | 0.9122698 | 0.3534583 | 0.999175987 |
| ENSMUSG00000044499 | Hs3st5 | protein_coding | 0.679712762 | 3.457253235 | 0.9121886 | 0.3534791 | 0.999175987 |
| ENSMUSG00000071662 | Polr2g | protein_coding | 0.32543402 | 8.576597452 | 0.9119734 | 0.3535344 | 0.999175987 |
| ENSMUSG00000092083 | Kcnb2 | protein_coding | 0.649233207 | 3.41738996 | 0.9119067 | 0.3535515 | 0.999175987 |
| ENSMUSG00000059866 | Tnip2 | protein_coding | -0.879672773 | 2.952295257 | 0.9109375 | 0.3538007 | 0.999175987 |
| ENSMUSG00000024943 | Smc5 | protein_coding | -0.640394229 | 4.320390629 | 0.9107795 | 0.3538413 | 0.999175987 |
| ENSMUSG00000031285 | Dcx | protein_coding | -0.673614517 | 3.778773421 | 0.9105833 | 0.3538918 | 0.999175987 |
| ENSMUSG00000021951 | Eef1akmt1 | protein_coding | 0.261858155 | 7.540472218 | 0.9101793 | 0.3539957 | 0.999175987 |
| ENSMUSG00000035486 | Plk5 | protein_coding | 0.75368253 | 3.165917379 | 0.9100897 | 0.3540188 | 0.999175987 |
| ENSMUSG00000035944 | Ttc38 | protein_coding | -0.68768518 | 3.406208913 | 0.9094501 | 0.3541835 | 0.999175987 |
| ENSMUSG00000053398 | Phgdh | protein_coding | -0.418244558 | 5.139658463 | 0.9092839 | 0.3542263 | 0.999175987 |
| ENSMUSG00000006423 | C330007P06Rik | protein_coding | -0.515371288 | 4.447876819 | 0.909016 | 0.3542953 | 0.999175987 |
| ENSMUSG00000021012 | Zc3h14 | protein_coding | -0.515137218 | 4.454841109 | 0.9089458 | 0.3543134 | 0.999175987 |
| ENSMUSG00000030161 | Gabarapl1 | protein_coding | 0.312021977 | 10.32629476 | 0.9085274 | 0.3544213 | 0.999175987 |
| ENSMUSG00000041115 | Iqsec2 | protein_coding | -0.520035436 | 4.473498805 | 0.9084159 | 0.35445 | 0.999175987 |
| ENSMUSG00000000486 | 1-Sep | protein_coding | 0.816782457 | 2.941187693 | 0.9078801 | 0.3545882 | 0.999175987 |
| ENSMUSG00000027763 | Mbnl1 | protein_coding | -0.348185759 | 6.597685114 | 0.9073449 | 0.3547263 | 0.999175987 |
| ENSMUSG00000061482 | Hist1h4d | protein_coding | -0.526400045 | 3.765522318 | 0.9062516 | 0.3550087 | 0.999175987 |
| ENSMUSG00000018841 | Rad51d | protein_coding | 0.437768762 | 5.734623597 | 0.9061119 | 0.3550448 | 0.999175987 |
| ENSMUSG00000037692 | Ahdc1 | protein_coding | -0.665347999 | 3.881753752 | 0.9056299 | 0.3551695 | 0.999175987 |
| ENSMUSG00000048310 | Pskh1 | protein_coding | 0.889576496 | 2.831746232 | 0.9048579 | 0.3553692 | 0.999175987 |
| ENSMUSG00000069539 | Scyl2 | protein_coding | 0.860249417 | 2.866869818 | 0.9045968 | 0.3554368 | 0.999175987 |
| ENSMUSG00000034731 | Dgkh | protein_coding | -0.70605188 | 3.239338009 | 0.9037126 | 0.3556658 | 0.999175987 |
| ENSMUSG00000035828 | Pim3 | protein_coding | -0.465048055 | 5.045713111 | 0.9033906 | 0.3557493 | 0.999175987 |
| ENSMUSG00000021357 | Exoc2 | protein_coding | -0.569182911 | 4.732963836 | 0.9029151 | 0.3558725 | 0.999175987 |
| ENSMUSG00000062328 | Rpl17 | protein_coding | 0.664302388 | 3.274014848 | 0.9019173 | 0.3561315 | 0.999175987 |
| ENSMUSG00000036918 | Ttc7 | protein_coding | -1.169388121 | 2.377443965 | 0.9018907 | 0.3561384 | 0.999175987 |
| ENSMUSG00000041642 | Kif21b | protein_coding | 0.665296841 | 3.805694586 | 0.9017022 | 0.3561873 | 0.999175987 |
| ENSMUSG00000002984 | Tomm40 | protein_coding | 0.528326783 | 4.313692104 | 0.901684 | 0.3561921 | 0.999175987 |
| ENSMUSG00000031885 | Cbfb | protein_coding | -0.502943132 | 4.202384283 | 0.9015254 | 0.3562333 | 0.999175987 |
| ENSMUSG00000050875 | A730017C20Rik | protein_coding | 0.348753534 | 8.826962204 | 0.901407 | 0.356264 | 0.999175987 |
| ENSMUSG00000030347 | D6Wsu163e | protein_coding | -0.383353246 | 5.091092312 | 0.9009977 | 0.3563703 | 0.999175987 |
| ENSMUSG00000041608 | Entpd3 | protein_coding | -0.757769375 | 3.918803749 | 0.8993056 | 0.3568331 | 0.999175987 |

| ensembl_gene_id | external_gene_name | gene_biotype | log2 (fold change) | log2 (counts per million) | F-value | p-value | False discovery rate |
|---|---|---|---|---|---|---|---|
| ENSMUSG00000002781 | Tmem143 | protein_coding | 0.601643046 | 3.594656534 | 0.8991939 | 0.3568396 | 0.999175987 |
| ENSMUSG00000038637 | Lrrc56 | protein_coding | -0.963036805 | 2.744091276 | 0.8991127 | 0.3568607 | 0.999175987 |
| ENSMUSG00000029545 | Acads | protein_coding | 0.725734624 | 3.275808124 | 0.8988028 | 0.3569414 | 0.999175987 |
| ENSMUSG00000062116 | Zfp954 | protein_coding | -0.523473426 | 5.134182017 | 0.8986224 | 0.3569884 | 0.999175987 |
| ENSMUSG00000091537 | Tma7 | protein_coding | 0.28199461 | 7.239049652 | 0.898475 | 0.3570268 | 0.999175987 |
| ENSMUSG00000059436 | Max | protein_coding | -0.560499437 | 5.952105046 | 0.9016155 | 0.3570869 | 0.999175987 |
| ENSMUSG00000038836 | Agbl3 | protein_coding | -0.403950936 | 5.249206097 | 0.8977297 | 0.3572211 | 0.999175987 |
| ENSMUSG00000017707 | Serinc3 | protein_coding | 0.364633575 | 6.07841394 | 0.8974797 | 0.3572863 | 0.999175987 |
| ENSMUSG00000033595 | Lgi3 | protein_coding | 0.367366735 | 5.970040407 | 0.8967833 | 0.357468 | 0.999175987 |
| ENSMUSG00000022750 | Klhl22 | protein_coding | 0.325988799 | 6.821613233 | 0.8961566 | 0.3576317 | 0.999175987 |
| ENSMUSG00000030595 | Nfkbib | protein_coding | 0.451924522 | 5.707432319 | 0.8956966 | 0.3577519 | 0.999175987 |
| ENSMUSG00000060032 | H2afj | protein_coding | 0.821196458 | 2.916554437 | 0.894908 | 0.3579581 | 0.999175987 |
| ENSMUSG00000046658 | Zfp316 | protein_coding | -0.915138381 | 2.786277545 | 0.8939161 | 0.3582177 | 0.999175987 |
| ENSMUSG00000031608 | Galnt7 | protein_coding | -0.776831517 | 3.425388949 | 0.8935774 | 0.3583064 | 0.999175987 |
| ENSMUSG00000066568 | Lsm14a | protein_coding | -0.35838575 | 5.43454583 | 0.893498 | 0.3583272 | 0.999175987 |
| ENSMUSG00000055319 | Sec23ip | protein_coding | -0.44780581 | 5.61426494 | 0.8931183 | 0.3584267 | 0.999175987 |
| ENSMUSG00000026288 | Inpp5d | protein_coding | -1.638960433 | 2.104741917 | 0.8928662 | 0.3585153 | 0.999175987 |
| ENSMUSG00000022494 | Shisa9 | protein_coding | 0.458499676 | 4.665970214 | 0.8927405 | 0.3585257 | 0.999175987 |
| ENSMUSG00000051855 | Mest | protein_coding | -0.729813265 | 3.702198442 | 0.891871 | 0.3587538 | 0.999175987 |
| ENSMUSG00000027472 | Pdrg1 | protein_coding | -0.28969111 | 7.819654886 | 0.8909738 | 0.3589893 | 0.999175987 |
| ENSMUSG00000026577 | Blzf1 | protein_coding | -0.431981511 | 5.141870769 | 0.890726 | 0.3590544 | 0.999175987 |
| ENSMUSG00000027589 | Pcmtd2 | protein_coding | -0.39788398 | 5.765213064 | 0.8906671 | 0.3590699 | 0.999175987 |
| ENSMUSG00000022752 | Tomm70a | protein_coding | -0.292387373 | 7.013920706 | 0.8905163 | 0.3591095 | 0.999175987 |
| ENSMUSG00000038388 | Mpp6 | protein_coding | -0.471754546 | 4.397720151 | 0.8904395 | 0.3591297 | 0.999175987 |
| ENSMUSG00000037152 | Ndufc1 | protein_coding | 0.346779896 | 9.412866579 | 0.8903777 | 0.3591459 | 0.999175987 |
| ENSMUSG00000038150 | Ormdl3 | protein_coding | 0.336939805 | 6.097584382 | 0.8903348 | 0.3591572 | 0.999175987 |
| ENSMUSG00000022889 | Mrpl39 | protein_coding | -0.361239354 | 6.207912801 | 0.8899669 | 0.3592539 | 0.999175987 |
| ENSMUSG00000026275 | Ppp1r7 | protein_coding | -0.285274597 | 7.815914867 | 0.8896694 | 0.3593322 | 0.999175987 |
| ENSMUSG00000078137 | Ankrd63 | protein_coding | -0.578477342 | 4.836188436 | 0.8894968 | 0.3594 | 0.999175987 |
| ENSMUSG00000037656 | Slc20a2 | protein_coding | -0.507091898 | 4.918344931 | 0.8886731 | 0.3595944 | 0.999175987 |
| ENSMUSG00000051527 | Usp29 | protein_coding | -0.419390369 | 5.338877038 | 0.8885353 | 0.3596306 | 0.999175987 |
| ENSMUSG00000059878 | Zfp422 | protein_coding | 0.434826015 | 5.255728432 | 0.8882975 | 0.3596933 | 0.999175987 |
| ENSMUSG00000034620 | Tmem5 | protein_coding | 0.470878445 | 4.559196807 | 0.8879109 | 0.3597951 | 0.999175987 |
| ENSMUSG00000011658 | Fuz | protein_coding | 0.962531341 | 2.572785091 | 0.8876519 | 0.3598634 | 0.999175987 |
| ENSMUSG00000064068 | Mtx1 | protein_coding | 0.325480736 | 5.985579184 | 0.8873403 | 0.3599455 | 0.999175987 |
| ENSMUSG00000001962 | Fam50a | protein_coding | -0.682177815 | 4.370050922 | 0.8873734 | 0.3599593 | 0.999175987 |
| ENSMUSG00000029438 | Bcl7a | protein_coding | -0.466504462 | 5.314470997 | 0.8872729 | 0.3599633 | 0.999175987 |
| ENSMUSG00000026102 | Inpp1 | protein_coding | -0.362088985 | 5.806926255 | 0.8872722 | 0.3599635 | 0.999175987 |
| ENSMUSG00000097820 | E530011L22Rik | antisense_RNA | 1.037152361 | 2.661232199 | 0.8870669 | 0.3600176 | 0.999175987 |
| ENSMUSG00000028187 | Rpf1 | protein_coding | 0.478565137 | 4.458672323 | 0.8870609 | 0.3600192 | 0.999175987 |
| ENSMUSG00000022897 | Dyrk1a | protein_coding | 0.358807666 | 6.246967347 | 0.8865872 | 0.3601442 | 0.999175987 |
| ENSMUSG00000021779 | Thrb | protein_coding | -0.494822173 | 5.798525101 | 0.8860594 | 0.360306 | 0.999175987 |
| ENSMUSG00000074220 | Zfp382 | protein_coding | -0.71641611 | 3.186077791 | 0.8852864 | 0.3604877 | 0.999175987 |
| ENSMUSG00000025607 | Copg2 | protein_coding | -0.373118176 | 7.752464664 | 0.884756 | 0.3606503 | 0.999175987 |
| ENSMUSG00000024963 | Dnajc4 | protein_coding | 0.376583123 | 6.0058377 | 0.8846064 | 0.3606675 | 0.999175987 |
| ENSMUSG00000001506 | Col1a1 | protein_coding | -1.479331467 | 2.015746299 | 0.8842048 | 0.3607737 | 0.999175987 |
| ENSMUSG00000002550 | Uck1 | protein_coding | 0.684617247 | 3.433031781 | 0.8841545 | 0.360787 | 0.999175987 |
| ENSMUSG00000043257 | Pigv | protein_coding | -0.940171212 | 2.814847498 | 0.8837968 | 0.3608817 | 0.999175987 |
| ENSMUSG00000048921 | Zfp689 | protein_coding | 0.629049523 | 3.831255787 | 0.883341 | 0.3610023 | 0.999175987 |
| ENSMUSG00000021090 | Lrrc9 | protein_coding | -1.066670423 | 2.667987494 | 0.8828254 | 0.3611389 | 0.999175987 |
| ENSMUSG00000024045 | Akap8 | protein_coding | -0.441555251 | 4.802518512 | 0.8818289 | 0.3614031 | 0.999175987 |
| ENSMUSG00000026975 | Dph7 | protein_coding | 0.323657665 | 6.119075733 | 0.8816184 | 0.3614589 | 0.999175987 |
| ENSMUSG00000084808 | 9430091E24Rik | processed_transcript | -0.624245436 | 3.72286899 | 0.8815028 | 0.3614896 | 0.999175987 |
| ENSMUSG00000020737 | Jpt1 | protein_coding | 0.277542472 | 8.50463391 | 0.8813643 | 0.3615263 | 0.999175987 |
| ENSMUSG00000036943 | Rab8b | protein_coding | -0.497290569 | 4.885630103 | 0.880648 | 0.3617165 | 0.999175987 |
| ENSMUSG00000042719 | Naa25 | protein_coding | 0.672437611 | 3.837378938 | 0.8803861 | 0.3617861 | 0.999175987 |
| ENSMUSG00000031762 | Mt2 | protein_coding | -0.628820073 | 5.28709843 | 0.8829416 | 0.3619758 | 0.999175987 |
| ENSMUSG00000019188 | H13 | protein_coding | 0.568661141 | 5.09278234 | 0.8795772 | 0.3620234 | 0.999175987 |
| ENSMUSG00000040907 | Atp1a3 | protein_coding | 0.278565871 | 9.92749935 | 0.8785173 | 0.362283 | 0.999175987 |
| ENSMUSG00000024661 | Fth1 | protein_coding | 1.45679815 | 12.1728887 | 0.9348565 | 0.3623227 | 0.999175987 |
| ENSMUSG00000003657 | Calb2 | protein_coding | -0.460212328 | 10.09150668 | 0.8805219 | 0.3625637 | 0.999175987 |
| ENSMUSG00000023118 | Sympk | protein_coding | 0.53216755 | 4.058834445 | 0.8773748 | 0.3625873 | 0.999175987 |
| ENSMUSG00000034263 | Ints14 | protein_coding | 0.482664686 | 4.562746553 | 0.8772806 | 0.3626124 | 0.999175987 |
| ENSMUSG00000074733 | Zfp950 | protein_coding | -0.414001578 | 5.169887919 | 0.8772713 | 0.3626149 | 0.999175987 |
| ENSMUSG00000047671 | Tctex1d4 | protein_coding | 0.762716371 | 3.345862456 | 0.8770454 | 0.3626751 | 0.999175987 |
| ENSMUSG00000034981 | Parm1 | protein_coding | -0.476302653 | 6.183718437 | 0.8777842 | 0.3627466 | 0.999175987 |
| ENSMUSG00000074165 | Zfp788 | protein_coding | 0.522103385 | 4.507406213 | 0.8763525 | 0.3628599 | 0.999175987 |
| ENSMUSG00000027109 | Sp3 | protein_coding | -0.44976353 | 4.960542373 | 0.8754744 | 0.3630943 | 0.999175987 |
| ENSMUSG00000030788 | Rnf141 | protein_coding | -0.40026644 | 5.494603547 | 0.8754407 | 0.3631033 | 0.999175987 |
| ENSMUSG00000048106 | 4632415L05Rik | lincRNA | -0.665515076 | 3.111521231 | 0.8751431 | 0.3631828 | 0.999175987 |
| ENSMUSG00000085881 | Gm15912 | processed_transcript | 0.640478185 | 3.367241813 | 0.8751278 | 0.3631869 | 0.999175987 |
| ENSMUSG00000035203 | Epn1 | protein_coding | 0.359558813 | 6.099293271 | 0.8738592 | 0.363526 | 0.999175987 |
| ENSMUSG00000047394 | Odf3b | protein_coding | 0.724816558 | 3.508796897 | 0.873729 | 0.3635609 | 0.999175987 |
| ENSMUSG00000022390 | Zc3h7b | protein_coding | -0.474641116 | 5.210253924 | 0.8736211 | 0.3635897 | 0.999175987 |
| ENSMUSG00000030505 | Prmt3 | protein_coding | 0.632924813 | 3.326721743 | 0.8736133 | 0.3635918 | 0.999175987 |
| ENSMUSG00000039058 | Ak5 | protein_coding | 0.393941645 | 5.474999154 | 0.8731281 | 0.3637217 | 0.999175987 |
| ENSMUSG00000019802 | Sec63 | protein_coding | -0.57485876 | 4.528129988 | 0.8729341 | 0.3637736 | 0.999175987 |
| ENSMUSG00000028992 | Nmnat1 | protein_coding | -0.604317057 | 3.606100699 | 0.8728842 | 0.363787 | 0.999175987 |

| ensembl_gene_id | external_gene_name | gene_biotype | log2 (fold change) | log2 (counts per million) | F-value | p-value | False discovery rate |
|---|---|---|---|---|---|---|---|
| ENSMUSG00000037771 | Slc32a1 | protein_coding | 0.741173838 | 4.616713118 | 0.876035 | 0.3638088 | 0.999175987 |
| ENSMUSG00000028657 | Ppt1 | protein_coding | 0.290530533 | 7.447321557 | 0.8719112 | 0.3640477 | 0.999175987 |
| ENSMUSG00000061740 | Cyp2d22 | protein_coding | -0.713948566 | 3.458521513 | 0.871694 | 0.3641059 | 0.999175987 |
| ENSMUSG00000037443 | Cep85 | protein_coding | -0.609960351 | 3.550220314 | 0.8707406 | 0.3643617 | 0.999175987 |
| ENSMUSG00000042705 | Commd10 | protein_coding | 0.284603823 | 6.755597794 | 0.8698811 | 0.3645925 | 0.999175987 |
| ENSMUSG00000033909 | Usp36 | protein_coding | -0.48932226 | 4.481393143 | 0.8696815 | 0.3646462 | 0.999175987 |
| ENSMUSG00000029703 | Lrwd1 | protein_coding | -0.737738595 | 3.433185979 | 0.869349 | 0.3647355 | 0.999175987 |
| ENSMUSG00000024642 | Tle4 | protein_coding | -0.434261045 | 5.585379741 | 0.868312 | 0.3650144 | 0.999175987 |
| ENSMUSG00000022142 | Nup155 | protein_coding | -0.741074062 | 2.915053981 | 0.8678764 | 0.3651317 | 0.999175987 |
| ENSMUSG00000033022 | Cdo1 | protein_coding | 0.393838459 | 5.595285487 | 0.8676986 | 0.3651796 | 0.999175987 |
| ENSMUSG00000056201 | Cfl1 | protein_coding | 0.278049337 | 10.34545569 | 0.8669259 | 0.3653878 | 0.999175987 |
| ENSMUSG00000031130 | Brs3 | protein_coding | 1.381148149 | 3.223265056 | 0.8748669 | 0.3654176 | 0.999175987 |
| ENSMUSG00000064302 | Clasp1 | protein_coding | 0.437306366 | 5.440027815 | 0.8667632 | 0.3654316 | 0.999175987 |
| ENSMUSG00000031090 | Nadsyn1 | protein_coding | -0.984689519 | 2.574717907 | 0.8665581 | 0.3654869 | 0.999175987 |
| ENSMUSG00000021194 | Chga | protein_coding | 0.298791215 | 7.63615483 | 0.866553 | 0.3654883 | 0.999175987 |
| ENSMUSG00000060397 | Zfp128 | protein_coding | 0.822653316 | 3.274873164 | 0.8657404 | 0.3657075 | 0.999175987 |
| ENSMUSG00000001783 | Rtcb | protein_coding | 0.37321391 | 6.885349732 | 0.865294 | 0.3658502 | 0.999175987 |
| ENSMUSG00000109836 | Gm45884 | TEC | -0.575379831 | 3.596234696 | 0.8648161 | 0.365957 | 0.999175987 |
| ENSMUSG00000032252 | Glce | protein_coding | 0.481966795 | 4.339791961 | 0.8646959 | 0.3659895 | 0.999175987 |
| ENSMUSG00000037990 | Sh3rf3 | protein_coding | -0.557229229 | 3.771848816 | 0.8638279 | 0.3662241 | 0.999175987 |
| ENSMUSG00000052676 | Zmat1 | protein_coding | 0.541771061 | 4.490497069 | 0.8636758 | 0.3662653 | 0.999175987 |
| ENSMUSG00000003778 | Brd8 | protein_coding | -0.371597637 | 6.983895678 | 0.8637505 | 0.3662672 | 0.999175987 |
| ENSMUSG00000086968 | 4933431E20Rik | antisense_RNA | -0.618934774 | 4.024833679 | 0.863442 | 0.3663285 | 0.999175987 |
| ENSMUSG00000037343 | Taf2 | protein_coding | 0.564124081 | 3.868903352 | 0.861437 | 0.3668716 | 0.999175987 |
| ENSMUSG00000066877 | Nck2 | protein_coding | -0.562953267 | 3.486586342 | 0.8610016 | 0.3669896 | 0.999175987 |
| ENSMUSG00000050017 | Pitpnb | protein_coding | -0.337311611 | 6.832640067 | 0.8605994 | 0.3670988 | 0.999175987 |
| ENSMUSG00000036845 | Lin37 | protein_coding | 0.28565846 | 6.935440291 | 0.8605619 | 0.367109 | 0.999175987 |
| ENSMUSG00000005886 | Ncoa2 | protein_coding | -0.50256758 | 4.780771823 | 0.8604508 | 0.3671391 | 0.999175987 |
| ENSMUSG00000021376 | Tpmt | protein_coding | -0.50377076 | 3.970889341 | 0.8603461 | 0.3671675 | 0.999175987 |
| ENSMUSG00000030092 | Cntn6 | protein_coding | -0.557396711 | 4.10731142 | 0.859898 | 0.3672892 | 0.999175987 |
| ENSMUSG00000047155 | Cyp4x1 | protein_coding | 0.591741303 | 3.539105573 | 0.8592715 | 0.3674594 | 0.999175987 |
| ENSMUSG00000027243 | Harbi1 | protein_coding | 0.645999565 | 4.173488341 | 0.859184 | 0.3674832 | 0.999175987 |
| ENSMUSG00000028312 | Smc2 | protein_coding | -0.959327564 | 2.554614109 | 0.8585523 | 0.3676549 | 0.999175987 |
| ENSMUSG00000009640 | Dmap1 | protein_coding | 0.350675894 | 5.696855416 | 0.8583554 | 0.3677085 | 0.999175987 |
| ENSMUSG00000031647 | Mfap3l | protein_coding | -0.552590228 | 3.545629981 | 0.8577995 | 0.3678598 | 0.999175987 |
| ENSMUSG00000020402 | Vdac1 | protein_coding | -0.278894517 | 9.386725854 | 0.8577609 | 0.3678703 | 0.999175987 |
| ENSMUSG00000022474 | Pmm1 | protein_coding | 0.254626205 | 8.903386654 | 0.8577429 | 0.3678752 | 0.999175987 |
| ENSMUSG00000002190 | Clgn | protein_coding | -1.058034534 | 2.905563367 | 0.8573837 | 0.367995 | 0.999175987 |
| ENSMUSG00000032263 | Bckdhb | protein_coding | 0.530727543 | 3.918022141 | 0.8564398 | 0.3682301 | 0.999175987 |
| ENSMUSG00000025066 | Sfr1 | protein_coding | 0.253589165 | 8.415339589 | 0.8558437 | 0.3683927 | 0.999175987 |
| ENSMUSG00000074622 | Mafb | protein_coding | -1.076999528 | 2.802937455 | 0.8550881 | 0.3685989 | 0.999175987 |
| ENSMUSG00000005871 | Apc | protein_coding | -0.433244039 | 5.403525281 | 0.8547792 | 0.3686833 | 0.999175987 |
| ENSMUSG00000033377 | Palmd | protein_coding | 0.790995235 | 3.561440039 | 0.854735 | 0.3687174 | 0.999175987 |
| ENSMUSG00000027714 | Exosc9 | protein_coding | -0.304601982 | 6.244827269 | 0.8544723 | 0.3687671 | 0.999175987 |
| ENSMUSG00000028015 | Ctso | protein_coding | 0.568866286 | 4.782366102 | 0.8544297 | 0.3688007 | 0.999175987 |
| ENSMUSG00000004394 | Tmed4 | protein_coding | 0.274255761 | 8.130152195 | 0.8539075 | 0.3689214 | 0.999175987 |
| ENSMUSG00000032523 | Hhatl | protein_coding | -1.129053398 | 3.133468773 | 0.8568088 | 0.3689838 | 0.999175987 |
| ENSMUSG00000037697 | Ddhd1 | protein_coding | -1.019260378 | 2.963253179 | 0.8535651 | 0.3690371 | 0.999175987 |
| ENSMUSG00000037049 | Smpd1 | protein_coding | 0.323175311 | 6.9351457 | 0.8531464 | 0.3691296 | 0.999175987 |
| ENSMUSG00000003444 | Med29 | protein_coding | 0.505867414 | 3.908901613 | 0.8528449 | 0.3692121 | 0.999175987 |
| ENSMUSG00000045482 | Trrap | protein_coding | -0.586948579 | 4.645502751 | 0.8522928 | 0.3693852 | 0.999175987 |
| ENSMUSG00000017760 | Ctsa | protein_coding | -0.268069412 | 6.797332357 | 0.8520043 | 0.3694422 | 0.999175987 |
| ENSMUSG00000010936 | Vac14 | protein_coding | 0.4675195 | 4.58232797 | 0.8516867 | 0.3695292 | 0.999175987 |
| ENSMUSG00000020790 | Ankfy1 | protein_coding | -0.534395242 | 4.178651392 | 0.8515807 | 0.3695583 | 0.999175987 |
| ENSMUSG00000019894 | Slc6a15 | protein_coding | -0.377927401 | 5.604783337 | 0.8513824 | 0.3696126 | 0.999175987 |
| ENSMUSG00000021660 | Btf3 | protein_coding | 0.33320072 | 6.743615765 | 0.8511497 | 0.3696764 | 0.999175987 |
| ENSMUSG00000021482 | Aaed1 | protein_coding | -0.521417229 | 4.010025099 | 0.850861 | 0.3697556 | 0.999175987 |
| ENSMUSG00000049739 | Zfp646 | protein_coding | -0.680328855 | 3.184393989 | 0.850653 | 0.3698127 | 0.999175987 |
| ENSMUSG00000068391 | Chrac1 | protein_coding | -0.31527888 | 6.829943269 | 0.8505199 | 0.3698492 | 0.999175987 |
| ENSMUSG00000066798 | Zbtb6 | protein_coding | -0.617005015 | 3.804663878 | 0.8501463 | 0.3699517 | 0.999175987 |
| ENSMUSG00000029757 | Dync1i1 | protein_coding | 0.354738092 | 7.483071199 | 0.8500052 | 0.3699904 | 0.999175987 |
| ENSMUSG00000059851 | Kmt5c | protein_coding | -0.828067516 | 3.019809196 | 0.8497322 | 0.3700654 | 0.999175987 |
| ENSMUSG00000064061 | Dzip3 | protein_coding | 0.344391788 | 5.567912203 | 0.8496857 | 0.3700782 | 0.999175987 |
| ENSMUSG00000056999 | Ide | protein_coding | -0.424066414 | 4.665093056 | 0.8489739 | 0.3702737 | 0.999175987 |
| ENSMUSG00000037499 | Nenf | protein_coding | 0.369168676 | 5.674159859 | 0.8488549 | 0.3703064 | 0.999175987 |
| ENSMUSG00000059119 | Nap1l4 | protein_coding | 0.38671298 | 8.400587276 | 0.8488922 | 0.3703181 | 0.999175987 |
| ENSMUSG00000020755 | Sap30bp | protein_coding | -0.260622086 | 6.926630139 | 0.8483519 | 0.3704447 | 0.999175987 |
| ENSMUSG00000017314 | Mpp2 | protein_coding | 0.338361556 | 6.632723407 | 0.8482694 | 0.3704674 | 0.999175987 |
| ENSMUSG00000022436 | Sh3bp1 | protein_coding | 0.291066797 | 7.128496643 | 0.8479901 | 0.3705442 | 0.999175987 |
| ENSMUSG00000037110 | Ralgapa2 | protein_coding | -1.021961241 | 2.806873281 | 0.8479685 | 0.3705502 | 0.999175987 |
| ENSMUSG00000041323 | Ak7 | protein_coding | -0.953319586 | 2.577050978 | 0.8477872 | 0.3706001 | 0.999175987 |
| ENSMUSG00000021245 | Mlh3 | protein_coding | -0.646947834 | 3.920570449 | 0.8473455 | 0.3707217 | 0.999175987 |
| ENSMUSG00000040102 | Klhl42 | protein_coding | 0.593391732 | 4.246095242 | 0.8471359 | 0.3707794 | 0.999175987 |
| ENSMUSG00000044906 | 4930503L19Rik | protein_coding | 1.059113263 | 2.687705148 | 0.8468135 | 0.3708682 | 0.999175987 |
| ENSMUSG00000020829 | Slc46a1 | protein_coding | 0.375920826 | 5.153076633 | 0.8464507 | 0.3709681 | 0.999175987 |
| ENSMUSG00000028224 | Nbn | protein_coding | -0.695147388 | 3.964025293 | 0.8464535 | 0.3709893 | 0.999175987 |
| ENSMUSG00000038608 | Dock10 | protein_coding | -0.563201114 | 4.563913217 | 0.8463594 | 0.3709933 | 0.999175987 |
| ENSMUSG00000021710 | Nln | protein_coding | 0.511140546 | 4.26161093 | 0.8454703 | 0.3712385 | 0.999175987 |

| ensembl_gene_id | external_gene_name | gene_biotype | log2 (fold change) | log2 (counts per million) | F-value | p-value | False discovery rate |
|---|---|---|---|---|---|---|---|
| ENSMUSG00000037386 | Rims2 | protein_coding | -0.298461965 | 6.12669985 | 0.8452988 | 0.3712858 | 0.999175987 |
| ENSMUSG00000021713 | Ppwd1 | protein_coding | -1.263279177 | 2.292183636 | 0.8441622 | 0.3715996 | 0.999175987 |
| ENSMUSG00000026280 | Atg4b | protein_coding | -0.333532369 | 6.247778244 | 0.8441196 | 0.3716114 | 0.999175987 |
| ENSMUSG00000060212 | Pcnx2 | protein_coding | -0.63992364 | 3.502275679 | 0.8440473 | 0.3716314 | 0.999175987 |
| ENSMUSG00000028035 | Dnajb4 | protein_coding | -0.32724263 | 6.584486295 | 0.8436833 | 0.371732 | 0.999175987 |
| ENSMUSG00000047139 | Cd24a | protein_coding | -0.56524961 | 4.013841585 | 0.8433975 | 0.371811 | 0.999175987 |
| ENSMUSG00000022255 | Mtdh | protein_coding | -0.504731276 | 4.957280925 | 0.8430638 | 0.3719033 | 0.999175987 |
| ENSMUSG00000109284 | B230311B06Rik | lincRNA | -0.695136048 | 3.309155098 | 0.8430309 | 0.3719124 | 0.999175987 |
| ENSMUSG00000010914 | Pdhx | protein_coding | 0.319859127 | 6.463653855 | 0.8429064 | 0.3719468 | 0.999175987 |
| ENSMUSG00000019297 | Nop9 | protein_coding | 0.938397883 | 2.89386203 | 0.8425326 | 0.3720503 | 0.999175987 |
| ENSMUSG00000039706 | Ldb2 | protein_coding | -0.332740391 | 6.280817788 | 0.8424435 | 0.3720749 | 0.999175987 |
| ENSMUSG00000021234 | Fam161b | protein_coding | -0.718237281 | 3.909700963 | 0.84252 | 0.3720756 | 0.999175987 |
| ENSMUSG00000034593 | Myo5a | protein_coding | -0.362119477 | 7.076444363 | 0.8423486 | 0.3721231 | 0.999175987 |
| ENSMUSG00000078578 | Ube2d3 | protein_coding | 0.259238945 | 7.32672066 | 0.8418814 | 0.3722306 | 0.999175987 |
| ENSMUSG00000005312 | Ubqln1 | protein_coding | 0.263231549 | 7.206045527 | 0.8416886 | 0.372284 | 0.999175987 |
| ENSMUSG00000038398 | Upf3a | protein_coding | -0.49933977 | 4.345430702 | 0.8413942 | 0.3723656 | 0.999175987 |
| ENSMUSG00000011096 | Akt1s1 | protein_coding | -0.859473046 | 2.861234585 | 0.841368 | 0.3723728 | 0.999175987 |
| ENSMUSG00000023806 | Rsph3b | protein_coding | 0.750636423 | 2.821497084 | 0.8412495 | 0.3724057 | 0.999175987 |
| ENSMUSG00000024501 | Dpysl3 | protein_coding | -0.410265264 | 6.205575626 | 0.8411267 | 0.3724615 | 0.999175987 |
| ENSMUSG00000024831 | Ighmbp2 | protein_coding | 0.891564599 | 3.065338584 | 0.840077 | 0.3727309 | 0.999175987 |
| ENSMUSG00000001445 | Mrpl10 | protein_coding | 0.333557308 | 6.93874724 | 0.8398352 | 0.372798 | 0.999175987 |
| ENSMUSG00000036898 | Zfp157 | protein_coding | -0.480988014 | 4.376131062 | 0.8396508 | 0.3728492 | 0.999175987 |
| ENSMUSG00000002345 | Borcs8 | protein_coding | 0.31507048 | 7.178621538 | 0.8392837 | 0.3729511 | 0.999175987 |
| ENSMUSG00000026317 | Cln8 | protein_coding | 0.4656682 | 5.354004225 | 0.8390309 | 0.3730214 | 0.999175987 |
| ENSMUSG00000006418 | Rnf114 | protein_coding | 0.300189418 | 7.368095382 | 0.8389428 | 0.3730458 | 0.999175987 |
| ENSMUSG00000020372 | Rack1 | protein_coding | 0.291987147 | 8.901453793 | 0.8385008 | 0.3731687 | 0.999175987 |
| ENSMUSG00000026558 | Uck2 | protein_coding | -0.596812138 | 3.764361791 | 0.8379574 | 0.3733198 | 0.999175987 |
| ENSMUSG00000030424 | Zfp939 | transcribed_unproces | -0.68542946 | 3.067086335 | 0.8377317 | 0.3733826 | 0.999175987 |
| ENSMUSG00000034898 | Filip1 | protein_coding | -0.538634533 | 4.020894749 | 0.8367533 | 0.373655 | 0.999175987 |
| ENSMUSG00000030322 | Mbd4 | protein_coding | 0.72775334 | 3.104928882 | 0.8359009 | 0.3738925 | 0.999175987 |
| ENSMUSG00000041939 | Mvk | protein_coding | 0.436110993 | 5.485362946 | 0.8355685 | 0.3739852 | 0.999175987 |
| ENSMUSG00000056900 | Usp13 | protein_coding | 0.551714889 | 4.009411994 | 0.8355367 | 0.3739941 | 0.999175987 |
| ENSMUSG00000009406 | Elk1 | protein_coding | 0.512930603 | 4.138443608 | 0.8348271 | 0.3741921 | 0.999175987 |
| ENSMUSG00000029066 | Mrpl20 | protein_coding | 0.296762389 | 8.825360776 | 0.8345751 | 0.3742624 | 0.999175987 |
| ENSMUSG00000021704 | Mtx3 | protein_coding | -0.473785841 | 4.285084892 | 0.8333596 | 0.3746021 | 0.999175987 |
| ENSMUSG00000028677 | Rnf220 | protein_coding | 0.27762873 | 6.594261474 | 0.8332491 | 0.374633 | 0.999175987 |
| ENSMUSG00000006931 | P3h4 | protein_coding | -0.548794702 | 3.879283003 | 0.8331534 | 0.3746597 | 0.999175987 |
| ENSMUSG00000030323 | Ift122 | protein_coding | -0.459613324 | 4.801334896 | 0.8319992 | 0.3749827 | 0.999175987 |
| ENSMUSG00000015002 | Efr3a | protein_coding | -0.331873771 | 5.889607461 | 0.8319905 | 0.3749851 | 0.999175987 |
| ENSMUSG00000018923 | Med11 | protein_coding | -0.408273313 | 5.137953717 | 0.8319744 | 0.3749897 | 0.999175987 |
| ENSMUSG00000026917 | Wdr5 | protein_coding | 0.619594182 | 3.894247691 | 0.8319467 | 0.3749974 | 0.999175987 |
| ENSMUSG00000055850 | Rnf181 | protein_coding | -0.257728955 | 7.940958936 | 0.8318827 | 0.3750153 | 0.999175987 |
| ENSMUSG00000028832 | Stmn1 | protein_coding | -0.278213291 | 7.60688748 | 0.8312149 | 0.3752024 | 0.999175987 |
| ENSMUSG00000041769 | Ppp2r2d | protein_coding | -0.292152586 | 6.312592525 | 0.8311844 | 0.375211 | 0.999175987 |
| ENSMUSG00000032370 | Lactb | protein_coding | -0.775677032 | 2.970459917 | 0.8311123 | 0.3752312 | 0.999175987 |
| ENSMUSG00000031226 | Pbdc1 | protein_coding | 0.287233898 | 6.387891772 | 0.831023 | 0.3752562 | 0.999175987 |
| ENSMUSG00000026049 | Tex30 | protein_coding | -0.477815457 | 4.010276521 | 0.8301707 | 0.3754952 | 0.999175987 |
| ENSMUSG00000042747 | Krtcap2 | protein_coding | 0.304470514 | 7.698861476 | 0.8298859 | 0.3755752 | 0.999175987 |
| ENSMUSG00000062169 | Cnih4 | protein_coding | 0.358209356 | 5.796278403 | 0.8296627 | 0.3756378 | 0.999175987 |
| ENSMUSG00000103649 | Gm37768 | antisense_RNA | -0.876881903 | 2.671521566 | 0.8291212 | 0.3757899 | 0.999175987 |
| ENSMUSG00000021112 | Mpp5 | protein_coding | -0.328348383 | 5.843263303 | 0.8283218 | 0.3760145 | 0.999175987 |
| ENSMUSG00000020374 | Rasgef1c | protein_coding | -0.701018371 | 3.537564496 | 0.8279187 | 0.3761279 | 0.999175987 |
| ENSMUSG00000000305 | Cdh4 | protein_coding | 0.370218974 | 5.535970234 | 0.8273891 | 0.3762769 | 0.999175987 |
| ENSMUSG00000004446 | Bid | protein_coding | 0.452022849 | 5.242118399 | 0.8273763 | 0.3762805 | 0.999175987 |
| ENSMUSG00000033883 | D3Ertd254e | protein_coding | -0.495401016 | 4.246837968 | 0.8272006 | 0.37633 | 0.999175987 |
| ENSMUSG00000027438 | Napb | protein_coding | 0.329035976 | 8.827453936 | 0.8264773 | 0.3765337 | 0.999175987 |
| ENSMUSG00000032131 | Abcg4 | protein_coding | -0.408641913 | 5.149193701 | 0.8261222 | 0.3766337 | 0.999175987 |
| ENSMUSG00000019790 | Stxbp5 | protein_coding | -0.400909811 | 5.111077448 | 0.825673 | 0.3767604 | 0.999175987 |
| ENSMUSG00000024158 | Hagh | protein_coding | 0.275927798 | 8.304436477 | 0.8255393 | 0.3767981 | 0.999175987 |
| ENSMUSG00000020928 | Higd1b | protein_coding | 0.86225922 | 2.886518888 | 0.8253159 | 0.3768611 | 0.999175987 |
| ENSMUSG00000020216 | Jsrp1 | protein_coding | 0.745507398 | 3.231822087 | 0.8243625 | 0.3771302 | 0.999175987 |
| ENSMUSG00000075232 | Amd1 | protein_coding | 0.472318231 | 3.991844883 | 0.8243183 | 0.3771427 | 0.999175987 |
| ENSMUSG00000036966 | Spryd3 | protein_coding | 0.354225473 | 8.680516596 | 0.8240768 | 0.3772109 | 0.999175987 |
| ENSMUSG00000020340 | Cyfip2 | protein_coding | 0.31206785 | 8.355776164 | 0.8234884 | 0.3773771 | 0.999175987 |
| ENSMUSG00000021133 | Susd6 | protein_coding | -0.660864649 | 3.394380395 | 0.8233515 | 0.3774158 | 0.999175987 |
| ENSMUSG00000060771 | Tsga10 | protein_coding | -0.773064444 | 3.247839513 | 0.8233331 | 0.377421 | 0.999175987 |
| ENSMUSG00000036188 | Ankmy2 | protein_coding | -0.310531895 | 6.939125546 | 0.8232847 | 0.3774347 | 0.999175987 |
| ENSMUSG00000075227 | Znhit2 | protein_coding | 0.626693104 | 3.470282843 | 0.8232224 | 0.3774523 | 0.999175987 |
| ENSMUSG00000053141 | Ptprt | protein_coding | -0.524315696 | 4.442839227 | 0.8230503 | 0.377501 | 0.999175987 |
| ENSMUSG00000003208 | Ccdc94 | protein_coding | 0.701633822 | 3.964551078 | 0.8231193 | 0.3775031 | 0.999175987 |
| ENSMUSG00000024767 | Otub1 | protein_coding | 0.325515963 | 10.83118066 | 0.8228332 | 0.3775624 | 0.999175987 |
| ENSMUSG00000003131 | Pafah1b2 | protein_coding | -0.286777264 | 7.544336209 | 0.8222043 | 0.3777404 | 0.999175987 |
| ENSMUSG00000026851 | BC005624 | protein_coding | 0.271268793 | 7.926438961 | 0.8222019 | 0.3777411 | 0.999175987 |
| ENSMUSG00000035234 | Abraxas1 | protein_coding | 0.807046407 | 2.869221752 | 0.8219657 | 0.3778079 | 0.999175987 |
| ENSMUSG00000034462 | Pkd2 | protein_coding | -0.803189638 | 2.742760216 | 0.8211883 | 0.3780281 | 0.999175987 |
| ENSMUSG00000008305 | Tle1 | protein_coding | -0.45406551 | 4.754083591 | 0.8211578 | 0.3780368 | 0.999175987 |
| ENSMUSG00000037788 | Vopp1 | protein_coding | -0.498685139 | 4.910781996 | 0.8208413 | 0.3781265 | 0.999175987 |
| ENSMUSG00000021240 | Abcd4 | protein_coding | 0.900381581 | 2.905909648 | 0.8194771 | 0.3785136 | 0.999175987 |

| ensembl_gene_id | external_gene_name | gene_biotype | log2 (fold change) | log2 (counts per million) | F-value | p-value | False discovery rate |
|---|---|---|---|---|---|---|---|
| ENSMUSG00000022849 | Hspbap1 | protein_coding | -0.847629635 | 3.208849033 | 0.8194053 | 0.3785339 | 0.999175987 |
| ENSMUSG00000030697 | Ppp4c | protein_coding | -0.343889231 | 5.388606369 | 0.8188628 | 0.378688 | 0.999175987 |
| ENSMUSG00000036264 | Fstl4 | protein_coding | -0.785472561 | 3.592194445 | 0.8186281 | 0.3787763 | 0.999175987 |
| ENSMUSG00000020954 | Strn3 | protein_coding | -0.417977937 | 5.765313245 | 0.8168391 | 0.3792637 | 0.999175987 |
| ENSMUSG00000036977 | Anapc10 | protein_coding | 0.424688698 | 5.966676204 | 0.8162424 | 0.3794552 | 0.999175987 |
| ENSMUSG00000042348 | Arl15 | protein_coding | 0.645149884 | 3.510940519 | 0.8158714 | 0.3795394 | 0.999175987 |
| ENSMUSG00000001525 | Tubb5 | protein_coding | 0.364988483 | 10.28704004 | 0.8155036 | 0.3796443 | 0.999175987 |
| ENSMUSG00000024413 | Npc1 | protein_coding | -0.514753867 | 4.208555159 | 0.8154917 | 0.3796477 | 0.999175987 |
| ENSMUSG00000018547 | Pip4k2b | protein_coding | 0.521355274 | 4.264914796 | 0.8153715 | 0.379682 | 0.999175987 |
| ENSMUSG00000063919 | Srrm4 | protein_coding | -0.509579939 | 4.491326494 | 0.8149063 | 0.3798147 | 0.999175987 |
| ENSMUSG00000003680 | Taf6l | protein_coding | 0.601768899 | 4.353004743 | 0.8147473 | 0.3798601 | 0.999175987 |
| ENSMUSG00000016382 | Pls3 | protein_coding | -0.311781161 | 6.281365613 | 0.8147328 | 0.3798642 | 0.999175987 |
| ENSMUSG00000028300 | 3110043O21Rik | protein_coding | 0.427854526 | 5.315404608 | 0.8143212 | 0.3799817 | 0.999175987 |
| ENSMUSG00000058486 | Wdr91 | protein_coding | 0.610329802 | 4.671450756 | 0.8142204 | 0.380032 | 0.999175987 |
| ENSMUSG00000094483 | Purb | protein_coding | -0.265140588 | 8.418996873 | 0.8139677 | 0.3800827 | 0.999175987 |
| ENSMUSG00000030683 | Sez6l2 | protein_coding | 0.279481896 | 7.401941601 | 0.8136981 | 0.3801597 | 0.999175987 |
| ENSMUSG00000019831 | Wasf1 | protein_coding | -0.283016583 | 7.054750933 | 0.8136255 | 0.3801805 | 0.999175987 |
| ENSMUSG00000022365 | Derl1 | protein_coding | -0.350623695 | 5.782097291 | 0.8134677 | 0.3802256 | 0.999175987 |
| ENSMUSG00000031027 | Stk33 | protein_coding | -0.731303036 | 3.030573786 | 0.8131836 | 0.3803068 | 0.999175987 |
| ENSMUSG00000096370 | Gm21992 | protein_coding | -0.763626128 | 3.226450449 | 0.8131285 | 0.3803226 | 0.999175987 |
| ENSMUSG00000025785 | Exosc7 | protein_coding | -0.287572248 | 7.102316384 | 0.8128668 | 0.3803974 | 0.999175987 |
| ENSMUSG00000085208 | Brip1os | antisense_RNA | -0.893820746 | 2.826125638 | 0.8117457 | 0.3807183 | 0.999175987 |
| ENSMUSG00000046879 | Irgm1 | protein_coding | 0.68967429 | 4.267996404 | 0.8116006 | 0.3808377 | 0.999175987 |
| ENSMUSG00000019467 | Arhgef25 | protein_coding | -0.45846119 | 4.862343665 | 0.811246 | 0.3808614 | 0.999175987 |
| ENSMUSG00000050199 | Lgr4 | protein_coding | 0.432236861 | 4.807990148 | 0.8111479 | 0.3808895 | 0.999175987 |
| ENSMUSG00000029697 | Fezf1 | protein_coding | -1.440100446 | 3.055115251 | 0.8172835 | 0.3809698 | 0.999175987 |
| ENSMUSG00000064246 | Chil1 | protein_coding | -0.593978617 | 4.459438099 | 0.8107933 | 0.3810126 | 0.999175987 |
| ENSMUSG00000038546 | Ranbp9 | protein_coding | -0.576188363 | 3.615199335 | 0.8103368 | 0.3811221 | 0.999175987 |
| ENSMUSG00000055541 | Lair1 | protein_coding | -0.727356327 | 3.076907307 | 0.8102563 | 0.3811452 | 0.999175987 |
| ENSMUSG00000025920 | Stau2 | protein_coding | -0.253968449 | 7.064248822 | 0.8102428 | 0.3811491 | 0.999175987 |
| ENSMUSG00000062203 | Gspt1 | protein_coding | -0.778037724 | 2.725209974 | 0.8099321 | 0.3812382 | 0.999175987 |
| ENSMUSG00000113722 | AC140477.3 | lincRNA | 0.422032294 | 4.754180436 | 0.8098971 | 0.3812483 | 0.999175987 |
| ENSMUSG00000024072 | Yipf4 | protein_coding | -0.427817733 | 5.55137118 | 0.8096452 | 0.3813206 | 0.999175987 |
| ENSMUSG00000029992 | Gfpt1 | protein_coding | -0.462455306 | 5.066774403 | 0.8095146 | 0.3813581 | 0.999175987 |
| ENSMUSG00000032383 | Ppib | protein_coding | 0.312828401 | 6.18592516 | 0.8090765 | 0.3814839 | 0.999175987 |
| ENSMUSG00000023960 | Enpp5 | protein_coding | -0.2410358 | 7.834927433 | 0.809038 | 0.3814949 | 0.999175987 |
| ENSMUSG00000039682 | Lap3 | protein_coding | -0.311472792 | 6.696552077 | 0.8089805 | 0.3815115 | 0.999175987 |
| ENSMUSG00000021285 | Ppp1r13b | protein_coding | -0.667667125 | 3.25431394 | 0.8088456 | 0.3815502 | 0.999175987 |
| ENSMUSG00000047388 | Atmin | protein_coding | 0.346846974 | 5.551702336 | 0.8085809 | 0.3816263 | 0.999175987 |
| ENSMUSG00000079598 | Clec2l | protein_coding | -0.448971978 | 4.576287823 | 0.8078317 | 0.3818417 | 0.999175987 |
| ENSMUSG00000033029 | 1700088E04Rik | protein_coding | 0.552138587 | 3.823155226 | 0.8076811 | 0.3818851 | 0.999175987 |
| ENSMUSG00000037214 | Thap1 | protein_coding | -0.495099496 | 3.795572481 | 0.806503 | 0.3822242 | 0.999175987 |
| ENSMUSG00000024222 | Fkbp5 | protein_coding | -0.544441031 | 4.536240414 | 0.8060675 | 0.3823497 | 0.999175987 |
| ENSMUSG00000041014 | Nrg3 | protein_coding | -0.470867561 | 4.681865863 | 0.8057105 | 0.3824526 | 0.999175987 |
| ENSMUSG00000034926 | Dhcr24 | protein_coding | 0.749687639 | 3.241174384 | 0.8056568 | 0.3824681 | 0.999175987 |
| ENSMUSG00000023991 | Foxp4 | protein_coding | 0.667594008 | 3.535904737 | 0.8053327 | 0.3825616 | 0.999175987 |
| ENSMUSG00000035226 | Rims4 | protein_coding | -0.445823803 | 5.01318547 | 0.8051939 | 0.3826016 | 0.999175987 |
| ENSMUSG00000006304 | Arpc2 | protein_coding | -0.261740987 | 7.838878671 | 0.8051141 | 0.3826247 | 0.999175987 |
| ENSMUSG00000050244 | Heatr1 | protein_coding | 0.811094518 | 2.870058153 | 0.803952 | 0.3829602 | 0.999175987 |
| ENSMUSG00000048895 | Cdk5r1 | protein_coding | -0.285197348 | 6.889457101 | 0.8039283 | 0.3829671 | 0.999175987 |
| ENSMUSG00000041684 | Bivm | protein_coding | -0.457969838 | 4.219318617 | 0.8037946 | 0.3830057 | 0.999175987 |
| ENSMUSG00000053091 | Lins1 | protein_coding | 0.726818558 | 3.081821021 | 0.8037414 | 0.3830211 | 0.999175987 |
| ENSMUSG00000034449 | Dhrs11 | protein_coding | 0.689559703 | 3.101245341 | 0.8033586 | 0.3831317 | 0.999175987 |
| ENSMUSG00000015575 | Atp6v0e | protein_coding | -0.38779879 | 7.084785121 | 0.8028979 | 0.3833424 | 0.999175987 |
| ENSMUSG00000026421 | Csrp1 | protein_coding | 0.283209664 | 8.291073707 | 0.8024938 | 0.3833819 | 0.999175987 |
| ENSMUSG00000025373 | Rnf41 | protein_coding | 0.30502272 | 7.783121997 | 0.8024187 | 0.3834036 | 0.999175987 |
| ENSMUSG00000027332 | Ivd | protein_coding | -0.403317534 | 6.87274443 | 0.802772 | 0.3834156 | 0.999175987 |
| ENSMUSG00000036218 | Pdzrn4 | protein_coding | 0.531140264 | 3.917302898 | 0.8021585 | 0.3834789 | 0.999175987 |
| ENSMUSG00000079339 | Ifit1bl1 | protein_coding | 1.728281185 | 2.273117401 | 0.8042558 | 0.3836986 | 0.999175987 |
| ENSMUSG00000039463 | Slc9a8 | protein_coding | -0.768253975 | 4.368778682 | 0.8049172 | 0.3837448 | 0.999175987 |
| ENSMUSG00000027479 | Mapre1 | protein_coding | -0.268404342 | 7.318382081 | 0.8003808 | 0.3839941 | 0.999175987 |
| ENSMUSG00000069072 | Slc7a14 | protein_coding | -0.319737559 | 6.089734951 | 0.800255 | 0.3840306 | 0.999175987 |
| ENSMUSG00000069049 | Eif2s3y | protein_coding | -0.47023754 | 5.038632738 | 0.8002296 | 0.384038 | 0.999175987 |
| ENSMUSG00000022314 | Rad21 | protein_coding | -0.359116221 | 5.920039703 | 0.8000472 | 0.3840909 | 0.999175987 |
| ENSMUSG00000035898 | Uba6 | protein_coding | -0.599291452 | 4.259359949 | 0.7993384 | 0.3842967 | 0.999175987 |
| ENSMUSG00000020098 | Pcbd1 | protein_coding | 0.269633155 | 7.694197222 | 0.7983742 | 0.3845769 | 0.999175987 |
| ENSMUSG00000054920 | Klhl5 | protein_coding | -0.430136171 | 4.89914094 | 0.7982773 | 0.384605 | 0.999175987 |
| ENSMUSG00000024182 | Axin1 | protein_coding | -0.464939561 | 4.05334865 | 0.7969173 | 0.3850008 | 0.999175987 |
| ENSMUSG00000029121 | Crmp1 | protein_coding | 0.557478215 | 4.814668844 | 0.7969532 | 0.3850116 | 0.999175987 |
| ENSMUSG00000013523 | Bcas1 | protein_coding | -0.550468219 | 5.245615683 | 0.796783 | 0.385117 | 0.999175987 |
| ENSMUSG00000006021 | Kptn | protein_coding | 0.511535066 | 5.224893488 | 0.7962597 | 0.3852136 | 0.999175987 |
| ENSMUSG00000050321 | Neto1 | protein_coding | -0.519736004 | 4.702651805 | 0.7961679 | 0.3852191 | 0.999175987 |
| ENSMUSG00000030905 | Crym | protein_coding | -0.537320834 | 7.711231282 | 0.8029972 | 0.385302 | 0.999175987 |
| ENSMUSG00000026229 | Psmd1 | protein_coding | 0.305978344 | 7.239032462 | 0.7956608 | 0.385367 | 0.999175987 |
| ENSMUSG00000036214 | Znrd1as | protein_coding | -0.599559773 | 3.733411621 | 0.7954999 | 0.3854139 | 0.999175987 |
| ENSMUSG00000046818 | Ddit4l | protein_coding | -0.495728599 | 4.730396204 | 0.7946793 | 0.3856534 | 0.999175987 |
| ENSMUSG00000030047 | Arhgap25 | protein_coding | -1.192966916 | 2.611281715 | 0.7944564 | 0.3857397 | 0.999175987 |
| ENSMUSG00000027962 | Vcam1 | protein_coding | -0.711689937 | 3.937435322 | 0.794294 | 0.3857871 | 0.999175987 |

| ensembl_gene_id | external_gene_name | gene_biotype | log2 (fold change) | log2 (counts per million) | F-value | p-value | False discovery rate |
|---|---|---|---|---|---|---|---|
| ENSMUSG00000032078 | Zpr1 | protein_coding | 0.550540037 | 4.798499595 | 0.7937948 | 0.385933 | 0.999175987 |
| ENSMUSG00000039195 | 1110008P14Rik | protein_coding | 0.537757829 | 4.218693784 | 0.7931893 | 0.3860888 | 0.999175987 |
| ENSMUSG00000087687 | Pet100 | protein_coding | 0.381372465 | 6.789267763 | 0.7931465 | 0.3861225 | 0.999175987 |
| ENSMUSG00000039630 | Hnrnpu | protein_coding | -0.243350808 | 7.5492458 | 0.7929849 | 0.3861486 | 0.999175987 |
| ENSMUSG00000020430 | Pes1 | protein_coding | 0.463071733 | 5.057633856 | 0.7928253 | 0.3861953 | 0.999175987 |
| ENSMUSG00000029499 | Pxmp2 | protein_coding | 0.590849667 | 3.719381021 | 0.7922327 | 0.3863687 | 0.999175987 |
| ENSMUSG00000086316 | 2210013O21Rik | protein_coding | 0.237453251 | 7.655400986 | 0.7922038 | 0.3863772 | 0.999175987 |
| ENSMUSG00000044042 | Fmn1 | protein_coding | -0.605562554 | 3.170494221 | 0.7919124 | 0.3864625 | 0.999175987 |
| ENSMUSG00000035024 | Ncapd3 | protein_coding | -0.598905842 | 3.666744051 | 0.7914907 | 0.3865861 | 0.999175987 |
| ENSMUSG00000053297 | AI854703 | protein_coding | 0.40487771 | 5.33239503 | 0.7914554 | 0.3865964 | 0.999175987 |
| ENSMUSG00000034412 | Tbc1d10a | protein_coding | -0.712446797 | 3.277671923 | 0.79142 | 0.3866068 | 0.999175987 |
| ENSMUSG00000071379 | Hpcal1 | protein_coding | 0.486610052 | 7.646691765 | 0.7975059 | 0.3866384 | 0.999175987 |
| ENSMUSG00000038844 | Kif16b | protein_coding | -0.721410712 | 3.107738631 | 0.7909769 | 0.3867367 | 0.999175987 |
| ENSMUSG00000025466 | Fuom | protein_coding | 0.307780173 | 7.227118524 | 0.7908425 | 0.3867761 | 0.999175987 |
| ENSMUSG00000069520 | Tmem19 | protein_coding | -0.347888443 | 5.323552314 | 0.7908019 | 0.386788 | 0.999175987 |
| ENSMUSG00000022489 | Pde1b | protein_coding | -0.293634738 | 7.601957761 | 0.7907695 | 0.3867975 | 0.999175987 |
| ENSMUSG00000030045 | Mrpl19 | protein_coding | 0.256285321 | 7.065750676 | 0.7906079 | 0.3868449 | 0.999175987 |
| ENSMUSG00000035681 | Kcnc2 | protein_coding | -0.450297701 | 5.261467745 | 0.7901362 | 0.3869833 | 0.999175987 |
| ENSMUSG00000032478 | Nme6 | protein_coding | 0.371178749 | 5.070254869 | 0.79007 | 0.3870027 | 0.999175987 |
| ENSMUSG00000058546 | Rpl23a | protein_coding | 0.629306825 | 3.615805118 | 0.7899587 | 0.3870354 | 0.999175987 |
| ENSMUSG00000054387 | Mdm4 | protein_coding | -0.348830519 | 5.099620466 | 0.7895997 | 0.3871408 | 0.999175987 |
| ENSMUSG00000045973 | Slc25a51 | protein_coding | 0.330077853 | 6.216469553 | 0.7894001 | 0.3871994 | 0.999175987 |
| ENSMUSG00000039275 | Foxk2 | protein_coding | -0.42457083 | 5.098792257 | 0.7885632 | 0.3874453 | 0.999175987 |
| ENSMUSG00000048796 | Cyb561d1 | protein_coding | 0.64913881 | 3.55124083 | 0.7883998 | 0.3874934 | 0.999175987 |
| ENSMUSG00000037236 | Matr3 | protein_coding | -0.274126066 | 8.047291825 | 0.7881988 | 0.3875525 | 0.999175987 |
| ENSMUSG00000031953 | Tmem170 | protein_coding | 0.552903309 | 4.277935296 | 0.788081 | 0.3875871 | 0.999175987 |
| ENSMUSG00000049232 | Tigd2 | protein_coding | 0.615142308 | 4.020976715 | 0.7880443 | 0.3875979 | 0.999175987 |
| ENSMUSG00000071669 | Snx29 | protein_coding | 0.651272242 | 3.290606917 | 0.7877573 | 0.3876824 | 0.999175987 |
| ENSMUSG00000029348 | Asphd2 | protein_coding | -0.367871265 | 5.911058479 | 0.7876812 | 0.3877048 | 0.999175987 |
| ENSMUSG00000034295 | Fhod3 | protein_coding | -0.68355638 | 2.940591565 | 0.787273 | 0.387825 | 0.999175987 |
| ENSMUSG00000032281 | Acsbg1 | protein_coding | -0.303617472 | 7.206270198 | 0.7868214 | 0.387958 | 0.999175987 |
| ENSMUSG00000025104 | Hdgfl3 | protein_coding | -0.342553846 | 6.422532832 | 0.7866114 | 0.3880198 | 0.999175987 |
| ENSMUSG00000029312 | Klhl8 | protein_coding | 0.447458816 | 4.255544535 | 0.7856049 | 0.3883166 | 0.999175987 |
| ENSMUSG00000061315 | Naca | protein_coding | 0.312054442 | 6.815709084 | 0.7854837 | 0.3883523 | 0.999175987 |
| ENSMUSG00000023967 | Mrps18a | protein_coding | 0.349446899 | 5.128822715 | 0.7853131 | 0.3884027 | 0.999175987 |
| ENSMUSG00000003545 | Fosb | protein_coding | 0.717824072 | 3.305642416 | 0.7848517 | 0.3885389 | 0.999175987 |
| ENSMUSG00000027993 | Trim2 | protein_coding | -0.259245506 | 7.573465912 | 0.7840382 | 0.3887792 | 0.999175987 |
| ENSMUSG00000029478 | Ncor2 | protein_coding | -0.587660078 | 3.755960441 | 0.7839345 | 0.3888098 | 0.999175987 |
| ENSMUSG00000047379 | B4gat1 | protein_coding | 0.360331572 | 7.438646218 | 0.7835449 | 0.388946 | 0.999175987 |
| ENSMUSG00000009739 | Pou6f1 | protein_coding | 0.427832624 | 4.692505443 | 0.7834031 | 0.3889669 | 0.999175987 |
| ENSMUSG00000037369 | Kdm6a | protein_coding | -0.581741071 | 3.870583074 | 0.7830635 | 0.3890674 | 0.999175987 |
| ENSMUSG00000032423 | Syncrip | protein_coding | 0.588738068 | 3.30915469 | 0.7826097 | 0.3892017 | 0.999175987 |
| ENSMUSG00000027533 | Fabp5 | protein_coding | 0.271006315 | 6.901610174 | 0.7826082 | 0.3892021 | 0.999175987 |
| ENSMUSG00000029616 | Erp29 | protein_coding | 0.259829075 | 7.669565449 | 0.7820235 | 0.3893753 | 0.999175987 |
| ENSMUSG00000038722 | Bud31 | protein_coding | 0.325718154 | 6.785825901 | 0.7807966 | 0.389739 | 0.999175987 |
| ENSMUSG00000021265 | Slc25a29 | protein_coding | 0.65370694 | 3.224824225 | 0.7804578 | 0.3898395 | 0.999175987 |
| ENSMUSG00000033066 | Gas7 | protein_coding | -0.402655287 | 6.236110823 | 0.7804859 | 0.3898521 | 0.999175987 |
| ENSMUSG00000027765 | P2ry1 | protein_coding | 0.70194431 | 3.143338903 | 0.7803374 | 0.3898752 | 0.999175987 |
| ENSMUSG00000019303 | Psmc3ip | protein_coding | -0.539682767 | 3.827689081 | 0.7797912 | 0.3900374 | 0.999175987 |
| ENSMUSG00000028226 | Mmp16 | protein_coding | -0.540255434 | 4.054769991 | 0.779681 | 0.3900701 | 0.999175987 |
| ENSMUSG00000024038 | Ndufv3 | protein_coding | 0.292531716 | 9.184277111 | 0.7794743 | 0.3901315 | 0.999175987 |
| ENSMUSG00000023249 | Parp3 | protein_coding | -0.806615633 | 2.909312316 | 0.7793136 | 0.3901792 | 0.999175987 |
| ENSMUSG00000049265 | Kcnk3 | protein_coding | 0.435962056 | 4.463893453 | 0.7786458 | 0.3903777 | 0.999175987 |
| ENSMUSG00000029213 | Commd8 | protein_coding | 0.287121103 | 7.193414177 | 0.7781317 | 0.3905307 | 0.999175987 |
| ENSMUSG00000002346 | Slc25a42 | protein_coding | 0.597042891 | 4.156457264 | 0.7780712 | 0.3905486 | 0.999175987 |
| ENSMUSG00000028646 | Rragc | protein_coding | -0.348862292 | 5.912414531 | 0.7778047 | 0.390628 | 0.999175987 |
| ENSMUSG00000044148 | 1810030O07Rik | protein_coding | 0.352450215 | 5.633291775 | 0.7776304 | 0.3906799 | 0.999175987 |
| ENSMUSG00000026627 | Tmem206 | protein_coding | -0.554962759 | 4.50691482 | 0.7771985 | 0.3908085 | 0.999175987 |
| ENSMUSG00000034290 | Nek9 | protein_coding | 0.612577473 | 3.854974297 | 0.7763338 | 0.3910662 | 0.999175987 |
| ENSMUSG00000022555 | Dgat1 | protein_coding | 0.685080128 | 3.529617965 | 0.776053 | 0.3911499 | 0.999175987 |
| ENSMUSG00000021978 | Extl3 | protein_coding | 0.42633776 | 5.404115986 | 0.7760194 | 0.3911599 | 0.999175987 |
| ENSMUSG00000022091 | Sorbs3 | protein_coding | 0.408817463 | 4.802148404 | 0.7754364 | 0.3913339 | 0.999175987 |
| ENSMUSG00000004018 | Fancl | protein_coding | 0.438016923 | 4.841292438 | 0.7752672 | 0.3913844 | 0.999175987 |
| ENSMUSG00000029861 | Fam131b | protein_coding | -0.623618176 | 3.633777531 | 0.775026 | 0.3914564 | 0.999175987 |
| ENSMUSG00000002428 | Hltf | protein_coding | -0.439703398 | 4.522616814 | 0.7750085 | 0.3914616 | 0.999175987 |
| ENSMUSG00000042390 | Gatad2b | protein_coding | -0.341239352 | 5.723837878 | 0.7745471 | 0.3915994 | 0.999175987 |
| ENSMUSG00000052310 | Slc39a1 | protein_coding | 0.653649725 | 4.916541823 | 0.7763847 | 0.3918617 | 0.999175987 |
| ENSMUSG00000024293 | Esco1 | protein_coding | -0.416733417 | 4.417822002 | 0.7732972 | 0.3919731 | 0.999175987 |
| ENSMUSG00000063063 | Ctnna2 | protein_coding | -0.291301983 | 6.555486821 | 0.7731718 | 0.3920106 | 0.999175987 |
| ENSMUSG00000022765 | Snap29 | protein_coding | -0.499778374 | 4.01884437 | 0.7731343 | 0.3920219 | 0.999175987 |
| ENSMUSG00000039410 | Prdm16 | protein_coding | 0.447101001 | 5.45472691 | 0.7728703 | 0.3921009 | 0.999175987 |
| ENSMUSG00000051166 | Eml5 | protein_coding | -0.521203032 | 3.710090778 | 0.7727029 | 0.392151 | 0.999175987 |
| ENSMUSG00000071072 | Ptges3 | protein_coding | -0.251015572 | 8.537608677 | 0.7725538 | 0.3921956 | 0.999175987 |
| ENSMUSG00000032062 | 2310030G06Rik | protein_coding | -0.670408842 | 3.466498892 | 0.7723303 | 0.3922626 | 0.999175987 |
| ENSMUSG00000006705 | Pknox1 | protein_coding | 0.488297144 | 4.007590986 | 0.7721262 | 0.3923237 | 0.999175987 |
| ENSMUSG00000038011 | Dnah10 | protein_coding | 0.861174189 | 2.859504059 | 0.772108 | 0.3923292 | 0.999175987 |
| ENSMUSG00000036422 | Pcdh8 | protein_coding | -0.560897831 | 4.455041384 | 0.7720262 | 0.3923537 | 0.999175987 |
| ENSMUSG00000026058 | Khdrbs2 | protein_coding | -0.399467496 | 4.980876755 | 0.7718382 | 0.39241 | 0.999175987 |

| ensembl_gene_id | external_gene_name | gene_biotype | log2 (fold change) | log2 (counts per million) | F-value | p-value | False discovery rate |
|---|---|---|---|---|---|---|---|
| ENSMUSG00000039483 | Asb6 | protein_coding | 0.355453966 | 6.025324549 | 0.7717366 | 0.3924405 | 0.999175987 |
| ENSMUSG00000049800 | Sertad2 | protein_coding | 0.492884332 | 3.710481192 | 0.7716499 | 0.3924665 | 0.999175987 |
| ENSMUSG00000044835 | Ankrd45 | protein_coding | -0.379957442 | 5.516952363 | 0.7706983 | 0.3927519 | 0.999175987 |
| ENSMUSG00000031403 | Dkc1 | protein_coding | -0.332566944 | 5.864226478 | 0.770303 | 0.3928705 | 0.999175987 |
| ENSMUSG00000068617 | Efcab1 | protein_coding | -0.350277653 | 5.490461706 | 0.7701555 | 0.3929148 | 0.999175987 |
| ENSMUSG00000048546 | Tob2 | protein_coding | -0.386422478 | 4.887642111 | 0.7698955 | 0.3929929 | 0.999175987 |
| ENSMUSG00000004892 | Bcan | protein_coding | -0.569513209 | 5.297750323 | 0.7716585 | 0.3930231 | 0.999175987 |
| ENSMUSG00000032744 | Heyl | protein_coding | -0.667712042 | 3.977557591 | 0.7689425 | 0.3933002 | 0.999175987 |
| ENSMUSG00000020713 | Gh | protein_coding | -1.026021648 | 3.159073707 | 0.7705215 | 0.3933365 | 0.999175987 |
| ENSMUSG00000049100 | Pcdh10 | protein_coding | -0.548976297 | 5.010605996 | 0.7685907 | 0.393406 | 0.999175987 |
| ENSMUSG00000054252 | Fgfr3 | protein_coding | -0.476986624 | 4.760512676 | 0.7681945 | 0.3935044 | 0.999175987 |
| ENSMUSG00000015484 | Fam163a | protein_coding | 0.793338514 | 3.346538733 | 0.7681094 | 0.39353 | 0.999175987 |
| ENSMUSG00000040415 | Dtx3 | protein_coding | 0.234933357 | 7.562950715 | 0.767886 | 0.3935972 | 0.999175987 |
| ENSMUSG00000021824 | Ap3m1 | protein_coding | -0.385585657 | 4.904119096 | 0.767833 | 0.3936132 | 0.999175987 |
| ENSMUSG00000060261 | Gtf2i | protein_coding | -0.334784632 | 6.664395869 | 0.7675967 | 0.3936843 | 0.999175987 |
| ENSMUSG00000079442 | St6galnac4 | protein_coding | 0.5514668 | 4.861569362 | 0.7674961 | 0.3937354 | 0.999175987 |
| ENSMUSG00000017417 | Plxdc1 | protein_coding | 0.517179057 | 3.993092198 | 0.7672331 | 0.3937939 | 0.999175987 |
| ENSMUSG00000026202 | Tuba4a | protein_coding | 0.363873872 | 9.099061636 | 0.7666214 | 0.3939991 | 0.999175987 |
| ENSMUSG00000045534 | Kcna5 | protein_coding | -0.582367069 | 3.796053901 | 0.7660897 | 0.3941386 | 0.999175987 |
| ENSMUSG00000005514 | Por | protein_coding | 0.373811891 | 6.626504381 | 0.7660156 | 0.3941817 | 0.999175987 |
| ENSMUSG00000002006 | Pdzd4 | protein_coding | -0.53123542 | 4.355802485 | 0.7659082 | 0.3941934 | 0.999175987 |
| ENSMUSG00000021830 | Txndc16 | protein_coding | -0.362456543 | 5.510028045 | 0.7658884 | 0.3941994 | 0.999175987 |
| ENSMUSG00000015461 | Atf6b | protein_coding | 0.609116715 | 4.014220969 | 0.7647607 | 0.3945399 | 0.999175987 |
| ENSMUSG00000029452 | Tmem116 | protein_coding | 0.643967883 | 3.387420825 | 0.7646397 | 0.3945765 | 0.999175987 |
| ENSMUSG00000035314 | Gdpd5 | protein_coding | 0.546352352 | 3.631621812 | 0.7646245 | 0.3945811 | 0.999175987 |
| ENSMUSG00000037610 | Kcnmb2 | protein_coding | 0.954869292 | 2.971614265 | 0.7645885 | 0.3946127 | 0.999175987 |
| ENSMUSG00000036819 | Jmjd4 | protein_coding | 0.487841014 | 4.267717668 | 0.7644673 | 0.3946286 | 0.999175987 |
| ENSMUSG00000006818 | Sod2 | protein_coding | 0.277081039 | 7.601455699 | 0.7633394 | 0.3949697 | 0.999175987 |
| ENSMUSG00000027737 | Slc7a11 | protein_coding | 0.734191962 | 3.218321286 | 0.7618465 | 0.395422 | 0.999175987 |
| ENSMUSG00000081225 | Cyp2j12 | protein_coding | -0.388835531 | 4.902637011 | 0.7614388 | 0.3955456 | 0.999175987 |
| ENSMUSG00000001700 | Gramd3 | protein_coding | -0.649765439 | 3.769375656 | 0.7612866 | 0.3955918 | 0.999175987 |
| ENSMUSG00000026933 | Camsap1 | protein_coding | -0.386793175 | 5.571069479 | 0.7612225 | 0.3956112 | 0.999175987 |
| ENSMUSG00000041488 | Stx3 | protein_coding | -0.506471278 | 3.858314056 | 0.7609465 | 0.395695 | 0.999175987 |
| ENSMUSG00000032571 | Pik3r4 | protein_coding | 0.560933609 | 4.210568491 | 0.7606462 | 0.3957862 | 0.999175987 |
| ENSMUSG00000028430 | Nol6 | protein_coding | 0.357024011 | 6.081971905 | 0.7604052 | 0.3958593 | 0.999175987 |
| ENSMUSG00000039901 | 9130011E15Rik | protein_coding | 0.593758287 | 3.936812485 | 0.7598318 | 0.3960335 | 0.999175987 |
| ENSMUSG00000031442 | Mcf2l | protein_coding | -0.431607728 | 4.428412457 | 0.7597986 | 0.3960436 | 0.999175987 |
| ENSMUSG00000019494 | Cops6 | protein_coding | 0.241673328 | 9.482982842 | 0.7597657 | 0.3960536 | 0.999175987 |
| ENSMUSG00000012429 | Mplkip | protein_coding | 0.889126066 | 2.538806212 | 0.7596741 | 0.3960815 | 0.999175987 |
| ENSMUSG00000019977 | Hbs1l | protein_coding | 0.400502543 | 5.924672831 | 0.7594668 | 0.3961445 | 0.999175987 |
| ENSMUSG00000045083 | Lingo2 | protein_coding | -0.392612722 | 4.931962584 | 0.7590809 | 0.3962618 | 0.999175987 |
| ENSMUSG00000015697 | Setdb1 | protein_coding | -0.512413283 | 3.872200855 | 0.7588725 | 0.3963252 | 0.999175987 |
| ENSMUSG00000024480 | Ap3s1 | protein_coding | 0.588366153 | 3.781050752 | 0.7587968 | 0.3963483 | 0.999175987 |
| ENSMUSG00000057788 | Ddx49 | protein_coding | 0.484160717 | 5.217780257 | 0.7587815 | 0.3963529 | 0.999175987 |
| ENSMUSG00000072915 | Gm12258 | protein_coding | -0.589528334 | 3.798291442 | 0.7576644 | 0.3966931 | 0.999175987 |
| ENSMUSG00000073002 | Vamp5 | protein_coding | -0.758299567 | 3.080028733 | 0.7574361 | 0.3967626 | 0.999175987 |
| ENSMUSG00000022674 | Ube2v2 | protein_coding | -0.401827637 | 4.830762363 | 0.7572684 | 0.3968137 | 0.999175987 |
| ENSMUSG00000036466 | Megf11 | protein_coding | -0.434969277 | 4.842826329 | 0.7572181 | 0.3968291 | 0.999175987 |
| ENSMUSG00000003549 | Ercc1 | protein_coding | 0.286751748 | 6.386600643 | 0.756308 | 0.3971067 | 0.999175987 |
| ENSMUSG00000021124 | Vti1b | protein_coding | 0.578701985 | 8.008059592 | 0.7679051 | 0.3971321 | 0.999175987 |
| ENSMUSG00000061979 | Rcc1l | protein_coding | -0.95325796 | 3.091089863 | 0.7560636 | 0.3972019 | 0.999175987 |
| ENSMUSG00000046785 | Epm2aip1 | protein_coding | -0.292647933 | 6.769392348 | 0.7553276 | 0.397406 | 0.999175987 |
| ENSMUSG00000073684 | Faap20 | protein_coding | -0.323703446 | 5.802245919 | 0.7551929 | 0.3974471 | 0.999175987 |
| ENSMUSG00000021719 | Rgs7bp | protein_coding | -0.251786294 | 7.127353161 | 0.7551263 | 0.3974675 | 0.999175987 |
| ENSMUSG00000020048 | Hsp90b1 | protein_coding | -0.24588784 | 7.071487555 | 0.7550331 | 0.397496 | 0.999175987 |
| ENSMUSG00000020572 | Nampt | protein_coding | -0.358972936 | 4.97681015 | 0.7548998 | 0.3975367 | 0.999175987 |
| ENSMUSG00000000552 | Zfp385a | protein_coding | 0.527428938 | 4.803536637 | 0.7547969 | 0.3975682 | 0.999175987 |
| ENSMUSG00000017969 | Ptgis | protein_coding | 0.969615903 | 2.848786097 | 0.7543532 | 0.3977039 | 0.999175987 |
| ENSMUSG00000045211 | Nudt18 | protein_coding | -0.373841402 | 5.721261591 | 0.7541902 | 0.3977537 | 0.999175987 |
| ENSMUSG00000097503 | 3110045C21Rik | antisense_RNA | 0.962746095 | 2.590778942 | 0.7540645 | 0.3977922 | 0.999175987 |
| ENSMUSG00000086370 | Ftx | lincRNA | -0.696545505 | 3.516053268 | 0.7538369 | 0.3978618 | 0.999175987 |
| ENSMUSG00000091636 | Akain1 | protein_coding | -1.18301168 | 2.482868168 | 0.7538009 | 0.3978728 | 0.999175987 |
| ENSMUSG00000036815 | Dpp10 | protein_coding | 0.339299142 | 6.009901135 | 0.7536494 | 0.3979192 | 0.999175987 |
| ENSMUSG00000051978 | Erich1 | protein_coding | 0.497594898 | 4.202474229 | 0.752772 | 0.3981879 | 0.999175987 |
| ENSMUSG00000034108 | Ccs | protein_coding | -0.29808986 | 6.176152604 | 0.752559 | 0.3982532 | 0.999175987 |
| ENSMUSG00000029686 | Cul1 | protein_coding | -0.300732907 | 6.502790169 | 0.7520515 | 0.3984088 | 0.999175987 |
| ENSMUSG00000018849 | Wwc1 | protein_coding | -0.397731772 | 5.078103555 | 0.7519807 | 0.3984305 | 0.999175987 |
| ENSMUSG00000024486 | Hbegf | protein_coding | 0.464480409 | 4.366426274 | 0.7515911 | 0.39855 | 0.999175987 |
| ENSMUSG00000055067 | Smyd3 | protein_coding | -0.499169454 | 6.563992212 | 0.7567849 | 0.3985885 | 0.999175987 |
| ENSMUSG00000049950 | Rpp38 | protein_coding | 0.341738427 | 5.107521382 | 0.7513571 | 0.3986219 | 0.999175987 |
| ENSMUSG00000109865 | Hspa14 | protein_coding | -0.346172475 | 5.770359006 | 0.751325 | 0.3986317 | 0.999175987 |
| ENSMUSG00000048279 | Sacs | protein_coding | -0.533435377 | 4.283864982 | 0.7511762 | 0.3986774 | 0.999175987 |
| ENSMUSG00000019897 | Ccdc59 | protein_coding | -0.2635339 | 7.036182763 | 0.7506068 | 0.3988522 | 0.999175987 |
| ENSMUSG00000033063 | Cntnap3 | protein_coding | -0.896943629 | 2.680127865 | 0.7499961 | 0.3990399 | 0.999175987 |
| ENSMUSG00000035086 | Becn1 | protein_coding | 0.57191445 | 3.339431092 | 0.748938 | 0.3993654 | 0.999175987 |
| ENSMUSG00000036613 | Tssc1 | protein_coding | 0.350370009 | 6.756816031 | 0.7484118 | 0.3995479 | 0.999175987 |
| ENSMUSG00000061589 | Dot1l | protein_coding | -0.603542899 | 3.989008239 | 0.7482101 | 0.3995895 | 0.999175987 |
| ENSMUSG00000045854 | Lyrm2 | protein_coding | 0.361376627 | 6.16688225 | 0.7475908 | 0.3997804 | 0.999175987 |

| ensembl_gene_id | external_gene_name | gene_biotype | log2 (fold change) | log2 (counts per million) | F-value | p-value | False discovery rate |
|---|---|---|---|---|---|---|---|
| ENSMUSG00000057322 | Rpl38 | protein_coding | 0.462177107 | 4.578530249 | 0.7474211 | 0.3998327 | 0.999175987 |
| ENSMUSG00000026131 | Dst | protein_coding | -0.444985722 | 5.014893336 | 0.7474059 | 0.3998374 | 0.999175987 |
| ENSMUSG00000046324 | Ermp1 | protein_coding | -0.583228054 | 4.23663807 | 0.7471146 | 0.3999273 | 0.999175987 |
| ENSMUSG00000038286 | Bphl | protein_coding | 0.250360768 | 6.518229383 | 0.7470759 | 0.3999392 | 0.999175987 |
| ENSMUSG00000007850 | Hnrnph1 | protein_coding | -0.273751219 | 6.886565233 | 0.7467679 | 0.4000342 | 0.999175987 |
| ENSMUSG00000066510 | Ankdd1a | protein_coding | -0.778305627 | 3.148513677 | 0.7467168 | 0.40005 | 0.999175987 |
| ENSMUSG00000026096 | Osgepl1 | protein_coding | -0.454885591 | 4.626322671 | 0.746653 | 0.4000697 | 0.999175987 |
| ENSMUSG00000036006 | Ripor2 | protein_coding | -0.413947972 | 5.540538924 | 0.7466454 | 0.400072 | 0.999175987 |
| ENSMUSG00000028843 | Sh3bgrl3 | protein_coding | 0.269978724 | 8.961784637 | 0.7465725 | 0.4000945 | 0.999175987 |
| ENSMUSG00000039952 | Dag1 | protein_coding | -0.41261658 | 4.91240035 | 0.7462046 | 0.4002081 | 0.999175987 |
| ENSMUSG00000050505 | Pcdh20 | protein_coding | 0.535975086 | 3.923139704 | 0.7460379 | 0.4002596 | 0.999175987 |
| ENSMUSG00000025511 | Tspan4 | protein_coding | 0.518532297 | 4.134808784 | 0.7450384 | 0.4005685 | 0.999175987 |
| ENSMUSG00000022517 | Mgrn1 | protein_coding | 0.596851997 | 6.009870723 | 0.7510918 | 0.4006475 | 0.999175987 |
| ENSMUSG00000042751 | Nmnat2 | protein_coding | -0.439112166 | 7.270128218 | 0.748704 | 0.4006831 | 0.999175987 |
| ENSMUSG00000018334 | Ksr1 | protein_coding | -0.655127489 | 3.4061664 | 0.7445852 | 0.4007087 | 0.999175987 |
| ENSMUSG00000023911 | Flywch2 | protein_coding | 0.325088048 | 5.777949659 | 0.7439931 | 0.4008919 | 0.999175987 |
| ENSMUSG00000060601 | Nr1h2 | protein_coding | 0.332934644 | 5.922656333 | 0.7436535 | 0.4009971 | 0.999175987 |
| ENSMUSG00000034216 | Vps18 | protein_coding | 0.576228514 | 5.153911227 | 0.7453141 | 0.4010321 | 0.999175987 |
| ENSMUSG00000024404 | Riok3 | protein_coding | -0.483695731 | 4.928221499 | 0.7424736 | 0.4013628 | 0.999175987 |
| ENSMUSG00000025041 | Nt5c2 | protein_coding | 0.371755109 | 5.858135852 | 0.7417807 | 0.4015778 | 0.999175987 |
| ENSMUSG00000020189 | Osbpl8 | protein_coding | -0.377113547 | 5.708584989 | 0.7415549 | 0.4016479 | 0.999175987 |
| ENSMUSG00000021147 | Wdr37 | protein_coding | -0.319449842 | 6.485592369 | 0.7414476 | 0.4016812 | 0.999175987 |
| ENSMUSG00000004393 | Ddx56 | protein_coding | 0.328188137 | 7.161439461 | 0.7402845 | 0.4020426 | 0.999175987 |
| ENSMUSG00000032086 | Bace1 | protein_coding | -0.357397082 | 5.419567206 | 0.7402387 | 0.4020569 | 0.999175987 |
| ENSMUSG00000042734 | Ttc9 | protein_coding | -0.403388813 | 5.301513792 | 0.7402196 | 0.4020628 | 0.999175987 |
| ENSMUSG00000026932 | Nacc2 | protein_coding | 0.374330458 | 6.10164232 | 0.7402007 | 0.4020687 | 0.999175987 |
| ENSMUSG00000027253 | Lrp4 | protein_coding | -0.656457606 | 4.078801821 | 0.7402304 | 0.4020798 | 0.999175987 |
| ENSMUSG00000026672 | Optn | protein_coding | -0.595668615 | 3.886804943 | 0.7400689 | 0.4021097 | 0.999175987 |
| ENSMUSG00000021495 | Fam193b | protein_coding | -0.788583059 | 3.323254284 | 0.740015 | 0.4021383 | 0.999175987 |
| ENSMUSG00000031683 | Lsm6 | protein_coding | -0.304147582 | 7.894789187 | 0.7398423 | 0.4021802 | 0.999175987 |
| ENSMUSG00000018401 | Mtmr4 | protein_coding | -0.401065293 | 5.241823159 | 0.7393067 | 0.4023469 | 0.999175987 |
| ENSMUSG00000026283 | Ing5 | protein_coding | -0.401739613 | 4.677544366 | 0.7381195 | 0.4027167 | 0.999175987 |
| ENSMUSG00000034243 | Golgb1 | protein_coding | -0.377528094 | 5.276950784 | 0.737815 | 0.4028117 | 0.999175987 |
| ENSMUSG00000025917 | Cops5 | protein_coding | -0.276877091 | 7.789224821 | 0.7377652 | 0.4028272 | 0.999175987 |
| ENSMUSG00000057530 | Ece1 | protein_coding | -0.365892971 | 5.311178642 | 0.7376893 | 0.4028509 | 0.999175987 |
| ENSMUSG00000024211 | Grm8 | protein_coding | -0.737024352 | 4.150109232 | 0.7398229 | 0.402976 | 0.999175987 |
| ENSMUSG00000042292 | Mkl1 | protein_coding | -0.418763645 | 4.91846856 | 0.7372306 | 0.402994 | 0.999175987 |
| ENSMUSG00000024064 | Galnt14 | protein_coding | 0.486948274 | 3.847379419 | 0.7371053 | 0.4030331 | 0.999175987 |
| ENSMUSG00000020648 | Dus4l | protein_coding | 0.87159515 | 2.939779055 | 0.737041 | 0.4030532 | 0.999175987 |
| ENSMUSG00000025959 | Klf7 | protein_coding | -0.343431048 | 5.180760094 | 0.736729 | 0.4031506 | 0.999175987 |
| ENSMUSG00000031879 | Fam96b | protein_coding | 0.310429741 | 6.989638524 | 0.7365763 | 0.4031983 | 0.999175987 |
| ENSMUSG00000026003 | Acadl | protein_coding | -0.455619539 | 5.020147576 | 0.7365715 | 0.4031998 | 0.999175987 |
| ENSMUSG00000097048 | 1600020E01Rik | antisense_RNA | 0.578704839 | 4.654607397 | 0.7365091 | 0.4032396 | 0.999175987 |
| ENSMUSG00000022376 | Adcy8 | protein_coding | 0.704954299 | 2.998032649 | 0.7362013 | 0.4033154 | 0.999175987 |
| ENSMUSG00000034714 | Ttyh2 | protein_coding | -0.390537214 | 7.227593125 | 0.737725 | 0.4033588 | 0.999175987 |
| ENSMUSG00000043668 | Tox3 | protein_coding | -0.620483415 | 3.709354237 | 0.7356414 | 0.4034904 | 0.999175987 |
| ENSMUSG00000025425 | St8sia5 | protein_coding | -0.90167995 | 2.602638481 | 0.7356021 | 0.4035027 | 0.999175987 |
| ENSMUSG00000022391 | Rangap1 | protein_coding | 0.279808419 | 7.905234947 | 0.7355253 | 0.4035267 | 0.999175987 |
| ENSMUSG00000024944 | Arl2 | protein_coding | 0.246620382 | 9.161043313 | 0.7354279 | 0.4035572 | 0.999175987 |
| ENSMUSG00000075478 | Slitrk1 | protein_coding | 0.456450817 | 4.949897252 | 0.7351747 | 0.4036364 | 0.999175987 |
| ENSMUSG00000038618 | Rassf7 | protein_coding | 0.723866544 | 3.373185711 | 0.7345586 | 0.4038292 | 0.999175987 |
| ENSMUSG00000042558 | Adprhl2 | protein_coding | 0.577516727 | 3.937292687 | 0.7344205 | 0.4038725 | 0.999175987 |
| ENSMUSG00000042857 | Gm9776 | bidirectional_promote | 0.819454389 | 2.515001981 | 0.7337573 | 0.4040802 | 0.999175987 |
| ENSMUSG00000108456 | 4732496C06Rik | lincRNA | -0.650812115 | 3.32660542 | 0.7333095 | 0.4042206 | 0.999175987 |
| ENSMUSG00000026185 | Igfbp5 | protein_coding | -0.432800446 | 5.404350889 | 0.733267 | 0.4042339 | 0.999175987 |
| ENSMUSG00000004285 | Atp6v1f | protein_coding | 0.263211482 | 9.236162641 | 0.7330089 | 0.4043149 | 0.999175987 |
| ENSMUSG00000022018 | Rgcc | protein_coding | -0.833685238 | 3.124808967 | 0.7326687 | 0.4044216 | 0.999175987 |
| ENSMUSG00000034781 | Gna11 | protein_coding | 0.31217663 | 8.147263377 | 0.7320094 | 0.4046285 | 0.999175987 |
| ENSMUSG00000044502 | Bod1 | protein_coding | -0.419871076 | 5.512528293 | 0.7319693 | 0.4046411 | 0.999175987 |
| ENSMUSG00000025921 | Rdh10 | protein_coding | -0.834860041 | 2.745578129 | 0.7316731 | 0.4047342 | 0.999175987 |
| ENSMUSG00000039887 | Alg14 | protein_coding | -0.315814988 | 5.90807092 | 0.7315811 | 0.4047631 | 0.999175987 |
| ENSMUSG00000032621 | Srek1 | protein_coding | -0.382249188 | 5.54780392 | 0.7315046 | 0.4047871 | 0.999175987 |
| ENSMUSG00000035847 | Ids | protein_coding | 0.295755948 | 6.718583316 | 0.7314539 | 0.404803 | 0.999175987 |
| ENSMUSG00000087380 | 2210408F21Rik | processed_transcript | -0.345978399 | 4.930598638 | 0.7312482 | 0.4048677 | 0.999175987 |
| ENSMUSG00000036067 | Slc2a6 | protein_coding | 0.551600139 | 4.741101066 | 0.7311627 | 0.4049148 | 0.999175987 |
| ENSMUSG00000040564 | Apoc1 | protein_coding | -0.673224743 | 3.600949766 | 0.7310776 | 0.4049213 | 0.999175987 |
| ENSMUSG00000003868 | Ruvbl2 | protein_coding | 0.367978884 | 6.01352007 | 0.7310118 | 0.404942 | 0.999175987 |
| ENSMUSG00000028557 | Rnf11 | protein_coding | 0.241743287 | 7.771785049 | 0.7307488 | 0.4050247 | 0.999175987 |
| ENSMUSG00000059897 | Zfp930 | protein_coding | 0.602345889 | 3.991080406 | 0.7299545 | 0.4052746 | 0.999175987 |
| ENSMUSG00000029416 | Slc15a4 | protein_coding | 0.457373228 | 4.24000187 | 0.7298693 | 0.4053015 | 0.999175987 |
| ENSMUSG00000021996 | Esd | protein_coding | 0.249811737 | 8.145446247 | 0.7297566 | 0.405337 | 0.999175987 |
| ENSMUSG00000024695 | Zfp91 | protein_coding | -0.315551669 | 5.647609796 | 0.7295797 | 0.4053926 | 0.999175987 |
| ENSMUSG00000008167 | Fbxw9 | protein_coding | 0.551368154 | 4.700894323 | 0.7296074 | 0.4054042 | 0.999175987 |
| ENSMUSG00000020953 | Coch | protein_coding | -0.556426841 | 3.620251044 | 0.7284077 | 0.405762 | 0.999175987 |
| ENSMUSG00000021276 | Cinp | protein_coding | 0.261042465 | 6.88534505 | 0.7282461 | 0.405813 | 0.999175987 |
| ENSMUSG00000026239 | Pde6d | protein_coding | -0.369463972 | 4.990558208 | 0.7279609 | 0.405903 | 0.999175987 |
| ENSMUSG00000031534 | Smim19 | protein_coding | -0.435156933 | 4.746452526 | 0.7278556 | 0.4059362 | 0.999175987 |
| ENSMUSG00000001052 | Sec24b | protein_coding | -0.3716802 | 4.881880454 | 0.7274958 | 0.4060498 | 0.999175987 |

| ensembl_gene_id | external_gene_name | gene_biotype | log2 (fold change) | log2 (counts per million) | F-value | p-value | False discovery rate |
|---|---|---|---|---|---|---|---|
| ENSMUSG00000047466 | 8030462N17Rik | protein_coding | 0.647737688 | 3.532953595 | 0.7268741 | 0.4062461 | 0.999175987 |
| ENSMUSG00000031516 | Dctn6 | protein_coding | 0.26705618 | 8.032111938 | 0.726837 | 0.4062579 | 0.999175987 |
| ENSMUSG00000075273 | Ttc30b | protein_coding | -0.424122599 | 4.086097015 | 0.7266243 | 0.4063251 | 0.999175987 |
| ENSMUSG00000002846 | Timmdc1 | protein_coding | 0.51212171 | 5.077553258 | 0.7266173 | 0.4063475 | 0.999175987 |
| ENSMUSG00000033184 | Tmed7 | protein_coding | -0.44742028 | 5.589289033 | 0.7263163 | 0.4064426 | 0.999175987 |
| ENSMUSG00000069255 | Dusp22 | protein_coding | -0.39935503 | 5.260943692 | 0.7259635 | 0.406534 | 0.999175987 |
| ENSMUSG00000038446 | Cdc40 | protein_coding | -0.490943307 | 5.089583073 | 0.7258544 | 0.4065887 | 0.999175987 |
| ENSMUSG00000021313 | Ryr2 | protein_coding | -0.661391258 | 4.29272841 | 0.7264796 | 0.4066135 | 0.999175987 |
| ENSMUSG00000066306 | Numa1 | protein_coding | 0.428502344 | 4.956374096 | 0.7255939 | 0.4066509 | 0.999175987 |
| ENSMUSG00000045095 | Magi1 | protein_coding | 0.312506091 | 6.208144879 | 0.7248559 | 0.4068846 | 0.999175987 |
| ENSMUSG00000026639 | Lamb3 | protein_coding | -0.82665704 | 3.199071581 | 0.7247195 | 0.4069278 | 0.999175987 |
| ENSMUSG00000066026 | Dhrs3 | protein_coding | -0.383869738 | 5.994403766 | 0.7246538 | 0.4069486 | 0.999175987 |
| ENSMUSG00000074218 | Cox7a1 | protein_coding | 0.48554862 | 3.850723237 | 0.7246056 | 0.4069639 | 0.999175987 |
| ENSMUSG00000021994 | Wnt5a | protein_coding | 0.621505198 | 3.22810331 | 0.7243889 | 0.4070325 | 0.999175987 |
| ENSMUSG00000027848 | Olfml3 | protein_coding | -0.486848956 | 4.462495399 | 0.7243885 | 0.4070327 | 0.999175987 |
| ENSMUSG00000021846 | Peli2 | protein_coding | -0.49042789 | 3.828429506 | 0.7242563 | 0.4070746 | 0.999175987 |
| ENSMUSG00000098108 | Gm27008 | antisense_RNA | 0.74232373 | 3.144980135 | 0.7239012 | 0.4071871 | 0.999175987 |
| ENSMUSG00000057455 | Rit2 | protein_coding | -0.287860253 | 9.683594199 | 0.7237534 | 0.407234 | 0.999175987 |
| ENSMUSG00000042410 | Agps | protein_coding | -0.445516324 | 4.159991566 | 0.7234797 | 0.4073208 | 0.999175987 |
| ENSMUSG00000049672 | Zbtb14 | protein_coding | 0.614205573 | 3.195792007 | 0.7233895 | 0.4073494 | 0.999175987 |
| ENSMUSG00000059495 | Arhgef12 | protein_coding | -0.348518594 | 5.470293133 | 0.723234 | 0.4073988 | 0.999175987 |
| ENSMUSG00000053286 | Trmt1l | protein_coding | -0.56778674 | 3.963026734 | 0.7231875 | 0.4074135 | 0.999175987 |
| ENSMUSG00000025178 | Pi4k2a | protein_coding | 0.356775134 | 6.30671777 | 0.7227538 | 0.4075512 | 0.999175987 |
| ENSMUSG00000048481 | Mypop | protein_coding | -0.465056807 | 3.951406015 | 0.7226845 | 0.4075732 | 0.999175987 |
| ENSMUSG00000079610 | Ankrd39 | protein_coding | 0.513040915 | 3.616515507 | 0.7226334 | 0.4075894 | 0.999175987 |
| ENSMUSG00000097545 | Mir124a-1hg | lincRNA | -0.626472757 | 3.619828558 | 0.7224767 | 0.4076392 | 0.999175987 |
| ENSMUSG00000036534 | Slc38a7 | protein_coding | 0.711628465 | 3.731879295 | 0.7220929 | 0.4077813 | 0.999175987 |
| ENSMUSG00000027797 | Dclk1 | protein_coding | -0.266905459 | 7.263284849 | 0.721317 | 0.4080078 | 0.999175987 |
| ENSMUSG00000037617 | Spag1 | protein_coding | -0.995202341 | 2.575763389 | 0.7206803 | 0.4082104 | 0.999175987 |
| ENSMUSG00000026197 | Zfand2b | protein_coding | -0.329424908 | 5.663000998 | 0.7200893 | 0.4083986 | 0.999175987 |
| ENSMUSG00000063531 | Sema3e | protein_coding | -0.749928598 | 2.783433438 | 0.7197819 | 0.4084966 | 0.999175987 |
| ENSMUSG00000005566 | Trim28 | protein_coding | 0.266464562 | 6.502968316 | 0.7193492 | 0.4086345 | 0.999175987 |
| ENSMUSG00000033055 | Ankrd54 | protein_coding | 0.549025286 | 3.488553445 | 0.7191974 | 0.4086829 | 0.999175987 |
| ENSMUSG00000066613 | Zfp932 | protein_coding | -0.497439147 | 3.590341987 | 0.7178639 | 0.4091084 | 0.999175987 |
| ENSMUSG00000036099 | Vezt | protein_coding | -0.439867335 | 5.092441276 | 0.7175826 | 0.4091983 | 0.999175987 |
| ENSMUSG00000097537 | 2610020C07Rik | lincRNA | -0.408130882 | 4.705334267 | 0.717473 | 0.4092333 | 0.999175987 |
| ENSMUSG00000027194 | Ttc17 | protein_coding | -0.464937891 | 3.704452578 | 0.7173572 | 0.4092703 | 0.999175987 |
| ENSMUSG00000043456 | Zfp536 | protein_coding | 0.651664921 | 3.772374222 | 0.7172844 | 0.4092936 | 0.999175987 |
| ENSMUSG00000071074 | Yipf3 | protein_coding | 0.410917095 | 5.260315035 | 0.7168216 | 0.4094416 | 0.999175987 |
| ENSMUSG00000034189 | Hsdl1 | protein_coding | -0.318341735 | 6.774531571 | 0.7163323 | 0.4095981 | 0.999175987 |
| ENSMUSG00000020889 | Nr1d1 | protein_coding | -0.40184378 | 6.574850309 | 0.717376 | 0.4096087 | 0.999175987 |
| ENSMUSG00000026788 | Zbtb43 | protein_coding | -0.421191965 | 4.339041362 | 0.7160889 | 0.409676 | 0.999175987 |
| ENSMUSG00000042350 | Arel1 | protein_coding | 0.523163142 | 4.131847776 | 0.7159815 | 0.4097104 | 0.999175987 |
| ENSMUSG00000071324 | Armc2 | protein_coding | -0.884009696 | 2.970125851 | 0.7156071 | 0.4098303 | 0.999175987 |
| ENSMUSG00000003038 | Hmgn2 | protein_coding | 0.335193639 | 5.561308851 | 0.7155934 | 0.4098346 | 0.999175987 |
| ENSMUSG00000025937 | Lactb2 | protein_coding | -0.581179832 | 3.577857389 | 0.7155839 | 0.4098377 | 0.999175987 |
| ENSMUSG00000037022 | Mmaa | protein_coding | -0.42079501 | 4.865514758 | 0.7150362 | 0.4100132 | 0.999175987 |
| ENSMUSG00000042688 | Mapk6 | protein_coding | -0.468820118 | 5.407855678 | 0.7146685 | 0.410151 | 0.999175987 |
| ENSMUSG00000073147 | 5031425E22Rik | lincRNA | -0.648153627 | 3.38855646 | 0.7142723 | 0.4102581 | 0.999175987 |
| ENSMUSG00000032030 | Cul5 | protein_coding | -0.351519738 | 5.90373286 | 0.713744 | 0.4104276 | 0.999175987 |
| ENSMUSG00000023079 | Gtf2ird1 | protein_coding | -0.497953622 | 4.057280364 | 0.7129254 | 0.4106905 | 0.999175987 |
| ENSMUSG00000020368 | Canx | protein_coding | -0.233077374 | 9.236388523 | 0.7124943 | 0.4108291 | 0.999175987 |
| ENSMUSG00000028826 | Tmem57 | protein_coding | -0.40336438 | 5.475240805 | 0.7121569 | 0.4109375 | 0.999175987 |
| ENSMUSG00000067787 | Blcap | protein_coding | 0.26320488 | 8.873666099 | 0.7120484 | 0.4109724 | 0.999175987 |
| ENSMUSG00000031963 | Bmper | protein_coding | -0.763954849 | 2.732481153 | 0.7118912 | 0.411023 | 0.999175987 |
| ENSMUSG00000024524 | Gnal | protein_coding | 0.292235416 | 7.129695087 | 0.7116809 | 0.4110907 | 0.999175987 |
| ENSMUSG00000021840 | Mapk1ip1l | protein_coding | 0.469293597 | 3.850109201 | 0.7114863 | 0.4111533 | 0.999175987 |
| ENSMUSG00000029414 | Kntc1 | protein_coding | -1.045644919 | 2.226839101 | 0.7114249 | 0.4111731 | 0.999175987 |
| ENSMUSG00000005533 | Igf1r | protein_coding | -0.481072615 | 4.398673351 | 0.7113781 | 0.4111881 | 0.999175987 |
| ENSMUSG00000031010 | Usp9x | protein_coding | -0.442624637 | 4.380907038 | 0.7103096 | 0.4115323 | 0.999175987 |
| ENSMUSG00000037103 | Dcaf15 | protein_coding | 0.744830618 | 3.497198755 | 0.709974 | 0.4116604 | 0.999175987 |
| ENSMUSG00000032586 | Traip | protein_coding | 0.41082395 | 4.521045935 | 0.7097428 | 0.4117151 | 0.999175987 |
| ENSMUSG00000029096 | Htra3 | protein_coding | 0.562637722 | 3.500425552 | 0.7096733 | 0.4117375 | 0.999175987 |
| ENSMUSG00000040363 | Bcor | protein_coding | 0.376777673 | 4.932756 | 0.7095959 | 0.4117624 | 0.999175987 |
| ENSMUSG00000027300 | Ubox5 | protein_coding | 0.432506992 | 4.192564187 | 0.7091747 | 0.4118984 | 0.999175987 |
| ENSMUSG00000042115 | Klhdc8a | protein_coding | 0.423306207 | 5.107572488 | 0.7089106 | 0.4119836 | 0.999175987 |
| ENSMUSG00000022962 | Gart | protein_coding | -0.506058512 | 4.633510336 | 0.7084722 | 0.4121252 | 0.999175987 |
| ENSMUSG00000024193 | Phf1 | protein_coding | 0.494490046 | 4.191752132 | 0.7082705 | 0.4121904 | 0.999175987 |
| ENSMUSG00000000056 | Narf | protein_coding | -0.344972888 | 5.785371926 | 0.7076932 | 0.412377 | 0.999175987 |
| ENSMUSG00000055761 | Nkain3 | protein_coding | 0.63835373 | 3.530744899 | 0.7072779 | 0.4125113 | 0.999175987 |
| ENSMUSG00000002546 | Golga2 | protein_coding | -0.399954411 | 5.61340806 | 0.7070826 | 0.4125745 | 0.999175987 |
| ENSMUSG00000024830 | Rps6kb2 | protein_coding | -0.989404423 | 3.145732886 | 0.7087025 | 0.4125979 | 0.999175987 |
| ENSMUSG00000052033 | Pfdn4 | protein_coding | 0.335315773 | 7.429777655 | 0.7058318 | 0.4129994 | 0.999175987 |
| ENSMUSG00000020801 | Med31 | protein_coding | 0.351116665 | 6.786437743 | 0.7058219 | 0.4130026 | 0.999175987 |
| ENSMUSG00000062054 | Iah1 | protein_coding | 0.347995432 | 5.522841567 | 0.7056555 | 0.4130367 | 0.999175987 |
| ENSMUSG00000021047 | Nova1 | protein_coding | -0.303088433 | 6.313334427 | 0.7053546 | 0.4131343 | 0.999175987 |
| ENSMUSG00000037784 | Dzip1l | protein_coding | -0.951996033 | 2.5623487 | 0.7053303 | 0.4131421 | 0.999175987 |
| ENSMUSG00000021185 | 9030617O03Rik | protein_coding | -0.694017102 | 4.068951454 | 0.7061995 | 0.4132376 | 0.999175987 |

| ensembl_gene_id | external_gene_name | gene_biotype | log2 (fold change) | log2 (counts per million) | F-value | p-value | False discovery rate |
|---|---|---|---|---|---|---|---|
| ENSMUSG00000041594 | Tmtc4 | protein_coding | 1.015617158 | 2.540256851 | 0.7044378 | 0.4134317 | 0.999175987 |
| ENSMUSG00000097867 | Lppos | lincRNA | -0.788584612 | 2.724646276 | 0.7043698 | 0.4134538 | 0.999175987 |
| ENSMUSG00000056629 | Fkbp2 | protein_coding | 0.285187052 | 9.108188008 | 0.7041707 | 0.4135185 | 0.999175987 |
| ENSMUSG00000046668 | Cxxc5 | protein_coding | -0.338494667 | 5.562815778 | 0.7036907 | 0.4136743 | 0.999175987 |
| ENSMUSG00000030342 | Cd9 | protein_coding | -0.34079959 | 6.841465186 | 0.7032465 | 0.4138386 | 0.999175987 |
| ENSMUSG00000048148 | Nwd1 | protein_coding | 0.598160704 | 3.57062 | 0.7029367 | 0.4139195 | 0.999175987 |
| ENSMUSG00000030657 | Xylt1 | protein_coding | -0.768665896 | 3.242131493 | 0.7028314 | 0.4139537 | 0.999175987 |
| ENSMUSG00000029502 | Golga3 | protein_coding | 0.39025569 | 5.084133763 | 0.7027213 | 0.4139895 | 0.999175987 |
| ENSMUSG00000022476 | Polr3h | protein_coding | 0.314617617 | 6.335432108 | 0.7027195 | 0.4139901 | 0.999175987 |
| ENSMUSG00000020910 | Adprm | protein_coding | 0.344035292 | 5.80374054 | 0.7023946 | 0.4140958 | 0.999175987 |
| ENSMUSG00000028945 | Rheb | protein_coding | -0.260188862 | 6.803813429 | 0.7020436 | 0.4142101 | 0.999175987 |
| ENSMUSG00000029603 | Dtx1 | protein_coding | -0.374035121 | 5.30201011 | 0.7014354 | 0.4144082 | 0.999175987 |
| ENSMUSG00000056050 | Mia3 | protein_coding | -0.341911533 | 5.465371818 | 0.7013673 | 0.4144304 | 0.999175987 |
| ENSMUSG00000006763 | Saal1 | protein_coding | 0.446679195 | 4.029950953 | 0.7013087 | 0.4144495 | 0.999175987 |
| ENSMUSG00000044098 | Rsbn1 | protein_coding | -0.747278736 | 2.860981529 | 0.7011734 | 0.4144936 | 0.999175987 |
| ENSMUSG00000028517 | Plpp3 | protein_coding | -0.425992618 | 6.071146533 | 0.7008934 | 0.4146568 | 0.999175987 |
| ENSMUSG00000061654 | Spry3 | protein_coding | 0.647617247 | 3.475000625 | 0.7001524 | 0.4148265 | 0.999175987 |
| ENSMUSG00000022338 | Eny2 | protein_coding | -0.262564941 | 7.138504758 | 0.700152 | 0.4148267 | 0.999175987 |
| ENSMUSG00000024976 | Shoc2 | protein_coding | -0.353479262 | 8.430827714 | 0.7001323 | 0.4148529 | 0.999175987 |
| ENSMUSG00000024654 | Asrgl1 | protein_coding | -0.247450529 | 7.648444656 | 0.6996909 | 0.4149772 | 0.999175987 |
| ENSMUSG00000027351 | Spred1 | protein_coding | -0.28339124 | 6.353192015 | 0.6996829 | 0.4149798 | 0.999175987 |
| ENSMUSG00000016619 | Nup50 | protein_coding | -0.427818662 | 4.860087363 | 0.6995228 | 0.4150321 | 0.999175987 |
| ENSMUSG00000030008 | Pradc1 | protein_coding | 0.70005425 | 3.403218735 | 0.6992252 | 0.4151293 | 0.999175987 |
| ENSMUSG00000044562 | Rasip1 | protein_coding | 0.85463305 | 3.036528528 | 0.6986726 | 0.4153099 | 0.999175987 |
| ENSMUSG00000068749 | Psma5 | protein_coding | -0.242874685 | 6.83374403 | 0.6985525 | 0.4153492 | 0.999175987 |
| ENSMUSG00000024740 | Ddb1 | protein_coding | -0.383245876 | 7.069296325 | 0.700079 | 0.4153549 | 0.999175987 |
| ENSMUSG00000078572 | Ndufaf8 | protein_coding | 0.341562021 | 6.474336998 | 0.6984359 | 0.4153873 | 0.999175987 |
| ENSMUSG00000108776 | Gm45169 | lincRNA | -0.622055247 | 3.493116215 | 0.6970426 | 0.4158434 | 0.999175987 |
| ENSMUSG00000066151 | Fkbp15 | protein_coding | -0.394579863 | 4.607863867 | 0.6968745 | 0.4158984 | 0.999175987 |
| ENSMUSG00000029994 | Anxa4 | protein_coding | -0.806806744 | 2.78056413 | 0.6967979 | 0.4159235 | 0.999175987 |
| ENSMUSG00000030204 | Ddx47 | protein_coding | 0.283426383 | 6.951213676 | 0.6967327 | 0.4159449 | 0.999175987 |
| ENSMUSG00000041798 | Gck | protein_coding | -0.488419616 | 5.389062002 | 0.6967472 | 0.4159599 | 0.999175987 |
| ENSMUSG00000000884 | Gnb1l | protein_coding | 0.515041322 | 5.353469977 | 0.6969822 | 0.4160998 | 0.999175987 |
| ENSMUSG00000021930 | Spryd7 | protein_coding | 0.245484711 | 6.699836475 | 0.695412 | 0.416378 | 0.999175987 |
| ENSMUSG00000038990 | Cables2 | protein_coding | -0.58644357 | 3.377384987 | 0.6953566 | 0.4163962 | 0.999175987 |
| ENSMUSG00000029547 | Ints1 | protein_coding | -0.61051091 | 4.190595333 | 0.694977 | 0.4165406 | 0.999175987 |
| ENSMUSG00000022881 | Rfc4 | protein_coding | -0.729064369 | 4.263347673 | 0.6978306 | 0.4165615 | 0.999175987 |
| ENSMUSG00000051331 | Cacna1c | protein_coding | -0.611581599 | 3.874655236 | 0.6945158 | 0.4166724 | 0.999175987 |
| ENSMUSG00000036298 | Slc2a13 | protein_coding | -0.438049432 | 5.510649288 | 0.6945713 | 0.4166738 | 0.999175987 |
| ENSMUSG00000028403 | Zdhhc21 | protein_coding | -0.45513837 | 4.489614356 | 0.6944512 | 0.4166936 | 0.999175987 |
| ENSMUSG00000030869 | Ndufab1 | protein_coding | 0.435287173 | 4.057246265 | 0.6941757 | 0.4167841 | 0.999175987 |
| ENSMUSG00000022774 | Ncbp2 | protein_coding | 0.231368103 | 7.268332187 | 0.6934516 | 0.4170223 | 0.999175987 |
| ENSMUSG00000032232 | Cgnl1 | protein_coding | -1.316480775 | 1.933576761 | 0.6933953 | 0.4170408 | 0.999175987 |
| ENSMUSG00000032350 | Gclc | protein_coding | -0.36326257 | 5.5127045 | 0.6933878 | 0.4170433 | 0.999175987 |
| ENSMUSG00000041164 | Zmiz2 | protein_coding | 0.293955362 | 6.669697262 | 0.692934 | 0.4171927 | 0.999175987 |
| ENSMUSG00000078566 | Bnip3 | protein_coding | -0.366657695 | 5.546691871 | 0.692887 | 0.4172081 | 0.999175987 |
| ENSMUSG00000036528 | Ppfibp2 | protein_coding | -0.661132412 | 3.651401463 | 0.6923198 | 0.4173949 | 0.999175987 |
| ENSMUSG00000039219 | Arid4b | protein_coding | -0.331585521 | 5.786670601 | 0.6919514 | 0.4175163 | 0.999175987 |
| ENSMUSG00000037852 | Cpe | protein_coding | -0.321557261 | 10.16661404 | 0.6919512 | 0.4175164 | 0.999175987 |
| ENSMUSG00000022721 | Trmt2a | protein_coding | 0.472149235 | 5.02992924 | 0.6915855 | 0.417637 | 0.999175987 |
| ENSMUSG00000029708 | Gcc1 | protein_coding | -0.485014289 | 3.788800149 | 0.6914862 | 0.4176698 | 0.999175987 |
| ENSMUSG00000033721 | Vav3 | protein_coding | -0.642399543 | 3.443028302 | 0.6912536 | 0.4177465 | 0.999175987 |
| ENSMUSG00000022905 | Kpna1 | protein_coding | -0.281211815 | 6.03260121 | 0.6911002 | 0.4177971 | 0.999175987 |
| ENSMUSG00000006333 | Rps9 | protein_coding | 0.271628142 | 9.87165058 | 0.6907836 | 0.4179016 | 0.999175987 |
| ENSMUSG00000045875 | Adra1a | protein_coding | -0.586919146 | 3.405582033 | 0.6907107 | 0.4179256 | 0.999175987 |
| ENSMUSG00000016346 | Kcnq2 | protein_coding | 0.293038527 | 5.984644413 | 0.6903287 | 0.4180518 | 0.999175987 |
| ENSMUSG00000020326 | Ccng1 | protein_coding | 0.41013725 | 4.812654868 | 0.69021 | 0.418091 | 0.999175987 |
| ENSMUSG00000041261 | Car8 | protein_coding | -0.512423299 | 4.349591531 | 0.6899232 | 0.4181858 | 0.999175987 |
| ENSMUSG00000032397 | Tipin | protein_coding | 0.33488213 | 6.106471395 | 0.6893971 | 0.4183597 | 0.999175987 |
| ENSMUSG00000057637 | Prdm2 | protein_coding | -0.386795669 | 5.321888145 | 0.6888981 | 0.4185248 | 0.999175987 |
| ENSMUSG00000046447 | Camk2n1 | protein_coding | -0.279246896 | 8.814693818 | 0.6888605 | 0.4185372 | 0.999175987 |
| ENSMUSG00000056215 | Lrguk | protein_coding | 1.038038099 | 2.904019546 | 0.6872094 | 0.4191037 | 0.999175987 |
| ENSMUSG00000020687 | Cdc27 | protein_coding | -0.340180956 | 5.098841015 | 0.6870919 | 0.4191231 | 0.999175987 |
| ENSMUSG00000108348 | Gm42372 | protein_coding | 0.295991935 | 6.623810946 | 0.6868407 | 0.4192064 | 0.999175987 |
| ENSMUSG00000013822 | Elof1 | protein_coding | 0.237360566 | 8.163650204 | 0.6856526 | 0.4196008 | 0.999175987 |
| ENSMUSG00000023755 | Rhebl1 | protein_coding | -0.645177155 | 4.079893955 | 0.6851734 | 0.4197796 | 0.999175987 |
| ENSMUSG00000016921 | Srsf6 | protein_coding | -0.444011513 | 5.095380755 | 0.6851061 | 0.4197824 | 0.999175987 |
| ENSMUSG00000024766 | Lipo3 | protein_coding | -0.623577582 | 3.2692085 | 0.6848242 | 0.4198761 | 0.999175987 |
| ENSMUSG00000090641 | Zfp712 | protein_coding | 0.902887914 | 2.588525085 | 0.6848239 | 0.4198762 | 0.999175987 |
| ENSMUSG00000066042 | Med18 | protein_coding | 0.402973739 | 4.473760807 | 0.6846381 | 0.419938 | 0.999175987 |
| ENSMUSG00000027079 | Clp1 | protein_coding | -0.536031465 | 4.119925071 | 0.6846049 | 0.4199491 | 0.999175987 |
| ENSMUSG00000020847 | Rph3al | protein_coding | 0.653162423 | 3.492750588 | 0.6845823 | 0.4199566 | 0.999175987 |
| ENSMUSG00000063087 | Gm10125 | processed_transcript | 0.857351163 | 2.386144775 | 0.6844683 | 0.4199945 | 0.999175987 |
| ENSMUSG00000022214 | Dcaf11 | protein_coding | 0.40029478 | 5.684362016 | 0.6840781 | 0.4201244 | 0.999175987 |
| ENSMUSG00000056204 | Pgpep1 | protein_coding | -0.38088285 | 5.433848232 | 0.6838212 | 0.4202099 | 0.999175987 |
| ENSMUSG00000036854 | Hspb6 | protein_coding | 0.413938079 | 5.283787309 | 0.68354 | 0.4203035 | 0.999175987 |
| ENSMUSG00000020778 | Ten1 | protein_coding | 0.242550026 | 7.058388038 | 0.683191 | 0.4204198 | 0.999175987 |
| ENSMUSG00000071064 | Zfp827 | protein_coding | -0.434623582 | 4.348245981 | 0.6830911 | 0.4204531 | 0.999175987 |

| ensembl_gene_id | external_gene_name | gene_biotype | log2 (fold change) | log2 (counts per million) | F-value | p-value | False discovery rate |
|---|---|---|---|---|---|---|---|
| ENSMUSG00000031246 | Sh3bgrl | protein_coding | 0.32596236 | 7.149897123 | 0.6831022 | 0.4204689 | 0.999175987 |
| ENSMUSG00000025492 | Ifitm3 | protein_coding | -0.490659652 | 4.155131299 | 0.6828401 | 0.4205368 | 0.999175987 |
| ENSMUSG00000028410 | Dnaja1 | protein_coding | -0.233758297 | 7.796491924 | 0.682755 | 0.4205651 | 0.999175987 |
| ENSMUSG00000038544 | Inip | protein_coding | -0.346560537 | 5.42137329 | 0.6826644 | 0.4205954 | 0.999175987 |
| ENSMUSG00000030265 | Kras | protein_coding | -0.374403078 | 5.529993642 | 0.6822795 | 0.4207237 | 0.999175987 |
| ENSMUSG00000036502 | Tmem255a | protein_coding | 0.31751726 | 6.151645705 | 0.6819975 | 0.4208178 | 0.999175987 |
| ENSMUSG00000028567 | Txndc12 | protein_coding | 0.289521088 | 6.104744527 | 0.6819603 | 0.4208302 | 0.999175987 |
| ENSMUSG00000022607 | Ptk2 | protein_coding | -0.332526436 | 5.103287679 | 0.6817388 | 0.4209042 | 0.999175987 |
| ENSMUSG00000024862 | Klc2 | protein_coding | 0.475827589 | 5.55964312 | 0.6812758 | 0.4211336 | 0.999175987 |
| ENSMUSG00000028649 | Macf1 | protein_coding | -0.490030254 | 5.171982175 | 0.6811054 | 0.4211352 | 0.999175987 |
| ENSMUSG00000037270 | 4932438A13Rik | protein_coding | -0.498021406 | 4.134086467 | 0.6803178 | 0.4213789 | 0.999175987 |
| ENSMUSG00000032060 | Cryab | protein_coding | -0.278485733 | 9.198418857 | 0.680052 | 0.4214679 | 0.999175987 |
| ENSMUSG00000009090 | Ap1b1 | protein_coding | 0.326612754 | 7.029784393 | 0.6798339 | 0.4215603 | 0.999175987 |
| ENSMUSG00000000787 | Ddx3x | protein_coding | -0.273363591 | 6.582524005 | 0.6796756 | 0.4215938 | 0.999175987 |
| ENSMUSG00000031483 | Erlin2 | protein_coding | 0.369671816 | 5.141166293 | 0.6794369 | 0.4216737 | 0.999175987 |
| ENSMUSG00000020669 | Sh3yl1 | protein_coding | 0.565305553 | 4.384485981 | 0.6785486 | 0.4219907 | 0.999175987 |
| ENSMUSG00000004642 | Slbp | protein_coding | 0.33402965 | 6.246675971 | 0.6784799 | 0.4219943 | 0.999175987 |
| ENSMUSG00000035642 | Aamdc | protein_coding | 0.277578311 | 7.16647457 | 0.677856 | 0.4222035 | 0.999175987 |
| ENSMUSG00000007777 | 0610009B22Rik | protein_coding | 0.297145155 | 7.420469813 | 0.6774313 | 0.422346 | 0.999175987 |
| ENSMUSG00000046480 | Scn4b | protein_coding | -0.481755129 | 5.135063588 | 0.6774592 | 0.4223561 | 0.999175987 |
| ENSMUSG00000042354 | Gnl3 | protein_coding | -0.366004219 | 5.427918584 | 0.6769304 | 0.4225142 | 0.999175987 |
| ENSMUSG00000040018 | Cox15 | protein_coding | 0.603909176 | 3.750012357 | 0.6768777 | 0.4225319 | 0.999175987 |
| ENSMUSG00000060988 | Galnt13 | protein_coding | -0.385385197 | 5.405734543 | 0.6768168 | 0.4225523 | 0.999175987 |
| ENSMUSG00000011960 | Ccnt1 | protein_coding | 0.361819242 | 5.296576263 | 0.6768011 | 0.4225576 | 0.999175987 |
| ENSMUSG00000026434 | Nucks1 | protein_coding | -0.269542161 | 7.003684231 | 0.676732 | 0.4225808 | 0.999175987 |
| ENSMUSG00000004508 | Gab2 | protein_coding | 0.46803804 | 3.938016817 | 0.6766613 | 0.4226046 | 0.999175987 |
| ENSMUSG00000030613 | Ccdc90b | protein_coding | -0.355333769 | 5.437788353 | 0.6765095 | 0.4226556 | 0.999175987 |
| ENSMUSG00000059474 | Mbtd1 | protein_coding | -0.521881787 | 3.747940993 | 0.6763678 | 0.4227032 | 0.999175987 |
| ENSMUSG00000055421 | Pcdh9 | protein_coding | -0.274278416 | 6.98252191 | 0.6761909 | 0.4227626 | 0.999175987 |
| ENSMUSG00000038827 | Fam206a | protein_coding | 0.384375206 | 4.517711697 | 0.6759457 | 0.4228451 | 0.999175987 |
| ENSMUSG00000020027 | Socs2 | protein_coding | 0.58396175 | 3.119462797 | 0.6759005 | 0.4228603 | 0.999175987 |
| ENSMUSG00000069020 | Urm1 | protein_coding | 0.246362357 | 6.34336436 | 0.675736 | 0.4229156 | 0.999175987 |
| ENSMUSG00000030284 | Creld1 | protein_coding | 0.316961665 | 6.220781924 | 0.6741811 | 0.4234391 | 0.999175987 |
| ENSMUSG00000002395 | Use1 | protein_coding | 0.262152057 | 8.642709291 | 0.6739534 | 0.4235158 | 0.999175987 |
| ENSMUSG00000030500 | Slc17a6 | protein_coding | -1.210074607 | 6.462160859 | 0.7123298 | 0.4235384 | 0.999175987 |
| ENSMUSG00000015476 | Prrt1 | protein_coding | 0.33581916 | 5.824490711 | 0.6729609 | 0.4238506 | 0.999175987 |
| ENSMUSG00000037426 | Depdc5 | protein_coding | -0.443834236 | 4.586188718 | 0.6724148 | 0.424035 | 0.999175987 |
| ENSMUSG00000046688 | Tifa | protein_coding | 0.925719045 | 2.772496425 | 0.6719476 | 0.4241928 | 0.999175987 |
| ENSMUSG00000027879 | Sec22b | protein_coding | -0.253260685 | 7.907287196 | 0.6717356 | 0.4242645 | 0.999175987 |
| ENSMUSG00000024998 | Plce1 | protein_coding | -0.650746889 | 2.957736556 | 0.6699832 | 0.4248575 | 0.999175987 |
| ENSMUSG00000049489 | Fam58b | protein_coding | 0.391575812 | 4.903408993 | 0.669783 | 0.4249253 | 0.999175987 |
| ENSMUSG00000027313 | Chac1 | protein_coding | 0.451043004 | 4.619900276 | 0.6696984 | 0.424954 | 0.999175987 |
| ENSMUSG00000046997 | Spsb4 | protein_coding | -0.910972413 | 2.451580769 | 0.6692281 | 0.4251134 | 0.999175987 |
| ENSMUSG00000040097 | Flywch1 | protein_coding | 0.763888106 | 7.261020424 | 0.6901013 | 0.4251205 | 0.999175987 |
| ENSMUSG00000034908 | Sidt2 | protein_coding | 0.444503162 | 4.407146496 | 0.6690979 | 0.4251575 | 0.999175987 |
| ENSMUSG00000040481 | Bptf | protein_coding | -0.31747149 | 6.101804355 | 0.6686714 | 0.4253022 | 0.999175987 |
| ENSMUSG00000022420 | Dnal4 | protein_coding | 0.309853305 | 6.838531098 | 0.6684545 | 0.4253758 | 0.999175987 |
| ENSMUSG00000057719 | Sh3rf2 | protein_coding | -0.663220069 | 3.397565118 | 0.6679237 | 0.4255561 | 0.999175987 |
| ENSMUSG00000022323 | Rida | protein_coding | -0.332802794 | 7.338296693 | 0.6677284 | 0.4256417 | 0.999175987 |
| ENSMUSG00000021432 | Slc35b3 | protein_coding | 0.686608588 | 3.149178668 | 0.6667863 | 0.4259427 | 0.999175987 |
| ENSMUSG00000061838 | Suclg2 | protein_coding | -0.686316027 | 3.038395466 | 0.6663633 | 0.4260866 | 0.999175987 |
| ENSMUSG00000024854 | Pold4 | protein_coding | -0.333940922 | 5.691467525 | 0.6661789 | 0.4261493 | 0.999175987 |
| ENSMUSG00000027774 | Gfm1 | protein_coding | -0.403453189 | 4.742264326 | 0.6660282 | 0.4262007 | 0.999175987 |
| ENSMUSG00000058569 | Tmed9 | protein_coding | -0.400185119 | 8.118184108 | 0.6682104 | 0.4262529 | 0.999175987 |
| ENSMUSG00000058325 | Dock1 | protein_coding | -0.486130178 | 4.34361608 | 0.6655802 | 0.4263532 | 0.999175987 |
| ENSMUSG00000062866 | Phactr2 | protein_coding | -0.352839921 | 5.45711007 | 0.6644001 | 0.4267555 | 0.999175987 |
| ENSMUSG00000002983 | Relb | protein_coding | 0.875682882 | 2.645367851 | 0.6643516 | 0.4267721 | 0.999175987 |
| ENSMUSG00000022180 | Slc7a8 | protein_coding | -0.373827011 | 4.673576006 | 0.6641945 | 0.4268257 | 0.999175987 |
| ENSMUSG00000052253 | Zfp622 | protein_coding | 0.428990031 | 5.262507283 | 0.6641264 | 0.4268489 | 0.999175987 |
| ENSMUSG00000025351 | Cd63 | protein_coding | -0.389196343 | 6.778323389 | 0.6656728 | 0.4268534 | 0.999175987 |
| ENSMUSG00000036894 | Rap2b | protein_coding | 0.359457921 | 6.864916024 | 0.6641497 | 0.4269108 | 0.999175987 |
| ENSMUSG00000046330 | Rpl37a | protein_coding | 0.421566465 | 7.908050191 | 0.6669915 | 0.4270419 | 0.999175987 |
| ENSMUSG00000038773 | Kdm3b | protein_coding | 0.425071144 | 3.911557916 | 0.663349 | 0.4271143 | 0.999175987 |
| ENSMUSG00000029561 | Oasl2 | protein_coding | 1.610994179 | 3.05310636 | 0.6769319 | 0.427167 | 0.999175987 |
| ENSMUSG00000028420 | Tmem38b | protein_coding | 0.711832231 | 3.341440637 | 0.6628821 | 0.4272739 | 0.999175987 |
| ENSMUSG00000055639 | Dach1 | protein_coding | -0.480455202 | 3.902452405 | 0.6624845 | 0.4274098 | 0.999175987 |
| ENSMUSG00000027997 | Casp6 | protein_coding | -0.753101188 | 2.845442278 | 0.6621983 | 0.4275077 | 0.999175987 |
| ENSMUSG00000032667 | Pon2 | protein_coding | -0.387802671 | 5.017661309 | 0.6618828 | 0.4276157 | 0.999175987 |
| ENSMUSG00000017291 | Taok1 | protein_coding | -0.454181718 | 3.751936543 | 0.6617052 | 0.4276765 | 0.999175987 |
| ENSMUSG00000029436 | Mmp17 | protein_coding | -0.38029761 | 5.137459336 | 0.6612401 | 0.4278357 | 0.999175987 |
| ENSMUSG00000021832 | Psmc6 | protein_coding | 0.247734523 | 7.89661194 | 0.660881 | 0.4279588 | 0.999175987 |
| ENSMUSG00000005716 | Pvalb | protein_coding | -0.457773518 | 7.802006038 | 0.6655988 | 0.4279858 | 0.999175987 |
| ENSMUSG00000004771 | Rab11a | protein_coding | -0.231627105 | 6.915055833 | 0.6606887 | 0.4280247 | 0.999175987 |
| ENSMUSG00000008140 | Emc10 | protein_coding | 0.349444128 | 5.708759003 | 0.6606118 | 0.428051 | 0.999175987 |
| ENSMUSG00000015755 | Tab2 | protein_coding | -0.438989806 | 5.182467895 | 0.6606062 | 0.4280529 | 0.999175987 |
| ENSMUSG00000042807 | Hecw2 | protein_coding | -0.478118086 | 4.125532092 | 0.6603601 | 0.4281373 | 0.999175987 |
| ENSMUSG00000037531 | Mrpl47 | protein_coding | 0.342657861 | 5.937371966 | 0.6601353 | 0.4282144 | 0.999175987 |
| ENSMUSG00000022013 | Dnajc15 | protein_coding | 0.263970616 | 7.804727818 | 0.6599378 | 0.4282822 | 0.999175987 |

| ensembl_gene_id | external_gene_name | gene_biotype | log2 (fold change) | log2 (counts per million) | F-value | p-value | False discovery rate |
|---|---|---|---|---|---|---|---|
| ENSMUSG00000028629 | Exo5 | protein_coding | 0.733766094 | 4.006934492 | 0.6619049 | 0.428351 | 0.999175987 |
| ENSMUSG00000055923 | Aasdh | protein_coding | -0.709316083 | 3.069737384 | 0.6596917 | 0.4283666 | 0.999175987 |
| ENSMUSG00000045180 | Shroom2 | protein_coding | 0.321685495 | 6.313908682 | 0.6596679 | 0.4283748 | 0.999175987 |
| ENSMUSG00000047832 | Cdca4 | protein_coding | 0.64706726 | 3.904284938 | 0.6597107 | 0.4283792 | 0.999175987 |
| ENSMUSG00000040055 | Gjb6 | protein_coding | -0.444951207 | 5.585771799 | 0.659395 | 0.4284876 | 0.999175987 |
| ENSMUSG00000037221 | Mospd3 | protein_coding | 0.736715393 | 4.650157983 | 0.6640735 | 0.4285038 | 0.999175987 |
| ENSMUSG00000037541 | Shank2 | protein_coding | -0.455086148 | 4.28895981 | 0.6589693 | 0.4286147 | 0.999175987 |
| ENSMUSG00000093483 | AA465934 | processed_transcript | -0.595346084 | 3.409305018 | 0.6586774 | 0.4287149 | 0.999175987 |
| ENSMUSG00000097789 | Gm2115 | protein_coding | -0.577601554 | 3.277335339 | 0.6586081 | 0.4287388 | 0.999175987 |
| ENSMUSG00000057469 | E2f6 | protein_coding | -0.471772092 | 4.490085816 | 0.6585503 | 0.4287586 | 0.999175987 |
| ENSMUSG00000001418 | Glmp | protein_coding | 0.285670437 | 6.461224664 | 0.6581695 | 0.4288895 | 0.999175987 |
| ENSMUSG00000006673 | Qrich1 | protein_coding | -0.287242315 | 5.913338784 | 0.6580015 | 0.4289473 | 0.999175987 |
| ENSMUSG00000030127 | Cops7a | protein_coding | 0.239136022 | 8.366976052 | 0.6572918 | 0.4291915 | 0.999175987 |
| ENSMUSG00000032101 | Ddx25 | protein_coding | -0.269958983 | 7.373943566 | 0.657044 | 0.4292768 | 0.999175987 |
| ENSMUSG00000071054 | Safb | protein_coding | 0.413524685 | 4.510427583 | 0.6566934 | 0.4293976 | 0.999175987 |
| ENSMUSG00000034308 | Sdr42e1 | protein_coding | -0.744356092 | 3.166429921 | 0.6565328 | 0.4294529 | 0.999175987 |
| ENSMUSG00000107585 | 3300002P13Rik | lincRNA | 0.785560262 | 2.696054502 | 0.6564873 | 0.4294686 | 0.999175987 |
| ENSMUSG00000022521 | Crebbp | protein_coding | -0.308377131 | 7.026471536 | 0.6561466 | 0.429586 | 0.999175987 |
| ENSMUSG00000021520 | Uqcrb | protein_coding | 0.334229931 | 8.483374154 | 0.655556 | 0.4298088 | 0.999175987 |
| ENSMUSG00000019947 | Arid5b | protein_coding | -0.37795761 | 5.545585943 | 0.6550361 | 0.4299692 | 0.999175987 |
| ENSMUSG00000019874 | Fabp7 | protein_coding | -0.517960047 | 6.935996241 | 0.6614523 | 0.4300774 | 0.999175987 |
| ENSMUSG00000039130 | Zc3hc1 | protein_coding | -0.308033892 | 6.442612538 | 0.6546119 | 0.4301156 | 0.999175987 |
| ENSMUSG00000020832 | Eral1 | protein_coding | 0.402756611 | 5.595521897 | 0.6545032 | 0.4301532 | 0.999175987 |
| ENSMUSG00000024037 | Wdr4 | protein_coding | -0.36655687 | 4.752591358 | 0.6540489 | 0.4303102 | 0.999175987 |
| ENSMUSG00000031137 | Fgf13 | protein_coding | 0.346511199 | 6.550759424 | 0.6539142 | 0.4303758 | 0.999175987 |
| ENSMUSG00000026142 | Rhbdd1 | protein_coding | -1.111772185 | 2.721468663 | 0.6540095 | 0.430393 | 0.999175987 |
| ENSMUSG00000035770 | Dync1li2 | protein_coding | -0.284683259 | 6.71919187 | 0.653628 | 0.4304557 | 0.999175987 |
| ENSMUSG00000038712 | Mindy1 | protein_coding | 0.486361249 | 4.8086224 | 0.6535455 | 0.4304842 | 0.999175987 |
| ENSMUSG00000079477 | Rab7 | protein_coding | 0.351199512 | 6.059395973 | 0.6532897 | 0.4305728 | 0.999175987 |
| ENSMUSG00000060534 | Dcc | protein_coding | -0.682657472 | 3.846926997 | 0.6534683 | 0.4305801 | 0.999175987 |
| ENSMUSG00000021036 | Sptlc2 | protein_coding | -0.559814177 | 3.764679548 | 0.6532627 | 0.4305821 | 0.999175987 |
| ENSMUSG00000034931 | Dhx8 | protein_coding | -0.434418124 | 4.399326705 | 0.6528623 | 0.4307207 | 0.999175987 |
| ENSMUSG00000006732 | Mettl1 | protein_coding | 0.29949171 | 6.02174802 | 0.65265 | 0.4307942 | 0.999175987 |
| ENSMUSG00000033102 | Cdc14b | protein_coding | 0.54343246 | 3.484622043 | 0.6524128 | 0.4308763 | 0.999175987 |
| ENSMUSG00000070644 | Etnk2 | protein_coding | 0.835525121 | 2.743692198 | 0.6523644 | 0.4308931 | 0.999175987 |
| ENSMUSG00000018634 | Crhr1 | protein_coding | 0.810111507 | 2.681756768 | 0.6519277 | 0.4310444 | 0.999175987 |
| ENSMUSG00000022836 | Mylk | protein_coding | 0.868971532 | 2.678548687 | 0.6518156 | 0.4310833 | 0.999175987 |
| ENSMUSG00000055239 | Kcmf1 | protein_coding | -0.356326541 | 5.772593006 | 0.6514053 | 0.4312255 | 0.999175987 |
| ENSMUSG00000028127 | Abcd3 | protein_coding | 0.365787953 | 5.816527553 | 0.6508919 | 0.4314037 | 0.999175987 |
| ENSMUSG00000028958 | Tmub1 | protein_coding | 0.264837485 | 6.454854854 | 0.6508235 | 0.4314274 | 0.999175987 |
| ENSMUSG00000024067 | Dpy30 | protein_coding | 0.265788525 | 8.230608169 | 0.6502727 | 0.4316186 | 0.999175987 |
| ENSMUSG00000028343 | Erp44 | protein_coding | -0.31661265 | 5.414849164 | 0.6502273 | 0.4316344 | 0.999175987 |
| ENSMUSG00000027660 | Skil | protein_coding | -0.268325575 | 6.513864919 | 0.6494882 | 0.4318913 | 0.999175987 |
| ENSMUSG00000034893 | Cog3 | protein_coding | -0.988457884 | 3.031055226 | 0.6505231 | 0.4319164 | 0.999175987 |
| ENSMUSG00000028039 | Efna3 | protein_coding | 0.47979476 | 4.154770139 | 0.6492471 | 0.4319751 | 0.999175987 |
| ENSMUSG00000003500 | Impdh1 | protein_coding | 0.567448449 | 4.04516045 | 0.6492416 | 0.431977 | 0.999175987 |
| ENSMUSG00000045639 | Zfp629 | protein_coding | 0.437117885 | 4.085104144 | 0.6491592 | 0.4320057 | 0.999175987 |
| ENSMUSG00000030493 | Faap24 | protein_coding | 0.477205826 | 4.212610145 | 0.6474573 | 0.4325983 | 0.999175987 |
| ENSMUSG00000014077 | Chp1 | protein_coding | 0.283829682 | 7.876562314 | 0.6473531 | 0.4326346 | 0.999175987 |
| ENSMUSG00000025019 | Lcor | protein_coding | 0.41866814 | 4.180898007 | 0.6472697 | 0.4326637 | 0.999175987 |
| ENSMUSG00000041570 | Camsap2 | protein_coding | -0.575279536 | 3.322878241 | 0.6470912 | 0.432726 | 0.999175987 |
| ENSMUSG00000091476 | Gm16432 | protein_coding | 0.804987484 | 2.639064719 | 0.6465695 | 0.432908 | 0.999175987 |
| ENSMUSG00000018347 | Zkscan6 | protein_coding | -0.930614444 | 3.488809905 | 0.6498505 | 0.4329378 | 0.999175987 |
| ENSMUSG00000031732 | Phlpp2 | protein_coding | 0.462658646 | 4.01577232 | 0.6461855 | 0.433042 | 0.999175987 |
| ENSMUSG00000024007 | Ppil1 | protein_coding | 0.263512084 | 6.774868429 | 0.645842 | 0.433162 | 0.999175987 |
| ENSMUSG00000093930 | Hmgcs1 | protein_coding | -0.301599595 | 7.080878806 | 0.6457976 | 0.4331775 | 0.999175987 |
| ENSMUSG00000009292 | Trpm2 | protein_coding | 0.627945614 | 3.85111753 | 0.6455817 | 0.4332718 | 0.999175987 |
| ENSMUSG00000053453 | Thoc7 | protein_coding | 0.321950211 | 7.306993074 | 0.6453237 | 0.433362 | 0.999175987 |
| ENSMUSG00000049657 | Zbtb5 | protein_coding | -0.70609995 | 3.629371052 | 0.6447841 | 0.4335507 | 0.999175987 |
| ENSMUSG00000001089 | Luzp1 | protein_coding | -0.3723879 | 5.210286883 | 0.6446912 | 0.4335643 | 0.999175987 |
| ENSMUSG00000061032 | Rrp1 | protein_coding | 0.286485612 | 8.565164267 | 0.6445946 | 0.4335981 | 0.999175987 |
| ENSMUSG00000004233 | Wars2 | protein_coding | -0.684196422 | 3.088499615 | 0.643488 | 0.4339855 | 0.999175987 |
| ENSMUSG00000018589 | Glra2 | protein_coding | 0.413623736 | 5.258017311 | 0.6430418 | 0.4341419 | 0.999175987 |
| ENSMUSG00000037458 | Azin1 | protein_coding | -0.258673193 | 6.855944283 | 0.6428326 | 0.4342153 | 0.999175987 |
| ENSMUSG00000042178 | Armc5 | protein_coding | -0.74488237 | 3.018035505 | 0.6426802 | 0.4342687 | 0.999175987 |
| ENSMUSG00000035351 | Nup37 | protein_coding | 0.471520106 | 4.096007473 | 0.641878 | 0.4345502 | 0.999175987 |
| ENSMUSG00000024800 | Rpp30 | protein_coding | 0.272131119 | 6.361405398 | 0.6418301 | 0.4345671 | 0.999175987 |
| ENSMUSG00000022749 | Tbc1d23 | protein_coding | -0.520722299 | 3.55848293 | 0.6414041 | 0.4347167 | 0.999175987 |
| ENSMUSG00000020284 | 1810043G02Rik | protein_coding | 0.291473378 | 6.869623216 | 0.6410957 | 0.434825 | 0.999175987 |
| ENSMUSG00000029440 | Psmd9 | protein_coding | 0.27966953 | 6.265659844 | 0.6402445 | 0.4351244 | 0.999175987 |
| ENSMUSG00000101585 | 1600010M07Rik | antisense_RNA | -0.778948335 | 2.830697685 | 0.6401003 | 0.4351751 | 0.999175987 |
| ENSMUSG00000041530 | Ago1 | protein_coding | -0.429966278 | 4.381906915 | 0.6389279 | 0.435588 | 0.999175987 |
| ENSMUSG00000032047 | Acat1 | protein_coding | -0.242310546 | 8.079841106 | 0.6387027 | 0.4356674 | 0.999175987 |
| ENSMUSG00000039217 | Il18 | protein_coding | -0.448189199 | 5.980826172 | 0.6406231 | 0.4356762 | 0.999175987 |
| ENSMUSG00000020321 | Mdh1 | protein_coding | -0.256926142 | 11.90593286 | 0.6381471 | 0.4358633 | 0.999175987 |
| ENSMUSG00000057880 | Abat | protein_coding | -0.262746154 | 8.308179823 | 0.6380417 | 0.4359005 | 0.999175987 |
| ENSMUSG00000097413 | A830052D11Rik | lincRNA | -0.82941047 | 2.599959074 | 0.6377175 | 0.4360149 | 0.999175987 |
| ENSMUSG00000052459 | Atp6v1a | protein_coding | -0.283764749 | 8.941268298 | 0.6374717 | 0.4361017 | 0.999175987 |

| ensembl_gene_id | external_gene_name | gene_biotype | log2 (fold change) | log2 (counts per million) | F-value | p-value | False discovery rate |
|---|---|---|---|---|---|---|---|
| ENSMUSG00000034111 | Tmed8 | protein_coding | -0.411494311 | 4.609132762 | 0.6373781 | 0.4361348 | 0.999175987 |
| ENSMUSG00000061665 | Cd2ap | protein_coding | 0.88356622 | 2.505229017 | 0.6369771 | 0.4362764 | 0.999175987 |
| ENSMUSG00000026342 | Slc35f5 | protein_coding | 0.794808783 | 3.268655608 | 0.6368821 | 0.4363287 | 0.999175987 |
| ENSMUSG00000054942 | Miga1 | protein_coding | -0.324332833 | 5.453316848 | 0.6366424 | 0.4363947 | 0.999175987 |
| ENSMUSG00000041815 | Poldip3 | protein_coding | -0.245050154 | 8.061233683 | 0.6363366 | 0.4365028 | 0.999175987 |
| ENSMUSG00000058835 | Abi1 | protein_coding | -0.251492783 | 6.933388157 | 0.6362977 | 0.4365166 | 0.999175987 |
| ENSMUSG00000038178 | Slc43a2 | protein_coding | 0.433543235 | 4.133100289 | 0.636284 | 0.4365214 | 0.999175987 |
| ENSMUSG00000024346 | Pfdn1 | protein_coding | 0.218020089 | 9.040204816 | 0.6359036 | 0.436656 | 0.999175987 |
| ENSMUSG00000039501 | Znfx1 | protein_coding | -0.502820448 | 3.843822906 | 0.6354164 | 0.4368284 | 0.999175987 |
| ENSMUSG00000022130 | Tgds | protein_coding | -0.894495014 | 2.404967611 | 0.635371 | 0.4368445 | 0.999175987 |
| ENSMUSG00000037936 | Scarb1 | protein_coding | -0.454703185 | 4.02383914 | 0.6353041 | 0.4368682 | 0.999175987 |
| ENSMUSG00000021540 | Smad5 | protein_coding | -0.365332142 | 4.475325274 | 0.6352456 | 0.4368889 | 0.999175987 |
| ENSMUSG00000048004 | Tmem196 | protein_coding | -0.496724554 | 4.160727152 | 0.6350273 | 0.4369662 | 0.999175987 |
| ENSMUSG00000030243 | Recql | protein_coding | 0.554234146 | 3.821508345 | 0.634644 | 0.437102 | 0.999175987 |
| ENSMUSG00000022856 | Tmem41a | protein_coding | 0.392118766 | 4.42514257 | 0.6346372 | 0.4371044 | 0.999175987 |
| ENSMUSG00000016554 | Eif3d | protein_coding | 0.246514322 | 6.399336327 | 0.6342751 | 0.4372328 | 0.999175987 |
| ENSMUSG00000026227 | 2810459M11Rik | protein_coding | -1.048030605 | 2.526631939 | 0.6339539 | 0.4373467 | 0.999175987 |
| ENSMUSG00000070737 | Tmem35b | protein_coding | -0.633905284 | 2.970141968 | 0.6339179 | 0.4373595 | 0.999175987 |
| ENSMUSG00000005125 | Ndrg1 | protein_coding | -0.346505005 | 5.389788109 | 0.6337042 | 0.4374353 | 0.999175987 |
| ENSMUSG00000063019 | Manbal | protein_coding | 0.243112715 | 7.243091372 | 0.6336299 | 0.4374617 | 0.999175987 |
| ENSMUSG00000022956 | Atp5o | protein_coding | 0.226846725 | 8.53452819 | 0.6333806 | 0.4375502 | 0.999175987 |
| ENSMUSG00000021326 | Trim27 | protein_coding | -0.5327325 | 3.769012894 | 0.6332499 | 0.4375966 | 0.999175987 |
| ENSMUSG00000021791 | Dydc2 | protein_coding | -0.694377125 | 2.879453552 | 0.6327658 | 0.4377685 | 0.999175987 |
| ENSMUSG00000018899 | Irf1 | protein_coding | -0.822278906 | 2.824912173 | 0.6325963 | 0.4378287 | 0.999175987 |
| ENSMUSG00000027805 | Pfn2 | protein_coding | 0.273625751 | 10.00691878 | 0.6325362 | 0.4378501 | 0.999175987 |
| ENSMUSG00000019952 | Poc1b | protein_coding | -0.907814578 | 2.66438283 | 0.6324022 | 0.4378977 | 0.999175987 |
| ENSMUSG00000056602 | Fry | protein_coding | -0.325416918 | 5.901134414 | 0.6318908 | 0.4380795 | 0.999175987 |
| ENSMUSG00000038576 | Susd4 | protein_coding | 0.273723388 | 7.183129061 | 0.631351 | 0.4382716 | 0.999175987 |
| ENSMUSG00000027291 | Vps39 | protein_coding | 0.449196666 | 5.778991377 | 0.6319824 | 0.4383999 | 0.999175987 |
| ENSMUSG00000022781 | Pak2 | protein_coding | -0.371144535 | 5.053092836 | 0.630883 | 0.4384382 | 0.999175987 |
| ENSMUSG00000003199 | Mpnd | protein_coding | 0.334780028 | 5.730802558 | 0.6308749 | 0.4384411 | 0.999175987 |
| ENSMUSG00000022412 | Mief1 | protein_coding | -0.44300068 | 5.00375469 | 0.6303205 | 0.4386386 | 0.999175987 |
| ENSMUSG00000097121 | D130020L05Rik | antisense_RNA | 0.489907757 | 3.731361309 | 0.6302914 | 0.438649 | 0.999175987 |
| ENSMUSG00000015094 | Npdc1 | protein_coding | 0.342088508 | 6.540215449 | 0.6301479 | 0.4387188 | 0.999175987 |
| ENSMUSG00000006058 | Snf8 | protein_coding | 0.387597884 | 5.101828562 | 0.629881 | 0.4387953 | 0.999175987 |
| ENSMUSG00000051029 | Serpinb1b | protein_coding | 0.399100272 | 4.661446212 | 0.6297523 | 0.4388412 | 0.999175987 |
| ENSMUSG00000026155 | Smap1 | protein_coding | -0.536304843 | 3.407709961 | 0.6297139 | 0.4388549 | 0.999175987 |
| ENSMUSG00000071660 | Ttc9c | protein_coding | -0.28751106 | 5.814119013 | 0.6295324 | 0.4389197 | 0.999175987 |
| ENSMUSG00000025232 | Hexa | protein_coding | -0.338450183 | 5.996006021 | 0.6295121 | 0.4389269 | 0.999175987 |
| ENSMUSG00000061232 | H2-K1 | protein_coding | -0.676715944 | 3.521797605 | 0.6294724 | 0.4389597 | 0.999175987 |
| ENSMUSG00000002055 | Spag5 | protein_coding | 1.122033571 | 2.194848976 | 0.6293148 | 0.4389973 | 0.999175987 |
| ENSMUSG00000014504 | Srp19 | protein_coding | 0.283017831 | 8.272780773 | 0.6291104 | 0.4390703 | 0.999175987 |
| ENSMUSG00000002103 | Acp2 | protein_coding | 0.596176296 | 4.573478707 | 0.6303149 | 0.4391164 | 0.999175987 |
| ENSMUSG00000051435 | Fhad1 | protein_coding | -0.316124605 | 5.190793183 | 0.6286387 | 0.4392387 | 0.999175987 |
| ENSMUSG00000025855 | Prkar1b | protein_coding | 0.367428132 | 10.12034065 | 0.62889 | 0.4392488 | 0.999175987 |
| ENSMUSG00000031078 | Cttn | protein_coding | 0.356180762 | 6.768797952 | 0.6281502 | 0.4394809 | 0.999175987 |
| ENSMUSG00000020231 | Dip2a | protein_coding | -0.439602762 | 4.17885028 | 0.6279607 | 0.439481 | 0.999175987 |
| ENSMUSG00000029701 | Rbm28 | protein_coding | -0.375310623 | 5.289942024 | 0.6278678 | 0.4395142 | 0.999175987 |
| ENSMUSG00000021193 | Pitrm1 | protein_coding | 0.477713071 | 4.532985356 | 0.627786 | 0.4395435 | 0.999175987 |
| ENSMUSG00000036098 | Myrf | protein_coding | -0.42424235 | 4.412695229 | 0.6276098 | 0.4396064 | 0.999175987 |
| ENSMUSG00000007050 | Lsm2 | protein_coding | 0.593585253 | 3.413094897 | 0.6275081 | 0.4396428 | 0.999175987 |
| ENSMUSG00000042766 | Trim46 | protein_coding | 0.295530634 | 5.439437472 | 0.6274584 | 0.4396606 | 0.999175987 |
| ENSMUSG00000060314 | Zfp941 | protein_coding | -0.288619561 | 6.292525636 | 0.6273387 | 0.4397034 | 0.999175987 |
| ENSMUSG00000033991 | Ttc37 | protein_coding | -0.61510851 | 3.353452265 | 0.6271716 | 0.4397632 | 0.999175987 |
| ENSMUSG00000023031 | Cela1 | protein_coding | 0.632403002 | 3.943138773 | 0.6271811 | 0.4397784 | 0.999175987 |
| ENSMUSG00000097207 | 6030443J06Rik | lincRNA | -0.599533716 | 3.104366737 | 0.6266087 | 0.4399648 | 0.999175987 |
| ENSMUSG00000035824 | Tk2 | protein_coding | 0.385842175 | 5.365178531 | 0.6266005 | 0.4399677 | 0.999175987 |
| ENSMUSG00000073771 | Btbd19 | protein_coding | -0.631877231 | 3.129784981 | 0.6263587 | 0.4400543 | 0.999175987 |
| ENSMUSG00000015843 | Rxrg | protein_coding | 0.255369169 | 6.702866231 | 0.6260509 | 0.4401646 | 0.999175987 |
| ENSMUSG00000040713 | Creg1 | protein_coding | -0.483839147 | 4.165173706 | 0.6252539 | 0.4404504 | 0.999175987 |
| ENSMUSG00000046861 | Hectd3 | protein_coding | -0.479421997 | 4.242507124 | 0.6250696 | 0.4405165 | 0.999175987 |
| ENSMUSG00000030307 | Slc6a11 | protein_coding | -0.368982627 | 6.060838409 | 0.6249706 | 0.4405706 | 0.999175987 |
| ENSMUSG00000007836 | Hnrnpa0 | protein_coding | -0.278021085 | 7.269052776 | 0.624708 | 0.4406463 | 0.999175987 |
| ENSMUSG00000024824 | Rad9a | protein_coding | 0.444803279 | 4.462150438 | 0.6246245 | 0.4406763 | 0.999175987 |
| ENSMUSG00000024962 | Vegfb | protein_coding | 0.362142168 | 5.333725805 | 0.6246062 | 0.4406829 | 0.999175987 |
| ENSMUSG00000035376 | Hacd2 | protein_coding | -0.50225424 | 3.720250957 | 0.6245494 | 0.4407033 | 0.999175987 |
| ENSMUSG00000027087 | Itgav | protein_coding | -0.629819776 | 5.549576828 | 0.6294872 | 0.4407435 | 0.999175987 |
| ENSMUSG00000039041 | Adrm1 | protein_coding | 0.431533174 | 4.483912201 | 0.6242339 | 0.4408166 | 0.999175987 |
| ENSMUSG00000035032 | Nek11 | protein_coding | -0.86111415 | 2.447111599 | 0.6239176 | 0.4409303 | 0.999175987 |
| ENSMUSG00000048304 | Slitrk3 | protein_coding | -0.468972158 | 4.373265654 | 0.6236168 | 0.4410384 | 0.999175987 |
| ENSMUSG00000039968 | Rsbn1l | protein_coding | -0.552851025 | 3.902598808 | 0.6234623 | 0.441094 | 0.999175987 |
| ENSMUSG00000022193 | Psmb5 | protein_coding | 0.235029519 | 6.843764382 | 0.6233742 | 0.4411256 | 0.999175987 |
| ENSMUSG00000037434 | Slc30a1 | protein_coding | 0.578653711 | 3.463613346 | 0.62314 | 0.4412099 | 0.999175987 |
| ENSMUSG00000019232 | Etnppl | protein_coding | -0.315317057 | 5.211151314 | 0.6230406 | 0.4412457 | 0.999175987 |
| ENSMUSG00000026674 | Ddr2 | protein_coding | -0.66810381 | 2.860090175 | 0.6228952 | 0.441298 | 0.999175987 |
| ENSMUSG00000026080 | Chst10 | protein_coding | 0.309115858 | 5.767554448 | 0.6225448 | 0.4414241 | 0.999175987 |
| ENSMUSG00000045761 | Togaram2 | protein_coding | -1.000639772 | 2.528370165 | 0.6224548 | 0.4414565 | 0.999175987 |
| ENSMUSG00000020366 | Mapk9 | protein_coding | 0.247020056 | 6.710022446 | 0.6224133 | 0.4414714 | 0.999175987 |

| ensembl_gene_id | external_gene_name | gene_biotype | log2 (fold change) | log2 (counts per million) | F-value | p-value | False discovery rate |
|---|---|---|---|---|---|---|---|
| ENSMUSG00000029403 | Cdkl2 | protein_coding | 0.346054399 | 5.228739217 | 0.6219875 | 0.4416248 | 0.999175987 |
| ENSMUSG00000035342 | Lzts2 | protein_coding | 0.778697552 | 3.030487529 | 0.6215641 | 0.4417774 | 0.999175987 |
| ENSMUSG00000047221 | Fam185a | protein_coding | -0.673110945 | 3.551582404 | 0.6215823 | 0.4417894 | 0.999175987 |
| ENSMUSG00000021681 | Aggf1 | protein_coding | -0.377887805 | 4.869183155 | 0.6211401 | 0.4419303 | 0.999175987 |
| ENSMUSG00000027076 | Timm10 | protein_coding | 0.281849531 | 7.283647256 | 0.6210747 | 0.4419539 | 0.999175987 |
| ENSMUSG00000038594 | Cep85l | protein_coding | -0.830256337 | 3.084781103 | 0.6206664 | 0.4421197 | 0.999175987 |
| ENSMUSG00000031157 | Pqbp1 | protein_coding | 0.253140397 | 7.908240987 | 0.6201514 | 0.4422872 | 0.999175987 |
| ENSMUSG00000015806 | Qdpr | protein_coding | -0.248360505 | 6.779004221 | 0.6199264 | 0.4423685 | 0.999175987 |
| ENSMUSG00000026333 | Gin1 | protein_coding | 0.490110132 | 4.444129451 | 0.6199178 | 0.4423716 | 0.999175987 |
| ENSMUSG00000050627 | Gpd1l | protein_coding | -0.386897618 | 5.678833813 | 0.6191916 | 0.4426341 | 0.999175987 |
| ENSMUSG00000032312 | Csk | protein_coding | 0.458527968 | 4.611753996 | 0.6191166 | 0.4426612 | 0.999175987 |
| ENSMUSG00000036606 | Plxnb2 | protein_coding | 0.58847503 | 3.938736052 | 0.6188788 | 0.4427657 | 0.999175987 |
| ENSMUSG00000030329 | Pianp | protein_coding | 0.30637694 | 6.795804784 | 0.618777 | 0.4427841 | 0.999175987 |
| ENSMUSG00000020211 | Sf3a2 | protein_coding | 0.476028297 | 3.820183956 | 0.6185771 | 0.4428564 | 0.999175987 |
| ENSMUSG00000097587 | 4930578M01Rik | lincRNA | -0.61678848 | 3.936053073 | 0.618608 | 0.4428637 | 0.999175987 |
| ENSMUSG00000004056 | Akt2 | protein_coding | 0.353866626 | 4.845874673 | 0.6183749 | 0.4429296 | 0.999175987 |
| ENSMUSG00000035126 | Wdr78 | protein_coding | -0.877788957 | 2.817560419 | 0.6178373 | 0.4431243 | 0.999175987 |
| ENSMUSG00000019772 | Vip | protein_coding | 0.604354776 | 3.329568435 | 0.6177988 | 0.4431382 | 0.999175987 |
| ENSMUSG00000003873 | Bax | protein_coding | 0.259745375 | 6.257187138 | 0.6177059 | 0.4431719 | 0.999175987 |
| ENSMUSG00000029469 | Ift81 | protein_coding | -0.352612366 | 5.347854405 | 0.6176454 | 0.4431938 | 0.999175987 |
| ENSMUSG00000060572 | Mfap2 | protein_coding | 0.474913547 | 4.044965662 | 0.6173821 | 0.4432893 | 0.999175987 |
| ENSMUSG00000000326 | Comt | protein_coding | 0.237983032 | 7.220056283 | 0.6171822 | 0.4433618 | 0.999175987 |
| ENSMUSG00000022324 | Matn2 | protein_coding | -0.637635287 | 3.743421757 | 0.6171856 | 0.443379 | 0.999175987 |
| ENSMUSG00000020792 | Exoc7 | protein_coding | -0.304807841 | 7.062563813 | 0.6169159 | 0.4434584 | 0.999175987 |
| ENSMUSG00000028247 | Coq3 | protein_coding | -0.310870397 | 6.189198552 | 0.6167852 | 0.4435057 | 0.999175987 |
| ENSMUSG00000023905 | Tnfrsf12a | protein_coding | 0.503135332 | 4.519644428 | 0.616659 | 0.4435515 | 0.999175987 |
| ENSMUSG00000071424 | Grid2 | protein_coding | 0.40030901 | 4.632350797 | 0.616568 | 0.4435846 | 0.999175987 |
| ENSMUSG00000022641 | Bbx | protein_coding | -0.374249384 | 5.438121776 | 0.6164652 | 0.4436219 | 0.999175987 |
| ENSMUSG00000067071 | Hes6 | protein_coding | -0.547789513 | 3.233856932 | 0.6157621 | 0.4438772 | 0.999175987 |
| ENSMUSG00000028145 | Them4 | protein_coding | 0.284864087 | 6.586980391 | 0.6151235 | 0.4441093 | 0.999175987 |
| ENSMUSG00000054455 | Vapb | protein_coding | -0.340211756 | 5.438720093 | 0.6150923 | 0.4441206 | 0.999175987 |
| ENSMUSG00000024579 | Pcyox1l | protein_coding | -0.454819722 | 3.862839148 | 0.6149964 | 0.4441555 | 0.999175987 |
| ENSMUSG00000032415 | Ube2cbp | protein_coding | -0.512542325 | 3.39040437 | 0.614923 | 0.4441822 | 0.999175987 |
| ENSMUSG00000036435 | Exoc1 | protein_coding | -0.376220002 | 5.289982686 | 0.6145397 | 0.4443216 | 0.999175987 |
| ENSMUSG00000049555 | Tmie | protein_coding | 0.461506345 | 4.130788794 | 0.6144936 | 0.4443384 | 0.999175987 |
| ENSMUSG00000038425 | Poli | protein_coding | -0.568941338 | 3.520332659 | 0.6141875 | 0.4444498 | 0.999175987 |
| ENSMUSG00000032340 | Neo1 | protein_coding | -0.426229852 | 4.574868656 | 0.6135147 | 0.4446949 | 0.999175987 |
| ENSMUSG00000035173 | Ccdc186 | protein_coding | -0.372981365 | 5.177513361 | 0.6128492 | 0.4449374 | 0.999175987 |
| ENSMUSG00000045064 | Zc2hc1c | protein_coding | -0.706416218 | 3.334567697 | 0.6124601 | 0.4450794 | 0.999175987 |
| ENSMUSG00000062519 | Zfp398 | protein_coding | -0.548448819 | 3.625922666 | 0.6120149 | 0.4452419 | 0.999175987 |
| ENSMUSG00000021040 | Slirp | protein_coding | 0.40968518 | 4.494074905 | 0.6117418 | 0.4453416 | 0.999175987 |
| ENSMUSG00000027384 | Ndufaf5 | protein_coding | 0.243662831 | 7.220808295 | 0.6113387 | 0.4454889 | 0.999175987 |
| ENSMUSG00000061981 | Flot2 | protein_coding | 0.282665727 | 6.29793369 | 0.6111557 | 0.4455558 | 0.999175987 |
| ENSMUSG00000030096 | Slc6a6 | protein_coding | 0.317939401 | 5.537140466 | 0.6111155 | 0.4455705 | 0.999175987 |
| ENSMUSG00000054426 | A930005H10Rik | processed_transcript | 0.362656737 | 4.744549992 | 0.6108088 | 0.4456826 | 0.999175987 |
| ENSMUSG00000028882 | Ppp1r8 | protein_coding | -0.313412629 | 5.707958089 | 0.6103791 | 0.4458397 | 0.999175987 |
| ENSMUSG00000005656 | Snx6 | protein_coding | -0.254201379 | 7.146636293 | 0.6103164 | 0.4458627 | 0.999175987 |
| ENSMUSG00000030207 | Fam234b | protein_coding | 0.336952618 | 6.781779912 | 0.610282 | 0.4458936 | 0.999175987 |
| ENSMUSG00000045932 | Ifit2 | protein_coding | 0.583041781 | 3.092937271 | 0.6098417 | 0.4460365 | 0.999175987 |
| ENSMUSG00000020604 | Arsg | protein_coding | -0.527665664 | 4.808082423 | 0.6096118 | 0.4461872 | 0.999175987 |
| ENSMUSG00000039461 | Tcta | protein_coding | 0.293731097 | 6.791598511 | 0.6092891 | 0.4462389 | 0.999175987 |
| ENSMUSG00000056679 | Gpr173 | protein_coding | -0.739541679 | 2.789059851 | 0.6082769 | 0.44661 | 0.999175987 |
| ENSMUSG00000062937 | Mtap | protein_coding | -0.375376024 | 5.135717786 | 0.608276 | 0.4466103 | 0.999175987 |
| ENSMUSG00000032648 | Pygm | protein_coding | -0.468304567 | 4.634923413 | 0.6080856 | 0.4466802 | 0.999175987 |
| ENSMUSG00000048240 | Gng7 | protein_coding | -0.302710062 | 8.62343048 | 0.6075343 | 0.4468826 | 0.999175987 |
| ENSMUSG00000000869 | Il4 | protein_coding | -0.430773655 | 4.629032743 | 0.6075281 | 0.4468849 | 0.999175987 |
| ENSMUSG00000039148 | Sart1 | protein_coding | 0.520944457 | 4.561525235 | 0.6066561 | 0.4472236 | 0.999175987 |
| ENSMUSG00000031642 | Sh3rf1 | protein_coding | 0.327294856 | 4.948869836 | 0.6065278 | 0.4472526 | 0.999175987 |
| ENSMUSG00000028706 | Nsun4 | protein_coding | 0.480875266 | 4.674578843 | 0.6064498 | 0.4472812 | 0.999175987 |
| ENSMUSG00000078908 | Mon1b | protein_coding | 0.932966186 | 2.834909832 | 0.6063049 | 0.4473528 | 0.999175987 |
| ENSMUSG00000020591 | Ntsr2 | protein_coding | -0.555887466 | 5.596747364 | 0.60935 | 0.4476599 | 0.999175987 |
| ENSMUSG00000001150 | Mcm3ap | protein_coding | -0.454178142 | 3.73704569 | 0.6048136 | 0.4478837 | 0.999175987 |
| ENSMUSG00000040048 | Ndufb10 | protein_coding | 0.247840462 | 10.18834066 | 0.6047649 | 0.4479017 | 0.999175987 |
| ENSMUSG00000039904 | Gpr37 | protein_coding | -0.333013893 | 6.394693251 | 0.6047515 | 0.4479172 | 0.999175987 |
| ENSMUSG00000042111 | Ccdc115 | protein_coding | 0.32413371 | 5.416773609 | 0.6047092 | 0.4479222 | 0.999175987 |
| ENSMUSG00000040151 | Hs2st1 | protein_coding | 0.431831493 | 4.851491335 | 0.6042572 | 0.4480889 | 0.999175987 |
| ENSMUSG00000034793 | G6pc3 | protein_coding | 0.338849629 | 6.389852792 | 0.6037696 | 0.448287 | 0.999175987 |
| ENSMUSG00000031634 | Ufsp2 | protein_coding | 0.220510657 | 7.256195566 | 0.6031731 | 0.4484891 | 0.999175987 |
| ENSMUSG00000061479 | Snrpa | protein_coding | 0.34397136 | 6.147516646 | 0.6031398 | 0.4485014 | 0.999175987 |
| ENSMUSG00000032402 | Smad3 | protein_coding | -0.452820963 | 4.130400917 | 0.6028834 | 0.4485961 | 0.999175987 |
| ENSMUSG00000029168 | Dpysl5 | protein_coding | -0.435992363 | 4.853573106 | 0.6028826 | 0.4485964 | 0.999175987 |
| ENSMUSG00000073155 | 1810058I24Rik | lincRNA | 0.228087701 | 6.7691686 | 0.6028269 | 0.448617 | 0.999175987 |
| ENSMUSG00000049225 | Pdp1 | protein_coding | -0.278461023 | 7.125775586 | 0.6027181 | 0.4486573 | 0.999175987 |
| ENSMUSG00000027956 | Tmem144 | protein_coding | -0.532197122 | 3.501868317 | 0.60256 | 0.4487157 | 0.999175987 |
| ENSMUSG00000013089 | Etv5 | protein_coding | -0.427844311 | 4.202614693 | 0.6014997 | 0.449108 | 0.999175987 |
| ENSMUSG00000069495 | Epc2 | protein_coding | -0.427146783 | 4.693634474 | 0.6014792 | 0.4491156 | 0.999175987 |
| ENSMUSG00000073664 | Nbeal1 | protein_coding | -0.547528869 | 3.613402452 | 0.601443 | 0.449129 | 0.999175987 |
| ENSMUSG00000029153 | Ociad2 | protein_coding | -0.207597184 | 7.775326845 | 0.6013545 | 0.4491618 | 0.999175987 |

| ensembl_gene_id | external_gene_name | gene_biotype | log2 (fold change) | log2 (counts per million) | F-value | p-value | False discovery rate |
|---|---|---|---|---|---|---|---|
| ENSMUSG00000030990 | Pgap2 | protein_coding | -0.361789874 | 4.838311353 | 0.6011679 | 0.4492309 | 0.999175987 |
| ENSMUSG00000025212 | Sfxn3 | protein_coding | 0.314092564 | 7.358371596 | 0.6008919 | 0.4493513 | 0.999175987 |
| ENSMUSG00000024011 | Pi16 | protein_coding | -0.888431883 | 3.090579411 | 0.6010344 | 0.4493673 | 0.999175987 |
| ENSMUSG00000090069 | E430024P14Rik | antisense_RNA | -0.839808511 | 3.360192038 | 0.6024497 | 0.4494182 | 0.999175987 |
| ENSMUSG00000025027 | Xpnpep1 | protein_coding | -0.317708984 | 6.097822581 | 0.6002033 | 0.4495884 | 0.999175987 |
| ENSMUSG00000044676 | Zfp612 | protein_coding | 0.605776589 | 3.313462669 | 0.5999303 | 0.4496897 | 0.999175987 |
| ENSMUSG00000045573 | Penk | protein_coding | 0.394498962 | 10.40961684 | 0.6017487 | 0.44972 | 0.999175987 |
| ENSMUSG00000003948 | Mmd | protein_coding | 0.440441123 | 6.024027836 | 0.6015839 | 0.4497809 | 0.999175987 |
| ENSMUSG00000033717 | Adra2a | protein_coding | -0.617173278 | 3.305469818 | 0.5995704 | 0.4498233 | 0.999175987 |
| ENSMUSG00000031453 | Rasa3 | protein_coding | 0.361754069 | 4.952652906 | 0.5995083 | 0.4498464 | 0.999175987 |
| ENSMUSG00000029128 | Rab28 | protein_coding | 0.296428999 | 6.833190454 | 0.5989234 | 0.4500636 | 0.999175987 |
| ENSMUSG00000027998 | Plrg1 | protein_coding | -0.315861974 | 6.257843665 | 0.5988053 | 0.4501075 | 0.999175987 |
| ENSMUSG00000057068 | Fam47e | protein_coding | -0.750418794 | 3.11634576 | 0.5985426 | 0.4502051 | 0.999175987 |
| ENSMUSG00000029127 | Zbtb49 | protein_coding | 1.374854107 | 2.185977151 | 0.5986369 | 0.4502359 | 0.999175987 |
| ENSMUSG00000004561 | Mettl17 | protein_coding | 0.407205188 | 4.410173875 | 0.5983189 | 0.4502883 | 0.999175987 |
| ENSMUSG00000038034 | Igsf8 | protein_coding | 0.276140479 | 5.971416614 | 0.5981527 | 0.4503501 | 0.999175987 |
| ENSMUSG00000030541 | Idh2 | protein_coding | 0.36091013 | 4.56323447 | 0.5977382 | 0.4505043 | 0.999175987 |
| ENSMUSG00000037730 | Mynn | protein_coding | -0.417240823 | 4.006506073 | 0.5975112 | 0.4505888 | 0.999175987 |
| ENSMUSG00000042010 | Acacb | protein_coding | 0.598836606 | 4.116900543 | 0.5974327 | 0.4506361 | 0.999175987 |
| ENSMUSG00000040111 | Gramd1b | protein_coding | 0.316954855 | 5.352504123 | 0.5973061 | 0.4506651 | 0.999175987 |
| ENSMUSG00000028514 | Usp24 | protein_coding | -0.688330466 | 3.687224854 | 0.5971917 | 0.4507735 | 0.999175987 |
| ENSMUSG00000032228 | Tcf12 | protein_coding | -0.396814301 | 4.905642339 | 0.5964879 | 0.45097 | 0.999175987 |
| ENSMUSG00000066235 | Pomgnt2 | protein_coding | 0.373768124 | 5.394538667 | 0.5964768 | 0.4509741 | 0.999175987 |
| ENSMUSG00000026809 | Spaca9 | protein_coding | -0.797580191 | 2.80355454 | 0.5962304 | 0.451066 | 0.999175987 |
| ENSMUSG00000042628 | Zfyve1 | protein_coding | 0.381496499 | 4.594394296 | 0.5958519 | 0.4512072 | 0.999175987 |
| ENSMUSG00000030216 | Wbp11 | protein_coding | 0.256366588 | 6.936768788 | 0.5946992 | 0.4516376 | 0.999175987 |
| ENSMUSG00000032115 | Hyou1 | protein_coding | -0.548733868 | 4.211311351 | 0.5938287 | 0.451981 | 0.999175987 |
| ENSMUSG00000029819 | Npy | protein_coding | -0.515707578 | 6.897415766 | 0.6014395 | 0.4519856 | 0.999175987 |
| ENSMUSG00000053192 | Mllt11 | protein_coding | 0.338742468 | 6.47669126 | 0.593667 | 0.4520415 | 0.999175987 |
| ENSMUSG00000033763 | Mtss1l | protein_coding | -0.30060315 | 5.913357951 | 0.593486 | 0.4520913 | 0.999175987 |
| ENSMUSG00000042406 | Atf4 | protein_coding | -0.244585791 | 7.274473083 | 0.5932895 | 0.4521648 | 0.999175987 |
| ENSMUSG00000066829 | Zfp810 | protein_coding | -0.681667351 | 3.844884453 | 0.5941915 | 0.452193 | 0.999175987 |
| ENSMUSG00000036323 | Srp72 | protein_coding | -0.22570754 | 7.571869453 | 0.5921236 | 0.4526016 | 0.999175987 |
| ENSMUSG00000022462 | Slc38a2 | protein_coding | -0.327553315 | 5.304103256 | 0.5915874 | 0.4528028 | 0.999175987 |
| ENSMUSG00000021186 | Fbln5 | protein_coding | 0.977395348 | 2.269903079 | 0.5912332 | 0.4529357 | 0.999175987 |
| ENSMUSG00000035686 | Thrsp | protein_coding | -0.386337177 | 6.131810158 | 0.5911687 | 0.4530254 | 0.999175987 |
| ENSMUSG00000022696 | Sidt1 | protein_coding | -0.630072411 | 3.069529831 | 0.5906077 | 0.4531707 | 0.999175987 |
| ENSMUSG00000019826 | Zbtb24 | protein_coding | 0.438850536 | 3.807397015 | 0.5905153 | 0.4532054 | 0.999175987 |
| ENSMUSG00000046240 | Hepacam | protein_coding | -0.510226656 | 4.693450718 | 0.5895917 | 0.4535707 | 0.999175987 |
| ENSMUSG00000029233 | Srd5a3 | protein_coding | -0.613487938 | 3.477859204 | 0.5894839 | 0.4535933 | 0.999175987 |
| ENSMUSG00000031827 | Cotl1 | protein_coding | 0.260699391 | 5.986020748 | 0.5891752 | 0.4537095 | 0.999175987 |
| ENSMUSG00000032564 | Cpne4 | protein_coding | -0.356015641 | 5.771340579 | 0.5889587 | 0.453791 | 0.999175987 |
| ENSMUSG00000048078 | Tenm4 | protein_coding | 0.476815446 | 4.19453274 | 0.5886165 | 0.4539199 | 0.999175987 |
| ENSMUSG00000006403 | Adamts4 | protein_coding | -0.631894835 | 3.873262184 | 0.5885117 | 0.4539773 | 0.999175987 |
| ENSMUSG00000014592 | Camta1 | protein_coding | -0.240439986 | 7.884971833 | 0.5882699 | 0.4540505 | 0.999175987 |
| ENSMUSG00000041617 | Ccdc74a | protein_coding | -0.552906008 | 3.585730457 | 0.5882128 | 0.454072 | 0.999175987 |
| ENSMUSG00000031432 | Prps1 | protein_coding | -0.217779489 | 7.358218619 | 0.5880979 | 0.4541153 | 0.999175987 |
| ENSMUSG00000027220 | Syt13 | protein_coding | 0.253798548 | 6.202347776 | 0.5878764 | 0.4541989 | 0.999175987 |
| ENSMUSG00000035933 | Cog5 | protein_coding | 0.607073928 | 3.09656944 | 0.5877548 | 0.4542447 | 0.999175987 |
| ENSMUSG00000024462 | Gabbr1 | protein_coding | -0.276080333 | 7.898759309 | 0.58688 | 0.4545749 | 0.999175987 |
| ENSMUSG00000029397 | Rchy1 | protein_coding | 0.260385792 | 6.608216687 | 0.5867965 | 0.4546064 | 0.999175987 |
| ENSMUSG00000037736 | Limch1 | protein_coding | -0.296319084 | 5.94337844 | 0.5867914 | 0.4546084 | 0.999175987 |
| ENSMUSG00000042156 | Dzip1 | protein_coding | -0.36709336 | 5.309869513 | 0.5861196 | 0.4548622 | 0.999175987 |
| ENSMUSG00000019820 | Utrn | protein_coding | -0.353961535 | 5.03436791 | 0.5859825 | 0.4549141 | 0.999175987 |
| ENSMUSG00000054708 | Ankrd24 | protein_coding | 0.493424117 | 3.893537642 | 0.5856835 | 0.4550272 | 0.999175987 |
| ENSMUSG00000025272 | Tro | protein_coding | -0.303070934 | 7.082353565 | 0.5857241 | 0.4550297 | 0.999175987 |
| ENSMUSG00000031513 | Leprotl1 | protein_coding | -0.336615351 | 6.383759213 | 0.585488 | 0.455119 | 0.999175987 |
| ENSMUSG00000035441 | Myo1d | protein_coding | -0.390341934 | 4.208263697 | 0.5854082 | 0.4551313 | 0.999175987 |
| ENSMUSG00000004071 | Cdip1 | protein_coding | 0.286224641 | 10.04724437 | 0.585239 | 0.4551953 | 0.999175987 |
| ENSMUSG00000058440 | Nrf1 | protein_coding | -0.342608078 | 4.588800568 | 0.585117 | 0.4552415 | 0.999175987 |
| ENSMUSG00000030304 | Ergic2 | protein_coding | -0.24753378 | 6.167755877 | 0.5848364 | 0.4553478 | 0.999175987 |
| ENSMUSG00000023044 | Csad | protein_coding | -0.493803372 | 4.287033396 | 0.5848228 | 0.4553529 | 0.999175987 |
| ENSMUSG00000081534 | Slc48a1 | protein_coding | 0.300569806 | 7.330296116 | 0.5844516 | 0.4555114 | 0.999175987 |
| ENSMUSG00000024286 | Ccny | protein_coding | 0.456887248 | 5.07855017 | 0.5840688 | 0.4556565 | 0.999175987 |
| ENSMUSG00000024170 | Telo2 | protein_coding | 0.465669499 | 4.060941039 | 0.5837225 | 0.45577 | 0.999175987 |
| ENSMUSG00000040618 | Pck2 | protein_coding | -0.357382585 | 5.48517705 | 0.5834076 | 0.4558894 | 0.999175987 |
| ENSMUSG00000058441 | Panx2 | protein_coding | -0.452582292 | 4.069168426 | 0.5827947 | 0.4561221 | 0.999175987 |
| ENSMUSG00000020090 | Npffr1 | protein_coding | 0.623313567 | 3.041290467 | 0.5826488 | 0.4561775 | 0.999175987 |
| ENSMUSG00000044071 | Fam19a2 | protein_coding | -0.420290756 | 4.660714796 | 0.5823565 | 0.4562886 | 0.999175987 |
| ENSMUSG00000021975 | Ints9 | protein_coding | 0.419809324 | 4.656545547 | 0.5818261 | 0.4564902 | 0.999175987 |
| ENSMUSG00000018537 | Pcgf2 | protein_coding | 0.559800487 | 3.225708111 | 0.5812949 | 0.4566923 | 0.999175987 |
| ENSMUSG00000040720 | Virma | protein_coding | -0.413463773 | 4.250629407 | 0.5809507 | 0.4568233 | 0.999175987 |
| ENSMUSG00000043432 | Leng9 | protein_coding | 0.406040008 | 4.373271908 | 0.5808024 | 0.4568798 | 0.999175987 |
| ENSMUSG00000097536 | 2610037D02Rik | lincRNA | -0.858247397 | 2.550039944 | 0.5803305 | 0.4570595 | 0.999175987 |
| ENSMUSG00000050538 | B230217C12Rik | protein_coding | 0.219894552 | 7.527299184 | 0.5800988 | 0.4571478 | 0.999175987 |
| ENSMUSG00000020873 | Slc35b1 | protein_coding | 0.224207713 | 7.164479832 | 0.5798177 | 0.457255 | 0.999175987 |
| ENSMUSG00000109336 | Samd4b | protein_coding | -0.394892946 | 5.297529667 | 0.5796627 | 0.4573141 | 0.999175987 |
| ENSMUSG00000023999 | Kif6 | protein_coding | -0.930394187 | 2.428414314 | 0.5796291 | 0.4573269 | 0.999175987 |

| ensembl_gene_id | external_gene_name | gene_biotype | log2 (fold change) | log2 (counts per million) | F-value | p-value | False discovery rate |
|---|---|---|---|---|---|---|---|
| ENSMUSG00000020691 | Mettl2 | protein_coding | 0.498935331 | 4.728498721 | 0.5793944 | 0.4574343 | 0.999175987 |
| ENSMUSG00000003348 | Mob3a | protein_coding | 0.869991701 | 2.797709961 | 0.5793288 | 0.4574415 | 0.999175987 |
| ENSMUSG00000048661 | Lemd3 | protein_coding | 0.435869869 | 3.832823905 | 0.578931 | 0.4575933 | 0.999175987 |
| ENSMUSG00000060924 | Csmd1 | protein_coding | -0.533614785 | 4.672874283 | 0.5790428 | 0.4576338 | 0.999175987 |
| ENSMUSG00000026594 | Ralgps2 | protein_coding | 0.381993405 | 4.878117885 | 0.5785567 | 0.4577363 | 0.999175987 |
| ENSMUSG00000003955 | Fam162a | protein_coding | 0.324984504 | 6.252638151 | 0.5780395 | 0.4579339 | 0.999175987 |
| ENSMUSG00000020333 | Acsl6 | protein_coding | -0.350994266 | 5.723656483 | 0.5779716 | 0.4579599 | 0.999175987 |
| ENSMUSG00000026819 | Slc25a25 | protein_coding | 0.281492319 | 5.904882908 | 0.5778868 | 0.4579923 | 0.999175987 |
| ENSMUSG00000019027 | Dnah1 | protein_coding | -0.434169213 | 5.188644309 | 0.5771563 | 0.4582895 | 0.999175987 |
| ENSMUSG00000020639 | Pfn4 | protein_coding | 0.460561321 | 4.264502937 | 0.5770909 | 0.4582968 | 0.999175987 |
| ENSMUSG00000063600 | Egfem1 | protein_coding | 0.490084948 | 3.503946583 | 0.577031 | 0.4583197 | 0.999175987 |
| ENSMUSG00000015342 | Xk | protein_coding | -1.046505971 | 3.146942023 | 0.5804634 | 0.4584132 | 0.999175987 |
| ENSMUSG00000023034 | Nr4a1 | protein_coding | -0.294890503 | 5.870254993 | 0.5766723 | 0.4584571 | 0.999175987 |
| ENSMUSG00000060803 | Gstp1 | protein_coding | 0.218588796 | 8.774563866 | 0.5762715 | 0.4586106 | 0.999175987 |
| ENSMUSG00000021481 | Zfp346 | protein_coding | -0.416874848 | 4.227970794 | 0.5762308 | 0.4586262 | 0.999175987 |
| ENSMUSG00000040687 | Madd | protein_coding | -0.296388647 | 5.786761892 | 0.5761152 | 0.4586705 | 0.999175987 |
| ENSMUSG00000090122 | Kcne1l | protein_coding | -0.986219481 | 2.519501753 | 0.5759358 | 0.4587393 | 0.999175987 |
| ENSMUSG00000038312 | Edem2 | protein_coding | -0.603855401 | 4.458289532 | 0.5776308 | 0.4587782 | 0.999175987 |
| ENSMUSG00000031661 | Nkd1 | protein_coding | 0.98183938 | 2.463176117 | 0.5755917 | 0.4588712 | 0.999175987 |
| ENSMUSG00000032590 | Apeh | protein_coding | 0.42062921 | 4.439069016 | 0.5754277 | 0.4589341 | 0.999175987 |
| ENSMUSG00000025321 | Itgb8 | protein_coding | 0.59684243 | 3.993392886 | 0.5745279 | 0.4592973 | 0.999175987 |
| ENSMUSG00000029228 | Lnx1 | protein_coding | 0.52792571 | 4.27486884 | 0.5744917 | 0.4593112 | 0.999175987 |
| ENSMUSG00000018427 | Ypel2 | protein_coding | -0.232855204 | 6.565486051 | 0.5742425 | 0.4593892 | 0.999175987 |
| ENSMUSG00000022199 | Slc22a17 | protein_coding | 0.250938422 | 8.546739703 | 0.5742243 | 0.4593962 | 0.999175987 |
| ENSMUSG00000084957 | Bbip1 | protein_coding | 0.236637252 | 7.193835222 | 0.5740466 | 0.4594645 | 0.999175987 |
| ENSMUSG00000029359 | Tesc | protein_coding | -0.471822003 | 4.607347057 | 0.5734039 | 0.4597117 | 0.999175987 |
| ENSMUSG00000021340 | Gpld1 | protein_coding | 0.512215842 | 3.527496448 | 0.5732404 | 0.4597746 | 0.999175987 |
| ENSMUSG00000026014 | Raph1 | protein_coding | 0.341312867 | 4.442371162 | 0.5731386 | 0.4598137 | 0.999175987 |
| ENSMUSG00000018417 | Myo1b | protein_coding | -0.419776956 | 3.79058272 | 0.5729366 | 0.4598915 | 0.999175987 |
| ENSMUSG00000043301 | Kcnj6 | protein_coding | -0.652584235 | 2.906168264 | 0.5728393 | 0.459929 | 0.999175987 |
| ENSMUSG00000032116 | Stt3a | protein_coding | -0.289856106 | 5.841603953 | 0.572777 | 0.4599529 | 0.999175987 |
| ENSMUSG00000069763 | Tmem100 | protein_coding | -0.423607528 | 5.158979621 | 0.5723598 | 0.4601136 | 0.999175987 |
| ENSMUSG00000034586 | Hid1 | protein_coding | 0.340278972 | 6.134864464 | 0.5722452 | 0.4601578 | 0.999175987 |
| ENSMUSG00000071547 | Nt5dc2 | protein_coding | 0.49225883 | 3.340278492 | 0.5722362 | 0.4601612 | 0.999175987 |
| ENSMUSG00000022000 | Zc3h13 | protein_coding | -0.335419297 | 5.432840269 | 0.5710205 | 0.4606301 | 0.999175987 |
| ENSMUSG00000051113 | Fam71e1 | protein_coding | 0.771494767 | 2.910561462 | 0.5709716 | 0.460649 | 0.999175987 |
| ENSMUSG00000040234 | Tm7sf3 | protein_coding | -0.38857174 | 4.603531011 | 0.5708774 | 0.4606853 | 0.999175987 |
| ENSMUSG00000009876 | Cox4i2 | protein_coding | -0.764719077 | 2.777996292 | 0.5706377 | 0.4607779 | 0.999175987 |
| ENSMUSG00000047935 | Gm5607 | sense_overlapping | -0.352098276 | 4.563505048 | 0.5701936 | 0.4609495 | 0.999175987 |
| ENSMUSG00000031398 | Plxna3 | protein_coding | 0.712341282 | 2.688513231 | 0.569981 | 0.4610316 | 0.999175987 |
| ENSMUSG00000047428 | Dlk2 | protein_coding | 0.623803114 | 3.196595907 | 0.5699785 | 0.4610326 | 0.999175987 |
| ENSMUSG00000098649 | Gm27199 | antisense_RNA | -0.659443667 | 2.751579585 | 0.569934 | 0.4610498 | 0.999175987 |
| ENSMUSG00000026234 | Ncl | protein_coding | -0.237348179 | 8.209735329 | 0.569812 | 0.461097 | 0.999175987 |
| ENSMUSG00000021595 | Nsun2 | protein_coding | -0.34923579 | 5.686086394 | 0.5696793 | 0.4611483 | 0.999175987 |
| ENSMUSG00000041774 | Ydjc | protein_coding | 0.620726151 | 3.276588869 | 0.5696306 | 0.4611671 | 0.999175987 |
| ENSMUSG00000028411 | Aptx | protein_coding | 0.234265415 | 6.507321975 | 0.5696256 | 0.4611691 | 0.999175987 |
| ENSMUSG00000030421 | Uri1 | protein_coding | -0.333036508 | 4.919120289 | 0.5695897 | 0.4611829 | 0.999175987 |
| ENSMUSG00000024376 | Epb41l4a | protein_coding | 0.566686849 | 3.386500114 | 0.56955 | 0.4611983 | 0.999175987 |
| ENSMUSG00000005804 | Bloc1s6 | protein_coding | 0.405433879 | 6.412023216 | 0.5712094 | 0.4612409 | 0.999175987 |
| ENSMUSG00000031585 | Gtf2e2 | protein_coding | 0.262963869 | 5.868931169 | 0.5693073 | 0.4612922 | 0.999175987 |
| ENSMUSG00000015748 | Prpf3 | protein_coding | 0.451610444 | 4.036123094 | 0.5688706 | 0.4614612 | 0.999175987 |
| ENSMUSG00000006736 | Tspan31 | protein_coding | 0.236798473 | 6.733065879 | 0.5687443 | 0.4615101 | 0.999175987 |
| ENSMUSG00000033216 | Eefsec | protein_coding | 0.452085586 | 4.607461669 | 0.5686199 | 0.4615583 | 0.999175987 |
| ENSMUSG00000027864 | Ptgfrn | protein_coding | -0.508280522 | 4.140524915 | 0.5685918 | 0.4615692 | 0.999175987 |
| ENSMUSG00000025402 | Nab2 | protein_coding | 0.555639122 | 3.529868613 | 0.5685695 | 0.4615778 | 0.999175987 |
| ENSMUSG00000053062 | Jam2 | protein_coding | -0.413795375 | 3.995873277 | 0.5677463 | 0.4618968 | 0.999175987 |
| ENSMUSG00000033389 | Arhgap44 | protein_coding | 0.357902382 | 8.415104961 | 0.5691901 | 0.4620199 | 0.999175987 |
| ENSMUSG00000001891 | Ugp2 | protein_coding | -0.240268728 | 6.183373685 | 0.567238 | 0.462094 | 0.999175987 |
| ENSMUSG00000011114 | Tbrg1 | protein_coding | -0.268365952 | 7.01306031 | 0.5671871 | 0.4621137 | 0.999175987 |
| ENSMUSG00000049323 | Smcr8 | protein_coding | -0.509609107 | 3.611316607 | 0.5670337 | 0.4621733 | 0.999175987 |
| ENSMUSG00000022973 | Synj1 | protein_coding | -0.281961842 | 7.54218916 | 0.5670269 | 0.4621759 | 0.999175987 |
| ENSMUSG00000019173 | Rab5c | protein_coding | -0.367107989 | 5.459684264 | 0.5668319 | 0.4622516 | 0.999175987 |
| ENSMUSG00000037640 | Zfp60 | protein_coding | -0.393133564 | 4.904706894 | 0.5667082 | 0.4622997 | 0.999175987 |
| ENSMUSG00000005148 | Klf5 | protein_coding | 0.534943708 | 3.456679139 | 0.5667036 | 0.4623014 | 0.999175987 |
| ENSMUSG00000067365 | Tmem128 | protein_coding | 0.277232874 | 6.726846509 | 0.5666766 | 0.4623119 | 0.999175987 |
| ENSMUSG00000003273 | Car11 | protein_coding | 0.378831525 | 5.660026022 | 0.5666293 | 0.4623303 | 0.999175987 |
| ENSMUSG00000020827 | Mink1 | protein_coding | 0.334581723 | 4.869979965 | 0.5663102 | 0.4624542 | 0.999175987 |
| ENSMUSG00000028470 | Hint2 | protein_coding | 0.501034009 | 4.938380281 | 0.5664478 | 0.4624646 | 0.999175987 |
| ENSMUSG00000020743 | Mif4gd | protein_coding | -0.329260747 | 5.246410867 | 0.5657902 | 0.4626564 | 0.999175987 |
| ENSMUSG00000022680 | Pdxdc1 | protein_coding | 0.34308576 | 5.315056667 | 0.5657592 | 0.4626684 | 0.999175987 |
| ENSMUSG00000029458 | Brap | protein_coding | -0.338885634 | 4.55708519 | 0.5653587 | 0.4628242 | 0.999175987 |
| ENSMUSG00000031574 | Star | protein_coding | -0.665001552 | 2.702912723 | 0.5650957 | 0.4629265 | 0.999175987 |
| ENSMUSG00000005470 | Asf1b | protein_coding | 1.431431952 | 2.029709115 | 0.56518 | 0.4629575 | 0.999175987 |
| ENSMUSG00000019470 | Xab2 | protein_coding | 0.447723981 | 5.777824972 | 0.5663343 | 0.4631252 | 0.999175987 |
| ENSMUSG00000033931 | Rbm34 | protein_coding | -0.366891244 | 4.615637206 | 0.5645479 | 0.4631398 | 0.999175987 |
| ENSMUSG00000029504 | Ddx51 | protein_coding | -1.217356483 | 2.463090803 | 0.5653468 | 0.4631843 | 0.999175987 |
| ENSMUSG00000032355 | Mlip | protein_coding | 0.432776319 | 4.229628769 | 0.564249 | 0.4632562 | 0.999175987 |
| ENSMUSG00000020883 | Fbxl20 | protein_coding | 0.445292453 | 3.978598993 | 0.5641068 | 0.4633117 | 0.999175987 |

| ensembl_gene_id | external_gene_name | gene_biotype | log2 (fold change) | log2 (counts per million) | F-value | p-value | False discovery rate |
|---|---|---|---|---|---|---|---|
| ENSMUSG00000025499 | Hras | protein_coding | 0.295682652 | 6.458936108 | 0.5639649 | 0.4633669 | 0.999175987 |
| ENSMUSG00000026640 | Plxna2 | protein_coding | -0.386891737 | 4.949512398 | 0.5639634 | 0.4633675 | 0.999175987 |
| ENSMUSG00000043388 | Tmem130 | protein_coding | 0.264980418 | 9.615115595 | 0.5639293 | 0.4633808 | 0.999175987 |
| ENSMUSG00000022346 | Myc | protein_coding | -0.704637496 | 3.014558506 | 0.5636531 | 0.4634885 | 0.999175987 |
| ENSMUSG00000025269 | Apex2 | protein_coding | 0.586869872 | 3.326395185 | 0.5634703 | 0.4635598 | 0.999175987 |
| ENSMUSG00000032046 | Abhd12 | protein_coding | -0.239576979 | 6.967606994 | 0.5631321 | 0.4636918 | 0.999175987 |
| ENSMUSG00000027674 | Pex5l | protein_coding | -0.545341091 | 3.225459184 | 0.5630022 | 0.4637425 | 0.999175987 |
| ENSMUSG00000035671 | Zswim4 | protein_coding | 0.421298334 | 3.825529101 | 0.5628491 | 0.4638023 | 0.999175987 |
| ENSMUSG00000030032 | Wdr54 | protein_coding | 0.297465513 | 5.463490421 | 0.562509 | 0.4639351 | 0.999175987 |
| ENSMUSG00000023019 | Gpd1 | protein_coding | -0.485202611 | 3.716180756 | 0.5623167 | 0.4640102 | 0.999175987 |
| ENSMUSG00000059552 | Trp53 | protein_coding | 0.319899468 | 5.633688644 | 0.562055 | 0.4641125 | 0.999175987 |
| ENSMUSG00000029482 | Aacs | protein_coding | -0.432643201 | 4.888742531 | 0.5619351 | 0.4641593 | 0.999175987 |
| ENSMUSG00000020467 | Efemp1 | protein_coding | -0.597478858 | 3.055751695 | 0.5618531 | 0.4641914 | 0.999175987 |
| ENSMUSG00000041670 | Rims1 | protein_coding | -0.261743536 | 6.013708503 | 0.5613427 | 0.464391 | 0.999175987 |
| ENSMUSG00000034848 | Ttc21b | protein_coding | -0.773283241 | 2.747969089 | 0.5612883 | 0.4644123 | 0.999175987 |
| ENSMUSG00000038539 | Atf5 | protein_coding | -0.772871149 | 2.495895301 | 0.5608893 | 0.4645685 | 0.999175987 |
| ENSMUSG00000052336 | Cx3cr1 | protein_coding | 0.492879955 | 4.189887564 | 0.5608416 | 0.4645871 | 0.999175987 |
| ENSMUSG00000044550 | Tceal3 | protein_coding | 0.203352527 | 7.206967085 | 0.560512 | 0.4647162 | 0.999175987 |
| ENSMUSG00000020719 | Ddx5 | protein_coding | -0.23216388 | 8.230626734 | 0.5601301 | 0.4648658 | 0.999175987 |
| ENSMUSG00000034557 | Zfyve9 | protein_coding | 0.311937897 | 5.23574843 | 0.5599752 | 0.4649266 | 0.999175987 |
| ENSMUSG00000026721 | Rabgap1l | protein_coding | -0.261189238 | 6.152632248 | 0.5594593 | 0.4651288 | 0.999175987 |
| ENSMUSG00000008734 | Gprc5b | protein_coding | -0.439452842 | 4.896268762 | 0.5591351 | 0.4652561 | 0.999175987 |
| ENSMUSG00000022378 | Fam49b | protein_coding | 0.231206812 | 6.70146057 | 0.5591106 | 0.4652657 | 0.999175987 |
| ENSMUSG00000024112 | Cacna1h | protein_coding | 0.921909815 | 2.669602741 | 0.5580563 | 0.4656798 | 0.999175987 |
| ENSMUSG00000028937 | Acot7 | protein_coding | 0.299329799 | 10.6617693 | 0.5578214 | 0.4657721 | 0.999175987 |
| ENSMUSG00000018932 | Map2k3 | protein_coding | -0.616453726 | 2.856853344 | 0.5574128 | 0.4659328 | 0.999175987 |
| ENSMUSG00000050373 | Snx21 | protein_coding | -0.411089854 | 4.355608397 | 0.5574107 | 0.4659337 | 0.999175987 |
| ENSMUSG00000039716 | Dock3 | protein_coding | -0.445249606 | 3.952430827 | 0.557307 | 0.4659744 | 0.999175987 |
| ENSMUSG00000021945 | Zmym2 | protein_coding | -0.322813367 | 5.437061708 | 0.5570516 | 0.466075 | 0.999175987 |
| ENSMUSG00000025858 | Get4 | protein_coding | -0.425470141 | 3.690348332 | 0.5568556 | 0.4661521 | 0.999175987 |
| ENSMUSG00000090266 | Mettl23 | protein_coding | 0.437524623 | 4.816072364 | 0.5566637 | 0.4662277 | 0.999175987 |
| ENSMUSG00000020149 | Rab1a | protein_coding | 0.250562282 | 8.128892063 | 0.556458 | 0.4663087 | 0.999175987 |
| ENSMUSG00000022557 | Bop1 | protein_coding | 0.40537259 | 5.784828613 | 0.5562155 | 0.4664674 | 0.999175987 |
| ENSMUSG00000028004 | Npy2r | protein_coding | 0.66037516 | 4.064201765 | 0.5575514 | 0.4665526 | 0.999175987 |
| ENSMUSG00000039745 | Htatip2 | protein_coding | 0.443058672 | 4.621718336 | 0.5558055 | 0.4665659 | 0.999175987 |
| ENSMUSG00000032540 | Abhd5 | protein_coding | -0.472521702 | 3.705300625 | 0.5558032 | 0.4665668 | 0.999175987 |
| ENSMUSG00000039656 | Rxrb | protein_coding | 0.334095243 | 5.001536529 | 0.5556047 | 0.4666451 | 0.999175987 |
| ENSMUSG00000035383 | Pmch | protein_coding | -1.003669654 | 6.988678681 | 0.584676 | 0.4667523 | 0.999175987 |
| ENSMUSG00000025880 | Smad7 | protein_coding | 0.545714189 | 3.443489442 | 0.5551977 | 0.4668056 | 0.999175987 |
| ENSMUSG00000026360 | Rgs2 | protein_coding | 0.3302676 | 7.852127661 | 0.5558286 | 0.466885 | 0.999175987 |
| ENSMUSG00000029554 | Mad1l1 | protein_coding | 0.304903983 | 5.278598499 | 0.5548768 | 0.4669323 | 0.999175987 |
| ENSMUSG00000039252 | Lgi2 | protein_coding | -0.352723316 | 4.613793116 | 0.5548313 | 0.4669503 | 0.999175987 |
| ENSMUSG00000073563 | Csnk1g3 | protein_coding | 0.321020257 | 5.131053326 | 0.554466 | 0.4670945 | 0.999175987 |
| ENSMUSG00000033006 | Sox10 | protein_coding | -0.368826808 | 5.169216814 | 0.5544063 | 0.4671181 | 0.999175987 |
| ENSMUSG00000031214 | Ophn1 | protein_coding | -0.374209392 | 4.328522202 | 0.5543782 | 0.4671292 | 0.999175987 |
| ENSMUSG00000031239 | Itm2a | protein_coding | 0.32424776 | 5.464426864 | 0.5543467 | 0.4671417 | 0.999175987 |
| ENSMUSG00000045466 | Zfp956 | protein_coding | -0.470026601 | 4.374954297 | 0.5537263 | 0.4673869 | 0.999175987 |
| ENSMUSG00000058715 | Fcer1g | protein_coding | -0.51914548 | 4.652242436 | 0.5533537 | 0.4675973 | 0.999175987 |
| ENSMUSG00000052609 | Plekhg3 | protein_coding | -0.527032149 | 3.484572256 | 0.5531628 | 0.4676099 | 0.999175987 |
| ENSMUSG00000064065 | Ipcef1 | protein_coding | 0.675216822 | 2.627359366 | 0.5528437 | 0.4677362 | 0.999175987 |
| ENSMUSG00000061288 | Taok3 | protein_coding | 0.265529788 | 7.069856673 | 0.5527309 | 0.4677809 | 0.999175987 |
| ENSMUSG00000025410 | Dctn2 | protein_coding | 0.281248627 | 8.007037595 | 0.5527126 | 0.4677881 | 0.999175987 |
| ENSMUSG00000045962 | Wnk1 | protein_coding | -0.318993131 | 5.624088413 | 0.5526335 | 0.4678194 | 0.999175987 |
| ENSMUSG00000032463 | Faim | protein_coding | 0.362617321 | 4.670042742 | 0.5525963 | 0.4678342 | 0.999175987 |
| ENSMUSG00000001440 | Kpnb1 | protein_coding | -0.250700838 | 6.058698675 | 0.5524777 | 0.4678812 | 0.999175987 |
| ENSMUSG00000005371 | Fbxo11 | protein_coding | -0.24882936 | 6.474362649 | 0.5524347 | 0.4678982 | 0.999175987 |
| ENSMUSG00000008763 | Man1a2 | protein_coding | -0.362849235 | 5.403876338 | 0.5520321 | 0.4680578 | 0.999175987 |
| ENSMUSG00000030987 | Stim1 | protein_coding | 0.283151393 | 5.982933369 | 0.5517253 | 0.4681794 | 0.999175987 |
| ENSMUSG00000026848 | Tor1b | protein_coding | -0.405452224 | 4.511419025 | 0.5515576 | 0.4682459 | 0.999175987 |
| ENSMUSG00000041552 | Ptchd1 | protein_coding | -0.515666285 | 3.196184494 | 0.5509942 | 0.4684695 | 0.999175987 |
| ENSMUSG00000029592 | Usp30 | protein_coding | -0.449084695 | 4.847948825 | 0.5509759 | 0.468494 | 0.999175987 |
| ENSMUSG00000026585 | Kifap3 | protein_coding | -0.237300683 | 7.567748367 | 0.5508719 | 0.4685181 | 0.999175987 |
| ENSMUSG00000050088 | 1600012H06Rik | protein_coding | 1.112779932 | 2.783557498 | 0.552534 | 0.4685297 | 0.999175987 |
| ENSMUSG00000000560 | Gabra2 | protein_coding | -0.4745147 | 6.153070811 | 0.5543501 | 0.4686188 | 0.999175987 |
| ENSMUSG00000056306 | Sertm1 | protein_coding | 0.572359037 | 3.222958381 | 0.5506152 | 0.46862 | 0.999175987 |
| ENSMUSG00000025354 | Dnajc14 | protein_coding | 0.404684467 | 4.999085832 | 0.550526 | 0.4686554 | 0.999175987 |
| ENSMUSG00000032834 | Pwp2 | protein_coding | -0.502024074 | 4.081071624 | 0.5504998 | 0.4686658 | 0.999175987 |
| ENSMUSG00000030826 | Bcat2 | protein_coding | 0.444636183 | 4.651427735 | 0.550185 | 0.4687909 | 0.999175987 |
| ENSMUSG00000021589 | Rhobtb3 | protein_coding | -0.47607716 | 3.888652789 | 0.5500874 | 0.4688297 | 0.999175987 |
| ENSMUSG00000075410 | Prcd | protein_coding | 0.800555307 | 3.463079104 | 0.5516326 | 0.4688864 | 0.999175987 |
| ENSMUSG00000004895 | Prcc | protein_coding | -0.438814121 | 3.576970569 | 0.5497644 | 0.4689582 | 0.999175987 |
| ENSMUSG00000029623 | Pdap1 | protein_coding | -0.226190012 | 6.886533944 | 0.549288 | 0.4691477 | 0.999175987 |
| ENSMUSG00000074001 | Klhl40 | protein_coding | -1.030519148 | 2.161219232 | 0.5491246 | 0.4692127 | 0.999175987 |
| ENSMUSG00000039976 | Tbc1d16 | protein_coding | -0.378464266 | 4.131890637 | 0.5490012 | 0.4692619 | 0.999175987 |
| ENSMUSG00000024033 | Rsph1 | protein_coding | -0.283953846 | 5.553946355 | 0.5485108 | 0.4694572 | 0.999175987 |
| ENSMUSG00000074247 | Dda1 | protein_coding | 0.349212187 | 5.191936078 | 0.5481815 | 0.4695884 | 0.999175987 |
| ENSMUSG00000024491 | Rbm27 | protein_coding | 0.329952474 | 5.31306942 | 0.5478209 | 0.4697321 | 0.999175987 |
| ENSMUSG00000068151 | A230006K03Rik | lincRNA | 0.370652537 | 4.432775736 | 0.5477051 | 0.4697783 | 0.999175987 |

| ensembl_gene_id | external_gene_name | gene_biotype | log2 (fold change) | log2 (counts per million) | F-value | p-value | False discovery rate |
|---|---|---|---|---|---|---|---|
| ENSMUSG00000027433 | Xrn2 | protein_coding | -0.281244152 | 5.935419471 | 0.547535 | 0.4698462 | 0.999175987 |
| ENSMUSG00000029430 | Ran | protein_coding | 0.235039405 | 8.036845059 | 0.5466641 | 0.4701939 | 0.999175987 |
| ENSMUSG00000033671 | Cep350 | protein_coding | -0.701448798 | 2.879776363 | 0.546497 | 0.4702606 | 0.999175987 |
| ENSMUSG00000022820 | Ndufb4 | protein_coding | 0.309243613 | 5.958522135 | 0.5461947 | 0.4703814 | 0.999175987 |
| ENSMUSG00000031872 | Bean1 | protein_coding | 0.719608237 | 2.534409631 | 0.5459827 | 0.4704662 | 0.999175987 |
| ENSMUSG00000029198 | Grpel1 | protein_coding | 0.220918527 | 8.453275361 | 0.5459077 | 0.4704962 | 0.999175987 |
| ENSMUSG00000010362 | Rdm1 | protein_coding | 0.39823088 | 5.33457854 | 0.5456706 | 0.470591 | 0.999175987 |
| ENSMUSG00000022111 | Uchl3 | protein_coding | 0.269855515 | 6.159675216 | 0.5456697 | 0.4705914 | 0.999175987 |
| ENSMUSG00000028617 | Lrrc42 | protein_coding | 0.331296312 | 4.923167692 | 0.5456189 | 0.4706117 | 0.999175987 |
| ENSMUSG00000001729 | Akt1 | protein_coding | 0.222423822 | 6.455986 | 0.5452178 | 0.4707722 | 0.999175987 |
| ENSMUSG00000040649 | Rimklb | protein_coding | -0.293327201 | 6.130343826 | 0.5449596 | 0.4708756 | 0.999175987 |
| ENSMUSG00000049106 | Dcaf5 | protein_coding | 0.27694006 | 5.500459631 | 0.5447231 | 0.4709703 | 0.999175987 |
| ENSMUSG00000034936 | Arl4d | protein_coding | -0.63584209 | 4.098473613 | 0.5461648 | 0.4710598 | 0.999175987 |
| ENSMUSG00000022890 | Atp5j | protein_coding | 0.253947981 | 10.19202003 | 0.5437941 | 0.4713427 | 0.999175987 |
| ENSMUSG00000060568 | Fam78b | protein_coding | -0.533208753 | 3.485171774 | 0.5435607 | 0.4714363 | 0.999175987 |
| ENSMUSG00000021720 | Rnf180 | protein_coding | 0.359582493 | 4.500126184 | 0.5433861 | 0.4715064 | 0.999175987 |
| ENSMUSG00000021895 | Arhgef3 | protein_coding | -0.318724254 | 5.025681311 | 0.5433713 | 0.4715123 | 0.999175987 |
| ENSMUSG00000026248 | Mrpl44 | protein_coding | 0.228167988 | 7.060938854 | 0.5426646 | 0.4717961 | 0.999175987 |
| ENSMUSG00000097527 | 1700112J16Rik | lincRNA | -0.7042078 | 2.740116355 | 0.5426448 | 0.4718041 | 0.999175987 |
| ENSMUSG00000043207 | Zmpste24 | protein_coding | 0.382546627 | 4.388046085 | 0.5424058 | 0.4719001 | 0.999175987 |
| ENSMUSG00000019433 | Gipc1 | protein_coding | 0.313986778 | 5.693327097 | 0.5422613 | 0.4719582 | 0.999175987 |
| ENSMUSG00000110156 | Gm42067 | antisense_RNA | 0.539144811 | 3.177236938 | 0.5421666 | 0.4719963 | 0.999175987 |
| ENSMUSG00000022426 | Josd1 | protein_coding | 0.38180288 | 5.076976247 | 0.5421206 | 0.4720148 | 0.999175987 |
| ENSMUSG00000033295 | Ptprf | protein_coding | 0.274509576 | 5.670309118 | 0.5416138 | 0.4722186 | 0.999175987 |
| ENSMUSG00000075467 | Dnlz | protein_coding | 0.466363516 | 4.734856396 | 0.5415838 | 0.4722478 | 0.999175987 |
| ENSMUSG00000032076 | Cadm1 | protein_coding | -0.229810916 | 7.275656087 | 0.5413531 | 0.4723235 | 0.999175987 |
| ENSMUSG00000030733 | Sh2b1 | protein_coding | -0.323539033 | 4.85364938 | 0.541243 | 0.4723678 | 0.999175987 |
| ENSMUSG00000023828 | Slc22a3 | protein_coding | -0.779206116 | 2.622116416 | 0.5412076 | 0.4723821 | 0.999175987 |
| ENSMUSG00000024640 | Psat1 | protein_coding | -0.200389019 | 7.37906699 | 0.5409978 | 0.4724665 | 0.999175987 |
| ENSMUSG00000039191 | Rbpj | protein_coding | -0.521056342 | 3.246021187 | 0.5404804 | 0.472675 | 0.999175987 |
| ENSMUSG00000031622 | Sin3b | protein_coding | 0.216838292 | 6.505434759 | 0.5404283 | 0.472696 | 0.999175987 |
| ENSMUSG00000038002 | Cramp1l | protein_coding | 0.52681696 | 3.327462723 | 0.5404207 | 0.472699 | 0.999175987 |
| ENSMUSG00000021928 | Ebpl | protein_coding | -0.351325104 | 4.421422157 | 0.540111 | 0.4728239 | 0.999175987 |
| ENSMUSG00000028847 | Trappc3 | protein_coding | 0.203145081 | 7.886045481 | 0.5400498 | 0.4728486 | 0.999175987 |
| ENSMUSG00000043629 | 1700019D03Rik | protein_coding | -0.488369936 | 3.571831769 | 0.5400016 | 0.472868 | 0.999175987 |
| ENSMUSG00000053181 | A830005F24Rik | protein_coding | -0.687576786 | 3.006694887 | 0.5397324 | 0.4729766 | 0.999175987 |
| ENSMUSG00000045160 | Bola3 | protein_coding | -0.399595641 | 4.126265745 | 0.539603 | 0.4730288 | 0.999175987 |
| ENSMUSG00000034994 | Eef2 | protein_coding | 0.272493054 | 10.00423559 | 0.53955 | 0.4730502 | 0.999175987 |
| ENSMUSG00000029512 | Ulk1 | protein_coding | -0.335135357 | 4.964379479 | 0.5391706 | 0.4732034 | 0.999175987 |
| ENSMUSG00000020812 | 1810032O08Rik | protein_coding | 0.648120763 | 2.780781175 | 0.5388981 | 0.4733134 | 0.999175987 |
| ENSMUSG00000030741 | Spns1 | protein_coding | 0.333550839 | 5.835921161 | 0.5382013 | 0.473595 | 0.999175987 |
| ENSMUSG00000026017 | Carf | protein_coding | -0.465028475 | 4.410506606 | 0.5373605 | 0.4739352 | 0.999175987 |
| ENSMUSG00000026775 | Yme1l1 | protein_coding | 0.305185076 | 6.837194338 | 0.5372897 | 0.4739809 | 0.999175987 |
| ENSMUSG00000032291 | Crabp1 | protein_coding | -0.457056605 | 4.617059342 | 0.5371908 | 0.4740039 | 0.999175987 |
| ENSMUSG00000040888 | Gfer | protein_coding | 0.338007533 | 4.47667064 | 0.5369315 | 0.4741089 | 0.999175987 |
| ENSMUSG00000062151 | Unc13c | protein_coding | -0.35637715 | 6.355882941 | 0.5368577 | 0.4742301 | 0.999175987 |
| ENSMUSG00000097119 | B230354K17Rik | lincRNA | 0.55466511 | 3.440706363 | 0.536242 | 0.4743884 | 0.999175987 |
| ENSMUSG00000024400 | Wdr33 | protein_coding | -0.246740096 | 6.93051735 | 0.5362419 | 0.4743884 | 0.999175987 |
| ENSMUSG00000016024 | Lbp | protein_coding | -0.934190496 | 2.970601539 | 0.5363913 | 0.4746495 | 0.999175987 |
| ENSMUSG00000020955 | Ap4s1 | protein_coding | 0.295587709 | 5.522627673 | 0.5351975 | 0.4748122 | 0.999175987 |
| ENSMUSG00000030291 | Med21 | protein_coding | 0.249493199 | 7.272504161 | 0.5351589 | 0.4748279 | 0.999175987 |
| ENSMUSG00000094626 | Cecr6 | protein_coding | 0.73173625 | 3.036081441 | 0.5348186 | 0.4749661 | 0.999175987 |
| ENSMUSG00000039069 | Mtg2 | protein_coding | -0.401246178 | 4.653762963 | 0.5345638 | 0.4750697 | 0.999175987 |
| ENSMUSG00000048249 | Crebrf | protein_coding | -0.236773358 | 6.30893955 | 0.5341004 | 0.4752581 | 0.999175987 |
| ENSMUSG00000007610 | Gtpbp3 | protein_coding | -0.624430943 | 2.835083554 | 0.5337436 | 0.4754033 | 0.999175987 |
| ENSMUSG00000019961 | Tmpo | protein_coding | -0.339442085 | 4.458552505 | 0.533548 | 0.4754829 | 0.999175987 |
| ENSMUSG00000042312 | S100a13 | protein_coding | -0.414191024 | 6.416420582 | 0.53604 | 0.4755035 | 0.999175987 |
| ENSMUSG00000030218 | Mgp | protein_coding | 0.956032931 | 2.11718742 | 0.5328137 | 0.475782 | 0.999175987 |
| ENSMUSG00000035329 | Fbxo33 | protein_coding | 0.325496023 | 5.444145011 | 0.5327692 | 0.4758001 | 0.999175987 |
| ENSMUSG00000031323 | Dmrtc1a | protein_coding | -0.507613064 | 3.489503687 | 0.5327541 | 0.4758062 | 0.999175987 |
| ENSMUSG00000010376 | Nedd8 | protein_coding | 0.23958273 | 9.396075068 | 0.5324425 | 0.4759333 | 0.999175987 |
| ENSMUSG00000057315 | Arhgap24 | protein_coding | -0.58117285 | 4.943632688 | 0.5348329 | 0.4759922 | 0.999175987 |
| ENSMUSG00000068732 | Tmem167b | protein_coding | -0.382346696 | 5.617912655 | 0.5321179 | 0.4760826 | 0.999175987 |
| ENSMUSG00000030579 | Tyrobp | protein_coding | -0.500325115 | 5.019160979 | 0.5331373 | 0.4760828 | 0.999175987 |
| ENSMUSG00000097326 | A330048O09Rik | lincRNA | -0.629741832 | 3.431106651 | 0.5317506 | 0.4762156 | 0.999175987 |
| ENSMUSG00000021068 | Nin | protein_coding | 0.551165067 | 3.182937395 | 0.5314937 | 0.4763204 | 0.999175987 |
| ENSMUSG00000072704 | Smim10l1 | protein_coding | -0.304761675 | 8.020245595 | 0.5313778 | 0.4763847 | 0.999175987 |
| ENSMUSG00000031924 | Cyb5b | protein_coding | -0.220288475 | 6.754261924 | 0.530965 | 0.4765363 | 0.999175987 |
| ENSMUSG00000021363 | Mak | protein_coding | -1.130395013 | 2.430673105 | 0.5309941 | 0.4765414 | 0.999175987 |
| ENSMUSG00000024855 | Pacs1 | protein_coding | 0.320213424 | 5.570134098 | 0.530903 | 0.4765617 | 0.999175987 |
| ENSMUSG00000020844 | Nxn | protein_coding | -0.462499037 | 3.816429731 | 0.5300152 | 0.4769247 | 0.999175987 |
| ENSMUSG00000035007 | Rundc1 | protein_coding | -0.48117939 | 4.464754984 | 0.5296392 | 0.4770955 | 0.999175987 |
| ENSMUSG00000104960 | Snhg8 | processed_transcript | 0.416894606 | 6.492113512 | 0.5323019 | 0.4771897 | 0.999175987 |
| ENSMUSG00000022964 | Tmem50b | protein_coding | 0.270810251 | 7.304724141 | 0.5293143 | 0.4772117 | 0.999175987 |
| ENSMUSG00000004936 | Map2k1 | protein_coding | -0.252664968 | 7.707290744 | 0.5290082 | 0.4773371 | 0.999175987 |
| ENSMUSG00000042532 | Golga7b | protein_coding | 0.279850542 | 7.965838073 | 0.5288135 | 0.4774168 | 0.999175987 |
| ENSMUSG00000023852 | Chd1 | protein_coding | -0.211986356 | 6.615970631 | 0.5287993 | 0.4774227 | 0.999175987 |
| ENSMUSG00000020599 | Rgs9 | protein_coding | -0.267746508 | 6.232323517 | 0.5281072 | 0.4777065 | 0.999175987 |

| ensembl_gene_id | external_gene_name | gene_biotype | log2 (fold change) | log2 (counts per million) | F-value | p-value | False discovery rate |
|---|---|---|---|---|---|---|---|
| ENSMUSG00000026384 | Ptpn4 | protein_coding | -0.278019386 | 5.596980002 | 0.5276971 | 0.4778748 | 0.999175987 |
| ENSMUSG00000043079 | Synpo | protein_coding | 0.325702936 | 5.277590938 | 0.5275728 | 0.4779259 | 0.999175987 |
| ENSMUSG00000024073 | Birc6 | protein_coding | -0.323463001 | 5.592319642 | 0.527566 | 0.4779287 | 0.999175987 |
| ENSMUSG00000041654 | Slc39a11 | protein_coding | 0.487847574 | 4.16824922 | 0.5275381 | 0.4779401 | 0.999175987 |
| ENSMUSG00000061983 | Rps12 | protein_coding | 0.542353767 | 3.176514267 | 0.5272231 | 0.4780695 | 0.999175987 |
| ENSMUSG00000015087 | Rabl6 | protein_coding | -0.233968742 | 6.476375965 | 0.5266887 | 0.4782891 | 0.999175987 |
| ENSMUSG00000033739 | Fkbpl | protein_coding | -0.340056577 | 5.200513763 | 0.5264969 | 0.478368 | 0.999175987 |
| ENSMUSG00000040054 | Baz2a | protein_coding | -0.433332789 | 4.379592687 | 0.5263414 | 0.478432 | 0.999175987 |
| ENSMUSG00000022307 | Oxr1 | protein_coding | -0.318083008 | 7.482523217 | 0.5267796 | 0.4784472 | 0.999175987 |
| ENSMUSG00000003346 | Abhd17a | protein_coding | 0.406267175 | 4.933499331 | 0.5262231 | 0.4784806 | 0.999175987 |
| ENSMUSG00000032965 | Ift57 | protein_coding | -0.62976968 | 3.069249186 | 0.5262082 | 0.4784868 | 0.999175987 |
| ENSMUSG00000025956 | Mettl21a | protein_coding | -0.357048721 | 4.838935925 | 0.5260336 | 0.4785586 | 0.999175987 |
| ENSMUSG00000019715 | Gle1 | protein_coding | 0.36455165 | 6.066302344 | 0.5260732 | 0.4785592 | 0.999175987 |
| ENSMUSG00000025198 | Erlin1 | protein_coding | 0.492325261 | 3.890880688 | 0.5259827 | 0.4785796 | 0.999175987 |
| ENSMUSG00000018040 | Rrp7a | protein_coding | 0.267593769 | 6.211056727 | 0.5259424 | 0.4785962 | 0.999175987 |
| ENSMUSG00000090399 | Gm38399 | lincRNA | -0.629759527 | 2.784710787 | 0.5253783 | 0.4788285 | 0.999175987 |
| ENSMUSG00000068740 | Celsr2 | protein_coding | 0.339747203 | 5.933846653 | 0.5253394 | 0.4788445 | 0.999175987 |
| ENSMUSG00000025702 | 8-Mar | protein_coding | -0.465906186 | 4.250494278 | 0.5253157 | 0.4788543 | 0.999175987 |
| ENSMUSG00000025571 | Tnrc6c | protein_coding | -0.283021771 | 5.659112533 | 0.5244858 | 0.4791964 | 0.999175987 |
| ENSMUSG00000040123 | Zmym5 | protein_coding | -0.302406628 | 5.017706106 | 0.5243838 | 0.4792384 | 0.999175987 |
| ENSMUSG00000002930 | Ppp1r17 | protein_coding | -0.758663662 | 3.752463583 | 0.5268674 | 0.4792387 | 0.999175987 |
| ENSMUSG00000040455 | Usp45 | protein_coding | -0.377193185 | 4.613921663 | 0.5243586 | 0.4792489 | 0.999175987 |
| ENSMUSG00000029642 | Polr1d | protein_coding | 0.235879046 | 6.929829932 | 0.5237123 | 0.4795156 | 0.999175987 |
| ENSMUSG00000031951 | Tmem231 | protein_coding | 0.423118465 | 4.565835644 | 0.5234678 | 0.4796166 | 0.999175987 |
| ENSMUSG00000029385 | Ccng2 | protein_coding | -0.415554425 | 4.500731803 | 0.5227005 | 0.4799338 | 0.999175987 |
| ENSMUSG00000019794 | Katna1 | protein_coding | 0.415125284 | 3.812878857 | 0.5222775 | 0.4801088 | 0.999175987 |
| ENSMUSG00000074519 | Zfp971 | protein_coding | 0.521182623 | 3.113243149 | 0.5222649 | 0.480114 | 0.999175987 |
| ENSMUSG00000051065 | Mb21d2 | protein_coding | 0.340621234 | 4.829982531 | 0.5222328 | 0.4801272 | 0.999175987 |
| ENSMUSG00000036955 | Kif1bp | protein_coding | -0.269340678 | 6.236492011 | 0.5220381 | 0.4802078 | 0.999175987 |
| ENSMUSG00000016510 | Mtif3 | protein_coding | -0.317834894 | 5.049693734 | 0.5220138 | 0.4802179 | 0.999175987 |
| ENSMUSG00000025156 | Gps1 | protein_coding | 0.264292921 | 5.477462297 | 0.5220026 | 0.4802225 | 0.999175987 |
| ENSMUSG00000004044 | Cavin1 | protein_coding | -0.305921641 | 5.39230093 | 0.5218562 | 0.4802831 | 0.999175987 |
| ENSMUSG00000021257 | Angel1 | protein_coding | -0.710122748 | 3.090701824 | 0.5217421 | 0.4803304 | 0.999175987 |
| ENSMUSG00000068329 | Htra2 | protein_coding | 0.379929211 | 4.739776655 | 0.5216533 | 0.4803672 | 0.999175987 |
| ENSMUSG00000033970 | Rfc3 | protein_coding | -0.308796996 | 4.936916151 | 0.5213186 | 0.4805058 | 0.999175987 |
| ENSMUSG00000005267 | Zfp287 | protein_coding | -0.507881545 | 3.255660461 | 0.5211717 | 0.4805667 | 0.999175987 |
| ENSMUSG00000101493 | 2810405F17Rik | lincRNA | -0.73831718 | 2.5205655 | 0.5209526 | 0.4806576 | 0.999175987 |
| ENSMUSG00000061950 | Ppp4r1 | protein_coding | 0.59603563 | 3.309485181 | 0.5208784 | 0.4806883 | 0.999175987 |
| ENSMUSG00000038280 | Ostm1 | protein_coding | 0.361274336 | 4.833108994 | 0.5208113 | 0.4807162 | 0.999175987 |
| ENSMUSG00000044991 | 1110034G24Rik | protein_coding | 0.334322231 | 5.267753978 | 0.5205673 | 0.4808174 | 0.999175987 |
| ENSMUSG00000028838 | Extl1 | protein_coding | 0.351278687 | 4.424538236 | 0.5204407 | 0.4808699 | 0.999175987 |
| ENSMUSG00000017831 | Rab5a | protein_coding | -0.417538495 | 5.016698875 | 0.5202505 | 0.4809489 | 0.999175987 |
| ENSMUSG00000078161 | Erich3 | protein_coding | -0.47156215 | 4.031065691 | 0.52001 | 0.4810487 | 0.999175987 |
| ENSMUSG00000046093 | Hpcal4 | protein_coding | 0.223923127 | 8.117955572 | 0.5199155 | 0.4810879 | 0.999175987 |
| ENSMUSG00000020679 | Hnf1b | protein_coding | 0.522936807 | 3.596697642 | 0.5197851 | 0.4811421 | 0.999175987 |
| ENSMUSG00000003360 | Ddx23 | protein_coding | -0.465250675 | 3.709420481 | 0.5196391 | 0.4812027 | 0.999175987 |
| ENSMUSG00000048495 | Tyw5 | protein_coding | 0.331647577 | 4.875694732 | 0.5195634 | 0.4812342 | 0.999175987 |
| ENSMUSG00000034390 | Cmip | protein_coding | 0.311753751 | 7.457598778 | 0.5191374 | 0.4814912 | 0.999175987 |
| ENSMUSG00000022523 | Fgf12 | protein_coding | 0.218508799 | 7.554931054 | 0.5189133 | 0.4815044 | 0.999175987 |
| ENSMUSG00000028441 | 1110017D15Rik | protein_coding | -0.290970701 | 5.39075987 | 0.5185469 | 0.4816569 | 0.999175987 |
| ENSMUSG00000020635 | Fkbp1b | protein_coding | 0.407607888 | 4.312844413 | 0.5184466 | 0.4816986 | 0.999175987 |
| ENSMUSG00000020077 | Srgn | protein_coding | -0.765963382 | 2.586494069 | 0.5181129 | 0.4818375 | 0.999175987 |
| ENSMUSG00000038365 | Fbxo25 | protein_coding | 0.277181031 | 6.579606532 | 0.5180577 | 0.4818605 | 0.999175987 |
| ENSMUSG00000026880 | Stom | protein_coding | 0.746057071 | 2.608445582 | 0.5178551 | 0.4819449 | 0.999175987 |
| ENSMUSG00000022525 | Hrasls | protein_coding | 0.289529949 | 6.23717462 | 0.5177299 | 0.481997 | 0.999175987 |
| ENSMUSG00000021361 | Tmem14c | protein_coding | 0.256406958 | 6.955817858 | 0.5176781 | 0.4820186 | 0.999175987 |
| ENSMUSG00000097638 | Carlr | lincRNA | 1.077046039 | 2.306244157 | 0.5172898 | 0.4821805 | 0.999175987 |
| ENSMUSG00000040365 | Trim41 | protein_coding | -0.494222562 | 3.930646536 | 0.5163363 | 0.4825782 | 0.999175987 |
| ENSMUSG00000020142 | Slc1a4 | protein_coding | 0.37124685 | 4.882143168 | 0.5161181 | 0.4826693 | 0.999175987 |
| ENSMUSG00000024212 | Mllt1 | protein_coding | -0.444753472 | 4.066042031 | 0.5160235 | 0.4827089 | 0.999175987 |
| ENSMUSG00000028333 | Anp32b | protein_coding | -0.333583705 | 5.070327518 | 0.5159843 | 0.4827252 | 0.999175987 |
| ENSMUSG00000097102 | 2310069G16Rik | lincRNA | 0.656909793 | 3.187040995 | 0.5159176 | 0.4827531 | 0.999175987 |
| ENSMUSG00000030519 | Apba2 | protein_coding | 0.350926074 | 5.802000429 | 0.5159187 | 0.4827693 | 0.999175987 |
| ENSMUSG00000092310 | Gm20509 | protein_coding | -0.522589154 | 3.420100164 | 0.5144841 | 0.4833525 | 0.999175987 |
| ENSMUSG00000030757 | Zkscan2 | protein_coding | -0.753308 | 2.533081149 | 0.5139941 | 0.4835577 | 0.999175987 |
| ENSMUSG00000025898 | Cwf19l2 | protein_coding | -0.408178053 | 4.25217616 | 0.5136437 | 0.4837046 | 0.999175987 |
| ENSMUSG00000026885 | Ttll11 | protein_coding | 0.399211314 | 4.24246534 | 0.5132599 | 0.4838655 | 0.999175987 |
| ENSMUSG00000021474 | Sfxn1 | protein_coding | 0.203063708 | 7.44795253 | 0.5132428 | 0.4838727 | 0.999175987 |
| ENSMUSG00000029752 | Asns | protein_coding | -0.375926481 | 8.118845226 | 0.5157552 | 0.4839031 | 0.999175987 |
| ENSMUSG00000025478 | Dpysl4 | protein_coding | 0.372198652 | 5.052584115 | 0.5131121 | 0.4839275 | 0.999175987 |
| ENSMUSG00000039983 | Ccdc32 | protein_coding | 0.217223573 | 7.137188915 | 0.5125556 | 0.4841611 | 0.999175987 |
| ENSMUSG00000020840 | Blmh | protein_coding | 0.229395061 | 7.288699791 | 0.512218 | 0.4843028 | 0.999175987 |
| ENSMUSG00000031980 | Agt | protein_coding | -0.28515625 | 6.828260512 | 0.5120808 | 0.4843605 | 0.999175987 |
| ENSMUSG00000094786 | Gm14403 | protein_coding | 0.509678931 | 3.592279995 | 0.5119007 | 0.4844362 | 0.999175987 |
| ENSMUSG00000023873 | 1700010I14Rik | protein_coding | 0.81710221 | 2.972098165 | 0.5119271 | 0.4844416 | 0.999175987 |
| ENSMUSG00000037262 | Kin | protein_coding | -0.361214438 | 4.991138795 | 0.511697 | 0.4845217 | 0.999175987 |
| ENSMUSG00000033684 | Qsox1 | protein_coding | -0.381428523 | 5.093274121 | 0.5116835 | 0.4845274 | 0.999175987 |
| ENSMUSG00000071037 | Camkmt | protein_coding | -0.427207212 | 4.431726802 | 0.511462 | 0.4846206 | 0.999175987 |

| ensembl_gene_id | external_gene_name | gene_biotype | log2 (fold change) | log2 (counts per million) | F-value | p-value | False discovery rate |
|---|---|---|---|---|---|---|---|
| ENSMUSG00000019808 | Adat2 | protein_coding | -0.582206471 | 2.972440111 | 0.5112759 | 0.4846988 | 0.999175987 |
| ENSMUSG00000029178 | Klf3 | protein_coding | -0.492477681 | 3.876269742 | 0.5109905 | 0.4848189 | 0.999175987 |
| ENSMUSG00000025979 | Mob4 | protein_coding | -0.217772957 | 7.801513554 | 0.510557 | 0.4850014 | 0.999175987 |
| ENSMUSG00000061689 | Dlgap4 | protein_coding | 0.243944089 | 6.687834921 | 0.5102684 | 0.485123 | 0.999175987 |
| ENSMUSG00000041147 | Brca2 | protein_coding | -0.562482114 | 3.318215813 | 0.5102246 | 0.4851414 | 0.999175987 |
| ENSMUSG00000071645 | Tut1 | protein_coding | 0.349681926 | 5.181577807 | 0.5100895 | 0.4851983 | 0.999175987 |
| ENSMUSG00000026516 | Nvl | protein_coding | -0.348876074 | 5.279776214 | 0.5100133 | 0.4852304 | 0.999175987 |
| ENSMUSG00000029640 | Usp12 | protein_coding | 0.597924111 | 3.012577258 | 0.5099682 | 0.4852494 | 0.999175987 |
| ENSMUSG00000025162 | Csnk1d | protein_coding | 0.264314889 | 6.384389758 | 0.5098391 | 0.4853039 | 0.999175987 |
| ENSMUSG00000040532 | Abhd11 | protein_coding | 0.239843525 | 6.511618318 | 0.5095494 | 0.485426 | 0.999175987 |
| ENSMUSG00000022160 | Mettl3 | protein_coding | -0.486376641 | 3.93605508 | 0.5084792 | 0.4858777 | 0.999175987 |
| ENSMUSG00000033355 | Rtp4 | protein_coding | 0.581921466 | 3.126391149 | 0.5083195 | 0.4859451 | 0.999175987 |
| ENSMUSG00000029126 | Nsg1 | protein_coding | 0.217613072 | 9.756698452 | 0.5082059 | 0.4859931 | 0.999175987 |
| ENSMUSG00000042472 | Zfp410 | protein_coding | -0.323151689 | 5.151333987 | 0.507653 | 0.4862269 | 0.999175987 |
| ENSMUSG00000041035 | Gm17018 | protein_coding | 0.209901451 | 8.814747378 | 0.5076518 | 0.4862274 | 0.999175987 |
| ENSMUSG00000029439 | Sfswap | protein_coding | -0.523066593 | 3.823565253 | 0.507486 | 0.4862975 | 0.999175987 |
| ENSMUSG00000034768 | Asb16 | protein_coding | 0.592520285 | 2.949665346 | 0.5073046 | 0.4863743 | 0.999175987 |
| ENSMUSG00000051223 | Bzw1 | protein_coding | -0.22502432 | 7.448089155 | 0.506936 | 0.4865303 | 0.999175987 |
| ENSMUSG00000025857 | Dnaaf5 | protein_coding | -1.763174825 | 1.77534841 | 0.5083643 | 0.4865658 | 0.999175987 |
| ENSMUSG00000027680 | Fxr1 | protein_coding | -0.361709473 | 5.052254019 | 0.5067774 | 0.4865974 | 0.999175987 |
| ENSMUSG00000033845 | Mrpl15 | protein_coding | 0.199380203 | 7.412366499 | 0.5063024 | 0.4867987 | 0.999175987 |
| ENSMUSG00000038122 | Tbc1d32 | protein_coding | 0.687171589 | 2.883341405 | 0.5061663 | 0.4868563 | 0.999175987 |
| ENSMUSG00000049957 | Ccdc137 | protein_coding | 0.315843639 | 4.829238784 | 0.5060891 | 0.4868891 | 0.999175987 |
| ENSMUSG00000071265 | 1700086L19Rik | transcribed_unitary_p | 0.273743372 | 5.636011337 | 0.5057902 | 0.4870158 | 0.999175987 |
| ENSMUSG00000042439 | Zfp532 | protein_coding | -0.325499407 | 4.906140566 | 0.5055072 | 0.4871358 | 0.999175987 |
| ENSMUSG00000029070 | Mxra8 | protein_coding | -1.305864409 | 3.850094941 | 0.5195196 | 0.4872141 | 0.999175987 |
| ENSMUSG00000037492 | Zmat4 | protein_coding | -0.383207411 | 5.848493678 | 0.5053038 | 0.4872969 | 0.999175987 |
| ENSMUSG00000029246 | Ppat | protein_coding | 0.55249363 | 4.337523247 | 0.5058366 | 0.4873086 | 0.999175987 |
| ENSMUSG00000062397 | Zfp706 | protein_coding | 0.226571232 | 8.195450523 | 0.5047192 | 0.4874704 | 0.999175987 |
| ENSMUSG00000096472 | Cdkn2d | protein_coding | 0.198078773 | 8.239729479 | 0.5046078 | 0.4875177 | 0.999175987 |
| ENSMUSG00000028416 | Bag1 | protein_coding | 0.248515909 | 6.204112354 | 0.5045699 | 0.4875338 | 0.999175987 |
| ENSMUSG00000009630 | Ppp2cb | protein_coding | -0.256456681 | 6.220185394 | 0.5044758 | 0.4875738 | 0.999175987 |
| ENSMUSG00000036057 | Ptpn23 | protein_coding | 0.469360758 | 3.875969448 | 0.5044017 | 0.4876053 | 0.999175987 |
| ENSMUSG00000013465 | Nelfb | protein_coding | 0.240869396 | 5.997050314 | 0.5040371 | 0.4877603 | 0.999175987 |
| ENSMUSG00000021929 | Kpna3 | protein_coding | 0.24031694 | 6.483431306 | 0.5037797 | 0.4878698 | 0.999175987 |
| ENSMUSG00000019966 | Kitl | protein_coding | -0.461937965 | 4.71695056 | 0.5035562 | 0.4879813 | 0.999175987 |
| ENSMUSG00000050022 | Amz1 | protein_coding | 0.362053182 | 4.858056728 | 0.5032948 | 0.4880761 | 0.999175987 |
| ENSMUSG00000058589 | Anks1b | protein_coding | -0.191464271 | 7.130867999 | 0.5027463 | 0.4883097 | 0.999175987 |
| ENSMUSG00000024302 | Dtna | protein_coding | -0.233271377 | 6.816373177 | 0.5026854 | 0.4883356 | 0.999175987 |
| ENSMUSG00000045257 | Morn2 | protein_coding | 0.252624707 | 5.872086206 | 0.5017557 | 0.4887321 | 0.999175987 |
| ENSMUSG00000031169 | Porcn | protein_coding | -0.375493933 | 5.106679883 | 0.5011849 | 0.4889757 | 0.999175987 |
| ENSMUSG00000031778 | Cx3cl1 | protein_coding | -0.227935775 | 7.131790328 | 0.5006341 | 0.4892111 | 0.999175987 |
| ENSMUSG00000054733 | Msra | protein_coding | 0.289742277 | 5.175527977 | 0.5005954 | 0.4892276 | 0.999175987 |
| ENSMUSG00000026312 | Cdh7 | protein_coding | -0.367941648 | 4.282803202 | 0.5004456 | 0.4892917 | 0.999175987 |
| ENSMUSG00000071337 | Tia1 | protein_coding | -0.464398313 | 3.8261691 | 0.5001511 | 0.4894176 | 0.999175987 |
| ENSMUSG00000056214 | Pard6g | protein_coding | 0.587055204 | 3.154507109 | 0.4999196 | 0.4895166 | 0.999175987 |
| ENSMUSG00000001053 | N4bp3 | protein_coding | 0.778782416 | 2.452162379 | 0.4995524 | 0.4896738 | 0.999175987 |
| ENSMUSG00000058587 | Tmod3 | protein_coding | 0.230797816 | 6.482394661 | 0.4993633 | 0.4897548 | 0.999175987 |
| ENSMUSG00000040612 | Ildr2 | protein_coding | -0.341086176 | 5.880273793 | 0.4993197 | 0.4897898 | 0.999175987 |
| ENSMUSG00000086629 | 2810403D21Rik | processed_transcript | -0.529799666 | 3.35964998 | 0.4990146 | 0.4899041 | 0.999175987 |
| ENSMUSG00000021991 | Cacna2d3 | protein_coding | -0.389998696 | 5.559323087 | 0.4987676 | 0.4900263 | 0.999175987 |
| ENSMUSG00000075705 | Msrb1 | protein_coding | 0.317842057 | 4.944493655 | 0.4985048 | 0.4901227 | 0.999175987 |
| ENSMUSG00000026986 | Hnmt | protein_coding | -0.215580353 | 6.539923914 | 0.4980614 | 0.4903128 | 0.999175987 |
| ENSMUSG00000035078 | Mtmr9 | protein_coding | -0.28745077 | 5.870127254 | 0.4980281 | 0.4903271 | 0.999175987 |
| ENSMUSG00000038215 | Cep44 | protein_coding | -0.650618755 | 3.199055763 | 0.4980202 | 0.4903305 | 0.999175987 |
| ENSMUSG00000020121 | Srgap1 | protein_coding | -0.469755961 | 4.487032466 | 0.4976528 | 0.4905045 | 0.999175987 |
| ENSMUSG00000054814 | Usp46 | protein_coding | 0.292069526 | 6.444932736 | 0.4975128 | 0.4905483 | 0.999175987 |
| ENSMUSG00000030727 | Rabep2 | protein_coding | -0.531780443 | 3.876506452 | 0.4973873 | 0.4906178 | 0.999175987 |
| ENSMUSG00000039218 | Srrm2 | protein_coding | -0.281232222 | 6.124599516 | 0.497211 | 0.490678 | 0.999175987 |
| ENSMUSG00000038260 | Trpm4 | protein_coding | 0.367620081 | 4.247393398 | 0.4969314 | 0.4907981 | 0.999175987 |
| ENSMUSG00000032481 | Smarcc1 | protein_coding | 0.2898876 | 5.557874425 | 0.4968376 | 0.4908384 | 0.999175987 |
| ENSMUSG00000030609 | Aen | protein_coding | 0.460777831 | 4.770893489 | 0.4968286 | 0.4908586 | 0.999175987 |
| ENSMUSG00000045625 | Pigz | protein_coding | 0.343211079 | 4.635725031 | 0.4967792 | 0.4908635 | 0.999175987 |
| ENSMUSG00000092607 | Scnm1 | protein_coding | 0.221930645 | 7.329508744 | 0.4967512 | 0.4908756 | 0.999175987 |
| ENSMUSG00000020457 | Drg1 | protein_coding | -0.212583458 | 8.483408787 | 0.496619 | 0.4909324 | 0.999175987 |
| ENSMUSG00000036572 | Upf3b | protein_coding | -0.287625187 | 5.476546156 | 0.4963209 | 0.4910607 | 0.999175987 |
| ENSMUSG00000031754 | Nudt21 | protein_coding | -0.375744727 | 3.847981517 | 0.4962783 | 0.491079 | 0.999175987 |
| ENSMUSG00000020134 | Peli1 | protein_coding | -0.344022584 | 5.027350715 | 0.4956565 | 0.4913466 | 0.999175987 |
| ENSMUSG00000038970 | Lmtk2 | protein_coding | -0.396306571 | 4.285647152 | 0.495633 | 0.4913568 | 0.999175987 |
| ENSMUSG00000021236 | Entpd5 | protein_coding | -0.371990179 | 4.612089594 | 0.4955045 | 0.4914121 | 0.999175987 |
| ENSMUSG00000057230 | Aak1 | protein_coding | -0.221651047 | 6.234859702 | 0.4953436 | 0.4914814 | 0.999175987 |
| ENSMUSG00000036646 | Man1b1 | protein_coding | 0.297126121 | 5.165876527 | 0.4950837 | 0.4915934 | 0.999175987 |
| ENSMUSG00000049090 | Zadh2 | protein_coding | -0.415870915 | 3.667105702 | 0.4948549 | 0.4916921 | 0.999175987 |
| ENSMUSG00000028578 | Caap1 | protein_coding | -0.849097008 | 2.471576403 | 0.4948374 | 0.4916996 | 0.999175987 |
| ENSMUSG00000013419 | Zfp651 | protein_coding | -0.417275847 | 4.179712255 | 0.4942392 | 0.4919577 | 0.999175987 |
| ENSMUSG00000025008 | Tctn3 | protein_coding | -0.768543632 | 3.740737521 | 0.4969745 | 0.4920646 | 0.999175987 |
| ENSMUSG00000070394 | Tmem256 | protein_coding | 0.289588971 | 8.328333018 | 0.4939952 | 0.4920792 | 0.999175987 |
| ENSMUSG00000031565 | Fgfr1 | protein_coding | -0.290647984 | 5.45452829 | 0.493915 | 0.4920976 | 0.999175987 |

| ensembl_gene_id | external_gene_name | gene_biotype | log2 (fold change) | log2 (counts per million) | F-value | p-value | False discovery rate |
|---|---|---|---|---|---|---|---|
| ENSMUSG00000038845 | Phb | protein_coding | 0.286257137 | 6.715993829 | 0.4939224 | 0.4921107 | 0.999175987 |
| ENSMUSG00000000627 | Sema4f | protein_coding | 0.422194171 | 4.304261615 | 0.4934356 | 0.4923047 | 0.999175987 |
| ENSMUSG00000022305 | Lrp12 | protein_coding | 0.454944892 | 4.229442977 | 0.4931346 | 0.4924348 | 0.999175987 |
| ENSMUSG00000029345 | Tfip11 | protein_coding | 0.485220984 | 4.892480515 | 0.4938742 | 0.4924443 | 0.999175987 |
| ENSMUSG00000020522 | Mfap3 | protein_coding | 0.367897636 | 4.986215384 | 0.4930675 | 0.4924638 | 0.999175987 |
| ENSMUSG00000039738 | Slx4 | protein_coding | 0.630118084 | 2.92523349 | 0.4929292 | 0.4925236 | 0.999175987 |
| ENSMUSG00000032816 | Igdcc4 | protein_coding | 0.56748826 | 3.276347996 | 0.492819 | 0.4925713 | 0.999175987 |
| ENSMUSG00000029777 | Gars | protein_coding | -0.243852592 | 6.473768453 | 0.492541 | 0.4926916 | 0.999175987 |
| ENSMUSG00000001473 | Tubb6 | protein_coding | -0.695399852 | 2.699897975 | 0.4924635 | 0.4927251 | 0.999175987 |
| ENSMUSG00000037493 | Cib2 | protein_coding | 0.297070202 | 5.405799481 | 0.4924428 | 0.4927341 | 0.999175987 |
| ENSMUSG00000002014 | Ssr4 | protein_coding | 0.269425519 | 7.614212691 | 0.4923187 | 0.4927878 | 0.999175987 |
| ENSMUSG00000059248 | 9-Sep | protein_coding | 0.508578216 | 5.15528298 | 0.4940065 | 0.4928675 | 0.999175987 |
| ENSMUSG00000038400 | Pmepa1 | protein_coding | -0.366966365 | 4.561801788 | 0.4921272 | 0.4928707 | 0.999175987 |
| ENSMUSG00000026463 | Atp2b4 | protein_coding | -0.260326893 | 6.025280533 | 0.4913893 | 0.4931904 | 0.999175987 |
| ENSMUSG00000029279 | Brdt | protein_coding | 0.483216422 | 3.734904963 | 0.4912624 | 0.4932454 | 0.999175987 |
| ENSMUSG00000028140 | Mrpl9 | protein_coding | -0.232621805 | 6.042274355 | 0.4911748 | 0.4932834 | 0.999175987 |
| ENSMUSG00000025408 | Ddit3 | protein_coding | -0.257957303 | 6.742839347 | 0.4908131 | 0.4934403 | 0.999175987 |
| ENSMUSG00000053411 | Cbx7 | protein_coding | -0.301086327 | 5.132280649 | 0.490527 | 0.4935644 | 0.999175987 |
| ENSMUSG00000000409 | Lck | protein_coding | 0.554046127 | 3.777898361 | 0.490442 | 0.4936175 | 0.999175987 |
| ENSMUSG00000047879 | Usp14 | protein_coding | -0.254760876 | 7.679212796 | 0.4902671 | 0.4936772 | 0.999175987 |
| ENSMUSG00000024101 | Washc1 | protein_coding | 0.305853735 | 5.043493892 | 0.4901402 | 0.4937323 | 0.999175987 |
| ENSMUSG00000020895 | Tmem107 | protein_coding | 0.287589155 | 5.722713389 | 0.4901348 | 0.4937347 | 0.999175987 |
| ENSMUSG00000051951 | Xkr4 | protein_coding | -0.583100048 | 3.610802665 | 0.4900545 | 0.4937857 | 0.999175987 |
| ENSMUSG00000031922 | Cep57 | protein_coding | -0.281525705 | 5.132438851 | 0.4900133 | 0.4937875 | 0.999175987 |
| ENSMUSG00000091864 | Gm17102 | lincRNA | -0.567399283 | 3.367061646 | 0.4897678 | 0.4938941 | 0.999175987 |
| ENSMUSG00000040972 | Igsf21 | protein_coding | -0.565541189 | 4.150300322 | 0.4904504 | 0.4939032 | 0.999175987 |
| ENSMUSG00000042595 | Fam199x | protein_coding | -0.480004332 | 3.526595474 | 0.4897338 | 0.4939089 | 0.999175987 |
| ENSMUSG00000048234 | Rnf149 | protein_coding | -0.634417182 | 3.818176469 | 0.4905026 | 0.4940261 | 0.999175987 |
| ENSMUSG00000015733 | Capza2 | protein_coding | -0.205788142 | 9.262856819 | 0.4893954 | 0.494056 | 0.999175987 |
| ENSMUSG00000027551 | Zfp64 | protein_coding | 0.38368846 | 5.124007071 | 0.4893781 | 0.4940635 | 0.999175987 |
| ENSMUSG00000020108 | Ddit4 | protein_coding | -0.576600841 | 3.351009333 | 0.4893454 | 0.4940777 | 0.999175987 |
| ENSMUSG00000020436 | Gabrg2 | protein_coding | 0.388945623 | 3.993808384 | 0.4893389 | 0.4940806 | 0.999175987 |
| ENSMUSG00000003355 | Fkbp11 | protein_coding | -0.553483017 | 3.242435879 | 0.4886103 | 0.4943976 | 0.999175987 |
| ENSMUSG00000032562 | Gnai2 | protein_coding | 0.225627575 | 7.278666785 | 0.4884628 | 0.4944618 | 0.999175987 |
| ENSMUSG00000057778 | Cyb5d2 | protein_coding | 0.403775373 | 4.542652472 | 0.4871131 | 0.4950501 | 0.999175987 |
| ENSMUSG00000029068 | Ccnl2 | protein_coding | 0.410879484 | 3.8551514 | 0.4868781 | 0.4951527 | 0.999175987 |
| ENSMUSG00000019699 | Akt3 | protein_coding | 0.311884335 | 5.479204504 | 0.4862538 | 0.4954253 | 0.999175987 |
| ENSMUSG00000024132 | Eci1 | protein_coding | 0.41421407 | 5.003862725 | 0.4861165 | 0.4955014 | 0.999175987 |
| ENSMUSG00000002326 | Gmpr2 | protein_coding | -0.32786698 | 5.289476233 | 0.4860333 | 0.4955217 | 0.999175987 |
| ENSMUSG00000064367 | mt-Nd5 | protein_coding | 0.335361116 | 9.814911841 | 0.487026 | 0.4955405 | 0.999175987 |
| ENSMUSG00000026705 | Klhl20 | protein_coding | -0.371023923 | 3.967137319 | 0.4858837 | 0.4955871 | 0.999175987 |
| ENSMUSG00000079330 | Lemd1 | protein_coding | -0.518657372 | 3.620997287 | 0.4851789 | 0.4958954 | 0.999175987 |
| ENSMUSG00000024006 | Stk38 | protein_coding | -0.437741683 | 3.52532164 | 0.4851303 | 0.4959166 | 0.999175987 |
| ENSMUSG00000002733 | Plekha3 | protein_coding | 0.343404461 | 4.711575466 | 0.4848565 | 0.4960365 | 0.999175987 |
| ENSMUSG00000020536 | Llgl1 | protein_coding | 0.349022943 | 4.550861631 | 0.4843844 | 0.4962433 | 0.999175987 |
| ENSMUSG00000038495 | Otud7b | protein_coding | -0.272507078 | 5.380455749 | 0.4839869 | 0.4964175 | 0.999175987 |
| ENSMUSG00000008153 | Clstn3 | protein_coding | 0.234642753 | 8.246138266 | 0.4830581 | 0.496825 | 0.999175987 |
| ENSMUSG00000066735 | Vkorc1l1 | protein_coding | -0.335331316 | 4.973310593 | 0.4829778 | 0.4968603 | 0.999175987 |
| ENSMUSG00000037416 | Dmxl1 | protein_coding | -0.468616238 | 3.915605645 | 0.4829742 | 0.4968619 | 0.999175987 |
| ENSMUSG00000051674 | Dcun1d4 | protein_coding | -0.261313667 | 6.531076334 | 0.4828556 | 0.496914 | 0.999175987 |
| ENSMUSG00000095139 | Pou3f2 | protein_coding | 0.321911739 | 4.942613641 | 0.4828539 | 0.4969147 | 0.999175987 |
| ENSMUSG00000027678 | Ncoa3 | protein_coding | 0.407264999 | 4.86053662 | 0.4819934 | 0.4972929 | 0.999175987 |
| ENSMUSG00000022635 | Zcrb1 | protein_coding | -0.315091333 | 5.522701848 | 0.4814464 | 0.4975336 | 0.999175987 |
| ENSMUSG00000014980 | Tsen15 | protein_coding | -0.308815592 | 4.869317829 | 0.4814459 | 0.4975338 | 0.999175987 |
| ENSMUSG00000086040 | Wipf3 | protein_coding | -0.677642332 | 2.898000175 | 0.4813988 | 0.4975545 | 0.999175987 |
| ENSMUSG00000048787 | Dcun1d3 | protein_coding | -0.440784343 | 3.568965369 | 0.4813507 | 0.4975757 | 0.999175987 |
| ENSMUSG00000021785 | Ngly1 | protein_coding | -0.383300915 | 4.401940673 | 0.4812753 | 0.4976089 | 0.999175987 |
| ENSMUSG00000036372 | Tmem258 | protein_coding | 0.324522664 | 7.304822578 | 0.4819216 | 0.4979448 | 0.999175987 |
| ENSMUSG00000017615 | Tnfaip1 | protein_coding | -0.291937235 | 6.625073356 | 0.4803991 | 0.498011 | 0.999175987 |
| ENSMUSG00000024873 | Cnih2 | protein_coding | 0.372900202 | 5.668024471 | 0.4803346 | 0.4980394 | 0.999175987 |
| ENSMUSG00000032388 | Spg21 | protein_coding | -0.310681507 | 5.239548365 | 0.4799471 | 0.4981943 | 0.999175987 |
| ENSMUSG00000023089 | Ndufa5 | protein_coding | 0.313672225 | 10.15552811 | 0.4797078 | 0.4983579 | 0.999175987 |
| ENSMUSG00000035829 | Ppp1r26 | protein_coding | 0.444712646 | 3.6414229 | 0.4795163 | 0.4983845 | 0.999175987 |
| ENSMUSG00000030000 | Add2 | protein_coding | 0.232770268 | 7.219291826 | 0.4795074 | 0.4983884 | 0.999175987 |
| ENSMUSG00000002477 | Snrpd1 | protein_coding | 0.258572835 | 7.826227714 | 0.4794242 | 0.4984251 | 0.999175987 |
| ENSMUSG00000024947 | Men1 | protein_coding | 0.313098473 | 5.013387401 | 0.479252 | 0.4985012 | 0.999175987 |
| ENSMUSG00000022707 | Gbe1 | protein_coding | -0.498998751 | 3.4421343 | 0.479031 | 0.4985988 | 0.999175987 |
| ENSMUSG00000068039 | Tcp1 | protein_coding | 0.236136812 | 8.356202542 | 0.478831 | 0.4986872 | 0.999175987 |
| ENSMUSG00000016541 | Atxn10 | protein_coding | 0.255198354 | 9.478745114 | 0.4788025 | 0.4986998 | 0.999175987 |
| ENSMUSG00000032846 | Zswim6 | processed_transcript | -0.59510798 | 3.842680476 | 0.4788796 | 0.4987512 | 0.999175987 |
| ENSMUSG00000059336 | Slc14a1 | protein_coding | -0.763297504 | 2.821442669 | 0.478359 | 0.4988959 | 0.999175987 |
| ENSMUSG00000026437 | Cdk18 | protein_coding | -0.405175262 | 5.932591511 | 0.4797015 | 0.4989213 | 0.999175987 |
| ENSMUSG00000031242 | 2610002M06Rik | protein_coding | -0.253299004 | 5.652742903 | 0.4779763 | 0.4990652 | 0.999175987 |
| ENSMUSG00000025475 | Adgra1 | protein_coding | 0.29017726 | 6.688221718 | 0.4780089 | 0.4990667 | 0.999175987 |
| ENSMUSG00000063253 | Scoc | protein_coding | -0.216396682 | 7.516493739 | 0.4777683 | 0.4991573 | 0.999175987 |
| ENSMUSG00000015222 | Map2 | protein_coding | -0.198518591 | 8.291809547 | 0.4776546 | 0.4992076 | 0.999175987 |
| ENSMUSG00000028973 | Abcb8 | protein_coding | -0.250203706 | 6.129467242 | 0.4773383 | 0.4993477 | 0.999175987 |
| ENSMUSG00000030087 | Klf15 | protein_coding | -0.509258068 | 3.61185396 | 0.4772809 | 0.4993731 | 0.999175987 |

| ensembl_gene_id | external_gene_name | gene_biotype | log2 (fold change) | log2 (counts per million) | F-value | p-value | False discovery rate |
|---|---|---|---|---|---|---|---|
| ENSMUSG00000024587 | Nars | protein_coding | -0.248351253 | 8.572052115 | 0.4771595 | 0.4994269 | 0.999175987 |
| ENSMUSG00000044533 | Rps2 | protein_coding | 0.383304041 | 4.659442267 | 0.4771363 | 0.4994372 | 0.999175987 |
| ENSMUSG00000053581 | Zfand2a | protein_coding | 0.328862725 | 6.114160358 | 0.4770707 | 0.4994822 | 0.999175987 |
| ENSMUSG00000071172 | Srsf3 | protein_coding | -0.215889807 | 7.133198226 | 0.4769919 | 0.4995012 | 0.999175987 |
| ENSMUSG00000049532 | Sall2 | protein_coding | -0.267044109 | 5.735768004 | 0.4767919 | 0.4995898 | 0.999175987 |
| ENSMUSG00000030815 | Phkg2 | protein_coding | -0.288001771 | 5.202607546 | 0.4761905 | 0.4998566 | 0.999175987 |
| ENSMUSG00000048562 | Sp8 | protein_coding | -0.397663159 | 5.007978714 | 0.4761033 | 0.4998953 | 0.999175987 |
| ENSMUSG00000028158 | Mttp | protein_coding | 0.477156918 | 3.913784523 | 0.4760365 | 0.499925 | 0.999175987 |
| ENSMUSG00000029596 | Sdsl | protein_coding | 0.6984867 | 2.705739573 | 0.4760016 | 0.4999405 | 0.999175987 |
| ENSMUSG00000039220 | Ppp1r10 | protein_coding | -0.36880088 | 5.239057029 | 0.4755506 | 0.5001408 | 0.999175987 |
| ENSMUSG00000028610 | Dmrtb1 | protein_coding | -0.83045994 | 3.322500982 | 0.4776849 | 0.5001613 | 0.999175987 |
| ENSMUSG00000061477 | Rps7 | protein_coding | 0.294665088 | 5.938776465 | 0.4746049 | 0.5005612 | 0.999175987 |
| ENSMUSG00000028184 | Adgrl2 | protein_coding | -0.334587582 | 4.233915542 | 0.4745884 | 0.5005686 | 0.999175987 |
| ENSMUSG00000059439 | Bcas3 | protein_coding | -0.422186541 | 4.684346565 | 0.4745465 | 0.5005872 | 0.999175987 |
| ENSMUSG00000049717 | Lig4 | protein_coding | 0.635680788 | 3.23115119 | 0.473894 | 0.5008777 | 0.999175987 |
| ENSMUSG00000039542 | Ncam1 | protein_coding | -0.240851353 | 8.750398292 | 0.4737206 | 0.5009549 | 0.999175987 |
| ENSMUSG00000041840 | Haus1 | protein_coding | -0.23494807 | 5.787045799 | 0.4734924 | 0.5010566 | 0.999175987 |
| ENSMUSG00000021282 | Eif5 | protein_coding | -0.20266306 | 6.971456499 | 0.4734106 | 0.5010931 | 0.999175987 |
| ENSMUSG00000017144 | Rnd3 | protein_coding | -0.344059136 | 5.151207108 | 0.4732235 | 0.5011765 | 0.999175987 |
| ENSMUSG00000054021 | Sirt5 | protein_coding | 0.331463 | 5.581060247 | 0.4730137 | 0.50127 | 0.999175987 |
| ENSMUSG00000025200 | Cwf19l1 | protein_coding | 0.35552367 | 4.413403784 | 0.472679 | 0.5014194 | 0.999175987 |
| ENSMUSG00000028568 | Btf3l4 | protein_coding | 0.312189937 | 4.661104131 | 0.4723235 | 0.501578 | 0.999175987 |
| ENSMUSG00000027933 | Ints3 | protein_coding | 0.339093667 | 4.900079861 | 0.4721414 | 0.5016594 | 0.999175987 |
| ENSMUSG00000029202 | Pds5a | protein_coding | 0.452858966 | 3.485613207 | 0.4720581 | 0.5016966 | 0.999175987 |
| ENSMUSG00000027067 | Ssrp1 | protein_coding | -0.237936465 | 6.496944194 | 0.4718486 | 0.5017902 | 0.999175987 |
| ENSMUSG00000013663 | Pten | protein_coding | 0.30821892 | 5.377594495 | 0.4714747 | 0.5019573 | 0.999175987 |
| ENSMUSG00000019775 | Rgs17 | protein_coding | -0.227275507 | 7.274097657 | 0.4710928 | 0.5021281 | 0.999175987 |
| ENSMUSG00000033732 | Sf3b3 | protein_coding | 0.298071675 | 5.440594887 | 0.4704907 | 0.5023976 | 0.999175987 |
| ENSMUSG00000038305 | Spats2l | protein_coding | -0.403622996 | 4.865048663 | 0.4700266 | 0.5026055 | 0.999175987 |
| ENSMUSG00000064293 | Cntn4 | protein_coding | -0.36300114 | 4.833247032 | 0.4699753 | 0.5026285 | 0.999175987 |
| ENSMUSG00000021270 | Hsp90aa1 | protein_coding | -0.209890005 | 7.648787366 | 0.469824 | 0.5026963 | 0.999175987 |
| ENSMUSG00000036686 | Cc2d1a | protein_coding | 0.410394355 | 4.848330308 | 0.469599 | 0.5027972 | 0.999175987 |
| ENSMUSG00000016487 | Ppfibp1 | protein_coding | -0.543924906 | 3.142430266 | 0.469519 | 0.502833 | 0.999175987 |
| ENSMUSG00000072623 | Zfp9 | protein_coding | -0.510396534 | 4.1590308 | 0.4691649 | 0.5030077 | 0.999175987 |
| ENSMUSG00000058488 | Kl | protein_coding | -1.270848916 | 3.713946412 | 0.4798833 | 0.5031492 | 0.999175987 |
| ENSMUSG00000061411 | Nol4l | protein_coding | -0.35617074 | 4.668333996 | 0.4684459 | 0.5033147 | 0.999175987 |
| ENSMUSG00000028309 | Rnf20 | protein_coding | -0.283836405 | 5.622701641 | 0.4680192 | 0.5035065 | 0.999175987 |
| ENSMUSG00000086075 | Gm15728 | lincRNA | 0.558547674 | 3.171824099 | 0.4676782 | 0.5036599 | 0.999175987 |
| ENSMUSG00000014355 | Anapc1 | protein_coding | 0.415363161 | 4.217643733 | 0.4676711 | 0.503663 | 0.999175987 |
| ENSMUSG00000027574 | Nkain4 | protein_coding | 0.426870933 | 6.610725985 | 0.4710331 | 0.5036871 | 0.999175987 |
| ENSMUSG00000031711 | Zfp330 | protein_coding | -0.227987589 | 5.953053569 | 0.4673095 | 0.5038257 | 0.999175987 |
| ENSMUSG00000022306 | Zfpm2 | protein_coding | 0.78610752 | 2.911789258 | 0.4673043 | 0.5038438 | 0.999175987 |
| ENSMUSG00000038764 | Ptpn3 | protein_coding | -0.610540323 | 4.327253601 | 0.4693658 | 0.5038591 | 0.999175987 |
| ENSMUSG00000019055 | Plod1 | protein_coding | 0.442732846 | 3.98051833 | 0.4671473 | 0.5038987 | 0.999175987 |
| ENSMUSG00000015289 | Lage3 | protein_coding | -0.297934825 | 5.787581657 | 0.4671424 | 0.5039009 | 0.999175987 |
| ENSMUSG00000026867 | Gapvd1 | protein_coding | -0.271788546 | 5.793790567 | 0.4670955 | 0.5039221 | 0.999175987 |
| ENSMUSG00000078427 | Sarnp | protein_coding | -0.248677673 | 6.551563415 | 0.4669088 | 0.5040061 | 0.999175987 |
| ENSMUSG00000036632 | Alg5 | protein_coding | -0.369865021 | 5.020217067 | 0.4668531 | 0.5040312 | 0.999175987 |
| ENSMUSG00000020087 | Tysnd1 | protein_coding | -0.883786205 | 2.574259294 | 0.4668516 | 0.5040319 | 0.999175987 |
| ENSMUSG00000063172 | Hspb11 | protein_coding | -0.2588305 | 5.895855498 | 0.4666168 | 0.5041376 | 0.999175987 |
| ENSMUSG00000055240 | Zfp101 | protein_coding | 0.596452521 | 3.021592044 | 0.4664987 | 0.5041909 | 0.999175987 |
| ENSMUSG00000026173 | Plcd4 | protein_coding | 0.475627911 | 3.898973072 | 0.4664309 | 0.5042214 | 0.999175987 |
| ENSMUSG00000018565 | Elp5 | protein_coding | 0.199777288 | 8.18573126 | 0.4663073 | 0.5042772 | 0.999175987 |
| ENSMUSG00000057894 | Zfp329 | protein_coding | -0.327529735 | 5.127169057 | 0.4661626 | 0.5043424 | 0.999175987 |
| ENSMUSG00000038085 | Cnbd2 | protein_coding | 0.485219495 | 3.309772347 | 0.4655673 | 0.5046109 | 0.999175987 |
| ENSMUSG00000020364 | Zfp354a | protein_coding | -0.546490448 | 3.351531826 | 0.4654199 | 0.5046774 | 0.999175987 |
| ENSMUSG00000049339 | Retreg2 | protein_coding | 0.233422065 | 6.910979518 | 0.4654158 | 0.5046793 | 0.999175987 |
| ENSMUSG00000047617 | Paxx | protein_coding | 0.202987244 | 7.046361588 | 0.465397 | 0.5046878 | 0.999175987 |
| ENSMUSG00000046671 | Mtfr1l | protein_coding | 0.205681898 | 7.657374727 | 0.4650496 | 0.5048446 | 0.999175987 |
| ENSMUSG00000034832 | Tet3 | protein_coding | 0.349091274 | 4.60386671 | 0.4648132 | 0.5049514 | 0.999175987 |
| ENSMUSG00000021685 | Otp | protein_coding | -0.687565977 | 2.728168457 | 0.4647294 | 0.5049893 | 0.999175987 |
| ENSMUSG00000020946 | Gosr2 | protein_coding | 0.249477173 | 6.890727503 | 0.464675 | 0.5050139 | 0.999175987 |
| ENSMUSG00000028089 | Chd1l | protein_coding | 0.584995053 | 3.109815379 | 0.4646423 | 0.5050287 | 0.999175987 |
| ENSMUSG00000004460 | Dnajb11 | protein_coding | 0.324811414 | 4.93697652 | 0.4644132 | 0.5051322 | 0.999175987 |
| ENSMUSG00000022814 | Umps | protein_coding | 0.343175477 | 5.473291853 | 0.4642427 | 0.5052093 | 0.999175987 |
| ENSMUSG00000059323 | Tonsl | protein_coding | 0.198208526 | 9.574008816 | 0.464188 | 0.5052341 | 0.999175987 |
| ENSMUSG00000039629 | Strip2 | protein_coding | -0.453334923 | 4.260741772 | 0.4637923 | 0.5054131 | 0.999175987 |
| ENSMUSG00000018379 | Srsf1 | protein_coding | -0.249867083 | 5.664583714 | 0.4637593 | 0.505428 | 0.999175987 |
| ENSMUSG00000090112 | Shprh | protein_coding | -0.406591907 | 4.266955647 | 0.4637245 | 0.5054438 | 0.999175987 |
| ENSMUSG00000027342 | Pcna | protein_coding | -0.276897621 | 5.227734481 | 0.4635402 | 0.5055272 | 0.999175987 |
| ENSMUSG00000068686 | Cd59b | protein_coding | 0.609331117 | 2.836200096 | 0.4635255 | 0.5055339 | 0.999175987 |
| ENSMUSG00000030720 | Cln3 | protein_coding | 0.466651924 | 4.129359323 | 0.4633675 | 0.5056054 | 0.999175987 |
| ENSMUSG00000006191 | Cdkal1 | protein_coding | -0.352370946 | 4.641648167 | 0.463077 | 0.505737 | 0.999175987 |
| ENSMUSG00000047710 | Champ1 | protein_coding | 0.366985977 | 4.904989309 | 0.4628728 | 0.5058296 | 0.999175987 |
| ENSMUSG00000026289 | Atg16l1 | protein_coding | 0.422255941 | 4.945084035 | 0.4621907 | 0.5061545 | 0.999175987 |
| ENSMUSG00000042303 | Sgsm3 | protein_coding | 0.312896414 | 6.572978962 | 0.4622548 | 0.5061666 | 0.999175987 |
| ENSMUSG00000022940 | Pigp | protein_coding | -0.299489289 | 5.775707716 | 0.4620754 | 0.5061912 | 0.999175987 |
| ENSMUSG00000062949 | Atp11c | protein_coding | 0.657724953 | 2.800095287 | 0.4617676 | 0.5063309 | 0.999175987 |

| ensembl_gene_id | external_gene_name | gene_biotype | log2 (fold change) | log2 (counts per million) | F-value | p-value | False discovery rate |
|---|---|---|---|---|---|---|---|
| ENSMUSG00000028896 | Rcc1 | protein_coding | 0.548260284 | 3.877977304 | 0.4616748 | 0.5063887 | 0.999175987 |
| ENSMUSG00000064345 | mt-Nd2 | protein_coding | 0.348761416 | 10.31322046 | 0.4627755 | 0.5064796 | 0.999175987 |
| ENSMUSG00000032878 | Ccdc85a | protein_coding | -0.279820713 | 5.289002769 | 0.4611234 | 0.5066236 | 0.999175987 |
| ENSMUSG00000020385 | Clk4 | protein_coding | 0.302507902 | 5.68838557 | 0.4609504 | 0.5067022 | 0.999175987 |
| ENSMUSG00000039988 | Ankrd13c | protein_coding | -0.248848365 | 5.982011039 | 0.4605847 | 0.5068685 | 0.999175987 |
| ENSMUSG00000021585 | Cast | protein_coding | -0.546305446 | 3.004113261 | 0.460546 | 0.5068861 | 0.999175987 |
| ENSMUSG00000044367 | Slc16a13 | protein_coding | 1.154750967 | 2.677996523 | 0.4621483 | 0.5071069 | 0.999175987 |
| ENSMUSG00000051067 | Lingo3 | protein_coding | -0.578164819 | 3.708246974 | 0.4600236 | 0.5071395 | 0.999175987 |
| ENSMUSG00000028484 | Psip1 | protein_coding | 0.318839255 | 4.976152638 | 0.45931 | 0.507449 | 0.999175987 |
| ENSMUSG00000017677 | Wsb1 | protein_coding | 0.418828779 | 4.565200691 | 0.459208 | 0.5074955 | 0.999175987 |
| ENSMUSG00000028760 | Eif4g3 | protein_coding | 0.282454487 | 5.792021111 | 0.4588145 | 0.5076749 | 0.999175987 |
| ENSMUSG00000014496 | Ankrd28 | protein_coding | -0.460505024 | 3.551917331 | 0.4588 | 0.5076816 | 0.999175987 |
| ENSMUSG00000114362 | AC124433.1 | antisense_RNA | -0.940617778 | 2.15005337 | 0.4586929 | 0.5077305 | 0.999175987 |
| ENSMUSG00000017288 | Vps53 | protein_coding | 0.307853144 | 6.081862636 | 0.4584993 | 0.5078188 | 0.999175987 |
| ENSMUSG00000022841 | Ap2m1 | protein_coding | 0.250794073 | 7.793836002 | 0.4583191 | 0.5079011 | 0.999175987 |
| ENSMUSG00000071691 | Gm960 | protein_coding | -0.914005515 | 2.546245427 | 0.457839 | 0.5081359 | 0.999175987 |
| ENSMUSG00000008393 | Carhsp1 | protein_coding | -0.230163033 | 6.747320037 | 0.4577687 | 0.5081525 | 0.999175987 |
| ENSMUSG00000039713 | Plekhg5 | protein_coding | 0.371181913 | 4.772621883 | 0.4576442 | 0.5082094 | 0.999175987 |
| ENSMUSG00000000753 | Serpinf1 | protein_coding | -0.453728474 | 4.379507595 | 0.4573393 | 0.5083643 | 0.999175987 |
| ENSMUSG00000027954 | Efna1 | protein_coding | 0.638672946 | 3.056885066 | 0.4572836 | 0.5083742 | 0.999175987 |
| ENSMUSG00000042305 | Tmem183a | protein_coding | 0.291878542 | 5.834486723 | 0.4569103 | 0.508545 | 0.999175987 |
| ENSMUSG00000051721 | Wdcp | protein_coding | -0.470887248 | 4.719304903 | 0.4568631 | 0.5086498 | 0.999175987 |
| ENSMUSG00000070880 | Gad1 | protein_coding | 0.214599261 | 8.867156089 | 0.4566269 | 0.5086747 | 0.999175987 |
| ENSMUSG00000048330 | Ric3 | protein_coding | 0.335323478 | 5.306662517 | 0.4565308 | 0.5087187 | 0.999175987 |
| ENSMUSG00000003421 | Nosip | protein_coding | 0.201928521 | 7.377059727 | 0.4563186 | 0.5088159 | 0.999175987 |
| ENSMUSG00000020962 | Gtf2a1 | protein_coding | -0.261048646 | 5.943670029 | 0.4562029 | 0.5088689 | 0.999175987 |
| ENSMUSG00000028799 | Zfp362 | protein_coding | -0.533417094 | 2.955070288 | 0.4560951 | 0.5089183 | 0.999175987 |
| ENSMUSG00000021466 | Ptch1 | protein_coding | 0.364483051 | 4.973170299 | 0.4558243 | 0.5090424 | 0.999175987 |
| ENSMUSG00000035413 | Tmem98 | protein_coding | 0.283771934 | 5.736134457 | 0.454767 | 0.5095275 | 0.999175987 |
| ENSMUSG00000027035 | Cers6 | protein_coding | 0.361817296 | 4.366479279 | 0.45462 | 0.509595 | 0.999175987 |
| ENSMUSG00000042607 | Asb4 | protein_coding | 0.481128145 | 4.45846655 | 0.4540869 | 0.5098554 | 0.999175987 |
| ENSMUSG00000040043 | Rbms2 | protein_coding | -0.43781588 | 3.612148986 | 0.4537594 | 0.5099905 | 0.999175987 |
| ENSMUSG00000052763 | Zfp212 | protein_coding | 0.478835129 | 3.652756293 | 0.4536044 | 0.5100618 | 0.999175987 |
| ENSMUSG00000033998 | Kcnk1 | protein_coding | 0.514255494 | 3.67952757 | 0.4535512 | 0.5100863 | 0.999175987 |
| ENSMUSG00000034591 | Slc41a2 | protein_coding | 0.37372138 | 3.769568938 | 0.4535468 | 0.5100883 | 0.999175987 |
| ENSMUSG00000054199 | Gon4l | protein_coding | -0.560391251 | 4.01507341 | 0.4535826 | 0.5102577 | 0.999175987 |
| ENSMUSG00000014786 | Slc9a5 | protein_coding | 0.485724996 | 3.415460126 | 0.4531348 | 0.5102779 | 0.999175987 |
| ENSMUSG00000017670 | Elmo2 | protein_coding | 0.27800103 | 6.908676478 | 0.4529855 | 0.5103621 | 0.999175987 |
| ENSMUSG00000028191 | Bcl10 | protein_coding | 0.294756878 | 4.911238581 | 0.452887 | 0.5103921 | 0.999175987 |
| ENSMUSG00000032869 | Psmf1 | protein_coding | -0.282384586 | 6.785877124 | 0.452824 | 0.5104365 | 0.999175987 |
| ENSMUSG00000028899 | Taf12 | protein_coding | -0.276249535 | 6.018048433 | 0.4526088 | 0.5105202 | 0.999175987 |
| ENSMUSG00000030521 | Mphosph10 | protein_coding | -0.316360191 | 4.697835945 | 0.452591 | 0.5105284 | 0.999175987 |
| ENSMUSG00000021745 | Ptprg | protein_coding | -0.444060792 | 4.195560502 | 0.4524644 | 0.5105868 | 0.999175987 |
| ENSMUSG00000028962 | Slc4a2 | protein_coding | 0.478553024 | 3.489299007 | 0.4524213 | 0.5106066 | 0.999175987 |
| ENSMUSG00000027207 | Galk2 | protein_coding | -0.28437867 | 6.042863571 | 0.4522647 | 0.5106788 | 0.999175987 |
| ENSMUSG00000014850 | Msh3 | protein_coding | -0.421351415 | 3.962347534 | 0.4520619 | 0.5107723 | 0.999175987 |
| ENSMUSG00000027984 | Hadh | protein_coding | -0.268387803 | 5.869148006 | 0.4516481 | 0.5109633 | 0.999175987 |
| ENSMUSG00000039959 | Hip1 | protein_coding | -0.317602634 | 4.775215907 | 0.4515602 | 0.5110038 | 0.999175987 |
| ENSMUSG00000024008 | Cpne5 | protein_coding | 0.286153876 | 7.202802253 | 0.4515752 | 0.5110123 | 0.999175987 |
| ENSMUSG00000022757 | Tfg | protein_coding | 0.296817225 | 6.460969884 | 0.4513566 | 0.5111132 | 0.999175987 |
| ENSMUSG00000061436 | Hipk2 | protein_coding | -0.493199624 | 4.56739696 | 0.451598 | 0.5112777 | 0.999175987 |
| ENSMUSG00000092368 | A930015D03Rik | antisense_RNA | 0.594895008 | 2.822723527 | 0.4508001 | 0.5113549 | 0.999175987 |
| ENSMUSG00000023909 | Paqr4 | protein_coding | 0.395717395 | 4.688858335 | 0.4507012 | 0.5114006 | 0.999175987 |
| ENSMUSG00000040928 | S100pbp | protein_coding | -0.286614477 | 5.027817843 | 0.4506915 | 0.5114051 | 0.999175987 |
| ENSMUSG00000036594 | H2-Aa | protein_coding | -1.054040545 | 2.156284574 | 0.4503058 | 0.5115834 | 0.999175987 |
| ENSMUSG00000082536 | Gm13456 | processed_pseudoge | 0.741441874 | 2.286606122 | 0.4502164 | 0.5116248 | 0.999175987 |
| ENSMUSG00000023262 | Acy1 | protein_coding | 0.365590687 | 4.934477265 | 0.4501277 | 0.5116658 | 0.999175987 |
| ENSMUSG00000034601 | 2700049A03Rik | protein_coding | -0.752294357 | 2.973556882 | 0.4500259 | 0.5117283 | 0.999175987 |
| ENSMUSG00000025875 | Tspan17 | protein_coding | 0.574583902 | 3.463116363 | 0.4499192 | 0.5117623 | 0.999175987 |
| ENSMUSG00000078786 | BC024978 | protein_coding | -0.396230999 | 4.375174023 | 0.4497941 | 0.5118202 | 0.999175987 |
| ENSMUSG00000021607 | Mrpl36 | protein_coding | 0.181231824 | 7.686975598 | 0.4496054 | 0.5119075 | 0.999175987 |
| ENSMUSG00000108414 | Snhg1 | processed_transcript | 0.330305117 | 4.599979832 | 0.4488883 | 0.5122398 | 0.999175987 |
| ENSMUSG00000022948 | Setd4 | protein_coding | 0.762809562 | 3.210991503 | 0.4495274 | 0.5123359 | 0.999175987 |
| ENSMUSG00000053166 | Cdh22 | protein_coding | -0.670279402 | 2.911345034 | 0.4486777 | 0.5123374 | 0.999175987 |
| ENSMUSG00000029836 | Cbx3 | protein_coding | 0.393481973 | 3.558430742 | 0.4486147 | 0.5123667 | 0.999175987 |
| ENSMUSG00000041609 | Bicdl1 | protein_coding | 0.430180692 | 4.482857563 | 0.4486122 | 0.5123678 | 0.999175987 |
| ENSMUSG00000022019 | Tdrd3 | protein_coding | -0.315657989 | 4.73038923 | 0.4485284 | 0.5124067 | 0.999175987 |
| ENSMUSG00000027430 | Dtd1 | protein_coding | 0.185173198 | 8.941783108 | 0.4483866 | 0.5124725 | 0.999175987 |
| ENSMUSG00000002524 | Puf60 | protein_coding | 0.205957479 | 8.566875765 | 0.4483531 | 0.512488 | 0.999175987 |
| ENSMUSG00000026728 | Vim | protein_coding | -0.441939655 | 4.265749196 | 0.4482839 | 0.5125201 | 0.999175987 |
| ENSMUSG00000027303 | Ptpra | protein_coding | 0.324805928 | 4.260738267 | 0.4476957 | 0.5127932 | 0.999175987 |
| ENSMUSG00000026811 | St6galnac6 | protein_coding | 0.247919057 | 5.915697612 | 0.4474724 | 0.512897 | 0.999175987 |
| ENSMUSG00000020846 | Rflnb | protein_coding | -0.469308334 | 3.672663658 | 0.4474604 | 0.5129025 | 0.999175987 |
| ENSMUSG00000021716 | Srek1ip1 | protein_coding | -0.245307287 | 6.026478648 | 0.4471328 | 0.5130548 | 0.999175987 |
| ENSMUSG00000026181 | Ppm1f | protein_coding | 0.317633921 | 5.832316627 | 0.4469744 | 0.5131285 | 0.999175987 |
| ENSMUSG00000034467 | Dynlrb2 | protein_coding | -0.445746174 | 4.88891651 | 0.4468005 | 0.513265 | 0.999175987 |
| ENSMUSG00000034194 | R3hcc1 | protein_coding | -0.286402632 | 5.675439554 | 0.4464902 | 0.5133537 | 0.999175987 |
| ENSMUSG00000028907 | Utp11 | protein_coding | 0.219435019 | 6.560606227 | 0.446357 | 0.5134157 | 0.999175987 |

| ensembl_gene_id | external_gene_name | gene_biotype | log2 (fold change) | log2 (counts per million) | F-value | p-value | False discovery rate |
|---|---|---|---|---|---|---|---|
| ENSMUSG00000025151 | Maged1 | protein_coding | 0.298126104 | 10.0312578 | 0.446368 | 0.5134259 | 0.999175987 |
| ENSMUSG00000029223 | Uchl1 | protein_coding | 0.200655167 | 10.33781782 | 0.4461684 | 0.5135035 | 0.999175987 |
| ENSMUSG00000020868 | Xylt2 | protein_coding | -0.407736692 | 3.705169652 | 0.4461671 | 0.5135041 | 0.999175987 |
| ENSMUSG00000028540 | Dph2 | protein_coding | 0.398768743 | 4.539993622 | 0.4460735 | 0.5135477 | 0.999175987 |
| ENSMUSG00000039210 | Gpatch2 | protein_coding | -0.786812073 | 2.641426006 | 0.4460175 | 0.5135738 | 0.999175987 |
| ENSMUSG00000019261 | Map1s | protein_coding | 0.513831517 | 4.671903465 | 0.4471873 | 0.5136235 | 0.999175987 |
| ENSMUSG00000003420 | Fcgrt | protein_coding | -0.558666997 | 3.467224619 | 0.4457577 | 0.5136948 | 0.999175987 |
| ENSMUSG00000047603 | Zfp235 | protein_coding | 0.311949782 | 4.396018757 | 0.4456621 | 0.5137394 | 0.999175987 |
| ENSMUSG00000020570 | Sypl | protein_coding | 0.240281266 | 6.263794843 | 0.4456425 | 0.5137485 | 0.999175987 |
| ENSMUSG00000004768 | Rab23 | protein_coding | -0.439912167 | 4.504457909 | 0.4455873 | 0.5137895 | 0.999175987 |
| ENSMUSG00000050732 | Vamp8 | protein_coding | -0.44981229 | 5.692453053 | 0.4473326 | 0.5138931 | 0.999175987 |
| ENSMUSG00000022394 | L3mbtl2 | protein_coding | 0.455967785 | 4.542254689 | 0.4452679 | 0.5139384 | 0.999175987 |
| ENSMUSG00000024392 | Bag6 | protein_coding | 0.300465825 | 5.578006025 | 0.4451518 | 0.5139773 | 0.999175987 |
| ENSMUSG00000063787 | Chchd1 | protein_coding | 0.230345943 | 8.251797352 | 0.4448529 | 0.5141168 | 0.999175987 |
| ENSMUSG00000039230 | Tbcd | protein_coding | -0.321392248 | 4.638417579 | 0.4447316 | 0.5141734 | 0.999175987 |
| ENSMUSG00000071604 | Fam189a2 | protein_coding | 0.728262234 | 2.759313938 | 0.4446402 | 0.514216 | 0.999175987 |
| ENSMUSG00000002102 | Psmc3 | protein_coding | -0.188716093 | 8.549169289 | 0.4445642 | 0.5142515 | 0.999175987 |
| ENSMUSG00000028334 | Nans | protein_coding | 0.445857058 | 3.834587374 | 0.4443177 | 0.5143666 | 0.999175987 |
| ENSMUSG00000032890 | Rims3 | protein_coding | 0.382243238 | 7.353189369 | 0.4474086 | 0.5143777 | 0.999175987 |
| ENSMUSG00000069670 | Nkain2 | protein_coding | -0.391946162 | 5.140302414 | 0.4439979 | 0.5145313 | 0.999175987 |
| ENSMUSG00000020752 | Recql5 | protein_coding | 0.46754576 | 3.326451752 | 0.4437022 | 0.5146542 | 0.999175987 |
| ENSMUSG00000031950 | Gabarapl2 | protein_coding | -0.20637472 | 6.722212429 | 0.4433807 | 0.5148046 | 0.999175987 |
| ENSMUSG00000060882 | Kcnd2 | protein_coding | 0.213350231 | 6.631503232 | 0.4433033 | 0.5148408 | 0.999175987 |
| ENSMUSG00000031785 | Adgrg1 | protein_coding | -0.240089654 | 6.230589893 | 0.4424309 | 0.5152493 | 0.999175987 |
| ENSMUSG00000026240 | Cops7b | protein_coding | 0.333962006 | 4.814157823 | 0.4423348 | 0.5152943 | 0.999175987 |
| ENSMUSG00000025525 | Apool | protein_coding | -0.417527651 | 4.103545178 | 0.4420119 | 0.5154456 | 0.999175987 |
| ENSMUSG00000071860 | 2900055J20Rik | protein_coding | 0.425773718 | 3.835069793 | 0.4418985 | 0.5154988 | 0.999175987 |
| ENSMUSG00000013921 | Clip3 | protein_coding | 0.239357387 | 8.694531453 | 0.4417538 | 0.5155667 | 0.999175987 |
| ENSMUSG00000030545 | Pex11a | protein_coding | -0.445022217 | 4.157662437 | 0.4414426 | 0.5157127 | 0.999175987 |
| ENSMUSG00000022216 | Psme1 | protein_coding | 0.213714689 | 7.421260387 | 0.4413395 | 0.515761 | 0.999175987 |
| ENSMUSG00000030035 | Wbp1 | protein_coding | 0.420654262 | 4.942696081 | 0.4412494 | 0.5158185 | 0.999175987 |
| ENSMUSG00000027520 | Zdbf2 | protein_coding | -0.392725367 | 4.992381259 | 0.4408739 | 0.5159949 | 0.999175987 |
| ENSMUSG00000044700 | Tmem201 | protein_coding | -0.352925605 | 4.577270721 | 0.4402463 | 0.5162746 | 0.999175987 |
| ENSMUSG00000041625 | Ggact | protein_coding | -0.290928303 | 5.141992694 | 0.4400597 | 0.5163624 | 0.999175987 |
| ENSMUSG00000027965 | Olfm3 | protein_coding | 0.348644683 | 4.733912521 | 0.4397569 | 0.5165048 | 0.999175987 |
| ENSMUSG00000027618 | Nfs1 | protein_coding | 0.33368426 | 4.773280756 | 0.4389978 | 0.5168623 | 0.999175987 |
| ENSMUSG00000029575 | Mmab | protein_coding | 0.308519151 | 5.092512116 | 0.4389306 | 0.516894 | 0.999175987 |
| ENSMUSG00000035399 | Oser1 | protein_coding | 0.355347235 | 3.785850373 | 0.4389026 | 0.5169071 | 0.999175987 |
| ENSMUSG00000003423 | Pih1d1 | protein_coding | 0.309721032 | 5.332552352 | 0.4382706 | 0.5172052 | 0.999175987 |
| ENSMUSG00000045215 | Asxl3 | protein_coding | -0.488701303 | 3.130231723 | 0.4382688 | 0.517206 | 0.999175987 |
| ENSMUSG00000033981 | Gria2 | protein_coding | -0.234294946 | 8.100468188 | 0.4375231 | 0.517558 | 0.999175987 |
| ENSMUSG00000008136 | Fhl2 | protein_coding | 0.709220013 | 3.259917399 | 0.4368026 | 0.5180485 | 0.999175987 |
| ENSMUSG00000050910 | Cdr2l | protein_coding | 0.35229929 | 5.033813653 | 0.4364583 | 0.5180614 | 0.999175987 |
| ENSMUSG00000020614 | Fam20a | protein_coding | 0.806416659 | 2.756788209 | 0.4362684 | 0.5181664 | 0.999175987 |
| ENSMUSG00000020794 | Ube2g1 | protein_coding | -0.251501634 | 5.654516639 | 0.4360049 | 0.518276 | 0.999175987 |
| ENSMUSG00000024544 | Ldlrad4 | protein_coding | 0.489354316 | 4.545320812 | 0.4364064 | 0.5183715 | 0.999175987 |
| ENSMUSG00000004843 | Chmp2b | protein_coding | -0.296845017 | 5.617063188 | 0.435745 | 0.5183991 | 0.999175987 |
| ENSMUSG00000039206 | Daglb | protein_coding | 0.298467808 | 4.974757951 | 0.4356375 | 0.51845 | 0.999175987 |
| ENSMUSG00000051730 | Mettl5 | protein_coding | -0.221187132 | 6.12775311 | 0.4355195 | 0.518506 | 0.999175987 |
| ENSMUSG00000039067 | Psmd7 | protein_coding | 0.183116343 | 8.601291436 | 0.4353837 | 0.5185703 | 0.999175987 |
| ENSMUSG00000028136 | Snx27 | protein_coding | -0.238087756 | 6.71847641 | 0.4353495 | 0.5185865 | 0.999175987 |
| ENSMUSG00000024867 | Pip5k1b | protein_coding | -0.28965281 | 5.233252831 | 0.4353254 | 0.5185979 | 0.999175987 |
| ENSMUSG00000028821 | Syf2 | protein_coding | -0.219684577 | 7.347891965 | 0.4351301 | 0.5186905 | 0.999175987 |
| ENSMUSG00000022139 | Mbnl2 | protein_coding | -0.218998254 | 7.573170751 | 0.4347826 | 0.5188554 | 0.999175987 |
| ENSMUSG00000024511 | Rab27b | protein_coding | -0.30482542 | 6.756997328 | 0.4352828 | 0.5188742 | 0.999175987 |
| ENSMUSG00000009863 | Sdhb | protein_coding | 0.207574544 | 10.22007054 | 0.4346848 | 0.5189018 | 0.999175987 |
| ENSMUSG00000036192 | Rorb | protein_coding | -0.579033746 | 5.784684724 | 0.4409006 | 0.5189599 | 0.999175987 |
| ENSMUSG00000058706 | 0610030E20Rik | protein_coding | 0.618227762 | 2.85645559 | 0.4342035 | 0.5191303 | 0.999175987 |
| ENSMUSG00000071650 | Ganab | protein_coding | 0.23004246 | 5.829963717 | 0.4339456 | 0.5192528 | 0.999175987 |
| ENSMUSG00000069206 | Zfp874a | protein_coding | 0.435615309 | 3.615797756 | 0.4336281 | 0.5194037 | 0.999175987 |
| ENSMUSG00000031431 | Tsc22d3 | protein_coding | -0.208703642 | 8.258900288 | 0.4335191 | 0.5194555 | 0.999175987 |
| ENSMUSG00000021767 | Kat6b | protein_coding | 0.363156901 | 4.30740422 | 0.4330457 | 0.5196807 | 0.999175987 |
| ENSMUSG00000053317 | Sec61b | protein_coding | 0.208747954 | 6.311453429 | 0.432829 | 0.5197839 | 0.999175987 |
| ENSMUSG00000103428 | Gm20754 | lincRNA | 0.879508617 | 2.397947304 | 0.4323913 | 0.5199923 | 0.999175987 |
| ENSMUSG00000027419 | Pcsk2 | protein_coding | -0.218119 | 6.944093578 | 0.4315351 | 0.5204005 | 0.999175987 |
| ENSMUSG00000026027 | Stradb | protein_coding | -0.343343297 | 4.655099218 | 0.4313715 | 0.5204786 | 0.999175987 |
| ENSMUSG00000072653 | Zfp783 | transcribed_unproces | 0.713071791 | 2.885958419 | 0.4312424 | 0.5205402 | 0.999175987 |
| ENSMUSG00000052684 | Jun | protein_coding | -0.454322403 | 3.651238088 | 0.4308638 | 0.5207209 | 0.999175987 |
| ENSMUSG00000033054 | Npat | protein_coding | -0.348634768 | 4.384894328 | 0.4308474 | 0.5207288 | 0.999175987 |
| ENSMUSG00000054400 | Cklf | protein_coding | -0.674449676 | 2.540702746 | 0.4308027 | 0.5207501 | 0.999175987 |
| ENSMUSG00000018377 | Vezf1 | protein_coding | -0.288050272 | 5.292392694 | 0.4307919 | 0.5207553 | 0.999175987 |
| ENSMUSG00000039601 | Rcan2 | protein_coding | -0.27982061 | 8.189909291 | 0.4307275 | 0.5208244 | 0.999175987 |
| ENSMUSG00000057766 | Ankrd29 | protein_coding | 0.269356393 | 5.195263539 | 0.4304339 | 0.5209263 | 0.999175987 |
| ENSMUSG00000021738 | Atxn7 | protein_coding | -0.458587177 | 3.312715562 | 0.4301573 | 0.5210585 | 0.999175987 |
| ENSMUSG00000043881 | Kbtbd7 | protein_coding | -0.350094943 | 4.593530467 | 0.4297776 | 0.5212402 | 0.999175987 |
| ENSMUSG00000024154 | Gtf2a1l | protein_coding | -0.726282755 | 2.657533945 | 0.4294857 | 0.5213799 | 0.999175987 |
| ENSMUSG00000020359 | Phykpl | protein_coding | 0.417460807 | 3.634499591 | 0.4289915 | 0.5216166 | 0.999175987 |
| ENSMUSG00000054893 | Zfp667 | protein_coding | 0.293619923 | 5.479913346 | 0.4288845 | 0.5216678 | 0.999175987 |

| ensembl_gene_id | external_gene_name | gene_biotype | log2 (fold change) | log2 (counts per million) | F-value | p-value | False discovery rate |
|---|---|---|---|---|---|---|---|
| ENSMUSG00000028165 | Cisd2 | protein_coding | 0.232335659 | 6.175455156 | 0.42888 | 0.52167 | 0.999175987 |
| ENSMUSG00000042079 | Hnrnpf | protein_coding | -0.502768061 | 2.968149836 | 0.4288437 | 0.5216874 | 0.999175987 |
| ENSMUSG00000045038 | Prkce | protein_coding | -0.20884382 | 7.599534377 | 0.4285646 | 0.5218212 | 0.999175987 |
| ENSMUSG00000039536 | Stau1 | protein_coding | -0.291059416 | 5.958777813 | 0.4284047 | 0.5218978 | 0.999175987 |
| ENSMUSG00000047003 | Zfp41 | protein_coding | 0.847408805 | 2.686236001 | 0.4274852 | 0.5223541 | 0.999175987 |
| ENSMUSG00000032497 | Lrrfip2 | protein_coding | -0.309873462 | 4.629103 | 0.4273508 | 0.5224038 | 0.999175987 |
| ENSMUSG00000021559 | Dapk1 | protein_coding | -0.274065929 | 5.818781129 | 0.4270574 | 0.5225448 | 0.999175987 |
| ENSMUSG00000025437 | Usp33 | protein_coding | 0.571457138 | 3.952541438 | 0.4278434 | 0.5225484 | 0.999175987 |
| ENSMUSG00000033083 | Tbc1d4 | protein_coding | -0.484651762 | 3.271941174 | 0.4268854 | 0.5226274 | 0.999175987 |
| ENSMUSG00000040667 | Nup88 | protein_coding | -0.273324344 | 6.057076029 | 0.4268764 | 0.5226318 | 0.999175987 |
| ENSMUSG00000021615 | Xrcc4 | protein_coding | 0.379961102 | 4.192460649 | 0.4267328 | 0.5227008 | 0.999175987 |
| ENSMUSG00000070047 | Fat1 | protein_coding | -0.508652505 | 3.598370895 | 0.4265202 | 0.5228031 | 0.999175987 |
| ENSMUSG00000041215 | Yeats2 | protein_coding | -0.537918472 | 5.218110279 | 0.4289749 | 0.5229004 | 0.999175987 |
| ENSMUSG00000019889 | Ptprk | protein_coding | -0.527986983 | 3.102611883 | 0.4260587 | 0.5230252 | 0.999175987 |
| ENSMUSG00000092060 | Bend4 | protein_coding | 0.470416603 | 4.913103167 | 0.427205 | 0.5230524 | 0.999175987 |
| ENSMUSG00000021028 | Mbip | protein_coding | -0.339090579 | 4.987960437 | 0.4258215 | 0.5231395 | 0.999175987 |
| ENSMUSG00000031021 | Tmem9b | protein_coding | -0.243333846 | 5.948648697 | 0.4257802 | 0.5231593 | 0.999175987 |
| ENSMUSG00000020409 | Slu7 | protein_coding | -0.217979392 | 7.116265078 | 0.4255804 | 0.5232556 | 0.999175987 |
| ENSMUSG00000040430 | Pitpnc1 | protein_coding | 0.243498881 | 7.311017842 | 0.4254568 | 0.5233152 | 0.999175987 |
| ENSMUSG00000028580 | Pum1 | protein_coding | 0.292963996 | 6.789635868 | 0.4254248 | 0.5233847 | 0.999175987 |
| ENSMUSG00000100599 | 1700120C14Rik | processed_transcript | -0.674215554 | 2.57138685 | 0.4252584 | 0.5234108 | 0.999175987 |
| ENSMUSG00000028645 | Slc2a1 | protein_coding | 0.502009577 | 3.207892489 | 0.4250288 | 0.5235215 | 0.999175987 |
| ENSMUSG00000006395 | Hyi | polymorphic_pseudog | -0.538356964 | 2.861671572 | 0.4248223 | 0.5236212 | 0.999175987 |
| ENSMUSG00000001323 | Srr | protein_coding | 0.287505601 | 4.668064243 | 0.4248152 | 0.5236246 | 0.999175987 |
| ENSMUSG00000030704 | Rab6a | protein_coding | -0.187898246 | 8.306047026 | 0.4245652 | 0.5237452 | 0.999175987 |
| ENSMUSG00000042428 | Mgat3 | protein_coding | 0.261271454 | 5.993891387 | 0.424556 | 0.5237497 | 0.999175987 |
| ENSMUSG00000052151 | Plpp2 | protein_coding | 0.328344391 | 4.438587461 | 0.4245419 | 0.5237565 | 0.999175987 |
| ENSMUSG00000062040 | Zfp27 | protein_coding | 0.402799699 | 4.00792207 | 0.4241665 | 0.5239378 | 0.999175987 |
| ENSMUSG00000030423 | Pop4 | protein_coding | 0.250383666 | 6.037266014 | 0.4240454 | 0.5239962 | 0.999175987 |
| ENSMUSG00000003865 | Gys1 | protein_coding | -0.405389091 | 4.209137295 | 0.4240307 | 0.5240034 | 0.999175987 |
| ENSMUSG00000021938 | Pspc1 | protein_coding | -0.372395314 | 4.338091377 | 0.4237118 | 0.5241575 | 0.999175987 |
| ENSMUSG00000109088 | Gm44593 | lincRNA | -0.600149776 | 2.679908493 | 0.423472 | 0.5242734 | 0.999175987 |
| ENSMUSG00000032468 | Armc8 | protein_coding | 0.251054303 | 6.763155801 | 0.4234367 | 0.5242905 | 0.999175987 |
| ENSMUSG00000056310 | Tyw1 | protein_coding | 0.344200459 | 4.485944164 | 0.4232585 | 0.5243767 | 0.999175987 |
| ENSMUSG00000038354 | Ankrd35 | protein_coding | 0.784208703 | 2.39288361 | 0.4229126 | 0.5245441 | 0.999175987 |
| ENSMUSG00000053310 | Nrgn | protein_coding | 0.246211796 | 7.189110252 | 0.4227946 | 0.5246012 | 0.999175987 |
| ENSMUSG00000067924 | Cxx1b | protein_coding | 0.272509626 | 4.928086095 | 0.4226691 | 0.524662 | 0.999175987 |
| ENSMUSG00000000594 | Gm2a | protein_coding | 0.213034071 | 7.111728144 | 0.422583 | 0.5247037 | 0.999175987 |
| ENSMUSG00000040033 | Stat2 | protein_coding | -0.714520522 | 2.603208494 | 0.4224304 | 0.5247776 | 0.999175987 |
| ENSMUSG00000029911 | Ssbp1 | protein_coding | 0.242405018 | 6.047762266 | 0.4221844 | 0.5248968 | 0.999175987 |
| ENSMUSG00000036873 | 2410004B18Rik | protein_coding | 0.415245177 | 4.247626815 | 0.4219491 | 0.5250109 | 0.999175987 |
| ENSMUSG00000048874 | Phf3 | protein_coding | -0.285855922 | 5.110300952 | 0.4219331 | 0.5250186 | 0.999175987 |
| ENSMUSG00000057969 | Sema3b | protein_coding | 1.124356552 | 2.483645804 | 0.4228036 | 0.525172 | 0.999175987 |
| ENSMUSG00000024856 | Cdk2ap2 | protein_coding | 0.250509742 | 5.554639604 | 0.4215576 | 0.5252007 | 0.999175987 |
| ENSMUSG00000020523 | Fam114a2 | protein_coding | 0.243581116 | 6.023009542 | 0.4213354 | 0.5253086 | 0.999175987 |
| ENSMUSG00000031393 | Mecp2 | protein_coding | 0.259898893 | 5.871023054 | 0.4213118 | 0.52532 | 0.999175987 |
| ENSMUSG00000113170 | AC132344.1 | lincRNA | 0.368414172 | 3.751730224 | 0.421235 | 0.5253573 | 0.999175987 |
| ENSMUSG00000031592 | Pcm1 | protein_coding | -0.314668278 | 5.501152417 | 0.421231 | 0.5253593 | 0.999175987 |
| ENSMUSG00000030447 | Cyfip1 | protein_coding | -0.294217169 | 5.532259004 | 0.4211173 | 0.5254145 | 0.999175987 |
| ENSMUSG00000087006 | Gm13889 | protein_coding | 0.43269733 | 3.586865845 | 0.4211106 | 0.5254177 | 0.999175987 |
| ENSMUSG00000002679 | Med6 | protein_coding | -0.283209781 | 5.727165433 | 0.4210784 | 0.5254333 | 0.999175987 |
| ENSMUSG00000008690 | Ncaph2 | protein_coding | 0.262579656 | 7.207697295 | 0.4208742 | 0.5255473 | 0.999175987 |
| ENSMUSG00000037887 | Dusp8 | protein_coding | 0.252681743 | 8.228678214 | 0.420619 | 0.5256713 | 0.999175987 |
| ENSMUSG00000032215 | Rsl24d1 | protein_coding | 0.206365484 | 6.40011498 | 0.4205288 | 0.5257003 | 0.999175987 |
| ENSMUSG00000037622 | Wdtc1 | protein_coding | 0.318940952 | 4.902716224 | 0.4203873 | 0.5257691 | 0.999175987 |
| ENSMUSG00000021222 | Dcaf4 | protein_coding | 0.26630717 | 6.168689986 | 0.4203533 | 0.5257857 | 0.999175987 |
| ENSMUSG00000026662 | Sephs1 | protein_coding | -0.395317532 | 4.295783792 | 0.420329 | 0.5257975 | 0.999175987 |
| ENSMUSG00000026224 | 4933407L21Rik | protein_coding | -0.348307 | 4.27700674 | 0.4202497 | 0.525836 | 0.999175987 |
| ENSMUSG00000020659 | Cbll1 | protein_coding | 0.350359421 | 4.67701495 | 0.4202477 | 0.525837 | 0.999175987 |
| ENSMUSG00000031701 | Dnaja2 | protein_coding | 0.220804185 | 7.840731032 | 0.4201162 | 0.525901 | 0.999175987 |
| ENSMUSG00000005481 | Ddx39 | protein_coding | -0.324217056 | 5.050695147 | 0.4200153 | 0.52595 | 0.999175987 |
| ENSMUSG00000017721 | Pigt | protein_coding | 0.387085752 | 4.034348605 | 0.419622 | 0.5261415 | 0.999175987 |
| ENSMUSG00000045106 | Ccdc73 | protein_coding | 0.400181565 | 3.773213786 | 0.4196045 | 0.5261499 | 0.999175987 |
| ENSMUSG00000021900 | Btd | protein_coding | -0.373700977 | 4.185095894 | 0.4195627 | 0.5261703 | 0.999175987 |
| ENSMUSG00000032532 | Cck | protein_coding | -0.544438052 | 3.604041295 | 0.419294 | 0.5263159 | 0.999175987 |
| ENSMUSG00000029474 | Rnf34 | protein_coding | 0.242731466 | 6.829315571 | 0.4190144 | 0.5264374 | 0.999175987 |
| ENSMUSG00000028454 | Pigo | protein_coding | 0.566351659 | 3.200219405 | 0.4189627 | 0.5264626 | 0.999175987 |
| ENSMUSG00000002343 | Armc6 | protein_coding | 0.297446306 | 6.423998086 | 0.4185496 | 0.5266787 | 0.999175987 |
| ENSMUSG00000017692 | Rhbdl3 | protein_coding | 0.487682833 | 3.989578344 | 0.4183354 | 0.5267832 | 0.999175987 |
| ENSMUSG00000056459 | Zbtb25 | protein_coding | -0.467736253 | 3.542563016 | 0.4177029 | 0.5270771 | 0.999175987 |
| ENSMUSG00000033900 | Map9 | protein_coding | -0.24430434 | 6.439453988 | 0.4167223 | 0.5275565 | 0.999175987 |
| ENSMUSG00000021982 | Cdadc1 | protein_coding | 0.220600177 | 6.603513022 | 0.4167063 | 0.5275643 | 0.999175987 |
| ENSMUSG00000039159 | Ube2h | protein_coding | 0.259330558 | 6.566999314 | 0.4166442 | 0.5275947 | 0.999175987 |
| ENSMUSG00000021010 | Npas3 | protein_coding | -0.749287827 | 2.326806981 | 0.4163196 | 0.5277536 | 0.999175987 |
| ENSMUSG00000074579 | Lekr1 | protein_coding | -0.502351209 | 3.043518446 | 0.4161842 | 0.5278199 | 0.999175987 |
| ENSMUSG00000036896 | C1qc | protein_coding | -0.59448813 | 4.251901366 | 0.4179876 | 0.5278335 | 0.999175987 |
| ENSMUSG00000030539 | Sema4b | protein_coding | -0.55486377 | 2.720905013 | 0.4160975 | 0.5278623 | 0.999175987 |
| ENSMUSG00000026495 | Efcab2 | protein_coding | 0.294958392 | 5.377614684 | 0.4158916 | 0.5279632 | 0.999175987 |

| ensembl_gene_id | external_gene_name | gene_biotype | log2 (fold change) | log2 (counts per million) | F-value | p-value | False discovery rate |
|---|---|---|---|---|---|---|---|
| ENSMUSG00000035476 | Tab3 | protein_coding | 0.616843944 | 3.025920837 | 0.4156835 | 0.5280651 | 0.999175987 |
| ENSMUSG00000029174 | Tbc1d1 | protein_coding | -0.521781629 | 3.007960314 | 0.4153547 | 0.5282263 | 0.999175987 |
| ENSMUSG00000027180 | Fbxo3 | protein_coding | -0.214307473 | 7.1115003 | 0.4153527 | 0.5282273 | 0.999175987 |
| ENSMUSG00000057561 | Eif1a | protein_coding | 0.212090056 | 7.456750366 | 0.4153438 | 0.5282317 | 0.999175987 |
| ENSMUSG00000040549 | Ckap5 | protein_coding | -0.207214908 | 6.407334351 | 0.415077 | 0.5283625 | 0.999175987 |
| ENSMUSG00000040591 | 1110051M20Rik | protein_coding | 0.229766865 | 8.374392058 | 0.4149284 | 0.5284354 | 0.999175987 |
| ENSMUSG00000085558 | 4930412C18Rik | processed_transcript | -0.449928813 | 3.611819385 | 0.414866 | 0.5284661 | 0.999175987 |
| ENSMUSG00000022336 | Eif3e | protein_coding | 0.228574362 | 7.651434213 | 0.4147456 | 0.5285252 | 0.999175987 |
| ENSMUSG00000028878 | Fam76a | protein_coding | -0.295904167 | 5.551510911 | 0.4147002 | 0.5285475 | 0.999175987 |
| ENSMUSG00000045078 | Rnf216 | protein_coding | -0.326066669 | 5.031410019 | 0.4146285 | 0.5285827 | 0.999175987 |
| ENSMUSG00000017009 | Sdc4 | protein_coding | -0.284129015 | 7.306298795 | 0.4149501 | 0.5286013 | 0.999175987 |
| ENSMUSG00000037058 | Paip2 | protein_coding | 0.175618988 | 7.838311285 | 0.4144435 | 0.5286735 | 0.999175987 |
| ENSMUSG00000062014 | Gmfb | protein_coding | 0.24830243 | 7.448109602 | 0.4143856 | 0.5287105 | 0.999175987 |
| ENSMUSG00000022789 | Dnm1l | protein_coding | 0.203311825 | 7.672574536 | 0.4142378 | 0.5287746 | 0.999175987 |
| ENSMUSG00000071655 | Ubxn1 | protein_coding | -0.192331858 | 6.632327151 | 0.413901 | 0.5289401 | 0.999175987 |
| ENSMUSG00000055733 | Nap1l3 | protein_coding | -0.279897651 | 5.768470532 | 0.4136444 | 0.5290663 | 0.999175987 |
| ENSMUSG00000056708 | Ier5 | protein_coding | -0.574346522 | 3.078491672 | 0.4135628 | 0.5291064 | 0.999175987 |
| ENSMUSG00000034292 | Traf3ip1 | protein_coding | -0.46263847 | 3.585416351 | 0.4131372 | 0.5293158 | 0.999175987 |
| ENSMUSG00000001964 | Emd | protein_coding | -0.321389104 | 5.350165237 | 0.41313 | 0.5293193 | 0.999175987 |
| ENSMUSG00000071176 | Arhgef10 | protein_coding | -0.346335421 | 4.175205598 | 0.4126217 | 0.5295696 | 0.999175987 |
| ENSMUSG00000030717 | Nupr1 | protein_coding | -0.489425253 | 5.056776348 | 0.4142976 | 0.5296359 | 0.999175987 |
| ENSMUSG00000020230 | Prmt2 | protein_coding | 0.282059766 | 9.243598961 | 0.4125677 | 0.5296495 | 0.999175987 |
| ENSMUSG00000055737 | Ghr | protein_coding | 0.477744671 | 2.985612569 | 0.4123093 | 0.5297236 | 0.999175987 |
| ENSMUSG00000018446 | C1qbp | protein_coding | -0.30447326 | 5.541922467 | 0.4123064 | 0.529725 | 0.999175987 |
| ENSMUSG00000040811 | Eml2 | protein_coding | 0.284818978 | 5.45460871 | 0.4121312 | 0.5298114 | 0.999175987 |
| ENSMUSG00000024181 | Mrpl28 | protein_coding | 0.18793384 | 8.842048939 | 0.4118063 | 0.5299717 | 0.999175987 |
| ENSMUSG00000038894 | Irs2 | protein_coding | 0.301489221 | 5.25886737 | 0.4116718 | 0.5300381 | 0.999175987 |
| ENSMUSG00000035293 | G2e3 | protein_coding | -0.701237291 | 2.867473345 | 0.4116667 | 0.5300406 | 0.999175987 |
| ENSMUSG00000057375 | Yipf1 | protein_coding | 0.213181739 | 6.431579522 | 0.4116181 | 0.5300646 | 0.999175987 |
| ENSMUSG00000020526 | Znhit3 | protein_coding | 0.192208903 | 7.147062682 | 0.411326 | 0.5302087 | 0.999175987 |
| ENSMUSG00000063065 | Mapk3 | protein_coding | 0.181624953 | 7.28755616 | 0.4112615 | 0.5302406 | 0.999175987 |
| ENSMUSG00000034345 | Gtf2h5 | protein_coding | 0.188920394 | 8.783717357 | 0.4110903 | 0.5303252 | 0.999175987 |
| ENSMUSG00000072769 | Gm10419 | lincRNA | -0.218029581 | 6.107039761 | 0.4110173 | 0.5303612 | 0.999175987 |
| ENSMUSG00000033961 | Zfp446 | protein_coding | 0.452414417 | 3.355200966 | 0.4107335 | 0.5305015 | 0.999175987 |
| ENSMUSG00000025384 | Faap100 | protein_coding | 0.463461231 | 4.090612436 | 0.4106725 | 0.5305462 | 0.999175987 |
| ENSMUSG00000049755 | Zfp672 | protein_coding | 0.305418863 | 5.688238535 | 0.4104834 | 0.5306251 | 0.999175987 |
| ENSMUSG00000097451 | Rian | processed_transcript | 0.250202961 | 5.684275856 | 0.4099341 | 0.5308969 | 0.999175987 |
| ENSMUSG00000033615 | Cplx1 | protein_coding | 0.223918952 | 9.61942052 | 0.4097802 | 0.5309731 | 0.999175987 |
| ENSMUSG00000086555 | Gm13446 | processed_transcript | -0.66296542 | 3.558892642 | 0.4102617 | 0.5313 | 0.999175987 |
| ENSMUSG00000059713 | Rcan3 | protein_coding | -0.242125762 | 5.330877705 | 0.4090338 | 0.5313429 | 0.999175987 |
| ENSMUSG00000056216 | Cebpg | protein_coding | 0.301864255 | 4.363496584 | 0.4089712 | 0.5313739 | 0.999175987 |
| ENSMUSG00000021986 | Amer2 | protein_coding | -0.328122179 | 5.07212423 | 0.4088833 | 0.5314175 | 0.999175987 |
| ENSMUSG00000022095 | Fam160b2 | protein_coding | -0.37461421 | 3.768285915 | 0.4087646 | 0.5314764 | 0.999175987 |
| ENSMUSG00000030303 | Far2 | protein_coding | 0.516743612 | 4.74885854 | 0.4105509 | 0.5314778 | 0.999175987 |
| ENSMUSG00000027199 | Gatm | protein_coding | -0.232387193 | 8.833057624 | 0.4086538 | 0.5315313 | 0.999175987 |
| ENSMUSG00000006522 | Itih3 | protein_coding | -0.390787702 | 5.66703038 | 0.4097257 | 0.5315648 | 0.999175987 |
| ENSMUSG00000050973 | Gdpgp1 | protein_coding | -0.432264965 | 4.282659624 | 0.4083412 | 0.5316865 | 0.999175987 |
| ENSMUSG00000030059 | Tmf1 | protein_coding | -0.392266922 | 6.469920041 | 0.4106306 | 0.5317963 | 0.999175987 |
| ENSMUSG00000084898 | Gm12371 | lincRNA | 0.491346278 | 3.240525916 | 0.4080374 | 0.5318373 | 0.999175987 |
| ENSMUSG00000037857 | Nufip2 | protein_coding | -0.279157325 | 6.648928208 | 0.407774 | 0.5319827 | 0.999175987 |
| ENSMUSG00000015747 | Vps45 | protein_coding | 0.22039232 | 6.472370461 | 0.4077436 | 0.5319832 | 0.999175987 |
| ENSMUSG00000020288 | Ahsa2 | protein_coding | 0.449011336 | 3.798943059 | 0.407737 | 0.5319865 | 0.999175987 |
| ENSMUSG00000034105 | Tldc1 | protein_coding | 0.414219397 | 3.480613273 | 0.4075389 | 0.5320849 | 0.999175987 |
| ENSMUSG00000046836 | Brox | protein_coding | 0.303037699 | 4.588628176 | 0.4072457 | 0.5322307 | 0.999175987 |
| ENSMUSG00000038039 | Gcc2 | protein_coding | -0.239863003 | 5.948137453 | 0.4071888 | 0.532259 | 0.999175987 |
| ENSMUSG00000023353 | Agap3 | protein_coding | 0.234020209 | 5.872220697 | 0.4071122 | 0.5322971 | 0.999175987 |
| ENSMUSG00000047141 | Zfp654 | protein_coding | 0.396440365 | 3.831966821 | 0.407088 | 0.5323092 | 0.999175987 |
| ENSMUSG00000043857 | Mgat5b | protein_coding | 0.352877176 | 4.474945001 | 0.4069153 | 0.5323951 | 0.999175987 |
| ENSMUSG00000109006 | B230209E15Rik | lincRNA | 0.244769132 | 6.39717013 | 0.4062242 | 0.5327392 | 0.999175987 |
| ENSMUSG00000073678 | Pgap1 | protein_coding | -0.312394816 | 4.58660552 | 0.4060983 | 0.5328019 | 0.999175987 |
| ENSMUSG00000027404 | Snrpb | protein_coding | 0.223394194 | 7.354725503 | 0.4055352 | 0.5330826 | 0.999175987 |
| ENSMUSG00000040943 | Tet2 | protein_coding | -0.316442889 | 5.009119942 | 0.4045635 | 0.5335677 | 0.999175987 |
| ENSMUSG00000063358 | Mapk1 | protein_coding | -0.232633561 | 8.06731653 | 0.404549 | 0.533575 | 0.999175987 |
| ENSMUSG00000056342 | Usp34 | protein_coding | -0.243527809 | 5.862036858 | 0.4044667 | 0.5336161 | 0.999175987 |
| ENSMUSG00000048603 | Gm9828 | bidirectional_promote | 0.496169338 | 3.293833343 | 0.4042265 | 0.5337362 | 0.999175987 |
| ENSMUSG00000038279 | Nop2 | protein_coding | 0.346153279 | 5.250246065 | 0.4040419 | 0.5338285 | 0.999175987 |
| ENSMUSG00000050541 | Adra1b | protein_coding | 0.745860709 | 2.48256062 | 0.4040167 | 0.5338411 | 0.999175987 |
| ENSMUSG00000060475 | Wtap | protein_coding | 0.4867432 | 5.476082691 | 0.4063137 | 0.533931 | 0.999175987 |
| ENSMUSG00000027575 | Arfgap1 | protein_coding | 0.219265331 | 6.255567342 | 0.4035505 | 0.5340743 | 0.999175987 |
| ENSMUSG00000036620 | Mgat4b | protein_coding | 0.666967682 | 2.571468355 | 0.4033543 | 0.5341726 | 0.999175987 |
| ENSMUSG00000031723 | Txnl4b | protein_coding | 0.281284777 | 6.709429716 | 0.4033576 | 0.5341853 | 0.999175987 |
| ENSMUSG00000020532 | Acaca | protein_coding | -0.38918167 | 3.55394166 | 0.4033134 | 0.534193 | 0.999175987 |
| ENSMUSG00000029146 | Snx17 | protein_coding | 0.232299099 | 8.228083589 | 0.4032708 | 0.5342144 | 0.999175987 |
| ENSMUSG00000005846 | Rsl1d1 | protein_coding | 0.21614046 | 7.142042132 | 0.4030016 | 0.5343492 | 0.999175987 |
| ENSMUSG00000049946 | BC030500 | protein_coding | 0.336871076 | 5.154877365 | 0.4027537 | 0.5344735 | 0.999175987 |
| ENSMUSG00000028318 | Polr1e | protein_coding | -0.362739972 | 4.451103162 | 0.4026823 | 0.5345092 | 0.999175987 |
| ENSMUSG00000074909 | Ranbp6 | protein_coding | -0.336429871 | 5.08870337 | 0.4024 | 0.5346508 | 0.999175987 |
| ENSMUSG00000096225 | Lhx8 | protein_coding | 0.375473859 | 4.460769682 | 0.4023495 | 0.5346761 | 0.999175987 |

| ensembl_gene_id | external_gene_name | gene_biotype | log2 (fold change) | log2 (counts per million) | F-value | p-value | False discovery rate |
|---|---|---|---|---|---|---|---|
| ENSMUSG00000017400 | Stac2 | protein_coding | 0.42990502 | 4.566754954 | 0.4021262 | 0.5348026 | 0.999175987 |
| ENSMUSG00000032855 | Pkd1 | protein_coding | 0.624578165 | 2.826235212 | 0.4019605 | 0.5348713 | 0.999175987 |
| ENSMUSG00000028902 | Sf3a3 | protein_coding | -0.21048623 | 6.182383047 | 0.4017972 | 0.5349533 | 0.999175987 |
| ENSMUSG00000018965 | Ywhah | protein_coding | 0.183190922 | 8.259081373 | 0.4017827 | 0.5349606 | 0.999175987 |
| ENSMUSG00000027248 | Pdia3 | protein_coding | -0.245429941 | 6.219888738 | 0.4017102 | 0.534997 | 0.999175987 |
| ENSMUSG00000034353 | Ramp1 | protein_coding | 0.345541706 | 6.466903309 | 0.4027775 | 0.5351809 | 0.999175987 |
| ENSMUSG00000052299 | Ltn1 | protein_coding | -0.353416447 | 4.44658231 | 0.4007594 | 0.5354748 | 0.999175987 |
| ENSMUSG00000068823 | Csde1 | protein_coding | -0.259568117 | 8.718473416 | 0.4007447 | 0.5354966 | 0.999175987 |
| ENSMUSG00000035637 | Grhpr | protein_coding | 0.212773241 | 7.164007776 | 0.4004515 | 0.5356297 | 0.999175987 |
| ENSMUSG00000032777 | Gtf3c1 | protein_coding | 0.331645351 | 5.043845491 | 0.4003532 | 0.5356792 | 0.999175987 |
| ENSMUSG00000033781 | Asb13 | protein_coding | -0.240821147 | 5.697185403 | 0.4000647 | 0.5358245 | 0.999175987 |
| ENSMUSG00000037395 | Rcor3 | protein_coding | 0.433707013 | 3.439315693 | 0.3999282 | 0.5358932 | 0.999175987 |
| ENSMUSG00000036376 | Abt1 | protein_coding | 0.26055434 | 6.069733086 | 0.3995603 | 0.5360786 | 0.999175987 |
| ENSMUSG00000046814 | Gchfr | protein_coding | -0.434571471 | 3.452972226 | 0.3995541 | 0.5360817 | 0.999175987 |
| ENSMUSG00000044147 | Arf6 | protein_coding | 0.263402189 | 5.076528552 | 0.3994617 | 0.5361283 | 0.999175987 |
| ENSMUSG00000027177 | Hipk3 | protein_coding | -0.295038765 | 5.20658307 | 0.3994613 | 0.5361285 | 0.999175987 |
| ENSMUSG00000092203 | 1110038B12Rik | processed_transcript | 0.265027153 | 5.013220904 | 0.3993662 | 0.5361765 | 0.999175987 |
| ENSMUSG00000047181 | Samd14 | protein_coding | 0.27851103 | 5.095027515 | 0.3992602 | 0.5362299 | 0.999175987 |
| ENSMUSG00000035151 | Elmod2 | protein_coding | -0.714519278 | 2.45865367 | 0.3987075 | 0.5365088 | 0.999175987 |
| ENSMUSG00000032382 | Snx1 | protein_coding | -0.201520026 | 7.24547317 | 0.3985897 | 0.5365683 | 0.999175987 |
| ENSMUSG00000025377 | Tepsin | protein_coding | 0.566262229 | 2.691341674 | 0.3985075 | 0.5366098 | 0.999175987 |
| ENSMUSG00000036273 | Lrrk2 | protein_coding | 0.565314566 | 2.917905266 | 0.3979448 | 0.5368941 | 0.999175987 |
| ENSMUSG00000000861 | Bcl11a | protein_coding | 0.388277896 | 5.098905753 | 0.397968 | 0.5368967 | 0.999175987 |
| ENSMUSG00000028329 | Xpa | protein_coding | 0.397128272 | 3.471350559 | 0.3978982 | 0.5369177 | 0.999175987 |
| ENSMUSG00000035048 | Anapc13 | protein_coding | 0.26792783 | 7.441151497 | 0.3978294 | 0.5370045 | 0.999175987 |
| ENSMUSG00000045671 | Spred2 | protein_coding | -0.393434779 | 4.025440524 | 0.3975732 | 0.537082 | 0.999175987 |
| ENSMUSG00000024743 | Syt7 | protein_coding | 0.257081679 | 5.813303596 | 0.3975419 | 0.5370979 | 0.999175987 |
| ENSMUSG00000029780 | Nt5c3 | protein_coding | -0.290891355 | 5.587108197 | 0.3974366 | 0.5371511 | 0.999175987 |
| ENSMUSG00000031561 | Tenm3 | protein_coding | -0.46623521 | 3.863586045 | 0.3972919 | 0.5372244 | 0.999175987 |
| ENSMUSG00000004791 | Pgf | protein_coding | -0.346468729 | 4.251531182 | 0.3971509 | 0.5372957 | 0.999175987 |
| ENSMUSG00000083844 | Ube2d-ps | transcribed_unproces | 0.235598594 | 5.765932777 | 0.3971197 | 0.5373115 | 0.999175987 |
| ENSMUSG00000025911 | Adhfe1 | protein_coding | -0.387548723 | 3.611597344 | 0.3968192 | 0.5374637 | 0.999175987 |
| ENSMUSG00000068921 | Dap3 | protein_coding | -0.193265303 | 7.196531851 | 0.396429 | 0.5376614 | 0.999175987 |
| ENSMUSG00000022772 | Senp5 | protein_coding | 0.404160652 | 3.340804341 | 0.3962885 | 0.5377326 | 0.999175987 |
| ENSMUSG00000055745 | Ldoc1l | protein_coding | 0.265574046 | 5.336780888 | 0.396048 | 0.5378546 | 0.999175987 |
| ENSMUSG00000033007 | Asic4 | protein_coding | 0.954664296 | 2.312963814 | 0.3960617 | 0.5378619 | 0.999175987 |
| ENSMUSG00000002844 | Adprh | protein_coding | 0.209733649 | 7.485039426 | 0.395731 | 0.5380154 | 0.999175987 |
| ENSMUSG00000021629 | Slc30a5 | protein_coding | -0.349729993 | 4.547295748 | 0.3956684 | 0.5380472 | 0.999175987 |
| ENSMUSG00000031834 | Pik3r2 | protein_coding | 0.301189391 | 4.690326057 | 0.3956167 | 0.5380734 | 0.999175987 |
| ENSMUSG00000041362 | Shtn1 | protein_coding | -0.270226728 | 5.496170292 | 0.3955385 | 0.5381131 | 0.999175987 |
| ENSMUSG00000027905 | Ddx20 | protein_coding | -0.550440573 | 3.076885073 | 0.3955306 | 0.5381172 | 0.999175987 |
| ENSMUSG00000024527 | Afg3l2 | protein_coding | 0.294819856 | 5.199336318 | 0.3954944 | 0.5381355 | 0.999175987 |
| ENSMUSG00000030888 | Rrp8 | protein_coding | 0.377480095 | 4.165826031 | 0.3954933 | 0.5381361 | 0.999175987 |
| ENSMUSG00000056666 | Retsat | protein_coding | 0.437851345 | 3.650956387 | 0.3952918 | 0.5382384 | 0.999175987 |
| ENSMUSG00000038967 | Pdk2 | protein_coding | -0.279339349 | 6.877811759 | 0.3952691 | 0.5383018 | 0.999175987 |
| ENSMUSG00000011427 | Zfp790 | protein_coding | 0.481661589 | 3.195718129 | 0.3949678 | 0.538403 | 0.999175987 |
| ENSMUSG00000030287 | Itpr2 | protein_coding | 0.641122909 | 2.660606631 | 0.3949286 | 0.538423 | 0.999175987 |
| ENSMUSG00000024777 | Ppp2r5b | protein_coding | 0.231030629 | 5.645437289 | 0.3948305 | 0.5384728 | 0.999175987 |
| ENSMUSG00000047976 | Kcna1 | protein_coding | 0.352142457 | 4.522790484 | 0.3948038 | 0.5384864 | 0.999175987 |
| ENSMUSG00000097062 | Gm17586 | lincRNA | 0.43533815 | 3.253532886 | 0.3947715 | 0.5385028 | 0.999175987 |
| ENSMUSG00000020024 | Cep83 | protein_coding | -0.372514345 | 4.004862667 | 0.3946967 | 0.5385408 | 0.999175987 |
| ENSMUSG00000029804 | Herc3 | protein_coding | 0.24540635 | 5.997854704 | 0.3946898 | 0.5385444 | 0.999175987 |
| ENSMUSG00000026730 | Pter | protein_coding | -0.504417827 | 4.465772053 | 0.3956982 | 0.5385856 | 0.999175987 |
| ENSMUSG00000032280 | Tle3 | protein_coding | -0.560973024 | 3.152596638 | 0.3943296 | 0.5387275 | 0.999175987 |
| ENSMUSG00000064030 | Pym1 | protein_coding | 0.301096205 | 4.753457262 | 0.3938937 | 0.5389494 | 0.999175987 |
| ENSMUSG00000038816 | Ctnnal1 | protein_coding | -0.389732024 | 3.863867728 | 0.3937955 | 0.5389994 | 0.999175987 |
| ENSMUSG00000024969 | Mark2 | protein_coding | 0.247392299 | 5.771430586 | 0.3936715 | 0.5390625 | 0.999175987 |
| ENSMUSG00000026553 | Copa | protein_coding | 0.262981309 | 6.898510448 | 0.3936643 | 0.5390805 | 0.999175987 |
| ENSMUSG00000028948 | Nol9 | protein_coding | -0.50734685 | 3.071242924 | 0.3935669 | 0.5391158 | 0.999175987 |
| ENSMUSG00000045763 | Basp1 | protein_coding | -0.185534146 | 7.471415446 | 0.3935491 | 0.5391249 | 0.999175987 |
| ENSMUSG00000030180 | Kdm5a | protein_coding | 0.281184026 | 5.70130828 | 0.3931468 | 0.5393299 | 0.999175987 |
| ENSMUSG00000070705 | Eid2b | protein_coding | 0.416290394 | 3.510431073 | 0.3929408 | 0.539435 | 0.999175987 |
| ENSMUSG00000033365 | Ipo13 | protein_coding | -0.293141832 | 5.310807978 | 0.3928813 | 0.5394654 | 0.999175987 |
| ENSMUSG00000021066 | Atl1 | protein_coding | 0.266123204 | 6.57713241 | 0.3927503 | 0.5395464 | 0.999175987 |
| ENSMUSG00000037349 | Nudt22 | protein_coding | 0.285842244 | 4.836511979 | 0.3925317 | 0.5396437 | 0.999175987 |
| ENSMUSG00000029110 | Rnf4 | protein_coding | -0.22758777 | 6.827518734 | 0.3925119 | 0.5396539 | 0.999175987 |
| ENSMUSG00000031166 | Wdr13 | protein_coding | -0.204125019 | 5.967335504 | 0.3923253 | 0.5397491 | 0.999175987 |
| ENSMUSG00000022442 | Ttll1 | protein_coding | 0.282361284 | 7.433576324 | 0.3930341 | 0.5397503 | 0.999175987 |
| ENSMUSG00000041377 | Ninj2 | protein_coding | -0.356668593 | 4.6352863 | 0.3921487 | 0.5398393 | 0.999175987 |
| ENSMUSG00000033813 | Tcea1 | protein_coding | -0.303965437 | 4.076756085 | 0.3918753 | 0.539979 | 0.999175987 |
| ENSMUSG00000029464 | Gpn3 | protein_coding | 0.298841317 | 5.759942848 | 0.3915057 | 0.5401679 | 0.999175987 |
| ENSMUSG00000004994 | Ccdc130 | protein_coding | 0.272304034 | 5.078397919 | 0.3913133 | 0.5402663 | 0.999175987 |
| ENSMUSG00000039914 | Coq10a | protein_coding | -0.318039587 | 5.652001736 | 0.3912087 | 0.540334 | 0.999175987 |
| ENSMUSG00000067199 | Frat1 | protein_coding | 0.300074449 | 7.740331066 | 0.3922307 | 0.5403481 | 0.999175987 |
| ENSMUSG00000036529 | Sbf1 | protein_coding | 0.275217929 | 5.353329883 | 0.3911361 | 0.5403569 | 0.999175987 |
| ENSMUSG00000054675 | Tmem119 | protein_coding | 0.565400193 | 3.114579512 | 0.390586 | 0.5406386 | 0.999175987 |
| ENSMUSG00000039648 | Kyat1 | protein_coding | -0.419713814 | 4.543496185 | 0.3904859 | 0.5407041 | 0.999175987 |
| ENSMUSG00000029050 | Ski | protein_coding | -0.325974774 | 5.481192407 | 0.3901862 | 0.5408434 | 0.999175987 |

| ensembl_gene_id | external_gene_name | gene_biotype | log2 (fold change) | log2 (counts per million) | F-value | p-value | False discovery rate |
|---|---|---|---|---|---|---|---|
| ENSMUSG00000049922 | Slc35c1 | protein_coding | -0.390348295 | 4.063966979 | 0.3900328 | 0.5409221 | 0.999175987 |
| ENSMUSG00000001844 | Zdhhc4 | protein_coding | 0.255200192 | 5.724670165 | 0.3887481 | 0.5415815 | 0.999175987 |
| ENSMUSG00000062729 | Ppox | protein_coding | 0.421228666 | 4.797051836 | 0.3888136 | 0.5415993 | 0.999175987 |
| ENSMUSG00000002980 | Bcam | protein_coding | 0.734125117 | 3.430448586 | 0.3900226 | 0.5417875 | 0.999175987 |
| ENSMUSG00000021666 | Gfm2 | protein_coding | 0.412390116 | 3.871473383 | 0.3883412 | 0.5417907 | 0.999175987 |
| ENSMUSG00000036561 | Ppp6r2 | protein_coding | 0.365561146 | 4.045764098 | 0.3881143 | 0.5419074 | 0.999175987 |
| ENSMUSG00000020747 | Tmem94 | protein_coding | -0.291130058 | 4.770877502 | 0.3880801 | 0.541925 | 0.999175987 |
| ENSMUSG00000025260 | Hsd17b10 | protein_coding | 0.182141234 | 7.110727815 | 0.3880693 | 0.5419306 | 0.999175987 |
| ENSMUSG00000026273 | Mterf4 | protein_coding | -0.249613593 | 5.478050371 | 0.3879645 | 0.5419845 | 0.999175987 |
| ENSMUSG00000052713 | Zfp608 | protein_coding | -0.366726192 | 4.004855985 | 0.3879432 | 0.5419955 | 0.999175987 |
| ENSMUSG00000094152 | Slc6a16 | protein_coding | -0.449525387 | 3.775682246 | 0.3879377 | 0.5419983 | 0.999175987 |
| ENSMUSG00000070044 | Fam149a | protein_coding | 0.332744176 | 4.159628302 | 0.3878839 | 0.542026 | 0.999175987 |
| ENSMUSG00000002266 | Zim1 | protein_coding | 0.448940035 | 3.127123016 | 0.3877097 | 0.5421157 | 0.999175987 |
| ENSMUSG00000070306 | Ccdc153 | protein_coding | -0.312363788 | 5.855935236 | 0.3876532 | 0.5421589 | 0.999175987 |
| ENSMUSG00000029924 | Slc37a3 | protein_coding | 0.331746937 | 5.162331966 | 0.3871755 | 0.5423908 | 0.999175987 |
| ENSMUSG00000033047 | Eif3l | protein_coding | -0.196246594 | 7.38292893 | 0.3871504 | 0.5424038 | 0.999175987 |
| ENSMUSG00000030276 | Ttll3 | protein_coding | 0.714167437 | 2.692870825 | 0.3871086 | 0.5424254 | 0.999175987 |
| ENSMUSG00000054509 | Parp4 | protein_coding | -0.678948211 | 2.787169199 | 0.3870604 | 0.5424501 | 0.999175987 |
| ENSMUSG00000067367 | Lyar | protein_coding | 0.487069729 | 3.281575278 | 0.386951 | 0.5425066 | 0.999175987 |
| ENSMUSG00000031311 | Nono | protein_coding | -0.304436668 | 6.397071582 | 0.3872992 | 0.5425588 | 0.999175987 |
| ENSMUSG00000024974 | Smc3 | protein_coding | -0.266598932 | 5.852107548 | 0.3868027 | 0.542583 | 0.999175987 |
| ENSMUSG00000029366 | Dck | protein_coding | -0.324910279 | 4.599533717 | 0.3866599 | 0.5426567 | 0.999175987 |
| ENSMUSG00000030259 | Rassf8 | protein_coding | -0.403711645 | 3.946340098 | 0.3866271 | 0.5426736 | 0.999175987 |
| ENSMUSG00000026526 | Fh1 | protein_coding | 0.190606553 | 7.555256967 | 0.3865409 | 0.5427181 | 0.999175987 |
| ENSMUSG00000032575 | Manf | protein_coding | 0.237702202 | 6.02686512 | 0.3864378 | 0.5427713 | 0.999175987 |
| ENSMUSG00000074829 | 2010315B03Rik | protein_coding | 0.360526366 | 4.00988942 | 0.3863062 | 0.5428392 | 0.999175987 |
| ENSMUSG00000028995 | Fam126a | protein_coding | -0.494146739 | 3.874948119 | 0.3861969 | 0.5429097 | 0.999175987 |
| ENSMUSG00000022708 | Zbtb20 | protein_coding | -0.289427467 | 6.260583793 | 0.3861727 | 0.5429222 | 0.999175987 |
| ENSMUSG00000031925 | Maml2 | protein_coding | 0.365778067 | 4.363464793 | 0.3860894 | 0.5429511 | 0.999175987 |
| ENSMUSG00000020814 | Mxra7 | protein_coding | -0.37415838 | 3.90545563 | 0.3860505 | 0.5429712 | 0.999175987 |
| ENSMUSG00000030357 | Fkbp4 | protein_coding | 0.189446233 | 7.238306529 | 0.3860395 | 0.5429769 | 0.999175987 |
| ENSMUSG00000064264 | Zfp428 | protein_coding | 0.426738019 | 3.237660318 | 0.3858573 | 0.543071 | 0.999175987 |
| ENSMUSG00000107002 | 0610012G03Rik | protein_coding | 0.342060559 | 4.20281554 | 0.385835 | 0.5430825 | 0.999175987 |
| ENSMUSG00000020561 | Twistnb | protein_coding | -0.404439689 | 3.707681291 | 0.3857726 | 0.5431148 | 0.999175987 |
| ENSMUSG00000056413 | Adap1 | protein_coding | 0.254893826 | 6.48871669 | 0.3855671 | 0.5432209 | 0.999175987 |
| ENSMUSG00000025436 | Atp23 | protein_coding | 0.504529521 | 3.030248704 | 0.3855313 | 0.5432395 | 0.999175987 |
| ENSMUSG00000021902 | Phf7 | protein_coding | 0.955220095 | 2.481842382 | 0.3855365 | 0.5432727 | 0.999175987 |
| ENSMUSG00000020115 | Tbk1 | protein_coding | 0.321738041 | 4.763770872 | 0.3853115 | 0.5433531 | 0.999175987 |
| ENSMUSG00000022610 | Mapk12 | protein_coding | -0.801224961 | 2.362859331 | 0.3852965 | 0.5433609 | 0.999175987 |
| ENSMUSG00000015776 | Med22 | protein_coding | 0.200023637 | 6.323748204 | 0.385092 | 0.5434666 | 0.999175987 |
| ENSMUSG00000041408 | Wapl | protein_coding | -0.189766441 | 6.719766122 | 0.3850623 | 0.543482 | 0.999175987 |
| ENSMUSG00000022710 | Usp7 | protein_coding | -0.23035963 | 5.755904599 | 0.3850288 | 0.5434993 | 0.999175987 |
| ENSMUSG00000022201 | Zfr | protein_coding | -0.182825265 | 8.293012069 | 0.3848142 | 0.5436104 | 0.999175987 |
| ENSMUSG00000087056 | Gm14004 | processed_transcript | 0.340040617 | 5.122867176 | 0.3845206 | 0.5437624 | 0.999175987 |
| ENSMUSG00000041351 | Rap1gap | protein_coding | 0.277118855 | 6.109828099 | 0.3843913 | 0.5438294 | 0.999175987 |
| ENSMUSG00000034656 | Cacna1a | protein_coding | 0.519734913 | 3.40050763 | 0.3843352 | 0.5438584 | 0.999175987 |
| ENSMUSG00000028879 | Stx12 | protein_coding | -0.233876946 | 7.947623175 | 0.3843235 | 0.5438645 | 0.999175987 |
| ENSMUSG00000032615 | Nt5m | protein_coding | -0.226247889 | 6.081307107 | 0.3843163 | 0.5438682 | 0.999175987 |
| ENSMUSG00000002608 | Ccdc97 | protein_coding | 0.311894659 | 5.891737021 | 0.3843167 | 0.543882 | 0.999175987 |
| ENSMUSG00000028676 | Srsf10 | protein_coding | 0.237592343 | 5.481555705 | 0.3838812 | 0.5440937 | 0.999175987 |
| ENSMUSG00000025092 | Hspa12a | protein_coding | 0.221197906 | 5.960480157 | 0.3838303 | 0.5441201 | 0.999175987 |
| ENSMUSG00000039531 | Zufsp | protein_coding | 0.353185453 | 3.712014587 | 0.3836532 | 0.5442119 | 0.999175987 |
| ENSMUSG00000046032 | Snx12 | protein_coding | -0.18306964 | 6.618575788 | 0.3829597 | 0.5445719 | 0.999175987 |
| ENSMUSG00000034532 | Fbxo16 | protein_coding | 0.285070761 | 5.05428201 | 0.3828207 | 0.544644 | 0.999175987 |
| ENSMUSG00000027489 | Necab3 | protein_coding | 0.497853272 | 5.198566932 | 0.3850486 | 0.5446876 | 0.999175987 |
| ENSMUSG00000022967 | Ifnar1 | protein_coding | 0.372431601 | 3.699766087 | 0.3827104 | 0.5447013 | 0.999175987 |
| ENSMUSG00000025816 | Sec61a2 | protein_coding | 0.308682356 | 4.587601177 | 0.3826823 | 0.544716 | 0.999175987 |
| ENSMUSG00000031320 | Rps4x | protein_coding | 0.242100816 | 6.337535257 | 0.3824704 | 0.5448261 | 0.999175987 |
| ENSMUSG00000055436 | Srsf11 | protein_coding | -0.253830562 | 5.796884674 | 0.3824689 | 0.5448269 | 0.999175987 |
| ENSMUSG00000056167 | Cnot10 | protein_coding | 0.306322306 | 5.128757025 | 0.3824237 | 0.5448503 | 0.999175987 |
| ENSMUSG00000025089 | Gfra1 | protein_coding | -0.241858404 | 5.701463054 | 0.3823505 | 0.5448884 | 0.999175987 |
| ENSMUSG00000014763 | Fam120b | protein_coding | -0.224930982 | 6.325271253 | 0.3821944 | 0.5449696 | 0.999175987 |
| ENSMUSG00000038025 | Phf2 | protein_coding | 0.537606154 | 3.106562759 | 0.3820229 | 0.5450588 | 0.999175987 |
| ENSMUSG00000004633 | Chn2 | protein_coding | -0.701902368 | 2.262162866 | 0.3820175 | 0.5450616 | 0.999175987 |
| ENSMUSG00000056211 | R3hdm1 | protein_coding | -0.181510812 | 6.656809729 | 0.3818936 | 0.5451261 | 0.999175987 |
| ENSMUSG00000017412 | Cacnb4 | protein_coding | -0.230852954 | 5.601837244 | 0.3818055 | 0.5451719 | 0.999175987 |
| ENSMUSG00000074754 | Smim26 | protein_coding | 0.212067538 | 6.791577845 | 0.3817883 | 0.5451809 | 0.999175987 |
| ENSMUSG00000034903 | Cobll1 | protein_coding | -0.845323201 | 2.680774634 | 0.3815489 | 0.5453195 | 0.999175987 |
| ENSMUSG00000059213 | Ddn | protein_coding | -0.428887806 | 5.670490936 | 0.3833491 | 0.5454759 | 0.999175987 |
| ENSMUSG00000060240 | Cend1 | protein_coding | 0.234441803 | 9.860547497 | 0.381103 | 0.5455377 | 0.999175987 |
| ENSMUSG00000025228 | Actr1a | protein_coding | 0.299202469 | 7.759377898 | 0.3821437 | 0.5455384 | 0.999175987 |
| ENSMUSG00000002496 | Tsc2 | protein_coding | 0.343607358 | 4.776192968 | 0.3806622 | 0.5457675 | 0.999175987 |
| ENSMUSG00000047881 | Rell1 | protein_coding | -0.778024772 | 2.650813724 | 0.3806597 | 0.5457688 | 0.999175987 |
| ENSMUSG00000050312 | Nsun3 | protein_coding | -0.397417695 | 3.963659659 | 0.3802189 | 0.5459988 | 0.999175987 |
| ENSMUSG00000032666 | 1700025G04Rik | protein_coding | 0.255036611 | 5.294701351 | 0.3801477 | 0.546036 | 0.999175987 |
| ENSMUSG00000022400 | Rbx1 | protein_coding | 0.180677605 | 9.326508806 | 0.3801014 | 0.5460602 | 0.999175987 |
| ENSMUSG00000025558 | Dock9 | protein_coding | -0.39098443 | 4.382703992 | 0.3798861 | 0.5461726 | 0.999175987 |
| ENSMUSG00000018774 | Cd68 | protein_coding | 0.335549849 | 5.002781027 | 0.3798002 | 0.5462174 | 0.999175987 |

| ensembl_gene_id | external_gene_name | gene_biotype | log2 (fold change) | log2 (counts per million) | F-value | p-value | False discovery rate |
|---|---|---|---|---|---|---|---|
| ENSMUSG00000020585 | Laptm4a | protein_coding | -0.213350038 | 8.703104482 | 0.3795269 | 0.5463602 | 0.999175987 |
| ENSMUSG00000097729 | 2310015A10Rik | processed_transcript | 0.507669289 | 3.141519068 | 0.3795247 | 0.5463614 | 0.999175987 |
| ENSMUSG00000026490 | Cdc42bpa | protein_coding | -0.242201647 | 6.237104324 | 0.3793945 | 0.5464294 | 0.999175987 |
| ENSMUSG00000023988 | Bysl | protein_coding | 0.288094107 | 5.257597428 | 0.3789705 | 0.5466511 | 0.999175987 |
| ENSMUSG00000018196 | Glrx2 | protein_coding | 0.278389652 | 5.315577468 | 0.3788691 | 0.5467042 | 0.999175987 |
| ENSMUSG00000095041 | AC149090.1 | protein_coding | 0.366596906 | 4.889587051 | 0.3782851 | 0.5470099 | 0.999175987 |
| ENSMUSG00000030316 | Tamm41 | protein_coding | -0.428995023 | 3.63002157 | 0.3781719 | 0.5470692 | 0.999175987 |
| ENSMUSG00000037993 | Dhx38 | protein_coding | 0.400427533 | 4.389647886 | 0.3777971 | 0.5472656 | 0.999175987 |
| ENSMUSG00000037916 | Ndufv1 | protein_coding | 0.205541622 | 9.217131399 | 0.3777412 | 0.547295 | 0.999175987 |
| ENSMUSG00000041879 | Ipo9 | protein_coding | 0.234040184 | 6.119711116 | 0.3777185 | 0.5473068 | 0.999175987 |
| ENSMUSG00000069631 | Strada | protein_coding | -0.358375947 | 4.689361972 | 0.3777071 | 0.5473128 | 0.999175987 |
| ENSMUSG00000040599 | Mis12 | protein_coding | 0.324037829 | 4.846130486 | 0.3776208 | 0.5473581 | 0.999175987 |
| ENSMUSG00000051346 | Spryd4 | protein_coding | -0.280475281 | 4.964114323 | 0.377149 | 0.5476056 | 0.999175987 |
| ENSMUSG00000020566 | Atp6v1c2 | protein_coding | -0.426433665 | 3.341533126 | 0.3771309 | 0.5476151 | 0.999175987 |
| ENSMUSG00000040867 | Begain | protein_coding | 0.294553447 | 4.989020161 | 0.3771045 | 0.547629 | 0.999175987 |
| ENSMUSG00000100164 | 2610306M01Rik | bidirectional_promote | 0.258579336 | 5.269928062 | 0.3768288 | 0.5477738 | 0.999175987 |
| ENSMUSG00000075700 | Selenot | protein_coding | 0.269504547 | 5.747603591 | 0.3766344 | 0.5478758 | 0.999175987 |
| ENSMUSG00000005102 | Eif2ak4 | protein_coding | -0.425121828 | 3.634315011 | 0.3766039 | 0.5478919 | 0.999175987 |
| ENSMUSG00000038000 | Acd | protein_coding | 0.259060433 | 6.959643338 | 0.3765789 | 0.5479189 | 0.999175987 |
| ENSMUSG00000048001 | Hes5 | protein_coding | 0.66564188 | 2.560221355 | 0.3764689 | 0.5479628 | 0.999175987 |
| ENSMUSG00000034247 | Plekhm1 | protein_coding | -0.375868727 | 3.720036447 | 0.3763432 | 0.5480289 | 0.999175987 |
| ENSMUSG00000020684 | Rasl10b | protein_coding | -0.374195544 | 5.728973139 | 0.3771741 | 0.5481308 | 0.999175987 |
| ENSMUSG00000097911 | Gm26691 | antisense_RNA | -0.558937065 | 2.904787901 | 0.3758168 | 0.5483057 | 0.999175987 |
| ENSMUSG00000035601 | Trmt10b | protein_coding | -0.215501882 | 5.834775679 | 0.3757602 | 0.5483355 | 0.999175987 |
| ENSMUSG00000029020 | Mfn2 | protein_coding | 0.288107581 | 7.370764413 | 0.376603 | 0.5484303 | 0.999175987 |
| ENSMUSG00000078958 | Atp6ap1l | protein_coding | -0.449391433 | 3.491809677 | 0.3755625 | 0.5484396 | 0.999175987 |
| ENSMUSG00000037306 | Man1c1 | protein_coding | -0.482430942 | 3.687824388 | 0.3750199 | 0.5487254 | 0.999175987 |
| ENSMUSG00000040351 | Ankib1 | protein_coding | -0.327790343 | 4.844242296 | 0.3749941 | 0.548739 | 0.999175987 |
| ENSMUSG00000084883 | Ccdc85c | protein_coding | 0.413699413 | 5.217603897 | 0.3756002 | 0.548927 | 0.999175987 |
| ENSMUSG00000080727 | C920021L13Rik | antisense_RNA | 0.580583512 | 2.533672648 | 0.3744653 | 0.5490178 | 0.999175987 |
| ENSMUSG00000055991 | Zkscan5 | protein_coding | -0.41197629 | 3.547191066 | 0.3741292 | 0.5491951 | 0.999175987 |
| ENSMUSG00000029265 | Dr1 | protein_coding | -0.367310791 | 4.416622029 | 0.3741193 | 0.5492004 | 0.999175987 |
| ENSMUSG00000038671 | Arfrp1 | protein_coding | 0.193869245 | 6.881633397 | 0.374026 | 0.5492496 | 0.999175987 |
| ENSMUSG00000030701 | Plekhb1 | protein_coding | -0.216706684 | 9.101479327 | 0.3739055 | 0.5493132 | 0.999175987 |
| ENSMUSG00000053550 | Shisa7 | protein_coding | 0.345836725 | 4.438449316 | 0.3738684 | 0.5493328 | 0.999175987 |
| ENSMUSG00000020182 | Ddc | protein_coding | -0.25011 | 5.347214123 | 0.3738385 | 0.5493486 | 0.999175987 |
| ENSMUSG00000041781 | Cpsf2 | protein_coding | 0.232207597 | 5.601195972 | 0.373697 | 0.5494233 | 0.999175987 |
| ENSMUSG00000022827 | Rabl3 | protein_coding | 0.363341114 | 4.725457774 | 0.3735145 | 0.5495198 | 0.999175987 |
| ENSMUSG00000030223 | Ptpro | protein_coding | 0.267118995 | 4.842292511 | 0.3733258 | 0.5496195 | 0.999175987 |
| ENSMUSG00000027349 | Fam98b | protein_coding | -0.203819046 | 6.291079979 | 0.373189 | 0.5496918 | 0.999175987 |
| ENSMUSG00000018848 | Rars | protein_coding | -0.223185191 | 6.479045316 | 0.3731101 | 0.5497336 | 0.999175987 |
| ENSMUSG00000024357 | Sil1 | protein_coding | 0.227993662 | 6.185084574 | 0.3730618 | 0.5497591 | 0.999175987 |
| ENSMUSG00000040675 | Mthfd1l | protein_coding | 0.447257442 | 3.427437564 | 0.373007 | 0.5497881 | 0.999175987 |
| ENSMUSG00000023473 | Celsr3 | protein_coding | 0.648655475 | 2.799257655 | 0.3725524 | 0.5500286 | 0.999175987 |
| ENSMUSG00000059970 | Hspa2 | protein_coding | -0.337199233 | 5.644125734 | 0.3725746 | 0.5500307 | 0.999175987 |
| ENSMUSG00000026107 | Nabp1 | protein_coding | -0.425536382 | 3.476302538 | 0.3723875 | 0.550116 | 0.999175987 |
| ENSMUSG00000040681 | Hmgn1 | protein_coding | 0.294406909 | 4.808654613 | 0.3722209 | 0.5502042 | 0.999175987 |
| ENSMUSG00000020628 | Trappc12 | protein_coding | 0.202966507 | 6.182638544 | 0.3717858 | 0.5504348 | 0.999175987 |
| ENSMUSG00000036469 | 1-Mar | protein_coding | -0.252784145 | 4.910146209 | 0.3713142 | 0.5506849 | 0.999175987 |
| ENSMUSG00000050830 | Vwc2 | protein_coding | 0.48942261 | 3.167405937 | 0.3713041 | 0.5506902 | 0.999175987 |
| ENSMUSG00000026192 | Atic | protein_coding | 0.304000054 | 5.524916594 | 0.3712773 | 0.5507045 | 0.999175987 |
| ENSMUSG00000046572 | Zfp518b | protein_coding | -0.713065047 | 2.427074338 | 0.3712134 | 0.5507384 | 0.999175987 |
| ENSMUSG00000049470 | Aff4 | protein_coding | -0.215992455 | 6.631452802 | 0.3711471 | 0.5507736 | 0.999175987 |
| ENSMUSG00000018372 | Cep95 | protein_coding | 0.330799607 | 3.87349461 | 0.3709905 | 0.5508567 | 0.999175987 |
| ENSMUSG00000041607 | Mbp | protein_coding | -0.268525689 | 11.80476767 | 0.3709804 | 0.5508758 | 0.999175987 |
| ENSMUSG00000025817 | Nudt5 | protein_coding | -0.247590606 | 5.502467842 | 0.3708953 | 0.5509072 | 0.999175987 |
| ENSMUSG00000044018 | Mrpl50 | protein_coding | 0.198101754 | 7.294997489 | 0.3704353 | 0.5511516 | 0.999175987 |
| ENSMUSG00000034501 | Pcnx4 | protein_coding | 0.308285592 | 4.806627932 | 0.3703336 | 0.5512056 | 0.999175987 |
| ENSMUSG00000043671 | Dpy19l3 | protein_coding | 0.385095989 | 4.19884492 | 0.37016 | 0.551298 | 0.999175987 |
| ENSMUSG00000039270 | Megf9 | protein_coding | -0.266483361 | 5.528848811 | 0.3696588 | 0.5515646 | 0.999175987 |
| ENSMUSG00000035171 | 1110059E24Rik | protein_coding | 0.196403528 | 7.010336727 | 0.3696542 | 0.5515671 | 0.999175987 |
| ENSMUSG00000038740 | Mvb12b | protein_coding | 0.313503877 | 4.934710172 | 0.3695265 | 0.551635 | 0.999175987 |
| ENSMUSG00000034278 | Dnajc17 | protein_coding | 0.314162926 | 4.320435729 | 0.369526 | 0.5516353 | 0.999175987 |
| ENSMUSG00000028894 | Inpp5b | protein_coding | -0.353464841 | 4.148575402 | 0.3691097 | 0.551857 | 0.999175987 |
| ENSMUSG00000029712 | Actl6b | protein_coding | -0.225851925 | 6.161214538 | 0.368984 | 0.551924 | 0.999175987 |
| ENSMUSG00000037210 | Fam193a | protein_coding | 0.339253614 | 4.910194379 | 0.3685841 | 0.5521372 | 0.999175987 |
| ENSMUSG00000038611 | Phrf1 | protein_coding | 0.32594128 | 4.427450614 | 0.3684576 | 0.5522046 | 0.999175987 |
| ENSMUSG00000038453 | Srcin1 | protein_coding | -0.234385861 | 5.644276852 | 0.3684317 | 0.5522185 | 0.999175987 |
| ENSMUSG00000022747 | St3gal6 | protein_coding | -0.41579467 | 3.685599646 | 0.3684275 | 0.5522207 | 0.999175987 |
| ENSMUSG00000034758 | Tle6 | protein_coding | 1.029307156 | 2.579938696 | 0.3696899 | 0.5522326 | 0.999175987 |
| ENSMUSG00000042557 | Sin3a | protein_coding | -0.538763848 | 2.824291115 | 0.3680025 | 0.5524475 | 0.999175987 |
| ENSMUSG00000078816 | Prkcg | protein_coding | -0.309748496 | 7.171257509 | 0.3694886 | 0.5524889 | 0.999175987 |
| ENSMUSG00000042396 | Rbm7 | protein_coding | -0.238686083 | 5.8744885 | 0.3678733 | 0.5525165 | 0.999175987 |
| ENSMUSG00000036368 | Rmdn2 | protein_coding | -0.643210307 | 2.618125722 | 0.3678306 | 0.5525393 | 0.999175987 |
| ENSMUSG00000024197 | Plin3 | protein_coding | -0.297837396 | 5.200013871 | 0.3678064 | 0.5525523 | 0.999175987 |
| ENSMUSG00000034640 | Tiparp | protein_coding | -0.392798948 | 4.451759757 | 0.3674053 | 0.5527666 | 0.999175987 |
| ENSMUSG00000048264 | Dip2c | protein_coding | -0.273893823 | 4.975060271 | 0.3673521 | 0.552795 | 0.999175987 |
| ENSMUSG00000050323 | Ndufaf6 | protein_coding | -0.597795379 | 2.951008596 | 0.3673242 | 0.5528099 | 0.999175987 |

| ensembl_gene_id | external_gene_name | gene_biotype | log2 (fold change) | log2 (counts per million) | F-value | p-value | False discovery rate |
|---|---|---|---|---|---|---|---|
| ENSMUSG00000044968 | Napepld | protein_coding | -0.534733689 | 3.709934025 | 0.3674001 | 0.5528191 | 0.999175987 |
| ENSMUSG00000034429 | Zfp707 | protein_coding | 0.391040122 | 4.951956453 | 0.3670942 | 0.5529466 | 0.999175987 |
| ENSMUSG00000039176 | Polg | protein_coding | -0.751413637 | 2.443151353 | 0.3670014 | 0.5529825 | 0.999175987 |
| ENSMUSG00000052364 | B630019K06Rik | protein_coding | -0.266303927 | 5.775078088 | 0.366679 | 0.553155 | 0.999175987 |
| ENSMUSG00000020078 | Vps26a | protein_coding | -0.216668466 | 6.546155547 | 0.3666739 | 0.5531578 | 0.999175987 |
| ENSMUSG00000033157 | Abhd10 | protein_coding | 0.233027726 | 5.680328417 | 0.3666417 | 0.553175 | 0.999175987 |
| ENSMUSG00000069744 | Psmb3 | protein_coding | 0.199281492 | 7.452217863 | 0.3660424 | 0.553496 | 0.999175987 |
| ENSMUSG00000025759 | Mfsd8 | protein_coding | 0.765797073 | 2.851632995 | 0.3659338 | 0.5535678 | 0.999175987 |
| ENSMUSG00000085042 | Abhd11os | transcribed_unitary_p | -0.297773771 | 5.216477272 | 0.3658018 | 0.5536249 | 0.999175987 |
| ENSMUSG00000027203 | Dut | protein_coding | 0.306545566 | 4.373505902 | 0.3657081 | 0.5536752 | 0.999175987 |
| ENSMUSG00000032786 | Alas1 | protein_coding | -0.316787411 | 6.36808781 | 0.366481 | 0.5537905 | 0.999175987 |
| ENSMUSG00000024528 | Srfbp1 | protein_coding | -0.323876324 | 4.73445239 | 0.365456 | 0.5538104 | 0.999175987 |
| ENSMUSG00000020361 | Hspa4 | protein_coding | -0.253081491 | 5.412967386 | 0.3653262 | 0.55388 | 0.999175987 |
| ENSMUSG00000007783 | Cpt1c | protein_coding | 0.226571731 | 6.709707017 | 0.3651901 | 0.553953 | 0.999175987 |
| ENSMUSG00000001054 | Rmnd5b | protein_coding | 0.248252158 | 5.63977067 | 0.3650725 | 0.5540162 | 0.999175987 |
| ENSMUSG00000040557 | Mettl27 | protein_coding | 0.37760477 | 4.134411456 | 0.364632 | 0.5542527 | 0.999175987 |
| ENSMUSG00000044030 | Irf2bp1 | protein_coding | 0.306517759 | 4.346356885 | 0.3645485 | 0.5542976 | 0.999175987 |
| ENSMUSG00000066037 | Hnrnpr | protein_coding | -0.199659326 | 6.498009891 | 0.3643279 | 0.5544162 | 0.999175987 |
| ENSMUSG00000026353 | Ubxn4 | protein_coding | 0.214341559 | 5.908342931 | 0.3642449 | 0.5544608 | 0.999175987 |
| ENSMUSG00000074555 | Gm10714 | antisense_RNA | 0.658223487 | 2.85445708 | 0.3642097 | 0.5544798 | 0.999175987 |
| ENSMUSG00000034022 | Cpsf1 | protein_coding | -0.327853045 | 4.900114075 | 0.3642066 | 0.5544814 | 0.999175987 |
| ENSMUSG00000030335 | Mrpl51 | protein_coding | 0.176516959 | 6.91307832 | 0.3640005 | 0.5545923 | 0.999175987 |
| ENSMUSG00000014606 | Slc25a11 | protein_coding | 0.180616475 | 8.391474826 | 0.3635945 | 0.5548108 | 0.999175987 |
| ENSMUSG00000001521 | Tulp3 | protein_coding | 0.461870493 | 3.3495638 | 0.3635611 | 0.5548287 | 0.999175987 |
| ENSMUSG00000052397 | Ezr | protein_coding | -0.350007751 | 4.466363293 | 0.363448 | 0.5548897 | 0.999175987 |
| ENSMUSG00000038533 | Cbfa2t2 | protein_coding | 0.346101566 | 6.28692202 | 0.3649414 | 0.5549143 | 0.999175987 |
| ENSMUSG00000021076 | Actr10 | protein_coding | -0.187503032 | 8.860257403 | 0.3632192 | 0.5550129 | 0.999175987 |
| ENSMUSG00000019791 | Hint3 | protein_coding | 0.333535266 | 4.098868876 | 0.3630426 | 0.5551081 | 0.999175987 |
| ENSMUSG00000068696 | Gpr88 | protein_coding | -0.506400197 | 6.335451933 | 0.3695379 | 0.5551295 | 0.999175987 |
| ENSMUSG00000003161 | Sri | protein_coding | -0.173661289 | 7.586269207 | 0.3629148 | 0.5551769 | 0.999175987 |
| ENSMUSG00000037062 | Sh3glb1 | protein_coding | -0.249490866 | 5.881245522 | 0.3627947 | 0.5552417 | 0.999175987 |
| ENSMUSG00000045312 | Lhfpl2 | protein_coding | -0.791363807 | 2.521567076 | 0.3626892 | 0.5552985 | 0.999175987 |
| ENSMUSG00000063694 | Cycs | protein_coding | -0.252212033 | 5.154449995 | 0.3624973 | 0.5554021 | 0.999175987 |
| ENSMUSG00000043831 | Lysmd4 | protein_coding | 0.186253412 | 6.736433411 | 0.3624897 | 0.5554062 | 0.999175987 |
| ENSMUSG00000034156 | Tspoap1 | protein_coding | 0.373479813 | 4.054410639 | 0.3623889 | 0.5554606 | 0.999175987 |
| ENSMUSG00000030451 | Herc2 | protein_coding | 0.295084507 | 5.246005761 | 0.3619824 | 0.55568 | 0.999175987 |
| ENSMUSG00000054737 | Zfp182 | protein_coding | 0.635222526 | 2.357138798 | 0.3619094 | 0.5557194 | 0.999175987 |
| ENSMUSG00000047897 | Ripply2 | protein_coding | 0.499993523 | 2.911430618 | 0.3616655 | 0.5558512 | 0.999175987 |
| ENSMUSG00000029108 | Pcdh7 | protein_coding | -0.244911192 | 5.97182435 | 0.3615885 | 0.5558928 | 0.999175987 |
| ENSMUSG00000028790 | Khdrbs1 | protein_coding | -0.259980916 | 5.447666043 | 0.3614642 | 0.55596 | 0.999175987 |
| ENSMUSG00000062991 | Nrg1 | protein_coding | -0.252133567 | 5.108745726 | 0.3610598 | 0.5561786 | 0.999175987 |
| ENSMUSG00000030752 | Kdm8 | protein_coding | -0.651168281 | 2.7555095 | 0.3608037 | 0.5563172 | 0.999175987 |
| ENSMUSG00000028163 | Nfkb1 | protein_coding | 0.456343641 | 3.062743816 | 0.3607549 | 0.5563436 | 0.999175987 |
| ENSMUSG00000071753 | C230004F18Rik | processed_transcript | 0.311817023 | 4.836514358 | 0.3605117 | 0.5564753 | 0.999175987 |
| ENSMUSG00000039652 | Cpeb3 | protein_coding | 0.293011917 | 5.090136494 | 0.3604249 | 0.5565223 | 0.999175987 |
| ENSMUSG00000028954 | Nub1 | protein_coding | -0.220796524 | 6.897385071 | 0.3603672 | 0.5565535 | 0.999175987 |
| ENSMUSG00000009905 | Kdsr | protein_coding | -0.250605222 | 5.742732626 | 0.3602418 | 0.5566215 | 0.999175987 |
| ENSMUSG00000027339 | Rassf2 | protein_coding | -0.296929412 | 4.579458605 | 0.360127 | 0.5566837 | 0.999175987 |
| ENSMUSG00000021368 | Tbc1d7 | protein_coding | 0.200904754 | 6.389363671 | 0.359724 | 0.5569022 | 0.999175987 |
| ENSMUSG00000031371 | Haus7 | protein_coding | -0.209685477 | 5.661302932 | 0.3596093 | 0.5569644 | 0.999175987 |
| ENSMUSG00000059208 | Hnrnpm | protein_coding | 0.238049847 | 6.265211582 | 0.3593926 | 0.557082 | 0.999175987 |
| ENSMUSG00000019370 | Calm3 | protein_coding | 0.177395932 | 9.637650955 | 0.3592306 | 0.5571699 | 0.999175987 |
| ENSMUSG00000055296 | Tmem245 | protein_coding | -0.353922694 | 4.413912717 | 0.358998 | 0.5572962 | 0.999175987 |
| ENSMUSG00000022498 | Txndc11 | protein_coding | 0.501029418 | 2.905277887 | 0.3589612 | 0.5573161 | 0.999175987 |
| ENSMUSG00000028552 | Eps15 | protein_coding | -0.19634555 | 6.687543463 | 0.3588085 | 0.5573991 | 0.999175987 |
| ENSMUSG00000017221 | Psmd3 | protein_coding | 0.256690167 | 5.503592171 | 0.3587753 | 0.5574172 | 0.999175987 |
| ENSMUSG00000061718 | Ppp1r1b | protein_coding | 0.579537069 | 2.587541766 | 0.3586391 | 0.5574912 | 0.999175987 |
| ENSMUSG00000033707 | Lrrc24 | protein_coding | 0.380076369 | 4.087936264 | 0.3584146 | 0.5576132 | 0.999175987 |
| ENSMUSG00000039110 | Mycbpap | protein_coding | -0.557270527 | 2.801243388 | 0.3583525 | 0.557647 | 0.999175987 |
| ENSMUSG00000003814 | Calr | protein_coding | 0.181812665 | 8.158670102 | 0.3582121 | 0.5577233 | 0.999175987 |
| ENSMUSG00000053702 | Nebl | protein_coding | -0.266494658 | 4.805923661 | 0.357802 | 0.5579465 | 0.999175987 |
| ENSMUSG00000027425 | Kat14 | protein_coding | -0.30083115 | 4.035801704 | 0.3577865 | 0.5579549 | 0.999175987 |
| ENSMUSG00000057054 | Inca1 | protein_coding | 0.352079495 | 3.848387541 | 0.3573342 | 0.5582012 | 0.999175987 |
| ENSMUSG00000028252 | Ccnc | protein_coding | 0.22420761 | 7.047618896 | 0.3573088 | 0.5582151 | 0.999175987 |
| ENSMUSG00000028199 | Cryz | protein_coding | -0.314389813 | 5.182555888 | 0.3572452 | 0.5582497 | 0.999175987 |
| ENSMUSG00000048047 | Zbtb33 | protein_coding | -0.573778773 | 2.590932394 | 0.3567988 | 0.5584931 | 0.999175987 |
| ENSMUSG00000074797 | Itpa | protein_coding | -0.189455399 | 6.312542748 | 0.3567196 | 0.5585362 | 0.999175987 |
| ENSMUSG00000027406 | Idh3b | protein_coding | 0.185533121 | 9.161263292 | 0.3564422 | 0.5586876 | 0.999175987 |
| ENSMUSG00000046573 | Lyrm4 | protein_coding | 0.240354927 | 7.735868849 | 0.3562861 | 0.5587863 | 0.999175987 |
| ENSMUSG00000024953 | Prdx5 | protein_coding | 0.174078443 | 10.25842496 | 0.3562002 | 0.5588197 | 0.999175987 |
| ENSMUSG00000006435 | Neurl1a | protein_coding | -0.270374198 | 5.92054398 | 0.3558887 | 0.5589899 | 0.999175987 |
| ENSMUSG00000074971 | Fibin | protein_coding | 0.564346129 | 2.923401288 | 0.3557238 | 0.55908 | 0.999175987 |
| ENSMUSG00000021431 | Snrnp48 | protein_coding | -0.326836087 | 4.203297408 | 0.3556791 | 0.5591044 | 0.999175987 |
| ENSMUSG00000035509 | Fbxl21 | protein_coding | 0.358992716 | 3.861763145 | 0.3556081 | 0.5591432 | 0.999175987 |
| ENSMUSG00000073725 | Lmbrd1 | protein_coding | -0.19322051 | 6.696811724 | 0.355572 | 0.5591629 | 0.999175987 |
| ENSMUSG00000046020 | Pofut1 | protein_coding | -0.572499402 | 2.626413497 | 0.3555019 | 0.5592013 | 0.999175987 |
| ENSMUSG00000071252 | 2210408I21Rik | protein_coding | -0.283869798 | 6.947680985 | 0.3561259 | 0.5593494 | 0.999175987 |
| ENSMUSG00000037608 | Bclaf1 | protein_coding | -0.210931247 | 7.408676833 | 0.3550265 | 0.5594613 | 0.999175987 |

| ensembl_gene_id | external_gene_name | gene_biotype | log2 (fold change) | log2 (counts per million) | F-value | p-value | False discovery rate |
|---|---|---|---|---|---|---|---|
| ENSMUSG00000022231 | Sema5a | protein_coding | -0.34148884 | 4.641101121 | 0.3548058 | 0.5595821 | 0.999175987 |
| ENSMUSG00000002728 | Naa20 | protein_coding | 0.181260351 | 7.108867017 | 0.3547499 | 0.5596127 | 0.999175987 |
| ENSMUSG00000032034 | Kcnj5 | protein_coding | -0.66638052 | 2.643768098 | 0.3545431 | 0.559726 | 0.999175987 |
| ENSMUSG00000036667 | Tcaf1 | protein_coding | -0.218967237 | 7.530609626 | 0.354529 | 0.5597337 | 0.999175987 |
| ENSMUSG00000003354 | Ccdc65 | protein_coding | -0.360289149 | 4.403472492 | 0.3544551 | 0.5597742 | 0.999175987 |
| ENSMUSG00000066687 | Zbtb16 | protein_coding | -0.608593535 | 2.852946376 | 0.354407 | 0.5598005 | 0.999175987 |
| ENSMUSG00000045039 | Megf8 | protein_coding | -0.358290239 | 4.716612134 | 0.3543726 | 0.5598194 | 0.999175987 |
| ENSMUSG00000019920 | Lims1 | protein_coding | 0.317809457 | 4.735409188 | 0.3541306 | 0.559952 | 0.999175987 |
| ENSMUSG00000031938 | 4931406C07Rik | protein_coding | -0.431881568 | 3.821503406 | 0.3540446 | 0.5599992 | 0.999175987 |
| ENSMUSG00000029821 | Dfna5 | protein_coding | -0.284727234 | 7.575944082 | 0.3549247 | 0.5600368 | 0.999175987 |
| ENSMUSG00000030868 | Dctn5 | protein_coding | 0.21242259 | 5.84964316 | 0.3539111 | 0.5600724 | 0.999175987 |
| ENSMUSG00000014426 | Map3k4 | protein_coding | -0.340724106 | 4.223032953 | 0.3538833 | 0.5600876 | 0.999175987 |
| ENSMUSG00000052949 | Rnf157 | protein_coding | -0.178455092 | 6.718553286 | 0.3537962 | 0.5601354 | 0.999175987 |
| ENSMUSG00000024122 | Pdpk1 | protein_coding | -0.203916994 | 6.553150578 | 0.3536364 | 0.5602231 | 0.999175987 |
| ENSMUSG00000004902 | Slc25a18 | protein_coding | -0.241730875 | 6.351347819 | 0.3536178 | 0.5602332 | 0.999175987 |
| ENSMUSG00000097675 | 1700101I11Rik | processed_transcript | 0.515849873 | 2.873337155 | 0.3535765 | 0.5602559 | 0.999175987 |
| ENSMUSG00000018340 | Anxa6 | protein_coding | 0.229033954 | 6.697405451 | 0.3531157 | 0.5605089 | 0.999175987 |
| ENSMUSG00000039849 | Pcif1 | protein_coding | 0.205943238 | 6.315142522 | 0.3530166 | 0.5605634 | 0.999175987 |
| ENSMUSG00000027646 | Src | protein_coding | -0.382173656 | 4.171514877 | 0.3529491 | 0.5606004 | 0.999175987 |
| ENSMUSG00000024187 | Fam234a | protein_coding | 0.409918332 | 4.779985029 | 0.3523895 | 0.5609797 | 0.999175987 |
| ENSMUSG00000024896 | Minpp1 | protein_coding | 0.317478473 | 4.77867413 | 0.3521507 | 0.5610395 | 0.999175987 |
| ENSMUSG00000037148 | Arhgap10 | protein_coding | -0.725116048 | 2.855312343 | 0.3517878 | 0.5612525 | 0.999175987 |
| ENSMUSG00000027475 | Kif3b | protein_coding | 0.275906143 | 4.750649199 | 0.3516989 | 0.5612882 | 0.999175987 |
| ENSMUSG00000028341 | Nr4a3 | protein_coding | -0.564286606 | 3.210663218 | 0.3516891 | 0.5612936 | 0.999175987 |
| ENSMUSG00000033530 | Ttc7b | protein_coding | 0.422186101 | 4.516190074 | 0.351638 | 0.5613702 | 0.999175987 |
| ENSMUSG00000022892 | App | protein_coding | -0.192405294 | 9.150916813 | 0.3514761 | 0.5614109 | 0.999175987 |
| ENSMUSG00000042331 | Specc1 | protein_coding | -0.322624258 | 5.358759164 | 0.351144 | 0.5616073 | 0.999175987 |
| ENSMUSG00000040731 | Eif4h | protein_coding | 0.184775819 | 8.360380443 | 0.3510893 | 0.5616241 | 0.999175987 |
| ENSMUSG00000068798 | Rap1a | protein_coding | -0.259827553 | 4.606959822 | 0.3503759 | 0.5620178 | 0.999175987 |
| ENSMUSG00000041153 | Osgin2 | protein_coding | 0.468026325 | 3.325612548 | 0.3502111 | 0.5621088 | 0.999175987 |
| ENSMUSG00000111977 | AC132265.1 | bidirectional_promote | -0.516890874 | 3.175579365 | 0.3499853 | 0.5622336 | 0.999175987 |
| ENSMUSG00000038563 | Efl1 | protein_coding | -0.267849075 | 4.51451738 | 0.3497073 | 0.5623872 | 0.999175987 |
| ENSMUSG00000043252 | Tmem64 | protein_coding | 0.347874211 | 4.869118991 | 0.3494396 | 0.5625353 | 0.999175987 |
| ENSMUSG00000057816 | 1700007G11Rik | protein_coding | -0.356586653 | 3.541309499 | 0.3493025 | 0.5626112 | 0.999175987 |
| ENSMUSG00000021417 | Eci2 | protein_coding | -0.273632375 | 5.131709028 | 0.3491563 | 0.5626921 | 0.999175987 |
| ENSMUSG00000043929 | Klhl15 | protein_coding | 0.388991526 | 4.742716628 | 0.3489697 | 0.5628087 | 0.999175987 |
| ENSMUSG00000020483 | Dynll2 | protein_coding | -0.267001182 | 8.058738353 | 0.3490866 | 0.5630695 | 0.999175987 |
| ENSMUSG00000022570 | Tsta3 | protein_coding | 0.173331287 | 6.72415395 | 0.3484211 | 0.5630993 | 0.999175987 |
| ENSMUSG00000026893 | Gca | protein_coding | 0.354251408 | 4.339121022 | 0.3483471 | 0.5631404 | 0.999175987 |
| ENSMUSG00000091625 | Lsm5 | protein_coding | 0.485029181 | 3.039674698 | 0.3482228 | 0.5632093 | 0.999175987 |
| ENSMUSG00000006782 | Cnp | protein_coding | -0.258024702 | 9.811024305 | 0.348202 | 0.563234 | 0.999175987 |
| ENSMUSG00000025964 | Adam23 | protein_coding | 0.211663534 | 6.261421837 | 0.3481041 | 0.5632751 | 0.999175987 |
| ENSMUSG00000017778 | Cox7c | protein_coding | 0.357259883 | 3.971414894 | 0.3480401 | 0.5633106 | 0.999175987 |
| ENSMUSG00000046329 | Slc25a23 | protein_coding | -0.336191351 | 3.69950055 | 0.3479602 | 0.5633549 | 0.999175987 |
| ENSMUSG00000021686 | Ap3b1 | protein_coding | 0.301134343 | 4.754380161 | 0.3477884 | 0.5634502 | 0.999175987 |
| ENSMUSG00000030314 | Atg7 | protein_coding | -0.377918251 | 6.424224491 | 0.3501151 | 0.5634527 | 0.999175987 |
| ENSMUSG00000038187 | Btbd10 | protein_coding | -0.239576303 | 5.697058457 | 0.3477518 | 0.5634705 | 0.999175987 |
| ENSMUSG00000001017 | Chtop | protein_coding | -0.197898749 | 5.922652074 | 0.3475835 | 0.563564 | 0.999175987 |
| ENSMUSG00000046637 | Ttc34 | protein_coding | 0.751690785 | 2.504701623 | 0.3473413 | 0.5636985 | 0.999175987 |
| ENSMUSG00000046532 | Ar | protein_coding | 0.248108666 | 5.82960088 | 0.3470865 | 0.5638401 | 0.999175987 |
| ENSMUSG00000002233 | Rhoc | protein_coding | 0.399407008 | 4.492495405 | 0.3471023 | 0.5638445 | 0.999175987 |
| ENSMUSG00000049670 | Morn4 | protein_coding | 0.162773643 | 7.601645334 | 0.3468817 | 0.5639539 | 0.999175987 |
| ENSMUSG00000022300 | Dcaf13 | protein_coding | 0.26965297 | 6.142582573 | 0.346785 | 0.5640209 | 0.999175987 |
| ENSMUSG00000087148 | 1700030C14Rik | antisense_RNA | 0.720191964 | 2.385949082 | 0.346683 | 0.5640644 | 0.999175987 |
| ENSMUSG00000033885 | Pxk | protein_coding | 0.298586017 | 5.072323438 | 0.3466762 | 0.5640682 | 0.999175987 |
| ENSMUSG00000057982 | Zfp809 | protein_coding | 0.357884947 | 4.043580567 | 0.3466199 | 0.5640995 | 0.999175987 |
| ENSMUSG00000021242 | Npc2 | protein_coding | -0.181480526 | 7.42587383 | 0.3466014 | 0.5641098 | 0.999175987 |
| ENSMUSG00000029670 | Ing3 | protein_coding | -0.25531398 | 5.85921492 | 0.3465705 | 0.564127 | 0.999175987 |
| ENSMUSG00000073131 | Vma21 | protein_coding | -0.189891082 | 6.259907015 | 0.3463842 | 0.5642306 | 0.999175987 |
| ENSMUSG00000056608 | Chd9 | protein_coding | -0.274476242 | 5.372321809 | 0.3463528 | 0.5642481 | 0.999175987 |
| ENSMUSG00000028413 | B4galt1 | protein_coding | 0.493914528 | 2.837557655 | 0.3463164 | 0.5642684 | 0.999175987 |
| ENSMUSG00000026696 | Vamp4 | protein_coding | 0.209246368 | 7.902477043 | 0.3463081 | 0.564273 | 0.999175987 |
| ENSMUSG00000003402 | Prkcsh | protein_coding | 0.346551958 | 5.289178934 | 0.3463224 | 0.5642782 | 0.999175987 |
| ENSMUSG00000022556 | Hsf1 | protein_coding | 0.336465334 | 4.899558544 | 0.3462977 | 0.5642788 | 0.999175987 |
| ENSMUSG00000021518 | Ptdss1 | protein_coding | -0.409376456 | 4.259465503 | 0.3460161 | 0.5644488 | 0.999175987 |
| ENSMUSG00000034203 | Chchd4 | protein_coding | 0.171100515 | 7.239234446 | 0.3459394 | 0.5644783 | 0.999175987 |
| ENSMUSG00000036430 | Tbcc | protein_coding | 0.435535832 | 3.39072153 | 0.3457288 | 0.5645956 | 0.999175987 |
| ENSMUSG00000020102 | Slc16a7 | protein_coding | 0.448632089 | 3.21491113 | 0.3451415 | 0.5649231 | 0.999175987 |
| ENSMUSG00000033159 | Cnppd1 | protein_coding | 0.251369037 | 5.282652657 | 0.345095 | 0.564949 | 0.999175987 |
| ENSMUSG00000039824 | Myl6b | protein_coding | 0.483437977 | 2.879614197 | 0.3449542 | 0.5650276 | 0.999175987 |
| ENSMUSG00000040521 | Tsfm | protein_coding | 0.213726245 | 6.541211415 | 0.3447797 | 0.565125 | 0.999175987 |
| ENSMUSG00000041057 | Wdr43 | protein_coding | -0.239735857 | 5.112090577 | 0.3447536 | 0.5651396 | 0.999175987 |
| ENSMUSG00000063142 | Kcnma1 | protein_coding | -0.195341547 | 6.222946109 | 0.3447356 | 0.5651496 | 0.999175987 |
| ENSMUSG00000036180 | Gatad2a | protein_coding | -0.251960983 | 4.89311502 | 0.3444724 | 0.5652966 | 0.999175987 |
| ENSMUSG00000032297 | Celf6 | protein_coding | 0.297789997 | 5.126596465 | 0.3444054 | 0.565334 | 0.999175987 |
| ENSMUSG00000007415 | Gatad1 | protein_coding | 0.21465099 | 7.549900113 | 0.3442675 | 0.5654111 | 0.999175987 |
| ENSMUSG00000008226 | Scrn3 | protein_coding | -0.315041084 | 5.023483344 | 0.344118 | 0.5654946 | 0.999175987 |
| ENSMUSG00000026544 | Dusp23 | protein_coding | 0.404704055 | 3.664859156 | 0.3440704 | 0.5655213 | 0.999175987 |

| ensembl_gene_id | external_gene_name | gene_biotype | log2 (fold change) | log2 (counts per million) | F-value | p-value | False discovery rate |
|---|---|---|---|---|---|---|---|
| ENSMUSG00000039308 | Ndst2 | protein_coding | -0.733505488 | 2.597608436 | 0.3435812 | 0.5657949 | 0.999175987 |
| ENSMUSG00000015341 | Golga7 | protein_coding | 0.211609514 | 5.737273515 | 0.3433998 | 0.5658964 | 0.999175987 |
| ENSMUSG00000016637 | Ift27 | protein_coding | 0.193016505 | 7.178958989 | 0.3432207 | 0.5659967 | 0.999175987 |
| ENSMUSG00000074671 | Tspyl3 | protein_coding | 0.285374243 | 5.022841253 | 0.3425517 | 0.5663716 | 0.999175987 |
| ENSMUSG00000070802 | Pnmal2 | protein_coding | 0.207435612 | 8.621589177 | 0.3424993 | 0.566401 | 0.999175987 |
| ENSMUSG00000091475 | 2810468N07Rik | processed_transcript | 0.194042664 | 6.296432369 | 0.3423529 | 0.5664831 | 0.999175987 |
| ENSMUSG00000020638 | Cmpk2 | protein_coding | 0.242074669 | 5.485329651 | 0.3423313 | 0.5664952 | 0.999175987 |
| ENSMUSG00000002289 | Angptl4 | protein_coding | -0.59122638 | 2.480448562 | 0.3423164 | 0.5665035 | 0.999175987 |
| ENSMUSG00000022035 | Ccdc25 | protein_coding | -0.195002699 | 6.514769902 | 0.342214 | 0.566561 | 0.999175987 |
| ENSMUSG00000056116 | H2-T22 | protein_coding | 0.368075235 | 4.006138147 | 0.3421402 | 0.5666024 | 0.999175987 |
| ENSMUSG00000085385 | Snhg17 | processed_transcript | -0.689010958 | 2.346316515 | 0.3412456 | 0.5671049 | 0.999175987 |
| ENSMUSG00000021747 | 4930452B06Rik | protein_coding | 0.322738243 | 3.836289135 | 0.3410939 | 0.5671902 | 0.999175987 |
| ENSMUSG00000006276 | Eps15l1 | protein_coding | 0.246519838 | 5.558917699 | 0.3407213 | 0.5673999 | 0.999175987 |
| ENSMUSG00000026385 | Dbi | protein_coding | -0.291805241 | 9.643997402 | 0.3412072 | 0.5676346 | 0.999175987 |
| ENSMUSG00000027078 | Ube2l6 | protein_coding | 0.331934309 | 4.029431324 | 0.3397557 | 0.5679438 | 0.999175987 |
| ENSMUSG00000078716 | Tmem8b | protein_coding | 0.357182597 | 4.758628436 | 0.3396383 | 0.56801 | 0.999175987 |
| ENSMUSG00000031668 | Eif2ak3 | protein_coding | 0.566403465 | 3.672571185 | 0.3403979 | 0.5680893 | 0.999175987 |
| ENSMUSG00000055692 | Tmem191c | protein_coding | 0.250352269 | 5.673885743 | 0.3394972 | 0.5680896 | 0.999175987 |
| ENSMUSG00000047084 | Ngrn | protein_coding | 0.222295332 | 6.385211894 | 0.3394637 | 0.5681085 | 0.999175987 |
| ENSMUSG00000000532 | Acvr1b | protein_coding | 0.330927691 | 4.535971057 | 0.3391635 | 0.5682779 | 0.999175987 |
| ENSMUSG00000035776 | Cd99l2 | protein_coding | -0.165093906 | 7.23010765 | 0.3383581 | 0.5687329 | 0.999175987 |
| ENSMUSG00000027667 | Zfp639 | protein_coding | 0.248944984 | 5.600760563 | 0.3383139 | 0.5687579 | 0.999175987 |
| ENSMUSG00000066551 | Hmgb1 | protein_coding | 0.172090752 | 6.525799609 | 0.3383052 | 0.5687628 | 0.999175987 |
| ENSMUSG00000022339 | Ebag9 | protein_coding | -0.27698491 | 4.976404661 | 0.3381685 | 0.5688402 | 0.999175987 |
| ENSMUSG00000036537 | Rnf113a1 | protein_coding | 0.27964999 | 5.094406699 | 0.3378894 | 0.5689981 | 0.999175987 |
| ENSMUSG00000089872 | Rps6kc1 | protein_coding | 0.263508796 | 4.90180226 | 0.3378146 | 0.5690404 | 0.999175987 |
| ENSMUSG00000035093 | Secisbp2l | protein_coding | -0.234518638 | 6.285429562 | 0.3373609 | 0.5692974 | 0.999175987 |
| ENSMUSG00000032398 | Snapc5 | protein_coding | -0.220274577 | 6.535597671 | 0.3372572 | 0.5693561 | 0.999175987 |
| ENSMUSG00000021591 | Glrx | protein_coding | 0.208875139 | 6.682961607 | 0.3368184 | 0.5696049 | 0.999175987 |
| ENSMUSG00000053332 | Gas5 | processed_transcript | 0.296273991 | 10.45470529 | 0.3373212 | 0.5698246 | 0.999175987 |
| ENSMUSG00000040621 | Gemin8 | protein_coding | -0.323258615 | 3.708095508 | 0.336341 | 0.5698758 | 0.999175987 |
| ENSMUSG00000059447 | Hadhb | protein_coding | -0.319161191 | 3.844665674 | 0.3360597 | 0.5700355 | 0.999175987 |
| ENSMUSG00000023990 | Tfeb | protein_coding | -0.784225533 | 2.436470241 | 0.3359661 | 0.5700887 | 0.999175987 |
| ENSMUSG00000053436 | Mapk14 | protein_coding | -0.27307845 | 5.388833924 | 0.3359296 | 0.5701095 | 0.999175987 |
| ENSMUSG00000021590 | Spata9 | protein_coding | 0.68326793 | 2.518650741 | 0.3358422 | 0.5701591 | 0.999175987 |
| ENSMUSG00000018559 | Ctdnep1 | protein_coding | 0.254376164 | 5.025039211 | 0.335342 | 0.5704435 | 0.999175987 |
| ENSMUSG00000070565 | Rasal2 | protein_coding | 0.29209427 | 5.085976539 | 0.3350783 | 0.5705936 | 0.999175987 |
| ENSMUSG00000000804 | Usp32 | protein_coding | -0.215331573 | 5.593076024 | 0.3350596 | 0.5706042 | 0.999175987 |
| ENSMUSG00000106379 | Lhfpl3 | protein_coding | -0.213451928 | 6.027799078 | 0.3349948 | 0.5706411 | 0.999175987 |
| ENSMUSG00000071573 | Rnls | protein_coding | 0.417861308 | 3.221765086 | 0.3349397 | 0.5706725 | 0.999175987 |
| ENSMUSG00000041977 | Arhgef11 | protein_coding | 0.476898656 | 3.297365252 | 0.334589 | 0.5708722 | 0.999175987 |
| ENSMUSG00000042761 | Mrap2 | protein_coding | 0.259566922 | 5.658619907 | 0.3344741 | 0.5709377 | 0.999175987 |
| ENSMUSG00000085741 | 5430405H02Rik | processed_transcript | 0.408542101 | 3.345413118 | 0.3342807 | 0.5710479 | 0.999175987 |
| ENSMUSG00000026596 | Abl2 | protein_coding | -0.29436672 | 4.873128831 | 0.3341836 | 0.5711033 | 0.999175987 |
| ENSMUSG00000048307 | Ankrd46 | protein_coding | -0.217916902 | 8.673423792 | 0.3341299 | 0.5711339 | 0.999175987 |
| ENSMUSG00000036390 | Gadd45a | protein_coding | -0.488183403 | 2.947059194 | 0.3340965 | 0.5711529 | 0.999175987 |
| ENSMUSG00000035595 | 1600002K03Rik | protein_coding | 0.376754344 | 3.89745917 | 0.3340416 | 0.5711843 | 0.999175987 |
| ENSMUSG00000021728 | Emb | protein_coding | 0.255541308 | 5.713795227 | 0.3340223 | 0.5711953 | 0.999175987 |
| ENSMUSG00000010392 | Gosr1 | protein_coding | -0.309148366 | 5.264274629 | 0.3338236 | 0.5713086 | 0.999175987 |
| ENSMUSG00000028133 | Rwdd3 | protein_coding | -0.268639209 | 4.843181953 | 0.3333549 | 0.5715762 | 0.999175987 |
| ENSMUSG00000026510 | Trp53bp2 | protein_coding | -0.308046023 | 4.712471166 | 0.3332626 | 0.5716289 | 0.999175987 |
| ENSMUSG00000030838 | Ush1c | protein_coding | 0.958767481 | 2.411824723 | 0.3334845 | 0.5717466 | 0.999175987 |
| ENSMUSG00000032259 | Drd2 | protein_coding | 0.330197987 | 5.144395593 | 0.3327899 | 0.571912 | 0.999175987 |
| ENSMUSG00000057506 | Bloc1s2 | protein_coding | 0.240162049 | 5.078684327 | 0.3327298 | 0.5719335 | 0.999175987 |
| ENSMUSG00000048371 | Pdp2 | protein_coding | -0.390602471 | 5.615301896 | 0.3337543 | 0.5721342 | 0.999175987 |
| ENSMUSG00000036949 | Slc39a12 | protein_coding | -0.35185335 | 4.149159605 | 0.3319683 | 0.5723692 | 0.999175987 |
| ENSMUSG00000022685 | Parn | protein_coding | 0.290298051 | 4.910161498 | 0.3312616 | 0.5727743 | 0.999175987 |
| ENSMUSG00000028639 | Ybx1 | protein_coding | -0.29673055 | 5.303736514 | 0.331235 | 0.5727896 | 0.999175987 |
| ENSMUSG00000042385 | Gzmk | protein_coding | -0.802618284 | 2.587285795 | 0.3311296 | 0.5728629 | 0.999175987 |
| ENSMUSG00000028684 | Urod | protein_coding | 0.217331326 | 7.131386531 | 0.3310934 | 0.5728708 | 0.999175987 |
| ENSMUSG00000042032 | Mat2b | protein_coding | -0.171865446 | 7.830334238 | 0.3308268 | 0.5730238 | 0.999175987 |
| ENSMUSG00000026765 | Lypd6b | protein_coding | 0.316325519 | 4.229487079 | 0.330577 | 0.5731673 | 0.999175987 |
| ENSMUSG00000096210 | H1f0 | protein_coding | 0.202334823 | 5.834766493 | 0.3302504 | 0.5733549 | 0.999175987 |
| ENSMUSG00000050394 | Armcx6 | protein_coding | 0.4510558 | 3.473449672 | 0.3300972 | 0.573443 | 0.999175987 |
| ENSMUSG00000018858 | Mrpl58 | protein_coding | 0.165765538 | 7.702655499 | 0.3298855 | 0.5735647 | 0.999175987 |
| ENSMUSG00000023032 | Slc4a8 | protein_coding | 0.28025215 | 4.621661121 | 0.3298652 | 0.5735764 | 0.999175987 |
| ENSMUSG00000098923 | Tmem185b | protein_coding | -0.309000535 | 4.597714848 | 0.3298613 | 0.5735787 | 0.999175987 |
| ENSMUSG00000049439 | Cyp20a1 | protein_coding | 0.333856394 | 4.164389823 | 0.3296435 | 0.5737039 | 0.999175987 |
| ENSMUSG00000037573 | Tob1 | protein_coding | -0.234716194 | 5.356894332 | 0.3295531 | 0.5737559 | 0.999175987 |
| ENSMUSG00000041297 | Cdk13 | protein_coding | -0.394624933 | 4.689720122 | 0.3294362 | 0.57387 | 0.999175987 |
| ENSMUSG00000054619 | Mettl7a1 | protein_coding | -0.426289381 | 3.794749842 | 0.3289216 | 0.5741197 | 0.999175987 |
| ENSMUSG00000037085 | Trmt12 | protein_coding | 0.2654741 | 5.50215 | 0.3286229 | 0.5742919 | 0.999176 |
| ENSMUSG00000028431 | Ikbkap | protein_coding | 0.2420556 | 5.09381 | 0.3286004 | 0.5743048 | 0.999176 |
| ENSMUSG00000028468 | Rgp1 | protein_coding | -0.4252379 | 3.156965 | 0.3285139 | 0.5743548 | 0.999176 |
| ENSMUSG00000026087 | Mrpl30 | protein_coding | 0.1902608 | 8.558954 | 0.3283907 | 0.5744258 | 0.999176 |
| ENSMUSG00000026259 | Ngef | protein_coding | -0.3221452 | 6.559742 | 0.3296325 | 0.574542 | 0.999176 |
| ENSMUSG00000037032 | Apbb1 | protein_coding | 0.220495 | 9.163307 | 0.3280512 | 0.5746217 | 0.999176 |
| ENSMUSG00000020978 | Klhdc2 | protein_coding | 0.1854262 | 6.554161 | 0.3280084 | 0.5746465 | 0.999176 |

| ensembl_gene_id | external_gene_name | gene_biotype | log2 (fold change) | log2 (counts per million) | F-value | p-value | False discovery rate |
|---|---|---|---|---|---|---|---|
| ENSMUSG00000063446 | Plppr1 | protein_coding | -0.2525636 | 4.933773 | 0.3278721 | 0.5747252 | 0.999176 |
| ENSMUSG00000017057 | Il13ra1 | protein_coding | -0.4384722 | 3.208088 | 0.3277623 | 0.5747886 | 0.999176 |
| ENSMUSG00000027457 | Snph | protein_coding | -0.2145949 | 6.359652 | 0.3275783 | 0.5748949 | 0.999176 |
| ENSMUSG00000024844 | Banf1 | protein_coding | 0.205473 | 7.963875 | 0.3274685 | 0.5749584 | 0.999176 |
| ENSMUSG00000048782 | Insc | protein_coding | -0.6773761 | 2.653835 | 0.3274228 | 0.5749848 | 0.999176 |
| ENSMUSG00000043463 | Rab9b | protein_coding | -0.2517763 | 5.675281 | 0.3274217 | 0.5749854 | 0.999176 |
| ENSMUSG00000020491 | 2810021J22Rik | protein_coding | 0.3918154 | 3.81392 | 0.3273176 | 0.5750456 | 0.999176 |
| ENSMUSG00000027375 | Mal | protein_coding | -0.1978011 | 7.87449 | 0.3270698 | 0.575189 | 0.999176 |
| ENSMUSG00000038418 | Egr1 | protein_coding | -0.228466 | 6.746067 | 0.3270341 | 0.5752096 | 0.999176 |
| ENSMUSG00000020307 | Cdc34 | protein_coding | 0.3121909 | 4.500428 | 0.3268857 | 0.5752955 | 0.999176 |
| ENSMUSG00000024241 | Sos1 | protein_coding | -0.4085437 | 3.380004 | 0.3268634 | 0.5753084 | 0.999176 |
| ENSMUSG00000007891 | Ctsd | protein_coding | 0.209129 | 6.348899 | 0.3268481 | 0.5753172 | 0.999176 |
| ENSMUSG00000069089 | Cdk7 | protein_coding | -0.2431678 | 5.207456 | 0.3268084 | 0.5753402 | 0.999176 |
| ENSMUSG00000020432 | Tcn2 | protein_coding | -0.4077589 | 4.297761 | 0.326691 | 0.5754209 | 0.999176 |
| ENSMUSG00000102869 | 2900097C17Rik | lincRNA | 0.2196604 | 7.970443 | 0.3265806 | 0.5754721 | 0.999176 |
| ENSMUSG00000015837 | Sqstm1 | protein_coding | 0.1867089 | 8.745918 | 0.3264984 | 0.5755197 | 0.999176 |
| ENSMUSG00000055531 | Cpsf6 | protein_coding | -0.2397736 | 5.774154 | 0.3264895 | 0.5755248 | 0.999176 |
| ENSMUSG00000025076 | Casp7 | protein_coding | 0.7294585 | 2.317534 | 0.3259957 | 0.575811 | 0.999176 |
| ENSMUSG00000023286 | Ube2j2 | protein_coding | 0.2907147 | 4.369163 | 0.3259094 | 0.575861 | 0.999176 |
| ENSMUSG00000019814 | Ltv1 | protein_coding | -0.3039989 | 4.246042 | 0.3255753 | 0.5760548 | 0.999176 |
| ENSMUSG00000031523 | Dlc1 | protein_coding | -0.4435257 | 3.484914 | 0.3254082 | 0.5761518 | 0.999176 |
| ENSMUSG00000086119 | Gm2415 | antisense_RNA | -0.5608405 | 2.633323 | 0.325054 | 0.5763575 | 0.999176 |
| ENSMUSG00000032009 | Sesn3 | protein_coding | -0.235189 | 6.287417 | 0.3248229 | 0.5764917 | 0.999176 |
| ENSMUSG00000041245 | Wnk3 | protein_coding | 0.4835793 | 3.176384 | 0.324807 | 0.576501 | 0.999176 |
| ENSMUSG00000021910 | Nisch | protein_coding | 0.167725 | 7.446222 | 0.3247629 | 0.5765266 | 0.999176 |
| ENSMUSG00000043065 | Spice1 | protein_coding | -0.4261047 | 3.476078 | 0.3246682 | 0.5765817 | 0.999176 |
| ENSMUSG00000028453 | Fancg | protein_coding | -0.3266069 | 3.980523 | 0.324548 | 0.5766516 | 0.999176 |
| ENSMUSG00000021468 | Sptlc1 | protein_coding | 0.2695118 | 4.391983 | 0.3238855 | 0.5770371 | 0.999176 |
| ENSMUSG00000032602 | Slc25a20 | protein_coding | -0.2688922 | 4.862579 | 0.3236608 | 0.5771679 | 0.999176 |
| ENSMUSG00000046994 | Mars2 | protein_coding | -0.4899616 | 3.103362 | 0.3235965 | 0.5772054 | 0.999176 |
| ENSMUSG00000029602 | Rasal1 | protein_coding | 0.302018 | 5.027994 | 0.3233346 | 0.5773581 | 0.999176 |
| ENSMUSG00000017999 | Ddx27 | protein_coding | -0.3146442 | 4.486232 | 0.3232938 | 0.5773818 | 0.999176 |
| ENSMUSG00000070002 | Ell | protein_coding | -0.2690904 | 5.354229 | 0.3225487 | 0.5778166 | 0.999176 |
| ENSMUSG00000034488 | Edil3 | protein_coding | -0.1740554 | 6.477289 | 0.3220853 | 0.5780873 | 0.999176 |
| ENSMUSG00000074259 | Gramd2 | protein_coding | -0.7339377 | 2.559464 | 0.3216967 | 0.5783145 | 0.999176 |
| ENSMUSG00000023027 | Atf1 | protein_coding | -0.6102066 | 2.517144 | 0.321426 | 0.5784729 | 0.999176 |
| ENSMUSG00000028347 | Tmeff1 | protein_coding | -0.2245068 | 5.739402 | 0.321356 | 0.5785139 | 0.999176 |
| ENSMUSG00000047658 | Gal3st3 | protein_coding | 0.5261772 | 3.135644 | 0.3213038 | 0.5785444 | 0.999176 |
| ENSMUSG00000015474 | Ppt2 | protein_coding | -0.4349854 | 2.910201 | 0.3212718 | 0.5785631 | 0.999176 |
| ENSMUSG00000036202 | Rif1 | protein_coding | -0.3348977 | 4.189043 | 0.32127 | 0.5785642 | 0.999176 |
| ENSMUSG00000026807 | Ak8 | protein_coding | -0.5949298 | 3.527732 | 0.321997 | 0.5786303 | 0.999176 |
| ENSMUSG00000074698 | Csnk2a1 | protein_coding | -0.2264237 | 5.274515 | 0.3211319 | 0.5786451 | 0.999176 |
| ENSMUSG00000042962 | Gm5436 | processed_pseudoge | 0.3323623 | 3.503939 | 0.3209449 | 0.5787546 | 0.999176 |
| ENSMUSG00000038459 | Abhd17c | protein_coding | -0.3193641 | 4.349748 | 0.3203753 | 0.5790885 | 0.999176 |
| ENSMUSG00000058260 | Serpina9 | protein_coding | -0.3647156 | 4.249093 | 0.3203504 | 0.5791031 | 0.999176 |
| ENSMUSG00000046844 | Vat1l | protein_coding | 0.2043419 | 7.407665 | 0.320211 | 0.5791849 | 0.999176 |
| ENSMUSG00000007812 | Zfp655 | protein_coding | 0.2339752 | 5.639644 | 0.320192 | 0.5791961 | 0.999176 |
| ENSMUSG00000032087 | Dscaml1 | protein_coding | 0.5096202 | 3.560281 | 0.3202052 | 0.5792009 | 0.999176 |
| ENSMUSG00000046058 | Eid2 | protein_coding | -0.3035975 | 4.64022 | 0.3201129 | 0.5792425 | 0.999176 |
| ENSMUSG00000002881 | Nab1 | protein_coding | -0.335787 | 4.198577 | 0.3200622 | 0.5792722 | 0.999176 |
| ENSMUSG00000039257 | Vstm2b | protein_coding | 0.2427091 | 5.263092 | 0.3196851 | 0.5794937 | 0.999176 |
| ENSMUSG00000043716 | Rpl7 | protein_coding | 0.2095499 | 8.566012 | 0.3196444 | 0.5795176 | 0.999176 |
| ENSMUSG00000038174 | Fam126b | protein_coding | -0.1931233 | 6.424129 | 0.3194656 | 0.5796226 | 0.999176 |
| ENSMUSG00000031551 | Ido1 | protein_coding | 0.3748732 | 3.973522 | 0.3194347 | 0.5796408 | 0.999176 |
| ENSMUSG00000087259 | 2610035D17Rik | lincRNA | -0.2352989 | 5.366583 | 0.3194298 | 0.5796436 | 0.999176 |
| ENSMUSG00000056458 | Mok | protein_coding | 0.3558744 | 4.404014 | 0.3190682 | 0.5798562 | 0.999176 |
| ENSMUSG00000029802 | Abcg2 | protein_coding | 0.5100159 | 2.876854 | 0.3188565 | 0.5799808 | 0.999176 |
| ENSMUSG00000024975 | Pdcd4 | protein_coding | 0.3173835 | 7.436718 | 0.3205533 | 0.5800622 | 0.999176 |
| ENSMUSG00000030654 | Arl6ip1 | protein_coding | 0.1621285 | 10.0542 | 0.318533 | 0.5801712 | 0.999176 |
| ENSMUSG00000051355 | Commd1 | protein_coding | 0.3320931 | 3.530139 | 0.3185062 | 0.580187 | 0.999176 |
| ENSMUSG00000029022 | Miip | protein_coding | 0.371884 | 3.589944 | 0.3184332 | 0.58023 | 0.999176 |
| ENSMUSG00000090061 | Nwd2 | protein_coding | -0.6634516 | 2.857913 | 0.3182864 | 0.5803291 | 0.999176 |
| ENSMUSG00000051238 | Swsap1 | protein_coding | 0.4804629 | 2.909239 | 0.318197 | 0.5803692 | 0.999176 |
| ENSMUSG00000068373 | D430041D05Rik | protein_coding | -0.2863431 | 4.776379 | 0.3181752 | 0.580382 | 0.999176 |
| ENSMUSG00000064284 | Cdpf1 | protein_coding | 0.2199529 | 6.128266 | 0.3179425 | 0.5805191 | 0.999176 |
| ENSMUSG00000040734 | Ppp1r13l | protein_coding | -0.595691 | 2.76236 | 0.3178146 | 0.5805945 | 0.999176 |
| ENSMUSG00000041912 | Tdrkh | protein_coding | -0.2073409 | 5.926393 | 0.3172967 | 0.5809001 | 0.999176 |
| ENSMUSG00000031302 | Nlgn3 | protein_coding | -0.183199 | 6.648637 | 0.3172934 | 0.580902 | 0.999176 |
| ENSMUSG00000020166 | Cnot2 | protein_coding | -0.2385801 | 5.701456 | 0.3170909 | 0.5810216 | 0.999176 |
| ENSMUSG00000055254 | Ntrk2 | protein_coding | -0.1925415 | 7.51989 | 0.3170634 | 0.5810379 | 0.999176 |
| ENSMUSG00000004937 | Sgta | protein_coding | 0.2671654 | 8.722332 | 0.3178583 | 0.5810564 | 0.999176 |
| ENSMUSG00000030105 | Arl8b | protein_coding | 0.1856579 | 8.049808 | 0.3170313 | 0.5810568 | 0.999176 |
| ENSMUSG00000028134 | Ptbp2 | protein_coding | -0.216954 | 6.239388 | 0.3170221 | 0.5810622 | 0.999176 |
| ENSMUSG00000020656 | Grhl1 | protein_coding | 0.5132869 | 3.637464 | 0.316926 | 0.5811864 | 0.999176 |
| ENSMUSG00000108621 | Gm37494 | lincRNA | -0.4811695 | 2.913499 | 0.3165052 | 0.5813677 | 0.999176 |
| ENSMUSG00000029038 | Ssu72 | protein_coding | 0.1567295 | 7.92076 | 0.3164323 | 0.5814109 | 0.999176 |
| ENSMUSG00000043991 | Pura | protein_coding | -0.1903215 | 6.957728 | 0.3164042 | 0.5814275 | 0.999176 |
| ENSMUSG00000026790 | Odf2 | protein_coding | 0.2671498 | 5.846599 | 0.316158 | 0.5815731 | 0.999176 |

| ensembl_gene_id | external_gene_name | gene_biotype | log2 (fold change) | log2 (counts per million) | F-value | p-value | False discovery rate |
|---|---|---|---|---|---|---|---|
| ENSMUSG00000014313 | Cox6c | protein_coding | 0.2217791 | 10.12988 | 0.3157557 | 0.5818113 | 0.999176 |
| ENSMUSG00000027324 | Rpusd2 | protein_coding | -0.5139733 | 2.668538 | 0.3156604 | 0.5818678 | 0.999176 |
| ENSMUSG00000066324 | Impad1 | protein_coding | -0.162348 | 6.810192 | 0.3154587 | 0.5819873 | 0.999176 |
| ENSMUSG00000086429 | Gt(ROSA)26Sor | antisense_RNA | 0.3221654 | 5.287848 | 0.3153205 | 0.5820817 | 0.999176 |
| ENSMUSG00000058886 | Deaf1 | protein_coding | 0.3107178 | 4.746997 | 0.3152453 | 0.5821137 | 0.999176 |
| ENSMUSG00000049281 | Scn3b | protein_coding | -0.1795796 | 6.319145 | 0.3152405 | 0.5821166 | 0.999176 |
| ENSMUSG00000038876 | Rnf146 | protein_coding | 0.2548121 | 4.722159 | 0.3151245 | 0.5821854 | 0.999176 |
| ENSMUSG00000032356 | Rasgrf1 | protein_coding | -0.2149308 | 8.150317 | 0.3148973 | 0.5823202 | 0.999176 |
| ENSMUSG00000029518 | Rab35 | protein_coding | 0.2283154 | 5.597696 | 0.3148801 | 0.5823304 | 0.999176 |
| ENSMUSG00000001123 | Lgals9 | protein_coding | -0.4289364 | 3.411263 | 0.3147942 | 0.5823813 | 0.999176 |
| ENSMUSG00000058444 | Map2k5 | protein_coding | 0.2933877 | 5.2632 | 0.3145509 | 0.5825258 | 0.999176 |
| ENSMUSG00000019785 | Clvs2 | protein_coding | -0.2235805 | 5.983016 | 0.3145403 | 0.582532 | 0.999176 |
| ENSMUSG00000005069 | Pex5 | protein_coding | -0.2974976 | 4.408127 | 0.3144574 | 0.5825812 | 0.999176 |
| ENSMUSG00000056692 | D17Wsu92e | protein_coding | -0.1763617 | 6.607724 | 0.3142996 | 0.582675 | 0.999176 |
| ENSMUSG00000003200 | Sh3gl1 | protein_coding | 0.3638512 | 4.023898 | 0.3142576 | 0.5826999 | 0.999176 |
| ENSMUSG00000069208 | Zfp825 | protein_coding | 0.3483584 | 3.55451 | 0.3138874 | 0.58292 | 0.999176 |
| ENSMUSG00000005054 | Cstb | protein_coding | 0.1886894 | 8.03171 | 0.3136519 | 0.58306 | 0.999176 |
| ENSMUSG00000074968 | Ano3 | protein_coding | 0.3188393 | 5.341786 | 0.3136597 | 0.5830679 | 0.999176 |
| ENSMUSG00000026739 | Bmi1 | protein_coding | -0.302644 | 4.297099 | 0.3135194 | 0.5831389 | 0.999176 |
| ENSMUSG00000028211 | Trp53inp1 | protein_coding | -0.398115 | 3.281706 | 0.3134797 | 0.5831625 | 0.999176 |
| ENSMUSG00000056121 | Fez2 | protein_coding | -0.2584625 | 4.617113 | 0.3133826 | 0.5832203 | 0.999176 |
| ENSMUSG00000018548 | Trim37 | protein_coding | -0.167179 | 6.692544 | 0.3133639 | 0.5832314 | 0.999176 |
| ENSMUSG00000004945 | Tmem242 | protein_coding | 0.1713851 | 6.377803 | 0.3129428 | 0.5834822 | 0.999176 |
| ENSMUSG00000018042 | Cyb5r3 | protein_coding | 0.2042159 | 7.649355 | 0.3128069 | 0.5835631 | 0.999176 |
| ENSMUSG00000027254 | Map1a | protein_coding | 0.231817 | 7.385342 | 0.3127252 | 0.5836243 | 0.999176 |
| ENSMUSG00000068284 | Usf3 | protein_coding | -0.4113592 | 3.504777 | 0.3122362 | 0.5839035 | 0.999176 |
| ENSMUSG00000087150 | BC064078 | transcribed_unproces | 0.4746918 | 3.367492 | 0.3122115 | 0.5839182 | 0.999176 |
| ENSMUSG00000052133 | Sema5b | protein_coding | -0.5051523 | 2.685226 | 0.3120289 | 0.5840272 | 0.999176 |
| ENSMUSG00000064145 | Arih2 | protein_coding | -0.2719949 | 4.752227 | 0.3119955 | 0.5840471 | 0.999176 |
| ENSMUSG00000059518 | Znhit1 | protein_coding | 0.1743044 | 7.934591 | 0.3119779 | 0.5840576 | 0.999176 |
| ENSMUSG00000037526 | Atg14 | protein_coding | -0.3250853 | 4.759605 | 0.3116355 | 0.5842621 | 0.999176 |
| ENSMUSG00000027340 | Slc23a2 | protein_coding | 0.2694888 | 5.141563 | 0.311517 | 0.5843329 | 0.999176 |
| ENSMUSG00000024188 | Luc7l | protein_coding | -0.1786155 | 6.515031 | 0.3115108 | 0.5843366 | 0.999176 |
| ENSMUSG00000028671 | Gale | protein_coding | -0.3132561 | 4.816076 | 0.3114893 | 0.5843495 | 0.999176 |
| ENSMUSG00000050164 | Mchr1 | protein_coding | 0.3510622 | 4.577819 | 0.3114807 | 0.5843546 | 0.999176 |
| ENSMUSG00000028964 | Park7 | protein_coding | 0.15947 | 10.1878 | 0.311418 | 0.5843921 | 0.999176 |
| ENSMUSG00000027010 | Slc25a12 | protein_coding | 0.2671369 | 7.805494 | 0.3121462 | 0.5844398 | 0.999176 |
| ENSMUSG00000032216 | Nedd4 | protein_coding | -0.2008838 | 8.223026 | 0.3112989 | 0.5844633 | 0.999176 |
| ENSMUSG00000039033 | Tasp1 | protein_coding | 0.2539147 | 5.575886 | 0.3111948 | 0.5845256 | 0.999176 |
| ENSMUSG00000019054 | Fis1 | protein_coding | 0.1801348 | 10.28494 | 0.3111641 | 0.5845439 | 0.999176 |
| ENSMUSG00000027293 | Ehd4 | protein_coding | 0.3688261 | 3.606209 | 0.3101531 | 0.5851492 | 0.999176 |
| ENSMUSG00000019087 | Atp6ap1 | protein_coding | 0.1608983 | 9.103616 | 0.3096322 | 0.5854615 | 0.999176 |
| ENSMUSG00000019854 | Reps1 | protein_coding | 0.4244222 | 4.63947 | 0.3101422 | 0.5856365 | 0.999176 |
| ENSMUSG00000042613 | Pbxip1 | protein_coding | 0.3376908 | 3.968771 | 0.3092814 | 0.5856721 | 0.999176 |
| ENSMUSG00000042207 | Kdm5b | protein_coding | -0.2140314 | 5.486205 | 0.3090372 | 0.5858187 | 0.999176 |
| ENSMUSG00000037098 | Rab11fip3 | protein_coding | 0.2953305 | 4.664842 | 0.3089309 | 0.5858826 | 0.999176 |
| ENSMUSG00000108350 | Gm44950 | TEC | -0.3348887 | 4.493757 | 0.3088312 | 0.5859425 | 0.999176 |
| ENSMUSG00000029145 | Eif2b4 | protein_coding | 0.227133 | 5.925867 | 0.3086808 | 0.5860329 | 0.999176 |
| ENSMUSG00000040760 | Appl1 | protein_coding | -0.2137602 | 5.821679 | 0.308548 | 0.5861128 | 0.999176 |
| ENSMUSG00000030726 | Pold3 | protein_coding | -0.2996059 | 5.231761 | 0.3084237 | 0.5861875 | 0.999176 |
| ENSMUSG00000079658 | Eloc | protein_coding | 0.1705315 | 8.764213 | 0.30833 | 0.5862439 | 0.999176 |
| ENSMUSG00000004099 | Dnmt1 | protein_coding | -0.2803162 | 4.668234 | 0.3077951 | 0.5865659 | 0.999176 |
| ENSMUSG00000022453 | Naga | protein_coding | 0.274041 | 5.324072 | 0.3076965 | 0.5866253 | 0.999176 |
| ENSMUSG00000036046 | 5031439G07Rik | protein_coding | 0.2421415 | 5.875617 | 0.3076422 | 0.586658 | 0.999176 |
| ENSMUSG00000028745 | Capzb | protein_coding | 0.1689285 | 8.992086 | 0.3072701 | 0.5868823 | 0.999176 |
| ENSMUSG00000024290 | Rock1 | protein_coding | -0.39371 | 3.749513 | 0.3069828 | 0.5870556 | 0.999176 |
| ENSMUSG00000022237 | Ankrd33b | protein_coding | 0.5725922 | 3.099321 | 0.3069615 | 0.5870809 | 0.999176 |
| ENSMUSG00000028464 | Tpm2 | protein_coding | -0.3470227 | 4.452469 | 0.3069148 | 0.5870967 | 0.999176 |
| ENSMUSG00000020387 | Jade2 | protein_coding | 0.2161328 | 5.322139 | 0.3068878 | 0.587113 | 0.999176 |
| ENSMUSG00000015478 | Rnf5 | protein_coding | 0.1654465 | 6.738153 | 0.3068357 | 0.5871444 | 0.999176 |
| ENSMUSG00000012483 | Rpa3 | protein_coding | -0.3355874 | 4.794647 | 0.3068308 | 0.5871474 | 0.999176 |
| ENSMUSG00000062761 | Zfp512 | protein_coding | -0.2063382 | 5.651798 | 0.3064713 | 0.5873645 | 0.999176 |
| ENSMUSG00000058966 | Fam57b | protein_coding | 0.4126656 | 4.642709 | 0.3062584 | 0.5878742 | 0.999176 |
| ENSMUSG00000028883 | Sema3a | protein_coding | -0.4460226 | 3.199348 | 0.3055643 | 0.5879129 | 0.999176 |
| ENSMUSG00000014813 | Stc1 | protein_coding | -0.3859831 | 3.672507 | 0.3055117 | 0.5879447 | 0.999176 |
| ENSMUSG00000005378 | Wbscr22 | protein_coding | 0.1843223 | 6.300555 | 0.3054631 | 0.5879741 | 0.999176 |
| ENSMUSG00000004880 | Lbr | protein_coding | -0.3744254 | 3.517131 | 0.305463 | 0.5879742 | 0.999176 |
| ENSMUSG00000030814 | Bcl7c | protein_coding | 0.5233102 | 2.647911 | 0.3053766 | 0.5880265 | 0.999176 |
| ENSMUSG00000031765 | Mt1 | protein_coding | -0.3346597 | 10.16084 | 0.3071371 | 0.5880708 | 0.999176 |
| ENSMUSG00000003134 | Tbc1d8 | protein_coding | 0.3733084 | 3.701744 | 0.3050921 | 0.5881988 | 0.999176 |
| ENSMUSG00000026718 | Stam | protein_coding | 0.2109666 | 5.868324 | 0.3048922 | 0.58832 | 0.999176 |
| ENSMUSG00000073418 | C4b | protein_coding | 0.7807603 | 2.333341 | 0.3047201 | 0.5884243 | 0.999176 |
| ENSMUSG00000020022 | Ndufa12 | protein_coding | 0.1608889 | 8.523937 | 0.3047057 | 0.5884331 | 0.999176 |
| ENSMUSG00000044328 | Trp53i13 | protein_coding | -0.3421744 | 3.714643 | 0.304619 | 0.5884856 | 0.999176 |
| ENSMUSG00000020178 | Adora2a | protein_coding | -0.3821177 | 4.517631 | 0.3046188 | 0.588498 | 0.999176 |
| ENSMUSG00000037624 | Kcnk2 | protein_coding | 0.2618698 | 5.383508 | 0.3045441 | 0.588531 | 0.999176 |
| ENSMUSG00000025340 | Rabgef1 | protein_coding | 0.228157 | 5.840808 | 0.30429 | 0.5886852 | 0.999176 |
| ENSMUSG00000034310 | Tmem132d | protein_coding | -0.4619373 | 2.986347 | 0.3042088 | 0.5887345 | 0.999176 |

| ensembl_gene_id | external_gene_name | gene_biotype | log2 (fold change) | log2 (counts per million) | F-value | p-value | False discovery rate |
|---|---|---|---|---|---|---|---|
| ENSMUSG00000023832 | Acat2 | protein_coding | -0.2100427 | 6.774164 | 0.3041569 | 0.588766 | 0.999176 |
| ENSMUSG00000073481 | 2-Mar | protein_coding | -0.3343087 | 4.460535 | 0.3041 | 0.5888005 | 0.999176 |
| ENSMUSG00000032064 | Dixdc1 | protein_coding | 0.1769868 | 6.123348 | 0.3038361 | 0.5889608 | 0.999176 |
| ENSMUSG00000024970 | AI846148 | protein_coding | -0.3108864 | 4.005249 | 0.3037343 | 0.5890227 | 0.999176 |
| ENSMUSG00000030530 | Furin | protein_coding | -0.4647807 | 2.998263 | 0.3036832 | 0.5890537 | 0.999176 |
| ENSMUSG00000025085 | Ablim1 | protein_coding | 0.256911 | 5.591578 | 0.3036341 | 0.5890836 | 0.999176 |
| ENSMUSG00000000934 | Top1mt | protein_coding | 0.2689474 | 5.354627 | 0.3031411 | 0.5893834 | 0.999176 |
| ENSMUSG00000002222 | Rmnd5a | protein_coding | -0.2328606 | 5.427513 | 0.3028586 | 0.5895553 | 0.999176 |
| ENSMUSG00000086171 | Pcsk2os1 | antisense_RNA | 0.5589327 | 2.510343 | 0.3027538 | 0.5896191 | 0.999176 |
| ENSMUSG00000024530 | Prelid3a | protein_coding | 0.1900085 | 6.054082 | 0.3026101 | 0.5897067 | 0.999176 |
| ENSMUSG00000018921 | Pelp1 | protein_coding | 0.3290494 | 3.52467 | 0.3025606 | 0.5897368 | 0.999176 |
| ENSMUSG00000004865 | Srpk1 | protein_coding | -0.2065905 | 6.884929 | 0.3023543 | 0.5898625 | 0.999176 |
| ENSMUSG00000039954 | Stk32a | protein_coding | -0.5905087 | 3.783347 | 0.3035567 | 0.5900973 | 0.999176 |
| ENSMUSG00000018648 | Dusp14 | protein_coding | -0.1945473 | 6.258316 | 0.3018603 | 0.5901638 | 0.999176 |
| ENSMUSG00000046519 | Golph3l | protein_coding | -0.3207877 | 4.820096 | 0.301786 | 0.5902091 | 0.999176 |
| ENSMUSG00000038677 | Scube3 | protein_coding | -0.5834122 | 2.596709 | 0.3017274 | 0.5902448 | 0.999176 |
| ENSMUSG00000021775 | Nr1d2 | protein_coding | -0.2193376 | 5.451793 | 0.3015658 | 0.5903435 | 0.999176 |
| ENSMUSG00000036781 | Rps27l | protein_coding | 0.2251495 | 6.17998 | 0.3015605 | 0.5903467 | 0.999176 |
| ENSMUSG00000026778 | Prkcq | protein_coding | -0.4267608 | 3.361687 | 0.3013701 | 0.590463 | 0.999176 |
| ENSMUSG00000070576 | Mn1 | protein_coding | 0.2987877 | 8.317536 | 0.3028833 | 0.5905058 | 0.999176 |
| ENSMUSG00000005672 | Kit | protein_coding | 0.4625229 | 3.26723 | 0.30104 | 0.5906647 | 0.999176 |
| ENSMUSG00000034522 | Zfp395 | protein_coding | -0.3414053 | 3.499836 | 0.3008028 | 0.5908097 | 0.999176 |
| ENSMUSG00000001995 | Sipa1l2 | protein_coding | 0.396986 | 3.881672 | 0.3005727 | 0.5909504 | 0.999176 |
| ENSMUSG00000069094 | Pde7a | protein_coding | 0.4313156 | 3.208383 | 0.3004056 | 0.5910527 | 0.999176 |
| ENSMUSG00000085218 | BB218582 | processed_transcript | 0.4760484 | 2.97013 | 0.2997587 | 0.5914489 | 0.999176 |
| ENSMUSG00000038991 | Txndc5 | protein_coding | 0.2953814 | 5.050696 | 0.2997227 | 0.591471 | 0.999176 |
| ENSMUSG00000025933 | Tmem14a | protein_coding | 0.1821852 | 6.769582 | 0.2993714 | 0.5916864 | 0.999176 |
| ENSMUSG00000070319 | Eif3g | protein_coding | 0.1533406 | 7.844694 | 0.2987615 | 0.5920608 | 0.999176 |
| ENSMUSG00000028419 | Chmp5 | protein_coding | 0.1570147 | 8.829841 | 0.2986354 | 0.5921383 | 0.999176 |
| ENSMUSG00000028772 | Zcchc17 | protein_coding | 0.1483417 | 8.941094 | 0.2985185 | 0.5922101 | 0.999176 |
| ENSMUSG00000060424 | Pantr1 | lincRNA | 0.3684586 | 7.079746 | 0.3017232 | 0.592328 | 0.999176 |
| ENSMUSG00000035067 | Xkr6 | protein_coding | 0.6050778 | 2.614267 | 0.298092 | 0.5924724 | 0.999176 |
| ENSMUSG00000005973 | Rcn1 | protein_coding | -0.2661022 | 5.132738 | 0.2980371 | 0.5925062 | 0.999176 |
| ENSMUSG00000008429 | Herpud2 | protein_coding | -0.3431523 | 4.954699 | 0.2977794 | 0.5926769 | 0.999176 |
| ENSMUSG00000034610 | Zcchc11 | protein_coding | -0.3281882 | 4.18104 | 0.297514 | 0.5928282 | 0.999176 |
| ENSMUSG00000085084 | 4930570G19Rik | lincRNA | -0.2859808 | 4.099745 | 0.2973539 | 0.5929268 | 0.999176 |
| ENSMUSG00000044770 | Scml4 | protein_coding | -0.4118449 | 3.261107 | 0.2973264 | 0.5929438 | 0.999176 |
| ENSMUSG00000056899 | Immp2l | protein_coding | -0.3265095 | 4.525311 | 0.2972622 | 0.5929833 | 0.999176 |
| ENSMUSG00000019998 | Stx7 | protein_coding | 0.1727338 | 7.73029 | 0.2972426 | 0.5929954 | 0.999176 |
| ENSMUSG00000025551 | Fgf14 | protein_coding | -0.2789438 | 5.81416 | 0.2971802 | 0.593046 | 0.999176 |
| ENSMUSG00000025793 | Hgs | protein_coding | 0.2161747 | 5.683935 | 0.2971347 | 0.5930619 | 0.999176 |
| ENSMUSG00000030527 | Crtc3 | protein_coding | 0.5107004 | 3.106435 | 0.2971138 | 0.5930748 | 0.999176 |
| ENSMUSG00000027889 | Ampd2 | protein_coding | 0.2484207 | 6.126254 | 0.2970915 | 0.5931006 | 0.999176 |
| ENSMUSG00000028851 | Nudc | protein_coding | 0.3225935 | 5.935962 | 0.2977429 | 0.5931569 | 0.999176 |
| ENSMUSG00000095362 | Gm14325 | protein_coding | -0.2445891 | 5.721333 | 0.2968885 | 0.5932137 | 0.999176 |
| ENSMUSG00000031870 | Pgr | protein_coding | 0.2662851 | 4.161279 | 0.2967899 | 0.5932746 | 0.999176 |
| ENSMUSG00000021384 | Susd3 | protein_coding | -0.5581343 | 2.919735 | 0.2966378 | 0.5933684 | 0.999176 |
| ENSMUSG00000020111 | Micu1 | protein_coding | -0.1782327 | 7.008005 | 0.2963112 | 0.5935701 | 0.999176 |
| ENSMUSG00000039747 | Orai2 | protein_coding | -0.3152625 | 4.359077 | 0.2961161 | 0.5936906 | 0.999176 |
| ENSMUSG00000027361 | Gabpb1 | protein_coding | 0.3041134 | 4.011358 | 0.295041 | 0.5943557 | 0.999176 |
| ENSMUSG00000037960 | Card19 | protein_coding | -0.3095683 | 4.101817 | 0.2939055 | 0.5950597 | 0.999176 |
| ENSMUSG00000021610 | Clptm1l | protein_coding | -0.2731895 | 5.774737 | 0.2938701 | 0.5950938 | 0.999176 |
| ENSMUSG00000032867 | Fbxw8 | protein_coding | -0.5425227 | 2.483065 | 0.2933496 | 0.5954051 | 0.999176 |
| ENSMUSG00000025971 | Maip1 | protein_coding | -0.2353096 | 4.989493 | 0.293218 | 0.5954869 | 0.999176 |
| ENSMUSG00000063626 | Unc5d | protein_coding | -0.2480924 | 5.112168 | 0.2930123 | 0.5956149 | 0.999176 |
| ENSMUSG00000031934 | Panx1 | protein_coding | 0.3728217 | 3.856228 | 0.2930079 | 0.5956176 | 0.999176 |
| ENSMUSG00000039782 | Cpeb2 | protein_coding | -0.2577377 | 4.842611 | 0.2929187 | 0.5956731 | 0.999176 |
| ENSMUSG00000037523 | Mavs | protein_coding | 0.3537776 | 4.38039 | 0.2928701 | 0.5957034 | 0.999176 |
| ENSMUSG00000032244 | Fem1b | protein_coding | -0.1824527 | 6.056166 | 0.2928487 | 0.5957167 | 0.999176 |
| ENSMUSG00000042642 | Flad1 | protein_coding | -0.2150183 | 6.006309 | 0.2925931 | 0.5958758 | 0.999176 |
| ENSMUSG00000020893 | Per1 | protein_coding | -0.3257555 | 4.241101 | 0.2915039 | 0.5965549 | 0.999176 |
| ENSMUSG00000021958 | Pinx1 | protein_coding | -0.2616924 | 4.210914 | 0.2914676 | 0.5965775 | 0.999176 |
| ENSMUSG00000032743 | D430042O09Rik | protein_coding | 0.5453354 | 2.810748 | 0.2912613 | 0.5967064 | 0.999176 |
| ENSMUSG00000038760 | Trhr | protein_coding | 0.7501978 | 2.638989 | 0.2901366 | 0.5974217 | 0.999176 |
| ENSMUSG00000059820 | AU019823 | protein_coding | -0.3957804 | 3.350332 | 0.2899928 | 0.5974998 | 0.999176 |
| ENSMUSG00000024826 | Dpf2 | protein_coding | -0.1989094 | 6.602827 | 0.2897786 | 0.597634 | 0.999176 |
| ENSMUSG00000040797 | Iqsec3 | protein_coding | -0.2783572 | 5.413933 | 0.2896885 | 0.5976904 | 0.999176 |
| ENSMUSG00000025978 | Rftn2 | protein_coding | -0.2529619 | 4.749615 | 0.2896582 | 0.5977095 | 0.999176 |
| ENSMUSG00000021532 | Fastkd3 | protein_coding | 0.2616979 | 5.03449 | 0.289554 | 0.5977748 | 0.999176 |
| ENSMUSG00000048285 | Frmd6 | protein_coding | 0.5862099 | 2.624951 | 0.2894912 | 0.5978142 | 0.999176 |
| ENSMUSG00000045009 | Prrt3 | protein_coding | 0.532869 | 2.804284 | 0.2894638 | 0.5978313 | 0.999176 |
| ENSMUSG00000038982 | Bloc1s5 | protein_coding | 0.2103978 | 5.813263 | 0.2892097 | 0.5979907 | 0.999176 |
| ENSMUSG00000020386 | Sar1b | protein_coding | -0.1931955 | 7.789737 | 0.2890264 | 0.5981057 | 0.999176 |
| ENSMUSG00000036285 | Noa1 | protein_coding | -0.3902323 | 3.49092 | 0.2889523 | 0.5981523 | 0.999176 |
| ENSMUSG00000042650 | Alkbh5 | protein_coding | 0.3055848 | 6.623995 | 0.2903723 | 0.5982045 | 0.999176 |
| ENSMUSG00000049119 | Fam110b | protein_coding | 0.1785871 | 6.340096 | 0.2886014 | 0.5983727 | 0.999176 |
| ENSMUSG00000019362 | D8Ertd738e | protein_coding | 0.1534513 | 8.808685 | 0.2885078 | 0.5984315 | 0.999176 |
| ENSMUSG00000029624 | Ptcd1 | protein_coding | 0.3629445 | 3.945843 | 0.2882866 | 0.5985705 | 0.999176 |

| ensembl_gene_id | external_gene_name | gene_biotype | log2 (fold change) | log2 (counts per million) | F-value | p-value | False discovery rate |
|---|---|---|---|---|---|---|---|
| ENSMUSG00000039585 | Myo9a | protein_coding | 0.4994567 | 2.871055 | 0.2882235 | 0.5986102 | 0.999176 |
| ENSMUSG00000024909 | Efemp2 | protein_coding | -0.5797633 | 4.049245 | 0.2897493 | 0.5988098 | 0.999176 |
| ENSMUSG00000027577 | Chrna4 | protein_coding | 0.4759994 | 4.640889 | 0.2894739 | 0.5988416 | 0.999176 |
| ENSMUSG00000034282 | Evpl | protein_coding | -0.3723501 | 3.8195 | 0.2878448 | 0.5988485 | 0.999176 |
| ENSMUSG00000026213 | Stk11ip | protein_coding | 0.4332874 | 3.316188 | 0.2876209 | 0.5989894 | 0.999176 |
| ENSMUSG00000033854 | Kcnk10 | protein_coding | 0.4353842 | 3.331572 | 0.2874009 | 0.5991279 | 0.999176 |
| ENSMUSG00000002871 | Tpra1 | protein_coding | 0.2359421 | 5.906543 | 0.2873921 | 0.5991335 | 0.999176 |
| ENSMUSG00000035131 | Brinp3 | protein_coding | -0.2463685 | 5.172184 | 0.2873824 | 0.5991396 | 0.999176 |
| ENSMUSG00000027411 | Vps16 | protein_coding | 0.2917073 | 5.55216 | 0.2873386 | 0.5991791 | 0.999176 |
| ENSMUSG00000026773 | Pfkfb3 | protein_coding | 0.4317659 | 3.827403 | 0.2872384 | 0.5992422 | 0.999176 |
| ENSMUSG00000023456 | Tpi1 | protein_coding | 0.1506687 | 9.877387 | 0.2870552 | 0.5993459 | 0.999176 |
| ENSMUSG00000045996 | Polr2k | protein_coding | 0.2901328 | 4.235161 | 0.2868529 | 0.5994735 | 0.999176 |
| ENSMUSG00000018740 | Slc25a35 | protein_coding | -0.6085838 | 2.782726 | 0.2863668 | 0.5997802 | 0.999176 |
| ENSMUSG00000028465 | Tln1 | protein_coding | 0.2325358 | 5.298575 | 0.2861541 | 0.5999146 | 0.999176 |
| ENSMUSG00000042608 | Stk40 | protein_coding | -0.4029696 | 3.739329 | 0.2851934 | 0.6005222 | 0.999176 |
| ENSMUSG00000061371 | Zfp873 | protein_coding | -0.3300424 | 4.085819 | 0.2848595 | 0.6007337 | 0.999176 |
| ENSMUSG00000024319 | Vps52 | protein_coding | -0.2430713 | 5.761971 | 0.2847856 | 0.6007806 | 0.999176 |
| ENSMUSG00000019878 | Hsf2 | protein_coding | 0.3347296 | 5.061571 | 0.2847992 | 0.6007837 | 0.999176 |
| ENSMUSG00000039735 | Fnbp1l | protein_coding | -0.2442398 | 4.974654 | 0.2847486 | 0.600804 | 0.999176 |
| ENSMUSG00000058748 | Zfp958 | protein_coding | 0.3809746 | 3.542261 | 0.2846464 | 0.6008688 | 0.999176 |
| ENSMUSG00000004934 | Pias4 | protein_coding | 0.8149597 | 2.13481 | 0.2846411 | 0.6008722 | 0.999176 |
| ENSMUSG00000040877 | Wdr25 | protein_coding | 0.4875424 | 3.857687 | 0.2852832 | 0.6008982 | 0.999176 |
| ENSMUSG00000067369 | Trmt2b | protein_coding | 0.3314308 | 4.472098 | 0.2845045 | 0.6009588 | 0.999176 |
| ENSMUSG00000005698 | Ctcf | protein_coding | 0.2060987 | 6.059742 | 0.2838426 | 0.6013789 | 0.999176 |
| ENSMUSG00000028020 | Glrb | protein_coding | 0.1579617 | 6.764038 | 0.2836899 | 0.6014758 | 0.999176 |
| ENSMUSG00000020902 | Ntn1 | protein_coding | 0.6794514 | 2.661445 | 0.2835403 | 0.601582 | 0.999176 |
| ENSMUSG00000033610 | Pank1 | protein_coding | -0.3299663 | 3.737444 | 0.2834742 | 0.6016129 | 0.999176 |
| ENSMUSG00000060904 | Arl1 | protein_coding | 0.1596658 | 6.881057 | 0.2833563 | 0.6016879 | 0.999176 |
| ENSMUSG00000027650 | Tti1 | protein_coding | 0.3004491 | 4.096852 | 0.2830023 | 0.601913 | 0.999176 |
| ENSMUSG00000111080 | AC153899.1 | lincRNA | 0.2104824 | 5.911816 | 0.2829717 | 0.6019325 | 0.999176 |
| ENSMUSG00000024900 | Cpt1a | protein_coding | -0.7003439 | 2.519855 | 0.2827193 | 0.6020932 | 0.999176 |
| ENSMUSG00000021973 | Micu2 | protein_coding | -0.3040943 | 4.362806 | 0.2827097 | 0.6020993 | 0.999176 |
| ENSMUSG00000004788 | Eif2b2 | protein_coding | 0.1784709 | 6.234374 | 0.2825843 | 0.6021792 | 0.999176 |
| ENSMUSG00000068663 | Clec16a | protein_coding | -0.4156616 | 4.878213 | 0.2835945 | 0.6022521 | 0.999176 |
| ENSMUSG00000000378 | Ccm2 | protein_coding | 0.1871296 | 5.708089 | 0.2823229 | 0.6023457 | 0.999176 |
| ENSMUSG00000006717 | Acot13 | protein_coding | 0.1718716 | 8.428009 | 0.2823097 | 0.6023541 | 0.999176 |
| ENSMUSG00000040250 | Ints13 | protein_coding | 0.3346177 | 3.385165 | 0.2821226 | 0.6024734 | 0.999176 |
| ENSMUSG00000024537 | Psmg2 | protein_coding | 0.175028 | 6.137538 | 0.2816169 | 0.602796 | 0.999176 |
| ENSMUSG00000021248 | Tmed10 | protein_coding | -0.1693009 | 6.82657 | 0.2815522 | 0.6028374 | 0.999176 |
| ENSMUSG00000040669 | Phc1 | protein_coding | 0.3350488 | 4.31482 | 0.281482 | 0.6028822 | 0.999176 |
| ENSMUSG00000067873 | Htatsf1 | protein_coding | 0.2059988 | 5.837514 | 0.2813607 | 0.6029597 | 0.999176 |
| ENSMUSG00000035198 | Tubg1 | protein_coding | 0.2416339 | 6.6972 | 0.281437 | 0.6029658 | 0.999176 |
| ENSMUSG00000066392 | Nrxn3 | protein_coding | 0.1748726 | 7.907882 | 0.2812704 | 0.6030174 | 0.999176 |
| ENSMUSG00000060402 | Chst8 | protein_coding | 0.47414 | 3.486512 | 0.2811399 | 0.6031125 | 0.999176 |
| ENSMUSG00000026278 | Bok | protein_coding | -0.2252704 | 5.664163 | 0.2810395 | 0.6031649 | 0.999176 |
| ENSMUSG00000051910 | Sox6 | protein_coding | -0.4010083 | 3.748186 | 0.2809328 | 0.6032331 | 0.999176 |
| ENSMUSG00000057615 | Ldoc1 | protein_coding | -0.3447679 | 3.467957 | 0.2805422 | 0.603483 | 0.999176 |
| ENSMUSG00000042429 | Adora1 | protein_coding | 0.2543038 | 5.825502 | 0.2802497 | 0.6036702 | 0.999176 |
| ENSMUSG00000022109 | Med4 | protein_coding | -0.1844567 | 6.378645 | 0.280249 | 0.6036707 | 0.999176 |
| ENSMUSG00000035941 | Ibtk | protein_coding | -0.3606202 | 3.603984 | 0.2802149 | 0.6036925 | 0.999176 |
| ENSMUSG00000019066 | Rab3d | protein_coding | 0.7889647 | 2.520366 | 0.2796767 | 0.6040491 | 0.999176 |
| ENSMUSG00000027860 | Vangl1 | protein_coding | -0.6192888 | 3.138917 | 0.279934 | 0.6041708 | 0.999176 |
| ENSMUSG00000060073 | Psma3 | protein_coding | 0.1908853 | 6.125811 | 0.2794089 | 0.6042092 | 0.999176 |
| ENSMUSG00000028063 | Lmna | protein_coding | 0.2274891 | 5.09596 | 0.2793925 | 0.6042197 | 0.999176 |
| ENSMUSG00000061607 | Mdc1 | protein_coding | -0.7710587 | 2.205417 | 0.2793847 | 0.6042247 | 0.999176 |
| ENSMUSG00000045636 | Mtus1 | protein_coding | -0.2597957 | 5.259558 | 0.2792774 | 0.6042936 | 0.999176 |
| ENSMUSG00000046841 | Ckap4 | protein_coding | -0.2817661 | 4.205988 | 0.2787567 | 0.604628 | 0.999176 |
| ENSMUSG00000029550 | Sppl3 | protein_coding | 0.2660097 | 4.979014 | 0.2785592 | 0.6047549 | 0.999176 |
| ENSMUSG00000015363 | Trabd | protein_coding | -0.3038623 | 4.444488 | 0.2785532 | 0.6047588 | 0.999176 |
| ENSMUSG00000038352 | Arl5c | protein_coding | -0.4651055 | 4.748158 | 0.2800658 | 0.6047855 | 0.999176 |
| ENSMUSG00000036721 | Zscan12 | protein_coding | -0.3748118 | 3.671864 | 0.2783262 | 0.6049048 | 0.999176 |
| ENSMUSG00000069237 | Fam8a1 | protein_coding | 0.2309657 | 5.511208 | 0.2779382 | 0.6051544 | 0.999176 |
| ENSMUSG00000030917 | Tmem159 | protein_coding | -0.2072184 | 5.458398 | 0.2779239 | 0.6051636 | 0.999176 |
| ENSMUSG00000024789 | Jak2 | protein_coding | 0.3194127 | 4.004079 | 0.2779143 | 0.6051699 | 0.999176 |
| ENSMUSG00000021764 | Ndufs4 | protein_coding | 0.2191474 | 8.839225 | 0.277825 | 0.605239 | 0.999176 |
| ENSMUSG00000030691 | Fchsd2 | protein_coding | 0.4552449 | 3.146761 | 0.2774157 | 0.605491 | 0.999176 |
| ENSMUSG00000036940 | Kdm1a | protein_coding | 0.3765514 | 3.272369 | 0.2773284 | 0.6055473 | 0.999176 |
| ENSMUSG00000028010 | Gar1 | protein_coding | 0.6356173 | 2.31815 | 0.2771942 | 0.6056339 | 0.999176 |
| ENSMUSG00000030760 | Acer3 | protein_coding | -0.2238827 | 4.766273 | 0.2770366 | 0.6057355 | 0.999176 |
| ENSMUSG00000068917 | Clk2 | protein_coding | -0.509382 | 3.160121 | 0.2770155 | 0.6057492 | 0.999176 |
| ENSMUSG00000041020 | Map7d2 | protein_coding | 0.1420197 | 7.66433 | 0.2769664 | 0.6057808 | 0.999176 |
| ENSMUSG00000039154 | Shd | protein_coding | 0.2226159 | 6.023462 | 0.2769635 | 0.6057827 | 0.999176 |
| ENSMUSG00000052738 | Suclg1 | protein_coding | -0.1483023 | 9.129198 | 0.2767146 | 0.6059433 | 0.999176 |
| ENSMUSG00000042797 | Aqp11 | protein_coding | -0.324703 | 3.620436 | 0.2766752 | 0.6059688 | 0.999176 |
| ENSMUSG00000020381 | Mrnip | protein_coding | 0.2366443 | 5.218173 | 0.2764766 | 0.6060971 | 0.999176 |
| ENSMUSG00000021552 | Gkap1 | protein_coding | -0.3913797 | 3.370369 | 0.2764337 | 0.6061248 | 0.999176 |
| ENSMUSG00000024176 | Sox8 | protein_coding | -0.2872354 | 4.254497 | 0.276069 | 0.6063605 | 0.999176 |
| ENSMUSG00000052914 | Cyp2j6 | protein_coding | -0.3863113 | 4.464167 | 0.2760567 | 0.6064307 | 0.999176 |

| ensembl_gene_id | external_gene_name | gene_biotype | log2 (fold change) | log2 (counts per million) | F-value | p-value | False discovery rate |
|---|---|---|---|---|---|---|---|
| ENSMUSG00000032267 | Usp28 | protein_coding | 0.6375223 | 2.665603 | 0.2755632 | 0.6066878 | 0.999176 |
| ENSMUSG00000020038 | Cry1 | protein_coding | -0.4801375 | 3.987715 | 0.2762222 | 0.6067113 | 0.999176 |
| ENSMUSG00000014873 | Surf2 | protein_coding | -0.1941969 | 6.407624 | 0.2749382 | 0.6070927 | 0.999176 |
| ENSMUSG00000025939 | Ube2w | protein_coding | 0.1548945 | 8.205577 | 0.2749099 | 0.6071111 | 0.999176 |
| ENSMUSG00000058153 | Sez6l | protein_coding | -0.1867204 | 5.904706 | 0.2748737 | 0.6071345 | 0.999176 |
| ENSMUSG00000032560 | Dnajc13 | protein_coding | -0.3243533 | 4.10239 | 0.2746568 | 0.6072752 | 0.999176 |
| ENSMUSG00000025006 | Sorbs1 | protein_coding | -0.1848082 | 6.236332 | 0.2744951 | 0.6073801 | 0.999176 |
| ENSMUSG00000021848 | Otx2 | protein_coding | -0.3989955 | 3.240621 | 0.274434 | 0.6074198 | 0.999176 |
| ENSMUSG00000028744 | Pqlc2 | protein_coding | 0.304738 | 4.249229 | 0.2742246 | 0.6075557 | 0.999176 |
| ENSMUSG00000034157 | Cipc | protein_coding | -0.1942636 | 6.034007 | 0.2742231 | 0.6075567 | 0.999176 |
| ENSMUSG00000031343 | Gabra3 | protein_coding | 0.2350976 | 5.587914 | 0.2740961 | 0.6076392 | 0.999176 |
| ENSMUSG00000078235 | Fam43b | protein_coding | -0.4361949 | 2.797049 | 0.2737496 | 0.6078644 | 0.999176 |
| ENSMUSG00000021816 | Ppp3cb | protein_coding | 0.1686238 | 7.762421 | 0.2737288 | 0.6078779 | 0.999176 |
| ENSMUSG00000107283 | Mpv17 | protein_coding | 0.1652896 | 6.94856 | 0.2736871 | 0.607905 | 0.999176 |
| ENSMUSG00000022771 | Ppil2 | protein_coding | -0.1999335 | 6.472392 | 0.2735029 | 0.6080248 | 0.999176 |
| ENSMUSG00000060206 | Zfp462 | protein_coding | 0.2929317 | 4.066286 | 0.2734584 | 0.6080537 | 0.999176 |
| ENSMUSG00000034024 | Cct2 | protein_coding | 0.1584448 | 8.002044 | 0.2734456 | 0.6080621 | 0.999176 |
| ENSMUSG00000001436 | Slc19a1 | protein_coding | 0.4040492 | 4.700138 | 0.2740182 | 0.6081375 | 0.999176 |
| ENSMUSG00000070000 | Fcho1 | protein_coding | 0.3496037 | 3.832682 | 0.2731747 | 0.6082384 | 0.999176 |
| ENSMUSG00000064215 | Ifi27 | protein_coding | 0.1760038 | 6.399141 | 0.273061 | 0.6083125 | 0.999176 |
| ENSMUSG00000025169 | Ogfod3 | protein_coding | -0.3242481 | 4.034517 | 0.2727495 | 0.6085154 | 0.999176 |
| ENSMUSG00000029501 | Ankle2 | protein_coding | -0.2702856 | 4.398548 | 0.2725236 | 0.6086626 | 0.999176 |
| ENSMUSG00000041258 | Zfp236 | protein_coding | 0.2698788 | 3.922609 | 0.2722156 | 0.6088635 | 0.999176 |
| ENSMUSG00000071103 | 1700029J07Rik | protein_coding | 0.8192689 | 2.093149 | 0.2722043 | 0.6088709 | 0.999176 |
| ENSMUSG00000007867 | Ift43 | protein_coding | 0.2063253 | 5.788973 | 0.2721475 | 0.6089079 | 0.999176 |
| ENSMUSG00000079235 | Ccdc13 | protein_coding | 0.3643959 | 3.461573 | 0.2720973 | 0.6089407 | 0.999176 |
| ENSMUSG00000040289 | Hey1 | protein_coding | -0.2597867 | 4.576717 | 0.2719355 | 0.6090464 | 0.999176 |
| ENSMUSG00000070366 | Plpp4 | protein_coding | 0.341276 | 4.061904 | 0.2717898 | 0.6091415 | 0.999176 |
| ENSMUSG00000045991 | Onecut2 | protein_coding | 0.2547975 | 4.746783 | 0.2716746 | 0.6092168 | 0.999176 |
| ENSMUSG00000024592 | C330018D20Rik | protein_coding | 0.2838562 | 4.791126 | 0.2713219 | 0.6094473 | 0.999176 |
| ENSMUSG00000026991 | Pkp4 | protein_coding | -0.1479528 | 6.884406 | 0.2711988 | 0.6095278 | 0.999176 |
| ENSMUSG00000038456 | Dennd2a | protein_coding | 0.358377 | 3.903286 | 0.2711445 | 0.6095633 | 0.999176 |
| ENSMUSG00000012076 | Brms1l | protein_coding | -0.1985757 | 5.794558 | 0.2710185 | 0.6096457 | 0.999176 |
| ENSMUSG00000044528 | Tram1l1 | protein_coding | -0.1514771 | 6.654882 | 0.2709188 | 0.609711 | 0.999176 |
| ENSMUSG00000021578 | Ccdc127 | protein_coding | 0.1733243 | 6.335168 | 0.2708114 | 0.6097813 | 0.999176 |
| ENSMUSG00000075229 | Ccdc58 | protein_coding | 0.2549758 | 4.81492 | 0.2706712 | 0.6098731 | 0.999176 |
| ENSMUSG00000031955 | Bcar1 | protein_coding | 0.2946066 | 4.683054 | 0.2704897 | 0.6099919 | 0.999176 |
| ENSMUSG00000038055 | Dexi | protein_coding | -0.3005312 | 4.185808 | 0.270409 | 0.6100449 | 0.999176 |
| ENSMUSG00000104784 | Gm42984 | antisense_RNA | 0.5461716 | 2.662388 | 0.2701633 | 0.6102059 | 0.999176 |
| ENSMUSG00000033436 | Armcx2 | protein_coding | 0.1924036 | 6.195995 | 0.2694209 | 0.6106931 | 0.999176 |
| ENSMUSG00000030806 | Stx1b | protein_coding | -0.18737 | 8.032937 | 0.2692181 | 0.6108263 | 0.999176 |
| ENSMUSG00000020262 | Adarb1 | protein_coding | -0.2675721 | 5.996686 | 0.2691155 | 0.6109353 | 0.999176 |
| ENSMUSG00000089917 | Uckl1 | protein_coding | 0.2579352 | 5.065466 | 0.2689951 | 0.6109729 | 0.999176 |
| ENSMUSG00000035142 | Nubpl | protein_coding | 0.3876513 | 3.289785 | 0.2689888 | 0.6109771 | 0.999176 |
| ENSMUSG00000027742 | Cog6 | protein_coding | -0.2424775 | 5.242379 | 0.2688649 | 0.6110586 | 0.999176 |
| ENSMUSG00000029521 | Chek2 | protein_coding | 0.4011106 | 3.049747 | 0.2686578 | 0.6111948 | 0.999176 |
| ENSMUSG00000028691 | Prdx1 | protein_coding | -0.1699147 | 9.357497 | 0.2685599 | 0.6112592 | 0.999176 |
| ENSMUSG00000037971 | 1110032A03Rik | protein_coding | 0.1450559 | 7.593221 | 0.2684854 | 0.6113082 | 0.999176 |
| ENSMUSG00000019647 | Sema6a | protein_coding | -0.2832568 | 4.673342 | 0.2684237 | 0.6113489 | 0.999176 |
| ENSMUSG00000031605 | Klhl2 | protein_coding | -0.1929751 | 6.092563 | 0.2683001 | 0.6114303 | 0.999176 |
| ENSMUSG00000033697 | Arhgap39 | protein_coding | -0.5240171 | 2.727259 | 0.2682713 | 0.6114492 | 0.999176 |
| ENSMUSG00000049305 | Ccdc71 | protein_coding | 0.2497015 | 5.392133 | 0.2682292 | 0.6114769 | 0.999176 |
| ENSMUSG00000038762 | Abcf1 | protein_coding | 0.1944629 | 6.924859 | 0.2682147 | 0.6114865 | 0.999176 |
| ENSMUSG00000044221 | Grsf1 | protein_coding | 0.172399 | 6.971258 | 0.2679679 | 0.6116492 | 0.999176 |
| ENSMUSG00000050663 | Trhde | protein_coding | -0.611671 | 2.690122 | 0.2679169 | 0.6116827 | 0.999176 |
| ENSMUSG00000031310 | Zmym3 | protein_coding | -0.1646282 | 6.639066 | 0.2676771 | 0.6118409 | 0.999176 |
| ENSMUSG00000022763 | Aifm3 | protein_coding | -0.245259 | 5.13719 | 0.2675127 | 0.6119493 | 0.999176 |
| ENSMUSG00000050954 | Zfp169 | protein_coding | 0.5349744 | 2.607588 | 0.2674065 | 0.6120194 | 0.999176 |
| ENSMUSG00000029767 | Calu | protein_coding | -0.2302822 | 5.515482 | 0.2673155 | 0.6120795 | 0.999176 |
| ENSMUSG00000097486 | Gm17733 | bidirectional_promote | 0.8824783 | 2.426052 | 0.2677463 | 0.6122102 | 0.999176 |
| ENSMUSG00000016526 | Dyrk3 | protein_coding | 0.6541399 | 2.250771 | 0.2671072 | 0.612217 | 0.999176 |
| ENSMUSG00000032875 | Arhgef17 | protein_coding | 0.1700003 | 6.310172 | 0.2670718 | 0.6122404 | 0.999176 |
| ENSMUSG00000035840 | Lysmd3 | protein_coding | 0.3158092 | 3.988317 | 0.2670404 | 0.6122611 | 0.999176 |
| ENSMUSG00000037509 | Arhgef4 | protein_coding | 0.1959004 | 6.287495 | 0.2670381 | 0.6122626 | 0.999176 |
| ENSMUSG00000040759 | Cmtm5 | protein_coding | -0.2199107 | 7.823721 | 0.2666732 | 0.6125452 | 0.999176 |
| ENSMUSG00000053279 | Aldh1a1 | protein_coding | -0.3121746 | 4.881994 | 0.266612 | 0.6125557 | 0.999176 |
| ENSMUSG00000045608 | Dbx2 | protein_coding | -0.3309055 | 3.945594 | 0.2665283 | 0.6125996 | 0.999176 |
| ENSMUSG00000064179 | Tnnt1 | protein_coding | -0.5051145 | 4.643141 | 0.2679249 | 0.6127841 | 0.999176 |
| ENSMUSG00000021891 | Mettl6 | protein_coding | 0.2290461 | 5.307869 | 0.266208 | 0.6128115 | 0.999176 |
| ENSMUSG00000021662 | Arhgef28 | protein_coding | -0.3507239 | 4.535902 | 0.2658176 | 0.6130814 | 0.999176 |
| ENSMUSG00000028680 | Plk3 | protein_coding | -0.4094597 | 3.886735 | 0.2655507 | 0.6132583 | 0.999176 |
| ENSMUSG00000024663 | Rab3il1 | protein_coding | -0.3933153 | 3.605714 | 0.2652044 | 0.6134766 | 0.999176 |
| ENSMUSG00000030512 | Snrpa1 | protein_coding | -0.1527441 | 6.516636 | 0.2646109 | 0.6138707 | 0.999176 |
| ENSMUSG00000028126 | Pip5k1a | protein_coding | 0.2119304 | 5.848862 | 0.2645445 | 0.6139148 | 0.999176 |
| ENSMUSG00000000632 | Sez6 | protein_coding | -0.2622132 | 6.55971 | 0.2650315 | 0.6140306 | 0.999176 |
| ENSMUSG00000047731 | Wbp1l | protein_coding | 0.3526557 | 4.18123 | 0.2638049 | 0.6144068 | 0.999176 |
| ENSMUSG00000030872 | Gga2 | protein_coding | -0.4742377 | 3.592282 | 0.2638482 | 0.6144321 | 0.999176 |
| ENSMUSG00000043909 | Trp53bp1 | protein_coding | -0.2813591 | 4.85283 | 0.2635539 | 0.6145739 | 0.999176 |

| ensembl_gene_id | external_gene_name | gene_biotype | log2 (fold change) | log2 (counts per million) | F-value | p-value | False discovery rate |
|---|---|---|---|---|---|---|---|
| ENSMUSG00000026249 | Serpine2 | protein_coding | 0.2203623 | 7.864507 | 0.2635326 | 0.6146292 | 0.999176 |
| ENSMUSG00000036499 | Eea1 | protein_coding | -0.2809171 | 4.669998 | 0.2633901 | 0.6146831 | 0.999176 |
| ENSMUSG00000070348 | Ccnd1 | protein_coding | -0.6348835 | 2.568117 | 0.263207 | 0.6148051 | 0.999176 |
| ENSMUSG00000044442 | N6amt1 | protein_coding | -0.1552893 | 6.418231 | 0.2631946 | 0.6148134 | 0.999176 |
| ENSMUSG00000026035 | Ppil3 | protein_coding | 0.2086712 | 6.186644 | 0.2630848 | 0.6148866 | 0.999176 |
| ENSMUSG00000033713 | Foxn3 | protein_coding | -0.2197688 | 6.874132 | 0.2629393 | 0.6149949 | 0.999176 |
| ENSMUSG00000057614 | Gnai1 | protein_coding | -0.152924 | 6.974124 | 0.2624723 | 0.6152953 | 0.999176 |
| ENSMUSG00000017453 | Pipox | protein_coding | -0.5468544 | 2.536586 | 0.2623791 | 0.6153576 | 0.999176 |
| ENSMUSG00000061603 | Akap6 | protein_coding | -0.1969987 | 5.891526 | 0.2623698 | 0.6153638 | 0.999176 |
| ENSMUSG00000022822 | Abcc5 | protein_coding | -0.1684125 | 6.360322 | 0.2623268 | 0.6153925 | 0.999176 |
| ENSMUSG00000025538 | Sumf2 | protein_coding | 0.3103217 | 5.000446 | 0.2623051 | 0.6154183 | 0.999176 |
| ENSMUSG00000066043 | Phactr4 | protein_coding | 0.4636421 | 2.921264 | 0.2621248 | 0.6155276 | 0.999176 |
| ENSMUSG00000016349 | Eef1a2 | protein_coding | 0.2504785 | 5.49195 | 0.2620655 | 0.6155672 | 0.999176 |
| ENSMUSG00000029472 | Anapc5 | protein_coding | 0.1884809 | 7.048917 | 0.262038 | 0.6155856 | 0.999176 |
| ENSMUSG00000023868 | Pde10a | protein_coding | -0.243812 | 6.422343 | 0.2620896 | 0.6155921 | 0.999176 |
| ENSMUSG00000043289 | Mei4 | protein_coding | 0.6283597 | 2.376394 | 0.2619303 | 0.6156576 | 0.999176 |
| ENSMUSG00000030706 | Mrpl48 | protein_coding | 0.1532065 | 7.445085 | 0.2618028 | 0.6157429 | 0.999176 |
| ENSMUSG00000028249 | Sdcbp | protein_coding | -0.1564641 | 8.82141 | 0.2617052 | 0.6158082 | 0.999176 |
| ENSMUSG00000028466 | Creb3 | protein_coding | -0.159298 | 7.342911 | 0.2611844 | 0.6161569 | 0.999176 |
| ENSMUSG00000026269 | Rnpepl1 | protein_coding | 0.3402675 | 3.453165 | 0.2610075 | 0.6162755 | 0.999176 |
| ENSMUSG00000072258 | Taf1a | protein_coding | -0.2876238 | 4.182932 | 0.2607726 | 0.6164329 | 0.999176 |
| ENSMUSG00000046230 | Vps13a | protein_coding | -0.3187343 | 3.501452 | 0.2605931 | 0.6165533 | 0.999176 |
| ENSMUSG00000005687 | Bcas2 | protein_coding | 0.1606461 | 8.263036 | 0.2605888 | 0.6165562 | 0.999176 |
| ENSMUSG00000032077 | Bud13 | protein_coding | -0.419551 | 4.458012 | 0.2611221 | 0.6166342 | 0.999176 |
| ENSMUSG00000055447 | Cd47 | protein_coding | -0.1609247 | 7.192666 | 0.2601955 | 0.6168202 | 0.999176 |
| ENSMUSG00000096199 | Ptrhd1 | protein_coding | 0.2220111 | 5.253314 | 0.2599632 | 0.6169763 | 0.999176 |
| ENSMUSG00000046079 | Lrrc8d | protein_coding | -0.2454026 | 4.444515 | 0.2599401 | 0.6169918 | 0.999176 |
| ENSMUSG00000067148 | Polr1c | protein_coding | -0.1504275 | 6.988973 | 0.2599393 | 0.6169923 | 0.999176 |
| ENSMUSG00000025130 | P4hb | protein_coding | -0.1862819 | 6.470405 | 0.2594704 | 0.6173077 | 0.999176 |
| ENSMUSG00000020605 | Hs1bp3 | protein_coding | -0.3129262 | 3.979064 | 0.2593819 | 0.6173672 | 0.999176 |
| ENSMUSG00000070287 | Slc35g2 | protein_coding | 0.2313193 | 5.014596 | 0.2588959 | 0.6176945 | 0.999176 |
| ENSMUSG00000051062 | Fbll1 | protein_coding | 0.2391585 | 5.381324 | 0.2584619 | 0.617987 | 0.999176 |
| ENSMUSG00000063870 | Chd4 | protein_coding | -0.1721071 | 6.028504 | 0.2584318 | 0.6180073 | 0.999176 |
| ENSMUSG00000040536 | Necab1 | protein_coding | -0.2982642 | 5.675722 | 0.2587583 | 0.6180716 | 0.999176 |
| ENSMUSG00000031783 | Polr2c | protein_coding | 0.1749489 | 5.726039 | 0.2583 | 0.6180963 | 0.999176 |
| ENSMUSG00000061322 | Dnaic1 | protein_coding | 0.3351173 | 3.965383 | 0.2581325 | 0.6182093 | 0.999176 |
| ENSMUSG00000025940 | Tmem70 | protein_coding | -0.2414184 | 4.937733 | 0.2578975 | 0.6183679 | 0.999176 |
| ENSMUSG00000020176 | Grb10 | protein_coding | -0.21916 | 5.982099 | 0.2578827 | 0.6183779 | 0.999176 |
| ENSMUSG00000031111 | Igsf1 | protein_coding | 0.2232574 | 5.180803 | 0.2577024 | 0.6184997 | 0.999176 |
| ENSMUSG00000018572 | Phf23 | protein_coding | 0.1982047 | 5.843593 | 0.2576508 | 0.6185346 | 0.999176 |
| ENSMUSG00000036278 | Macrod1 | protein_coding | 0.2490049 | 5.257612 | 0.2576483 | 0.6185363 | 0.999176 |
| ENSMUSG00000025894 | Aasdhppt | protein_coding | 0.1822827 | 6.465649 | 0.2574625 | 0.6186619 | 0.999176 |
| ENSMUSG00000069662 | Marcks | protein_coding | -0.1435016 | 7.706302 | 0.2572601 | 0.6187987 | 0.999176 |
| ENSMUSG00000001707 | Eef1e1 | protein_coding | 0.167173 | 7.812688 | 0.2571612 | 0.6188657 | 0.999176 |
| ENSMUSG00000084159 | Gm12696 | processed_pseudoge | -0.4105231 | 2.892749 | 0.2571352 | 0.6188833 | 0.999176 |
| ENSMUSG00000070643 | Sox13 | protein_coding | -0.6035026 | 2.646484 | 0.2571118 | 0.6188991 | 0.999176 |
| ENSMUSG00000027495 | Fam210b | protein_coding | -0.2559827 | 4.9453 | 0.2570423 | 0.6189462 | 0.999176 |
| ENSMUSG00000060038 | Dhps | protein_coding | -0.2545777 | 5.142539 | 0.2569063 | 0.6190382 | 0.999176 |
| ENSMUSG00000040189 | Ccdc114 | protein_coding | -1.055292 | 2.034086 | 0.2575161 | 0.6190578 | 0.999176 |
| ENSMUSG00000030301 | Ccdc91 | protein_coding | 0.1964516 | 5.880554 | 0.2568434 | 0.6190808 | 0.999176 |
| ENSMUSG00000000563 | Atp5f1 | protein_coding | -0.1443725 | 9.728584 | 0.2566979 | 0.6191793 | 0.999176 |
| ENSMUSG00000031511 | Arhgef7 | protein_coding | -0.2477277 | 5.673301 | 0.2566442 | 0.6192157 | 0.999176 |
| ENSMUSG00000038296 | Galnt18 | protein_coding | -0.273795 | 4.651286 | 0.2564948 | 0.619317 | 0.999176 |
| ENSMUSG00000039183 | Nubp2 | protein_coding | 0.1552297 | 6.605297 | 0.2564938 | 0.6193177 | 0.999176 |
| ENSMUSG00000097915 | A330009N23Rik | lincRNA | -0.3871876 | 4.085278 | 0.2564329 | 0.61937 | 0.999176 |
| ENSMUSG00000057069 | Ero1lb | protein_coding | -0.658531 | 2.65998 | 0.2564166 | 0.6193811 | 0.999176 |
| ENSMUSG00000029551 | Psmg3 | protein_coding | 0.4469058 | 2.713556 | 0.2563957 | 0.6193842 | 0.999176 |
| ENSMUSG00000037843 | Vstm2l | protein_coding | -0.5044968 | 2.609806 | 0.2563034 | 0.6194467 | 0.999176 |
| ENSMUSG00000090698 | Apold1 | protein_coding | 0.4775543 | 3.368476 | 0.2562958 | 0.619463 | 0.999176 |
| ENSMUSG00000085181 | Gm12709 | antisense_RNA | -0.2336691 | 5.139537 | 0.2558209 | 0.6197741 | 0.999176 |
| ENSMUSG00000048978 | Nrsn1 | protein_coding | -0.1479098 | 9.176901 | 0.2557347 | 0.6198326 | 0.999176 |
| ENSMUSG00000089665 | Fcor | protein_coding | 0.2676757 | 4.812904 | 0.2556789 | 0.6198705 | 0.999176 |
| ENSMUSG00000020496 | Rnf187 | protein_coding | 0.1668263 | 8.020059 | 0.2556367 | 0.6198991 | 0.999176 |
| ENSMUSG00000027465 | Tbc1d20 | protein_coding | 0.2332581 | 5.170926 | 0.2554892 | 0.6199993 | 0.999176 |
| ENSMUSG00000028597 | Gpx7 | protein_coding | -0.3403079 | 3.864938 | 0.2553069 | 0.6201232 | 0.999176 |
| ENSMUSG00000026578 | Ccdc181 | protein_coding | -0.1950145 | 5.907735 | 0.2550999 | 0.620264 | 0.999176 |
| ENSMUSG00000062421 | Arf2 | protein_coding | -0.2046375 | 6.704695 | 0.2547423 | 0.6205073 | 0.999176 |
| ENSMUSG00000057388 | Mrpl18 | protein_coding | 0.1472673 | 8.058049 | 0.2546202 | 0.6205904 | 0.999176 |
| ENSMUSG00000038619 | Ensa | protein_coding | 0.1551429 | 9.758407 | 0.254471 | 0.620692 | 0.999176 |
| ENSMUSG00000038976 | Ppp1r9b | protein_coding | -0.2123614 | 7.021831 | 0.2544212 | 0.620737 | 0.999176 |
| ENSMUSG00000029625 | Cpsf4 | protein_coding | -0.4501683 | 3.032354 | 0.2543905 | 0.6207468 | 0.999176 |
| ENSMUSG00000067567 | Hdac8 | protein_coding | -0.4948147 | 3.922088 | 0.2552753 | 0.6208185 | 0.999176 |
| ENSMUSG00000024885 | Aldh3b1 | protein_coding | -0.3394274 | 3.921266 | 0.2542108 | 0.6208693 | 0.999176 |
| ENSMUSG00000031487 | Brf2 | protein_coding | -0.2322213 | 5.216124 | 0.2541336 | 0.6209219 | 0.999176 |
| ENSMUSG00000020519 | Sap30l | protein_coding | -0.4417433 | 2.941718 | 0.2541097 | 0.6209382 | 0.999176 |
| ENSMUSG00000028577 | Plaa | protein_coding | 0.2415085 | 4.639243 | 0.2540572 | 0.620974 | 0.999176 |
| ENSMUSG00000064363 | mt-Nd4 | protein_coding | 0.2491543 | 10.34043 | 0.2544275 | 0.6211091 | 0.999176 |
| ENSMUSG00000047712 | Ust | protein_coding | 0.4164337 | 3.075891 | 0.2537465 | 0.6211859 | 0.999176 |

| ensembl_gene_id | external_gene_name | gene_biotype | log2 (fold change) | log2 (counts per million) | F-value | p-value | False discovery rate |
|---|---|---|---|---|---|---|---|
| ENSMUSG00000059920 | 4930453N24Rik | protein_coding | -0.2564242 | 4.780918 | 0.2535096 | 0.6213476 | 0.999176 |
| ENSMUSG00000003849 | Nqo1 | protein_coding | -0.3732916 | 3.758895 | 0.2534941 | 0.6213582 | 0.999176 |
| ENSMUSG00000030662 | Ipo5 | protein_coding | -0.1866821 | 6.205584 | 0.253471 | 0.6213741 | 0.999176 |
| ENSMUSG00000036698 | Ago2 | protein_coding | -0.2093107 | 5.261315 | 0.2534165 | 0.6214112 | 0.999176 |
| ENSMUSG00000029185 | Fam114a1 | protein_coding | -0.7986293 | 2.987001 | 0.2548473 | 0.6214316 | 0.999176 |
| ENSMUSG00000074748 | Atxn7l3b | protein_coding | 0.1407484 | 7.701684 | 0.2531058 | 0.6216235 | 0.999176 |
| ENSMUSG00000032018 | Sc5d | protein_coding | 0.1586275 | 6.268769 | 0.2527666 | 0.6218555 | 0.999176 |
| ENSMUSG00000034800 | Zfp661 | protein_coding | -0.5467332 | 2.819067 | 0.2526428 | 0.6219401 | 0.999176 |
| ENSMUSG00000090110 | Cmc4 | protein_coding | -0.2889755 | 3.930517 | 0.2525956 | 0.6219725 | 0.999176 |
| ENSMUSG00000005609 | Ctr9 | protein_coding | -0.2274469 | 5.35963 | 0.2525068 | 0.6220333 | 0.999176 |
| ENSMUSG00000024666 | Tmem138 | protein_coding | -0.2510835 | 5.47453 | 0.2523808 | 0.6221195 | 0.999176 |
| ENSMUSG00000041623 | D11Wsu47e | protein_coding | 0.3526447 | 3.499928 | 0.2523552 | 0.622137 | 0.999176 |
| ENSMUSG00000020807 | 4933427D14Rik | protein_coding | 0.3215062 | 3.610853 | 0.252302 | 0.6221734 | 0.999176 |
| ENSMUSG00000043683 | Fem1a | protein_coding | -0.2572237 | 4.919768 | 0.2521898 | 0.6222503 | 0.999176 |
| ENSMUSG00000013698 | Pea15a | protein_coding | 0.1859279 | 10.33338 | 0.2519676 | 0.6224026 | 0.999176 |
| ENSMUSG00000027751 | Supt20 | protein_coding | 0.3242263 | 4.469264 | 0.2519411 | 0.6224207 | 0.999176 |
| ENSMUSG00000041930 | Fam222a | protein_coding | -0.6502743 | 2.589838 | 0.2518697 | 0.6224696 | 0.999176 |
| ENSMUSG00000032803 | Cdv3 | protein_coding | 0.19323 | 5.753626 | 0.2518645 | 0.6224732 | 0.999176 |
| ENSMUSG00000028910 | Mecr | protein_coding | -0.2439818 | 5.23956 | 0.2518365 | 0.6224924 | 0.999176 |
| ENSMUSG00000063632 | Sox11 | protein_coding | -0.1907336 | 6.041335 | 0.2518335 | 0.6224945 | 0.999176 |
| ENSMUSG00000091705 | H2-Q2 | protein_coding | 0.5416549 | 2.492497 | 0.2518142 | 0.6225077 | 0.999176 |
| ENSMUSG00000035051 | Dhx57 | protein_coding | -0.2792606 | 4.112709 | 0.251632 | 0.6226327 | 0.999176 |
| ENSMUSG00000058351 | Smim4 | protein_coding | 0.2674416 | 5.621327 | 0.2514058 | 0.6227989 | 0.999176 |
| ENSMUSG00000000085 | Scmh1 | protein_coding | -0.2319623 | 5.071774 | 0.2511735 | 0.6229474 | 0.999176 |
| ENSMUSG00000027628 | Aar2 | protein_coding | 0.1817411 | 6.484043 | 0.2507161 | 0.6232617 | 0.999176 |
| ENSMUSG00000031753 | Cog4 | protein_coding | -0.3442039 | 4.12655 | 0.2503425 | 0.6235187 | 0.999176 |
| ENSMUSG00000028978 | Nos3 | protein_coding | 0.38676 | 3.220517 | 0.249869 | 0.6238448 | 0.999176 |
| ENSMUSG00000040447 | Spns2 | protein_coding | 0.3907826 | 4.353384 | 0.2500901 | 0.6239722 | 0.999176 |
| ENSMUSG00000029103 | Lrpap1 | protein_coding | 0.1420245 | 9.146954 | 0.2495419 | 0.6240703 | 0.999176 |
| ENSMUSG00000052488 | Cherp | protein_coding | 0.4570409 | 4.638329 | 0.2508894 | 0.6240792 | 0.999176 |
| ENSMUSG00000041264 | Uspl1 | protein_coding | -0.2143574 | 5.374579 | 0.249166 | 0.6243296 | 0.999176 |
| ENSMUSG00000026158 | Ogfrl1 | protein_coding | -0.2022074 | 6.023285 | 0.2490919 | 0.6243807 | 0.999176 |
| ENSMUSG00000032549 | Rab6b | protein_coding | 0.1473321 | 8.764897 | 0.2490693 | 0.6243964 | 0.999176 |
| ENSMUSG00000084416 | Rpl10a-ps1 | processed_pseudoge | 0.3820671 | 2.88622 | 0.2488149 | 0.6245721 | 0.999176 |
| ENSMUSG00000024812 | Tjp2 | protein_coding | -0.3282615 | 3.754612 | 0.2487749 | 0.6245997 | 0.999176 |
| ENSMUSG00000040712 | Camta2 | protein_coding | 0.2774128 | 6.010244 | 0.2493367 | 0.6246102 | 0.999176 |
| ENSMUSG00000024137 | E4f1 | protein_coding | -0.4115964 | 3.098904 | 0.2487574 | 0.6246118 | 0.999176 |
| ENSMUSG00000026074 | Map4k4 | protein_coding | -0.2683624 | 4.142939 | 0.2487164 | 0.6246402 | 0.999176 |
| ENSMUSG00000024085 | Man2a1 | protein_coding | 0.7144766 | 2.203002 | 0.2487016 | 0.6246504 | 0.999176 |
| ENSMUSG00000026484 | Rnf2 | protein_coding | -0.2205194 | 5.037564 | 0.2485928 | 0.6247256 | 0.999176 |
| ENSMUSG00000027286 | Lrrc57 | protein_coding | -0.2035381 | 5.643363 | 0.2484852 | 0.6247999 | 0.999176 |
| ENSMUSG00000047945 | Marcksl1 | protein_coding | 0.2603634 | 4.95628 | 0.2481481 | 0.6250331 | 0.999176 |
| ENSMUSG00000035297 | Cops4 | protein_coding | -0.1427875 | 8.429447 | 0.2481437 | 0.6250362 | 0.999176 |
| ENSMUSG00000047777 | Phf13 | protein_coding | 0.321723 | 3.441622 | 0.2479769 | 0.6251517 | 0.999176 |
| ENSMUSG00000035189 | Ano4 | protein_coding | -0.2646638 | 4.021599 | 0.2476566 | 0.6253735 | 0.999176 |
| ENSMUSG00000061613 | U2af1 | protein_coding | 0.2508286 | 4.063219 | 0.2476204 | 0.6253986 | 0.999176 |
| ENSMUSG00000018433 | Nol11 | protein_coding | -0.238284 | 5.26625 | 0.2474751 | 0.6254993 | 0.999176 |
| ENSMUSG00000038152 | 5033430I15Rik | lincRNA | 0.451314 | 2.755846 | 0.2472388 | 0.6256632 | 0.999176 |
| ENSMUSG00000001025 | S100a6 | protein_coding | -0.2685342 | 5.360027 | 0.2469412 | 0.6258697 | 0.999176 |
| ENSMUSG00000036352 | Ubac1 | protein_coding | 0.2052625 | 5.215569 | 0.2469265 | 0.6258799 | 0.999176 |
| ENSMUSG00000020952 | Scfd1 | protein_coding | -0.1616959 | 6.245588 | 0.2466459 | 0.6260748 | 0.999176 |
| ENSMUSG00000018451 | 6330403K07Rik | lincRNA | 0.1554748 | 10.97421 | 0.2465988 | 0.6261075 | 0.999176 |
| ENSMUSG00000039450 | Dcxr | protein_coding | 0.3138393 | 4.156764 | 0.2465528 | 0.6261394 | 0.999176 |
| ENSMUSG00000052752 | Traf7 | protein_coding | -0.2122136 | 4.874099 | 0.246465 | 0.6262005 | 0.999176 |
| ENSMUSG00000033628 | Pik3c3 | protein_coding | -0.2231873 | 5.531721 | 0.2463758 | 0.6262625 | 0.999176 |
| ENSMUSG00000034566 | Atp5h | protein_coding | 0.1734351 | 8.487547 | 0.2463315 | 0.6262932 | 0.999176 |
| ENSMUSG00000041313 | Slc7a1 | protein_coding | 0.2939813 | 3.810195 | 0.246216 | 0.6263736 | 0.999176 |
| ENSMUSG00000011306 | Sugp1 | protein_coding | 0.2103373 | 6.264809 | 0.2461987 | 0.6263856 | 0.999176 |
| ENSMUSG00000030256 | Bhlhe41 | protein_coding | 0.4536318 | 7.981567 | 0.2517056 | 0.6265128 | 0.999176 |
| ENSMUSG00000068134 | Zfp120 | protein_coding | 0.2960928 | 4.443658 | 0.245628 | 0.6267828 | 0.999176 |
| ENSMUSG00000057409 | Zfp53 | protein_coding | 0.5936326 | 2.40912 | 0.2455877 | 0.6268109 | 0.999176 |
| ENSMUSG00000033863 | Klf9 | protein_coding | -0.1544607 | 6.174847 | 0.2455449 | 0.6268407 | 0.999176 |
| ENSMUSG00000025591 | Tma16 | protein_coding | 0.2432799 | 4.834137 | 0.2453606 | 0.6269691 | 0.999176 |
| ENSMUSG00000022359 | Wdyhv1 | protein_coding | -0.1779378 | 5.773108 | 0.2453113 | 0.6270035 | 0.999176 |
| ENSMUSG00000041685 | Fcho2 | protein_coding | -0.2928665 | 3.901869 | 0.2452977 | 0.627013 | 0.999176 |
| ENSMUSG00000003541 | Ier3 | protein_coding | -0.2150416 | 4.830211 | 0.2449231 | 0.6272742 | 0.999176 |
| ENSMUSG00000049624 | Slc17a5 | protein_coding | 0.2749688 | 4.188864 | 0.2446359 | 0.6274747 | 0.999176 |
| ENSMUSG00000033059 | Pygb | protein_coding | 0.2284036 | 6.971583 | 0.2450319 | 0.6274748 | 0.999176 |
| ENSMUSG00000042446 | Zmym4 | protein_coding | -0.5170064 | 2.574492 | 0.244503 | 0.6275675 | 0.999176 |
| ENSMUSG00000038141 | Tmem181a | protein_coding | -0.3074407 | 3.611116 | 0.2443313 | 0.6276875 | 0.999176 |
| ENSMUSG00000047554 | Tmem41b | protein_coding | -0.2468131 | 4.38952 | 0.2443278 | 0.62769 | 0.999176 |
| ENSMUSG00000051705 | Senp8 | protein_coding | -0.2722604 | 4.582202 | 0.2440221 | 0.6279036 | 0.999176 |
| ENSMUSG00000027833 | Shox2 | protein_coding | -0.9584204 | 4.859037 | 0.2542016 | 0.6279167 | 0.999176 |
| ENSMUSG00000035443 | Thyn1 | protein_coding | -0.1430249 | 7.727956 | 0.2437888 | 0.6280669 | 0.999176 |
| ENSMUSG00000020882 | Cacnb1 | protein_coding | 0.1811581 | 5.735647 | 0.243717 | 0.6281171 | 0.999176 |
| ENSMUSG00000025371 | Chmp6 | protein_coding | 0.2371181 | 5.972847 | 0.2436131 | 0.6282006 | 0.999176 |
| ENSMUSG00000021701 | Plk2 | protein_coding | -0.1550269 | 7.131696 | 0.2435534 | 0.6282317 | 0.999176 |
| ENSMUSG00000052373 | Mpp3 | protein_coding | 0.2574913 | 5.007329 | 0.2435099 | 0.6282621 | 0.999176 |

| ensembl_gene_id | external_gene_name | gene_biotype | log2 (fold change) | log2 (counts per million) | F-value | p-value | False discovery rate |
|---|---|---|---|---|---|---|---|
| ENSMUSG00000062190 | Lancl2 | protein_coding | 0.5556532 | 3.077569 | 0.2433139 | 0.6284102 | 0.999176 |
| ENSMUSG00000028007 | Snx7 | protein_coding | -0.3024806 | 3.934943 | 0.2432479 | 0.6284457 | 0.999176 |
| ENSMUSG00000001082 | Mfsd10 | protein_coding | 0.4092003 | 3.679602 | 0.2430673 | 0.628583 | 0.999176 |
| ENSMUSG00000054226 | Tprkb | protein_coding | 0.1386929 | 7.424712 | 0.2430334 | 0.628596 | 0.999176 |
| ENSMUSG00000036501 | Fam13b | protein_coding | -0.2952486 | 3.975044 | 0.2429274 | 0.6286703 | 0.999176 |
| ENSMUSG00000087497 | 2810001G20Rik | antisense_RNA | 0.2375164 | 4.972236 | 0.2428459 | 0.6287276 | 0.999176 |
| ENSMUSG00000020869 | Lrrc59 | protein_coding | 0.2073707 | 5.356799 | 0.2427411 | 0.6288011 | 0.999176 |
| ENSMUSG00000030612 | Mrpl46 | protein_coding | 0.1382685 | 7.796074 | 0.2427361 | 0.6288046 | 0.999176 |
| ENSMUSG00000079109 | Pms2 | protein_coding | -0.5696619 | 2.451916 | 0.242398 | 0.6290419 | 0.999176 |
| ENSMUSG00000026494 | Kif26b | protein_coding | -0.3641139 | 3.373506 | 0.2423648 | 0.6290652 | 0.999176 |
| ENSMUSG00000025139 | Tollip | protein_coding | -0.1879751 | 7.020218 | 0.242354 | 0.6290728 | 0.999176 |
| ENSMUSG00000004626 | Stxbp2 | protein_coding | -0.2874315 | 4.167827 | 0.2422775 | 0.6291265 | 0.999176 |
| ENSMUSG00000045176 | Borcs6 | protein_coding | -0.3251735 | 3.728894 | 0.2422686 | 0.6291328 | 0.999176 |
| ENSMUSG00000024083 | Pja2 | protein_coding | -0.1781829 | 9.105464 | 0.2421907 | 0.6291876 | 0.999176 |
| ENSMUSG00000023980 | Taf8 | protein_coding | -0.2129184 | 5.220399 | 0.2418185 | 0.6294492 | 0.999176 |
| ENSMUSG00000028291 | Akirin2 | protein_coding | -0.2164542 | 5.350127 | 0.2414065 | 0.6297392 | 0.999176 |
| ENSMUSG00000031189 | Aff2 | protein_coding | -0.4516082 | 2.7727 | 0.2412939 | 0.6298185 | 0.999176 |
| ENSMUSG00000022769 | Sdf2l1 | protein_coding | 0.4676599 | 2.50465 | 0.2412868 | 0.6298234 | 0.999176 |
| ENSMUSG00000067370 | B3galt4 | protein_coding | 0.4648396 | 3.638787 | 0.2416331 | 0.6298539 | 0.999176 |
| ENSMUSG00000031960 | Aars | protein_coding | 0.2017097 | 7.021415 | 0.2411995 | 0.6298957 | 0.999176 |
| ENSMUSG00000046854 | Pip5kl1 | protein_coding | 0.9320597 | 2.460729 | 0.2420687 | 0.629927 | 0.999176 |
| ENSMUSG00000040006 | Ginm1 | protein_coding | -0.2278754 | 5.331043 | 0.2411347 | 0.6299306 | 0.999176 |
| ENSMUSG00000028172 | Tacr3 | protein_coding | -0.4255922 | 2.731954 | 0.2411328 | 0.629932 | 0.999176 |
| ENSMUSG00000111088 | AC122428.3 | TEC | 0.4649287 | 3.354418 | 0.2410884 | 0.629974 | 0.999176 |
| ENSMUSG00000029587 | Zfp12 | protein_coding | 0.5736846 | 3.779491 | 0.242461 | 0.6300429 | 0.999176 |
| ENSMUSG00000054428 | Atpif1 | protein_coding | 0.1898629 | 9.284074 | 0.240891 | 0.6301024 | 0.999176 |
| ENSMUSG00000037754 | Ppp1r16b | protein_coding | -0.1954426 | 8.079456 | 0.2406236 | 0.6303017 | 0.999176 |
| ENSMUSG00000054469 | Lclat1 | protein_coding | -0.3762526 | 3.717781 | 0.2404167 | 0.630437 | 0.999176 |
| ENSMUSG00000020898 | Ctc1 | protein_coding | 0.2745082 | 4.549826 | 0.2403401 | 0.6304911 | 0.999176 |
| ENSMUSG00000022623 | Shank3 | protein_coding | -0.4228321 | 2.872524 | 0.2403094 | 0.6305128 | 0.999176 |
| ENSMUSG00000037649 | H2-DMa | protein_coding | 0.3100044 | 4.709291 | 0.2402188 | 0.6305874 | 0.999176 |
| ENSMUSG00000021427 | Ssr1 | protein_coding | -0.1607026 | 8.643972 | 0.2400988 | 0.6306615 | 0.999176 |
| ENSMUSG00000018481 | Appbp2 | protein_coding | -0.1480717 | 6.661614 | 0.2399831 | 0.6307433 | 0.999176 |
| ENSMUSG00000032299 | Commd4 | protein_coding | 0.1617171 | 7.67794 | 0.2399643 | 0.6307566 | 0.999176 |
| ENSMUSG00000027007 | Ssfa2 | protein_coding | -0.4703435 | 3.402115 | 0.2399344 | 0.6307884 | 0.999176 |
| ENSMUSG00000042133 | Ppig | protein_coding | -0.1829845 | 6.474704 | 0.2398936 | 0.6308066 | 0.999176 |
| ENSMUSG00000042632 | Pla2g6 | protein_coding | -0.2153292 | 5.294237 | 0.2397769 | 0.6308891 | 0.999176 |
| ENSMUSG00000002949 | Timm44 | protein_coding | 0.1954766 | 6.385037 | 0.2397529 | 0.630906 | 0.999176 |
| ENSMUSG00000038683 | Pak1ip1 | protein_coding | 0.1779546 | 7.238186 | 0.239677 | 0.6309597 | 0.999176 |
| ENSMUSG00000040249 | Lrp1 | protein_coding | -0.2154859 | 5.675659 | 0.2395512 | 0.6310487 | 0.999176 |
| ENSMUSG00000030711 | Sult1a1 | protein_coding | -0.4629502 | 3.215347 | 0.239509 | 0.6310786 | 0.999176 |
| ENSMUSG00000021576 | Pdcd6 | protein_coding | 0.1467945 | 7.915905 | 0.2394842 | 0.6310961 | 0.999176 |
| ENSMUSG00000042066 | Tmcc2 | protein_coding | 0.2034479 | 6.075979 | 0.2393723 | 0.6311753 | 0.999176 |
| ENSMUSG00000023938 | Aars2 | protein_coding | -0.5293137 | 2.54337 | 0.2393685 | 0.631178 | 0.999176 |
| ENSMUSG00000019923 | Zwint | protein_coding | 0.1824207 | 12.04741 | 0.2392812 | 0.6312398 | 0.999176 |
| ENSMUSG00000025582 | Nptx1 | protein_coding | -0.2586437 | 4.495207 | 0.2392558 | 0.6312578 | 0.999176 |
| ENSMUSG00000027882 | Stxbp3 | protein_coding | 0.251239 | 4.760749 | 0.239083 | 0.6313802 | 0.999176 |
| ENSMUSG00000022195 | 6030458C11Rik | protein_coding | 0.2898852 | 5.026615 | 0.2390551 | 0.6314107 | 0.999176 |
| ENSMUSG00000041801 | Phlda3 | protein_coding | 0.2111659 | 5.543786 | 0.2389944 | 0.631443 | 0.999176 |
| ENSMUSG00000057594 | Arl16 | protein_coding | 0.1531683 | 6.821007 | 0.2389778 | 0.6314547 | 0.999176 |
| ENSMUSG00000051586 | Mical3 | protein_coding | -0.3263564 | 4.129779 | 0.2389014 | 0.6315088 | 0.999176 |
| ENSMUSG00000026854 | Usp20 | protein_coding | 0.204587 | 5.976016 | 0.2386871 | 0.6316608 | 0.999176 |
| ENSMUSG00000048379 | Socs4 | protein_coding | -0.5007199 | 2.690754 | 0.238504 | 0.6317907 | 0.999176 |
| ENSMUSG00000020817 | Rabep1 | protein_coding | 0.166195 | 6.820638 | 0.2383024 | 0.6319337 | 0.999176 |
| ENSMUSG00000058392 | Rrp1b | protein_coding | 0.2342527 | 4.924291 | 0.2381387 | 0.6320499 | 0.999176 |
| ENSMUSG00000022075 | Rhobtb2 | protein_coding | -0.215631 | 5.279648 | 0.2381358 | 0.632052 | 0.999176 |
| ENSMUSG00000049940 | Pgrmc2 | protein_coding | -0.2078423 | 5.786279 | 0.2380639 | 0.6321031 | 0.999176 |
| ENSMUSG00000029291 | Rufy3 | protein_coding | -0.1555394 | 8.129277 | 0.2379532 | 0.6321817 | 0.999176 |
| ENSMUSG00000079045 | Prox1os | antisense_RNA | -0.3971342 | 4.162944 | 0.2381554 | 0.6322542 | 0.999176 |
| ENSMUSG00000029922 | Mkrn1 | protein_coding | 0.1535074 | 6.384668 | 0.2378249 | 0.6322729 | 0.999176 |
| ENSMUSG00000029028 | Lrrc47 | protein_coding | 0.4717536 | 2.925906 | 0.237765 | 0.6323154 | 0.999176 |
| ENSMUSG00000070939 | Tgfbrap1 | protein_coding | 0.2611513 | 4.076641 | 0.2377178 | 0.632349 | 0.999176 |
| ENSMUSG00000038879 | Nipal2 | protein_coding | -0.5672951 | 3.331697 | 0.2380749 | 0.6326002 | 0.999176 |
| ENSMUSG00000039512 | Uhrf1bp1 | protein_coding | 0.3627866 | 3.174498 | 0.2373177 | 0.6326336 | 0.999176 |
| ENSMUSG00000029326 | Enoph1 | protein_coding | -0.1460409 | 7.226128 | 0.2370055 | 0.6328559 | 0.999176 |
| ENSMUSG00000022023 | Wbp4 | protein_coding | 0.1404532 | 6.758043 | 0.2369927 | 0.632865 | 0.999176 |
| ENSMUSG00000031847 | 1700030J22Rik | protein_coding | 0.3141477 | 3.621339 | 0.2369813 | 0.6328731 | 0.999176 |
| ENSMUSG00000027176 | Cstf3 | protein_coding | 0.1697616 | 6.278475 | 0.2369804 | 0.6328738 | 0.999176 |
| ENSMUSG00000024030 | Abcg1 | protein_coding | -0.1819056 | 7.836224 | 0.236975 | 0.6328776 | 0.999176 |
| ENSMUSG00000029106 | Add1 | protein_coding | 0.1444338 | 7.534351 | 0.2369133 | 0.6329216 | 0.999176 |
| ENSMUSG00000029131 | Dnajb6 | protein_coding | 0.2549215 | 4.287202 | 0.2363544 | 0.63332 | 0.999176 |
| ENSMUSG00000042211 | Fbxo38 | protein_coding | 0.2026506 | 5.164586 | 0.2359771 | 0.6335894 | 0.999176 |
| ENSMUSG00000021428 | Riok1 | protein_coding | -0.2326611 | 4.878698 | 0.2356521 | 0.6338216 | 0.999176 |
| ENSMUSG00000037851 | Iars | protein_coding | -0.2596514 | 4.834015 | 0.2355107 | 0.6339227 | 0.999176 |
| ENSMUSG00000036061 | Smug1 | protein_coding | -0.3515555 | 3.632797 | 0.2354991 | 0.633931 | 0.999176 |
| ENSMUSG00000038733 | Wdr26 | protein_coding | -0.2254184 | 6.151535 | 0.2354431 | 0.6339816 | 0.999176 |
| ENSMUSG00000015488 | Cacfd1 | protein_coding | 0.2763567 | 4.855522 | 0.2353023 | 0.6340717 | 0.999176 |
| ENSMUSG00000015829 | Tnr | protein_coding | 0.3068826 | 3.979884 | 0.2352254 | 0.6341268 | 0.999176 |

| ensembl_gene_id | external_gene_name | gene_biotype | log2 (fold change) | log2 (counts per million) | F-value | p-value | False discovery rate |
|---|---|---|---|---|---|---|---|
| ENSMUSG00000025246 | Tbl1x | protein_coding | -0.1823172 | 6.155743 | 0.2351687 | 0.6341674 | 0.999176 |
| ENSMUSG00000036932 | Aifm1 | protein_coding | -0.2308918 | 5.357849 | 0.2350784 | 0.634232 | 0.999176 |
| ENSMUSG00000046341 | Gm11223 | processed_pseudoge | -0.3324549 | 3.200806 | 0.2350296 | 0.6342669 | 0.999176 |
| ENSMUSG00000033319 | Fem1c | protein_coding | -0.283926 | 3.986097 | 0.234984 | 0.6342996 | 0.999176 |
| ENSMUSG00000006715 | Gmnn | protein_coding | -0.3409092 | 4.321437 | 0.234915 | 0.6343596 | 0.999176 |
| ENSMUSG00000030417 | Pdcd5 | protein_coding | 0.3013456 | 4.304474 | 0.2348275 | 0.6344117 | 0.999176 |
| ENSMUSG00000018583 | G3bp1 | protein_coding | -0.1758448 | 6.057078 | 0.234774 | 0.63445 | 0.999176 |
| ENSMUSG00000033460 | Armcx1 | protein_coding | -0.1836861 | 6.736641 | 0.2347621 | 0.6344585 | 0.999176 |
| ENSMUSG00000040822 | 1700123O20Rik | protein_coding | -0.162028 | 6.609926 | 0.2345468 | 0.6346128 | 0.999176 |
| ENSMUSG00000002833 | Hdgfl2 | protein_coding | 0.2366642 | 5.880915 | 0.2343561 | 0.6347602 | 0.999176 |
| ENSMUSG00000026942 | Traf2 | protein_coding | -0.2622352 | 4.716629 | 0.2342942 | 0.634794 | 0.999176 |
| ENSMUSG00000072582 | Ptrh2 | protein_coding | -0.1756386 | 6.181313 | 0.2342742 | 0.6348083 | 0.999176 |
| ENSMUSG00000009535 | Rnmt | protein_coding | -0.1771532 | 6.789872 | 0.2340457 | 0.6349724 | 0.999176 |
| ENSMUSG00000109975 | Gm19196 | transcribed_unproces | 0.265386 | 3.871162 | 0.2335988 | 0.6352934 | 0.999176 |
| ENSMUSG00000024712 | Rfk | protein_coding | 0.1675322 | 6.43847 | 0.2333591 | 0.6354657 | 0.999176 |
| ENSMUSG00000045994 | B3gat1 | protein_coding | 0.2383746 | 4.885531 | 0.2331593 | 0.6356094 | 0.999176 |
| ENSMUSG00000002602 | Axl | protein_coding | -0.4349282 | 3.171211 | 0.2330545 | 0.6356848 | 0.999176 |
| ENSMUSG00000026356 | Dars | protein_coding | -0.2101616 | 6.447265 | 0.2329134 | 0.635797 | 0.999176 |
| ENSMUSG00000064373 | Selenop | protein_coding | -0.2018944 | 7.836043 | 0.2329001 | 0.6358066 | 0.999176 |
| ENSMUSG00000097068 | Gm26760 | lincRNA | 0.4773349 | 2.564849 | 0.2326045 | 0.636009 | 0.999176 |
| ENSMUSG00000071657 | Bscl2 | protein_coding | 0.1455331 | 7.871977 | 0.2325497 | 0.6360484 | 0.999176 |
| ENSMUSG00000036565 | Ttyh3 | protein_coding | 0.2936018 | 4.395065 | 0.2325457 | 0.6360513 | 0.999176 |
| ENSMUSG00000026457 | Adipor1 | protein_coding | -0.20016 | 5.534447 | 0.2318601 | 0.6365459 | 0.999176 |
| ENSMUSG00000032264 | Zw10 | protein_coding | 0.3134837 | 4.109129 | 0.2317877 | 0.6365982 | 0.999176 |
| ENSMUSG00000028609 | Magoh | protein_coding | -0.1831033 | 6.642206 | 0.2314953 | 0.6368095 | 0.999176 |
| ENSMUSG00000056222 | Spock1 | protein_coding | 0.1443995 | 6.523959 | 0.2314942 | 0.6368103 | 0.999176 |
| ENSMUSG00000002325 | Irf9 | protein_coding | 0.3492788 | 3.572885 | 0.2314423 | 0.6368478 | 0.999176 |
| ENSMUSG00000096188 | Cmtm4 | protein_coding | 0.2916509 | 4.718666 | 0.2311476 | 0.6370609 | 0.999176 |
| ENSMUSG00000001016 | Ilf2 | protein_coding | -0.1460101 | 7.841223 | 0.2311188 | 0.6370818 | 0.999176 |
| ENSMUSG00000028277 | Ube2j1 | protein_coding | 0.1872063 | 6.01649 | 0.2311043 | 0.6370922 | 0.999176 |
| ENSMUSG00000028101 | Pias3 | protein_coding | 0.3079203 | 5.43273 | 0.231609 | 0.6371115 | 0.999176 |
| ENSMUSG00000020261 | Slc36a1 | protein_coding | -0.4346086 | 3.689175 | 0.2311527 | 0.6371612 | 0.999176 |
| ENSMUSG00000028890 | Mtf1 | protein_coding | 0.214159 | 4.908269 | 0.2308886 | 0.6372484 | 0.999176 |
| ENSMUSG00000096935 | 1700113A16Rik | antisense_RNA | -0.5683738 | 2.436714 | 0.2307528 | 0.6373467 | 0.999176 |
| ENSMUSG00000035805 | Mlc1 | protein_coding | -0.3147002 | 6.046922 | 0.2319812 | 0.6373543 | 0.999176 |
| ENSMUSG00000019856 | Fam184a | protein_coding | -0.3143584 | 3.474339 | 0.2304765 | 0.6375469 | 0.999176 |
| ENSMUSG00000001924 | Uba1 | protein_coding | 0.1500714 | 7.770458 | 0.2302765 | 0.6376918 | 0.999176 |
| ENSMUSG00000052681 | Rap1b | protein_coding | -0.208845 | 5.520196 | 0.2302551 | 0.6377073 | 0.999176 |
| ENSMUSG00000036890 | Gtdc1 | protein_coding | -0.2725449 | 4.693489 | 0.2302444 | 0.6377151 | 0.999176 |
| ENSMUSG00000033596 | Rfwd3 | protein_coding | -0.3746599 | 3.150217 | 0.2302063 | 0.6377427 | 0.999176 |
| ENSMUSG00000029219 | Slc10a4 | protein_coding | 0.3181358 | 3.807946 | 0.2300333 | 0.6378683 | 0.999176 |
| ENSMUSG00000024617 | Camk2a | protein_coding | -0.1974135 | 7.991298 | 0.2299516 | 0.637938 | 0.999176 |
| ENSMUSG00000022947 | Cbr3 | protein_coding | -0.3911267 | 3.369841 | 0.2297387 | 0.6380821 | 0.999176 |
| ENSMUSG00000031422 | Morf4l2 | protein_coding | 0.166785 | 8.396678 | 0.2296622 | 0.6381377 | 0.999176 |
| ENSMUSG00000013622 | Atraid | protein_coding | 0.1669248 | 7.682198 | 0.2296607 | 0.6381388 | 0.999176 |
| ENSMUSG00000004187 | Kifc2 | protein_coding | 0.1798117 | 6.489651 | 0.2294948 | 0.6382593 | 0.999176 |
| ENSMUSG00000021391 | Cenpp | protein_coding | 0.3270304 | 3.797326 | 0.2291786 | 0.6384892 | 0.999176 |
| ENSMUSG00000034867 | Ankrd27 | protein_coding | 0.3058423 | 4.345784 | 0.228871 | 0.638713 | 0.999176 |
| ENSMUSG00000032309 | Fbxo22 | protein_coding | 0.2797059 | 4.513697 | 0.2288687 | 0.6387147 | 0.999176 |
| ENSMUSG00000057132 | Rpgrip1 | protein_coding | 0.3752342 | 3.430311 | 0.2285945 | 0.6389144 | 0.999176 |
| ENSMUSG00000037907 | Ankrd13b | protein_coding | -0.4017571 | 3.239422 | 0.228591 | 0.6389169 | 0.999176 |
| ENSMUSG00000025495 | Ptdss2 | protein_coding | -0.2429962 | 5.09732 | 0.2284123 | 0.6390471 | 0.999176 |
| ENSMUSG00000071654 | Uqcc3 | protein_coding | -0.1790208 | 6.871598 | 0.2283475 | 0.6390943 | 0.999176 |
| ENSMUSG00000031770 | Herpud1 | protein_coding | 0.1642758 | 6.778396 | 0.2282536 | 0.6391628 | 0.999176 |
| ENSMUSG00000050029 | Rap2c | protein_coding | 0.2655453 | 4.081773 | 0.228193 | 0.639207 | 0.999176 |
| ENSMUSG00000037475 | Thoc2 | protein_coding | -0.2709224 | 4.708719 | 0.2275114 | 0.6397045 | 0.999176 |
| ENSMUSG00000049690 | Nckap5 | protein_coding | 0.6405169 | 2.575472 | 0.2273234 | 0.6398522 | 0.999176 |
| ENSMUSG00000027365 | Trpm7 | protein_coding | -0.2766154 | 3.924431 | 0.2272666 | 0.6398834 | 0.999176 |
| ENSMUSG00000071302 | 2610044O15Rik8 | protein_coding | 0.2221466 | 4.95283 | 0.2272069 | 0.6399271 | 0.999176 |
| ENSMUSG00000042502 | Cd2bp2 | protein_coding | -0.1907987 | 6.020396 | 0.2270884 | 0.6400137 | 0.999176 |
| ENSMUSG00000042569 | Dhrs7b | protein_coding | 0.2017248 | 5.48156 | 0.2270834 | 0.6400174 | 0.999176 |
| ENSMUSG00000025220 | Mgea5 | protein_coding | -0.2614596 | 4.184251 | 0.2270682 | 0.6400285 | 0.999176 |
| ENSMUSG00000021250 | Fos | protein_coding | 0.2225411 | 6.54085 | 0.2269507 | 0.6401629 | 0.999176 |
| ENSMUSG00000022175 | Lrp10 | protein_coding | 0.3394682 | 3.736933 | 0.2267026 | 0.640296 | 0.999176 |
| ENSMUSG00000024622 | Hmgxb3 | protein_coding | 0.3769558 | 3.198519 | 0.2266262 | 0.640352 | 0.999176 |
| ENSMUSG00000029386 | Tctn2 | processed_transcript | -0.2250569 | 4.322254 | 0.2265412 | 0.6404143 | 0.999176 |
| ENSMUSG00000038888 | Ctu1 | protein_coding | 0.5217589 | 3.03594 | 0.2264239 | 0.6405106 | 0.999176 |
| ENSMUSG00000013787 | Ehmt2 | protein_coding | 0.2104803 | 5.583196 | 0.2261832 | 0.6406766 | 0.999176 |
| ENSMUSG00000040435 | Ppp1r15a | protein_coding | -0.282974 | 4.902967 | 0.2261576 | 0.6406954 | 0.999176 |
| ENSMUSG00000048485 | Zbtb8b | protein_coding | 0.4322737 | 2.654409 | 0.2261219 | 0.6407216 | 0.999176 |
| ENSMUSG00000055720 | Ubl7 | protein_coding | 0.1543916 | 7.952824 | 0.2261181 | 0.6407244 | 0.999176 |
| ENSMUSG00000021156 | Zmynd11 | protein_coding | -0.2614661 | 4.511812 | 0.2260657 | 0.6407628 | 0.999176 |
| ENSMUSG00000019843 | Fyn | protein_coding | 0.2337079 | 4.647894 | 0.2260576 | 0.6407687 | 0.999176 |
| ENSMUSG00000020181 | Nav3 | protein_coding | -0.2103391 | 5.391696 | 0.2258641 | 0.6409107 | 0.999176 |
| ENSMUSG00000001441 | Npepps | protein_coding | -0.1423772 | 6.845432 | 0.2257857 | 0.6409683 | 0.999176 |
| ENSMUSG00000024594 | Prrc1 | protein_coding | -0.6673442 | 2.087124 | 0.2257217 | 0.6410153 | 0.999176 |
| ENSMUSG00000063439 | B9d2 | protein_coding | 0.1447793 | 7.162229 | 0.2256474 | 0.6410698 | 0.999176 |
| ENSMUSG00000058975 | Kcnc1 | protein_coding | 0.2077193 | 5.324145 | 0.2255346 | 0.6411526 | 0.999176 |

| ensembl_gene_id | external_gene_name | gene_biotype | log2 (fold change) | log2 (counts per million) | F-value | p-value | False discovery rate |
|---|---|---|---|---|---|---|---|
| ENSMUSG00000034300 | Fam53c | protein_coding | -0.2343142 | 5.380149 | 0.2255222 | 0.6411617 | 0.999176 |
| ENSMUSG00000020923 | Ubtf | protein_coding | 0.1816247 | 6.328264 | 0.2254852 | 0.641189 | 0.999176 |
| ENSMUSG00000056268 | Dennd1b | protein_coding | -0.4938522 | 2.79048 | 0.2253888 | 0.6412598 | 0.999176 |
| ENSMUSG00000025892 | Gria4 | protein_coding | 0.2110599 | 5.401966 | 0.2247645 | 0.6417189 | 0.999176 |
| ENSMUSG00000020621 | Rdh14 | protein_coding | 0.2149258 | 5.32637 | 0.2247528 | 0.6417276 | 0.999176 |
| ENSMUSG00000021748 | Pdhb | protein_coding | -0.1293479 | 7.858429 | 0.2246844 | 0.6417779 | 0.999176 |
| ENSMUSG00000049807 | Arhgap23 | protein_coding | -0.2774175 | 4.266676 | 0.2246497 | 0.6418035 | 0.999176 |
| ENSMUSG00000036699 | Zcchc12 | protein_coding | 0.2840207 | 9.901316 | 0.2258954 | 0.6418174 | 0.999176 |
| ENSMUSG00000040722 | Scamp5 | protein_coding | 0.1386112 | 7.379338 | 0.2245811 | 0.6418539 | 0.999176 |
| ENSMUSG00000039178 | Tbc1d19 | protein_coding | -0.1784311 | 5.520886 | 0.2245544 | 0.6418737 | 0.999176 |
| ENSMUSG00000005204 | Senp3 | protein_coding | -0.2217739 | 5.074072 | 0.2241269 | 0.6421887 | 0.999176 |
| ENSMUSG00000047996 | Prrg1 | protein_coding | -0.3177076 | 3.350187 | 0.2240999 | 0.6422086 | 0.999176 |
| ENSMUSG00000052387 | Trpm3 | protein_coding | -0.4073957 | 4.387658 | 0.2247451 | 0.6422612 | 0.999176 |
| ENSMUSG00000023092 | Fhl1 | protein_coding | 0.1636315 | 7.233668 | 0.2238402 | 0.6424002 | 0.999176 |
| ENSMUSG00000020514 | Mrpl22 | protein_coding | 0.1649176 | 6.224654 | 0.2237161 | 0.6424918 | 0.999176 |
| ENSMUSG00000054910 | 4931415C17Rik | processed_transcript | -0.5512098 | 2.736321 | 0.223673 | 0.6425236 | 0.999176 |
| ENSMUSG00000028049 | Scamp3 | protein_coding | 0.1576891 | 6.211899 | 0.2235295 | 0.6426296 | 0.999176 |
| ENSMUSG00000048755 | Mcat | protein_coding | 0.4566863 | 3.814426 | 0.2236872 | 0.6429373 | 0.999176 |
| ENSMUSG00000060860 | Ube2s | protein_coding | 0.4576255 | 2.896767 | 0.223058 | 0.6429781 | 0.999176 |
| ENSMUSG00000059857 | Ntng1 | protein_coding | -0.2942182 | 5.235176 | 0.2230168 | 0.6430566 | 0.999176 |
| ENSMUSG00000037563 | Rps16 | protein_coding | 0.4039101 | 2.81622 | 0.2226748 | 0.6432616 | 0.999176 |
| ENSMUSG00000002017 | Fam98a | protein_coding | -0.2382942 | 4.176499 | 0.2225208 | 0.6433757 | 0.999176 |
| ENSMUSG00000028274 | Rngtt | protein_coding | 0.2493379 | 5.010766 | 0.2224982 | 0.6433925 | 0.999176 |
| ENSMUSG00000003233 | Dvl3 | protein_coding | 0.3160833 | 4.062869 | 0.2224492 | 0.6434287 | 0.999176 |
| ENSMUSG00000020328 | Nudcd2 | protein_coding | 0.185077 | 5.547238 | 0.2224105 | 0.6434574 | 0.999176 |
| ENSMUSG00000038257 | Glra3 | protein_coding | -0.3396884 | 3.159078 | 0.2223066 | 0.6435344 | 0.999176 |
| ENSMUSG00000027540 | Ptpn1 | protein_coding | 0.2493237 | 4.791763 | 0.2221572 | 0.6436451 | 0.999176 |
| ENSMUSG00000066406 | Akap13 | protein_coding | 0.390874 | 3.010877 | 0.222006 | 0.6437572 | 0.999176 |
| ENSMUSG00000042589 | Cux2 | protein_coding | 0.3571821 | 5.710128 | 0.2230493 | 0.6439813 | 0.999176 |
| ENSMUSG00000030279 | C2cd5 | protein_coding | 0.2317481 | 4.359382 | 0.2217007 | 0.6439838 | 0.999176 |
| ENSMUSG00000038271 | Iffo1 | protein_coding | -0.2918319 | 4.008016 | 0.2216867 | 0.6439942 | 0.999176 |
| ENSMUSG00000068015 | Lrch1 | protein_coding | -0.4330912 | 3.917522 | 0.2221848 | 0.644022 | 0.999176 |
| ENSMUSG00000053646 | Plxnb1 | protein_coding | 0.569693 | 3.351708 | 0.2224302 | 0.6440658 | 0.999176 |
| ENSMUSG00000026466 | Tor1aip1 | protein_coding | -0.2773566 | 3.687747 | 0.2215726 | 0.6440789 | 0.999176 |
| ENSMUSG00000039384 | Dusp10 | protein_coding | -0.3783553 | 4.084572 | 0.2216144 | 0.6441028 | 0.999176 |
| ENSMUSG00000018286 | Psmb6 | protein_coding | 0.1489939 | 9.771319 | 0.2215242 | 0.6441149 | 0.999176 |
| ENSMUSG00000015126 | Tsr3 | protein_coding | 0.2339617 | 4.581791 | 0.2212353 | 0.6443295 | 0.999176 |
| ENSMUSG00000031729 | Ist1 | protein_coding | -0.2469799 | 5.56111 | 0.2212399 | 0.6443363 | 0.999176 |
| ENSMUSG00000011257 | Pabpc4 | protein_coding | 0.3407543 | 3.198286 | 0.2211523 | 0.6443913 | 0.999176 |
| ENSMUSG00000032120 | C2cd2l | protein_coding | 0.1730246 | 6.011684 | 0.2210005 | 0.6445041 | 0.999176 |
| ENSMUSG00000018411 | Mapt | protein_coding | 0.1911512 | 8.897667 | 0.2209865 | 0.6445247 | 0.999176 |
| ENSMUSG00000033578 | Tmem35a | protein_coding | 0.1523456 | 6.739431 | 0.2202359 | 0.6450735 | 0.999176 |
| ENSMUSG00000097353 | A430046D13Rik | lincRNA | 0.271569 | 3.666363 | 0.2202157 | 0.6450885 | 0.999176 |
| ENSMUSG00000021065 | Fut8 | protein_coding | 0.2220554 | 4.914718 | 0.219949 | 0.6452874 | 0.999176 |
| ENSMUSG00000029420 | Rimbp2 | protein_coding | -0.3094967 | 3.871528 | 0.2198666 | 0.6453489 | 0.999176 |
| ENSMUSG00000066456 | Hmgn3 | protein_coding | -0.1812025 | 7.04536 | 0.2196188 | 0.6455339 | 0.999176 |
| ENSMUSG00000032042 | Srpr | protein_coding | 0.1736098 | 5.738268 | 0.2194701 | 0.6456449 | 0.999176 |
| ENSMUSG00000040035 | Disp2 | protein_coding | 0.245915 | 7.018399 | 0.2198369 | 0.6459899 | 0.999176 |
| ENSMUSG00000033061 | Resp18 | protein_coding | 0.1534792 | 9.264314 | 0.2189586 | 0.6460273 | 0.999176 |
| ENSMUSG00000002319 | Ipo4 | protein_coding | 0.2021271 | 6.218564 | 0.2185391 | 0.6463413 | 0.999176 |
| ENSMUSG00000033285 | Wdr3 | protein_coding | 0.6218655 | 2.434929 | 0.2184954 | 0.646374 | 0.999176 |
| ENSMUSG00000047067 | Dusp28 | protein_coding | 0.2543895 | 3.811465 | 0.2184711 | 0.6463923 | 0.999176 |
| ENSMUSG00000039018 | Mtg1 | protein_coding | 0.1776628 | 6.316392 | 0.2184579 | 0.6464021 | 0.999176 |
| ENSMUSG00000075590 | Nrbp2 | protein_coding | -0.3102496 | 4.648333 | 0.2183997 | 0.6464559 | 0.999176 |
| ENSMUSG00000031835 | Mbtps1 | protein_coding | 0.1975722 | 5.428852 | 0.218374 | 0.6464649 | 0.999176 |
| ENSMUSG00000037855 | Zfp365 | protein_coding | -0.1378814 | 7.027594 | 0.218328 | 0.6464995 | 0.999176 |
| ENSMUSG00000038437 | Mllt6 | protein_coding | 0.3444881 | 3.657008 | 0.2181598 | 0.6466255 | 0.999176 |
| ENSMUSG00000041236 | Vps41 | protein_coding | 0.1741393 | 7.767959 | 0.2178933 | 0.6468254 | 0.999176 |
| ENSMUSG00000023106 | Denr | protein_coding | 0.1408453 | 7.385994 | 0.2178807 | 0.6468348 | 0.999176 |
| ENSMUSG00000014418 | Hps5 | protein_coding | 0.4057213 | 3.081212 | 0.2176741 | 0.6469899 | 0.999176 |
| ENSMUSG00000032127 | Vps11 | protein_coding | -0.2341633 | 6.488605 | 0.218054 | 0.6470563 | 0.999176 |
| ENSMUSG00000059040 | Eno1b | protein_coding | -0.9070181 | 2.216287 | 0.2179442 | 0.6471802 | 0.999176 |
| ENSMUSG00000053819 | Camk2d | protein_coding | -0.1442763 | 7.235924 | 0.2174203 | 0.6471805 | 0.999176 |
| ENSMUSG00000058740 | Kcnt1 | protein_coding | 0.2911634 | 3.781216 | 0.2173846 | 0.6472073 | 0.999176 |
| ENSMUSG00000025588 | Nat1 | protein_coding | -0.5777034 | 2.428411 | 0.217343 | 0.6472386 | 0.999176 |
| ENSMUSG00000011148 | Adssl1 | protein_coding | -0.2116452 | 4.719642 | 0.2172346 | 0.6473201 | 0.999176 |
| ENSMUSG00000066643 | Wdr35 | protein_coding | 0.2198995 | 5.363478 | 0.2167497 | 0.6476848 | 0.999176 |
| ENSMUSG00000010175 | Prox1 | protein_coding | 0.3190986 | 4.41398 | 0.2166604 | 0.6477622 | 0.999176 |
| ENSMUSG00000024359 | Hspa9 | protein_coding | -0.1650023 | 6.97433 | 0.2165469 | 0.6478376 | 0.999176 |
| ENSMUSG00000000682 | Cd52 | protein_coding | -0.5400159 | 2.638471 | 0.2164162 | 0.647936 | 0.999176 |
| ENSMUSG00000033186 | Mzt1 | protein_coding | 0.1406559 | 7.298393 | 0.2161389 | 0.6481451 | 0.999176 |
| ENSMUSG00000027784 | Ppm1l | protein_coding | -0.1991895 | 5.153257 | 0.2160345 | 0.6482238 | 0.999176 |
| ENSMUSG00000027185 | Nat10 | protein_coding | 0.295554 | 3.557745 | 0.2160001 | 0.6482497 | 0.999176 |
| ENSMUSG00000079003 | Samd1 | protein_coding | 0.4164598 | 3.283322 | 0.2159109 | 0.648317 | 0.999176 |
| ENSMUSG00000025199 | Chuk | protein_coding | -0.4598016 | 3.326487 | 0.2156267 | 0.6485416 | 0.999176 |
| ENSMUSG00000029388 | Eif2b1 | protein_coding | 0.1366041 | 6.756538 | 0.2154285 | 0.6486813 | 0.999176 |
| ENSMUSG00000023912 | Slc25a27 | protein_coding | 0.1977309 | 4.990672 | 0.2153543 | 0.6487373 | 0.999176 |
| ENSMUSG00000047842 | Diras2 | protein_coding | -0.2240573 | 5.535113 | 0.2153018 | 0.648777 | 0.999176 |

| ensembl_gene_id | external_gene_name | gene_biotype | log2 (fold change) | log2 (counts per million) | F-value | p-value | False discovery rate |
|---|---|---|---|---|---|---|---|
| ENSMUSG00000022354 | Ndufb9 | protein_coding | 0.1811457 | 10.17307 | 0.2152486 | 0.6488173 | 0.999176 |
| ENSMUSG00000028373 | Astn2 | protein_coding | -0.2217536 | 5.382909 | 0.2152107 | 0.6488459 | 0.999176 |
| ENSMUSG00000022800 | Fyttd1 | protein_coding | 0.1642972 | 6.968083 | 0.2148318 | 0.6491325 | 0.999176 |
| ENSMUSG00000049122 | Frmd3 | protein_coding | 0.4311479 | 3.188286 | 0.2147581 | 0.6491883 | 0.999176 |
| ENSMUSG00000039100 | 6-Mar | protein_coding | -0.1770408 | 6.267039 | 0.2147258 | 0.6492128 | 0.999176 |
| ENSMUSG00000027281 | Slx4ip | protein_coding | 0.3020851 | 3.290613 | 0.2147094 | 0.6492252 | 0.999176 |
| ENSMUSG00000055932 | Fto | protein_coding | 0.2751745 | 8.114774 | 0.2159494 | 0.6492667 | 0.999176 |
| ENSMUSG00000050071 | Bex1 | protein_coding | 0.1415329 | 8.358076 | 0.2144864 | 0.649394 | 0.999176 |
| ENSMUSG00000027424 | Mgme1 | protein_coding | 0.2536525 | 3.799947 | 0.2144482 | 0.649423 | 0.999176 |
| ENSMUSG00000020515 | Cnot8 | protein_coding | 0.1760488 | 6.86437 | 0.2143776 | 0.6494765 | 0.999176 |
| ENSMUSG00000045464 | 2810002D19Rik | processed_transcript | 0.3376237 | 4.38772 | 0.2140717 | 0.6497309 | 0.999176 |
| ENSMUSG00000022265 | Ank | protein_coding | -0.1668284 | 6.57825 | 0.2139491 | 0.6498014 | 0.999176 |
| ENSMUSG00000045438 | Cox19 | protein_coding | 0.153211 | 7.156108 | 0.21393 | 0.6498159 | 0.999176 |
| ENSMUSG00000025173 | Wdr45b | protein_coding | 0.1479202 | 6.486144 | 0.2139086 | 0.6498322 | 0.999176 |
| ENSMUSG00000070436 | Serpinh1 | protein_coding | -0.3988972 | 3.617528 | 0.2137194 | 0.6499858 | 0.999176 |
| ENSMUSG00000052926 | Rnaseh2a | protein_coding | 0.1800531 | 5.930473 | 0.2135741 | 0.6500862 | 0.999176 |
| ENSMUSG00000025189 | Cnnm1 | protein_coding | 0.1628103 | 5.847749 | 0.2134079 | 0.6502124 | 0.999176 |
| ENSMUSG00000032754 | Slc8b1 | protein_coding | 0.4523474 | 2.605505 | 0.2131952 | 0.6503741 | 0.999176 |
| ENSMUSG00000024906 | Mus81 | protein_coding | 0.250043 | 4.973404 | 0.2130522 | 0.6504829 | 0.999176 |
| ENSMUSG00000042992 | Borcs5 | protein_coding | 0.2634868 | 5.352749 | 0.2127319 | 0.6507367 | 0.999176 |
| ENSMUSG00000020300 | Cpeb4 | protein_coding | -0.1955082 | 5.245711 | 0.212614 | 0.6508165 | 0.999176 |
| ENSMUSG00000024039 | Cbs | protein_coding | -0.2742131 | 4.681259 | 0.212589 | 0.6508355 | 0.999176 |
| ENSMUSG00000058093 | Zfp729b | protein_coding | 0.5234068 | 2.732909 | 0.2125123 | 0.6508939 | 0.999176 |
| ENSMUSG00000032456 | Nmnat3 | protein_coding | 0.4437535 | 2.80927 | 0.2124035 | 0.6509768 | 0.999176 |
| ENSMUSG00000096433 | Zfp994 | protein_coding | -0.5309995 | 2.366309 | 0.2121042 | 0.651205 | 0.999176 |
| ENSMUSG00000023921 | Mut | protein_coding | -0.2126039 | 5.101189 | 0.211885 | 0.6513723 | 0.999176 |
| ENSMUSG00000041736 | Tspo | protein_coding | -0.3233867 | 3.492472 | 0.2117795 | 0.6514528 | 0.999176 |
| ENSMUSG00000024308 | Tapbp | protein_coding | 0.2058095 | 4.960715 | 0.2115905 | 0.6515972 | 0.999176 |
| ENSMUSG00000022342 | Kcnv1 | protein_coding | 0.3693496 | 3.770553 | 0.2115865 | 0.6516102 | 0.999176 |
| ENSMUSG00000064115 | Cadm2 | protein_coding | -0.2128883 | 5.531232 | 0.2113242 | 0.6518007 | 0.999176 |
| ENSMUSG00000024647 | Cbln2 | protein_coding | -0.22795 | 4.755811 | 0.2113196 | 0.6518042 | 0.999176 |
| ENSMUSG00000041577 | Prelp | protein_coding | -0.5803673 | 2.359008 | 0.2112974 | 0.6518212 | 0.999176 |
| ENSMUSG00000040147 | Maob | protein_coding | -0.2838182 | 6.056154 | 0.212 | 0.6518908 | 0.999176 |
| ENSMUSG00000029723 | Tsc22d4 | protein_coding | 0.2514349 | 7.493527 | 0.2123049 | 0.6519038 | 0.999176 |
| ENSMUSG00000084289 | Gm6977 | processed_pseudoge | 0.5855797 | 3.235062 | 0.2116562 | 0.6520442 | 0.999176 |
| ENSMUSG00000003435 | Supt5 | protein_coding | 0.2567428 | 5.680562 | 0.2109407 | 0.6521302 | 0.999176 |
| ENSMUSG00000025364 | Pa2g4 | protein_coding | 0.1560992 | 7.354737 | 0.2106726 | 0.6522993 | 0.999176 |
| ENSMUSG00000040746 | Rnf167 | protein_coding | 0.1437275 | 7.602094 | 0.2106057 | 0.6523505 | 0.999176 |
| ENSMUSG00000046699 | Slitrk4 | protein_coding | -0.2324983 | 4.671662 | 0.2105963 | 0.6523578 | 0.999176 |
| ENSMUSG00000033439 | Trmt13 | protein_coding | -0.4195912 | 2.783158 | 0.2102358 | 0.6526341 | 0.999176 |
| ENSMUSG00000024816 | Frmd8 | protein_coding | 0.2639852 | 5.575738 | 0.2102155 | 0.6526961 | 0.999176 |
| ENSMUSG00000025510 | Cd151 | protein_coding | 0.3649299 | 3.943163 | 0.2101426 | 0.6527155 | 0.999176 |
| ENSMUSG00000031644 | Nek1 | protein_coding | -0.3717401 | 3.354983 | 0.2099239 | 0.6528734 | 0.999176 |
| ENSMUSG00000024268 | Celf4 | protein_coding | 0.190537 | 9.063432 | 0.2096535 | 0.653091 | 0.999176 |
| ENSMUSG00000030718 | Ppme1 | protein_coding | 0.2071343 | 7.347338 | 0.2098918 | 0.6531513 | 0.999176 |
| ENSMUSG00000037679 | Inf2 | protein_coding | -0.2365859 | 4.847573 | 0.2095221 | 0.653182 | 0.999176 |
| ENSMUSG00000052273 | Dnah3 | protein_coding | 0.4881552 | 2.341484 | 0.2094377 | 0.6532469 | 0.999176 |
| ENSMUSG00000025485 | Ric8a | protein_coding | 0.2020414 | 5.790533 | 0.209299 | 0.6533536 | 0.999176 |
| ENSMUSG00000022550 | Adck5 | protein_coding | 0.3168385 | 4.420545 | 0.2090189 | 0.6535789 | 0.999176 |
| ENSMUSG00000058064 | Gm10036 | protein_coding | -0.7749662 | 2.964324 | 0.2102262 | 0.6536054 | 0.999176 |
| ENSMUSG00000000171 | Sdhd | protein_coding | 0.1361755 | 9.071525 | 0.2087689 | 0.6537615 | 0.999176 |
| ENSMUSG00000026737 | Pip4k2a | protein_coding | 0.1685089 | 5.635831 | 0.2085413 | 0.6539369 | 0.999176 |
| ENSMUSG00000050668 | Gpatch11 | protein_coding | 0.1484354 | 6.136313 | 0.2085127 | 0.6539589 | 0.999176 |
| ENSMUSG00000024456 | Diaph1 | protein_coding | -0.2776428 | 4.603003 | 0.2084911 | 0.6539756 | 0.999176 |
| ENSMUSG00000021364 | Elovl2 | protein_coding | -0.6094631 | 2.667074 | 0.2084118 | 0.6540466 | 0.999176 |
| ENSMUSG00000073411 | H2-D1 | protein_coding | -0.3904938 | 4.646071 | 0.2091547 | 0.6540662 | 0.999176 |
| ENSMUSG00000020850 | Prpf8 | protein_coding | -0.15742 | 6.32823 | 0.2081919 | 0.6542063 | 0.999176 |
| ENSMUSG00000020053 | Igf1 | protein_coding | -0.6712273 | 2.511684 | 0.2081596 | 0.6542411 | 0.999176 |
| ENSMUSG00000048216 | Gpr85 | protein_coding | 0.182284 | 6.060611 | 0.2080746 | 0.6542968 | 0.999176 |
| ENSMUSG00000026688 | Mgst3 | protein_coding | -0.1316251 | 8.852649 | 0.2078795 | 0.6544474 | 0.999176 |
| ENSMUSG00000026749 | Nek6 | protein_coding | 0.2649774 | 4.33166 | 0.2078533 | 0.6544677 | 0.999176 |
| ENSMUSG00000057421 | Las1l | protein_coding | -0.1757957 | 5.820323 | 0.2078415 | 0.6544768 | 0.999176 |
| ENSMUSG00000020218 | Wif1 | protein_coding | 0.4453653 | 3.057694 | 0.207803 | 0.6545065 | 0.999176 |
| ENSMUSG00000032224 | Fam81a | protein_coding | -0.204144 | 8.374444 | 0.2080314 | 0.6545168 | 0.999176 |
| ENSMUSG00000031357 | Syap1 | protein_coding | -0.2986458 | 3.793192 | 0.2077163 | 0.6545735 | 0.999176 |
| ENSMUSG00000060288 | Ppih | protein_coding | 0.3406761 | 3.363588 | 0.2076609 | 0.6546163 | 0.999176 |
| ENSMUSG00000045822 | Zswim3 | protein_coding | -0.3194787 | 3.408236 | 0.2075329 | 0.6547152 | 0.999176 |
| ENSMUSG00000057716 | Tmem178b | protein_coding | -0.2098228 | 5.542624 | 0.2074903 | 0.6547481 | 0.999176 |
| ENSMUSG00000048486 | Fitm2 | protein_coding | -0.2503876 | 4.319731 | 0.2074395 | 0.6547875 | 0.999176 |
| ENSMUSG00000022131 | Gpr180 | protein_coding | -0.3106813 | 3.381276 | 0.2073193 | 0.6548804 | 0.999176 |
| ENSMUSG00000024109 | Nrxn1 | protein_coding | -0.1621653 | 8.156306 | 0.2072367 | 0.6549443 | 0.999176 |
| ENSMUSG00000040964 | Arhgef10l | protein_coding | -0.2876035 | 3.819511 | 0.2071314 | 0.6550258 | 0.999176 |
| ENSMUSG00000038916 | Soga3 | protein_coding | 0.221976 | 5.687259 | 0.2069141 | 0.655194 | 0.999176 |
| ENSMUSG00000039108 | Lsm14b | protein_coding | -0.2469905 | 4.354057 | 0.2066369 | 0.6554088 | 0.999176 |
| ENSMUSG00000031392 | Irak1 | protein_coding | -0.1916259 | 5.165498 | 0.2066227 | 0.6554198 | 0.999176 |
| ENSMUSG00000021773 | Comtd1 | protein_coding | 0.3820584 | 3.079362 | 0.2063379 | 0.6556406 | 0.999176 |
| ENSMUSG00000026603 | Smyd2 | protein_coding | -0.2921256 | 3.360309 | 0.2062862 | 0.6556807 | 0.999176 |
| ENSMUSG00000038759 | Nup205 | protein_coding | 0.3534367 | 3.095527 | 0.2062466 | 0.6557115 | 0.999176 |

| ensembl_gene_id | external_gene_name | gene_biotype | log2 (fold change) | log2 (counts per million) | F-value | p-value | False discovery rate |
|---|---|---|---|---|---|---|---|
| ENSMUSG00000021453 | Gadd45g | protein_coding | 0.231809 | 5.830834 | 0.2062284 | 0.6557354 | 0.999176 |
| ENSMUSG00000066196 | Spag8 | protein_coding | 0.3672053 | 3.189612 | 0.2062151 | 0.6557359 | 0.999176 |
| ENSMUSG00000036241 | Ube2r2 | protein_coding | -0.1390818 | 6.41431 | 0.2061144 | 0.655814 | 0.999176 |
| ENSMUSG00000048022 | Tmem229a | protein_coding | -0.2775482 | 4.868359 | 0.206107 | 0.6558296 | 0.999176 |
| ENSMUSG00000074505 | Fat3 | protein_coding | -0.2957152 | 4.060654 | 0.2059469 | 0.6559441 | 0.999176 |
| ENSMUSG00000032745 | Gpbp1 | protein_coding | -0.1750966 | 6.97776 | 0.2058499 | 0.6560195 | 0.999176 |
| ENSMUSG00000005893 | Nr2c2 | protein_coding | -0.2330137 | 4.94102 | 0.2057394 | 0.6561053 | 0.999176 |
| ENSMUSG00000036752 | Tubb4b | protein_coding | 0.1652944 | 7.66865 | 0.2056009 | 0.6562129 | 0.999176 |
| ENSMUSG00000052794 | 1700030K09Rik | protein_coding | 0.2727012 | 4.025646 | 0.205528 | 0.6562697 | 0.999176 |
| ENSMUSG00000085829 | Gm4285 | antisense_RNA | 0.5654417 | 2.39625 | 0.2054775 | 0.6563089 | 0.999176 |
| ENSMUSG00000054871 | Tmem158 | protein_coding | -0.3084792 | 4.795903 | 0.205493 | 0.6563325 | 0.999176 |
| ENSMUSG00000043572 | Pars2 | protein_coding | -0.2414561 | 3.973159 | 0.2053234 | 0.6564287 | 0.999176 |
| ENSMUSG00000032051 | Fdx1 | protein_coding | -0.3039609 | 3.433172 | 0.2050163 | 0.6566679 | 0.999176 |
| ENSMUSG00000049493 | Pls1 | protein_coding | -0.2821296 | 4.987409 | 0.2048727 | 0.6567894 | 0.999176 |
| ENSMUSG00000032336 | Nptn | protein_coding | 0.1364512 | 8.112727 | 0.2047537 | 0.6568724 | 0.999176 |
| ENSMUSG00000054792 | Klhl18 | protein_coding | 0.2250986 | 4.768381 | 0.2046729 | 0.6569354 | 0.999176 |
| ENSMUSG00000059013 | Sh2d3c | protein_coding | 0.2897992 | 4.324343 | 0.2046505 | 0.6569528 | 0.999176 |
| ENSMUSG00000086191 | Zfp652os | antisense_RNA | 0.3732799 | 2.93421 | 0.2046098 | 0.6569846 | 0.999176 |
| ENSMUSG00000038708 | Golga4 | protein_coding | -0.1826725 | 5.144765 | 0.204256 | 0.6572606 | 0.999176 |
| ENSMUSG00000051579 | Tceal8 | protein_coding | -0.1447136 | 7.46592 | 0.203986 | 0.6574714 | 0.999176 |
| ENSMUSG00000049728 | Zfp668 | protein_coding | -0.3046382 | 3.787904 | 0.2037862 | 0.6576276 | 0.999176 |
| ENSMUSG00000026031 | Cflar | protein_coding | -0.316374 | 3.42663 | 0.2037737 | 0.6576373 | 0.999176 |
| ENSMUSG00000022451 | Twf1 | protein_coding | -0.458905 | 2.529936 | 0.2036804 | 0.6577103 | 0.999176 |
| ENSMUSG00000028636 | Ppcs | protein_coding | 0.3526552 | 4.117835 | 0.2034575 | 0.65792 | 0.999176 |
| ENSMUSG00000036792 | Mbd5 | protein_coding | -0.2026396 | 4.797091 | 0.2033666 | 0.6579557 | 0.999176 |
| ENSMUSG00000055762 | Eef1d | protein_coding | 0.1454386 | 6.868658 | 0.203301 | 0.6580071 | 0.999176 |
| ENSMUSG00000055491 | Pprc1 | protein_coding | -0.3649072 | 2.946577 | 0.2031506 | 0.6581248 | 0.999176 |
| ENSMUSG00000036989 | Trim3 | protein_coding | 0.1627046 | 6.159598 | 0.2030728 | 0.6581857 | 0.999176 |
| ENSMUSG00000024853 | Sf3b2 | protein_coding | 0.1500711 | 9.008018 | 0.2027837 | 0.6584123 | 0.999176 |
| ENSMUSG00000035473 | Galm | protein_coding | -0.6432487 | 2.245935 | 0.2027779 | 0.6584168 | 0.999176 |
| ENSMUSG00000024044 | Epb41l3 | protein_coding | -0.169613 | 7.654101 | 0.2026593 | 0.6585098 | 0.999176 |
| ENSMUSG00000037977 | 6430571L13Rik | protein_coding | -0.3746006 | 3.498635 | 0.2025732 | 0.6585774 | 0.999176 |
| ENSMUSG00000035248 | Zcchc6 | protein_coding | -0.338946 | 3.076259 | 0.2025009 | 0.6586341 | 0.999176 |
| ENSMUSG00000052302 | Tbc1d30 | protein_coding | -0.1818328 | 5.205387 | 0.2024791 | 0.6586512 | 0.999176 |
| ENSMUSG00000042462 | Dctpp1 | protein_coding | 0.2138381 | 5.573849 | 0.2024252 | 0.6586935 | 0.999176 |
| ENSMUSG00000022263 | Trio | protein_coding | -0.1944304 | 5.44701 | 0.2023832 | 0.6587264 | 0.999176 |
| ENSMUSG00000043251 | Exoc3l | protein_coding | 0.6004447 | 2.066138 | 0.2021573 | 0.6589038 | 0.999176 |
| ENSMUSG00000000826 | Dnajc5 | protein_coding | -0.1396445 | 8.254323 | 0.2018553 | 0.6591411 | 0.999176 |
| ENSMUSG00000040843 | Tiprl | protein_coding | -0.1497215 | 7.344424 | 0.2018253 | 0.6591647 | 0.999176 |
| ENSMUSG00000026134 | Prim2 | protein_coding | -0.2235052 | 4.413398 | 0.2016072 | 0.6593362 | 0.999176 |
| ENSMUSG00000023861 | Mpc1 | protein_coding | -0.1417055 | 6.224462 | 0.2014526 | 0.6594579 | 0.999176 |
| ENSMUSG00000043531 | Sorcs1 | protein_coding | 0.2693129 | 3.709531 | 0.2013189 | 0.6595631 | 0.999176 |
| ENSMUSG00000037533 | Rapgef6 | protein_coding | 0.2173955 | 5.157853 | 0.2011938 | 0.6596617 | 0.999176 |
| ENSMUSG00000029918 | Mrps33 | protein_coding | 0.1720982 | 6.96422 | 0.2010854 | 0.6597471 | 0.999176 |
| ENSMUSG00000032897 | Nfyc | protein_coding | 0.2361333 | 4.118223 | 0.2010034 | 0.6598117 | 0.999176 |
| ENSMUSG00000024750 | Zfand5 | protein_coding | -0.1607877 | 6.01802 | 0.2007897 | 0.6599802 | 0.999176 |
| ENSMUSG00000029310 | Nudt9 | protein_coding | 0.1902085 | 5.614689 | 0.2007669 | 0.6599982 | 0.999176 |
| ENSMUSG00000063972 | Nr6a1 | protein_coding | -0.3792637 | 3.21914 | 0.2007127 | 0.6600409 | 0.999176 |
| ENSMUSG00000046111 | Cep295 | protein_coding | -0.3732802 | 2.984215 | 0.2006806 | 0.6600662 | 0.999176 |
| ENSMUSG00000021504 | B4galt7 | protein_coding | 0.2352598 | 4.868777 | 0.2005029 | 0.6602065 | 0.999176 |
| ENSMUSG00000022899 | Slc15a2 | protein_coding | -0.354418 | 3.89264 | 0.2005112 | 0.6602096 | 0.999176 |
| ENSMUSG00000004462 | Tbccd1 | protein_coding | -0.2821937 | 4.312858 | 0.2004689 | 0.6602333 | 0.999176 |
| ENSMUSG00000006476 | Nsmf | protein_coding | -0.2334674 | 6.35506 | 0.2003543 | 0.6606985 | 0.999176 |
| ENSMUSG00000068966 | Zbtb34 | protein_coding | 0.3940822 | 2.946998 | 0.1998125 | 0.660752 | 0.999176 |
| ENSMUSG00000014959 | Gorasp2 | protein_coding | 0.1386251 | 6.525025 | 0.1997122 | 0.6608313 | 0.999176 |
| ENSMUSG00000021302 | Ggps1 | protein_coding | 0.1358367 | 6.849266 | 0.1996824 | 0.6608549 | 0.999176 |
| ENSMUSG00000021096 | Ppm1a | protein_coding | -0.1195585 | 7.718985 | 0.1993795 | 0.6610947 | 0.999176 |
| ENSMUSG00000030638 | Sh3gl3 | protein_coding | 0.2310306 | 5.200237 | 0.1992153 | 0.6612248 | 0.999176 |
| ENSMUSG00000038582 | Pptc7 | protein_coding | 0.2517702 | 4.485504 | 0.1989568 | 0.6614298 | 0.999176 |
| ENSMUSG00000060981 | Hist1h4h | protein_coding | 0.535502 | 2.313796 | 0.198902 | 0.6614732 | 0.999176 |
| ENSMUSG00000028889 | Yrdc | protein_coding | -0.2811099 | 4.415055 | 0.1988286 | 0.6615314 | 0.999176 |
| ENSMUSG00000022103 | Gfra2 | protein_coding | 0.228285 | 5.329227 | 0.1988138 | 0.6615432 | 0.999176 |
| ENSMUSG00000066705 | Fxyd6 | protein_coding | 0.1389917 | 9.078928 | 0.1986988 | 0.6616345 | 0.999176 |
| ENSMUSG00000024664 | Fads3 | protein_coding | 0.358213 | 4.010319 | 0.1987277 | 0.6616465 | 0.999176 |
| ENSMUSG00000019877 | Serinc1 | protein_coding | 0.1243121 | 9.253247 | 0.1986834 | 0.6616466 | 0.999176 |
| ENSMUSG00000030556 | Lrrc28 | protein_coding | 0.1562882 | 5.877709 | 0.1986623 | 0.6616634 | 0.999176 |
| ENSMUSG00000041961 | Znrf3 | protein_coding | -0.2324223 | 5.062821 | 0.1985263 | 0.6617713 | 0.999176 |
| ENSMUSG00000048388 | Fam171b | protein_coding | -0.1391691 | 7.116365 | 0.1985091 | 0.661785 | 0.999176 |
| ENSMUSG00000030166 | Rad52 | protein_coding | 0.2146163 | 4.5797 | 0.1984684 | 0.6618174 | 0.999176 |
| ENSMUSG00000027706 | Sec62 | protein_coding | -0.1306981 | 7.909474 | 0.1983935 | 0.6618768 | 0.999176 |
| ENSMUSG00000020918 | Kat2a | protein_coding | 0.1812942 | 5.123733 | 0.1981669 | 0.6620569 | 0.999176 |
| ENSMUSG00000033124 | Atg9a | protein_coding | 0.4849954 | 3.263364 | 0.1983784 | 0.6620837 | 0.999176 |
| ENSMUSG00000023965 | Fbxl17 | protein_coding | 0.1802631 | 5.385806 | 0.1979321 | 0.6622437 | 0.999176 |
| ENSMUSG00000044340 | Phlpp1 | protein_coding | -0.1907855 | 5.578807 | 0.1978419 | 0.6623154 | 0.999176 |
| ENSMUSG00000027525 | Phactr3 | protein_coding | 0.1412263 | 6.714599 | 0.1973782 | 0.6626847 | 0.999176 |
| ENSMUSG00000038351 | Sgsm2 | protein_coding | -0.2184008 | 4.882328 | 0.1973752 | 0.6626871 | 0.999176 |
| ENSMUSG00000031808 | Slc27a1 | protein_coding | -0.2355193 | 5.635909 | 0.197362 | 0.6627072 | 0.999176 |
| ENSMUSG00000020271 | Fbxw11 | protein_coding | -0.1690422 | 6.205915 | 0.1973432 | 0.6627126 | 0.999176 |

| ensembl_gene_id | external_gene_name | gene_biotype | log2 (fold change) | log2 (counts per million) | F-value | p-value | False discovery rate |
|---|---|---|---|---|---|---|---|
| ENSMUSG00000041729 | Coro2b | protein_coding | -0.1767953 | 5.548722 | 0.1973248 | 0.6627272 | 0.999176 |
| ENSMUSG00000006134 | Crkl | protein_coding | -0.2392706 | 3.826481 | 0.1972541 | 0.6627835 | 0.999176 |
| ENSMUSG00000023391 | Dlx2 | protein_coding | 0.4271695 | 2.893844 | 0.197249 | 0.6627876 | 0.999176 |
| ENSMUSG00000038806 | Sde2 | protein_coding | 0.1429787 | 6.141264 | 0.1970672 | 0.6629326 | 0.999176 |
| ENSMUSG00000025147 | Mob2 | protein_coding | 0.1335556 | 6.615878 | 0.1969137 | 0.6630551 | 0.999176 |
| ENSMUSG00000025034 | Trim8 | protein_coding | -0.1823265 | 5.425124 | 0.1968889 | 0.6630749 | 0.999176 |
| ENSMUSG00000002413 | Braf | protein_coding | -0.1717472 | 6.16161 | 0.1968476 | 0.6631078 | 0.999176 |
| ENSMUSG00000029135 | Fosl2 | protein_coding | -0.3808659 | 3.461338 | 0.1966068 | 0.6633001 | 0.999176 |
| ENSMUSG00000042535 | Gtpbp1 | protein_coding | -0.3004568 | 3.944895 | 0.1965355 | 0.663357 | 0.999176 |
| ENSMUSG00000038828 | Tmem214 | protein_coding | -0.2331358 | 4.820999 | 0.1962387 | 0.6635943 | 0.999176 |
| ENSMUSG00000053925 | Gm9929 | processed_transcript | 0.3925349 | 2.965742 | 0.1962246 | 0.6636056 | 0.999176 |
| ENSMUSG00000022792 | Yars2 | protein_coding | 0.2567701 | 3.83504 | 0.1962244 | 0.6636058 | 0.999176 |
| ENSMUSG00000040028 | Elavl1 | protein_coding | -0.1739178 | 5.450016 | 0.1959695 | 0.6638097 | 0.999176 |
| ENSMUSG00000054256 | Msi1 | protein_coding | -0.3022113 | 4.022097 | 0.195805 | 0.6639413 | 0.999176 |
| ENSMUSG00000052727 | Map1b | protein_coding | -0.3060236 | 9.563345 | 0.1978409 | 0.6639524 | 0.999176 |
| ENSMUSG00000034269 | Setd5 | protein_coding | -0.2245717 | 4.314935 | 0.1957842 | 0.663958 | 0.999176 |
| ENSMUSG00000047123 | Ticam1 | protein_coding | -0.4937371 | 2.958279 | 0.1955133 | 0.6641846 | 0.999176 |
| ENSMUSG00000031796 | Cfap20 | protein_coding | 0.2092407 | 6.06102 | 0.1953158 | 0.6643429 | 0.999176 |
| ENSMUSG00000028496 | Mllt3 | protein_coding | -0.1841949 | 5.335619 | 0.1949186 | 0.6646521 | 0.999176 |
| ENSMUSG00000025613 | Cct8 | protein_coding | -0.1761414 | 8.639491 | 0.1948416 | 0.6647234 | 0.999176 |
| ENSMUSG00000022536 | Glyr1 | protein_coding | 0.1641082 | 7.648162 | 0.1947811 | 0.6647625 | 0.999176 |
| ENSMUSG00000025207 | Sema4g | protein_coding | -0.4206363 | 3.063198 | 0.194665 | 0.6648558 | 0.999176 |
| ENSMUSG00000022031 | Elp3 | protein_coding | 0.2007485 | 6.756834 | 0.1946736 | 0.6648999 | 0.999176 |
| ENSMUSG00000006526 | Tmem110 | protein_coding | -0.3822828 | 3.326176 | 0.1946033 | 0.6649054 | 0.999176 |
| ENSMUSG00000033342 | Plppr5 | protein_coding | 0.2147972 | 4.639042 | 0.1945844 | 0.6649205 | 0.999176 |
| ENSMUSG00000021149 | Gtpbp4 | protein_coding | -0.1519478 | 6.593417 | 0.1945547 | 0.6649445 | 0.999176 |
| ENSMUSG00000025878 | Uimc1 | protein_coding | -0.222478 | 4.440031 | 0.1944929 | 0.6649941 | 0.999176 |
| ENSMUSG00000039958 | Etfbkmt | protein_coding | -0.2743124 | 4.189351 | 0.1944575 | 0.6650226 | 0.999176 |
| ENSMUSG00000090841 | Myl6 | protein_coding | 0.1375589 | 7.729845 | 0.1943551 | 0.6651049 | 0.999176 |
| ENSMUSG00000015120 | Ube2i | protein_coding | 0.2931599 | 3.638053 | 0.1943156 | 0.6651367 | 0.999176 |
| ENSMUSG00000030630 | Fah | protein_coding | 0.1868988 | 5.376408 | 0.1942664 | 0.6651763 | 0.999176 |
| ENSMUSG00000019777 | Hdac2 | protein_coding | -0.2286802 | 5.824625 | 0.1942332 | 0.6652125 | 0.999176 |
| ENSMUSG00000021592 | Arsk | protein_coding | -0.3967436 | 3.16525 | 0.1942198 | 0.6652138 | 0.999176 |
| ENSMUSG00000023460 | Rab12 | protein_coding | 0.169051 | 5.846957 | 0.1941803 | 0.6652455 | 0.999176 |
| ENSMUSG00000033577 | Myo6 | protein_coding | -0.1949795 | 5.153348 | 0.1936711 | 0.6656557 | 0.999176 |
| ENSMUSG00000010517 | Faf1 | protein_coding | 0.2249847 | 5.091484 | 0.1929302 | 0.6662535 | 0.999176 |
| ENSMUSG00000022855 | Senp2 | protein_coding | -0.1604153 | 6.140583 | 0.1928556 | 0.6663138 | 0.999176 |
| ENSMUSG00000021130 | Galnt16 | protein_coding | -0.1527027 | 6.390037 | 0.1928403 | 0.6663262 | 0.999176 |
| ENSMUSG00000046791 | Riox1 | protein_coding | -0.3992546 | 2.938539 | 0.1928274 | 0.6663366 | 0.999176 |
| ENSMUSG00000020224 | Llph | protein_coding | 0.2021998 | 4.444462 | 0.1927943 | 0.6663634 | 0.999176 |
| ENSMUSG00000061360 | Phf5a | protein_coding | 0.1652602 | 6.590205 | 0.1925416 | 0.6665677 | 0.999176 |
| ENSMUSG00000056770 | Setd3 | protein_coding | -0.1404206 | 6.514828 | 0.1925291 | 0.6665778 | 0.999176 |
| ENSMUSG00000031930 | Wwp2 | protein_coding | -0.2361902 | 5.60701 | 0.1924064 | 0.6666865 | 0.999176 |
| ENSMUSG00000042660 | Wdr55 | protein_coding | 0.2005249 | 4.827519 | 0.1923685 | 0.6667077 | 0.999176 |
| ENSMUSG00000089911 | Mfsd14a | protein_coding | 0.265467 | 4.328371 | 0.1921953 | 0.666848 | 0.999176 |
| ENSMUSG00000032126 | Hmbs | protein_coding | -0.3093994 | 3.707314 | 0.1919705 | 0.66703 | 0.999176 |
| ENSMUSG00000039163 | Cmc1 | protein_coding | 0.2403405 | 4.134332 | 0.1919208 | 0.6670703 | 0.999176 |
| ENSMUSG00000032985 | 5730522E02Rik | protein_coding | 0.4125949 | 3.229163 | 0.1917764 | 0.6671874 | 0.999176 |
| ENSMUSG00000097290 | 1300002E11Rik | lincRNA | 0.2225782 | 4.732783 | 0.1916256 | 0.6673097 | 0.999176 |
| ENSMUSG00000020912 | Krt12 | protein_coding | -0.468237 | 2.457497 | 0.1915174 | 0.6673975 | 0.999176 |
| ENSMUSG00000060519 | Tor3a | protein_coding | -0.4837678 | 2.469767 | 0.1914919 | 0.6674181 | 0.999176 |
| ENSMUSG00000034889 | Cactin | protein_coding | 0.5092601 | 2.7493 | 0.19147 | 0.6674359 | 0.999176 |
| ENSMUSG00000053460 | Ggcx | protein_coding | 0.4569465 | 3.244691 | 0.1914093 | 0.6674946 | 0.999176 |
| ENSMUSG00000074582 | Arfgef2 | protein_coding | 0.3045579 | 4.268294 | 0.1913337 | 0.667556 | 0.999176 |
| ENSMUSG00000029629 | Phf14 | protein_coding | -0.2245521 | 4.886093 | 0.1913119 | 0.6675642 | 0.999176 |
| ENSMUSG00000033220 | Rac2 | protein_coding | -0.4677572 | 2.777567 | 0.1912773 | 0.6675924 | 0.999176 |
| ENSMUSG00000038406 | Scaf1 | protein_coding | 0.2384391 | 4.706982 | 0.1910338 | 0.6677902 | 0.999176 |
| ENSMUSG00000035478 | Mbd3 | protein_coding | -0.2271501 | 5.413148 | 0.1909981 | 0.6678192 | 0.999176 |
| ENSMUSG00000054640 | Slc8a1 | protein_coding | -0.347233 | 3.279024 | 0.1907685 | 0.6680058 | 0.999176 |
| ENSMUSG00000048109 | Rbm15 | protein_coding | 0.3189004 | 5.00797 | 0.1910914 | 0.6681083 | 0.999176 |
| ENSMUSG00000039096 | Rsad1 | protein_coding | 0.39581 | 3.080677 | 0.1905811 | 0.6681584 | 0.999176 |
| ENSMUSG00000048029 | Eno4 | protein_coding | -0.3541407 | 3.208913 | 0.1905349 | 0.6681959 | 0.999176 |
| ENSMUSG00000028759 | Hp1bp3 | protein_coding | 0.152275 | 7.591972 | 0.1904388 | 0.6682742 | 0.999176 |
| ENSMUSG00000002052 | Supt6 | protein_coding | -0.1645029 | 6.119792 | 0.1902213 | 0.6684514 | 0.999176 |
| ENSMUSG00000024603 | Dctn4 | protein_coding | 0.2100757 | 7.585918 | 0.1906649 | 0.6684546 | 0.999176 |
| ENSMUSG00000054766 | Set | protein_coding | -0.1618414 | 8.405428 | 0.1901604 | 0.6685009 | 0.999176 |
| ENSMUSG00000073609 | D2hgdh | protein_coding | -0.3769911 | 3.018182 | 0.1900517 | 0.6685896 | 0.999176 |
| ENSMUSG00000028798 | Eif3i | protein_coding | 0.1272388 | 9.126721 | 0.1899917 | 0.6686385 | 0.999176 |
| ENSMUSG00000060675 | Pla2g16 | protein_coding | -0.2100185 | 6.167451 | 0.1899778 | 0.6686592 | 0.999176 |
| ENSMUSG00000038007 | Acer2 | protein_coding | 0.2339457 | 4.297018 | 0.1899218 | 0.6686955 | 0.999176 |
| ENSMUSG00000007815 | Rhoa | protein_coding | 0.1906226 | 5.81972 | 0.189917 | 0.6686994 | 0.999176 |
| ENSMUSG00000037958 | Nsrp1 | protein_coding | -0.172553 | 6.11231 | 0.1894288 | 0.6690979 | 0.999176 |
| ENSMUSG00000001420 | Tmem79 | protein_coding | -0.4212433 | 2.657697 | 0.1891131 | 0.6693559 | 0.999176 |
| ENSMUSG00000036291 | Ap5m1 | protein_coding | -0.2820449 | 3.986148 | 0.1890986 | 0.6693677 | 0.999176 |
| ENSMUSG00000041915 | Ammecr1l | protein_coding | 0.3271216 | 3.297962 | 0.1890897 | 0.669375 | 0.999176 |
| ENSMUSG00000029245 | Epha5 | protein_coding | -0.2119905 | 5.230723 | 0.1889399 | 0.6694976 | 0.999176 |
| ENSMUSG00000066900 | Suds3 | protein_coding | 0.1666302 | 5.49123 | 0.1887303 | 0.6696691 | 0.999176 |
| ENSMUSG00000072949 | Acot1 | protein_coding | -0.3883006 | 2.791619 | 0.1887253 | 0.6696732 | 0.999176 |

| ensembl_gene_id | external_gene_name | gene_biotype | log2 (fold change) | log2 (counts per million) | F-value | p-value | False discovery rate |
|---|---|---|---|---|---|---|---|
| ENSMUSG00000025893 | Kbtbd3 | protein_coding | 0.1951947 | 5.278128 | 0.1886595 | 0.669727 | 0.999176 |
| ENSMUSG00000032462 | Pik3cb | protein_coding | -0.4008664 | 3.05208 | 0.1886244 | 0.6697557 | 0.999176 |
| ENSMUSG00000021877 | Arf4 | protein_coding | 0.1537039 | 8.805973 | 0.1884892 | 0.6698665 | 0.999176 |
| ENSMUSG00000035152 | Ap2b1 | protein_coding | -0.1570481 | 6.570604 | 0.1884725 | 0.6698802 | 0.999176 |
| ENSMUSG00000085971 | Gm15411 | processed_transcript | 0.5270451 | 2.348571 | 0.1884189 | 0.6699241 | 0.999176 |
| ENSMUSG00000020859 | Spag9 | protein_coding | -0.1479659 | 6.745141 | 0.1882942 | 0.6700263 | 0.999176 |
| ENSMUSG00000079614 | Seh1l | protein_coding | -0.1601426 | 6.686796 | 0.1882646 | 0.6700506 | 0.999176 |
| ENSMUSG00000000308 | Ckmt1 | protein_coding | 0.1854773 | 8.958998 | 0.1882576 | 0.6700656 | 0.999176 |
| ENSMUSG00000042492 | Tbc1d10b | protein_coding | 0.1748924 | 5.264163 | 0.1882424 | 0.6700688 | 0.999176 |
| ENSMUSG00000036570 | Fxyd1 | protein_coding | -0.274704 | 6.278205 | 0.1889523 | 0.6702852 | 0.999176 |
| ENSMUSG00000022791 | Tnk2 | protein_coding | -0.2121935 | 5.128254 | 0.1878103 | 0.6704233 | 0.999176 |
| ENSMUSG00000006932 | Ctnnb1 | protein_coding | -0.3461699 | 3.323877 | 0.1877317 | 0.6704878 | 0.999176 |
| ENSMUSG00000069227 | Gprin1 | protein_coding | -0.2549585 | 3.988466 | 0.1876839 | 0.6705271 | 0.999176 |
| ENSMUSG00000066571 | 4931406P16Rik | protein_coding | 0.2481821 | 3.750216 | 0.1875354 | 0.6706491 | 0.999176 |
| ENSMUSG00000031917 | Nip7 | protein_coding | -0.158618 | 6.92253 | 0.1874791 | 0.6706953 | 0.999176 |
| ENSMUSG00000024908 | Ppp6r3 | protein_coding | -0.2356522 | 4.469791 | 0.1873296 | 0.6708183 | 0.999176 |
| ENSMUSG00000031167 | Rbm3 | protein_coding | 0.2177046 | 4.087314 | 0.1872202 | 0.6709083 | 0.999176 |
| ENSMUSG00000059482 | 2610301B20Rik | protein_coding | -0.1379743 | 6.793203 | 0.1872143 | 0.6709131 | 0.999176 |
| ENSMUSG00000037418 | Best1 | protein_coding | -0.5063289 | 2.896601 | 0.1869465 | 0.6711428 | 0.999176 |
| ENSMUSG00000029669 | Tspan12 | protein_coding | -0.1910631 | 6.187879 | 0.1869298 | 0.6711472 | 0.999176 |
| ENSMUSG00000074607 | Tox2 | protein_coding | 0.2825131 | 5.99781 | 0.1877326 | 0.6712439 | 0.999176 |
| ENSMUSG00000050390 | C77080 | protein_coding | -0.2596125 | 4.3369 | 0.1866404 | 0.6713856 | 0.999176 |
| ENSMUSG00000047242 | Taf9b | protein_coding | -0.1676284 | 6.562037 | 0.1865948 | 0.6714232 | 0.999176 |
| ENSMUSG00000074345 | Tnfaip8l3 | protein_coding | -0.3673387 | 3.454021 | 0.1864973 | 0.6715036 | 0.999176 |
| ENSMUSG00000062931 | Zfp938 | protein_coding | -0.1350055 | 6.3488 | 0.1864932 | 0.6715069 | 0.999176 |
| ENSMUSG00000005262 | Ufd1 | protein_coding | 0.1677082 | 7.572505 | 0.1864738 | 0.6715322 | 0.999176 |
| ENSMUSG00000001424 | Snd1 | protein_coding | 0.1698225 | 5.712354 | 0.1864144 | 0.6715719 | 0.999176 |
| ENSMUSG00000032911 | Cspg4 | protein_coding | 0.3549558 | 3.823511 | 0.1860138 | 0.6719118 | 0.999176 |
| ENSMUSG00000040767 | Snrnp25 | protein_coding | 0.1414286 | 6.893031 | 0.185992 | 0.6719205 | 0.999176 |
| ENSMUSG00000046909 | Tefm | protein_coding | -0.1924304 | 5.180163 | 0.1859863 | 0.6719252 | 0.999176 |
| ENSMUSG00000030894 | Tpp1 | protein_coding | -0.203464 | 6.126024 | 0.1858807 | 0.6720217 | 0.999176 |
| ENSMUSG00000037013 | Ss18 | protein_coding | -0.2715754 | 4.372799 | 0.1857802 | 0.6720955 | 0.999176 |
| ENSMUSG00000033658 | Ddx19b | protein_coding | 0.3432958 | 2.951968 | 0.1855719 | 0.6722677 | 0.999176 |
| ENSMUSG00000032112 | Trappc4 | protein_coding | 0.1177051 | 8.584497 | 0.1854956 | 0.6723308 | 0.999176 |
| ENSMUSG00000037355 | Uvssa | protein_coding | -0.4176516 | 2.934808 | 0.1854198 | 0.6723935 | 0.999176 |
| ENSMUSG00000030669 | Calca | protein_coding | -0.6443722 | 3.089485 | 0.1863745 | 0.6723965 | 0.999176 |
| ENSMUSG00000000876 | Pxmp4 | protein_coding | 0.1771546 | 5.150634 | 0.1853354 | 0.6724633 | 0.999176 |
| ENSMUSG00000009207 | Lnpk | protein_coding | -0.1763712 | 5.53237 | 0.1853255 | 0.6724715 | 0.999176 |
| ENSMUSG00000021007 | Spata7 | protein_coding | 0.2010162 | 5.025591 | 0.1851721 | 0.6725985 | 0.999176 |
| ENSMUSG00000002455 | Prpf6 | protein_coding | -0.1949373 | 6.141371 | 0.1851443 | 0.6726308 | 0.999176 |
| ENSMUSG00000009418 | Nav1 | protein_coding | 0.1759314 | 5.623536 | 0.1850068 | 0.6727354 | 0.999176 |
| ENSMUSG00000022574 | Naprt | protein_coding | 0.3864645 | 3.163064 | 0.1849342 | 0.6727956 | 0.999176 |
| ENSMUSG00000028932 | Psmc2 | protein_coding | -0.1333323 | 8.499906 | 0.1848697 | 0.672849 | 0.999176 |
| ENSMUSG00000024165 | Hn1l | protein_coding | 0.2558757 | 4.286298 | 0.1848425 | 0.6728716 | 0.999176 |
| ENSMUSG00000022212 | Cpne6 | protein_coding | -0.2185693 | 7.179004 | 0.1854934 | 0.672895 | 0.999176 |
| ENSMUSG00000022179 | 4931414P19Rik | protein_coding | -0.4558515 | 2.714418 | 0.1847978 | 0.6729086 | 0.999176 |
| ENSMUSG00000097785 | B230217O12Rik | lincRNA | 0.211695 | 5.528692 | 0.1847731 | 0.6729291 | 0.999176 |
| ENSMUSG00000026926 | Pmpca | protein_coding | -0.2065955 | 6.246171 | 0.1847693 | 0.6729415 | 0.999176 |
| ENSMUSG00000057522 | Spop | protein_coding | -0.1543888 | 8.272857 | 0.1846251 | 0.6730518 | 0.999176 |
| ENSMUSG00000032349 | Elovl5 | protein_coding | 0.1570197 | 5.954479 | 0.1846034 | 0.6730699 | 0.999176 |
| ENSMUSG00000032551 | 1110059G10Rik | protein_coding | -0.1755964 | 5.850312 | 0.1845667 | 0.6731003 | 0.999176 |
| ENSMUSG00000069581 | Tspear | protein_coding | 0.4464792 | 2.714406 | 0.1845017 | 0.6731542 | 0.999176 |
| ENSMUSG00000022403 | St13 | protein_coding | -0.1554472 | 9.582884 | 0.1844326 | 0.6732116 | 0.999176 |
| ENSMUSG00000021890 | Eaf1 | protein_coding | 0.3560003 | 3.042197 | 0.1843408 | 0.6732878 | 0.999176 |
| ENSMUSG00000029422 | Rsrc2 | protein_coding | -0.1408969 | 7.293159 | 0.1842917 | 0.6733285 | 0.999176 |
| ENSMUSG00000074918 | Inafm2 | protein_coding | 0.1643525 | 7.812709 | 0.1842037 | 0.6734016 | 0.999176 |
| ENSMUSG00000031617 | Tmem184c | protein_coding | -0.2402071 | 4.827263 | 0.1839455 | 0.6736162 | 0.999176 |
| ENSMUSG00000063931 | Pepd | protein_coding | -0.1842027 | 6.764993 | 0.1839074 | 0.673657 | 0.999176 |
| ENSMUSG00000029726 | Mepce | protein_coding | -0.1805068 | 5.396212 | 0.1838288 | 0.6737132 | 0.999176 |
| ENSMUSG00000022040 | Ephx2 | protein_coding | -0.1781428 | 4.957748 | 0.1836756 | 0.6738406 | 0.999176 |
| ENSMUSG00000002910 | Arrdc2 | protein_coding | -0.4222987 | 2.68252 | 0.1836355 | 0.673874 | 0.999176 |
| ENSMUSG00000006998 | Psmd2 | protein_coding | 0.1682475 | 7.721677 | 0.1835863 | 0.6739242 | 0.999176 |
| ENSMUSG00000024410 | 3110002H16Rik | protein_coding | -0.192358 | 5.443925 | 0.1835568 | 0.6739395 | 0.999176 |
| ENSMUSG00000034192 | Lsm3 | protein_coding | -0.2106971 | 6.577786 | 0.1838711 | 0.6739595 | 0.999176 |
| ENSMUSG00000000194 | Gpr107 | protein_coding | -0.2397988 | 4.487791 | 0.1833865 | 0.6740813 | 0.999176 |
| ENSMUSG00000055675 | Kbtbd11 | protein_coding | -0.2647336 | 5.211613 | 0.183387 | 0.6741143 | 0.999176 |
| ENSMUSG00000051373 | Plpp7 | protein_coding | -0.2032085 | 4.706038 | 0.1832415 | 0.6742021 | 0.999176 |
| ENSMUSG00000028842 | Ago3 | protein_coding | 0.2839734 | 4.372174 | 0.1830496 | 0.674362 | 0.999176 |
| ENSMUSG00000055320 | Tead1 | protein_coding | -0.249372 | 3.646695 | 0.1828387 | 0.6745379 | 0.999176 |
| ENSMUSG00000029009 | Mthfr | protein_coding | -0.4272287 | 3.157893 | 0.1828237 | 0.6745596 | 0.999176 |
| ENSMUSG00000026172 | Bcs1l | protein_coding | -0.1637742 | 6.012627 | 0.1825685 | 0.6747635 | 0.999176 |
| ENSMUSG00000022895 | Ets2 | protein_coding | 0.2427348 | 5.444192 | 0.1825736 | 0.6747684 | 0.999176 |
| ENSMUSG00000039671 | Zmynd8 | protein_coding | 0.1338322 | 6.581414 | 0.1824941 | 0.6748256 | 0.999176 |
| ENSMUSG00000021266 | Wars | protein_coding | 0.2558415 | 7.014787 | 0.1835215 | 0.674862 | 0.999176 |
| ENSMUSG00000052331 | Ankrd44 | protein_coding | -0.4920549 | 2.45703 | 0.1822525 | 0.6750274 | 0.999176 |
| ENSMUSG00000022698 | Naa50 | protein_coding | -0.1439912 | 5.95297 | 0.1821964 | 0.6750743 | 0.999176 |
| ENSMUSG00000046152 | Fut10 | protein_coding | -0.3719307 | 2.851395 | 0.1821057 | 0.6751502 | 0.999176 |
| ENSMUSG00000017264 | Exosc10 | protein_coding | 0.2142693 | 5.261205 | 0.1819991 | 0.6752393 | 0.999176 |

| ensembl_gene_id | external_gene_name | gene_biotype | log2 (fold change) | log2 (counts per million) | F-value | p-value | False discovery rate |
|---|---|---|---|---|---|---|---|
| ENSMUSG00000021692 | Dimt1 | protein_coding | -0.2330737 | 3.968309 | 0.1819142 | 0.6753103 | 0.999176 |
| ENSMUSG00000050890 | Pdik1l | protein_coding | -0.2366183 | 4.280959 | 0.1818377 | 0.6753744 | 0.999176 |
| ENSMUSG00000039263 | Npepl1 | protein_coding | -0.7564682 | 2.256624 | 0.1812081 | 0.6760754 | 0.999176 |
| ENSMUSG00000003452 | Bicd1 | protein_coding | 0.1655771 | 5.506908 | 0.1806967 | 0.6763312 | 0.999176 |
| ENSMUSG00000097055 | Gm4419 | lincRNA | 0.2938347 | 3.336072 | 0.1806083 | 0.6764055 | 0.999176 |
| ENSMUSG00000039533 | Mmd2 | protein_coding | -0.1776801 | 7.96871 | 0.1804556 | 0.676543 | 0.999176 |
| ENSMUSG00000036282 | Naa30 | protein_coding | 0.2553803 | 4.880532 | 0.1802041 | 0.6767545 | 0.999176 |
| ENSMUSG00000074358 | Ccdc61 | protein_coding | 0.278529 | 3.424313 | 0.1801933 | 0.6767546 | 0.999176 |
| ENSMUSG00000025381 | Cnpy2 | protein_coding | 0.1266269 | 7.272331 | 0.1800904 | 0.6768411 | 0.999176 |
| ENSMUSG00000090247 | Bloc1s1 | protein_coding | 0.1882991 | 5.050698 | 0.1800478 | 0.676877 | 0.999176 |
| ENSMUSG00000028560 | Usp1 | protein_coding | -0.3471091 | 3.026843 | 0.1800344 | 0.6768883 | 0.999176 |
| ENSMUSG00000026839 | Upp2 | protein_coding | 0.2490492 | 4.290427 | 0.1799587 | 0.676952 | 0.999176 |
| ENSMUSG00000039199 | Zdhhc1 | protein_coding | 0.3963638 | 2.754887 | 0.1798556 | 0.6770389 | 0.999176 |
| ENSMUSG00000066880 | Zfp617 | protein_coding | 0.2207241 | 3.954047 | 0.1798295 | 0.6770609 | 0.999176 |
| ENSMUSG00000021418 | Rpp40 | protein_coding | 0.2583457 | 3.772606 | 0.1797797 | 0.6771029 | 0.999176 |
| ENSMUSG00000050074 | Spink8 | protein_coding | 0.3802911 | 3.449042 | 0.1795936 | 0.6772688 | 0.999176 |
| ENSMUSG00000033543 | Gtf2a2 | protein_coding | -0.13626 | 7.381251 | 0.1795409 | 0.6773042 | 0.999176 |
| ENSMUSG00000029580 | Actb | protein_coding | 0.2308351 | 12.7409 | 0.1799238 | 0.6773127 | 0.999176 |
| ENSMUSG00000062963 | Ufc1 | protein_coding | 0.1276353 | 8.43534 | 0.1794579 | 0.6773742 | 0.999176 |
| ENSMUSG00000009731 | Kcnd1 | protein_coding | 0.6213786 | 2.365476 | 0.1792542 | 0.6775551 | 0.999176 |
| ENSMUSG00000040446 | Rprd1a | protein_coding | 0.2687753 | 3.754154 | 0.1792369 | 0.6775607 | 0.999176 |
| ENSMUSG00000026433 | Rab29 | protein_coding | -0.4049536 | 3.270457 | 0.1791928 | 0.677607 | 0.999176 |
| ENSMUSG00000063801 | Ap3s2 | protein_coding | -0.1343508 | 6.881999 | 0.1790556 | 0.6777138 | 0.999176 |
| ENSMUSG00000015217 | Hmgb3 | protein_coding | -0.2444809 | 3.751086 | 0.1790027 | 0.6777585 | 0.999176 |
| ENSMUSG00000038622 | Med30 | protein_coding | 0.2891788 | 3.868645 | 0.1787727 | 0.6779529 | 0.999176 |
| ENSMUSG00000004637 | Wwox | protein_coding | 0.1967294 | 6.345005 | 0.1787368 | 0.6779923 | 0.999176 |
| ENSMUSG00000060739 | Nsa2 | protein_coding | 0.2576751 | 3.599786 | 0.1787213 | 0.6779963 | 0.999176 |
| ENSMUSG00000055024 | Ep300 | protein_coding | 0.2958443 | 3.593335 | 0.1787185 | 0.6779987 | 0.999176 |
| ENSMUSG00000026511 | Srp9 | protein_coding | 0.1222642 | 7.039766 | 0.1786674 | 0.6780419 | 0.999176 |
| ENSMUSG00000022257 | Laptm4b | protein_coding | -0.1576376 | 6.147959 | 0.1783569 | 0.6783047 | 0.999176 |
| ENSMUSG00000060427 | Zfp868 | protein_coding | -0.2814681 | 4.786684 | 0.1783526 | 0.6783173 | 0.999176 |
| ENSMUSG00000043702 | Pde12 | protein_coding | 0.4858763 | 2.914669 | 0.1783156 | 0.6783487 | 0.999176 |
| ENSMUSG00000084403 | Rps15a-ps8 | processed_pseudoge | -1.038554 | 1.126235 | 0.178288 | 0.678363 | 0.999176 |
| ENSMUSG00000026051 | 1500015O10Rik | protein_coding | 0.4380382 | 3.884914 | 0.1785994 | 0.6784036 | 0.999176 |
| ENSMUSG00000003184 | Irf3 | protein_coding | -0.3344764 | 3.25301 | 0.1780879 | 0.6785325 | 0.999176 |
| ENSMUSG00000032178 | Ilf3 | protein_coding | 0.2359233 | 7.480326 | 0.1787825 | 0.6787179 | 0.999176 |
| ENSMUSG00000028407 | Toporsos | protein_coding | 0.272798 | 3.603595 | 0.1778005 | 0.6787762 | 0.999176 |
| ENSMUSG00000046753 | Ccdc66 | protein_coding | -0.3388989 | 3.093185 | 0.1777924 | 0.6787831 | 0.999176 |
| ENSMUSG00000052533 | Nup188 | protein_coding | -0.468869 | 2.716645 | 0.1777725 | 0.6788 | 0.999176 |
| ENSMUSG00000042425 | Frmpd3 | protein_coding | -0.4193838 | 2.928075 | 0.1777581 | 0.6788122 | 0.999176 |
| ENSMUSG00000022313 | Utp23 | protein_coding | -0.2463688 | 4.466096 | 0.1776999 | 0.6788616 | 0.999176 |
| ENSMUSG00000001774 | Chordc1 | protein_coding | 0.1260731 | 6.496847 | 0.1775496 | 0.6789891 | 0.999176 |
| ENSMUSG00000074892 | B3galt5 | protein_coding | -0.1996336 | 4.952349 | 0.1774217 | 0.6790977 | 0.999176 |
| ENSMUSG00000022253 | Nadk2 | protein_coding | 0.2117805 | 4.544091 | 0.1773981 | 0.6791178 | 0.999176 |
| ENSMUSG00000068762 | Gstm6 | protein_coding | 0.1503427 | 6.187683 | 0.1773172 | 0.6791865 | 0.999176 |
| ENSMUSG00000021413 | Prpf4b | protein_coding | -0.1833044 | 5.7847 | 0.1772618 | 0.6792336 | 0.999176 |
| ENSMUSG00000047213 | Ythdf3 | protein_coding | 0.2243116 | 4.409674 | 0.1771033 | 0.6793684 | 0.999176 |
| ENSMUSG00000059554 | Ccdc28a | protein_coding | 0.1466849 | 6.973734 | 0.1770731 | 0.679394 | 0.999176 |
| ENSMUSG00000025962 | Fastkd2 | protein_coding | 0.20247 | 5.107571 | 0.1770655 | 0.6794005 | 0.999176 |
| ENSMUSG00000051890 | Klhdc1 | protein_coding | 0.2126309 | 4.471963 | 0.1767998 | 0.6796265 | 0.999176 |
| ENSMUSG00000028901 | Gmeb1 | protein_coding | 0.2701506 | 4.08369 | 0.1767811 | 0.6796424 | 0.999176 |
| ENSMUSG00000003929 | Zfp81 | protein_coding | -0.3275292 | 3.661235 | 0.176196 | 0.6801409 | 0.999176 |
| ENSMUSG00000020435 | Osbp2 | protein_coding | 0.3018673 | 3.854302 | 0.1759978 | 0.68031 | 0.999176 |
| ENSMUSG00000029328 | Hnrnpdl | protein_coding | -0.1264716 | 6.699063 | 0.1759278 | 0.6803698 | 0.999176 |
| ENSMUSG00000020828 | Pld2 | protein_coding | -0.2823547 | 3.411783 | 0.1757018 | 0.6805628 | 0.999176 |
| ENSMUSG00000031618 | Nr3c2 | protein_coding | -0.2670337 | 4.117893 | 0.1756806 | 0.6805808 | 0.999176 |
| ENSMUSG00000027350 | Chgb | protein_coding | 0.1319698 | 7.54377 | 0.1756803 | 0.6805811 | 0.999176 |
| ENSMUSG00000062232 | Rapgef2 | protein_coding | -0.1911636 | 5.343581 | 0.1756733 | 0.6805871 | 0.999176 |
| ENSMUSG00000024269 | Tpgs2 | protein_coding | 0.1299515 | 6.61225 | 0.1754977 | 0.6807372 | 0.999176 |
| ENSMUSG00000024436 | Mrps18b | protein_coding | -0.203594 | 4.956158 | 0.1753466 | 0.6808664 | 0.999176 |
| ENSMUSG00000032598 | Nckipsd | protein_coding | -0.1683252 | 5.747789 | 0.1751704 | 0.6810171 | 0.999176 |
| ENSMUSG00000075324 | Fign | protein_coding | -0.3254083 | 4.297662 | 0.1746615 | 0.6816211 | 0.999176 |
| ENSMUSG00000035397 | Klf16 | protein_coding | 0.1912436 | 5.288355 | 0.1744017 | 0.6816759 | 0.999176 |
| ENSMUSG00000035878 | Hykk | protein_coding | -0.6525636 | 2.69386 | 0.1746773 | 0.6817865 | 0.999176 |
| ENSMUSG00000020086 | H2afy2 | protein_coding | 0.4654524 | 2.575478 | 0.1741762 | 0.6818694 | 0.999176 |
| ENSMUSG00000063873 | Slc24a3 | protein_coding | -0.1821847 | 6.203921 | 0.1741541 | 0.6818884 | 0.999176 |
| ENSMUSG00000034544 | Rsrc1 | protein_coding | -0.1499597 | 6.519232 | 0.174067 | 0.6819631 | 0.999176 |
| ENSMUSG00000029415 | Sdad1 | protein_coding | -0.2737158 | 4.236614 | 0.1740483 | 0.6819792 | 0.999176 |
| ENSMUSG00000014075 | Tctex1d2 | protein_coding | 0.1331313 | 6.552979 | 0.173947 | 0.6820663 | 0.999176 |
| ENSMUSG00000014232 | Cluap1 | protein_coding | -0.1797398 | 4.900014 | 0.1737107 | 0.6822694 | 0.999176 |
| ENSMUSG00000041130 | Zfp598 | protein_coding | 0.3740608 | 3.231571 | 0.1734264 | 0.682514 | 0.999176 |
| ENSMUSG00000109554 | A230057D06Rik | lincRNA | -0.5274689 | 2.696357 | 0.1733509 | 0.682588 | 0.999176 |
| ENSMUSG00000032306 | Mpi | protein_coding | 0.1408472 | 7.010687 | 0.1732819 | 0.6826385 | 0.999176 |
| ENSMUSG00000099696 | 2900052N01Rik | lincRNA | -0.1624266 | 5.699291 | 0.1732608 | 0.6826567 | 0.999176 |
| ENSMUSG00000030846 | Tial1 | protein_coding | 0.1563726 | 6.111303 | 0.1729644 | 0.682912 | 0.999176 |
| ENSMUSG00000026999 | Nup35 | protein_coding | 0.2110837 | 4.376576 | 0.1728813 | 0.6829837 | 0.999176 |
| ENSMUSG00000062376 | Borcs7 | protein_coding | 0.1211963 | 7.115717 | 0.1728758 | 0.6829885 | 0.999176 |
| ENSMUSG00000097717 | 1700066J03Rik | lincRNA | 0.4039539 | 3.003968 | 0.1727208 | 0.6831223 | 0.999176 |

| ensembl_gene_id | external_gene_name | gene_biotype | log2 (fold change) | log2 (counts per million) | F-value | p-value | False discovery rate |
|---|---|---|---|---|---|---|---|
| ENSMUSG00000027560 | Dok5 | protein_coding | 0.1721902 | 5.700823 | 0.1726208 | 0.6832085 | 0.999176 |
| ENSMUSG00000030393 | Zik1 | protein_coding | -0.3002745 | 3.691285 | 0.1725473 | 0.683272 | 0.999176 |
| ENSMUSG00000052921 | Arhgef15 | protein_coding | 0.1635384 | 6.334669 | 0.1724713 | 0.6833376 | 0.999176 |
| ENSMUSG00000037287 | Tbcel | protein_coding | -0.2115231 | 4.496772 | 0.1720289 | 0.68372 | 0.999176 |
| ENSMUSG00000013275 | Slc41a1 | protein_coding | 0.2117783 | 4.441277 | 0.1719965 | 0.6837481 | 0.999176 |
| ENSMUSG00000022191 | Drosha | protein_coding | -0.1549966 | 6.414153 | 0.1719942 | 0.68375 | 0.999176 |
| ENSMUSG00000039913 | Pak7 | protein_coding | -0.1927389 | 4.788729 | 0.1719696 | 0.6837713 | 0.999176 |
| ENSMUSG00000047187 | Rab2a | protein_coding | -0.117103 | 8.097975 | 0.1718194 | 0.6839013 | 0.999176 |
| ENSMUSG00000019832 | Rab32 | protein_coding | -0.7535191 | 1.917971 | 0.1717038 | 0.6840102 | 0.999176 |
| ENSMUSG00000064368 | mt-Nd6 | protein_coding | 0.2144599 | 8.052756 | 0.1722637 | 0.6840566 | 0.999176 |
| ENSMUSG00000020335 | Zfp354b | protein_coding | -0.3121517 | 3.462588 | 0.1716251 | 0.6840695 | 0.999176 |
| ENSMUSG00000022639 | Dubr | lincRNA | -0.3180712 | 3.784583 | 0.1715162 | 0.6841639 | 0.999176 |
| ENSMUSG00000058672 | Tubb2a | protein_coding | 0.156093 | 9.43981 | 0.1714254 | 0.6842426 | 0.999176 |
| ENSMUSG00000039917 | Rhbdd2 | protein_coding | -0.3505436 | 6.52721 | 0.1744139 | 0.6842432 | 0.999176 |
| ENSMUSG00000019837 | Gtf3c6 | protein_coding | -0.1431298 | 6.84451 | 0.171393 | 0.6842707 | 0.999176 |
| ENSMUSG00000024613 | Tcof1 | protein_coding | 0.2258673 | 4.449652 | 0.1713901 | 0.6842732 | 0.999176 |
| ENSMUSG00000037740 | Mrps26 | protein_coding | 0.1817499 | 4.850607 | 0.1713184 | 0.6843353 | 0.999176 |
| ENSMUSG00000035984 | Nme5 | protein_coding | 0.1538648 | 6.20005 | 0.1711851 | 0.684451 | 0.999176 |
| ENSMUSG00000006575 | Rundc3a | protein_coding | 0.1551496 | 6.00834 | 0.1710883 | 0.684535 | 0.999176 |
| ENSMUSG00000098112 | Bin2 | protein_coding | -0.6008731 | 2.315691 | 0.1708571 | 0.6847357 | 0.999176 |
| ENSMUSG00000023236 | Scg5 | protein_coding | -0.1357889 | 9.418818 | 0.1708527 | 0.6847394 | 0.999176 |
| ENSMUSG00000043448 | Gjc2 | protein_coding | -0.5176994 | 2.313015 | 0.170803 | 0.6847827 | 0.999176 |
| ENSMUSG00000049411 | Tmem241 | protein_coding | -0.3161546 | 4.270905 | 0.1707604 | 0.684862 | 0.999176 |
| ENSMUSG00000040174 | Alkbh3 | protein_coding | 0.3567985 | 2.893481 | 0.1704275 | 0.6851091 | 0.999176 |
| ENSMUSG00000047412 | Zbtb44 | protein_coding | -0.3114201 | 3.245285 | 0.1703737 | 0.6851559 | 0.999176 |
| ENSMUSG00000032386 | Trip4 | protein_coding | -0.1793812 | 4.950811 | 0.1703543 | 0.6851727 | 0.999176 |
| ENSMUSG00000000355 | Mcts1 | protein_coding | 0.1299317 | 7.797103 | 0.1703492 | 0.6851771 | 0.999176 |
| ENSMUSG00000058301 | Upf1 | protein_coding | 0.2134765 | 4.106843 | 0.1702745 | 0.6852421 | 0.999176 |
| ENSMUSG00000005505 | Kbtbd4 | protein_coding | 0.2893158 | 3.95557 | 0.1702021 | 0.6853052 | 0.999176 |
| ENSMUSG00000058297 | Spock2 | protein_coding | -0.122602 | 7.721449 | 0.1700772 | 0.6854139 | 0.999176 |
| ENSMUSG00000025372 | Baiap2 | protein_coding | -0.1473619 | 7.550733 | 0.1700716 | 0.6854188 | 0.999176 |
| ENSMUSG00000026520 | Pycr2 | protein_coding | -0.2577279 | 5.07548 | 0.1700451 | 0.6854506 | 0.999176 |
| ENSMUSG00000037017 | Zscan21 | protein_coding | 0.2360328 | 4.9613 | 0.1699661 | 0.6855106 | 0.999176 |
| ENSMUSG00000063052 | Lrrc40 | protein_coding | -0.1996215 | 5.17975 | 0.1699509 | 0.685524 | 0.999176 |
| ENSMUSG00000041415 | Dicer1 | protein_coding | 0.2463786 | 4.065796 | 0.1698911 | 0.685576 | 0.999176 |
| ENSMUSG00000026858 | Miga2 | protein_coding | 0.1809653 | 5.634868 | 0.1698411 | 0.6856196 | 0.999176 |
| ENSMUSG00000031750 | Il34 | protein_coding | 0.1770246 | 5.729123 | 0.1696953 | 0.6857467 | 0.999176 |
| ENSMUSG00000040213 | Kyat3 | protein_coding | -0.3261659 | 3.134047 | 0.1696114 | 0.6858199 | 0.999176 |
| ENSMUSG00000025157 | Zdhhc16 | protein_coding | -0.243689 | 3.534691 | 0.1695088 | 0.6859094 | 0.999176 |
| ENSMUSG00000025350 | Rdh5 | protein_coding | -0.4406995 | 3.235797 | 0.1694133 | 0.6860247 | 0.999176 |
| ENSMUSG00000050891 | Tatdn1 | protein_coding | -0.163875 | 5.328397 | 0.1693526 | 0.6860457 | 0.999176 |
| ENSMUSG00000006307 | Kmt2b | protein_coding | -0.395397 | 3.089273 | 0.1691234 | 0.6862459 | 0.999176 |
| ENSMUSG00000090386 | Mir99ahg | processed_transcript | -0.2617639 | 4.000903 | 0.1690142 | 0.6863413 | 0.999176 |
| ENSMUSG00000028782 | Adgrb2 | protein_coding | -0.329099 | 3.140052 | 0.1690047 | 0.6863496 | 0.999176 |
| ENSMUSG00000074576 | Mocs3 | protein_coding | 0.5210979 | 2.22411 | 0.1689863 | 0.6863657 | 0.999176 |
| ENSMUSG00000041544 | Disp3 | protein_coding | -0.3531504 | 3.256826 | 0.1689671 | 0.6863825 | 0.999176 |
| ENSMUSG00000001127 | Araf | protein_coding | -0.1193981 | 9.716487 | 0.1688264 | 0.6865055 | 0.999176 |
| ENSMUSG00000023826 | Park2 | protein_coding | 0.2073902 | 4.804067 | 0.1687911 | 0.6865364 | 0.999176 |
| ENSMUSG00000027273 | Snap25 | protein_coding | 0.1709623 | 12.17103 | 0.1687107 | 0.6866068 | 0.999176 |
| ENSMUSG00000074575 | Kcng1 | protein_coding | -0.3017138 | 3.10265 | 0.1686164 | 0.6866892 | 0.999176 |
| ENSMUSG00000074457 | S100a16 | protein_coding | -0.1541085 | 8.358843 | 0.1683887 | 0.6868886 | 0.999176 |
| ENSMUSG00000061474 | Mrps36 | protein_coding | 0.1997978 | 4.525805 | 0.16832 | 0.6869489 | 0.999176 |
| ENSMUSG00000019969 | Psen1 | protein_coding | 0.166436 | 5.820382 | 0.1683184 | 0.6869503 | 0.999176 |
| ENSMUSG00000033106 | Slc7a6os | protein_coding | -0.3258416 | 3.552646 | 0.1682989 | 0.6869673 | 0.999176 |
| ENSMUSG00000060376 | Bckdha | protein_coding | 0.3167587 | 3.830388 | 0.1679971 | 0.6872407 | 0.999176 |
| ENSMUSG00000027804 | Ppid | protein_coding | 0.1177426 | 6.992843 | 0.167887 | 0.6873285 | 0.999176 |
| ENSMUSG00000001419 | Mef2d | protein_coding | -0.1509633 | 5.675215 | 0.1677362 | 0.6874609 | 0.999176 |
| ENSMUSG00000022048 | Dpysl2 | protein_coding | -0.2260796 | 11.92064 | 0.1680049 | 0.6875646 | 0.999176 |
| ENSMUSG00000031012 | Cask | protein_coding | -0.2096149 | 7.302078 | 0.1680459 | 0.6877213 | 0.999176 |
| ENSMUSG00000022329 | Stk3 | protein_coding | 0.2674315 | 4.112229 | 0.1674262 | 0.6877332 | 0.999176 |
| ENSMUSG00000021645 | Smn1 | protein_coding | -0.1634773 | 5.462921 | 0.1672378 | 0.6878989 | 0.999176 |
| ENSMUSG00000021275 | Tecpr2 | protein_coding | -0.2222258 | 4.203638 | 0.1671819 | 0.687948 | 0.999176 |
| ENSMUSG00000069835 | Sat2 | protein_coding | 0.161696 | 5.743221 | 0.166798 | 0.6882859 | 0.999176 |
| ENSMUSG00000050822 | Slc29a4 | protein_coding | 0.2889505 | 3.279898 | 0.1667816 | 0.6883004 | 0.999176 |
| ENSMUSG00000056305 | Usp39 | protein_coding | 0.2002262 | 4.208385 | 0.1667677 | 0.6883126 | 0.999176 |
| ENSMUSG00000078813 | Leng1 | protein_coding | 0.4060493 | 2.817331 | 0.1667515 | 0.688327 | 0.999176 |
| ENSMUSG00000041426 | Hibch | protein_coding | -0.1850393 | 5.204313 | 0.1666372 | 0.6884277 | 0.999176 |
| ENSMUSG00000026856 | Dolpp1 | protein_coding | 0.2004478 | 4.741128 | 0.1665424 | 0.6885112 | 0.999176 |
| ENSMUSG00000055567 | Unc80 | protein_coding | -0.1492865 | 6.175451 | 0.1665371 | 0.6885158 | 0.999176 |
| ENSMUSG00000001376 | Vps50 | protein_coding | 0.2069787 | 5.090107 | 0.1664253 | 0.6886145 | 0.999176 |
| ENSMUSG00000052298 | Cdc42se2 | protein_coding | 0.1613387 | 7.224808 | 0.166281 | 0.6887505 | 0.999176 |
| ENSMUSG00000046201 | Scaf8 | protein_coding | -0.2274197 | 4.444221 | 0.166262 | 0.6887585 | 0.999176 |
| ENSMUSG00000072437 | Nanos1 | protein_coding | -0.2110011 | 4.553238 | 0.1662478 | 0.6887711 | 0.999176 |
| ENSMUSG00000074781 | Ube2n | protein_coding | 0.1406189 | 6.876333 | 0.1661592 | 0.6888493 | 0.999176 |
| ENSMUSG00000047044 | D030056L22Rik | protein_coding | -0.2422401 | 3.731754 | 0.1661496 | 0.6888578 | 0.999176 |
| ENSMUSG00000027612 | Mmp24 | protein_coding | 0.3392127 | 3.063833 | 0.1661217 | 0.6888824 | 0.999176 |
| ENSMUSG00000009575 | Cbx5 | protein_coding | -0.1150772 | 8.581273 | 0.16598 | 0.6890075 | 0.999176 |
| ENSMUSG00000043061 | Tmem18 | protein_coding | -0.1991715 | 4.516317 | 0.16596 | 0.6890252 | 0.999176 |

| ensembl_gene_id | external_gene_name | gene_biotype | log2 (fold change) | log2 (counts per million) | F-value | p-value | False discovery rate |
|---|---|---|---|---|---|---|---|
| ENSMUSG00000037242 | Clic4 | protein_coding | 0.2054892 | 5.378088 | 0.1658826 | 0.6890936 | 0.999176 |
| ENSMUSG00000043496 | Tril | protein_coding | -0.3698454 | 4.010766 | 0.1662633 | 0.689095 | 0.999176 |
| ENSMUSG00000027522 | Stx16 | protein_coding | -0.2323488 | 5.559586 | 0.1658873 | 0.6891211 | 0.999176 |
| ENSMUSG00000022562 | Oplah | protein_coding | 0.4157818 | 2.77596 | 0.165823 | 0.6891463 | 0.999176 |
| ENSMUSG00000023952 | Gtpbp2 | protein_coding | 0.2353429 | 4.165418 | 0.1657368 | 0.6892225 | 0.999176 |
| ENSMUSG00000021103 | Mnat1 | protein_coding | 0.1382428 | 6.011421 | 0.1656184 | 0.6893272 | 0.999176 |
| ENSMUSG00000031993 | Snx19 | protein_coding | 0.1904889 | 5.03412 | 0.1656102 | 0.6893344 | 0.999176 |
| ENSMUSG00000037503 | Fam168b | protein_coding | 0.1330894 | 6.432526 | 0.1656022 | 0.6893415 | 0.999176 |
| ENSMUSG00000028976 | Slc2a5 | protein_coding | -0.6489344 | 2.785864 | 0.1661168 | 0.6894155 | 0.999176 |
| ENSMUSG00000032184 | Lysmd2 | protein_coding | -0.1291484 | 6.451402 | 0.1654929 | 0.6894382 | 0.999176 |
| ENSMUSG00000001138 | Cnnm3 | protein_coding | -0.2513638 | 4.023338 | 0.1651013 | 0.689785 | 0.999176 |
| ENSMUSG00000052040 | Klf13 | protein_coding | 0.1668909 | 8.3196 | 0.1649098 | 0.6899634 | 0.999176 |
| ENSMUSG00000057236 | Rbbp4 | protein_coding | 0.1606233 | 7.282346 | 0.1647811 | 0.6900776 | 0.999176 |
| ENSMUSG00000007987 | Ift22 | protein_coding | 0.1152529 | 8.388343 | 0.1647524 | 0.6900944 | 0.999176 |
| ENSMUSG00000041353 | Tmem29 | protein_coding | 0.1300646 | 6.878533 | 0.1647445 | 0.6901014 | 0.999176 |
| ENSMUSG00000021661 | Ankra2 | protein_coding | 0.1633442 | 5.411819 | 0.1647379 | 0.6901073 | 0.999176 |
| ENSMUSG00000083282 | Ctsf | protein_coding | -0.2158115 | 5.385651 | 0.1646742 | 0.6901638 | 0.999176 |
| ENSMUSG00000029716 | Tfr2 | protein_coding | 0.2541551 | 4.001578 | 0.1644469 | 0.6903656 | 0.999176 |
| ENSMUSG00000039599 | Fam149b | protein_coding | 0.3034003 | 3.162414 | 0.1642114 | 0.6905749 | 0.999176 |
| ENSMUSG00000039128 | Cdc123 | protein_coding | 0.128873 | 8.754662 | 0.1638571 | 0.6908901 | 0.999176 |
| ENSMUSG00000033352 | Map2k4 | protein_coding | -0.1257064 | 7.402422 | 0.163773 | 0.690965 | 0.999176 |
| ENSMUSG00000054793 | Cadm4 | protein_coding | 0.2925516 | 4.443615 | 0.1636888 | 0.6910713 | 0.999176 |
| ENSMUSG00000043885 | Slc36a4 | protein_coding | 0.1930017 | 5.179743 | 0.1636438 | 0.69108 | 0.999176 |
| ENSMUSG00000031197 | Vbp1 | protein_coding | -0.1165892 | 7.867598 | 0.1636163 | 0.6911045 | 0.999176 |
| ENSMUSG00000097311 | Gm26871 | processed_transcript | 0.4772888 | 2.800489 | 0.1635435 | 0.6911693 | 0.999176 |
| ENSMUSG00000035653 | Lrfn5 | protein_coding | -0.1934206 | 5.506145 | 0.1634889 | 0.691218 | 0.999176 |
| ENSMUSG00000021178 | Psmc1 | protein_coding | -0.1171637 | 9.082673 | 0.163418 | 0.6912812 | 0.999176 |
| ENSMUSG00000032012 | Nectin1 | protein_coding | 0.2761669 | 4.533619 | 0.1633935 | 0.6913117 | 0.999176 |
| ENSMUSG00000021018 | Polr2h | protein_coding | -0.1384841 | 6.125717 | 0.1632406 | 0.6914394 | 0.999176 |
| ENSMUSG00000031176 | Dynlt3 | protein_coding | 0.1232726 | 8.581355 | 0.1631637 | 0.691508 | 0.999176 |
| ENSMUSG00000027285 | Haus2 | protein_coding | 0.1498135 | 6.04553 | 0.1629042 | 0.6917397 | 0.999176 |
| ENSMUSG00000056258 | Kcnq3 | protein_coding | 0.2200717 | 5.110849 | 0.1627486 | 0.6918787 | 0.999176 |
| ENSMUSG00000031536 | Polb | protein_coding | -0.1092639 | 7.048116 | 0.1627429 | 0.6918838 | 0.999176 |
| ENSMUSG00000021508 | Cxcl14 | protein_coding | -0.2905842 | 4.651295 | 0.1628053 | 0.6919828 | 0.999176 |
| ENSMUSG00000016918 | Sulf1 | protein_coding | -0.3550224 | 3.857128 | 0.162601 | 0.6921454 | 0.999176 |
| ENSMUSG00000054091 | 1810037I17Rik | protein_coding | -0.1515851 | 7.637371 | 0.1621847 | 0.6923832 | 0.999176 |
| ENSMUSG00000024759 | Atl3 | protein_coding | -0.3441222 | 3.070248 | 0.1620924 | 0.6924659 | 0.999176 |
| ENSMUSG00000025532 | Crcp | protein_coding | 0.1255249 | 7.105217 | 0.1620873 | 0.6924704 | 0.999176 |
| ENSMUSG00000004127 | Trmt10a | protein_coding | 0.2166196 | 4.116831 | 0.1620593 | 0.6924954 | 0.999176 |
| ENSMUSG00000030822 | Prr14 | protein_coding | 0.2078591 | 5.126155 | 0.1619783 | 0.6925681 | 0.999176 |
| ENSMUSG00000037709 | Fam13a | protein_coding | -0.2431644 | 3.768904 | 0.1617835 | 0.6927427 | 0.999176 |
| ENSMUSG00000040997 | Abhd4 | protein_coding | -0.1314851 | 6.797734 | 0.1616591 | 0.6928543 | 0.999176 |
| ENSMUSG00000028161 | Ppp3ca | protein_coding | 0.1149203 | 9.571254 | 0.1615886 | 0.6929175 | 0.999176 |
| ENSMUSG00000014245 | Pigl | protein_coding | 0.3677128 | 2.891857 | 0.1613324 | 0.6931476 | 0.999176 |
| ENSMUSG00000005936 | Kctd20 | protein_coding | -0.185772 | 5.24216 | 0.1612375 | 0.6932328 | 0.999176 |
| ENSMUSG00000038612 | Mcl1 | protein_coding | -0.2389661 | 4.875636 | 0.161235 | 0.6932351 | 0.999176 |
| ENSMUSG00000031161 | Hdac6 | protein_coding | 0.2273722 | 4.879868 | 0.1611684 | 0.6932949 | 0.999176 |
| ENSMUSG00000036104 | Rab3gap1 | protein_coding | -0.1654415 | 6.155083 | 0.1611645 | 0.6932985 | 0.999176 |
| ENSMUSG00000038718 | Pbx3 | protein_coding | 0.2029822 | 5.394241 | 0.1611431 | 0.6933177 | 0.999176 |
| ENSMUSG00000041319 | Thoc6 | protein_coding | 0.2936944 | 4.222496 | 0.1610659 | 0.6933956 | 0.999176 |
| ENSMUSG00000034341 | Wbp2 | protein_coding | 0.168756 | 10.24062 | 0.1610008 | 0.6934542 | 0.999176 |
| ENSMUSG00000015542 | Nat9 | protein_coding | 0.1693333 | 5.966384 | 0.1609572 | 0.6934848 | 0.999176 |
| ENSMUSG00000038780 | Smurf1 | protein_coding | 0.2187642 | 3.930852 | 0.1608258 | 0.693603 | 0.999176 |
| ENSMUSG00000027985 | Lef1 | protein_coding | -0.650338 | 3.205573 | 0.1623285 | 0.6936063 | 0.999176 |
| ENSMUSG00000036112 | Metap2 | protein_coding | 0.1356662 | 7.48961 | 0.160711 | 0.6937064 | 0.999176 |
| ENSMUSG00000021476 | Habp4 | protein_coding | 0.14102 | 6.948834 | 0.1606079 | 0.6937992 | 0.999176 |
| ENSMUSG00000057113 | Npm1 | protein_coding | 0.1192799 | 8.151726 | 0.1604467 | 0.6939445 | 0.999176 |
| ENSMUSG00000016940 | Kctd2 | protein_coding | -0.1909993 | 5.365807 | 0.1604201 | 0.6939684 | 0.999176 |
| ENSMUSG00000027108 | Ola1 | protein_coding | 0.1201402 | 7.789074 | 0.1603526 | 0.6940292 | 0.999176 |
| ENSMUSG00000086943 | 4732414G09Rik | lincRNA | 0.5685092 | 2.822619 | 0.1607023 | 0.6940454 | 0.999176 |
| ENSMUSG00000059146 | Ntrk3 | protein_coding | -0.1441517 | 6.11167 | 0.1601767 | 0.6941879 | 0.999176 |
| ENSMUSG00000021252 | Erg28 | protein_coding | -0.1289432 | 7.380018 | 0.1600229 | 0.6943267 | 0.999176 |
| ENSMUSG00000024242 | Map4k3 | protein_coding | -0.2022379 | 5.042346 | 0.1599778 | 0.6943673 | 0.999176 |
| ENSMUSG00000024384 | Iws1 | protein_coding | -0.2505435 | 3.589956 | 0.1598278 | 0.6945027 | 0.999176 |
| ENSMUSG00000042042 | Csgalnact2 | protein_coding | -0.4012765 | 2.890966 | 0.1598115 | 0.6945175 | 0.999176 |
| ENSMUSG00000032531 | Amotl2 | protein_coding | -0.388813 | 3.205997 | 0.1595521 | 0.6947604 | 0.999176 |
| ENSMUSG00000070476 | Fam217b | protein_coding | -0.1404336 | 5.854552 | 0.1593862 | 0.6949019 | 0.999176 |
| ENSMUSG00000024079 | Eif2ak2 | protein_coding | -0.7958708 | 1.681538 | 0.159301 | 0.6949875 | 0.999176 |
| ENSMUSG00000056486 | Chn1 | protein_coding | 0.1141269 | 7.212606 | 0.1590315 | 0.695223 | 0.999176 |
| ENSMUSG00000030270 | Cpne9 | protein_coding | 0.4374881 | 5.478852 | 0.1618445 | 0.6952256 | 0.999176 |
| ENSMUSG00000073415 | Gm10501 | antisense_RNA | -0.3616626 | 2.689804 | 0.1587123 | 0.6955124 | 0.999176 |
| ENSMUSG00000031918 | Mtmr2 | protein_coding | -0.1862092 | 5.617266 | 0.1587003 | 0.6955232 | 0.999176 |
| ENSMUSG00000022545 | Ercc4 | protein_coding | 0.3526537 | 4.237129 | 0.1591157 | 0.6955706 | 0.999176 |
| ENSMUSG00000109398 | Gm3854 | protein_coding | -0.3845597 | 3.106398 | 0.15848 | 0.6957231 | 0.999176 |
| ENSMUSG00000049659 | Aftph | protein_coding | 0.1583 | 5.669653 | 0.1583007 | 0.6958859 | 0.999176 |
| ENSMUSG00000041216 | Clvs1 | protein_coding | 0.2330202 | 4.457796 | 0.1581695 | 0.6960051 | 0.999176 |
| ENSMUSG00000004364 | Cul3 | protein_coding | 0.1289861 | 6.451955 | 0.1580095 | 0.6961506 | 0.999176 |
| ENSMUSG00000048915 | Efna5 | protein_coding | -0.2404877 | 6.53037 | 0.1588037 | 0.6961535 | 0.999176 |

| ensembl_gene_id | external_gene_name | gene_biotype | log2 (fold change) | log2 (counts per million) | F-value | p-value | False discovery rate |
|---|---|---|---|---|---|---|---|
| ENSMUSG00000028778 | Hcrtr1 | protein_coding | 0.8149537 | 3.40245 | 0.161251 | 0.6962799 | 0.999176 |
| ENSMUSG00000044934 | Zfp367 | protein_coding | -0.2077347 | 4.72127 | 0.1576953 | 0.6964364 | 0.999176 |
| ENSMUSG00000052139 | Babam2 | protein_coding | 0.1389737 | 6.603074 | 0.157677 | 0.6964531 | 0.999176 |
| ENSMUSG00000027739 | Rab33b | protein_coding | -0.1628763 | 5.633893 | 0.157605 | 0.6965186 | 0.999176 |
| ENSMUSG00000078862 | Gm14326 | protein_coding | 0.223829 | 4.483002 | 0.1574896 | 0.6966237 | 0.999176 |
| ENSMUSG00000033940 | Brk1 | protein_coding | 0.111598 | 9.750061 | 0.1573928 | 0.6967119 | 0.999176 |
| ENSMUSG00000093606 | B130034C11Rik | processed_transcript | 0.4301148 | 2.838126 | 0.1570933 | 0.6969849 | 0.999176 |
| ENSMUSG00000023330 | Dtwd1 | protein_coding | 0.1486901 | 5.508982 | 0.1570756 | 0.6970011 | 0.999176 |
| ENSMUSG00000039043 | Arpin | protein_coding | -0.2021244 | 5.129652 | 0.1568539 | 0.6972035 | 0.999176 |
| ENSMUSG00000008384 | Sertad1 | protein_coding | 0.1941273 | 4.17892 | 0.156829 | 0.6972263 | 0.999176 |
| ENSMUSG00000022528 | Hes1 | protein_coding | -0.3258651 | 2.960253 | 0.1568029 | 0.69725 | 0.999176 |
| ENSMUSG00000042605 | Atxn2 | protein_coding | 0.1487328 | 5.778145 | 0.1567959 | 0.6972564 | 0.999176 |
| ENSMUSG00000026188 | Tmem169 | protein_coding | 0.2262469 | 3.928873 | 0.1566074 | 0.6974287 | 0.999176 |
| ENSMUSG00000037010 | Apln | protein_coding | -0.2986246 | 3.299784 | 0.1565234 | 0.6975055 | 0.999176 |
| ENSMUSG00000039982 | Dtx4 | protein_coding | 0.2204845 | 4.140135 | 0.1565021 | 0.6975249 | 0.999176 |
| ENSMUSG00000074896 | Ifit3 | protein_coding | -0.3133792 | 3.128248 | 0.1564946 | 0.6975317 | 0.999176 |
| ENSMUSG00000085925 | Rtl1 | protein_coding | -0.2503646 | 4.657097 | 0.1564555 | 0.6975759 | 0.999176 |
| ENSMUSG00000024644 | Cndp2 | protein_coding | -0.1920836 | 6.333925 | 0.1564531 | 0.69761 | 0.999176 |
| ENSMUSG00000023935 | Spats1 | protein_coding | -0.3634435 | 2.793313 | 0.1563963 | 0.6976217 | 0.999176 |
| ENSMUSG00000034354 | Mtmr3 | protein_coding | 0.2096942 | 5.133219 | 0.1561432 | 0.6978533 | 0.999176 |
| ENSMUSG00000025503 | Taldo1 | protein_coding | -0.1103952 | 8.388558 | 0.1561356 | 0.6978602 | 0.999176 |
| ENSMUSG00000021279 | Cdc42bpb | protein_coding | -0.2799999 | 3.217432 | 0.1558416 | 0.6981296 | 0.999176 |
| ENSMUSG00000034647 | Ankrd12 | protein_coding | -0.1834435 | 6.328543 | 0.155846 | 0.698134 | 0.999176 |
| ENSMUSG00000027890 | Gstm4 | protein_coding | 0.172694 | 4.802565 | 0.1555937 | 0.6983569 | 0.999176 |
| ENSMUSG00000027245 | Hypk | protein_coding | 0.2160177 | 4.01193 | 0.1554992 | 0.6984437 | 0.999176 |
| ENSMUSG00000057134 | Ado | protein_coding | -0.1640428 | 8.137692 | 0.1553716 | 0.6985691 | 0.999176 |
| ENSMUSG00000027298 | Tyro3 | protein_coding | -0.1820159 | 5.381614 | 0.155329 | 0.6985999 | 0.999176 |
| ENSMUSG00000034473 | Sec22a | protein_coding | 0.1319135 | 6.196041 | 0.1552077 | 0.6987113 | 0.999176 |
| ENSMUSG00000032381 | Fam96a | protein_coding | 0.129467 | 6.727894 | 0.1550096 | 0.6988933 | 0.999176 |
| ENSMUSG00000004591 | Pkn2 | protein_coding | 0.2743861 | 3.453411 | 0.1547842 | 0.6991007 | 0.999176 |
| ENSMUSG00000032434 | Cmtm6 | protein_coding | -0.4202154 | 2.968209 | 0.1546707 | 0.6992052 | 0.999176 |
| ENSMUSG00000022983 | Scaf4 | protein_coding | 0.4077493 | 3.146398 | 0.1545877 | 0.69929 | 0.999176 |
| ENSMUSG00000036249 | Rbm43 | protein_coding | 0.2022685 | 5.194965 | 0.1545515 | 0.6993149 | 0.999176 |
| ENSMUSG00000039611 | Tmem246 | protein_coding | 0.1469832 | 6.775407 | 0.1544793 | 0.6993814 | 0.999176 |
| ENSMUSG00000022200 | Golph3 | protein_coding | 0.1762162 | 5.640856 | 0.1544395 | 0.6994181 | 0.999176 |
| ENSMUSG00000041669 | Prima1 | protein_coding | 0.4550396 | 2.634618 | 0.1543953 | 0.6994588 | 0.999176 |
| ENSMUSG00000039563 | 2210406O10Rik | lincRNA | -0.2830378 | 3.185292 | 0.1542402 | 0.6996018 | 0.999176 |
| ENSMUSG00000056856 | Jakmip3 | protein_coding | 0.2143322 | 4.05951 | 0.1542089 | 0.6996307 | 0.999176 |
| ENSMUSG00000021395 | Spin1 | protein_coding | -0.1264643 | 7.491725 | 0.1541904 | 0.6996477 | 0.999176 |
| ENSMUSG00000029920 | Smarcad1 | protein_coding | -0.2572182 | 3.781083 | 0.1541811 | 0.6996563 | 0.999176 |
| ENSMUSG00000037685 | Atp8a1 | protein_coding | -0.2671846 | 3.50241 | 0.1541554 | 0.69968 | 0.999176 |
| ENSMUSG00000042787 | Exog | protein_coding | 0.222688 | 5.02664 | 0.1538942 | 0.699921 | 0.999176 |
| ENSMUSG00000063450 | Syne2 | protein_coding | -0.3750139 | 3.309267 | 0.1538705 | 0.6999513 | 0.999176 |
| ENSMUSG00000001507 | Itga3 | protein_coding | 0.2503463 | 3.834342 | 0.1537978 | 0.7000101 | 0.999176 |
| ENSMUSG00000031325 | 4930519F16Rik | transcribed_unitary_p | -0.40001 | 2.588531 | 0.1537827 | 0.700024 | 0.999176 |
| ENSMUSG00000087269 | D330023K18Rik | processed_transcript | 0.1604174 | 5.379798 | 0.1537752 | 0.7000309 | 0.999176 |
| ENSMUSG00000033735 | Spr | protein_coding | 0.2503496 | 5.109773 | 0.1538073 | 0.7000316 | 0.999176 |
| ENSMUSG00000010660 | Plcd1 | protein_coding | 0.4492538 | 3.059502 | 0.1538 | 0.7000384 | 0.999176 |
| ENSMUSG00000029735 | Tpk1 | protein_coding | -0.5314467 | 2.615277 | 0.1536973 | 0.7001113 | 0.999176 |
| ENSMUSG00000070570 | Slc17a7 | protein_coding | 0.5064214 | 2.764083 | 0.153691 | 0.7001171 | 0.999176 |
| ENSMUSG00000008090 | Fgfrl1 | protein_coding | -0.4870899 | 2.626197 | 0.1535844 | 0.7002073 | 0.999176 |
| ENSMUSG00000019726 | Lyst | protein_coding | 0.2406629 | 3.969897 | 0.1535494 | 0.7002396 | 0.999176 |
| ENSMUSG00000097709 | 2810429I04Rik | lincRNA | -0.2858367 | 3.47789 | 0.1534844 | 0.7002997 | 0.999176 |
| ENSMUSG00000021948 | Prkcd | protein_coding | -0.2834171 | 5.889248 | 0.1543231 | 0.7003362 | 0.999176 |
| ENSMUSG00000038738 | Shank1 | protein_coding | 0.2059588 | 5.506 | 0.1534243 | 0.7003636 | 0.999176 |
| ENSMUSG00000060450 | Rnf14 | protein_coding | 0.1624295 | 8.835925 | 0.1534039 | 0.7003825 | 0.999176 |
| ENSMUSG00000022120 | Rnf219 | protein_coding | -0.2373871 | 4.36941 | 0.1532527 | 0.700514 | 0.999176 |
| ENSMUSG00000027596 | a | protein_coding | -0.3549884 | 2.687726 | 0.1530979 | 0.7006574 | 0.999176 |
| ENSMUSG00000014074 | Rnf168 | protein_coding | -0.2018533 | 4.749605 | 0.1530535 | 0.7006985 | 0.999176 |
| ENSMUSG00000025451 | Paip1 | protein_coding | -0.2344523 | 8.314838 | 0.15381 | 0.7008082 | 0.999176 |
| ENSMUSG00000009470 | Tnpo1 | protein_coding | -0.2126457 | 4.766316 | 0.1527692 | 0.700962 | 0.999176 |
| ENSMUSG00000039670 | Oxld1 | protein_coding | 0.4174487 | 2.546772 | 0.1525699 | 0.7011469 | 0.999176 |
| ENSMUSG00000031278 | Acsl4 | protein_coding | -0.1556093 | 6.224603 | 0.152186 | 0.7015034 | 0.999176 |
| ENSMUSG00000018707 | Dync1h1 | protein_coding | -0.1913318 | 5.922824 | 0.1521448 | 0.7015499 | 0.999176 |
| ENSMUSG00000031813 | Mvb12a | protein_coding | 0.2169161 | 4.721394 | 0.1521174 | 0.7015671 | 0.999176 |
| ENSMUSG00000044986 | Tst | protein_coding | -0.1892809 | 6.522074 | 0.152054 | 0.7017941 | 0.999176 |
| ENSMUSG00000044912 | Syt16 | protein_coding | -0.193835 | 4.870033 | 0.1517157 | 0.7019408 | 0.999176 |
| ENSMUSG00000060279 | Ap2a1 | protein_coding | 0.4491558 | 4.119981 | 0.1527865 | 0.7020384 | 0.999176 |
| ENSMUSG00000025134 | Alyref | protein_coding | -0.2096796 | 4.548844 | 0.1515103 | 0.7021321 | 0.999176 |
| ENSMUSG00000025781 | Atp5c1 | protein_coding | 0.1139349 | 10.33953 | 0.1514541 | 0.7021845 | 0.999176 |
| ENSMUSG00000097039 | Pvt1 | lincRNA | -0.1859727 | 4.786443 | 0.1514455 | 0.7021925 | 0.999176 |
| ENSMUSG00000036678 | Aaas | protein_coding | -0.1883847 | 5.155665 | 0.1513789 | 0.7022545 | 0.999176 |
| ENSMUSG00000020069 | Hnrnph3 | protein_coding | 0.2184841 | 4.443108 | 0.1510895 | 0.7025245 | 0.999176 |
| ENSMUSG00000030134 | Rasgef1a | protein_coding | -0.1565733 | 7.583744 | 0.1510097 | 0.7026072 | 0.999176 |
| ENSMUSG00000079557 | 2-Mar | protein_coding | 0.1725759 | 5.59789 | 0.1509538 | 0.7026511 | 0.999176 |
| ENSMUSG00000026960 | Arl6ip6 | protein_coding | 0.2350129 | 3.486996 | 0.1509364 | 0.7026674 | 0.999176 |
| ENSMUSG00000018846 | Pank3 | protein_coding | -0.1825926 | 4.737048 | 0.1509267 | 0.7026765 | 0.999176 |
| ENSMUSG00000035232 | Pdk3 | protein_coding | -0.3584573 | 4.387157 | 0.1515514 | 0.7027465 | 0.999176 |

| ensembl_gene_id | external_gene_name | gene_biotype | log2 (fold change) | log2 (counts per million) | F-value | p-value | False discovery rate |
|---|---|---|---|---|---|---|---|
| ENSMUSG00000043940 | Wdfy3 | protein_coding | -0.1923952 | 5.725793 | 0.1508513 | 0.7027469 | 0.999176 |
| ENSMUSG00000073424 | Cyp4f15 | protein_coding | -0.6269605 | 2.390839 | 0.1508787 | 0.7027534 | 0.999176 |
| ENSMUSG00000020864 | Ankrd40 | protein_coding | 0.1533508 | 5.976956 | 0.1507214 | 0.7028682 | 0.999176 |
| ENSMUSG00000020894 | Vamp2 | protein_coding | 0.1194852 | 9.219984 | 0.1506085 | 0.7029738 | 0.999176 |
| ENSMUSG00000043300 | B3galnt1 | protein_coding | 0.1449314 | 6.205818 | 0.1505509 | 0.7030276 | 0.999176 |
| ENSMUSG00000040136 | Abcc8 | protein_coding | 0.800261 | 2.080395 | 0.1508349 | 0.7030819 | 0.999176 |
| ENSMUSG00000060923 | Acyp2 | protein_coding | -0.1136426 | 7.248511 | 0.1504365 | 0.7031346 | 0.999176 |
| ENSMUSG00000002043 | Trappc6a | protein_coding | -0.1753419 | 4.975157 | 0.150267 | 0.7032932 | 0.999176 |
| ENSMUSG00000034432 | Cops8 | protein_coding | 0.1212248 | 8.78802 | 0.1502539 | 0.7033055 | 0.999176 |
| ENSMUSG00000001383 | Zmat2 | protein_coding | 0.1221584 | 9.115223 | 0.1502502 | 0.703309 | 0.999176 |
| ENSMUSG00000027942 | 4933434E20Rik | protein_coding | 0.1094788 | 8.603961 | 0.150212 | 0.7033448 | 0.999176 |
| ENSMUSG00000100837 | 1700063D05Rik | lincRNA | 0.2758897 | 3.53966 | 0.1501183 | 0.7034325 | 0.999176 |
| ENSMUSG00000002908 | Kcnn1 | protein_coding | 0.4742399 | 2.593229 | 0.1500521 | 0.7034945 | 0.999176 |
| ENSMUSG00000027423 | Snx5 | protein_coding | -0.1397599 | 6.114144 | 0.1500094 | 0.7035345 | 0.999176 |
| ENSMUSG00000004110 | Cacna1e | protein_coding | -0.2162166 | 4.923583 | 0.1499899 | 0.7035528 | 0.999176 |
| ENSMUSG00000042873 | Lhfpl4 | protein_coding | -0.1334404 | 6.044645 | 0.1499647 | 0.7035764 | 0.999176 |
| ENSMUSG00000041949 | Tango6 | protein_coding | 0.2256585 | 3.555739 | 0.149946 | 0.7035939 | 0.999176 |
| ENSMUSG00000009079 | Ewsr1 | protein_coding | -0.1481558 | 5.93226 | 0.1498618 | 0.7036729 | 0.999176 |
| ENSMUSG00000041936 | Agrn | protein_coding | 0.2229111 | 4.31919 | 0.1498374 | 0.7036958 | 0.999176 |
| ENSMUSG00000021301 | Hecw1 | protein_coding | -0.1760916 | 5.21215 | 0.149662 | 0.7038603 | 0.999176 |
| ENSMUSG00000021573 | Tppp | protein_coding | -0.1211971 | 7.734587 | 0.1496175 | 0.7039021 | 0.999176 |
| ENSMUSG00000001240 | Ramp2 | protein_coding | -0.3986926 | 2.702734 | 0.1496019 | 0.7039167 | 0.999176 |
| ENSMUSG00000032570 | Atp2c1 | protein_coding | -0.1802657 | 5.012945 | 0.1494212 | 0.7040863 | 0.999176 |
| ENSMUSG00000030536 | Iqgap1 | protein_coding | -0.3174134 | 3.601172 | 0.1492628 | 0.7042352 | 0.999176 |
| ENSMUSG00000004931 | Apba3 | protein_coding | -0.3658459 | 2.610797 | 0.1492545 | 0.7042429 | 0.999176 |
| ENSMUSG00000033014 | Trim33 | protein_coding | -0.1789821 | 4.77645 | 0.1492123 | 0.7042826 | 0.999176 |
| ENSMUSG00000021131 | Erh | protein_coding | 0.2334334 | 3.646676 | 0.1491967 | 0.7042973 | 0.999176 |
| ENSMUSG00000028800 | Hdac1 | protein_coding | 0.2233633 | 3.739194 | 0.1488117 | 0.7046595 | 0.999176 |
| ENSMUSG00000030898 | Cckbr | protein_coding | -0.3544413 | 3.17254 | 0.1487277 | 0.7047386 | 0.999176 |
| ENSMUSG00000001467 | Cyp51 | protein_coding | -0.153768 | 5.708777 | 0.1486844 | 0.7047794 | 0.999176 |
| ENSMUSG00000026032 | Ndufb3 | protein_coding | 0.1573313 | 8.401086 | 0.1486733 | 0.704798 | 0.999176 |
| ENSMUSG00000031107 | Rbmx2 | protein_coding | -0.1828315 | 4.255341 | 0.1485873 | 0.7048709 | 0.999176 |
| ENSMUSG00000027634 | Ndrg3 | protein_coding | -0.1679396 | 9.40626 | 0.1485437 | 0.7049201 | 0.999176 |
| ENSMUSG00000029722 | Agfg2 | protein_coding | -0.1783097 | 5.019407 | 0.1483909 | 0.705056 | 0.999176 |
| ENSMUSG00000022350 | Washc5 | protein_coding | 0.1885749 | 4.438826 | 0.1483191 | 0.7051237 | 0.999176 |
| ENSMUSG00000040785 | Ttc3 | protein_coding | -0.1209859 | 9.858795 | 0.1481983 | 0.7052377 | 0.999176 |
| ENSMUSG00000005667 | Mthfd2 | protein_coding | 0.1678998 | 5.701857 | 0.1479657 | 0.7054572 | 0.999176 |
| ENSMUSG00000040423 | Rc3h1 | protein_coding | -0.1688394 | 4.84348 | 0.1478329 | 0.7055827 | 0.999176 |
| ENSMUSG00000030058 | Copg1 | protein_coding | 0.1493751 | 7.934758 | 0.1477443 | 0.7056746 | 0.999176 |
| ENSMUSG00000027977 | Ndst3 | protein_coding | 0.3780582 | 2.657613 | 0.1477313 | 0.7056788 | 0.999176 |
| ENSMUSG00000045598 | Zfp553 | protein_coding | 0.237933 | 3.911544 | 0.1476916 | 0.7057163 | 0.999176 |
| ENSMUSG00000049044 | Rapgef4 | protein_coding | -0.1788053 | 6.176588 | 0.1476524 | 0.7057615 | 0.999176 |
| ENSMUSG00000034993 | Vat1 | protein_coding | 0.1454146 | 6.200072 | 0.147622 | 0.7057822 | 0.999176 |
| ENSMUSG00000048170 | Mcmbp | protein_coding | -0.1681917 | 5.869127 | 0.1475352 | 0.7058642 | 0.999176 |
| ENSMUSG00000047246 | Hist1h2be | protein_coding | -0.3500564 | 2.832654 | 0.1474781 | 0.7059183 | 0.999176 |
| ENSMUSG00000020818 | Mfsd11 | protein_coding | -0.1602814 | 5.25114 | 0.1474672 | 0.7059286 | 0.999176 |
| ENSMUSG00000020525 | Ppm1d | protein_coding | 0.2918777 | 3.35679 | 0.1473786 | 0.7060125 | 0.999176 |
| ENSMUSG00000004631 | Sgce | protein_coding | 0.1360507 | 5.822566 | 0.147291 | 0.7060954 | 0.999176 |
| ENSMUSG00000031633 | Slc25a4 | protein_coding | 0.1193552 | 11.73554 | 0.1472734 | 0.706112 | 0.999176 |
| ENSMUSG00000027201 | Myef2 | protein_coding | -0.3734171 | 2.690353 | 0.147231 | 0.7061522 | 0.999176 |
| ENSMUSG00000036391 | Sec24a | protein_coding | -0.2398796 | 3.856373 | 0.1471937 | 0.7061876 | 0.999176 |
| ENSMUSG00000063200 | Nol7 | protein_coding | 0.1476497 | 6.244344 | 0.1471211 | 0.7062564 | 0.999176 |
| ENSMUSG00000052981 | Ube2ql1 | protein_coding | 0.1327295 | 6.859346 | 0.147023 | 0.7063494 | 0.999176 |
| ENSMUSG00000087461 | C230014O12Rik | lincRNA | -0.3673485 | 2.858043 | 0.1469632 | 0.706406 | 0.999176 |
| ENSMUSG00000041124 | Msantd4 | protein_coding | -0.1391455 | 6.795587 | 0.146852 | 0.7065115 | 0.999176 |
| ENSMUSG00000046096 | BC030336 | protein_coding | -0.1535805 | 5.853572 | 0.1468283 | 0.706534 | 0.999176 |
| ENSMUSG00000043091 | Tuba1c | protein_coding | -0.8364149 | 1.936353 | 0.1470607 | 0.706629 | 0.999176 |
| ENSMUSG00000063275 | Hacd1 | protein_coding | -0.4104668 | 2.469024 | 0.1465516 | 0.7067967 | 0.999176 |
| ENSMUSG00000042197 | Zfp451 | protein_coding | 0.1494965 | 5.442189 | 0.1465444 | 0.7068035 | 0.999176 |
| ENSMUSG00000026821 | Ralgds | protein_coding | -0.1703606 | 5.58292 | 0.1463097 | 0.7070265 | 0.999176 |
| ENSMUSG00000037325 | Bbs7 | protein_coding | 0.1952465 | 4.537572 | 0.1462281 | 0.7071041 | 0.999176 |
| ENSMUSG00000031446 | Cul4a | protein_coding | -0.2377104 | 4.612522 | 0.1461064 | 0.7072199 | 0.999176 |
| ENSMUSG00000033790 | Tubgcp5 | protein_coding | -0.1742825 | 4.515937 | 0.1460583 | 0.7072657 | 0.999176 |
| ENSMUSG00000047030 | Spata2 | protein_coding | 0.235435 | 4.479293 | 0.1459712 | 0.7073486 | 0.999176 |
| ENSMUSG00000062198 | 2700097O09Rik | protein_coding | -0.202532 | 4.295032 | 0.1459616 | 0.7073577 | 0.999176 |
| ENSMUSG00000022108 | Itm2b | protein_coding | -0.1131436 | 8.615978 | 0.1458895 | 0.7074263 | 0.999176 |
| ENSMUSG00000019461 | Plscr3 | protein_coding | 0.2020488 | 4.880977 | 0.145872 | 0.7074431 | 0.999176 |
| ENSMUSG00000026361 | Cdc73 | protein_coding | -0.17936 | 4.876864 | 0.1457413 | 0.7075675 | 0.999176 |
| ENSMUSG00000026771 | Spopl | protein_coding | -0.2725609 | 3.182406 | 0.1454579 | 0.7078377 | 0.999176 |
| ENSMUSG00000057715 | A830018L16Rik | protein_coding | -0.1902584 | 4.93703 | 0.14536 | 0.7079311 | 0.999176 |
| ENSMUSG00000023923 | Tbc1d5 | protein_coding | -0.1990521 | 4.134017 | 0.1451047 | 0.7081749 | 0.999176 |
| ENSMUSG00000018470 | Kcnab3 | protein_coding | 0.2334378 | 3.594479 | 0.1450571 | 0.7082204 | 0.999176 |
| ENSMUSG00000061086 | Myl4 | protein_coding | -0.3246072 | 2.909886 | 0.144989 | 0.7082854 | 0.999176 |
| ENSMUSG00000031627 | Irf2 | protein_coding | -0.1558915 | 4.944027 | 0.1449044 | 0.7083663 | 0.999176 |
| ENSMUSG00000086938 | 4930481A15Rik | antisense_RNA | 0.2046769 | 3.896317 | 0.1448348 | 0.7084328 | 0.999176 |
| ENSMUSG00000056429 | Tgoln1 | protein_coding | -0.1155909 | 6.55358 | 0.1447272 | 0.7085357 | 0.999176 |
| ENSMUSG00000008333 | Snrpb2 | protein_coding | -0.1471343 | 6.805208 | 0.1446246 | 0.708634 | 0.999176 |
| ENSMUSG00000002957 | Ap2a2 | protein_coding | 0.1535683 | 7.263748 | 0.1445903 | 0.7086748 | 0.999176 |

| ensembl_gene_id | external_gene_name | gene_biotype | log2 (fold change) | log2 (counts per million) | F-value | p-value | False discovery rate |
|---|---|---|---|---|---|---|---|
| ENSMUSG00000021852 | Slc35f4 | protein_coding | -0.2504044 | 3.695873 | 0.1444612 | 0.7087904 | 0.999176 |
| ENSMUSG00000022965 | Ifngr2 | protein_coding | 0.1599141 | 5.187981 | 0.144379 | 0.7088691 | 0.999176 |
| ENSMUSG00000028328 | Tmod1 | protein_coding | 0.192283 | 4.726566 | 0.1442968 | 0.7089479 | 0.999176 |
| ENSMUSG00000029993 | Nfu1 | protein_coding | 0.1104077 | 8.380903 | 0.1441453 | 0.7090931 | 0.999176 |
| ENSMUSG00000049313 | Sorl1 | protein_coding | -0.1361721 | 6.405108 | 0.1440556 | 0.7091792 | 0.999176 |
| ENSMUSG00000025357 | Dgka | protein_coding | -0.4493412 | 2.571707 | 0.1438644 | 0.7093626 | 0.999176 |
| ENSMUSG00000037204 | Atg101 | protein_coding | -0.1115103 | 6.702083 | 0.1438144 | 0.7094106 | 0.999176 |
| ENSMUSG00000037259 | Dzank1 | protein_coding | -0.2319913 | 4.315431 | 0.1437842 | 0.7094396 | 0.999176 |
| ENSMUSG00000030811 | Fbxl19 | protein_coding | 0.5572517 | 2.807103 | 0.1440817 | 0.709466 | 0.999176 |
| ENSMUSG00000054237 | Fra10ac1 | protein_coding | 0.1538977 | 5.39657 | 0.143657 | 0.7095618 | 0.999176 |
| ENSMUSG00000020949 | Fkbp3 | protein_coding | 0.1151698 | 9.46343 | 0.1435854 | 0.7096306 | 0.999176 |
| ENSMUSG00000040550 | Otud6b | protein_coding | -0.1667832 | 5.995032 | 0.1434836 | 0.7097285 | 0.999176 |
| ENSMUSG00000022240 | Ctnnd2 | protein_coding | -0.1116576 | 7.273597 | 0.1434148 | 0.7097946 | 0.999176 |
| ENSMUSG00000048878 | Hexim1 | protein_coding | -0.1874515 | 4.585484 | 0.1433628 | 0.7098447 | 0.999176 |
| ENSMUSG00000026987 | Baz2b | protein_coding | -0.1652546 | 4.862876 | 0.1433273 | 0.7098788 | 0.999176 |
| ENSMUSG00000018574 | Acadvl | protein_coding | -0.1948528 | 5.426074 | 0.1433097 | 0.7098958 | 0.999176 |
| ENSMUSG00000032328 | Tmem30a | protein_coding | -0.1075842 | 7.184056 | 0.1433052 | 0.7099001 | 0.999176 |
| ENSMUSG00000025950 | Idh1 | protein_coding | -0.1187637 | 6.523686 | 0.1431049 | 0.7100928 | 0.999176 |
| ENSMUSG00000034158 | Lrrc58 | protein_coding | -0.1731026 | 5.653377 | 0.1429782 | 0.7102149 | 0.999176 |
| ENSMUSG00000061458 | Nol10 | protein_coding | 0.2134131 | 4.249978 | 0.1426814 | 0.710501 | 0.999176 |
| ENSMUSG00000059659 | Gm10069 | antisense_RNA | 0.3154678 | 3.082249 | 0.1426195 | 0.7105608 | 0.999176 |
| ENSMUSG00000022558 | Mroh1 | protein_coding | -0.1957 | 4.88403 | 0.1425995 | 0.71058 | 0.999176 |
| ENSMUSG00000071454 | Dtnb | protein_coding | 0.1632314 | 4.995226 | 0.1425994 | 0.7105801 | 0.999176 |
| ENSMUSG00000034336 | Ina | protein_coding | 0.1123403 | 7.552056 | 0.1425388 | 0.7106386 | 0.999176 |
| ENSMUSG00000051159 | Cited1 | protein_coding | -0.2310574 | 5.49966 | 0.1427259 | 0.7106623 | 0.999176 |
| ENSMUSG00000071359 | Tbpl1 | protein_coding | -0.1479261 | 6.377243 | 0.1424419 | 0.7107321 | 0.999176 |
| ENSMUSG00000028849 | Map7d1 | protein_coding | -0.1256102 | 6.582568 | 0.1420638 | 0.7110974 | 0.999176 |
| ENSMUSG00000021215 | Net1 | protein_coding | -0.3655165 | 2.936039 | 0.1419973 | 0.7111618 | 0.999176 |
| ENSMUSG00000026637 | Traf5 | protein_coding | 0.2952029 | 2.904135 | 0.1419409 | 0.7112163 | 0.999176 |
| ENSMUSG00000070284 | Gmppb | protein_coding | 0.2104217 | 4.677332 | 0.1419088 | 0.7112474 | 0.999176 |
| ENSMUSG00000034543 | Morc2a | protein_coding | 0.2110503 | 3.987447 | 0.1418513 | 0.711303 | 0.999176 |
| ENSMUSG00000032966 | Fkbp1a | protein_coding | -0.1166461 | 9.154511 | 0.1417313 | 0.7114192 | 0.999176 |
| ENSMUSG00000006369 | Fbln1 | protein_coding | 0.3408083 | 3.155323 | 0.1416952 | 0.7114542 | 0.999176 |
| ENSMUSG00000032011 | Thy1 | protein_coding | 0.1476842 | 7.286845 | 0.1416047 | 0.7115498 | 0.999176 |
| ENSMUSG00000023809 | Rps6ka2 | protein_coding | -0.2264826 | 4.225924 | 0.1413435 | 0.711795 | 0.999176 |
| ENSMUSG00000029249 | Rest | protein_coding | -0.4891749 | 2.127345 | 0.1413337 | 0.7118045 | 0.999176 |
| ENSMUSG00000062906 | Hdac10 | protein_coding | 0.404415 | 3.334555 | 0.1415199 | 0.7118376 | 0.999176 |
| ENSMUSG00000056851 | Pcbp2 | protein_coding | 0.111846 | 6.502847 | 0.1412882 | 0.7118486 | 0.999176 |
| ENSMUSG00000030677 | Kif22 | protein_coding | 0.2407872 | 3.465538 | 0.1411491 | 0.7119836 | 0.999176 |
| ENSMUSG00000021218 | Gdi2 | protein_coding | -0.1307654 | 8.659816 | 0.14111 | 0.7120216 | 0.999176 |
| ENSMUSG00000014195 | Dnajc7 | protein_coding | -0.1230826 | 6.780053 | 0.1409618 | 0.7121655 | 0.999176 |
| ENSMUSG00000030302 | Atp2b2 | protein_coding | -0.1255708 | 8.150042 | 0.1409136 | 0.7122122 | 0.999176 |
| ENSMUSG00000037938 | Chchd5 | protein_coding | 0.1174555 | 6.292169 | 0.1408912 | 0.712234 | 0.999176 |
| ENSMUSG00000026614 | Slc30a10 | protein_coding | -0.3476658 | 3.19946 | 0.1407151 | 0.7124052 | 0.999176 |
| ENSMUSG00000020917 | Acly | protein_coding | 0.1487191 | 7.114004 | 0.1406633 | 0.7124635 | 0.999176 |
| ENSMUSG00000031024 | St5 | protein_coding | -0.2924676 | 3.050118 | 0.1405843 | 0.7125324 | 0.999176 |
| ENSMUSG00000020250 | Txnrd1 | protein_coding | -0.150642 | 5.659791 | 0.1405452 | 0.7125704 | 0.999176 |
| ENSMUSG00000025375 | Aatk | protein_coding | -0.1669824 | 5.730584 | 0.1405215 | 0.7125935 | 0.999176 |
| ENSMUSG00000094910 | D430019H16Rik | protein_coding | -0.203996 | 5.818414 | 0.1405568 | 0.7125971 | 0.999176 |
| ENSMUSG00000058656 | Samd12 | protein_coding | 0.24219 | 4.076291 | 0.1405052 | 0.7126094 | 0.999176 |
| ENSMUSG00000056427 | Slit3 | protein_coding | 0.3304433 | 2.882208 | 0.1404959 | 0.7126184 | 0.999176 |
| ENSMUSG00000039633 | Lonrf1 | protein_coding | 0.2913824 | 3.347394 | 0.140472 | 0.7126417 | 0.999176 |
| ENSMUSG00000026806 | Ddx31 | protein_coding | 0.3826944 | 2.812605 | 0.140015 | 0.7130869 | 0.999176 |
| ENSMUSG00000065979 | Cpped1 | protein_coding | 0.148025 | 6.910962 | 0.1398999 | 0.7131991 | 0.999176 |
| ENSMUSG00000103034 | Gm8797 | protein_coding | -0.280703 | 2.947512 | 0.139767 | 0.7133288 | 0.999176 |
| ENSMUSG00000028354 | Fmn2 | protein_coding | 0.2101656 | 4.40803 | 0.1396383 | 0.7134545 | 0.999176 |
| ENSMUSG00000029684 | Wasl | protein_coding | 0.1313871 | 6.066732 | 0.1395792 | 0.7135122 | 0.999176 |
| ENSMUSG00000029221 | Slc30a9 | protein_coding | 0.1193221 | 6.804376 | 0.13948 | 0.7136091 | 0.999176 |
| ENSMUSG00000006360 | Crip1 | protein_coding | -0.2802536 | 4.025109 | 0.1394612 | 0.7136354 | 0.999176 |
| ENSMUSG00000031862 | Atp13a1 | protein_coding | 0.3005163 | 3.36037 | 0.1394484 | 0.71364 | 0.999176 |
| ENSMUSG00000062352 | Itgb1bp1 | protein_coding | 0.1472463 | 6.48329 | 0.1393301 | 0.7137556 | 0.999176 |
| ENSMUSG00000029649 | Pomp | protein_coding | 0.1202645 | 9.627769 | 0.1393228 | 0.7137627 | 0.999176 |
| ENSMUSG00000052395 | Rft1 | protein_coding | -0.1914331 | 4.127595 | 0.1392951 | 0.7137898 | 0.999176 |
| ENSMUSG00000022507 | 1810013L24Rik | protein_coding | -0.1144618 | 6.407993 | 0.139269 | 0.7138154 | 0.999176 |
| ENSMUSG00000023845 | Lnpep | protein_coding | 0.4899737 | 2.367203 | 0.1392429 | 0.7138409 | 0.999176 |
| ENSMUSG00000039519 | Cyp7b1 | protein_coding | -0.2931844 | 3.486739 | 0.1392199 | 0.7138634 | 0.999176 |
| ENSMUSG00000004789 | Dlst | protein_coding | -0.1383538 | 6.924176 | 0.1391817 | 0.7139008 | 0.999176 |
| ENSMUSG00000044876 | Zfp444 | protein_coding | 0.1897146 | 5.12086 | 0.1391203 | 0.7139609 | 0.999176 |
| ENSMUSG00000036550 | Cnot1 | protein_coding | 0.1919792 | 4.58406 | 0.1390704 | 0.7140097 | 0.999176 |
| ENSMUSG00000093460 | Six3os1 | processed_transcript | -0.2832764 | 3.604495 | 0.1388326 | 0.7142426 | 0.999176 |
| ENSMUSG00000062202 | Btbd9 | protein_coding | 0.1763683 | 5.808402 | 0.1384832 | 0.7145852 | 0.999176 |
| ENSMUSG00000021057 | Akap5 | protein_coding | -0.131083 | 6.766826 | 0.1384044 | 0.7146625 | 0.999176 |
| ENSMUSG00000108532 | Gm32647 | lincRNA | 0.4632319 | 2.491067 | 0.1383929 | 0.7146739 | 0.999176 |
| ENSMUSG00000026723 | Trdmt1 | protein_coding | 0.2938796 | 3.159474 | 0.1383843 | 0.7146822 | 0.999176 |
| ENSMUSG00000024430 | Cabyr | protein_coding | -0.412416 | 2.451007 | 0.1383422 | 0.7147236 | 0.999176 |
| ENSMUSG00000030057 | Cnbp | protein_coding | -0.1089012 | 10.42447 | 0.1382287 | 0.7148351 | 0.999176 |
| ENSMUSG00000085793 | Lin52 | protein_coding | -0.1443975 | 5.724162 | 0.1381482 | 0.7149142 | 0.999176 |
| ENSMUSG00000033326 | Kdm4a | protein_coding | -0.3938161 | 2.537096 | 0.1380358 | 0.7150246 | 0.999176 |

| ensembl_gene_id | external_gene_name | gene_biotype | log2 (fold change) | log2 (counts per million) | F-value | p-value | False discovery rate |
|---|---|---|---|---|---|---|---|
| ENSMUSG00000036315 | Znrd1 | protein_coding | 0.1582842 | 6.192883 | 0.1378699 | 0.7151878 | 0.999176 |
| ENSMUSG00000066113 | Adamtsl1 | protein_coding | 0.4602029 | 3.038575 | 0.138069 | 0.7151925 | 0.999176 |
| ENSMUSG00000010066 | Cacna2d2 | protein_coding | -0.2213716 | 4.557097 | 0.1378514 | 0.7152059 | 0.999176 |
| ENSMUSG00000020088 | Sar1a | protein_coding | 0.1120904 | 7.259951 | 0.1377458 | 0.7153099 | 0.999176 |
| ENSMUSG00000031979 | Cog2 | protein_coding | -0.1908215 | 4.659814 | 0.1374096 | 0.715641 | 0.999176 |
| ENSMUSG00000057093 | Zfp607b | protein_coding | 0.2659652 | 3.589036 | 0.1373617 | 0.7156883 | 0.999176 |
| ENSMUSG00000050721 | Plekho2 | protein_coding | -0.2073433 | 4.310671 | 0.1373555 | 0.7156943 | 0.999176 |
| ENSMUSG00000038005 | Hpf1 | protein_coding | 0.1244079 | 5.994375 | 0.1373186 | 0.7157307 | 0.999176 |
| ENSMUSG00000018395 | Kif3a | protein_coding | -0.1477347 | 7.687994 | 0.1372831 | 0.7157735 | 0.999176 |
| ENSMUSG00000045658 | Pid1 | protein_coding | 0.1907202 | 4.859348 | 0.1371998 | 0.7158479 | 0.999176 |
| ENSMUSG00000021262 | Evl | protein_coding | 0.2687145 | 3.886539 | 0.1371185 | 0.7159281 | 0.999176 |
| ENSMUSG00000037737 | Actrt3 | protein_coding | -0.5218864 | 2.315385 | 0.1371137 | 0.7159328 | 0.999176 |
| ENSMUSG00000059263 | Usp47 | protein_coding | 0.1496853 | 5.505801 | 0.1369325 | 0.7161117 | 0.999176 |
| ENSMUSG00000066152 | Slc31a2 | protein_coding | -0.2190266 | 3.622977 | 0.1368538 | 0.7161895 | 0.999176 |
| ENSMUSG00000042404 | Dennd4b | protein_coding | 0.1889778 | 4.202061 | 0.1368179 | 0.7162249 | 0.999176 |
| ENSMUSG00000037553 | Zdhhc18 | protein_coding | -0.2299391 | 4.081427 | 0.1367646 | 0.7162776 | 0.999176 |
| ENSMUSG00000002778 | Kdelr1 | protein_coding | 0.1892806 | 5.216956 | 0.1367624 | 0.7162798 | 0.999176 |
| ENSMUSG00000025465 | Echs1 | protein_coding | -0.1020213 | 7.770651 | 0.1367624 | 0.7162798 | 0.999176 |
| ENSMUSG00000021180 | Rps6ka5 | protein_coding | -0.2840826 | 4.180149 | 0.1366879 | 0.7163734 | 0.999176 |
| ENSMUSG00000032525 | Nktr | protein_coding | 0.2639455 | 4.031667 | 0.1366602 | 0.7163809 | 0.999176 |
| ENSMUSG00000002329 | Mdp1 | protein_coding | -0.103894 | 7.037462 | 0.1365729 | 0.7164671 | 0.999176 |
| ENSMUSG00000010136 | Pifo | protein_coding | -0.1883338 | 4.485324 | 0.1365641 | 0.7164759 | 0.999176 |
| ENSMUSG00000048376 | F2r | protein_coding | -0.3743606 | 2.762144 | 0.1364945 | 0.7165447 | 0.999176 |
| ENSMUSG00000060126 | Tpt1 | protein_coding | 0.2372246 | 3.675309 | 0.1364595 | 0.7165793 | 0.999176 |
| ENSMUSG00000027822 | Slc33a1 | protein_coding | -0.1959901 | 4.249301 | 0.1364369 | 0.7166017 | 0.999176 |
| ENSMUSG00000044715 | Gskip | protein_coding | -0.1074064 | 7.505913 | 0.136303 | 0.7167343 | 0.999176 |
| ENSMUSG00000025231 | Sufu | protein_coding | -0.3182144 | 3.201262 | 0.1362401 | 0.7167965 | 0.999176 |
| ENSMUSG00000032420 | Nt5e | protein_coding | 0.3898567 | 2.641089 | 0.1362357 | 0.7168009 | 0.999176 |
| ENSMUSG00000019960 | Dusp6 | protein_coding | 0.1754178 | 5.189858 | 0.1362163 | 0.7168201 | 0.999176 |
| ENSMUSG00000043733 | Ptpn11 | protein_coding | -0.187152 | 5.782792 | 0.1362089 | 0.7168352 | 0.999176 |
| ENSMUSG00000050332 | Amer1 | protein_coding | -0.3316121 | 3.496552 | 0.1359269 | 0.7171148 | 0.999176 |
| ENSMUSG00000058192 | Zfp846 | protein_coding | -0.2387212 | 4.151515 | 0.1357748 | 0.7172579 | 0.999176 |
| ENSMUSG00000043154 | Ppp2r3a | protein_coding | -0.1884244 | 4.875236 | 0.1357082 | 0.7173239 | 0.999176 |
| ENSMUSG00000030088 | Aldh1l1 | protein_coding | -0.2622838 | 4.311707 | 0.1354492 | 0.717589 | 0.999176 |
| ENSMUSG00000031099 | Smarca1 | protein_coding | 0.1741018 | 5.353965 | 0.1351725 | 0.7178563 | 0.999176 |
| ENSMUSG00000022995 | Enah | protein_coding | 0.1188176 | 6.376214 | 0.1351542 | 0.7178745 | 0.999176 |
| ENSMUSG00000003604 | Aven | protein_coding | -0.2405049 | 4.14355 | 0.1349649 | 0.7180629 | 0.999176 |
| ENSMUSG00000056917 | Sipa1 | protein_coding | -0.2438321 | 4.245049 | 0.134659 | 0.7183678 | 0.999176 |
| ENSMUSG00000032883 | Acsl3 | protein_coding | -0.1601304 | 5.964221 | 0.13454 | 0.7184864 | 0.999176 |
| ENSMUSG00000020477 | Mrps24 | protein_coding | 0.2077501 | 4.177255 | 0.134458 | 0.7185682 | 0.999176 |
| ENSMUSG00000034006 | Pqlc1 | protein_coding | 0.186538 | 5.558147 | 0.134424 | 0.7186022 | 0.999176 |
| ENSMUSG00000031095 | Cul4b | protein_coding | -0.2504164 | 3.859404 | 0.1343509 | 0.7186752 | 0.999176 |
| ENSMUSG00000033906 | Zdhhc15 | protein_coding | -0.4485362 | 2.513277 | 0.1338777 | 0.7191481 | 0.999176 |
| ENSMUSG00000032599 | Ip6k2 | protein_coding | 0.1159936 | 6.790141 | 0.1336626 | 0.7193634 | 0.999176 |
| ENSMUSG00000029916 | Agk | protein_coding | -0.157524 | 4.775771 | 0.1336119 | 0.7194142 | 0.999176 |
| ENSMUSG00000033768 | Nrxn2 | protein_coding | 0.1805742 | 4.840511 | 0.1335851 | 0.719441 | 0.999176 |
| ENSMUSG00000000120 | Ngfr | protein_coding | -0.3348165 | 2.713912 | 0.1335681 | 0.719458 | 0.999176 |
| ENSMUSG00000024985 | Tcf7l2 | protein_coding | -0.522018 | 8.165222 | 0.1384065 | 0.7196706 | 0.999176 |
| ENSMUSG00000029167 | Ppargc1a | protein_coding | -0.1554391 | 5.318008 | 0.1332143 | 0.7198127 | 0.999176 |
| ENSMUSG00000040374 | Pex2 | protein_coding | -0.1173702 | 6.584902 | 0.1330657 | 0.7199619 | 0.999176 |
| ENSMUSG00000026643 | Nmt2 | protein_coding | -0.1494761 | 8.632762 | 0.1330357 | 0.7199996 | 0.999176 |
| ENSMUSG00000024251 | Thada | protein_coding | 0.2513339 | 3.259668 | 0.1329542 | 0.7200737 | 0.999176 |
| ENSMUSG00000020802 | Ube2o | protein_coding | 0.1678808 | 4.962172 | 0.1328296 | 0.7201989 | 0.999176 |
| ENSMUSG00000027550 | Lrrcc1 | protein_coding | -0.4076808 | 4.236048 | 0.1337744 | 0.7202241 | 0.999176 |
| ENSMUSG00000042745 | Id1 | protein_coding | -0.2207041 | 3.894868 | 0.1328022 | 0.7202265 | 0.999176 |
| ENSMUSG00000054770 | Kctd18 | protein_coding | 0.2795822 | 3.477737 | 0.1327959 | 0.7202328 | 0.999176 |
| ENSMUSG00000034071 | Zfp551 | protein_coding | 0.5996899 | 2.569406 | 0.1329 | 0.7204265 | 0.999176 |
| ENSMUSG00000037470 | Uggt1 | protein_coding | -0.2164542 | 4.094917 | 0.1323635 | 0.7206678 | 0.999176 |
| ENSMUSG00000029319 | Coq2 | protein_coding | -0.1635877 | 4.903824 | 0.1323491 | 0.7206823 | 0.999176 |
| ENSMUSG00000018326 | Ywhab | protein_coding | 0.1013152 | 9.025884 | 0.1321684 | 0.7208643 | 0.999176 |
| ENSMUSG00000072494 | Ppp1r3e | protein_coding | 0.2702671 | 3.247804 | 0.1321684 | 0.7208644 | 0.999176 |
| ENSMUSG00000027500 | Stmn2 | protein_coding | -0.1070573 | 10.26119 | 0.1321083 | 0.7209249 | 0.999176 |
| ENSMUSG00000002741 | Ykt6 | protein_coding | -0.1759159 | 5.749169 | 0.1320703 | 0.7209632 | 0.999176 |
| ENSMUSG00000062867 | Impdh2 | protein_coding | 0.2137763 | 4.062558 | 0.1320388 | 0.720995 | 0.999176 |
| ENSMUSG00000042709 | Atpaf2 | protein_coding | -0.1556882 | 5.953941 | 0.1317658 | 0.7212704 | 0.999176 |
| ENSMUSG00000026150 | Mff | protein_coding | -0.1256256 | 7.330525 | 0.1317637 | 0.7212725 | 0.999176 |
| ENSMUSG00000035437 | Rabgap1 | protein_coding | -0.3063831 | 3.378998 | 0.1317101 | 0.7213266 | 0.999176 |
| ENSMUSG00000023932 | Cdc5l | protein_coding | -0.1446289 | 5.338713 | 0.1316437 | 0.7213937 | 0.999176 |
| ENSMUSG00000029089 | 5730480H06Rik | protein_coding | 0.2754208 | 3.863149 | 0.1313914 | 0.7216563 | 0.999176 |
| ENSMUSG00000028655 | Mfsd2a | protein_coding | 0.6362028 | 2.405621 | 0.1313577 | 0.7219791 | 0.999176 |
| ENSMUSG00000032470 | Mras | protein_coding | 0.1172472 | 6.058891 | 0.1309292 | 0.7221165 | 0.999176 |
| ENSMUSG00000021731 | Mrps30 | protein_coding | 0.2282967 | 4.13705 | 0.1307119 | 0.7223367 | 0.999176 |
| ENSMUSG00000003810 | Mast2 | protein_coding | -0.1457335 | 5.41546 | 0.1306936 | 0.7223554 | 0.999176 |
| ENSMUSG00000009894 | Snap47 | protein_coding | 0.1525057 | 10.21677 | 0.130676 | 0.7223808 | 0.999176 |
| ENSMUSG00000000325 | Arvcf | protein_coding | -0.2302953 | 3.429181 | 0.1305248 | 0.7225266 | 0.999176 |
| ENSMUSG00000038467 | Chmp4b | protein_coding | -0.1688991 | 5.568378 | 0.1305218 | 0.7225296 | 0.999176 |
| ENSMUSG00000053929 | Cyhr1 | protein_coding | 0.1075352 | 6.914207 | 0.1302848 | 0.7227703 | 0.999176 |
| ENSMUSG00000038473 | Nos1ap | protein_coding | 0.1476948 | 5.935122 | 0.1301243 | 0.7229335 | 0.999176 |

| ensembl_gene_id | external_gene_name | gene_biotype | log2 (fold change) | log2 (counts per million) | F-value | p-value | False discovery rate |
|---|---|---|---|---|---|---|---|
| ENSMUSG00000019951 | Uhrf1bp1l | protein_coding | 0.146432 | 5.28349 | 0.1300854 | 0.722973 | 0.999176 |
| ENSMUSG00000048747 | E130114P18Rik | protein_coding | 0.2505233 | 4.640998 | 0.1300196 | 0.7230594 | 0.999176 |
| ENSMUSG00000035953 | Tmem55b | protein_coding | 0.1601618 | 5.558918 | 0.1299794 | 0.7230808 | 0.999176 |
| ENSMUSG00000026335 | Pam | protein_coding | 0.1399451 | 6.194626 | 0.1298887 | 0.7231731 | 0.999176 |
| ENSMUSG00000040297 | Suco | protein_coding | -0.1761806 | 4.478072 | 0.1298182 | 0.7232449 | 0.999176 |
| ENSMUSG00000045983 | Eif4g1 | protein_coding | 0.1587207 | 5.209677 | 0.1297665 | 0.7232975 | 0.999176 |
| ENSMUSG00000045691 | Thtpa | protein_coding | -0.2200557 | 5.335795 | 0.129778 | 0.7233265 | 0.999176 |
| ENSMUSG00000042699 | Dhx9 | protein_coding | -0.1940715 | 4.60899 | 0.1294121 | 0.7236587 | 0.999176 |
| ENSMUSG00000097908 | 4933404O12Rik | lincRNA | 0.2873055 | 3.217325 | 0.1292937 | 0.7237794 | 0.999176 |
| ENSMUSG00000022634 | Yaf2 | protein_coding | -0.1157464 | 6.893969 | 0.1292248 | 0.7238498 | 0.999176 |
| ENSMUSG00000026576 | Atp1b1 | protein_coding | -0.1356719 | 8.629833 | 0.1292246 | 0.72385 | 0.999176 |
| ENSMUSG00000022464 | Slc38a4 | protein_coding | 0.3818173 | 2.634928 | 0.1289819 | 0.7240979 | 0.999176 |
| ENSMUSG00000004069 | Dnaja3 | protein_coding | -0.1345886 | 6.564819 | 0.1289046 | 0.7241769 | 0.999176 |
| ENSMUSG00000101930 | Gm5441 | antisense_RNA | 0.1205331 | 6.242236 | 0.1288864 | 0.7241954 | 0.999176 |
| ENSMUSG00000027296 | Itpka | protein_coding | 0.247902 | 5.082096 | 0.1291484 | 0.7242035 | 0.999176 |
| ENSMUSG00000021111 | Papola | protein_coding | 0.1379825 | 6.108188 | 0.1288726 | 0.7242096 | 0.999176 |
| ENSMUSG00000020308 | Tpgs1 | protein_coding | -0.3066253 | 4.572861 | 0.1293145 | 0.7242187 | 0.999176 |
| ENSMUSG00000028194 | Ddah1 | protein_coding | -0.1341248 | 7.196138 | 0.1287631 | 0.7243216 | 0.999176 |
| ENSMUSG00000029196 | Tada2b | protein_coding | -0.2375138 | 3.571372 | 0.128726 | 0.7243595 | 0.999176 |
| ENSMUSG00000030435 | U2af2 | protein_coding | -0.2382315 | 3.44095 | 0.1285474 | 0.7245422 | 0.999176 |
| ENSMUSG00000025980 | Hspd1 | protein_coding | -0.1158714 | 7.043086 | 0.1285461 | 0.7245436 | 0.999176 |
| ENSMUSG00000024846 | Cst6 | protein_coding | 0.1873994 | 4.547372 | 0.1285064 | 0.7245842 | 0.999176 |
| ENSMUSG00000050989 | Selenon | protein_coding | -0.3108299 | 3.328667 | 0.1283178 | 0.7247775 | 0.999176 |
| ENSMUSG00000060743 | H3f3a | protein_coding | -0.1905917 | 3.826425 | 0.1282927 | 0.7248032 | 0.999176 |
| ENSMUSG00000021051 | Ppp2r5e | protein_coding | 0.1290498 | 5.915104 | 0.1281835 | 0.7249151 | 0.999176 |
| ENSMUSG00000061024 | Rrs1 | protein_coding | -0.2427744 | 3.855579 | 0.1279094 | 0.7251964 | 0.999176 |
| ENSMUSG00000050272 | Dscam | protein_coding | 0.1777622 | 4.946494 | 0.1278802 | 0.7252264 | 0.999176 |
| ENSMUSG00000007564 | Ppp2r1a | protein_coding | 0.1281068 | 9.864952 | 0.1275981 | 0.7255162 | 0.999176 |
| ENSMUSG00000021770 | Samd8 | protein_coding | 0.1627218 | 5.08333 | 0.127537 | 0.7255791 | 0.999176 |
| ENSMUSG00000075020 | Mir670hg | lincRNA | -0.2604681 | 4.707948 | 0.1276972 | 0.7256064 | 0.999176 |
| ENSMUSG00000000346 | Dazap2 | protein_coding | -0.1096846 | 8.57023 | 0.1274229 | 0.7256964 | 0.999176 |
| ENSMUSG00000013701 | Timm23 | protein_coding | -0.2616955 | 3.061887 | 0.1273483 | 0.7257732 | 0.999176 |
| ENSMUSG00000053838 | Nudcd3 | protein_coding | 0.146651 | 7.133424 | 0.127234 | 0.7258983 | 0.999176 |
| ENSMUSG00000056476 | Med12l | protein_coding | 0.2561413 | 3.301636 | 0.1271225 | 0.7260057 | 0.999176 |
| ENSMUSG00000032187 | Smarca4 | protein_coding | 0.1632421 | 5.949081 | 0.1270869 | 0.7260424 | 0.999176 |
| ENSMUSG00000025024 | Smndc1 | protein_coding | -0.1781645 | 4.371693 | 0.126849 | 0.7262877 | 0.999176 |
| ENSMUSG00000113701 | AC121583.1 | transcribed_processe | -0.1769637 | 4.360973 | 0.1267498 | 0.72639 | 0.999176 |
| ENSMUSG00000061451 | Tmem151a | protein_coding | 0.1468107 | 5.821231 | 0.1267063 | 0.7264349 | 0.999176 |
| ENSMUSG00000036167 | Pphln1 | protein_coding | -0.1592499 | 5.291291 | 0.1266217 | 0.7265222 | 0.999176 |
| ENSMUSG00000031104 | Rab33a | protein_coding | 0.1191061 | 6.802207 | 0.126531 | 0.726616 | 0.999176 |
| ENSMUSG00000031568 | Rwdd4a | protein_coding | -0.1666839 | 4.719291 | 0.1265181 | 0.7266293 | 0.999176 |
| ENSMUSG00000057182 | Scn3a | protein_coding | 0.1405944 | 5.406071 | 0.1262781 | 0.7268773 | 0.999176 |
| ENSMUSG00000005625 | Psmd4 | protein_coding | 0.1060842 | 8.64351 | 0.1261611 | 0.7269983 | 0.999176 |
| ENSMUSG00000030614 | Tmem126b | protein_coding | 0.1147007 | 7.484927 | 0.1260966 | 0.7270652 | 0.999176 |
| ENSMUSG00000023043 | Krt18 | protein_coding | -0.3610512 | 3.282091 | 0.1259877 | 0.7271854 | 0.999176 |
| ENSMUSG00000037235 | Mxd4 | protein_coding | -0.2070751 | 4.331217 | 0.1259114 | 0.7272568 | 0.999176 |
| ENSMUSG00000028293 | Slc35a1 | protein_coding | 0.1566699 | 5.190767 | 0.1256921 | 0.7274842 | 0.999176 |
| ENSMUSG00000036160 | Surf6 | protein_coding | 0.2196456 | 4.498405 | 0.1254944 | 0.7276893 | 0.999176 |
| ENSMUSG00000021694 | Ercc8 | protein_coding | -0.2353951 | 4.122049 | 0.1250077 | 0.728195 | 0.999176 |
| ENSMUSG00000000581 | C1d | protein_coding | 0.1211956 | 9.17731 | 0.1249653 | 0.728239 | 0.999176 |
| ENSMUSG00000029290 | Zfp326 | protein_coding | -0.1926377 | 4.10124 | 0.1249209 | 0.7282852 | 0.999176 |
| ENSMUSG00000040029 | Ipo8 | protein_coding | -0.1501295 | 5.019495 | 0.1245671 | 0.7286537 | 0.999176 |
| ENSMUSG00000081552 | Gm12328 | processed_pseudoge | 0.3208737 | 2.681846 | 0.1244254 | 0.7288014 | 0.999176 |
| ENSMUSG00000029392 | Rilpl1 | protein_coding | -0.1977106 | 4.592945 | 0.1243601 | 0.7288695 | 0.999176 |
| ENSMUSG00000031948 | Kars | protein_coding | 0.1516341 | 7.945595 | 0.1243854 | 0.7288701 | 0.999176 |
| ENSMUSG00000041375 | Ccdc9 | protein_coding | 0.2726638 | 3.067043 | 0.1243255 | 0.7289056 | 0.999176 |
| ENSMUSG00000038160 | Atg5 | protein_coding | -0.1098336 | 6.381313 | 0.1242754 | 0.7289579 | 0.999176 |
| ENSMUSG00000032714 | Syde1 | protein_coding | -0.4938364 | 2.675543 | 0.1242948 | 0.7289646 | 0.999176 |
| ENSMUSG00000026657 | Frmd4a | protein_coding | -0.1657026 | 4.795458 | 0.124226 | 0.7290094 | 0.999176 |
| ENSMUSG00000048920 | Fkrp | protein_coding | -0.1671645 | 4.622353 | 0.1242019 | 0.7290346 | 0.999176 |
| ENSMUSG00000048175 | Asb8 | protein_coding | 0.1263745 | 7.244738 | 0.1241997 | 0.7290369 | 0.999176 |
| ENSMUSG00000022620 | Arsa | protein_coding | -0.1990266 | 4.693227 | 0.1241477 | 0.7290912 | 0.999176 |
| ENSMUSG00000030714 | Sgf29 | protein_coding | -0.1640873 | 5.825383 | 0.1241189 | 0.7291212 | 0.999176 |
| ENSMUSG00000052798 | Nup107 | protein_coding | -0.1753715 | 4.67763 | 0.123937 | 0.7293113 | 0.999176 |
| ENSMUSG00000034509 | Mad2l1bp | protein_coding | -0.1504496 | 5.073676 | 0.1238773 | 0.7293737 | 0.999176 |
| ENSMUSG00000060671 | Atp8b2 | protein_coding | -0.2488639 | 3.471157 | 0.1238416 | 0.729411 | 0.999176 |
| ENSMUSG00000074461 | Gm10699 | TEC | -0.1237694 | 5.997096 | 0.1237416 | 0.7295156 | 0.999176 |
| ENSMUSG00000031059 | Ndufb11 | protein_coding | 0.1279678 | 9.367675 | 0.123675 | 0.7295853 | 0.999176 |
| ENSMUSG00000053870 | Fpgt | protein_coding | 0.2689595 | 3.953646 | 0.123502 | 0.7297738 | 0.999176 |
| ENSMUSG00000025137 | Pcyt2 | protein_coding | 0.1289639 | 7.272872 | 0.1233306 | 0.7299459 | 0.999176 |
| ENSMUSG00000028656 | Cap1 | protein_coding | -0.139342 | 6.506581 | 0.1233299 | 0.7299467 | 0.999176 |
| ENSMUSG00000055044 | Pdlim1 | protein_coding | -0.3652089 | 2.578696 | 0.1232727 | 0.7300067 | 0.999176 |
| ENSMUSG00000097195 | Snhg5 | processed_transcript | 0.1939766 | 4.857033 | 0.1232212 | 0.7300607 | 0.999176 |
| ENSMUSG00000021254 | Gpatch2l | protein_coding | -0.2935736 | 3.772627 | 0.1231741 | 0.7301174 | 0.999176 |
| ENSMUSG00000018995 | Nars2 | protein_coding | 0.2275619 | 3.68733 | 0.1231439 | 0.7301418 | 0.999176 |
| ENSMUSG00000030751 | Psma1 | protein_coding | -0.1067416 | 8.507515 | 0.122997 | 0.7302959 | 0.999176 |
| ENSMUSG00000057531 | Dtnbp1 | protein_coding | 0.1247238 | 6.867281 | 0.1229164 | 0.7303806 | 0.999176 |
| ENSMUSG00000024533 | Spire1 | protein_coding | -0.1546431 | 5.694342 | 0.1228332 | 0.7304679 | 0.999176 |

| ensembl_gene_id | external_gene_name | gene_biotype | log2 (fold change) | log2 (counts per million) | F-value | p-value | False discovery rate |
|---|---|---|---|---|---|---|---|
| ENSMUSG00000003527 | Dgcr14 | protein_coding | 0.154417 | 5.29142 | 0.1228071 | 0.7304954 | 0.999176 |
| ENSMUSG00000042148 | Cox10 | protein_coding | -0.1488753 | 5.255918 | 0.1226407 | 0.7306702 | 0.999176 |
| ENSMUSG00000021192 | Golga5 | protein_coding | -0.2154888 | 3.686893 | 0.122597 | 0.7307161 | 0.999176 |
| ENSMUSG00000018651 | Tada2a | protein_coding | 0.1950727 | 5.035906 | 0.122567 | 0.7307477 | 0.999176 |
| ENSMUSG00000037151 | Lrrc20 | protein_coding | 0.1867538 | 4.799613 | 0.1224091 | 0.7309139 | 0.999176 |
| ENSMUSG00000001334 | Fndc5 | protein_coding | -0.1736653 | 5.763481 | 0.1223966 | 0.7309344 | 0.999176 |
| ENSMUSG00000002731 | Prkra | protein_coding | -0.1250236 | 6.252493 | 0.1223434 | 0.7309831 | 0.999176 |
| ENSMUSG00000021327 | Zkscan3 | protein_coding | 0.2131365 | 3.967107 | 0.122329 | 0.7309982 | 0.999176 |
| ENSMUSG00000037275 | Gemin5 | protein_coding | -0.2480838 | 3.516746 | 0.1222316 | 0.7311008 | 0.999176 |
| ENSMUSG00000022682 | Rrn3 | protein_coding | 0.1982572 | 7.475948 | 0.1227066 | 0.7312627 | 0.999176 |
| ENSMUSG00000047719 | Ubiad1 | protein_coding | 0.2546468 | 3.207248 | 0.1220305 | 0.7313128 | 0.999176 |
| ENSMUSG00000035713 | Usp35 | protein_coding | -0.2649982 | 3.237843 | 0.1218836 | 0.7314677 | 0.999176 |
| ENSMUSG00000039725 | Trp53rka | protein_coding | 0.3355514 | 3.216719 | 0.121881 | 0.7314704 | 0.999176 |
| ENSMUSG00000029576 | Radil | protein_coding | 0.2646465 | 3.178133 | 0.1215953 | 0.731772 | 0.999176 |
| ENSMUSG00000024068 | Spast | protein_coding | 0.2079844 | 4.623685 | 0.121509 | 0.7318632 | 0.999176 |
| ENSMUSG00000049421 | Zfp260 | protein_coding | 0.1561545 | 5.497328 | 0.1213731 | 0.7320069 | 0.999176 |
| ENSMUSG00000028488 | Sh3gl2 | protein_coding | 0.115726 | 7.615827 | 0.1212429 | 0.7321446 | 0.999176 |
| ENSMUSG00000033545 | Znrf1 | protein_coding | -0.1484087 | 5.488763 | 0.1211032 | 0.7322926 | 0.999176 |
| ENSMUSG00000020549 | Elac2 | protein_coding | 0.234274 | 3.621125 | 0.1210936 | 0.7323027 | 0.999176 |
| ENSMUSG00000049091 | Sephs2 | protein_coding | 0.21506 | 5.725464 | 0.1213242 | 0.7323423 | 0.999176 |
| ENSMUSG00000045333 | Zfp423 | protein_coding | -0.2032566 | 4.549716 | 0.1210329 | 0.732367 | 0.999176 |
| ENSMUSG00000018378 | Cuedc1 | protein_coding | 0.2105677 | 3.943363 | 0.1208574 | 0.732553 | 0.999176 |
| ENSMUSG00000027534 | Snx16 | protein_coding | -0.3012701 | 3.75528 | 0.1208037 | 0.7326365 | 0.999176 |
| ENSMUSG00000037361 | Sf3b6 | protein_coding | -0.1180221 | 8.375933 | 0.1206944 | 0.7327259 | 0.999176 |
| ENSMUSG00000093738 | AI606473 | processed_transcript | 0.2932094 | 3.792607 | 0.120672 | 0.732757 | 0.999176 |
| ENSMUSG00000075284 | Wipf1 | protein_coding | -0.2596 | 3.142378 | 0.1206493 | 0.7327738 | 0.999176 |
| ENSMUSG00000024261 | Syt4 | protein_coding | -0.1097464 | 8.374893 | 0.120606 | 0.7328197 | 0.999176 |
| ENSMUSG00000027994 | Mcub | protein_coding | 0.3137434 | 3.540386 | 0.1206107 | 0.732822 | 0.999176 |
| ENSMUSG00000009394 | Syn2 | protein_coding | 0.1046492 | 8.464177 | 0.1205391 | 0.7328907 | 0.999176 |
| ENSMUSG00000034940 | Synrg | protein_coding | -0.1252456 | 6.096106 | 0.1204242 | 0.7330127 | 0.999176 |
| ENSMUSG00000021013 | Ttc8 | protein_coding | -0.144728 | 5.003376 | 0.1202975 | 0.7331474 | 0.999176 |
| ENSMUSG00000062270 | Morf4l1 | protein_coding | -0.1123235 | 6.593493 | 0.1201466 | 0.7333078 | 0.999176 |
| ENSMUSG00000006699 | Cdc42 | protein_coding | -0.09579156 | 9.861276 | 0.1200889 | 0.7333692 | 0.999176 |
| ENSMUSG00000030532 | Hddc3 | protein_coding | 0.2164387 | 5.144441 | 0.1200517 | 0.7334274 | 0.999176 |
| ENSMUSG00000025856 | Pdgfa | protein_coding | 0.1476613 | 5.019665 | 0.119922 | 0.7335468 | 0.999176 |
| ENSMUSG00000053329 | D10Jhu81e | protein_coding | 0.1560558 | 5.636115 | 0.1198924 | 0.7335783 | 0.999176 |
| ENSMUSG00000031691 | Tnpo2 | protein_coding | 0.3005139 | 3.370183 | 0.1197013 | 0.7337819 | 0.999176 |
| ENSMUSG00000038290 | Smg6 | protein_coding | -0.1726117 | 4.756265 | 0.1196818 | 0.7338027 | 0.999176 |
| ENSMUSG00000018666 | Cbx1 | protein_coding | 0.1110003 | 6.193501 | 0.1195308 | 0.7339638 | 0.999176 |
| ENSMUSG00000042408 | Zmym6 | protein_coding | 0.3064306 | 2.984924 | 0.119454 | 0.7340457 | 0.999176 |
| ENSMUSG00000015846 | Rxra | protein_coding | 0.1515111 | 4.583473 | 0.1192429 | 0.7342712 | 0.999176 |
| ENSMUSG00000037519 | Ppfia1 | protein_coding | 0.2218012 | 3.816122 | 0.1191844 | 0.7343337 | 0.999176 |
| ENSMUSG00000038344 | Txlng | protein_coding | -0.2386603 | 3.688827 | 0.1191325 | 0.7343891 | 0.999176 |
| ENSMUSG00000081051 | Gm15427 | processed_pseudoge | 0.281432 | 2.959743 | 0.1189574 | 0.7345763 | 0.999176 |
| ENSMUSG00000054309 | Cpsf3 | protein_coding | -0.1508489 | 6.587729 | 0.1189263 | 0.7346168 | 0.999176 |
| ENSMUSG00000074170 | Plekhf1 | protein_coding | -0.2651259 | 3.880265 | 0.1189035 | 0.7346412 | 0.999176 |
| ENSMUSG00000030598 | Fbxo17 | protein_coding | 0.2292482 | 4.047246 | 0.1187979 | 0.7347469 | 0.999176 |
| ENSMUSG00000087620 | 5330434G04Rik | lincRNA | 0.206887 | 4.666726 | 0.1187839 | 0.734762 | 0.999176 |
| ENSMUSG00000055409 | Nell1 | protein_coding | -0.1168652 | 5.86049 | 0.1186476 | 0.7349079 | 0.999176 |
| ENSMUSG00000008682 | Rpl10 | protein_coding | -0.2633985 | 3.033878 | 0.11858 | 0.7349803 | 0.999176 |
| ENSMUSG00000025404 | R3hdm2 | protein_coding | 0.1294136 | 5.633321 | 0.1184878 | 0.7350791 | 0.999176 |
| ENSMUSG00000024360 | Etf1 | protein_coding | -0.1247703 | 6.051335 | 0.11847 | 0.7350982 | 0.999176 |
| ENSMUSG00000033569 | Adgrb3 | protein_coding | -0.1696367 | 5.96116 | 0.1184317 | 0.7351465 | 0.999176 |
| ENSMUSG00000063568 | Jazf1 | protein_coding | 0.159201 | 4.730185 | 0.1182139 | 0.735373 | 0.999176 |
| ENSMUSG00000007594 | Hapln4 | protein_coding | 0.1680252 | 5.143696 | 0.1181669 | 0.7354234 | 0.999176 |
| ENSMUSG00000060843 | Ctnna3 | protein_coding | -0.2734469 | 2.978928 | 0.1176106 | 0.7360215 | 0.999176 |
| ENSMUSG00000040396 | Abhd13 | protein_coding | -0.2115338 | 3.696865 | 0.117568 | 0.7360674 | 0.999176 |
| ENSMUSG00000041986 | Elmod1 | protein_coding | -0.189921 | 6.195305 | 0.1177726 | 0.7361263 | 0.999176 |
| ENSMUSG00000073293 | Nudt10 | protein_coding | -0.1880715 | 5.579234 | 0.1175051 | 0.7361423 | 0.999176 |
| ENSMUSG00000020611 | Gna13 | protein_coding | -0.1496183 | 5.400921 | 0.1174493 | 0.7361953 | 0.999176 |
| ENSMUSG00000056579 | Tug1 | processed_transcript | -0.1852964 | 6.521282 | 0.1177757 | 0.7362124 | 0.999176 |
| ENSMUSG00000017390 | Aldoc | protein_coding | -0.1094932 | 10.98812 | 0.117196 | 0.7364683 | 0.999176 |
| ENSMUSG00000059602 | Syn3 | protein_coding | 0.2516952 | 3.615109 | 0.1171251 | 0.7365448 | 0.999176 |
| ENSMUSG00000032109 | Nlrx1 | protein_coding | 0.5176374 | 2.261596 | 0.1170343 | 0.7366427 | 0.999176 |
| ENSMUSG00000031451 | Gas6 | protein_coding | -0.3150455 | 2.816423 | 0.1170162 | 0.7366623 | 0.999176 |
| ENSMUSG00000028909 | Ptpru | protein_coding | -0.2879872 | 3.507828 | 0.1167992 | 0.7368967 | 0.999176 |
| ENSMUSG00000041638 | Gcn1l1 | protein_coding | -0.1755125 | 4.073845 | 0.1167808 | 0.7369166 | 0.999176 |
| ENSMUSG00000030801 | Kat8 | protein_coding | -0.1542226 | 4.772854 | 0.1167294 | 0.7369722 | 0.999176 |
| ENSMUSG00000029661 | Col1a2 | protein_coding | 0.3885733 | 3.187938 | 0.1167903 | 0.7370516 | 0.999176 |
| ENSMUSG00000010097 | Nxf1 | protein_coding | 0.09380267 | 7.710549 | 0.1164999 | 0.7372204 | 0.999176 |
| ENSMUSG00000096257 | Ccer2 | protein_coding | -0.3673937 | 2.987225 | 0.1164926 | 0.7372355 | 0.999176 |
| ENSMUSG00000026077 | Npas2 | protein_coding | -0.2018981 | 3.989063 | 0.1164557 | 0.7372682 | 0.999176 |
| ENSMUSG00000113216 | AL590503.1 | lincRNA | 0.2285695 | 4.160543 | 0.1164333 | 0.7372925 | 0.999176 |
| ENSMUSG00000085139 | A730046J19Rik | protein_coding | 0.437675 | 2.27924 | 0.116301 | 0.7374358 | 0.999176 |
| ENSMUSG00000060260 | Pwwp2b | protein_coding | -0.2220631 | 4.463805 | 0.1162415 | 0.7375002 | 0.999176 |
| ENSMUSG00000099689 | Zfp383 | protein_coding | 0.2529955 | 3.063736 | 0.1162162 | 0.7375276 | 0.999176 |
| ENSMUSG00000059981 | Taok2 | protein_coding | -0.1942223 | 4.662938 | 0.1161764 | 0.7375708 | 0.999176 |
| ENSMUSG00000036810 | Cnep1r1 | protein_coding | -0.151007 | 5.715661 | 0.1161131 | 0.7376394 | 0.999176 |

| ensembl_gene_id | external_gene_name | gene_biotype | log2 (fold change) | log2 (counts per million) | F-value | p-value | False discovery rate |
|---|---|---|---|---|---|---|---|
| ENSMUSG00000026279 | Thap4 | protein_coding | 0.1482632 | 4.762738 | 0.1160626 | 0.7376942 | 0.999176 |
| ENSMUSG00000042680 | Garem1 | protein_coding | -0.2306262 | 3.877893 | 0.116044 | 0.7377143 | 0.999176 |
| ENSMUSG00000041268 | Dmxl2 | protein_coding | -0.143967 | 5.706292 | 0.1160246 | 0.7377354 | 0.999176 |
| ENSMUSG00000025885 | Myo5b | protein_coding | 0.1856548 | 4.374766 | 0.1160119 | 0.7377492 | 0.999176 |
| ENSMUSG00000073633 | Fbxo36 | protein_coding | 0.1840729 | 4.715544 | 0.1159965 | 0.7377659 | 0.999176 |
| ENSMUSG00000051695 | Pcbp1 | protein_coding | -0.09860938 | 6.761843 | 0.1159372 | 0.7378302 | 0.999176 |
| ENSMUSG00000100975 | Gm28875 | lincRNA | -0.3459082 | 3.55139 | 0.116137 | 0.7378809 | 0.999176 |
| ENSMUSG00000030095 | Tmem43 | protein_coding | -0.1347872 | 5.636489 | 0.1158145 | 0.7379635 | 0.999176 |
| ENSMUSG00000027397 | Slc20a1 | protein_coding | -0.2946627 | 2.862421 | 0.1156228 | 0.7381717 | 0.999176 |
| ENSMUSG00000038374 | Rbm8a | protein_coding | 0.1019534 | 7.190097 | 0.1155981 | 0.7381986 | 0.999176 |
| ENSMUSG00000044408 | Sptssa | protein_coding | 0.1146565 | 6.727465 | 0.1155528 | 0.7382478 | 0.999176 |
| ENSMUSG00000063714 | Sp3os | antisense_RNA | 0.230674 | 3.530542 | 0.115471 | 0.7383367 | 0.999176 |
| ENSMUSG00000038368 | Focad | protein_coding | -0.3035858 | 2.994303 | 0.1154344 | 0.7383765 | 0.999176 |
| ENSMUSG00000049985 | Ankrd55 | protein_coding | 0.2633177 | 3.601224 | 0.1153548 | 0.7384632 | 0.999176 |
| ENSMUSG00000093436 | Gm20646 | sense_overlapping | -0.2086923 | 3.643241 | 0.1152639 | 0.7385621 | 0.999176 |
| ENSMUSG00000026034 | Clk1 | protein_coding | -0.1916523 | 5.304603 | 0.1151848 | 0.7386554 | 0.999176 |
| ENSMUSG00000034912 | Mdga2 | protein_coding | 0.1949482 | 5.501504 | 0.115125 | 0.7387205 | 0.999176 |
| ENSMUSG00000067336 | Bmpr2 | protein_coding | -0.1326338 | 5.50909 | 0.114914 | 0.7389434 | 0.999176 |
| ENSMUSG00000097164 | Cep83os | bidirectional_promote | -0.1488498 | 5.223949 | 0.1147946 | 0.7390737 | 0.999176 |
| ENSMUSG00000026655 | Fam107b | protein_coding | 0.423126 | 3.145437 | 0.1150237 | 0.7390994 | 0.999176 |
| ENSMUSG00000045969 | Ing1 | protein_coding | -0.1258297 | 5.778529 | 0.1146176 | 0.7392669 | 0.999176 |
| ENSMUSG00000030272 | Camk1 | protein_coding | 0.2019359 | 3.713427 | 0.1143303 | 0.7395809 | 0.999176 |
| ENSMUSG00000007570 | Fance | protein_coding | -0.3682805 | 2.876004 | 0.1142114 | 0.739711 | 0.999176 |
| ENSMUSG00000034653 | Ythdc2 | protein_coding | -0.2388747 | 3.328496 | 0.1141626 | 0.7397644 | 0.999176 |
| ENSMUSG00000032609 | Klhdc8b | protein_coding | 0.1570873 | 5.787174 | 0.1140373 | 0.7399016 | 0.999176 |
| ENSMUSG00000031479 | Vps36 | protein_coding | 0.1452151 | 5.319831 | 0.1139659 | 0.7399798 | 0.999176 |
| ENSMUSG00000032410 | Xrn1 | protein_coding | 0.2080569 | 4.021214 | 0.1139568 | 0.7399897 | 0.999176 |
| ENSMUSG00000015882 | Lcorl | protein_coding | 0.2738603 | 3.036229 | 0.1138531 | 0.7401035 | 0.999176 |
| ENSMUSG00000020375 | Rufy1 | protein_coding | 0.1539819 | 4.966513 | 0.1137813 | 0.7401821 | 0.999176 |
| ENSMUSG00000044950 | Pwwp2a | protein_coding | -0.1852842 | 4.588339 | 0.1137599 | 0.7402057 | 0.999176 |
| ENSMUSG00000038848 | Ythdf1 | protein_coding | 0.1772973 | 5.274372 | 0.1135341 | 0.7404535 | 0.999176 |
| ENSMUSG00000054648 | Zfp869 | protein_coding | 0.2433013 | 3.365948 | 0.1134951 | 0.7404962 | 0.999176 |
| ENSMUSG00000023206 | Il15ra | protein_coding | 0.2603386 | 3.39111 | 0.1134398 | 0.7405571 | 0.999176 |
| ENSMUSG00000091896 | Ube2d2a | protein_coding | -0.1302144 | 5.745542 | 0.1134216 | 0.740577 | 0.999176 |
| ENSMUSG00000034648 | Lrrn1 | protein_coding | -0.2019915 | 4.623744 | 0.1133042 | 0.740706 | 0.999176 |
| ENSMUSG00000043445 | Pgp | protein_coding | 0.1712141 | 4.355717 | 0.1131686 | 0.7408552 | 0.999176 |
| ENSMUSG00000042487 | Leo1 | protein_coding | -0.1676049 | 6.183719 | 0.1131577 | 0.7408927 | 0.999176 |
| ENSMUSG00000023192 | Grm2 | protein_coding | 0.3462332 | 3.081457 | 0.112905 | 0.7411453 | 0.999176 |
| ENSMUSG00000015852 | Fcrls | protein_coding | -0.4036647 | 3.534349 | 0.1131799 | 0.7413989 | 0.999176 |
| ENSMUSG00000032026 | Rexo2 | protein_coding | 0.1303515 | 5.948169 | 0.1123157 | 0.7417954 | 0.999176 |
| ENSMUSG00000022054 | Nefm | protein_coding | -0.2052384 | 7.01896 | 0.1129282 | 0.7418029 | 0.999176 |
| ENSMUSG00000021224 | Numb | protein_coding | 0.2038309 | 4.281775 | 0.1123009 | 0.7418117 | 0.999176 |
| ENSMUSG00000021144 | Mta1 | protein_coding | 0.1839143 | 5.067801 | 0.1122074 | 0.7419151 | 0.999176 |
| ENSMUSG00000052605 | Isoc2b | protein_coding | 0.1927256 | 4.50165 | 0.1121173 | 0.7420146 | 0.999176 |
| ENSMUSG00000006215 | Zbtb17 | protein_coding | 0.2495776 | 4.625696 | 0.1122794 | 0.7420237 | 0.999176 |
| ENSMUSG00000019853 | Hebp2 | protein_coding | -0.1845138 | 3.909299 | 0.1120762 | 0.7420601 | 0.999176 |
| ENSMUSG00000051853 | Arf3 | protein_coding | -0.1082649 | 9.245092 | 0.112038 | 0.7421023 | 0.999176 |
| ENSMUSG00000067194 | Eif1ax | protein_coding | 0.1024447 | 7.481457 | 0.1120331 | 0.7421078 | 0.999176 |
| ENSMUSG00000099964 | Gm29514 | antisense_RNA | -0.1874432 | 3.670834 | 0.112029 | 0.7421123 | 0.999176 |
| ENSMUSG00000027878 | Notch2 | protein_coding | 0.2864263 | 3.242618 | 0.1120102 | 0.7421331 | 0.999176 |
| ENSMUSG00000048120 | Entpd1 | protein_coding | 0.2238728 | 3.685527 | 0.1118749 | 0.7422829 | 0.999176 |
| ENSMUSG00000021550 | 2210016F16Rik | protein_coding | 0.2309947 | 3.754677 | 0.1117618 | 0.7424081 | 0.999176 |
| ENSMUSG00000090125 | Pou3f1 | protein_coding | 0.2080886 | 3.97771 | 0.1117473 | 0.7424242 | 0.999176 |
| ENSMUSG00000046562 | Unc119b | protein_coding | -0.2830955 | 3.136479 | 0.1115554 | 0.7426368 | 0.999176 |
| ENSMUSG00000035274 | Tpbg | protein_coding | -0.3075494 | 3.203808 | 0.1115513 | 0.7426414 | 0.999176 |
| ENSMUSG00000037166 | Ppp1r14a | protein_coding | 0.2456612 | 3.985494 | 0.1115198 | 0.7426833 | 0.999176 |
| ENSMUSG00000021792 | Fam213a | protein_coding | -0.1161418 | 7.844559 | 0.1113606 | 0.742853 | 0.999176 |
| ENSMUSG00000031703 | Itfg1 | protein_coding | -0.1422704 | 6.881238 | 0.1113523 | 0.7428691 | 0.999176 |
| ENSMUSG00000019139 | Isyna1 | protein_coding | 0.2022693 | 5.384045 | 0.1113528 | 0.742899 | 0.999176 |
| ENSMUSG00000087478 | 4930506C21Rik | processed_transcript | -0.338792 | 2.649937 | 0.111188 | 0.7430446 | 0.999176 |
| ENSMUSG00000019738 | Polr2i | protein_coding | 0.1297605 | 6.638654 | 0.1111541 | 0.7430823 | 0.999176 |
| ENSMUSG00000021577 | Sdha | protein_coding | -0.1457524 | 9.626183 | 0.1111099 | 0.7431383 | 0.999176 |
| ENSMUSG00000039768 | Dnajc11 | protein_coding | 0.1631978 | 6.270258 | 0.1111134 | 0.7431529 | 0.999176 |
| ENSMUSG00000021140 | Pcnx | protein_coding | 0.2514714 | 3.713047 | 0.1110167 | 0.743235 | 0.999176 |
| ENSMUSG00000022456 | 3-Sep | protein_coding | 0.1045809 | 8.375689 | 0.1110048 | 0.7432482 | 0.999176 |
| ENSMUSG00000020075 | Ddx21 | protein_coding | -0.1893903 | 3.940671 | 0.1108441 | 0.743427 | 0.999176 |
| ENSMUSG00000067578 | Cbln4 | protein_coding | 0.22989 | 4.364327 | 0.1105892 | 0.7437178 | 0.999176 |
| ENSMUSG00000016253 | Nelfcd | protein_coding | 0.1482988 | 6.811903 | 0.1105485 | 0.7437816 | 0.999176 |
| ENSMUSG00000021947 | Cryl1 | protein_coding | -0.1390095 | 5.011776 | 0.1104187 | 0.7439009 | 0.999176 |
| ENSMUSG00000025428 | Atp5a1 | protein_coding | 0.130822 | 10.55176 | 0.1103105 | 0.7440216 | 0.999176 |
| ENSMUSG00000066438 | Plekhd1 | protein_coding | -0.3747003 | 2.449474 | 0.1102807 | 0.7440549 | 0.999176 |
| ENSMUSG00000049148 | Plcxd3 | protein_coding | 0.167031 | 5.864057 | 0.1102636 | 0.7440809 | 0.999176 |
| ENSMUSG00000015597 | Zfp318 | protein_coding | -0.166864 | 4.926959 | 0.1101611 | 0.7441884 | 0.999176 |
| ENSMUSG00000023015 | Racgap1 | protein_coding | -0.2068898 | 3.922597 | 0.1099744 | 0.744397 | 0.999176 |
| ENSMUSG00000084215 | Gm11814 | processed_pseudoge | -0.2940438 | 3.05476 | 0.1098702 | 0.7445135 | 0.999176 |
| ENSMUSG00000050619 | Zscan29 | protein_coding | 0.3495084 | 2.682584 | 0.1098354 | 0.7445524 | 0.999176 |
| ENSMUSG00000032607 | Amt | protein_coding | 0.2582434 | 5.327477 | 0.1103622 | 0.7445569 | 0.999176 |
| ENSMUSG00000037957 | Wdr20 | protein_coding | -0.1898664 | 4.071078 | 0.1098239 | 0.7445653 | 0.999176 |

| ensembl_gene_id | external_gene_name | gene_biotype | log2 (fold change) | log2 (counts per million) | F-value | p-value | False discovery rate |
|---|---|---|---|---|---|---|---|
| ENSMUSG00000038860 | Garnl3 | protein_coding | 0.1725704 | 5.181698 | 0.1098202 | 0.7445694 | 0.999176 |
| ENSMUSG00000002332 | Dhrs1 | protein_coding | 0.1034443 | 7.310267 | 0.1096424 | 0.7447684 | 0.999176 |
| ENSMUSG00000021990 | Spata13 | protein_coding | 0.2122364 | 4.088362 | 0.1094414 | 0.7449936 | 0.999176 |
| ENSMUSG00000020589 | Fam49a | protein_coding | -0.2645823 | 3.251559 | 0.1093694 | 0.7450743 | 0.999176 |
| ENSMUSG00000038112 | AW551984 | protein_coding | -0.1605584 | 6.598811 | 0.1093445 | 0.7452787 | 0.999176 |
| ENSMUSG00000030621 | Me3 | protein_coding | 0.2126265 | 4.224413 | 0.1091051 | 0.7453709 | 0.999176 |
| ENSMUSG00000008200 | Fnbp4 | protein_coding | -0.1886354 | 3.868927 | 0.1090382 | 0.7454459 | 0.999176 |
| ENSMUSG00000021730 | Hcn1 | protein_coding | -0.1557566 | 4.651421 | 0.1089942 | 0.7454953 | 0.999176 |
| ENSMUSG00000003464 | Pex19 | protein_coding | -0.1194257 | 7.878206 | 0.108942 | 0.745554 | 0.999176 |
| ENSMUSG00000025812 | Pard3 | protein_coding | -0.1588228 | 4.74977 | 0.1087636 | 0.7457545 | 0.999176 |
| ENSMUSG00000034789 | Rab24 | protein_coding | 0.206792 | 4.431595 | 0.1086549 | 0.7458769 | 0.999176 |
| ENSMUSG00000072980 | Oip5 | protein_coding | 0.2344446 | 3.608399 | 0.1086286 | 0.7459065 | 0.999176 |
| ENSMUSG00000025035 | Arl3 | protein_coding | 0.1024395 | 9.134159 | 0.1085295 | 0.746018 | 0.999176 |
| ENSMUSG00000078485 | Plekhn1 | protein_coding | -0.4175248 | 3.134278 | 0.1087316 | 0.7461051 | 0.999176 |
| ENSMUSG00000026959 | Grin1 | protein_coding | 0.3345116 | 2.545413 | 0.1082892 | 0.7462888 | 0.999176 |
| ENSMUSG00000034842 | Art3 | protein_coding | -0.4695185 | 2.48698 | 0.1082254 | 0.7463675 | 0.999176 |
| ENSMUSG00000057572 | Zbtb8os | protein_coding | 0.1680537 | 4.871847 | 0.1081995 | 0.7463899 | 0.999176 |
| ENSMUSG00000035245 | Eogt | protein_coding | -0.2969508 | 2.911415 | 0.1081724 | 0.7464205 | 0.999176 |
| ENSMUSG00000026632 | Tatdn3 | protein_coding | 0.1713622 | 4.434897 | 0.1080126 | 0.7466009 | 0.999176 |
| ENSMUSG00000050565 | Tor1aip2 | protein_coding | 0.1076869 | 6.521708 | 0.1080097 | 0.7466041 | 0.999176 |
| ENSMUSG00000036009 | Mettl25 | protein_coding | -0.266353 | 2.9951 | 0.1079988 | 0.7466165 | 0.999176 |
| ENSMUSG00000066892 | Fbxl12 | protein_coding | -0.2251092 | 3.706845 | 0.1079484 | 0.7466734 | 0.999176 |
| ENSMUSG00000075266 | Cenpw | protein_coding | -0.3090025 | 3.134142 | 0.1079462 | 0.7466758 | 0.999176 |
| ENSMUSG00000026425 | Srgap2 | protein_coding | -0.1334897 | 5.390911 | 0.1078076 | 0.7468324 | 0.999176 |
| ENSMUSG00000046204 | Pnma2 | protein_coding | -0.1119684 | 7.583794 | 0.1077916 | 0.7468505 | 0.999176 |
| ENSMUSG00000057649 | Brd9 | protein_coding | 0.1345286 | 5.955179 | 0.1077223 | 0.7469288 | 0.999176 |
| ENSMUSG00000018102 | Hist1h2bc | protein_coding | -0.1880219 | 5.968763 | 0.1079433 | 0.7469453 | 0.999176 |
| ENSMUSG00000025762 | Larp1b | protein_coding | 0.2715293 | 3.278689 | 0.1076555 | 0.7470044 | 0.999176 |
| ENSMUSG00000026944 | Abca2 | protein_coding | -0.1402076 | 6.357413 | 0.1076526 | 0.7470076 | 0.999176 |
| ENSMUSG00000048965 | Mrgpre | protein_coding | 0.1901796 | 4.437605 | 0.1076466 | 0.7470144 | 0.999176 |
| ENSMUSG00000031954 | Cfdp1 | protein_coding | 0.08986124 | 8.6815 | 0.1076383 | 0.7470239 | 0.999176 |
| ENSMUSG00000025474 | Tubgcp2 | protein_coding | 0.1649385 | 5.413787 | 0.1076156 | 0.7470495 | 0.999176 |
| ENSMUSG00000035284 | Vps13c | protein_coding | -0.1735848 | 4.709764 | 0.1073436 | 0.7473575 | 0.999176 |
| ENSMUSG00000033712 | Ccar2 | protein_coding | 0.16708 | 5.487467 | 0.1072748 | 0.7474354 | 0.999176 |
| ENSMUSG00000044317 | Gpr4 | protein_coding | -0.4106367 | 2.413927 | 0.1071932 | 0.747528 | 0.999176 |
| ENSMUSG00000031337 | Mtm1 | protein_coding | 0.3804354 | 2.849706 | 0.1070816 | 0.7476545 | 0.999176 |
| ENSMUSG00000000167 | Pih1d2 | protein_coding | 0.1969428 | 3.671754 | 0.1070679 | 0.74767 | 0.999176 |
| ENSMUSG00000034226 | Rhov | protein_coding | 0.3554152 | 2.758435 | 0.1070515 | 0.7476886 | 0.999176 |
| ENSMUSG00000031290 | Lrch2 | protein_coding | 0.1795831 | 3.809746 | 0.1069005 | 0.74786 | 0.999176 |
| ENSMUSG00000035597 | Prpf39 | protein_coding | -0.18321 | 4.118614 | 0.1067282 | 0.7480558 | 0.999176 |
| ENSMUSG00000073096 | Lrrc61 | protein_coding | -0.2256395 | 4.166504 | 0.1067177 | 0.7480678 | 0.999176 |
| ENSMUSG00000004897 | Hdgf | protein_coding | -0.09449987 | 7.340818 | 0.1067017 | 0.7480859 | 0.999176 |
| ENSMUSG00000083695 | Rnf138rt1 | protein_coding | 0.2760627 | 3.793234 | 0.1067047 | 0.7480893 | 0.999176 |
| ENSMUSG00000028391 | Wdr31 | protein_coding | -0.1975994 | 5.189648 | 0.1066981 | 0.7480969 | 0.999176 |
| ENSMUSG00000074064 | Mlycd | protein_coding | -0.2014698 | 4.413334 | 0.1066123 | 0.7481875 | 0.999176 |
| ENSMUSG00000028582 | Cc2d1b | protein_coding | -0.2584054 | 3.403676 | 0.1065509 | 0.7482574 | 0.999176 |
| ENSMUSG00000060733 | Ipmk | protein_coding | -0.1773462 | 3.940778 | 0.106492 | 0.7483244 | 0.999176 |
| ENSMUSG00000034312 | Iqsec1 | protein_coding | 0.1510277 | 5.851192 | 0.1064143 | 0.7484129 | 0.999176 |
| ENSMUSG00000029059 | Fam213b | protein_coding | 0.1290734 | 9.193709 | 0.1063523 | 0.7484834 | 0.999176 |
| ENSMUSG00000030486 | Zfp108 | protein_coding | -0.3471194 | 2.501826 | 0.1062418 | 0.7486093 | 0.999176 |
| ENSMUSG00000026176 | Ctdsp1 | protein_coding | 0.2443563 | 3.061701 | 0.1060947 | 0.748777 | 0.999176 |
| ENSMUSG00000021712 | Trim23 | protein_coding | 0.17933 | 5.553012 | 0.1059221 | 0.7489807 | 0.999176 |
| ENSMUSG00000028218 | Fam92a | protein_coding | 0.1182233 | 5.899351 | 0.1057027 | 0.7492245 | 0.999176 |
| ENSMUSG00000028456 | Unc13b | protein_coding | -0.1282774 | 5.591984 | 0.1055627 | 0.7493845 | 0.999176 |
| ENSMUSG00000032405 | Pias1 | protein_coding | 0.1224527 | 7.09337 | 0.1053638 | 0.749612 | 0.999176 |
| ENSMUSG00000049092 | Gpr137c | protein_coding | 0.309628 | 2.982284 | 0.1052389 | 0.7497551 | 0.999176 |
| ENSMUSG00000024780 | Cdc37l1 | protein_coding | -0.1026889 | 6.340182 | 0.1051832 | 0.7498189 | 0.999176 |
| ENSMUSG00000003345 | Csnk1g2 | protein_coding | 0.1756377 | 5.019597 | 0.1051313 | 0.7498784 | 0.999176 |
| ENSMUSG00000019948 | Actr6 | protein_coding | -0.1100477 | 6.39779 | 0.1050757 | 0.7499421 | 0.999176 |
| ENSMUSG00000042529 | Kcnj12 | protein_coding | -0.2602682 | 3.044336 | 0.10486 | 0.7501895 | 0.999176 |
| ENSMUSG00000026116 | Tmem131 | protein_coding | 0.1881281 | 3.967844 | 0.1048312 | 0.7502226 | 0.999176 |
| ENSMUSG00000030341 | Tnfrsf1a | protein_coding | 0.3450451 | 2.560832 | 0.1048016 | 0.7502567 | 0.999176 |
| ENSMUSG00000032353 | Tmed3 | protein_coding | 0.09720678 | 7.049319 | 0.104795 | 0.7502642 | 0.999176 |
| ENSMUSG00000008540 | Mgst1 | protein_coding | -0.2806464 | 5.354485 | 0.1053278 | 0.750456 | 0.999176 |
| ENSMUSG00000000538 | Tom1l2 | protein_coding | -0.1288322 | 7.247091 | 0.1046218 | 0.7504699 | 0.999176 |
| ENSMUSG00000041341 | Atg2b | protein_coding | 0.2029394 | 3.948558 | 0.1046067 | 0.7504805 | 0.999176 |
| ENSMUSG00000010057 | Nprl2 | protein_coding | -0.1331198 | 6.552188 | 0.1044962 | 0.7506075 | 0.999176 |
| ENSMUSG00000032475 | Nck1 | protein_coding | -0.1359218 | 5.32682 | 0.1044207 | 0.7506944 | 0.999176 |
| ENSMUSG00000047649 | Cd3eap | protein_coding | 0.1203994 | 6.083497 | 0.1044037 | 0.7507139 | 0.999176 |
| ENSMUSG00000051864 | Tbc1d22a | protein_coding | 0.1757566 | 4.965718 | 0.1042072 | 0.7509402 | 0.999176 |
| ENSMUSG00000070697 | Utp3 | protein_coding | 0.1016595 | 6.400541 | 0.1041123 | 0.7510495 | 0.999176 |
| ENSMUSG00000017897 | Eya2 | protein_coding | 0.232732 | 3.059167 | 0.1039007 | 0.7512935 | 0.999176 |
| ENSMUSG00000027673 | Ndufb5 | protein_coding | 0.09965975 | 9.868636 | 0.1038554 | 0.7513457 | 0.999176 |
| ENSMUSG00000078515 | Ddi2 | protein_coding | -0.1485586 | 5.409686 | 0.103823 | 0.7513831 | 0.999176 |
| ENSMUSG00000029229 | Chic2 | protein_coding | 0.2229881 | 3.422372 | 0.103722 | 0.7514997 | 0.999176 |
| ENSMUSG00000058291 | Zfp68 | protein_coding | 0.1315108 | 5.270613 | 0.1036134 | 0.7516251 | 0.999176 |
| ENSMUSG00000087178 | A230056P14Rik | processed_transcript | 0.3038408 | 3.420153 | 0.1035418 | 0.7517146 | 0.999176 |
| ENSMUSG00000023104 | Rfc2 | protein_coding | -0.1034086 | 6.38448 | 0.1034397 | 0.7518259 | 0.999176 |

| ensembl_gene_id | external_gene_name | gene_biotype | log2 (fold change) | log2 (counts per million) | F-value | p-value | False discovery rate |
|---|---|---|---|---|---|---|---|
| ENSMUSG00000023915 | Tnfrsf21 | protein_coding | -0.1306986 | 5.548326 | 0.1034343 | 0.7518323 | 0.999176 |
| ENSMUSG00000030652 | Coq7 | protein_coding | 0.08744028 | 7.843275 | 0.1034249 | 0.7518431 | 0.999176 |
| ENSMUSG00000047766 | Lrrc49 | protein_coding | 0.1231238 | 6.572947 | 0.1033742 | 0.7519017 | 0.999176 |
| ENSMUSG00000034059 | Ypel4 | protein_coding | 0.1094638 | 6.631334 | 0.1032583 | 0.7520359 | 0.999176 |
| ENSMUSG00000039414 | Heatr5b | protein_coding | 0.2184573 | 3.549098 | 0.1030764 | 0.7522465 | 0.999176 |
| ENSMUSG00000030226 | Lmo3 | protein_coding | -0.1026894 | 7.101001 | 0.1029642 | 0.7523765 | 0.999176 |
| ENSMUSG00000054808 | Actn4 | protein_coding | -0.1806102 | 4.526153 | 0.1029177 | 0.7524305 | 0.999176 |
| ENSMUSG00000032601 | Prkar2a | protein_coding | 0.1090574 | 5.871923 | 0.1028728 | 0.7524826 | 0.999176 |
| ENSMUSG00000026123 | Plekhb2 | protein_coding | 0.185288 | 9.104808 | 0.103235 | 0.7526347 | 0.999176 |
| ENSMUSG00000006169 | Clint1 | protein_coding | -0.147392 | 4.742446 | 0.1027042 | 0.7526782 | 0.999176 |
| ENSMUSG00000035949 | Fbxw2 | protein_coding | -0.1866523 | 5.108968 | 0.102683 | 0.7527095 | 0.999176 |
| ENSMUSG00000038403 | Hfe2 | protein_coding | -0.3122699 | 2.767877 | 0.102591 | 0.7528097 | 0.999176 |
| ENSMUSG00000097881 | Celrr | lincRNA | -0.386048 | 2.872963 | 0.1025796 | 0.7528297 | 0.999176 |
| ENSMUSG00000074657 | Kif5a | protein_coding | -0.1278857 | 8.897046 | 0.102439 | 0.7529931 | 0.999176 |
| ENSMUSG00000061244 | Exoc5 | protein_coding | -0.1518435 | 4.7149 | 0.1023909 | 0.7530423 | 0.999176 |
| ENSMUSG00000020423 | Btg2 | protein_coding | -0.2626657 | 5.9536 | 0.1033668 | 0.7530541 | 0.999176 |
| ENSMUSG00000036327 | Qsox2 | protein_coding | 0.3272168 | 2.993633 | 0.1023525 | 0.753087 | 0.999176 |
| ENSMUSG00000001802 | Lrp3 | protein_coding | 0.193305 | 4.149815 | 0.1022129 | 0.7532494 | 0.999176 |
| ENSMUSG00000020657 | Dnajc27 | protein_coding | 0.0969164 | 6.523434 | 0.102201 | 0.7532633 | 0.999176 |
| ENSMUSG00000070780 | Rbm47 | protein_coding | -0.4405569 | 2.40204 | 0.102103 | 0.7533774 | 0.999176 |
| ENSMUSG00000040697 | Dnajc16 | protein_coding | -0.2249537 | 4.041503 | 0.1020721 | 0.7534135 | 0.999176 |
| ENSMUSG00000038705 | Gmeb2 | protein_coding | 0.2308257 | 3.403363 | 0.1020082 | 0.7534879 | 0.999176 |
| ENSMUSG00000061524 | Zic2 | protein_coding | -0.3322344 | 2.582368 | 0.1019067 | 0.7536062 | 0.999176 |
| ENSMUSG00000014349 | Ube2z | protein_coding | -0.11189 | 6.026996 | 0.1018987 | 0.7536156 | 0.999176 |
| ENSMUSG00000044763 | Trmt10c | protein_coding | -0.1782664 | 4.636812 | 0.1017286 | 0.753814 | 0.999176 |
| ENSMUSG00000062929 | Cfl2 | protein_coding | 0.09262393 | 7.973811 | 0.1016627 | 0.753891 | 0.999176 |
| ENSMUSG00000030089 | Slc41a3 | protein_coding | -0.2593177 | 3.496802 | 0.1014998 | 0.7540813 | 0.999176 |
| ENSMUSG00000019864 | Rtn4ip1 | protein_coding | -0.1707193 | 4.815927 | 0.1013211 | 0.7542902 | 0.999176 |
| ENSMUSG00000000340 | Dbt | protein_coding | -0.2698599 | 3.982134 | 0.1012633 | 0.7544838 | 0.999176 |
| ENSMUSG00000029600 | Rita1 | protein_coding | 0.1253465 | 5.501721 | 0.1010211 | 0.7546415 | 0.999176 |
| ENSMUSG00000048058 | Ldlrad3 | protein_coding | -0.2766183 | 3.027808 | 0.1009973 | 0.7546693 | 0.999176 |
| ENSMUSG00000029657 | Hsph1 | protein_coding | -0.1005222 | 7.818311 | 0.1009526 | 0.7547217 | 0.999176 |
| ENSMUSG00000034826 | Nup54 | protein_coding | 0.2442469 | 3.960241 | 0.1008698 | 0.7548254 | 0.999176 |
| ENSMUSG00000032908 | Sgpp2 | protein_coding | 0.5673257 | 2.288441 | 0.1009974 | 0.7548383 | 0.999176 |
| ENSMUSG00000038212 | Mfsd14b | protein_coding | 0.178793 | 4.628873 | 0.1007762 | 0.7549286 | 0.999176 |
| ENSMUSG00000027499 | Pkia | protein_coding | -0.100035 | 7.198927 | 0.1007507 | 0.7549586 | 0.999176 |
| ENSMUSG00000021809 | Nudt13 | protein_coding | 0.1945822 | 4.202713 | 0.1006621 | 0.7550626 | 0.999176 |
| ENSMUSG00000027584 | Oprl1 | protein_coding | 0.126786 | 6.424173 | 0.1005068 | 0.7552449 | 0.999176 |
| ENSMUSG00000027649 | Ctnnbl1 | protein_coding | 0.1289598 | 6.22898 | 0.100472 | 0.7552858 | 0.999176 |
| ENSMUSG00000033955 | Tnks1bp1 | protein_coding | 0.3713923 | 2.644913 | 0.1004382 | 0.7553255 | 0.999176 |
| ENSMUSG00000038132 | Rbm24 | protein_coding | 0.1757564 | 4.232803 | 0.1004264 | 0.7553394 | 0.999176 |
| ENSMUSG00000020745 | Pafah1b1 | protein_coding | -0.09806206 | 7.459928 | 0.1004081 | 0.7553609 | 0.999176 |
| ENSMUSG00000055725 | Paqr3 | protein_coding | -0.2970083 | 5.3312 | 0.1011702 | 0.7555974 | 0.999176 |
| ENSMUSG00000022770 | Dlg1 | protein_coding | -0.1327591 | 5.375226 | 0.1001701 | 0.755641 | 0.999176 |
| ENSMUSG00000053519 | Kcnip1 | protein_coding | 0.1124619 | 6.198908 | 0.1000998 | 0.7557237 | 0.999176 |
| ENSMUSG00000042167 | Papd4 | protein_coding | 0.2344879 | 3.182694 | 0.1000363 | 0.7557985 | 0.999176 |
| ENSMUSG00000003031 | Cdkn1b | protein_coding | 0.1767937 | 5.14848 | 0.0999065 | 0.7559515 | 0.999176 |
| ENSMUSG00000039478 | Micu3 | protein_coding | 0.1621964 | 8.724197 | 0.1002054 | 0.7560003 | 0.999176 |
| ENSMUSG00000071661 | Zbtb3 | protein_coding | 0.2586849 | 3.124409 | 0.0998574 | 0.7560094 | 0.999176 |
| ENSMUSG00000043059 | Zfp513 | protein_coding | -0.4059337 | 2.252912 | 0.0997612 | 0.7561229 | 0.999176 |
| ENSMUSG00000041423 | Paqr6 | protein_coding | -0.1911183 | 3.762613 | 0.0995834 | 0.7563328 | 0.999176 |
| ENSMUSG00000030979 | Uros | protein_coding | 0.1033619 | 6.304832 | 0.0995641 | 0.7563556 | 0.999176 |
| ENSMUSG00000047866 | Lonp2 | protein_coding | -0.1105496 | 6.782646 | 0.0995295 | 0.7563964 | 0.999176 |
| ENSMUSG00000040040 | Ift88 | protein_coding | 0.189001 | 4.567561 | 0.0995272 | 0.7563991 | 0.999176 |
| ENSMUSG00000008318 | Relt | protein_coding | 0.2794319 | 3.40533 | 0.0994852 | 0.7564488 | 0.999176 |
| ENSMUSG00000056185 | Snx32 | protein_coding | 0.09920018 | 7.503684 | 0.0994555 | 0.7564839 | 0.999176 |
| ENSMUSG00000025026 | Add3 | protein_coding | -0.1165651 | 6.071689 | 0.0994393 | 0.7565031 | 0.999176 |
| ENSMUSG00000032199 | Polr2m | protein_coding | -0.09851171 | 6.886651 | 0.0994283 | 0.756516 | 0.999176 |
| ENSMUSG00000001300 | Efnb2 | protein_coding | -0.2062465 | 4.524014 | 0.099296 | 0.7566791 | 0.999176 |
| ENSMUSG00000022270 | Retreg1 | protein_coding | -0.1172742 | 5.980789 | 0.0992427 | 0.7567355 | 0.999176 |
| ENSMUSG00000021203 | Otub2 | protein_coding | 0.3270759 | 3.546583 | 0.0994216 | 0.7567786 | 0.999176 |
| ENSMUSG00000032217 | Rnf111 | protein_coding | -0.1586088 | 4.463265 | 0.0991919 | 0.7567956 | 0.999176 |
| ENSMUSG00000028884 | Rpa2 | protein_coding | 0.1541055 | 5.220255 | 0.0989708 | 0.7570576 | 0.999176 |
| ENSMUSG00000025277 | Abhd6 | protein_coding | 0.1175227 | 6.217602 | 0.0989488 | 0.7570836 | 0.999176 |
| ENSMUSG00000063849 | Ppcdc | protein_coding | 0.1313053 | 5.543888 | 0.0988834 | 0.7571612 | 0.999176 |
| ENSMUSG00000027655 | Dhx35 | protein_coding | 0.2140479 | 4.104663 | 0.0988662 | 0.7571816 | 0.999176 |
| ENSMUSG00000022223 | Sdr39u1 | protein_coding | 0.09071426 | 8.470249 | 0.0988227 | 0.7572331 | 0.999176 |
| ENSMUSG00000024776 | Stambpl1 | protein_coding | -0.1446661 | 4.764129 | 0.0987702 | 0.7572954 | 0.999176 |
| ENSMUSG00000049526 | Tmem202 | protein_coding | 0.4365972 | 2.358322 | 0.0986044 | 0.7574922 | 0.999176 |
| ENSMUSG00000024145 | Pigf | protein_coding | -0.1785104 | 5.052367 | 0.0985668 | 0.7575368 | 0.999176 |
| ENSMUSG00000025722 | Wdr73 | protein_coding | -0.1253096 | 5.634726 | 0.0984952 | 0.757622 | 0.999176 |
| ENSMUSG00000096141 | Dnah7a | protein_coding | -0.1755683 | 3.695229 | 0.0984169 | 0.757715 | 0.999176 |
| ENSMUSG00000024382 | Ercc3 | protein_coding | 0.1619339 | 4.834173 | 0.0983386 | 0.7578081 | 0.999176 |
| ENSMUSG00000021116 | Eif2s1 | protein_coding | -0.09123042 | 6.639159 | 0.0983255 | 0.7578237 | 0.999176 |
| ENSMUSG00000058267 | Mrps14 | protein_coding | -0.09421746 | 8.757601 | 0.0982992 | 0.7578549 | 0.999176 |
| ENSMUSG00000001151 | Pcnt | protein_coding | -0.274689 | 3.436727 | 0.0981823 | 0.7580006 | 0.999176 |
| ENSMUSG00000021831 | Ero1l | protein_coding | -0.309199 | 3.134149 | 0.0981492 | 0.7580335 | 0.999176 |
| ENSMUSG00000034570 | Inpp5j | protein_coding | 0.2522151 | 3.112371 | 0.0979574 | 0.7582619 | 0.999176 |

| ensembl_gene_id | external_gene_name | gene_biotype | log2 (fold change) | log2 (counts per million) | F-value | p-value | False discovery rate |
|---|---|---|---|---|---|---|---|
| ENSMUSG00000001098 | Kctd10 | protein_coding | -0.2784432 | 3.615845 | 0.0979226 | 0.7583099 | 0.999176 |
| ENSMUSG00000041380 | Htr2c | protein_coding | -0.1255614 | 5.4926 | 0.0979122 | 0.7583159 | 0.999176 |
| ENSMUSG00000090291 | Lrrc10b | protein_coding | 0.2091355 | 4.112852 | 0.0978663 | 0.7583706 | 0.999176 |
| ENSMUSG00000029634 | Rnf6 | protein_coding | 0.1385517 | 7.052887 | 0.0977788 | 0.7585099 | 0.999176 |
| ENSMUSG00000011877 | Git1 | protein_coding | 0.09723899 | 6.426263 | 0.0977275 | 0.7585361 | 0.999176 |
| ENSMUSG00000075376 | Rc3h2 | protein_coding | -0.1833795 | 5.503473 | 0.0977169 | 0.7585792 | 0.999176 |
| ENSMUSG00000026904 | Slc4a10 | protein_coding | -0.1409897 | 5.330997 | 0.0974937 | 0.7588153 | 0.999176 |
| ENSMUSG00000044952 | Kctd21 | protein_coding | 0.1954689 | 4.021925 | 0.0974667 | 0.7588476 | 0.999176 |
| ENSMUSG00000031748 | Gnao1 | protein_coding | 0.09879989 | 8.517778 | 0.0973527 | 0.7589839 | 0.999176 |
| ENSMUSG00000078941 | Ak6 | protein_coding | 0.1343253 | 5.524307 | 0.0971405 | 0.7592378 | 0.999176 |
| ENSMUSG00000050730 | Arhgap42 | protein_coding | 0.4796422 | 2.211126 | 0.0971106 | 0.7592736 | 0.999176 |
| ENSMUSG00000056091 | St3gal5 | protein_coding | -0.1510197 | 4.890994 | 0.0970083 | 0.7593962 | 0.999176 |
| ENSMUSG00000048701 | Ccdc6 | protein_coding | -0.1007458 | 6.963462 | 0.0969776 | 0.759433 | 0.999176 |
| ENSMUSG00000010290 | AI597479 | protein_coding | 0.1512055 | 5.439774 | 0.0968756 | 0.7595552 | 0.999176 |
| ENSMUSG00000075701 | Selenos | protein_coding | -0.1244483 | 7.051739 | 0.096684 | 0.7597916 | 0.999176 |
| ENSMUSG00000026489 | Coq8a | protein_coding | -0.1432188 | 5.611938 | 0.0966447 | 0.7598324 | 0.999176 |
| ENSMUSG00000108112 | Gm45193 | processed_transcript | -0.2168874 | 3.974443 | 0.0966319 | 0.7598477 | 0.999176 |
| ENSMUSG00000071659 | Hnrnpul2 | protein_coding | -0.136022 | 5.875006 | 0.0965507 | 0.7599452 | 0.999176 |
| ENSMUSG00000024091 | Vapa | protein_coding | -0.08527265 | 8.549084 | 0.0965363 | 0.7599625 | 0.999176 |
| ENSMUSG00000003809 | Gcdh | protein_coding | -0.1810924 | 4.636543 | 0.0963989 | 0.7601277 | 0.999176 |
| ENSMUSG00000070639 | Lrrc8b | protein_coding | -0.1637399 | 4.292065 | 0.0963431 | 0.7601949 | 0.999176 |
| ENSMUSG00000008855 | Hdac5 | protein_coding | 0.1483981 | 5.346066 | 0.0962855 | 0.7602641 | 0.999176 |
| ENSMUSG00000032517 | Mobp | protein_coding | -0.09553654 | 7.823792 | 0.0962126 | 0.7603519 | 0.999176 |
| ENSMUSG00000038807 | Rap1gap2 | protein_coding | -0.1491277 | 5.432079 | 0.0962069 | 0.7603587 | 0.999176 |
| ENSMUSG00000038302 | Afg1l | protein_coding | -0.1710017 | 4.729017 | 0.0961032 | 0.7604836 | 0.999176 |
| ENSMUSG00000042496 | Prdm10 | protein_coding | -0.2402099 | 3.933504 | 0.0960514 | 0.7605524 | 0.999176 |
| ENSMUSG00000027259 | Adal | protein_coding | -0.1648935 | 4.295735 | 0.0960268 | 0.7605756 | 0.999176 |
| ENSMUSG00000019986 | Ahi1 | protein_coding | -0.1188815 | 8.777982 | 0.0960199 | 0.760584 | 0.999176 |
| ENSMUSG00000110185 | Igip | protein_coding | -0.1547329 | 5.364581 | 0.0958962 | 0.760733 | 0.999176 |
| ENSMUSG00000047909 | Ankrd16 | protein_coding | -0.3460462 | 2.792144 | 0.0957832 | 0.7608693 | 0.999176 |
| ENSMUSG00000027642 | Rpn2 | protein_coding | -0.08226491 | 7.653254 | 0.0956501 | 0.76103 | 0.999176 |
| ENSMUSG00000026189 | Pecr | protein_coding | -0.2681838 | 2.841674 | 0.0955606 | 0.7611381 | 0.999176 |
| ENSMUSG00000007872 | Id3 | protein_coding | -0.09939891 | 7.392787 | 0.0954672 | 0.761251 | 0.999176 |
| ENSMUSG00000020803 | Txndc17 | protein_coding | 0.08520757 | 7.832674 | 0.0954585 | 0.7612615 | 0.999176 |
| ENSMUSG00000040560 | Wdr7 | protein_coding | -0.1106396 | 6.034944 | 0.0954487 | 0.7612734 | 0.999176 |
| ENSMUSG00000050144 | Slc25a44 | protein_coding | 0.1027931 | 7.042172 | 0.0954147 | 0.7613145 | 0.999176 |
| ENSMUSG00000079036 | Alkbh1 | protein_coding | -0.1565329 | 4.201293 | 0.0953686 | 0.7613702 | 0.999176 |
| ENSMUSG00000045757 | Zfp764 | protein_coding | -0.5223862 | 2.13166 | 0.0953579 | 0.7613897 | 0.999176 |
| ENSMUSG00000030138 | Bms1 | protein_coding | -0.2209391 | 3.655077 | 0.0953193 | 0.7614299 | 0.999176 |
| ENSMUSG00000051335 | Gfod1 | protein_coding | -0.1792052 | 3.964177 | 0.0952444 | 0.7615206 | 0.999176 |
| ENSMUSG00000005150 | Wdr83 | protein_coding | 0.117112 | 6.399766 | 0.0950967 | 0.7616994 | 0.999176 |
| ENSMUSG00000021870 | Slmap | protein_coding | -0.1565368 | 5.716525 | 0.0950957 | 0.761707 | 0.999176 |
| ENSMUSG00000025352 | Gdf11 | protein_coding | -0.1729845 | 4.577488 | 0.0949769 | 0.7618446 | 0.999176 |
| ENSMUSG00000027011 | Ube2e3 | protein_coding | -0.08742119 | 6.868104 | 0.0949379 | 0.7618919 | 0.999176 |
| ENSMUSG00000025280 | Polr3a | protein_coding | 0.2030185 | 3.709209 | 0.0947972 | 0.7620626 | 0.999176 |
| ENSMUSG00000034858 | Fam214a | protein_coding | 0.169656 | 4.271266 | 0.0947148 | 0.7621626 | 0.999176 |
| ENSMUSG00000042507 | Elmsan1 | protein_coding | 0.2760761 | 2.772001 | 0.0945705 | 0.7623379 | 0.999176 |
| ENSMUSG00000008301 | Phax | protein_coding | -0.08275615 | 7.65944 | 0.094529 | 0.7623883 | 0.999176 |
| ENSMUSG00000028893 | Sesn2 | protein_coding | 0.2525463 | 3.422479 | 0.0943459 | 0.7626109 | 0.999176 |
| ENSMUSG00000029119 | Man2b2 | protein_coding | 0.2294232 | 3.495649 | 0.0941053 | 0.7629039 | 0.999176 |
| ENSMUSG00000054459 | Vsnl1 | protein_coding | 0.1071798 | 10.53234 | 0.0938489 | 0.7632166 | 0.999176 |
| ENSMUSG00000014932 | Yes1 | protein_coding | -0.3452156 | 2.407524 | 0.0937488 | 0.7633388 | 0.999176 |
| ENSMUSG00000021732 | Fgf10 | protein_coding | -0.3412277 | 2.872667 | 0.0936757 | 0.7634281 | 0.999176 |
| ENSMUSG00000019996 | Map7 | protein_coding | -0.2096147 | 3.207849 | 0.093625 | 0.76349 | 0.999176 |
| ENSMUSG00000028563 | Tm2d1 | protein_coding | 0.1864939 | 4.406927 | 0.0935888 | 0.7635342 | 0.999176 |
| ENSMUSG00000028397 | Kdm4c | protein_coding | -0.1720025 | 4.255248 | 0.0935149 | 0.7636246 | 0.999176 |
| ENSMUSG00000040725 | Hnrnpul1 | protein_coding | -0.1598208 | 5.585243 | 0.0932352 | 0.7639732 | 0.999176 |
| ENSMUSG00000025580 | Eif4a3 | protein_coding | -0.09716875 | 6.742675 | 0.0932189 | 0.7639869 | 0.999176 |
| ENSMUSG00000074994 | Qser1 | protein_coding | 0.3454364 | 2.574662 | 0.0931507 | 0.7640703 | 0.999176 |
| ENSMUSG00000025332 | Kdm5c | protein_coding | -0.1667465 | 4.416504 | 0.0930986 | 0.7641342 | 0.999176 |
| ENSMUSG00000052305 | Hbb-bs | protein_coding | -0.1608586 | 5.942453 | 0.0931016 | 0.7641536 | 0.999176 |
| ENSMUSG00000020843 | Timm22 | protein_coding | 0.1090217 | 7.608336 | 0.0929283 | 0.7643431 | 0.999176 |
| ENSMUSG00000039372 | 4-Mar | protein_coding | -0.209035 | 3.590618 | 0.0927678 | 0.7645401 | 0.999176 |
| ENSMUSG00000006641 | Slc5a6 | protein_coding | 0.1641343 | 4.429236 | 0.0927277 | 0.7645893 | 0.999176 |
| ENSMUSG00000112964 | CT009718.1 | lincRNA | -0.2414145 | 3.165645 | 0.0927266 | 0.7645906 | 0.999176 |
| ENSMUSG00000005045 | Chd5 | protein_coding | -0.1187315 | 6.187594 | 0.0926635 | 0.7646682 | 0.999176 |
| ENSMUSG00000064127 | Med14 | protein_coding | 0.1763053 | 4.923459 | 0.0923551 | 0.7650476 | 0.999176 |
| ENSMUSG00000029998 | Pcyox1 | protein_coding | 0.1072251 | 6.16802 | 0.092163 | 0.7652842 | 0.999176 |
| ENSMUSG00000058230 | Arhgap35 | protein_coding | -0.1265455 | 6.11753 | 0.0919394 | 0.76556 | 0.999176 |
| ENSMUSG00000020458 | Rtn4 | protein_coding | 0.1025677 | 8.419616 | 0.0919368 | 0.7655632 | 0.999176 |
| ENSMUSG00000032536 | Trak1 | protein_coding | 0.11454 | 5.859127 | 0.0917534 | 0.7657896 | 0.999176 |
| ENSMUSG00000029189 | Sel1l3 | protein_coding | 0.2847799 | 2.904287 | 0.0915754 | 0.7660096 | 0.999176 |
| ENSMUSG00000054728 | Phactr1 | protein_coding | -0.08799708 | 8.711496 | 0.0915723 | 0.7660135 | 0.999176 |
| ENSMUSG00000025159 | Mms19 | protein_coding | 0.1743597 | 4.342493 | 0.0914864 | 0.7661198 | 0.999176 |
| ENSMUSG00000074793 | Hspa12b | protein_coding | 0.2701951 | 2.950966 | 0.0914832 | 0.7661237 | 0.999176 |
| ENSMUSG00000043262 | Uevld | protein_coding | -0.3004478 | 2.707248 | 0.0913785 | 0.7662534 | 0.999176 |
| ENSMUSG00000032435 | Dync1li1 | protein_coding | -0.1493683 | 7.157449 | 0.0915841 | 0.7662712 | 0.999176 |
| ENSMUSG00000021549 | Rasa1 | protein_coding | -0.184278 | 3.628781 | 0.0913304 | 0.766313 | 0.999176 |

| ensembl_gene_id | external_gene_name | gene_biotype | log2 (fold change) | log2 (counts per million) | F-value | p-value | False discovery rate |
|---|---|---|---|---|---|---|---|
| ENSMUSG00000064043 | Trerf1 | processed_transcript | -0.1504218 | 4.211829 | 0.0913077 | 0.7663411 | 0.999176 |
| ENSMUSG00000011154 | Cfap161 | protein_coding | -0.165992 | 4.412086 | 0.0912322 | 0.7664347 | 0.999176 |
| ENSMUSG00000027827 | Kcnab1 | protein_coding | -0.09276532 | 6.617754 | 0.091215 | 0.7664559 | 0.999176 |
| ENSMUSG00000030137 | Tuba8 | protein_coding | -0.2947019 | 3.065757 | 0.0911521 | 0.766534 | 0.999176 |
| ENSMUSG00000066637 | Ttc32 | protein_coding | 0.3184485 | 2.762092 | 0.0910788 | 0.7666248 | 0.999176 |
| ENSMUSG00000003153 | Slc2a3 | protein_coding | 0.08989605 | 6.362209 | 0.0909728 | 0.7667564 | 0.999176 |
| ENSMUSG00000044636 | Csrnp2 | protein_coding | -0.186851 | 4.041868 | 0.0909335 | 0.7668051 | 0.999176 |
| ENSMUSG00000064225 | Paqr9 | protein_coding | 0.186612 | 4.614825 | 0.0908524 | 0.7669059 | 0.999176 |
| ENSMUSG00000000416 | Cttnbp2 | protein_coding | -0.1596531 | 5.046223 | 0.0908084 | 0.7669605 | 0.999176 |
| ENSMUSG00000020538 | Srebf1 | protein_coding | -0.2005622 | 4.174964 | 0.0907744 | 0.7670028 | 0.999176 |
| ENSMUSG00000015354 | Pcolce2 | protein_coding | 0.2925852 | 2.68756 | 0.0907728 | 0.7670048 | 0.999176 |
| ENSMUSG00000005198 | Polr2a | protein_coding | -0.1986055 | 4.727635 | 0.0907481 | 0.7670418 | 0.999176 |
| ENSMUSG00000028990 | Lzic | protein_coding | 0.1810466 | 4.126036 | 0.0907397 | 0.767046 | 0.999176 |
| ENSMUSG00000029363 | Rfc5 | protein_coding | -0.2961096 | 2.780363 | 0.0906818 | 0.7671179 | 0.999176 |
| ENSMUSG00000020922 | Lsm12 | protein_coding | 0.1281698 | 6.04997 | 0.0906146 | 0.7672015 | 0.999176 |
| ENSMUSG00000001380 | Hars | protein_coding | 0.1002591 | 8.238367 | 0.0905983 | 0.7672218 | 0.999176 |
| ENSMUSG00000017428 | Psmd11 | protein_coding | -0.09038118 | 7.521195 | 0.0905607 | 0.7672686 | 0.999176 |
| ENSMUSG00000029433 | Diablo | protein_coding | 0.1258368 | 4.820748 | 0.0905216 | 0.7673172 | 0.999176 |
| ENSMUSG00000028637 | Ccdc30 | protein_coding | 0.1284767 | 5.308539 | 0.090473 | 0.7673777 | 0.999176 |
| ENSMUSG00000028683 | Eif2b3 | protein_coding | 0.1103038 | 6.287356 | 0.0904598 | 0.7673941 | 0.999176 |
| ENSMUSG00000045410 | Akr1e1 | protein_coding | -0.09738742 | 6.841923 | 0.0903908 | 0.7674801 | 0.999176 |
| ENSMUSG00000040466 | Blvrb | protein_coding | 0.1163576 | 5.167729 | 0.0903754 | 0.7674993 | 0.999176 |
| ENSMUSG00000028137 | Celf3 | protein_coding | 0.1573223 | 5.279092 | 0.0903304 | 0.7675554 | 0.999176 |
| ENSMUSG00000025742 | Prps2 | protein_coding | 0.1512592 | 4.834751 | 0.0903123 | 0.7675779 | 0.999176 |
| ENSMUSG00000031532 | Saraf | protein_coding | -0.1133484 | 7.23265 | 0.0901181 | 0.7678201 | 0.999176 |
| ENSMUSG00000043923 | Ccdc84 | protein_coding | -0.4095411 | 2.271908 | 0.0901097 | 0.7678306 | 0.999176 |
| ENSMUSG00000062444 | Ap3b2 | protein_coding | 0.1204033 | 6.661807 | 0.0900491 | 0.7679062 | 0.999176 |
| ENSMUSG00000000301 | Pemt | protein_coding | -0.2171586 | 3.320569 | 0.0897558 | 0.7682727 | 0.999176 |
| ENSMUSG00000044349 | Snhg11 | protein_coding | -0.1430841 | 7.403342 | 0.089942 | 0.7682813 | 0.999176 |
| ENSMUSG00000032757 | Bet1 | protein_coding | -0.09021008 | 6.576748 | 0.0896885 | 0.7683568 | 0.999176 |
| ENSMUSG00000060181 | Slc35e3 | protein_coding | -0.2077919 | 3.131692 | 0.0896712 | 0.7683785 | 0.999176 |
| ENSMUSG00000034042 | Gpbp1l1 | protein_coding | -0.1921071 | 3.696996 | 0.0894275 | 0.7686837 | 0.999176 |
| ENSMUSG00000041120 | Nbl1 | protein_coding | -0.1907107 | 3.953416 | 0.0894178 | 0.7686958 | 0.999176 |
| ENSMUSG00000038564 | Ift172 | protein_coding | 0.2228478 | 3.500283 | 0.0893475 | 0.7687839 | 0.999176 |
| ENSMUSG00000026502 | Desi2 | protein_coding | -0.1602406 | 4.263984 | 0.089327 | 0.7688097 | 0.999176 |
| ENSMUSG00000020647 | Ncoa1 | protein_coding | 0.1597387 | 6.32571 | 0.0894551 | 0.7688897 | 0.999176 |
| ENSMUSG00000029599 | Ddx54 | protein_coding | 0.2366477 | 3.394294 | 0.0891704 | 0.7690061 | 0.999176 |
| ENSMUSG00000039470 | Zdhhc2 | protein_coding | -0.1429051 | 5.225885 | 0.089169 | 0.7690077 | 0.999176 |
| ENSMUSG00000029608 | Rph3a | protein_coding | -0.1411376 | 7.193579 | 0.0893503 | 0.7690209 | 0.999176 |
| ENSMUSG00000031224 | Magee2 | protein_coding | 0.1658036 | 5.034566 | 0.0891448 | 0.7690381 | 0.999176 |
| ENSMUSG00000038349 | Plcl1 | protein_coding | 0.1363903 | 4.898905 | 0.0891004 | 0.7690939 | 0.999176 |
| ENSMUSG00000020903 | Stx8 | protein_coding | -0.08855351 | 6.872107 | 0.0890592 | 0.7691456 | 0.999176 |
| ENSMUSG00000009076 | Zmat5 | protein_coding | 0.1200672 | 5.897945 | 0.0890376 | 0.7691728 | 0.999176 |
| ENSMUSG00000024966 | Stip1 | protein_coding | -0.1168148 | 8.728755 | 0.0890196 | 0.7691953 | 0.999176 |
| ENSMUSG00000032507 | Fbxl2 | protein_coding | -0.1502321 | 5.618177 | 0.0889606 | 0.7692695 | 0.999176 |
| ENSMUSG00000034055 | Phka1 | protein_coding | -0.2997057 | 2.741177 | 0.0889558 | 0.7692755 | 0.999176 |
| ENSMUSG00000016495 | Plgrkt | protein_coding | -0.1361425 | 5.452037 | 0.0887904 | 0.7694835 | 0.999176 |
| ENSMUSG00000027257 | Pacsin3 | protein_coding | -0.1813402 | 4.267206 | 0.0885976 | 0.7697262 | 0.999176 |
| ENSMUSG00000047414 | Flrt2 | protein_coding | 0.164396 | 4.531359 | 0.0885947 | 0.7697298 | 0.999176 |
| ENSMUSG00000052920 | Prkg1 | protein_coding | 0.2317044 | 3.423851 | 0.0885845 | 0.7697426 | 0.999176 |
| ENSMUSG00000021114 | Atp6v1d | protein_coding | 0.08536891 | 9.903446 | 0.088561 | 0.7697723 | 0.999176 |
| ENSMUSG00000027104 | Atf2 | protein_coding | -0.107308 | 7.127111 | 0.0884503 | 0.7699118 | 0.999176 |
| ENSMUSG00000014846 | Tppp3 | protein_coding | 0.1101197 | 8.447402 | 0.0884209 | 0.7699488 | 0.999176 |
| ENSMUSG00000060716 | Plekhh1 | protein_coding | 0.2608538 | 3.439005 | 0.0884101 | 0.7699686 | 0.999176 |
| ENSMUSG00000039683 | Sdk1 | protein_coding | -0.3616224 | 2.43321 | 0.0883895 | 0.7699884 | 0.999176 |
| ENSMUSG00000032307 | Ube2q2 | protein_coding | -0.1198199 | 5.230685 | 0.0883169 | 0.77008 | 0.999176 |
| ENSMUSG00000018412 | Kansl1 | protein_coding | -0.1430674 | 5.409291 | 0.0882551 | 0.7701581 | 0.999176 |
| ENSMUSG00000019795 | Pcmt1 | protein_coding | 0.0909787 | 8.737573 | 0.0882071 | 0.7702186 | 0.999176 |
| ENSMUSG00000040209 | Zfp704 | protein_coding | -0.2809475 | 2.930958 | 0.0881992 | 0.7702285 | 0.999176 |
| ENSMUSG00000022992 | Kansl2 | protein_coding | 0.2706241 | 2.796807 | 0.0881043 | 0.7703485 | 0.999176 |
| ENSMUSG00000023030 | Slc11a2 | protein_coding | 0.1876701 | 4.58606 | 0.0878892 | 0.7706204 | 0.999176 |
| ENSMUSG00000020736 | Nt5c | protein_coding | 0.08188691 | 7.343854 | 0.0878163 | 0.7707126 | 0.999176 |
| ENSMUSG00000023284 | Zfp605 | protein_coding | -0.2186733 | 3.455144 | 0.0877791 | 0.7707597 | 0.999176 |
| ENSMUSG00000052928 | Ctif | protein_coding | 0.1425401 | 4.956431 | 0.0876891 | 0.7708737 | 0.999176 |
| ENSMUSG00000021596 | Mctp1 | protein_coding | -0.1665201 | 6.109102 | 0.0878128 | 0.7709553 | 0.999176 |
| ENSMUSG00000027669 | Gnb4 | protein_coding | 0.10262 | 6.130793 | 0.0874212 | 0.7712132 | 0.999176 |
| ENSMUSG00000094002 | Gm9866 | lincRNA | 0.09157712 | 6.629774 | 0.0872875 | 0.771383 | 0.999176 |
| ENSMUSG00000024163 | Mapk8ip3 | protein_coding | 0.08595155 | 7.311189 | 0.0872458 | 0.7714359 | 0.999176 |
| ENSMUSG00000042155 | Klhl23 | protein_coding | 0.1346844 | 5.279083 | 0.0870737 | 0.7716546 | 0.999176 |
| ENSMUSG00000046417 | Lrrc75a | protein_coding | 0.3529577 | 2.392586 | 0.0870682 | 0.7716616 | 0.999176 |
| ENSMUSG00000034673 | Pbx2 | protein_coding | 0.1516428 | 4.721256 | 0.0869949 | 0.7717548 | 0.999176 |
| ENSMUSG00000041592 | Sdk2 | protein_coding | 0.2582144 | 3.169685 | 0.0869819 | 0.7717713 | 0.999176 |
| ENSMUSG00000022132 | Cldn10 | protein_coding | -0.223938 | 5.794281 | 0.0874337 | 0.7717925 | 0.999176 |
| ENSMUSG00000029179 | Zcchc4 | protein_coding | 0.2344835 | 3.609632 | 0.0869648 | 0.7717931 | 0.999176 |
| ENSMUSG00000017376 | Nlk | protein_coding | 0.1286696 | 5.430242 | 0.0868475 | 0.7719424 | 0.999176 |
| ENSMUSG00000027950 | Chrnb2 | protein_coding | 0.1574759 | 4.510799 | 0.0866318 | 0.7722172 | 0.999176 |
| ENSMUSG00000014859 | E2f4 | protein_coding | 0.1892306 | 3.896729 | 0.0866196 | 0.7722328 | 0.999176 |
| ENSMUSG00000026404 | Ddx59 | protein_coding | -0.2337776 | 2.920961 | 0.0866185 | 0.7722342 | 0.999176 |

| ensembl_gene_id | external_gene_name | gene_biotype | log2 (fold change) | log2 (counts per million) | F-value | p-value | False discovery rate |
|---|---|---|---|---|---|---|---|
| ENSMUSG00000028651 | Ppie | protein_coding | 0.1366424 | 4.584049 | 0.0865624 | 0.7723057 | 0.999176 |
| ENSMUSG00000021807 | 2700060E02Rik | protein_coding | -0.0965817 | 6.484399 | 0.0863479 | 0.7725795 | 0.999176 |
| ENSMUSG00000032932 | Hspa13 | protein_coding | -0.1245722 | 5.451744 | 0.0862087 | 0.7727573 | 0.999176 |
| ENSMUSG00000035834 | Polr3g | protein_coding | -0.4482361 | 1.97997 | 0.086175 | 0.7728005 | 0.999176 |
| ENSMUSG00000000282 | Mnt | protein_coding | 0.1450887 | 4.597142 | 0.0859602 | 0.7730753 | 0.999176 |
| ENSMUSG00000020180 | Snrpd3 | protein_coding | 0.09871398 | 8.64158 | 0.0858763 | 0.7731827 | 0.999176 |
| ENSMUSG00000032251 | Irak1bp1 | protein_coding | 0.1414167 | 4.895446 | 0.0856954 | 0.7734146 | 0.999176 |
| ENSMUSG00000030002 | Dusp11 | protein_coding | -0.1683891 | 3.916513 | 0.085603 | 0.7735332 | 0.999176 |
| ENSMUSG00000020590 | Snx13 | protein_coding | -0.1396033 | 4.74667 | 0.085551 | 0.7736 | 0.999176 |
| ENSMUSG00000032633 | Flcn | protein_coding | -0.1656863 | 5.481049 | 0.0855323 | 0.77363 | 0.999176 |
| ENSMUSG00000031988 | Vps26b | protein_coding | -0.1282925 | 5.714869 | 0.0854452 | 0.7737357 | 0.999176 |
| ENSMUSG00000091264 | Smim13 | protein_coding | 0.08981201 | 6.418625 | 0.0852245 | 0.7740195 | 0.999176 |
| ENSMUSG00000021919 | Chat | protein_coding | -0.2544534 | 3.470664 | 0.0851997 | 0.7740575 | 0.999176 |
| ENSMUSG00000023156 | Rpp14 | protein_coding | 0.08949378 | 6.519071 | 0.0851851 | 0.7740702 | 0.999176 |
| ENSMUSG00000054074 | Skida1 | protein_coding | 0.3031244 | 2.6732 | 0.0850983 | 0.7741819 | 0.999176 |
| ENSMUSG00000003848 | Nob1 | protein_coding | -0.1180292 | 6.283997 | 0.0849144 | 0.7744189 | 0.999176 |
| ENSMUSG00000112639 | AC134329.1 | lincRNA | -0.2420446 | 2.83498 | 0.0848229 | 0.7745367 | 0.999176 |
| ENSMUSG00000019302 | Atp6v0a1 | protein_coding | 0.0996764 | 7.714074 | 0.084813 | 0.7745495 | 0.999176 |
| ENSMUSG00000073557 | Ppp1r12b | protein_coding | 0.3081086 | 2.649361 | 0.0848112 | 0.7745519 | 0.999176 |
| ENSMUSG00000020456 | Ogdh | protein_coding | 0.1226192 | 8.134128 | 0.0848081 | 0.774562 | 0.999176 |
| ENSMUSG00000027422 | Rrbp1 | protein_coding | 0.1687487 | 3.93115 | 0.0847196 | 0.7746701 | 0.999176 |
| ENSMUSG00000026036 | Nif3l1 | protein_coding | 0.1461145 | 6.001363 | 0.0845883 | 0.7748457 | 0.999176 |
| ENSMUSG00000019852 | Arfgef3 | protein_coding | -0.1180527 | 5.896841 | 0.0845441 | 0.7748967 | 0.999176 |
| ENSMUSG00000067825 | Pex26 | protein_coding | 0.2102191 | 3.49403 | 0.0843942 | 0.7750905 | 0.999176 |
| ENSMUSG00000074182 | Znhit6 | protein_coding | 0.1319339 | 4.740682 | 0.084275 | 0.7752446 | 0.999176 |
| ENSMUSG00000027894 | Slc6a17 | protein_coding | -0.1073243 | 6.12109 | 0.0842102 | 0.7753286 | 0.999176 |
| ENSMUSG00000044792 | Isca1 | protein_coding | 0.09530104 | 6.796536 | 0.0841293 | 0.7754334 | 0.999176 |
| ENSMUSG00000049536 | Tceal1 | protein_coding | -0.07944771 | 7.57697 | 0.084036 | 0.7755543 | 0.999176 |
| ENSMUSG00000023827 | Agpat4 | protein_coding | -0.09877268 | 7.118019 | 0.0840076 | 0.7755911 | 0.999176 |
| ENSMUSG00000029769 | Ccdc136 | protein_coding | 0.1452407 | 6.416345 | 0.0840434 | 0.7756979 | 0.999176 |
| ENSMUSG00000020780 | Srp68 | protein_coding | -0.09563463 | 6.48061 | 0.0838956 | 0.7757363 | 0.999176 |
| ENSMUSG00000043987 | Cep164 | protein_coding | 0.2712025 | 2.793294 | 0.083879 | 0.7757579 | 0.999176 |
| ENSMUSG00000054405 | Dnajc8 | protein_coding | -0.07822519 | 8.553718 | 0.0838699 | 0.7757696 | 0.999176 |
| ENSMUSG00000056755 | Grm7 | protein_coding | -0.1409757 | 5.387432 | 0.0836898 | 0.7760036 | 0.999176 |
| ENSMUSG00000027333 | Smox | protein_coding | -0.1346926 | 4.904103 | 0.0836607 | 0.7760414 | 0.999176 |
| ENSMUSG00000025266 | Gnl3l | protein_coding | 0.1699486 | 8.742299 | 0.0839271 | 0.7762777 | 0.999176 |
| ENSMUSG00000027552 | E2f5 | protein_coding | -0.2211906 | 2.977907 | 0.0833899 | 0.7763935 | 0.999176 |
| ENSMUSG00000039449 | Prpf18 | protein_coding | -0.1293872 | 5.607846 | 0.0832461 | 0.7765808 | 0.999176 |
| ENSMUSG00000025986 | Slc39a10 | protein_coding | -0.1022637 | 5.661117 | 0.0831363 | 0.7767239 | 0.999176 |
| ENSMUSG00000024921 | Smarca2 | protein_coding | -0.1058886 | 6.061484 | 0.0831224 | 0.776742 | 0.999176 |
| ENSMUSG00000044857 | Lemd2 | protein_coding | -0.245169 | 3.49921 | 0.0829754 | 0.7769338 | 0.999176 |
| ENSMUSG00000028696 | Ipp | protein_coding | -0.2277584 | 3.869325 | 0.0829351 | 0.7769925 | 0.999176 |
| ENSMUSG00000021699 | Pde4d | protein_coding | -0.1104564 | 5.435247 | 0.0828424 | 0.7771075 | 0.999176 |
| ENSMUSG00000035790 | Cep19 | protein_coding | -0.08314467 | 7.533054 | 0.0828232 | 0.7771327 | 0.999176 |
| ENSMUSG00000041459 | Tardbp | protein_coding | 0.1001768 | 6.1373 | 0.0827555 | 0.7772211 | 0.999176 |
| ENSMUSG00000022661 | Cd200 | protein_coding | -0.1000198 | 8.070795 | 0.0822744 | 0.7778509 | 0.999176 |
| ENSMUSG00000038102 | Trappc11 | protein_coding | -0.1337856 | 4.883764 | 0.0821879 | 0.7779645 | 0.999176 |
| ENSMUSG00000030077 | Chl1 | protein_coding | -0.111178 | 6.308067 | 0.082092 | 0.7780903 | 0.999176 |
| ENSMUSG00000055065 | Ddx17 | protein_coding | 0.1332045 | 5.50697 | 0.082021 | 0.7781835 | 0.999176 |
| ENSMUSG00000061306 | Slc38a10 | protein_coding | -0.1629242 | 5.193891 | 0.0817314 | 0.7785702 | 0.999176 |
| ENSMUSG00000034275 | Igsf9b | protein_coding | 0.1778542 | 4.741213 | 0.0816735 | 0.7786465 | 0.999176 |
| ENSMUSG00000037712 | Fermt2 | protein_coding | 0.1232422 | 5.541512 | 0.0816317 | 0.7786956 | 0.999176 |
| ENSMUSG00000027447 | Cst3 | protein_coding | -0.1766049 | 11.15738 | 0.0819525 | 0.7787787 | 0.999176 |
| ENSMUSG00000039530 | Tusc3 | protein_coding | 0.1141733 | 7.370832 | 0.0814741 | 0.7789091 | 0.999176 |
| ENSMUSG00000025429 | Pstpip2 | protein_coding | -0.3116624 | 2.547899 | 0.0814317 | 0.7789591 | 0.999176 |
| ENSMUSG00000044496 | 2510039O18Rik | protein_coding | -0.2482392 | 3.529021 | 0.0812639 | 0.7791864 | 0.999176 |
| ENSMUSG00000020092 | Pald1 | protein_coding | -0.3660212 | 2.379015 | 0.0812461 | 0.7792041 | 0.999176 |
| ENSMUSG00000025413 | Ttc4 | protein_coding | -0.1257574 | 5.924255 | 0.0810731 | 0.7794325 | 0.999176 |
| ENSMUSG00000043384 | Gprasp1 | protein_coding | 0.255446 | 3.130955 | 0.081058 | 0.7794526 | 0.999176 |
| ENSMUSG00000006412 | Pfdn2 | protein_coding | 0.09532986 | 8.392454 | 0.0809972 | 0.7795329 | 0.999176 |
| ENSMUSG00000031790 | Mmp15 | protein_coding | 0.1712976 | 3.531036 | 0.0809556 | 0.779588 | 0.999176 |
| ENSMUSG00000066189 | Cacng3 | protein_coding | 0.08384358 | 7.43677 | 0.0808943 | 0.7796691 | 0.999176 |
| ENSMUSG00000040452 | Cdh12 | protein_coding | -0.2594135 | 3.246182 | 0.0805514 | 0.7801233 | 0.999176 |
| ENSMUSG00000003269 | Cyth2 | protein_coding | -0.1427446 | 5.170669 | 0.0805242 | 0.7801595 | 0.999176 |
| ENSMUSG00000024454 | Hdac3 | protein_coding | -0.09133889 | 7.052944 | 0.0804609 | 0.7802435 | 0.999176 |
| ENSMUSG00000078670 | Fam174b | protein_coding | -0.2537606 | 3.171189 | 0.0803761 | 0.7803561 | 0.999176 |
| ENSMUSG00000022842 | Ece2 | protein_coding | -0.1320328 | 5.36422 | 0.0803554 | 0.7803836 | 0.999176 |
| ENSMUSG00000055917 | Zfp277 | protein_coding | -0.1641203 | 5.231259 | 0.0803569 | 0.7803875 | 0.999176 |
| ENSMUSG00000057363 | Uxs1 | protein_coding | -0.1904901 | 3.824973 | 0.0803495 | 0.7803914 | 0.999176 |
| ENSMUSG00000004947 | Dtx2 | protein_coding | 0.285121 | 2.572194 | 0.0802663 | 0.7805019 | 0.999176 |
| ENSMUSG00000004896 | Rrnad1 | protein_coding | 0.178638 | 3.883038 | 0.0801901 | 0.7806032 | 0.999176 |
| ENSMUSG00000031987 | Egln1 | protein_coding | 0.119608 | 5.548783 | 0.0800577 | 0.7807793 | 0.999176 |
| ENSMUSG00000097277 | 2900076A07Rik | lincRNA | 0.1665942 | 3.85274 | 0.0800392 | 0.7808041 | 0.999176 |
| ENSMUSG00000004270 | Lpcat3 | protein_coding | 0.1886808 | 4.037494 | 0.0800062 | 0.7808479 | 0.999176 |
| ENSMUSG00000068706 | Gm10250 | protein_coding | 0.190044 | 3.267852 | 0.0799588 | 0.7809111 | 0.999176 |
| ENSMUSG00000027255 | Arfgap2 | protein_coding | 0.1712613 | 4.61517 | 0.0798409 | 0.7810681 | 0.999176 |
| ENSMUSG00000019373 | Cops3 | protein_coding | -0.07784038 | 7.15497 | 0.0798387 | 0.781071 | 0.999176 |
| ENSMUSG00000001674 | Ddx18 | protein_coding | 0.1413726 | 4.228747 | 0.0798259 | 0.7810881 | 0.999176 |

| ensembl_gene_id | external_gene_name | gene_biotype | log2 (fold change) | log2 (counts per million) | F-value | p-value | False discovery rate |
|---|---|---|---|---|---|---|---|
| ENSMUSG00000022636 | Alcam | protein_coding | 0.1095404 | 7.059555 | 0.079819 | 0.7810974 | 0.999176 |
| ENSMUSG00000046750 | Selenov | protein_coding | 0.1394025 | 4.165584 | 0.079752 | 0.7811867 | 0.999176 |
| ENSMUSG00000039697 | Ncoa7 | protein_coding | 0.09780084 | 5.96673 | 0.0796956 | 0.7812618 | 0.999176 |
| ENSMUSG00000074890 | Lcmt2 | protein_coding | 0.2586783 | 3.090682 | 0.0796781 | 0.7812852 | 0.999176 |
| ENSMUSG00000027274 | Mkks | protein_coding | 0.1054069 | 7.441327 | 0.0796274 | 0.7813529 | 0.999176 |
| ENSMUSG00000067629 | Syngap1 | protein_coding | 0.1793137 | 3.426344 | 0.079592 | 0.7814002 | 0.999176 |
| ENSMUSG00000034098 | Fstl5 | protein_coding | -0.1260791 | 5.253126 | 0.0793449 | 0.7817304 | 0.999176 |
| ENSMUSG00000040990 | Sh3kbp1 | protein_coding | 0.1087403 | 5.563579 | 0.079189 | 0.781939 | 0.999176 |
| ENSMUSG00000024099 | Ndufv2 | protein_coding | 0.08898121 | 10.34246 | 0.0790928 | 0.7820679 | 0.999176 |
| ENSMUSG00000034265 | Zdhhc14 | protein_coding | 0.1763542 | 4.272817 | 0.0790905 | 0.7820709 | 0.999176 |
| ENSMUSG00000029461 | Fam168a | protein_coding | 0.1078731 | 7.503392 | 0.0790305 | 0.7821512 | 0.999176 |
| ENSMUSG00000051615 | Rap2a | protein_coding | 0.11567 | 5.783771 | 0.0788581 | 0.7823825 | 0.999176 |
| ENSMUSG00000026277 | Stk25 | protein_coding | 0.09277116 | 7.586586 | 0.0788296 | 0.7824208 | 0.999176 |
| ENSMUSG00000074886 | Grk6 | protein_coding | -0.1423808 | 4.896833 | 0.0787573 | 0.7825178 | 0.999176 |
| ENSMUSG00000033416 | Gucd1 | protein_coding | -0.2715058 | 3.277746 | 0.0787111 | 0.7825857 | 0.999176 |
| ENSMUSG00000058076 | Sdhc | protein_coding | 0.08400667 | 9.022961 | 0.0786706 | 0.7826342 | 0.999176 |
| ENSMUSG00000051736 | Fam229b | protein_coding | -0.1143272 | 5.95975 | 0.0785744 | 0.7827635 | 0.999176 |
| ENSMUSG00000022414 | Tab1 | protein_coding | -0.1709285 | 5.193363 | 0.0785449 | 0.782809 | 0.999176 |
| ENSMUSG00000086401 | Gm15559 | antisense_RNA | 0.2973616 | 3.268404 | 0.0785213 | 0.782866 | 0.999176 |
| ENSMUSG00000024150 | Mcfd2 | protein_coding | -0.08490372 | 7.672832 | 0.0784799 | 0.7828906 | 0.999176 |
| ENSMUSG00000037257 | Aagab | protein_coding | -0.1244502 | 6.616531 | 0.0784092 | 0.7829916 | 0.999176 |
| ENSMUSG00000003934 | Efnb3 | protein_coding | -0.09167517 | 7.187374 | 0.0783133 | 0.7831149 | 0.999176 |
| ENSMUSG00000042225 | Ammecr1 | protein_coding | -0.19691 | 3.249758 | 0.0783051 | 0.783126 | 0.999176 |
| ENSMUSG00000052087 | Rgs14 | protein_coding | 0.134078 | 5.188153 | 0.0782785 | 0.7831618 | 0.999176 |
| ENSMUSG00000001627 | Ifrd1 | protein_coding | -0.2296306 | 3.542411 | 0.0782004 | 0.783267 | 0.999176 |
| ENSMUSG00000027405 | Nop56 | protein_coding | 0.0989126 | 7.276843 | 0.0781959 | 0.7832731 | 0.999176 |
| ENSMUSG00000044772 | Sntn | protein_coding | 0.2174133 | 3.33697 | 0.0780908 | 0.7834148 | 0.999176 |
| ENSMUSG00000055302 | Mrfap1 | protein_coding | 0.0820731 | 7.938451 | 0.0780389 | 0.7834849 | 0.999176 |
| ENSMUSG00000024570 | Rbfa | protein_coding | -0.190981 | 3.664244 | 0.0780005 | 0.7835366 | 0.999176 |
| ENSMUSG00000028530 | Jak1 | protein_coding | -0.08750502 | 6.6457 | 0.0779185 | 0.7836474 | 0.999176 |
| ENSMUSG00000026575 | Nme7 | protein_coding | -0.1535706 | 4.734404 | 0.0779088 | 0.7836605 | 0.999176 |
| ENSMUSG00000036377 | C530008M17Rik | protein_coding | -0.1437458 | 4.964827 | 0.0778011 | 0.783806 | 0.999176 |
| ENSMUSG00000037428 | Vgf | protein_coding | 0.1862233 | 4.032652 | 0.0776296 | 0.7840379 | 0.999176 |
| ENSMUSG00000031858 | Mau2 | protein_coding | 0.1410835 | 5.019563 | 0.0775788 | 0.7841067 | 0.999176 |
| ENSMUSG00000047342 | Zfp286 | protein_coding | 0.1286273 | 5.013948 | 0.0775401 | 0.784159 | 0.999176 |
| ENSMUSG00000040543 | Pitpnm3 | protein_coding | 0.180247 | 3.960727 | 0.0774415 | 0.7842926 | 0.999176 |
| ENSMUSG00000022843 | Clcn2 | protein_coding | -0.3256254 | 2.632605 | 0.0772325 | 0.784576 | 0.999176 |
| ENSMUSG00000048332 | Lhfp | protein_coding | -0.2344911 | 3.587935 | 0.0772066 | 0.784617 | 0.999176 |
| ENSMUSG00000047907 | Tshz2 | protein_coding | 0.1396908 | 4.067112 | 0.0769815 | 0.7849171 | 0.999176 |
| ENSMUSG00000054934 | Kcnmb4 | protein_coding | 0.3040412 | 2.524913 | 0.0769282 | 0.7849895 | 0.999176 |
| ENSMUSG00000028149 | Rap1gds1 | protein_coding | 0.08281884 | 7.472949 | 0.076787 | 0.7851816 | 0.999176 |
| ENSMUSG00000027086 | Fastkd1 | protein_coding | 0.1469696 | 4.208332 | 0.0767675 | 0.7852081 | 0.999176 |
| ENSMUSG00000002578 | Ikzf4 | protein_coding | -0.1533611 | 4.382671 | 0.0767422 | 0.7852426 | 0.999176 |
| ENSMUSG00000019822 | Smpd2 | protein_coding | -0.1763884 | 4.065078 | 0.0766012 | 0.7854347 | 0.999176 |
| ENSMUSG00000038534 | Osbpl7 | protein_coding | -0.1348266 | 4.381384 | 0.0765603 | 0.7854904 | 0.999176 |
| ENSMUSG00000020921 | Tmem101 | protein_coding | -0.1113523 | 5.629055 | 0.076543 | 0.7855141 | 0.999176 |
| ENSMUSG00000006273 | Atp6v1b2 | protein_coding | 0.09413082 | 9.50591 | 0.0764216 | 0.7856797 | 0.999176 |
| ENSMUSG00000030805 | Stx4a | protein_coding | -0.1064866 | 6.13173 | 0.0763989 | 0.7857106 | 0.999176 |
| ENSMUSG00000031429 | Psmd10 | protein_coding | -0.0887777 | 6.949199 | 0.0761372 | 0.7860682 | 0.999176 |
| ENSMUSG00000027498 | Cstf1 | protein_coding | -0.1173541 | 5.412005 | 0.0760791 | 0.7861477 | 0.999176 |
| ENSMUSG00000047539 | Fbxo28 | protein_coding | -0.1279957 | 4.946112 | 0.075994 | 0.7862642 | 0.999176 |
| ENSMUSG00000039770 | Ypel5 | protein_coding | -0.077723 | 8.954683 | 0.0759192 | 0.7863666 | 0.999176 |
| ENSMUSG00000028445 | Enho | protein_coding | -0.1246817 | 6.246349 | 0.0758748 | 0.7864331 | 0.999176 |
| ENSMUSG00000039809 | Gabbr2 | protein_coding | -0.1185293 | 5.83455 | 0.0757802 | 0.7865571 | 0.999176 |
| ENSMUSG00000021054 | Sgpp1 | protein_coding | 0.1337419 | 4.598487 | 0.0756415 | 0.7867473 | 0.999176 |
| ENSMUSG00000032322 | Pstpip1 | protein_coding | 0.3488565 | 2.260252 | 0.0756219 | 0.7867743 | 0.999176 |
| ENSMUSG00000035062 | Zc4h2 | protein_coding | 0.1161701 | 5.682404 | 0.0755823 | 0.7868286 | 0.999176 |
| ENSMUSG00000039234 | Sec24d | protein_coding | -0.2179004 | 3.237169 | 0.0755531 | 0.7868687 | 0.999176 |
| ENSMUSG00000048271 | Rbm33 | protein_coding | 0.1150547 | 5.378287 | 0.0755326 | 0.7868969 | 0.999176 |
| ENSMUSG00000056023 | Gm9989 | processed_transcript | 0.2137777 | 3.63811 | 0.0755174 | 0.7869178 | 0.999176 |
| ENSMUSG00000027168 | Pax6 | protein_coding | -0.1703396 | 3.913722 | 0.0755156 | 0.7869202 | 0.999176 |
| ENSMUSG00000020100 | Slc29a3 | protein_coding | -0.2309622 | 3.130166 | 0.0755143 | 0.7869219 | 0.999176 |
| ENSMUSG00000026309 | Ilkap | protein_coding | 0.1266503 | 5.086444 | 0.0754636 | 0.7869917 | 0.999176 |
| ENSMUSG00000030224 | Strap | protein_coding | 0.0860237 | 6.523885 | 0.0754325 | 0.7870344 | 0.999176 |
| ENSMUSG00000053768 | Chchd3 | protein_coding | 0.07558198 | 6.995114 | 0.0753835 | 0.7871018 | 0.999176 |
| ENSMUSG00000041836 | Ptpre | protein_coding | 0.149757 | 4.376135 | 0.0753126 | 0.7871993 | 0.999176 |
| ENSMUSG00000020528 | Prpsap2 | protein_coding | -0.1302799 | 5.112066 | 0.075312 | 0.7872001 | 0.999176 |
| ENSMUSG00000029049 | Morn1 | protein_coding | 0.22013 | 3.800028 | 0.0752738 | 0.7872582 | 0.999176 |
| ENSMUSG00000024875 | Yif1a | protein_coding | 0.09500283 | 6.495636 | 0.0752229 | 0.7873226 | 0.999176 |
| ENSMUSG00000052214 | Opa3 | protein_coding | 0.08793125 | 6.439299 | 0.07515 | 0.787423 | 0.999176 |
| ENSMUSG00000030980 | Knop1 | protein_coding | 0.1078725 | 5.433076 | 0.0751265 | 0.7874554 | 0.999176 |
| ENSMUSG00000031201 | Brcc3 | protein_coding | 0.09554173 | 7.292193 | 0.0749951 | 0.7876365 | 0.999176 |
| ENSMUSG00000024899 | Papss2 | protein_coding | -0.1895122 | 3.466976 | 0.0748213 | 0.7878763 | 0.999176 |
| ENSMUSG00000037408 | Cnnm4 | protein_coding | -0.1554678 | 5.394827 | 0.0747165 | 0.7880267 | 0.999176 |
| ENSMUSG00000031385 | Plxnb3 | protein_coding | 0.2844698 | 2.818135 | 0.0746778 | 0.7880745 | 0.999176 |
| ENSMUSG00000022515 | Anks3 | protein_coding | 0.1747765 | 4.97173 | 0.0746307 | 0.7881452 | 0.999176 |
| ENSMUSG00000026088 | Mitd1 | protein_coding | -0.1568436 | 4.400588 | 0.0745598 | 0.7882377 | 0.999176 |
| ENSMUSG00000036948 | BC037034 | protein_coding | 0.1765352 | 3.583442 | 0.0745532 | 0.7882467 | 0.999176 |

| ensembl_gene_id | external_gene_name | gene_biotype | log2 (fold change) | log2 (counts per million) | F-value | p-value | False discovery rate |
|---|---|---|---|---|---|---|---|
| ENSMUSG00000049176 | Frmpd4 | protein_coding | -0.1239365 | 5.270505 | 0.074419 | 0.7884324 | 0.999176 |
| ENSMUSG00000033579 | Fa2h | protein_coding | -0.1346664 | 4.900478 | 0.0743836 | 0.7884815 | 0.999176 |
| ENSMUSG00000031314 | Taf1 | protein_coding | -0.1344689 | 5.394955 | 0.0743491 | 0.7885293 | 0.999176 |
| ENSMUSG00000045374 | Wdr81 | protein_coding | 0.2742023 | 2.921892 | 0.0743301 | 0.7885556 | 0.999176 |
| ENSMUSG00000020283 | Pex13 | protein_coding | -0.150362 | 4.641363 | 0.0742801 | 0.7886249 | 0.999176 |
| ENSMUSG00000015085 | Entpd2 | protein_coding | -0.2291646 | 2.870822 | 0.0742239 | 0.7887027 | 0.999176 |
| ENSMUSG00000074923 | Pak6 | protein_coding | 0.1317759 | 4.581499 | 0.0740397 | 0.7889583 | 0.999176 |
| ENSMUSG00000023010 | Tmbim6 | protein_coding | 0.0891212 | 9.694523 | 0.0739845 | 0.789035 | 0.999176 |
| ENSMUSG00000027428 | Rbbp9 | protein_coding | -0.1140261 | 6.297843 | 0.0738631 | 0.7892037 | 0.999176 |
| ENSMUSG00000078794 | Dact3 | protein_coding | -0.1459573 | 4.872785 | 0.0738458 | 0.7892277 | 0.999176 |
| ENSMUSG00000022401 | Xpnpep3 | protein_coding | -0.1926262 | 3.690002 | 0.0738394 | 0.7892367 | 0.999176 |
| ENSMUSG00000022151 | Ttc33 | protein_coding | 0.1241423 | 4.688553 | 0.0735735 | 0.7896067 | 0.999176 |
| ENSMUSG00000001576 | Ergic1 | protein_coding | 0.1169514 | 5.837601 | 0.0735011 | 0.7897076 | 0.999176 |
| ENSMUSG00000054423 | Cadps | protein_coding | -0.08113726 | 7.090859 | 0.073492 | 0.7897203 | 0.999176 |
| ENSMUSG00000053093 | Myh7 | protein_coding | -0.2267948 | 2.92177 | 0.0734097 | 0.7898349 | 0.999176 |
| ENSMUSG00000042045 | Sln | protein_coding | -0.2057706 | 6.315764 | 0.0740165 | 0.789947 | 0.999176 |
| ENSMUSG00000054720 | Lrrc8c | protein_coding | -0.1528463 | 3.937073 | 0.0733223 | 0.789957 | 0.999176 |
| ENSMUSG00000028865 | Cd164l2 | protein_coding | -0.1464691 | 3.901544 | 0.0732344 | 0.7900797 | 0.999176 |
| ENSMUSG00000018189 | Uchl5 | protein_coding | 0.08104866 | 6.939611 | 0.0731732 | 0.7901652 | 0.999176 |
| ENSMUSG00000055305 | Zfp93 | protein_coding | -0.2654033 | 2.68119 | 0.0731628 | 0.7901796 | 0.999176 |
| ENSMUSG00000071341 | Egr4 | protein_coding | -0.4318563 | 1.664415 | 0.0730673 | 0.7903132 | 0.999176 |
| ENSMUSG00000064037 | Gpn1 | protein_coding | -0.08224855 | 6.995837 | 0.0729586 | 0.7904653 | 0.999176 |
| ENSMUSG00000028233 | Tgs1 | protein_coding | 0.101959 | 5.482857 | 0.0729498 | 0.7904775 | 0.999176 |
| ENSMUSG00000039704 | Lmbrd2 | protein_coding | -0.3144539 | 2.339369 | 0.0729403 | 0.7904908 | 0.999176 |
| ENSMUSG00000031245 | Hmgn5 | protein_coding | 0.1920951 | 3.447119 | 0.0728765 | 0.7905802 | 0.999176 |
| ENSMUSG00000038175 | Mylip | protein_coding | 0.2154458 | 3.224945 | 0.07284 | 0.7906313 | 0.999176 |
| ENSMUSG00000017734 | Dbndd2 | protein_coding | -0.1241534 | 5.176144 | 0.0728079 | 0.7906762 | 0.999176 |
| ENSMUSG00000021709 | Erbin | protein_coding | 0.138147 | 4.967908 | 0.0727839 | 0.7907098 | 0.999176 |
| ENSMUSG00000067818 | Myl9 | protein_coding | -0.1413376 | 4.619615 | 0.0727346 | 0.790779 | 0.999176 |
| ENSMUSG00000024993 | Fam45a | protein_coding | -0.09465165 | 6.807849 | 0.072698 | 0.7908302 | 0.999176 |
| ENSMUSG00000049811 | Fam161a | protein_coding | -0.2193663 | 2.959913 | 0.0726846 | 0.7908492 | 0.999176 |
| ENSMUSG00000030268 | Bcat1 | protein_coding | -0.1174838 | 7.583349 | 0.0726985 | 0.7908725 | 0.999176 |
| ENSMUSG00000024381 | Bin1 | protein_coding | 0.1038304 | 6.261509 | 0.0726555 | 0.7908898 | 0.999176 |
| ENSMUSG00000032316 | Clk3 | protein_coding | 0.1189735 | 5.25566 | 0.0726415 | 0.7909095 | 0.999176 |
| ENSMUSG00000020664 | Dld | protein_coding | -0.1078649 | 7.390086 | 0.0726281 | 0.7909338 | 0.999176 |
| ENSMUSG00000006390 | Elovl1 | protein_coding | -0.1290921 | 4.990622 | 0.0725239 | 0.7910745 | 0.999176 |
| ENSMUSG00000059772 | Slx1b | protein_coding | 0.143144 | 4.103154 | 0.0725196 | 0.7910806 | 0.999176 |
| ENSMUSG00000036779 | Papd5 | protein_coding | 0.1510179 | 4.076825 | 0.072496 | 0.7911137 | 0.999176 |
| ENSMUSG00000020899 | Pfas | protein_coding | 0.1295899 | 5.876813 | 0.0724543 | 0.7911778 | 0.999176 |
| ENSMUSG00000034403 | Pja1 | protein_coding | 0.1170807 | 8.745548 | 0.0723902 | 0.7912825 | 0.999176 |
| ENSMUSG00000041037 | Irgq | protein_coding | -0.1322644 | 4.537449 | 0.0722428 | 0.7914696 | 0.999176 |
| ENSMUSG00000027305 | Ndufaf1 | protein_coding | 0.09039977 | 5.910435 | 0.0721368 | 0.7916188 | 0.999176 |
| ENSMUSG00000032449 | Slc25a36 | protein_coding | 0.1040624 | 5.766422 | 0.0720706 | 0.791712 | 0.999176 |
| ENSMUSG00000003418 | St8sia6 | protein_coding | -0.2861749 | 2.527976 | 0.0720035 | 0.7918066 | 0.999176 |
| ENSMUSG00000030738 | Eif3c | protein_coding | -0.09208943 | 7.789053 | 0.0720008 | 0.7918103 | 0.999176 |
| ENSMUSG00000062309 | Rpp25 | protein_coding | -0.1429187 | 4.690937 | 0.0719786 | 0.7918416 | 0.999176 |
| ENSMUSG00000010755 | Cars | protein_coding | 0.1654289 | 5.871904 | 0.0720835 | 0.7919697 | 0.999176 |
| ENSMUSG00000034617 | Mtrr | protein_coding | -0.3003697 | 3.455731 | 0.0719682 | 0.7921392 | 0.999176 |
| ENSMUSG00000041703 | Zic5 | protein_coding | -0.2243043 | 3.725062 | 0.0716495 | 0.7923117 | 0.999176 |
| ENSMUSG00000019143 | Hars2 | protein_coding | -0.1449967 | 4.390408 | 0.0716136 | 0.7923568 | 0.999176 |
| ENSMUSG00000021375 | Kif13a | protein_coding | 0.1802983 | 3.691016 | 0.0716032 | 0.7923716 | 0.999176 |
| ENSMUSG00000027875 | Hmgcs2 | protein_coding | 0.1396884 | 4.074244 | 0.0715062 | 0.7925088 | 0.999176 |
| ENSMUSG00000030509 | Asb7 | protein_coding | -0.1381108 | 4.074378 | 0.0714988 | 0.7925192 | 0.999176 |
| ENSMUSG00000090946 | Ccdc71l | protein_coding | 0.2949273 | 2.466316 | 0.0714057 | 0.7926509 | 0.999176 |
| ENSMUSG00000015652 | Steap1 | protein_coding | 0.60691 | 2.662124 | 0.0718816 | 0.7927187 | 0.999176 |
| ENSMUSG00000028837 | Psmb2 | protein_coding | 0.07792266 | 8.876894 | 0.0712602 | 0.7928571 | 0.999176 |
| ENSMUSG00000029247 | Paics | protein_coding | -0.07896766 | 7.582639 | 0.0711137 | 0.7930648 | 0.999176 |
| ENSMUSG00000047495 | Dlgap2 | protein_coding | -0.1426338 | 4.648678 | 0.071047 | 0.7931594 | 0.999176 |
| ENSMUSG00000040385 | Ppp1ca | protein_coding | 0.07748427 | 9.192561 | 0.071016 | 0.7932035 | 0.999176 |
| ENSMUSG00000037933 | Bicd2 | protein_coding | 0.1351037 | 4.460973 | 0.071003 | 0.7932219 | 0.999176 |
| ENSMUSG00000026478 | Lamc1 | protein_coding | 0.2150861 | 3.02326 | 0.0709672 | 0.7932728 | 0.999176 |
| ENSMUSG00000109156 | Gm45194 | protein_coding | 0.1739217 | 4.03125 | 0.0709423 | 0.7933081 | 0.999176 |
| ENSMUSG00000050447 | Lypd6 | protein_coding | -0.1505014 | 4.566478 | 0.0709343 | 0.7933195 | 0.999176 |
| ENSMUSG00000021939 | Ctsb | protein_coding | 0.08207465 | 10.06289 | 0.0709006 | 0.7933674 | 0.999176 |
| ENSMUSG00000029681 | Bcl7b | protein_coding | 0.1047713 | 6.484288 | 0.0708687 | 0.7934127 | 0.999176 |
| ENSMUSG00000064349 | mt-Tc | Mt_tRNA | 0.2259724 | 2.982255 | 0.0708512 | 0.7934376 | 0.999176 |
| ENSMUSG00000001173 | Ocrl | protein_coding | -0.1618848 | 5.090428 | 0.070794 | 0.7935244 | 0.999176 |
| ENSMUSG00000028756 | Pink1 | protein_coding | 0.09141481 | 7.559031 | 0.070687 | 0.7936712 | 0.999176 |
| ENSMUSG00000024104 | Washc2 | protein_coding | -0.1417055 | 5.126618 | 0.0705764 | 0.7938287 | 0.999176 |
| ENSMUSG00000026812 | Tsc1 | protein_coding | -0.1712003 | 3.868228 | 0.0705424 | 0.7938771 | 0.999176 |
| ENSMUSG00000041658 | Rragb | protein_coding | 0.3101514 | 5.21502 | 0.0715992 | 0.7940082 | 0.999176 |
| ENSMUSG00000024477 | Pggt1b | protein_coding | -0.1783366 | 3.324418 | 0.0703724 | 0.7941196 | 0.999176 |
| ENSMUSG00000045795 | Whamm | protein_coding | -0.2863471 | 2.505883 | 0.0703376 | 0.7941691 | 0.999176 |
| ENSMUSG00000050796 | B3galt6 | protein_coding | 0.1859779 | 3.461346 | 0.0702873 | 0.794241 | 0.999176 |
| ENSMUSG00000012535 | Tnpo3 | protein_coding | -0.1526678 | 5.407096 | 0.07025 | 0.7942997 | 0.999176 |
| ENSMUSG00000041084 | Ostc | protein_coding | 0.07519409 | 7.486781 | 0.0701321 | 0.7944628 | 0.999176 |
| ENSMUSG00000034032 | Rpap1 | protein_coding | 0.1875737 | 3.352134 | 0.069969 | 0.794696 | 0.999176 |
| ENSMUSG00000031517 | Gpm6a | protein_coding | -0.0764897 | 9.910279 | 0.0699664 | 0.7946997 | 0.999176 |

| ensembl_gene_id | external_gene_name | gene_biotype | log2 (fold change) | log2 (counts per million) | F-value | p-value | False discovery rate |
|---|---|---|---|---|---|---|---|
| ENSMUSG00000034066 | Farp2 | protein_coding | -0.2254619 | 2.859175 | 0.0698756 | 0.7948298 | 0.999176 |
| ENSMUSG00000006651 | Aplp1 | protein_coding | 0.1102731 | 9.291884 | 0.0697136 | 0.7950673 | 0.999176 |
| ENSMUSG00000006024 | Napa | protein_coding | 0.08693346 | 9.942773 | 0.0696735 | 0.7951194 | 0.999176 |
| ENSMUSG00000038383 | Pigu | protein_coding | 0.1263706 | 5.496662 | 0.0695454 | 0.7953033 | 0.999176 |
| ENSMUSG00000031344 | Gabrq | protein_coding | -0.1700661 | 4.170745 | 0.0692898 | 0.7956707 | 0.999176 |
| ENSMUSG00000032705 | Exd2 | protein_coding | 0.1456121 | 4.414391 | 0.0692101 | 0.7957854 | 0.999176 |
| ENSMUSG00000032294 | Pkm | protein_coding | 0.1160154 | 10.22861 | 0.0691737 | 0.7958431 | 0.999176 |
| ENSMUSG00000041695 | Kcnj2 | protein_coding | 0.4220999 | 2.23646 | 0.0691577 | 0.7958662 | 0.999176 |
| ENSMUSG00000034201 | Gas2l1 | protein_coding | -0.2203816 | 3.767729 | 0.0690348 | 0.7960433 | 0.999176 |
| ENSMUSG00000026824 | Kcnj3 | protein_coding | -0.1224483 | 4.637257 | 0.0690156 | 0.7960656 | 0.999176 |
| ENSMUSG00000021373 | Cap2 | protein_coding | -0.08886181 | 7.06883 | 0.0689996 | 0.7960886 | 0.999176 |
| ENSMUSG00000019868 | Vta1 | protein_coding | 0.08064536 | 6.910587 | 0.068728 | 0.7964806 | 0.999176 |
| ENSMUSG00000047921 | Trappc9 | protein_coding | 0.1175692 | 6.268041 | 0.0687145 | 0.7965056 | 0.999176 |
| ENSMUSG00000024782 | Ak3 | protein_coding | 0.220166 | 2.964821 | 0.0687064 | 0.7965119 | 0.999176 |
| ENSMUSG00000023007 | Prpf40b | protein_coding | -0.1430488 | 5.256868 | 0.0686045 | 0.7966592 | 0.999176 |
| ENSMUSG00000040265 | Dnm3 | protein_coding | 0.1173998 | 5.663832 | 0.0685913 | 0.7966783 | 0.999176 |
| ENSMUSG00000066838 | Zfp772 | protein_coding | 0.2127068 | 3.21986 | 0.0685557 | 0.7967299 | 0.999176 |
| ENSMUSG00000042694 | Stn1 | protein_coding | 0.1235419 | 4.440789 | 0.0684139 | 0.7969352 | 0.999176 |
| ENSMUSG00000024539 | Ptpn2 | protein_coding | -0.1328094 | 4.713151 | 0.0682831 | 0.7971247 | 0.999176 |
| ENSMUSG00000006463 | Zdhhc24 | protein_coding | 0.1700175 | 3.984993 | 0.0682734 | 0.7971388 | 0.999176 |
| ENSMUSG00000045287 | Rtn4rl1 | protein_coding | -0.1495336 | 3.9353 | 0.0682315 | 0.7971996 | 0.999176 |
| ENSMUSG00000063406 | Tmed5 | protein_coding | -0.1304932 | 5.217523 | 0.0681916 | 0.7972575 | 0.999176 |
| ENSMUSG00000047786 | Lix1 | protein_coding | -0.1288683 | 5.070151 | 0.0681203 | 0.7973609 | 0.999176 |
| ENSMUSG00000026894 | Morn5 | protein_coding | 0.2149355 | 3.188589 | 0.0681147 | 0.7973691 | 0.999176 |
| ENSMUSG00000025825 | Iscu | protein_coding | -0.1196275 | 4.795017 | 0.0681086 | 0.7973779 | 0.999176 |
| ENSMUSG00000030583 | Sipa1l3 | protein_coding | -0.2199892 | 3.080284 | 0.0680861 | 0.7974106 | 0.999176 |
| ENSMUSG00000027671 | Actl6a | protein_coding | -0.1392658 | 4.085813 | 0.068068 | 0.7974369 | 0.999176 |
| ENSMUSG00000052031 | Tagap1 | protein_coding | 0.126608 | 5.618157 | 0.0679699 | 0.7975795 | 0.999176 |
| ENSMUSG00000020388 | Pdlim4 | protein_coding | -0.1940822 | 3.714482 | 0.067831 | 0.7977815 | 0.999176 |
| ENSMUSG00000016150 | Tenm1 | protein_coding | -0.1480199 | 4.381688 | 0.0678184 | 0.7977998 | 0.999176 |
| ENSMUSG00000048707 | Tprn | protein_coding | 0.1483548 | 4.150385 | 0.0677997 | 0.797827 | 0.999176 |
| ENSMUSG00000046500 | Fam19a4 | protein_coding | -0.305904 | 2.344135 | 0.0677319 | 0.7979257 | 0.999176 |
| ENSMUSG00000056673 | Kdm5d | protein_coding | 0.2120514 | 2.893253 | 0.0677077 | 0.7979611 | 0.999176 |
| ENSMUSG00000042670 | Immp1l | protein_coding | -0.1075958 | 5.770079 | 0.0676561 | 0.7980362 | 0.999176 |
| ENSMUSG00000027676 | Ccdc39 | protein_coding | -0.2276333 | 3.229859 | 0.0676532 | 0.7980404 | 0.999176 |
| ENSMUSG00000032301 | Psma4 | protein_coding | -0.08773533 | 7.577675 | 0.0675819 | 0.7981443 | 0.999176 |
| ENSMUSG00000000399 | Ndufa9 | protein_coding | 0.07482171 | 9.013706 | 0.0675554 | 0.798183 | 0.999176 |
| ENSMUSG00000005034 | Prkacb | protein_coding | 0.08924193 | 8.785419 | 0.0675428 | 0.7982013 | 0.999176 |
| ENSMUSG00000028613 | Lrp8 | protein_coding | 0.1746737 | 4.297032 | 0.0675293 | 0.7982211 | 0.999176 |
| ENSMUSG00000028788 | Ptp4a2 | protein_coding | -0.09404812 | 8.424714 | 0.0674345 | 0.7983594 | 0.999176 |
| ENSMUSG00000014599 | Csf1 | protein_coding | 0.2093716 | 2.841161 | 0.0673902 | 0.7984241 | 0.999176 |
| ENSMUSG00000000441 | Raf1 | protein_coding | -0.2500149 | 2.610805 | 0.0673896 | 0.7984249 | 0.999176 |
| ENSMUSG00000040842 | Szrd1 | protein_coding | -0.1186148 | 5.368235 | 0.0673471 | 0.798487 | 0.999176 |
| ENSMUSG00000059540 | Tcea2 | protein_coding | -0.164255 | 4.568438 | 0.0672287 | 0.7986601 | 0.999176 |
| ENSMUSG00000032913 | Lrig2 | protein_coding | -0.1402524 | 3.96009 | 0.0671728 | 0.7987418 | 0.999176 |
| ENSMUSG00000036452 | Arhgap26 | protein_coding | 0.1130962 | 5.249432 | 0.0670963 | 0.7988538 | 0.999176 |
| ENSMUSG00000030739 | Myh14 | protein_coding | 0.1792539 | 3.342815 | 0.06708 | 0.7988776 | 0.999176 |
| ENSMUSG00000029047 | Pex10 | protein_coding | 0.1129011 | 4.967392 | 0.0670748 | 0.7988852 | 0.999176 |
| ENSMUSG00000045896 | Paip2b | protein_coding | 0.09840253 | 5.696697 | 0.0670537 | 0.7989161 | 0.999176 |
| ENSMUSG00000030091 | Nup210 | protein_coding | -0.2781702 | 2.789626 | 0.067004 | 0.7989889 | 0.999176 |
| ENSMUSG00000002320 | Tm9sf1 | protein_coding | 0.1943166 | 3.63793 | 0.0669847 | 0.7990173 | 0.999176 |
| ENSMUSG00000022602 | Arc | protein_coding | 0.1411698 | 5.510501 | 0.0669773 | 0.7990334 | 0.999176 |
| ENSMUSG00000102252 | Snrpn | protein_coding | -0.1116335 | 5.158851 | 0.0669081 | 0.7991294 | 0.999176 |
| ENSMUSG00000021226 | Acot2 | protein_coding | 0.1452955 | 5.08056 | 0.0668334 | 0.799239 | 0.999176 |
| ENSMUSG00000047694 | Yipf6 | protein_coding | -0.1675099 | 4.256792 | 0.0668258 | 0.7992501 | 0.999176 |
| ENSMUSG00000085885 | Gm11906 | processed_transcript | -0.1335236 | 4.577047 | 0.0668058 | 0.7992796 | 0.999176 |
| ENSMUSG00000071451 | Psmg4 | protein_coding | 0.1166527 | 5.430962 | 0.0667403 | 0.7993757 | 0.999176 |
| ENSMUSG00000021884 | Hacl1 | protein_coding | 0.1781381 | 3.644692 | 0.0666563 | 0.799499 | 0.999176 |
| ENSMUSG00000023046 | Igfbp6 | protein_coding | 0.1651906 | 4.68971 | 0.0666118 | 0.7995696 | 0.999176 |
| ENSMUSG00000029162 | Khk | protein_coding | -0.1036898 | 5.107388 | 0.0665437 | 0.7996645 | 0.999176 |
| ENSMUSG00000046679 | C87436 | protein_coding | -0.1401298 | 4.824338 | 0.0665425 | 0.7996662 | 0.999176 |
| ENSMUSG00000026766 | Mmadhc | protein_coding | -0.08711782 | 6.389746 | 0.0663511 | 0.7999479 | 0.999176 |
| ENSMUSG00000038975 | Rabggtb | protein_coding | 0.06915397 | 7.224132 | 0.0662702 | 0.800067 | 0.999176 |
| ENSMUSG00000019179 | Mdh2 | protein_coding | 0.09014978 | 10.78112 | 0.066231 | 0.8001248 | 0.999176 |
| ENSMUSG00000078970 | Wdr92 | protein_coding | 0.1588595 | 4.645181 | 0.0661945 | 0.8001786 | 0.999176 |
| ENSMUSG00000038630 | Zkscan16 | protein_coding | -0.2194659 | 2.95242 | 0.066193 | 0.8001808 | 0.999176 |
| ENSMUSG00000024792 | Zfpl1 | protein_coding | 0.2138167 | 3.797925 | 0.0661711 | 0.8002185 | 0.999176 |
| ENSMUSG00000032177 | Pde4a | protein_coding | -0.08017655 | 6.70643 | 0.066044 | 0.8004006 | 0.999176 |
| ENSMUSG00000025809 | Itgb1 | protein_coding | 0.1466768 | 4.43776 | 0.0660271 | 0.8004256 | 0.999176 |
| ENSMUSG00000021264 | Yy1 | protein_coding | -0.1497943 | 4.865081 | 0.0659393 | 0.8005553 | 0.999176 |
| ENSMUSG00000057367 | Birc2 | protein_coding | -0.132092 | 5.390642 | 0.0659014 | 0.8006113 | 0.999176 |
| ENSMUSG00000083674 | Zfp133-ps | transcribed_unitary_p | -0.2259362 | 3.335927 | 0.0658226 | 0.8007278 | 0.999176 |
| ENSMUSG00000051351 | Zfp46 | protein_coding | 0.1774493 | 3.5209 | 0.0657523 | 0.8008318 | 0.999176 |
| ENSMUSG00000041650 | Pcca | protein_coding | 0.1509384 | 4.350312 | 0.0657181 | 0.8008824 | 0.999176 |
| ENSMUSG00000053754 | Chd8 | protein_coding | 0.2250628 | 3.055712 | 0.0657179 | 0.8008827 | 0.999176 |
| ENSMUSG00000033161 | Atp1a1 | protein_coding | -0.1433664 | 4.929642 | 0.0655447 | 0.8011392 | 0.999176 |
| ENSMUSG00000039717 | Ralyl | protein_coding | 0.07715011 | 7.948292 | 0.0655443 | 0.8011397 | 0.999176 |
| ENSMUSG00000025521 | Tmem192 | protein_coding | -0.1641468 | 3.808438 | 0.0655404 | 0.8011455 | 0.999176 |

| ensembl_gene_id | external_gene_name | gene_biotype | log2 (fold change) | log2 (counts per million) | F-value | p-value | False discovery rate |
|---|---|---|---|---|---|---|---|
| ENSMUSG00000022571 | Pycrl | protein_coding | -0.08335217 | 6.757808 | 0.0654576 | 0.8012684 | 0.999176 |
| ENSMUSG00000028920 | Fbxo42 | protein_coding | 0.1563854 | 3.880096 | 0.0654302 | 0.8013089 | 0.999176 |
| ENSMUSG00000036591 | Arhgap21 | protein_coding | -0.1534013 | 4.502984 | 0.0654252 | 0.8013164 | 0.999176 |
| ENSMUSG00000037072 | Selenof | protein_coding | -0.08680768 | 8.841279 | 0.0653859 | 0.8013748 | 0.999176 |
| ENSMUSG00000040339 | Fam102b | protein_coding | -0.09622411 | 5.639958 | 0.065325 | 0.8014651 | 0.999176 |
| ENSMUSG00000079108 | Srp54b | protein_coding | -0.1826888 | 3.340424 | 0.0653194 | 0.8014734 | 0.999176 |
| ENSMUSG00000029486 | Mrpl1 | protein_coding | 0.108149 | 5.919402 | 0.0653052 | 0.8014945 | 0.999176 |
| ENSMUSG00000037111 | Setd7 | protein_coding | 0.1145085 | 4.984668 | 0.0652927 | 0.8015131 | 0.999176 |
| ENSMUSG00000035783 | Acta2 | protein_coding | 0.2684526 | 3.19598 | 0.0651455 | 0.8017371 | 0.999176 |
| ENSMUSG00000029120 | Ppp2r2c | protein_coding | -0.07473195 | 7.304229 | 0.0651385 | 0.8017423 | 0.999176 |
| ENSMUSG00000029636 | Wasf3 | protein_coding | 0.0999912 | 5.556638 | 0.0651304 | 0.8017543 | 0.999176 |
| ENSMUSG00000030695 | Aldoa | protein_coding | 0.08701496 | 11.99141 | 0.0650474 | 0.8018779 | 0.999176 |
| ENSMUSG00000073643 | Wdfy1 | protein_coding | -0.3112026 | 4.899426 | 0.0660964 | 0.8018867 | 0.999176 |
| ENSMUSG00000026623 | Lpgat1 | protein_coding | -0.08073856 | 6.424591 | 0.0650339 | 0.8018979 | 0.999176 |
| ENSMUSG00000030986 | Dhx32 | protein_coding | 0.1302709 | 4.597175 | 0.0649943 | 0.8019568 | 0.999176 |
| ENSMUSG00000032119 | Hinfp | protein_coding | 0.2366347 | 3.263631 | 0.0648772 | 0.8021313 | 0.999176 |
| ENSMUSG00000062075 | Lmnb2 | protein_coding | 0.2059562 | 3.29539 | 0.0648406 | 0.8021858 | 0.999176 |
| ENSMUSG00000017132 | Cyth1 | protein_coding | 0.09829864 | 5.502106 | 0.0648202 | 0.8022163 | 0.999176 |
| ENSMUSG00000071856 | Mcc | protein_coding | -0.3836526 | 2.394947 | 0.0647779 | 0.8022846 | 0.999176 |
| ENSMUSG00000021044 | Adck1 | protein_coding | -0.12747 | 5.485597 | 0.0646736 | 0.8024349 | 0.999176 |
| ENSMUSG00000039953 | Clstn1 | protein_coding | -0.07912459 | 7.959737 | 0.0645869 | 0.8025645 | 0.999176 |
| ENSMUSG00000056073 | Grik2 | protein_coding | -0.1144872 | 5.420069 | 0.0645437 | 0.802629 | 0.999176 |
| ENSMUSG00000059890 | Ube4a | protein_coding | -0.1527711 | 3.818743 | 0.0645346 | 0.8026426 | 0.999176 |
| ENSMUSG00000042035 | Igsf3 | protein_coding | 0.1135697 | 5.476602 | 0.0644844 | 0.8027176 | 0.999176 |
| ENSMUSG00000040412 | 5330417C22Rik | protein_coding | -0.1388523 | 4.990528 | 0.0642938 | 0.8030028 | 0.999176 |
| ENSMUSG00000032285 | Dnaja4 | protein_coding | -0.2331516 | 2.787859 | 0.0641813 | 0.8031713 | 0.999176 |
| ENSMUSG00000052144 | Ppp4r2 | protein_coding | 0.1119324 | 4.914874 | 0.0639792 | 0.8034746 | 0.999176 |
| ENSMUSG00000032905 | Atg12 | protein_coding | -0.07359021 | 6.552263 | 0.0639735 | 0.8034832 | 0.999176 |
| ENSMUSG00000039286 | Fndc3b | protein_coding | 0.160462 | 3.988064 | 0.0639442 | 0.8035271 | 0.999176 |
| ENSMUSG00000031065 | Cdk16 | protein_coding | 0.102977 | 6.001277 | 0.0638721 | 0.8036355 | 0.999176 |
| ENSMUSG00000029771 | Irf5 | protein_coding | 0.2441644 | 2.8811 | 0.0638225 | 0.80371 | 0.999176 |
| ENSMUSG00000026442 | Nfasc | protein_coding | 0.08614231 | 5.774131 | 0.063815 | 0.8037213 | 0.999176 |
| ENSMUSG00000036639 | Nudt1 | protein_coding | 0.1261654 | 4.102397 | 0.0637851 | 0.8037663 | 0.999176 |
| ENSMUSG00000024383 | Map3k2 | protein_coding | -0.1576592 | 3.762127 | 0.0637578 | 0.8038073 | 0.999176 |
| ENSMUSG00000039318 | Rab3gap2 | protein_coding | 0.1629948 | 4.163688 | 0.0637384 | 0.8038365 | 0.999176 |
| ENSMUSG00000022960 | Donson | protein_coding | -0.4040422 | 2.715215 | 0.0638436 | 0.8039447 | 0.999176 |
| ENSMUSG00000072964 | Bhlhb9 | protein_coding | -0.09897366 | 5.937865 | 0.0636411 | 0.803983 | 0.999176 |
| ENSMUSG00000030235 | Slco1c1 | protein_coding | 0.1686426 | 3.677615 | 0.0634635 | 0.8042506 | 0.999176 |
| ENSMUSG00000031969 | Acad8 | protein_coding | 0.111884 | 5.460269 | 0.0633847 | 0.8043695 | 0.999176 |
| ENSMUSG00000043461 | Sptssb | protein_coding | -0.2471897 | 2.800324 | 0.0632418 | 0.8045852 | 0.999176 |
| ENSMUSG00000015501 | Hivep2 | protein_coding | -0.2709954 | 2.894832 | 0.0632007 | 0.8046474 | 0.999176 |
| ENSMUSG00000040451 | Sgms1 | protein_coding | 0.1439977 | 4.122359 | 0.0631525 | 0.8047202 | 0.999176 |
| ENSMUSG00000022122 | Ednrb | protein_coding | 0.1098832 | 6.100862 | 0.0630492 | 0.8048764 | 0.999176 |
| ENSMUSG00000053070 | 9230110C19Rik | protein_coding | 0.1453452 | 4.620341 | 0.0630191 | 0.804922 | 0.999176 |
| ENSMUSG00000029016 | Clcn6 | protein_coding | 0.09859606 | 5.548645 | 0.0629873 | 0.8049701 | 0.999176 |
| ENSMUSG00000040747 | Cd53 | protein_coding | -0.1718146 | 4.078831 | 0.0629759 | 0.8049874 | 0.999176 |
| ENSMUSG00000024955 | Esrra | protein_coding | -0.2925488 | 2.598698 | 0.0629139 | 0.8050813 | 0.999176 |
| ENSMUSG00000022468 | Endou | protein_coding | -0.4203545 | 3.049066 | 0.0630913 | 0.8054337 | 0.999176 |
| ENSMUSG00000050751 | Pgbd5 | protein_coding | 0.0822564 | 6.575469 | 0.0626402 | 0.8054964 | 0.999176 |
| ENSMUSG00000021936 | Mapk8 | protein_coding | -0.1046594 | 5.316795 | 0.0625993 | 0.8055585 | 0.999176 |
| ENSMUSG00000054679 | Srsf12 | protein_coding | -0.1454904 | 5.326645 | 0.0625986 | 0.8055647 | 0.999176 |
| ENSMUSG00000064343 | mt-Tq | Mt_tRNA | 0.2609444 | 2.419131 | 0.0625854 | 0.8055796 | 0.999176 |
| ENSMUSG00000038128 | Camk4 | protein_coding | -0.1132512 | 6.331167 | 0.0624081 | 0.8058543 | 0.999176 |
| ENSMUSG00000023571 | C1qtnf12 | protein_coding | -0.2459934 | 3.022559 | 0.0623058 | 0.806005 | 0.999176 |
| ENSMUSG00000031875 | Cmtm3 | protein_coding | 0.1810963 | 3.327982 | 0.0622846 | 0.8060373 | 0.999176 |
| ENSMUSG00000063229 | Ldha | protein_coding | 0.07422931 | 8.555127 | 0.0622512 | 0.8060882 | 0.999176 |
| ENSMUSG00000040490 | Lrfn2 | protein_coding | -0.1963181 | 2.951924 | 0.0622478 | 0.8060933 | 0.999176 |
| ENSMUSG00000029176 | Anapc4 | protein_coding | -0.0905048 | 6.037862 | 0.0621775 | 0.8062005 | 0.999176 |
| ENSMUSG00000030282 | Cmas | protein_coding | 0.07244523 | 6.744705 | 0.0621632 | 0.8062222 | 0.999176 |
| ENSMUSG00000031266 | Gla | protein_coding | 0.1862886 | 3.42668 | 0.0621525 | 0.8062386 | 0.999176 |
| ENSMUSG00000049878 | Rlf | protein_coding | 0.15982 | 3.754234 | 0.0620961 | 0.8063246 | 0.999176 |
| ENSMUSG00000034107 | Ano7 | protein_coding | -0.3014216 | 2.574093 | 0.0620633 | 0.8063746 | 0.999176 |
| ENSMUSG00000054263 | Lifr | protein_coding | 0.1732731 | 3.79113 | 0.0620516 | 0.8063925 | 0.999176 |
| ENSMUSG00000018339 | Gpx3 | protein_coding | 0.09251401 | 9.533676 | 0.0620178 | 0.806444 | 0.999176 |
| ENSMUSG00000020516 | Rps6kb1 | protein_coding | 0.1220524 | 5.715168 | 0.0620051 | 0.8064635 | 0.999176 |
| ENSMUSG00000027835 | Pdcd10 | protein_coding | 0.09031449 | 6.234613 | 0.0618534 | 0.8066953 | 0.999176 |
| ENSMUSG00000031558 | Slit2 | protein_coding | -0.182784 | 3.494477 | 0.0618144 | 0.8067548 | 0.999176 |
| ENSMUSG00000021061 | Sptb | protein_coding | -0.1928465 | 3.921842 | 0.0616759 | 0.8069718 | 0.999176 |
| ENSMUSG00000028719 | Cmpk1 | protein_coding | -0.08473688 | 7.302322 | 0.0615893 | 0.8070994 | 0.999176 |
| ENSMUSG00000030884 | Uqcrc2 | protein_coding | -0.07551652 | 8.887544 | 0.0614593 | 0.8072987 | 0.999176 |
| ENSMUSG00000052534 | Pbx1 | protein_coding | 0.07990774 | 7.052924 | 0.0614457 | 0.8073197 | 0.999176 |
| ENSMUSG00000053046 | Brsk2 | protein_coding | 0.1075694 | 5.330846 | 0.061372 | 0.8074327 | 0.999176 |
| ENSMUSG00000028634 | Hivep3 | protein_coding | 0.1487401 | 4.620867 | 0.0613211 | 0.8075109 | 0.999176 |
| ENSMUSG00000052906 | Ubxn8 | protein_coding | -0.0925682 | 6.051718 | 0.0612938 | 0.8075528 | 0.999176 |
| ENSMUSG00000024097 | Srsf7 | protein_coding | 0.08199915 | 8.150589 | 0.0612378 | 0.8076389 | 0.999176 |
| ENSMUSG00000039661 | Dusp26 | protein_coding | -0.07838009 | 7.594853 | 0.0611935 | 0.8077069 | 0.999176 |
| ENSMUSG00000059734 | Ndufs8 | protein_coding | 0.07736643 | 9.673631 | 0.0610434 | 0.807938 | 0.999176 |
| ENSMUSG00000043872 | Zmym1 | protein_coding | -0.1837679 | 3.116851 | 0.0609799 | 0.8080357 | 0.999176 |

| ensembl_gene_id | external_gene_name | gene_biotype | log2 (fold change) | log2 (counts per million) | F-value | p-value | False discovery rate |
|---|---|---|---|---|---|---|---|
| ENSMUSG00000019971 | Cep290 | protein_coding | -0.1875336 | 3.45879 | 0.0608666 | 0.8082103 | 0.999176 |
| ENSMUSG00000040738 | Ints8 | protein_coding | -0.1669417 | 3.724447 | 0.060722 | 0.8084334 | 0.999176 |
| ENSMUSG00000054477 | Kcnn2 | protein_coding | -0.1476132 | 3.612488 | 0.0606476 | 0.8085484 | 0.999176 |
| ENSMUSG00000053617 | Sh3pxd2a | protein_coding | -0.2437646 | 3.092813 | 0.0606386 | 0.8085622 | 0.999176 |
| ENSMUSG00000026496 | Parp1 | protein_coding | 0.1018619 | 5.257757 | 0.0606321 | 0.8085723 | 0.999176 |
| ENSMUSG00000000142 | Axin2 | protein_coding | -0.1776876 | 3.53976 | 0.0605983 | 0.8086245 | 0.999176 |
| ENSMUSG00000036427 | Gpi1 | protein_coding | 0.103431 | 9.013078 | 0.0605595 | 0.8086895 | 0.999176 |
| ENSMUSG00000031594 | Fgl1 | protein_coding | -0.2678934 | 2.721859 | 0.0604372 | 0.8088737 | 0.999176 |
| ENSMUSG00000075486 | Commd6 | protein_coding | -0.09326263 | 7.794029 | 0.0604118 | 0.8089129 | 0.999176 |
| ENSMUSG00000078307 | AI593442 | protein_coding | 0.1202855 | 8.096052 | 0.0605176 | 0.8089572 | 0.999176 |
| ENSMUSG00000025586 | Cpeb1 | protein_coding | 0.1517942 | 5.172912 | 0.0601257 | 0.8093692 | 0.999176 |
| ENSMUSG00000027746 | Ufm1 | protein_coding | -0.08703118 | 6.165844 | 0.0601074 | 0.8093849 | 0.999176 |
| ENSMUSG00000024558 | Mapk4 | protein_coding | -0.1165905 | 4.869485 | 0.0600857 | 0.8094185 | 0.999176 |
| ENSMUSG00000028139 | Riiad1 | protein_coding | -0.1154978 | 5.223337 | 0.0599592 | 0.809615 | 0.999176 |
| ENSMUSG00000050052 | Tdrp | protein_coding | -0.1351528 | 4.529503 | 0.0599576 | 0.8096176 | 0.999176 |
| ENSMUSG00000026626 | Ppp2r5a | protein_coding | -0.1465803 | 4.445671 | 0.0599465 | 0.8096347 | 0.999176 |
| ENSMUSG00000025421 | Hdhd2 | protein_coding | 0.07024418 | 8.431282 | 0.0599175 | 0.8096799 | 0.999176 |
| ENSMUSG00000075415 | Fnbp1 | protein_coding | -0.0924788 | 6.990679 | 0.0598853 | 0.80973 | 0.999176 |
| ENSMUSG00000041765 | Ubac2 | protein_coding | 0.1031068 | 5.337075 | 0.0598292 | 0.8098173 | 0.999176 |
| ENSMUSG00000003923 | Tfam | protein_coding | 0.09797851 | 5.862559 | 0.0598019 | 0.8098596 | 0.999176 |
| ENSMUSG00000028034 | Fubp1 | protein_coding | -0.08896892 | 6.71912 | 0.0597097 | 0.8100032 | 0.999176 |
| ENSMUSG00000018507 | Trpv2 | protein_coding | 0.1827082 | 4.6955 | 0.0598264 | 0.8100041 | 0.999176 |
| ENSMUSG00000044150 | A830080D01Rik | protein_coding | 0.2131712 | 3.460707 | 0.0595955 | 0.8101813 | 0.999176 |
| ENSMUSG00000036062 | Phf24 | protein_coding | -0.08431432 | 6.335359 | 0.05957 | 0.810221 | 0.999176 |
| ENSMUSG00000039431 | Mtmr7 | protein_coding | -0.09457604 | 6.083458 | 0.0594407 | 0.8104228 | 0.999176 |
| ENSMUSG00000032870 | Smap2 | protein_coding | 0.06553877 | 7.112394 | 0.0593881 | 0.8105049 | 0.999176 |
| ENSMUSG00000030647 | Ndufc2 | protein_coding | 0.09000867 | 9.492655 | 0.0592481 | 0.8107238 | 0.999176 |
| ENSMUSG00000020225 | Tmbim4 | protein_coding | 0.07964881 | 6.932846 | 0.059195 | 0.8108069 | 0.999176 |
| ENSMUSG00000031497 | Tnfsf13b | protein_coding | 0.2206594 | 3.010661 | 0.0591692 | 0.8108473 | 0.999176 |
| ENSMUSG00000000439 | Mkrn2 | protein_coding | -0.09884417 | 5.914693 | 0.0591091 | 0.8109414 | 0.999176 |
| ENSMUSG00000024665 | Fads2 | protein_coding | -0.1443191 | 4.030559 | 0.0590875 | 0.8109752 | 0.999176 |
| ENSMUSG00000040586 | Ofd1 | protein_coding | 0.2518587 | 2.848027 | 0.0590305 | 0.8110645 | 0.999176 |
| ENSMUSG00000031068 | Glrx3 | protein_coding | 0.07535222 | 6.392049 | 0.0589533 | 0.8111855 | 0.999176 |
| ENSMUSG00000020034 | Tcp11l2 | protein_coding | 0.1546974 | 5.714037 | 0.0590997 | 0.8112045 | 0.999176 |
| ENSMUSG00000089759 | 3632454L22Rik | processed_transcript | -0.2411594 | 2.614672 | 0.0589132 | 0.8112484 | 0.999176 |
| ENSMUSG00000030747 | Dgat2 | protein_coding | 0.1033534 | 5.811964 | 0.0588978 | 0.8112726 | 0.999176 |
| ENSMUSG00000026504 | Sdccag8 | protein_coding | 0.165649 | 4.755366 | 0.0588619 | 0.811347 | 0.999176 |
| ENSMUSG00000032850 | Rnft2 | protein_coding | 0.1298425 | 4.516266 | 0.058772 | 0.8114702 | 0.999176 |
| ENSMUSG00000039686 | Zer1 | protein_coding | 0.1757314 | 5.816097 | 0.0590215 | 0.8115617 | 0.999176 |
| ENSMUSG00000034317 | Trim59 | protein_coding | 0.1921325 | 4.038573 | 0.0587137 | 0.8115667 | 0.999176 |
| ENSMUSG00000020739 | Nup85 | protein_coding | 0.1188335 | 5.327087 | 0.0586421 | 0.8116743 | 0.999176 |
| ENSMUSG00000037905 | Bri3bp | protein_coding | 0.09088284 | 5.625456 | 0.0584556 | 0.8119679 | 0.999176 |
| ENSMUSG00000026135 | Zfp142 | protein_coding | -0.3001964 | 2.531789 | 0.0584504 | 0.8119761 | 0.999176 |
| ENSMUSG00000056158 | Car10 | protein_coding | -0.1282896 | 6.295306 | 0.0583016 | 0.8124024 | 0.999176 |
| ENSMUSG00000042423 | Fbrs | protein_coding | -0.1768818 | 3.257938 | 0.0581765 | 0.8124081 | 0.999176 |
| ENSMUSG00000038742 | Angptl6 | protein_coding | 0.1580787 | 4.581477 | 0.0581781 | 0.8124106 | 0.999176 |
| ENSMUSG00000020642 | Rnf144a | protein_coding | -0.1690455 | 3.501786 | 0.0581537 | 0.8124442 | 0.999176 |
| ENSMUSG00000024431 | Nr3c1 | protein_coding | -0.0758609 | 6.420683 | 0.0580519 | 0.812605 | 0.999176 |
| ENSMUSG00000022197 | Pdzd2 | protein_coding | -0.1729687 | 3.694721 | 0.0580115 | 0.8126689 | 0.999176 |
| ENSMUSG00000030098 | Grip2 | protein_coding | 0.2283617 | 3.227024 | 0.0580084 | 0.8126739 | 0.999176 |
| ENSMUSG00000061132 | Blnk | protein_coding | 0.4610436 | 2.085736 | 0.0580404 | 0.8127165 | 0.999176 |
| ENSMUSG00000001847 | Rac1 | protein_coding | -0.07742179 | 6.536388 | 0.0579197 | 0.8128142 | 0.999176 |
| ENSMUSG00000042763 | Maneal | protein_coding | -0.1425561 | 4.41117 | 0.0578901 | 0.812861 | 0.999176 |
| ENSMUSG00000039879 | Heca | protein_coding | -0.1358804 | 3.829113 | 0.0578812 | 0.8128751 | 0.999176 |
| ENSMUSG00000031007 | Atp6ap2 | protein_coding | -0.06972541 | 8.573246 | 0.0578242 | 0.8129654 | 0.999176 |
| ENSMUSG00000058174 | Gm5148 | protein_coding | 0.1286082 | 4.361022 | 0.0577661 | 0.8130575 | 0.999176 |
| ENSMUSG00000026563 | Tada1 | protein_coding | 0.2426641 | 2.606274 | 0.0576509 | 0.8132402 | 0.999176 |
| ENSMUSG00000028538 | St3gal3 | protein_coding | -0.1771125 | 3.241666 | 0.0576475 | 0.8132456 | 0.999176 |
| ENSMUSG00000020704 | Asic2 | protein_coding | -0.08269921 | 5.867011 | 0.0576161 | 0.8132956 | 0.999176 |
| ENSMUSG00000034575 | Papd7 | protein_coding | -0.1158392 | 4.694228 | 0.0575808 | 0.8133515 | 0.999176 |
| ENSMUSG00000025016 | Tm9sf3 | protein_coding | -0.09874368 | 6.079951 | 0.0575752 | 0.8133605 | 0.999176 |
| ENSMUSG00000026782 | Abi2 | protein_coding | -0.07703507 | 7.192862 | 0.0575524 | 0.8133967 | 0.999176 |
| ENSMUSG00000019945 | 1700040L02Rik | protein_coding | -0.1592803 | 3.569374 | 0.0574591 | 0.813545 | 0.999176 |
| ENSMUSG00000024513 | Mbd2 | protein_coding | -0.1504421 | 3.827741 | 0.0572254 | 0.8139169 | 0.999176 |
| ENSMUSG00000057835 | Zfp119a | protein_coding | -0.2293684 | 3.001433 | 0.0572036 | 0.8139516 | 0.999176 |
| ENSMUSG00000031391 | L1cam | protein_coding | -0.09020204 | 5.602604 | 0.0569983 | 0.8142791 | 0.999176 |
| ENSMUSG00000044783 | Hjurp | protein_coding | -0.1140085 | 4.757925 | 0.0569975 | 0.8142803 | 0.999176 |
| ENSMUSG00000029064 | Gnb1 | protein_coding | 0.1211737 | 12.19785 | 0.0570355 | 0.8142902 | 0.999176 |
| ENSMUSG00000031482 | Slc25a15 | protein_coding | -0.1750599 | 3.020939 | 0.0569467 | 0.8143615 | 0.999176 |
| ENSMUSG00000000197 | Nalcn | protein_coding | -0.1195836 | 5.58775 | 0.0569063 | 0.814426 | 0.999176 |
| ENSMUSG00000025102 | 3110040N11Rik | protein_coding | -0.08430896 | 6.076094 | 0.0569018 | 0.8144332 | 0.999176 |
| ENSMUSG00000006586 | Runx1t1 | protein_coding | -0.1169218 | 6.322734 | 0.056833 | 0.8145693 | 0.999176 |
| ENSMUSG00000050079 | Rspry1 | protein_coding | 0.114507 | 4.87343 | 0.0567155 | 0.8147312 | 0.999176 |
| ENSMUSG00000040723 | Rcsd1 | protein_coding | 0.2635103 | 2.563252 | 0.0566758 | 0.8147949 | 0.999176 |
| ENSMUSG00000027184 | Caprin1 | protein_coding | -0.08355932 | 6.502733 | 0.056658 | 0.8148233 | 0.999176 |
| ENSMUSG00000058594 | Fbxo18 | protein_coding | -0.09001354 | 5.709384 | 0.0565714 | 0.8149621 | 0.999176 |
| ENSMUSG00000049858 | Suox | protein_coding | -0.1883607 | 3.829617 | 0.056536 | 0.8150236 | 0.999176 |
| ENSMUSG00000020198 | Ap3d1 | protein_coding | -0.09103405 | 6.565677 | 0.0564982 | 0.8150795 | 0.999176 |

| ensembl_gene_id | external_gene_name | gene_biotype | log2 (fold change) | log2 (counts per million) | F-value | p-value | False discovery rate |
|---|---|---|---|---|---|---|---|
| ENSMUSG00000024140 | Epas1 | protein_coding | -0.1534872 | 4.29406 | 0.0564125 | 0.815217 | 0.999176 |
| ENSMUSG00000032174 | Icam5 | protein_coding | -0.3145277 | 2.171268 | 0.0563993 | 0.8152381 | 0.999176 |
| ENSMUSG00000021311 | Mtr | protein_coding | -0.2020229 | 3.017225 | 0.0563583 | 0.8153041 | 0.999176 |
| ENSMUSG00000004096 | Cwc15 | protein_coding | 0.07834531 | 8.159173 | 0.0563511 | 0.8153155 | 0.999176 |
| ENSMUSG00000033653 | Vps8 | protein_coding | 0.1358196 | 4.79667 | 0.0563223 | 0.8153619 | 0.999176 |
| ENSMUSG00000032295 | Man2c1 | protein_coding | 0.1297949 | 5.04176 | 0.056297 | 0.8154025 | 0.999176 |
| ENSMUSG00000023791 | Pigx | protein_coding | -0.1711264 | 4.053542 | 0.0562366 | 0.8155045 | 0.999176 |
| ENSMUSG00000103183 | Gm37090 | TEC | -0.3740436 | 2.36524 | 0.0561942 | 0.8155726 | 0.999176 |
| ENSMUSG00000028207 | Asph | protein_coding | -0.1053995 | 5.442789 | 0.056067 | 0.8157725 | 0.999176 |
| ENSMUSG00000037190 | Cyb561d2 | protein_coding | 0.08606899 | 6.107874 | 0.0560595 | 0.8157847 | 0.999176 |
| ENSMUSG00000042271 | Nxt2 | protein_coding | -0.09697919 | 6.680657 | 0.0559746 | 0.8159262 | 0.999176 |
| ENSMUSG00000064036 | Mro | protein_coding | -0.09329731 | 5.294121 | 0.0558989 | 0.8160435 | 0.999176 |
| ENSMUSG00000042073 | Abhd14b | protein_coding | 0.1632402 | 3.539142 | 0.0558288 | 0.8161566 | 0.999176 |
| ENSMUSG00000020546 | Stxbp4 | protein_coding | 0.1494346 | 4.780325 | 0.0558054 | 0.8161993 | 0.999176 |
| ENSMUSG00000028521 | Slc35d1 | protein_coding | 0.1713297 | 3.083162 | 0.0557899 | 0.8162195 | 0.999176 |
| ENSMUSG00000040354 | Mars | protein_coding | -0.1443702 | 4.800314 | 0.0557596 | 0.8162733 | 0.999176 |
| ENSMUSG00000021639 | Gtf2h2 | protein_coding | -0.08822284 | 5.981032 | 0.0557 | 0.8163648 | 0.999176 |
| ENSMUSG00000020929 | Eftud2 | protein_coding | -0.1096049 | 5.697167 | 0.0555865 | 0.8165483 | 0.999176 |
| ENSMUSG00000030103 | Bhlhe40 | protein_coding | -0.1013619 | 6.30507 | 0.0554735 | 0.8167312 | 0.999176 |
| ENSMUSG00000028469 | Npr2 | protein_coding | 0.1824885 | 3.538516 | 0.055116 | 0.8173112 | 0.999176 |
| ENSMUSG00000075296 | Aldh3b2 | protein_coding | -0.1087709 | 5.345022 | 0.0550453 | 0.8174261 | 0.999176 |
| ENSMUSG00000073102 | Drc1 | protein_coding | 0.1096222 | 4.88221 | 0.0549914 | 0.8175139 | 0.999176 |
| ENSMUSG00000024990 | Rbp4 | protein_coding | 0.2221248 | 2.894015 | 0.0549618 | 0.817562 | 0.999176 |
| ENSMUSG00000036040 | Adamtsl2 | protein_coding | -0.1619204 | 3.447616 | 0.0548936 | 0.8176731 | 0.999176 |
| ENSMUSG00000026784 | Pdss1 | protein_coding | 0.1707143 | 3.362815 | 0.0548771 | 0.8177 | 0.999176 |
| ENSMUSG00000022508 | Bcl6 | protein_coding | 0.2066613 | 3.048045 | 0.0548494 | 0.8177451 | 0.999176 |
| ENSMUSG00000059742 | Kcnh7 | protein_coding | -0.131615 | 3.864898 | 0.0548482 | 0.817747 | 0.999176 |
| ENSMUSG00000068040 | Tm9sf4 | protein_coding | 0.08333527 | 6.085283 | 0.0548251 | 0.8177847 | 0.999176 |
| ENSMUSG00000062591 | Tubb4a | protein_coding | 0.1110294 | 11.07819 | 0.054757 | 0.8179132 | 0.999176 |
| ENSMUSG00000014905 | Dnajb9 | protein_coding | 0.116678 | 5.484626 | 0.0547446 | 0.817916 | 0.999176 |
| ENSMUSG00000072872 | Rybp | protein_coding | -0.2342565 | 2.674842 | 0.054698 | 0.8179919 | 0.999176 |
| ENSMUSG00000079414 | Gm11110 | protein_coding | 0.28086 | 2.61234 | 0.0546067 | 0.8181411 | 0.999176 |
| ENSMUSG00000021514 | Zfp369 | protein_coding | -0.1805039 | 3.534013 | 0.05459 | 0.8181684 | 0.999176 |
| ENSMUSG00000055334 | Snupn | protein_coding | -0.09933837 | 5.270604 | 0.0545064 | 0.8183049 | 0.999176 |
| ENSMUSG00000020891 | Alox8 | protein_coding | 0.2720523 | 2.964018 | 0.0544641 | 0.8183788 | 0.999176 |
| ENSMUSG00000062127 | Cttnbp2nl | protein_coding | -0.1400575 | 3.730858 | 0.0544507 | 0.8183961 | 0.999176 |
| ENSMUSG00000027091 | Zc3h15 | protein_coding | 0.0929191 | 6.253886 | 0.0544365 | 0.8184193 | 0.999176 |
| ENSMUSG00000024534 | Sncaip | protein_coding | -0.2733046 | 2.571522 | 0.0544272 | 0.8184345 | 0.999176 |
| ENSMUSG00000068876 | Cgn | protein_coding | -0.1737019 | 3.525451 | 0.0543312 | 0.8185916 | 0.999176 |
| ENSMUSG00000032118 | Fez1 | protein_coding | -0.08391935 | 9.195684 | 0.0543136 | 0.8186204 | 0.999176 |
| ENSMUSG00000031781 | Ciapin1 | protein_coding | 0.07263563 | 7.512321 | 0.0543081 | 0.8186294 | 0.999176 |
| ENSMUSG00000052593 | Adam17 | protein_coding | -0.3012636 | 2.509099 | 0.0542858 | 0.818666 | 0.999176 |
| ENSMUSG00000031399 | Fam3a | protein_coding | 0.1018105 | 5.11049 | 0.0541683 | 0.8188585 | 0.999176 |
| ENSMUSG00000028668 | Eloa | protein_coding | -0.183663 | 3.051228 | 0.0541485 | 0.8188911 | 0.999176 |
| ENSMUSG00000035629 | Rubcn | protein_coding | -0.114499 | 5.172054 | 0.054131 | 0.8189199 | 0.999176 |
| ENSMUSG00000029148 | Nrbp1 | protein_coding | -0.1020042 | 6.827885 | 0.05404 | 0.819074 | 0.999176 |
| ENSMUSG00000033799 | Fam208b | protein_coding | -0.2869959 | 2.470024 | 0.0539573 | 0.8192051 | 0.999176 |
| ENSMUSG00000001542 | Ell2 | protein_coding | -0.1644847 | 3.625463 | 0.0539215 | 0.8192639 | 0.999176 |
| ENSMUSG00000032579 | Hemk1 | protein_coding | -0.1222504 | 4.990889 | 0.0538816 | 0.8193295 | 0.999176 |
| ENSMUSG00000022848 | Dirc2 | protein_coding | 0.1120658 | 5.275796 | 0.0538578 | 0.8193687 | 0.999176 |
| ENSMUSG00000021470 | Ercc6l2 | protein_coding | 0.1620656 | 3.592633 | 0.0538364 | 0.819404 | 0.999176 |
| ENSMUSG00000008976 | Gabpa | protein_coding | -0.1824784 | 2.888752 | 0.0538034 | 0.8194582 | 0.999176 |
| ENSMUSG00000002265 | Peg3 | protein_coding | -0.08318481 | 8.47597 | 0.0537998 | 0.8194641 | 0.999176 |
| ENSMUSG00000020173 | Cobl | protein_coding | -0.1141863 | 4.979196 | 0.0537943 | 0.8194731 | 0.999176 |
| ENSMUSG00000001034 | Mapk7 | protein_coding | -0.2562014 | 2.897732 | 0.0537804 | 0.8194961 | 0.999176 |
| ENSMUSG00000024146 | Cript | protein_coding | -0.07548767 | 8.259973 | 0.0537377 | 0.8195665 | 0.999176 |
| ENSMUSG00000015189 | Casd1 | protein_coding | 0.09712487 | 5.565797 | 0.0536809 | 0.8196601 | 0.999176 |
| ENSMUSG00000007880 | Arid1a | protein_coding | 0.1108584 | 6.022103 | 0.0536612 | 0.8196973 | 0.999176 |
| ENSMUSG00000021901 | Bap1 | protein_coding | -0.06841628 | 6.542754 | 0.0535956 | 0.8198006 | 0.999176 |
| ENSMUSG00000049866 | Arl4c | protein_coding | 0.09855033 | 5.539892 | 0.0534399 | 0.8200577 | 0.999176 |
| ENSMUSG00000067424 | Zfp563 | protein_coding | 0.1354423 | 3.802605 | 0.0534098 | 0.8201074 | 0.999176 |
| ENSMUSG00000029004 | Kmt2e | protein_coding | -0.09506166 | 6.199021 | 0.0532686 | 0.8203409 | 0.999176 |
| ENSMUSG00000030245 | Golt1b | protein_coding | -0.08101029 | 7.497291 | 0.0532241 | 0.8204146 | 0.999176 |
| ENSMUSG00000044033 | Ccdc141 | protein_coding | -0.2104515 | 2.953889 | 0.0532016 | 0.8204518 | 0.999176 |
| ENSMUSG00000023945 | Slc5a7 | protein_coding | -0.168511 | 3.343748 | 0.0531116 | 0.8206009 | 0.999176 |
| ENSMUSG00000031537 | Ikbkb | protein_coding | -0.110992 | 4.460226 | 0.0531016 | 0.8206176 | 0.999176 |
| ENSMUSG00000002825 | Qtrt1 | protein_coding | -0.1001909 | 5.198671 | 0.0530948 | 0.8206288 | 0.999176 |
| ENSMUSG00000026199 | Ankzf1 | protein_coding | -0.1470017 | 3.976779 | 0.0530574 | 0.8206908 | 0.999176 |
| ENSMUSG00000035493 | Tgfbi | protein_coding | -0.2851946 | 2.223096 | 0.0530378 | 0.8207233 | 0.999176 |
| ENSMUSG00000030145 | Zfp248 | protein_coding | 0.1315292 | 4.432423 | 0.0529982 | 0.8207891 | 0.999176 |
| ENSMUSG00000022114 | Spry2 | protein_coding | 0.1834817 | 6.703076 | 0.0536092 | 0.8209996 | 0.999176 |
| ENSMUSG00000074194 | Zfp791 | protein_coding | 0.163176 | 3.226863 | 0.0528368 | 0.8210571 | 0.999176 |
| ENSMUSG00000015711 | Prune1 | protein_coding | 0.134271 | 5.363997 | 0.0527533 | 0.8212006 | 0.999176 |
| ENSMUSG00000037706 | Cd81 | protein_coding | -0.07340899 | 9.865788 | 0.0525882 | 0.8214708 | 0.999176 |
| ENSMUSG00000049892 | Rasd1 | protein_coding | -0.1112 | 6.061518 | 0.0525892 | 0.8214738 | 0.999176 |
| ENSMUSG00000005299 | Letm1 | protein_coding | -0.09489983 | 5.787352 | 0.0524507 | 0.8217 | 0.999176 |
| ENSMUSG00000021241 | Isca2 | protein_coding | -0.06171084 | 8.298252 | 0.0524085 | 0.8217704 | 0.999176 |
| ENSMUSG00000036622 | Atp13a2 | protein_coding | -0.1223455 | 5.209432 | 0.0523065 | 0.8219408 | 0.999176 |

| ensembl_gene_id | external_gene_name | gene_biotype | log2 (fold change) | log2 (counts per million) | F-value | p-value | False discovery rate |
|---|---|---|---|---|---|---|---|
| ENSMUSG00000112280 | AC153962.3 | processed_transcript | -0.1369753 | 3.952618 | 0.0523036 | 0.8219456 | 0.999176 |
| ENSMUSG00000043639 | Rbm20 | protein_coding | 0.2465622 | 2.429036 | 0.0522957 | 0.8219588 | 0.999176 |
| ENSMUSG00000024983 | Vti1a | protein_coding | 0.1187537 | 5.623949 | 0.0521879 | 0.8221437 | 0.999176 |
| ENSMUSG00000032239 | Rp9 | protein_coding | 0.1074027 | 4.929226 | 0.0521833 | 0.8221467 | 0.999176 |
| ENSMUSG00000043192 | Gm1840 | processed_pseudoge | 0.3259517 | 2.559595 | 0.0521314 | 0.8222383 | 0.999176 |
| ENSMUSG00000024172 | St6gal2 | protein_coding | 0.2465436 | 2.881336 | 0.0520993 | 0.8222873 | 0.999176 |
| ENSMUSG00000026889 | Rbm18 | protein_coding | 0.07153628 | 6.793041 | 0.0520799 | 0.8223198 | 0.999176 |
| ENSMUSG00000021693 | Kif2a | protein_coding | -0.08040073 | 5.858301 | 0.0520703 | 0.8223358 | 0.999176 |
| ENSMUSG00000031349 | Nsdhl | protein_coding | 0.1386378 | 5.791567 | 0.0520866 | 0.8224893 | 0.999176 |
| ENSMUSG00000044122 | Proca1 | protein_coding | -0.1371506 | 3.871431 | 0.0519733 | 0.8224985 | 0.999176 |
| ENSMUSG00000074794 | Arrdc3 | protein_coding | -0.0898625 | 5.529297 | 0.0517066 | 0.8229461 | 0.999176 |
| ENSMUSG00000033499 | Larp4b | protein_coding | -0.07882538 | 6.176355 | 0.0515796 | 0.8231597 | 0.999176 |
| ENSMUSG00000026341 | Actr3 | protein_coding | 0.07362514 | 6.223189 | 0.0515745 | 0.8231683 | 0.999176 |
| ENSMUSG00000044017 | Adgrd1 | protein_coding | 0.1709975 | 2.958359 | 0.0515254 | 0.823251 | 0.999176 |
| ENSMUSG00000005103 | Wdr1 | protein_coding | 0.1064773 | 7.835163 | 0.0516275 | 0.823259 | 0.999176 |
| ENSMUSG00000064370 | mt-Cytb | protein_coding | 0.1291283 | 12.19261 | 0.0516426 | 0.8232852 | 0.999176 |
| ENSMUSG00000015357 | Clpx | protein_coding | 0.09759481 | 5.129758 | 0.0514898 | 0.823311 | 0.999176 |
| ENSMUSG00000038902 | Pogz | protein_coding | -0.1150867 | 4.852381 | 0.0514456 | 0.8233855 | 0.999176 |
| ENSMUSG00000055022 | Cntn1 | protein_coding | -0.0651332 | 7.891555 | 0.0513804 | 0.8234955 | 0.999176 |
| ENSMUSG00000032308 | Ulk3 | protein_coding | 0.2266392 | 3.059244 | 0.0513664 | 0.8235189 | 0.999176 |
| ENSMUSG00000002010 | Idh3g | protein_coding | -0.06628559 | 8.415721 | 0.0513393 | 0.8235648 | 0.999176 |
| ENSMUSG00000021696 | Elovl7 | protein_coding | -0.1809978 | 3.896188 | 0.0512429 | 0.823732 | 0.999176 |
| ENSMUSG00000042590 | Ipo11 | protein_coding | 0.160782 | 3.436435 | 0.0510851 | 0.8239942 | 0.999176 |
| ENSMUSG00000031562 | Dctd | protein_coding | -0.1502904 | 4.074162 | 0.0509849 | 0.8241639 | 0.999176 |
| ENSMUSG00000021669 | Col4a3bp | protein_coding | -0.1019728 | 4.648174 | 0.0509074 | 0.8242952 | 0.999176 |
| ENSMUSG00000022280 | Rnf19a | protein_coding | -0.1254878 | 4.079641 | 0.050904 | 0.8243009 | 0.999176 |
| ENSMUSG00000030518 | Fam189a1 | protein_coding | -0.1771407 | 4.028151 | 0.0508829 | 0.8243413 | 0.999176 |
| ENSMUSG00000039354 | Smarcal1 | protein_coding | -0.1468269 | 4.604777 | 0.0508453 | 0.824405 | 0.999176 |
| ENSMUSG00000052656 | Rnf103 | protein_coding | 0.174256 | 4.638592 | 0.050677 | 0.8248644 | 0.999176 |
| ENSMUSG00000020732 | Rab37 | protein_coding | -0.2991483 | 4.520753 | 0.051292 | 0.8250132 | 0.999176 |
| ENSMUSG00000039852 | Rere | protein_coding | 0.08762975 | 6.268422 | 0.0504613 | 0.8250531 | 0.999176 |
| ENSMUSG00000043019 | Edem3 | protein_coding | 0.1671344 | 3.210635 | 0.050389 | 0.8251761 | 0.999176 |
| ENSMUSG00000030753 | Thap12 | protein_coding | -0.1033455 | 4.885945 | 0.0502649 | 0.8253879 | 0.999176 |
| ENSMUSG00000029875 | Ccdc184 | protein_coding | -0.11275 | 6.120348 | 0.050226 | 0.8254709 | 0.999176 |
| ENSMUSG00000041141 | Pnmal1 | protein_coding | -0.09065617 | 7.956656 | 0.0501893 | 0.8255214 | 0.999176 |
| ENSMUSG00000015023 | Ddx19a | protein_coding | -0.1840275 | 2.818227 | 0.0501579 | 0.8255705 | 0.999176 |
| ENSMUSG00000029227 | Fip1l1 | protein_coding | -0.1521524 | 4.152034 | 0.0501124 | 0.8256482 | 0.999176 |
| ENSMUSG00000058655 | Eif4b | protein_coding | 0.09913689 | 7.164427 | 0.0501349 | 0.8256627 | 0.999176 |
| ENSMUSG00000007817 | Zmiz1 | protein_coding | -0.09487582 | 5.827892 | 0.050064 | 0.825731 | 0.999176 |
| ENSMUSG00000085071 | Gm14066 | processed_transcript | -0.2139353 | 3.106832 | 0.0499418 | 0.8259401 | 0.999176 |
| ENSMUSG00000032547 | Ryk | protein_coding | -0.1671569 | 3.831006 | 0.0498872 | 0.8260335 | 0.999176 |
| ENSMUSG00000020740 | Gga3 | protein_coding | -0.09265507 | 5.90623 | 0.0498313 | 0.8261294 | 0.999176 |
| ENSMUSG00000073987 | Ggh | protein_coding | -0.1068678 | 5.054664 | 0.0498194 | 0.8261497 | 0.999176 |
| ENSMUSG00000021171 | Esyt2 | protein_coding | 0.1193729 | 4.758011 | 0.0498111 | 0.826164 | 0.999176 |
| ENSMUSG00000110080 | Gm6145 | processed_transcript | 0.2739238 | 3.087889 | 0.0498728 | 0.8262241 | 0.999176 |
| ENSMUSG00000074682 | Zcchc3 | protein_coding | -0.156943 | 3.928975 | 0.0497502 | 0.8262684 | 0.999176 |
| ENSMUSG00000051344 | Plekhm3 | protein_coding | -0.1135546 | 4.396974 | 0.0496127 | 0.8265045 | 0.999176 |
| ENSMUSG00000042050 | Wdr60 | protein_coding | 0.1459215 | 3.910344 | 0.0495592 | 0.8265965 | 0.999176 |
| ENSMUSG00000028708 | Mknk1 | protein_coding | 0.116723 | 4.546988 | 0.0495552 | 0.8266033 | 0.999176 |
| ENSMUSG00000051401 | Kctd16 | protein_coding | 0.1989437 | 2.779443 | 0.0495331 | 0.8266413 | 0.999176 |
| ENSMUSG00000023033 | Scn8a | protein_coding | -0.09476715 | 5.789346 | 0.0495088 | 0.826683 | 0.999176 |
| ENSMUSG00000021771 | Vdac2 | protein_coding | -0.068984 | 9.130915 | 0.0494785 | 0.8267352 | 0.999176 |
| ENSMUSG00000026426 | Arl8a | protein_coding | 0.07298711 | 6.998272 | 0.0494652 | 0.8267581 | 0.999176 |
| ENSMUSG00000003644 | Rps6ka1 | protein_coding | 0.1123877 | 4.109013 | 0.0493466 | 0.8269624 | 0.999176 |
| ENSMUSG00000040761 | Spen | protein_coding | 0.1605754 | 4.045662 | 0.0493489 | 0.8269628 | 0.999176 |
| ENSMUSG00000050931 | Sgms2 | protein_coding | 0.3178304 | 2.269272 | 0.0493204 | 0.8270074 | 0.999176 |
| ENSMUSG00000052712 | BC004004 | protein_coding | 0.06358172 | 6.97332 | 0.0492121 | 0.8271942 | 0.999176 |
| ENSMUSG00000051177 | Plcb1 | protein_coding | -0.1577184 | 3.797052 | 0.0491067 | 0.8273763 | 0.999176 |
| ENSMUSG00000034765 | Dusp5 | protein_coding | 0.2508082 | 2.49791 | 0.0490971 | 0.8273927 | 0.999176 |
| ENSMUSG00000029465 | Arpc3 | protein_coding | 0.08452505 | 8.65012 | 0.0490931 | 0.8273996 | 0.999176 |
| ENSMUSG00000029125 | Stx18 | protein_coding | -0.0924168 | 5.479895 | 0.0489688 | 0.8276146 | 0.999176 |
| ENSMUSG00000008435 | Rdh13 | protein_coding | 0.08444165 | 6.406502 | 0.0488636 | 0.8277967 | 0.999176 |
| ENSMUSG00000024271 | Elp2 | protein_coding | -0.08769869 | 6.92205 | 0.0488406 | 0.8278408 | 0.999176 |
| ENSMUSG00000039552 | Rsph4a | protein_coding | -0.2522717 | 2.769765 | 0.0488314 | 0.8278525 | 0.999176 |
| ENSMUSG00000039254 | Pomt1 | protein_coding | 0.1991304 | 2.894868 | 0.0487956 | 0.8279145 | 0.999176 |
| ENSMUSG00000030432 | Rpl28 | protein_coding | 0.07249418 | 6.389848 | 0.0487821 | 0.8279379 | 0.999176 |
| ENSMUSG00000027849 | Syt6 | protein_coding | -0.1613927 | 4.318855 | 0.048768 | 0.8279668 | 0.999176 |
| ENSMUSG00000027806 | Tsc22d2 | protein_coding | -0.1139253 | 4.615469 | 0.0487181 | 0.8280488 | 0.999176 |
| ENSMUSG00000087075 | A230065H16Rik | protein_coding | -0.08314649 | 6.340547 | 0.0486736 | 0.8281261 | 0.999176 |
| ENSMUSG00000041712 | Ubr7 | protein_coding | -0.1151932 | 4.846411 | 0.0486515 | 0.8281644 | 0.999176 |
| ENSMUSG00000113679 | AC159633.2 | processed_transcript | -0.1451299 | 3.571614 | 0.0486036 | 0.8282475 | 0.999176 |
| ENSMUSG00000038072 | Galnt11 | protein_coding | -0.1525515 | 3.71815 | 0.0485957 | 0.8282614 | 0.999176 |
| ENSMUSG00000018363 | Smurf2 | protein_coding | -0.1209938 | 4.56087 | 0.0485938 | 0.8282645 | 0.999176 |
| ENSMUSG00000097287 | D130017N08Rik | lincRNA | -0.1378143 | 3.860943 | 0.0485662 | 0.8283126 | 0.999176 |
| ENSMUSG00000036087 | Slain2 | protein_coding | -0.2226303 | 3.484801 | 0.0485187 | 0.8285097 | 0.999176 |
| ENSMUSG00000041396 | Mettl18 | protein_coding | -0.1093494 | 4.638652 | 0.0484191 | 0.8285684 | 0.999176 |
| ENSMUSG00000018476 | Kdm6b | protein_coding | 0.168046 | 5.499899 | 0.0485876 | 0.8286589 | 0.999176 |
| ENSMUSG00000036450 | Hif1an | protein_coding | -0.1326269 | 4.210287 | 0.0482084 | 0.8289355 | 0.999176 |

| ensembl_gene_id | external_gene_name | gene_biotype | log2 (fold change) | log2 (counts per million) | F-value | p-value | False discovery rate |
|---|---|---|---|---|---|---|---|
| ENSMUSG00000014867 | Surf4 | protein_coding | 0.09624624 | 5.453203 | 0.0481652 | 0.8290108 | 0.999176 |
| ENSMUSG00000102674 | 8030442B05Rik | lincRNA | -0.1808665 | 3.225546 | 0.0481347 | 0.8290639 | 0.999176 |
| ENSMUSG00000029156 | Sgcb | protein_coding | 0.1073226 | 5.265247 | 0.0481192 | 0.8290911 | 0.999176 |
| ENSMUSG00000068739 | Sars | protein_coding | 0.07978291 | 8.568066 | 0.0480609 | 0.829193 | 0.999176 |
| ENSMUSG00000047036 | Zfp445 | protein_coding | 0.1115662 | 4.849598 | 0.0480467 | 0.8292178 | 0.999176 |
| ENSMUSG00000036983 | Tfb1m | protein_coding | 0.1379781 | 3.668715 | 0.0480123 | 0.8292779 | 0.999176 |
| ENSMUSG00000033009 | Ogfod1 | protein_coding | -0.07667545 | 6.537882 | 0.0479973 | 0.829304 | 0.999176 |
| ENSMUSG00000025037 | Maoa | protein_coding | -0.09413974 | 5.565644 | 0.0479256 | 0.8294294 | 0.999176 |
| ENSMUSG00000025786 | Zdhhc3 | protein_coding | -0.1209779 | 5.3499 | 0.0479242 | 0.8294363 | 0.999176 |
| ENSMUSG00000037104 | Socs5 | protein_coding | -0.08104272 | 5.97034 | 0.0478969 | 0.8294796 | 0.999176 |
| ENSMUSG00000097166 | 9330179D12Rik | antisense_RNA | -0.1674389 | 3.657326 | 0.0478853 | 0.8294999 | 0.999176 |
| ENSMUSG00000019189 | Rnf145 | protein_coding | 0.1062696 | 5.142569 | 0.0478831 | 0.8295038 | 0.999176 |
| ENSMUSG00000005442 | Cic | protein_coding | -0.128261 | 3.706909 | 0.0478733 | 0.829521 | 0.999176 |
| ENSMUSG00000027006 | Dnajc10 | protein_coding | -0.07415257 | 5.867636 | 0.047789 | 0.8296686 | 0.999176 |
| ENSMUSG00000019810 | Fuca2 | protein_coding | 0.0854791 | 6.321854 | 0.0477225 | 0.8297852 | 0.999176 |
| ENSMUSG00000019916 | P4ha1 | protein_coding | 0.1221293 | 4.435432 | 0.0476826 | 0.8298551 | 0.999176 |
| ENSMUSG00000035325 | Sec31a | protein_coding | -0.08551499 | 6.048448 | 0.047654 | 0.8299053 | 0.999176 |
| ENSMUSG00000097383 | 1500026H17Rik | bidirectional_promote | 0.1190875 | 3.728712 | 0.0475947 | 0.8300095 | 0.999176 |
| ENSMUSG00000035845 | Alg12 | protein_coding | 0.154546 | 3.806612 | 0.0474793 | 0.8302123 | 0.999176 |
| ENSMUSG00000004347 | Pde1c | protein_coding | -0.1102342 | 5.082863 | 0.0474777 | 0.830215 | 0.999176 |
| ENSMUSG00000020705 | Ddx42 | protein_coding | 0.08102588 | 5.811683 | 0.0474068 | 0.8303398 | 0.999176 |
| ENSMUSG00000025795 | Rassf3 | protein_coding | 0.1897521 | 2.979774 | 0.0473318 | 0.8304718 | 0.999176 |
| ENSMUSG00000022100 | Xpo7 | protein_coding | -0.09047234 | 5.314992 | 0.0473074 | 0.8305147 | 0.999176 |
| ENSMUSG00000022024 | Sugt1 | protein_coding | -0.06252145 | 7.235566 | 0.0472799 | 0.8305631 | 0.999176 |
| ENSMUSG00000025268 | Maged2 | protein_coding | -0.09413198 | 7.775686 | 0.0471555 | 0.8308062 | 0.999176 |
| ENSMUSG00000028248 | Pnisr | protein_coding | -0.08866941 | 5.828569 | 0.0471061 | 0.8308698 | 0.999176 |
| ENSMUSG00000036887 | C1qa | protein_coding | -0.1760665 | 3.80681 | 0.0468894 | 0.8312572 | 0.999176 |
| ENSMUSG00000017210 | Med24 | protein_coding | -0.1137435 | 5.606036 | 0.0468794 | 0.8312748 | 0.999176 |
| ENSMUSG00000056771 | Gm10010 | lincRNA | 0.3089464 | 2.306701 | 0.0467476 | 0.8315039 | 0.999176 |
| ENSMUSG00000024535 | Snx24 | protein_coding | -0.1302863 | 4.484463 | 0.0467318 | 0.831532 | 0.999176 |
| ENSMUSG00000036817 | Sun1 | protein_coding | 0.1261383 | 5.080491 | 0.0465914 | 0.8317854 | 0.999176 |
| ENSMUSG00000050587 | Lrrc4c | protein_coding | -0.06285553 | 6.84102 | 0.0465743 | 0.8318114 | 0.999176 |
| ENSMUSG00000001260 | Gabrg1 | protein_coding | -0.07756094 | 6.075152 | 0.0464374 | 0.8320548 | 0.999176 |
| ENSMUSG00000034274 | Thoc5 | protein_coding | -0.09972252 | 5.329992 | 0.0462645 | 0.8323626 | 0.999176 |
| ENSMUSG00000036306 | Lzts1 | protein_coding | 0.07145397 | 8.608718 | 0.0461953 | 0.832486 | 0.999176 |
| ENSMUSG00000006906 | Stambp | protein_coding | -0.07391289 | 6.173513 | 0.0461889 | 0.8324974 | 0.999176 |
| ENSMUSG00000003438 | Timm50 | protein_coding | -0.1049426 | 4.759467 | 0.0461231 | 0.8326148 | 0.999176 |
| ENSMUSG00000091512 | Lamtor3 | protein_coding | 0.07203558 | 6.80698 | 0.0460484 | 0.8327484 | 0.999176 |
| ENSMUSG00000049932 | H2afx | protein_coding | -0.2164504 | 2.595392 | 0.0459342 | 0.8329524 | 0.999176 |
| ENSMUSG00000097616 | 1110019D14Rik | lincRNA | 0.1702862 | 3.245475 | 0.0459307 | 0.8329587 | 0.999176 |
| ENSMUSG00000010067 | Rassf1 | protein_coding | -0.108287 | 4.84378 | 0.0458257 | 0.8331467 | 0.999176 |
| ENSMUSG00000025854 | Fam20c | protein_coding | 0.1469311 | 4.759858 | 0.0458277 | 0.8331639 | 0.999176 |
| ENSMUSG00000010086 | Rnf112 | protein_coding | -0.1308936 | 4.615825 | 0.0457936 | 0.8332041 | 0.999176 |
| ENSMUSG00000062580 | Timm17a | protein_coding | 0.05874586 | 9.022383 | 0.0456193 | 0.8335169 | 0.999176 |
| ENSMUSG00000020460 | Rps27a | protein_coding | -0.1282545 | 3.584484 | 0.0455945 | 0.8335614 | 0.999176 |
| ENSMUSG00000013997 | Nit1 | protein_coding | -0.08156168 | 5.735274 | 0.0455451 | 0.8336501 | 0.999176 |
| ENSMUSG00000097574 | C920006O11Rik | lincRNA | -0.1293518 | 3.671111 | 0.0455377 | 0.8336634 | 0.999176 |
| ENSMUSG00000037126 | Psd | protein_coding | -0.1034847 | 5.103726 | 0.0453583 | 0.8339861 | 0.999176 |
| ENSMUSG00000034486 | Gbx2 | protein_coding | 0.259241 | 3.344052 | 0.0455011 | 0.8339932 | 0.999176 |
| ENSMUSG00000024258 | Polr2d | protein_coding | -0.06294835 | 7.113924 | 0.0452449 | 0.8341906 | 0.999176 |
| ENSMUSG00000022505 | Emp2 | protein_coding | -0.1924831 | 3.382256 | 0.0452123 | 0.8342536 | 0.999176 |
| ENSMUSG00000034973 | Dopey1 | protein_coding | -0.1687685 | 3.857142 | 0.0452068 | 0.8342636 | 0.999176 |
| ENSMUSG00000068479 | Mfap1a | protein_coding | 0.1099341 | 4.009951 | 0.0451376 | 0.8343841 | 0.999176 |
| ENSMUSG00000020057 | Dram1 | protein_coding | -0.1110886 | 4.043758 | 0.0451212 | 0.8344138 | 0.999176 |
| ENSMUSG00000022044 | Stmn4 | protein_coding | -0.08045871 | 8.719824 | 0.0451195 | 0.8344167 | 0.999176 |
| ENSMUSG00000015932 | Dstn | protein_coding | -0.06553835 | 8.557412 | 0.0451101 | 0.8344339 | 0.999176 |
| ENSMUSG00000039323 | Igfbp2 | protein_coding | 0.1994479 | 3.213649 | 0.0450411 | 0.8345584 | 0.999176 |
| ENSMUSG00000051989 | Smim11 | protein_coding | 0.1060987 | 5.259884 | 0.0450314 | 0.834576 | 0.999176 |
| ENSMUSG00000068184 | Ndufaf2 | protein_coding | -0.0747002 | 7.347231 | 0.0450274 | 0.8345833 | 0.999176 |
| ENSMUSG00000019558 | Slc6a8 | protein_coding | -0.09220412 | 5.170337 | 0.0449665 | 0.8346933 | 0.999176 |
| ENSMUSG00000020297 | Nsg2 | protein_coding | -0.05975249 | 9.10325 | 0.0449371 | 0.8347466 | 0.999176 |
| ENSMUSG00000002280 | Narfl | protein_coding | 0.1555146 | 3.72307 | 0.0449071 | 0.8348009 | 0.999176 |
| ENSMUSG00000023944 | Hsp90ab1 | protein_coding | -0.07423372 | 11.18772 | 0.0448387 | 0.8349248 | 0.999176 |
| ENSMUSG00000097059 | Fam120aos | lincRNA | -0.1901626 | 2.730748 | 0.0447259 | 0.8351293 | 0.999176 |
| ENSMUSG00000027828 | Ssr3 | protein_coding | -0.0563313 | 7.518891 | 0.0446572 | 0.8352539 | 0.999176 |
| ENSMUSG00000028952 | Zbtb48 | protein_coding | -0.1229541 | 4.685662 | 0.044627 | 0.8353088 | 0.999176 |
| ENSMUSG00000044881 | Coa4 | protein_coding | -0.1049605 | 4.985126 | 0.0445419 | 0.8354633 | 0.999176 |
| ENSMUSG00000063445 | Nmral1 | protein_coding | 0.07335493 | 6.160853 | 0.0445388 | 0.835469 | 0.999176 |
| ENSMUSG00000003762 | Coq8b | protein_coding | 0.1951555 | 2.662125 | 0.0444857 | 0.8355657 | 0.999176 |
| ENSMUSG00000020708 | Psmc5 | protein_coding | 0.06453284 | 9.994708 | 0.044462 | 0.8356087 | 0.999176 |
| ENSMUSG00000034518 | Hmgxb4 | protein_coding | -0.1134829 | 3.850291 | 0.0444425 | 0.8356443 | 0.999176 |
| ENSMUSG00000054027 | Nt5dc3 | protein_coding | -0.1181985 | 4.951661 | 0.0443928 | 0.8357347 | 0.999176 |
| ENSMUSG00000031728 | Zfp821 | protein_coding | -0.1202421 | 4.634155 | 0.0443569 | 0.8358001 | 0.999176 |
| ENSMUSG00000061808 | Ttr | protein_coding | 0.313584 | 11.33229 | 0.0458428 | 0.8358953 | 0.999176 |
| ENSMUSG00000000915 | Hip1r | protein_coding | -0.1285905 | 4.062825 | 0.044288 | 0.8359257 | 0.999176 |
| ENSMUSG00000031093 | Dock11 | protein_coding | -0.2692767 | 2.307063 | 0.0442861 | 0.8359292 | 0.999176 |
| ENSMUSG00000075595 | Zfp652 | protein_coding | -0.1054972 | 4.281857 | 0.0442628 | 0.8359717 | 0.999176 |
| ENSMUSG00000046897 | Zfp740 | protein_coding | 0.09414544 | 4.84519 | 0.0441762 | 0.8361297 | 0.999176 |

| ensembl_gene_id | external_gene_name | gene_biotype | log2 (fold change) | log2 (counts per million) | F-value | p-value | False discovery rate |
|---|---|---|---|---|---|---|---|
| ENSMUSG00000004562 | Arhgef40 | protein_coding | 0.126104 | 5.044574 | 0.0440925 | 0.8362869 | 0.999176 |
| ENSMUSG00000024560 | Cxxc1 | protein_coding | 0.1566717 | 4.553512 | 0.0441001 | 0.8363558 | 0.999176 |
| ENSMUSG00000006289 | Osgep | protein_coding | 0.1223883 | 4.895725 | 0.0440085 | 0.8364362 | 0.999176 |
| ENSMUSG00000027188 | Pamr1 | protein_coding | -0.2023677 | 2.718553 | 0.043885 | 0.8366623 | 0.999176 |
| ENSMUSG00000027200 | Sema6d | protein_coding | -0.09896594 | 4.817092 | 0.0438058 | 0.8368075 | 0.999176 |
| ENSMUSG00000041679 | Lrrc29 | protein_coding | 0.1894013 | 2.848384 | 0.0437815 | 0.8368521 | 0.999176 |
| ENSMUSG00000044674 | Fzd1 | protein_coding | 0.2704383 | 2.389092 | 0.0437124 | 0.8369789 | 0.999176 |
| ENSMUSG00000021687 | Scamp1 | protein_coding | -0.06037121 | 7.292855 | 0.0437079 | 0.8369872 | 0.999176 |
| ENSMUSG00000026471 | Mr1 | protein_coding | -0.4141883 | 2.113181 | 0.043787 | 0.8370047 | 0.999176 |
| ENSMUSG00000022787 | Wdr53 | protein_coding | -0.09791367 | 4.781929 | 0.0436668 | 0.8370626 | 0.999176 |
| ENSMUSG00000026220 | Slc16a14 | protein_coding | -0.2759904 | 3.227649 | 0.0438358 | 0.8370888 | 0.999176 |
| ENSMUSG00000051403 | Ppp1r37 | protein_coding | -0.09974433 | 4.556806 | 0.0436423 | 0.8371077 | 0.999176 |
| ENSMUSG00000064120 | Mocs1 | protein_coding | 0.2822018 | 2.373373 | 0.0436296 | 0.837131 | 0.999176 |
| ENSMUSG00000045294 | Insig1 | protein_coding | -0.09345403 | 5.877428 | 0.0435487 | 0.8372799 | 0.999176 |
| ENSMUSG00000073838 | Tufm | protein_coding | 0.1221516 | 3.580336 | 0.0435364 | 0.8373024 | 0.999176 |
| ENSMUSG00000029560 | Snx8 | protein_coding | 0.1423759 | 4.573281 | 0.0434517 | 0.8374627 | 0.999176 |
| ENSMUSG00000020070 | Rufy2 | protein_coding | 0.1000496 | 5.062097 | 0.0432935 | 0.83775 | 0.999176 |
| ENSMUSG00000022150 | Dab2 | protein_coding | -0.1696431 | 3.048865 | 0.0431931 | 0.8379355 | 0.999176 |
| ENSMUSG00000008575 | Nfib | protein_coding | -0.1051725 | 5.048399 | 0.0431765 | 0.837966 | 0.999176 |
| ENSMUSG00000039737 | Prkrip1 | protein_coding | -0.07341021 | 6.355338 | 0.0431485 | 0.8380178 | 0.999176 |
| ENSMUSG00000054099 | Slc25a40 | protein_coding | -0.1500399 | 3.363761 | 0.0430952 | 0.8381164 | 0.999176 |
| ENSMUSG00000045259 | Klhdc9 | protein_coding | 0.1139376 | 4.511998 | 0.0430653 | 0.8381718 | 0.999176 |
| ENSMUSG00000043510 | Hscb | protein_coding | -0.1082913 | 4.868453 | 0.0430356 | 0.8382267 | 0.999176 |
| ENSMUSG00000038861 | Pi4kb | protein_coding | 0.1427832 | 3.52774 | 0.0429988 | 0.8382949 | 0.999176 |
| ENSMUSG00000046550 | Spin2c | protein_coding | 0.1005188 | 5.370067 | 0.042883 | 0.8385094 | 0.999176 |
| ENSMUSG00000059173 | Pde1a | protein_coding | -0.07739098 | 6.814878 | 0.0428752 | 0.8385239 | 0.999176 |
| ENSMUSG00000021557 | Agtpbp1 | protein_coding | -0.07610893 | 6.052023 | 0.0428529 | 0.8385654 | 0.999176 |
| ENSMUSG00000022808 | Snx4 | protein_coding | -0.09121026 | 5.947934 | 0.0428464 | 0.8385773 | 0.999176 |
| ENSMUSG00000029063 | Nadk | protein_coding | 0.09788915 | 4.830288 | 0.0428241 | 0.8386187 | 0.999176 |
| ENSMUSG00000027865 | Gdap2 | protein_coding | -0.1156026 | 5.846193 | 0.0428621 | 0.8386335 | 0.999176 |
| ENSMUSG00000029869 | Ephb6 | protein_coding | 0.1410327 | 4.249871 | 0.0428155 | 0.8386347 | 0.999176 |
| ENSMUSG00000024096 | Ralbp1 | protein_coding | 0.07624576 | 5.912336 | 0.0427981 | 0.838667 | 0.999176 |
| ENSMUSG00000073207 | Ccdc160 | protein_coding | 0.1736129 | 3.255507 | 0.0427692 | 0.8387206 | 0.999176 |
| ENSMUSG00000026608 | Kctd3 | protein_coding | -0.1167099 | 4.79177 | 0.0427335 | 0.8387869 | 0.999176 |
| ENSMUSG00000022807 | Osbpl11 | protein_coding | -0.1350429 | 3.436729 | 0.0427159 | 0.8388197 | 0.999176 |
| ENSMUSG00000097868 | A330069E16Rik | lincRNA | -0.144014 | 4.657656 | 0.0426924 | 0.8388786 | 0.999176 |
| ENSMUSG00000063382 | Bcl9l | protein_coding | 0.1415408 | 4.306736 | 0.0426376 | 0.8389695 | 0.999176 |
| ENSMUSG00000048483 | Zdhhc22 | protein_coding | -0.1518873 | 7.562365 | 0.0432141 | 0.8389764 | 0.999176 |
| ENSMUSG00000028949 | Smarcd3 | protein_coding | 0.12659 | 3.796073 | 0.0426033 | 0.839029 | 0.999176 |
| ENSMUSG00000073478 | D730003I15Rik | lincRNA | 0.1321759 | 4.288159 | 0.0425921 | 0.8390499 | 0.999176 |
| ENSMUSG00000025766 | D3Ertd751e | protein_coding | -0.1185019 | 4.628733 | 0.0424011 | 0.8394058 | 0.999176 |
| ENSMUSG00000054611 | Kdm2a | protein_coding | -0.09211998 | 4.94849 | 0.042399 | 0.8394097 | 0.999176 |
| ENSMUSG00000030870 | Ubfd1 | protein_coding | 0.06736636 | 6.262235 | 0.0423805 | 0.8394443 | 0.999176 |
| ENSMUSG00000037007 | Zfp113 | protein_coding | -0.1557094 | 3.32483 | 0.0422917 | 0.83961 | 0.999176 |
| ENSMUSG00000021905 | Dph3 | protein_coding | -0.07718081 | 6.193024 | 0.0422747 | 0.8396417 | 0.999176 |
| ENSMUSG00000041298 | Katnal1 | protein_coding | -0.07964859 | 6.237263 | 0.0420814 | 0.8400034 | 0.999176 |
| ENSMUSG00000066640 | Fbxl18 | protein_coding | -0.1230092 | 4.1088 | 0.0420622 | 0.8400394 | 0.999176 |
| ENSMUSG00000059361 | Nrsn2 | protein_coding | -0.1493352 | 3.125861 | 0.0420436 | 0.8400742 | 0.999176 |
| ENSMUSG00000062944 | 9130023H24Rik | protein_coding | -0.1467088 | 3.946583 | 0.0419647 | 0.8402221 | 0.999176 |
| ENSMUSG00000038615 | Nfe2l1 | protein_coding | -0.05963607 | 6.872198 | 0.0419402 | 0.840268 | 0.999176 |
| ENSMUSG00000013495 | Tmem175 | protein_coding | -0.1021947 | 5.834362 | 0.0418951 | 0.8403568 | 0.999176 |
| ENSMUSG00000021048 | Mthfd1 | protein_coding | -0.1132577 | 4.127595 | 0.0418506 | 0.8404363 | 0.999176 |
| ENSMUSG00000029705 | Cux1 | protein_coding | -0.07868015 | 6.038084 | 0.0418481 | 0.840441 | 0.999176 |
| ENSMUSG00000015143 | Actn1 | protein_coding | 0.09373035 | 4.509325 | 0.0418434 | 0.8404498 | 0.999176 |
| ENSMUSG00000058741 | Prr19 | protein_coding | -0.2929526 | 2.448788 | 0.0418245 | 0.8404895 | 0.999176 |
| ENSMUSG00000060657 | Marf1 | protein_coding | 0.1855907 | 7.25331 | 0.0425683 | 0.8405946 | 0.999176 |
| ENSMUSG00000086607 | 4930511M06Rik | processed_transcript | 0.1675794 | 3.102547 | 0.0417079 | 0.8407045 | 0.999176 |
| ENSMUSG00000042364 | Snx18 | protein_coding | 0.1466263 | 3.823215 | 0.0416939 | 0.8407308 | 0.999176 |
| ENSMUSG00000027883 | Gpsm2 | protein_coding | -0.1510885 | 2.962966 | 0.0416827 | 0.840752 | 0.999176 |
| ENSMUSG00000037278 | Tmem97 | protein_coding | 0.09750765 | 4.261965 | 0.0416122 | 0.8408847 | 0.999176 |
| ENSMUSG00000022817 | Itgb5 | protein_coding | -0.1860315 | 2.879385 | 0.0415944 | 0.8409183 | 0.999176 |
| ENSMUSG00000030779 | Rbbp6 | protein_coding | 0.07573791 | 5.982878 | 0.0415337 | 0.8410328 | 0.999176 |
| ENSMUSG00000071866 | Ppia | protein_coding | 0.05967068 | 7.158663 | 0.0415333 | 0.8410336 | 0.999176 |
| ENSMUSG00000036120 | Rfxank | protein_coding | -0.09598842 | 4.926668 | 0.0414969 | 0.841102 | 0.999176 |
| ENSMUSG00000038366 | Lasp1 | protein_coding | -0.09624152 | 5.692131 | 0.0413961 | 0.8412924 | 0.999176 |
| ENSMUSG00000039105 | Atp6v1g1 | protein_coding | -0.05709825 | 8.41965 | 0.0413273 | 0.8414224 | 0.999176 |
| ENSMUSG00000022106 | Rcbtb2 | protein_coding | -0.1707946 | 3.564056 | 0.0413059 | 0.841467 | 0.999176 |
| ENSMUSG00000020412 | Ascc2 | protein_coding | -0.1395826 | 3.282316 | 0.0413037 | 0.8414671 | 0.999176 |
| ENSMUSG00000029371 | Cxcl5 | protein_coding | -0.3470541 | 2.469771 | 0.0413842 | 0.8414752 | 0.999176 |
| ENSMUSG00000032121 | Tmem218 | protein_coding | -0.08412169 | 5.52412 | 0.0412594 | 0.8415508 | 0.999176 |
| ENSMUSG00000031985 | Gnpat | protein_coding | -0.09444327 | 5.414486 | 0.0412344 | 0.8415982 | 0.999176 |
| ENSMUSG00000040479 | Dgkz | protein_coding | -0.08557102 | 6.007074 | 0.0412148 | 0.8416353 | 0.999176 |
| ENSMUSG00000000711 | Rab5b | protein_coding | 0.10443 | 6.431024 | 0.0412888 | 0.8416554 | 0.999176 |
| ENSMUSG00000022377 | Asap1 | protein_coding | -0.09676781 | 5.303139 | 0.0411735 | 0.8417136 | 0.999176 |
| ENSMUSG00000019876 | Pkib | protein_coding | 0.08894209 | 4.94944 | 0.0411642 | 0.8417312 | 0.999176 |
| ENSMUSG00000024561 | Mbd1 | protein_coding | 0.09864819 | 5.181179 | 0.0410807 | 0.8418893 | 0.999176 |
| ENSMUSG00000029462 | Vps29 | protein_coding | 0.0590878 | 9.414166 | 0.040995 | 0.8420521 | 0.999176 |
| ENSMUSG00000021235 | Coq6 | protein_coding | -0.1075608 | 5.401148 | 0.040965 | 0.8421089 | 0.999176 |

| ensembl_gene_id | external_gene_name | gene_biotype | log2 (fold change) | log2 (counts per million) | F-value | p-value | False discovery rate |
|---|---|---|---|---|---|---|---|
| ENSMUSG00000024583 | Txnl1 | protein_coding | -0.06285271 | 7.176357 | 0.0408649 | 0.8422993 | 0.999176 |
| ENSMUSG00000040666 | Sh3bgr | protein_coding | -0.2923842 | 2.521521 | 0.0408615 | 0.8423098 | 0.999176 |
| ENSMUSG00000037363 | Letm2 | protein_coding | -0.1483042 | 3.508788 | 0.0408422 | 0.8423424 | 0.999176 |
| ENSMUSG00000047238 | Mageh1 | protein_coding | -0.06757258 | 6.19135 | 0.0407967 | 0.8424291 | 0.999176 |
| ENSMUSG00000010048 | Ifrd2 | protein_coding | 0.1745553 | 3.343852 | 0.0407721 | 0.8424759 | 0.999176 |
| ENSMUSG00000053931 | Cnn3 | protein_coding | 0.1171625 | 4.961678 | 0.0407662 | 0.8424871 | 0.999176 |
| ENSMUSG00000041235 | Chd7 | protein_coding | -0.1294581 | 3.645944 | 0.0407235 | 0.8425684 | 0.999176 |
| ENSMUSG00000020190 | Mknk2 | protein_coding | 0.1358406 | 3.544431 | 0.0406964 | 0.8426201 | 0.999176 |
| ENSMUSG00000005873 | Reep5 | protein_coding | 0.0600819 | 8.580468 | 0.0406543 | 0.8427003 | 0.999176 |
| ENSMUSG00000032750 | Gab3 | protein_coding | -0.1403454 | 3.593346 | 0.0406341 | 0.8427389 | 0.999176 |
| ENSMUSG00000032679 | Cd59a | protein_coding | -0.09654035 | 5.657192 | 0.0406177 | 0.8427701 | 0.999176 |
| ENSMUSG00000022553 | Maf1 | protein_coding | 0.1182627 | 5.636313 | 0.0406222 | 0.8428446 | 0.999176 |
| ENSMUSG00000029621 | Arpc1a | protein_coding | -0.05476716 | 9.187905 | 0.0405554 | 0.8428891 | 0.999176 |
| ENSMUSG00000022022 | Mtrf1 | protein_coding | 0.1206496 | 4.82886 | 0.0405441 | 0.8429107 | 0.999176 |
| ENSMUSG00000064289 | Tank | protein_coding | -0.08185615 | 5.599823 | 0.0404807 | 0.8430318 | 0.999176 |
| ENSMUSG00000049422 | Chchd10 | protein_coding | 0.09927473 | 5.888722 | 0.0403881 | 0.843213 | 0.999176 |
| ENSMUSG00000005580 | Adcy9 | protein_coding | 0.1549463 | 4.012219 | 0.0403231 | 0.8433375 | 0.999176 |
| ENSMUSG00000057691 | Zfp746 | protein_coding | 0.1234549 | 3.760092 | 0.0403155 | 0.8433479 | 0.999176 |
| ENSMUSG00000024201 | Kdm4b | protein_coding | -0.1625434 | 4.745666 | 0.0402539 | 0.8437136 | 0.999176 |
| ENSMUSG00000029402 | Snrnp35 | protein_coding | -0.09312036 | 5.151248 | 0.0399574 | 0.8440354 | 0.999176 |
| ENSMUSG00000041231 | Ublcp1 | protein_coding | -0.07471516 | 5.847101 | 0.0397249 | 0.8444833 | 0.999176 |
| ENSMUSG00000021772 | Nkiras1 | protein_coding | -0.05519581 | 8.25803 | 0.0396561 | 0.8446162 | 0.999176 |
| ENSMUSG00000036432 | Siah2 | protein_coding | -0.1528243 | 3.735799 | 0.039593 | 0.8447382 | 0.999176 |
| ENSMUSG00000021286 | Zfyve21 | protein_coding | 0.09589518 | 5.297217 | 0.0395375 | 0.8448456 | 0.999176 |
| ENSMUSG00000079157 | Fam155a | protein_coding | -0.07616546 | 5.533616 | 0.0394479 | 0.8450191 | 0.999176 |
| ENSMUSG00000020268 | Lyrm7 | protein_coding | 0.07060325 | 5.825075 | 0.0393707 | 0.8451686 | 0.999176 |
| ENSMUSG00000052942 | Glis3 | protein_coding | 0.1436269 | 3.356678 | 0.0393512 | 0.8452064 | 0.999176 |
| ENSMUSG00000019254 | Ppp1r12c | protein_coding | 0.1019068 | 4.829505 | 0.0393382 | 0.8452317 | 0.999176 |
| ENSMUSG00000024556 | Me2 | protein_coding | 0.1102604 | 4.00399 | 0.0393162 | 0.8452743 | 0.999176 |
| ENSMUSG00000028221 | Tmem55a | protein_coding | -0.05780355 | 7.03247 | 0.0392791 | 0.8453463 | 0.999176 |
| ENSMUSG00000034613 | Ppm1h | protein_coding | 0.1176515 | 7.551319 | 0.0394272 | 0.8454506 | 0.999176 |
| ENSMUSG00000034190 | Chmp7 | protein_coding | 0.08237793 | 6.459668 | 0.0391631 | 0.8455718 | 0.999176 |
| ENSMUSG00000029389 | Ddx55 | protein_coding | 0.08485504 | 5.187898 | 0.0391232 | 0.8456494 | 0.999176 |
| ENSMUSG00000015291 | Gdi1 | protein_coding | 0.09436251 | 10.42006 | 0.039125 | 0.8457218 | 0.999176 |
| ENSMUSG00000039676 | Capsl | protein_coding | 0.1441306 | 4.066368 | 0.0390687 | 0.8457595 | 0.999176 |
| ENSMUSG00000027247 | Arhgap1 | protein_coding | -0.09142592 | 6.717357 | 0.0389904 | 0.8459656 | 0.999176 |
| ENSMUSG00000021494 | Ddx41 | protein_coding | -0.08357843 | 5.333515 | 0.0388677 | 0.8461472 | 0.999176 |
| ENSMUSG00000053289 | Ddx10 | protein_coding | 0.1035817 | 5.026413 | 0.0388638 | 0.8461548 | 0.999176 |
| ENSMUSG00000031696 | Vps35 | protein_coding | -0.05731271 | 7.691568 | 0.0388356 | 0.8462099 | 0.999176 |
| ENSMUSG00000021819 | Zswim8 | protein_coding | -0.1007863 | 4.343797 | 0.0387841 | 0.8463105 | 0.999176 |
| ENSMUSG00000026849 | Tor1a | protein_coding | 0.09701168 | 5.71609 | 0.0386588 | 0.8465557 | 0.999176 |
| ENSMUSG00000060962 | Dmkn | protein_coding | -0.1506192 | 4.473782 | 0.038679 | 0.8466181 | 0.999176 |
| ENSMUSG00000036941 | Elac1 | protein_coding | -0.1006436 | 4.921003 | 0.0385372 | 0.8467939 | 0.999176 |
| ENSMUSG00000021684 | Pde8b | protein_coding | 0.09724562 | 5.163452 | 0.0385366 | 0.8467951 | 0.999176 |
| ENSMUSG00000024601 | Isoc1 | protein_coding | 0.09915088 | 5.166928 | 0.0384787 | 0.8469086 | 0.999176 |
| ENSMUSG00000087128 | Gm12655 | processed_transcript | 0.2123692 | 2.679447 | 0.0384768 | 0.8469125 | 0.999176 |
| ENSMUSG00000040502 | 9-Mar | protein_coding | -0.1549821 | 3.218846 | 0.03847 | 0.8469257 | 0.999176 |
| ENSMUSG00000046947 | Adck2 | protein_coding | -0.1983058 | 3.163244 | 0.0384186 | 0.8470307 | 0.999176 |
| ENSMUSG00000040236 | Trappc5 | protein_coding | 0.06501186 | 6.273591 | 0.0384143 | 0.847035 | 0.999176 |
| ENSMUSG00000108644 | Gm44920 | lincRNA | 0.2282885 | 2.522787 | 0.0383312 | 0.8471984 | 0.999176 |
| ENSMUSG00000014668 | Chfr | protein_coding | 0.2786332 | 2.161271 | 0.0382987 | 0.8472622 | 0.999176 |
| ENSMUSG00000048281 | Dleu7 | protein_coding | 0.1154569 | 3.777677 | 0.0381812 | 0.8474937 | 0.999176 |
| ENSMUSG00000031556 | Tm2d2 | protein_coding | 0.06552454 | 7.927169 | 0.0380806 | 0.847692 | 0.999176 |
| ENSMUSG00000032262 | Elovl4 | protein_coding | -0.07088961 | 5.962058 | 0.0379879 | 0.8478749 | 0.999176 |
| ENSMUSG00000004151 | Etv1 | protein_coding | -0.08333226 | 5.695037 | 0.0379524 | 0.8479452 | 0.999176 |
| ENSMUSG00000021963 | Sap18 | protein_coding | 0.05681725 | 6.985811 | 0.037919 | 0.8480111 | 0.999176 |
| ENSMUSG00000047248 | C2cd3 | protein_coding | -0.2824424 | 2.391019 | 0.0378506 | 0.8481505 | 0.999176 |
| ENSMUSG00000032773 | Chrm1 | protein_coding | -0.1053751 | 5.403171 | 0.0377968 | 0.848257 | 0.999176 |
| ENSMUSG00000045092 | S1pr1 | protein_coding | -0.09583989 | 5.733063 | 0.0377931 | 0.8482603 | 0.999176 |
| ENSMUSG00000025369 | Smarcc2 | protein_coding | -0.06767377 | 6.432758 | 0.0377821 | 0.8482823 | 0.999176 |
| ENSMUSG00000110277 | Gm45871 | protein_coding | -0.1572919 | 3.264618 | 0.037764 | 0.8483181 | 0.999176 |
| ENSMUSG00000027488 | Snta1 | protein_coding | 0.1024795 | 4.29758 | 0.0377442 | 0.8483574 | 0.999176 |
| ENSMUSG00000049556 | Lingo1 | protein_coding | 0.08860373 | 5.534299 | 0.0377176 | 0.84841 | 0.999176 |
| ENSMUSG00000034327 | Kctd9 | protein_coding | -0.116465 | 3.862374 | 0.0376542 | 0.8485358 | 0.999176 |
| ENSMUSG00000033444 | Specc1l | protein_coding | 0.09351385 | 4.588919 | 0.037552 | 0.8487387 | 0.999176 |
| ENSMUSG00000031153 | Gripap1 | protein_coding | -0.05967657 | 7.663268 | 0.0375339 | 0.8487747 | 0.999176 |
| ENSMUSG00000054034 | Tceal5 | protein_coding | -0.06276882 | 6.457638 | 0.0375225 | 0.8487974 | 0.999176 |
| ENSMUSG00000039826 | Trub2 | protein_coding | 0.1272951 | 5.013049 | 0.0374806 | 0.8489018 | 0.999176 |
| ENSMUSG00000020956 | Dtd2 | protein_coding | 0.1527976 | 5.506275 | 0.0376106 | 0.8490043 | 0.999176 |
| ENSMUSG00000015112 | Slc25a13 | protein_coding | -0.1549765 | 3.432725 | 0.0374165 | 0.8490082 | 0.999176 |
| ENSMUSG00000024949 | Sf1 | protein_coding | -0.06860614 | 5.964314 | 0.0373102 | 0.8492201 | 0.999176 |
| ENSMUSG00000058624 | Gda | protein_coding | -0.06983332 | 6.490384 | 0.0371514 | 0.8495372 | 0.999176 |
| ENSMUSG00000042572 | Ube2q1 | protein_coding | 0.08025883 | 5.737949 | 0.0371305 | 0.8495788 | 0.999176 |
| ENSMUSG00000030611 | Mrps11 | protein_coding | 0.05884359 | 7.327359 | 0.0371277 | 0.8495845 | 0.999176 |
| ENSMUSG00000031839 | Hsbp1 | protein_coding | 0.06276997 | 10.62593 | 0.0370421 | 0.8497558 | 0.999176 |
| ENSMUSG00000054408 | Spcs3 | protein_coding | -0.07074552 | 5.892873 | 0.0370064 | 0.8498271 | 0.999176 |
| ENSMUSG00000001175 | Calm1 | protein_coding | 0.06191815 | 12.85395 | 0.0369872 | 0.8498657 | 0.999176 |
| ENSMUSG00000031731 | Ap1g1 | protein_coding | 0.155744 | 3.130153 | 0.0369805 | 0.8498791 | 0.999176 |

| ensembl_gene_id | external_gene_name | gene_biotype | log2 (fold change) | log2 (counts per million) | F-value | p-value | False discovery rate |
|---|---|---|---|---|---|---|---|
| ENSMUSG00000022125 | Cln5 | protein_coding | -0.1451624 | 3.400347 | 0.0369454 | 0.8499494 | 0.999176 |
| ENSMUSG00000020085 | Aifm2 | protein_coding | 0.1384492 | 4.330429 | 0.0369422 | 0.8499597 | 0.999176 |
| ENSMUSG00000020184 | Mdm2 | protein_coding | -0.09110476 | 4.968489 | 0.0369079 | 0.8500247 | 0.999176 |
| ENSMUSG00000027195 | Hsd17b12 | protein_coding | -0.05467191 | 7.43842 | 0.0368928 | 0.850055 | 0.999176 |
| ENSMUSG00000039860 | Srrm3 | protein_coding | 0.1557522 | 2.993256 | 0.0367147 | 0.8504126 | 0.999176 |
| ENSMUSG00000051726 | Kcnf1 | protein_coding | 0.1229017 | 4.39067 | 0.0366728 | 0.8504968 | 0.999176 |
| ENSMUSG00000020671 | Rab10 | protein_coding | 0.1586163 | 2.972998 | 0.0366612 | 0.8505202 | 0.999176 |
| ENSMUSG00000055204 | Ankrd17 | protein_coding | -0.06593768 | 6.665546 | 0.0365025 | 0.85084 | 0.999176 |
| ENSMUSG00000032044 | Rpusd4 | protein_coding | 0.103553 | 4.453726 | 0.0363719 | 0.8511036 | 0.999176 |
| ENSMUSG00000044134 | Fam109a | protein_coding | -0.2887962 | 2.104216 | 0.0363565 | 0.8511346 | 0.999176 |
| ENSMUSG00000097219 | Gm26551 | lincRNA | 0.1262353 | 3.711355 | 0.0363299 | 0.8511885 | 0.999176 |
| ENSMUSG00000062488 | Ifit3b | protein_coding | -0.1883638 | 2.687332 | 0.0362976 | 0.8512537 | 0.999176 |
| ENSMUSG00000007338 | Mrpl49 | protein_coding | 0.05660883 | 7.222372 | 0.0362889 | 0.8512715 | 0.999176 |
| ENSMUSG00000021871 | Pnp | protein_coding | 0.1679719 | 2.8071 | 0.0361709 | 0.8515103 | 0.999176 |
| ENSMUSG00000027374 | Mrps5 | protein_coding | -0.07099105 | 5.855083 | 0.0361635 | 0.8515252 | 0.999176 |
| ENSMUSG00000020866 | Cacna1g | protein_coding | 0.1686147 | 3.129771 | 0.0360871 | 0.8516803 | 0.999176 |
| ENSMUSG00000019295 | Tmem129 | protein_coding | -0.08752823 | 5.344032 | 0.0360305 | 0.8517952 | 0.999176 |
| ENSMUSG00000018821 | Avpi1 | protein_coding | 0.06293329 | 7.044228 | 0.0360267 | 0.8518029 | 0.999176 |
| ENSMUSG00000031352 | Hccs | protein_coding | 0.115514 | 4.604802 | 0.0360188 | 0.8518188 | 0.999176 |
| ENSMUSG00000028167 | Bdh2 | protein_coding | -0.1244272 | 3.684437 | 0.0360186 | 0.8518194 | 0.999176 |
| ENSMUSG00000042105 | Inpp5f | protein_coding | 0.07376222 | 8.659098 | 0.0359782 | 0.8519014 | 0.999176 |
| ENSMUSG00000099034 | 2810039B14Rik | transcribed_unitary_p | 0.1375855 | 3.229646 | 0.0359651 | 0.8519279 | 0.999176 |
| ENSMUSG00000054484 | Tmem62 | protein_coding | -0.112237 | 4.323852 | 0.0359276 | 0.8520043 | 0.999176 |
| ENSMUSG00000048832 | Vps37c | protein_coding | -0.1123686 | 4.872612 | 0.0359219 | 0.8520159 | 0.999176 |
| ENSMUSG00000006127 | Inpp5k | protein_coding | -0.09500034 | 5.897337 | 0.0359044 | 0.8520553 | 0.999176 |
| ENSMUSG00000032279 | Idh3a | protein_coding | -0.06513912 | 9.358749 | 0.0358711 | 0.8521193 | 0.999176 |
| ENSMUSG00000047789 | Slc38a9 | protein_coding | 0.1040265 | 3.84426 | 0.0358682 | 0.8521251 | 0.999176 |
| ENSMUSG00000001985 | Grik3 | protein_coding | -0.09663631 | 4.438518 | 0.0358619 | 0.8521379 | 0.999176 |
| ENSMUSG00000024095 | Hnrnpll | protein_coding | 0.1006503 | 5.037192 | 0.0358566 | 0.8521487 | 0.999176 |
| ENSMUSG00000078700 | D030028A08Rik | processed_transcript | -0.2119766 | 2.384145 | 0.0358186 | 0.852226 | 0.999176 |
| ENSMUSG00000068874 | Selenbp1 | protein_coding | -0.1218189 | 3.323188 | 0.0357575 | 0.8523505 | 0.999176 |
| ENSMUSG00000046707 | Csnk2a2 | protein_coding | -0.0794743 | 5.413038 | 0.0357089 | 0.8524497 | 0.999176 |
| ENSMUSG00000028771 | Ptpn12 | protein_coding | -0.1306896 | 4.087179 | 0.0356271 | 0.8526167 | 0.999176 |
| ENSMUSG00000015766 | Eps8 | protein_coding | 0.1154156 | 4.559608 | 0.0355356 | 0.8528036 | 0.999176 |
| ENSMUSG00000026004 | Kansl1l | protein_coding | 0.1577152 | 3.449601 | 0.0355019 | 0.8528727 | 0.999176 |
| ENSMUSG00000035529 | Prdm4 | protein_coding | 0.1342253 | 3.940585 | 0.0353882 | 0.8531056 | 0.999176 |
| ENSMUSG00000000339 | Rtca | protein_coding | -0.1258205 | 4.546336 | 0.0353232 | 0.8532425 | 0.999176 |
| ENSMUSG00000020037 | Rfx4 | protein_coding | -0.1023581 | 3.874908 | 0.0353016 | 0.8532831 | 0.999176 |
| ENSMUSG00000047797 | Gjb1 | protein_coding | -0.1102332 | 6.266397 | 0.0353816 | 0.8533508 | 0.999176 |
| ENSMUSG00000021987 | Mtmr6 | protein_coding | -0.07343158 | 6.883621 | 0.0351834 | 0.8535258 | 0.999176 |
| ENSMUSG00000035247 | Hectd1 | protein_coding | -0.1155677 | 4.236072 | 0.0351735 | 0.8535463 | 0.999176 |
| ENSMUSG00000038274 | Fau | protein_coding | -0.09880658 | 4.187083 | 0.0351419 | 0.8536112 | 0.999176 |
| ENSMUSG00000024259 | Slc25a46 | protein_coding | -0.0763793 | 5.508648 | 0.0350947 | 0.8537084 | 0.999176 |
| ENSMUSG00000020706 | Ftsj3 | protein_coding | 0.1135999 | 4.824402 | 0.0350719 | 0.8537591 | 0.999176 |
| ENSMUSG00000055540 | Epha6 | protein_coding | 0.1334992 | 3.574034 | 0.0350629 | 0.8537739 | 0.999176 |
| ENSMUSG00000021189 | Atxn3 | protein_coding | -0.122575 | 3.618118 | 0.0350106 | 0.8538816 | 0.999176 |
| ENSMUSG00000027566 | Psma7 | protein_coding | 0.05995047 | 9.496866 | 0.0350061 | 0.8538909 | 0.999176 |
| ENSMUSG00000038949 | Cnst | protein_coding | 0.118255 | 4.301464 | 0.0349719 | 0.8539615 | 0.999176 |
| ENSMUSG00000017176 | Nt5c3b | protein_coding | -0.05848164 | 6.755126 | 0.0349588 | 0.8539885 | 0.999176 |
| ENSMUSG00000051306 | Usp42 | protein_coding | 0.1088855 | 4.038454 | 0.0349541 | 0.8539981 | 0.999176 |
| ENSMUSG00000039328 | Rnf122 | protein_coding | -0.1051325 | 4.06267 | 0.0348579 | 0.8541967 | 0.999176 |
| ENSMUSG00000038506 | Dcun1d2 | protein_coding | -0.08716744 | 5.279137 | 0.0348106 | 0.8542944 | 0.999176 |
| ENSMUSG00000033983 | Coil | protein_coding | -0.09481048 | 4.528481 | 0.0347928 | 0.8543311 | 0.999176 |
| ENSMUSG00000028709 | Mob3c | protein_coding | -0.2835045 | 2.685665 | 0.0348484 | 0.854363 | 0.999176 |
| ENSMUSG00000064351 | mt-Co1 | protein_coding | -0.1086608 | 11.50948 | 0.0348511 | 0.8544448 | 0.999176 |
| ENSMUSG00000038520 | Tbc1d17 | protein_coding | -0.07327825 | 6.289338 | 0.0347322 | 0.8544567 | 0.999176 |
| ENSMUSG00000087200 | Lrp8os3 | bidirectional_promote | -0.1240949 | 3.219259 | 0.0346244 | 0.8546798 | 0.999176 |
| ENSMUSG00000057897 | Camk2b | protein_coding | 0.05834452 | 8.455445 | 0.0346109 | 0.8547078 | 0.999176 |
| ENSMUSG00000045136 | Tubb2b | protein_coding | 0.07343134 | 5.423478 | 0.0345753 | 0.8547818 | 0.999176 |
| ENSMUSG00000026238 | Ptma | protein_coding | 0.06096688 | 6.344462 | 0.0345285 | 0.8548787 | 0.999176 |
| ENSMUSG00000071648 | Rom1 | protein_coding | -0.1429291 | 3.346983 | 0.0344856 | 0.8549679 | 0.999176 |
| ENSMUSG00000021190 | Lgmn | protein_coding | 0.06788391 | 7.624437 | 0.0344288 | 0.8550859 | 0.999176 |
| ENSMUSG00000036968 | Cnpy4 | protein_coding | -0.09007055 | 5.693023 | 0.0343676 | 0.8552132 | 0.999176 |
| ENSMUSG00000001366 | Fbxo9 | protein_coding | 0.05450226 | 6.406147 | 0.0343373 | 0.8552763 | 0.999176 |
| ENSMUSG00000025867 | Cplx2 | protein_coding | 0.05756462 | 8.226537 | 0.0341592 | 0.8556477 | 0.999176 |
| ENSMUSG00000020849 | Ywhae | protein_coding | -0.06180593 | 11.16071 | 0.0341242 | 0.8557206 | 0.999176 |
| ENSMUSG00000048905 | 4930539E08Rik | protein_coding | 0.2256997 | 2.498641 | 0.0340842 | 0.8558044 | 0.999176 |
| ENSMUSG00000073639 | Rab18 | protein_coding | -0.06100383 | 8.258164 | 0.0340741 | 0.8558254 | 0.999176 |
| ENSMUSG00000030222 | Rerg | protein_coding | 0.07418399 | 5.623907 | 0.0340675 | 0.8558392 | 0.999176 |
| ENSMUSG00000041773 | Enc1 | protein_coding | 0.0723969 | 8.466463 | 0.0340122 | 0.8559586 | 0.999176 |
| ENSMUSG00000032373 | Car12 | protein_coding | 0.1977645 | 3.89923 | 0.0341484 | 0.8559657 | 0.999176 |
| ENSMUSG00000020097 | Sgpl1 | protein_coding | 0.1535207 | 3.309907 | 0.0339824 | 0.8560172 | 0.999176 |
| ENSMUSG00000031610 | Scrg1 | protein_coding | -0.1855697 | 5.703642 | 0.0344248 | 0.8561786 | 0.999176 |
| ENSMUSG00000031539 | Ap3m2 | protein_coding | 0.07608616 | 7.416457 | 0.0338766 | 0.8562425 | 0.999176 |
| ENSMUSG00000030110 | Ret | protein_coding | 0.1220079 | 3.503713 | 0.0338484 | 0.856298 | 0.999176 |
| ENSMUSG00000071723 | Gspt2 | protein_coding | -0.07635444 | 5.012476 | 0.0338084 | 0.8563819 | 0.999176 |
| ENSMUSG00000021366 | Hivep1 | protein_coding | -0.2137516 | 2.754576 | 0.0337566 | 0.8564907 | 0.999176 |
| ENSMUSG00000037373 | Ctbp1 | protein_coding | 0.06258626 | 6.682707 | 0.0336823 | 0.8566469 | 0.999176 |

| ensembl_gene_id | external_gene_name | gene_biotype | log2 (fold change) | log2 (counts per million) | F-value | p-value | False discovery rate |
|---|---|---|---|---|---|---|---|
| ENSMUSG00000061533 | Cep128 | protein_coding | -0.1722354 | 3.424717 | 0.0336818 | 0.8566516 | 0.999176 |
| ENSMUSG00000008035 | Mid1ip1 | protein_coding | -0.08308059 | 6.306995 | 0.0336568 | 0.8567042 | 0.999176 |
| ENSMUSG00000061755 | Bod1l | protein_coding | -0.09166484 | 5.390227 | 0.033273 | 0.8575103 | 0.999176 |
| ENSMUSG00000096910 | Zfp955b | protein_coding | 0.1776666 | 2.486901 | 0.0332468 | 0.8575657 | 0.999176 |
| ENSMUSG00000027509 | Rae1 | protein_coding | 0.08853273 | 4.866388 | 0.0331246 | 0.8578247 | 0.999176 |
| ENSMUSG00000036356 | Csgalnact1 | protein_coding | 0.1517797 | 3.439312 | 0.033117 | 0.8578406 | 0.999176 |
| ENSMUSG00000020594 | Pum2 | protein_coding | 0.05609221 | 7.253719 | 0.0330025 | 0.8580839 | 0.999176 |
| ENSMUSG00000028546 | Elavl4 | protein_coding | 0.07177242 | 7.370551 | 0.0329852 | 0.8581242 | 0.999176 |
| ENSMUSG00000014361 | Mertk | protein_coding | -0.2027209 | 2.70796 | 0.0329742 | 0.8581441 | 0.999176 |
| ENSMUSG00000061759 | Armt1 | protein_coding | 0.08884753 | 4.666926 | 0.0327705 | 0.8585777 | 0.999176 |
| ENSMUSG00000048076 | Arf1 | protein_coding | 0.05831655 | 9.201644 | 0.0327282 | 0.8586679 | 0.999176 |
| ENSMUSG00000026021 | Sumo1 | protein_coding | 0.05486998 | 8.260689 | 0.0327151 | 0.8586959 | 0.999176 |
| ENSMUSG00000032763 | Ilvbl | protein_coding | 0.1054598 | 5.377961 | 0.0326499 | 0.8588389 | 0.999176 |
| ENSMUSG00000078877 | Gm14295 | protein_coding | 0.1132494 | 3.695255 | 0.0326218 | 0.8588953 | 0.999176 |
| ENSMUSG00000034780 | B3galt1 | protein_coding | 0.07565123 | 5.905182 | 0.0324951 | 0.8591665 | 0.999176 |
| ENSMUSG00000032342 | Mto1 | protein_coding | 0.1141822 | 3.611282 | 0.0324473 | 0.8592688 | 0.999176 |
| ENSMUSG00000038145 | Snrk | protein_coding | -0.09985121 | 4.663586 | 0.0324473 | 0.859269 | 0.999176 |
| ENSMUSG00000004383 | Large1 | protein_coding | -0.07443388 | 5.054282 | 0.0324387 | 0.8592872 | 0.999176 |
| ENSMUSG00000028016 | Ints12 | protein_coding | 0.08130365 | 4.737708 | 0.0323229 | 0.8595359 | 0.999176 |
| ENSMUSG00000027272 | Ubr1 | protein_coding | 0.0994749 | 3.851007 | 0.0322399 | 0.8597142 | 0.999176 |
| ENSMUSG00000024423 | Impact | protein_coding | -0.0791541 | 10.99091 | 0.0320952 | 0.8600295 | 0.999176 |
| ENSMUSG00000019906 | Lin7a | protein_coding | -0.07128128 | 6.307667 | 0.0320901 | 0.8600368 | 0.999176 |
| ENSMUSG00000074925 | Ptar1 | protein_coding | 0.2993495 | 2.381256 | 0.0321037 | 0.8600761 | 0.999176 |
| ENSMUSG00000037089 | Slc35b2 | protein_coding | -0.09272658 | 5.014499 | 0.0320654 | 0.8600901 | 0.999176 |
| ENSMUSG00000079316 | Rab9 | protein_coding | -0.06457855 | 6.864755 | 0.0320168 | 0.8601951 | 0.999176 |
| ENSMUSG00000041203 | Trir | protein_coding | 0.09349052 | 4.336468 | 0.0319084 | 0.8604292 | 0.999176 |
| ENSMUSG00000042506 | Usp22 | protein_coding | 0.07456237 | 8.288348 | 0.0319028 | 0.8604449 | 0.999176 |
| ENSMUSG00000025221 | Kcnip2 | protein_coding | 0.06312105 | 6.323219 | 0.0318723 | 0.8605072 | 0.999176 |
| ENSMUSG00000032479 | Map4 | protein_coding | -0.0686131 | 8.692961 | 0.0317829 | 0.8607008 | 0.999176 |
| ENSMUSG00000018765 | Fxr2 | protein_coding | -0.0701295 | 5.496279 | 0.0317797 | 0.8607077 | 0.999176 |
| ENSMUSG00000002129 | Sf3a1 | protein_coding | 0.07494374 | 6.106237 | 0.0317594 | 0.8607518 | 0.999176 |
| ENSMUSG00000025050 | Pcgf6 | protein_coding | -0.1933937 | 2.536866 | 0.0317534 | 0.8607649 | 0.999176 |
| ENSMUSG00000021037 | Ahsa1 | protein_coding | -0.05306738 | 7.846548 | 0.0316864 | 0.8609101 | 0.999176 |
| ENSMUSG00000041272 | Tox | protein_coding | 0.1667461 | 6.058984 | 0.0321787 | 0.8609201 | 0.999176 |
| ENSMUSG00000070498 | Tmem132b | protein_coding | 0.08440301 | 4.839907 | 0.0316575 | 0.860973 | 0.999176 |
| ENSMUSG00000025213 | Kazald1 | protein_coding | 0.1647776 | 2.824612 | 0.0316253 | 0.8610429 | 0.999176 |
| ENSMUSG00000018217 | Pmp22 | protein_coding | -0.08542681 | 5.840557 | 0.0315652 | 0.8611734 | 0.999176 |
| ENSMUSG00000022865 | Cxadr | protein_coding | -0.1438868 | 2.89959 | 0.0315299 | 0.8612503 | 0.999176 |
| ENSMUSG00000038014 | Fam120a | protein_coding | -0.08444814 | 5.576354 | 0.0315172 | 0.8612779 | 0.999176 |
| ENSMUSG00000046138 | 9930021J03Rik | protein_coding | -0.09283434 | 5.84891 | 0.0314834 | 0.8613551 | 0.999176 |
| ENSMUSG00000038650 | Rnh1 | protein_coding | 0.06582674 | 7.089744 | 0.0313978 | 0.861538 | 0.999176 |
| ENSMUSG00000026767 | Mindy3 | protein_coding | 0.06163448 | 6.240347 | 0.0313753 | 0.861587 | 0.999176 |
| ENSMUSG00000037525 | Bcdin3d | protein_coding | -0.1309166 | 4.013762 | 0.0313664 | 0.86161 | 0.999176 |
| ENSMUSG00000024487 | Yipf5 | protein_coding | -0.06609608 | 5.843377 | 0.0313276 | 0.8616912 | 0.999176 |
| ENSMUSG00000056004 | 9330182L06Rik | protein_coding | -0.08195575 | 5.064103 | 0.0313148 | 0.8617191 | 0.999176 |
| ENSMUSG00000022387 | Brd1 | protein_coding | -0.08802858 | 4.94356 | 0.0312749 | 0.8618062 | 0.999176 |
| ENSMUSG00000029863 | Casp2 | protein_coding | 0.1513472 | 2.862351 | 0.0311543 | 0.8620701 | 0.999176 |
| ENSMUSG00000028869 | Gnl2 | protein_coding | -0.1145525 | 5.825994 | 0.0311888 | 0.8622117 | 0.999176 |
| ENSMUSG00000089832 | Shkbp1 | protein_coding | -0.2296982 | 2.761606 | 0.0310845 | 0.8622264 | 0.999176 |
| ENSMUSG00000001983 | Taco1 | protein_coding | -0.1196167 | 3.448332 | 0.0309985 | 0.8624115 | 0.999176 |
| ENSMUSG00000041438 | Utp4 | protein_coding | 0.09225292 | 5.843236 | 0.0309907 | 0.8624322 | 0.999176 |
| ENSMUSG00000025648 | Pfkfb4 | protein_coding | -0.1439677 | 3.353676 | 0.0309566 | 0.8625036 | 0.999176 |
| ENSMUSG00000031060 | Rbm10 | protein_coding | 0.09428767 | 4.31473 | 0.0309363 | 0.8625481 | 0.999176 |
| ENSMUSG00000043323 | Fbrsl1 | protein_coding | 0.1271212 | 3.680549 | 0.0309147 | 0.8625956 | 0.999176 |
| ENSMUSG00000029234 | Tmem165 | protein_coding | -0.1487091 | 3.647535 | 0.0309066 | 0.8626171 | 0.999176 |
| ENSMUSG00000090093 | Gm14399 | protein_coding | -0.1696952 | 2.701888 | 0.03088 | 0.8626719 | 0.999176 |
| ENSMUSG00000028100 | Nudt17 | protein_coding | -0.1197604 | 4.79388 | 0.0308684 | 0.8627144 | 0.999176 |
| ENSMUSG00000028618 | Tmem59 | protein_coding | 0.05170426 | 9.205034 | 0.0308287 | 0.8627849 | 0.999176 |
| ENSMUSG00000036707 | Cab39 | protein_coding | 0.06770596 | 6.547629 | 0.030781 | 0.8628898 | 0.999176 |
| ENSMUSG00000032939 | Nup93 | protein_coding | -0.08759246 | 4.417205 | 0.030772 | 0.8629097 | 0.999176 |
| ENSMUSG00000004221 | Ikbkg | protein_coding | -0.0967465 | 5.073525 | 0.0307675 | 0.8629196 | 0.999176 |
| ENSMUSG00000038335 | Tsr1 | protein_coding | -0.08277062 | 5.146566 | 0.0307532 | 0.8629512 | 0.999176 |
| ENSMUSG00000011254 | Thg1l | protein_coding | -0.08571031 | 5.01098 | 0.0307518 | 0.8629542 | 0.999176 |
| ENSMUSG00000084843 | B230312C02Rik | lincRNA | 0.1379903 | 3.027622 | 0.0307004 | 0.8630675 | 0.999176 |
| ENSMUSG00000049971 | Glt1d1 | protein_coding | 0.143817 | 3.186405 | 0.0306723 | 0.8631297 | 0.999176 |
| ENSMUSG00000046768 | Rhoj | protein_coding | 0.1809974 | 2.680213 | 0.0306529 | 0.8631723 | 0.999176 |
| ENSMUSG00000038984 | Tspyl5 | protein_coding | -0.07299279 | 6.739904 | 0.0306031 | 0.8632859 | 0.999176 |
| ENSMUSG00000029455 | Aldh2 | protein_coding | 0.07750809 | 5.057758 | 0.0305589 | 0.86338 | 0.999176 |
| ENSMUSG00000026761 | Orc4 | protein_coding | -0.06196136 | 6.233258 | 0.0305321 | 0.8634395 | 0.999176 |
| ENSMUSG00000040420 | Cdh18 | protein_coding | 0.1088051 | 3.766926 | 0.0305084 | 0.8634919 | 0.999176 |
| ENSMUSG00000035666 | Gtf3c4 | protein_coding | -0.1178479 | 3.779221 | 0.0305074 | 0.863494 | 0.999176 |
| ENSMUSG00000032329 | Hmg20a | protein_coding | 0.06921771 | 6.190942 | 0.0304473 | 0.863627 | 0.999176 |
| ENSMUSG00000036095 | Dgkb | protein_coding | 0.06614239 | 7.206414 | 0.0303888 | 0.8637567 | 0.999176 |
| ENSMUSG00000027951 | Adar | protein_coding | -0.1982021 | 3.419892 | 0.0304456 | 0.8638115 | 0.999176 |
| ENSMUSG00000025968 | Ndufs1 | protein_coding | -0.09155206 | 5.702322 | 0.0302989 | 0.8639598 | 0.999176 |
| ENSMUSG00000024245 | Tmem178 | protein_coding | -0.1732835 | 3.659489 | 0.0302449 | 0.8642127 | 0.999176 |
| ENSMUSG00000027900 | Dram2 | protein_coding | -0.07013441 | 5.679447 | 0.0301783 | 0.8642244 | 0.999176 |
| ENSMUSG00000073616 | Cops9 | protein_coding | 0.0738079 | 8.663055 | 0.03017 | 0.8642465 | 0.999176 |

| ensembl_gene_id | external_gene_name | gene_biotype | log2 (fold change) | log2 (counts per million) | F-value | p-value | False discovery rate |
|---|---|---|---|---|---|---|---|
| ENSMUSG00000104864 | Gm43655 | processed_pseudoge | 0.1409411 | 2.811837 | 0.0301231 | 0.8643474 | 0.999176 |
| ENSMUSG00000041460 | Cacna2d4 | protein_coding | 0.1912025 | 2.942425 | 0.0301159 | 0.8643635 | 0.999176 |
| ENSMUSG00000022283 | Pabpc1 | protein_coding | 0.07446251 | 5.356367 | 0.0300922 | 0.8644161 | 0.999176 |
| ENSMUSG00000027099 | Mtx2 | protein_coding | 0.08481269 | 5.076598 | 0.0300819 | 0.8644391 | 0.999176 |
| ENSMUSG00000042671 | Rgs8 | protein_coding | -0.076882 | 6.430974 | 0.0300262 | 0.8645669 | 0.999176 |
| ENSMUSG00000070699 | Sars2 | protein_coding | -0.1136774 | 4.124512 | 0.0299097 | 0.8648235 | 0.999176 |
| ENSMUSG00000063273 | Naa15 | protein_coding | 0.08604493 | 5.168553 | 0.029885 | 0.8648788 | 0.999176 |
| ENSMUSG00000053641 | Dennd4a | protein_coding | -0.09346042 | 4.413776 | 0.0298673 | 0.8649186 | 0.999176 |
| ENSMUSG00000089889 | 0610040B10Rik | antisense_RNA | -0.1107362 | 3.573181 | 0.0297773 | 0.86512 | 0.999176 |
| ENSMUSG00000004267 | Eno2 | protein_coding | -0.069494 | 10.2375 | 0.0297639 | 0.86515 | 0.999176 |
| ENSMUSG00000032788 | Pdxk | protein_coding | 0.09669875 | 7.679197 | 0.0298961 | 0.8651522 | 0.999176 |
| ENSMUSG00000043311 | D17H6S53E | protein_coding | 0.1345325 | 5.399066 | 0.0298784 | 0.8651918 | 0.999176 |
| ENSMUSG00000027030 | Stk39 | protein_coding | 0.06969766 | 5.963511 | 0.0297434 | 0.865196 | 0.999176 |
| ENSMUSG00000000134 | Tfe3 | protein_coding | -0.08782285 | 4.709438 | 0.0297405 | 0.8652025 | 0.999176 |
| ENSMUSG00000037653 | Kctd8 | protein_coding | -0.1165368 | 3.702297 | 0.0297024 | 0.8652879 | 0.999176 |
| ENSMUSG00000027012 | Dync1i2 | protein_coding | 0.05191207 | 7.194548 | 0.0296978 | 0.8652984 | 0.999176 |
| ENSMUSG00000037196 | Pacrg | protein_coding | -0.05731707 | 6.01575 | 0.0296916 | 0.8653122 | 0.999176 |
| ENSMUSG00000030275 | Etnk1 | protein_coding | 0.2073137 | 2.811777 | 0.0296534 | 0.865401 | 0.999176 |
| ENSMUSG00000041046 | Ramp3 | protein_coding | -0.1067921 | 6.002881 | 0.0296327 | 0.865656 | 0.999176 |
| ENSMUSG00000047260 | Emc6 | protein_coding | 0.09714143 | 4.534436 | 0.0295148 | 0.8657097 | 0.999176 |
| ENSMUSG00000049881 | 2810025M15Rik | transcribed_processe | -0.06563854 | 5.804135 | 0.0294984 | 0.8657467 | 0.999176 |
| ENSMUSG00000035227 | Spcs2 | protein_coding | 0.05626221 | 8.275561 | 0.0294815 | 0.8657845 | 0.999176 |
| ENSMUSG00000026827 | Gpd2 | protein_coding | 0.05526513 | 6.542161 | 0.0294779 | 0.8657927 | 0.999176 |
| ENSMUSG00000031066 | Usp11 | protein_coding | 0.07877236 | 7.700349 | 0.0294848 | 0.8658937 | 0.999176 |
| ENSMUSG00000040128 | Pnrc1 | protein_coding | -0.06516267 | 5.564301 | 0.0294241 | 0.8659139 | 0.999176 |
| ENSMUSG00000029860 | Zyx | protein_coding | -0.09323468 | 4.171575 | 0.0294237 | 0.8659148 | 0.999176 |
| ENSMUSG00000046434 | Hnrnpa1 | protein_coding | 0.09223314 | 3.845622 | 0.0293906 | 0.8659896 | 0.999176 |
| ENSMUSG00000038240 | Pdss2 | protein_coding | 0.0795771 | 4.928223 | 0.0293824 | 0.866008 | 0.999176 |
| ENSMUSG00000022094 | Slc39a14 | protein_coding | -0.1556545 | 3.064126 | 0.0293371 | 0.8661104 | 0.999176 |
| ENSMUSG00000035212 | Leprot | protein_coding | -0.08157243 | 5.555849 | 0.0293336 | 0.8661182 | 0.999176 |
| ENSMUSG00000000295 | Hddc2 | protein_coding | 0.06095128 | 7.10108 | 0.0292246 | 0.8663647 | 0.999176 |
| ENSMUSG00000034211 | Mrps17 | protein_coding | 0.05245632 | 7.895572 | 0.0292167 | 0.8663825 | 0.999176 |
| ENSMUSG00000024902 | Mrpl11 | protein_coding | 0.04985696 | 7.861114 | 0.0291865 | 0.8664509 | 0.999176 |
| ENSMUSG00000024213 | Nudt3 | protein_coding | 0.05391953 | 8.135176 | 0.0291809 | 0.8664635 | 0.999176 |
| ENSMUSG00000041794 | Myrip | protein_coding | -0.1180114 | 3.511553 | 0.0291634 | 0.8665032 | 0.999176 |
| ENSMUSG00000037447 | Arid5a | protein_coding | 0.1198153 | 4.065855 | 0.0291228 | 0.8665953 | 0.999176 |
| ENSMUSG00000107476 | Zfp862-ps | transcribed_unitary_p | -0.1543355 | 2.780248 | 0.0290987 | 0.8666498 | 0.999176 |
| ENSMUSG00000021619 | Atg10 | protein_coding | -0.05800028 | 5.973823 | 0.0290901 | 0.8666694 | 0.999176 |
| ENSMUSG00000009291 | Pttg1ip | protein_coding | -0.06263705 | 6.171832 | 0.0290546 | 0.8667498 | 0.999176 |
| ENSMUSG00000086272 | BC039966 | antisense_RNA | -0.1377546 | 3.834085 | 0.0290334 | 0.8668016 | 0.999176 |
| ENSMUSG00000020701 | Tmem132e | protein_coding | 0.13009 | 3.462793 | 0.0290157 | 0.8668381 | 0.999176 |
| ENSMUSG00000060121 | Gemin2 | protein_coding | 0.07779908 | 4.848683 | 0.0289984 | 0.8668775 | 0.999176 |
| ENSMUSG00000037892 | Pcdh18 | protein_coding | 0.129084 | 3.706613 | 0.0289611 | 0.8669622 | 0.999176 |
| ENSMUSG00000015247 | Nipsnap3b | protein_coding | -0.0616097 | 6.272603 | 0.0289254 | 0.8670434 | 0.999176 |
| ENSMUSG00000027165 | B230118H07Rik | protein_coding | 0.05594846 | 7.438964 | 0.0288818 | 0.8671428 | 0.999176 |
| ENSMUSG00000069171 | Nr2f1 | protein_coding | -0.08139811 | 4.58568 | 0.0288204 | 0.8672825 | 0.999176 |
| ENSMUSG00000053333 | Dis3l2 | protein_coding | -0.1566919 | 2.901809 | 0.0288072 | 0.8673126 | 0.999176 |
| ENSMUSG00000030298 | Sec13 | protein_coding | -0.05218023 | 7.532545 | 0.0287744 | 0.8673875 | 0.999176 |
| ENSMUSG00000043421 | Hilpda | protein_coding | 0.08014943 | 4.813176 | 0.0287278 | 0.8674939 | 0.999176 |
| ENSMUSG00000034379 | Wdr5b | protein_coding | 0.1298692 | 3.381514 | 0.0287022 | 0.8675522 | 0.999176 |
| ENSMUSG00000025337 | Sbds | protein_coding | -0.04683053 | 8.185857 | 0.0286857 | 0.86759 | 0.999176 |
| ENSMUSG00000028099 | Polr3c | protein_coding | 0.0917858 | 5.569448 | 0.0286498 | 0.8676754 | 0.999176 |
| ENSMUSG00000041278 | Ttc1 | protein_coding | 0.04578449 | 7.870328 | 0.0286373 | 0.8677005 | 0.999176 |
| ENSMUSG00000024240 | Epc1 | protein_coding | -0.08318495 | 4.111177 | 0.0286002 | 0.8677854 | 0.999176 |
| ENSMUSG00000056267 | Cep70 | protein_coding | -0.08693625 | 4.424573 | 0.028498 | 0.8680195 | 0.999176 |
| ENSMUSG00000079084 | Ccdc82 | protein_coding | 0.09785253 | 4.377176 | 0.0284155 | 0.8682088 | 0.999176 |
| ENSMUSG00000028603 | Scp2 | protein_coding | 0.145989 | 3.075464 | 0.0282451 | 0.8686006 | 0.999176 |
| ENSMUSG00000021113 | Snapc1 | protein_coding | -0.06372465 | 5.663202 | 0.0281464 | 0.8688281 | 0.999176 |
| ENSMUSG00000040918 | Slc19a2 | protein_coding | -0.1362793 | 3.250005 | 0.0280564 | 0.869036 | 0.999176 |
| ENSMUSG00000040003 | Magi2 | protein_coding | -0.06892076 | 5.226439 | 0.0280563 | 0.8690362 | 0.999176 |
| ENSMUSG00000018001 | Cyth3 | protein_coding | 0.1033008 | 3.947395 | 0.0280292 | 0.8690986 | 0.999176 |
| ENSMUSG00000028398 | Tmem261 | protein_coding | -0.05612175 | 6.682361 | 0.0279941 | 0.8691799 | 0.999176 |
| ENSMUSG00000062646 | Ganc | protein_coding | -0.09499633 | 3.615466 | 0.0279405 | 0.8693041 | 0.999176 |
| ENSMUSG00000038665 | Dgki | protein_coding | -0.1124569 | 3.62175 | 0.0279332 | 0.8693209 | 0.999176 |
| ENSMUSG00000064341 | mt-Nd1 | protein_coding | 0.08685413 | 12.02551 | 0.0279699 | 0.8693221 | 0.999176 |
| ENSMUSG00000020593 | Lpin1 | protein_coding | 0.0982448 | 4.675133 | 0.0279159 | 0.869361 | 0.999176 |
| ENSMUSG00000038822 | Hace1 | protein_coding | -0.09566092 | 4.195022 | 0.0279051 | 0.8693861 | 0.999176 |
| ENSMUSG00000062184 | Hs6st2 | protein_coding | -0.06914236 | 5.469407 | 0.0278562 | 0.8694993 | 0.999176 |
| ENSMUSG00000053293 | Pom121 | protein_coding | -0.1168636 | 4.121407 | 0.0278443 | 0.869527 | 0.999176 |
| ENSMUSG00000032612 | Usp4 | protein_coding | 0.05754739 | 6.156897 | 0.0277089 | 0.8698415 | 0.999176 |
| ENSMUSG00000015656 | Hspa8 | protein_coding | 0.08052258 | 8.910846 | 0.0276703 | 0.8700616 | 0.999176 |
| ENSMUSG00000007656 | Arpp19 | protein_coding | 0.06639725 | 9.503378 | 0.027487 | 0.8703587 | 0.999176 |
| ENSMUSG00000036197 | Gxylt1 | protein_coding | 0.1156567 | 3.207011 | 0.0274549 | 0.8704337 | 0.999176 |
| ENSMUSG00000087651 | 1500009L16Rik | protein_coding | -0.05398769 | 6.322883 | 0.0273095 | 0.8707738 | 0.999176 |
| ENSMUSG00000028848 | Gpn2 | protein_coding | -0.1493253 | 2.981743 | 0.0272476 | 0.8709191 | 0.999176 |
| ENSMUSG00000041895 | Wipi1 | protein_coding | -0.1442824 | 4.434203 | 0.0272815 | 0.8710421 | 0.999176 |
| ENSMUSG00000025236 | Adpgk | protein_coding | 0.1556762 | 3.119366 | 0.0271715 | 0.8710976 | 0.999176 |
| ENSMUSG00000091474 | 2610021A01Rik | protein_coding | 0.1157834 | 3.469323 | 0.0270981 | 0.8712701 | 0.999176 |

| ensembl_gene_id | external_gene_name | gene_biotype | log2 (fold change) | log2 (counts per million) | F-value | p-value | False discovery rate |
|---|---|---|---|---|---|---|---|
| ENSMUSG00000035064 | Eef2k | protein_coding | -0.09565667 | 3.847566 | 0.0270958 | 0.8712756 | 0.999176 |
| ENSMUSG00000028811 | Yars | protein_coding | -0.06744322 | 7.068007 | 0.0270774 | 0.8713222 | 0.999176 |
| ENSMUSG00000046613 | Vwa5b2 | protein_coding | 0.1520894 | 2.903224 | 0.0270476 | 0.8713892 | 0.999176 |
| ENSMUSG00000025240 | Sacm1l | protein_coding | -0.15616 | 3.773464 | 0.0271001 | 0.8713945 | 0.999176 |
| ENSMUSG00000029651 | Mtus2 | protein_coding | 0.06246619 | 5.408452 | 0.0269959 | 0.8715108 | 0.999176 |
| ENSMUSG00000029763 | Exoc4 | protein_coding | 0.06610634 | 5.708464 | 0.0269463 | 0.8716279 | 0.999176 |
| ENSMUSG00000049882 | Vcpkmt | protein_coding | -0.1359191 | 2.982139 | 0.0269061 | 0.8717228 | 0.999176 |
| ENSMUSG00000017466 | Timp2 | protein_coding | -0.0488584 | 8.165905 | 0.0269022 | 0.8717319 | 0.999176 |
| ENSMUSG00000079242 | C730034F03Rik | protein_coding | 0.1575412 | 3.111918 | 0.0268839 | 0.8717751 | 0.999176 |
| ENSMUSG00000058446 | Znrf2 | protein_coding | -0.1044226 | 3.726275 | 0.0268638 | 0.8718228 | 0.999176 |
| ENSMUSG00000020151 | Ptprr | protein_coding | -0.07209857 | 4.876246 | 0.0268534 | 0.8718472 | 0.999176 |
| ENSMUSG00000037455 | Slc18b1 | protein_coding | 0.1276703 | 3.006307 | 0.0268371 | 0.8718858 | 0.999176 |
| ENSMUSG00000020453 | Patz1 | protein_coding | 0.1257423 | 3.463946 | 0.0267949 | 0.8719855 | 0.999176 |
| ENSMUSG00000010663 | Fads1 | protein_coding | -0.06293766 | 5.969592 | 0.0267745 | 0.8720339 | 0.999176 |
| ENSMUSG00000022092 | Ppp3cc | protein_coding | 0.1414985 | 3.041517 | 0.0267536 | 0.8720834 | 0.999176 |
| ENSMUSG00000047013 | Fbxo41 | protein_coding | -0.08674844 | 5.23185 | 0.0266761 | 0.8722671 | 0.999176 |
| ENSMUSG00000020277 | Pfkl | protein_coding | 0.07855774 | 7.516622 | 0.0267058 | 0.8723245 | 0.999176 |
| ENSMUSG00000023336 | Wfdc1 | protein_coding | -0.2222412 | 2.318903 | 0.0266043 | 0.8724375 | 0.999176 |
| ENSMUSG00000037172 | E330009J07Rik | protein_coding | -0.08555409 | 4.419616 | 0.026566 | 0.8725284 | 0.999176 |
| ENSMUSG00000024479 | Mal2 | protein_coding | -0.06442045 | 5.853344 | 0.0265313 | 0.8726109 | 0.999176 |
| ENSMUSG00000033316 | Galnt9 | protein_coding | 0.09413949 | 4.225352 | 0.0264964 | 0.872694 | 0.999176 |
| ENSMUSG00000021750 | Fam107a | protein_coding | -0.06427059 | 6.542336 | 0.0264034 | 0.8729155 | 0.999176 |
| ENSMUSG00000041907 | Gpr45 | protein_coding | -0.1177307 | 3.999235 | 0.0263793 | 0.8729729 | 0.999176 |
| ENSMUSG00000044250 | Pced1b | protein_coding | -0.1329035 | 2.992363 | 0.0263755 | 0.872982 | 0.999176 |
| ENSMUSG00000055553 | Kxd1 | protein_coding | 0.04891603 | 7.517411 | 0.0263687 | 0.8729983 | 0.999176 |
| ENSMUSG00000046449 | C77370 | protein_coding | -0.08763102 | 4.756877 | 0.0262816 | 0.8732063 | 0.999176 |
| ENSMUSG00000017493 | Igfbp4 | protein_coding | -0.1262616 | 4.47855 | 0.0263261 | 0.8732135 | 0.999176 |
| ENSMUSG00000018449 | Rpain | protein_coding | 0.06361388 | 5.882815 | 0.0262757 | 0.8732203 | 0.999176 |
| ENSMUSG00000051413 | Plagl2 | protein_coding | -0.1254378 | 2.963118 | 0.0261953 | 0.8734127 | 0.999176 |
| ENSMUSG00000052557 | Gan | protein_coding | 0.115789 | 3.380615 | 0.0261872 | 0.8734321 | 0.999176 |
| ENSMUSG00000009070 | Rsph14 | protein_coding | -0.1307186 | 3.214989 | 0.0261459 | 0.8735311 | 0.999176 |
| ENSMUSG00000050556 | Kcnb1 | protein_coding | 0.08395323 | 5.253583 | 0.026054 | 0.8737514 | 0.999176 |
| ENSMUSG00000062995 | Ica1 | protein_coding | -0.0741798 | 6.464475 | 0.02599 | 0.8739084 | 0.999176 |
| ENSMUSG00000107872 | Gm44511 | protein_coding | -0.1335565 | 3.305189 | 0.0259743 | 0.873943 | 0.999176 |
| ENSMUSG00000029765 | Plxna4 | protein_coding | -0.0858652 | 4.416683 | 0.0259696 | 0.8739541 | 0.999176 |
| ENSMUSG00000086877 | A230072C01Rik | processed_transcript | 0.07785817 | 4.659432 | 0.0259483 | 0.8740055 | 0.999176 |
| ENSMUSG00000024312 | Wdr46 | protein_coding | 0.1505805 | 2.90048 | 0.0259137 | 0.8740886 | 0.999176 |
| ENSMUSG00000054693 | Adam10 | protein_coding | 0.05616633 | 5.802093 | 0.0259048 | 0.8741101 | 0.999176 |
| ENSMUSG00000027809 | Etfdh | protein_coding | 0.07883125 | 4.954824 | 0.0258984 | 0.8741254 | 0.999176 |
| ENSMUSG00000021483 | Cdk20 | protein_coding | -0.07888668 | 4.943557 | 0.0258726 | 0.8741876 | 0.999176 |
| ENSMUSG00000048537 | Phldb1 | protein_coding | 0.07613892 | 4.84687 | 0.0258684 | 0.8741978 | 0.999176 |
| ENSMUSG00000033486 | Catsper2 | protein_coding | 0.1218931 | 3.225605 | 0.0258188 | 0.8743173 | 0.999176 |
| ENSMUSG00000031353 | Rbbp7 | protein_coding | -0.05468603 | 7.023522 | 0.0257925 | 0.8743806 | 0.999176 |
| ENSMUSG00000026154 | Sdhaf4 | protein_coding | 0.05830651 | 8.414364 | 0.0257822 | 0.8744055 | 0.999176 |
| ENSMUSG00000044864 | Ankrd50 | protein_coding | 0.08838488 | 4.326761 | 0.0257552 | 0.8744709 | 0.999176 |
| ENSMUSG00000039285 | Azi2 | protein_coding | 0.06977834 | 5.542732 | 0.0257379 | 0.8745126 | 0.999176 |
| ENSMUSG00000028029 | Aimp1 | protein_coding | 0.05737086 | 7.77312 | 0.0257285 | 0.8745353 | 0.999176 |
| ENSMUSG00000032606 | Nicn1 | protein_coding | -0.05205269 | 8.336556 | 0.0257252 | 0.8745433 | 0.999176 |
| ENSMUSG00000046182 | Gsg1l | protein_coding | -0.1256651 | 3.75918 | 0.0256974 | 0.8746137 | 0.999176 |
| ENSMUSG00000039804 | Ncoa5 | protein_coding | -0.08903394 | 4.500782 | 0.0256937 | 0.8746195 | 0.999176 |
| ENSMUSG00000030876 | Mettl9 | protein_coding | -0.08235725 | 5.042005 | 0.0256919 | 0.8746237 | 0.999176 |
| ENSMUSG00000031660 | Brd7 | protein_coding | -0.07070037 | 5.766306 | 0.0256839 | 0.874643 | 0.999176 |
| ENSMUSG00000051232 | Tmem199 | protein_coding | 0.04808427 | 7.900209 | 0.0256409 | 0.874747 | 0.999176 |
| ENSMUSG00000032097 | Ddx6 | protein_coding | -0.0496648 | 6.223176 | 0.025617 | 0.8748049 | 0.999176 |
| ENSMUSG00000014856 | Tmem208 | protein_coding | 0.05499912 | 6.816027 | 0.0255845 | 0.8748837 | 0.999176 |
| ENSMUSG00000074093 | Svip | protein_coding | 0.06516339 | 6.117589 | 0.0255351 | 0.8750034 | 0.999176 |
| ENSMUSG00000037197 | Rbm17 | protein_coding | -0.07194958 | 4.756585 | 0.0254894 | 0.8751143 | 0.999176 |
| ENSMUSG00000063160 | Numbl | protein_coding | 0.110947 | 3.218412 | 0.0254786 | 0.8751406 | 0.999176 |
| ENSMUSG00000057551 | Zfp317 | protein_coding | -0.08935993 | 4.910414 | 0.0254415 | 0.8752307 | 0.999176 |
| ENSMUSG00000069184 | Zfp72 | protein_coding | -0.1294612 | 3.149836 | 0.0254045 | 0.8753208 | 0.999176 |
| ENSMUSG00000034807 | Colgalt1 | protein_coding | -0.1271729 | 3.792657 | 0.0253324 | 0.8754993 | 0.999176 |
| ENSMUSG00000009073 | Nf2 | protein_coding | -0.07700396 | 4.799165 | 0.0252783 | 0.875628 | 0.999176 |
| ENSMUSG00000048578 | Mlec | protein_coding | 0.06290556 | 5.514728 | 0.0252383 | 0.8757256 | 0.999176 |
| ENSMUSG00000071234 | Syndig1l | protein_coding | -0.08915026 | 5.554383 | 0.0251967 | 0.8758304 | 0.999176 |
| ENSMUSG00000040688 | Tbl3 | protein_coding | -0.0794203 | 5.372885 | 0.0251863 | 0.8758524 | 0.999176 |
| ENSMUSG00000057278 | Snrpg | protein_coding | 0.08552018 | 5.204688 | 0.025147 | 0.8759486 | 0.999176 |
| ENSMUSG00000027651 | Rprd1b | protein_coding | 0.09204399 | 4.07536 | 0.0251363 | 0.8759747 | 0.999176 |
| ENSMUSG00000022861 | Dgkg | protein_coding | -0.05367096 | 6.621568 | 0.0251145 | 0.876028 | 0.999176 |
| ENSMUSG00000052137 | Rbm12b2 | protein_coding | 0.1340897 | 3.454938 | 0.0251083 | 0.8760431 | 0.999176 |
| ENSMUSG00000032002 | Dcun1d5 | protein_coding | 0.05589109 | 5.856713 | 0.0250238 | 0.87625 | 0.999176 |
| ENSMUSG00000026094 | Stk17b | protein_coding | 0.27664 | 2.255065 | 0.0249911 | 0.8763475 | 0.999176 |
| ENSMUSG00000004798 | Ulk2 | protein_coding | -0.07689444 | 5.441128 | 0.024975 | 0.8763697 | 0.999176 |
| ENSMUSG00000036879 | Phkb | protein_coding | -0.07836634 | 4.60745 | 0.0249284 | 0.876484 | 0.999176 |
| ENSMUSG00000027206 | Cops2 | protein_coding | 0.05579984 | 7.228815 | 0.0249097 | 0.87653 | 0.999176 |
| ENSMUSG00000036257 | Pnpla8 | protein_coding | 0.05248786 | 6.179466 | 0.0248802 | 0.8766025 | 0.999176 |
| ENSMUSG00000032625 | Thsd7a | protein_coding | -0.1234336 | 4.466603 | 0.0249241 | 0.8766107 | 0.999176 |
| ENSMUSG00000029250 | Polr2b | protein_coding | -0.06502048 | 5.613263 | 0.0248709 | 0.8766253 | 0.999176 |
| ENSMUSG00000003863 | Ppfia3 | protein_coding | 0.07379428 | 4.668905 | 0.0248561 | 0.8766618 | 0.999176 |

| ensembl_gene_id | external_gene_name | gene_biotype | log2 (fold change) | log2 (counts per million) | F-value | p-value | False discovery rate |
|---|---|---|---|---|---|---|---|
| ENSMUSG00000022010 | Tsc22d1 | protein_coding | 0.0470105 | 9.219451 | 0.0248543 | 0.8766661 | 0.999176 |
| ENSMUSG00000018678 | Sp2 | protein_coding | 0.2099745 | 2.48432 | 0.0247891 | 0.8768268 | 0.999176 |
| ENSMUSG00000033478 | Fam160b1 | protein_coding | -0.09813974 | 3.699283 | 0.0247496 | 0.876924 | 0.999176 |
| ENSMUSG00000021209 | Ppp4r4 | protein_coding | 0.08616341 | 4.206088 | 0.0247322 | 0.8769669 | 0.999176 |
| ENSMUSG00000038602 | Slc35f1 | protein_coding | -0.06928919 | 5.437663 | 0.0246601 | 0.8771448 | 0.999176 |
| ENSMUSG00000041598 | Cdc42ep4 | protein_coding | -0.08321163 | 4.979978 | 0.0245611 | 0.8773895 | 0.999176 |
| ENSMUSG00000074771 | Ankef1 | protein_coding | -0.1642444 | 2.771985 | 0.0244346 | 0.8777029 | 0.999176 |
| ENSMUSG00000021000 | Ctage5 | protein_coding | -0.09759453 | 4.251313 | 0.0244272 | 0.8777213 | 0.999176 |
| ENSMUSG00000095105 | Edaradd | protein_coding | 0.3142374 | 1.826958 | 0.024423 | 0.8777349 | 0.999176 |
| ENSMUSG00000023992 | Trem2 | protein_coding | -0.09778688 | 4.01697 | 0.0243693 | 0.8778651 | 0.999176 |
| ENSMUSG00000074088 | Snrnp40 | protein_coding | 0.0797317 | 5.159838 | 0.0242843 | 0.8780765 | 0.999176 |
| ENSMUSG00000055866 | Per2 | protein_coding | -0.1479267 | 3.069343 | 0.0242611 | 0.8781342 | 0.999176 |
| ENSMUSG00000024070 | Prkd3 | protein_coding | 0.104512 | 3.494451 | 0.0242588 | 0.8781401 | 0.999176 |
| ENSMUSG00000003813 | Rad23a | protein_coding | 0.04881749 | 8.108862 | 0.024256 | 0.8781468 | 0.999176 |
| ENSMUSG00000021196 | Pfkp | protein_coding | -0.06531127 | 7.67635 | 0.0242216 | 0.8782358 | 0.999176 |
| ENSMUSG00000080316 | Spaca6 | protein_coding | -0.09772905 | 3.488998 | 0.024186 | 0.8783213 | 0.999176 |
| ENSMUSG00000036980 | Taf6 | protein_coding | 0.09124896 | 5.72807 | 0.0241885 | 0.8784026 | 0.999176 |
| ENSMUSG00000022677 | Fopnl | protein_coding | -0.04720838 | 8.132747 | 0.0241133 | 0.8785028 | 0.999176 |
| ENSMUSG00000026610 | Esrrg | protein_coding | 0.08352308 | 4.177014 | 0.0240891 | 0.8785633 | 0.999176 |
| ENSMUSG00000034285 | Nipsnap1 | protein_coding | -0.04358229 | 8.165716 | 0.0240734 | 0.8786024 | 0.999176 |
| ENSMUSG00000037253 | Mex3c | protein_coding | -0.1108868 | 3.9381 | 0.0240705 | 0.8786098 | 0.999176 |
| ENSMUSG00000033214 | Slitrk5 | protein_coding | -0.07201893 | 4.334733 | 0.0240138 | 0.8787515 | 0.999176 |
| ENSMUSG00000025630 | Hprt | protein_coding | -0.04467673 | 9.881868 | 0.0239832 | 0.8788283 | 0.999176 |
| ENSMUSG00000029782 | Tmem209 | protein_coding | 0.09548275 | 3.896533 | 0.0239674 | 0.8788678 | 0.999176 |
| ENSMUSG00000020948 | Klhl28 | protein_coding | -0.06690727 | 4.843609 | 0.0239498 | 0.878912 | 0.999176 |
| ENSMUSG00000039579 | Grin3a | protein_coding | 0.05848018 | 5.533032 | 0.0239178 | 0.8789922 | 0.999176 |
| ENSMUSG00000024927 | Rela | protein_coding | -0.07872748 | 5.165239 | 0.0238146 | 0.8792514 | 0.999176 |
| ENSMUSG00000035268 | Pkig | protein_coding | 0.04929715 | 7.141991 | 0.0237981 | 0.8792929 | 0.999176 |
| ENSMUSG00000022292 | Rrm2b | protein_coding | -0.07555919 | 4.879641 | 0.0237898 | 0.8793137 | 0.999176 |
| ENSMUSG00000086350 | B230369F24Rik | antisense_RNA | -0.09564795 | 3.605259 | 0.0237815 | 0.8793345 | 0.999176 |
| ENSMUSG00000021273 | Fdft1 | protein_coding | -0.05409356 | 7.190816 | 0.023708 | 0.8795195 | 0.999176 |
| ENSMUSG00000027227 | Sord | protein_coding | 0.08139873 | 5.580487 | 0.0235966 | 0.8798037 | 0.999176 |
| ENSMUSG00000027341 | Tmem230 | protein_coding | 0.07869416 | 4.332293 | 0.0235794 | 0.8798441 | 0.999176 |
| ENSMUSG00000031935 | Med17 | protein_coding | -0.112056 | 3.988972 | 0.0235625 | 0.8798867 | 0.999176 |
| ENSMUSG00000018474 | Chd3 | protein_coding | -0.04875277 | 6.673163 | 0.0235522 | 0.8799128 | 0.999176 |
| ENSMUSG00000038299 | Wdr36 | protein_coding | 0.08973503 | 4.820469 | 0.0234876 | 0.8800762 | 0.999176 |
| ENSMUSG00000019505 | Ubb | protein_coding | -0.05030046 | 10.89795 | 0.0234354 | 0.8802085 | 0.999176 |
| ENSMUSG00000025083 | Afap1l2 | protein_coding | -0.1949315 | 2.378909 | 0.023284 | 0.880593 | 0.999176 |
| ENSMUSG00000026797 | Stxbp1 | protein_coding | 0.07035306 | 9.183461 | 0.023308 | 0.8805943 | 0.999176 |
| ENSMUSG00000060152 | Pop5 | protein_coding | 0.07208861 | 5.033105 | 0.0231437 | 0.8809504 | 0.999176 |
| ENSMUSG00000008206 | Cers4 | protein_coding | -0.06219665 | 5.832784 | 0.0231091 | 0.8810385 | 0.999176 |
| ENSMUSG00000045613 | Chrm2 | protein_coding | -0.0824911 | 4.448618 | 0.0230958 | 0.8810724 | 0.999176 |
| ENSMUSG00000039427 | Alg1 | protein_coding | -0.1233461 | 3.429039 | 0.0230617 | 0.8811597 | 0.999176 |
| ENSMUSG00000024352 | Spata24 | protein_coding | 0.08224485 | 4.421206 | 0.0230574 | 0.8811706 | 0.999176 |
| ENSMUSG00000031659 | Adcy7 | protein_coding | 0.1185499 | 3.286394 | 0.0230131 | 0.881284 | 0.999176 |
| ENSMUSG00000075703 | Selenoi | protein_coding | -0.09223387 | 5.201008 | 0.022957 | 0.8814306 | 0.999176 |
| ENSMUSG00000021243 | Fcf1 | protein_coding | 0.07457933 | 5.818225 | 0.022936 | 0.8814844 | 0.999176 |
| ENSMUSG00000112390 | CT025654.1 | processed_transcript | 0.1363979 | 3.127734 | 0.0228573 | 0.8816833 | 0.999176 |
| ENSMUSG00000005362 | Crbn | protein_coding | 0.06353168 | 6.15556 | 0.0228158 | 0.8817897 | 0.999176 |
| ENSMUSG00000078695 | Cisd3 | protein_coding | 0.04519411 | 6.448157 | 0.0228115 | 0.8818009 | 0.999176 |
| ENSMUSG00000002221 | Paxip1 | protein_coding | -0.08197941 | 4.563425 | 0.0227061 | 0.8820721 | 0.999176 |
| ENSMUSG00000026888 | Grb14 | protein_coding | -0.06791847 | 6.202079 | 0.0227066 | 0.8820738 | 0.999176 |
| ENSMUSG00000028439 | Fam219a | protein_coding | 0.1411963 | 2.820742 | 0.0226886 | 0.8821172 | 0.999176 |
| ENSMUSG00000028381 | Ugcg | protein_coding | 0.06259519 | 5.413771 | 0.0226751 | 0.882152 | 0.999176 |
| ENSMUSG00000026153 | Fam135a | protein_coding | 0.1451334 | 3.403982 | 0.0225882 | 0.8823793 | 0.999176 |
| ENSMUSG00000020482 | Ccdc117 | protein_coding | -0.120785 | 3.228346 | 0.0225791 | 0.8823998 | 0.999176 |
| ENSMUSG00000039000 | Ube3c | protein_coding | -0.06294226 | 5.459231 | 0.0225607 | 0.8824473 | 0.999176 |
| ENSMUSG00000022559 | Fbxl6 | protein_coding | -0.1216291 | 3.018727 | 0.0225441 | 0.88249 | 0.999176 |
| ENSMUSG00000038205 | Prkab2 | protein_coding | -0.1699182 | 2.740696 | 0.0224881 | 0.882635 | 0.999176 |
| ENSMUSG00000033720 | Sfxn5 | protein_coding | -0.07571365 | 8.774875 | 0.0225058 | 0.8827401 | 0.999176 |
| ENSMUSG00000053347 | Zfp943 | protein_coding | -0.125242 | 4.009395 | 0.0224493 | 0.8827926 | 0.999176 |
| ENSMUSG00000038233 | Fam198a | protein_coding | -0.2091095 | 2.170744 | 0.0223865 | 0.8828983 | 0.999176 |
| ENSMUSG00000087143 | A830082K12Rik | antisense_RNA | -0.09061932 | 3.857802 | 0.0223781 | 0.8829202 | 0.999176 |
| ENSMUSG00000025745 | Hadha | protein_coding | -0.06312937 | 5.641656 | 0.0223612 | 0.882964 | 0.999176 |
| ENSMUSG00000023805 | Synj2 | protein_coding | 0.076118 | 4.969784 | 0.0223332 | 0.8830366 | 0.999176 |
| ENSMUSG00000039828 | Wdr70 | protein_coding | 0.05734704 | 5.70049 | 0.0223017 | 0.8831186 | 0.999176 |
| ENSMUSG00000046010 | Zfp830 | protein_coding | -0.07527539 | 4.747411 | 0.0222153 | 0.8833433 | 0.999176 |
| ENSMUSG00000024588 | Fech | protein_coding | 0.08226841 | 5.324262 | 0.0222048 | 0.8833739 | 0.999176 |
| ENSMUSG00000050063 | Klk6 | protein_coding | -0.1235084 | 3.952359 | 0.0221778 | 0.8834573 | 0.999176 |
| ENSMUSG00000023942 | Slc29a1 | protein_coding | -0.08021242 | 4.982933 | 0.0221396 | 0.8835407 | 0.999176 |
| ENSMUSG00000013076 | Amotl1 | protein_coding | -0.1120048 | 7.00135 | 0.0224185 | 0.8835857 | 0.999176 |
| ENSMUSG00000032841 | Prr5l | protein_coding | 0.08316511 | 5.449711 | 0.0220822 | 0.8836938 | 0.999176 |
| ENSMUSG00000032220 | Myo1e | protein_coding | -0.1378538 | 2.891452 | 0.0220238 | 0.8838434 | 0.999176 |
| ENSMUSG00000028801 | Stpg1 | protein_coding | 0.1228963 | 3.122919 | 0.0219757 | 0.8839692 | 0.999176 |
| ENSMUSG00000005374 | Tbl2 | protein_coding | -0.1007766 | 3.460875 | 0.0218692 | 0.8842487 | 0.999176 |
| ENSMUSG00000026825 | Dnm1 | protein_coding | -0.05314969 | 6.683177 | 0.0218606 | 0.8842712 | 0.999176 |
| ENSMUSG00000031657 | Heatr3 | protein_coding | -0.08624405 | 4.264893 | 0.0218204 | 0.8843769 | 0.999176 |
| ENSMUSG00000004113 | Cacna1b | protein_coding | -0.0816097 | 4.851316 | 0.0217797 | 0.884484 | 0.999176 |

| ensembl_gene_id | external_gene_name | gene_biotype | log2 (fold change) | log2 (counts per million) | F-value | p-value | False discovery rate |
|---|---|---|---|---|---|---|---|
| ENSMUSG00000065954 | Tacc1 | protein_coding | 0.05361804 | 6.421679 | 0.021774 | 0.8844989 | 0.999176 |
| ENSMUSG00000027395 | Polr1b | protein_coding | -0.1580185 | 2.592148 | 0.0217725 | 0.8845029 | 0.999176 |
| ENSMUSG00000027187 | Cat | protein_coding | -0.06947001 | 5.842333 | 0.0215951 | 0.8849708 | 0.999176 |
| ENSMUSG00000047844 | Bex4 | protein_coding | -0.05981537 | 5.169282 | 0.0214847 | 0.8852629 | 0.999176 |
| ENSMUSG00000032968 | Inha | protein_coding | -0.1686069 | 2.437308 | 0.021445 | 0.8853682 | 0.999176 |
| ENSMUSG00000033808 | Tmem87a | protein_coding | 0.154627 | 2.727547 | 0.0214181 | 0.8854396 | 0.999176 |
| ENSMUSG00000027630 | Tbl1xr1 | protein_coding | -0.06315599 | 5.024175 | 0.0213876 | 0.8855207 | 0.999176 |
| ENSMUSG00000038250 | Usp38 | protein_coding | 0.1318645 | 4.033245 | 0.021427 | 0.8855302 | 0.999176 |
| ENSMUSG00000027944 | Hax1 | protein_coding | -0.0549639 | 5.327219 | 0.0212719 | 0.8858282 | 0.999176 |
| ENSMUSG00000028541 | B4galt2 | protein_coding | -0.09449353 | 3.494569 | 0.0212709 | 0.8858309 | 0.999176 |
| ENSMUSG00000033364 | Usp37 | protein_coding | -0.09620041 | 5.195302 | 0.0212571 | 0.8859427 | 0.999176 |
| ENSMUSG00000038909 | Kat7 | protein_coding | 0.06438555 | 6.342928 | 0.0212217 | 0.8859651 | 0.999176 |
| ENSMUSG00000028573 | Fggy | protein_coding | 0.06898485 | 5.243829 | 0.0212054 | 0.8860056 | 0.999176 |
| ENSMUSG00000002799 | Jag2 | protein_coding | 0.1685712 | 2.957579 | 0.021204 | 0.8860122 | 0.999176 |
| ENSMUSG00000052446 | Zfp961 | protein_coding | -0.08212909 | 4.061899 | 0.0211669 | 0.8861083 | 0.999176 |
| ENSMUSG00000044795 | Cyb5d1 | protein_coding | 0.1418801 | 3.322606 | 0.0211024 | 0.8862836 | 0.999176 |
| ENSMUSG00000020074 | Ccar1 | protein_coding | -0.06122944 | 5.314642 | 0.0209462 | 0.8866991 | 0.999176 |
| ENSMUSG00000070462 | Tlnrd1 | protein_coding | 0.09241764 | 4.426053 | 0.0208641 | 0.8869199 | 0.999176 |
| ENSMUSG00000040464 | Gtpbp10 | protein_coding | -0.1391556 | 2.572768 | 0.0208376 | 0.8869912 | 0.999176 |
| ENSMUSG00000020994 | Pnn | protein_coding | -0.06037505 | 5.97914 | 0.0208081 | 0.8870707 | 0.999176 |
| ENSMUSG00000030407 | Qpctl | protein_coding | -0.06725738 | 4.971527 | 0.0207854 | 0.8871317 | 0.999176 |
| ENSMUSG00000037286 | Stag1 | protein_coding | -0.06473402 | 5.854712 | 0.0207672 | 0.8871808 | 0.999176 |
| ENSMUSG00000020160 | Meis1 | protein_coding | 0.158968 | 3.539786 | 0.0207737 | 0.8873396 | 0.999176 |
| ENSMUSG00000055760 | Gemin6 | protein_coding | 0.09134604 | 4.423543 | 0.0207079 | 0.8873408 | 0.999176 |
| ENSMUSG00000036885 | Arhgef26 | protein_coding | 0.1127846 | 3.349895 | 0.020671 | 0.8874406 | 0.999176 |
| ENSMUSG00000030816 | Rnf40 | protein_coding | -0.06723829 | 4.378119 | 0.0206465 | 0.8875069 | 0.999176 |
| ENSMUSG00000006599 | Gtf2h1 | protein_coding | -0.07824433 | 4.521942 | 0.0206144 | 0.8875936 | 0.999176 |
| ENSMUSG00000105891 | A230001M10Rik | lincRNA | 0.09919663 | 3.815658 | 0.0205715 | 0.8877098 | 0.999176 |
| ENSMUSG00000038658 | Ric1 | protein_coding | 0.09807674 | 4.409328 | 0.0205396 | 0.8877991 | 0.999176 |
| ENSMUSG00000028062 | Lamtor2 | protein_coding | 0.05027538 | 9.130326 | 0.0205112 | 0.8878732 | 0.999176 |
| ENSMUSG00000020485 | Supt4a | protein_coding | -0.04365449 | 6.644403 | 0.0204585 | 0.8880165 | 0.999176 |
| ENSMUSG00000031932 | Gpr83 | protein_coding | 0.07149371 | 5.762852 | 0.0204028 | 0.8881708 | 0.999176 |
| ENSMUSG00000029878 | Dbpht2 | lincRNA | 0.08236898 | 8.515574 | 0.0204828 | 0.8882142 | 0.999176 |
| ENSMUSG00000020541 | Tom1l1 | protein_coding | -0.05956941 | 5.201903 | 0.0203431 | 0.8883305 | 0.999176 |
| ENSMUSG00000000605 | Clcn4 | protein_coding | -0.04360414 | 7.351839 | 0.0202625 | 0.8885504 | 0.999176 |
| ENSMUSG00000056941 | Commd7 | protein_coding | -0.0650615 | 6.256937 | 0.020241 | 0.888612 | 0.999176 |
| ENSMUSG00000032724 | Abtb2 | protein_coding | 0.1699801 | 2.47082 | 0.020221 | 0.8886635 | 0.999176 |
| ENSMUSG00000041852 | Tcf20 | protein_coding | 0.06824311 | 5.79242 | 0.0201696 | 0.8888044 | 0.999176 |
| ENSMUSG00000051451 | Crebzf | protein_coding | -0.1436208 | 2.589739 | 0.0201356 | 0.8888972 | 0.999176 |
| ENSMUSG00000042616 | Oscp1 | protein_coding | 0.05207533 | 6.296014 | 0.0201107 | 0.8889656 | 0.999176 |
| ENSMUSG00000050737 | Ptges | protein_coding | -0.1328768 | 2.854793 | 0.0201106 | 0.8889659 | 0.999176 |
| ENSMUSG00000028030 | Tbck | protein_coding | -0.08083384 | 6.406016 | 0.0201692 | 0.888979 | 0.999176 |
| ENSMUSG00000048799 | Cep120 | protein_coding | -0.06779875 | 4.95278 | 0.0200916 | 0.889018 | 0.999176 |
| ENSMUSG00000049134 | Nrap | protein_coding | -0.3542347 | 1.73894 | 0.0201317 | 0.8890259 | 0.999176 |
| ENSMUSG00000037896 | Rcor1 | protein_coding | 0.09136719 | 4.042405 | 0.0200602 | 0.8891041 | 0.999176 |
| ENSMUSG00000070687 | Htr1d | protein_coding | -0.2153178 | 2.709893 | 0.0200785 | 0.8891645 | 0.999176 |
| ENSMUSG00000024997 | Prdx3 | protein_coding | -0.0390244 | 8.346245 | 0.0200301 | 0.8891867 | 0.999176 |
| ENSMUSG00000019828 | Grm1 | protein_coding | -0.05778161 | 5.464319 | 0.0200123 | 0.8892357 | 0.999176 |
| ENSMUSG00000028933 | Xrcc2 | protein_coding | 0.09586916 | 3.257362 | 0.0199799 | 0.8893248 | 0.999176 |
| ENSMUSG00000040482 | Dxo | protein_coding | 0.0771965 | 4.449984 | 0.0199746 | 0.8893392 | 0.999176 |
| ENSMUSG00000085067 | Gm15631 | lincRNA | -0.07985472 | 5.229904 | 0.0199739 | 0.8893441 | 0.999176 |
| ENSMUSG00000024190 | Dusp1 | protein_coding | 0.07772305 | 4.473586 | 0.0199066 | 0.8895264 | 0.999176 |
| ENSMUSG00000018398 | 8-Sep | protein_coding | -0.0402688 | 6.969691 | 0.0198899 | 0.8895724 | 0.999176 |
| ENSMUSG00000020776 | Fbf1 | protein_coding | -0.08878889 | 3.362885 | 0.0198882 | 0.889577 | 0.999176 |
| ENSMUSG00000029034 | Ints11 | protein_coding | -0.1204839 | 3.329846 | 0.0198667 | 0.8896365 | 0.999176 |
| ENSMUSG00000052102 | Gnpda1 | protein_coding | -0.0691828 | 5.444722 | 0.0198597 | 0.8896557 | 0.999176 |
| ENSMUSG00000027244 | Atg13 | protein_coding | 0.05754424 | 6.631336 | 0.0198338 | 0.8897273 | 0.999176 |
| ENSMUSG00000054855 | Rnd1 | protein_coding | -0.07236103 | 5.994944 | 0.0198336 | 0.889738 | 0.999176 |
| ENSMUSG00000048814 | Lonrf2 | protein_coding | -0.0643387 | 6.85701 | 0.0197863 | 0.8899092 | 0.999176 |
| ENSMUSG00000036198 | Arhgap36 | protein_coding | -0.06494486 | 5.900012 | 0.0197199 | 0.890042 | 0.999176 |
| ENSMUSG00000020697 | Lig3 | protein_coding | 0.1373684 | 2.791082 | 0.0196575 | 0.8902149 | 0.999176 |
| ENSMUSG00000020009 | Ifngr1 | protein_coding | -0.08030645 | 4.016043 | 0.0196566 | 0.8902175 | 0.999176 |
| ENSMUSG00000040850 | Psme4 | protein_coding | 0.07177599 | 4.122706 | 0.0196461 | 0.8902467 | 0.999176 |
| ENSMUSG00000032482 | Cspg5 | protein_coding | 0.05411037 | 7.199515 | 0.019642 | 0.890258 | 0.999176 |
| ENSMUSG00000003033 | Ap1m1 | protein_coding | -0.05947623 | 7.141287 | 0.019628 | 0.8902997 | 0.999176 |
| ENSMUSG00000031787 | Katnb1 | protein_coding | 0.07273788 | 6.12597 | 0.0196087 | 0.8904038 | 0.999176 |
| ENSMUSG00000013736 | Trnt1 | protein_coding | 0.0627064 | 6.412083 | 0.0195556 | 0.8905008 | 0.999176 |
| ENSMUSG00000020362 | Cnot6 | protein_coding | -0.06729267 | 5.369454 | 0.0195306 | 0.8905674 | 0.999176 |
| ENSMUSG00000024317 | Rnf138 | protein_coding | -0.07015686 | 4.51356 | 0.0195254 | 0.8905821 | 0.999176 |
| ENSMUSG00000038069 | Cdkn2aip | protein_coding | -0.1034346 | 3.533201 | 0.0194433 | 0.8908106 | 0.999176 |
| ENSMUSG00000050945 | Zfp438 | protein_coding | -0.1437923 | 3.362616 | 0.0194347 | 0.8908448 | 0.999176 |
| ENSMUSG00000041757 | Plekha6 | protein_coding | 0.07130642 | 4.590887 | 0.0193741 | 0.8910037 | 0.999176 |
| ENSMUSG00000074024 | 4632427E13Rik | bidirectional_promote | -0.2411526 | 2.528839 | 0.0194146 | 0.8910305 | 0.999176 |
| ENSMUSG00000036957 | Lrfn3 | protein_coding | -0.1730557 | 2.342208 | 0.0193595 | 0.8910446 | 0.999176 |
| ENSMUSG00000036270 | Edc4 | protein_coding | -0.07500064 | 4.747503 | 0.0192453 | 0.8913644 | 0.999176 |
| ENSMUSG00000038252 | Ncapd2 | protein_coding | -0.192713 | 2.557168 | 0.0192227 | 0.8914304 | 0.999176 |
| ENSMUSG00000033916 | Chmp2a | protein_coding | 0.03828546 | 9.030814 | 0.0192094 | 0.8914649 | 0.999176 |
| ENSMUSG00000020720 | Psmd12 | protein_coding | 0.04185081 | 8.112983 | 0.0192059 | 0.8914747 | 0.999176 |

| ensembl_gene_id | external_gene_name | gene_biotype | log2 (fold change) | log2 (counts per million) | F-value | p-value | False discovery rate |
|---|---|---|---|---|---|---|---|
| ENSMUSG00000024277 | Mapre2 | protein_coding | 0.05146444 | 8.384898 | 0.0191853 | 0.8915326 | 0.999176 |
| ENSMUSG00000034951 | Cog7 | protein_coding | 0.06559417 | 5.961421 | 0.0191438 | 0.891652 | 0.999176 |
| ENSMUSG00000001056 | Nhp2 | protein_coding | 0.04728313 | 7.596108 | 0.0189861 | 0.8920933 | 0.999176 |
| ENSMUSG00000035762 | Tmem161b | protein_coding | -0.06602221 | 4.604567 | 0.0189736 | 0.8921286 | 0.999176 |
| ENSMUSG00000022788 | Fgd4 | protein_coding | 0.07578701 | 3.779366 | 0.0189606 | 0.8921654 | 0.999176 |
| ENSMUSG00000028423 | Nfx1 | protein_coding | 0.05519901 | 6.440802 | 0.0189254 | 0.8922649 | 0.999176 |
| ENSMUSG00000029238 | Clock | protein_coding | -0.06677713 | 5.392086 | 0.018921 | 0.8922772 | 0.999176 |
| ENSMUSG00000029538 | Srsf9 | protein_coding | -0.0669408 | 5.304815 | 0.0188765 | 0.8924031 | 0.999176 |
| ENSMUSG00000023272 | Creld2 | protein_coding | -0.1413755 | 3.296779 | 0.0188568 | 0.8924689 | 0.999176 |
| ENSMUSG00000006494 | Pdk1 | protein_coding | 0.09280592 | 4.046963 | 0.018837 | 0.8925151 | 0.999176 |
| ENSMUSG00000053134 | Supt7l | protein_coding | 0.06673432 | 4.772758 | 0.0187695 | 0.8927066 | 0.999176 |
| ENSMUSG00000040865 | Ino80d | protein_coding | -0.08364453 | 3.5589 | 0.0187646 | 0.8927206 | 0.999176 |
| ENSMUSG00000034621 | Gpatch8 | protein_coding | -0.05818119 | 5.727575 | 0.0187609 | 0.892731 | 0.999176 |
| ENSMUSG00000003234 | Abcf3 | protein_coding | -0.05855825 | 7.064598 | 0.0187259 | 0.8928332 | 0.999176 |
| ENSMUSG00000026349 | Ccnt2 | protein_coding | -0.06964031 | 4.920342 | 0.0187107 | 0.8928737 | 0.999176 |
| ENSMUSG00000038042 | Ptpdc1 | protein_coding | -0.085283 | 4.040816 | 0.0186948 | 0.8929189 | 0.999176 |
| ENSMUSG00000042684 | Npl | protein_coding | -0.1249116 | 3.115437 | 0.0186561 | 0.8930292 | 0.999176 |
| ENSMUSG00000032582 | Rbm6 | protein_coding | 0.07020802 | 4.699736 | 0.0186503 | 0.8930456 | 0.999176 |
| ENSMUSG00000048756 | Foxo3 | protein_coding | -0.06829786 | 5.828438 | 0.0186049 | 0.8931776 | 0.999176 |
| ENSMUSG00000029761 | Cald1 | protein_coding | -0.1067419 | 4.789122 | 0.0186344 | 0.8932279 | 0.999176 |
| ENSMUSG00000054863 | Fam19a5 | protein_coding | 0.04933278 | 6.303145 | 0.0185795 | 0.8932474 | 0.999176 |
| ENSMUSG00000030101 | Sumf1 | protein_coding | -0.1032233 | 4.039993 | 0.0185573 | 0.8933136 | 0.999176 |
| ENSMUSG00000049516 | Spty2d1 | protein_coding | 0.0813924 | 3.873396 | 0.0184427 | 0.8936387 | 0.999176 |
| ENSMUSG00000007659 | Bcl2l1 | protein_coding | -0.04236286 | 8.134873 | 0.0183687 | 0.8938507 | 0.999176 |
| ENSMUSG00000056014 | A430033K04Rik | protein_coding | 0.07935396 | 4.071843 | 0.0183136 | 0.8940092 | 0.999176 |
| ENSMUSG00000033793 | Atp6v1h | protein_coding | 0.04753969 | 8.77314 | 0.0182081 | 0.8943127 | 0.999176 |
| ENSMUSG00000028331 | Trmo | protein_coding | -0.1835874 | 3.101889 | 0.0182577 | 0.8943352 | 0.999176 |
| ENSMUSG00000051246 | Msantd1 | protein_coding | 0.114079 | 3.377605 | 0.0181772 | 0.8944019 | 0.999176 |
| ENSMUSG00000027710 | Acad9 | protein_coding | -0.1429191 | 3.779285 | 0.0182397 | 0.8944097 | 0.999176 |
| ENSMUSG00000055963 | Triqk | protein_coding | 0.04620534 | 5.89038 | 0.0181661 | 0.894434 | 0.999176 |
| ENSMUSG00000031374 | Zfp92 | protein_coding | -0.1278122 | 2.641579 | 0.0181533 | 0.8944711 | 0.999176 |
| ENSMUSG00000048960 | Prex2 | protein_coding | -0.1729368 | 3.109216 | 0.0181884 | 0.8944851 | 0.999176 |
| ENSMUSG00000048458 | Fam212b | protein_coding | -0.0768618 | 5.08737 | 0.0181233 | 0.8945578 | 0.999176 |
| ENSMUSG00000039307 | Hexdc | protein_coding | 0.08487768 | 3.888877 | 0.0181196 | 0.8945683 | 0.999176 |
| ENSMUSG00000038127 | Ccdc50 | protein_coding | -0.04972525 | 5.518561 | 0.0181027 | 0.8946171 | 0.999176 |
| ENSMUSG00000048503 | Tmem136 | protein_coding | -0.08200872 | 3.529242 | 0.0180615 | 0.8947364 | 0.999176 |
| ENSMUSG00000026209 | Dnpep | protein_coding | 0.04740885 | 6.566438 | 0.0180483 | 0.8947746 | 0.999176 |
| ENSMUSG00000029610 | Aimp2 | protein_coding | -0.04298659 | 6.109982 | 0.018021 | 0.8948539 | 0.999176 |
| ENSMUSG00000028830 | AU040320 | protein_coding | -0.05691629 | 5.325685 | 0.0180197 | 0.8948575 | 0.999176 |
| ENSMUSG00000031959 | Wdr59 | protein_coding | 0.08247268 | 3.814673 | 0.0179771 | 0.8949812 | 0.999176 |
| ENSMUSG00000040270 | Bach2 | protein_coding | -0.1036833 | 3.307613 | 0.0179604 | 0.8950295 | 0.999176 |
| ENSMUSG00000095845 | Gm5741 | protein_coding | -0.5983509 | 2.704893 | 0.0184306 | 0.8954138 | 0.999176 |
| ENSMUSG00000018509 | Cenpv | protein_coding | 0.07175516 | 4.041841 | 0.0178223 | 0.8954314 | 0.999176 |
| ENSMUSG00000042063 | Zfp386 | protein_coding | 0.06529683 | 5.182604 | 0.0177724 | 0.8955771 | 0.999176 |
| ENSMUSG00000015759 | Cnih1 | protein_coding | -0.04094829 | 7.541103 | 0.017764 | 0.8956017 | 0.999176 |
| ENSMUSG00000097156 | Gm3764 | lincRNA | -0.06437312 | 5.037532 | 0.0177194 | 0.8957317 | 0.999176 |
| ENSMUSG00000073468 | Sft2d1 | protein_coding | 0.06753815 | 4.717821 | 0.0176902 | 0.8958172 | 0.999176 |
| ENSMUSG00000014039 | Prdm15 | protein_coding | -0.1554554 | 2.373502 | 0.0176678 | 0.8958828 | 0.999176 |
| ENSMUSG00000003458 | Ncstn | protein_coding | -0.06042825 | 5.951255 | 0.0176661 | 0.8958877 | 0.999176 |
| ENSMUSG00000022860 | Chodl | protein_coding | 0.1165264 | 3.315417 | 0.0175928 | 0.8961025 | 0.999176 |
| ENSMUSG00000025060 | Slk | protein_coding | 0.08634896 | 4.682045 | 0.0175838 | 0.8961318 | 0.999176 |
| ENSMUSG00000050428 | Fbxo46 | protein_coding | -0.06150483 | 4.897243 | 0.0175751 | 0.8961546 | 0.999176 |
| ENSMUSG00000020196 | Cabin1 | protein_coding | -0.07771592 | 5.004643 | 0.0175636 | 0.896191 | 0.999176 |
| ENSMUSG00000009647 | Mcu | protein_coding | 0.04775611 | 6.260622 | 0.0175589 | 0.8962023 | 0.999176 |
| ENSMUSG00000031402 | Mpp1 | protein_coding | -0.05483711 | 6.297692 | 0.0175504 | 0.896227 | 0.999176 |
| ENSMUSG00000068747 | Sort1 | protein_coding | 0.06264008 | 5.841446 | 0.0175289 | 0.8962902 | 0.999176 |
| ENSMUSG00000031843 | Mphosph6 | protein_coding | 0.05250221 | 6.02101 | 0.0175069 | 0.896355 | 0.999176 |
| ENSMUSG00000022247 | Brix1 | protein_coding | 0.04129382 | 6.667334 | 0.017471 | 0.8964608 | 0.999176 |
| ENSMUSG00000037001 | Zfp39 | protein_coding | -0.08460427 | 3.344581 | 0.0174607 | 0.896491 | 0.999176 |
| ENSMUSG00000059669 | Taf1b | protein_coding | -0.06996844 | 4.718519 | 0.0174563 | 0.8965041 | 0.999176 |
| ENSMUSG00000032359 | Ctsh | protein_coding | 0.1007279 | 3.698744 | 0.0174149 | 0.8966259 | 0.999176 |
| ENSMUSG00000040570 | Rundc3b | protein_coding | 0.06458564 | 5.015461 | 0.0173915 | 0.8966952 | 0.999176 |
| ENSMUSG00000010095 | Slc3a2 | protein_coding | -0.06715501 | 5.734787 | 0.0173783 | 0.8967367 | 0.999176 |
| ENSMUSG00000024370 | Cdc23 | protein_coding | -0.07573415 | 5.487269 | 0.0173576 | 0.896798 | 0.999176 |
| ENSMUSG00000022724 | Riox2 | protein_coding | -0.1023867 | 3.831923 | 0.0173455 | 0.8968335 | 0.999176 |
| ENSMUSG00000039963 | Ccdc40 | protein_coding | -0.09843399 | 3.778706 | 0.0173178 | 0.896913 | 0.999176 |
| ENSMUSG00000032637 | Atxn2l | protein_coding | 0.09975109 | 3.237501 | 0.0173078 | 0.8969423 | 0.999176 |
| ENSMUSG00000029405 | G3bp2 | protein_coding | 0.05479018 | 7.275277 | 0.0172484 | 0.897121 | 0.999176 |
| ENSMUSG00000032412 | Atp1b3 | protein_coding | 0.05727801 | 6.144788 | 0.0172421 | 0.8971372 | 0.999176 |
| ENSMUSG00000036835 | Psenen | protein_coding | 0.05549219 | 4.977597 | 0.0172391 | 0.8971458 | 0.999176 |
| ENSMUSG00000001280 | Sp1 | protein_coding | -0.08235767 | 3.948195 | 0.017207 | 0.8972411 | 0.999176 |
| ENSMUSG00000029581 | Fscn1 | protein_coding | 0.04964926 | 5.563898 | 0.0171915 | 0.8972871 | 0.999176 |
| ENSMUSG00000064339 | mt-Rnr2 | Mt_rRNA | -0.08604671 | 13.836 | 0.0172425 | 0.8973442 | 0.999176 |
| ENSMUSG00000017631 | Abr | protein_coding | 0.04254529 | 7.151394 | 0.0171485 | 0.8974148 | 0.999176 |
| ENSMUSG00000037949 | Ano10 | protein_coding | 0.08332065 | 4.049037 | 0.0171431 | 0.8974311 | 0.999176 |
| ENSMUSG00000023147 | Wrb | protein_coding | 0.0373648 | 8.827525 | 0.0171209 | 0.897497 | 0.999176 |
| ENSMUSG00000042834 | Nrep | protein_coding | 0.04874338 | 6.433059 | 0.0171005 | 0.8975577 | 0.999176 |
| ENSMUSG00000068114 | Ccdc134 | protein_coding | 0.1096398 | 3.203838 | 0.0170966 | 0.8975695 | 0.999176 |

| ensembl_gene_id | external_gene_name | gene_biotype | log2 (fold change) | log2 (counts per million) | F-value | p-value | False discovery rate |
|---|---|---|---|---|---|---|---|
| ENSMUSG00000061650 | Med9 | protein_coding | 0.07778726 | 4.350181 | 0.0170849 | 0.8976044 | 0.999176 |
| ENSMUSG00000026514 | Cnih3 | protein_coding | -0.06742832 | 4.499759 | 0.0170801 | 0.8976187 | 0.999176 |
| ENSMUSG00000025512 | Chid1 | protein_coding | -0.04426274 | 6.302928 | 0.0170663 | 0.8976597 | 0.999176 |
| ENSMUSG00000047714 | Ppp1r2 | protein_coding | 0.05990176 | 5.611116 | 0.0170478 | 0.8977147 | 0.999176 |
| ENSMUSG00000019809 | Pex3 | protein_coding | 0.06696548 | 5.017499 | 0.0170161 | 0.8978093 | 0.999176 |
| ENSMUSG00000024799 | Tm7sf2 | protein_coding | -0.08682434 | 4.114726 | 0.0169906 | 0.8978854 | 0.999176 |
| ENSMUSG00000007617 | Homer1 | protein_coding | -0.05803645 | 5.92905 | 0.0169661 | 0.8979587 | 0.999176 |
| ENSMUSG00000010721 | Lmbr1 | protein_coding | -0.05715345 | 4.769063 | 0.0169644 | 0.8979637 | 0.999176 |
| ENSMUSG00000038256 | Bcl9 | protein_coding | 0.08173441 | 4.243007 | 0.016892 | 0.8981804 | 0.999176 |
| ENSMUSG00000013707 | Tnfaip8l2 | protein_coding | -0.1562498 | 3.013897 | 0.0168903 | 0.8982016 | 0.999176 |
| ENSMUSG00000044224 | Dnajc21 | protein_coding | -0.06736622 | 4.34374 | 0.0168718 | 0.8982411 | 0.999176 |
| ENSMUSG00000025224 | Gbf1 | protein_coding | -0.05833734 | 5.734217 | 0.016846 | 0.8983185 | 0.999176 |
| ENSMUSG00000036155 | Mgat5 | protein_coding | -0.0880336 | 3.778047 | 0.016838 | 0.8983424 | 0.999176 |
| ENSMUSG00000109776 | Gm30400 | processed_transcript | -0.1530177 | 2.474284 | 0.0165995 | 0.8990607 | 0.999176 |
| ENSMUSG00000061313 | Ddhd2 | protein_coding | -0.07694531 | 7.095675 | 0.0166426 | 0.8991821 | 0.999176 |
| ENSMUSG00000005610 | Eif4g2 | protein_coding | -0.04874139 | 8.797452 | 0.0165511 | 0.8992071 | 0.999176 |
| ENSMUSG00000034574 | Daam1 | protein_coding | -0.06826233 | 4.869052 | 0.0164858 | 0.899405 | 0.999176 |
| ENSMUSG00000042523 | Dnal1 | protein_coding | -0.05851518 | 5.155331 | 0.0164854 | 0.8994063 | 0.999176 |
| ENSMUSG00000026566 | Mpzl1 | protein_coding | -0.1111938 | 3.369693 | 0.0164609 | 0.8994806 | 0.999176 |
| ENSMUSG00000016833 | Mrps18c | protein_coding | -0.05510811 | 7.215217 | 0.0164496 | 0.8995242 | 0.999176 |
| ENSMUSG00000022759 | Lrrc74b | protein_coding | -0.09959347 | 3.165219 | 0.0164372 | 0.8995525 | 0.999176 |
| ENSMUSG00000038422 | Hdhd3 | protein_coding | 0.06137772 | 5.091884 | 0.0163835 | 0.8997156 | 0.999176 |
| ENSMUSG00000033773 | Rpap2 | protein_coding | -0.08543405 | 4.518198 | 0.0163407 | 0.8998487 | 0.999176 |
| ENSMUSG00000004565 | Pnpla6 | protein_coding | -0.0633672 | 4.932264 | 0.0161831 | 0.9003275 | 0.999176 |
| ENSMUSG00000020023 | Tmcc3 | protein_coding | 0.1791649 | 2.391542 | 0.0161267 | 0.9005003 | 0.999176 |
| ENSMUSG00000022864 | D16Ertd472e | protein_coding | -0.1583035 | 2.559572 | 0.0161229 | 0.9005121 | 0.999176 |
| ENSMUSG00000024260 | Sap130 | protein_coding | 0.08629509 | 3.683428 | 0.0160979 | 0.9005886 | 0.999176 |
| ENSMUSG00000005417 | Mprip | protein_coding | -0.05599283 | 5.350902 | 0.0160819 | 0.9006377 | 0.999176 |
| ENSMUSG00000063235 | Ptpmt1 | processed_transcript | 0.07219157 | 3.817351 | 0.0160669 | 0.900684 | 0.999176 |
| ENSMUSG00000026427 | Eif2d | protein_coding | 0.05540527 | 5.48917 | 0.0159932 | 0.9009106 | 0.999176 |
| ENSMUSG00000028789 | Azin2 | protein_coding | -0.06379794 | 4.471869 | 0.0159578 | 0.9010198 | 0.999176 |
| ENSMUSG00000045284 | Dcaf12l1 | protein_coding | 0.04513651 | 6.420948 | 0.0159568 | 0.9010227 | 0.999176 |
| ENSMUSG00000072875 | Gpr27 | protein_coding | -0.07872424 | 4.238632 | 0.0159098 | 0.9011679 | 0.999176 |
| ENSMUSG00000064202 | 4430402I18Rik | protein_coding | 0.1115482 | 3.057453 | 0.0158871 | 0.9012381 | 0.999176 |
| ENSMUSG00000027810 | Eif2a | protein_coding | 0.03784066 | 6.400074 | 0.0158594 | 0.9013237 | 0.999176 |
| ENSMUSG00000030772 | Dkk3 | protein_coding | 0.06783049 | 5.609323 | 0.0158122 | 0.9014725 | 0.999176 |
| ENSMUSG00000020079 | Supv3l1 | protein_coding | 0.1262995 | 3.067614 | 0.0157778 | 0.9015763 | 0.999176 |
| ENSMUSG00000015536 | Mocs2 | protein_coding | -0.03604414 | 7.941275 | 0.0157543 | 0.9016494 | 0.999176 |
| ENSMUSG00000026988 | Wdsub1 | protein_coding | 0.08191552 | 3.950591 | 0.0157286 | 0.9017291 | 0.999176 |
| ENSMUSG00000022477 | Aco2 | protein_coding | 0.04287118 | 8.493534 | 0.0157056 | 0.9018005 | 0.999176 |
| ENSMUSG00000070372 | Capza1 | protein_coding | -0.07605218 | 3.747599 | 0.015683 | 0.9018709 | 0.999176 |
| ENSMUSG00000014602 | Kif1a | protein_coding | 0.04002658 | 8.85731 | 0.0156626 | 0.9019344 | 0.999176 |
| ENSMUSG00000040387 | Klhl32 | protein_coding | -0.06476471 | 4.658008 | 0.0156299 | 0.9020362 | 0.999176 |
| ENSMUSG00000037536 | Fbxo34 | protein_coding | -0.07559987 | 3.509492 | 0.0155707 | 0.9022208 | 0.999176 |
| ENSMUSG00000071398 | 2410004P03Rik | protein_coding | -0.09582477 | 3.660535 | 0.0155445 | 0.9023028 | 0.999176 |
| ENSMUSG00000051483 | Cbr1 | protein_coding | -0.04479902 | 7.257001 | 0.0155413 | 0.9023129 | 0.999176 |
| ENSMUSG00000038502 | Ptov1 | protein_coding | -0.04906761 | 6.606393 | 0.0155335 | 0.9023372 | 0.999176 |
| ENSMUSG00000041028 | Ghitm | protein_coding | -0.039332 | 9.508313 | 0.0154168 | 0.9027027 | 0.999176 |
| ENSMUSG00000038013 | Wipf2 | protein_coding | -0.04103729 | 6.46317 | 0.015376 | 0.9028308 | 0.999176 |
| ENSMUSG00000013662 | Atad1 | protein_coding | -0.043798 | 6.835618 | 0.0153277 | 0.9029829 | 0.999176 |
| ENSMUSG00000026018 | Ica1l | protein_coding | 0.0592171 | 4.956794 | 0.0153209 | 0.9030043 | 0.999176 |
| ENSMUSG00000031355 | Arhgap6 | protein_coding | 0.06271283 | 4.359166 | 0.0153205 | 0.9030055 | 0.999176 |
| ENSMUSG00000036371 | Serbp1 | protein_coding | -0.03620146 | 7.425136 | 0.0153142 | 0.9030251 | 0.999176 |
| ENSMUSG00000016257 | Prelid3b | protein_coding | -0.03799814 | 7.019083 | 0.015242 | 0.9032528 | 0.999176 |
| ENSMUSG00000047446 | Arl4a | protein_coding | -0.06453765 | 6.549602 | 0.0152465 | 0.9033348 | 0.999176 |
| ENSMUSG00000031840 | Rab3a | protein_coding | -0.06277287 | 9.649739 | 0.0152476 | 0.9033373 | 0.999176 |
| ENSMUSG00000002210 | Smg9 | protein_coding | -0.0673538 | 4.213804 | 0.0152019 | 0.9033796 | 0.999176 |
| ENSMUSG00000005510 | Ndufs3 | protein_coding | 0.03352915 | 7.416448 | 0.0151786 | 0.9034532 | 0.999176 |
| ENSMUSG00000039253 | Fn3krp | protein_coding | -0.1401784 | 3.490452 | 0.0152176 | 0.9034804 | 0.999176 |
| ENSMUSG00000001158 | Snrnp27 | protein_coding | 0.04501301 | 8.0721 | 0.0151647 | 0.9034972 | 0.999176 |
| ENSMUSG00000105843 | Gm19439 | TEC | -0.1476881 | 4.624204 | 0.0153588 | 0.9035008 | 0.999176 |
| ENSMUSG00000052544 | St6galnac3 | protein_coding | -0.0786171 | 3.529796 | 0.0151626 | 0.9035038 | 0.999176 |
| ENSMUSG00000032058 | Ppp2r1b | protein_coding | -0.0646429 | 4.656753 | 0.0151565 | 0.9035231 | 0.999176 |
| ENSMUSG00000038957 | Edc3 | protein_coding | -0.06003487 | 5.546286 | 0.0151533 | 0.9035334 | 0.999176 |
| ENSMUSG00000048163 | Selplg | protein_coding | -0.08305712 | 4.155606 | 0.0151368 | 0.9035857 | 0.999176 |
| ENSMUSG00000026305 | Lrrfip1 | protein_coding | 0.06591196 | 4.908103 | 0.0151229 | 0.9036296 | 0.999176 |
| ENSMUSG00000008348 | Ubc | protein_coding | 0.04584403 | 8.069522 | 0.0151165 | 0.90365 | 0.999176 |
| ENSMUSG00000062296 | Trank1 | protein_coding | 0.07701168 | 4.826616 | 0.0150953 | 0.9037196 | 0.999176 |
| ENSMUSG00000054162 | Spock3 | protein_coding | 0.03727343 | 7.329592 | 0.0150168 | 0.9039666 | 0.999176 |
| ENSMUSG00000021385 | Ippk | protein_coding | 0.07528455 | 4.175137 | 0.0149983 | 0.9040253 | 0.999176 |
| ENSMUSG00000021097 | Clmn | protein_coding | 0.05834121 | 5.483942 | 0.0149981 | 0.9040259 | 0.999176 |
| ENSMUSG00000016200 | Syt14 | protein_coding | 0.09188268 | 3.445937 | 0.0149713 | 0.9041112 | 0.999176 |
| ENSMUSG00000084885 | 3010001F23Rik | processed_transcript | 0.103495 | 2.823728 | 0.0149243 | 0.904261 | 0.999176 |
| ENSMUSG00000031590 | Frg1 | protein_coding | -0.03805257 | 7.061393 | 0.0149194 | 0.9042768 | 0.999176 |
| ENSMUSG00000003721 | Insig2 | protein_coding | -0.03614803 | 6.989022 | 0.0148876 | 0.9043784 | 0.999176 |
| ENSMUSG00000016528 | Mapkapk2 | protein_coding | -0.1378628 | 2.80941 | 0.0148393 | 0.9045326 | 0.999176 |
| ENSMUSG00000035158 | Mitf | protein_coding | 0.1386708 | 2.72306 | 0.0148029 | 0.9046493 | 0.999176 |
| ENSMUSG00000062373 | Tmem65 | protein_coding | 0.04958483 | 5.907265 | 0.0147964 | 0.9046701 | 0.999176 |

| ensembl_gene_id | external_gene_name | gene_biotype | log2 (fold change) | log2 (counts per million) | F-value | p-value | False discovery rate |
|---|---|---|---|---|---|---|---|
| ENSMUSG00000007039 | Ddah2 | protein_coding | -0.05740091 | 4.694999 | 0.0147906 | 0.9046887 | 0.999176 |
| ENSMUSG00000054676 | 1600014C10Rik | protein_coding | -0.06281227 | 4.517362 | 0.0146653 | 0.9050911 | 0.999176 |
| ENSMUSG00000031697 | Orc6 | protein_coding | -0.0516605 | 6.865655 | 0.0146627 | 0.9051019 | 0.999176 |
| ENSMUSG00000031295 | Phka2 | protein_coding | -0.113888 | 3.009511 | 0.014651 | 0.9051373 | 0.999176 |
| ENSMUSG00000056124 | B4galt6 | protein_coding | -0.04018363 | 6.352547 | 0.0146485 | 0.9051452 | 0.999176 |
| ENSMUSG00000020205 | Phlda1 | protein_coding | -0.07343973 | 4.042855 | 0.0145234 | 0.905549 | 0.999176 |
| ENSMUSG00000028693 | Nasp | protein_coding | -0.06631185 | 4.348576 | 0.0145179 | 0.9055669 | 0.999176 |
| ENSMUSG00000085882 | 2610507I01Rik | processed_transcript | 0.093785 | 3.233353 | 0.0144805 | 0.905688 | 0.999176 |
| ENSMUSG00000035049 | Rrp12 | protein_coding | -0.06366984 | 4.31385 | 0.0144748 | 0.9057066 | 0.999176 |
| ENSMUSG00000079056 | Kcnip3 | protein_coding | 0.03773969 | 6.305152 | 0.0144734 | 0.905711 | 0.999176 |
| ENSMUSG00000015668 | Pdzd11 | protein_coding | -0.03430923 | 7.534399 | 0.0144591 | 0.9057572 | 0.999176 |
| ENSMUSG00000066621 | Tecpr1 | protein_coding | -0.05800187 | 5.740775 | 0.0144544 | 0.9057727 | 0.999176 |
| ENSMUSG00000034818 | Celf5 | protein_coding | -0.04084274 | 6.515982 | 0.0144476 | 0.9057947 | 0.999176 |
| ENSMUSG00000001105 | Ift20 | protein_coding | 0.03339352 | 7.772655 | 0.0144441 | 0.9058059 | 0.999176 |
| ENSMUSG00000031786 | Drc7 | protein_coding | -0.1430229 | 3.185636 | 0.0144049 | 0.906021 | 0.999176 |
| ENSMUSG00000021268 | Meg3 | lincRNA | -0.08936949 | 7.508684 | 0.0144862 | 0.9062891 | 0.999176 |
| ENSMUSG00000094707 | A830019P07Rik | lincRNA | -0.1479615 | 2.449637 | 0.0142872 | 0.9063163 | 0.999176 |
| ENSMUSG00000033126 | Ybey | protein_coding | -0.07460677 | 4.195007 | 0.0142634 | 0.906394 | 0.999176 |
| ENSMUSG00000022096 | Hr | protein_coding | 0.1139349 | 2.578029 | 0.0140882 | 0.9069677 | 0.999176 |
| ENSMUSG00000040383 | Aqr | protein_coding | 0.0651631 | 4.519893 | 0.0140458 | 0.9071071 | 0.999176 |
| ENSMUSG00000021400 | Wrnip1 | protein_coding | -0.05301371 | 5.408688 | 0.013967 | 0.9073668 | 0.999176 |
| ENSMUSG00000074282 | Zfp94 | protein_coding | 0.05628866 | 4.398317 | 0.0139524 | 0.907415 | 0.999176 |
| ENSMUSG00000108608 | Gm6916 | transcribed_processe | 0.3577988 | 2.57295 | 0.0142017 | 0.9074512 | 0.999176 |
| ENSMUSG00000037722 | Gnpnat1 | protein_coding | 0.05937056 | 4.307557 | 0.0139394 | 0.907458 | 0.999176 |
| ENSMUSG00000029169 | Dhx15 | protein_coding | -0.05068842 | 6.032587 | 0.0138154 | 0.9078686 | 0.999176 |
| ENSMUSG00000026245 | Farsb | protein_coding | -0.03986049 | 7.306 | 0.013803 | 0.9079097 | 0.999176 |
| ENSMUSG00000067028 | Cntnap5b | protein_coding | -0.1024291 | 2.916094 | 0.0137917 | 0.9079471 | 0.999176 |
| ENSMUSG00000039050 | Osbpl2 | protein_coding | 0.04625087 | 5.793113 | 0.0137764 | 0.9079981 | 0.999176 |
| ENSMUSG00000038569 | Rad9b | protein_coding | 0.1633583 | 2.652886 | 0.0137381 | 0.9081278 | 0.999176 |
| ENSMUSG00000047473 | Zfp30 | protein_coding | -0.0545783 | 4.606517 | 0.0136442 | 0.9084385 | 0.999176 |
| ENSMUSG00000031913 | Vps4a | protein_coding | -0.07781708 | 5.448353 | 0.0136592 | 0.9084793 | 0.999176 |
| ENSMUSG00000034235 | Usp54 | protein_coding | -0.1408435 | 2.469394 | 0.0136023 | 0.9085786 | 0.999176 |
| ENSMUSG00000031628 | Casp3 | protein_coding | -0.1067865 | 3.475954 | 0.0135934 | 0.9086104 | 0.999176 |
| ENSMUSG00000046807 | Lrrc75b | protein_coding | 0.06703332 | 4.251037 | 0.0135411 | 0.9087835 | 0.999176 |
| ENSMUSG00000020660 | Pomc | protein_coding | 0.07228973 | 4.148241 | 0.0135296 | 0.9088219 | 0.999176 |
| ENSMUSG00000032324 | Tspan3 | protein_coding | -0.04723883 | 7.434399 | 0.013481 | 0.9089873 | 0.999176 |
| ENSMUSG00000045767 | B230219D22Rik | protein_coding | 0.04446292 | 5.908005 | 0.0134301 | 0.9091562 | 0.999176 |
| ENSMUSG00000058388 | Phtf1 | protein_coding | -0.07407267 | 4.011756 | 0.0134251 | 0.9091731 | 0.999176 |
| ENSMUSG00000068099 | 1500009C09Rik | protein_coding | 0.04159797 | 5.961929 | 0.013352 | 0.9094194 | 0.999176 |
| ENSMUSG00000019899 | Lama2 | protein_coding | 0.1091831 | 2.772552 | 0.0133356 | 0.9094749 | 0.999176 |
| ENSMUSG00000028977 | Casz1 | protein_coding | 0.06964413 | 3.652476 | 0.0133212 | 0.9095234 | 0.999176 |
| ENSMUSG00000029470 | P2rx4 | protein_coding | -0.09774865 | 3.457032 | 0.0133109 | 0.9095583 | 0.999176 |
| ENSMUSG00000038936 | Sccpdh | protein_coding | -0.03912429 | 7.11729 | 0.0133017 | 0.9095894 | 0.999176 |
| ENSMUSG00000022159 | Rab2b | protein_coding | 0.04803199 | 4.849299 | 0.0132415 | 0.9097934 | 0.999176 |
| ENSMUSG00000039908 | Slc26a11 | protein_coding | 0.09130454 | 3.558136 | 0.0132003 | 0.9099329 | 0.999176 |
| ENSMUSG00000025969 | Nrp2 | protein_coding | 0.06816931 | 4.422069 | 0.0131957 | 0.9099486 | 0.999176 |
| ENSMUSG00000019710 | Mrpl24 | protein_coding | 0.03568172 | 6.704733 | 0.0131805 | 0.9100003 | 0.999176 |
| ENSMUSG00000051185 | Fam174a | protein_coding | 0.05233414 | 5.226009 | 0.013159 | 0.9100733 | 0.999176 |
| ENSMUSG00000001794 | Capns1 | protein_coding | 0.03374554 | 7.233773 | 0.0130993 | 0.9102768 | 0.999176 |
| ENSMUSG00000033430 | Terf2ip | protein_coding | -0.04909904 | 5.577778 | 0.0130738 | 0.9103638 | 0.999176 |
| ENSMUSG00000039375 | Wdr17 | protein_coding | 0.06509629 | 4.796423 | 0.013059 | 0.9104142 | 0.999176 |
| ENSMUSG00000041890 | Git2 | protein_coding | -0.07425856 | 4.123003 | 0.0130508 | 0.9104423 | 0.999176 |
| ENSMUSG00000036908 | Unc93b1 | protein_coding | 0.1338619 | 3.136844 | 0.0130663 | 0.9104474 | 0.999176 |
| ENSMUSG00000113314 | AC122013.1 | lincRNA | -0.1277046 | 2.498443 | 0.0129974 | 0.9106248 | 0.999176 |
| ENSMUSG00000097466 | D430036J16Rik | antisense_RNA | 0.09058406 | 3.323471 | 0.0129879 | 0.9106572 | 0.999176 |
| ENSMUSG00000026111 | Unc50 | protein_coding | 0.04641991 | 5.589463 | 0.0129831 | 0.9106736 | 0.999176 |
| ENSMUSG00000076432 | Ywhaq | protein_coding | -0.03094865 | 8.082582 | 0.0129418 | 0.9108151 | 0.999176 |
| ENSMUSG00000022419 | Deptor | protein_coding | -0.08539886 | 4.525747 | 0.0129412 | 0.9108637 | 0.999176 |
| ENSMUSG00000038143 | Stox2 | protein_coding | 0.04703389 | 6.042122 | 0.0128503 | 0.9111296 | 0.999176 |
| ENSMUSG00000015149 | Sirt2 | protein_coding | -0.03268573 | 8.342603 | 0.0128185 | 0.9112393 | 0.999176 |
| ENSMUSG00000059921 | Unc5c | protein_coding | 0.07508366 | 4.057898 | 0.012816 | 0.9112479 | 0.999176 |
| ENSMUSG00000061393 | Acvr2b | protein_coding | 0.1160943 | 2.642841 | 0.0127597 | 0.9114422 | 0.999176 |
| ENSMUSG00000022828 | Gtf2e1 | protein_coding | -0.0698107 | 3.880362 | 0.0127513 | 0.9114713 | 0.999176 |
| ENSMUSG00000038537 | Mc3r | protein_coding | -0.1611601 | 2.441414 | 0.0127151 | 0.9115985 | 0.999176 |
| ENSMUSG00000022554 | Hgh1 | protein_coding | -0.1049608 | 2.63937 | 0.0126443 | 0.9118416 | 0.999176 |
| ENSMUSG00000044365 | Cxxc4 | protein_coding | -0.05078374 | 5.37961 | 0.0126361 | 0.9118701 | 0.999176 |
| ENSMUSG00000026792 | Lrsam1 | protein_coding | -0.07938409 | 4.490933 | 0.012624 | 0.9119145 | 0.999176 |
| ENSMUSG00000040424 | Hipk4 | protein_coding | -0.05878537 | 4.448475 | 0.0126091 | 0.9119638 | 0.999176 |
| ENSMUSG00000062961 | Ccdc177 | protein_coding | 0.07254802 | 3.767734 | 0.0126079 | 0.9119679 | 0.999176 |
| ENSMUSG00000044629 | Cnrip1 | protein_coding | 0.0316848 | 7.542334 | 0.0125991 | 0.9119985 | 0.999176 |
| ENSMUSG00000034729 | Mrps10 | protein_coding | 0.03822889 | 6.322368 | 0.0125986 | 0.9120003 | 0.999176 |
| ENSMUSG00000004846 | Plod3 | protein_coding | -0.08386586 | 3.690383 | 0.0125792 | 0.9120678 | 0.999176 |
| ENSMUSG00000027854 | Sike1 | protein_coding | -0.07490186 | 4.124326 | 0.0125487 | 0.9121742 | 0.999176 |
| ENSMUSG00000067925 | Cxx1a | protein_coding | -0.06277197 | 3.974048 | 0.0124304 | 0.9125871 | 0.999176 |
| ENSMUSG00000034453 | Polr3b | protein_coding | -0.07331362 | 3.459292 | 0.0124166 | 0.9126355 | 0.999176 |
| ENSMUSG00000034035 | Ccdc17 | protein_coding | -0.1167121 | 2.517725 | 0.0124138 | 0.9126452 | 0.999176 |
| ENSMUSG00000038020 | Rapgefl1 | protein_coding | 0.05755069 | 5.041302 | 0.0124101 | 0.9126582 | 0.999176 |
| ENSMUSG00000041923 | Nol4 | protein_coding | 0.03941482 | 6.112689 | 0.0123745 | 0.9127831 | 0.999176 |

| ensembl_gene_id | external_gene_name | gene_biotype | log2 (fold change) | log2 (counts per million) | F-value | p-value | False discovery rate |
|---|---|---|---|---|---|---|---|
| ENSMUSG00000040044 | Orc3 | protein_coding | -0.04928942 | 6.00196 | 0.0123611 | 0.9128303 | 0.999176 |
| ENSMUSG00000021725 | Parp8 | protein_coding | -0.07752752 | 3.485293 | 0.012343 | 0.9128938 | 0.999176 |
| ENSMUSG00000049571 | Cfap46 | protein_coding | 0.100133 | 3.510505 | 0.0123415 | 0.9129012 | 0.999176 |
| ENSMUSG00000025509 | Pnpla2 | protein_coding | -0.07245491 | 5.141812 | 0.0123164 | 0.9130292 | 0.999176 |
| ENSMUSG00000041777 | Cir1 | protein_coding | 0.08385283 | 2.987627 | 0.0122957 | 0.9130601 | 0.999176 |
| ENSMUSG00000022684 | Bfar | protein_coding | -0.04301721 | 5.79158 | 0.0122884 | 0.9130857 | 0.999176 |
| ENSMUSG00000050192 | Eif5a2 | protein_coding | 0.0448196 | 5.390785 | 0.0122836 | 0.9131028 | 0.999176 |
| ENSMUSG00000066647 | Gm5113 | protein_coding | 0.04852854 | 4.778625 | 0.0122421 | 0.913249 | 0.999176 |
| ENSMUSG00000025314 | Ptprj | protein_coding | -0.1039033 | 2.738238 | 0.0122385 | 0.9132618 | 0.999176 |
| ENSMUSG00000045348 | Nyap1 | protein_coding | 0.09079025 | 3.371112 | 0.0122277 | 0.9132999 | 0.999176 |
| ENSMUSG00000042662 | Dusp15 | protein_coding | 0.09049088 | 2.943083 | 0.0121728 | 0.9134937 | 0.999176 |
| ENSMUSG00000039753 | Fbxl5 | protein_coding | 0.03407556 | 6.427889 | 0.0121601 | 0.9135388 | 0.999176 |
| ENSMUSG00000027940 | Tpm3 | protein_coding | -0.03356183 | 6.752328 | 0.0121546 | 0.9135582 | 0.999176 |
| ENSMUSG00000004360 | 9330159F19Rik | protein_coding | -0.07229421 | 4.996475 | 0.0121087 | 0.9137328 | 0.999176 |
| ENSMUSG00000092035 | Peg10 | protein_coding | -0.05299524 | 4.614871 | 0.012088 | 0.9137942 | 0.999176 |
| ENSMUSG00000044600 | Smim7 | protein_coding | 0.02982729 | 8.103929 | 0.0120242 | 0.914021 | 0.999176 |
| ENSMUSG00000027239 | Mdk | protein_coding | -0.09357919 | 4.884047 | 0.012027 | 0.9141988 | 0.999176 |
| ENSMUSG00000041817 | Fam169a | protein_coding | -0.05229572 | 5.870143 | 0.011967 | 0.9142251 | 0.999176 |
| ENSMUSG00000051234 | Rnf7 | protein_coding | -0.03010311 | 7.855072 | 0.0119652 | 0.9142313 | 0.999176 |
| ENSMUSG00000016664 | Pacsin2 | protein_coding | 0.05469651 | 4.843475 | 0.0119609 | 0.9142469 | 0.999176 |
| ENSMUSG00000036644 | Tbc1d9b | protein_coding | 0.05129794 | 5.502997 | 0.0119535 | 0.9142734 | 0.999176 |
| ENSMUSG00000039047 | Pigk | protein_coding | 0.04027576 | 6.55221 | 0.01194 | 0.9143216 | 0.999176 |
| ENSMUSG00000029432 | Nipsnap2 | protein_coding | 0.04561193 | 8.17946 | 0.0119317 | 0.9143533 | 0.999176 |
| ENSMUSG00000015671 | Psma2 | protein_coding | 0.03646257 | 8.831763 | 0.0119265 | 0.9143698 | 0.999176 |
| ENSMUSG00000026670 | Uap1 | protein_coding | -0.05066015 | 5.067554 | 0.0119257 | 0.9143726 | 0.999176 |
| ENSMUSG00000003731 | Kpna6 | protein_coding | 0.05098071 | 4.824788 | 0.0119156 | 0.9144087 | 0.999176 |
| ENSMUSG00000024294 | Mib1 | protein_coding | -0.04336163 | 6.045811 | 0.0118756 | 0.9145517 | 0.999176 |
| ENSMUSG00000008859 | Rala | protein_coding | -0.04444887 | 5.3935 | 0.0118603 | 0.9146068 | 0.999176 |
| ENSMUSG00000005882 | Uqcc1 | protein_coding | 0.03885387 | 7.093361 | 0.0117928 | 0.914849 | 0.999176 |
| ENSMUSG00000020741 | Cluh | protein_coding | 0.0454236 | 5.834446 | 0.0117642 | 0.9149517 | 0.999176 |
| ENSMUSG00000075033 | Nxpe3 | protein_coding | 0.08307451 | 3.370713 | 0.0117597 | 0.9149682 | 0.999176 |
| ENSMUSG00000029270 | Fam69a | protein_coding | -0.04356591 | 5.543228 | 0.0117543 | 0.9149874 | 0.999176 |
| ENSMUSG00000058503 | Fam133b | protein_coding | 0.03911371 | 6.062375 | 0.0117266 | 0.9150872 | 0.999176 |
| ENSMUSG00000028758 | Kif17 | protein_coding | 0.05418935 | 4.78534 | 0.0117262 | 0.915089 | 0.999176 |
| ENSMUSG00000032492 | Pth1r | protein_coding | -0.08444662 | 3.535234 | 0.0117126 | 0.9151379 | 0.999176 |
| ENSMUSG00000067242 | Lgi1 | protein_coding | 0.05705241 | 4.060192 | 0.0116514 | 0.915359 | 0.999176 |
| ENSMUSG00000060992 | Copz1 | protein_coding | -0.03291919 | 8.602942 | 0.0116221 | 0.9154649 | 0.999176 |
| ENSMUSG00000030397 | Mark4 | protein_coding | 0.0601787 | 4.220878 | 0.011617 | 0.9154836 | 0.999176 |
| ENSMUSG00000051652 | Lrrc3 | protein_coding | -0.131363 | 3.470694 | 0.0116394 | 0.915587 | 0.999176 |
| ENSMUSG00000073889 | Il11ra1 | protein_coding | 0.06007601 | 3.906026 | 0.0115812 | 0.9156132 | 0.999176 |
| ENSMUSG00000061111 | Mcrip1 | protein_coding | 0.04487526 | 7.096391 | 0.0115379 | 0.9157725 | 0.999176 |
| ENSMUSG00000026113 | Inpp4a | protein_coding | -0.05919163 | 9.065526 | 0.0115563 | 0.9158736 | 0.999176 |
| ENSMUSG00000039473 | Ubn1 | protein_coding | -0.05950004 | 4.132009 | 0.0114791 | 0.9159847 | 0.999176 |
| ENSMUSG00000090223 | Pcp4 | protein_coding | 0.04871985 | 8.449283 | 0.0114361 | 0.9161962 | 0.999176 |
| ENSMUSG00000028803 | Nipal3 | protein_coding | -0.04835979 | 5.249474 | 0.0114191 | 0.9162036 | 0.999176 |
| ENSMUSG00000046157 | Tmem229b | protein_coding | 0.07418648 | 3.999142 | 0.0114142 | 0.9162213 | 0.999176 |
| ENSMUSG00000075012 | Fjx1 | protein_coding | -0.09853687 | 3.278787 | 0.0113924 | 0.9163013 | 0.999176 |
| ENSMUSG00000066760 | Psg16 | protein_coding | -0.140201 | 2.924307 | 0.0113958 | 0.9163461 | 0.999176 |
| ENSMUSG00000026204 | Ptprn | protein_coding | -0.03715795 | 7.339597 | 0.0113263 | 0.9165434 | 0.999176 |
| ENSMUSG00000023055 | Calcoco1 | protein_coding | -0.04481158 | 6.086909 | 0.0112892 | 0.9166797 | 0.999176 |
| ENSMUSG00000063410 | Stk24 | protein_coding | 0.05789417 | 3.879405 | 0.0112322 | 0.9168894 | 0.999176 |
| ENSMUSG00000022353 | Mtss1 | protein_coding | -0.04289773 | 5.152988 | 0.0112051 | 0.9169891 | 0.999176 |
| ENSMUSG00000028672 | Hmgcl | protein_coding | 0.03438623 | 7.215861 | 0.0112049 | 0.91699 | 0.999176 |
| ENSMUSG00000041468 | Gpr12 | protein_coding | -0.05642042 | 4.750508 | 0.0111541 | 0.9171775 | 0.999176 |
| ENSMUSG00000030560 | Ctsc | protein_coding | 0.1400149 | 2.630697 | 0.0111014 | 0.9173748 | 0.999176 |
| ENSMUSG00000019891 | Dcbld1 | protein_coding | -0.1217408 | 2.688547 | 0.0110929 | 0.9174044 | 0.999176 |
| ENSMUSG00000031848 | Lsm4 | protein_coding | -0.0959135 | 4.547719 | 0.0111306 | 0.9174316 | 0.999176 |
| ENSMUSG00000030232 | Aebp2 | protein_coding | 0.04670967 | 5.211686 | 0.0110517 | 0.9175572 | 0.999176 |
| ENSMUSG00000022744 | Cldnd1 | protein_coding | 0.03109113 | 7.721319 | 0.0110441 | 0.9175856 | 0.999176 |
| ENSMUSG00000019124 | Scrn1 | protein_coding | -0.03977007 | 7.737757 | 0.0110162 | 0.9176892 | 0.999176 |
| ENSMUSG00000026117 | Zap70 | protein_coding | 0.1654374 | 2.329439 | 0.0109711 | 0.9178594 | 0.999176 |
| ENSMUSG00000036438 | Calm2 | protein_coding | 0.03799441 | 12.20622 | 0.0109675 | 0.9178708 | 0.999176 |
| ENSMUSG00000025578 | Cbx8 | protein_coding | -0.08293682 | 3.142789 | 0.0109512 | 0.9179315 | 0.999176 |
| ENSMUSG00000036299 | BC031181 | protein_coding | 0.03113482 | 9.008097 | 0.0108668 | 0.9182471 | 0.999176 |
| ENSMUSG00000025261 | Huwe1 | protein_coding | -0.03666691 | 5.898106 | 0.0108371 | 0.9183584 | 0.999176 |
| ENSMUSG00000039990 | Edrf1 | protein_coding | -0.08992046 | 3.26626 | 0.0108349 | 0.9183669 | 0.999176 |
| ENSMUSG00000023075 | Akirin1 | protein_coding | 0.06253655 | 4.188818 | 0.0108323 | 0.9183764 | 0.999176 |
| ENSMUSG00000055629 | B4galnt4 | protein_coding | -0.0436547 | 5.69011 | 0.0108105 | 0.9184582 | 0.999176 |
| ENSMUSG00000026618 | Iars2 | protein_coding | -0.0752729 | 3.85648 | 0.0107862 | 0.9185496 | 0.999176 |
| ENSMUSG00000002804 | Nudt14 | protein_coding | -0.03880992 | 5.569466 | 0.0106678 | 0.9189962 | 0.999176 |
| ENSMUSG00000031958 | Ldhd | protein_coding | -0.08251054 | 4.177067 | 0.0106318 | 0.9191717 | 0.999176 |
| ENSMUSG00000042121 | Ssh1 | protein_coding | -0.04901104 | 4.392451 | 0.0105944 | 0.9192744 | 0.999176 |
| ENSMUSG00000020650 | Bcap29 | protein_coding | -0.03737746 | 6.32447 | 0.0105712 | 0.9193623 | 0.999176 |
| ENSMUSG00000055805 | Fmnl1 | protein_coding | 0.05812882 | 4.428709 | 0.0105581 | 0.9194123 | 0.999176 |
| ENSMUSG00000042046 | Dstyk | protein_coding | 0.04395857 | 5.169483 | 0.0105109 | 0.9195919 | 0.999176 |
| ENSMUSG00000084349 | Rpl3-ps1 | processed_pseudoge | -0.08345898 | 3.37498 | 0.0104735 | 0.9197345 | 0.999176 |
| ENSMUSG00000000794 | Kcnn3 | protein_coding | -0.08010364 | 3.752233 | 0.010441 | 0.9198608 | 0.999176 |
| ENSMUSG00000006378 | Gcat | protein_coding | -0.05134714 | 5.257506 | 0.0104228 | 0.9199283 | 0.999176 |

| ensembl_gene_id | external_gene_name | gene_biotype | log2 (fold change) | log2 (counts per million) | F-value | p-value | False discovery rate |
|---|---|---|---|---|---|---|---|
| ENSMUSG00000026254 | Eif4e2 | protein_coding | -0.03536745 | 6.461111 | 0.0103792 | 0.9200953 | 0.999176 |
| ENSMUSG00000034037 | Fgd5 | protein_coding | 0.07874687 | 3.056743 | 0.0103518 | 0.9202006 | 0.999176 |
| ENSMUSG00000000959 | Oxa1l | protein_coding | -0.03749909 | 6.370403 | 0.010345 | 0.9202266 | 0.999176 |
| ENSMUSG00000027932 | Slc27a3 | protein_coding | 0.1176698 | 2.496548 | 0.0102923 | 0.9204295 | 0.999176 |
| ENSMUSG00000029787 | Avl9 | protein_coding | 0.05098953 | 4.490577 | 0.0102872 | 0.9204489 | 0.999176 |
| ENSMUSG00000031776 | Arl2bp | protein_coding | 0.03256037 | 6.444304 | 0.0102732 | 0.9205031 | 0.999176 |
| ENSMUSG00000039795 | Zfand1 | protein_coding | -0.04142884 | 5.231061 | 0.010243 | 0.9206194 | 0.999176 |
| ENSMUSG00000028757 | Ddost | protein_coding | -0.05214547 | 5.732806 | 0.0102127 | 0.9207386 | 0.999176 |
| ENSMUSG00000041133 | Smc1a | protein_coding | -0.04181849 | 6.176883 | 0.010203 | 0.9207743 | 0.999176 |
| ENSMUSG00000025403 | Shmt2 | protein_coding | 0.06813344 | 4.196182 | 0.0101958 | 0.9208021 | 0.999176 |
| ENSMUSG00000019763 | Rmnd1 | protein_coding | -0.05321423 | 3.994863 | 0.0101769 | 0.9208752 | 0.999176 |
| ENSMUSG00000029407 | Uso1 | protein_coding | -0.04261856 | 5.557245 | 0.0101645 | 0.9209232 | 0.999176 |
| ENSMUSG00000024991 | Eif3a | protein_coding | -0.04568673 | 6.49309 | 0.0101606 | 0.9209404 | 0.999176 |
| ENSMUSG00000003992 | Ssbp2 | protein_coding | 0.02878181 | 7.282676 | 0.0100816 | 0.9212451 | 0.999176 |
| ENSMUSG00000018697 | Aatf | protein_coding | -0.07717099 | 3.967412 | 0.0100819 | 0.9212461 | 0.999176 |
| ENSMUSG00000048349 | Pou4f1 | protein_coding | -0.3547098 | 1.427937 | 0.0101355 | 0.9212673 | 0.999176 |
| ENSMUSG00000033174 | Mgll | protein_coding | -0.04068262 | 6.84298 | 0.0100751 | 0.9212724 | 0.999176 |
| ENSMUSG00000020900 | Myh10 | protein_coding | 0.04087718 | 6.443935 | 0.0100447 | 0.921389 | 0.999176 |
| ENSMUSG00000019518 | Ap4m1 | protein_coding | -0.08640163 | 2.850539 | 0.0100298 | 0.9214469 | 0.999176 |
| ENSMUSG00000096974 | Gm26881 | antisense_RNA | -0.07932991 | 2.940712 | 0.0100072 | 0.9215352 | 0.999176 |
| ENSMUSG00000028988 | Ctnnbip1 | protein_coding | 0.07437619 | 3.844295 | 0.0100014 | 0.9215578 | 0.999176 |
| ENSMUSG00000020627 | Klhl29 | protein_coding | -0.06533843 | 4.320473 | 0.009999 | 0.9215673 | 0.999176 |
| ENSMUSG00000019210 | Atp6v1e1 | protein_coding | -0.02856138 | 10.11011 | 0.0099883 | 0.9216089 | 0.999176 |
| ENSMUSG00000076928 | Trac | TR_C_gene | 0.1458539 | 2.46587 | 0.0099823 | 0.9216345 | 0.999176 |
| ENSMUSG00000050812 | AI314180 | protein_coding | -0.04255507 | 5.528442 | 0.0099556 | 0.9217373 | 0.999176 |
| ENSMUSG00000063524 | Eno1 | protein_coding | 0.03811563 | 8.641938 | 0.009926 | 0.9218532 | 0.999176 |
| ENSMUSG00000050103 | Agmo | protein_coding | 0.09955616 | 2.957131 | 0.0098754 | 0.9220518 | 0.999176 |
| ENSMUSG00000018287 | Spag7 | protein_coding | 0.0268088 | 8.922909 | 0.0098625 | 0.9221025 | 0.999176 |
| ENSMUSG00000029413 | Naaa | protein_coding | -0.06480958 | 4.870786 | 0.0098594 | 0.9221168 | 0.999176 |
| ENSMUSG00000047514 | Tspyl1 | protein_coding | -0.03474674 | 8.789174 | 0.0097999 | 0.9223494 | 0.999176 |
| ENSMUSG00000025235 | Bbs4 | protein_coding | -0.04604073 | 6.460609 | 0.0097539 | 0.9225384 | 0.999176 |
| ENSMUSG00000047648 | Fbxo30 | protein_coding | 0.09112829 | 2.810722 | 0.0097399 | 0.9225864 | 0.999176 |
| ENSMUSG00000040276 | Pacsin1 | protein_coding | -0.03207118 | 7.277581 | 0.0097375 | 0.922596 | 0.999176 |
| ENSMUSG00000024474 | Ik | protein_coding | 0.03551148 | 7.511914 | 0.0097164 | 0.9226796 | 0.999176 |
| ENSMUSG00000037234 | Hook3 | protein_coding | -0.05853566 | 4.477801 | 0.0097154 | 0.9226835 | 0.999176 |
| ENSMUSG00000012640 | Zfp715 | protein_coding | -0.06469655 | 3.863644 | 0.0096727 | 0.9228531 | 0.999176 |
| ENSMUSG00000029161 | Cgref1 | protein_coding | 0.0467487 | 5.114377 | 0.0096081 | 0.9231104 | 0.999176 |
| ENSMUSG00000023169 | Slc38a1 | protein_coding | 0.03263256 | 7.064916 | 0.0096041 | 0.9231265 | 0.999176 |
| ENSMUSG00000029071 | Dvl1 | protein_coding | -0.0512647 | 4.183195 | 0.0095886 | 0.923188 | 0.999176 |
| ENSMUSG00000048490 | Nrip1 | protein_coding | 0.05160376 | 4.388132 | 0.0095817 | 0.9232159 | 0.999176 |
| ENSMUSG00000028243 | Ubxn2b | protein_coding | -0.04097529 | 5.753516 | 0.009581 | 0.9232185 | 0.999176 |
| ENSMUSG00000052419 | 2610001J05Rik | protein_coding | -0.02975748 | 7.768716 | 0.00954 | 0.9233823 | 0.999176 |
| ENSMUSG00000026470 | Stx6 | protein_coding | 0.04331167 | 5.73493 | 0.0095216 | 0.9234562 | 0.999176 |
| ENSMUSG00000020089 | Ppa1 | protein_coding | 0.03487661 | 7.898839 | 0.0095034 | 0.9235289 | 0.999176 |
| ENSMUSG00000024050 | Wiz | protein_coding | 0.06565267 | 3.622252 | 0.0094917 | 0.923576 | 0.999176 |
| ENSMUSG00000036093 | Arl5a | protein_coding | 0.04876469 | 4.645275 | 0.0094611 | 0.9236987 | 0.999176 |
| ENSMUSG00000031921 | Terf2 | protein_coding | 0.04715506 | 4.78876 | 0.009449 | 0.9237476 | 0.999176 |
| ENSMUSG00000026644 | Acbd7 | protein_coding | -0.07294525 | 3.201098 | 0.0094455 | 0.9237615 | 0.999176 |
| ENSMUSG00000045404 | Kcnk13 | protein_coding | 0.09036079 | 3.372933 | 0.0094411 | 0.9237812 | 0.999176 |
| ENSMUSG00000030079 | Ruvbl1 | protein_coding | 0.04435409 | 5.052224 | 0.0094379 | 0.923792 | 0.999176 |
| ENSMUSG00000021668 | Polk | protein_coding | -0.07442156 | 3.087683 | 0.0094224 | 0.9238545 | 0.999176 |
| ENSMUSG00000024078 | Ttc27 | protein_coding | -0.07742191 | 4.069331 | 0.0093418 | 0.924189 | 0.999176 |
| ENSMUSG00000033966 | Cdkl4 | protein_coding | -0.06103585 | 4.235279 | 0.0093354 | 0.9242056 | 0.999176 |
| ENSMUSG00000045210 | Vcpip1 | protein_coding | -0.03727642 | 5.468058 | 0.0092723 | 0.9244614 | 0.999176 |
| ENSMUSG00000024941 | Scyl1 | protein_coding | -0.04009451 | 5.491645 | 0.009255 | 0.9245317 | 0.999176 |
| ENSMUSG00000029406 | Pitpnm2 | protein_coding | 0.04542139 | 5.065492 | 0.0092534 | 0.9245383 | 0.999176 |
| ENSMUSG00000068742 | Cry2 | protein_coding | 0.03861955 | 5.848385 | 0.0092371 | 0.9246045 | 0.999176 |
| ENSMUSG00000097736 | 9530059O14Rik | lincRNA | 0.1492789 | 2.537023 | 0.0091914 | 0.9247925 | 0.999176 |
| ENSMUSG00000084799 | Ino80dos | lincRNA | -0.07998983 | 2.857562 | 0.0091772 | 0.9248488 | 0.999176 |
| ENSMUSG00000002342 | Tmem161a | protein_coding | 0.05503719 | 4.850548 | 0.0091628 | 0.9249074 | 0.999176 |
| ENSMUSG00000066415 | Msl2 | protein_coding | -0.06117912 | 4.015006 | 0.0091467 | 0.9249731 | 0.999176 |
| ENSMUSG00000024253 | Dync2li1 | protein_coding | -0.03504089 | 5.827767 | 0.0091456 | 0.9249777 | 0.999176 |
| ENSMUSG00000021314 | Amph | protein_coding | 0.04704706 | 7.151885 | 0.0091424 | 0.9250697 | 0.999176 |
| ENSMUSG00000026519 | Tmem63a | protein_coding | 0.05770663 | 4.576867 | 0.0091123 | 0.9251138 | 0.999176 |
| ENSMUSG00000034168 | Irf2bpl | protein_coding | 0.04736249 | 4.404039 | 0.0090852 | 0.9252252 | 0.999176 |
| ENSMUSG00000097875 | Gm26704 | lincRNA | 0.06550861 | 3.844734 | 0.0090776 | 0.9252561 | 0.999176 |
| ENSMUSG00000029480 | Dhx37 | protein_coding | 0.06584142 | 3.67094 | 0.0090643 | 0.925311 | 0.999176 |
| ENSMUSG00000025791 | Pgm2 | protein_coding | -0.04087687 | 7.022991 | 0.009061 | 0.9253263 | 0.999176 |
| ENSMUSG00000024270 | Slc39a6 | protein_coding | -0.05000011 | 5.133147 | 0.0090575 | 0.9253385 | 0.999176 |
| ENSMUSG00000070282 | 3000002C10Rik | transcribed_processe | 0.1300774 | 2.154411 | 0.0090218 | 0.9254857 | 0.999176 |
| ENSMUSG00000038485 | Socs7 | protein_coding | -0.0427144 | 5.37011 | 0.0090181 | 0.9255009 | 0.999176 |
| ENSMUSG00000048154 | Kmt2d | protein_coding | -0.05160479 | 6.139164 | 0.0090081 | 0.9256115 | 0.999176 |
| ENSMUSG00000022110 | Sucla2 | protein_coding | 0.0319948 | 8.265718 | 0.0089868 | 0.9256299 | 0.999176 |
| ENSMUSG00000020681 | Ace | protein_coding | -0.08198011 | 3.361367 | 0.0089769 | 0.9256704 | 0.999176 |
| ENSMUSG00000029434 | Vps33a | protein_coding | 0.04784483 | 5.761476 | 0.0089765 | 0.9256741 | 0.999176 |
| ENSMUSG00000054945 | Gm9958 | protein_coding | 0.1787857 | 1.942478 | 0.008971 | 0.9256951 | 0.999176 |
| ENSMUSG00000048355 | Arxes1 | protein_coding | -0.05113501 | 4.788125 | 0.0089019 | 0.925981 | 0.999176 |
| ENSMUSG00000018425 | Dhx40 | protein_coding | -0.0440045 | 4.628552 | 0.0088696 | 0.9261148 | 0.999176 |

| ensembl_gene_id | external_gene_name | gene_biotype | log2 (fold change) | log2 (counts per million) | F-value | p-value | False discovery rate |
|---|---|---|---|---|---|---|---|
| ENSMUSG00000038079 | Tmem237 | protein_coding | 0.04060328 | 6.257935 | 0.008838 | 0.9262462 | 0.999176 |
| ENSMUSG00000052428 | Tmco1 | protein_coding | -0.03125224 | 7.076711 | 0.0088081 | 0.9263706 | 0.999176 |
| ENSMUSG00000028426 | Rad23b | protein_coding | -0.0366969 | 5.858985 | 0.008808 | 0.9263709 | 0.999176 |
| ENSMUSG00000031538 | Plat | protein_coding | -0.05900857 | 4.335393 | 0.008806 | 0.9263794 | 0.999176 |
| ENSMUSG00000034324 | Tmem132c | protein_coding | -0.1019266 | 2.717999 | 0.0088025 | 0.9263938 | 0.999176 |
| ENSMUSG00000046727 | Cystm1 | protein_coding | 0.02776837 | 8.856224 | 0.0088019 | 0.9263963 | 0.999176 |
| ENSMUSG00000020993 | Trappc6b | protein_coding | 0.0390242 | 5.564164 | 0.0087951 | 0.9264248 | 0.999176 |
| ENSMUSG00000028152 | Tspan5 | protein_coding | 0.03133712 | 6.044116 | 0.0087754 | 0.9265071 | 0.999176 |
| ENSMUSG00000079508 | Apoo | protein_coding | 0.04507796 | 4.398445 | 0.0087397 | 0.926656 | 0.999176 |
| ENSMUSG00000038482 | Tfdp1 | protein_coding | -0.04997932 | 4.65646 | 0.0087396 | 0.9266567 | 0.999176 |
| ENSMUSG00000028926 | Cdk14 | protein_coding | -0.04403612 | 7.358195 | 0.0087546 | 0.9266664 | 0.999176 |
| ENSMUSG00000079317 | Trappc2 | protein_coding | -0.02859788 | 6.535423 | 0.0087251 | 0.9267174 | 0.999176 |
| ENSMUSG00000028524 | Sgip1 | protein_coding | 0.04316967 | 4.884116 | 0.0087117 | 0.9267734 | 0.999176 |
| ENSMUSG00000031447 | Lamp1 | protein_coding | 0.06201053 | 3.552952 | 0.008709 | 0.9267848 | 0.999176 |
| ENSMUSG00000087598 | Zfp111 | protein_coding | -0.07817776 | 2.73204 | 0.0086899 | 0.9268647 | 0.999176 |
| ENSMUSG00000004207 | Psap | protein_coding | -0.03075503 | 9.011553 | 0.008681 | 0.926902 | 0.999176 |
| ENSMUSG00000043153 | Ispd | protein_coding | -0.05561555 | 4.227465 | 0.0086752 | 0.9269263 | 0.999176 |
| ENSMUSG00000031545 | Gpat4 | protein_coding | 0.03937299 | 6.547038 | 0.0086295 | 0.9271205 | 0.999176 |
| ENSMUSG00000052512 | Nav2 | protein_coding | 0.04712324 | 5.384332 | 0.0086194 | 0.9271611 | 0.999176 |
| ENSMUSG00000006235 | Epor | protein_coding | -0.07554829 | 2.937805 | 0.0085948 | 0.9272649 | 0.999176 |
| ENSMUSG00000025782 | Taf3 | protein_coding | 0.04303407 | 4.814054 | 0.008588 | 0.9272937 | 0.999176 |
| ENSMUSG00000063018 | 2010204K13Rik | processed_transcript | 0.03263512 | 7.865833 | 0.008566 | 0.9273864 | 0.999176 |
| ENSMUSG00000032040 | Dcps | protein_coding | 0.04630458 | 4.857427 | 0.0085305 | 0.9275365 | 0.999176 |
| ENSMUSG00000029212 | Gabrb1 | protein_coding | -0.0333224 | 6.368172 | 0.0085079 | 0.9276322 | 0.999176 |
| ENSMUSG00000040782 | Rfwd2 | protein_coding | 0.07853845 | 3.50021 | 0.0084749 | 0.9277723 | 0.999176 |
| ENSMUSG00000050914 | Ankrd37 | protein_coding | 0.04081517 | 4.687637 | 0.0084337 | 0.9279475 | 0.999176 |
| ENSMUSG00000020056 | Washc3 | protein_coding | 0.03406341 | 6.423033 | 0.0083774 | 0.9281877 | 0.999176 |
| ENSMUSG00000019929 | Dcn | protein_coding | 0.06813855 | 3.742445 | 0.0083459 | 0.9283225 | 0.999176 |
| ENSMUSG00000031841 | Cdh13 | protein_coding | 0.0251757 | 7.328179 | 0.0083356 | 0.9283667 | 0.999176 |
| ENSMUSG00000039787 | Cercam | protein_coding | -0.1365491 | 3.018023 | 0.0083539 | 0.9284444 | 0.999176 |
| ENSMUSG00000023072 | Cep89 | protein_coding | 0.06134405 | 3.758035 | 0.0082764 | 0.928621 | 0.999176 |
| ENSMUSG00000018796 | Acsl1 | protein_coding | -0.0490297 | 4.888774 | 0.0082666 | 0.9286628 | 0.999176 |
| ENSMUSG00000029267 | Mtf2 | protein_coding | 0.04478842 | 4.569289 | 0.0082432 | 0.9287639 | 0.999176 |
| ENSMUSG00000051048 | P4ha3 | protein_coding | 0.1093537 | 2.603269 | 0.0082327 | 0.9288089 | 0.999176 |
| ENSMUSG00000087403 | Kantr | protein_coding | 0.0396194 | 6.665899 | 0.0081854 | 0.9290149 | 0.999176 |
| ENSMUSG00000041959 | S100a10 | protein_coding | 0.03116001 | 6.685286 | 0.0081454 | 0.9291863 | 0.999176 |
| ENSMUSG00000070371 | Prss36 | protein_coding | 0.06108069 | 4.542792 | 0.0081207 | 0.9292953 | 0.999176 |
| ENSMUSG00000021916 | Glt8d1 | protein_coding | 0.05986392 | 4.044384 | 0.0081161 | 0.9293133 | 0.999176 |
| ENSMUSG00000020334 | Slc22a4 | protein_coding | 0.07545301 | 3.121546 | 0.0080921 | 0.9294176 | 0.999176 |
| ENSMUSG00000001120 | Pcbp3 | protein_coding | 0.06007432 | 6.02875 | 0.0081034 | 0.9295225 | 0.999176 |
| ENSMUSG00000047284 | Neurl4 | protein_coding | -0.05013441 | 5.238738 | 0.0080483 | 0.9296102 | 0.999176 |
| ENSMUSG00000022351 | Sqle | protein_coding | -0.04119227 | 5.667981 | 0.0080204 | 0.9297304 | 0.999176 |
| ENSMUSG00000035575 | Utp6 | protein_coding | -0.04628332 | 4.648041 | 0.0080137 | 0.9297595 | 0.999176 |
| ENSMUSG00000030795 | Fus | protein_coding | 0.03953604 | 5.029174 | 0.0079475 | 0.9300496 | 0.999176 |
| ENSMUSG00000001416 | Cct3 | protein_coding | -0.0289818 | 7.87536 | 0.0079408 | 0.9300789 | 0.999176 |
| ENSMUSG00000021906 | Oxnad1 | protein_coding | -0.04564734 | 4.883129 | 0.0078794 | 0.9303488 | 0.999176 |
| ENSMUSG00000059486 | Kbtbd2 | protein_coding | -0.03726681 | 5.815665 | 0.00784 | 0.9305229 | 0.999176 |
| ENSMUSG00000027708 | Dcun1d1 | protein_coding | 0.03734453 | 5.159537 | 0.0078157 | 0.9306304 | 0.999176 |
| ENSMUSG00000024758 | Rtn3 | protein_coding | 0.0302953 | 8.667424 | 0.0078133 | 0.9306409 | 0.999176 |
| ENSMUSG00000029376 | Mthfd2l | protein_coding | -0.04465086 | 4.601838 | 0.0077775 | 0.9307997 | 0.999176 |
| ENSMUSG00000064325 | Hhip | protein_coding | 0.08286446 | 2.593914 | 0.0077629 | 0.9308645 | 0.999176 |
| ENSMUSG00000073236 | 2500004C02Rik | processed_transcript | -0.0436406 | 4.325523 | 0.0077403 | 0.9309648 | 0.999176 |
| ENSMUSG00000054277 | Arfgap3 | protein_coding | -0.04185157 | 5.491943 | 0.0077387 | 0.9309717 | 0.999176 |
| ENSMUSG00000017929 | B4galt5 | protein_coding | 0.0414454 | 4.772614 | 0.0077089 | 0.9311047 | 0.999176 |
| ENSMUSG00000050854 | Tmem125 | protein_coding | 0.0710557 | 3.590189 | 0.0077072 | 0.9311122 | 0.999176 |
| ENSMUSG00000002900 | Lamb1 | protein_coding | 0.06872548 | 3.348845 | 0.0077063 | 0.9311161 | 0.999176 |
| ENSMUSG00000029173 | Sepsecs | protein_coding | -0.08713933 | 3.513727 | 0.0077043 | 0.9311699 | 0.999176 |
| ENSMUSG00000024217 | Snrpc | protein_coding | -0.04296662 | 5.049866 | 0.0076826 | 0.9312217 | 0.999176 |
| ENSMUSG00000020305 | Asb3 | protein_coding | 0.03969404 | 5.649299 | 0.0076 | 0.9315916 | 0.999176 |
| ENSMUSG00000026820 | Ptges2 | protein_coding | 0.03623403 | 5.297392 | 0.0075942 | 0.9316177 | 0.999176 |
| ENSMUSG00000024639 | Gnaq | protein_coding | 0.03598466 | 8.272938 | 0.0075837 | 0.9316665 | 0.999176 |
| ENSMUSG00000024369 | Nelfe | protein_coding | 0.03209801 | 6.009581 | 0.0075779 | 0.9316909 | 0.999176 |
| ENSMUSG00000005087 | Cd44 | protein_coding | 0.09350592 | 2.696449 | 0.0075681 | 0.9317349 | 0.999176 |
| ENSMUSG00000020721 | Helz | protein_coding | -0.03345939 | 5.362375 | 0.0075587 | 0.931777 | 0.999176 |
| ENSMUSG00000044646 | Zbtb7c | protein_coding | 0.08483216 | 2.786384 | 0.0075496 | 0.9318184 | 0.999176 |
| ENSMUSG00000030835 | Nomo1 | protein_coding | 0.0307811 | 6.283509 | 0.0075249 | 0.9319295 | 0.999176 |
| ENSMUSG00000031889 | D230025D16Rik | protein_coding | 0.04716961 | 4.334105 | 0.0074983 | 0.9320496 | 0.999176 |
| ENSMUSG00000003746 | Man1a | protein_coding | -0.04392426 | 4.613703 | 0.0074872 | 0.9320998 | 0.999176 |
| ENSMUSG00000018442 | Derl2 | protein_coding | -0.0350279 | 5.580025 | 0.0074763 | 0.9321491 | 0.999176 |
| ENSMUSG00000063239 | Grm4 | protein_coding | 0.0438089 | 4.528026 | 0.0074564 | 0.9322394 | 0.999176 |
| ENSMUSG00000049721 | Gal3st1 | protein_coding | 0.0757528 | 3.604891 | 0.0074555 | 0.9322451 | 0.999176 |
| ENSMUSG00000092094 | Zfp804b | protein_coding | -0.07504511 | 2.697702 | 0.0074505 | 0.932266 | 0.999176 |
| ENSMUSG00000002625 | Akap8l | protein_coding | -0.03446914 | 5.114808 | 0.0074323 | 0.9323486 | 0.999176 |
| ENSMUSG00000054280 | Prr14l | protein_coding | -0.0711278 | 3.099167 | 0.0074086 | 0.9324563 | 0.999176 |
| ENSMUSG00000028796 | Phc2 | protein_coding | 0.03575414 | 5.428603 | 0.007373 | 0.9326183 | 0.999176 |
| ENSMUSG00000030761 | Myo7a | protein_coding | 0.09771056 | 2.559451 | 0.0073674 | 0.9326437 | 0.999176 |
| ENSMUSG00000028479 | Gne | protein_coding | -0.04150618 | 4.514314 | 0.0073611 | 0.9326726 | 0.999176 |
| ENSMUSG00000022358 | Fbxo32 | protein_coding | -0.06701331 | 3.737421 | 0.0073595 | 0.9326797 | 0.999176 |

| ensembl_gene_id | external_gene_name | gene_biotype | log2 (fold change) | log2 (counts per million) | F-value | p-value | False discovery rate |
|---|---|---|---|---|---|---|---|
| ENSMUSG00000034751 | Mast4 | protein_coding | -0.05805393 | 3.656852 | 0.0073578 | 0.9326875 | 0.999176 |
| ENSMUSG00000049658 | Bdp1 | protein_coding | -0.05456993 | 3.75333 | 0.007349 | 0.9327277 | 0.999176 |
| ENSMUSG00000020329 | Polrmt | protein_coding | -0.07841611 | 3.309973 | 0.0072748 | 0.9330671 | 0.999176 |
| ENSMUSG00000022621 | Rabl2 | protein_coding | 0.03460078 | 6.451841 | 0.0072645 | 0.9331148 | 0.999176 |
| ENSMUSG00000064138 | Fam172a | protein_coding | 0.06331262 | 3.102864 | 0.0072492 | 0.9331847 | 0.999176 |
| ENSMUSG00000027455 | Nsfl1c | protein_coding | 0.02620741 | 7.593697 | 0.0072226 | 0.9333071 | 0.999176 |
| ENSMUSG00000039623 | Ap5z1 | protein_coding | 0.06996127 | 3.859693 | 0.0071891 | 0.9334636 | 0.999176 |
| ENSMUSG00000036902 | Neto2 | protein_coding | 0.04850928 | 4.898348 | 0.0071776 | 0.933515 | 0.999176 |
| ENSMUSG00000005161 | Prdx2 | protein_coding | 0.02351581 | 9.485122 | 0.0071704 | 0.933548 | 0.999176 |
| ENSMUSG00000059005 | Hnrnpa3 | protein_coding | -0.0431292 | 3.88491 | 0.0071525 | 0.9336309 | 0.999176 |
| ENSMUSG00000040459 | Arglu1 | protein_coding | -0.03175673 | 7.490945 | 0.0071461 | 0.9336606 | 0.999176 |
| ENSMUSG00000039458 | Mtmr12 | protein_coding | 0.05677406 | 3.255889 | 0.0070776 | 0.9339786 | 0.999176 |
| ENSMUSG00000020694 | Tlk2 | protein_coding | -0.03535333 | 5.276387 | 0.0070717 | 0.934006 | 0.999176 |
| ENSMUSG00000043140 | Tmem186 | protein_coding | 0.04712076 | 5.17311 | 0.0070319 | 0.9341914 | 0.999176 |
| ENSMUSG00000021179 | Nrde2 | protein_coding | -0.05715788 | 4.271588 | 0.0070155 | 0.9342679 | 0.999176 |
| ENSMUSG00000035960 | Apex1 | protein_coding | 0.03357565 | 5.558135 | 0.0070034 | 0.9343248 | 0.999176 |
| ENSMUSG00000039835 | Nhsl1 | protein_coding | 0.06561682 | 3.785744 | 0.0069921 | 0.9343789 | 0.999176 |
| ENSMUSG00000071719 | Tmem28 | protein_coding | 0.05074181 | 3.903333 | 0.0069872 | 0.9344006 | 0.999176 |
| ENSMUSG00000003154 | Foxj2 | protein_coding | 0.05225946 | 3.932938 | 0.0069654 | 0.9345027 | 0.999176 |
| ENSMUSG00000034723 | Tmx4 | protein_coding | -0.02570014 | 7.659285 | 0.006954 | 0.934556 | 0.999176 |
| ENSMUSG00000004032 | Gstm5 | protein_coding | -0.03024352 | 9.655987 | 0.0068855 | 0.9348781 | 0.999176 |
| ENSMUSG00000048351 | Coa7 | protein_coding | -0.0304607 | 5.885506 | 0.0068834 | 0.9348884 | 0.999176 |
| ENSMUSG00000020475 | Pgam2 | protein_coding | -0.04208682 | 4.34265 | 0.0068684 | 0.9349589 | 0.999176 |
| ENSMUSG00000054874 | Pcnx3 | protein_coding | 0.09469098 | 2.791372 | 0.0068669 | 0.9349664 | 0.999176 |
| ENSMUSG00000037594 | BC022687 | protein_coding | -0.06385621 | 3.430829 | 0.0068616 | 0.9349911 | 0.999176 |
| ENSMUSG00000027597 | Ahcy | protein_coding | -0.05045802 | 3.452462 | 0.0068405 | 0.9350908 | 0.999176 |
| ENSMUSG00000026554 | Dcaf8 | protein_coding | -0.0342388 | 7.101565 | 0.0068318 | 0.935134 | 0.999176 |
| ENSMUSG00000094344 | Gm11942 | processed_pseudoge | 0.1917823 | 1.97259 | 0.0068169 | 0.9352812 | 0.999176 |
| ENSMUSG00000024130 | Abca3 | protein_coding | 0.07487211 | 2.720833 | 0.0067711 | 0.9354204 | 0.999176 |
| ENSMUSG00000063480 | Snu13 | protein_coding | -0.03349235 | 5.062152 | 0.0067124 | 0.9356999 | 0.999176 |
| ENSMUSG00000030061 | Uba3 | protein_coding | -0.02658679 | 6.785816 | 0.0067109 | 0.9357074 | 0.999176 |
| ENSMUSG00000038323 | 1700066M21Rik | protein_coding | -0.07372825 | 3.20764 | 0.006626 | 0.9361144 | 0.999176 |
| ENSMUSG00000040848 | Sft2d2 | protein_coding | 0.05585957 | 3.875666 | 0.0066114 | 0.9361846 | 0.999176 |
| ENSMUSG00000028549 | Itgb3bp | protein_coding | 0.04041618 | 4.194755 | 0.0066084 | 0.9361993 | 0.999176 |
| ENSMUSG00000026620 | Mark1 | protein_coding | 0.0419815 | 4.913049 | 0.006575 | 0.9363603 | 0.999176 |
| ENSMUSG00000057914 | Cacnb2 | protein_coding | 0.04781898 | 4.629845 | 0.0065454 | 0.936503 | 0.999176 |
| ENSMUSG00000004054 | Map3k11 | protein_coding | 0.08280991 | 2.518704 | 0.0065424 | 0.9365178 | 0.999176 |
| ENSMUSG00000061130 | Ppm1b | protein_coding | -0.03433446 | 5.232651 | 0.0065414 | 0.9365227 | 0.999176 |
| ENSMUSG00000000983 | Wfdc18 | protein_coding | -0.05912759 | 3.370804 | 0.0065148 | 0.9366514 | 0.999176 |
| ENSMUSG00000028633 | Ctps | protein_coding | 0.03532111 | 5.140252 | 0.0065072 | 0.9366882 | 0.999176 |
| ENSMUSG00000059991 | Nptx2 | protein_coding | 0.06322171 | 3.168592 | 0.0064863 | 0.9367899 | 0.999176 |
| ENSMUSG00000027456 | Sdcbp2 | protein_coding | 0.08295129 | 3.053528 | 0.0064093 | 0.9371655 | 0.999176 |
| ENSMUSG00000047126 | Cltc | protein_coding | -0.02314298 | 8.080627 | 0.0064061 | 0.9371808 | 0.999176 |
| ENSMUSG00000061080 | Lsamp | protein_coding | -0.0215043 | 7.720072 | 0.0063978 | 0.9372215 | 0.999176 |
| ENSMUSG00000028351 | Brinp1 | protein_coding | -0.02739541 | 6.494578 | 0.0063737 | 0.9373394 | 0.999176 |
| ENSMUSG00000032565 | Nudt16 | protein_coding | -0.04183271 | 4.087896 | 0.0063614 | 0.9373999 | 0.999176 |
| ENSMUSG00000003068 | Stk11 | protein_coding | -0.03735473 | 5.192737 | 0.0063466 | 0.9374726 | 0.999176 |
| ENSMUSG00000053119 | Chmp3 | protein_coding | 0.02480922 | 7.855505 | 0.0063296 | 0.9375561 | 0.999176 |
| ENSMUSG00000024151 | Msh2 | protein_coding | -0.04862157 | 5.128753 | 0.0063278 | 0.937567 | 0.999176 |
| ENSMUSG00000003135 | Cnot11 | protein_coding | 0.05938555 | 2.972213 | 0.0063273 | 0.9375676 | 0.999176 |
| ENSMUSG00000078546 | Zfp995 | protein_coding | -0.06899693 | 2.66709 | 0.006322 | 0.9375936 | 0.999176 |
| ENSMUSG00000035621 | Midn | protein_coding | 0.05955592 | 2.965152 | 0.0063106 | 0.9376501 | 0.999176 |
| ENSMUSG00000022434 | Fam118a | protein_coding | -0.03449984 | 5.011475 | 0.0062931 | 0.9377363 | 0.999176 |
| ENSMUSG00000031578 | Mak16 | protein_coding | 0.03892335 | 4.675582 | 0.0062901 | 0.9377512 | 0.999176 |
| ENSMUSG00000053774 | Ubxn7 | protein_coding | 0.06712795 | 3.20918 | 0.0062758 | 0.9378219 | 0.999176 |
| ENSMUSG00000022415 | Syngr1 | protein_coding | 0.02149543 | 7.596962 | 0.0062601 | 0.9378991 | 0.999176 |
| ENSMUSG00000025935 | Tram1 | protein_coding | -0.03821127 | 5.064605 | 0.0062566 | 0.9379168 | 0.999176 |
| ENSMUSG00000055835 | Zfp1 | protein_coding | -0.045501 | 4.897413 | 0.0062555 | 0.9379223 | 0.999176 |
| ENSMUSG00000072825 | Cep170b | protein_coding | 0.03020635 | 6.618849 | 0.0062467 | 0.9379656 | 0.999176 |
| ENSMUSG00000021911 | Parg | protein_coding | -0.04223684 | 4.457585 | 0.0062448 | 0.9379749 | 0.999176 |
| ENSMUSG00000098332 | 2310009A05Rik | processed_transcript | -0.03725396 | 5.400427 | 0.0062361 | 0.9380183 | 0.999176 |
| ENSMUSG00000110444 | Gm10033 | transcribed_unproces | 0.08012883 | 3.567409 | 0.0062351 | 0.9380843 | 0.999176 |
| ENSMUSG00000001785 | Pwp1 | protein_coding | -0.02998151 | 5.304626 | 0.0062176 | 0.9381099 | 0.999176 |
| ENSMUSG00000091478 | Gm10039 | processed_pseudoge | 0.05982004 | 2.898953 | 0.0062054 | 0.9381706 | 0.999176 |
| ENSMUSG00000052026 | Slc6a7 | protein_coding | 0.05352246 | 3.676003 | 0.0062002 | 0.9381965 | 0.999176 |
| ENSMUSG00000027288 | Zfp106 | protein_coding | 0.03135575 | 5.455992 | 0.0061997 | 0.9381987 | 0.999176 |
| ENSMUSG00000033170 | Card10 | protein_coding | -0.1478811 | 2.435957 | 0.0061933 | 0.9383263 | 0.999176 |
| ENSMUSG00000052613 | Pcdh15 | protein_coding | -0.04838521 | 4.05166 | 0.0061648 | 0.9383728 | 0.999176 |
| ENSMUSG00000035168 | Tanc1 | protein_coding | -0.04794294 | 3.707262 | 0.006157 | 0.9384118 | 0.999176 |
| ENSMUSG00000058881 | Zfp516 | protein_coding | -0.08930073 | 2.882451 | 0.0061496 | 0.9384502 | 0.999176 |
| ENSMUSG00000040483 | Xaf1 | protein_coding | 0.0434979 | 4.266146 | 0.0061492 | 0.9384509 | 0.999176 |
| ENSMUSG00000038811 | Gngt2 | protein_coding | -0.07575903 | 2.841497 | 0.0061359 | 0.9385171 | 0.999176 |
| ENSMUSG00000007908 | Hmgcll1 | protein_coding | 0.04714751 | 4.293572 | 0.0061129 | 0.9386321 | 0.999176 |
| ENSMUSG00000034755 | Pcdh11x | protein_coding | -0.0355088 | 5.168562 | 0.0060962 | 0.938716 | 0.999176 |
| ENSMUSG00000055489 | Ano5 | protein_coding | -0.06888615 | 4.588167 | 0.0060943 | 0.9388487 | 0.999176 |
| ENSMUSG00000030982 | 9030624J02Rik | protein_coding | 0.04283707 | 6.165526 | 0.0060115 | 0.9392024 | 0.999176 |
| ENSMUSG00000050846 | Zfp623 | protein_coding | 0.0405978 | 4.333015 | 0.005955 | 0.9394284 | 0.999176 |
| ENSMUSG00000020677 | Ddx52 | protein_coding | -0.02936772 | 5.73686 | 0.0058954 | 0.9397316 | 0.999176 |

| ensembl_gene_id | external_gene_name | gene_biotype | log2 (fold change) | log2 (counts per million) | F-value | p-value | False discovery rate |
|---|---|---|---|---|---|---|---|
| ENSMUSG00000029826 | Zc3hav1 | protein_coding | -0.05248045 | 3.92524 | 0.0058932 | 0.9397429 | 0.999176 |
| ENSMUSG00000036712 | Cyld | protein_coding | -0.02484254 | 6.61124 | 0.0058836 | 0.9397918 | 0.999176 |
| ENSMUSG00000021671 | Poc5 | protein_coding | -0.03898311 | 4.47209 | 0.0058785 | 0.9398179 | 0.999176 |
| ENSMUSG00000045576 | St7l | protein_coding | -0.04305857 | 4.455684 | 0.0058776 | 0.9398223 | 0.999176 |
| ENSMUSG00000039131 | Gipc2 | protein_coding | -0.04963901 | 3.787852 | 0.0058391 | 0.9400194 | 0.999176 |
| ENSMUSG00000021754 | Map3k1 | protein_coding | 0.04760158 | 3.794129 | 0.0058069 | 0.9401848 | 0.999176 |
| ENSMUSG00000044566 | Cage1 | protein_coding | 0.05180907 | 3.494822 | 0.0057841 | 0.9403018 | 0.999176 |
| ENSMUSG00000020171 | Yeats4 | protein_coding | -0.0442595 | 4.308758 | 0.005776 | 0.9403437 | 0.999176 |
| ENSMUSG00000031901 | Dus2 | protein_coding | 0.03520518 | 5.148644 | 0.0056898 | 0.9407898 | 0.999176 |
| ENSMUSG00000026313 | Hdac4 | protein_coding | -0.03230466 | 5.850927 | 0.0056743 | 0.9408703 | 0.999176 |
| ENSMUSG00000004085 | Map3k20 | protein_coding | -0.07572993 | 3.118671 | 0.0056709 | 0.9408881 | 0.999176 |
| ENSMUSG00000031672 | Got2 | protein_coding | 0.03287129 | 7.037328 | 0.0056472 | 0.9410165 | 0.999176 |
| ENSMUSG00000039157 | Fam102a | protein_coding | 0.02502314 | 6.538389 | 0.0056341 | 0.9410797 | 0.999176 |
| ENSMUSG00000022533 | Atp13a3 | protein_coding | -0.04378001 | 4.077992 | 0.0055648 | 0.9414422 | 0.999176 |
| ENSMUSG00000079481 | Nhsl2 | protein_coding | -0.03484566 | 5.128428 | 0.0055558 | 0.9414895 | 0.999176 |
| ENSMUSG00000024127 | Prepl | protein_coding | 0.0675642 | 7.106367 | 0.0056261 | 0.9416573 | 0.999176 |
| ENSMUSG00000070426 | Rnf121 | protein_coding | 0.02598474 | 5.735342 | 0.0055208 | 0.9416739 | 0.999176 |
| ENSMUSG00000028664 | Ephb2 | protein_coding | -0.06488347 | 2.859947 | 0.0055128 | 0.9417159 | 0.999176 |
| ENSMUSG00000025278 | Flnb | protein_coding | -0.07642988 | 3.310573 | 0.0055081 | 0.9417488 | 0.999176 |
| ENSMUSG00000040282 | BC052040 | protein_coding | -0.03950869 | 4.305138 | 0.00545 | 0.9420485 | 0.999176 |
| ENSMUSG00000048897 | Zfp710 | protein_coding | 0.03817872 | 4.031453 | 0.0054395 | 0.9421039 | 0.999176 |
| ENSMUSG00000051391 | Ywhag | protein_coding | 0.02881379 | 9.147746 | 0.0053688 | 0.9424809 | 0.999176 |
| ENSMUSG00000029613 | Eif2ak1 | protein_coding | 0.03560302 | 4.672484 | 0.0053548 | 0.9425555 | 0.999176 |
| ENSMUSG00000032314 | Etfa | protein_coding | 0.02113982 | 6.758427 | 0.0053117 | 0.9427869 | 0.999176 |
| ENSMUSG00000009549 | Srp14 | protein_coding | -0.02362017 | 9.519047 | 0.0053077 | 0.9428085 | 0.999176 |
| ENSMUSG00000040390 | Map3k10 | protein_coding | -0.07863087 | 3.157233 | 0.0052683 | 0.9430221 | 0.999176 |
| ENSMUSG00000089687 | Rab42 | protein_coding | 0.06638817 | 2.882036 | 0.0052674 | 0.9430253 | 0.999176 |
| ENSMUSG00000015944 | Gatsl2 | protein_coding | 0.02403292 | 6.85164 | 0.0052589 | 0.9430717 | 0.999176 |
| ENSMUSG00000056749 | Nfil3 | protein_coding | 0.05582071 | 3.514817 | 0.0052344 | 0.9432042 | 0.999176 |
| ENSMUSG00000031749 | St3gal2 | protein_coding | 0.03164959 | 5.079419 | 0.0052279 | 0.9432391 | 0.999176 |
| ENSMUSG00000024667 | Tmem216 | protein_coding | -0.03048194 | 5.05072 | 0.0051884 | 0.9434539 | 0.999176 |
| ENSMUSG00000085396 | Firre | processed_transcript | -0.05105983 | 3.688051 | 0.00518 | 0.9434994 | 0.999176 |
| ENSMUSG00000104283 | Gm37459 | antisense_RNA | -0.03897656 | 4.559866 | 0.0051774 | 0.9435135 | 0.999176 |
| ENSMUSG00000026110 | Mgat4a | protein_coding | -0.03892451 | 4.36464 | 0.005168 | 0.9435646 | 0.999176 |
| ENSMUSG00000079179 | Rab10os | antisense_RNA | 0.0316656 | 6.31361 | 0.0051667 | 0.9435732 | 0.999176 |
| ENSMUSG00000034064 | Poglut1 | protein_coding | -0.02914914 | 6.366769 | 0.0051629 | 0.9435924 | 0.999176 |
| ENSMUSG00000006800 | Sulf2 | protein_coding | -0.03491967 | 5.070675 | 0.0051418 | 0.9437077 | 0.999176 |
| ENSMUSG00000044847 | Lsm11 | protein_coding | -0.0628386 | 3.172878 | 0.0051256 | 0.9437965 | 0.999176 |
| ENSMUSG00000072235 | Tuba1a | protein_coding | -0.02724541 | 9.916995 | 0.0051076 | 0.9438946 | 0.999176 |
| ENSMUSG00000026663 | Atf6 | protein_coding | -0.03753487 | 4.068018 | 0.0051032 | 0.9439191 | 0.999176 |
| ENSMUSG00000034109 | Golim4 | protein_coding | -0.09409171 | 2.425925 | 0.0050927 | 0.9439769 | 0.999176 |
| ENSMUSG00000078861 | Zfp931 | protein_coding | 0.04298076 | 5.300697 | 0.0050816 | 0.9440411 | 0.999176 |
| ENSMUSG00000043542 | Zc2hc1a | protein_coding | 0.02154795 | 7.16744 | 0.0050643 | 0.9441325 | 0.999176 |
| ENSMUSG00000030678 | Maz | protein_coding | 0.04090782 | 4.644659 | 0.0050486 | 0.9442194 | 0.999176 |
| ENSMUSG00000035735 | Dagla | protein_coding | -0.05056002 | 3.672132 | 0.0050449 | 0.9442395 | 0.999176 |
| ENSMUSG00000035469 | Rcbtb1 | protein_coding | -0.03606375 | 5.131448 | 0.005015 | 0.9444049 | 0.999176 |
| ENSMUSG00000045031 | Cetn4 | protein_coding | 0.0424015 | 3.838305 | 0.0050009 | 0.9444831 | 0.999176 |
| ENSMUSG00000005357 | Slc1a6 | protein_coding | 0.04179863 | 4.159537 | 0.0049923 | 0.9445308 | 0.999176 |
| ENSMUSG00000027852 | Nras | protein_coding | 0.0200638 | 7.041033 | 0.0049916 | 0.9445347 | 0.999176 |
| ENSMUSG00000024516 | Sec11c | protein_coding | 0.02830451 | 7.690143 | 0.0049837 | 0.9445796 | 0.999176 |
| ENSMUSG00000033419 | Snap91 | protein_coding | 0.02242948 | 7.467942 | 0.0049592 | 0.9447147 | 0.999176 |
| ENSMUSG00000024646 | Cyb5a | protein_coding | -0.02500881 | 8.284304 | 0.0049333 | 0.9448589 | 0.999176 |
| ENSMUSG00000040112 | Mrps35 | protein_coding | 0.04103443 | 4.382661 | 0.0049296 | 0.9448793 | 0.999176 |
| ENSMUSG00000051811 | Cox6b2 | protein_coding | -0.03906222 | 4.008247 | 0.0048883 | 0.9451105 | 0.999176 |
| ENSMUSG00000090626 | Tex9 | protein_coding | 0.04687249 | 3.860998 | 0.0048879 | 0.9451128 | 0.999176 |
| ENSMUSG00000015889 | Lta4h | protein_coding | 0.02537369 | 6.22151 | 0.0048826 | 0.9451422 | 0.999176 |
| ENSMUSG00000024081 | Cebpz | protein_coding | 0.03145073 | 5.211814 | 0.0048587 | 0.9452763 | 0.999176 |
| ENSMUSG00000018143 | Mafk | protein_coding | -0.06034813 | 3.372681 | 0.0048444 | 0.9453573 | 0.999176 |
| ENSMUSG00000002058 | Unc119 | protein_coding | 0.03818808 | 4.628577 | 0.0048362 | 0.9454032 | 0.999176 |
| ENSMUSG00000032265 | Fam46a | protein_coding | 0.04172418 | 3.586635 | 0.0048313 | 0.9454307 | 0.999176 |
| ENSMUSG00000042826 | Fgf11 | protein_coding | 0.05024718 | 3.445099 | 0.0048308 | 0.9454337 | 0.999176 |
| ENSMUSG00000015568 | Lpl | protein_coding | -0.05409767 | 3.603553 | 0.0048259 | 0.9454614 | 0.999176 |
| ENSMUSG00000033933 | Vhl | protein_coding | -0.03454539 | 5.045703 | 0.0047873 | 0.9456797 | 0.999176 |
| ENSMUSG00000023328 | Ache | protein_coding | 0.02339307 | 6.236612 | 0.0047845 | 0.9456953 | 0.999176 |
| ENSMUSG00000051343 | Rab11fip5 | protein_coding | -0.03096874 | 5.378459 | 0.0047734 | 0.9457582 | 0.999176 |
| ENSMUSG00000052160 | Pld4 | protein_coding | -0.06265931 | 3.550459 | 0.0047501 | 0.945892 | 0.999176 |
| ENSMUSG00000038295 | Atg9b | protein_coding | -0.03985928 | 4.819288 | 0.0047304 | 0.9460025 | 0.999176 |
| ENSMUSG00000031029 | Eif3f | protein_coding | 0.02677312 | 6.45489 | 0.0047173 | 0.9460772 | 0.999176 |
| ENSMUSG00000028443 | Nudt2 | protein_coding | 0.02216142 | 6.671103 | 0.0046829 | 0.9462738 | 0.999176 |
| ENSMUSG00000020986 | Sec23a | protein_coding | 0.04949725 | 4.691122 | 0.0046855 | 0.9462872 | 0.999176 |
| ENSMUSG00000031378 | Abcd1 | protein_coding | 0.07090881 | 2.850826 | 0.0046742 | 0.9463237 | 0.999176 |
| ENSMUSG00000032518 | Rpsa | protein_coding | 0.04094182 | 3.711441 | 0.0046697 | 0.9463499 | 0.999176 |
| ENSMUSG00000040818 | Dennd6a | protein_coding | -0.04104571 | 3.729117 | 0.0046605 | 0.9464027 | 0.999176 |
| ENSMUSG00000069917 | Hba-a2 | protein_coding | -0.04942571 | 3.810028 | 0.0046502 | 0.9464616 | 0.999176 |
| ENSMUSG00000027193 | Api5 | protein_coding | -0.0253856 | 5.813266 | 0.0046327 | 0.9465623 | 0.999176 |
| ENSMUSG00000036555 | Iqce | protein_coding | 0.04297574 | 4.140155 | 0.0046216 | 0.9466261 | 0.999176 |
| ENSMUSG00000097310 | A930038B10Rik | lincRNA | -0.08651075 | 2.32461 | 0.0046025 | 0.9467366 | 0.999176 |
| ENSMUSG00000039057 | Myo16 | protein_coding | 0.0695518 | 2.929518 | 0.0045972 | 0.9467671 | 0.999176 |

| ensembl_gene_id | external_gene_name | gene_biotype | log2 (fold change) | log2 (counts per million) | F-value | p-value | False discovery rate |
|---|---|---|---|---|---|---|---|
| ENSMUSG00000030823 | 9130019O22Rik | protein_coding | -0.1090277 | 1.986 | 0.0045749 | 0.9468964 | 0.999176 |
| ENSMUSG00000020630 | Rnaseh1 | protein_coding | -0.03812038 | 4.392954 | 0.004569 | 0.9469306 | 0.999176 |
| ENSMUSG00000034321 | Exosc1 | protein_coding | 0.02643105 | 6.375685 | 0.0045565 | 0.9470029 | 0.999176 |
| ENSMUSG00000090015 | Gm15446 | protein_coding | 0.06597134 | 3.140717 | 0.0045365 | 0.947119 | 0.999176 |
| ENSMUSG00000006711 | D130043K22Rik | protein_coding | 0.05681355 | 2.919162 | 0.0045298 | 0.9471579 | 0.999176 |
| ENSMUSG00000023067 | Cdkn1a | protein_coding | -0.05994991 | 4.338982 | 0.0045231 | 0.9472787 | 0.999176 |
| ENSMUSG00000040724 | Kcna2 | protein_coding | -0.03230231 | 5.230898 | 0.0044886 | 0.9473987 | 0.999176 |
| ENSMUSG00000051790 | Nlgn2 | protein_coding | -0.02394252 | 6.043379 | 0.004485 | 0.9474195 | 0.999176 |
| ENSMUSG00000031155 | Pim2 | protein_coding | 0.04611438 | 5.437067 | 0.0044957 | 0.9474255 | 0.999176 |
| ENSMUSG00000097571 | Jpx | lincRNA | -0.06662977 | 2.721412 | 0.0044645 | 0.9475398 | 0.999176 |
| ENSMUSG00000009995 | Taz | protein_coding | 0.03330468 | 4.485729 | 0.0044218 | 0.9477908 | 0.999176 |
| ENSMUSG00000109321 | A330076H08Rik | processed_transcript | 0.0411277 | 3.772798 | 0.0043956 | 0.9479458 | 0.999176 |
| ENSMUSG00000019872 | Smpdl3a | protein_coding | -0.03521024 | 4.45535 | 0.0043475 | 0.9482307 | 0.999176 |
| ENSMUSG00000029647 | Pan3 | protein_coding | 0.03346712 | 4.615411 | 0.0043474 | 0.9482315 | 0.999176 |
| ENSMUSG00000038026 | Kcnj9 | protein_coding | -0.05275437 | 4.98654 | 0.0043609 | 0.9482504 | 0.999176 |
| ENSMUSG00000033728 | Lrrc14 | protein_coding | 0.03634359 | 4.191297 | 0.0043437 | 0.9482533 | 0.999176 |
| ENSMUSG00000047635 | 2810006K23Rik | protein_coding | 0.03031907 | 5.227096 | 0.0043383 | 0.9482852 | 0.999176 |
| ENSMUSG00000032020 | Ubash3b | protein_coding | -0.03987692 | 6.093082 | 0.004349 | 0.9483018 | 0.999176 |
| ENSMUSG00000001942 | Siae | protein_coding | 0.03395676 | 4.62853 | 0.0043088 | 0.9484611 | 0.999176 |
| ENSMUSG00000025823 | Pdia4 | protein_coding | -0.0246966 | 5.920512 | 0.0042991 | 0.948519 | 0.999176 |
| ENSMUSG00000045435 | Tmem60 | protein_coding | -0.02500125 | 7.072074 | 0.0042889 | 0.9485804 | 0.999176 |
| ENSMUSG00000033565 | Rbfox2 | protein_coding | 0.02018043 | 6.285509 | 0.0042799 | 0.9486342 | 0.999176 |
| ENSMUSG00000028917 | Plekhm2 | protein_coding | 0.05031831 | 3.915621 | 0.0042729 | 0.9486774 | 0.999176 |
| ENSMUSG00000029578 | Wipi2 | protein_coding | -0.02055156 | 6.482686 | 0.0042509 | 0.9488084 | 0.999176 |
| ENSMUSG00000020170 | Frs2 | protein_coding | -0.03431228 | 4.047673 | 0.0042331 | 0.948915 | 0.999176 |
| ENSMUSG00000095930 | Nim1k | protein_coding | -0.02725226 | 5.445213 | 0.0042301 | 0.9489333 | 0.999176 |
| ENSMUSG00000049164 | Zfp518a | protein_coding | -0.05261506 | 2.828007 | 0.0042131 | 0.9490358 | 0.999176 |
| ENSMUSG00000032180 | Tmed1 | protein_coding | 0.02571589 | 5.381311 | 0.0042008 | 0.9491104 | 0.999176 |
| ENSMUSG00000024892 | Pcx | protein_coding | -0.02670469 | 5.608768 | 0.0041906 | 0.9491723 | 0.999176 |
| ENSMUSG00000037818 | Abhd18 | protein_coding | 0.04952838 | 3.236697 | 0.0041804 | 0.9492338 | 0.999176 |
| ENSMUSG00000054079 | Utp18 | protein_coding | 0.03307861 | 4.102257 | 0.0041726 | 0.9492809 | 0.999176 |
| ENSMUSG00000030257 | Srgap3 | protein_coding | 0.02468087 | 6.169413 | 0.0041433 | 0.9494595 | 0.999176 |
| ENSMUSG00000021706 | Zfyve16 | protein_coding | 0.04711689 | 3.227398 | 0.0041265 | 0.949562 | 0.999176 |
| ENSMUSG00000007646 | Rad51c | protein_coding | -0.05704777 | 3.20479 | 0.0041104 | 0.9496598 | 0.999176 |
| ENSMUSG00000073702 | Rpl31 | protein_coding | 0.03744455 | 3.938346 | 0.0040635 | 0.9499476 | 0.999176 |
| ENSMUSG00000048794 | Cfap100 | protein_coding | 0.07310923 | 2.308253 | 0.0040552 | 0.949999 | 0.999176 |
| ENSMUSG00000025133 | Ints4 | protein_coding | -0.03407061 | 4.854162 | 0.0040545 | 0.9500032 | 0.999176 |
| ENSMUSG00000002812 | Flii | protein_coding | -0.03716178 | 5.044575 | 0.0040435 | 0.950072 | 0.999176 |
| ENSMUSG00000040389 | Wdr47 | protein_coding | 0.0238172 | 6.1562 | 0.0040331 | 0.9501347 | 0.999176 |
| ENSMUSG00000024276 | Zfp397 | protein_coding | -0.03987766 | 4.089526 | 0.0040319 | 0.9501423 | 0.999176 |
| ENSMUSG00000028854 | Slc9a1 | protein_coding | -0.03399521 | 4.56503 | 0.0040079 | 0.9502908 | 0.999176 |
| ENSMUSG00000030082 | Sec61a1 | protein_coding | 0.027624 | 4.915063 | 0.0040075 | 0.9502932 | 0.999176 |
| ENSMUSG00000063904 | 4930533O14Rik | protein_coding | 0.03772522 | 5.120789 | 0.0040052 | 0.9503089 | 0.999176 |
| ENSMUSG00000068250 | Amn1 | protein_coding | 0.02590652 | 5.656344 | 0.0040006 | 0.9503359 | 0.999176 |
| ENSMUSG00000043241 | Upf2 | protein_coding | 0.0341929 | 5.165862 | 0.0039902 | 0.9504008 | 0.999176 |
| ENSMUSG00000000184 | Ccnd2 | protein_coding | -0.03029313 | 5.165602 | 0.0039729 | 0.9505078 | 0.999176 |
| ENSMUSG00000038429 | Usp5 | protein_coding | 0.03252837 | 8.40786 | 0.003983 | 0.9505217 | 0.999176 |
| ENSMUSG00000004319 | Clcn3 | protein_coding | -0.01846703 | 7.199468 | 0.003937 | 0.9507318 | 0.999176 |
| ENSMUSG00000028128 | F3 | protein_coding | -0.03576215 | 5.787862 | 0.0039038 | 0.9509717 | 0.999176 |
| ENSMUSG00000025190 | Got1 | protein_coding | -0.02558531 | 9.60611 | 0.0038752 | 0.951121 | 0.999176 |
| ENSMUSG00000034551 | Hdx | protein_coding | -0.09092679 | 2.241437 | 0.0038713 | 0.9511441 | 0.999176 |
| ENSMUSG00000041762 | Gpr155 | protein_coding | -0.042103 | 4.032146 | 0.0038638 | 0.9511914 | 0.999176 |
| ENSMUSG00000048807 | Slc35e4 | protein_coding | -0.03569753 | 4.387236 | 0.0038612 | 0.9512078 | 0.999176 |
| ENSMUSG00000035011 | Zbtb7a | protein_coding | 0.03038044 | 5.645805 | 0.003852 | 0.951266 | 0.999176 |
| ENSMUSG00000032477 | Cdc25a | protein_coding | -0.06227639 | 3.205353 | 0.0038485 | 0.9512892 | 0.999176 |
| ENSMUSG00000063804 | Lin28b | protein_coding | 0.0483608 | 3.228588 | 0.0038452 | 0.9513088 | 0.999176 |
| ENSMUSG00000005501 | Usp40 | protein_coding | 0.05199692 | 2.889454 | 0.0038394 | 0.9513455 | 0.999176 |
| ENSMUSG00000033429 | Mcee | protein_coding | 0.02250523 | 6.651822 | 0.0038374 | 0.951358 | 0.999176 |
| ENSMUSG00000032246 | Calml4 | protein_coding | 0.05366025 | 3.876237 | 0.0038267 | 0.9514272 | 0.999176 |
| ENSMUSG00000052962 | Mrpl35 | protein_coding | -0.01678687 | 7.399186 | 0.0037821 | 0.9517097 | 0.999176 |
| ENSMUSG00000017686 | Rhot1 | protein_coding | 0.02737832 | 5.268119 | 0.0037701 | 0.9517858 | 0.999176 |
| ENSMUSG00000036840 | Siah1a | protein_coding | 0.03850934 | 3.974478 | 0.003762 | 0.9518379 | 0.999176 |
| ENSMUSG00000028138 | Adh5 | protein_coding | 0.01781238 | 7.589361 | 0.0037518 | 0.9519029 | 0.999176 |
| ENSMUSG00000059939 | 9430015G10Rik | protein_coding | 0.03734127 | 3.558765 | 0.0037495 | 0.9519179 | 0.999176 |
| ENSMUSG00000037060 | Cavin3 | protein_coding | -0.0268431 | 5.077329 | 0.0037487 | 0.9519231 | 0.999176 |
| ENSMUSG00000028675 | Pnrc2 | protein_coding | 0.02706955 | 6.454868 | 0.0037341 | 0.9520178 | 0.999176 |
| ENSMUSG00000063236 | 1110038F14Rik | protein_coding | -0.02539536 | 5.6278 | 0.0037189 | 0.9521142 | 0.999176 |
| ENSMUSG00000034850 | Tmem127 | protein_coding | 0.02961359 | 6.472458 | 0.0037126 | 0.9521775 | 0.999176 |
| ENSMUSG00000046709 | Mapk10 | protein_coding | 0.02171154 | 8.463617 | 0.0036859 | 0.9523266 | 0.999176 |
| ENSMUSG00000015672 | Mrpl32 | protein_coding | 0.02301139 | 6.614622 | 0.0036784 | 0.9523751 | 0.999176 |
| ENSMUSG00000028078 | Dclk2 | protein_coding | -0.02559337 | 4.942542 | 0.003662 | 0.9524814 | 0.999176 |
| ENSMUSG00000073982 | Rhog | protein_coding | 0.02498971 | 6.224358 | 0.0036328 | 0.9526707 | 0.999176 |
| ENSMUSG00000035215 | Lsm7 | protein_coding | 0.03673931 | 3.81251 | 0.0036294 | 0.9526931 | 0.999176 |
| ENSMUSG00000028532 | Cachd1 | protein_coding | -0.03923129 | 3.756837 | 0.0036226 | 0.952737 | 0.999176 |
| ENSMUSG00000033454 | Zbtb1 | protein_coding | 0.06083837 | 2.644817 | 0.0036134 | 0.9527975 | 0.999176 |
| ENSMUSG00000024897 | Apba1 | protein_coding | 0.02635543 | 5.352297 | 0.0036109 | 0.9528135 | 0.999176 |
| ENSMUSG00000050379 | 6-Sep | protein_coding | 0.02827307 | 6.389172 | 0.0036042 | 0.9528606 | 0.999176 |
| ENSMUSG00000038510 | Rpf2 | protein_coding | -0.02031126 | 6.323683 | 0.0035901 | 0.9529495 | 0.999176 |

| ensembl_gene_id | external_gene_name | gene_biotype | log2 (fold change) | log2 (counts per million) | F-value | p-value | False discovery rate |
|---|---|---|---|---|---|---|---|
| ENSMUSG00000042682 | Selenok | protein_coding | 0.02127666 | 9.326717 | 0.0035574 | 0.9531641 | 0.999176 |
| ENSMUSG00000032498 | Mlh1 | protein_coding | 0.05332402 | 2.910645 | 0.0035491 | 0.9532188 | 0.999176 |
| ENSMUSG00000037217 | Syn1 | protein_coding | 0.02194825 | 6.267628 | 0.0035465 | 0.9532357 | 0.999176 |
| ENSMUSG00000027502 | Rtfdc1 | protein_coding | -0.01967933 | 7.356181 | 0.0035206 | 0.9534068 | 0.999176 |
| ENSMUSG00000085316 | D330050G23Rik | lincRNA | 0.07609937 | 2.556195 | 0.0034995 | 0.9535463 | 0.999176 |
| ENSMUSG00000027665 | Pik3ca | protein_coding | -0.02371857 | 5.507951 | 0.0034465 | 0.953899 | 0.999176 |
| ENSMUSG00000024077 | Strn | protein_coding | 0.02051608 | 6.419243 | 0.0034413 | 0.9539338 | 0.999176 |
| ENSMUSG00000057666 | Gapdh | protein_coding | -0.03196402 | 4.429048 | 0.0034237 | 0.9540514 | 0.999176 |
| ENSMUSG00000025486 | Sirt3 | protein_coding | -0.01874827 | 7.720723 | 0.0034131 | 0.9541227 | 0.999176 |
| ENSMUSG00000026439 | Rbbp5 | protein_coding | 0.02650891 | 5.23177 | 0.0034106 | 0.9541392 | 0.999176 |
| ENSMUSG00000032641 | Gpr19 | protein_coding | -0.0208134 | 5.7969 | 0.0033893 | 0.9542827 | 0.999176 |
| ENSMUSG00000025138 | Sirt7 | protein_coding | -0.05180189 | 3.093682 | 0.0033677 | 0.9544283 | 0.999176 |
| ENSMUSG00000019883 | Echdc1 | protein_coding | 0.03720599 | 3.896635 | 0.0033658 | 0.9544416 | 0.999176 |
| ENSMUSG00000020255 | D10Wsu102e | protein_coding | -0.02979629 | 5.333206 | 0.0033512 | 0.9545402 | 0.999176 |
| ENSMUSG00000062778 | Chia1 | protein_coding | -0.09100437 | 2.15256 | 0.003349 | 0.9545551 | 0.999176 |
| ENSMUSG00000036975 | Tmem177 | protein_coding | -0.0380983 | 3.831928 | 0.003328 | 0.9546972 | 0.999176 |
| ENSMUSG00000073680 | Tmem88b | protein_coding | 0.02896944 | 5.823505 | 0.0033281 | 0.9546982 | 0.999176 |
| ENSMUSG00000024878 | Cbwd1 | protein_coding | 0.02586584 | 4.853471 | 0.0032929 | 0.9549366 | 0.999176 |
| ENSMUSG00000025545 | Clybl | protein_coding | -0.017265 | 7.084662 | 0.0032735 | 0.9550694 | 0.999176 |
| ENSMUSG00000040414 | Slc25a28 | protein_coding | -0.03611789 | 4.194077 | 0.0032717 | 0.9550819 | 0.999176 |
| ENSMUSG00000038267 | Slc22a23 | protein_coding | -0.02763433 | 4.568708 | 0.0032698 | 0.9550951 | 0.999176 |
| ENSMUSG00000035495 | Tstd2 | protein_coding | 0.03968604 | 3.222865 | 0.0032684 | 0.9551047 | 0.999176 |
| ENSMUSG00000024218 | Taf11 | protein_coding | -0.01714996 | 6.628796 | 0.0032677 | 0.9551092 | 0.999176 |
| ENSMUSG00000018861 | Fdxr | protein_coding | 0.03582046 | 3.581502 | 0.0032605 | 0.9551585 | 0.999176 |
| ENSMUSG00000028991 | Mtor | protein_coding | -0.02913263 | 4.714339 | 0.0032586 | 0.9551718 | 0.999176 |
| ENSMUSG00000023723 | Mrps23 | protein_coding | 0.01774478 | 6.937091 | 0.003254 | 0.9552033 | 0.999176 |
| ENSMUSG00000002279 | Lmf1 | protein_coding | 0.03461633 | 4.875749 | 0.0032539 | 0.9552054 | 0.999176 |
| ENSMUSG00000055013 | Agap1 | protein_coding | -0.01977068 | 6.38573 | 0.0032514 | 0.9552214 | 0.999176 |
| ENSMUSG00000049804 | Armcx4 | protein_coding | 0.04087749 | 4.416622 | 0.0032455 | 0.9552627 | 0.999176 |
| ENSMUSG00000038784 | Cnot4 | protein_coding | -0.0223852 | 6.426545 | 0.0032392 | 0.955305 | 0.999176 |
| ENSMUSG00000017843 | Ppp2r5c | protein_coding | -0.01856891 | 7.854451 | 0.0032373 | 0.9553181 | 0.999176 |
| ENSMUSG00000008822 | Acyp1 | protein_coding | 0.01867681 | 6.363257 | 0.0032171 | 0.9554578 | 0.999176 |
| ENSMUSG00000042804 | Gpr153 | protein_coding | -0.0368476 | 3.871507 | 0.0031911 | 0.9556383 | 0.999176 |
| ENSMUSG00000030029 | Lrig1 | protein_coding | 0.04230216 | 3.127579 | 0.0031882 | 0.9556582 | 0.999176 |
| ENSMUSG00000034377 | Tulp4 | protein_coding | -0.02119653 | 5.511358 | 0.0031832 | 0.9556931 | 0.999176 |
| ENSMUSG00000020990 | Cdkl1 | protein_coding | 0.03093884 | 4.588057 | 0.0031824 | 0.9556982 | 0.999176 |
| ENSMUSG00000023572 | Ccndbp1 | protein_coding | -0.01750406 | 7.556442 | 0.0031823 | 0.955699 | 0.999176 |
| ENSMUSG00000020032 | Nuak1 | protein_coding | 0.02873763 | 4.065983 | 0.0031394 | 0.9559984 | 0.999176 |
| ENSMUSG00000033904 | Ccp110 | protein_coding | -0.02214759 | 5.276147 | 0.0031325 | 0.9560471 | 0.999176 |
| ENSMUSG00000032555 | Topbp1 | protein_coding | 0.07365497 | 2.651248 | 0.0031039 | 0.9562489 | 0.999176 |
| ENSMUSG00000019732 | Calr3 | protein_coding | -0.07267123 | 3.324885 | 0.0031081 | 0.9563127 | 0.999176 |
| ENSMUSG00000060681 | Slc9a6 | protein_coding | -0.02089172 | 5.876186 | 0.0030785 | 0.9564271 | 0.999176 |
| ENSMUSG00000031299 | Pdha1 | protein_coding | -0.03289817 | 7.285463 | 0.0030901 | 0.9564524 | 0.999176 |
| ENSMUSG00000005907 | Pex1 | protein_coding | 0.04947096 | 3.524649 | 0.0030531 | 0.9566076 | 0.999176 |
| ENSMUSG00000062822 | 4833420G17Rik | protein_coding | 0.02506901 | 5.39347 | 0.0030274 | 0.9567899 | 0.999176 |
| ENSMUSG00000021537 | Cetn3 | protein_coding | -0.02171146 | 7.856869 | 0.0030259 | 0.9568012 | 0.999176 |
| ENSMUSG00000029776 | Hibadh | protein_coding | -0.02132788 | 6.256325 | 0.0030206 | 0.9568377 | 0.999176 |
| ENSMUSG00000021548 | Ccnh | protein_coding | -0.01670293 | 7.141782 | 0.0030135 | 0.9568884 | 0.999176 |
| ENSMUSG00000055782 | Abcd2 | protein_coding | -0.0421427 | 3.920591 | 0.0029955 | 0.9570185 | 0.999176 |
| ENSMUSG00000022672 | Prkdc | protein_coding | -0.07270602 | 2.643443 | 0.0029947 | 0.9570243 | 0.999176 |
| ENSMUSG00000084910 | C630043F03Rik | processed_transcript | -0.0350851 | 3.895248 | 0.0029899 | 0.957058 | 0.999176 |
| ENSMUSG00000018428 | Akap1 | protein_coding | -0.04037528 | 3.861081 | 0.002987 | 0.9570784 | 0.999176 |
| ENSMUSG00000032657 | Fam189b | protein_coding | 0.03328516 | 3.935315 | 0.0029809 | 0.9571221 | 0.999176 |
| ENSMUSG00000054667 | Irs4 | protein_coding | -0.03190519 | 4.720076 | 0.002965 | 0.9572371 | 0.999176 |
| ENSMUSG00000025743 | Sdc3 | protein_coding | 0.0394274 | 3.22556 | 0.0029456 | 0.9573768 | 0.999176 |
| ENSMUSG00000028467 | Gba2 | protein_coding | -0.02137378 | 5.388188 | 0.0029397 | 0.9574194 | 0.999176 |
| ENSMUSG00000022664 | Slc35a5 | protein_coding | 0.02244896 | 5.115607 | 0.0029358 | 0.9574477 | 0.999176 |
| ENSMUSG00000079055 | Slc8a3 | protein_coding | 0.03079124 | 4.08651 | 0.0029085 | 0.9576456 | 0.999176 |
| ENSMUSG00000051934 | Spats2 | protein_coding | -0.02449684 | 4.648541 | 0.0028942 | 0.9577496 | 0.999176 |
| ENSMUSG00000039936 | Pik3cd | protein_coding | -0.05701759 | 2.828825 | 0.0028455 | 0.9581065 | 0.999176 |
| ENSMUSG00000029287 | Tgfbr3 | protein_coding | 0.06685349 | 2.92535 | 0.0028394 | 0.9581553 | 0.999176 |
| ENSMUSG00000028614 | Ndc1 | protein_coding | 0.05383471 | 3.068111 | 0.0028256 | 0.958253 | 0.999176 |
| ENSMUSG00000031072 | Oraov1 | protein_coding | 0.02204787 | 5.31626 | 0.0028176 | 0.9583122 | 0.999176 |
| ENSMUSG00000066357 | Wdr6 | protein_coding | 0.04375389 | 7.794394 | 0.0028335 | 0.9585233 | 0.999176 |
| ENSMUSG00000019888 | Mgat4c | protein_coding | 0.03057458 | 4.655285 | 0.0027763 | 0.9586183 | 0.999176 |
| ENSMUSG00000078584 | AU022252 | protein_coding | 0.02470034 | 4.762003 | 0.0027687 | 0.9586755 | 0.999176 |
| ENSMUSG00000060950 | Trmt61a | protein_coding | -0.03640103 | 3.441034 | 0.0027636 | 0.9587131 | 0.999176 |
| ENSMUSG00000051224 | Tceanc | protein_coding | -0.05422898 | 2.642795 | 0.0027617 | 0.9587272 | 0.999176 |
| ENSMUSG00000001569 | Nom1 | protein_coding | -0.02642409 | 4.537748 | 0.0027607 | 0.9587347 | 0.999176 |
| ENSMUSG00000024982 | Zdhhc6 | protein_coding | 0.02274629 | 4.825079 | 0.0027407 | 0.9588844 | 0.999176 |
| ENSMUSG00000055897 | Ppp4r1l-ps | transcribed_unproces | 0.03626538 | 3.477519 | 0.0027384 | 0.9589019 | 0.999176 |
| ENSMUSG00000016256 | Ctsz | protein_coding | -0.02254265 | 6.410511 | 0.0027284 | 0.9589776 | 0.999176 |
| ENSMUSG00000036580 | Spg20 | protein_coding | -0.02339674 | 5.385816 | 0.0027175 | 0.9590586 | 0.999176 |
| ENSMUSG00000100618 | Gm29595 | lincRNA | -0.05193757 | 2.59172 | 0.0027172 | 0.9590606 | 0.999176 |
| ENSMUSG00000079427 | Mthfsl | protein_coding | 0.02855135 | 3.962724 | 0.0027092 | 0.9591209 | 0.999176 |
| ENSMUSG00000025616 | Usp16 | protein_coding | -0.02453637 | 4.813608 | 0.0026819 | 0.9593271 | 0.999176 |
| ENSMUSG00000029650 | Slc46a3 | protein_coding | 0.04691132 | 2.794713 | 0.0026757 | 0.9593748 | 0.999176 |
| ENSMUSG00000049511 | Htr1b | protein_coding | 0.03972208 | 3.790007 | 0.0026581 | 0.9595079 | 0.999176 |

| ensembl_gene_id | external_gene_name | gene_biotype | log2 (fold change) | log2 (counts per million) | F-value | p-value | False discovery rate |
|---|---|---|---|---|---|---|---|
| ENSMUSG00000017548 | Suz12 | protein_coding | -0.02551013 | 4.468042 | 0.0026519 | 0.9595551 | 0.999176 |
| ENSMUSG00000035967 | Ints6l | protein_coding | 0.05000422 | 2.801393 | 0.0026428 | 0.9596247 | 0.999176 |
| ENSMUSG00000008604 | Ubqln4 | protein_coding | -0.01988042 | 5.605717 | 0.0026414 | 0.9596357 | 0.999176 |
| ENSMUSG00000079197 | Psme2 | protein_coding | 0.0168191 | 6.05263 | 0.0026046 | 0.959917 | 0.999176 |
| ENSMUSG00000026452 | Syt2 | protein_coding | 0.0266834 | 4.272955 | 0.0025822 | 0.9600895 | 0.999176 |
| ENSMUSG00000064372 | mt-Tp | Mt_tRNA | 0.03246195 | 4.066601 | 0.0025738 | 0.9601546 | 0.999176 |
| ENSMUSG00000059401 | Mamld1 | protein_coding | 0.02818868 | 4.822807 | 0.0025384 | 0.9604297 | 0.999176 |
| ENSMUSG00000071533 | Pcnp | protein_coding | -0.01484038 | 8.028674 | 0.0025369 | 0.9604416 | 0.999176 |
| ENSMUSG00000056076 | Eif3b | protein_coding | -0.01891019 | 5.889642 | 0.0025286 | 0.9605061 | 0.999176 |
| ENSMUSG00000091002 | Tcerg1l | protein_coding | 0.03087268 | 4.286746 | 0.0025184 | 0.9605854 | 0.999176 |
| ENSMUSG00000037465 | Klf10 | protein_coding | 0.02981295 | 3.896626 | 0.0025076 | 0.9606703 | 0.999176 |
| ENSMUSG00000032504 | Pdcd6ip | protein_coding | 0.02392628 | 5.101561 | 0.0024963 | 0.9607584 | 0.999176 |
| ENSMUSG00000020099 | Unc5b | protein_coding | 0.03022037 | 4.237964 | 0.0024931 | 0.9607835 | 0.999176 |
| ENSMUSG00000090523 | Gypc | protein_coding | -0.07874668 | 1.789288 | 0.0024745 | 0.9609302 | 0.999176 |
| ENSMUSG00000026103 | Gls | protein_coding | -0.02150441 | 6.054019 | 0.0024486 | 0.9611351 | 0.999176 |
| ENSMUSG00000036850 | Mrpl41 | protein_coding | 0.01383128 | 8.315884 | 0.0024276 | 0.9613017 | 0.999176 |
| ENSMUSG00000017639 | Rab11fip4 | protein_coding | 0.02128949 | 7.063124 | 0.0024254 | 0.9613235 | 0.999176 |
| ENSMUSG00000021541 | Trpc7 | protein_coding | -0.02283353 | 5.123096 | 0.0024034 | 0.9614949 | 0.999176 |
| ENSMUSG00000038244 | Mical2 | protein_coding | 0.03830088 | 3.840898 | 0.0023987 | 0.9615339 | 0.999176 |
| ENSMUSG00000014550 | Rbsn | protein_coding | -0.0217662 | 5.928369 | 0.0023791 | 0.9616906 | 0.999176 |
| ENSMUSG00000024347 | Psd2 | protein_coding | 0.03413785 | 4.892577 | 0.0023793 | 0.9617134 | 0.999176 |
| ENSMUSG00000072946 | Ptgr2 | protein_coding | -0.02652724 | 5.058398 | 0.0023616 | 0.961831 | 0.999176 |
| ENSMUSG00000059142 | Zfp945 | protein_coding | -0.02439055 | 4.555714 | 0.0023585 | 0.9618563 | 0.999176 |
| ENSMUSG00000038525 | Armc10 | protein_coding | 0.03770052 | 3.275491 | 0.0023551 | 0.9618839 | 0.999176 |
| ENSMUSG00000041528 | Rnf123 | protein_coding | -0.06673634 | 2.638792 | 0.0023539 | 0.9618943 | 0.999176 |
| ENSMUSG00000056962 | Jmjd6 | protein_coding | -0.01966845 | 5.364288 | 0.0023513 | 0.9619146 | 0.999176 |
| ENSMUSG00000040302 | Rbm48 | protein_coding | -0.02416075 | 4.467891 | 0.0023443 | 0.9619709 | 0.999176 |
| ENSMUSG00000032652 | Crebl2 | protein_coding | 0.02350997 | 7.062361 | 0.0023477 | 0.9619812 | 0.999176 |
| ENSMUSG00000029641 | Rasl11a | protein_coding | -0.04238691 | 3.948291 | 0.0023352 | 0.9620773 | 0.999176 |
| ENSMUSG00000023021 | Cers5 | protein_coding | -0.02265285 | 5.106722 | 0.0023146 | 0.9622128 | 0.999176 |
| ENSMUSG00000046962 | Zbtb21 | protein_coding | -0.02031512 | 5.212693 | 0.0023069 | 0.9622752 | 0.999176 |
| ENSMUSG00000044627 | Swi5 | protein_coding | -0.01707402 | 9.426211 | 0.0022849 | 0.9624559 | 0.999176 |
| ENSMUSG00000028114 | Mettl14 | protein_coding | -0.02525779 | 4.797595 | 0.0022629 | 0.9626368 | 0.999176 |
| ENSMUSG00000044285 | Gm1821 | processed_pseudoge | -0.07365121 | 5.447086 | 0.0023309 | 0.9626403 | 0.999176 |
| ENSMUSG00000038646 | Fam103a1 | protein_coding | -0.01474372 | 6.473152 | 0.0022551 | 0.9627007 | 0.999176 |
| ENSMUSG00000054976 | Nyap2 | protein_coding | -0.02445877 | 4.313599 | 0.0022428 | 0.9628025 | 0.999176 |
| ENSMUSG00000041702 | Btbd7 | protein_coding | -0.03190476 | 3.842871 | 0.0022294 | 0.9629138 | 0.999176 |
| ENSMUSG00000031791 | Tmem38a | protein_coding | 0.01369752 | 7.106816 | 0.0022068 | 0.9631024 | 0.999176 |
| ENSMUSG00000028822 | Tmem50a | protein_coding | 0.0134537 | 8.468054 | 0.0021918 | 0.9632282 | 0.999176 |
| ENSMUSG00000070738 | Dgkd | protein_coding | 0.02767596 | 4.513298 | 0.002175 | 0.9633687 | 0.999176 |
| ENSMUSG00000017421 | Zfp207 | protein_coding | 0.0153786 | 7.026282 | 0.002169 | 0.9634192 | 0.999176 |
| ENSMUSG00000025575 | Cant1 | protein_coding | -0.02763195 | 4.199755 | 0.0021686 | 0.9634231 | 0.999176 |
| ENSMUSG00000034187 | Nsf | protein_coding | 0.02373295 | 9.831543 | 0.0021455 | 0.963656 | 0.999176 |
| ENSMUSG00000021957 | Tkt | protein_coding | -0.01818559 | 8.534921 | 0.0021348 | 0.9637097 | 0.999176 |
| ENSMUSG00000029017 | Pmpcb | protein_coding | 0.01399085 | 7.100039 | 0.0021057 | 0.9639571 | 0.999176 |
| ENSMUSG00000026456 | Cyb5r1 | protein_coding | 0.02169023 | 5.561723 | 0.0020989 | 0.964015 | 0.999176 |
| ENSMUSG00000025279 | Dnase1l3 | protein_coding | -0.07681505 | 1.9818 | 0.0020914 | 0.9640793 | 0.999176 |
| ENSMUSG00000020524 | Gria1 | protein_coding | -0.01367396 | 8.109252 | 0.0020912 | 0.9640814 | 0.999176 |
| ENSMUSG00000027263 | Tubgcp4 | protein_coding | 0.03616625 | 3.016042 | 0.002083 | 0.9641515 | 0.999176 |
| ENSMUSG00000032059 | Alg9 | protein_coding | -0.02707342 | 4.222902 | 0.0020791 | 0.9641852 | 0.999176 |
| ENSMUSG00000024891 | Slc29a2 | protein_coding | 0.02468218 | 4.347253 | 0.0020752 | 0.964219 | 0.999176 |
| ENSMUSG00000073940 | Hbb-bt | protein_coding | -0.03421019 | 3.790203 | 0.0020689 | 0.9642728 | 0.999176 |
| ENSMUSG00000066129 | Kndc1 | protein_coding | -0.02404906 | 4.972632 | 0.0020456 | 0.9644748 | 0.999176 |
| ENSMUSG00000036766 | Dner | protein_coding | 0.01831587 | 6.807641 | 0.0020368 | 0.9645516 | 0.999176 |
| ENSMUSG00000044155 | Lsm8 | protein_coding | 0.01962724 | 6.069493 | 0.0020308 | 0.9646033 | 0.999176 |
| ENSMUSG00000020376 | Rnf130 | protein_coding | 0.02956634 | 3.97007 | 0.0020262 | 0.9646436 | 0.999176 |
| ENSMUSG00000058298 | Mcm9 | protein_coding | -0.03909937 | 2.917679 | 0.0020212 | 0.9646868 | 0.999176 |
| ENSMUSG00000028793 | Rnf19b | protein_coding | -0.02152112 | 4.540441 | 0.0020141 | 0.9647489 | 0.999176 |
| ENSMUSG00000037353 | Letmd1 | protein_coding | 0.02016386 | 5.638733 | 0.0020083 | 0.9647994 | 0.999176 |
| ENSMUSG00000069919 | Hba-a1 | protein_coding | -0.03165623 | 4.43709 | 0.0020049 | 0.9648306 | 0.999176 |
| ENSMUSG00000044791 | Setd2 | protein_coding | -0.02637508 | 4.536649 | 0.0020017 | 0.9648575 | 0.999176 |
| ENSMUSG00000051154 | Commd3 | protein_coding | -0.01327593 | 8.214268 | 0.0019869 | 0.9649876 | 0.999176 |
| ENSMUSG00000037868 | Egr2 | protein_coding | 0.04618627 | 2.595433 | 0.0019805 | 0.9650438 | 0.999176 |
| ENSMUSG00000024169 | Ift140 | protein_coding | -0.03698621 | 2.989982 | 0.0019682 | 0.9651525 | 0.999176 |
| ENSMUSG00000031158 | Timm17b | protein_coding | -0.01612147 | 6.777739 | 0.0019485 | 0.9653274 | 0.999176 |
| ENSMUSG00000030584 | Dpf1 | protein_coding | -0.02250012 | 4.907281 | 0.0019382 | 0.9654191 | 0.999176 |
| ENSMUSG00000018239 | Zcchc10 | protein_coding | 0.0177806 | 5.560306 | 0.0019234 | 0.9655517 | 0.999176 |
| ENSMUSG00000057147 | Dph6 | protein_coding | -0.01784783 | 5.277549 | 0.0019152 | 0.9656249 | 0.999176 |
| ENSMUSG00000034930 | Rtkn | protein_coding | 0.02208716 | 4.703299 | 0.0019107 | 0.9656653 | 0.999176 |
| ENSMUSG00000004698 | Hdac9 | protein_coding | 0.0202563 | 5.084762 | 0.0018953 | 0.9658036 | 0.999176 |
| ENSMUSG00000048706 | Lurap1l | protein_coding | 0.02302033 | 4.247336 | 0.0018944 | 0.9658113 | 0.999176 |
| ENSMUSG00000038065 | Mturn | protein_coding | -0.02085118 | 6.05953 | 0.0018943 | 0.9658132 | 0.999176 |
| ENSMUSG00000025184 | R3hcc1l | protein_coding | 0.02817481 | 3.570688 | 0.0018897 | 0.9658543 | 0.999176 |
| ENSMUSG00000022526 | Zfp251 | protein_coding | -0.02233798 | 4.270325 | 0.0018559 | 0.9661611 | 0.999176 |
| ENSMUSG00000021484 | Lman2 | protein_coding | -0.02067347 | 5.33373 | 0.001853 | 0.9661875 | 0.999176 |
| ENSMUSG00000022205 | Sub1 | protein_coding | 0.01447321 | 10.23803 | 0.0018472 | 0.9662397 | 0.999176 |
| ENSMUSG00000090546 | Cdr1 | protein_coding | 0.03622289 | 4.856865 | 0.001851 | 0.9662754 | 0.999176 |
| ENSMUSG00000026890 | Lhx6 | protein_coding | -0.01993938 | 6.00367 | 0.0018167 | 0.9665209 | 0.999176 |

| ensembl_gene_id | external_gene_name | gene_biotype | log2 (fold change) | log2 (counts per million) | F-value | p-value | False discovery rate |
|---|---|---|---|---|---|---|---|
| ENSMUSG00000042104 | Uggt2 | protein_coding | -0.02802828 | 4.359632 | 0.001803 | 0.9666465 | 0.999176 |
| ENSMUSG00000046441 | Cmtr2 | protein_coding | -0.040914 | 3.056757 | 0.0018012 | 0.9666626 | 0.999176 |
| ENSMUSG00000036086 | Zranb3 | protein_coding | -0.04757143 | 2.837932 | 0.0017987 | 0.9666856 | 0.999176 |
| ENSMUSG00000039840 | Epg5 | protein_coding | 0.04050916 | 3.388539 | 0.001782 | 0.9668416 | 0.999176 |
| ENSMUSG00000058571 | Gpc6 | protein_coding | 0.03171591 | 3.164254 | 0.0017745 | 0.9669106 | 0.999176 |
| ENSMUSG00000028550 | Atg4c | protein_coding | -0.01826994 | 5.355859 | 0.0017733 | 0.9669217 | 0.999176 |
| ENSMUSG00000002489 | Tiam1 | protein_coding | -0.01876983 | 5.244388 | 0.0017722 | 0.9669325 | 0.999176 |
| ENSMUSG00000050299 | Gm9843 | protein_coding | 0.02019936 | 4.564892 | 0.0017678 | 0.9669736 | 0.999176 |
| ENSMUSG00000035045 | Zc3h12b | protein_coding | 0.02117033 | 4.206359 | 0.0017672 | 0.9669791 | 0.999176 |
| ENSMUSG00000022037 | Clu | protein_coding | 0.01544013 | 9.993372 | 0.0017486 | 0.967153 | 0.999176 |
| ENSMUSG00000030469 | Zfp719 | protein_coding | 0.02718494 | 3.742662 | 0.0017398 | 0.9672352 | 0.999176 |
| ENSMUSG00000029471 | Camkk2 | protein_coding | -0.02180629 | 4.662696 | 0.0017339 | 0.9672912 | 0.999176 |
| ENSMUSG00000064105 | Cnnm2 | protein_coding | 0.03366381 | 3.696486 | 0.0017333 | 0.9672973 | 0.999176 |
| ENSMUSG00000019577 | Pdk4 | protein_coding | -0.04698736 | 2.415017 | 0.0017221 | 0.9674026 | 0.999176 |
| ENSMUSG00000061911 | Myt1l | protein_coding | -0.01193582 | 7.934149 | 0.0017157 | 0.9674634 | 0.999176 |
| ENSMUSG00000012117 | Dhdds | protein_coding | 0.01537263 | 6.15706 | 0.0017029 | 0.9675847 | 0.999176 |
| ENSMUSG00000045871 | Slitrk6 | protein_coding | -0.0522534 | 2.800353 | 0.0016993 | 0.9676196 | 0.999176 |
| ENSMUSG00000111336 | AC160338.2 | processed_pseudoge | -0.03993165 | 2.495704 | 0.0016959 | 0.9676512 | 0.999176 |
| ENSMUSG00000005338 | Cadm3 | protein_coding | 0.01759144 | 5.578297 | 0.0016905 | 0.967703 | 0.999176 |
| ENSMUSG00000028108 | Ecm1 | protein_coding | 0.04373501 | 2.62974 | 0.0016824 | 0.9677799 | 0.999176 |
| ENSMUSG00000038866 | Zcchc2 | protein_coding | 0.03171056 | 2.898669 | 0.0016624 | 0.9679724 | 0.999176 |
| ENSMUSG00000037643 | Prkci | protein_coding | -0.02821386 | 4.427018 | 0.0016563 | 0.9680318 | 0.999176 |
| ENSMUSG00000030271 | Ogg1 | protein_coding | -0.0225546 | 3.966878 | 0.0016535 | 0.9680579 | 0.999176 |
| ENSMUSG00000027367 | Stard7 | protein_coding | -0.01240395 | 6.499195 | 0.0016417 | 0.9681726 | 0.999176 |
| ENSMUSG00000050473 | Slc35d3 | protein_coding | -0.03633461 | 3.052098 | 0.0016005 | 0.9685737 | 0.999176 |
| ENSMUSG00000024283 | Wac | protein_coding | 0.01514357 | 5.655655 | 0.0015976 | 0.9686018 | 0.999176 |
| ENSMUSG00000085389 | 1700003M07Rik | lincRNA | -0.02820647 | 3.252358 | 0.0015908 | 0.9686691 | 0.999176 |
| ENSMUSG00000100417 | Gm28578 | lincRNA | -0.04871843 | 2.375551 | 0.0015732 | 0.9688429 | 0.999176 |
| ENSMUSG00000054604 | Cggbp1 | protein_coding | -0.01453114 | 5.712912 | 0.0015337 | 0.9692358 | 0.999176 |
| ENSMUSG00000031379 | Pir | protein_coding | 0.02969845 | 3.804556 | 0.0015121 | 0.9694538 | 0.999176 |
| ENSMUSG00000025202 | Scd3 | protein_coding | 0.02265316 | 4.335583 | 0.0014941 | 0.9696359 | 0.999176 |
| ENSMUSG00000031928 | Mre11a | protein_coding | 0.02878816 | 3.140149 | 0.001467 | 0.9699127 | 0.999176 |
| ENSMUSG00000035585 | Tsen34 | protein_coding | 0.02248443 | 4.467189 | 0.0014631 | 0.9699519 | 0.999176 |
| ENSMUSG00000051518 | Rps19bp1 | protein_coding | -0.02219977 | 4.065044 | 0.0014626 | 0.9699575 | 0.999176 |
| ENSMUSG00000026509 | Capn2 | protein_coding | -0.01952313 | 4.854153 | 0.0014586 | 0.9699985 | 0.999176 |
| ENSMUSG00000039194 | Rlbp1 | protein_coding | 0.029019 | 4.133638 | 0.001455 | 0.9700366 | 0.999176 |
| ENSMUSG00000047613 | A430005L14Rik | protein_coding | -0.01053503 | 7.307186 | 0.0014462 | 0.9701258 | 0.999176 |
| ENSMUSG00000068735 | Trp53i11 | protein_coding | -0.01604144 | 5.63439 | 0.0014458 | 0.97013 | 0.999176 |
| ENSMUSG00000019432 | Ddx39b | protein_coding | -0.01227134 | 7.952589 | 0.0014359 | 0.9702325 | 0.999176 |
| ENSMUSG00000032680 | 6820408C15Rik | protein_coding | 0.02640909 | 4.21574 | 0.0014339 | 0.9702545 | 0.999176 |
| ENSMUSG00000029199 | Lias | protein_coding | -0.01223851 | 6.878862 | 0.0014276 | 0.9703193 | 0.999176 |
| ENSMUSG00000000881 | Dlg3 | protein_coding | -0.01618371 | 5.349494 | 0.0014214 | 0.9703836 | 0.999176 |
| ENSMUSG00000003039 | Fam32a | protein_coding | 0.01049648 | 8.117999 | 0.0014078 | 0.970525 | 0.999176 |
| ENSMUSG00000004268 | Emg1 | protein_coding | -0.01078963 | 7.510536 | 0.0014043 | 0.9705616 | 0.999176 |
| ENSMUSG00000036636 | Clcn7 | protein_coding | -0.02130252 | 4.562534 | 0.0014009 | 0.9705978 | 0.999176 |
| ENSMUSG00000059423 | Zfp933 | protein_coding | 0.03295463 | 3.282108 | 0.0013882 | 0.970731 | 0.999176 |
| ENSMUSG00000031360 | Ctps2 | protein_coding | 0.02016135 | 5.768637 | 0.0013859 | 0.9707578 | 0.999176 |
| ENSMUSG00000032261 | Sh3bgrl2 | protein_coding | 0.02321012 | 5.20158 | 0.0013783 | 0.9708387 | 0.999176 |
| ENSMUSG00000040859 | Bsdc1 | protein_coding | 0.01287412 | 5.986133 | 0.0013694 | 0.9709299 | 0.999176 |
| ENSMUSG00000058056 | Palld | protein_coding | -0.03066523 | 3.232378 | 0.0013682 | 0.9709431 | 0.999176 |
| ENSMUSG00000002997 | Prkar2b | protein_coding | -0.01490251 | 6.045391 | 0.001365 | 0.9709767 | 0.999176 |
| ENSMUSG00000036672 | Cenpt | protein_coding | 0.01725831 | 5.315053 | 0.0013632 | 0.9709958 | 0.999176 |
| ENSMUSG00000074825 | Itpripl1 | protein_coding | 0.03026501 | 3.798788 | 0.0013599 | 0.9710312 | 0.999176 |
| ENSMUSG00000026455 | Klhl12 | protein_coding | -0.02349589 | 4.628528 | 0.0013504 | 0.9711327 | 0.999176 |
| ENSMUSG00000018008 | Cyth4 | protein_coding | -0.02281408 | 3.659424 | 0.0013489 | 0.9711486 | 0.999176 |
| ENSMUSG00000020723 | Cacng4 | protein_coding | -0.01753587 | 5.752117 | 0.0013475 | 0.971163 | 0.999176 |
| ENSMUSG00000029387 | Gtf2h3 | protein_coding | 0.01614021 | 5.419839 | 0.0013407 | 0.9712356 | 0.999176 |
| ENSMUSG00000026956 | Uap1l1 | protein_coding | -0.02790044 | 3.614291 | 0.0013406 | 0.9712369 | 0.999176 |
| ENSMUSG00000020640 | Itsn2 | protein_coding | 0.02158163 | 3.894198 | 0.0013296 | 0.9713557 | 0.999176 |
| ENSMUSG00000026853 | Crat | protein_coding | 0.019136 | 4.648622 | 0.0013292 | 0.9713593 | 0.999176 |
| ENSMUSG00000100691 | 2010320M18Rik | lincRNA | -0.01322148 | 5.567277 | 0.0013261 | 0.9713935 | 0.999176 |
| ENSMUSG00000040520 | Manea | protein_coding | 0.02094769 | 4.031957 | 0.0012961 | 0.9717178 | 0.999176 |
| ENSMUSG00000004567 | Mcoln1 | protein_coding | 0.01964732 | 4.725138 | 0.0012961 | 0.9717179 | 0.999176 |
| ENSMUSG00000024335 | Brd2 | protein_coding | 0.01145874 | 6.404588 | 0.0012945 | 0.9717358 | 0.999176 |
| ENSMUSG00000059923 | Grb2 | protein_coding | -0.01006275 | 6.953409 | 0.001292 | 0.9717627 | 0.999176 |
| ENSMUSG00000112343 | AC154734.1 | processed_transcript | 0.04771158 | 2.490998 | 0.0012877 | 0.97181 | 0.999176 |
| ENSMUSG00000020612 | Prkar1a | protein_coding | 0.01820475 | 9.700402 | 0.0012878 | 0.9718362 | 0.999176 |
| ENSMUSG00000030750 | Nsmce1 | protein_coding | -0.01207129 | 6.353034 | 0.0012836 | 0.9718547 | 0.999176 |
| ENSMUSG00000040648 | Ppip5k2 | protein_coding | 0.01785722 | 4.174445 | 0.0012797 | 0.9718981 | 0.999176 |
| ENSMUSG00000056260 | Lrif1 | protein_coding | -0.01538051 | 5.034576 | 0.0012715 | 0.9719873 | 0.999176 |
| ENSMUSG00000031730 | Dhodh | protein_coding | 0.01329517 | 6.121564 | 0.001271 | 0.9719931 | 0.999176 |
| ENSMUSG00000001305 | Rrp15 | protein_coding | 0.01924249 | 4.420876 | 0.0012669 | 0.9720387 | 0.999176 |
| ENSMUSG00000032537 | Ephb1 | protein_coding | 0.03870602 | 2.574719 | 0.0012358 | 0.972384 | 0.999176 |
| ENSMUSG00000027508 | Pag1 | protein_coding | 0.02919531 | 3.019349 | 0.0012351 | 0.9723912 | 0.999176 |
| ENSMUSG00000038070 | Cntln | protein_coding | -0.02479252 | 3.697883 | 0.0012267 | 0.9724857 | 0.999176 |
| ENSMUSG00000079017 | Ifi27l2a | protein_coding | -0.03932183 | 2.489333 | 0.0012115 | 0.9726563 | 0.999176 |
| ENSMUSG00000041645 | Ddx24 | protein_coding | 0.01113509 | 8.638056 | 0.0012104 | 0.972669 | 0.999176 |
| ENSMUSG00000027569 | Mrgbp | protein_coding | -0.02891984 | 3.122265 | 0.0012072 | 0.9727051 | 0.999176 |

| ensembl_gene_id | external_gene_name | gene_biotype | log2 (fold change) | log2 (counts per million) | F-value | p-value | False discovery rate |
|---|---|---|---|---|---|---|---|
| ENSMUSG00000031774 | Fam192a | protein_coding | 0.01536995 | 6.846179 | 0.0011956 | 0.9728384 | 0.999176 |
| ENSMUSG00000061666 | Gdpd1 | protein_coding | 0.01032727 | 6.787748 | 0.0011838 | 0.972971 | 0.999176 |
| ENSMUSG00000038324 | Trpc4ap | protein_coding | -0.01078048 | 6.560717 | 0.0011816 | 0.9729959 | 0.999176 |
| ENSMUSG00000014547 | Wdfy2 | protein_coding | -0.02827239 | 3.165944 | 0.0011597 | 0.9732466 | 0.999176 |
| ENSMUSG00000020607 | Fam84a | protein_coding | -0.0107267 | 7.303704 | 0.001138 | 0.9734983 | 0.999176 |
| ENSMUSG00000000088 | Cox5a | protein_coding | -0.01552841 | 4.608528 | 0.0011236 | 0.973667 | 0.999176 |
| ENSMUSG00000061887 | Ssbp3 | protein_coding | 0.01316347 | 8.011994 | 0.0011177 | 0.9737368 | 0.999176 |
| ENSMUSG00000022829 | Stxbp5l | protein_coding | -0.02093253 | 3.902236 | 0.0011133 | 0.9737882 | 0.999176 |
| ENSMUSG00000022812 | Gsk3b | protein_coding | 0.01894835 | 9.531559 | 0.0011128 | 0.9738497 | 0.999176 |
| ENSMUSG00000003062 | Stard3nl | protein_coding | 0.009858541 | 7.038506 | 0.0010887 | 0.9740784 | 0.999176 |
| ENSMUSG00000064337 | mt-Rnr1 | Mt_rRNA | 0.02349745 | 10.86487 | 0.0010783 | 0.974314 | 0.999176 |
| ENSMUSG00000035439 | Haus8 | protein_coding | 0.01411056 | 5.555559 | 0.0010534 | 0.9745016 | 0.999176 |
| ENSMUSG00000057858 | Fam204a | protein_coding | 0.01423084 | 5.679382 | 0.0010426 | 0.9746332 | 0.999176 |
| ENSMUSG00000097213 | Gm26745 | lincRNA | -0.02924467 | 2.703867 | 0.0010333 | 0.9747464 | 0.999176 |
| ENSMUSG00000017754 | Pltp | protein_coding | -0.01840884 | 4.931605 | 0.0010284 | 0.9748062 | 0.999176 |
| ENSMUSG00000028607 | Cpt2 | protein_coding | -0.02074651 | 3.913189 | 0.0010196 | 0.974915 | 0.999176 |
| ENSMUSG00000071266 | Zfp946 | protein_coding | 0.02990073 | 2.83478 | 0.0010176 | 0.9749394 | 0.999176 |
| ENSMUSG00000050840 | Cdh20 | protein_coding | 0.03713 | 2.865321 | 0.001016 | 0.9749588 | 0.999176 |
| ENSMUSG00000046873 | Mbtps2 | protein_coding | -0.01988031 | 3.778127 | 0.0009984 | 0.9751766 | 0.999176 |
| ENSMUSG00000036611 | Eepd1 | protein_coding | 0.02188917 | 3.164771 | 0.0009926 | 0.9752483 | 0.999176 |
| ENSMUSG00000015165 | Hnrnpl | protein_coding | -0.01071554 | 6.560718 | 0.0009865 | 0.9753254 | 0.999176 |
| ENSMUSG00000033417 | Cacul1 | protein_coding | -0.01452606 | 5.058587 | 0.0009848 | 0.9753458 | 0.999176 |
| ENSMUSG00000001687 | Ubl3 | protein_coding | 0.00922123 | 8.338305 | 0.0009807 | 0.9753971 | 0.999176 |
| ENSMUSG00000039844 | Rapgef1 | protein_coding | 0.01667664 | 4.591279 | 0.0009779 | 0.9754322 | 0.999176 |
| ENSMUSG00000079020 | Slc45a4 | protein_coding | -0.02243351 | 3.704989 | 0.0009771 | 0.9754421 | 0.999176 |
| ENSMUSG00000021115 | Vrk1 | protein_coding | -0.01270416 | 5.201208 | 0.0009721 | 0.9755061 | 0.999176 |
| ENSMUSG00000020175 | Rab36 | protein_coding | -0.01256086 | 6.417358 | 0.0009674 | 0.975565 | 0.999176 |
| ENSMUSG00000014402 | Tsg101 | protein_coding | -0.009186034 | 6.732789 | 0.0009655 | 0.9755887 | 0.999176 |
| ENSMUSG00000028654 | Mycl | protein_coding | -0.02820987 | 2.975843 | 0.000956 | 0.9757097 | 0.999176 |
| ENSMUSG00000031652 | N4bp1 | protein_coding | 0.01237835 | 5.833231 | 0.0009401 | 0.9759119 | 0.999176 |
| ENSMUSG00000047146 | Tet1 | protein_coding | -0.01928301 | 3.973923 | 0.0009385 | 0.9759323 | 0.999176 |
| ENSMUSG00000020183 | Cpm | protein_coding | 0.02103558 | 3.680921 | 0.0009286 | 0.9760591 | 0.999176 |
| ENSMUSG00000050310 | Rictor | protein_coding | 0.01513736 | 4.044007 | 0.0009199 | 0.9761725 | 0.999176 |
| ENSMUSG00000027808 | Serp1 | protein_coding | -0.01660572 | 4.652541 | 0.0009177 | 0.9762005 | 0.999176 |
| ENSMUSG00000085156 | Snhg15 | processed_transcript | 0.02236447 | 3.023525 | 0.0008809 | 0.9766828 | 0.999176 |
| ENSMUSG00000059796 | Eif4a1 | protein_coding | -0.009479787 | 6.153011 | 0.0008804 | 0.9766894 | 0.999176 |
| ENSMUSG00000061751 | Kalrn | protein_coding | -0.01209945 | 5.52477 | 0.0008767 | 0.9767375 | 0.999176 |
| ENSMUSG00000027573 | Gid8 | protein_coding | 0.01285599 | 5.507733 | 0.0008751 | 0.9767594 | 0.999176 |
| ENSMUSG00000040710 | St8sia4 | protein_coding | -0.03249928 | 2.98006 | 0.0008714 | 0.9768087 | 0.999176 |
| ENSMUSG00000046793 | Gpr61 | protein_coding | 0.01665885 | 4.355468 | 0.0008712 | 0.9768112 | 0.999176 |
| ENSMUSG00000044352 | Sowaha | protein_coding | -0.01916601 | 5.361426 | 0.0008704 | 0.9768429 | 0.999176 |
| ENSMUSG00000057177 | Gsk3a | protein_coding | 0.01275905 | 4.855786 | 0.0008686 | 0.9768461 | 0.999176 |
| ENSMUSG00000021460 | Auh | protein_coding | 0.01333093 | 5.498864 | 0.0008632 | 0.9769181 | 0.999176 |
| ENSMUSG00000057604 | Lmcd1 | protein_coding | -0.02394059 | 3.76095 | 0.0008614 | 0.9769421 | 0.999176 |
| ENSMUSG00000023345 | Poc1a | protein_coding | -0.01844002 | 4.073827 | 0.0008528 | 0.9770572 | 0.999176 |
| ENSMUSG00000044748 | Defb1 | protein_coding | -0.04729684 | 3.007935 | 0.0008518 | 0.9771276 | 0.999176 |
| ENSMUSG00000070814 | 6330408A02Rik | protein_coding | 0.02509356 | 3.603769 | 0.0008366 | 0.9772768 | 0.999176 |
| ENSMUSG00000008892 | Vdac3 | protein_coding | 0.008085308 | 7.591641 | 0.0008337 | 0.9773158 | 0.999176 |
| ENSMUSG00000038542 | Pcid2 | protein_coding | -0.02354342 | 3.686813 | 0.0008335 | 0.9773182 | 0.999176 |
| ENSMUSG00000050240 | Hic2 | protein_coding | 0.03494133 | 3.345082 | 0.0008222 | 0.9775071 | 0.999176 |
| ENSMUSG00000029311 | Hsd17b11 | protein_coding | -0.02501352 | 3.207451 | 0.0008116 | 0.9775293 | 0.999176 |
| ENSMUSG00000056598 | Drc3 | protein_coding | 0.01837593 | 3.995195 | 0.0008069 | 0.9776827 | 0.999176 |
| ENSMUSG00000001229 | Dpp9 | protein_coding | 0.01461068 | 5.095201 | 0.0008063 | 0.9776912 | 0.999176 |
| ENSMUSG00000020537 | Drg2 | protein_coding | 0.01114241 | 7.094896 | 0.0007961 | 0.977834 | 0.999176 |
| ENSMUSG00000020610 | Amz2 | protein_coding | -0.009989341 | 6.095369 | 0.0007856 | 0.9779793 | 0.999176 |
| ENSMUSG00000098659 | 1110015O18Rik | lincRNA | 0.02778436 | 2.736696 | 0.0007801 | 0.9780562 | 0.999176 |
| ENSMUSG00000026241 | Nppc | protein_coding | -0.02198068 | 3.26284 | 0.0007733 | 0.9781532 | 0.999176 |
| ENSMUSG00000037791 | Phf12 | protein_coding | 0.01681748 | 4.091899 | 0.0007709 | 0.9781859 | 0.999176 |
| ENSMUSG00000004366 | Sst | protein_coding | -0.0120571 | 8.917731 | 0.0007678 | 0.978233 | 0.999176 |
| ENSMUSG00000026311 | Asb1 | protein_coding | -0.01494454 | 4.529768 | 0.0007658 | 0.9782593 | 0.999176 |
| ENSMUSG00000014769 | Psmb1 | protein_coding | 0.007917241 | 10.0284 | 0.0007554 | 0.9784069 | 0.999176 |
| ENSMUSG00000037664 | Cdkn1c | protein_coding | -0.03570712 | 2.32899 | 0.0007523 | 0.9784519 | 0.999176 |
| ENSMUSG00000032940 | Rbm11 | protein_coding | -0.01508641 | 4.260309 | 0.000748 | 0.9785135 | 0.999176 |
| ENSMUSG00000014748 | Tex261 | protein_coding | 0.01292945 | 5.957895 | 0.0007461 | 0.9785408 | 0.999176 |
| ENSMUSG00000027427 | Polr3f | protein_coding | -0.01474476 | 4.529544 | 0.0007426 | 0.9785904 | 0.999176 |
| ENSMUSG00000020427 | Igfbp3 | protein_coding | -0.02643883 | 2.704675 | 0.0007415 | 0.9786064 | 0.999176 |
| ENSMUSG00000030060 | Hmces | protein_coding | -0.0133925 | 5.224205 | 0.0007297 | 0.9787771 | 0.999176 |
| ENSMUSG00000047098 | Rnf31 | protein_coding | -0.01795042 | 4.120448 | 0.0007297 | 0.9787776 | 0.999176 |
| ENSMUSG00000059995 | Atxn7l3 | protein_coding | 0.01166814 | 5.429082 | 0.0007287 | 0.9787915 | 0.999176 |
| ENSMUSG00000030515 | Tarsl2 | protein_coding | 0.02203384 | 2.914048 | 0.0007277 | 0.9788069 | 0.999176 |
| ENSMUSG00000042606 | Hirip3 | protein_coding | -0.01489778 | 4.380832 | 0.0007226 | 0.9788813 | 0.999176 |
| ENSMUSG00000027692 | Tnik | protein_coding | -0.01498806 | 5.675198 | 0.0007211 | 0.978905 | 0.999176 |
| ENSMUSG00000031174 | Rpgr | protein_coding | -0.01267279 | 4.631533 | 0.0006994 | 0.9792223 | 0.999176 |
| ENSMUSG00000038135 | Crygn | protein_coding | -0.02351815 | 3.045605 | 0.0006963 | 0.9792682 | 0.999176 |
| ENSMUSG00000029426 | Scarb2 | protein_coding | 0.01240971 | 5.566328 | 0.0006853 | 0.9794327 | 0.999176 |
| ENSMUSG00000029207 | Apbb2 | protein_coding | 0.00873405 | 6.354913 | 0.0006812 | 0.9794937 | 0.999176 |
| ENSMUSG00000029516 | Cit | protein_coding | -0.01562816 | 8.525516 | 0.0006718 | 0.9796866 | 0.999176 |
| ENSMUSG00000018171 | Vmp1 | protein_coding | 0.007741984 | 7.624051 | 0.0006526 | 0.9799298 | 0.999176 |

| ensembl_gene_id | external_gene_name | gene_biotype | log2 (fold change) | log2 (counts per million) | F-value | p-value | False discovery rate |
|---|---|---|---|---|---|---|---|
| ENSMUSG00000108030 | 9530062K07Rik | bidirectional_promote | -0.03121922 | 3.071295 | 0.0006508 | 0.9799739 | 0.999176 |
| ENSMUSG00000039233 | Tbce | protein_coding | 0.01344307 | 5.077394 | 0.0006389 | 0.9801409 | 0.999176 |
| ENSMUSG00000029401 | Rilpl2 | protein_coding | 0.01529846 | 4.513482 | 0.0006388 | 0.9801435 | 0.999176 |
| ENSMUSG00000024350 | Dnajc18 | protein_coding | 0.009677349 | 7.567937 | 0.0006366 | 0.9801774 | 0.999176 |
| ENSMUSG00000045193 | Cirbp | protein_coding | -0.007594611 | 9.040231 | 0.0006327 | 0.9802372 | 0.999176 |
| ENSMUSG00000055041 | Commd5 | protein_coding | 0.00898142 | 6.244817 | 0.0006286 | 0.9803023 | 0.999176 |
| ENSMUSG00000023966 | Rsph9 | protein_coding | 0.007592782 | 6.664529 | 0.0006145 | 0.9805241 | 0.999176 |
| ENSMUSG00000026750 | Psmb7 | protein_coding | 0.006895874 | 8.167253 | 0.0006108 | 0.9805819 | 0.999176 |
| ENSMUSG00000030774 | Pak1 | protein_coding | -0.008694163 | 8.523615 | 0.0006081 | 0.9806259 | 0.999176 |
| ENSMUSG00000034152 | Exoc3 | protein_coding | 0.007963429 | 7.219471 | 0.0006065 | 0.980651 | 0.999176 |
| ENSMUSG00000069895 | Atxn1l | protein_coding | -0.02158682 | 3.812985 | 0.0006048 | 0.9806876 | 0.999176 |
| ENSMUSG00000039126 | Prune2 | protein_coding | -0.01732273 | 8.868721 | 0.0006089 | 0.9807107 | 0.999176 |
| ENSMUSG00000039703 | Nploc4 | protein_coding | -0.01083423 | 4.976777 | 0.0005864 | 0.9809742 | 0.999176 |
| ENSMUSG00000015653 | Steap2 | protein_coding | 0.01513103 | 3.443828 | 0.0005807 | 0.9810672 | 0.999176 |
| ENSMUSG00000060477 | Irak2 | protein_coding | 0.01598096 | 3.333829 | 0.0005623 | 0.9813693 | 0.999176 |
| ENSMUSG00000002015 | Bcap31 | protein_coding | -0.007275396 | 7.856845 | 0.0005583 | 0.9814358 | 0.999176 |
| ENSMUSG00000025040 | Fundc1 | protein_coding | 0.007639787 | 7.658821 | 0.0005539 | 0.9815084 | 0.999176 |
| ENSMUSG00000020646 | Mboat2 | protein_coding | 0.01143477 | 4.425745 | 0.0005518 | 0.9815441 | 0.999176 |
| ENSMUSG00000054836 | Elp6 | protein_coding | -0.009012825 | 5.750769 | 0.0005471 | 0.9816223 | 0.999176 |
| ENSMUSG00000028437 | Ubap1 | protein_coding | 0.009888074 | 5.993347 | 0.0005444 | 0.9816675 | 0.999176 |
| ENSMUSG00000044337 | Ackr3 | protein_coding | -0.03024664 | 2.35736 | 0.0005408 | 0.9817294 | 0.999176 |
| ENSMUSG00000003279 | Dlgap1 | protein_coding | 0.007907469 | 6.176424 | 0.0005254 | 0.9819916 | 0.999176 |
| ENSMUSG00000036598 | Ccdc113 | protein_coding | 0.01133782 | 4.425322 | 0.0005252 | 0.9819945 | 0.999176 |
| ENSMUSG00000074649 | BC029722 | protein_coding | -0.01453497 | 4.08322 | 0.000524 | 0.9820148 | 0.999176 |
| ENSMUSG00000033392 | Clasp2 | protein_coding | -0.01005332 | 7.111265 | 0.0005223 | 0.9820467 | 0.999176 |
| ENSMUSG00000039428 | Tmem135 | protein_coding | 0.01004309 | 5.142708 | 0.0005097 | 0.9822618 | 0.999176 |
| ENSMUSG00000019718 | L3hypdh | protein_coding | 0.01612737 | 3.288056 | 0.000503 | 0.9823792 | 0.999176 |
| ENSMUSG00000022234 | Cct5 | protein_coding | -0.007570738 | 9.14548 | 0.0004987 | 0.9824542 | 0.999176 |
| ENSMUSG00000017715 | Pgs1 | protein_coding | -0.01155548 | 4.039492 | 0.0004958 | 0.9825056 | 0.999176 |
| ENSMUSG00000037260 | Hgsnat | protein_coding | 0.01153604 | 4.476159 | 0.0004602 | 0.9831461 | 0.999176 |
| ENSMUSG00000019338 | Zfp687 | protein_coding | 0.01224878 | 3.918077 | 0.0004478 | 0.9833735 | 0.999176 |
| ENSMUSG00000037280 | Galnt6 | protein_coding | -0.01355188 | 3.473774 | 0.0004461 | 0.9834052 | 0.999176 |
| ENSMUSG00000035392 | Dennd1a | protein_coding | 0.01020494 | 4.580423 | 0.0004381 | 0.9835556 | 0.999176 |
| ENSMUSG00000001666 | Ddt | protein_coding | 0.008659267 | 5.151267 | 0.0004376 | 0.9835637 | 0.999176 |
| ENSMUSG00000022913 | Psmg1 | protein_coding | -0.009740049 | 4.951692 | 0.000436 | 0.9835942 | 0.999176 |
| ENSMUSG00000018459 | Slc13a3 | protein_coding | -0.02567545 | 3.046342 | 0.0004348 | 0.9836289 | 0.999176 |
| ENSMUSG00000079038 | D130040H23Rik | protein_coding | -0.02271946 | 2.546525 | 0.0004331 | 0.983648 | 0.999176 |
| ENSMUSG00000022186 | Oxct1 | protein_coding | 0.006119416 | 8.154561 | 0.0004326 | 0.9836589 | 0.999176 |
| ENSMUSG00000079434 | Neu2 | protein_coding | -0.01802171 | 3.211015 | 0.0004258 | 0.9837874 | 0.999176 |
| ENSMUSG00000024131 | Slc3a1 | protein_coding | -0.006608219 | 7.649142 | 0.0004239 | 0.9838243 | 0.999176 |
| ENSMUSG00000038116 | Phf20 | protein_coding | -0.00641986 | 6.675524 | 0.0004113 | 0.984066 | 0.999176 |
| ENSMUSG00000051768 | Xrcc1 | protein_coding | -0.01067089 | 4.455173 | 0.0004098 | 0.9840947 | 0.999176 |
| ENSMUSG00000025925 | Terf1 | protein_coding | -0.01289963 | 3.790795 | 0.0004053 | 0.9841831 | 0.999176 |
| ENSMUSG00000030120 | Mlf2 | protein_coding | -0.00533839 | 7.984126 | 0.0004005 | 0.9842766 | 0.999176 |
| ENSMUSG00000022375 | Lrrc6 | protein_coding | 0.01751315 | 3.852469 | 0.0003916 | 0.9844627 | 0.999176 |
| ENSMUSG00000034255 | Arhgap27 | protein_coding | -0.009015242 | 5.421061 | 0.0003857 | 0.9845693 | 0.999176 |
| ENSMUSG00000031386 | Hcfc1 | protein_coding | -0.01125199 | 5.197956 | 0.0003819 | 0.9846463 | 0.999176 |
| ENSMUSG00000025192 | Entpd7 | protein_coding | 0.01462956 | 3.783402 | 0.0003753 | 0.9847796 | 0.999176 |
| ENSMUSG00000022722 | Arl6 | protein_coding | -0.006570694 | 7.395387 | 0.0003691 | 0.9849046 | 0.999176 |
| ENSMUSG00000019471 | Cdc37 | protein_coding | 0.005221235 | 7.155076 | 0.0003647 | 0.9849962 | 0.999176 |
| ENSMUSG00000020988 | L2hgdh | protein_coding | 0.0117079 | 3.62081 | 0.0003641 | 0.9850072 | 0.999176 |
| ENSMUSG00000001911 | Nfix | protein_coding | 0.008615472 | 5.602618 | 0.0003625 | 0.9850403 | 0.999176 |
| ENSMUSG00000046567 | 4930430F08Rik | protein_coding | -0.009632948 | 4.367057 | 0.0003623 | 0.9850438 | 0.999176 |
| ENSMUSG00000025872 | Thoc3 | protein_coding | 0.008915674 | 5.108572 | 0.0003584 | 0.9851263 | 0.999176 |
| ENSMUSG00000021217 | Tshz3 | protein_coding | -0.01436356 | 2.952394 | 0.0003506 | 0.9852879 | 0.999176 |
| ENSMUSG00000025237 | Parp6 | protein_coding | 0.006639265 | 5.991047 | 0.0003489 | 0.9853236 | 0.999176 |
| ENSMUSG00000031751 | Amfr | protein_coding | 0.008304074 | 6.103462 | 0.0003445 | 0.9854165 | 0.999176 |
| ENSMUSG00000028703 | Lrrc41 | protein_coding | 0.01276803 | 3.908572 | 0.0003435 | 0.9854379 | 0.999176 |
| ENSMUSG00000033373 | Fntb | protein_coding | -0.007680678 | 5.852447 | 0.0003278 | 0.9857754 | 0.999176 |
| ENSMUSG00000040596 | Pogk | protein_coding | -0.009167538 | 4.689285 | 0.0003245 | 0.9858473 | 0.999176 |
| ENSMUSG00000020358 | Hnrnpab | protein_coding | 0.01716879 | 2.762757 | 0.0003199 | 0.9859477 | 0.999176 |
| ENSMUSG00000035206 | Sppl2b | protein_coding | -0.01111998 | 3.960929 | 0.0003179 | 0.9859901 | 0.999176 |
| ENSMUSG00000023452 | Pisd | protein_coding | -0.006470002 | 5.562064 | 0.0003139 | 0.9860795 | 0.999176 |
| ENSMUSG00000014503 | Pkd2l2 | protein_coding | -0.00657965 | 5.54669 | 0.0003112 | 0.9861392 | 0.999176 |
| ENSMUSG00000029056 | Pank4 | protein_coding | 0.008230863 | 4.982484 | 0.0003016 | 0.9863538 | 0.999176 |
| ENSMUSG00000022141 | Nipbl | protein_coding | 0.01255908 | 4.41639 | 0.000301 | 0.9863687 | 0.999176 |
| ENSMUSG00000029447 | Cct6a | processed_transcript | 0.005227063 | 8.111928 | 0.0002971 | 0.9864573 | 0.999176 |
| ENSMUSG00000073755 | 5730409E04Rik | protein_coding | -0.006889686 | 5.339607 | 0.0002945 | 0.9865168 | 0.999176 |
| ENSMUSG00000017781 | Pitpna | protein_coding | 0.004993971 | 7.228052 | 0.0002942 | 0.9865236 | 0.999176 |
| ENSMUSG00000027820 | Mme | protein_coding | 0.02098764 | 2.654254 | 0.0002919 | 0.9865771 | 0.999176 |
| ENSMUSG00000030188 | Magohb | protein_coding | 0.008181766 | 5.418425 | 0.0002887 | 0.9866508 | 0.999176 |
| ENSMUSG00000011158 | Brf1 | protein_coding | -0.01048392 | 3.943898 | 0.0002815 | 0.9868174 | 0.999176 |
| ENSMUSG00000031916 | Cog8 | protein_coding | -0.0190218 | 2.43214 | 0.0002783 | 0.9868923 | 0.999176 |
| ENSMUSG00000026930 | Gpsm1 | protein_coding | 0.006527326 | 6.004158 | 0.0002761 | 0.9869441 | 0.999176 |
| ENSMUSG00000030007 | Cct7 | protein_coding | -0.006175282 | 9.448791 | 0.0002717 | 0.9870496 | 0.999176 |
| ENSMUSG00000055371 | Stam2 | protein_coding | 0.008303688 | 4.296967 | 0.0002602 | 0.9873259 | 0.999176 |
| ENSMUSG00000048429 | Timm29 | protein_coding | -0.007647431 | 5.196734 | 0.0002588 | 0.9873611 | 0.999176 |
| ENSMUSG00000034748 | Sirt6 | protein_coding | 0.01002777 | 4.765305 | 0.0002553 | 0.9874456 | 0.999176 |

| ensembl_gene_id | external_gene_name | gene_biotype | log2 (fold change) | log2 (counts per million) | F-value | p-value | False discovery rate |
|---|---|---|---|---|---|---|---|
| ENSMUSG00000043587 | Pxylp1 | protein_coding | -0.01009996 | 3.931387 | 0.0002506 | 0.9875624 | 0.999176 |
| ENSMUSG00000022443 | Myh9 | protein_coding | 0.01034776 | 3.503121 | 0.000245 | 0.9877004 | 0.999176 |
| ENSMUSG00000090063 | Dlx6os1 | antisense_RNA | 0.01111699 | 4.559342 | 0.0002439 | 0.987731 | 0.999176 |
| ENSMUSG00000097416 | Gm26670 | antisense_RNA | -0.01560081 | 2.35226 | 0.0002405 | 0.9878146 | 0.999176 |
| ENSMUSG00000044813 | Shb | protein_coding | 0.01492025 | 3.022881 | 0.0002323 | 0.9880255 | 0.999176 |
| ENSMUSG00000031631 | Cfap97 | protein_coding | 0.006518111 | 5.248264 | 0.0002319 | 0.9880348 | 0.999176 |
| ENSMUSG00000028032 | Papss1 | protein_coding | 0.004555057 | 6.70691 | 0.0002285 | 0.9881239 | 0.999176 |
| ENSMUSG00000015850 | Adamtsl4 | protein_coding | 0.01856431 | 2.532289 | 0.0002226 | 0.9882763 | 0.999176 |
| ENSMUSG00000033021 | Gmppa | protein_coding | -0.007112146 | 5.686106 | 0.0002197 | 0.9883532 | 0.999176 |
| ENSMUSG00000028700 | Pomgnt1 | protein_coding | -0.007388206 | 5.107725 | 0.0002158 | 0.9884576 | 0.999176 |
| ENSMUSG00000027198 | Ext2 | protein_coding | 0.009853098 | 3.945682 | 0.0002158 | 0.9884578 | 0.999176 |
| ENSMUSG00000021597 | Slf1 | protein_coding | 0.008262134 | 4.400162 | 0.0002146 | 0.988491 | 0.999176 |
| ENSMUSG00000026946 | Nmi | protein_coding | -0.01066649 | 3.253389 | 0.0002131 | 0.9885305 | 0.999176 |
| ENSMUSG00000036097 | Slf2 | protein_coding | -0.006480395 | 5.11073 | 0.0002108 | 0.9885927 | 0.999176 |
| ENSMUSG00000026458 | Ppfia4 | protein_coding | -0.006548812 | 5.51051 | 0.0002059 | 0.9887265 | 0.999176 |
| ENSMUSG00000000976 | Heatr6 | protein_coding | -0.007187193 | 4.608351 | 0.0001957 | 0.989008 | 0.999176 |
| ENSMUSG00000030603 | Psmc4 | protein_coding | 0.004041695 | 8.493986 | 0.0001945 | 0.9890411 | 0.999176 |
| ENSMUSG00000022241 | Tars | protein_coding | -0.006546991 | 5.555736 | 0.0001876 | 0.9892372 | 0.999176 |
| ENSMUSG00000003518 | Dusp3 | protein_coding | -0.005927688 | 8.317146 | 0.000183 | 0.9893713 | 0.999176 |
| ENSMUSG00000041193 | Pla2g5 | protein_coding | 0.01384766 | 2.690347 | 0.0001822 | 0.989395 | 0.999176 |
| ENSMUSG00000023963 | Cyp39a1 | protein_coding | -0.02148669 | 2.887793 | 0.0001771 | 0.9895653 | 0.999176 |
| ENSMUSG00000031576 | Kcnu1 | protein_coding | 0.01627213 | 2.930506 | 0.0001706 | 0.989738 | 0.999176 |
| ENSMUSG00000060703 | Cd302 | protein_coding | -0.006941005 | 5.575287 | 0.0001655 | 0.9898935 | 0.999176 |
| ENSMUSG00000019803 | Nr2e1 | protein_coding | 0.01560707 | 2.888782 | 0.0001652 | 0.9899007 | 0.999176 |
| ENSMUSG00000016831 | Tox4 | protein_coding | -0.005273818 | 6.124848 | 0.0001649 | 0.9899097 | 0.999176 |
| ENSMUSG00000028452 | Vcp | protein_coding | 0.01227438 | 2.751293 | 0.0001592 | 0.9900856 | 0.999176 |
| ENSMUSG00000029304 | Spp1 | protein_coding | 0.009123312 | 4.246993 | 0.0001553 | 0.9902081 | 0.999176 |
| ENSMUSG00000021023 | 1110008L16Rik | protein_coding | -0.009236164 | 3.348714 | 0.0001537 | 0.9902604 | 0.999176 |
| ENSMUSG00000040537 | Adam22 | protein_coding | 0.004353863 | 6.220759 | 0.0001527 | 0.9902918 | 0.999176 |
| ENSMUSG00000021423 | Ly86 | protein_coding | -0.007306163 | 4.544424 | 0.0001524 | 0.9903002 | 0.999176 |
| ENSMUSG00000037940 | Inpp4b | protein_coding | 0.007884296 | 4.371399 | 0.0001518 | 0.9903176 | 0.999176 |
| ENSMUSG00000021134 | Srsf5 | protein_coding | -0.003860239 | 7.734244 | 0.0001515 | 0.9903282 | 0.999176 |
| ENSMUSG00000029674 | Limk1 | protein_coding | -0.01373437 | 2.789344 | 0.0001478 | 0.9904489 | 0.999176 |
| ENSMUSG00000038702 | Dsel | protein_coding | -0.01005893 | 3.507318 | 0.0001473 | 0.9904649 | 0.999176 |
| ENSMUSG00000027002 | Nckap1 | protein_coding | 0.003980413 | 7.916297 | 0.0001466 | 0.990488 | 0.999176 |
| ENSMUSG00000060227 | Casc4 | protein_coding | 0.005003859 | 5.34359 | 0.0001438 | 0.990579 | 0.999176 |
| ENSMUSG00000032594 | Ip6k1 | protein_coding | 0.003784349 | 7.034764 | 0.000142 | 0.9906379 | 0.999176 |
| ENSMUSG00000026918 | Brd3 | protein_coding | 0.003847372 | 6.624802 | 0.0001368 | 0.9908101 | 0.999176 |
| ENSMUSG00000052833 | Sae1 | protein_coding | 0.004630923 | 7.901328 | 0.0001311 | 0.9910044 | 0.999176 |
| ENSMUSG00000031533 | Mrps31 | protein_coding | -0.005238305 | 6.396049 | 0.0001309 | 0.9910116 | 0.999176 |
| ENSMUSG00000000759 | Tubgcp3 | protein_coding | -0.005680356 | 4.82796 | 0.0001292 | 0.9910688 | 0.999176 |
| ENSMUSG00000113771 | AC160108.1 | antisense_RNA | -0.009876228 | 3.105544 | 0.0001275 | 0.9911286 | 0.999176 |
| ENSMUSG00000075271 | Ttc30a1 | protein_coding | -0.01113803 | 2.943197 | 0.0001267 | 0.9911553 | 0.999176 |
| ENSMUSG00000037872 | Ackr1 | protein_coding | 0.00593139 | 4.243581 | 0.0001227 | 0.9912974 | 0.999176 |
| ENSMUSG00000037570 | Mcrs1 | protein_coding | 0.0049422 | 4.744328 | 0.0001212 | 0.9913515 | 0.999176 |
| ENSMUSG00000037935 | Smarce1 | protein_coding | -0.004749153 | 5.526289 | 0.0001144 | 0.9915948 | 0.999176 |
| ENSMUSG00000013236 | Ptprs | protein_coding | -0.003778761 | 7.138867 | 0.0001106 | 0.991735 | 0.999176 |
| ENSMUSG00000028608 | 0610037L13Rik | protein_coding | -0.003198004 | 6.864864 | 0.000104 | 0.9919887 | 0.999176 |
| ENSMUSG00000022295 | Atp6v1c1 | protein_coding | -0.003205368 | 9.301729 | 0.0001035 | 0.9920056 | 0.999176 |
| ENSMUSG00000028132 | Tmem56 | protein_coding | 0.006566445 | 4.35704 | 0.0001026 | 0.9920426 | 0.999176 |
| ENSMUSG00000073062 | Zxdb | protein_coding | -0.007855567 | 3.790263 | 0.0001025 | 0.9920454 | 0.999176 |
| ENSMUSG00000030922 | Lyrm1 | protein_coding | 0.003586364 | 5.692454 | 0.0001025 | 0.9920456 | 0.999176 |
| ENSMUSG00000034040 | Wbscr17 | protein_coding | -0.005563477 | 4.346021 | 0.0001016 | 0.9920809 | 0.999176 |
| ENSMUSG00000067860 | Zic3 | protein_coding | 0.005466649 | 3.91092 | 9.87E-05 | 0.9921926 | 0.999176 |
| ENSMUSG00000029672 | Fam3c | protein_coding | 0.004034892 | 5.096168 | 9.69E-05 | 0.9922656 | 0.999176 |
| ENSMUSG00000079467 | Gm14966 | antisense_RNA | -0.008490752 | 1.889569 | 9.57E-05 | 0.9923137 | 0.999176 |
| ENSMUSG00000019792 | Trmt11 | protein_coding | -0.006422943 | 4.368418 | 9.18E-05 | 0.9924721 | 0.999176 |
| ENSMUSG00000026914 | Psmd14 | protein_coding | -0.003031427 | 8.344814 | 8.95E-05 | 0.9925657 | 0.999176 |
| ENSMUSG00000034163 | Zfc3h1 | protein_coding | 0.009387836 | 2.682464 | 8.83E-05 | 0.9926146 | 0.999176 |
| ENSMUSG00000022951 | Rcan1 | protein_coding | 0.003527836 | 5.367255 | 8.79E-05 | 0.9926334 | 0.999176 |
| ENSMUSG00000087197 | Gm13780 | lincRNA | -0.006917981 | 3.154261 | 8.74E-05 | 0.992654 | 0.999176 |
| ENSMUSG00000020925 | Ccdc43 | protein_coding | 0.003569512 | 5.857816 | 8.61E-05 | 0.9927099 | 0.999176 |
| ENSMUSG00000028229 | Rmdn1 | protein_coding | -0.004777989 | 4.287708 | 8.18E-05 | 0.9928922 | 0.999176 |
| ENSMUSG00000022678 | Nde1 | protein_coding | 0.005023209 | 5.028135 | 7.92E-05 | 0.993007 | 0.999176 |
| ENSMUSG00000028284 | Map3k7 | protein_coding | -0.005191648 | 3.835598 | 7.71E-05 | 0.9930993 | 0.999176 |
| ENSMUSG00000036564 | Ndrg4 | protein_coding | -0.004218015 | 11.22155 | 7.65E-05 | 0.9931289 | 0.999176 |
| ENSMUSG00000035557 | Krt17 | protein_coding | -0.007898592 | 2.761114 | 7.55E-05 | 0.9931738 | 0.999176 |
| ENSMUSG00000031505 | Naxd | protein_coding | -0.002851739 | 6.902995 | 7.52E-05 | 0.9931846 | 0.999176 |
| ENSMUSG00000021810 | Ecd | protein_coding | -0.003779976 | 6.14806 | 7.47E-05 | 0.9932101 | 0.999176 |
| ENSMUSG00000086851 | A430108G06Rik | processed_transcript | 0.01528131 | 2.553515 | 7.30E-05 | 0.9932955 | 0.999176 |
| ENSMUSG00000030688 | Stard10 | protein_coding | 0.004224854 | 4.304016 | 7.05E-05 | 0.9934036 | 0.999176 |
| ENSMUSG00000000131 | Xpo6 | protein_coding | 0.005473867 | 5.28173 | 6.68E-05 | 0.9935826 | 0.999176 |
| ENSMUSG00000091811 | Inafm1 | protein_coding | 0.004784052 | 4.222061 | 6.44E-05 | 0.9936967 | 0.999176 |
| ENSMUSG00000003037 | Rab8a | protein_coding | 0.002386743 | 7.49502 | 6.10E-05 | 0.993863 | 0.999176 |
| ENSMUSG00000057176 | Ccdc189 | protein_coding | -0.003618511 | 4.692951 | 6.01E-05 | 0.9939076 | 0.999176 |
| ENSMUSG00000079018 | Ly6c1 | protein_coding | -0.009155825 | 2.6312 | 5.66E-05 | 0.9940885 | 0.999176 |
| ENSMUSG00000022679 | Mpv17l | protein_coding | -0.003157569 | 5.775121 | 5.39E-05 | 0.9942327 | 0.999176 |
| ENSMUSG00000042604 | Kcna4 | protein_coding | 0.005211755 | 3.988613 | 5.32E-05 | 0.9942685 | 0.999176 |

| ensembl_gene_id | external_gene_name | gene_biotype | log2 (fold change) | log2 (counts per million) | F-value | p-value | False discovery rate |
|---|---|---|---|---|---|---|---|
| ENSMUSG00000024502 | Jakmip2 | protein_coding | -0.002511836 | 6.19856 | 5.20E-05 | 0.9943353 | 0.999176 |
| ENSMUSG00000038074 | Fkbp14 | protein_coding | 0.00511095 | 3.38283 | 5.14E-05 | 0.9943679 | 0.999176 |
| ENSMUSG00000028953 | Abcf2 | protein_coding | -0.004117201 | 6.907046 | 5.14E-05 | 0.9943792 | 0.999176 |
| ENSMUSG00000024614 | Tmx3 | protein_coding | 0.002806888 | 5.685585 | 5.01E-05 | 0.9944402 | 0.999176 |
| ENSMUSG00000087260 | Lamtor5 | protein_coding | -0.002198433 | 7.529657 | 4.29E-05 | 0.9948513 | 0.999176 |
| ENSMUSG00000066057 | Gm1976 | lincRNA | -0.005481826 | 3.030686 | 4.23E-05 | 0.9948925 | 0.999176 |
| ENSMUSG00000063894 | Zkscan8 | protein_coding | -0.004621748 | 3.195151 | 3.93E-05 | 0.9950741 | 0.999176 |
| ENSMUSG00000032305 | Fam219b | protein_coding | -0.003042011 | 5.79132 | 3.83E-05 | 0.9951367 | 0.999176 |
| ENSMUSG00000044231 | Nhlrc1 | protein_coding | -0.00413707 | 3.692156 | 3.82E-05 | 0.9951407 | 0.999176 |
| ENSMUSG00000020520 | Galnt10 | protein_coding | 0.004304801 | 3.456536 | 3.81E-05 | 0.9951528 | 0.999176 |
| ENSMUSG00000057265 | Bbof1 | protein_coding | -0.006966259 | 2.443926 | 3.74E-05 | 0.9951972 | 0.999176 |
| ENSMUSG00000019731 | Slc35e1 | protein_coding | -0.004054672 | 4.019406 | 3.70E-05 | 0.9952233 | 0.999176 |
| ENSMUSG00000022783 | Spag6l | protein_coding | 0.003933602 | 3.789404 | 3.64E-05 | 0.9952582 | 0.999176 |
| ENSMUSG00000026887 | Mrrf | protein_coding | -0.002577657 | 5.462209 | 3.61E-05 | 0.9952777 | 0.999176 |
| ENSMUSG00000005683 | Cs | protein_coding | -0.002487168 | 8.087842 | 3.48E-05 | 0.995363 | 0.999176 |
| ENSMUSG00000031527 | Eri1 | protein_coding | 0.00656455 | 2.766883 | 3.48E-05 | 0.995366 | 0.999176 |
| ENSMUSG00000039367 | Sec24c | protein_coding | 0.008818892 | 2.482692 | 3.43E-05 | 0.9953968 | 0.999176 |
| ENSMUSG00000028982 | Slc25a33 | protein_coding | 0.003696791 | 4.497905 | 3.30E-05 | 0.9954841 | 0.999176 |
| ENSMUSG00000052848 | C130026L21Rik | lincRNA | 0.002916134 | 4.471492 | 3.21E-05 | 0.9955475 | 0.999176 |
| ENSMUSG00000030663 | 1110004F10Rik | protein_coding | 0.001636855 | 8.634743 | 3.21E-05 | 0.9955513 | 0.999176 |
| ENSMUSG00000001901 | Kcnh6 | protein_coding | -0.005326083 | 2.875577 | 3.16E-05 | 0.9955863 | 0.999176 |
| ENSMUSG00000003526 | Prodh | protein_coding | -0.004008042 | 4.696585 | 3.08E-05 | 0.9956436 | 0.999176 |
| ENSMUSG00000041096 | Tspyl2 | protein_coding | -0.001878549 | 6.912719 | 2.99E-05 | 0.9957049 | 0.999176 |
| ENSMUSG00000037894 | H2afz | protein_coding | -0.002757611 | 3.996994 | 2.97E-05 | 0.9957189 | 0.999176 |
| ENSMUSG00000028583 | Pdpn | protein_coding | -0.003595378 | 4.183131 | 2.94E-05 | 0.9957371 | 0.999176 |
| ENSMUSG00000062031 | Pgghg | protein_coding | 0.008494405 | 2.076155 | 2.88E-05 | 0.995781 | 0.999176 |
| ENSMUSG00000030411 | Nova2 | protein_coding | 0.002474574 | 5.176793 | 2.87E-05 | 0.9957876 | 0.999176 |
| ENSMUSG00000009030 | Pdcl | protein_coding | -0.002404972 | 5.005015 | 2.80E-05 | 0.9958448 | 0.999176 |
| ENSMUSG00000026083 | Eif5b | protein_coding | 0.002313648 | 6.243742 | 2.79E-05 | 0.9958508 | 0.999176 |
| ENSMUSG00000020580 | Rock2 | protein_coding | -0.001746088 | 6.259631 | 2.71E-05 | 0.995911 | 0.999176 |
| ENSMUSG00000057895 | Zfp105 | protein_coding | -0.004770397 | 2.794362 | 2.60E-05 | 0.9959937 | 0.999176 |
| ENSMUSG00000024581 | Napg | protein_coding | 0.001596632 | 7.799077 | 2.41E-05 | 0.996145 | 0.999176 |
| ENSMUSG00000021024 | Psma6 | protein_coding | 0.001517325 | 9.087785 | 2.40E-05 | 0.9961524 | 0.999176 |
| ENSMUSG00000017307 | Acot8 | protein_coding | -0.002414677 | 5.065815 | 2.28E-05 | 0.9962452 | 0.999176 |
| ENSMUSG00000078622 | Ccdc47 | protein_coding | 0.002159901 | 5.70067 | 2.13E-05 | 0.9963715 | 0.999176 |
| ENSMUSG00000003824 | Syce2 | protein_coding | 0.003981212 | 3.052079 | 2.08E-05 | 0.9964156 | 0.999176 |
| ENSMUSG00000033880 | Lgals3bp | protein_coding | -0.003457033 | 4.716345 | 1.99E-05 | 0.9965002 | 0.999176 |
| ENSMUSG00000006342 | Susd2 | protein_coding | 0.003530157 | 4.020237 | 1.94E-05 | 0.9965347 | 0.999176 |
| ENSMUSG00000097145 | 9230114K14Rik | processed_transcript | 0.004844123 | 2.639944 | 1.76E-05 | 0.996702 | 0.999176 |
| ENSMUSG00000063646 | Jakmip1 | protein_coding | 0.00136199 | 5.848734 | 1.65E-05 | 0.9968068 | 0.999176 |
| ENSMUSG00000016559 | H3f3b | protein_coding | 0.001237413 | 9.584167 | 1.60E-05 | 0.996857 | 0.999176 |
| ENSMUSG00000027163 | Commd9 | protein_coding | -0.001332618 | 7.20849 | 1.31E-05 | 0.9971545 | 0.999176 |
| ENSMUSG00000033972 | Zfp944 | protein_coding | -0.001983209 | 3.843087 | 1.19E-05 | 0.9972861 | 0.999176 |
| ENSMUSG00000005483 | Dnajb1 | protein_coding | -0.001138864 | 6.68527 | 1.11E-05 | 0.9973858 | 0.999176 |
| ENSMUSG00000052337 | Immt | protein_coding | -0.001287779 | 7.21927 | 1.10E-05 | 0.997399 | 0.999176 |
| ENSMUSG00000047022 | Mipol1 | protein_coding | 0.001825321 | 4.213269 | 9.42E-06 | 0.9975886 | 0.999176 |
| ENSMUSG00000027357 | Crls1 | protein_coding | -0.001314806 | 4.835742 | 9.40E-06 | 0.9975903 | 0.999176 |
| ENSMUSG00000043259 | Fam13c | protein_coding | 0.00113979 | 5.533975 | 8.93E-06 | 0.9976522 | 0.999176 |
| ENSMUSG00000024754 | Tmem2 | protein_coding | 0.002594463 | 3.101772 | 8.87E-06 | 0.9976594 | 0.999176 |
| ENSMUSG00000004530 | Coro1c | protein_coding | 0.001623153 | 6.021975 | 8.87E-06 | 0.997661 | 0.999176 |
| ENSMUSG00000023170 | Gps2 | protein_coding | -0.001285752 | 4.892226 | 7.73E-06 | 0.997816 | 0.999176 |
| ENSMUSG00000022052 | Ppp2r2a | protein_coding | -0.001126621 | 6.583689 | 7.64E-06 | 0.9978286 | 0.999176 |
| ENSMUSG00000097554 | Gm26825 | lincRNA | 0.002336705 | 2.7009 | 7.12E-06 | 0.9979036 | 0.999176 |
| ENSMUSG00000066442 | Mthfs | protein_coding | -0.001980444 | 2.85745 | 6.69E-06 | 0.997968 | 0.999176 |
| ENSMUSG00000052105 | Mtcl1 | protein_coding | 0.001931582 | 3.993372 | 5.98E-06 | 0.9980778 | 0.9991929 |
| ENSMUSG00000030327 | Necap1 | protein_coding | -0.000876498 | 9.043498 | 4.80E-06 | 0.998278 | 0.9993003 |
| ENSMUSG00000028710 | Atpaf1 | protein_coding | 0.001587352 | 3.757951 | 4.25E-06 | 0.9983792 | 0.9993086 |
| ENSMUSG00000026864 | Hspa5 | protein_coding | -0.000538537 | 8.346771 | 3.76E-06 | 0.9984773 | 0.9993137 |
| ENSMUSG00000029171 | Pgm1 | protein_coding | -0.001202607 | 2.853407 | 3.20E-06 | 0.9985938 | 0.9993373 |
| ENSMUSG00000026565 | Pou2f1 | protein_coding | 0.001319838 | 3.145915 | 2.32E-06 | 0.9988038 | 0.9994544 |
| ENSMUSG00000066149 | Cdc26 | protein_coding | -0.000529065 | 5.822246 | 1.75E-06 | 0.9989599 | 0.9995177 |
| ENSMUSG00000023977 | Ubr2 | protein_coding | -0.000697967 | 3.830373 | 9.95E-07 | 0.9992164 | 0.9996166 |
| ENSMUSG00000027999 | Pla2g12a | protein_coding | -0.000409406 | 4.434527 | 9.24E-07 | 0.9992448 | 0.9996166 |
| ENSMUSG00000063808 | Gpatch1 | protein_coding | -0.000338567 | 5.479377 | 6.26E-07 | 0.9993785 | 0.9996574 |
| ENSMUSG00000016477 | E2f3 | protein_coding | -0.000381122 | 3.467306 | 3.27E-07 | 0.9995509 | 0.9997006 |
| ENSMUSG00000047230 | Cldn2 | protein_coding | -0.1860568 | 1.917465 | 2.13E-07 | 0.9996387 | 0.9997006 |
| ENSMUSG00000022012 | Enox1 | protein_coding | -0.000154512 | 4.283157 | 1.45E-07 | 0.9997006 | 0.9997006 |
